# Genetic control of KRAB-ZFP genes explains distal CpG-site methylation which associates with human disease phenotypes

**DOI:** 10.1101/2022.01.18.476742

**Authors:** Andrew D. Bretherick, Yanni Zeng, Rosie M. Walker, Caroline Hayward, David J. Porteous, Kathryn L. Evans, Andrew M. McIntosh, J. Kenneth Baillie, Chris Haley, Chris P. Ponting

## Abstract

Krüppel-associated box zinc finger proteins (KRAB-ZFPs) are the largest gene family of transcriptional regulators in higher vertebrates. We have developed a method for inferring these factors’ DNA binding-sites *in vivo*. We achieved this by combining the genome-wide association of methylation at 573,027 CpG-sites (meQTL) in blood and KRAB-ZFP RNA expression changes (eQTL). This method’s efficacy is clearly demonstrated by showing that the CpG-sites whose methylation is affected by KRAB-ZFP expression changes occur preferentially near KRAB-ZFP binding-sites in the HEK293T cell line. We found such an enrichment to be significant for 31% of factors tested. In addition, we found that binding-sites of many KRAB-ZFPs are significantly enriched (or sometimes depleted) in disease-associated CpG-sites. In total, 9% (11 of 125) of the traits tested, and 14% (32 of 222) of the factors (KRAB-ZFPs and TIF1-beta) tested, were enriched or depleted in one or more trait-factor pairing. Rheumatoid arthritis and human immunodeficiency virus (HIV) infection were associated with the largest number of KRAB-ZFP enrichments. There have been variable reports on the effects on HIV infection dynamics of transcription intermediary factor 1-beta (TIF1-beta, also known as KAP1 or TRIM28), a nuclear corepressor for KRAB-ZFPs. We provide evidence that KRAB-ZFP-independent effects of TIF1-beta are responsible for the decreased methylation of CpG-sites within its binding-sites that is observed in HIV infection. In conclusion, KRAB-ZFPs often affect CpG-site methylation within the proximity of their binding-sites, and CpG-sites within the binding-sites of KRAB-ZFPs are enriched for association with many traits, including rheumatoid arthritis and HIV infection.

## Introduction

Krüppel-associated box domain-containing zinc-finger proteins (KRAB-ZFPs) contain an N-terminal KRAB-domain and a C-terminal array of C2H2 zinc-finger domains. They are the largest gene family of transcriptional regulators in higher vertebrates (Ecco et al., 2017), with the human genome encoding approximately 350 (Imbeault et al., 2017). KRAB-ZFPs have been associated with the control of transposable elements and lay the foundation of many species-specific regulatory networks by transcriptionally repressing newly inserted transposable element sequences (Ecco et al., 2017; Wolf et al., 2020). In addition, a large minority do not associate with transposable elements but bind instead to promoters, simple repeats, and poly-zinc finger protein genes (Ecco et al., 2017).

KRAB-ZFPs have many physiological roles, including in development, cell-differentiation, and metabolism. For example, the binding of ZFP57 to a methylated hexanucleotide at imprinting- control regions results in the preservation of imprinting (Li et al., 2008; Quenneville et al., 2011; Strogantsev et al., 2015). TIF1-beta (TRIM28, also known as KAP1) binds to many KRAB-ZFPs and acts as a scaffold for a silencing complex with repressive effects on transcription. However, not all KRAB-ZFPs bind TIF1-beta; instead some associate with other protein types, including transcriptional activators (Schmitges et al., 2016).

In this study, we performed genome-wide association of methylation at 573,027 CpG-sites measured in whole blood from 4,101 individuals from the Generation Scotland cohort and identified KRAB-ZFPs as considerably over-represented amongst co-localising local- expression quantitative trait loci (eQTLs) and distal-methylation quantitative trait loci (meQTLs). We sought to determine whether local genetic control of KRAB-ZFPs influences distal CpG-site methylation, and whether KRAB-ZFPs are associated with CpG-sites whose methylation has previously been associated with human disease. To address these questions, the results from these 573,027 genome-wide association (GWA) studies were integrated with HEK293T cell line genome-wide binding data for hundreds of KRAB-ZFPs, and subsequently the binding-site data with the epigenome-wide associations of 125 human traits.

We first replicated a previous finding (Hop et al., 2020; Huan et al., 2019) that ZFP expression is commonly associated with altered methylation at distal CpG-sites. Next, we showed that, for nearly one-third of KRAB-ZFPs tested, these associations of CpG-site methylation and KRAB-ZFP expression occur preferentially at their binding-sites in HEK293T cells (Imbeault et al., 2017). As the relationship between KRAB-ZFP abundance and CpG-sites is a one-to-many relationship, this implicates these factors as causal modulators of CpG-site methylation. Our evidence that KRAB-ZFPs causally alter distal CpG-site methylation is strong. This is because we require a SNP to be both a local-eQTL for a KRAB-ZFP gene and a meQTL for a distal CpG-site. Furthermore, KRAB-ZFP binding is known to result in methylation change for nearby CpG-sites (Quenneville et al., 2012; Ying et al., 2015).

Finally, for traits such as rheumatoid arthritis and HIV infection, we demonstrated strong enrichment of trait-associated CpG-sites within the binding-sites of many KRAB-ZFPs. This builds upon previous evidence that rheumatoid arthritis alters CpG-site methylation at multiple sites (Min et al., 2021), and observations linking other autoimmune diseases (systemic lupus erythematosus in mice and multiple sclerosis in humans) and endogenous retrovirus (a type of transposable element) protein expression (Antony et al., 2011; Baudino et al., 2010). This study is unable to determine whether KRAB-ZFP-mediated changes in CpG-site methylation causally alter human disease risk, are downstream consequences, or simply represent divergent paths from a shared locus. Disease-associated DNA variants have previously been shown to have widespread distant effects on CpG-site methylation, potentially reflecting differential occupancy of distant DNA binding-sites by a locally regulated transcription factor (Bonder et al., 2017). However, evidence is accumulating that CpG-site methylation changes are causal for disease in only a minority of cases (Huan et al., 2019; Min et al., 2021). Nevertheless, even when not directly affecting disease incidence, such changes could be a predictive biomarker and/or modulate disease symptoms or progression.

## Results

For our initial analyses, we made use of SNPs that are both an eQTL to a local target (KRAB-ZFP / TIF1-beta) gene (SNP to transcriptional start site < 1Mb) and a meQTL to one or more distal CpG-sites. Henceforth, we refer to such a CpG-site as a CpG_eQTL_ and the eQTL’s target gene as factor *F_i_* (e.g. ZFP57). In this nomenclature CpG_eQTL_(*F_i_*) is a CpG-site that is significantly associated with a local-eQTL for the gene encoding factor *F_i_*. We refer to the SNP and CpG as ‘distal’ when they are on different chromosomes, or else are more than 10Mb apart.

### Trans-acting modifiers of CpG methylation

We considered DNA CpG-site methylation as a quantitative trait, with meQTLs containing SNPs linked to the genetic control of methylation. We ascertained meQTL through large-scale genome-wide association testing. A genome-wide association test was performed for each of 573,027 CpG-sites, measured in whole blood from 4,101 individuals (Methods). We fitted both technical and biological covariables, including estimated cell counts, in a mixed linear model prior to genome-wide association testing (Methods). To avoid over-counting meQTL associations in our results, we defined a non-redundant set of 497,877 CpGs (hereafter termed “independent” CpG-sites) among which no pair of CpG-sites within 10Mb of each other was correlated with an absolute Pearson’s correlation coefficient of more than 0.4 (Methods). Methylation for one-third (172,544; 35%) of these independent CpG-sites was significantly (p-value < 1x10^-13^) associated with one or more meQTL. A third of the independent CpG-sites (165,080; 33%) had at least one local (*cis*) acting meQTL. Moreover, 12,424 (2.5%) independent CpG-sites had at least one significant distal-meQTL. When the 172,544 Manhattan plots for these CpG-sites were ordered by their genomic location, stacked, and viewed from above, the composite plot appeared as in Figure 1.

**Figure 1:**
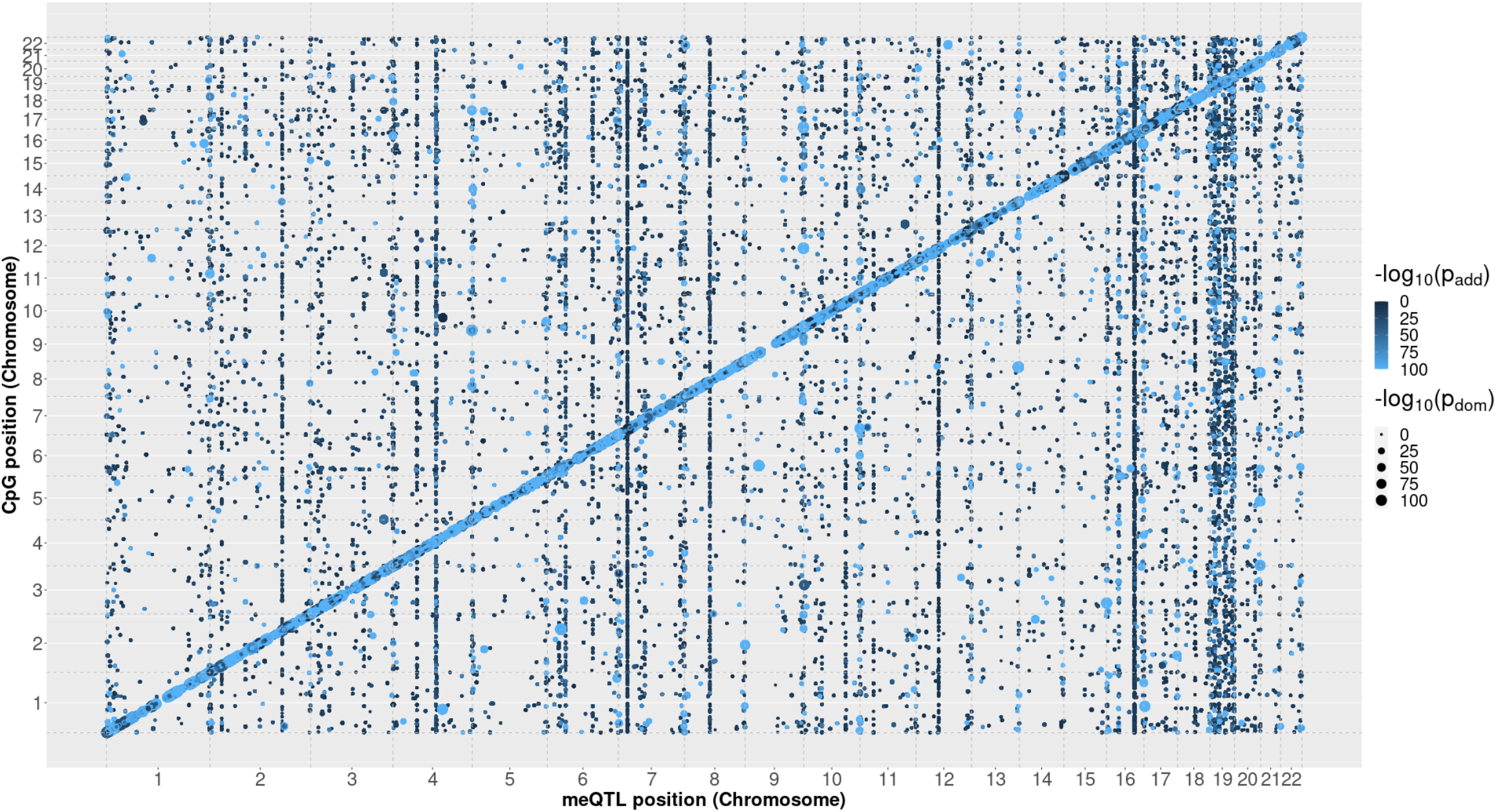
Genetic effects influencing CpG-site methylation commonly act locally (in cis; along the diagonal) but also distally, in trans. Genetic associations for 172,544 independent CpG-sites, stacked by genomic position, with an meQTL with a p-value of < 1x10^-13^. Colour indicates the strength of association of the additive component of the model. The size of the data point represents the strength of association of the dominance component. To aid visualisation, all p-values in this plot were capped at -log_10_(p-value) = 100.

Many meQTLs are associated with nearby CpG sites in *cis* (diagonal; Figure 1). In contrast, the vertical patterns in this plot illustrate meQTL ‘hub’ loci. These are associated with methylation levels at multiple CpG-sites across the genome, such as meQTL on chromosomes 2, 4, 6, 7, 8, 10, 12, 16 and, most notably, 19. Our hypothesis is that each hub locus lies in the chromosomal vicinity (i.e., is in *cis*) of a gene whose protein’s activity causes CpG-site methylation change elsewhere in the genome (i.e., in *trans*). An example of a meQTL hub locus is rs4473920 (chr7:21974610, within the *CDCA7L* gene) which had significant associations with 1,119 independent distal CpG-sites spread across all autosomes.

Figure 2 presents the numbers of distal associations of meQTL across the human genome and highlights the many loci that were associated with multiple independent CpG-sites. When a threshold of more than 50 independent CpG-sites is applied, we identified 20 such hub loci (Supplementary Table 1; Supplementary File 1), twice that of previous studies when similarly defined (Hop et al., 2020; Huan et al., 2019). These loci have previously been associated with transcriptional regulation, transcription factor expression (including KRAB-ZFPs), and active chromatin.

**Figure 2:**
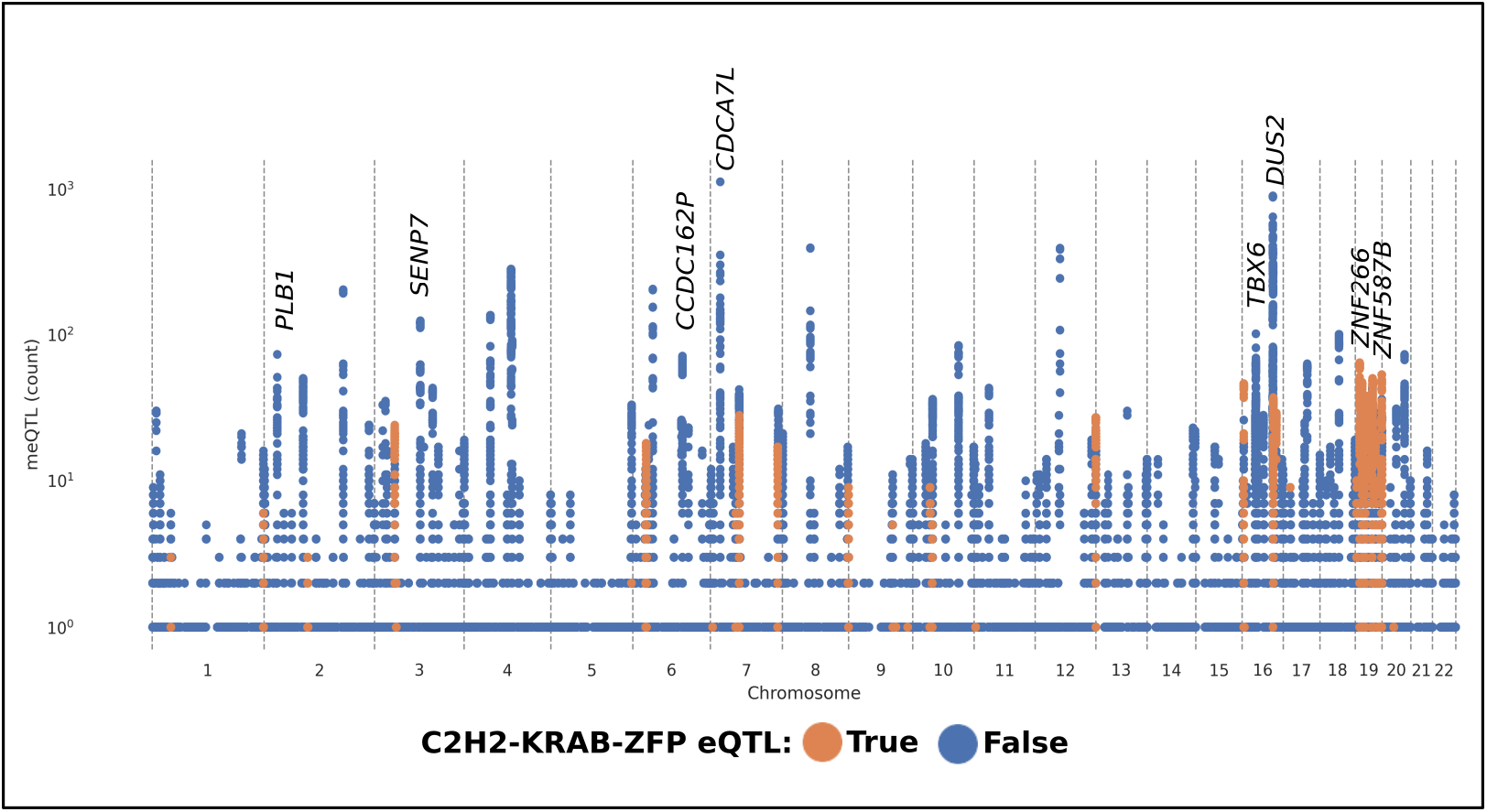
C2H2 KRAB-ZFP gene expression commonly affects methylation at multiple distal independent CpG-sites. meQTL hub loci influence CpG-site methylation at multiple locations at distal loci (in *trans*). The x-axis represents the chromosomal location of the meQTL SNP, and the y-axis indicates the number of significant (p-value < 1x10^-13^) independent CpG-site associations of the SNP, on a different chromosome, or else greater than 10Mb distant. eQTLs associated with a C2H2 KRAB-ZFP in whole blood are highlighted in orange. As can be seen, C2H2 KRAB-ZFPs are responsible for many hub loci present across the genome, in particular those located on chromosome 19. To aid comprehension, where available, gene labels have been added to the 20 hub loci: these relate to the most significant eQTL association of the lead-SNP of the peak in whole blood (Methods). Detailed plots of each locus are available in Supplementary File 1.

Among the distal-CpG_eQTL_ genes, zinc-finger proteins were considerably over-represented (2.4-fold enriched, p-value = 1x10^-12^; Supplementary Table 2; Supplementary File 2; Supplementary File 3). This enrichment is illustrated in Figure 2, where the eQTL of C2H2 KRAB-ZFPs are highlighted in orange. Given their large number, established biological importance, and known effects on transcriptional repression (Ecco et al., 2017; Imbeault et al., 2017), we chose to further determine the effects of KRAB-ZFPs on CpG-site methylation, as well as their downstream effects upon human traits.

Co-localisation of a local-eQTL of a KRAB-ZFP and a distal-meQTL implies a link between the target KRAB-ZFPs (*F_i_*) and the distal CpG-site. Of the 344 KRAB-ZFPs identified by Lambert et al. (Lambert et al., 2018), 121 were associated with one or more eQTL identified in whole blood within 1Mb of its transcription start-site (GTEx v7; GTEx Consortium, 2017). Of these 121, 85% (103) were associated via their eQTL with one or more independent distal-CpG_eQTL_. Furthermore, an eQTL of half of these (52%; 54 of 103) had 9 or more independent distal- CpG_eQTL_. This number of sites was chosen as it yields 5% of all SNPs associated with CpG-site methylation (22,166 SNPs were associated with 9 or more independent distal CpG-sites among 452,600 SNPs associated with one or more). This indicates that the eQTL SNPs of C2H2 KRAB-ZFPs commonly affect methylation at more CpG-sites in *trans-* than SNPs located elsewhere.

### KRAB-ZFPs modulate distal CpG-site methylation near to their binding-sites

We first investigated whether KRAB-ZFPs’ DNA binding-sites occur preferentially near to their distal-CpG_eQTL_. As discussed below, these binding sites were identified in the HEK293T cell line rather than in blood, yet this analysis was assisted by availability of KRAB-ZFPs’ genome-wide DNA-binding data via ChIP-exo (chromatin immunoprecipitation followed by exonuclease digestion) (Imbeault et al., 2017).

When considering all 222 *F_i_* factors with available ChIP-exo data (Imbeault et al., 2017), 33 KRAB-ZFP *F_i_*s had at least one distal-CpG_eQTL_(*F_i_*) located either within a binding-site or within 100bp of one. This number increased to 40 *F_i_* factors within 1kb (Figure 3). In total, there were 992 loci whose CpG methylation was sensitive to the expression and nearby binding activity of a KRAB-ZFP (Supplementary File 5 and Supplementary File 6).

**Figure 3:**
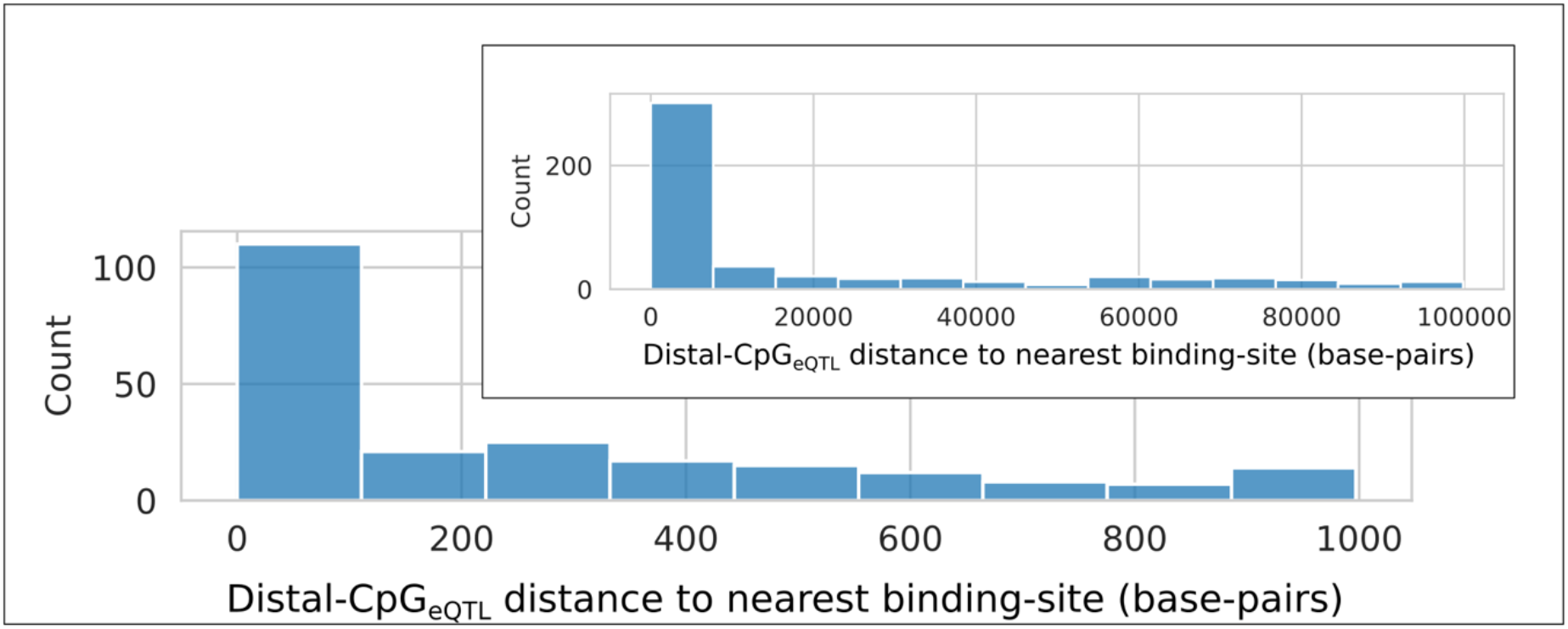
Distal-CpG_eQTL_(F_i_) represent a high-resolution method of binding-site prediction. Histogram of the distances to the closest binding-site of *F_i_* for all independent distal- CpG_eQTL_(*F_i_*). Limited to a distance of 0 to 1kb, with 0 to 100kb shown in the inset. A distance of 0 indicates that the distal-CpG_eQTL_ lies within a binding-site. Of the 222 factors, *F_i_*, for which ChIP-exo data was available ^2^, 86 had an eQTL in whole blood, and 72 had one or more distal- CpG_eQTL_.

We analysed these data for the 72 (32%; 72/222) factors *F_i_* that (a) had at least one locally-acting whole blood eQTL (GTEx Consortium, 2017), and (b) whose whole blood eQTL was also a distal meQTL for at least one CpG-site. For each factor *F_i_* we tested whether its experimental DNA binding-sites were enriched for distal-CpG_eQTL_(*F_i_*). Nearly one-third (31%; 22 of 72) of these KRAB-ZFPs showed such an enrichment (Bonferroni correction; p-value < 0.05/72; Figure 4). Of these 22 factors, ZFP57 had a replicate ChIP-exo data set available, and ZNF100 had two. All three replicate data sets also yielded significant enrichments (Bonferroni correction; p-value < 0.05/3; Supplementary File 4) with the expected direction-of-effect.

**Figure 4:**
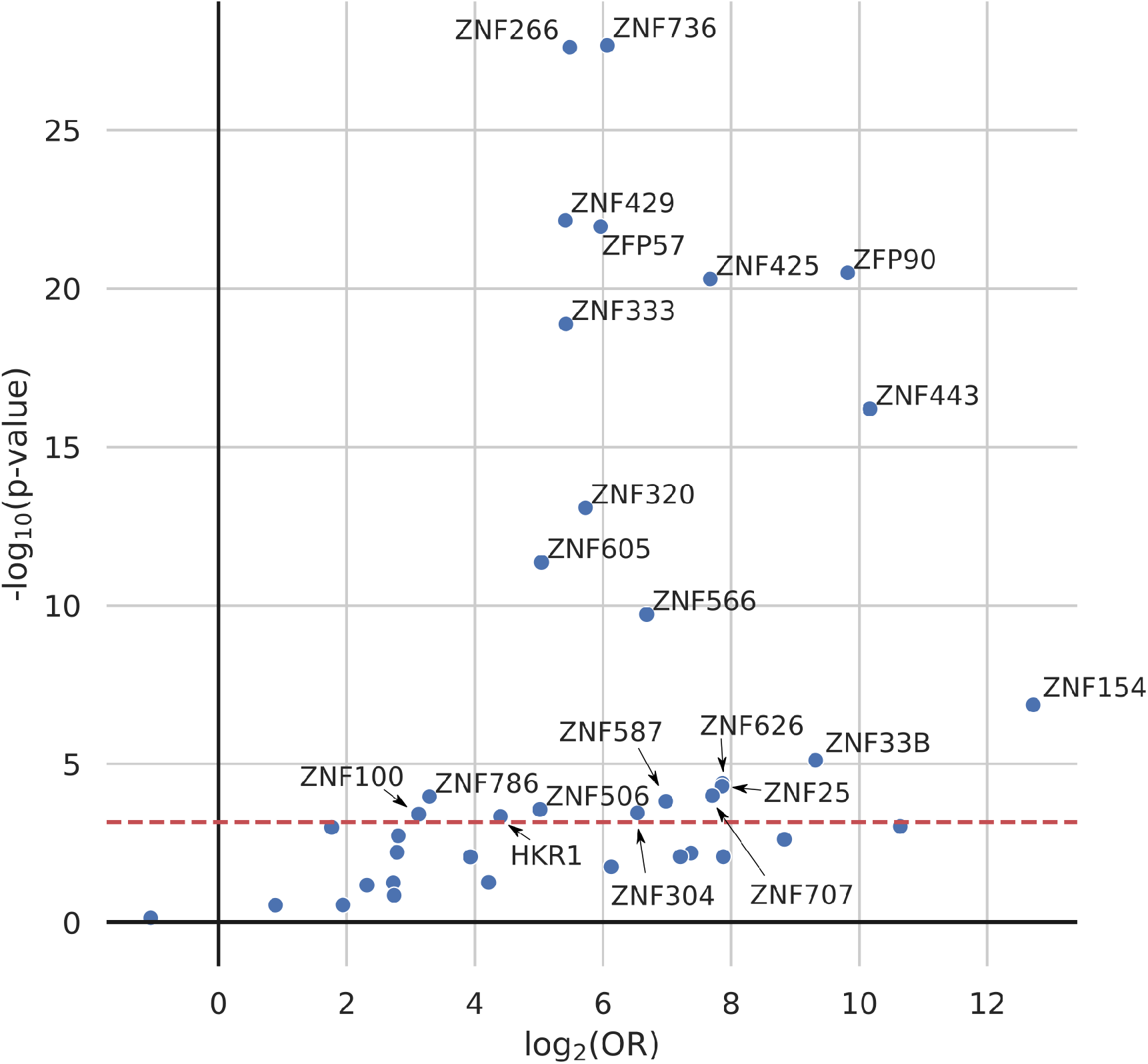
KRAB-ZFPs influence CpG-site methylation near their binding-sites. Volcano plot. Binding-sites for 22 KRAB-ZFPs occur preferentially near to differentially methylated CpG-sites. y-axis: -log_10_(p-value) of the enrichment of distal-CpG_eQTL_(*F_i_*) within factor *F_i_*‘s ChIP-exo defined binding-sites; x-axis: log_2_(OR) (log-base 2 odds-ratio) of distal- CpG_eQTL_(*F_i_*) within experimentally-defined binding-sites (± 1kb) of each KRAB-ZFP, *F_i_*. Significant KRAB-ZFPs are labelled by name. The horizontal dashed red line indicates the Bonferroni significance threshold (p-value < 0.05/72; there were no significant depletions). Enrichments of differentially methylated CpG-sites within the binding-sites of KRAB-ZFPs (with respect to KRAB-ZFP locally-acting eQTL) are provided in Supplementary File 4.

As an example, we present the *GNAS* locus in Figure 5. A local eQTL of ZFP57 (rs365052; on chromosome 6) is also a meQTL to multiple clustered CpG-sites on chromosome 20, proximal to ZFP57-binding sites lying within the promoter of *GNAS-AS1*: a paternally-imprinted anti- sense RNA transcript. These observations support ZFP57 as being a *trans-*acting modulator of methylation at the *GNAS-AS1* promoter. This prediction is supported by findings in patients with transient neonatal diabetes mellitus 1, who are known to carry disrupting mutations in *ZFP57* and to show hypomethylation of *GNAS-AS1* (Mackay et al., 2008, p. 57). It is notable that, both the eQTL of *ZFP57* (rs365052) and the *GNAS* locus are associated with body size (Plagge et al., 2004; Richardson et al., 2020).

**Figure 5:**
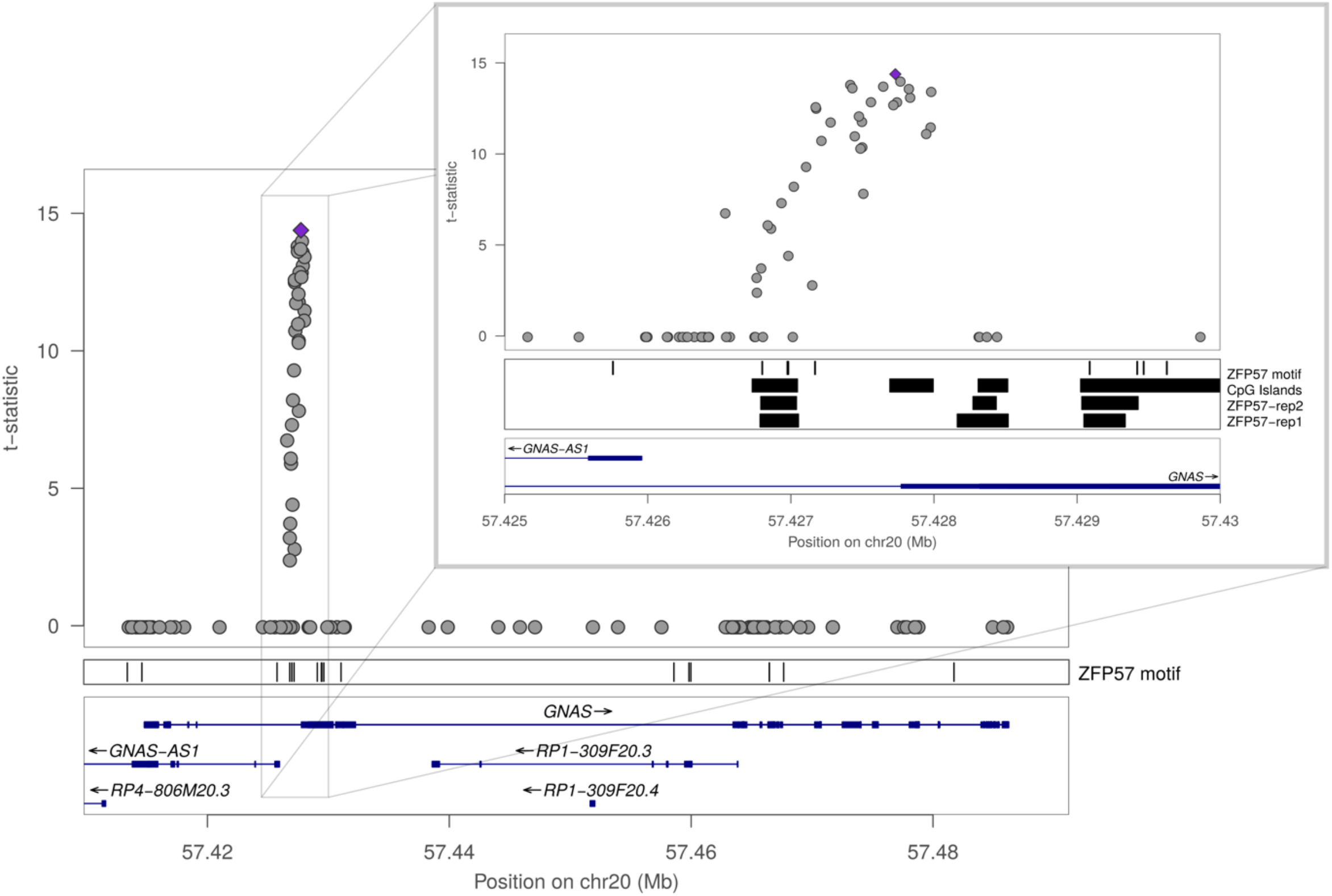
Genetic effects on ZFP57 expression alter CpG-site methylation across the GNAS locus. The effects of a local ZFP57 eQTL (rs365052) on distal CpG site methylation across the *GNAS* locus and, more specifically, within the promoter of *GNAS-AS1* (inset). *GNAS-AS1* is an imprinted RNA transcript that regulates the *GNAS* complex locus. Each point on the plot indicates a CpG-site (not a SNP). x-axis: chromosomal position of the CpG-site; y-axis: t- statistic of the additive component of the meQTL model. A positive t-statistic indicates that methylation of that CpG-site increases with increased expression of ZFP57 mRNA. The ZFP57 motif track indicates canonical ZFP57 binding motif locations: TGCCGC (Quenneville et al., 2011). **Inset**: The effect of the ZFP57 eQTL is greatest 1-2kb upstream of the *GNAS-AS1* transcriptional start site and within 1kb of ZFP57 binding sites defined by two replicates (“rep1”, “rep2”) of ZFP57 ChIP-exo sequence reads. CpG-sites without an association with rs365052 at a p-value of < 1x10^-3^ are recorded as having a t-statistic of zero on this plot because CpG-sites with p-value_model_ > 1x10^-3^ were discarded during processing. x-axis gaps thus indicate an absence of measured CpG-sites, not an absence of association. Additional plots for ZFP57, as well as those relating to all other KRAB-ZFPs for which ChIP-exo data and a suitable eQTL were available, can be found in Supplementary File 5.

Locating a KRAB-ZFP’s distal-CpG_eQTL_(*F_i_*) may thus represent a high-resolution approach to infer these molecules’ DNA binding locations *in vivo*, in particular when ChIP data in physiologically relevant cell lines is absent. In addition to the precise co-localisation of CpG_eQTL_(*F_i_*) with many KRAB-ZFP *F_i_*s’ known binding-sites, other CpG_eQTL_(*F_i_*) are situated away from known binding-sites. Approximately 85% of the independent CpG_eQTL_(*F_i_*) are located more than 1kb away from a binding-site of their respective factor *F_i_* (1,324 of 1,553). These loci may represent KRAB-ZFP binding-sites that are occupied *in vivo* but not in HEK293T cell- lines (Imbeault et al., 2017). To assist investigation of specific loci Supplementary File 5 contains plots of all CpG_eQTL_(*F_i_*)-containing genomic loci relevant to 64 KRAB-ZFPs for which ChIP-exo data was available (Imbeault et al., 2017); Supplementary File 6 contains plots for 32 KRAB-ZFPs for which ChIP-exo data is not currently available.

### Associating human traits with KRAB-ZFP–mediated modulation of CpG methylation

Next, we investigated whether CpG-site methylation within KRAB-ZFP binding-sites is associated with specific human traits. For this analysis we took advantage of discoveries from human epigenome-wide association studies (EWASs). These seek to define trait-associated CpG-site methylation variation by regressing CpG-site methylation status against a trait of interest, in a similar manner to a genome-wide association study.

We hypothesised that a factor *F_i_*, which alters CpG-site methylation within its binding-sites, could be associated with a trait if CpG-sites associated with this trait are unexpectedly abundant within *F_i_*’s binding-sites.

Two types of enrichment analyses were performed to minimise both biological biases (e.g., uneven distribution of CpG-sites across the genome, and correlation among CpG-sites) and technical biases (e.g., biased selection of CpG-sites for inclusion on the array). We first performed an analysis of individual CpG-sites, before completing a pair of analyses based upon CpG-site regions. Below we discuss each approach in turn, and then review consensus results.

### CpG-site based factor-trait enrichment analysis

For this analysis, we defined *F_i_*-associated CpG-sites as those lying within *F_i_’s* experimentally defined binding-site peaks. Enrichment analyses were performed for 29,500 factor-trait pairs: 221 unique KRAB-ZFPs and TIF1-beta (for which genome-wide binding-site data was available (Imbeault et al., 2017)), across 125 traits whose EWAS associations were acquired from the MRC-IEU EWAS catalog (http://www.ewascatalog.org/).

The distributions of binding-site sizes for 221 unique KRAB-ZFPs and TIF1-beta are shown in Figure 6A. CpG-sites were divided between those associated (or not) with trait in an EWAS, and further divided into those that fall within (or not) these factors’ binding-sites, and enrichment assessed. As an exemplar, consider ZNF202 and HIV infection as a factor-trait pair; ZNF202 was chosen as it was the factor most significantly associated with HIV in this analysis. ZNF202 had 10,804 binding-sites genome-wide whose mean size was 603 base-pairs (Figure 6B); HIV infection was associated by EWAS (p-value < 1x10^-7^) with 6,357 CpG-sites (Figure 6C). From the resulting 2x2 contingency table (Figure 6D) a Fisher’s exact test was performed. Of the 6,357 HIV infection-associated CpG-sites, 953 (15%) lay within ZNF202’s 10,804 binding-sites. Comparison to non-HIV associated CpG-sites yielded an odds-ratio of 2.1 and a Fisher’s exact test p-value of 1.5x10^-79^.

**Figure 6:**
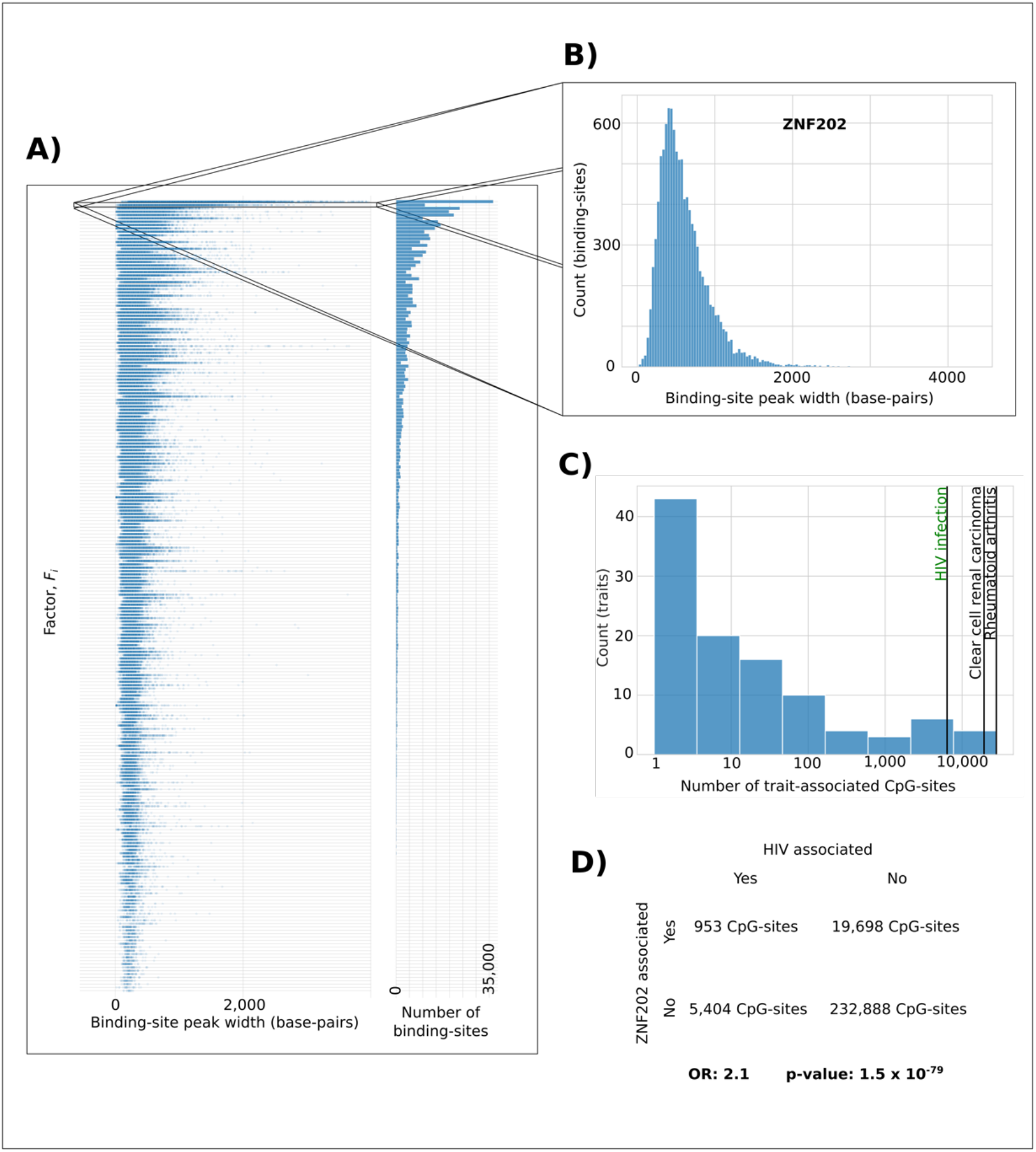
CpG-site based analyses. A) Left: Distributions of binding-site sizes (in base-pairs [bp]) identified in 236 ChIP-exo experiments, relating to 221 unique KRAB-ZFPs and TIF1-beta (Imbeault et al., 2017). Right: The number of binding-site peaks identified per factor. Factors are ordered by the summed size of their binding-sites (largest-to-smallest, top-to-bottom). The binding-site peak width distribution plot is truncated at 4,000bp to aid visualisation. B) Histogram of the binding-site peak widths of ZNF202. Of the 10,804 binding-sites identified, the mean size was 603bp. C) Numbers of CpG-sites associated with HIV infection (n = 6,357), clear cell renal carcinoma (n = 19,187), and rheumatoid arthritis (n = 27,725) marked as black vertical lines on a plot of the number of independent CpG-sites associated with each of the 125 traits examined (logarithmic scale) against the count of traits associated with this number of independent CpG sites. D) Contingency table of the independent CpG-sites (n = 258,943 on the Illumina 450k array) that are HIV associated (EWAS p-value < 1x10^-7^), or not, and those that fall within a binding-site, or not.

In total, we identified 332 unique significant (p-value < 0.05/29,500) factor-trait pairs. In order to obtain this number, we considered all available ChIP-exo experiments (n = 236; Supplementary File 8). This number was reduced to 318 significant (p-value < 0.05/29,500) factor-trait pairs when only a single ChIP-exo experiment per factor was considered (Figure 7). Of these 318 factor-trait pairs, 96 (30%) were enrichments and 222 (70%) were depletions, and they involved 74 unique factors and 20 unique traits.

**Figure 7:**
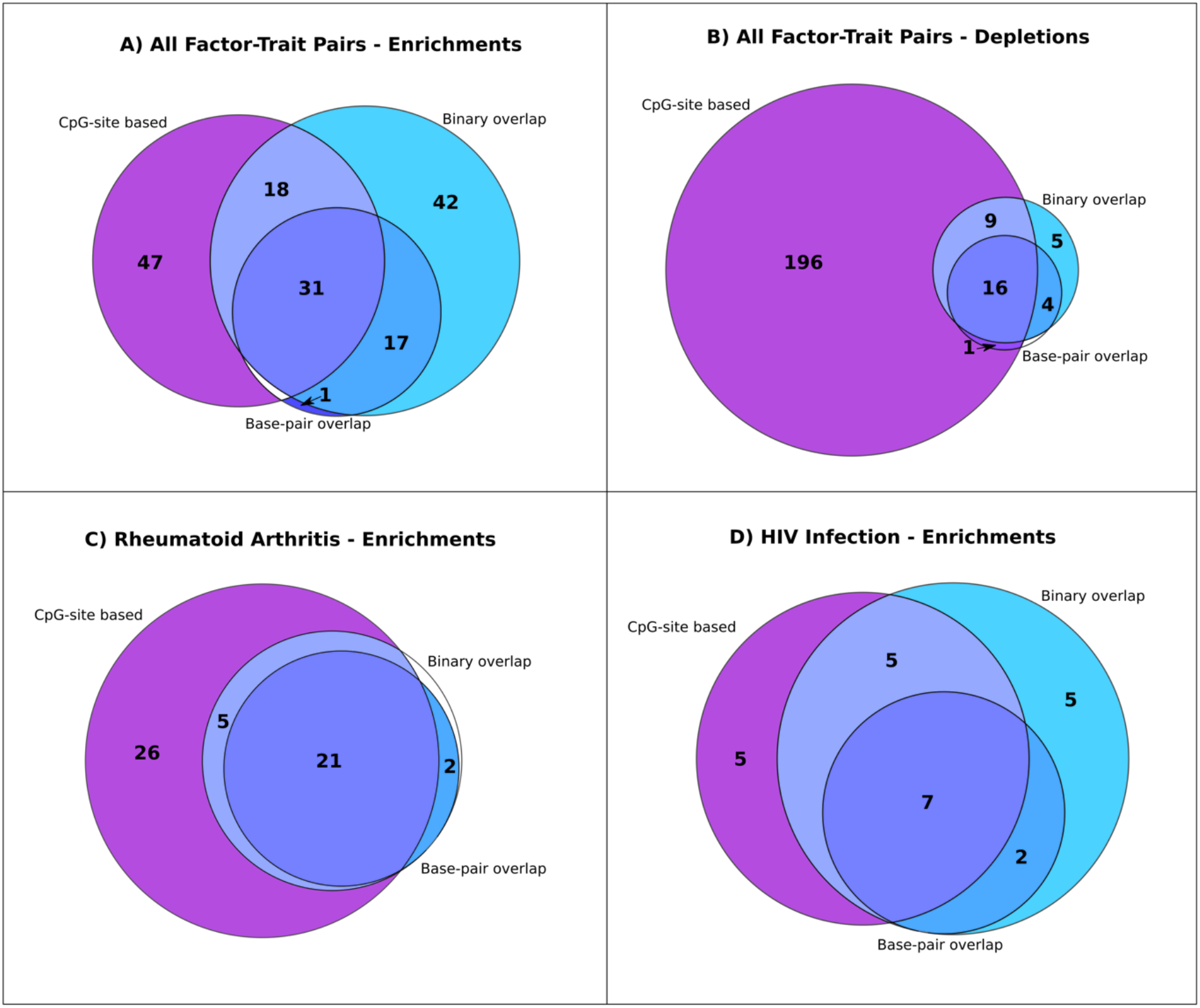
Consistency of significant findings between CpG-site and region-based analyses. Overlap of the factor-trait pairs with a significant A) enrichment, or B) depletion, of trait- associated CpG-sites within the factor’s binding-sites in one or more of the CpG-site (p-value < 0.05/29,500) and the two region-based analyses (empirical p-value ≤ 0.05/25,000); and overlap of all significantly enriched factors associated with C) rheumatoid arthritis, or D) HIV infection, across all three analyses.

Depletion implies fewer trait-associated CpG-sites within a factor’s binding-sites than elsewhere in the genome, whereas enrichment implies increased trait-associated CpG-sites within the factor’s binding-sites. Most of these significant enrichments cannot be explained by several trait-associated CpG-sites lying within a small number of factor binding-sites. When we limited, per factor, the number of EWAS hits within each binding-site to one, whilst leaving values in the other three quadrants of the contingency table unaltered (i.e., a conservative approach), 69% (75) of 108 significant enrichments in the initial analysis remained significant (Supplementary File 8).

### CpG-region based factor-trait enrichment analyses

We next undertook two region-based analyses (Supplementary Figure 1; Methods). In summary, ±10kb regions surrounding CpG sites on the array were merged if overlapping and then, per trait, were assigned as containing, or not, an EWAS-associated CpG site. Subsequently, again per trait, we tested whether such trait-linked CpG regions were enriched in a factor’s, *F_i_*’s, binding-sites. Enrichment was assessed against 500,000 random permutations matched by region-size, and CpG-site density. Permutations were generated by shuffling which CpG regions were considered to be trait-associated.

Region-binding site overlap was assessed both by 1) binary overlap: whether the region overlapped with a factor’s (*F_i_*’s) binding-sites, and 2) base-pair overlap: the number (*n*) of base-pairs in a region that lie within a factor’s, *F_i_*’s, binding-sites. The latter analysis is more conservative because a factor’s binding-sites tend to be much smaller than the CpG-site regions (mean 297 base-pairs vs 43,883 base-pairs, respectively).

These two analyses yielded 121 and 55 trait-factor pair enrichments, and 34 and 21 trait-factor pair depletions, for binary and base-pair overlap approaches, respectively (Supplementary File 9). We defined as significant those factor-trait pairs for which no permutation had a greater (enrichment), or lesser (depletion), overlap respectively.

### Ensemble consensus

The two types of test – CpG site- and region-based – for factor-trait pair CpG-site co-location are largely complementary because of how they account for biases. Nevertheless, results between the two methods were mostly concordant. Only a small minority of the 318 significant factor-trait pair associations in the site-based analysis yielded discordant (opposite, significant direction-of-effect) results in the region-based analyses: 4.7% (15/318) and 3.1% (10/318) in the binary interval and base-pair overlap analyses, respectively. Consequently, we took a consensus approach, considering factor-EWAS trait pairs that were significantly enriched (or depleted), with a consistent direction-of-effect, in all three analyses as being *high quality* predictions. Additionally, factor-trait pair predictions were assigned as being of *good quality* when significantly enriched (or depleted), with a consistent direction-of-effect, in the CpG site-based test and in one of the two region-based tests. This approach yielded 31 high quality (and 18 good quality) enriched factor-trait pairs, as well as 16 high quality (and 10 good quality) depleted factor-trait pairs (Figure 7).

In these analyses, an enriched factor-trait pair is one in which trait-associated CpG-methylation status is unusually common at the factor’s binding-sites.

Nine per cent (11 of 125) of traits, and 14% (32 of 222) of factors, were listed among these high- or good-quality predictions (Supplementary File 7, Supplementary File 8, Supplementary File 9). The three traits associated with most high- or good-quality predictions were: rheumatoid arthritis (n = 26 factors), clear cell renal carcinoma (n = 19 factors), and HIV infection (n = 12 factors). ZNF202 was the factor associated with the most traits (n = 7 traits) which included HIV infection and rheumatoid arthritis (both enrichments) and clear cell renal carcinoma (depletion). ZNF441 was associated with 5 traits, and ZNF519, ZNF783, ZNF263, ZNF273, and ZNF343 all with 4 traits. These results may be due, in part, to the statistical power afforded to these factors by the large number and size of the ChIP-exo binding-sites associated with them.

Clear cell renal carcinoma was the only trait associated with three or more high-quality depletions (Supplementary Table 3; Supplementary Table 6). Depletion implies fewer trait-associated CpG-sites within a factor’s binding-sites compared with elsewhere in the genome. This observation is consistent with clear cell renal carcinoma commonly being associated with a functional or actual loss of the von Hippel-Lindau (*VHL*) tumour suppressor gene that leads to substantial alterations to the DNA methylome (Robinson et al., 2018).

Depletion of trait-associated CpG-sites within a factor’s binding-sites does not directly implicate that factor in that trait. Conversely, enrichment implies increased trait-associated CpG-sites within the factor’s binding-sites and thus thereby directly implicates the factor. We therefore focus the subsequent discussion on enriched associations, of which the most numerous were for rheumatoid arthritis and HIV infection. For rheumatoid arthritis, 21 factors displayed enrichment across all three analyses; for HIV infection, 7 factors display enrichment across all the analyses (Supplementary Table 4, Supplementary Table 5).

### TIF1-beta

Among CpG-sites that were differentially methylated between cases and controls in rheumatoid arthritis or HIV infection, most factors were associated with increased CpG-site methylation within their binding-sites in cases compared to controls (Figure 8). TIF1-beta in HIV infection was a notable exception to this trend.

**Figure 8:**
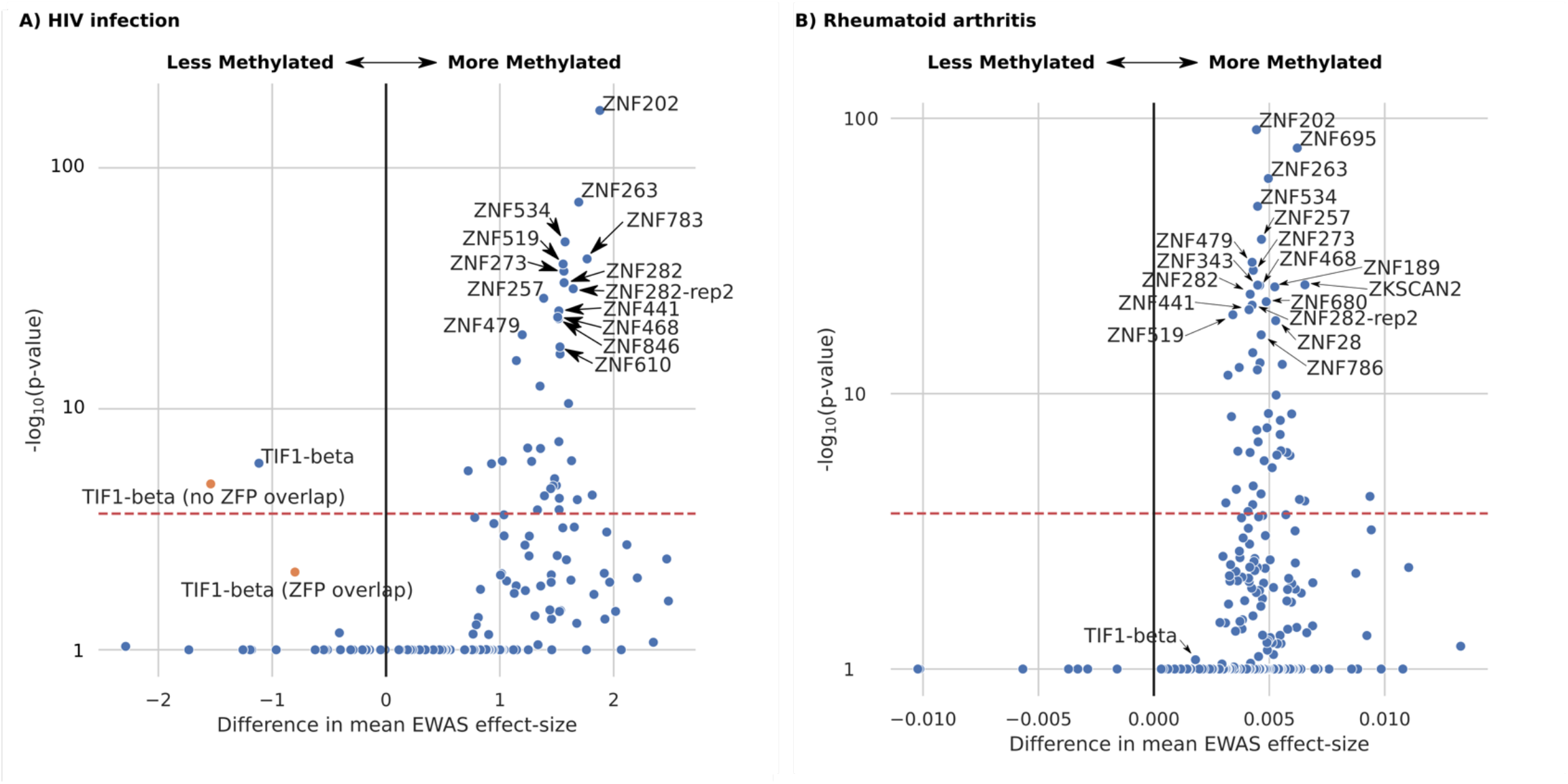
Direction-of-effect of HIV on factor binding-site CpG-site methylation. Mean difference in HIV EWAS effect-size estimate between independent CpG-sites located in the binding-sites of factor, *F_i_*, or not. Each panel can be likened to a volcano plot. Each point represents a specific factor, *F_i_*. Factors falling to the right of the vertical line x = 0 represent factors for which CpG-sites within their binding-sites are, on average, more methylated in HIV patients than in controls, and less methylated for factors falling to the left of the x = 0 line. The horizontal dashed red lines represent the significance threshold (p-value < 0.05/236; 236 ChIP-exo experiments) for the test comparing independent CpG-sites lying within or outside of the factor *F_i_*’s binding-sites. In order to aid presentation, the y-axis scale is logarithmic and has been clipped at a -log_10_(p-value) of 1, corresponding to a p-value of 0.1. For reasons of clarity, only TIF1-beta and significant factors (p-value < 0.05/236; 1,000 permutations; Methods) are labelled (Supplementary File 10). Panel A) is with respect to HIV infection; and A) to Rheumatoid arthritis. The orange points in A) are a subdivision of the TIF1-beta binding-sites, specifically those that overlap (or not) binding-sites of one or more measured ZFP (n = 221). The difference between the two values (the orange points) is significant (p-value = 0.03; two-tailed t-test; mean_ZFP-colocated_ -1.84, SD 1.02, n_CpGs_ 30; mean_not-ZFP-colocated_ -1.11, SD 1.53, n_CpGs_ 41). This analysis is robust to many biases in the location of TIF1-beta binding-sites.

TIF1-beta’s binding-sites tended to be less methylated amongst HIV infected individuals compared with controls (Figure 8; Supplementary File 10), a pattern not seen in rheumatoid arthritis (Supplementary File 11).

TIF1-beta is a nuclear corepressor for KRAB-ZFPs yet also has KRAB-ZFP-independent roles (Iyengar and Farnham, 2011). We were thus interested in whether TIF1-beta’s direction-of- effect on CpG-site methylation within its binding-sites varies according to whether or not a KRAB-ZFP also binds there. Indeed, co-binding (co-localisation of ChIP-exo binding-sites) of TIF1-beta with one or more of the 221 KRAB-ZFPs was associated with a significant increase (less negative; smaller magnitude) in average EWAS effect size (that is, more CpG-site methylation; Figure 8). This result implies that the decreased methylation of CpG-sites associated with HIV infection observed within TIF1-beta binding-sites may be consequent to its non-KRAB-ZFP associated effects. This is consistent with previous evidence that TIF1-beta has a KRAB-ZFP independent role in HIV proviral activation (Morton et al., 2019).

## Discussion

This study provides genome-wide association summary statistics of whole blood DNA methylation at 573,027 CpG-sites for a large single cohort of 4,101 individuals (DataShare DOI: XXX). The human genome contains many meQTL whose genetic variation controls methylation at multiple CpG-sites elsewhere in the genome. These loci have previously been shown to contain genes involved in transcriptional regulation, and to be enriched in KRAB-ZFPs (Hop et al., 2020; Huan et al., 2019). With respect to KRAB-ZFPs, we replicated this enrichment (Figure 2) then built upon it to reach two principal conclusions.

Firstly, KRAB-ZFPs often modulate CpG-site methylation at distal CpG-sites specifically within, or near to, their binding-sites (Figure 4). We demonstrated this for nearly one-third (22) of 72 KRAB-ZFPs tested. The identified KRAB-ZFP distal-CpG_eQTL_ loci are consistent with binding-sites experimentally identified in cultured cell lines (Imbeault et al., 2017) (Supplementary File 5) being predictive of their binding-sites in primary blood cells *in vivo*. The methodology is applicable to both factors for which ChIP-exo data is currently available (Supplementary File 5), and those where it is not (Supplementary File 6).

Secondly, using trait-associated CpG-sites from EWASs, we demonstrated that 11 human diseases or traits are associated with modulation of CpG-site methylation at binding-sites of KRAB-ZFP factors and/or their co-factor TIF1-beta. Rheumatoid arthritis and HIV infection were associated with the largest number of high-/good-quality enrichments in our analyses (26 and 12, respectively; Supplementary Table 4 and Supplementary Table 5).

There is compelling evidence that the observation that KRAB-ZFPs often modulate CpG-site methylation at distal CpG-sites specifically within, or near to, their binding-sites (Figure 4) is biologically important, and we demonstrate this with three examples. First, it was previously demonstrated that ZFP57, together with TIF1-beta, recognises a methylated hexanucleotide and affects chromatin and DNA CpG-site methylation at imprinting control regions in embryonic stem cells (Quenneville et al., 2011). We extend this observation by demonstrating that its effects are observable *in vivo*, in adult cells, and at concentration changes occurring within the general population.

Second, ZNF154 was associated with the largest odds-ratio of any of the KRAB-ZFPs (OR = 6727). This factor is a predicted tumour suppressor gene, whose expression is downregulated by promoter hypermethylation in nasopharyngeal carcinoma (Hu et al., 2017). Indeed, promoter hypermethylation of ZNF154 has shown promise as a pan-cancer biomarker (Miller et al., 2021). Here, we demonstrated that this molecule exerts effects on CpG-site methylation *in vivo* on chromosomes 2, 10, and 19 (Supplementary File 5).

Third, the factor most strongly associated (lowest p-value; p-value = 2.14x10^-28^) with altered methylation at its binding-sites, ZNF736, has been shown to be both hypermethylated and down-regulated in the pre-cancerous condition proliferative verrucous leucoplakia (Herreros- Pomares et al., 2021).

By linking KRAB-ZFPs and CpG-site methylation at multiple sites across the genome, and the association of CpG-sites within the binding-sites of these factors with human disease phenotypes, we have presented evidence that is consistent with KRAB-ZFPs being responsible either for the pathogenesis of specific disease phenotypes, or for their consequences. Specifically, this evidence is consistent with the hypothesis that either KRAB-ZFPs cause disease by altering CpG-site methylation, or that they are responsible for observed consequences of that disease, including alteration of the methylome. The extent to which altered CpG-site methylation is responsible for the pathological manifestations of disease remains unknown. For example, altered expression of some KRAB-ZFPs may predispose individuals to developing autoimmunity in rheumatoid arthritis; or, following HIV infection, altered expression of a specific KRAB-ZFP may control the extent to which a cell suppresses viral replication, and hence modulates viral load and progression to AIDS.

A major barrier to curing HIV-1 is viral latency (whereby the virus lies dormant within cells). KRAB-ZFPs have previously been implicated in the control of endogenous retroelements (Ecco et al., 2017) including endogenous retroviruses. HIV-1 is an exogenous retrovirus and may fall under similar control once integrated into nuclear DNA. ZNF282, for example, binds to the U5 repressive element of the human T cell leukaemia virus (HTLV) type I long terminal repeat (LTR) and exerts a strong repressive effect (Okumura et al., 1997). Indeed, the results we observe for the KRAB-ZFPs themselves in HIV are consistent with this classical view, specifically that there is an increase in DNA CpG-site methylation at their binding-sites associated with HIV infection. However, it should be noted that this situation is far from clear-cut because some KRAB-ZFPs, especially those containing SCAN and DUF3669 domains, in fact bind at promoters and do not recruit TIF1-beta (Ecco et al., 2017; Imbeault et al., 2017; Schmitges et al., 2016).

The only exception to the association of HIV with increased DNA CpG-site methylation within factors’ experimentally determined binding-sites was for TIF1-beta itself. TIF1-beta is required for silencing of endogenous and introduced retroviruses in ES cells and early embryos (Matsui et al., 2010; Rowe et al., 2010; Turelli et al., 2014) and has complex effects on HIV-1. These effects include inhibition of proviral integration (Allouch et al., 2011), contribution to ZBRK1 and ZNF10 mediated HIV-1 LTR repression (Nishitsuji et al., 2015, 2012), and recruitment of an inactive form of P-TEFb to many genes with a paused RNA polymerase II at their promoters, including the HIV-1 LTR (Kauzlaric et al., 2020; Ma et al., 2019; McNamara et al., 2016; Morton et al., 2019). In line with this complex picture, TIF1- beta has been reported by different groups to have differing effects upon HIV-1 latency and reactivation in different cell-types (Ait-Ammar et al., 2021; Ma et al., 2019; McNamara et al., 2016; Morton et al., 2019; Nishitsuji et al., 2015, 2012; Taura et al., 2019). We provide evidence that TIF1-beta’s KRAB-ZFP independent binding is associated with decreased local CpG-site methylation, consistent with a potential activating role.

Whilst it is not proven that rheumatoid arthritis is caused by an infectious agent, there has been much speculation with respect to its association with human endogenous retroviruses (Freimanis et al., 2010; Nelson et al., 2014; Reynier et al., 2009), which may undergo repression by KRAB-ZFPs. However, despite its unknown aetiology, evidence implicates epigenetic processes in the pathogenesis of rheumatoid arthritis. It has previously been reported that epigenetic changes may mediate the development of chronic inflammation by increasing the expression of pro-inflammatory cytokines, such as IL-1β, TNF-α, IL-6, and reactive oxygen species (Ciechomska et al., 2019).

The two traits, rheumatoid arthritis and HIV infection, with the largest number of high- and good-quality associations with KRAB-ZFPs are both associated with a symmetrical polyarthritis. It is thus tempting to speculate that KRAB-ZFPs may represent a unifying pathway, as cause of or consequent to, the musculoskeletal effects in both cases.

There are five main limitations of this study. First, it considers an incomplete coverage of all CpG-sites because it is limited to those present on the array. Second, whilst we have striven to correct the analyses for correlations between CpG-sites – by fitting biological and technical covariates, and limiting ourselves to an ‘independent’ set of CpG-sites – this correlation structure is highly variable and difficult to fully control for. Third, DNA CpG-site methylation has been estimated using whole blood, and whilst corrections for cell-type biases were made (Methods), these are never perfect, especially in conditions known to perturb cell-type frequency. This limitation may be of particular relevance to proteins affecting haematopoiesis such as ZNF589, ZNF268, ZNF300, and TIF1-beta (Santoni de Sio et al., 2012b, 2012a; Venturini et al., 2016; Xu et al., 2010; Zeng et al., 2012). There are also two limitations when using the CpG-sites within a factor *F_i_*’s binding-sites to link *F_i_* to trait. The first of these is that we make no claim as to whether modulation of the CpG-sites’ methylation is cause of or consequent to trait change. Secondly, we cannot completely exclude the possibility that our factor-trait links are confounded by another genomic annotation.

## Data Availability

For the purpose of open access, the author has applied a Creative Commons Attribution (CC BY) licence to any Author Accepted Manuscript version arising from this submission.

### GWAS results

GWAS summary statistics with a p-value < 1x10^-7^ are freely available from Edinburgh University DataShare DOI: XXX.

### Individual level data

Individual level data is available to bona fide researchers, subject to approval by the Generation Scotland data access committee: https://www.ed.ac.uk/generation-scotland/for-researchers/access.

## Methods

### GWAS

#### Samples

Generation Scotland: Scottish Family Health Study (GS) is a population- and family-based cohort from the Scottish population collected between 2006 and 2011 and has been described in detail elsewhere (Smith et al., 2013).

#### Genotypes

Genotyping and imputation of the Generation Scotland cohort has been described previously (Nagy et al., 2017), but are summarised here for completeness.

Samples for DNA extraction (blood, or occasionally saliva) were obtained at the time of recruitment. Genotyping was carried out using the HumanOmniExpressExome-8 v1.0 or v1.2 BeadChips with Infinium chemistry (Illumina). Genotypes were processed using the GenomeStudio Analysis software v2011.1 (Illumina) and called using BeadStudio-Gencall v3.0 (Illumina). The details of blood collection and DNA extraction are also provided elsewhere (Smith et al., 2006). Subsequent quality control steps removed individuals with < 98% call rate, SNPs with < 98% call rate, and SNPs with a Hardy-Weinberg equilibrium p-value < 1x10^-6^. After initial quality control, 604,858 genotyped autosomal SNPs remained. The genotyped data were imputed utilising the Sanger Imputation Service to the HRC panel v1.1. Data were pre-phased using SHAPEIT v2.r873 (Delaneau et al., 2013) and duohmm12 (O’Connell et al., 2014) and imputed with PBWT (Durbin, 2014).

#### Phenotypes

The genesis of the data, and its initial processing are described in Zeng et al. (Zeng et al., 2019) but are summarised here for completeness.

Whole blood genomic DNA (500ng) was treated with sodium bisulfite using the EZ-96 DNA Methylation Kit (Zymo Research) and DNA methylation was assessed using Illumina Infinium MethylationEPIC BeadChip technology (Illumina), as per the manufacturer’s instructions. The arrays were scanned using an Illumina HiScan scanner (Illumina) and initial inspection of array quality was carried out using Illumina GenomeStudio Analysis software v2011.1 (Illumina).

Quality control of the DNA methylation data was carried out before normalisation. The R package shinyMethyl (Fortin et al., 2014) was used for preliminary quality control. This quality control step removed 81 samples based on the following criteria: 1) overall array signal intensity and control probe performance outliers, 2) samples with a mismatch between recorded sex and predicted sex based on X and Y chromosome DNA methylation, and 3) genetic ethnic outliers for the cohort identified by principal component analysis (Patterson et al., 2006). Further quality control was performed using the ‘pfilter’ function in the R package watermelon (Pidsley et al., 2013). Samples were removed if ≥ 1% sites had a detection p-value of > 0.05. This removed 18 further samples. Finally, 5,101 samples remained for further processing. Before normalization, individual probe-sample pairs with a detection p-value of > 0.05 were removed. Normalization was performed using function ‘preprocessNoob’ in the R package minfi (Aryee et al., 2014).

In order to remove potential technical confounders, linear mixed modelling was used to pre-correct each probe. This model included the following fixed effects: top 50 principal components of control probe intensities (which explained 99% of variation in control probe intensities), appointment clinic centre, processing batch, year of the visit, and Sentrix position (position of the sample in Illumina slide); and the random effects: appointment date and Sentrix ID (Illumina slide). The model converged successfully for 712,595 sites, and the resultant residualised M-values were used as DNA methylation phenotypes in downstream analysis. For individual sites, outlier samples with residualised-M-values more than five interquartile ranges from the nearest quartile were removed.

Finally, we fit biological covariates to create residuals for GWA. Cell-type proportions were estimated for granulocytes, monocytes, B-lymphocytes, natural killer cells, CD4+ T-lymphocytes and CD8+ T- lymphocytes using the ‘estimateCellCounts’ function in R package minfi (Aryee et al., 2014).

The following mixed linear model was fit: Fixed effects) age, age^2^, gender, cell-type proportions for granulocytes, B-lymphocytes, natural killer cells, CD4+ T-lymphocytes and CD8+ T-lymphocytes, season of the visit, appointment time of the day, appointment day of the week; Random effects) genomic relationship matrices, G (genomic relationship matrix) and K (kinship relationship matrix), and three environmental relationship matrices, F (environmental matrix representing nuclear-family-member relationships), S (environmental matrix representing full-sibling relationships) and C (environmental matrix representing couple relationships). The resulting residuals were inverse rank transformed prior to GWA analysis in a simple linear model.

#### GWA model

A combined, additive and dominance, model was fit to the dataset (Equation 1). Individuals were randomly split into a discovery set of 4,101, and a replication set of 1,000 individuals. This model was fit to all CpG-sites for which the residualisation (above) had successfully completed (n = 573,027).

Following model fitting, SNPs were retained if (in both the full Discovery and Replication sets):

(1) Imputation information score of ≥ 0.9; 2) a minimum genotype count of ≥ 3 for each of the three possible genotypes: ‘AA’, ‘AB’, ‘BB’; and 3) HWE p-value ≥ 1x10^-6^.

#### Equation 1

Additive and dominant model.

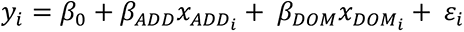

*y_i_* is the residualised methylation values (above); *β*_0_ is the intercept term; *β_ADD_* and *β_DOM_* are the additive and dominance effect-size estimates; *x_ADDi_* is, for individual *i*, the probability of the genotypes BB minus the probability of the genotypes AA: p(BB)-p(AA); *x_DOMi_* is, for individual *i*, the probability of genotypes AB: p(AB); *ε_i_* is the error term.

Data from the discovery set only are used in all subsequent analyses reported in this paper.

### CpG-site independence

CpG-sites present in the Infinium MethylationEPIC v1.0 B4 CpG manifest (https://emea.support.illumina.com/; accessed 27 Aug 2018) were limited to only those autosomal CpG-sites that converged and passed stringent quality control (as described in Zeng et al. (Zeng et al., 2019)). In order to obtain a set of pseudo-independent CpG-sites the following algorithm was executed:

1) Of all CpG-sites under consideration, an ‘active’ list was created.
2) A CpG was randomly removed from the active list and placed on the independent list.
3) Any other CpG-site (a) within 10Mb of the selected CpG, and (b) with a squared Pearson’s correlation coefficient (r^2^) ≥ 0.16 between their residuals having fit the two models described above (technical confounders and biological covariates; Phenotypes) and the selected CpG-site, was also removed from the active list.
4) The above steps were repeated until the active list was empty.

497,877 ‘independent’ CpG-sites were thus identified

### Molecular moieties underlying trans-meQTL – protein annotation

Using the independent CpG-sites defined above, for each SNP, we computed the number of CpG-sites that met the following criteria: 1) meQTL p-value < 1x10^-13^; and 2) Distally-acting: SNP and CpG on different chromosomes, or > 10Mb between them. We then determined the bounds of contiguous loci, that is, regions containing gaps of no larger than 100kb between SNPs associated with > 50 such CpG-sites. Taking the lead-SNP (that is, the SNP with the largest number of independent CpG-site associations), we identified the gene whose mRNA expression it was most significantly associated with in GTEx v7 whole blood. We then compared this list with the list of genes whose genetic instruments were associated with 50 or more CpG sites by Hop et al. Table S5 (Hop et al., 2020).

We define a CpG_eQTL_ as a CpG-site that passes two criteria: its methylation is significantly (meQTL p-value < 1x10^-13^) associated with a SNP, and this SNP is also a significant (per gene, GTEx v7) locally-acting eQTL, whose distance to the target gene’s transcriptional start site is within 1Mb. It is a distal-CpG_eQTL_ when the SNP and CpG-site are on different chromosomes, or have > 10Mb between them. To denote when the SNP in question is associated with the expression of a specific factor, we add this in brackets. So, CpG_eQTL_(*F_i_*) represents a CpG that is significantly associated with a locally-acting eQTL for factor *F_i_*, and distal-CpG_eQTL_(ZFP57) represents a CpG that is significantly associated with a locally-acting eQTL for ZFP57, and the CpG-site and SNP are either located on different chromosomes, or there are > 10Mb between them.

With respect to protein group enrichment among distal-CpG_eQTL_ genes, the set of protein coding genes (as per GENCODE19) associated with biallelic autosomal eQTL of all distal- CpG_eQTL_ in whole blood (GTEx v7) was identified. This set was compared against two background sets: 1) the full set of protein coding genes with ≥ 1 locally-acting eQTL in GTEx v7 from whole blood; and 2) the set of protein coding genes associated with a set of SNPs (of equivalent size to the number of distally-acting meQTL) selected at random from all significant (p-value < 1x10^-13^) meQTL. Following identification, the lists of Ensembl Gene IDs were annotated using the StringDB v11 (Szklarczyk et al., 2019) API (https://string-db.org/; accessed 08 Sep 2020), results were collated and an odds-ratio, and p-value (Fisher’s exact test) calculated. A strict Bonferroni correction was applied per category of annotation.

Plots under all peaks (p-value < 1x10^-13^) were created using a modified version of LocusZoom Standalone v1.4 (Pruim et al., 2010) (https://github.com/statgen/locuszoom-standalone; downloaded 29 Jun 2020). Where used, the track ‘CpG Islands’ is the UCSC CpG Islands Track ‘cpgIslandExt’ (http://genome-euro.ucsc.edu/).

To make Figure 2, the Ensembl Gene IDs of all C2H2 KRAB-ZFPs identified by Lambert et al. (Lambert et al., 2018) were used to query the eQTL from whole blood of GTEx v7. Lead-eQTL labels are those of the transcript with the smallest associated p-value in Whole Blood in GTEx v7.

### KRAB-ZFPs modulate DNA CpG-site methylation local to their binding-sites

Using the independent CpG set (497,877 CpG-sites), for each KRAB-ZFP (or TIF1-beta) factor *F_i_*, we identified the set of significant distal-CpG_eQTL_(*F_i_*) for all significant locally-acting (TSS ± 1Mb) eQTL identified in whole blood (per gene, GTEx v7) for factor *F_i_*.

The location of each CpG-site in the independent CpG set – within a ChIP-exo defined binding-site for factor *F_i_*, or not – was determined through comparison with the ChIP-exo data of Imbeault et al. (Imbeault et al., 2017) (GEO GSE78099; https://www.ncbi.nlm.nih.gov/geo/query/acc.cgi?acc=GSE78099; accessed 13 Mar 2020) ± 1kb of the called peaks.

A contingency table – within a binding-site (yes/no) vs. distal-CpG_eQTL_(*F_i_*) (yes/no) – was constructed, and odds-ratio and Fisher’s exact test p-value calculated.

Figure 5. With regard to the ZFP57 motif track, motif methylation was not considered. Gene annotation is as per GENCODE.

### EWAS trait-associated CpG-site enrichment within KRAB-ZFP binding-sites

Using EWAS studies conducted with the IlluminaHumanMethylation450 array from the EWAS catalog (http://www.ewascatalog.org/; accessed 21 Mar 2020; data version 03-07-2019), CpG-sites that were significantly trait-associated (p-value_EWAS_ < 1x10^-7^ (approximate

Bonferroni correction for 450,000 tests) were identified. Those CpG-sites present on both the IlluminaHumanMethylation450 and MethylationEPIC arrays (as per Illumina Infinium MethylationEPIC v1.0 B4 CpG Manifest; https://emea.support.illumina.com/; accessed 27 Aug 2018) and present within the independent CpG set (as defined above) were considered. For each trait *T_j_* (n = 125) and factor *F_i_* (n = 236; factor replicates considered separately) a contingency table was created – trait-associated CpG (yes/no) vs. within factor *F_i_* binding-site (yes/no) – and the odds-ratio and p-value (Fisher’s exact test) calculated. Factor *F_i_* ChIP-exo defined binding-sites were as per Imbeault et al. (Imbeault et al., 2017) (GEO GSE78099; https://www.ncbi.nlm.nih.gov/geo/query/acc.cgi?acc=GSE78099; accessed 13 Mar 2020).

For each trait, *T_j_*, the odds-ratio captures the enrichment of trait-associated CpG-sites within the binding-sites of each factor, *F_i_*. Depletion represents a lack of trait-associated CpG-sites within the binding-sites of factor *F_i_* when compared to CpG-sites located elsewhere in the genome. Therefore, enrichment putatively associates factor and trait, whereas depletion does not.

### Region based analysis

We undertook a region-based analyses, inspired by GAT (Heger et al., 2013), as illustrated in Supplementary Figure 1.

CpG-site ‘regions’ were created as follows: 1) A 10kb flank was added to either side of all CpG-site locations identified as 450k array (as used for the EWASs) CpG-sites in the Illumina Epic array manifest (v-1-0_B4; Supplementary Figure 1.1A-B), and 2) overlapping regions were combined into a single combined region (Supplementary Figure 1.1C). This produced a set of CpG-site regions where the maximum distance between adjacent CpG-sites was 20kb. Once formed, those regions that contained one or more trait-associated (i.e., EWAS CpG-to-trait association <1x10^-7^) CpG-sites were defined as trait-associated regions (Supplementary Figure 1.2). Factor binding-site overlap was assessed in two separate ways: 1) binary overlap (Supplementary Figure 1.3A), and 2) base-pair overlap (Supplementary Figure 1.3B).

When considering ‘base-pair overlap’ between trait-associated regions and non-trait-associated regions, the metric used for comparison was the number of base-pairs overlapped by a factor’s binding-sites compared with the number of base-pairs that were not. Whereas, when considering ‘binary overlap’, the metric for comparison was the number of trait-associated regions that contained any overlap with a factor’s binding-sites compared with the number that did not.

Enrichment was assessed against a permuted background. Null permutations were obtained by shuffling which regions were considered as EWAS-associated, matched for region-size, and region CpG-site density. Region-size and region CpG-site density were split into 20 evenly sized quantiles each (400 bins in total, duplicate bins merged) and samples obtained that preserved the frequency within each bin.

Finally, an empirical p-value was obtained by comparison to 500,000 permutations. In order to reduce the computational burden of permutations, filtering steps were undertaken at 1,000, 10,000, and 100,000 permutations, whereby those factor-trait pairs that were not more significant than these thresholds were excluded, and further permutations omitted.

### Direction of effect on methylation within binding-sites: TIF1-beta

Using those significantly (p-value_EWAS_ < 1x10^-7^) trait-associated (in whole blood) CpG-sites on the Illumina HumanMethylation450 array from Gross et al. (Gross et al., 2016) (HIV) and Liu et al. (Liu et al., 2013) (rheumatoid arthritis) in the EWAS catalog (http://www.ewascatalog.org/; accessed 21 Mar 2020; data version 03-07-2019; Supplementary File 12), we sought to compare the EWAS effect-size estimates of methylation of CpG-sites that are within factor *F_i_* binding-sites, with those that are not. Factor *F_i_* binding- sites were defined as per the ChIP-exo data of Imbeault et al. (Imbeault et al., 2017) (GEO GSE78099; https://www.ncbi.nlm.nih.gov/geo/query/acc.cgi?acc=GSE78099; accessed 13 Mar 2020). These findings cannot be ascribed trivially to CpG sites’ associations to trait because we only considered in these analyses CpG-sites that are trait associated.

We achieved this in two ways: 1) we compared the EWAS effect-size estimates of all trait-associated CpG-sites from the independent CpG set within the binding-sites of factor *F_i_* to all other trait-associated CpG-sites; and 2) a more conservative analysis: we compared the EWAS effect-size estimates of trait-associated CpG-sites within a given set of ChIP-exo defined binding sites (limited to a maximum of one, randomly chosen, CpG-site per binding-site) to an equivalently sized randomly chosen set from all other trait-associated CpG-sites not within a factor *F_i_* binding-site. This procedure was repeated 1,000 times, and the minimum and maximum odds-ratio and p-values reported.

### Ethics

All participants provided written informed consent. Ethical approval was provided by the East of Scotland Research Ethics Service committee on research ethics (REC reference: 15/ES/0040). Ethical approval for the GS:SFHS study was obtained from the Tayside Committee on Medical Research Ethics (on behalf of the National Health Service).

## Acknowledgements

- A debt of gratitude is owed to all the participants in all cohorts used, without whom this work would not have been possible.
- Genotyping of the GS samples was carried out by the Genetics Core Laboratory at the Wellcome Trust Clinical Research Facility, Edinburgh, Scotland.
- The Genotype-Tissue Expression (GTEx) Project was supported by the Common Fund of the Office of the Director of the National Institutes of Health, and by NCI, NHGRI, NHLBI, NIDA, NIMH, and NINDS.

## Supplementary Materials

**Supplementary Figure 1:**
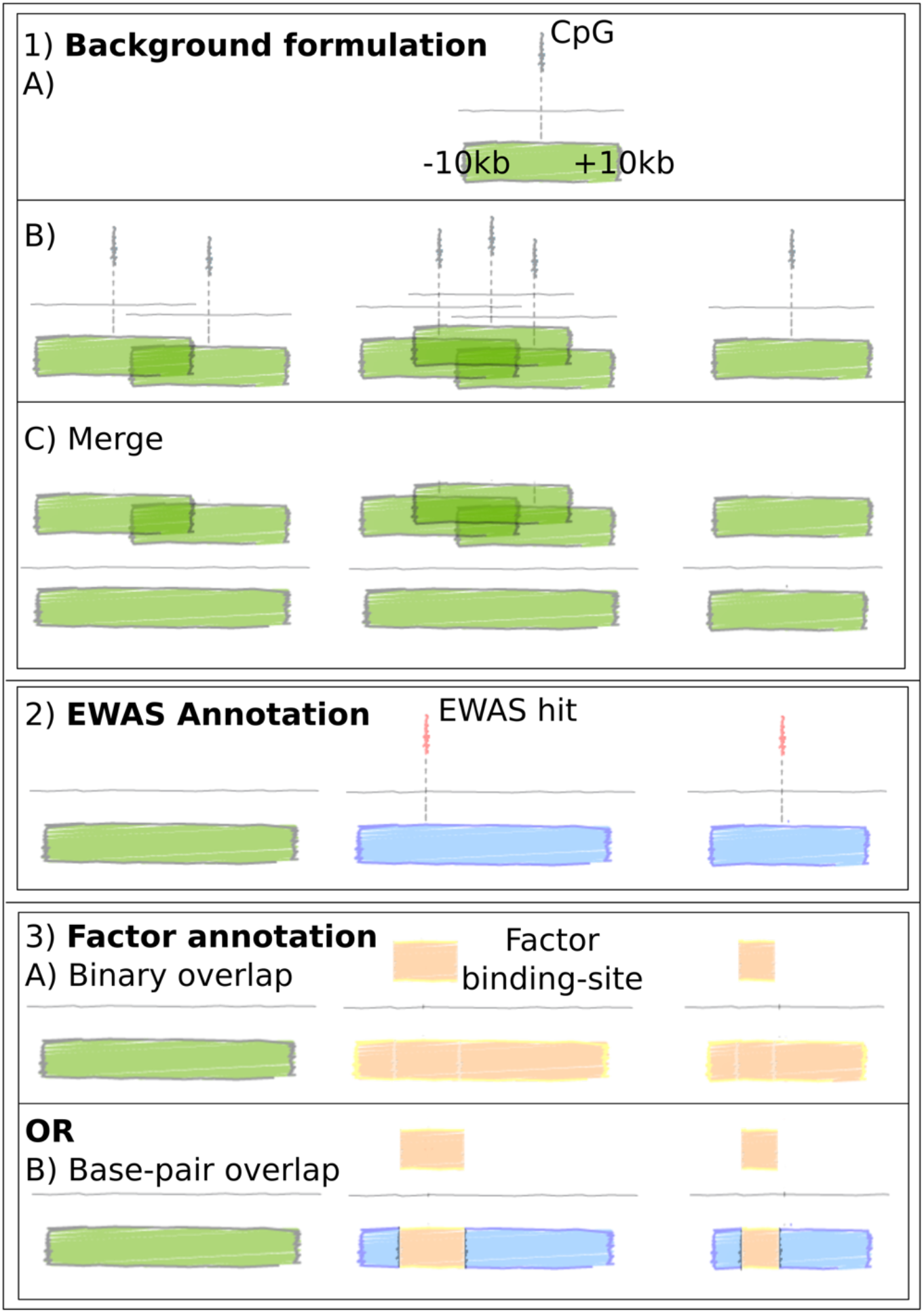
CpG-region based analysis. CpG-regions were defined as contiguous regions wherein the distance between adjacent CpG-sites was maximally 20kb. That is, a 10kb flank was added to each CpG-site (Supplementary Figure 1.1A), and then overlapping regions were merged (Supplementary Figure 1.1B,C). Once defined, these regions were identified as being EWAS-associated if they contained a significant (p-value < 1x10^-7^) EWAS hit (Supplementary Figure 1.2), or not. Region overlap was assessed in two ways: 1) binary assignment of a region as containing a factor binding-site (Supplementary Figure 1.3A), or 2) by calculating the base-pair overlap between the factor binding-site’s and the CpG-region’s (Supplementary Figure 1.3B). The observed overlap of EWAS and factor-associated CpG-regions was then compared to a permuted background set matched for region-sizes and CpG-site densities.

**Supplementary Table 1:**
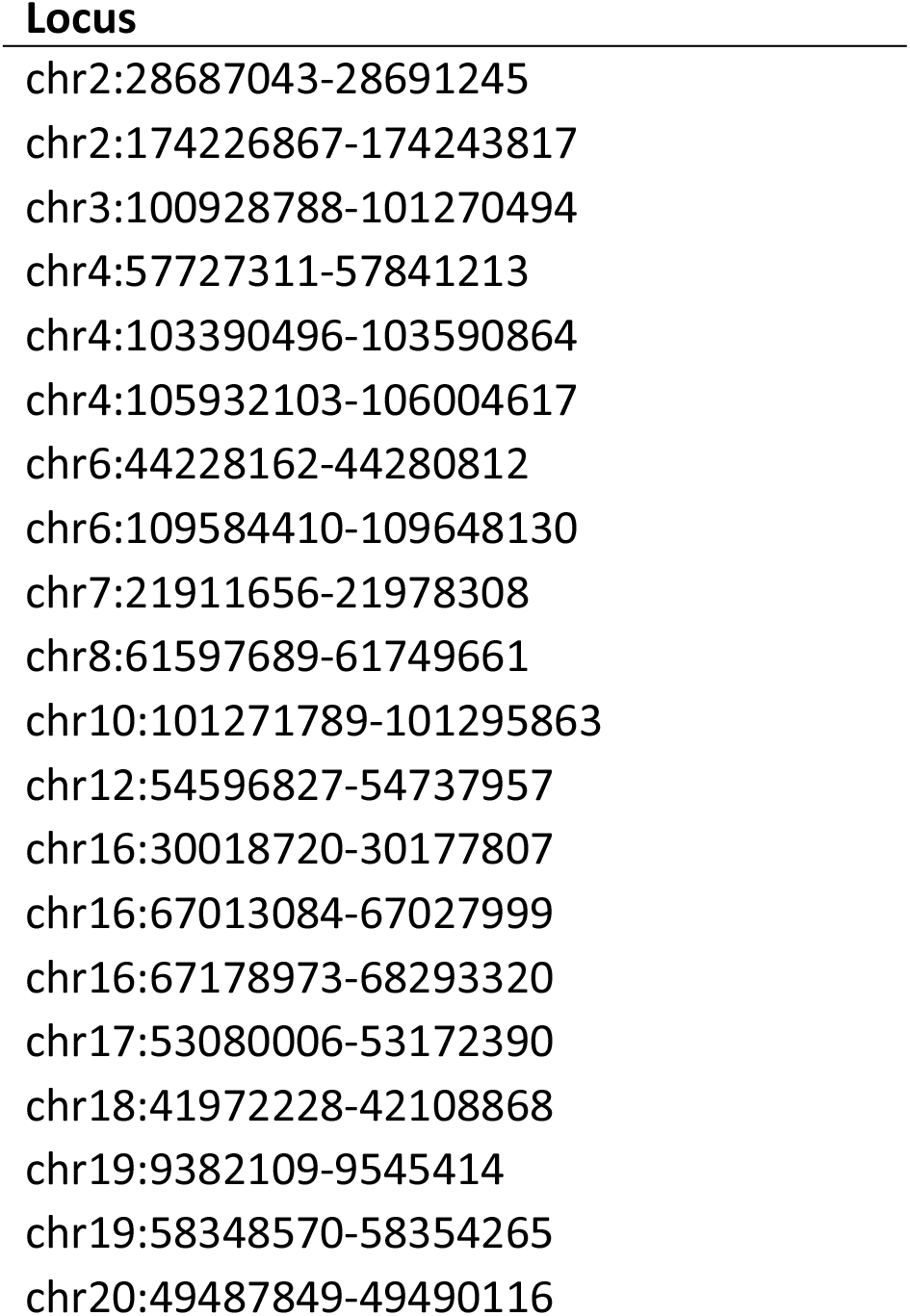
Distally-acting meQTL ‘hub’ loci, contiguous SNPs. Contiguous loci with < 100kb between SNPs associated with > 50 significant (p-value < 1x10^-13^) distal (SNP and CpG-site on different chromosomes, or > 10Mb between them) independent (Methods) CpG-site associations. Chromosome and position on GRCh37.

**Supplementary Table 2:**
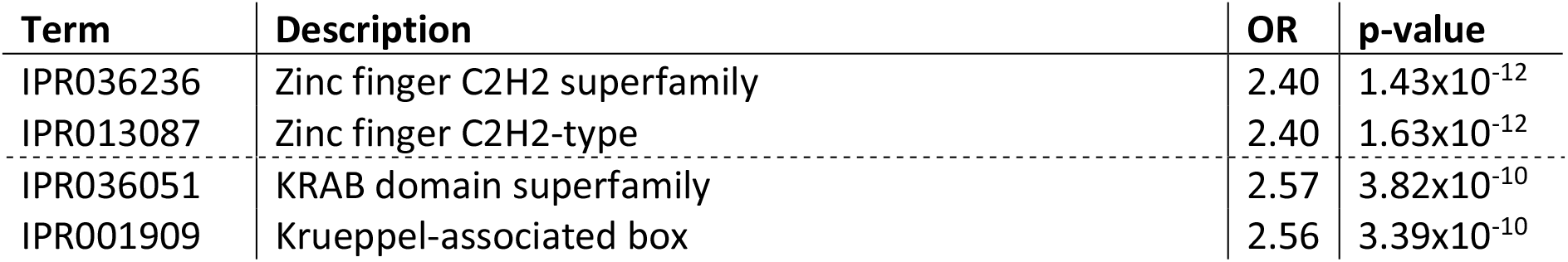
InterPro entries showing significant enrichment of distal-CpGeQTL genes. InterPro entries showing significant (Bonferroni correction, p-value < 0.05/6,790) enrichment of distal-CpG_eQTL_ genes, when compared to a random background (Methods). OR: odds-ratio of the enrichment; p-value: Fisher’s exact test p-value. The grouped rows are ‘Overlapping entries’. An InterPro entry is considered related to a homologous superfamily (i.e. overlapping) if its sequence matches overlap and either the Jaccard index (equivalent) or containment index (parent/child) of the matching sequence sets is greater than 0.75.

**Supplementary Table 3:** CpG-site individual- and region-based enrichment analysis – EWAS centric. Per trait, the number of factors (KRAB-ZFP factors, or TIF1-beta, *F_i_*) for which significant enrichment (or depletion) of trait-associated CpG-sites was identified within the binding-sites of *F_i_*. Significance defined as a p-value < 0.05/29,500 (236 ChIP-exo experiments x 125 EWAS traits) in the site-based analysis and an empirical p-value ≤ 0.05/25,000 in the region-based analyses. Enrichment (depletion) represents increased (decreased) variance explained by trait upon CpG-sites within the KRAB-ZFP or TIF1-beta binding-sites when compared to CpG-sites located elsewhere in the genome. Traits with no significant enrichments or depletions are not shown. ‘Site-based analysis’ is the analysis based upon individual independent CpG-sites; ‘Region-based analysis’ is that based upon contiguous regions with less that 20kb between CpG-sites (Supplementary Figure 1.1; Methods). ‘Binary overlap’ and ‘Base-pair overlap’ are the region-based analysis as shown in Supplementary Figure 1.3A, and 3B respectively (Methods). All ChIP-exo experimental runs were considered separately (222 unique factors; 236 ChIP-exo runs).

|  | CpG-site based analysis |  | Region-based analysis |  |  |  |
| --- | --- | --- | --- | --- | --- | --- |
|  | Enrichment | Depletion | Binary overlap |  | Base-pair overlap |  |
|  |  |  | Enrichment | Depletion | Enrichment | Depletion |
| Age | 1 | 4 | 1 | - | - | - |
| Age 4 vs. Age 0 | - | 23 | 3 | 1 | 2 | 1 |
| Ageing | - | 18 | - | 4 | - | - |
| Alcohol consumption per day | - | 1 | - | - | - | - |
| Arsenic exposure | 1 | - | - | - | - | - |
| Body mass index | - | 1 | 2 | - | 1 | - |
| C-reactive protein | - | - | 3 | - | - | - |
| Victim of Child abuse | 17 | - | 13 | - | 3 | - |
| Clear cell renal carcinoma | 1 | 53 | - | 22 | - | 17 |
| Fetal vs adult liver | - | 39 | 13 | 1 | 8 | 1 |
| Gestational age | - | 71 | 3 | 3 | 3 | 2 |
| HIV infection | 19 | - | 21 | - | 9 | - |
| Maternal smoking in pregnancy | - | 1 | 2 | - | - | - |
| Non-syndromic cleft lip and palate | 3 | - | 2 | - | - | - |
| Pancreatic ductal adenocarcinoma | 4 | 5 | 2 | 3 | - | - |
| Primary Sjogrens syndrome | - | 5 | 9 | - | 1 | - |
| Rheumatoid arthritis | 58 | 3 | 30 | - | 24 | - |
| Right vs. left colon adenomas | 2 | - | - | - | - | - |
| Sex | - | 10 | 6 | - | 2 | - |
| Smoking | - | 9 | 9 | - | 2 | - |
| Smoking pack-years | 2 | - | 1 | - | - | - |
| Tobacco smoking | - | - | 1 | - | - | - |

**Supplementary Table 4:**
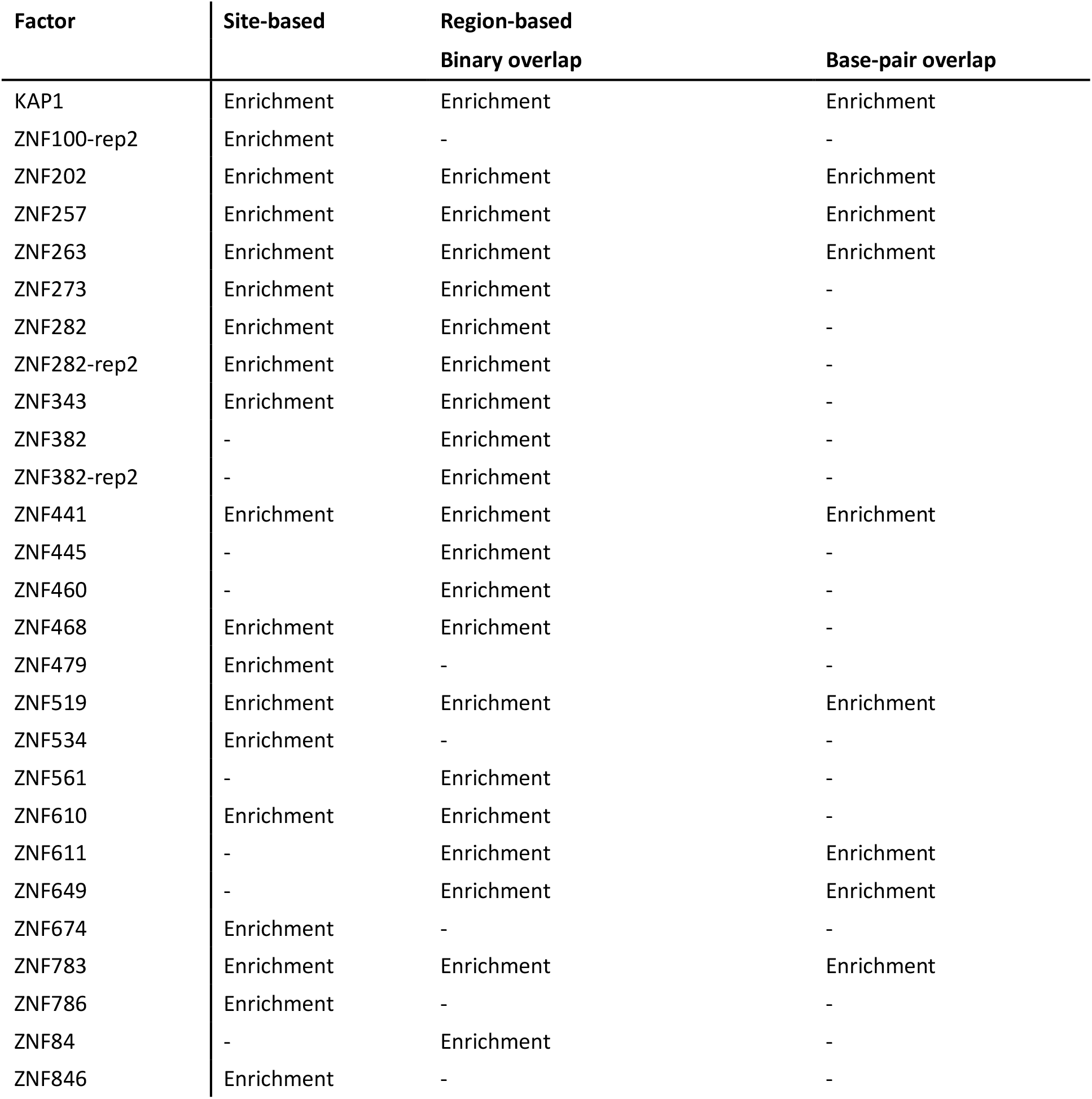
Significant enrichments or depletions of HIV infection associated CpG-sites within factor binding-sites. Factors, *F_i_*, within whose binding-sites there are significant enrichments or depletions of HIV infection associated CpG-sites, identified by EWAS. Significance defined as a p-value < 0.05/29,500 (236 ChIP-exo experiments x 125 EWAS traits) in the site-based analysis and an empirical p-value ≤ 0.05/25,000 in the region-based analyses. Enrichment (depletion) represents increased (decreased) variation explained by trait upon CpG-sites within the KRAB-ZFP or TIF1-beta binding-sites when compared to CpG-sites located elsewhere in the genome. Multiple ChIP-exo data sets for a single factor were considered individually. Where included in the table, traits or factors without significant enriched or depleted are indicated by ‘-‘. ‘Site-based analysis’ is the analysis based upon individual independent (Methods) CpG-sites; ‘Region-based analysis’ is that based upon contiguous regions with less that 20kb between CpG-sites (Supplementary Figure 1.1). ‘Binary overlap’ and ‘base-pair overlap’ are the region-based analysis as shown in Supplementary Figure 1.3A, and 3B respectively.

| Factor | Site-based | Region-based |  |
| --- | --- | --- | --- |
|  |  | Binary overlap | Base-pair overlap |
| KAP1 | Enrichment | Enrichment | Enrichment |
| ZNF100-rep2 | Enrichment | - | - |
| ZNF202 | Enrichment | Enrichment | Enrichment |
| ZNF257 | Enrichment | Enrichment | Enrichment |
| ZNF263 | Enrichment | Enrichment | Enrichment |
| ZNF273 | Enrichment | Enrichment | - |
| ZNF282 | Enrichment | Enrichment | - |
| ZNF282-rep2 | Enrichment | Enrichment | - |
| ZNF343 | Enrichment | Enrichment | - |
| ZNF382 | - | Enrichment | - |
| ZNF382-rep2 | - | Enrichment | - |
| ZNF441 | Enrichment | Enrichment | Enrichment |
| ZNF445 | - | Enrichment | - |
| ZNF460 | - | Enrichment | - |
| ZNF468 | Enrichment | Enrichment | - |
| ZNF479 | Enrichment | - | - |
| ZNF519 | Enrichment | Enrichment | Enrichment |
| ZNF534 | Enrichment | - | - |
| ZNF561 | - | Enrichment | - |
| ZNF610 | Enrichment | Enrichment | - |
| ZNF611 | - | Enrichment | Enrichment |
| ZNF649 | - | Enrichment | Enrichment |
| ZNF674 | Enrichment | - | - |
| ZNF783 | Enrichment | Enrichment | Enrichment |
| ZNF786 | Enrichment | - | - |
| ZNF84 | - | Enrichment | - |
| ZNF846 | Enrichment | - | - |

**Supplementary Table 5:**
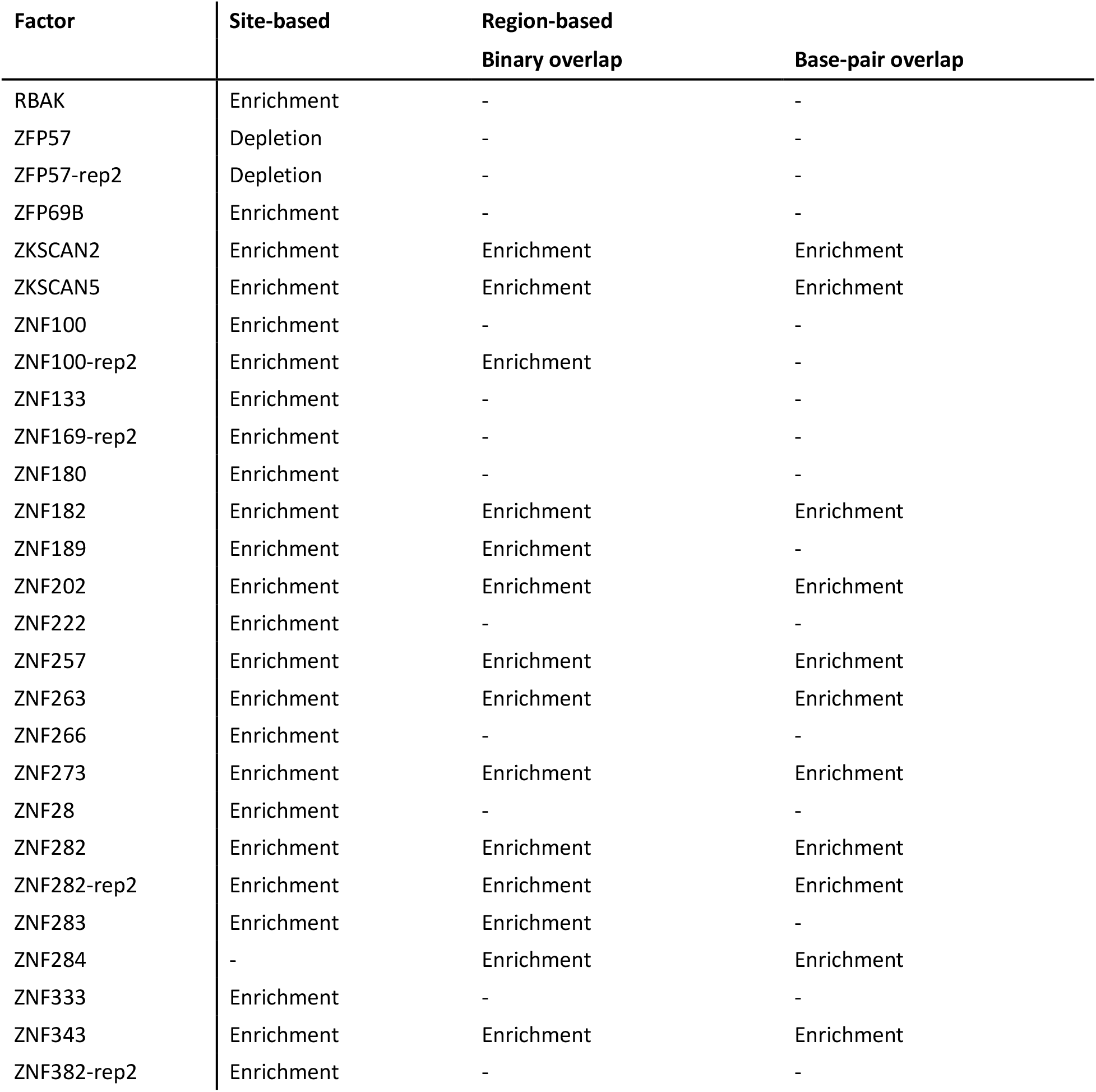
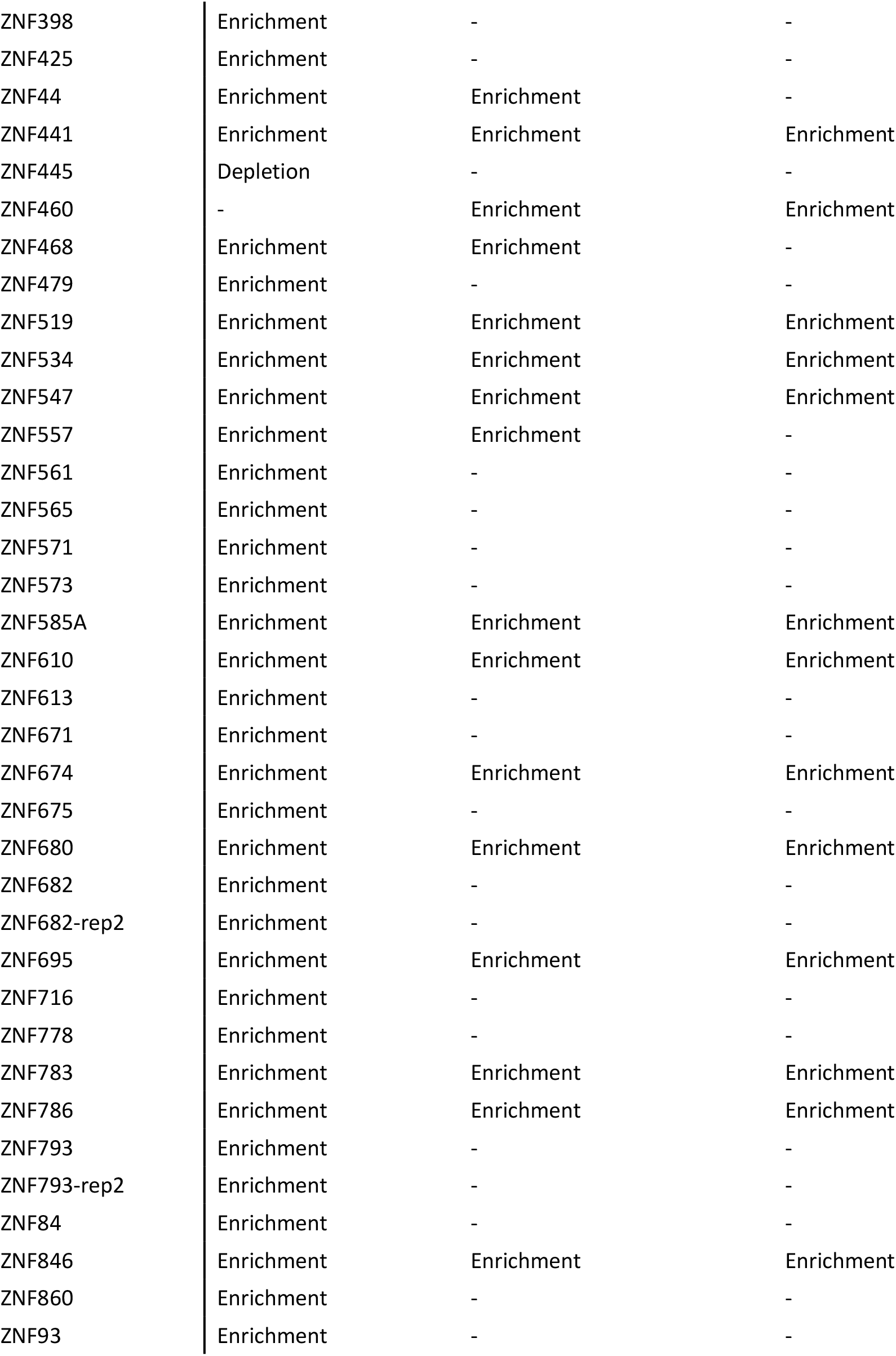
Significant enrichments or depletions of rheumatoid arthritis associated CpG-sites within factor binding-sites. Factors, *F_i_*, within whose binding-sites there were significant enrichments or depletions of rheumatoid arthritis associated CpG-sites, identified by EWAS. Significance defined as a p-value < 0.05/29,500 (236 ChIP-exo experiments x 125 EWAS traits) in the site-based analysis and an empirical p-value ≤ 0.05/25,000 in the region-based analyses. Enrichment (depletion) represents increased (decreased) variation explained by trait upon CpG-sites within the KRAB-ZFP or TIF1-beta binding-sites when compared to CpG-sites located elsewhere in the genome. Multiple ChIP-exo data sets for a single factor were considered individually. Where included in the table, traits or factors without significant enriched or depleted are indicated by ‘-‘. ‘Site-based analysis’ is the analysis based upon individual independent (Methods) CpG-sites; ‘Region-based analysis’ is that based upon contiguous regions with less that 20kb between CpG-sites (Supplementary Figure 1.1). ‘Binary overlap’ and ‘base-pair overlap’ are the region-based analysis as shown in Supplementary Figure 1.3A, and 3B respectively.

| Factor | Site-based | Region-based |  |
| --- | --- | --- | --- |
|  |  | Binary overlap | Base-pair overlap |
| RBAK | Enrichment | - | - |
| ZFP57 | Depletion | - | - |
| ZFP57-rep2 | Depletion | - | - |
| ZFP69B | Enrichment | - | - |
| ZKSCAN2 | Enrichment | Enrichment | Enrichment |
| ZKSCAN5 | Enrichment | Enrichment | Enrichment |
| ZNF100 | Enrichment | - | - |
| ZNF100-rep2 | Enrichment | Enrichment | - |
| ZNF133 | Enrichment | - | - |
| ZNF169-rep2 | Enrichment | - | - |
| ZNF180 | Enrichment | - | - |
| ZNF182 | Enrichment | Enrichment | Enrichment |
| ZNF189 | Enrichment | Enrichment | - |
| ZNF202 | Enrichment | Enrichment | Enrichment |
| ZNF222 | Enrichment | - | - |
| ZNF257 | Enrichment | Enrichment | Enrichment |
| ZNF263 | Enrichment | Enrichment | Enrichment |
| ZNF266 | Enrichment | - | - |
| ZNF273 | Enrichment | Enrichment | Enrichment |
| ZNF28 | Enrichment | - | - |
| ZNF282 | Enrichment | Enrichment | Enrichment |
| ZNF282-rep2 | Enrichment | Enrichment | Enrichment |
| ZNF283 | Enrichment | Enrichment | - |
| ZNF284 | - | Enrichment | Enrichment |
| ZNF333 | Enrichment | - | - |
| ZNF343 | Enrichment | Enrichment | Enrichment |
| ZNF382-rep2 | Enrichment | - | - |
| ZNF398 | Enrichment | - | - |
| ZNF425 | Enrichment | - | - |
| ZNF44 | Enrichment | Enrichment | - |
| ZNF441 | Enrichment | Enrichment | Enrichment |
| ZNF445 | Depletion | - | - |
| ZNF460 | - | Enrichment | Enrichment |
| ZNF468 | Enrichment | Enrichment | - |
| ZNF479 | Enrichment | - | - |
| ZNF519 | Enrichment | Enrichment | Enrichment |
| ZNF534 | Enrichment | Enrichment | Enrichment |
| ZNF547 | Enrichment | Enrichment | Enrichment |
| ZNF557 | Enrichment | Enrichment | - |
| ZNF561 | Enrichment | - | - |
| ZNF565 | Enrichment | - | - |
| ZNF571 | Enrichment | - | - |
| ZNF573 | Enrichment | - | - |
| ZNF585A | Enrichment | Enrichment | Enrichment |
| ZNF610 | Enrichment | Enrichment | Enrichment |
| ZNF613 | Enrichment | - | - |
| ZNF671 | Enrichment | - | - |
| ZNF674 | Enrichment | Enrichment | Enrichment |
| ZNF675 | Enrichment | - | - |
| ZNF680 | Enrichment | Enrichment | Enrichment |
| ZNF682 | Enrichment | - | - |
| ZNF682-rep2 | Enrichment | - | - |
| ZNF695 | Enrichment | Enrichment | Enrichment |
| ZNF716 | Enrichment | - | - |
| ZNF778 | Enrichment | - | - |
| ZNF783 | Enrichment | Enrichment | Enrichment |
| ZNF786 | Enrichment | Enrichment | Enrichment |
| ZNF793 | Enrichment | - | - |
| ZNF793-rep2 | Enrichment | - | - |
| ZNF84 | Enrichment | - | - |
| ZNF846 | Enrichment | Enrichment | Enrichment |
| ZNF860 | Enrichment | - | - |
| ZNF93 | Enrichment | - | - |

**Supplementary Table 6:** Significant enrichments or depletions of clear cell renal carcinoma associated CpG-sites within factor binding-sites. Factors, *F_i_*, within whose binding-sites there were significant enrichments or depletions of clear cell renal carcinoma associated CpG-sites, identified by EWAS. Significance defined as a p-value < 0.05/29,500 (236 ChIP-exo experiments x 125 EWAS traits) in the site-based analysis and an empirical p-value ≤ 0.05/25,000 in the region-based analyses. Enrichment (depletion) represents increased (decreased) variation explained by trait upon CpG-sites within the KRAB-ZFP or TIF1-beta binding-sites when compared to CpG-sites located elsewhere in the genome. Multiple ChIP-exo data sets for a single factor were considered individually. Where included in the table, traits or factors without significant enriched or depleted are indicated by ‘-‘. ‘Site-based analysis’ is the analysis based upon individual independent (Methods) CpG-sites; ‘Region-based analysis’ is that based upon contiguous regions with less that 20kb between CpG-sites (Supplementary Figure 1.1). ‘Binary overlap’ and ‘base-pair overlap’ are the region-based analysis as shown in Supplementary Figure 1.3A, and 3B respectively.

| Factor | Site-based | Region-based |  |
| --- | --- | --- | --- |
|  |  | Binary overlap | Base-pair overlap |
| KAP1 | - | Depletion | Depletion |
| ZFP57 | Enrichment | - | - |
| ZFP69B | Depletion | - | - |
| ZIM3 | - | Depletion | Depletion |
| ZKSCAN2 | Depletion | Depletion | Depletion |
| ZKSCAN5 | Depletion | Depletion | Depletion |
| ZNF100-rep2 | Depletion | - | - |
| ZNF141 | Depletion | - | - |
| ZNF180 | Depletion | - | - |
| ZNF189 | Depletion | Depletion | Depletion |
| ZNF202 | Depletion | Depletion | Depletion |
| ZNF257 | Depletion | Depletion | Depletion |
| ZNF263 | Depletion | Depletion | Depletion |
| ZNF273 | Depletion | Depletion | - |
| ZNF28 | Depletion | - | - |
| ZNF282 | Depletion | - | - |
| ZNF282-rep2 | Depletion | - | - |
| ZNF283 | Depletion | - | - |
| ZNF284 | - | Depletion | Depletion |
| ZNF333 | Depletion | - | - |
| ZNF343 | Depletion | Depletion | Depletion |
| ZNF382-rep2 | Depletion | - | - |
| ZNF383 | Depletion | - | - |
| ZNF425 | Depletion | - | - |
| ZNF44 | Depletion | - | - |
| ZNF441 | Depletion | Depletion | Depletion |
| ZNF460 | Depletion | Depletion | Depletion |
| ZNF468 | Depletion | - | - |
| ZNF479 | Depletion | - | - |
| ZNF519 | Depletion | Depletion | Depletion |
| ZNF534 | Depletion | Depletion | - |
| ZNF543 | Depletion | - | - |
| ZNF547 | Depletion | - | - |
| ZNF549-rep2 | Depletion | - | - |
| ZNF557 | Depletion | Depletion | - |
| ZNF561 | Depletion | - | - |
| ZNF565 | Depletion | - | - |
| ZNF566 | Depletion | - | - |
| ZNF571 | Depletion | - | - |
| ZNF585A | Depletion | Depletion | - |
| ZNF610 | Depletion | - | - |
| ZNF611 | Depletion | - | Depletion |
| ZNF613 | Depletion | - | - |
| ZNF671 | Depletion | - | - |
| ZNF674 | Depletion | - | - |
| ZNF675 | Depletion | - | - |
| ZNF680 | Depletion | - | - |
| ZNF682 | Depletion | - | - |
| ZNF682-rep2 | Depletion | - | - |
| ZNF695 | Depletion | Depletion | Depletion |
| ZNF765 | Depletion | - | - |
| ZNF766 | - | Depletion | - |
| ZNF778 | Depletion | - | - |
| ZNF783 | Depletion | Depletion | Depletion |
| ZNF786 | Depletion | Depletion | Depletion |
| ZNF84 | Depletion | - | - |
| ZNF846 | Depletion | Depletion | - |
| ZNF860 | Depletion | - | - |

## Supplementary File 1

LocusZoom (Pruim et al., 2010) plots of the regions underlying the loci detailed in Supplementary Table 1. x-axis: genomic position of the SNP; y-axis: count (in thousands) of the number of independent CpG-sites to which each SNP is significantly (p-value < 1x10^-13^) associated. Linkage disequilibrium is displayed with respect to the SNP with the largest number of independent CpG-sites associated with it. Plotted SNPs are only those that are associated with ≥ 1 significant distal independent CpG-sites.

## Supplementary File 2

Full results of the enrichment results of those genes under the control of a locally-acting eQTL affecting the methylation status of at least one CpG-site at a distal site compared to all genes for which a significant (as per GTEx v7) locally-acting eQTL was identified in GTEx.

Gzipped, tab-separated file. Columns:

- ’category’: source (Component: Cellular Component (GO), Function: Molecular Function (GO), InterPro: INTERPRO Protein Domains and Features, Keyword: UniProt Keywords, Pfam: PFAM Protein Domains, Process: Biological Process (GO), RCTM: Reactome Pathways, SMART: SMART Protein Domains).

- ‘term’: term from ‘category’.

- ’description’: verbose description of ‘term’.

- ’OR’: odds ratio.

- ’pval’: p-value of the odds ratio (‘OR’).

- ’signif’: is the odds ratio (OR) significantly different from 1 (True/False).

- ’n_category’: Number of tested ‘terms’ within ‘category’.

- ’number_of_genes_in_set_subset’: Number of genes in the set (those associated with the ‘term’ in question) present within the subset of interest (colocalising locally-acting eQTL and distally-acting meQTL).

- ’inputGenes_subset’: Ensembl gene IDs of those genes in ‘’number_of_genes_in_set_subset’

- ’preferredNames_subset’: StringDB preferred names of those genes in ’number_of_genes_in_set_subset’

- ’n_subset’: Number of genes in the input subset.

- ’number_of_genes_in_set_all’: Number of genes in the set (those associated with the ‘term’ in question) present within all genes with a significant (as per GTEx v7) locally-acting eQTL.

- ’inputGenes_all’: Ensembl gene IDs of those genes in ’number_of_genes_in_set_all’.

- ’preferredNames_all’: StringDB preferred names of those genes in ’number_of_genes_in_set_all’

- ’n_all’: Number of genes in GTEx (v7) with a significant (as per GTEx v7) locally-acting eQTL.

## Supplementary File 3

Enrichment results of those genes under the control of a locally-acting eQTL affecting the methylation status of at least one CpG-site at a distal site compared to an additional background. The additional background was identified as follows: a set of SNPs of equivalent size to the number of distally-acting meQTL was selected at random from all significant (p-value < 1x10^-13^) meQTL. Then, genes that had ≥ 1 colocalised significant (as per GTEx v7) locally-acting eQTL from GTEx with this set were identified and used as background.

Gzipped, tab-separated file. Columns:

- ‘term’: term from ‘category’.

- ’description’: verbose description of ‘term’.

- ’OR’: odds ratio.

- ’pval’: p-value of the odds ratio (‘OR’).

- ’signif’: is the odds ratio (OR) significantly different from 1 (True/False).

- ’n_category’: Number of tested ‘terms’ within ‘category’.

- ’number_of_genes_in_set_dist’: Number of genes in the set (those associated with the ‘term’ in question) present within the subset of interest (colocalising locally-acting eQTL and distally-acting meQTL).

- ’inputGenes_dist’: Ensembl gene IDs of those genes in ’number_of_genes_in_set_dist’

- ’preferredNames_dist’: StringDB preferred names of those genes in ’number_of_genes_in_set_dist’

- ’n_dist’: Number of genes in the input subset.

- ’number_of_genes_in_set_rand’: Number of genes in the set (those associated with the ‘term’ in question) present within the randomly selected background.

- ’inputGenes_rand’: Ensembl gene IDs of those genes in ’number_of_genes_in_set_rand’.

- ’preferredNames_rand’: StringDB preferred names of those genes in ’number_of_genes_in_set_rand’

- ’n_rand’: Number of genes in the randomly selected background.

## Supplementary File 4

Full results of enrichments of differentially methylated CpG-sites, with respect to KRAB-ZFP locally-acting (TSS +/- 1Mb) eQTL within its binding-sites +/- 1kb. Limited to the independent CpG set.

Tab-separated file. Columns:

- ‘ZFP’: The KRAB domain containing zinc finger protein the locally-acting eQTL with respect to which the CpG-sites are differentially methylated.

- ‘crosstab’: List [[a,b],[c,d]]:

`o` a: Number of distal CpG-sites significantly (Bonferroni correction; p-value threshold 1x10^-13^) affected by a locally-acting eQTL of the KRAB-ZFP within one of its binding-sites +/- 1kb.
`o` b: Number of CpG-sites not significantly (Bonferroni correction; p-value threshold 1x10^-13^) affected by a locally-acting eQTL of the KRAB-ZFP within one of its binding-sites +/- 1kb.
`o` c: Number of distal CpG-sites significantly (Bonferroni correction; p-value threshold 1x10^-13^) affected by a locally-acting eQTL of the KRAB-ZFP not within one of its binding-sites +/- 1kb.
`o` d: Number of CpG-sites not significantly (Bonferroni correction; p-value threshold 1x10^-13^) affected by a locally-acting eQTL of the KRAB-ZFP not within one of its binding-sites +/- 1kb.

- ‘OR’: The odds ratio of the crosstab.

- ‘p’: The p-value associated with the odds ratio.

## Supplementary File 5

‘CpGZoom’ of all significant (p-value < 1x10^-13^) distal-CpG_eQTL_ peaks. Plotting was iterated until all significant CpG-sites were included. Lead-CpG_eQTL_ ±25kb plotted in each case. A positive t-statistic (y-axis) indicates that CpG-site methylation (additive effect) goes up as the mRNA of the factor increases. The lead-eQTL for all C2H2 KRAB-ZFPs (Lambert et al., 2018) from GTEx (v7) Whole Blood, were included. Those C2H2 KRAB-ZFPs (Lambert et al., 2018) with ChIP-exo data (Imbeault et al., 2017) are included (690 plots; 64 factors).

CpGZoom (modified LocusZoom (Pruim et al., 2010)): each point on the plot indicates a CpG (not a SNP); the x-axis is genomic position; the y-axis is the t-statistic of the additive component of the meQTL model (note that a positive t-statistic indicates that methylation (additive effect) of that CpG increases with increased expression of the denoted KRAB-ZFP / TIF1-beta). Banners are ChIP-exo defined binding-sites (Imbeault et al., 2017). ‘cpgIslandExt’ is the UCSC CpG Islands Track of the same name (http://genome-euro.ucsc.edu/). Gene annotation is as per GENCODE, CpG-sites without an association with the lead-eQTL of the KRAB-ZFP / TIF1-beta at a p-model of <1x10^-3^ are recorded as a t-statistic of zero on this plot (results with a p-model > 1x10^-3^ not retained during processing due to storage limitations) – gaps therefore mean an absence of measured CpG-sites, not an absolute absence of association.

## Supplementary File 6

As per Supplementary File 5, but for the C2H2 KRAB-ZFPs (Lambert et al., 2018) without ChIP-exo data (Imbeault et al., 2017) (302 plots; 32 factors).

## Supplementary File 7

High- and good-quality trait-factor associations. A high-quality prediction is one that was found to be significant in all three analyses: site-based, region-based – binary overlap, and region-based – base-pair overlap. A good-quality prediction was one that was significant in the site-based and one of the region-based analyses. See Main Text for further detail.

Further detail of the results of the site- and region-based analyses can be found in Supplementary File 7 and Supplementary File 8, respectively. Significance defined as a p-value < 0.05/29,500 (236 ChIP-exo experiments x 125 EWAS traits) in the site-based analysis and an empirical p-value ≤ 0.05/25,000 in the region-based analyses.

## Supplementary File 8

ChIP-exo / EWAS enrichment results. Limited to the independent CpG-site set. Tab-separated file.

Columns:

- ’EWAS’: The EWAS study name from the EWAS catalog.

- ’ZFP’: The ZFP against which the ChIP-exo was performed.

- ’n_exoseq’: Number of ChIP-exo defined binding sites.

- ’crosstab_indep’: list [[a,b],[c,d]]

- a: Number of differentially methylated CpG-sites within a ChIP-exo defined binding-site.

- b: Number of not differentially methylated CpG-sites within a ChIP-exo defined binding-site.

- c: Number of differentially methylated CpG-sites not within a ChIP-exo defined binding-site.

- d: Number of not differentially methylated CpG-sites not within a ChIP-exo defined binding-site.

- a+b+c+d = Number of CpG-sites with in the manifest.

- a+b = Number of CpG-sites within a ChIP-exo defined binding site.

- a+c = Number of differentially methylated CpG-sites.

- ’OR_indep’: The odds ratio of the crosstab.

- ’p_indep’: Fisher’s exact test p-value of the crosstab.

- ’n_exoseq_cpg_indep’: Number of ChIP-exo defined binding sites with one or more CpG.

- ’n_exoseq_dms_indep’: Number of ChIP-exo defined binding sites with one or more differentially methylated CpG.

## Supplementary File 9

Significant (empirical p-value < 0.05/25,000) results of CpG-region analyses described above.

- bp_overlap: Number of base-pairs within EWAS associated regions overlapping a factor binding-site.

- bp: Number of base-pairs within EWAS associated regions.

- n_intervals_with_overlap: Number of EWAS associated regions containing an overlap with a factor binding-site.

- n_intervals: Number of EWAS associated regions.

- frac_bp_overlap: ’bp_overlap’/’bp’

- ewas: The trait under study with respect to the EWAS.

- zfp: The factor under study with respect to the binding-sites.

- nperms_gt: The number of permutations that were greater than the real data.

- nperms_eq: The number of permutations that were equal to the real data.

- nperms_lt: The number of permutations that were less than the real data.

- measure: The measure with which the real data was compared to the permutations.

## Supplementary File 10

Direction-of-effect on methylation amongst CpG-sites differentially methylated between HIV cases and controls, within and out with KRAB-ZFP and TIF1-beta ChIP-exo defined binding-sites. This analysis was undertaken using the independent set of CpGs, as described in the Methods.

- tissue: tissue the EWAS results were derived from.

- zfp: the KRAB-ZFP to which the ChIP-exo defined binding-sites correspond.

- n_exo: the number of ChIP-exo defined binding-sites with at least one differentially methylated CpG within it.

- n_cpg_exo: number of differentially methylated CpG-sites within a ChIP-exo defined binding-site.

- mean_exo: the mean beta value of the CpG-sites within a ChIP-exo defined binding-site.

- sd_exo: the standard deviation of ’mean_exo’.

- n_cpg_notexo: number of differentially methylated CpG-sites NOT within a ChIP-exo defined binding-site.

- mean_notexo: the mean beta value of the CpG-sites NOT within a ChIP-exo defined binding-site.

- sd_notexo: the standard deviation of ’mean_notexo’.

- t: t-statistic comparing ’mean_exo’ and ’mean_notexo’.

- pval: the p-value of ’t’.

- min_t_iter1000: the minimum t-statistic encountered in 1,000 iterations of the more conservative test (2), detailed above.

- max_t_iter1000: the maximum t-statistic encountered in 1,000 iterations of the more conservative test (2), detailed above.

- min_p_iter1000: the minimum p-value encountered in 1,000 iterations of the more conservative test (2), detailed above.

- max_p_iter1000: the maximum p-value encountered in 1,000 iterations of the more conservative test (2), detailed above.

## Supplementary File 11

‘df_ra_ewas_hits_binding_sites.tsv’

As per Supplementary File 10, but with respect to rheumatoid arthritis, rather than HIV infection.

## Supplementary File 12

EWAS catalog original sources’ list references.

‘ewas_catalog_sources.tsv’ Tab-separated file.

Columns:

- ‘Author’: the first author of the publication.

- ‘PMID’: the PubMed ID of the publication.

- ‘Trait’: the phenotype under study.

