## Supplementary File 1 for "Genetic control of KRAB-ZFP genes explains distal CpG-site methylation which associates with human disease phenotypes"

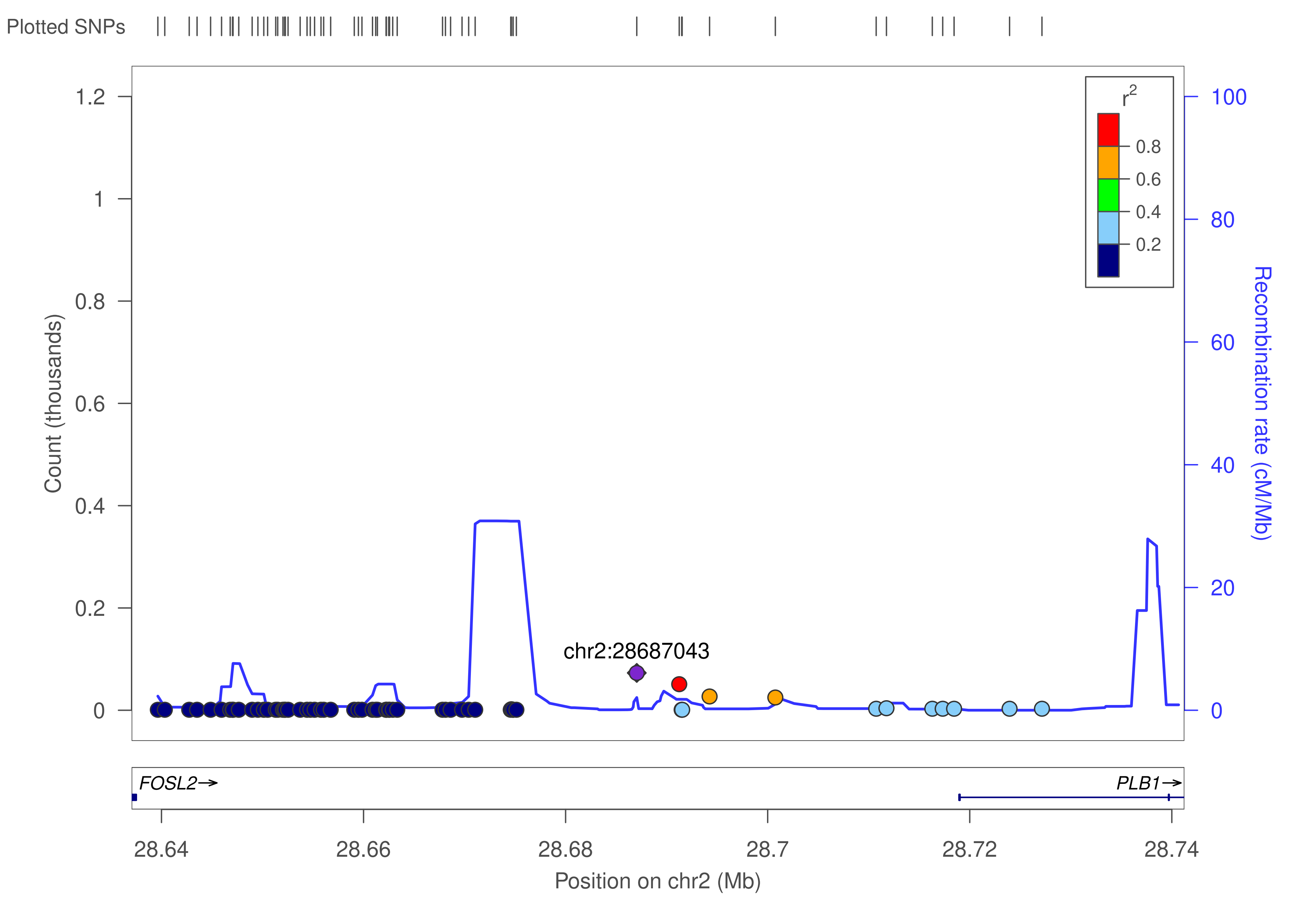

Plotted SNPs
Count (thousands)
1.2
1
0.8
0.6
0.4
0.2
0
Recombination rate (cM/Mb)
100
80
60
40
20
0
r2
0.8
0.6
0.4
0.2
chr2:28687043
FOSL2→
PLB1→
28.64
28.66
28.68
28.7
28.72
28.74
Position on chr2 (Mb)

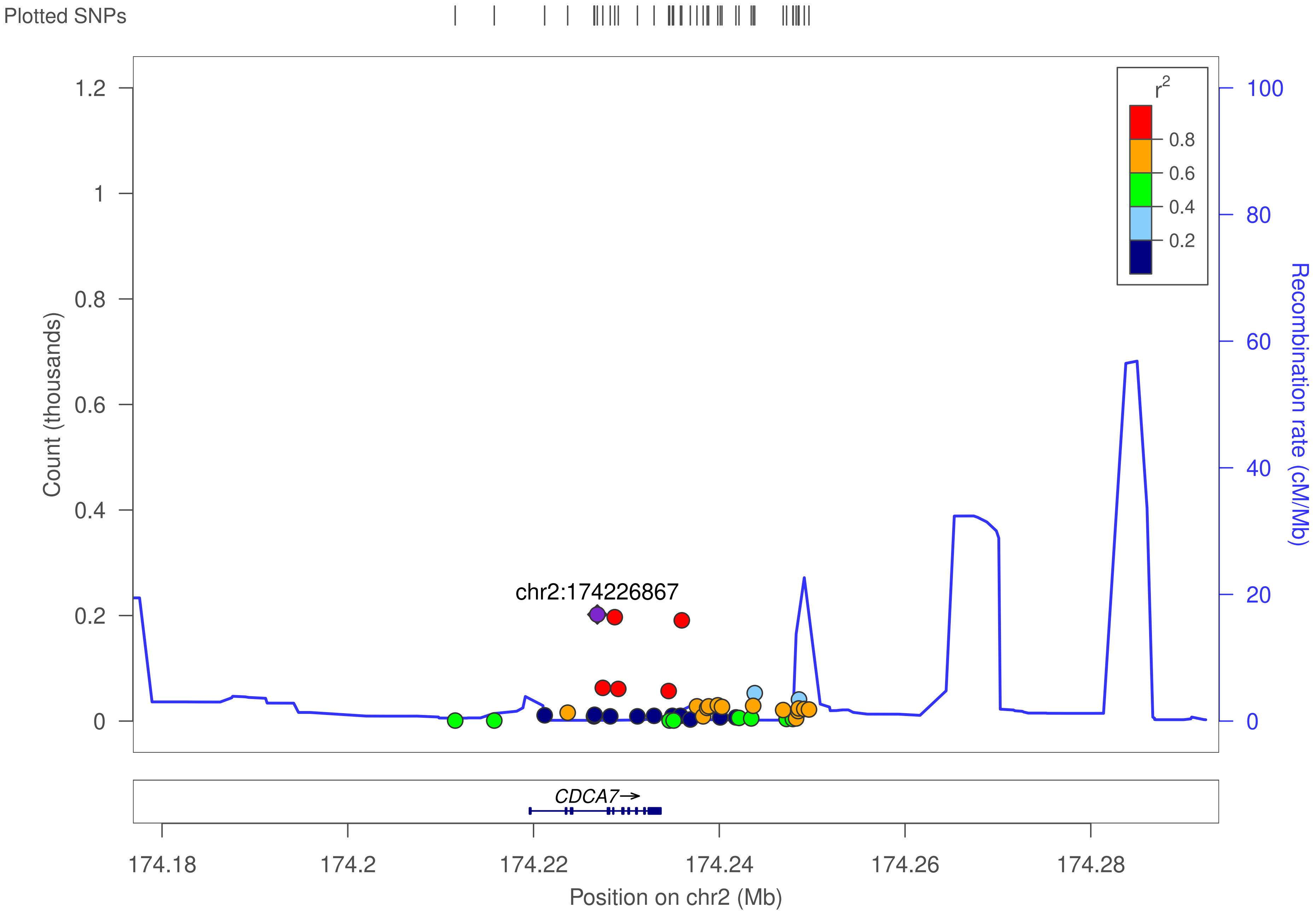
Plotted SNPs
r2
0.8
0.6
0.4
0.2
1.2
1
0.8
0.6
0.4
0.2
0
Count (thousands)
100
80
60
40
20
0
Recombination rate (cM/Mb)
chr2:174226867
CDCA7→
174.18
174.2
174.22
174.24
174.26
174.28
Position on chr2 (Mb)

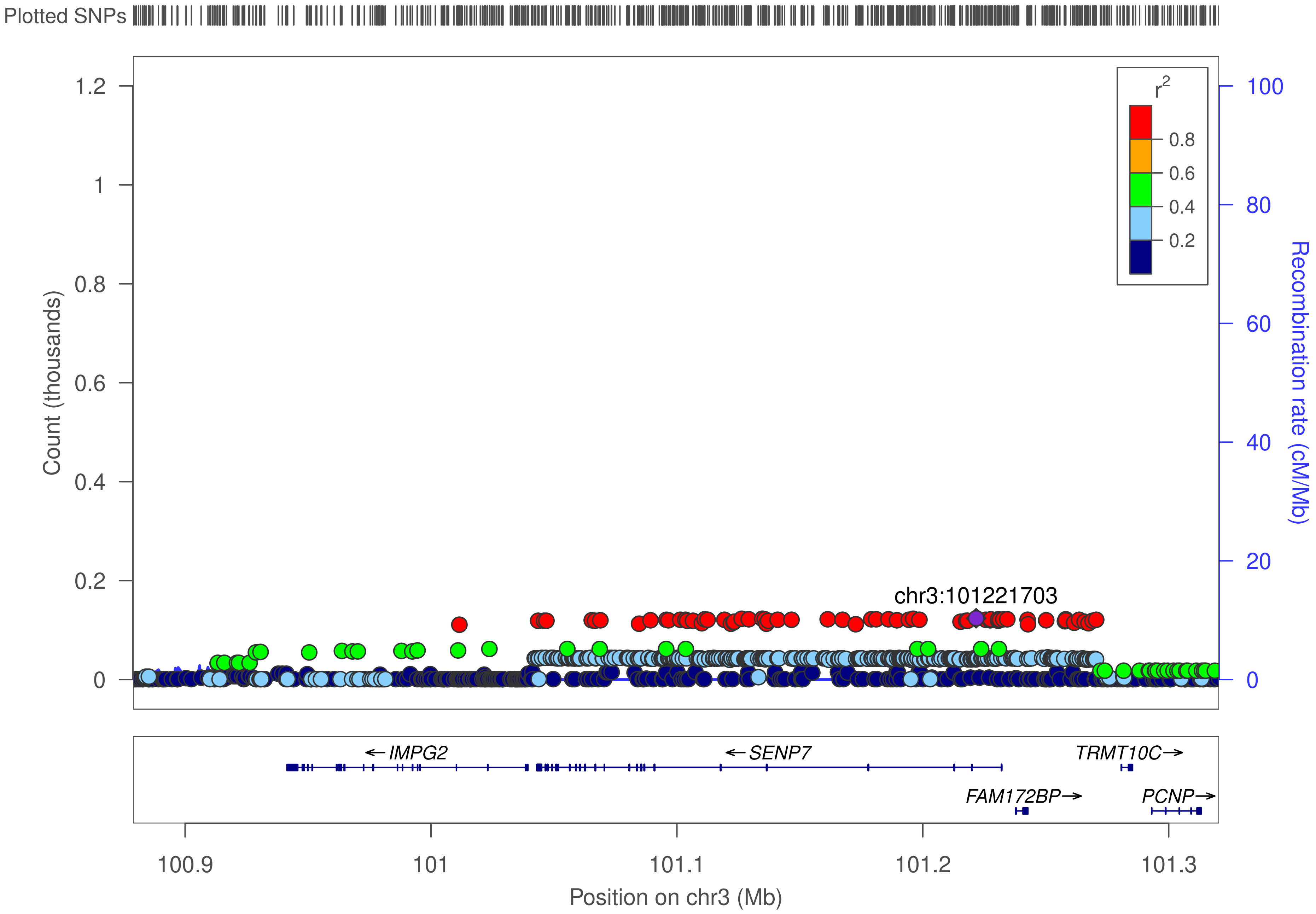

Plotted SNPs
Count (thousands)
1.2
1
0.8
0.6
0.4
0.2
0
r2
0.8
0.6
0.4
0.2
100
80
60
40
20
0
Recombination rate (cM/Mb)
chr3:101221703
← IMPG2
← SENP7
TRMT10C →
FAM172BP →
PCNP →
100.9
101
101.1
101.2
101.3
Position on chr3 (Mb)

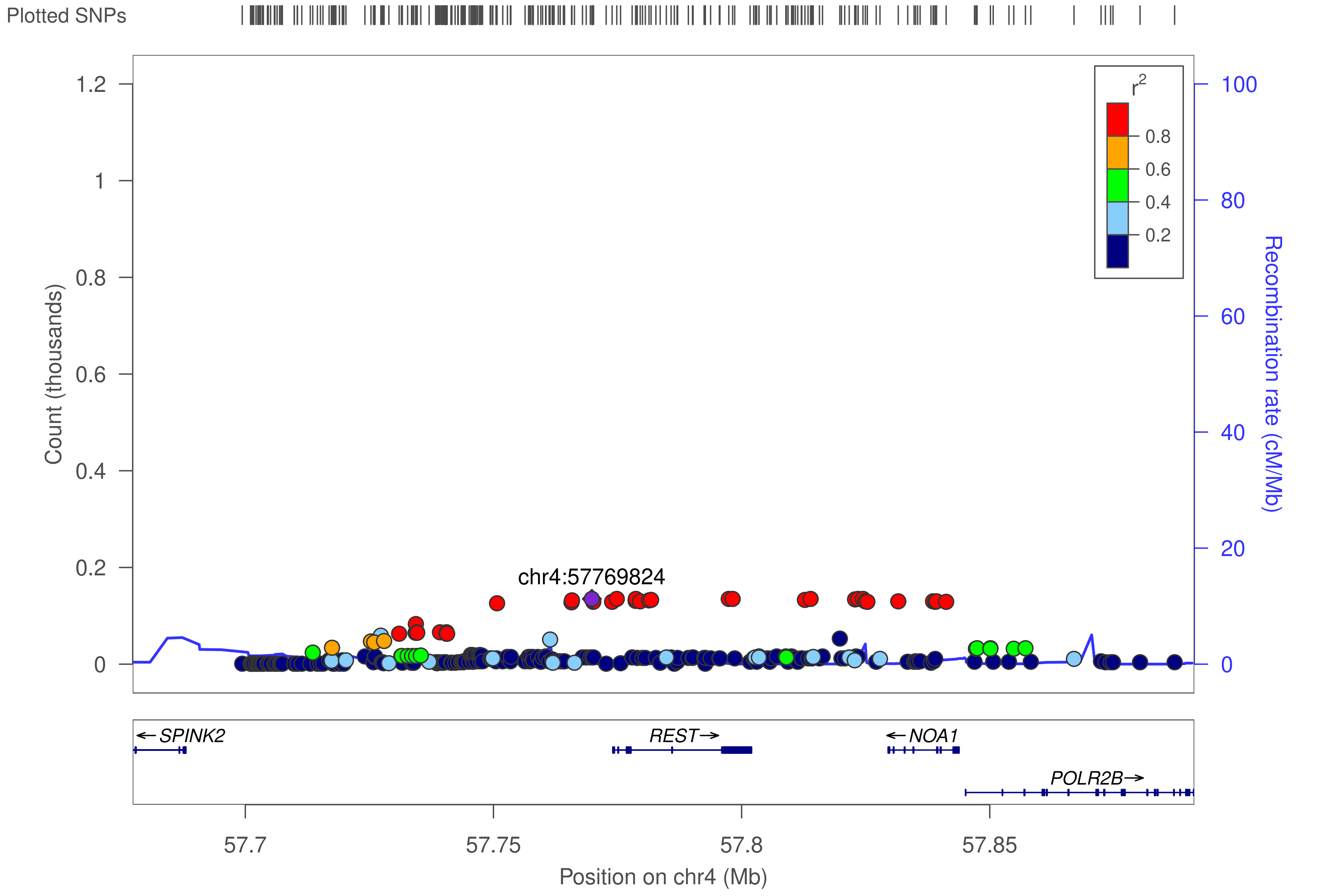

Plotted SNPs
Count (thousands)
1.2
1
0.8
0.6
0.4
0.2
0
r2
0.8
0.6
0.4
0.2
100
80
60
40
20
0
Recombination rate (cM/Mb)
chr4:57769824
← SPINK2
REST→
← NOA1
POLR2B→
57.7
57.75
57.8
57.85
Position on chr4 (Mb)

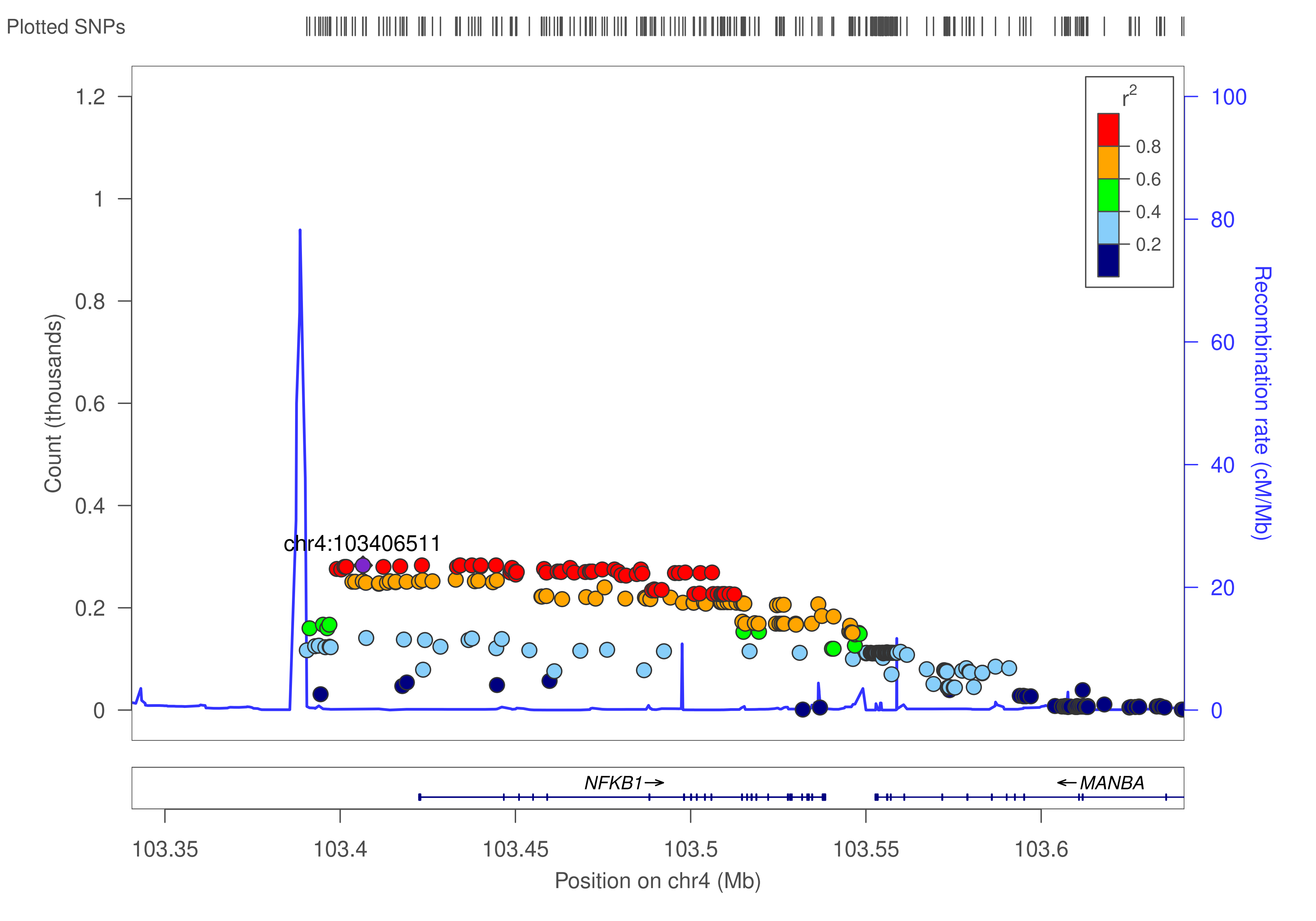

Plotted SNPs
chr4:103406511
Count (thousands)
0
0.2
0.4
0.6
0.8
1
1.2
Recombination rate (cM/Mb)
0
20
40
60
80
100
$r^2$
0.8
0.6
0.4
0.2
NFKB1→
←MANBA
103.35
103.4
103.45
103.5
103.55
103.6
Position on chr4 (Mb)

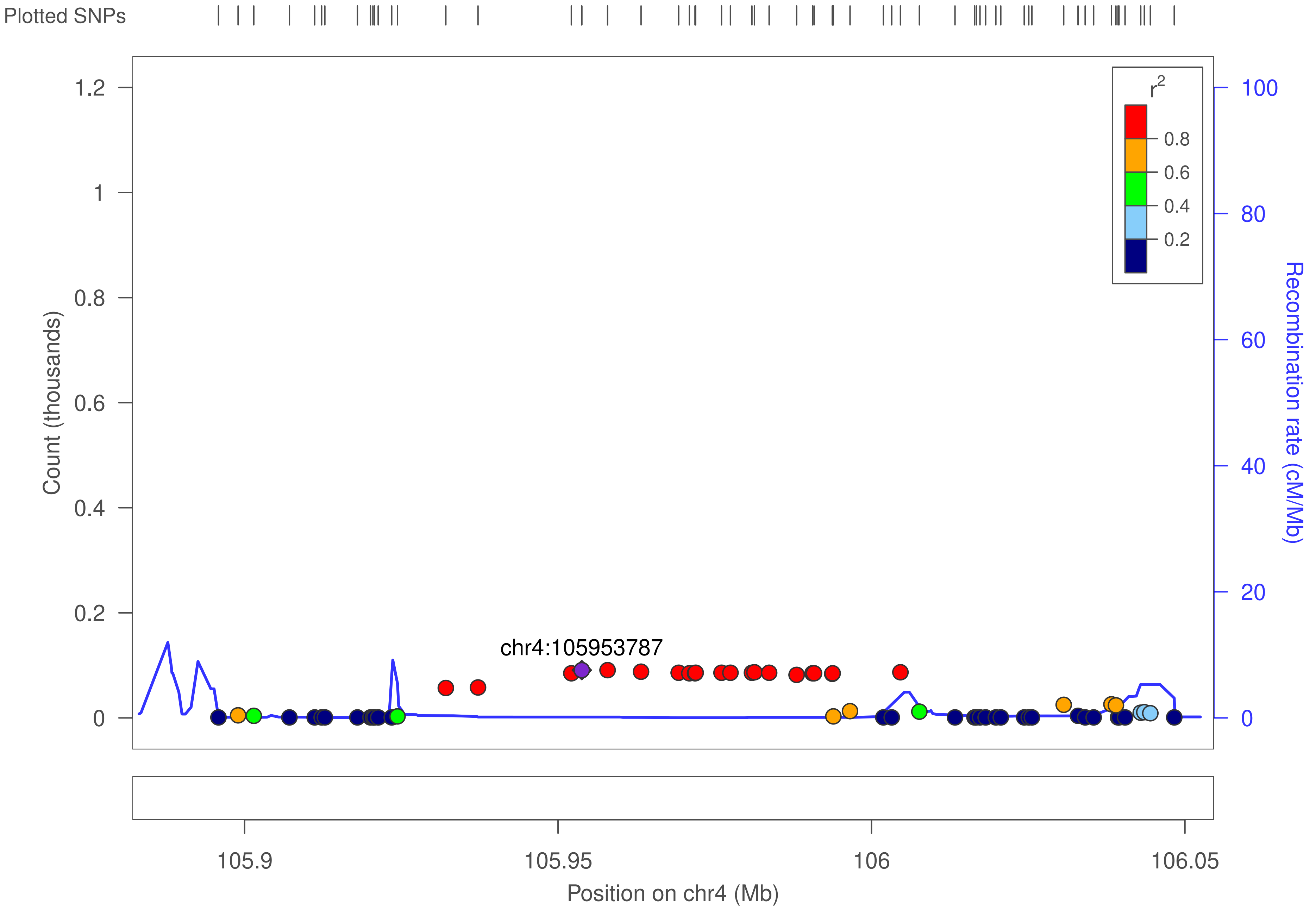

Plotted SNPs
Count (thousands)
1.2
1
0.8
0.6
0.4
0.2
0
chr4:105953787
r²
0.8
0.6
0.4
0.2
100
80
60
40
20
0
Recombination rate (cM/Mb)
105.9
105.95
106
106.05
Position on chr4 (Mb)

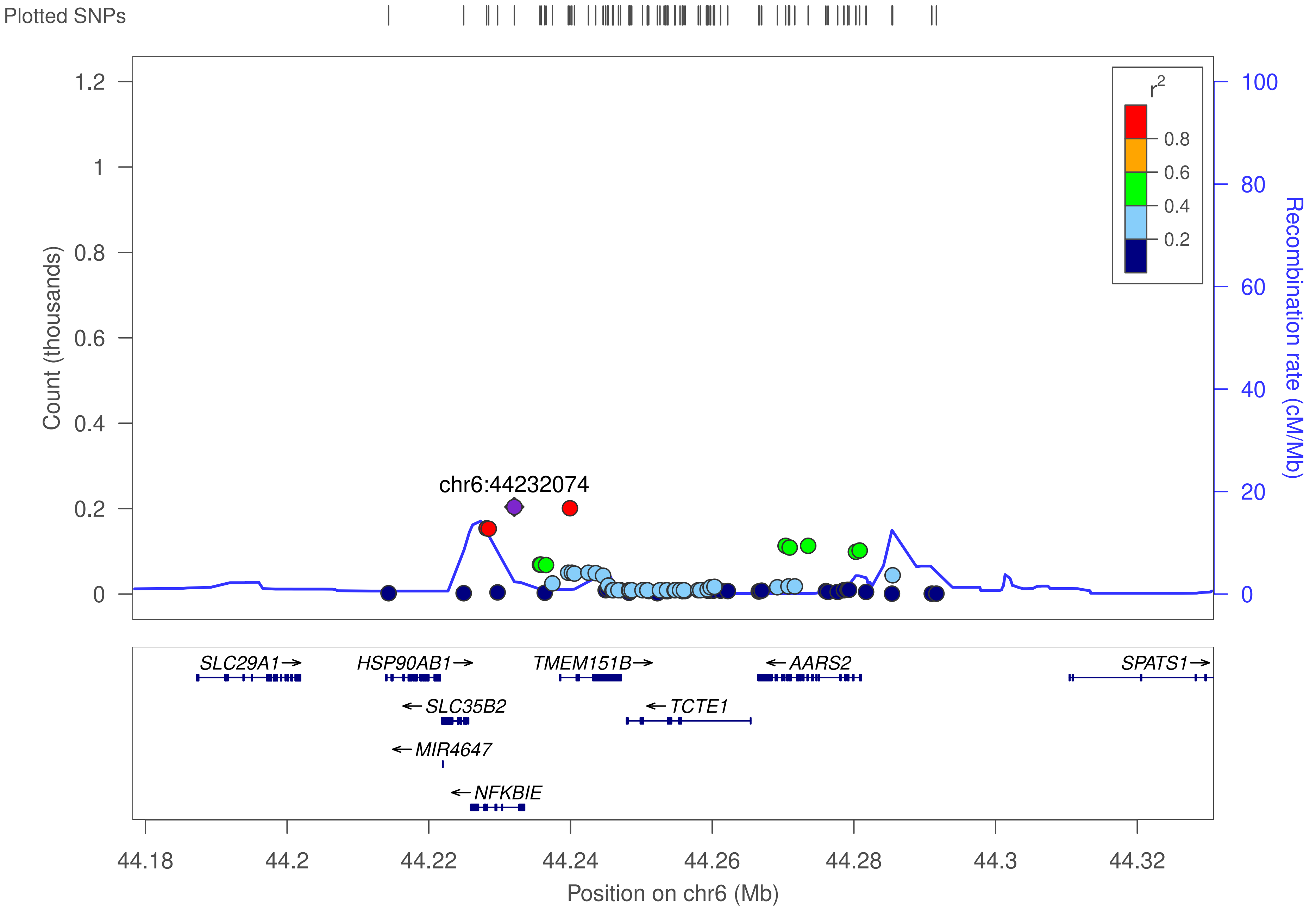

Plotted SNPs
Count (thousands)
1.2
1
0.8
0.6
0.4
0.2
0
chr6:44232074
r2
0.8
0.6
0.4
0.2
100
80
60
40
20
0
Recombination rate (cM/Mb)
SLC29A1→
HSP90AB1→
TMEM151B→
←AARS2
SPATS1→
←SLC35B2
←TCTE1
←MIR4647
←NFKBIE
44.18
44.2
44.22
44.24
44.26
44.28
44.3
44.32
Position on chr6 (Mb)

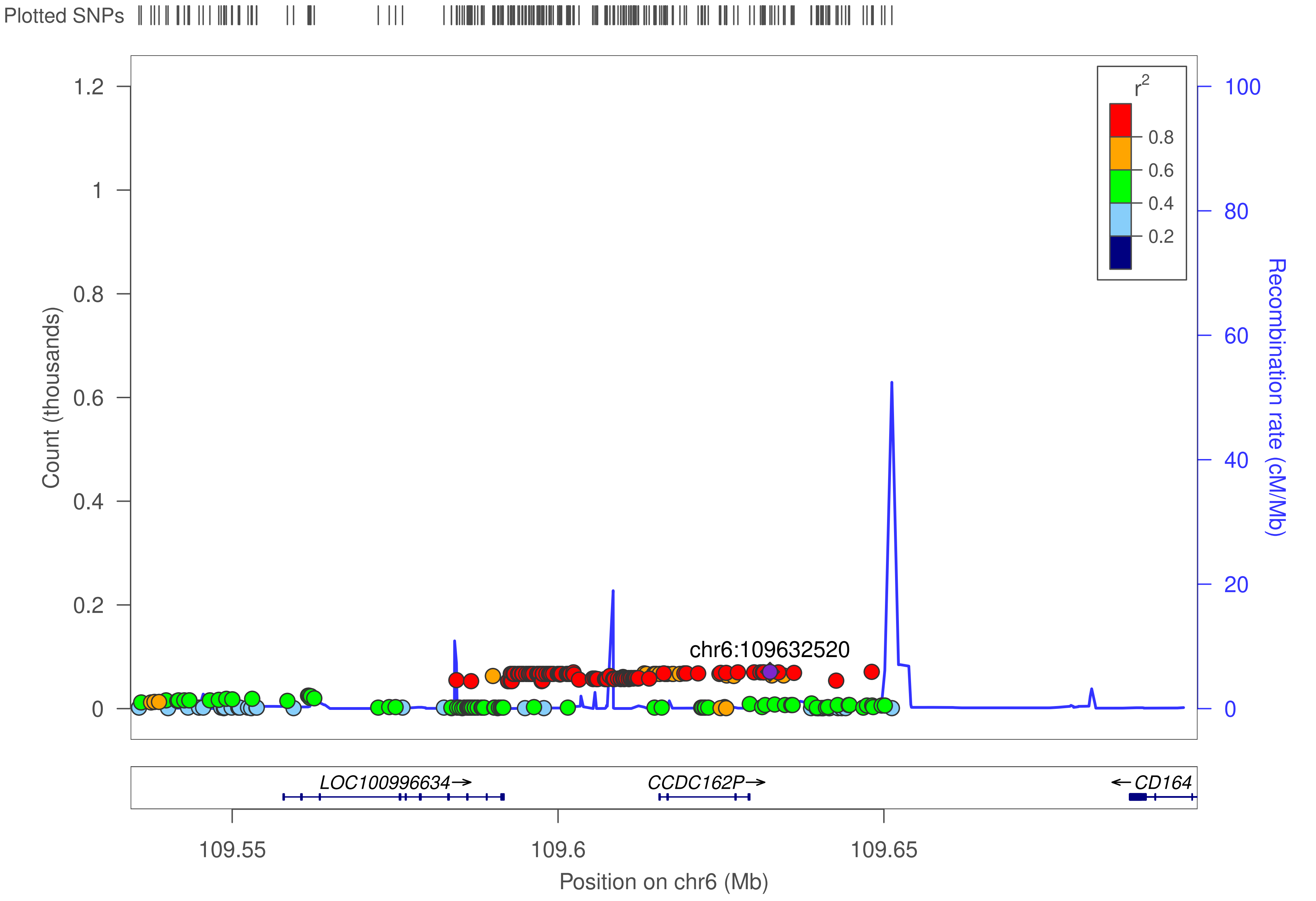

Plotted SNPs
Count (thousands)
1.2
1
0.8
0.6
0.4
0.2
0
r2
0.8
0.6
0.4
0.2
100
80
60
40
20
0
Recombination rate (cM/Mb)
chr6:109632520
LOC100996634→
CCDC162P→
←CD164
109.55
109.6
109.65
Position on chr6 (Mb)

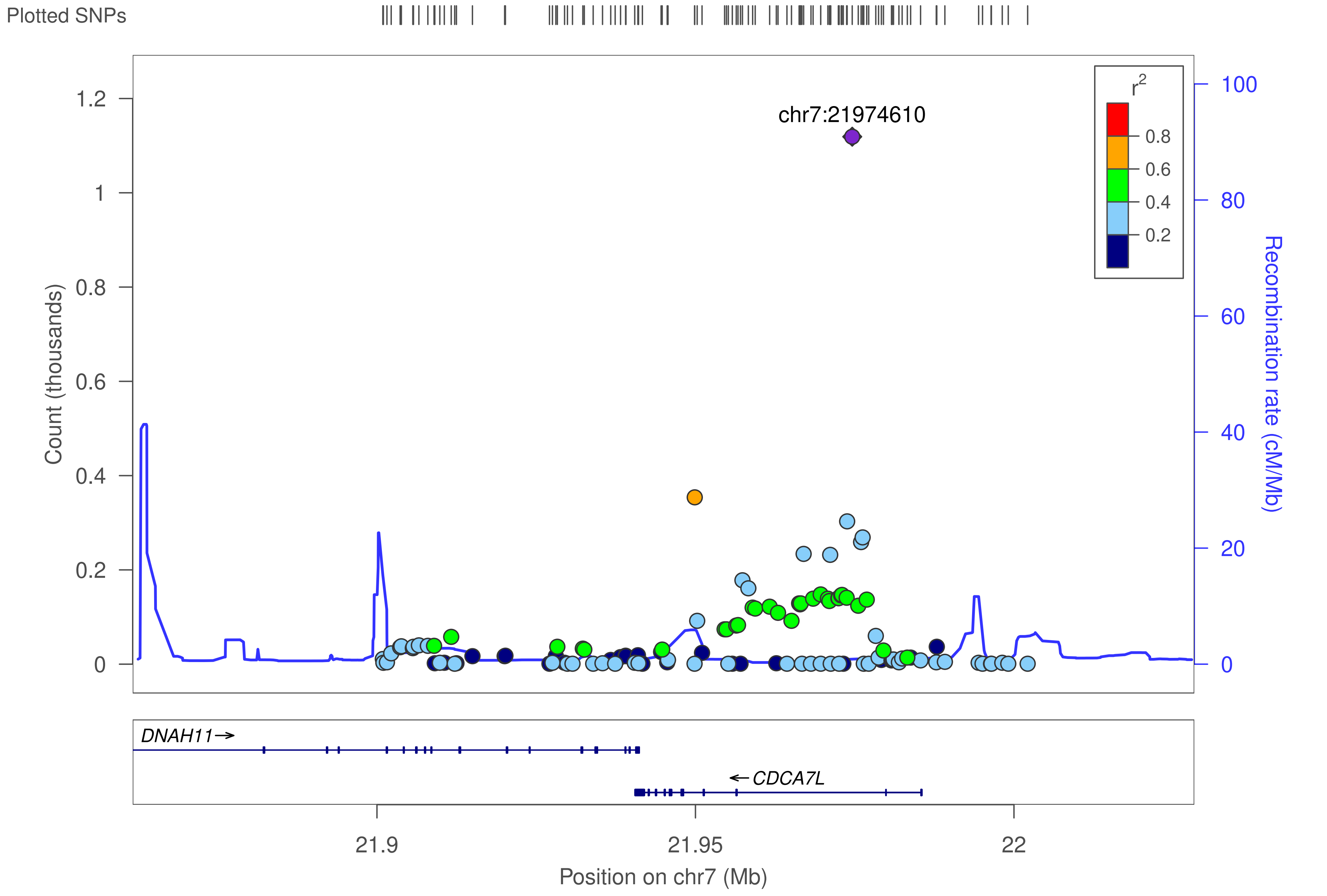
Plotted SNPs
chr7:21974610
r2
0.8
0.6
0.4
0.2
1.2
1
0.8
0.6
0.4
0.2
0
Count (thousands)
100
80
60
40
20
0
Recombination rate (cM/Mb)
DNAH11→
←CDCA7L
21.9
21.95
22
Position on chr7 (Mb)

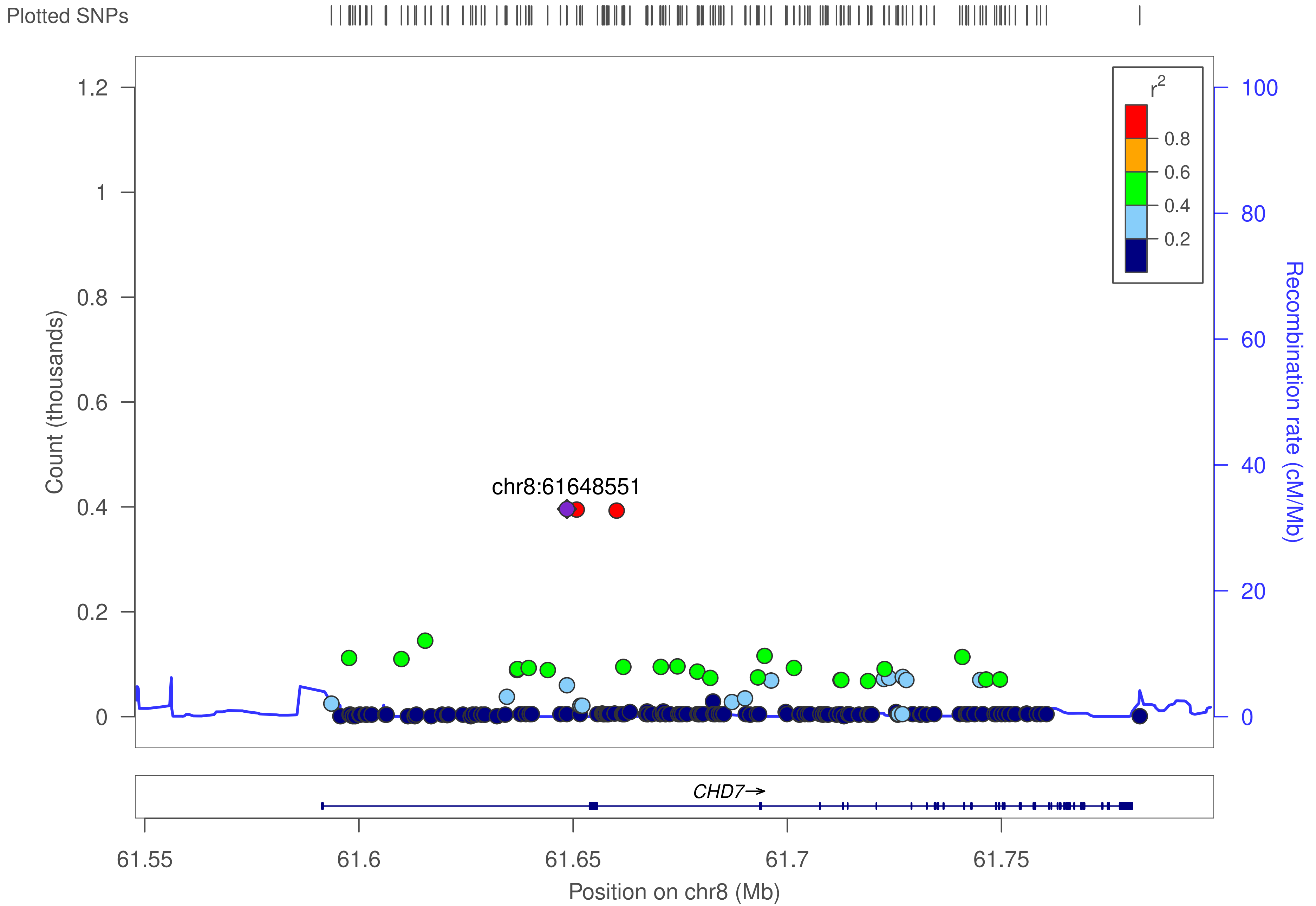

Plotted SNPs
Count (thousands)
1.2
1
0.8
0.6
0.4
0.2
0
chr8:61648551
r2
0.8
0.6
0.4
0.2
100
80
60
40
20
0
Recombination rate (cM/Mb)
CHD7→
61.55
61.6
61.65
61.7
61.75
Position on chr8 (Mb)

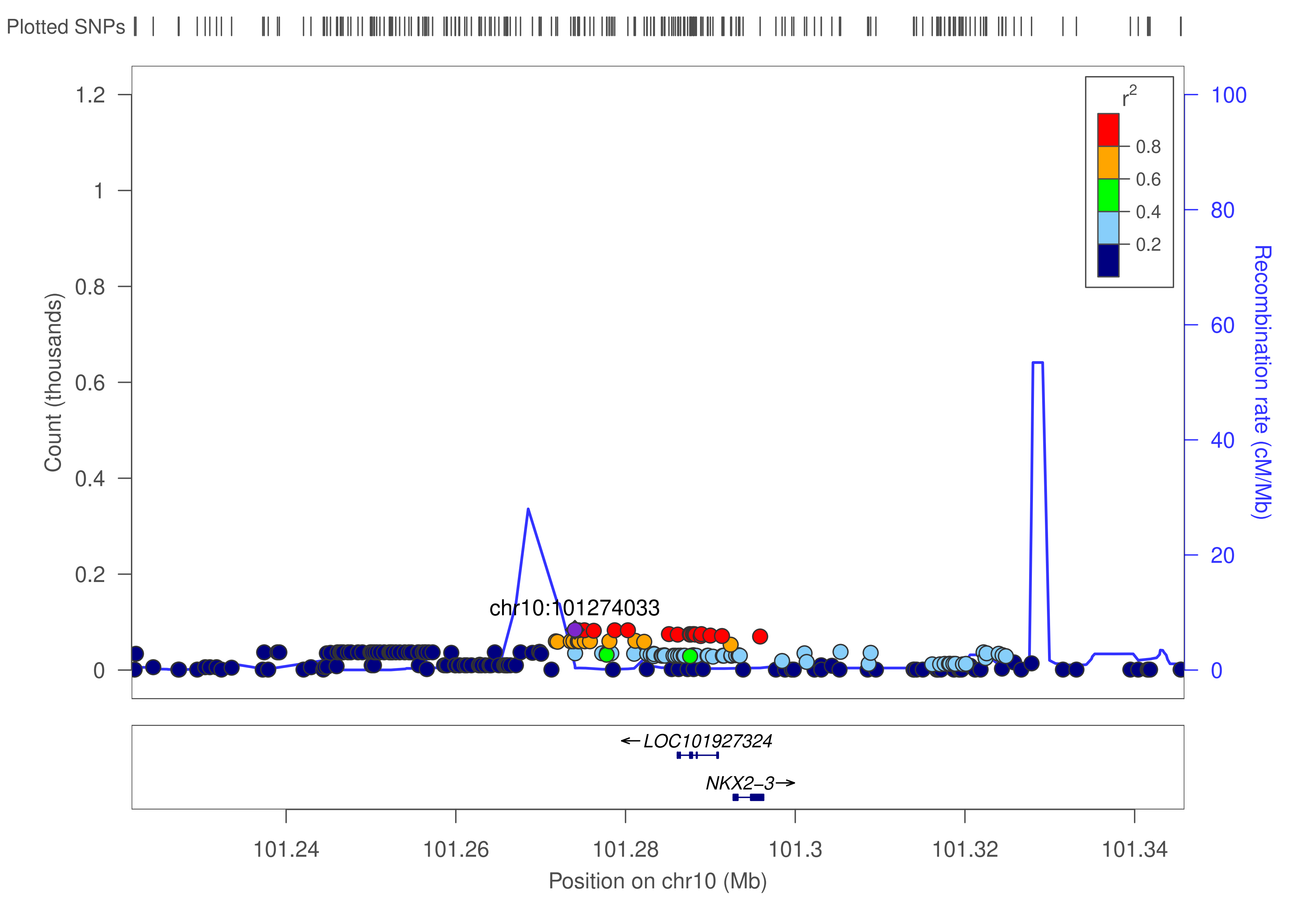

Plotted SNPs
Count (thousands)
1.2
1
0.8
0.6
0.4
0.2
0
chr10:101274033
r2
0.8
0.6
0.4
0.2
100
80
60
40
20
0
Recombination rate (cM/Mb)
LOC101927324
NKX2-3
101.24
101.26
101.28
101.3
101.32
101.34
Position on chr10 (Mb)

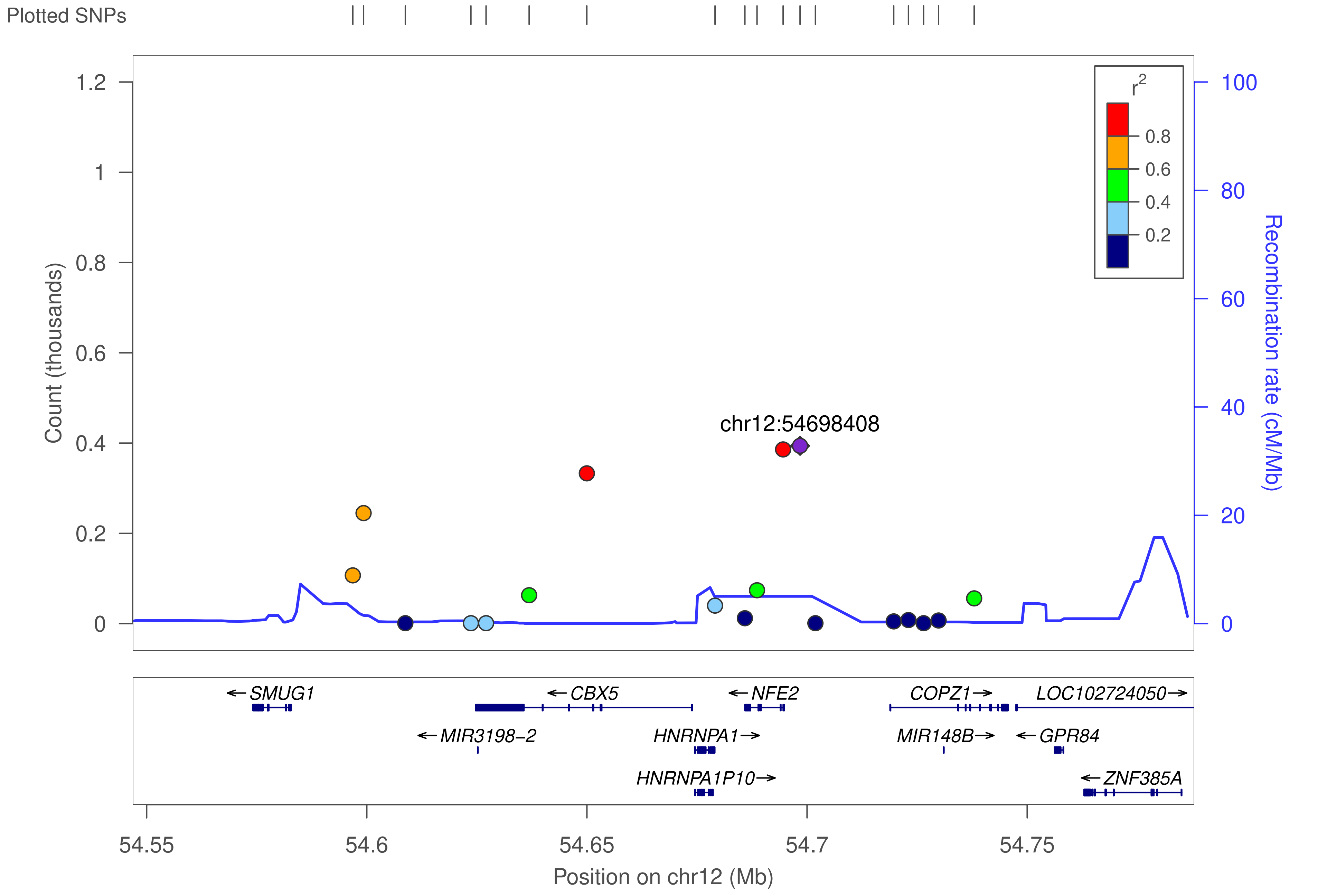

Plotted SNPs
Count (thousands)
1.2
1
0.8
0.6
0.4
0.2
0
Recombination rate (cM/Mb)
100
80
60
40
20
0
r²
0.8
0.6
0.4
0.2
chr12:54698408
←SMUG1
←CBX5
←NFE2
COPZ1→
LOC102724050→
←MIR3198−2
HNRNPA1→
MIR148B→
←GPR84
HNRNPA1P10→
←ZNF385A
54.55
54.6
54.65
54.7
54.75
Position on chr12 (Mb)

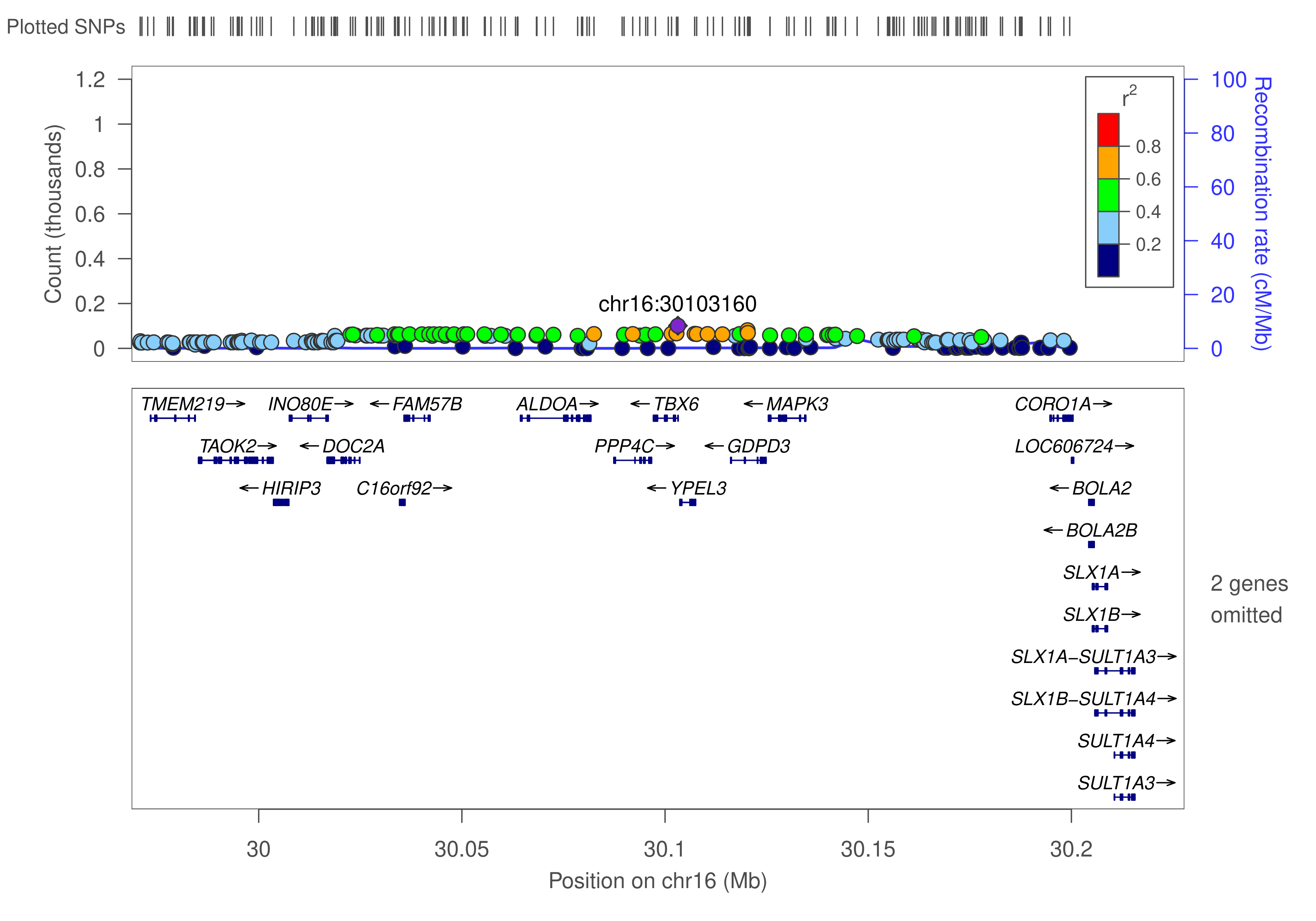

Plotted SNPs
Count (thousands)
1.2
1
0.8
0.6
0.4
0.2
0
chr16:30103160
r2
0.8
0.6
0.4
0.2
100
80
60
40
20
0
Recombination rate (cM/Mb)
TMEM219
INO80E
FAM57B
ALDOA
TBX6
MAPK3
CORO1A
TAOK2
DOC2A
PPP4C
GDPD3
LOC606724
HIRIP3
C16orf92
YPEL3
BOLA2
BOLA2B
SLX1A
SLX1B
SLX1A−SULT1A3
SLX1B−SULT1A4
SULT1A4
SULT1A3
2 genes omitted
30
30.05
30.1
30.15
30.2
Position on chr16 (Mb)

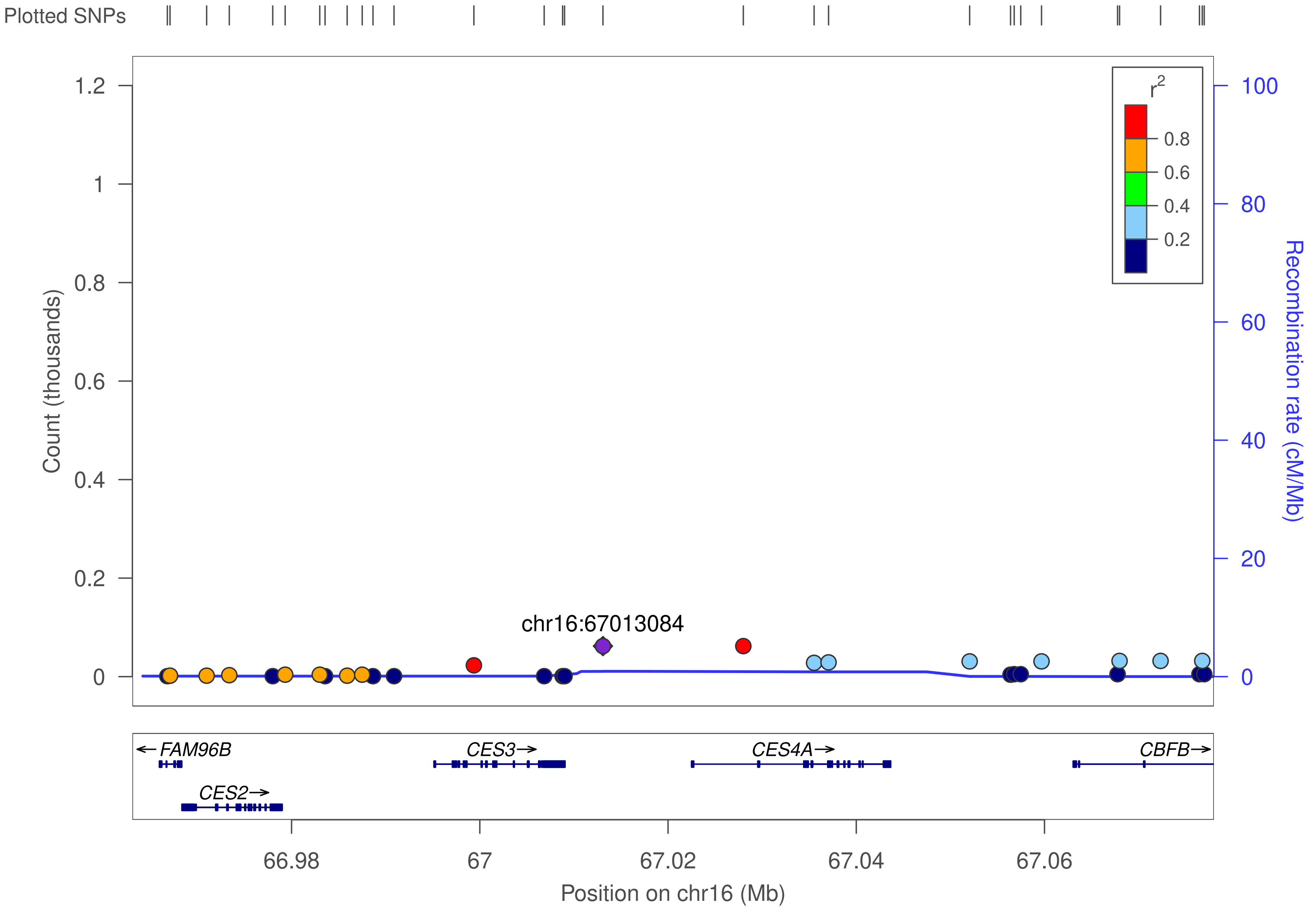

Plotted SNPs
Count (thousands)
1.2
1
0.8
0.6
0.4
0.2
0
r2
0.8
0.6
0.4
0.2
100
80
60
40
20
0
Recombination rate (cM/Mb)
chr16:67013084
FAM96B
CES3
CES4A
CBFB
CES2
66.98
67
67.02
67.04
67.06
Position on chr16 (Mb)

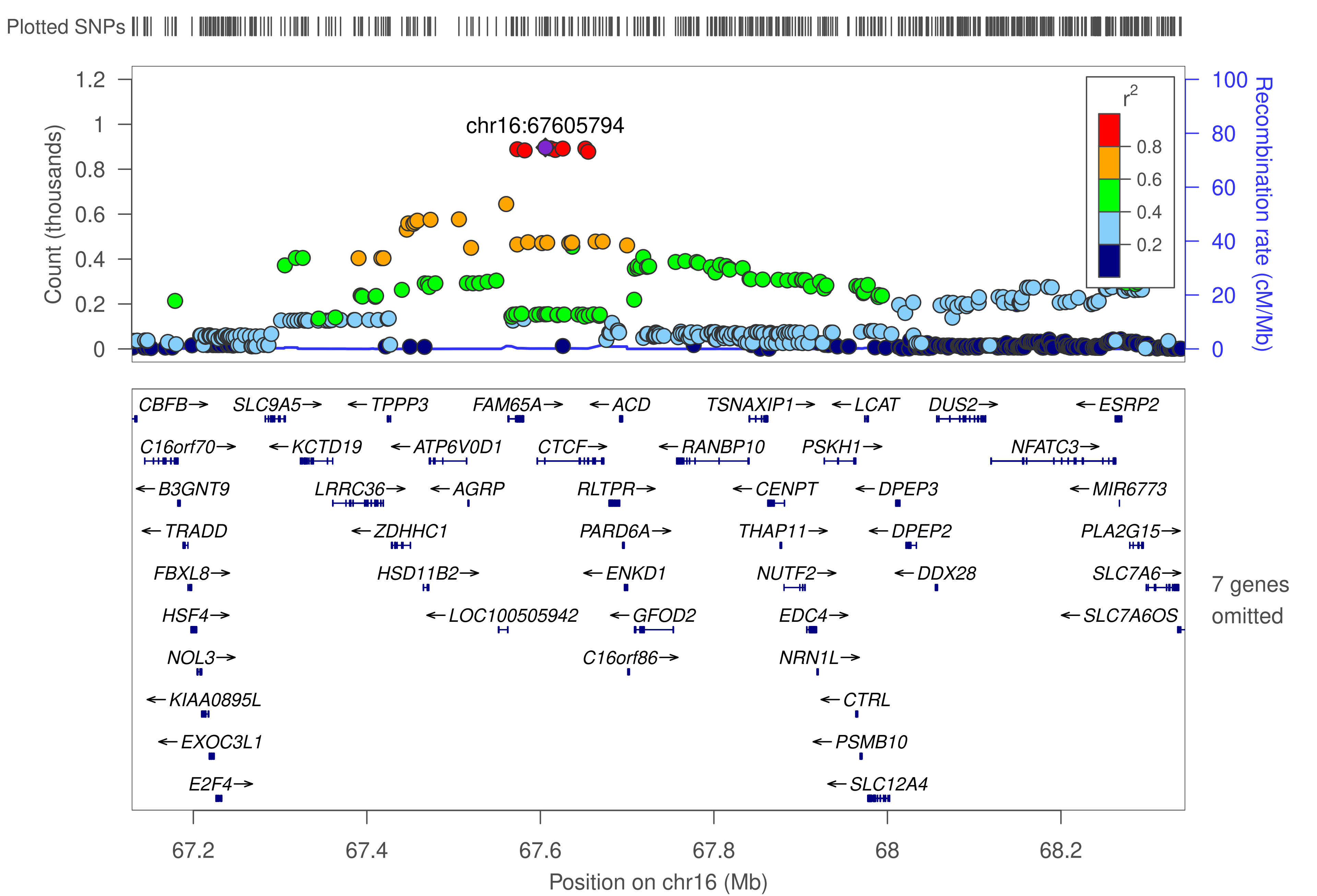

Plotted SNPs
chr16:67605794
Count (thousands)
Recombination rate (cM/Mb)
r2
0.8
0.6
0.4
0.2
CBFB→
SLC9A5→
←TPPP3
FAM65A→
←ACD
TSNAXIP1→
←LCAT
DUS2→
←ESRP2
C16orf70→
←KCTD19
←ATP6V0D1
CTCF→
←RANBP10
PSKH1→
NFATC3→
←B3GNT9
LRRC36→
←AGRP
RLTPR→
←CENPT
←DPEP3
←MIR6773
←TRADD
←ZDHHC1
PARD6A→
THAP11→
←DPEP2
PLA2G15→
FBXL8→
HSD11B2→
←ENKD1
NUTF2→
←DDX28
SLC7A6→
HSF4→
←LOC100505942
←GFOD2
EDC4→
←SLC7A6OS
NOL3→
C16orf86→
NRN1L→
←KIAA0895L
←CTRL
←EXOC3L1
←PSMB10
E2F4→
←SLC12A4
7 genes omitted
67.2
67.4
67.6
67.8
68
68.2
Position on chr16 (Mb)

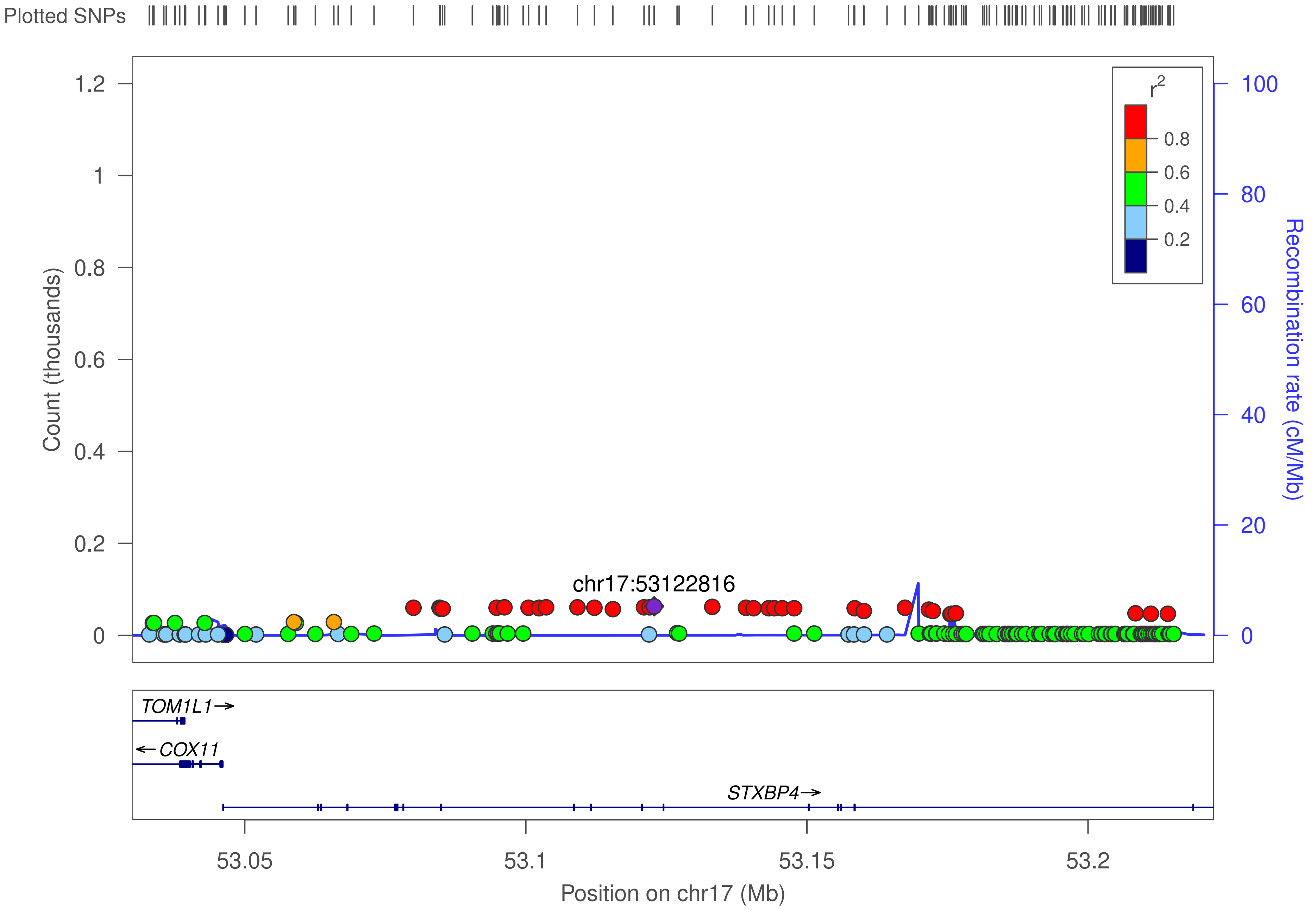

Plotted SNPs
Count (thousands)
1.2
1
0.8
0.6
0.4
0.2
0
r2
0.8
0.6
0.4
0.2
100
80
60
40
20
0
Recombination rate (cM/Mb)
chr17:53122816
TOM1L1→
←COX11
STXBP4→
53.05
53.1
53.15
53.2
Position on chr17 (Mb)

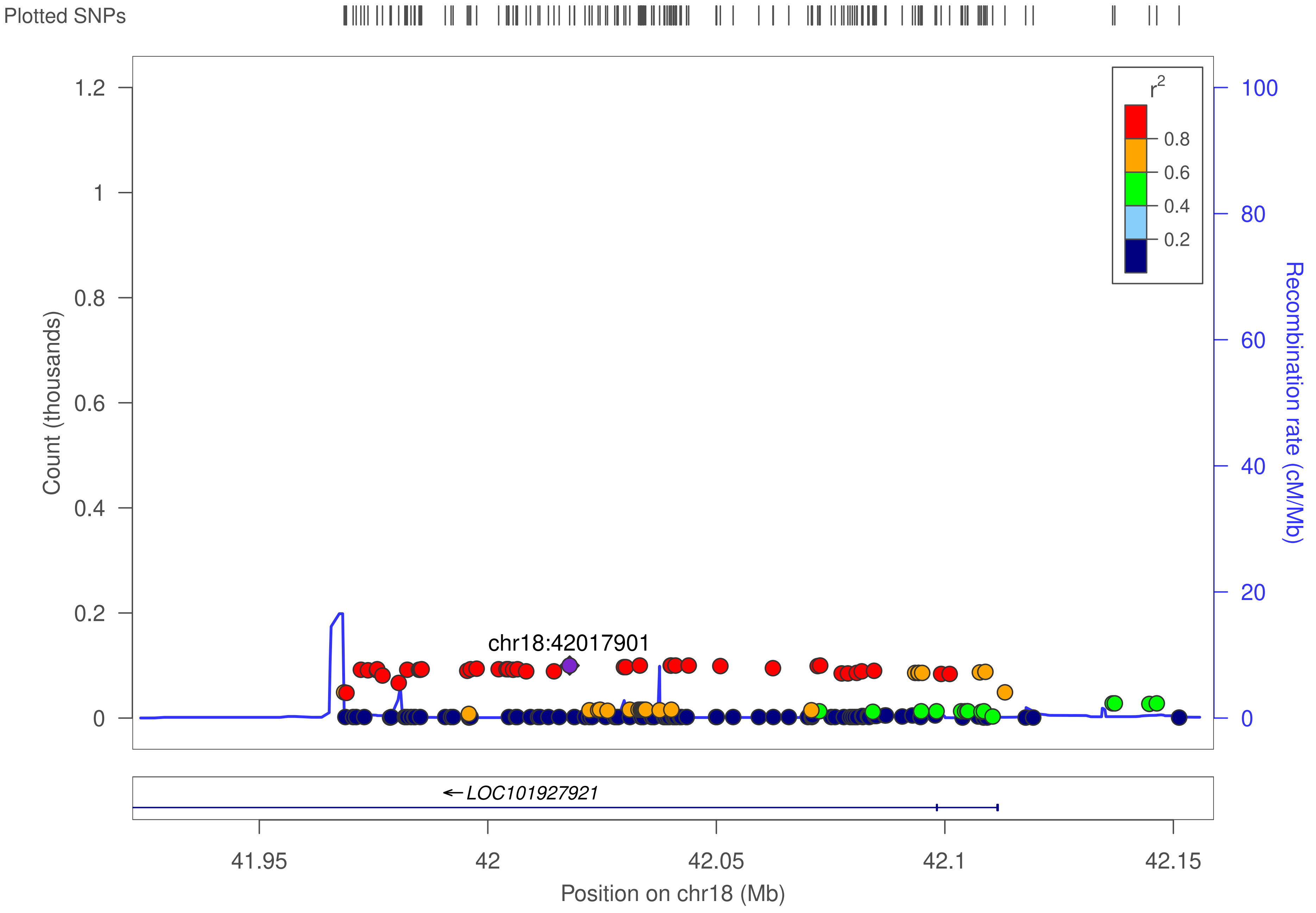
Plotted SNPs
Count (thousands)
1.2
1
0.8
0.6
0.4
0.2
0
chr18:42017901
r2
0.8
0.6
0.4
0.2
Recombination rate (cM/Mb)
100
80
60
40
20
0
← LOC101927921
41.95
42
42.05
42.1
42.15
Position on chr18 (Mb)

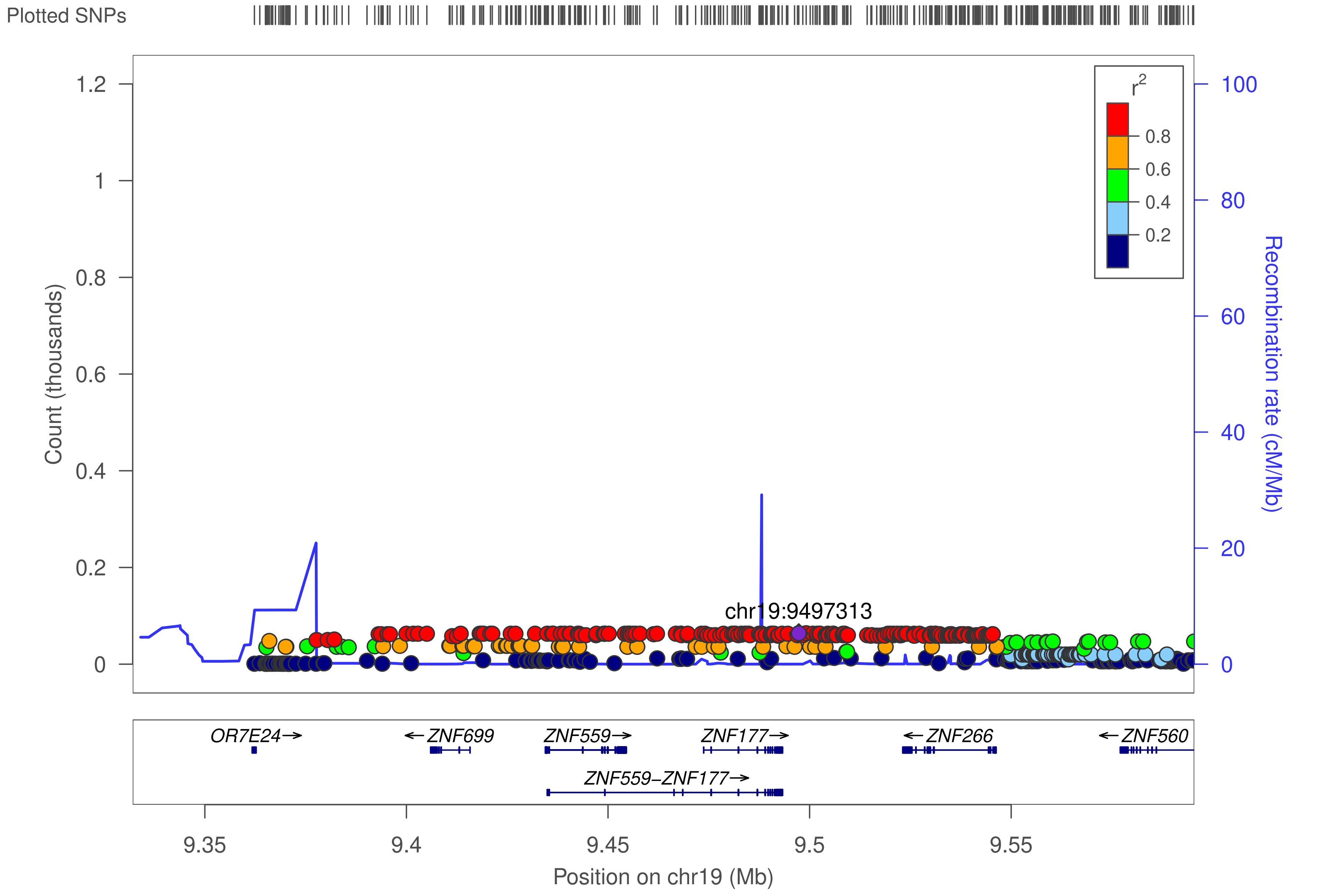

Plotted SNPs
Count (thousands)
1.2
1
0.8
0.6
0.4
0.2
0
Recombination rate (cM/Mb)
100
80
60
40
20
0
r²
0.8
0.6
0.4
0.2
chr19:9497313
OR7E24→
←ZNF699
ZNF559→
ZNF177→
←ZNF266
←ZNF560
ZNF559–ZNF177→
9.35
9.4
9.45
9.5
9.55
Position on chr19 (Mb)

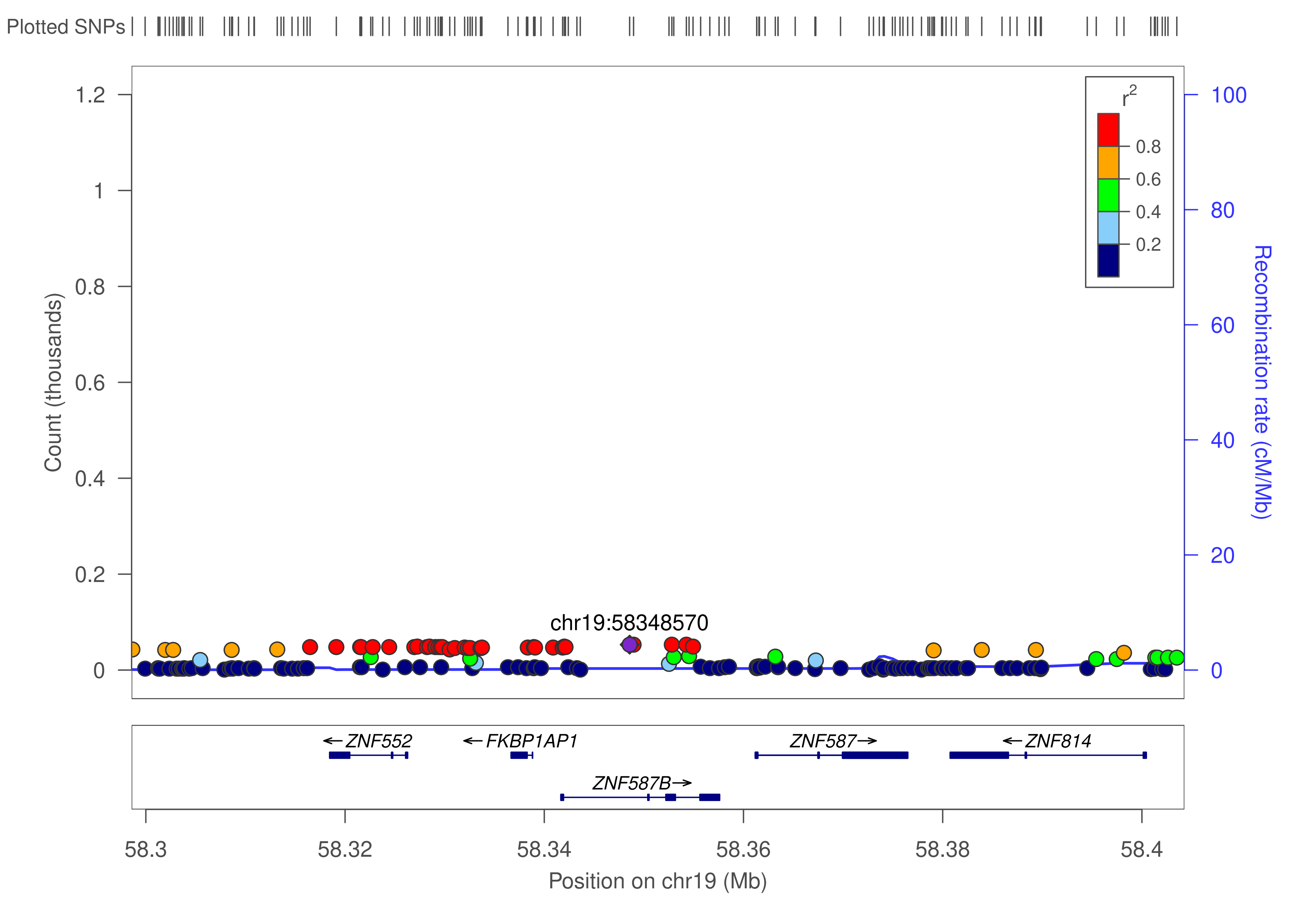

Plotted SNPs
Count (thousands)
1.2
1
0.8
0.6
0.4
0.2
0
Recombination rate (cM/Mb)
100
80
60
40
20
0
$r^2$
0.8
0.6
0.4
0.2
chr19:58348570
← ZNF552
← FKBP1AP1
ZNF587 →
← ZNF814
ZNF587B →
58.3
58.32
58.34
58.36
58.38
58.4
Position on chr19 (Mb)

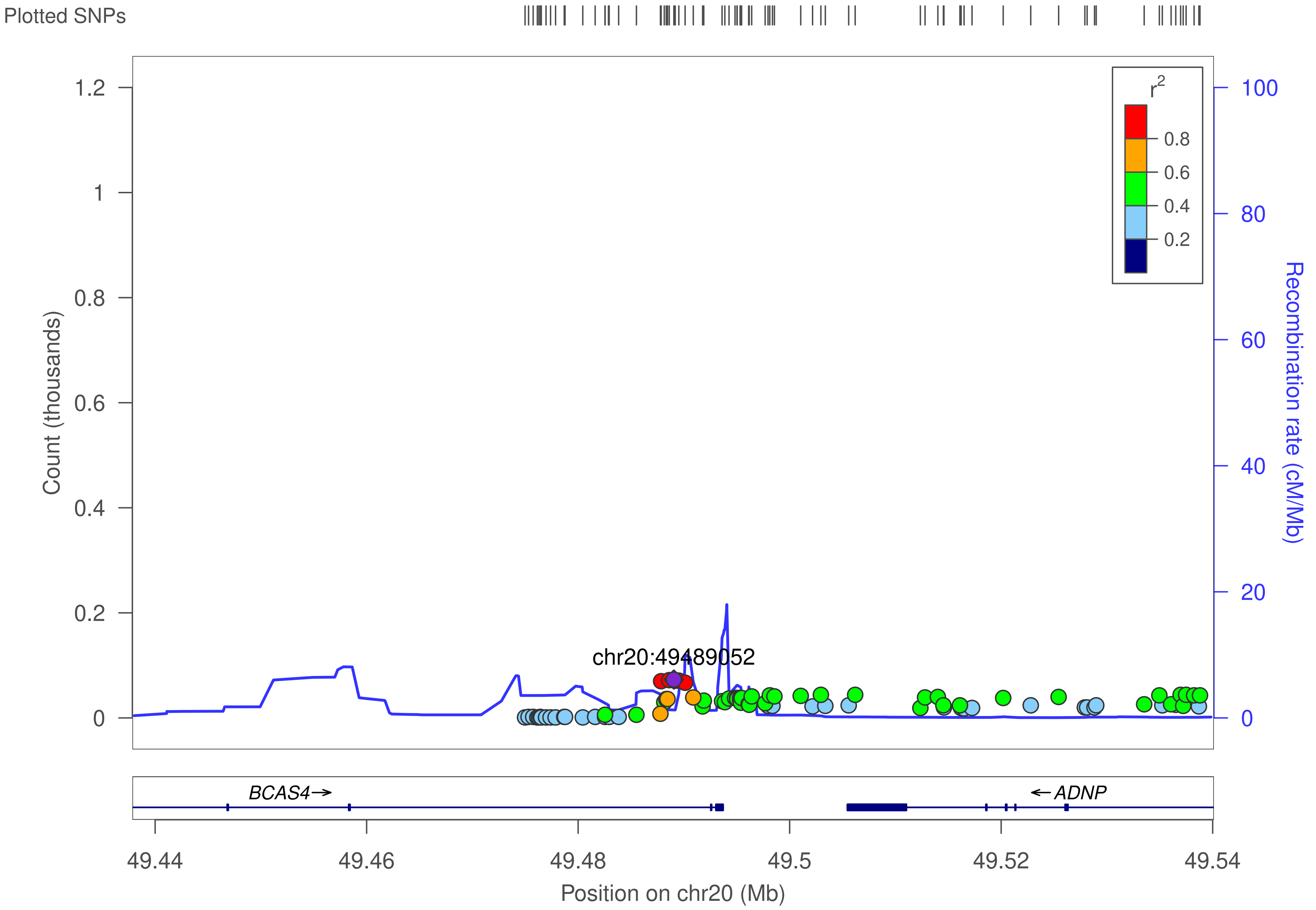

Plotted SNPs
Count (thousands)
1.2
1
0.8
0.6
0.4
0.2
0
r2
0.8
0.6
0.4
0.2
100
80
60
40
20
0
Recombination rate (cM/Mb)
chr20:49489052
BCAS4→
←ADNP
49.44
49.46
49.48
49.5
49.52
49.54
Position on chr20 (Mb)
