## Supplementary File 5 for "Genetic control of KRAB-ZFP genes explains distal CpG-site methylation which associates with human disease phenotypes"

### ZNF506

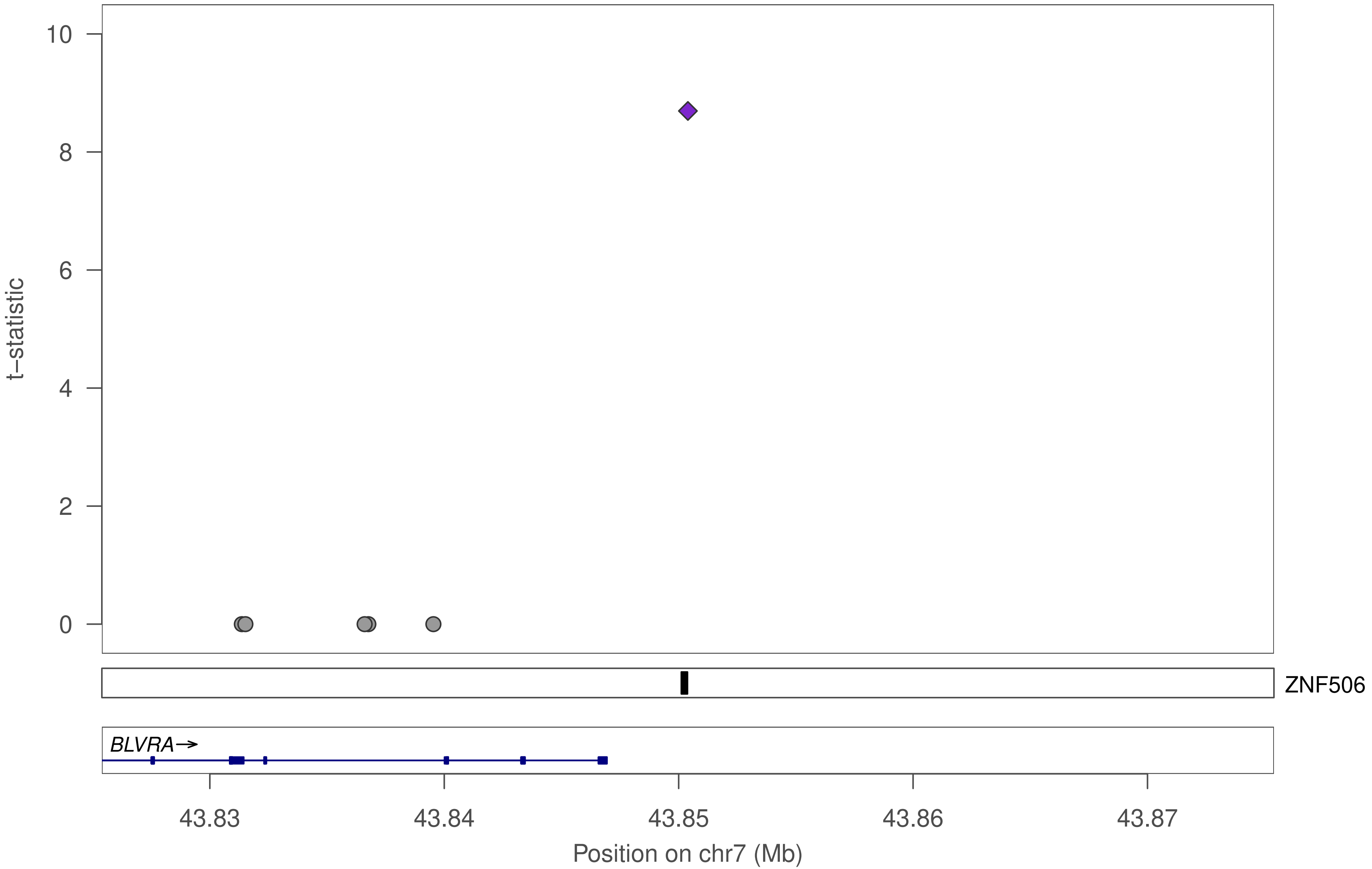

t–statistic
10
8
6
4
2
0
ZNF506
BLVRA→
43.83
43.84
43.85
43.86
43.87
Position on chr7 (Mb)

### ZNF506

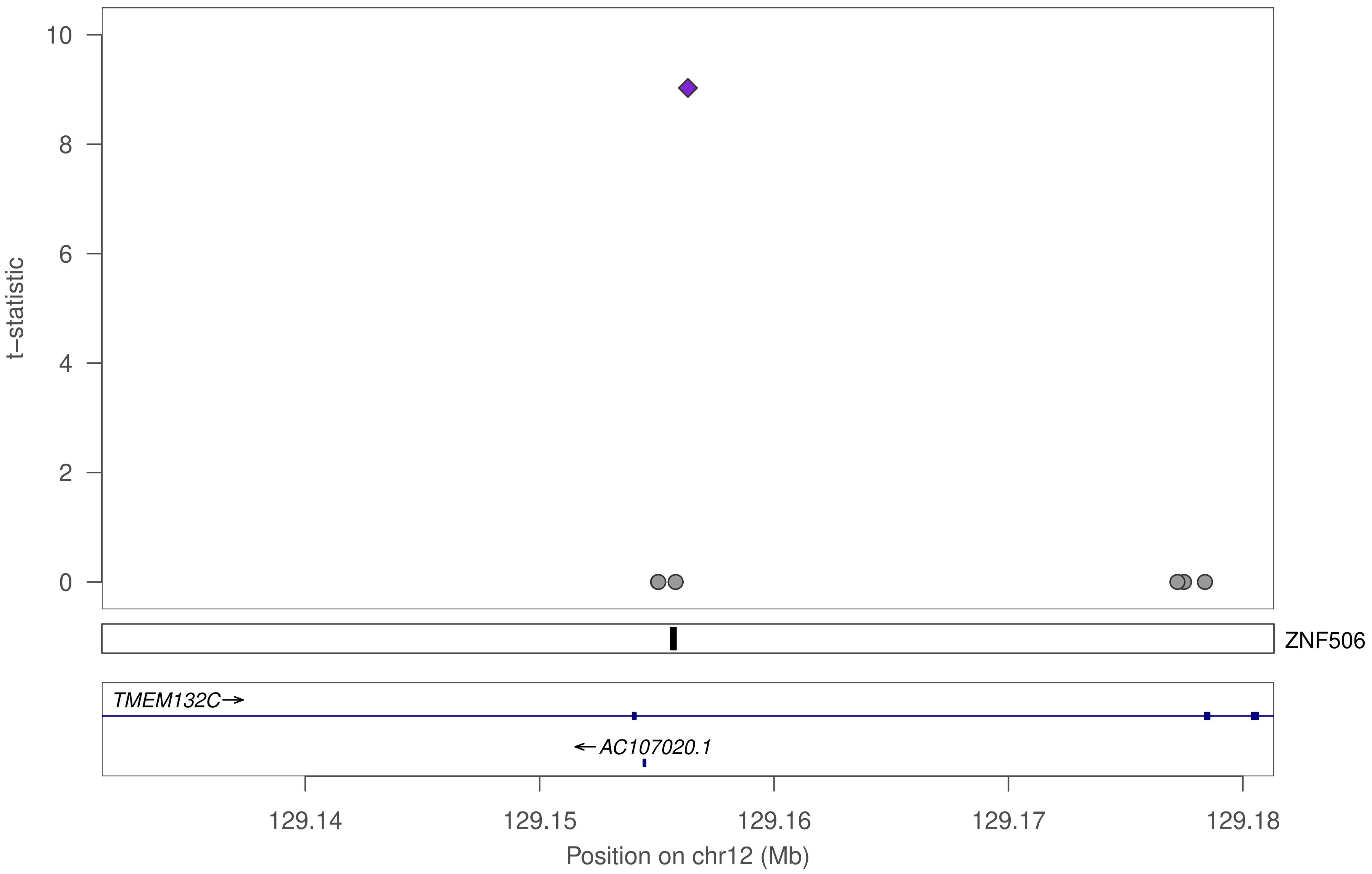

10
8
6
4
2
0
t−statistic
ZNF506
TMEM132C→
←AC107020.1
129.14
129.15
129.16
129.17
129.18
Position on chr12 (Mb)

### ZNF506

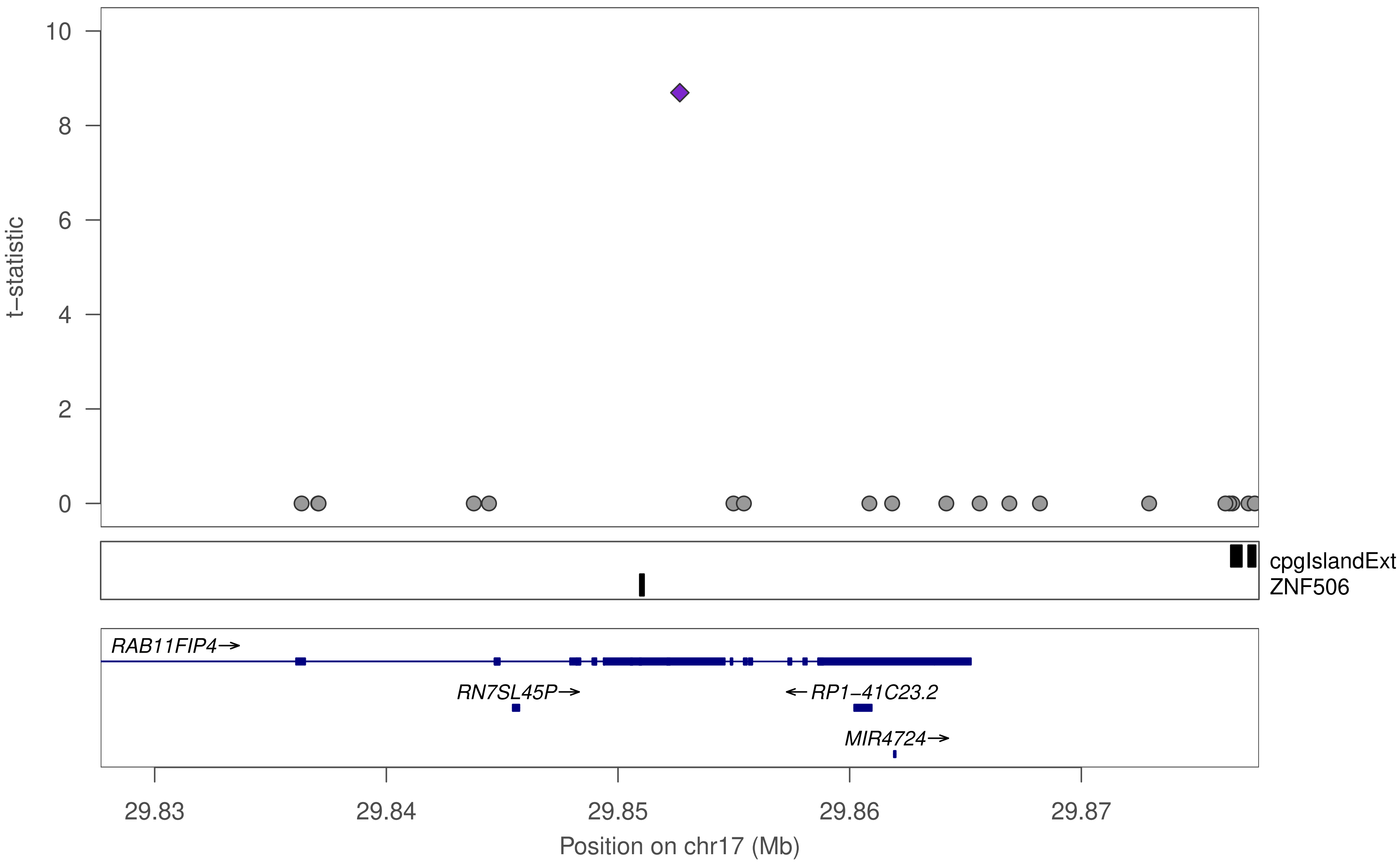

10
8
6
4
2
0
t−statistic
cpgIslandExt
ZNF506
RAB11FIP4→
RN7SL45P→
←RP1−41C23.2
MIR4724→
29.83
29.84
29.85
29.86
29.87
Position on chr17 (Mb)

### ZNF506

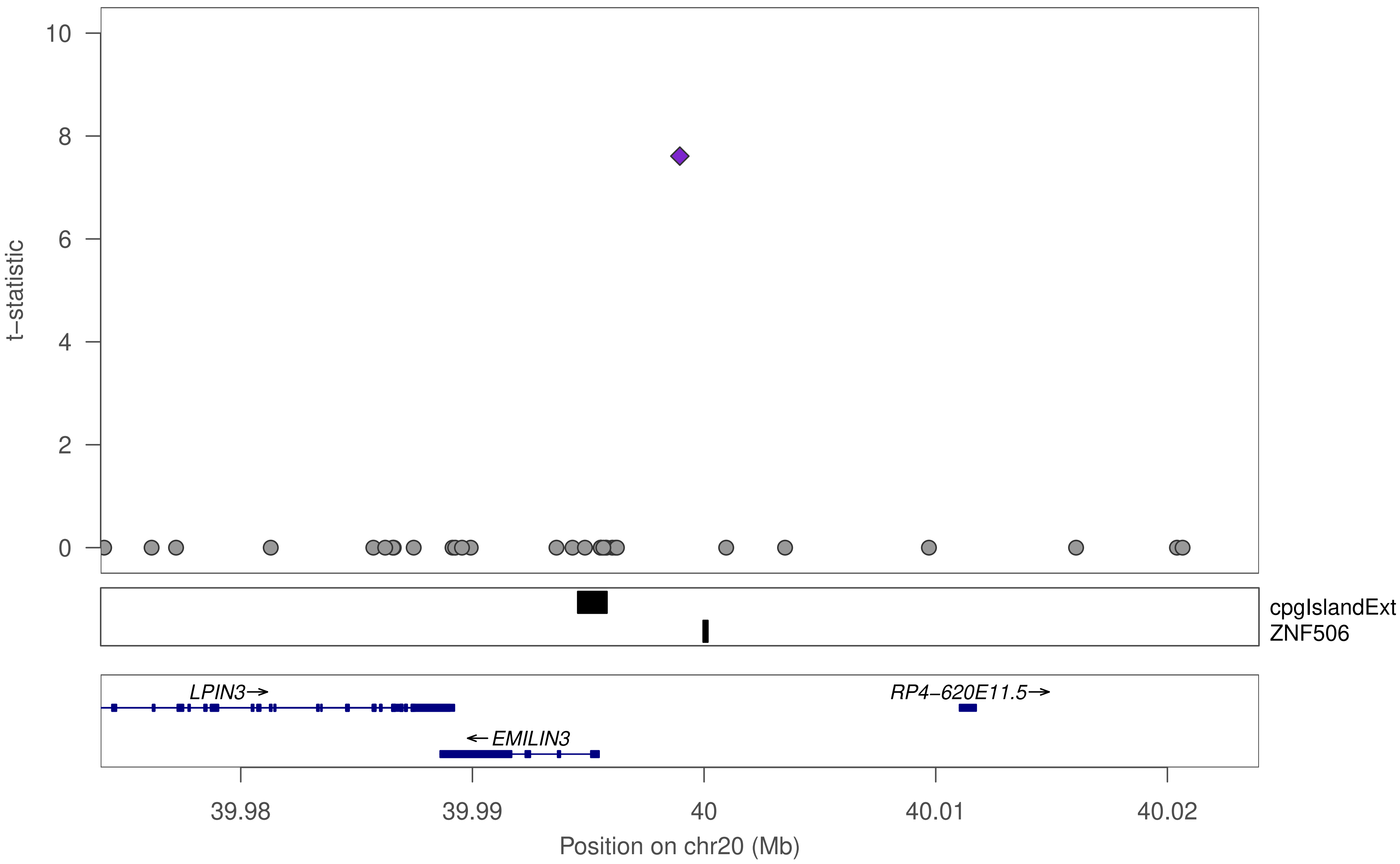

10
8
6
4
2
0
t−statistic
cpgIslandExt
ZNF506
LPIN3→
RP4−620E11.5→
←EMILIN3
39.98
39.99
40
40.01
40.02
Position on chr20 (Mb)

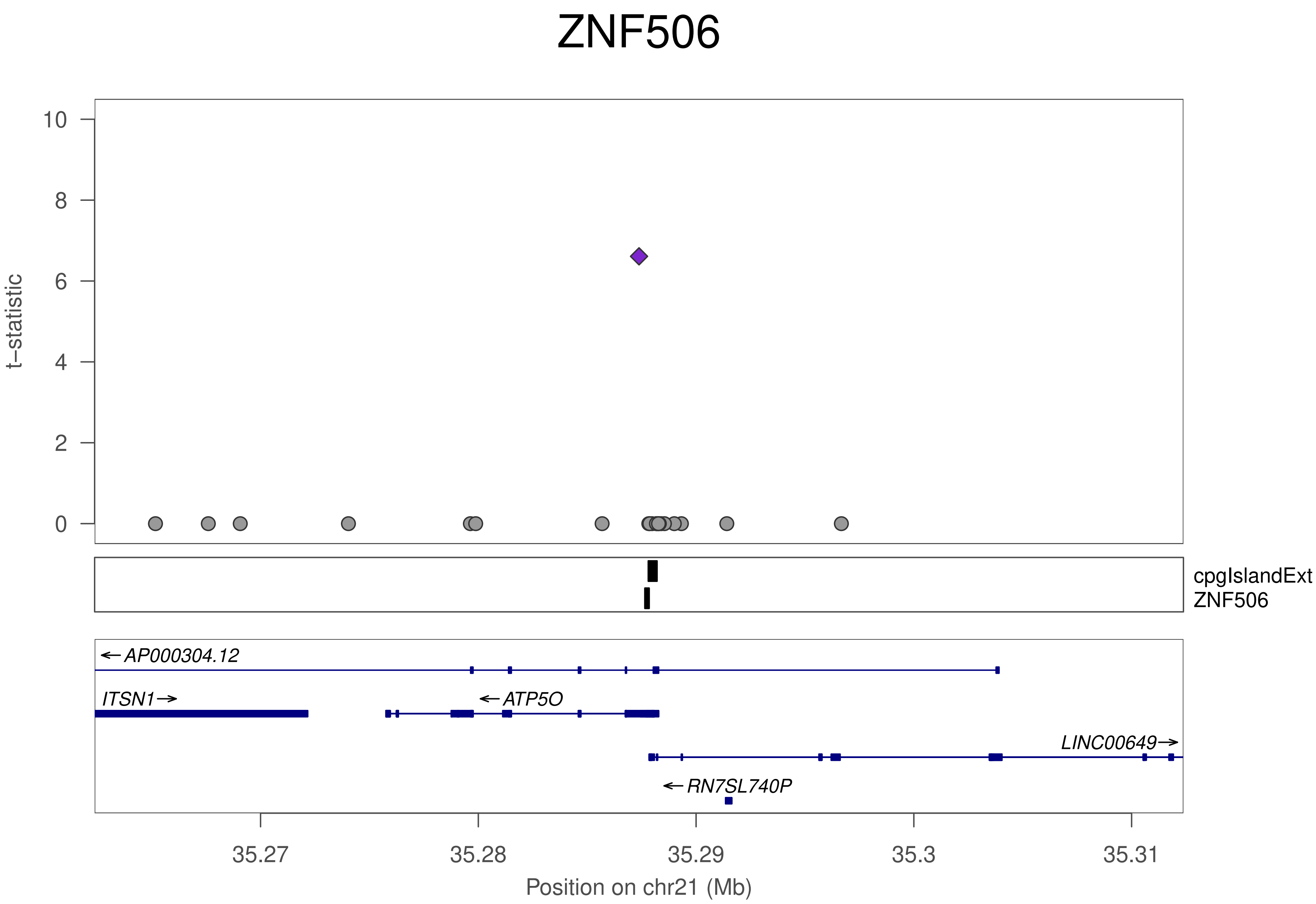

ZNF506
t−statistic
10
8
6
4
2
0
cpgIslandExt
ZNF506
←AP000304.12
ITSN1→
←ATP5O
LINC00649→
←RN7SL740P
35.27
35.28
35.29
35.3
35.31
Position on chr21 (Mb)

### ZNF430

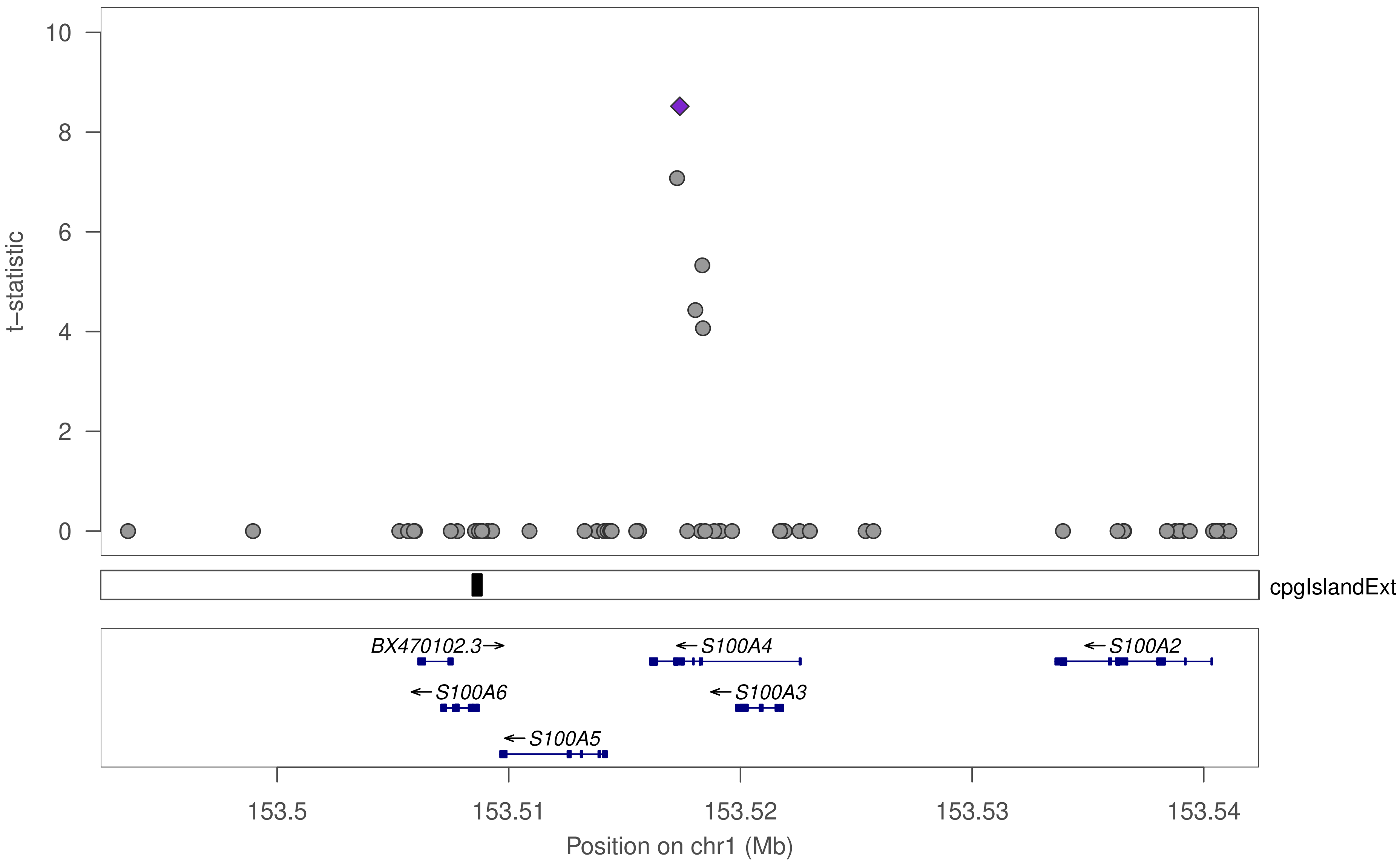

t–statistic
10
8
6
4
2
0
cpgIslandExt
BX470102.3→
←S100A4
←S100A2
←S100A6
←S100A3
←S100A5
153.5
153.51
153.52
153.53
153.54
Position on chr1 (Mb)

### ZNF430

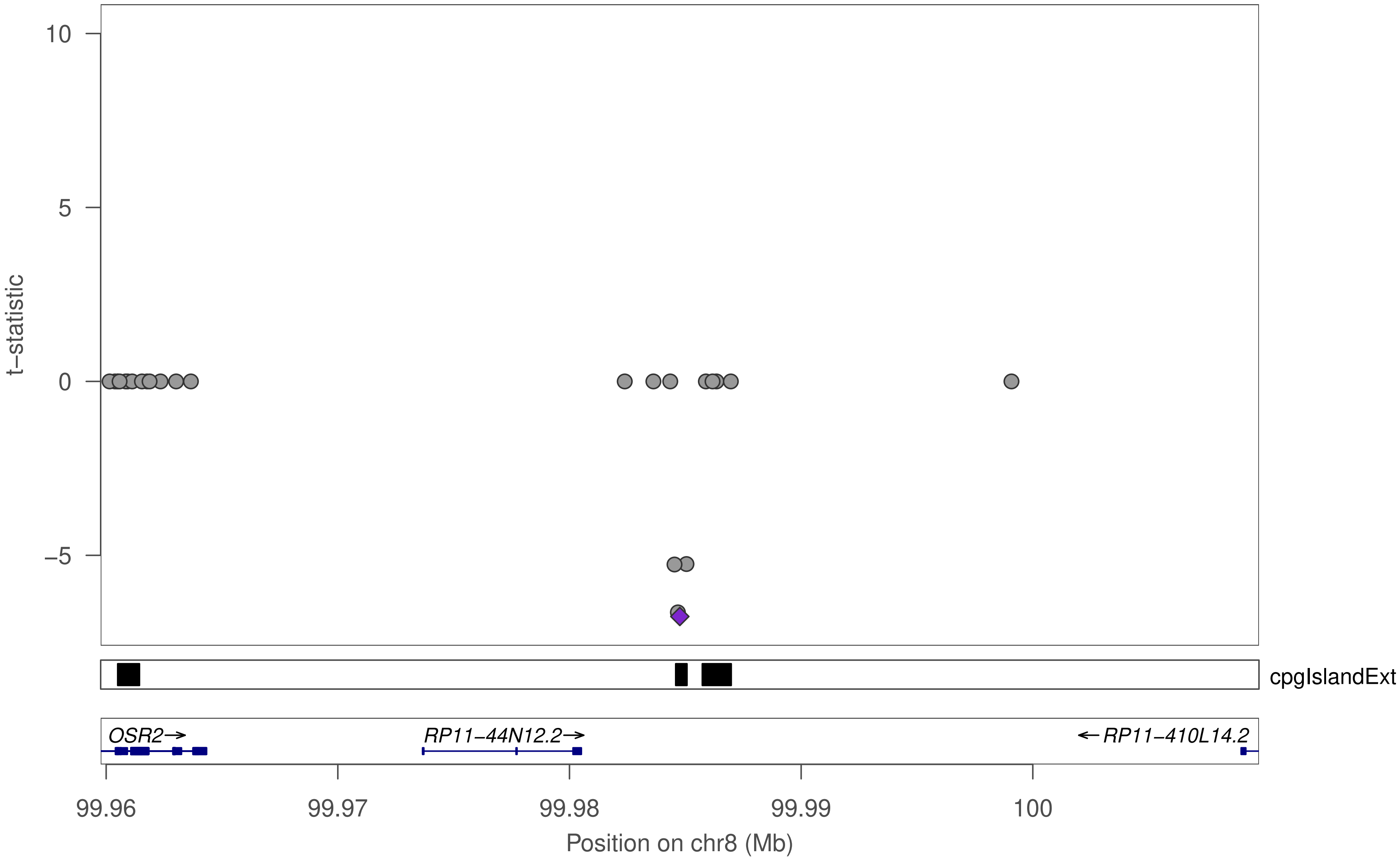

10
5
0
−5
t−statistic
cpgIslandExt
OSR2→
RP11−44N12.2→
←RP11−410L14.2
99.96
99.97
99.98
99.99
100
Position on chr8 (Mb)

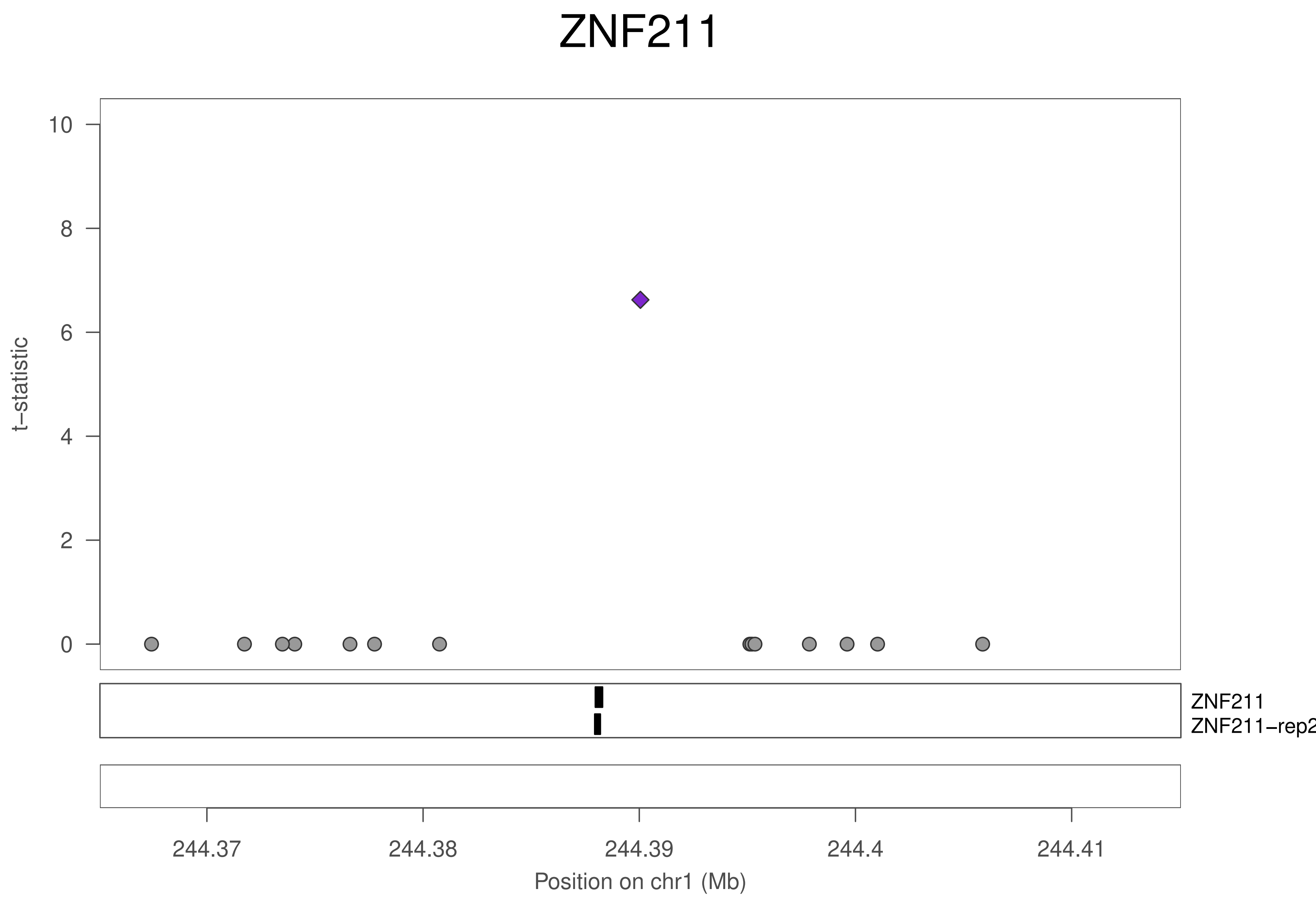

ZNF211
t−statistic
10
8
6
4
2
0
ZNF211
ZNF211−rep2
244.37
244.38
244.39
244.4
244.41
Position on chr1 (Mb)

### ZNF211

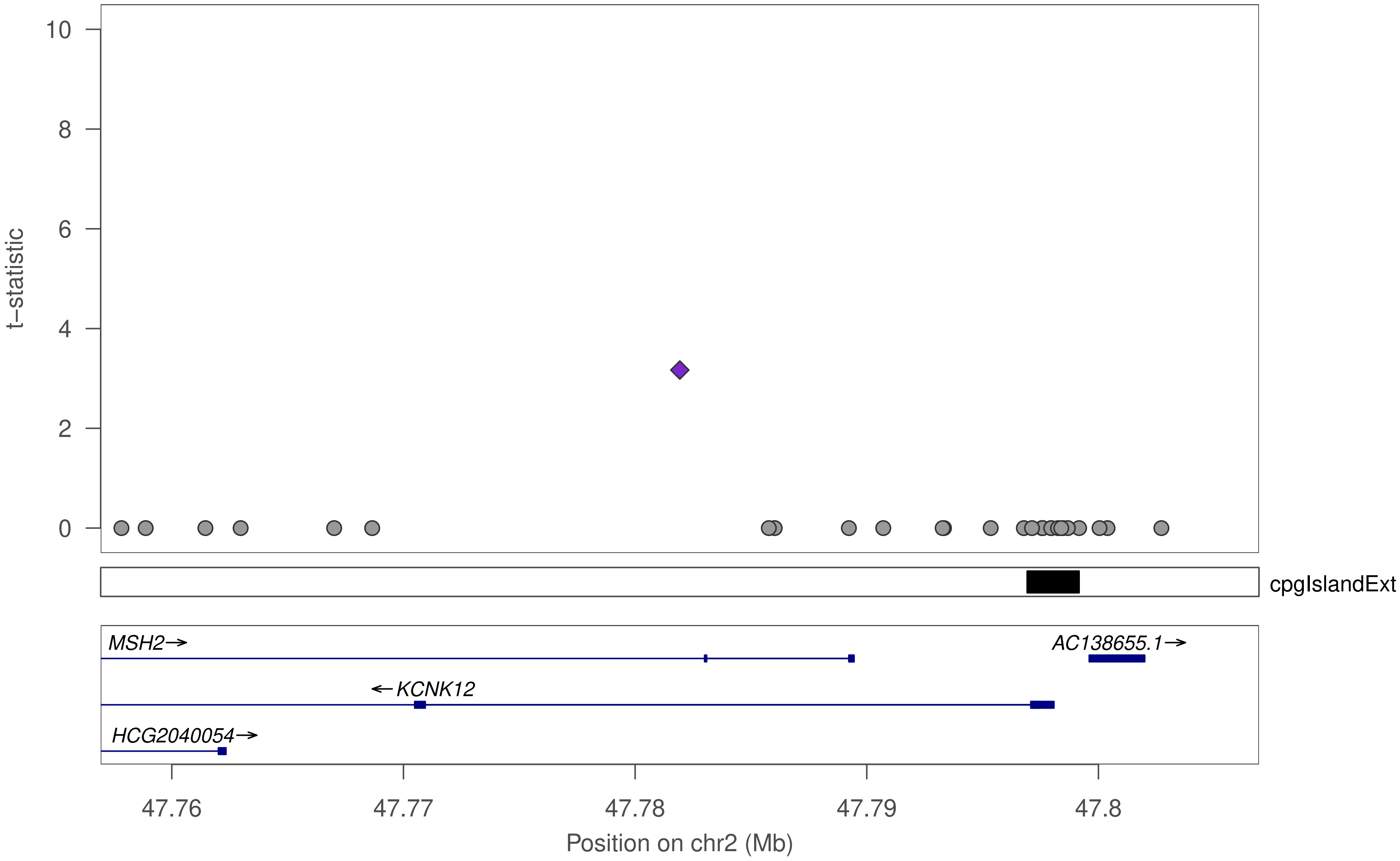

t–statistic
10
8
6
4
2
0
cpgIslandExt
MSH2→
AC138655.1→
←KCNK12
HCG2040054→
47.76
47.77
47.78
47.79
47.8
Position on chr2 (Mb)

### ZNF211

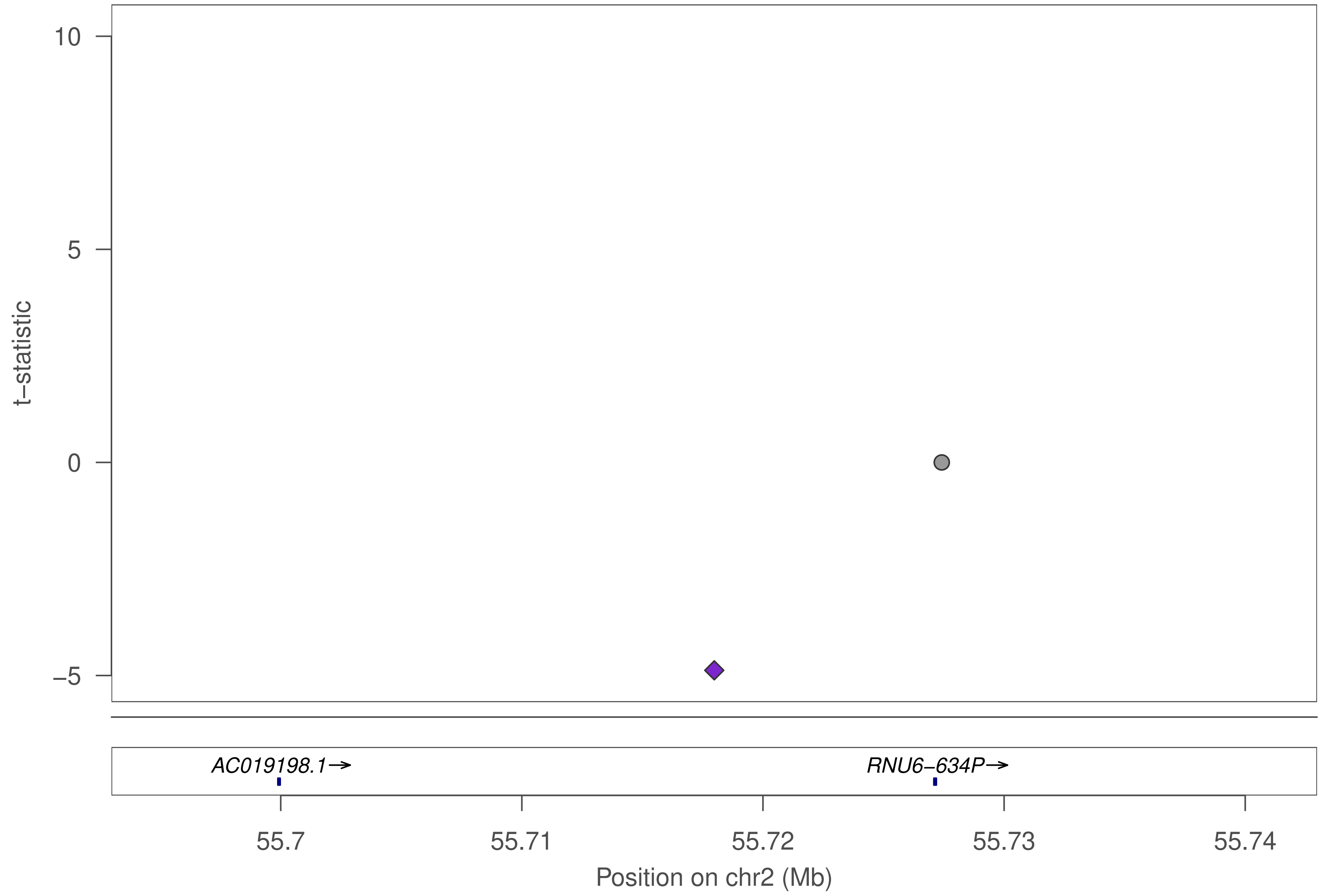

10
5
0
−5
t−statistic
AC019198.1→
RNU6−634P→
55.7
55.71
55.72
55.73
55.74
Position on chr2 (Mb)

### ZNF211

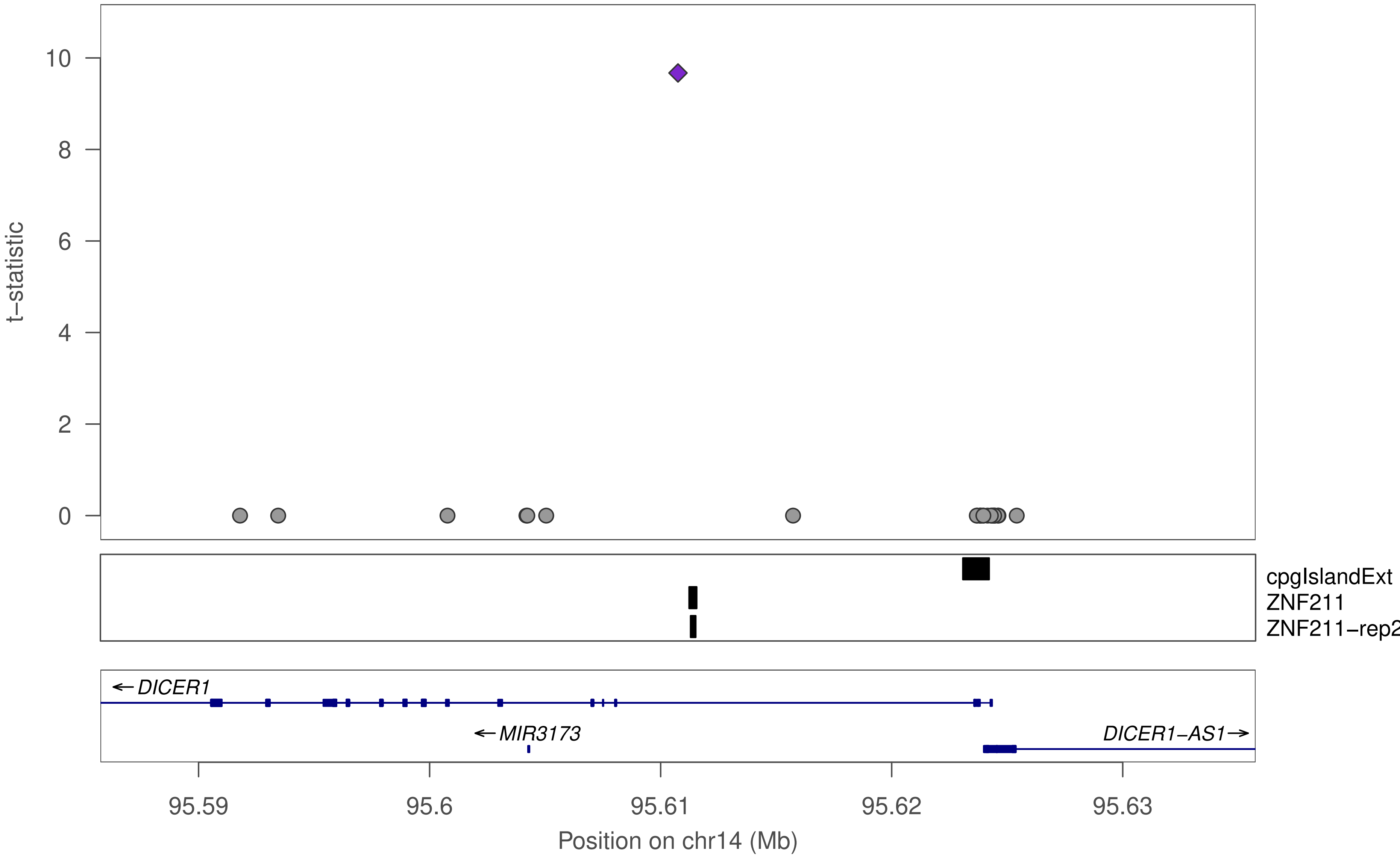

t−statistic
10
8
6
4
2
0
cpgIslandExt
ZNF211
ZNF211−rep2
←DICER1
←MIR3173
DICER1−AS1→
95.59
95.6
95.61
95.62
95.63
Position on chr14 (Mb)

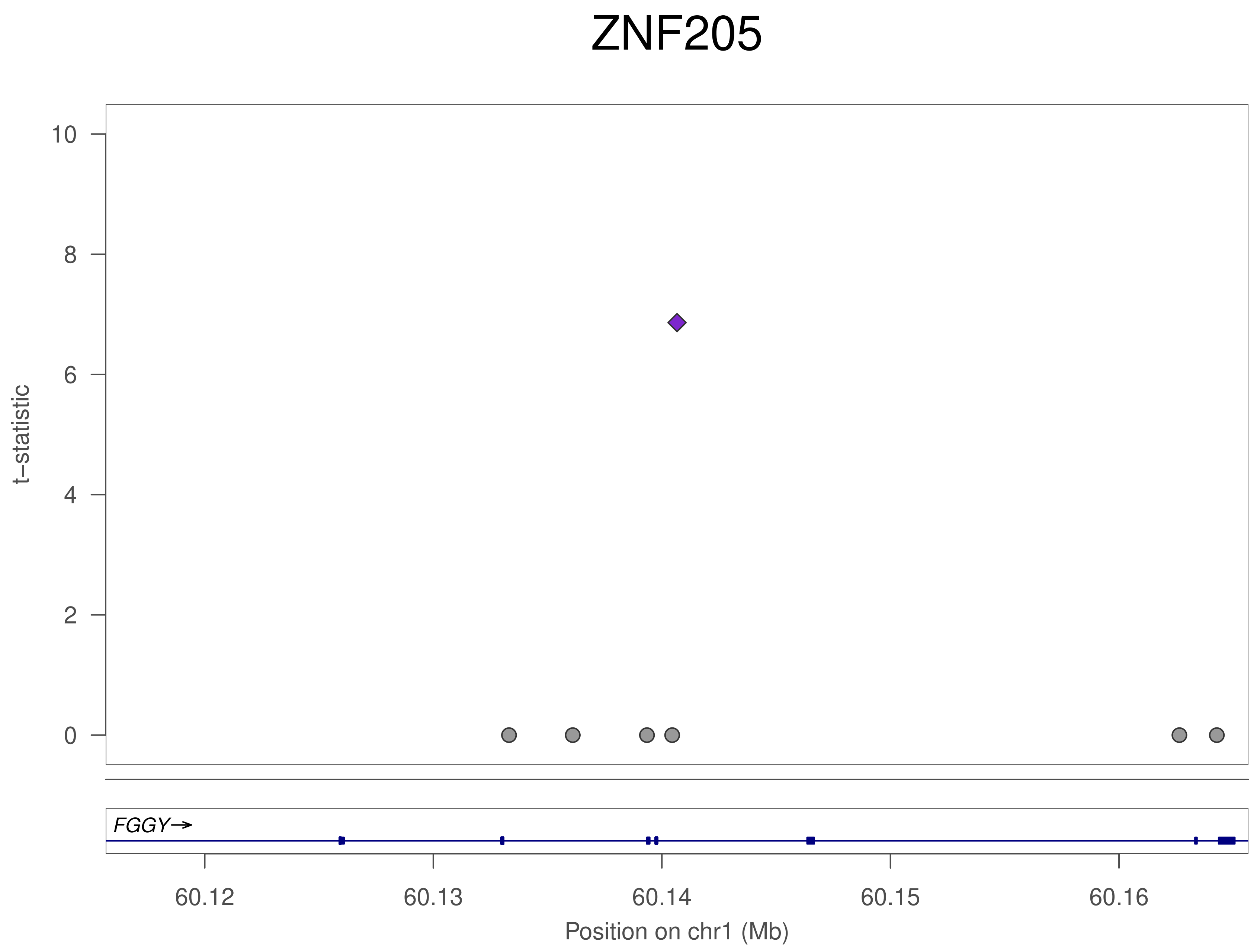
ZNF205
t−statistic
10
8
6
4
2
0
FGGY→
60.12
60.13
60.14
60.15
60.16
Position on chr1 (Mb)

### ZNF205

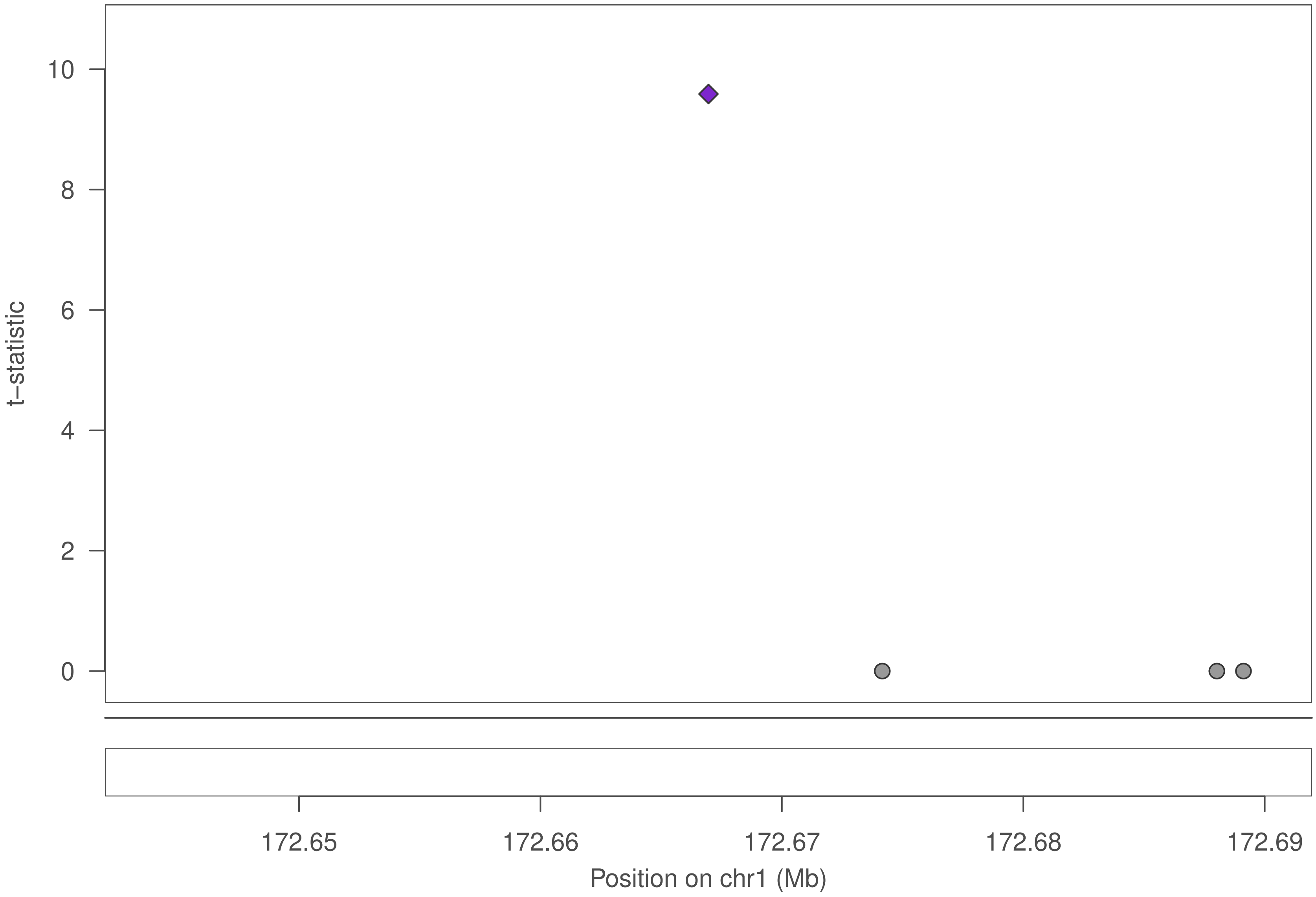

10
8
6
4
2
0
t−statistic
172.65
172.66
172.67
172.68
172.69
Position on chr1 (Mb)

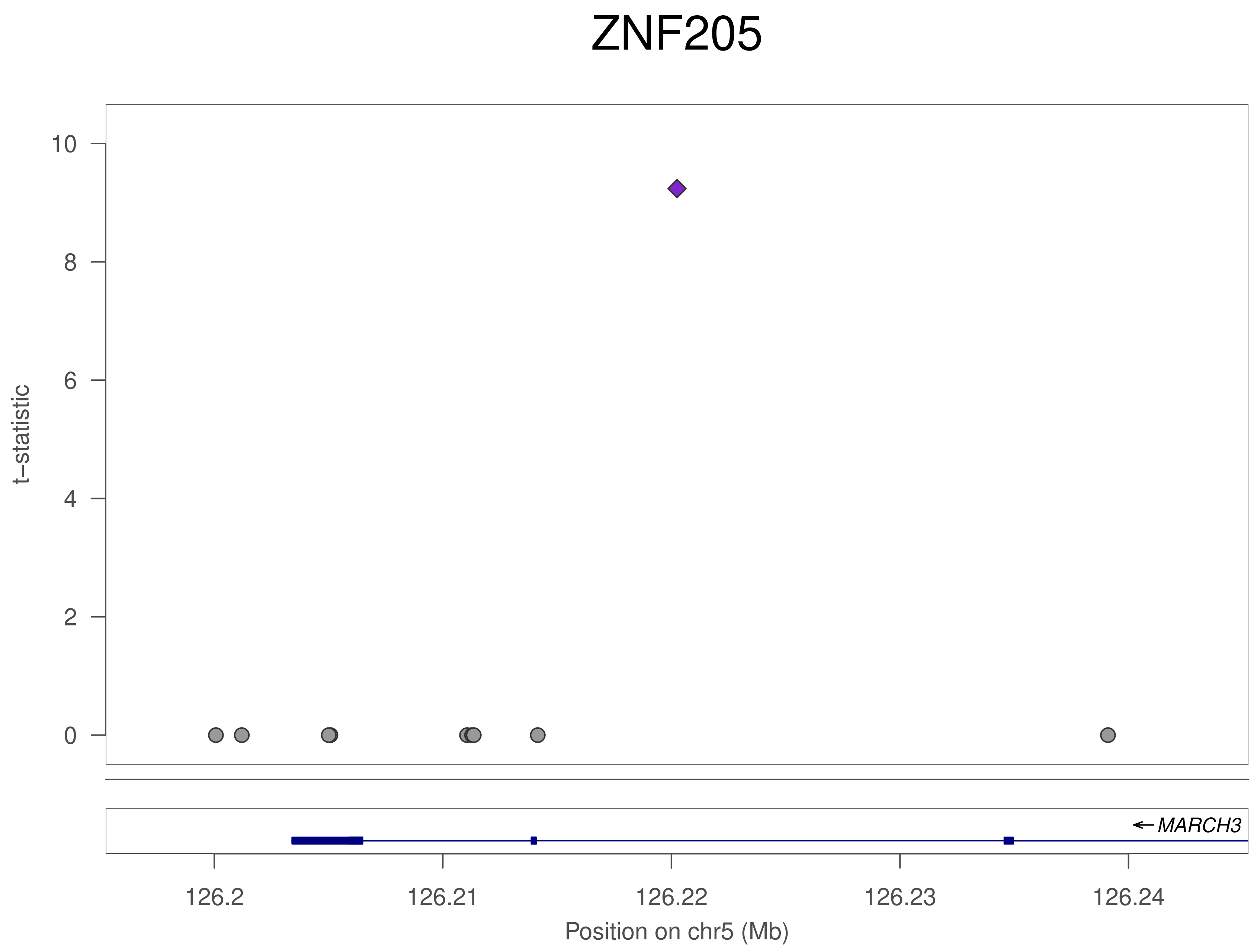

ZNF205
t–statistic
10
8
6
4
2
0
MARCH3
126.2
126.21
126.22
126.23
126.24
Position on chr5 (Mb)

### ZNF205

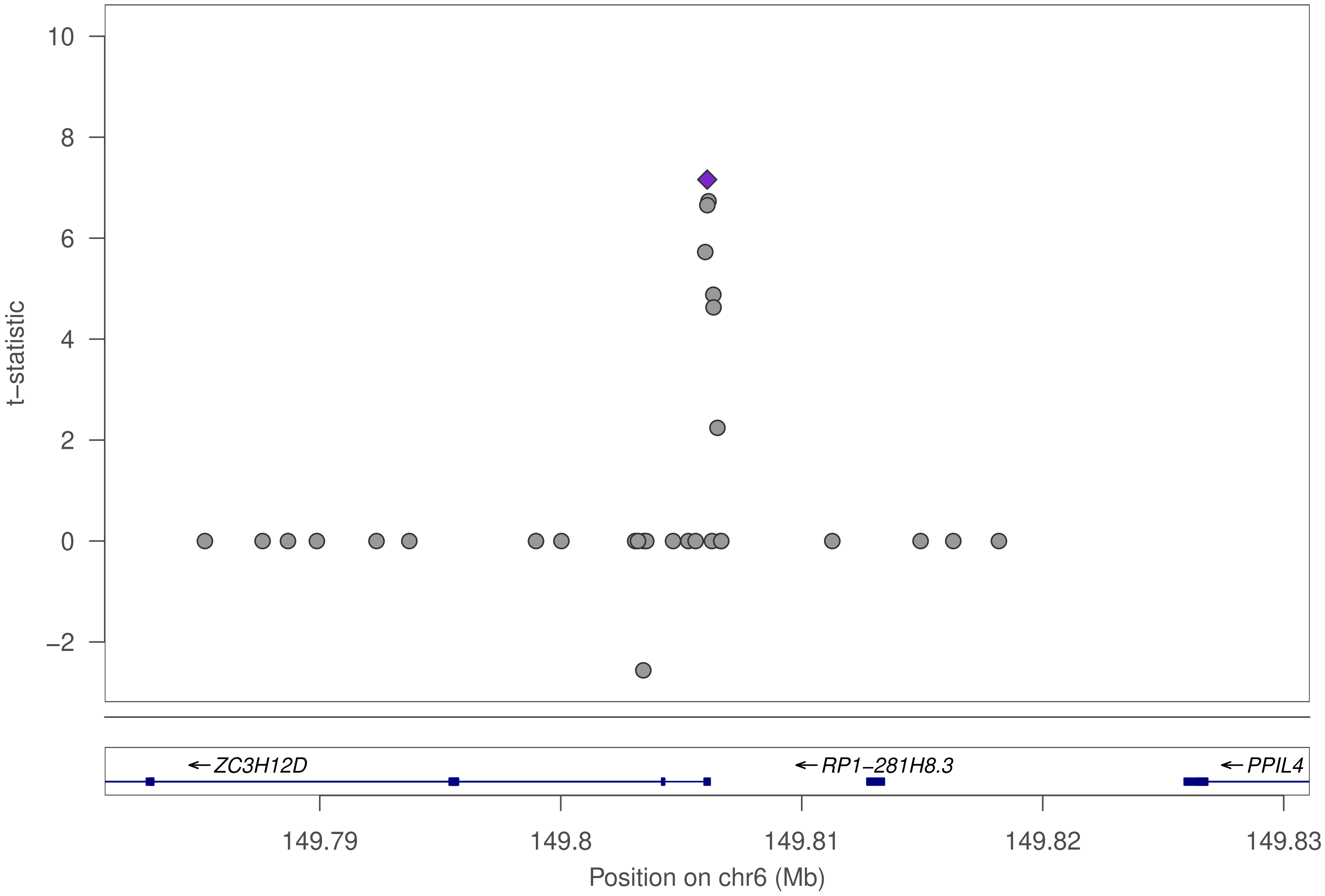

10
8
6
4
2
0
−2
t−statistic
←ZC3H12D
←RP1−281H8.3
←PPIL4
149.79
149.8
149.81
149.82
149.83
Position on chr6 (Mb)

### ZNF205

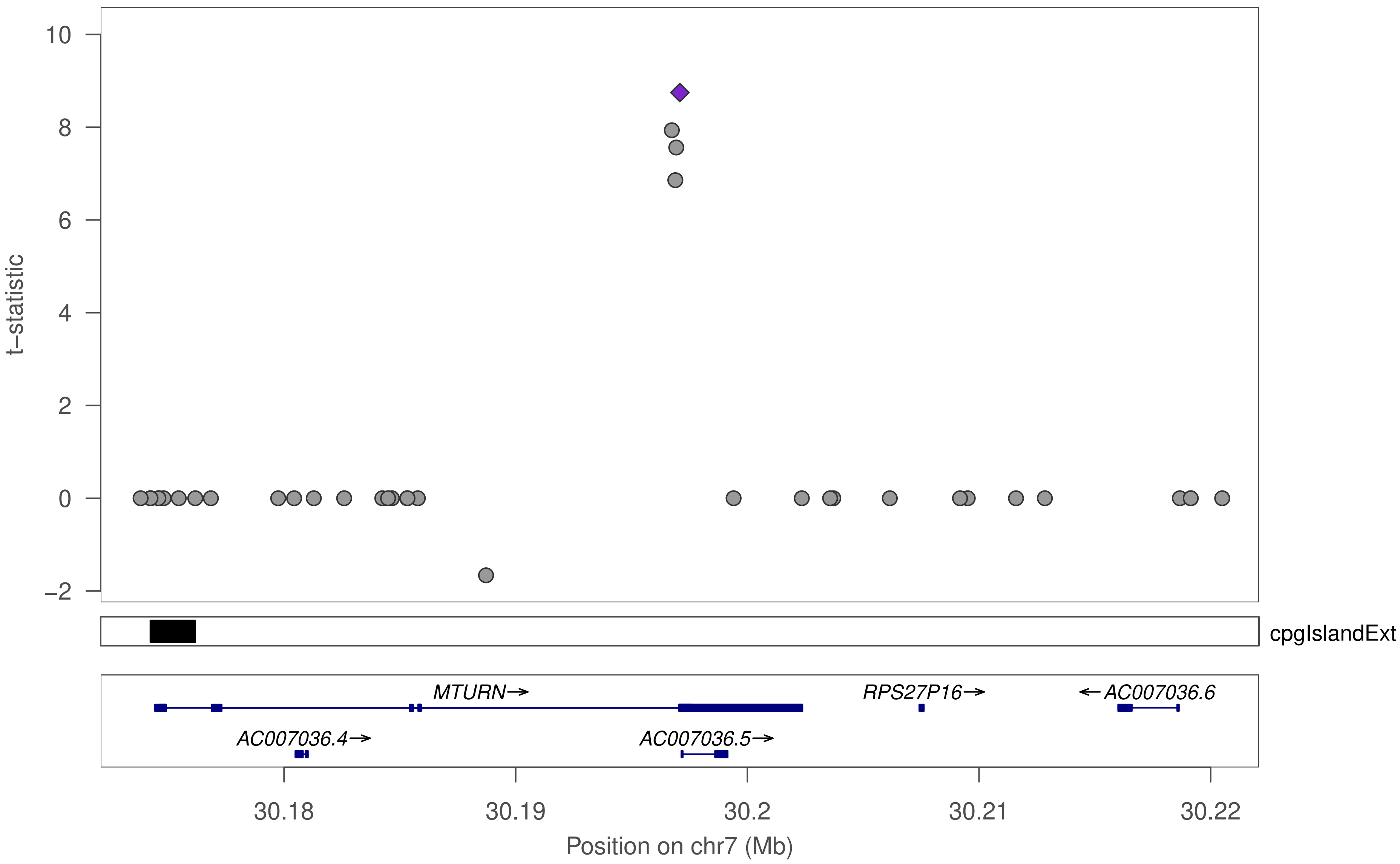

10
8
6
4
2
0
−2
t−statistic
cpgIslandExt
MTURN→
RPS27P16→
←AC007036.6
AC007036.4→
AC007036.5→
30.18
30.19
30.2
30.21
30.22
Position on chr7 (Mb)

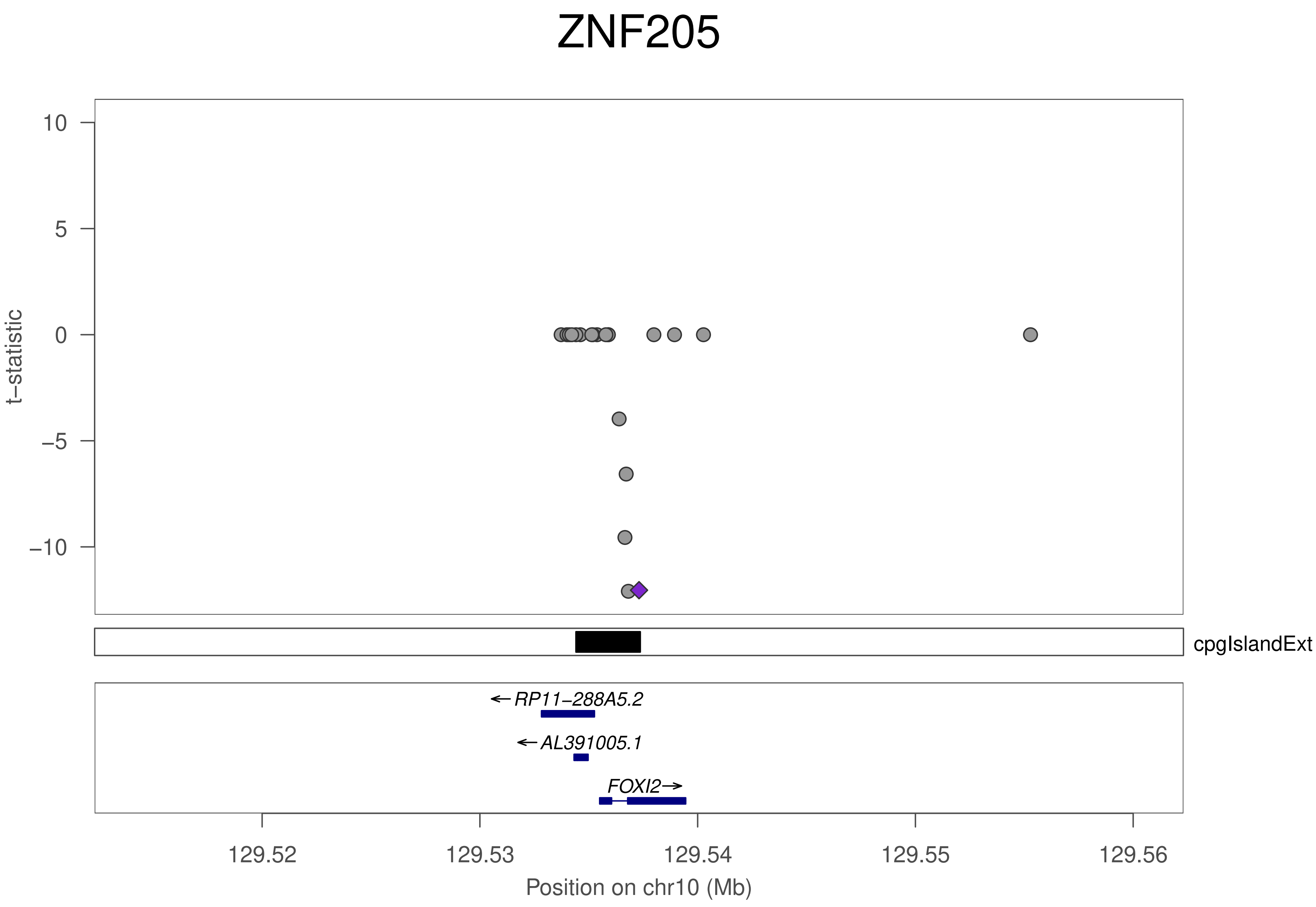

ZNF205
t−statistic
10
5
0
−5
−10
cpgIslandExt
RP11−288A5.2
AL391005.1
FOXI2
129.52
129.53
129.54
129.55
129.56
Position on chr10 (Mb)

ZNF205

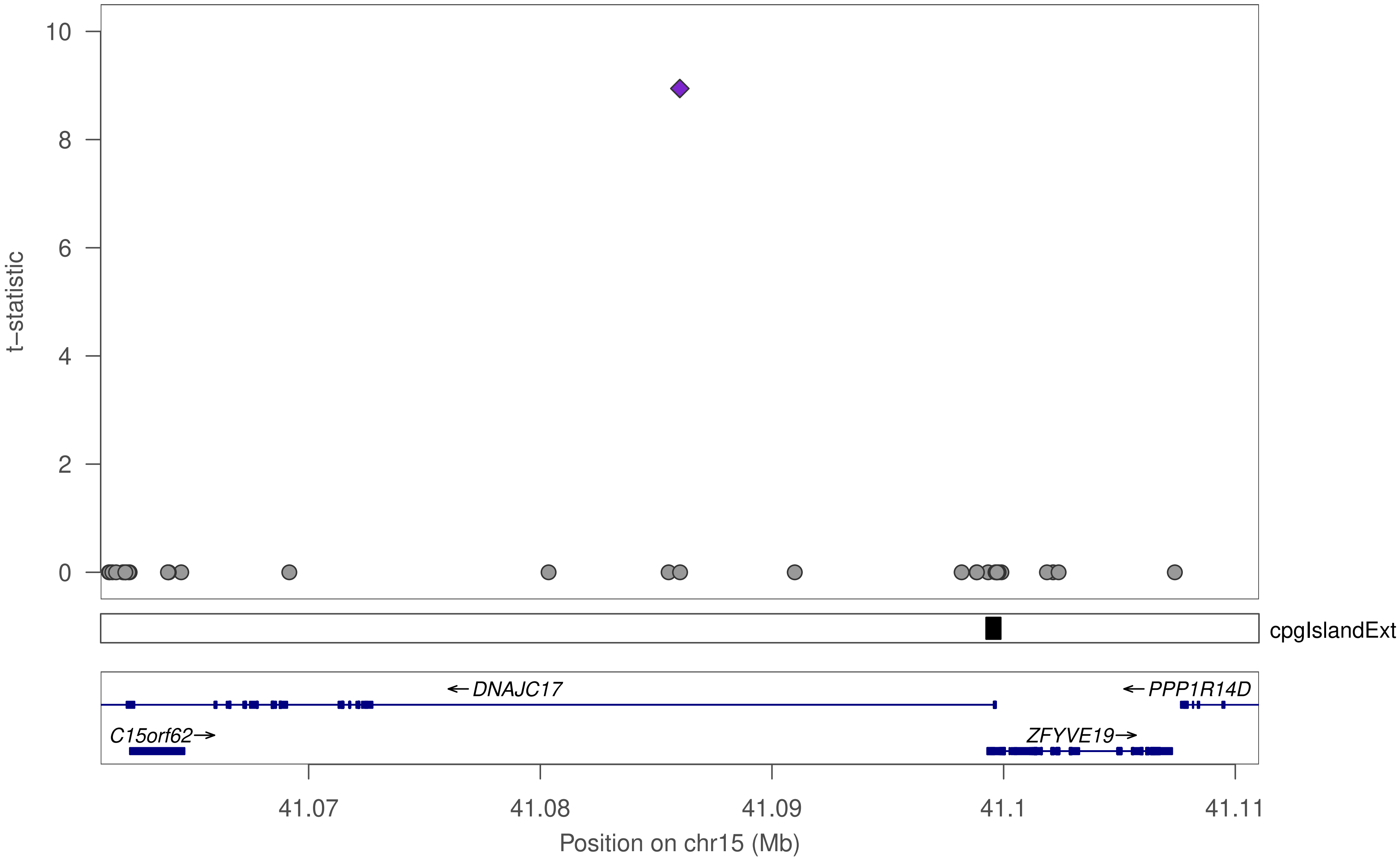

10
8
6
4
2
0
t−statistic
cpgIslandExt
←DNAJC17
←PPP1R14D
C15orf62→
ZFYVE19→
41.07
41.08
41.09
41.1
41.11
Position on chr15 (Mb)

### ZNF205

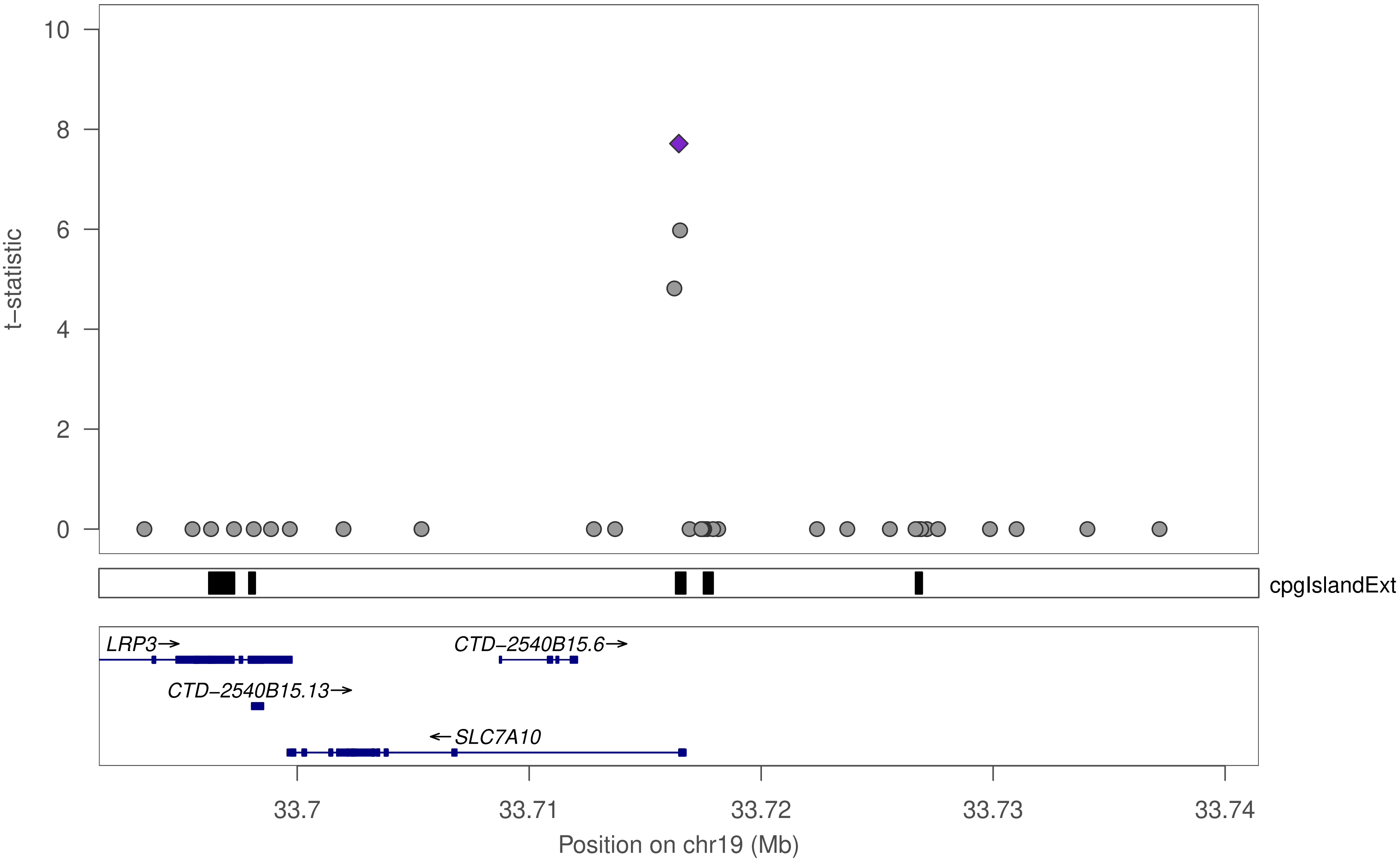

t−statistic
10
8
6
4
2
0
cpgIslandExt
LRP3→
CTD−2540B15.6→
CTD−2540B15.13→
←SLC7A10
33.7
33.71
33.72
33.73
33.74
Position on chr19 (Mb)

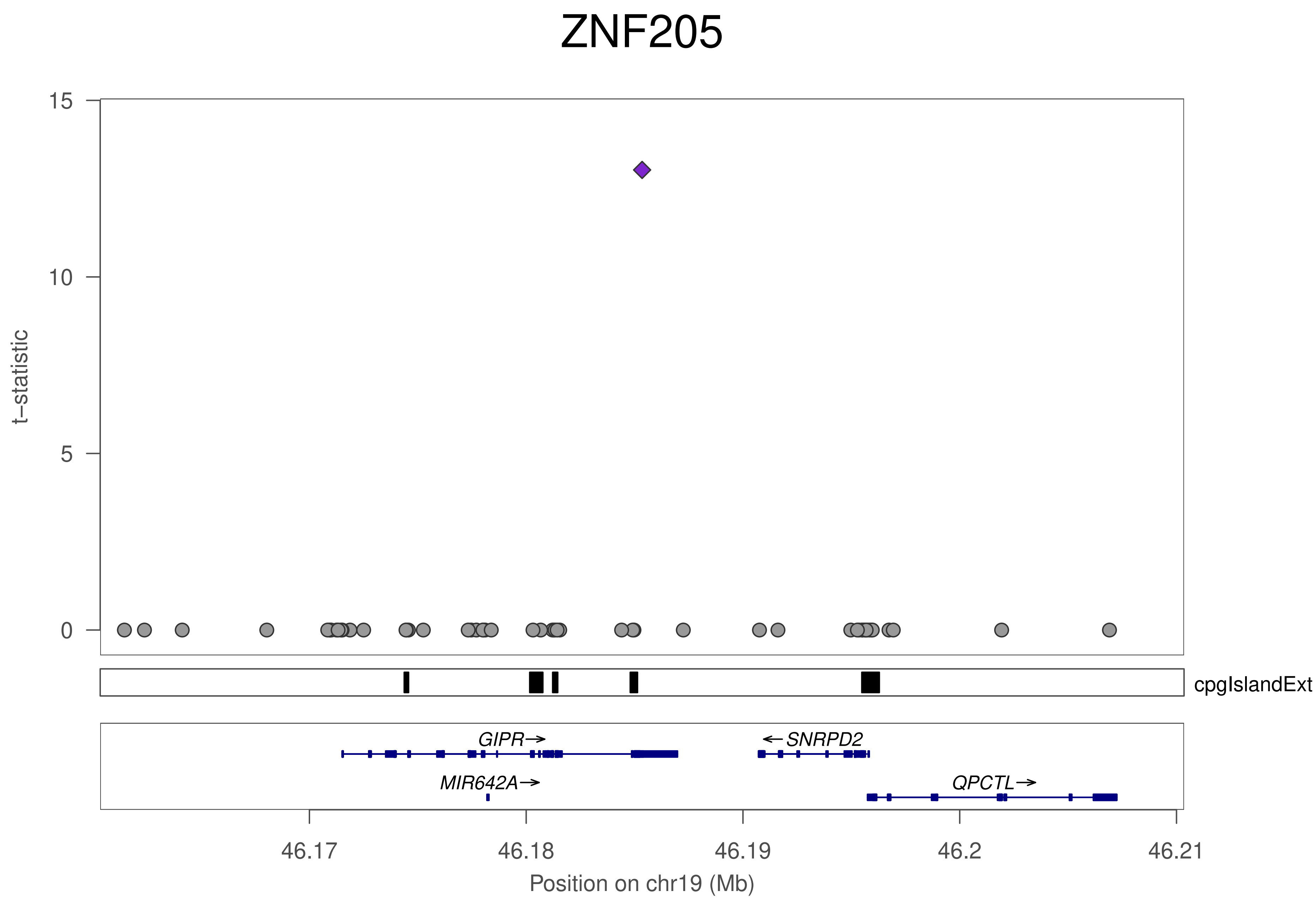
ZNF205
t−statistic
15
10
5
0
cpgIslandExt
GIPR→
←SNRPD2
MIR642A→
QPCTL→
46.17
46.18
46.19
46.2
46.21
Position on chr19 (Mb)

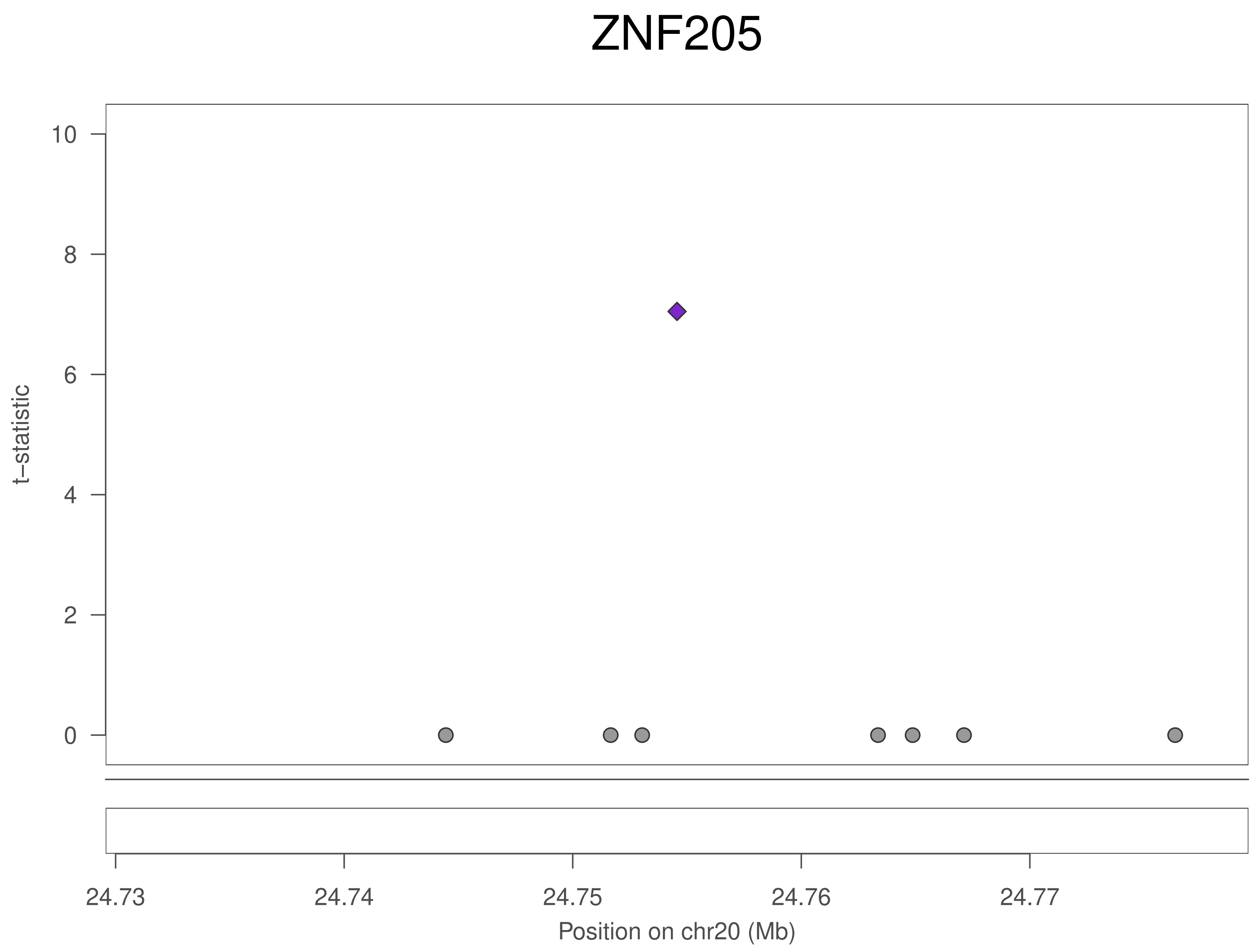

ZNF205
t–statistic
0
2
4
6
8
10
24.73
24.74
24.75
24.76
24.77
Position on chr20 (Mb)

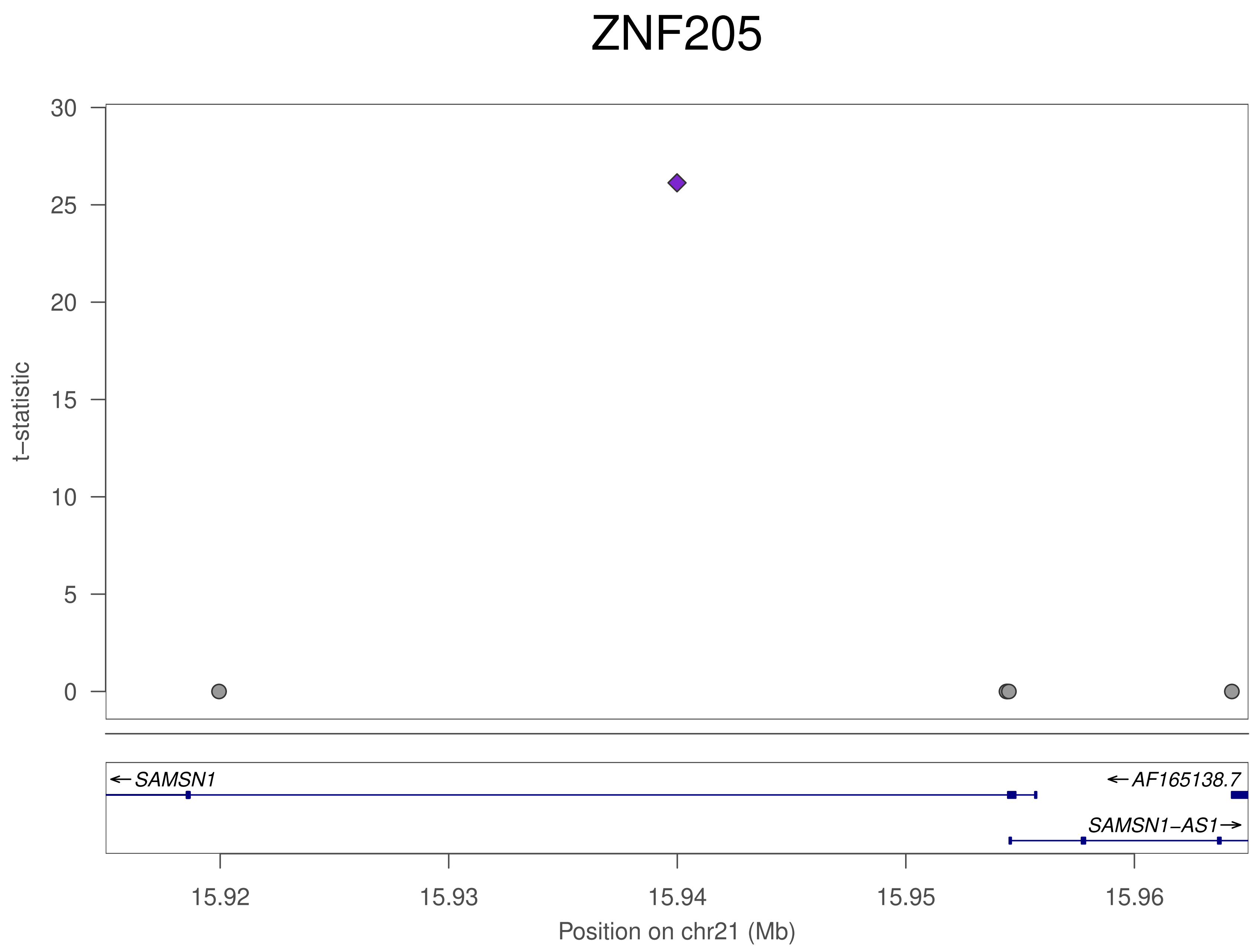
ZNF205
t−statistic
30
25
20
15
10
5
0
SAMSN1
AF165138.7
SAMSN1−AS1
15.92
15.93
15.94
15.95
15.96
Position on chr21 (Mb)

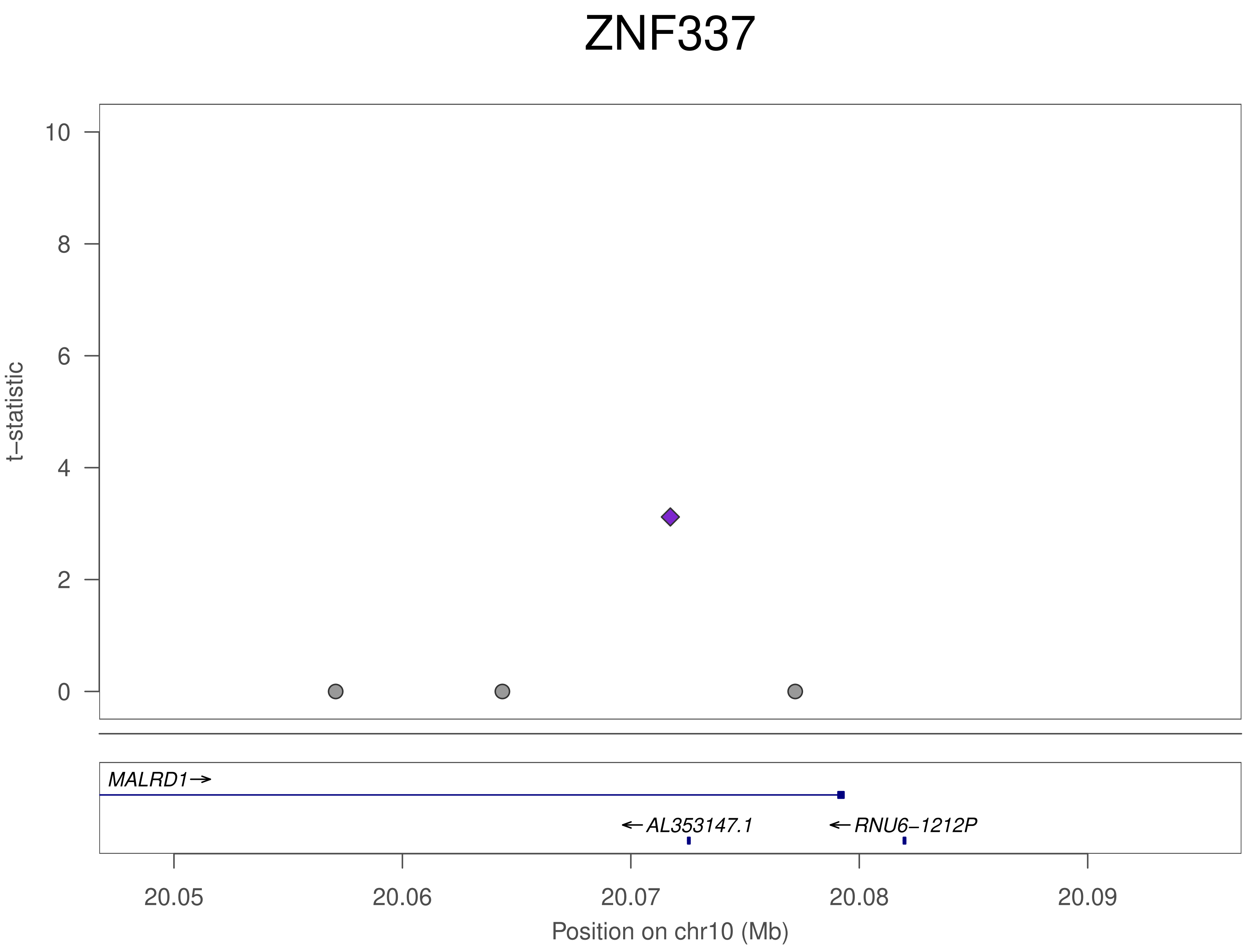
ZNF337
t−statistic
10
8
6
4
2
0
MALRD1→
←AL353147.1
←RNU6−1212P
20.05
20.06
20.07
20.08
20.09
Position on chr10 (Mb)

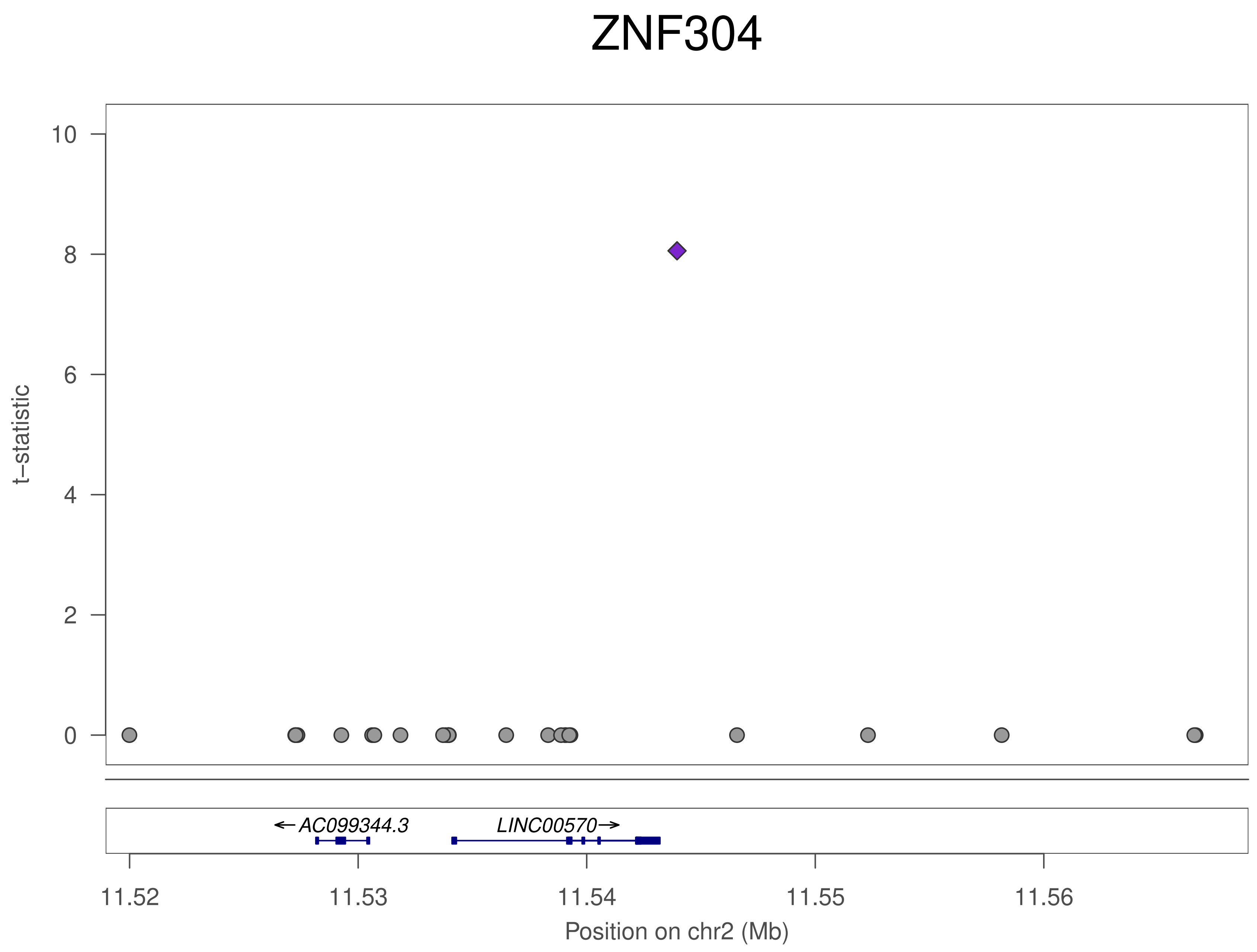
ZNF304
t–statistic
10
8
6
4
2
0
AC099344.3
LINC00570
11.52
11.53
11.54
11.55
11.56
Position on chr2 (Mb)

### ZNF304

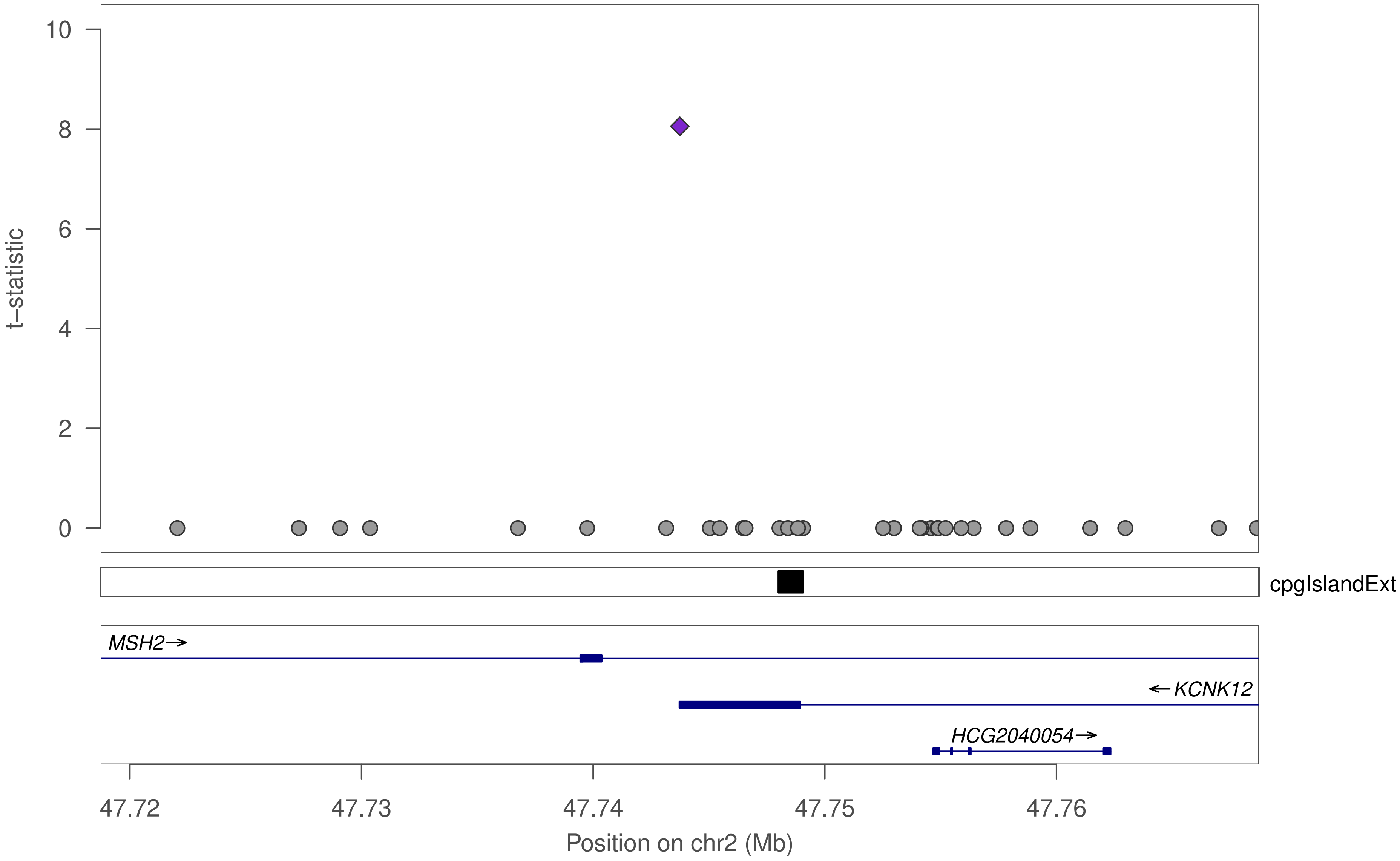

t–statistic
10
8
6
4
2
0
cpgIslandExt
MSH2→
←KCNK12
HCG2040054→
47.72
47.73
47.74
47.75
47.76
Position on chr2 (Mb)

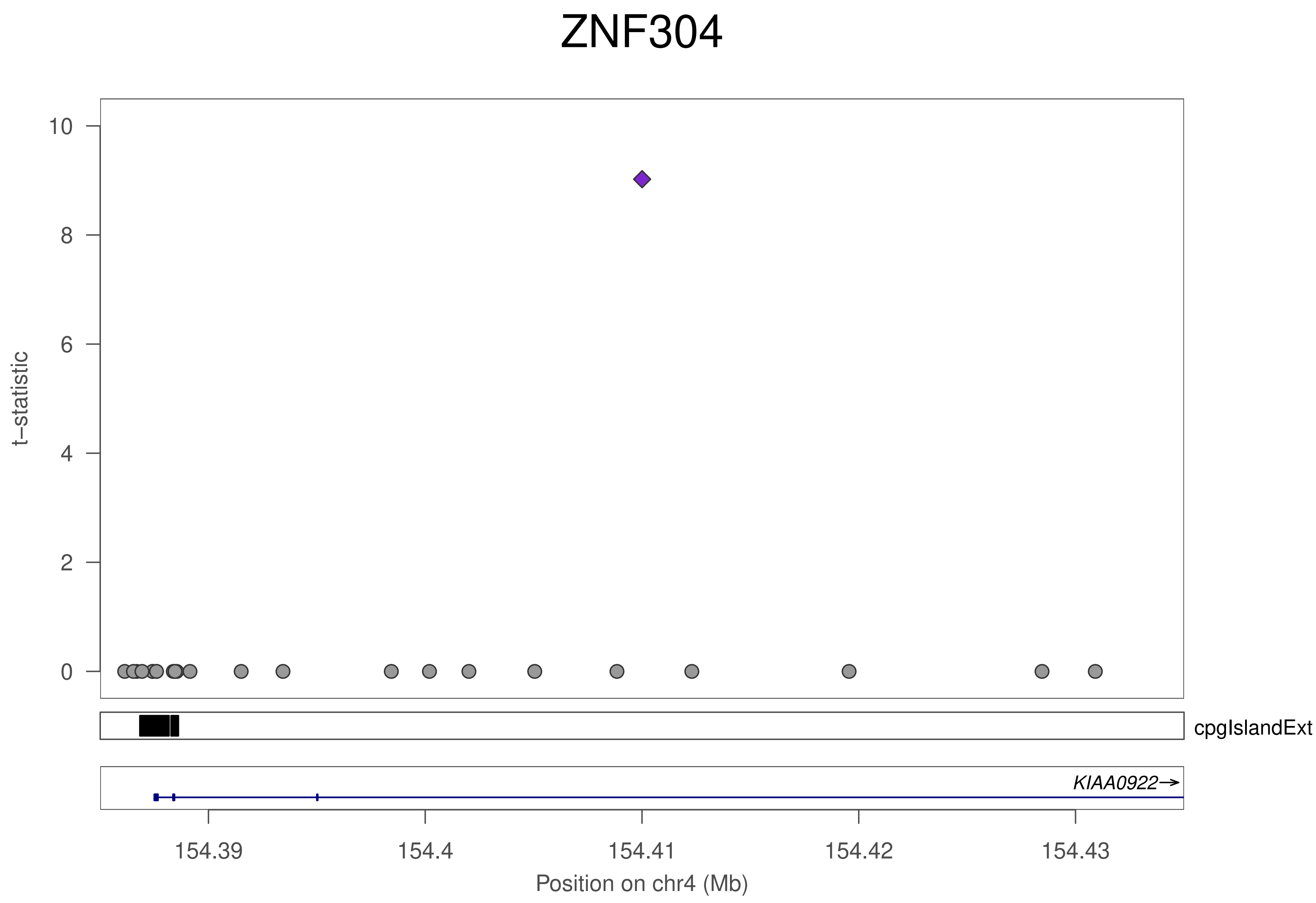
ZNF304
t−statistic
0
2
4
6
8
10
cpgIslandExt
KIAA0922→
154.39
154.4
154.41
154.42
154.43
Position on chr4 (Mb)

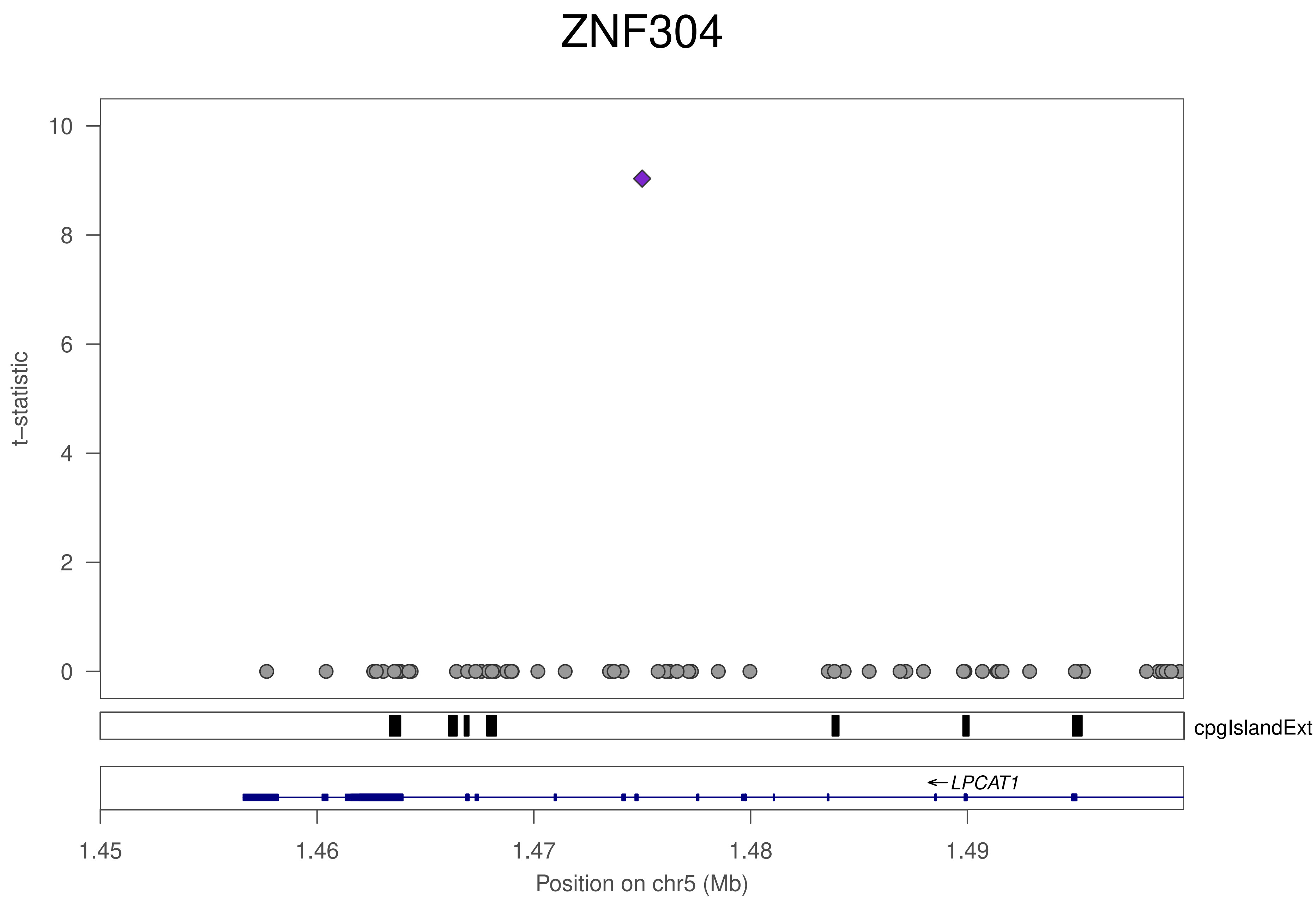
ZNF304
t−statistic
0
2
4
6
8
10
cpgIslandExt
←LPCAT1
1.45
1.46
1.47
1.48
1.49
Position on chr5 (Mb)

### ZNF304

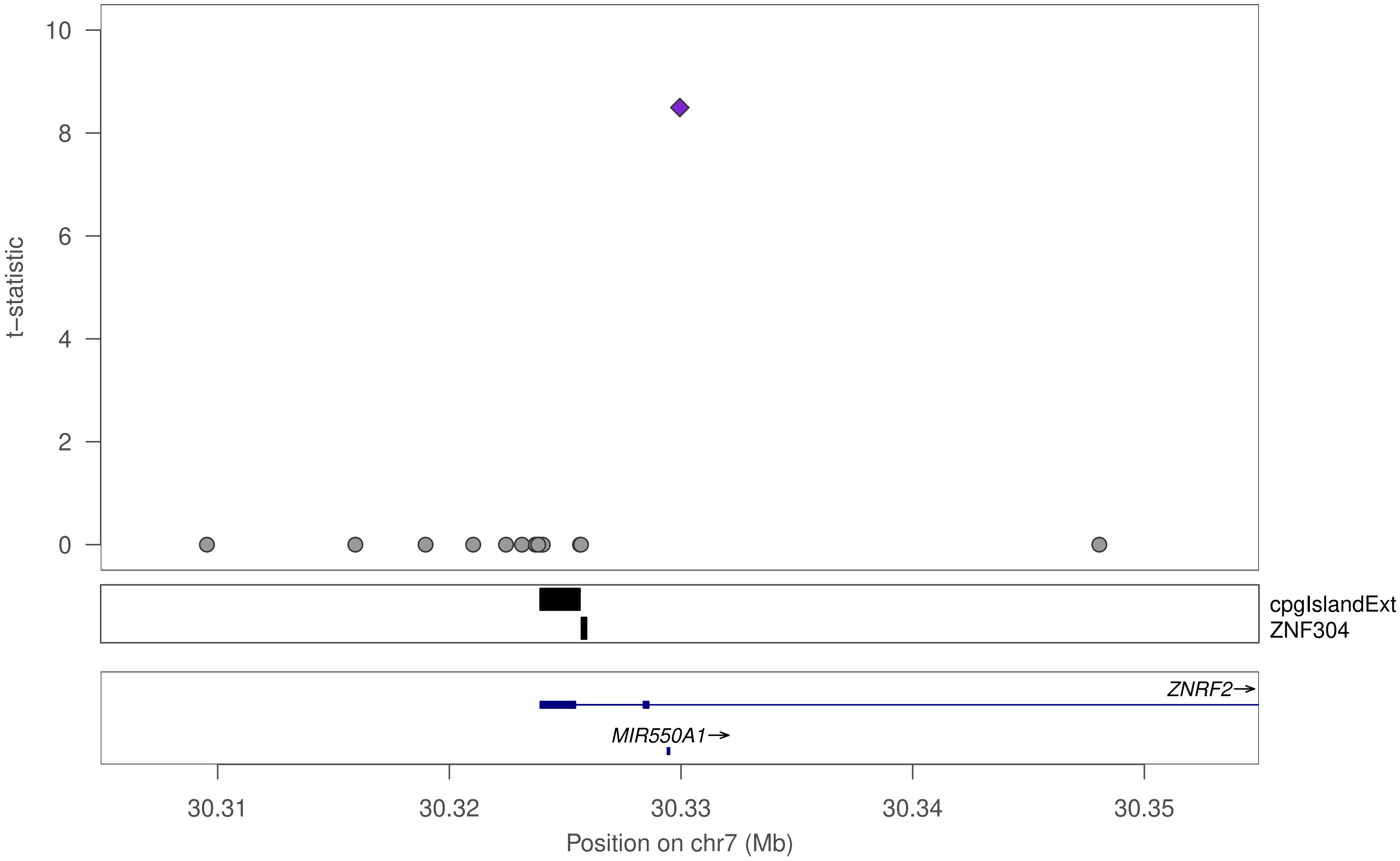

10
8
6
4
2
0
t–statistic
cpgIslandExt
ZNF304
ZNRF2→
MIR550A1→
30.31
30.32
30.33
30.34
30.35
Position on chr7 (Mb)

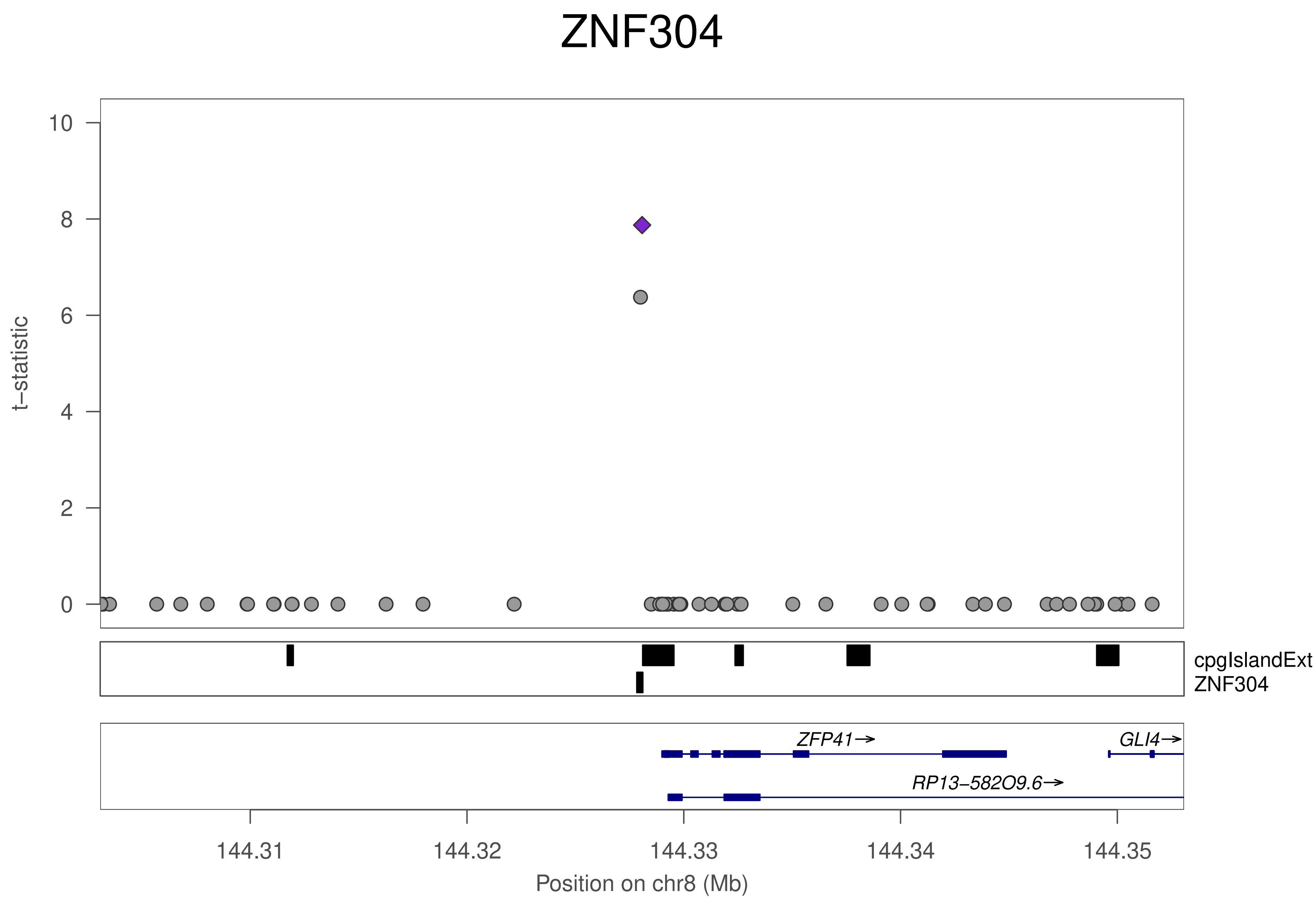

ZNF304
t−statistic
10
8
6
4
2
0
cpgIslandExt
ZNF304
ZFP41→
GLI4→
RP13−582O9.6→
144.31
144.32
144.33
144.34
144.35
Position on chr8 (Mb)

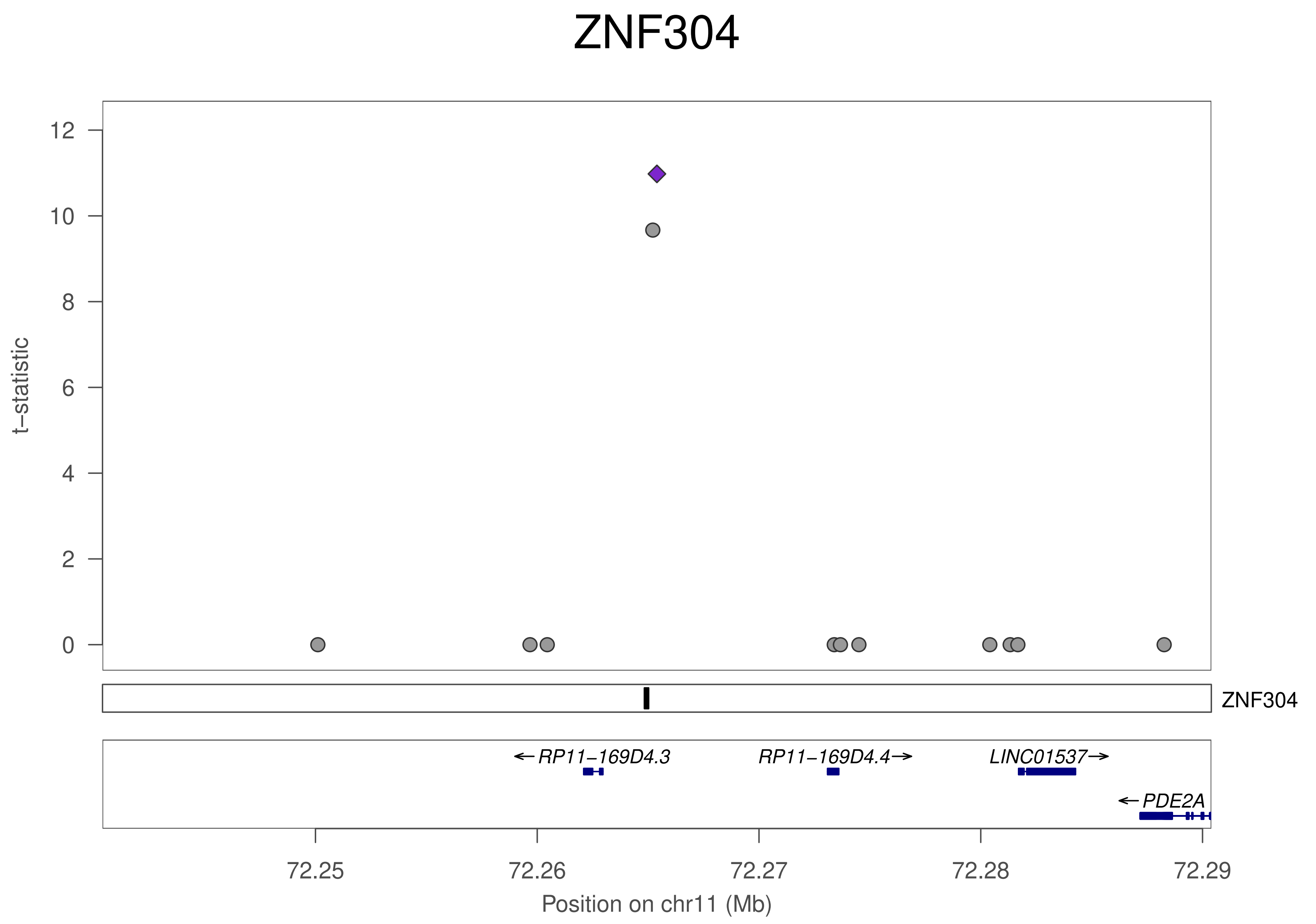
ZNF304
t−statistic
12
10
8
6
4
2
0
ZNF304
←RP11−169D4.3
RP11−169D4.4→
LINC01537→
←PDE2A
72.25
72.26
72.27
72.28
72.29
Position on chr11 (Mb)

### ZNF304

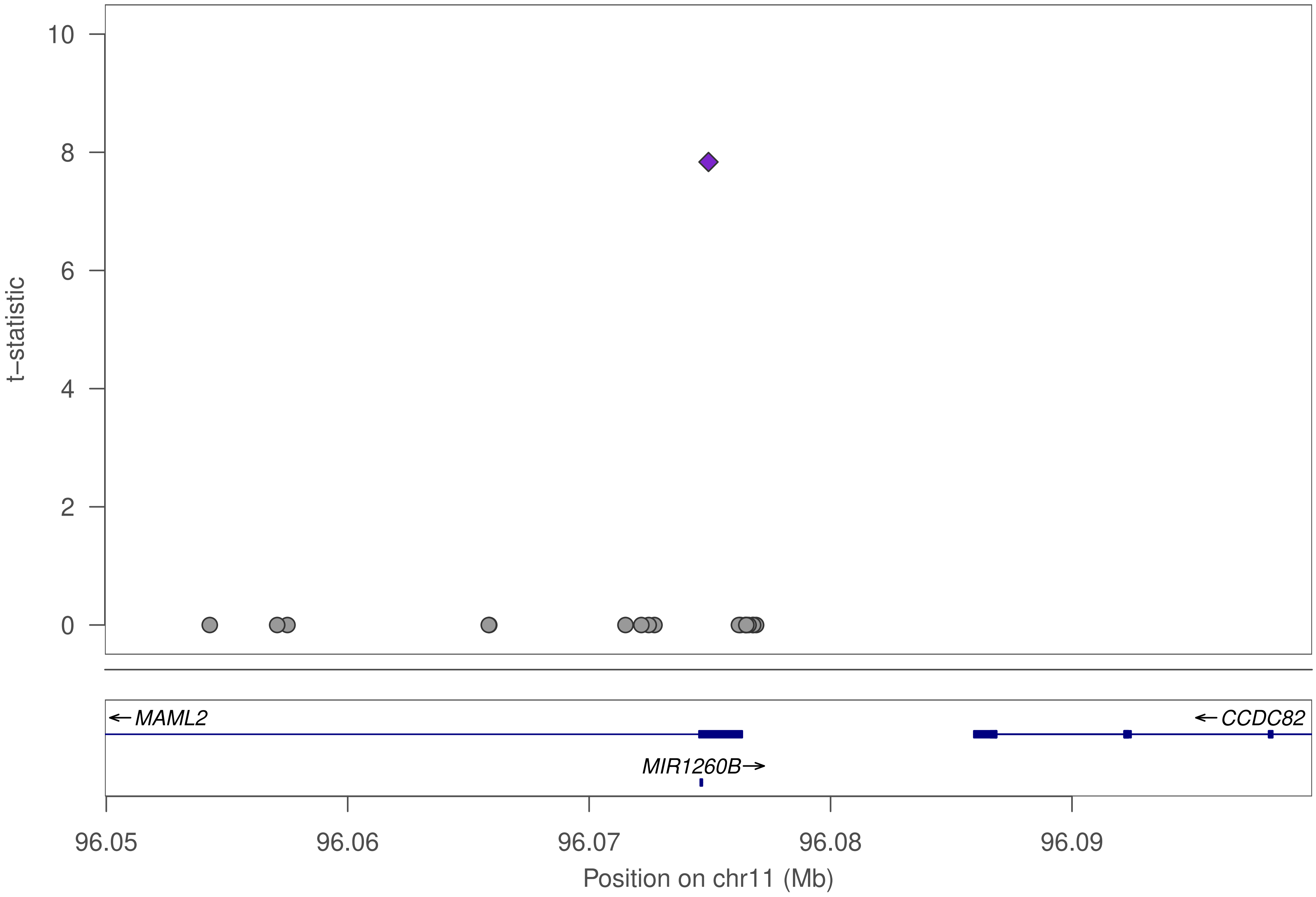

10
8
6
4
2
0
t–statistic
← MAML2
← CCDC82
MIR1260B→
96.05
96.06
96.07
96.08
96.09
Position on chr11 (Mb)

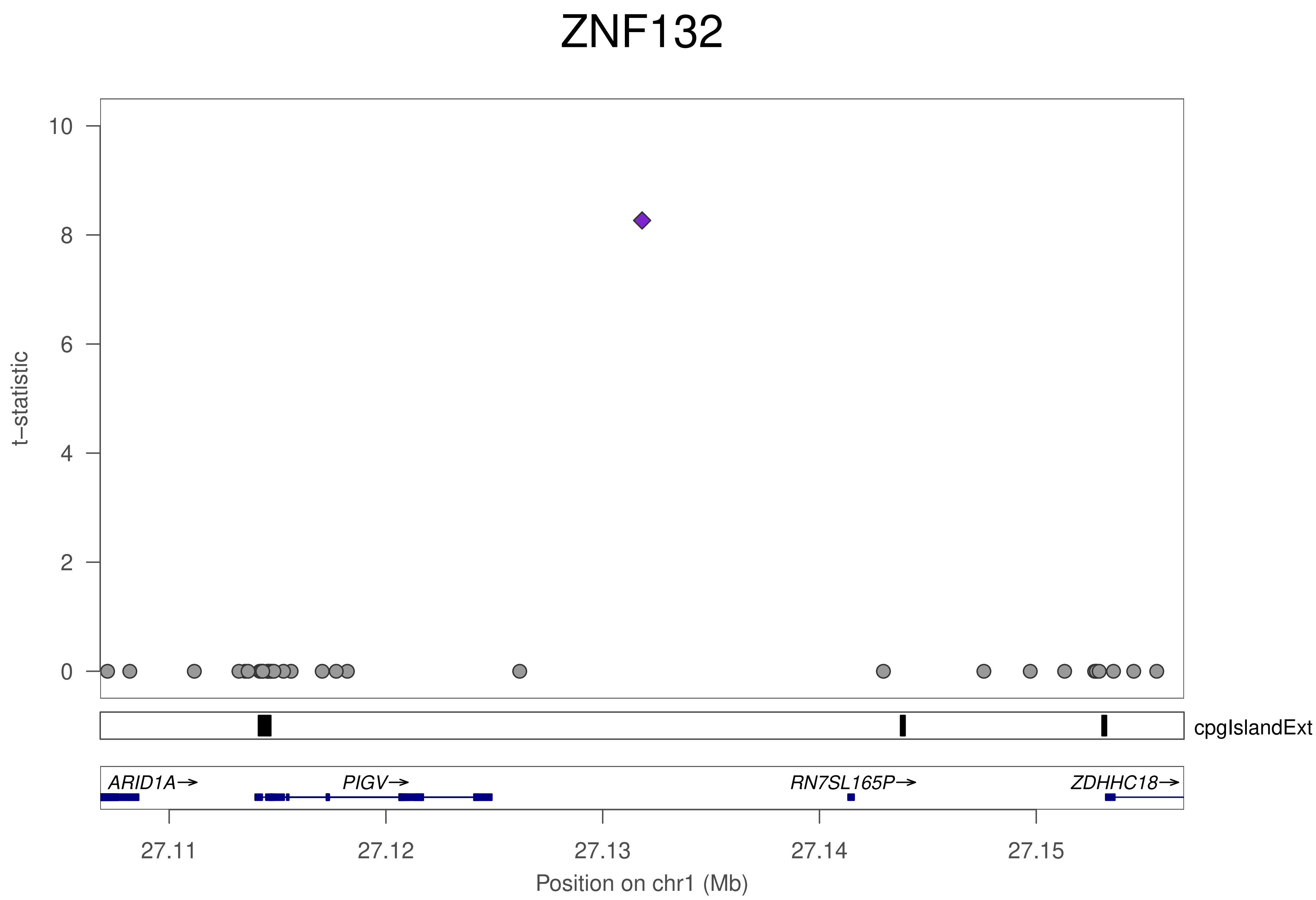
ZNF132
t−statistic
10
8
6
4
2
0
cpgIslandExt
ARID1A→
PIGV→
RN7SL165P→
ZDHHC18→
27.11
27.12
27.13
27.14
27.15
Position on chr1 (Mb)

### ZNF132

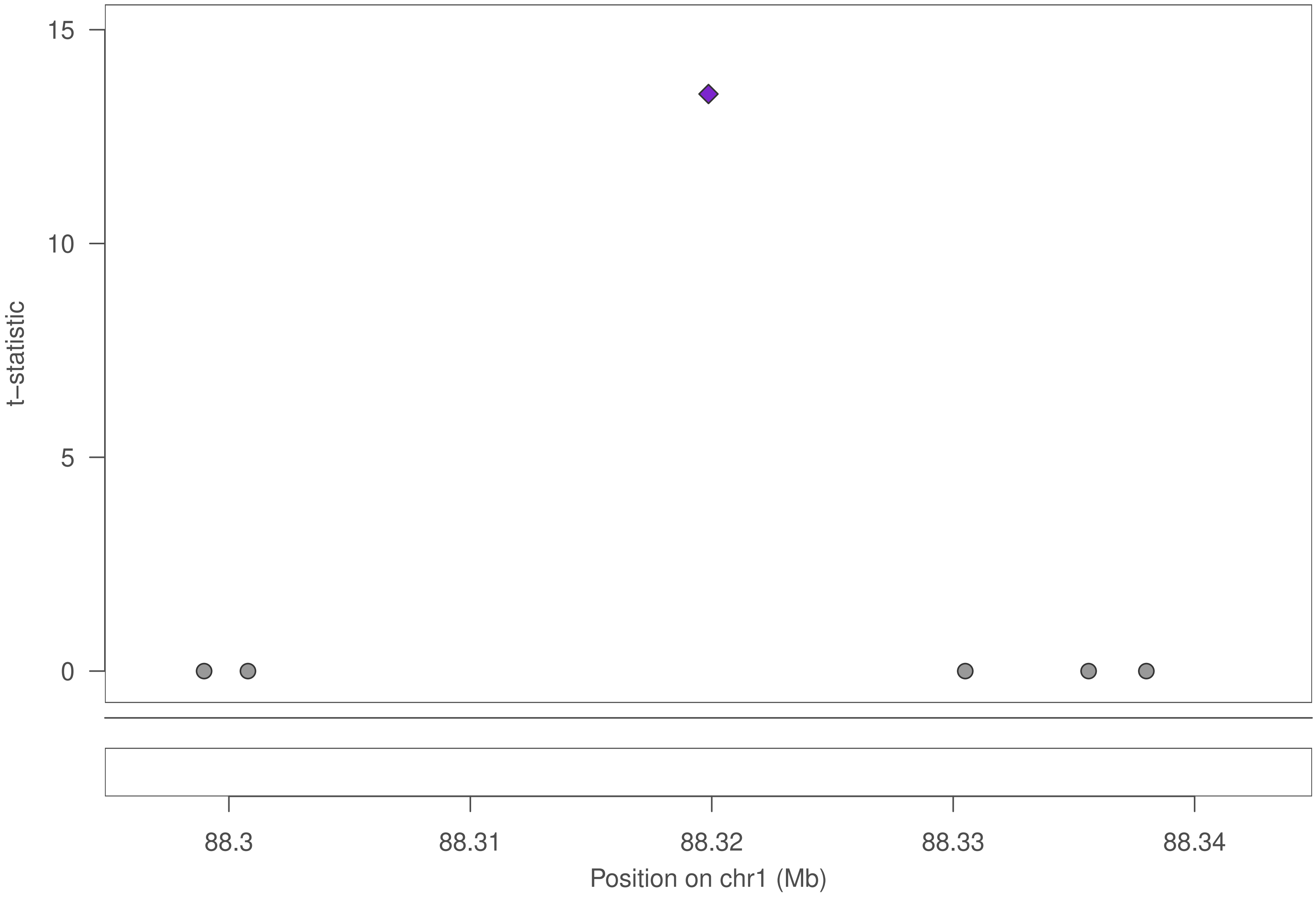

15
10
5
0
t–statistic
88.3
88.31
88.32
88.33
88.34
Position on chr1 (Mb)

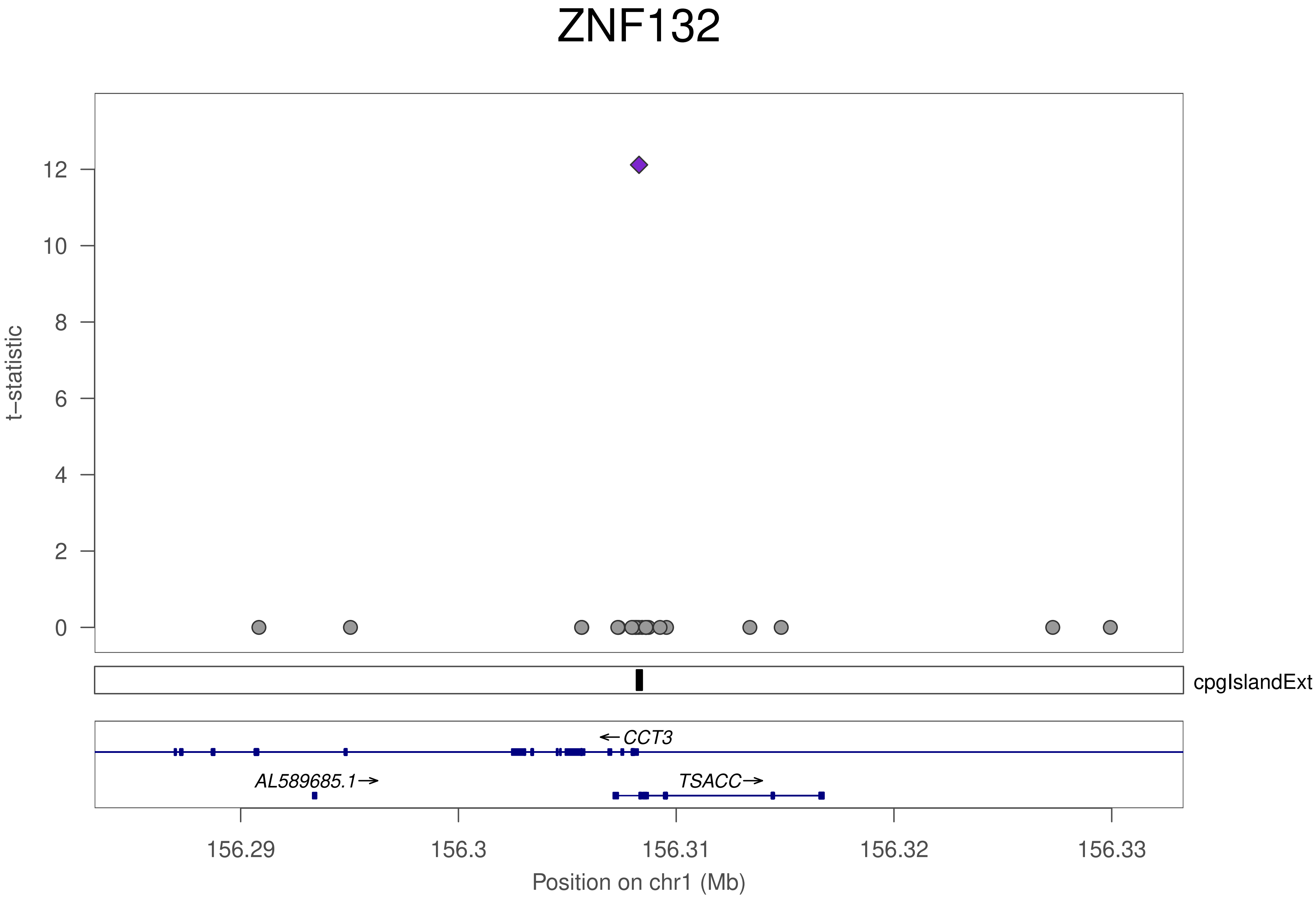
ZNF132
t−statistic
12
10
8
6
4
2
0
cpgIslandExt
←CCT3
AL589685.1→
TSACC→
156.29
156.3
156.31
156.32
156.33
Position on chr1 (Mb)

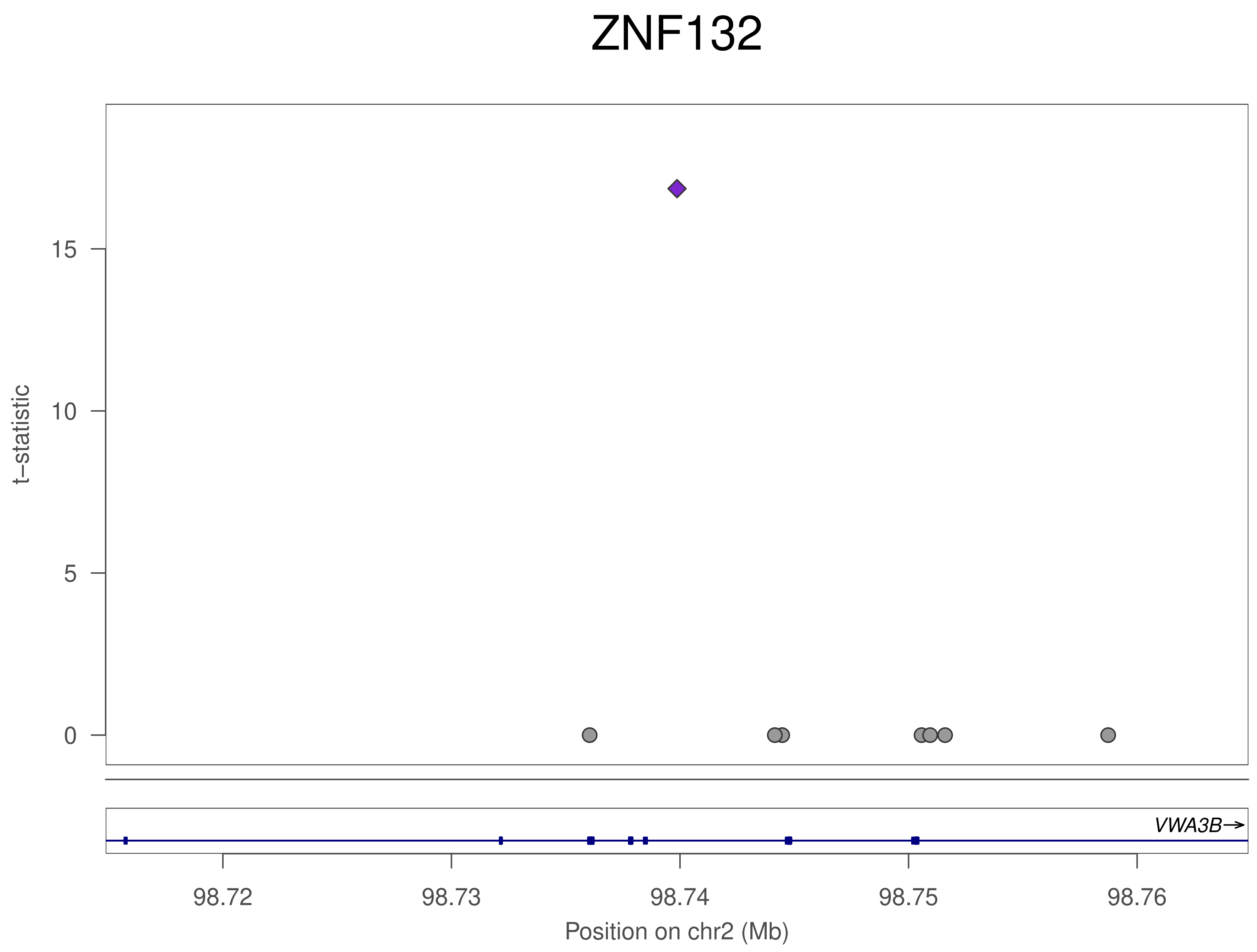
ZNF132
t–statistic
0
5
10
15
VWA3B→
98.72
98.73
98.74
98.75
98.76
Position on chr2 (Mb)

### ZNF132

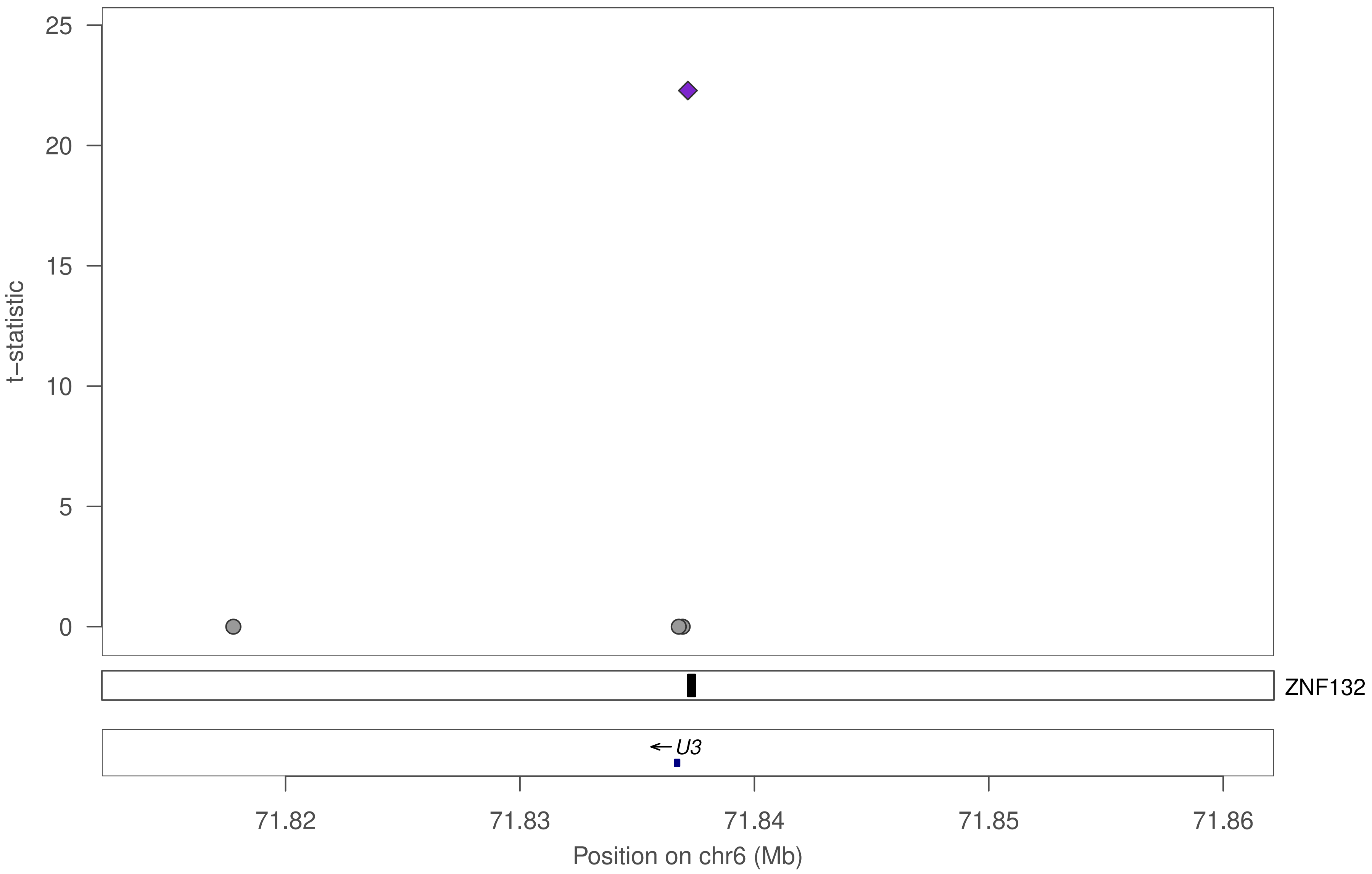

25
20
15
10
5
0
t−statistic
ZNF132
← U3
71.82
71.83
71.84
71.85
71.86
Position on chr6 (Mb)

### ZNF132

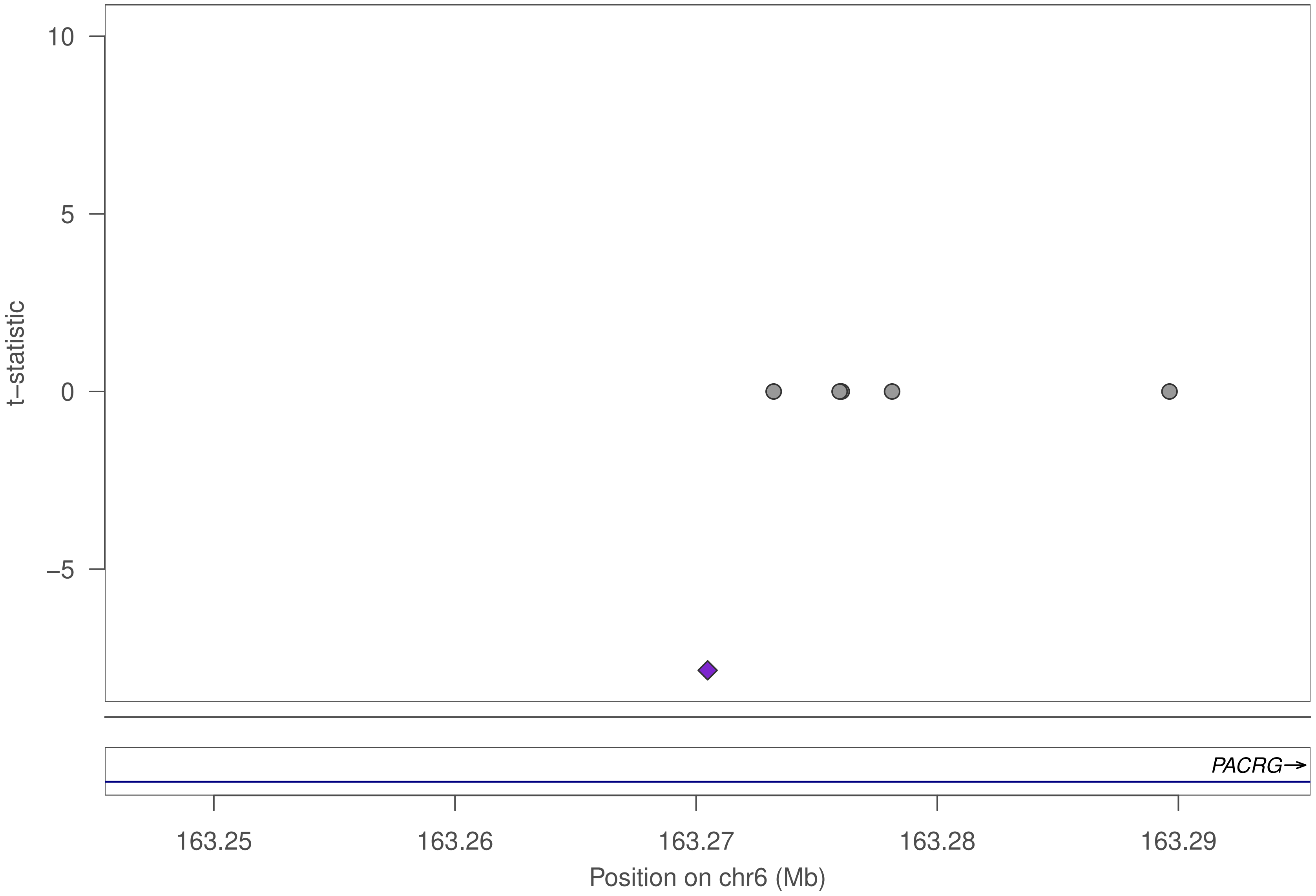

10
5
0
−5
t−statistic
PACRG→
163.25
163.26
163.27
163.28
163.29
Position on chr6 (Mb)

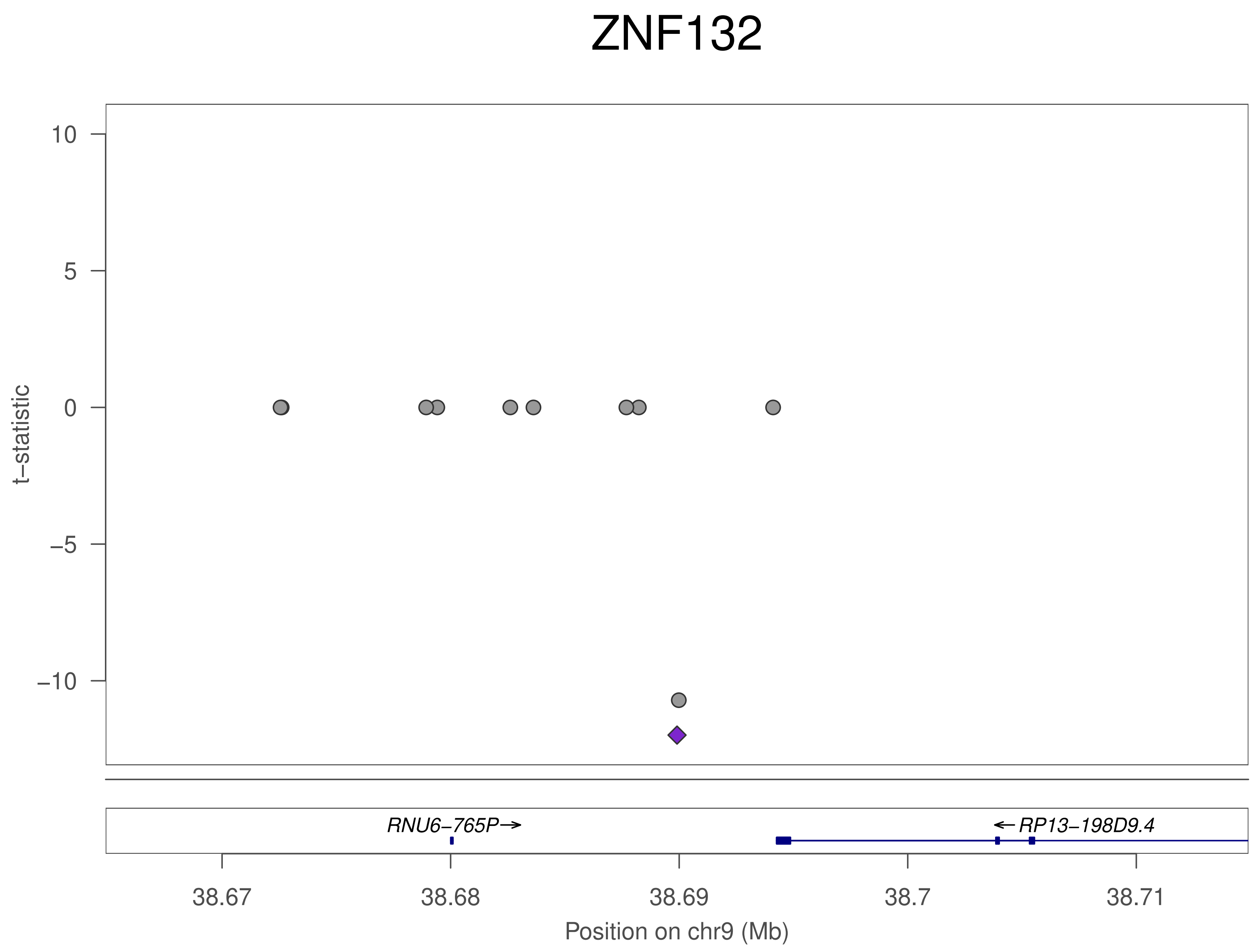
ZNF132
t−statistic
10
5
0
−5
−10
RNU6−765P→
←RP13−198D9.4
38.67
38.68
38.69
38.7
38.71
Position on chr9 (Mb)

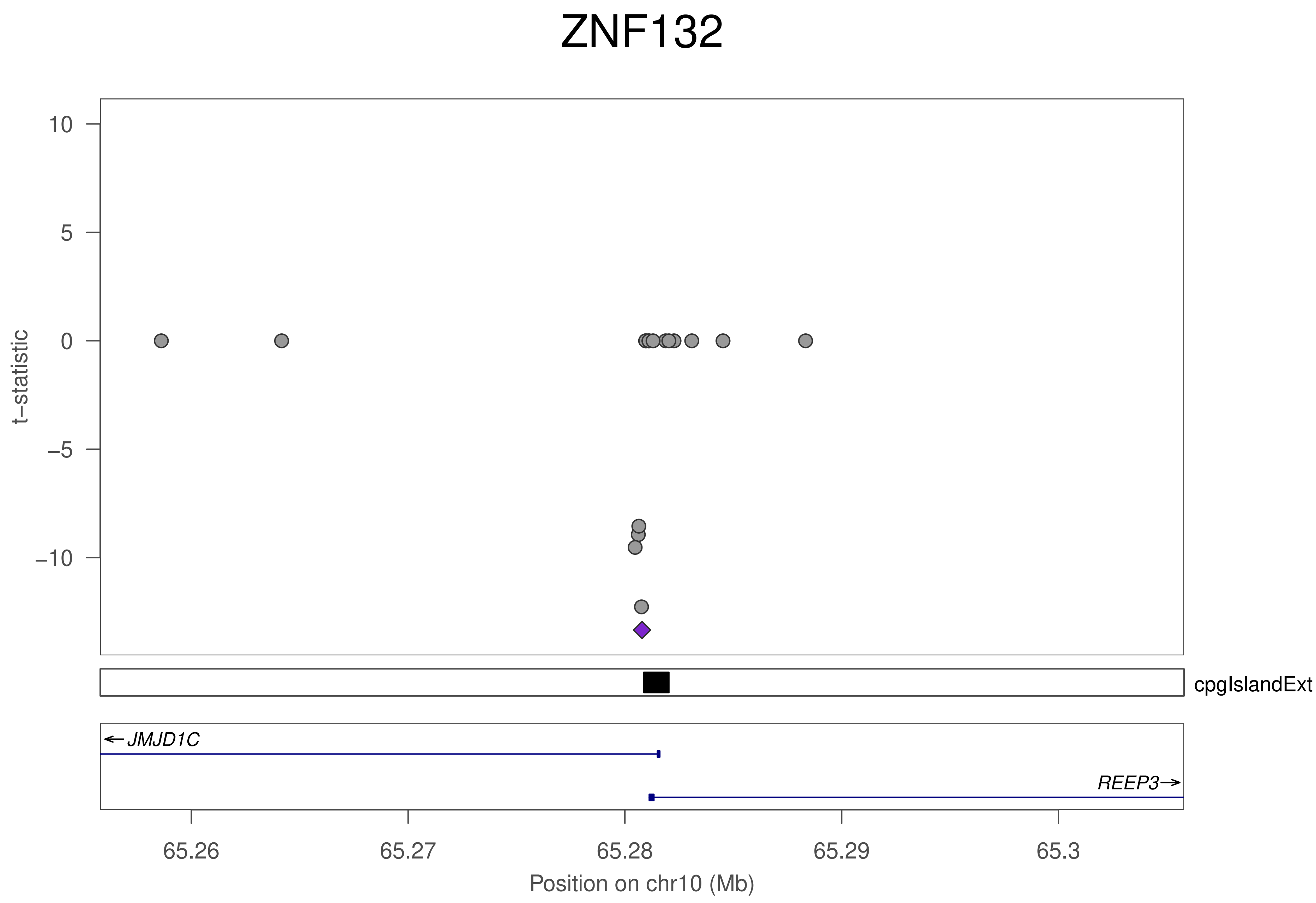
ZNF132
t−statistic
10
5
0
−5
−10
cpgIslandExt
← JMJD1C
REEP3→
65.26
65.27
65.28
65.29
65.3
Position on chr10 (Mb)

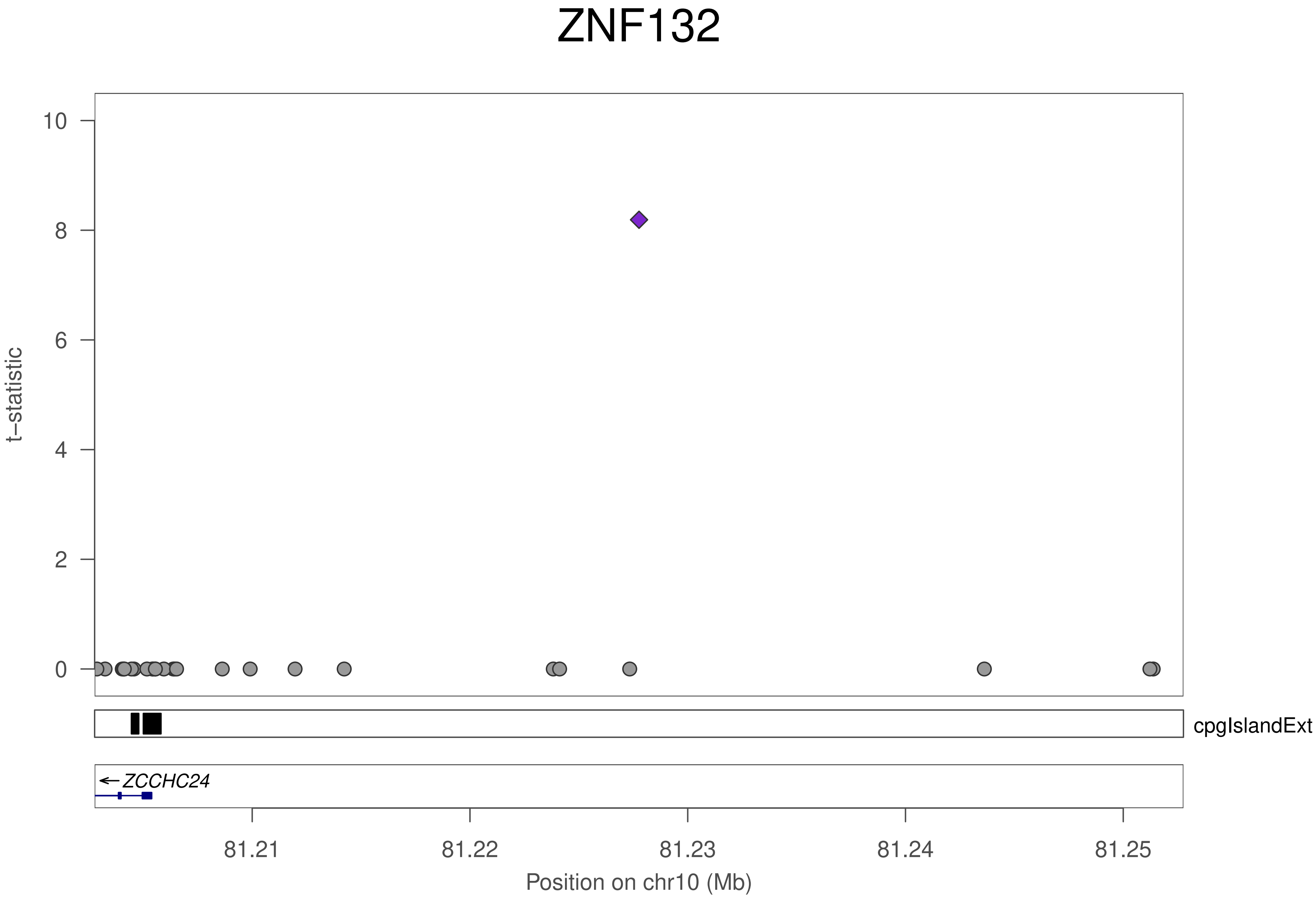

ZNF132
t–statistic
10
8
6
4
2
0
cpgIslandExt
ZCCHC24
81.21
81.22
81.23
81.24
81.25
Position on chr10 (Mb)

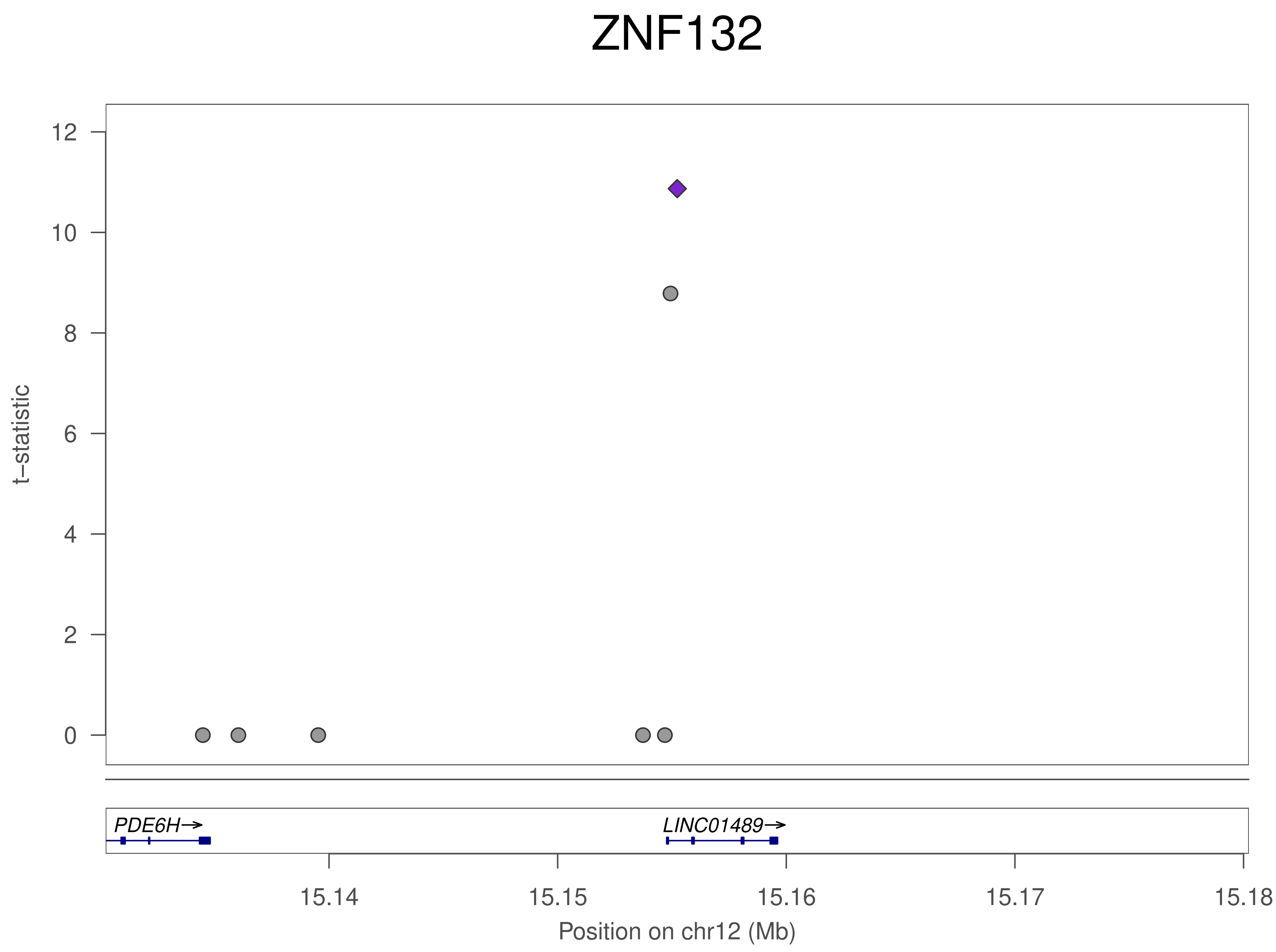
ZNF132
t–statistic
12
10
8
6
4
2
0
PDE6H→
LINC01489→
15.14
15.15
15.16
15.17
15.18
Position on chr12 (Mb)

### ZNF132

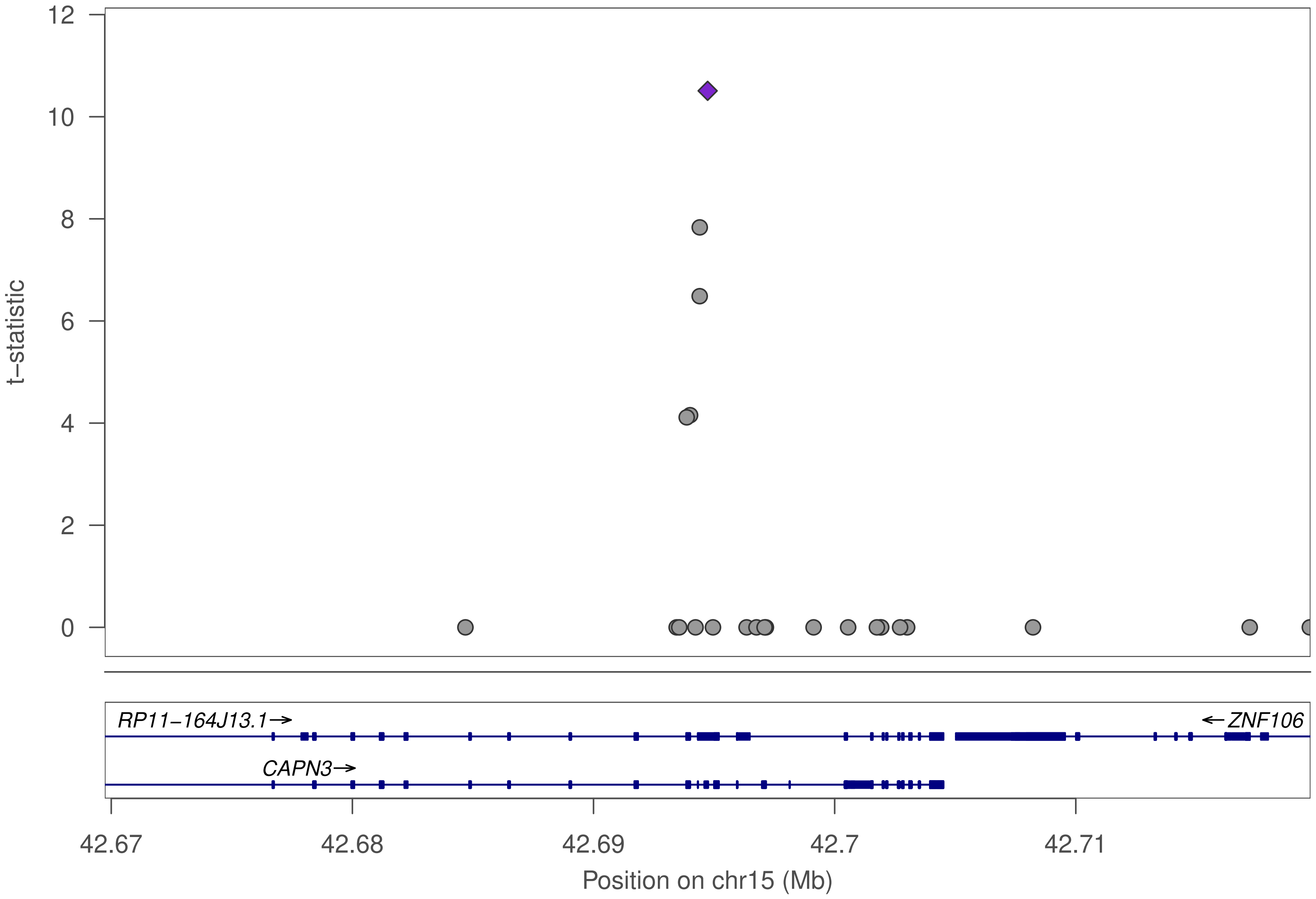

12
10
8
6
4
2
0
t−statistic
RP11−164J13.1→
←ZNF106
CAPN3→
42.67
42.68
42.69
42.7
42.71
Position on chr15 (Mb)

### ZNF132

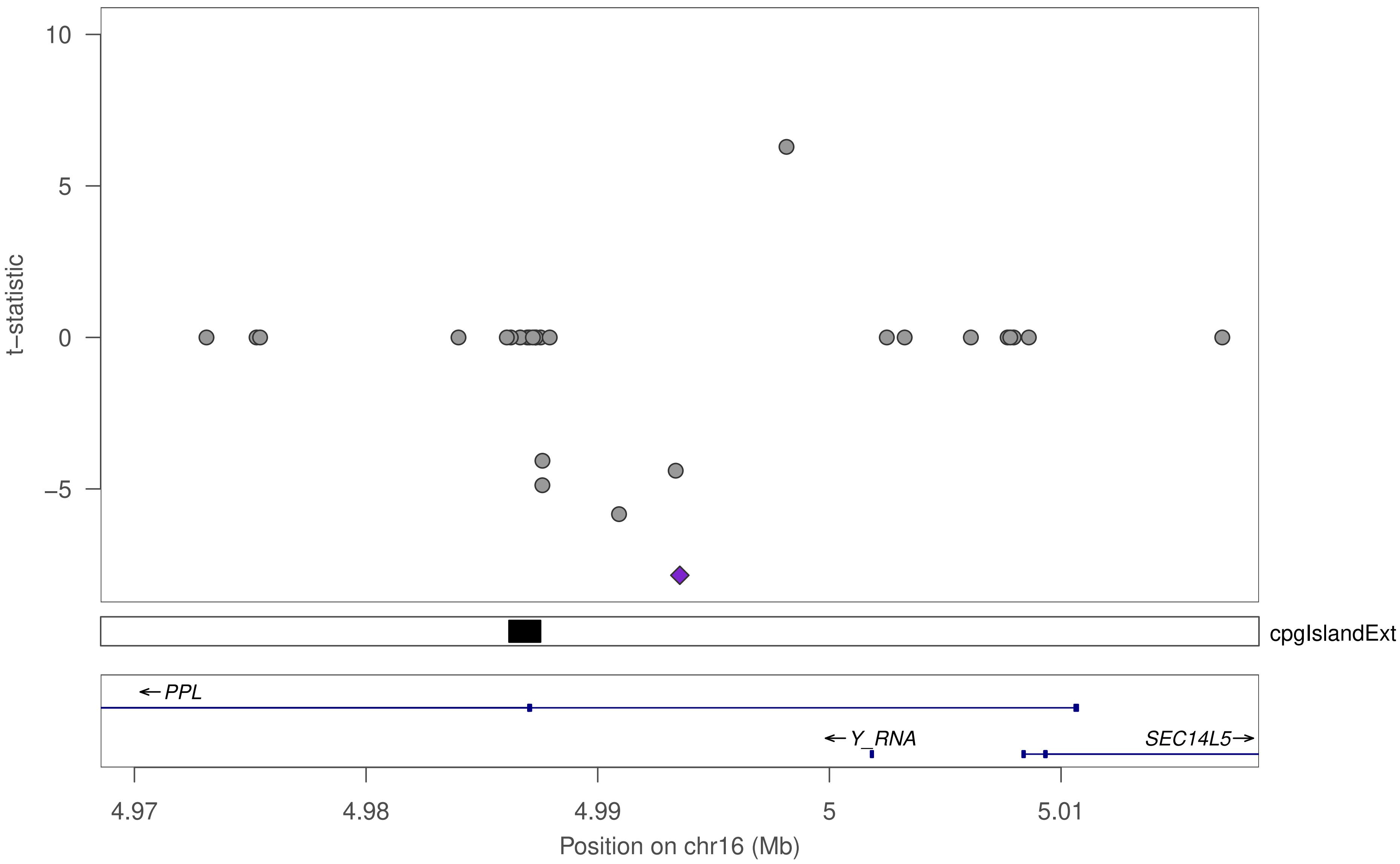

10
5
0
−5
t−statistic
cpgIslandExt
← PPL
← Y_RNA
SEC14L5→
4.97
4.98
4.99
5
5.01
Position on chr16 (Mb)

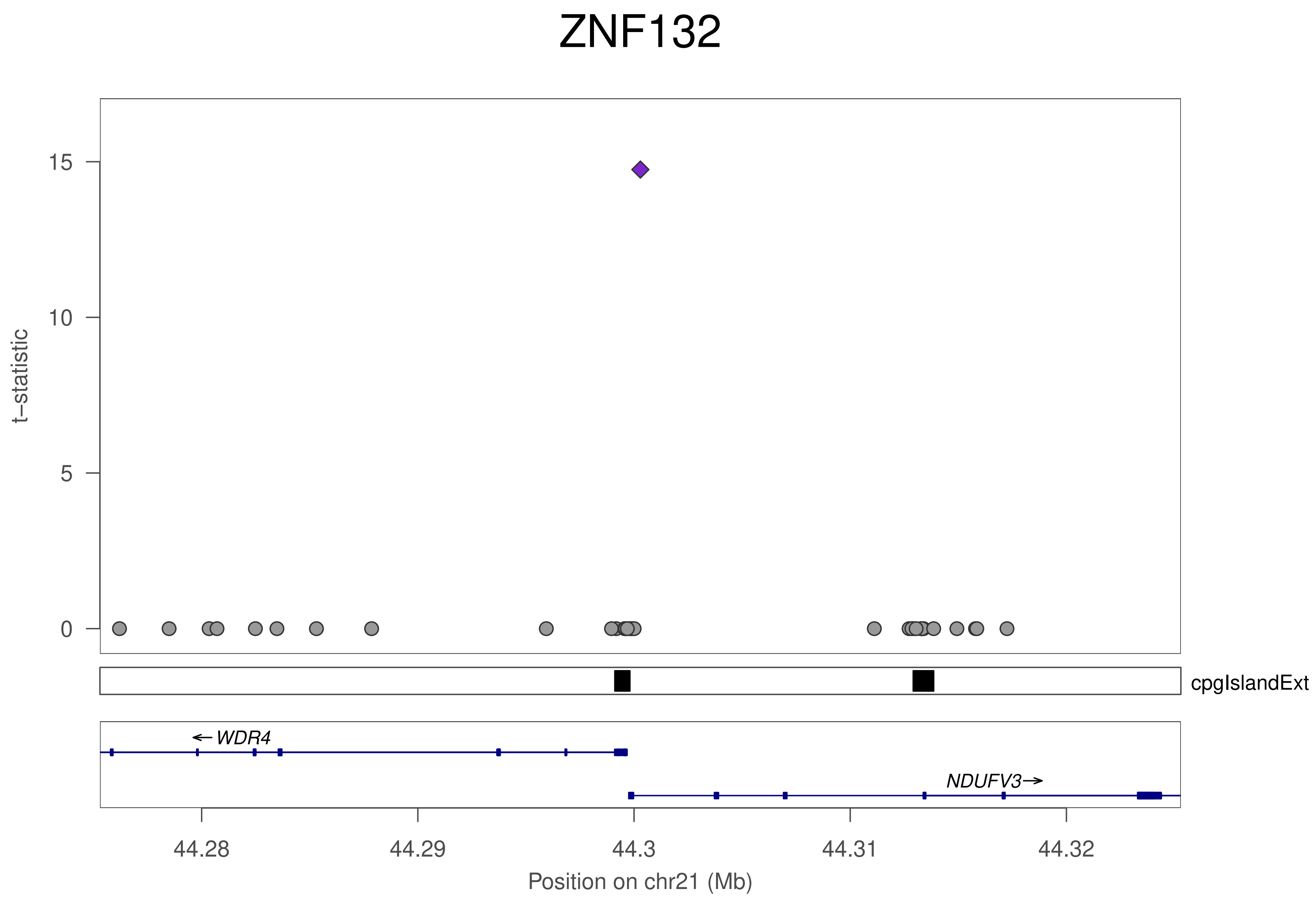
ZNF132
t−statistic
15
10
5
0
cpgIslandExt
←WDR4
NDUFV3→
44.28
44.29
44.3
44.31
44.32
Position on chr21 (Mb)

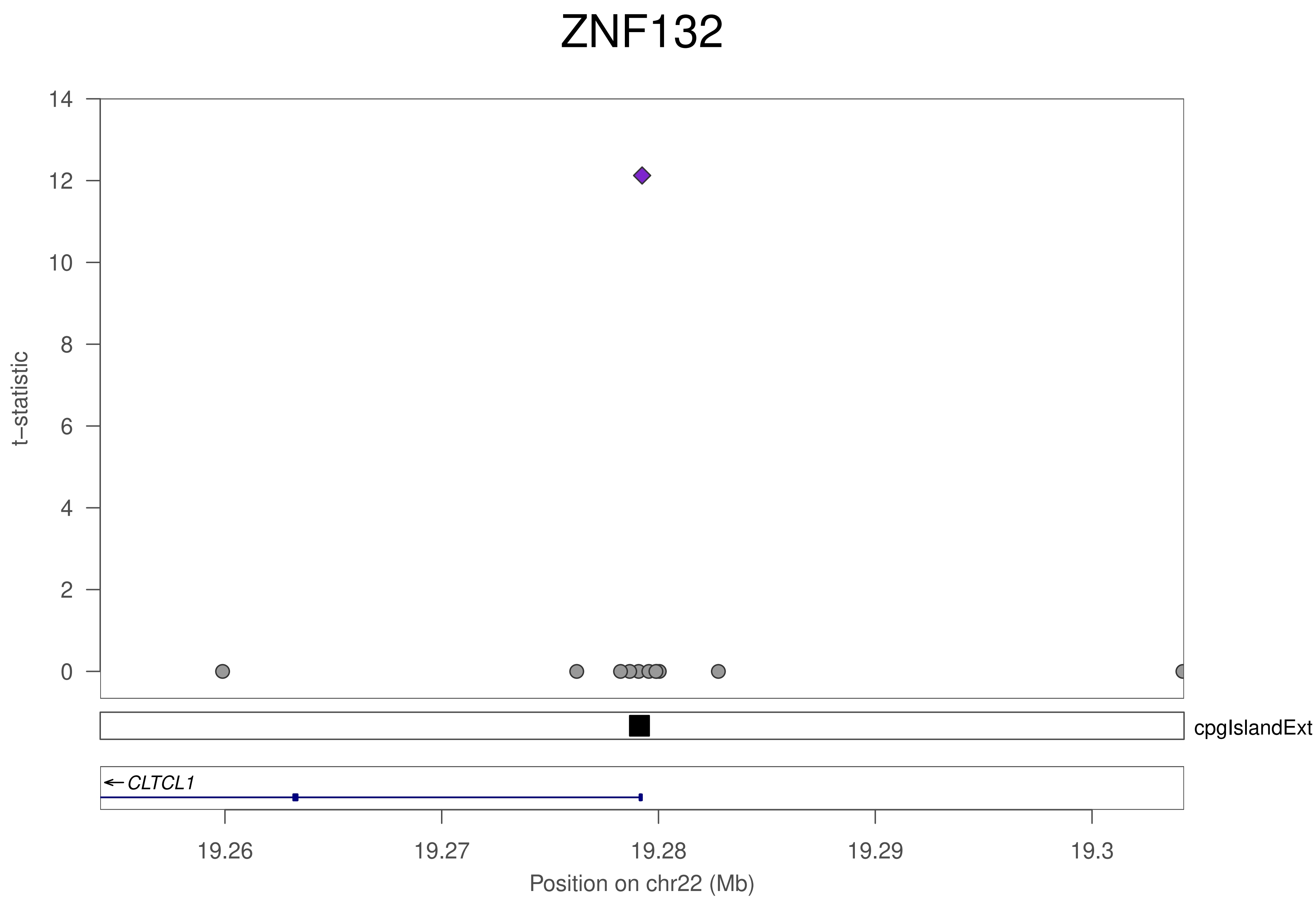
ZNF132
t−statistic
14
12
10
8
6
4
2
0
cpgIslandExt
← CLTCL1
19.26
19.27
19.28
19.29
19.3
Position on chr22 (Mb)

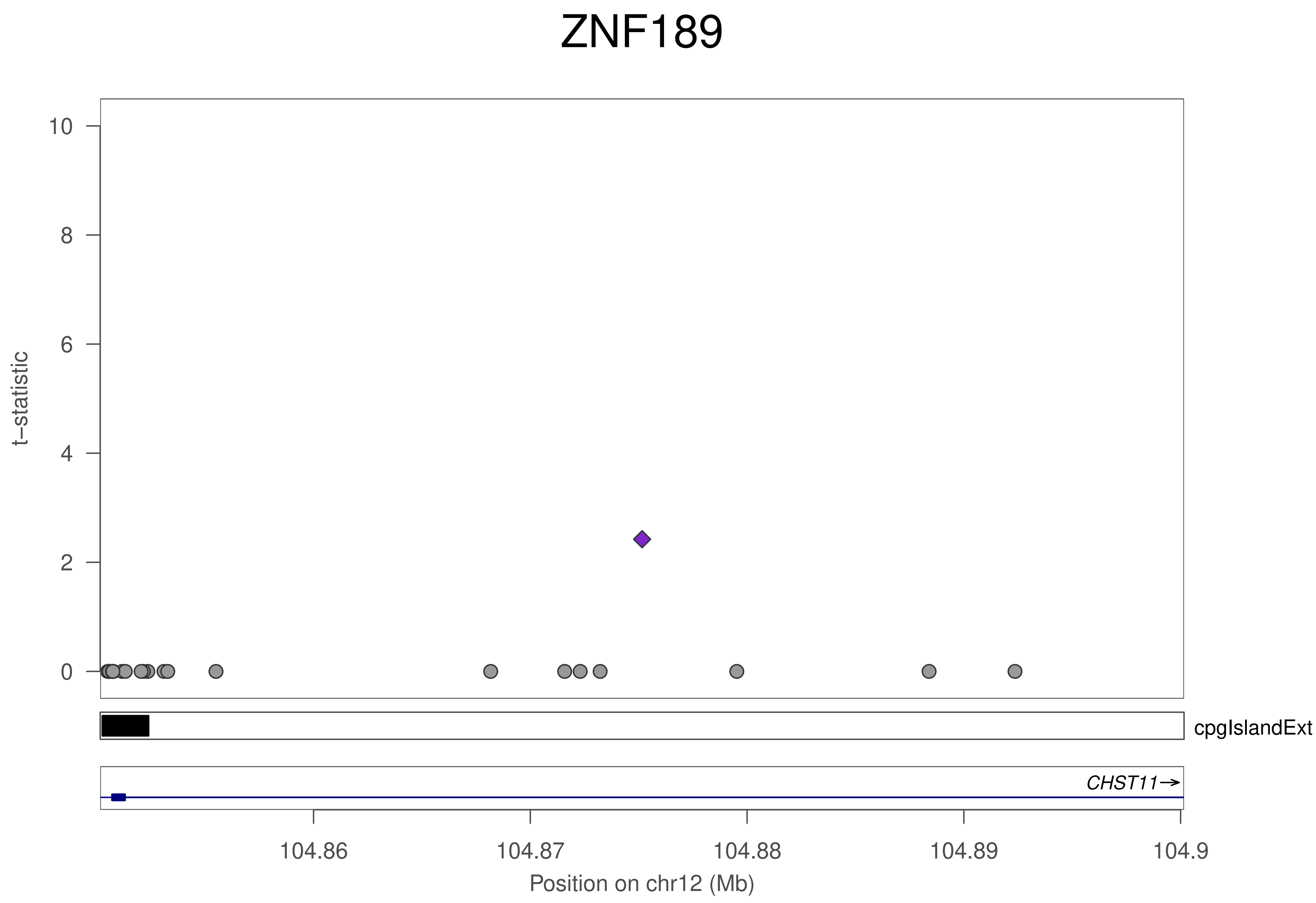
ZNF189
t–statistic
10
8
6
4
2
0
cpgIslandExt
CHST11
104.86
104.87
104.88
104.89
104.9
Position on chr12 (Mb)

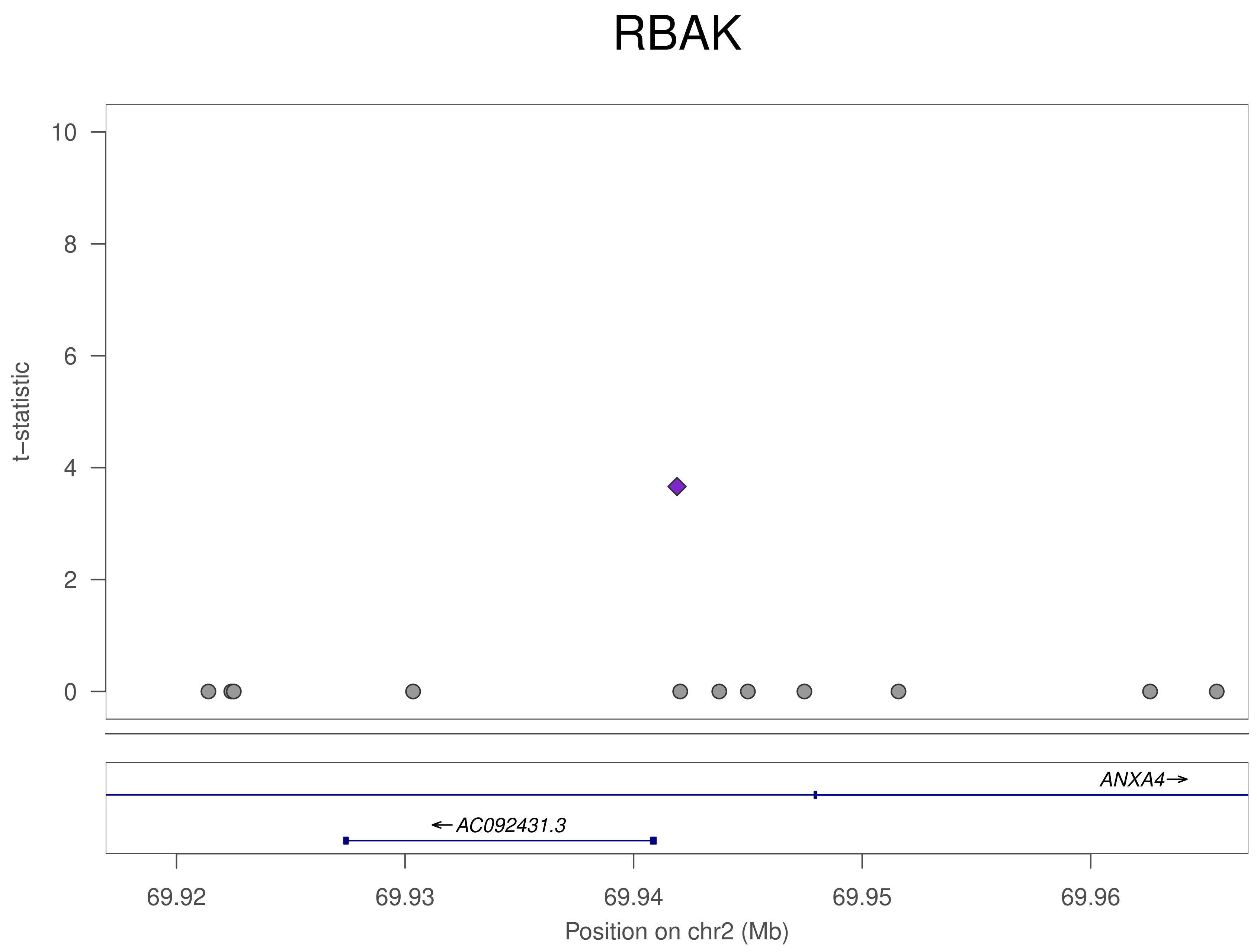
RBAK
t−statistic
10
8
6
4
2
0
ANXA4→
←AC092431.3
69.92
69.93
69.94
69.95
69.96
Position on chr2 (Mb)

### ZNF714

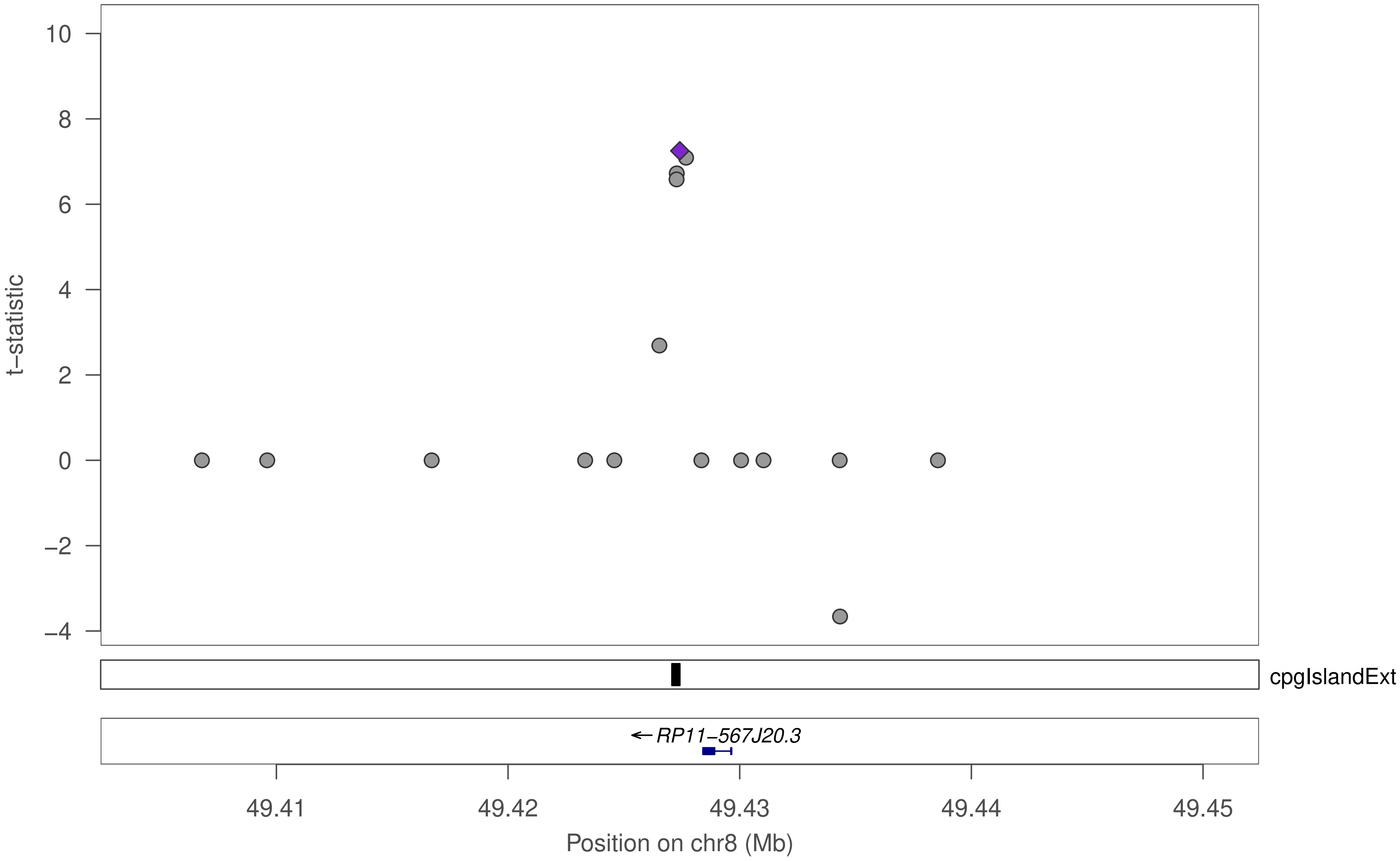

t–statistic
10
8
6
4
2
0
−2
−4
cpgIslandExt
← RP11−567J20.3
49.41
49.42
49.43
49.44
49.45
Position on chr8 (Mb)

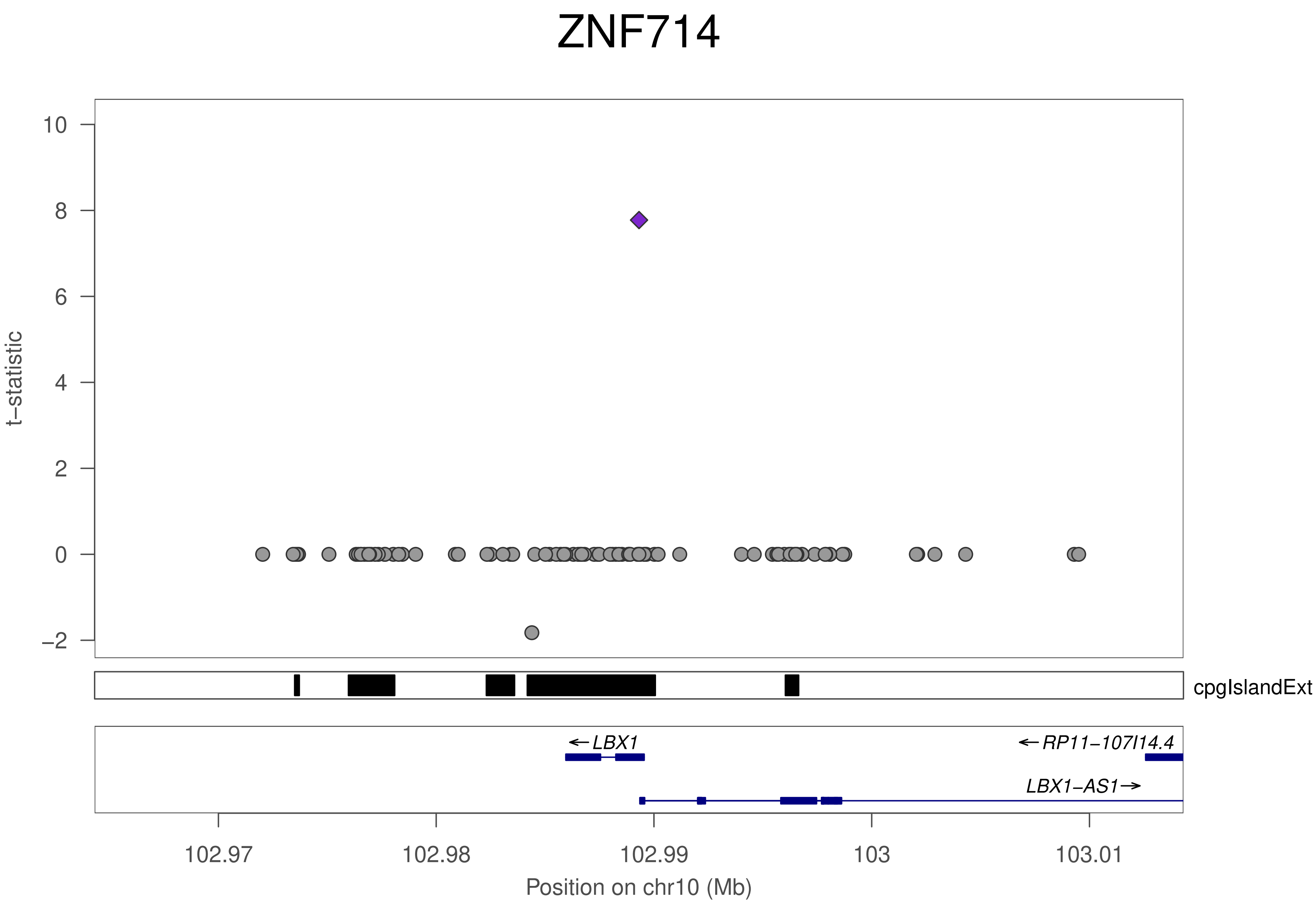
ZNF714
t−statistic
10
8
6
4
2
0
−2
cpgIslandExt
←LBX1
←RP11−107I14.4
LBX1−AS1→
102.97
102.98
102.99
103
103.01
Position on chr10 (Mb)

### ZNF714

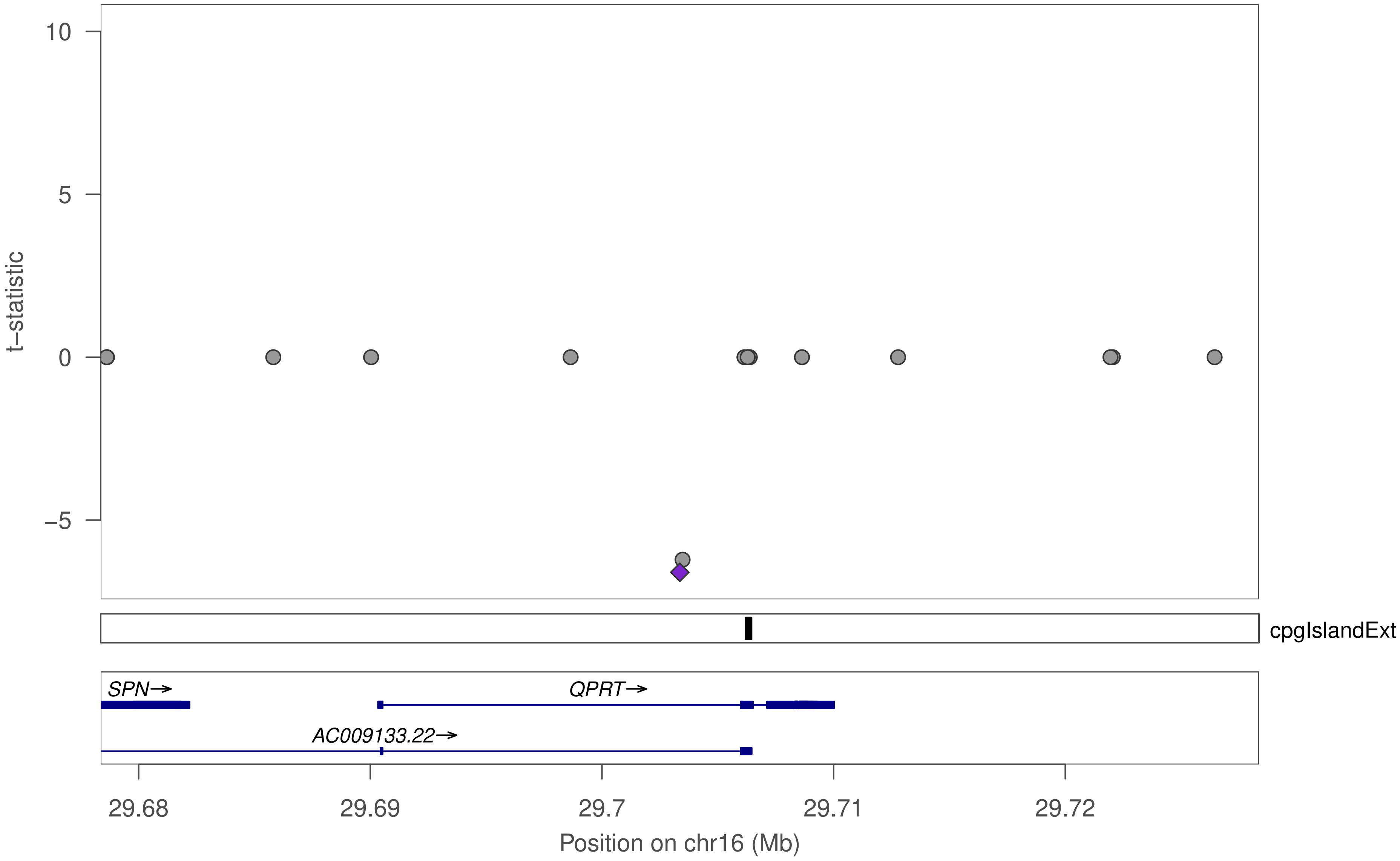

10
5
0
−5
t−statistic
cpgIslandExt
SPN→
QPRT→
AC009133.22→
29.68
29.69
29.7
29.71
29.72
Position on chr16 (Mb)

### ZNF333

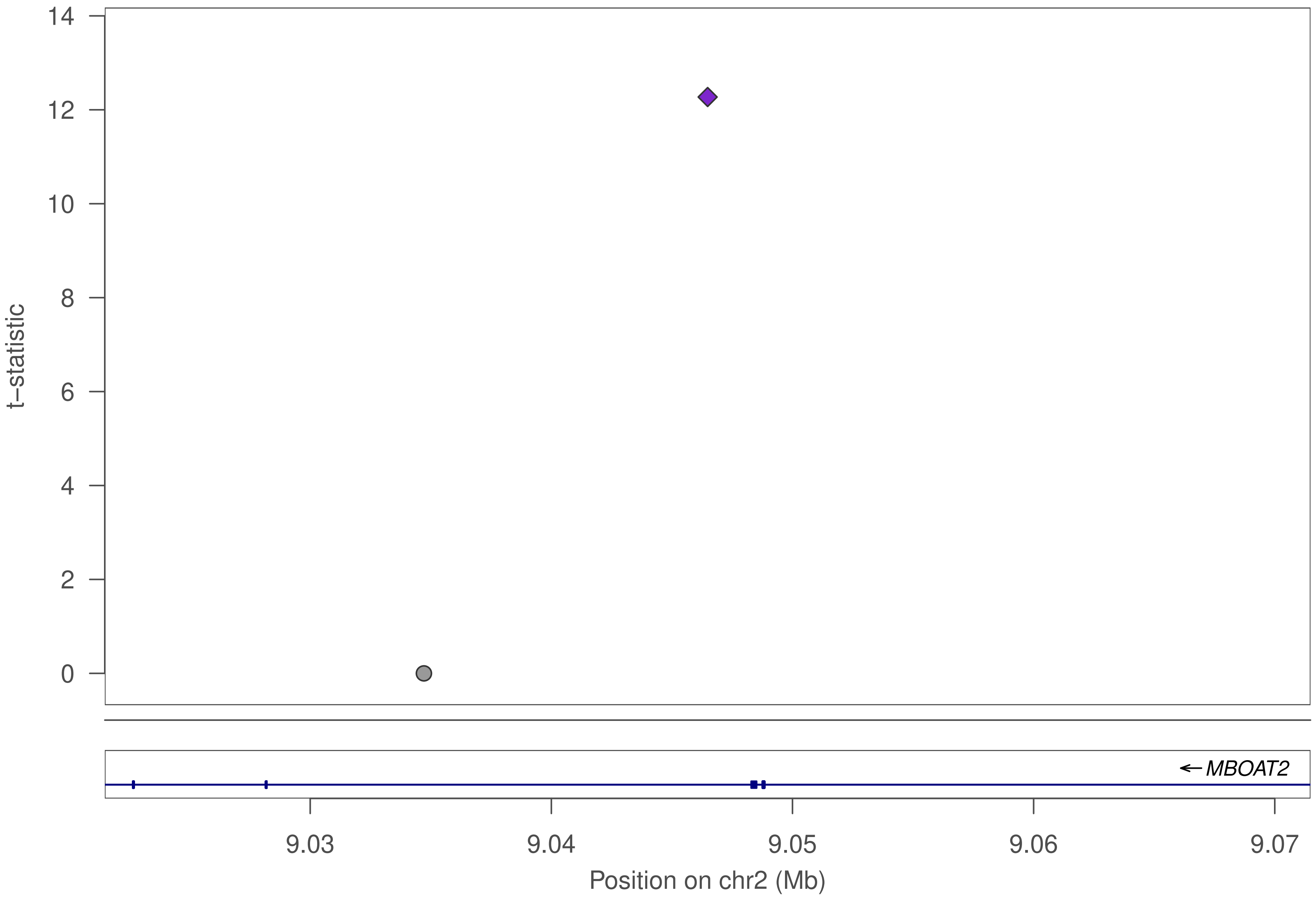

14
12
10
8
6
4
2
0
t–statistic
MBOAT2
9.03
9.04
9.05
9.06
9.07
Position on chr2 (Mb)

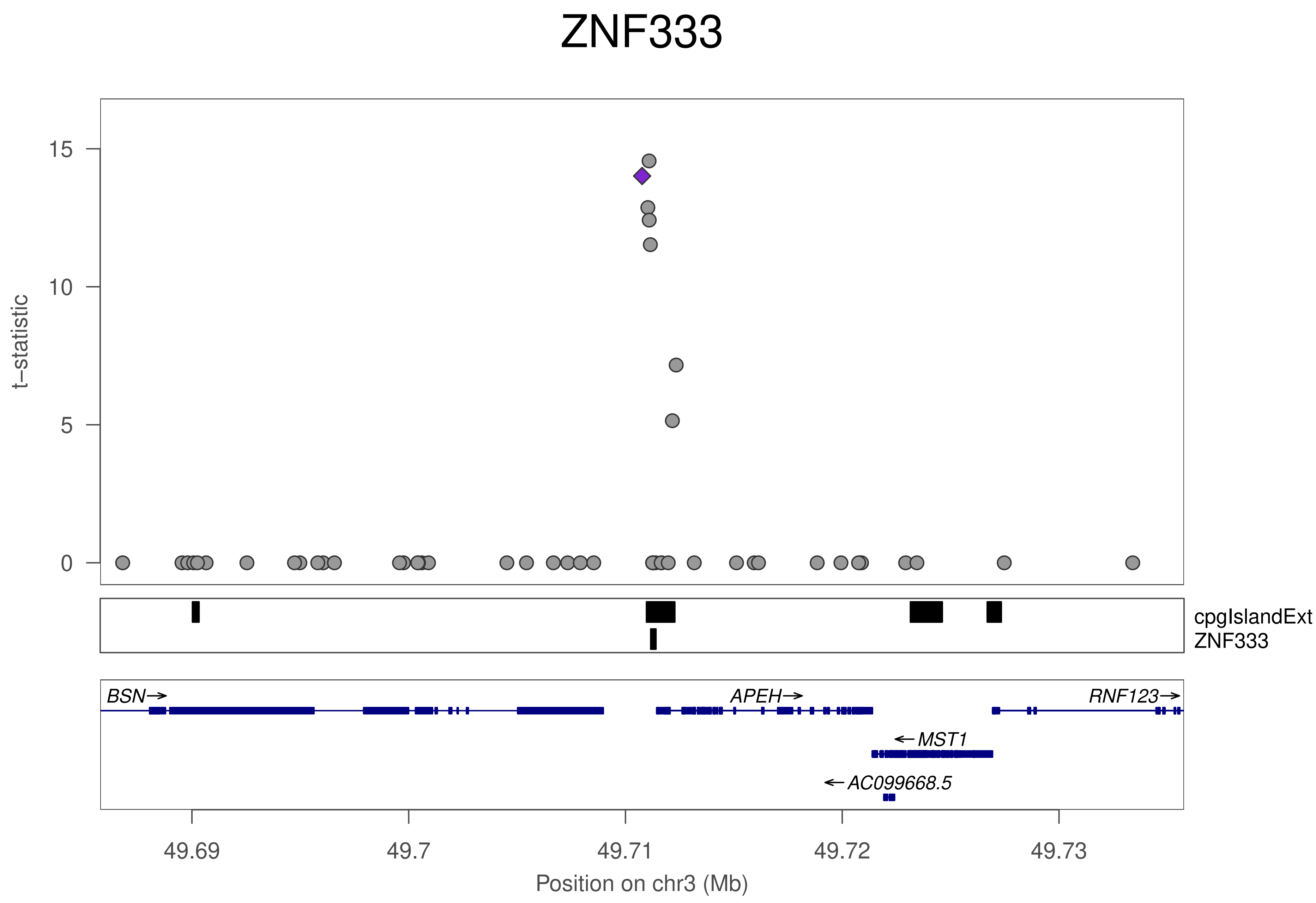
ZNF333
t−statistic
15
10
5
0
cpgIslandExt
ZNF333
BSN→
APEH→
RNF123→
←MST1
←AC099668.5
49.69
49.7
49.71
49.72
49.73
Position on chr3 (Mb)

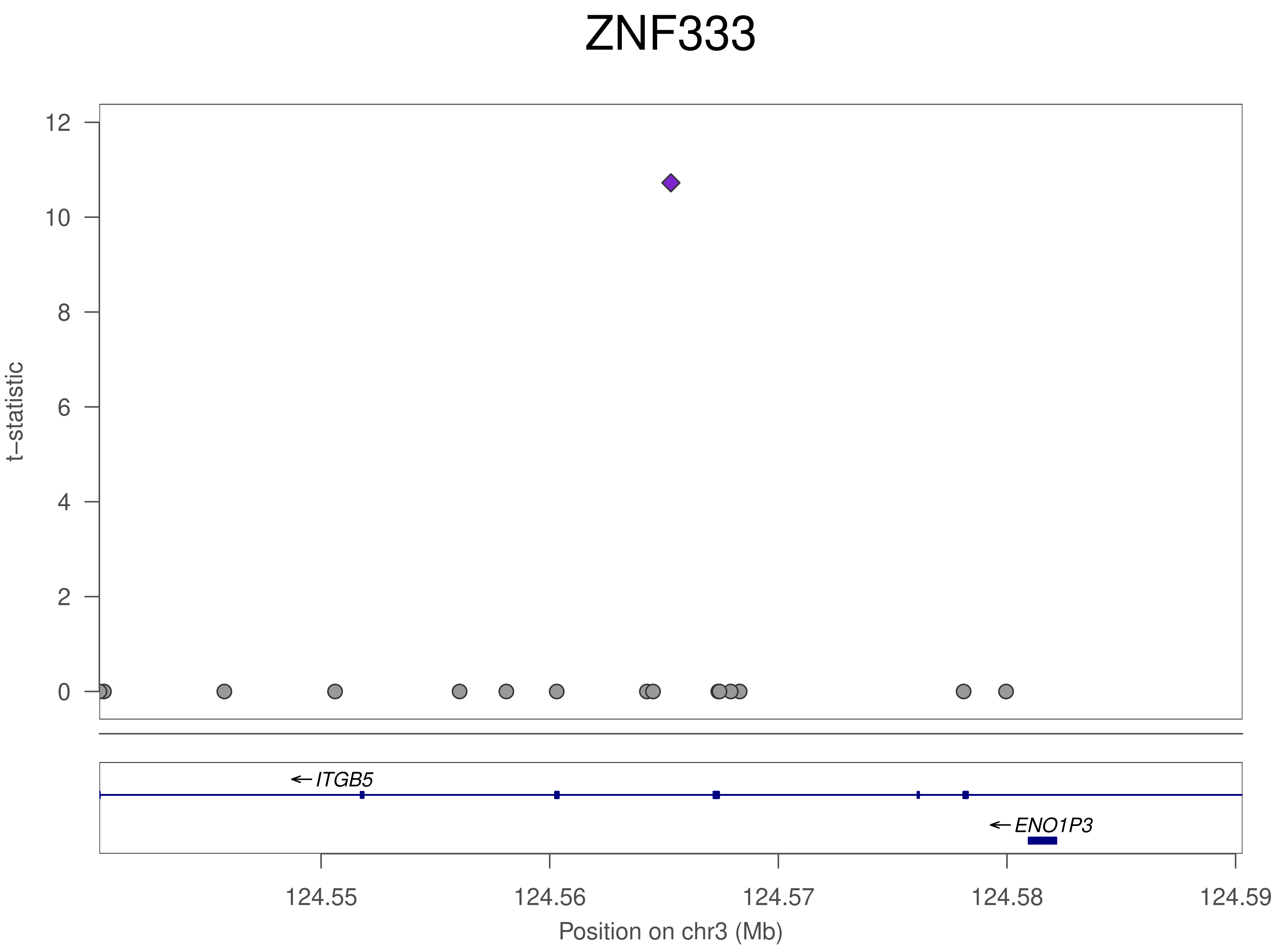
ZNF333
t−statistic
12
10
8
6
4
2
0
ITGB5
ENO1P3
124.55
124.56
124.57
124.58
124.59
Position on chr3 (Mb)

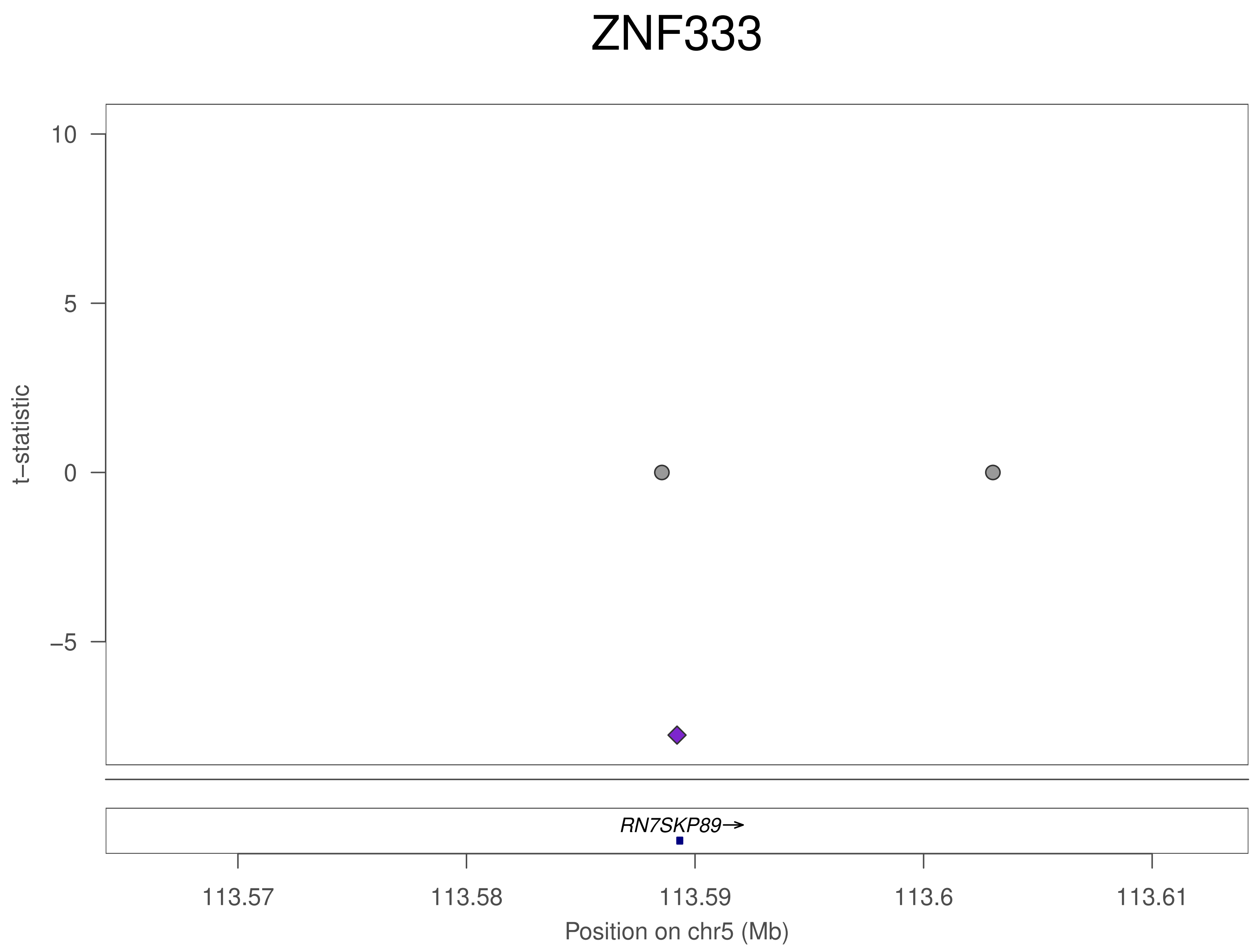
ZNF333
10
5
0
−5
t−statistic
RN7SKP89→
113.57
113.58
113.59
113.6
113.61
Position on chr5 (Mb)

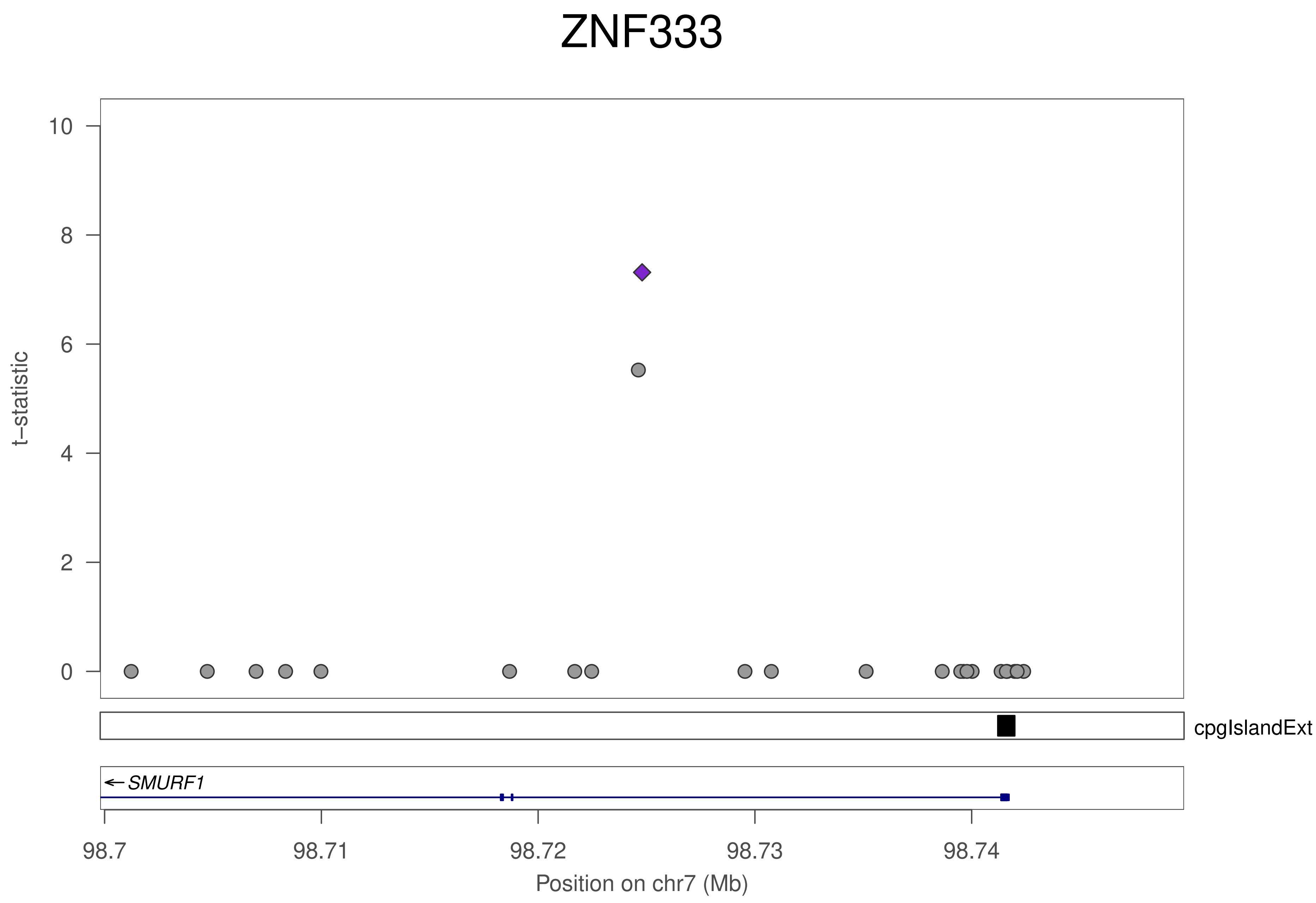

ZNF333
t–statistic
0
2
4
6
8
10
cpgIslandExt
SMURF1
98.7
98.71
98.72
98.73
98.74
Position on chr7 (Mb)

### ZNF333

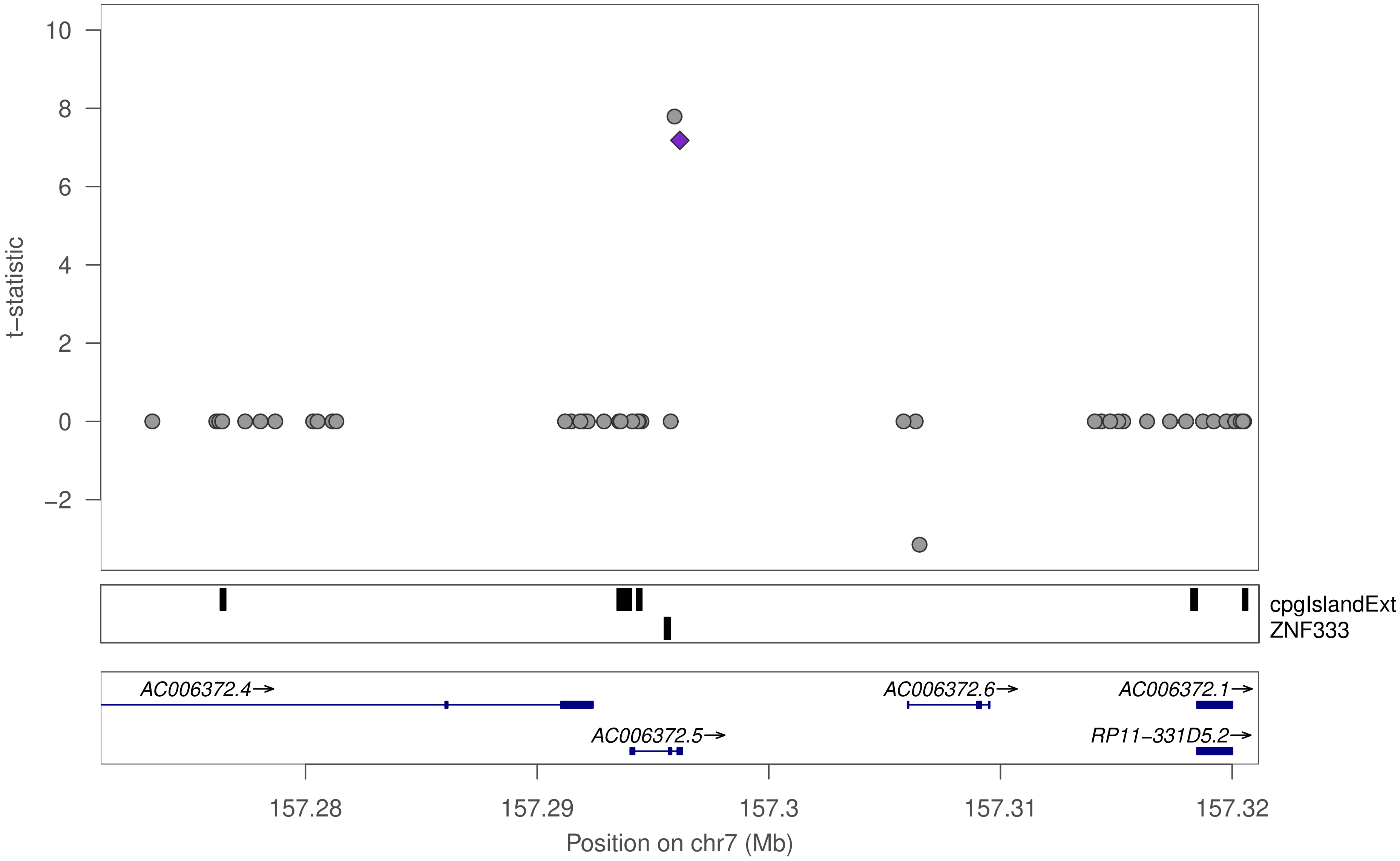

t−statistic
10
8
6
4
2
0
−2
cpgIslandExt
ZNF333
AC006372.4→
AC006372.6→
AC006372.1→
AC006372.5→
RP11−331D5.2→
157.28
157.29
157.3
157.31
157.32
Position on chr7 (Mb)

### ZNF333

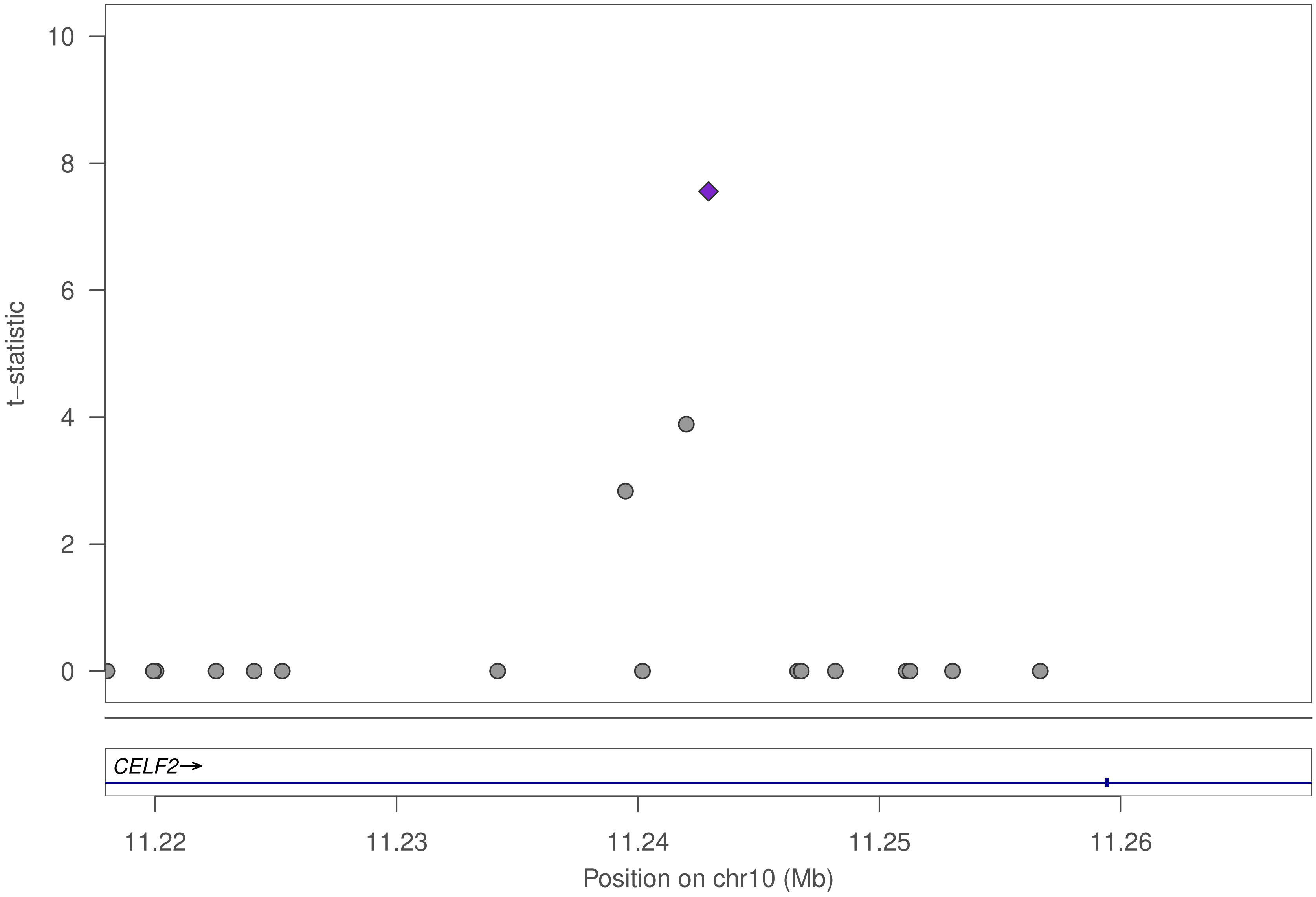

10
8
6
4
2
0
t−statistic
CELF2→
11.22
11.23
11.24
11.25
11.26
Position on chr10 (Mb)

### ZNF333

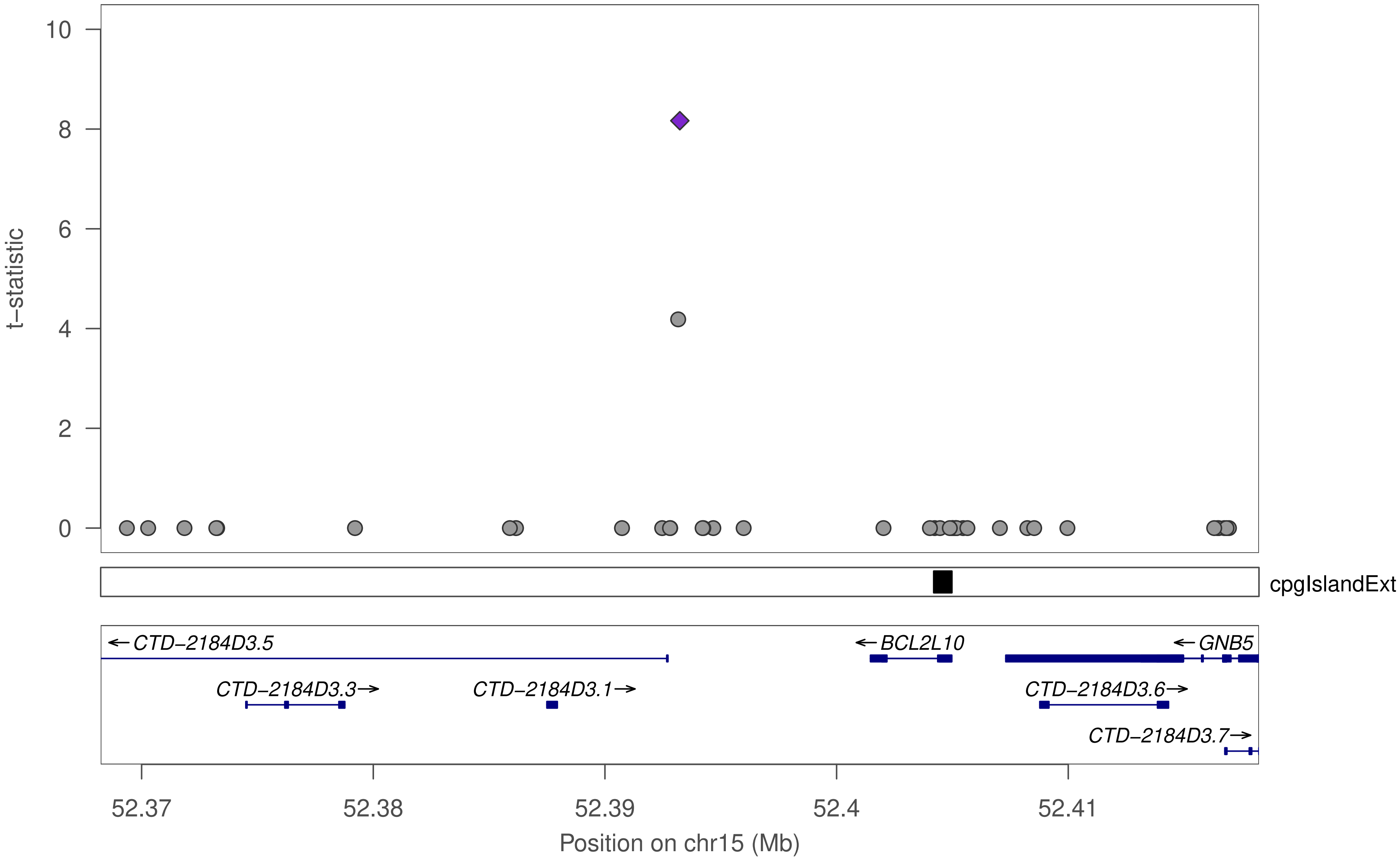

t−statistic
10
8
6
4
2
0
cpgIslandExt
←CTD−2184D3.5
←BCL2L10
←GNB5
CTD−2184D3.3→
CTD−2184D3.1→
CTD−2184D3.6→
CTD−2184D3.7→
52.37
52.38
52.39
52.4
52.41
Position on chr15 (Mb)

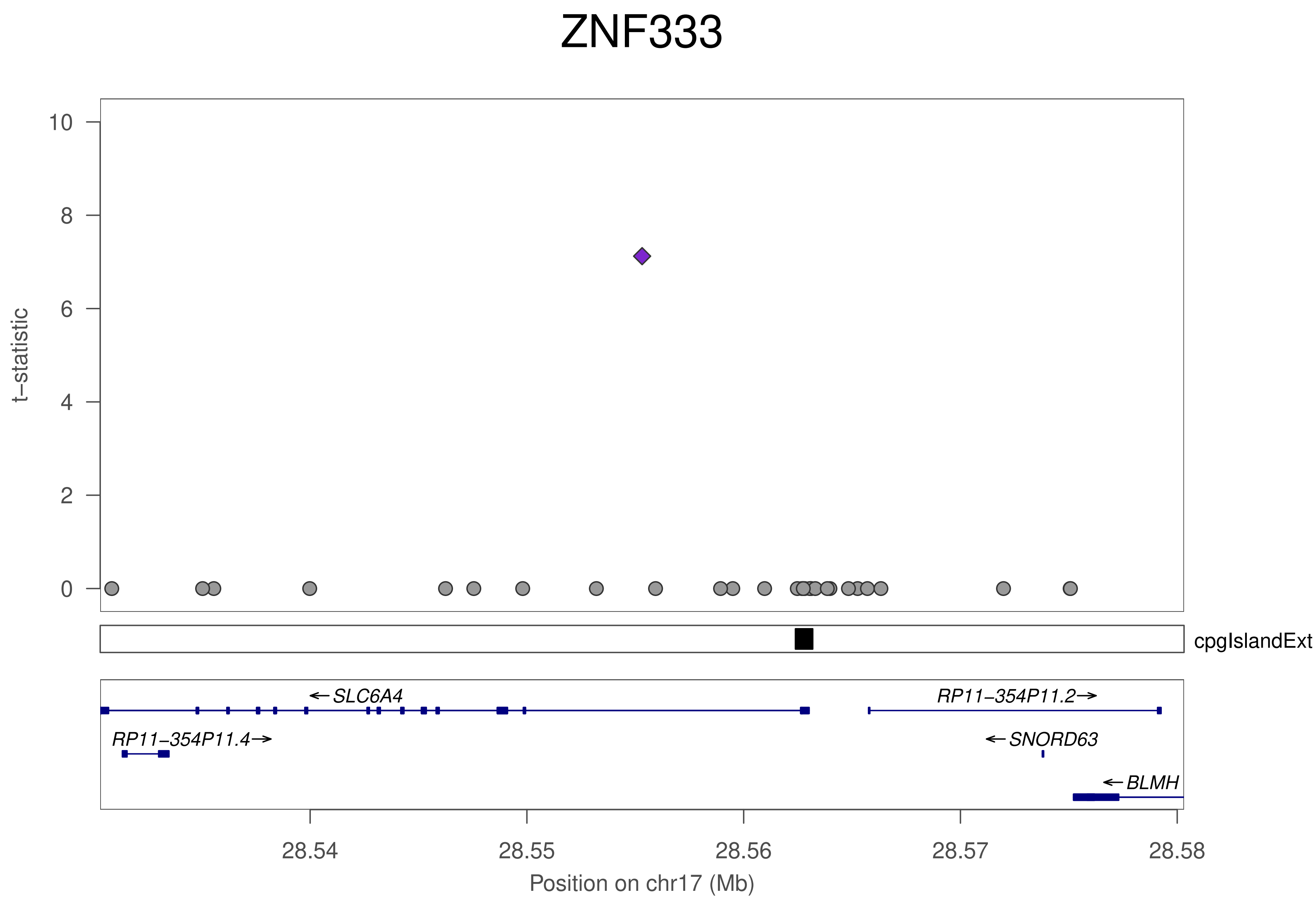
ZNF333
t−statistic
10
8
6
4
2
0
cpgIslandExt
←SLC6A4
RP11−354P11.2→
RP11−354P11.4→
←SNORD63
←BLMH
28.54
28.55
28.56
28.57
28.58
Position on chr17 (Mb)

### ZNF333

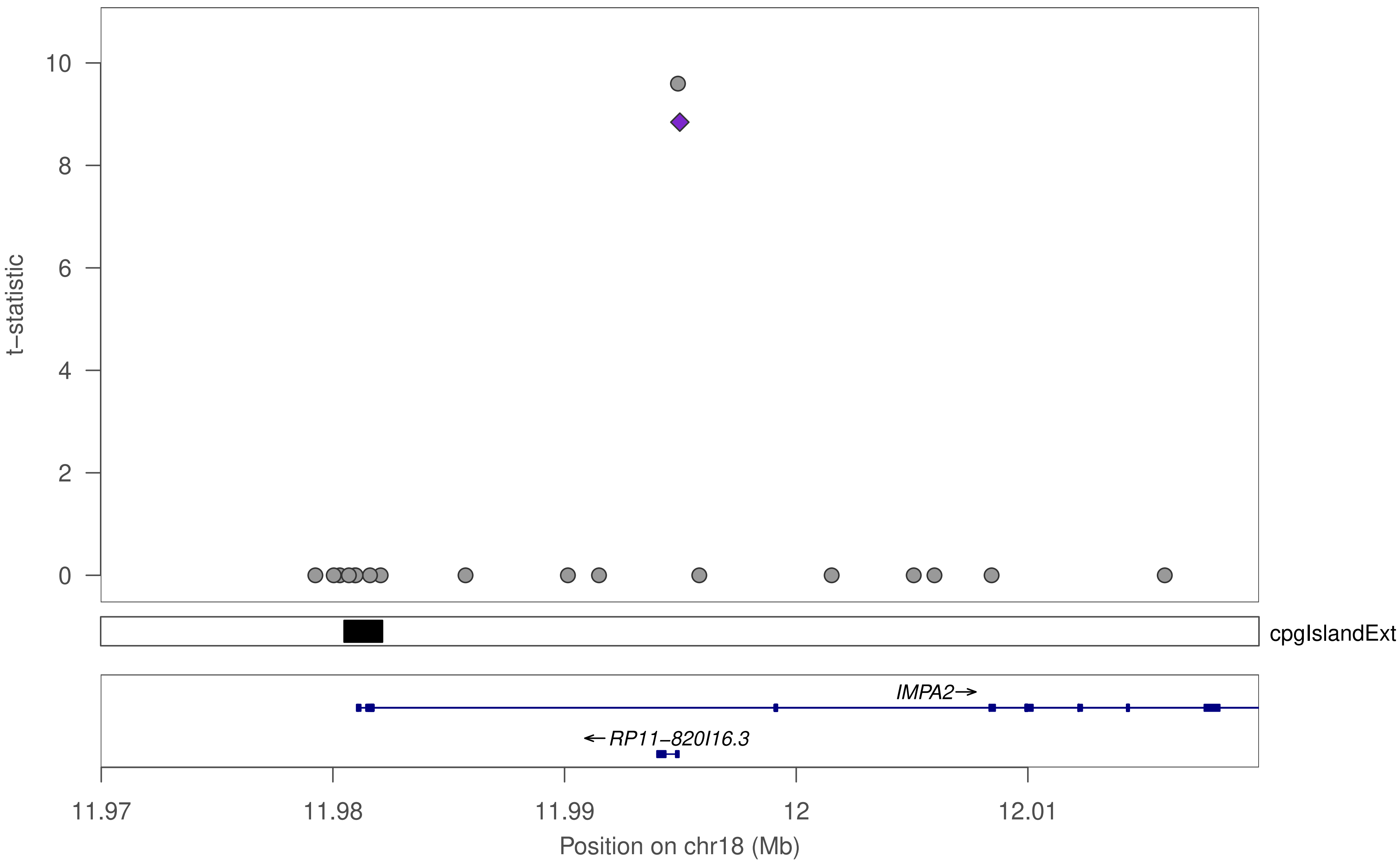

10
8
6
4
2
0
t–statistic
cpgIslandExt
IMPA2→
←RP11–820I16.3
11.97
11.98
11.99
12
12.01
Position on chr18 (Mb)

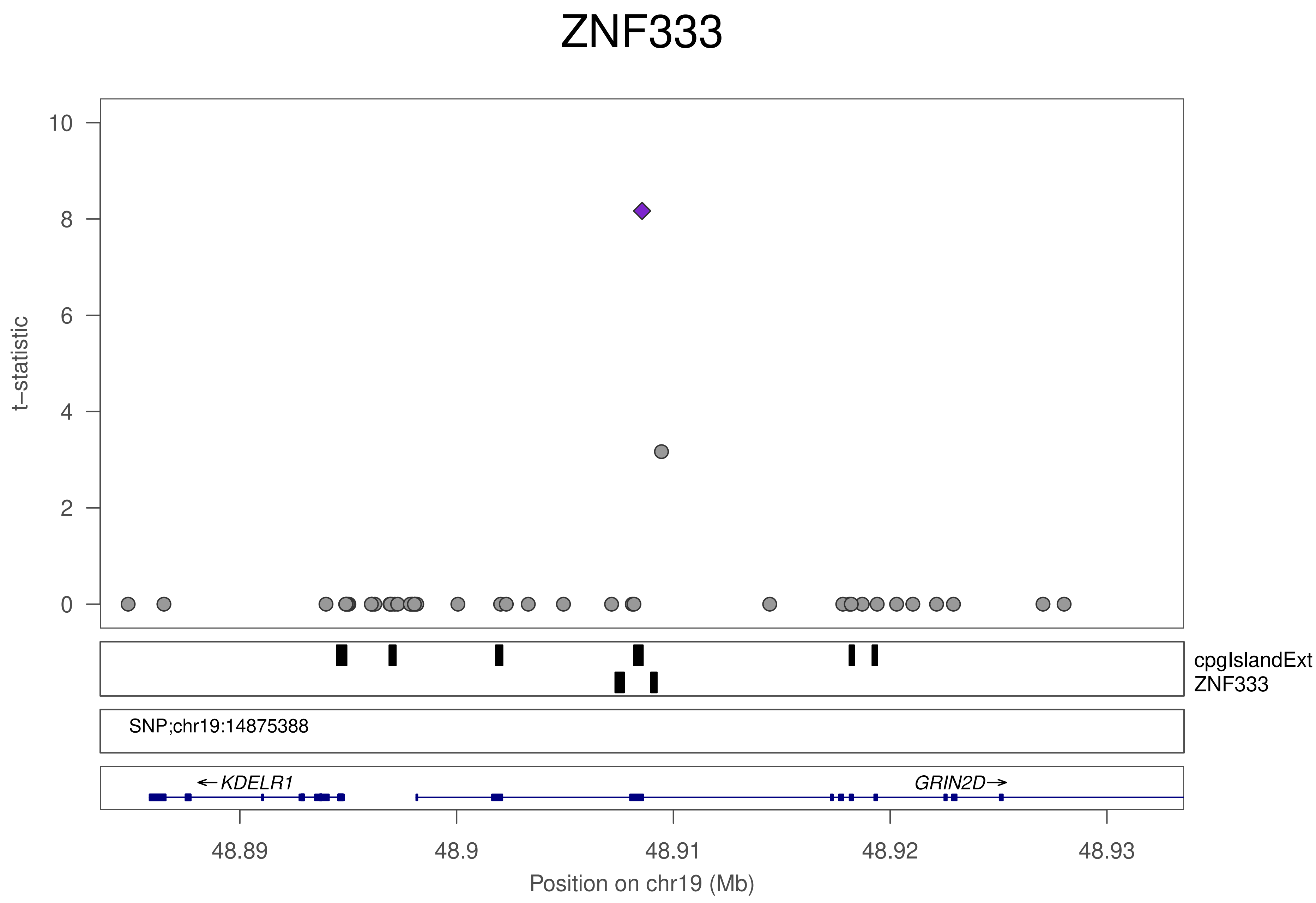
ZNF333
t−statistic
10
8
6
4
2
0
cpgIslandExt
ZNF333
SNP;chr19:14875388
←KDELR1
GRIN2D→
48.89
48.9
48.91
48.92
48.93
Position on chr19 (Mb)

### ZNF496

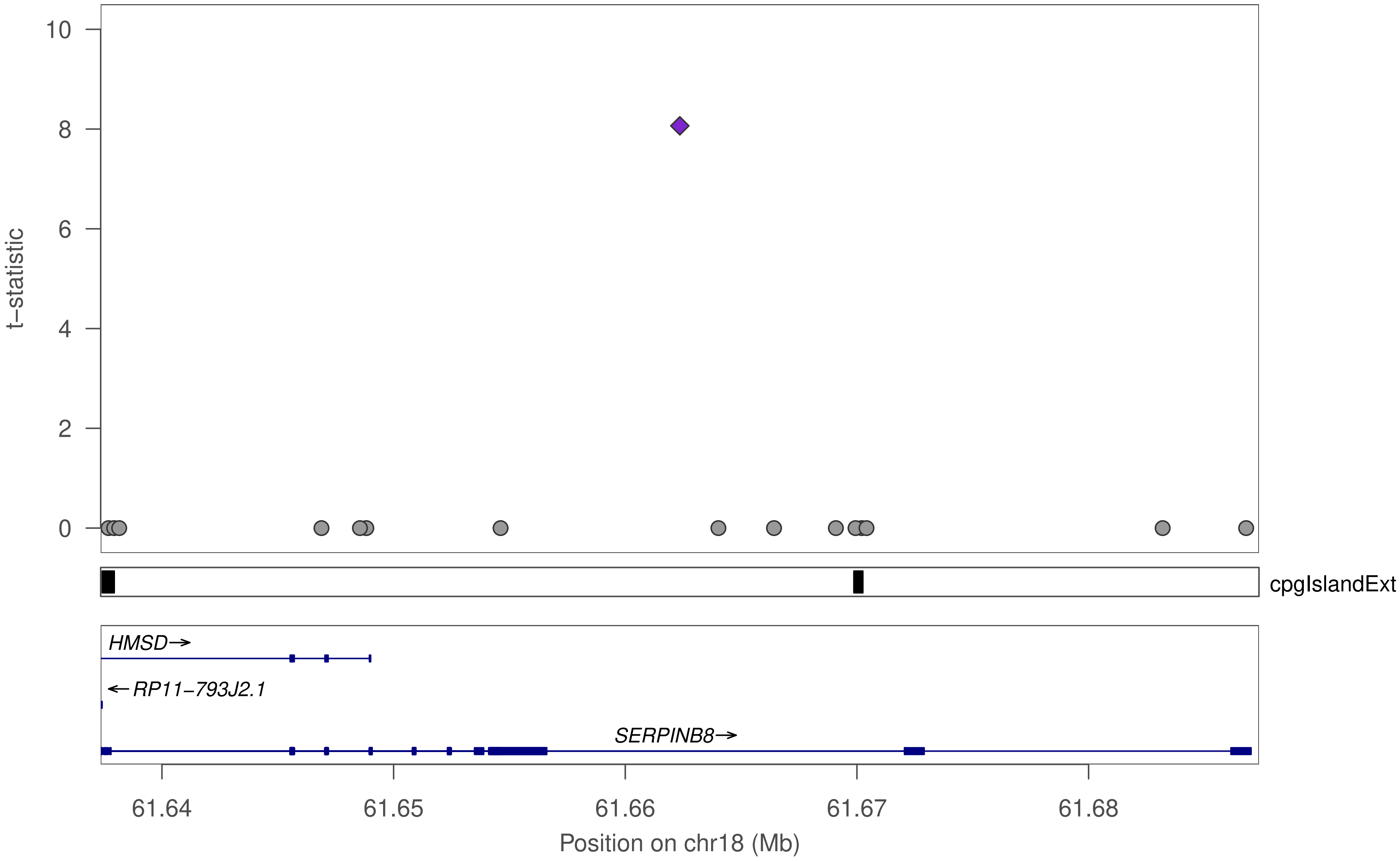

t−statistic
10
8
6
4
2
0
cpgIslandExt
HMSD→
←RP11−793J2.1
SERPINB8→
61.64
61.65
61.66
61.67
61.68
Position on chr18 (Mb)

### ZNF180

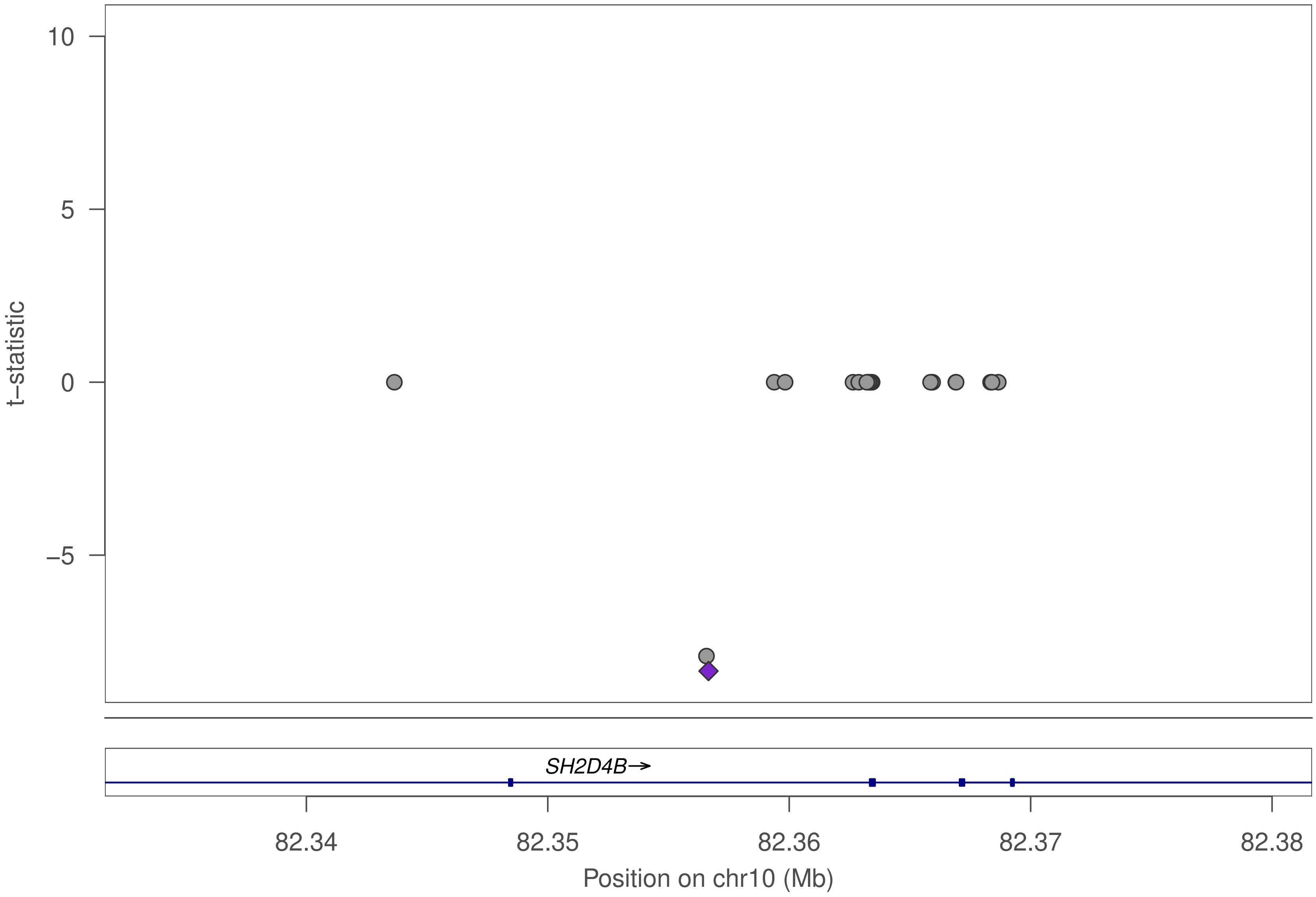

10
5
0
−5
t−statistic
SH2D4B→
82.34
82.35
82.36
82.37
82.38
Position on chr10 (Mb)

### ZNF180

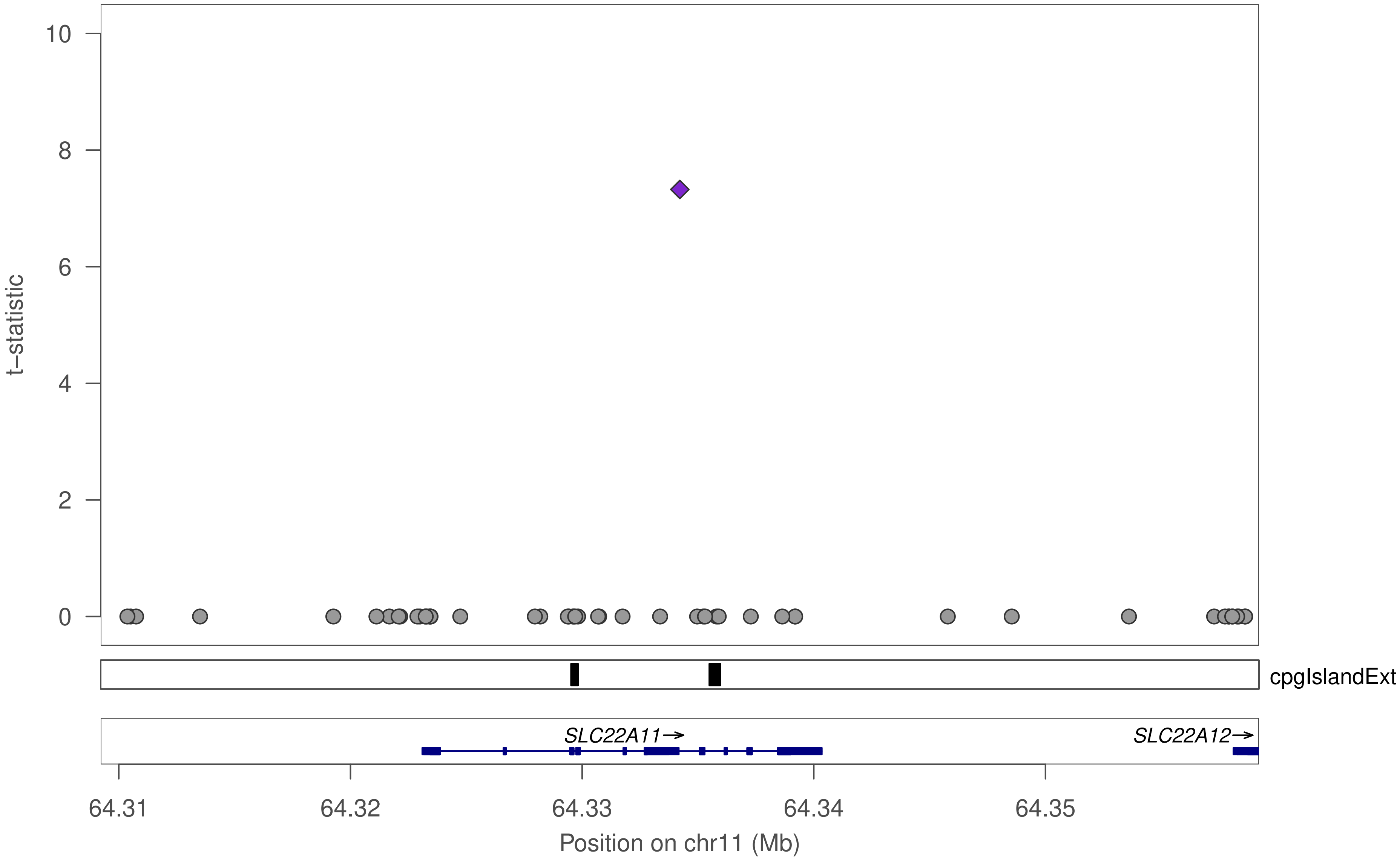

10
8
6
4
2
0
t–statistic
cpgIslandExt
SLC22A11→
SLC22A12→
64.31
64.32
64.33
64.34
64.35
Position on chr11 (Mb)

### ZNF180

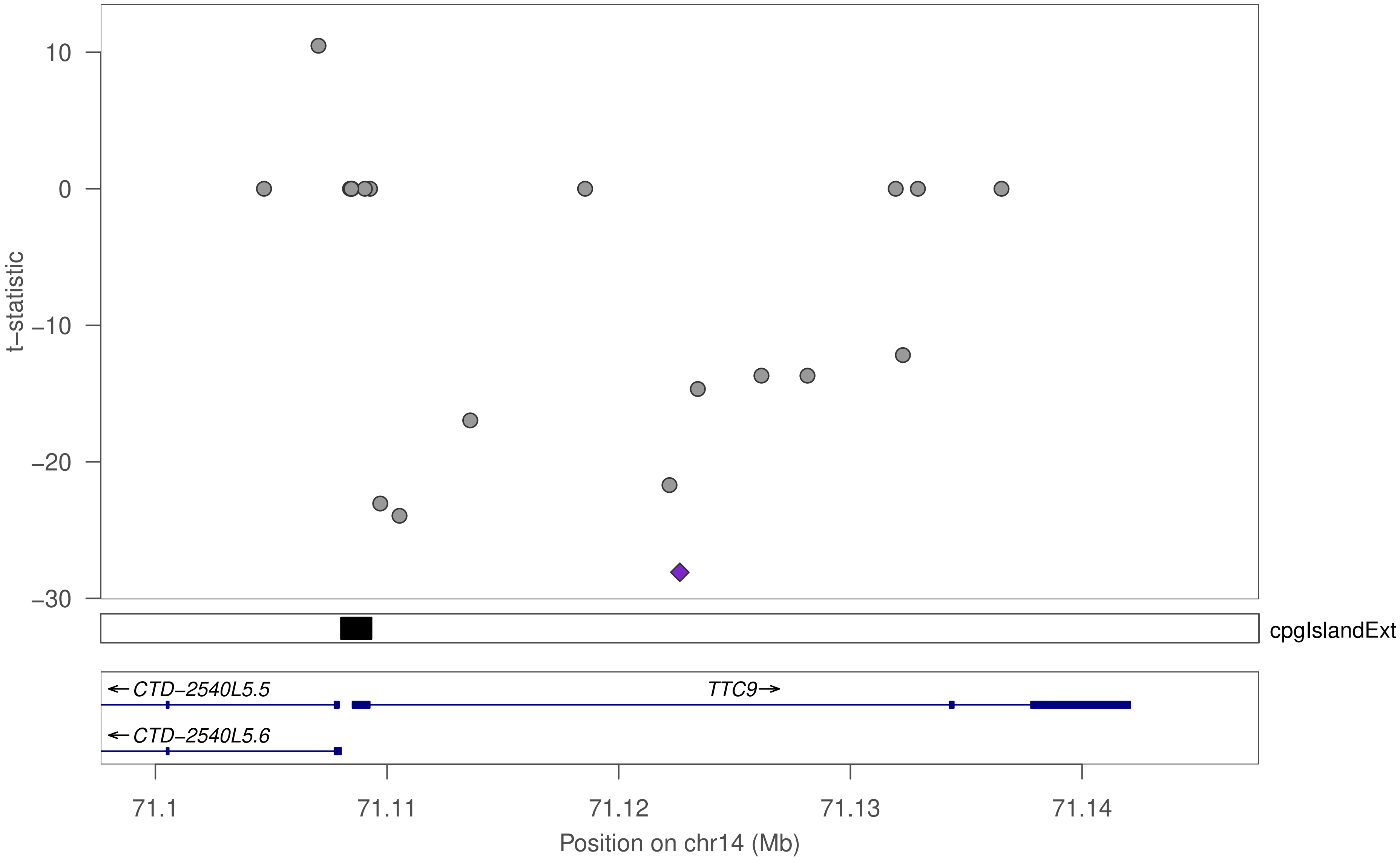

10
0
−10
−20
−30
t−statistic
cpgIslandExt
←CTD−2540L5.5
TTC9→
←CTD−2540L5.6
71.1
71.11
71.12
71.13
71.14
Position on chr14 (Mb)

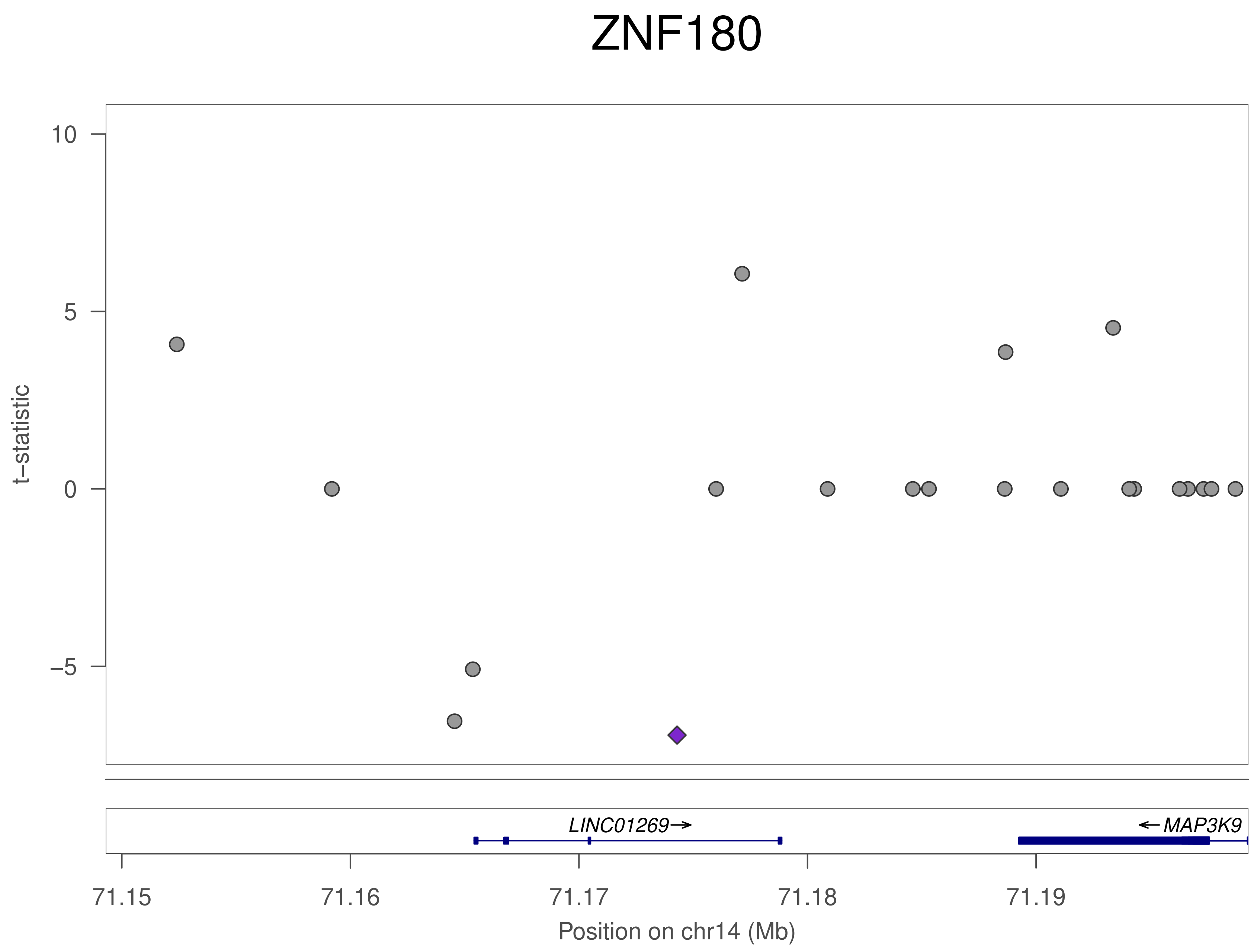
ZNF180
t–statistic
10
5
0
–5
LINC01269→
←MAP3K9
71.15
71.16
71.17
71.18
71.19
Position on chr14 (Mb)

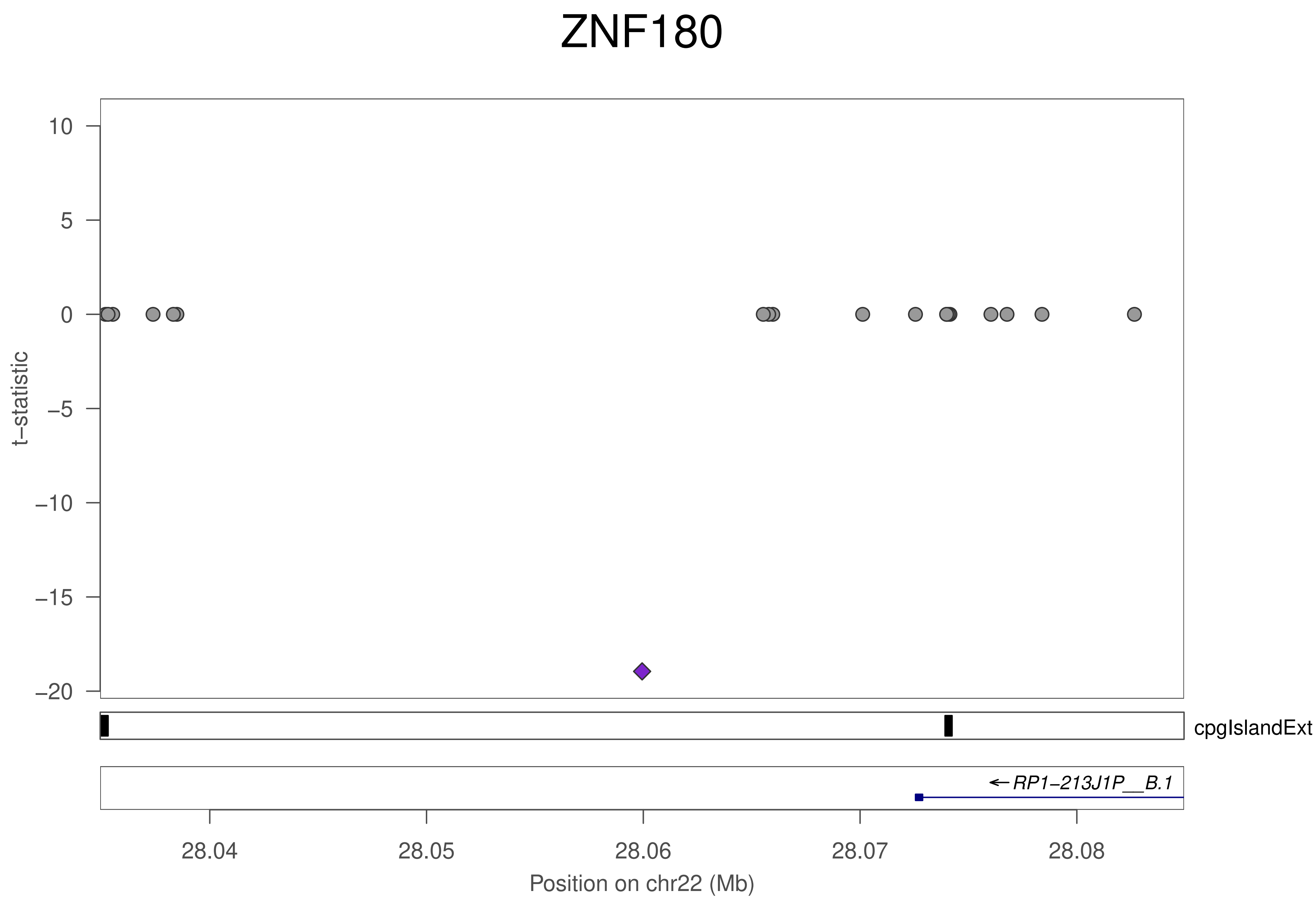
ZNF180
t−statistic
10
5
0
−5
−10
−15
−20
cpgIslandExt
← RP1−213J1P__B.1
28.04
28.05
28.06
28.07
28.08
Position on chr22 (Mb)

### ZNF528

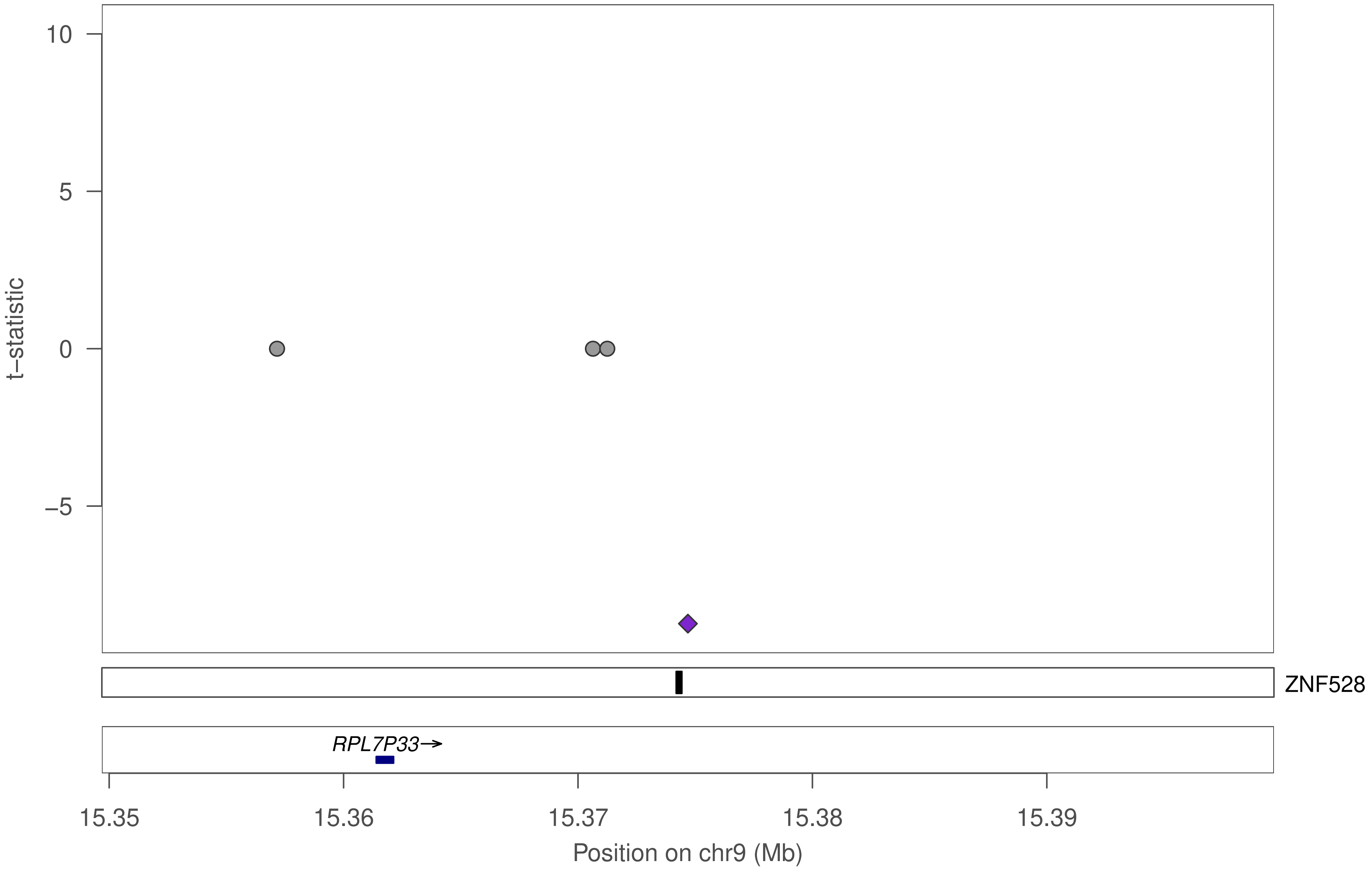

10
5
0
−5
t−statistic
ZNF528
RPL7P33→
15.35
15.36
15.37
15.38
15.39
Position on chr9 (Mb)

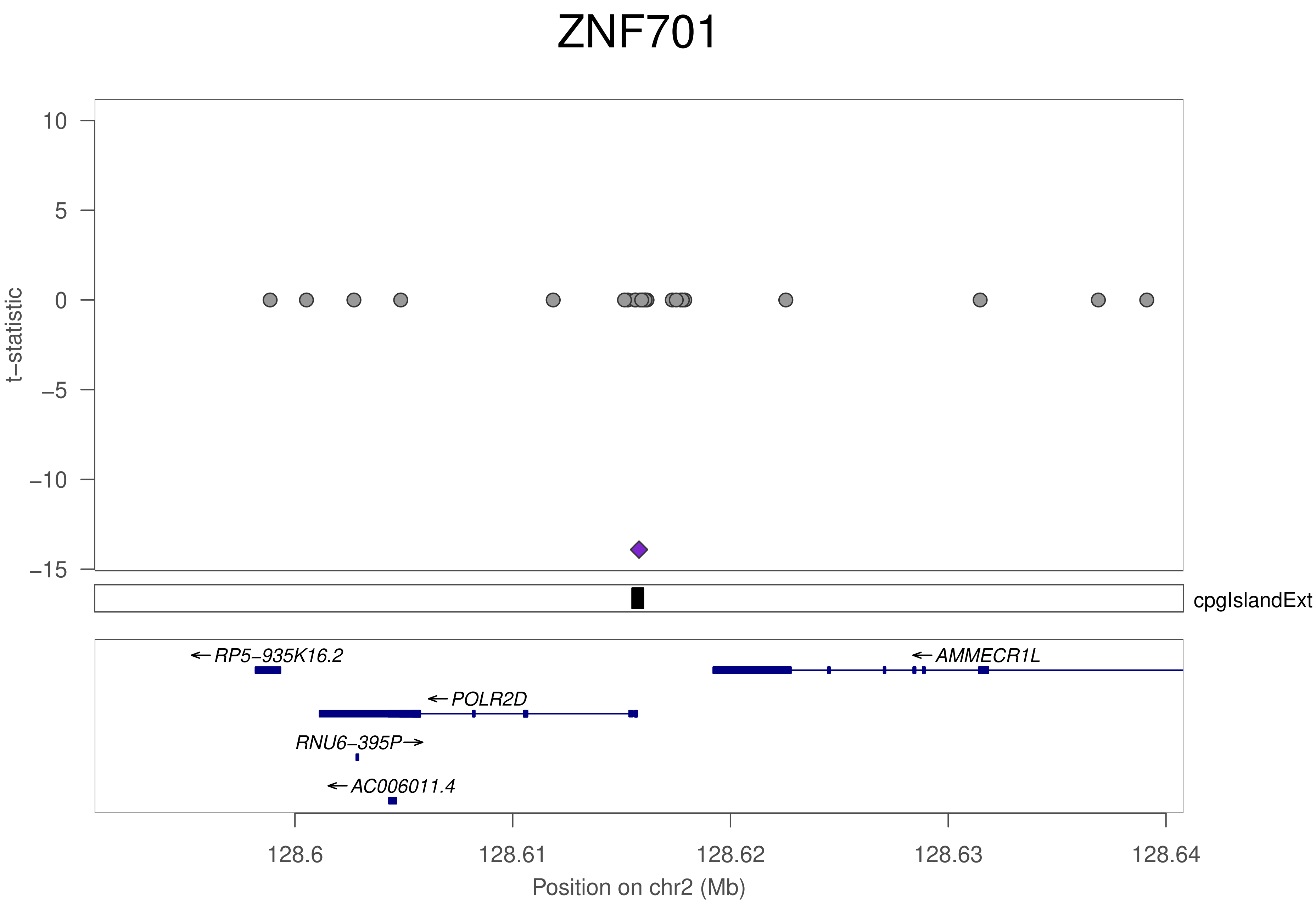
ZNF701
t–statistic
10
5
0
–5
–10
–15
cpgIslandExt
RP5–935K16.2
AMMECR1L
POLR2D
RNU6–395P
AC006011.4
128.6
128.61
128.62
128.63
128.64
Position on chr2 (Mb)

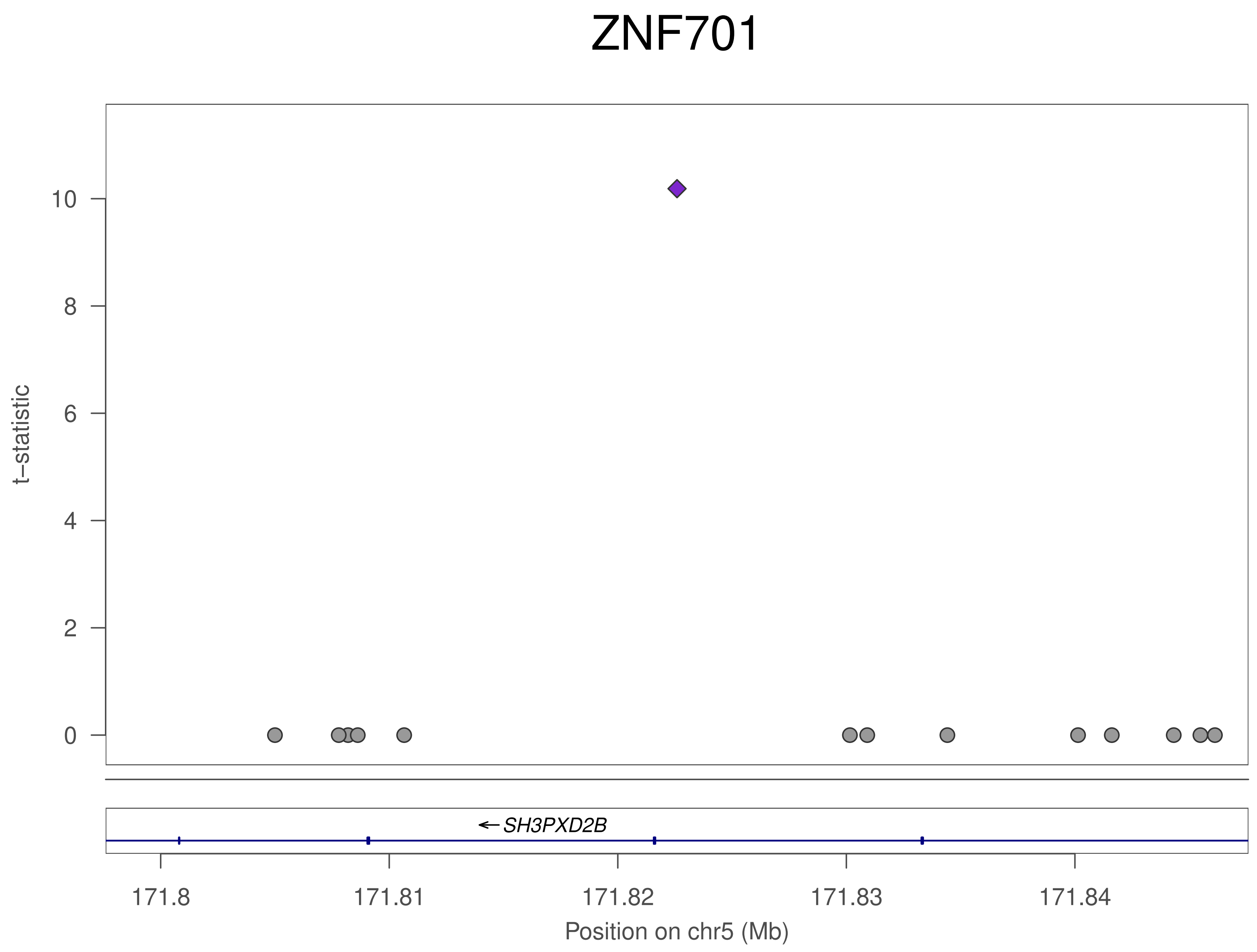
ZNF701
t−statistic
10
8
6
4
2
0
←SH3PXD2B
171.8
171.81
171.82
171.83
171.84
Position on chr5 (Mb)

ZNF701

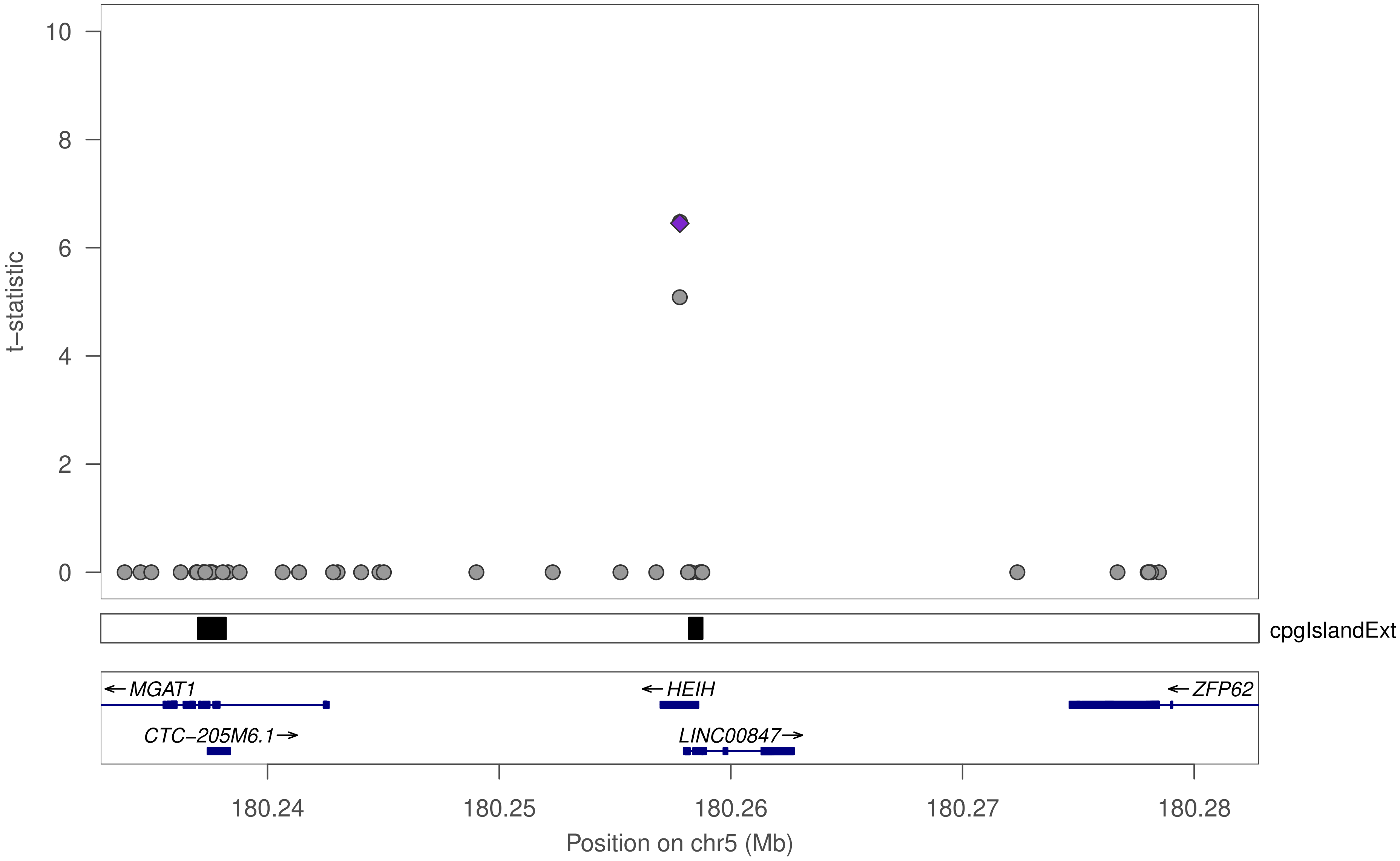

t−statistic
10
8
6
4
2
0
cpgIslandExt
← MGAT1
← HEIH
← ZFP62
CTC−205M6.1 →
LINC00847 →
180.24
180.25
180.26
180.27
180.28
Position on chr5 (Mb)

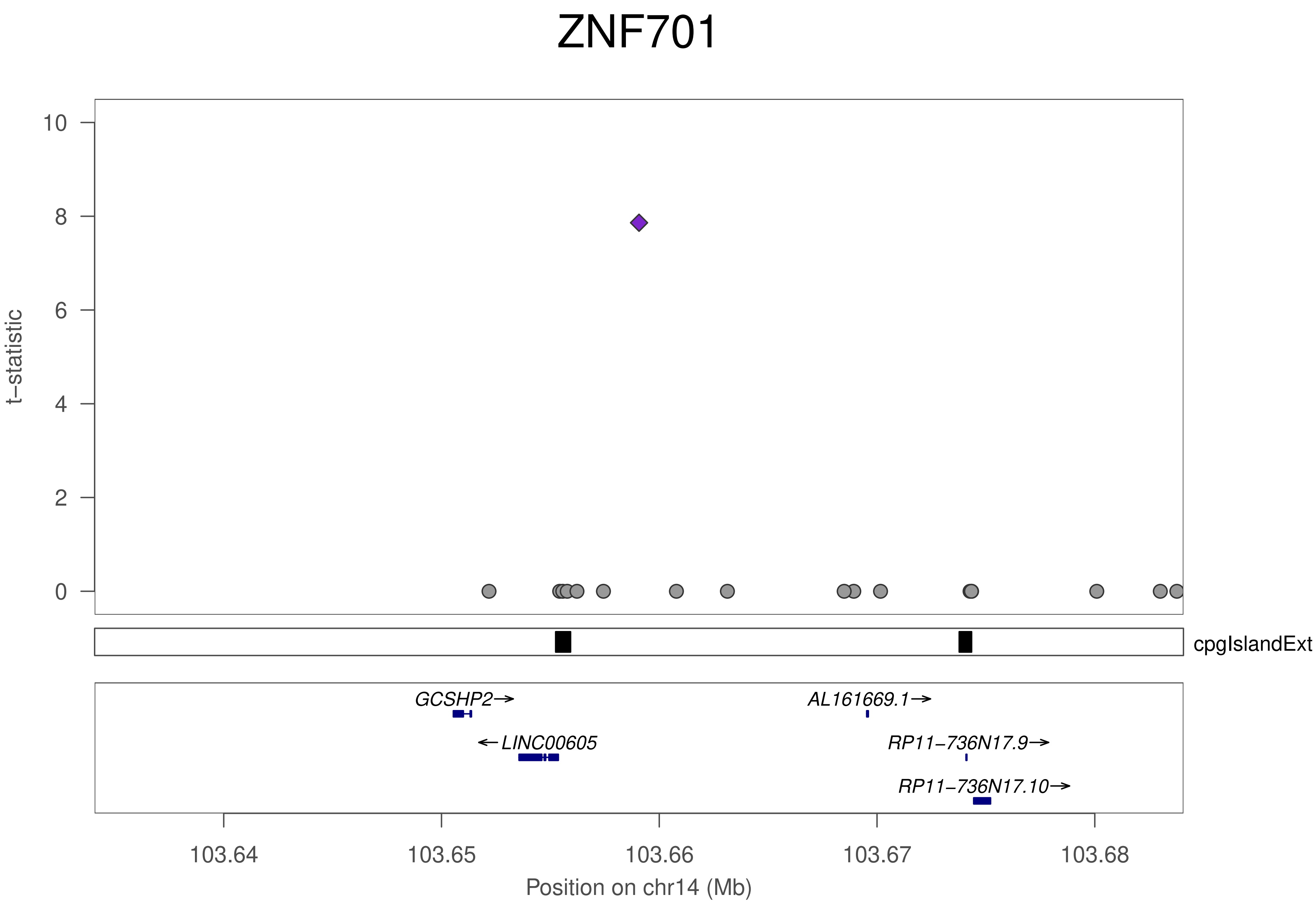

ZNF701
t−statistic
10
8
6
4
2
0
cpgIslandExt
GCSHP2→
←LINC00605
AL161669.1→
RP11−736N17.9→
RP11−736N17.10→
103.64
103.65
103.66
103.67
103.68
Position on chr14 (Mb)

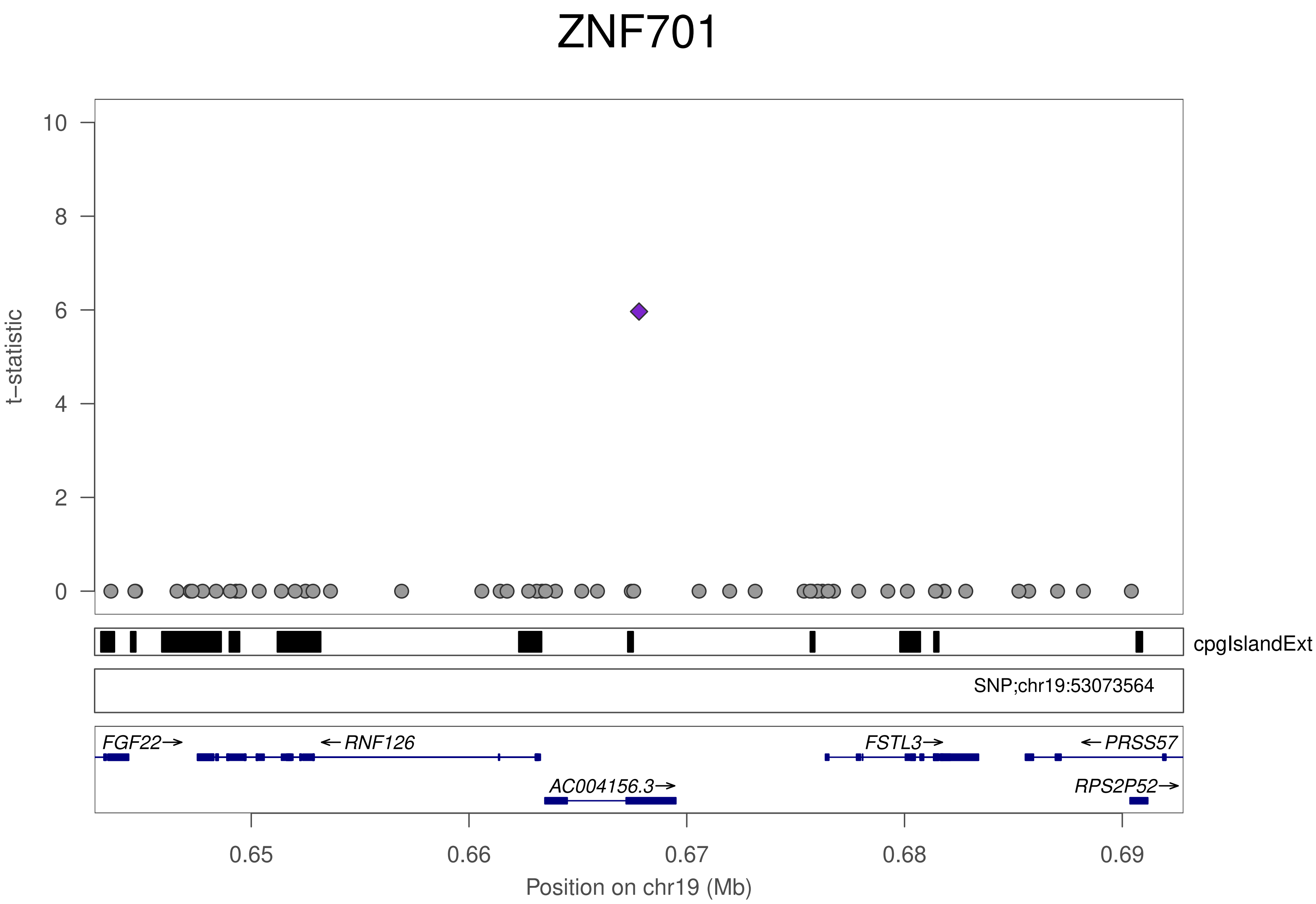

ZNF701
t–statistic
0
2
4
6
8
10
cpgIslandExt
SNP;chr19:53073564
FGF22→
←RNF126
FSTL3→
←PRSS57
AC004156.3→
RPS2P52→
0.65
0.66
0.67
0.68
0.69
Position on chr19 (Mb)

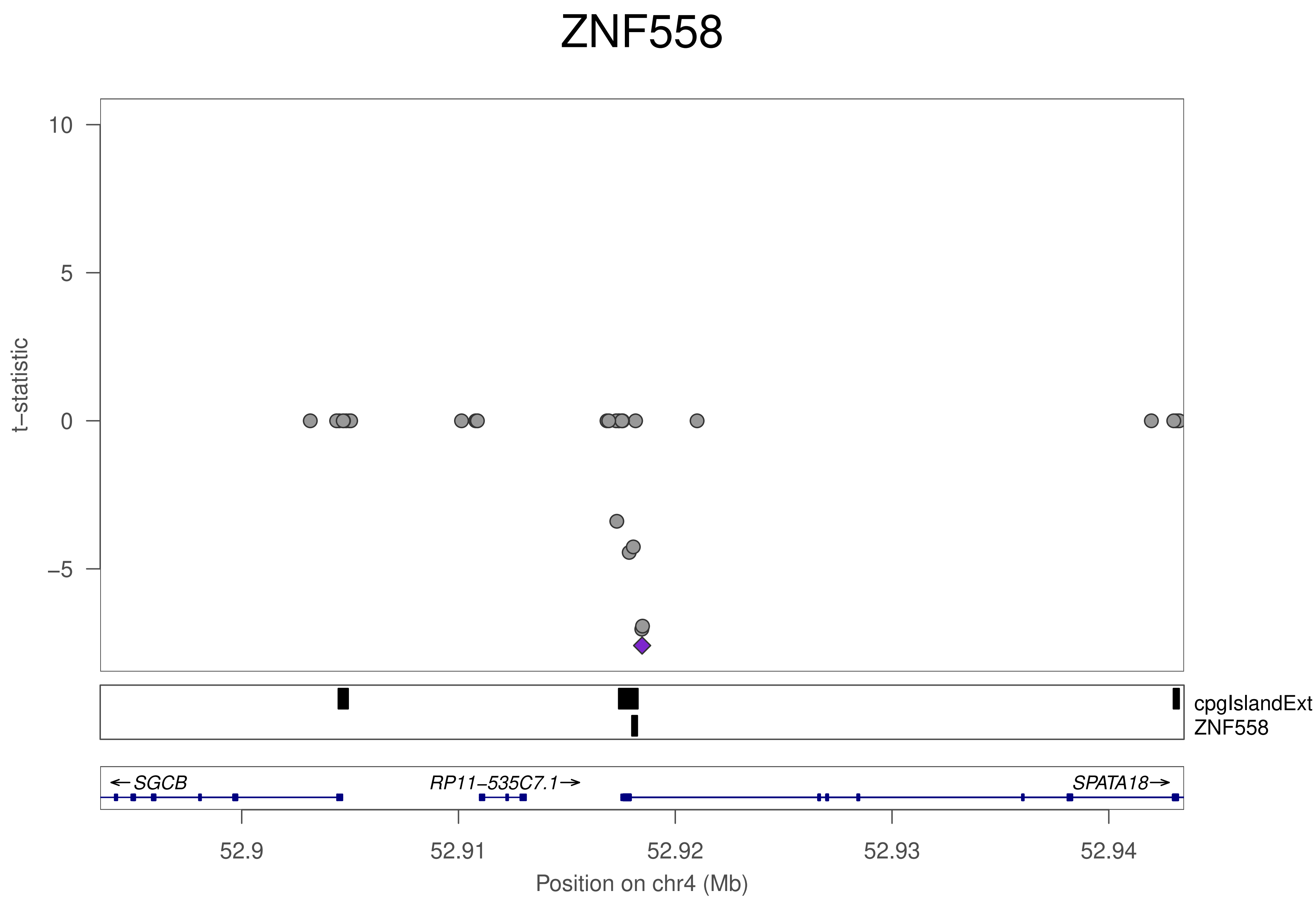

ZNF558
t−statistic
10
5
0
−5
cpgIslandExt
ZNF558
←SGCB
RP11−535C7.1→
SPATA18→
52.9
52.91
52.92
52.93
52.94
Position on chr4 (Mb)

### ZNF212

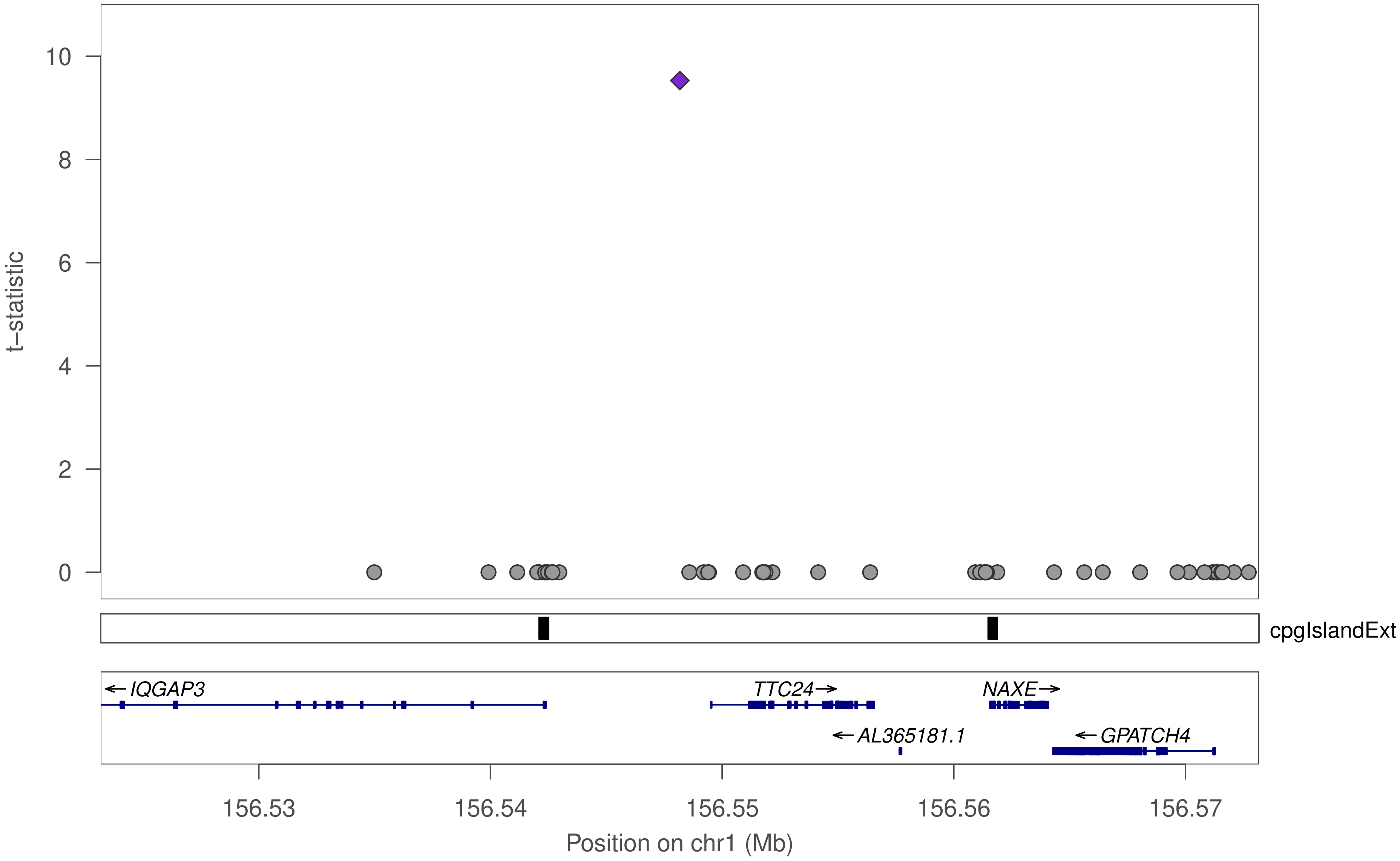

t−statistic
10
8
6
4
2
0
cpgIslandExt
IQGAP3
TTC24
NAXE
AL365181.1
GPATCH4
156.53
156.54
156.55
156.56
156.57
Position on chr1 (Mb)

### ZNF212

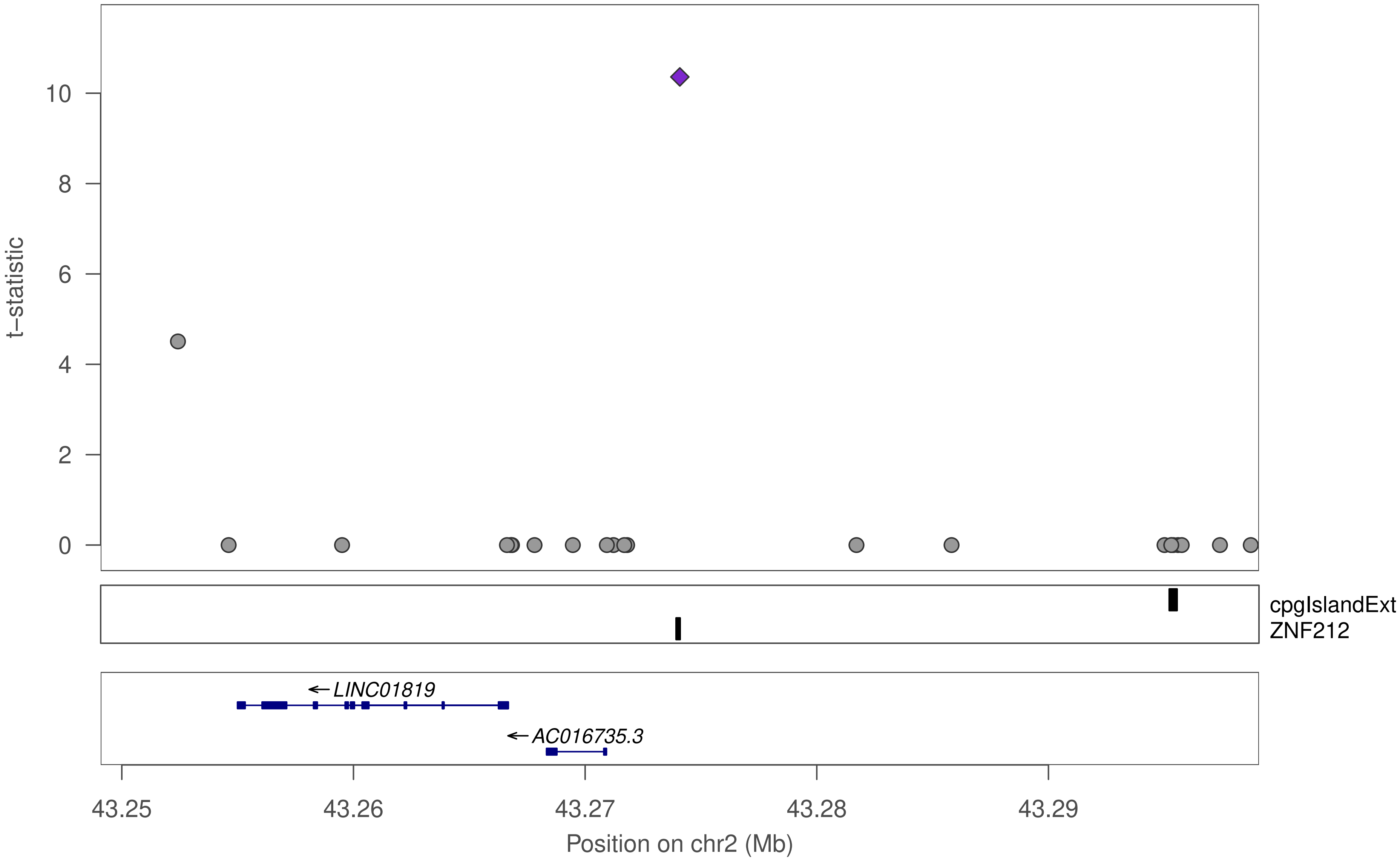
t–statistic
0
2
4
6
8
10
cpgIslandExt
ZNF212
LINC01819
AC016735.3
43.25
43.26
43.27
43.28
43.29
Position on chr2 (Mb)

### ZNF282

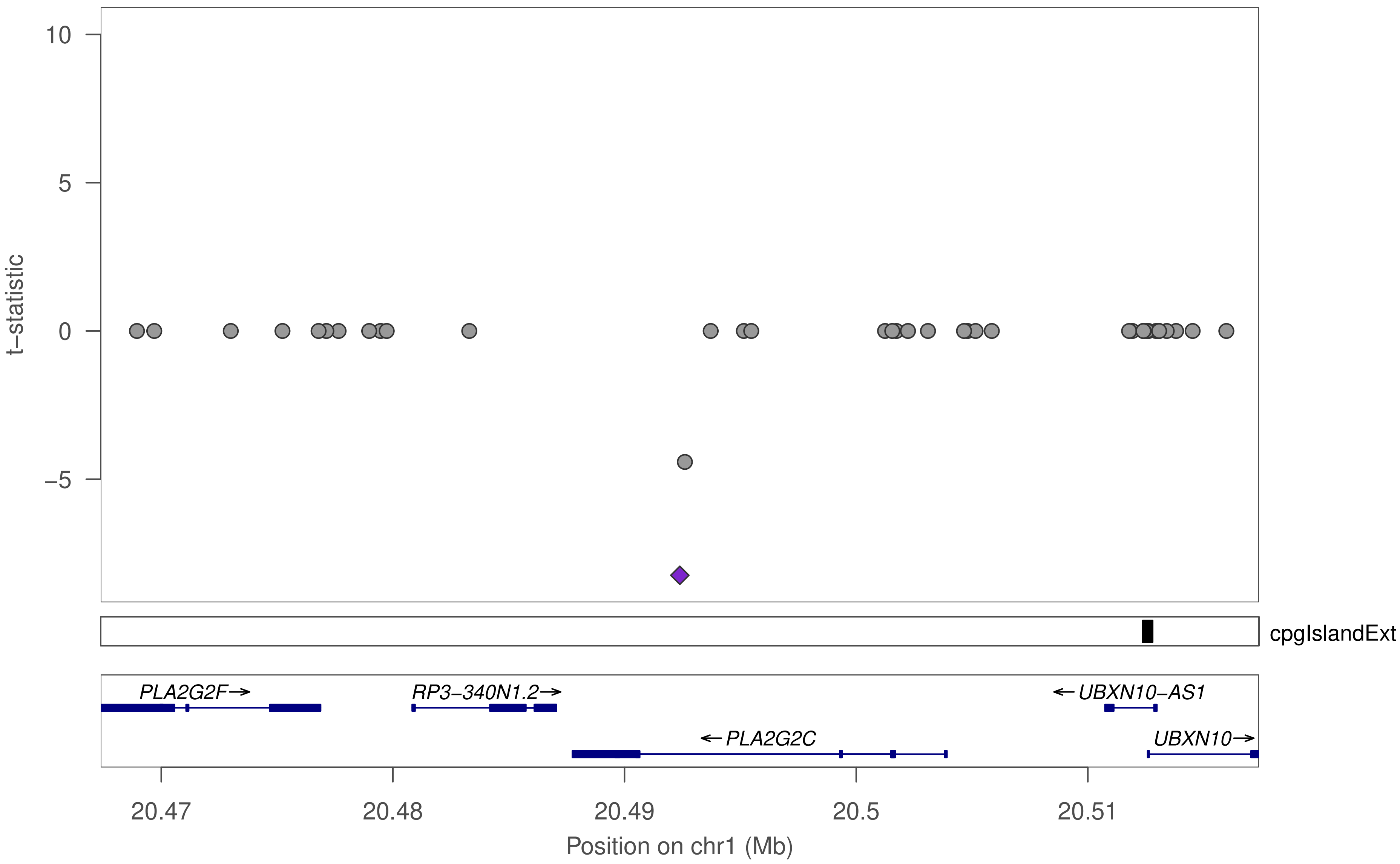

10
5
0
−5
t−statistic
cpgIslandExt
PLA2G2F→
RP3−340N1.2→
←UBXN10−AS1
←PLA2G2C
UBXN10→
20.47
20.48
20.49
20.5
20.51
Position on chr1 (Mb)

### ZNF282

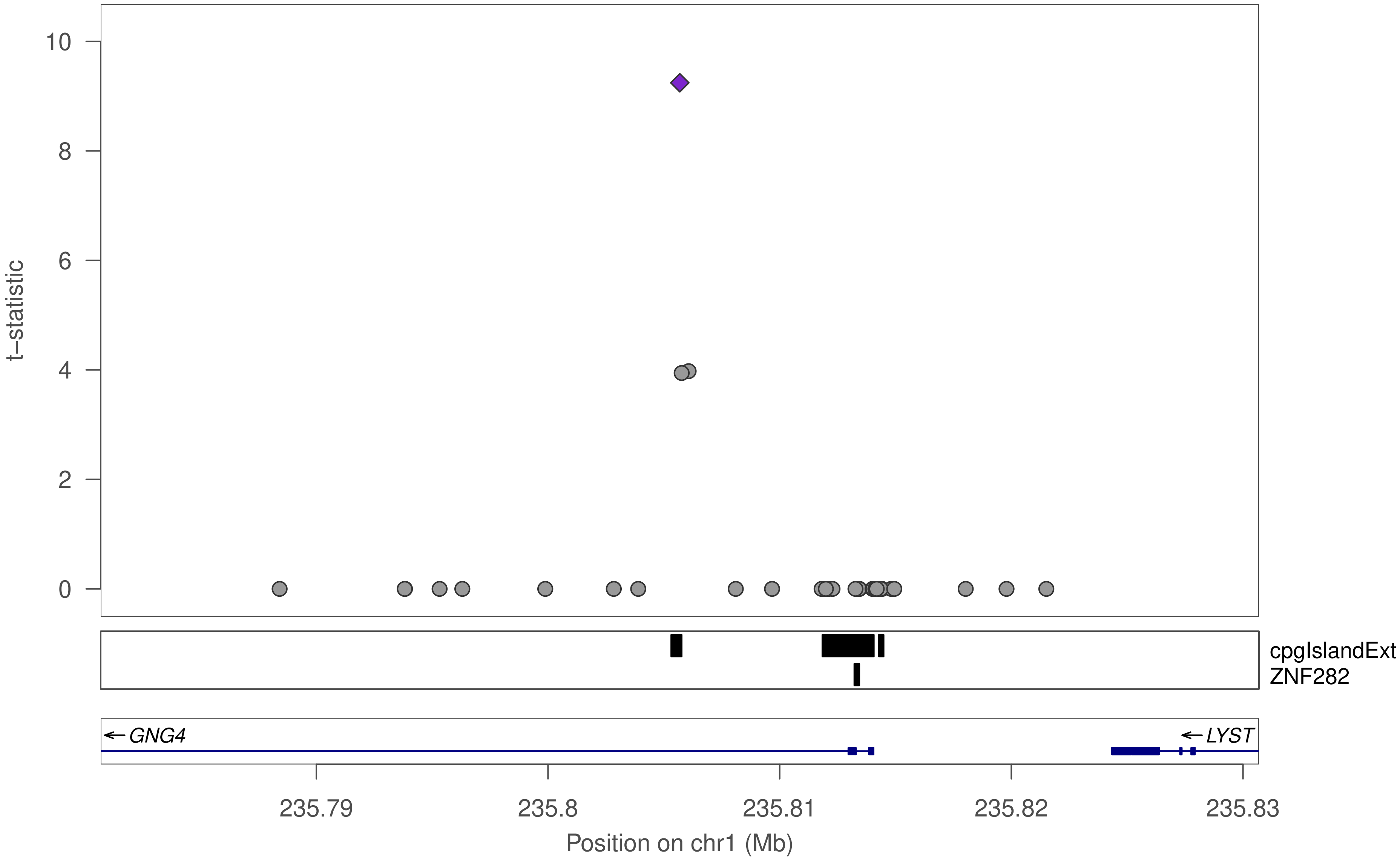

10
8
6
4
2
0
t−statistic
cpgIslandExt
ZNF282
GNG4
LYST
235.79
235.8
235.81
235.82
235.83
Position on chr1 (Mb)

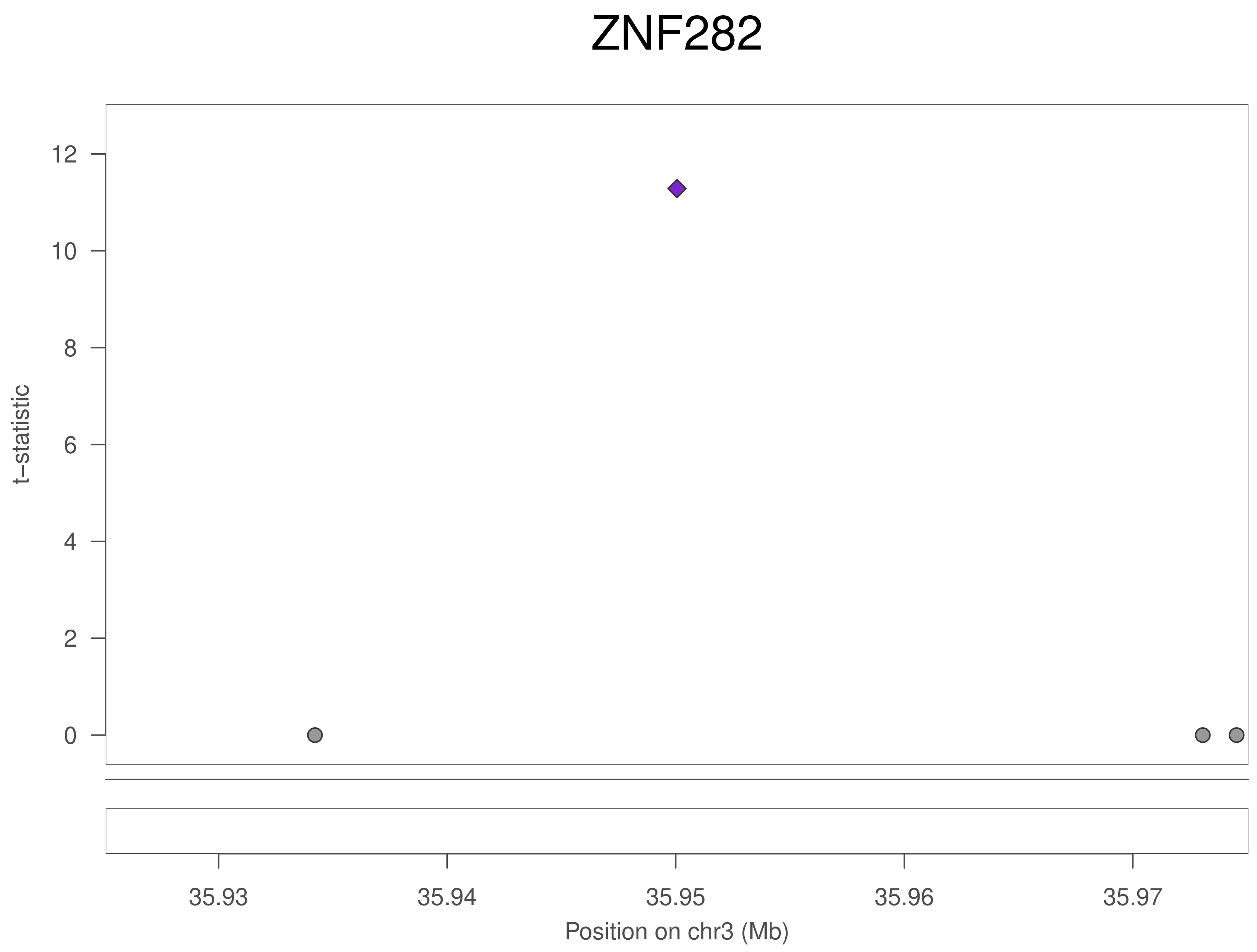

ZNF282
t−statistic
12
10
8
6
4
2
0
35.93
35.94
35.95
35.96
35.97
Position on chr3 (Mb)

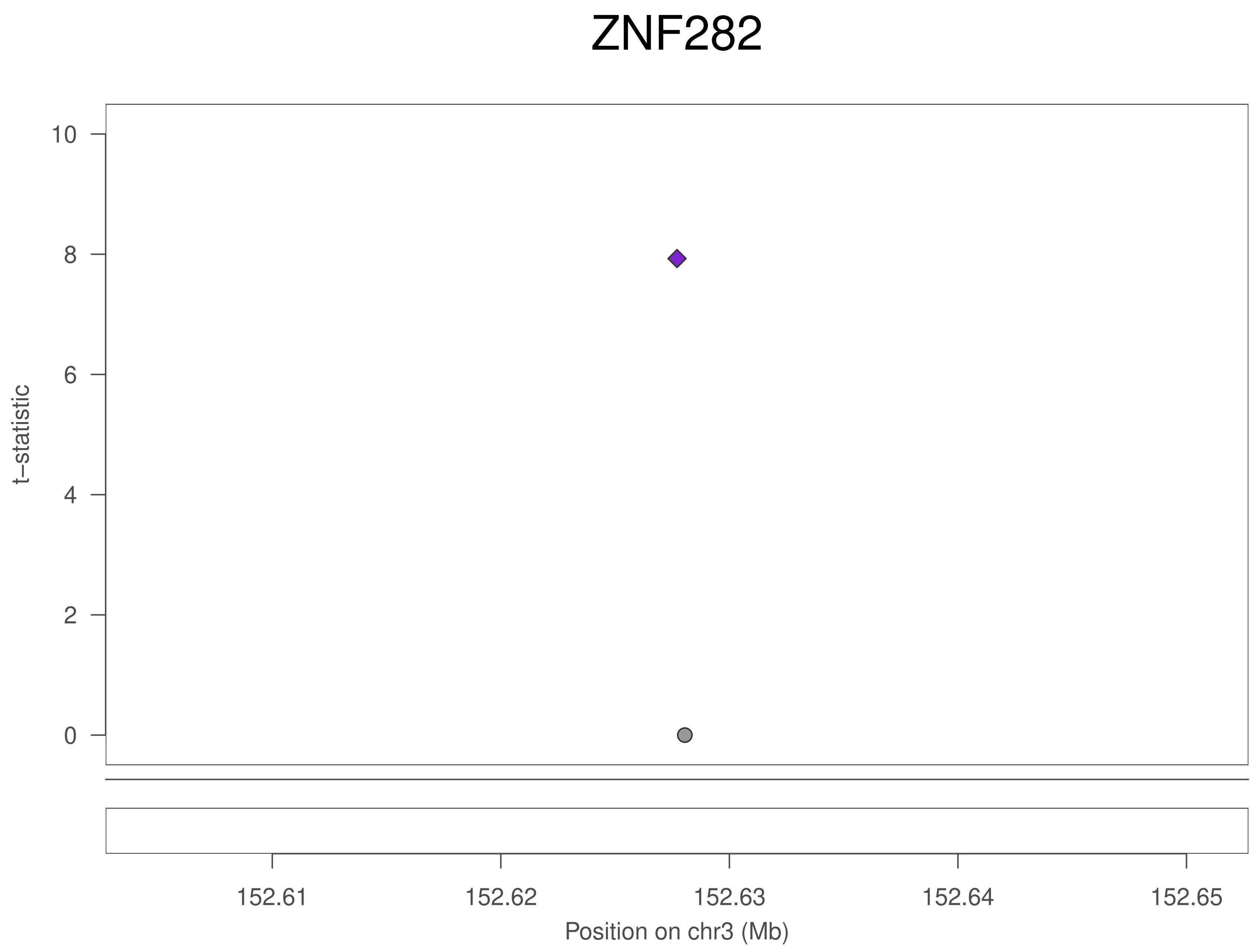
ZNF282
10
8
6
4
2
0
t−statistic
152.61
152.62
152.63
152.64
152.65
Position on chr3 (Mb)

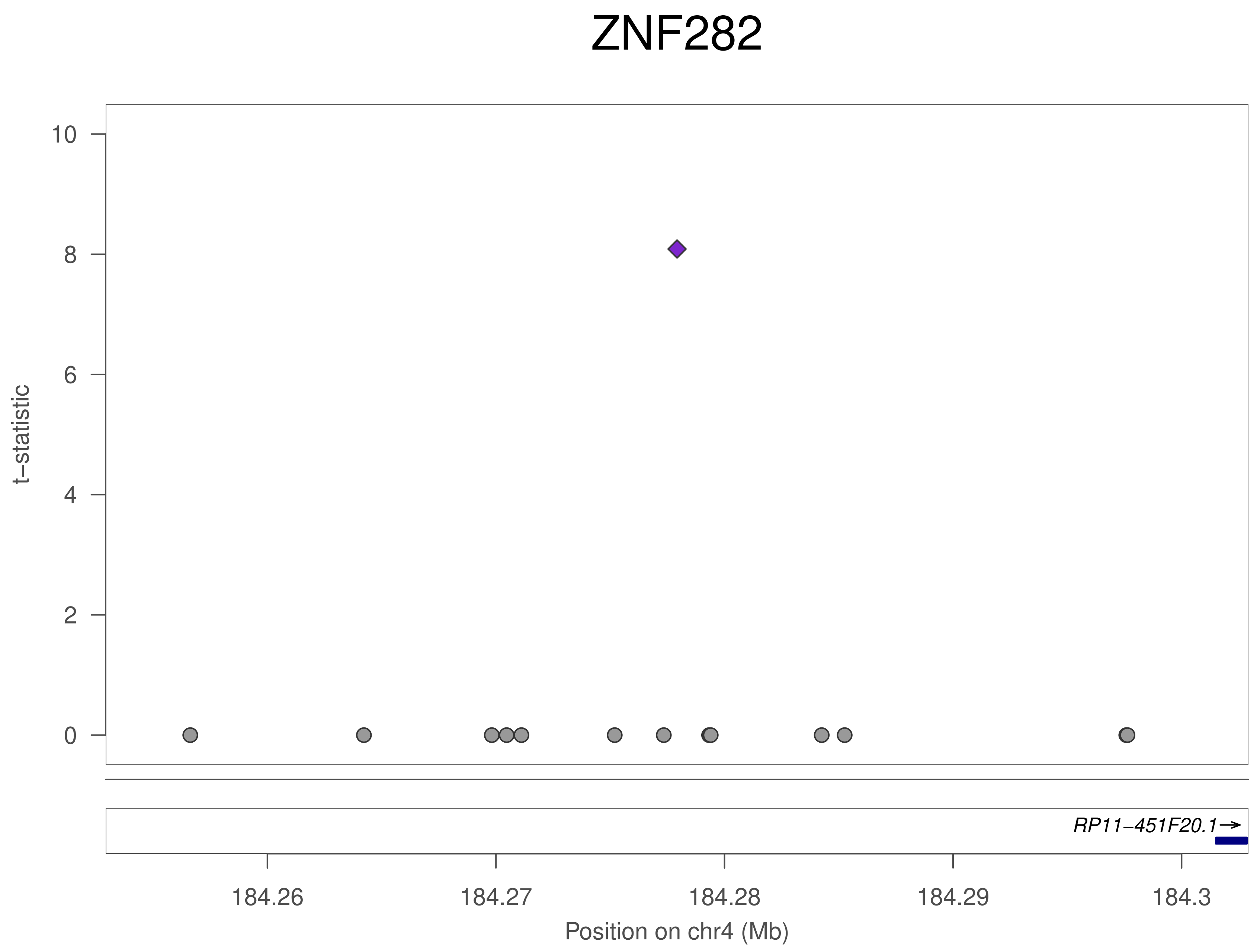
ZNF282
t–statistic
10
8
6
4
2
0
RP11–451F20.1→
184.26
184.27
184.28
184.29
184.3
Position on chr4 (Mb)

### ZNF282

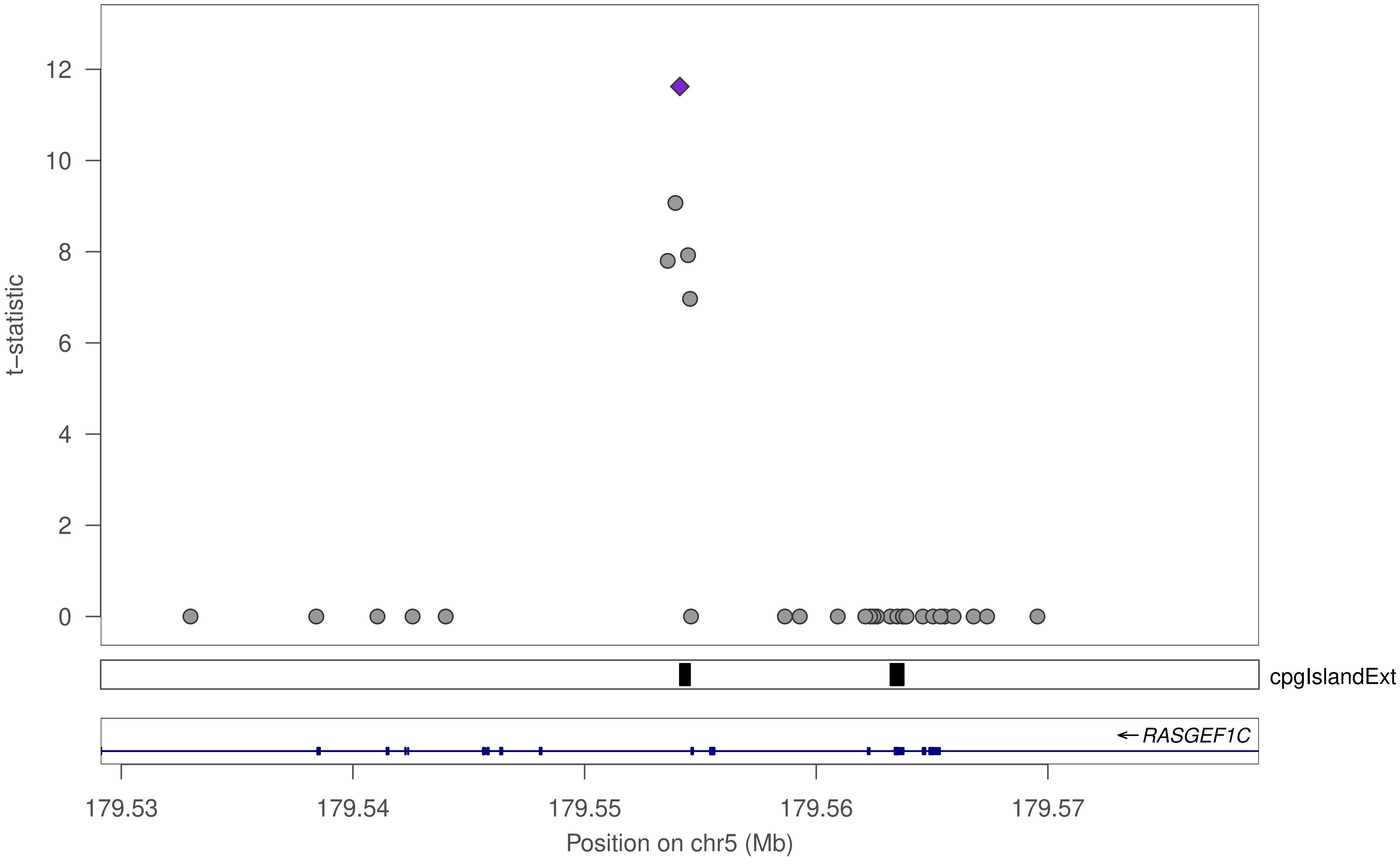

12
10
8
6
4
2
0
t−statistic
cpgIslandExt
← RASGEF1C
179.53
179.54
179.55
179.56
179.57
Position on chr5 (Mb)

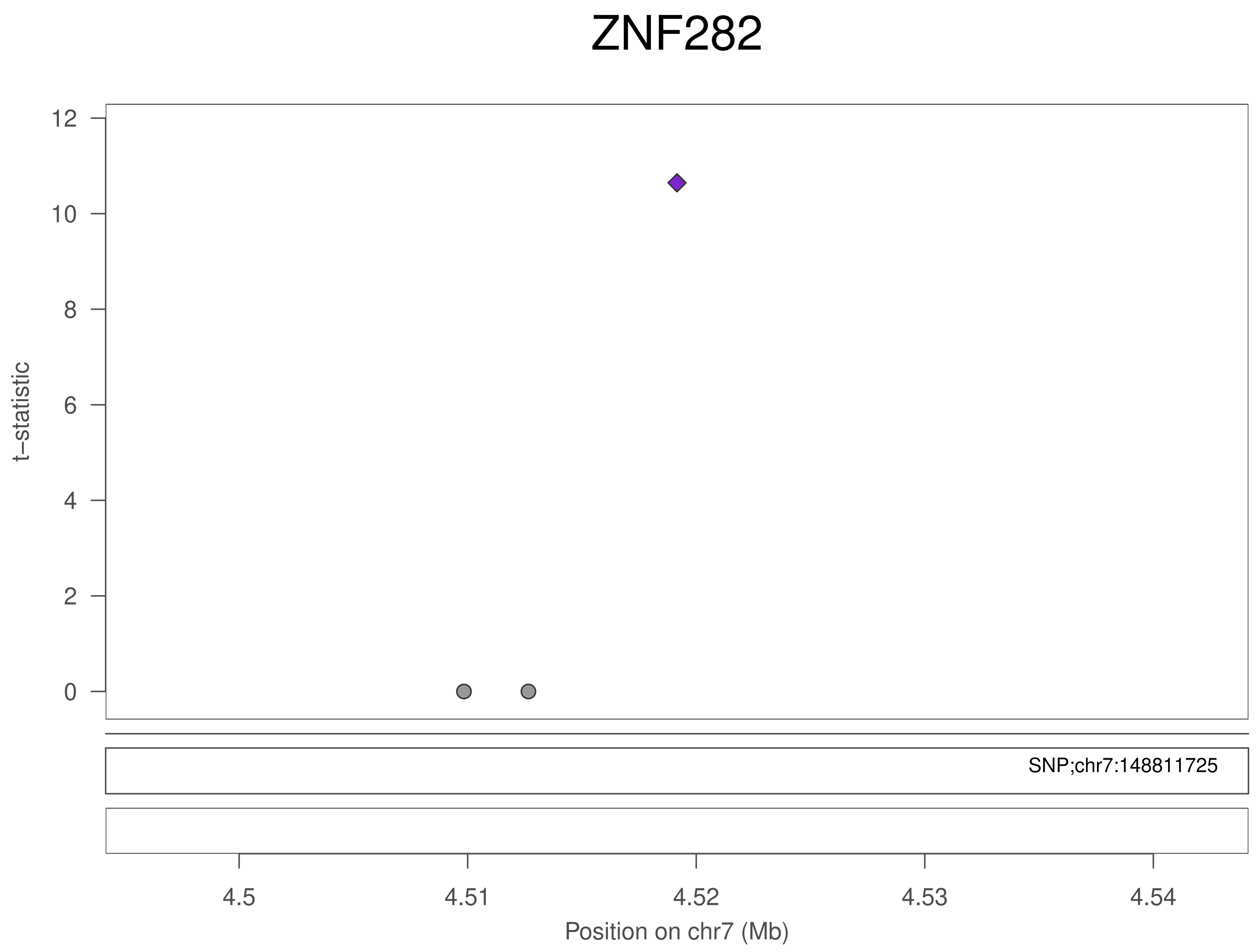
ZNF282
t−statistic
12
10
8
6
4
2
0
SNP;chr7:148811725
4.5
4.51
4.52
4.53
4.54
Position on chr7 (Mb)

### ZNF282

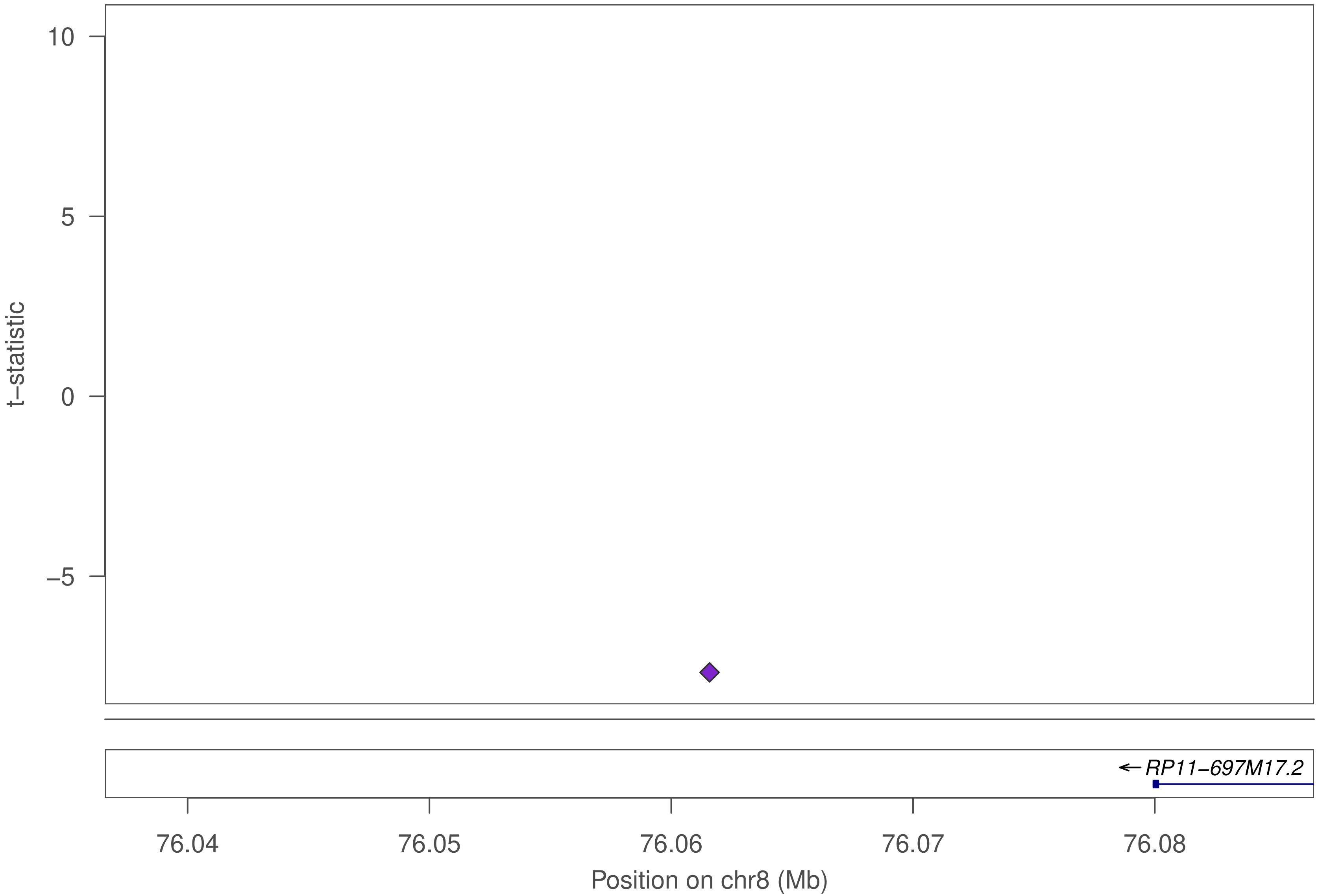

10
5
0
–5
t–statistic
RP11–697M17.2
76.04
76.05
76.06
76.07
76.08
Position on chr8 (Mb)

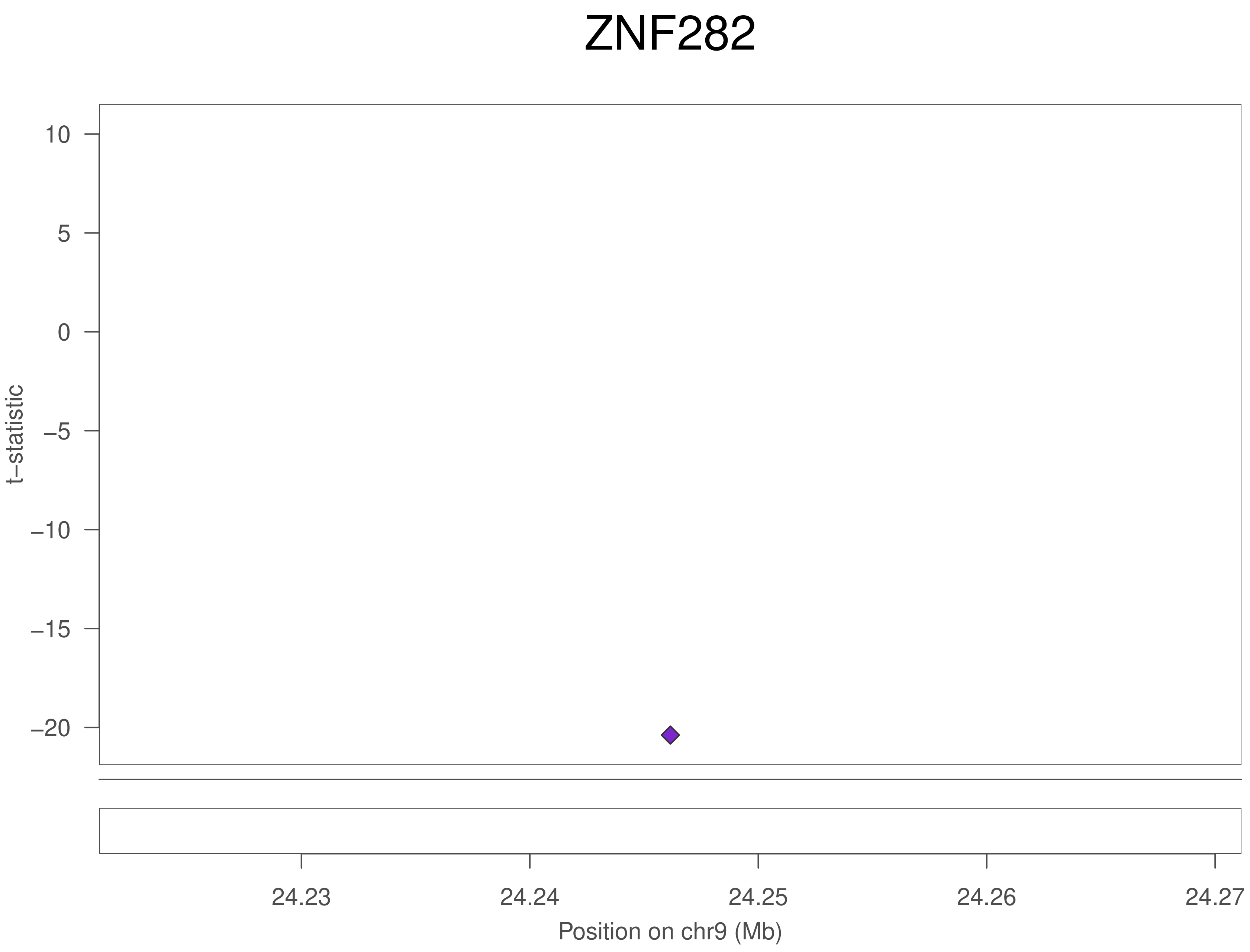
ZNF282
t–statistic
10
5
0
–5
–10
–15
–20
24.23
24.24
24.25
24.26
24.27
Position on chr9 (Mb)

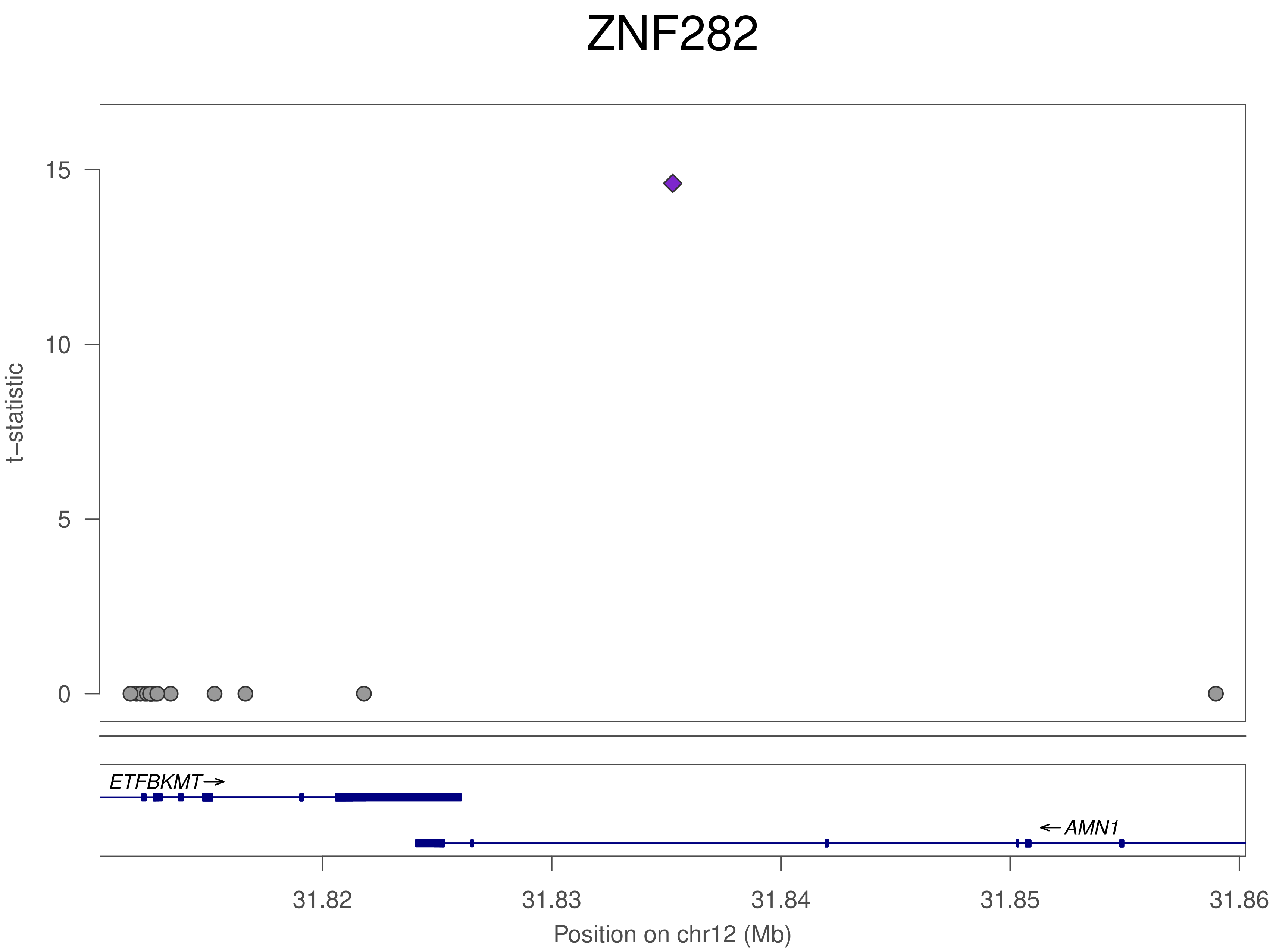
ZNF282
t−statistic
15
10
5
0
ETFBKMT
AMN1
31.82
31.83
31.84
31.85
31.86
Position on chr12 (Mb)

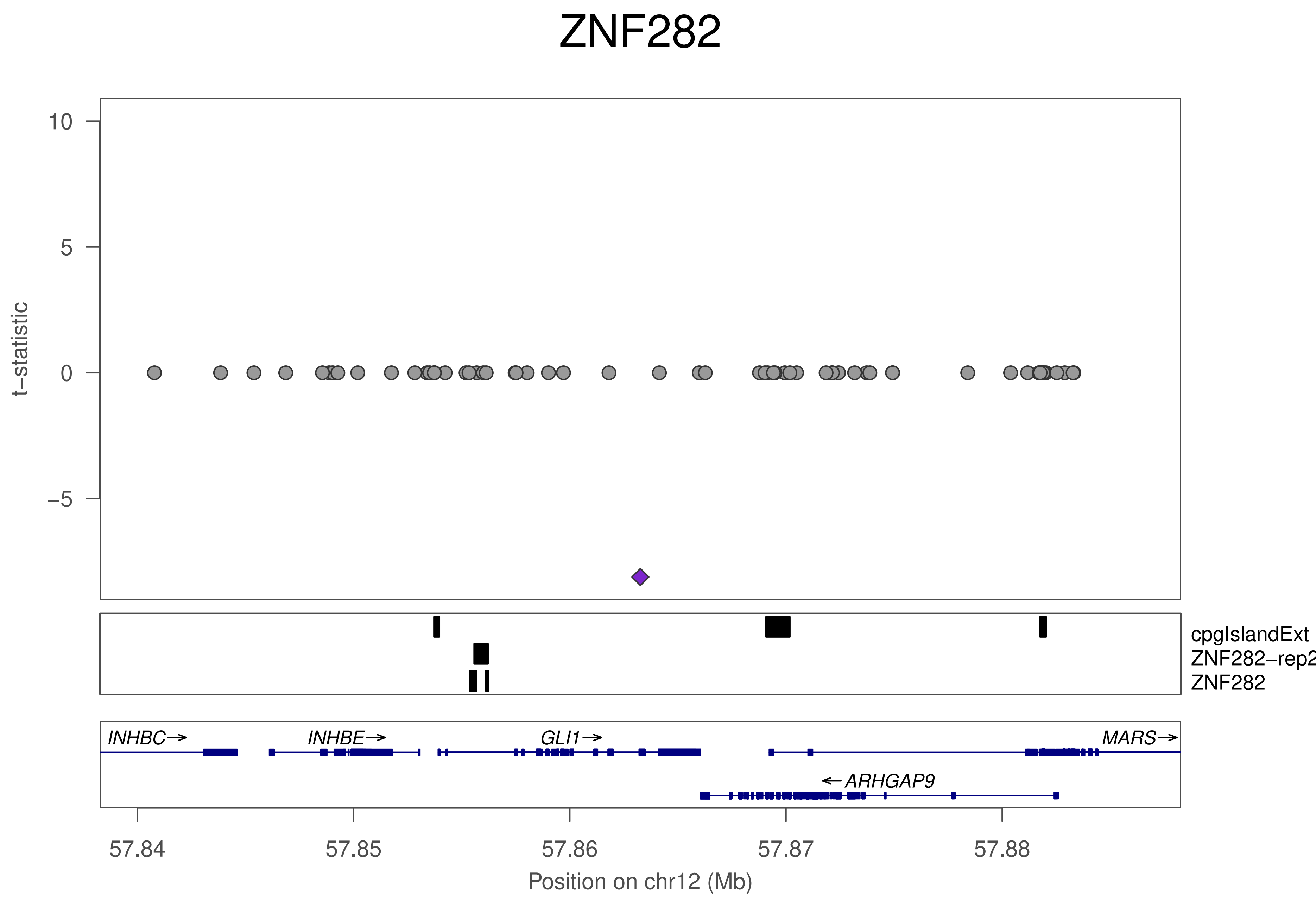
ZNF282
10
5
0
−5
t−statistic
cpgIslandExt
ZNF282−rep2
ZNF282
INHBC→
INHBE→
GLI1→
MARS→
←ARHGAP9
57.84
57.85
57.86
57.87
57.88
Position on chr12 (Mb)

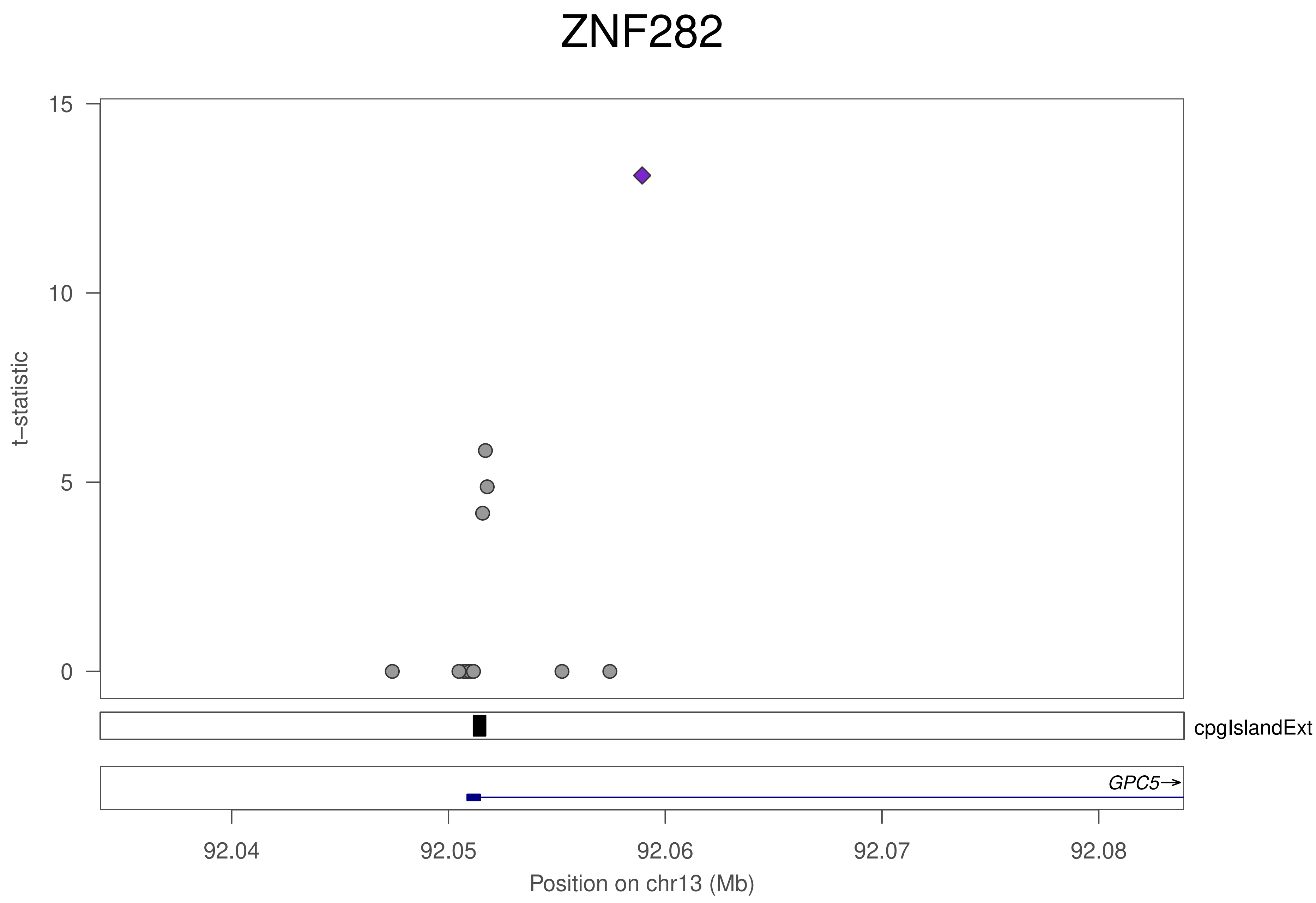
ZNF282
t−statistic
15
10
5
0
cpgIslandExt
GPC5→
92.04
92.05
92.06
92.07
92.08
Position on chr13 (Mb)

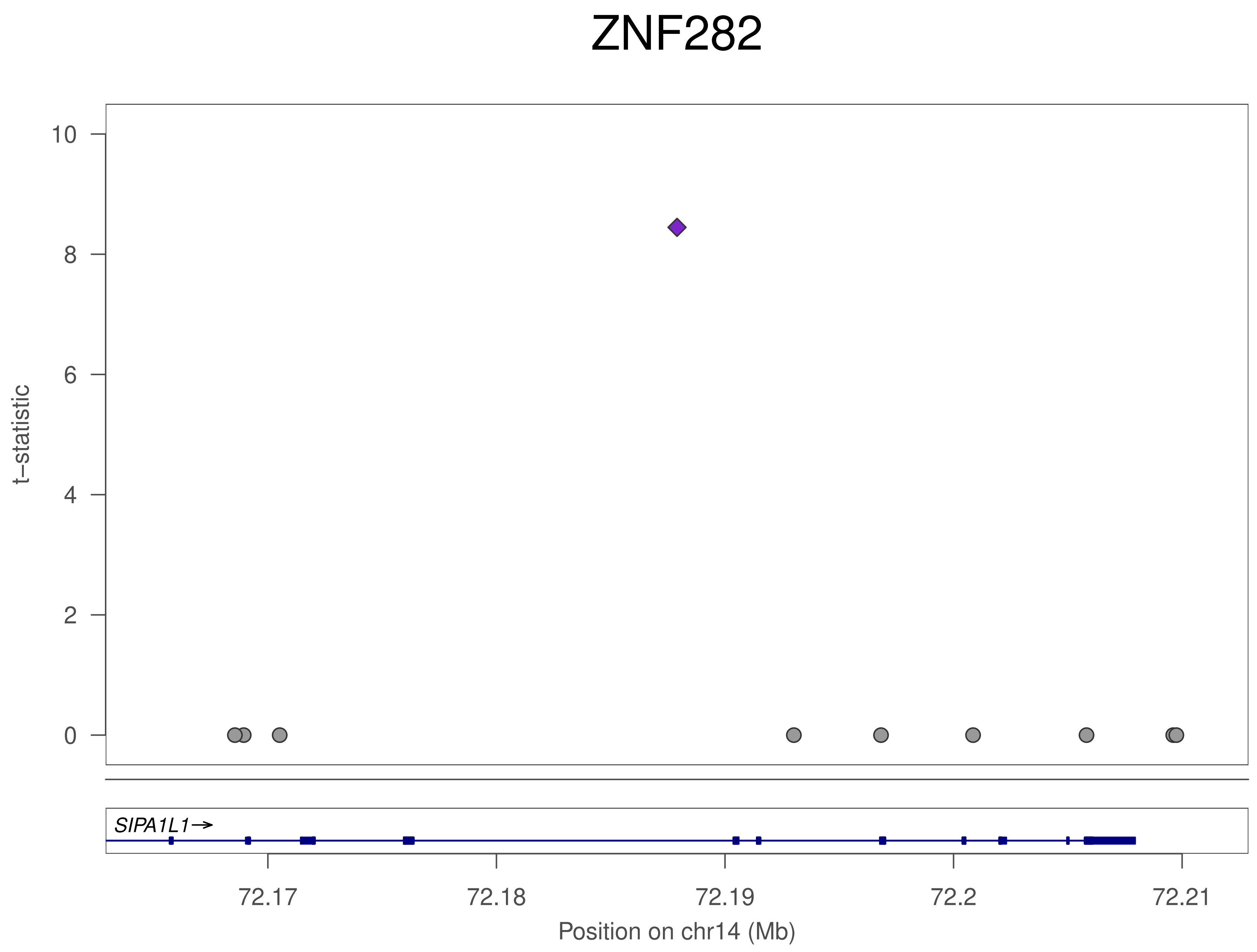

ZNF282
t–statistic
10
8
6
4
2
0
SIPA1L1→
72.17
72.18
72.19
72.2
72.21
Position on chr14 (Mb)

### ZNF282

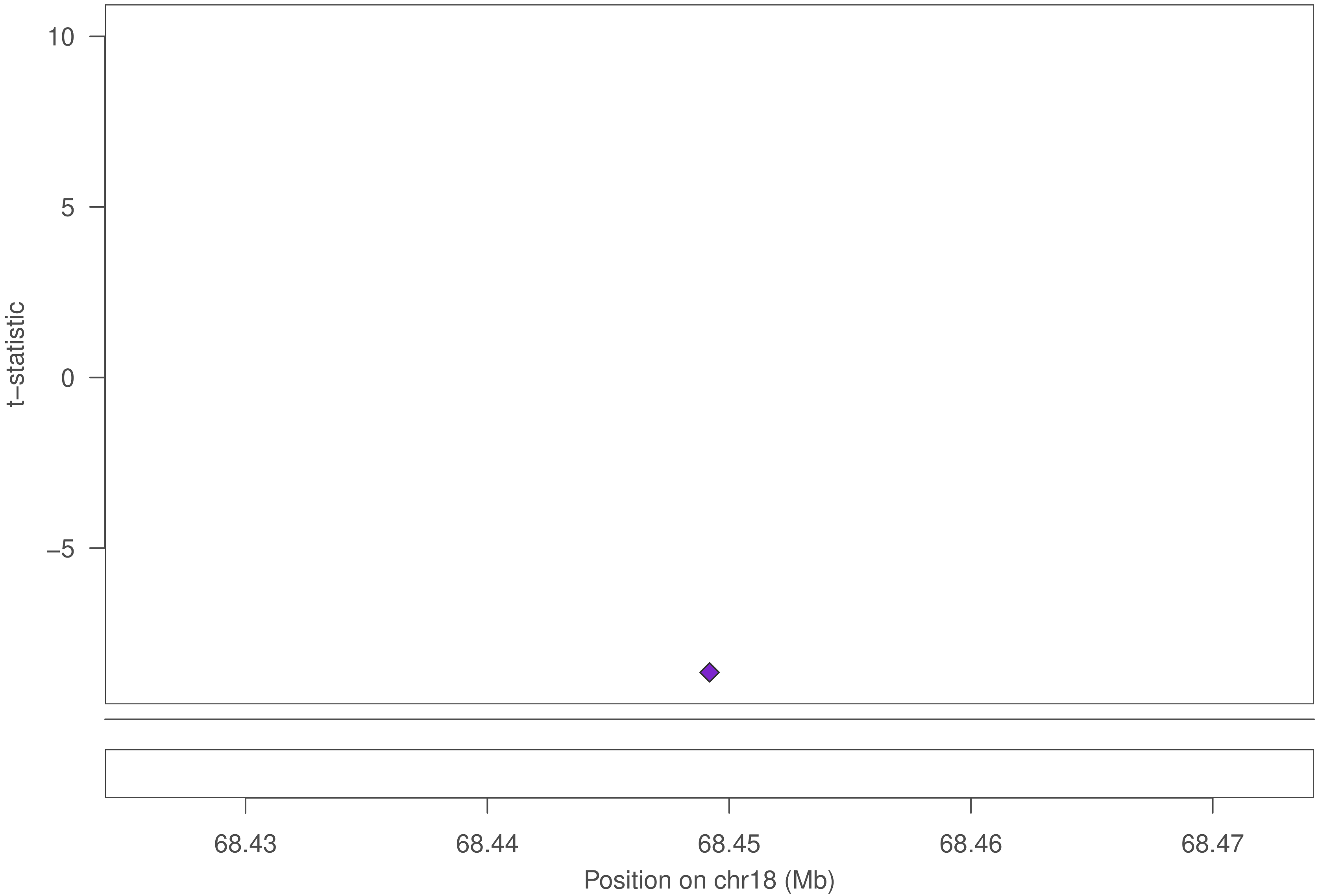

10
5
0
−5
t−statistic
68.43
68.44
68.45
68.46
68.47
Position on chr18 (Mb)

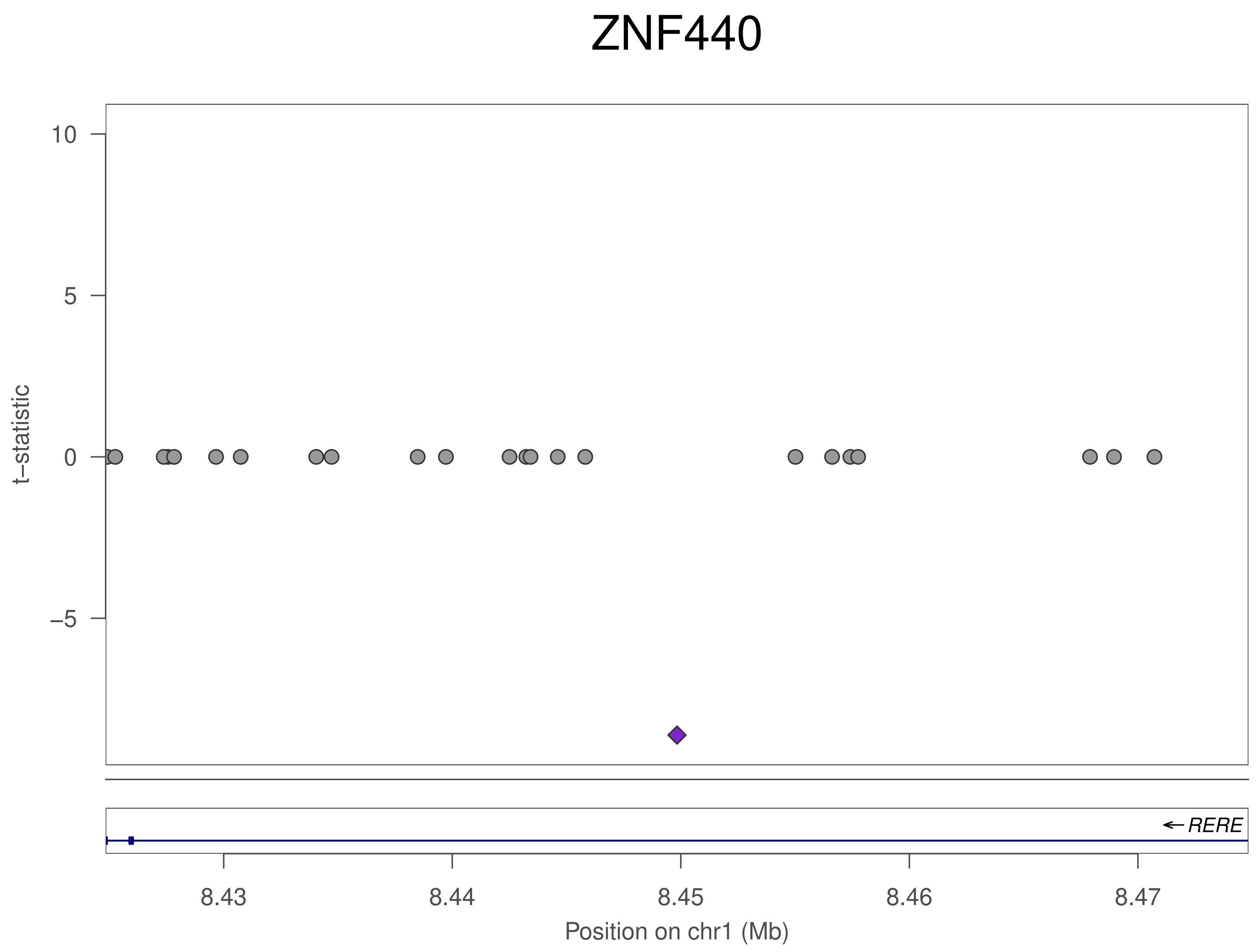

ZNF440
t–statistic
10
5
0
–5
RERE
8.43
8.44
8.45
8.46
8.47
Position on chr1 (Mb)

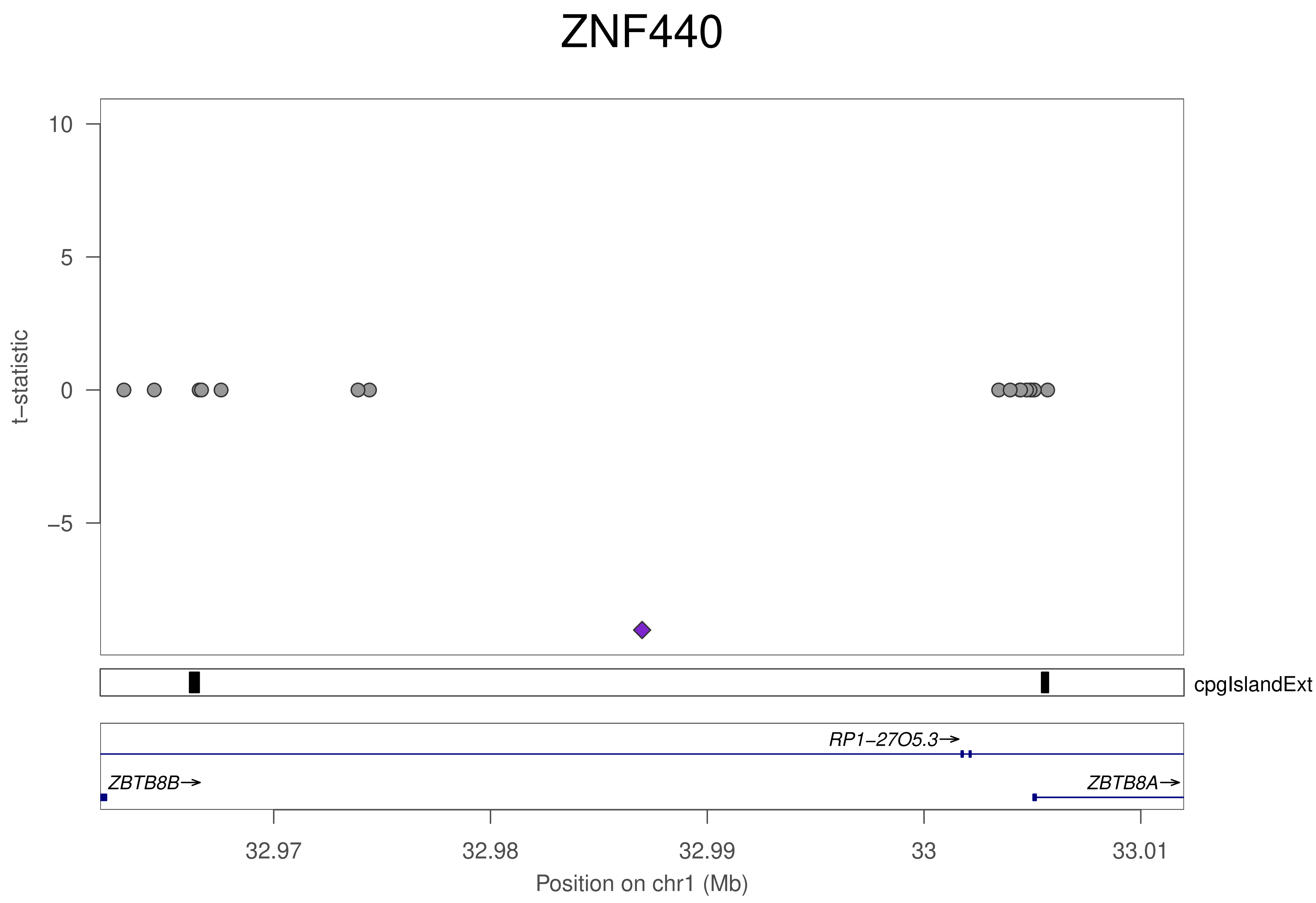

ZNF440
t−statistic
10
5
0
−5
cpgIslandExt
RP1−27O5.3→
ZBTB8B→
ZBTB8A→
32.97
32.98
32.99
33
33.01
Position on chr1 (Mb)

### ZNF440

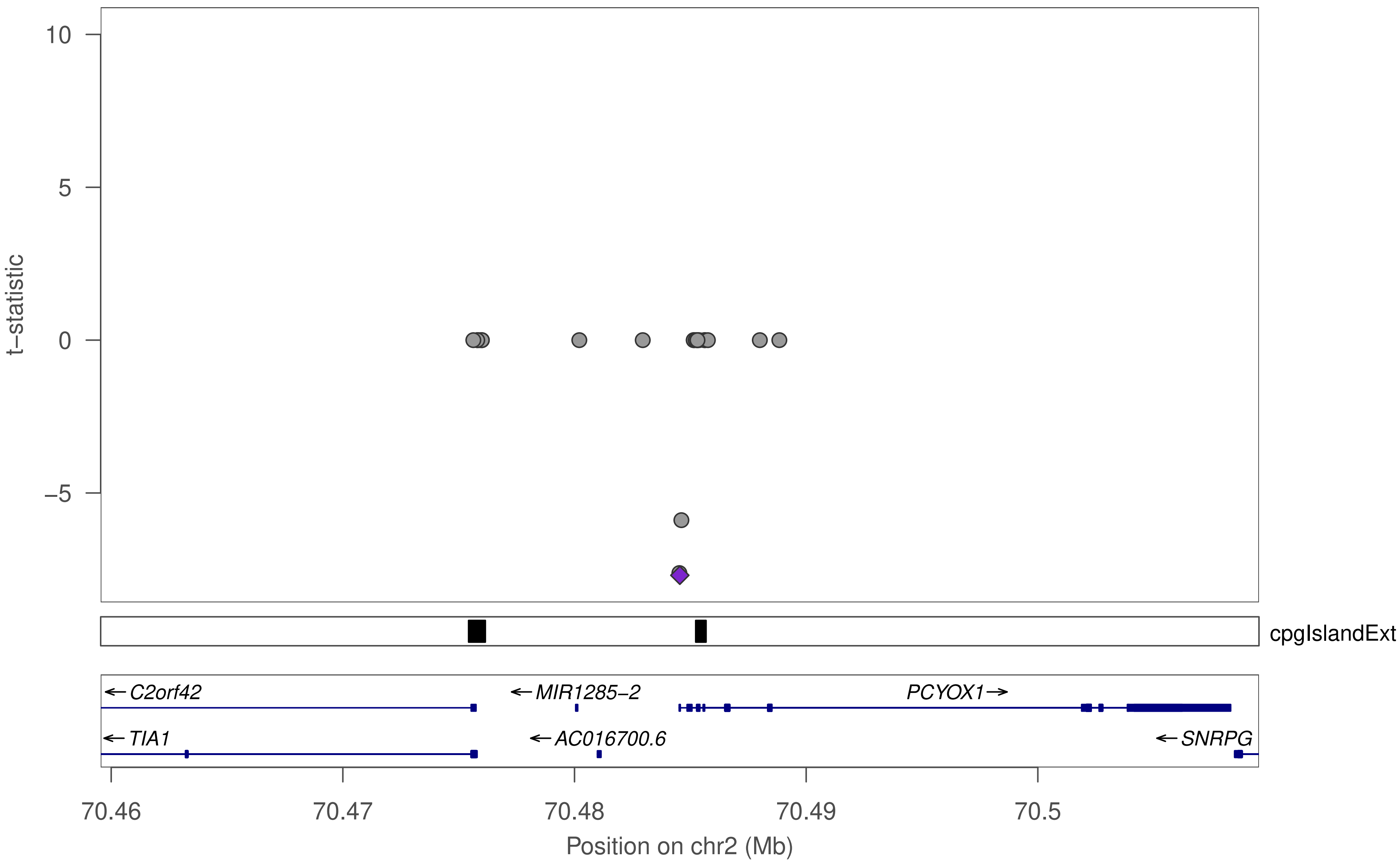

10
5
0
−5
t−statistic
cpgIslandExt
←C2orf42
←MIR1285−2
PCYOX1→
←TIA1
←AC016700.6
←SNRPG
70.46
70.47
70.48
70.49
70.5
Position on chr2 (Mb)

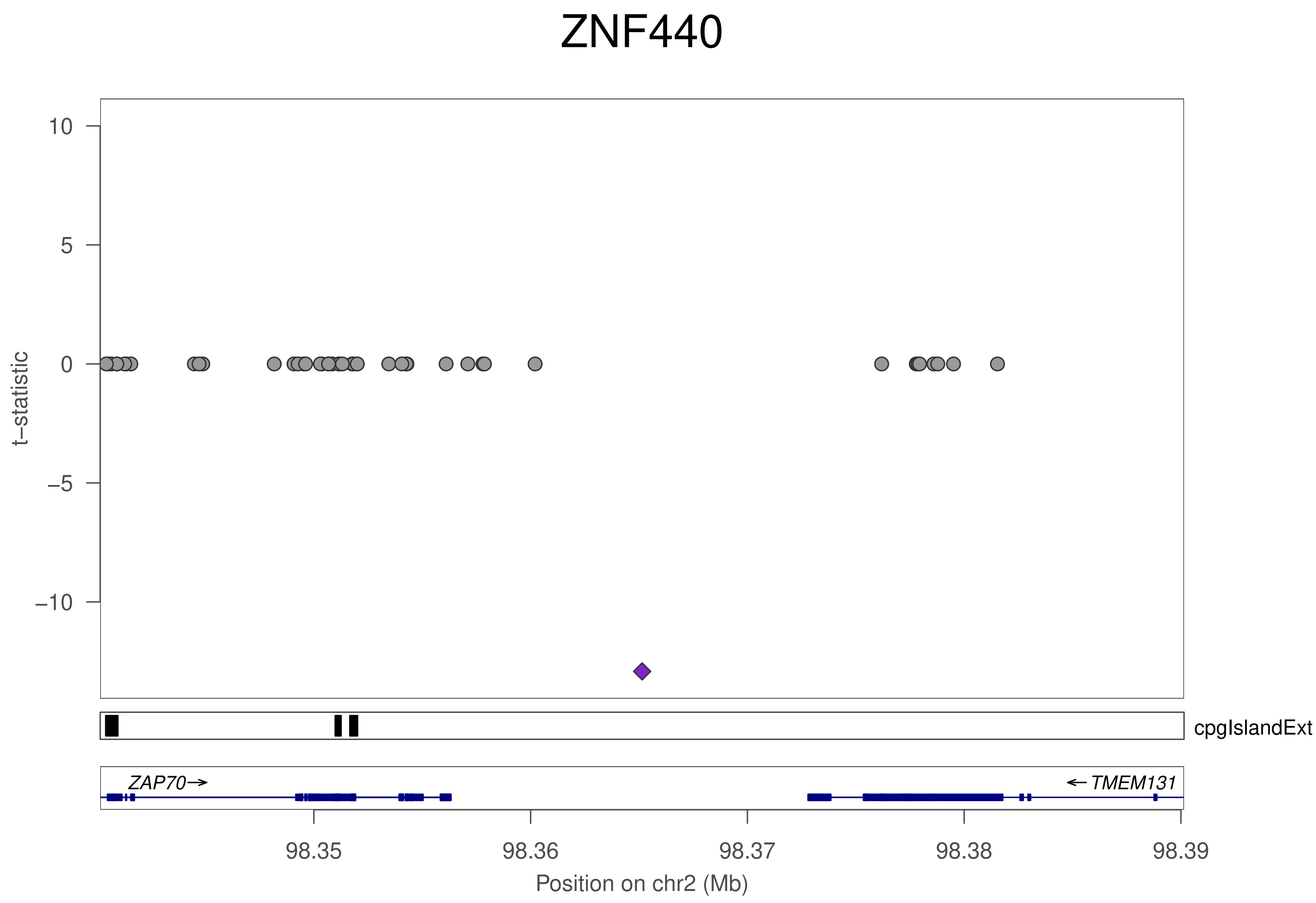
ZNF440
t−statistic
10
5
0
−5
−10
cpgIslandExt
ZAP70→
←TMEM131
98.35
98.36
98.37
98.38
98.39
Position on chr2 (Mb)

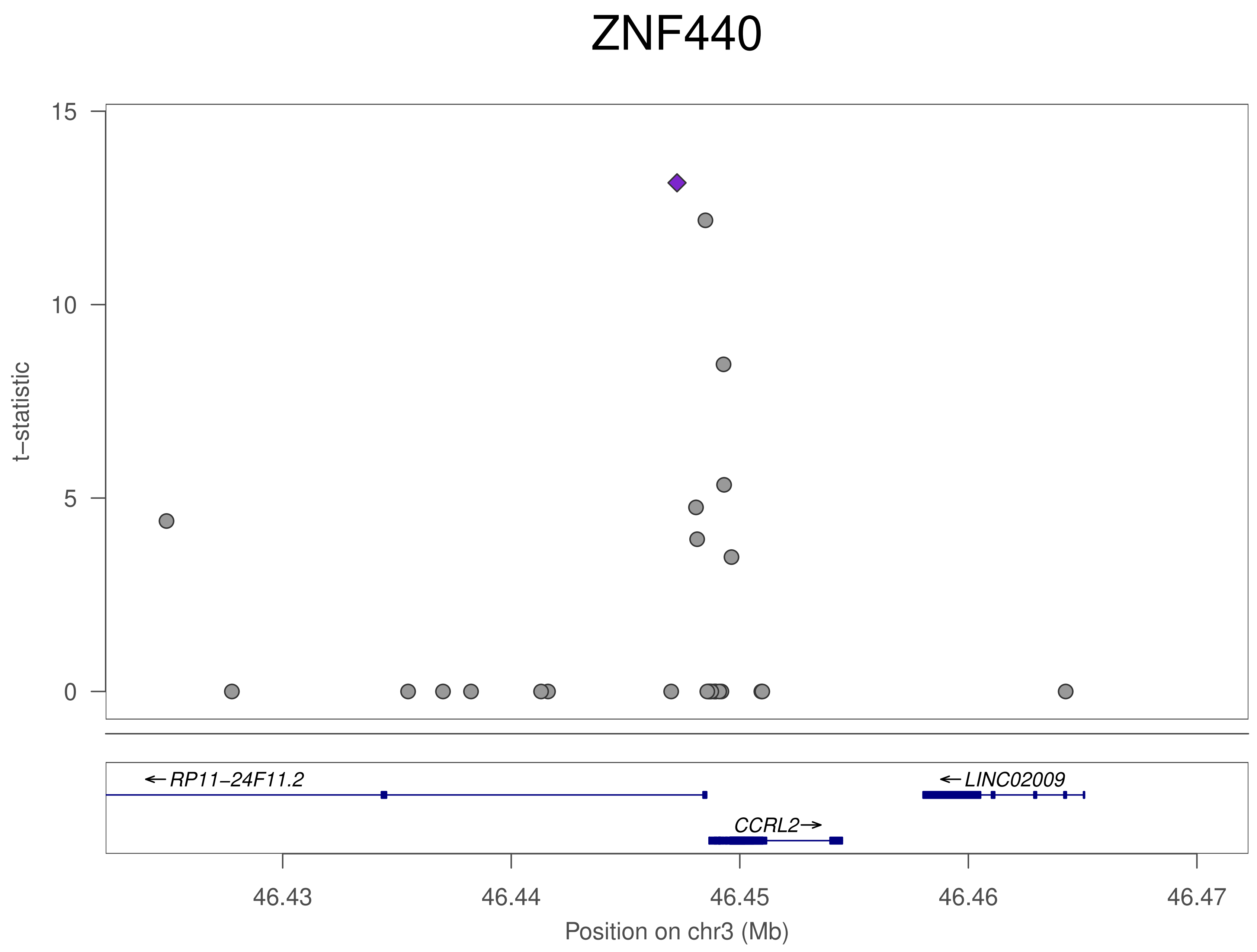
ZNF440
t−statistic
15
10
5
0
← RP11−24F11.2
← LINC02009
CCRL2→
46.43
46.44
46.45
46.46
46.47
Position on chr3 (Mb)

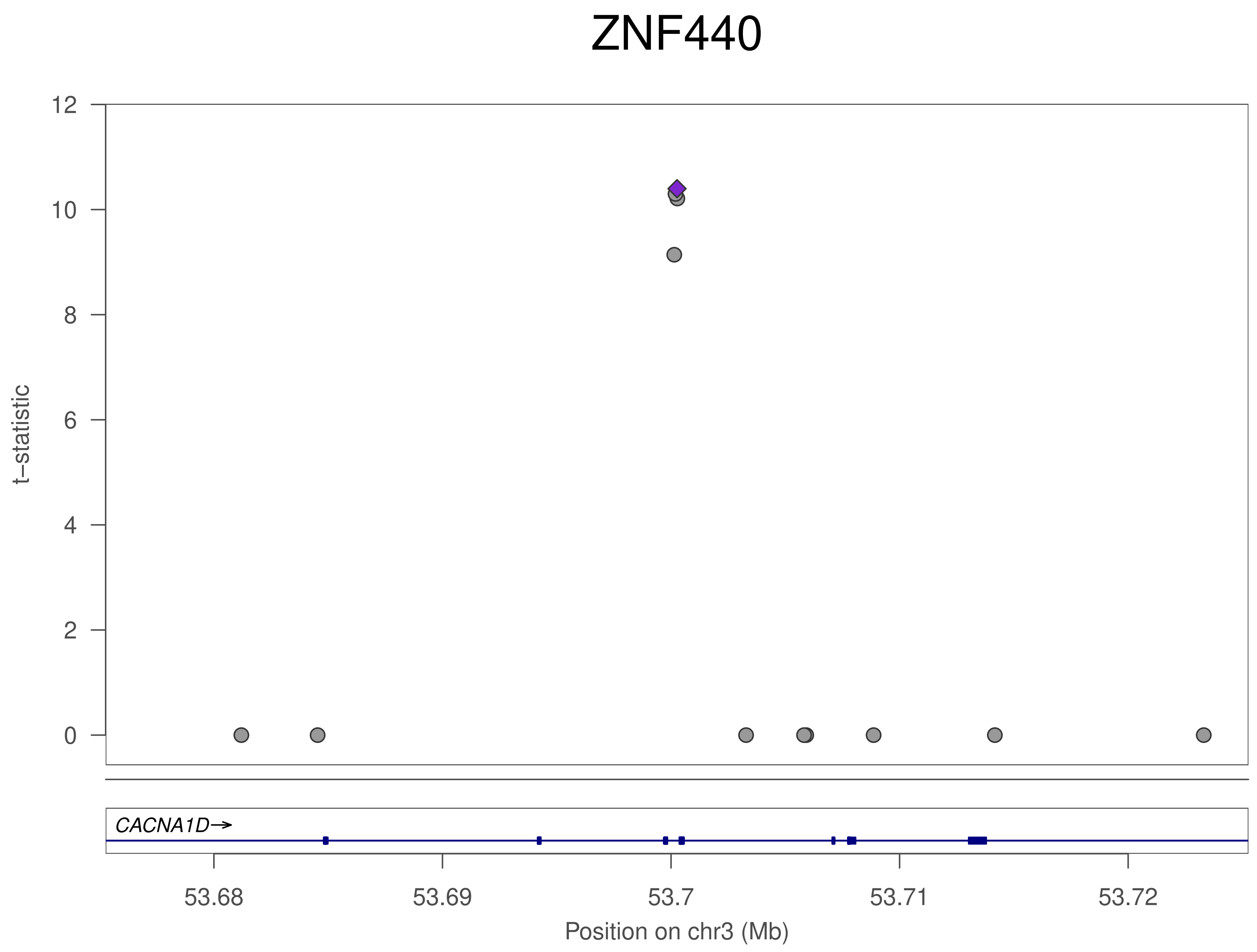
ZNF440
t−statistic
12
10
8
6
4
2
0
CACNA1D→
53.68
53.69
53.7
53.71
53.72
Position on chr3 (Mb)

### ZNF440

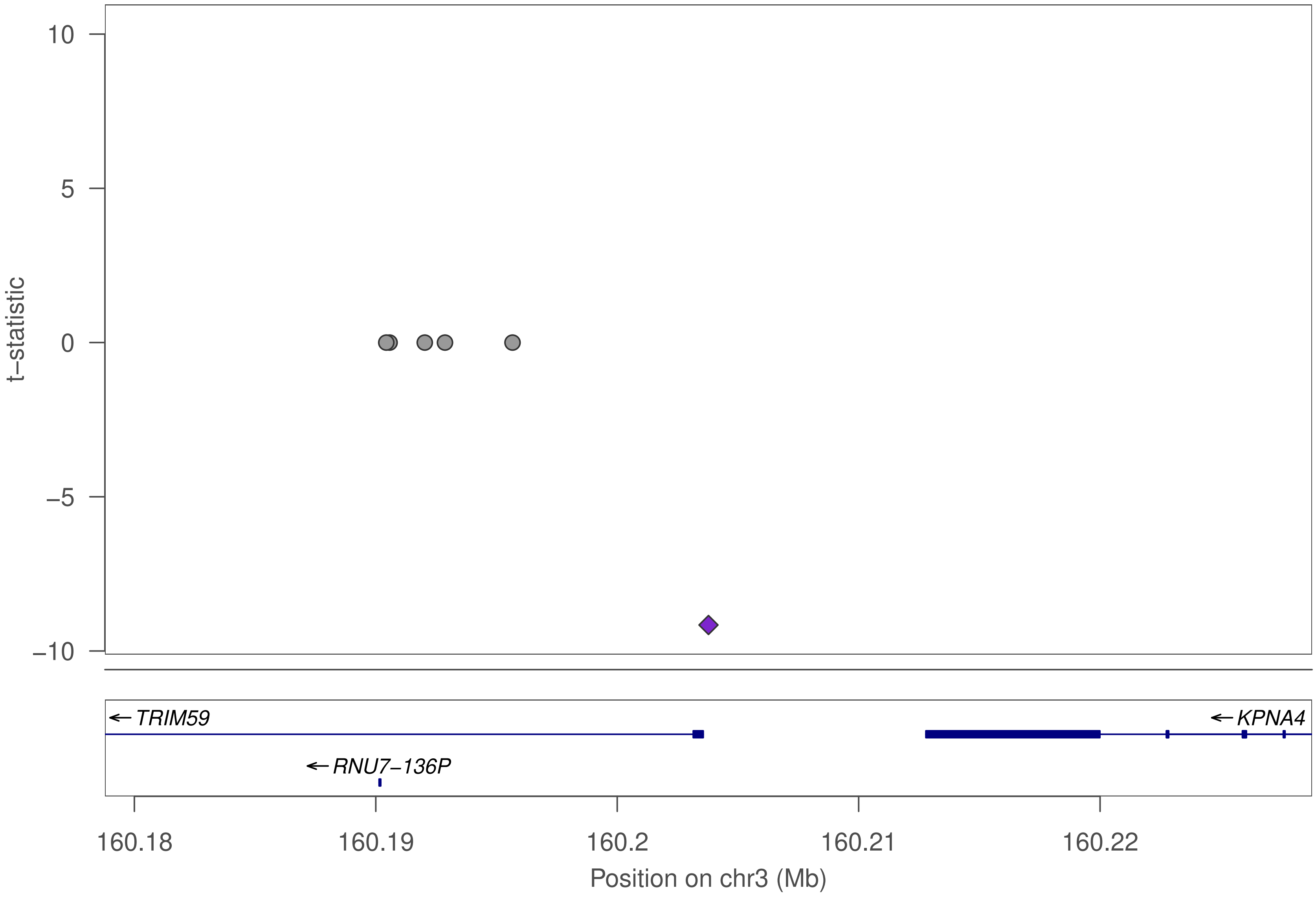

10
5
0
−5
−10
t−statistic
TRIM59
RNU7−136P
KPNA4
160.18
160.19
160.2
160.21
160.22
Position on chr3 (Mb)

### ZNF440

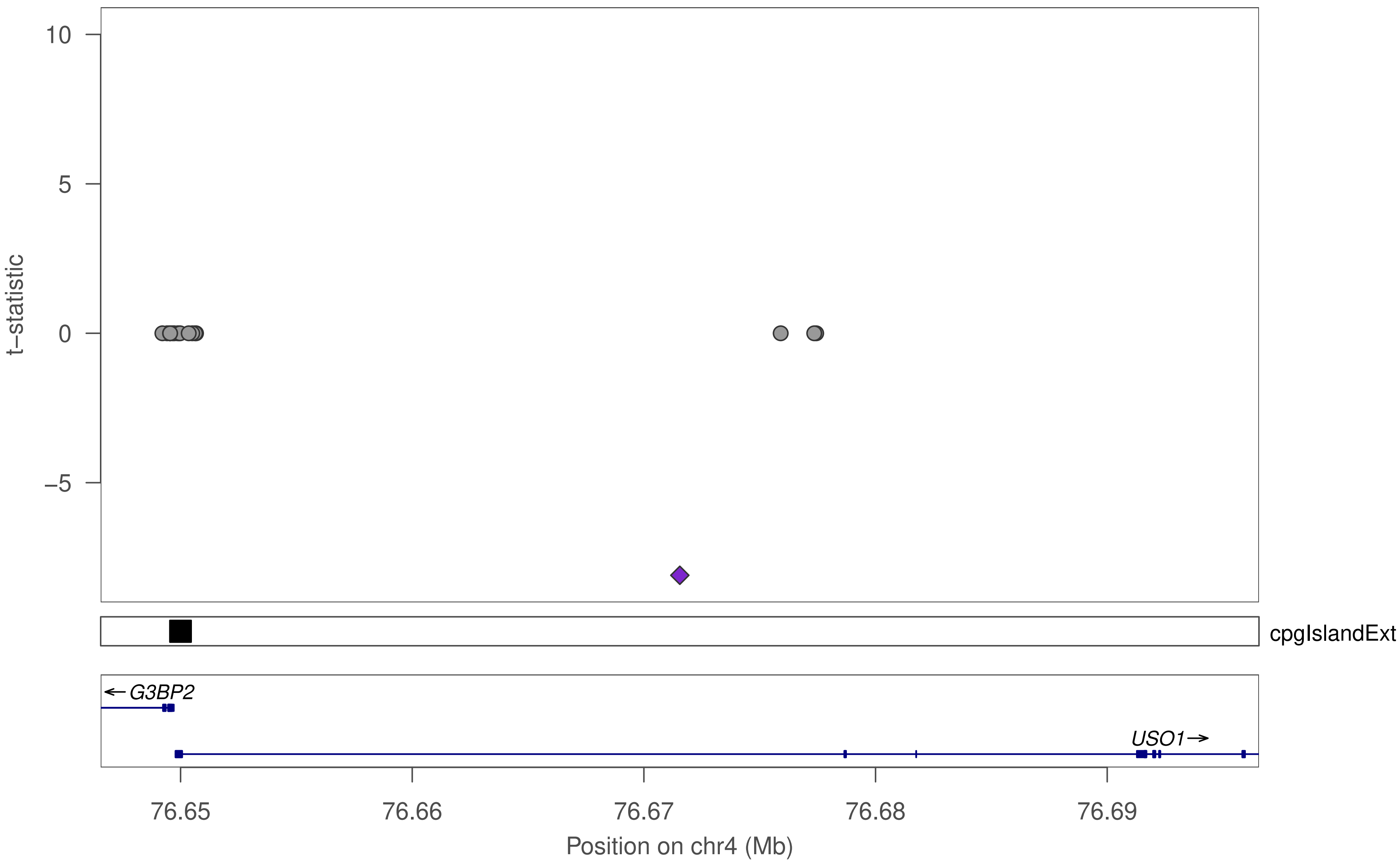

10
5
0
−5
t−statistic
cpgIslandExt
G3BP2
USO1
76.65
76.66
76.67
76.68
76.69
Position on chr4 (Mb)

### ZNF440

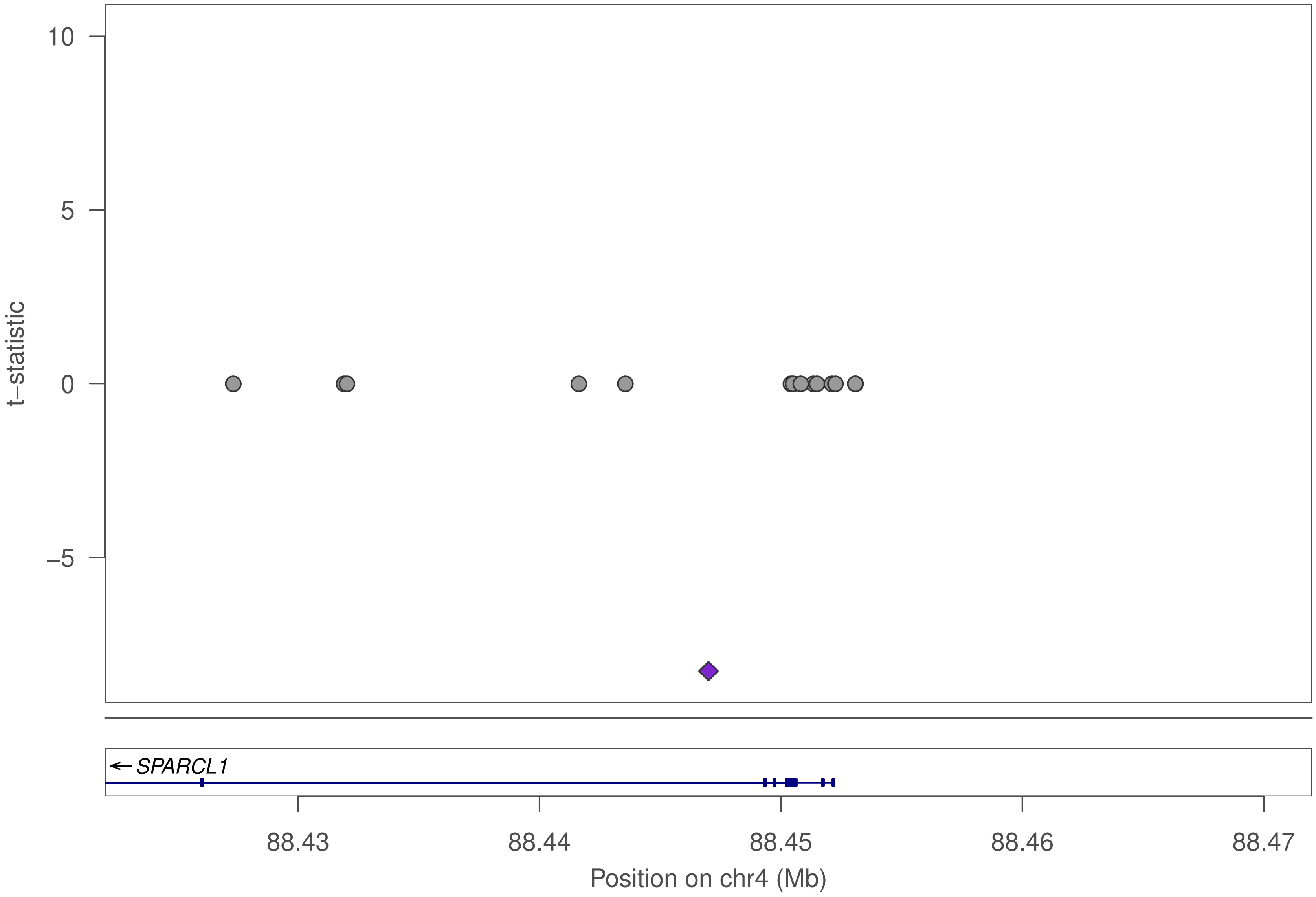

10
5
0
−5
t−statistic
SPARCL1
88.43
88.44
88.45
88.46
88.47
Position on chr4 (Mb)

### ZNF440

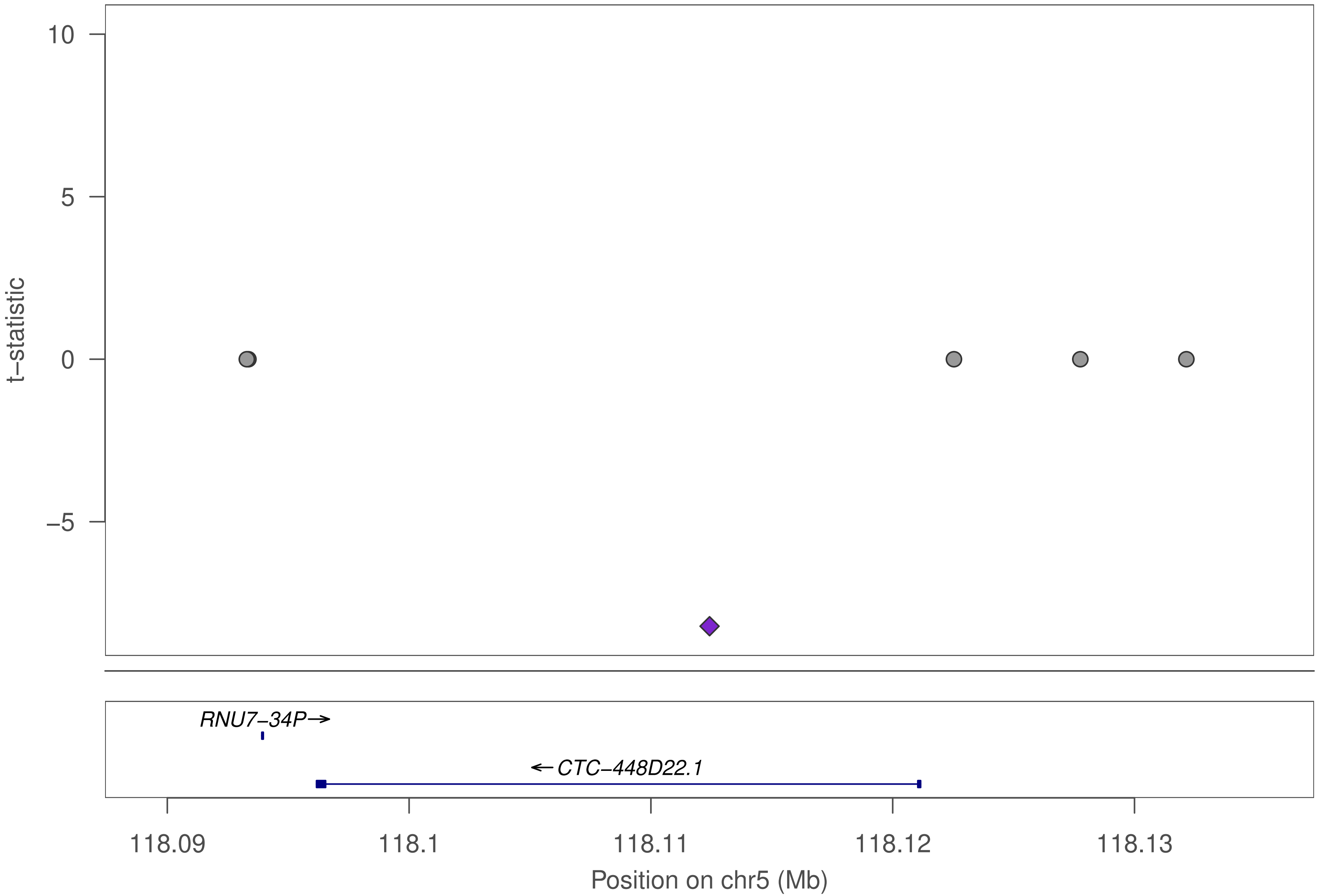

10
5
0
−5
t−statistic
RNU7−34P→
←CTC−448D22.1
118.09
118.1
118.11
118.12
118.13
Position on chr5 (Mb)

### ZNF440

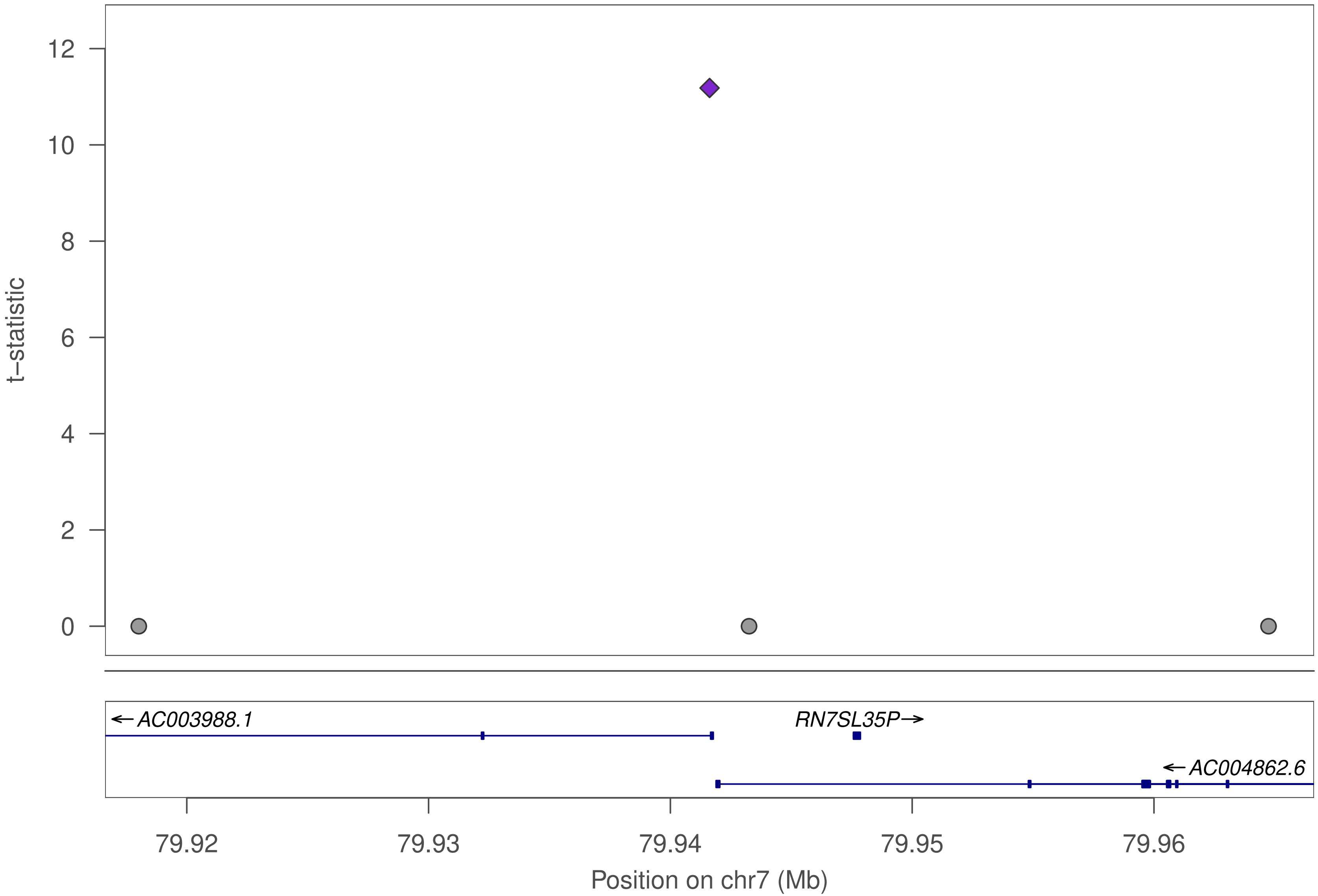

t–statistic
12
10
8
6
4
2
0
← AC003988.1
RN7SL35P→
← AC004862.6
79.92
79.93
79.94
79.95
79.96
Position on chr7 (Mb)

### ZNF440

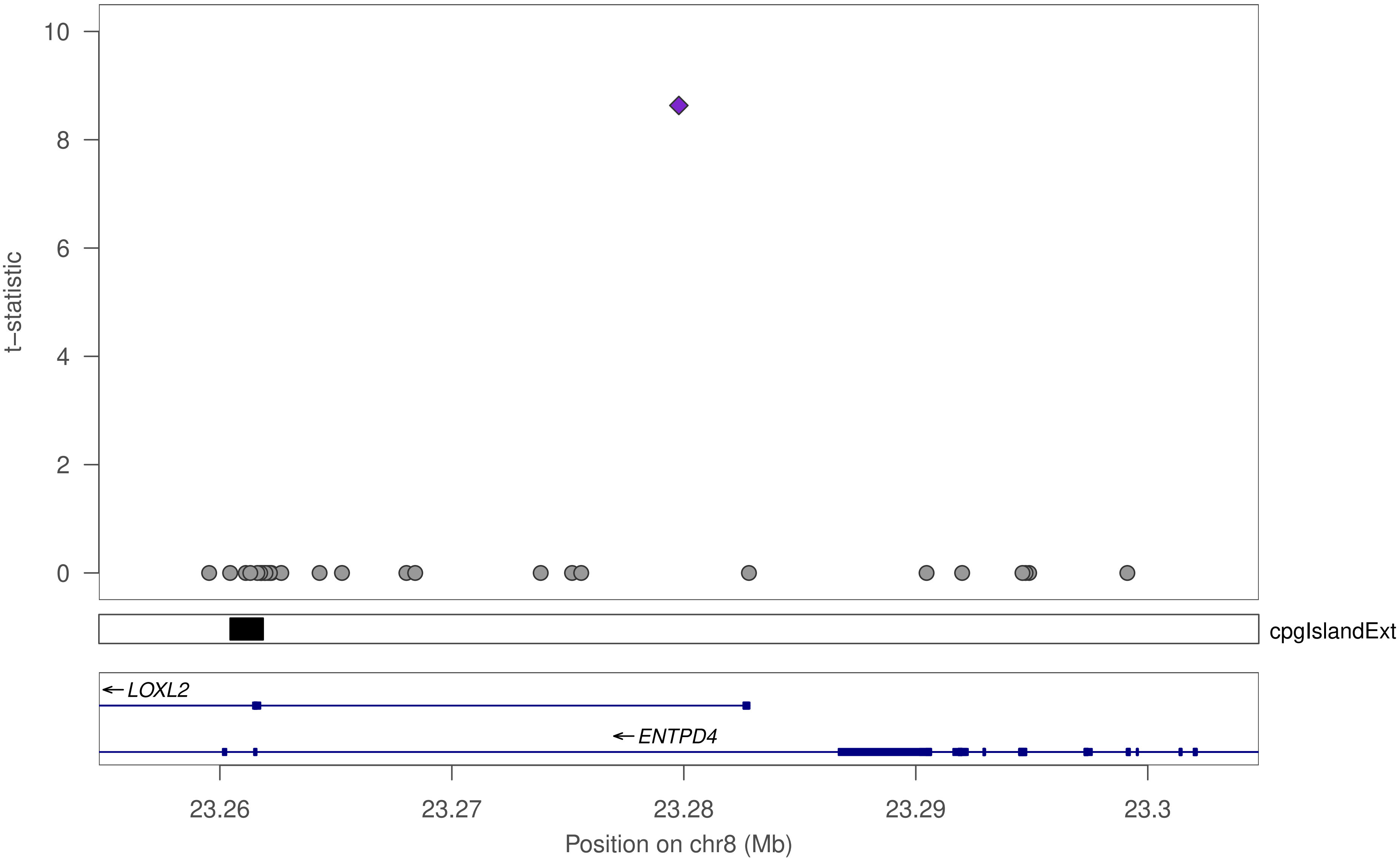

10
8
6
4
2
0
t–statistic
cpgIslandExt
LOXL2
ENTPD4
23.26
23.27
23.28
23.29
23.3
Position on chr8 (Mb)

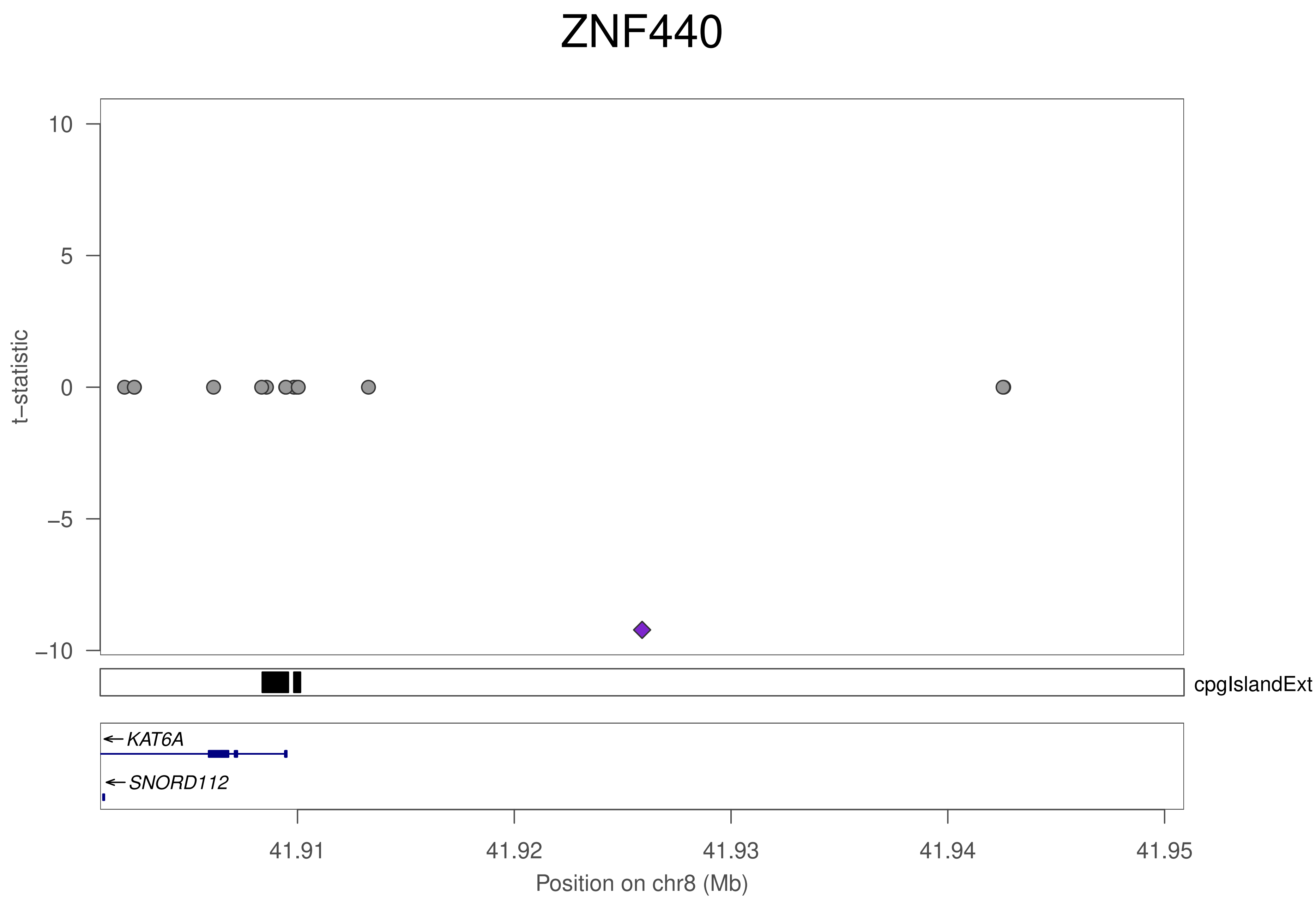

ZNF440
t−statistic
10
5
0
−5
−10
cpgIslandExt
KAT6A
SNORD112
41.91
41.92
41.93
41.94
41.95
Position on chr8 (Mb)

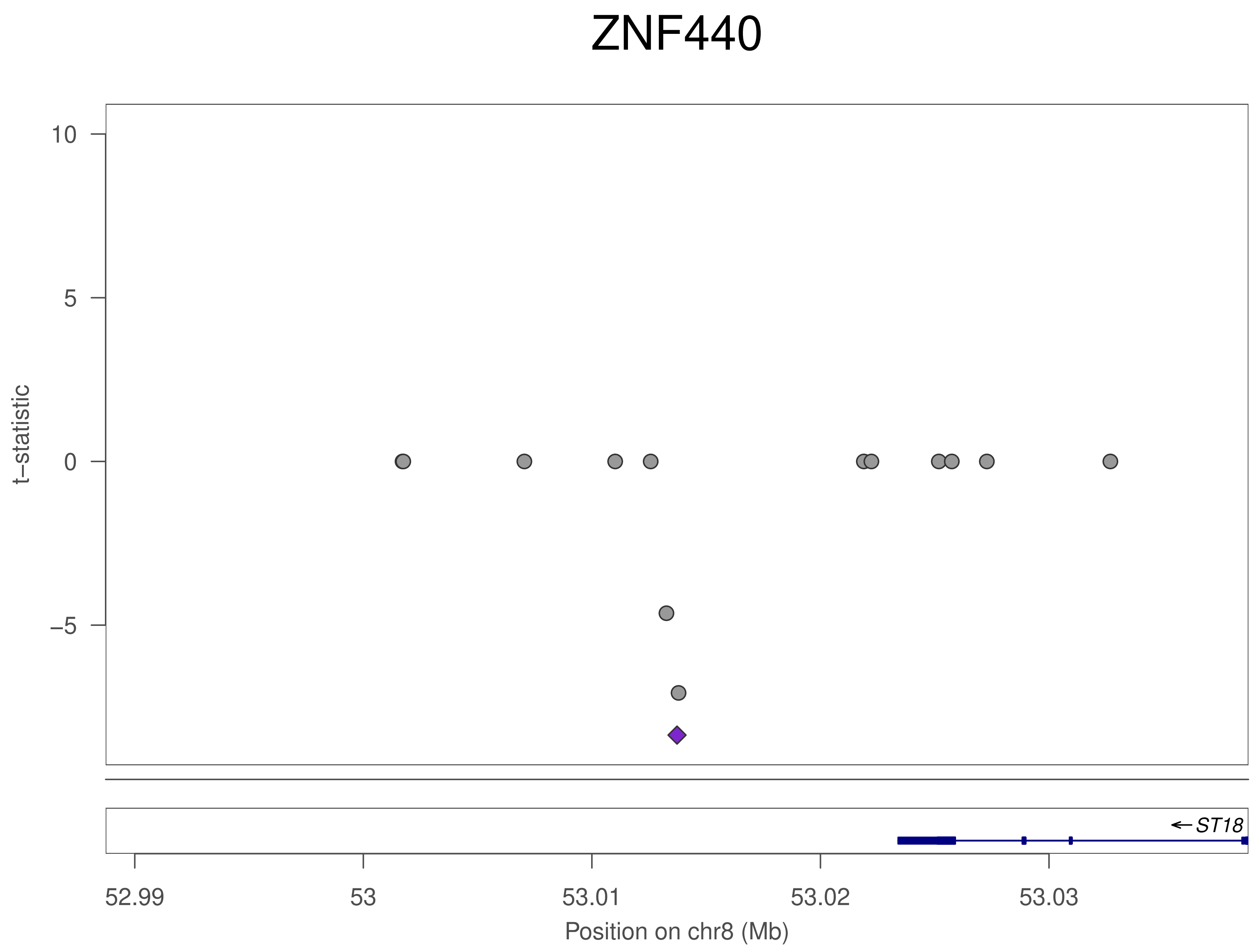
ZNF440
t–statistic
10
5
0
–5
ST18
52.99
53
53.01
53.02
53.03
Position on chr8 (Mb)

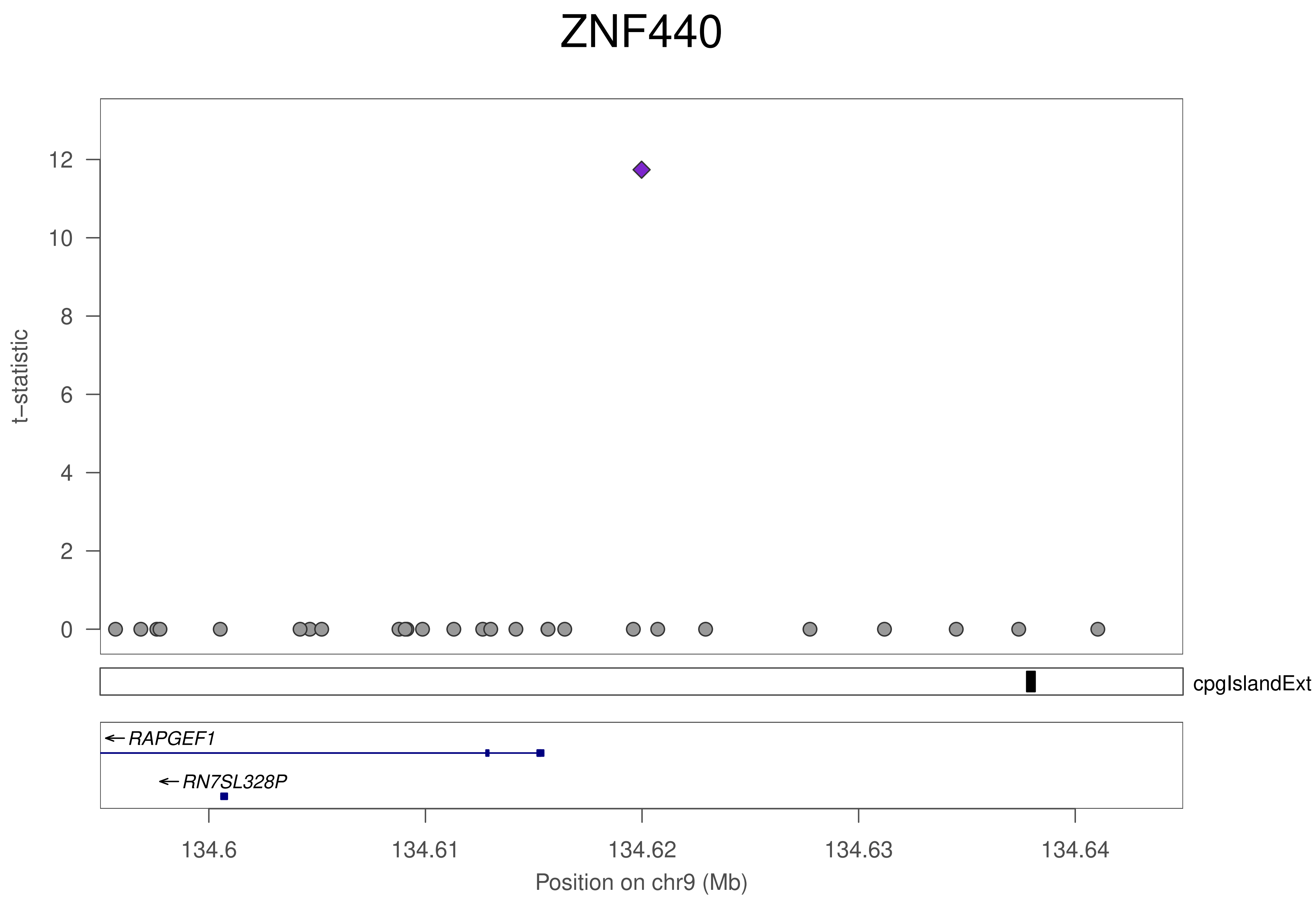

ZNF440
t–statistic
12
10
8
6
4
2
0
cpgIslandExt
RAPGEF1
RN7SL328P
134.6
134.61
134.62
134.63
134.64
Position on chr9 (Mb)

### ZNF440

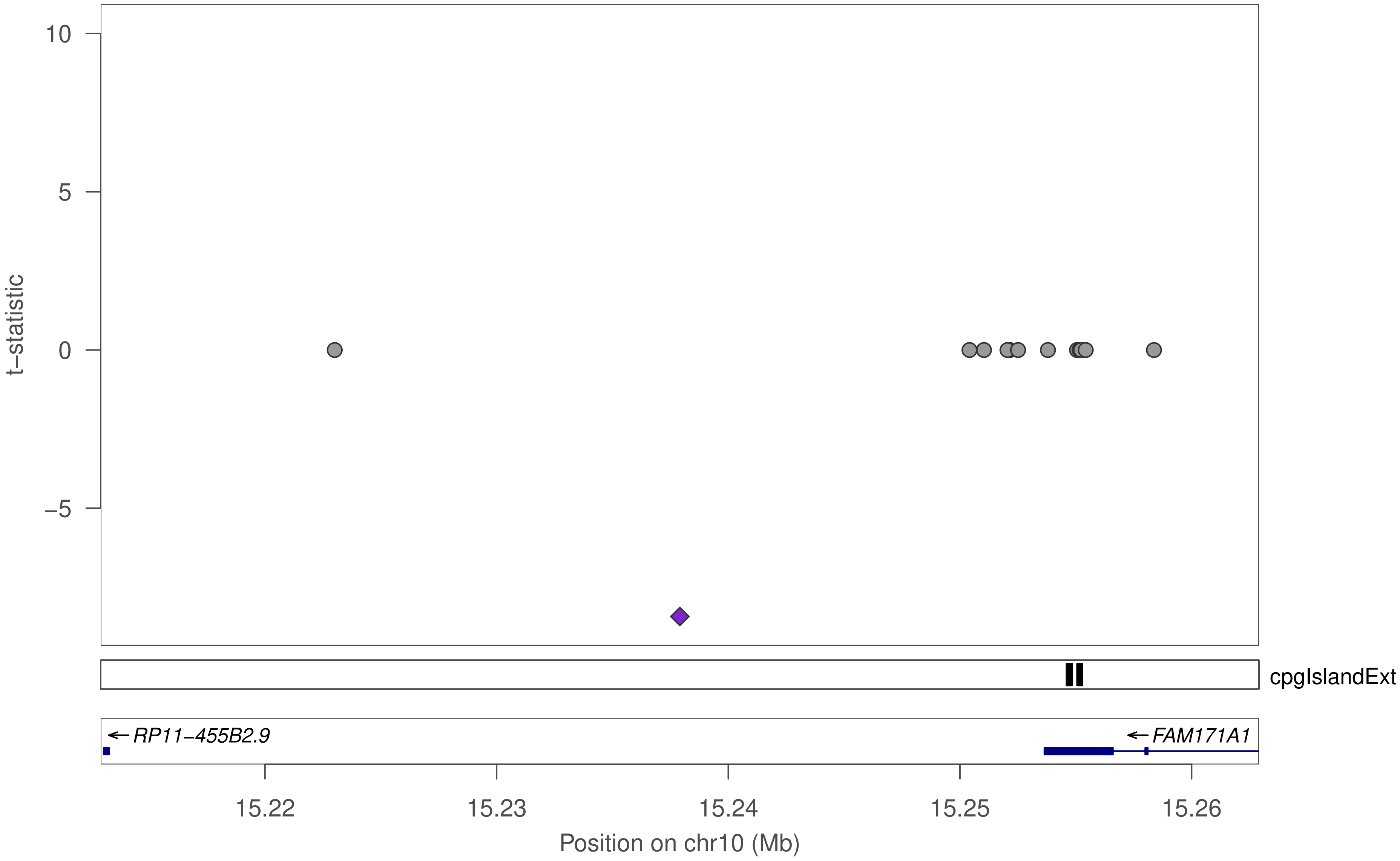

10
5
0
−5
t−statistic
cpgIslandExt
RP11−455B2.9
FAM171A1
15.22
15.23
15.24
15.25
15.26
Position on chr10 (Mb)

### ZNF440

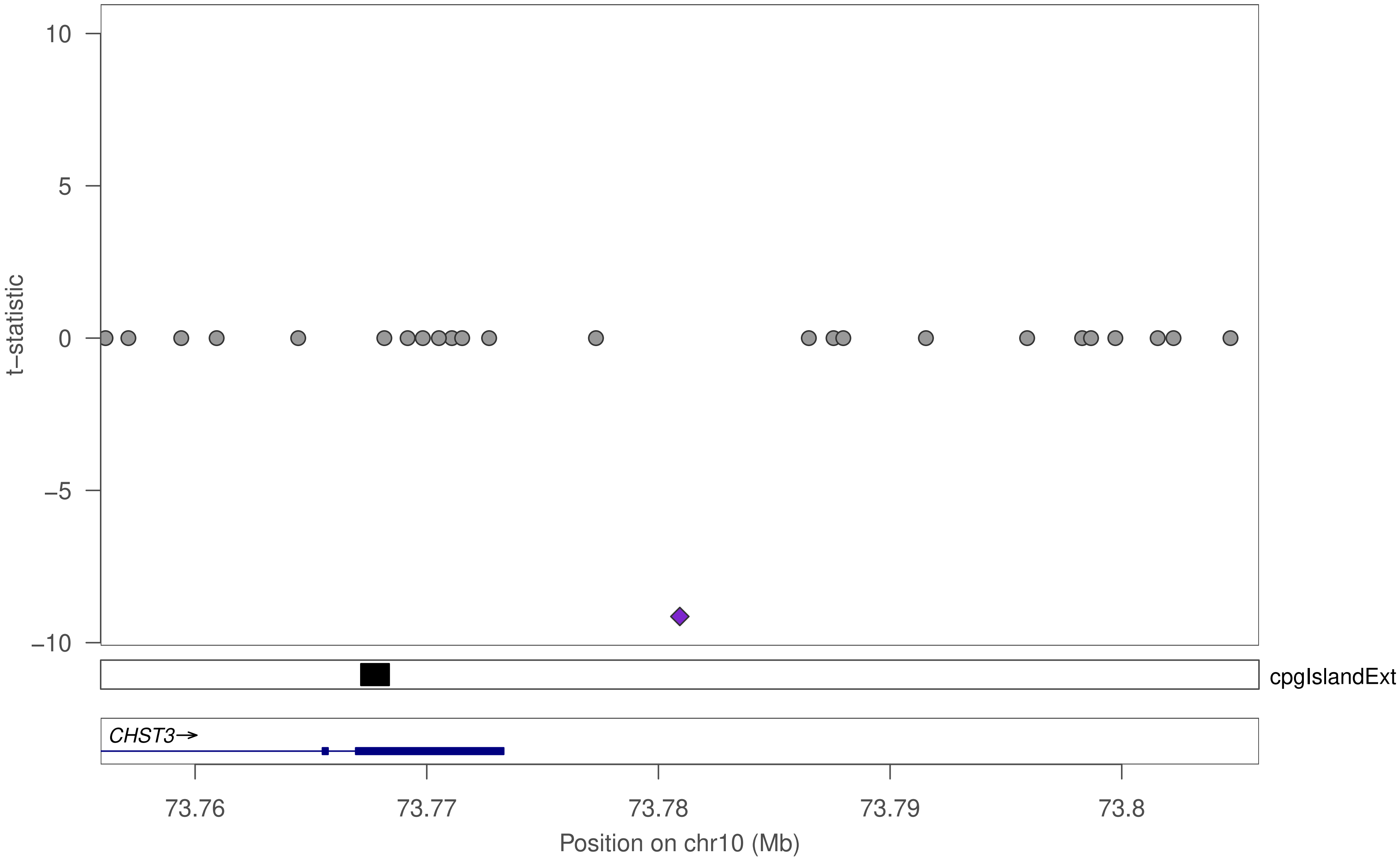

10
5
0
−5
−10
t−statistic
cpgIslandExt
CHST3→
73.76
73.77
73.78
73.79
73.8
Position on chr10 (Mb)

### ZNF440

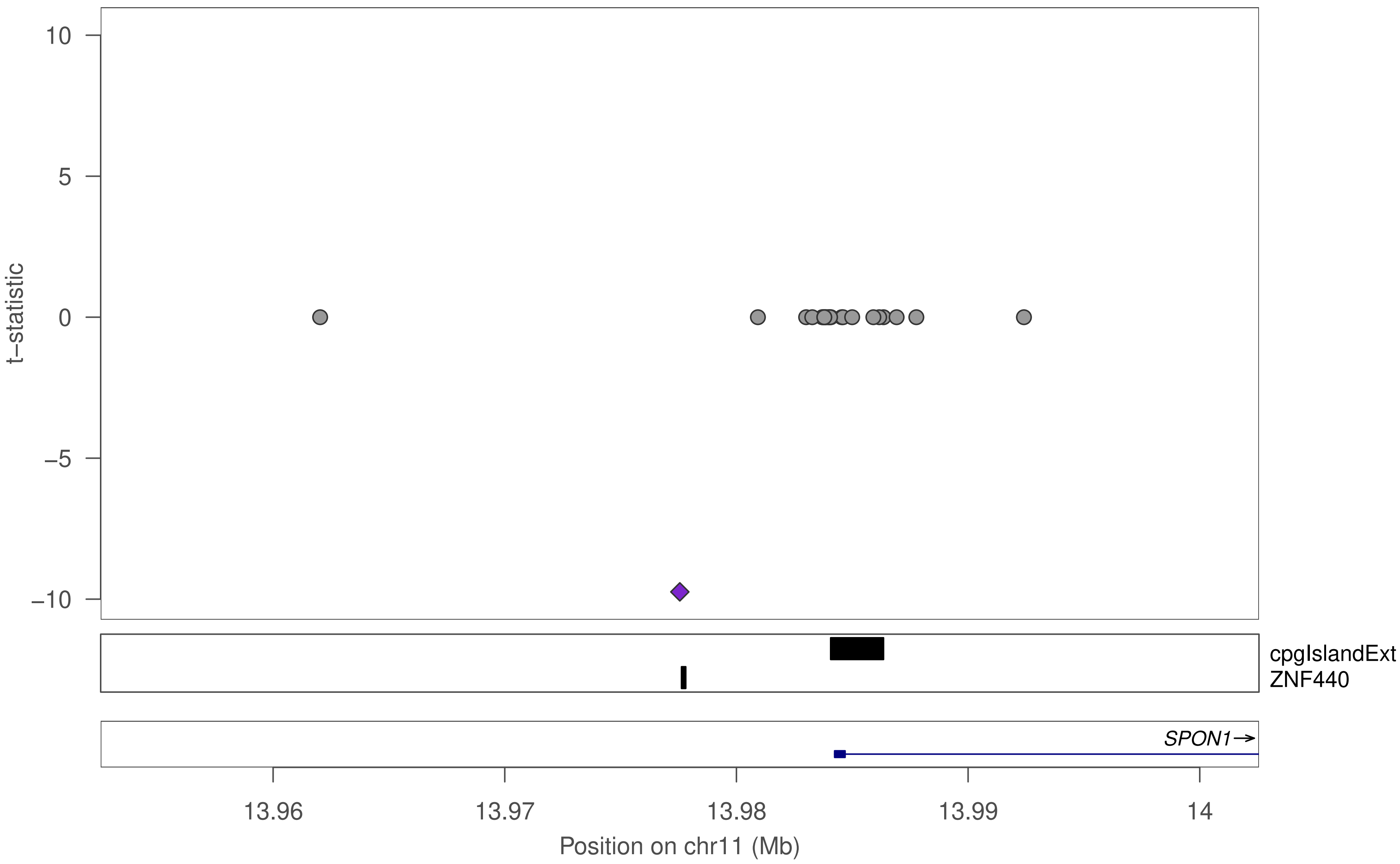

10
5
0
−5
−10
t−statistic
cpgIslandExt
ZNF440
SPON1→
13.96
13.97
13.98
13.99
14
Position on chr11 (Mb)

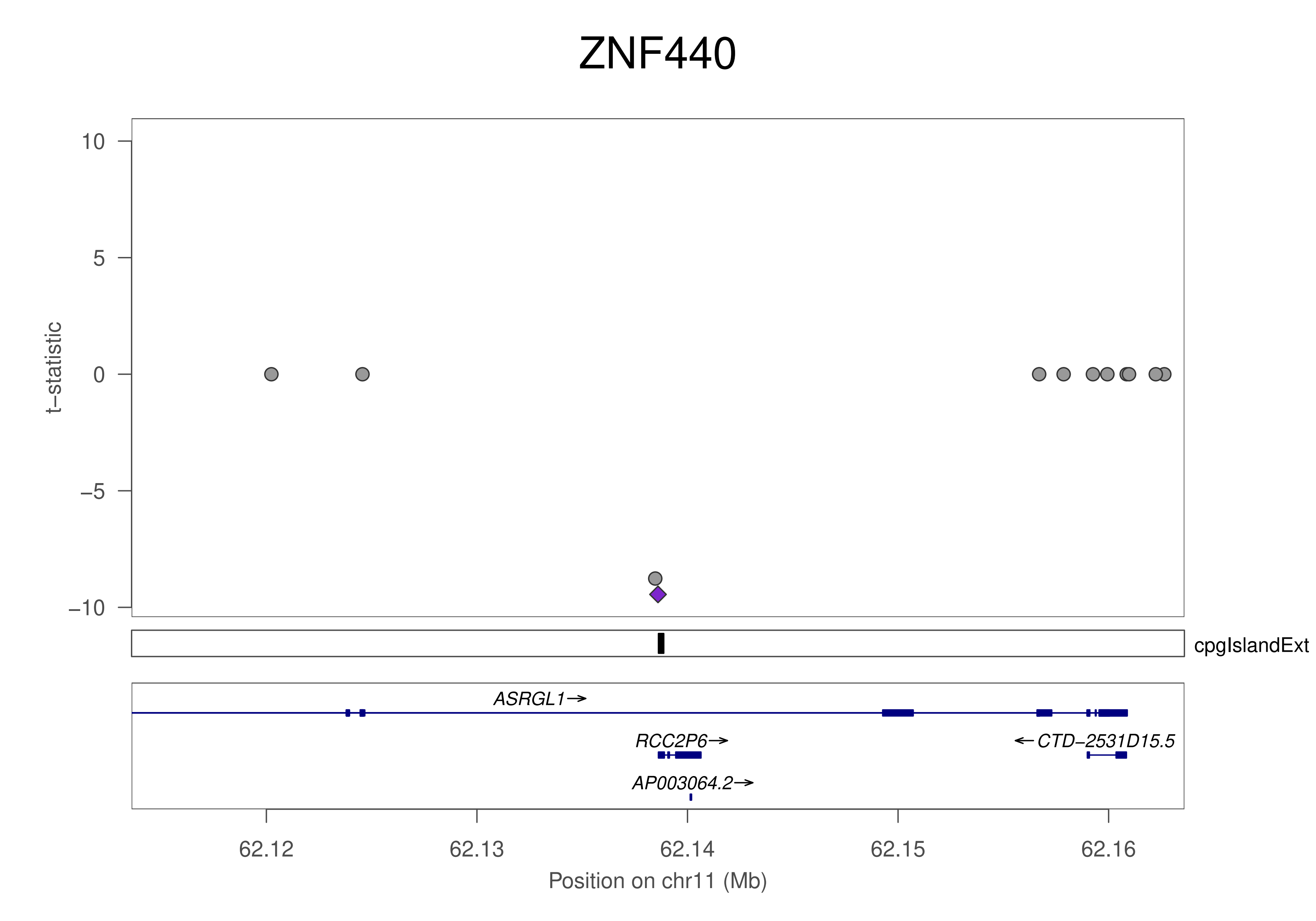

ZNF440
10
5
0
−5
−10
t−statistic
cpgIslandExt
ASRGL1→
RCC2P6→
←CTD−2531D15.5
AP003064.2→
62.12
62.13
62.14
62.15
62.16
Position on chr11 (Mb)

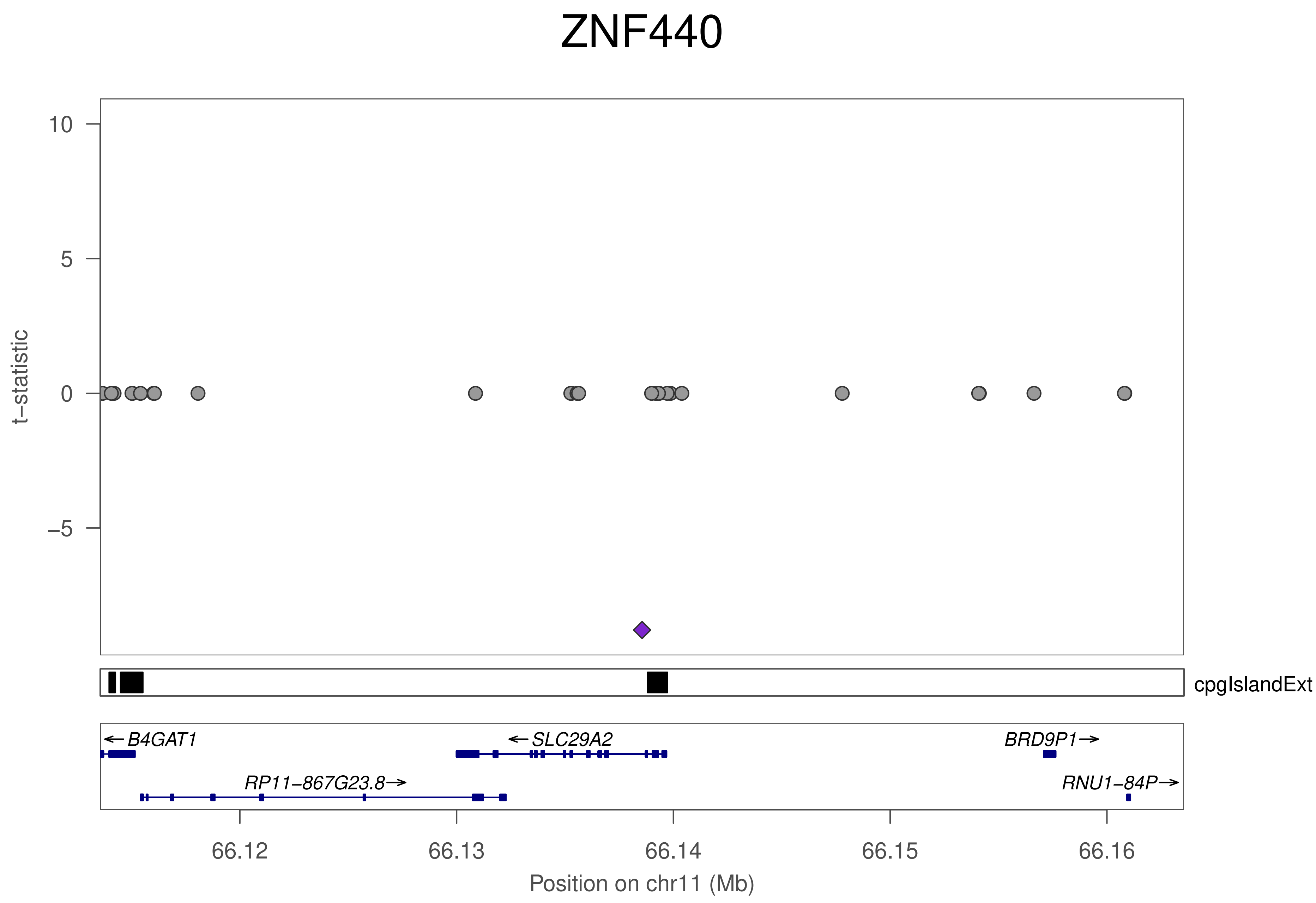
ZNF440
t–statistic
10
5
0
–5
cpgIslandExt
← B4GAT1
← SLC29A2
BRD9P1 →
RP11–867G23.8 →
RNU1–84P →
66.12
66.13
66.14
66.15
66.16
Position on chr11 (Mb)

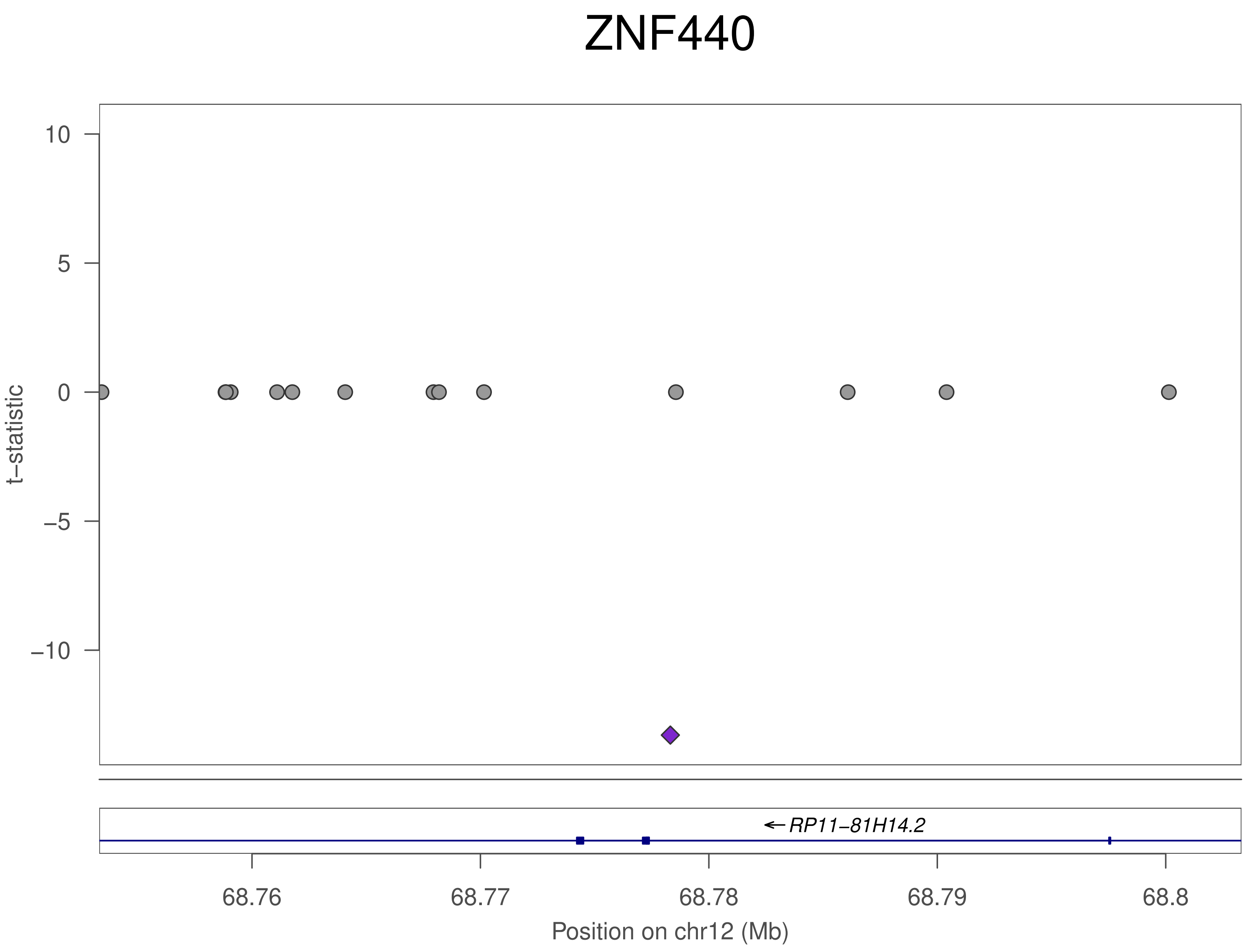

ZNF440
10
5
0
−5
−10
t−statistic
← RP11−81H14.2
68.76
68.77
68.78
68.79
68.8
Position on chr12 (Mb)

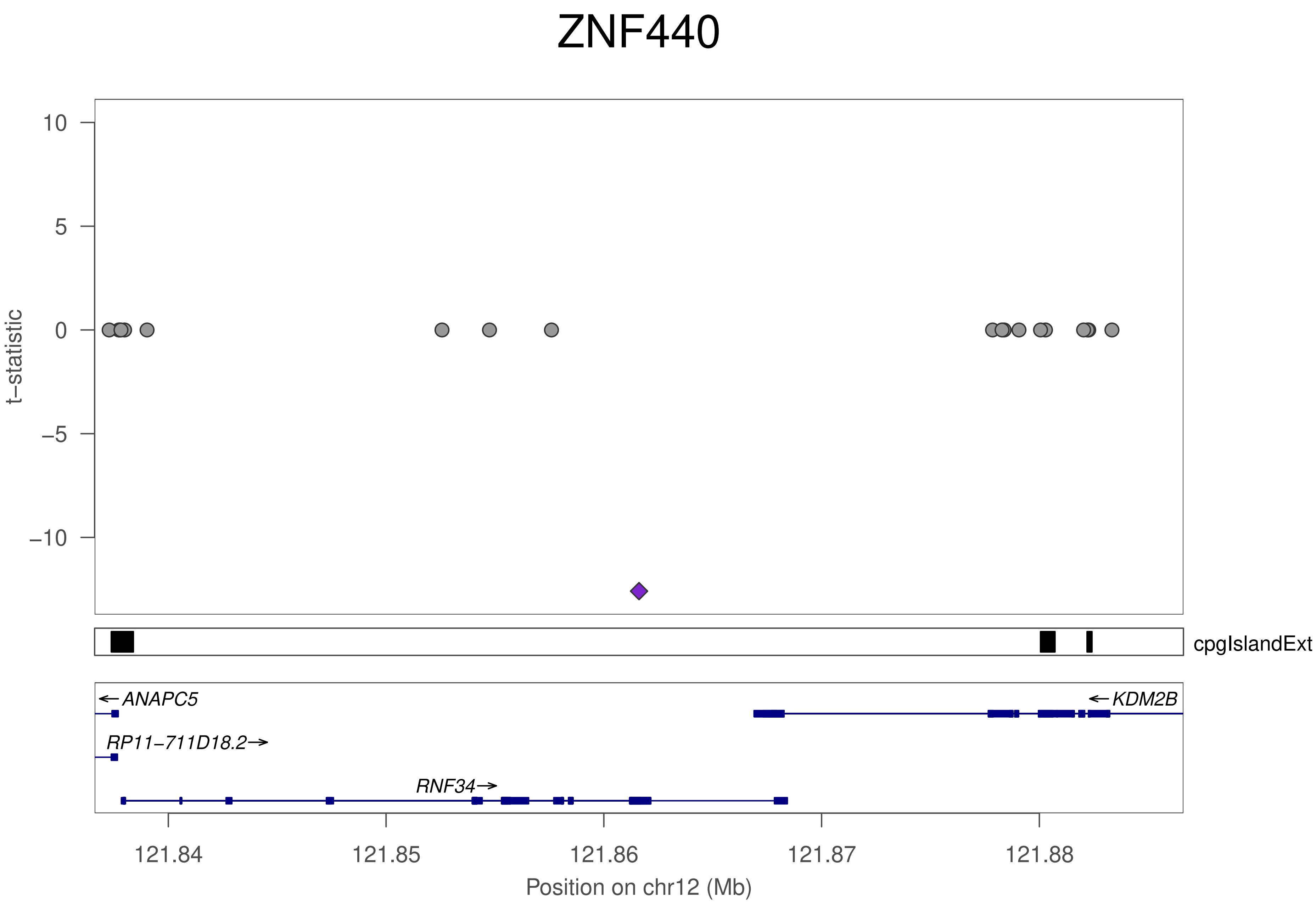

ZNF440
t−statistic
10
5
0
−5
−10
cpgIslandExt
ANAPC5
KDM2B
RP11−711D18.2
RNF34
121.84
121.85
121.86
121.87
121.88
Position on chr12 (Mb)

### ZNF440

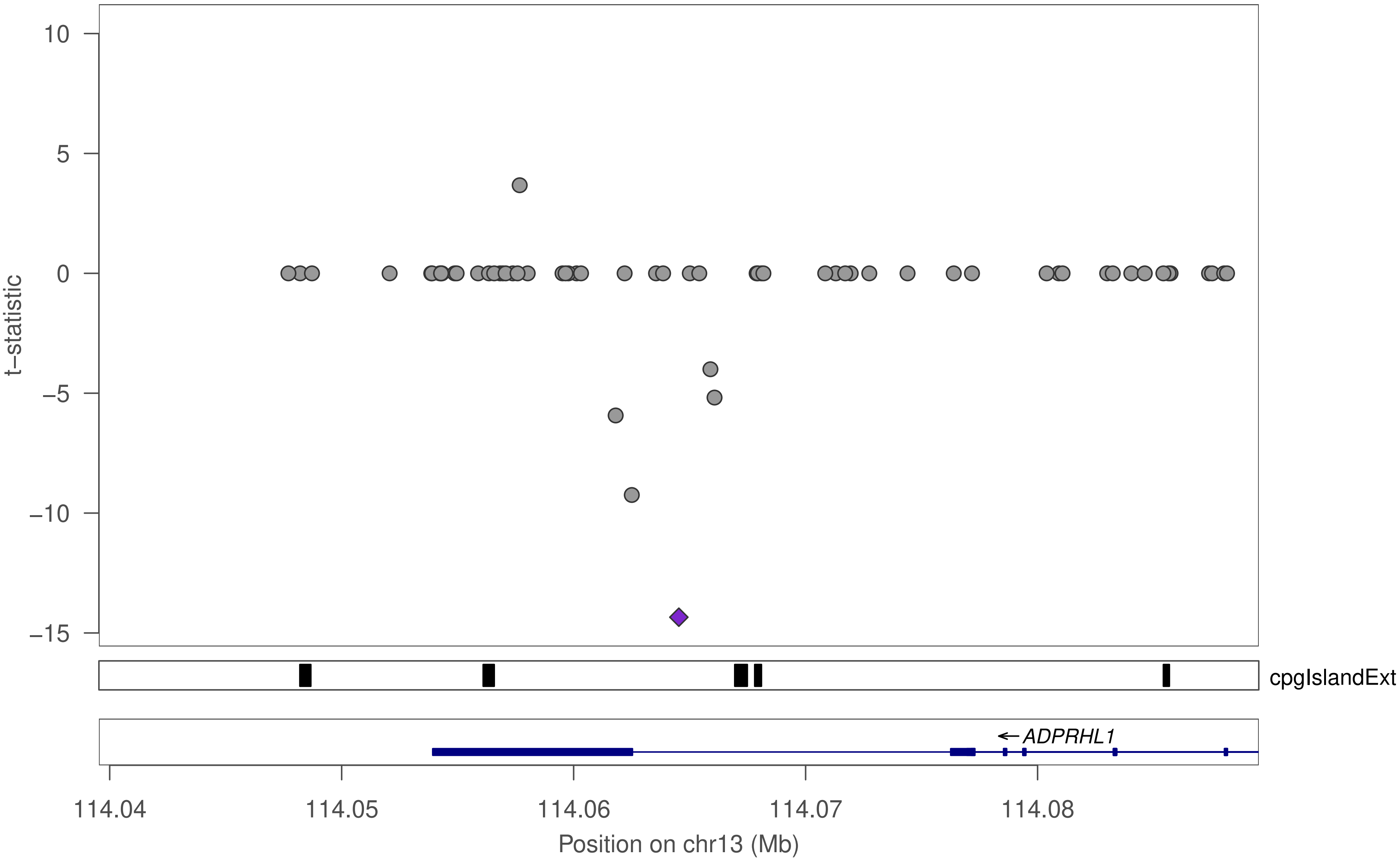

10
5
0
−5
−10
−15
t−statistic
cpgIslandExt
ADPRHL1
114.04
114.05
114.06
114.07
114.08
Position on chr13 (Mb)

### ZNF440

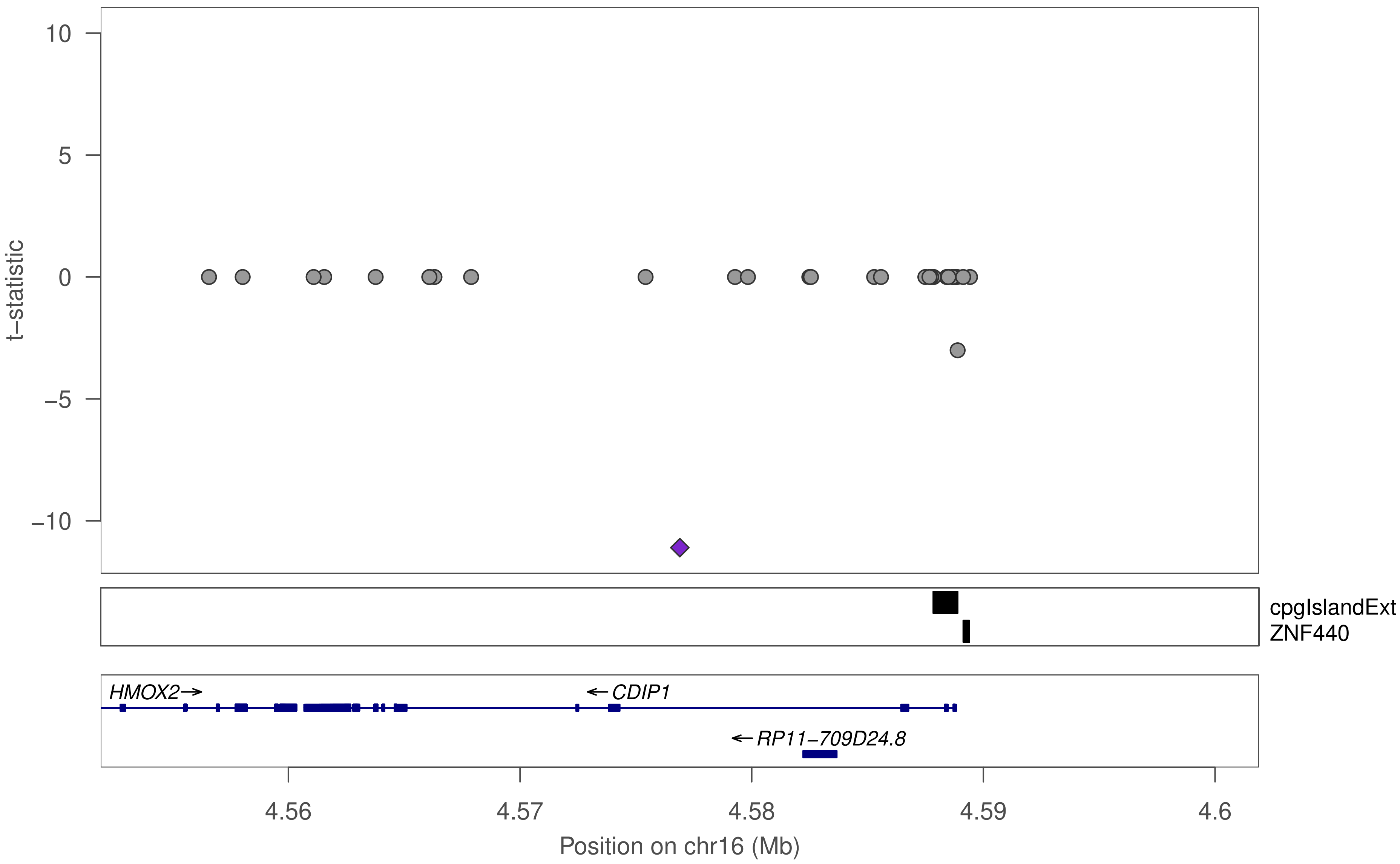

10
5
0
−5
−10
t−statistic
cpgIslandExt
ZNF440
HMOX2→
←CDIP1
←RP11−709D24.8
4.56
4.57
4.58
4.59
4.6
Position on chr16 (Mb)

### ZNF440

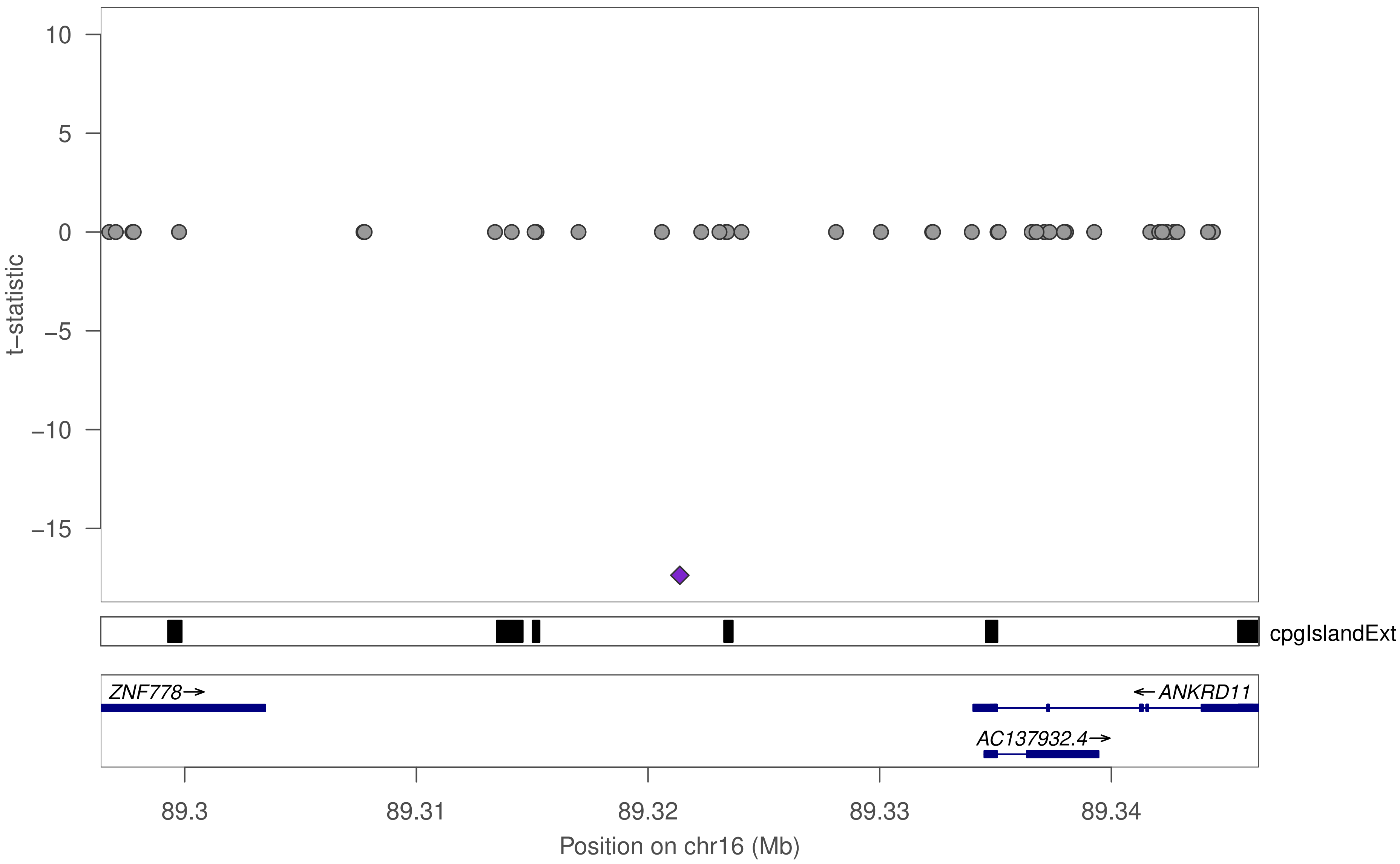

10
5
0
−5
−10
−15
t−statistic
cpgIslandExt
ZNF778→
←ANKRD11
AC137932.4→
89.3
89.31
89.32
89.33
89.34
Position on chr16 (Mb)

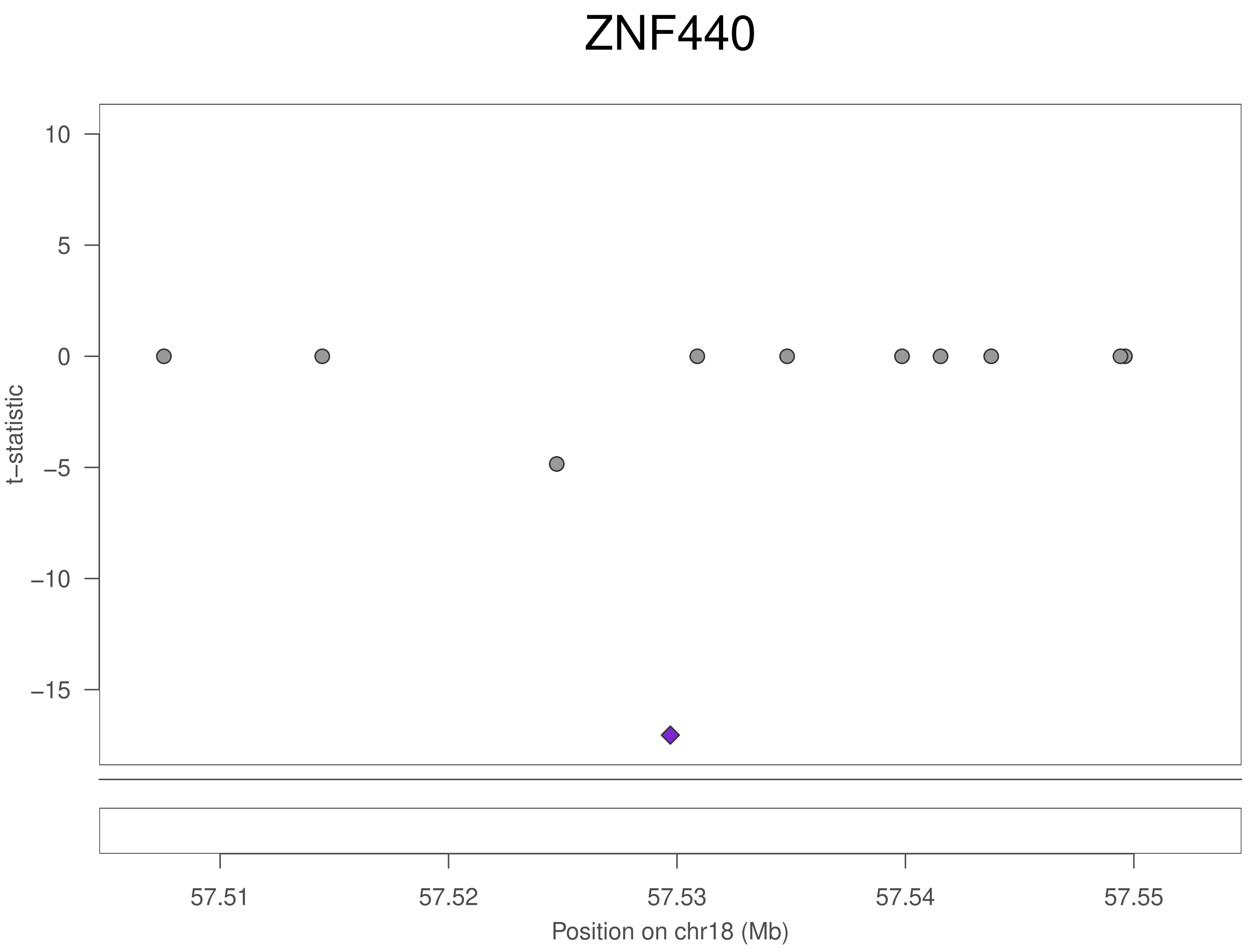
ZNF440
t–statistic
10
5
0
–5
–10
–15
57.51
57.52
57.53
57.54
57.55
Position on chr18 (Mb)

### ZNF440

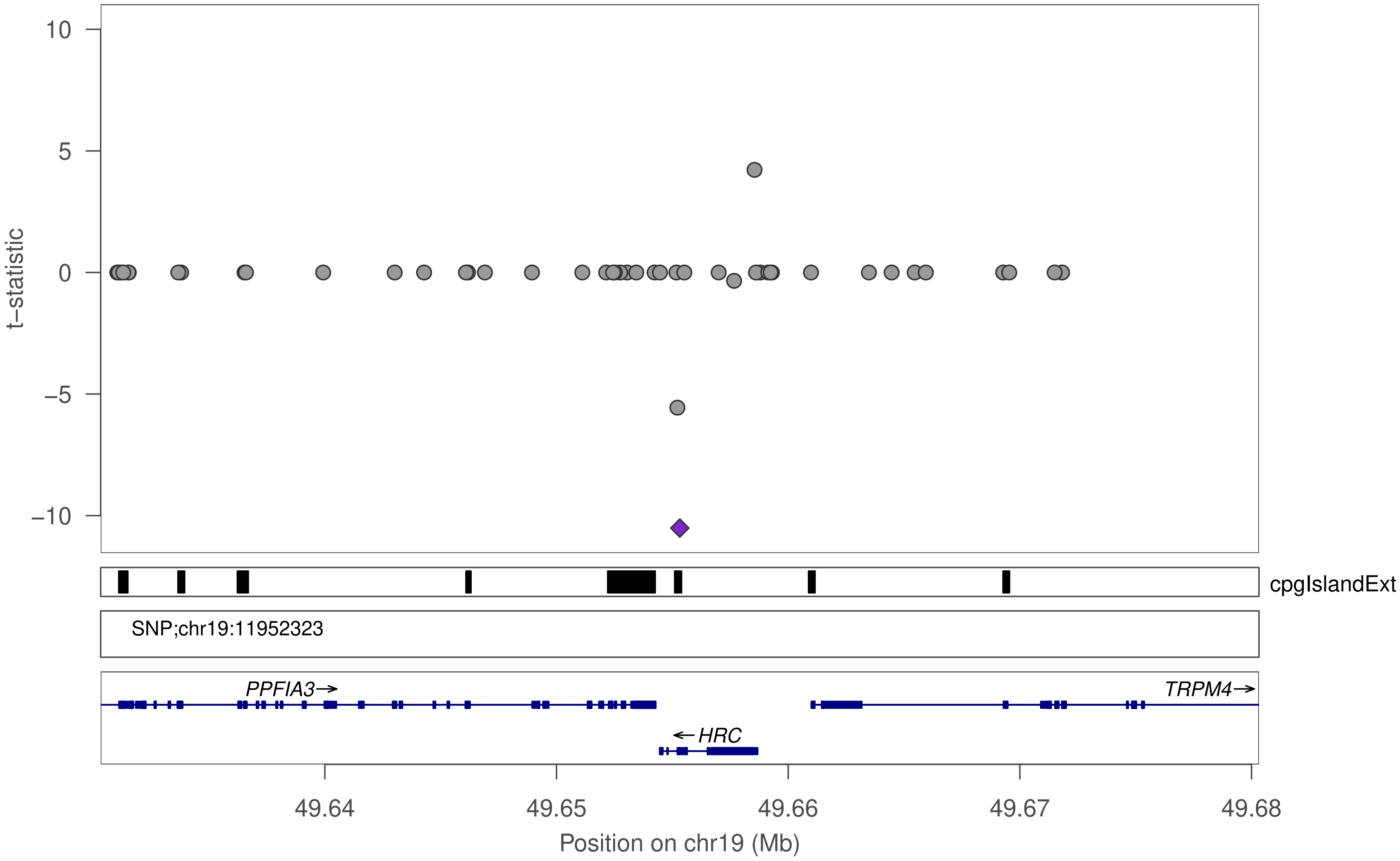

10
5
0
−5
−10
t−statistic
cpgIslandExt
SNP;chr19:11952323
PPFIA3→
TRPM4→
←HRC
49.64
49.65
49.66
49.67
49.68
Position on chr19 (Mb)

### ZNF440

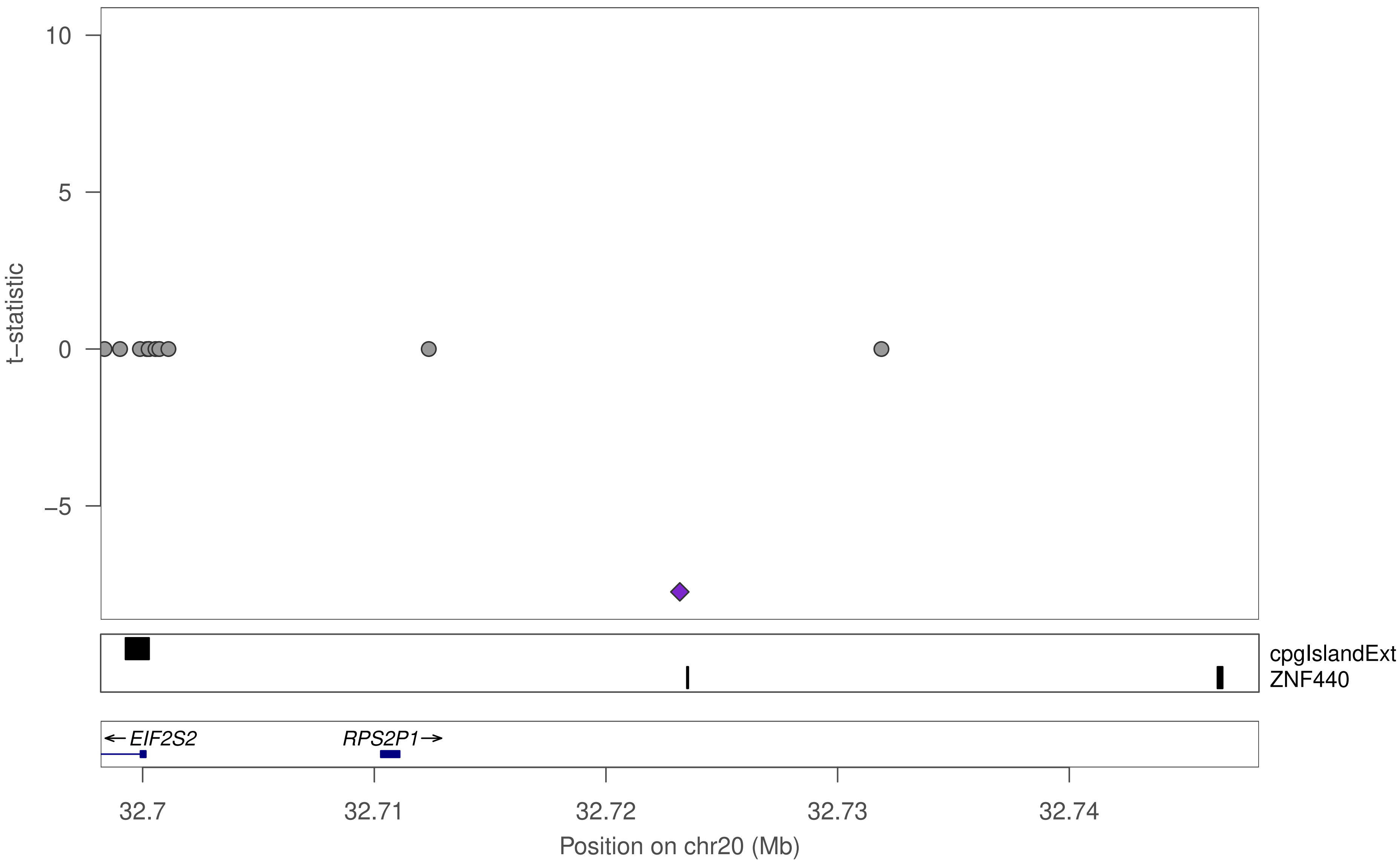

10
5
0
−5
t−statistic
cpgIslandExt
ZNF440
← EIF2S2
RPS2P1→
32.7
32.71
32.72
32.73
32.74
Position on chr20 (Mb)

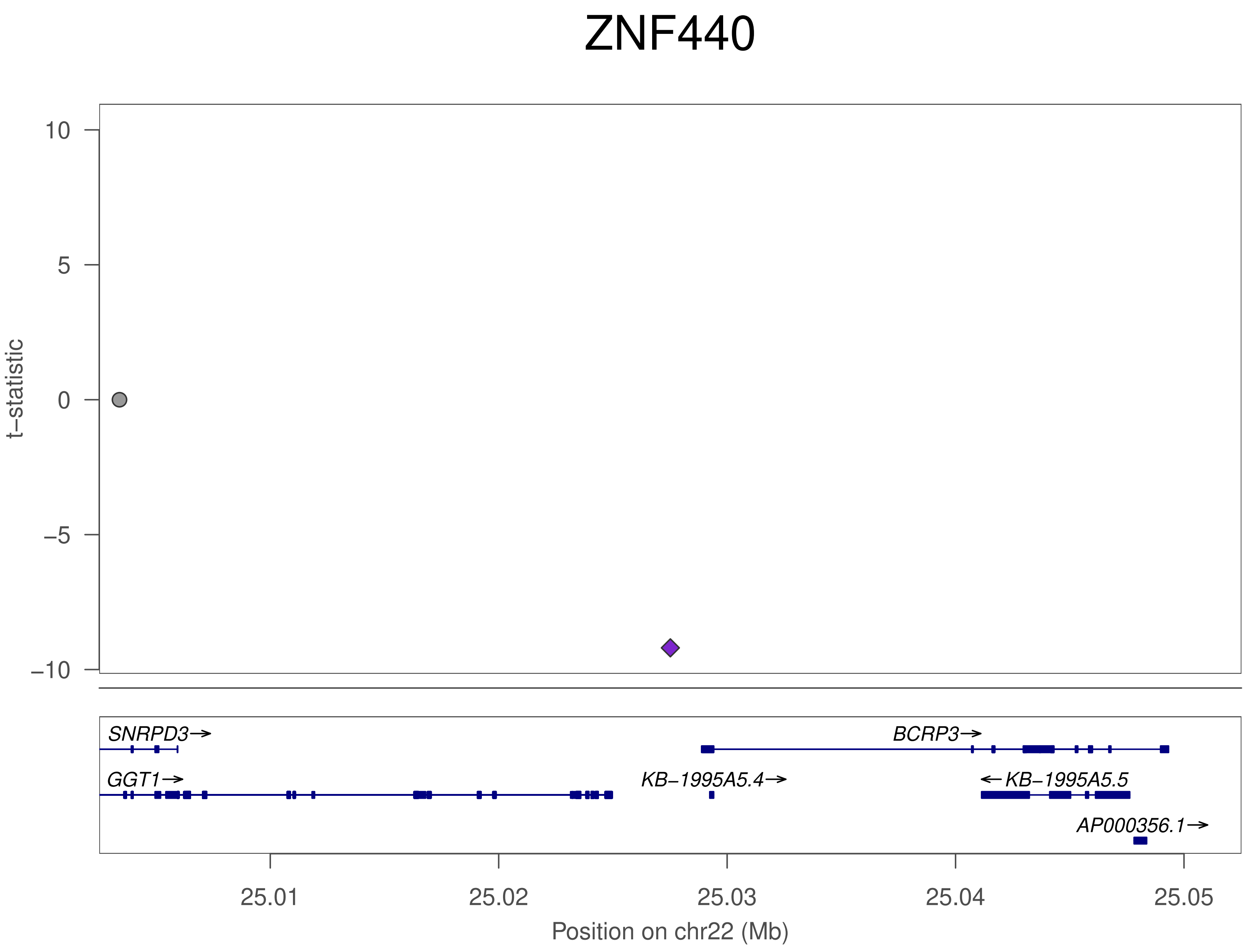
ZNF440
t−statistic
10
5
0
−5
−10
SNRPD3→
BCRP3→
GGT1→
KB−1995A5.4→
←KB−1995A5.5
AP000356.1→
25.01
25.02
25.03
25.04
25.05
Position on chr22 (Mb)

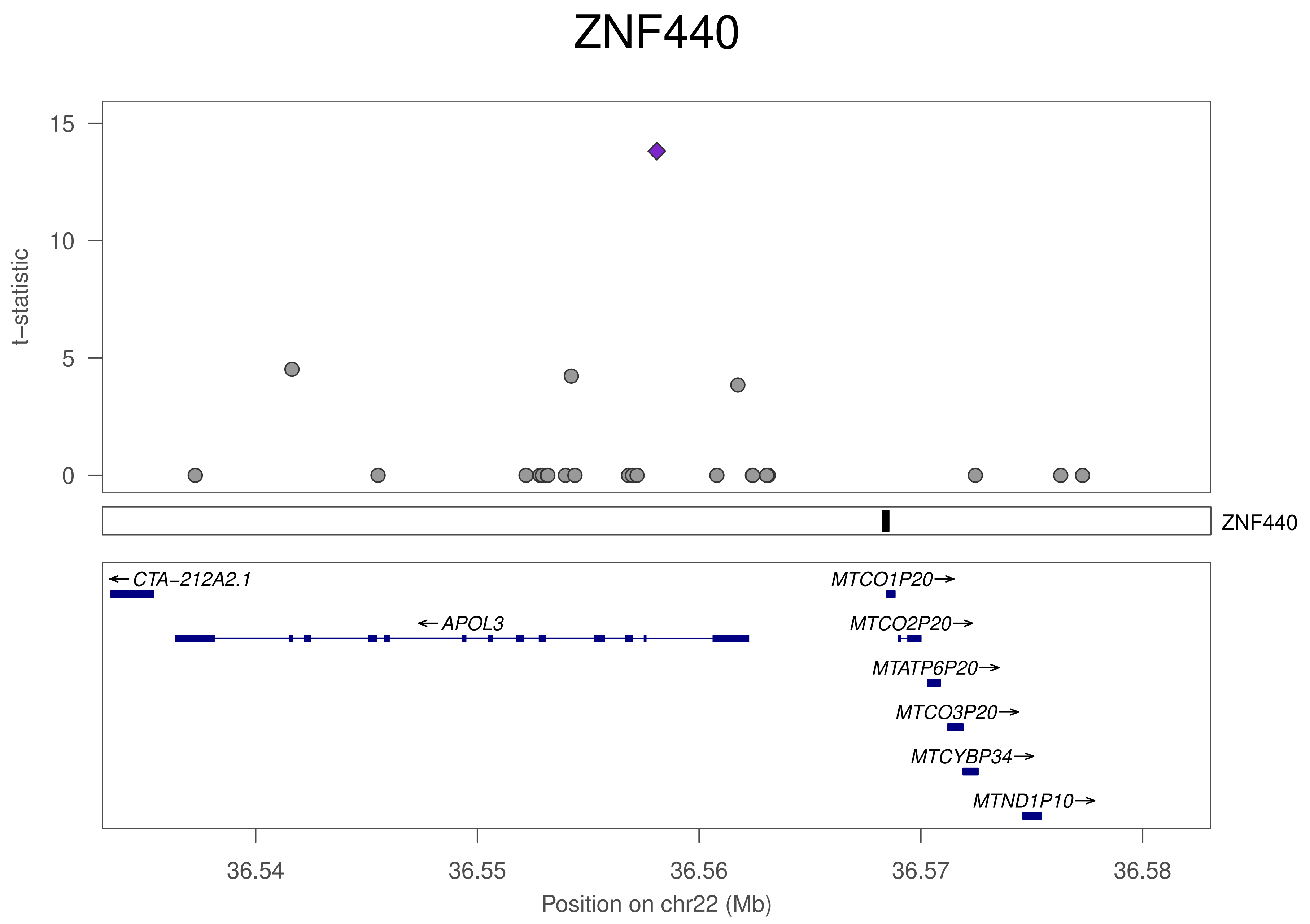
ZNF440
t−statistic
15
10
5
0
ZNF440
←CTA−212A2.1
←APOL3
MTCO1P20→
MTCO2P20→
MTATP6P20→
MTCO3P20→
MTCYBP34→
MTND1P10→
36.54
36.55
36.56
36.57
36.58
Position on chr22 (Mb)

### ZNF561

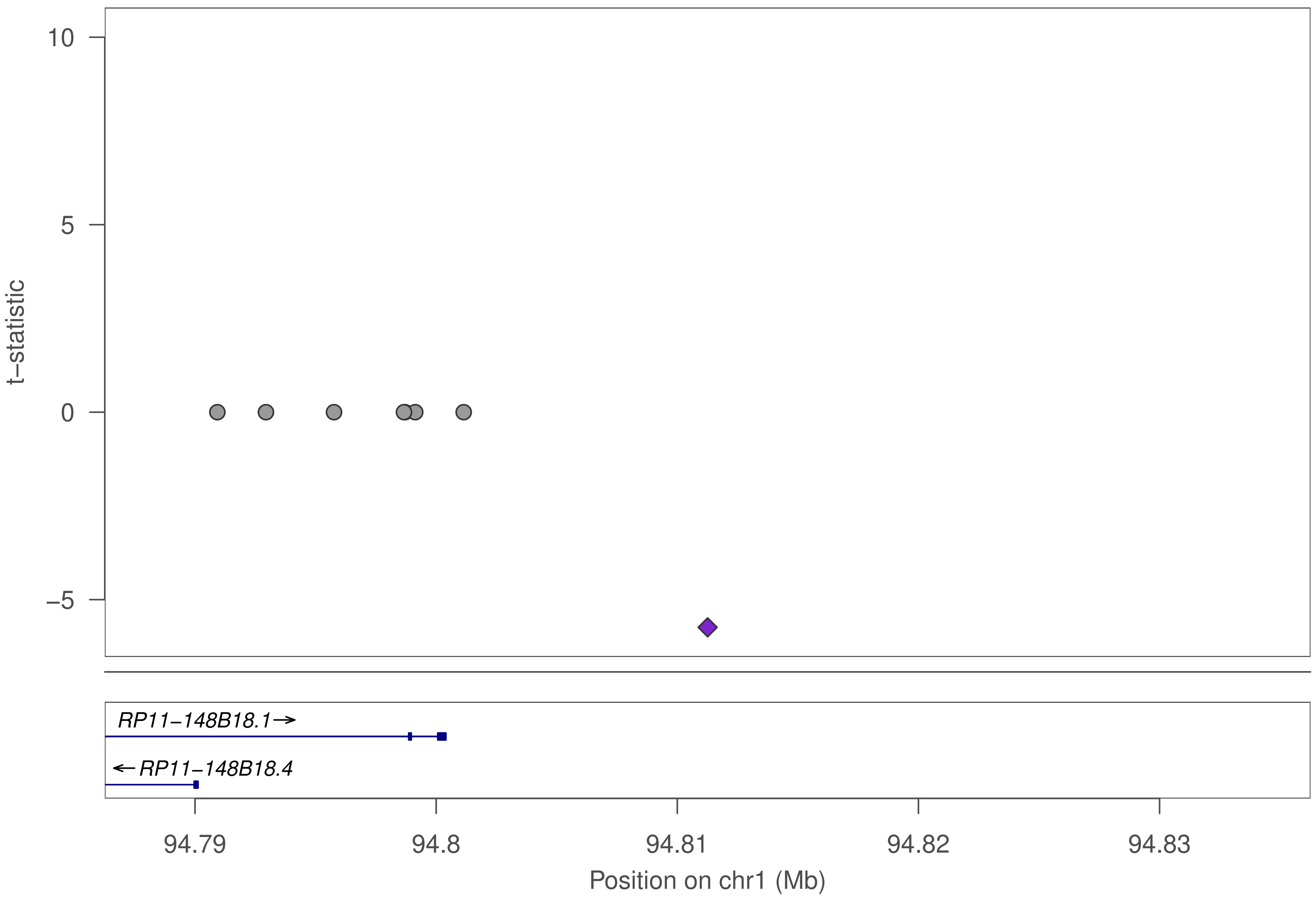

10
5
0
−5
t−statistic
RP11−148B18.1→
←RP11−148B18.4
94.79
94.8
94.81
94.82
94.83
Position on chr1 (Mb)

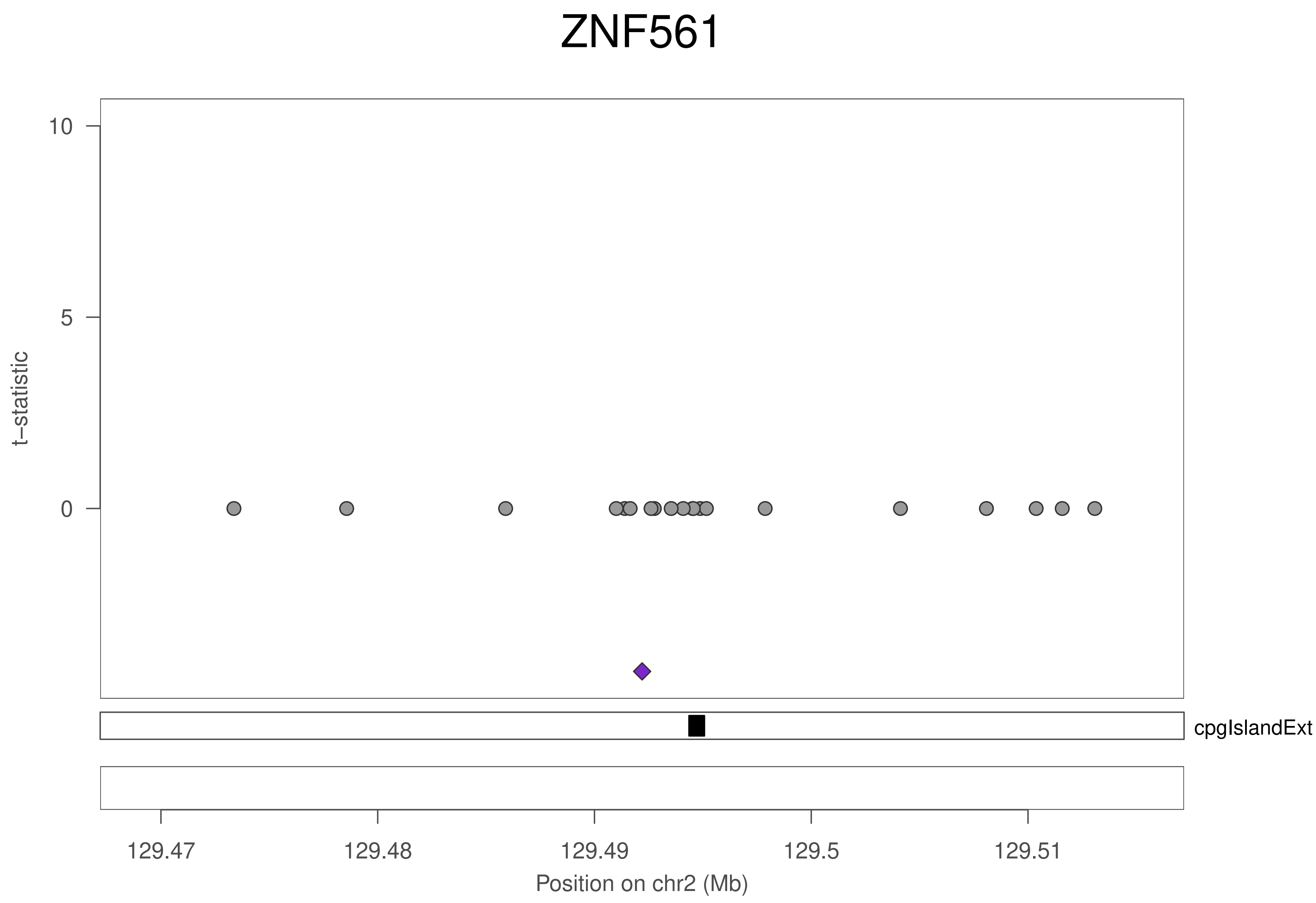

ZNF561
t−statistic
10
5
0
cpgIslandExt
129.47
129.48
129.49
129.5
129.51
Position on chr2 (Mb)

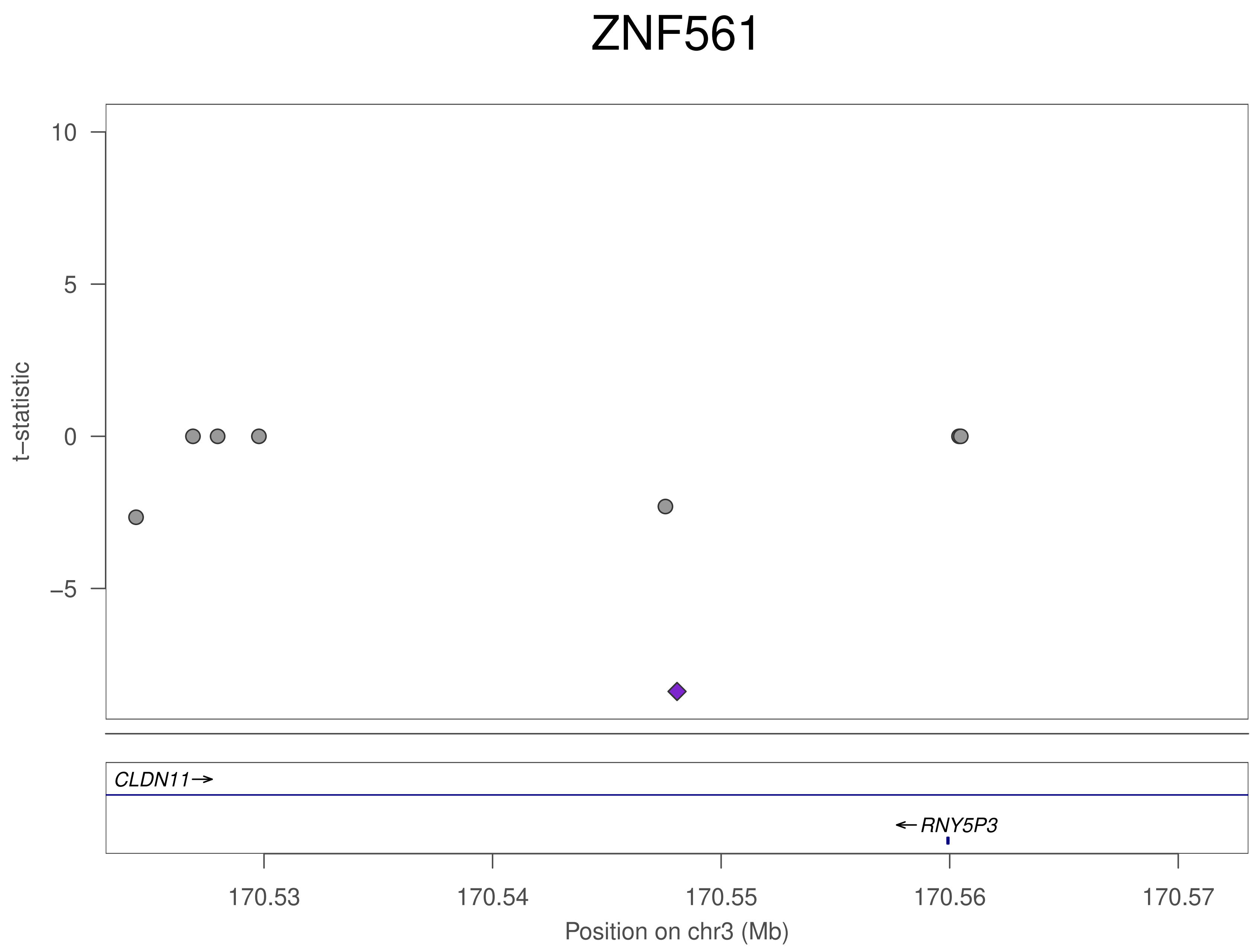
ZNF561
t−statistic
10
5
0
−5
CLDN11→
←RNY5P3
170.53
170.54
170.55
170.56
170.57
Position on chr3 (Mb)

### ZNF561

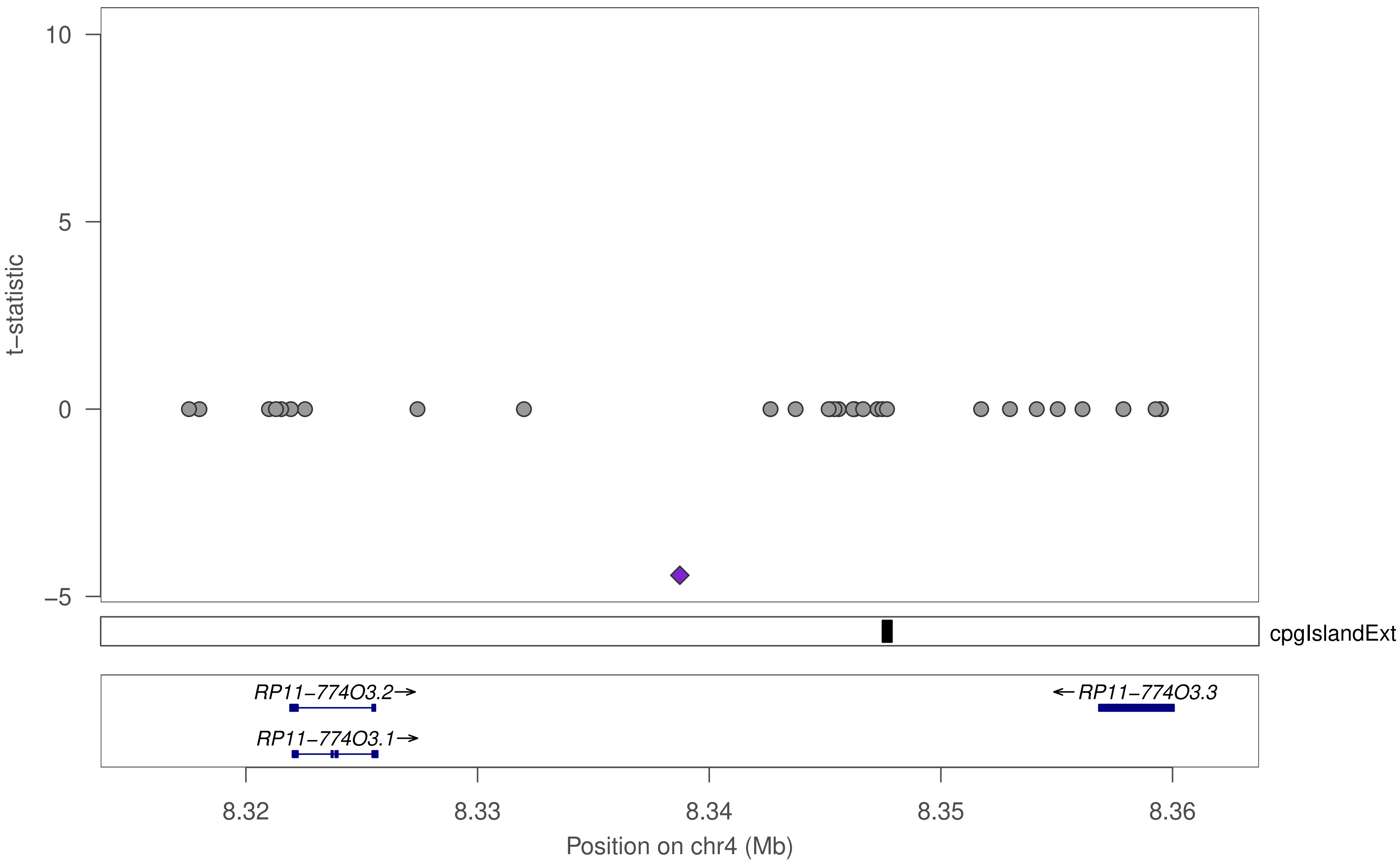

10
5
0
−5
t−statistic
cpgIslandExt
RP11−774O3.2→
RP11−774O3.1→
←RP11−774O3.3
8.32
8.33
8.34
8.35
8.36
Position on chr4 (Mb)

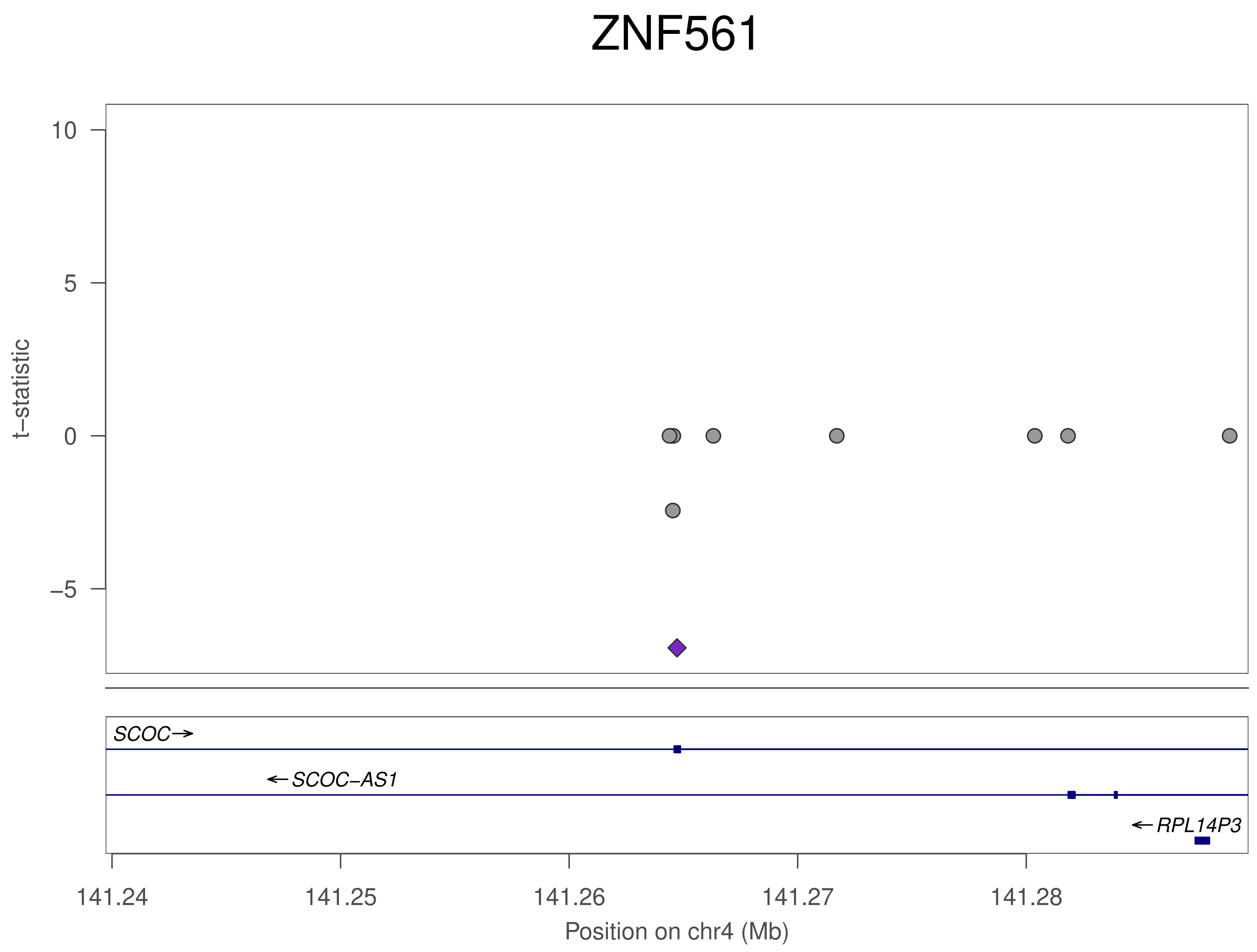

ZNF561
t−statistic
10
5
0
−5
SCOC→
←SCOC−AS1
←RPL14P3
141.24
141.25
141.26
141.27
141.28
Position on chr4 (Mb)

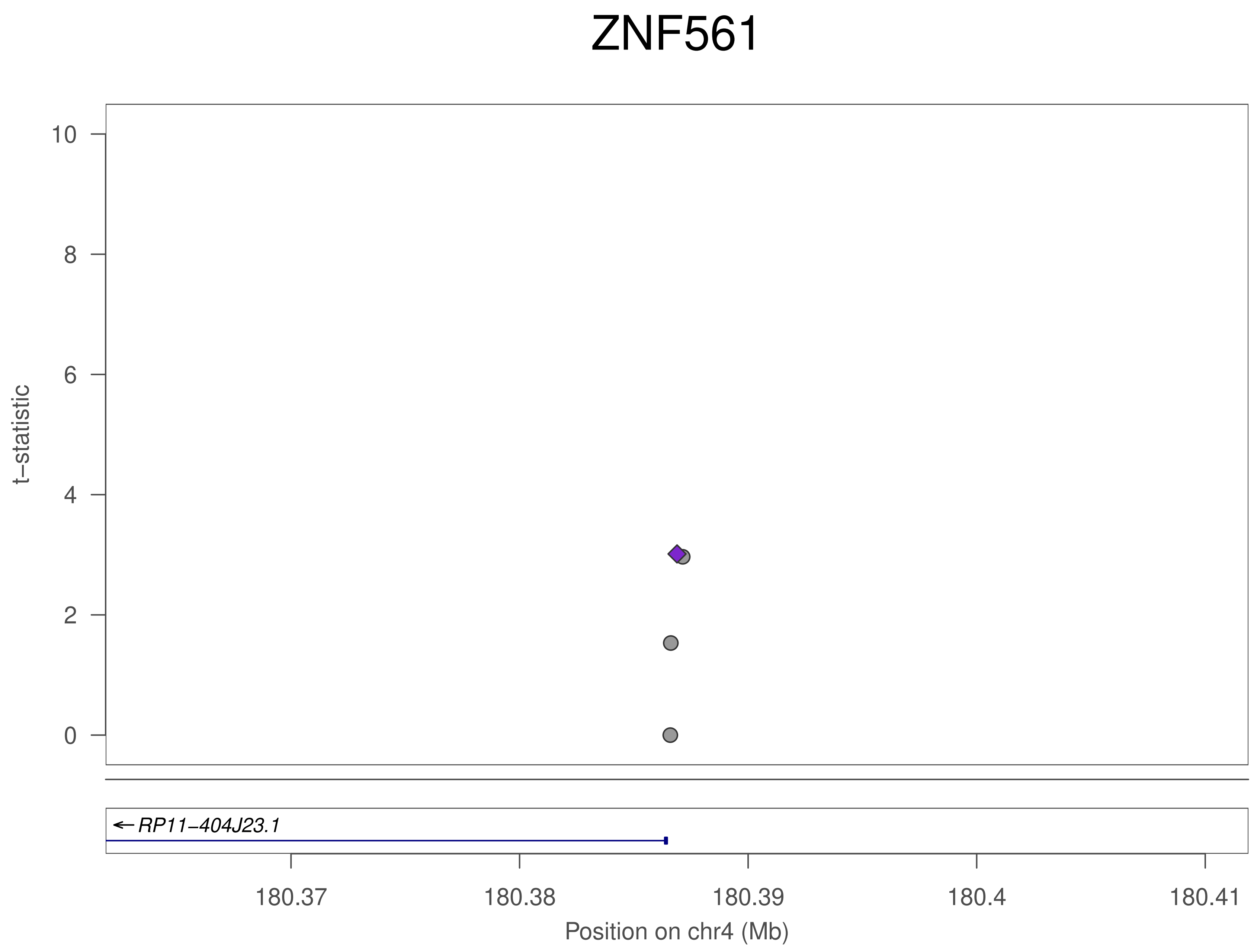
ZNF561
t−statistic
10
8
6
4
2
0
RP11−404J23.1
180.37
180.38
180.39
180.4
180.41
Position on chr4 (Mb)

### ZNF561

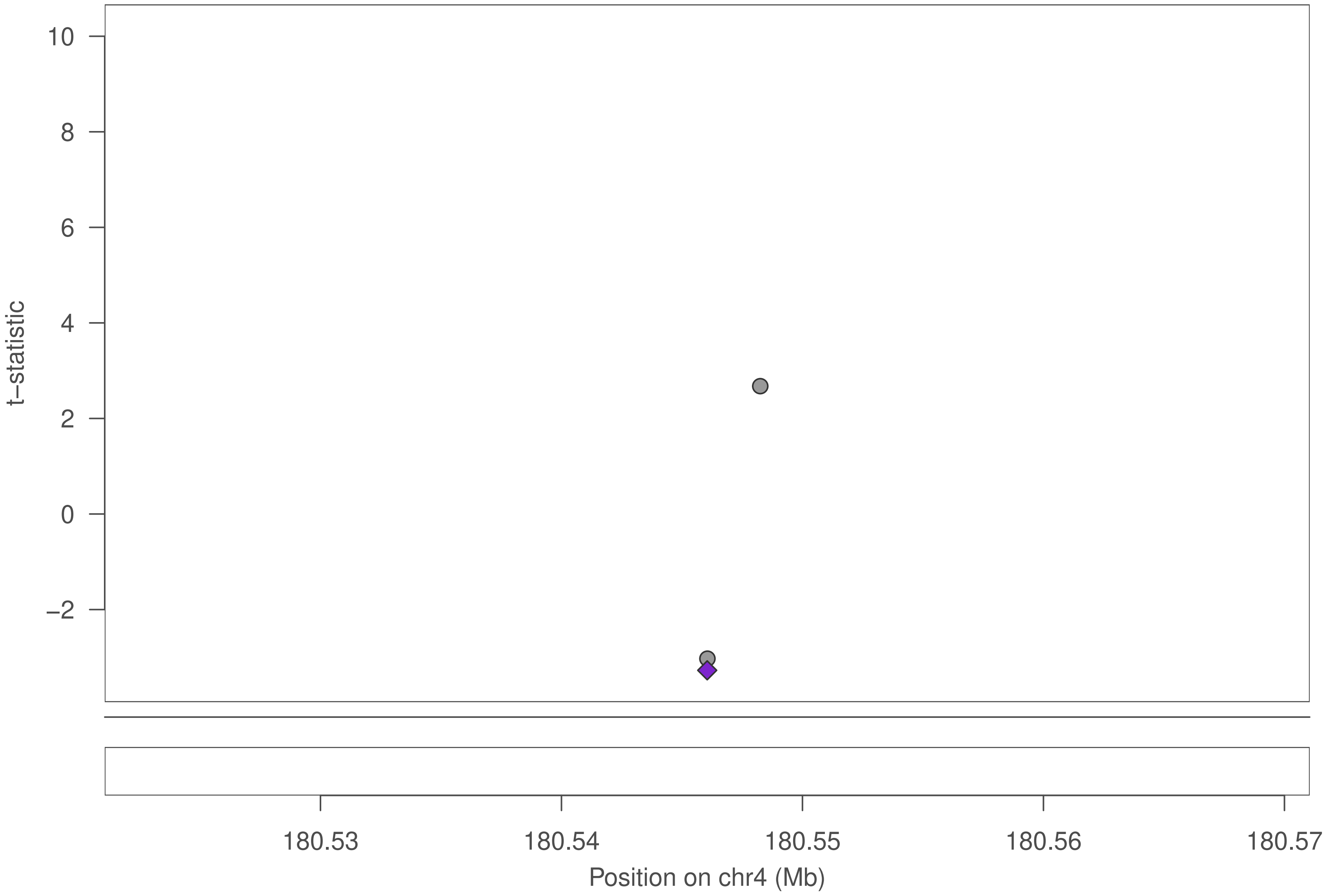
t–statistic
10
8
6
4
2
0
−2
180.53
180.54
180.55
180.56
180.57
Position on chr4 (Mb)

### ZNF561

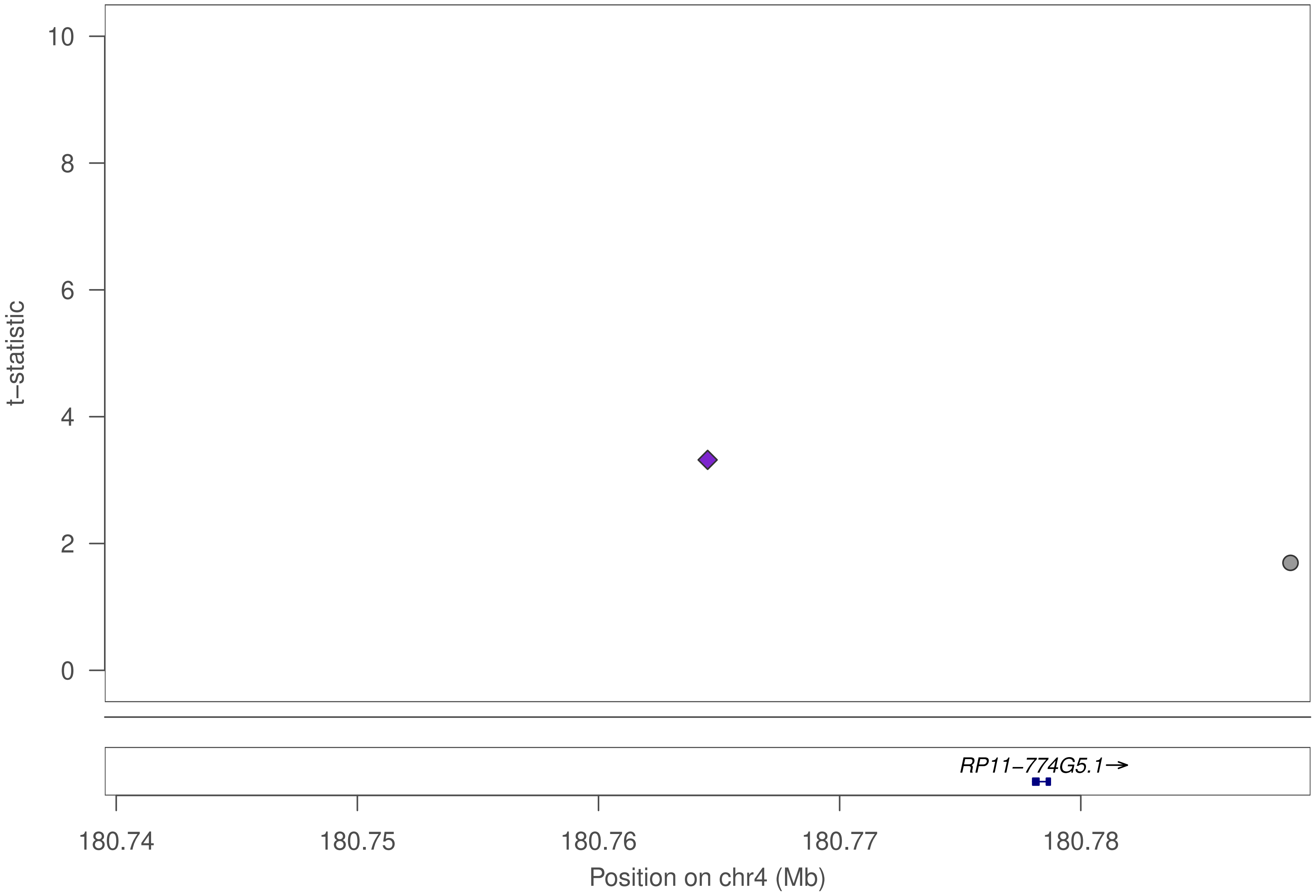

t−statistic
10
8
6
4
2
0
RP11−774G5.1→
180.74
180.75
180.76
180.77
180.78
Position on chr4 (Mb)

### ZNF561

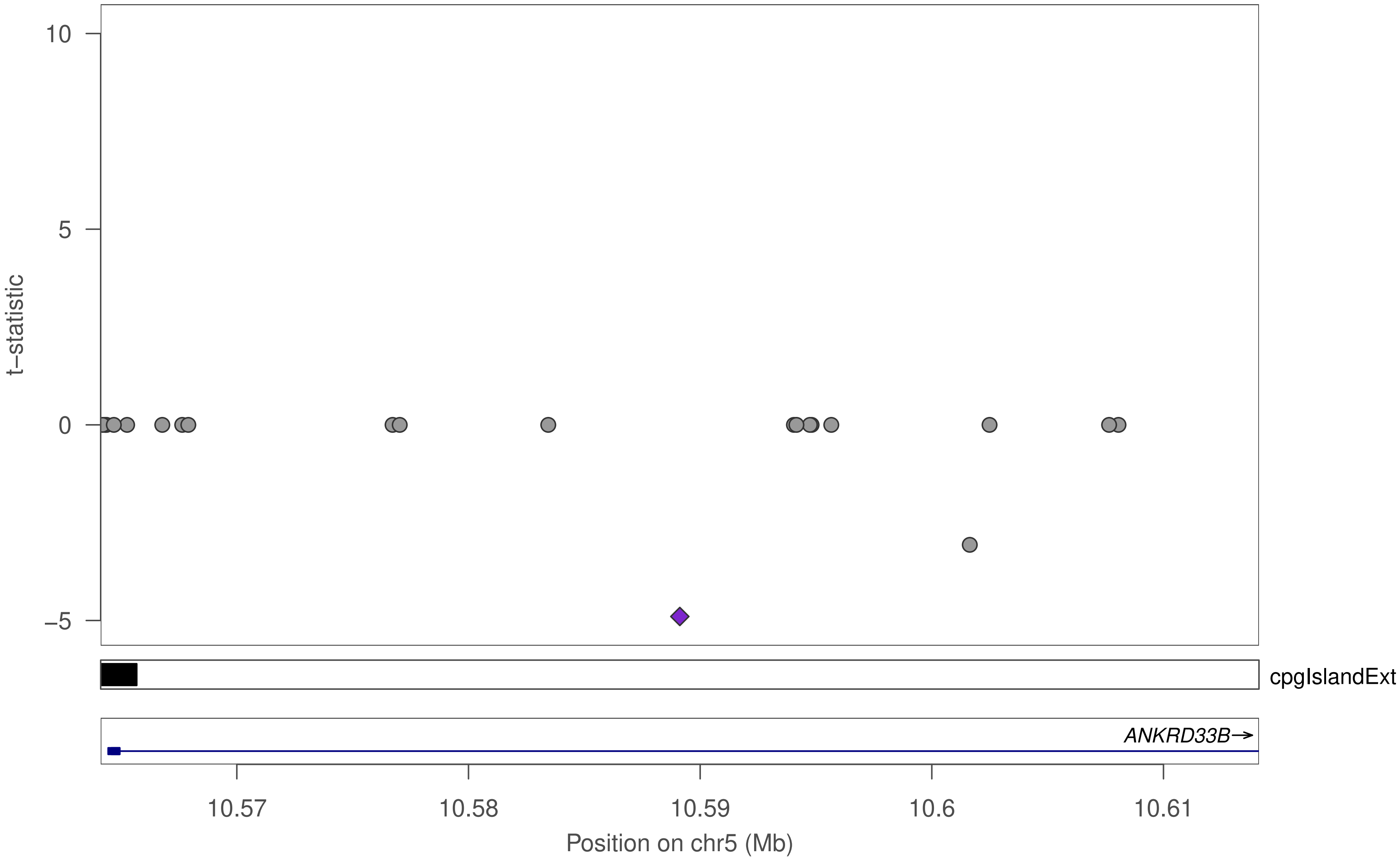

10
5
0
−5
t−statistic
cpgIslandExt
ANKRD33B→
10.57
10.58
10.59
10.6
10.61
Position on chr5 (Mb)

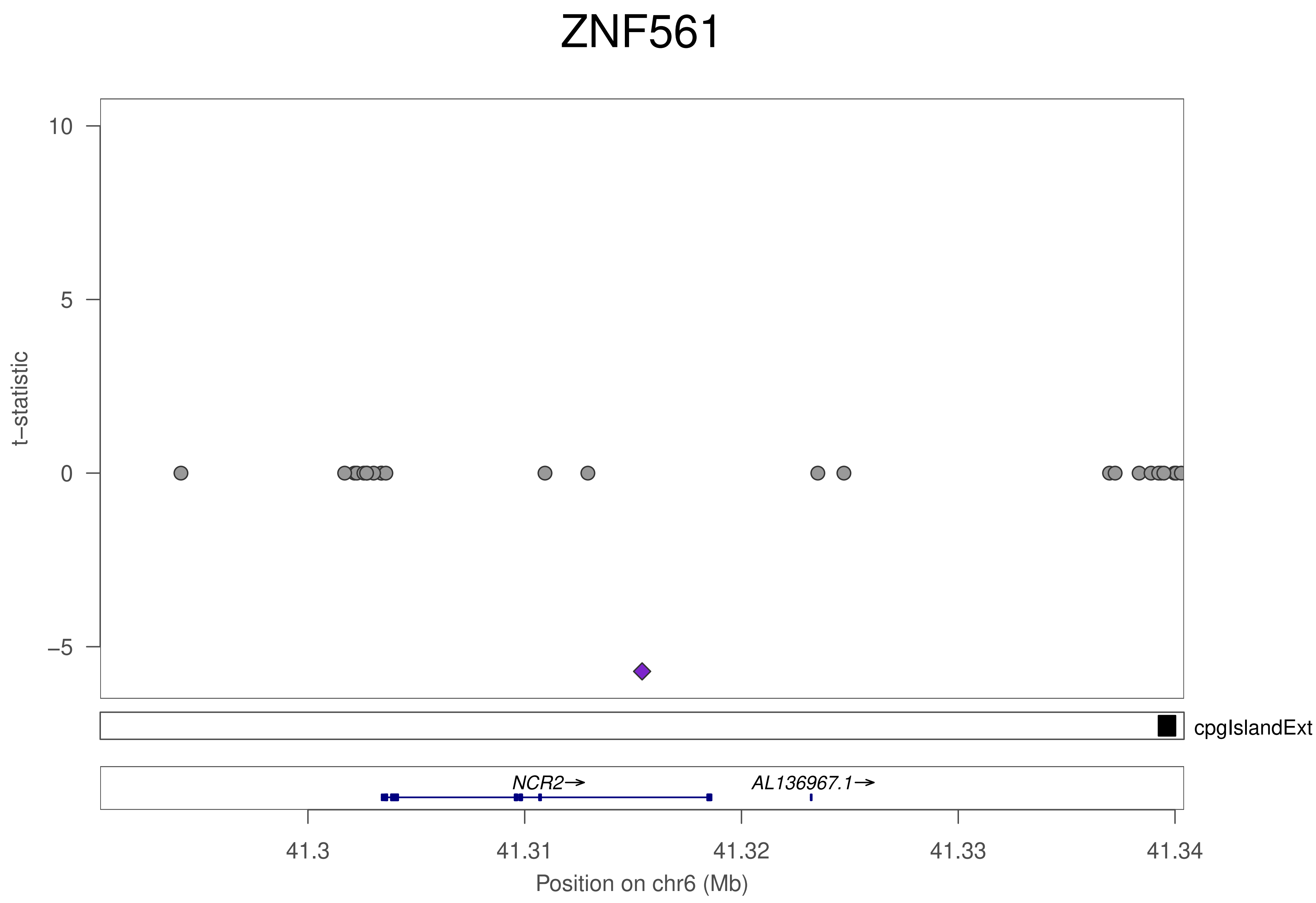
ZNF561
t–statistic
10
5
0
–5
cpgIslandExt
NCR2→
AL136967.1→
41.3
41.31
41.32
41.33
41.34
Position on chr6 (Mb)

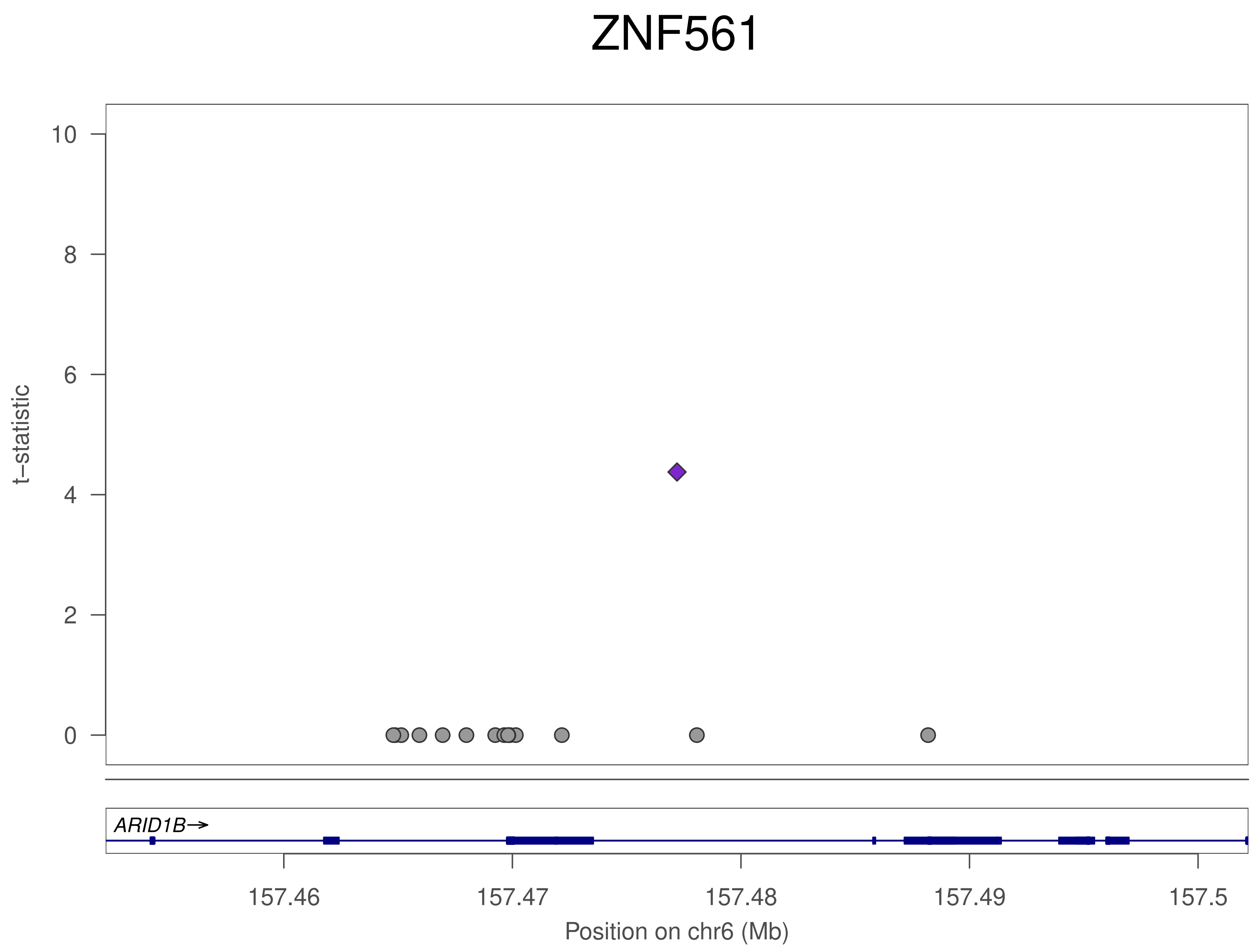

ZNF561
t−statistic
10
8
6
4
2
0
ARID1B→
157.46
157.47
157.48
157.49
157.5
Position on chr6 (Mb)

### ZNF561

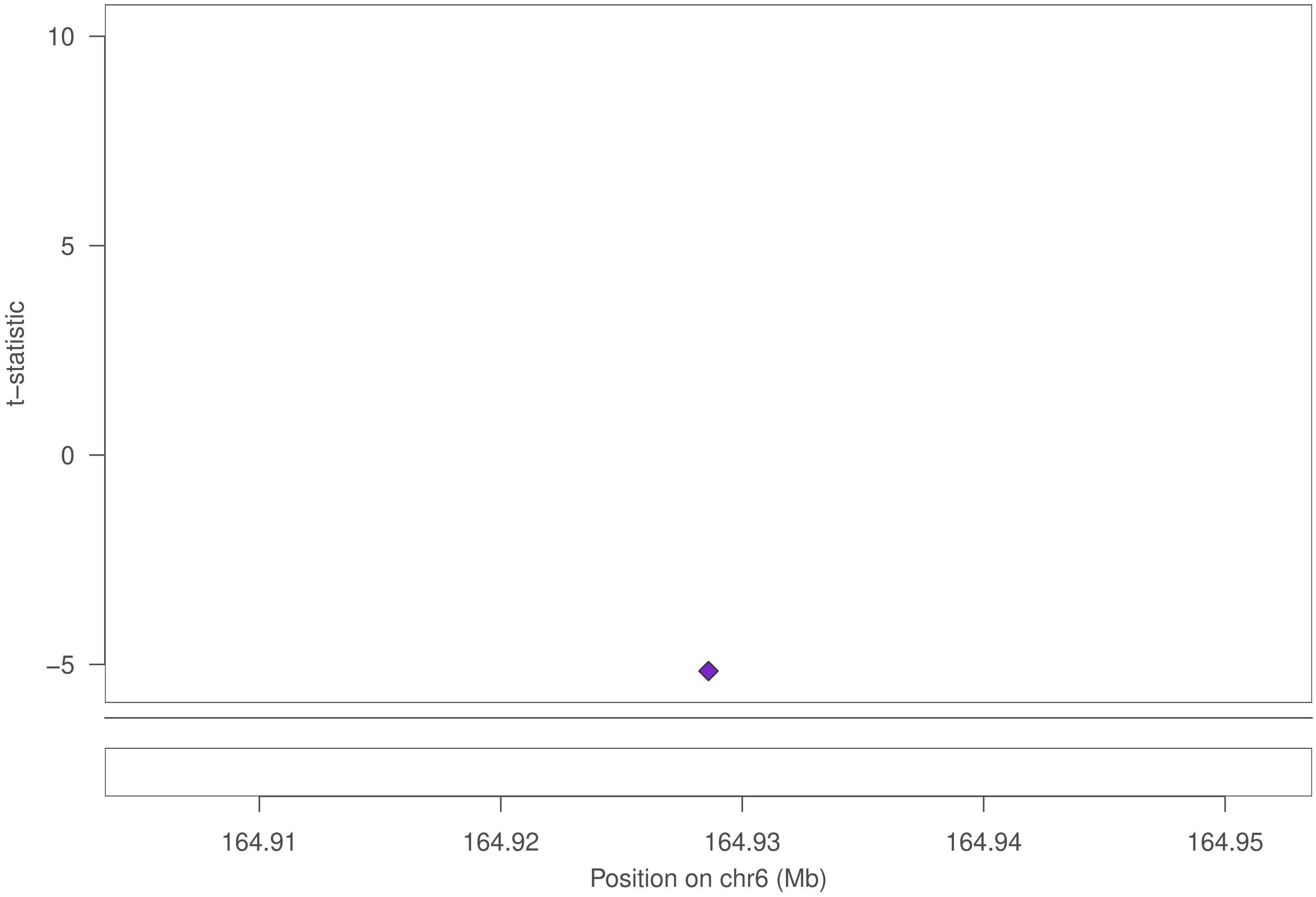
10
5
0
−5
t−statistic
164.91
164.92
164.93
164.94
164.95
Position on chr6 (Mb)

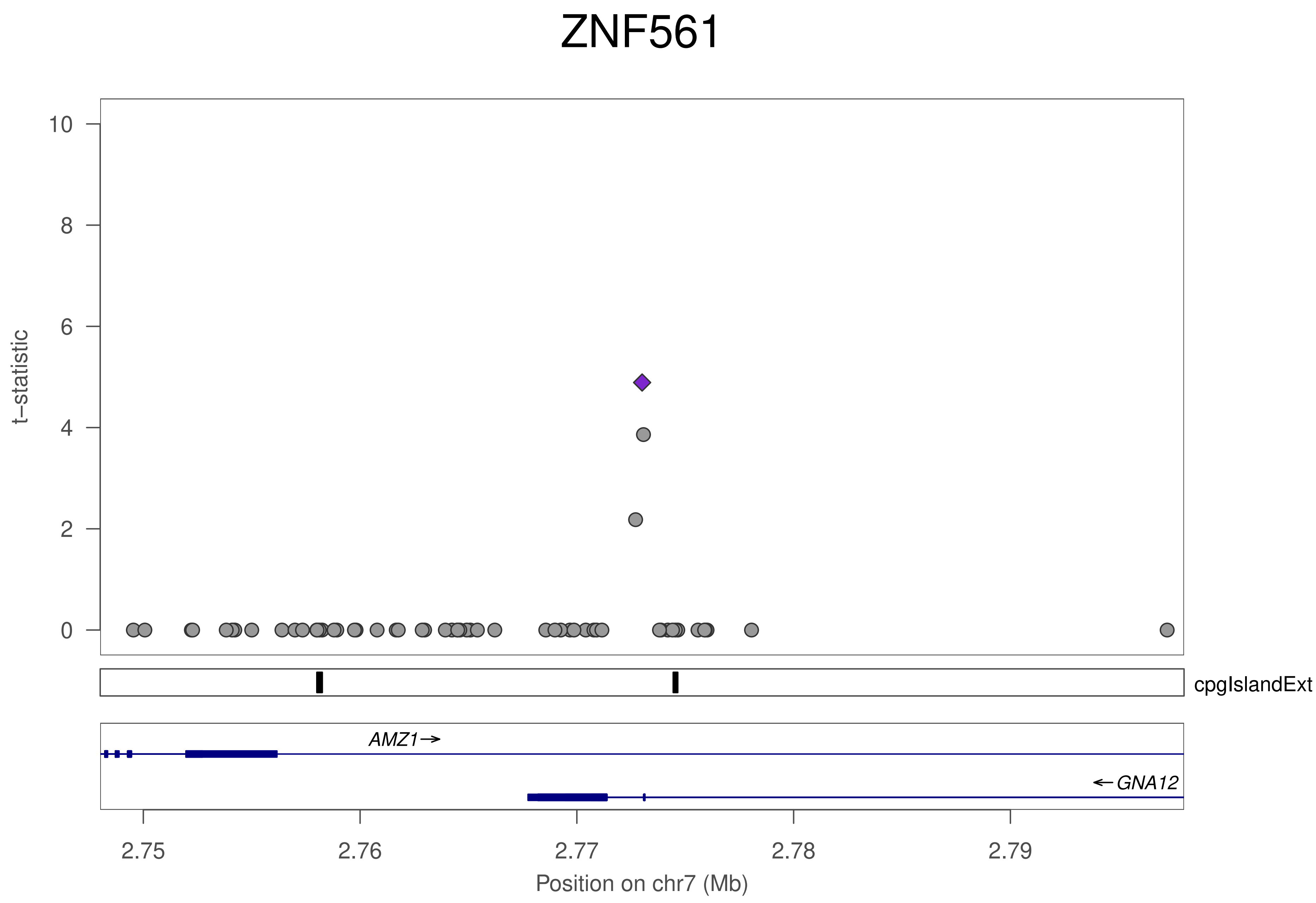

ZNF561
t−statistic
10
8
6
4
2
0
cpgIslandExt
AMZ1
GNA12
2.75
2.76
2.77
2.78
2.79
Position on chr7 (Mb)

### ZNF561

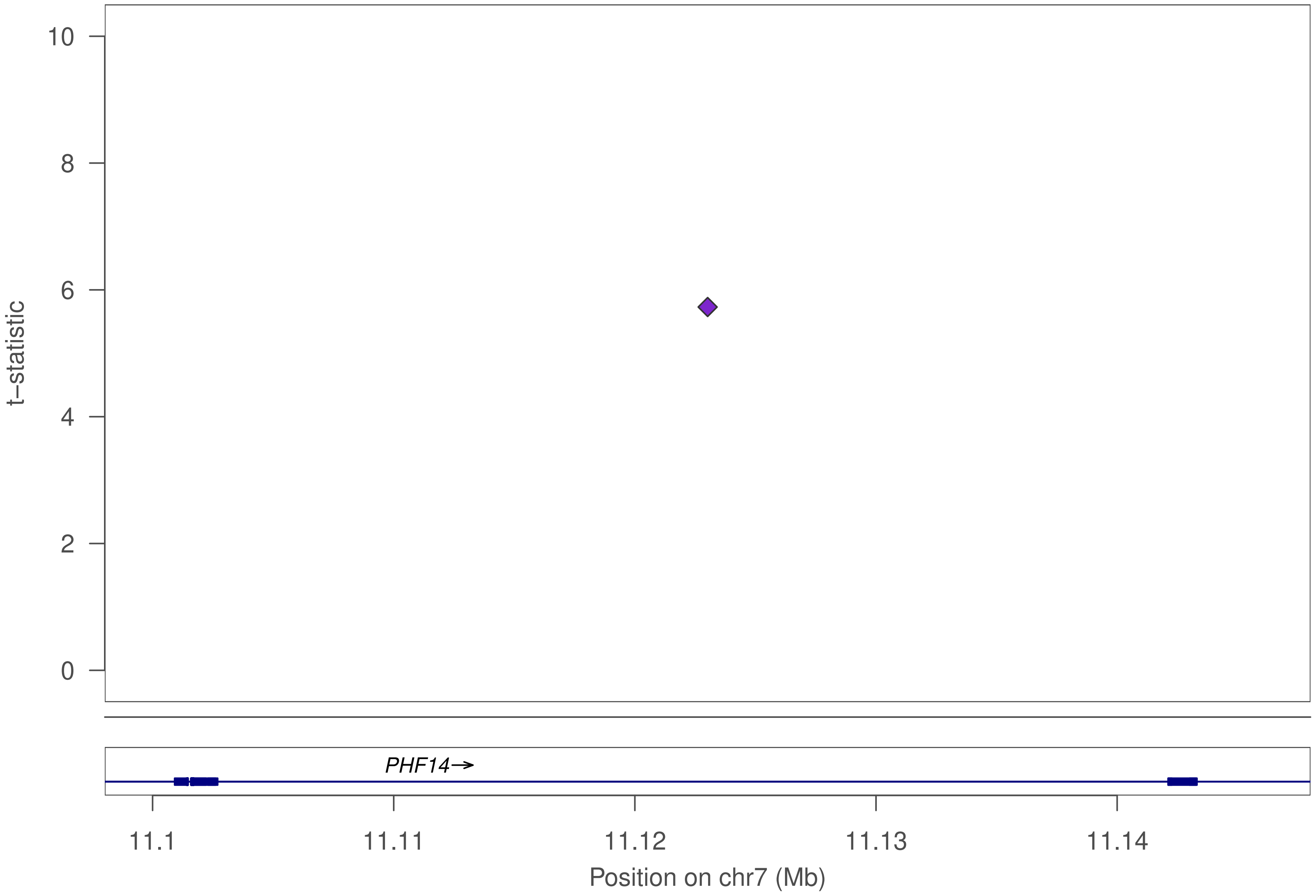
10
8
6
4
2
0
t−statistic
PHF14→
11.1
11.11
11.12
11.13
11.14
Position on chr7 (Mb)

### ZNF561

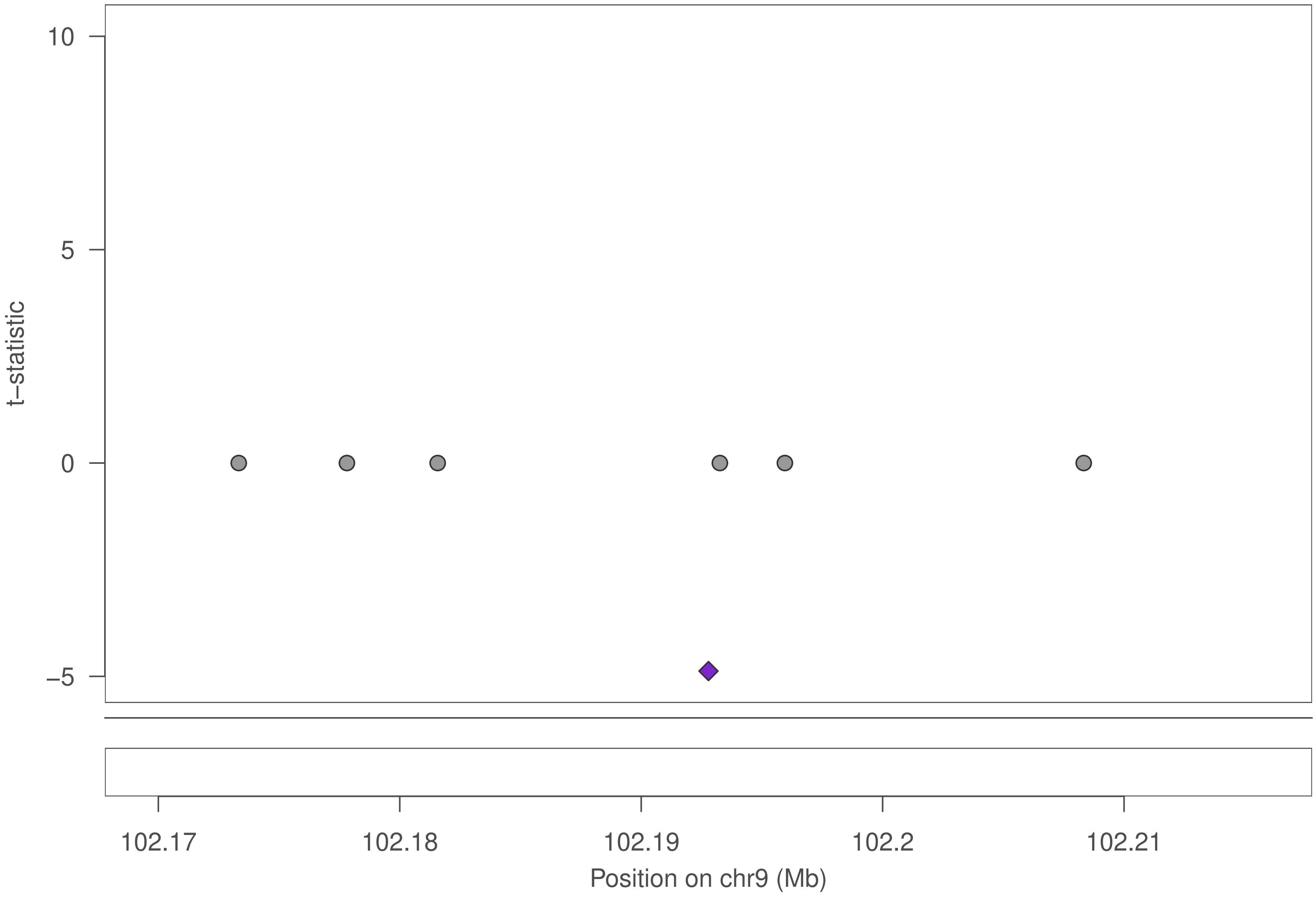

10
5
0
−5
t−statistic
102.17
102.18
102.19
102.2
102.21
Position on chr9 (Mb)

### ZNF561

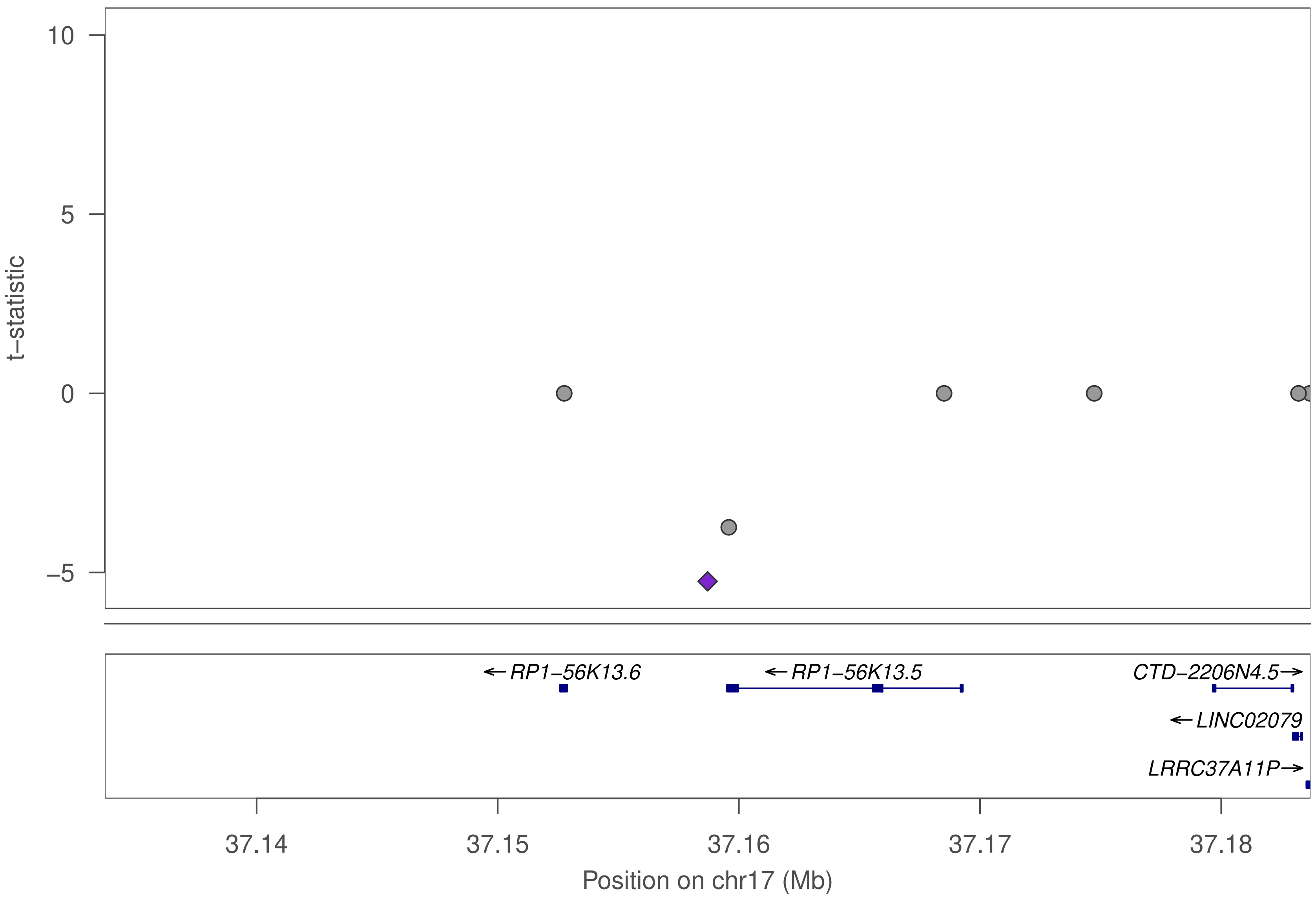

t−statistic
10
5
0
−5
←RP1−56K13.6
←RP1−56K13.5
CTD−2206N4.5→
←LINC02079
LRRC37A11P→
37.14
37.15
37.16
37.17
37.18
Position on chr17 (Mb)

### ZNF561

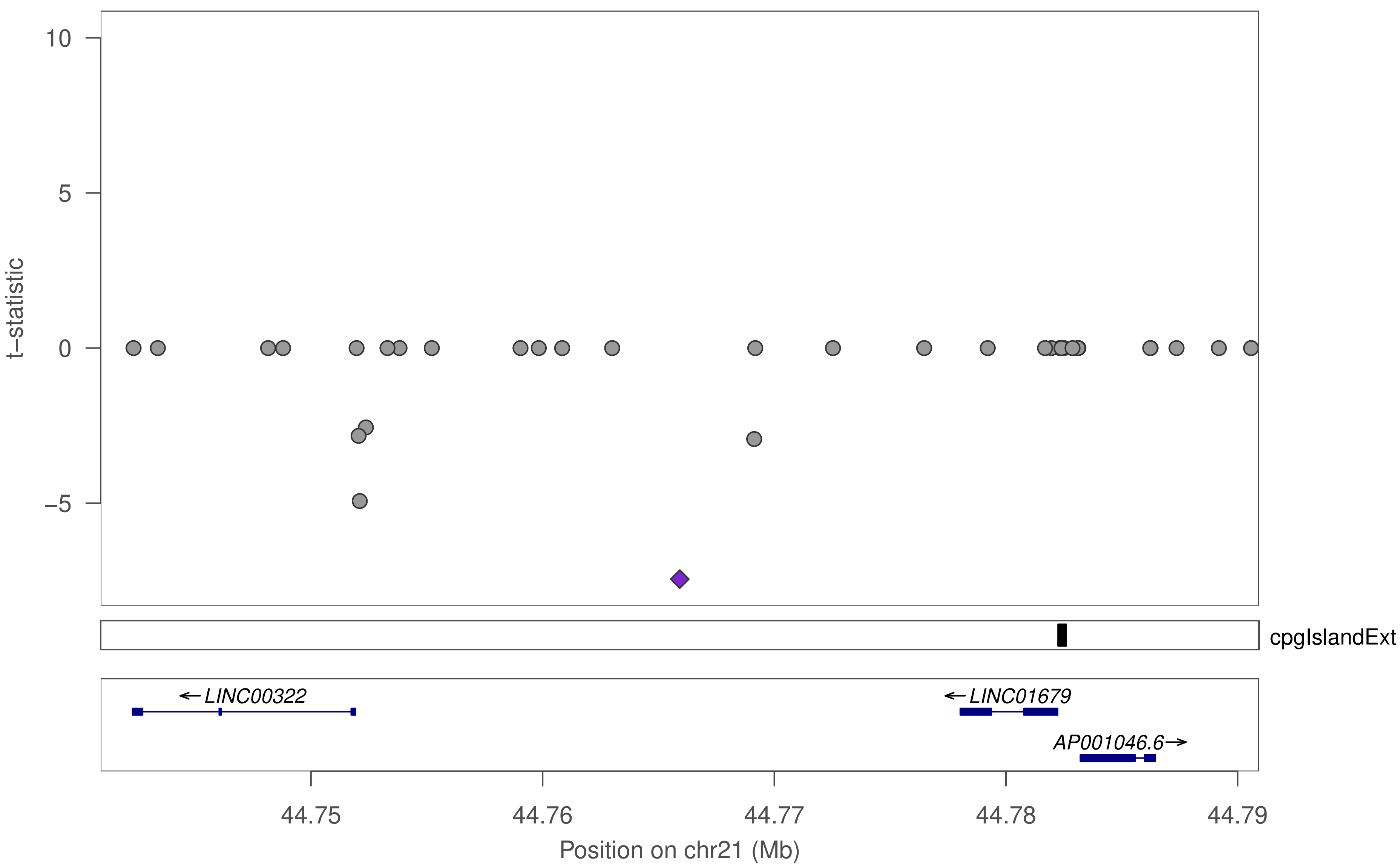

10
5
0
−5
t−statistic
cpgIslandExt
←LINC00322
←LINC01679
AP001046.6→
44.75
44.76
44.77
44.78
44.79
Position on chr21 (Mb)

### ZNF561

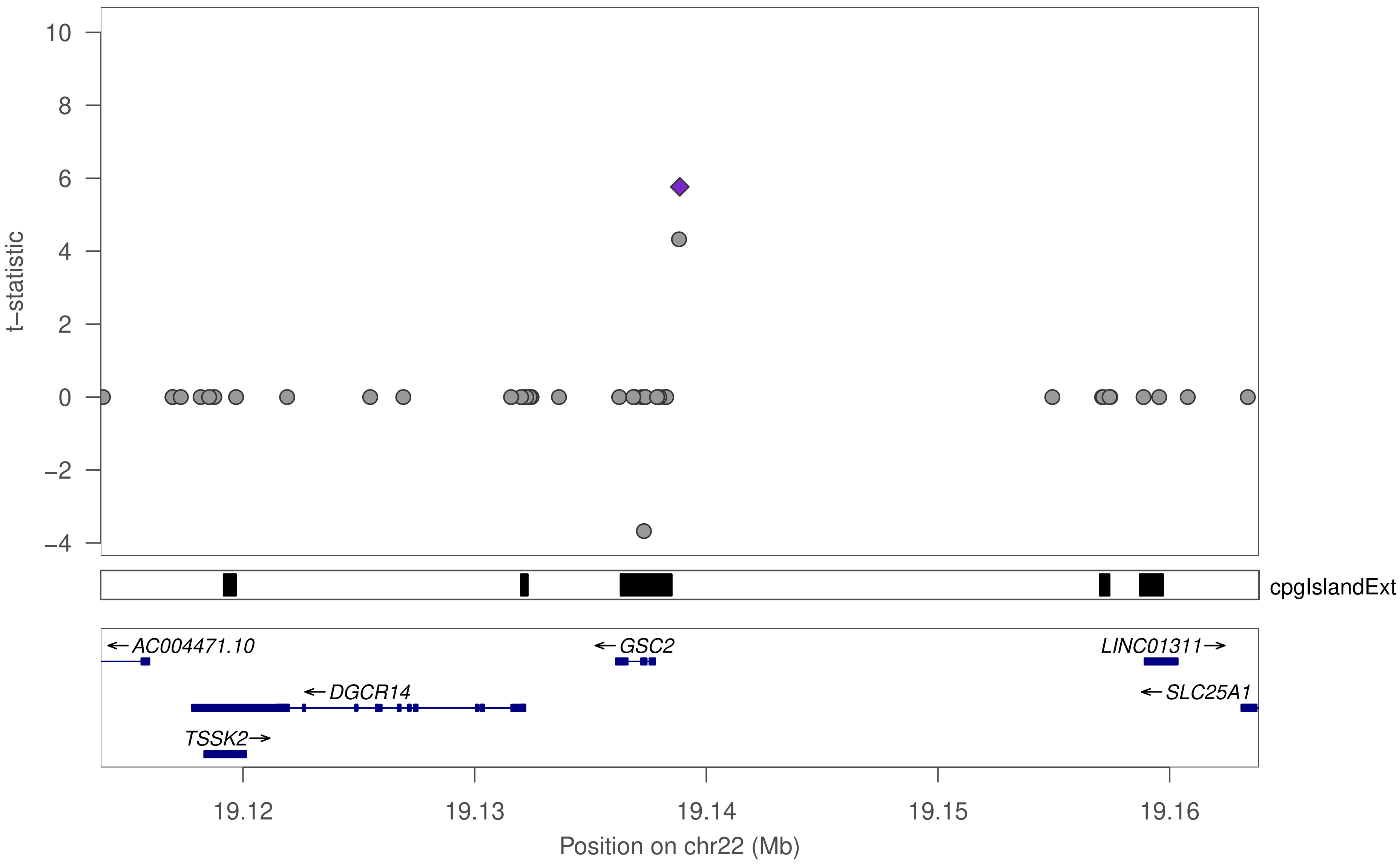

t−statistic
10
8
6
4
2
0
−2
−4
cpgIslandExt
AC004471.10
GSC2
LINC01311
DGCR14
SLC25A1
TSSK2
19.12
19.13
19.14
19.15
19.16
Position on chr22 (Mb)

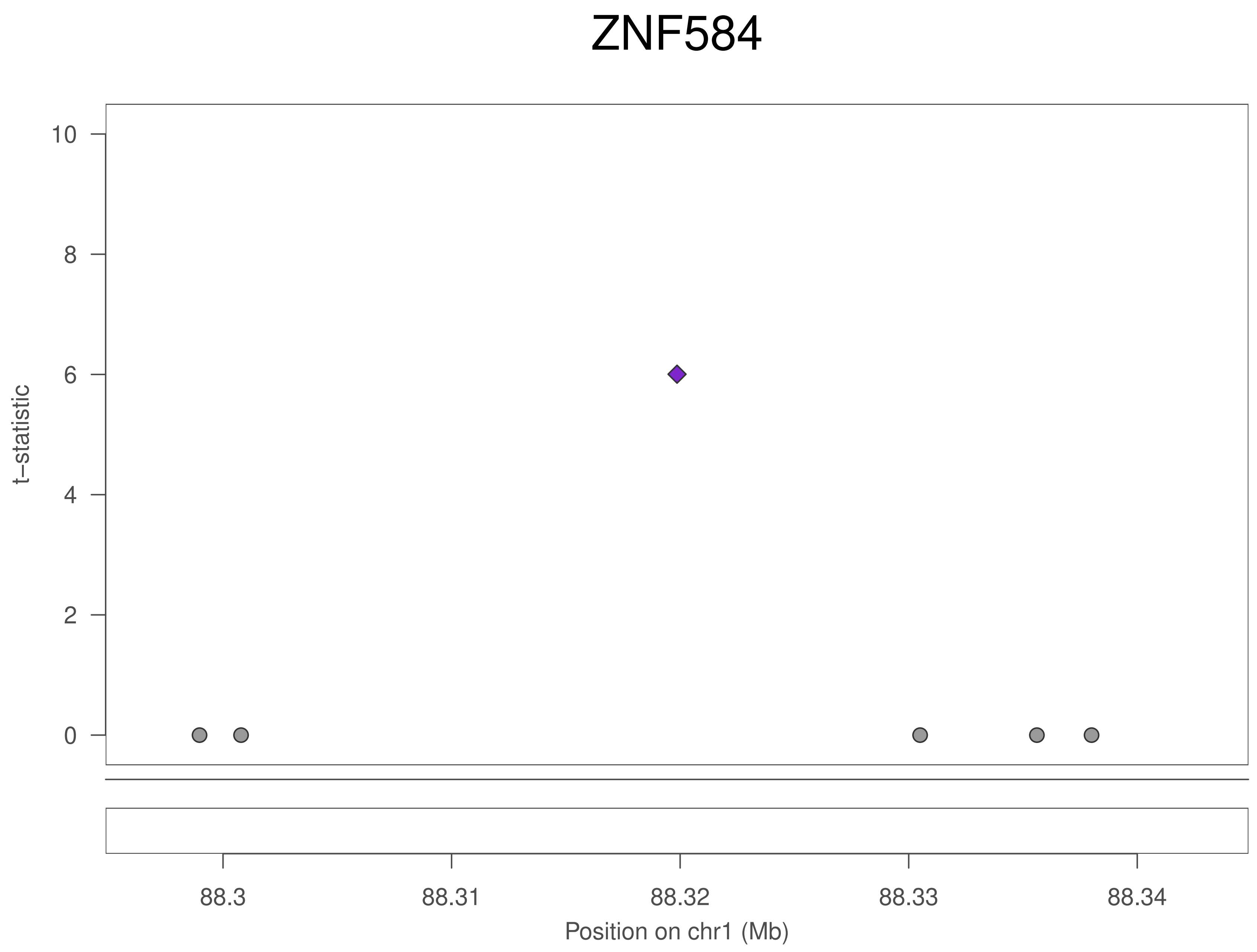

ZNF584
t–statistic
10
8
6
4
2
0
88.3
88.31
88.32
88.33
88.34
Position on chr1 (Mb)

### ZNF584

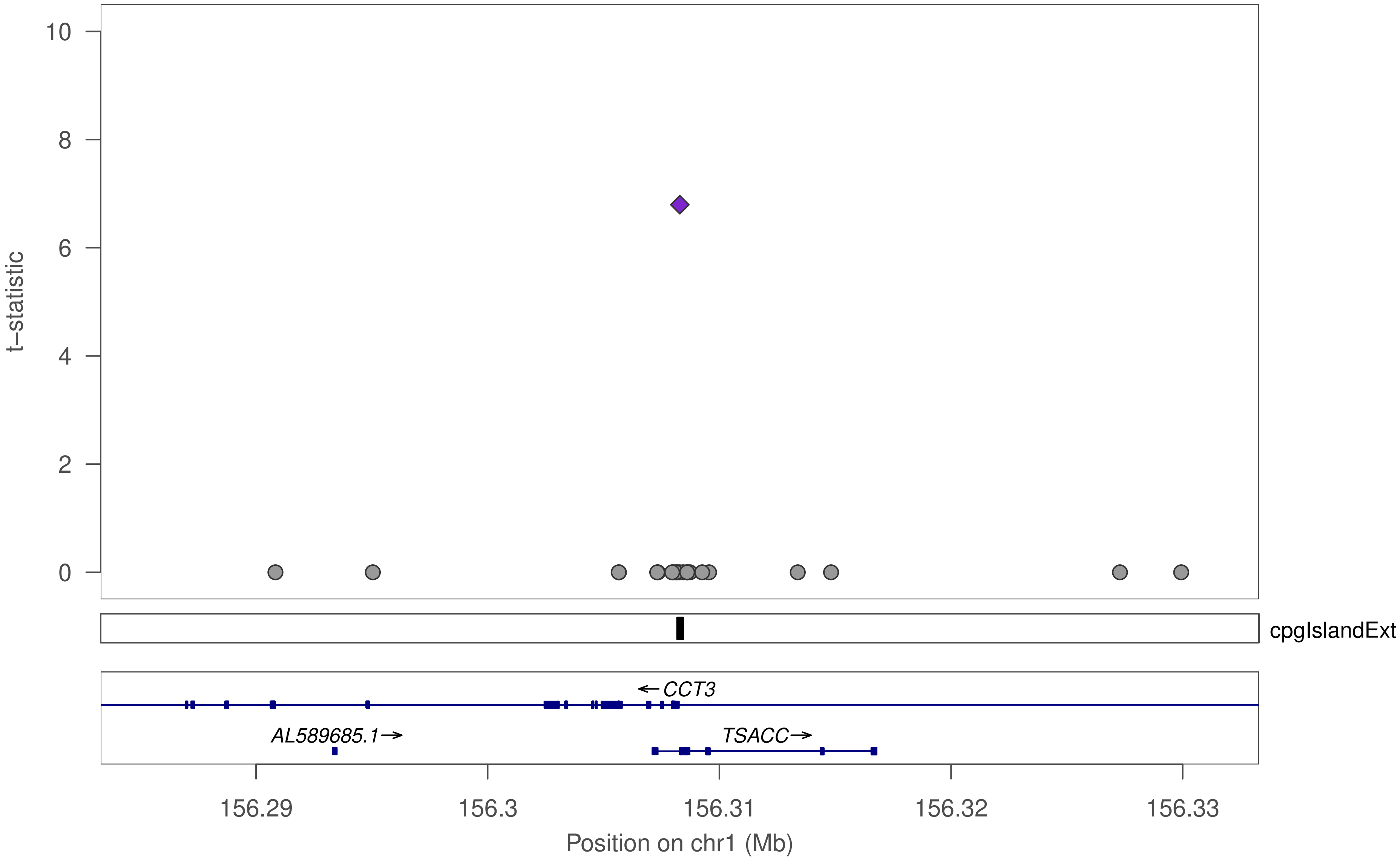

10
8
6
4
2
0
t−statistic
cpgIslandExt
←CCT3
AL589685.1→
TSACC→
156.29
156.3
156.31
156.32
156.33
Position on chr1 (Mb)

### ZNF584

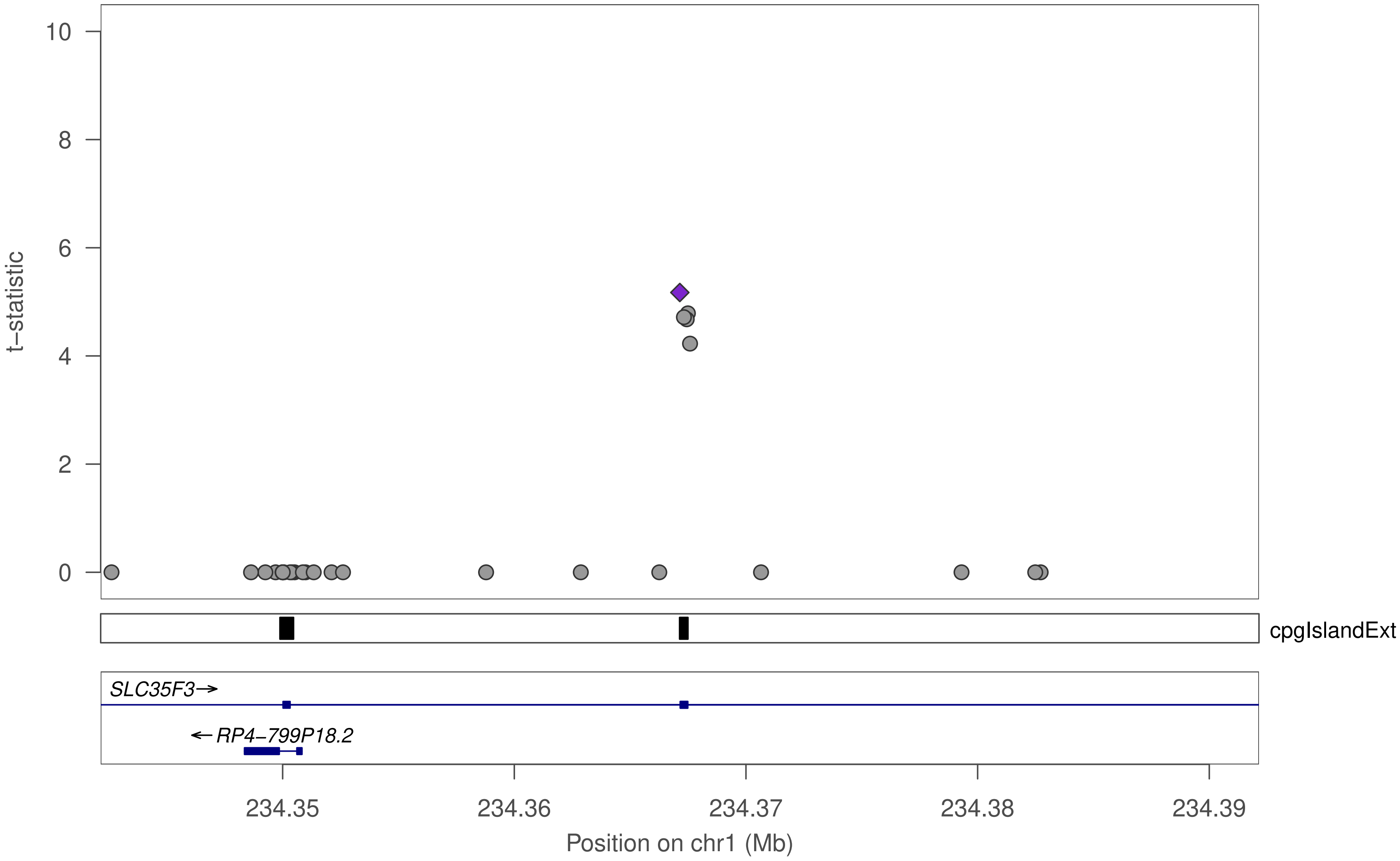

t−statistic
10
8
6
4
2
0
cpgIslandExt
SLC35F3→
←RP4−799P18.2
234.35
234.36
234.37
234.38
234.39
Position on chr1 (Mb)

### ZNF584

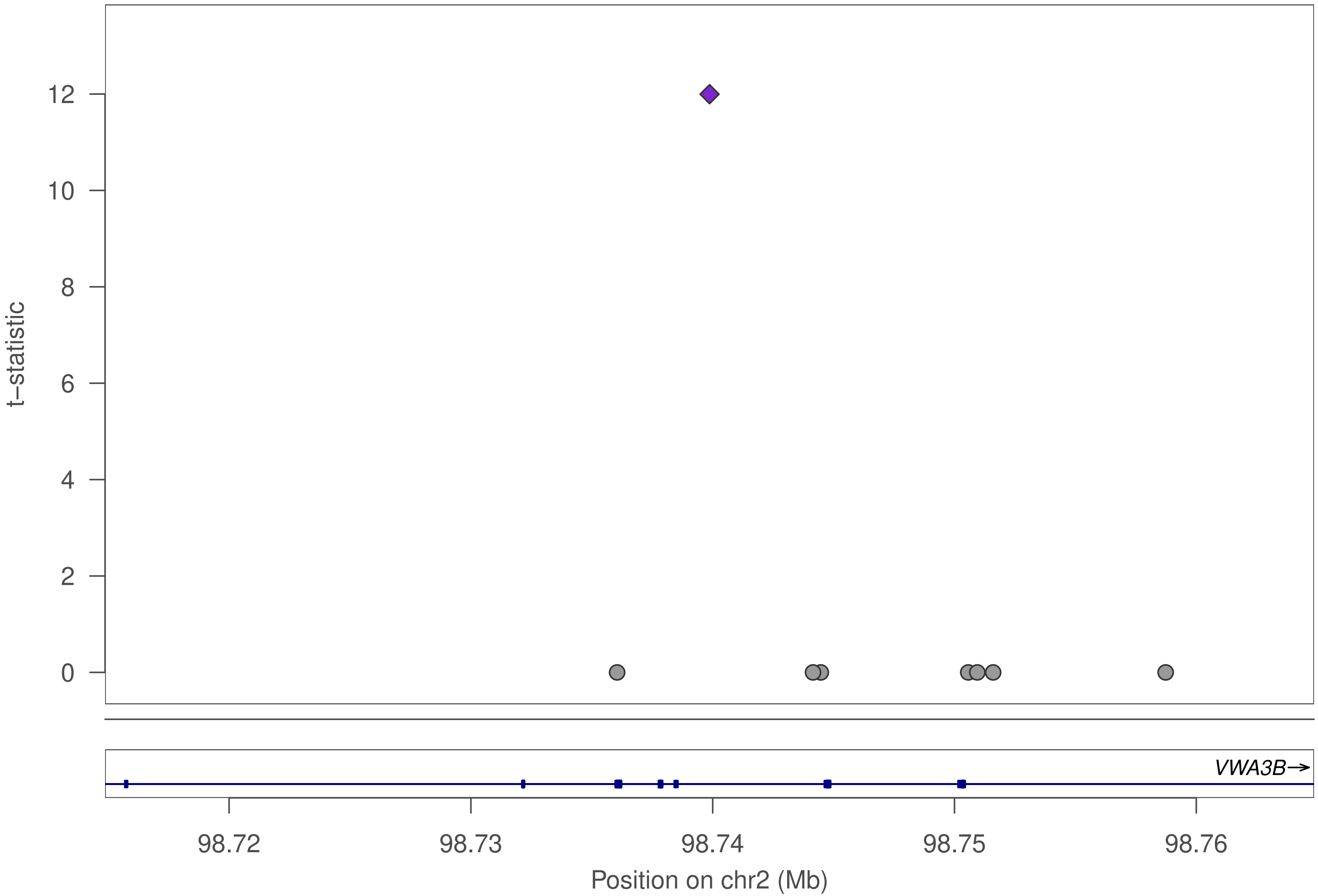

t–statistic
0
2
4
6
8
10
12
VWA3B→
98.72
98.73
98.74
98.75
98.76
Position on chr2 (Mb)

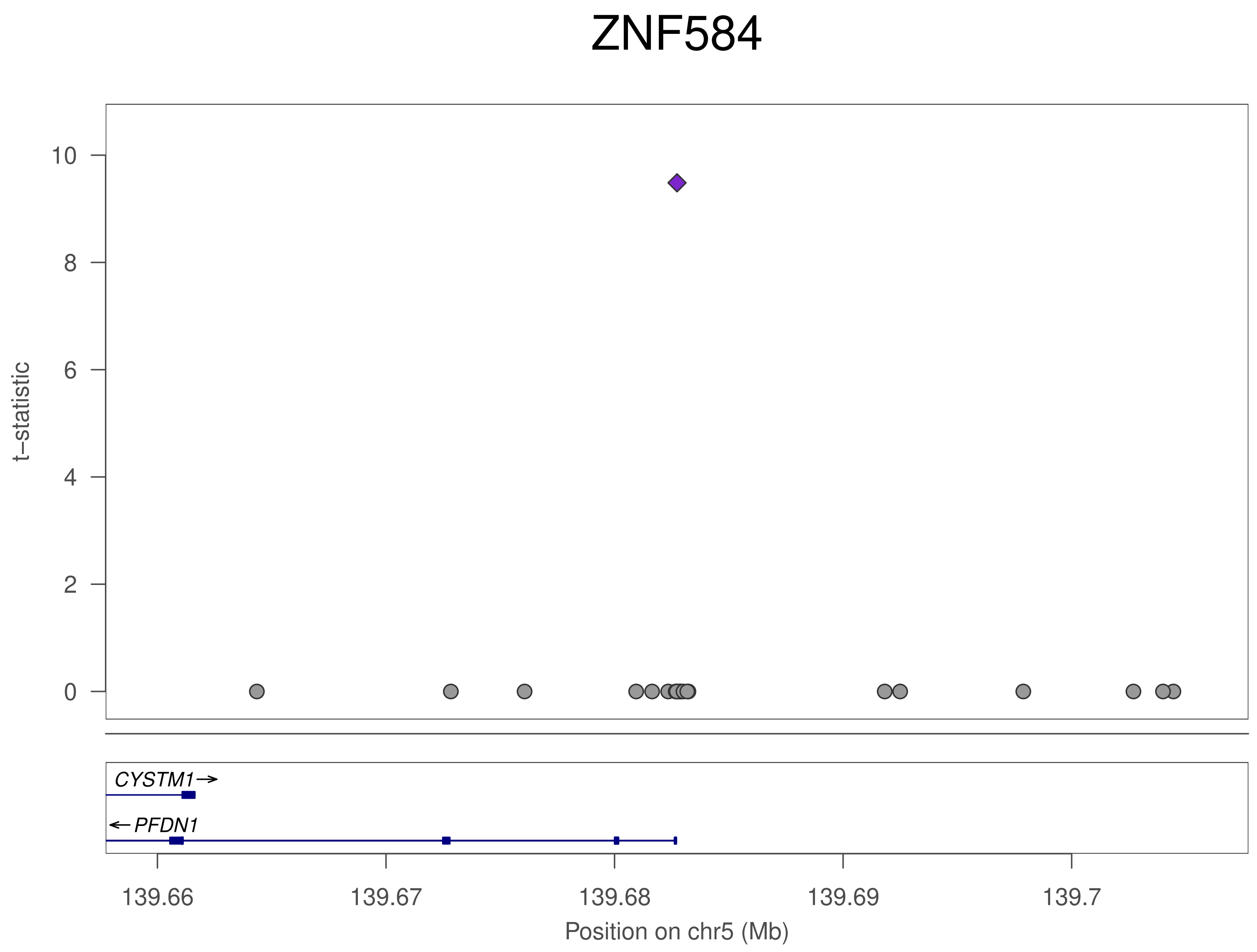
ZNF584
t−statistic
10
8
6
4
2
0
CYSTM1→
←PFDN1
139.66
139.67
139.68
139.69
139.7
Position on chr5 (Mb)

### ZNF584

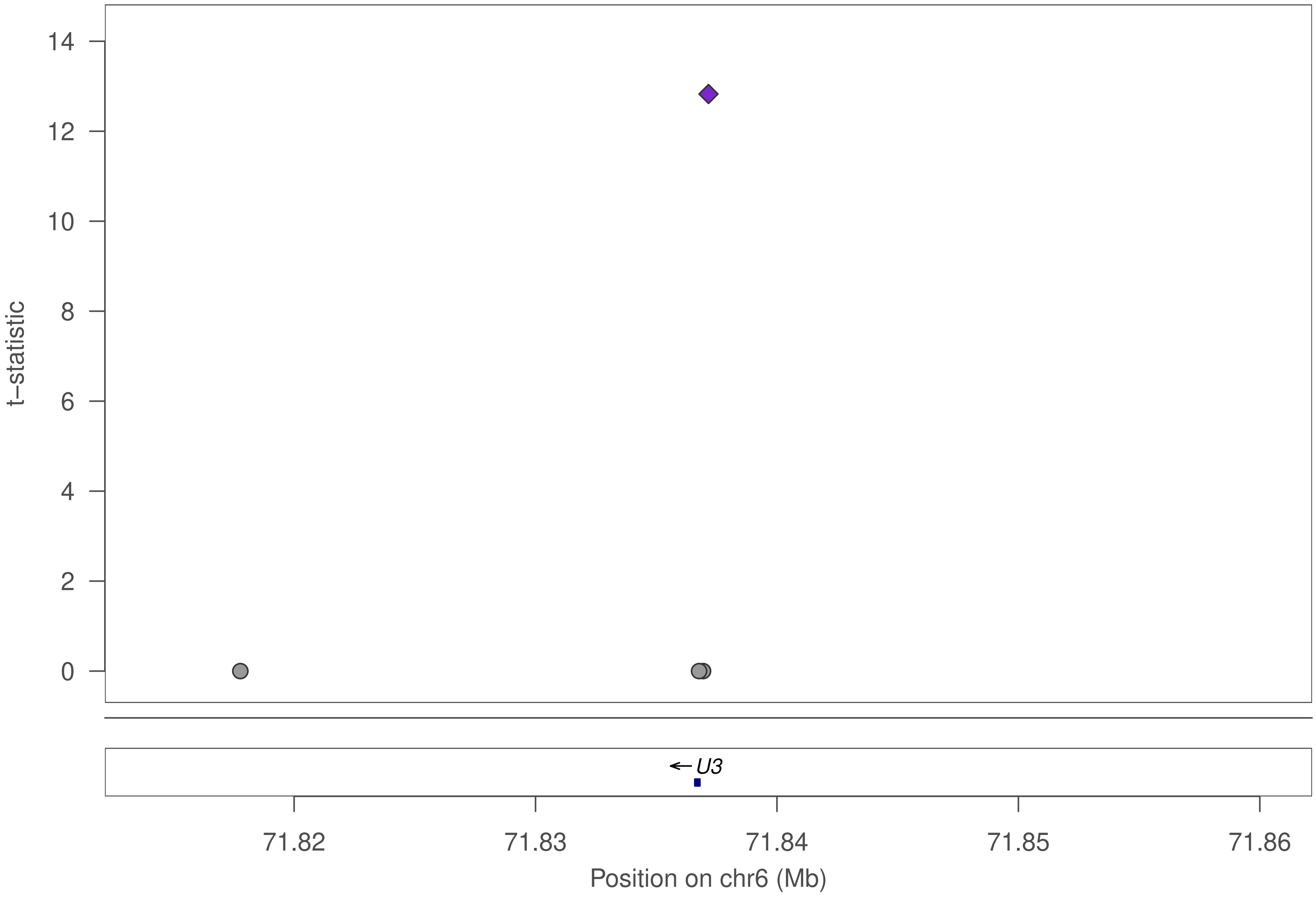

t–statistic
14
12
10
8
6
4
2
0
← U3
71.82
71.83
71.84
71.85
71.86
Position on chr6 (Mb)

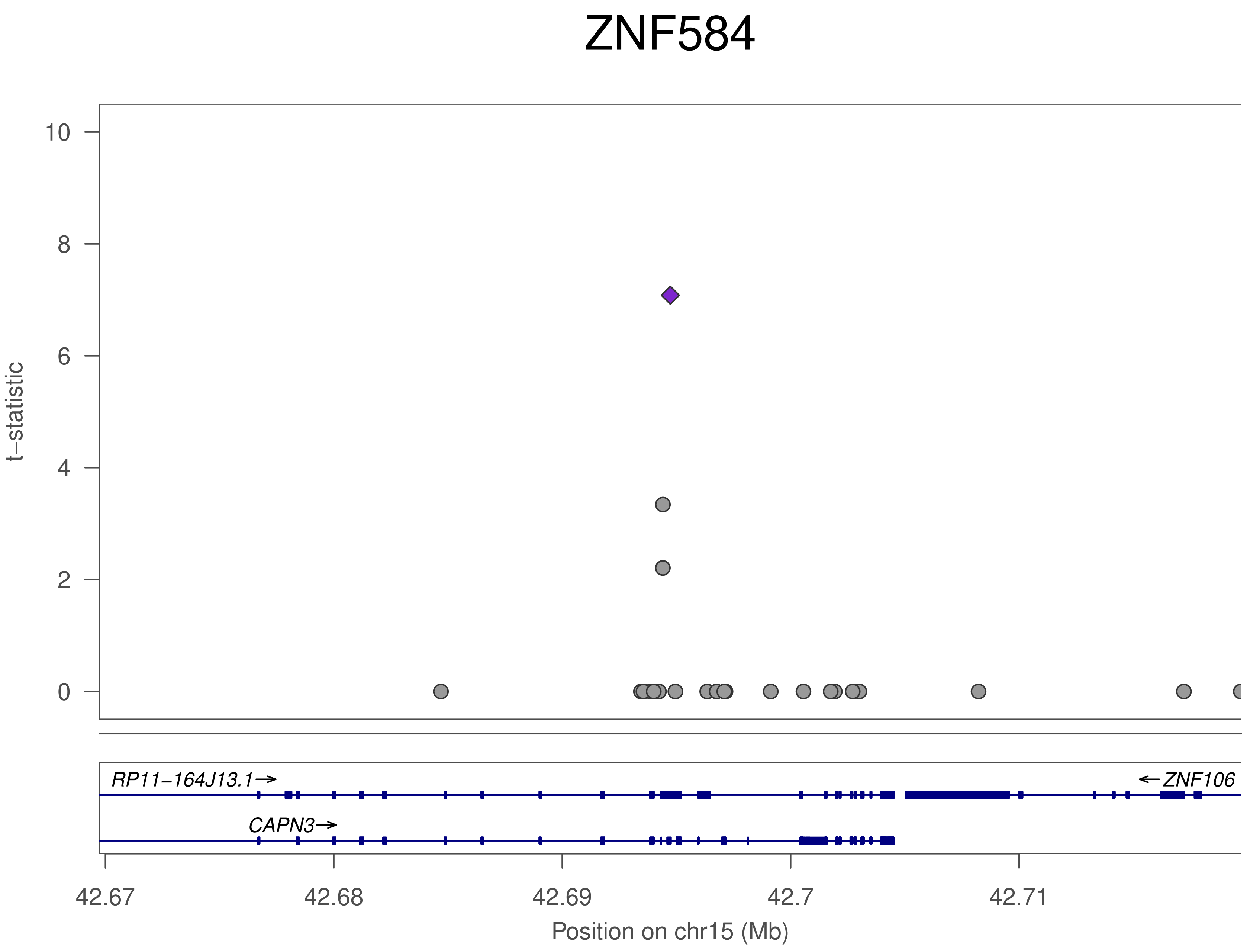
ZNF584
t−statistic
10
8
6
4
2
0
RP11−164J13.1→
←ZNF106
CAPN3→
42.67
42.68
42.69
42.7
42.71
Position on chr15 (Mb)

### ZNF584

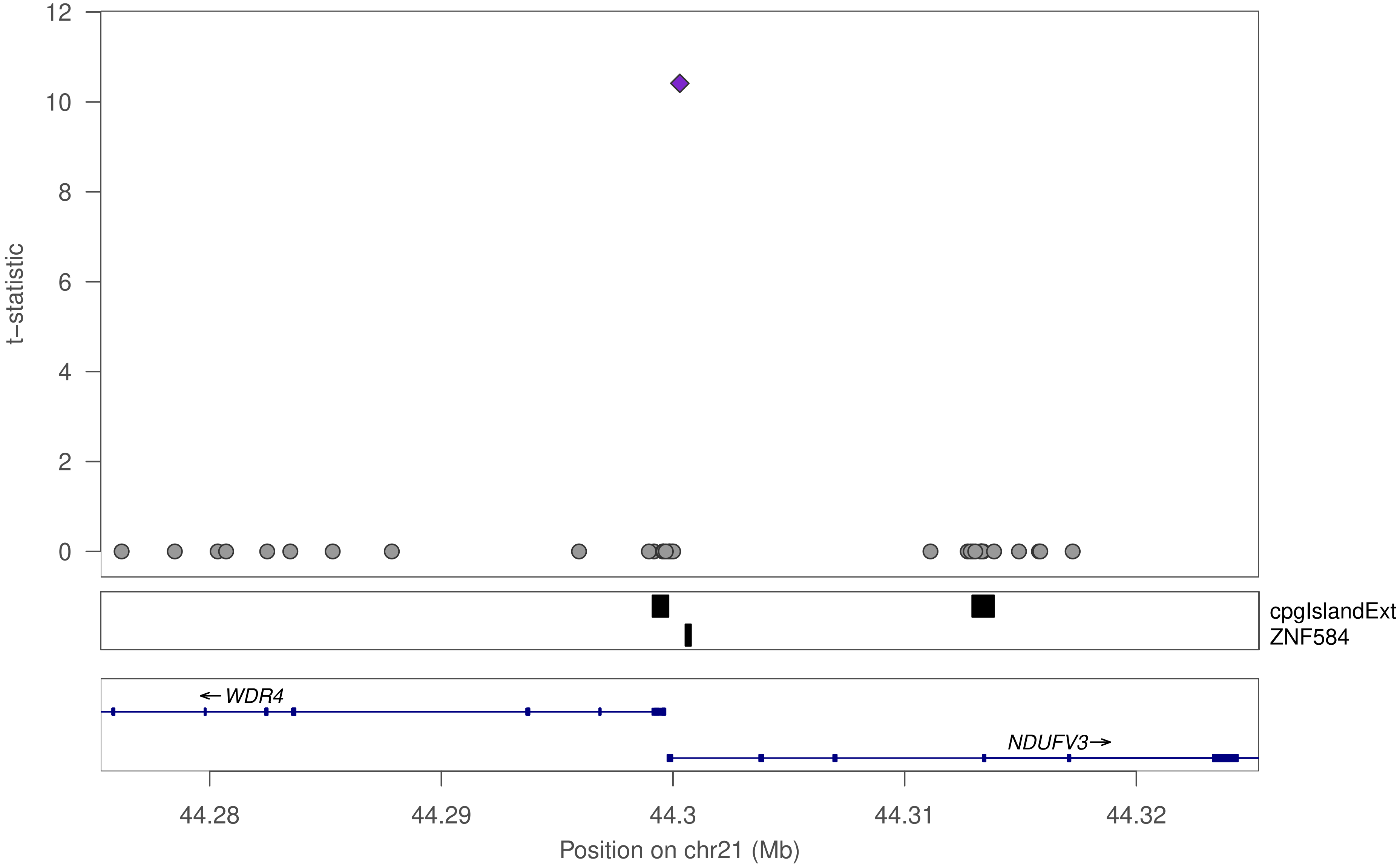

12
10
8
6
4
2
0
t−statistic
cpgIslandExt
ZNF584
← WDR4
NDUFV3→
44.28
44.29
44.3
44.31
44.32
Position on chr21 (Mb)

### ZNF584

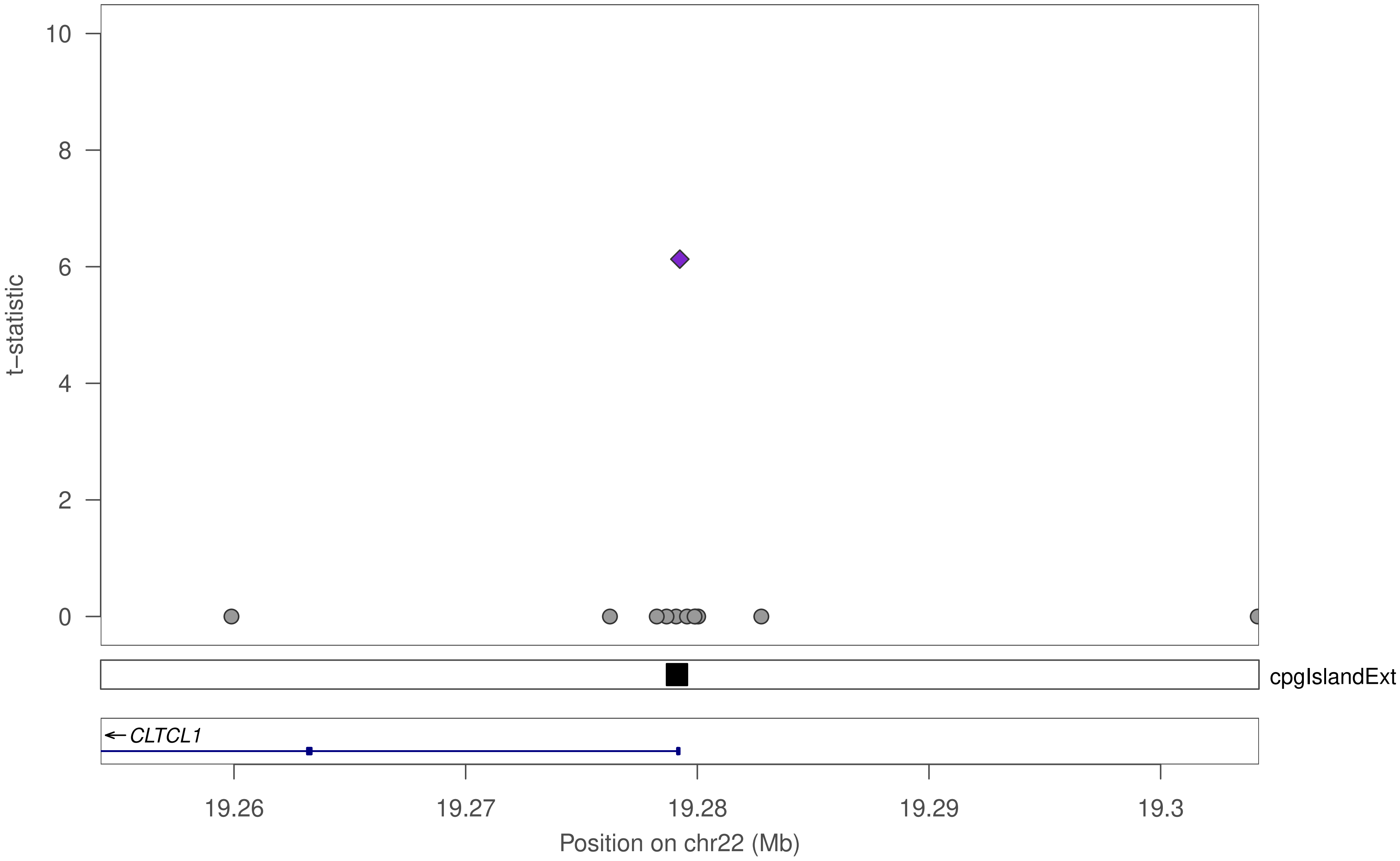

10
8
6
4
2
0
t–statistic
cpgIslandExt
← CLTCL1
19.26
19.27
19.28
19.29
19.3
Position on chr22 (Mb)

### ZNF680

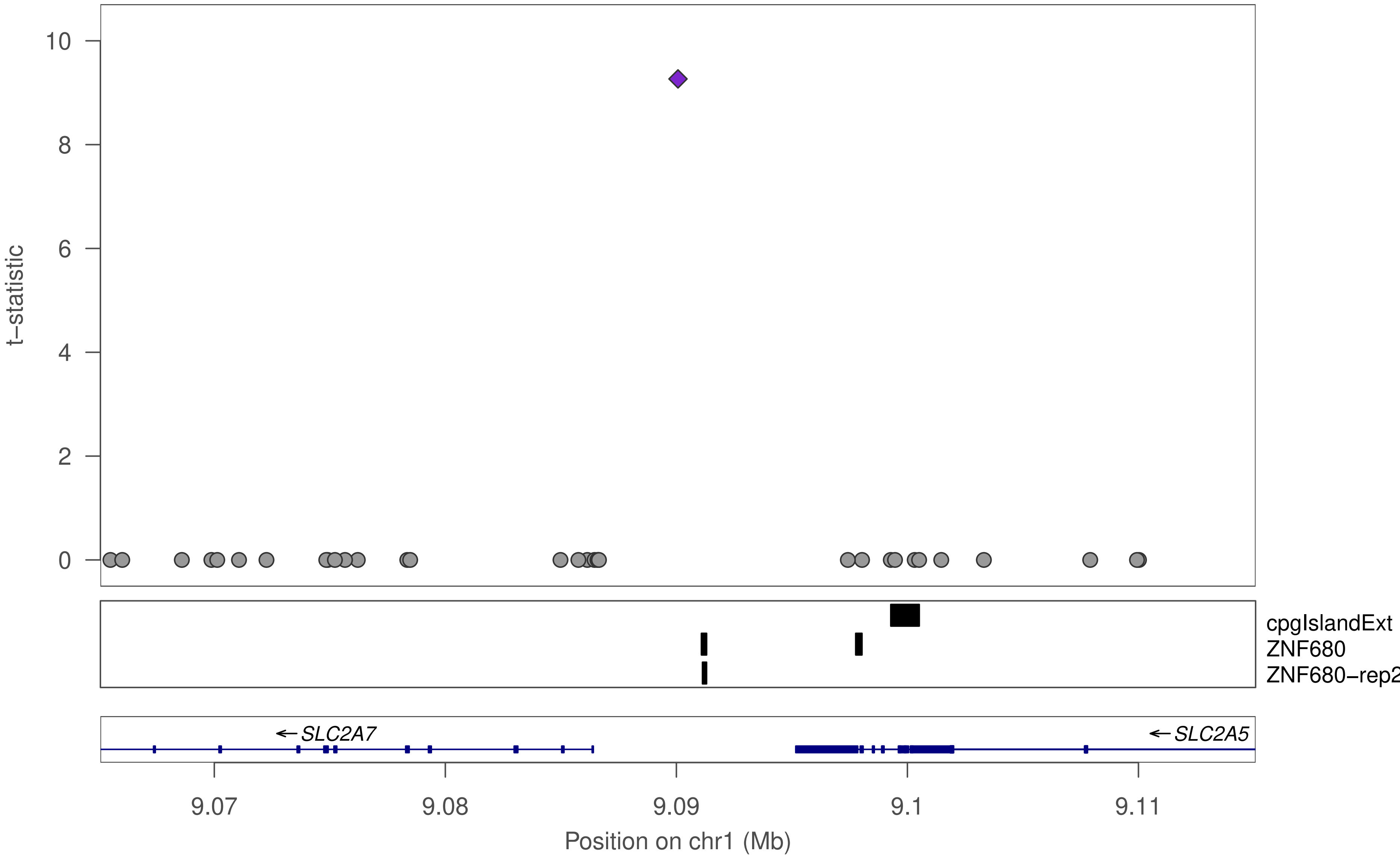

t−statistic
10
8
6
4
2
0
cpgIslandExt
ZNF680
ZNF680−rep2
SLC2A7
SLC2A5
9.07
9.08
9.09
9.1
9.11
Position on chr1 (Mb)

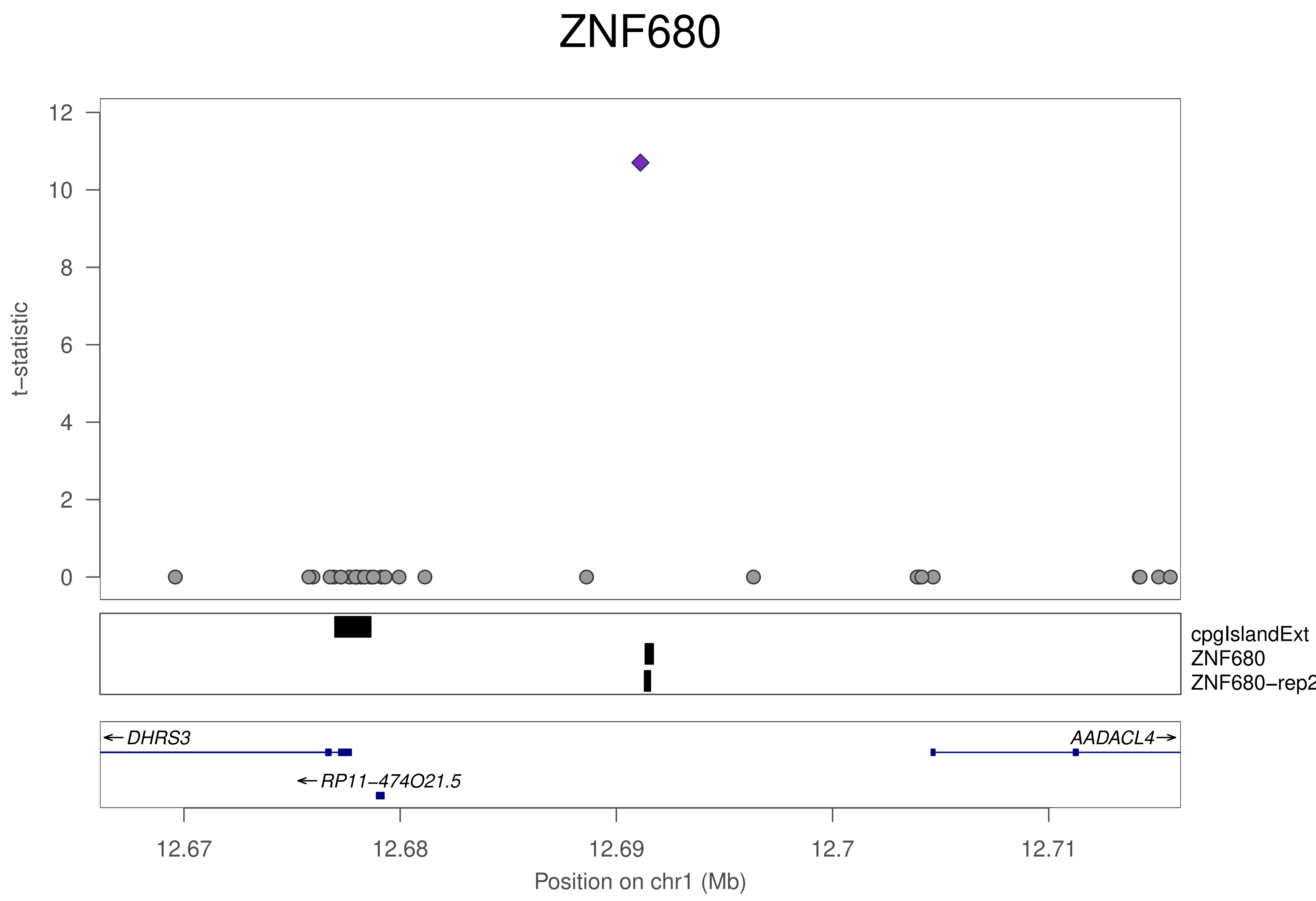
ZNF680
t−statistic
12
10
8
6
4
2
0
cpgIslandExt
ZNF680
ZNF680−rep2
DHRS3
AADACL4
RP11−474O21.5
12.67
12.68
12.69
12.7
12.71
Position on chr1 (Mb)

### ZNF680

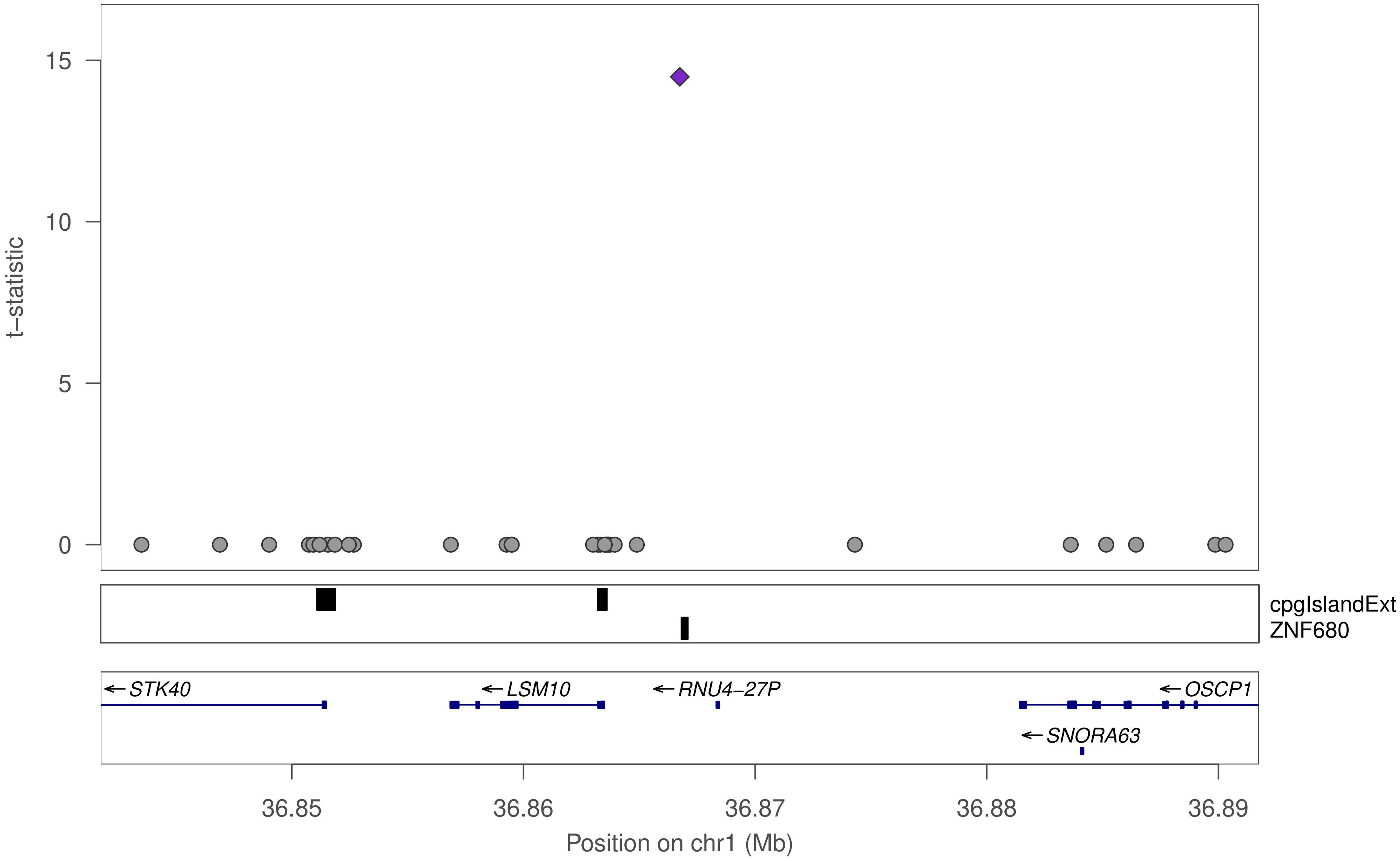

t−statistic
15
10
5
0
cpgIslandExt
ZNF680
STK40
LSM10
RNU4−27P
OSCP1
SNORA63
36.85
36.86
36.87
36.88
36.89
Position on chr1 (Mb)

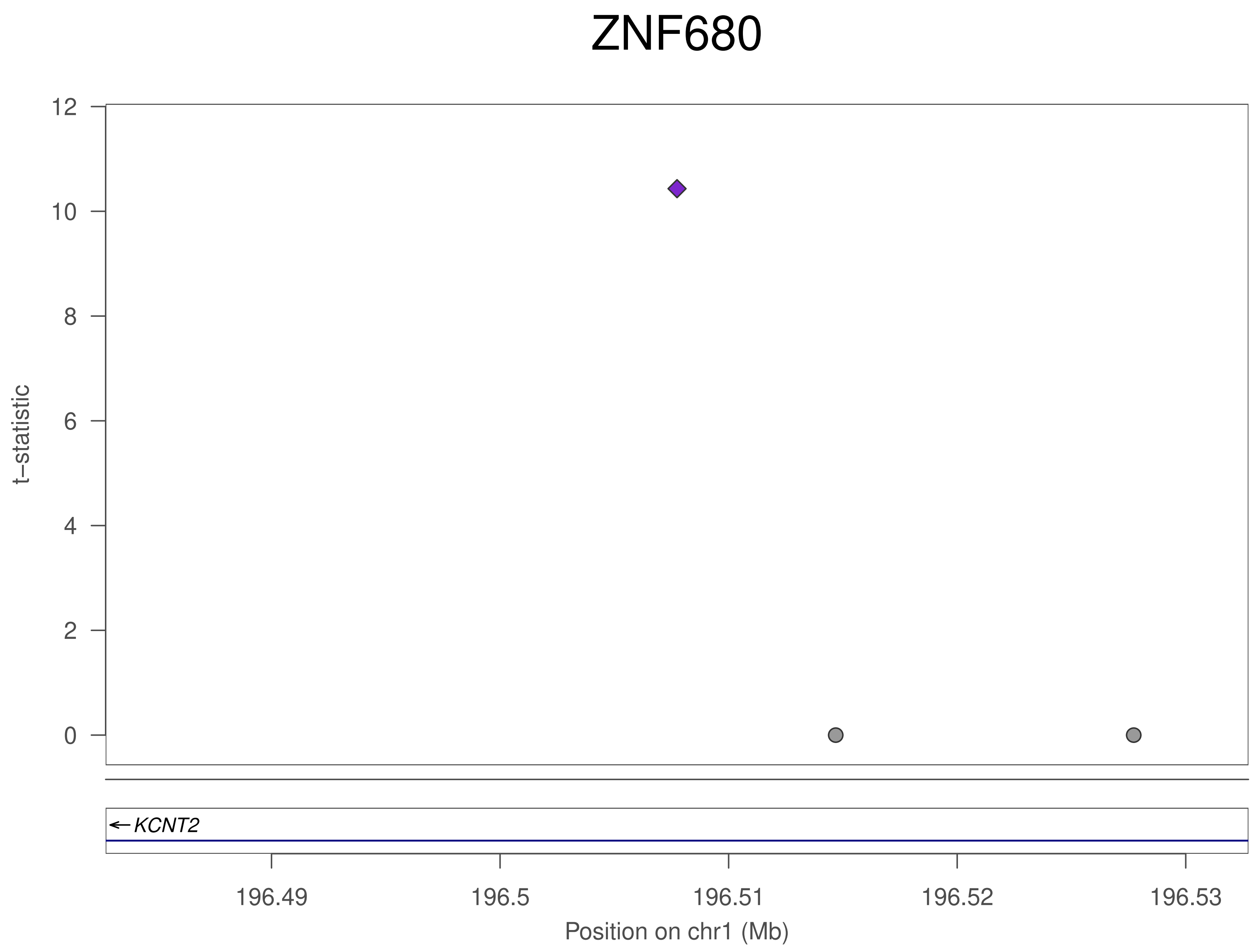
ZNF680
t–statistic
12
10
8
6
4
2
0
KCNT2
196.49
196.5
196.51
196.52
196.53
Position on chr1 (Mb)

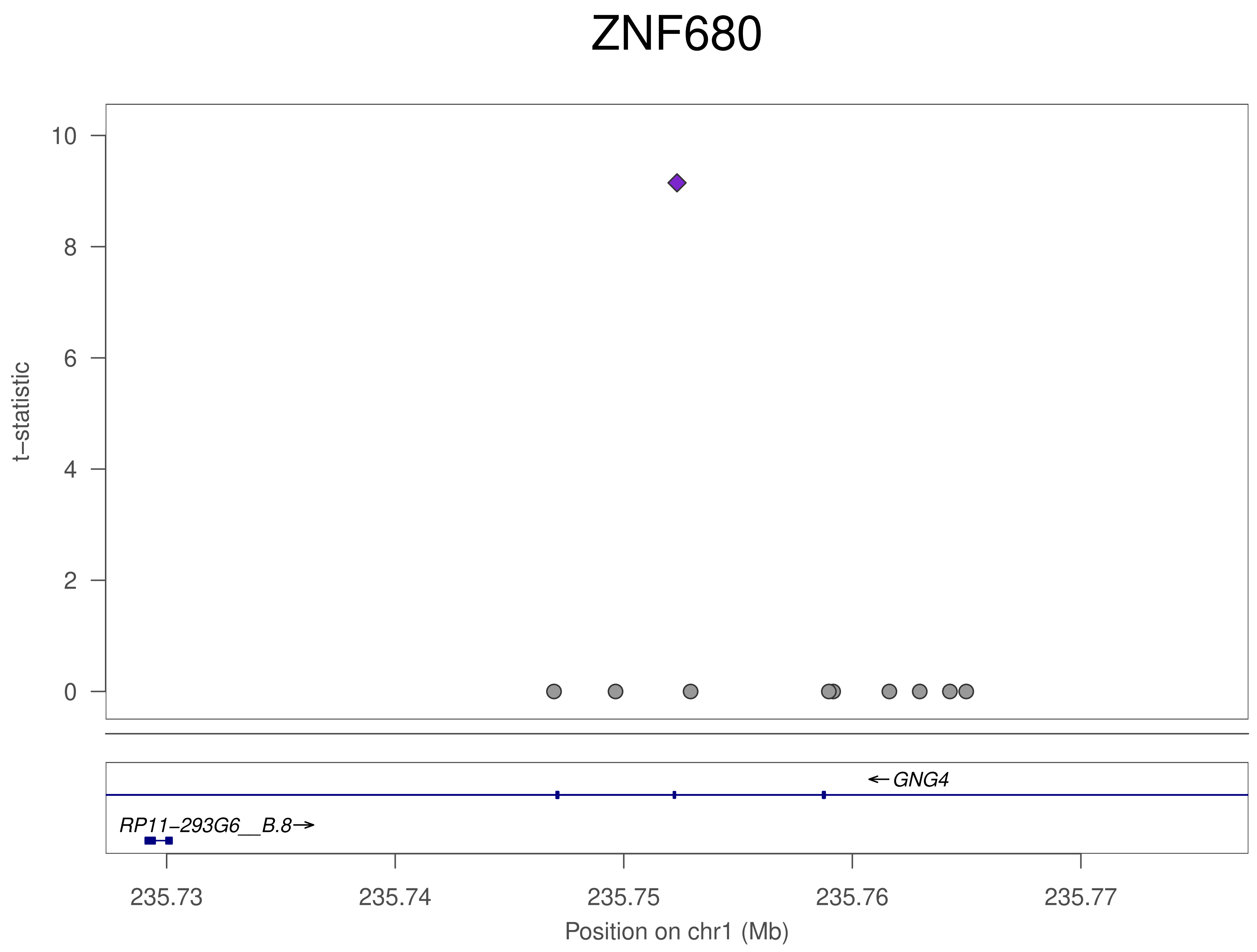
ZNF680
t−statistic
10
8
6
4
2
0
GNG4
RP11−293G6__B.8
235.73
235.74
235.75
235.76
235.77
Position on chr1 (Mb)

### ZNF680

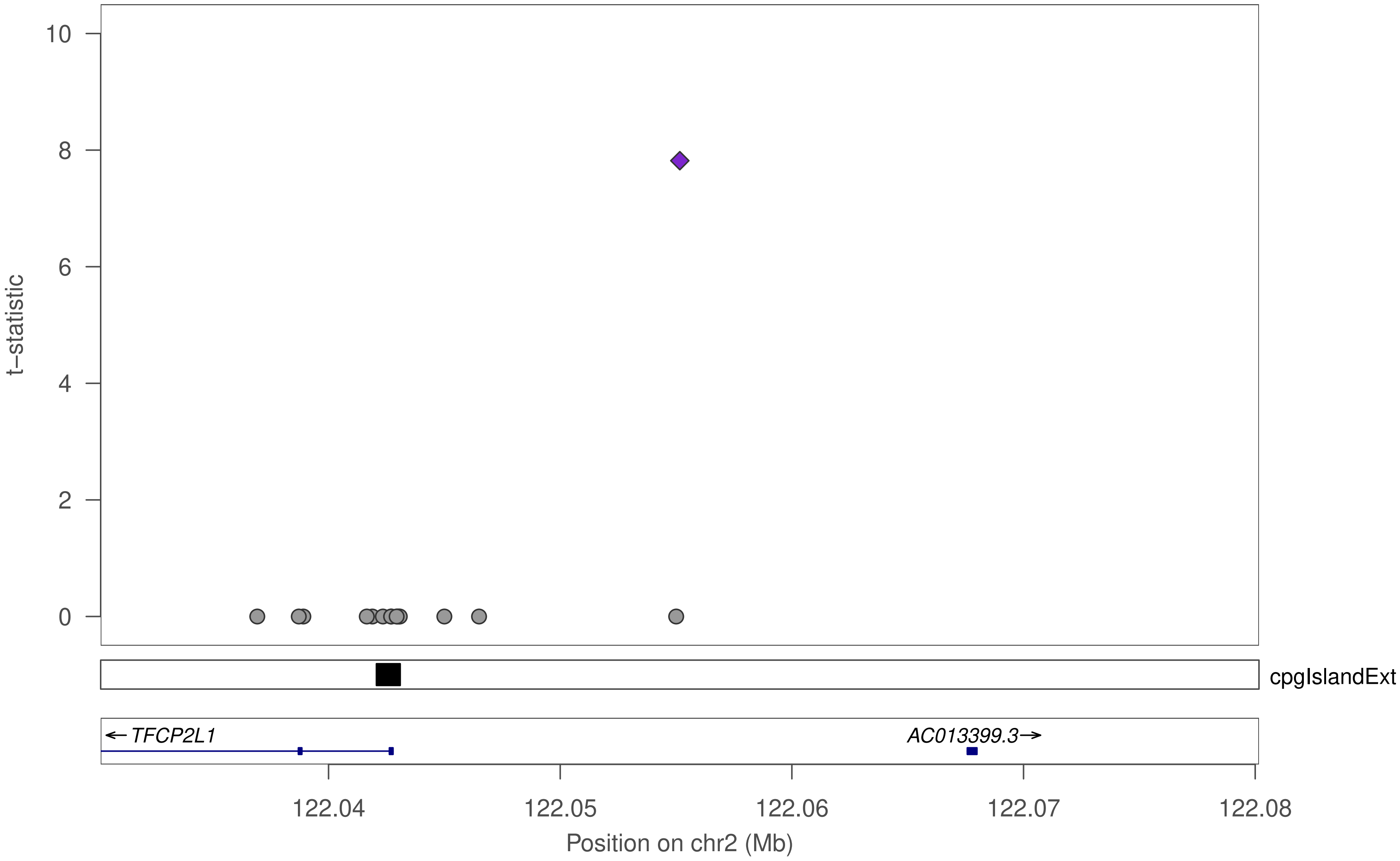

10
8
6
4
2
0
t−statistic
cpgIslandExt
← TFCP2L1
AC013399.3→
122.04
122.05
122.06
122.07
122.08
Position on chr2 (Mb)

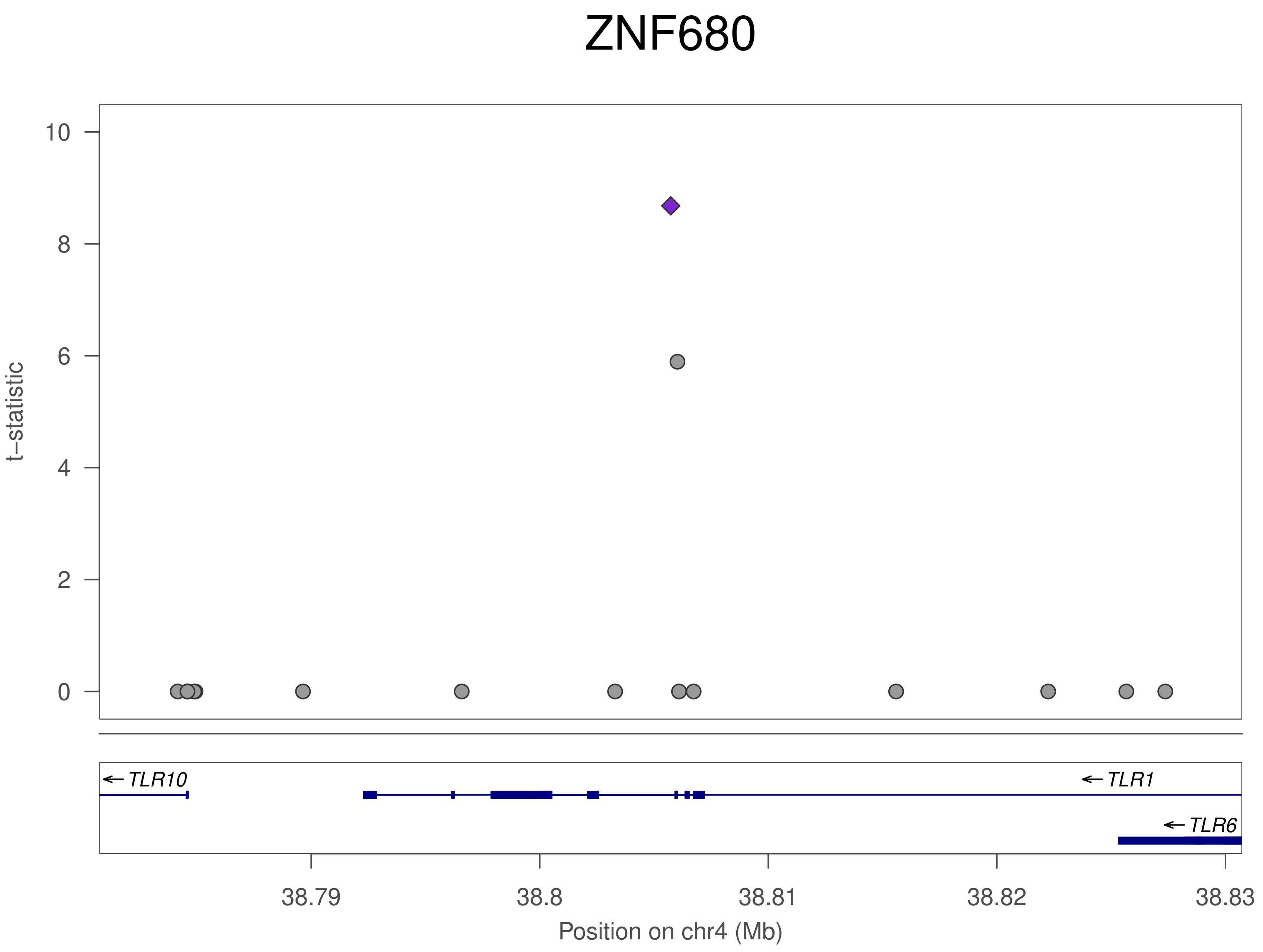

ZNF680
t–statistic
10
8
6
4
2
0
TLR10
TLR1
TLR6
38.79
38.8
38.81
38.82
38.83
Position on chr4 (Mb)

### ZNF680

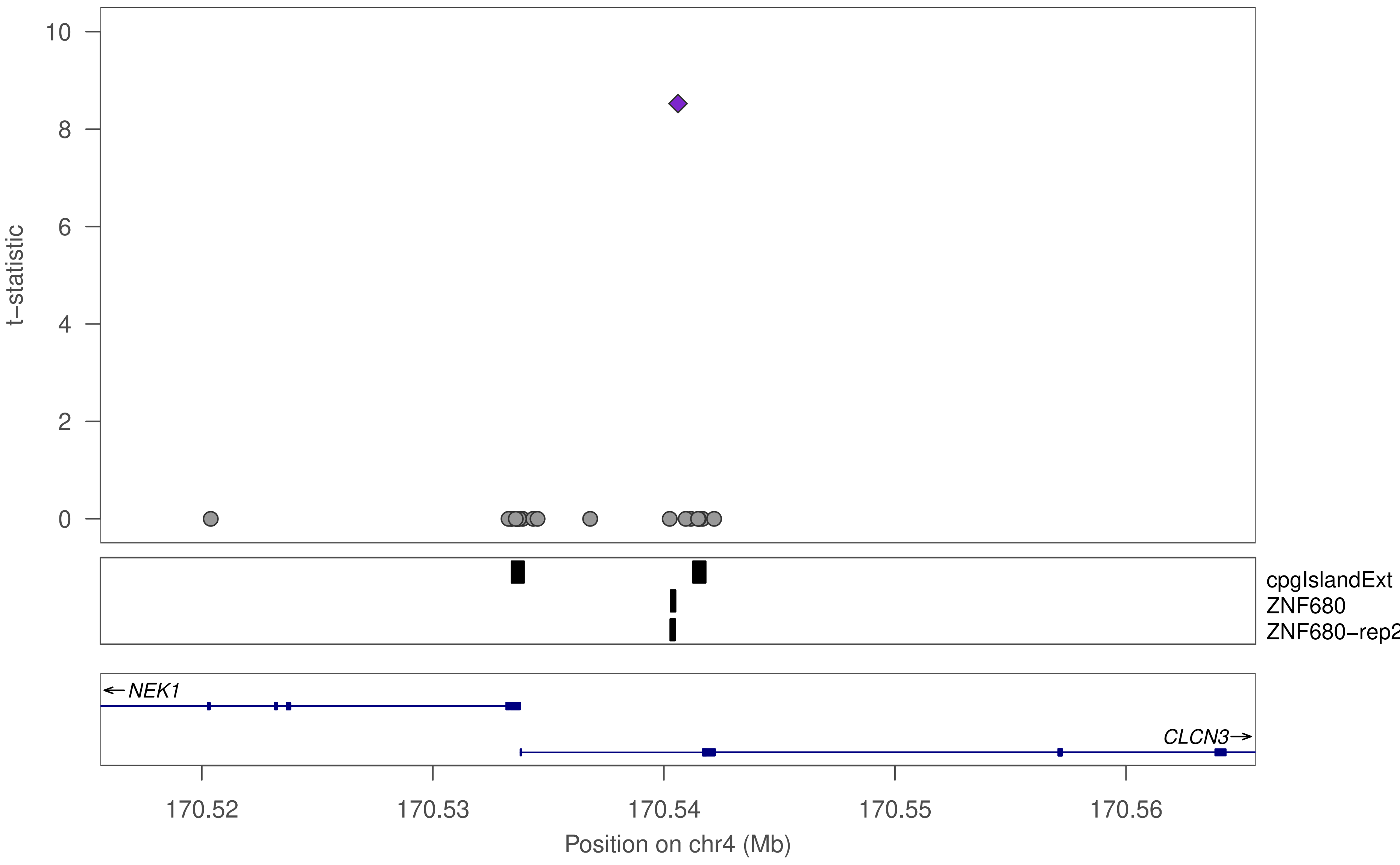

t−statistic
10
8
6
4
2
0
cpgIslandExt
ZNF680
ZNF680−rep2
NEK1
CLCN3
170.52
170.53
170.54
170.55
170.56
Position on chr4 (Mb)

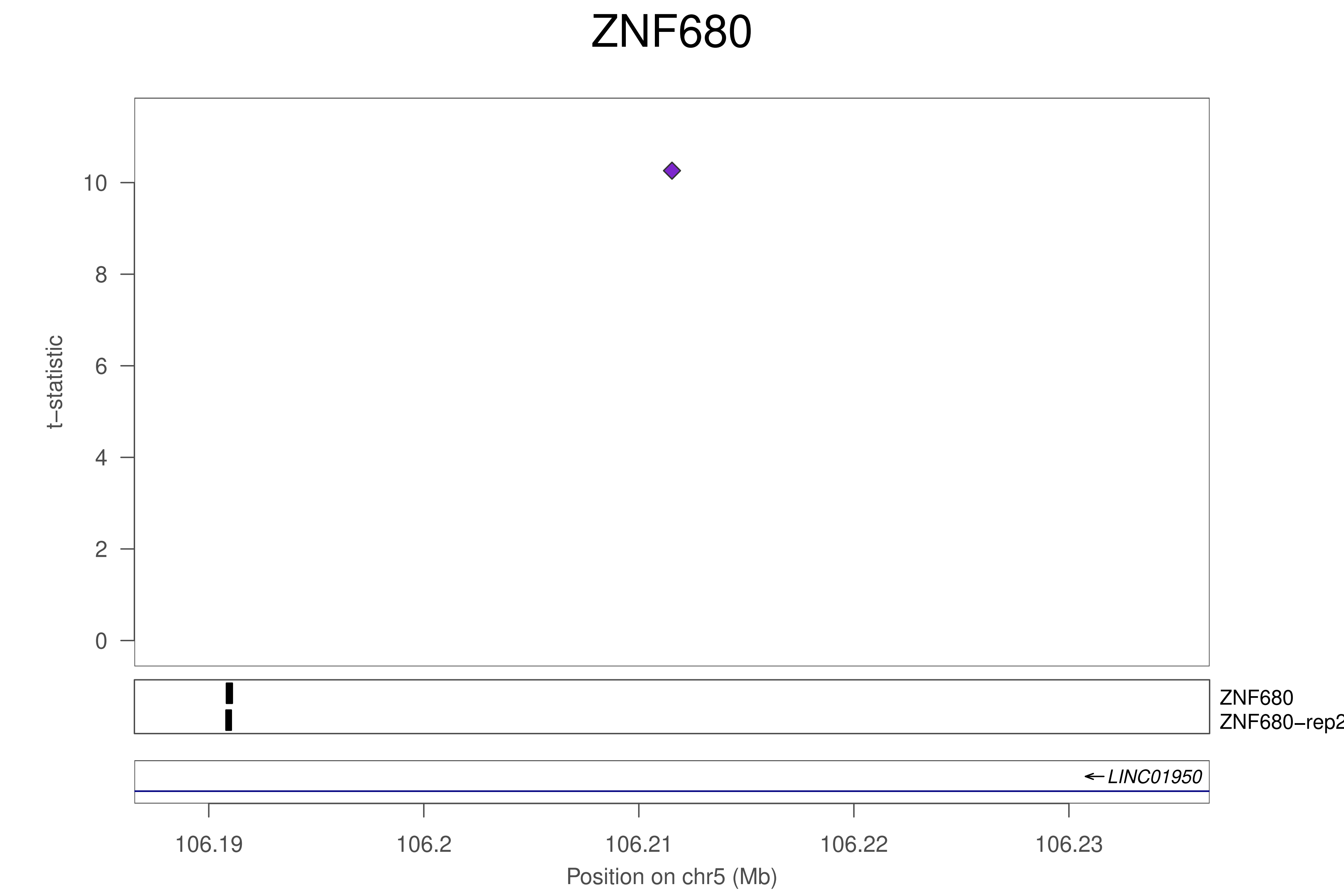

ZNF680
10
8
6
4
2
0
t−statistic
ZNF680
ZNF680−rep2
LINC01950
106.19
106.2
106.21
106.22
106.23
Position on chr5 (Mb)

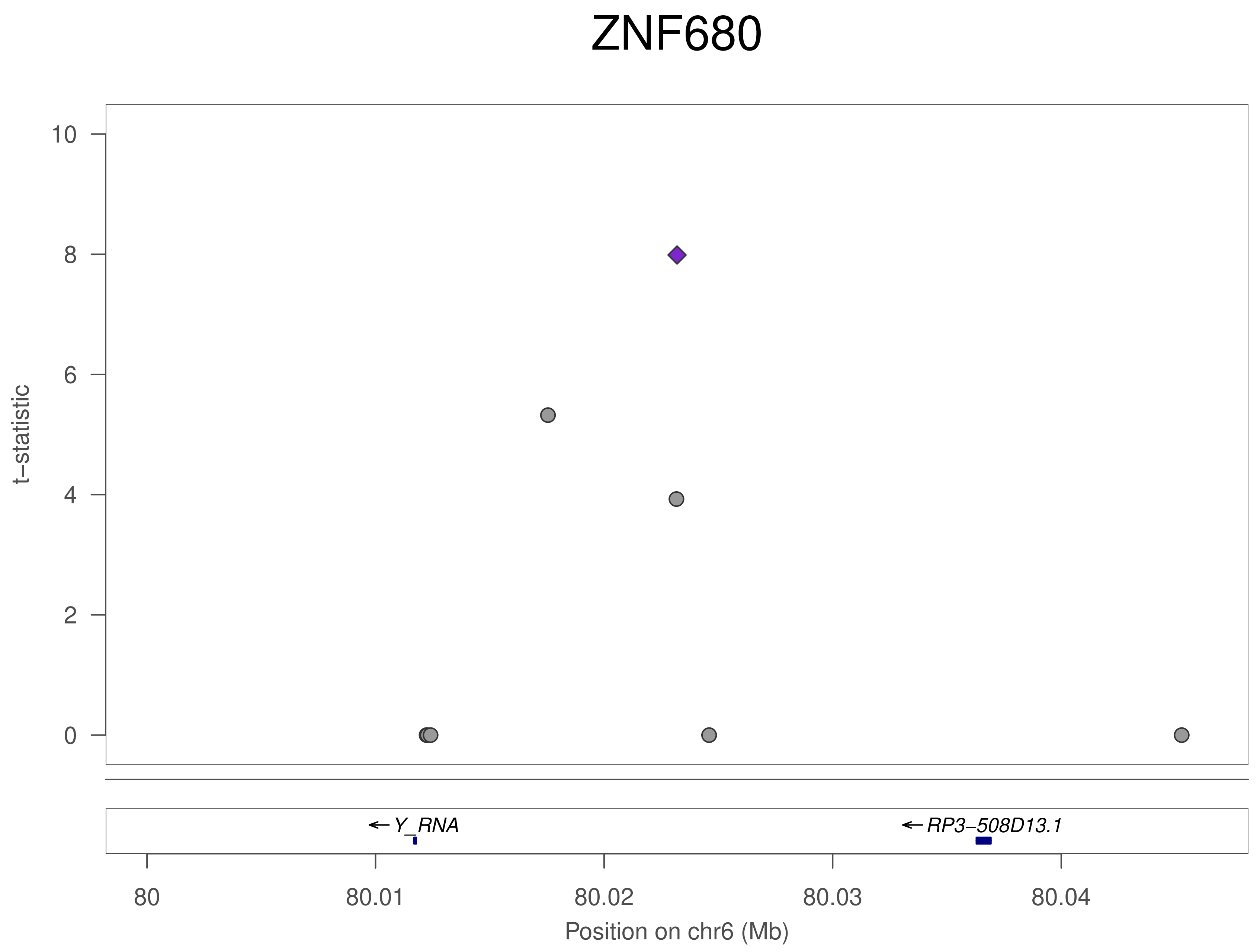

ZNF680
t−statistic
10
8
6
4
2
0
← Y_RNA
← RP3−508D13.1
80
80.01
80.02
80.03
80.04
Position on chr6 (Mb)

### ZNF680

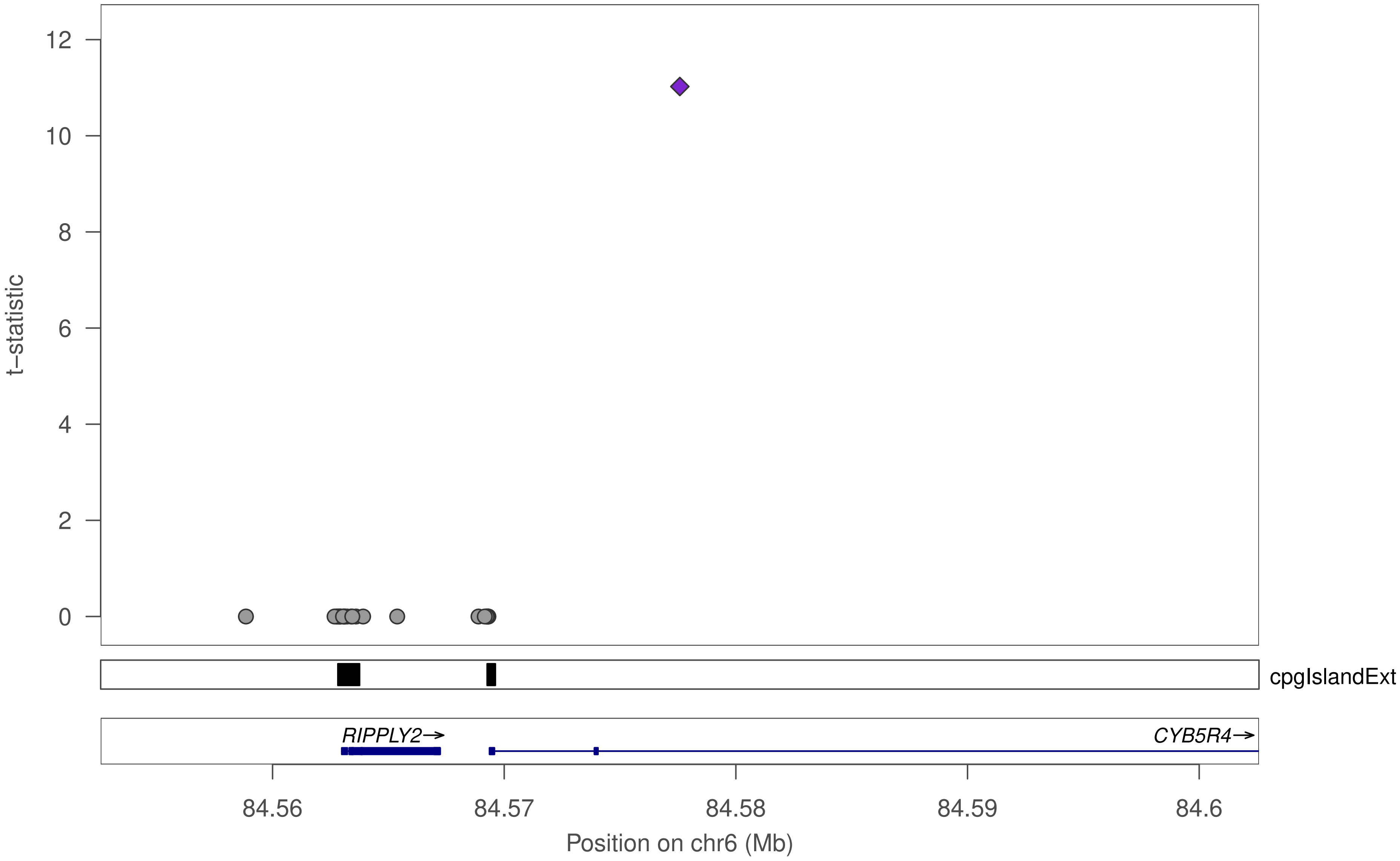

12
10
8
6
4
2
0
t–statistic
cpgIslandExt
RIPPLY2→
CYB5R4→
84.56
84.57
84.58
84.59
84.6
Position on chr6 (Mb)

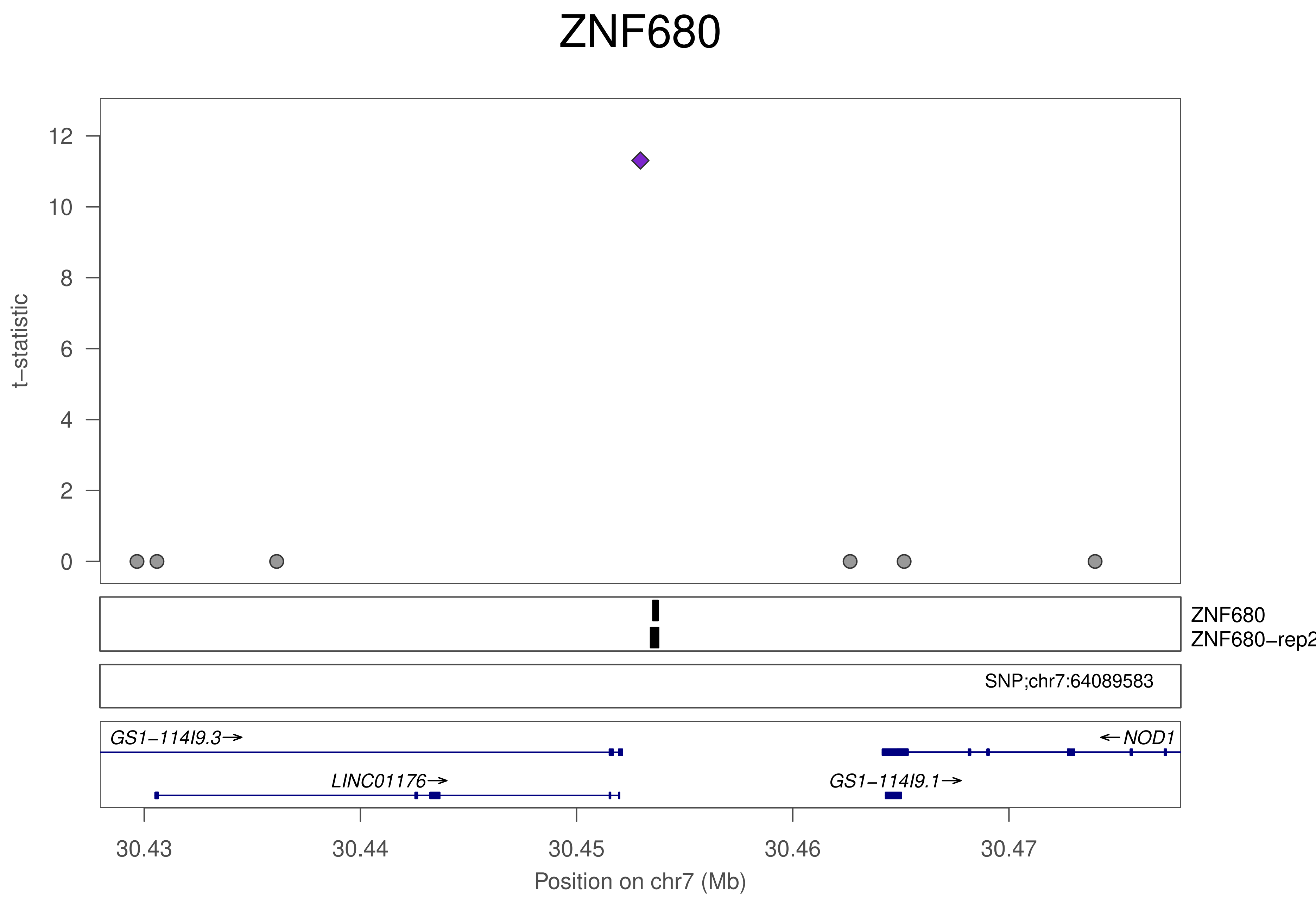

ZNF680
t−statistic
12
10
8
6
4
2
0
ZNF680
ZNF680−rep2
SNP;chr7:64089583
GS1−114I9.3→
←NOD1
LINC01176→
GS1−114I9.1→
30.43
30.44
30.45
30.46
30.47
Position on chr7 (Mb)

### ZNF680

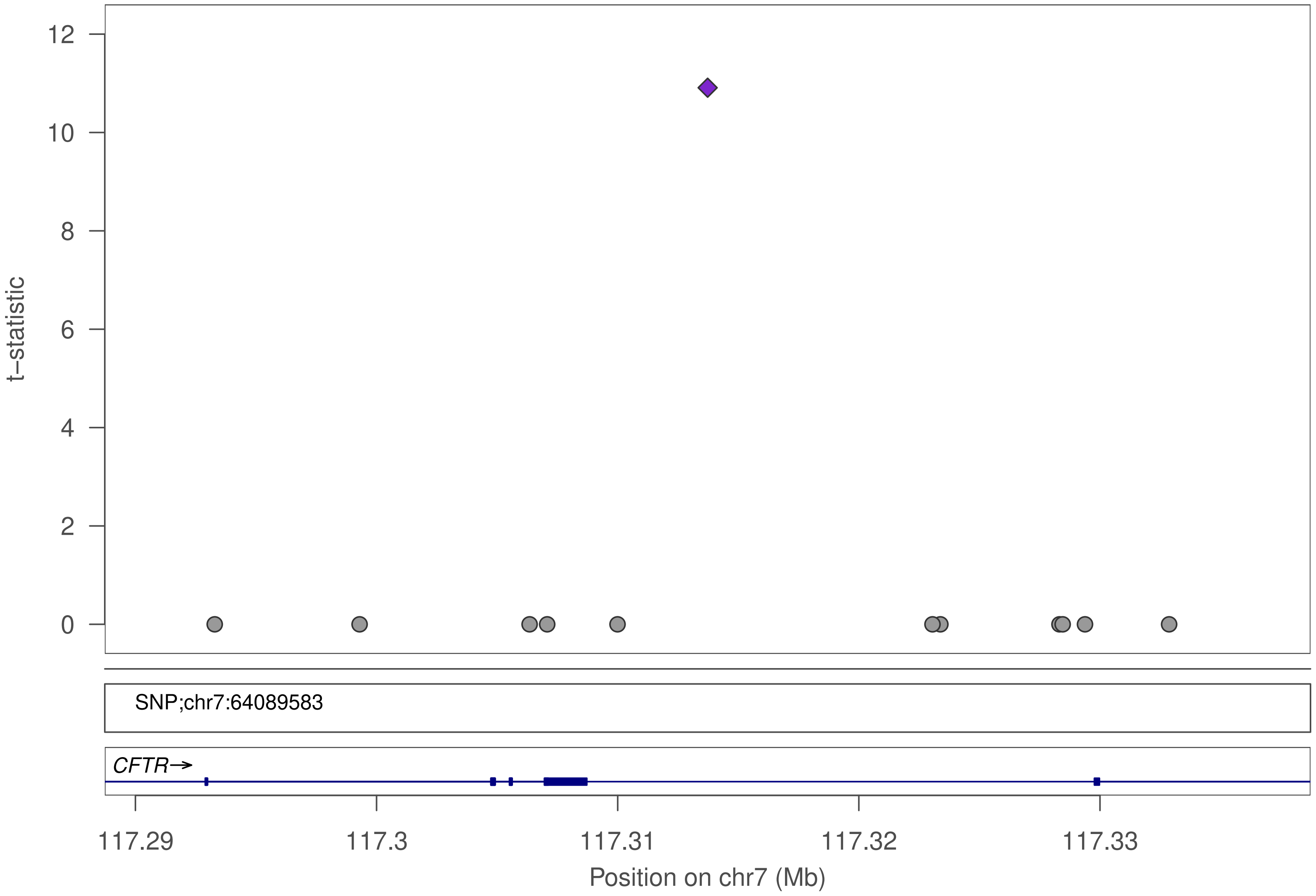

t–statistic
12
10
8
6
4
2
0
SNP;chr7:64089583
CFTR→
117.29
117.3
117.31
117.32
117.33
Position on chr7 (Mb)

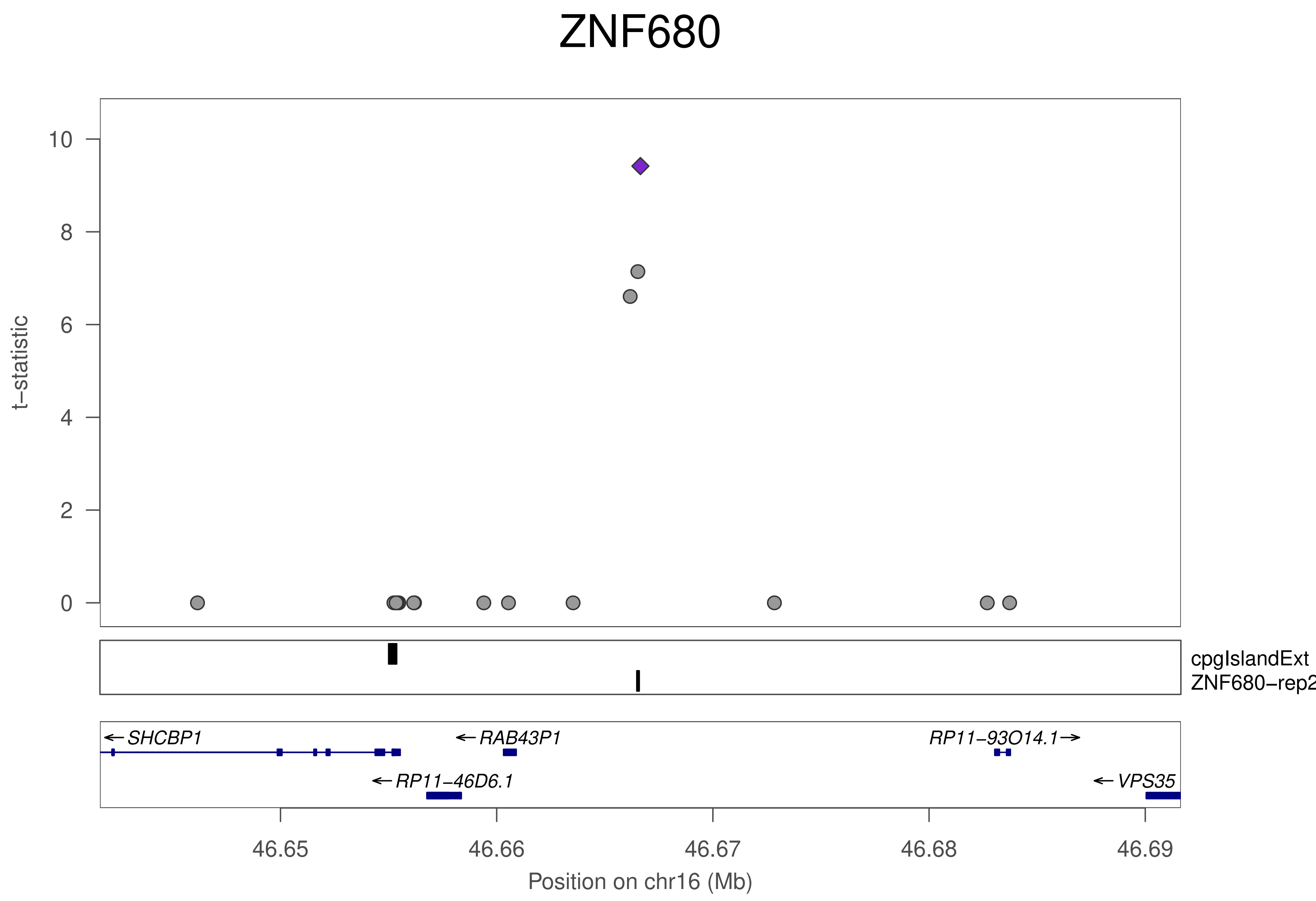

ZNF680
t−statistic
10
8
6
4
2
0
cpgIslandExt
ZNF680−rep2
SHCBP1
RAB43P1
RP11−93O14.1
RP11−46D6.1
VPS35
46.65
46.66
46.67
46.68
46.69
Position on chr16 (Mb)

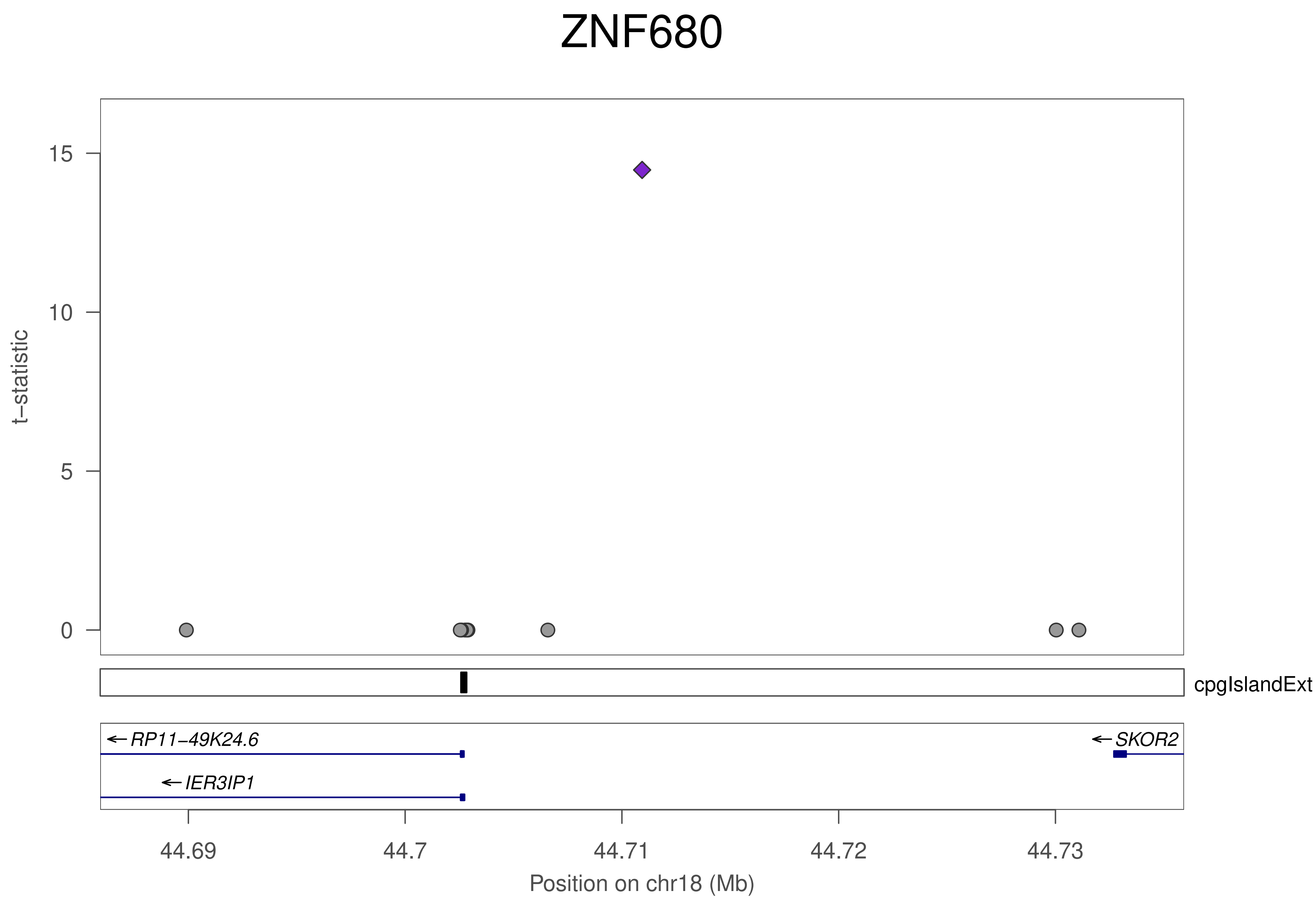
ZNF680
t−statistic
15
10
5
0
cpgIslandExt
← RP11−49K24.6
← IER3IP1
← SKOR2
44.69
44.7
44.71
44.72
44.73
Position on chr18 (Mb)

### ZNF680

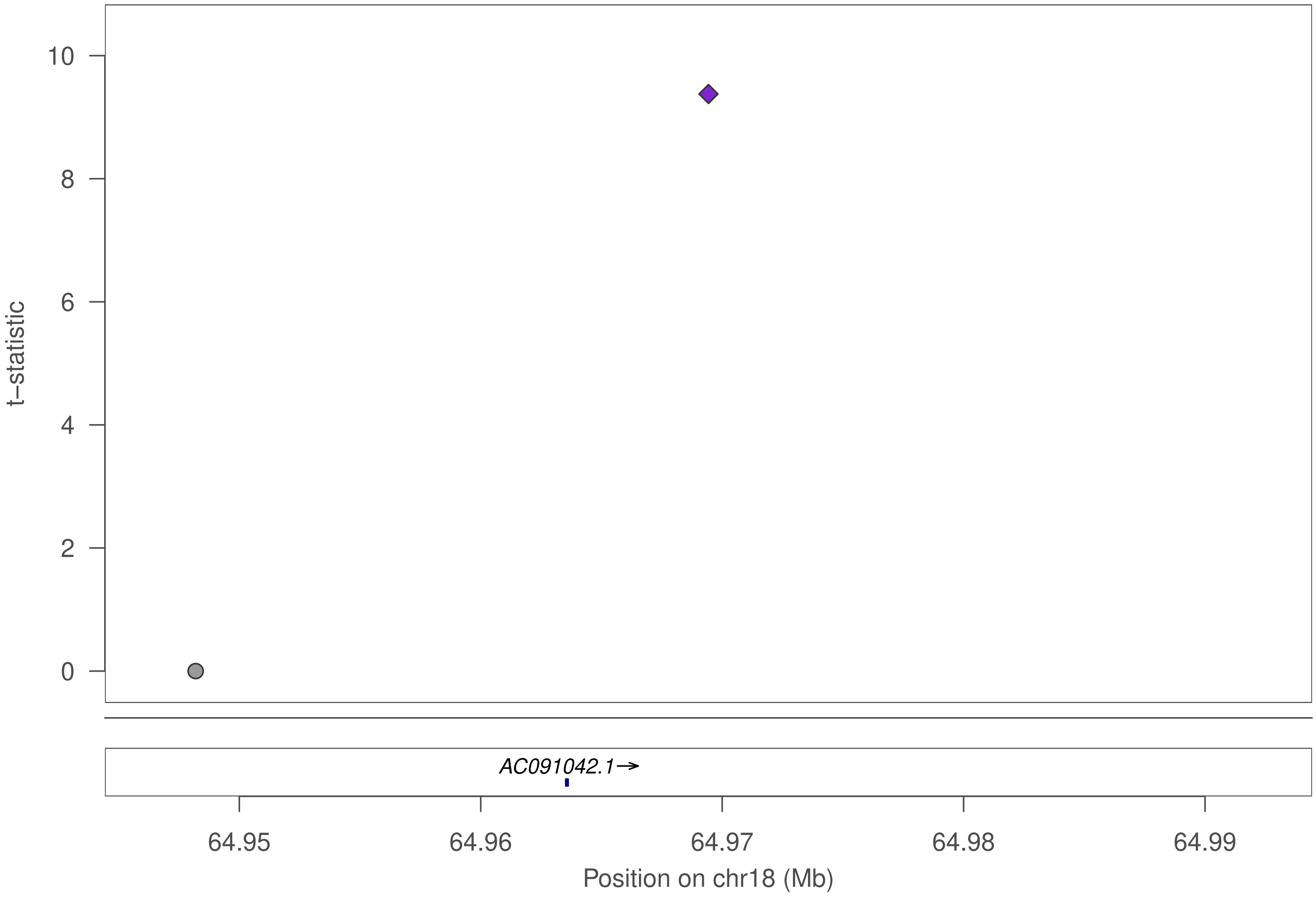

10
8
6
4
2
0
t−statistic
AC091042.1→
64.95
64.96
64.97
64.98
64.99
Position on chr18 (Mb)

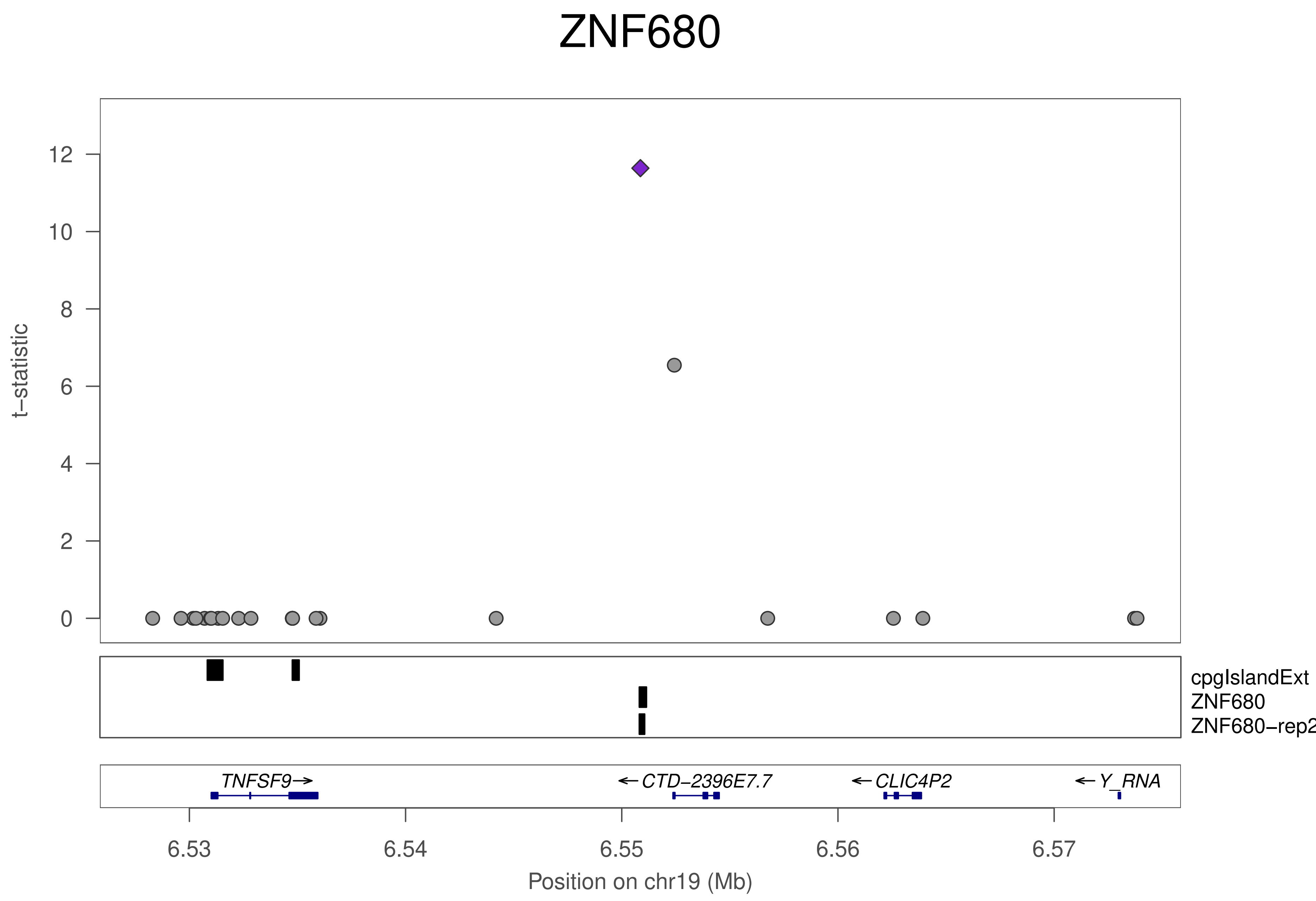
ZNF680
t−statistic
12
10
8
6
4
2
0
cpgIslandExt
ZNF680
ZNF680−rep2
TNFSF9→
← CTD−2396E7.7
← CLIC4P2
← Y_RNA
6.53
6.54
6.55
6.56
6.57
Position on chr19 (Mb)

### ZNF680

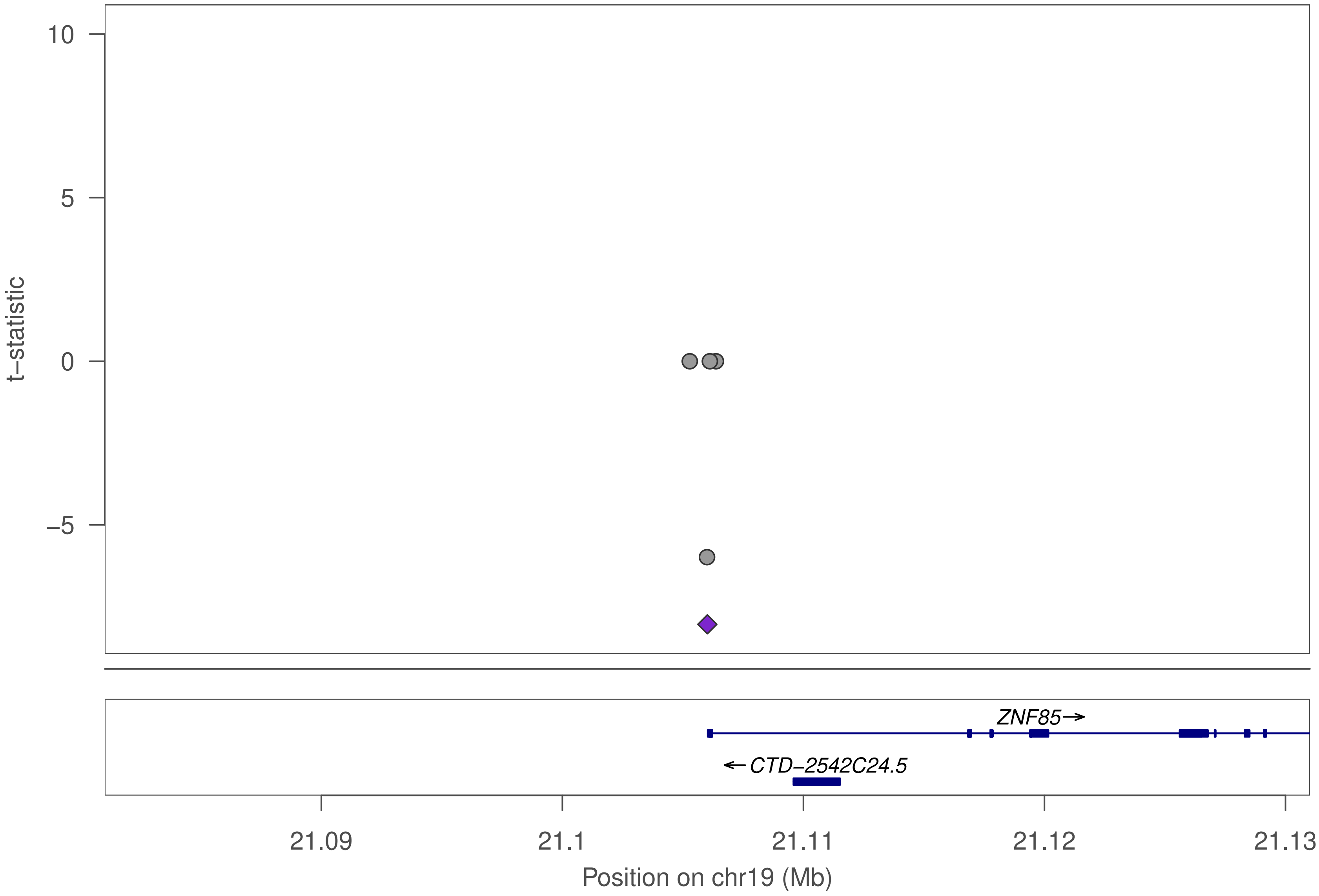

10
5
0
−5
t−statistic
ZNF85→
←CTD−2542C24.5
21.09
21.1
21.11
21.12
21.13
Position on chr19 (Mb)

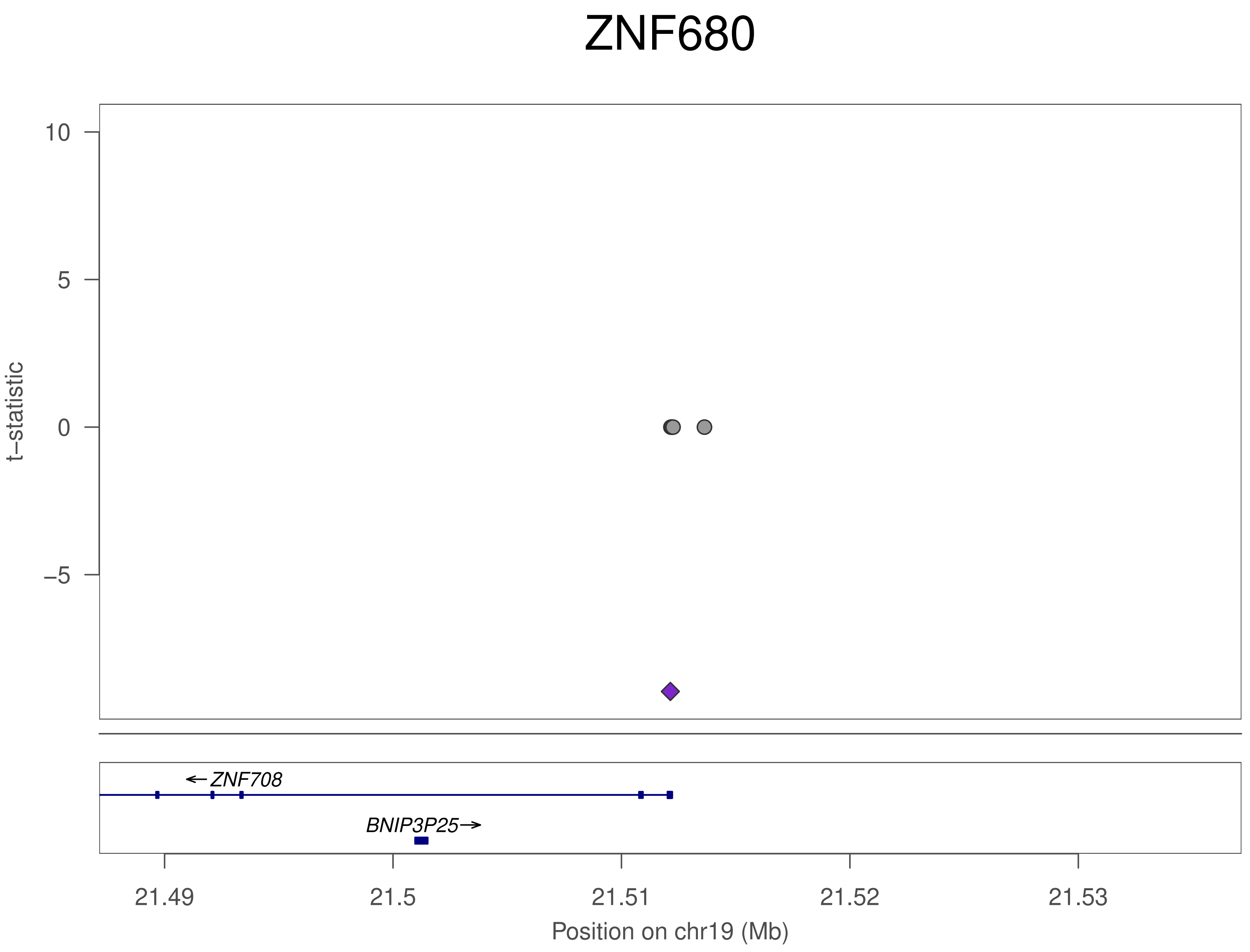
ZNF680
t−statistic
10
5
0
−5
ZNF708
BNIP3P25
21.49
21.5
21.51
21.52
21.53
Position on chr19 (Mb)

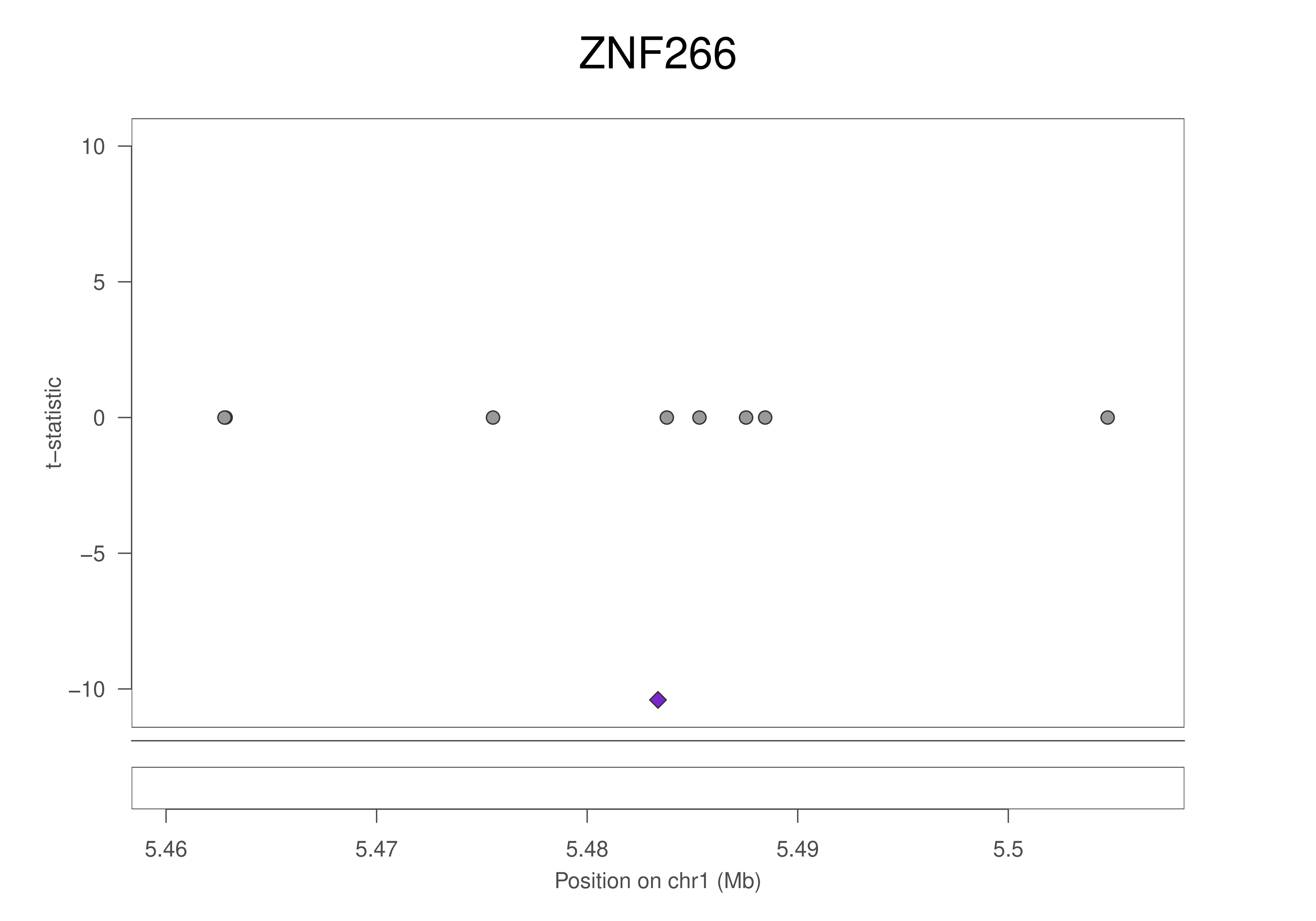
ZNF266
t−statistic
10
5
0
−5
−10
5.46
5.47
5.48
5.49
5.5
Position on chr1 (Mb)

### ZNF266

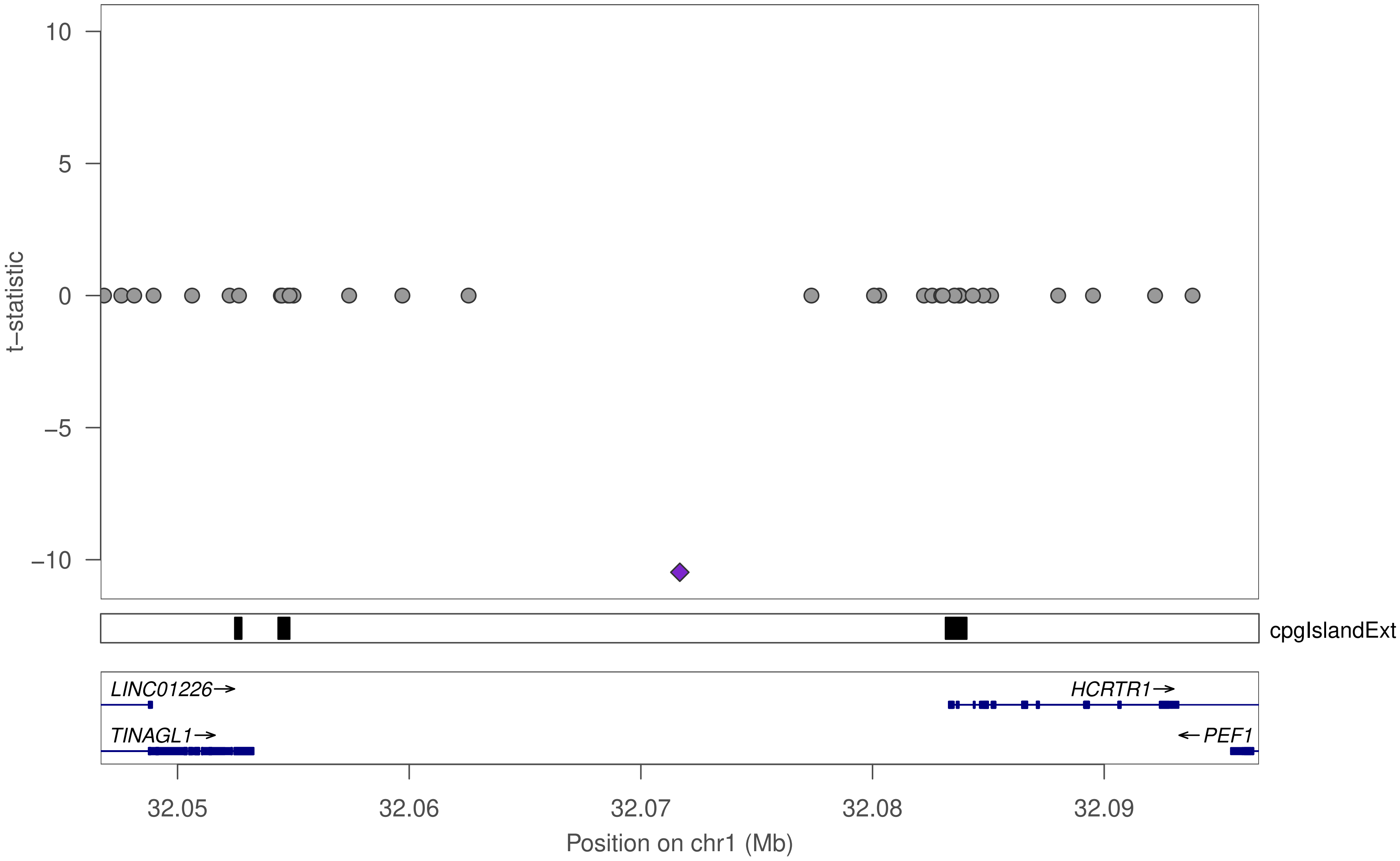

10
5
0
−5
−10
t−statistic
cpgIslandExt
LINC01226→
TINAGL1→
HCRTR1→
←PEF1
32.05
32.06
32.07
32.08
32.09
Position on chr1 (Mb)

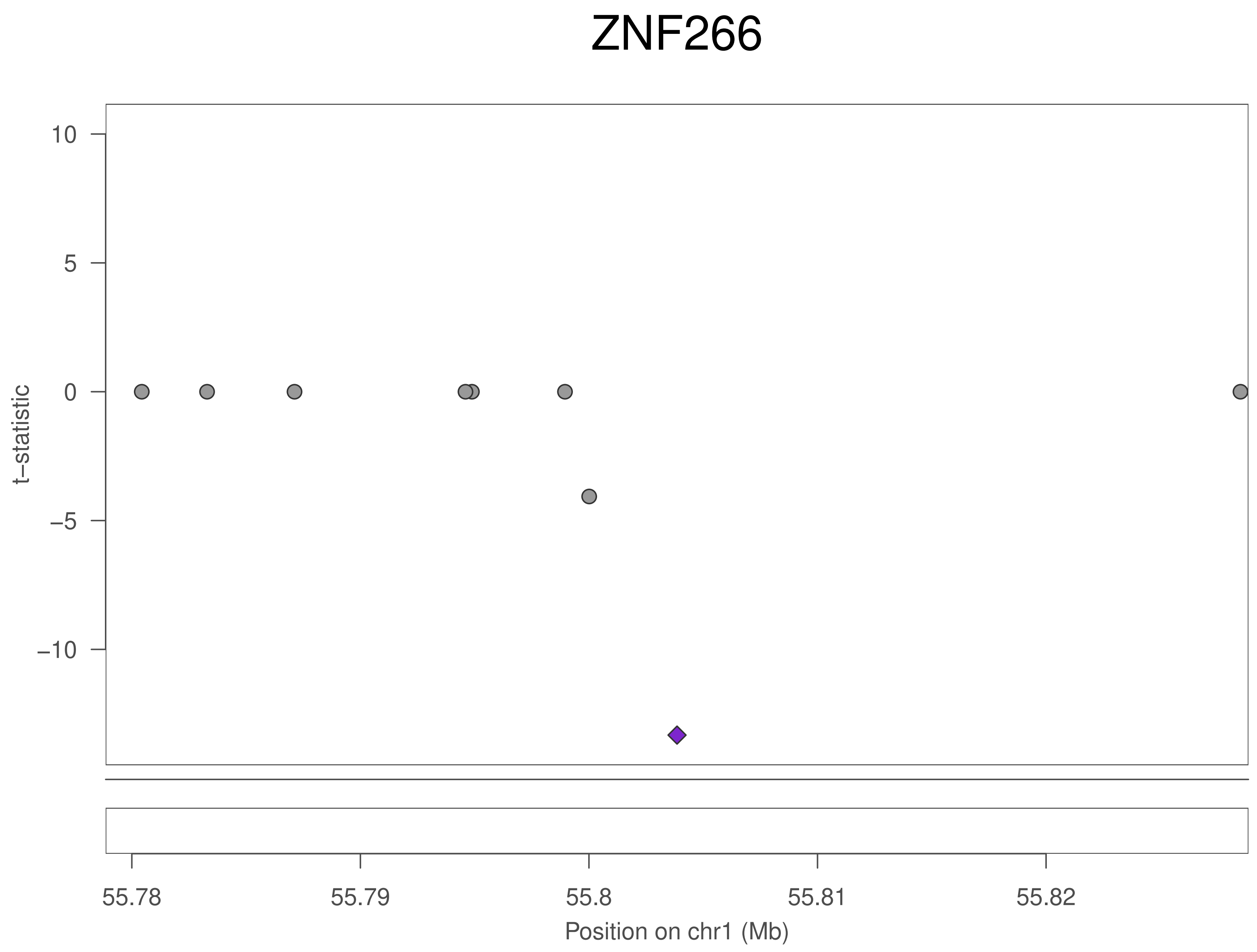
ZNF266
t–statistic
10
5
0
−5
−10
55.78
55.79
55.8
55.81
55.82
Position on chr1 (Mb)

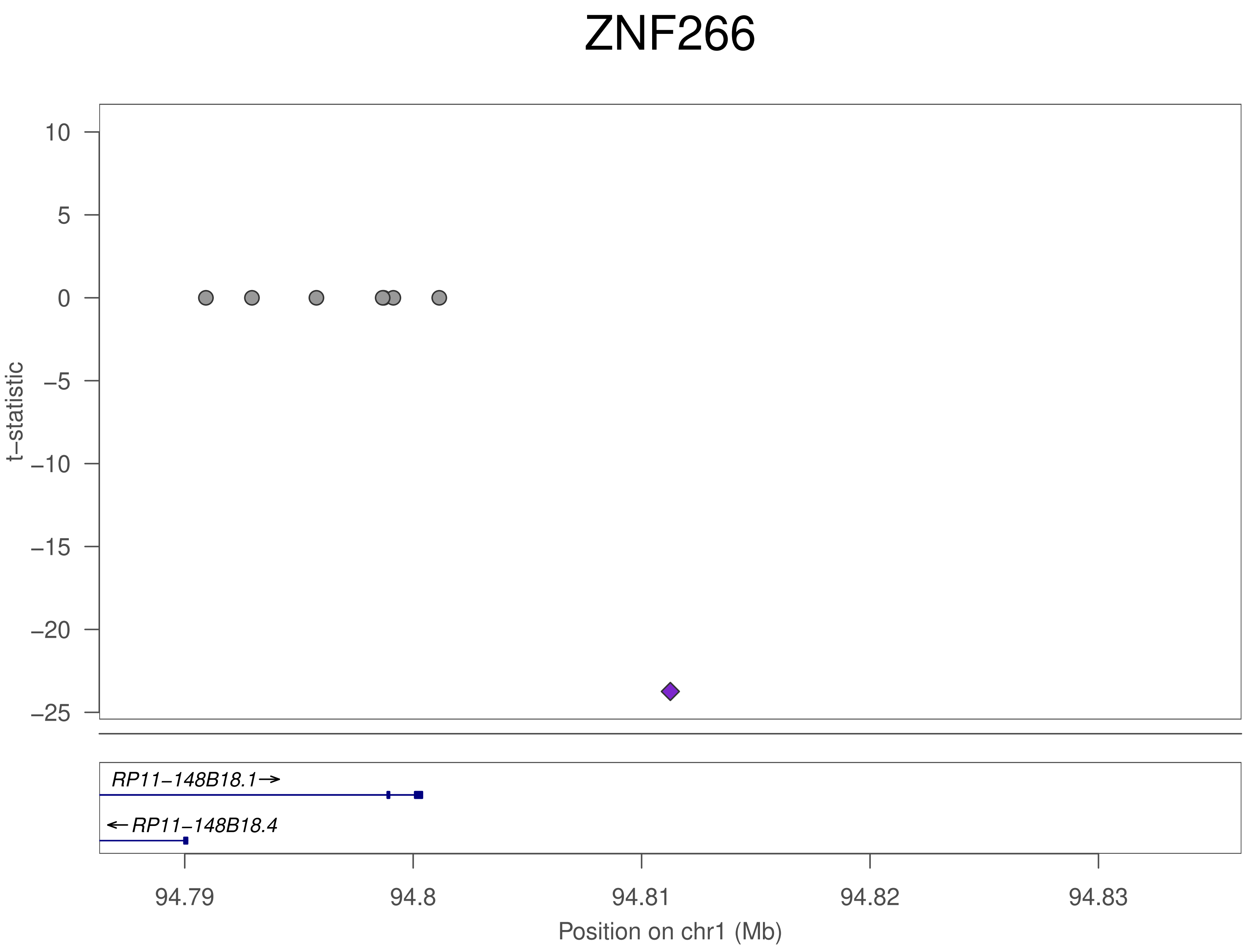
ZNF266
t−statistic
10
5
0
−5
−10
−15
−20
−25
RP11−148B18.1→
←RP11−148B18.4
94.79
94.8
94.81
94.82
94.83
Position on chr1 (Mb)

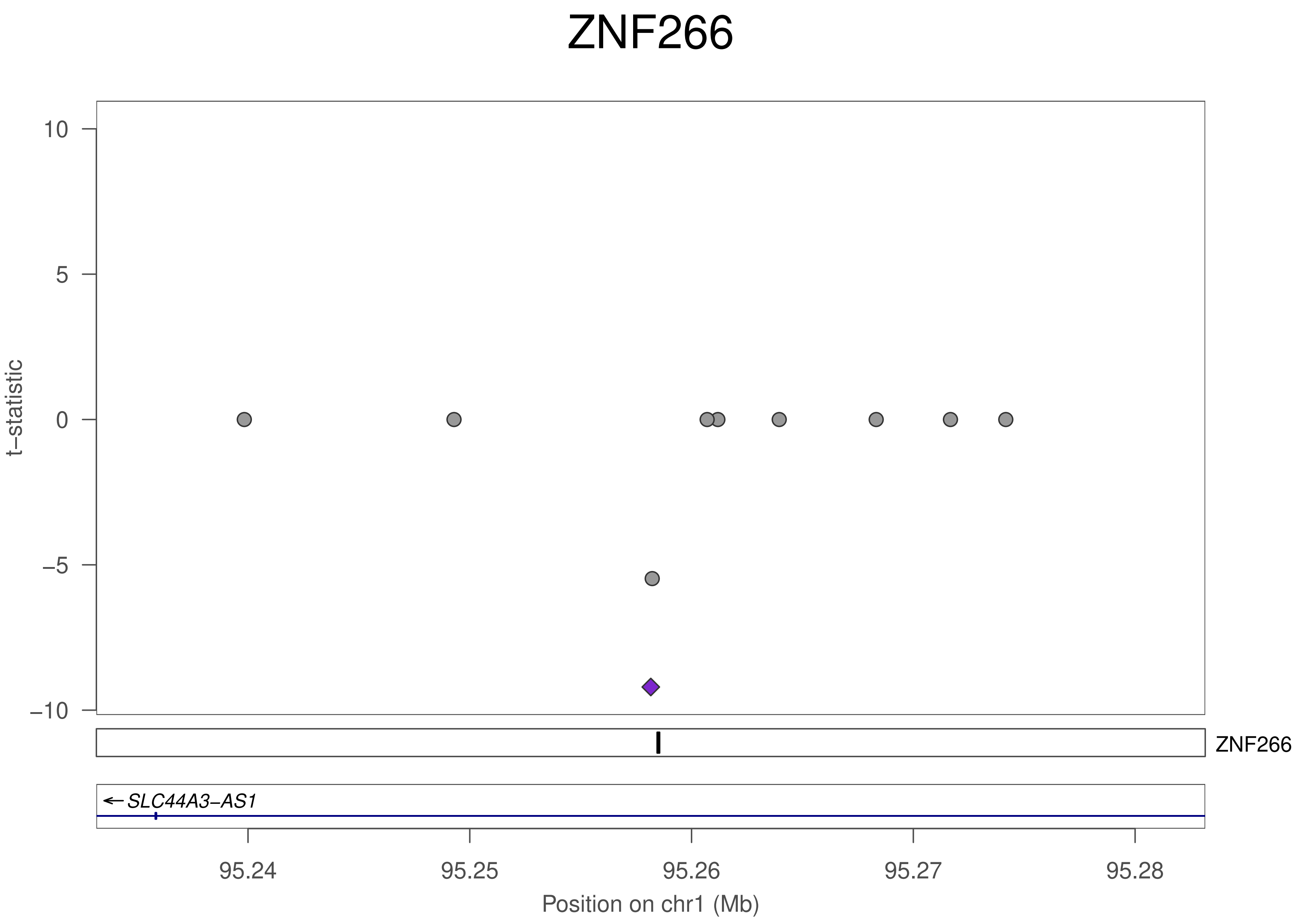
ZNF266
t−statistic
10
5
0
−5
−10
ZNF266
SLC44A3−AS1
95.24
95.25
95.26
95.27
95.28
Position on chr1 (Mb)

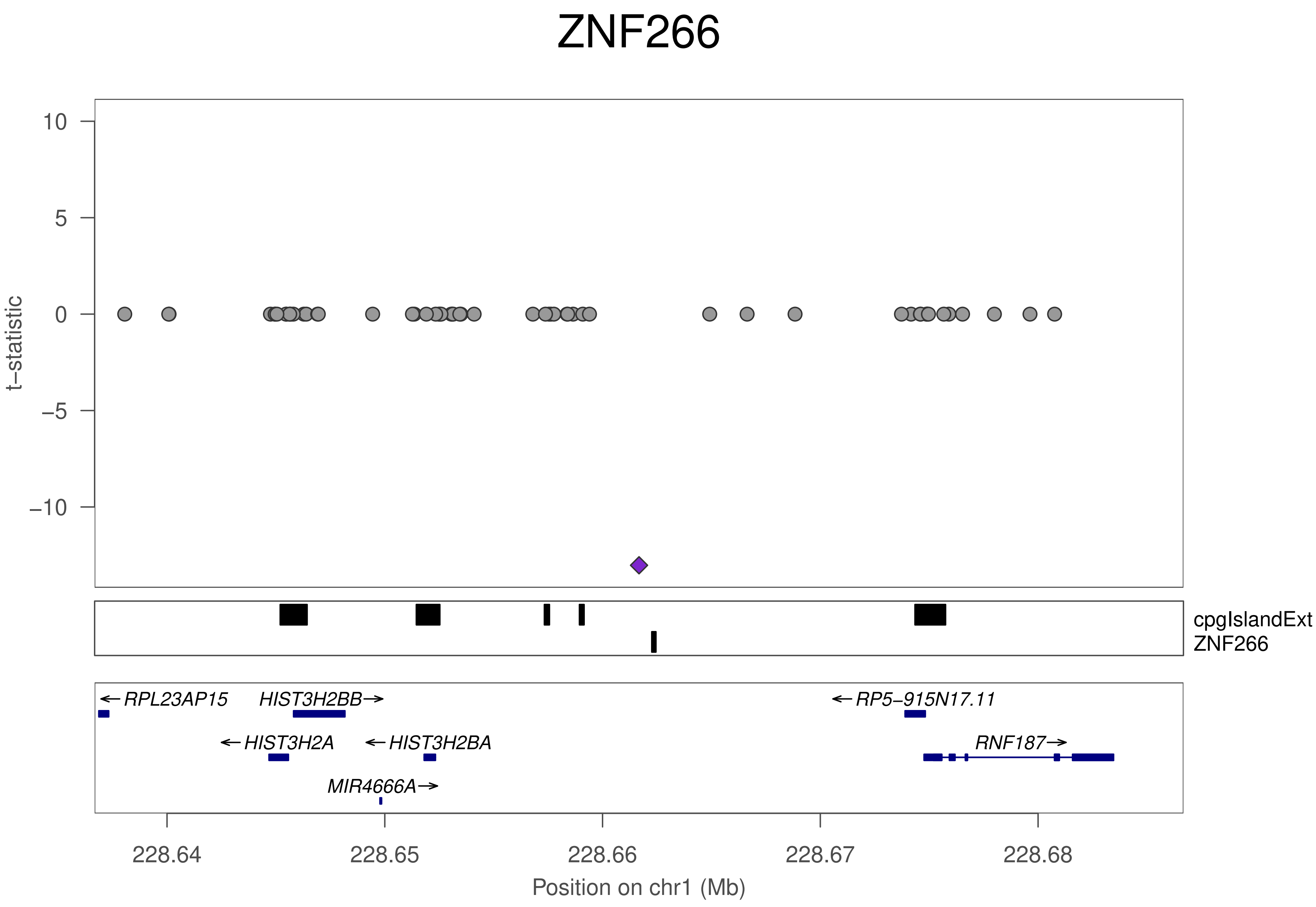

ZNF266
10
5
0
−5
−10
t−statistic
cpgIslandExt
ZNF266
←RPL23AP15
HIST3H2BB→
←RP5−915N17.11
←HIST3H2A
←HIST3H2BA
RNF187→
MIR4666A→
228.64
228.65
228.66
228.67
228.68
Position on chr1 (Mb)

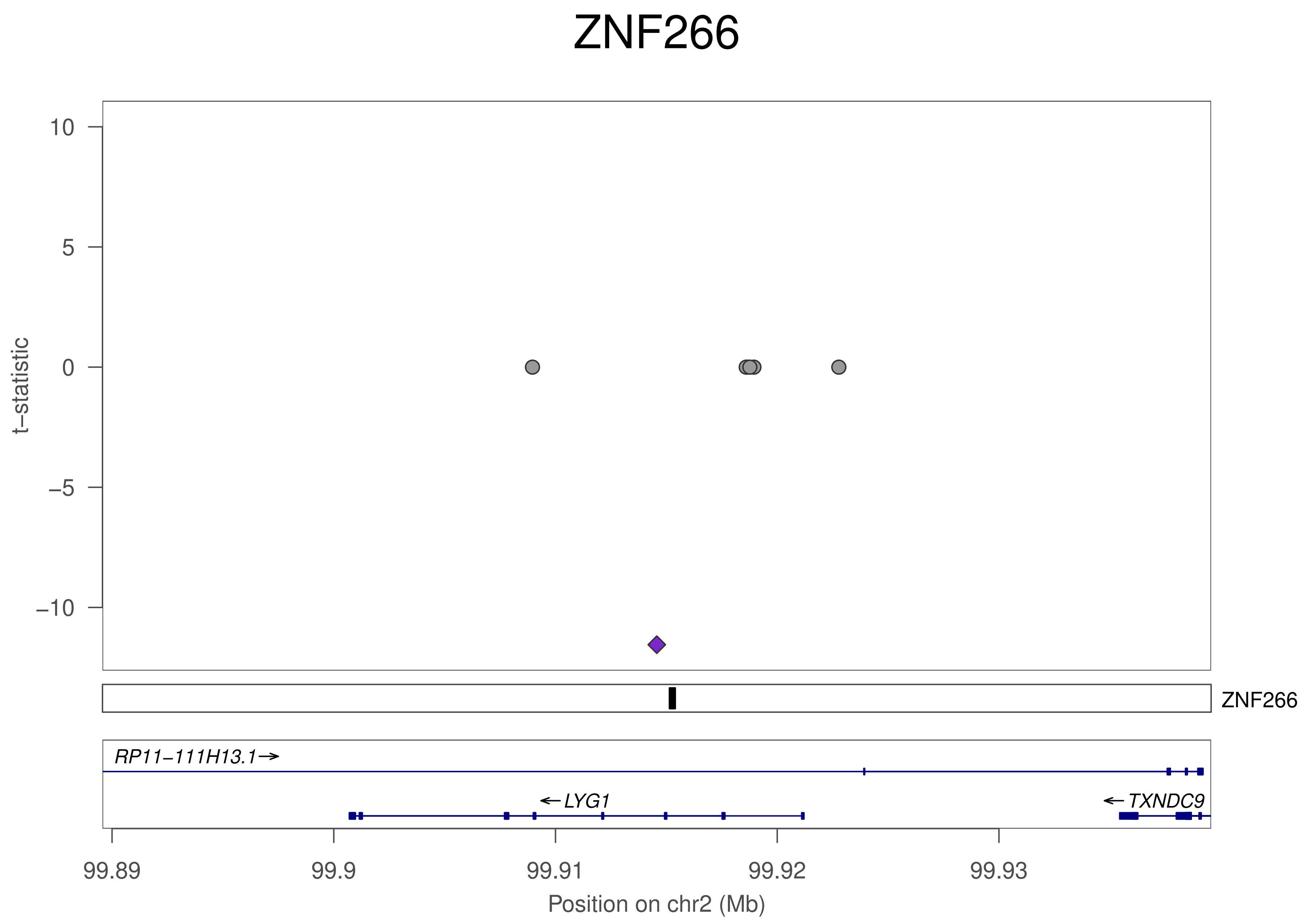

ZNF266
t−statistic
10
5
0
−5
−10
ZNF266
RP11−111H13.1→
←LYG1
←TXNDC9
99.89
99.9
99.91
99.92
99.93
Position on chr2 (Mb)

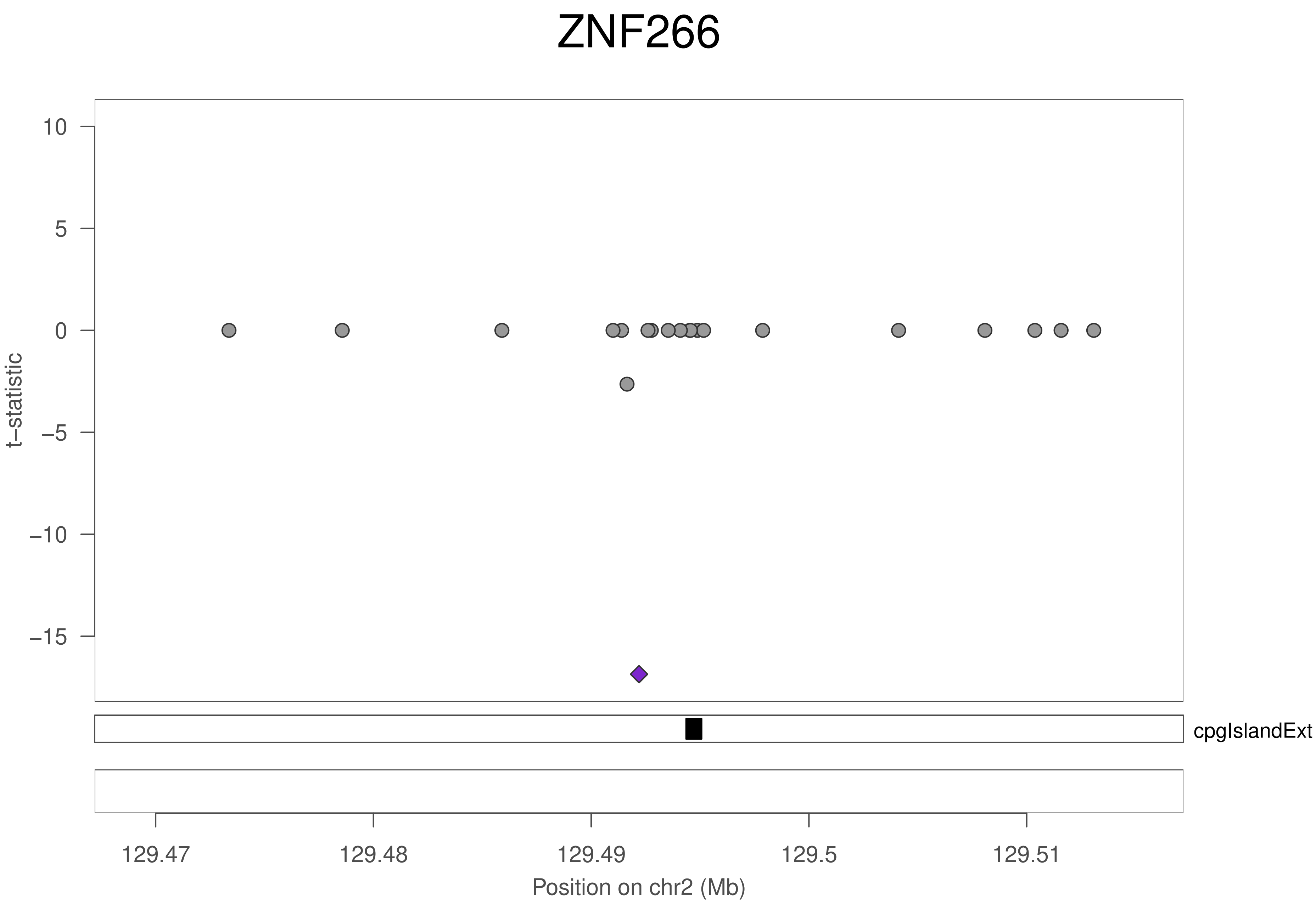

ZNF266
10
5
0
−5
−10
−15
t−statistic
cpgIslandExt
129.47
129.48
129.49
129.5
129.51
Position on chr2 (Mb)

### ZNF266

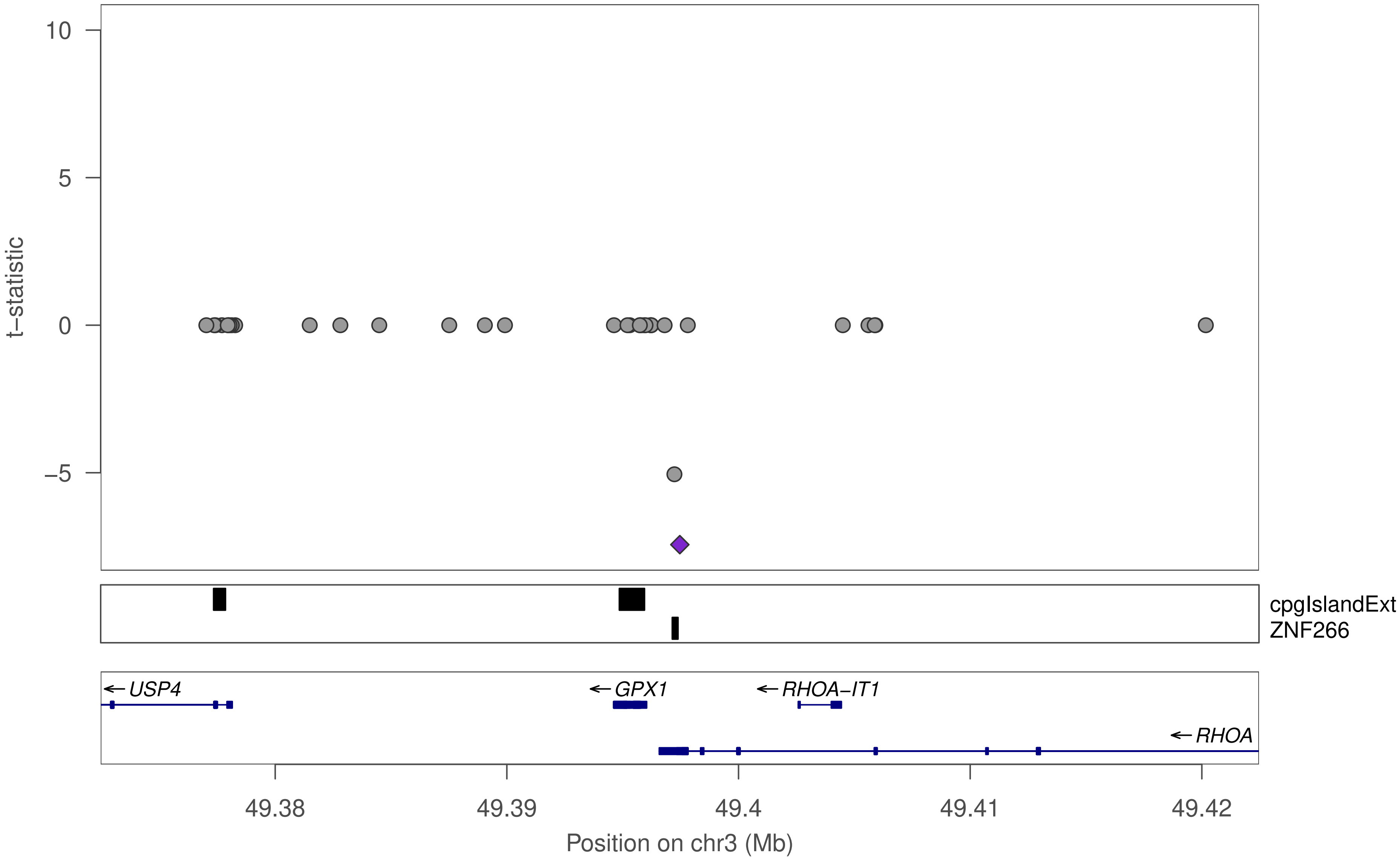
t−statistic
10
5
0
−5
cpgIslandExt
ZNF266
USP4
GPX1
RHOA−IT1
RHOA
49.38
49.39
49.4
49.41
49.42
Position on chr3 (Mb)

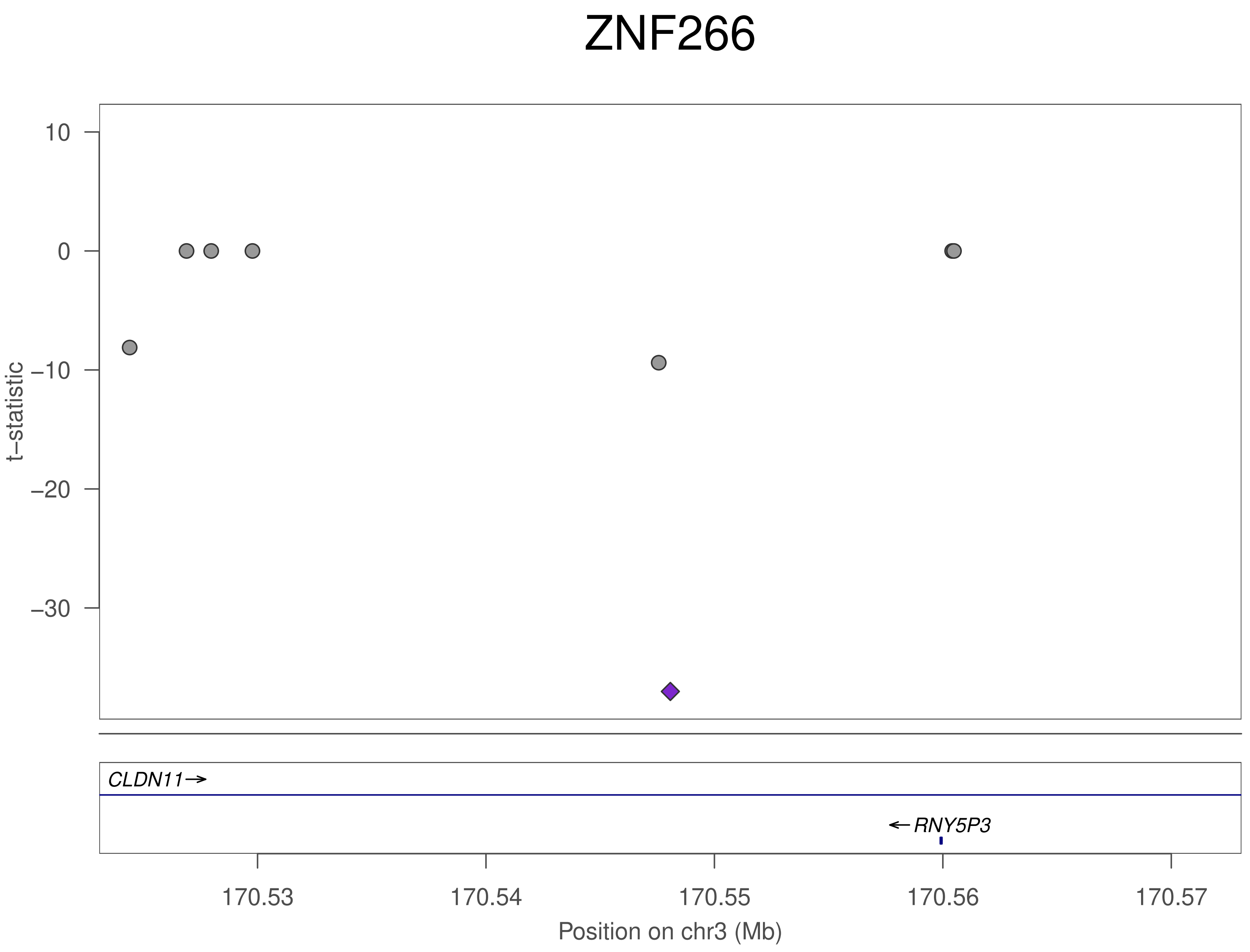
ZNF266
t–statistic
10
0
–10
–20
–30
CLDN11→
←RNY5P3
170.53
170.54
170.55
170.56
170.57
Position on chr3 (Mb)

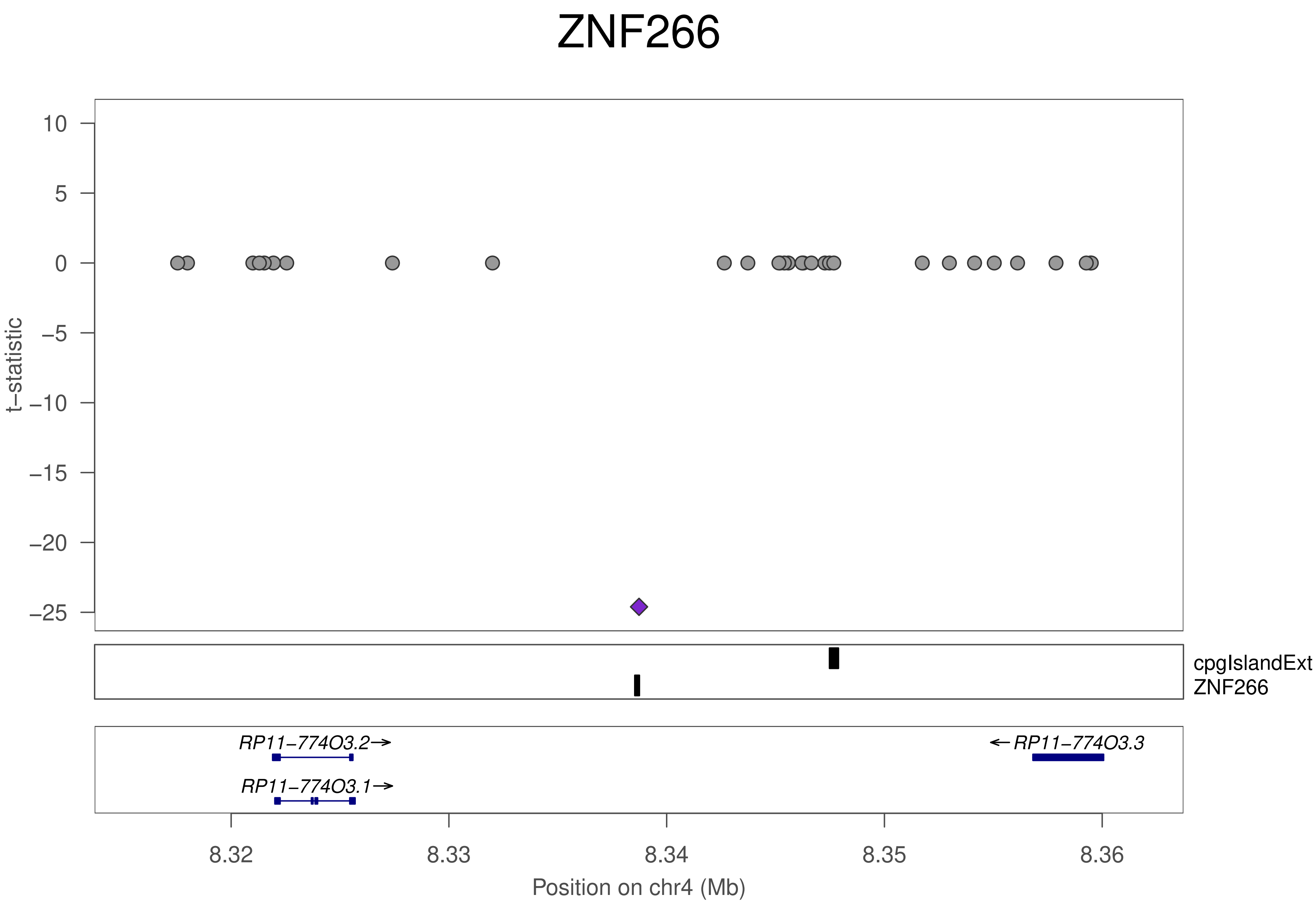
ZNF266
t−statistic
10
5
0
−5
−10
−15
−20
−25
cpgIslandExt
ZNF266
RP11−774O3.2→
RP11−774O3.1→
←RP11−774O3.3
8.32
8.33
8.34
8.35
8.36
Position on chr4 (Mb)

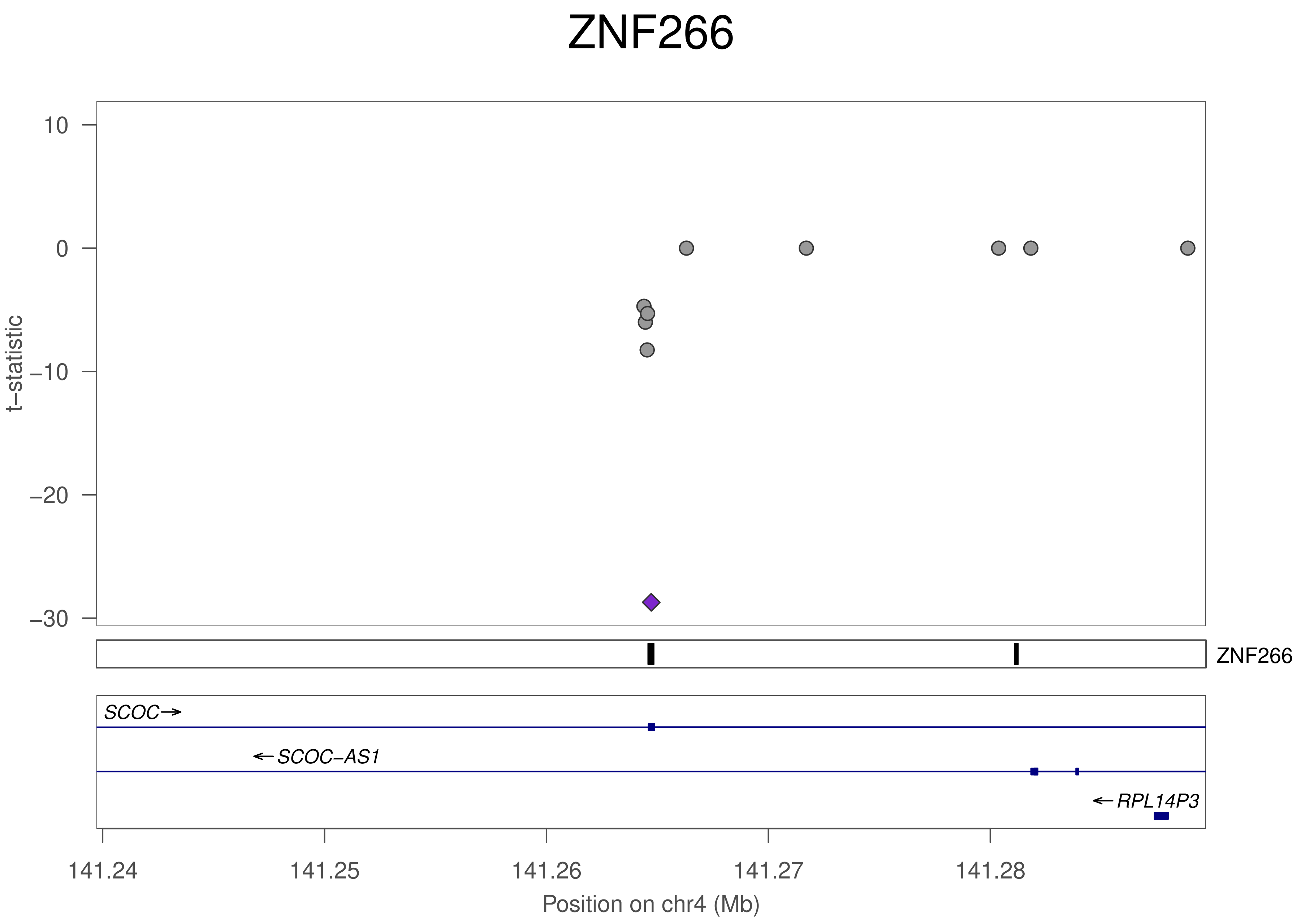
ZNF266
t−statistic
10
0
−10
−20
−30
ZNF266
SCOC
SCOC−AS1
RPL14P3
141.24
141.25
141.26
141.27
141.28
Position on chr4 (Mb)

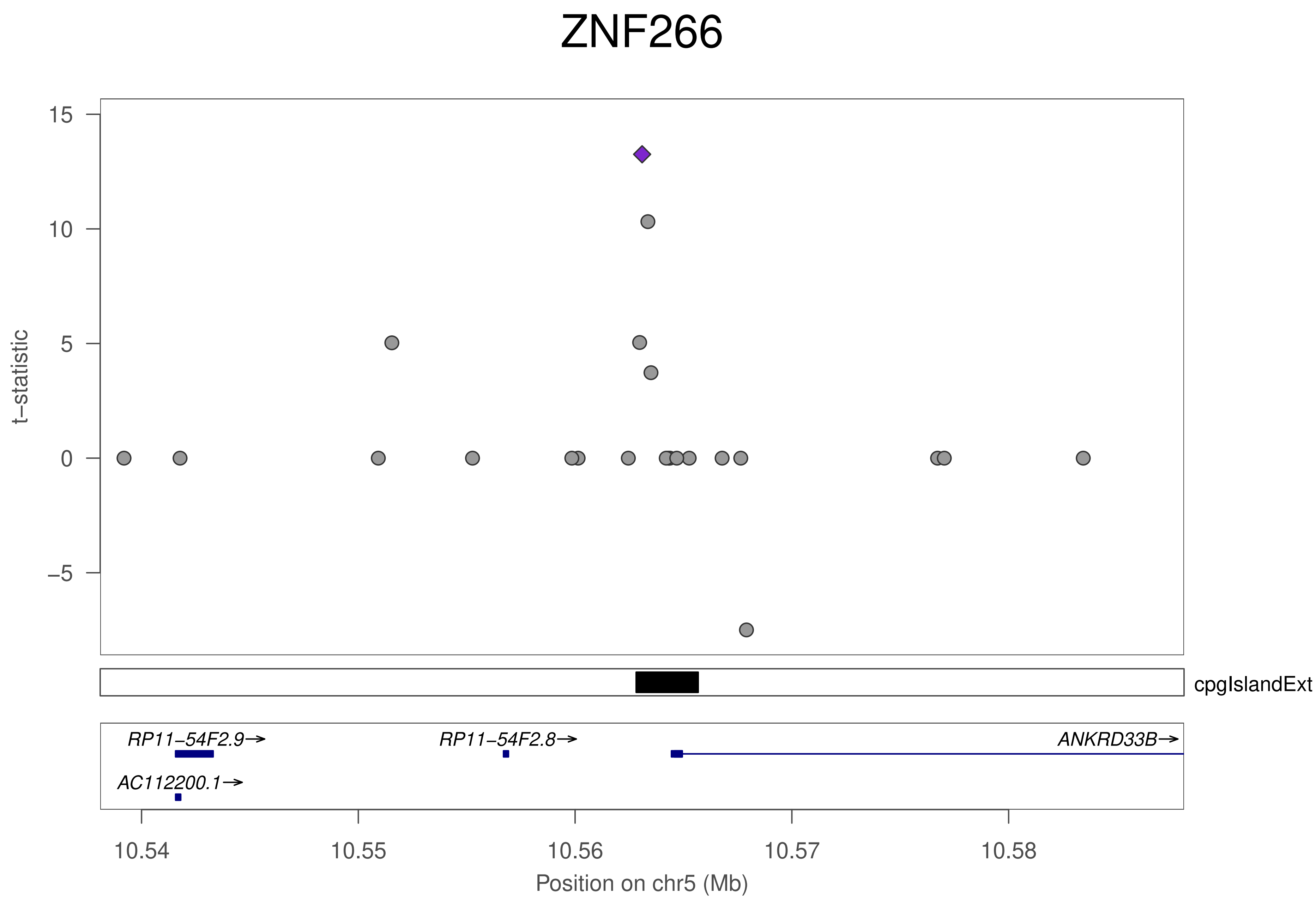
ZNF266
t−statistic
15
10
5
0
−5
cpgIslandExt
RP11−54F2.9→
RP11−54F2.8→
ANKRD33B→
AC112200.1→
10.54
10.55
10.56
10.57
10.58
Position on chr5 (Mb)

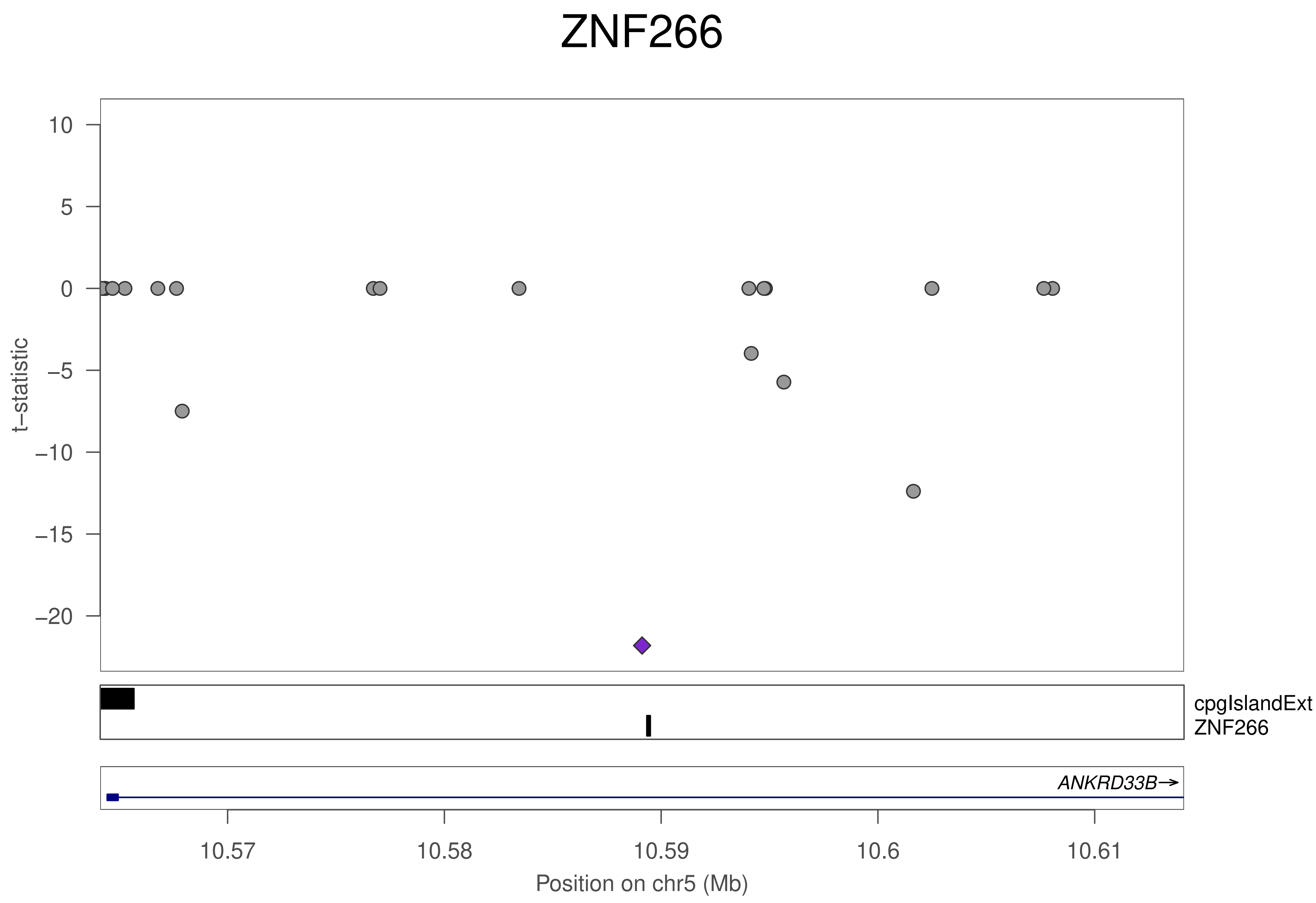
ZNF266
t−statistic
10
5
0
−5
−10
−15
−20
cpgIslandExt
ZNF266
ANKRD33B→
10.57
10.58
10.59
10.6
10.61
Position on chr5 (Mb)

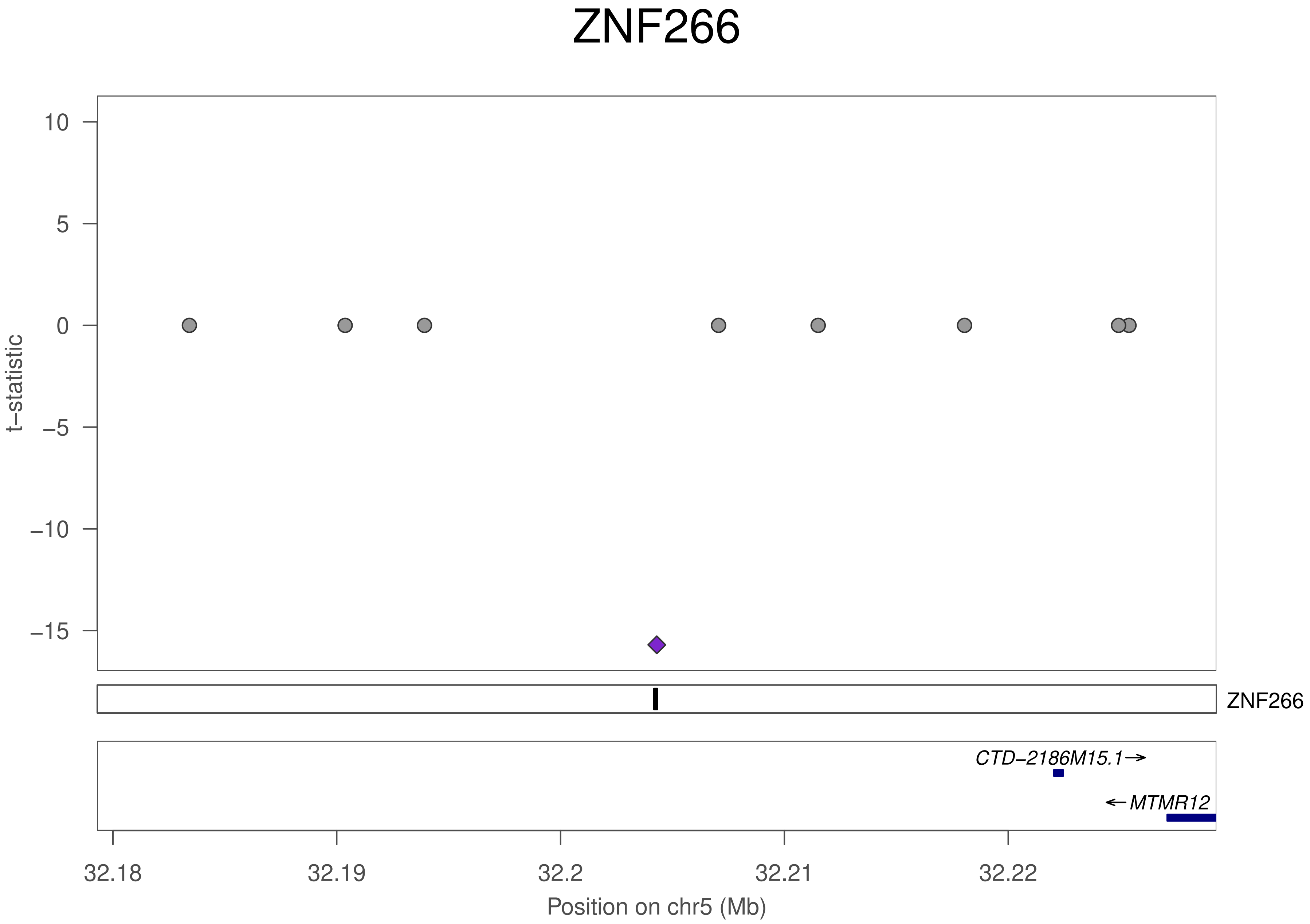
ZNF266
t−statistic
10
5
0
−5
−10
−15
ZNF266
CTD−2186M15.1→
←MTMR12
32.18
32.19
32.2
32.21
32.22
Position on chr5 (Mb)

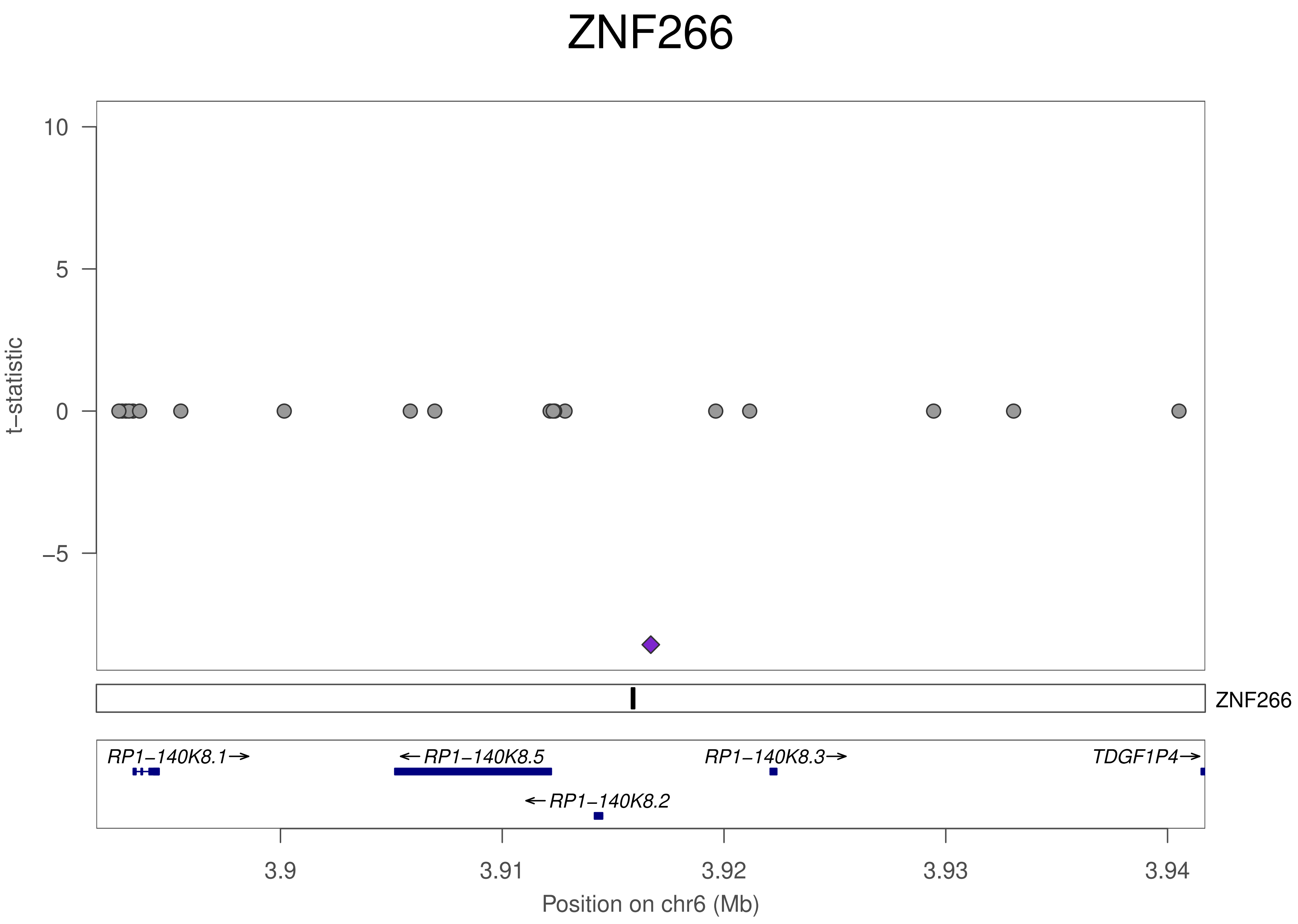
ZNF266
10
5
0
−5
t−statistic
ZNF266
RP1−140K8.1→
←RP1−140K8.5
RP1−140K8.3→
TDGF1P4→
←RP1−140K8.2
3.9
3.91
3.92
3.93
3.94
Position on chr6 (Mb)

### ZNF266

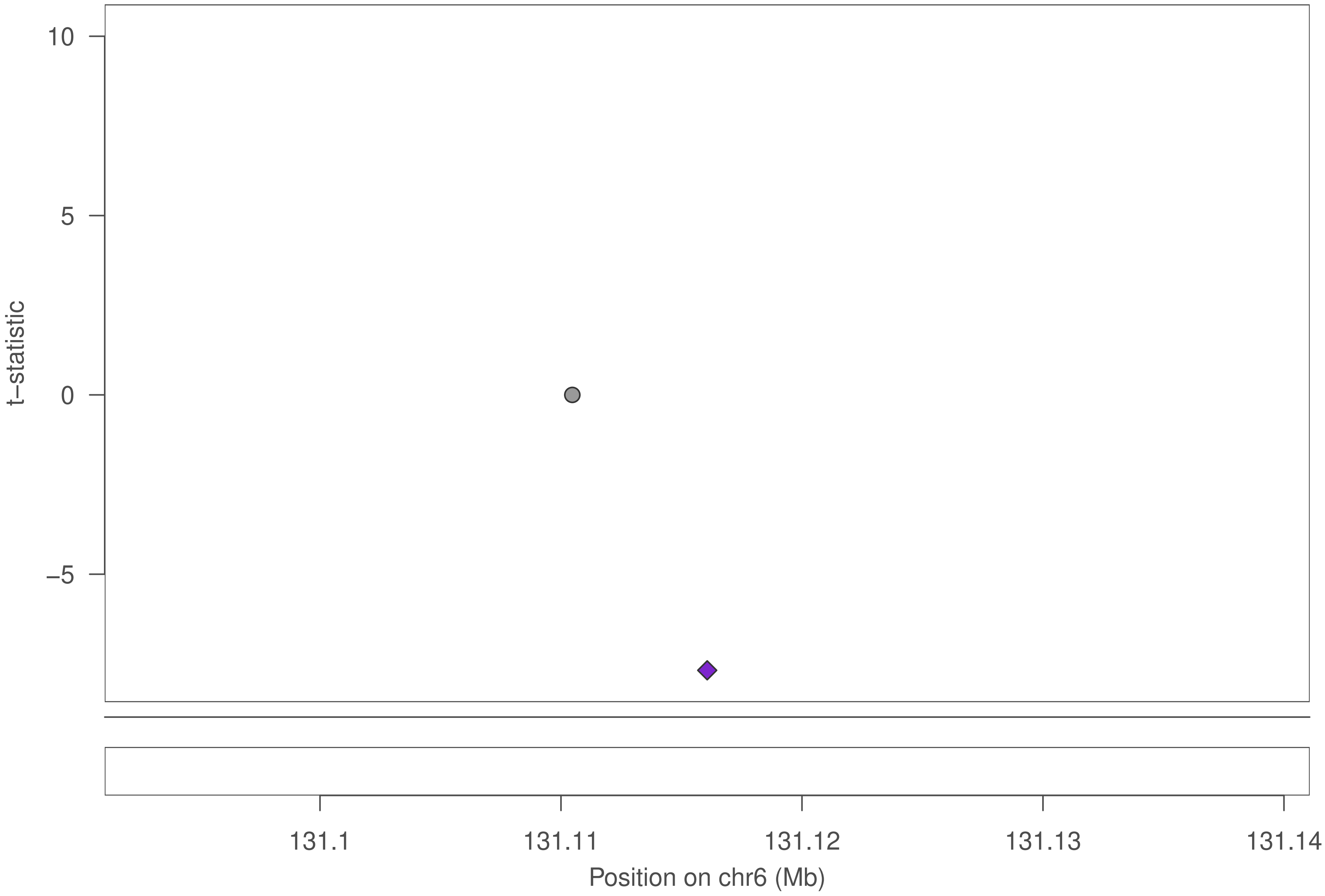

10
5
0
−5
t−statistic
131.1
131.11
131.12
131.13
131.14
Position on chr6 (Mb)

### ZNF266

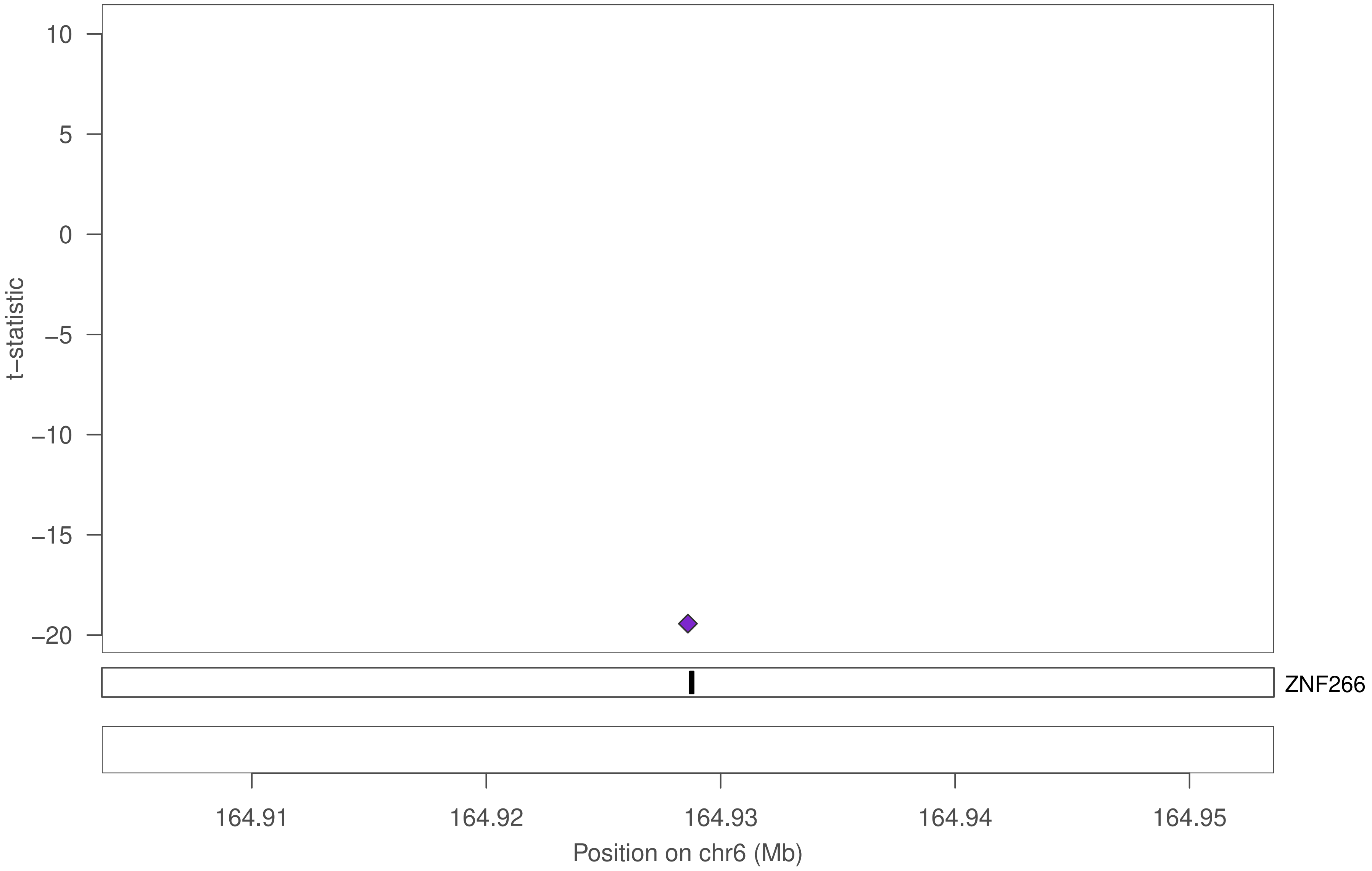

10
5
0
−5
−10
−15
−20
t−statistic
ZNF266
164.91
164.92
164.93
164.94
164.95
Position on chr6 (Mb)

### ZNF266

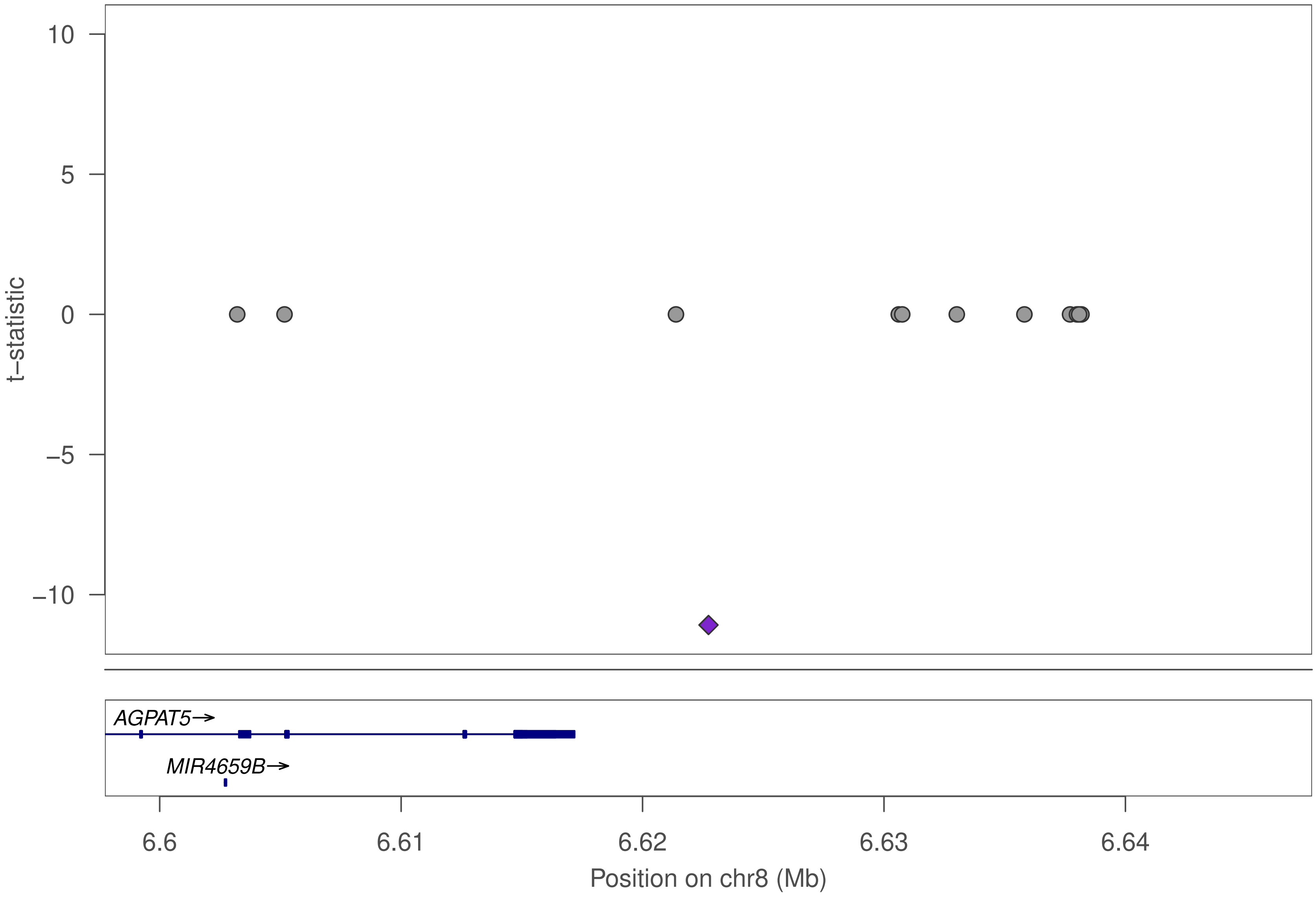

t−statistic
10
5
0
−5
−10
AGPAT5→
MIR4659B→
6.6
6.61
6.62
6.63
6.64
Position on chr8 (Mb)

### ZNF266

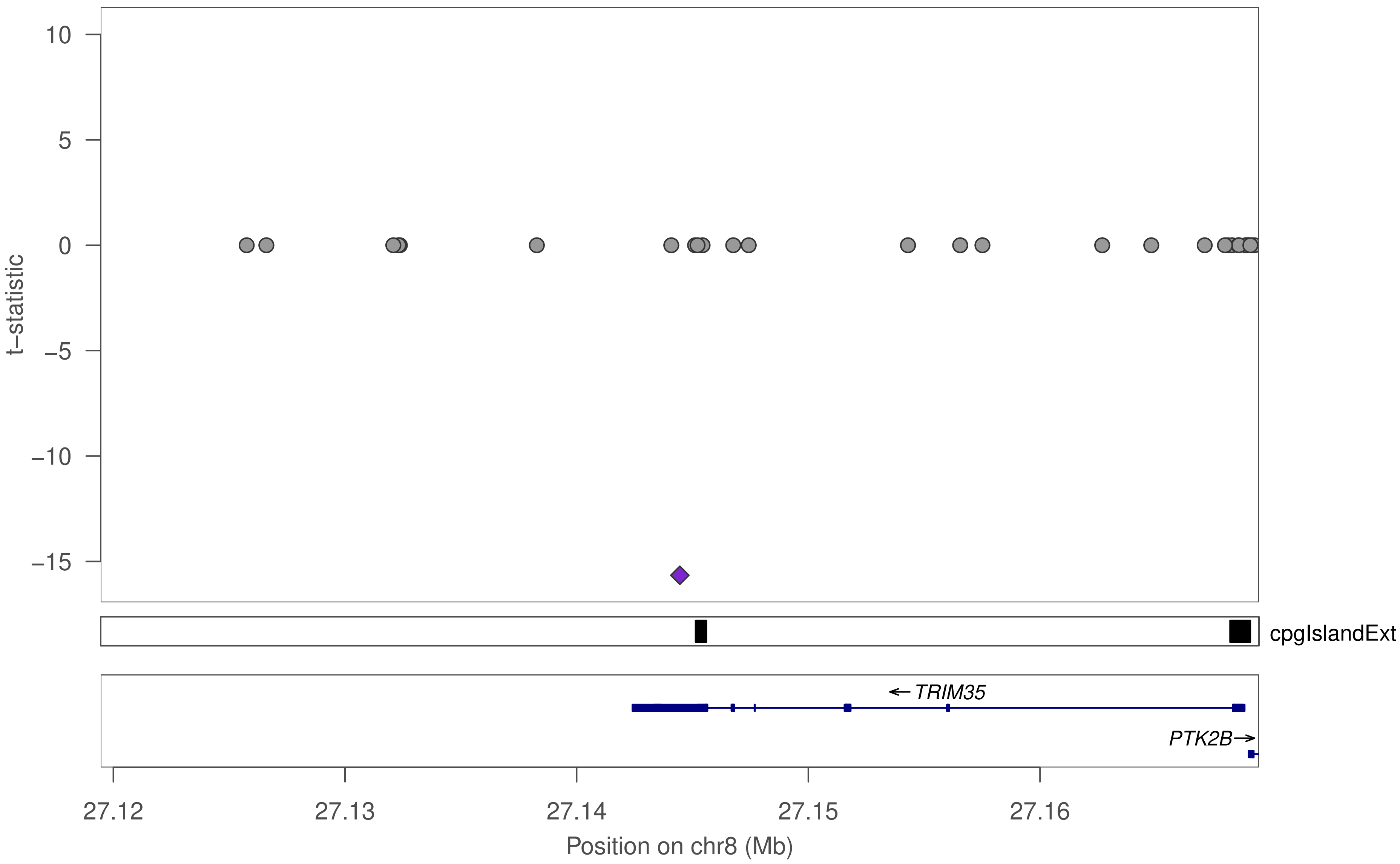

10
5
0
−5
−10
−15
t−statistic
cpgIslandExt
← TRIM35
PTK2B→
27.12
27.13
27.14
27.15
27.16
Position on chr8 (Mb)

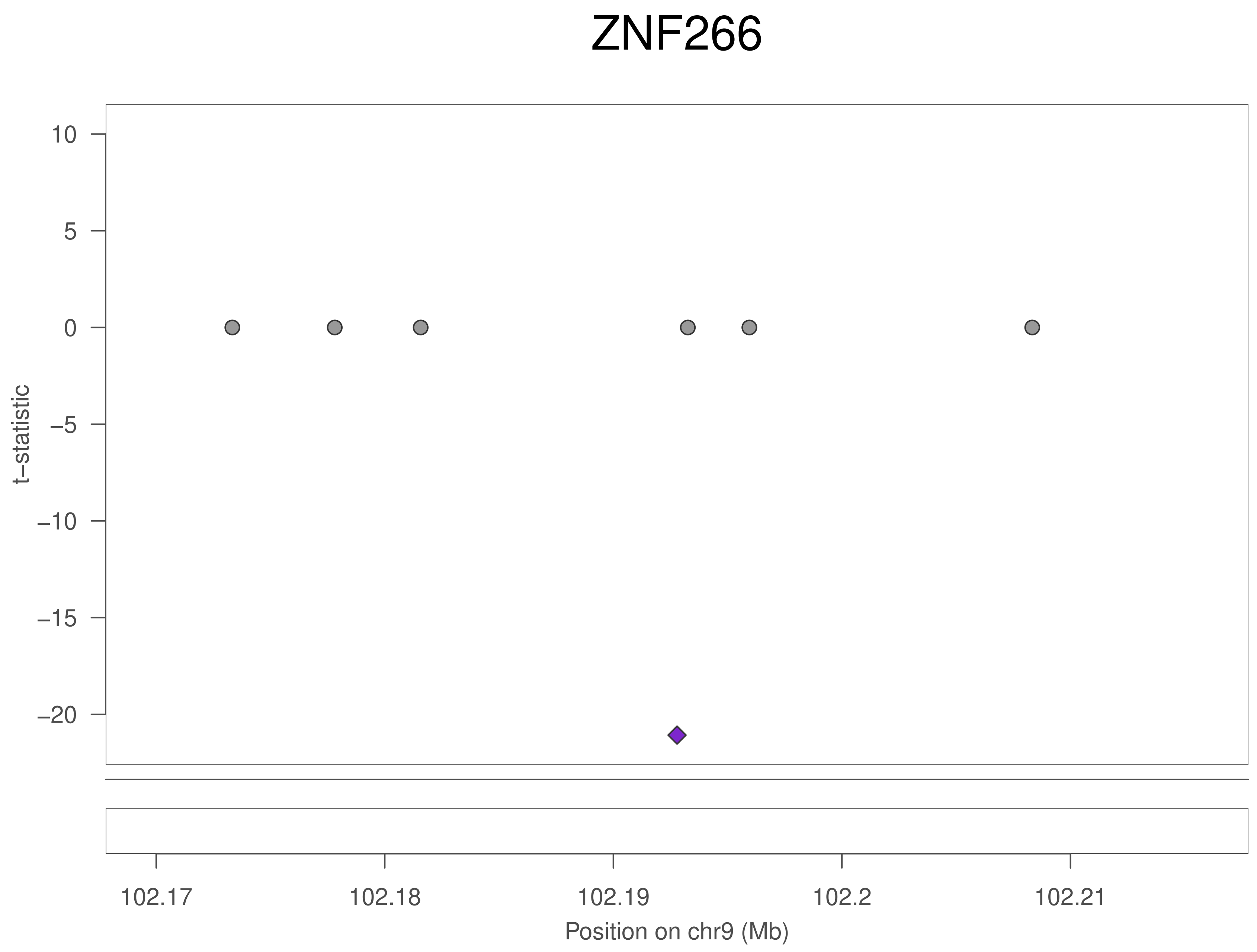
ZNF266
10
5
0
−5
−10
−15
−20
t−statistic
102.17
102.18
102.19
102.2
102.21
Position on chr9 (Mb)

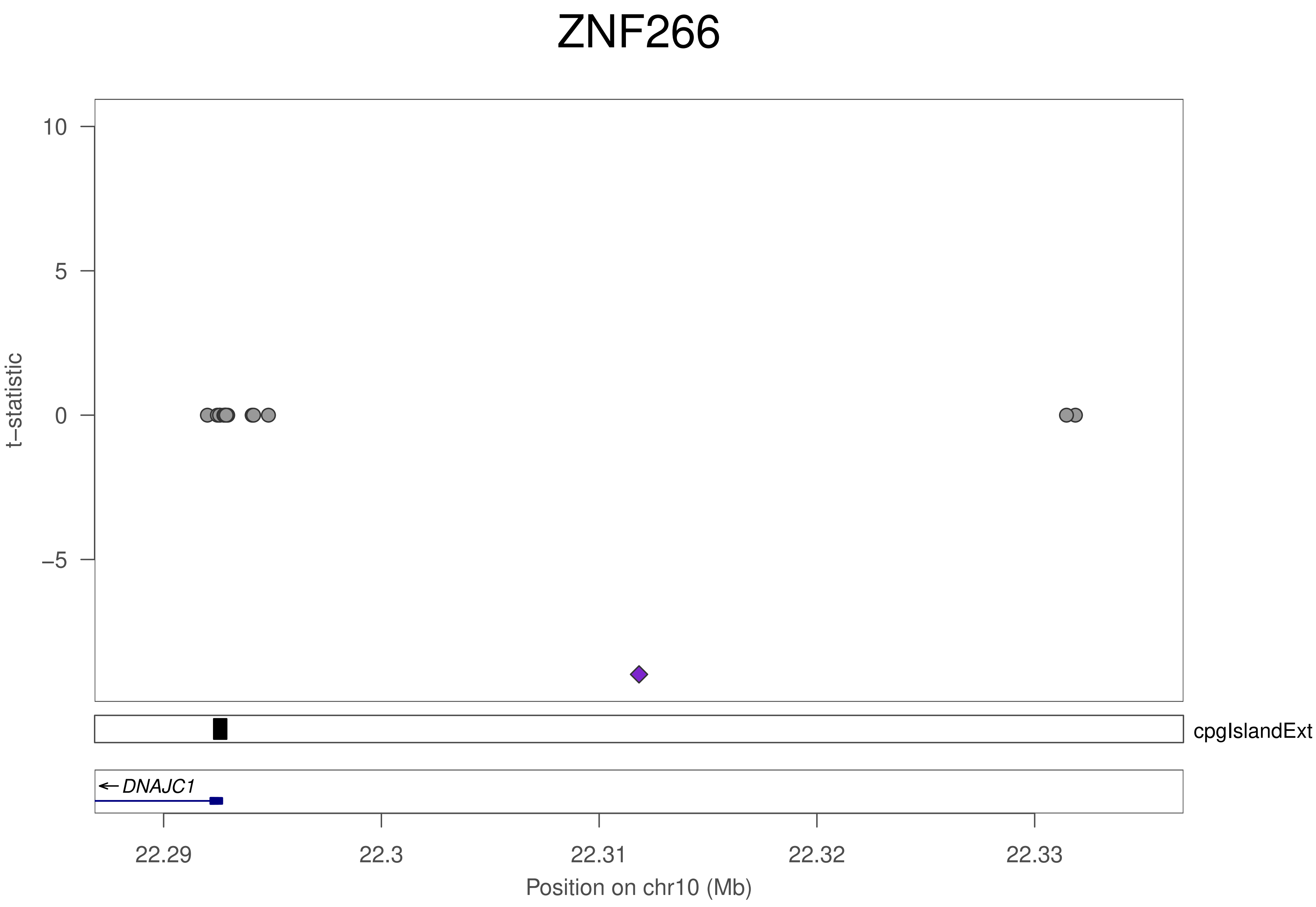

ZNF266
t−statistic
10
5
0
−5
cpgIslandExt
DNAJC1
22.29
22.3
22.31
22.32
22.33
Position on chr10 (Mb)

### ZNF266

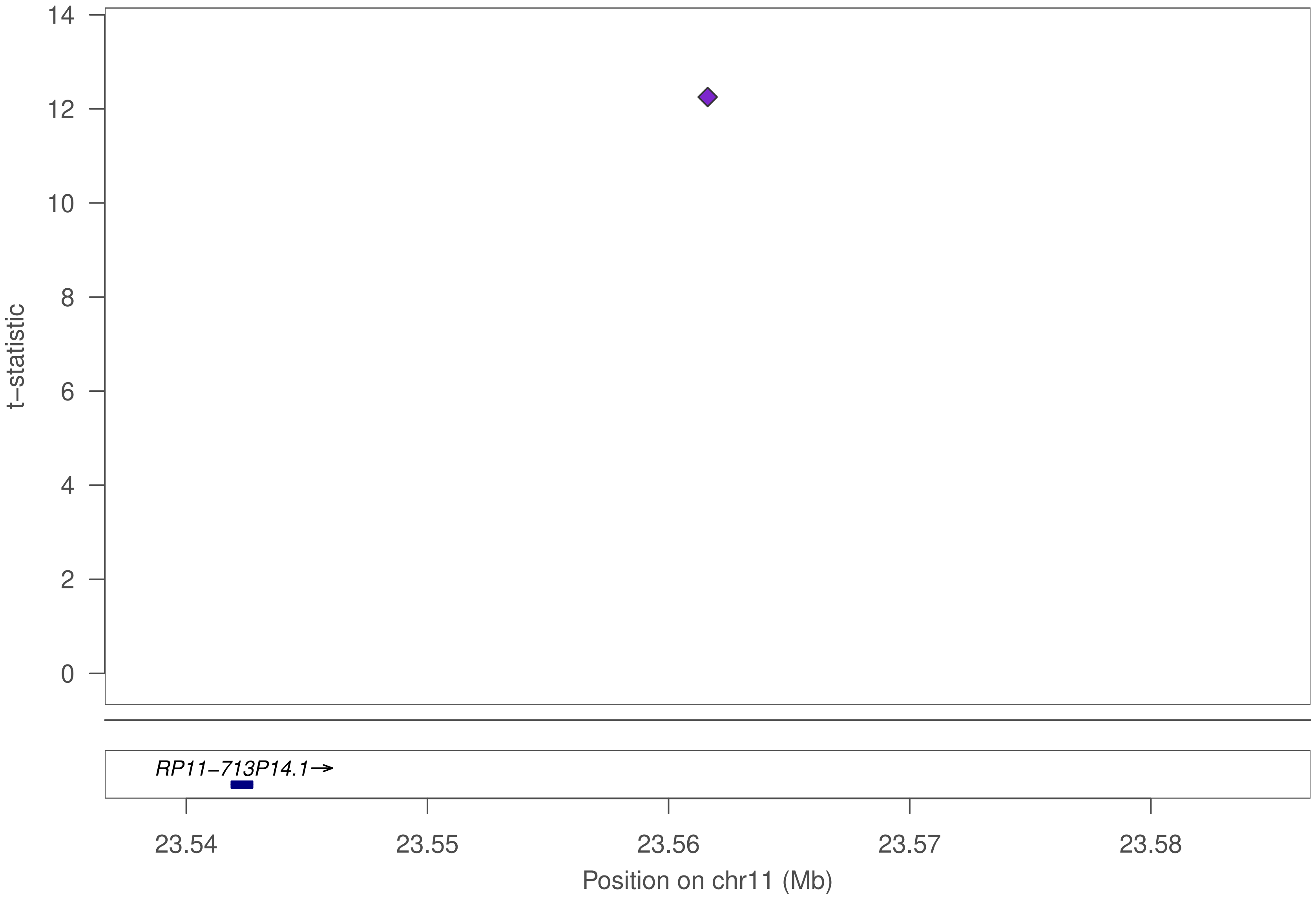

t–statistic
14
12
10
8
6
4
2
0
RP11−713P14.1→
23.54
23.55
23.56
23.57
23.58
Position on chr11 (Mb)

### ZNF266

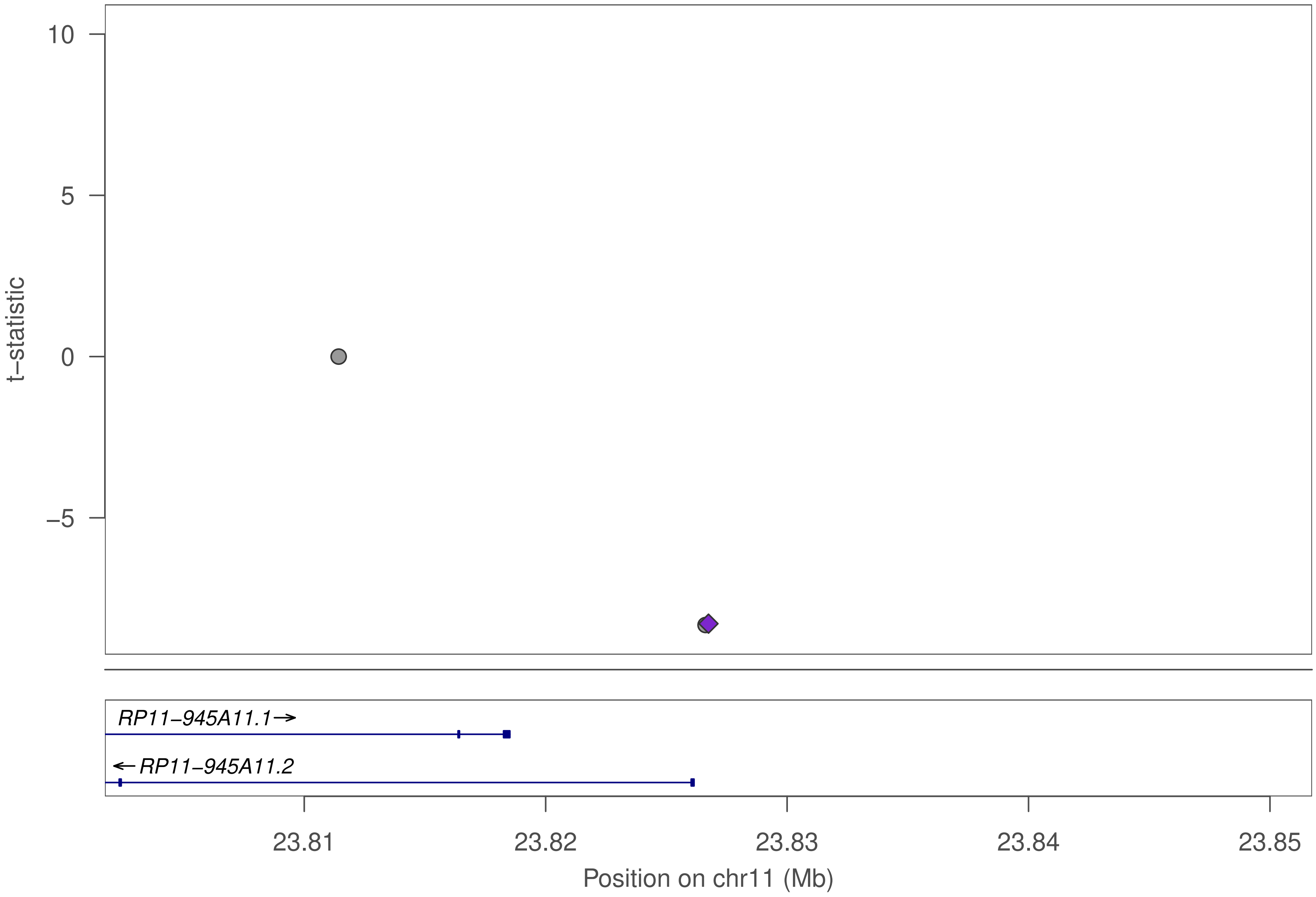

10
5
0
−5
t−statistic
RP11−945A11.1→
←RP11−945A11.2
23.81
23.82
23.83
23.84
23.85
Position on chr11 (Mb)

### ZNF266

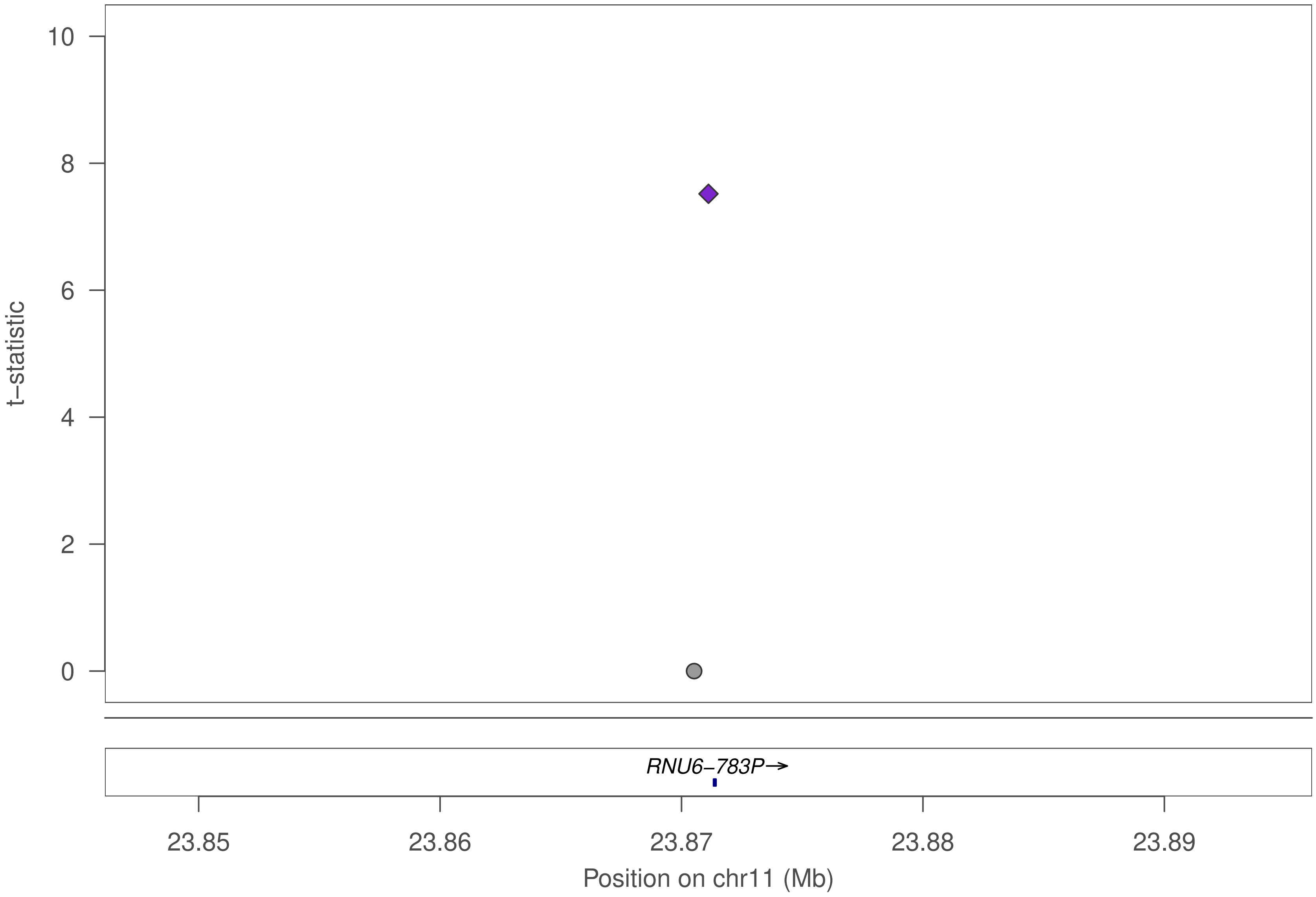

10
8
6
4
2
0
t−statistic
RNU6−783P→
23.85
23.86
23.87
23.88
23.89
Position on chr11 (Mb)

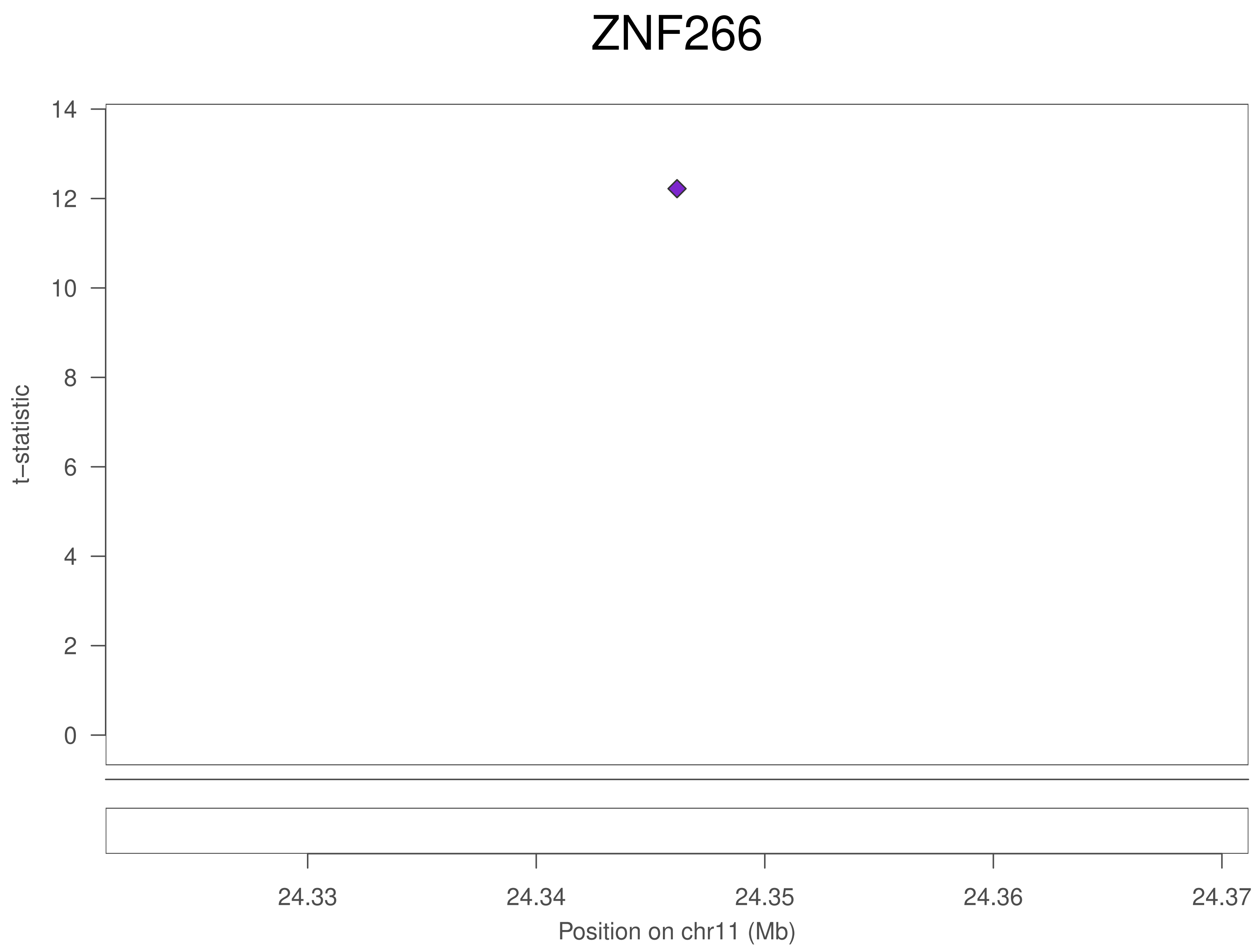

ZNF266
t−statistic
14
12
10
8
6
4
2
0
24.33
24.34
24.35
24.36
24.37
Position on chr11 (Mb)

### ZNF266

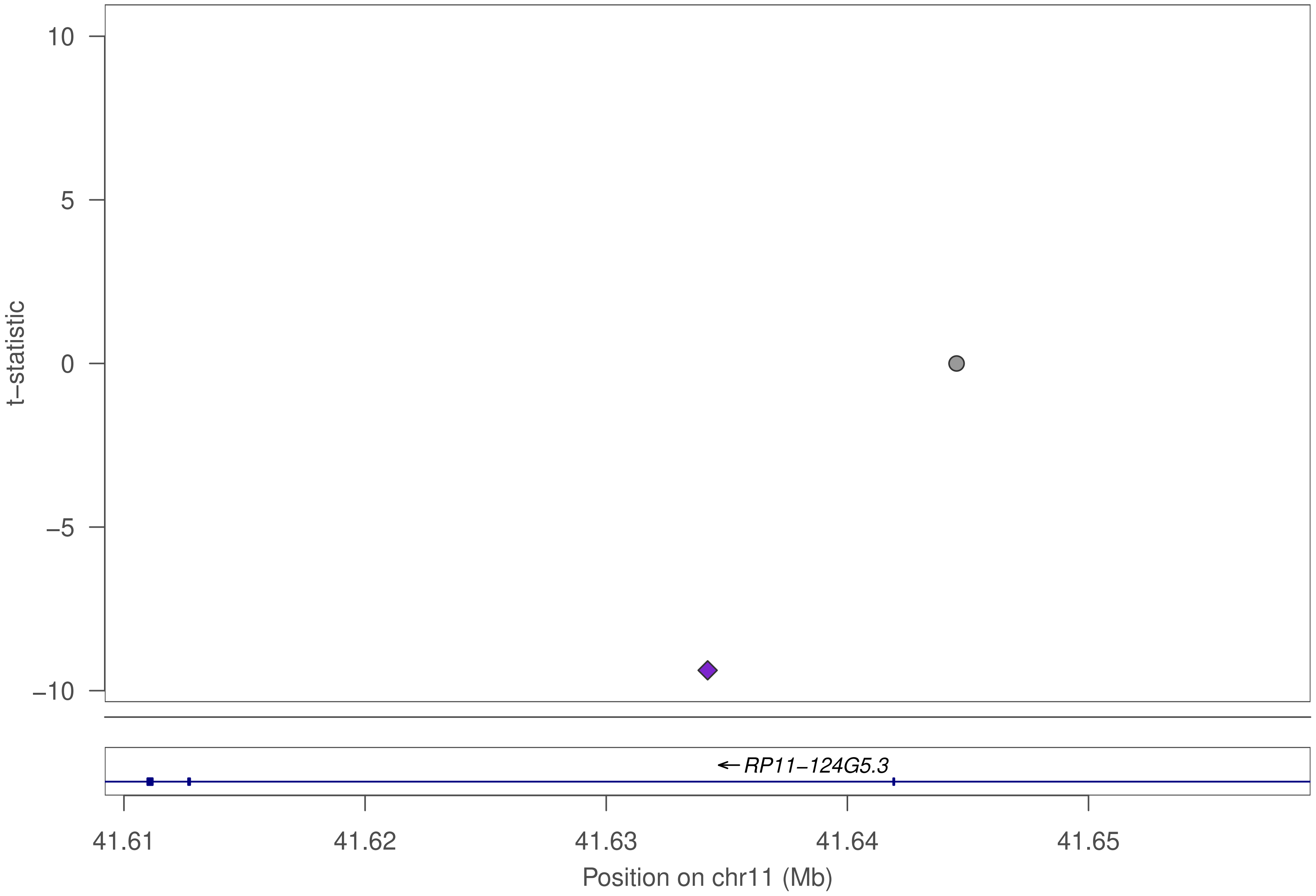

10
5
0
−5
−10
t−statistic
RP11−124G5.3
41.61
41.62
41.63
41.64
41.65
Position on chr11 (Mb)

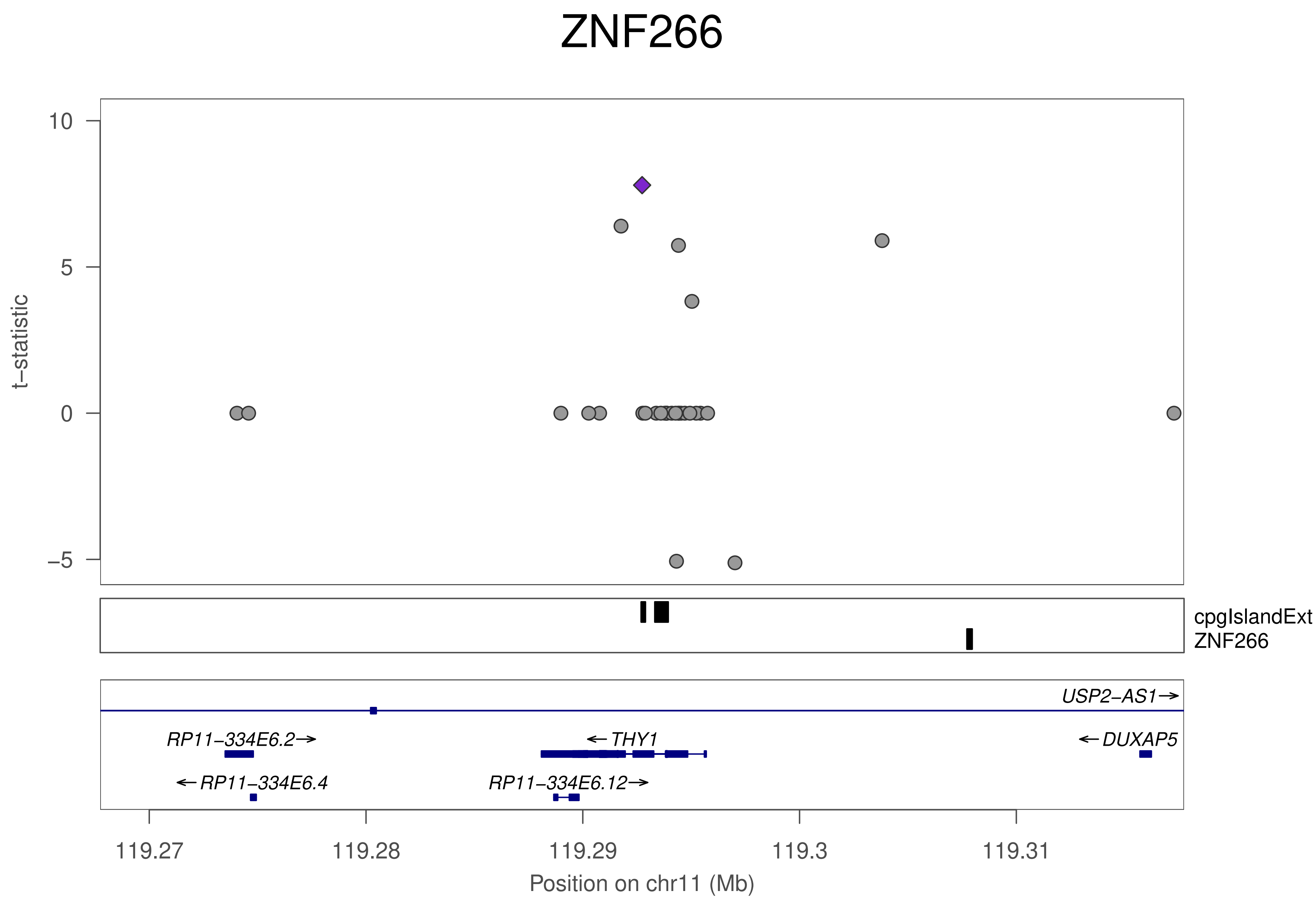

ZNF266
t–statistic
10
5
0
–5
cpgIslandExt
ZNF266
USP2–AS1
RP11–334E6.2
THY1
DUXAP5
RP11–334E6.4
RP11–334E6.12
119.27
119.28
119.29
119.3
119.31
Position on chr11 (Mb)

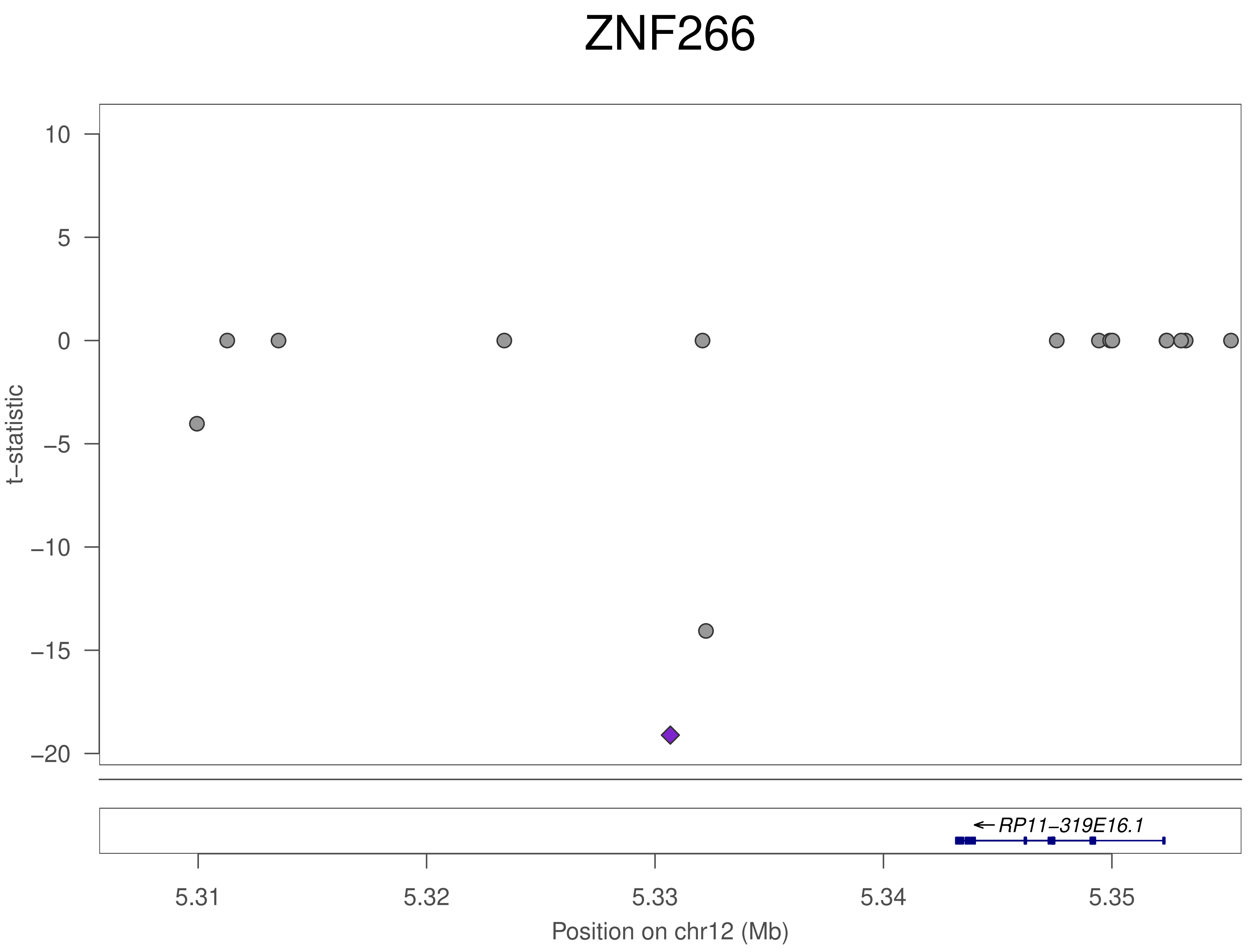

ZNF266
t–statistic
10
5
0
–5
–10
–15
–20
RP11–319E16.1
5.31
5.32
5.33
5.34
5.35
Position on chr12 (Mb)

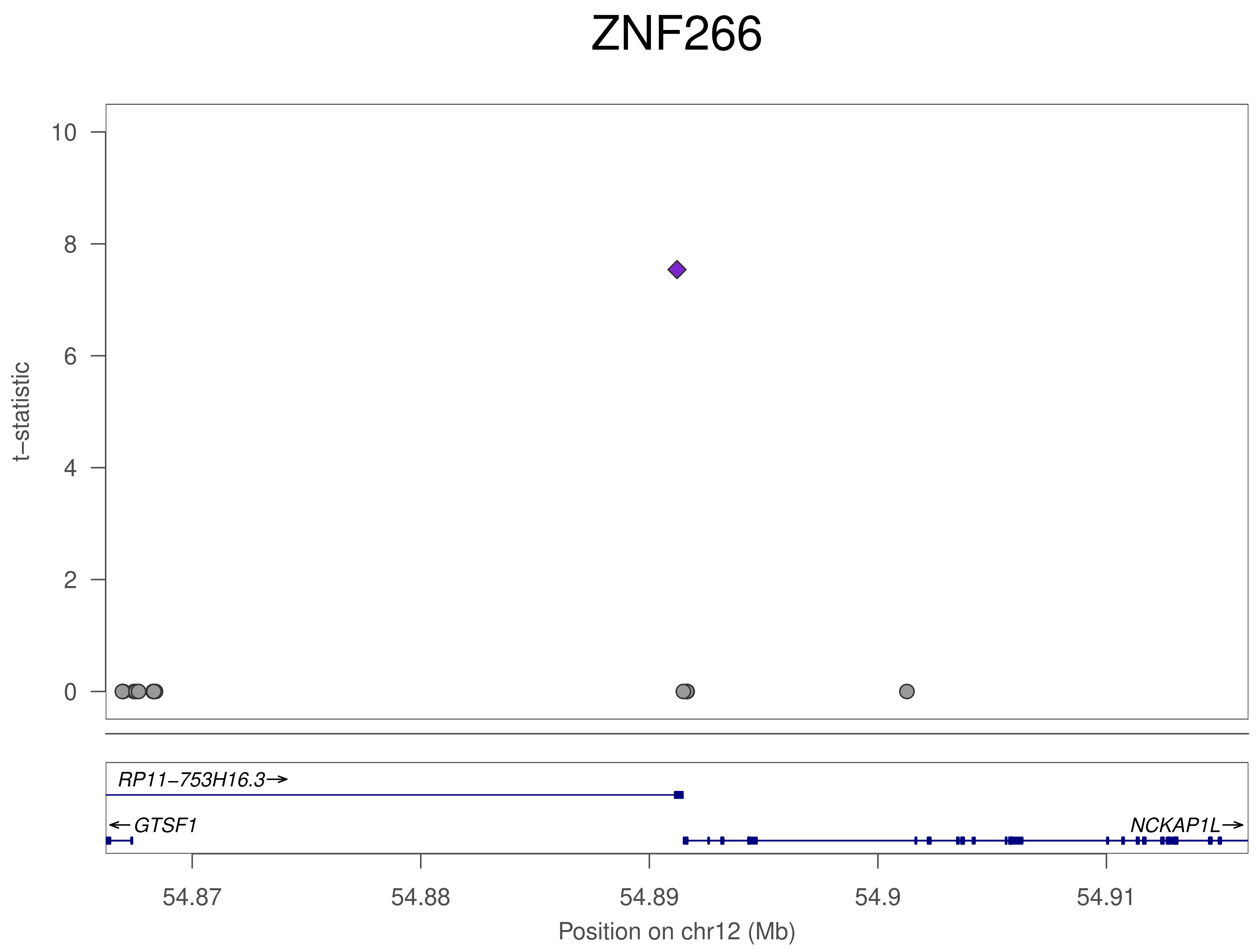
ZNF266
t−statistic
10
8
6
4
2
0
RP11−753H16.3→
← GTSF1
NCKAP1L→
54.87
54.88
54.89
54.9
54.91
Position on chr12 (Mb)

### ZNF266

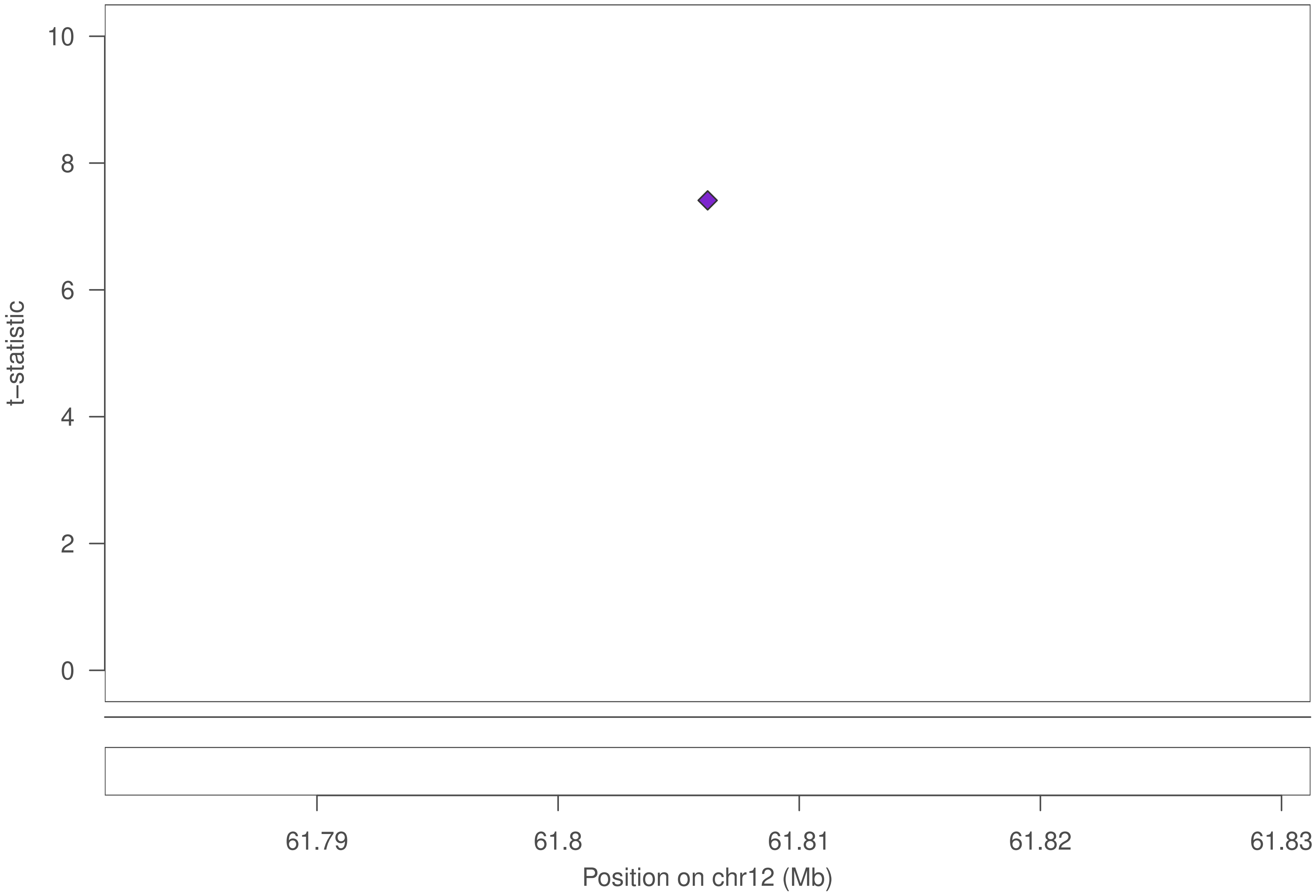

t–statistic
10
8
6
4
2
0
61.79
61.8
61.81
61.82
61.83
Position on chr12 (Mb)

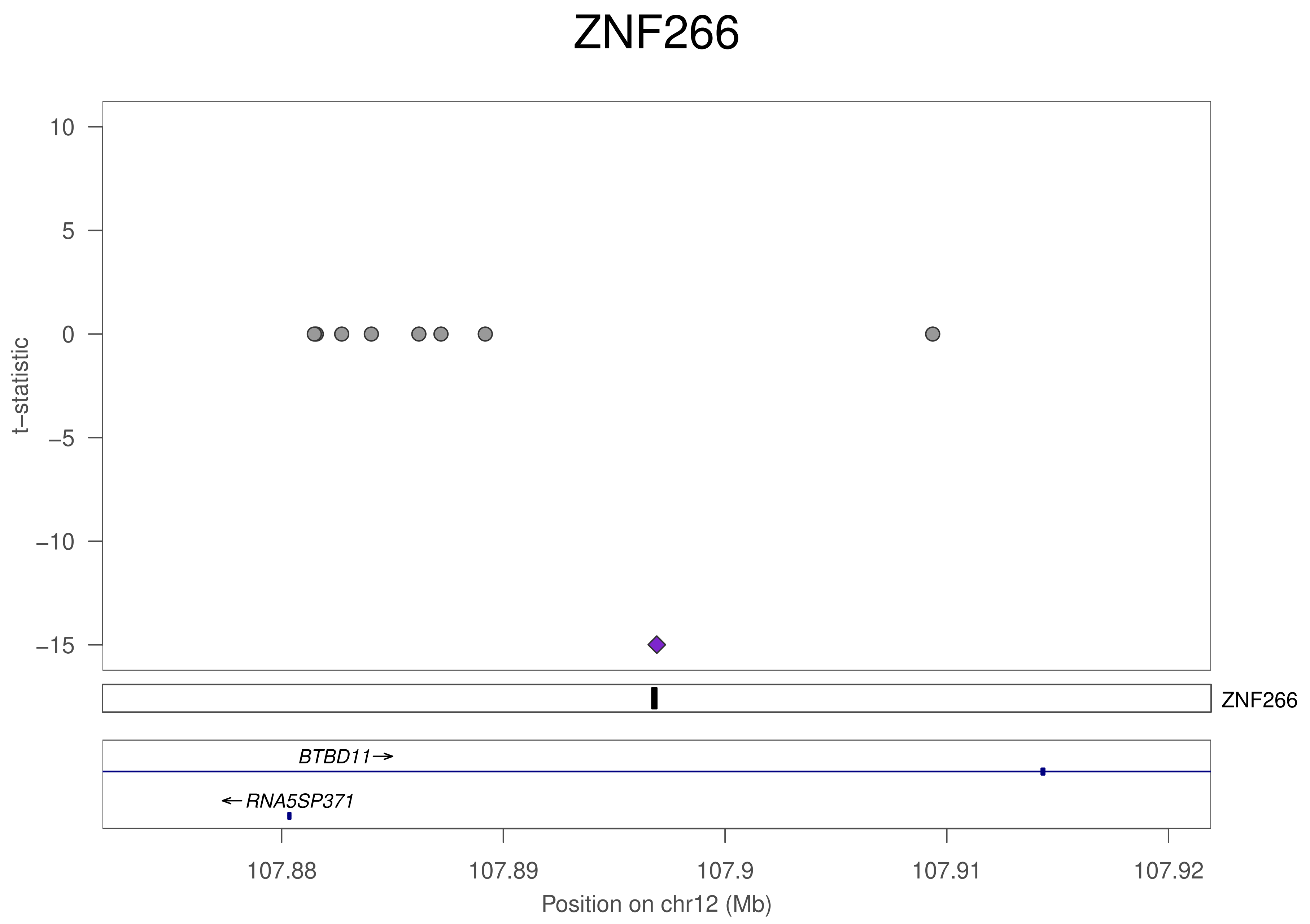
ZNF266
t−statistic
10
5
0
−5
−10
−15
ZNF266
BTBD11→
←RNA5SP371
107.88
107.89
107.9
107.91
107.92
Position on chr12 (Mb)

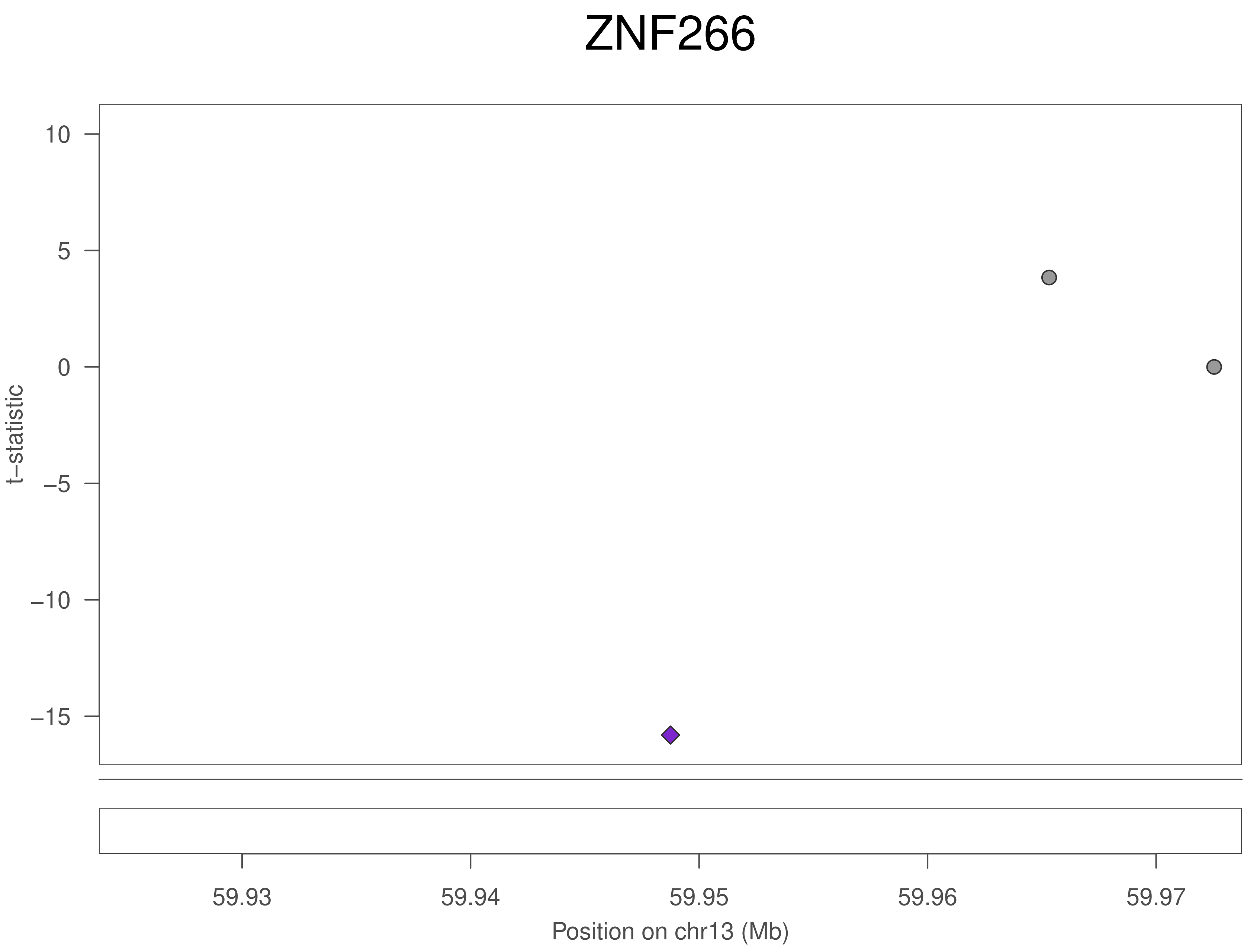
ZNF266
10
5
0
−5
−10
−15
t−statistic
59.93
59.94
59.95
59.96
59.97
Position on chr13 (Mb)

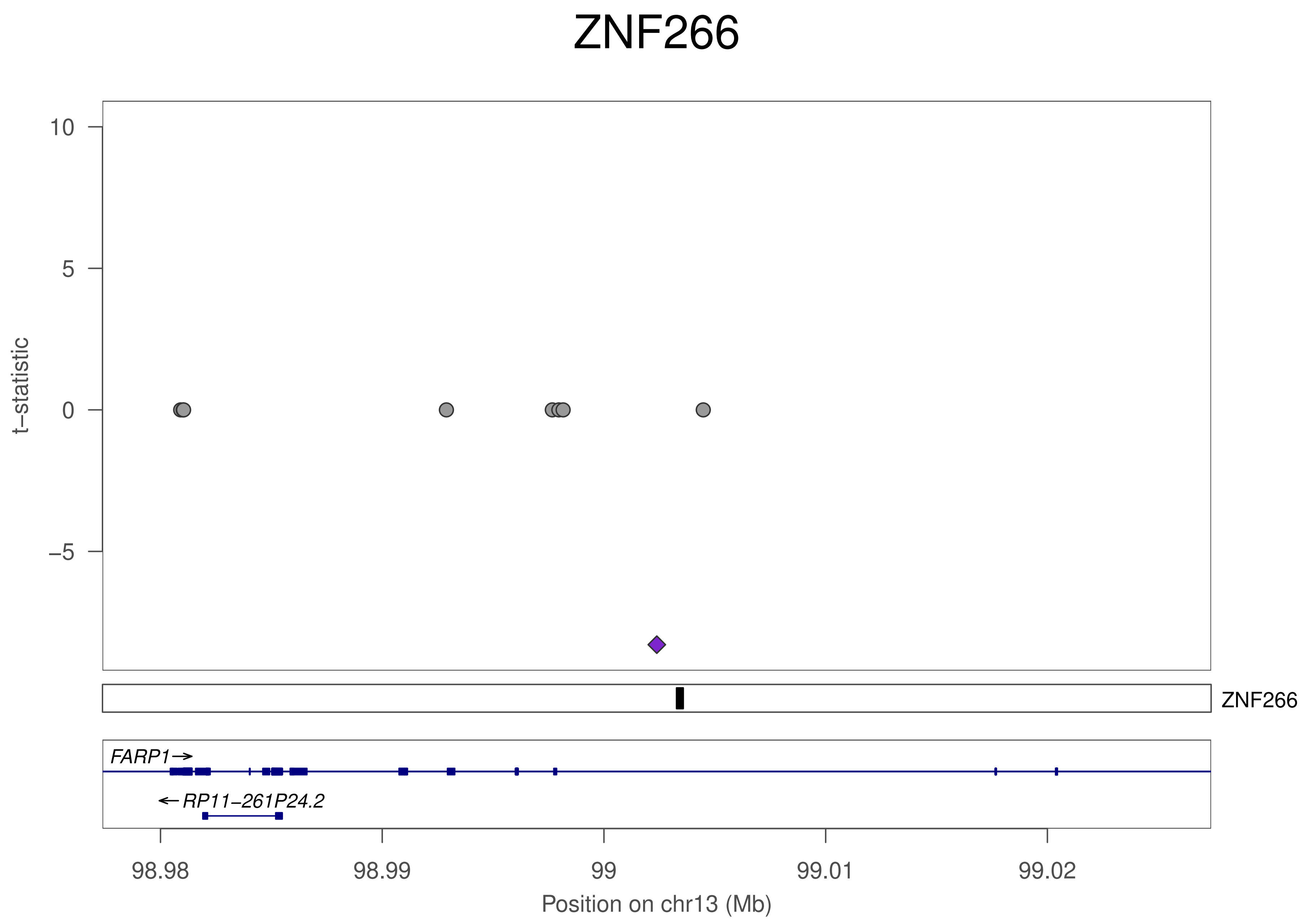
ZNF266
t−statistic
10
5
0
−5
ZNF266
FARP1→
←RP11−261P24.2
98.98
98.99
99
99.01
99.02
Position on chr13 (Mb)

### ZNF266

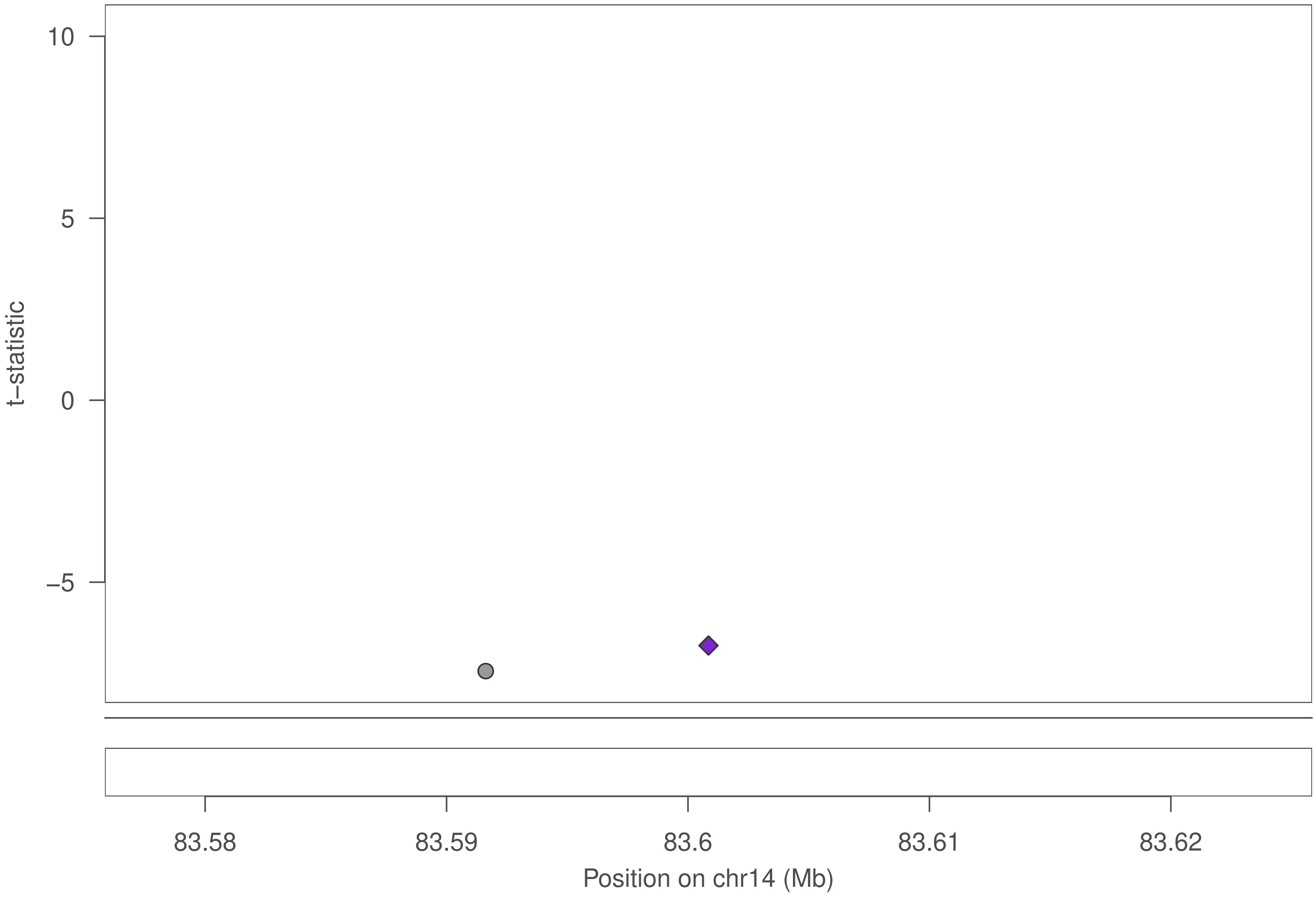

10
5
0
−5
t−statistic
83.58
83.59
83.6
83.61
83.62
Position on chr14 (Mb)

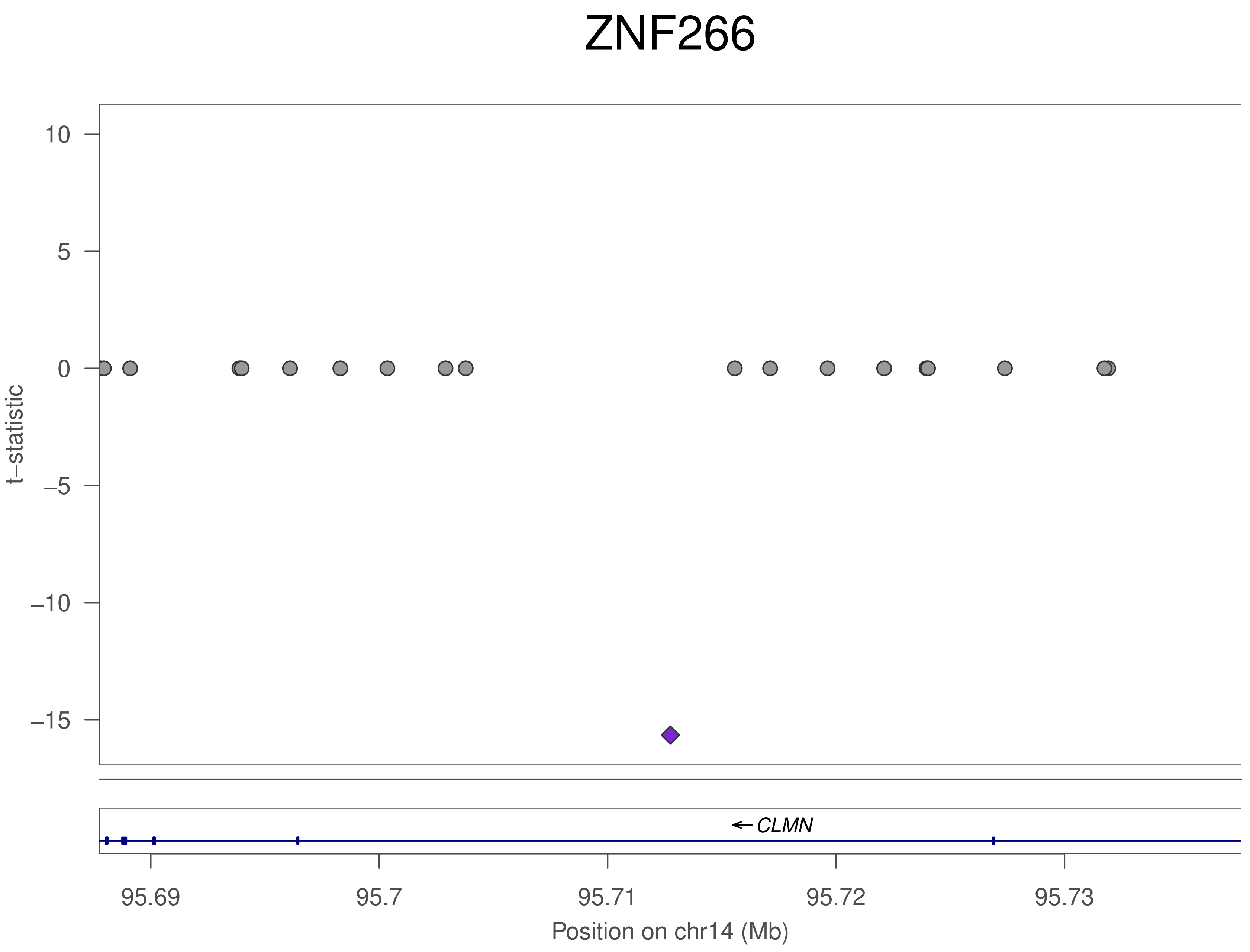
ZNF266
t–statistic
10
5
0
–5
–10
–15
CLMN
95.69
95.7
95.71
95.72
95.73
Position on chr14 (Mb)

### ZNF266

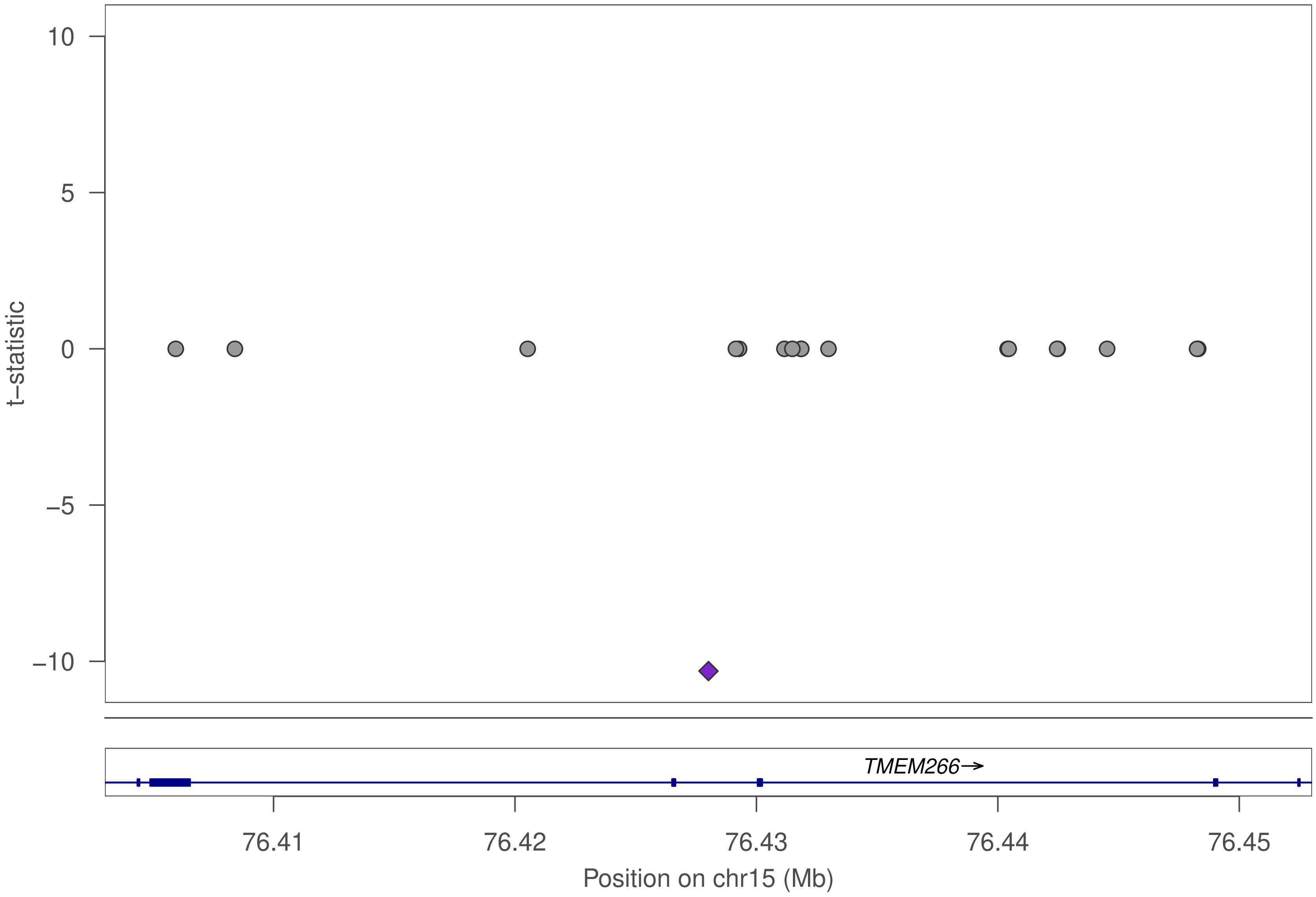

10
5
0
–5
–10
t–statistic
TMEM266→
76.41
76.42
76.43
76.44
76.45
Position on chr15 (Mb)

### ZNF266

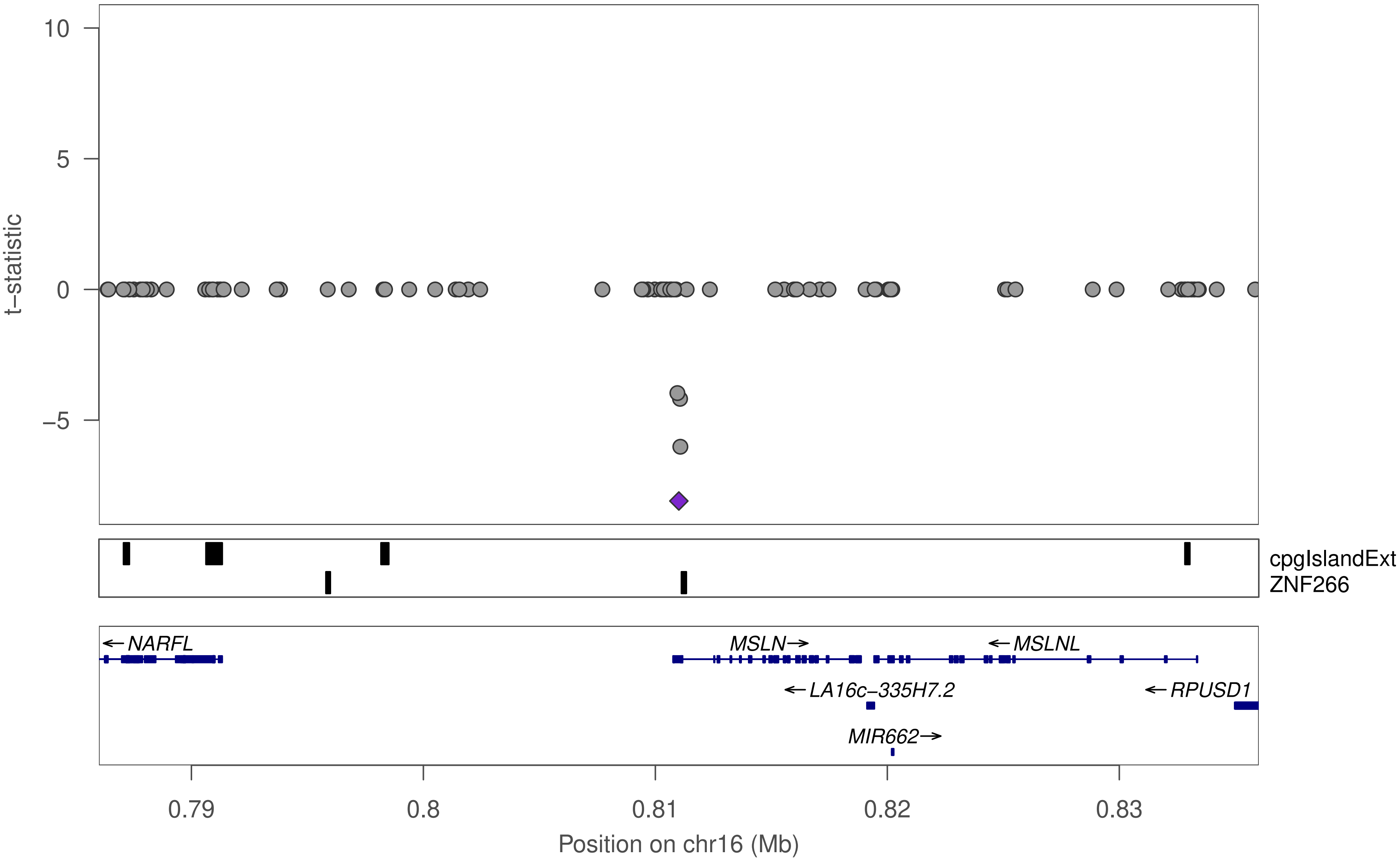

t−statistic
10
5
0
−5
cpgIslandExt
ZNF266
←NARFL
MSLN→
←MSLNL
←LA16c−335H7.2
←RPUSD1
MIR662→
0.79
0.8
0.81
0.82
0.83
Position on chr16 (Mb)

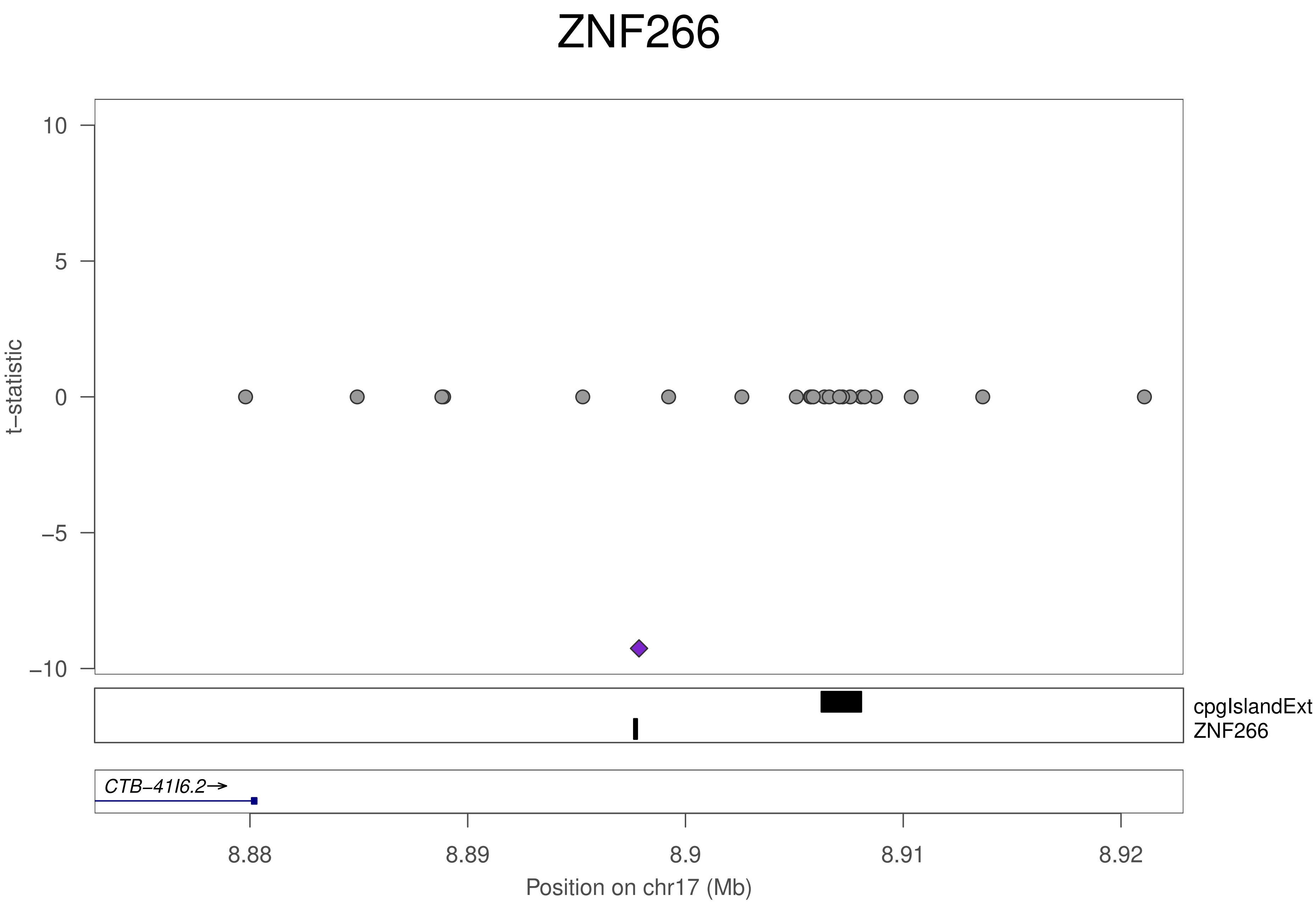

ZNF266
t−statistic
10
5
0
−5
−10
cpgIslandExt
ZNF266
CTB−41I6.2→
8.88
8.89
8.9
8.91
8.92
Position on chr17 (Mb)

### ZNF266

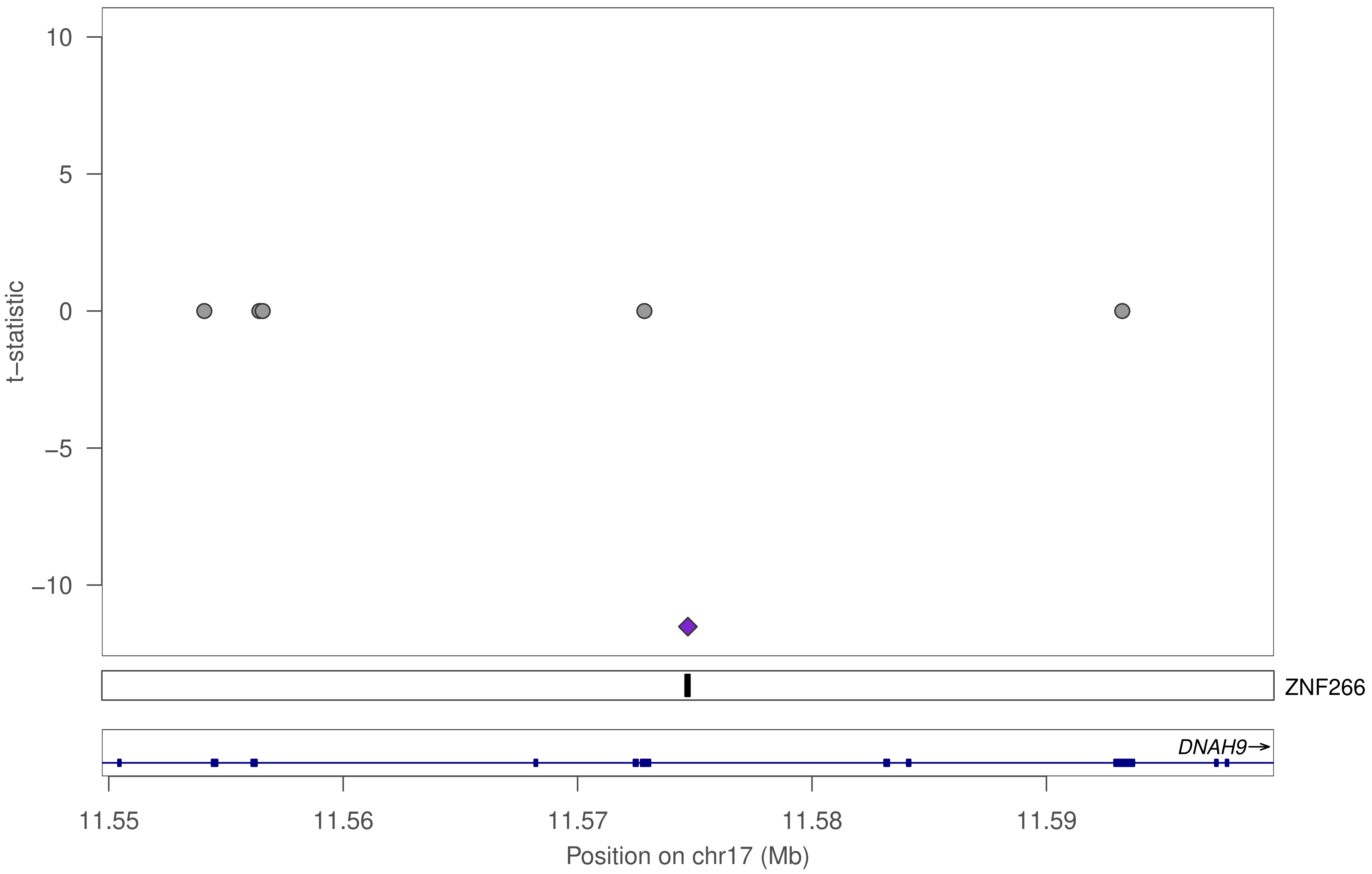
10
5
0
−5
−10
t−statistic
ZNF266
DNAH9→
11.55
11.56
11.57
11.58
11.59
Position on chr17 (Mb)

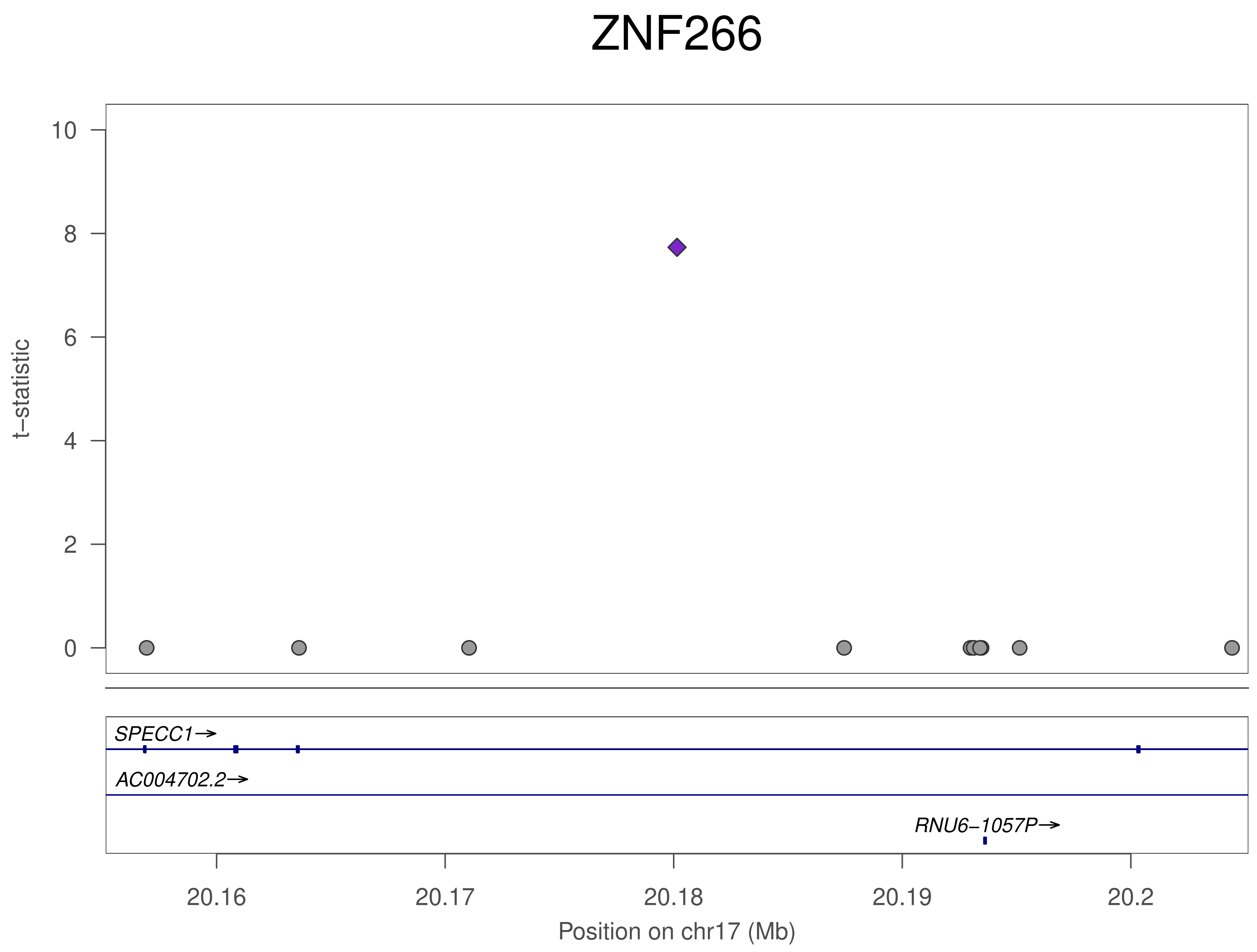
ZNF266
t−statistic
10
8
6
4
2
0
SPECC1→
AC004702.2→
RNU6−1057P→
20.16
20.17
20.18
20.19
20.2
Position on chr17 (Mb)

#### ZNF266

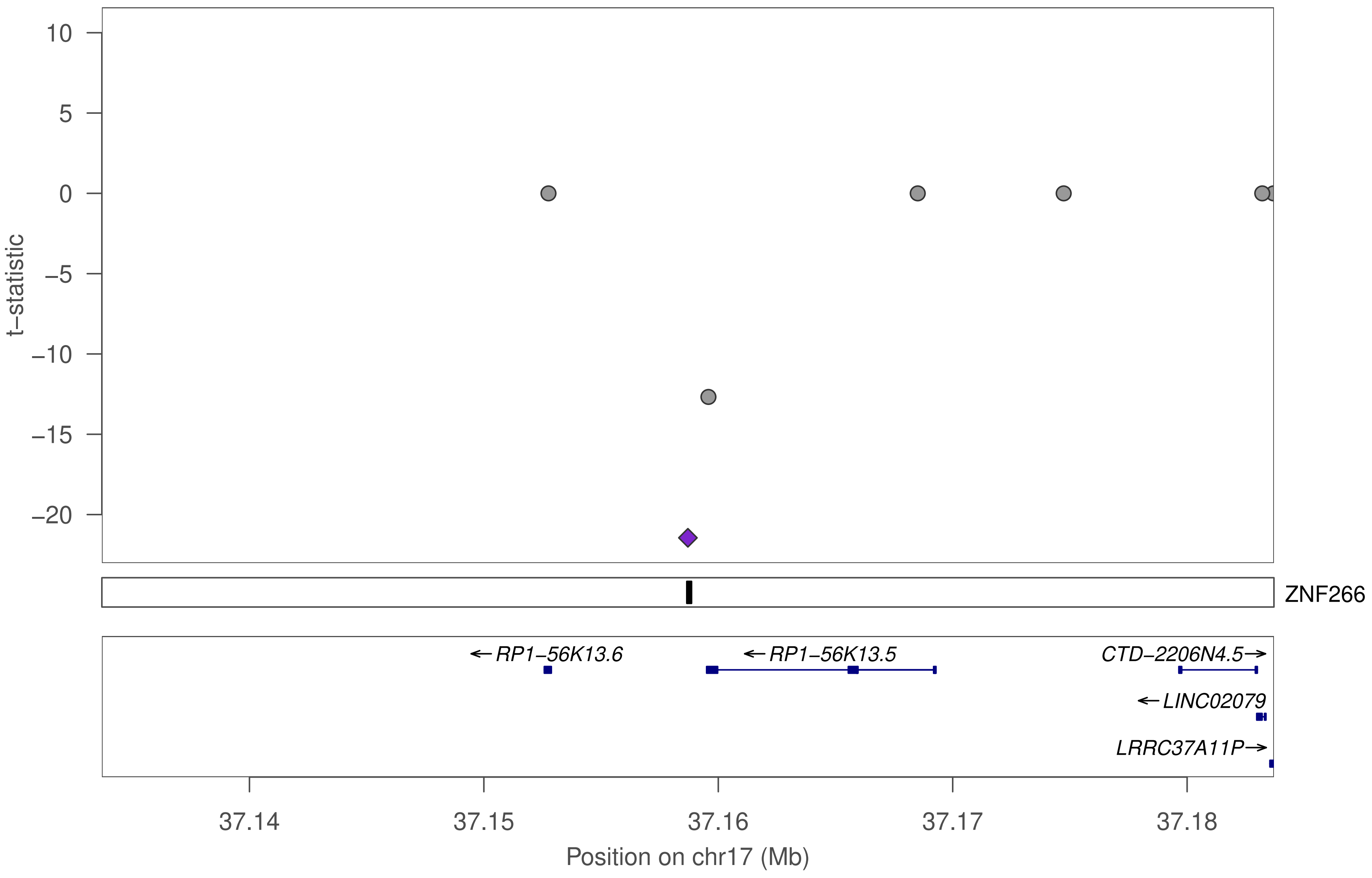

ZNF266
10
5
0
−5
−10
−15
−20
t−statistic
ZNF266
←RP1−56K13.6
←RP1−56K13.5
CTD−2206N4.5→
←LINC02079
LRRC37A11P→
37.14
37.15
37.16
37.17
37.18
Position on chr17 (Mb)

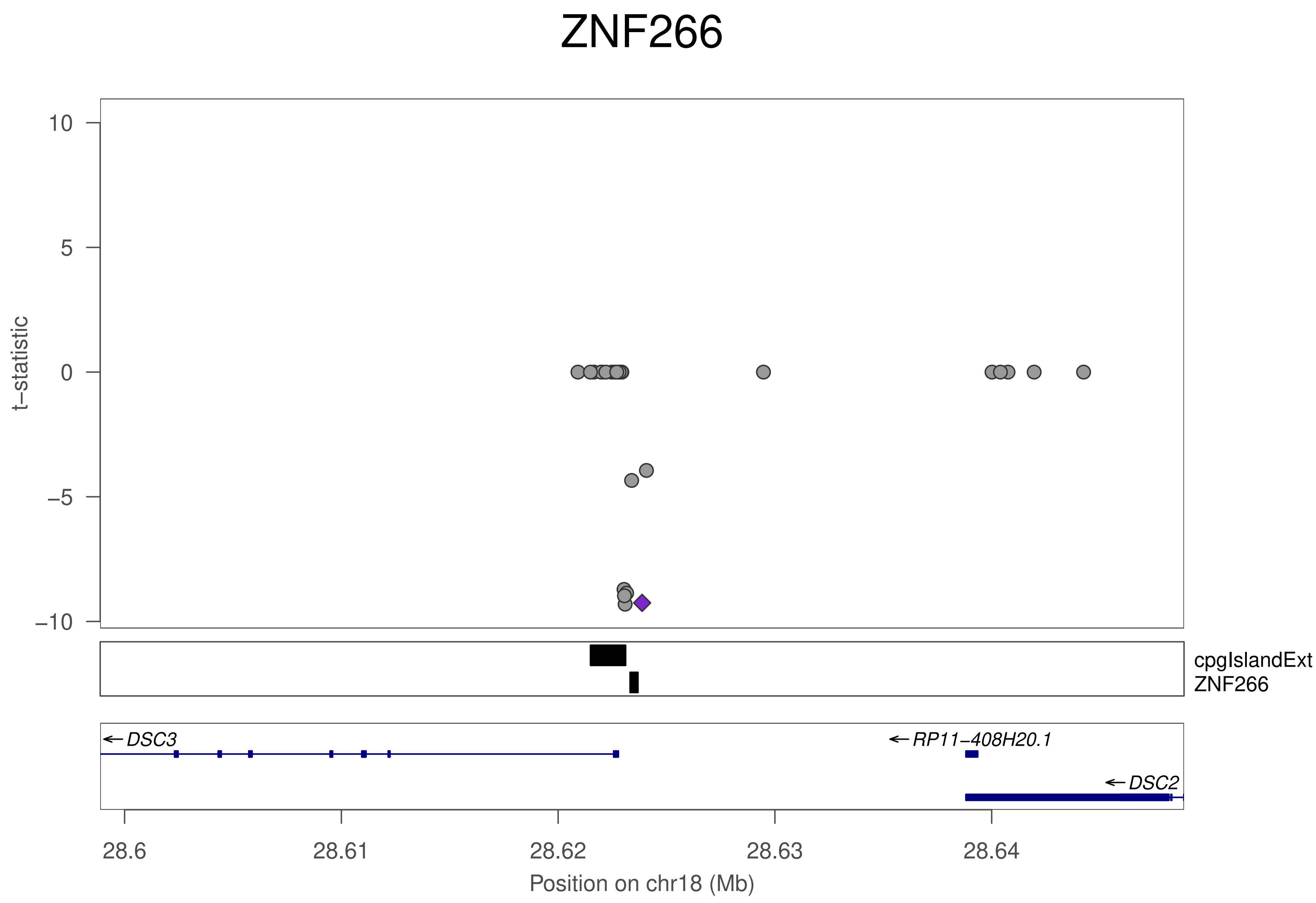
ZNF266
t−statistic
10
5
0
−5
−10
cpgIslandExt
ZNF266
DSC3
RP11−408H20.1
DSC2
28.6
28.61
28.62
28.63
28.64
Position on chr18 (Mb)

### ZNF266

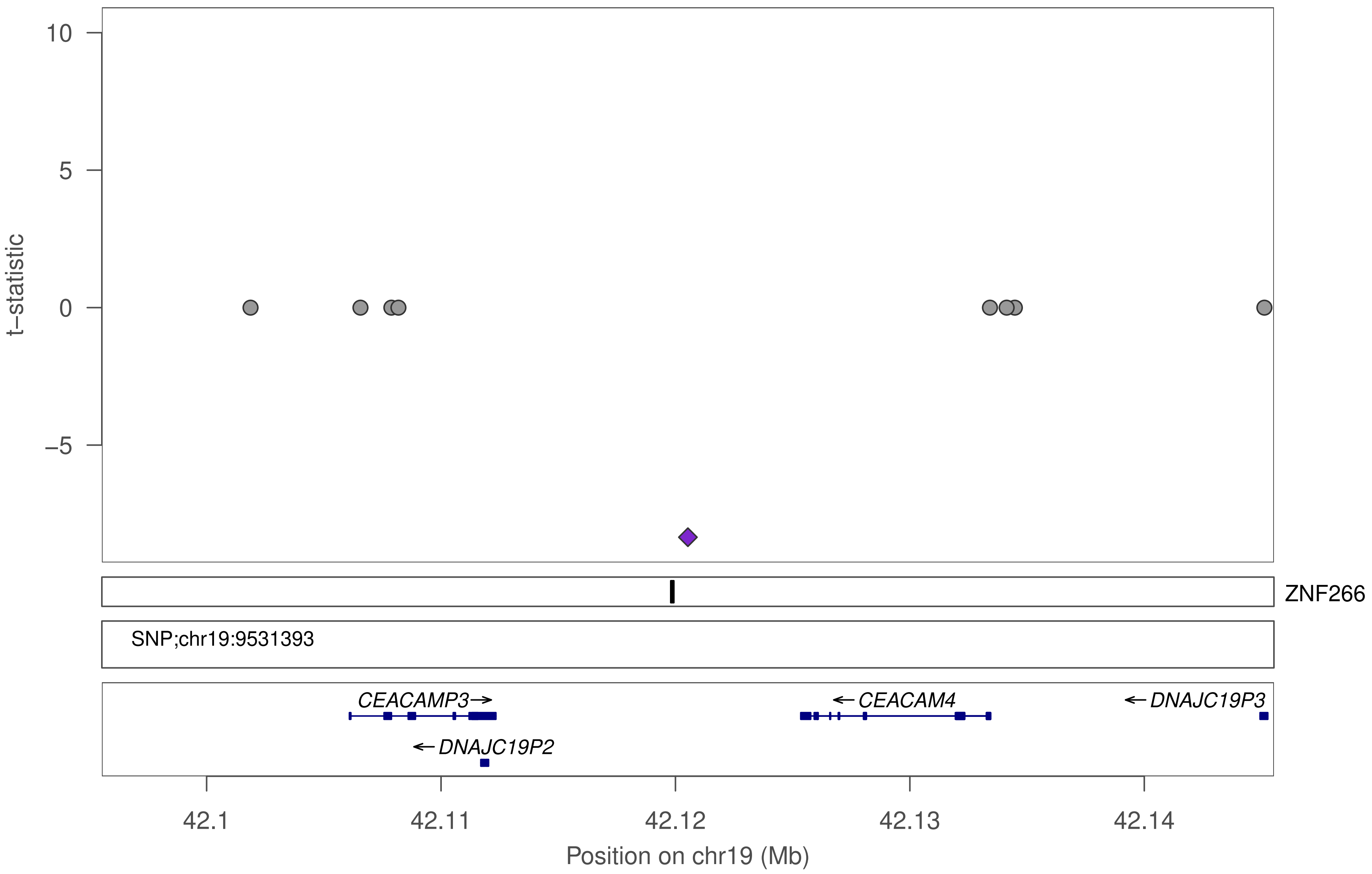

10
5
0
−5
t−statistic
ZNF266
SNP;chr19:9531393
CEACAMP3
CEACAM4
DNAJC19P3
DNAJC19P2
42.1
42.11
42.12
42.13
42.14
Position on chr19 (Mb)

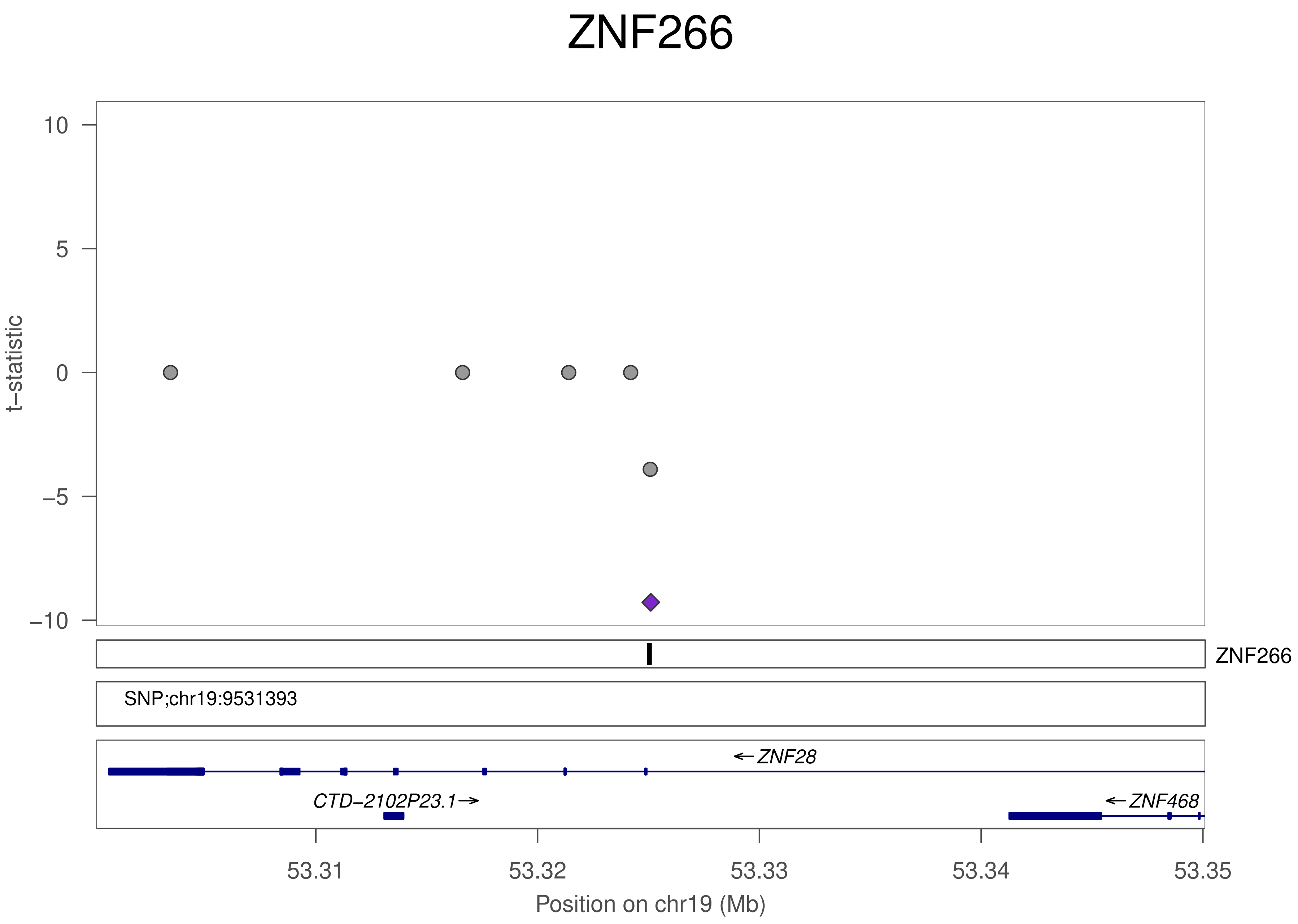
ZNF266
10
5
0
−5
−10
t−statistic
ZNF266
SNP;chr19:9531393
←ZNF28
CTD−2102P23.1→
←ZNF468
53.31
53.32
53.33
53.34
53.35
Position on chr19 (Mb)

### ZNF266

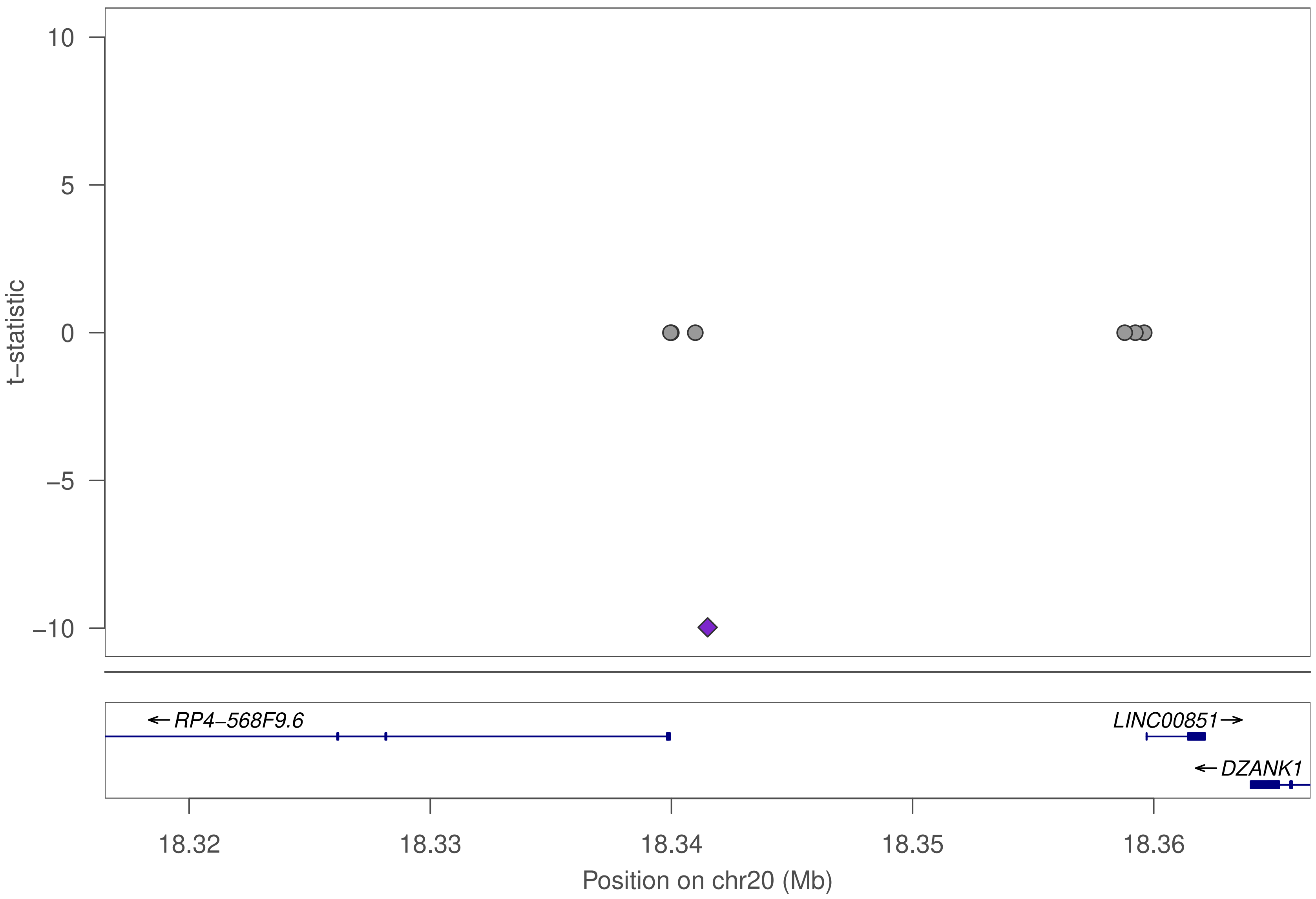

10
5
0
−5
−10
t−statistic
← RP4−568F9.6
LINC00851 →
← DZANK1
18.32
18.33
18.34
18.35
18.36
Position on chr20 (Mb)

### ZNF266

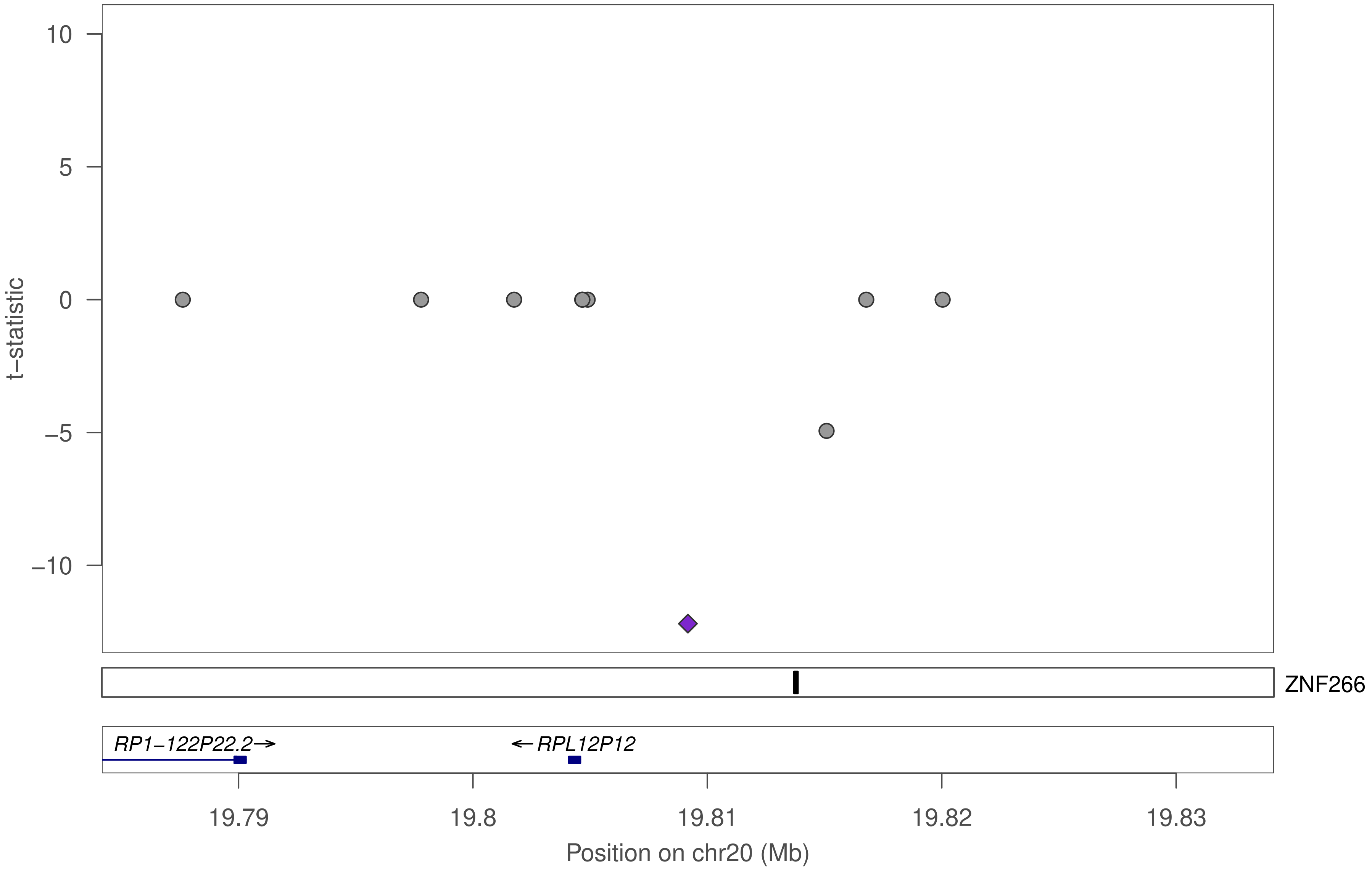

10
5
0
−5
−10
t−statistic
ZNF266
RP1−122P22.2→
←RPL12P12
19.79
19.8
19.81
19.82
19.83
Position on chr20 (Mb)

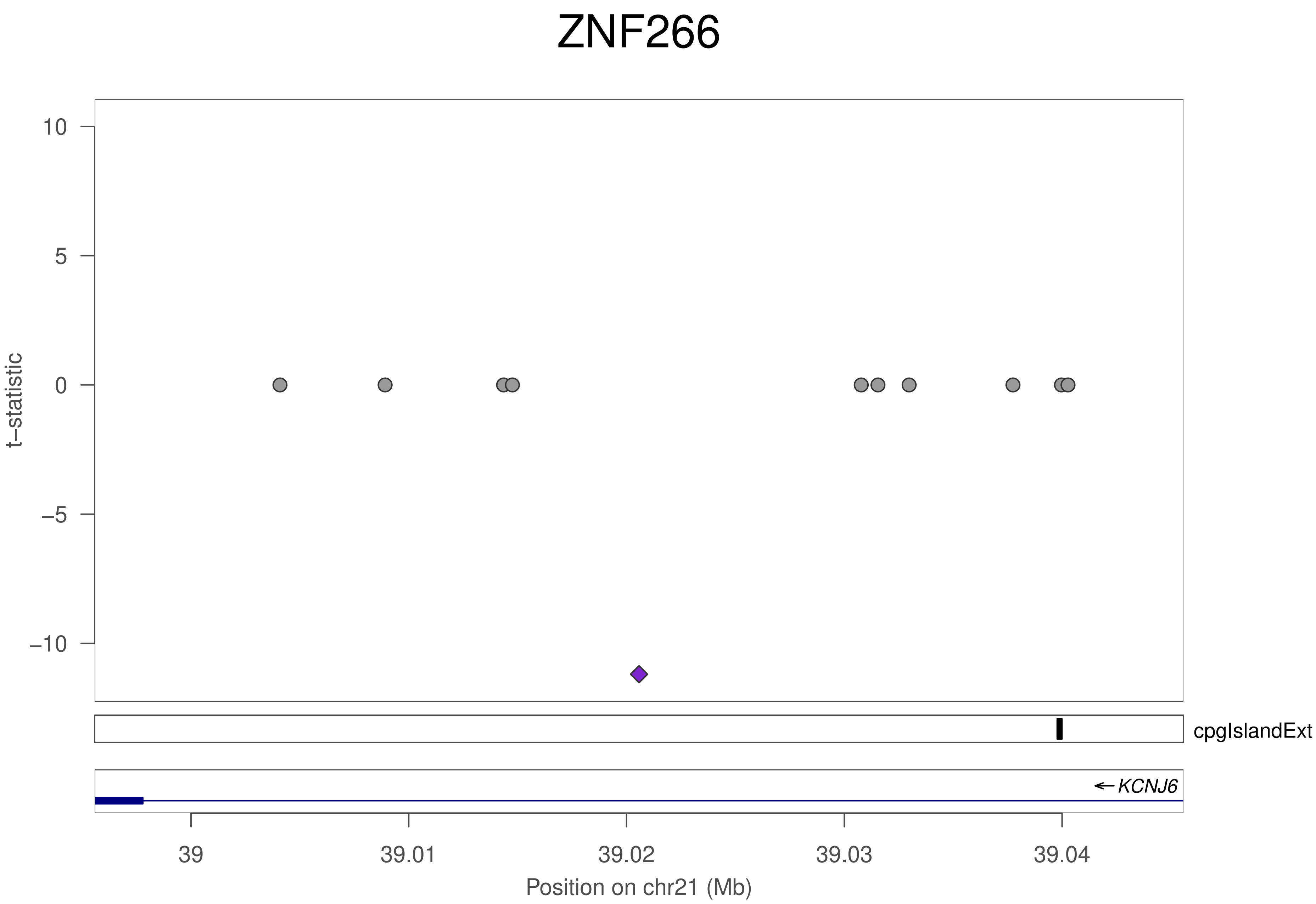
ZNF266
t–statistic
10
5
0
–5
–10
cpgIslandExt
KCNJ6
39
39.01
39.02
39.03
39.04
Position on chr21 (Mb)

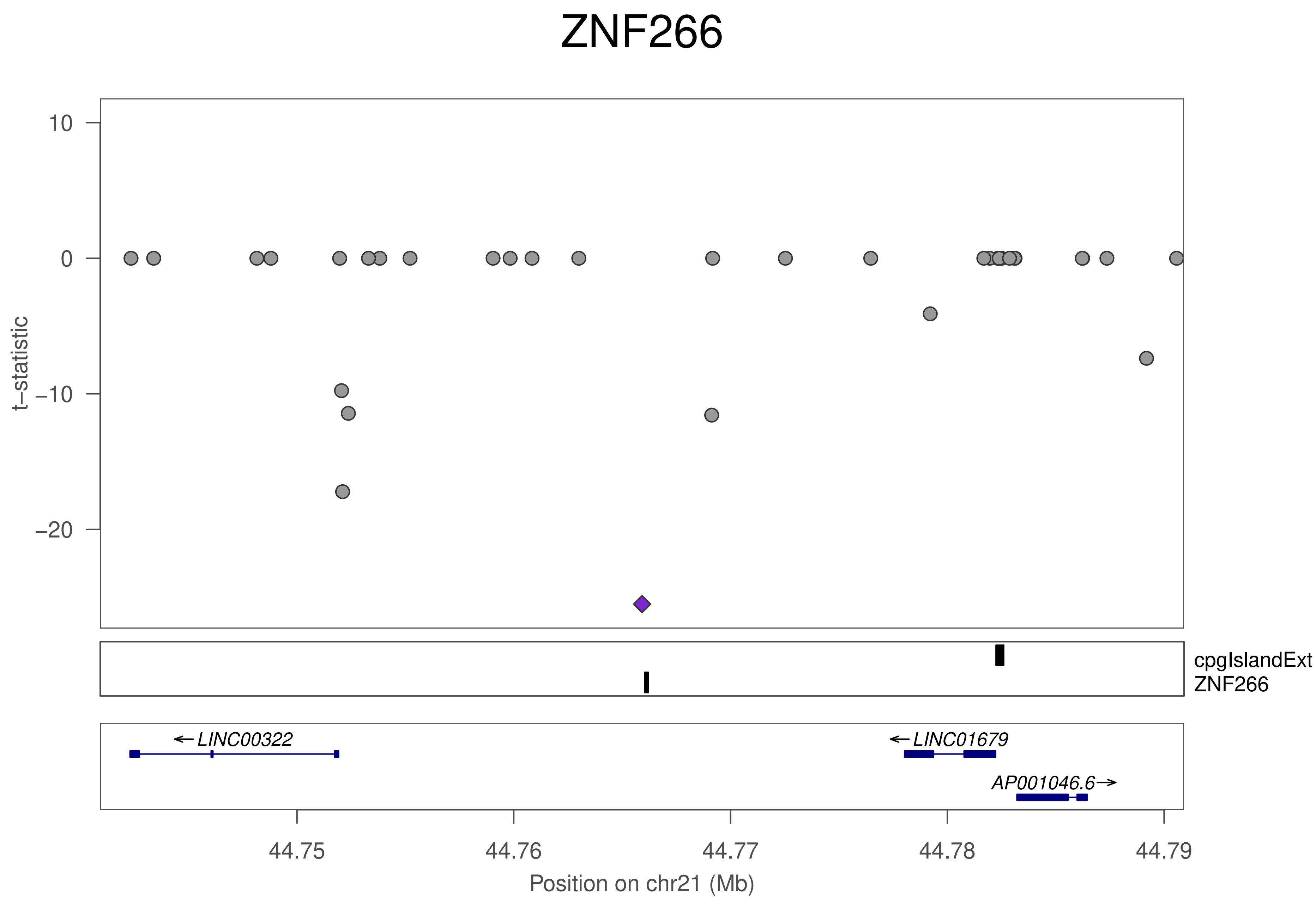
ZNF266
t−statistic
10
0
−10
−20
cpgIslandExt
ZNF266
LINC00322
LINC01679
AP001046.6
44.75
44.76
44.77
44.78
44.79
Position on chr21 (Mb)

### ZNF266

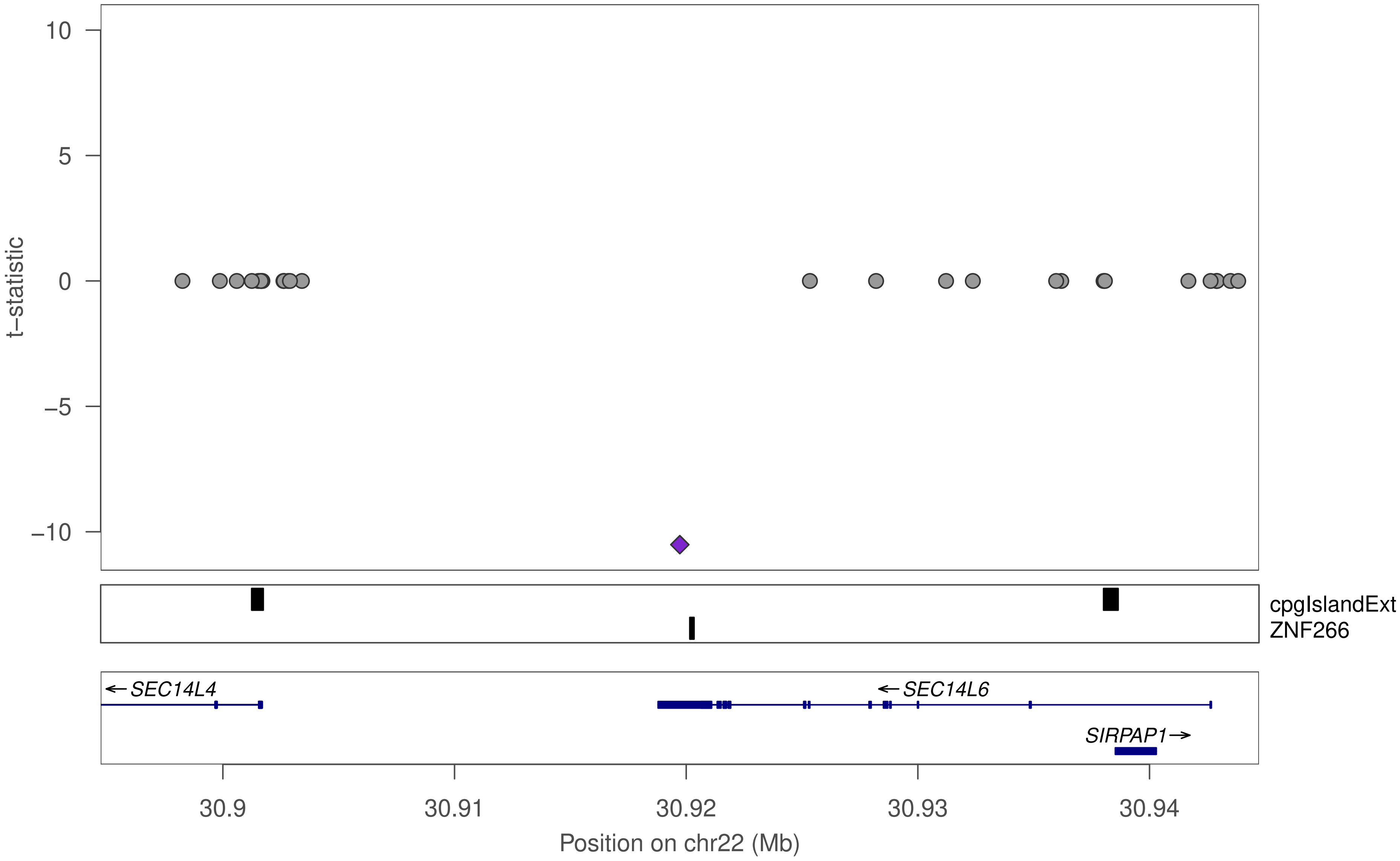

t–statistic
10
5
0
–5
–10
cpgIslandExt
ZNF266
SEC14L4
SEC14L6
SIRPAP1
30.9
30.91
30.92
30.93
30.94
Position on chr22 (Mb)

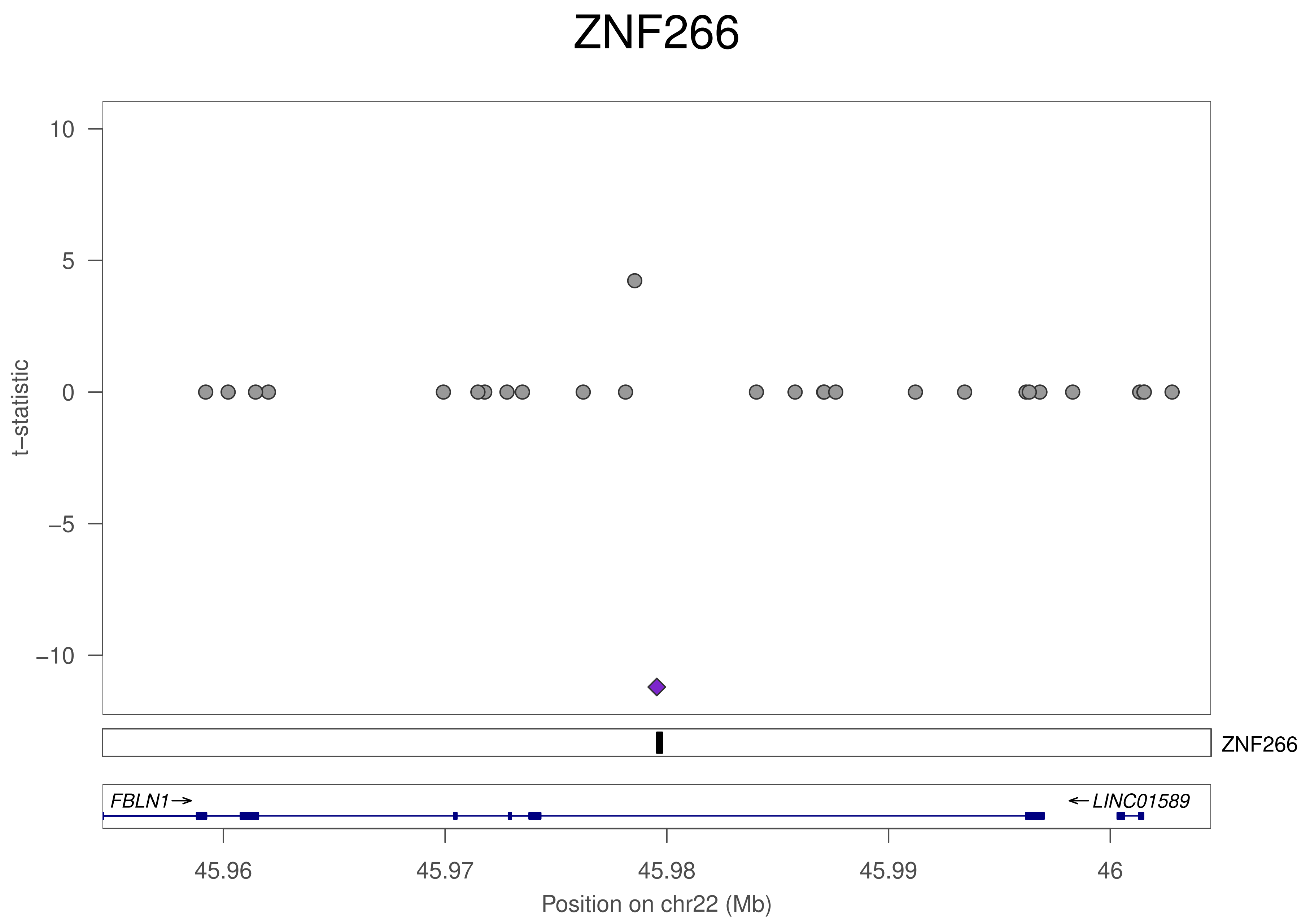
ZNF266
10
5
0
−5
−10
t−statistic
ZNF266
FBLN1→
←LINC01589
45.96
45.97
45.98
45.99
46
Position on chr22 (Mb)

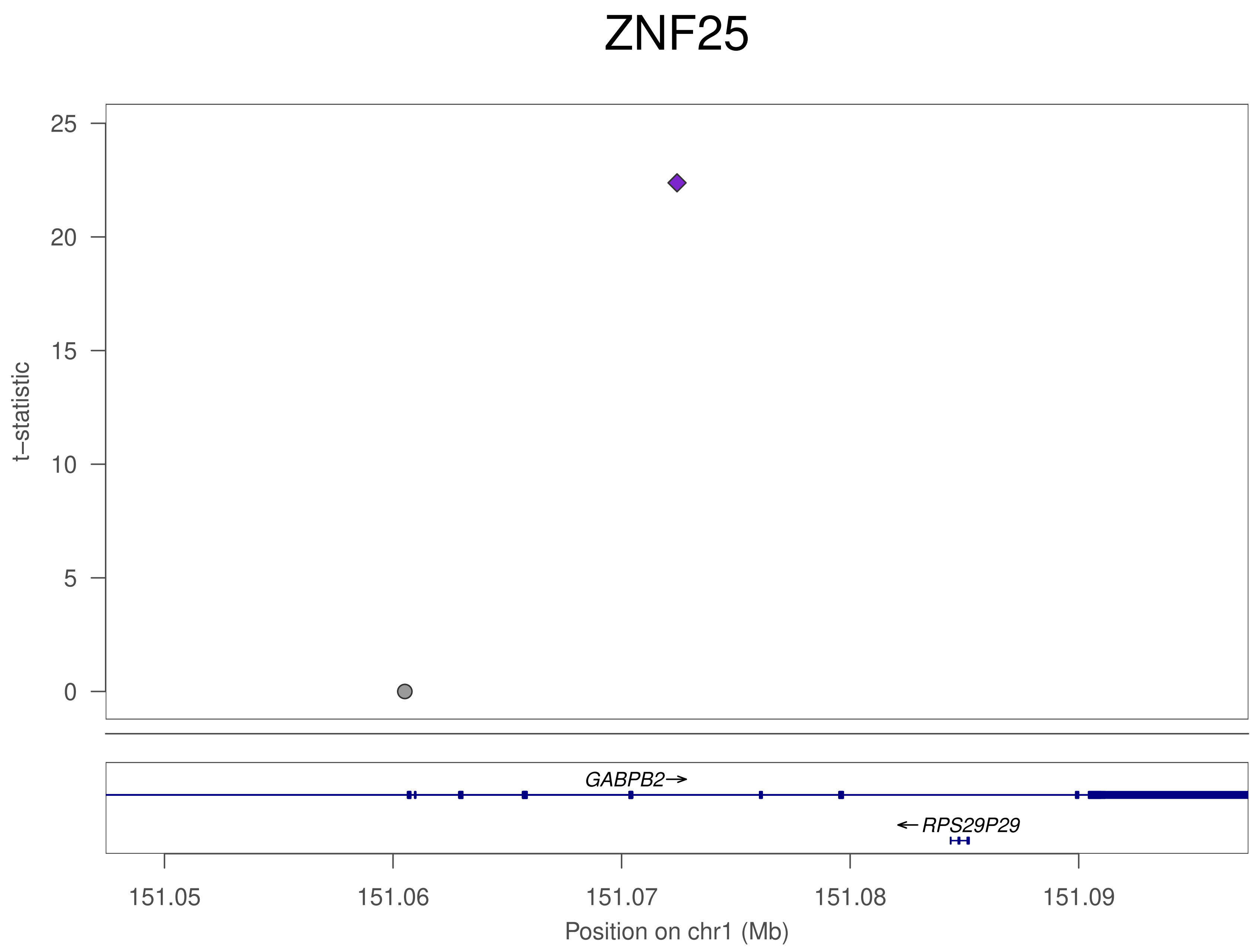

ZNF25
t−statistic
25
20
15
10
5
0
GABPB2→
←RPS29P29
151.05
151.06
151.07
151.08
151.09
Position on chr1 (Mb)

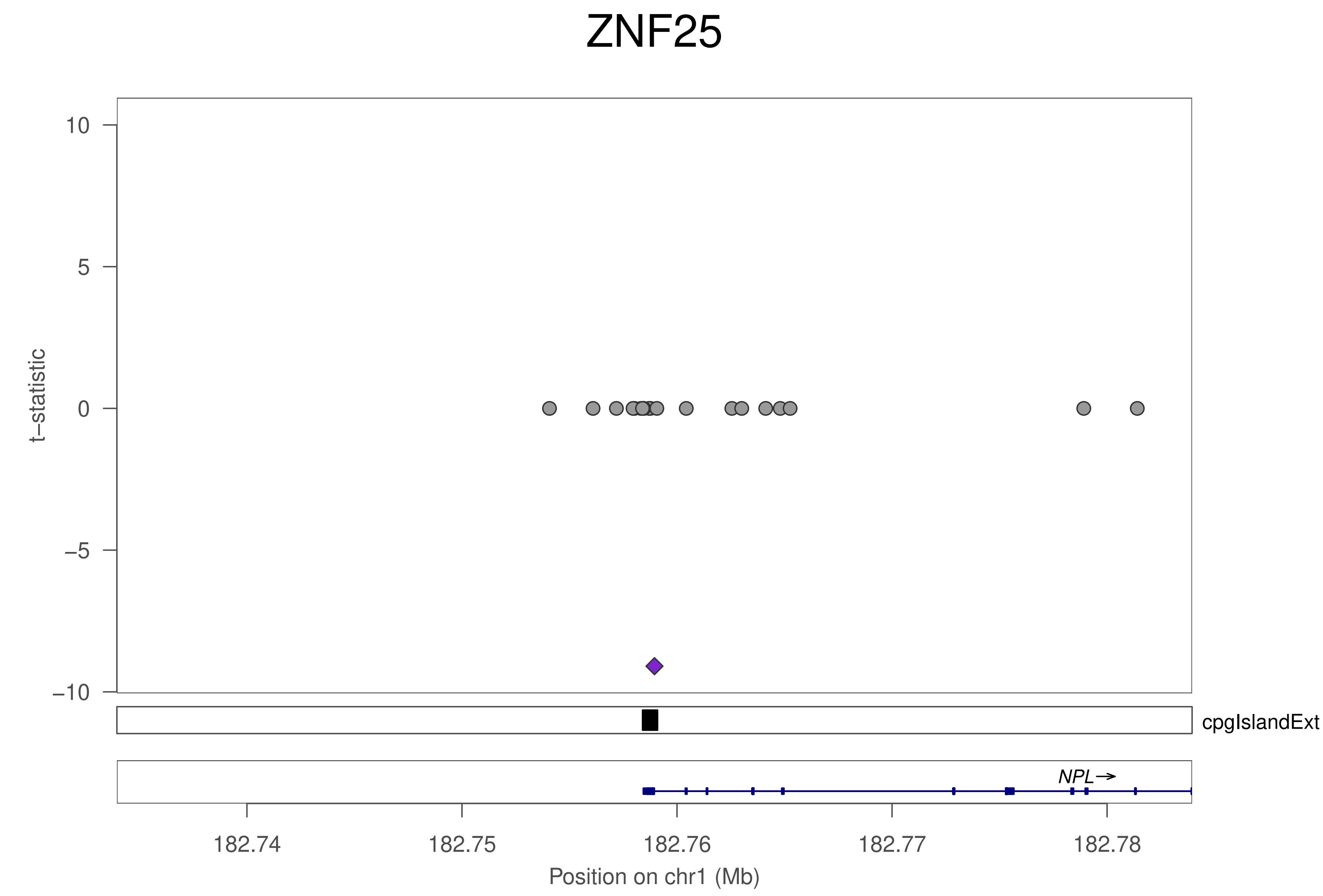

ZNF25
t–statistic
10
5
0
–5
–10
cpgIslandExt
NPL→
182.74
182.75
182.76
182.77
182.78
Position on chr1 (Mb)

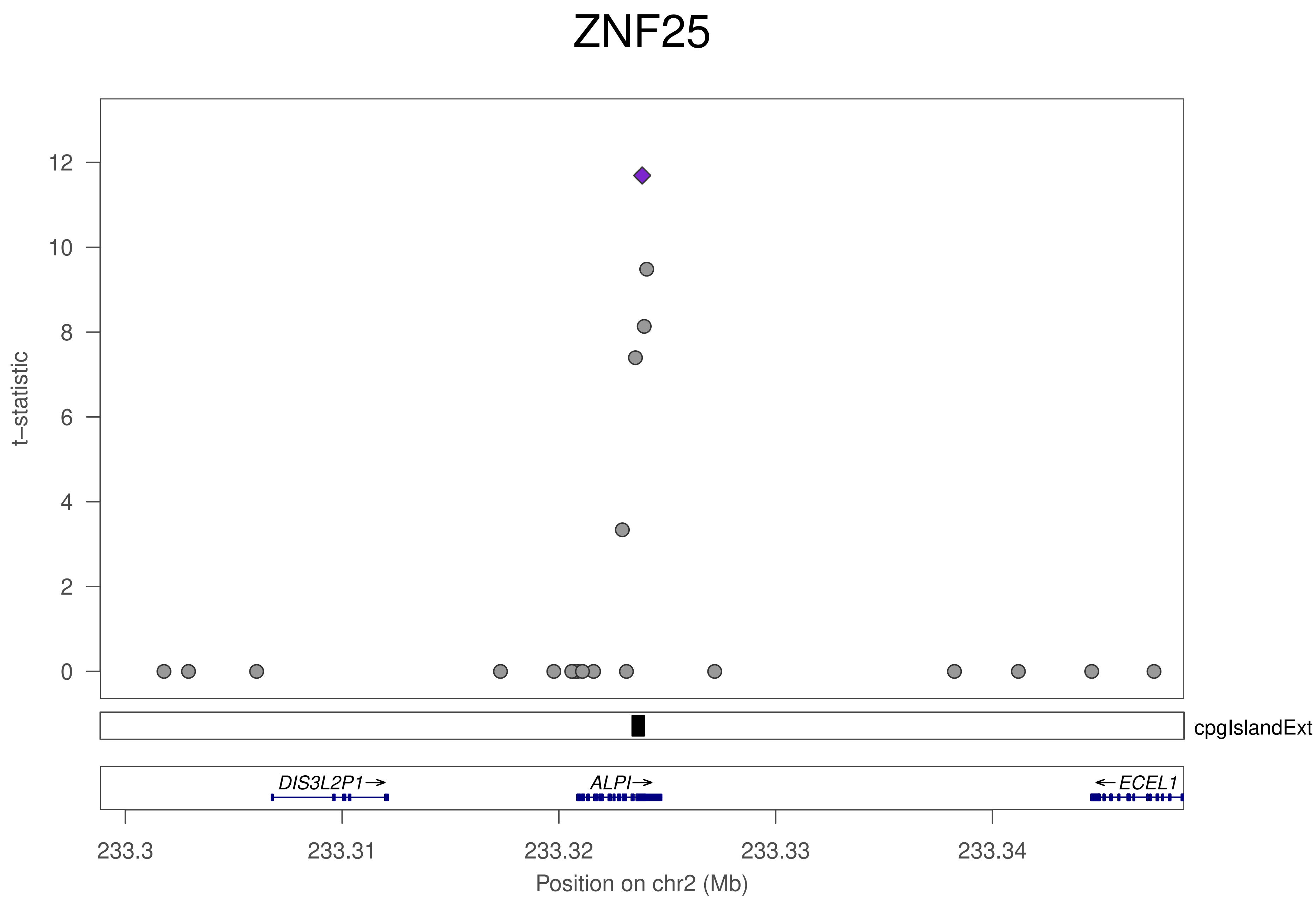
ZNF25
t−statistic
0
2
4
6
8
10
12
cpgIslandExt
DIS3L2P1→
ALPI→
←ECEL1
233.3
233.31
233.32
233.33
233.34
Position on chr2 (Mb)

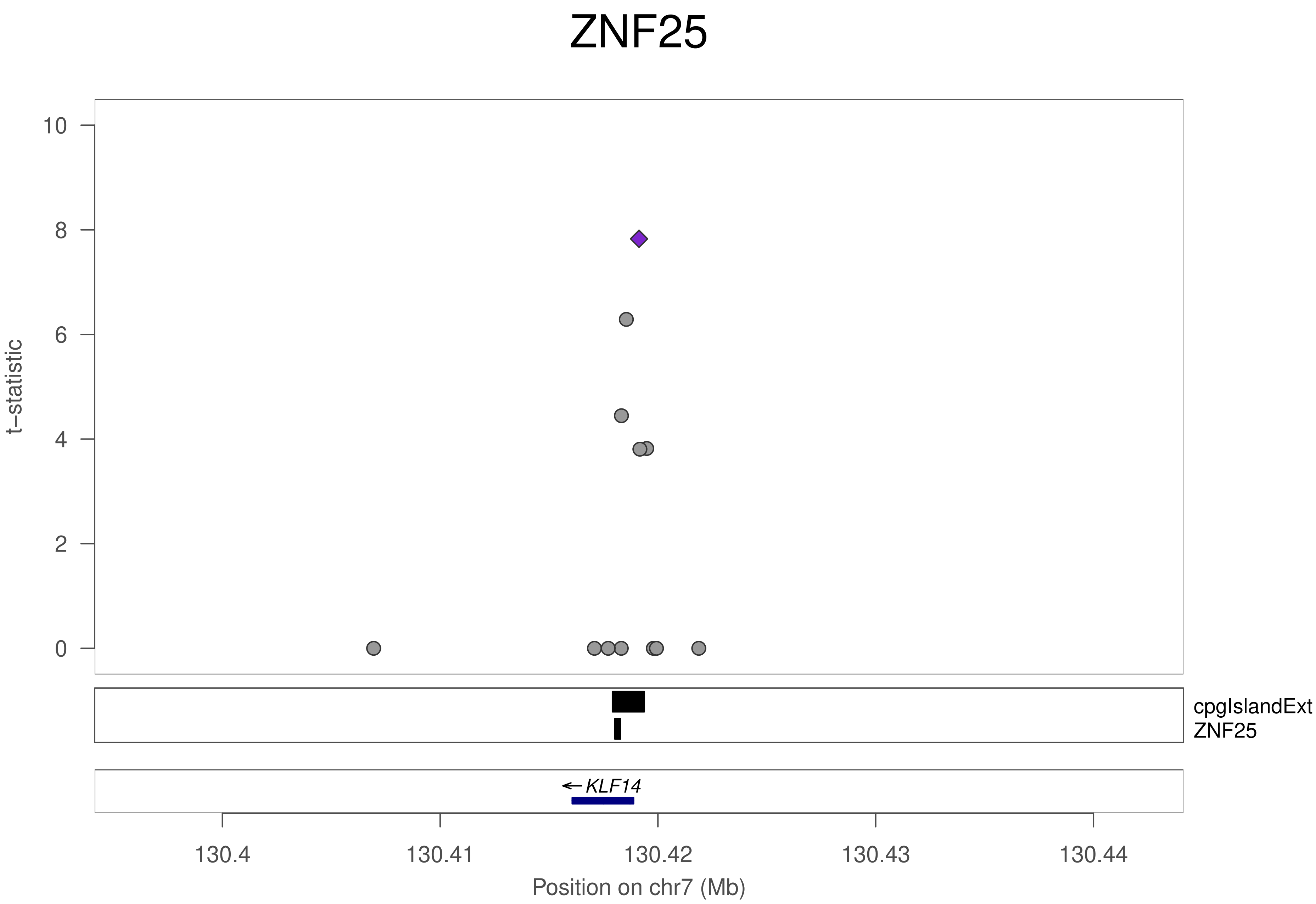
ZNF25
t−statistic
10
8
6
4
2
0
cpgIslandExt
ZNF25
←KLF14
130.4
130.41
130.42
130.43
130.44
Position on chr7 (Mb)

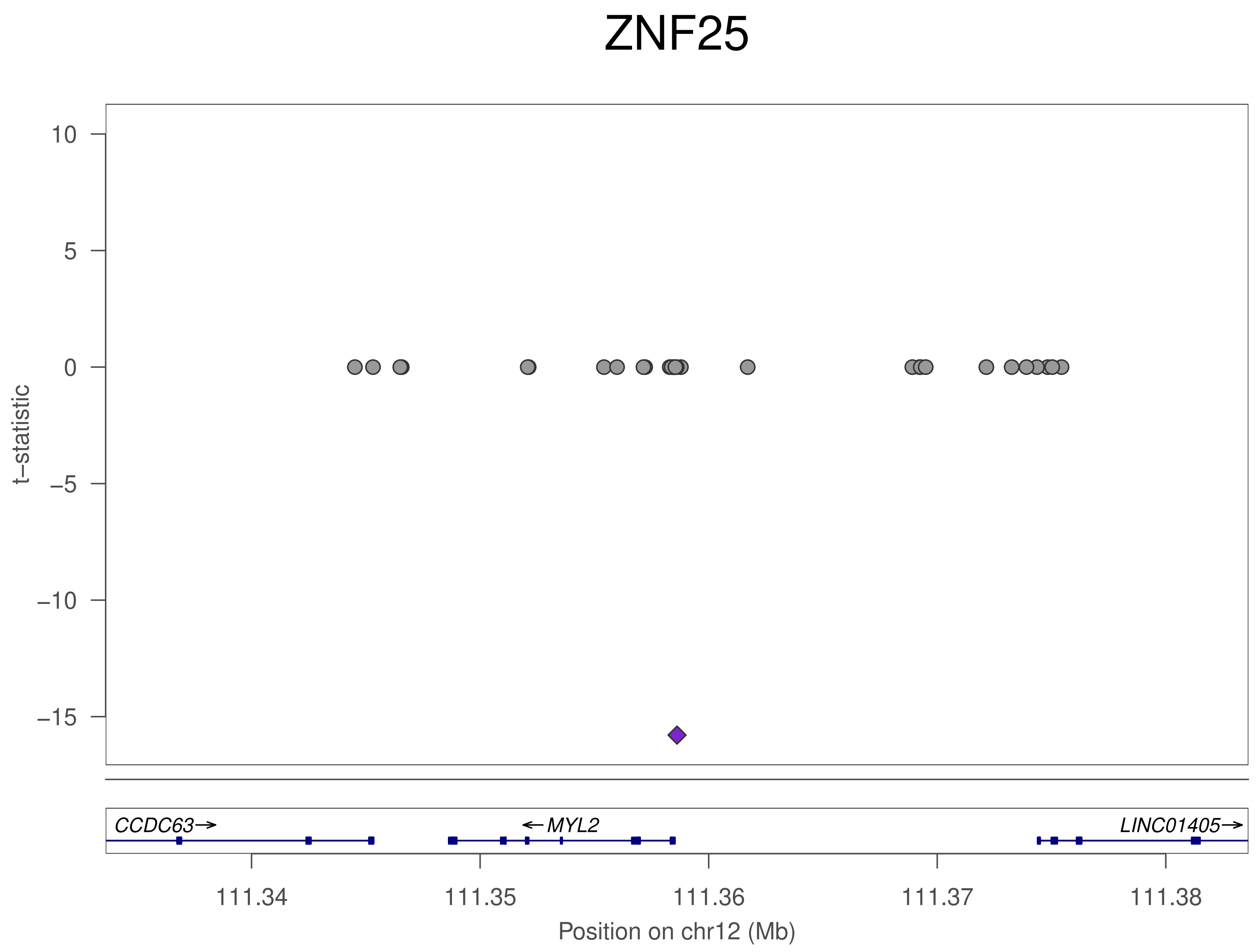

ZNF25
10
5
0
−5
−10
−15
t−statistic
CCDC63→
←MYL2
LINC01405→
111.34
111.35
111.36
111.37
111.38
Position on chr12 (Mb)

### ZNF25

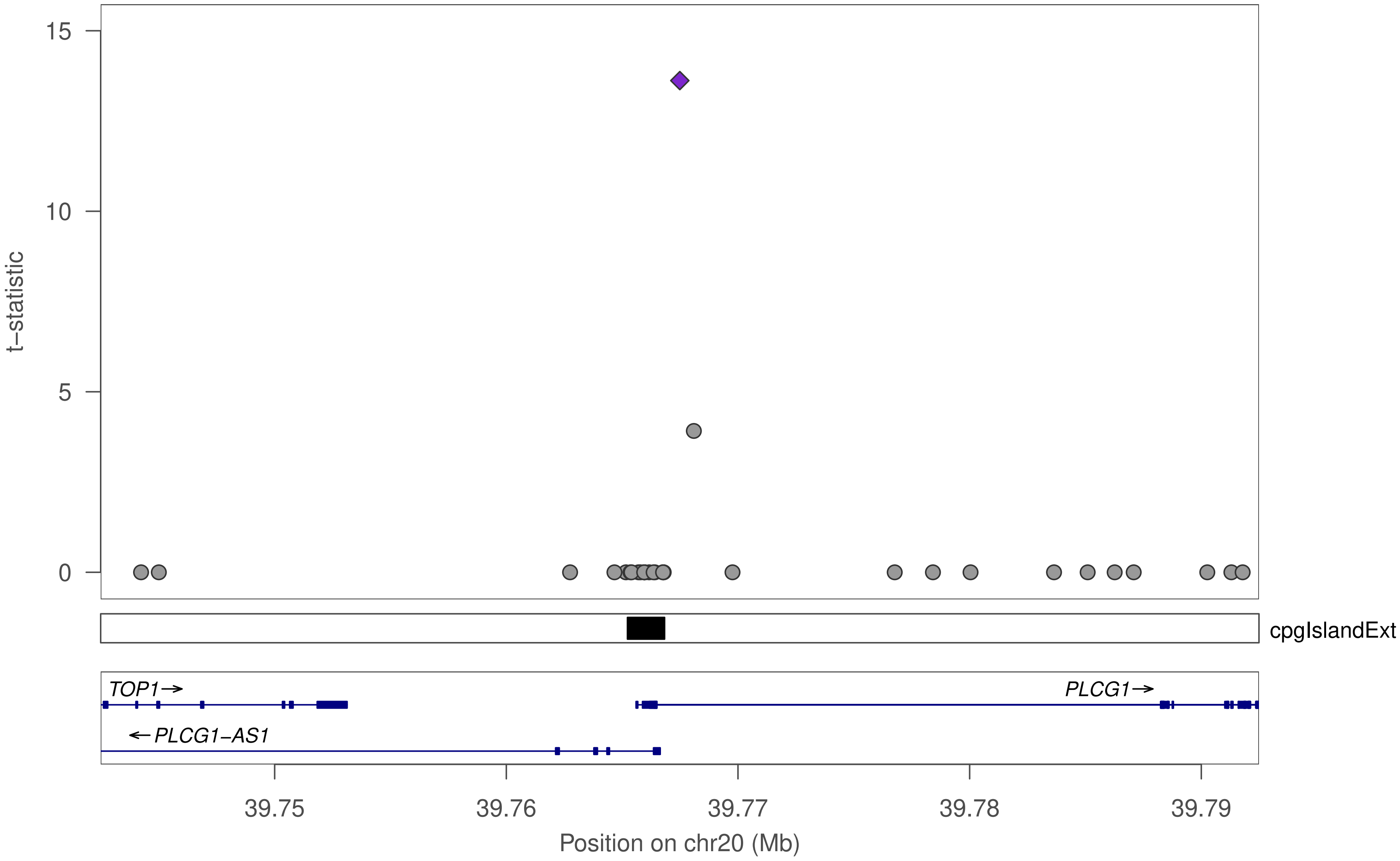

15
10
5
0
t−statistic
cpgIslandExt
TOP1→
PLCG1→
←PLCG1−AS1
39.75
39.76
39.77
39.78
39.79
Position on chr20 (Mb)

### ZNF169

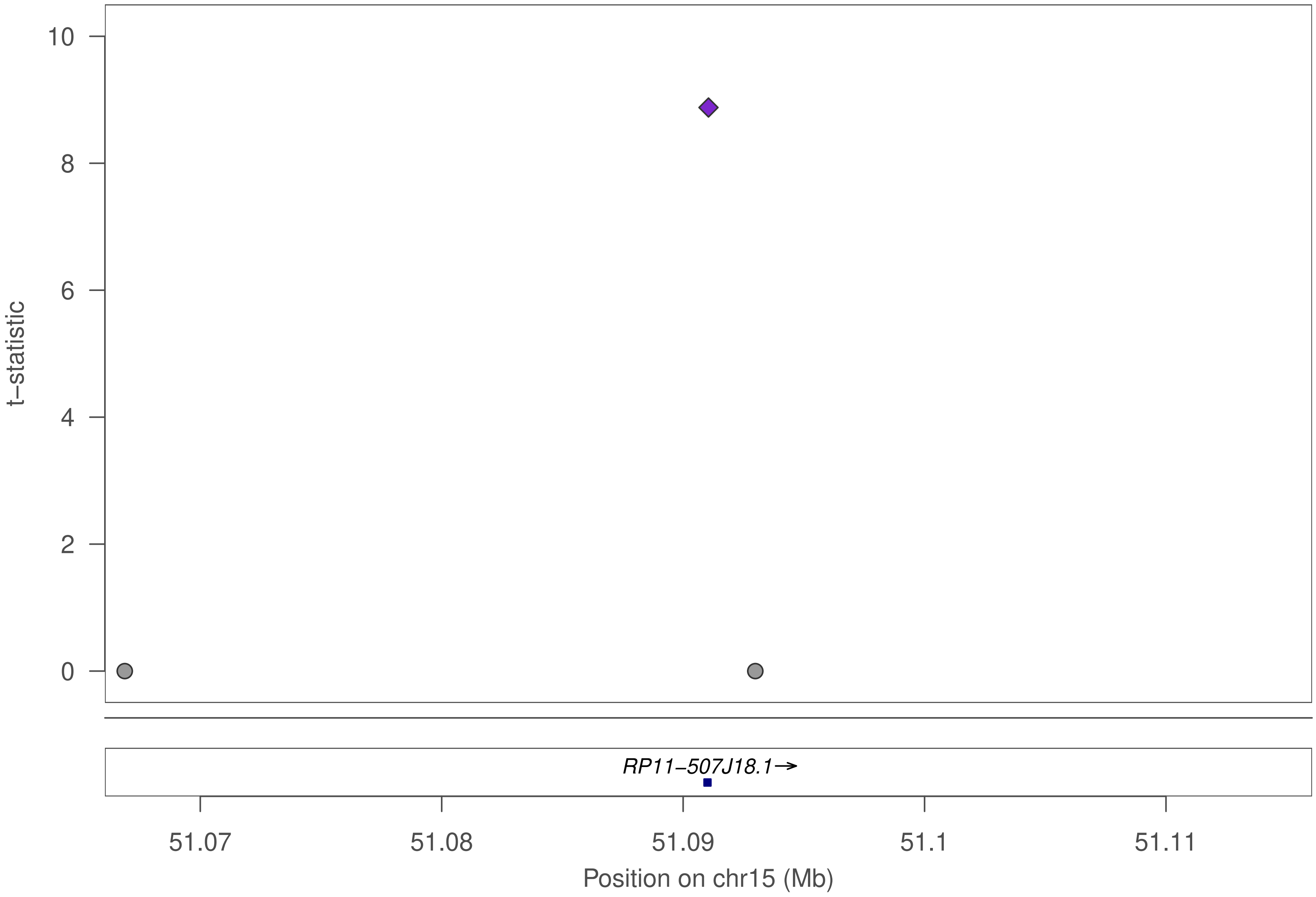

10
8
6
4
2
0
t−statistic
RP11−507J18.1→
51.07
51.08
51.09
51.1
51.11
Position on chr15 (Mb)

### ZNF613

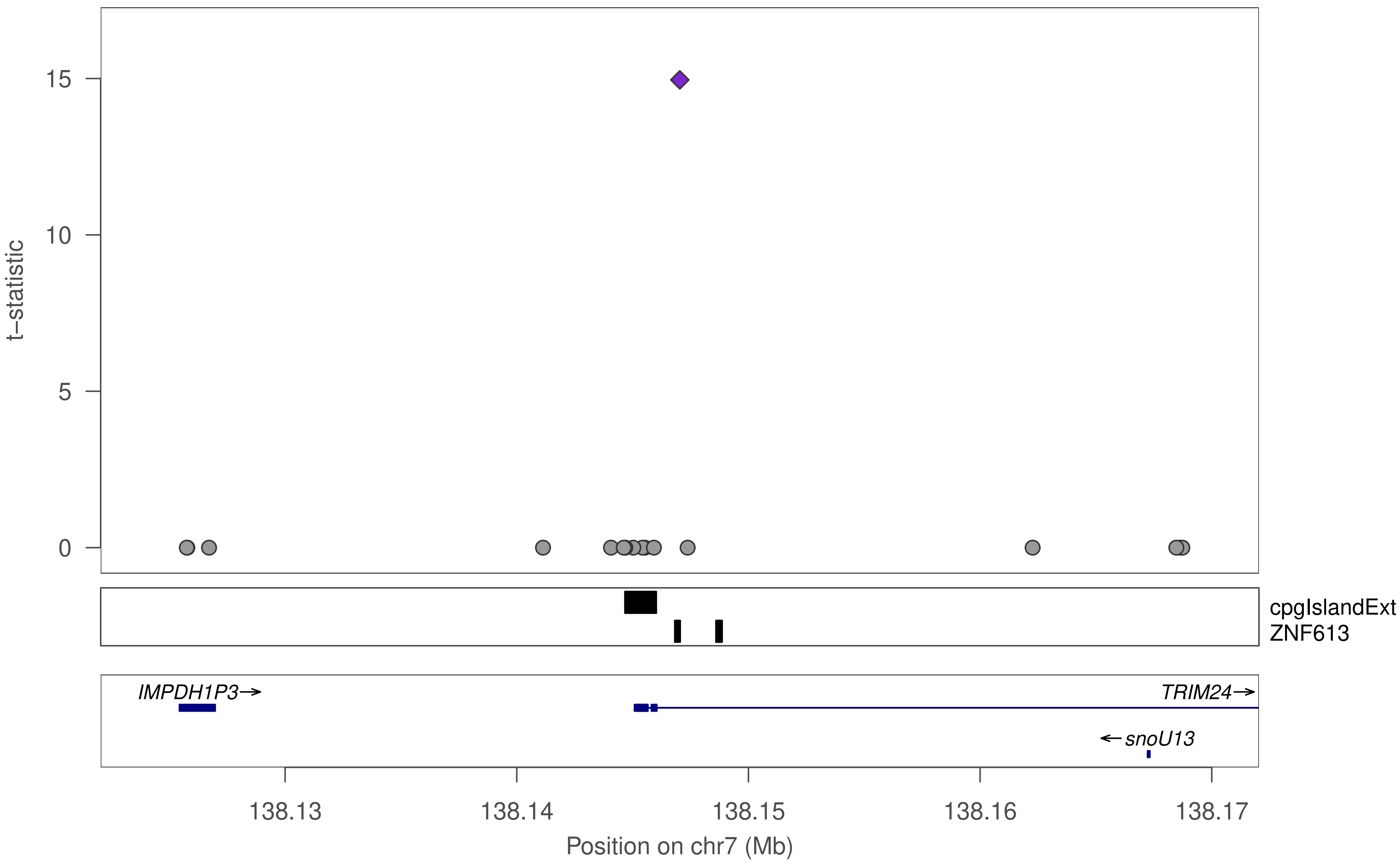

t−statistic
15
10
5
0
cpgIslandExt
ZNF613
IMPDH1P3
TRIM24
snoU13
138.13
138.14
138.15
138.16
138.17
Position on chr7 (Mb)

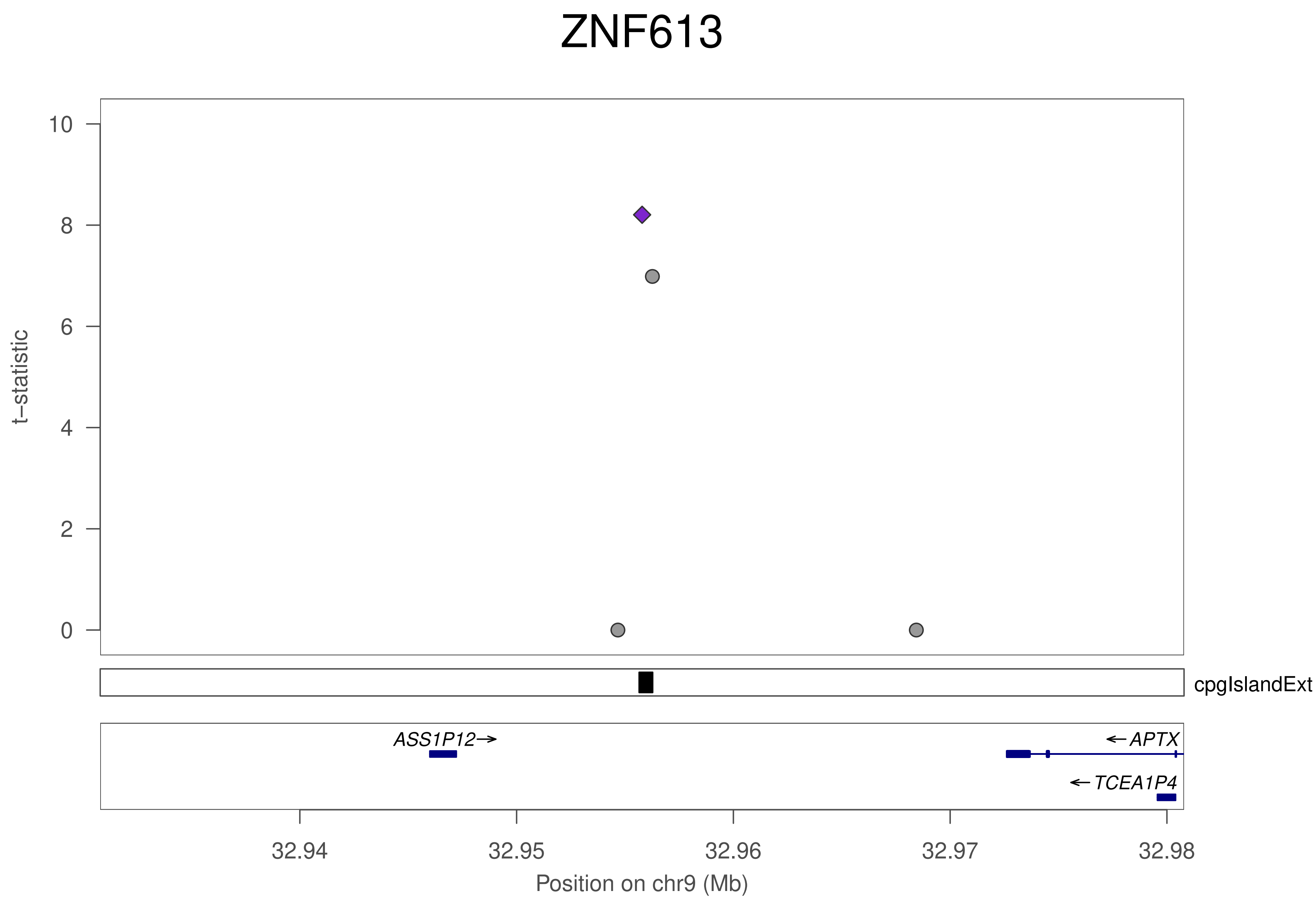
ZNF613
t−statistic
10
8
6
4
2
0
cpgIslandExt
ASS1P12→
←APTX
←TCEA1P4
32.94
32.95
32.96
32.97
32.98
Position on chr9 (Mb)

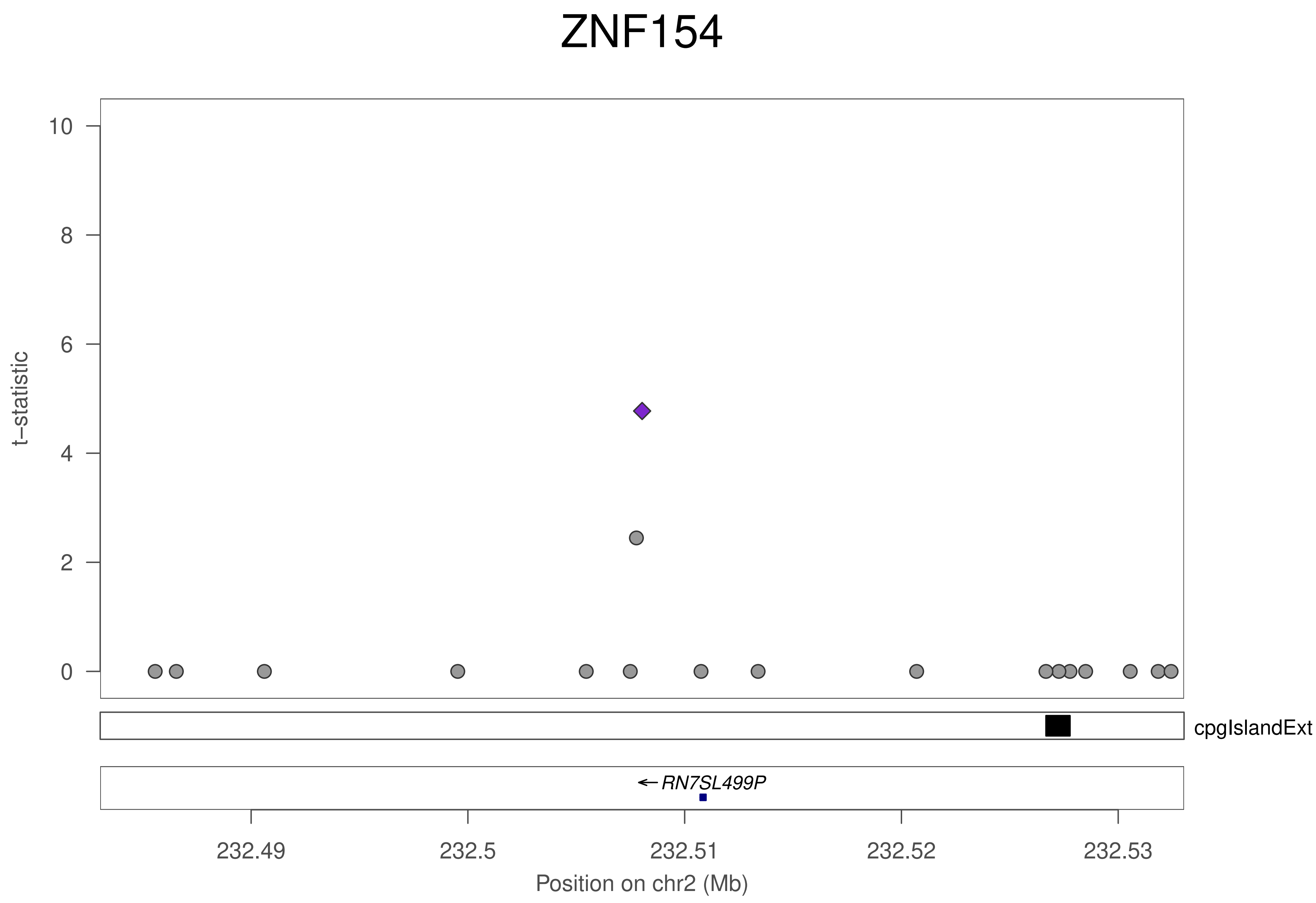

ZNF154
t–statistic
10
8
6
4
2
0
cpgIslandExt
RN7SL499P
232.49
232.5
232.51
232.52
232.53
Position on chr2 (Mb)

### ZNF154

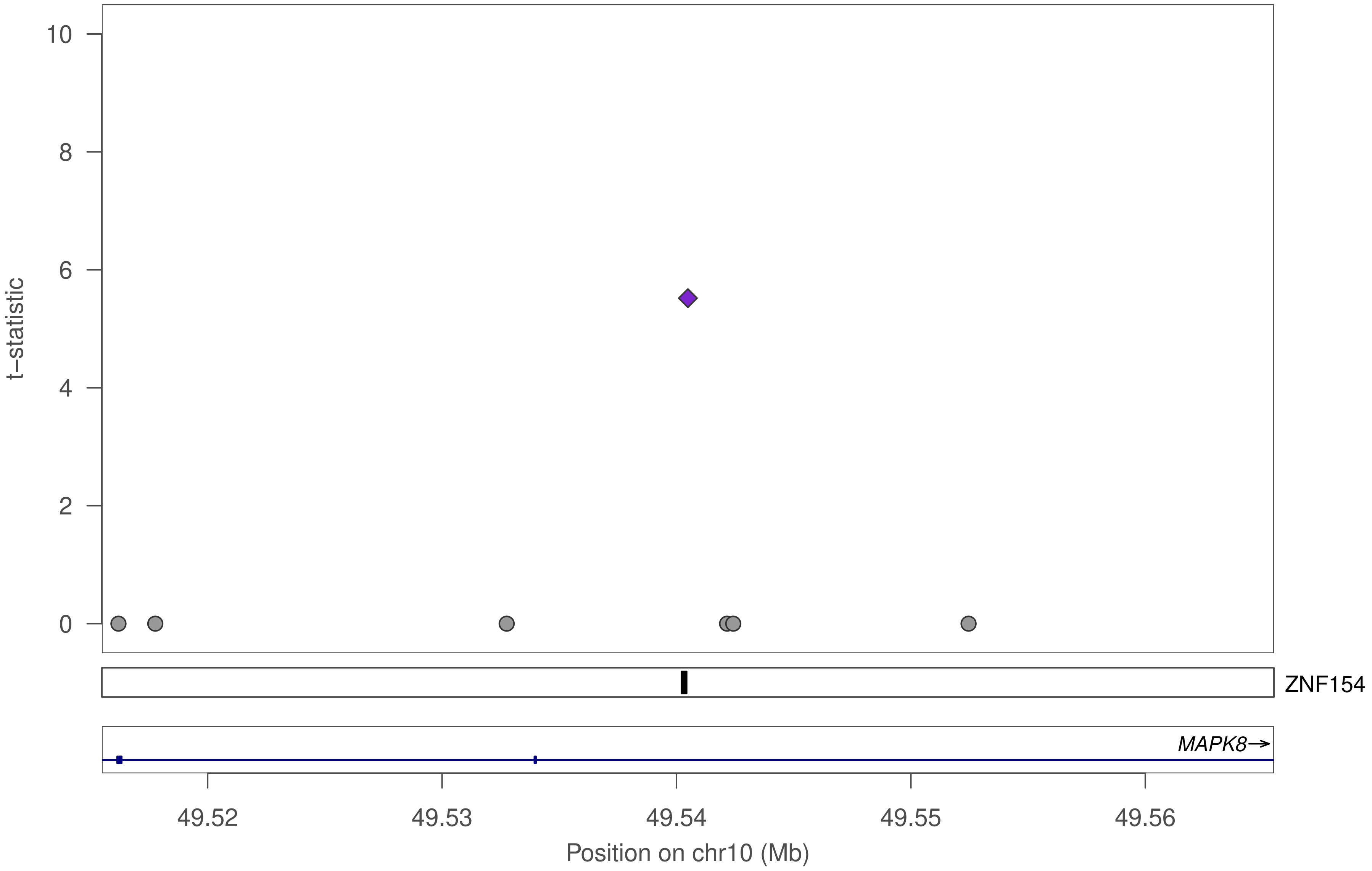

t−statistic
10
8
6
4
2
0
ZNF154
MAPK8→
49.52
49.53
49.54
49.55
49.56
Position on chr10 (Mb)

### ZNF154

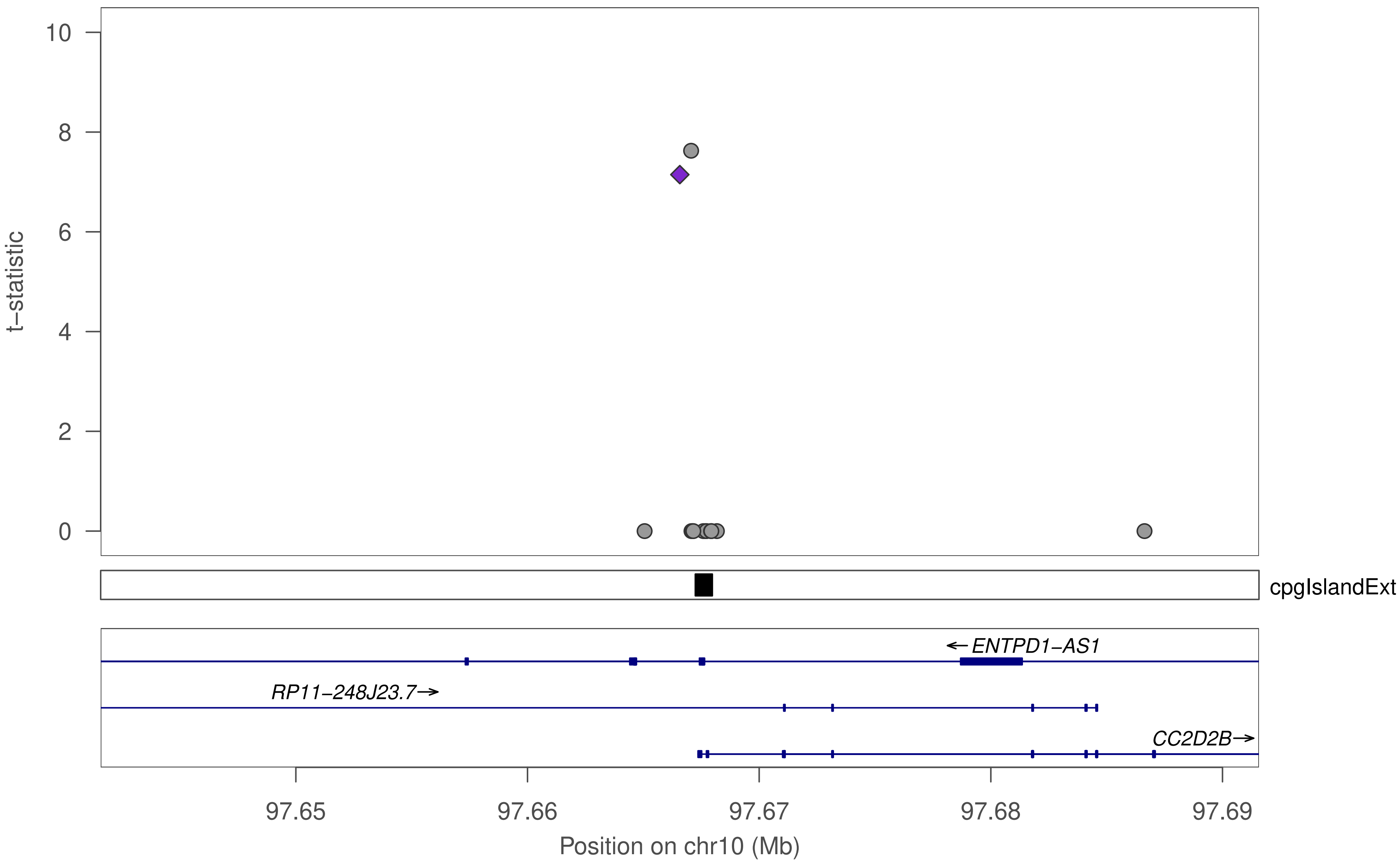

t−statistic
10
8
6
4
2
0
cpgIslandExt
←ENTPD1−AS1
RP11−248J23.7→
CC2D2B→
97.65
97.66
97.67
97.68
97.69
Position on chr10 (Mb)

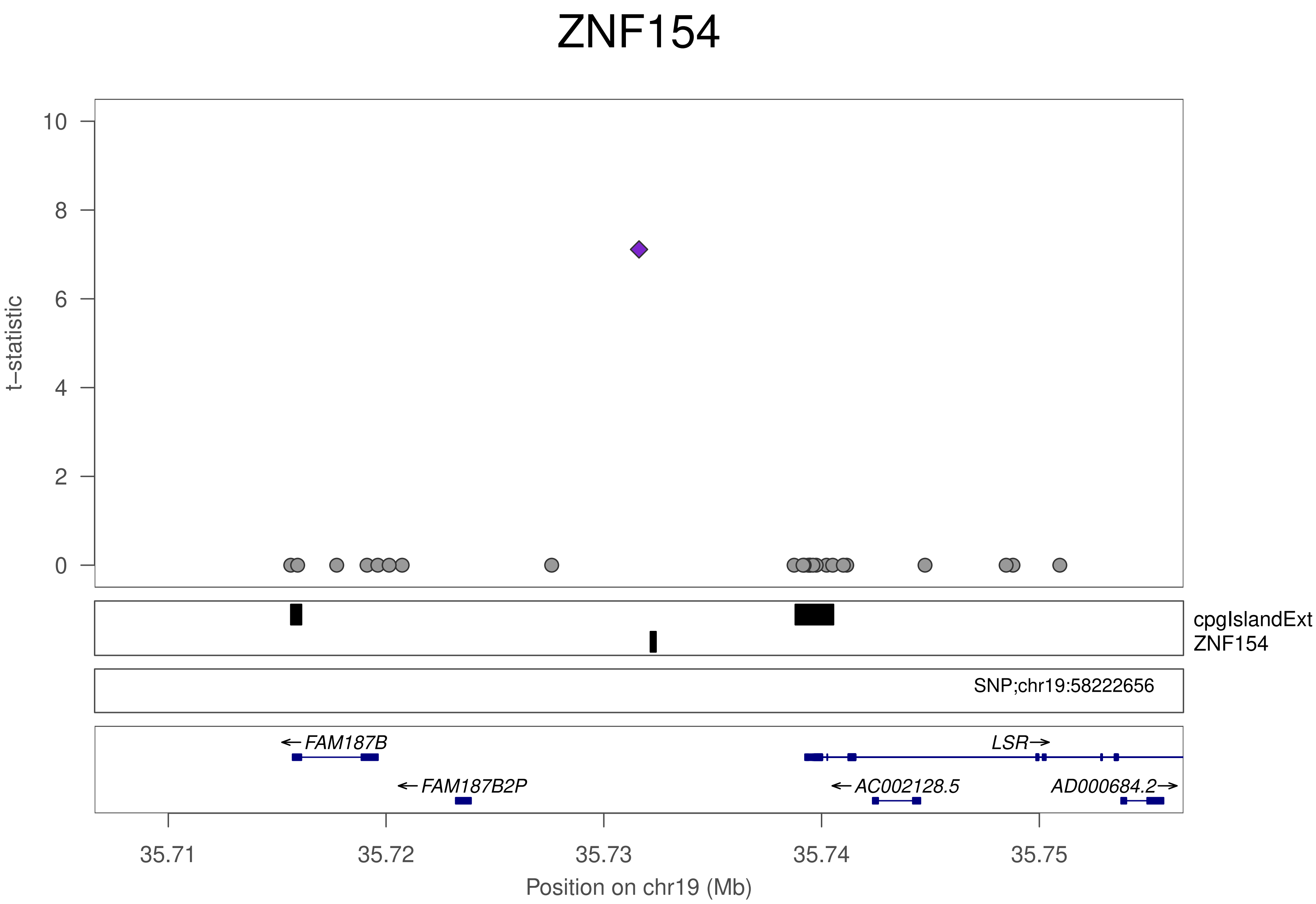
ZNF154
t−statistic
0
2
4
6
8
10
cpgIslandExt
ZNF154
SNP;chr19:58222656
←FAM187B
←FAM187B2P
LSR→
←AC002128.5
AD000684.2→
35.71
35.72
35.73
35.74
35.75
Position on chr19 (Mb)

### ZNF443

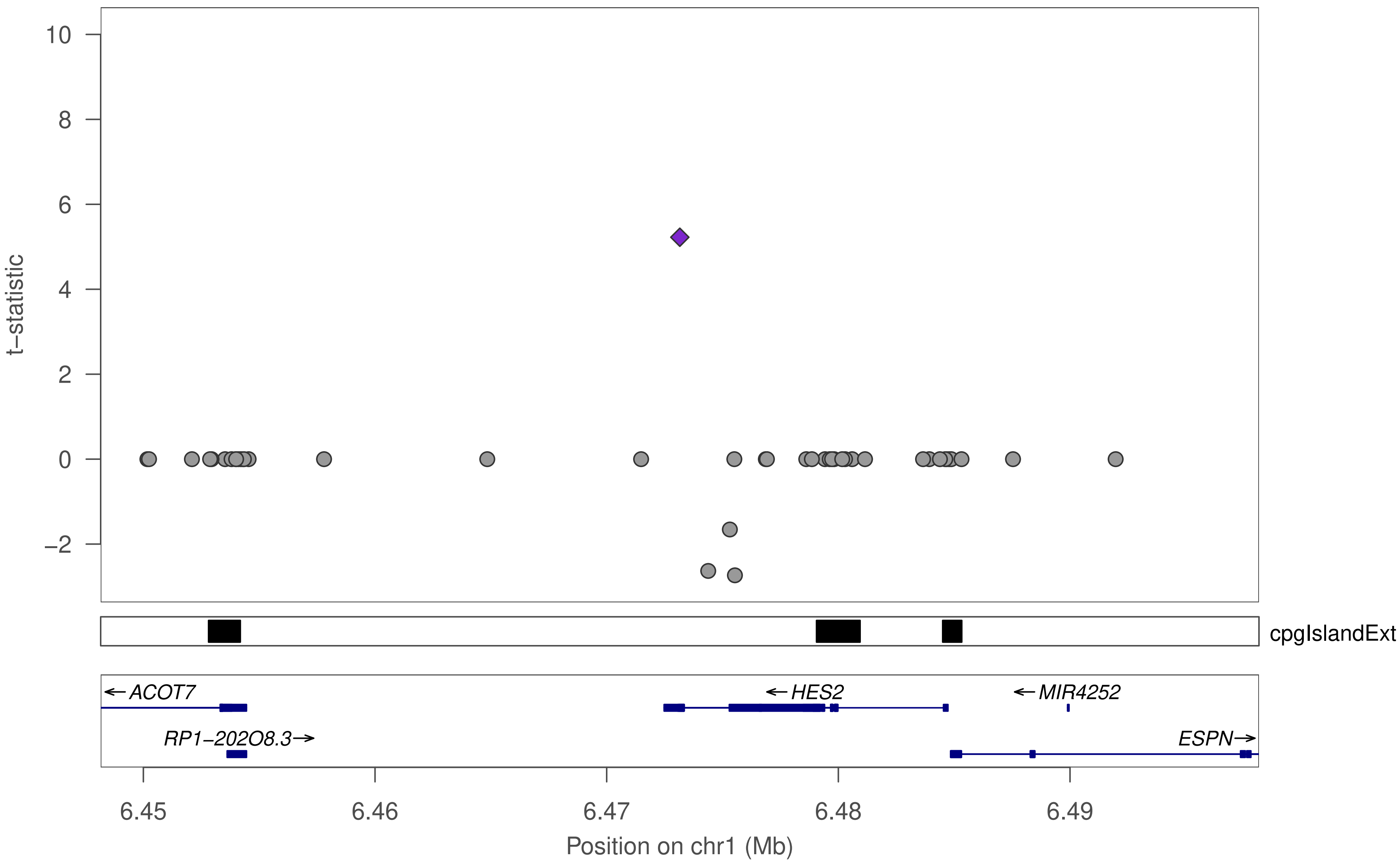

t−statistic
10
8
6
4
2
0
−2
cpgIslandExt
← ACOT7
← HES2
← MIR4252
RP1−202O8.3 →
ESPN →
6.45
6.46
6.47
6.48
6.49
Position on chr1 (Mb)

### ZNF443

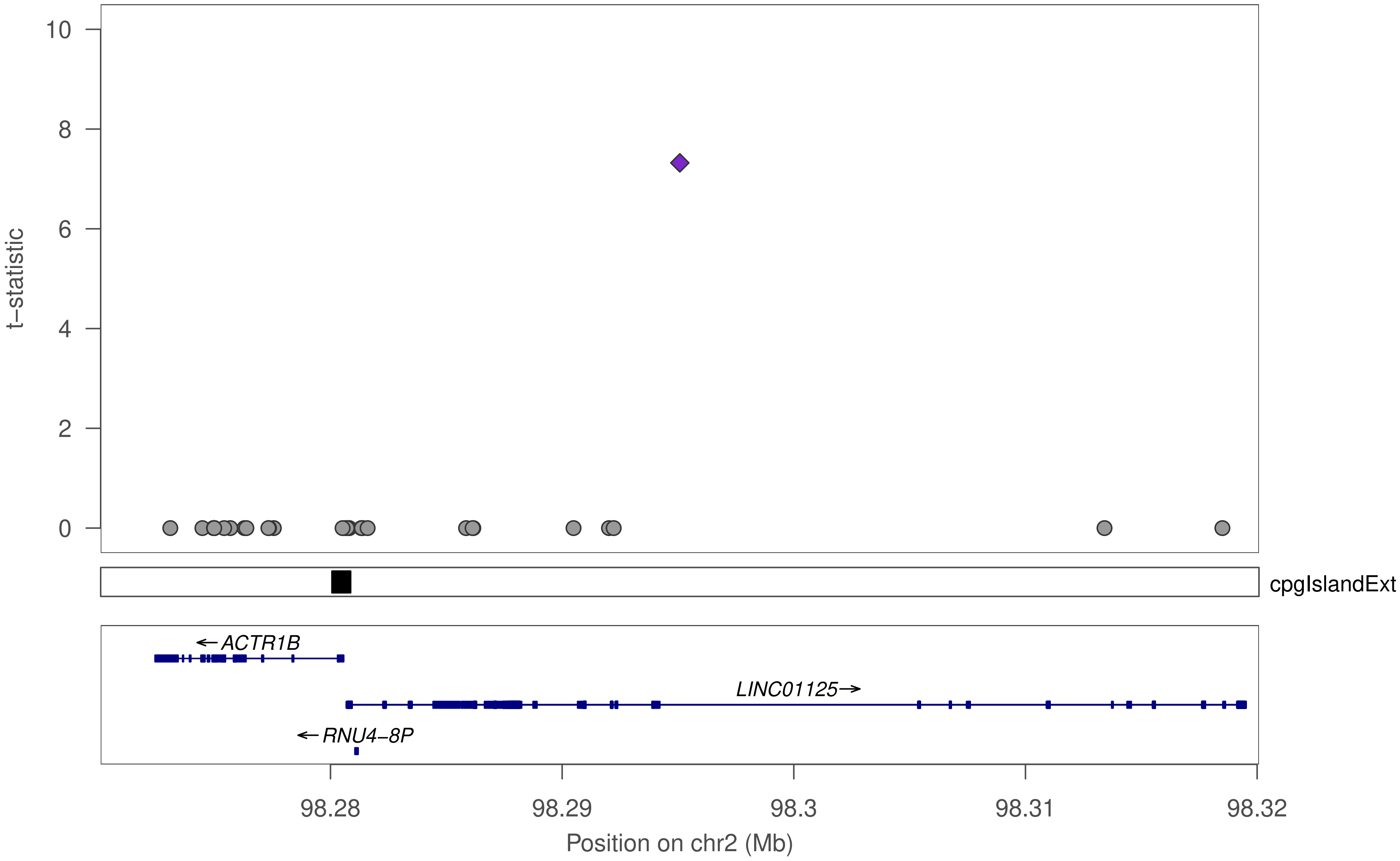

10
8
6
4
2
0
t−statistic
cpgIslandExt
←ACTR1B
LINC01125→
←RNU4−8P
98.28
98.29
98.3
98.31
98.32
Position on chr2 (Mb)

### ZNF443

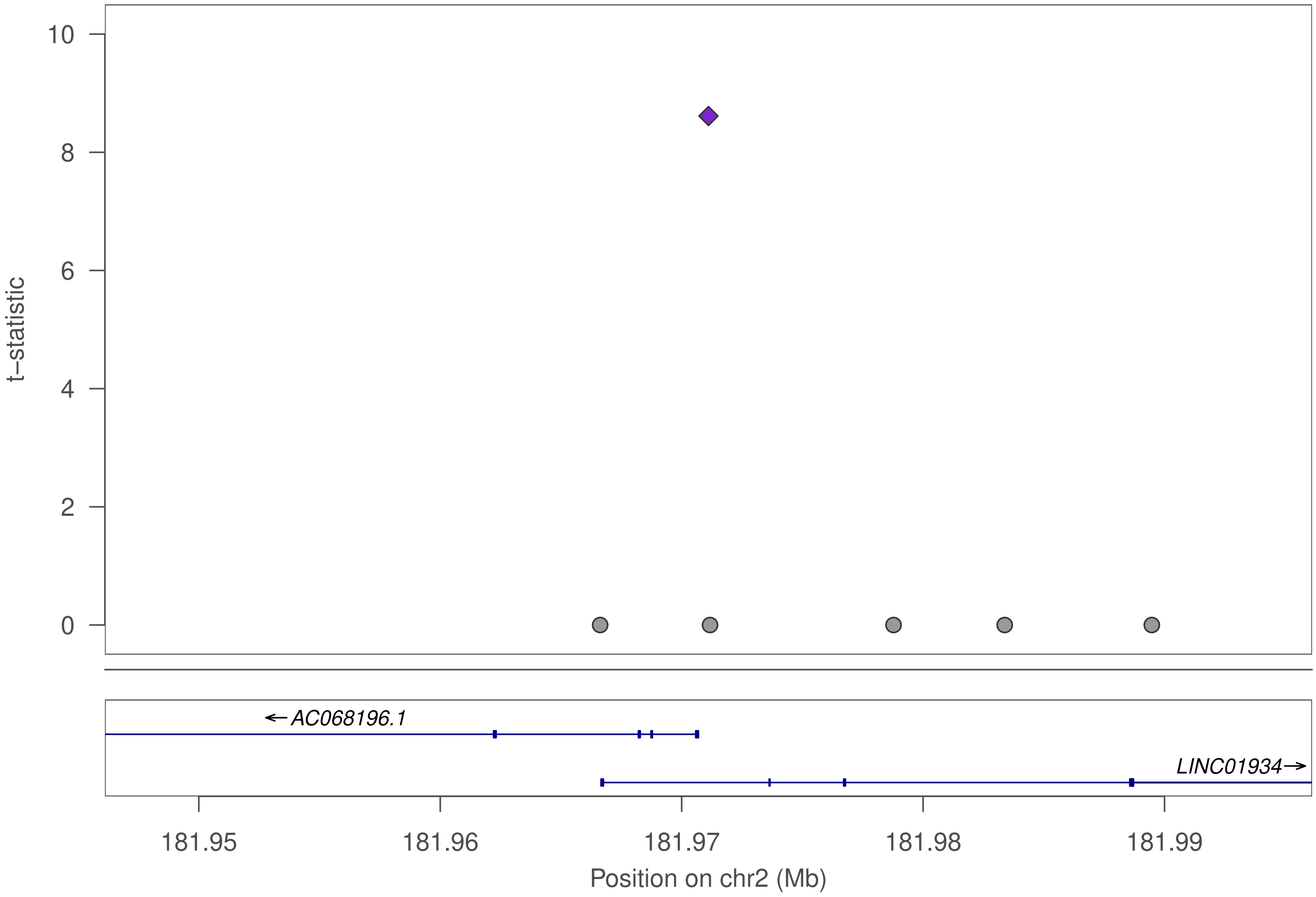

10
8
6
4
2
0
t−statistic
←AC068196.1
LINC01934→
181.95
181.96
181.97
181.98
181.99
Position on chr2 (Mb)

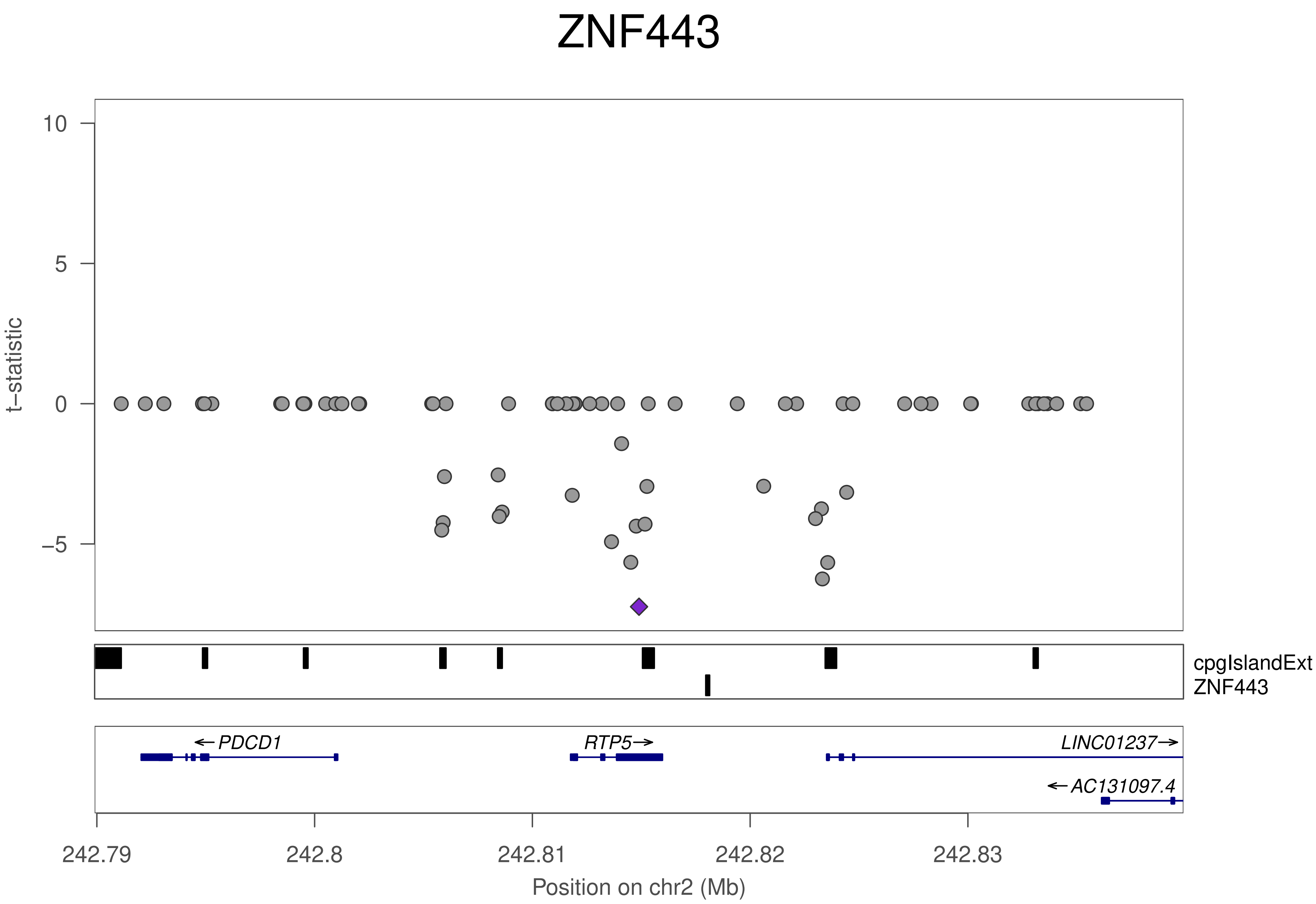
ZNF443
t−statistic
10
5
0
−5
cpgIslandExt
ZNF443
←PDCD1
RTP5→
LINC01237→
←AC131097.4
242.79
242.8
242.81
242.82
242.83
Position on chr2 (Mb)

### ZNF443

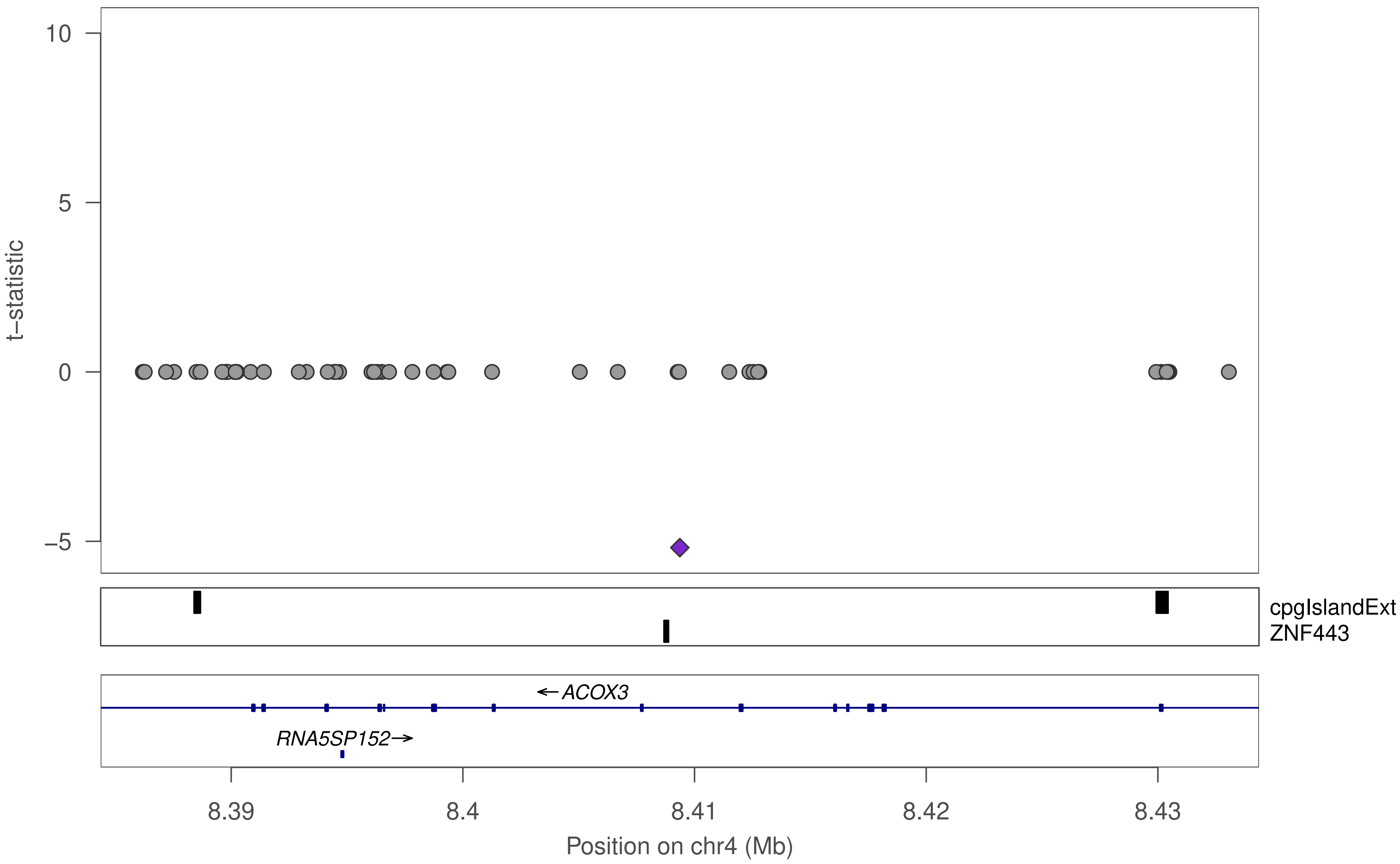

10
5
0
−5
t−statistic
cpgIslandExt
ZNF443
← ACOX3
RNA5SP152 →
8.39
8.4
8.41
8.42
8.43
Position on chr4 (Mb)

### ZNF443

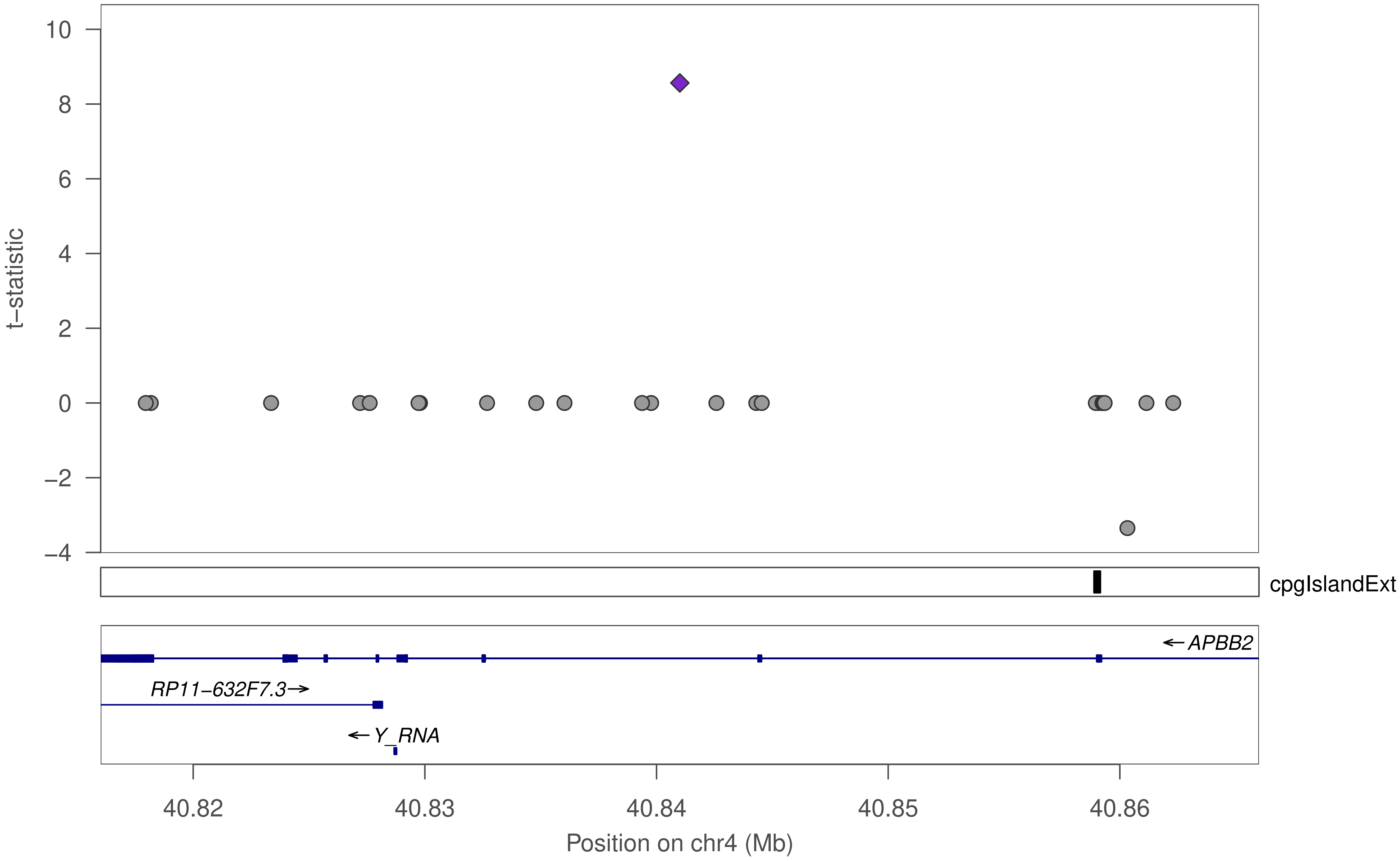

10
8
6
4
2
0
−2
−4
t−statistic
cpgIslandExt
← APBB2
RP11−632F7.3→
← Y_RNA
40.82
40.83
40.84
40.85
40.86
Position on chr4 (Mb)

### ZNF443

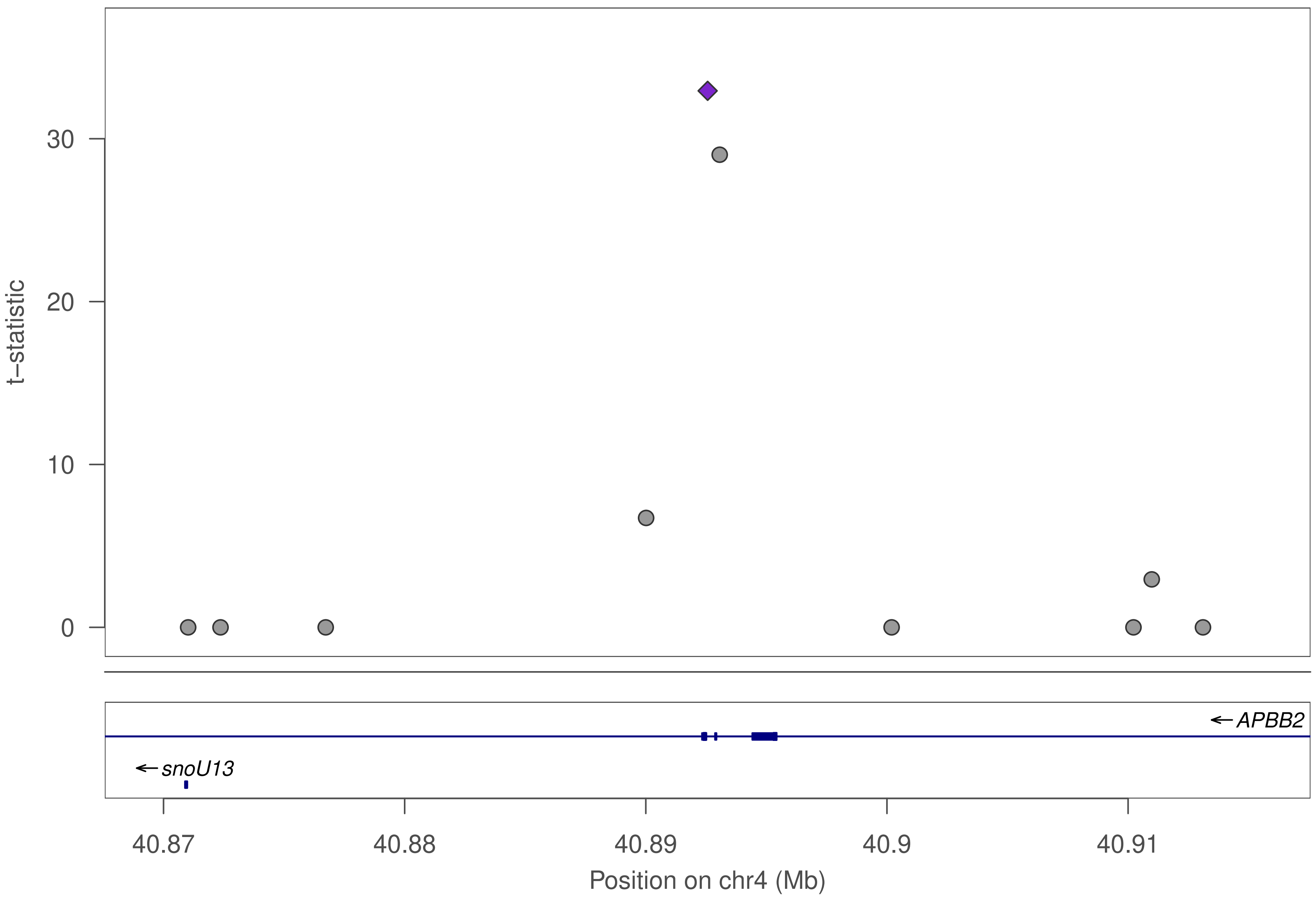

t–statistic
0
10
20
30
APBB2
snoU13
40.87
40.88
40.89
40.9
40.91
Position on chr4 (Mb)

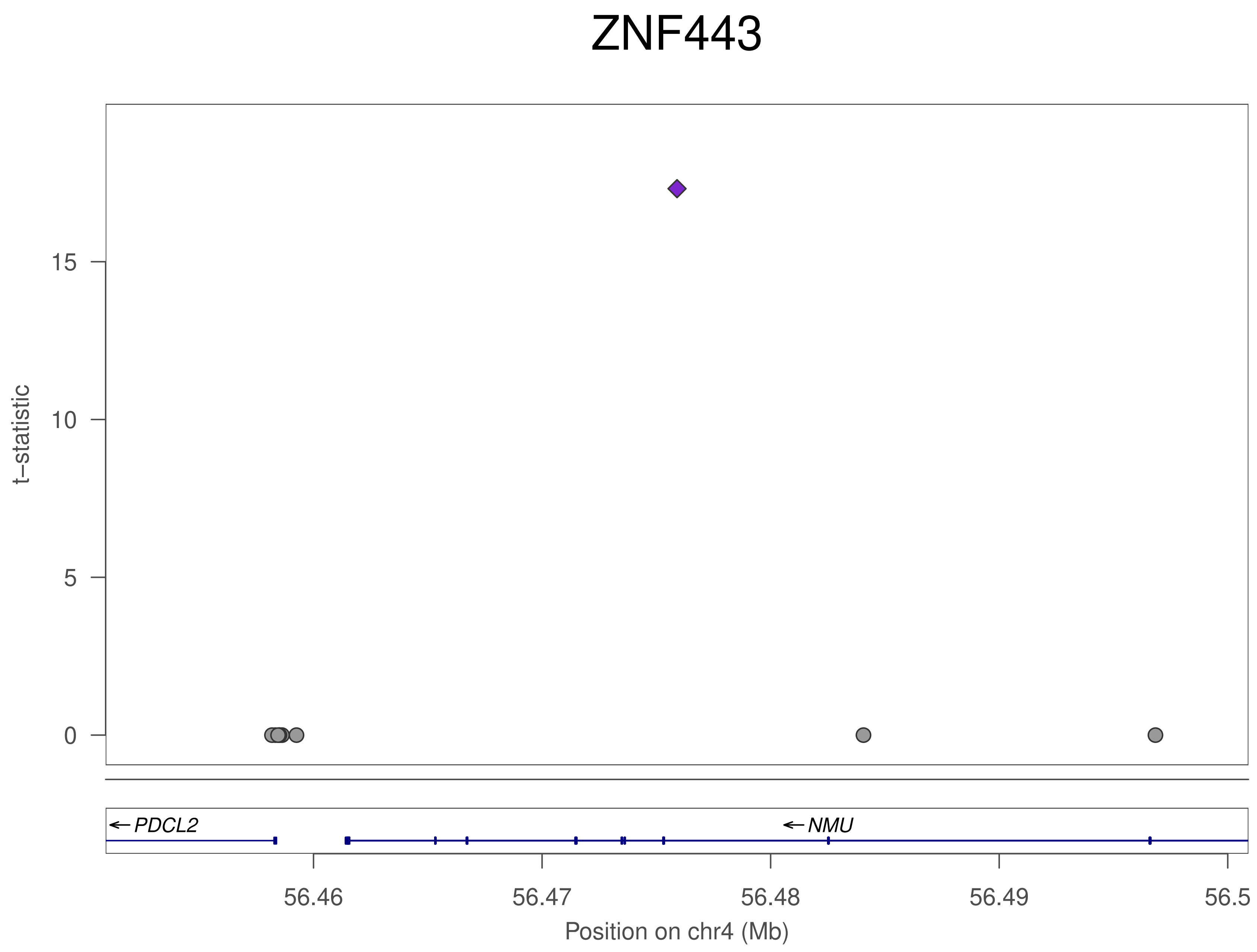
ZNF443
t–statistic
0
5
10
15
PDCL2
NMU
56.46
56.47
56.48
56.49
56.5
Position on chr4 (Mb)

### ZNF443

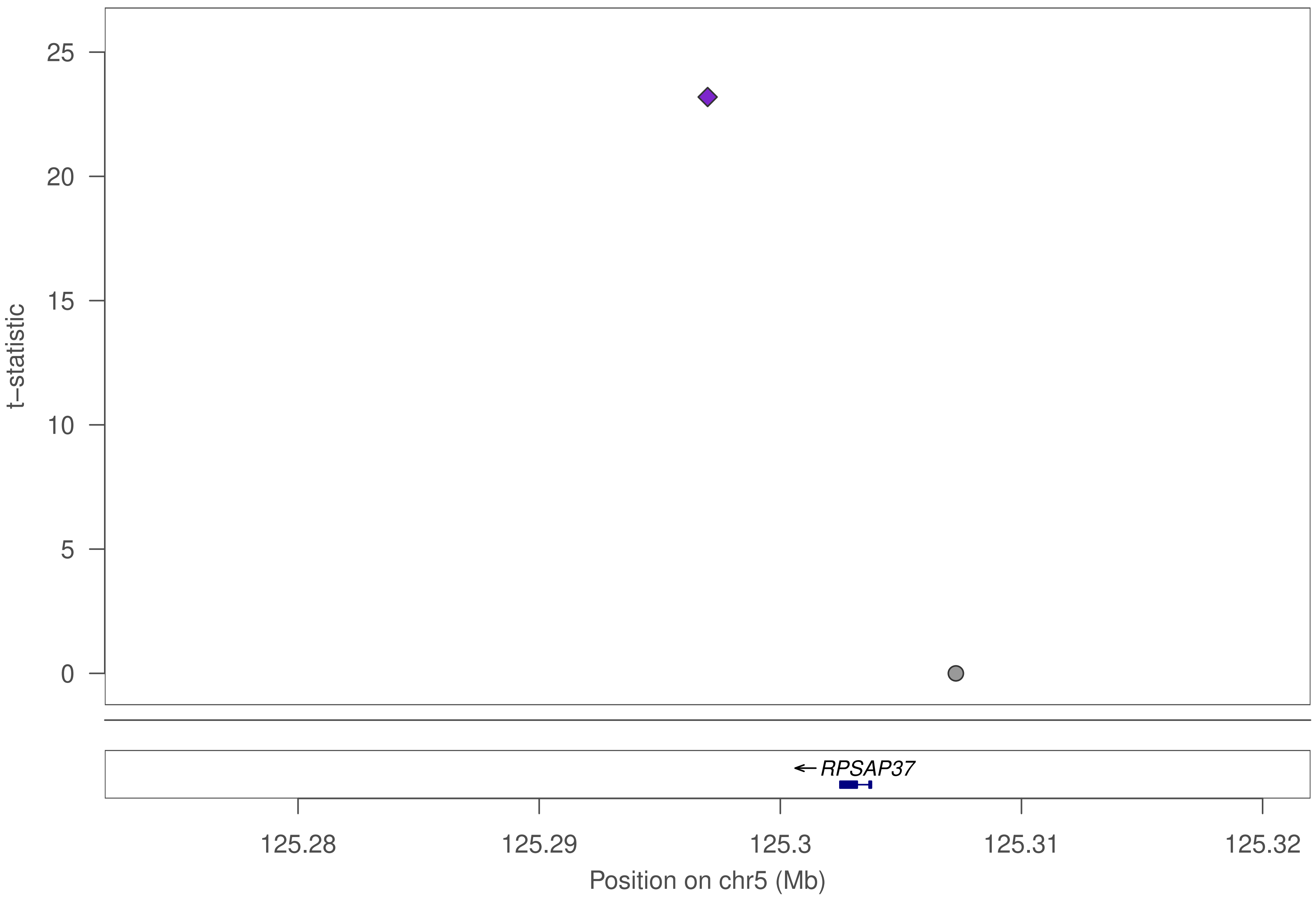

t–statistic
25
20
15
10
5
0
RPSAP37
125.28
125.29
125.3
125.31
125.32
Position on chr5 (Mb)

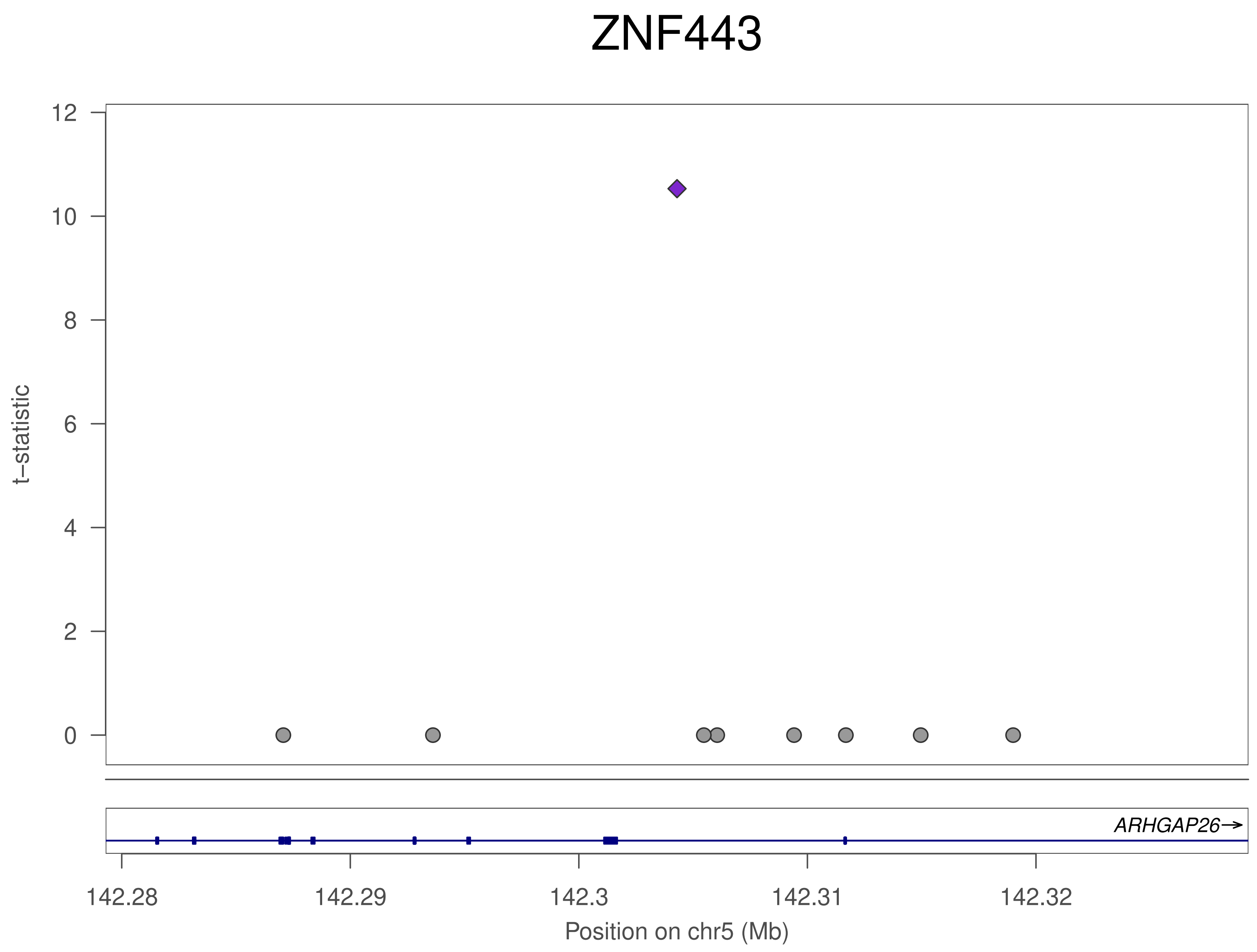

ZNF443
t−statistic
12
10
8
6
4
2
0
ARHGAP26→
142.28
142.29
142.3
142.31
142.32
Position on chr5 (Mb)

### ZNF443

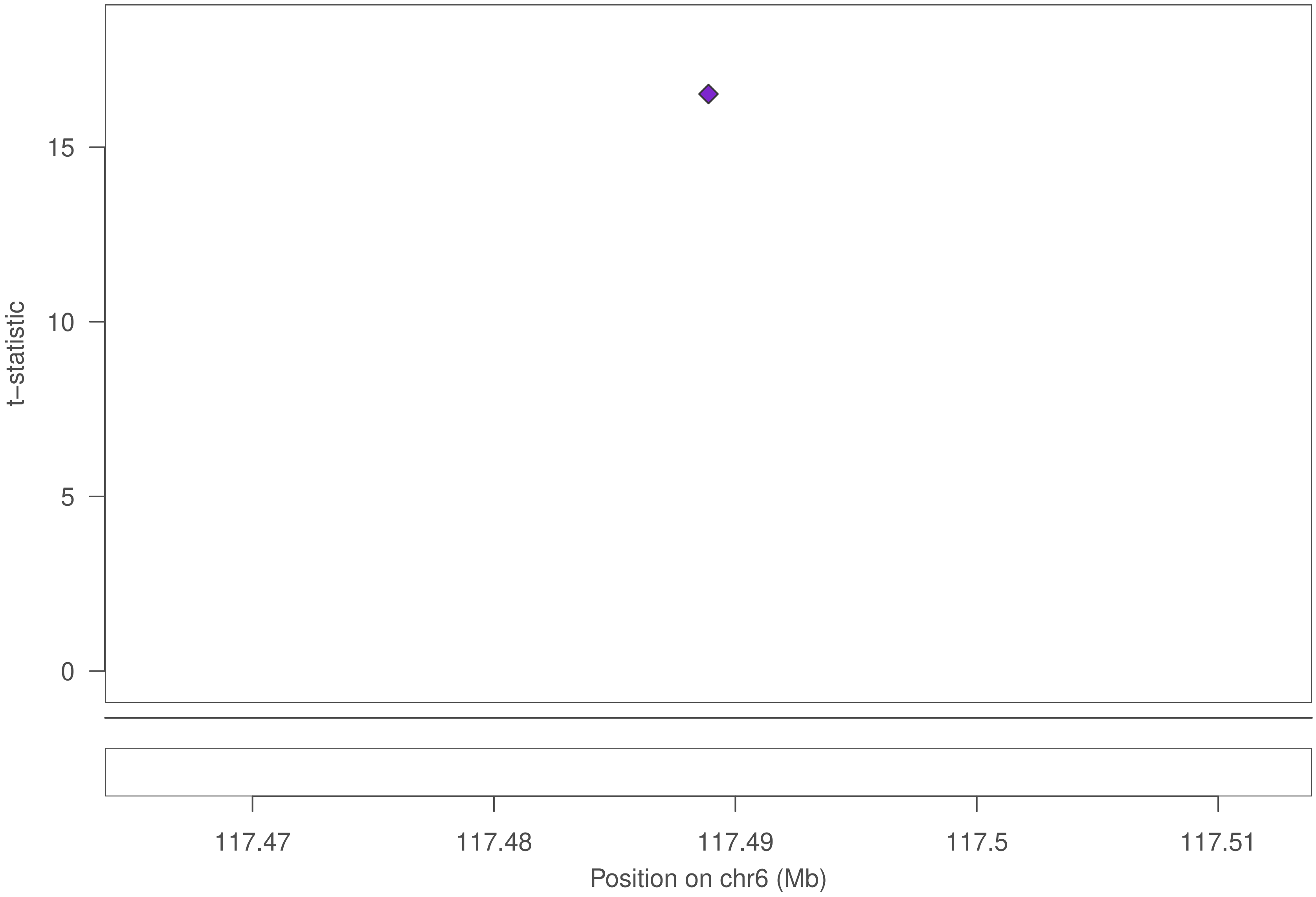
t–statistic
15
10
5
0
117.47
117.48
117.49
117.5
117.51
Position on chr6 (Mb)

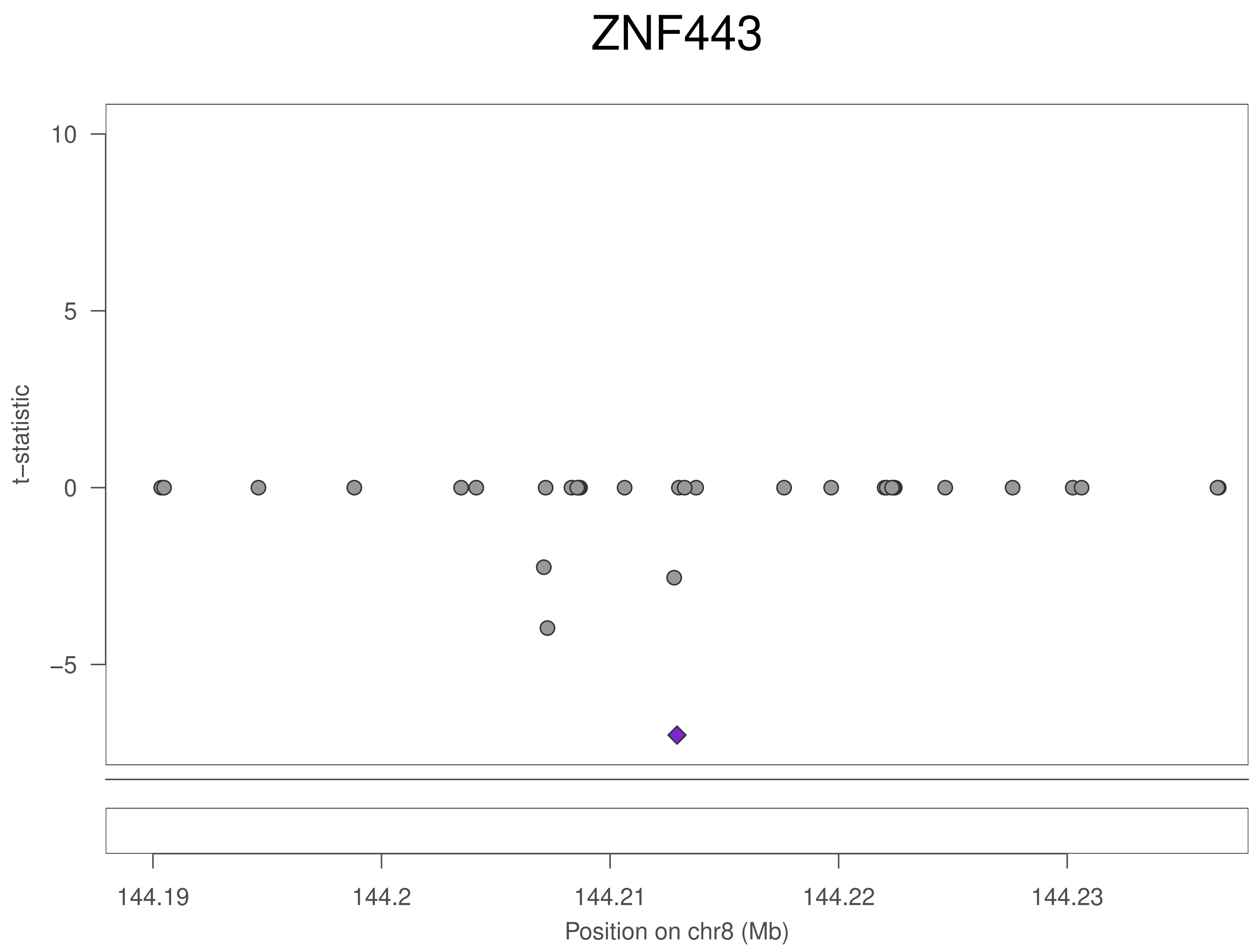
ZNF443
t–statistic
10
5
0
–5
144.19
144.2
144.21
144.22
144.23
Position on chr8 (Mb)

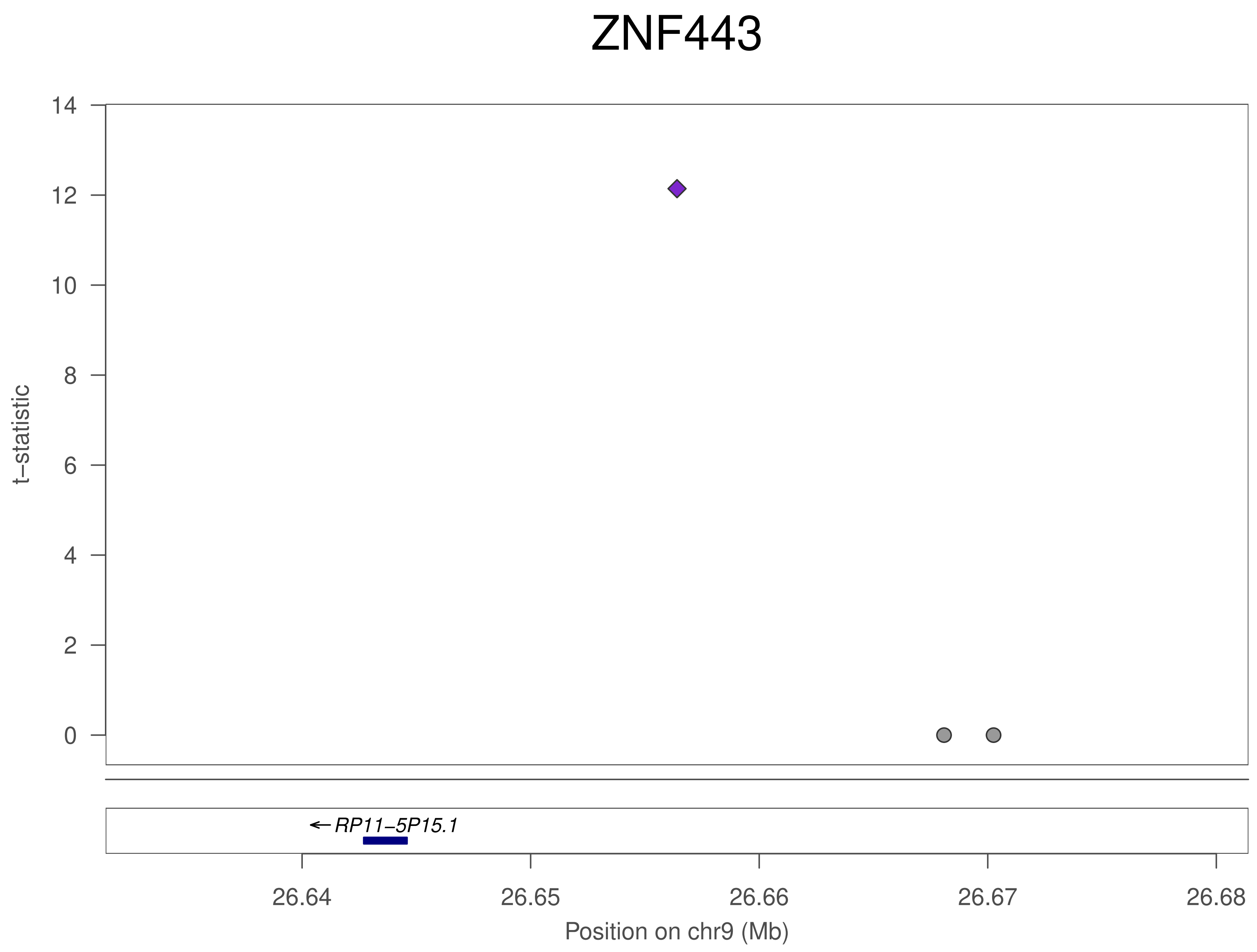
ZNF443
t–statistic
14
12
10
8
6
4
2
0
←RP11–5P15.1
26.64
26.65
26.66
26.67
26.68
Position on chr9 (Mb)

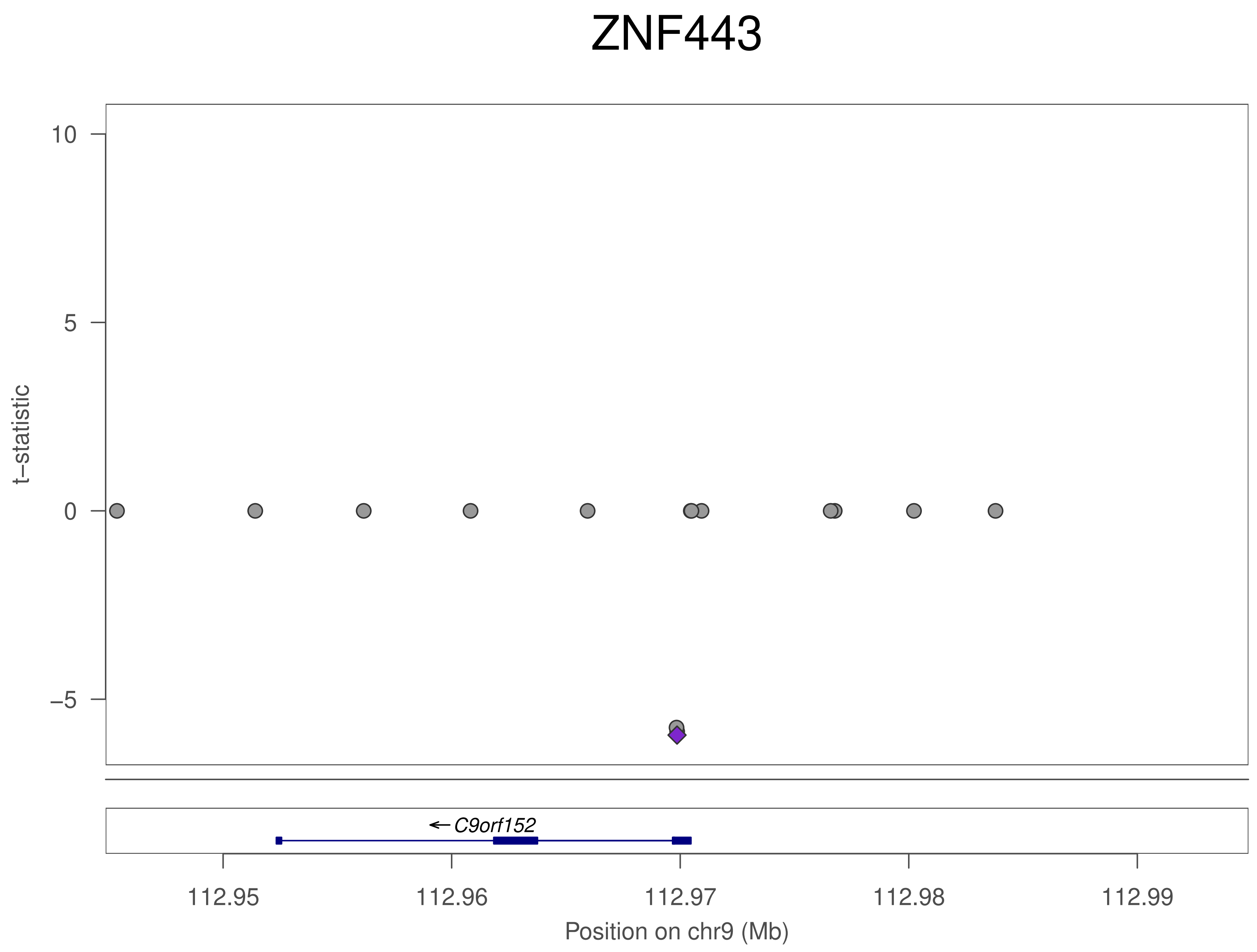

ZNF443
t–statistic
10
5
0
–5
C9orf152
112.95
112.96
112.97
112.98
112.99
Position on chr9 (Mb)

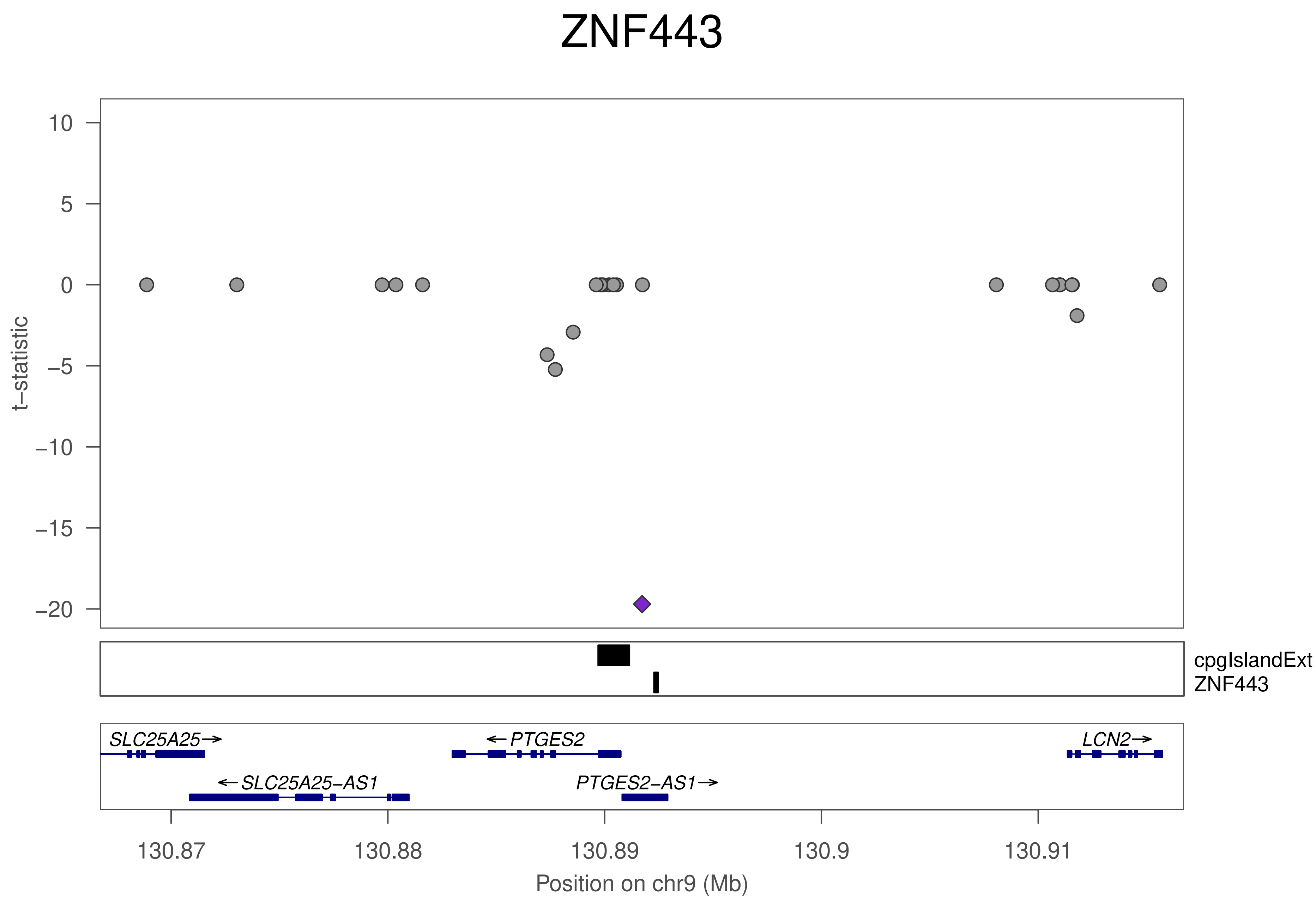
ZNF443
t−statistic
10
5
0
−5
−10
−15
−20
cpgIslandExt
ZNF443
SLC25A25→
←PTGES2
LCN2→
←SLC25A25−AS1
PTGES2−AS1→
130.87
130.88
130.89
130.9
130.91
Position on chr9 (Mb)

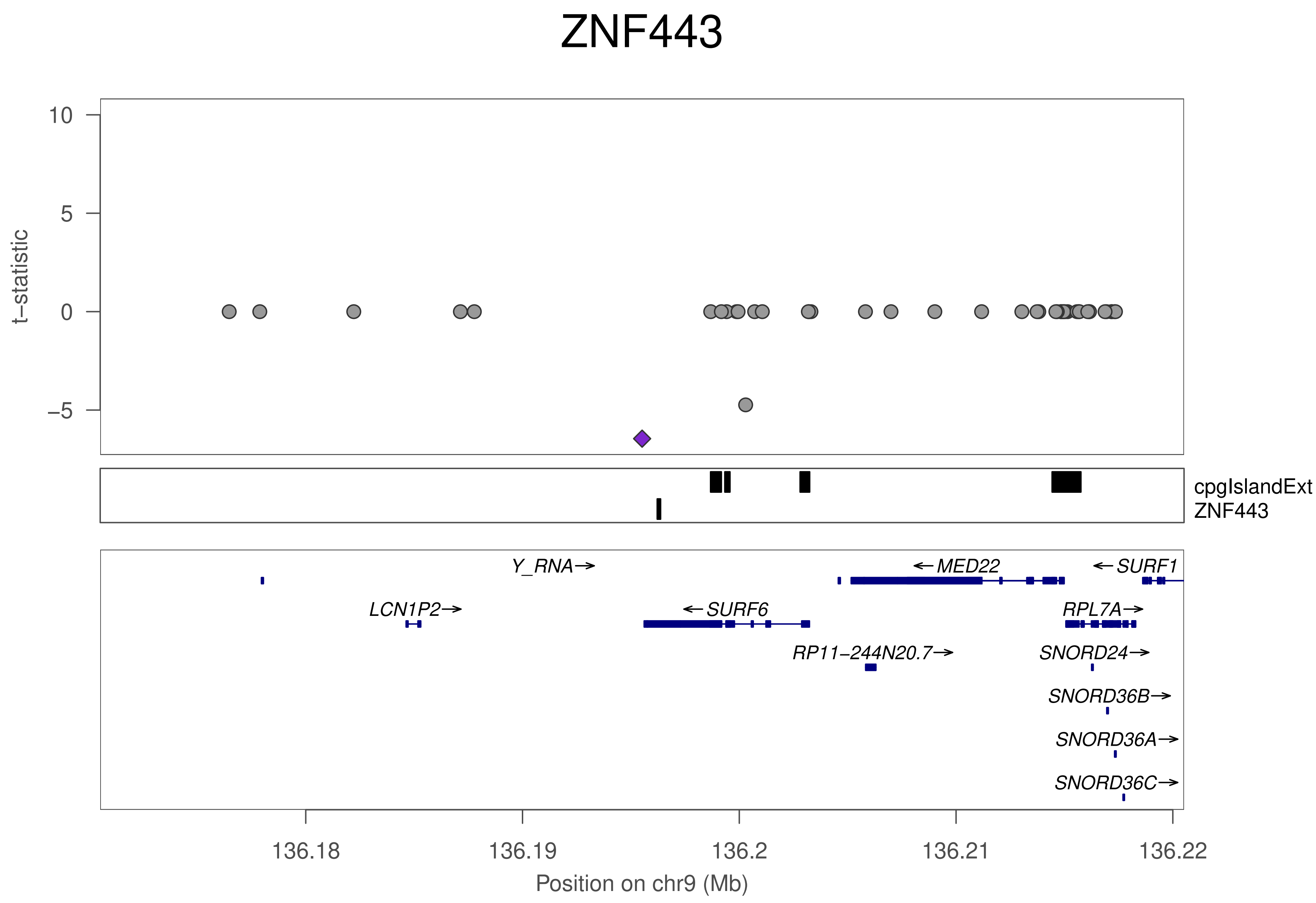
ZNF443
t−statistic
10
5
0
−5
cpgIslandExt
ZNF443
Y_RNA→
←MED22
←SURF1
LCN1P2→
←SURF6
RPL7A→
RP11−244N20.7→
SNORD24→
SNORD36B→
SNORD36A→
SNORD36C→
136.18
136.19
136.2
136.21
136.22
Position on chr9 (Mb)

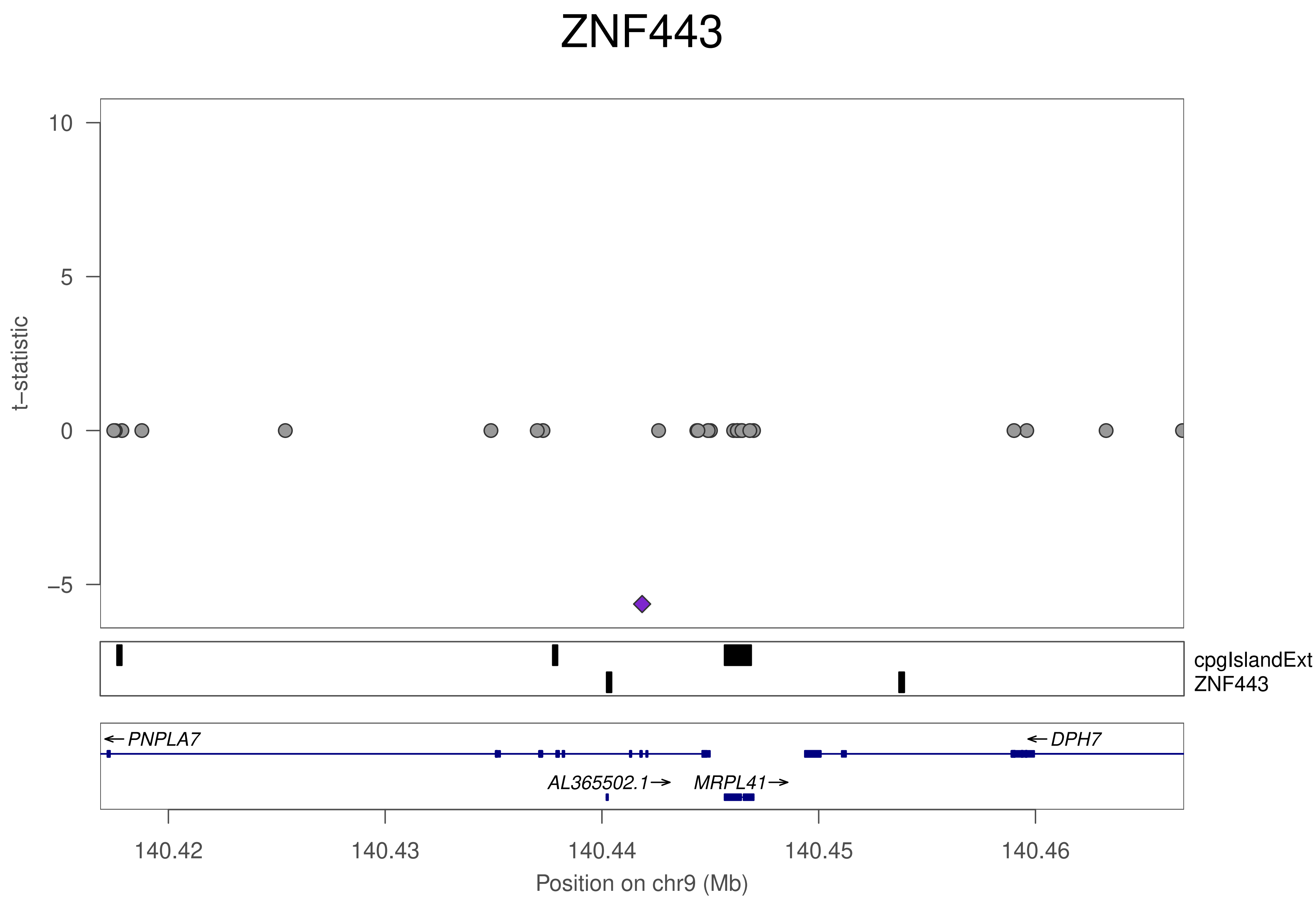
ZNF443
t–statistic
10
5
0
–5
cpgIslandExt
ZNF443
PNPLA7
DPH7
AL365502.1
MRPL41
140.42
140.43
140.44
140.45
140.46
Position on chr9 (Mb)

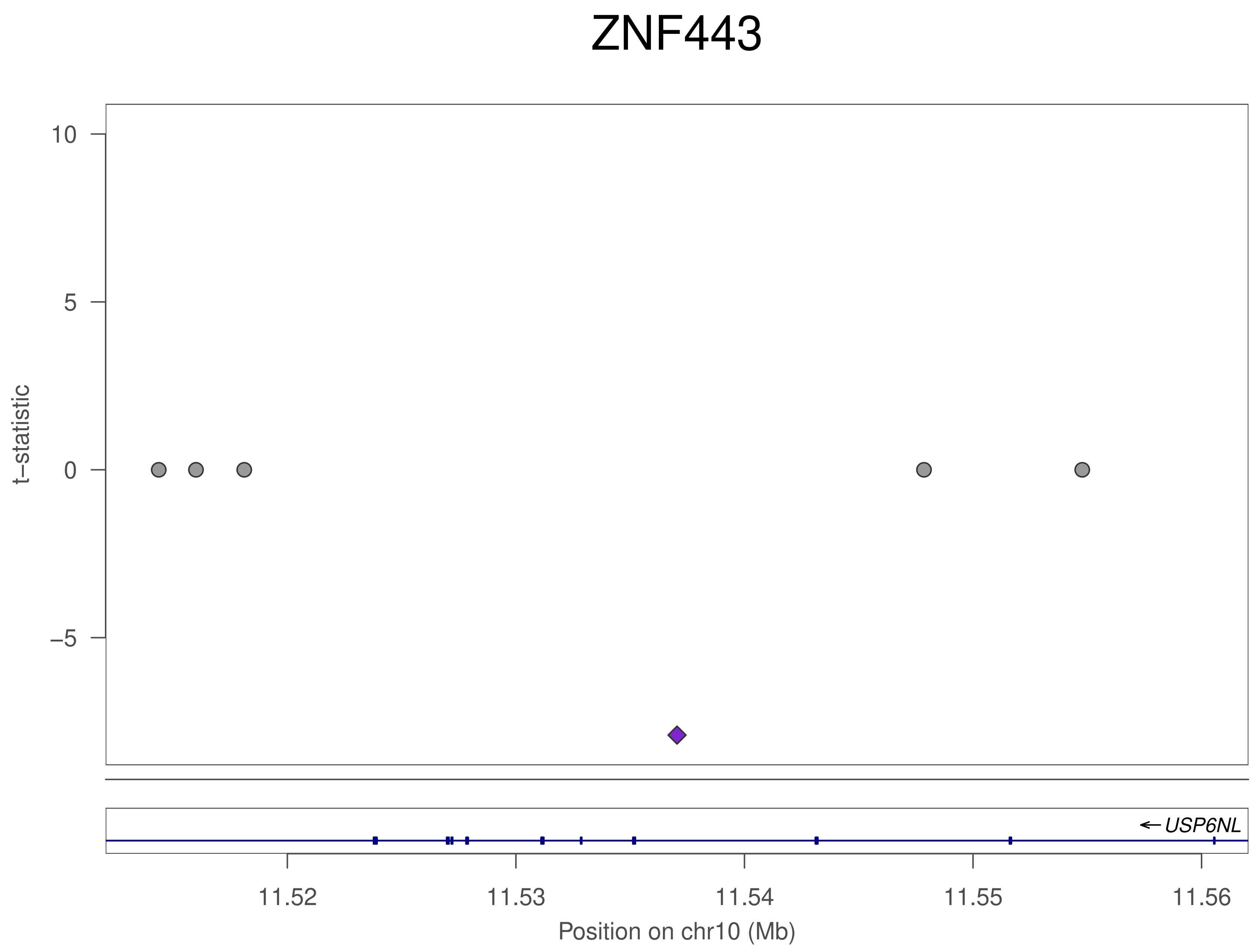

ZNF443
10
5
0
−5
t−statistic
USP6NL
11.52
11.53
11.54
11.55
11.56
Position on chr10 (Mb)

### ZNF443

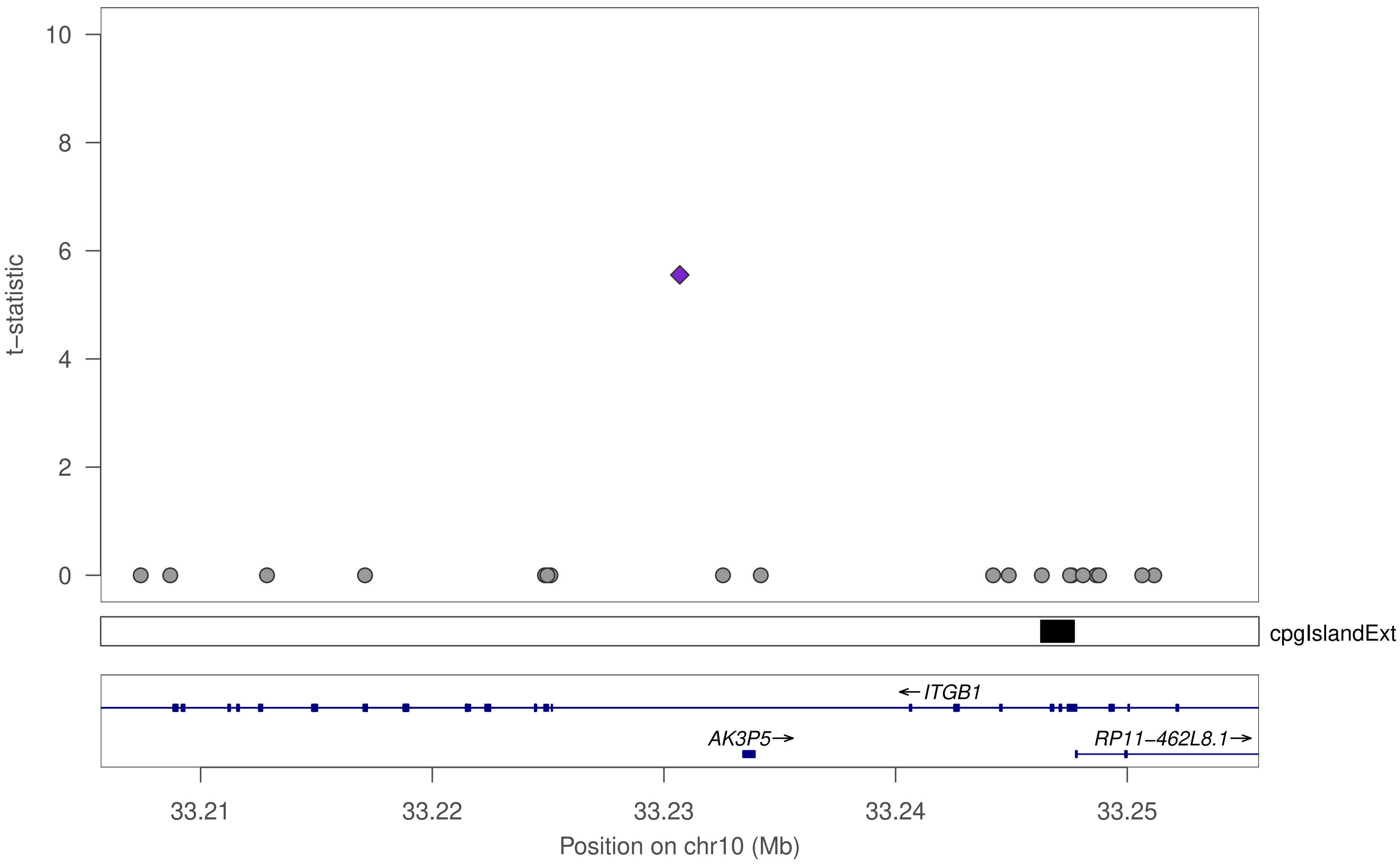

10
8
6
4
2
0
t−statistic
cpgIslandExt
←ITGB1
AK3P5→
RP11−462L8.1→
33.21
33.22
33.23
33.24
33.25
Position on chr10 (Mb)

### ZNF443

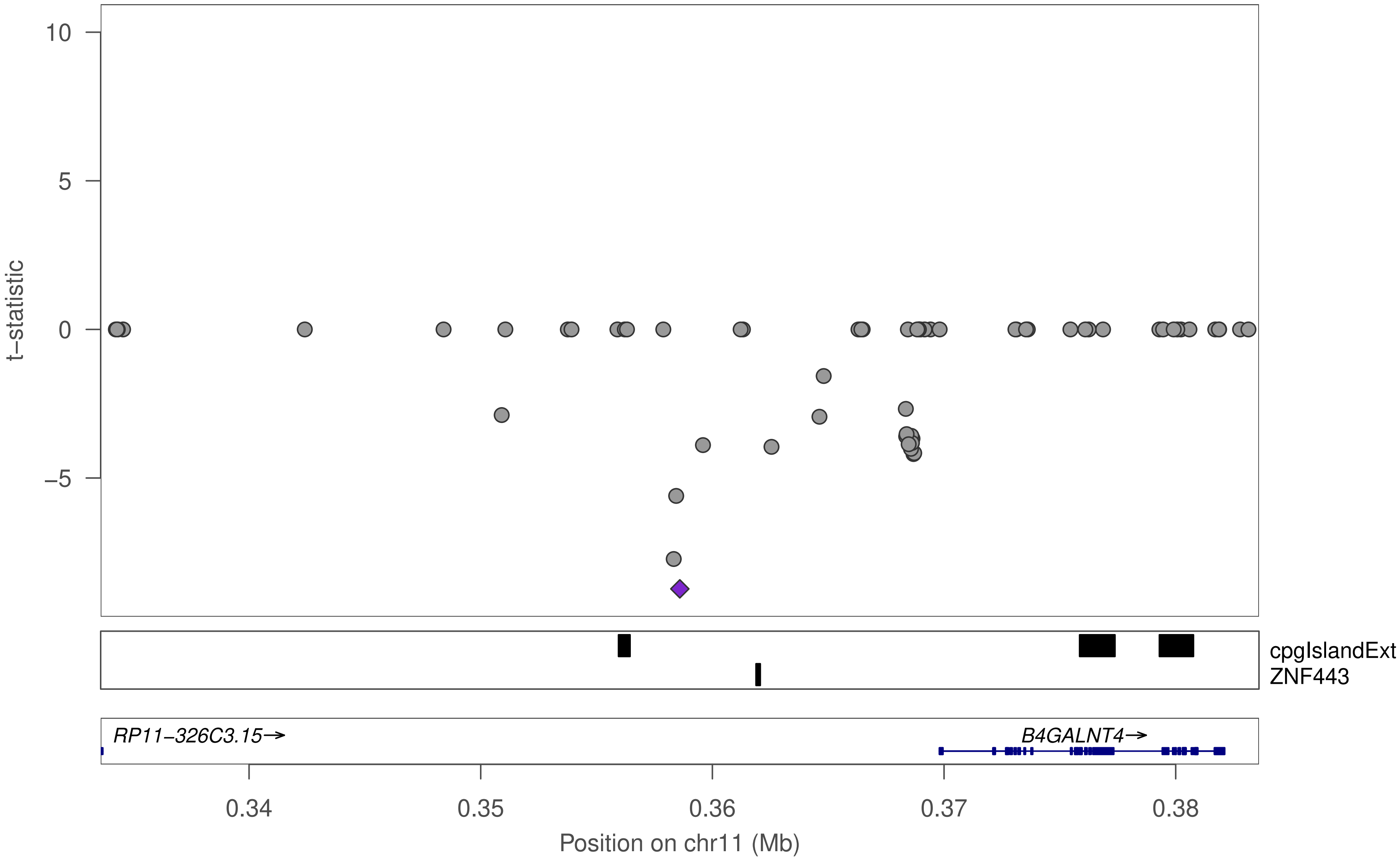

10
5
0
−5
t−statistic
cpgIslandExt
ZNF443
RP11−326C3.15→
B4GALNT4→
0.34
0.35
0.36
0.37
0.38
Position on chr11 (Mb)

### ZNF443

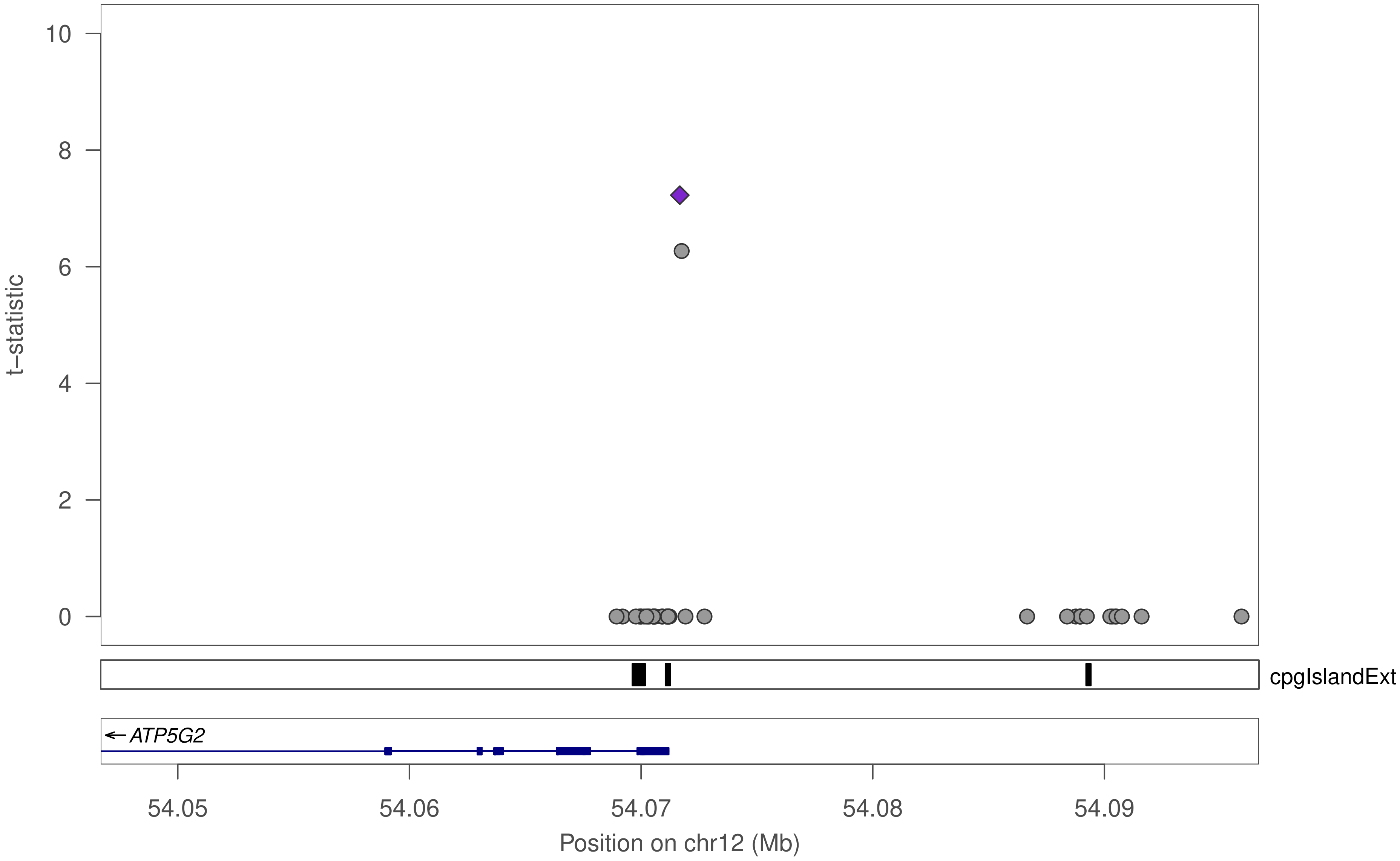

10
8
6
4
2
0
t−statistic
cpgIslandExt
ATP5G2
54.05
54.06
54.07
54.08
54.09
Position on chr12 (Mb)

### ZNF443

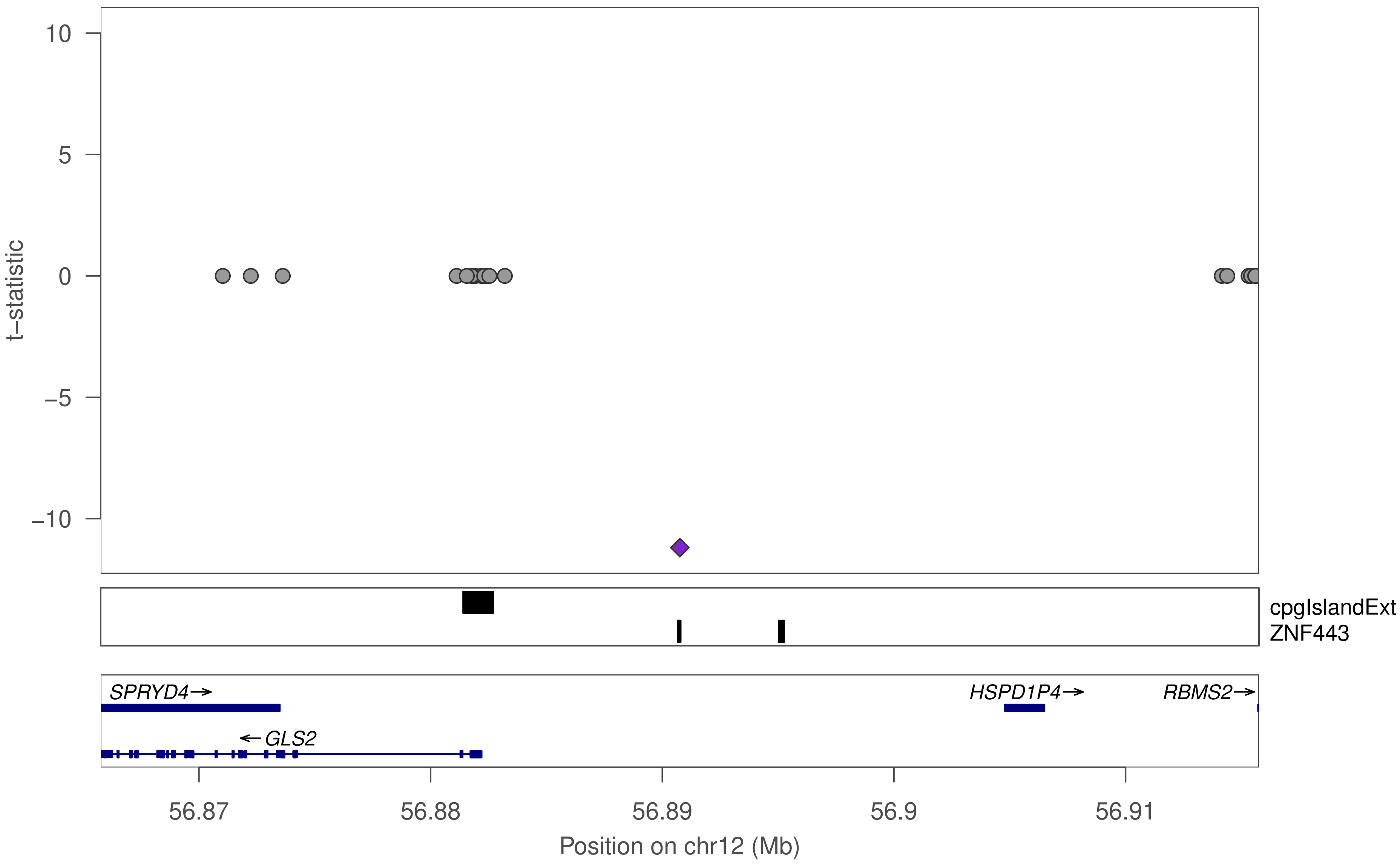

10
5
0
−5
−10
t−statistic
cpgIslandExt
ZNF443
SPRYD4→
←GLS2
HSPD1P4→
RBMS2→
56.87
56.88
56.89
56.9
56.91
Position on chr12 (Mb)

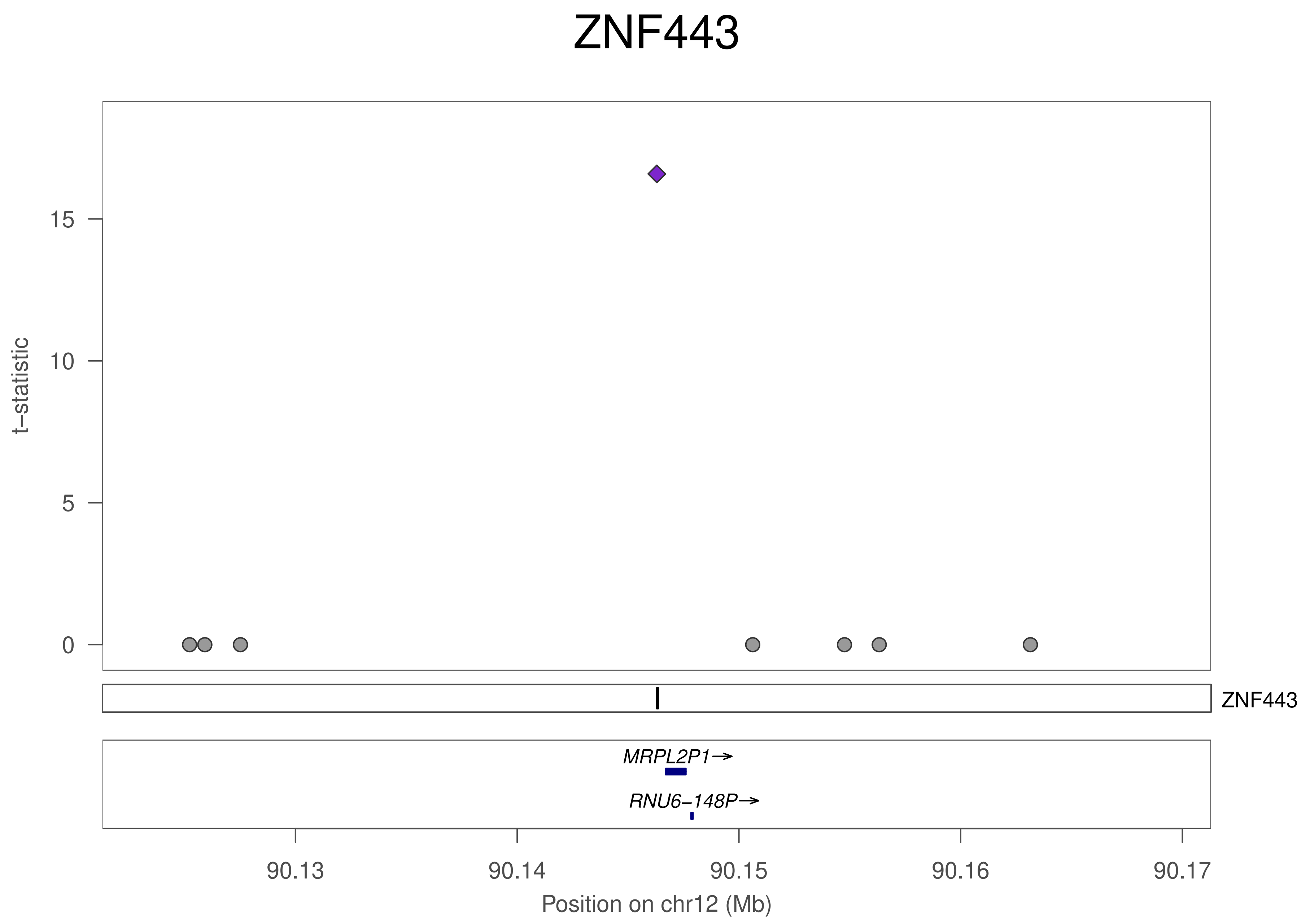

ZNF443
t−statistic
0
5
10
15
ZNF443
MRPL2P1→
RNU6−148P→
90.13
90.14
90.15
90.16
90.17
Position on chr12 (Mb)

### ZNF443

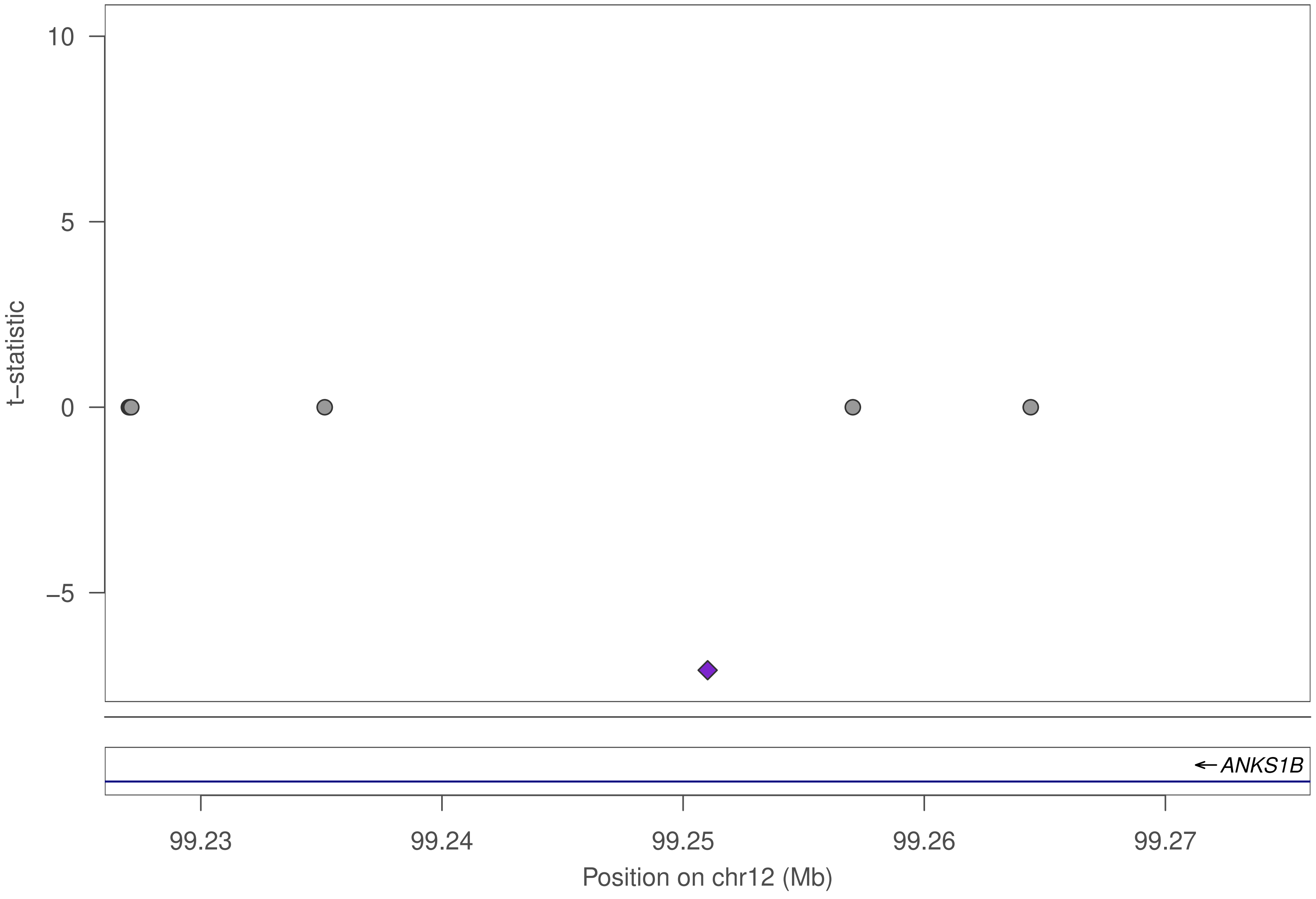
10
5
0
−5
t−statistic
← ANKS1B
99.23
99.24
99.25
99.26
99.27
Position on chr12 (Mb)

### ZNF443

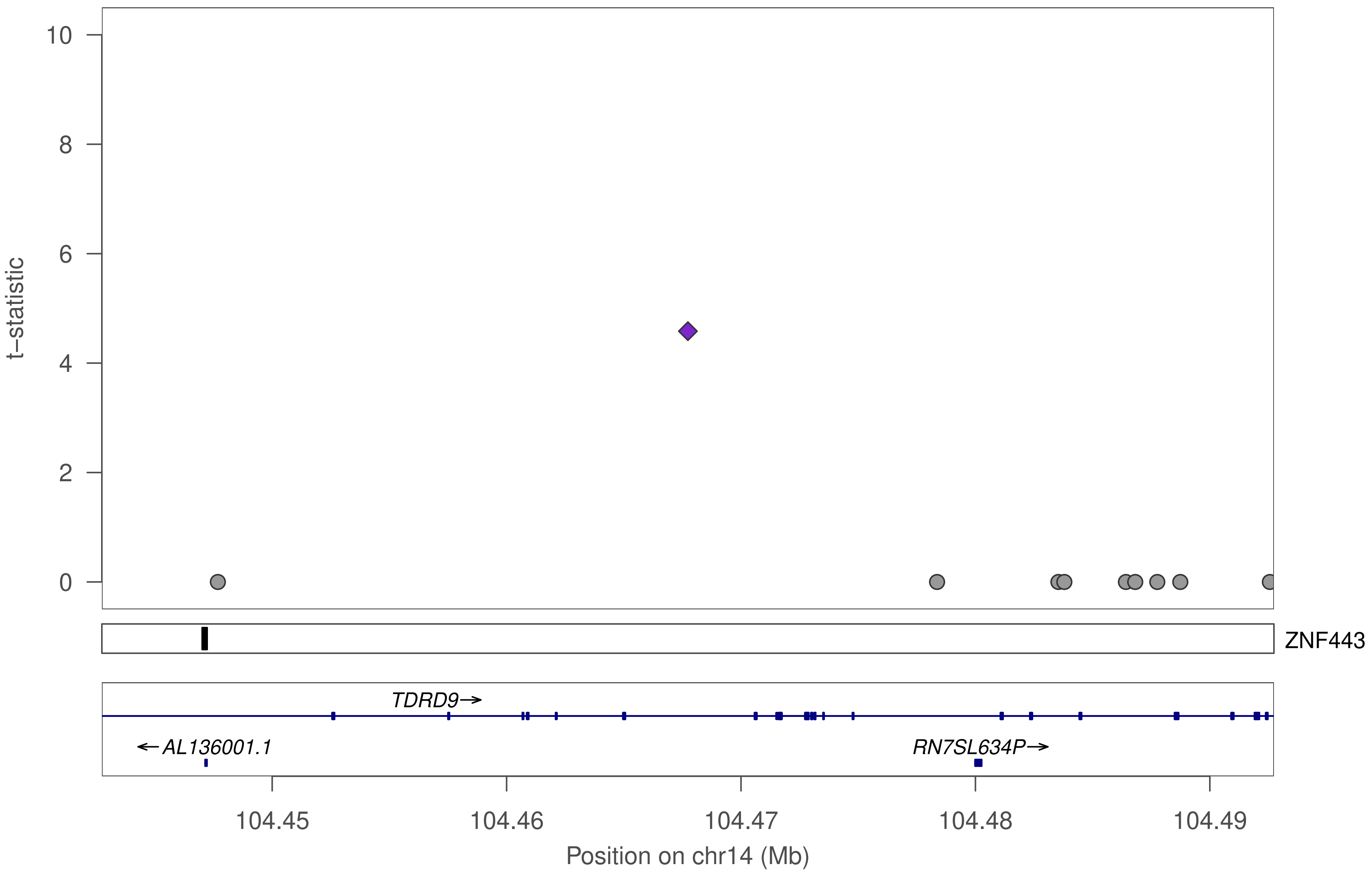

10
8
6
4
2
0
t−statistic
ZNF443
TDRD9→
←AL136001.1
RN7SL634P→
104.45
104.46
104.47
104.48
104.49
Position on chr14 (Mb)

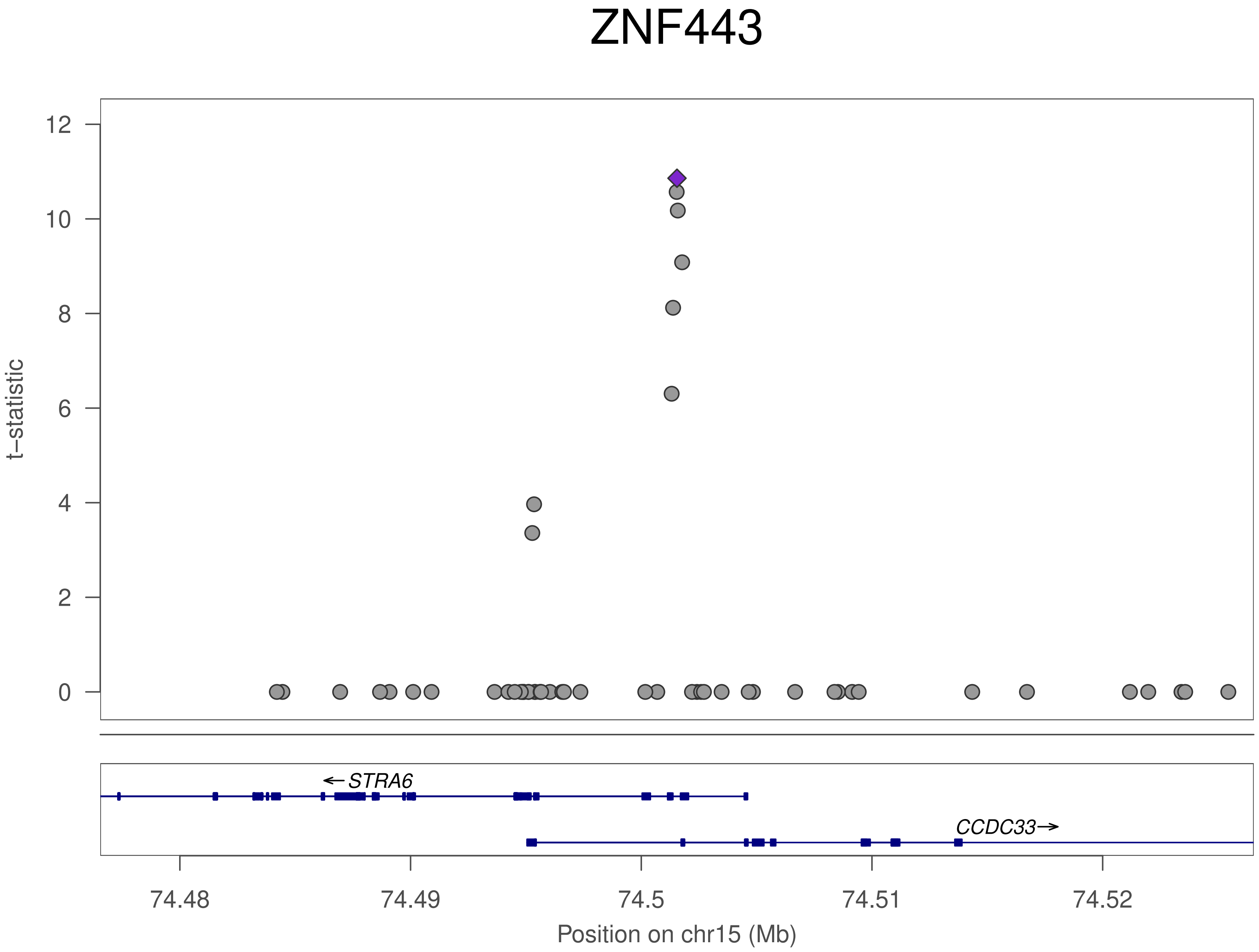

ZNF443
t−statistic
12
10
8
6
4
2
0
←STRA6
CCDC33→
74.48
74.49
74.5
74.51
74.52
Position on chr15 (Mb)

### ZNF443

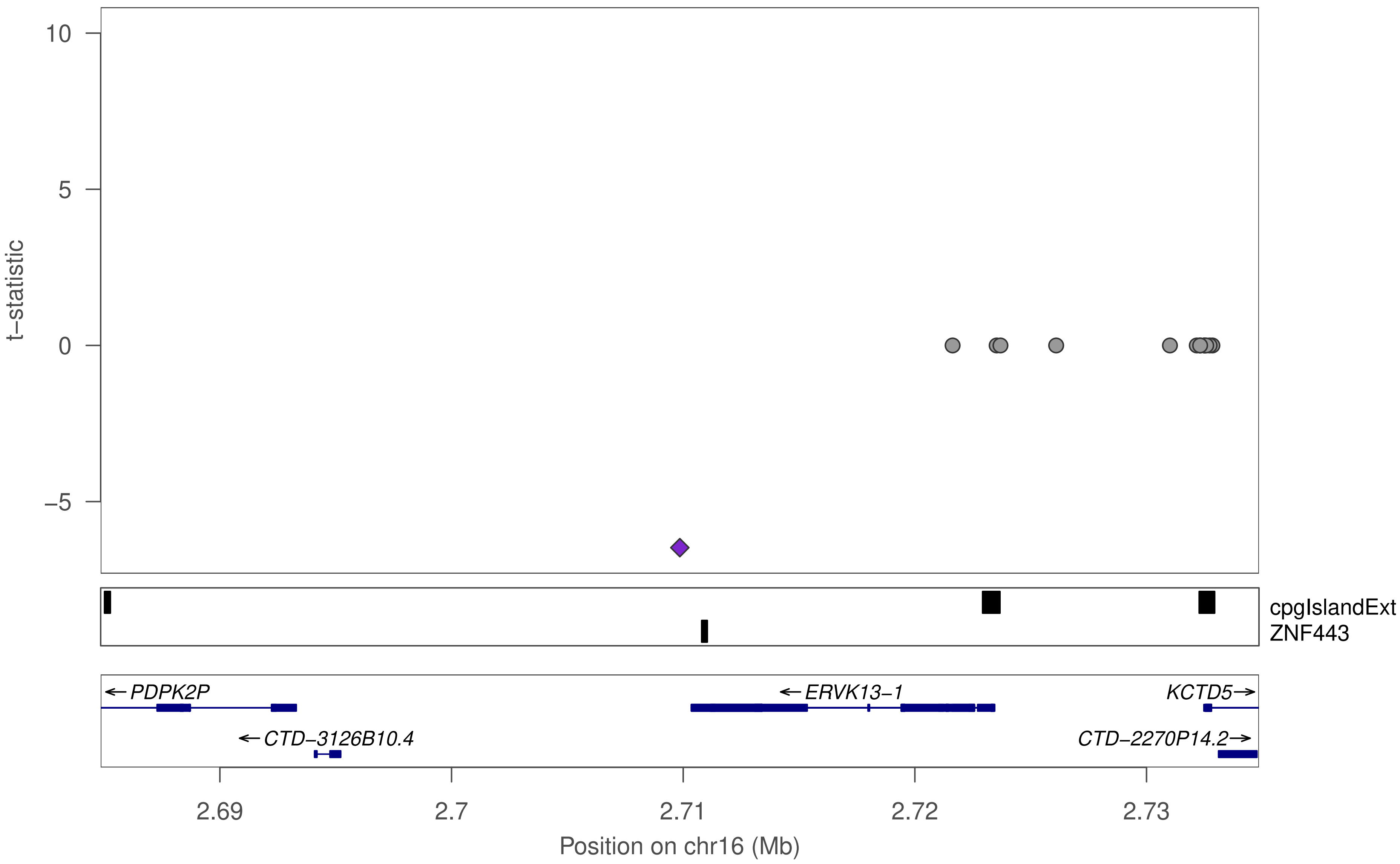

t−statistic
10
5
0
−5
cpgIslandExt
ZNF443
← PDPK2P
← ERVK13−1
KCTD5→
← CTD−3126B10.4
CTD−2270P14.2→
2.69
2.7
2.71
2.72
2.73
Position on chr16 (Mb)

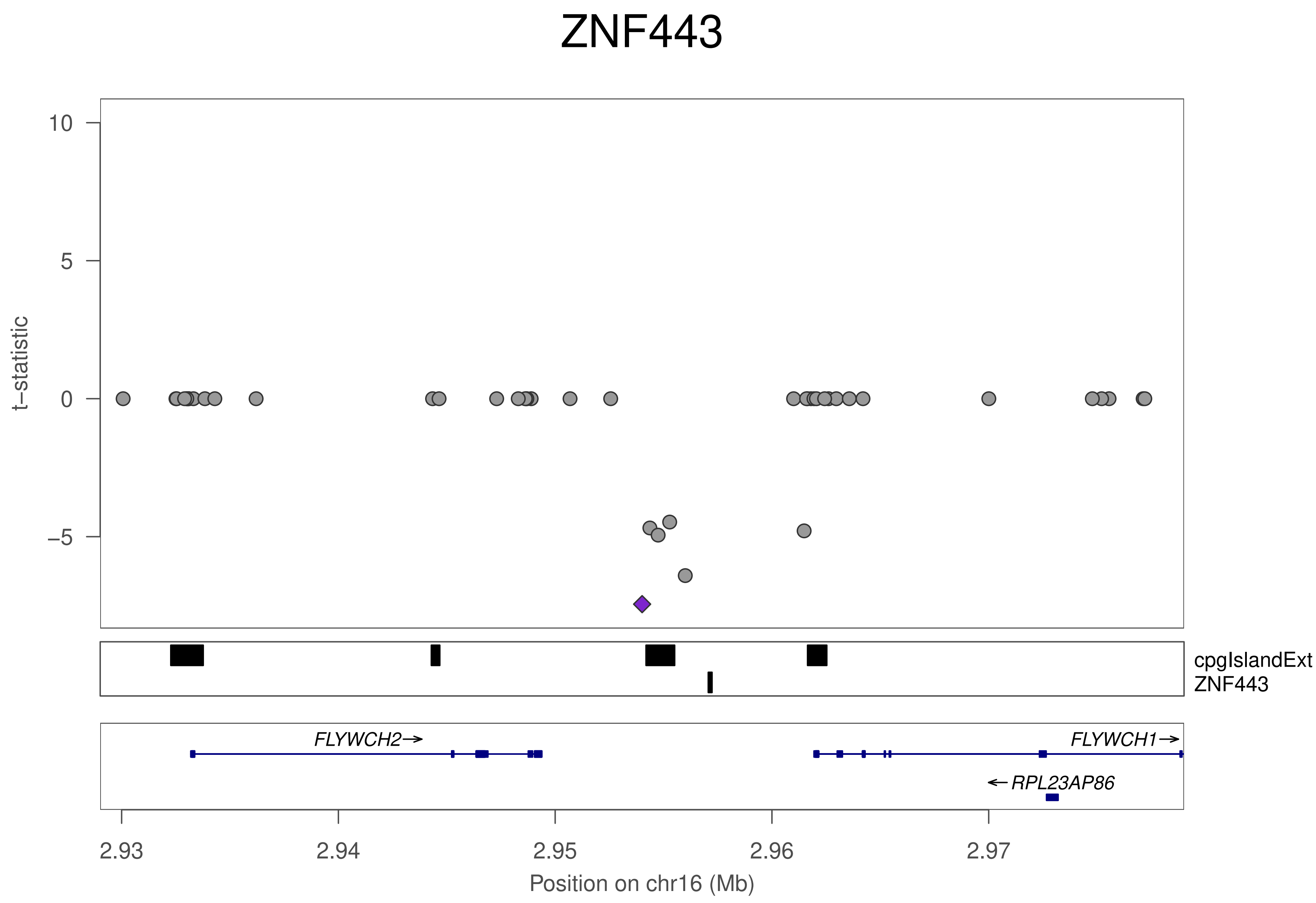
ZNF443
t−statistic
10
5
0
−5
cpgIslandExt
ZNF443
FLYWCH2→
FLYWCH1→
←RPL23AP86
2.93
2.94
2.95
2.96
2.97
Position on chr16 (Mb)

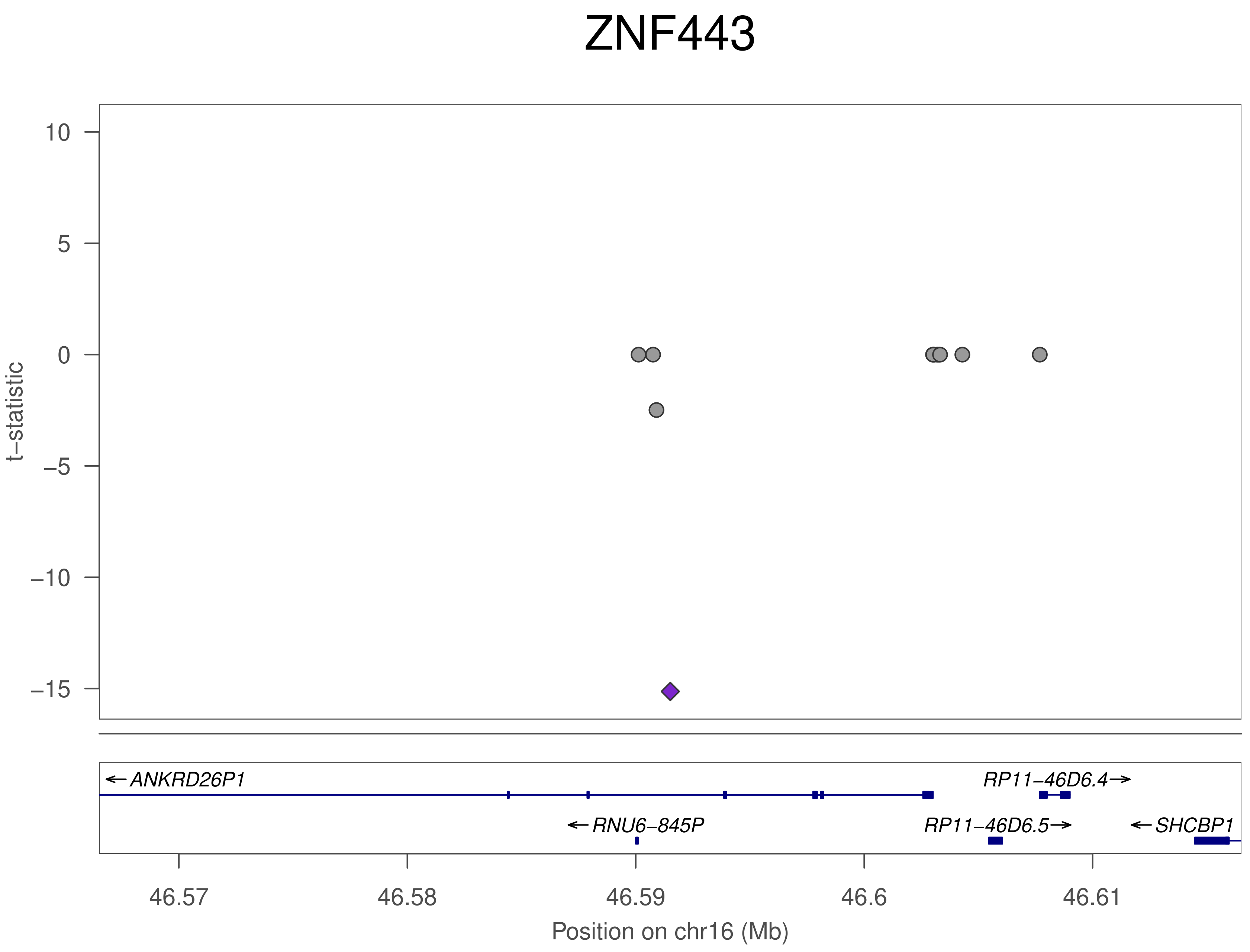

ZNF443
t−statistic
10
5
0
−5
−10
−15
← ANKRD26P1
RP11−46D6.4→
← RNU6−845P
RP11−46D6.5→
← SHCBP1
46.57
46.58
46.59
46.6
46.61
Position on chr16 (Mb)

### ZNF443

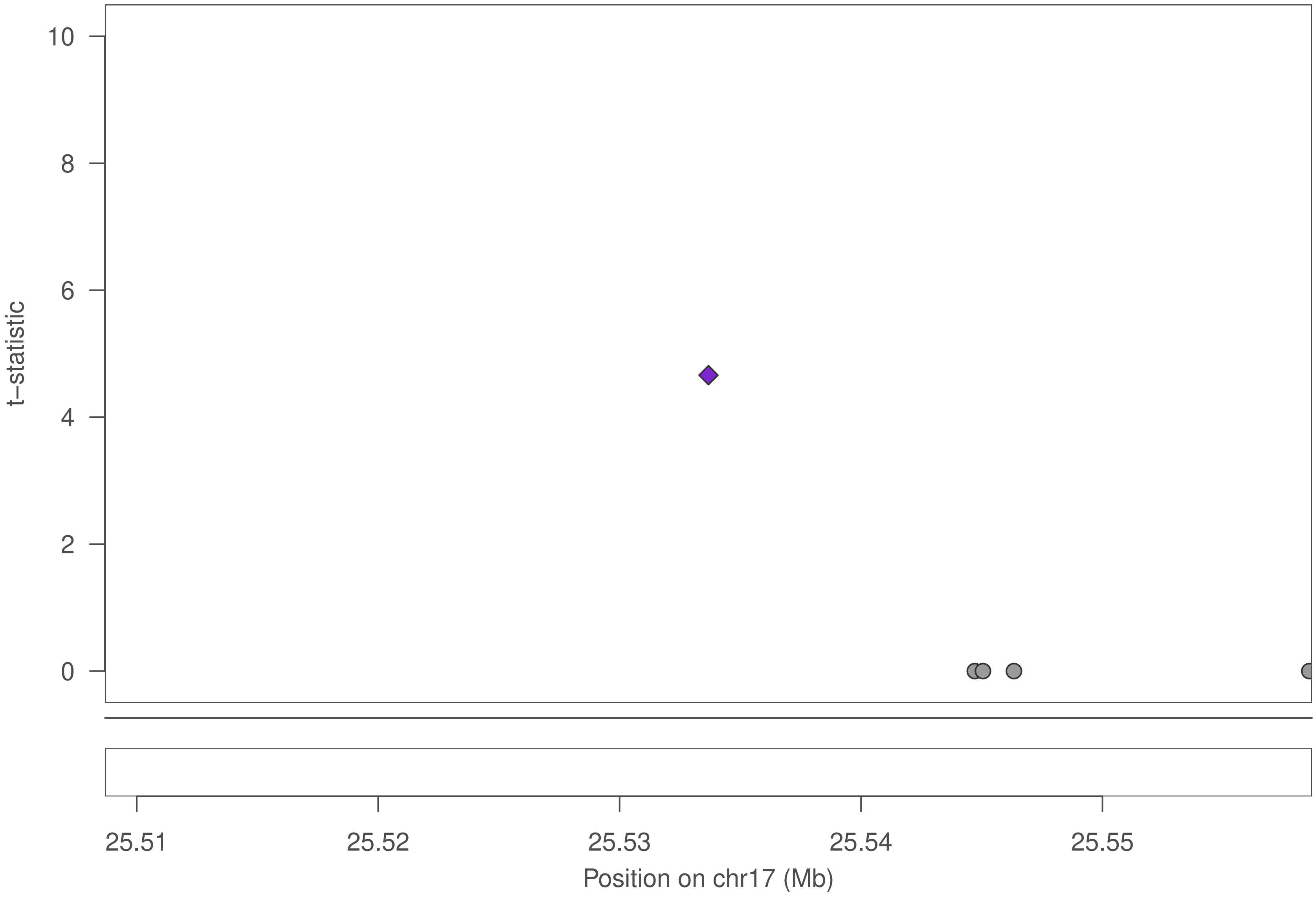

10
8
6
4
2
0
t−statistic
25.51
25.52
25.53
25.54
25.55
Position on chr17 (Mb)

### ZNF443

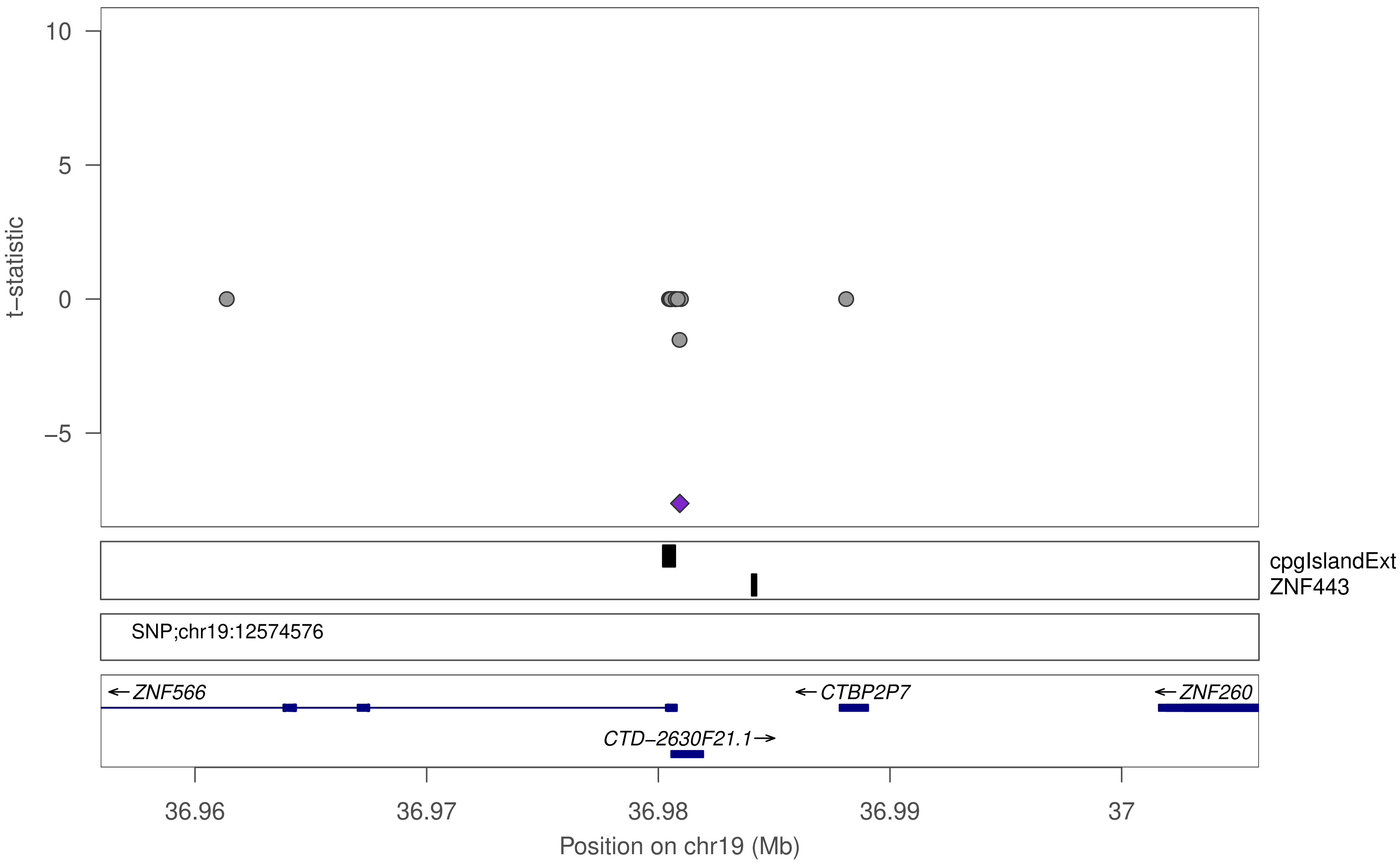

10
5
0
−5
t−statistic
cpgIslandExt
ZNF443
SNP;chr19:12574576
←ZNF566
←CTBP2P7
←ZNF260
CTD−2630F21.1→
36.96
36.97
36.98
36.99
37
Position on chr19 (Mb)

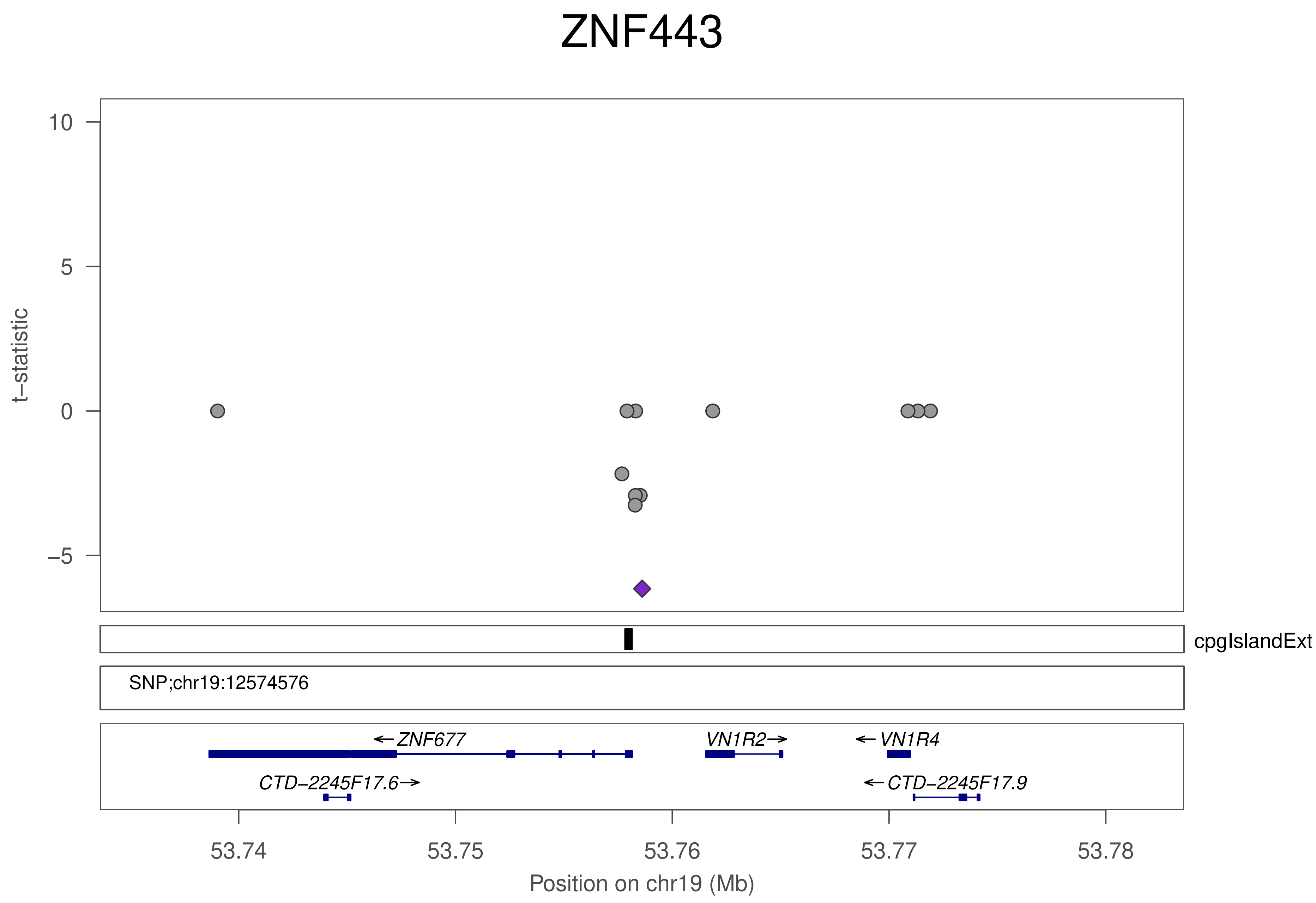
ZNF443
t−statistic
10
5
0
−5
cpgIslandExt
SNP;chr19:12574576
←ZNF677
VN1R2→
←VN1R4
CTD−2245F17.6→
←CTD−2245F17.9
53.74
53.75
53.76
53.77
53.78
Position on chr19 (Mb)

### ZNF443

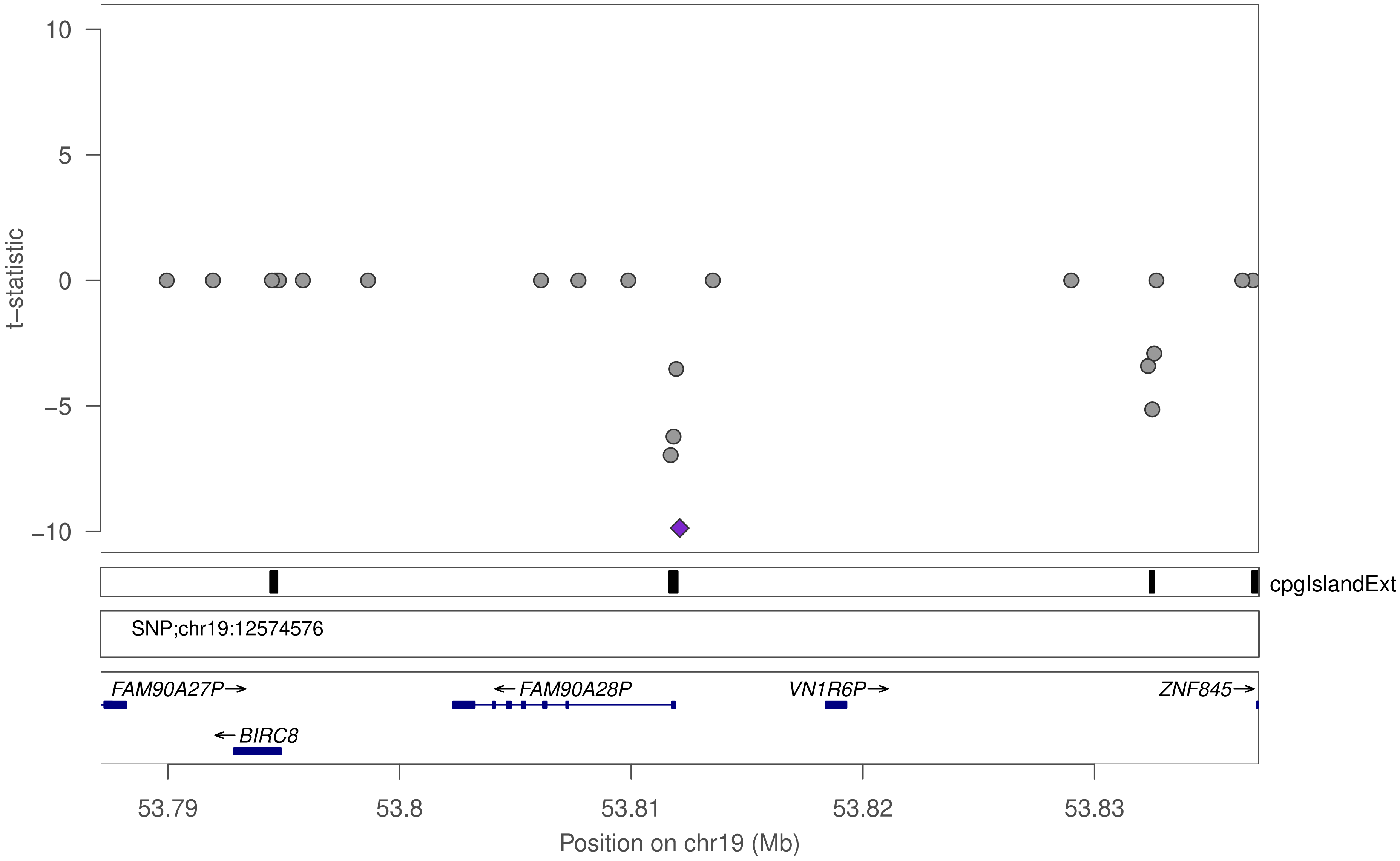

t−statistic
10
5
0
−5
−10
cpgIslandExt
SNP;chr19:12574576
FAM90A27P→
←FAM90A28P
VN1R6P→
ZNF845→
←BIRC8
53.79
53.8
53.81
53.82
53.83
Position on chr19 (Mb)

### ZNF443

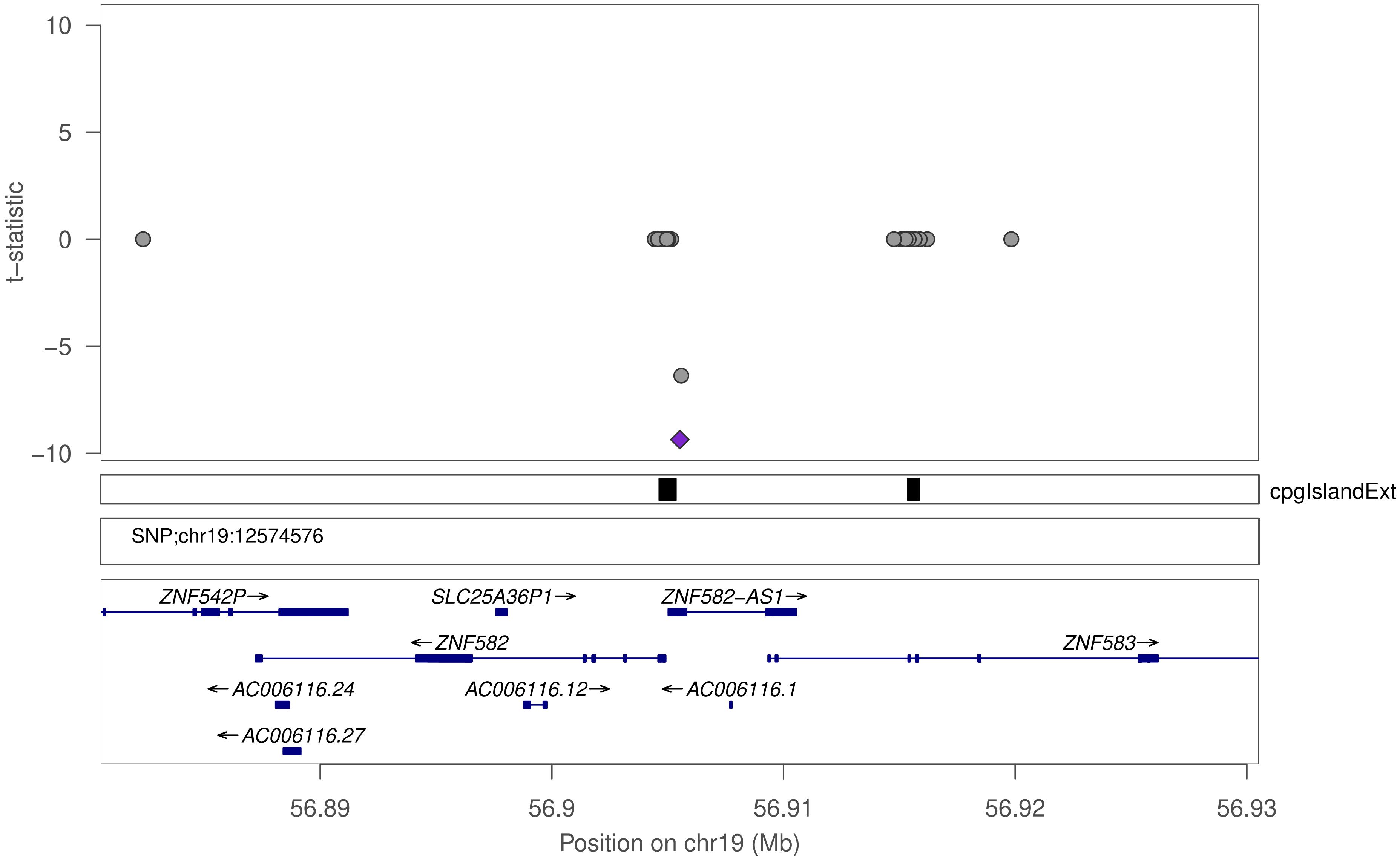
t–statistic
10
5
0
–5
–10
cpgIslandExt
SNP;chr19:12574576
ZNF542P→
SLC25A36P1→
ZNF582–AS1→
←ZNF582
ZNF583→
←AC006116.24
AC006116.12→
←AC006116.1
←AC006116.27
56.89
56.9
56.91
56.92
56.93
Position on chr19 (Mb)

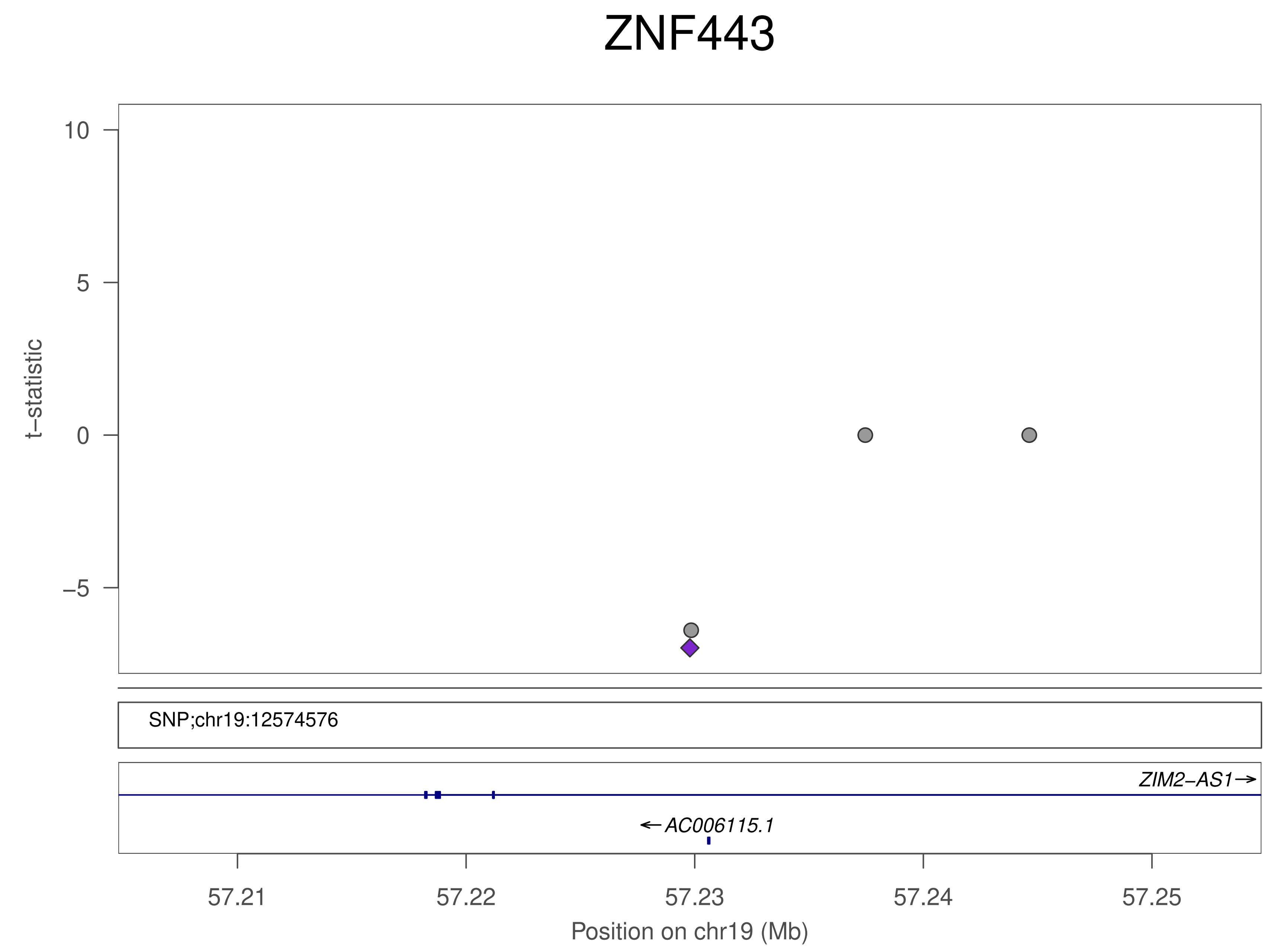
ZNF443
t−statistic
10
5
0
−5
SNP;chr19:12574576
ZIM2−AS1→
←AC006115.1
57.21
57.22
57.23
57.24
57.25
Position on chr19 (Mb)

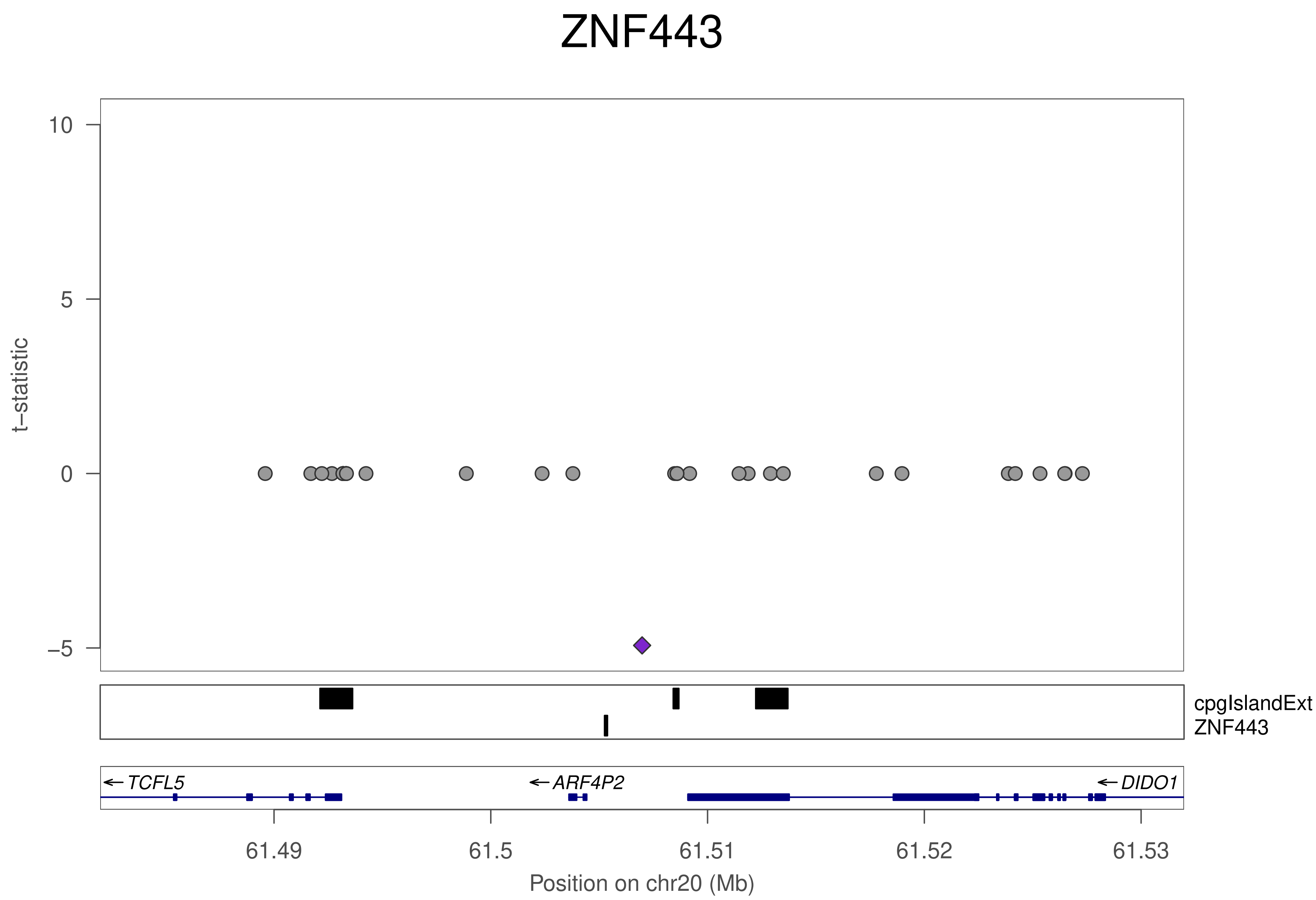

ZNF443
10
5
0
−5
t−statistic
cpgIslandExt
ZNF443
← TCFL5
← ARF4P2
← DIDO1
61.49
61.5
61.51
61.52
61.53
Position on chr20 (Mb)

### ZNF443

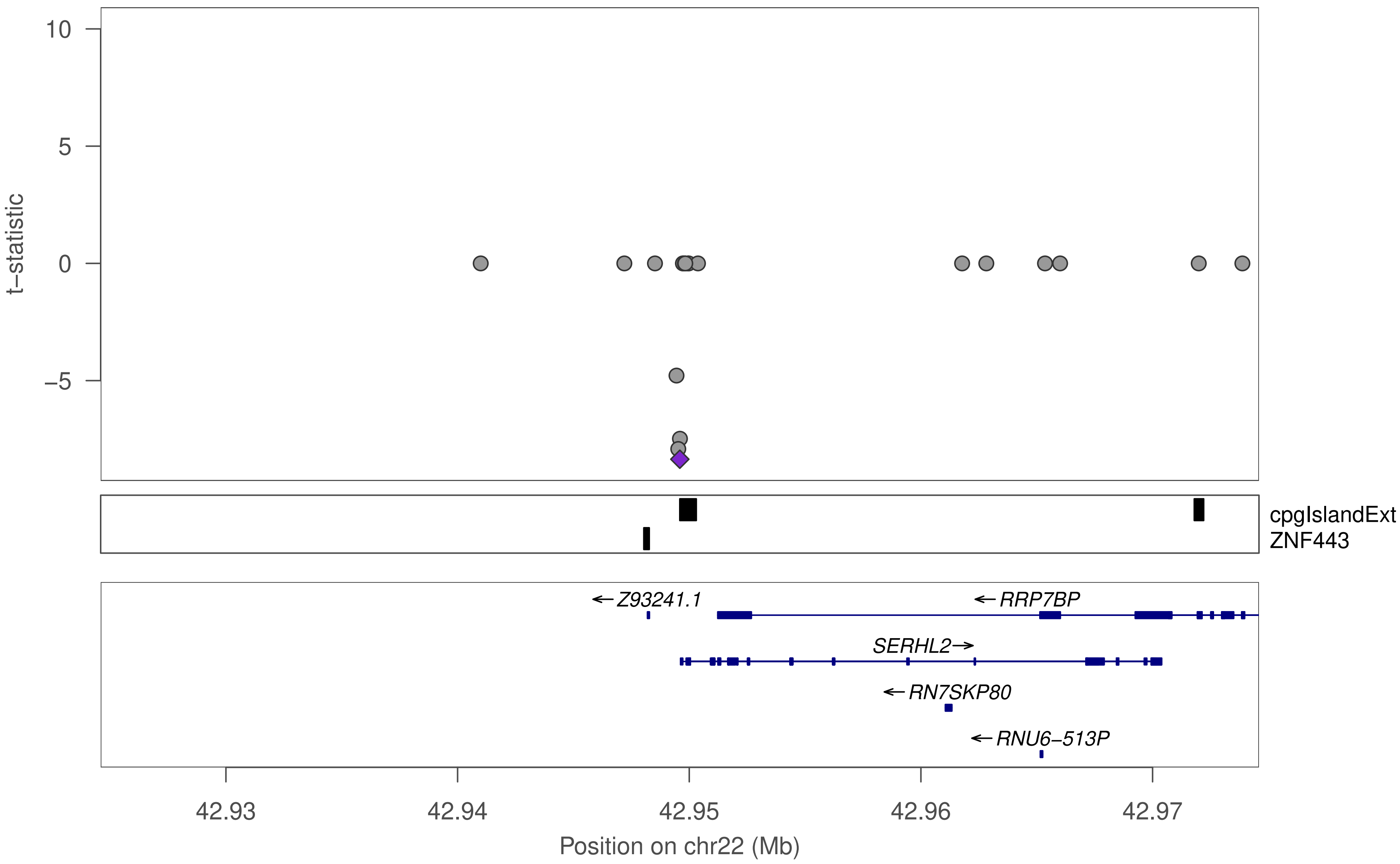

10
5
0
−5
t−statistic
cpgIslandExt
ZNF443
←Z93241.1
←RRP7BP
SERHL2→
←RN7SKP80
←RNU6−513P
42.93
42.94
42.95
42.96
42.97
Position on chr22 (Mb)

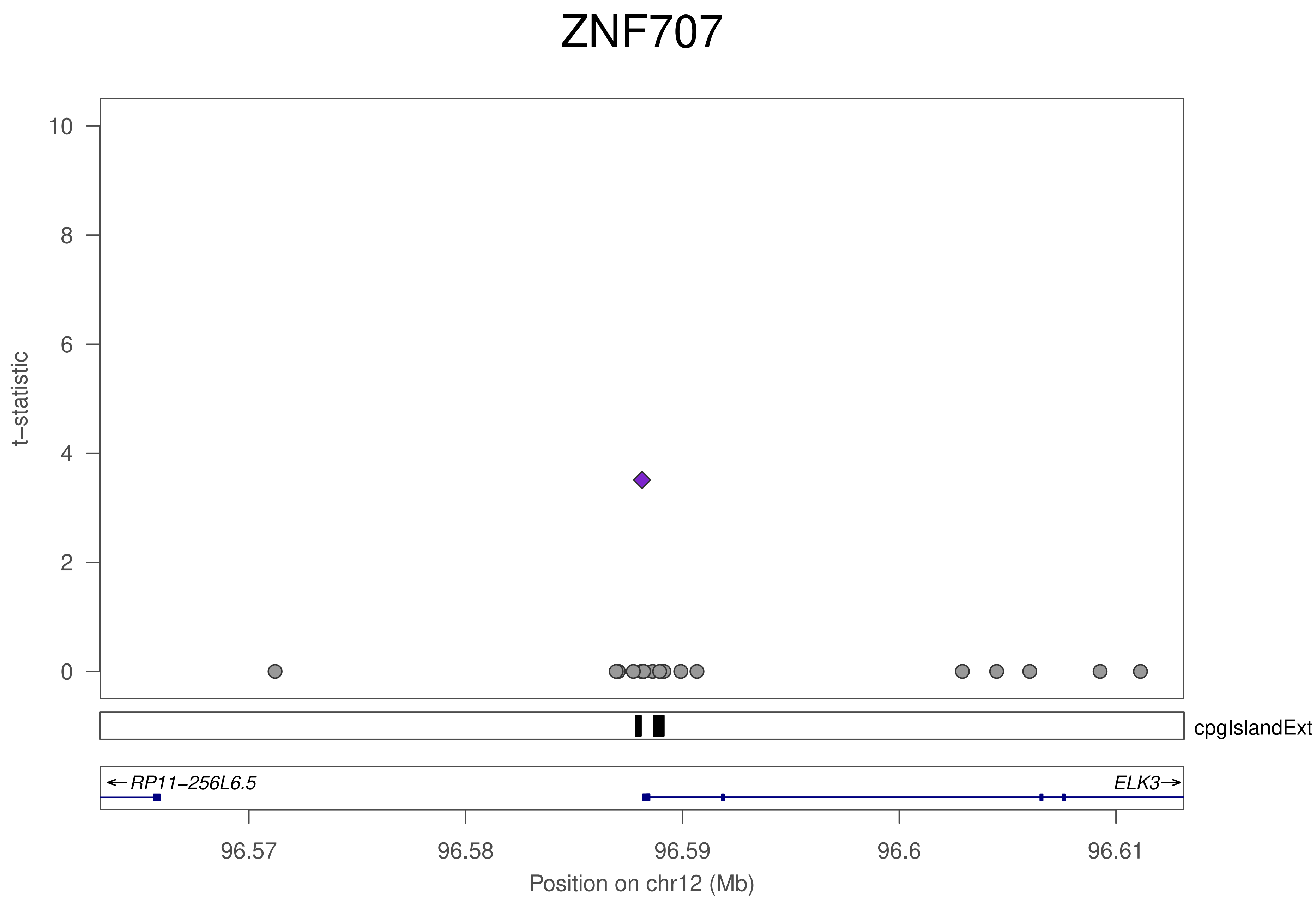
ZNF707
t–statistic
10
8
6
4
2
0
cpgIslandExt
←RP11–256L6.5
ELK3→
96.57
96.58
96.59
96.6
96.61
Position on chr12 (Mb)

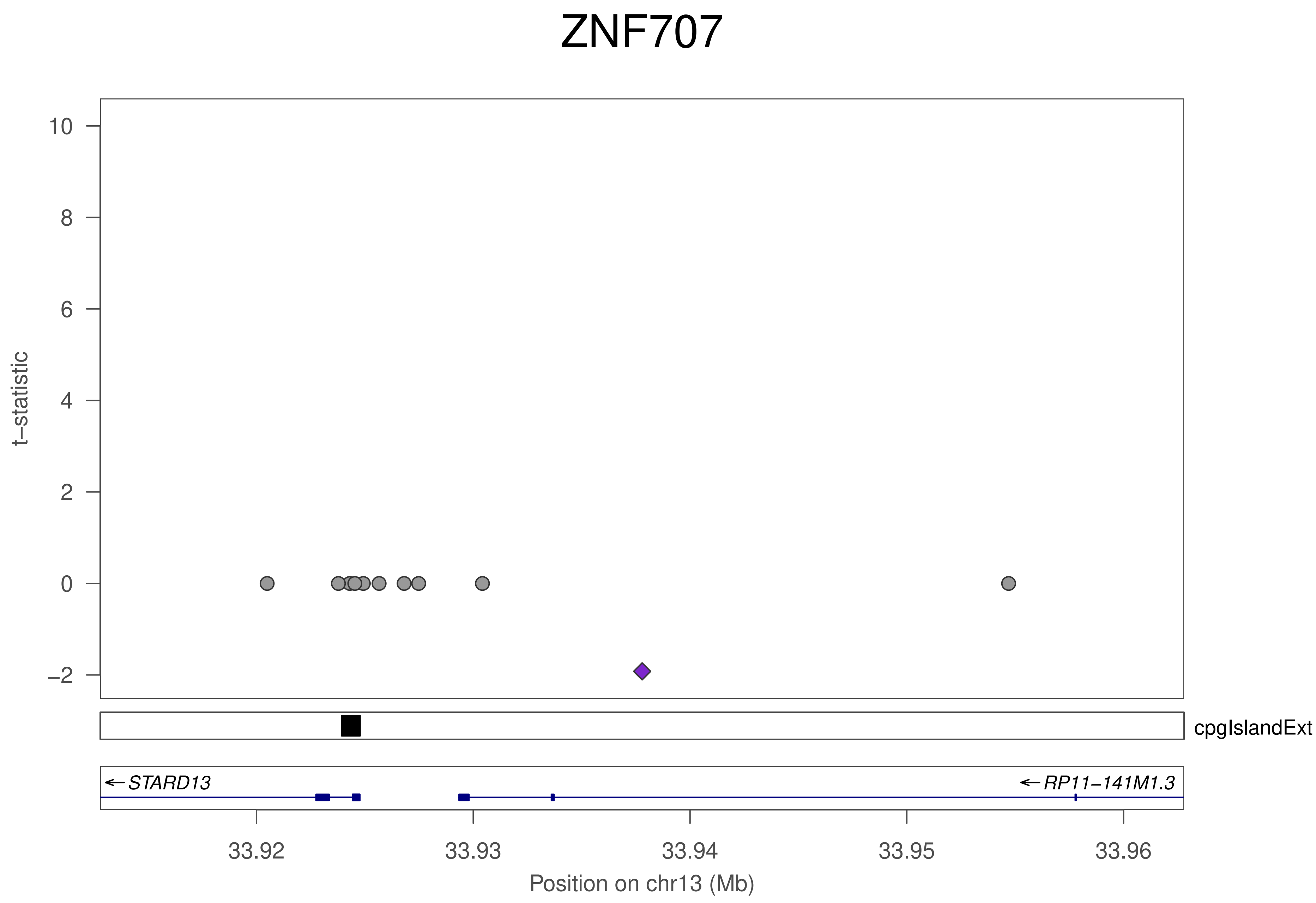

ZNF707
t−statistic
10
8
6
4
2
0
−2
cpgIslandExt
← STARD13
← RP11−141M1.3
33.92
33.93
33.94
33.95
33.96
Position on chr13 (Mb)

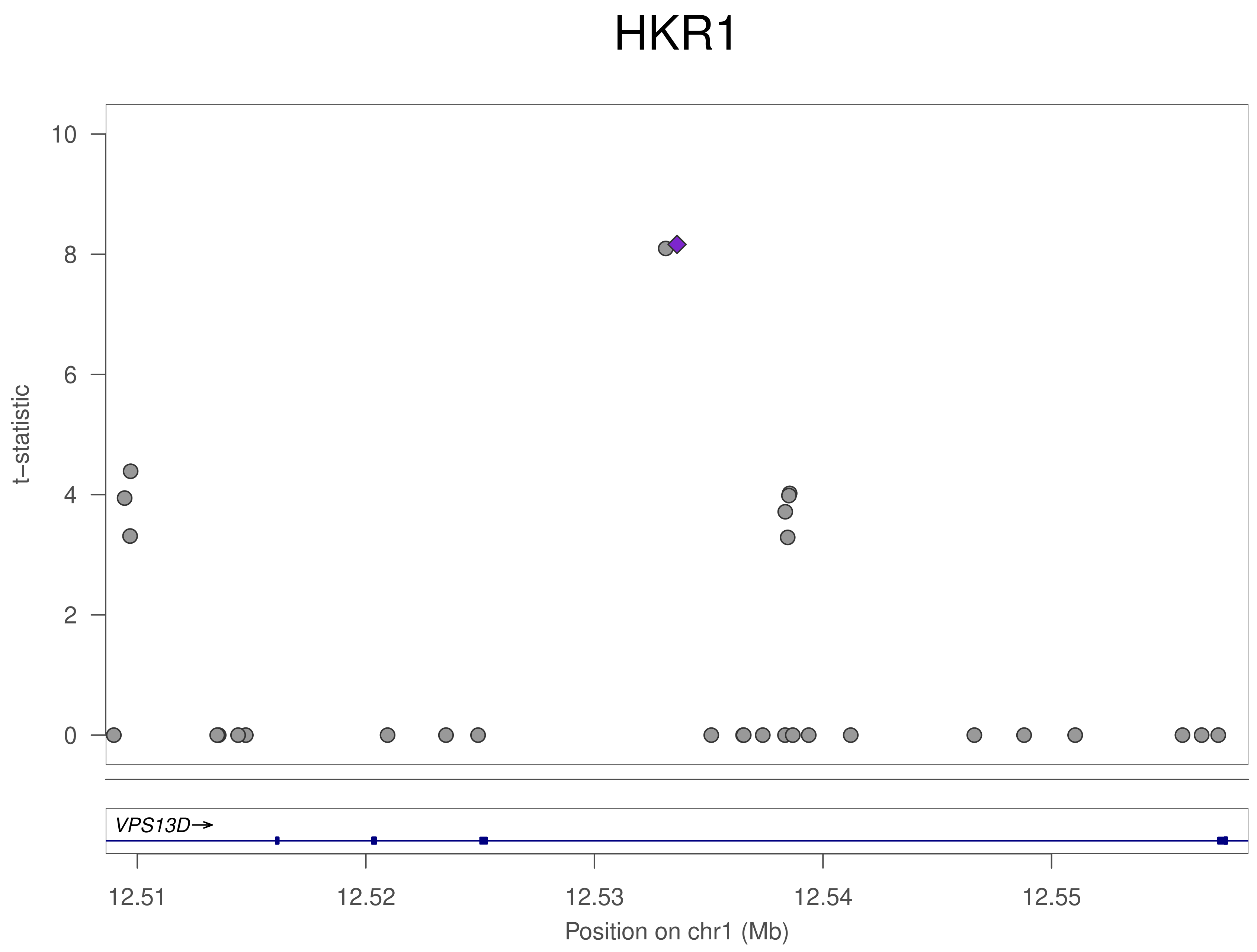

HKR1
t−statistic
10
8
6
4
2
0
VPS13D→
12.51
12.52
12.53
12.54
12.55
Position on chr1 (Mb)

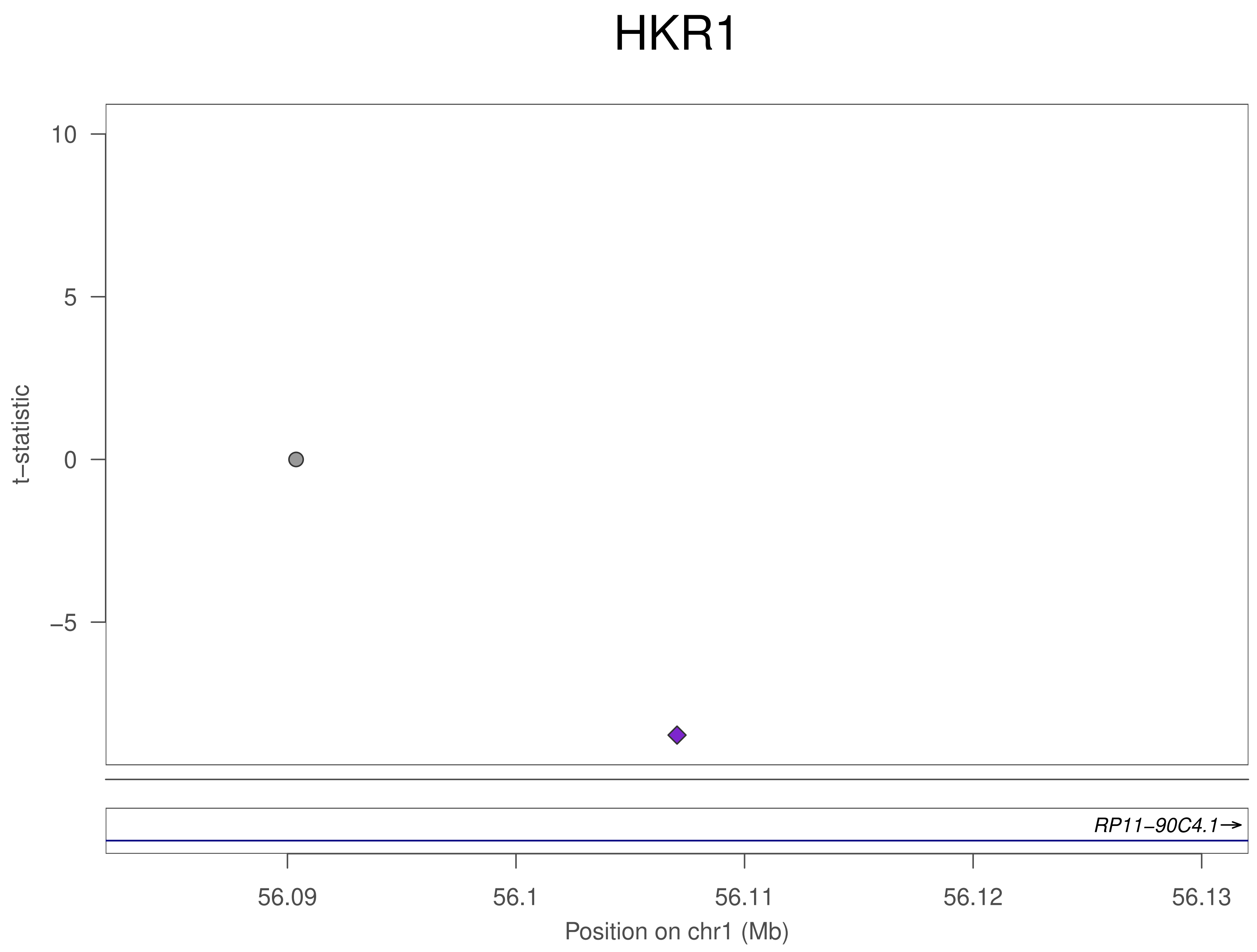

HKR1
10
5
0
–5
t–statistic
RP11–90C4.1→
56.09
56.1
56.11
56.12
56.13
Position on chr1 (Mb)

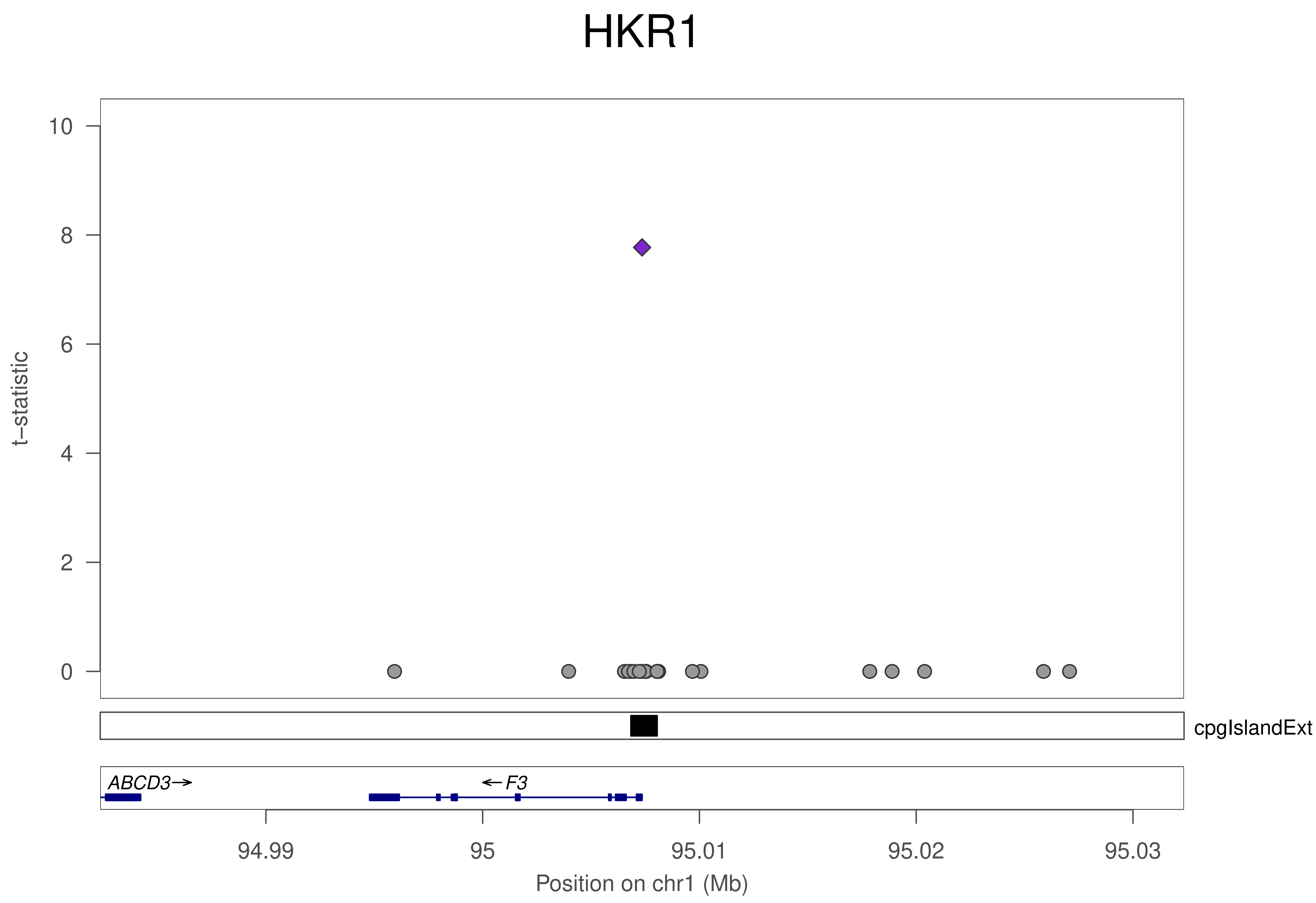
HKR1
t−statistic
10
8
6
4
2
0
cpgIslandExt
ABCD3→
←F3
94.99
95
95.01
95.02
95.03
Position on chr1 (Mb)

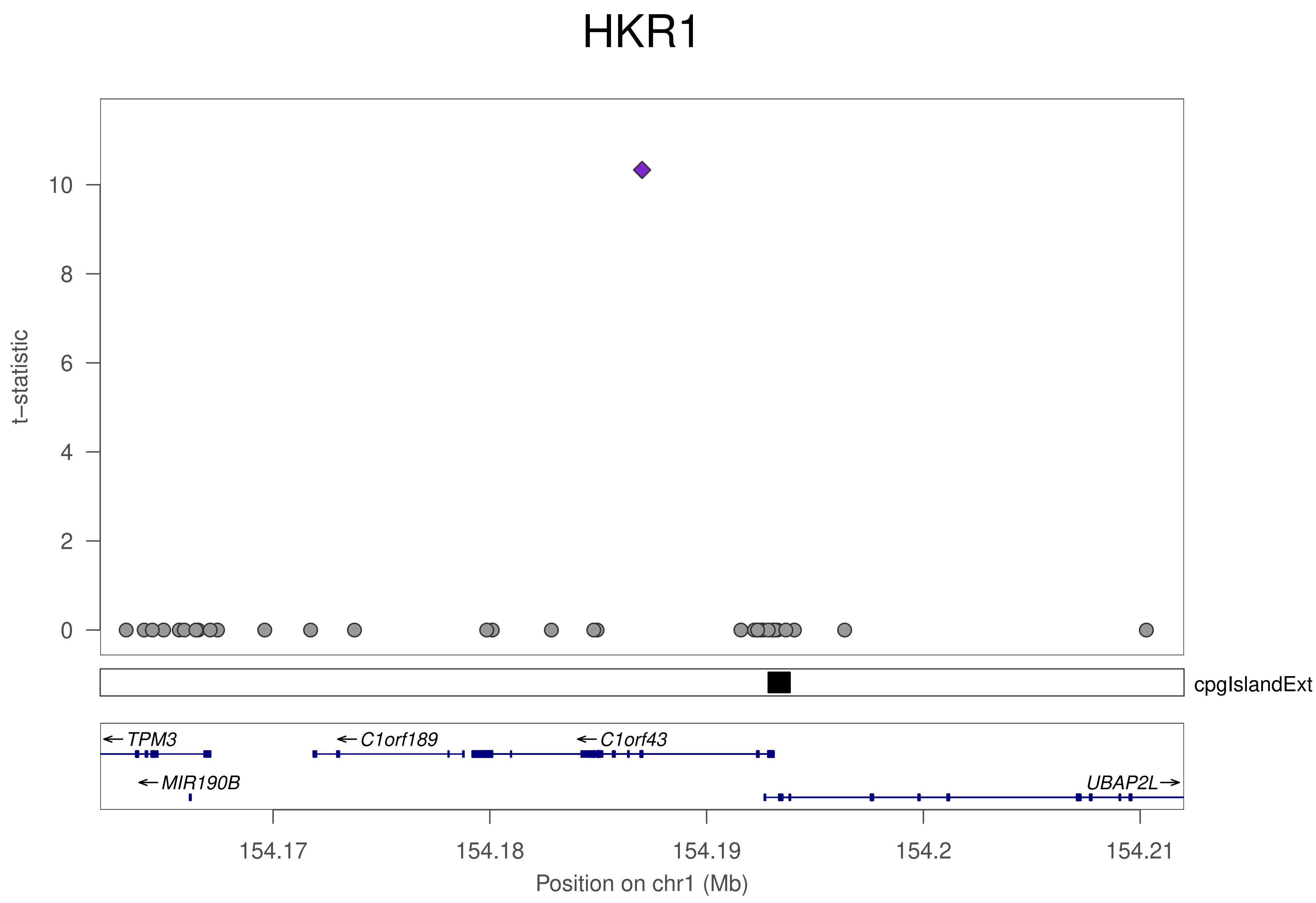
HKR1
t−statistic
0
2
4
6
8
10
cpgIslandExt
←TPM3
←C1orf189
←C1orf43
←MIR190B
UBAP2L→
154.17
154.18
154.19
154.2
154.21
Position on chr1 (Mb)

### HKR1

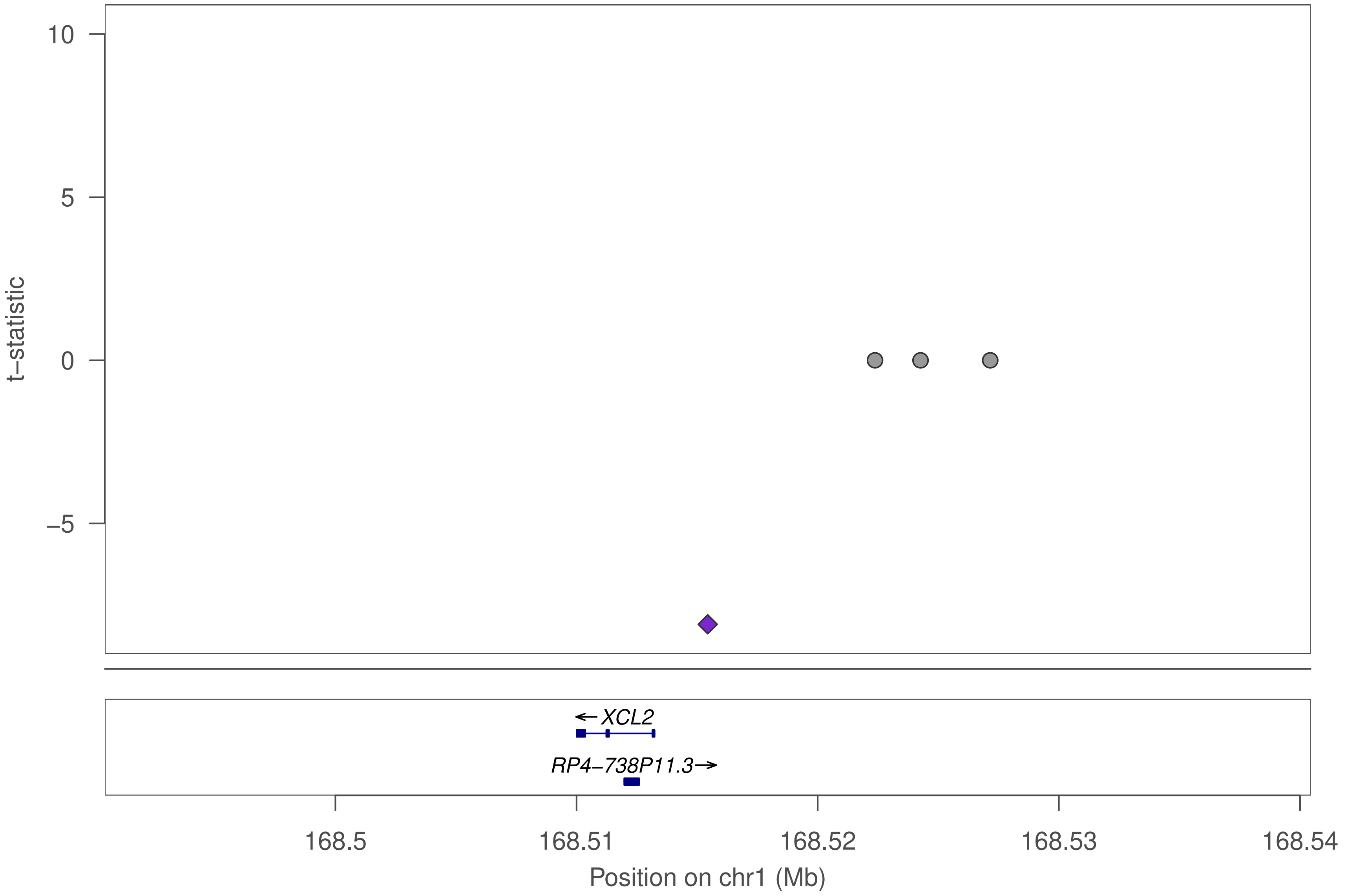

10
5
0
−5
t−statistic
←XCL2
RP4−738P11.3→
168.5
168.51
168.52
168.53
168.54
Position on chr1 (Mb)

### HKR1

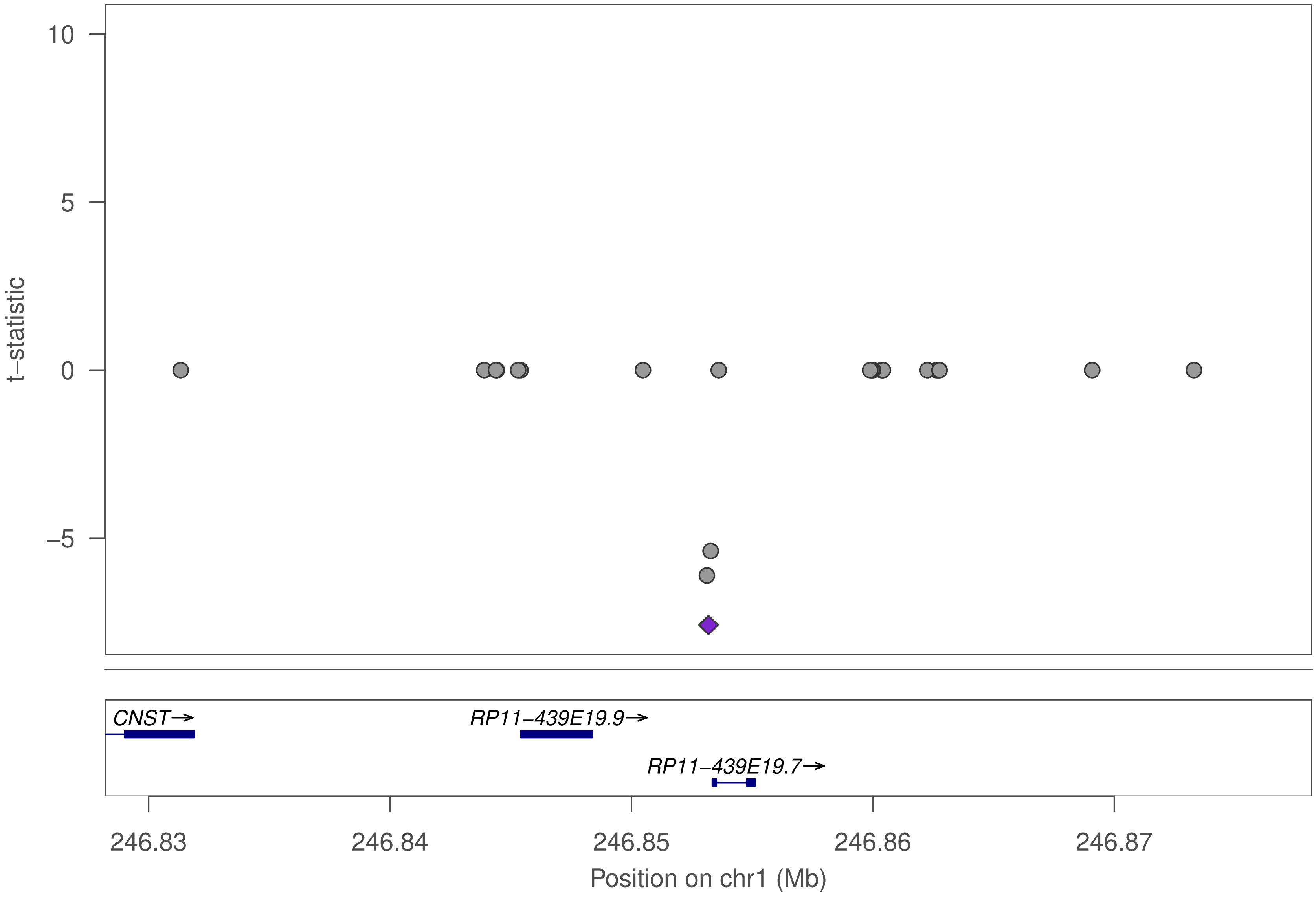

10
5
0
−5
t−statistic
CNST→
RP11−439E19.9→
RP11−439E19.7→
246.83
246.84
246.85
246.86
246.87
Position on chr1 (Mb)

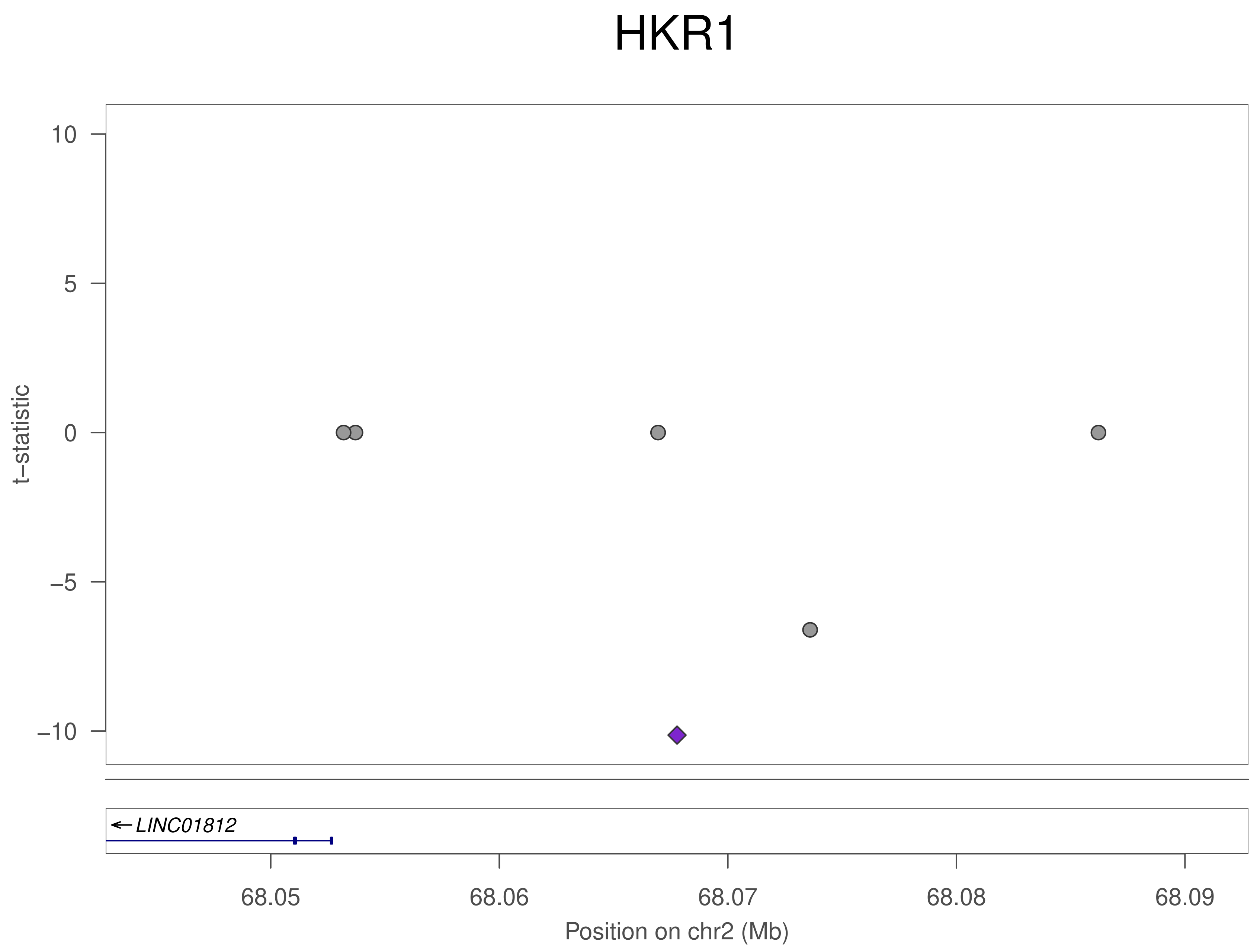

HKR1
10
5
0
−5
−10
t−statistic
LINC01812
68.05
68.06
68.07
68.08
68.09
Position on chr2 (Mb)

### HKR1

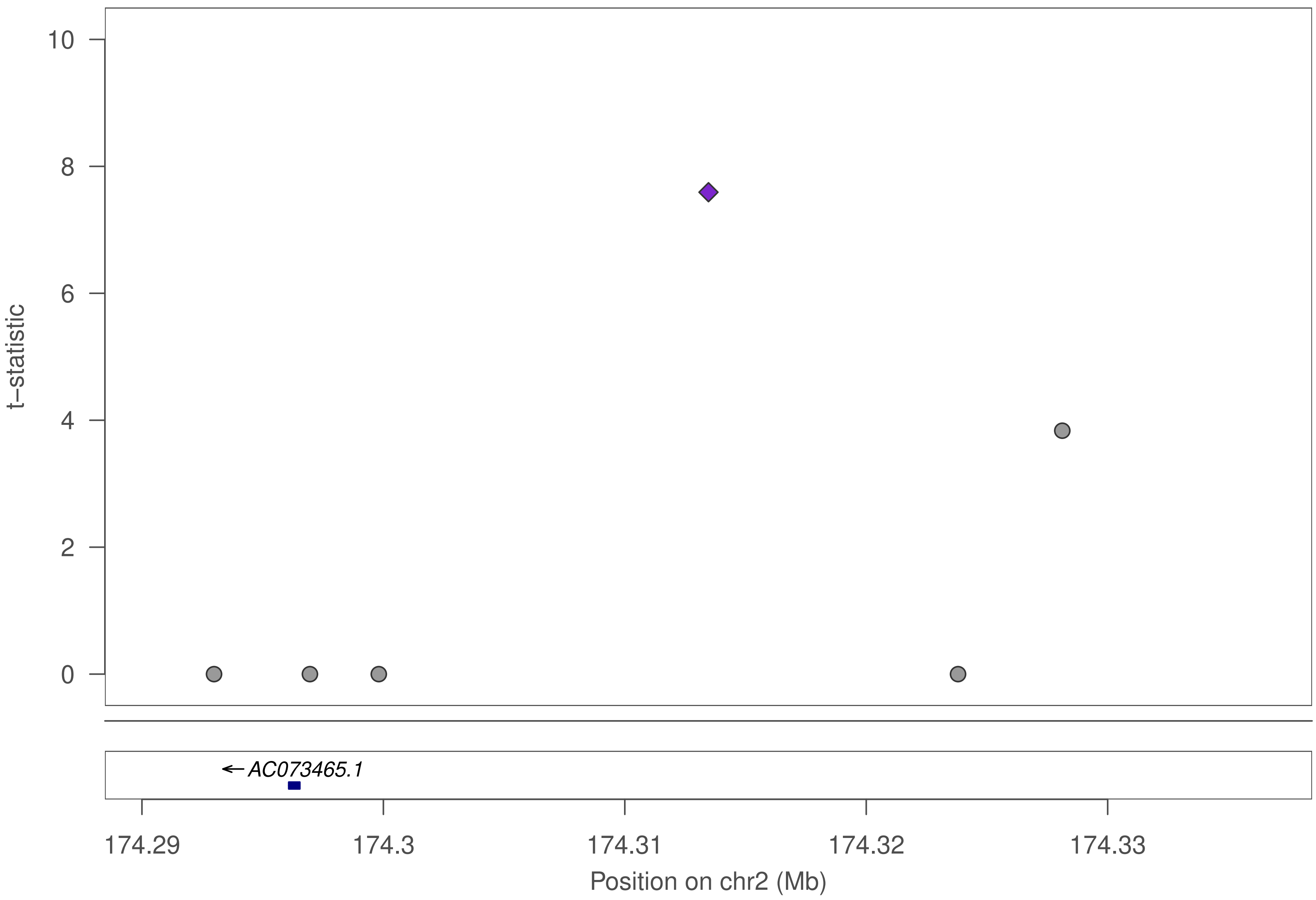

10
8
6
4
2
0
t−statistic
AC073465.1
174.29
174.3
174.31
174.32
174.33
Position on chr2 (Mb)

### HKR1

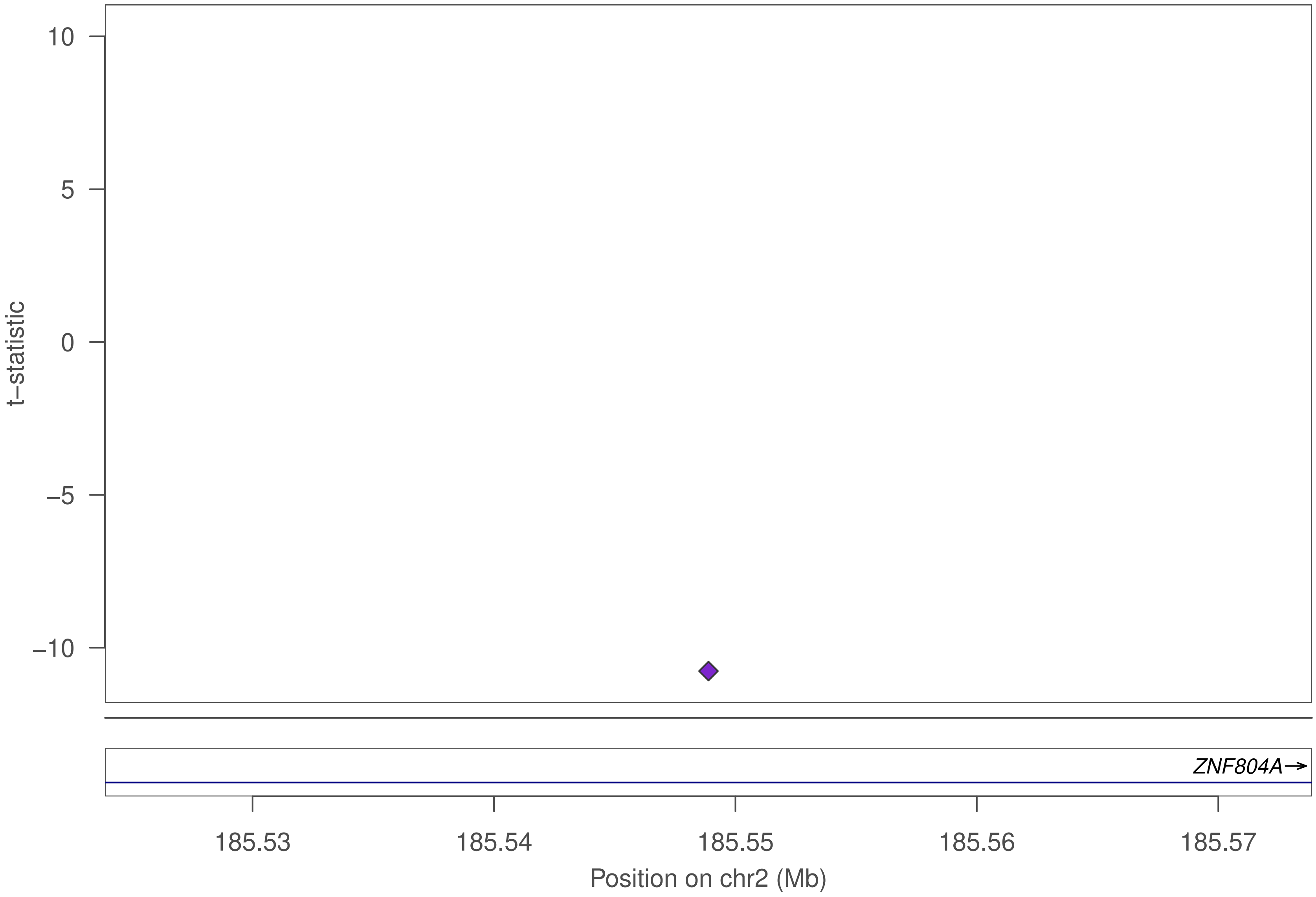

10
5
0
−5
−10
t−statistic
ZNF804A
185.53
185.54
185.55
185.56
185.57
Position on chr2 (Mb)

### HKR1

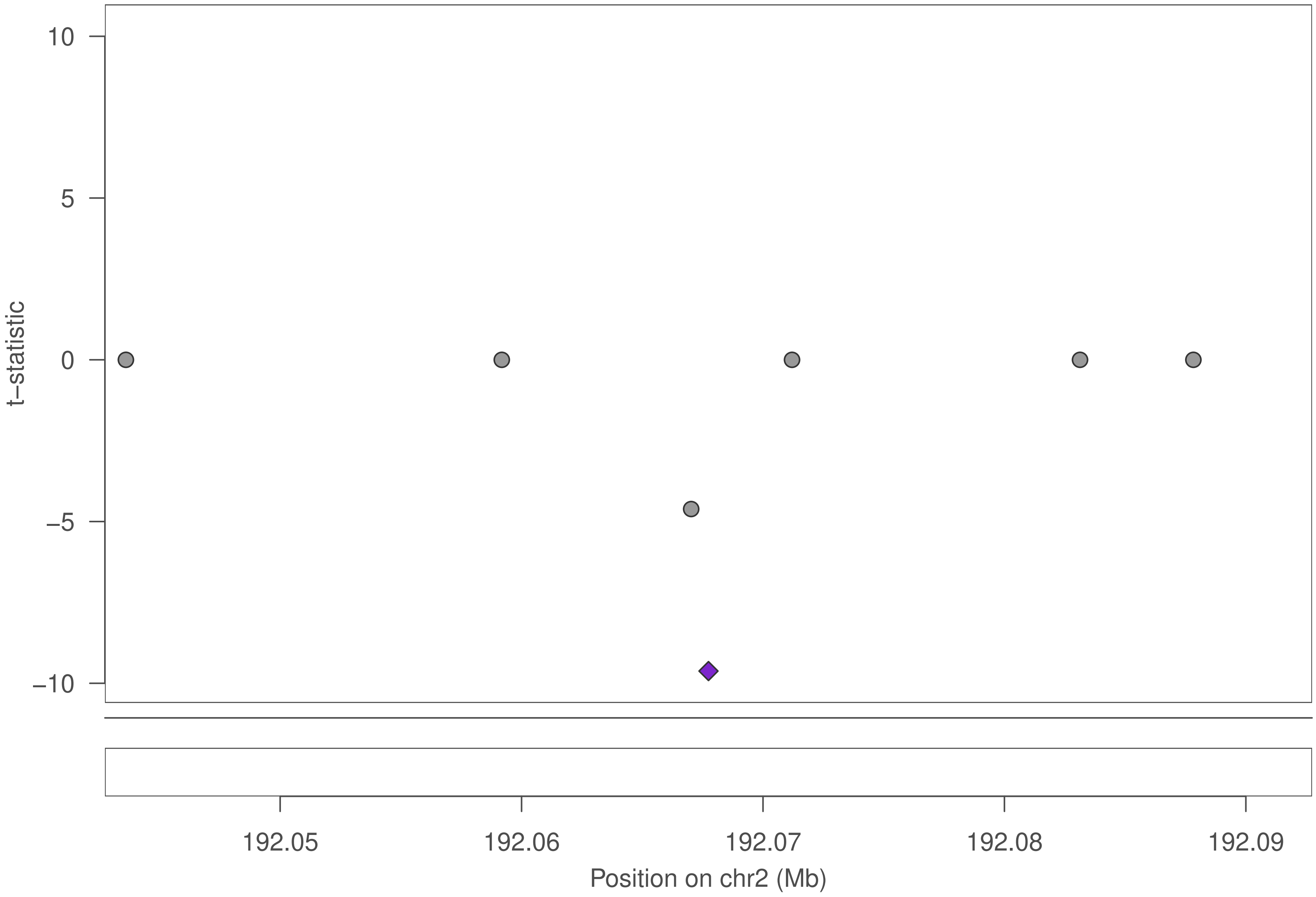

10
5
0
−5
−10
t−statistic
192.05
192.06
192.07
192.08
192.09
Position on chr2 (Mb)

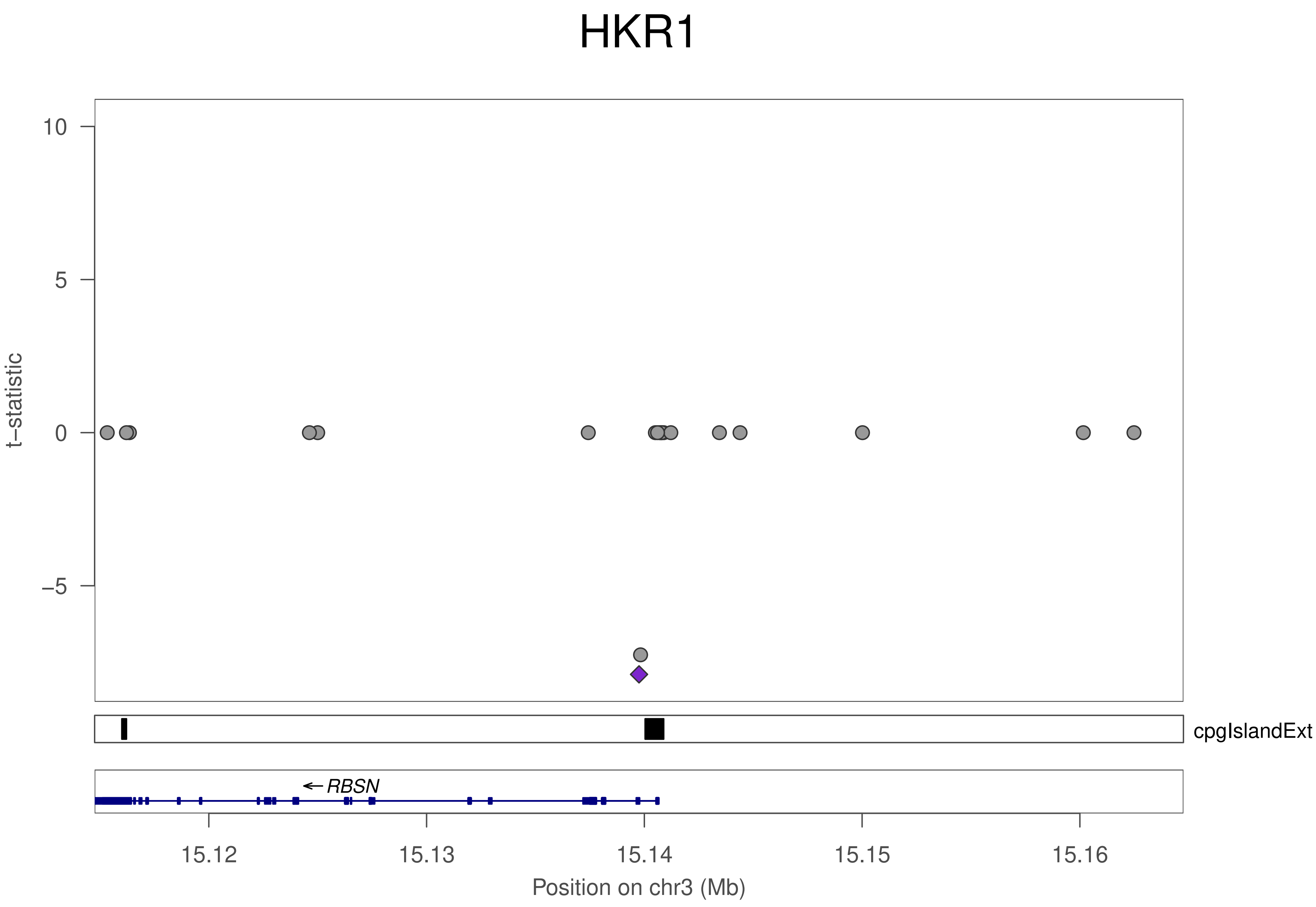

HKR1
t−statistic
10
5
0
−5
cpgIslandExt
← RBSN
15.12
15.13
15.14
15.15
15.16
Position on chr3 (Mb)

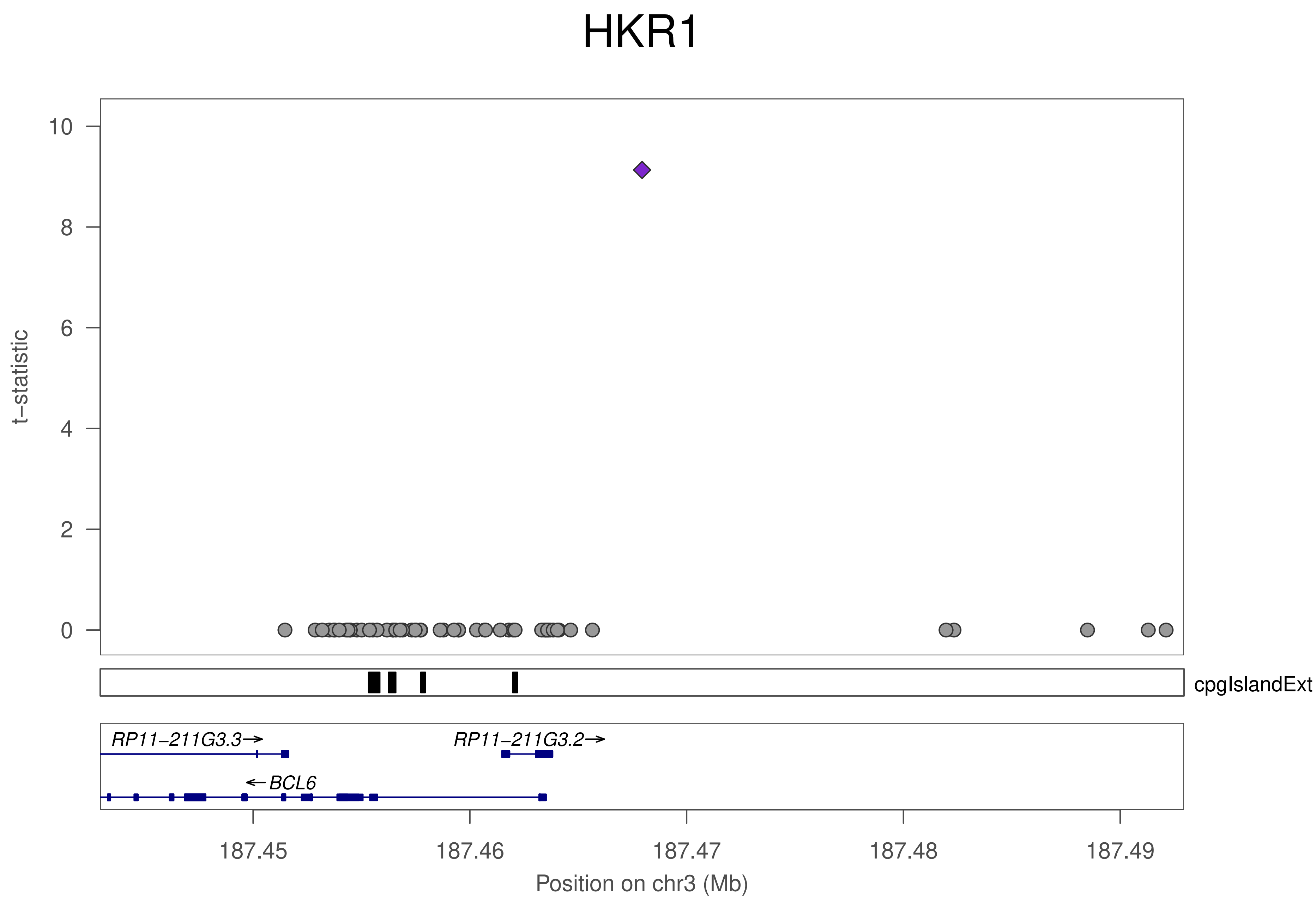

HKR1
t−statistic
0
2
4
6
8
10
cpgIslandExt
RP11−211G3.3→
RP11−211G3.2→
←BCL6
187.45
187.46
187.47
187.48
187.49
Position on chr3 (Mb)

### HKR1

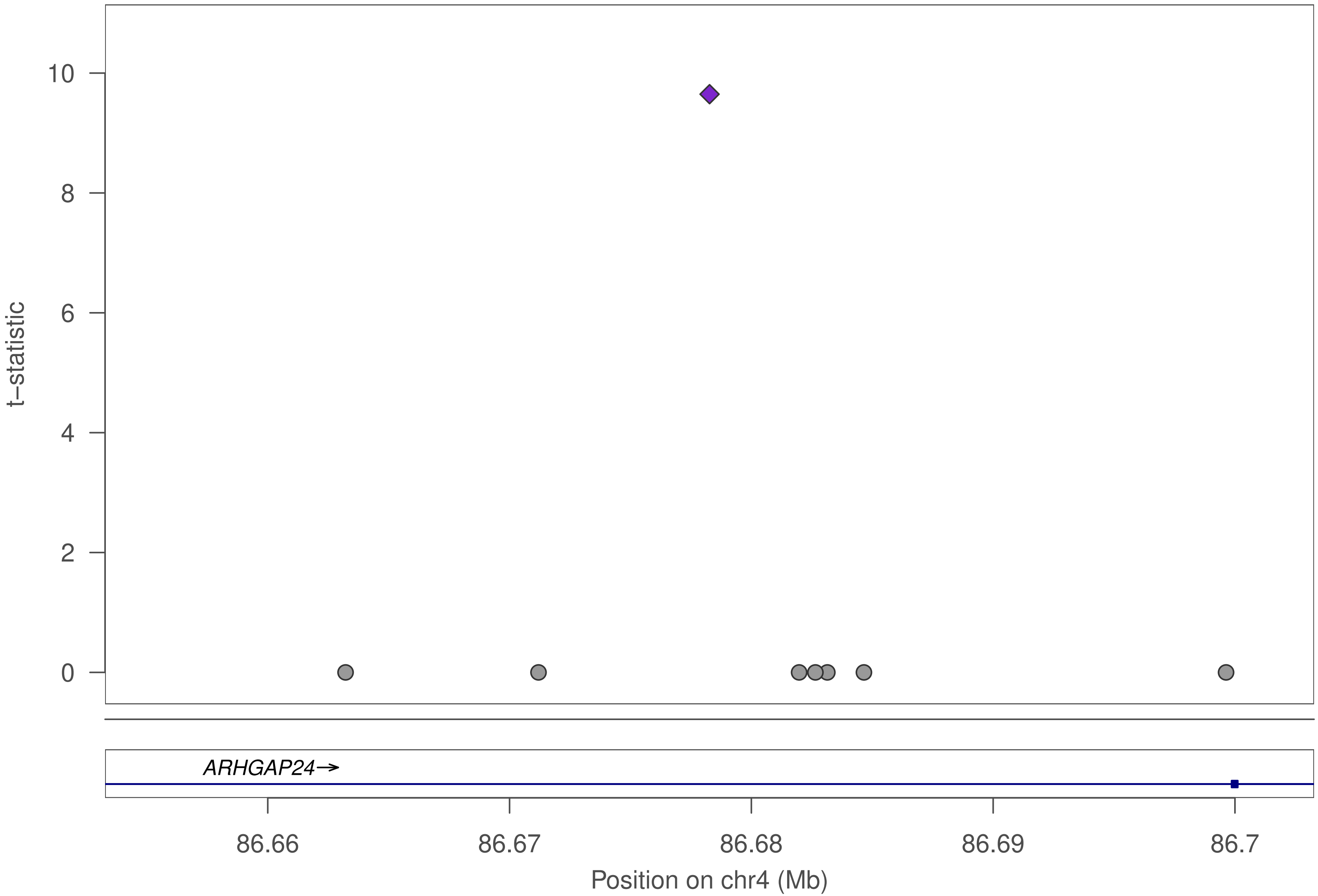

10
8
6
4
2
0
t–statistic
ARHGAP24→
86.66
86.67
86.68
86.69
86.7
Position on chr4 (Mb)

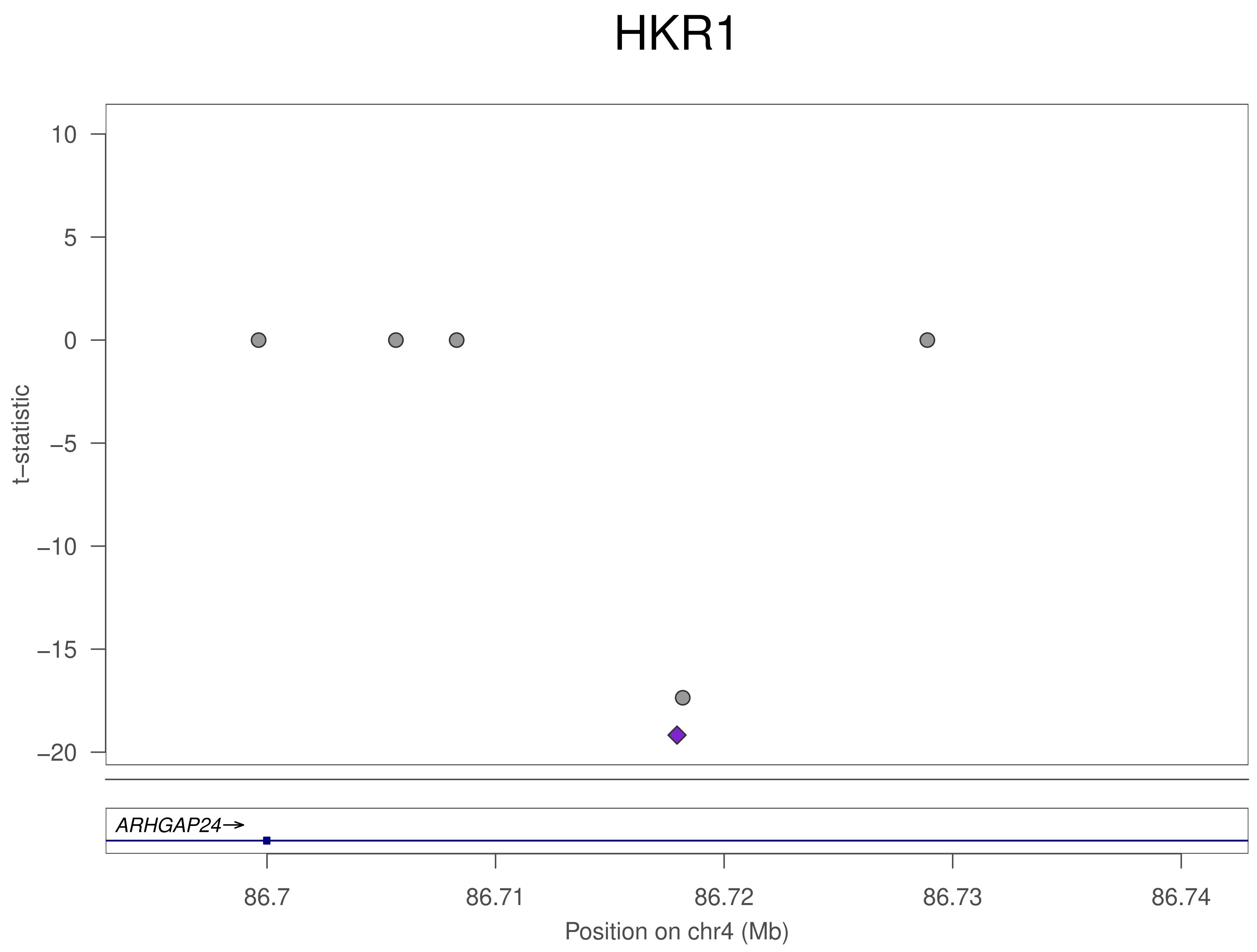
HKR1
10
5
0
−5
−10
−15
−20
t−statistic
ARHGAP24→
86.7
86.71
86.72
86.73
86.74
Position on chr4 (Mb)

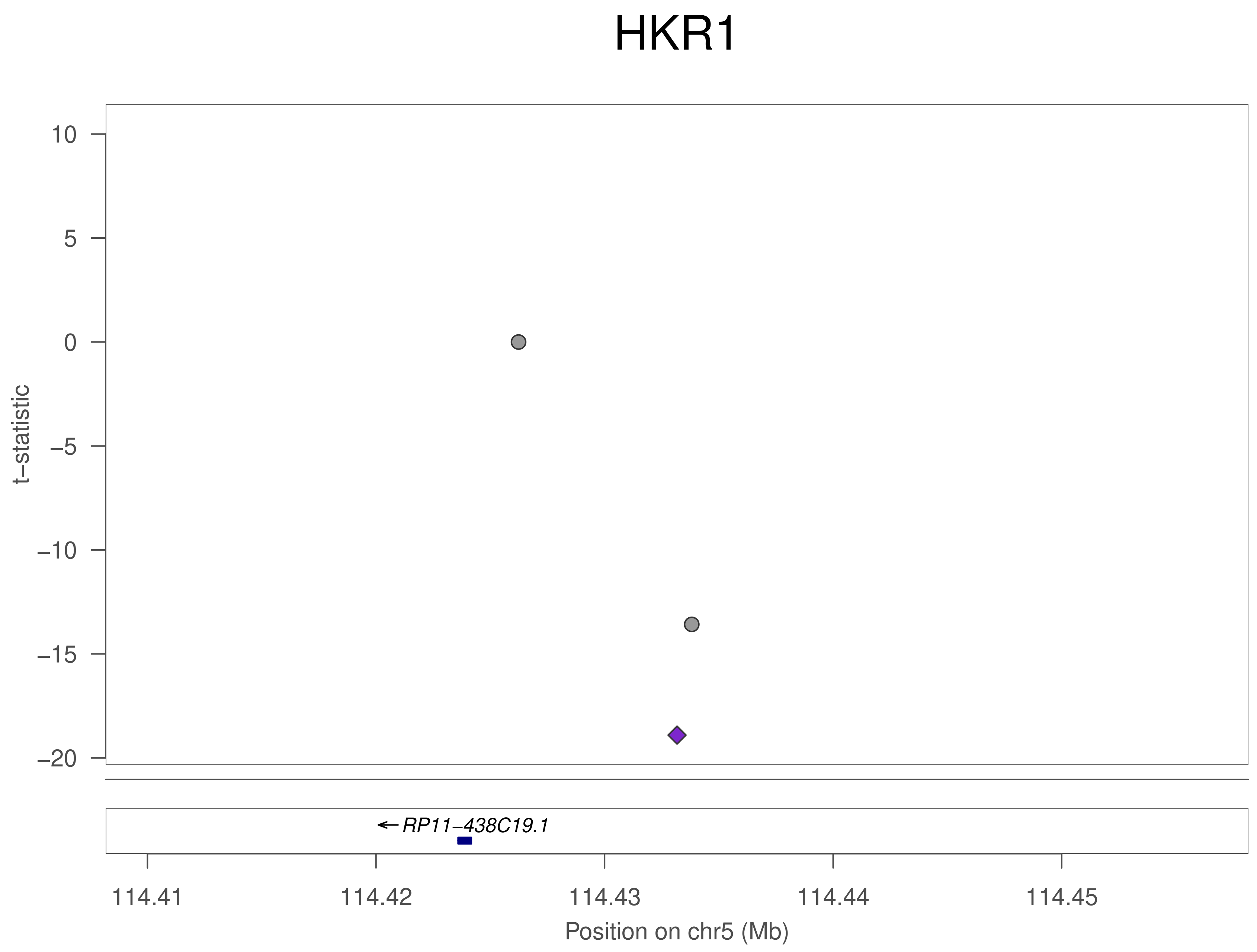

HKR1
10
5
0
−5
−10
−15
−20
t−statistic
RP11−438C19.1
114.41
114.42
114.43
114.44
114.45
Position on chr5 (Mb)

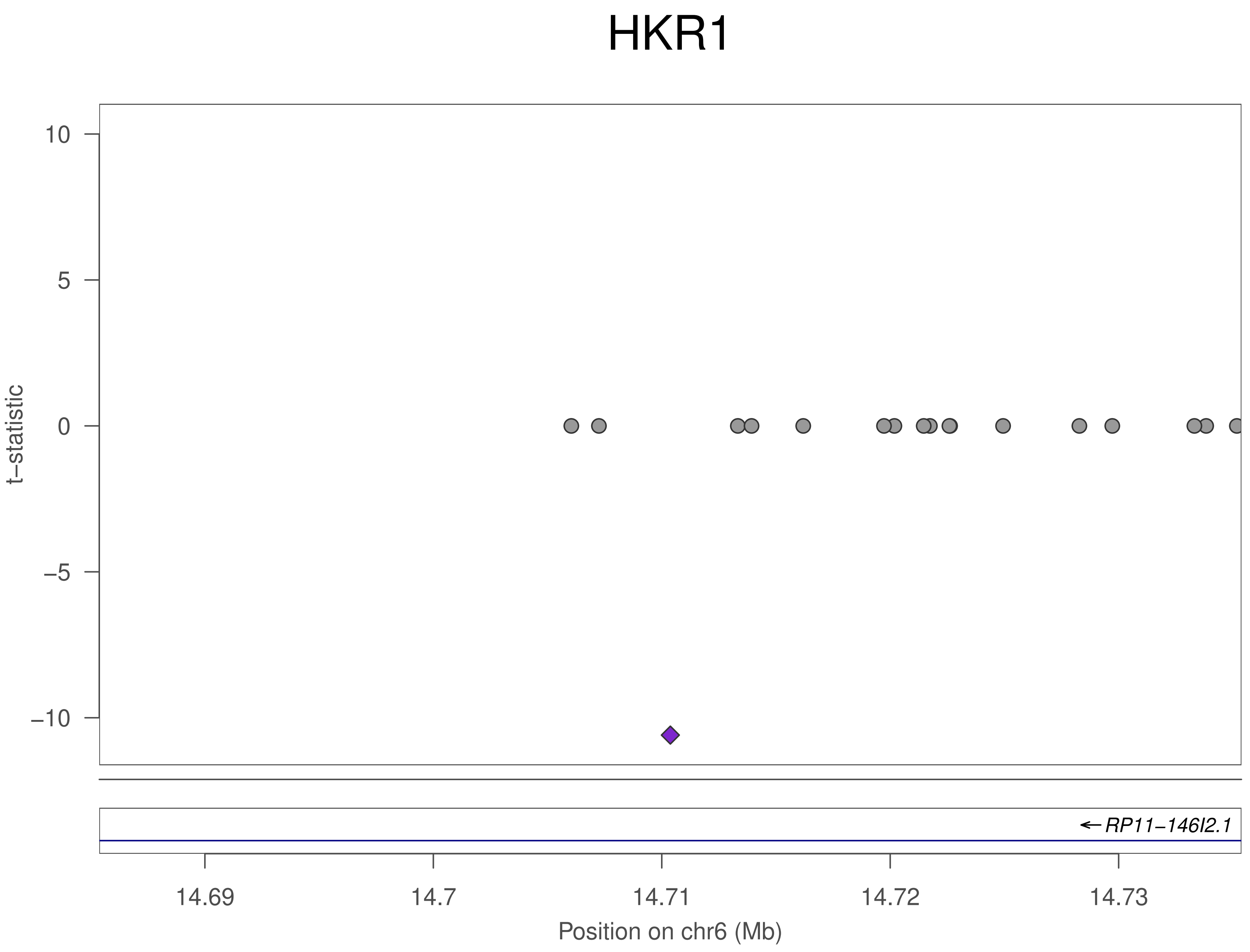

HKR1
10
5
0
−5
−10
t−statistic
RP11−146I2.1
14.69
14.7
14.71
14.72
14.73
Position on chr6 (Mb)

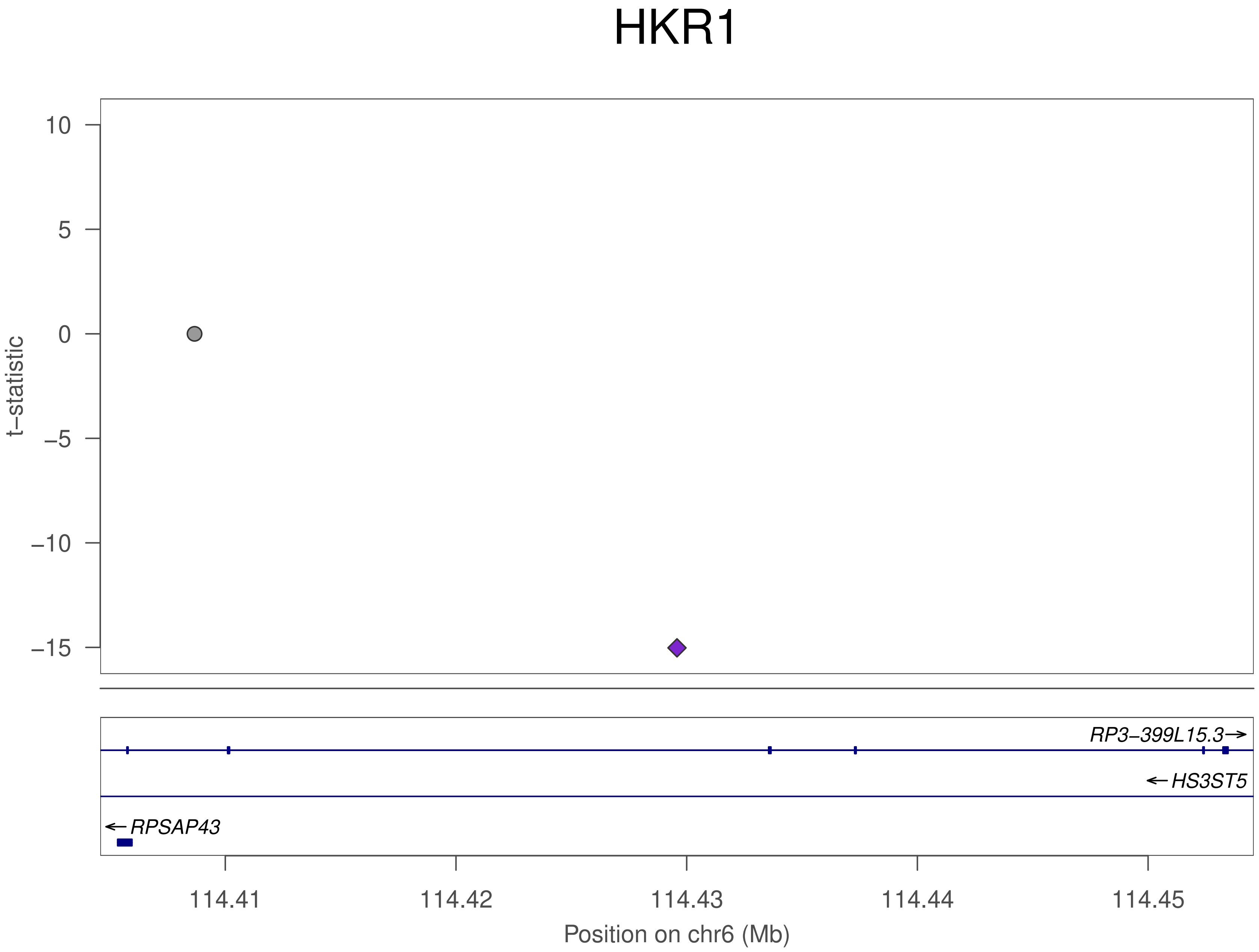
HKR1
t−statistic
10
5
0
−5
−10
−15
RP3−399L15.3
HS3ST5
RPSAP43
114.41
114.42
114.43
114.44
114.45
Position on chr6 (Mb)

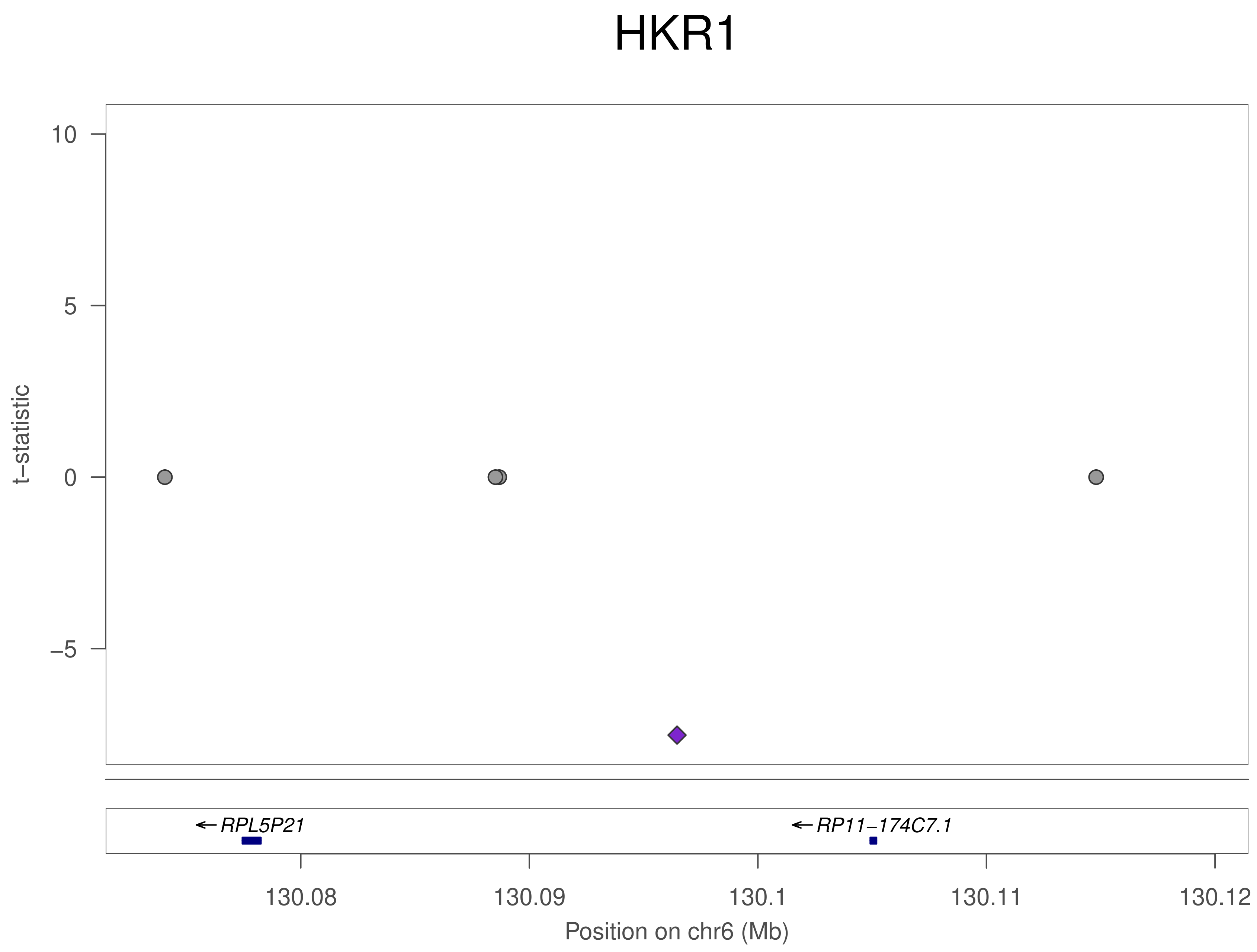
HKR1
t−statistic
10
5
0
−5
←RPL5P21
←RP11−174C7.1
130.08
130.09
130.1
130.11
130.12
Position on chr6 (Mb)

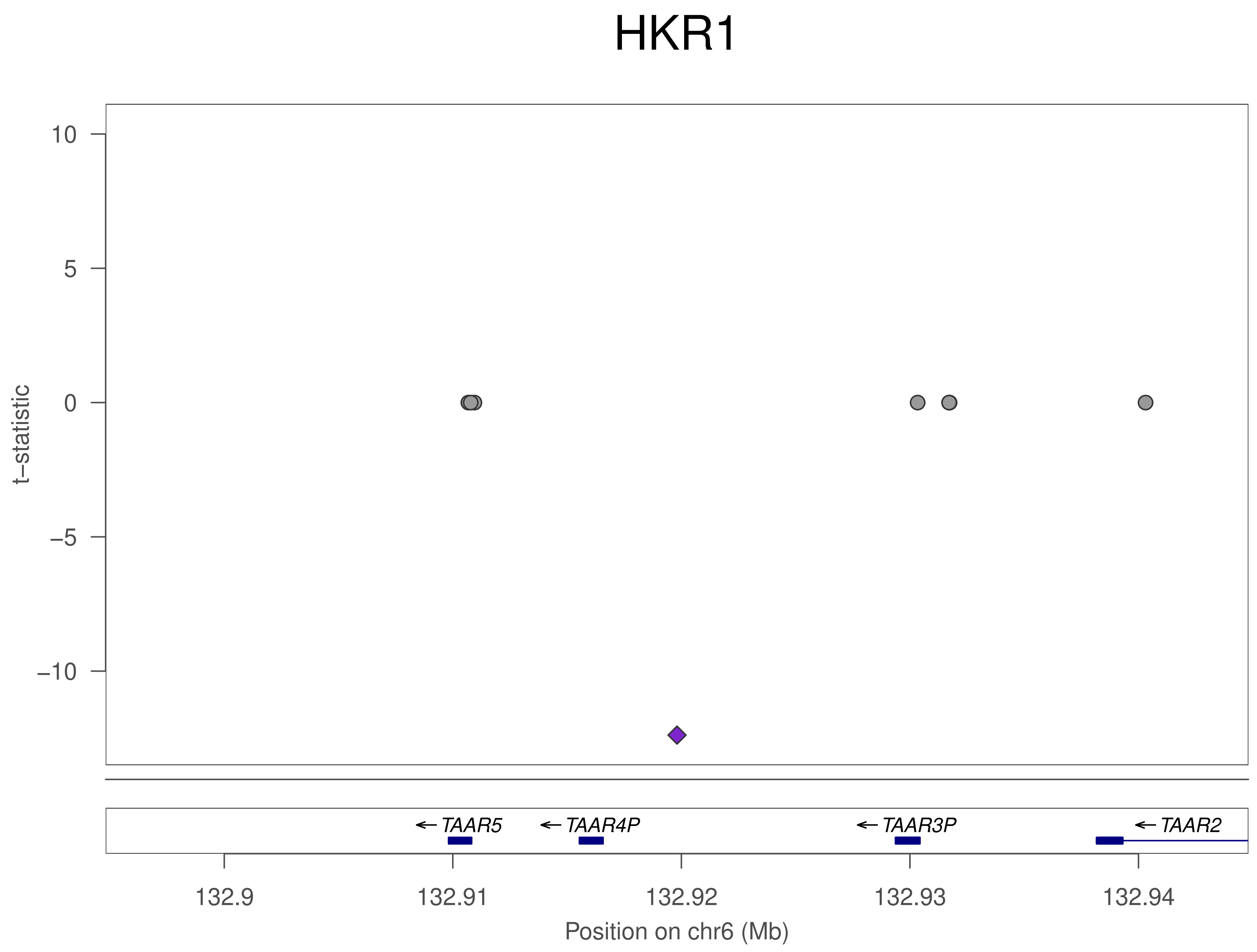

HKR1
10
5
0
−5
−10
t−statistic
TAAR5
TAAR4P
TAAR3P
TAAR2
132.9
132.91
132.92
132.93
132.94
Position on chr6 (Mb)

### HKR1

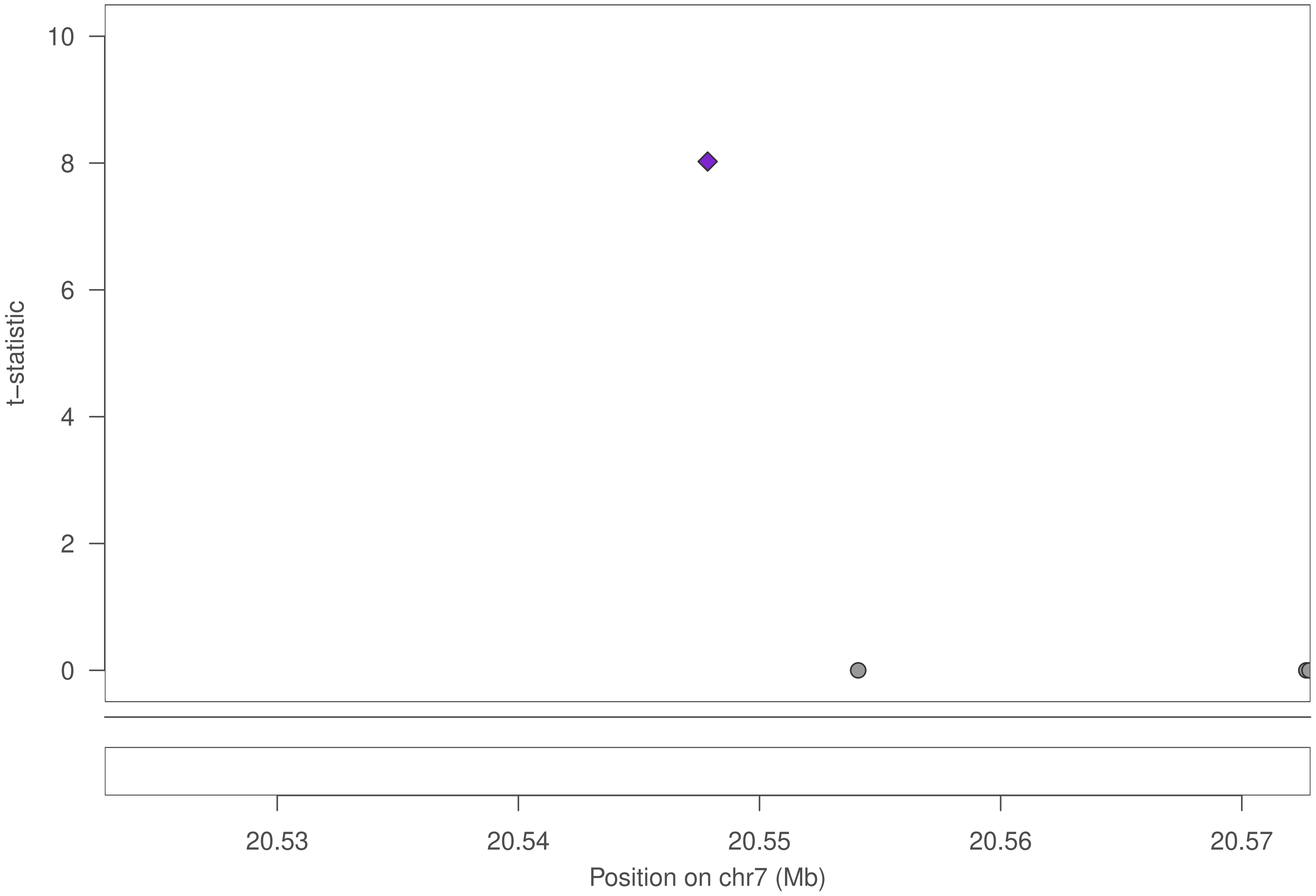

10
8
6
4
2
0
t−statistic
20.53
20.54
20.55
20.56
20.57
Position on chr7 (Mb)

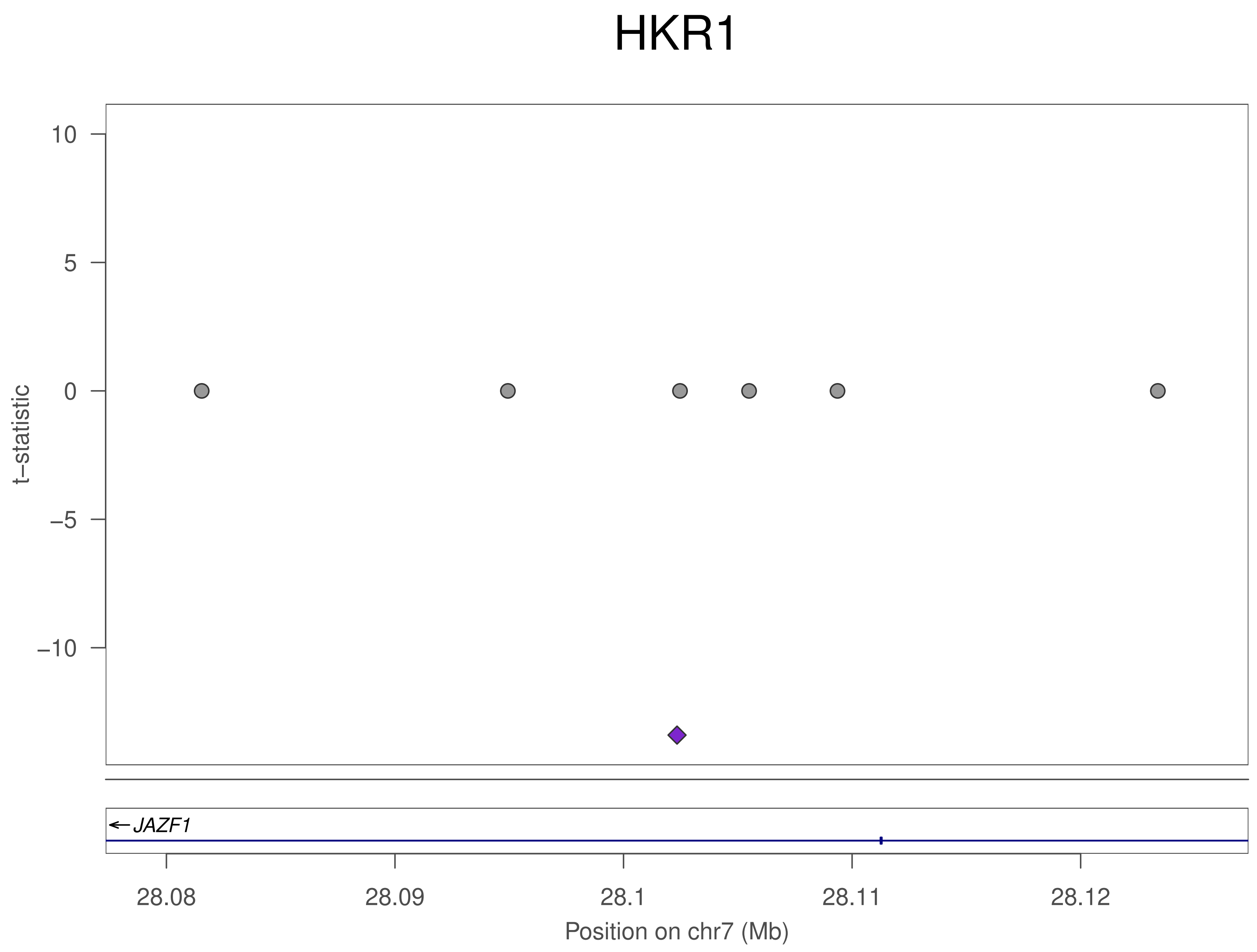

HKR1
t−statistic
10
5
0
−5
−10
JAZF1
28.08
28.09
28.1
28.11
28.12
Position on chr7 (Mb)

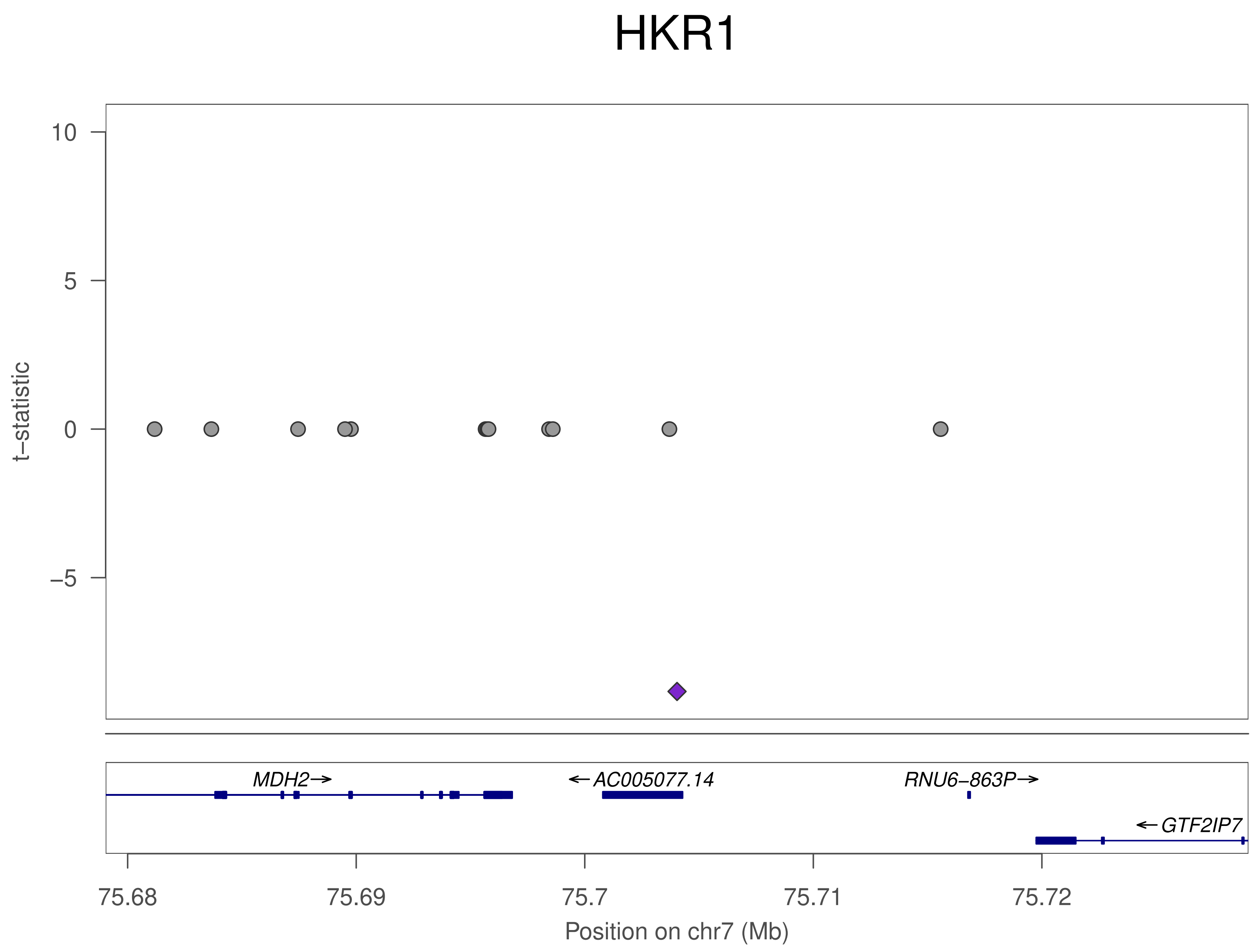
HKR1
t−statistic
10
5
0
−5
MDH2→
←AC005077.14
RNU6−863P→
←GTF2IP7
75.68
75.69
75.7
75.71
75.72
Position on chr7 (Mb)

### HKR1

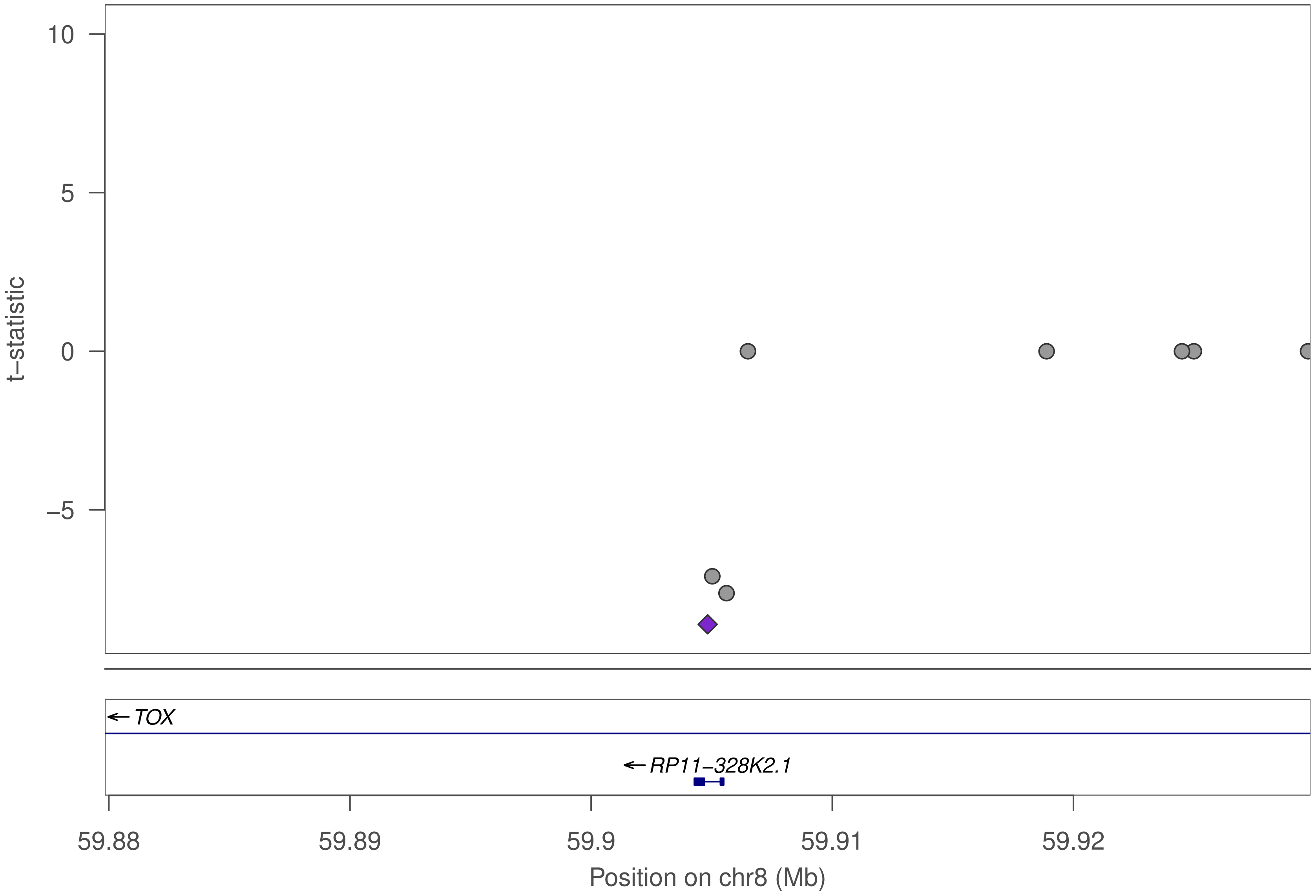

10
5
0
−5
t−statistic
TOX
RP11−328K2.1
59.88
59.89
59.9
59.91
59.92
Position on chr8 (Mb)

### HKR1

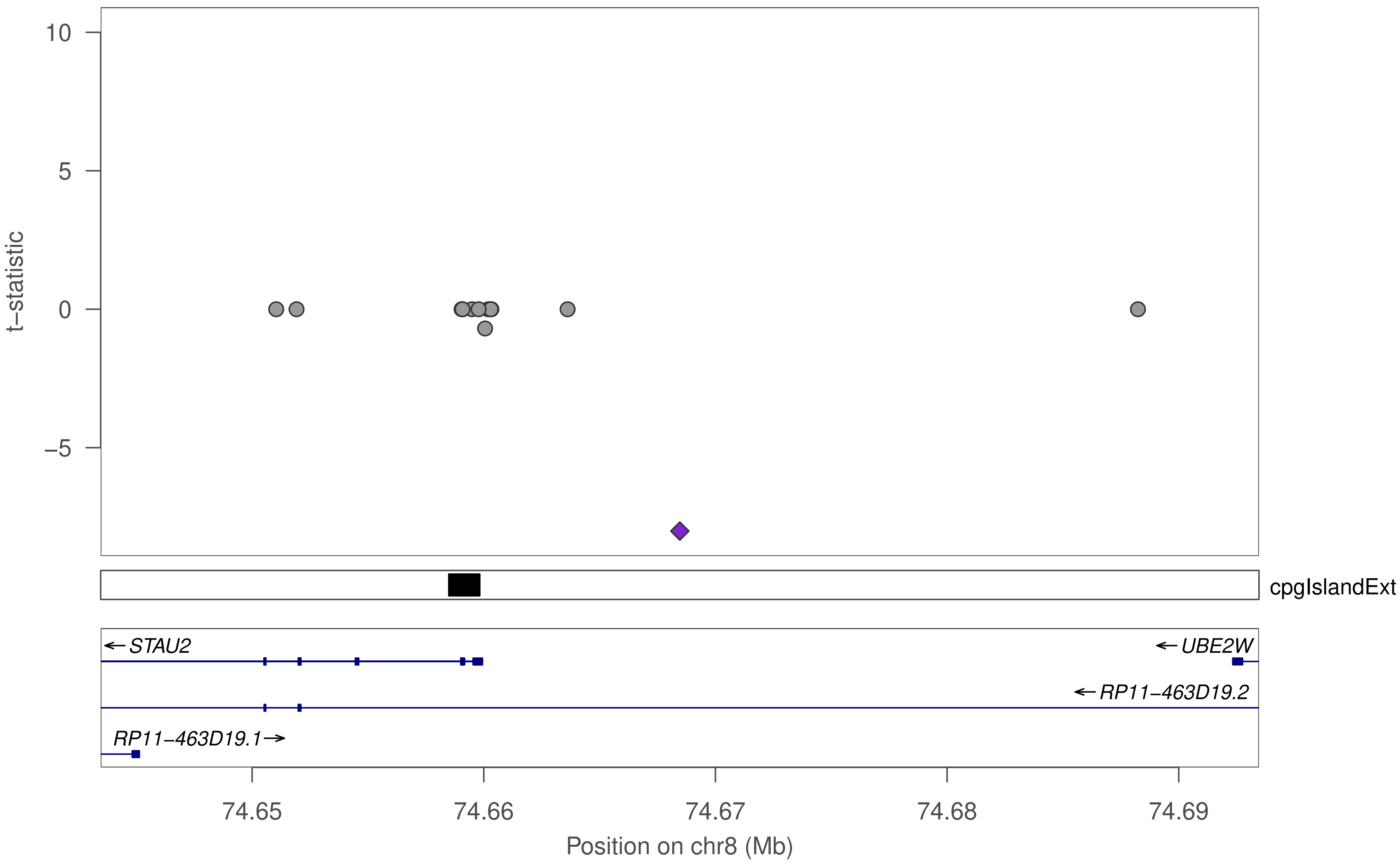

10
5
0
−5
t−statistic
cpgIslandExt
STAU2
UBE2W
RP11−463D19.2
RP11−463D19.1
74.65
74.66
74.67
74.68
74.69
Position on chr8 (Mb)

### HKR1

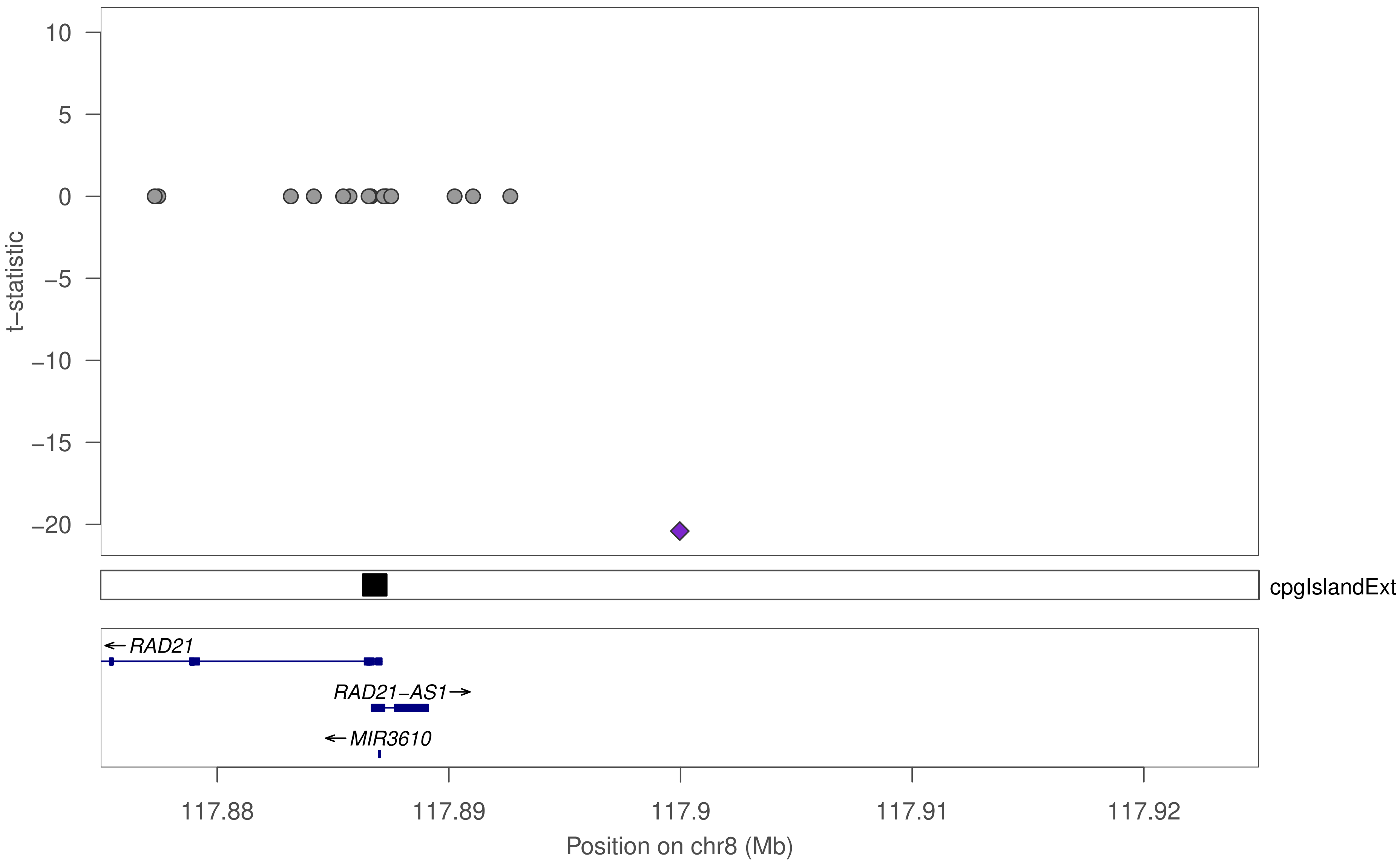

10
5
0
−5
−10
−15
−20
t−statistic
cpgIslandExt
RAD21
RAD21−AS1
MIR3610
117.88
117.89
117.9
117.91
117.92
Position on chr8 (Mb)

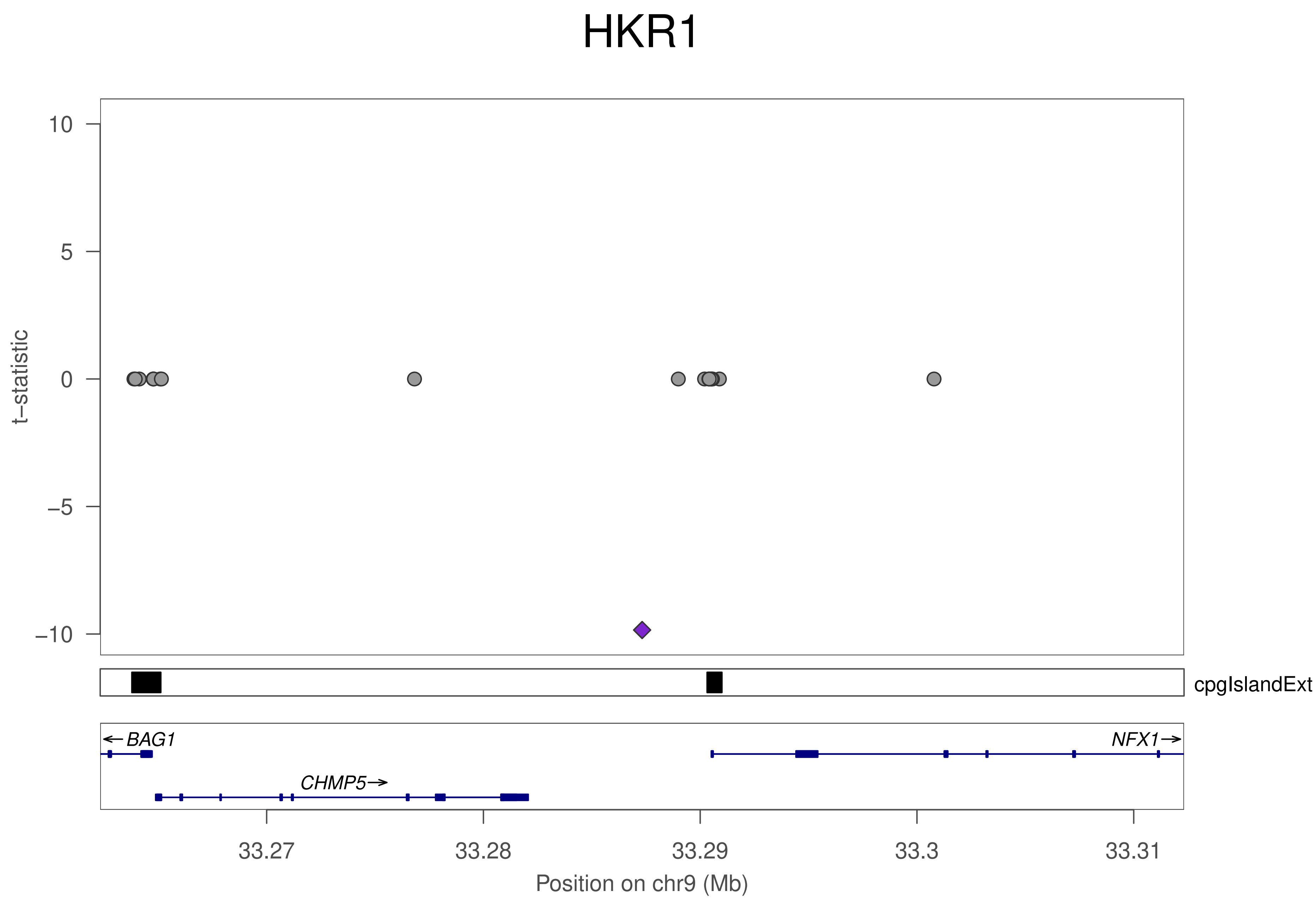
HKR1
t−statistic
10
5
0
−5
−10
cpgIslandExt
BAG1
NFX1
CHMP5
33.27
33.28
33.29
33.3
33.31
Position on chr9 (Mb)

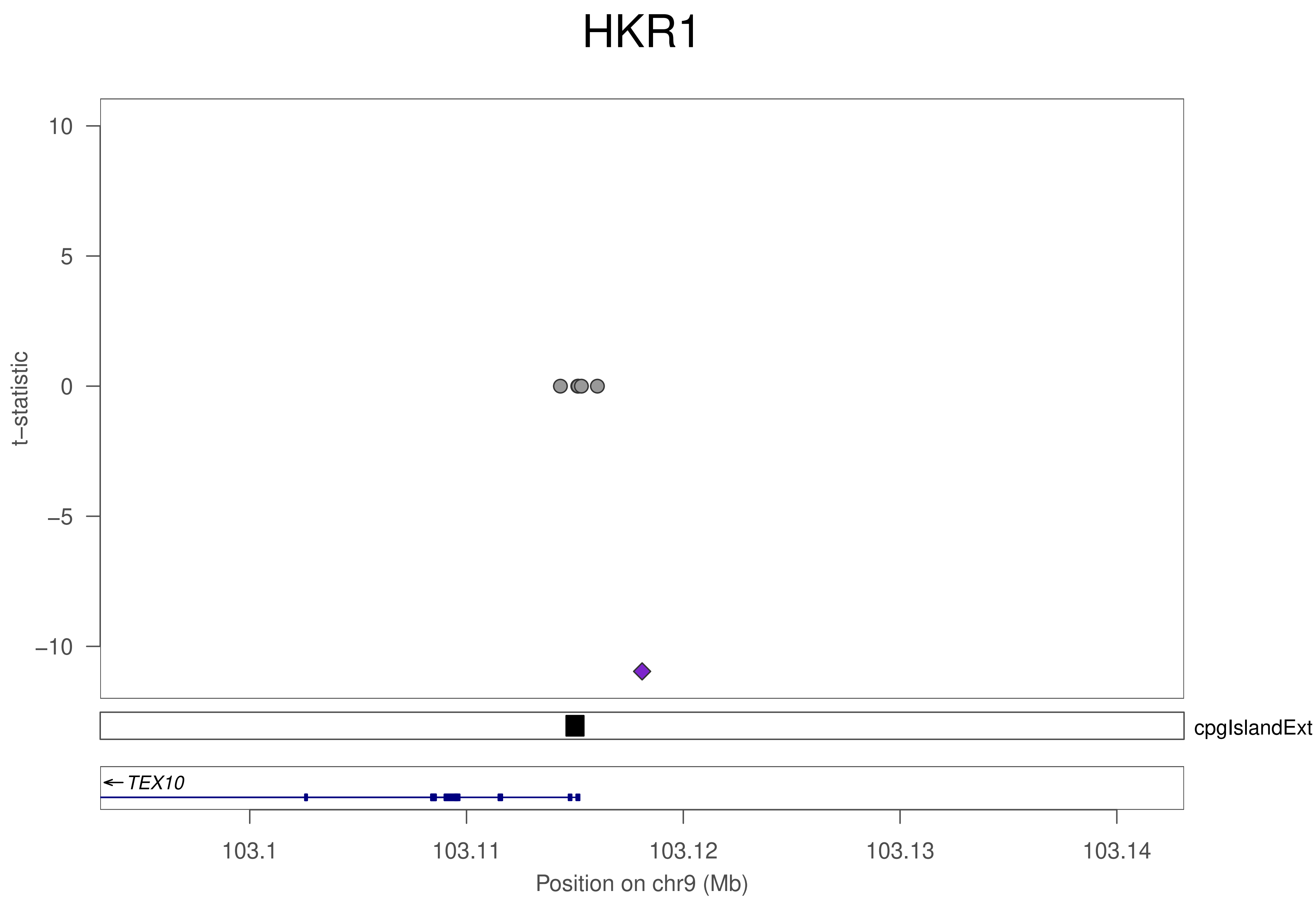
HKR1
t−statistic
10
5
0
−5
−10
cpgIslandExt
TEX10
103.1
103.11
103.12
103.13
103.14
Position on chr9 (Mb)

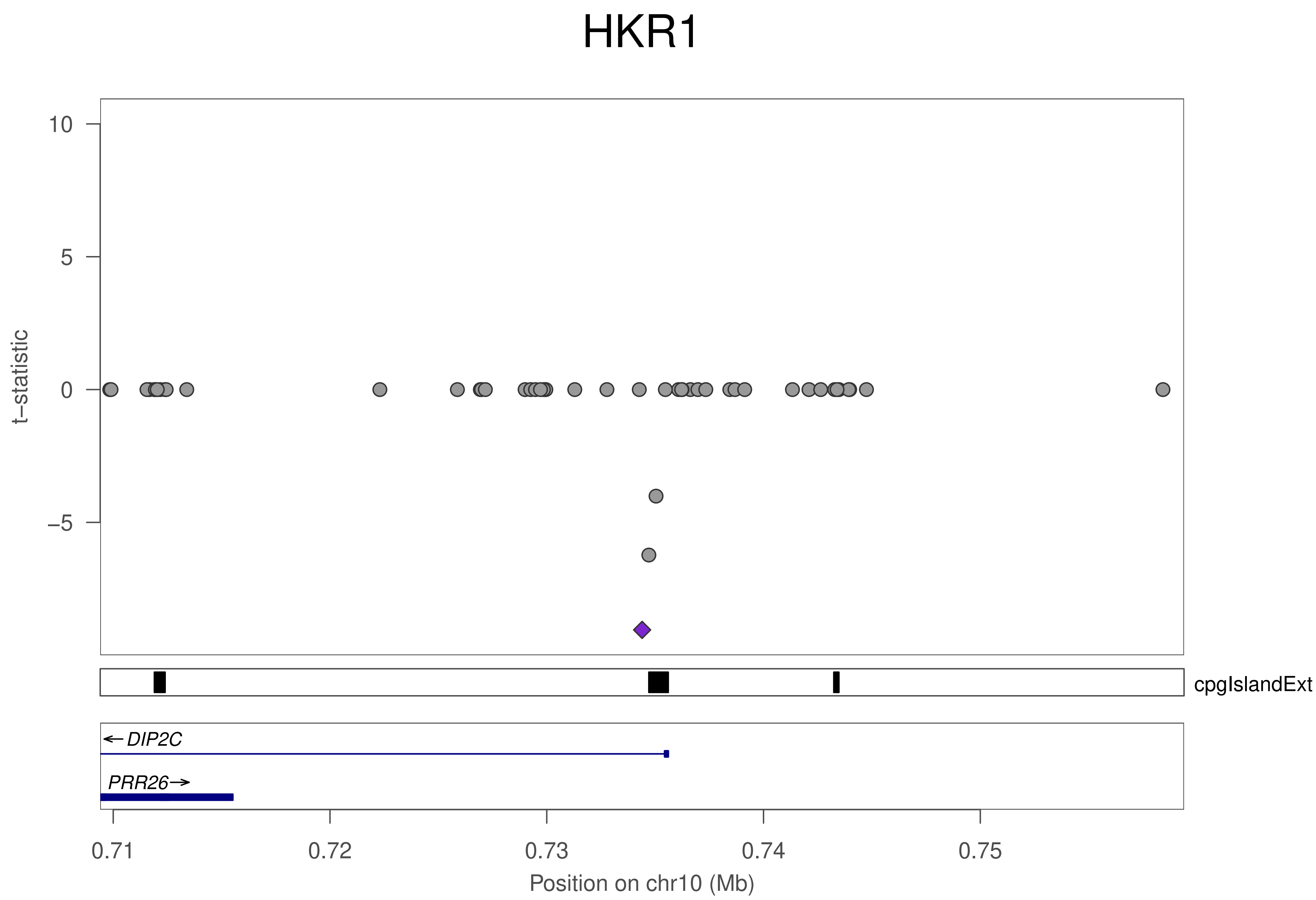
HKR1
10
5
0
−5
t−statistic
cpgIslandExt
DIP2C
PRR26
0.71
0.72
0.73
0.74
0.75
Position on chr10 (Mb)

### HKR1

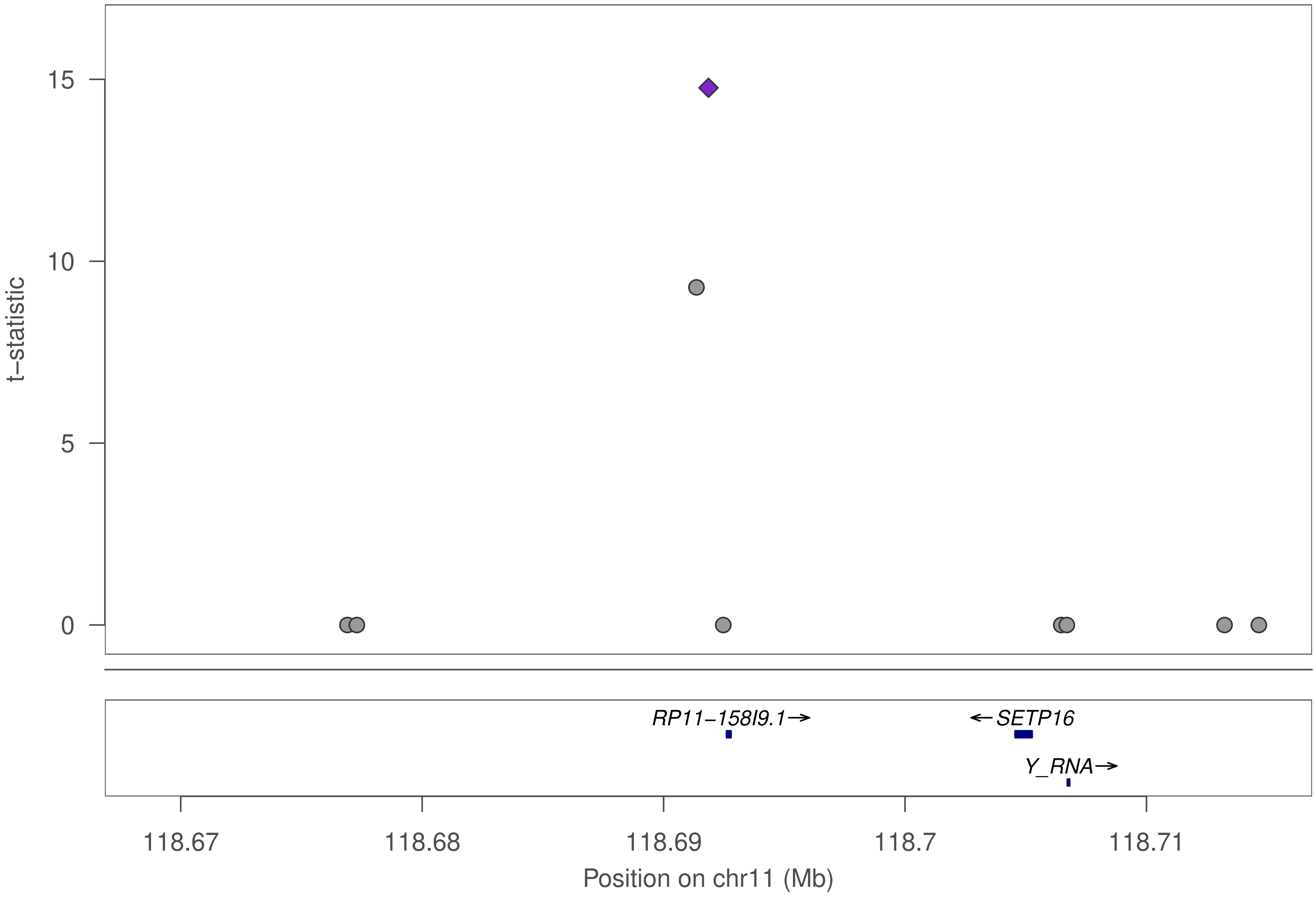

t−statistic
15
10
5
0
RP11−158I9.1→
←SETP16
Y_RNA→
118.67
118.68
118.69
118.7
118.71
Position on chr11 (Mb)

### HKR1

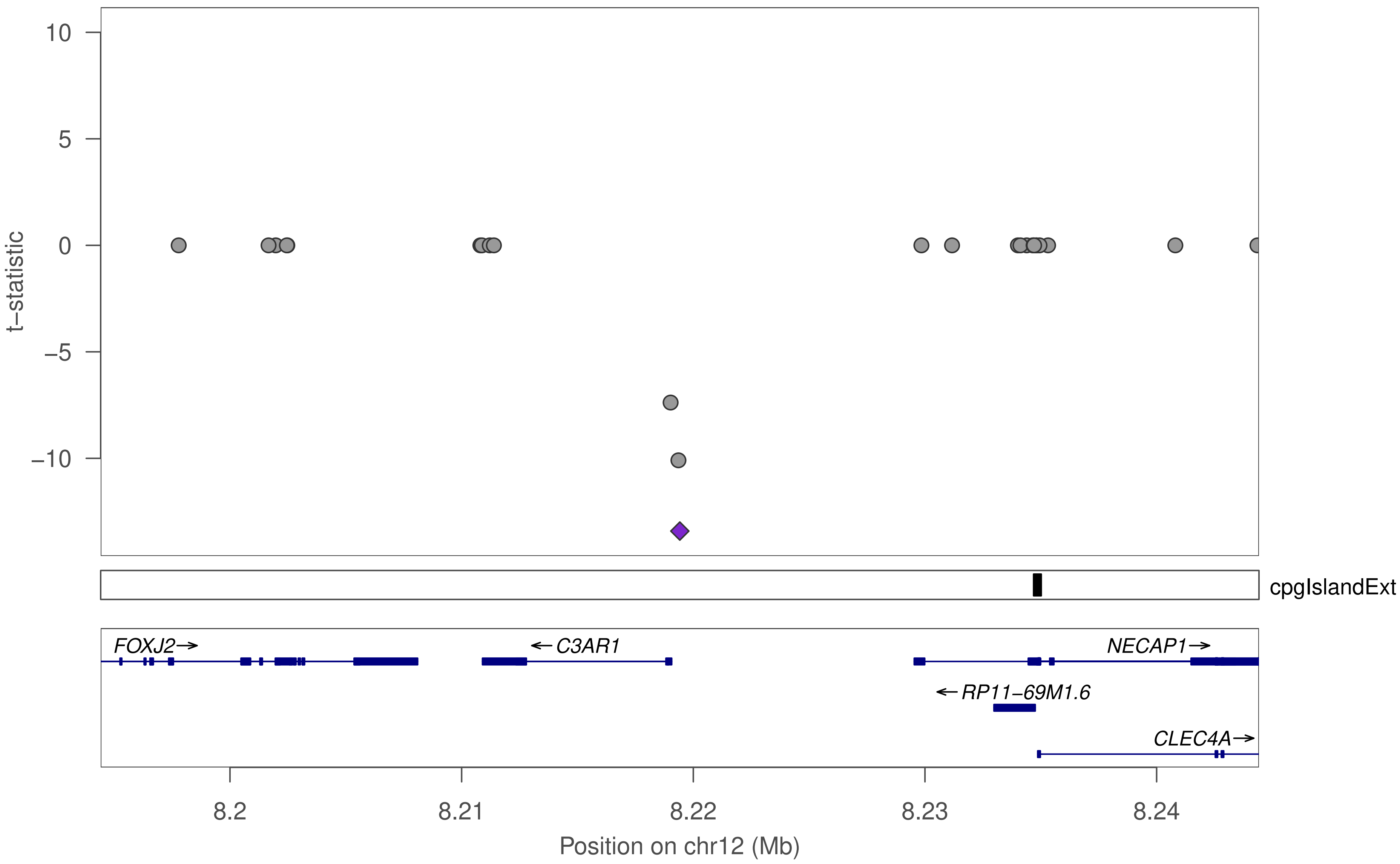

10
5
0
−5
−10
t−statistic
cpgIslandExt
FOXJ2→
←C3AR1
NECAP1→
←RP11−69M1.6
CLEC4A→
8.2
8.21
8.22
8.23
8.24
Position on chr12 (Mb)

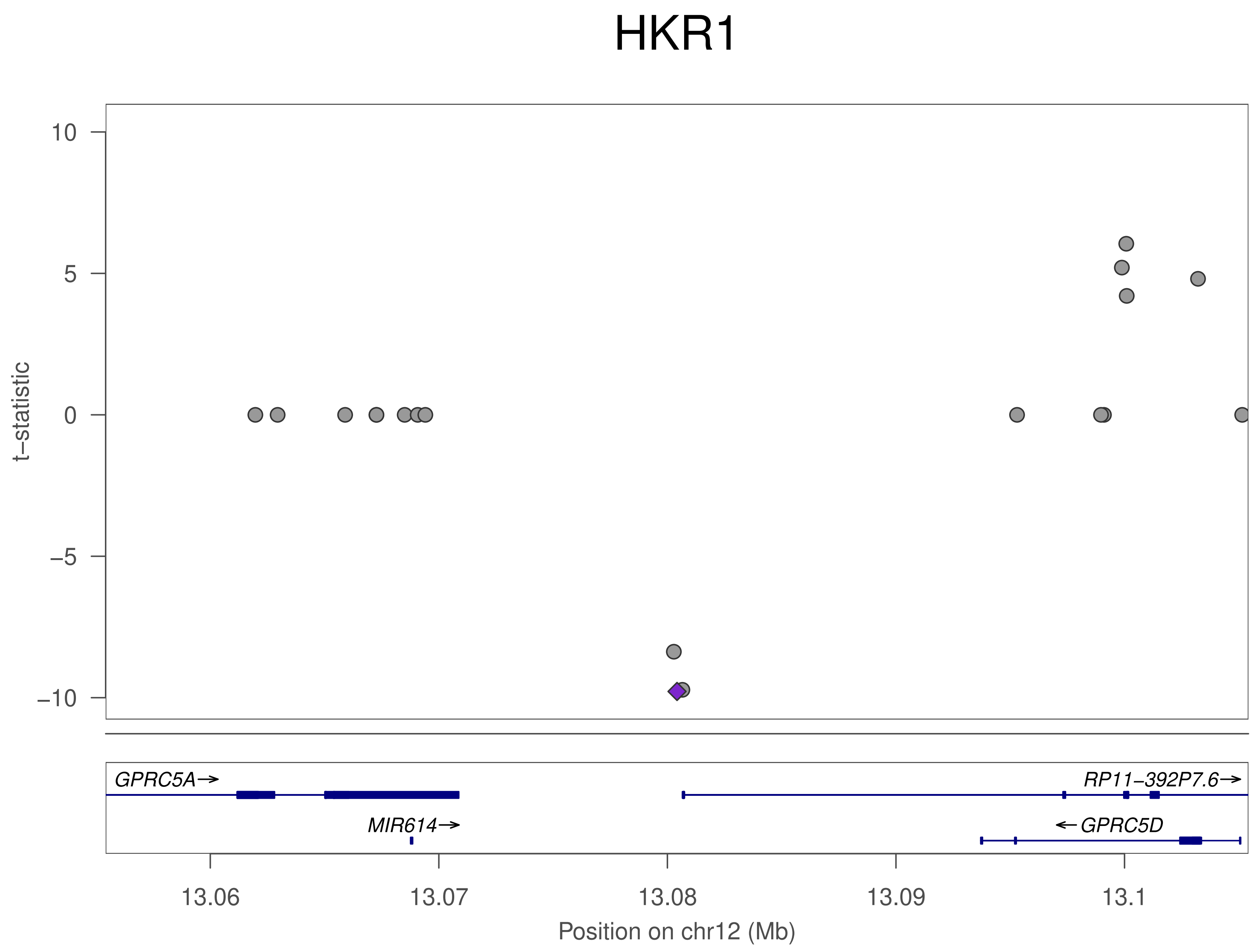
HKR1
t−statistic
10
5
0
−5
−10
GPRC5A→
MIR614→
RP11−392P7.6→
←GPRC5D
13.06
13.07
13.08
13.09
13.1
Position on chr12 (Mb)

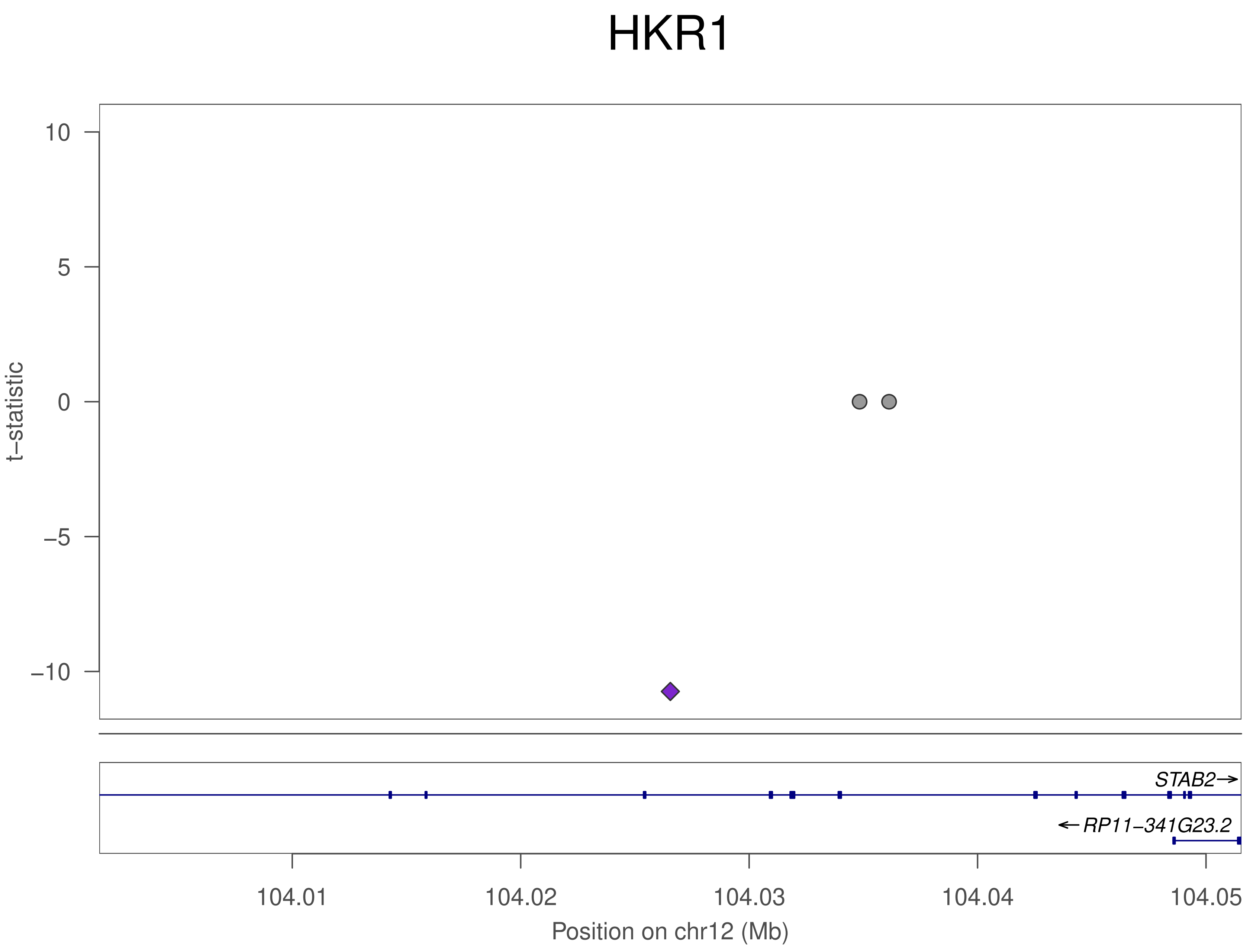
HKR1
t−statistic
10
5
0
−5
−10
STAB2→
←RP11−341G23.2
104.01
104.02
104.03
104.04
104.05
Position on chr12 (Mb)

### HKR1

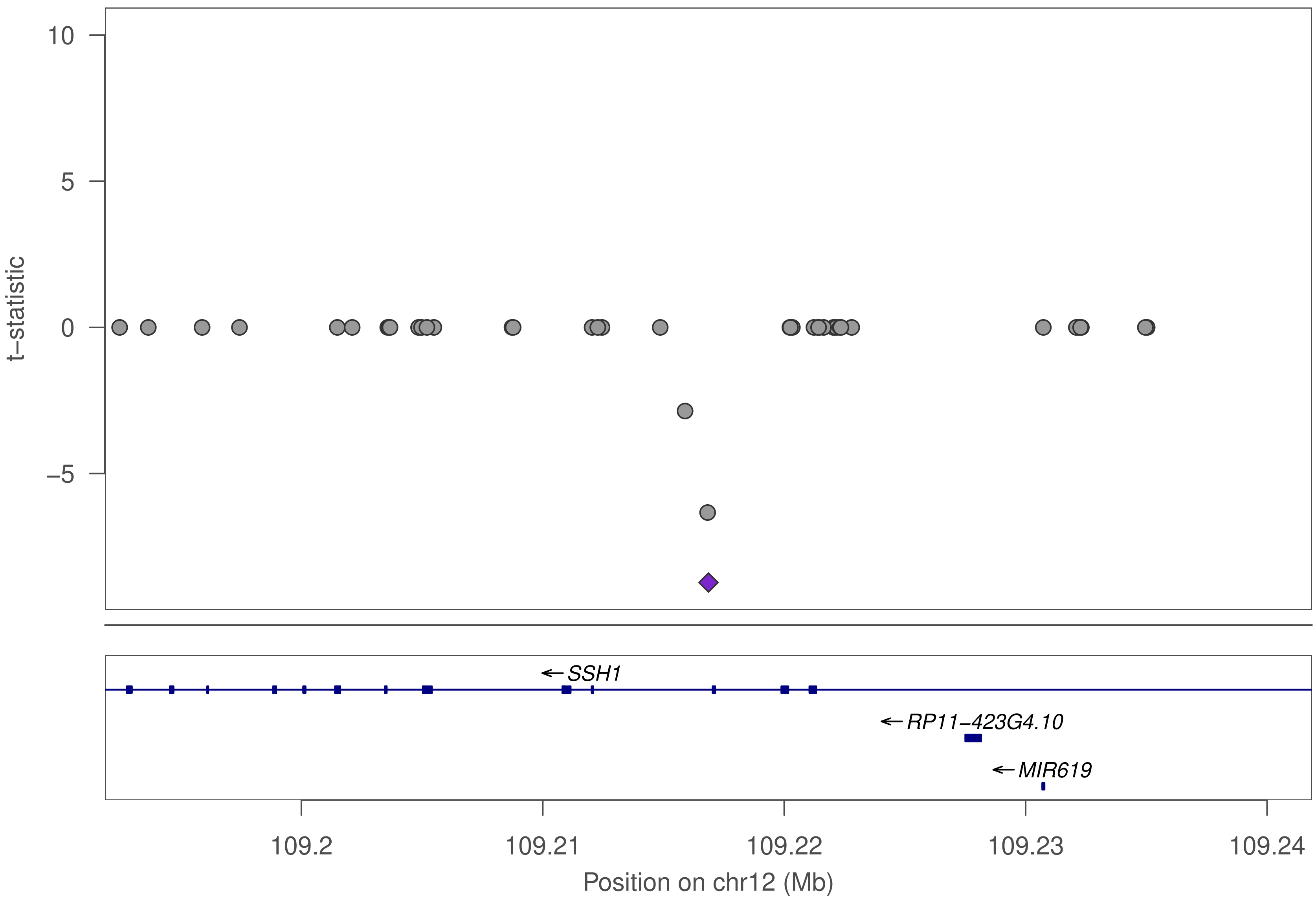
10
5
0
−5
t−statistic
SSH1
RP11−423G4.10
MIR619
109.2
109.21
109.22
109.23
109.24
Position on chr12 (Mb)

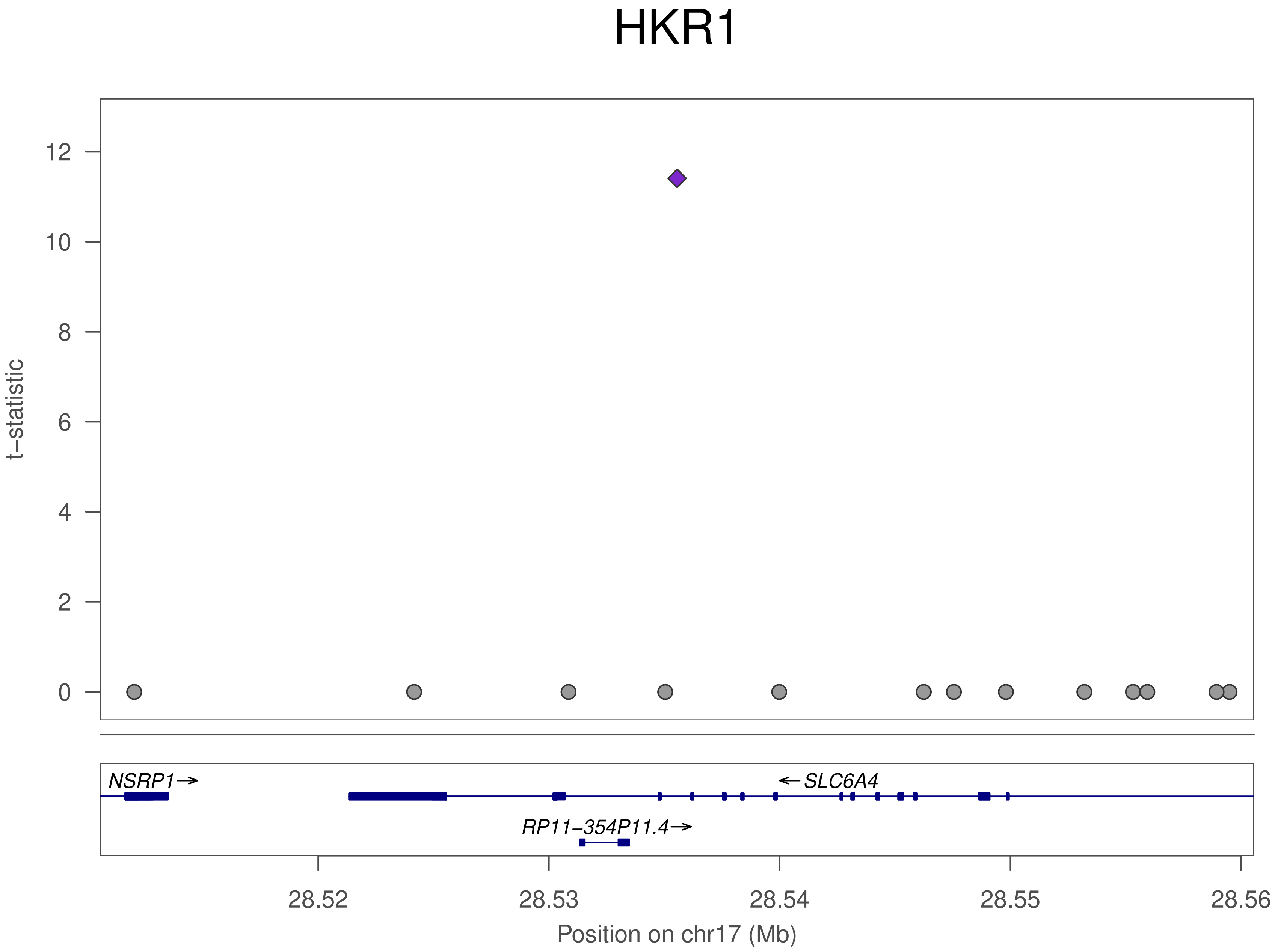

HKR1
t−statistic
12
10
8
6
4
2
0
NSRP1→
←SLC6A4
RP11−354P11.4→
28.52
28.53
28.54
28.55
28.56
Position on chr17 (Mb)

### HKR1

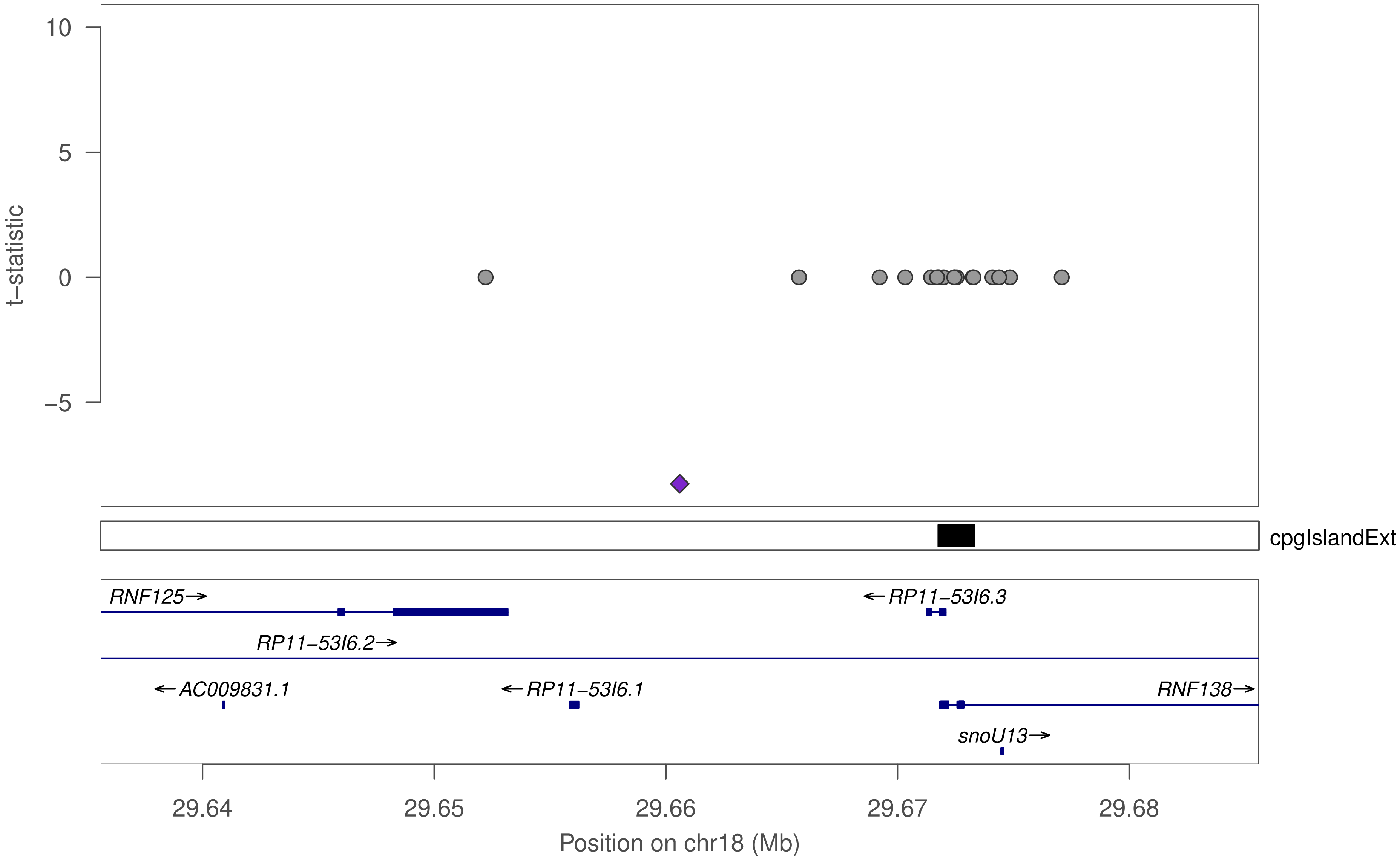

10
5
0
−5
t−statistic
cpgIslandExt
RNF125→
←RP11−53I6.3
RP11−53I6.2→
←AC009831.1
←RP11−53I6.1
RNF138→
snoU13→
29.64
29.65
29.66
29.67
29.68
Position on chr18 (Mb)

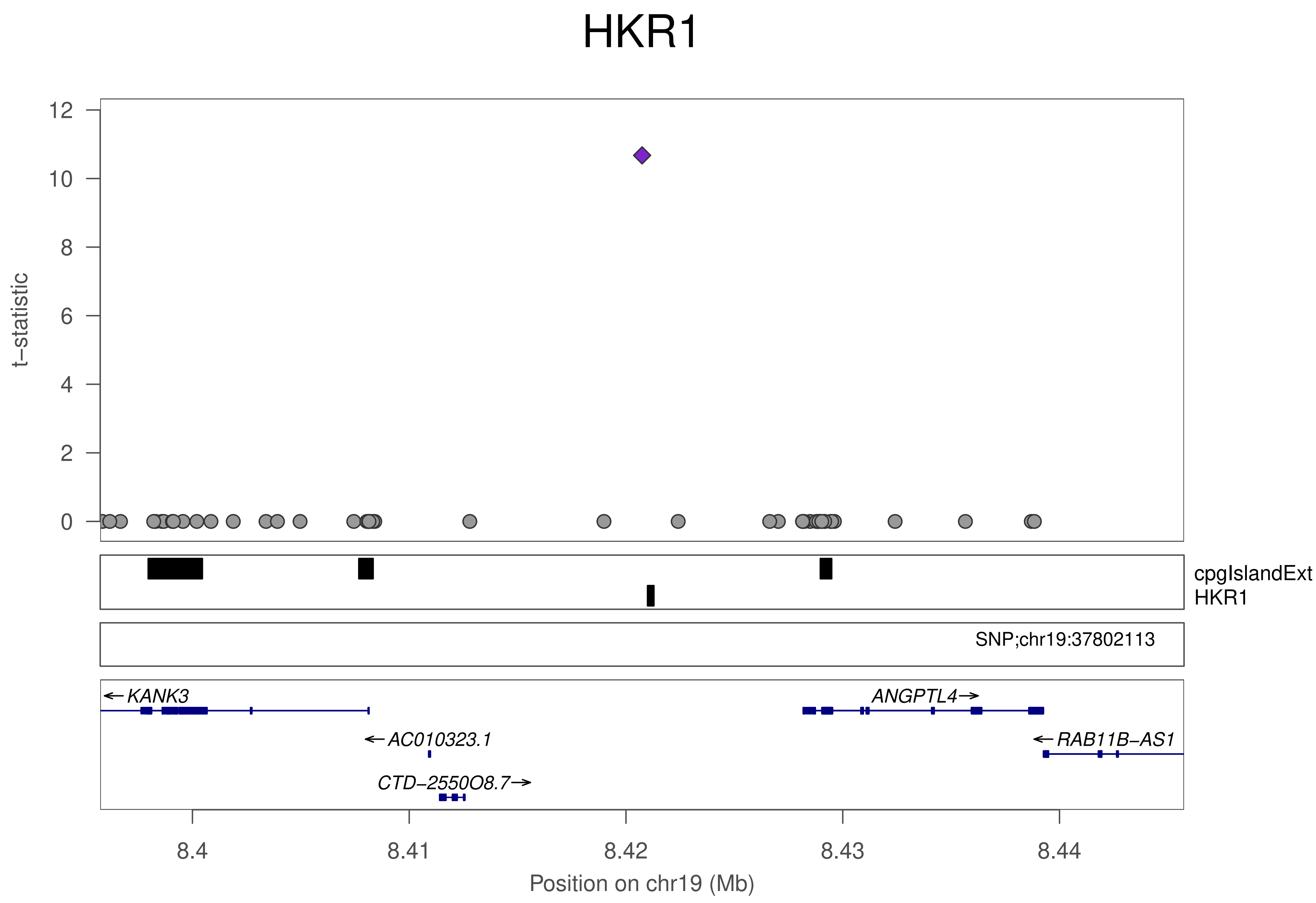

HKR1
t−statistic
12
10
8
6
4
2
0
cpgIslandExt
HKR1
SNP;chr19:37802113
KANK3
ANGPTL4
AC010323.1
RAB11B−AS1
CTD−2550O8.7
8.4
8.41
8.42
8.43
8.44
Position on chr19 (Mb)

### ZNF320

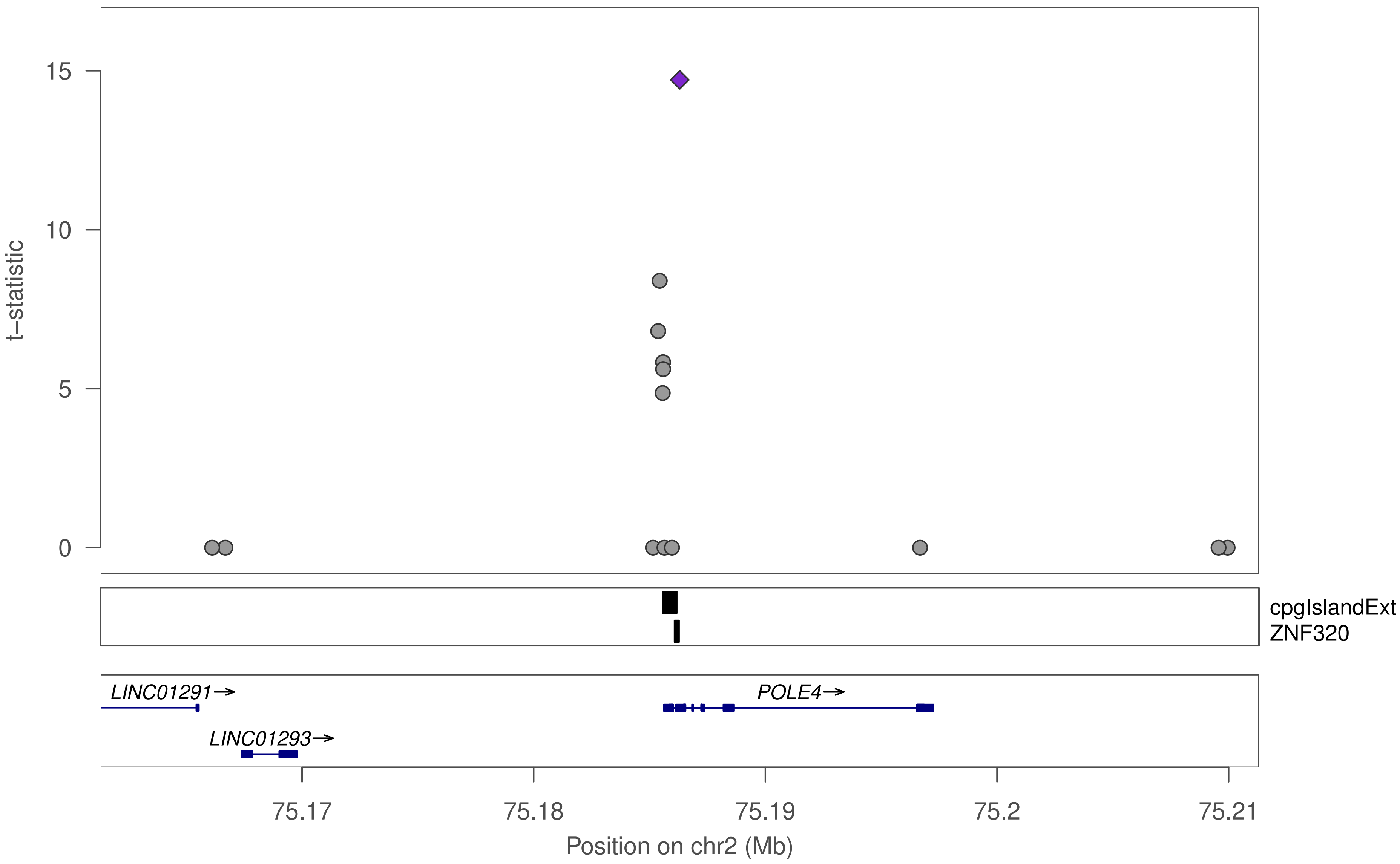
t−statistic
15
10
5
0
cpgIslandExt
ZNF320
LINC01291→
LINC01293→
POLE4→
75.17
75.18
75.19
75.2
75.21
Position on chr2 (Mb)

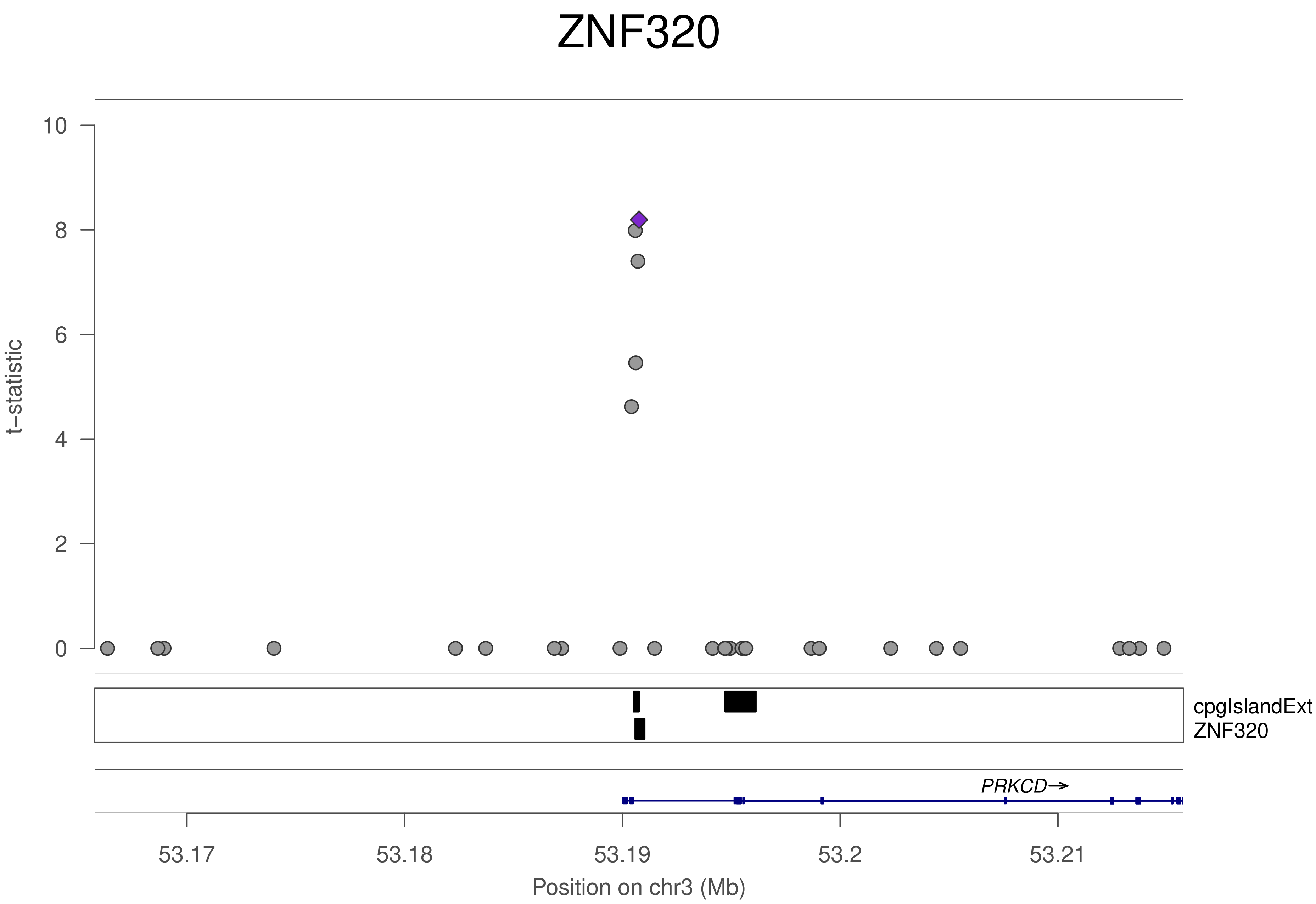

ZNF320
t−statistic
10
8
6
4
2
0
cpgIslandExt
ZNF320
PRKCD→
53.17
53.18
53.19
53.2
53.21
Position on chr3 (Mb)

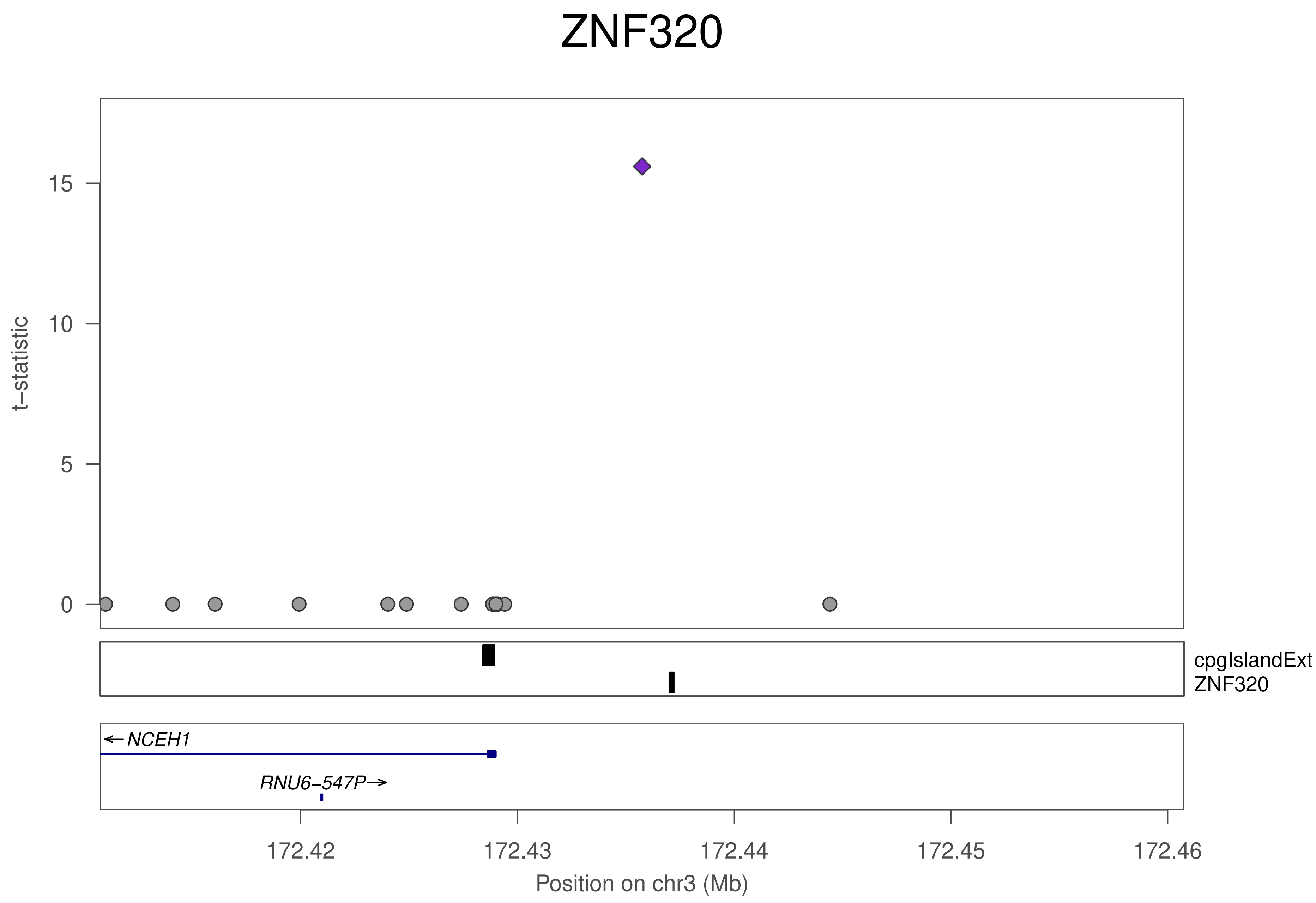

ZNF320
t−statistic
15
10
5
0
cpgIslandExt
ZNF320
NCEH1
RNU6−547P
172.42
172.43
172.44
172.45
172.46
Position on chr3 (Mb)

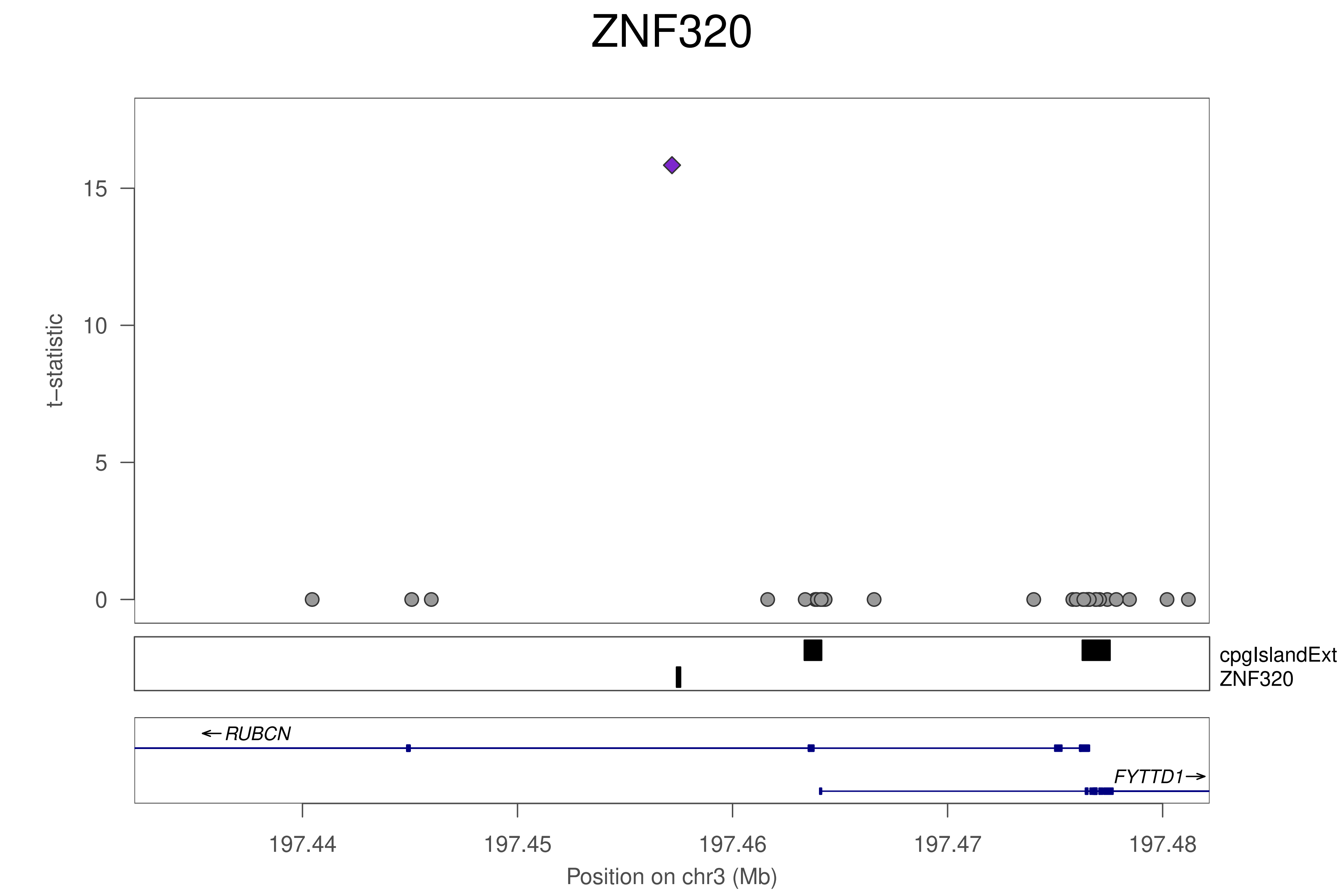
ZNF320
t–statistic
0
5
10
15
cpgIslandExt
ZNF320
RUBCN
FYTTD1
197.44
197.45
197.46
197.47
197.48
Position on chr3 (Mb)

### ZNF320

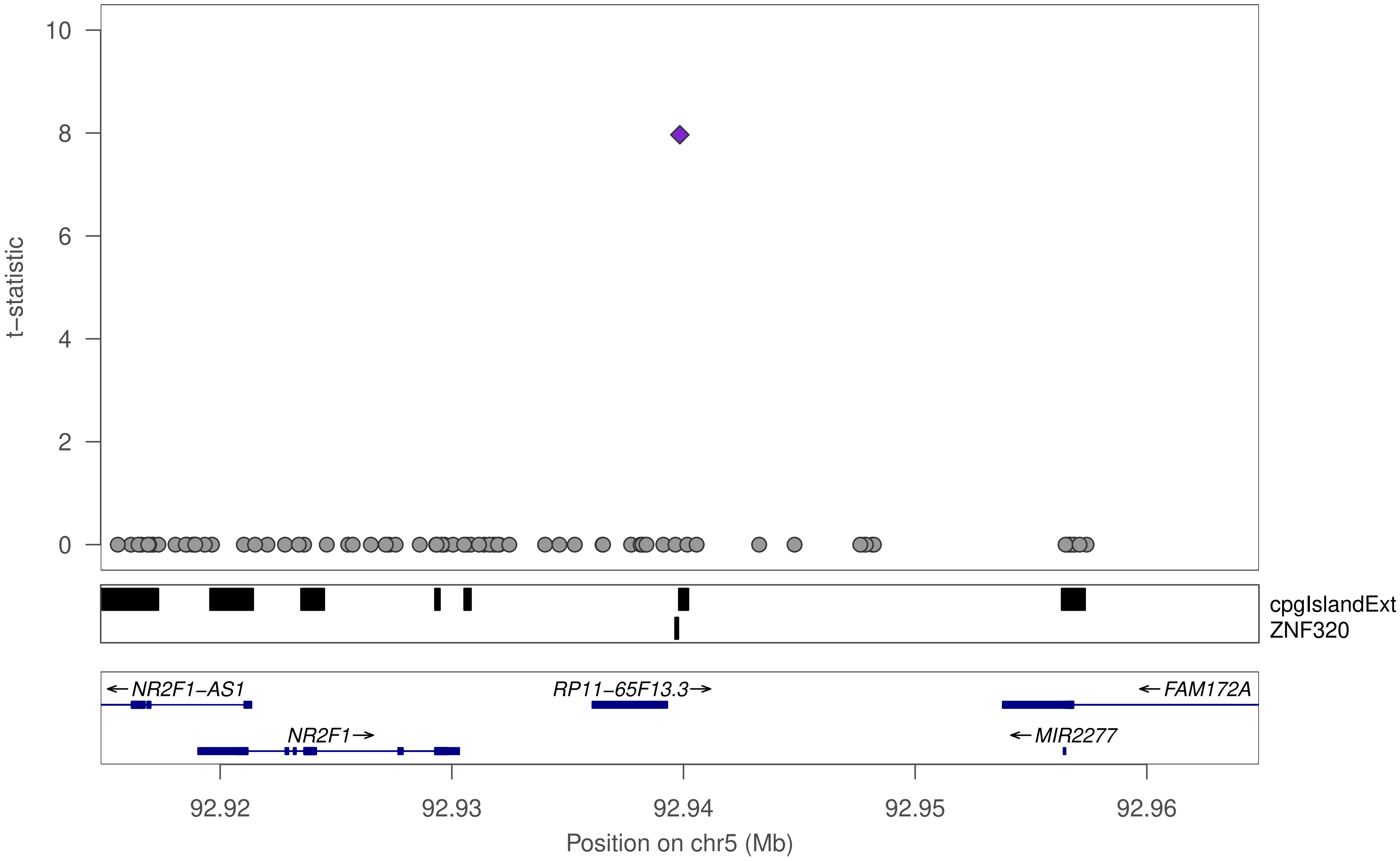

10
8
6
4
2
0
t−statistic
cpgIslandExt
ZNF320
←NR2F1−AS1
RP11−65F13.3→
←FAM172A
NR2F1→
←MIR2277
92.92
92.93
92.94
92.95
92.96
Position on chr5 (Mb)

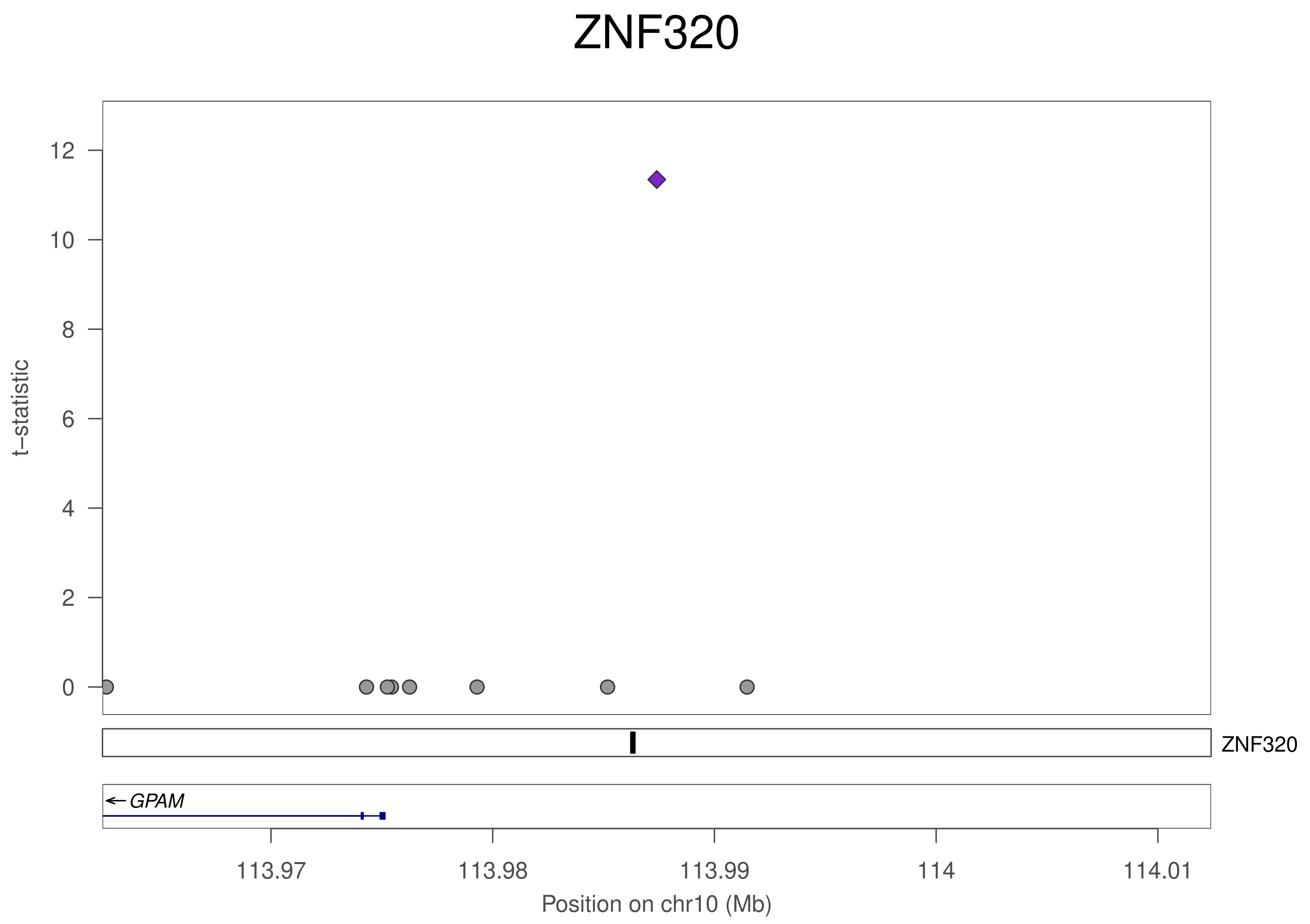

ZNF320
12
10
8
6
4
2
0
t–statistic
ZNF320
GPAM
113.97
113.98
113.99
114
114.01
Position on chr10 (Mb)

### ZNF320

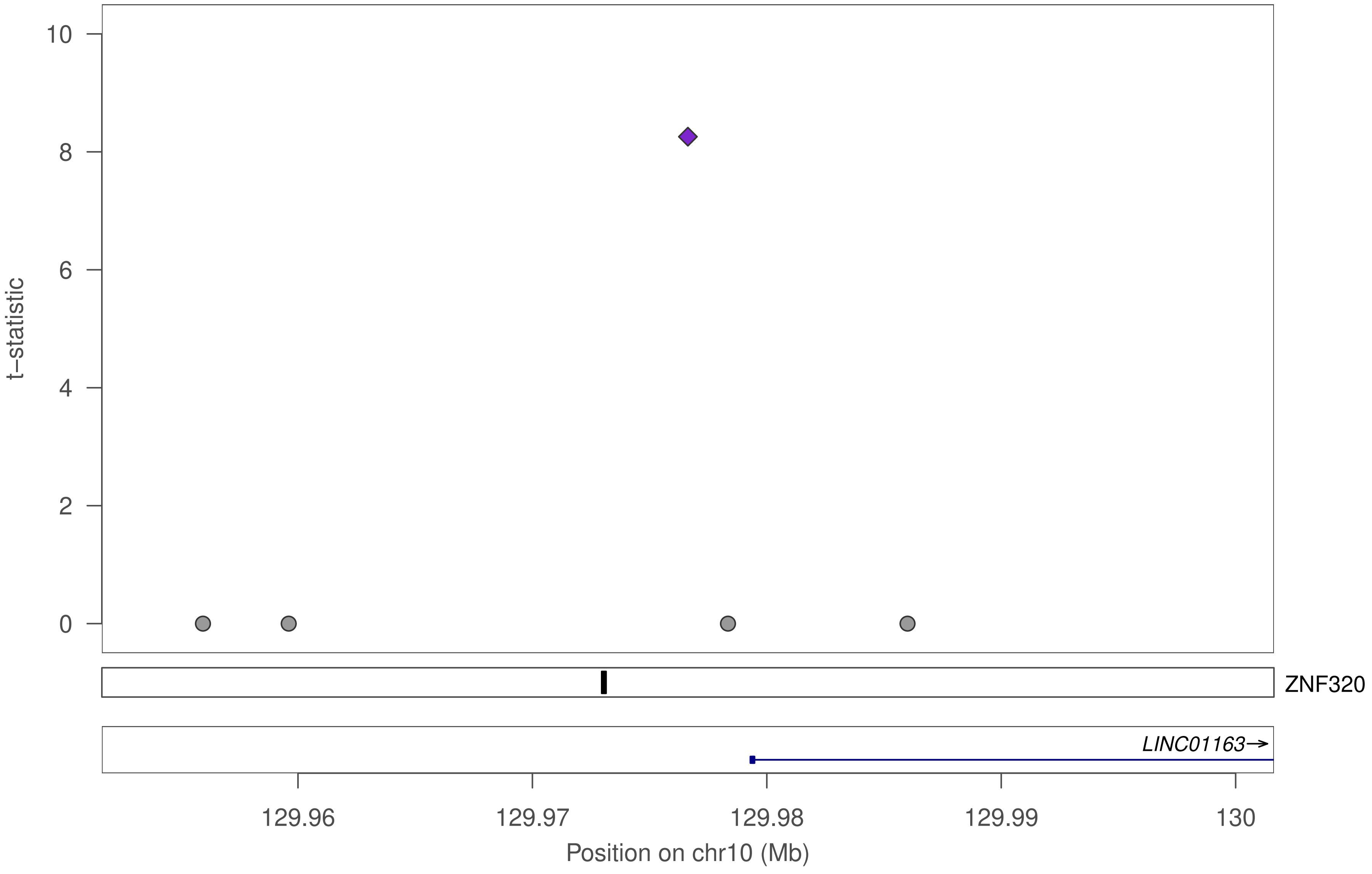

t−statistic
10
8
6
4
2
0
ZNF320
LINC01163→
129.96
129.97
129.98
129.99
130
Position on chr10 (Mb)

### ZNF320

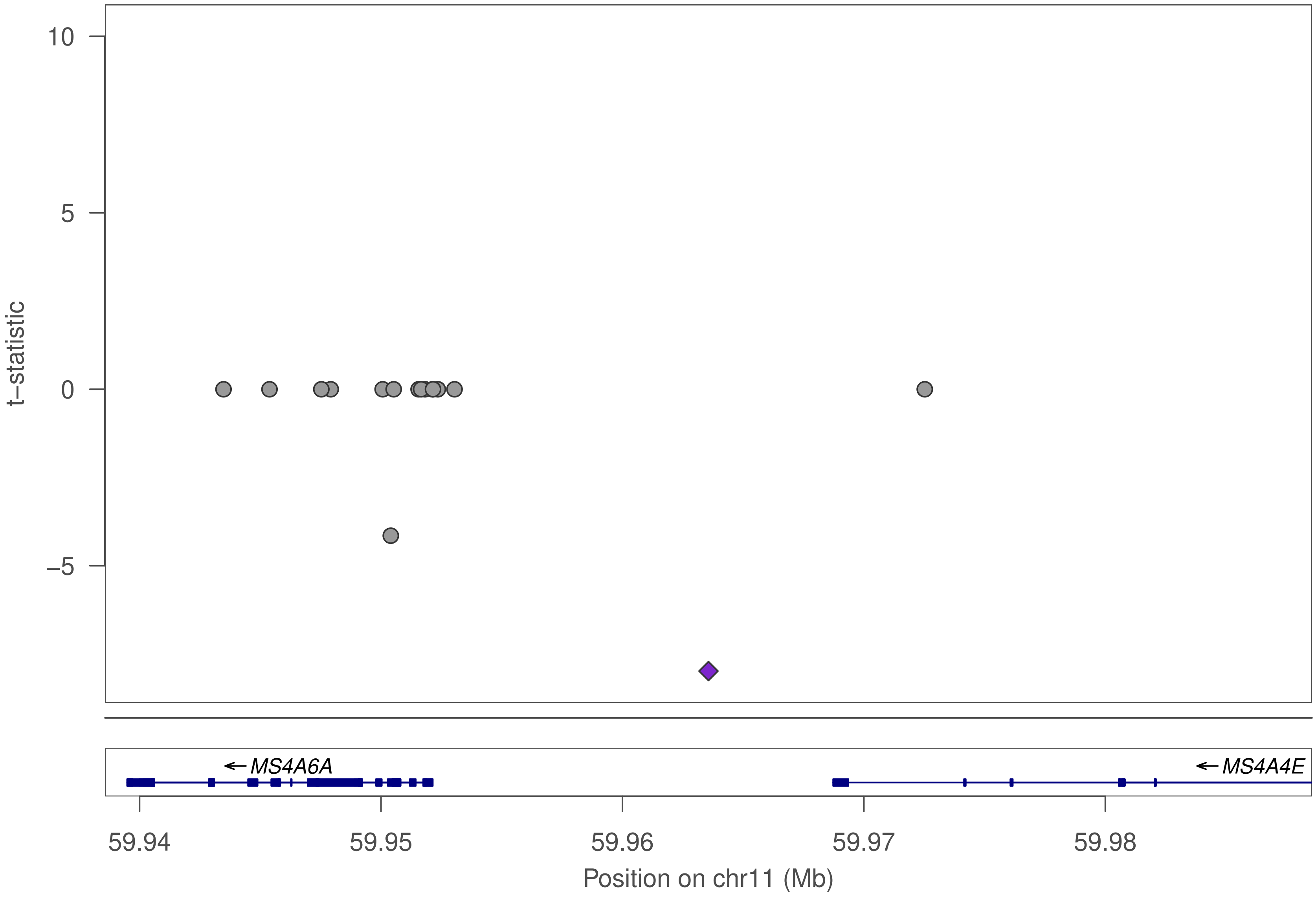

10
5
0
−5
t−statistic
MS4A6A
MS4A4E
59.94
59.95
59.96
59.97
59.98
Position on chr11 (Mb)

### ZNF320

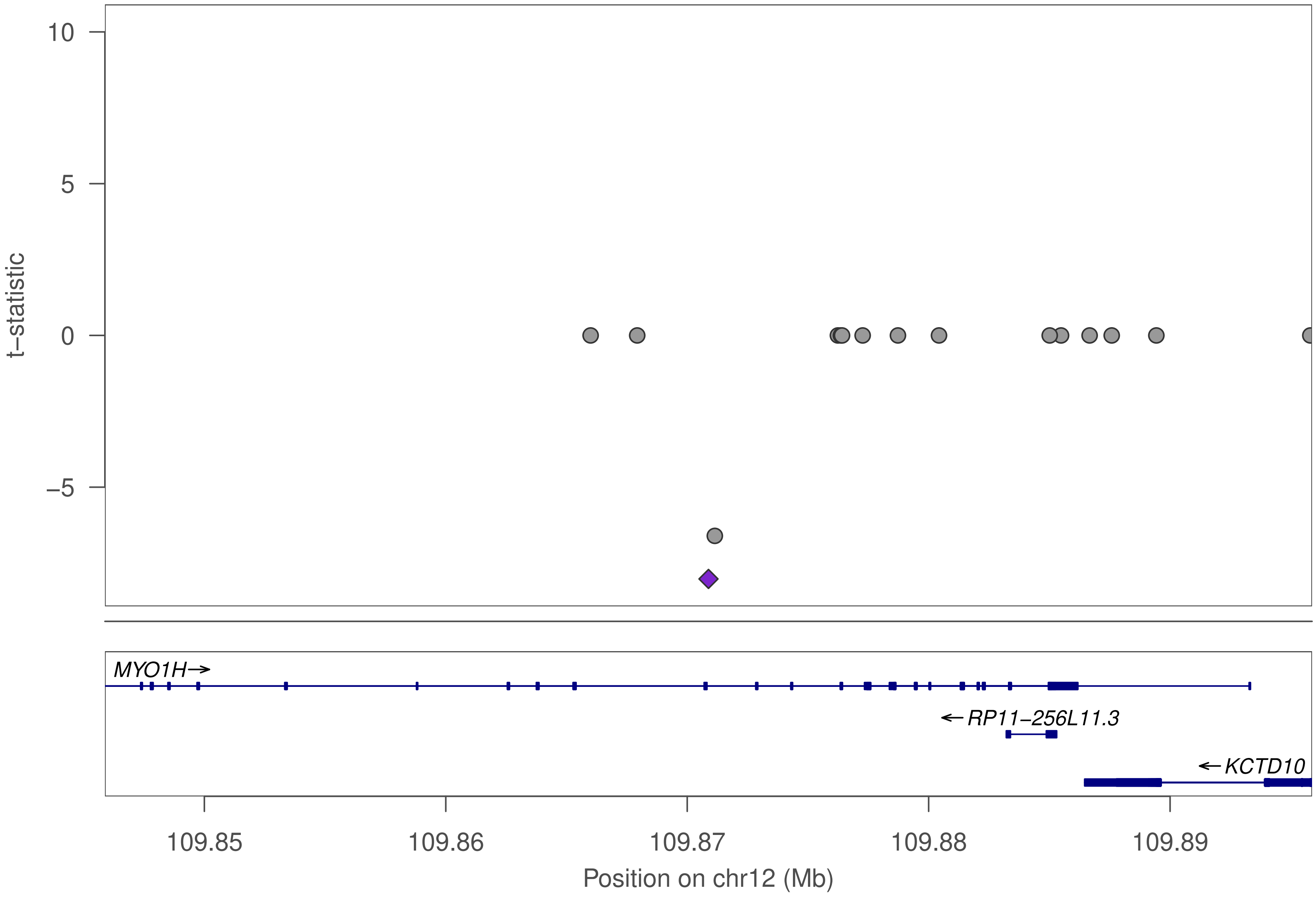

10
5
0
−5
t−statistic
MYO1H→
←RP11−256L11.3
←KCTD10
109.85
109.86
109.87
109.88
109.89
Position on chr12 (Mb)

### ZNF320

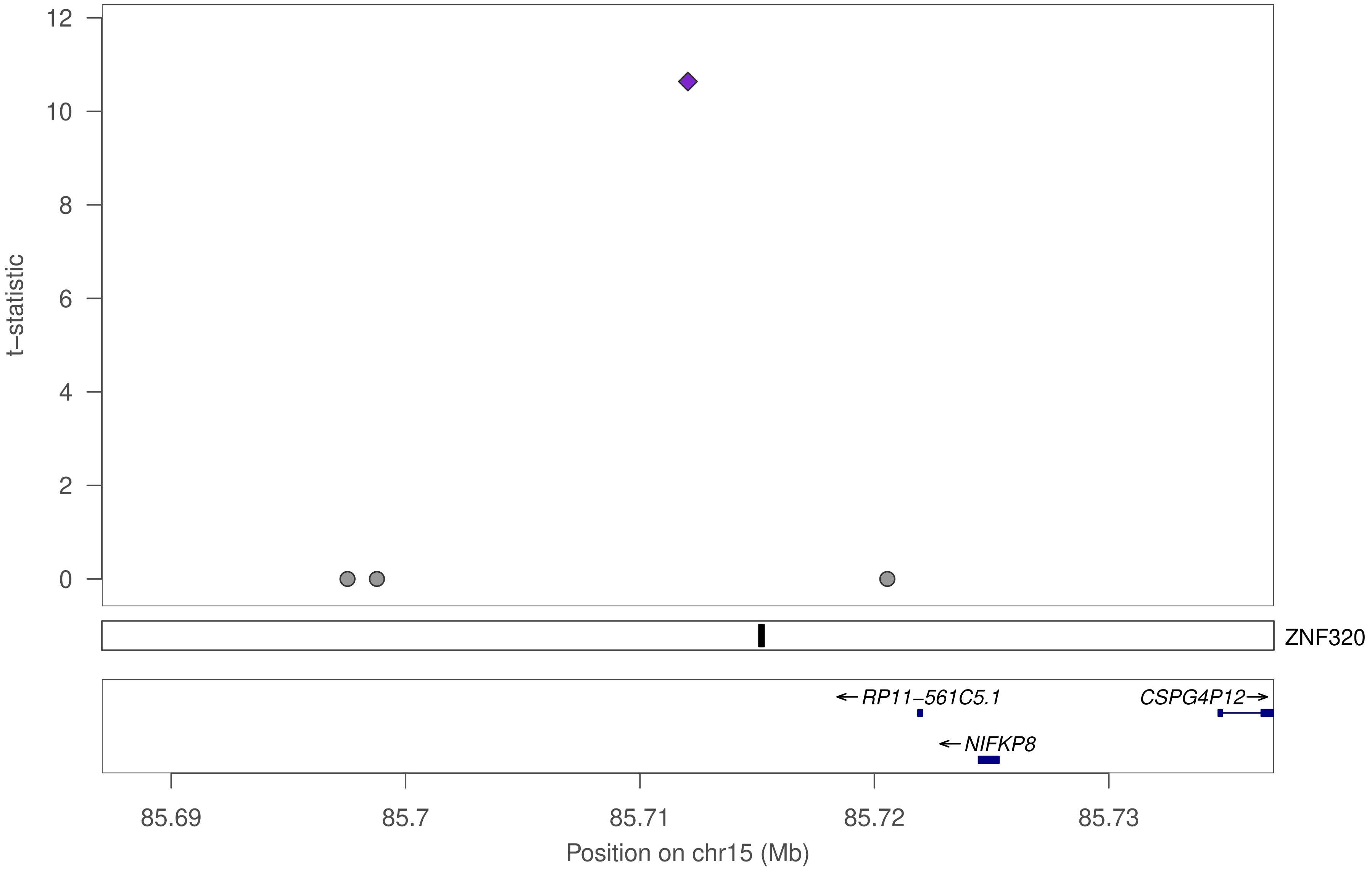

12
10
8
6
4
2
0
t−statistic
ZNF320
←RP11−561C5.1
CSPG4P12→
←NIFKP8
85.69
85.7
85.71
85.72
85.73
Position on chr15 (Mb)

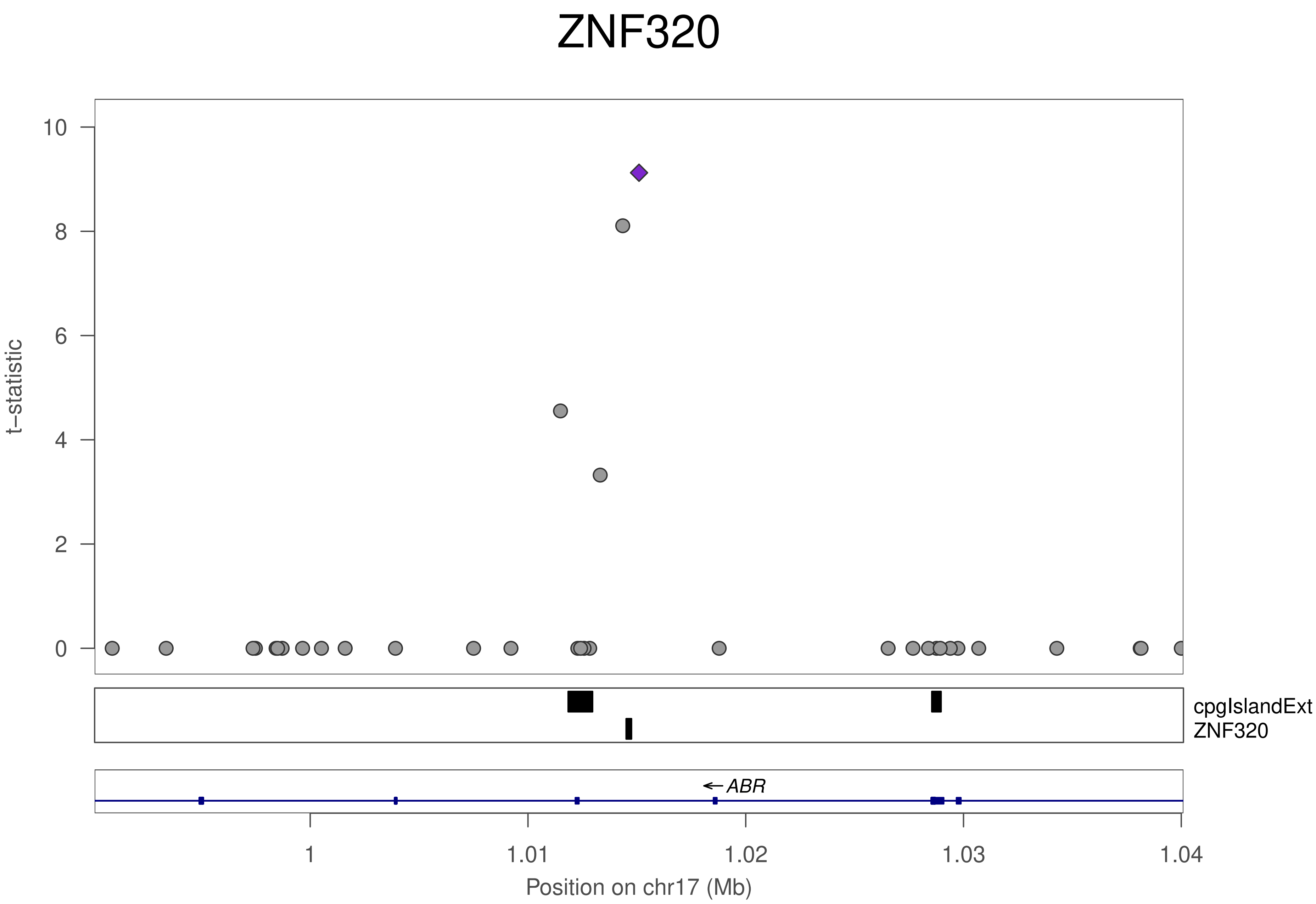
ZNF320
t−statistic
10
8
6
4
2
0
cpgIslandExt
ZNF320
← ABR
1
1.01
1.02
1.03
1.04
Position on chr17 (Mb)

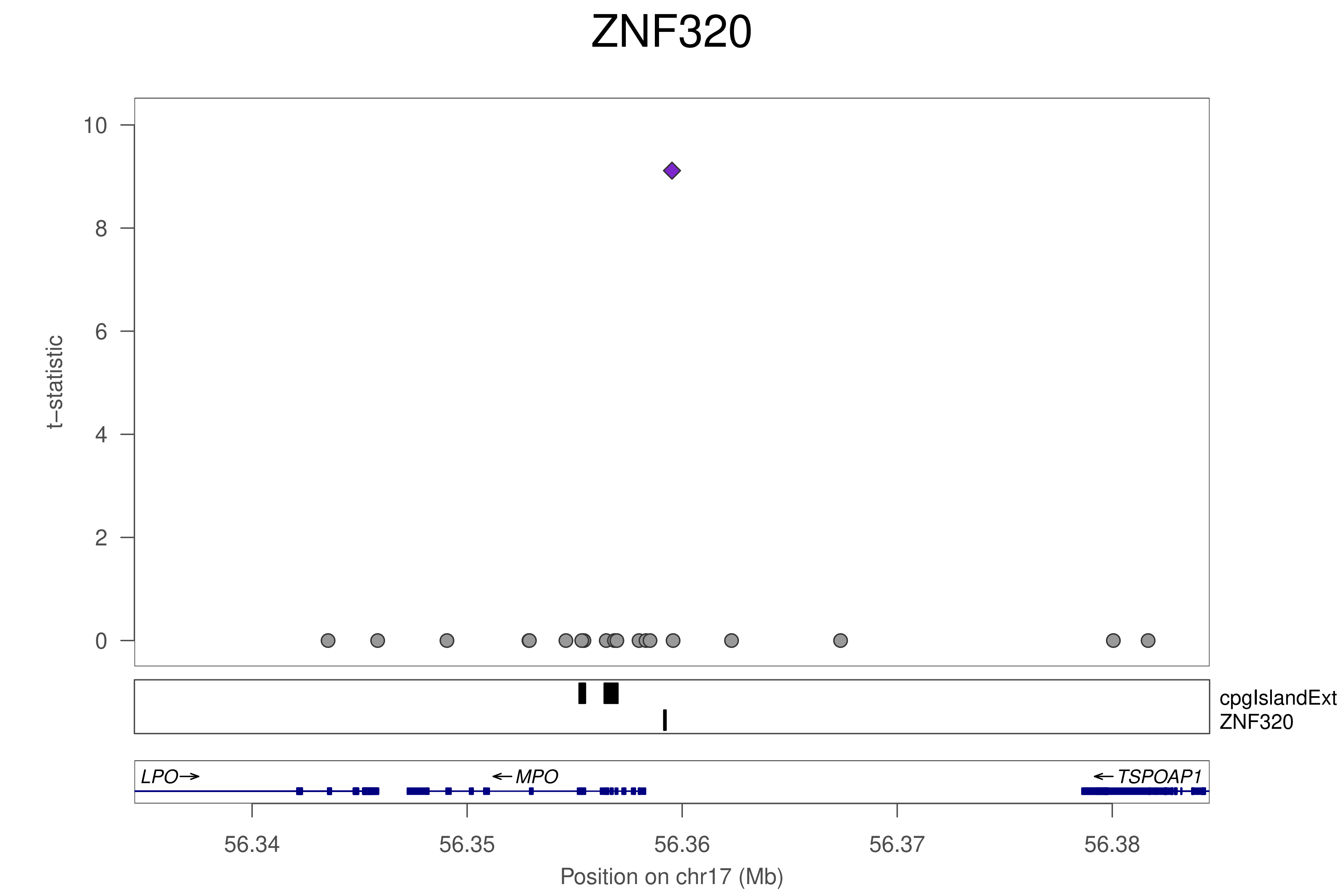

ZNF320
t−statistic
10
8
6
4
2
0
cpgIslandExt
ZNF320
LPO→
←MPO
←TSPOAP1
56.34
56.35
56.36
56.37
56.38
Position on chr17 (Mb)

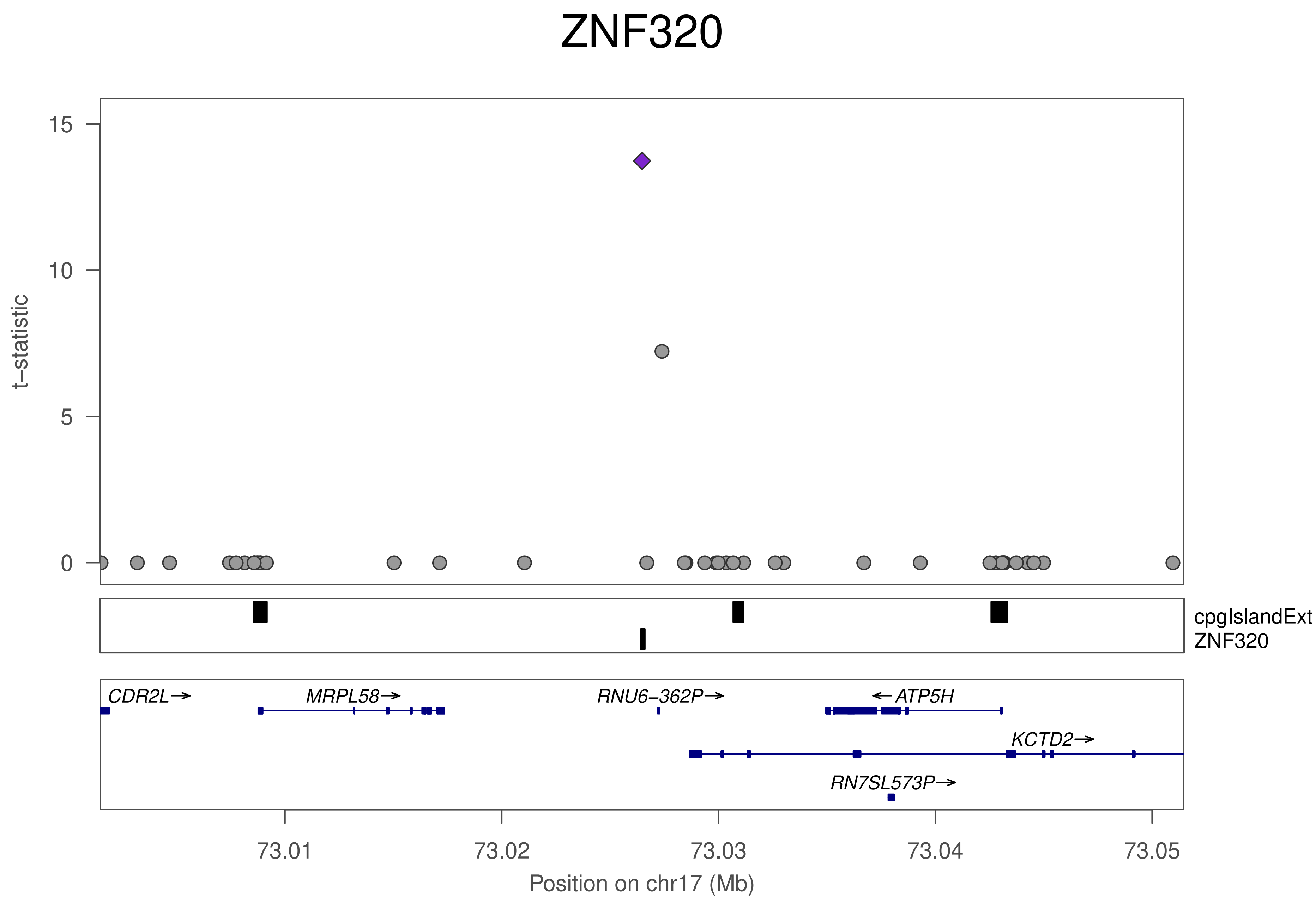

ZNF320
t−statistic
15
10
5
0
cpgIslandExt
ZNF320
CDR2L→
MRPL58→
RNU6−362P→
←ATP5H
KCTD2→
RN7SL573P→
73.01
73.02
73.03
73.04
73.05
Position on chr17 (Mb)

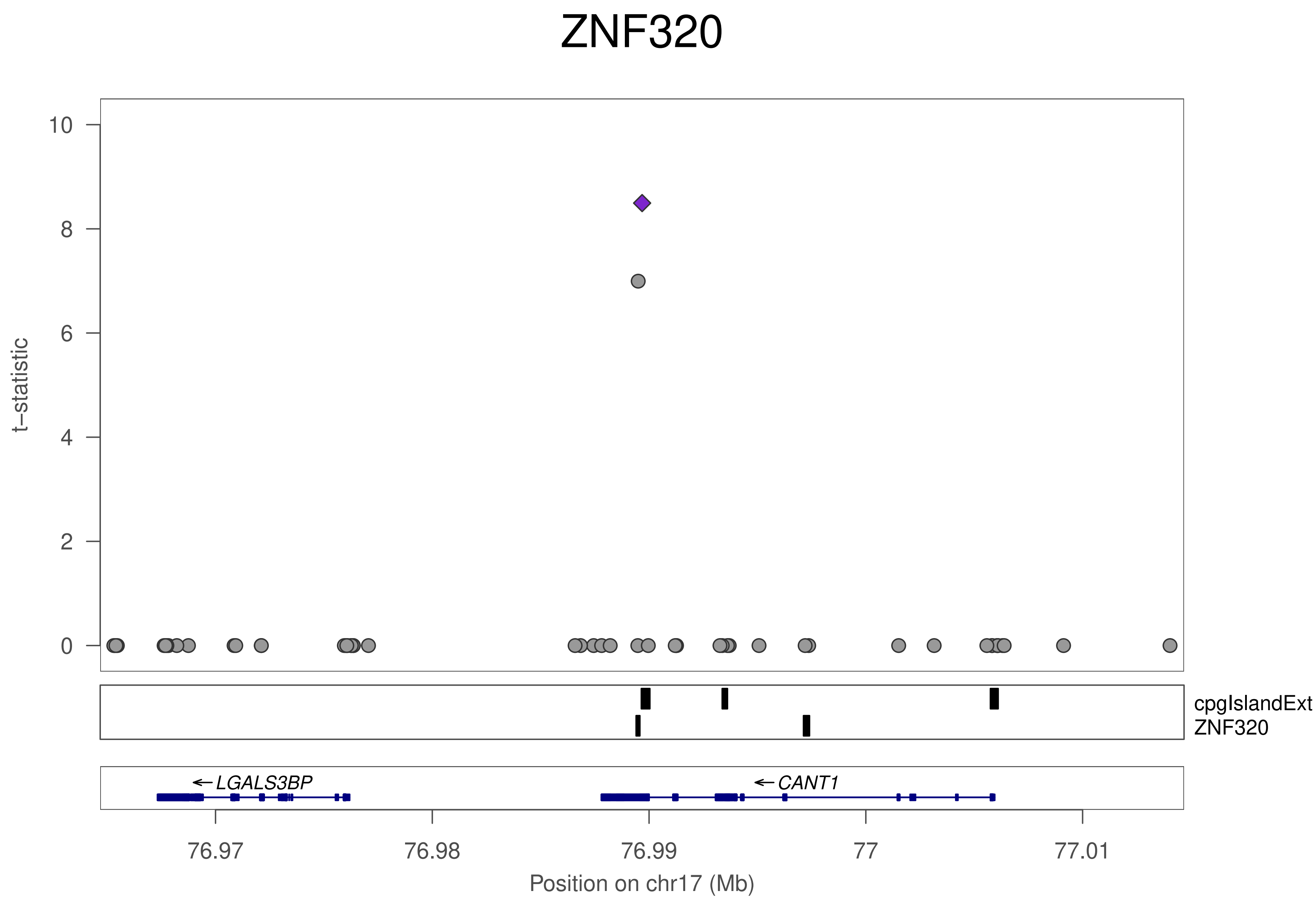
ZNF320
t−statistic
10
8
6
4
2
0
cpgIslandExt
ZNF320
←LGALS3BP
←CANT1
76.97
76.98
76.99
77
77.01
Position on chr17 (Mb)

ZNF320

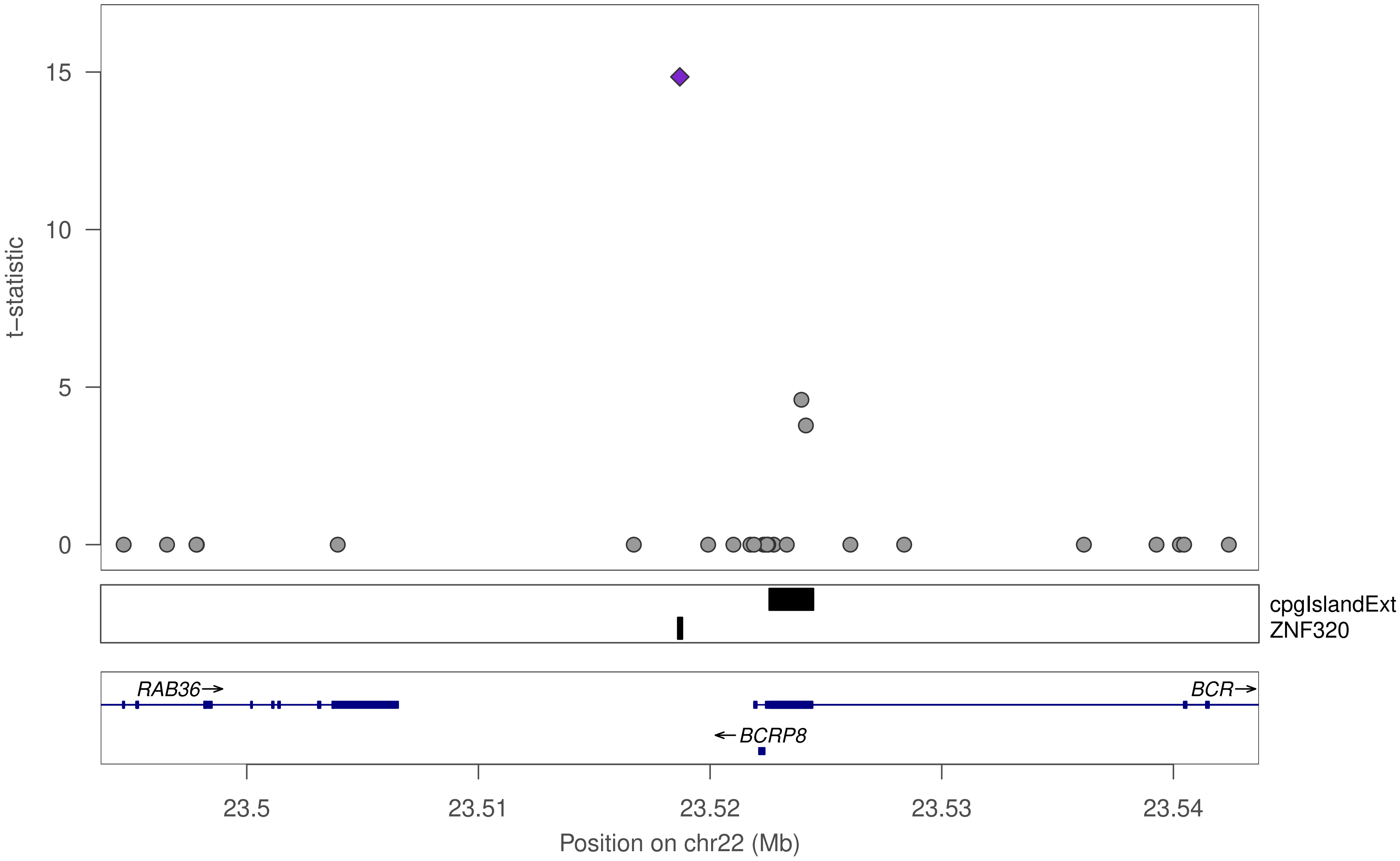

15
10
5
0
t−statistic
cpgIslandExt
ZNF320
RAB36→
BCR→
←BCRP8
23.5
23.51
23.52
23.53
23.54
Position on chr22 (Mb)

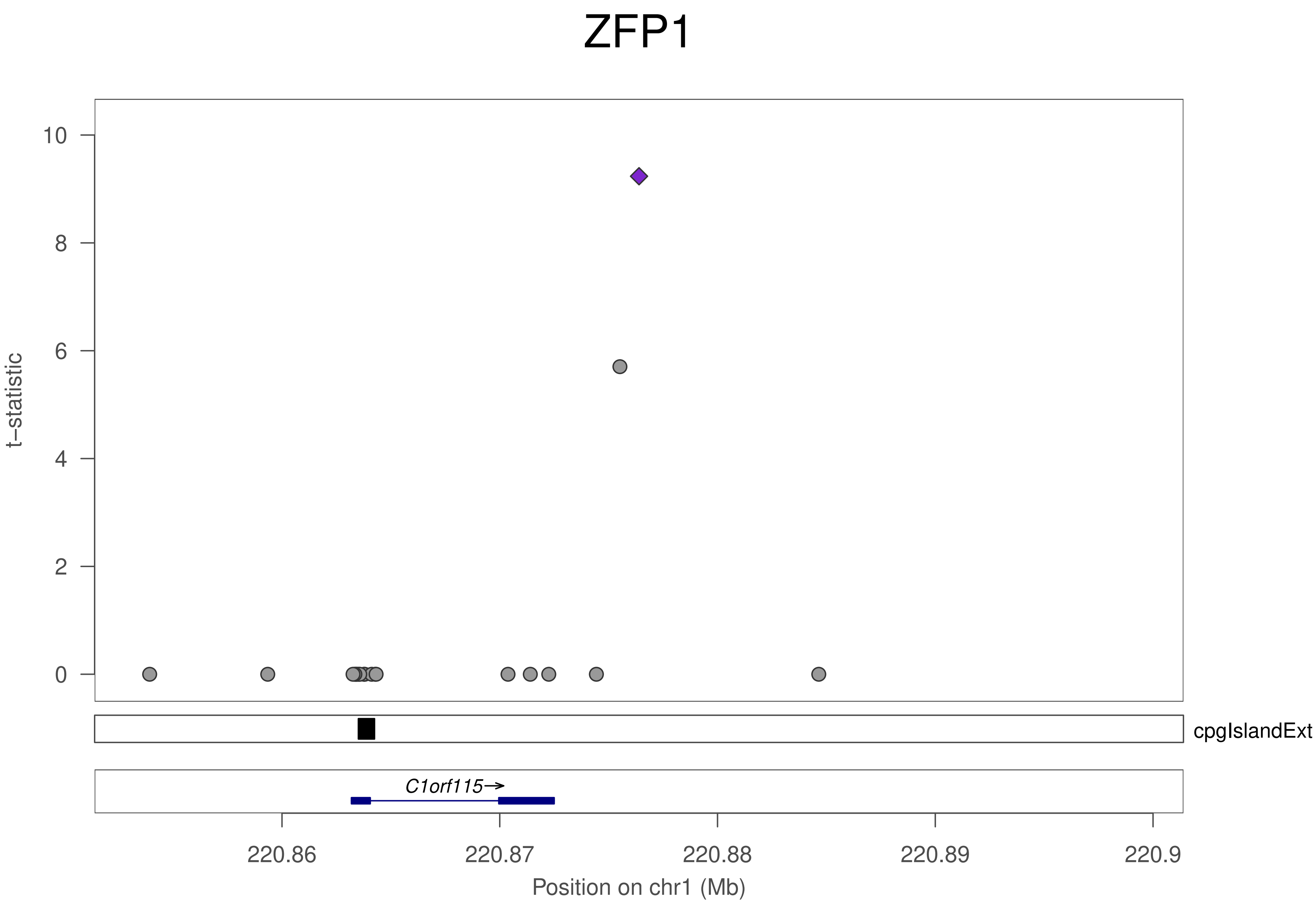
ZFP1
t–statistic
10
8
6
4
2
0
cpgIslandExt
C1orf115→
220.86
220.87
220.88
220.89
220.9
Position on chr1 (Mb)

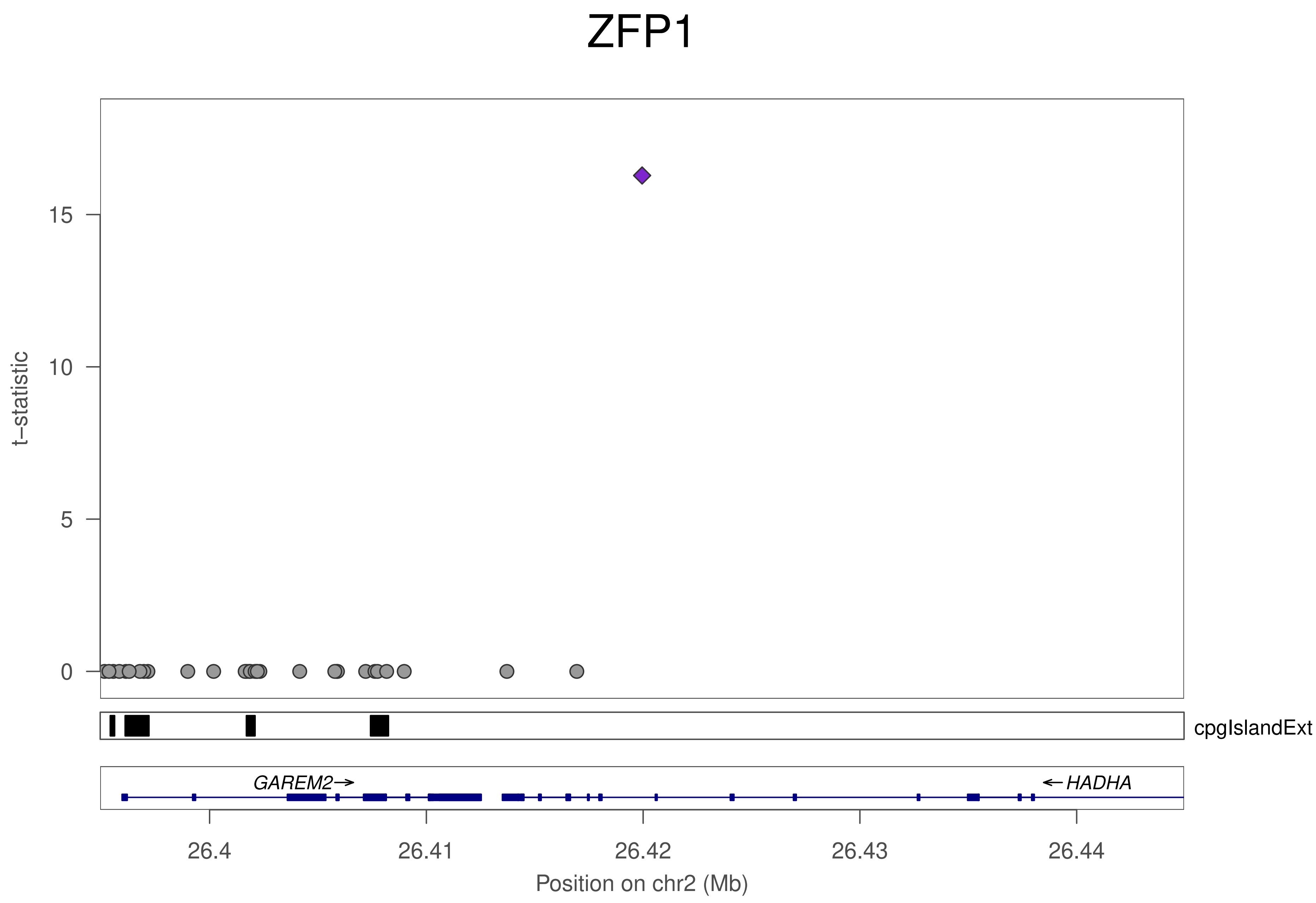

ZFP1
t−statistic
15
10
5
0
cpgIslandExt
GAREM2→
← HADHA
26.4
26.41
26.42
26.43
26.44
Position on chr2 (Mb)

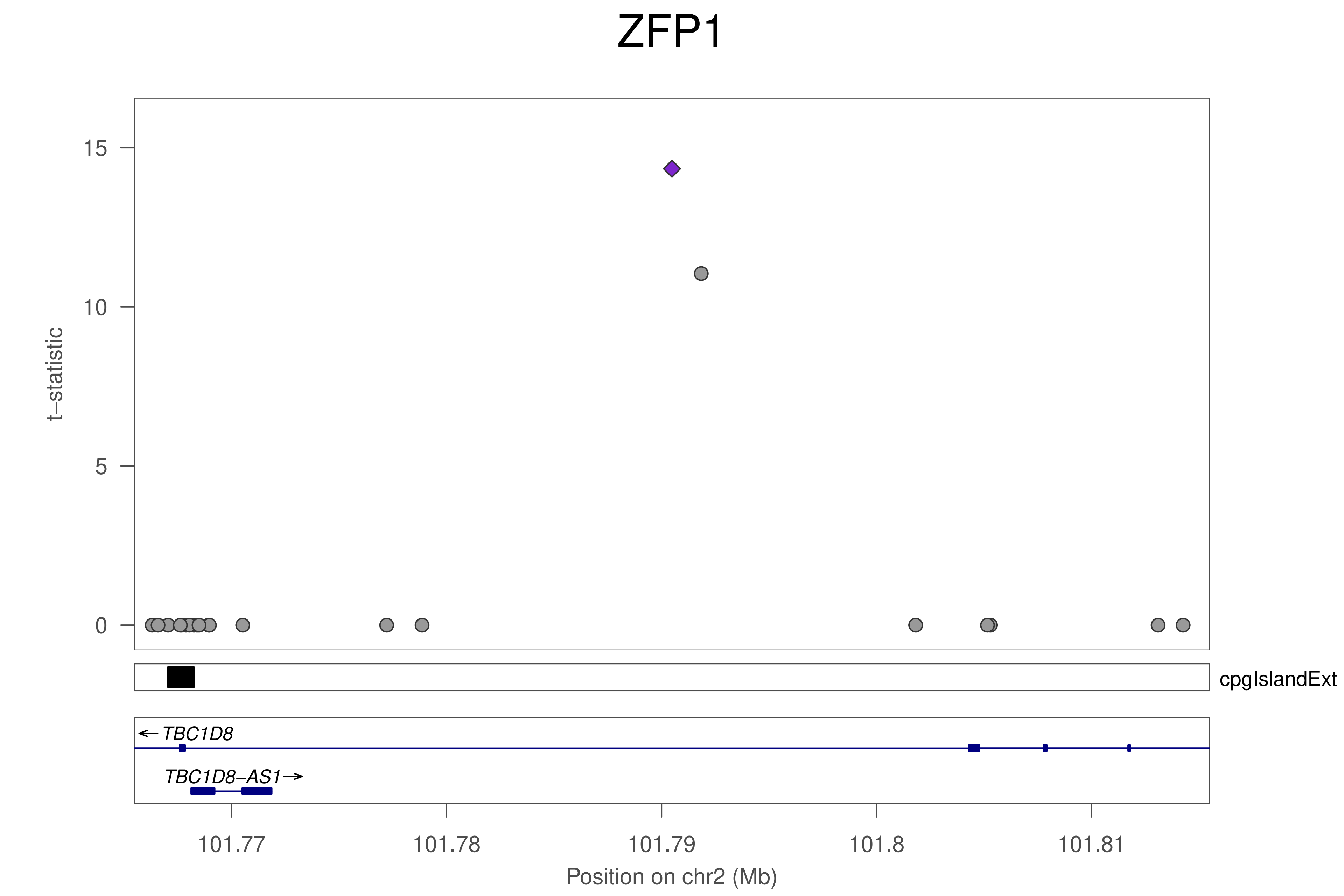
ZFP1
t−statistic
15
10
5
0
cpgIslandExt
← TBC1D8
TBC1D8−AS1→
101.77
101.78
101.79
101.8
101.81
Position on chr2 (Mb)

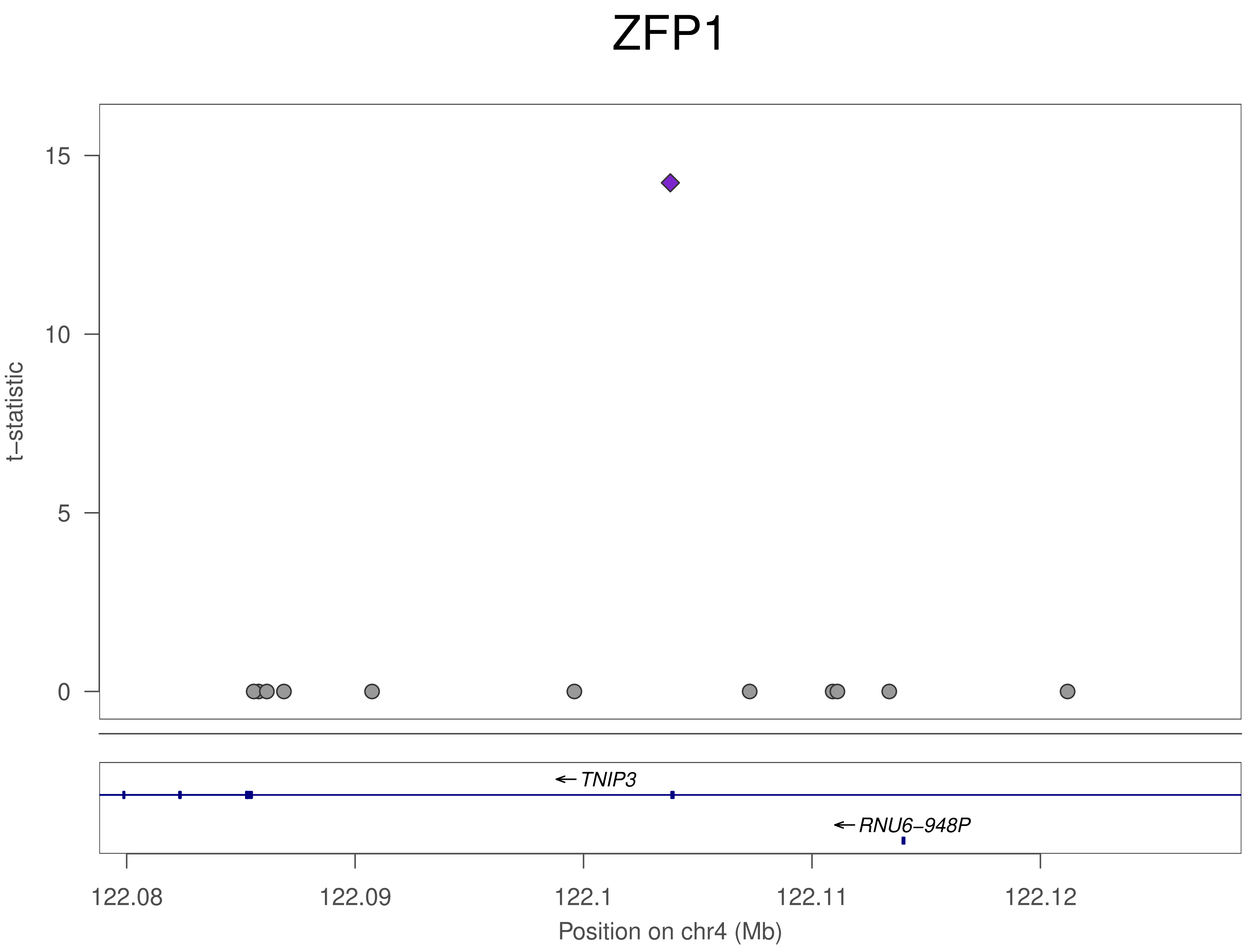

ZFP1
15
10
5
0
t−statistic
← TNIP3
← RNU6−948P
122.08
122.09
122.1
122.11
122.12
Position on chr4 (Mb)

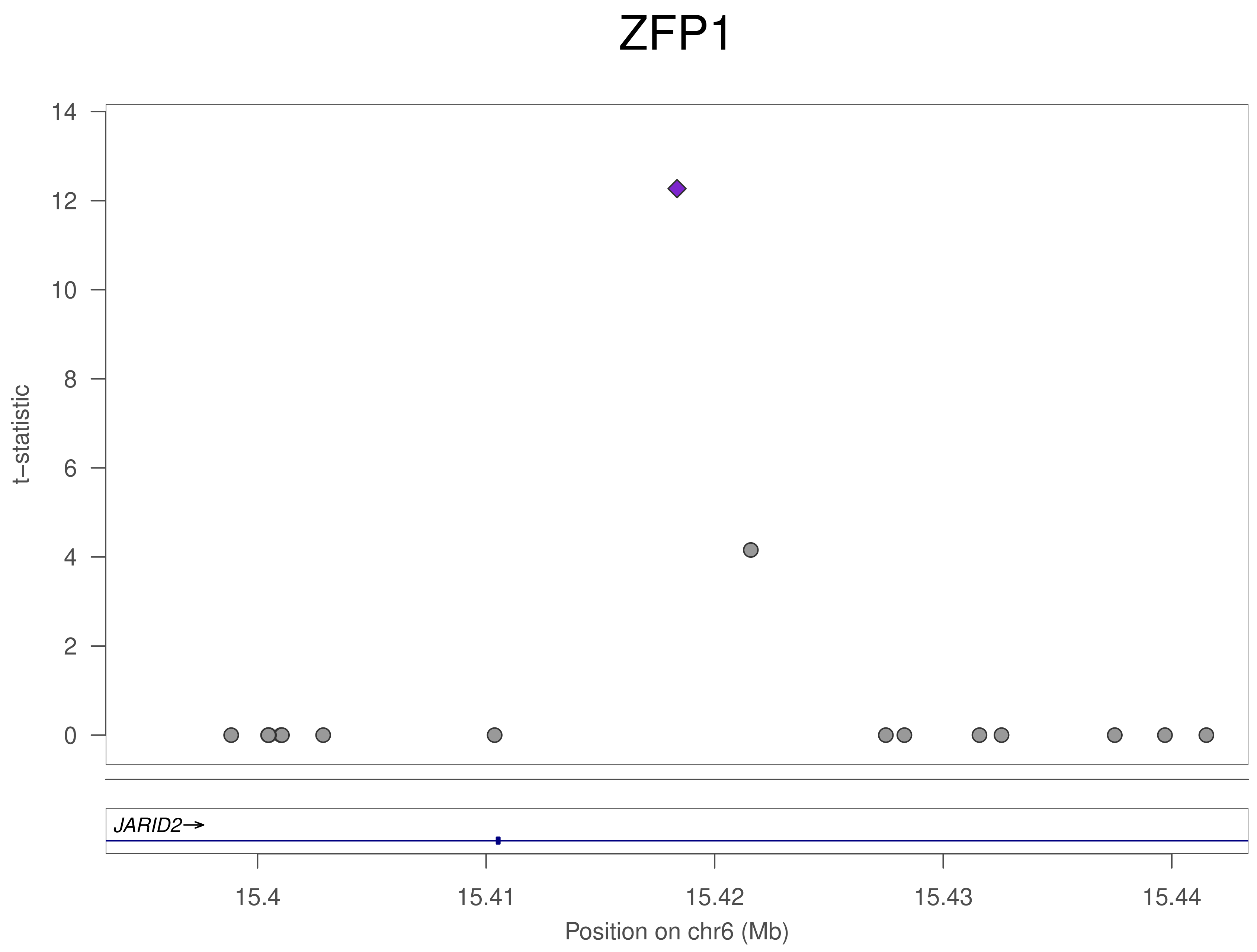

ZFP1
t−statistic
14
12
10
8
6
4
2
0
JARID2→
15.4
15.41
15.42
15.43
15.44
Position on chr6 (Mb)

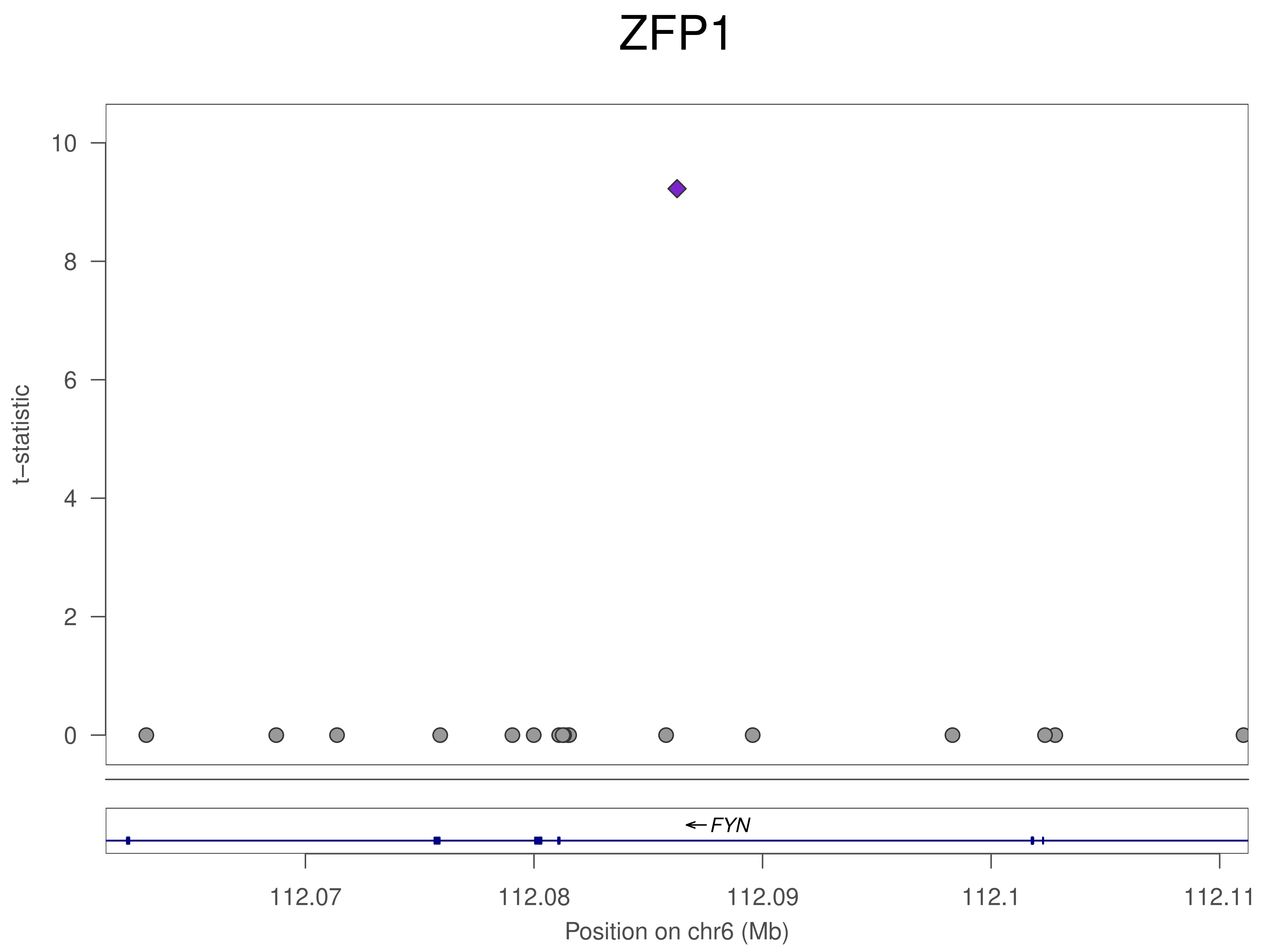
ZFP1
t–statistic
10
8
6
4
2
0
← FYN
112.07
112.08
112.09
112.1
112.11
Position on chr6 (Mb)

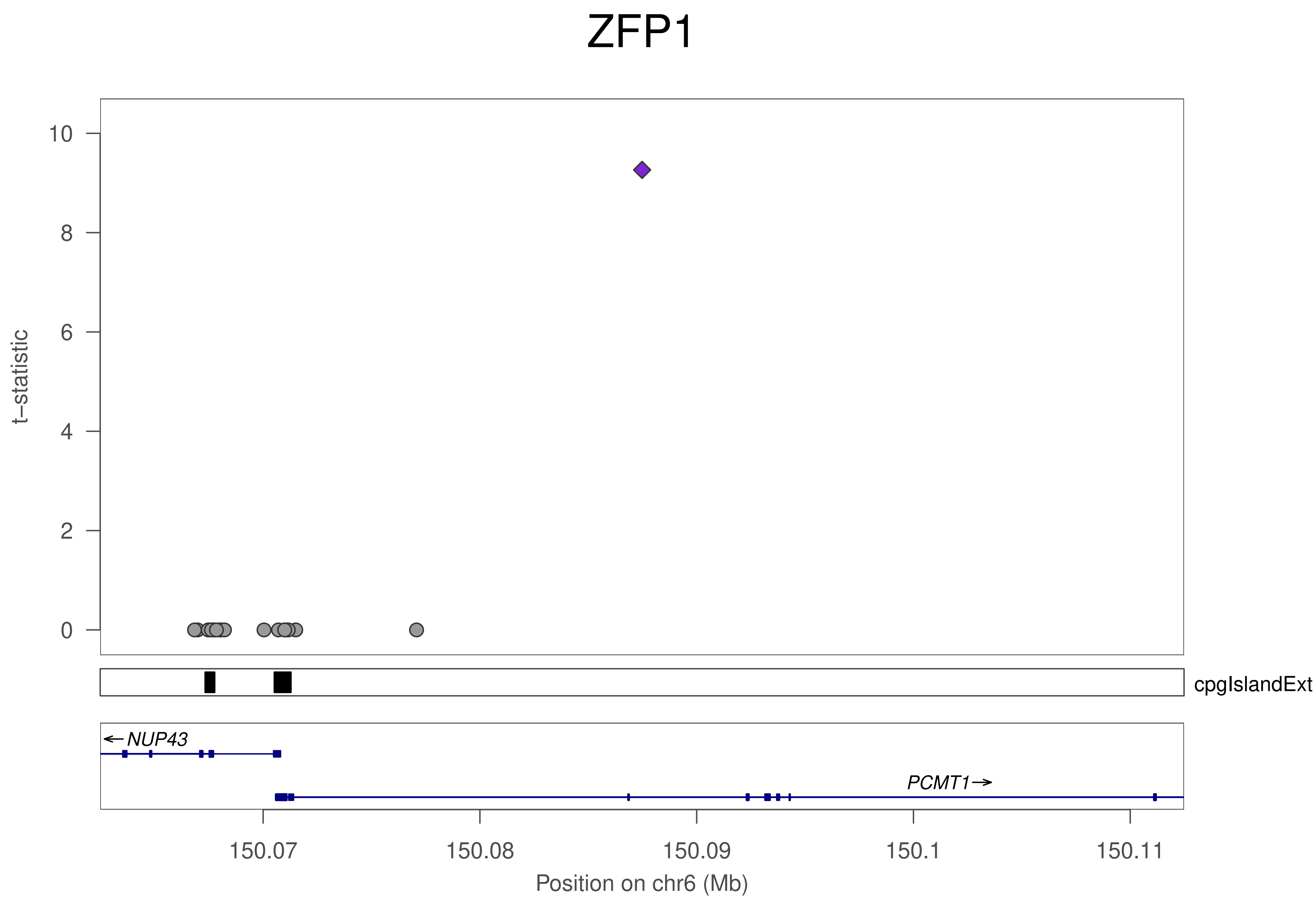

ZFP1
t–statistic
10
8
6
4
2
0
cpgIslandExt
NUP43
PCMT1
150.07
150.08
150.09
150.1
150.11
Position on chr6 (Mb)

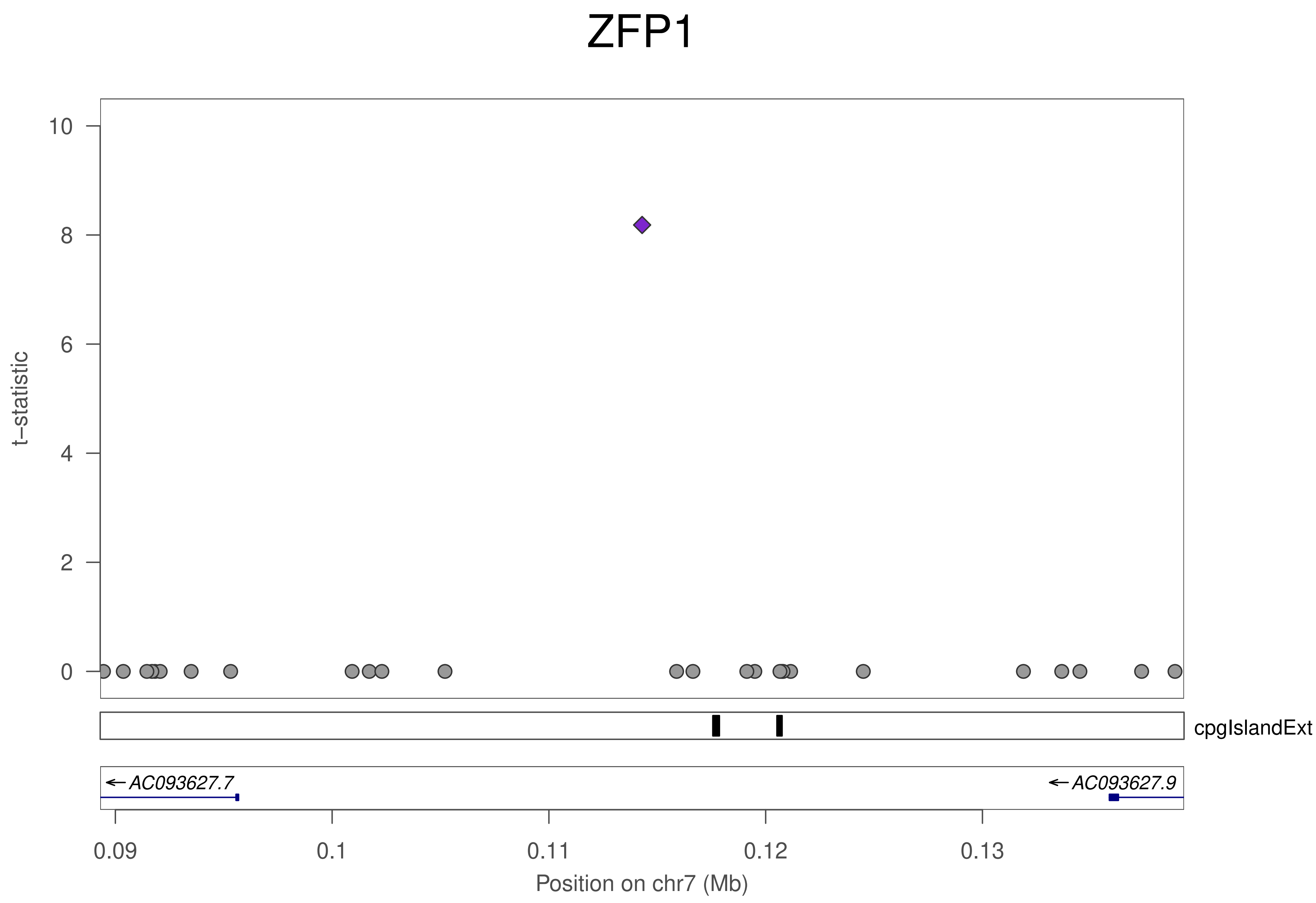
ZFP1
t–statistic
10
8
6
4
2
0
cpgIslandExt
← AC093627.7
← AC093627.9
0.09
0.1
0.11
0.12
0.13
Position on chr7 (Mb)

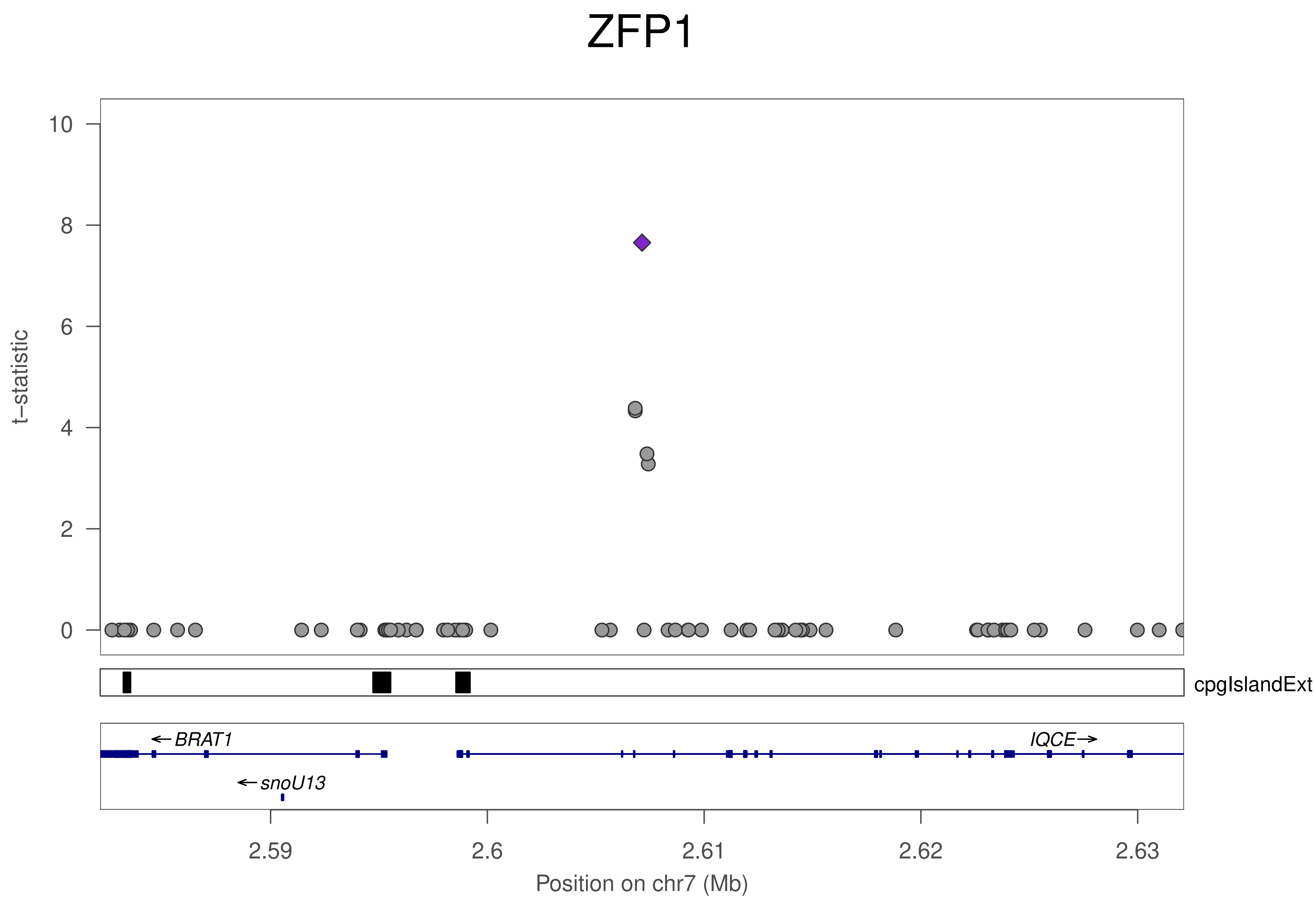

ZFP1
t−statistic
10
8
6
4
2
0
cpgIslandExt
←BRAT1
←snoU13
IQCE→
2.59
2.6
2.61
2.62
2.63
Position on chr7 (Mb)

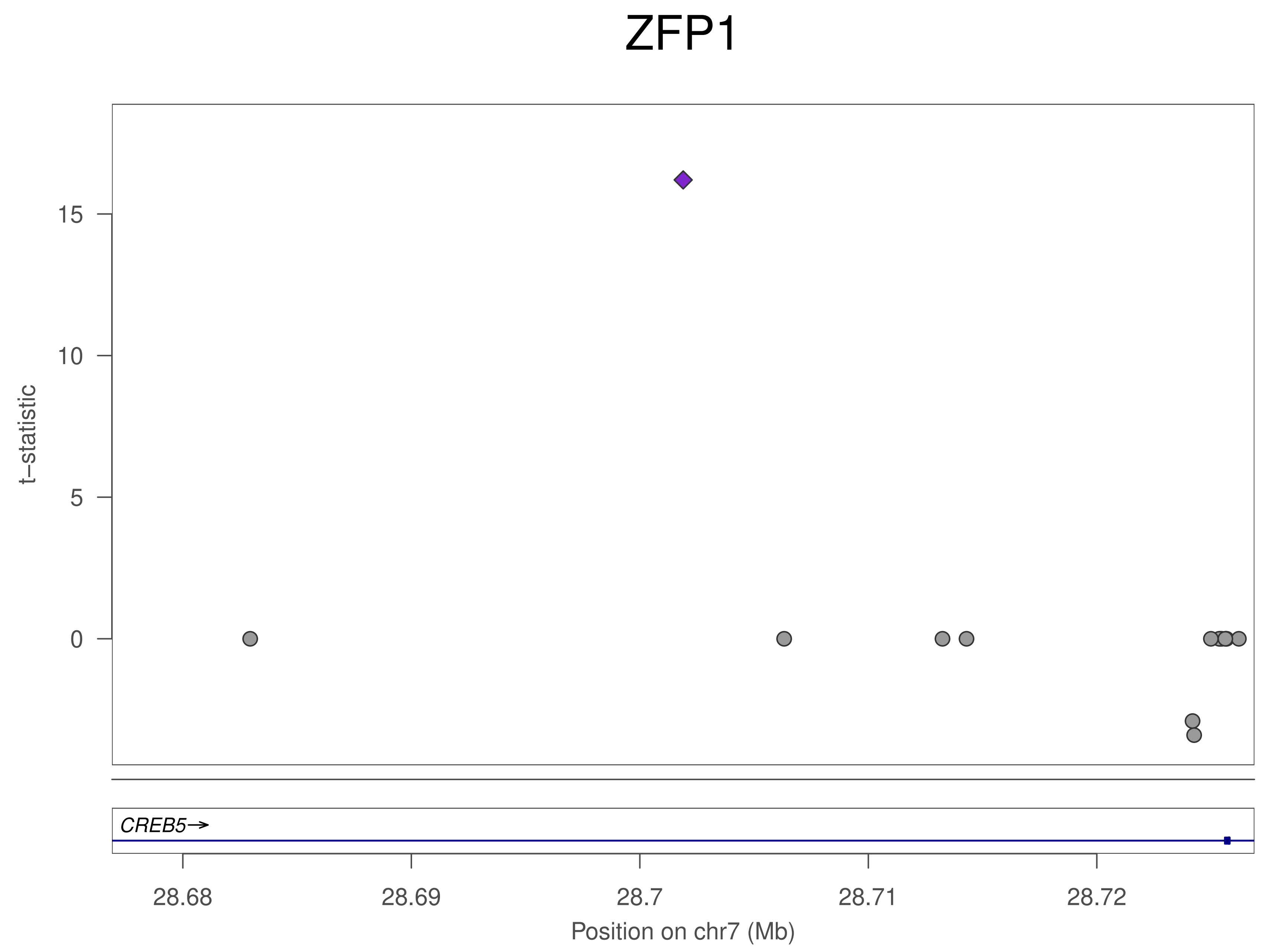
ZFP1
t–statistic
15
10
5
0
CREB5→
28.68
28.69
28.7
28.71
28.72
Position on chr7 (Mb)

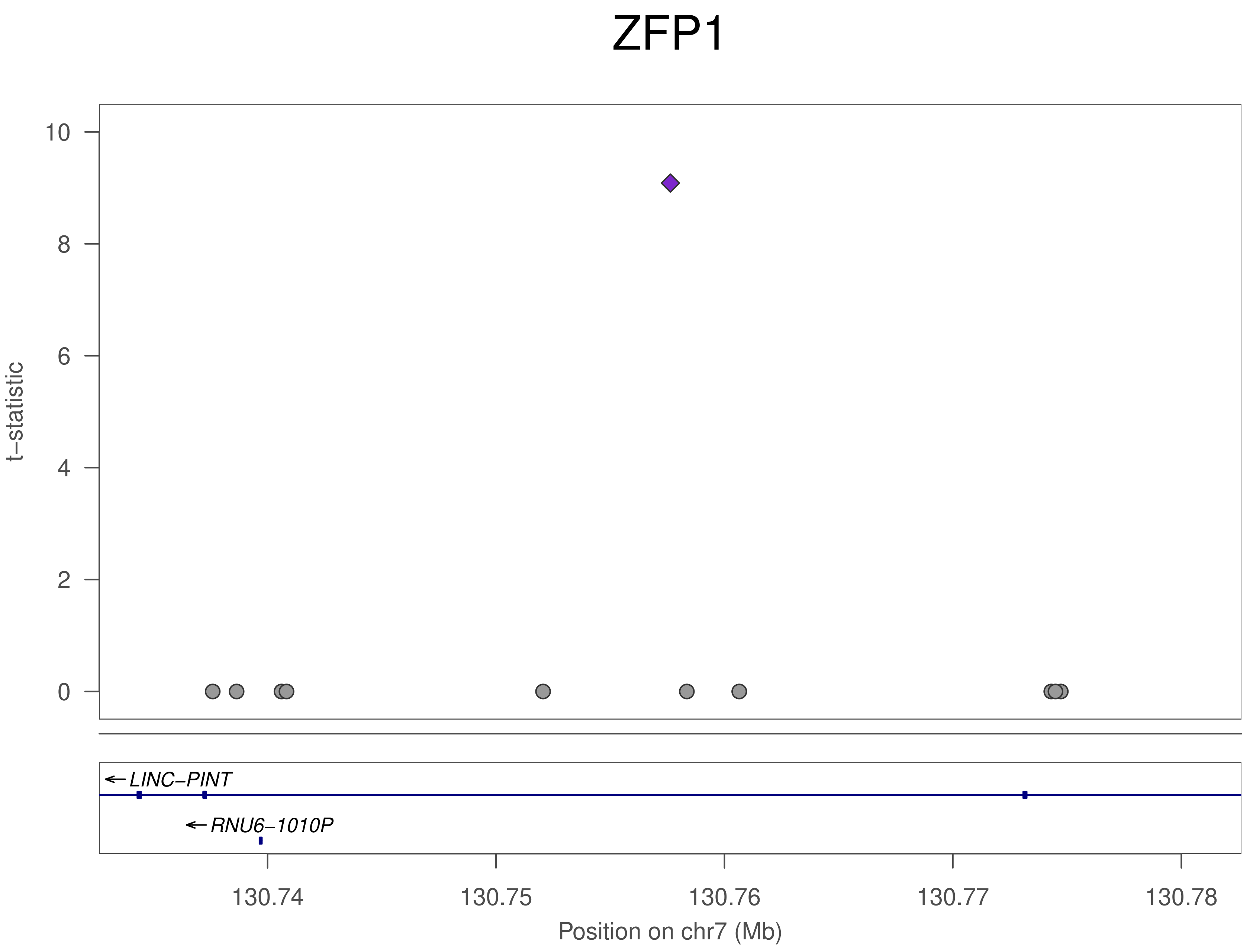
ZFP1
t−statistic
10
8
6
4
2
0
LINC−PINT
RNU6−1010P
130.74
130.75
130.76
130.77
130.78
Position on chr7 (Mb)

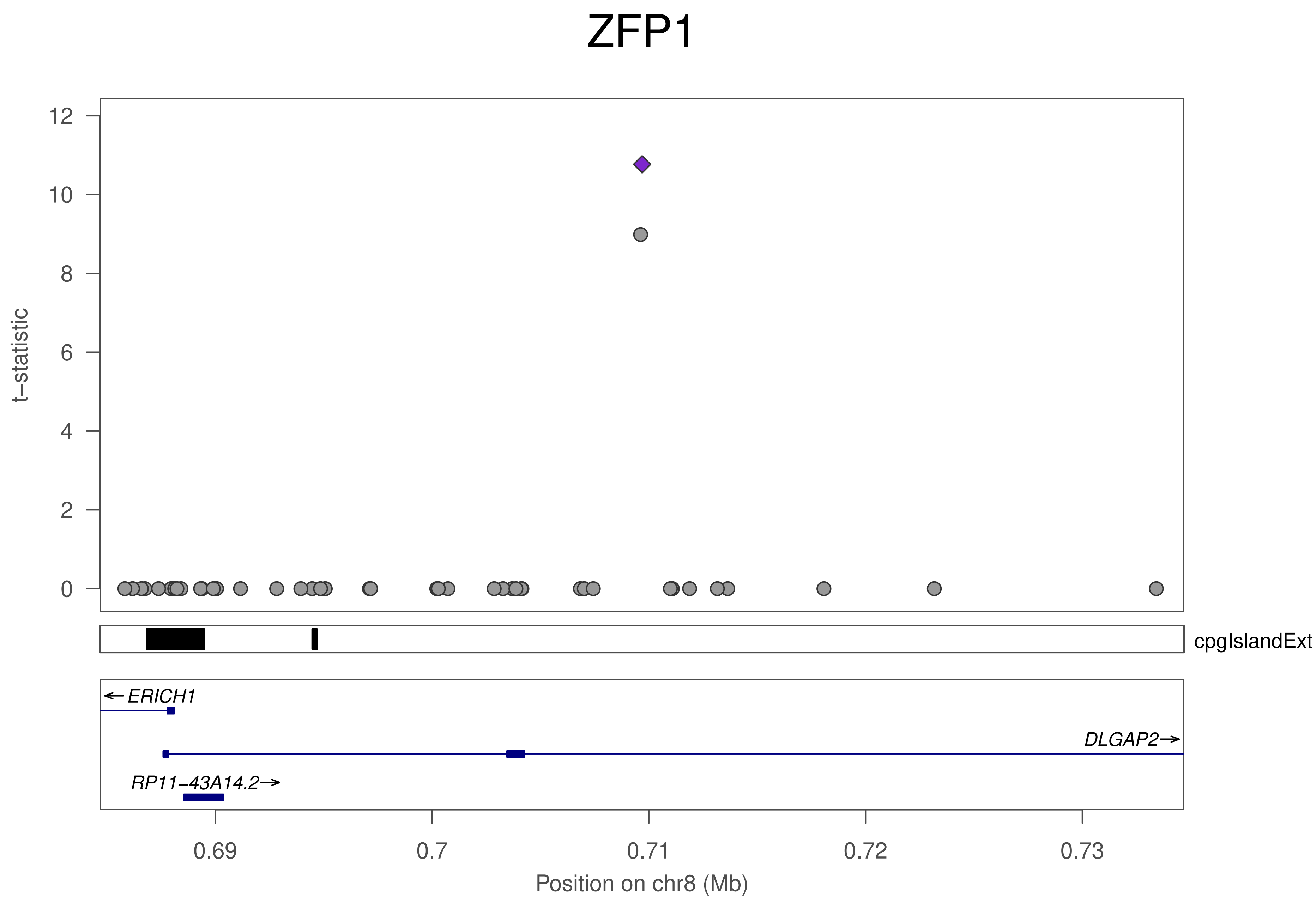
ZFP1
t−statistic
12
10
8
6
4
2
0
cpgIslandExt
←ERICH1
DLGAP2→
RP11−43A14.2→
0.69
0.7
0.71
0.72
0.73
Position on chr8 (Mb)

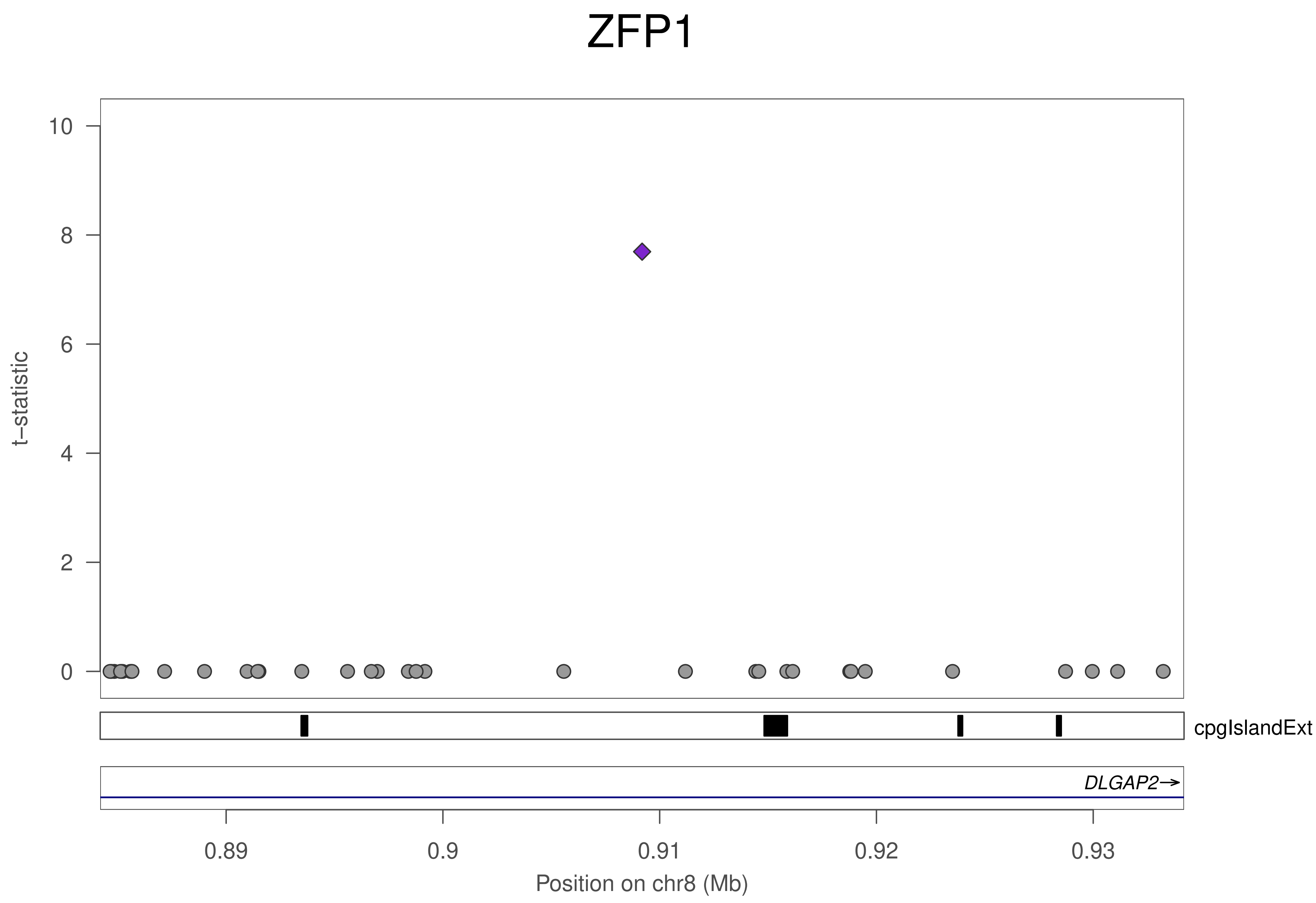

ZFP1
t−statistic
10
8
6
4
2
0
cpgIslandExt
DLGAP2→
0.89
0.9
0.91
0.92
0.93
Position on chr8 (Mb)

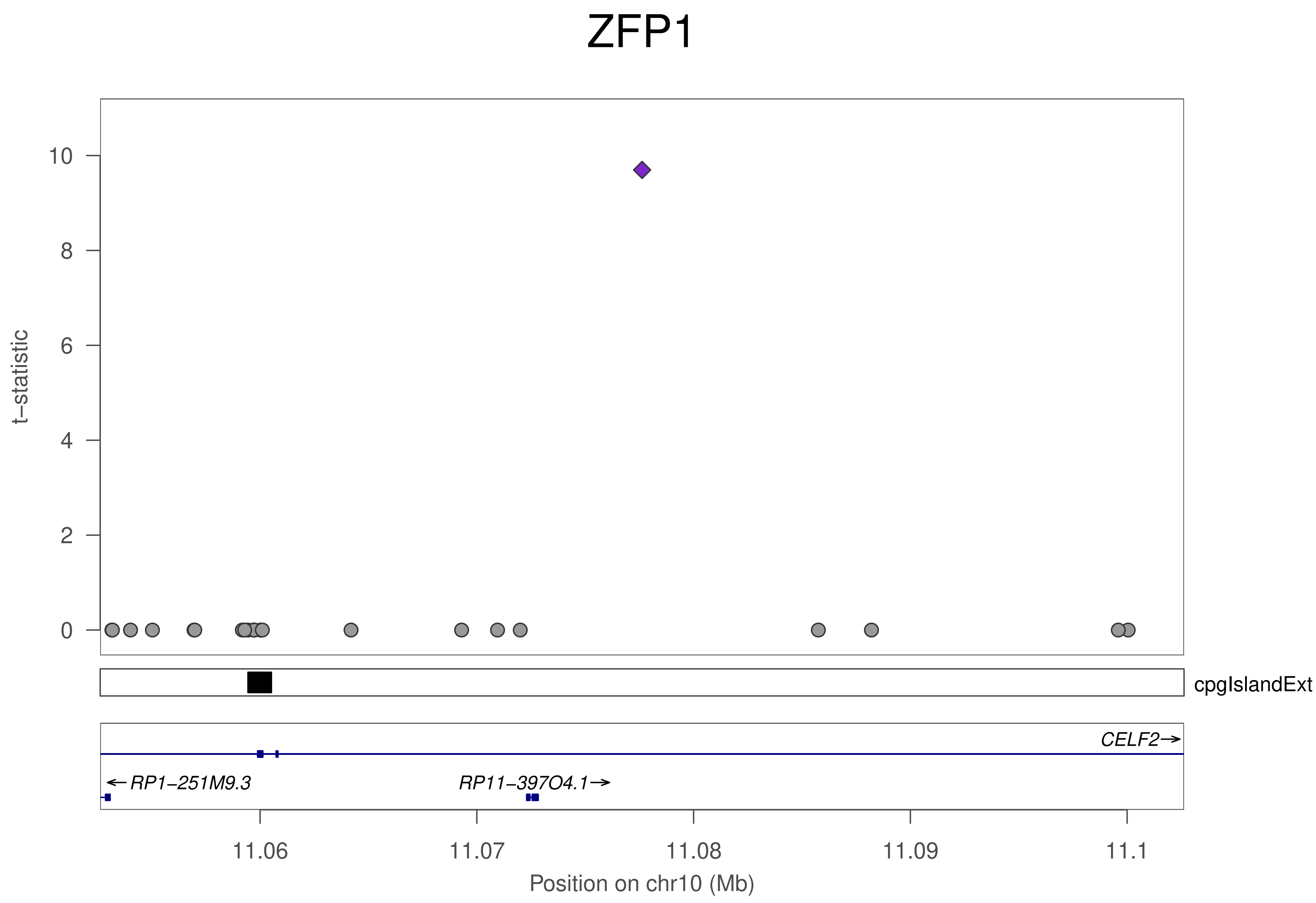
ZFP1
t–statistic
10
8
6
4
2
0
cpgIslandExt
CELF2→
←RP1–251M9.3
RP11–397O4.1→
11.06
11.07
11.08
11.09
11.1
Position on chr10 (Mb)

### ZFP1

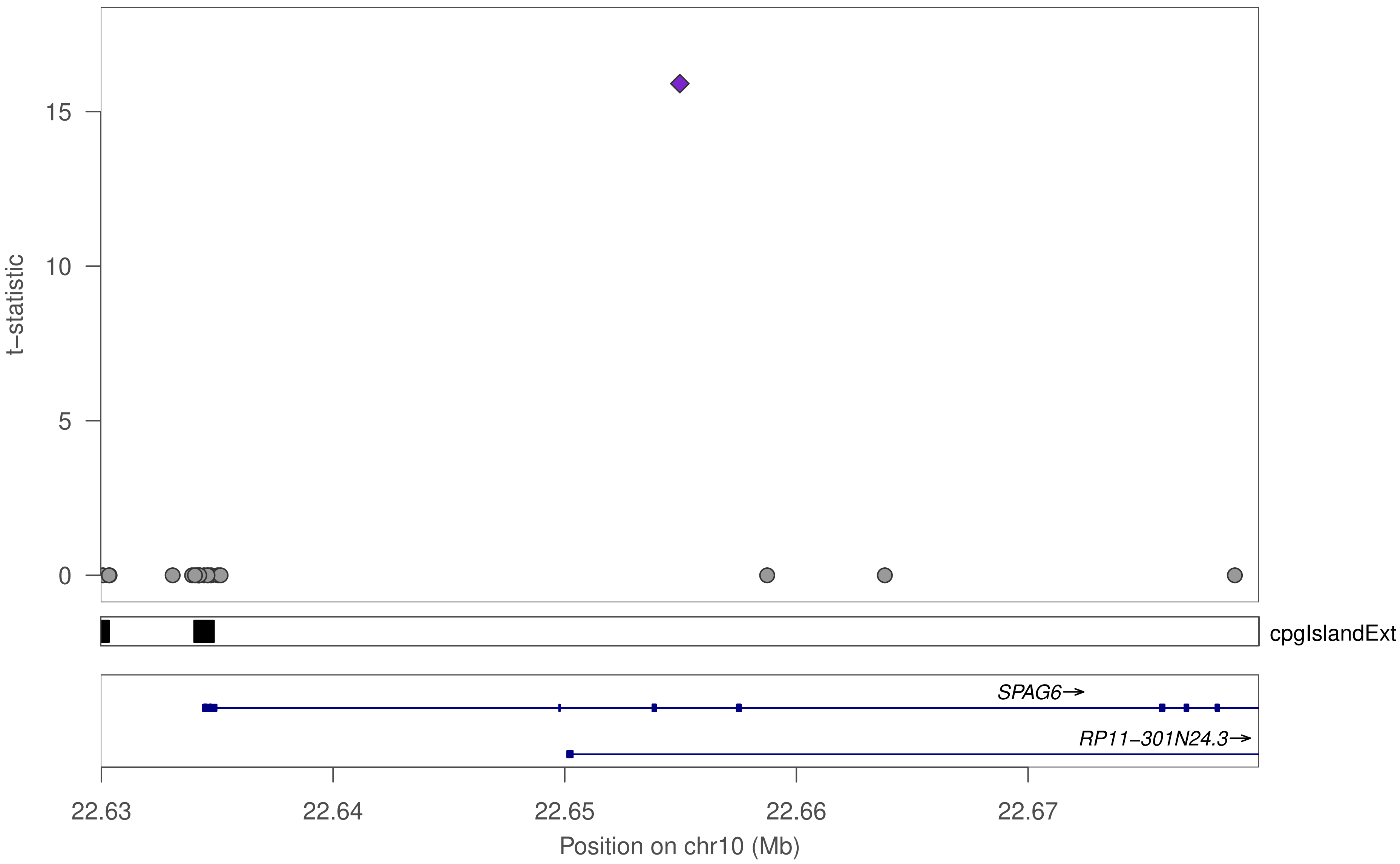

15
10
5
0
t−statistic
cpgIslandExt
SPAG6→
RP11−301N24.3→
22.63
22.64
22.65
22.66
22.67
Position on chr10 (Mb)

### ZFP1

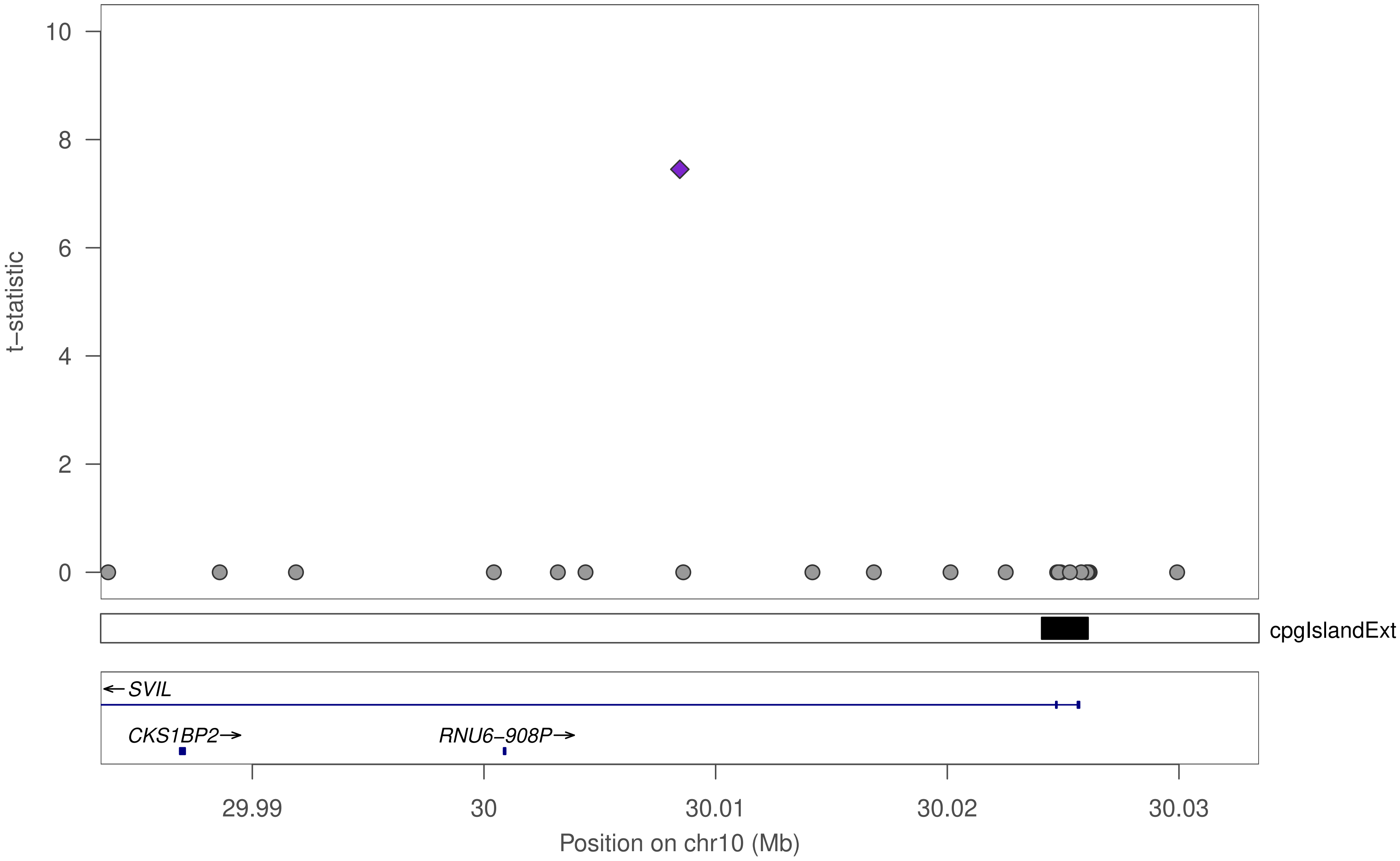

10
8
6
4
2
0
t−statistic
cpgIslandExt
SVIL
CKS1BP2
RNU6−908P
29.99
30
30.01
30.02
30.03
Position on chr10 (Mb)

### ZFP1

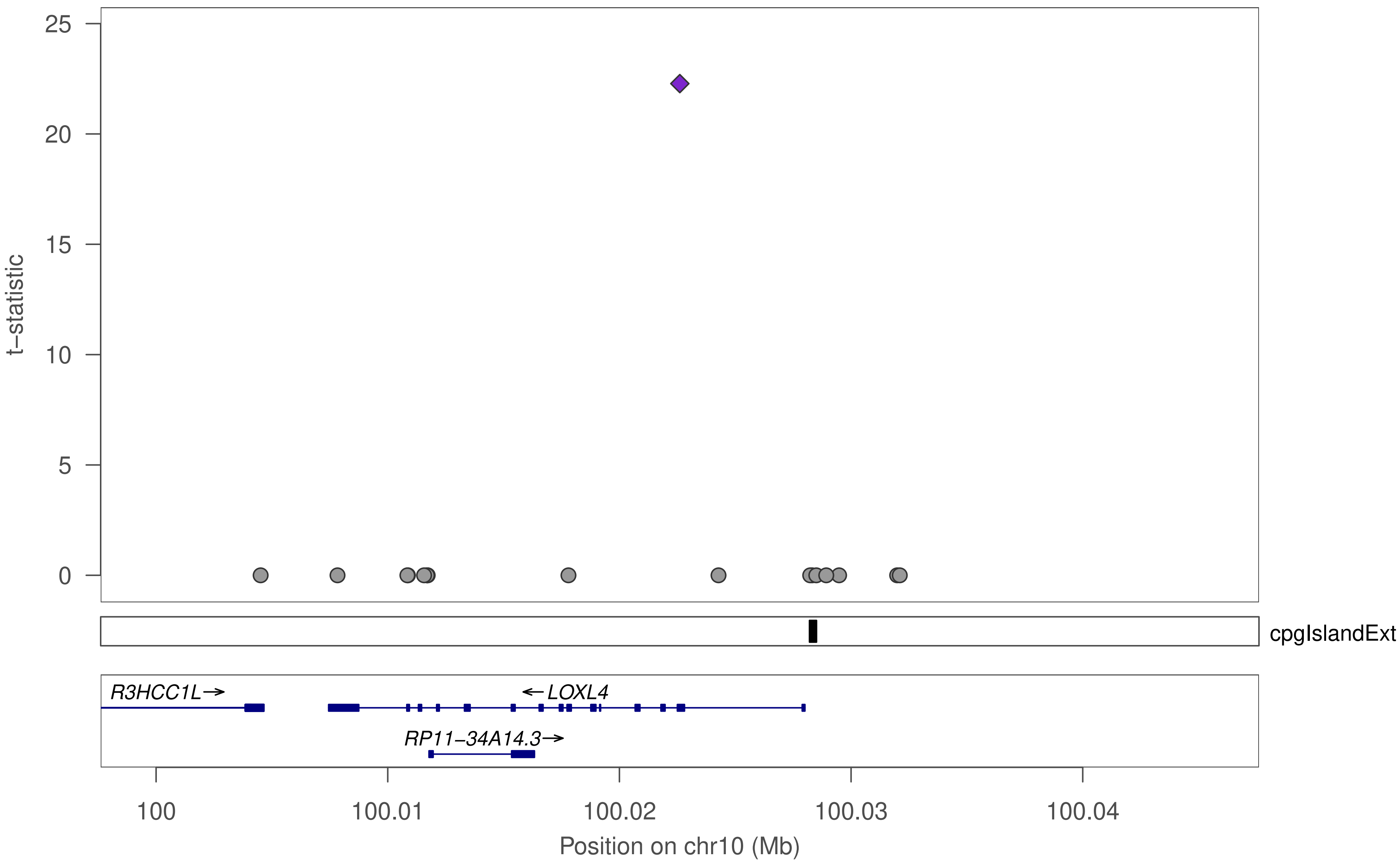

t−statistic
25
20
15
10
5
0
cpgIslandExt
R3HCC1L→
←LOXL4
RP11−34A14.3→
100
100.01
100.02
100.03
100.04
Position on chr10 (Mb)

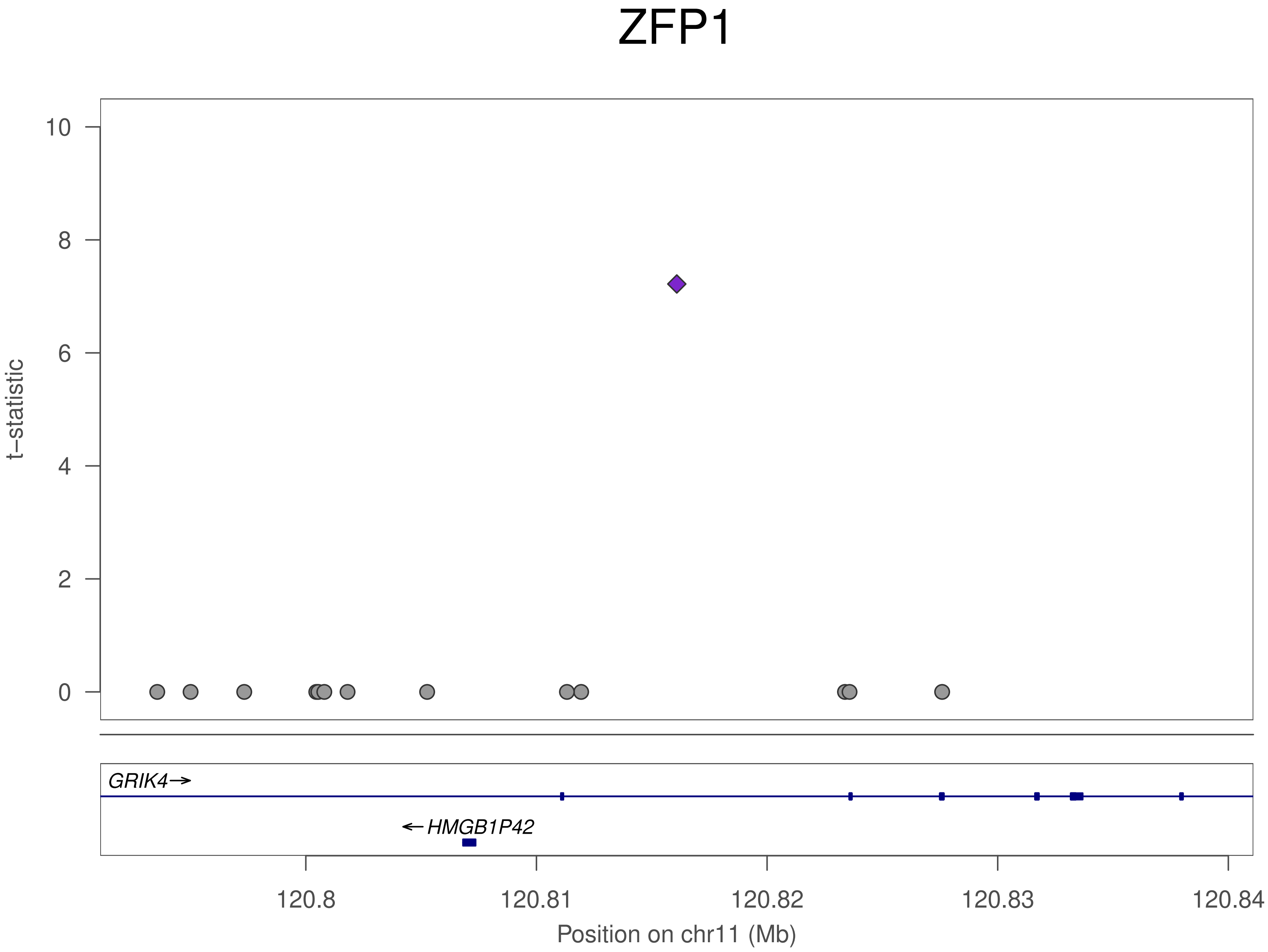

ZFP1
t–statistic
10
8
6
4
2
0
GRIK4
HMGB1P42
120.8
120.81
120.82
120.83
120.84
Position on chr11 (Mb)

### ZFP1

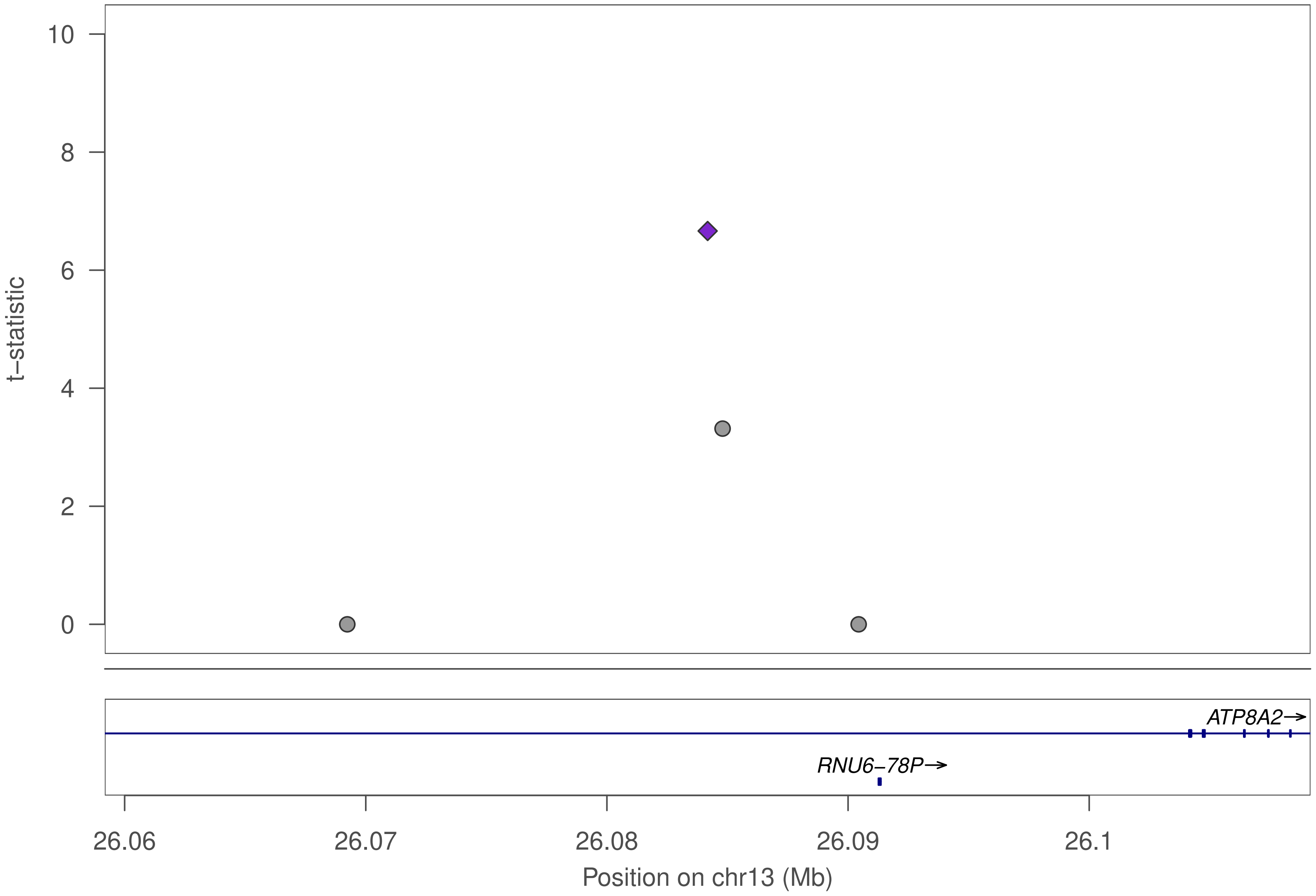

t−statistic
10
8
6
4
2
0
ATP8A2→
RNU6−78P→
26.06
26.07
26.08
26.09
26.1
Position on chr13 (Mb)

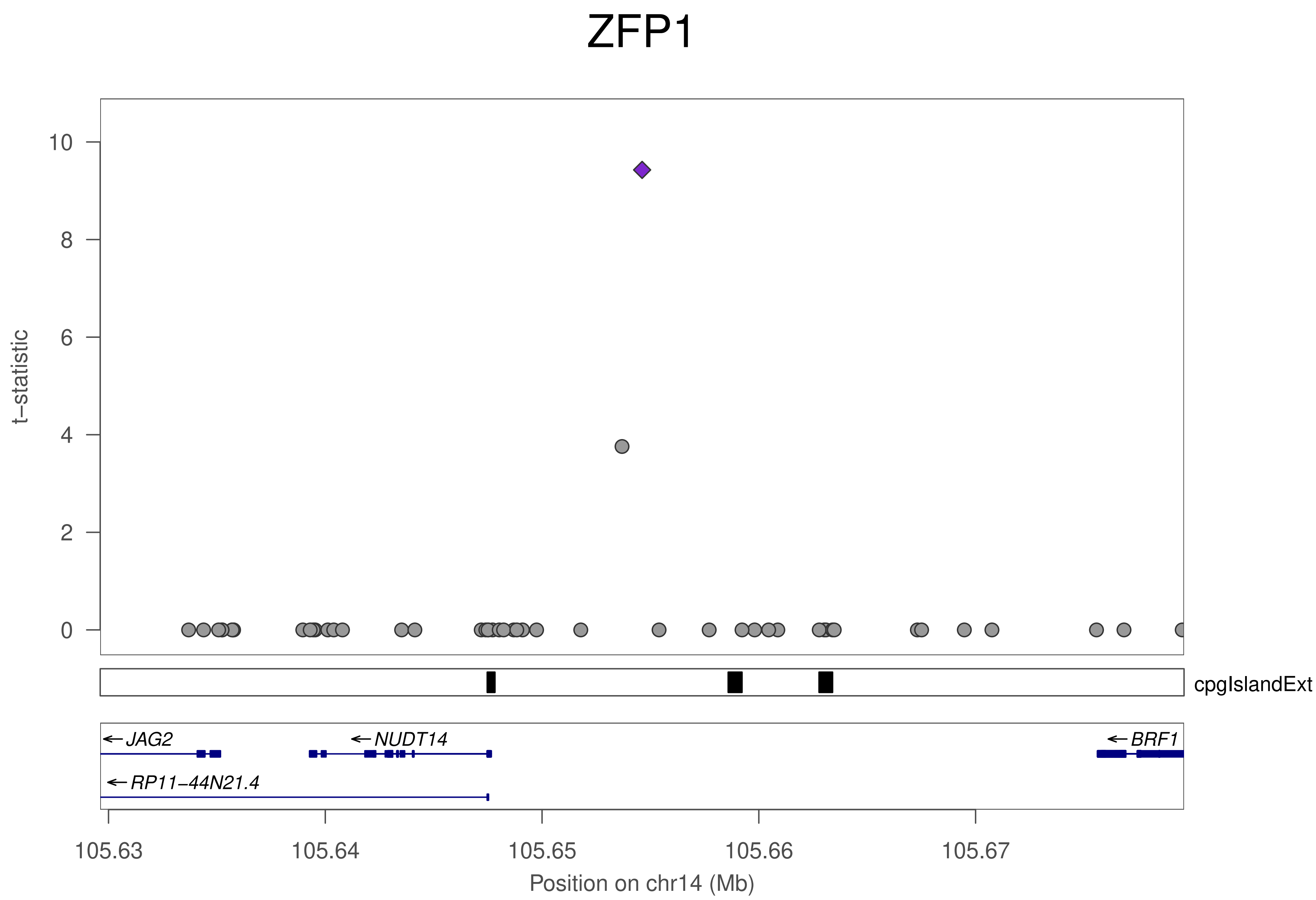

ZFP1
t–statistic
10
8
6
4
2
0
cpgIslandExt
JAG2
NUDT14
BRF1
RP11–44N21.4
105.63
105.64
105.65
105.66
105.67
Position on chr14 (Mb)

### ZFP1

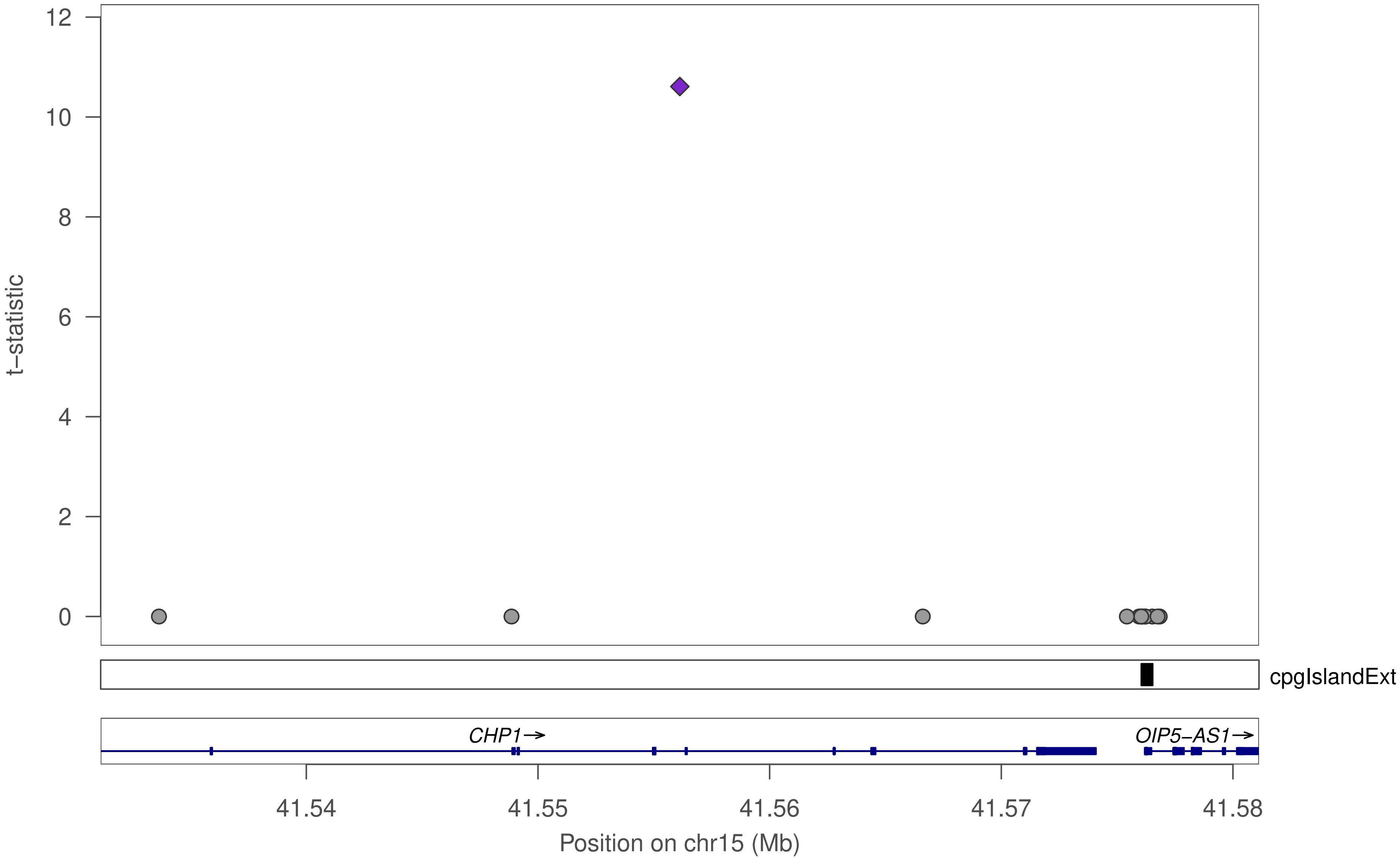
t−statistic
12
10
8
6
4
2
0
cpgIslandExt
CHP1→
OIP5−AS1→
41.54
41.55
41.56
41.57
41.58
Position on chr15 (Mb)

### ZFP1

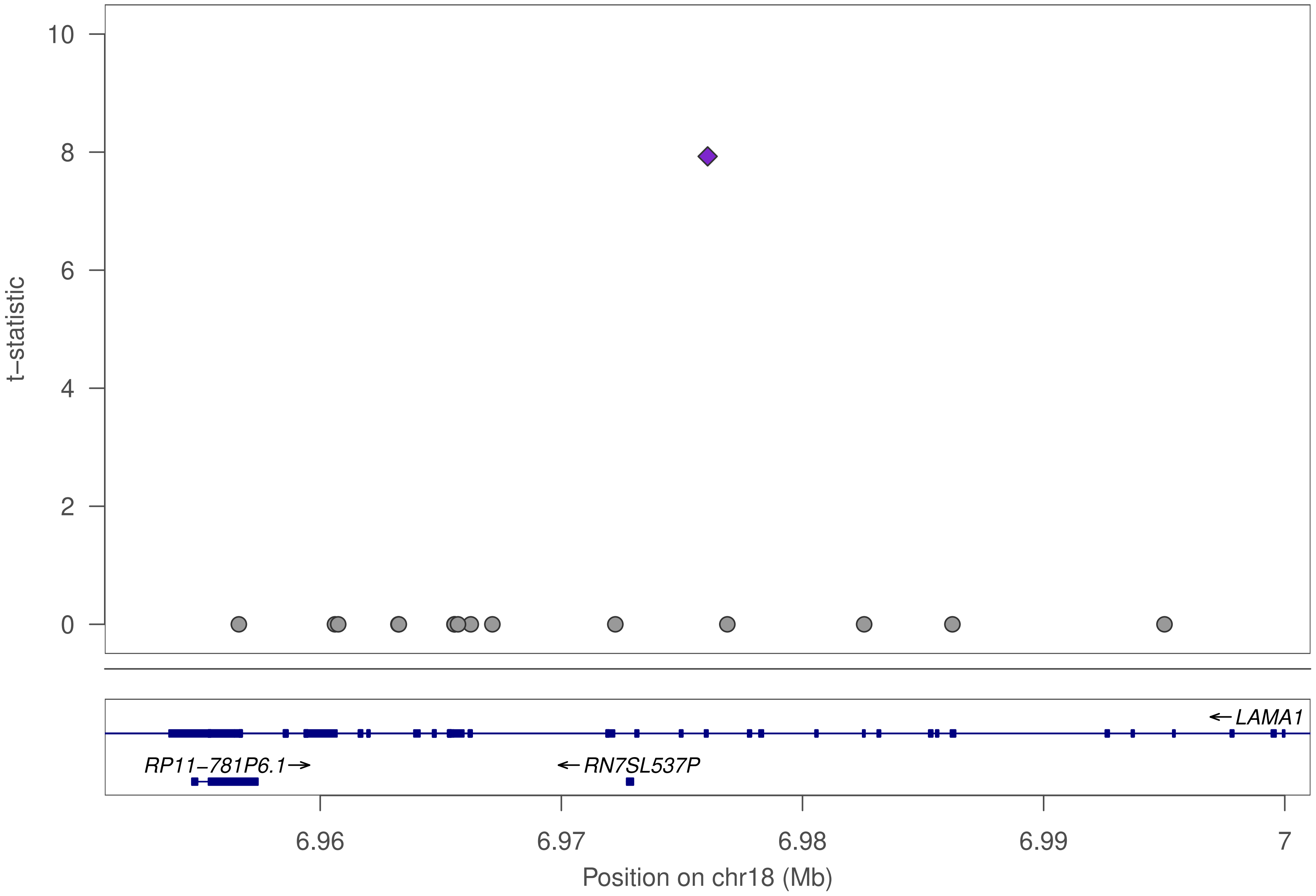

t–statistic
10
8
6
4
2
0
LAMA1
RP11–781P6.1
RN7SL537P
6.96
6.97
6.98
6.99
7
Position on chr18 (Mb)

### ZFP1

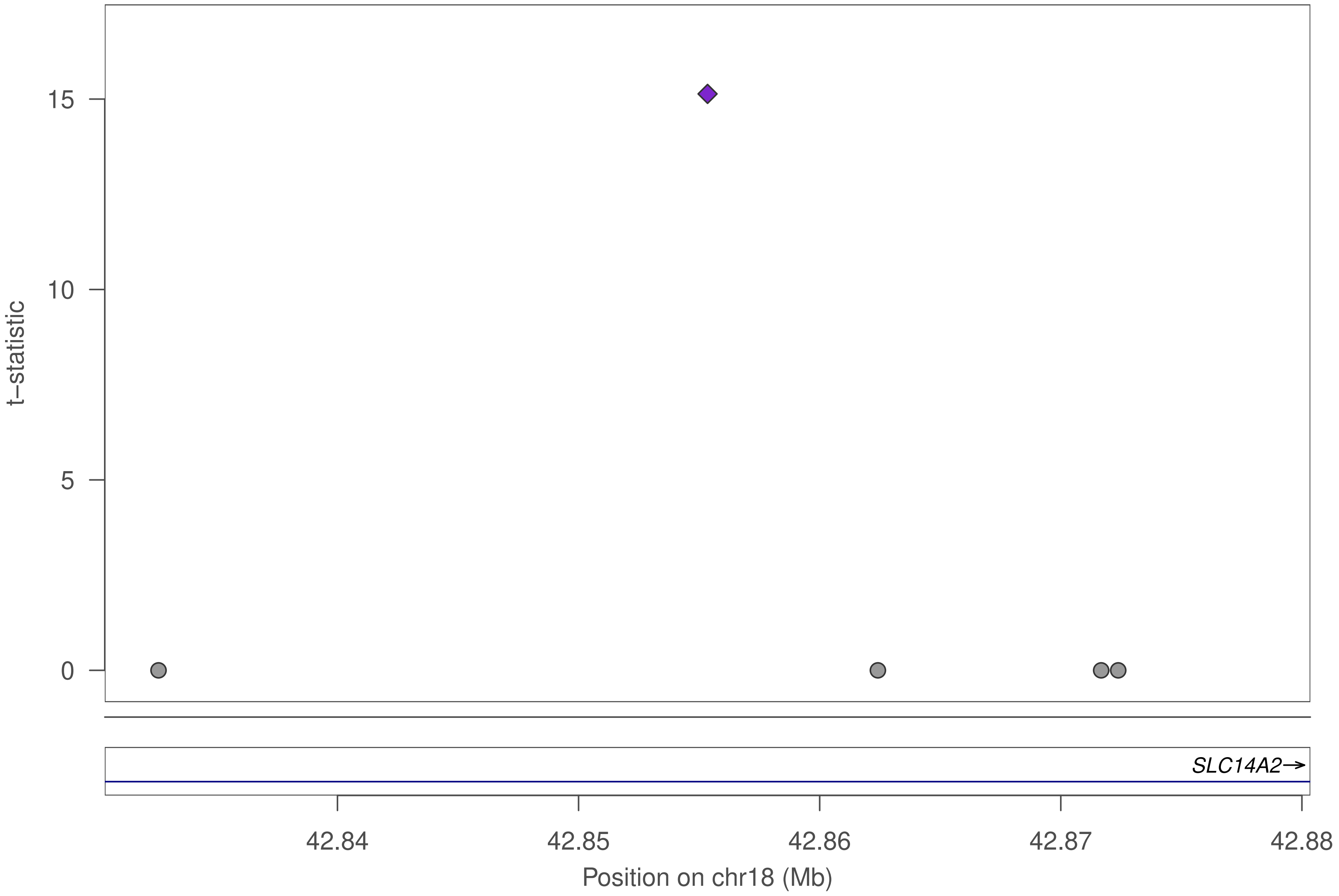

15
10
5
0
t–statistic
SLC14A2→
42.84
42.85
42.86
42.87
42.88
Position on chr18 (Mb)

ZFP1

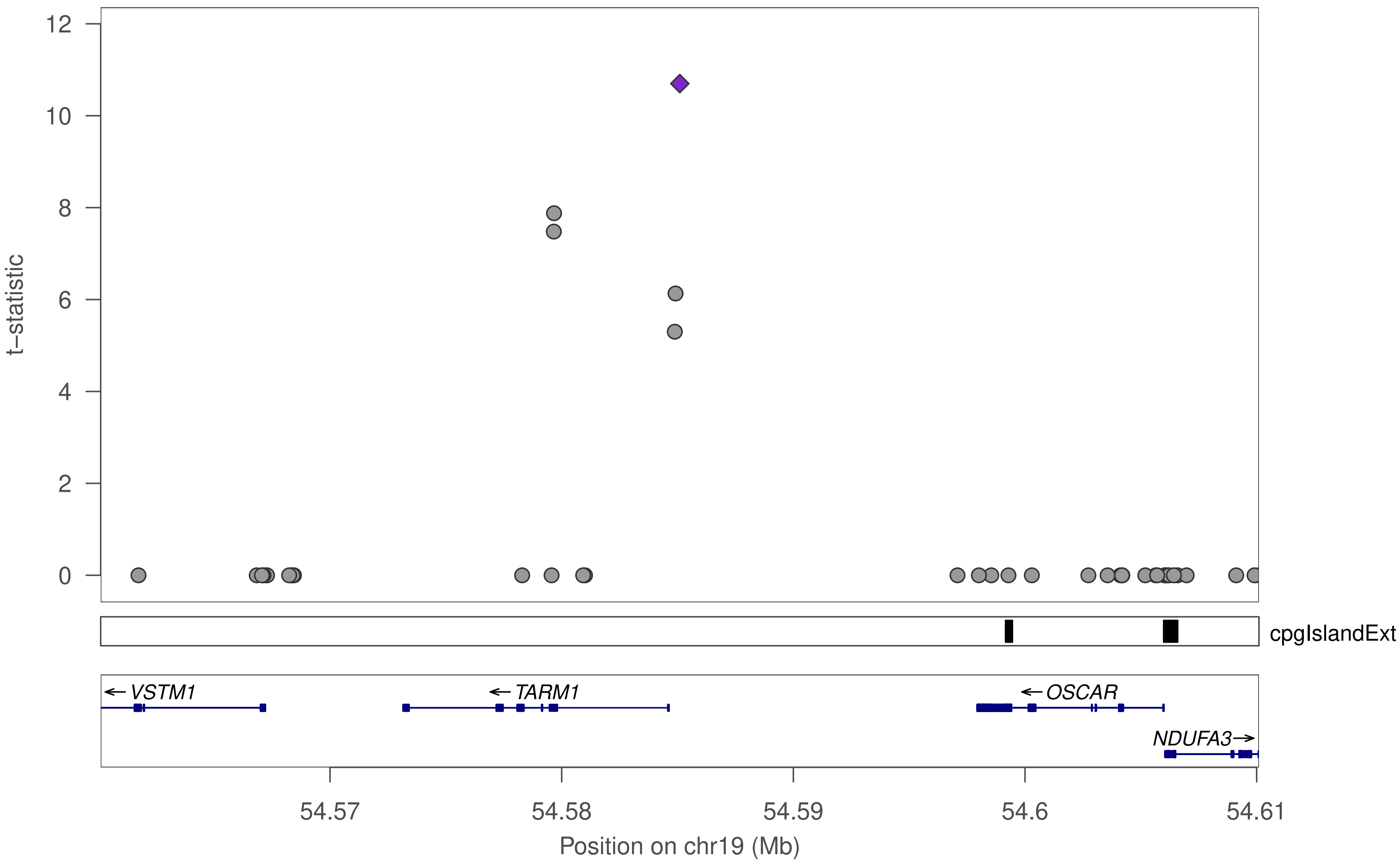

12
10
8
6
4
2
0
t−statistic
cpgIslandExt
← VSTM1
← TARM1
← OSCAR
NDUFA3→
54.57
54.58
54.59
54.6
54.61
Position on chr19 (Mb)

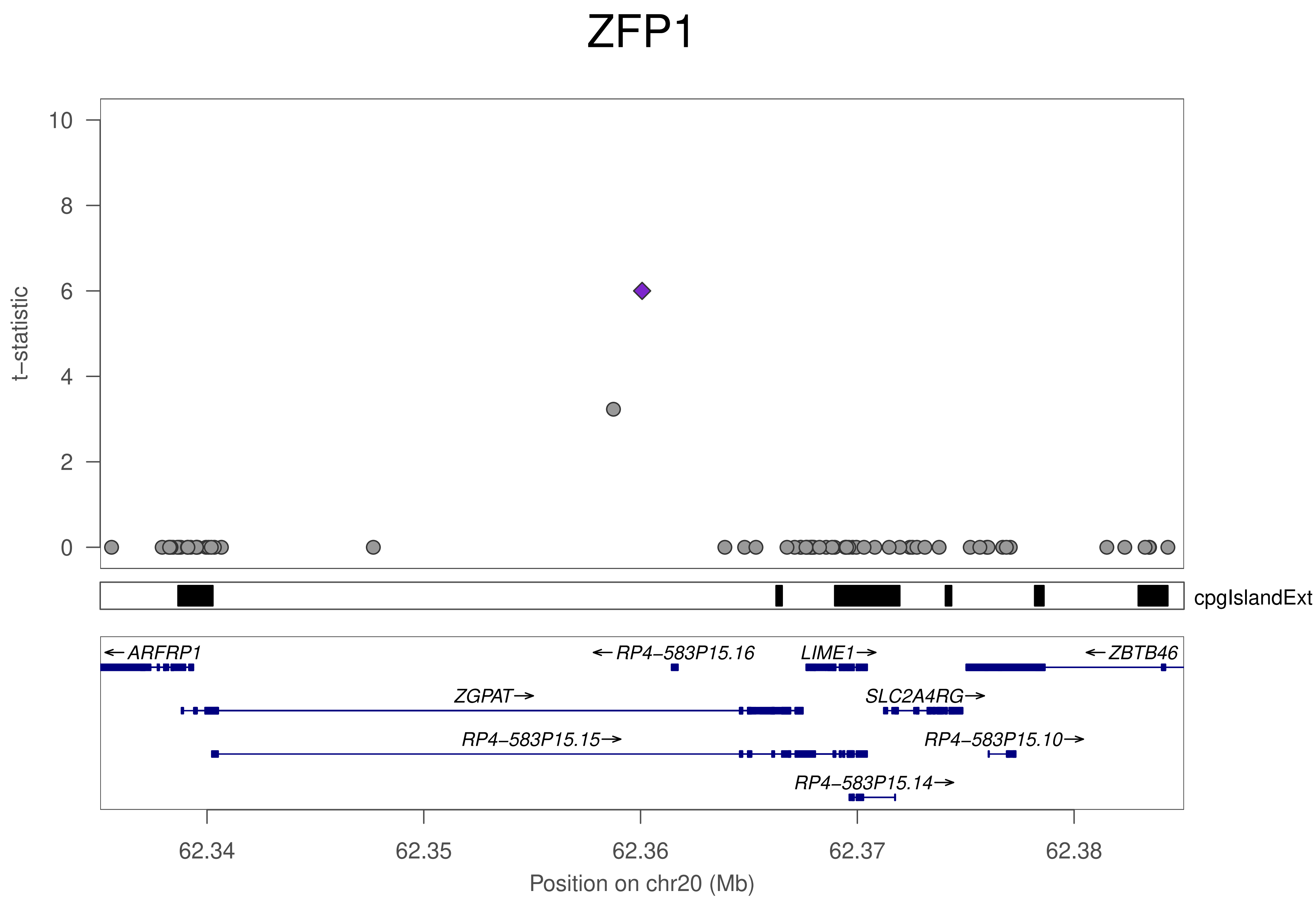

ZFP1
t−statistic
10
8
6
4
2
0
cpgIslandExt
←ARFRP1
←RP4−583P15.16
LIME1→
←ZBTB46
ZGPAT→
SLC2A4RG→
RP4−583P15.15→
RP4−583P15.10→
RP4−583P15.14→
62.34
62.35
62.36
62.37
62.38
Position on chr20 (Mb)

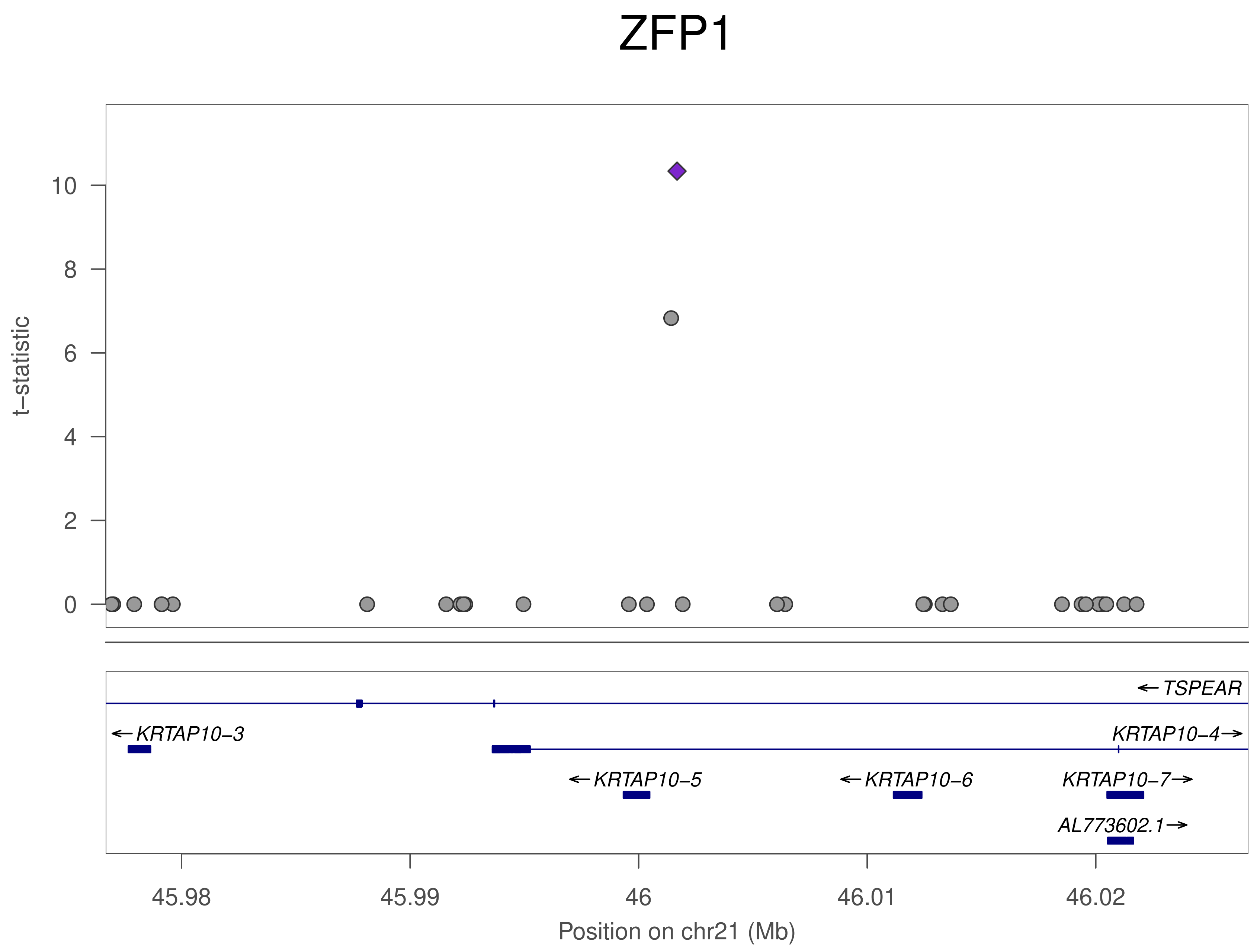

ZFP1
t−statistic
0
2
4
6
8
10
TSPEAR
KRTAP10−3
KRTAP10−4
KRTAP10−5
KRTAP10−6
KRTAP10−7
AL773602.1
45.98
45.99
46
46.01
46.02
Position on chr21 (Mb)

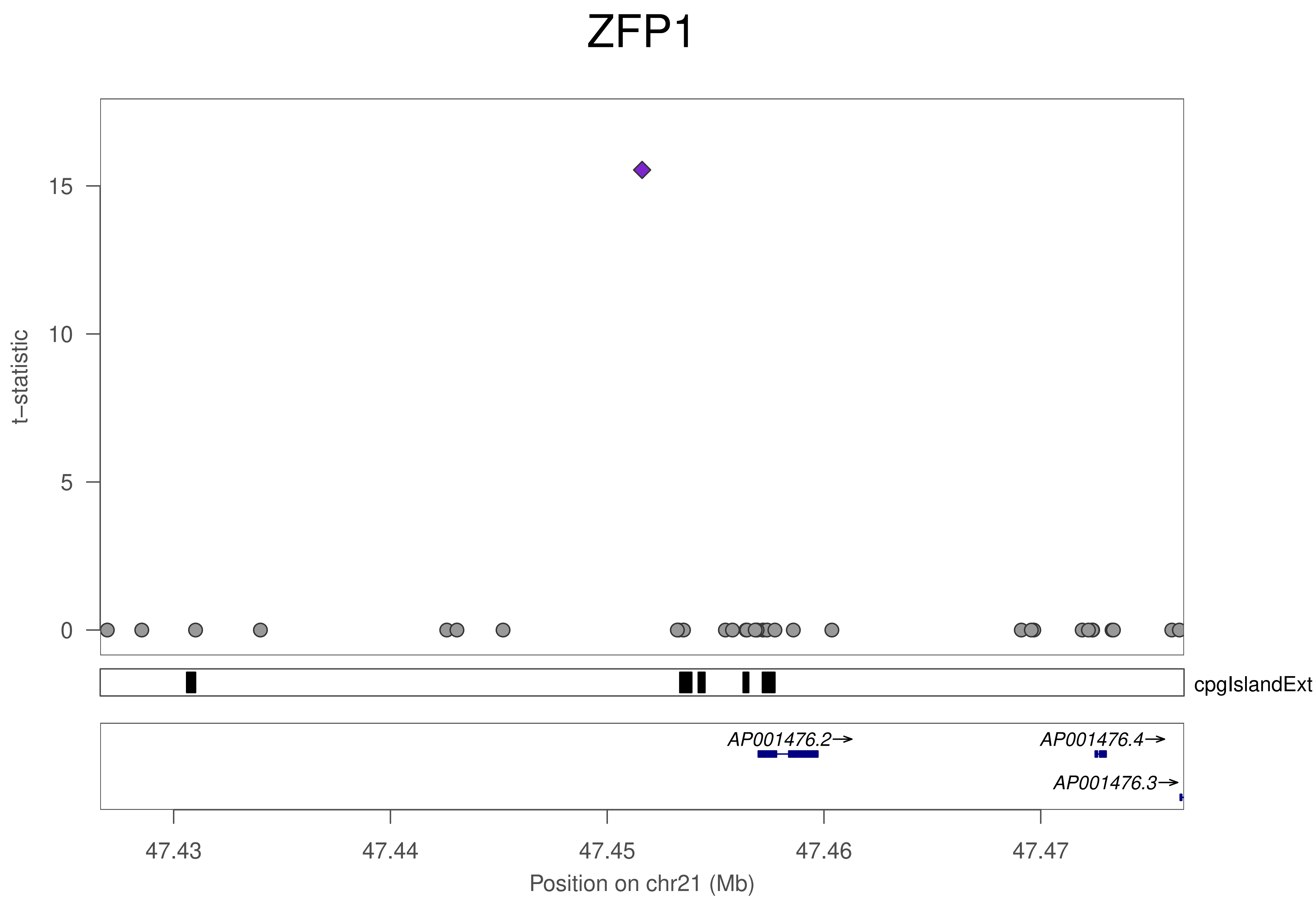

ZFP1
t−statistic
15
10
5
0
cpgIslandExt
AP001476.2→
AP001476.4→
AP001476.3→
47.43
47.44
47.45
47.46
47.47
Position on chr21 (Mb)

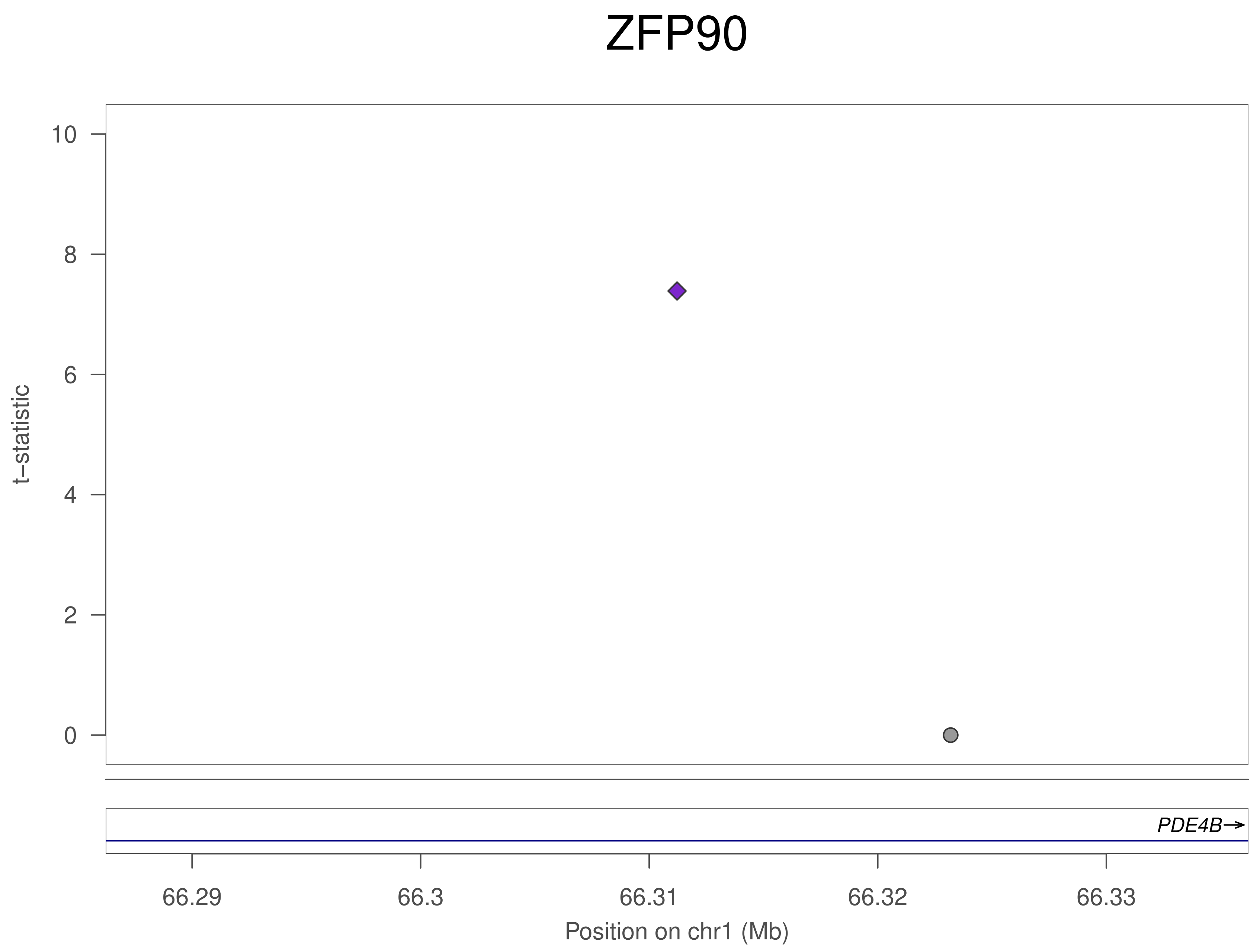

ZFP90
t−statistic
10
8
6
4
2
0
PDE4B→
66.29
66.3
66.31
66.32
66.33
Position on chr1 (Mb)

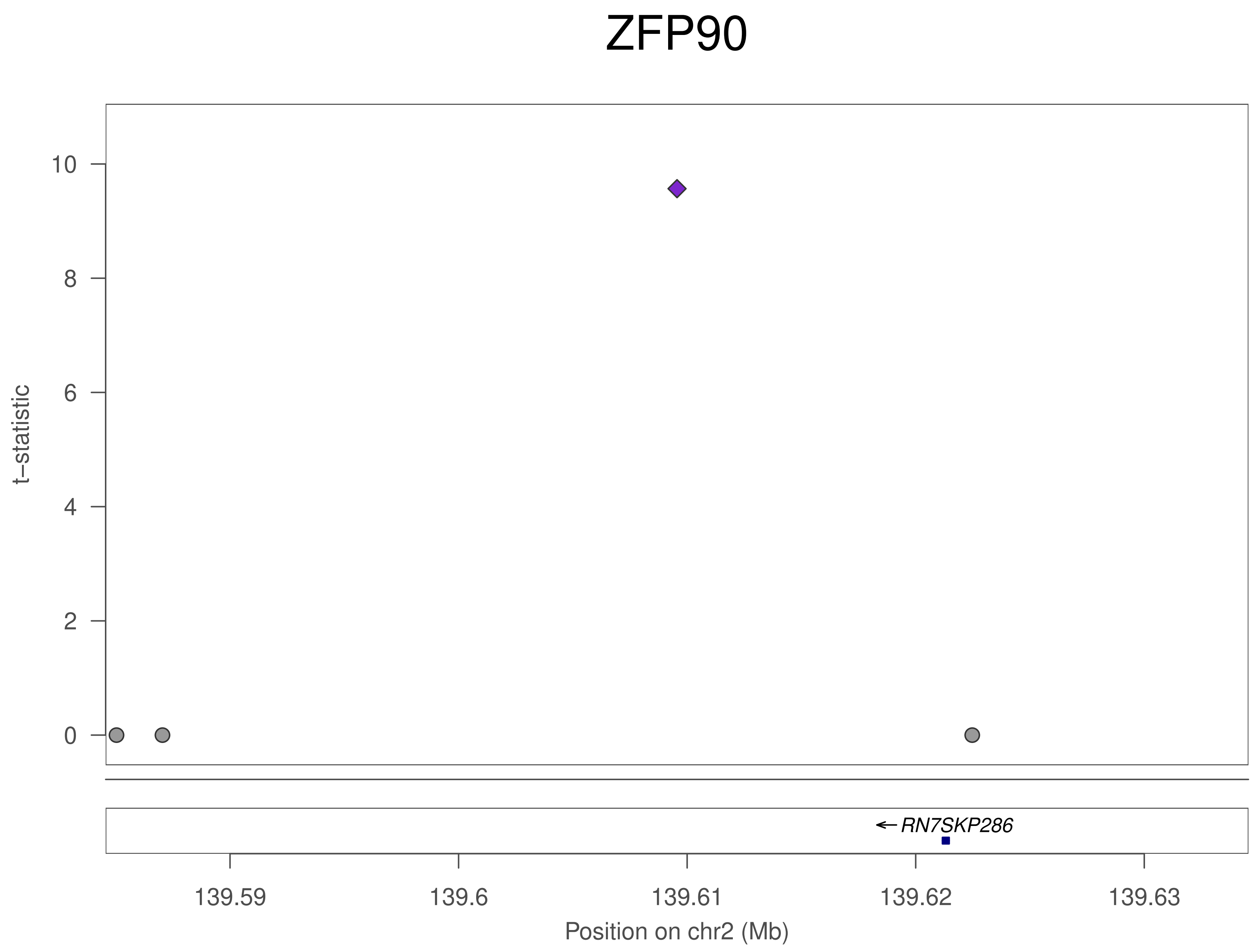

ZFP90
t–statistic
10
8
6
4
2
0
RN7SKP286
139.59
139.6
139.61
139.62
139.63
Position on chr2 (Mb)

### ZFP90

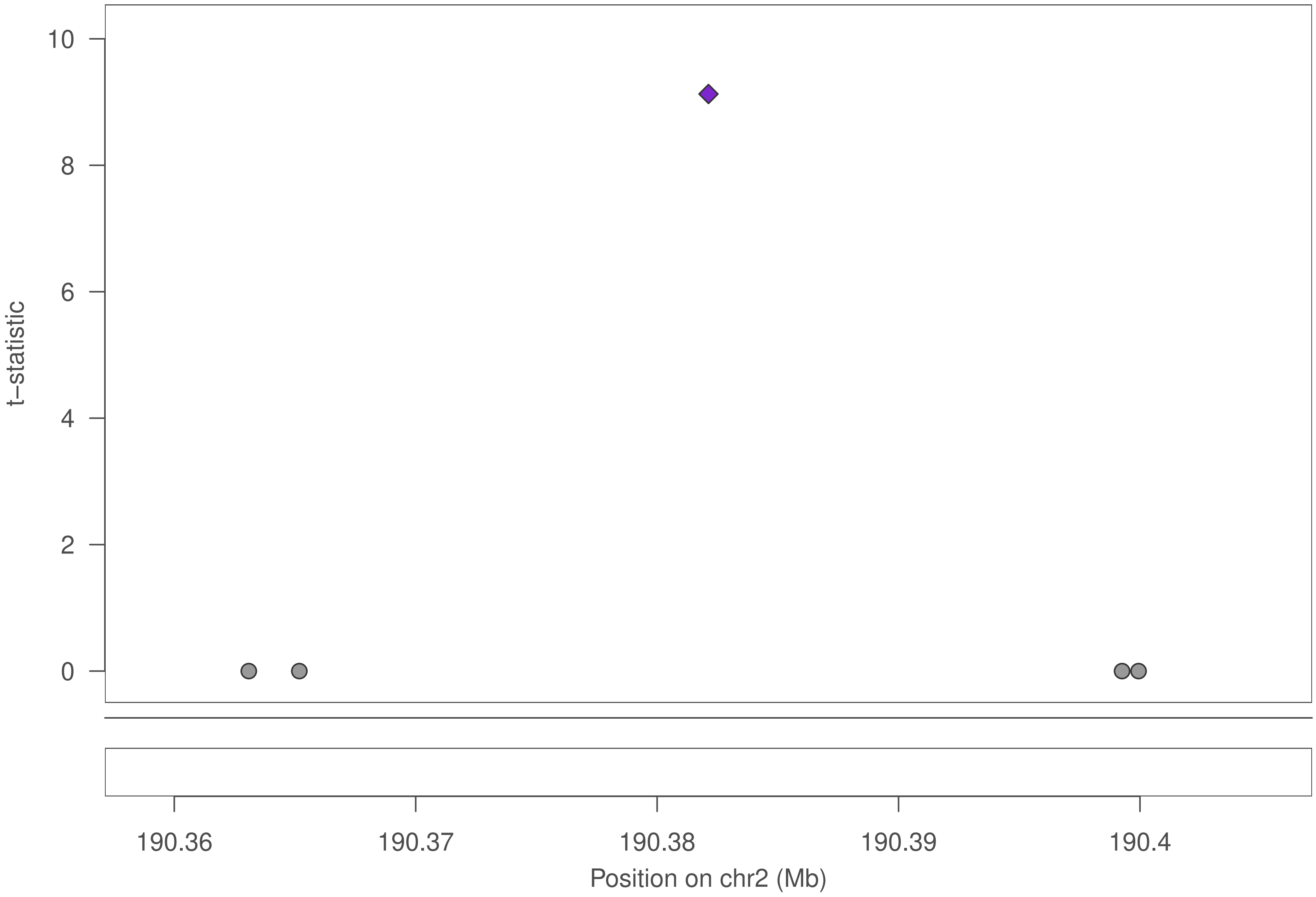

10
8
6
4
2
0
t−statistic
190.36
190.37
190.38
190.39
190.4
Position on chr2 (Mb)

### ZFP90

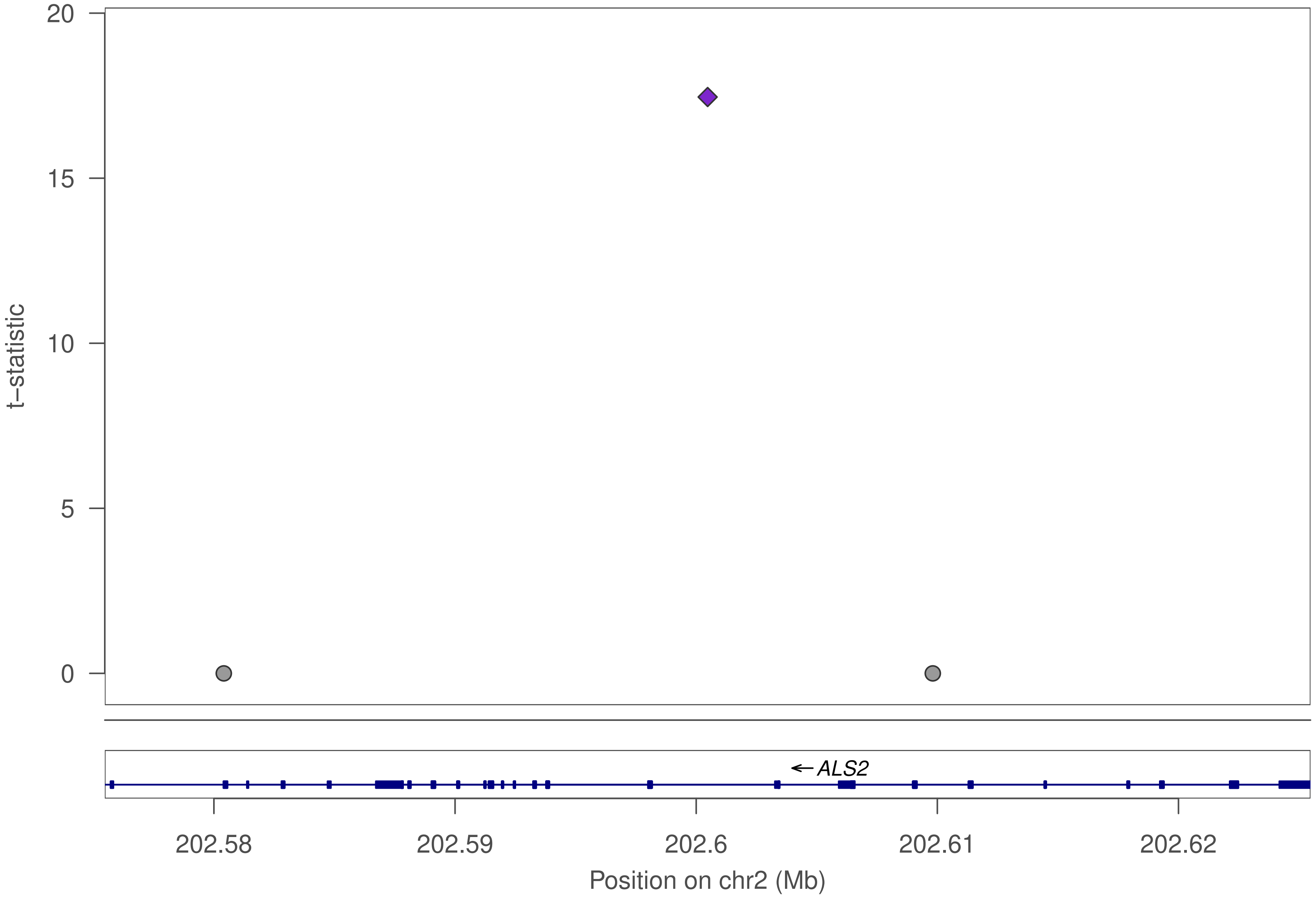

20
15
10
5
0
t−statistic
ALS2
202.58
202.59
202.6
202.61
202.62
Position on chr2 (Mb)

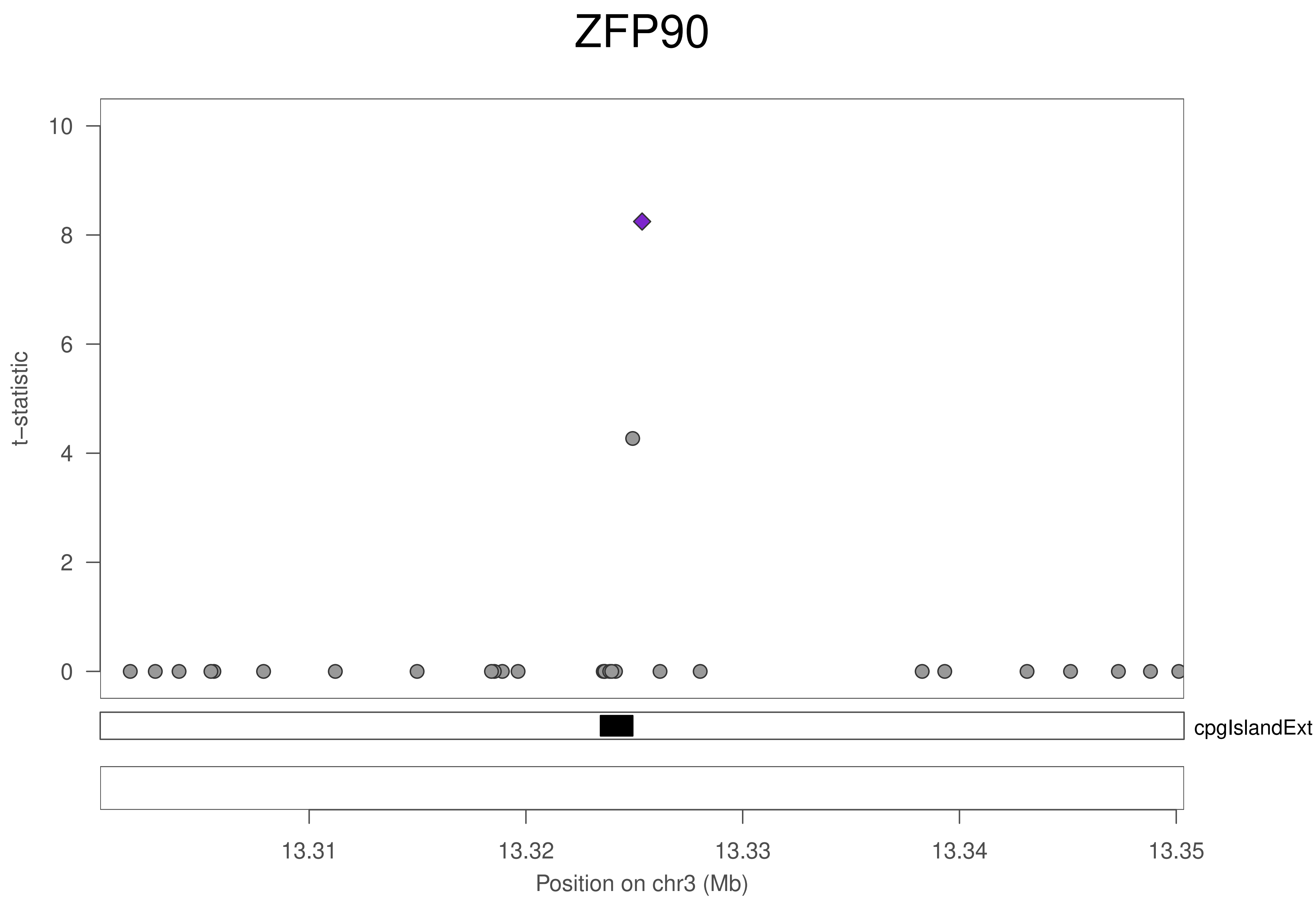
ZFP90
t−statistic
0
2
4
6
8
10
cpgIslandExt
13.31
13.32
13.33
13.34
13.35
Position on chr3 (Mb)

### ZFP90

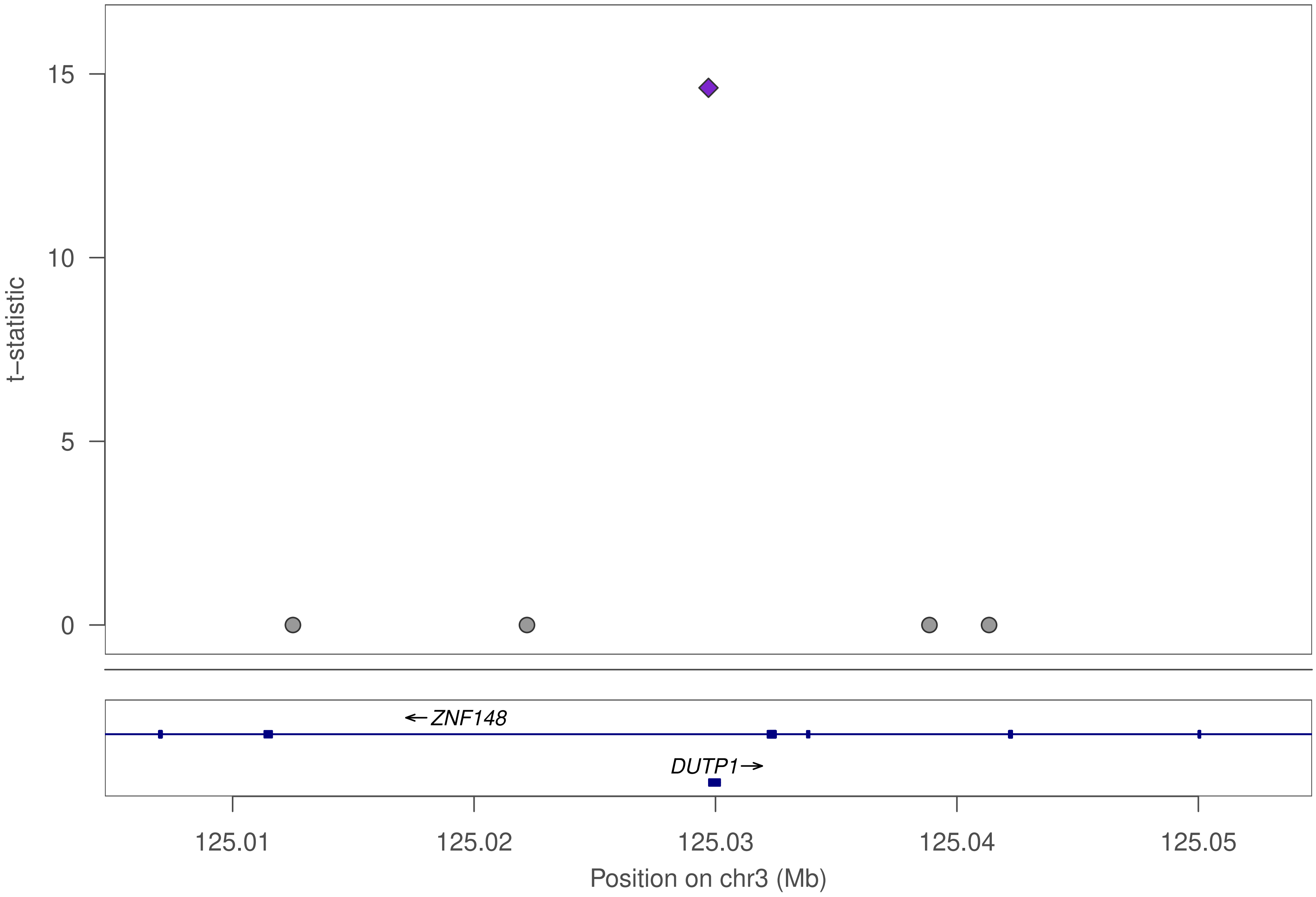

t–statistic
15
10
5
0
←ZNF148
DUTP1→
125.01
125.02
125.03
125.04
125.05
Position on chr3 (Mb)

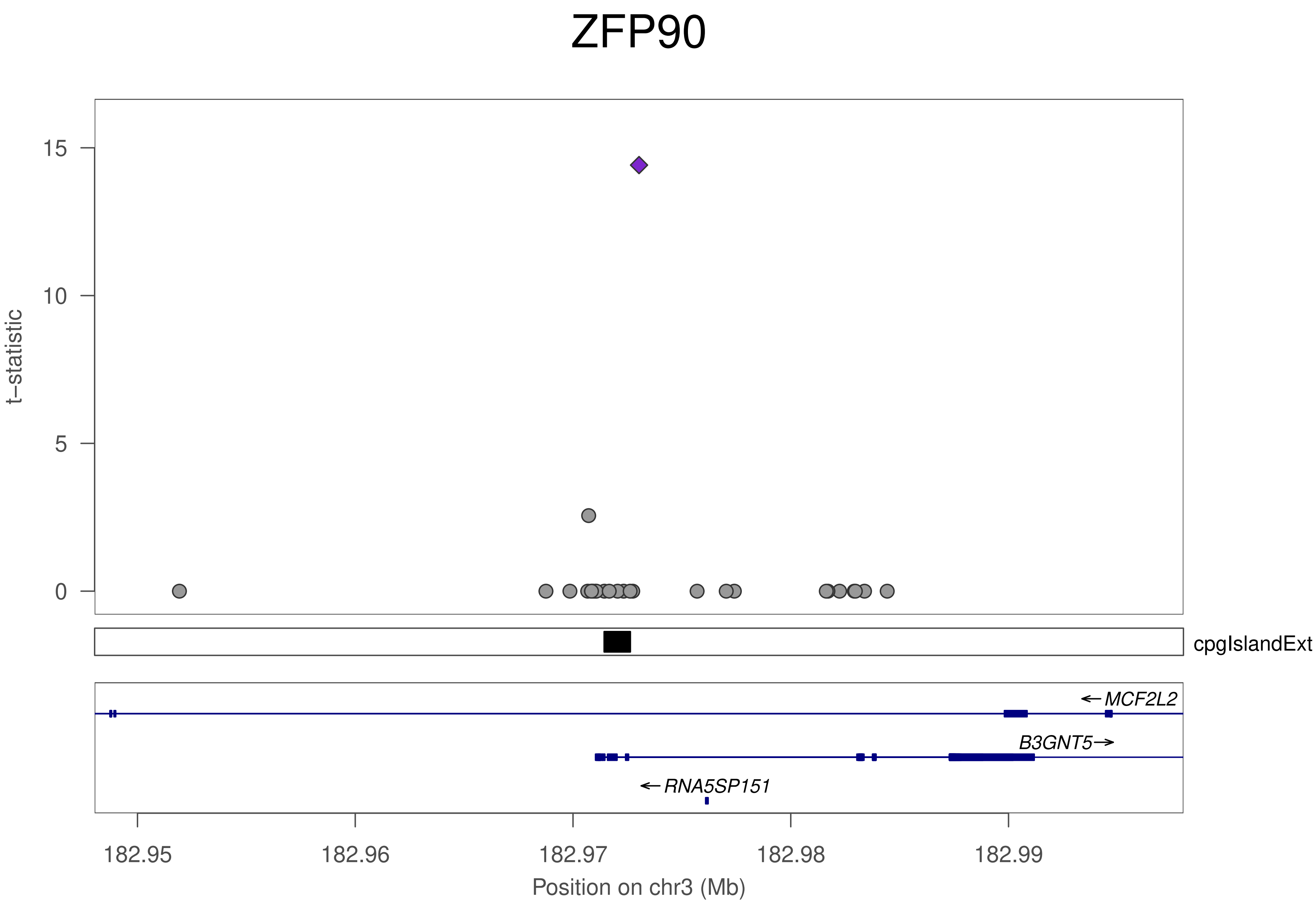

ZFP90
t−statistic
15
10
5
0
cpgIslandExt
MCF2L2
B3GNT5
RNA5SP151
182.95
182.96
182.97
182.98
182.99
Position on chr3 (Mb)

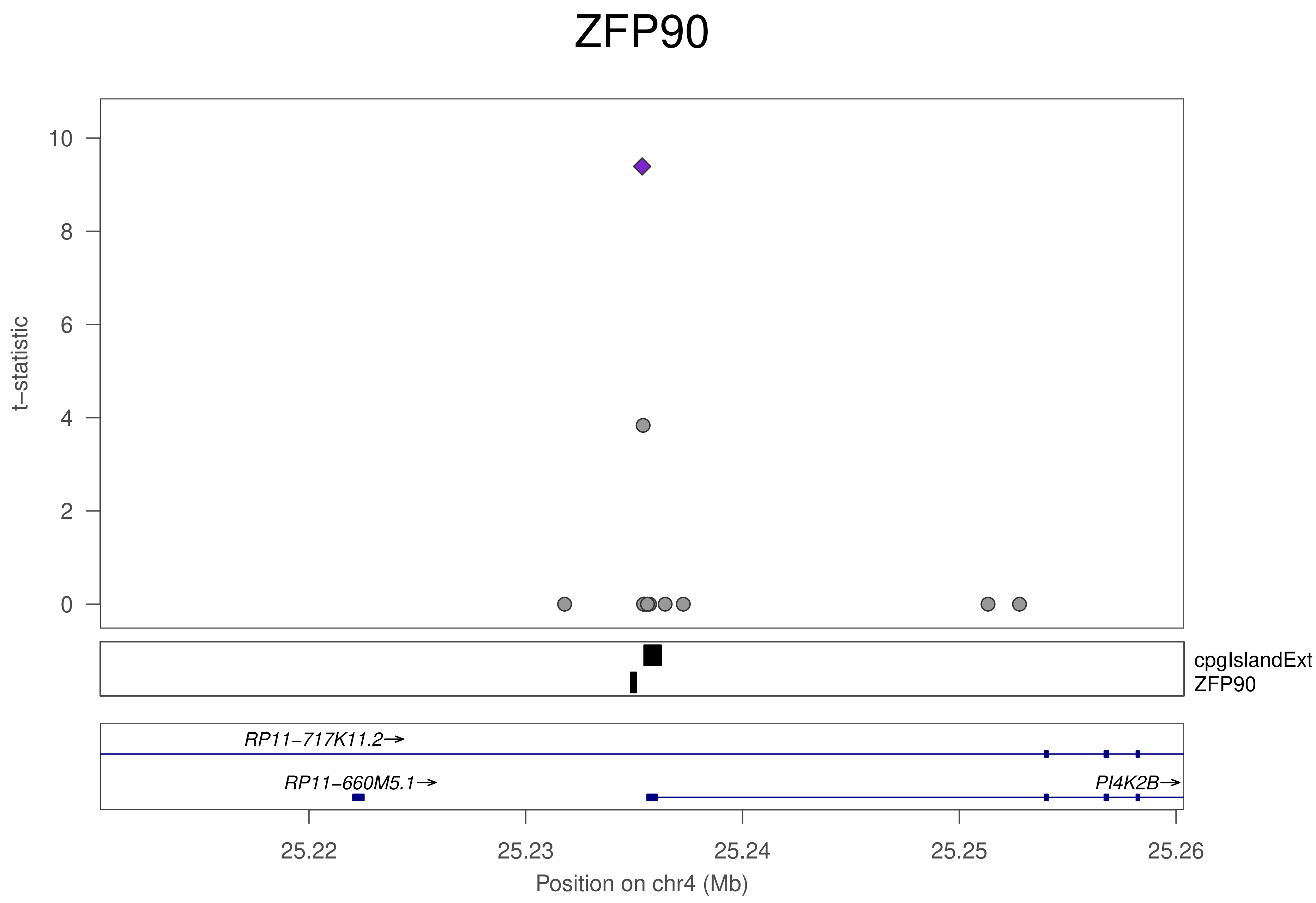
ZFP90
t−statistic
10
8
6
4
2
0
cpgIslandExt
ZFP90
RP11−717K11.2→
RP11−660M5.1→
PI4K2B→
25.22
25.23
25.24
25.25
25.26
Position on chr4 (Mb)

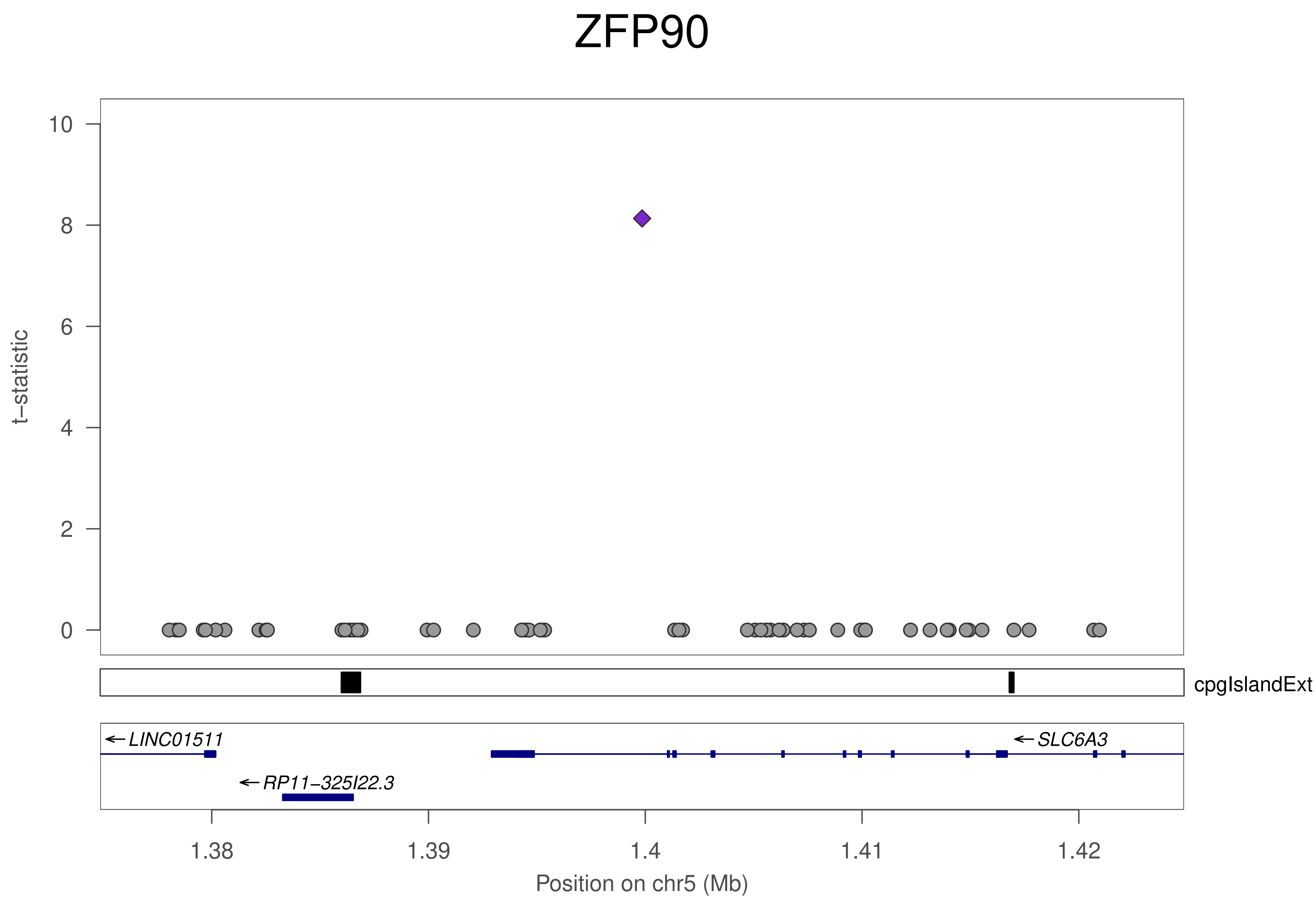

ZFP90
t−statistic
10
8
6
4
2
0
cpgIslandExt
LINC01511
RP11−325I22.3
SLC6A3
1.38
1.39
1.4
1.41
1.42
Position on chr5 (Mb)

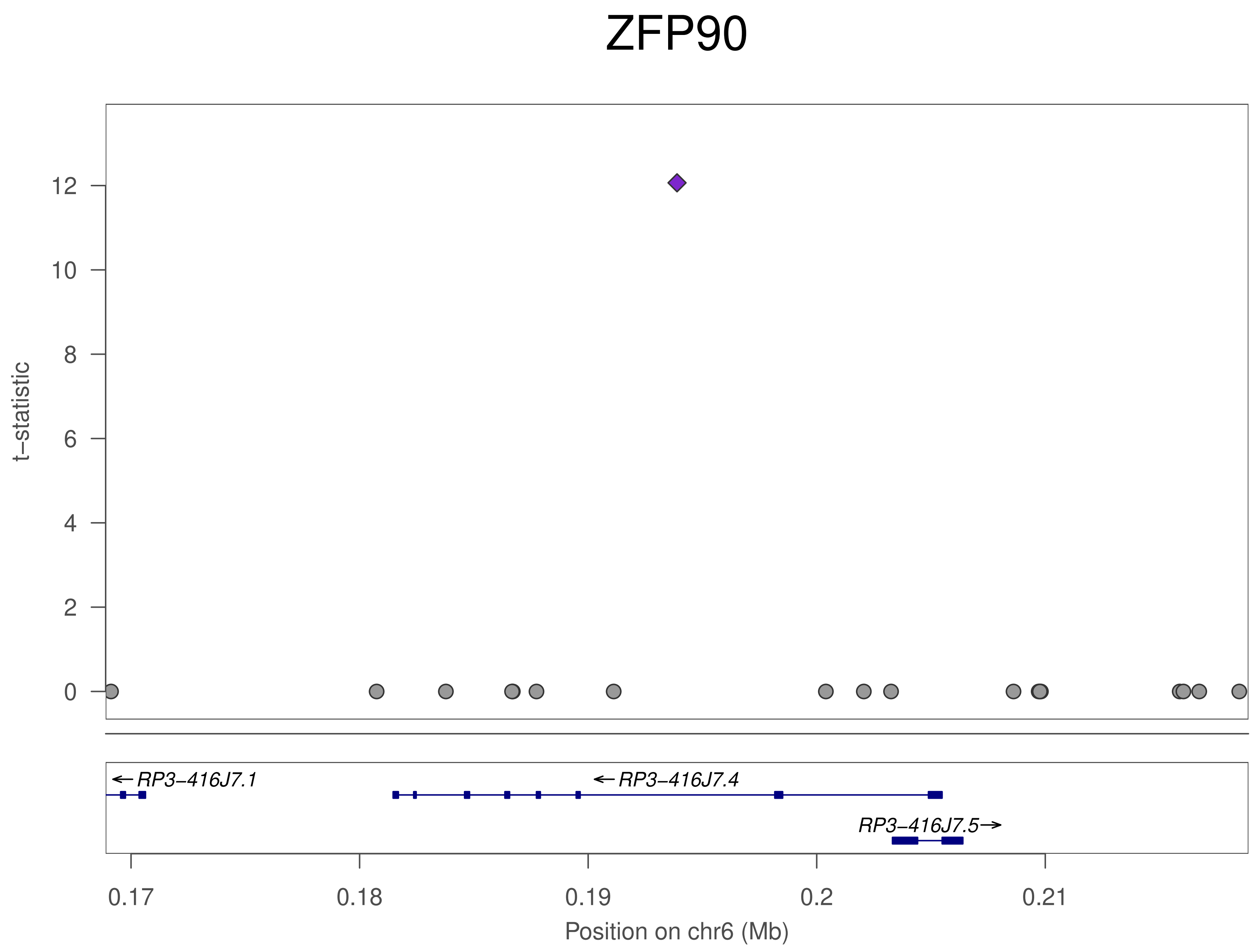

ZFP90
t–statistic
12
10
8
6
4
2
0
←RP3–416J7.1
←RP3–416J7.4
RP3–416J7.5→
0.17
0.18
0.19
0.2
0.21
Position on chr6 (Mb)

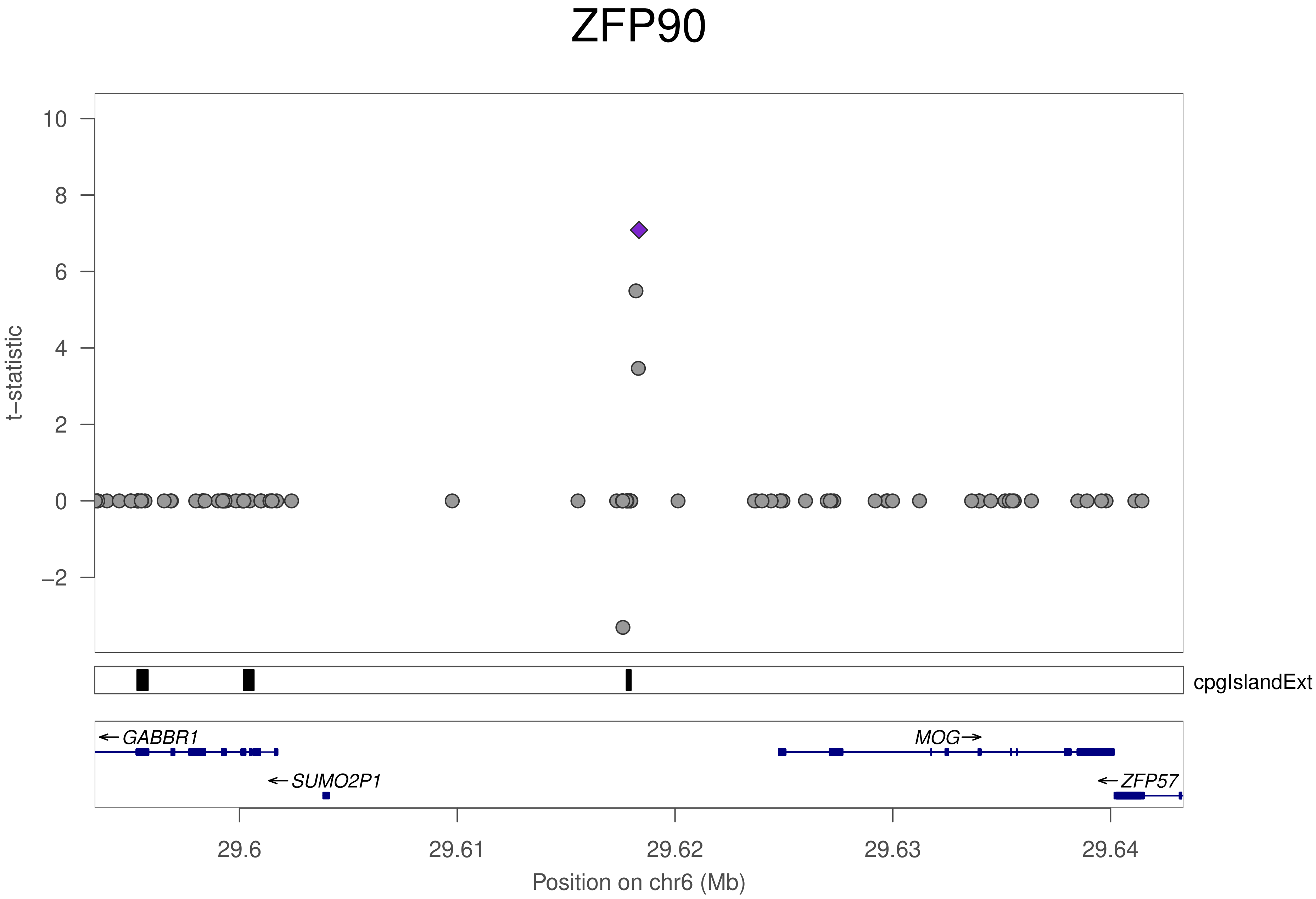
ZFP90
t−statistic
10
8
6
4
2
0
−2
cpgIslandExt
←GABBR1
MOG→
←SUMO2P1
←ZFP57
29.6
29.61
29.62
29.63
29.64
Position on chr6 (Mb)

### ZFP90

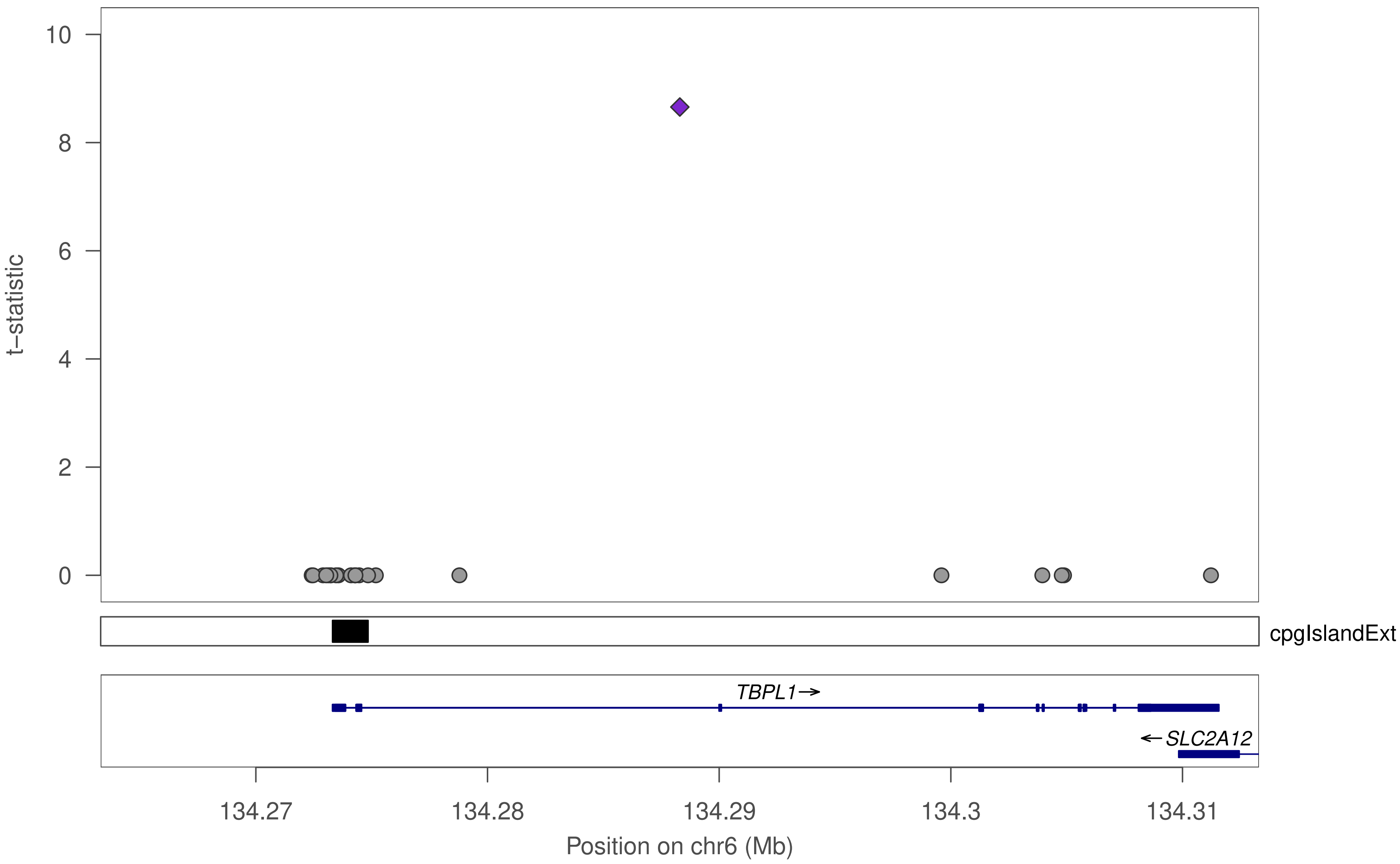

t−statistic
10
8
6
4
2
0
cpgIslandExt
TBPL1→
←SLC2A12
134.27
134.28
134.29
134.3
134.31
Position on chr6 (Mb)

### ZFP90

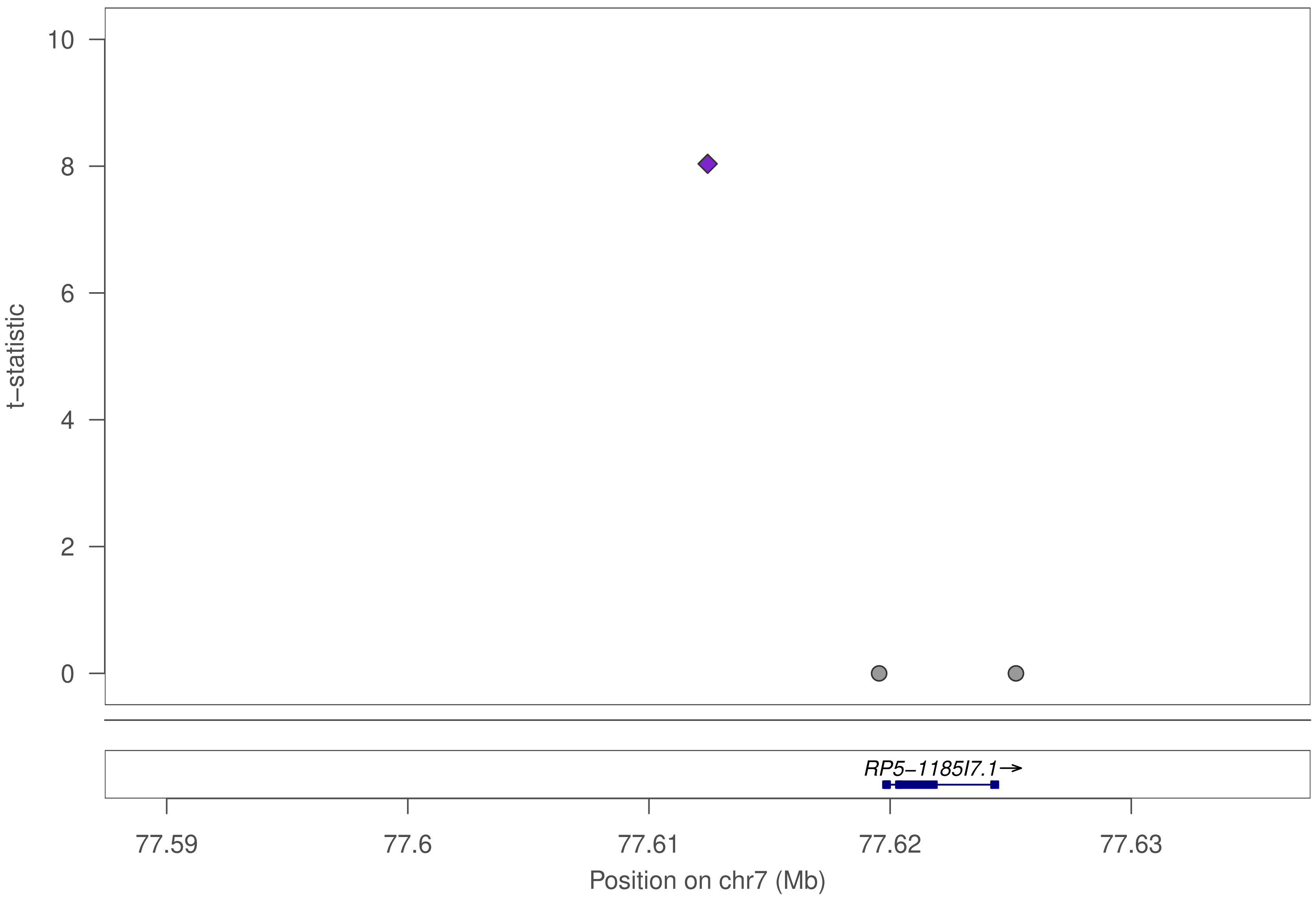
10
8
6
4
2
0
t−statistic
RP5−1185I7.1→
77.59
77.6
77.61
77.62
77.63
Position on chr7 (Mb)

### ZFP90

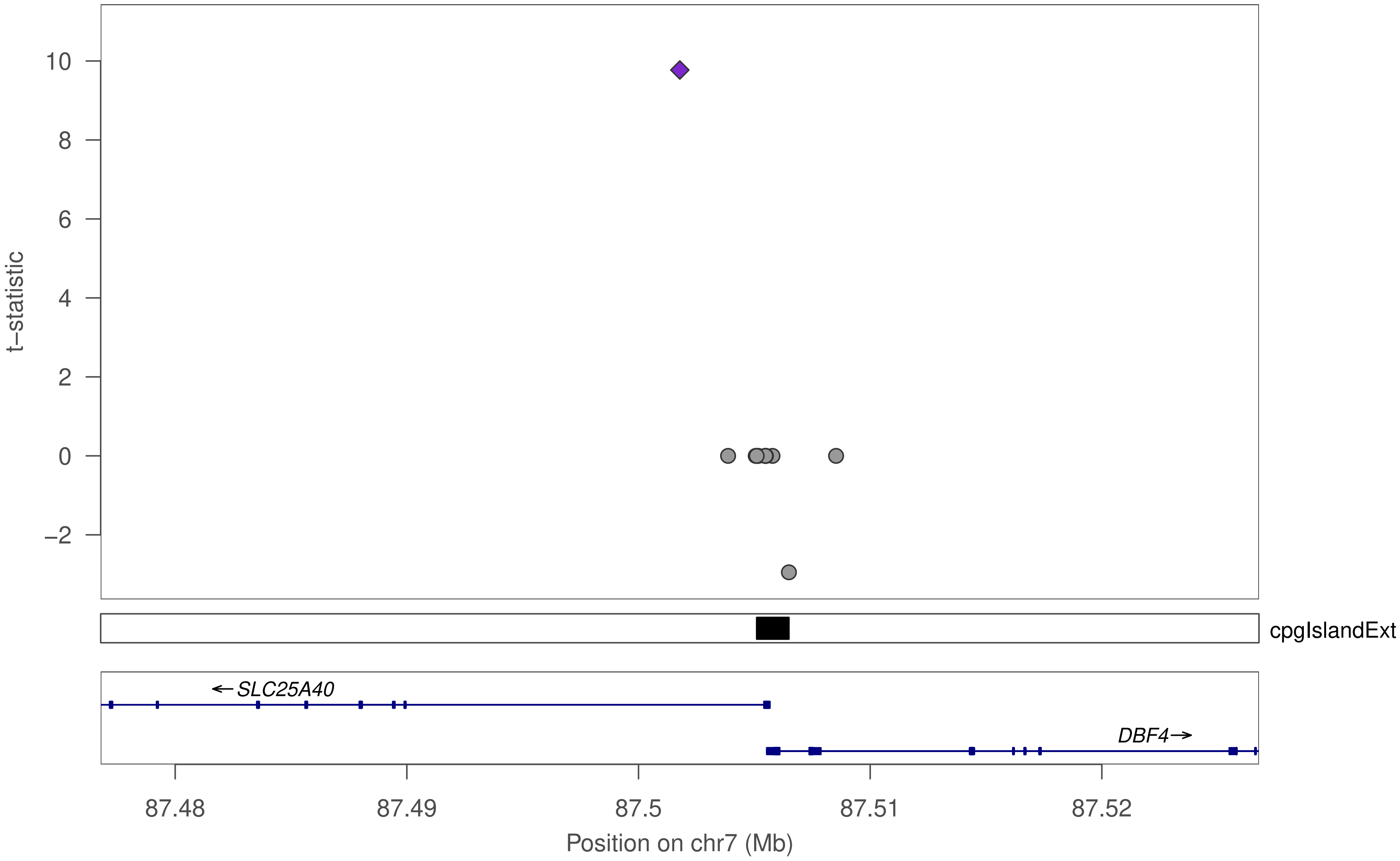

10
8
6
4
2
0
−2
t−statistic
cpgIslandExt
←SLC25A40
DBF4→
87.48
87.49
87.5
87.51
87.52
Position on chr7 (Mb)

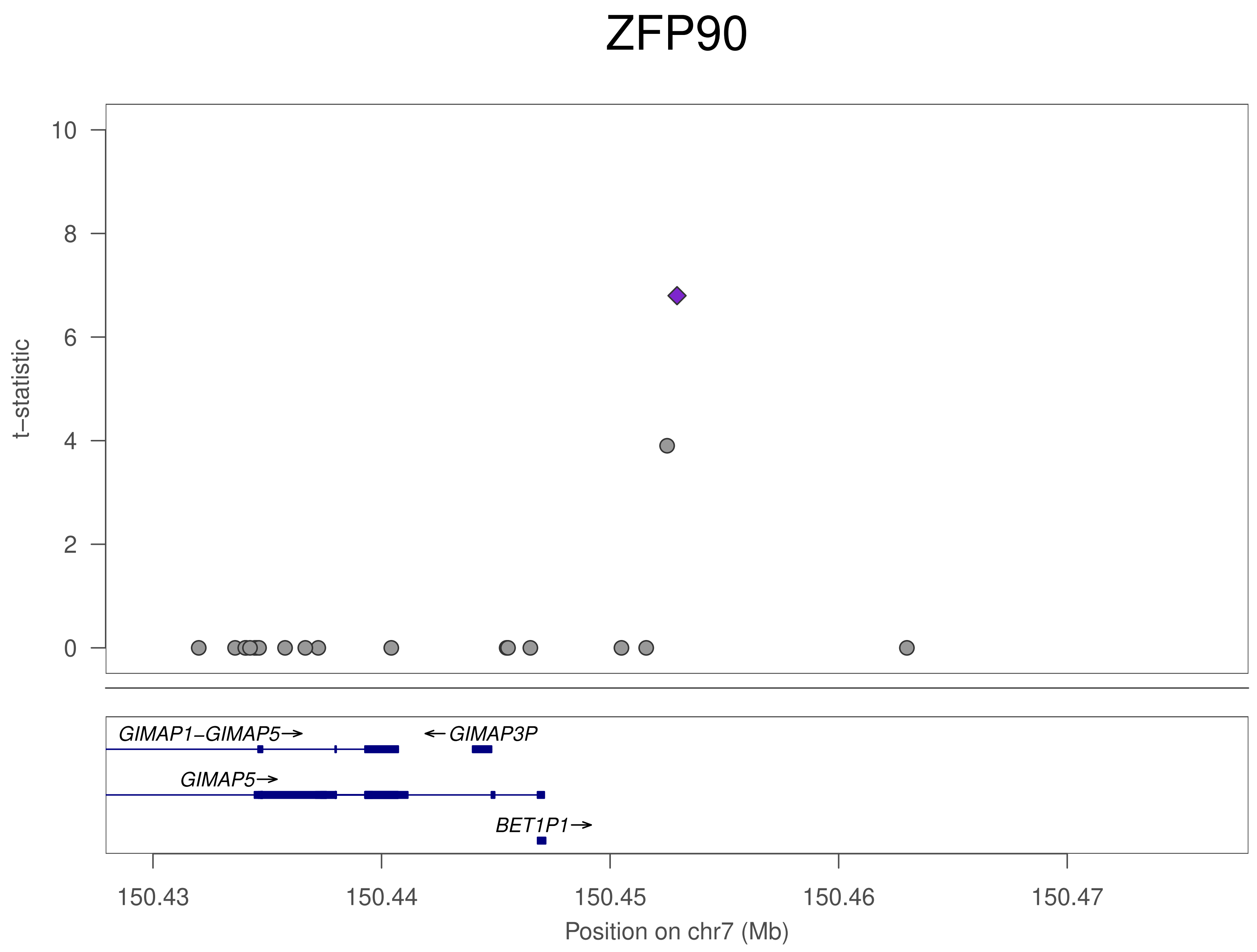
ZFP90
t−statistic
10
8
6
4
2
0
GIMAP1−GIMAP5→
←GIMAP3P
GIMAP5→
BET1P1→
150.43
150.44
150.45
150.46
150.47
Position on chr7 (Mb)

### ZFP90

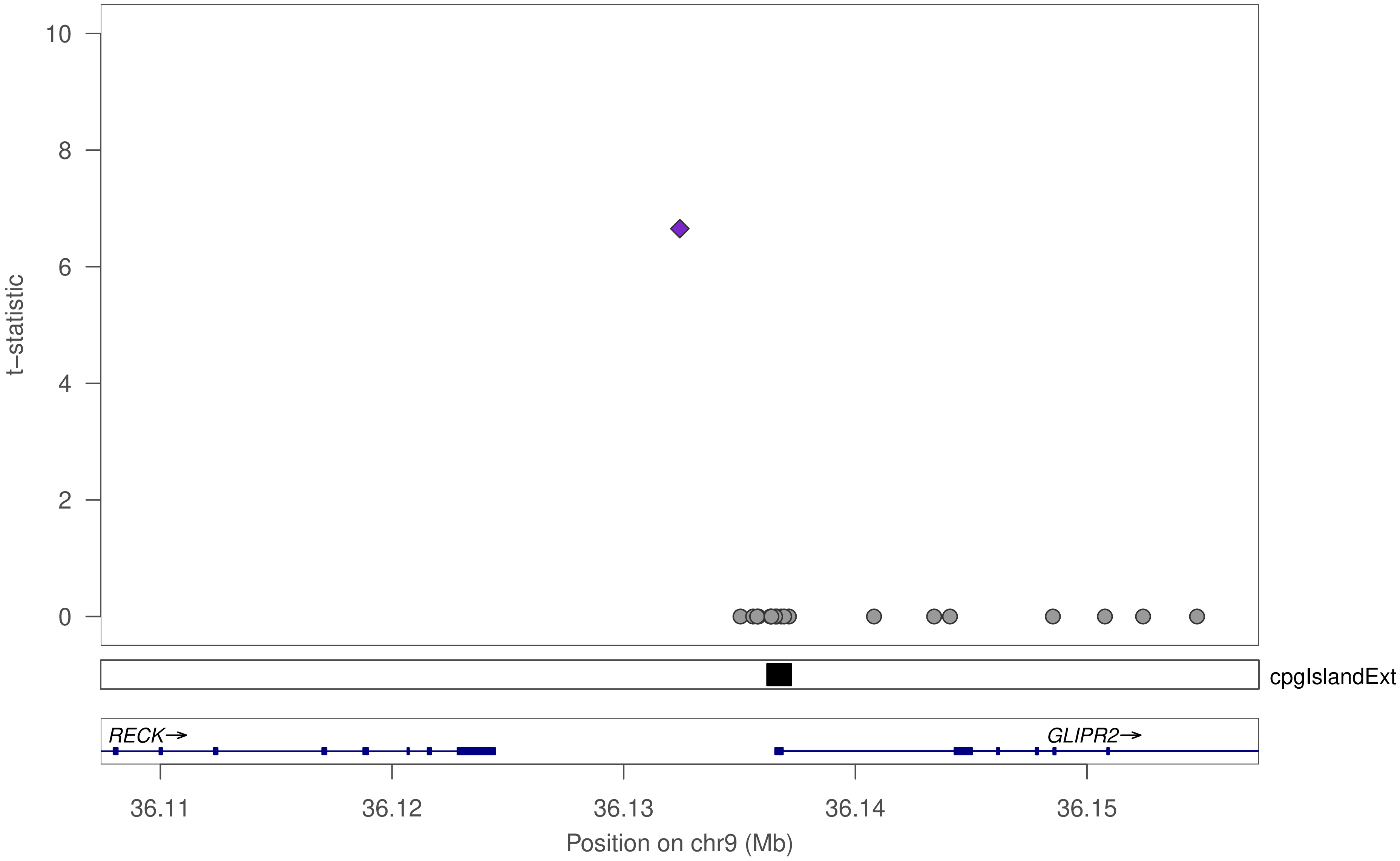

10
8
6
4
2
0
t–statistic
cpgIslandExt
RECK→
GLIPR2→
36.11
36.12
36.13
36.14
36.15
Position on chr9 (Mb)

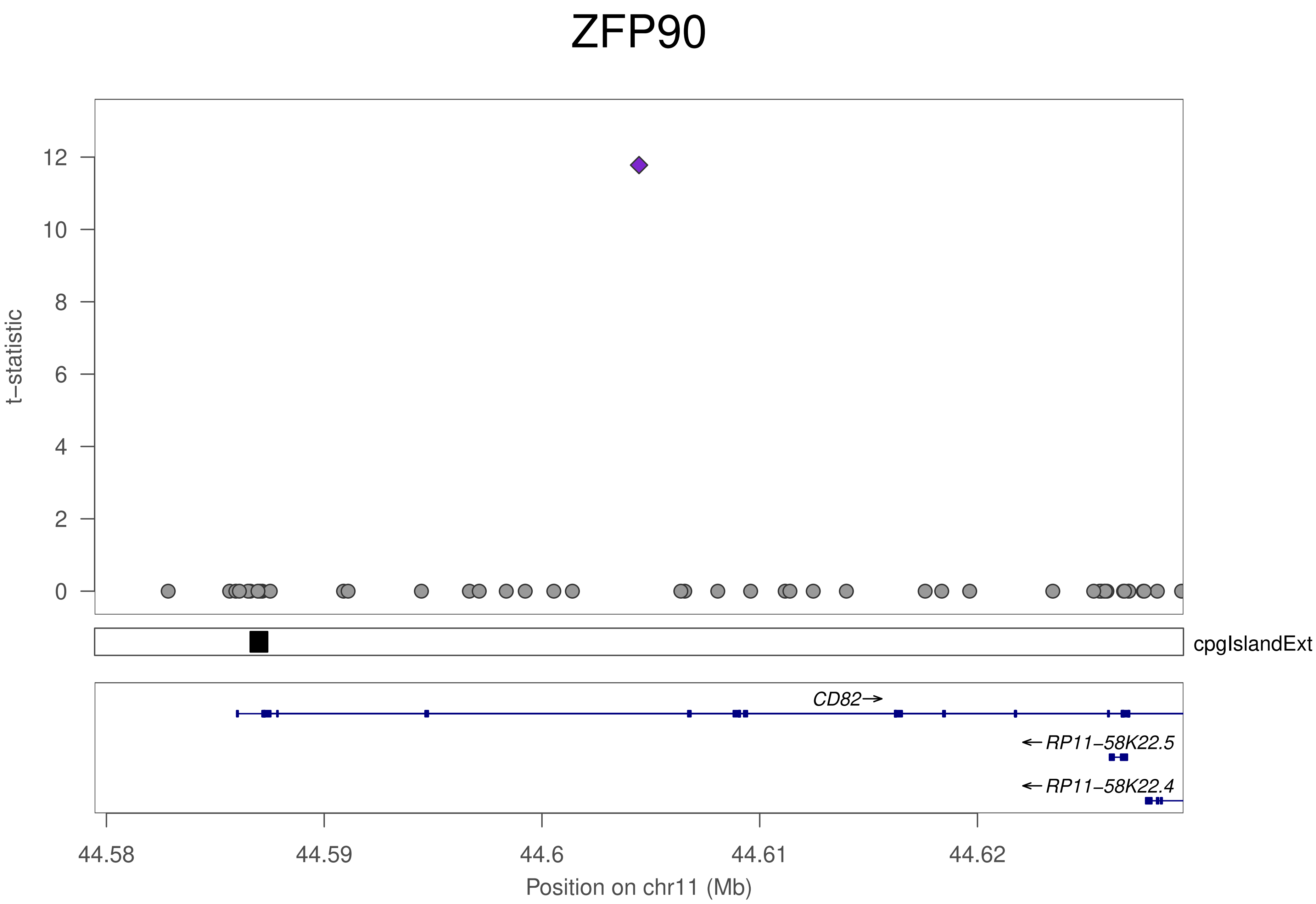

ZFP90
t–statistic
12
10
8
6
4
2
0
cpgIslandExt
CD82→
←RP11–58K22.5
←RP11–58K22.4
44.58
44.59
44.6
44.61
44.62
Position on chr11 (Mb)

### ZFP90

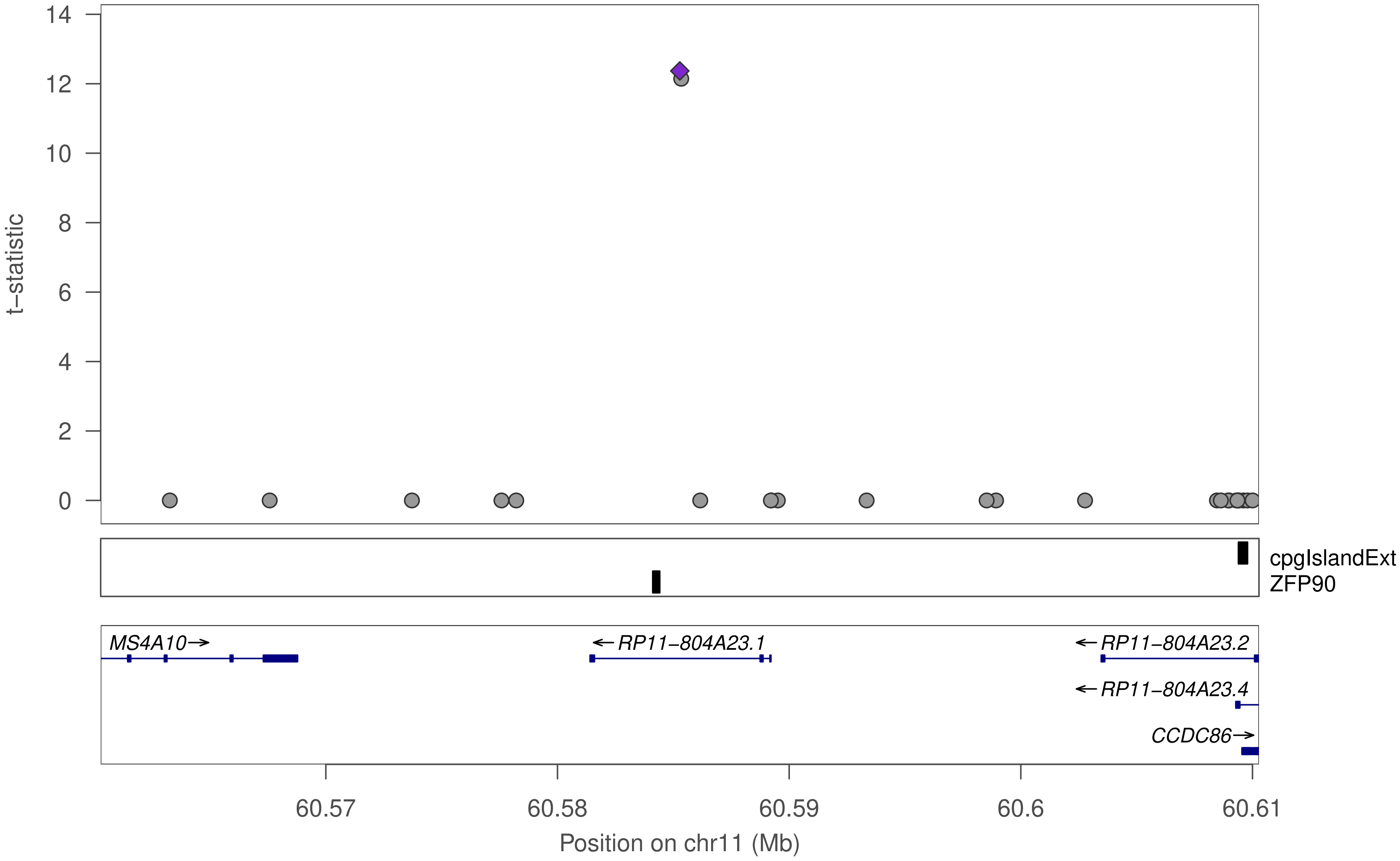

t−statistic
14
12
10
8
6
4
2
0
cpgIslandExt
ZFP90
MS4A10→
←RP11−804A23.1
←RP11−804A23.2
←RP11−804A23.4
CCDC86→
60.57
60.58
60.59
60.6
60.61
Position on chr11 (Mb)

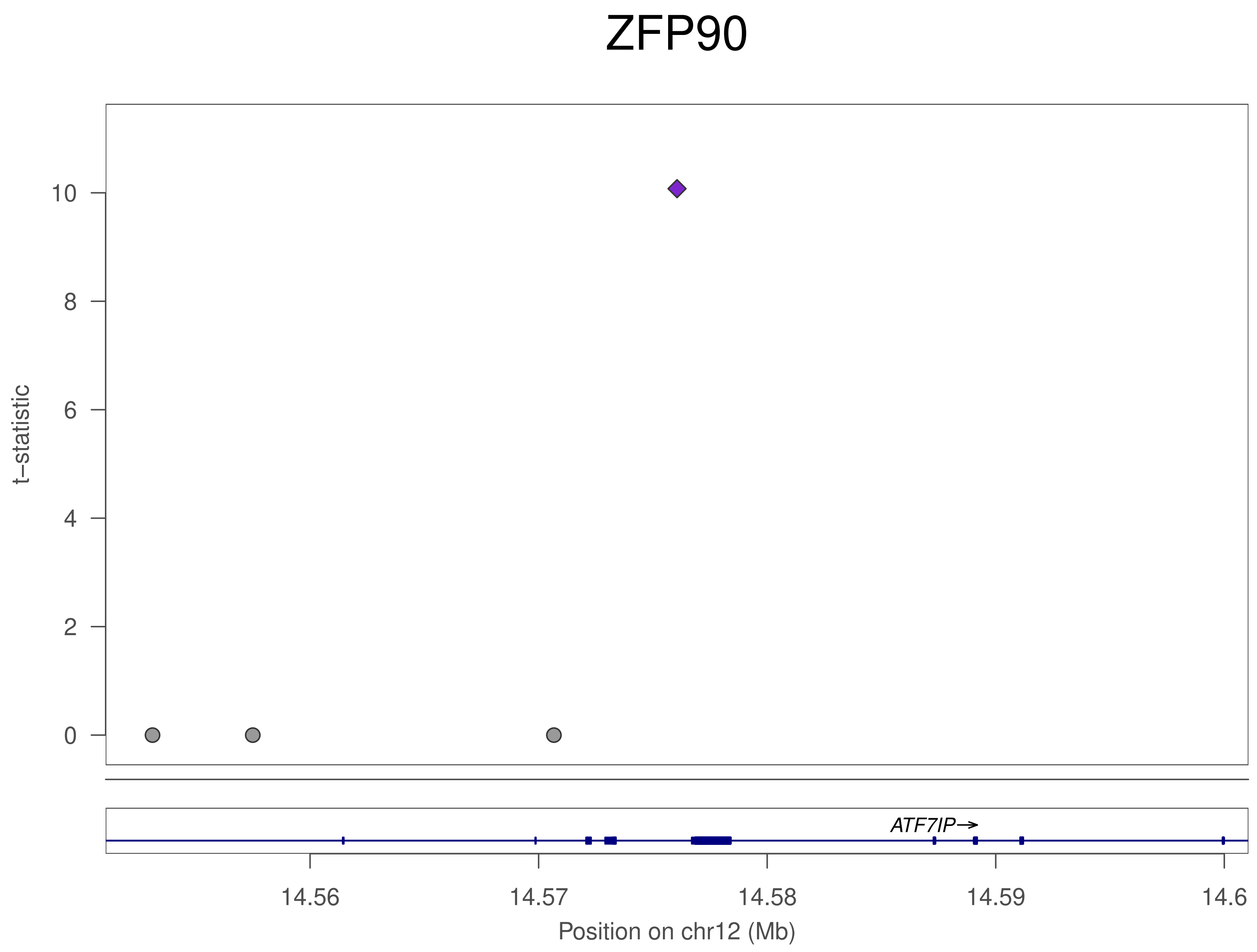
ZFP90
t−statistic
10
8
6
4
2
0
ATF7IP→
14.56
14.57
14.58
14.59
14.6
Position on chr12 (Mb)

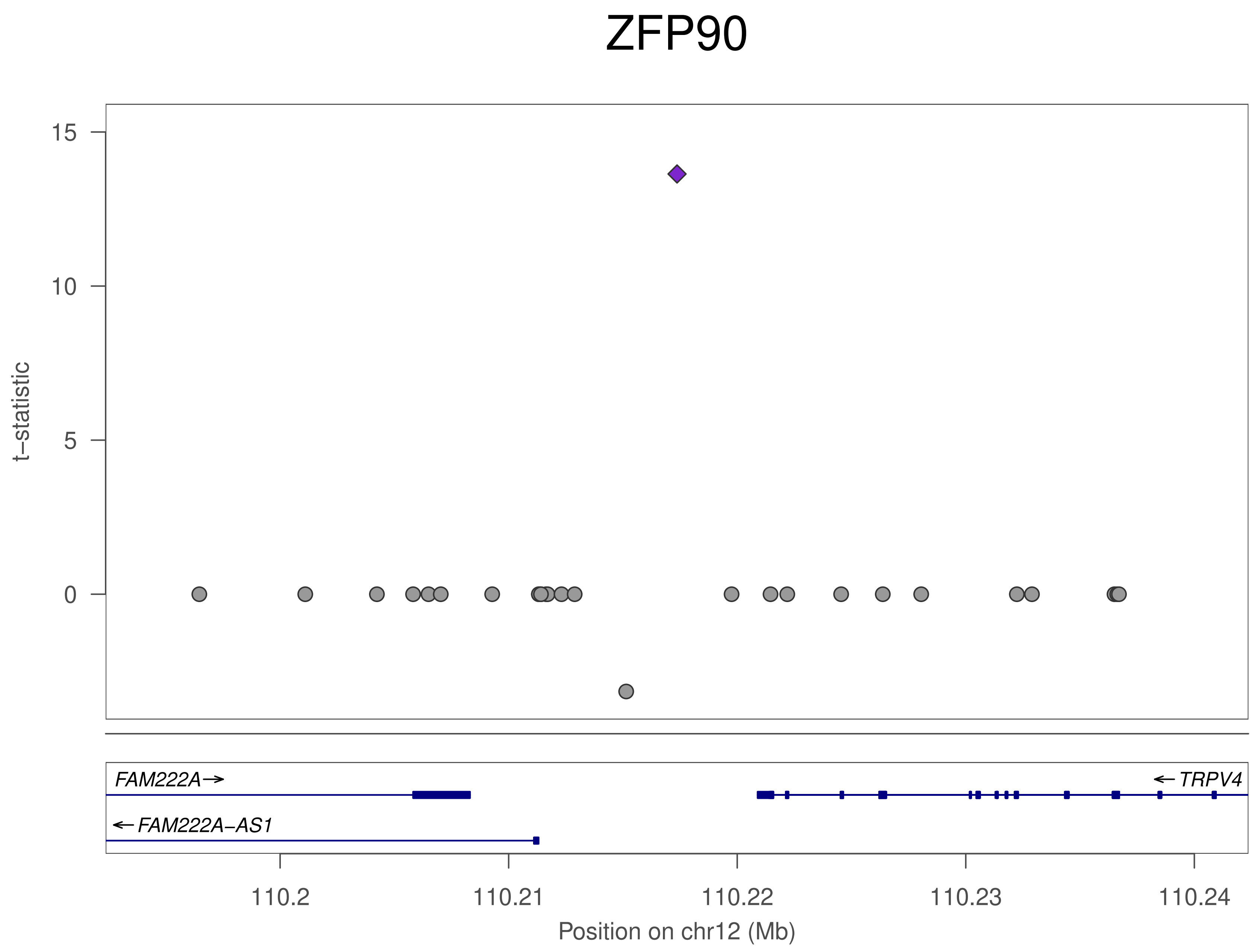
ZFP90
15
10
5
0
t−statistic
FAM222A→
←TRPV4
←FAM222A−AS1
110.2
110.21
110.22
110.23
110.24
Position on chr12 (Mb)

ZFP90

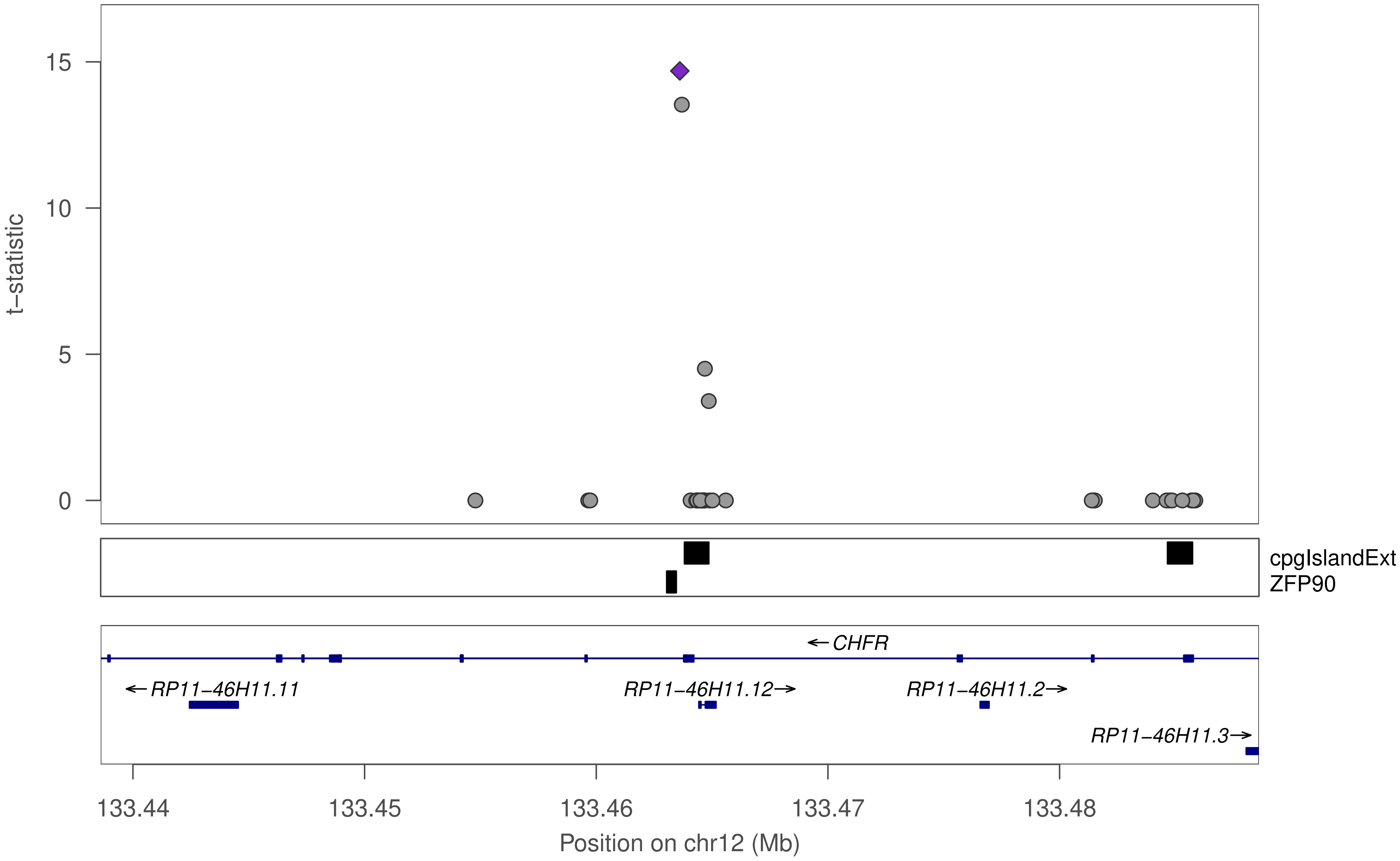

t−statistic
15
10
5
0
cpgIslandExt
ZFP90
←CHFR
←RP11−46H11.11
RP11−46H11.12→
RP11−46H11.2→
RP11−46H11.3→
133.44
133.45
133.46
133.47
133.48
Position on chr12 (Mb)

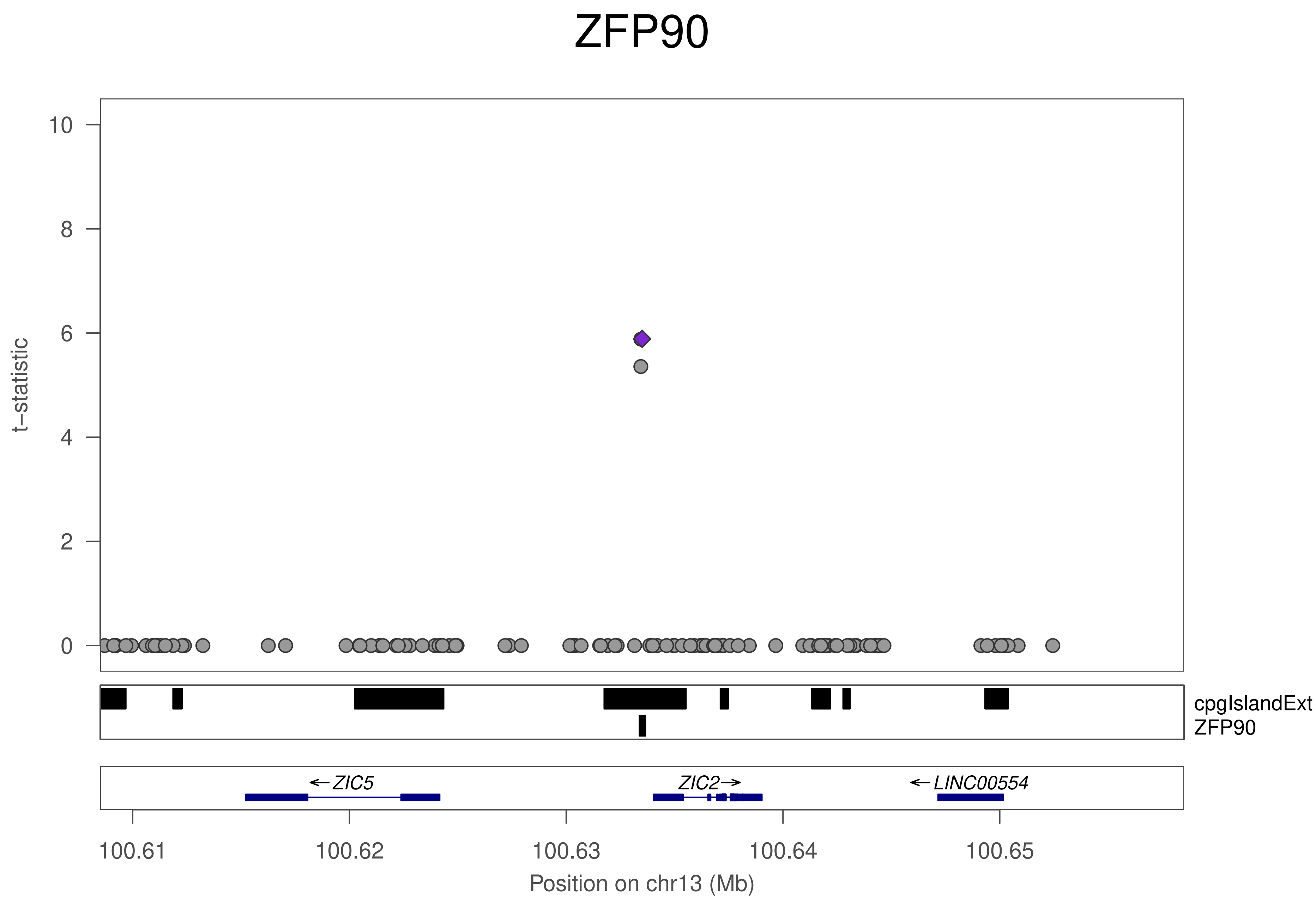

ZFP90
t−statistic
10
8
6
4
2
0
cpgIslandExt
ZFP90
←ZIC5
ZIC2→
←LINC00554
100.61
100.62
100.63
100.64
100.65
Position on chr13 (Mb)

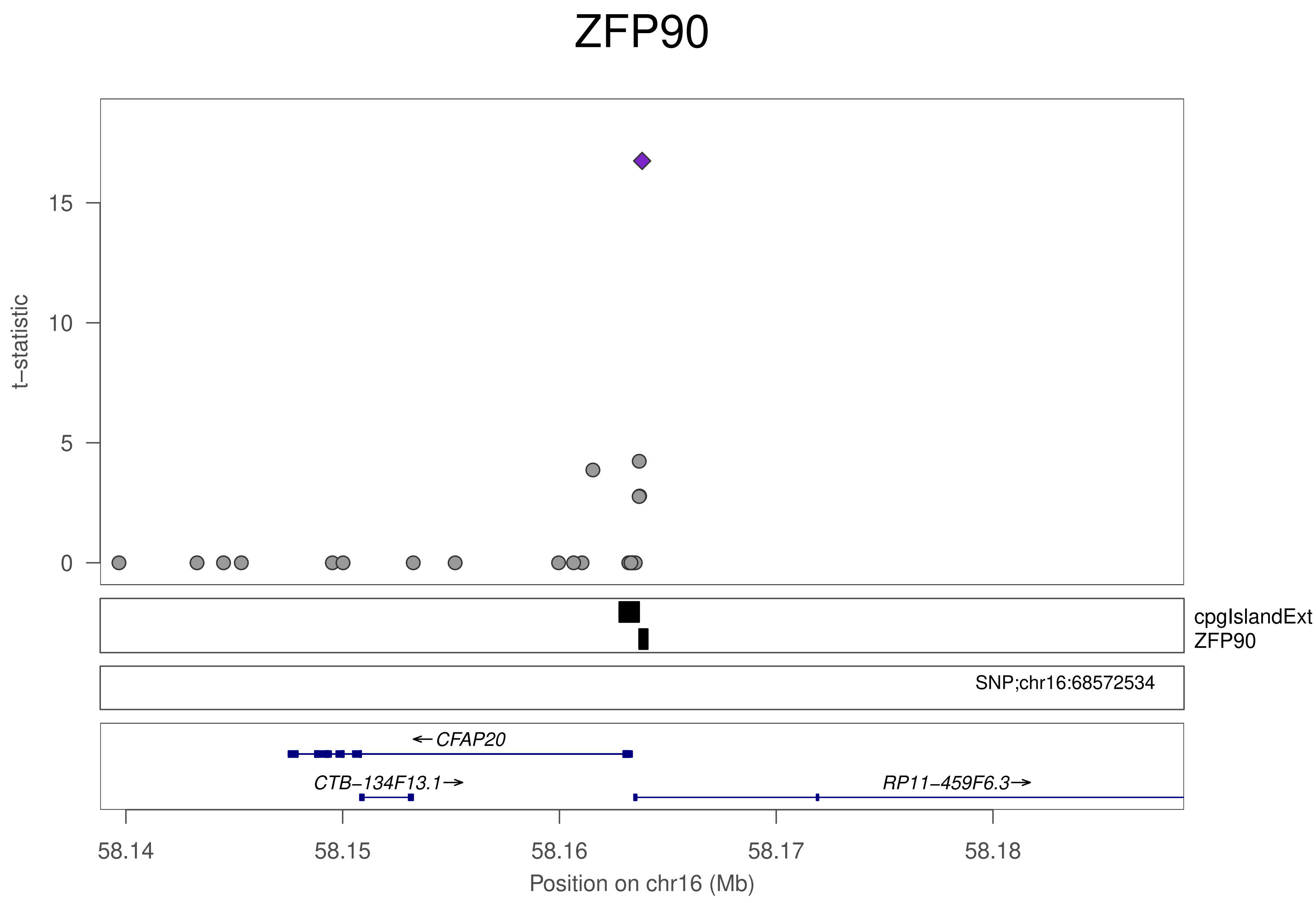
ZFP90
t−statistic
0
5
10
15
cpgIslandExt
ZFP90
SNP;chr16:68572534
←CFAP20
CTB−134F13.1→
RP11−459F6.3→
58.14
58.15
58.16
58.17
58.18
Position on chr16 (Mb)

### ZFP90

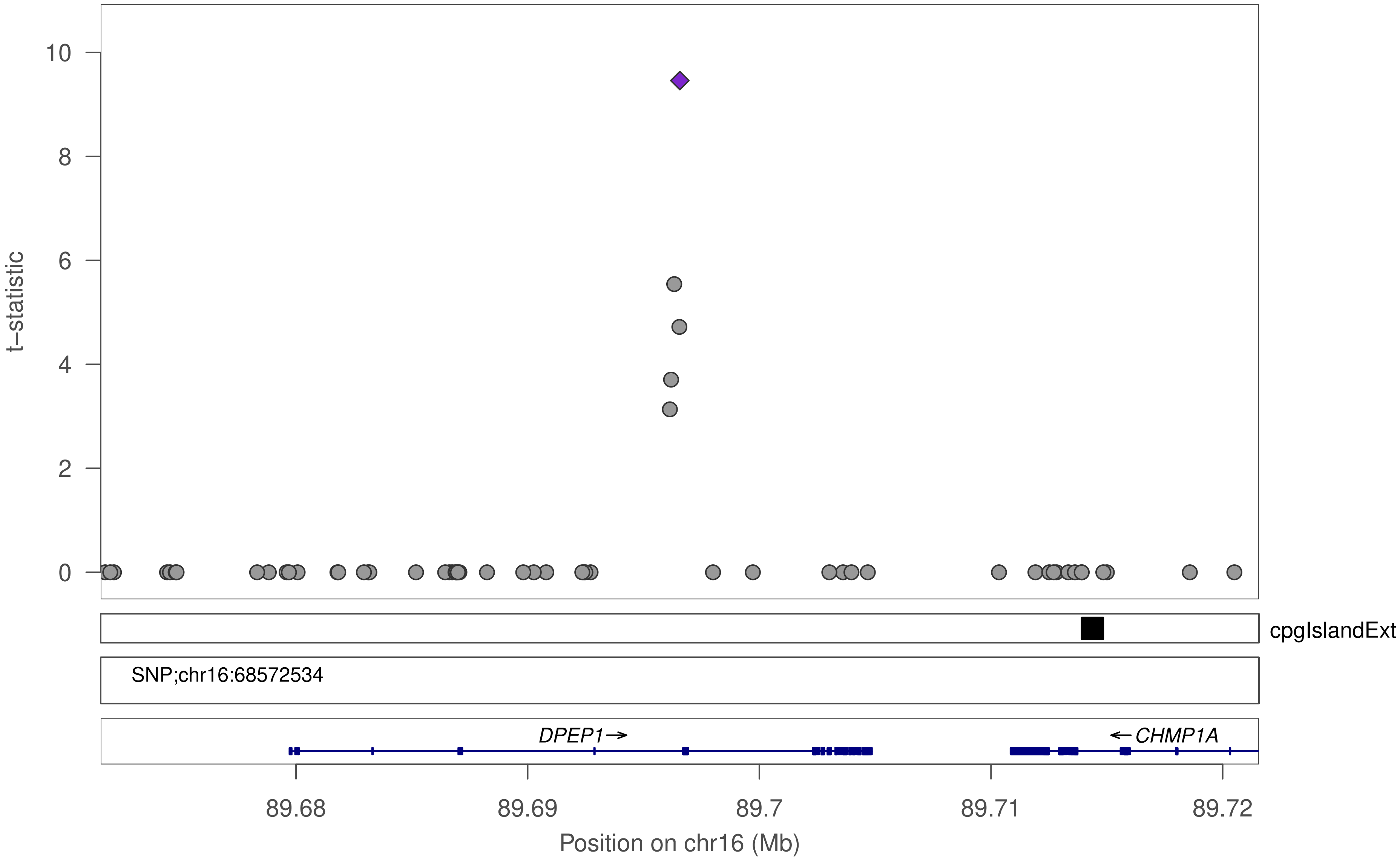

t–statistic
10
8
6
4
2
0
cpgIslandExt
SNP;chr16:68572534
DPEP1→
←CHMP1A
89.68
89.69
89.7
89.71
89.72
Position on chr16 (Mb)

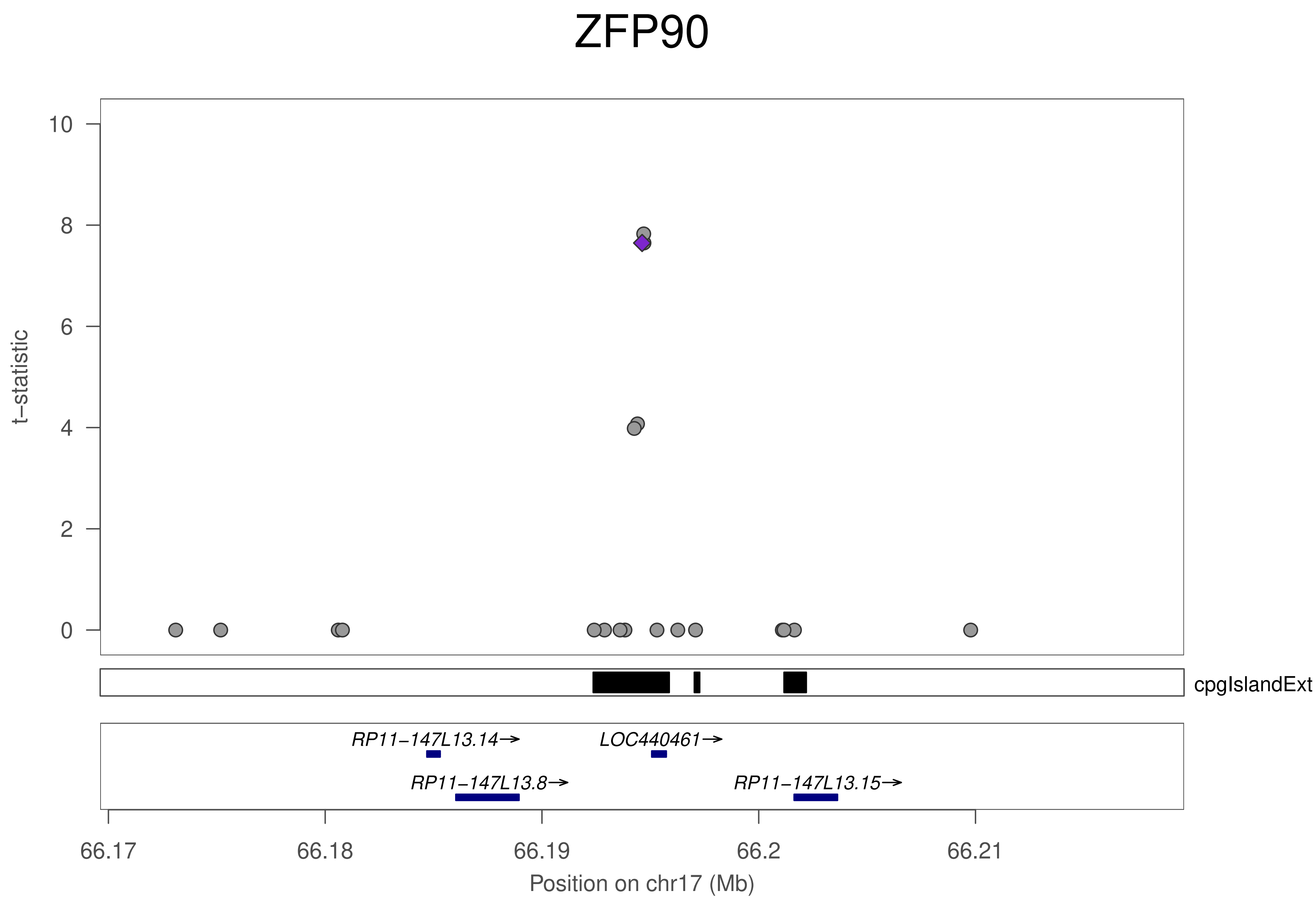
ZFP90
t−statistic
10
8
6
4
2
0
cpgIslandExt
RP11−147L13.14→
LOC440461→
RP11−147L13.8→
RP11−147L13.15→
66.17
66.18
66.19
66.2
66.21
Position on chr17 (Mb)

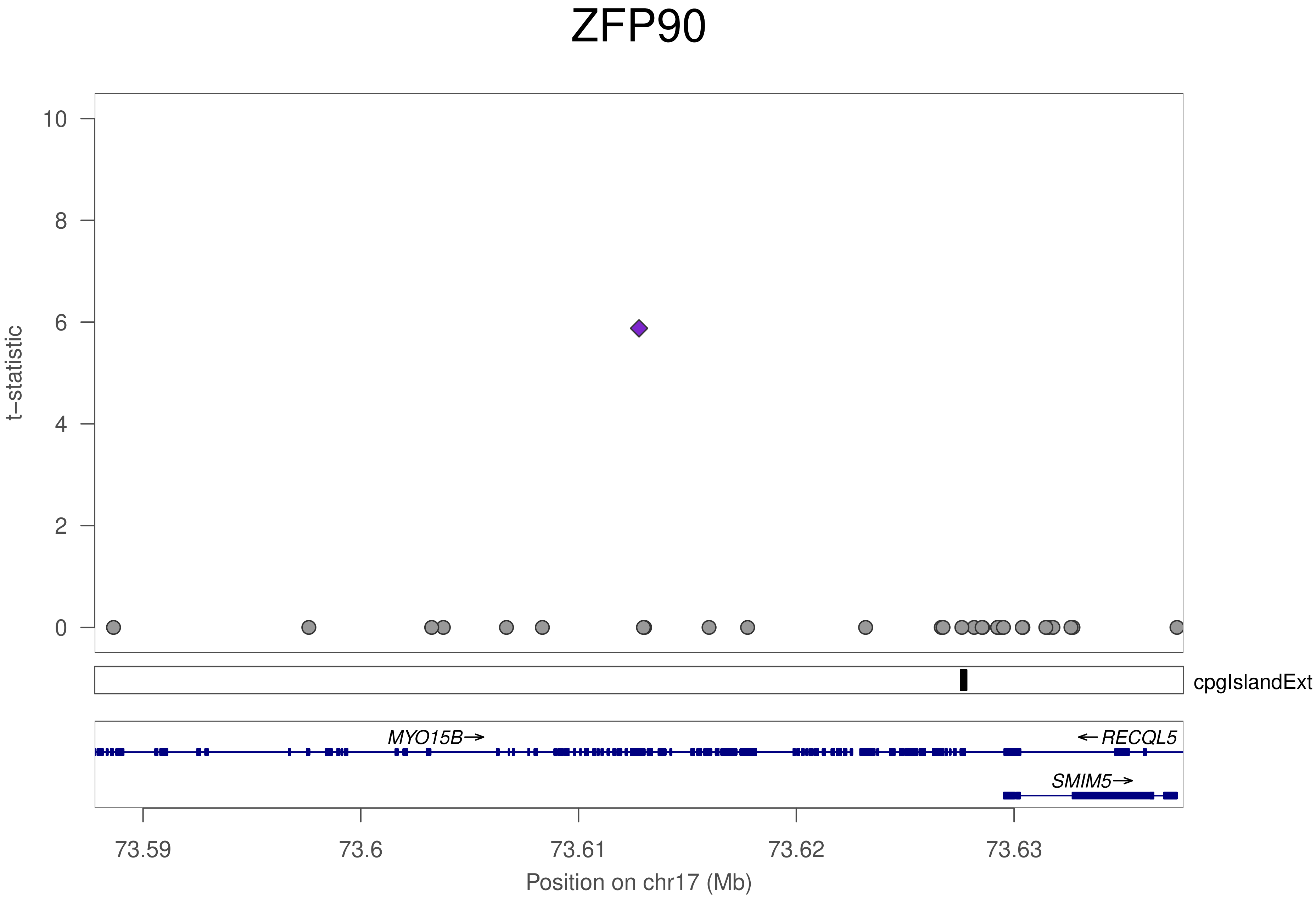

ZFP90
t−statistic
10
8
6
4
2
0
cpgIslandExt
MYO15B→
← RECQL5
SMIM5→
73.59
73.6
73.61
73.62
73.63
Position on chr17 (Mb)

### ZFP90

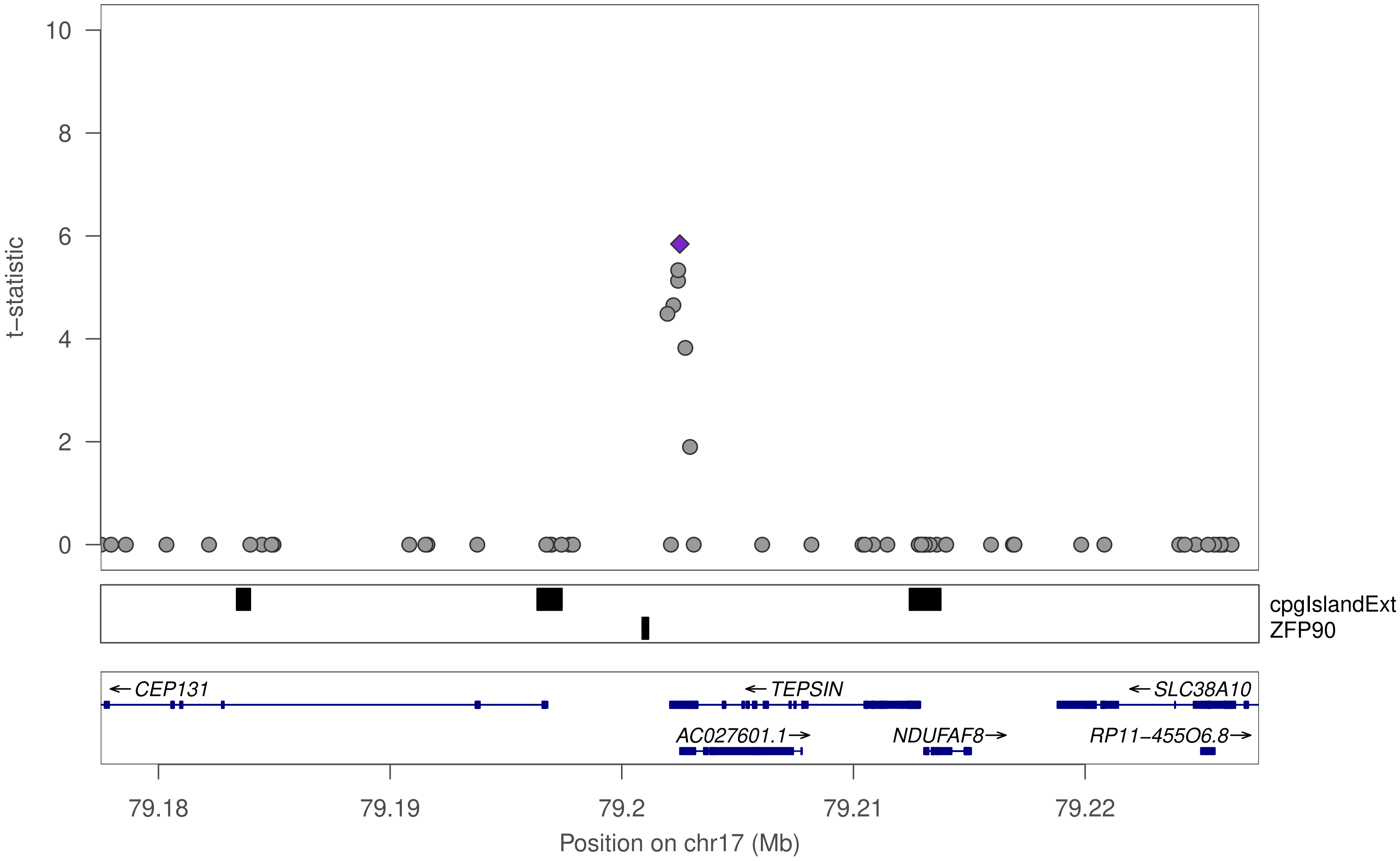

t−statistic
10
8
6
4
2
0
cpgIslandExt
ZFP90
← CEP131
← TEPSIN
← SLC38A10
AC027601.1→
NDUFAF8→
RP11−455O6.8→
79.18
79.19
79.2
79.21
79.22
Position on chr17 (Mb)

### ZFP90

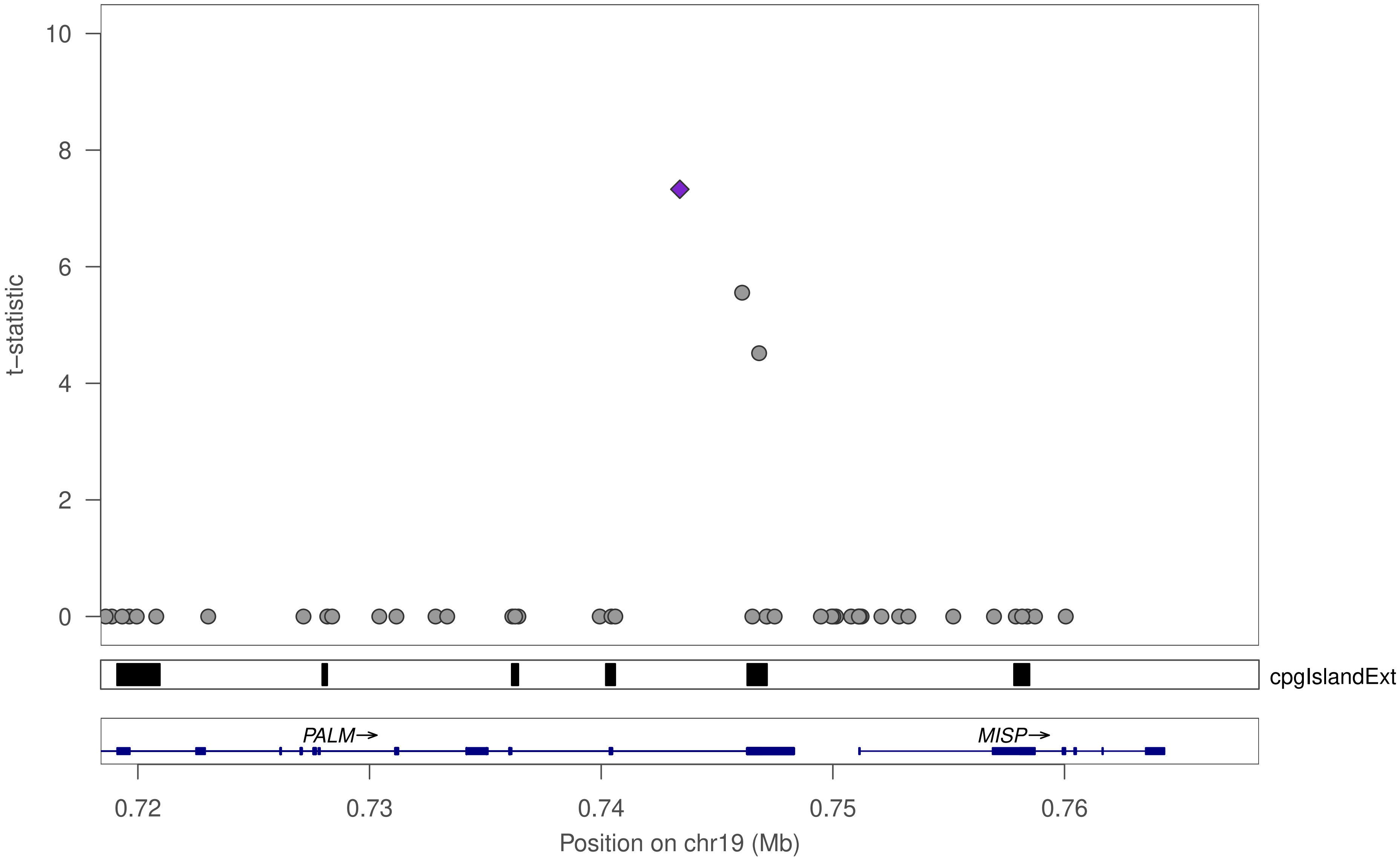

t−statistic
10
8
6
4
2
0
cpgIslandExt
PALM→
MISP→
0.72
0.73
0.74
0.75
0.76
Position on chr19 (Mb)

### ZFP90

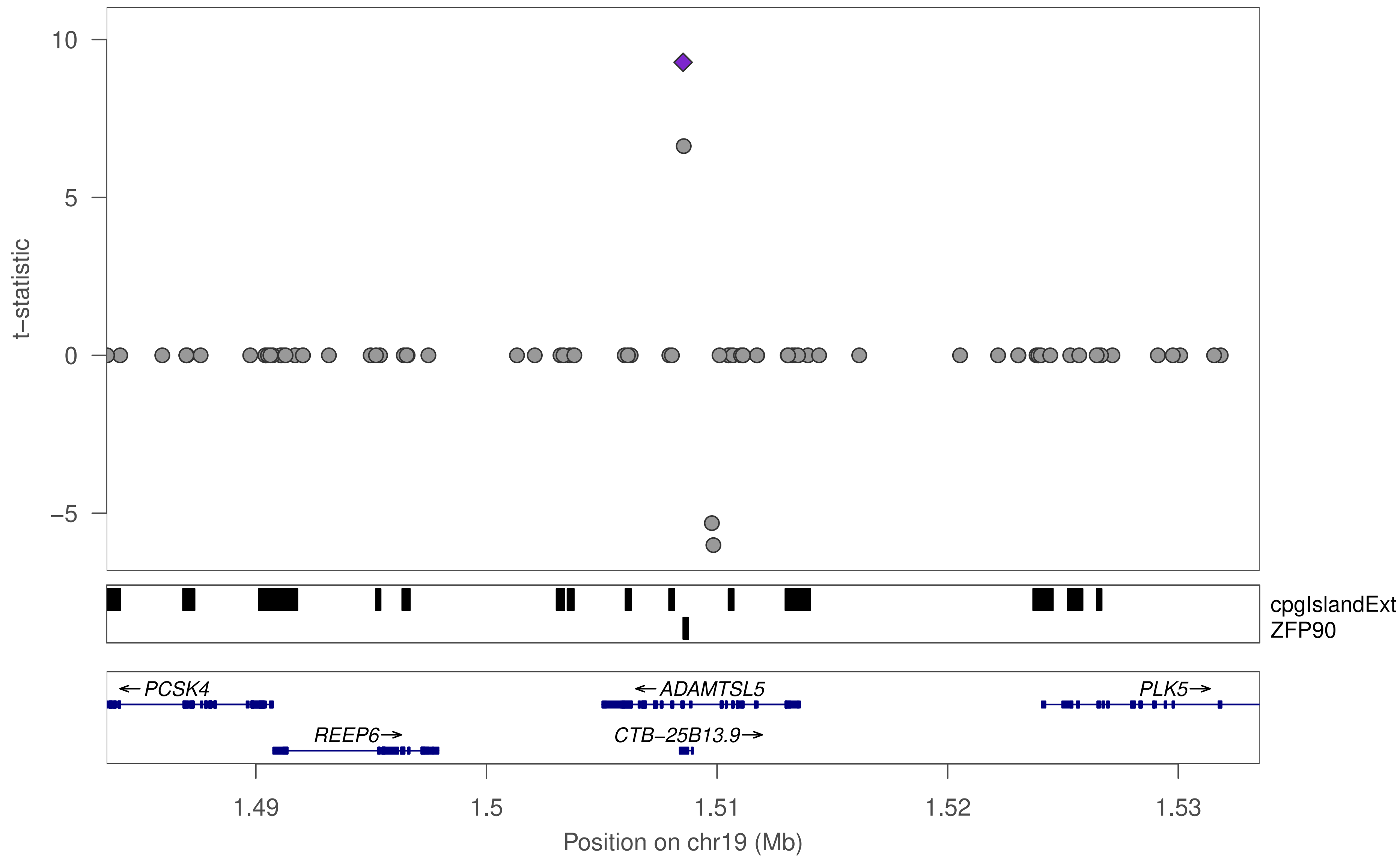

t−statistic
10
5
0
−5
cpgIslandExt
ZFP90
← PCSK4
← ADAMTSL5
PLK5→
REEP6→
CTB−25B13.9→
1.49
1.5
1.51
1.52
1.53
Position on chr19 (Mb)

### ZFP90

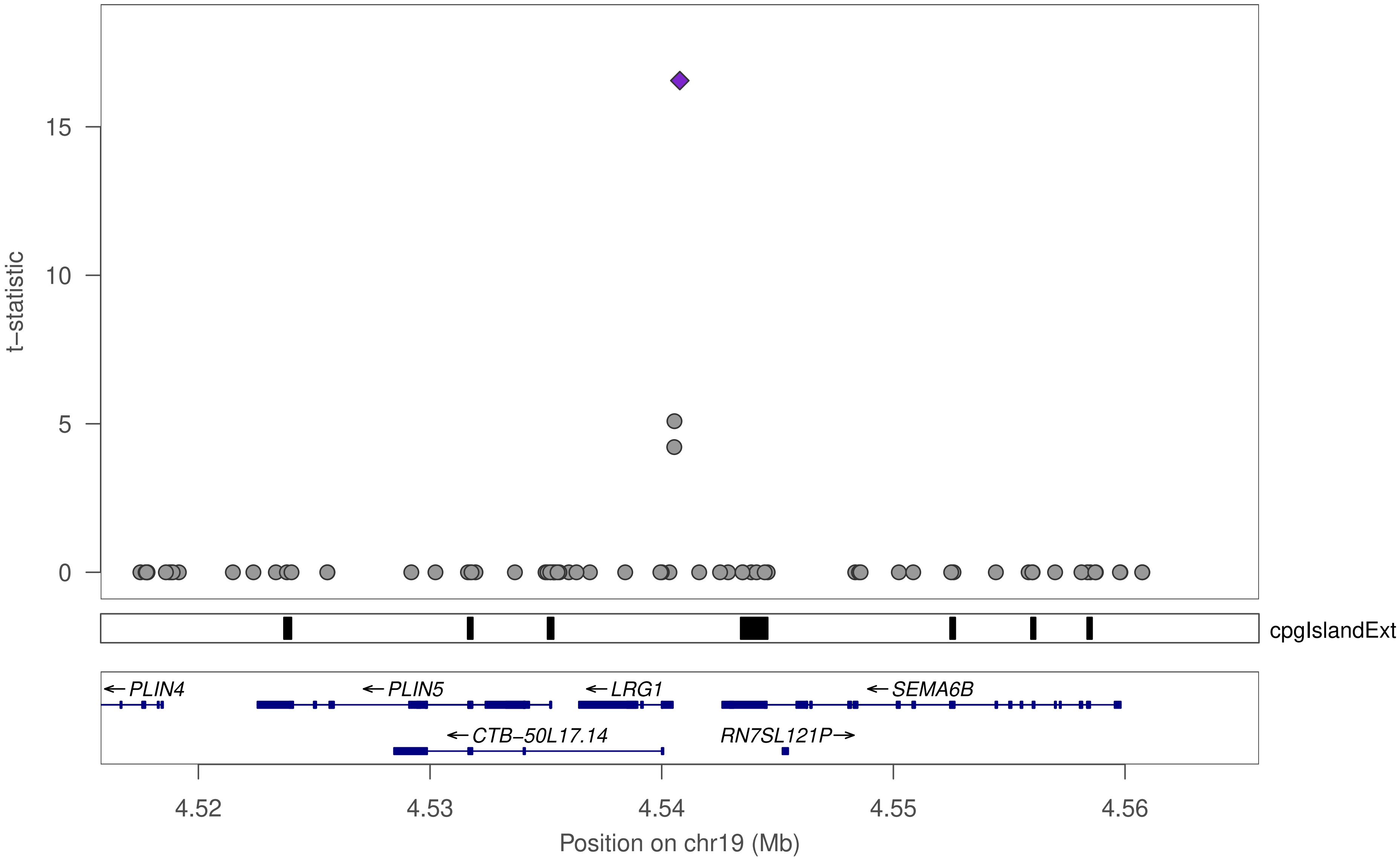

t–statistic
15
10
5
0
cpgIslandExt
←PLIN4
←PLIN5
←LRG1
←SEMA6B
←CTB–50L17.14
RN7SL121P→
4.52
4.53
4.54
4.55
4.56
Position on chr19 (Mb)

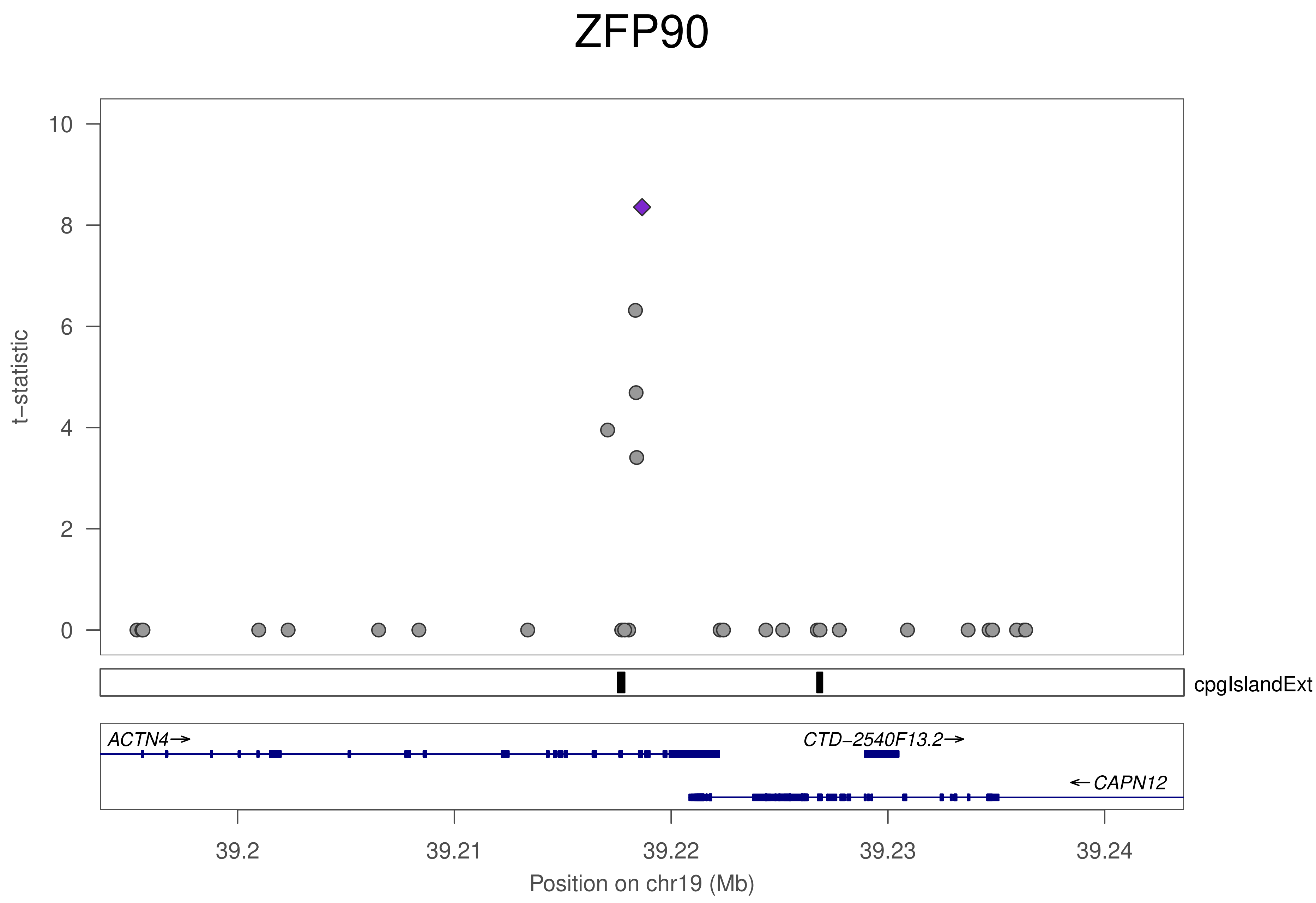

ZFP90
t−statistic
10
8
6
4
2
0
cpgIslandExt
ACTN4→
CTD−2540F13.2→
←CAPN12
39.2
39.21
39.22
39.23
39.24
Position on chr19 (Mb)

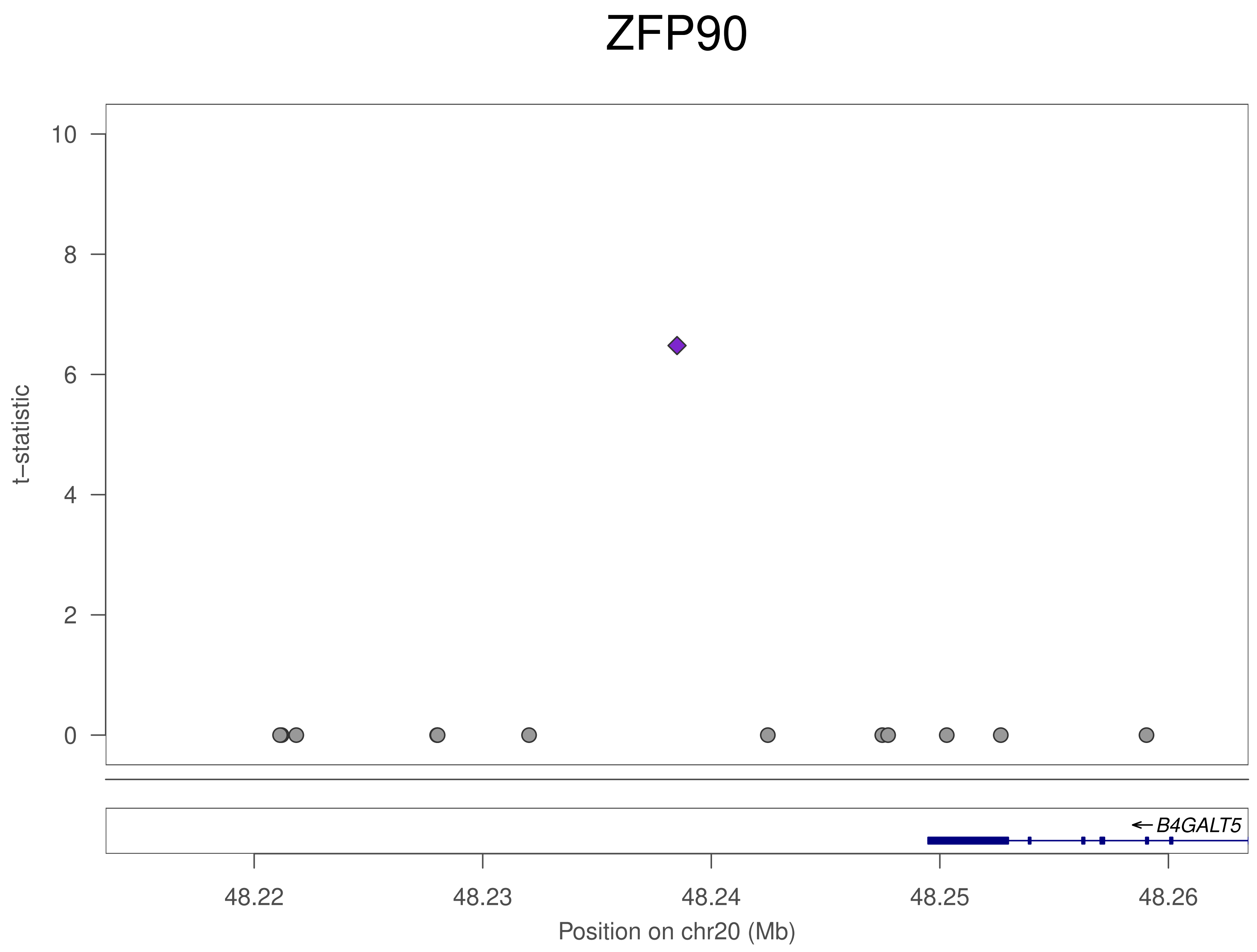
ZFP90
10
8
6
4
2
0
t−statistic
B4GALT5
48.22
48.23
48.24
48.25
48.26
Position on chr20 (Mb)

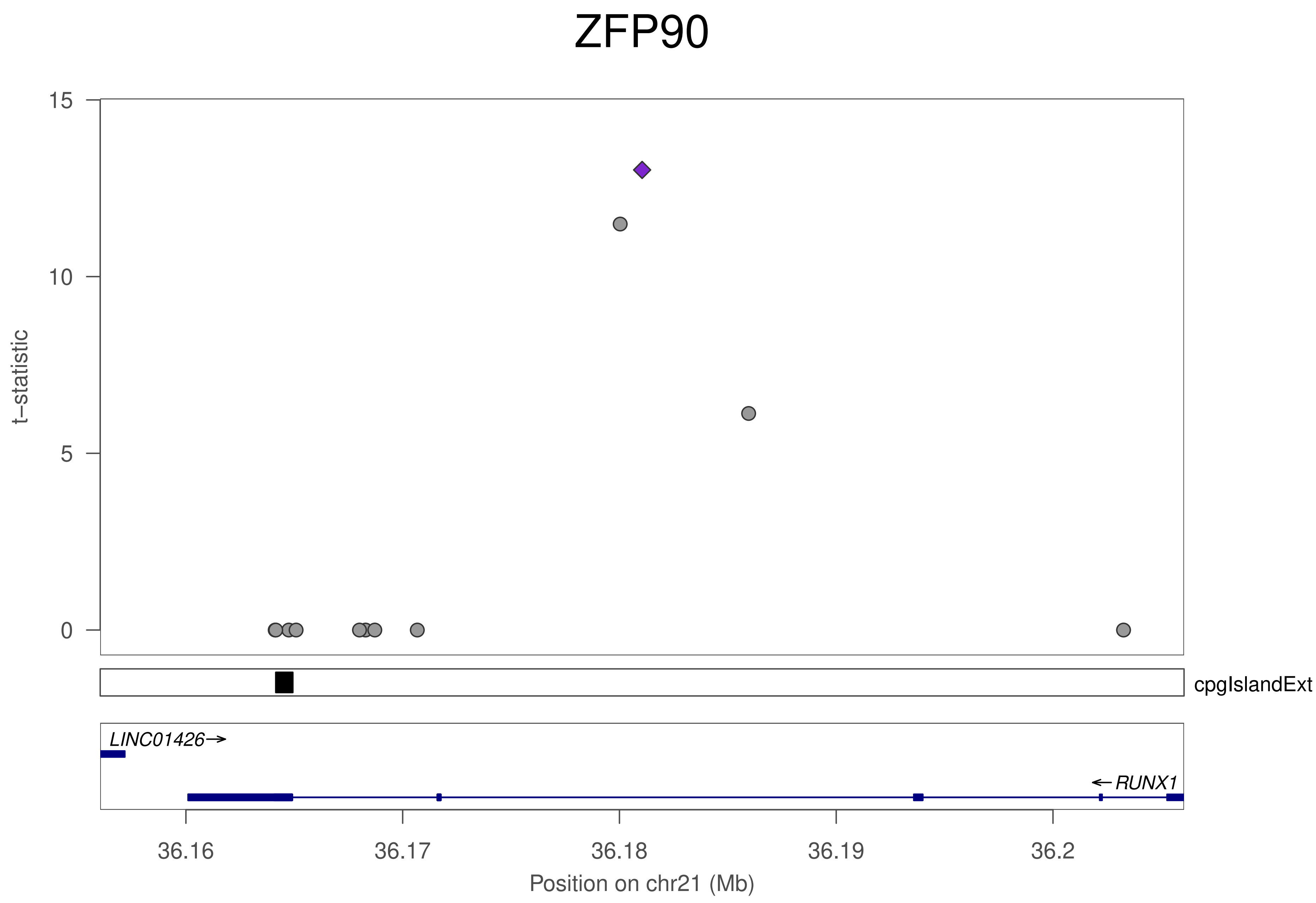
ZFP90
t−statistic
15
10
5
0
cpgIslandExt
LINC01426→
←RUNX1
36.16
36.17
36.18
36.19
36.2
Position on chr21 (Mb)

### ZNF566

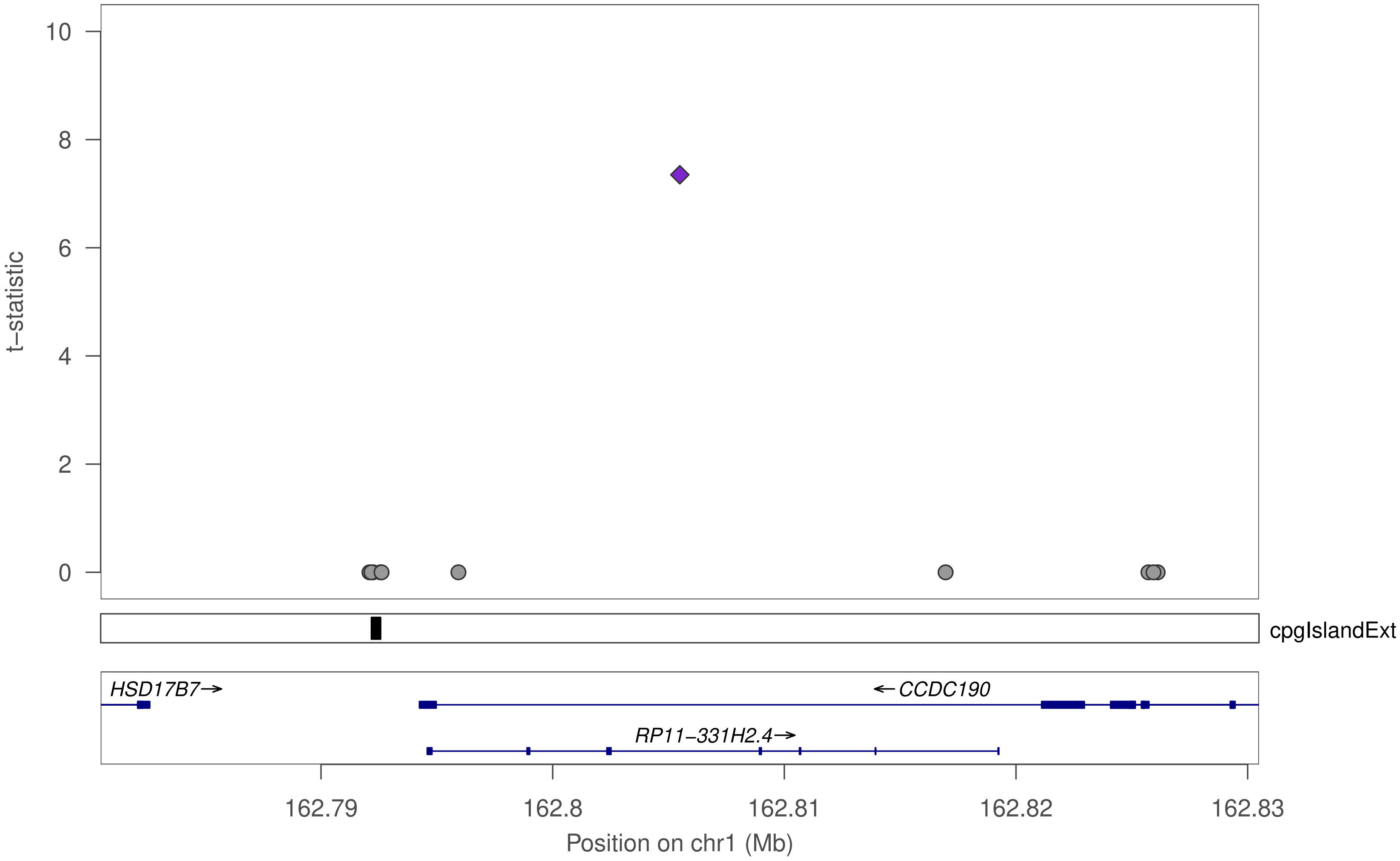

10
8
6
4
2
0
t−statistic
cpgIslandExt
HSD17B7→
←CCDC190
RP11−331H2.4→
162.79
162.8
162.81
162.82
162.83
Position on chr1 (Mb)

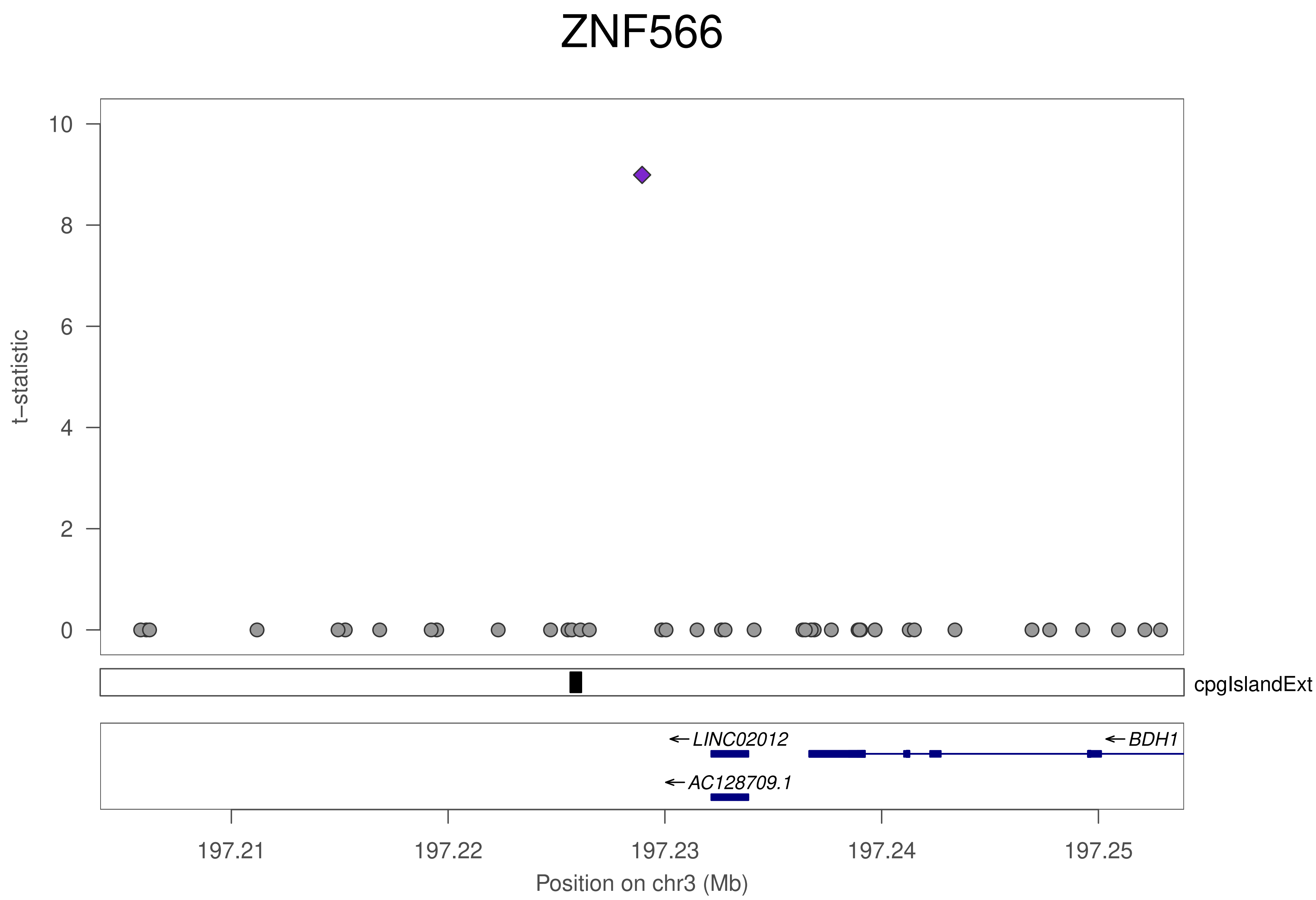
ZNF566
t−statistic
0
2
4
6
8
10
cpgIslandExt
←LINC02012
←BDH1
←AC128709.1
197.21
197.22
197.23
197.24
197.25
Position on chr3 (Mb)

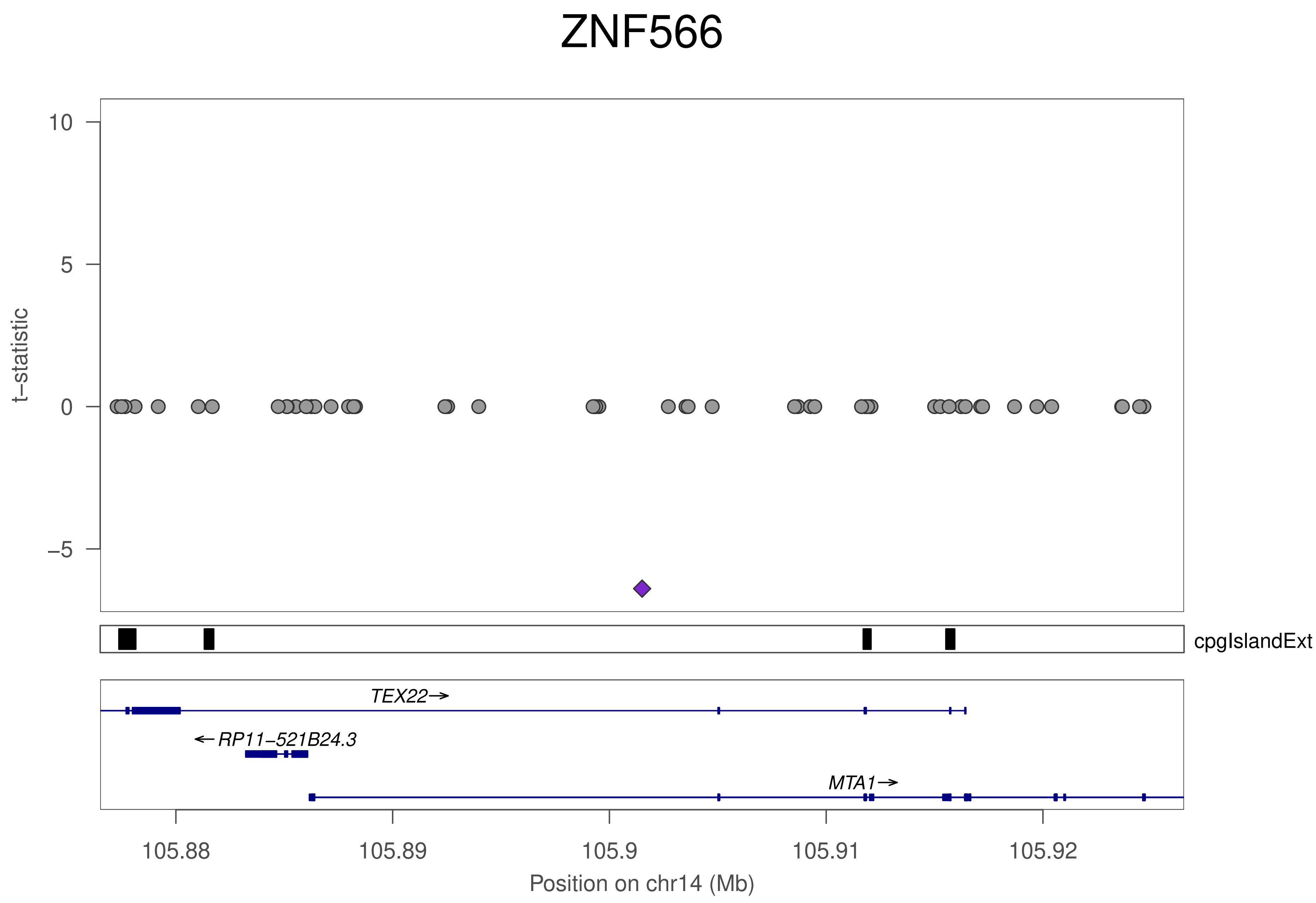

ZNF566
t−statistic
10
5
0
−5
cpgIslandExt
TEX22→
←RP11−521B24.3
MTA1→
105.88
105.89
105.9
105.91
105.92
Position on chr14 (Mb)

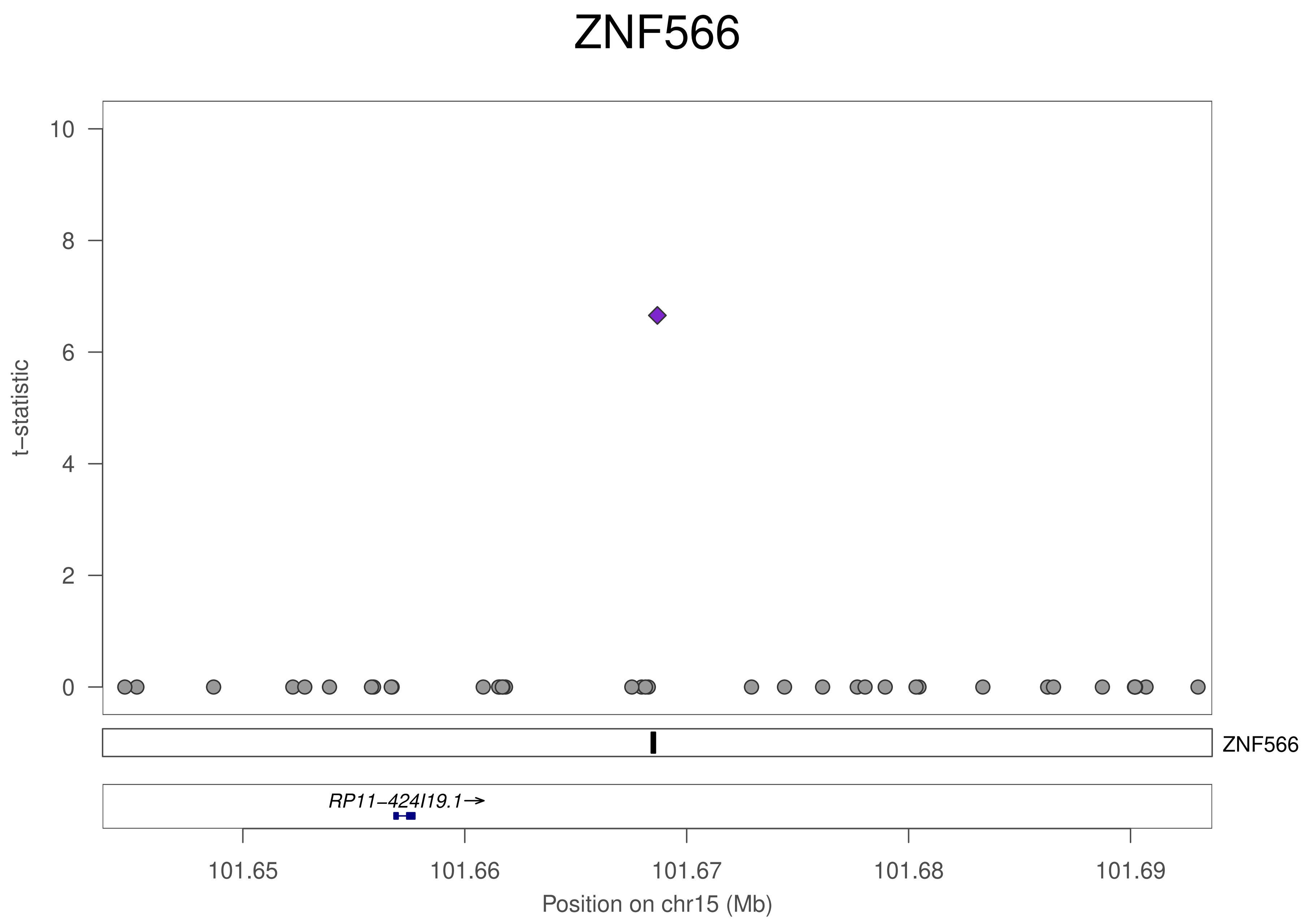
ZNF566
t−statistic
10
8
6
4
2
0
ZNF566
RP11−424I19.1→
101.65
101.66
101.67
101.68
101.69
Position on chr15 (Mb)

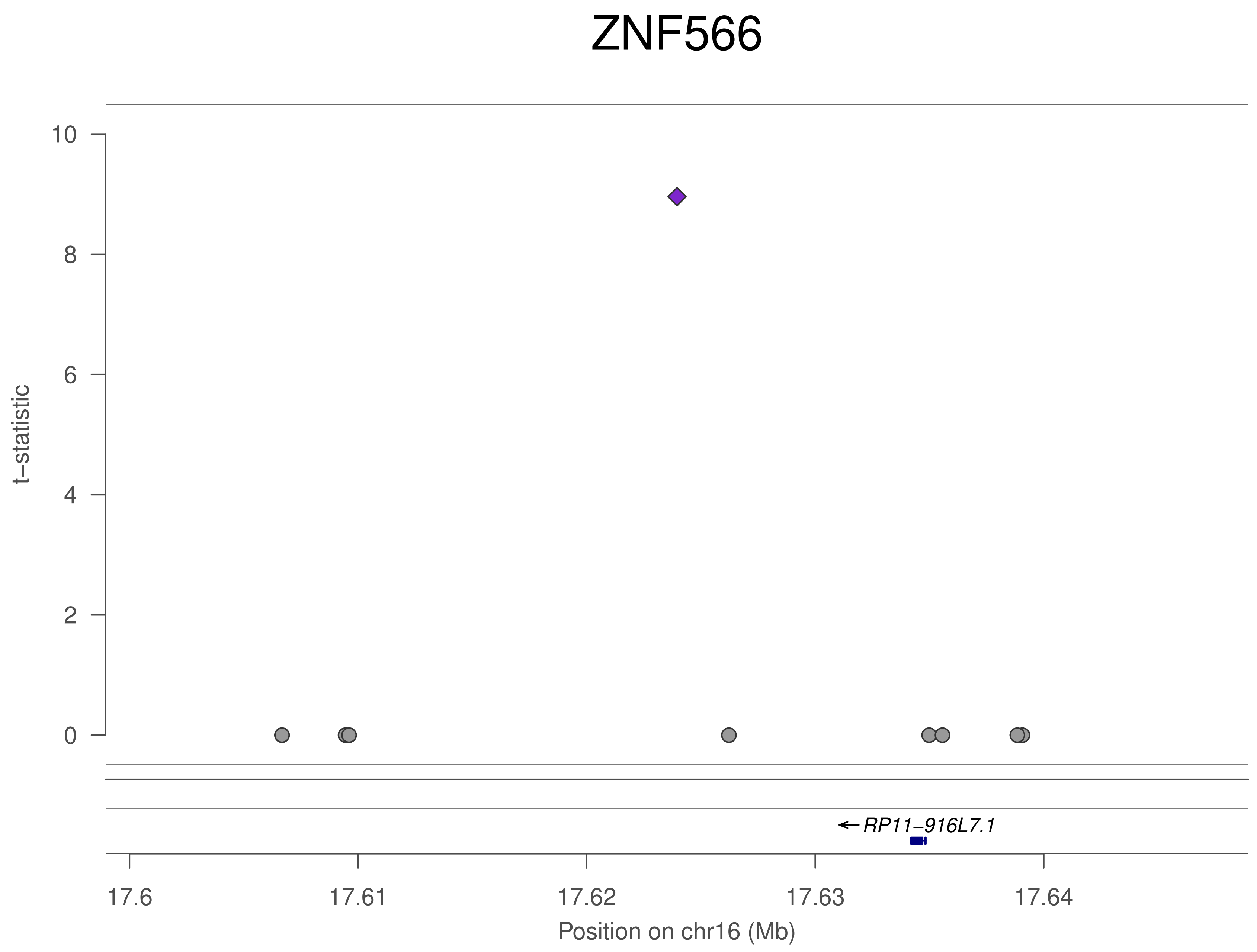
ZNF566
t–statistic
10
8
6
4
2
0
RP11–916L7.1
17.6
17.61
17.62
17.63
17.64
Position on chr16 (Mb)

### ZNF284

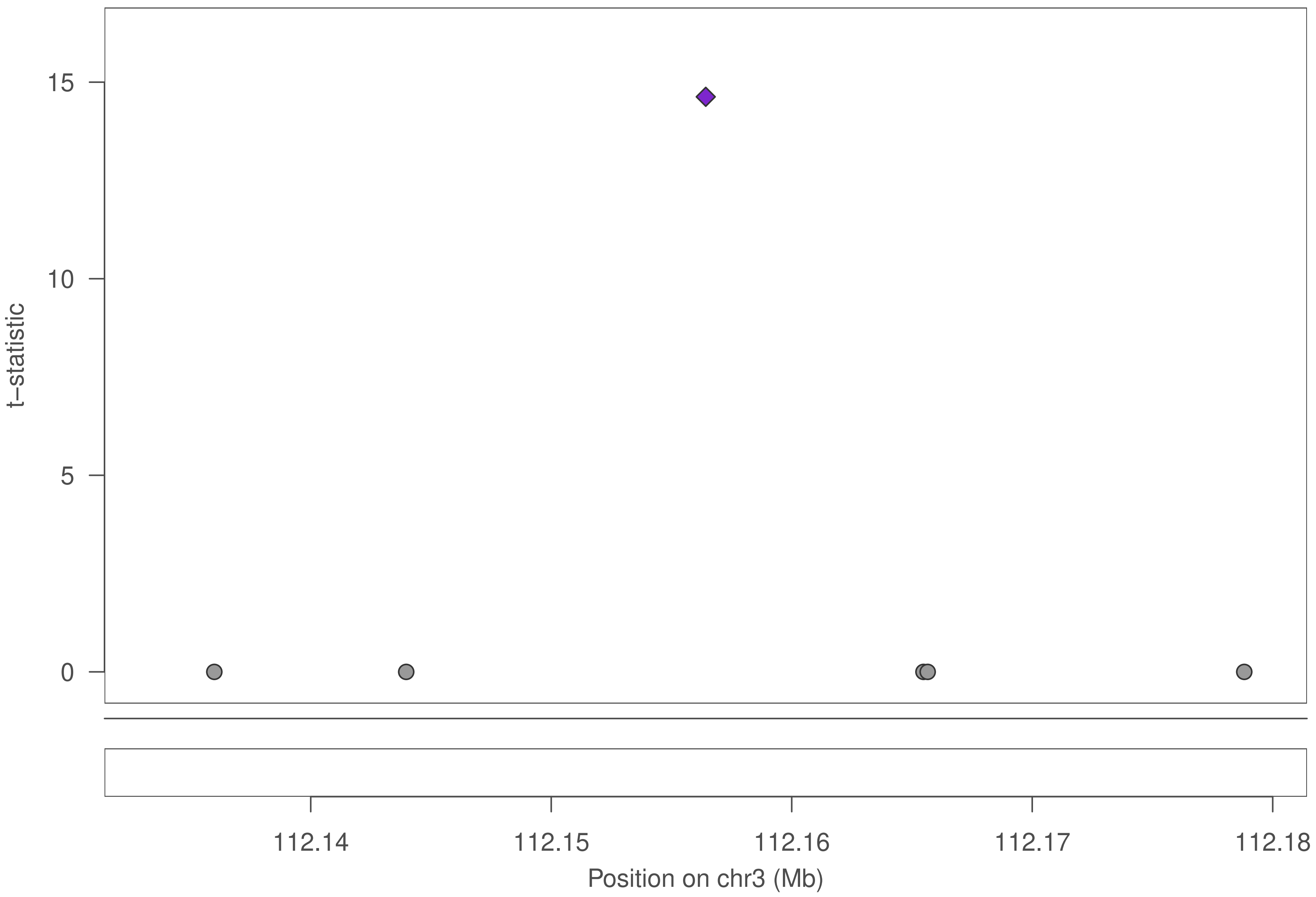

15
10
5
0
t−statistic
112.14
112.15
112.16
112.17
112.18
Position on chr3 (Mb)

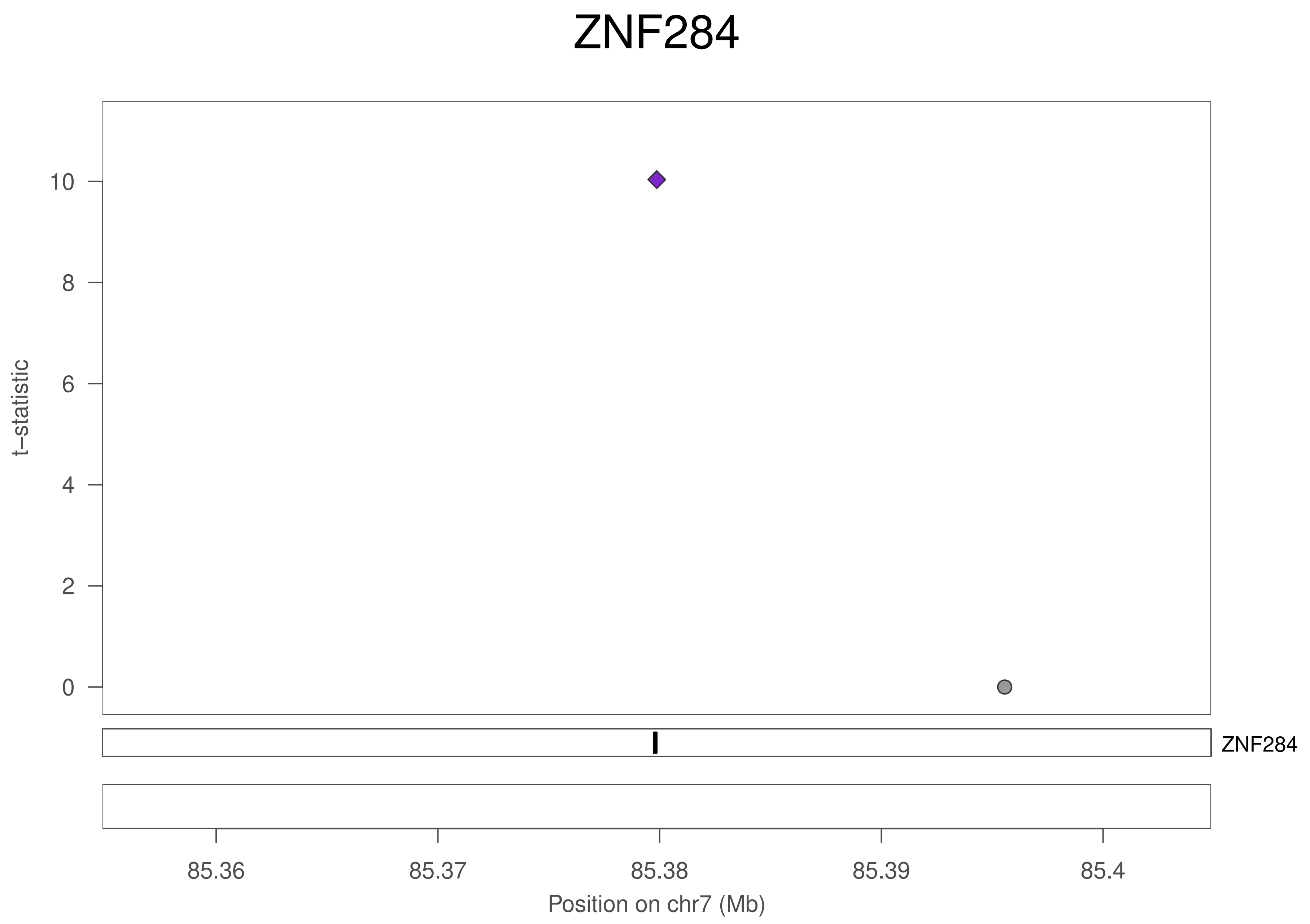

ZNF284
10
8
6
4
2
0
t−statistic
ZNF284
85.36
85.37
85.38
85.39
85.4
Position on chr7 (Mb)

### ZNF284

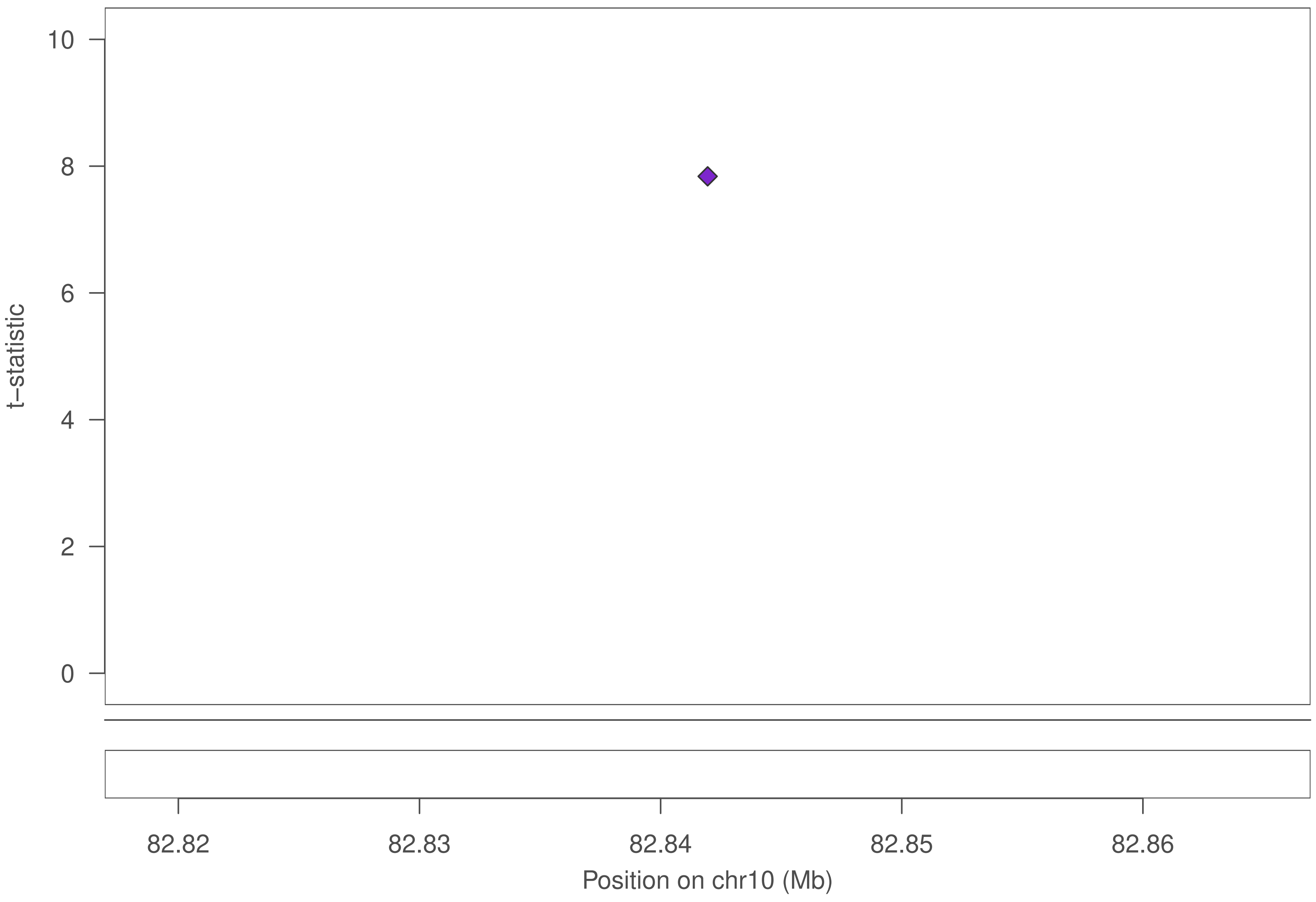

t−statistic
10
8
6
4
2
0
82.82
82.83
82.84
82.85
82.86
Position on chr10 (Mb)

### ZNF284

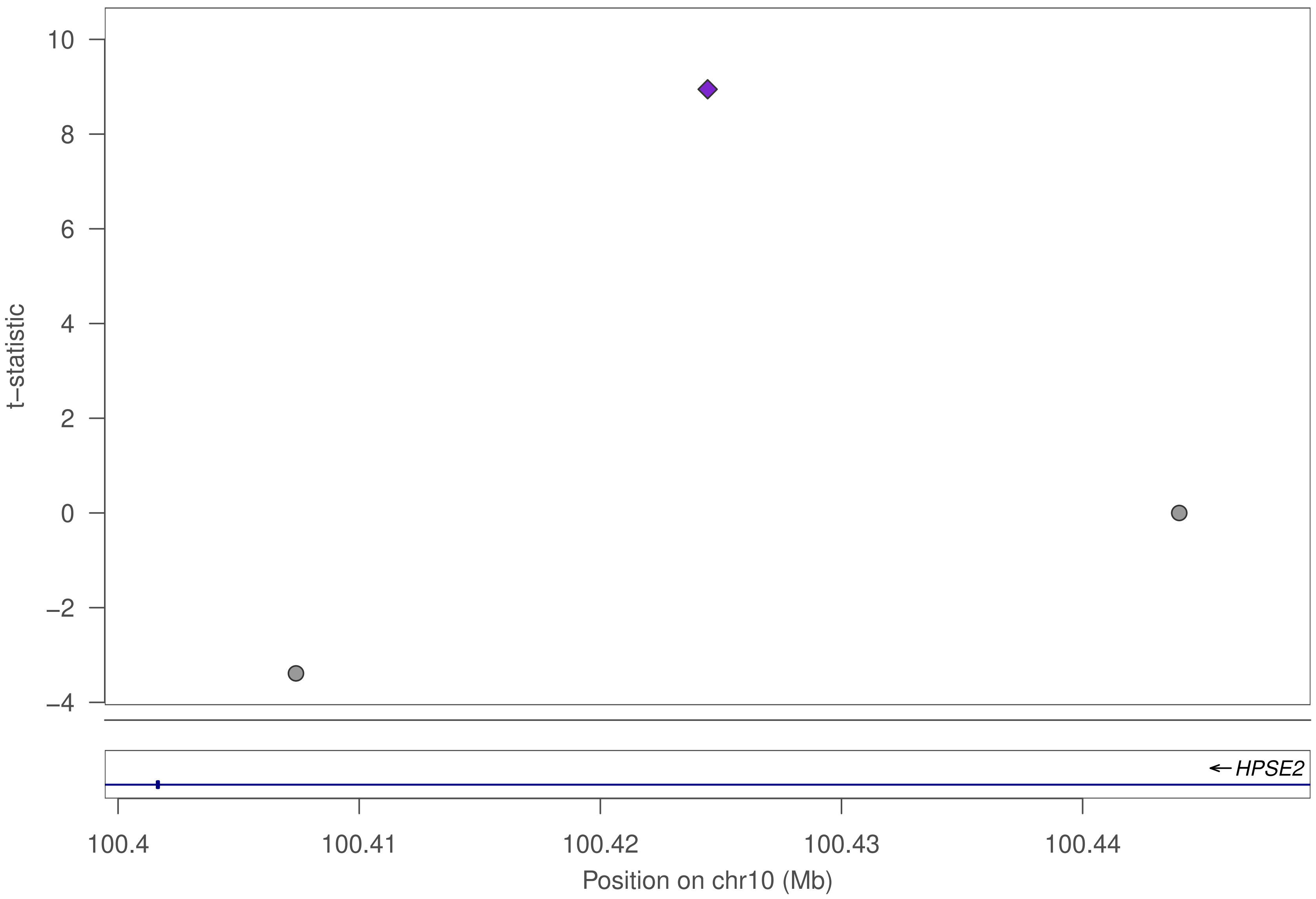

10
8
6
4
2
0
−2
−4
t−statistic
HPSE2
100.4
100.41
100.42
100.43
100.44
Position on chr10 (Mb)

### ZNF284

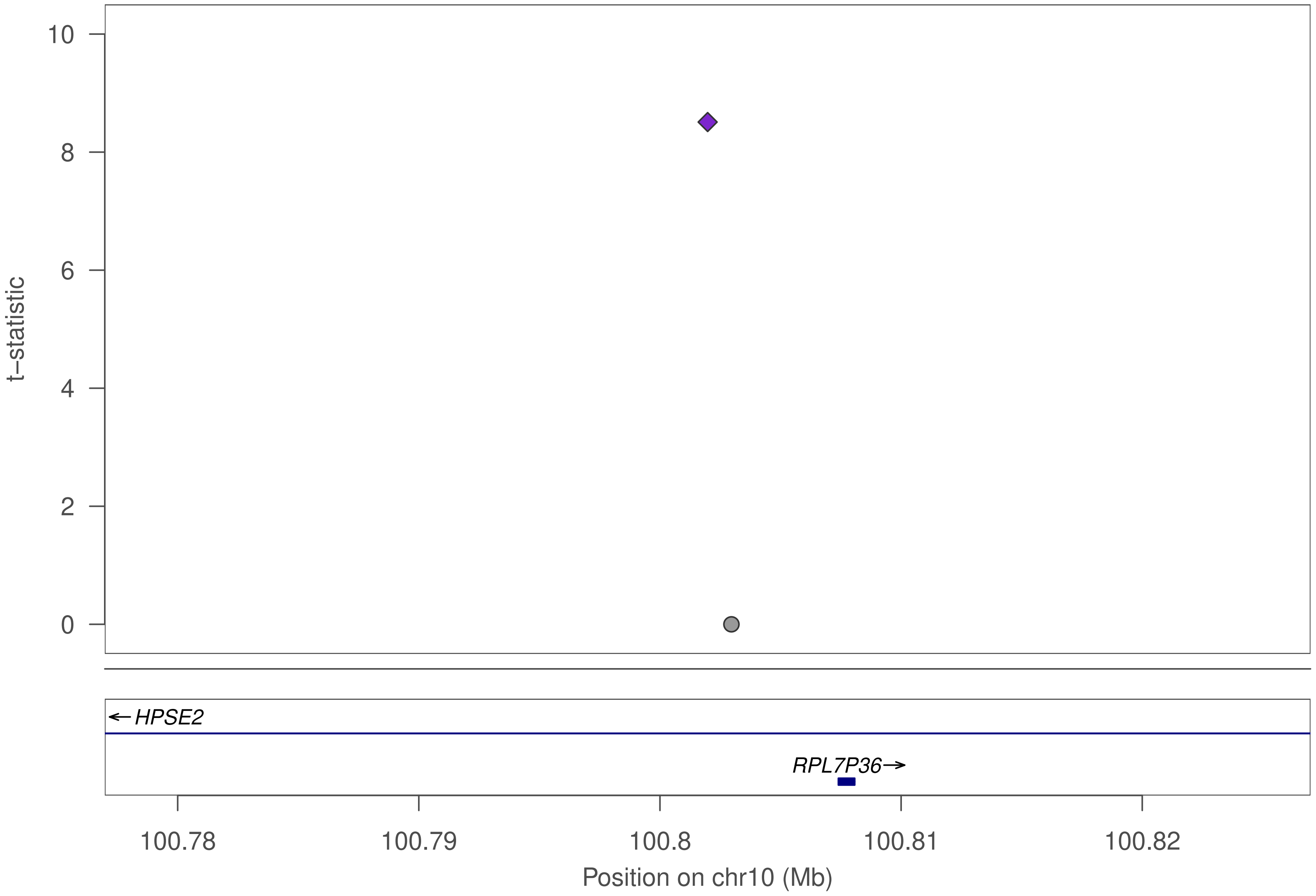

t–statistic
10
8
6
4
2
0
← HPSE2
RPL7P36→
100.78
100.79
100.8
100.81
100.82
Position on chr10 (Mb)

### ZNF749

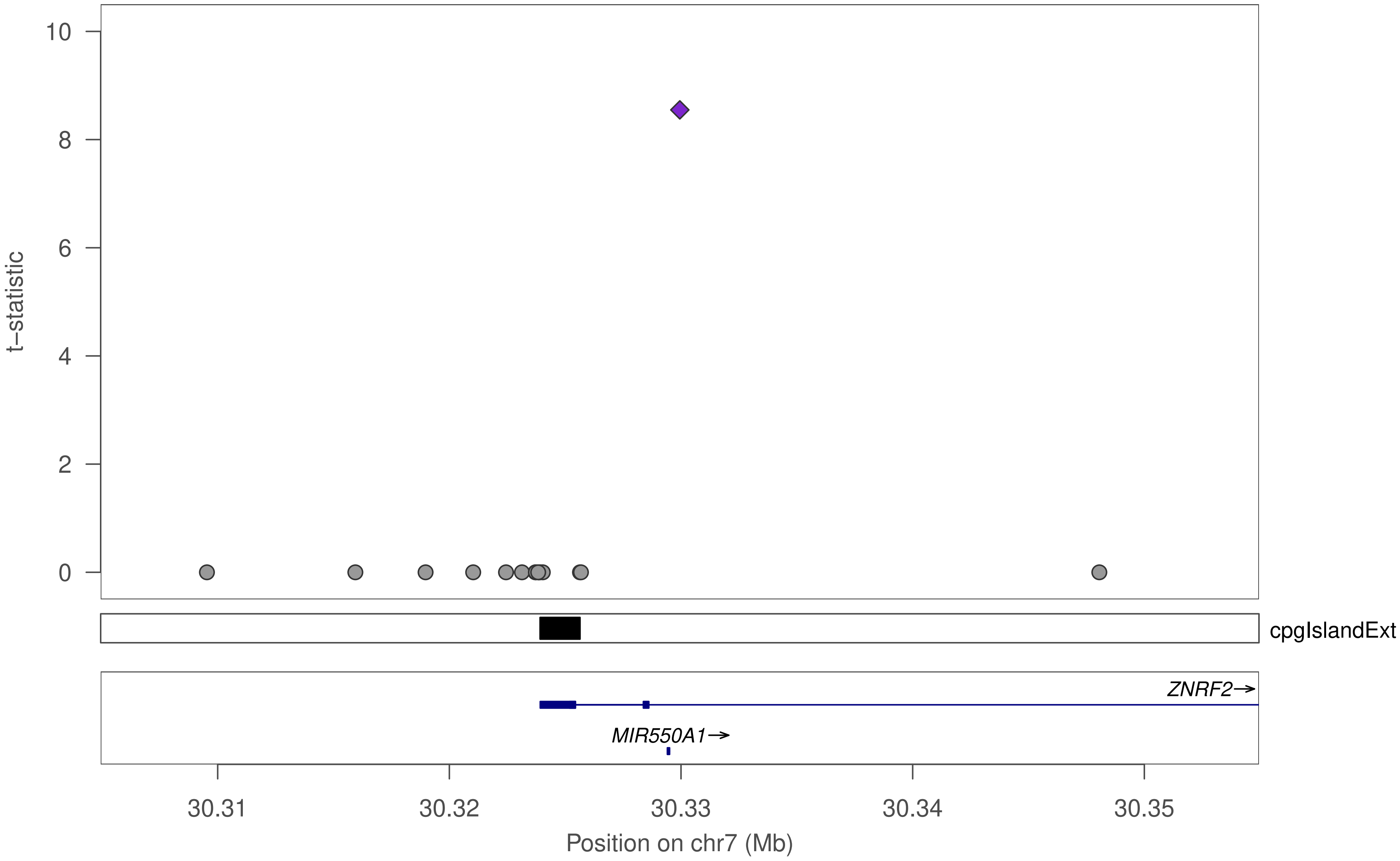

10
8
6
4
2
0
t−statistic
cpgIslandExt
ZNRF2→
MIR550A1→
30.31
30.32
30.33
30.34
30.35
Position on chr7 (Mb)

### ZNF749

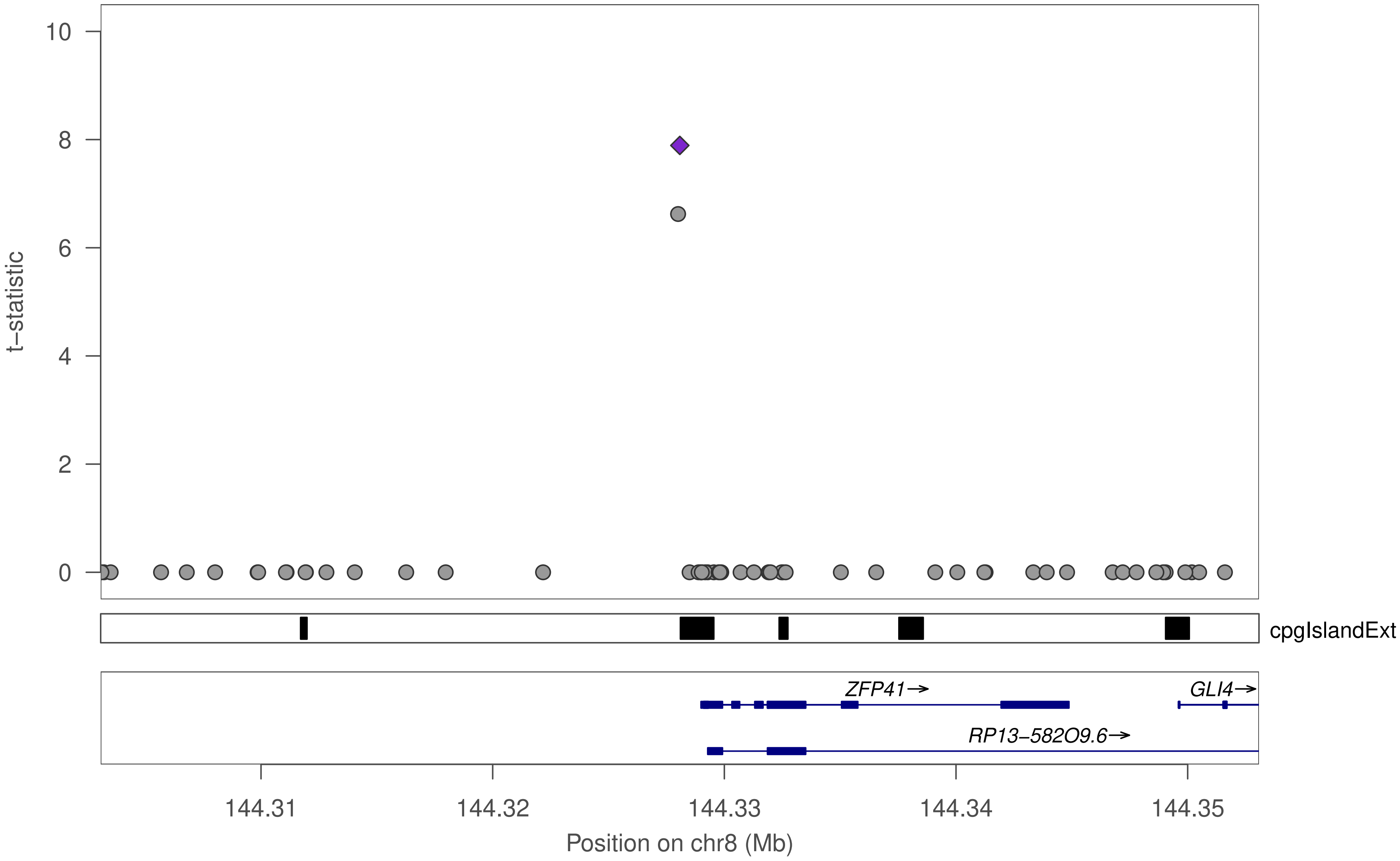

t–statistic
10
8
6
4
2
0
cpgIslandExt
ZFP41→
GLI4→
RP13–582O9.6→
144.31
144.32
144.33
144.34
144.35
Position on chr8 (Mb)

### ZNF749

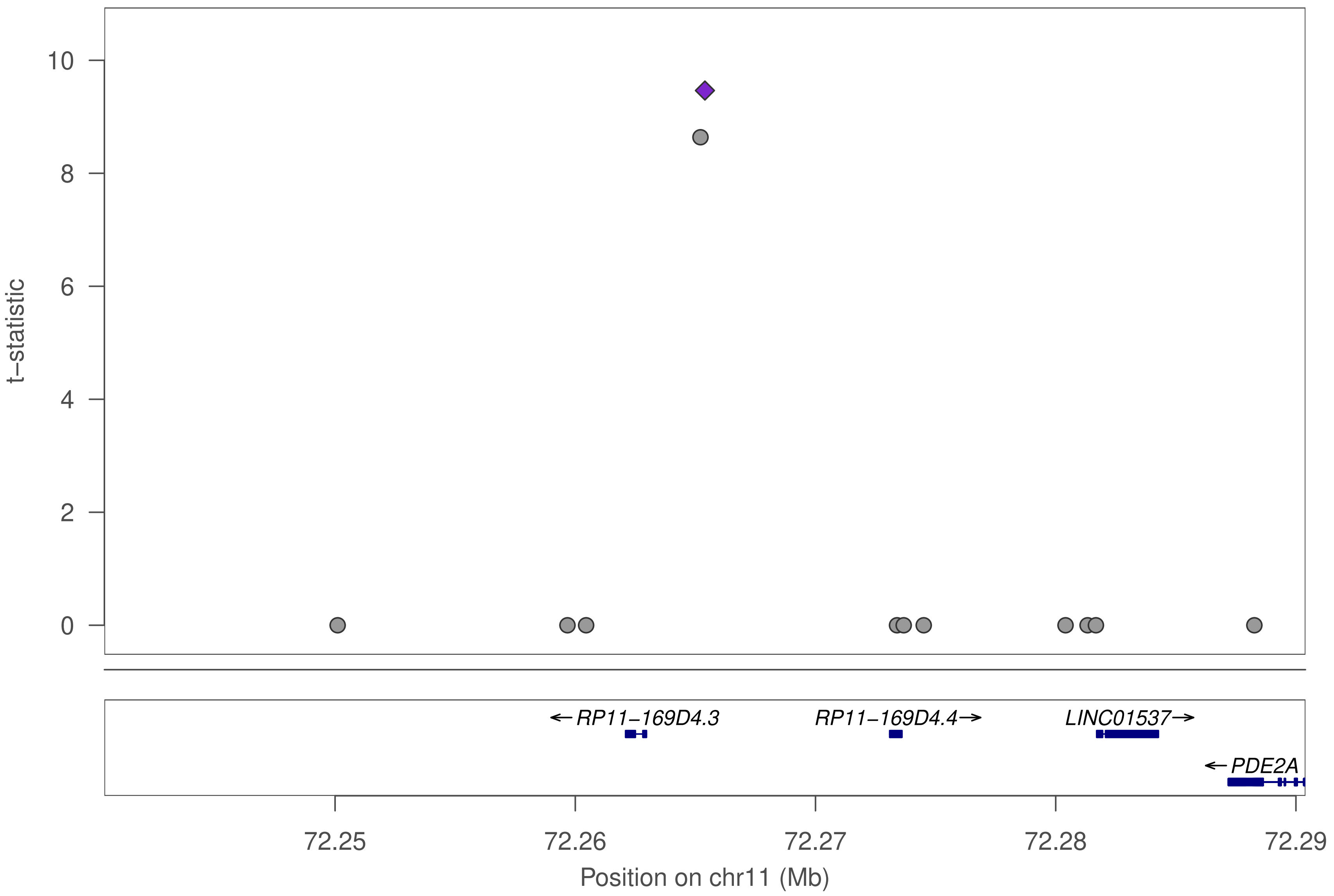

10
8
6
4
2
0
t−statistic
←RP11−169D4.3
RP11−169D4.4→
LINC01537→
←PDE2A
72.25
72.26
72.27
72.28
72.29
Position on chr11 (Mb)

### ZNF197

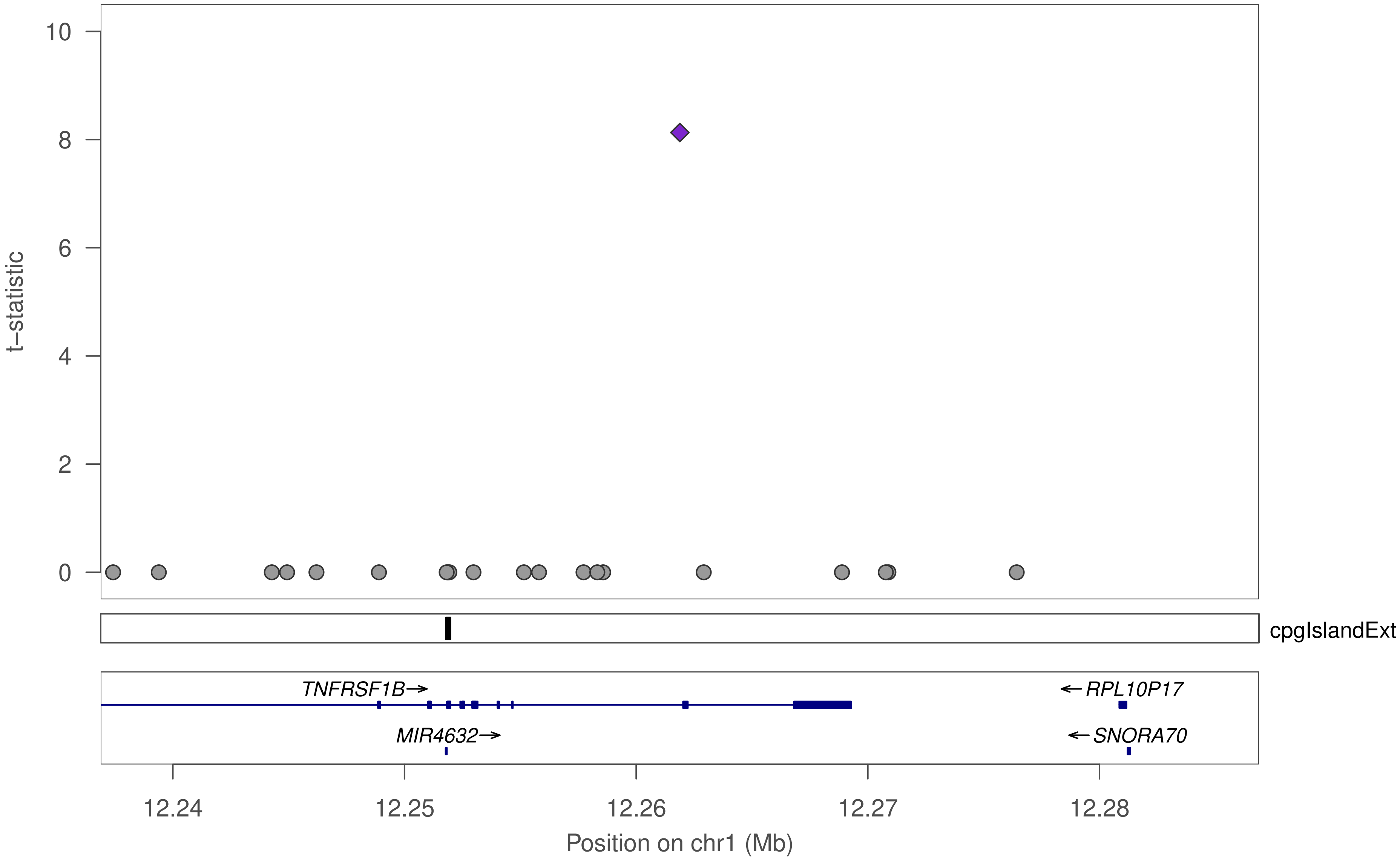
t−statistic
10
8
6
4
2
0
cpgIslandExt
TNFRSF1B→
←RPL10P17
MIR4632→
←SNORA70
12.24
12.25
12.26
12.27
12.28
Position on chr1 (Mb)

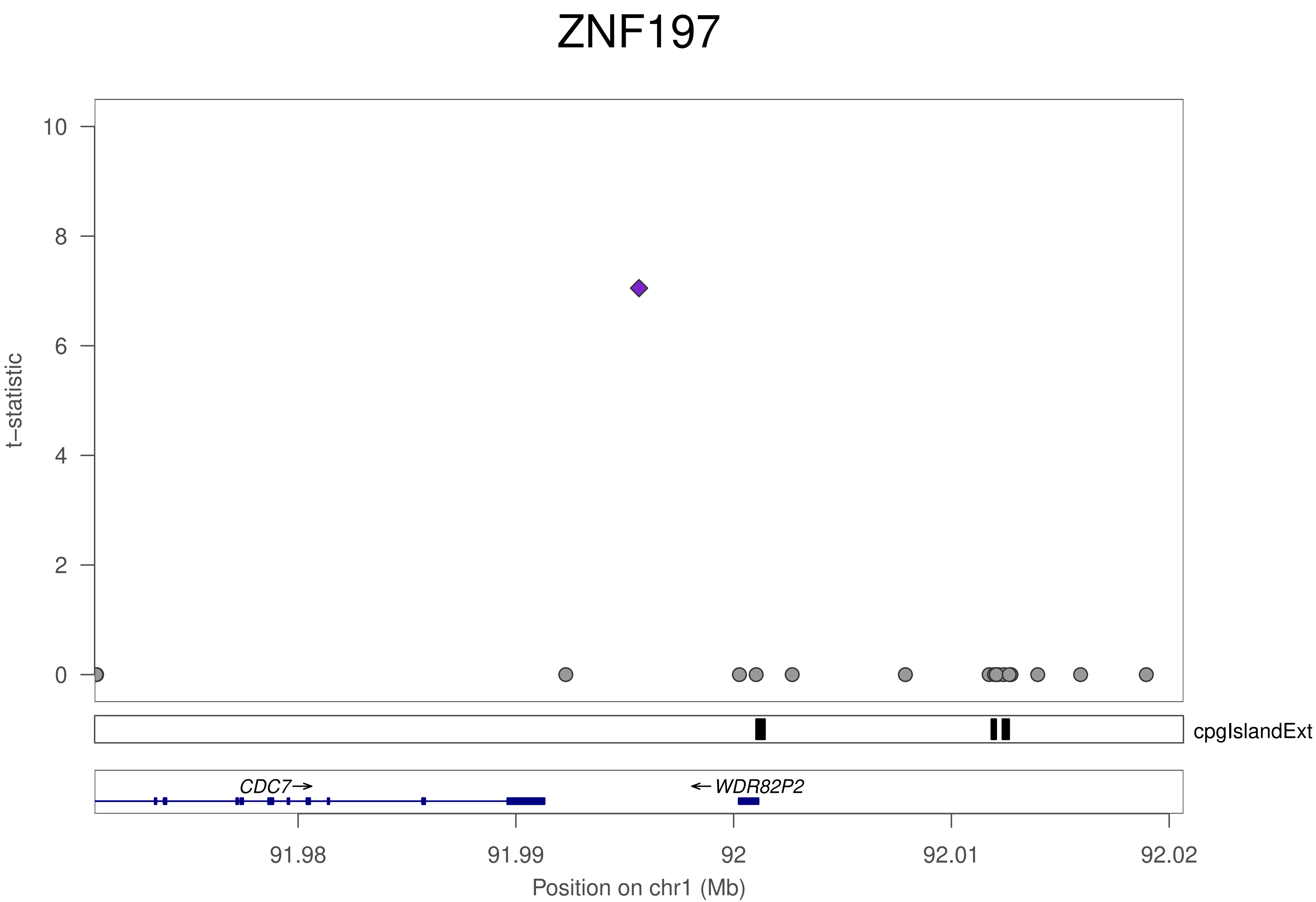
ZNF197
t−statistic
10
8
6
4
2
0
cpgIslandExt
CDC7→
←WDR82P2
91.98
91.99
92
92.01
92.02
Position on chr1 (Mb)

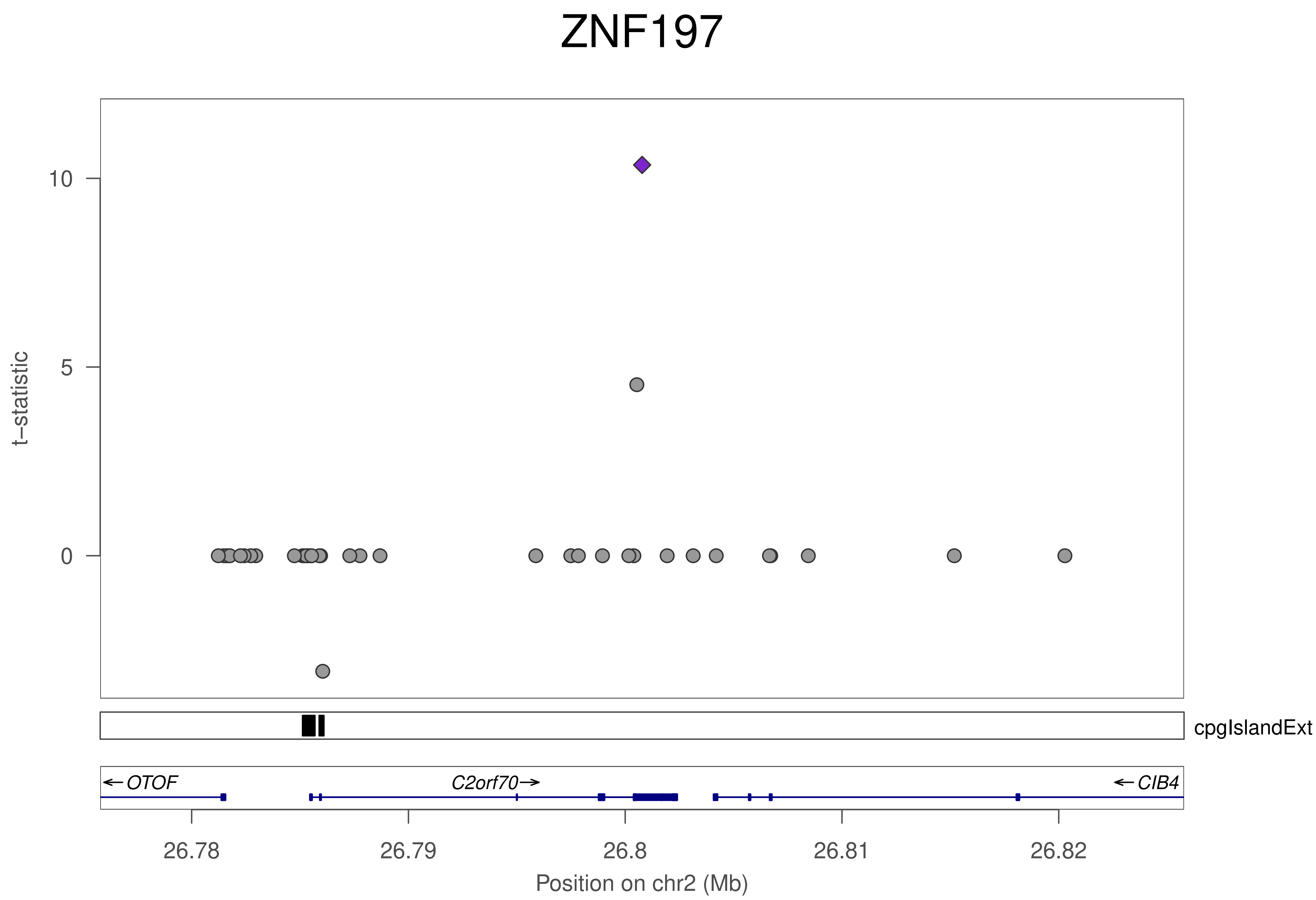

ZNF197
t–statistic
10
5
0
cpgIslandExt
← OTOF
C2orf70→
← CIB4
26.78
26.79
26.8
26.81
26.82
Position on chr2 (Mb)

### ZNF197

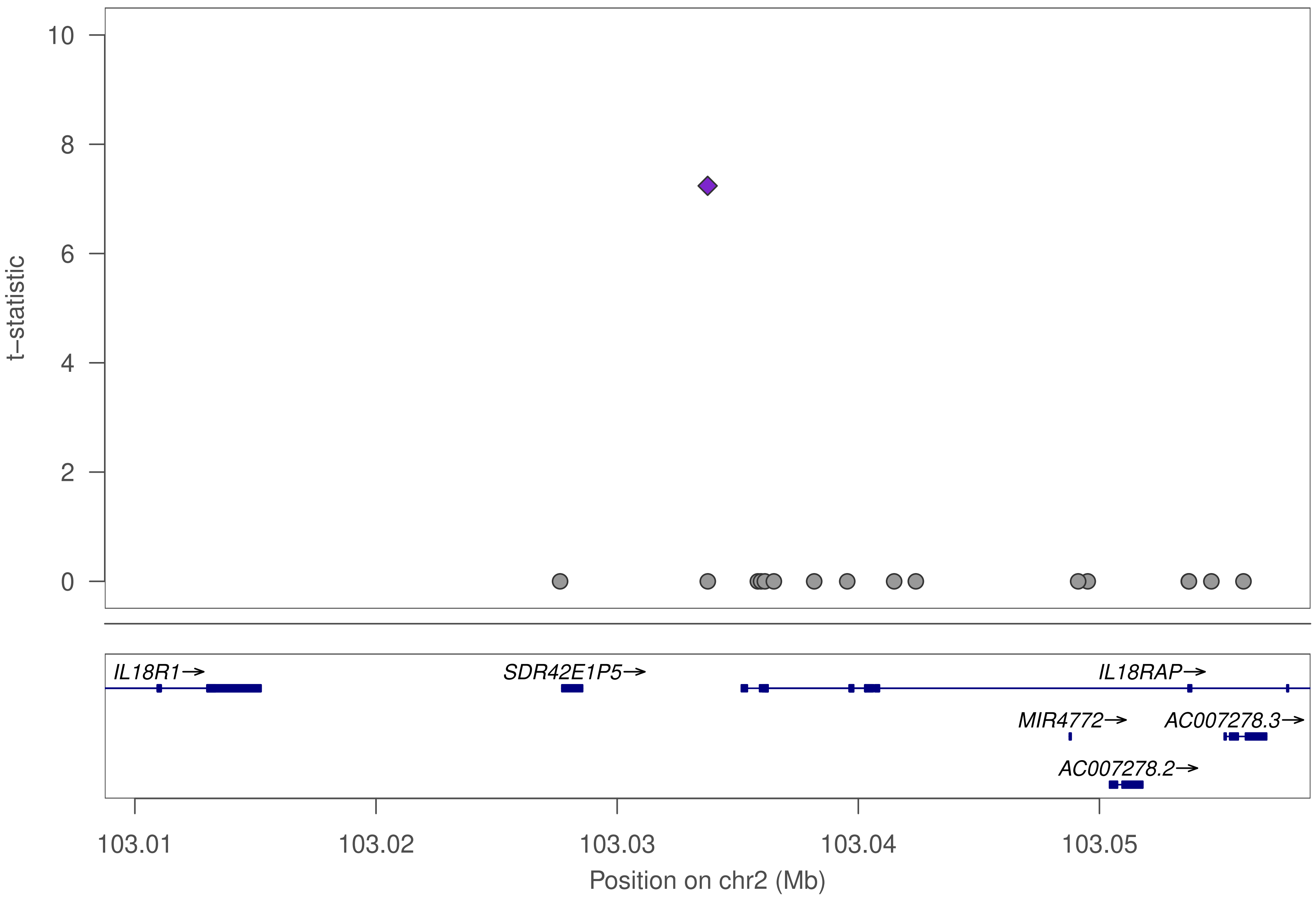

10
8
6
4
2
0
t–statistic
IL18R1→
SDR42E1P5→
IL18RAP→
MIR4772→
AC007278.3→
AC007278.2→
103.01
103.02
103.03
103.04
103.05
Position on chr2 (Mb)

### ZNF197

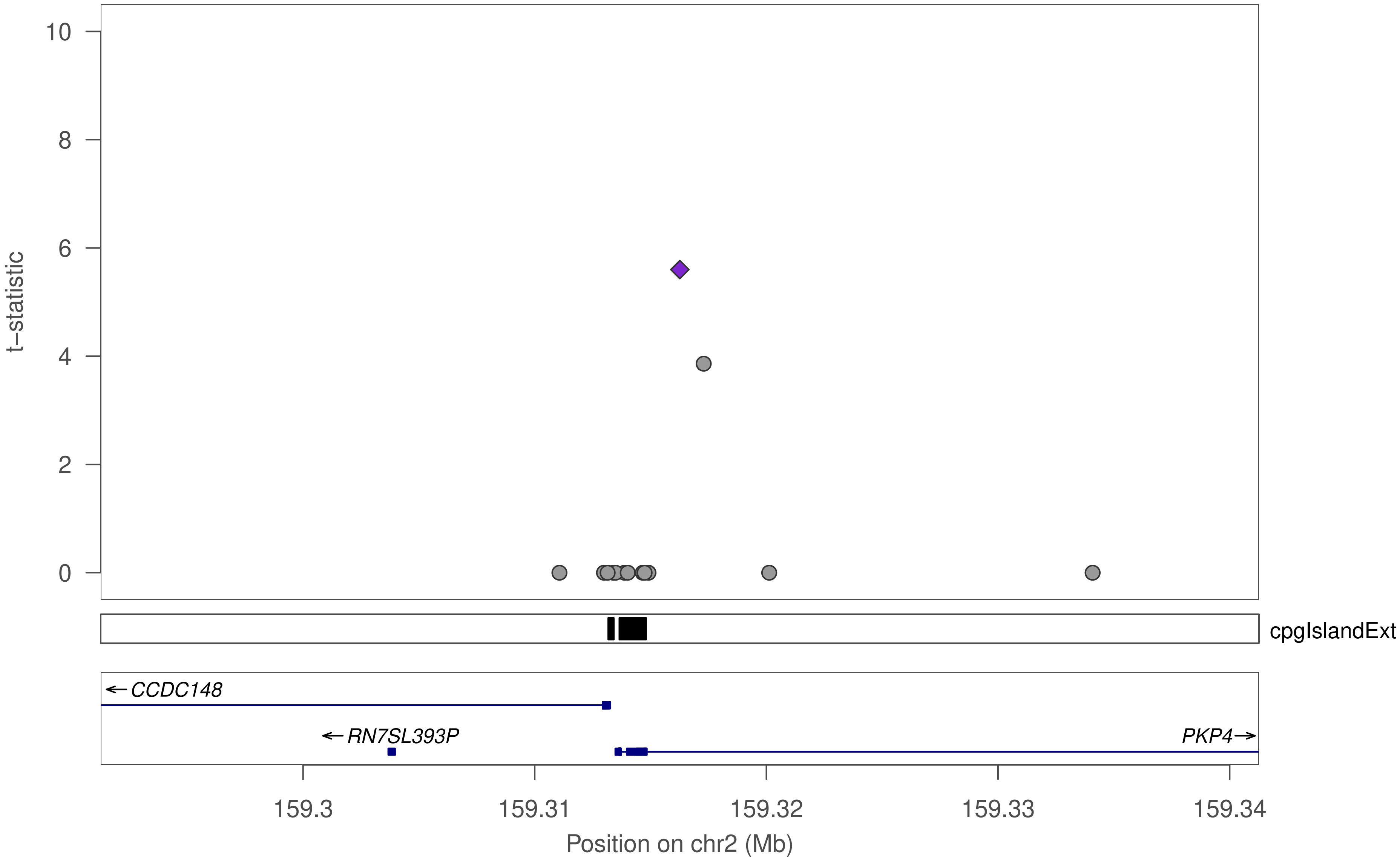

t–statistic
10
8
6
4
2
0
cpgIslandExt
←CCDC148
←RN7SL393P
PKP4→
159.3
159.31
159.32
159.33
159.34
Position on chr2 (Mb)

### ZNF197

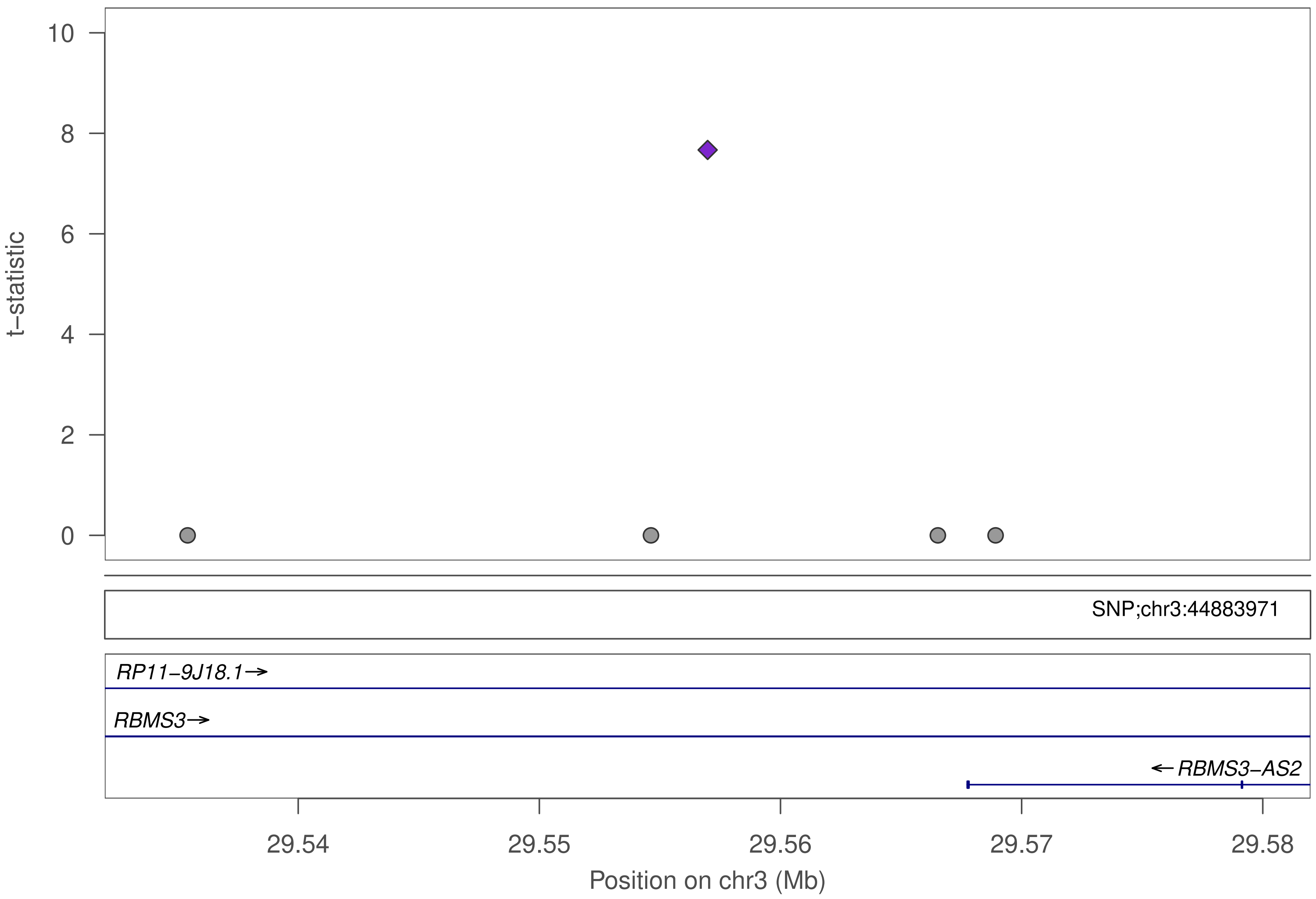

t−statistic
10
8
6
4
2
0
SNP;chr3:44883971
RP11−9J18.1→
RBMS3→
← RBMS3−AS2
29.54
29.55
29.56
29.57
29.58
Position on chr3 (Mb)

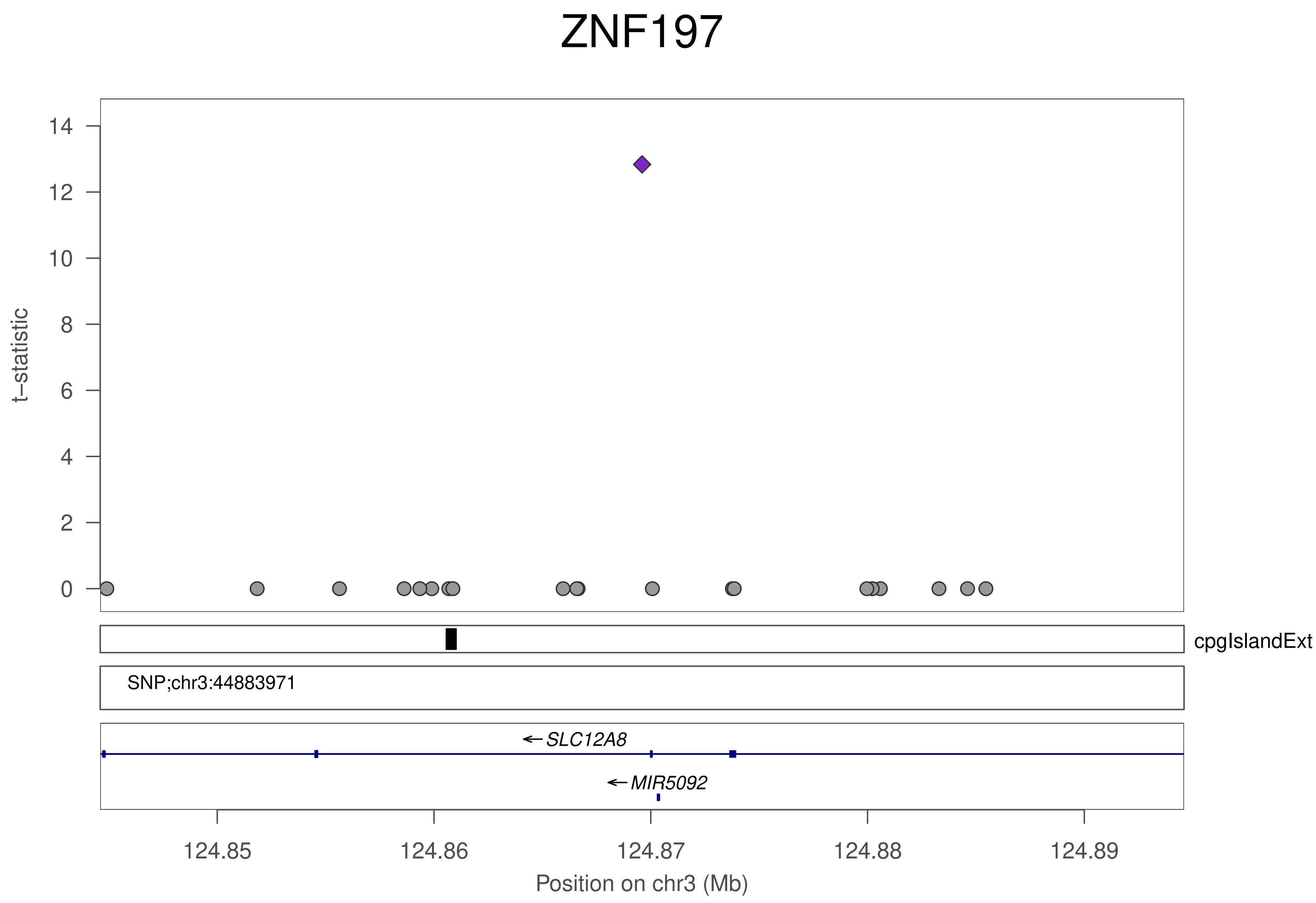

ZNF197
t–statistic
14
12
10
8
6
4
2
0
cpgIslandExt
SNP;chr3:44883971
←SLC12A8
←MIR5092
124.85
124.86
124.87
124.88
124.89
Position on chr3 (Mb)

### ZNF197

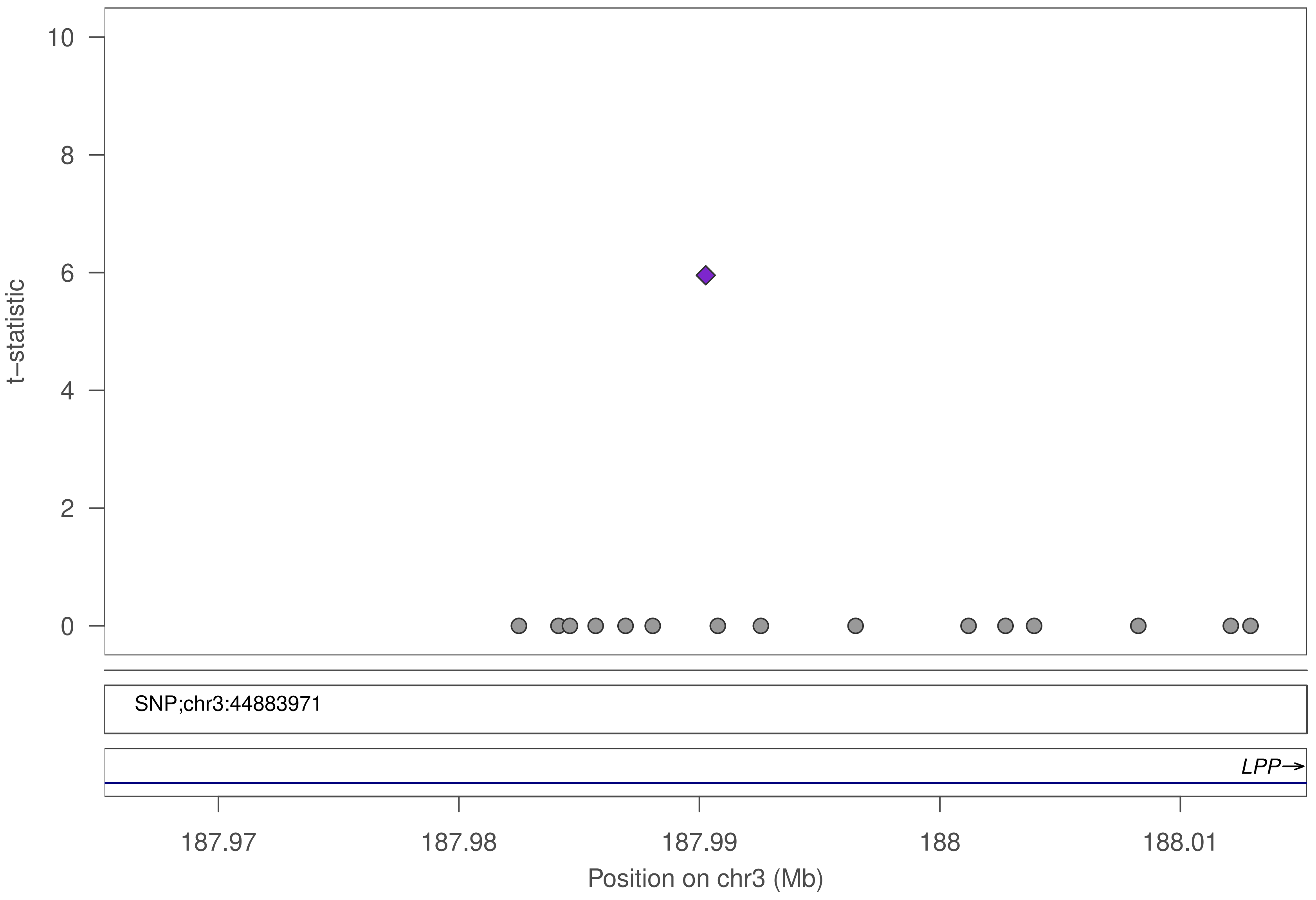

t–statistic
10
8
6
4
2
0
SNP;chr3:44883971
LPP→
187.97
187.98
187.99
188
188.01
Position on chr3 (Mb)

### ZNF197

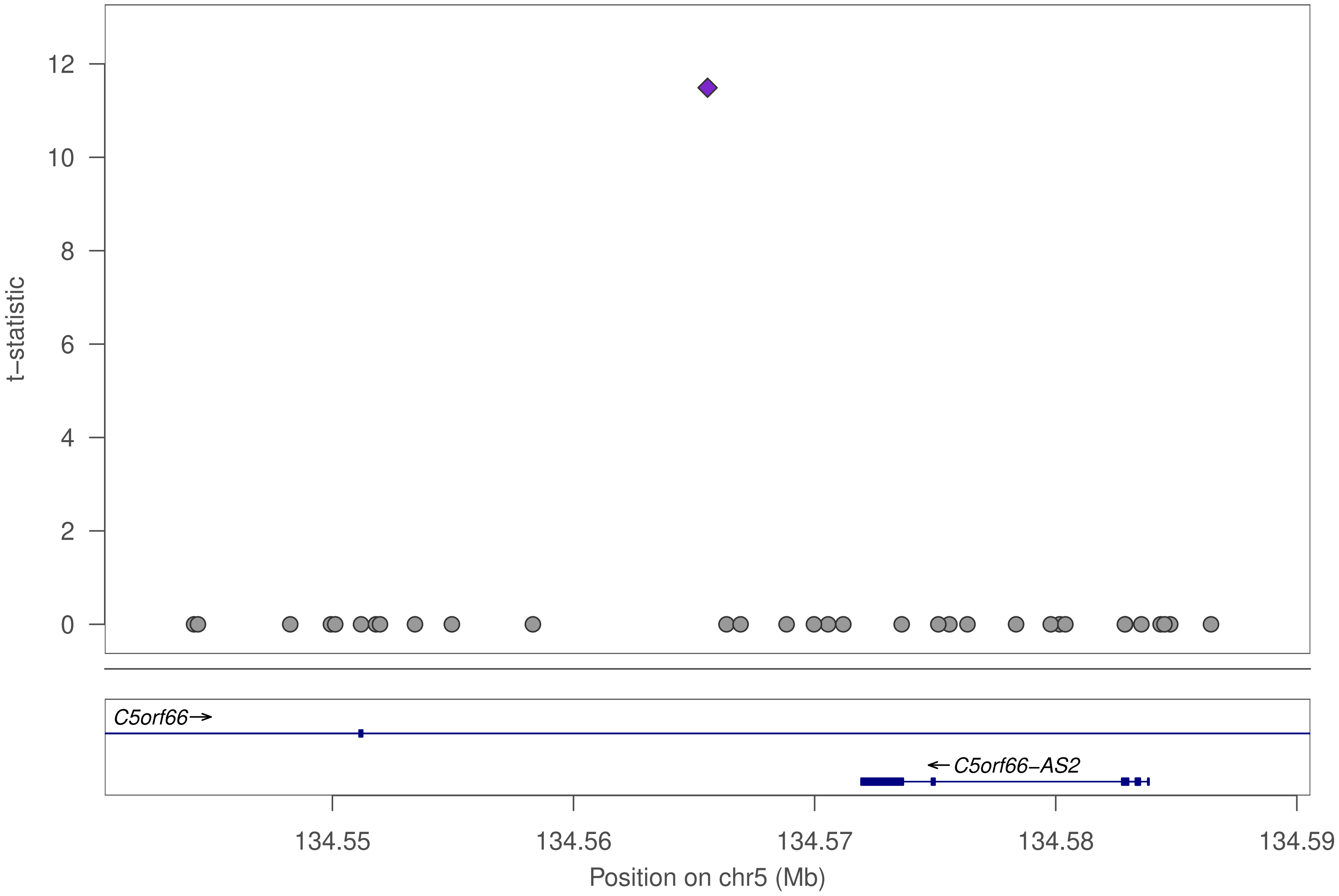

t–statistic
12
10
8
6
4
2
0
C5orf66→
←C5orf66–AS2
134.55
134.56
134.57
134.58
134.59
Position on chr5 (Mb)

### ZNF197

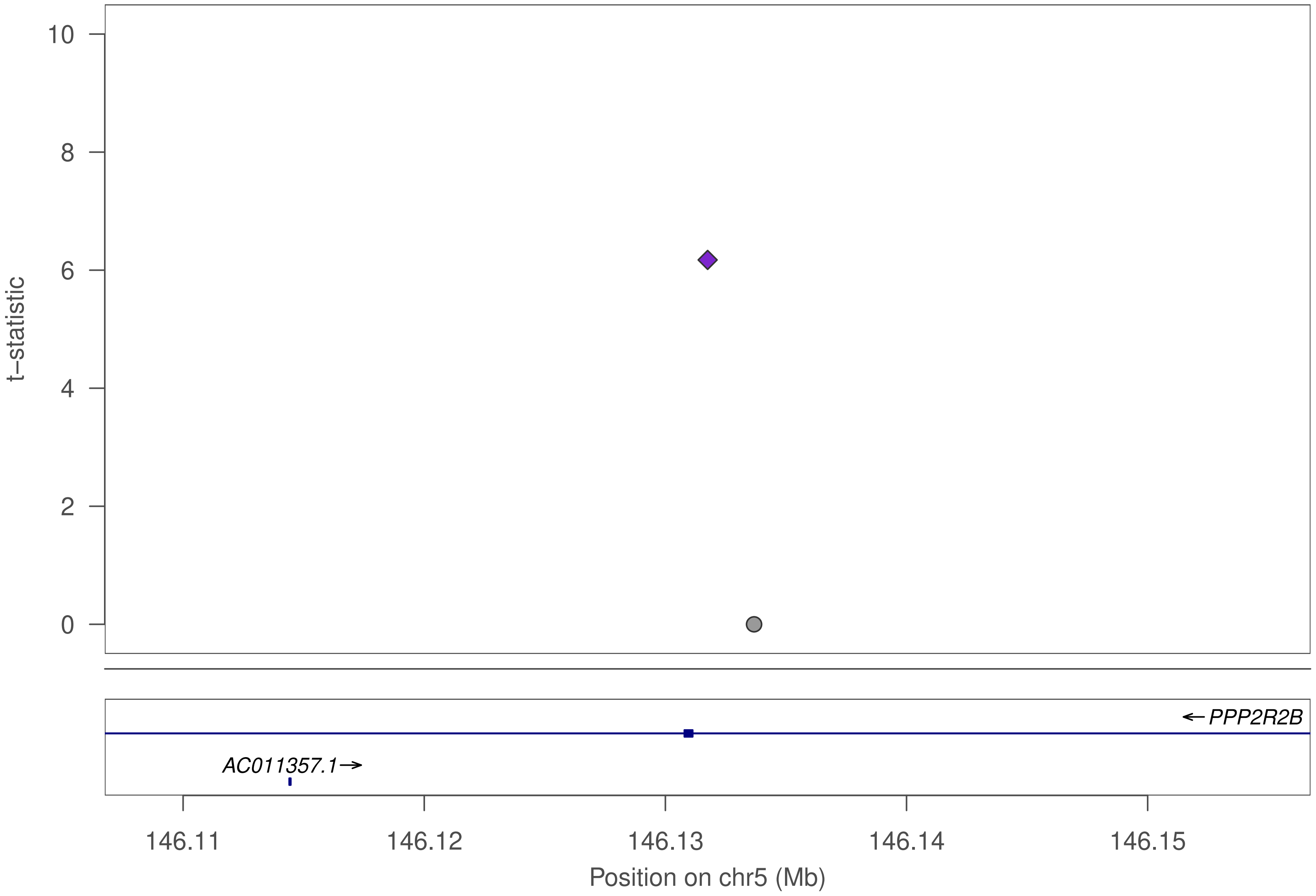

10
8
6
4
2
0
t−statistic
← PPP2R2B
AC011357.1→
146.11
146.12
146.13
146.14
146.15
Position on chr5 (Mb)

### ZNF197

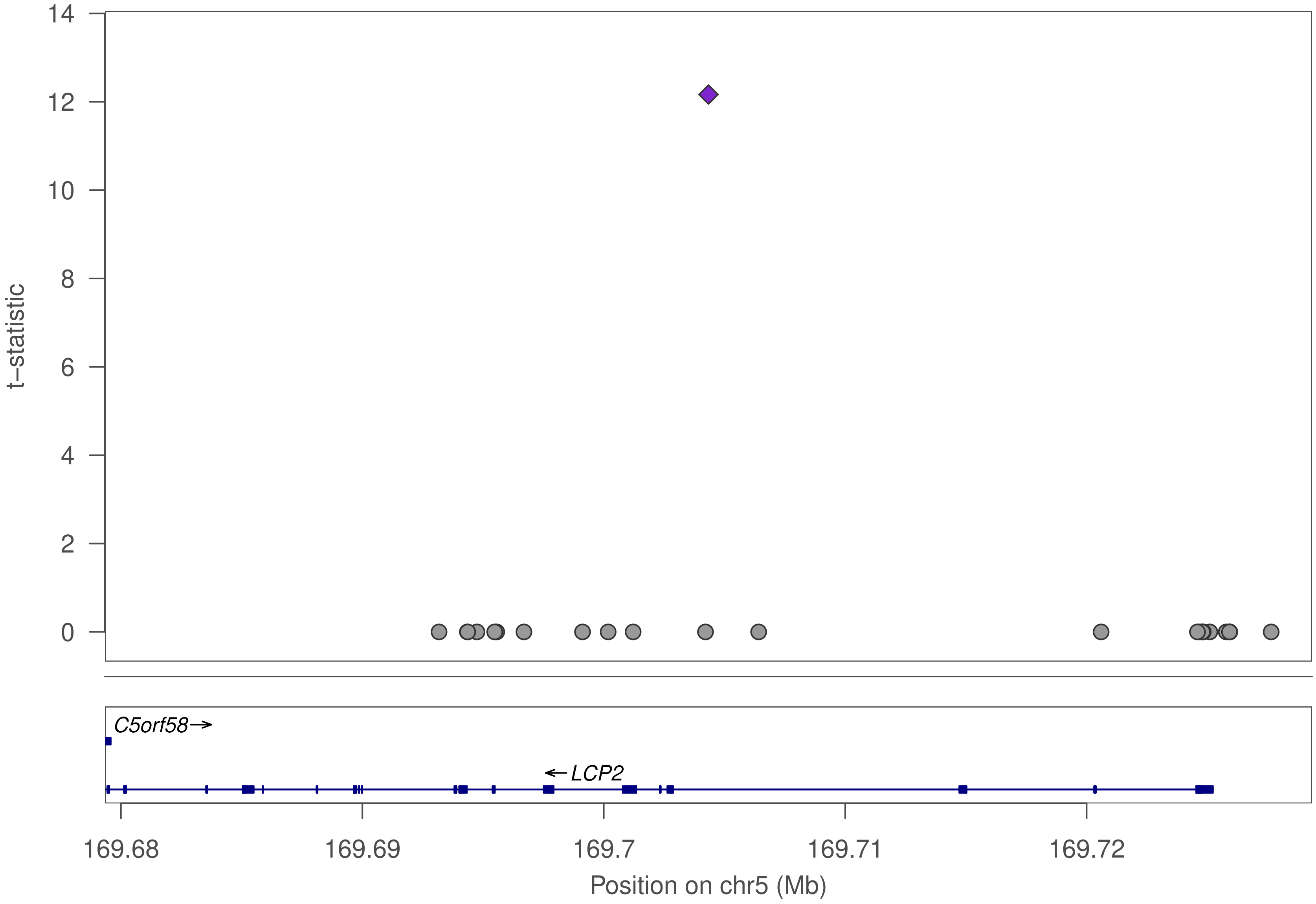

t−statistic
0
2
4
6
8
10
12
14
C5orf58→
←LCP2
169.68
169.69
169.7
169.71
169.72
Position on chr5 (Mb)

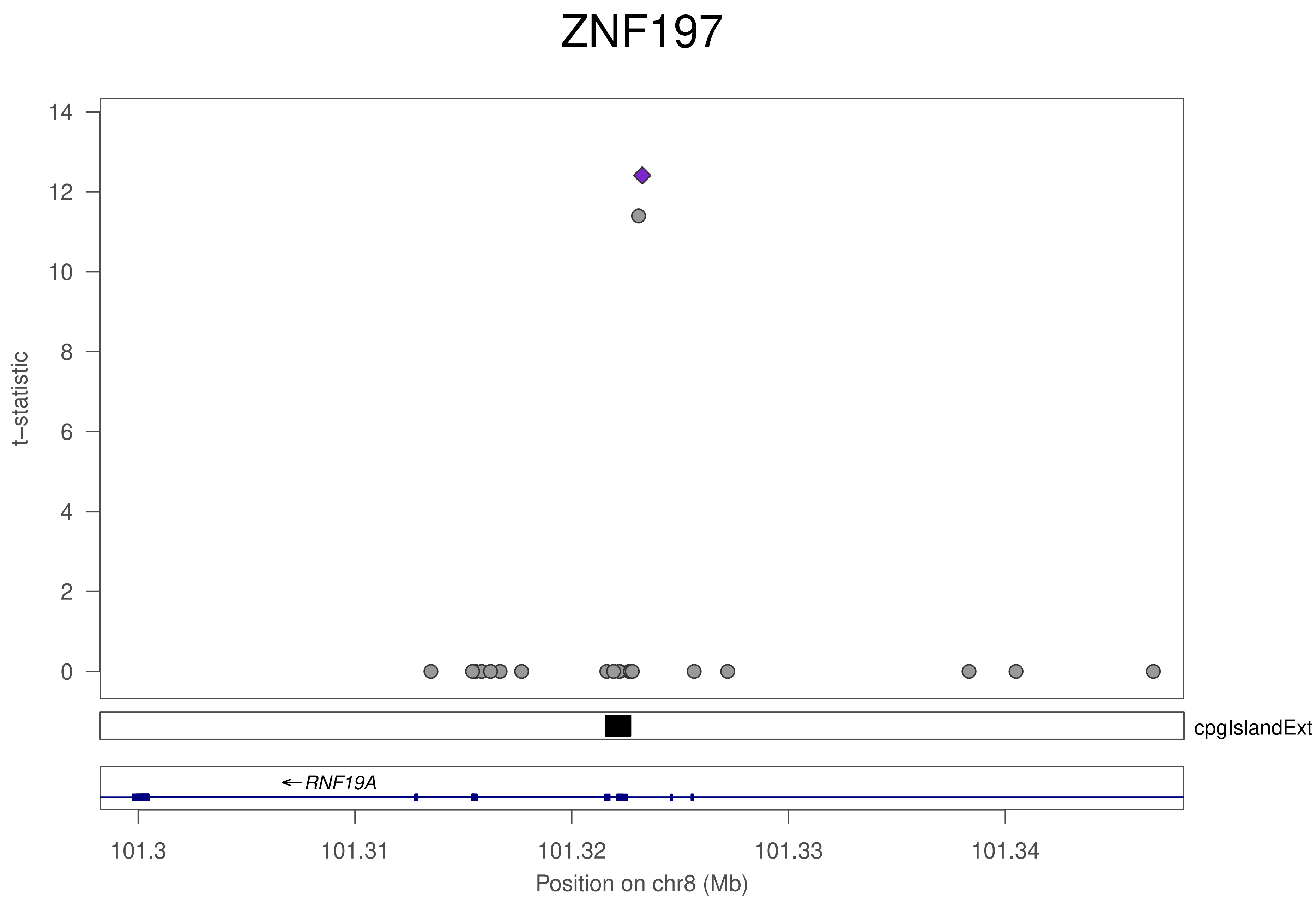
ZNF197
t−statistic
14
12
10
8
6
4
2
0
cpgIslandExt
← RNF19A
101.3
101.31
101.32
101.33
101.34
Position on chr8 (Mb)

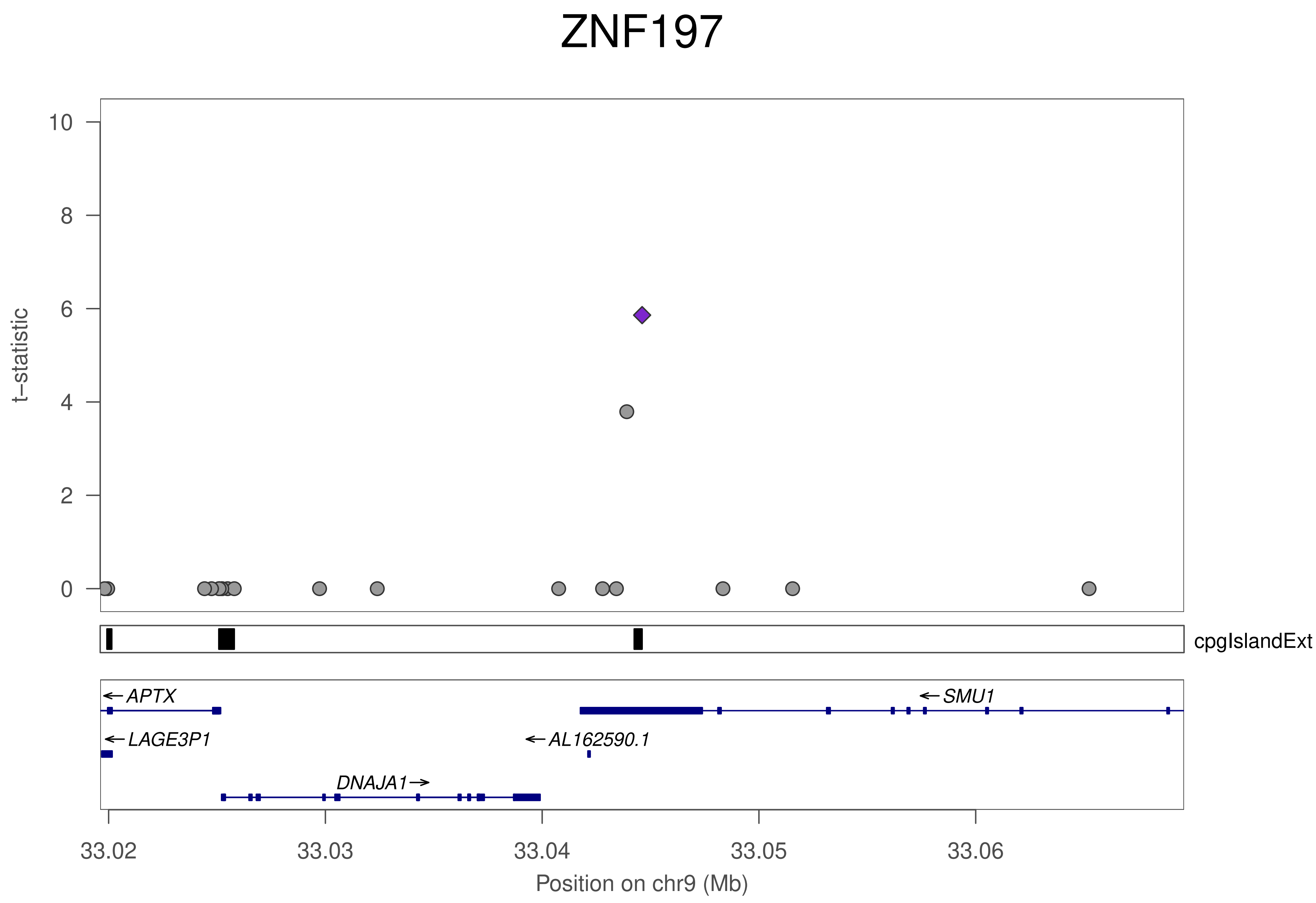
ZNF197
t−statistic
0
2
4
6
8
10
cpgIslandExt
APTX
SMU1
LAGE3P1
AL162590.1
DNAJA1
33.02
33.03
33.04
33.05
33.06
Position on chr9 (Mb)

### ZNF197

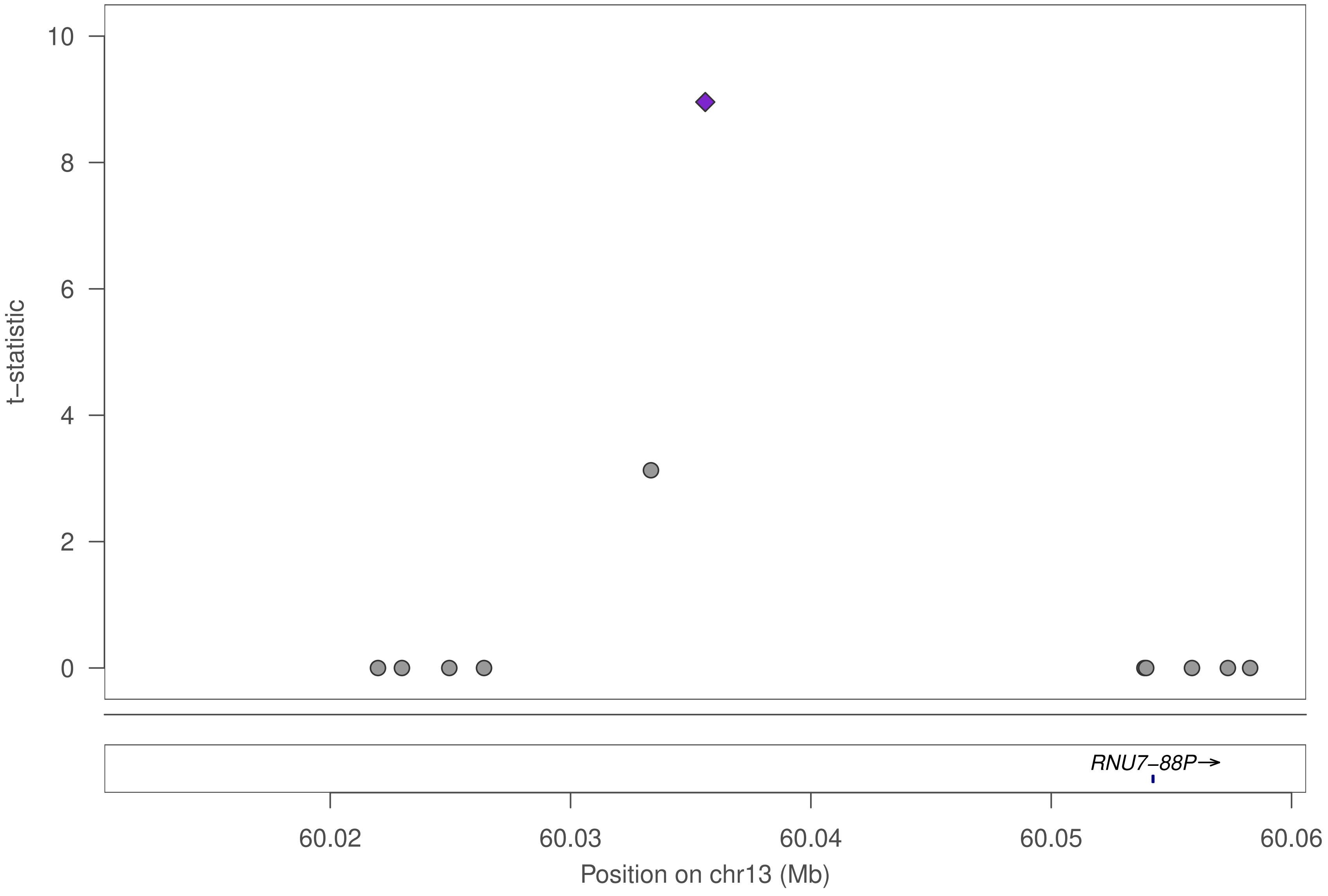

10
8
6
4
2
0
t−statistic
RNU7−88P→
60.02
60.03
60.04
60.05
60.06
Position on chr13 (Mb)

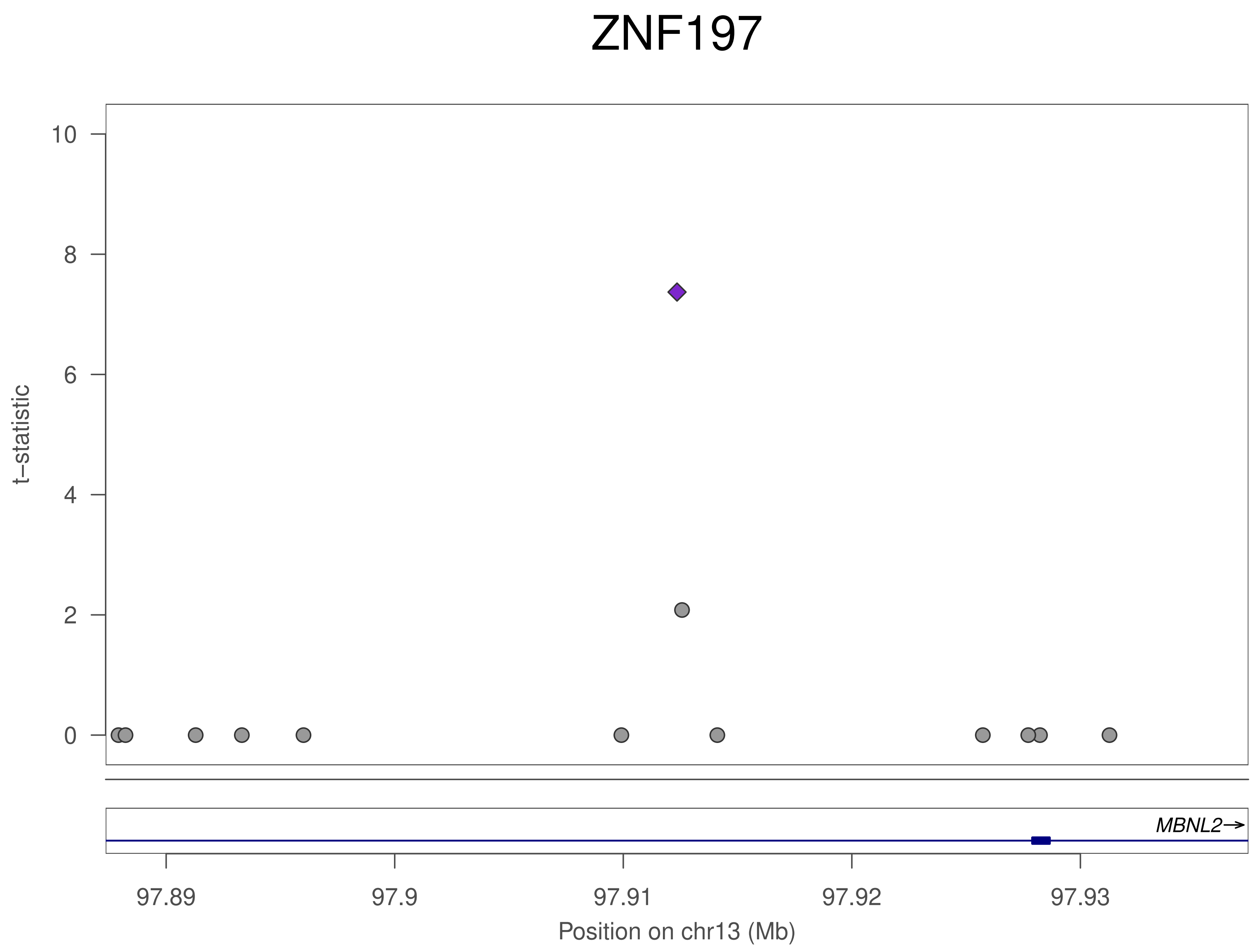
ZNF197
t−statistic
10
8
6
4
2
0
MBNL2→
97.89
97.9
97.91
97.92
97.93
Position on chr13 (Mb)

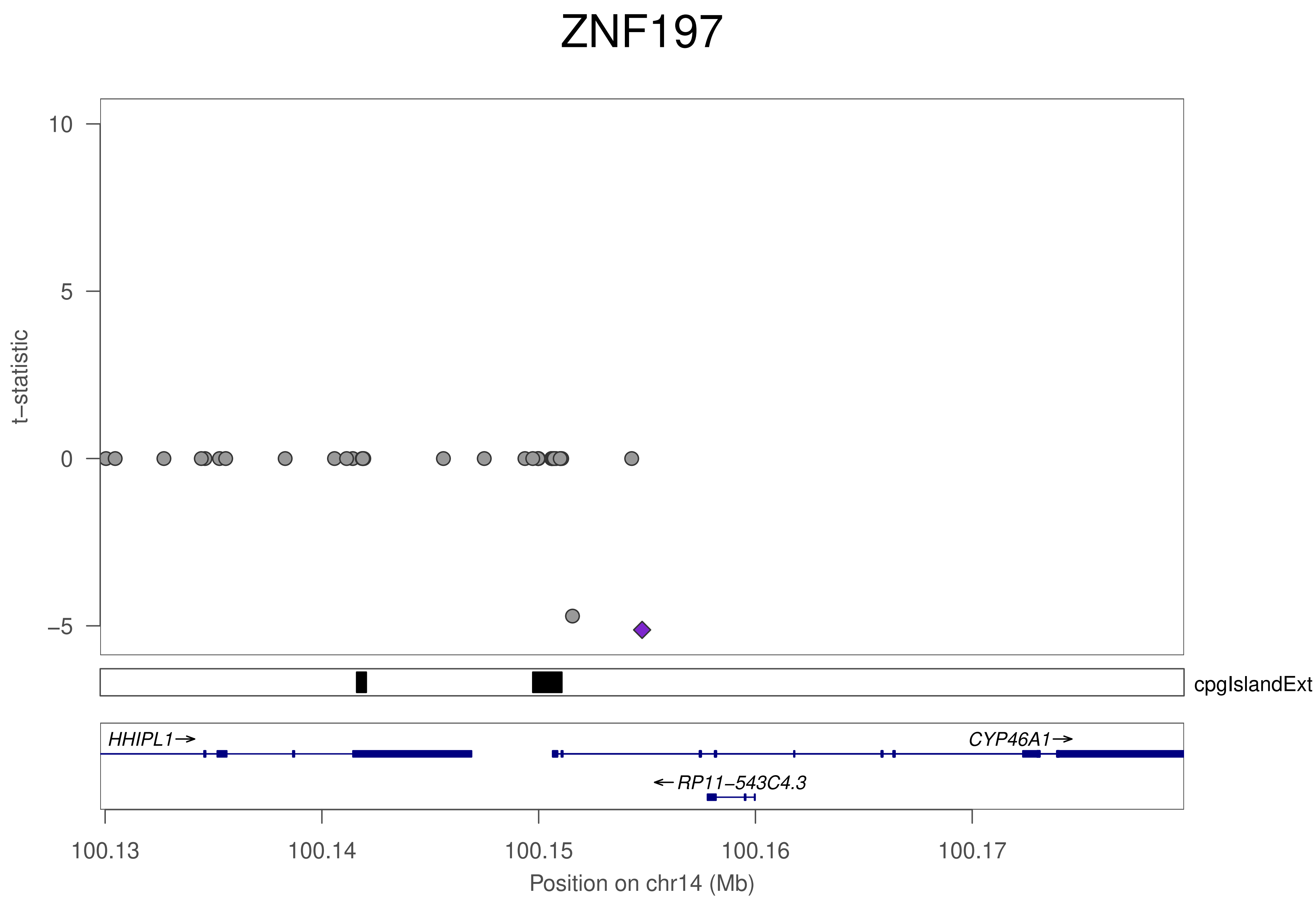
ZNF197
t−statistic
10
5
0
−5
cpgIslandExt
HHIPL1→
CYP46A1→
←RP11−543C4.3
100.13
100.14
100.15
100.16
100.17
Position on chr14 (Mb)

### ZNF197

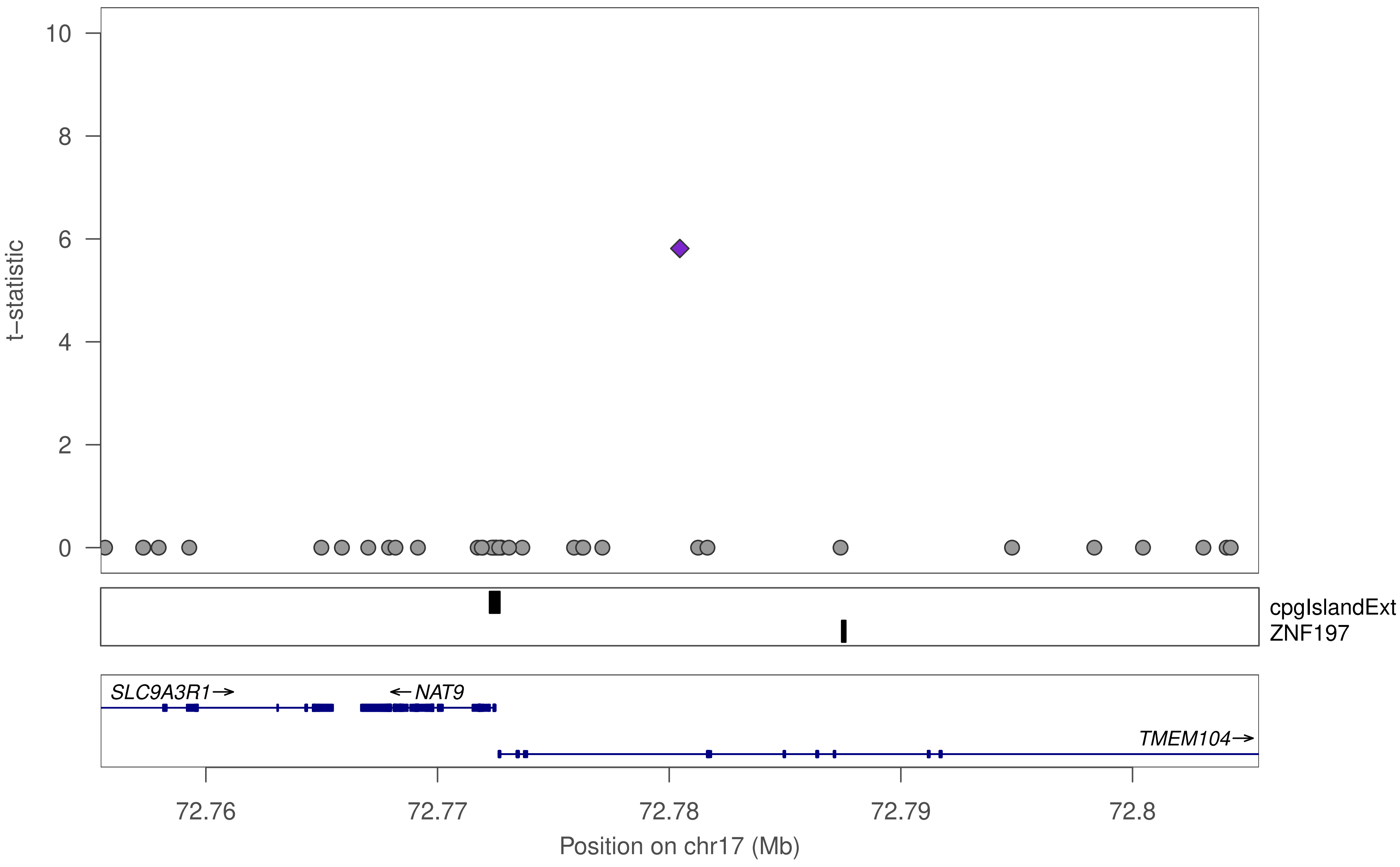

t−statistic
0
2
4
6
8
10
cpgIslandExt
ZNF197
SLC9A3R1→
←NAT9
TMEM104→
72.76
72.77
72.78
72.79
72.8
Position on chr17 (Mb)

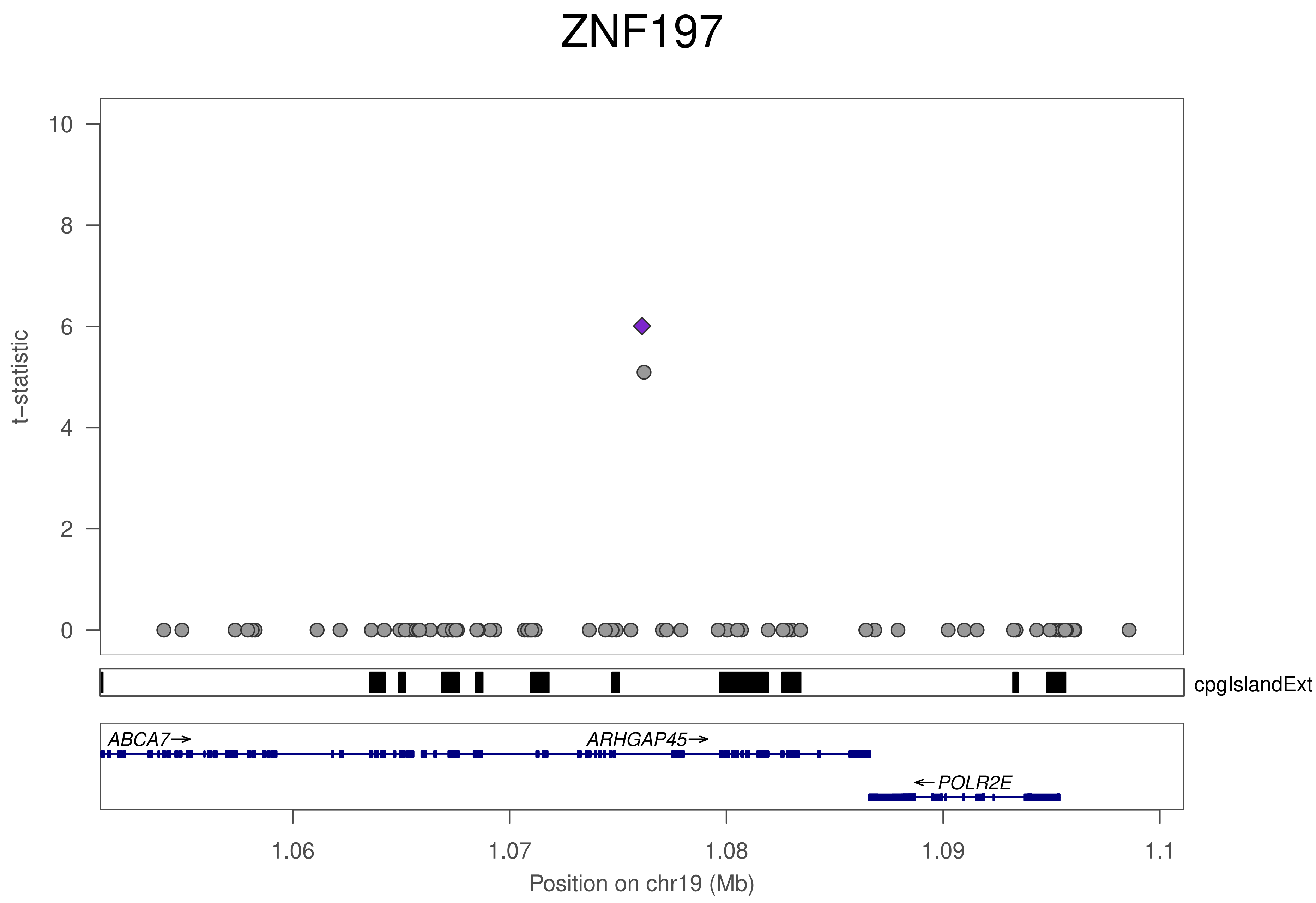
ZNF197
t−statistic
0
2
4
6
8
10
cpgIslandExt
ABCA7→
ARHGAP45→
←POLR2E
1.06
1.07
1.08
1.09
1.1
Position on chr19 (Mb)

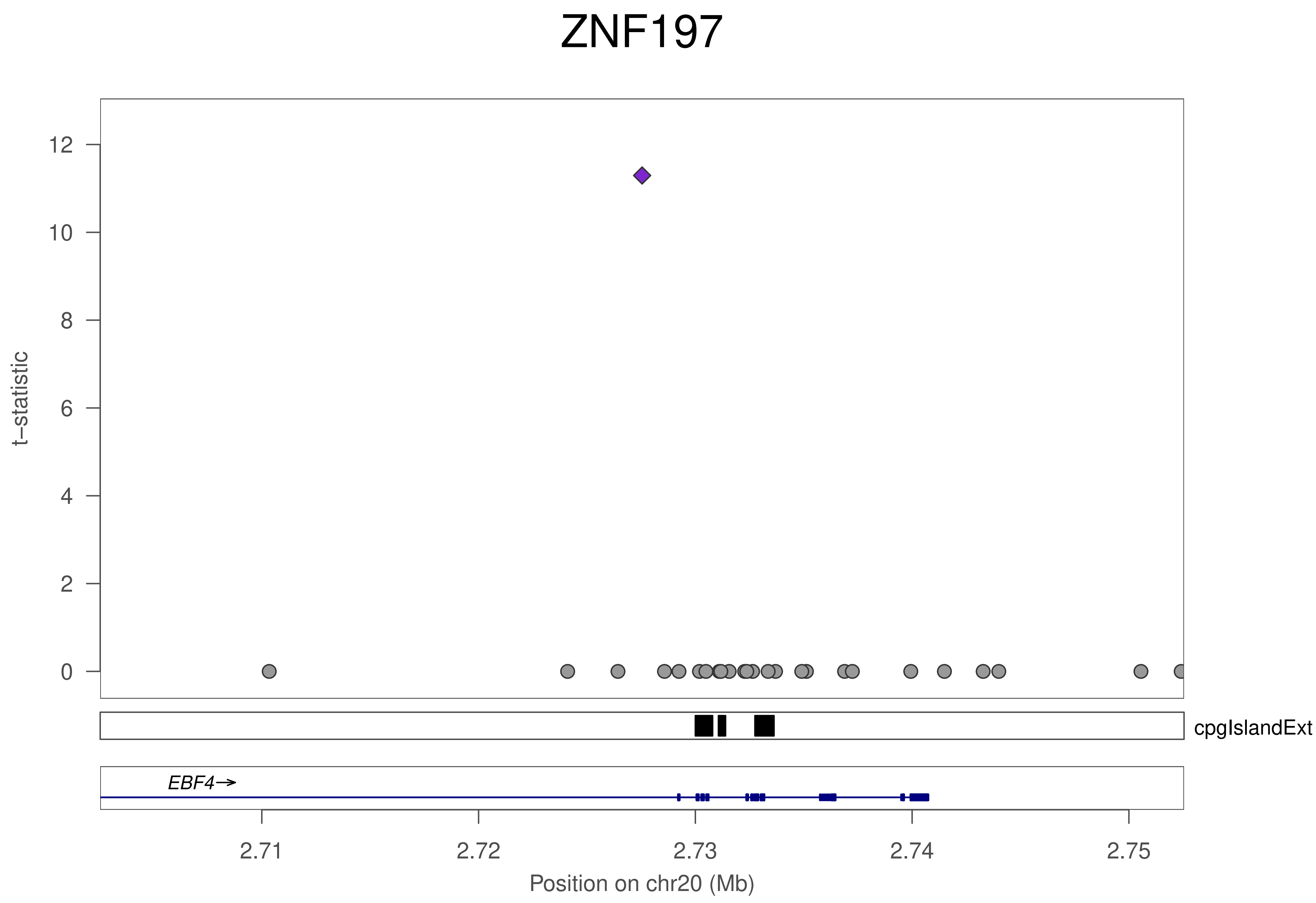

ZNF197
t–statistic
0
2
4
6
8
10
12
cpgIslandExt
EBF4→
2.71
2.72
2.73
2.74
2.75
Position on chr20 (Mb)

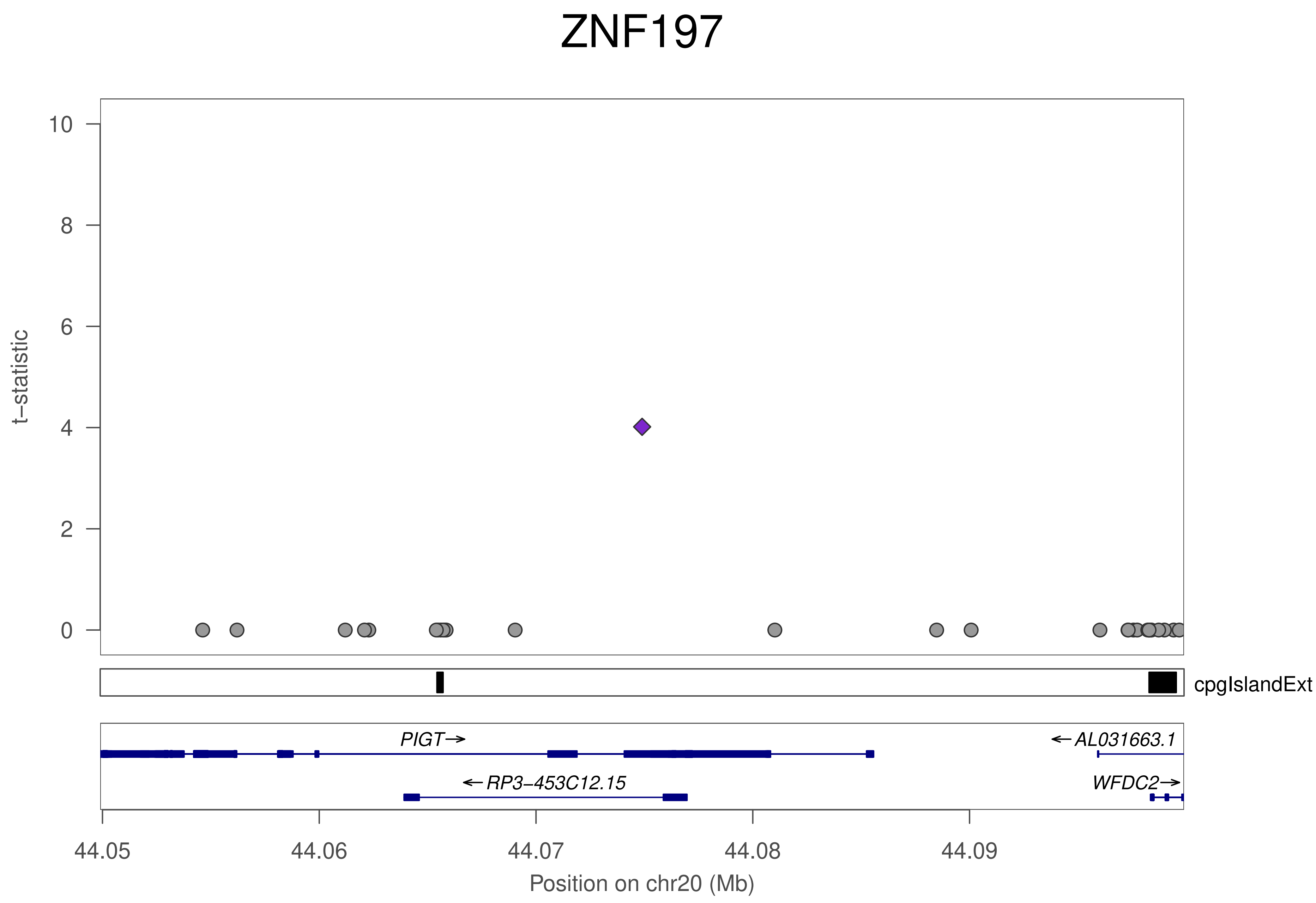

ZNF197
t−statistic
10
8
6
4
2
0
cpgIslandExt
PIGT→
←AL031663.1
←RP3−453C12.15
WFDC2→
44.05
44.06
44.07
44.08
44.09
Position on chr20 (Mb)

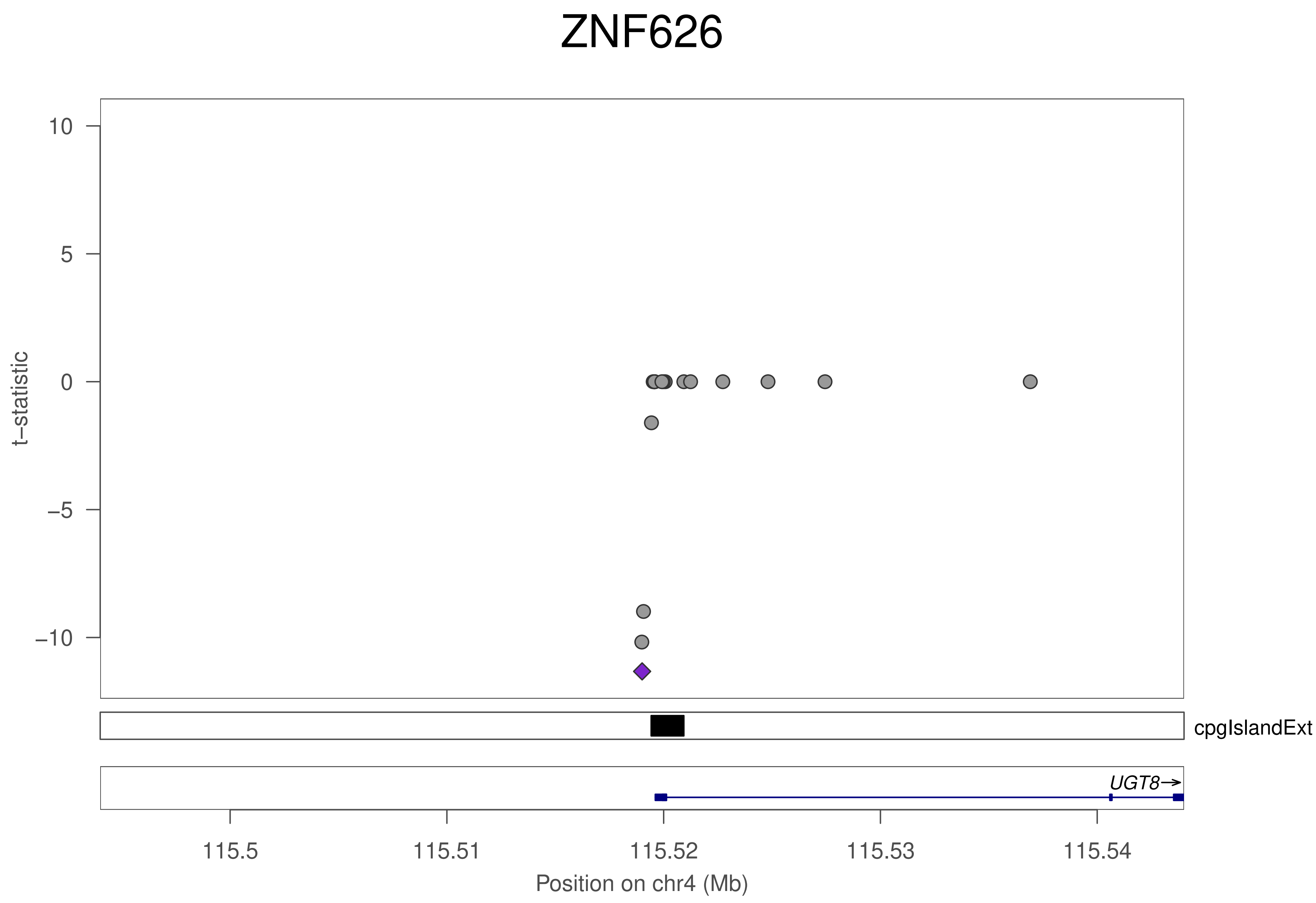

ZNF626
10
5
0
−5
−10
t−statistic
cpgIslandExt
UGT8→
115.5
115.51
115.52
115.53
115.54
Position on chr4 (Mb)

### ZNF626

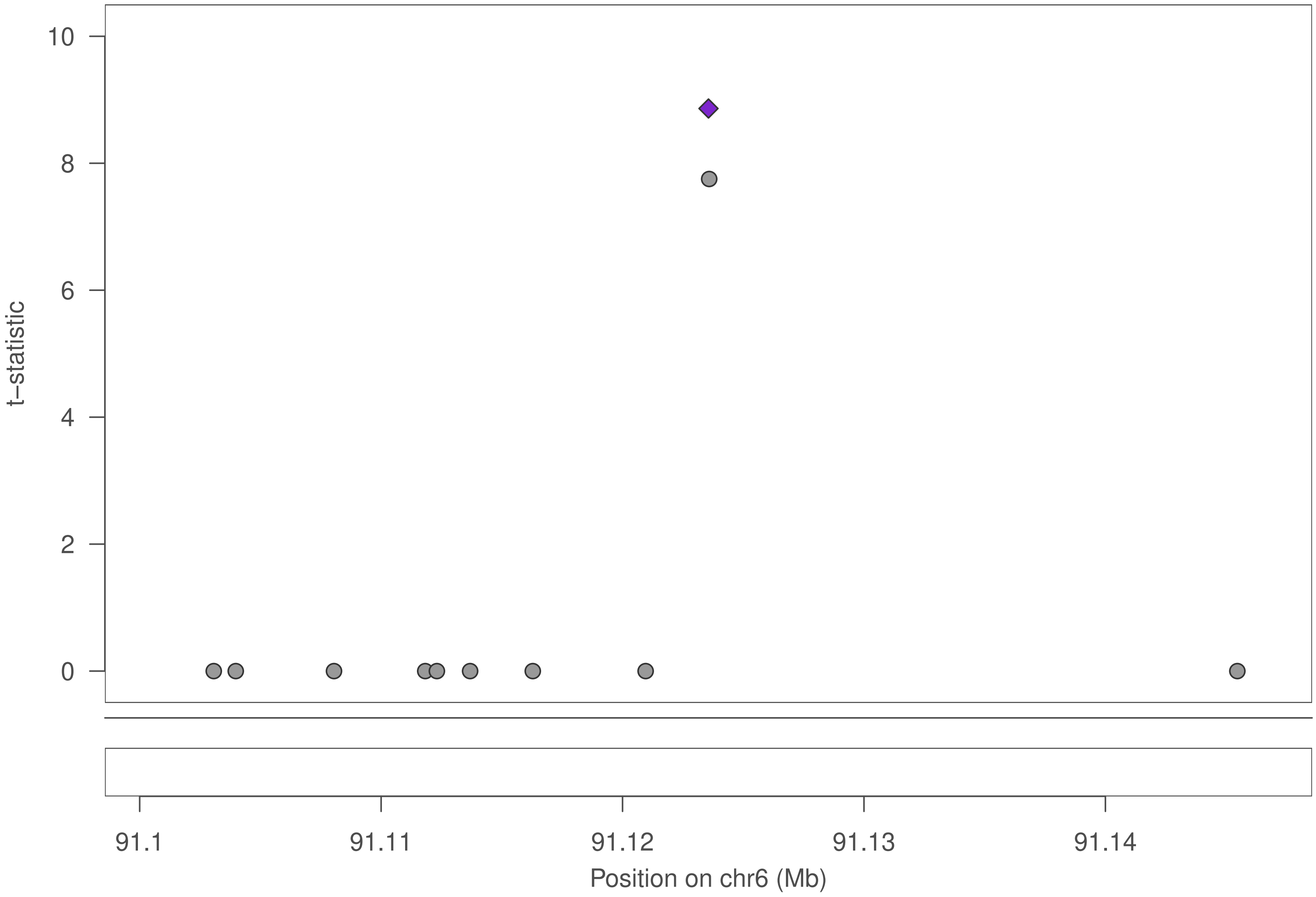

10
8
6
4
2
0
t−statistic
91.1
91.11
91.12
91.13
91.14
Position on chr6 (Mb)

### ZNF626

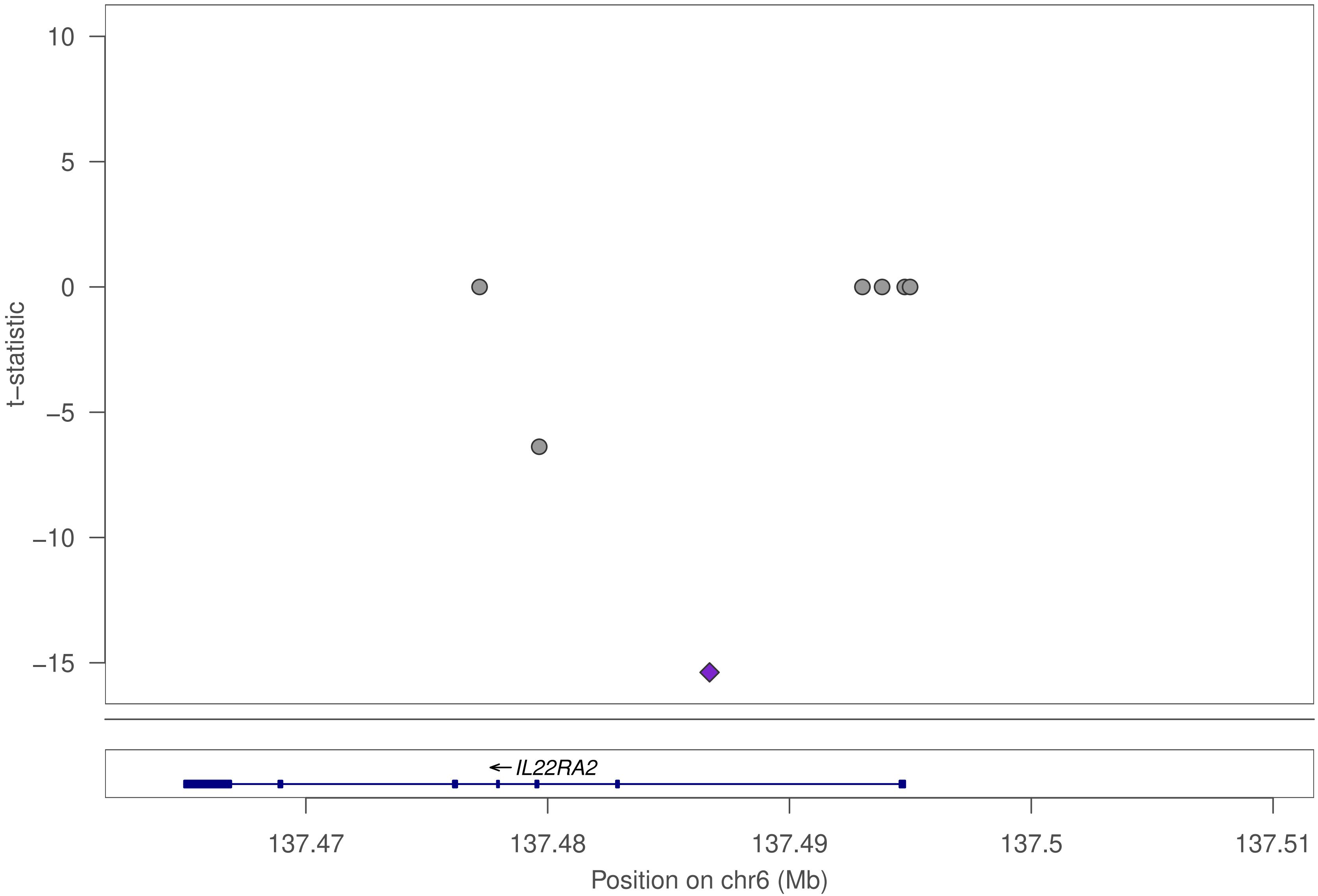

10
5
0
−5
−10
−15
t−statistic
IL22RA2
137.47
137.48
137.49
137.5
137.51
Position on chr6 (Mb)

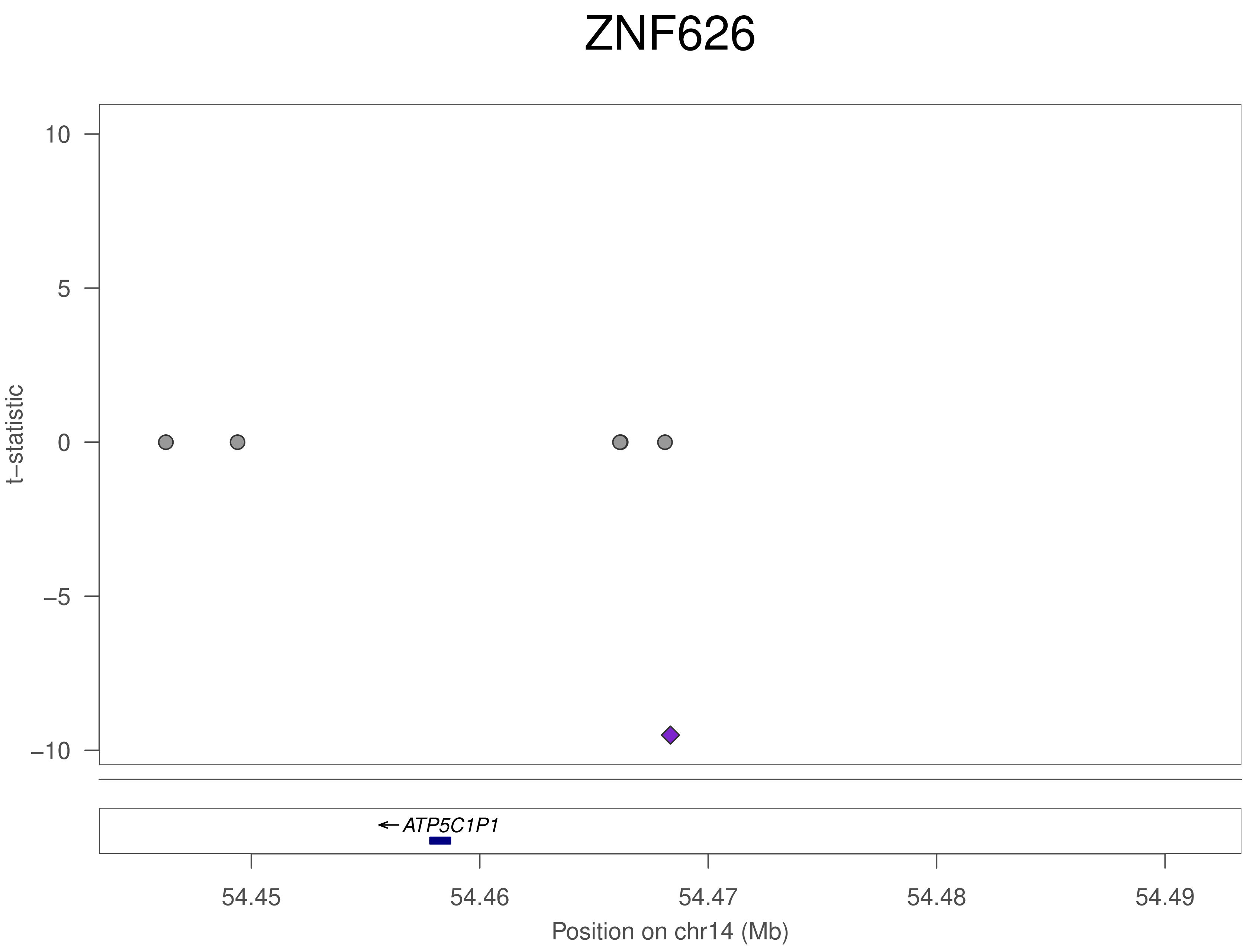
ZNF626
10
5
0
−5
−10
t−statistic
←ATP5C1P1
54.45
54.46
54.47
54.48
54.49
Position on chr14 (Mb)

### ZNF793

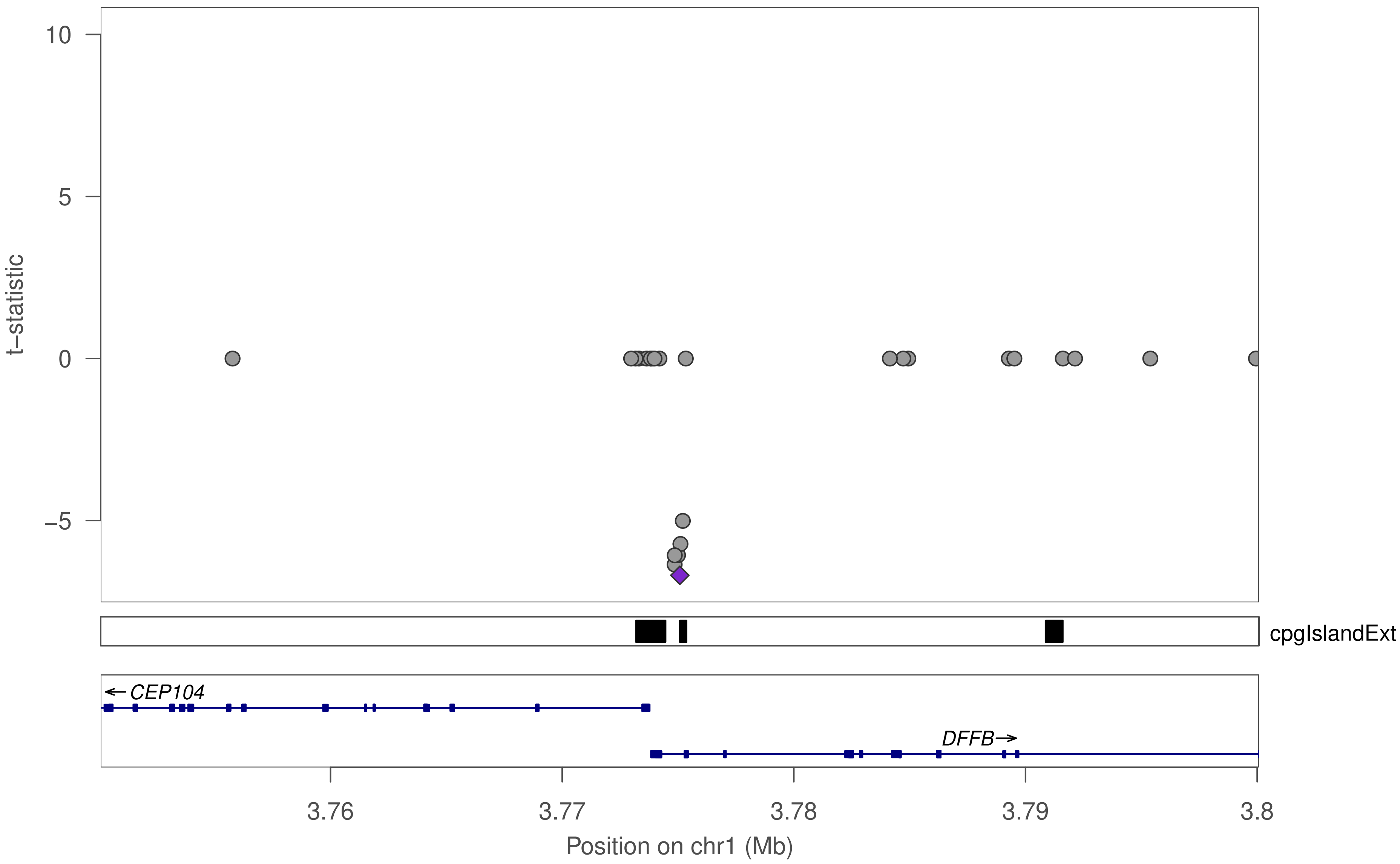

10
5
0
−5
t−statistic
cpgIslandExt
CEP104
DFFB
3.76
3.77
3.78
3.79
3.8
Position on chr1 (Mb)

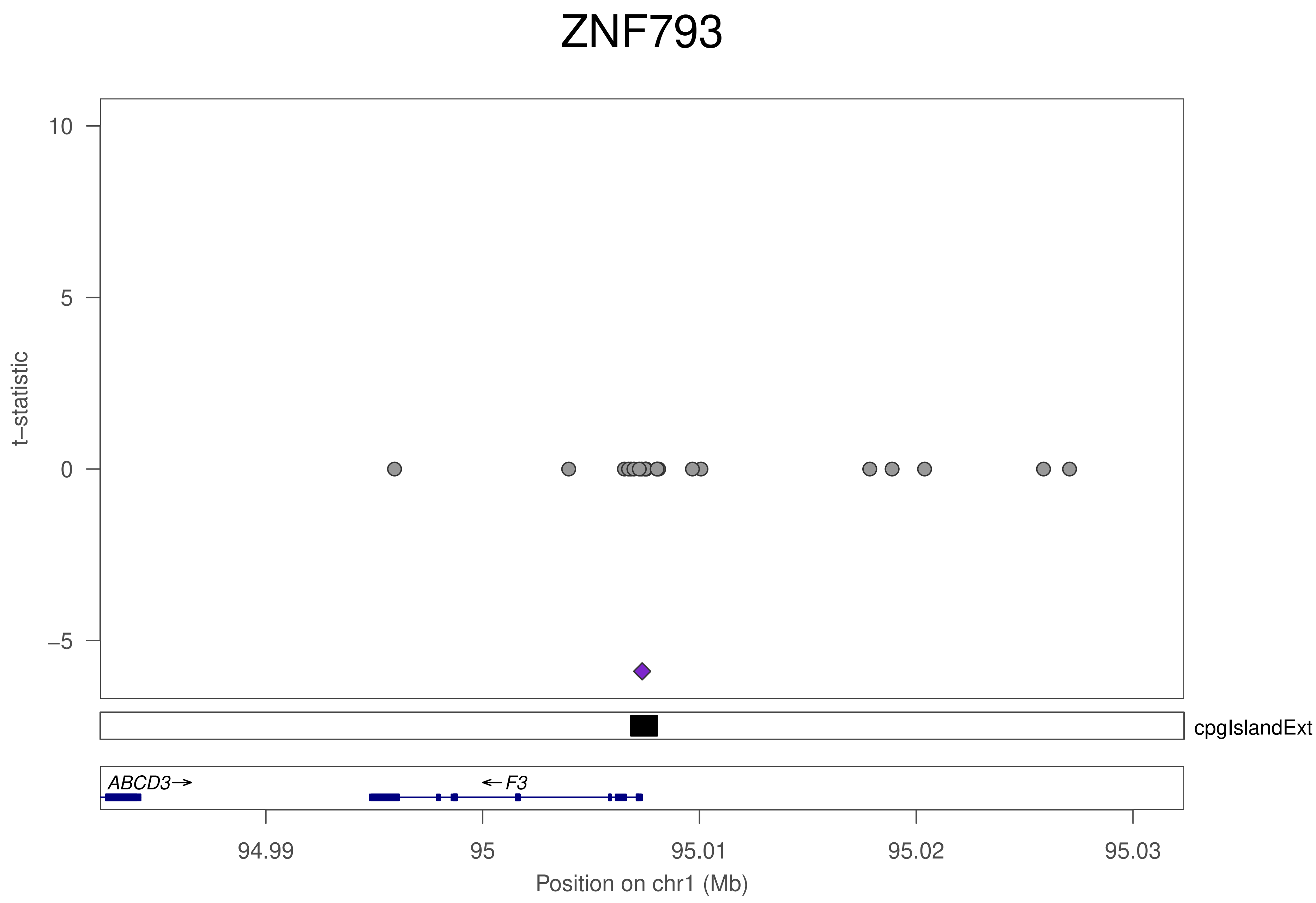
ZNF793
t–statistic
10
5
0
–5
cpgIslandExt
ABCD3→
←F3
94.99
95
95.01
95.02
95.03
Position on chr1 (Mb)

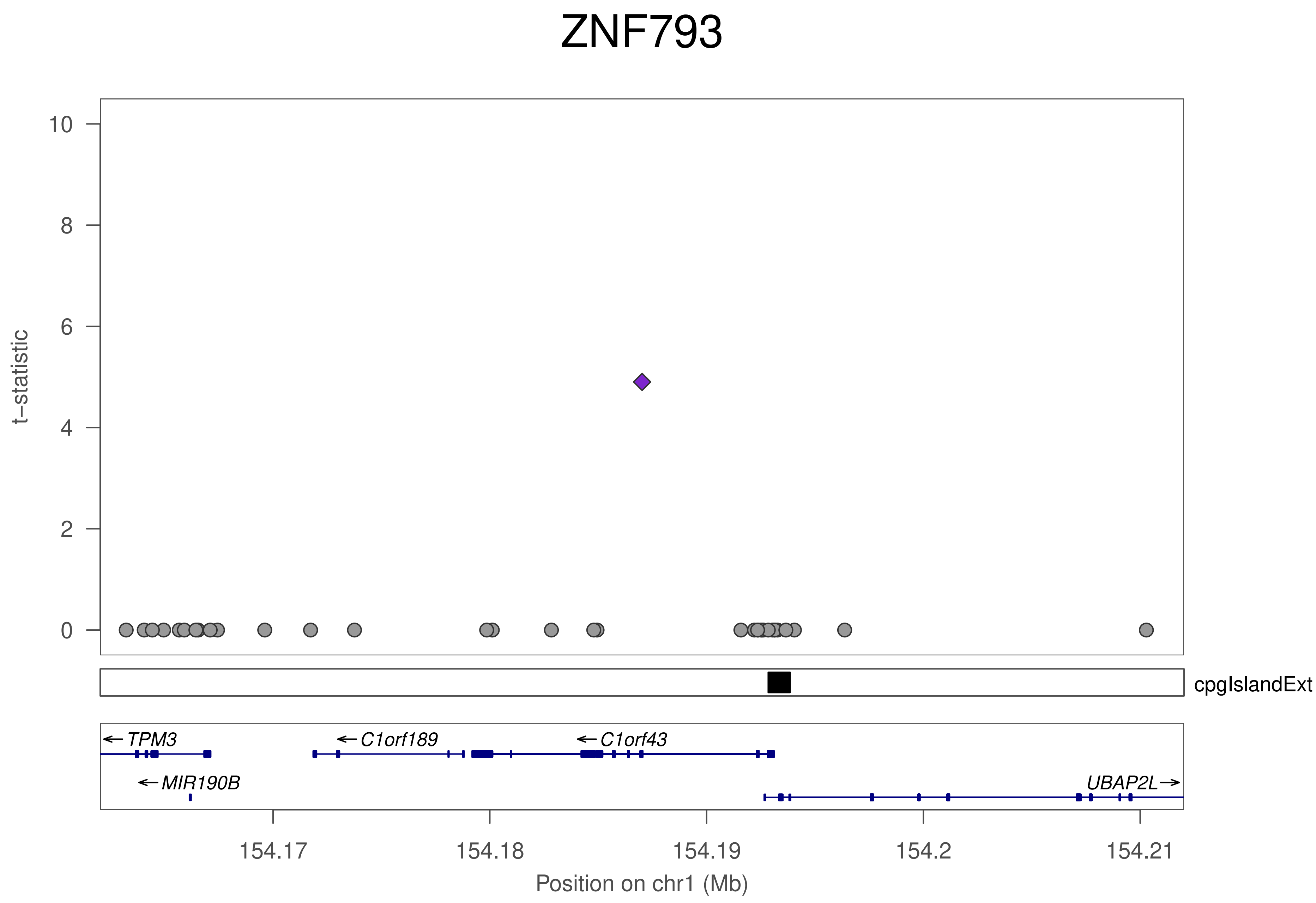

ZNF793
t−statistic
0
2
4
6
8
10
cpgIslandExt
TPM3
MIR190B
C1orf189
C1orf43
UBAP2L
154.17
154.18
154.19
154.2
154.21
Position on chr1 (Mb)

### ZNF793

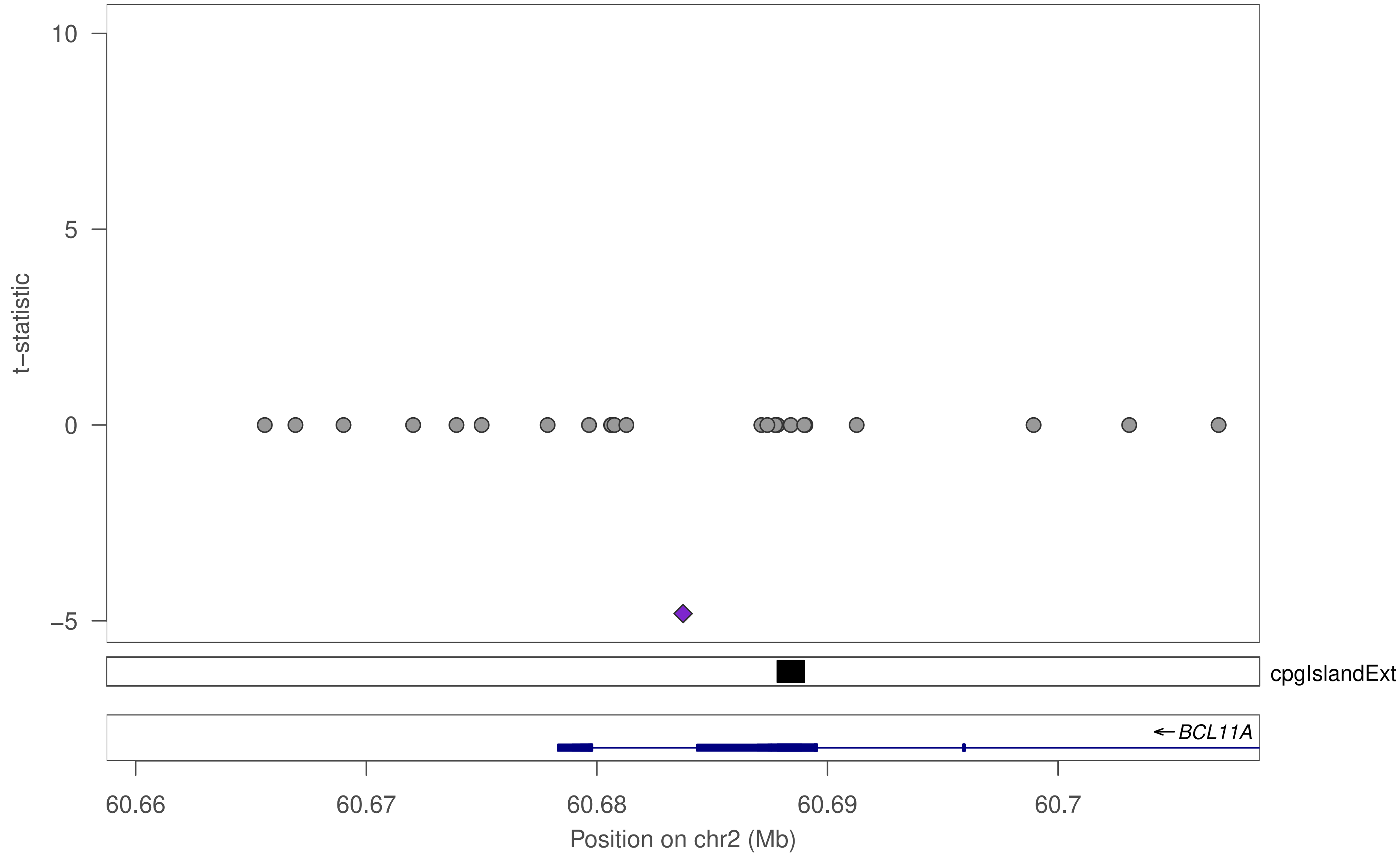

10
5
0
−5
t−statistic
cpgIslandExt
← BCL11A
60.66
60.67
60.68
60.69
60.7
Position on chr2 (Mb)

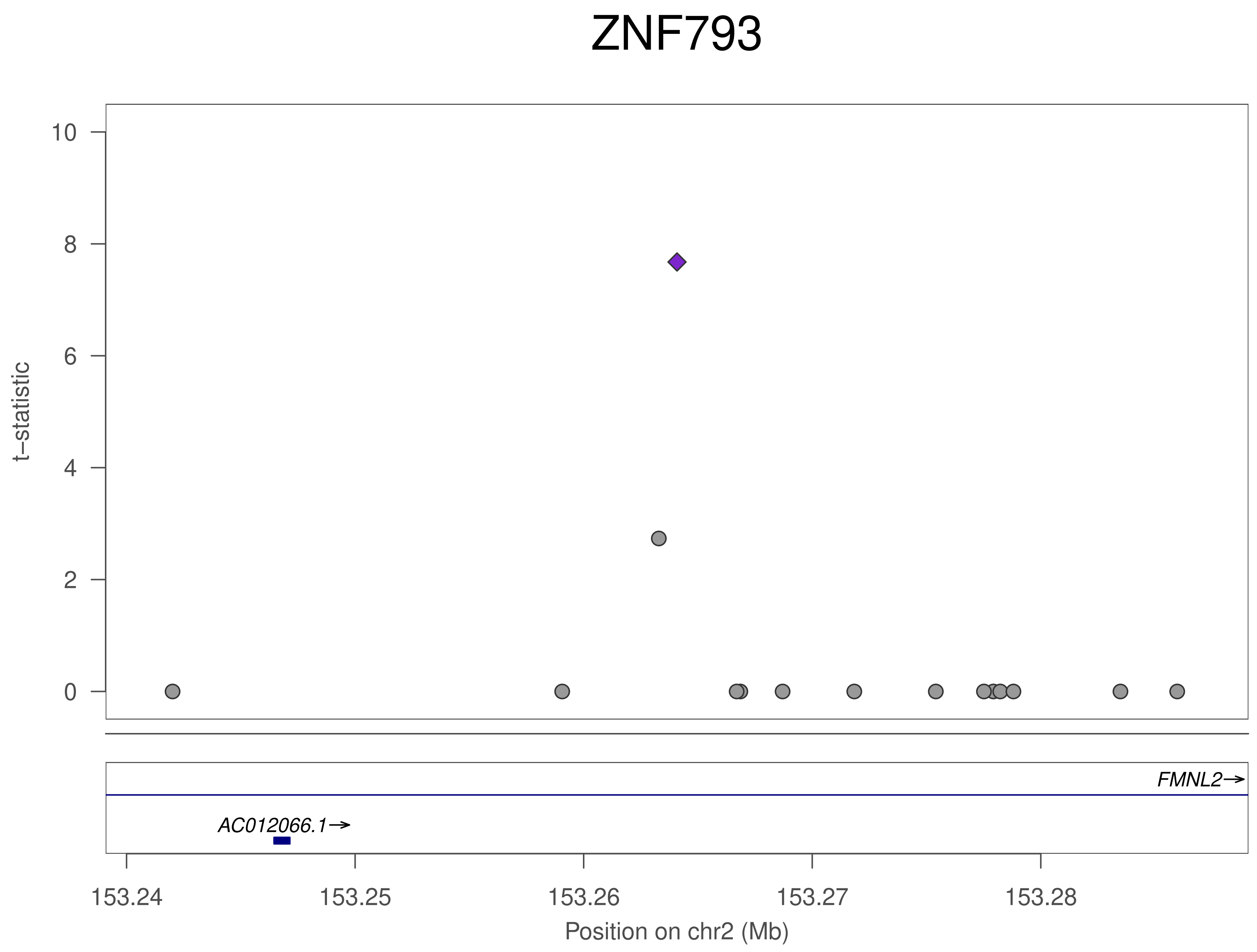
ZNF793
t–statistic
10
8
6
4
2
0
FMNL2→
AC012066.1→
153.24
153.25
153.26
153.27
153.28
Position on chr2 (Mb)

### ZNF793

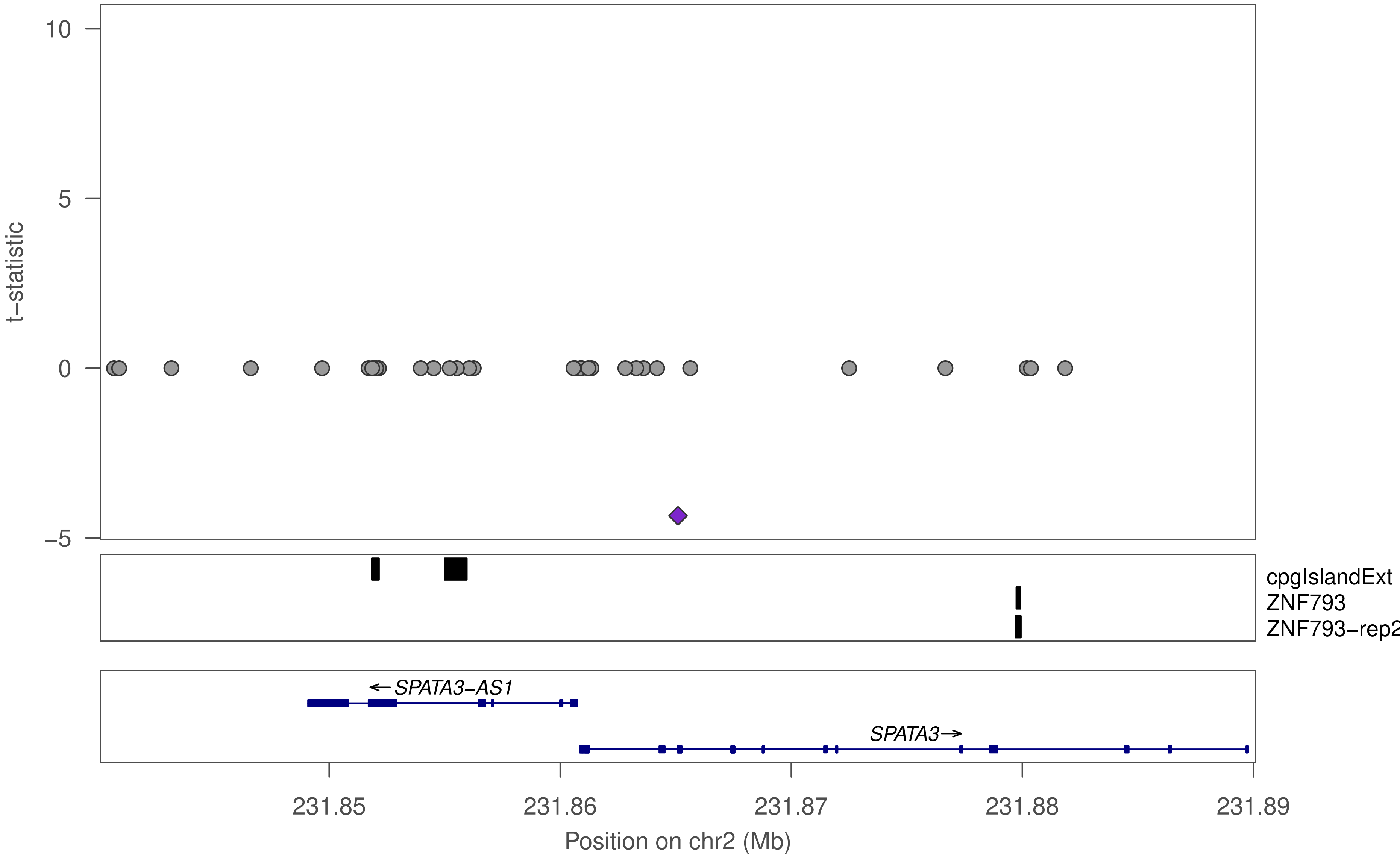

t−statistic
10
5
0
−5
cpgIslandExt
ZNF793
ZNF793−rep2
←SPATA3−AS1
SPATA3→
231.85
231.86
231.87
231.88
231.89
Position on chr2 (Mb)

### ZNF793

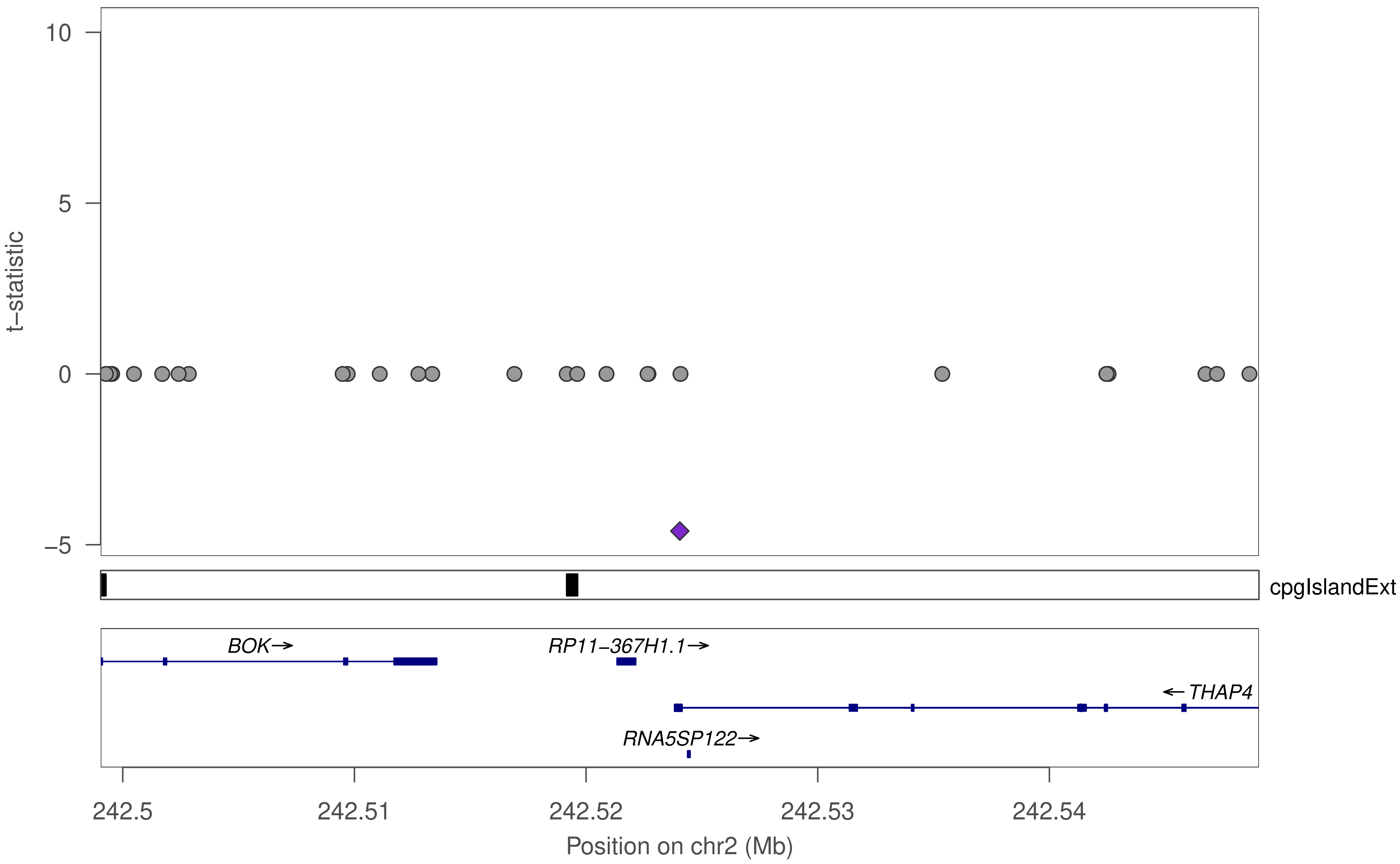

10
5
0
−5
t−statistic
cpgIslandExt
BOK→
RP11−367H1.1→
←THAP4
RNA5SP122→
242.5
242.51
242.52
242.53
242.54
Position on chr2 (Mb)

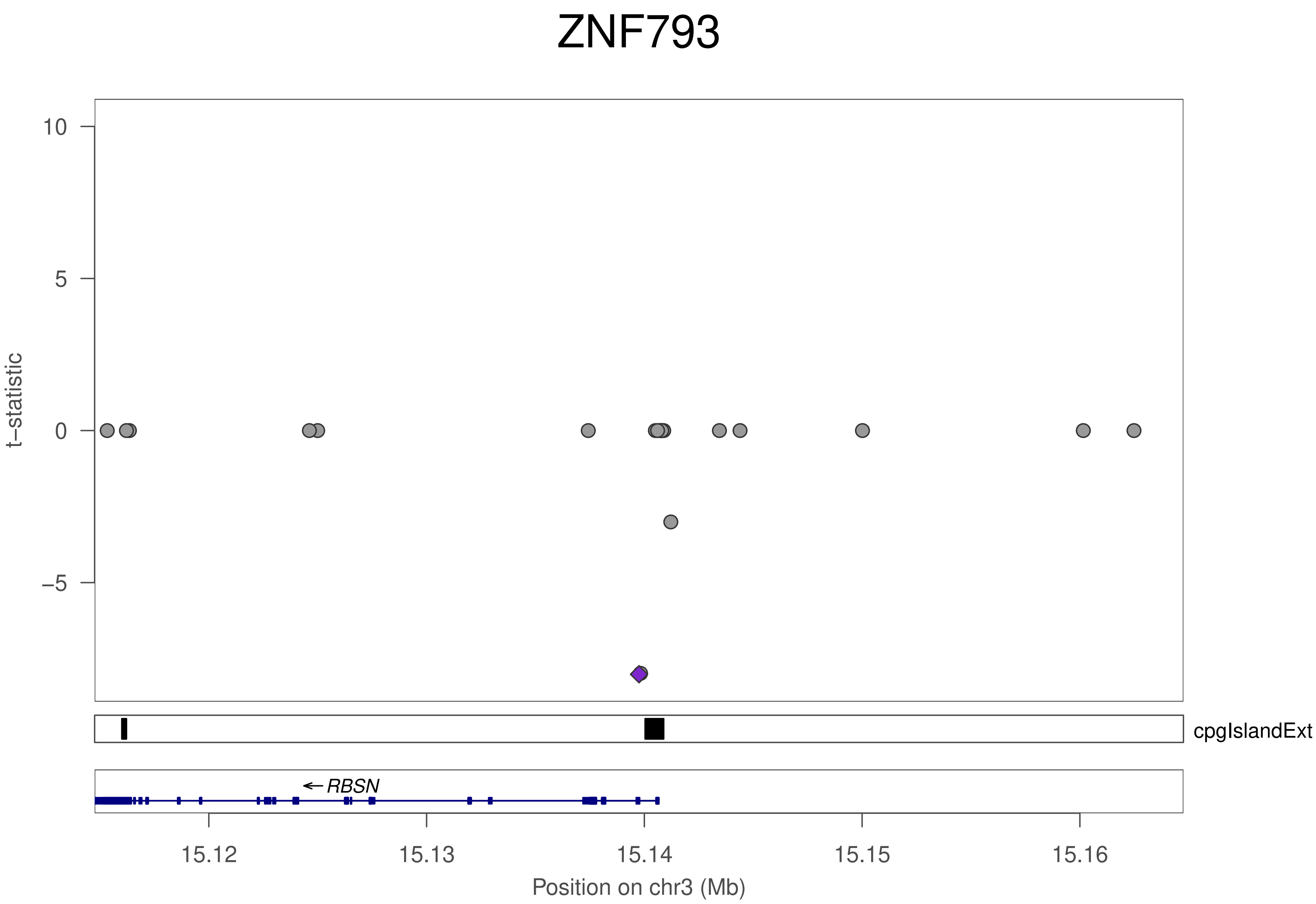
ZNF793
t−statistic
10
5
0
−5
cpgIslandExt
RBSN
15.12
15.13
15.14
15.15
15.16
Position on chr3 (Mb)

### ZNF793

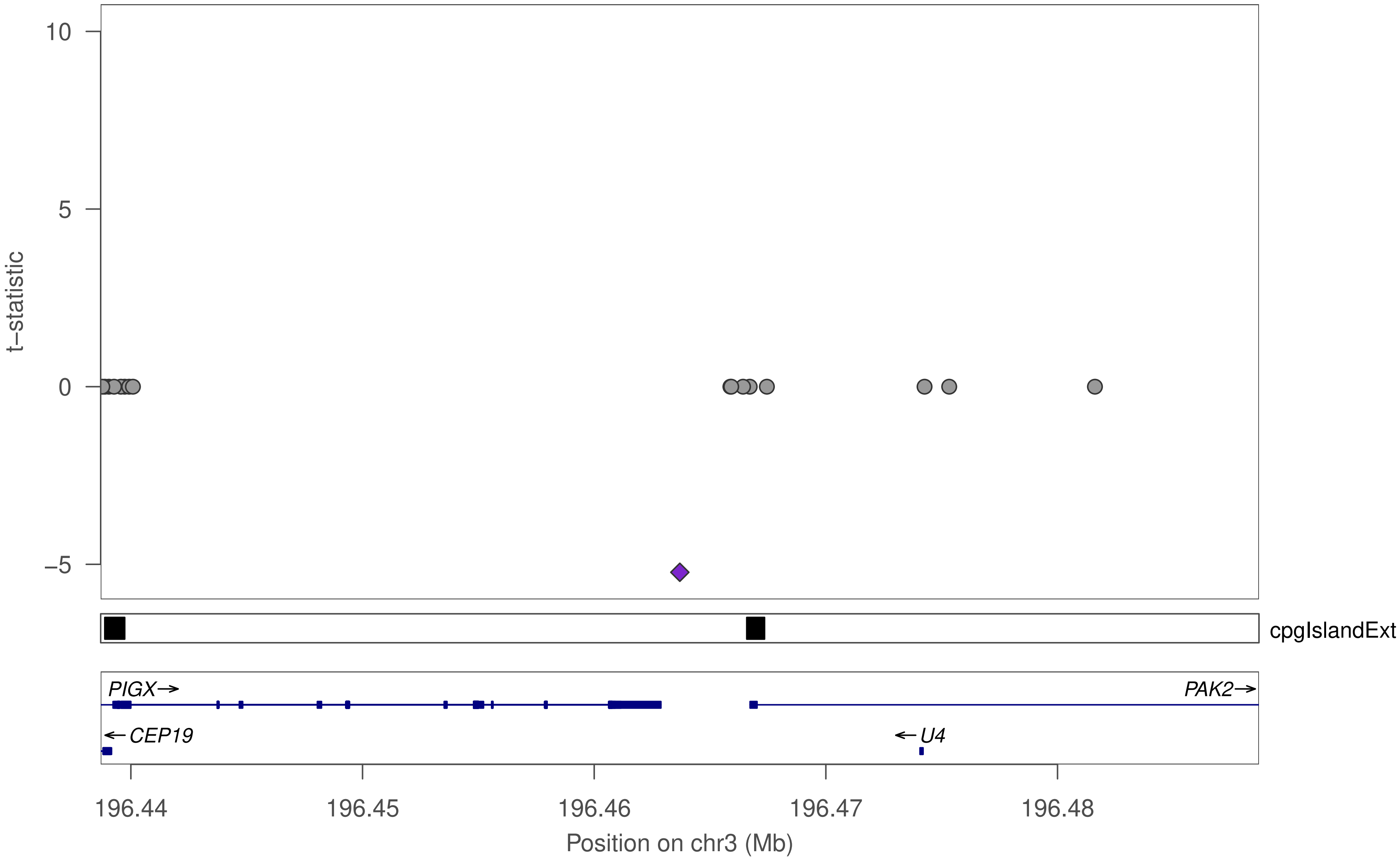
10
5
0
−5
t−statistic
cpgIslandExt
PIGX→
PAK2→
←CEP19
←U4
196.44
196.45
196.46
196.47
196.48
Position on chr3 (Mb)

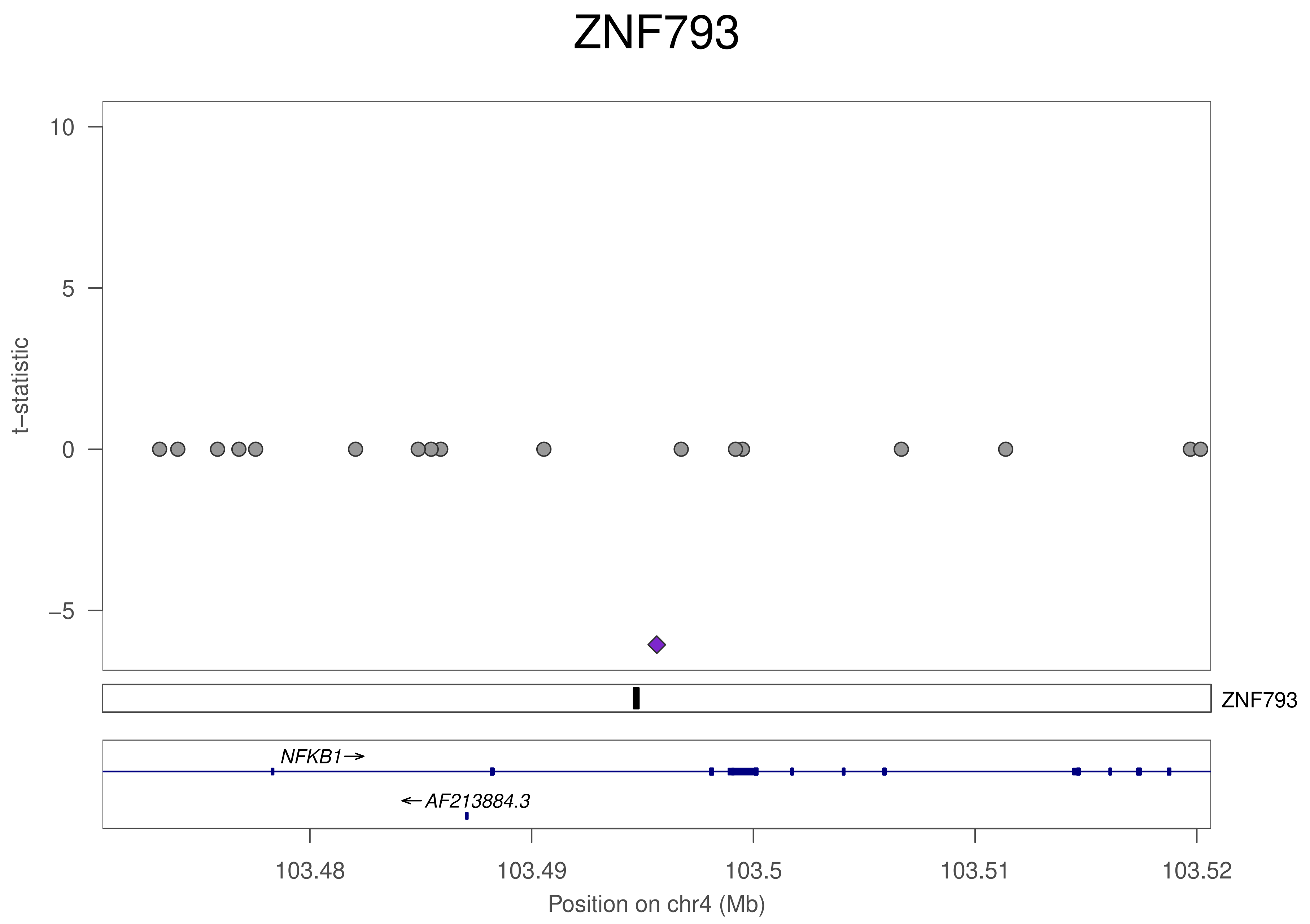
ZNF793
t−statistic
10
5
0
−5
ZNF793
NFKB1→
←AF213884.3
103.48
103.49
103.5
103.51
103.52
Position on chr4 (Mb)

### ZNF793

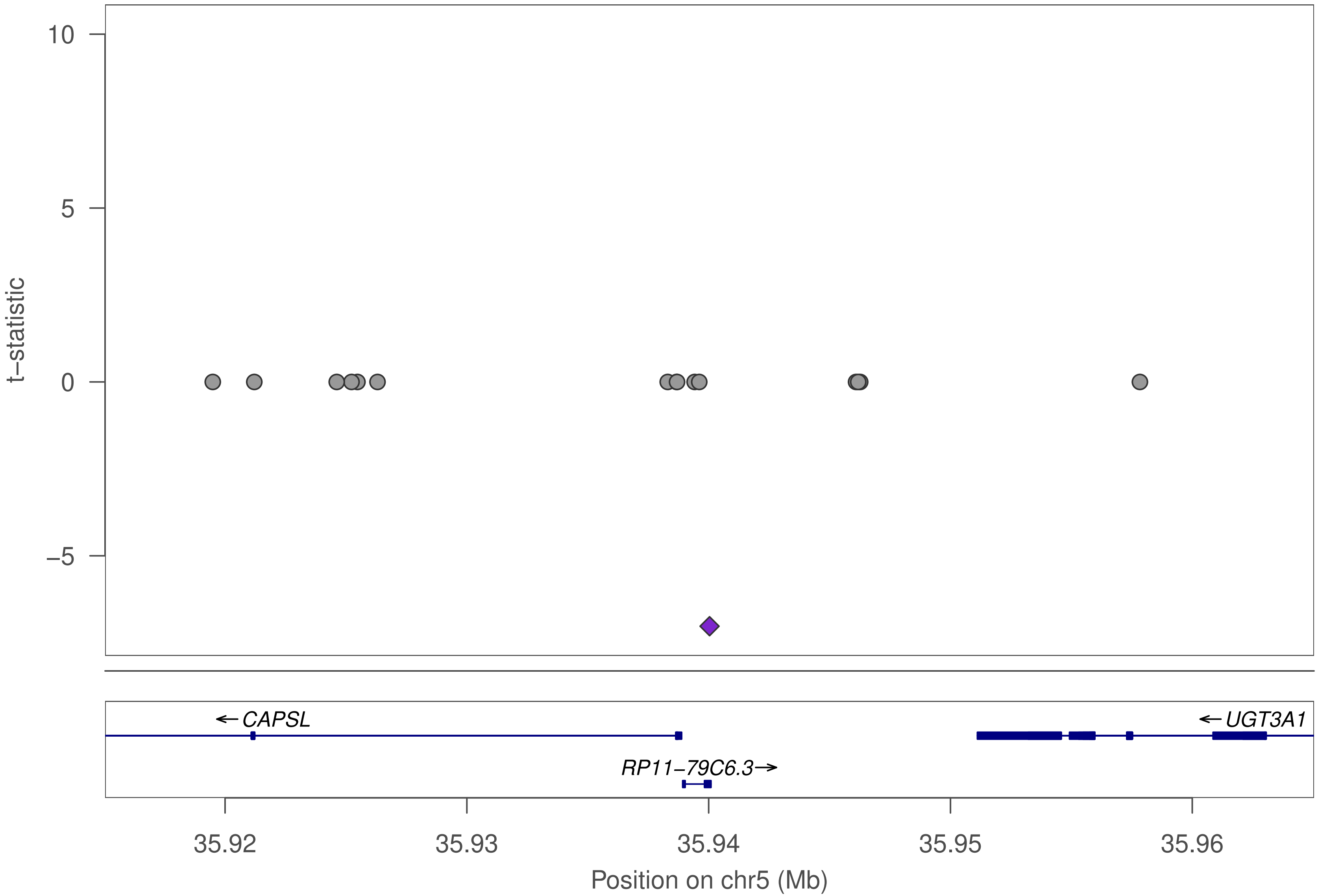

t−statistic
10
5
0
−5
←CAPSL
←UGT3A1
RP11−79C6.3→
35.92
35.93
35.94
35.95
35.96
Position on chr5 (Mb)

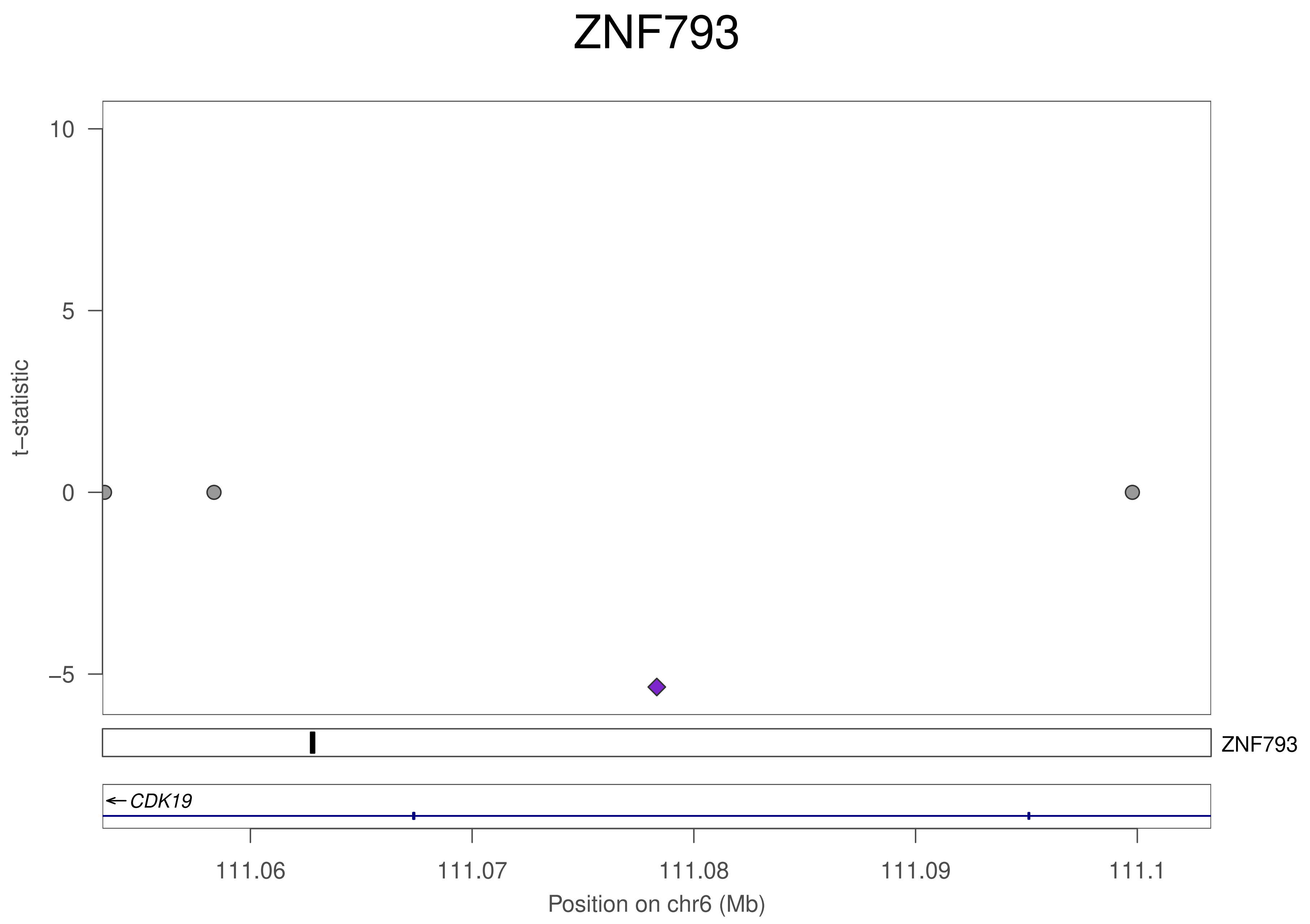

ZNF793
10
5
0
−5
t−statistic
ZNF793
CDK19
111.06
111.07
111.08
111.09
111.1
Position on chr6 (Mb)

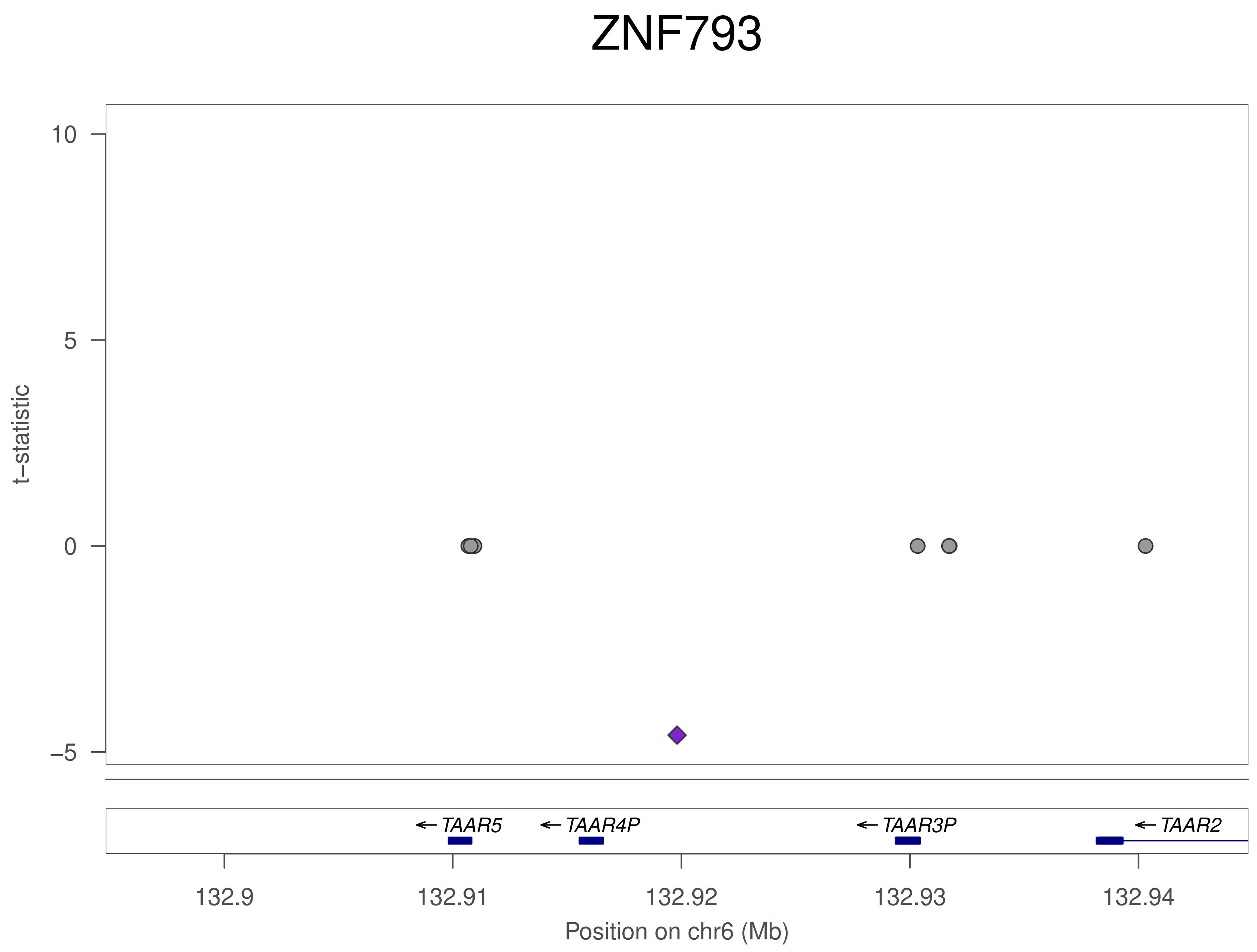
ZNF793
10
5
0
−5
t−statistic
TAAR5
TAAR4P
TAAR3P
TAAR2
132.9
132.91
132.92
132.93
132.94
Position on chr6 (Mb)

### ZNF793

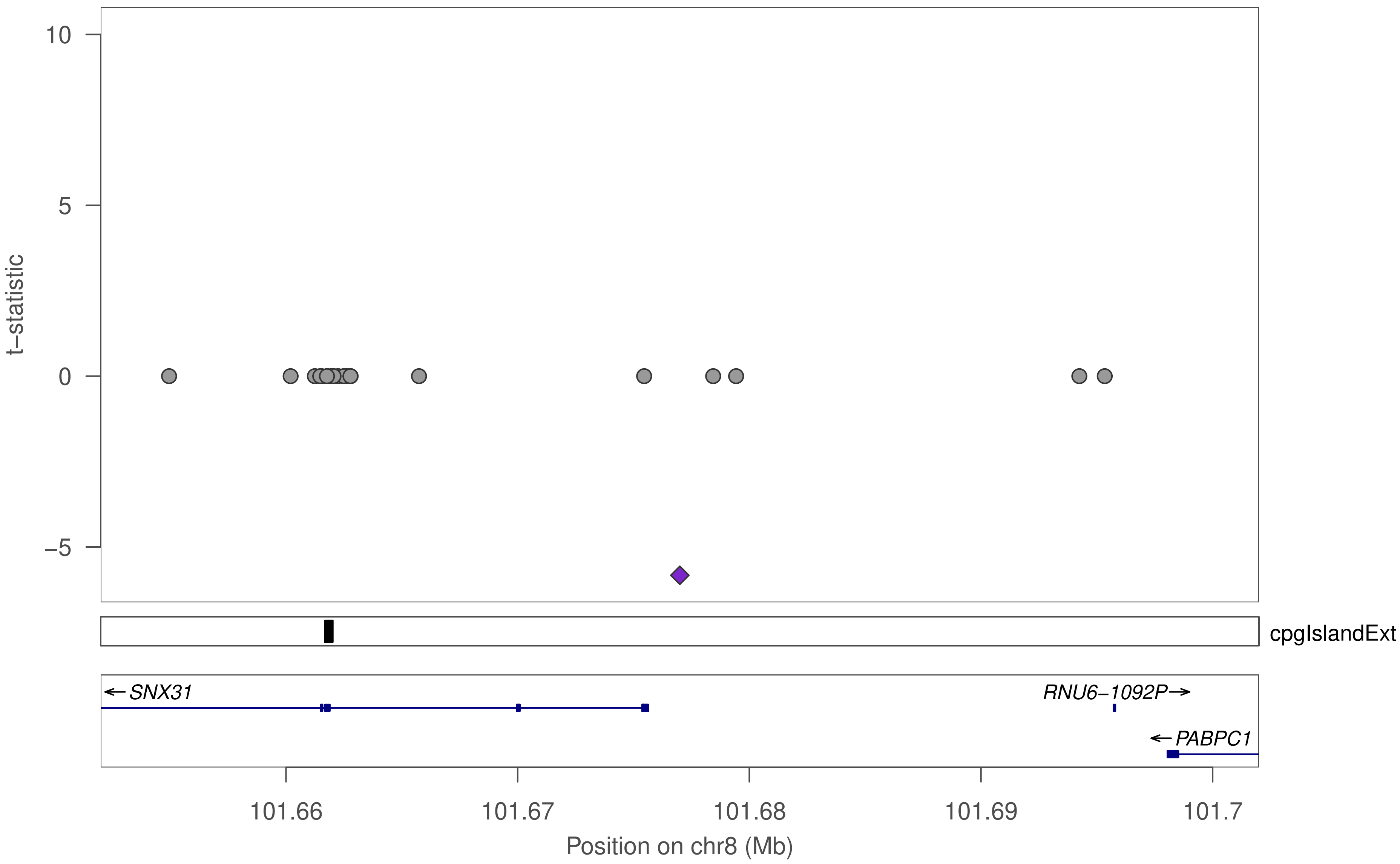

10
5
0
−5
t−statistic
cpgIslandExt
SNX31
RNU6−1092P
PABPC1
101.66
101.67
101.68
101.69
101.7
Position on chr8 (Mb)

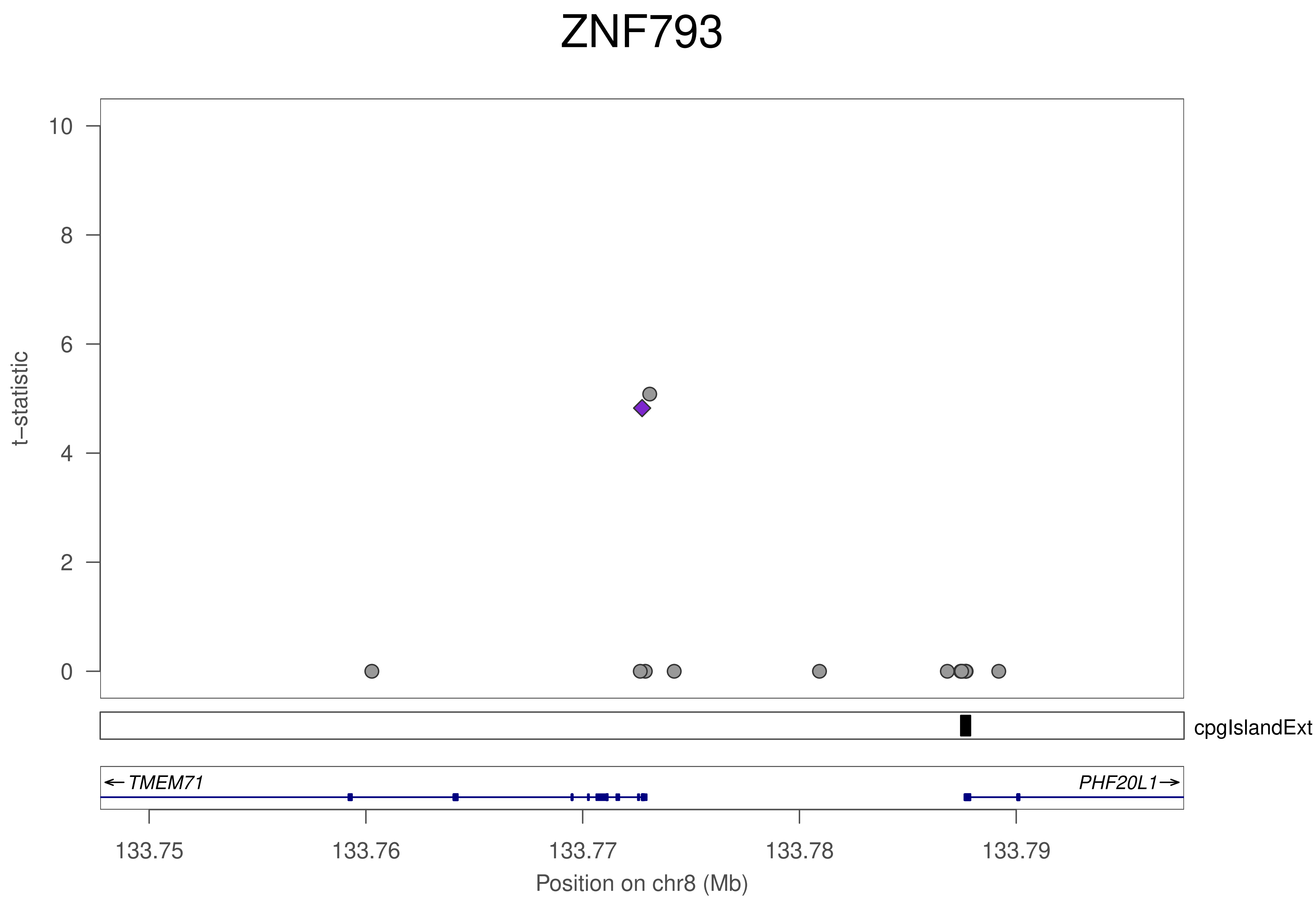
ZNF793
t−statistic
10
8
6
4
2
0
cpgIslandExt
← TMEM71
PHF20L1→
133.75
133.76
133.77
133.78
133.79
Position on chr8 (Mb)

### ZNF793

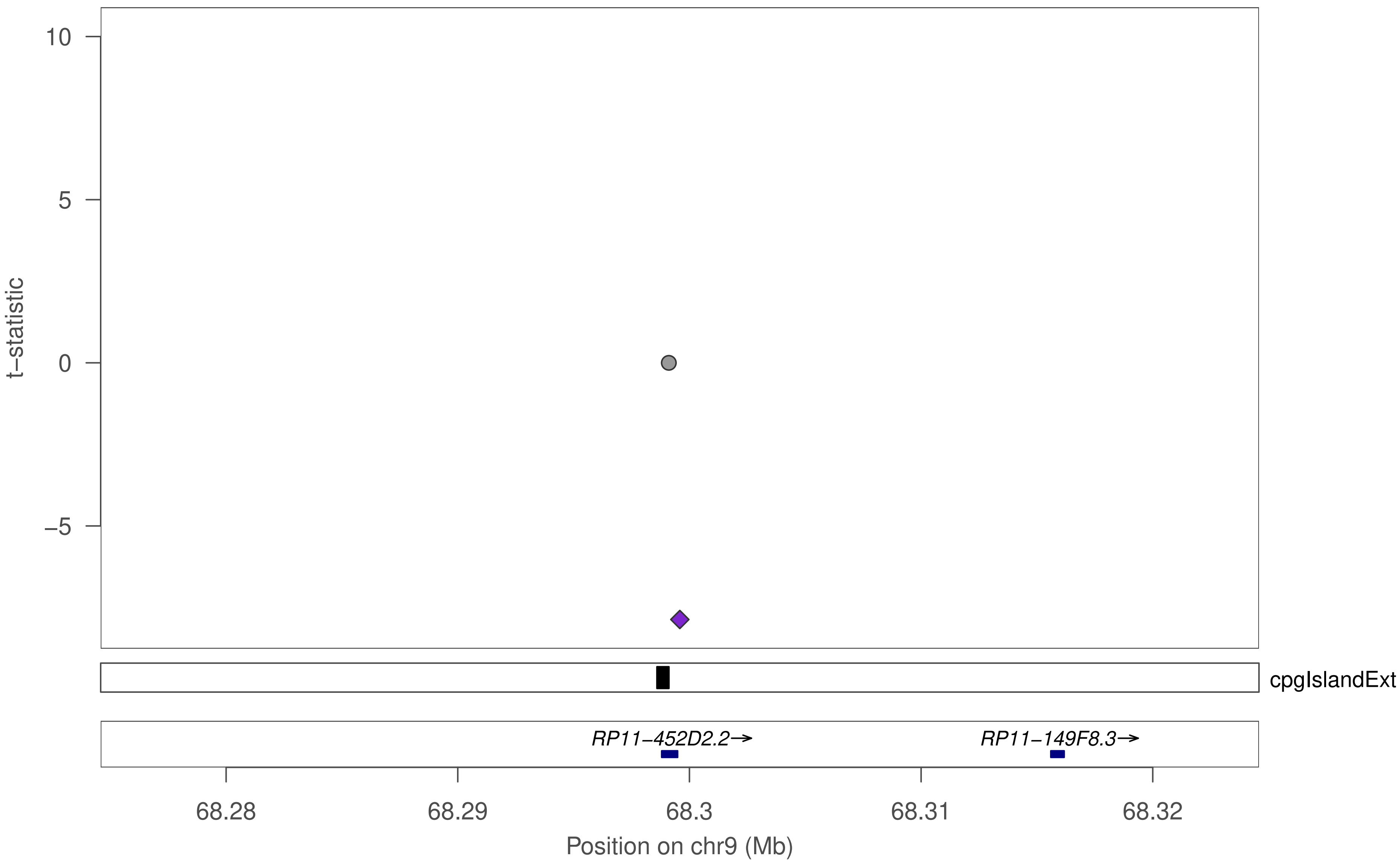

10
5
0
−5
t−statistic
cpgIslandExt
RP11−452D2.2→
RP11−149F8.3→
68.28
68.29
68.3
68.31
68.32
Position on chr9 (Mb)

### ZNF793

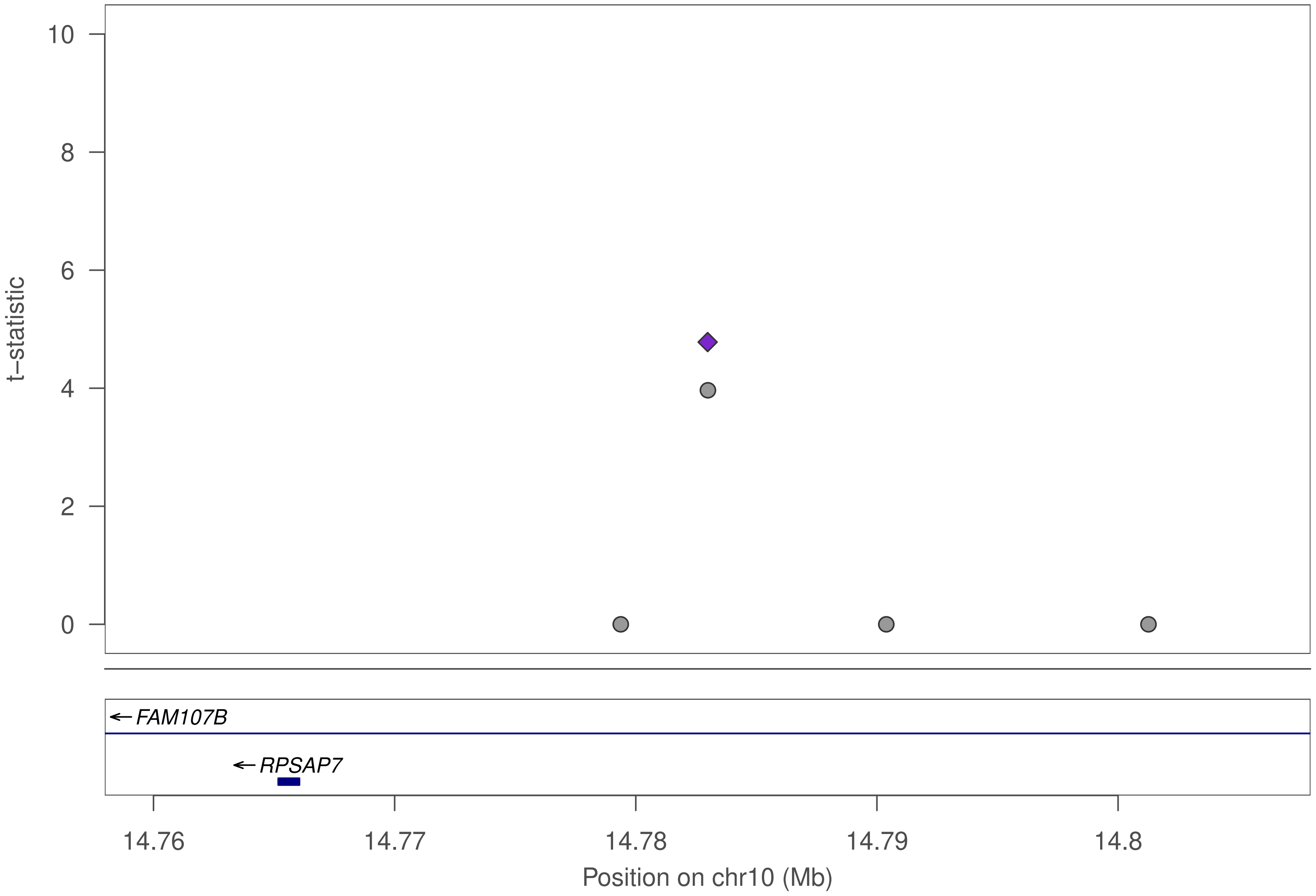

10
8
6
4
2
0
t−statistic
FAM107B
RPSAP7
14.76
14.77
14.78
14.79
14.8
Position on chr10 (Mb)

### ZNF793

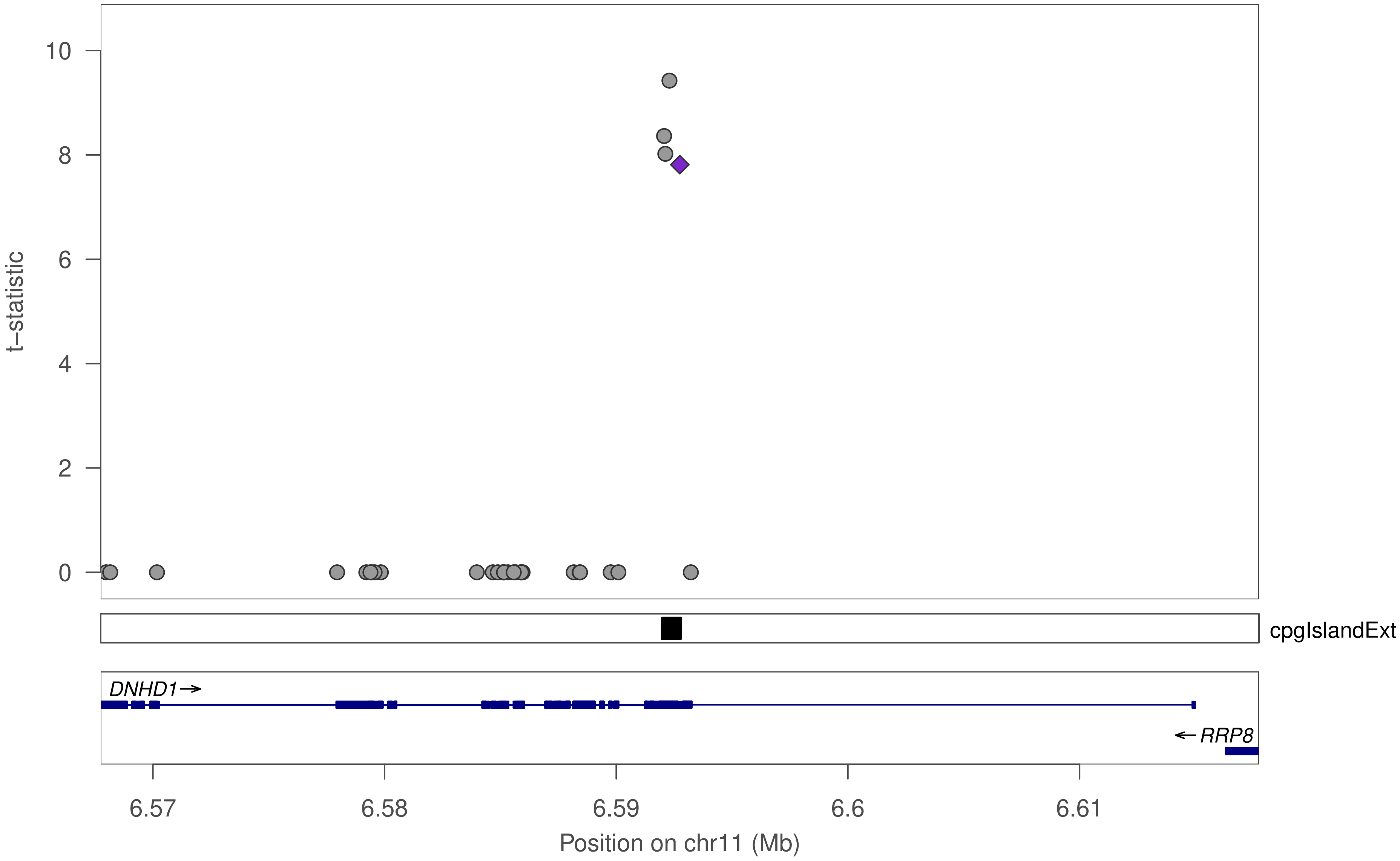

10
8
6
4
2
0
t–statistic
cpgIslandExt
DNHD1→
←RRP8
6.57
6.58
6.59
6.6
6.61
Position on chr11 (Mb)

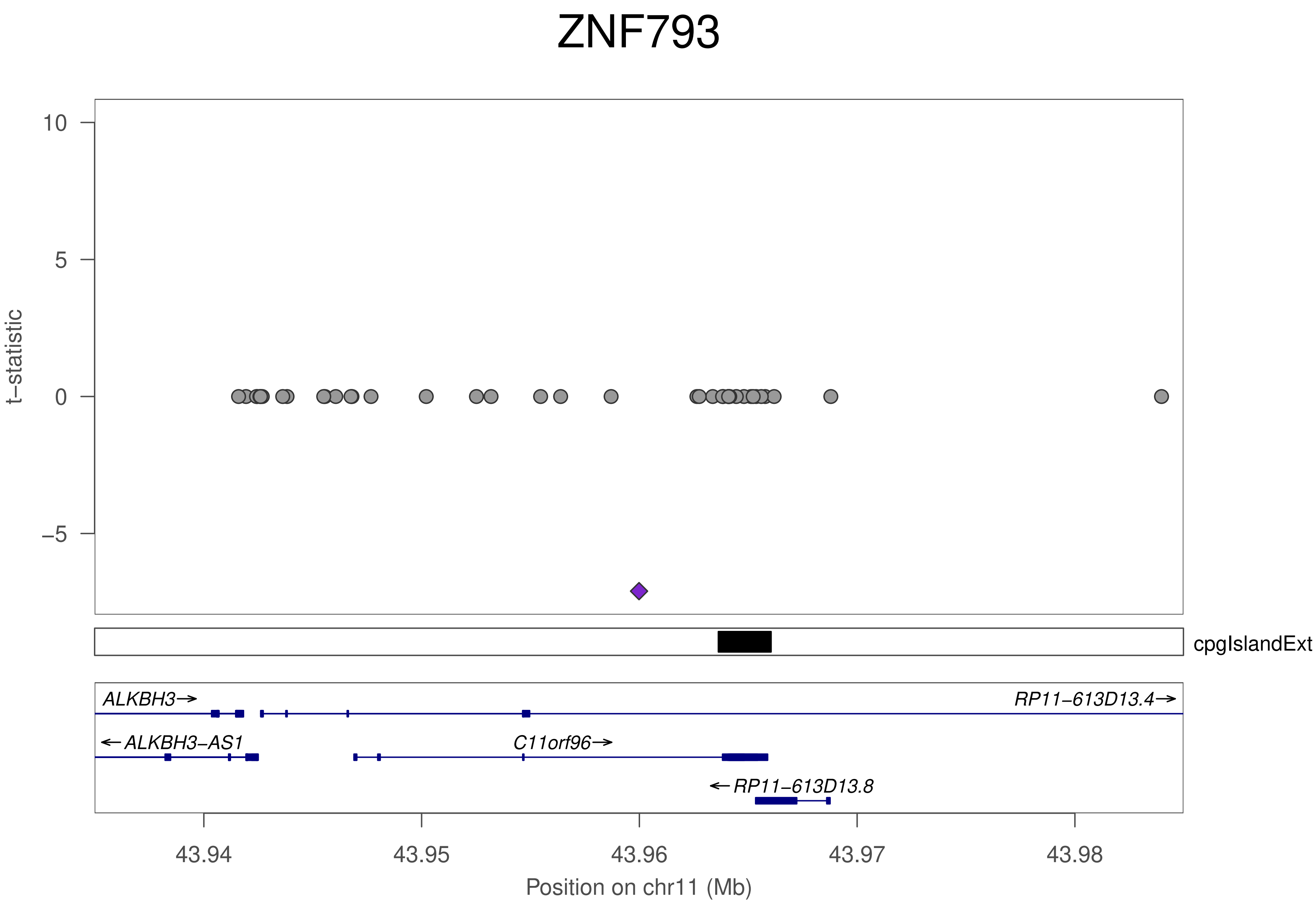
ZNF793
t−statistic
10
5
0
−5
cpgIslandExt
ALKBH3→
RP11−613D13.4→
←ALKBH3−AS1
C11orf96→
←RP11−613D13.8
43.94
43.95
43.96
43.97
43.98
Position on chr11 (Mb)

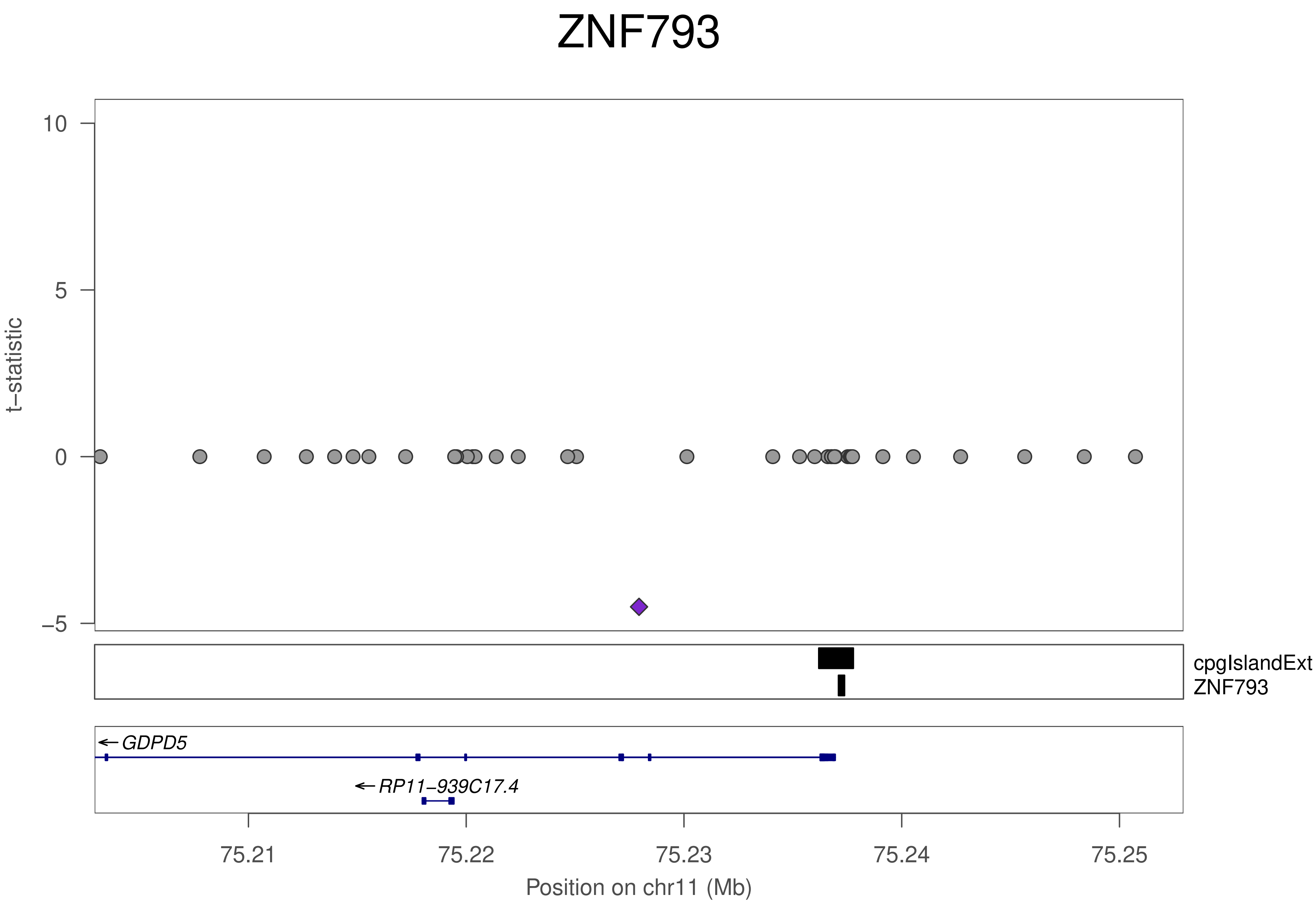

ZNF793
10
5
0
−5
t−statistic
cpgIslandExt
ZNF793
GDPD5
RP11−939C17.4
75.21
75.22
75.23
75.24
75.25
Position on chr11 (Mb)

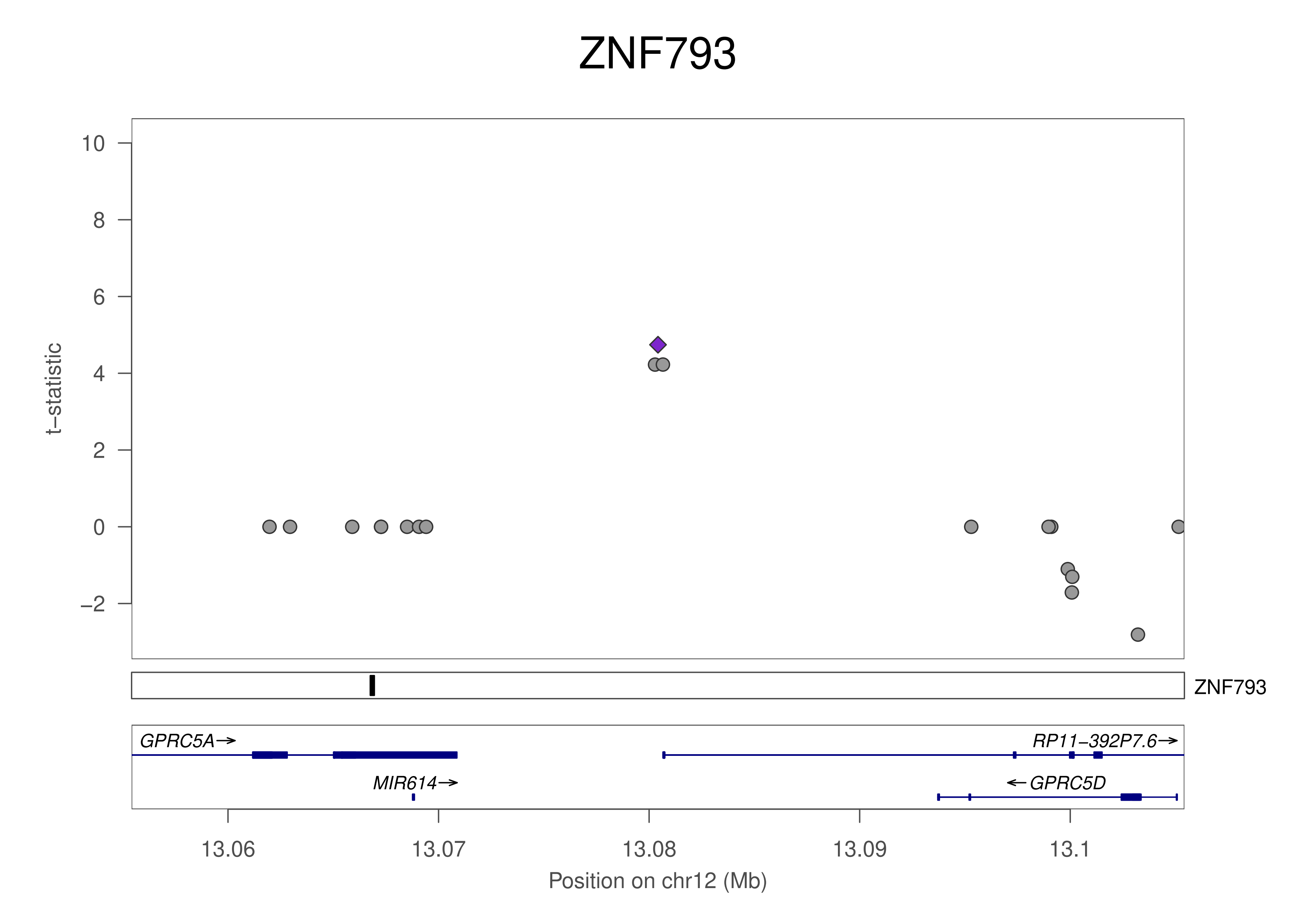

ZNF793
t−statistic
10
8
6
4
2
0
−2
ZNF793
GPRC5A→
RP11−392P7.6→
MIR614→
←GPRC5D
13.06
13.07
13.08
13.09
13.1
Position on chr12 (Mb)

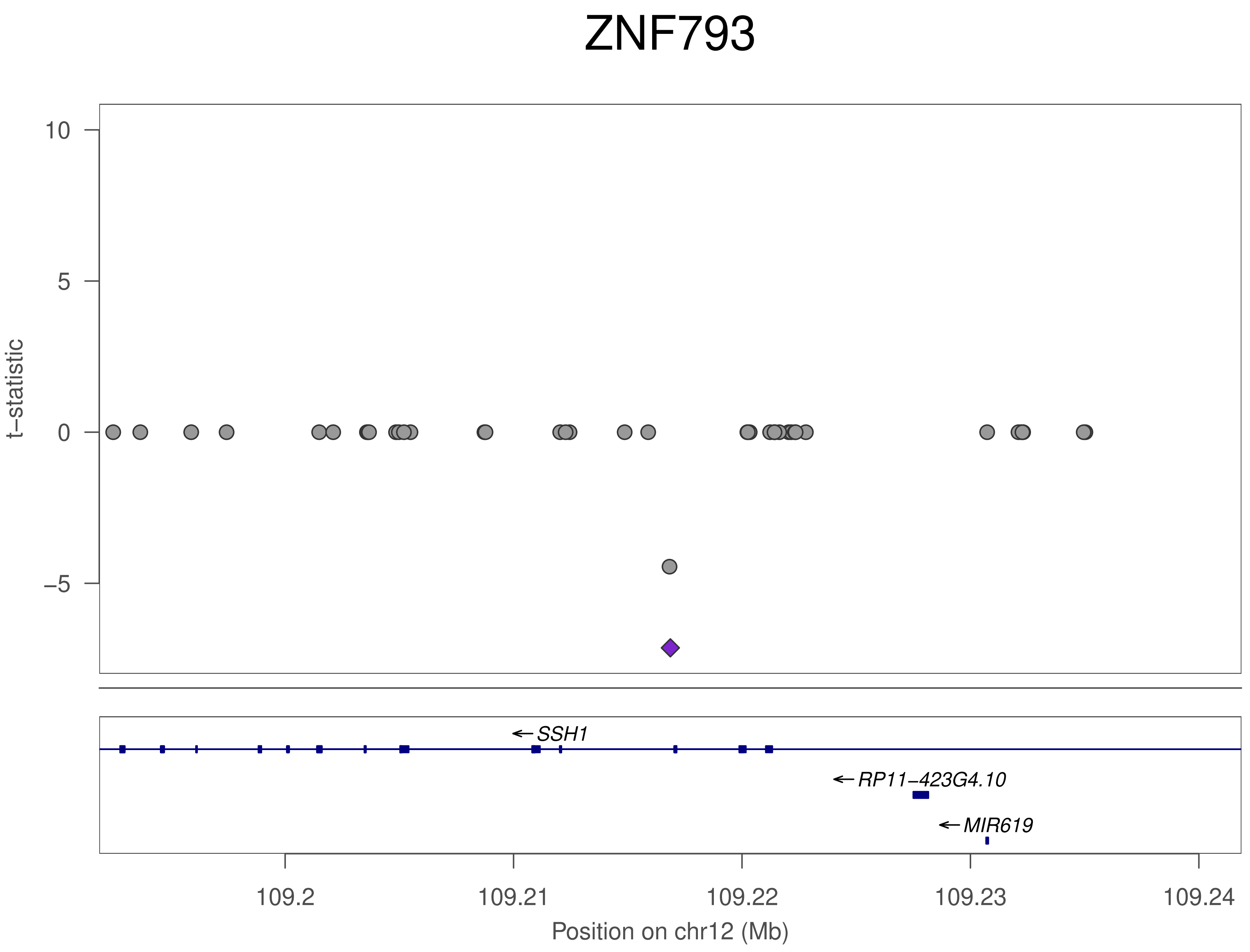

ZNF793
t−statistic
10
5
0
−5
SSH1
RP11−423G4.10
MIR619
109.2
109.21
109.22
109.23
109.24
Position on chr12 (Mb)

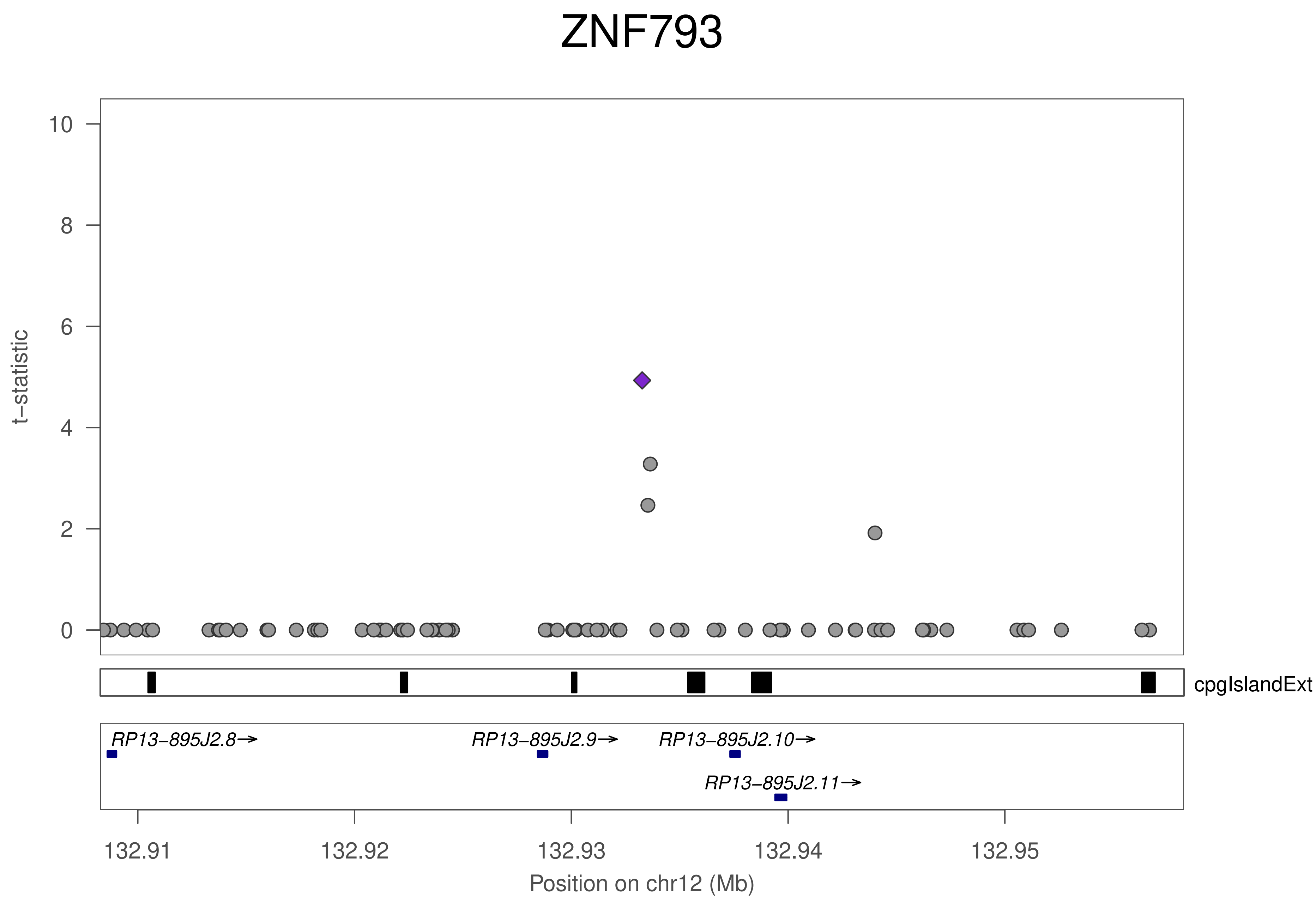

ZNF793
t−statistic
0
2
4
6
8
10
cpgIslandExt
RP13−895J2.8→
RP13−895J2.9→
RP13−895J2.10→
RP13−895J2.11→
132.91
132.92
132.93
132.94
132.95
Position on chr12 (Mb)

### ZNF793

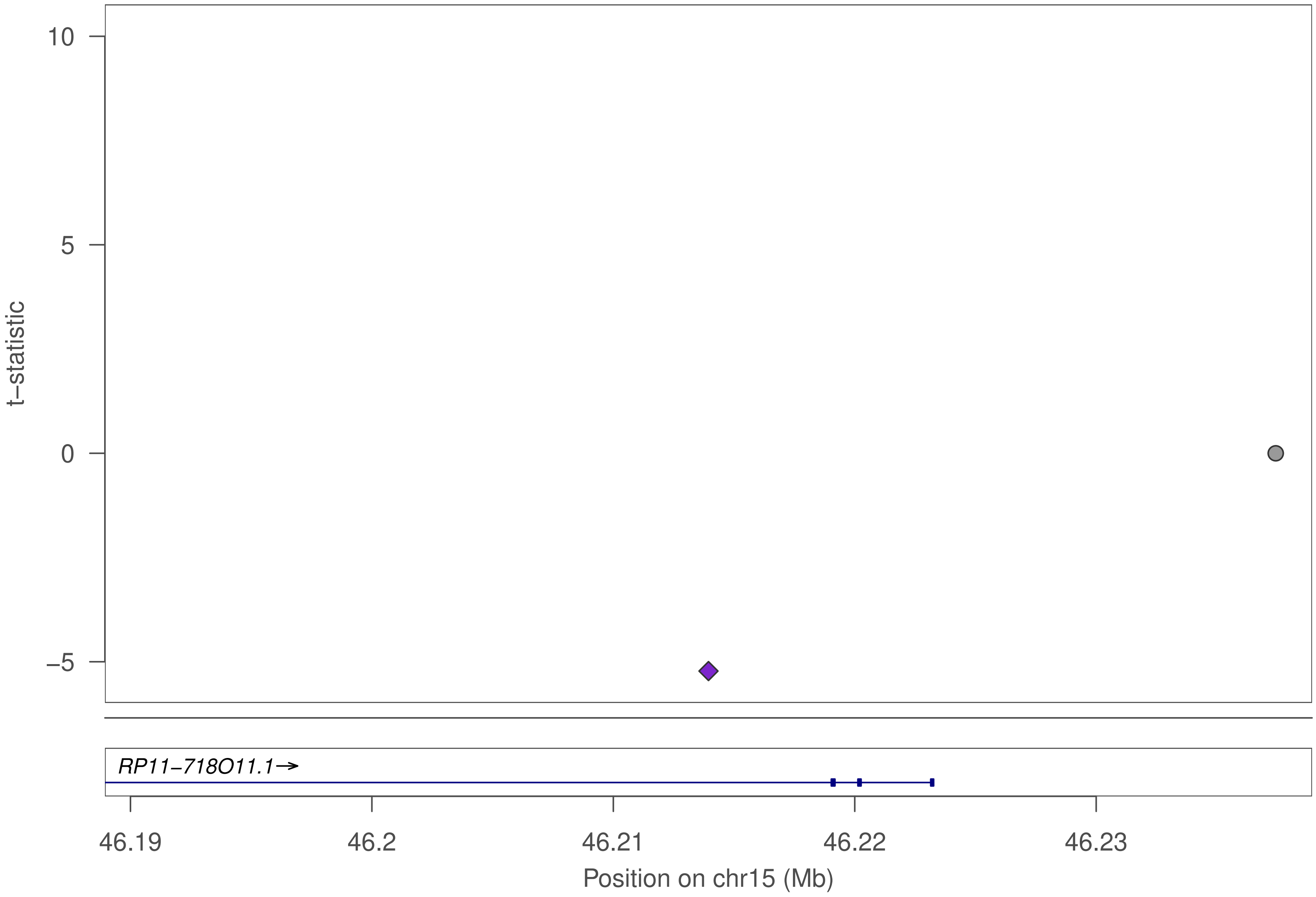

10
5
0
−5
t−statistic
RP11−718O11.1→
46.19
46.2
46.21
46.22
46.23
Position on chr15 (Mb)

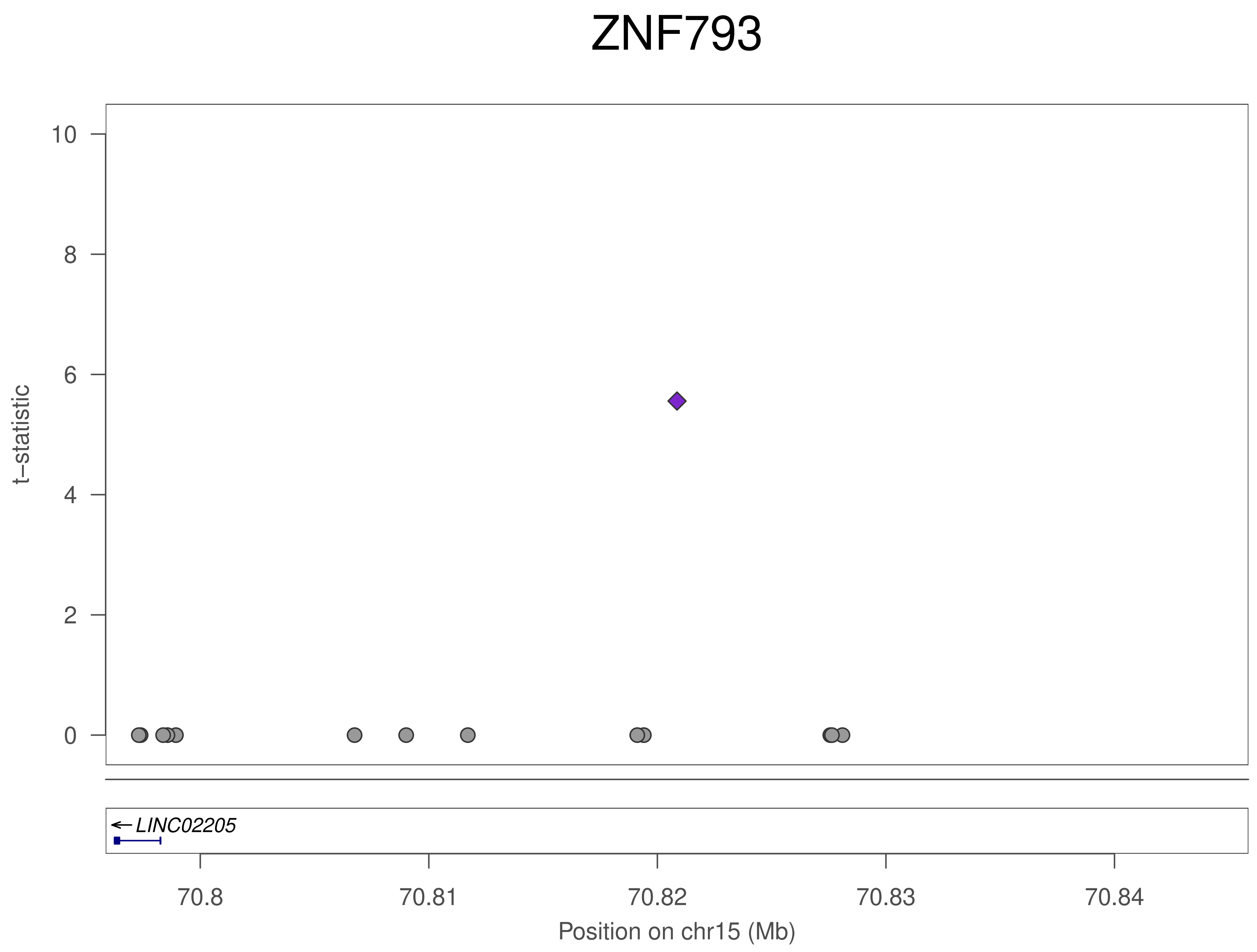
ZNF793
t–statistic
0
2
4
6
8
10
LINC02205
70.8
70.81
70.82
70.83
70.84
Position on chr15 (Mb)

### ZNF793

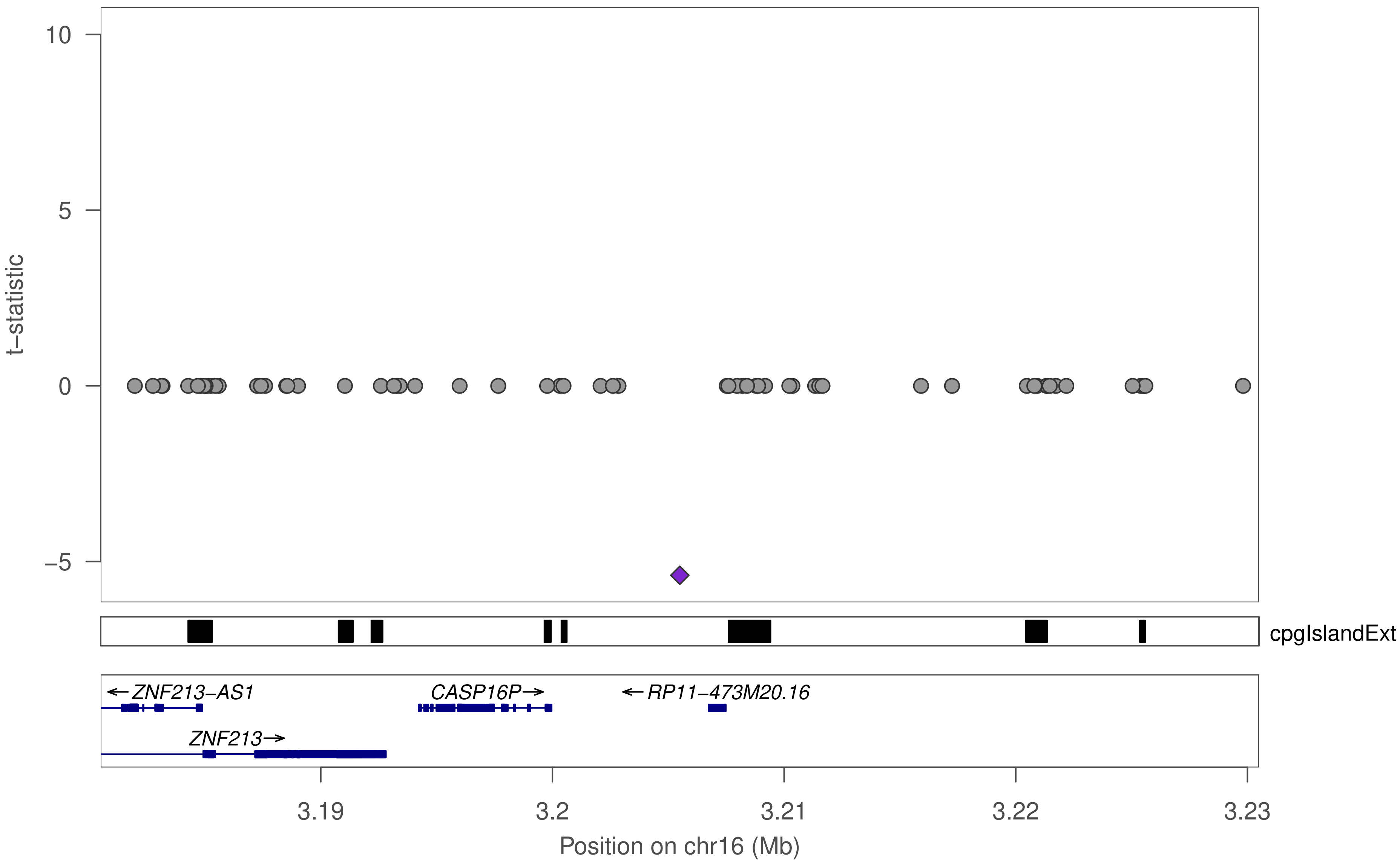

t−statistic
10
5
0
−5
cpgIslandExt
←ZNF213−AS1
CASP16P→
←RP11−473M20.16
ZNF213→
3.19
3.2
3.21
3.22
3.23
Position on chr16 (Mb)

### ZNF793

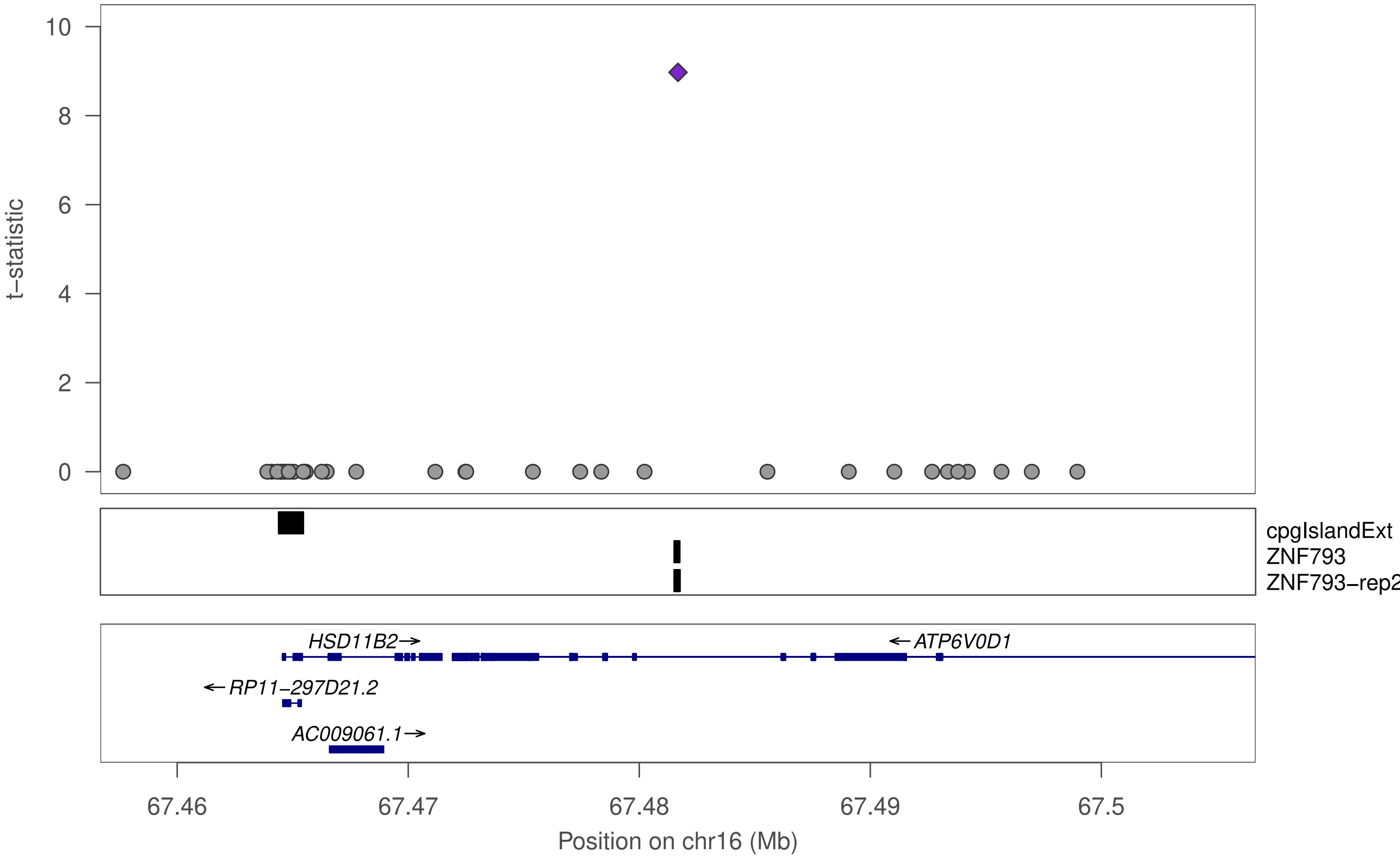

10
8
6
4
2
0
t−statistic
cpgIslandExt
ZNF793
ZNF793−rep2
HSD11B2→
←ATP6V0D1
←RP11−297D21.2
AC009061.1→
67.46
67.47
67.48
67.49
67.5
Position on chr16 (Mb)

### ZNF793

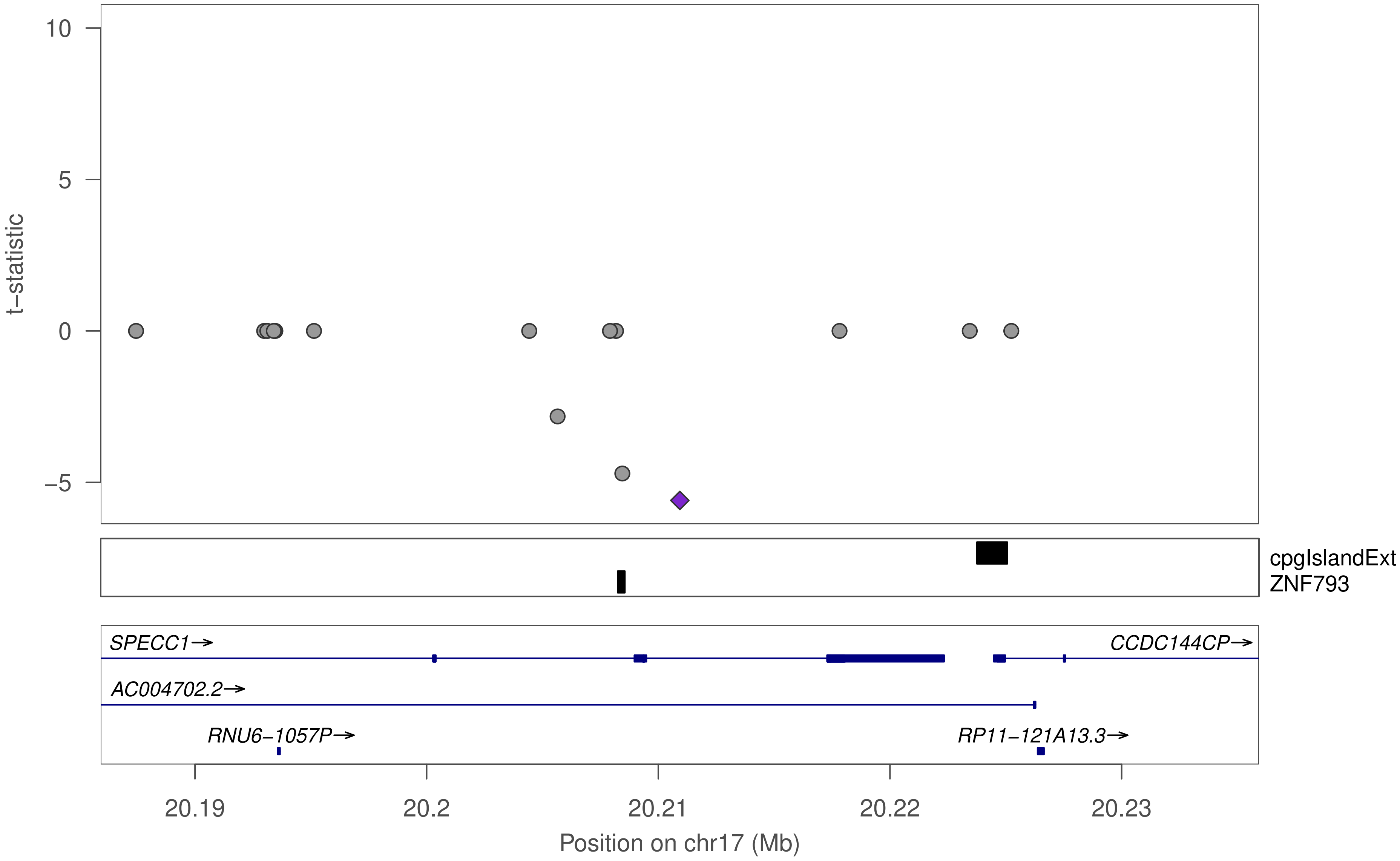

10
5
0
−5
t−statistic
cpgIslandExt
ZNF793
SPECC1→
CCDC144CP→
AC004702.2→
RNU6−1057P→
RP11−121A13.3→
20.19
20.2
20.21
20.22
20.23
Position on chr17 (Mb)

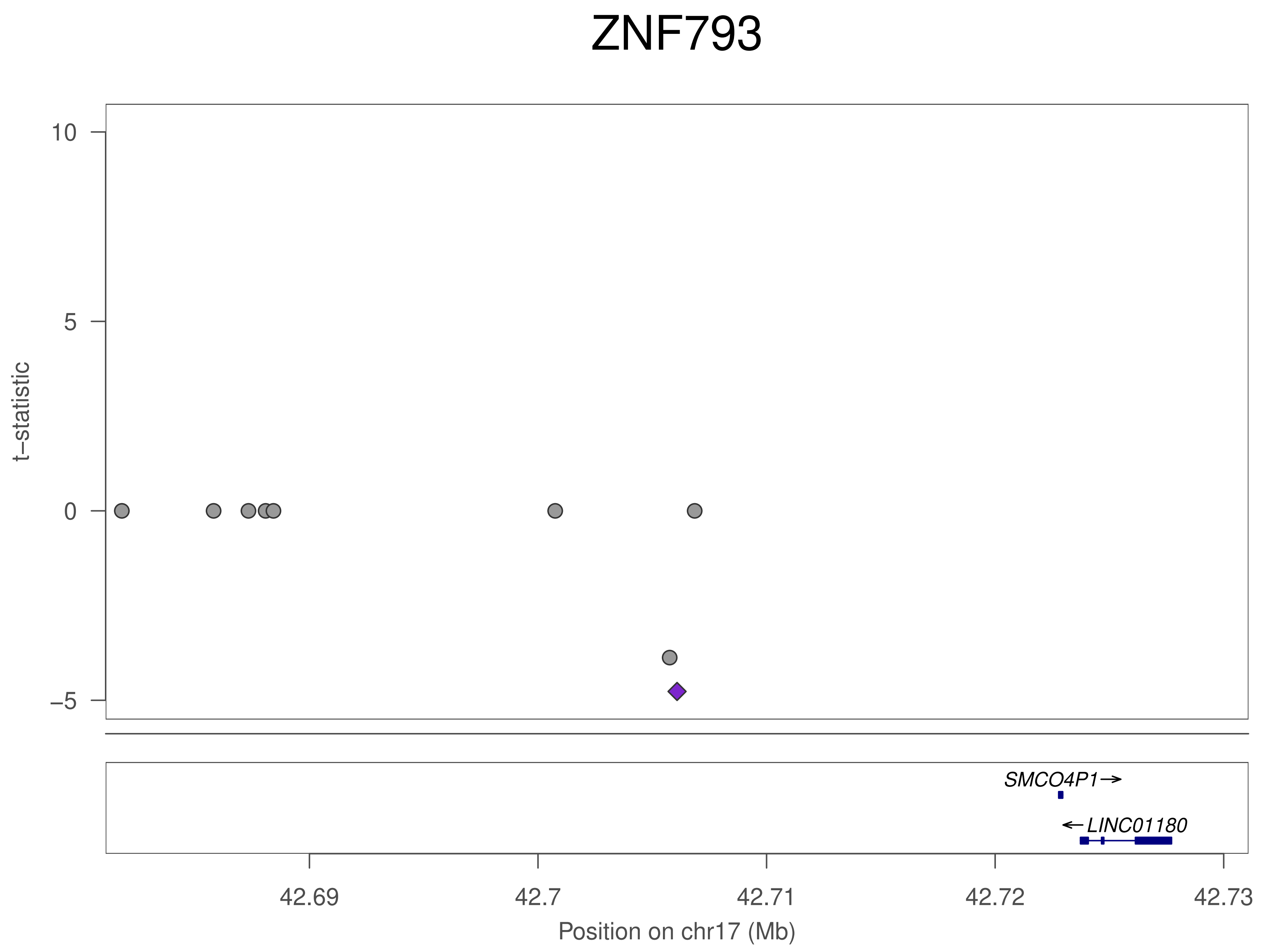

ZNF793
10
5
0
−5
t−statistic
SMCO4P1→
←LINC01180
42.69
42.7
42.71
42.72
42.73
Position on chr17 (Mb)

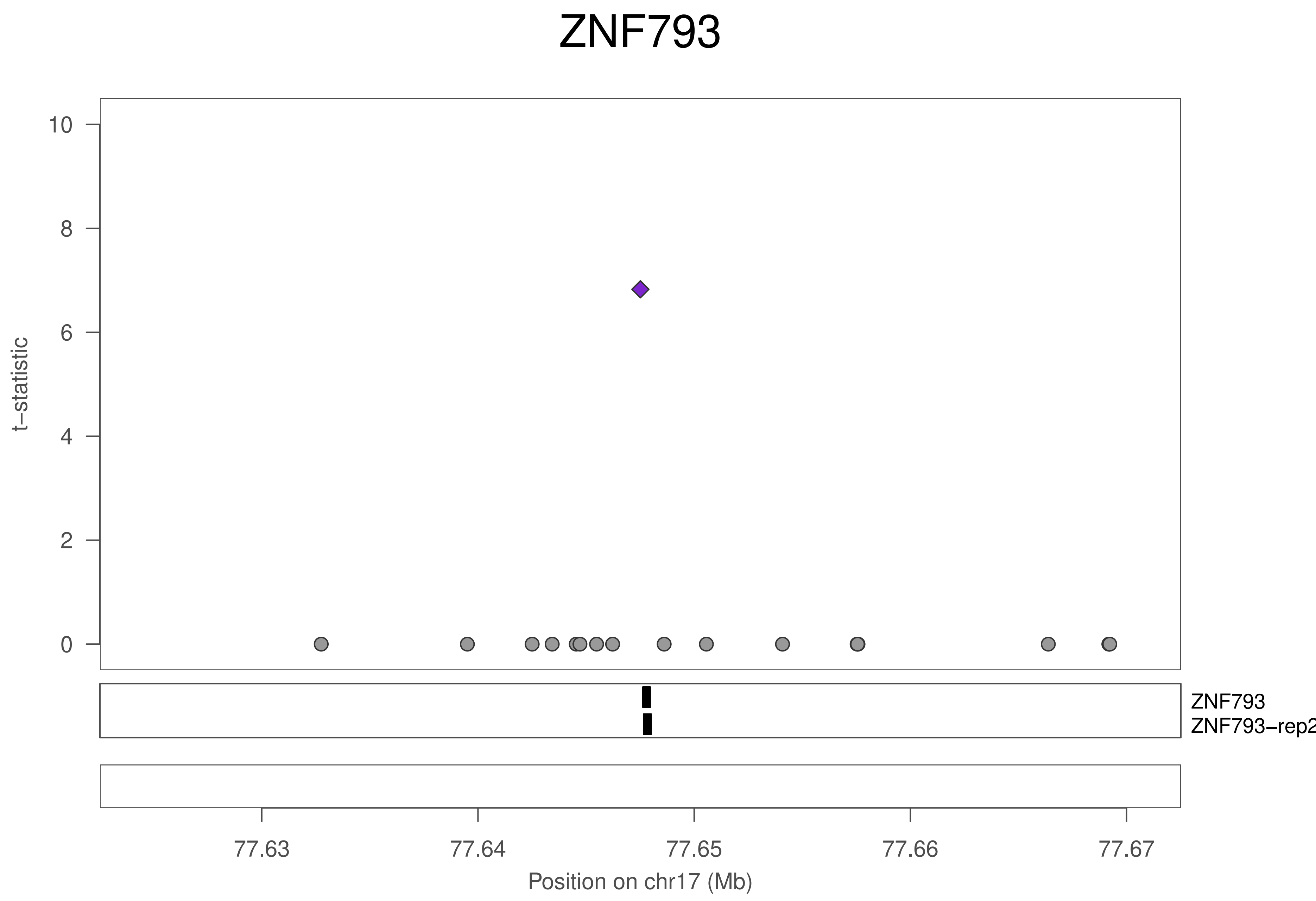
ZNF793
t−statistic
10
8
6
4
2
0
ZNF793
ZNF793−rep2
77.63
77.64
77.65
77.66
77.67
Position on chr17 (Mb)

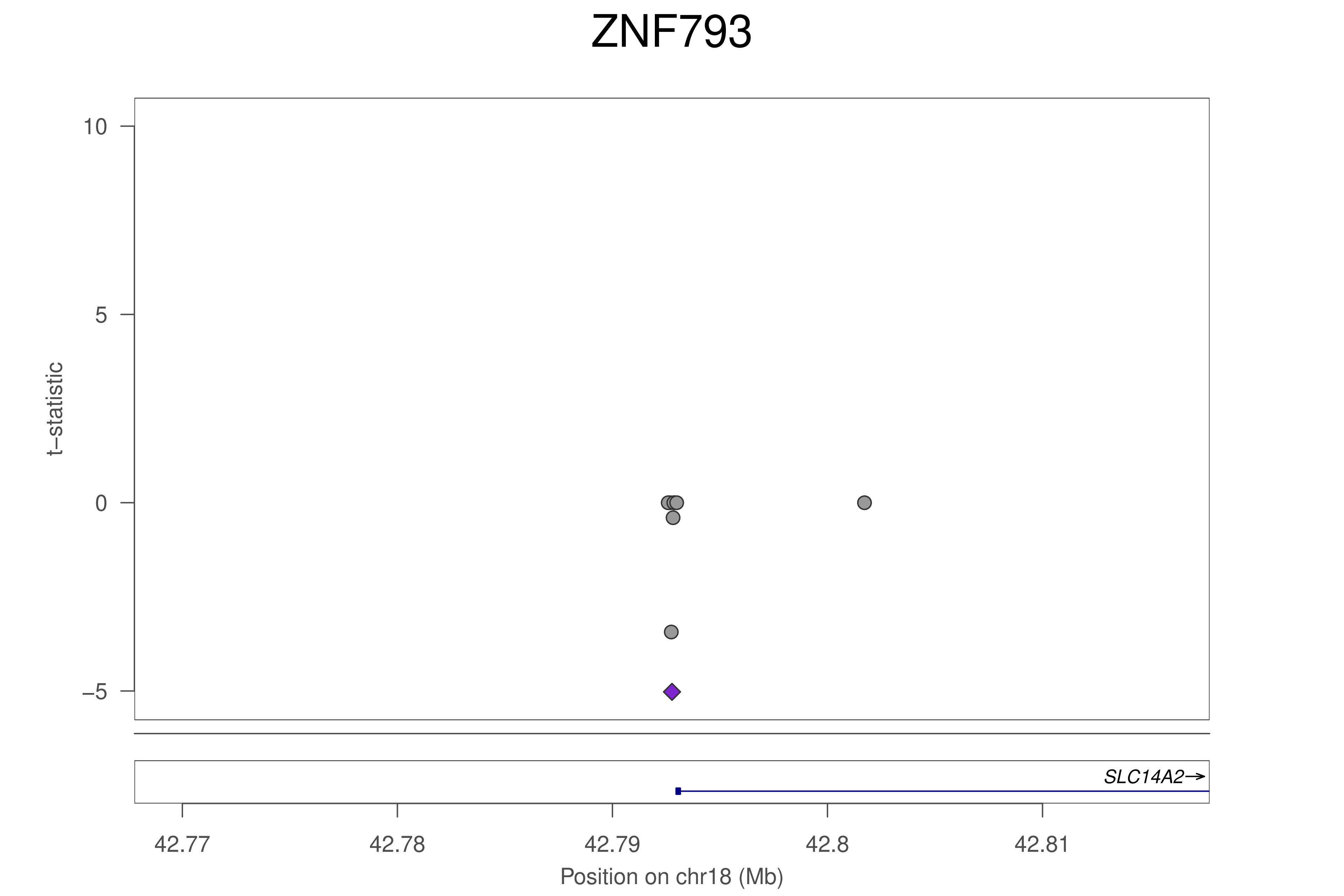

ZNF793
t−statistic
10
5
0
−5
SLC14A2→
42.77
42.78
42.79
42.8
42.81
Position on chr18 (Mb)

### ZNF793

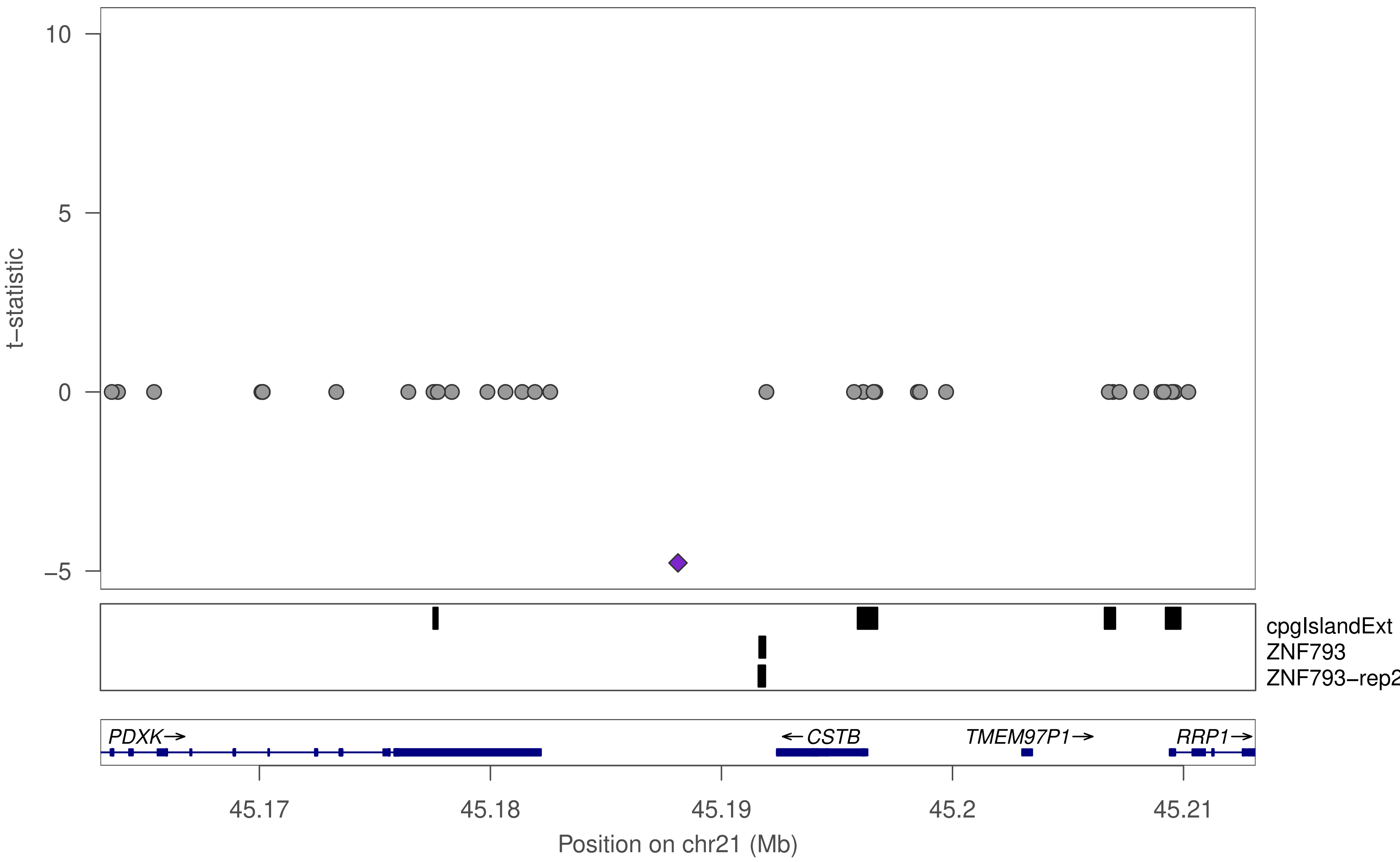

t−statistic
10
5
0
−5
cpgIslandExt
ZNF793
ZNF793−rep2
PDXK→
←CSTB
TMEM97P1→
RRP1→
45.17
45.18
45.19
45.2
45.21
Position on chr21 (Mb)

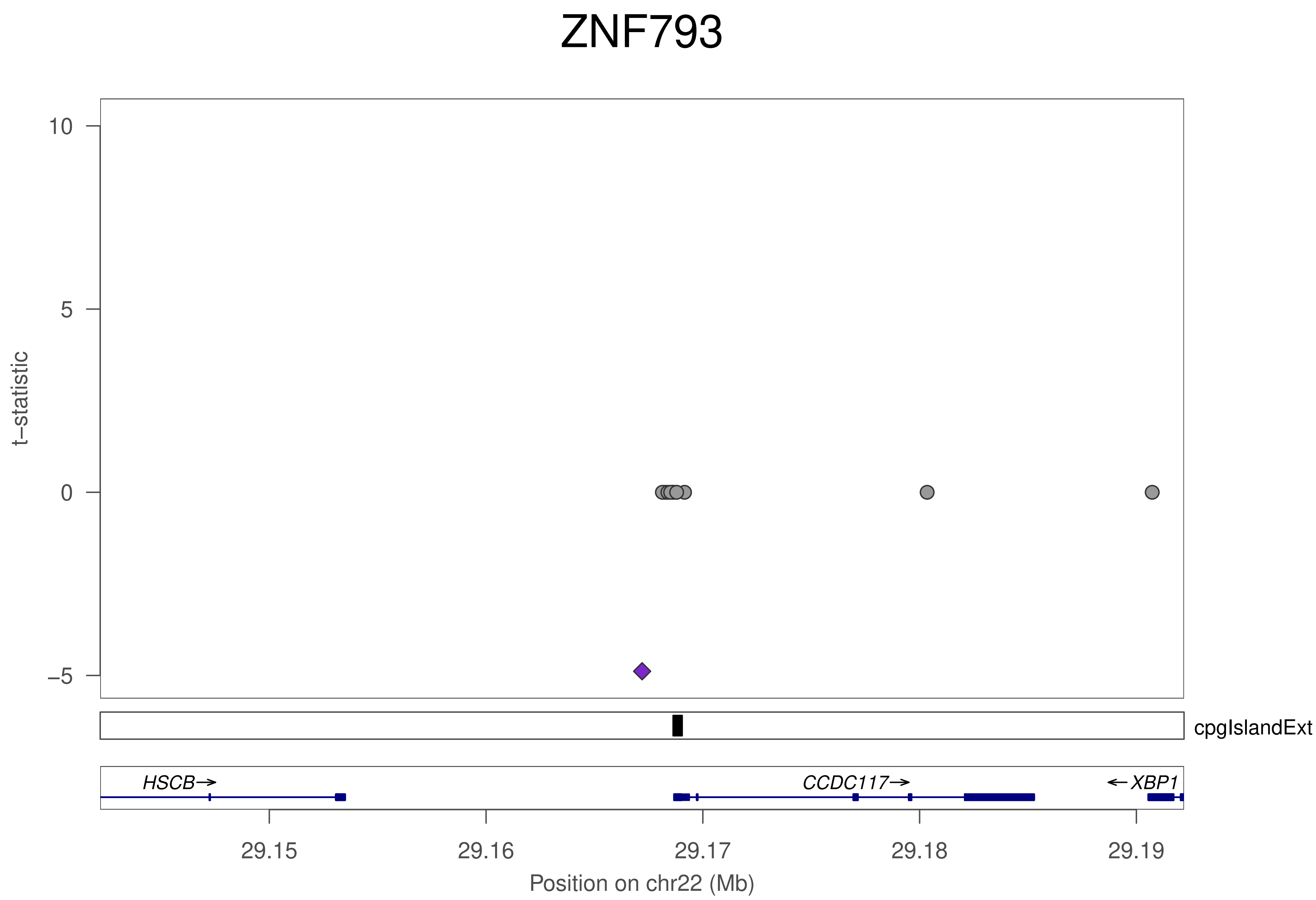
ZNF793
t−statistic
10
5
0
−5
cpgIslandExt
HSCB→
CCDC117→
←XBP1
29.15
29.16
29.17
29.18
29.19
Position on chr22 (Mb)

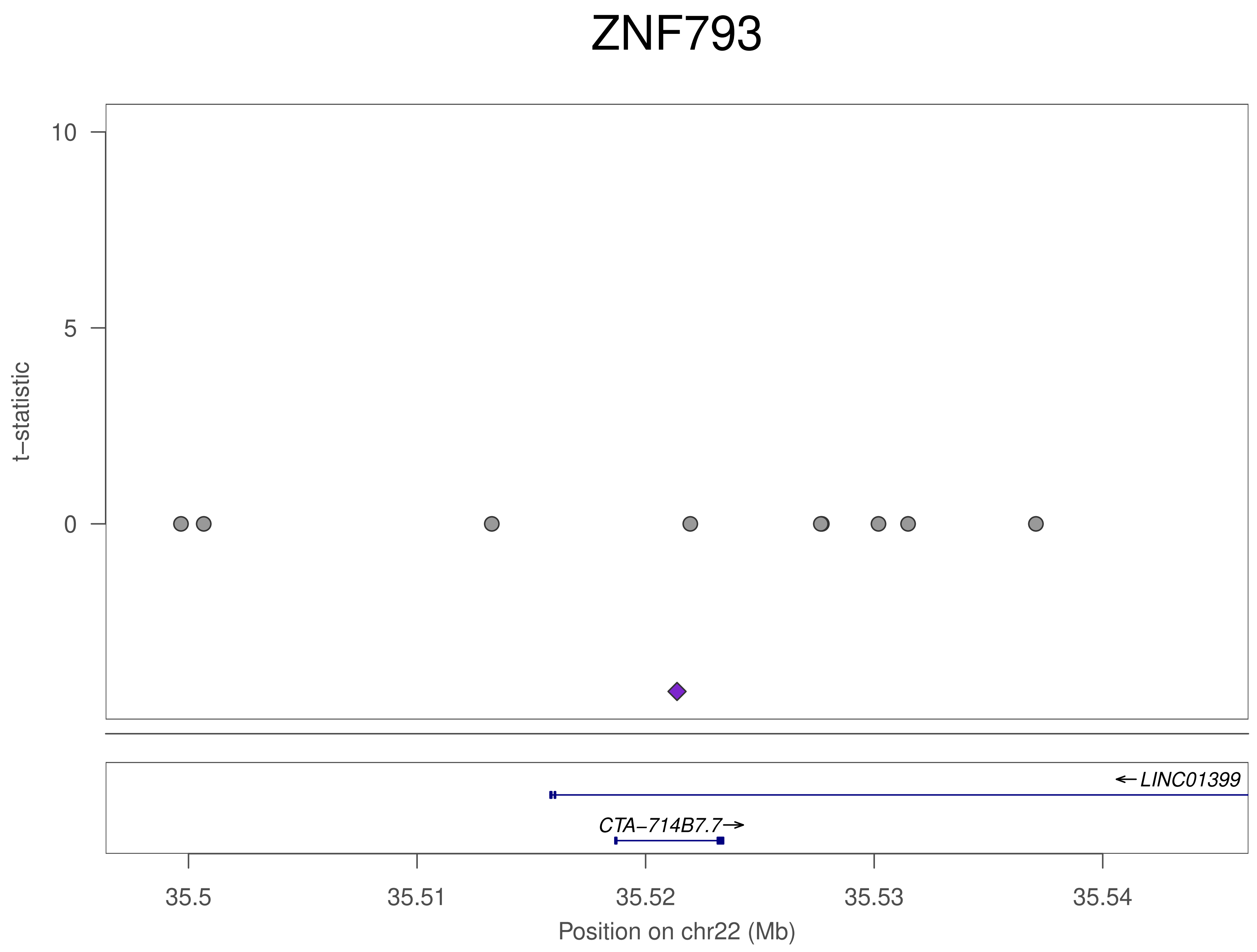
ZNF793
t−statistic
10
5
0
←LINC01399
CTA−714B7.7→
35.5
35.51
35.52
35.53
35.54
Position on chr22 (Mb)

### ZNF548

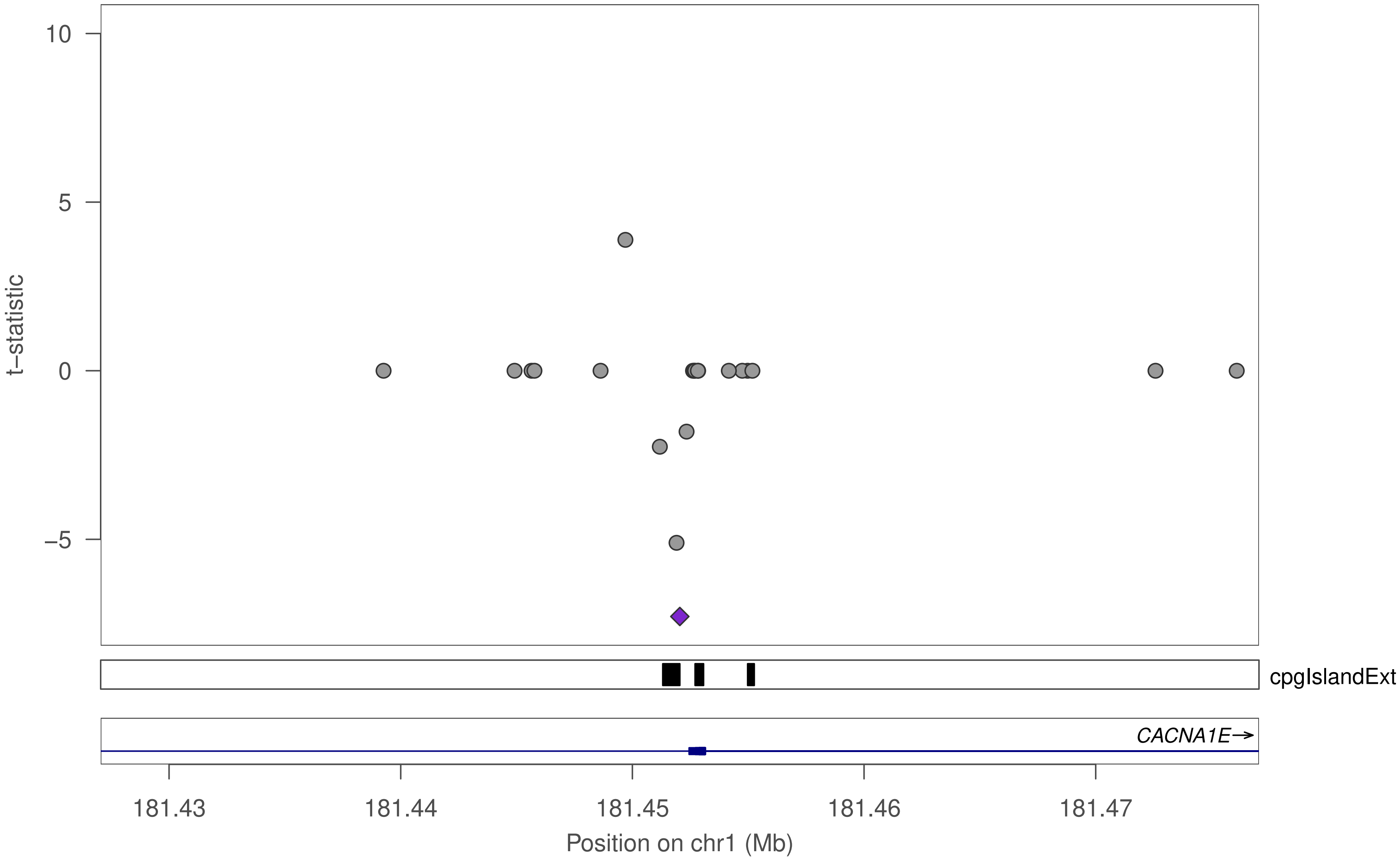

10
5
0
−5
t−statistic
cpgIslandExt
CACNA1E→
181.43
181.44
181.45
181.46
181.47
Position on chr1 (Mb)

### ZNF548

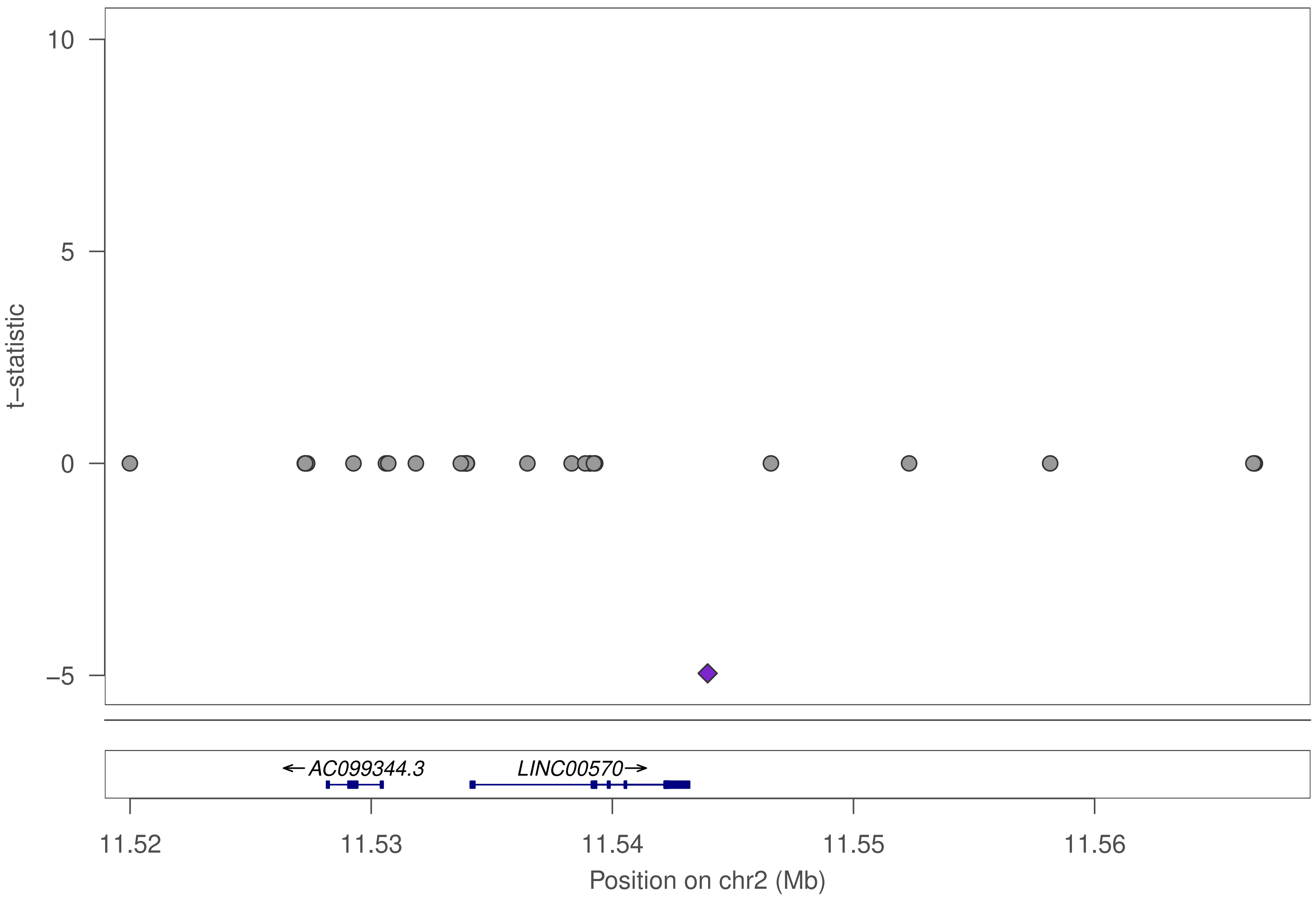

10
5
0
−5
t−statistic
←AC099344.3
LINC00570→
11.52
11.53
11.54
11.55
11.56
Position on chr2 (Mb)

### ZNF548

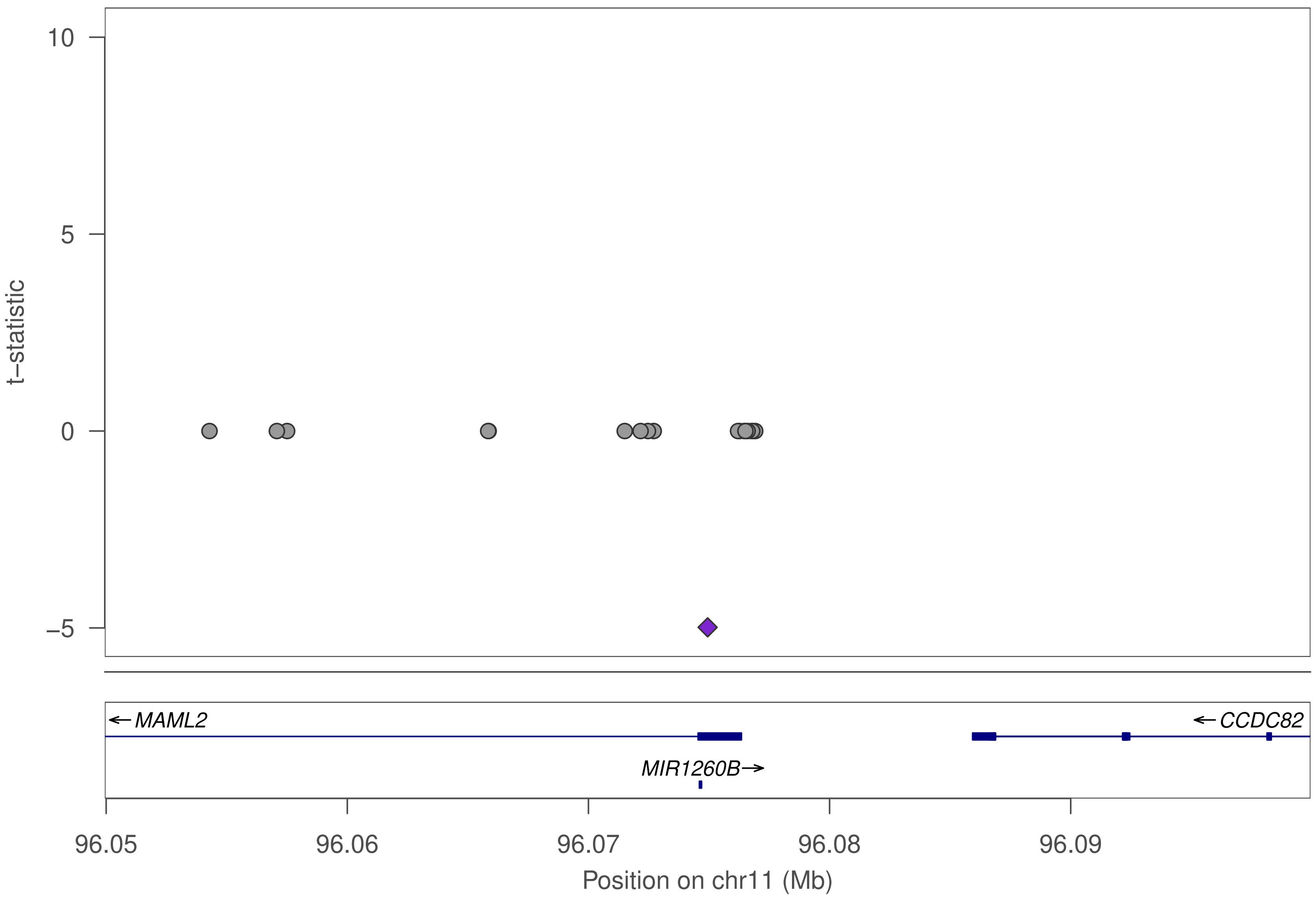

t−statistic
10
5
0
−5
MAML2
CCDC82
MIR1260B
96.05
96.06
96.07
96.08
96.09
Position on chr11 (Mb)

### ZNF548

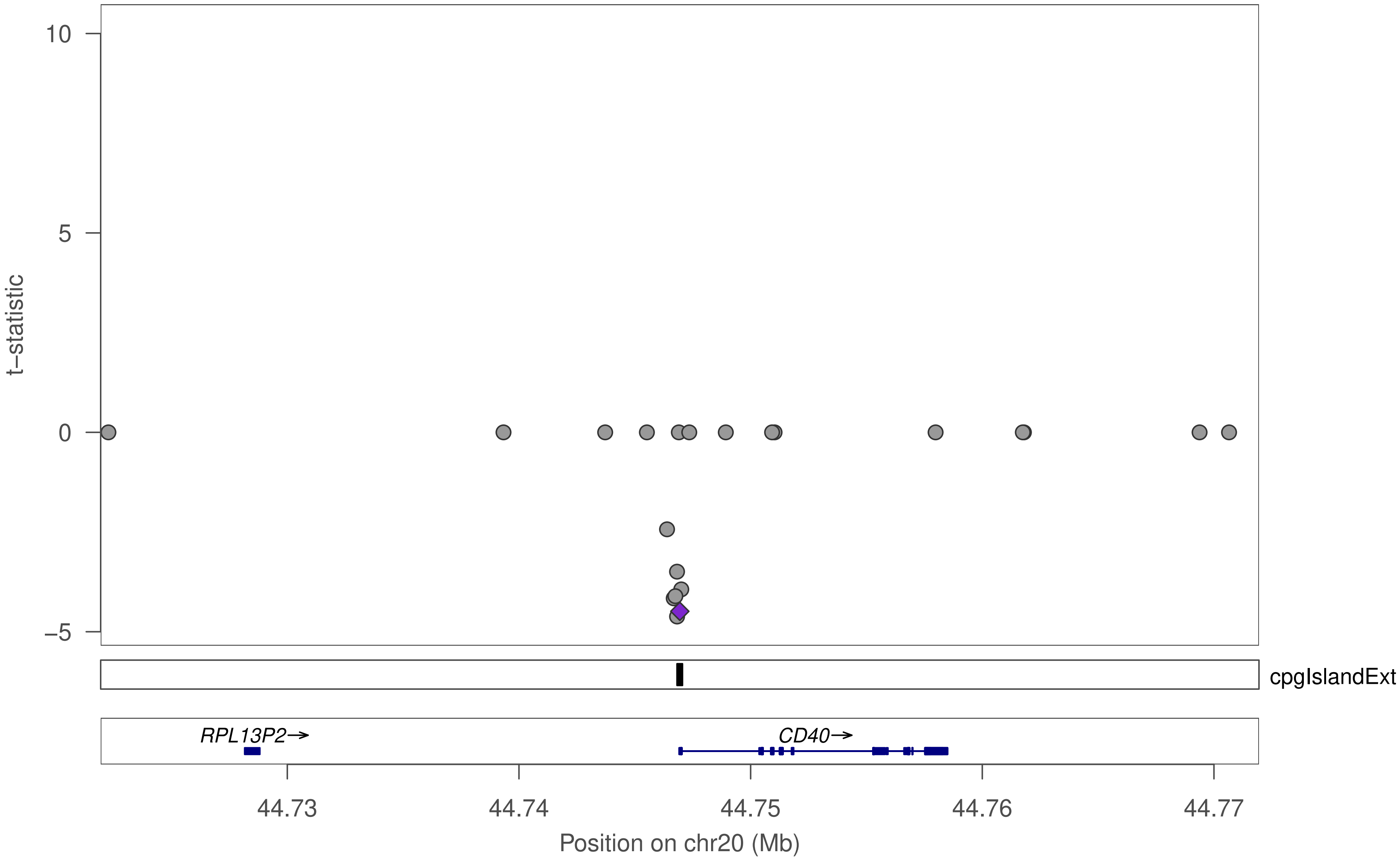

10
5
0
−5
t−statistic
cpgIslandExt
RPL13P2→
CD40→
44.73
44.74
44.75
44.76
44.77
Position on chr20 (Mb)

### ZNF548

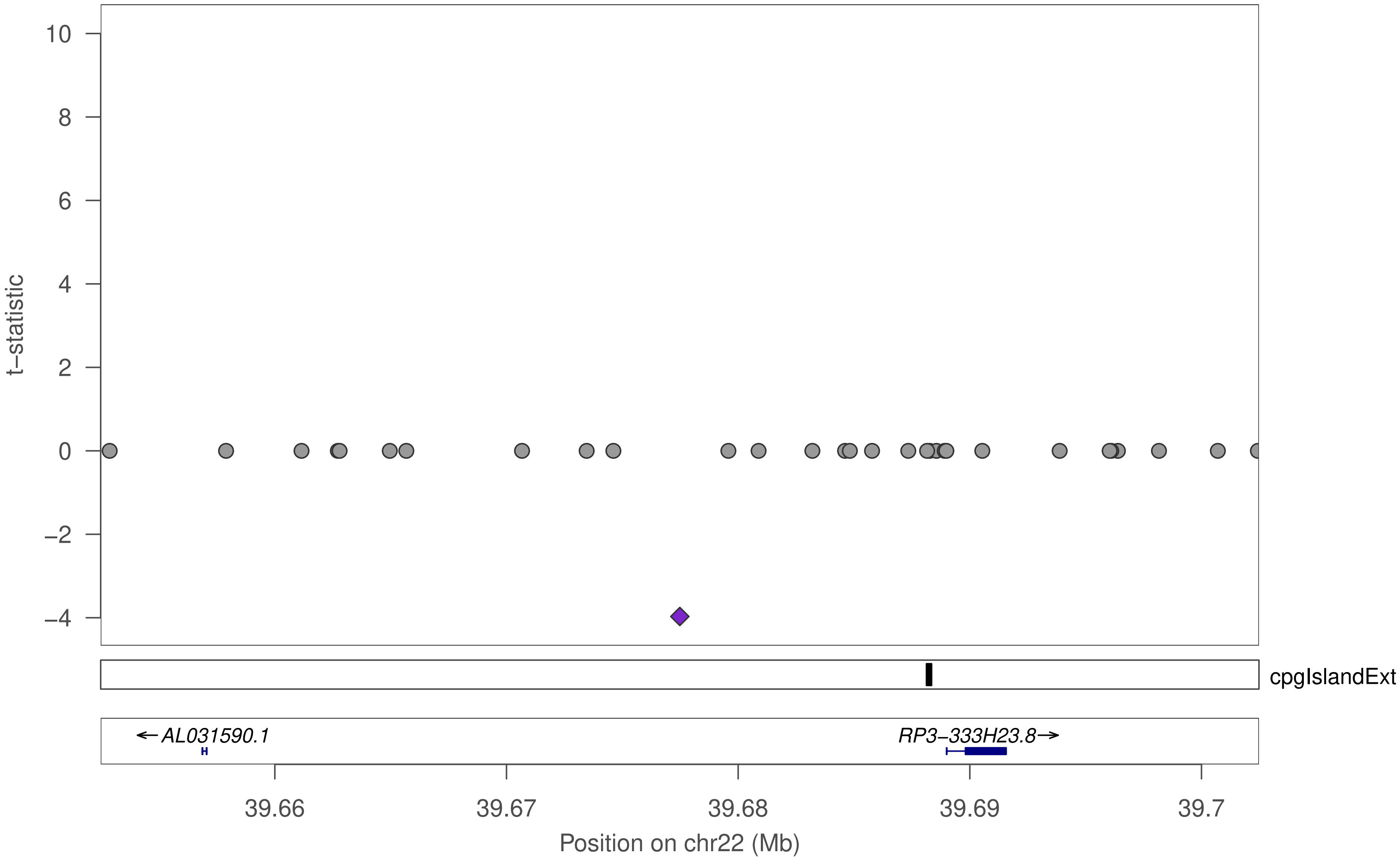

10
8
6
4
2
0
−2
−4
t−statistic
cpgIslandExt
←AL031590.1
RP3−333H23.8→
39.66
39.67
39.68
39.69
39.7
Position on chr22 (Mb)

### ZKSCAN3

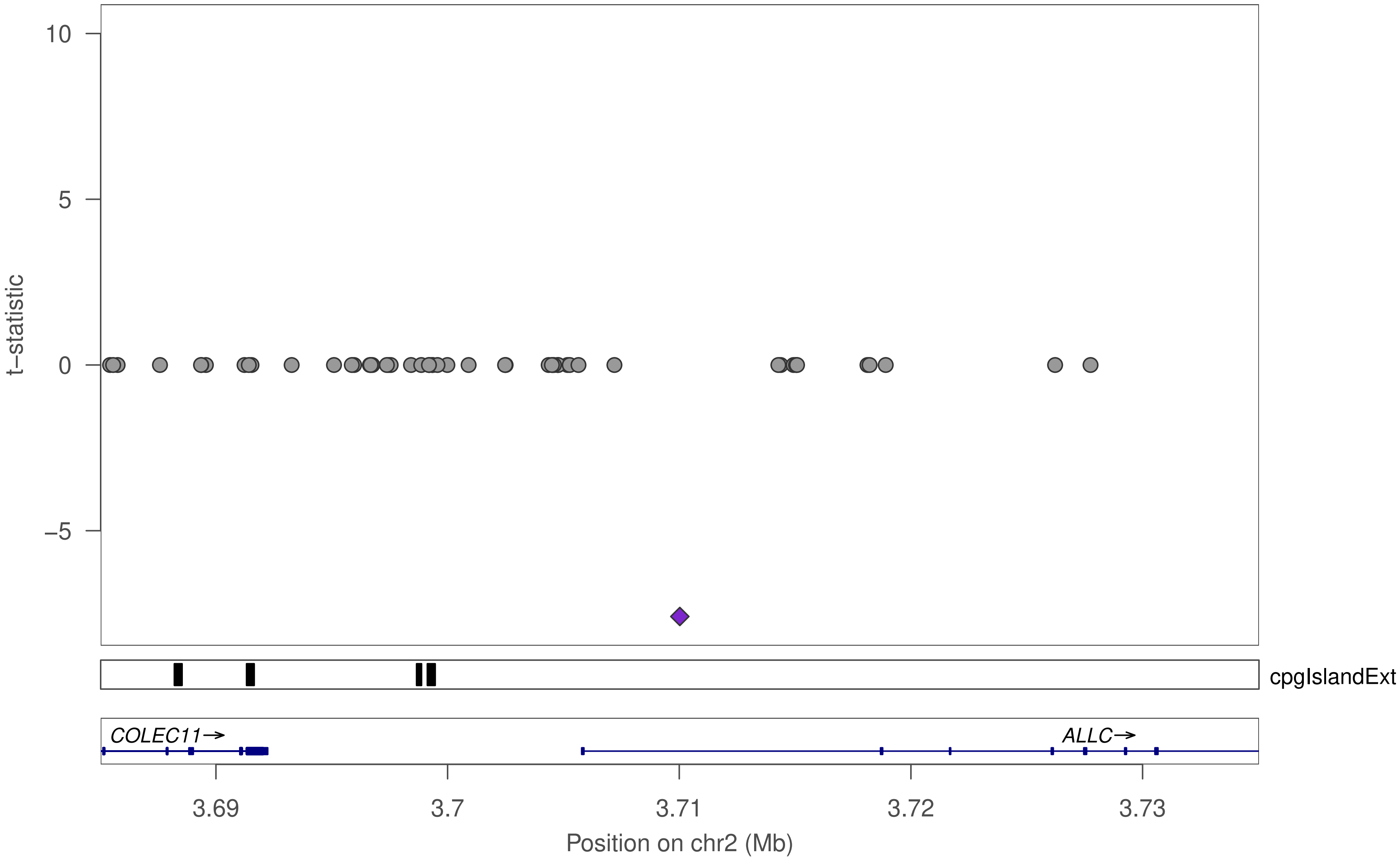

t–statistic
10
5
0
–5
cpgIslandExt
COLEC11→
ALLC→
3.69
3.7
3.71
3.72
3.73
Position on chr2 (Mb)

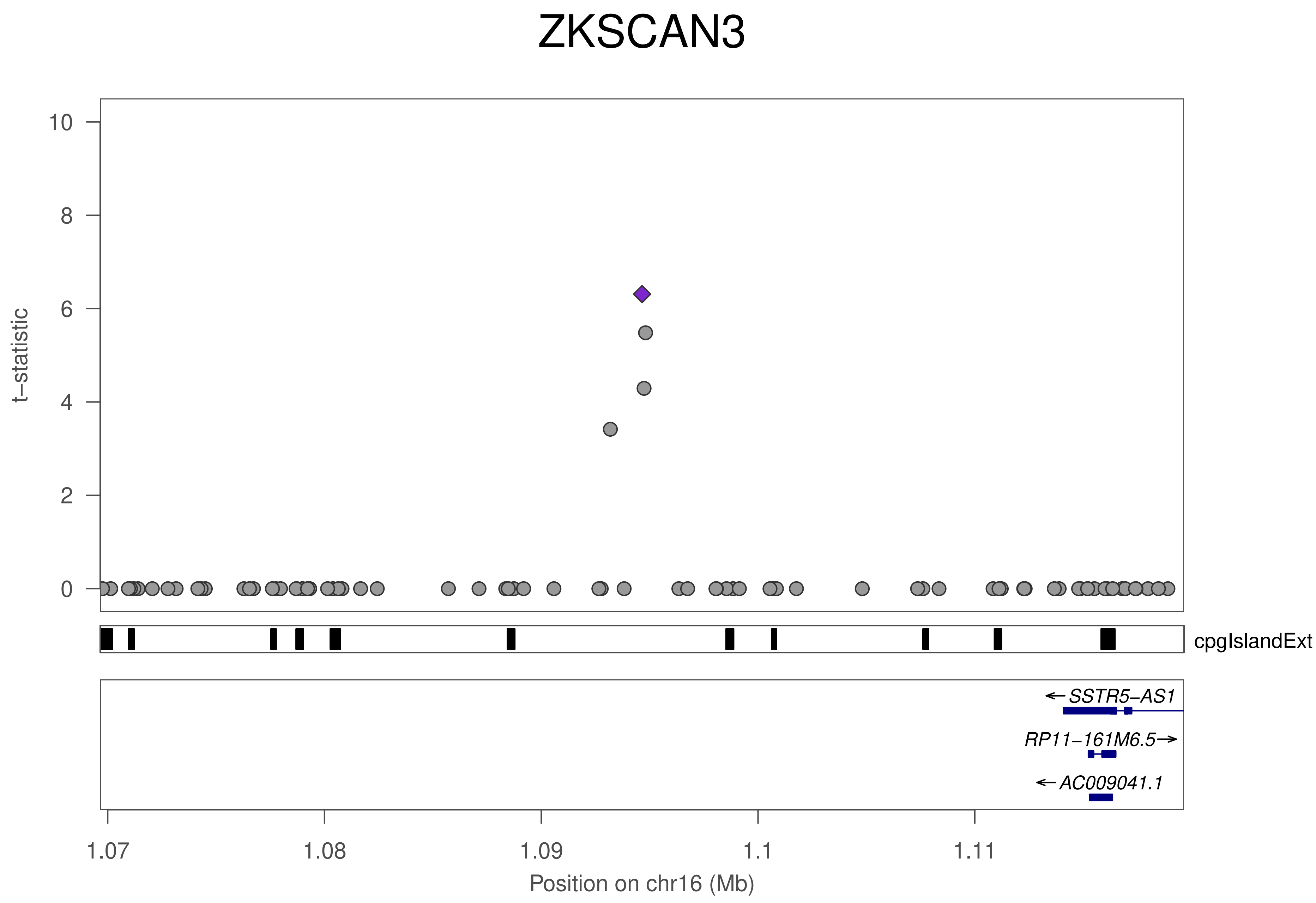

ZKSCAN3
t−statistic
0
2
4
6
8
10
cpgIslandExt
SSTR5−AS1
RP11−161M6.5
AC009041.1
1.07
1.08
1.09
1.1
1.11
Position on chr16 (Mb)

### ZNF79

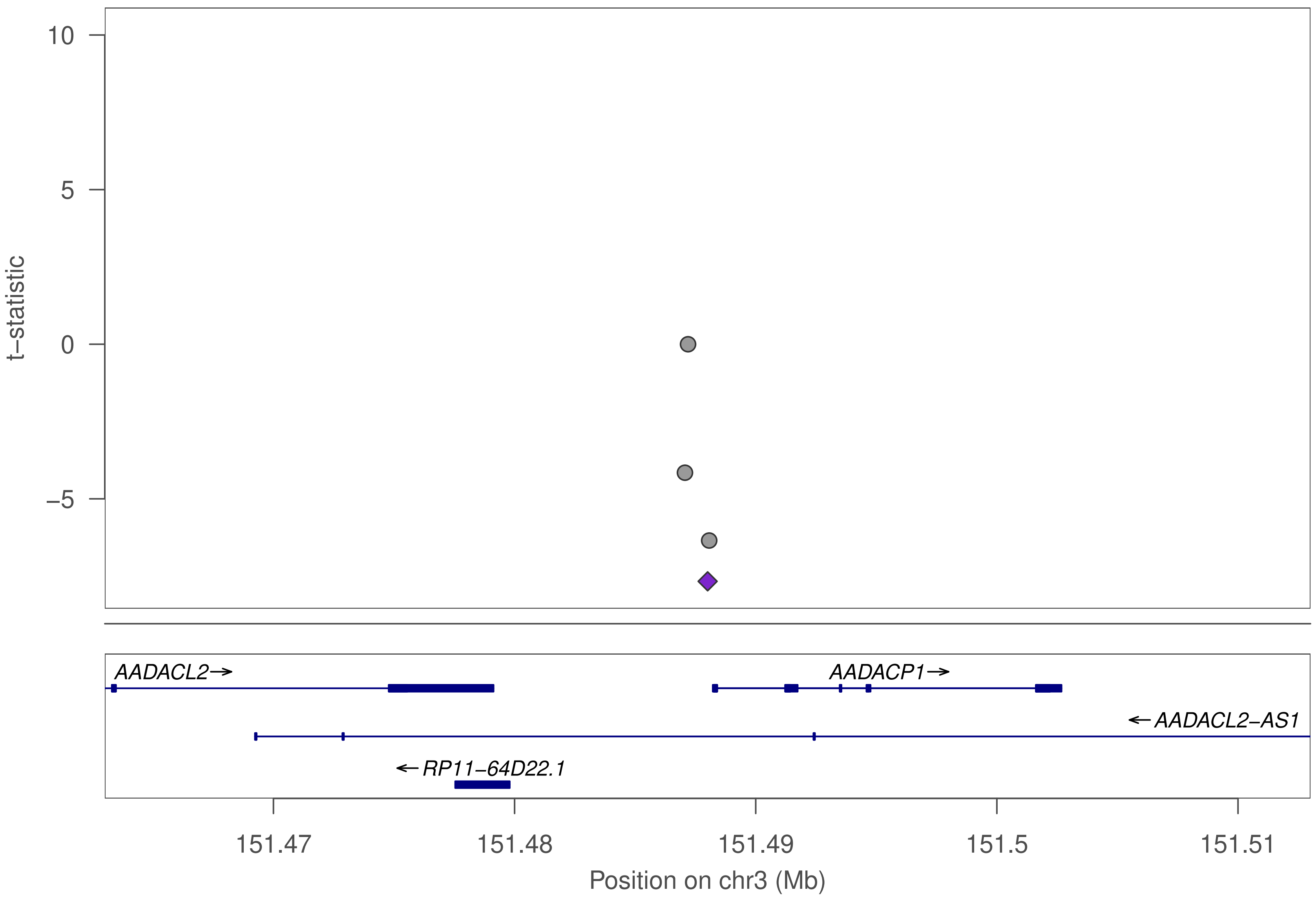
10
5
0
−5
t−statistic
AADACL2→
AADACP1→
←AADACL2−AS1
←RP11−64D22.1
151.47
151.48
151.49
151.5
151.51
Position on chr3 (Mb)

### ZNF605

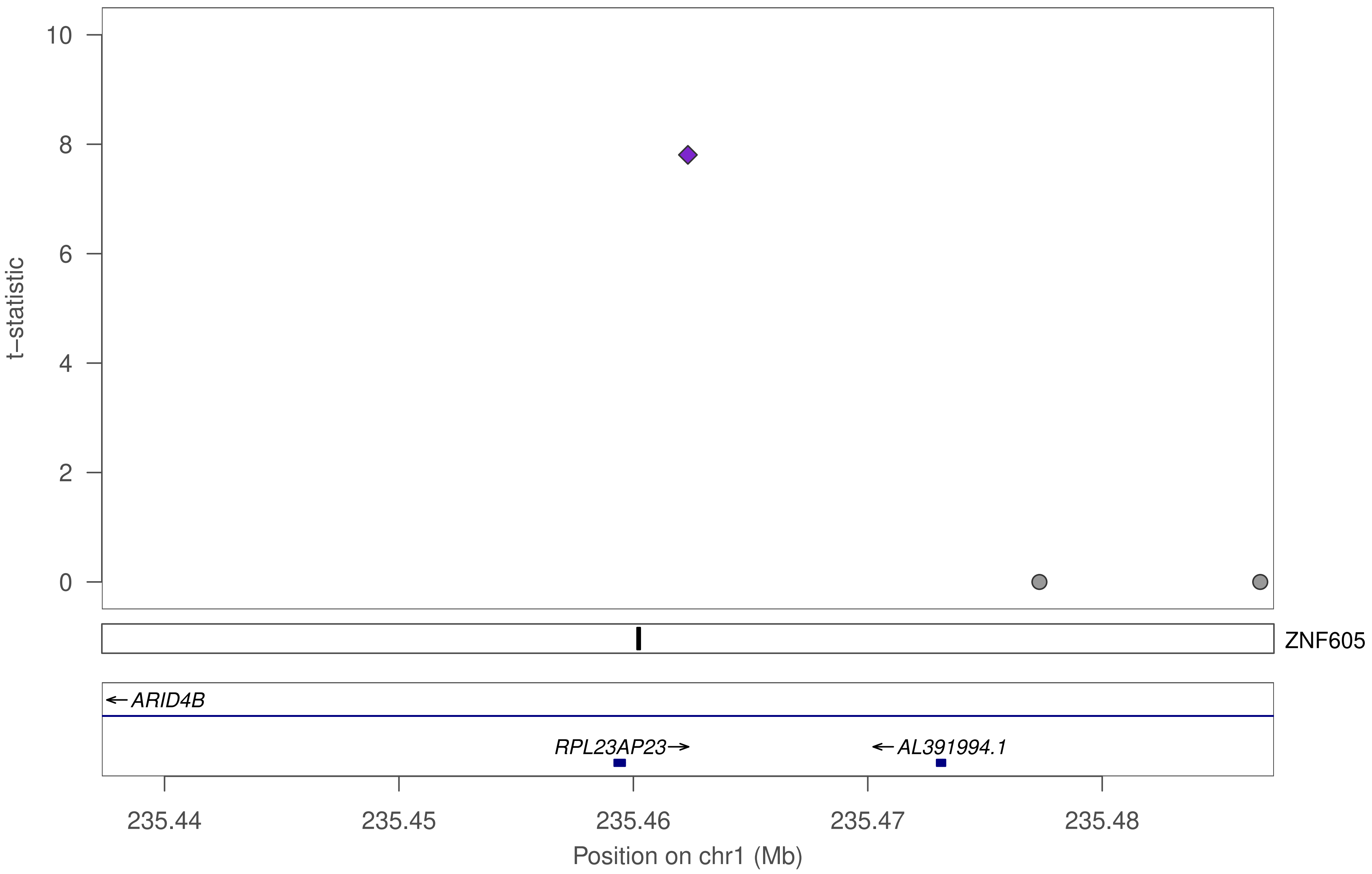

10
8
6
4
2
0
t−statistic
ZNF605
←ARID4B
RPL23AP23→
←AL391994.1
235.44
235.45
235.46
235.47
235.48
Position on chr1 (Mb)

### ZNF605

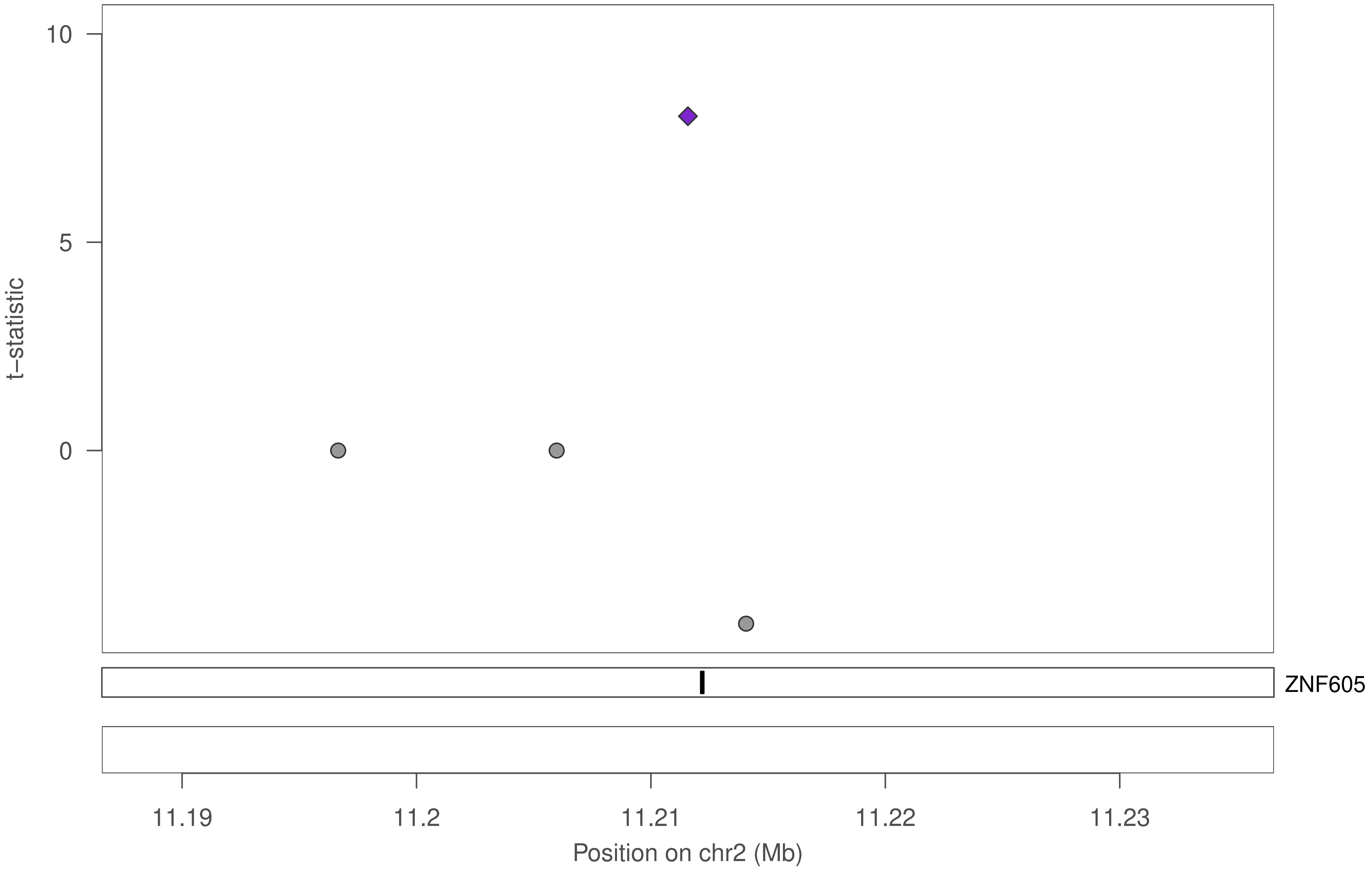

10
5
0
t−statistic
ZNF605
11.19
11.2
11.21
11.22
11.23
Position on chr2 (Mb)

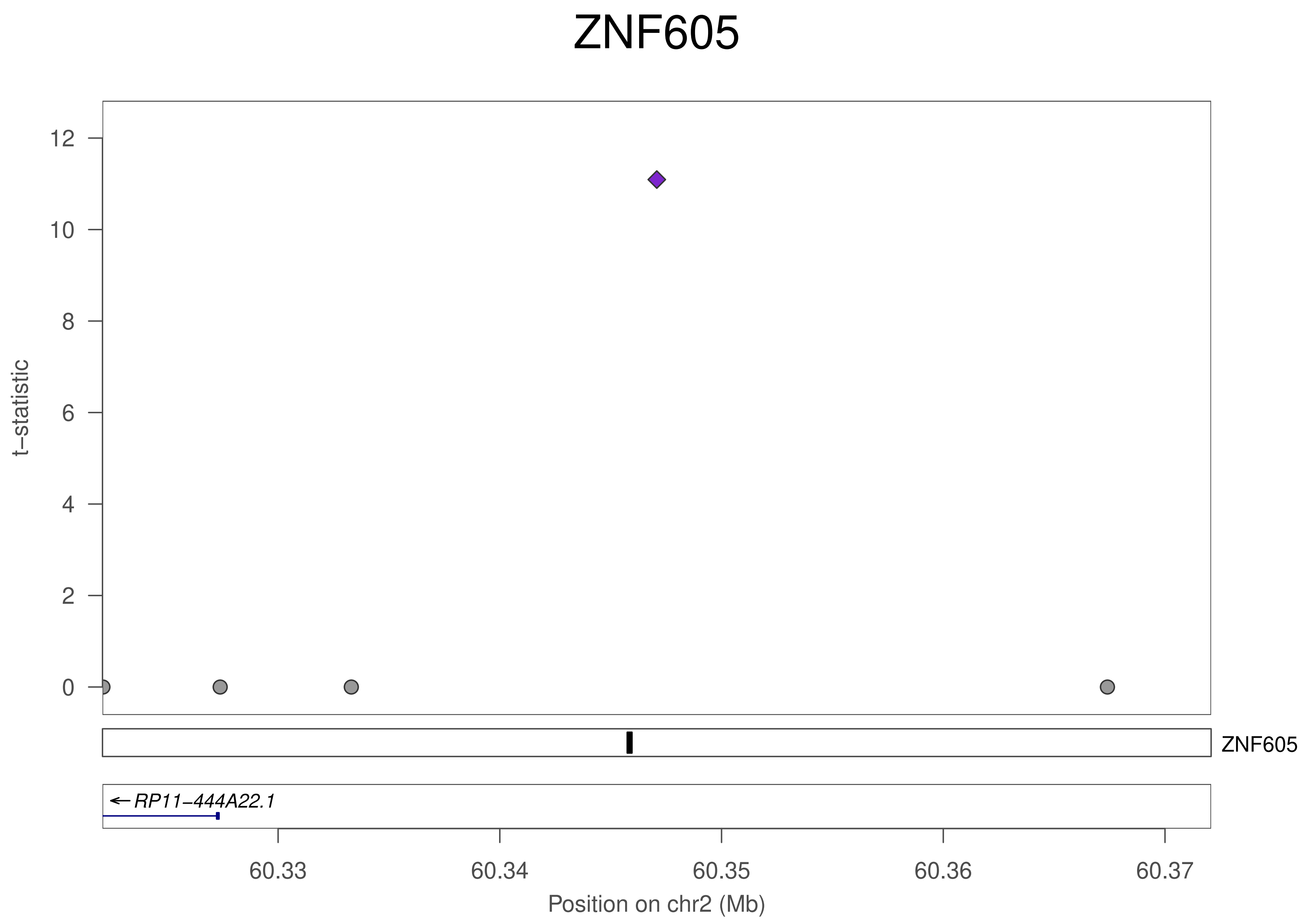

ZNF605
12
10
8
6
4
2
0
t−statistic
ZNF605
RP11−444A22.1
60.33
60.34
60.35
60.36
60.37
Position on chr2 (Mb)

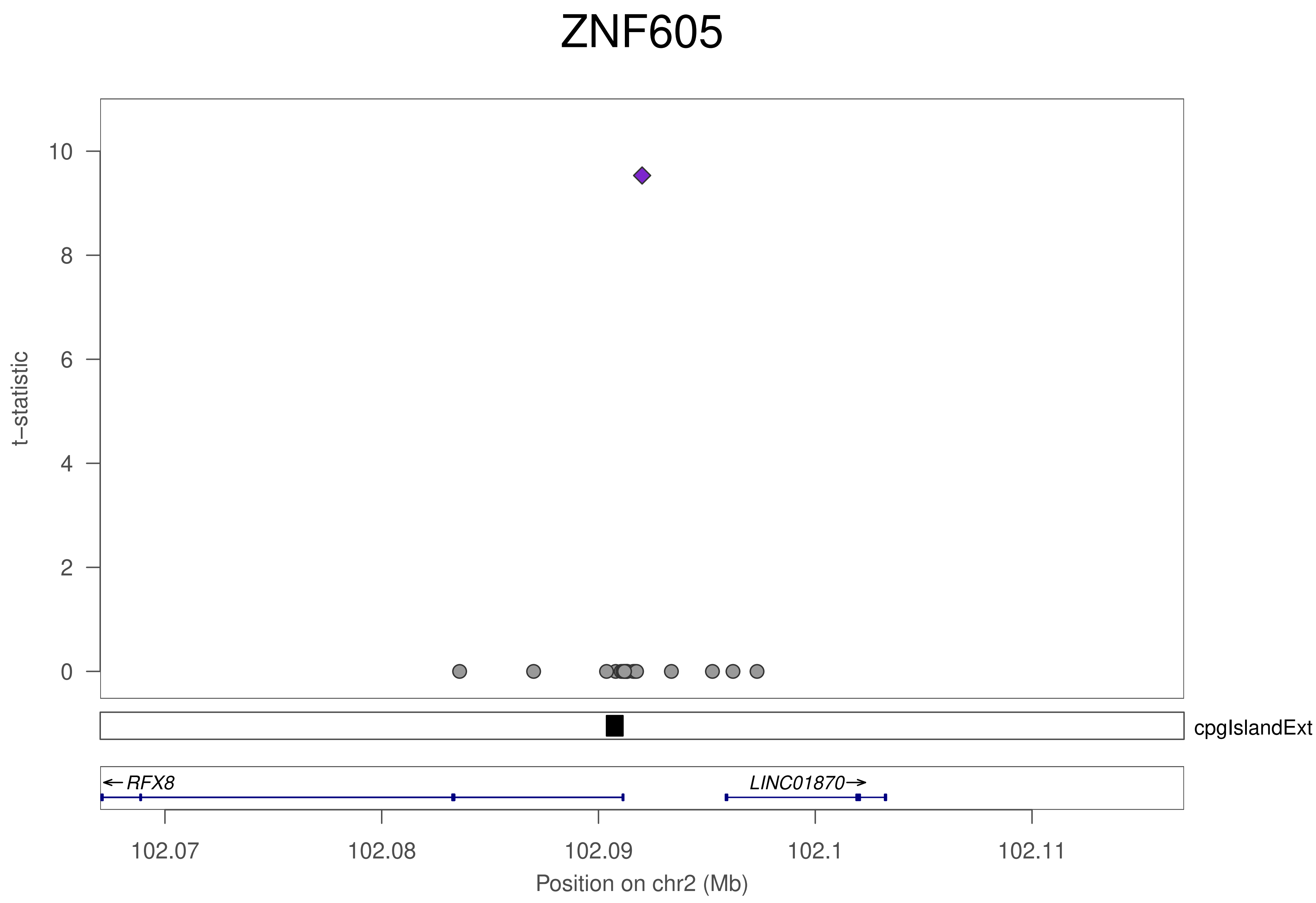
ZNF605
t–statistic
10
8
6
4
2
0
cpgIslandExt
←RFX8
LINC01870→
102.07
102.08
102.09
102.1
102.11
Position on chr2 (Mb)

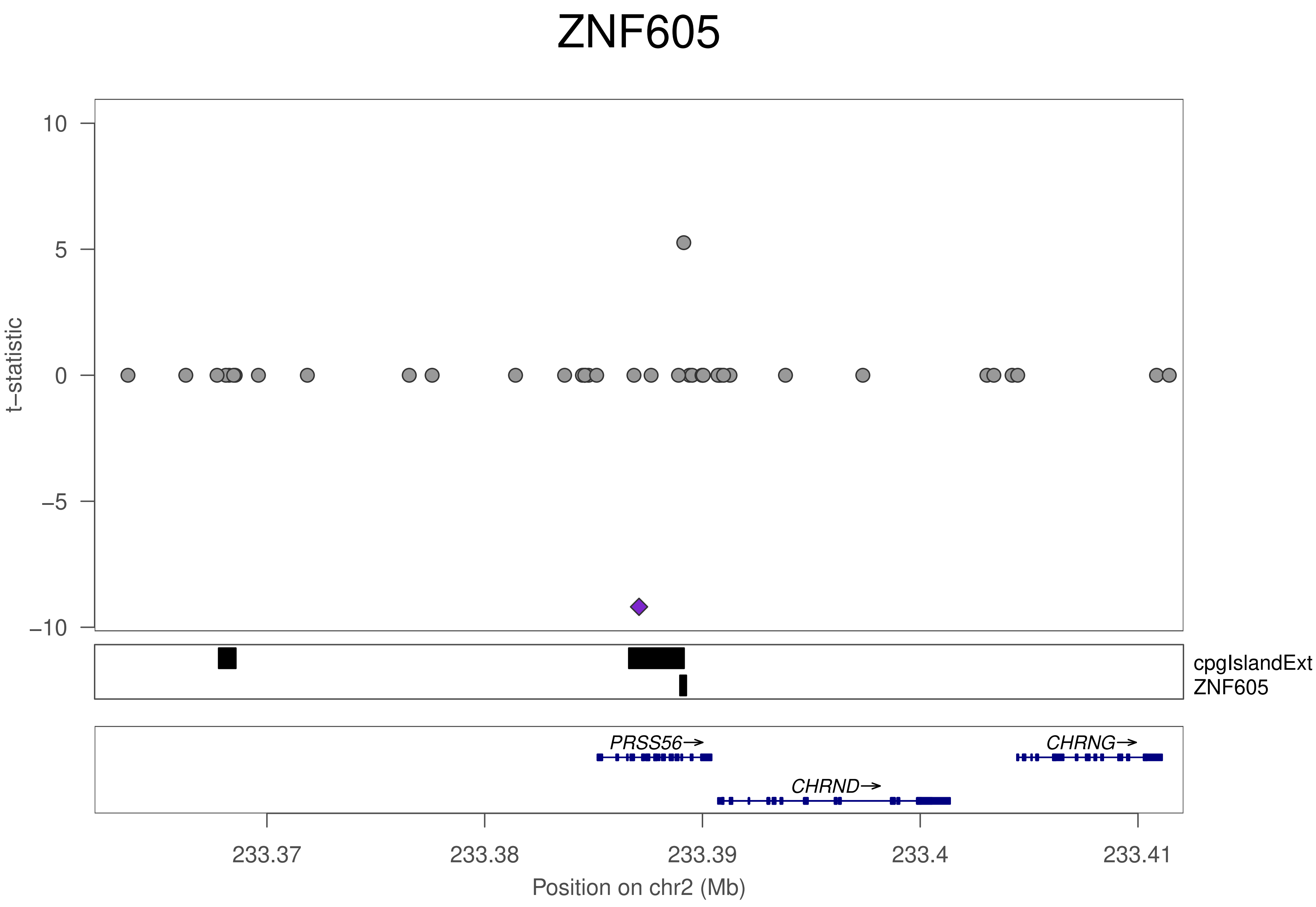
ZNF605
10
5
0
−5
−10
t−statistic
cpgIslandExt
ZNF605
PRSS56→
CHRNG→
CHRND→
233.37
233.38
233.39
233.4
233.41
Position on chr2 (Mb)

### ZNF605

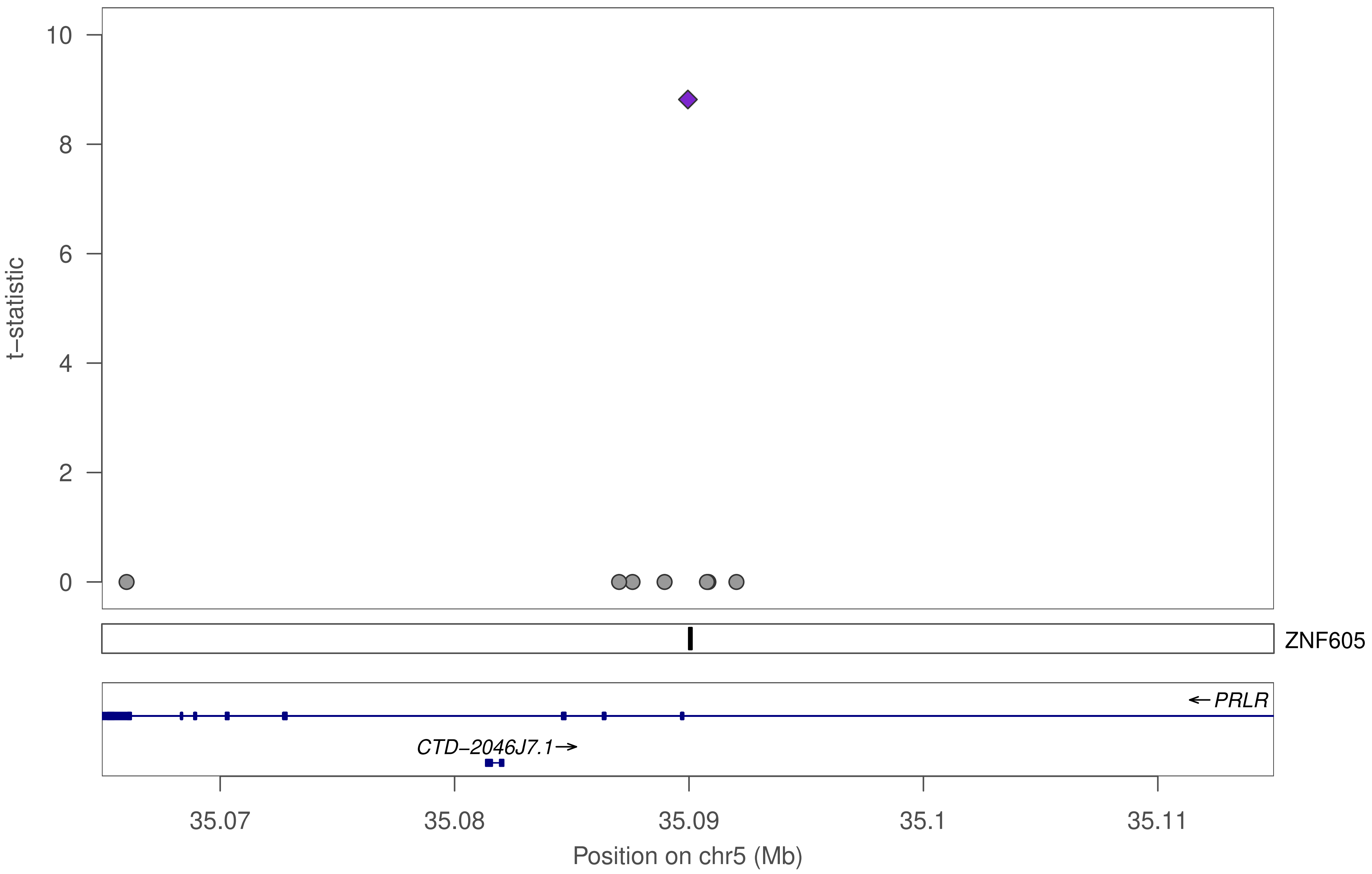

10
8
6
4
2
0
t−statistic
ZNF605
← PRLR
CTD−2046J7.1→
35.07
35.08
35.09
35.1
35.11
Position on chr5 (Mb)

### ZNF605

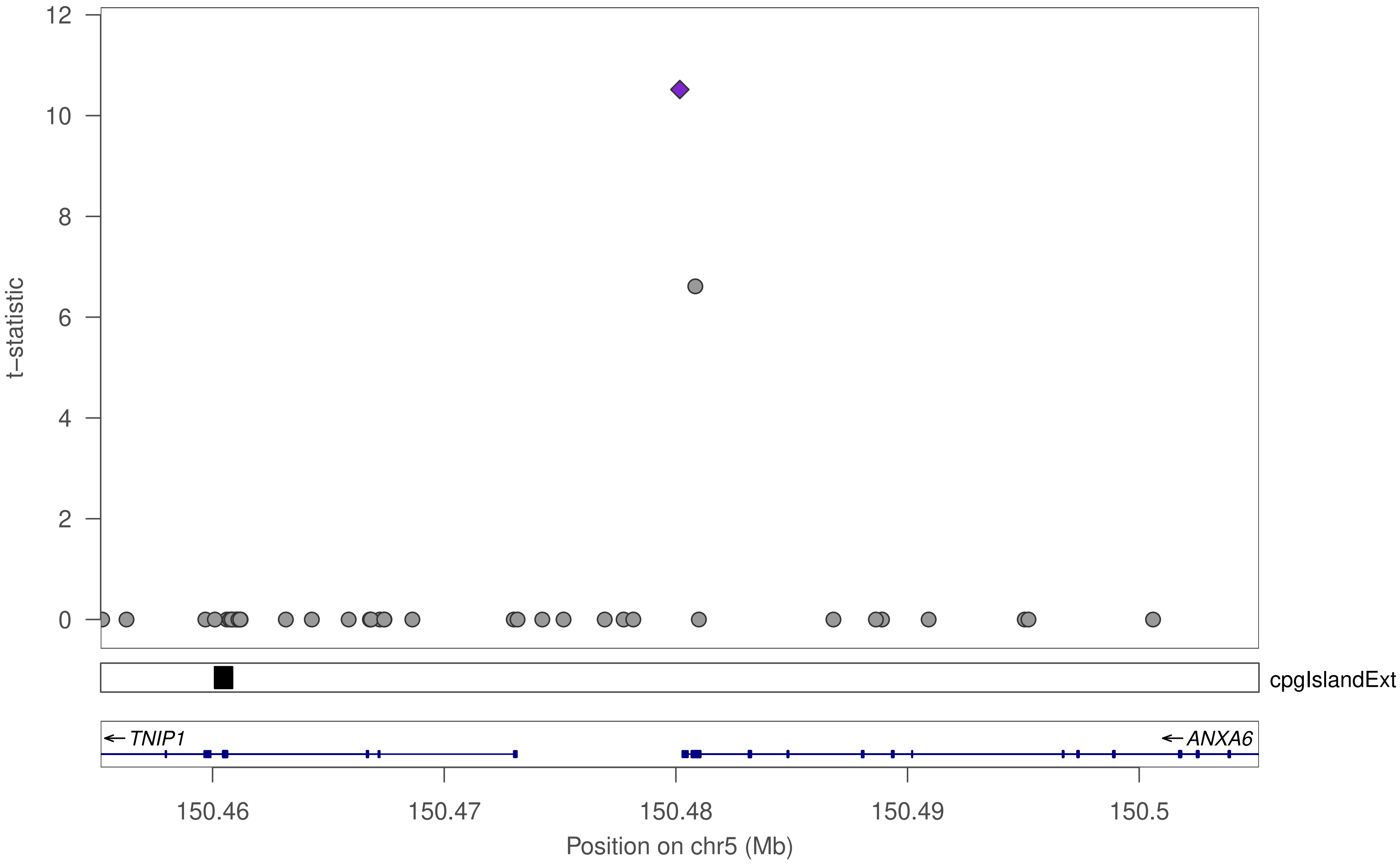

t–statistic
12
10
8
6
4
2
0
cpgIslandExt
TNIP1
ANXA6
150.46
150.47
150.48
150.49
150.5
Position on chr5 (Mb)

### ZNF605

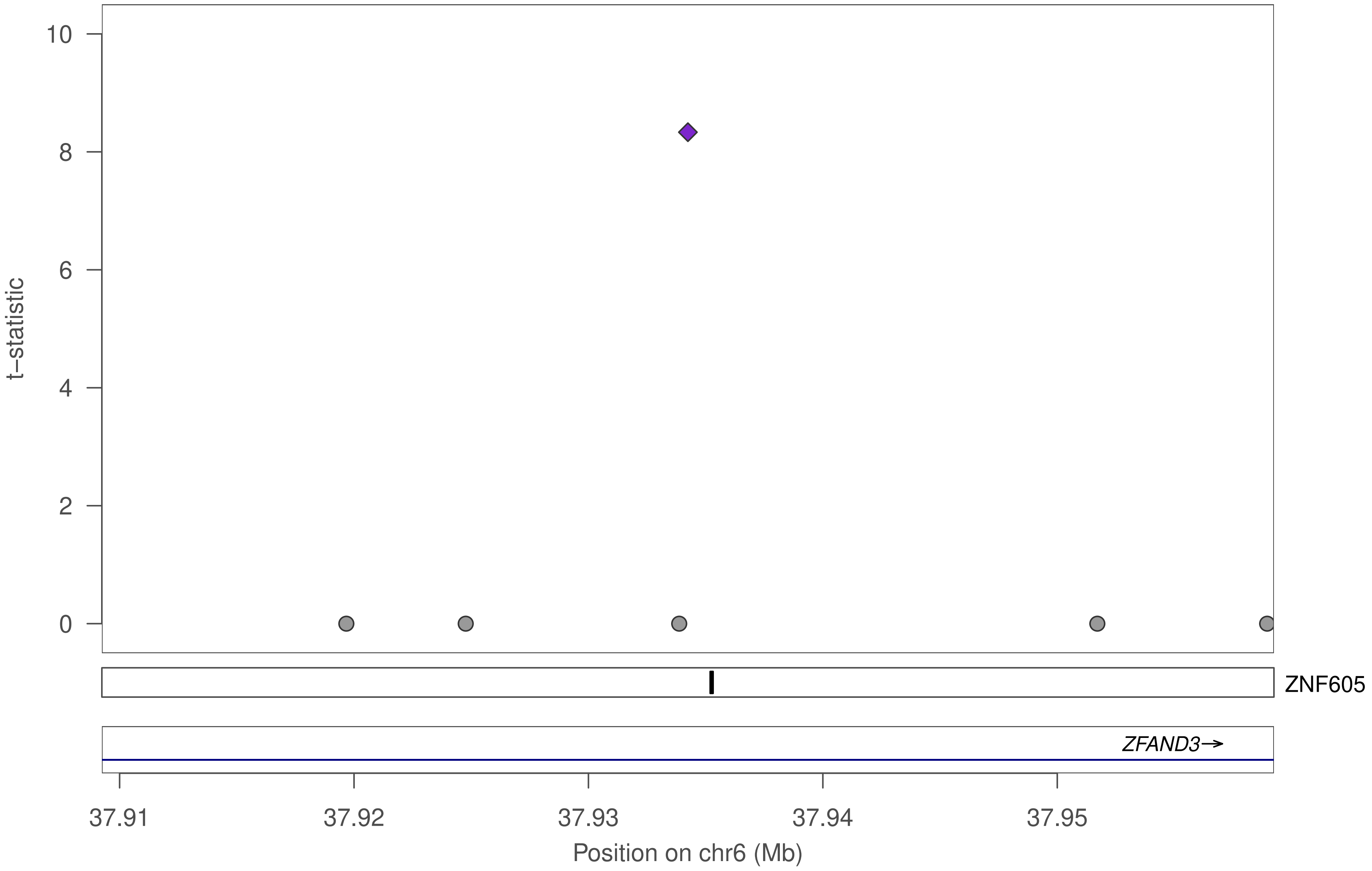

10
8
6
4
2
0
t–statistic
ZNF605
ZFAND3→
37.91
37.92
37.93
37.94
37.95
Position on chr6 (Mb)

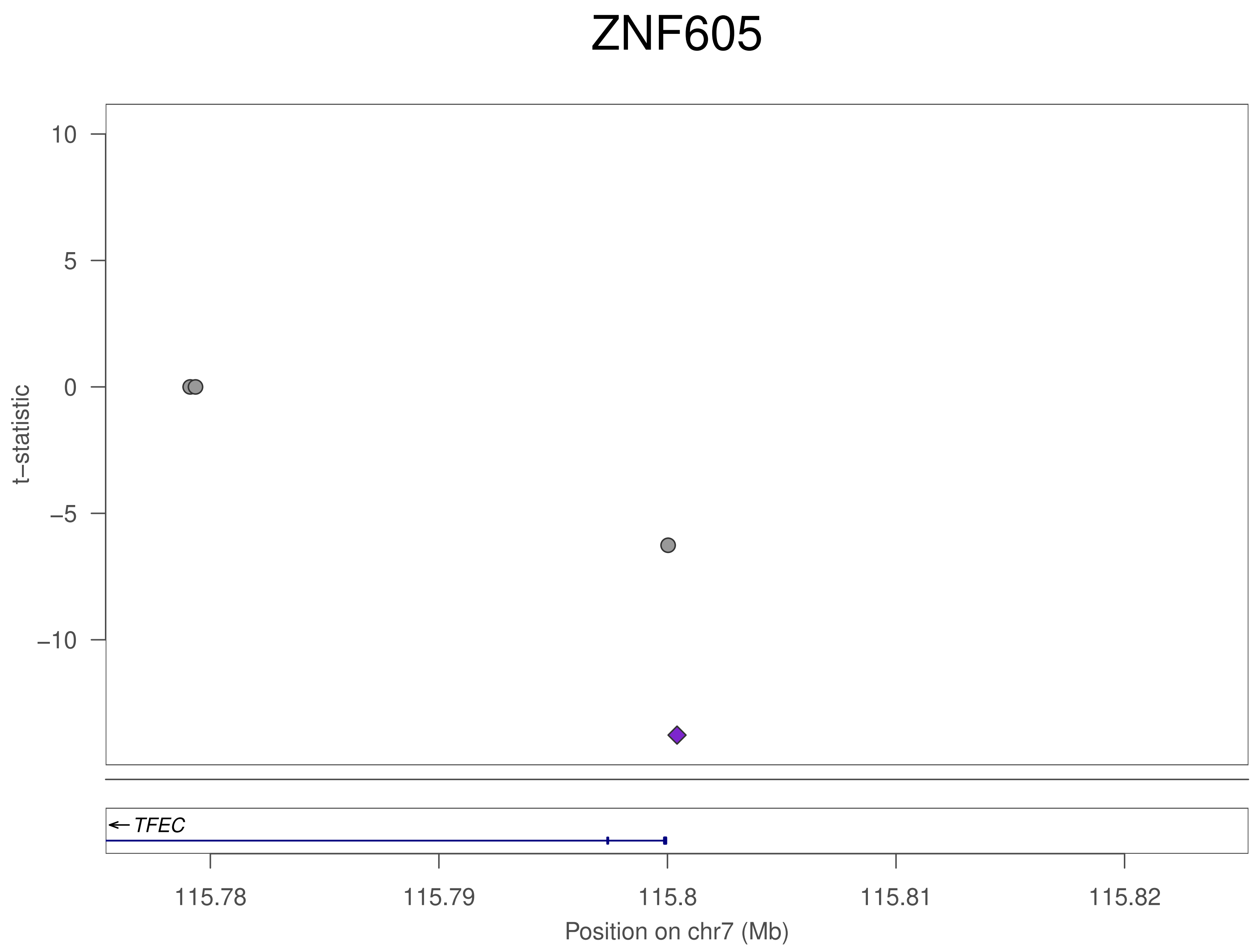
ZNF605
t−statistic
10
5
0
−5
−10
TFEC
115.78
115.79
115.8
115.81
115.82
Position on chr7 (Mb)

### ZNF605

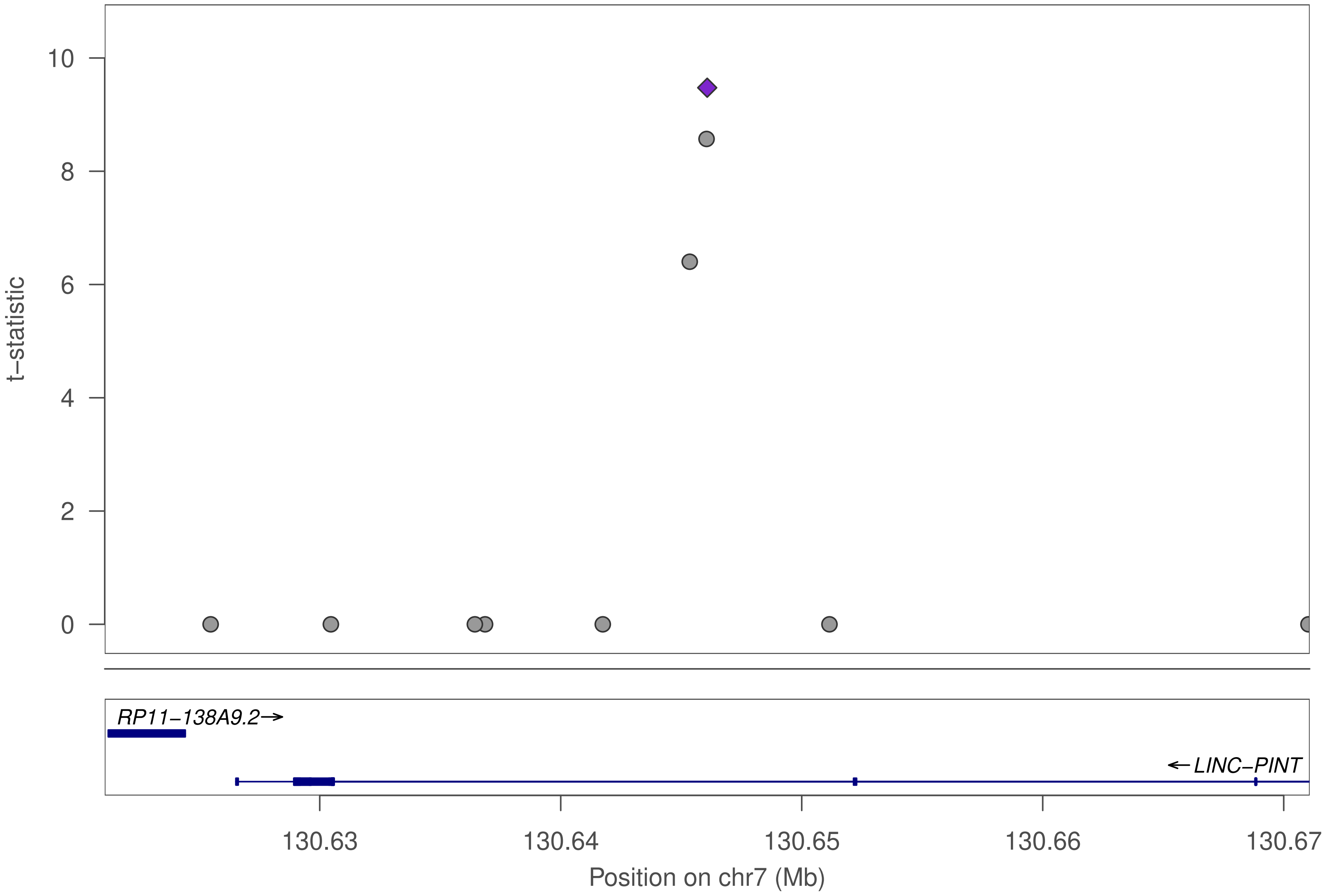

10
8
6
4
2
0
t−statistic
RP11−138A9.2→
←LINC−PINT
130.63
130.64
130.65
130.66
130.67
Position on chr7 (Mb)

### ZNF605

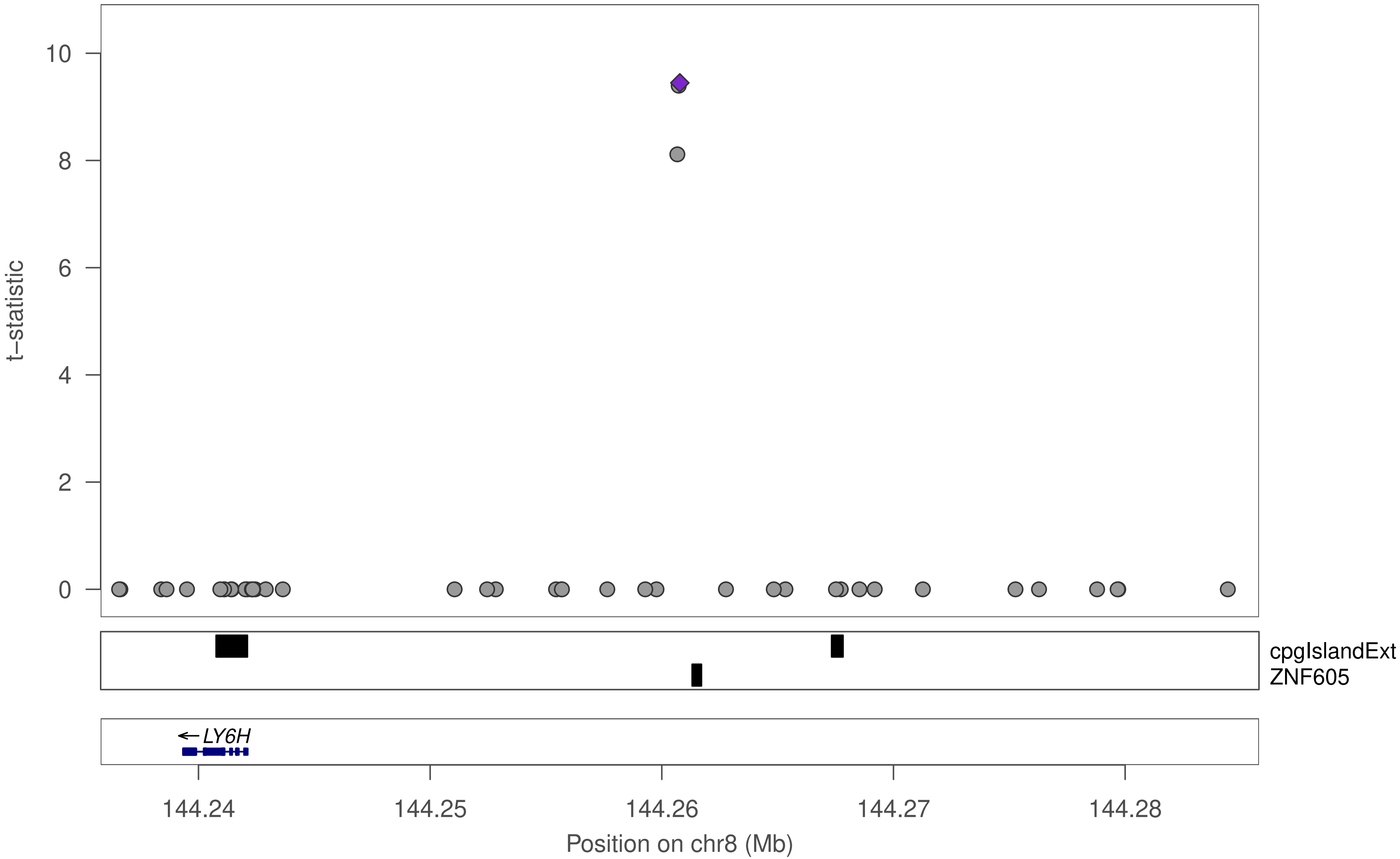

10
8
6
4
2
0
t−statistic
cpgIslandExt
ZNF605
LY6H
144.24
144.25
144.26
144.27
144.28
Position on chr8 (Mb)

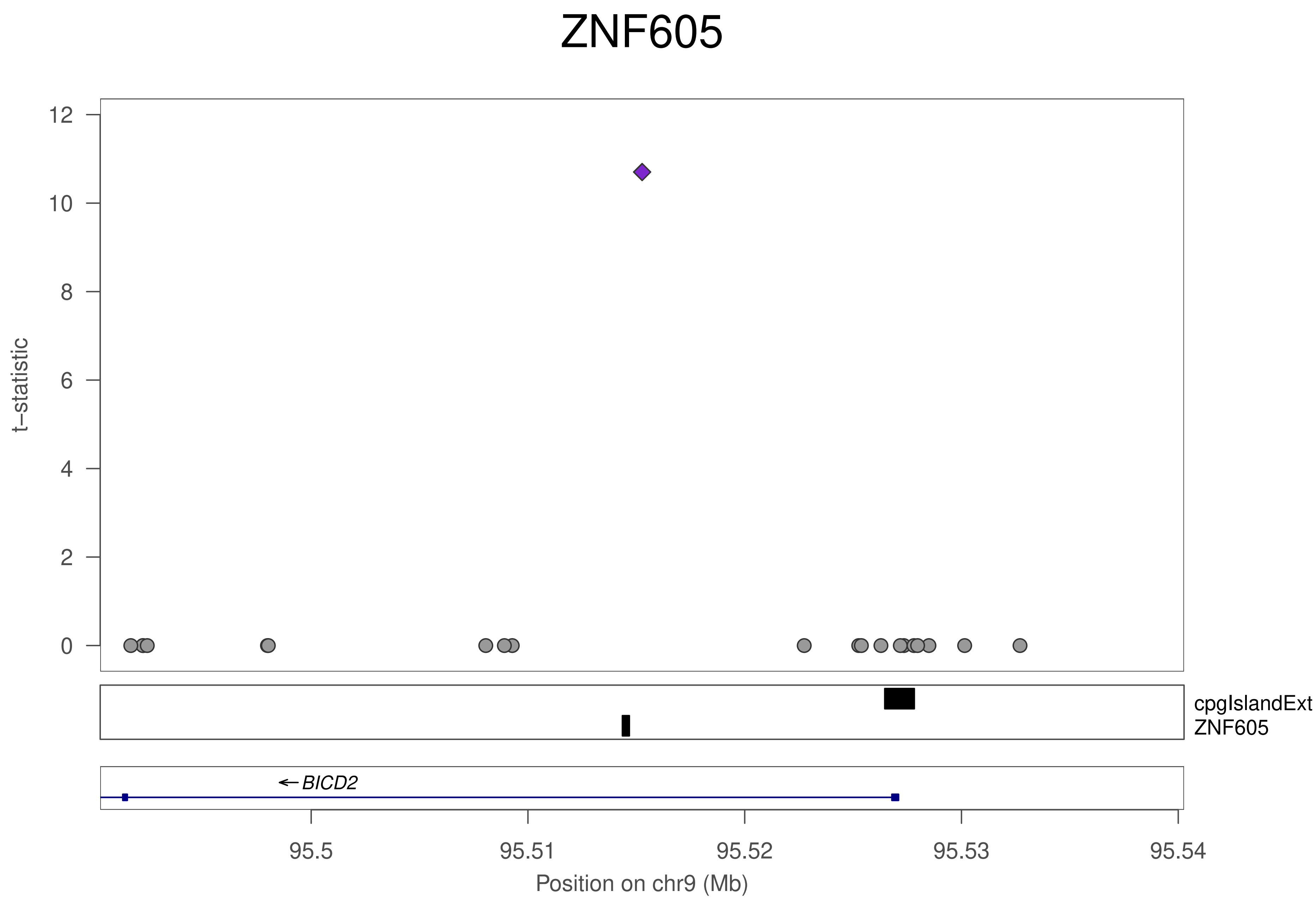

ZNF605
t−statistic
12
10
8
6
4
2
0
cpgIslandExt
ZNF605
BICD2
95.5
95.51
95.52
95.53
95.54
Position on chr9 (Mb)

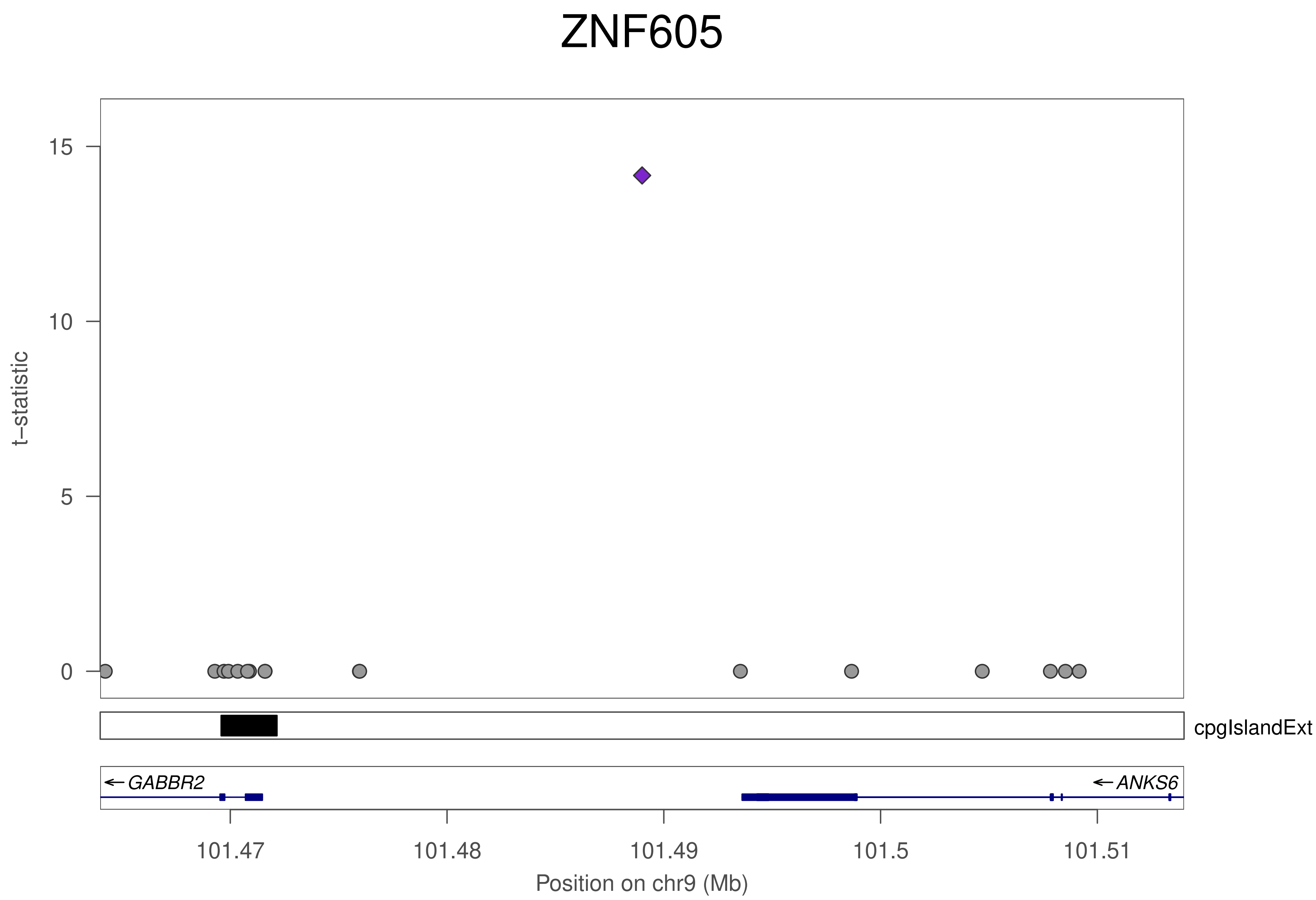
ZNF605
t–statistic
15
10
5
0
cpgIslandExt
GABBR2
ANKS6
101.47
101.48
101.49
101.5
101.51
Position on chr9 (Mb)

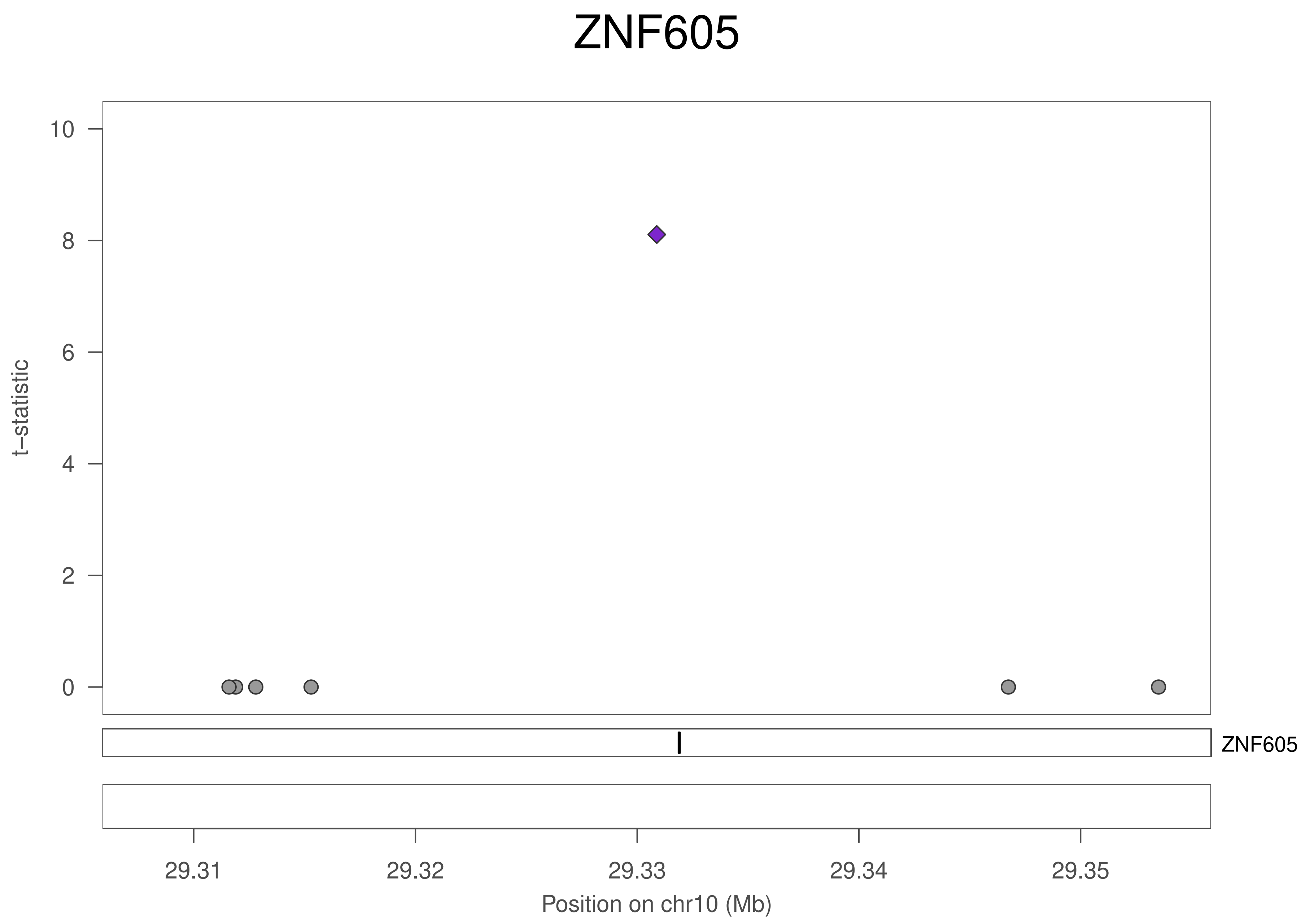
ZNF605
t–statistic
10
8
6
4
2
0
ZNF605
29.31
29.32
29.33
29.34
29.35
Position on chr10 (Mb)

### ZNF605

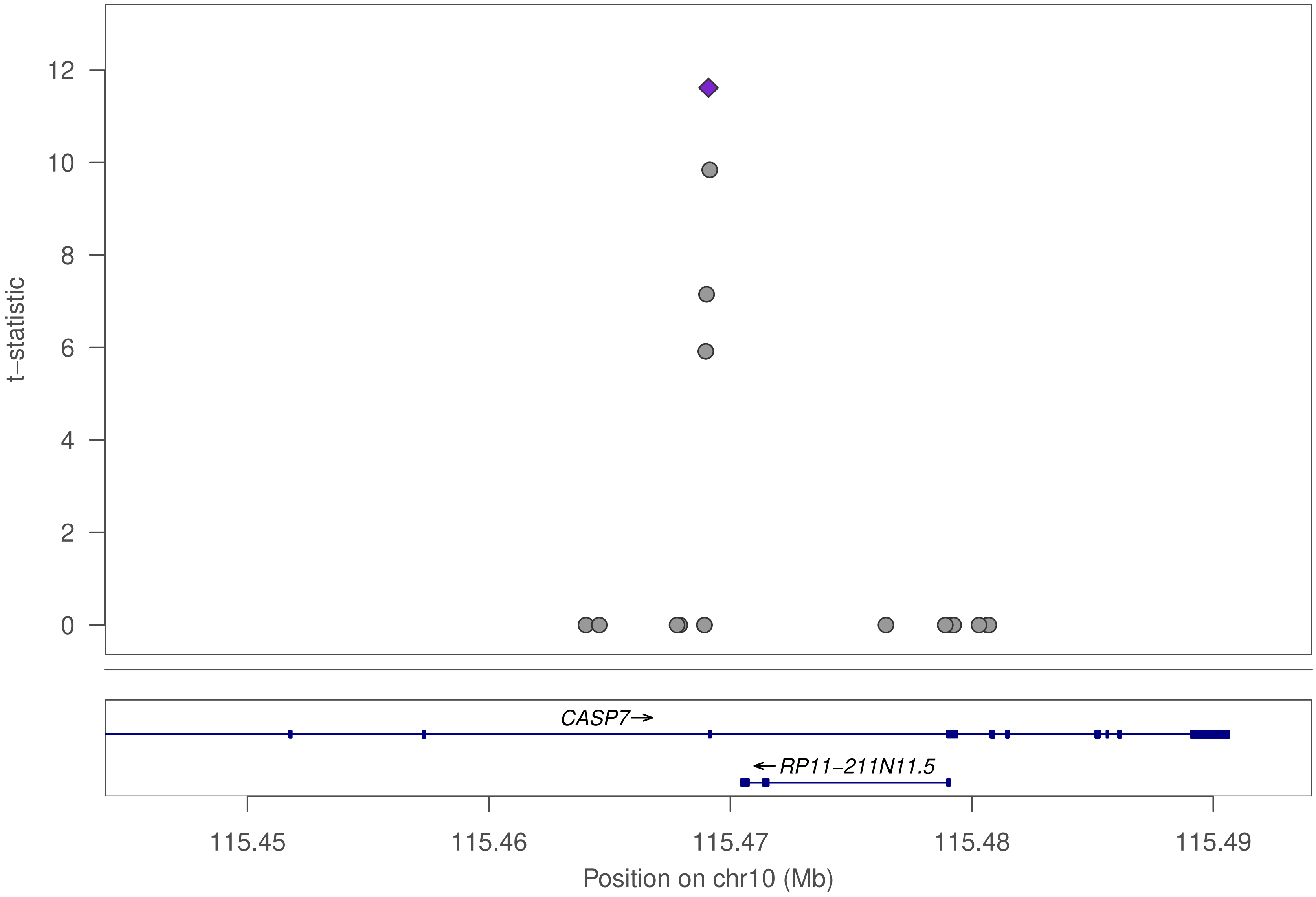

12
10
8
6
4
2
0
t−statistic
CASP7→
←RP11−211N11.5
115.45
115.46
115.47
115.48
115.49
Position on chr10 (Mb)

### ZNF605

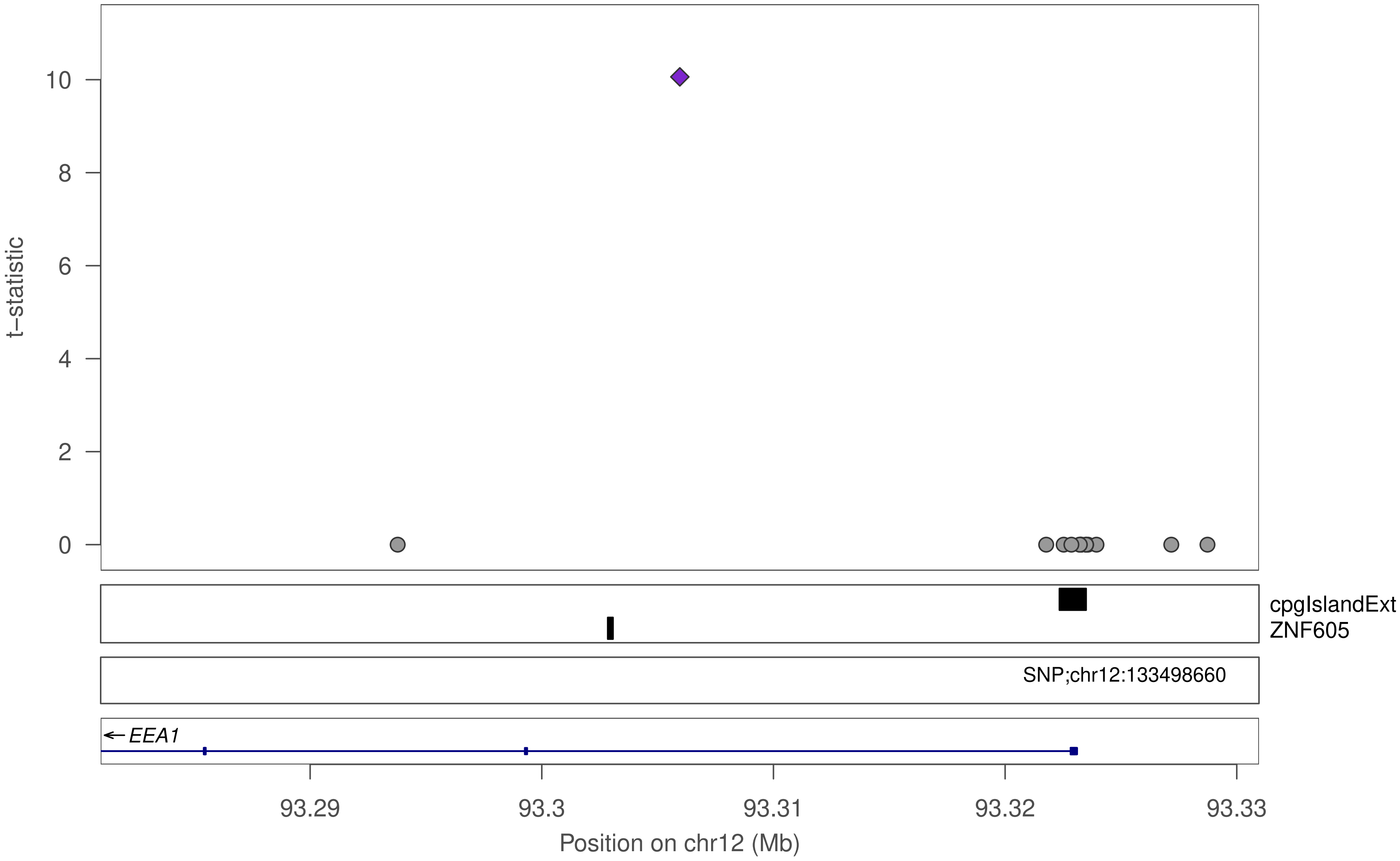

t–statistic
10
8
6
4
2
0
cpgIslandExt
ZNF605
SNP;chr12:133498660
EEA1
93.29
93.3
93.31
93.32
93.33
Position on chr12 (Mb)

### ZNF605

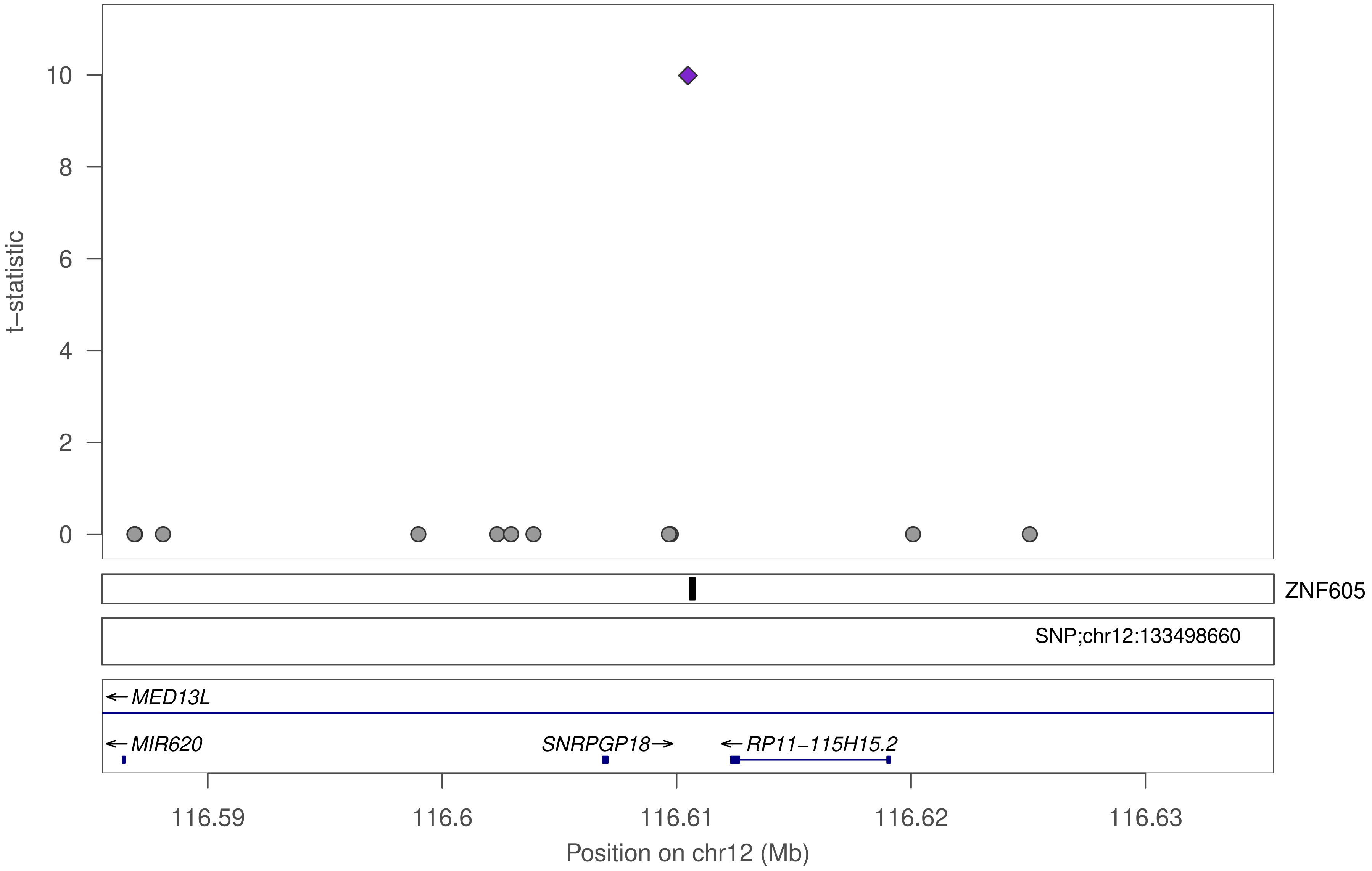

t−statistic
10
8
6
4
2
0
ZNF605
SNP;chr12:133498660
MED13L
MIR620
SNRPGP18
RP11−115H15.2
116.59
116.6
116.61
116.62
116.63
Position on chr12 (Mb)

### ZNF605

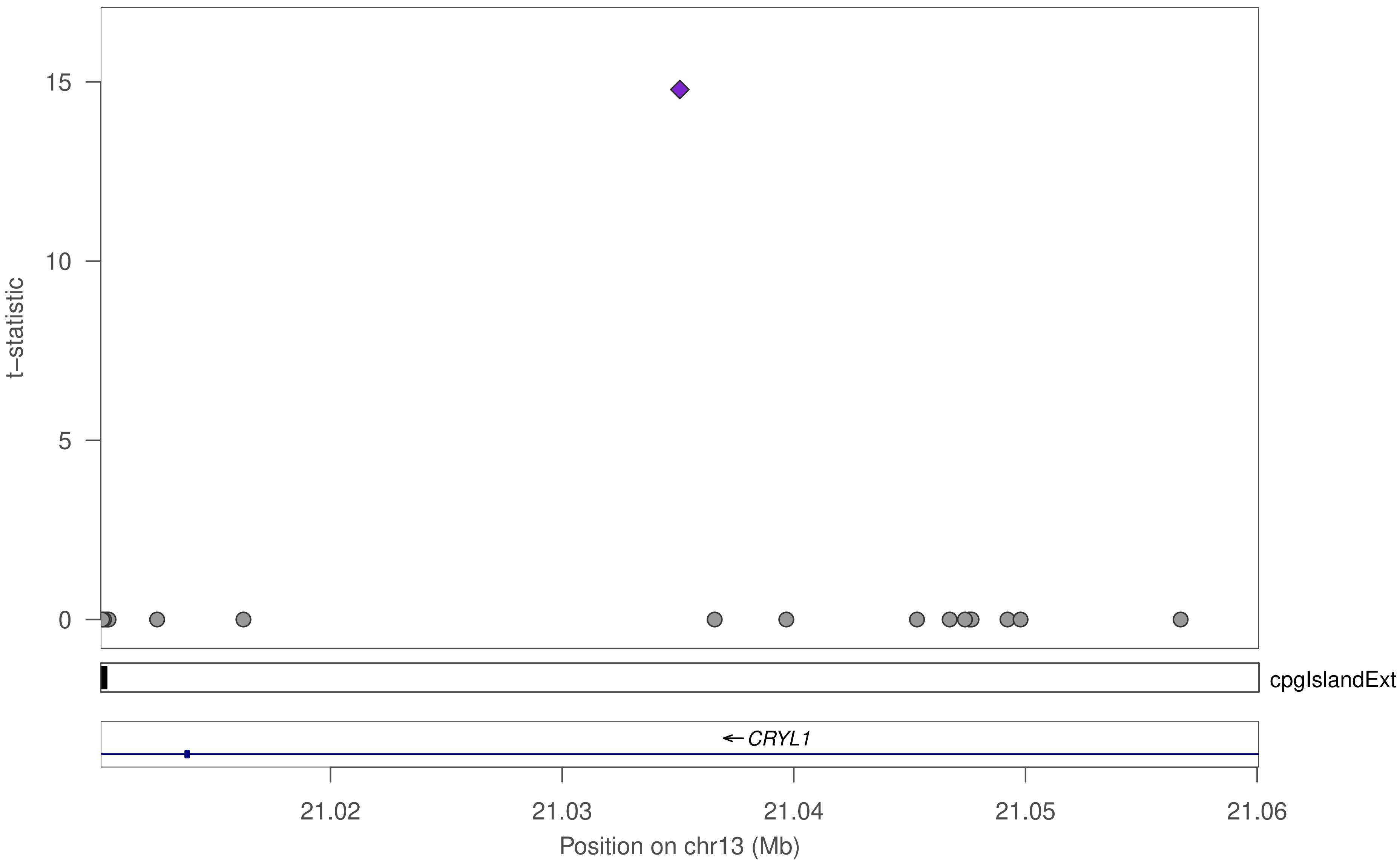

15
10
5
0
t−statistic
cpgIslandExt
← CRYL1
21.02
21.03
21.04
21.05
21.06
Position on chr13 (Mb)

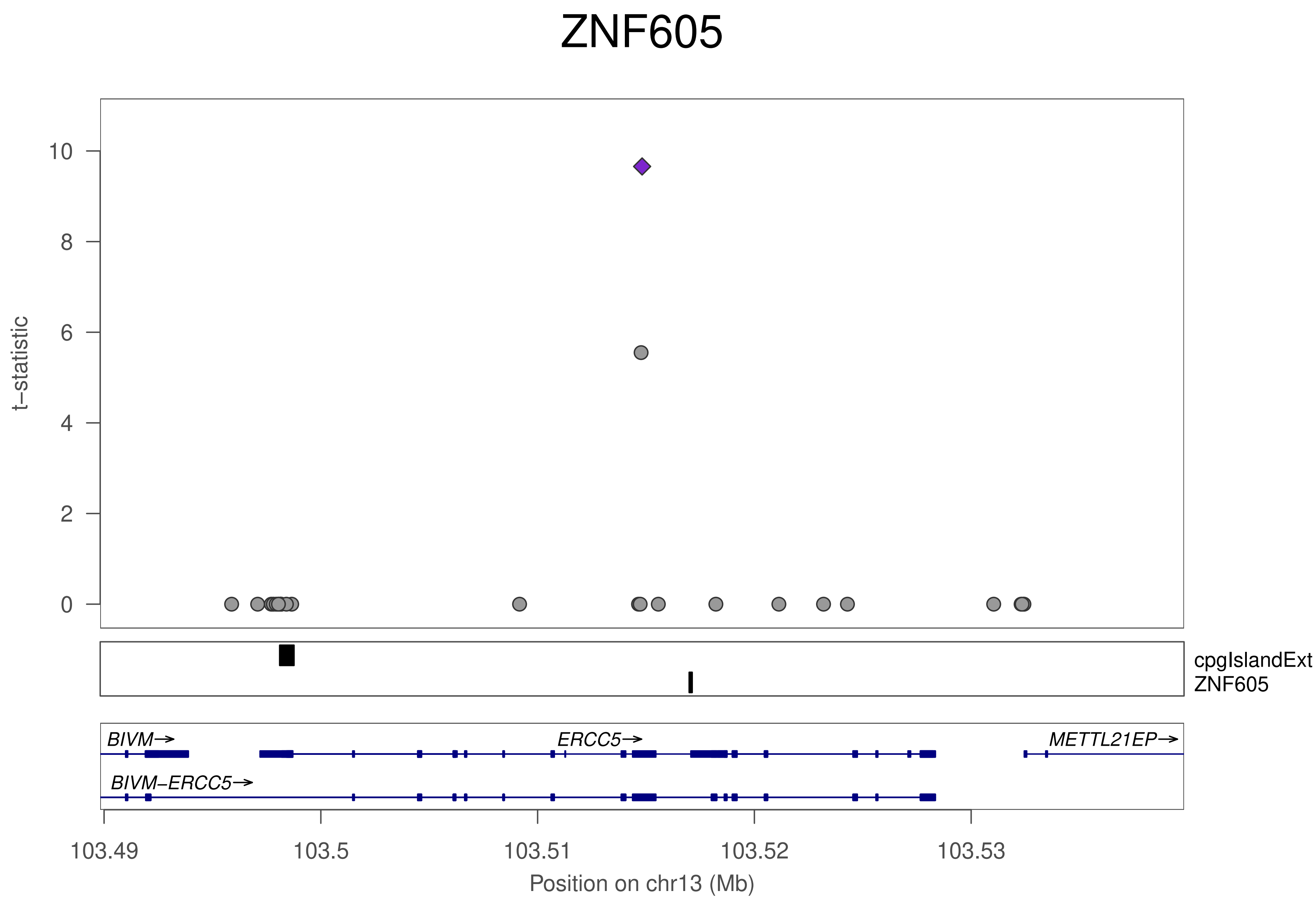
ZNF605
t−statistic
0
2
4
6
8
10
cpgIslandExt
ZNF605
BIVM→
ERCC5→
METTL21EP→
BIVM−ERCC5→
103.49
103.5
103.51
103.52
103.53
Position on chr13 (Mb)

### ZNF605

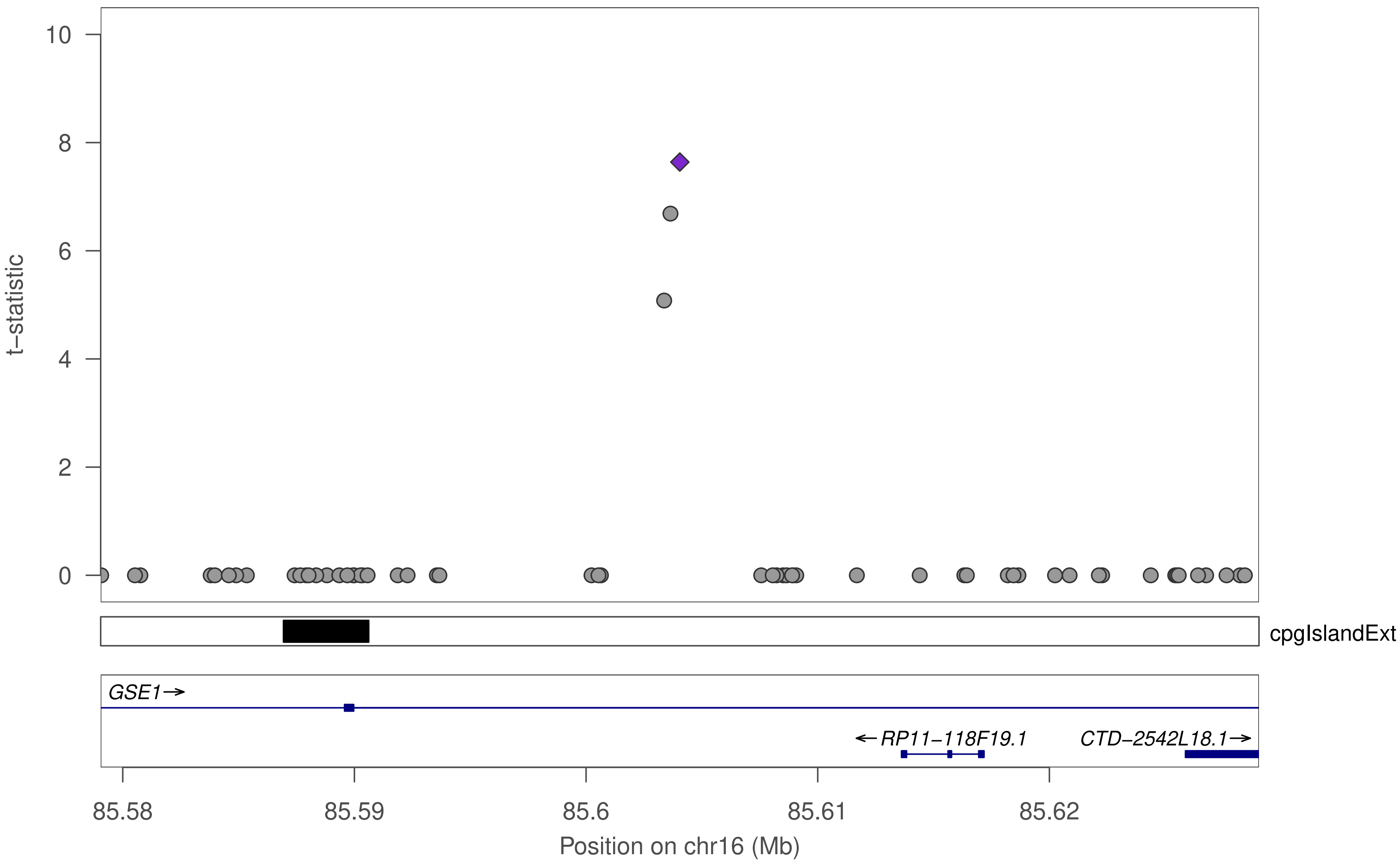

t−statistic
10
8
6
4
2
0
cpgIslandExt
GSE1→
←RP11−118F19.1
CTD−2542L18.1→
85.58
85.59
85.6
85.61
85.62
Position on chr16 (Mb)

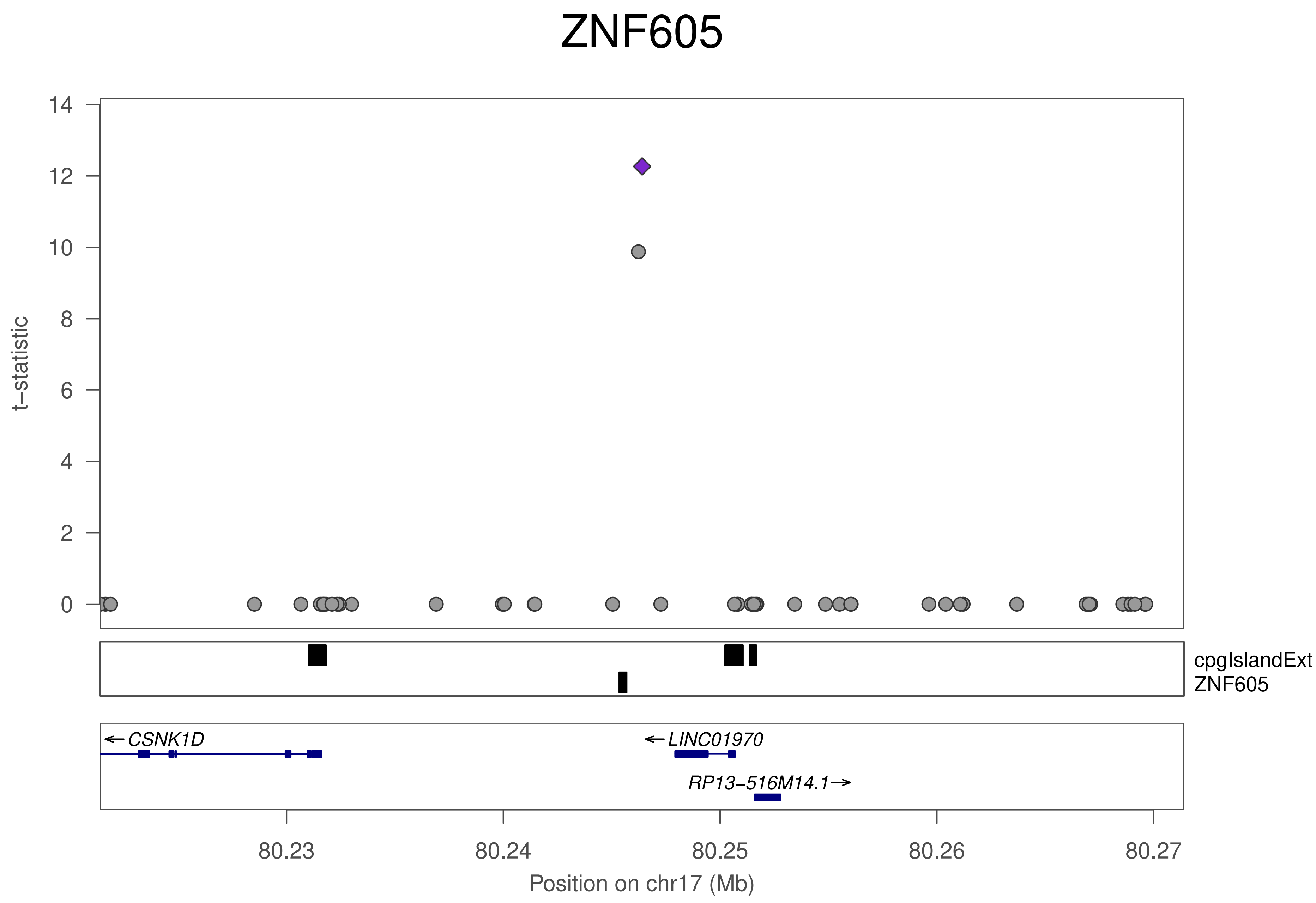

ZNF605
t–statistic
0
2
4
6
8
10
12
14
cpgIslandExt
ZNF605
← CSNK1D
← LINC01970
RP13–516M14.1 →
80.23
80.24
80.25
80.26
80.27
Position on chr17 (Mb)

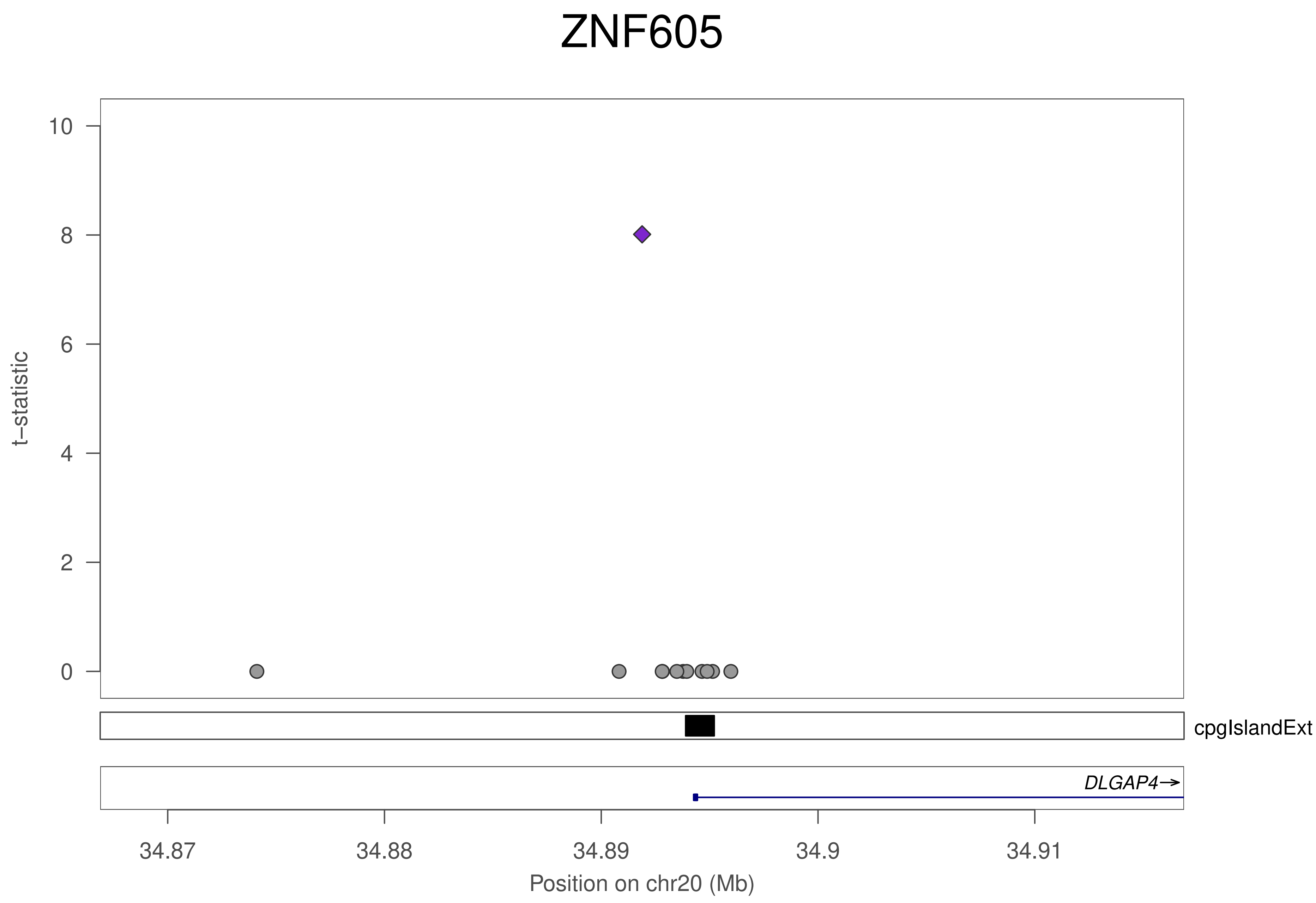
ZNF605
t−statistic
10
8
6
4
2
0
cpgIslandExt
DLGAP4→
34.87
34.88
34.89
34.9
34.91
Position on chr20 (Mb)

### ZNF605

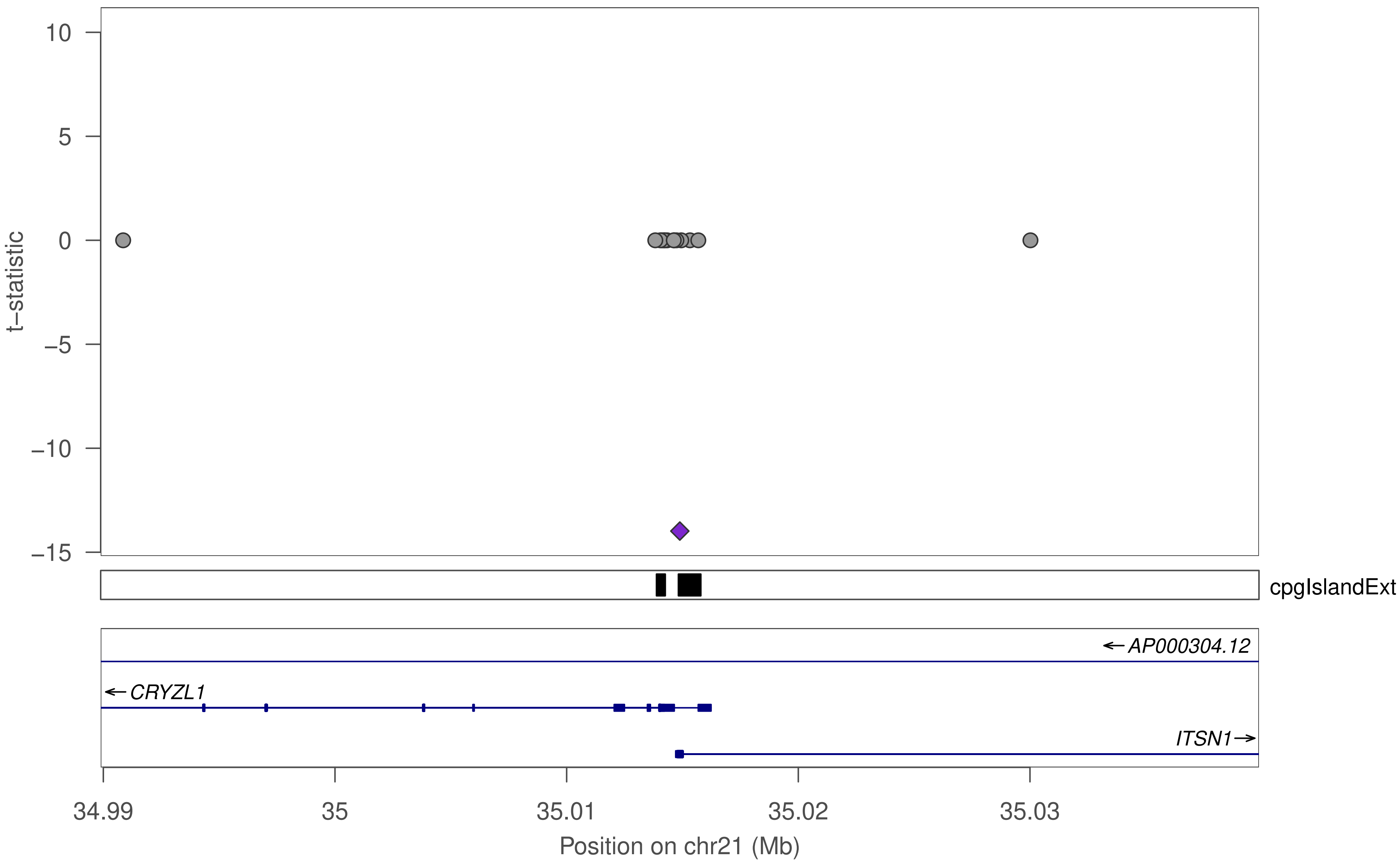
10
5
0
−5
−10
−15
t−statistic
cpgIslandExt
AP000304.12
CRYZL1
ITSN1
34.99
35
35.01
35.02
35.03
Position on chr21 (Mb)

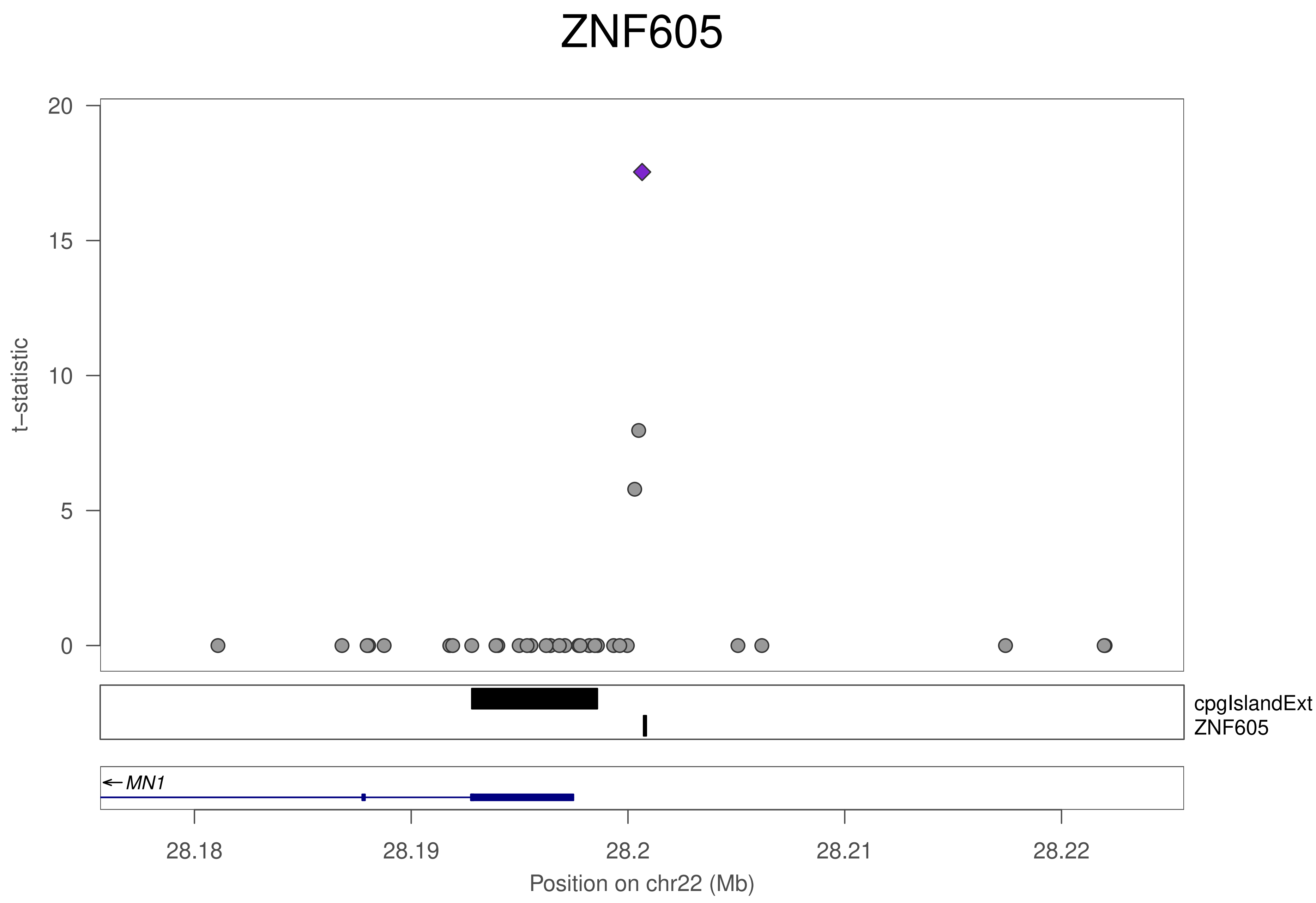

ZNF605
t−statistic
20
15
10
5
0
cpgIslandExt
ZNF605
← MN1
28.18
28.19
28.2
28.21
28.22
Position on chr22 (Mb)

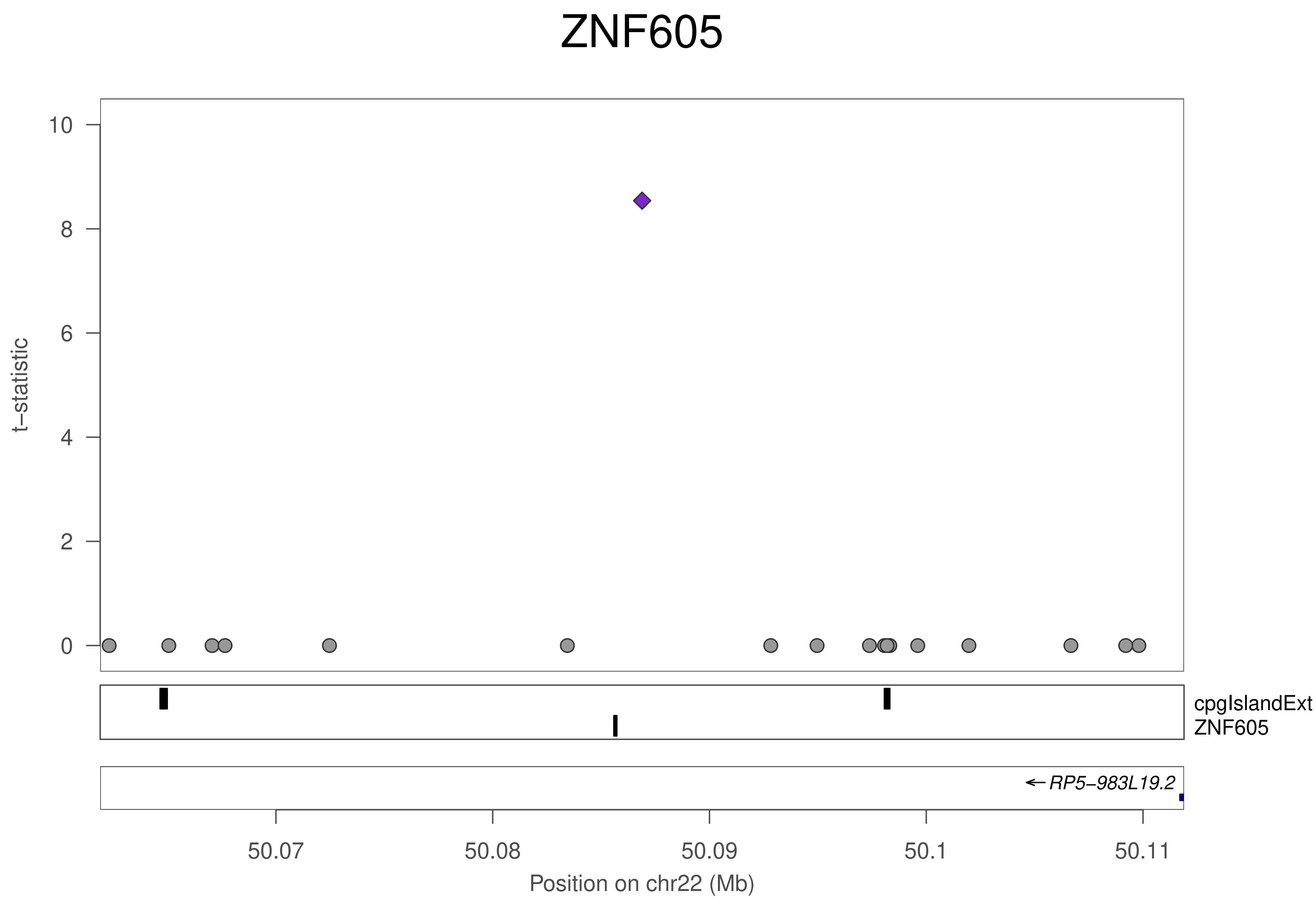
ZNF605
t−statistic
10
8
6
4
2
0
cpgIslandExt
ZNF605
← RP5−983L19.2
50.07
50.08
50.09
50.1
50.11
Position on chr22 (Mb)

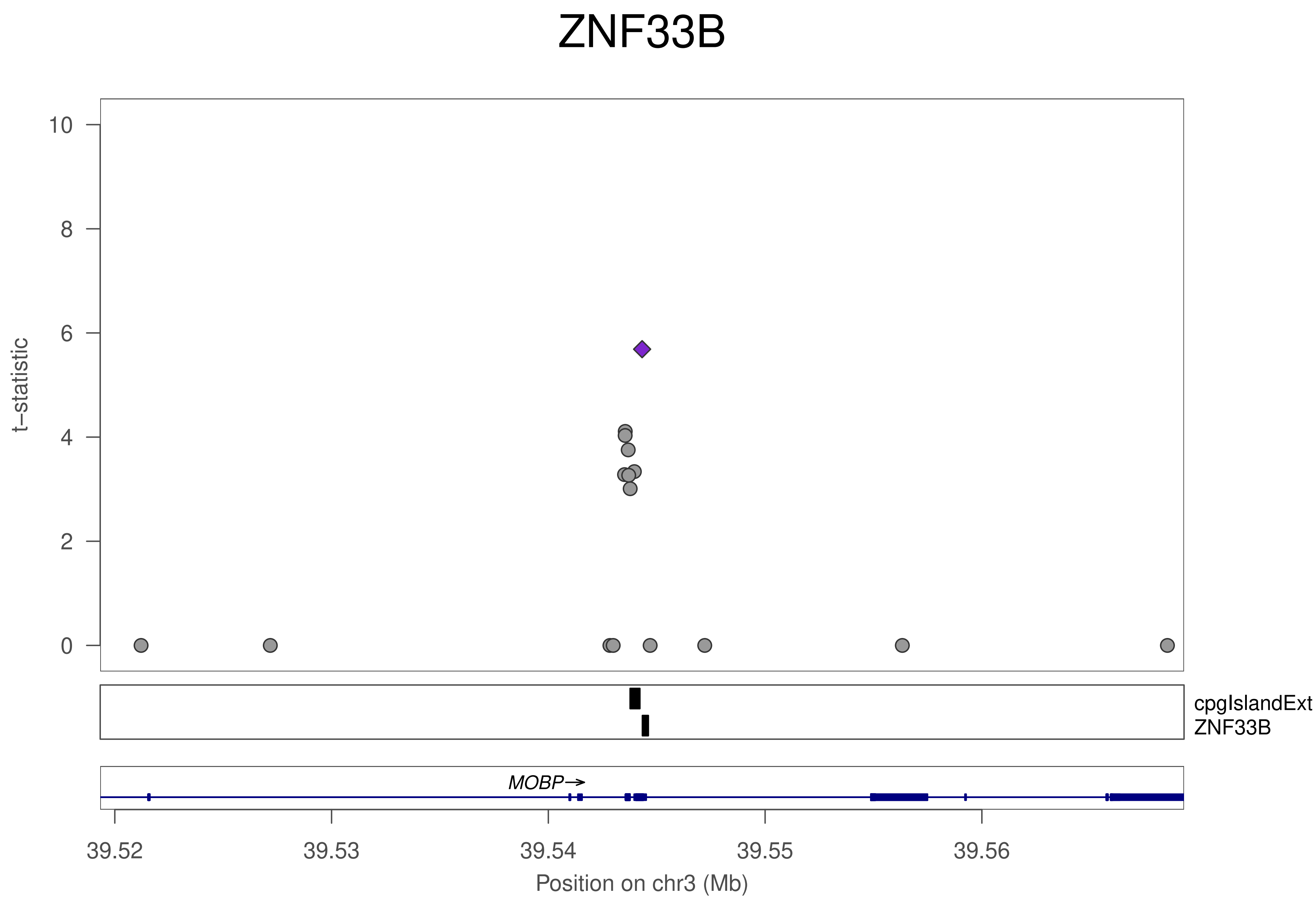
ZNF33B
t−statistic
10
8
6
4
2
0
cpgIslandExt
ZNF33B
MOBP→
39.52
39.53
39.54
39.55
39.56
Position on chr3 (Mb)

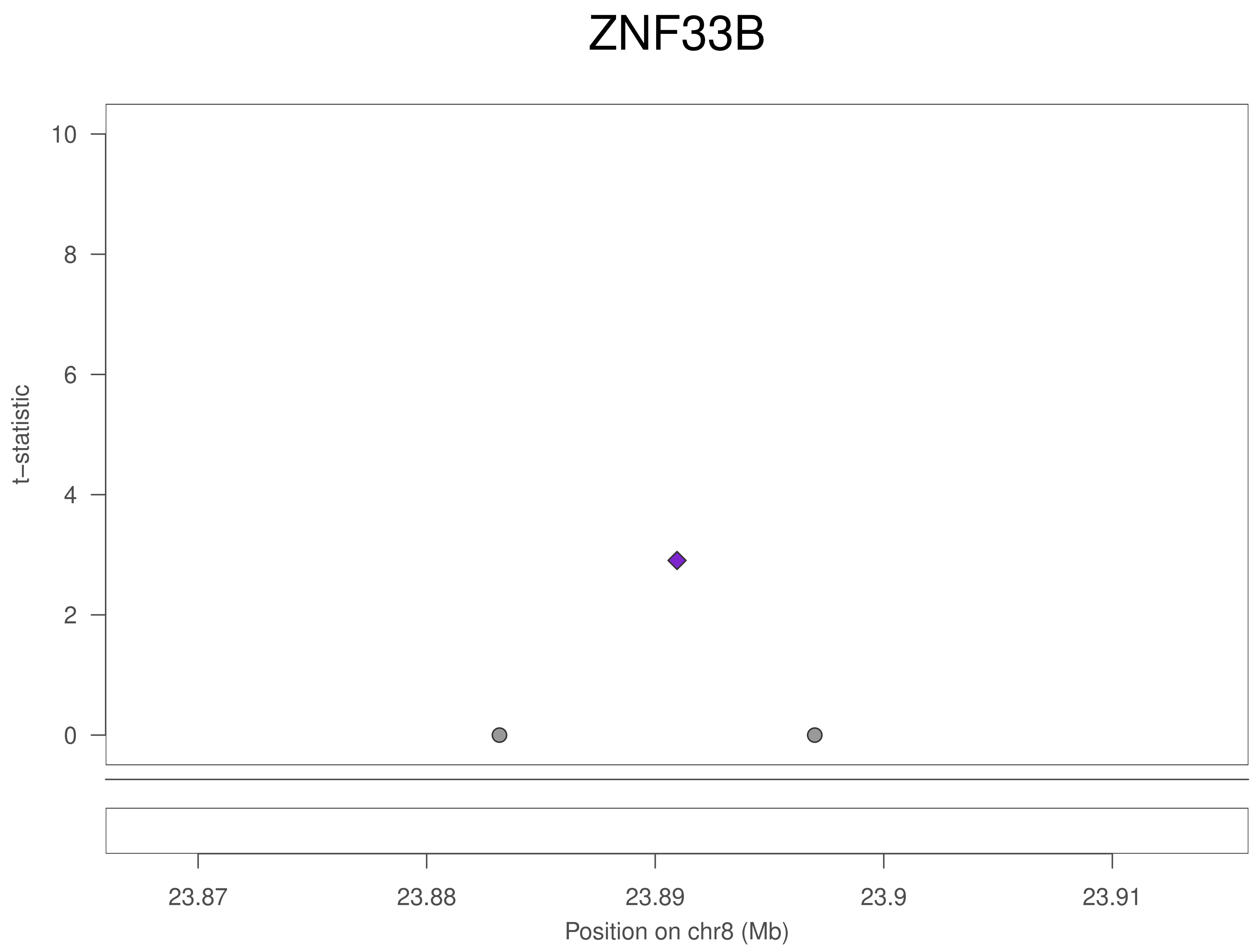
ZNF33B
t–statistic
10
8
6
4
2
0
23.87
23.88
23.89
23.9
23.91
Position on chr8 (Mb)

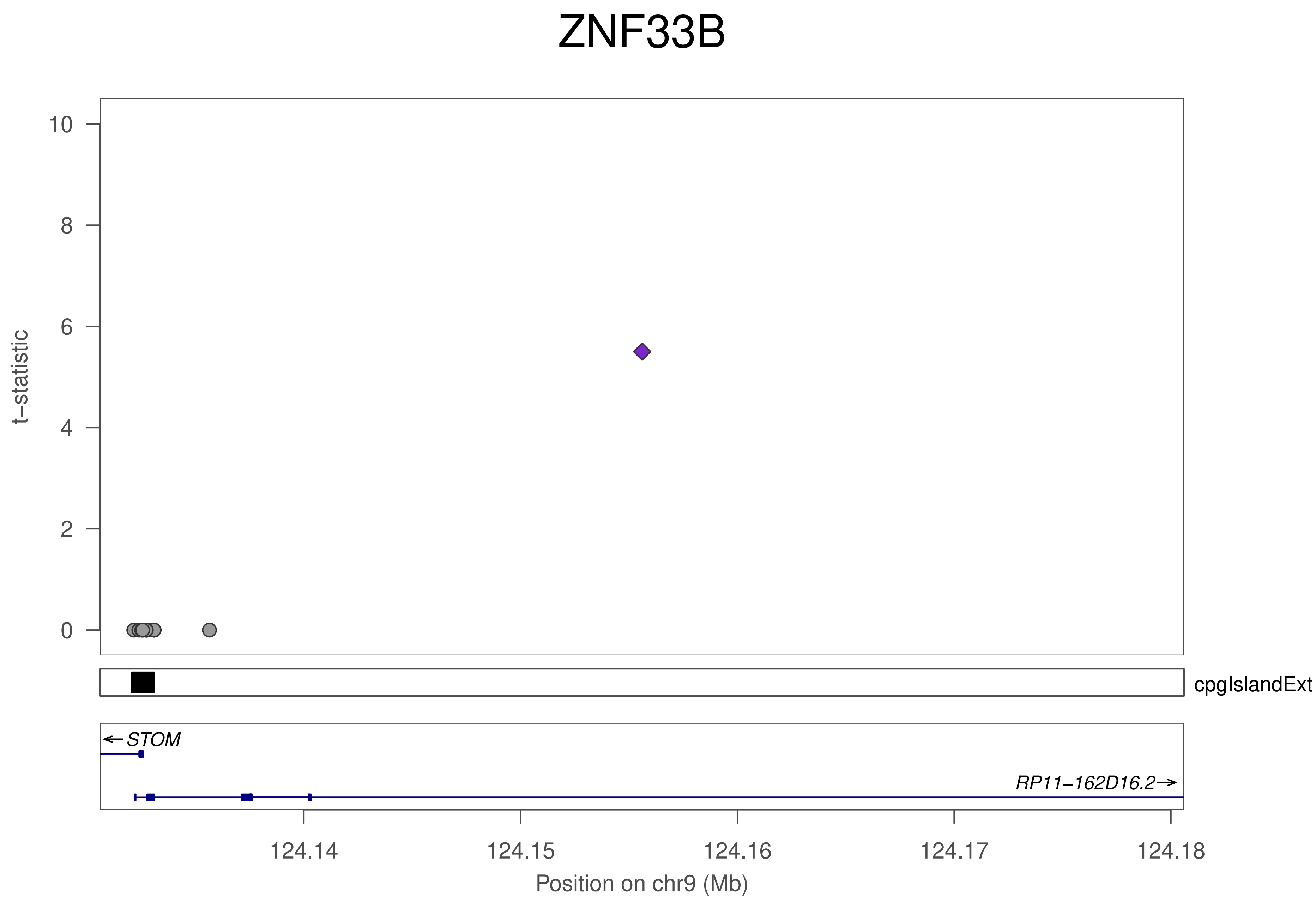
ZNF33B
t−statistic
10
8
6
4
2
0
cpgIslandExt
←STOM
RP11−162D16.2→
124.14
124.15
124.16
124.17
124.18
Position on chr9 (Mb)

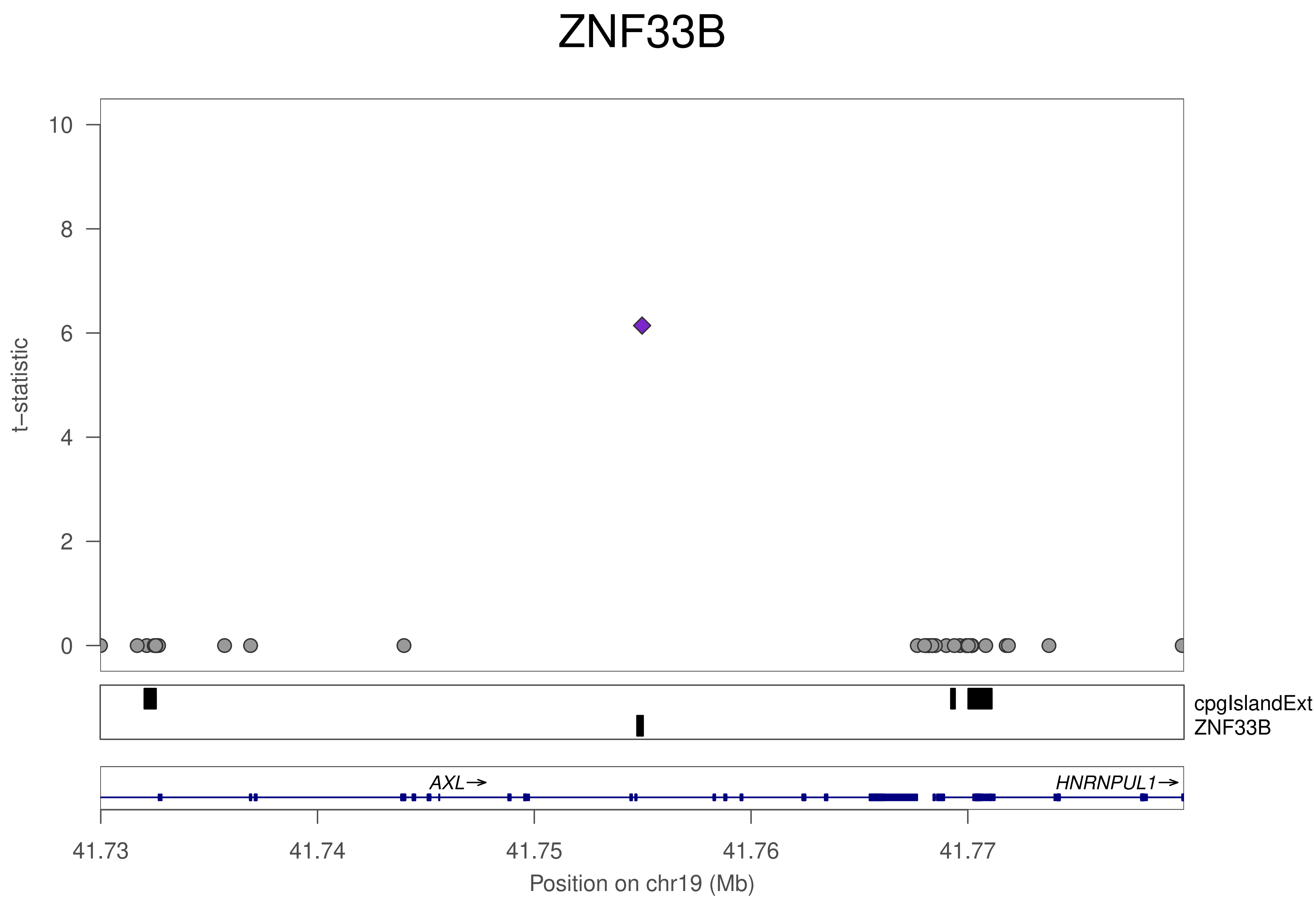
ZNF33B
t−statistic
10
8
6
4
2
0
cpgIslandExt
ZNF33B
AXL→
HNRNPUL1→
41.73
41.74
41.75
41.76
41.77
Position on chr19 (Mb)

### ZNF33B

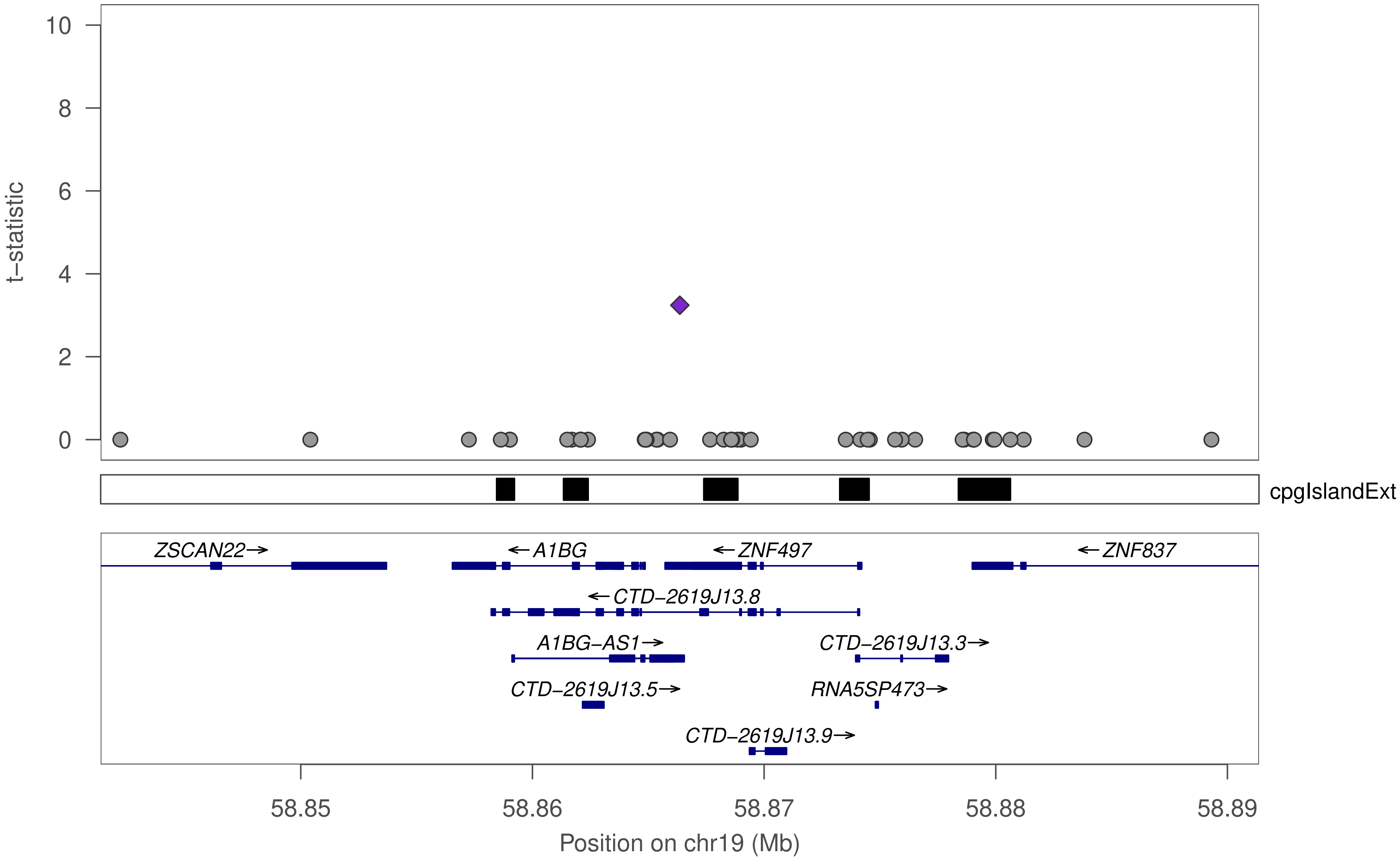

10
8
6
4
2
0
t−statistic
cpgIslandExt
ZSCAN22→
←A1BG
←ZNF497
←ZNF837
←CTD−2619J13.8
A1BG−AS1→
CTD−2619J13.3→
CTD−2619J13.5→
RNA5SP473→
CTD−2619J13.9→
58.85
58.86
58.87
58.88
58.89
Position on chr19 (Mb)

### ZNF431

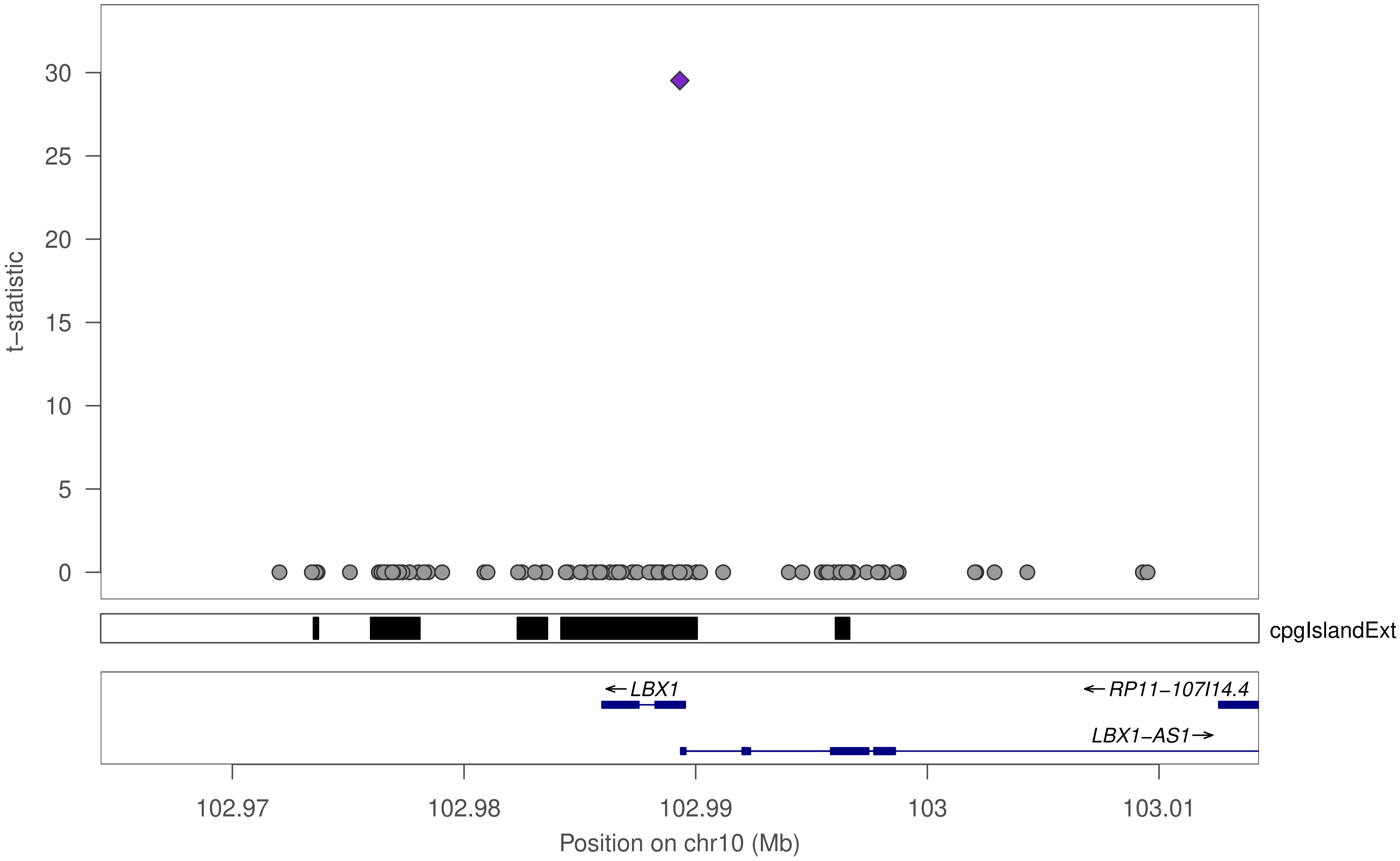

t−statistic
30
25
20
15
10
5
0
cpgIslandExt
←LBX1
←RP11−107I14.4
LBX1−AS1→
102.97
102.98
102.99
103
103.01
Position on chr10 (Mb)

### ZNF418

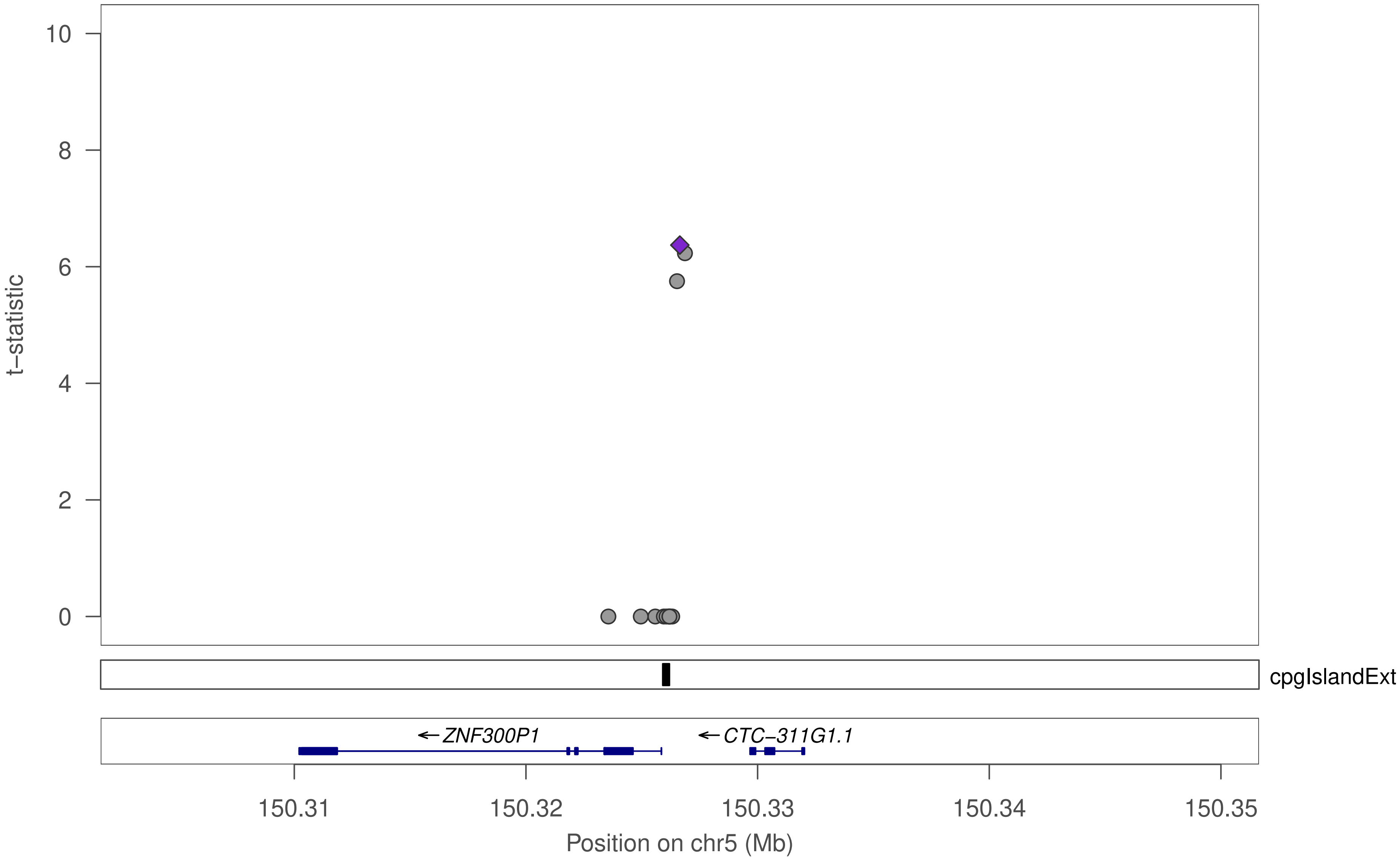

10
8
6
4
2
0
t−statistic
cpgIslandExt
←ZNF300P1
←CTC−311G1.1
150.31
150.32
150.33
150.34
150.35
Position on chr5 (Mb)

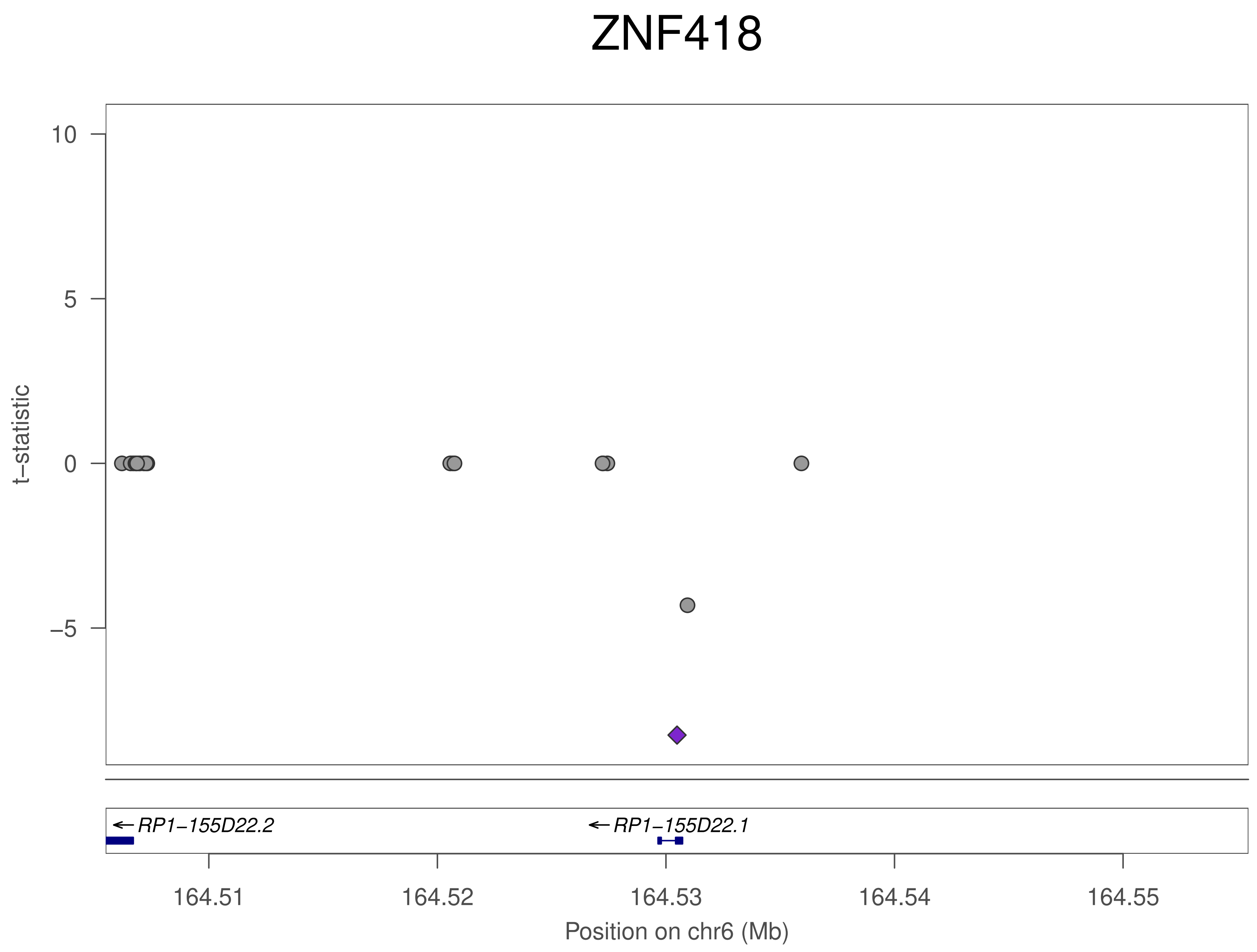
ZNF418
10
5
0
−5
t−statistic
RP1−155D22.2
RP1−155D22.1
164.51
164.52
164.53
164.54
164.55
Position on chr6 (Mb)

### ZNF418

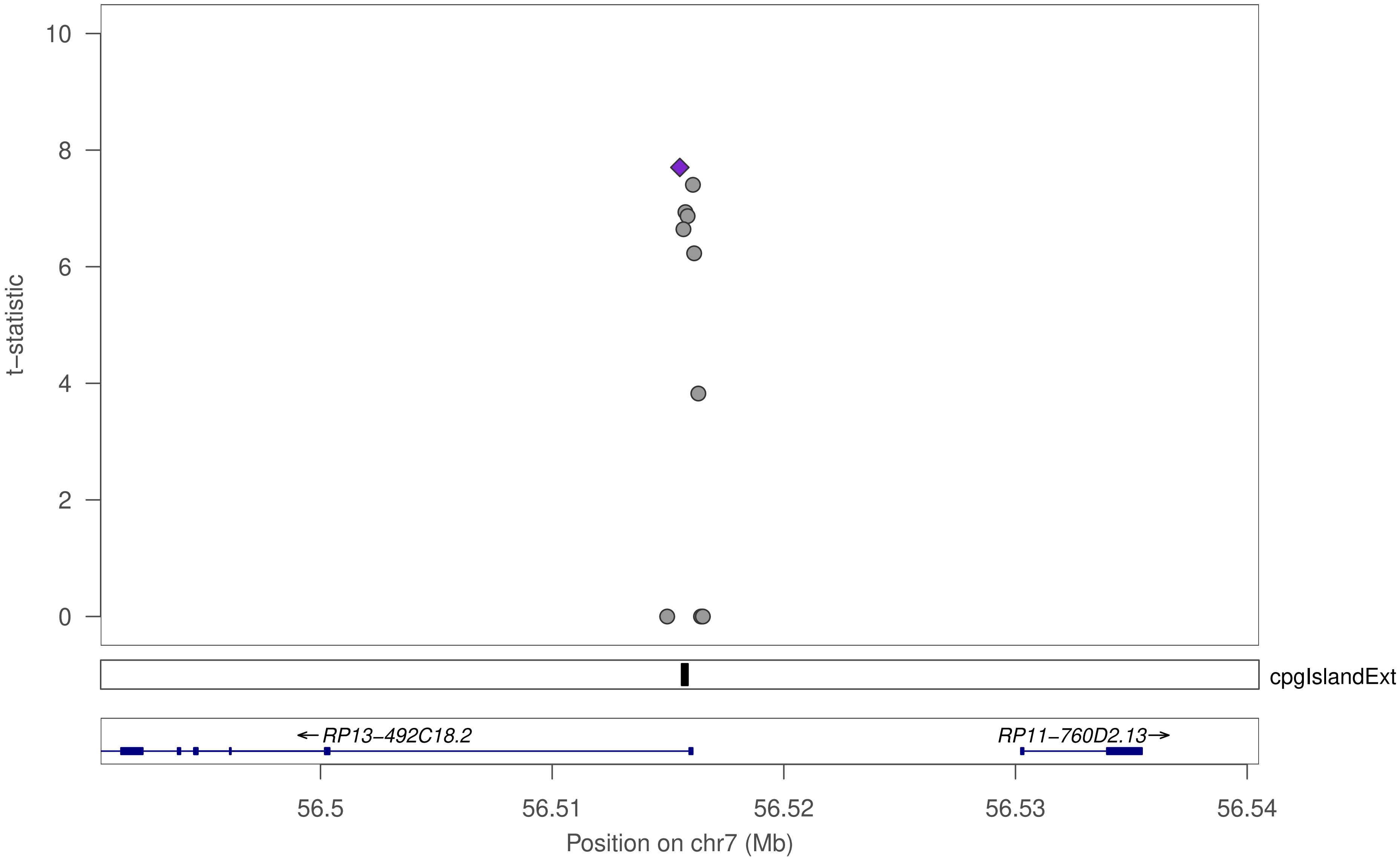

t−statistic
10
8
6
4
2
0
cpgIslandExt
←RP13−492C18.2
RP11−760D2.13→
56.5
56.51
56.52
56.53
56.54
Position on chr7 (Mb)

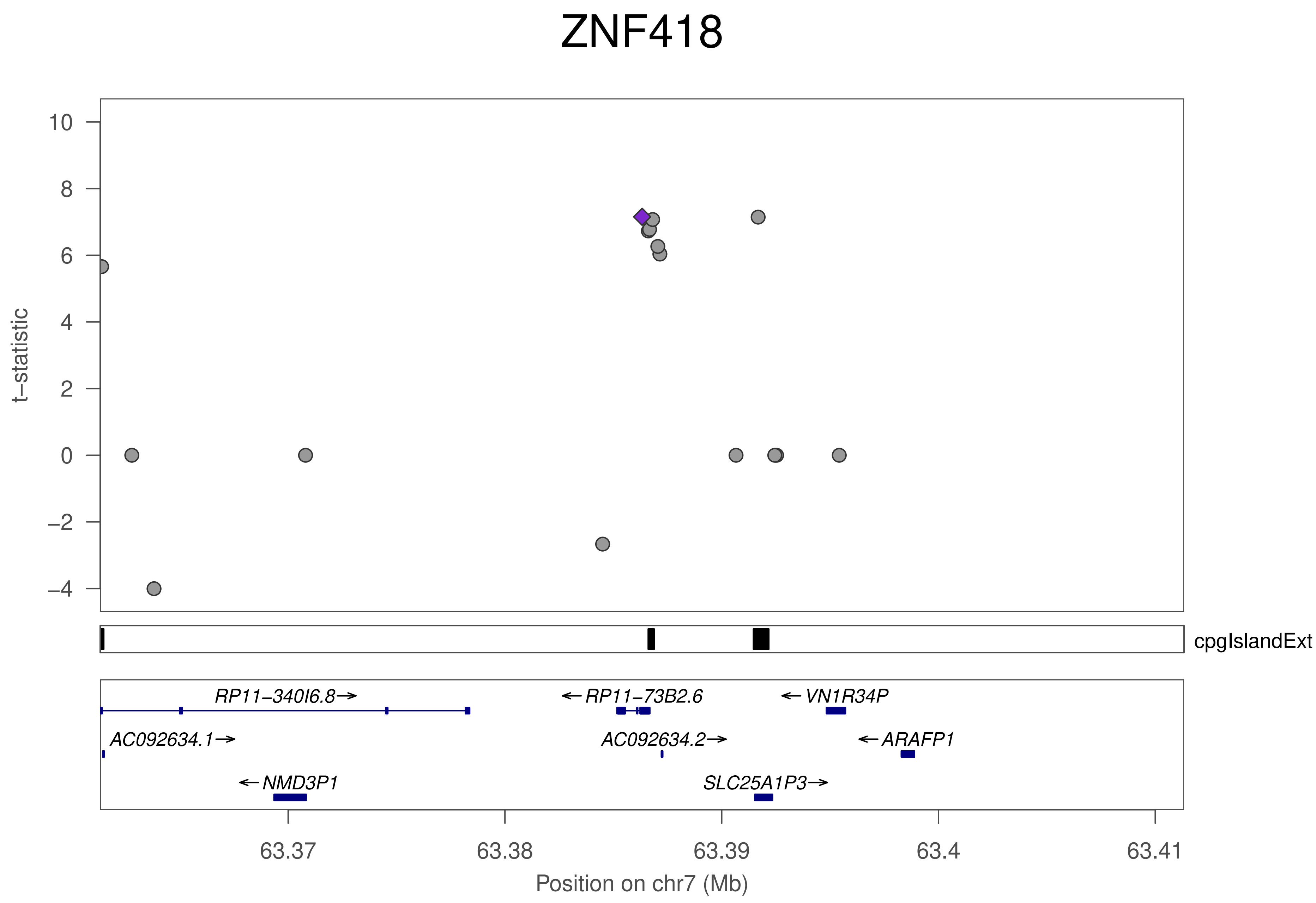

ZNF418
t−statistic
10
8
6
4
2
0
−2
−4
cpgIslandExt
RP11−340I6.8→
←RP11−73B2.6
←VN1R34P
AC092634.1→
AC092634.2→
←ARAFP1
←NMD3P1
SLC25A1P3→
63.37
63.38
63.39
63.4
63.41
Position on chr7 (Mb)

### ZNF418

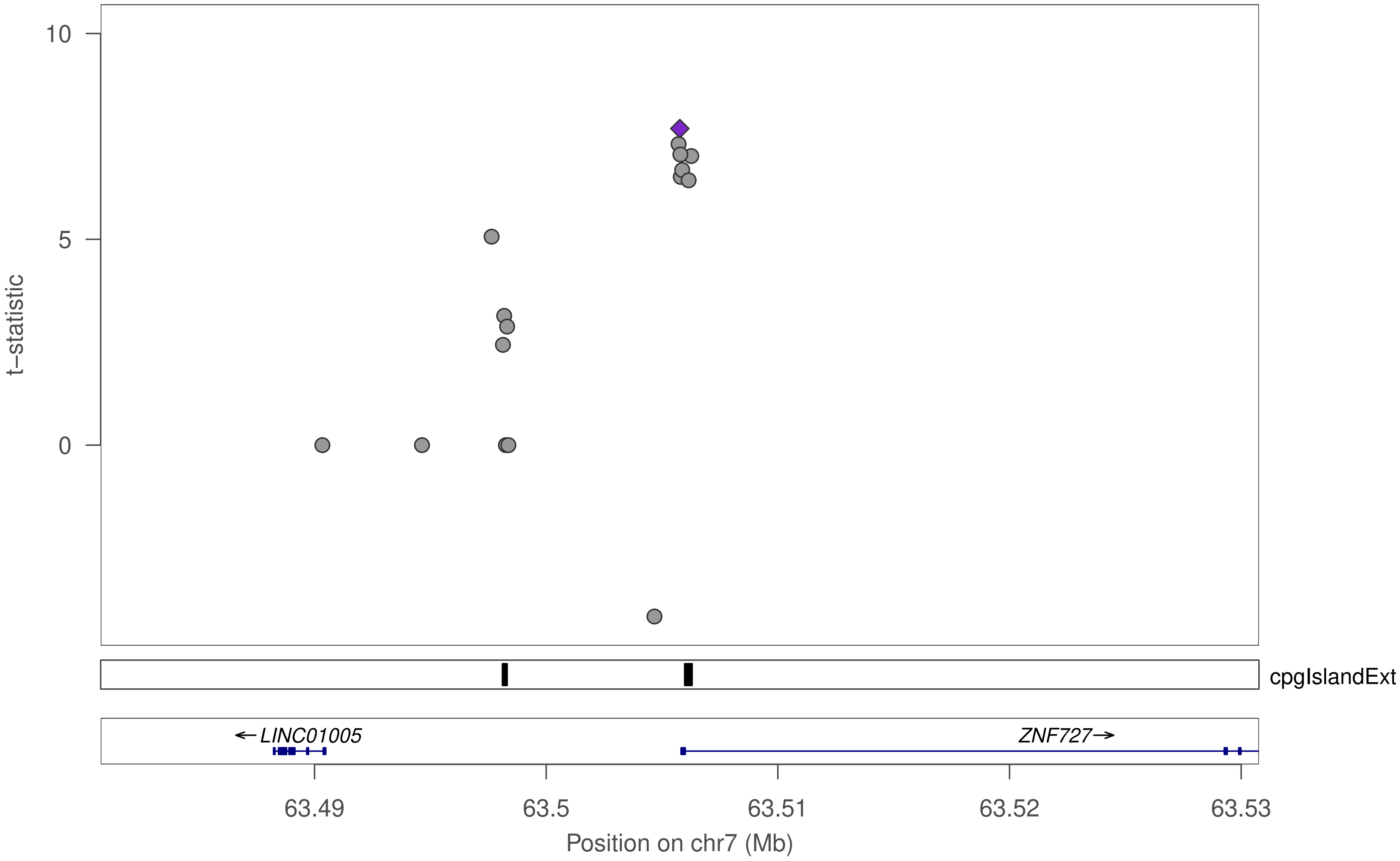

10
5
0
t−statistic
cpgIslandExt
←LINC01005
ZNF727→
63.49
63.5
63.51
63.52
63.53
Position on chr7 (Mb)

### ZNF418

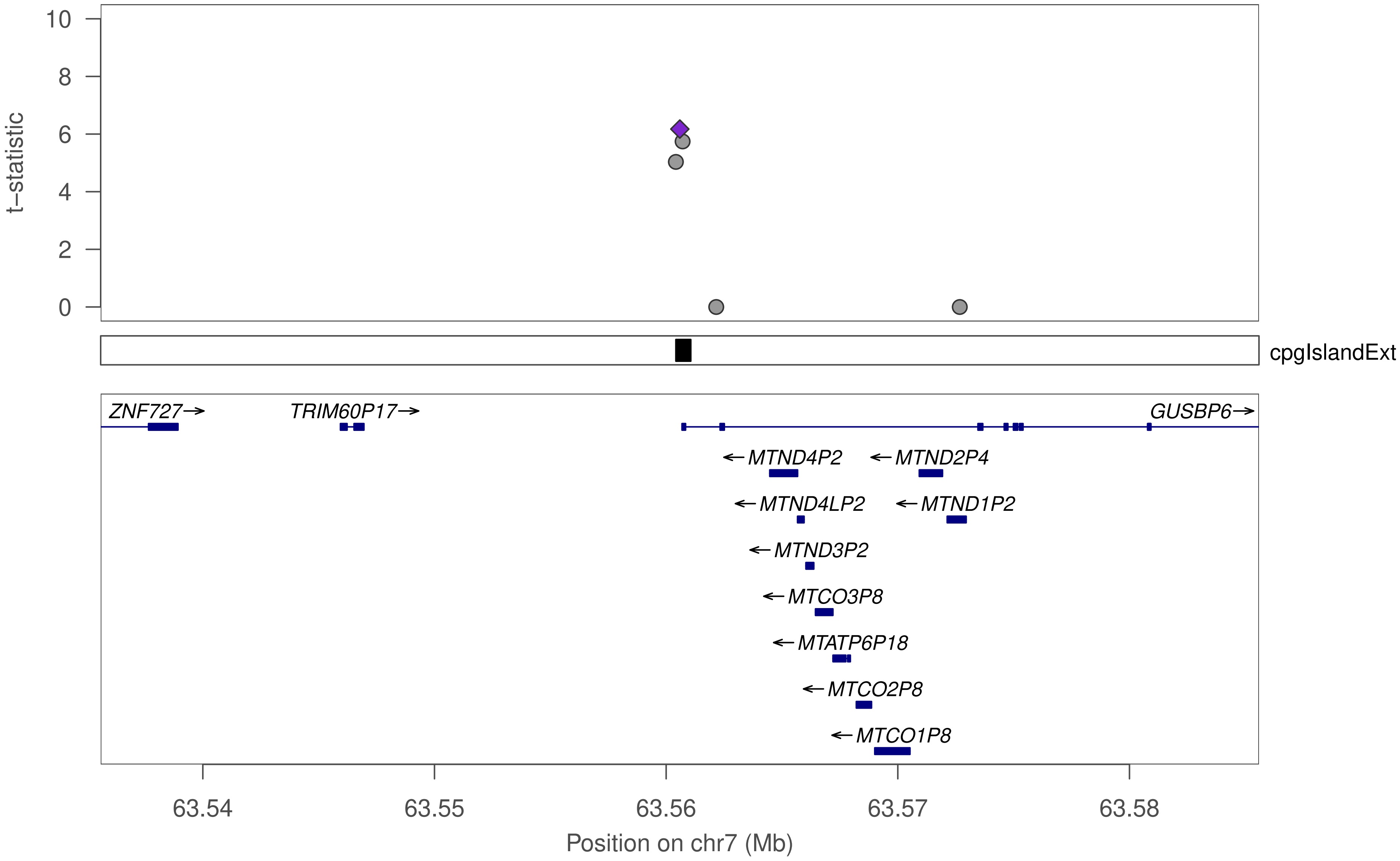

t–statistic
10
8
6
4
2
0
cpgIslandExt
ZNF727→
TRIM60P17→
GUSBP6→
←MTND4P2
←MTND2P4
←MTND4LP2
←MTND1P2
←MTND3P2
←MTCO3P8
←MTATP6P18
←MTCO2P8
←MTCO1P8
63.54
63.55
63.56
63.57
63.58
Position on chr7 (Mb)

### ZNF418

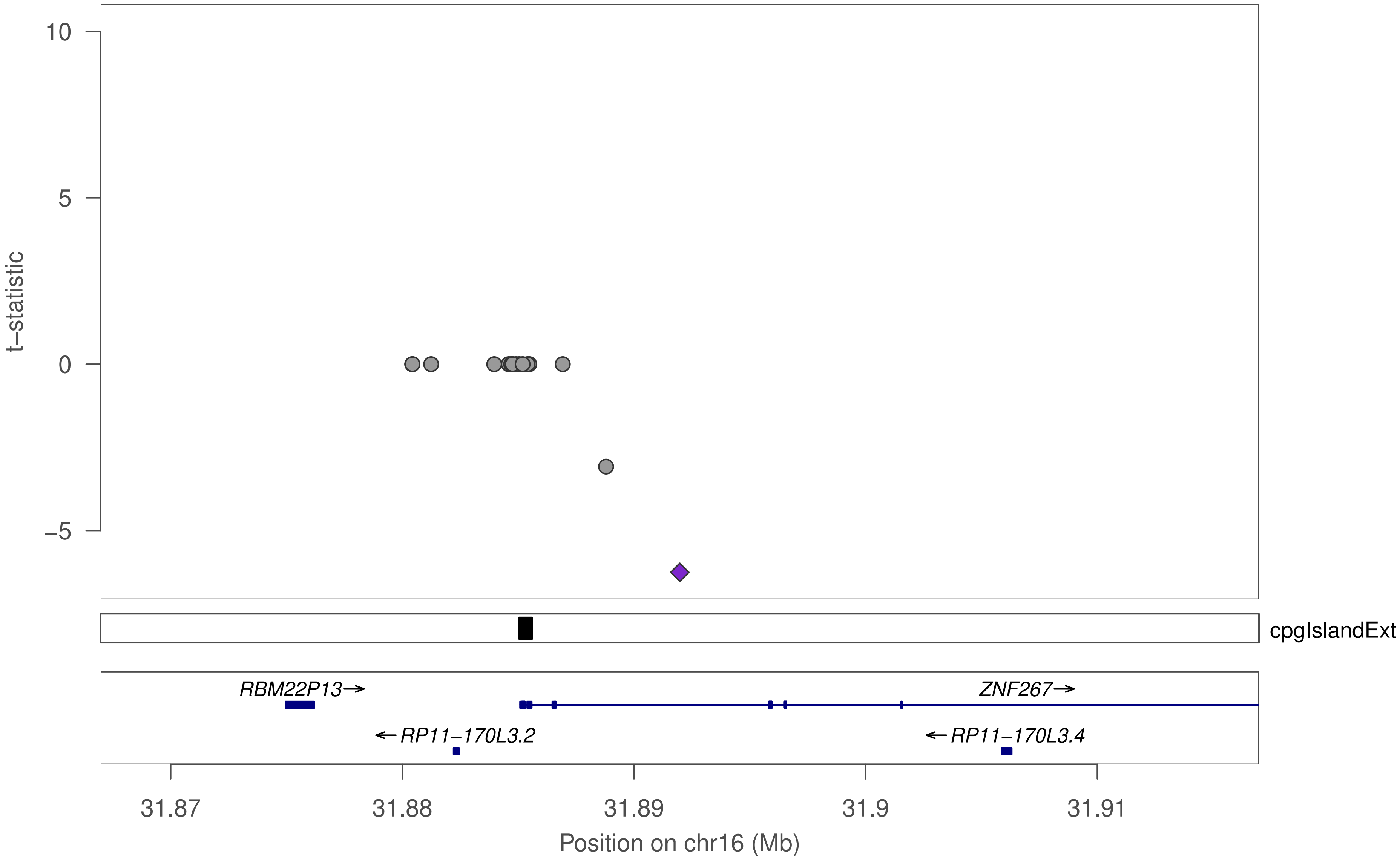

10
5
0
−5
t−statistic
cpgIslandExt
RBM22P13→
ZNF267→
←RP11−170L3.2
←RP11−170L3.4
31.87
31.88
31.89
31.9
31.91
Position on chr16 (Mb)

### ZNF418

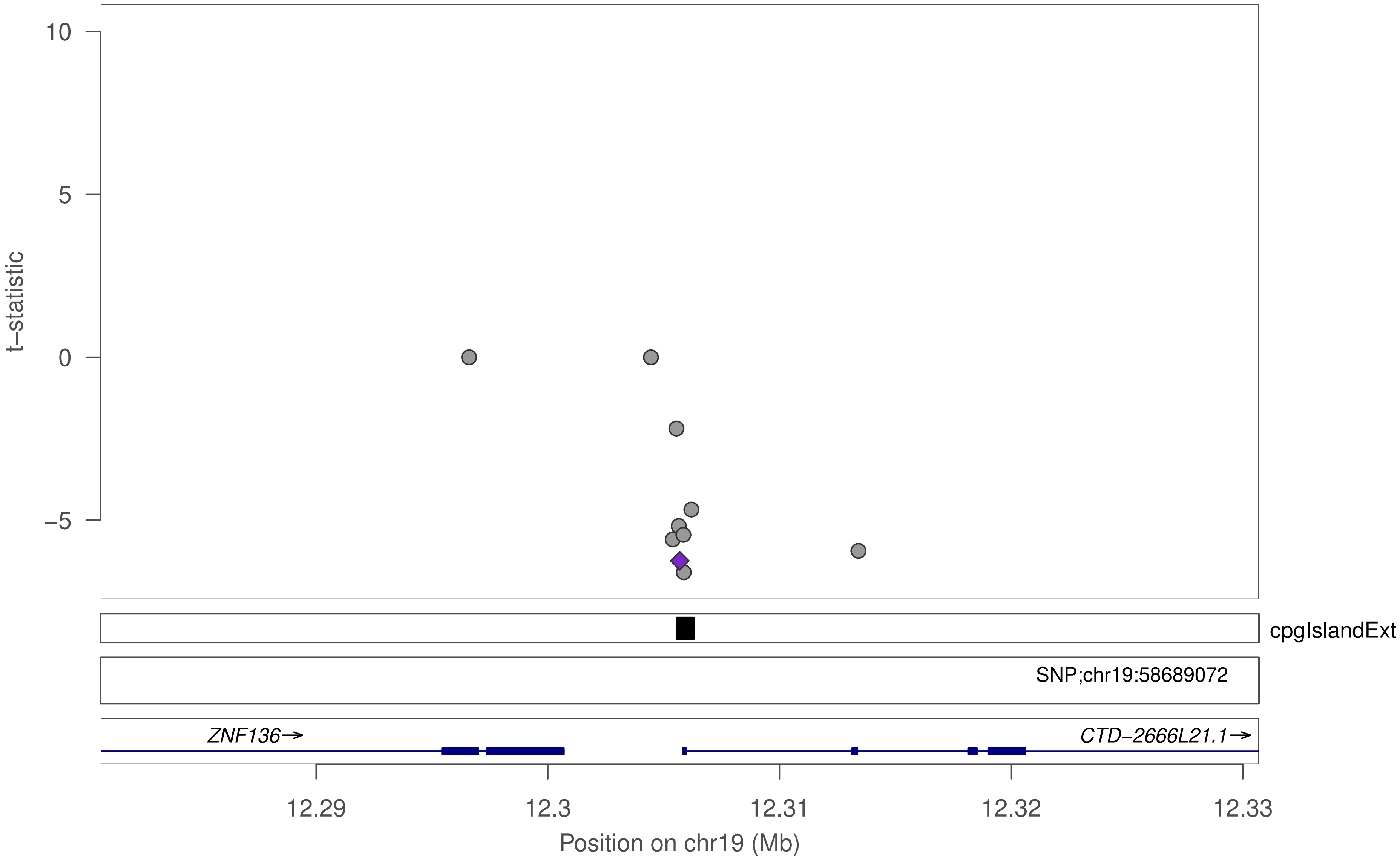

10
5
0
−5
t−statistic
cpgIslandExt
SNP;chr19:58689072
ZNF136→
CTD−2666L21.1→
12.29
12.3
12.31
12.32
12.33
Position on chr19 (Mb)

### ZNF418

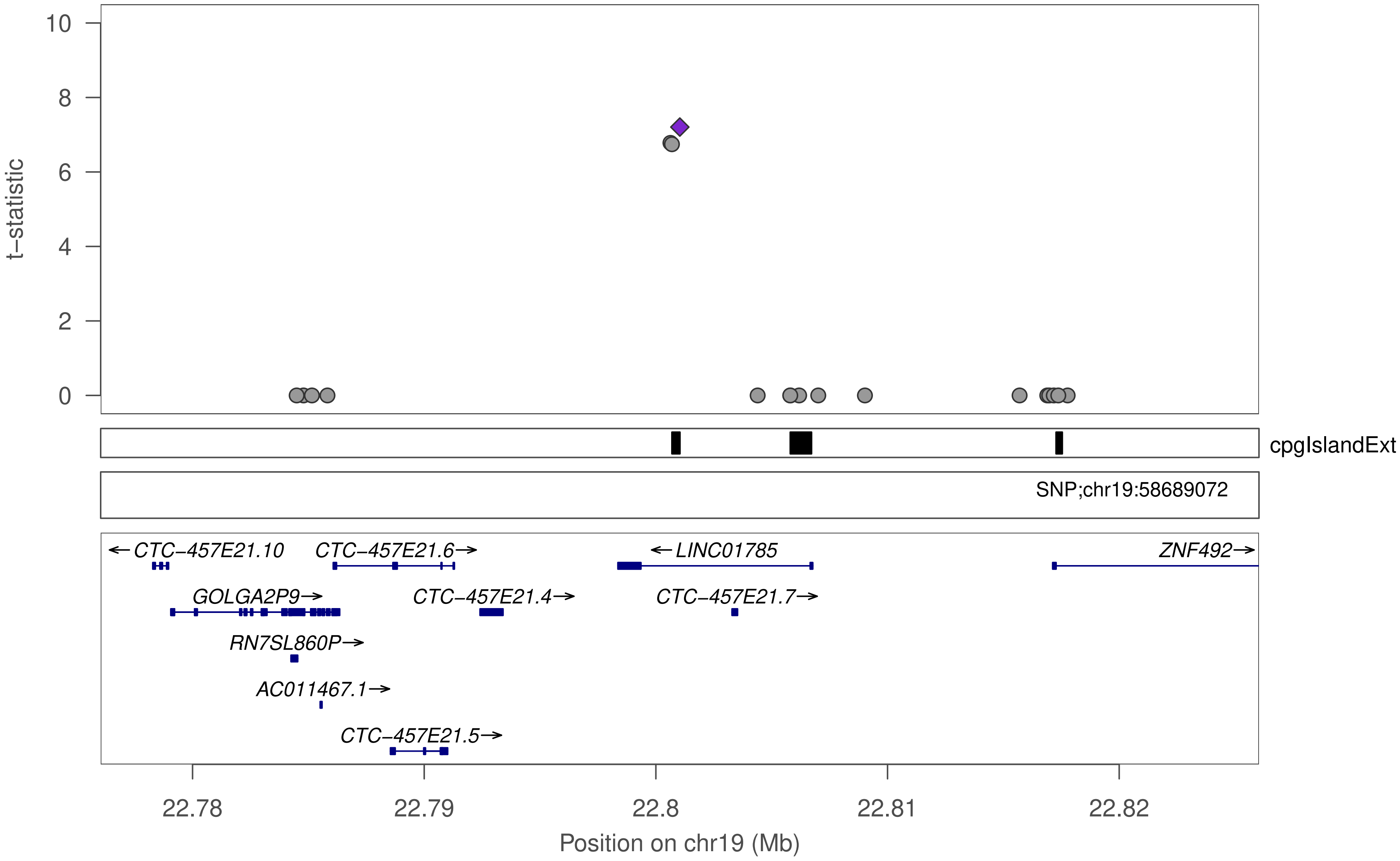
t–statistic
10
8
6
4
2
0
cpgIslandExt
SNP;chr19:58689072
←CTC–457E21.10
CTC–457E21.6→
←LINC01785
ZNF492→
GOLGA2P9→
CTC–457E21.4→
CTC–457E21.7→
RN7SL860P→
AC011467.1→
CTC–457E21.5→
22.78
22.79
22.8
22.81
22.82
Position on chr19 (Mb)

### ZNF418

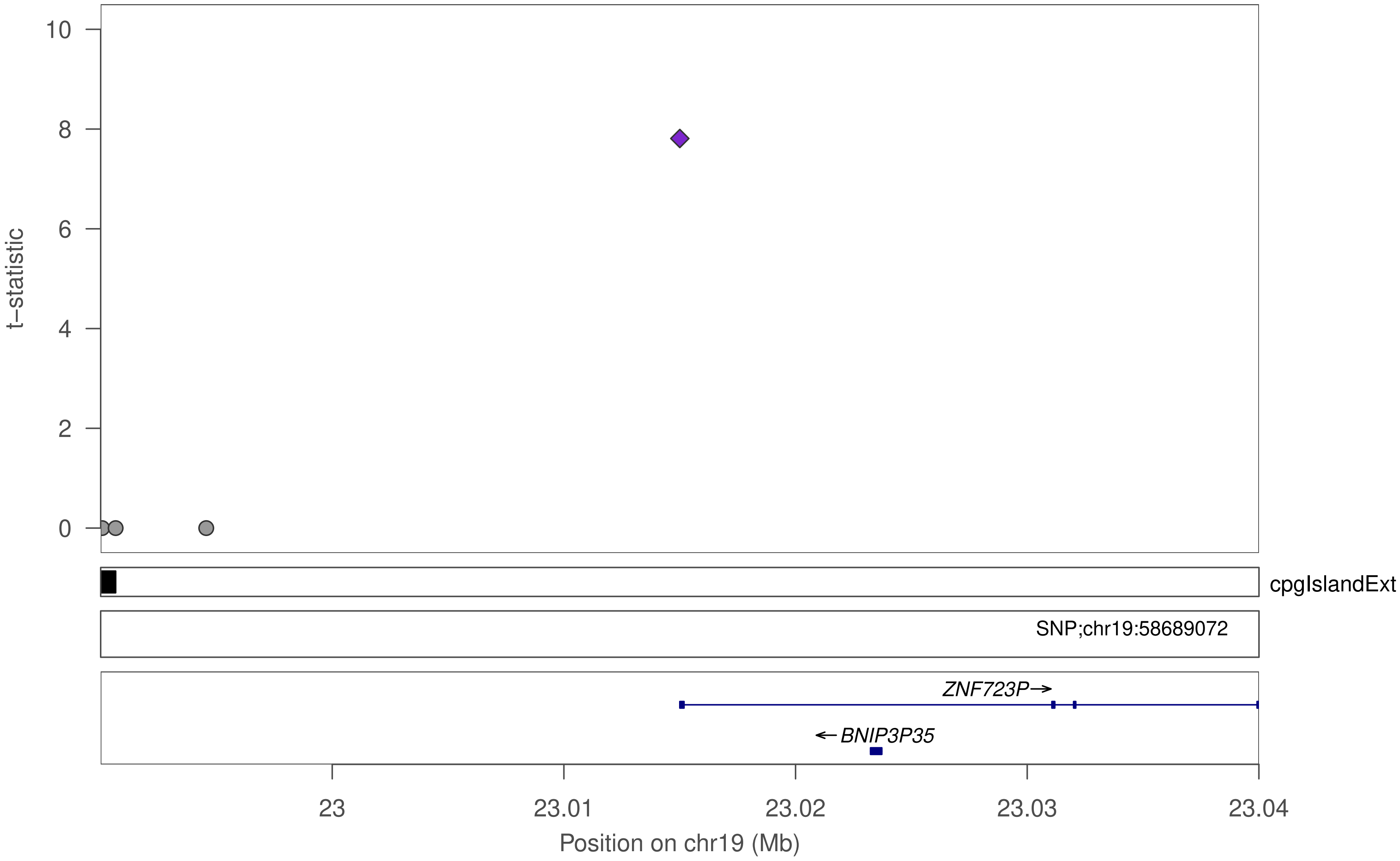

10
8
6
4
2
0
t−statistic
cpgIslandExt
SNP;chr19:58689072
ZNF723P→
←BNIP3P35
23
23.01
23.02
23.03
23.04
Position on chr19 (Mb)

### ZNF418

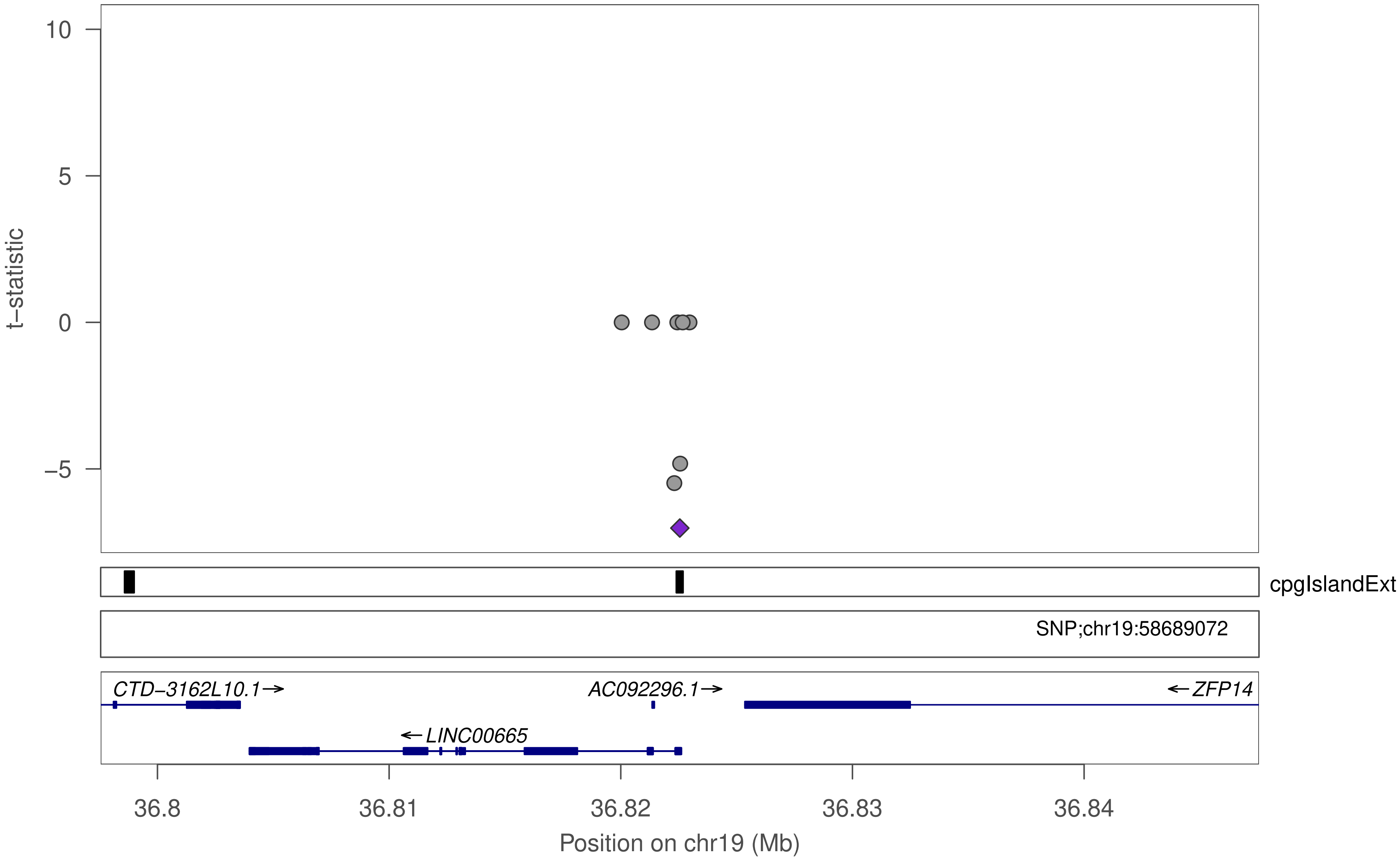

10
5
0
−5
t−statistic
cpgIslandExt
SNP;chr19:58689072
CTD−3162L10.1→
AC092296.1→
←ZFP14
←LINC00665
36.8
36.81
36.82
36.83
36.84
Position on chr19 (Mb)

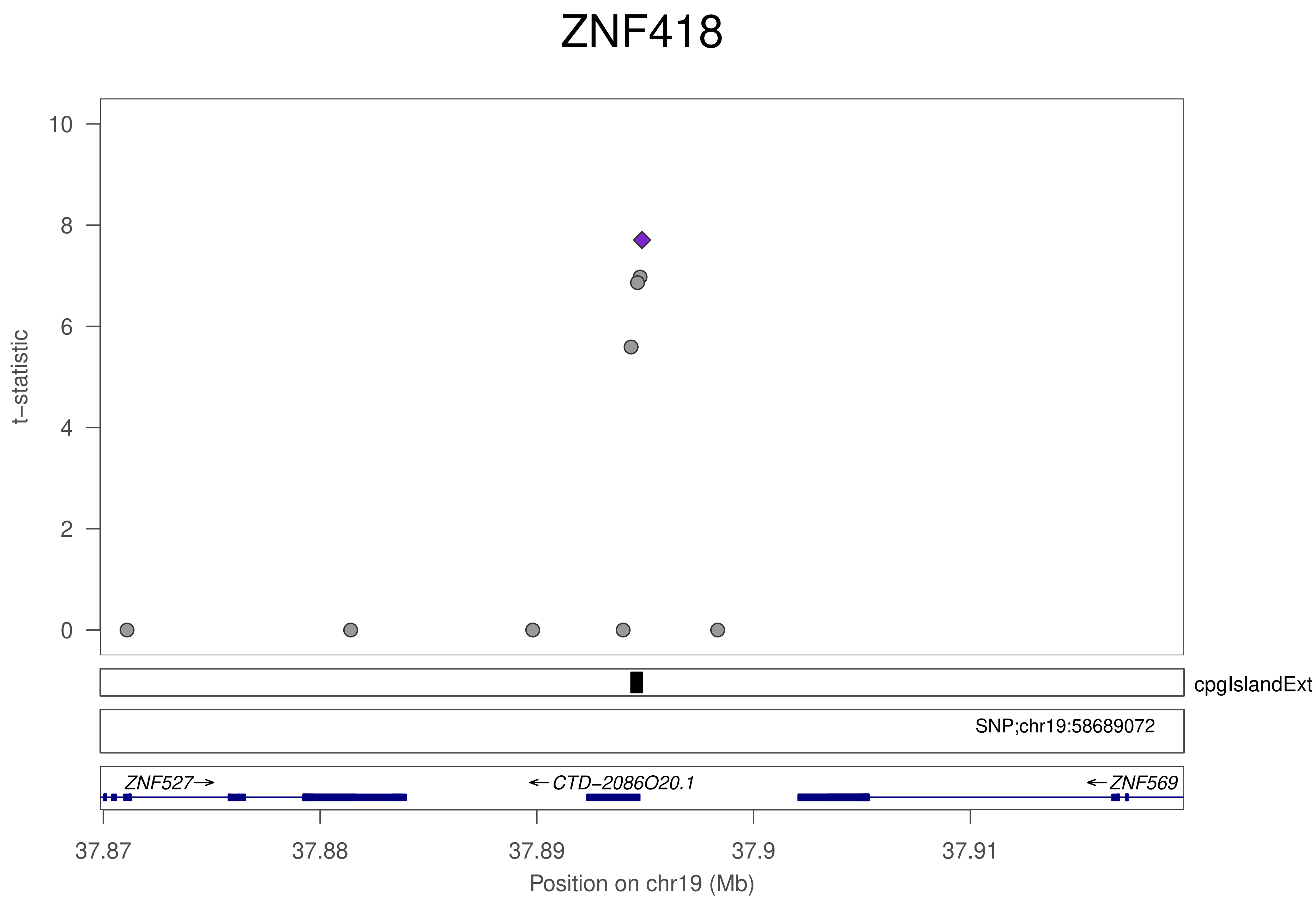
ZNF418
t−statistic
10
8
6
4
2
0
cpgIslandExt
SNP;chr19:58689072
ZNF527→
←CTD−2086O20.1
←ZNF569
37.87
37.88
37.89
37.9
37.91
Position on chr19 (Mb)

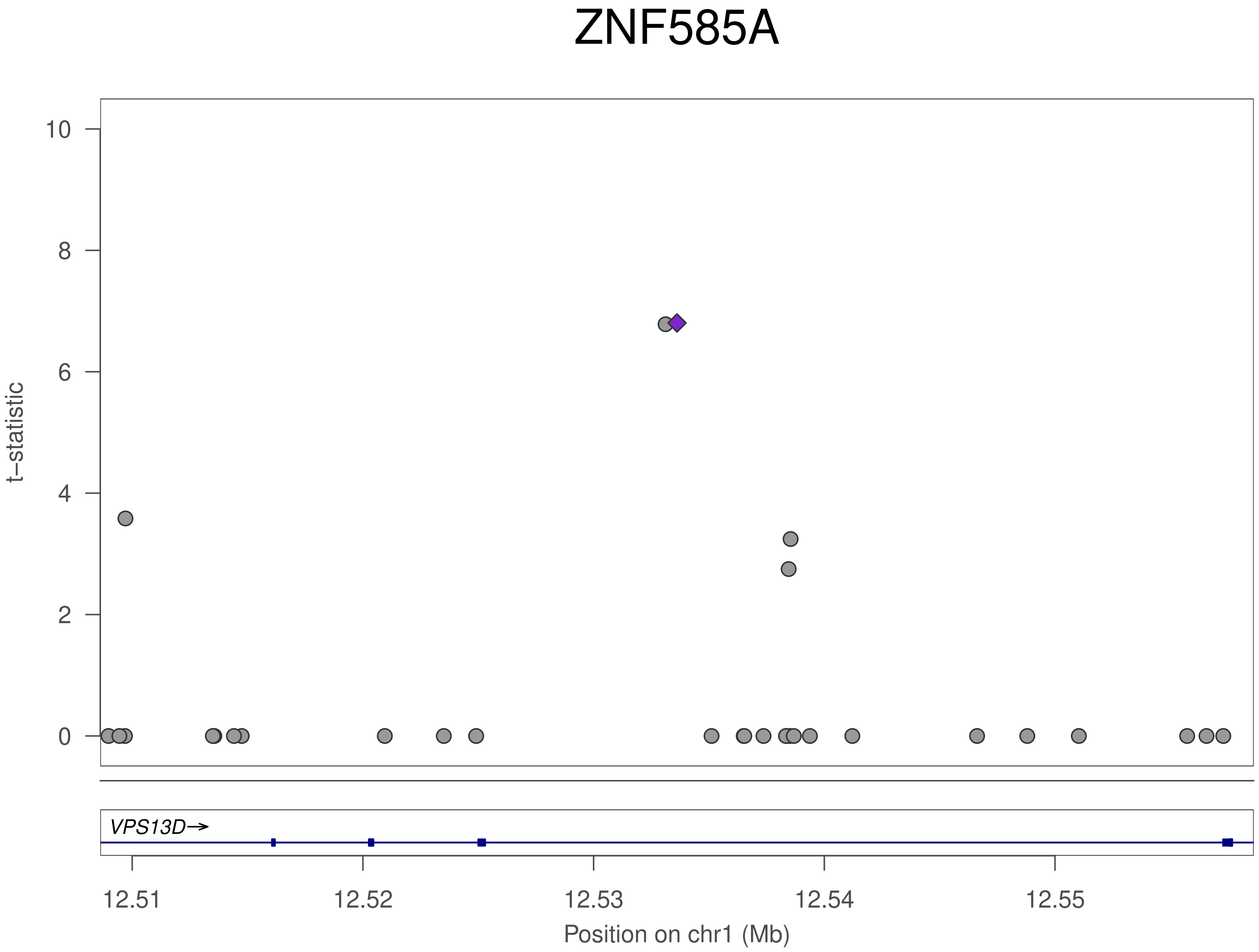
ZNF585A
t–statistic
10
8
6
4
2
0
VPS13D→
12.51
12.52
12.53
12.54
12.55
Position on chr1 (Mb)

### ZNF585A

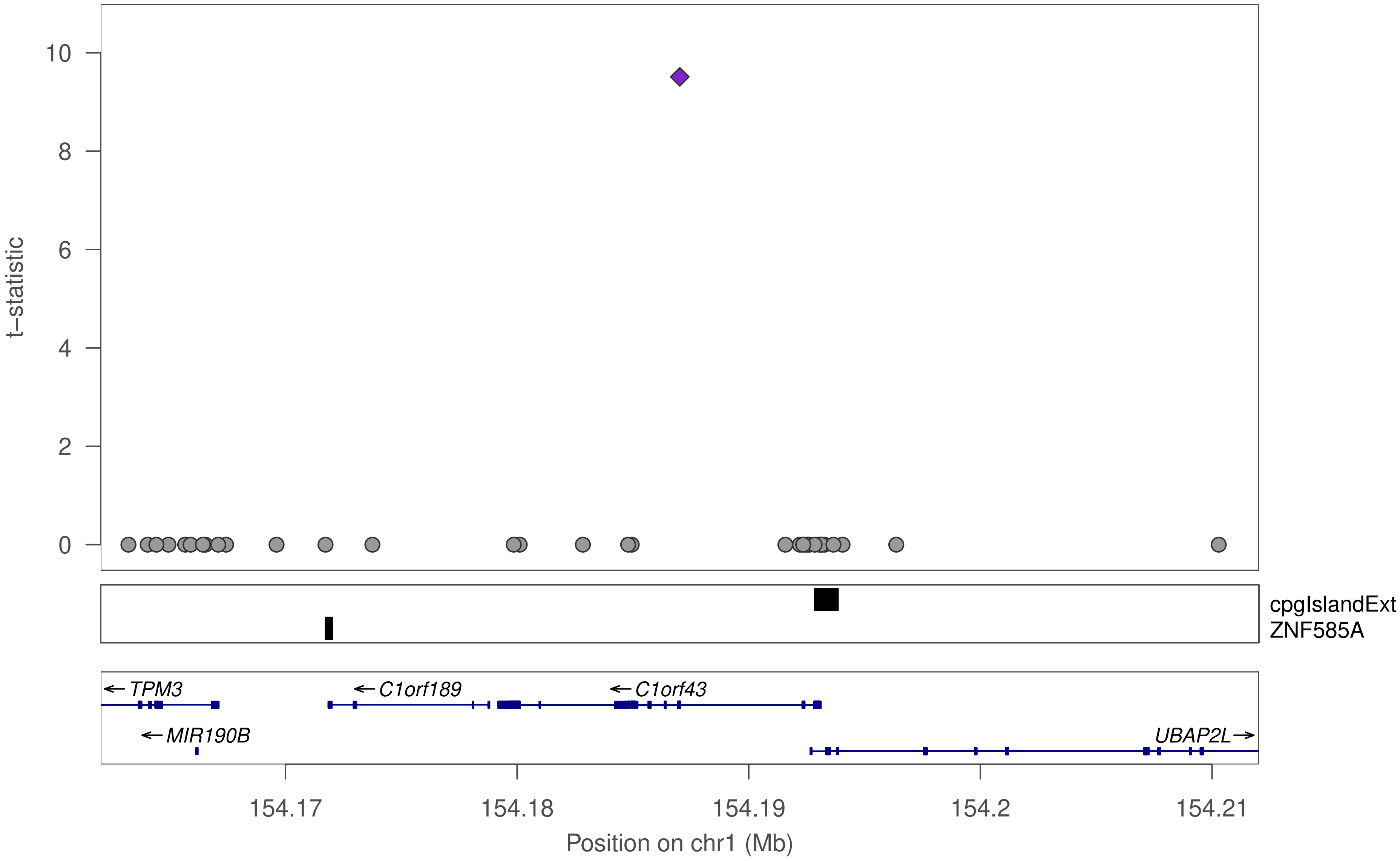

t–statistic
10
8
6
4
2
0
cpgIslandExt
ZNF585A
← TPM3
← C1orf189
← C1orf43
← MIR190B
UBAP2L →
154.17
154.18
154.19
154.2
154.21
Position on chr1 (Mb)

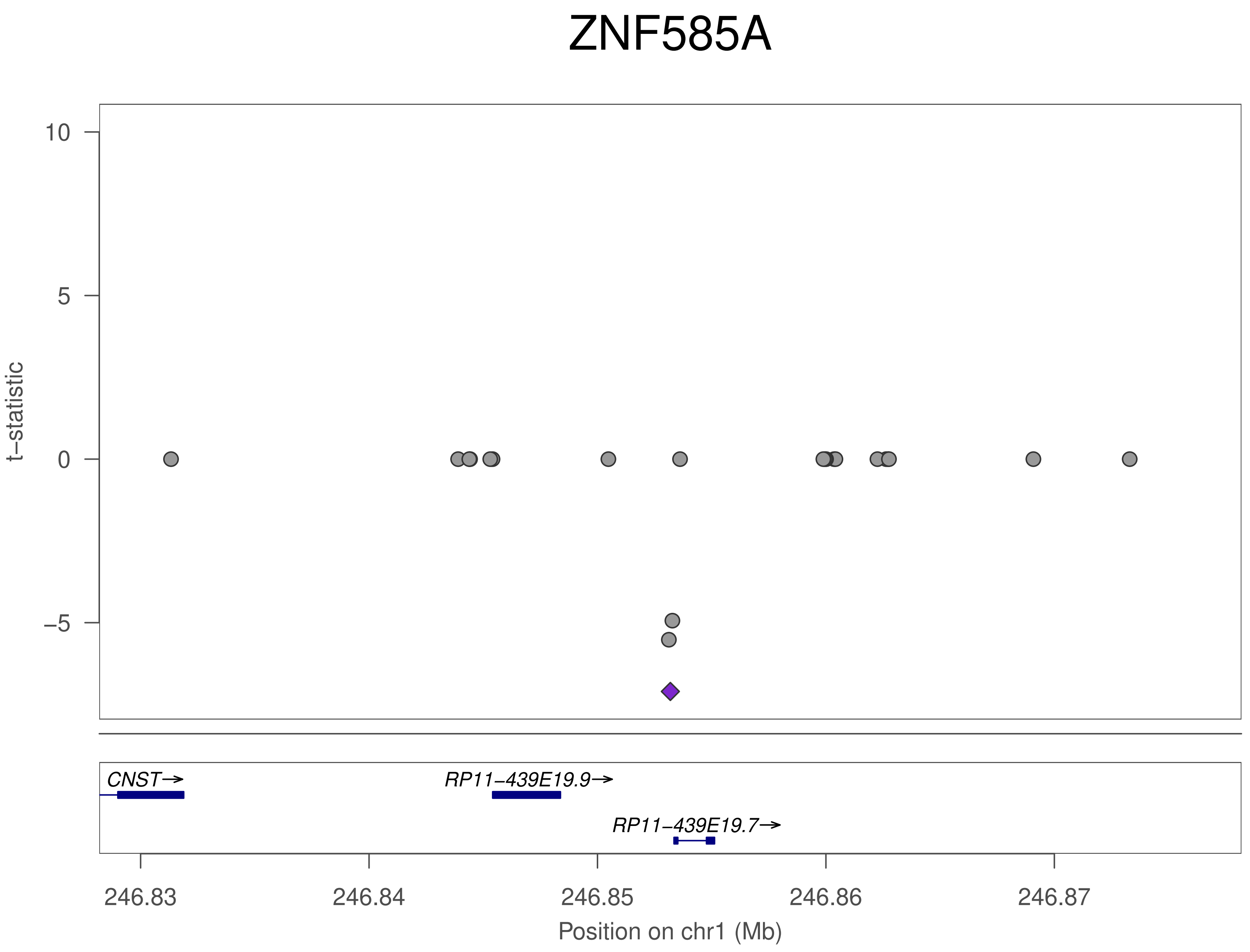

ZNF585A
10
5
0
−5
t−statistic
CNST→
RP11−439E19.9→
RP11−439E19.7→
246.83
246.84
246.85
246.86
246.87
Position on chr1 (Mb)

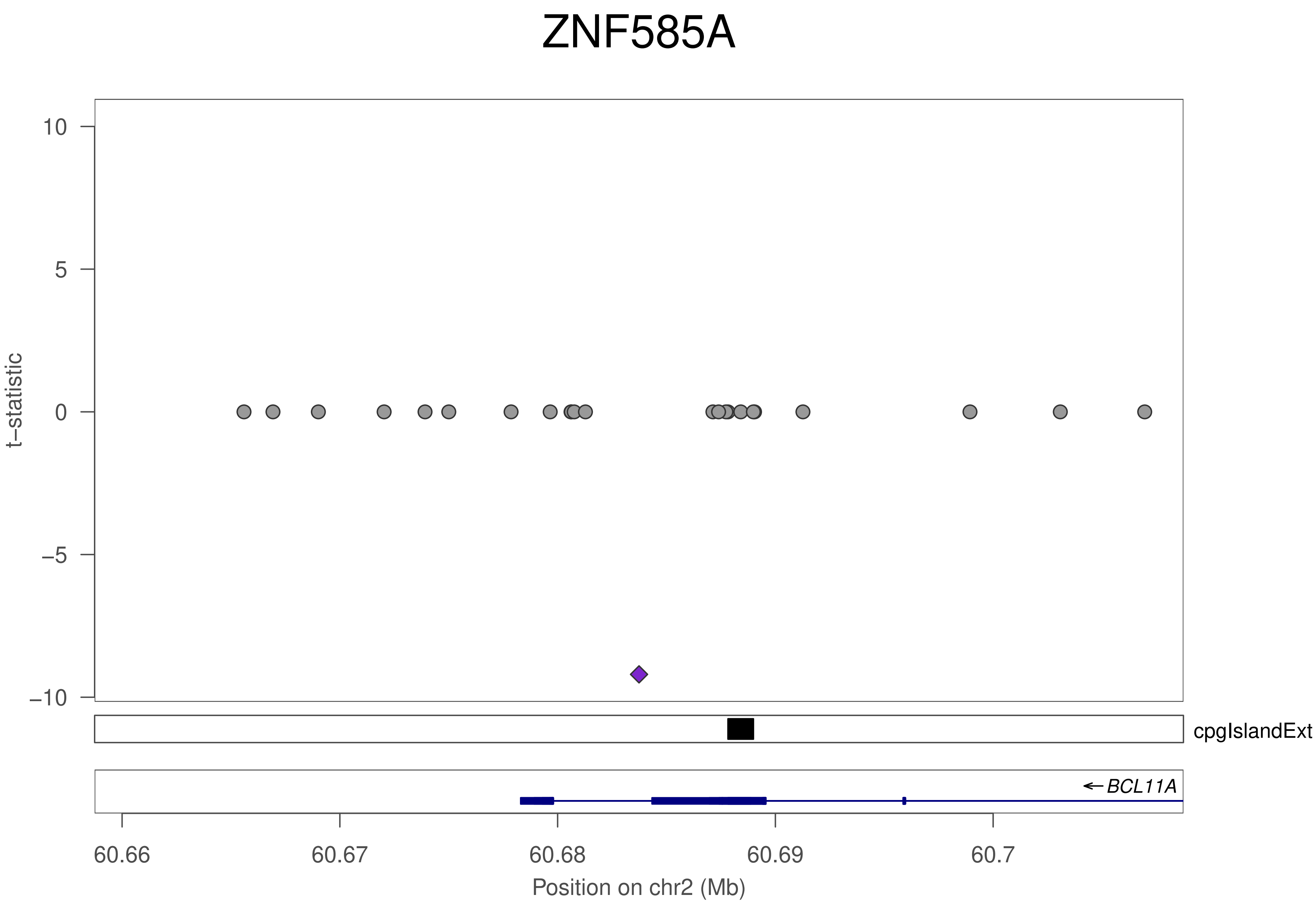
ZNF585A
t−statistic
10
5
0
−5
−10
cpgIslandExt
BCL11A
60.66
60.67
60.68
60.69
60.7
Position on chr2 (Mb)

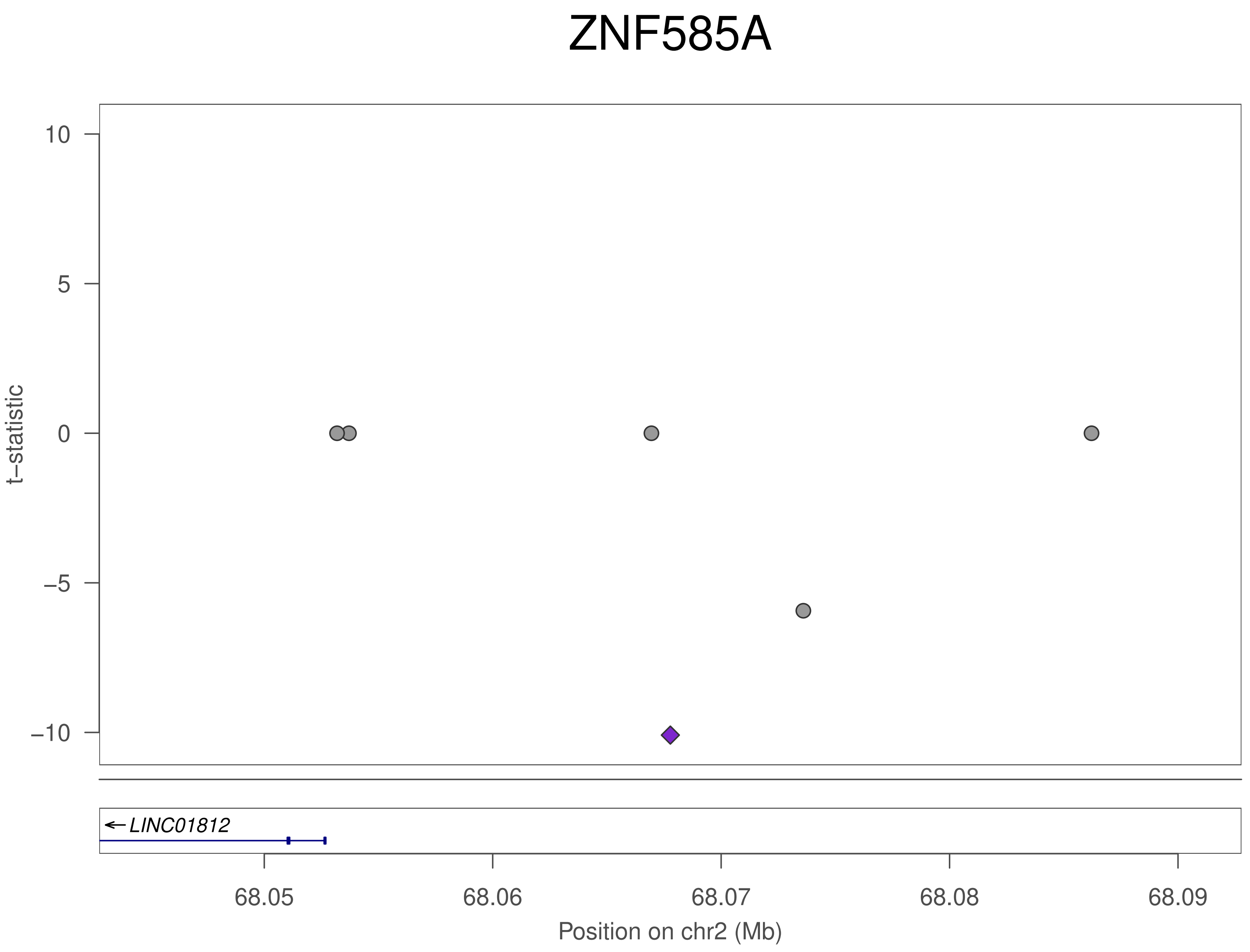

ZNF585A
t–statistic
10
5
0
–5
–10
LINC01812
68.05
68.06
68.07
68.08
68.09
Position on chr2 (Mb)

### ZNF585A

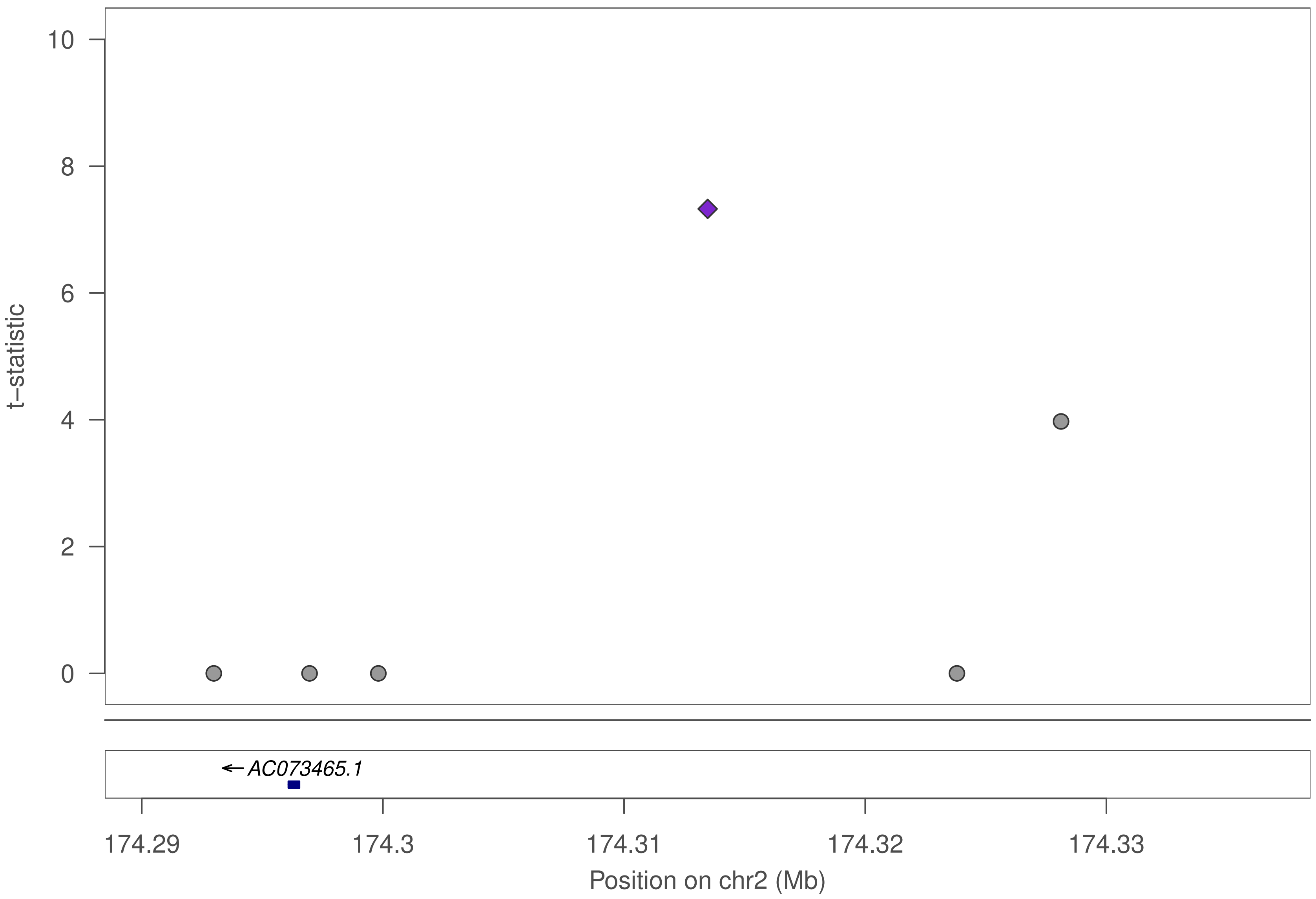

t–statistic
10
8
6
4
2
0
AC073465.1
174.29
174.3
174.31
174.32
174.33
Position on chr2 (Mb)

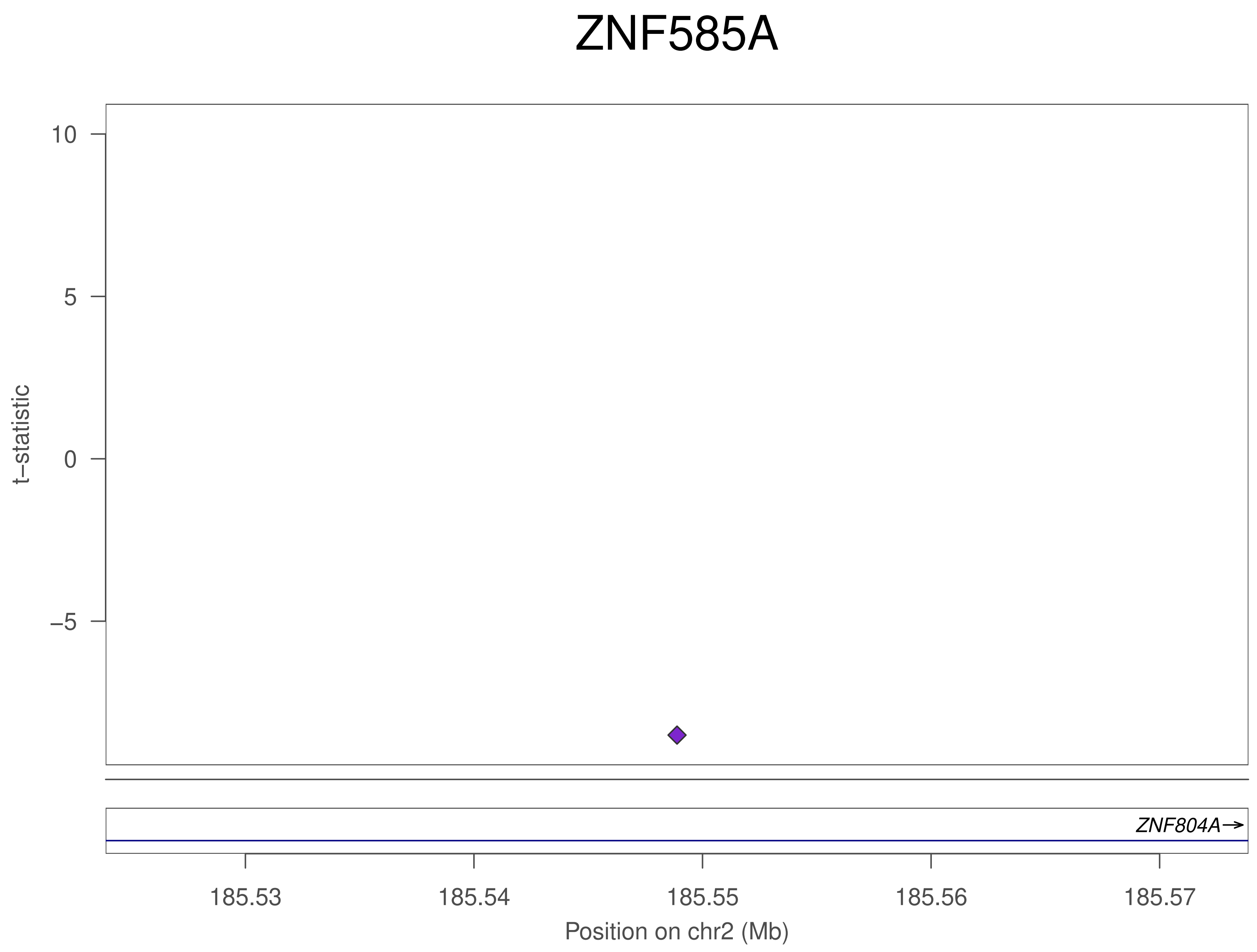
ZNF585A
t–statistic
10
5
0
–5
ZNF804A→
185.53
185.54
185.55
185.56
185.57
Position on chr2 (Mb)

### ZNF585A

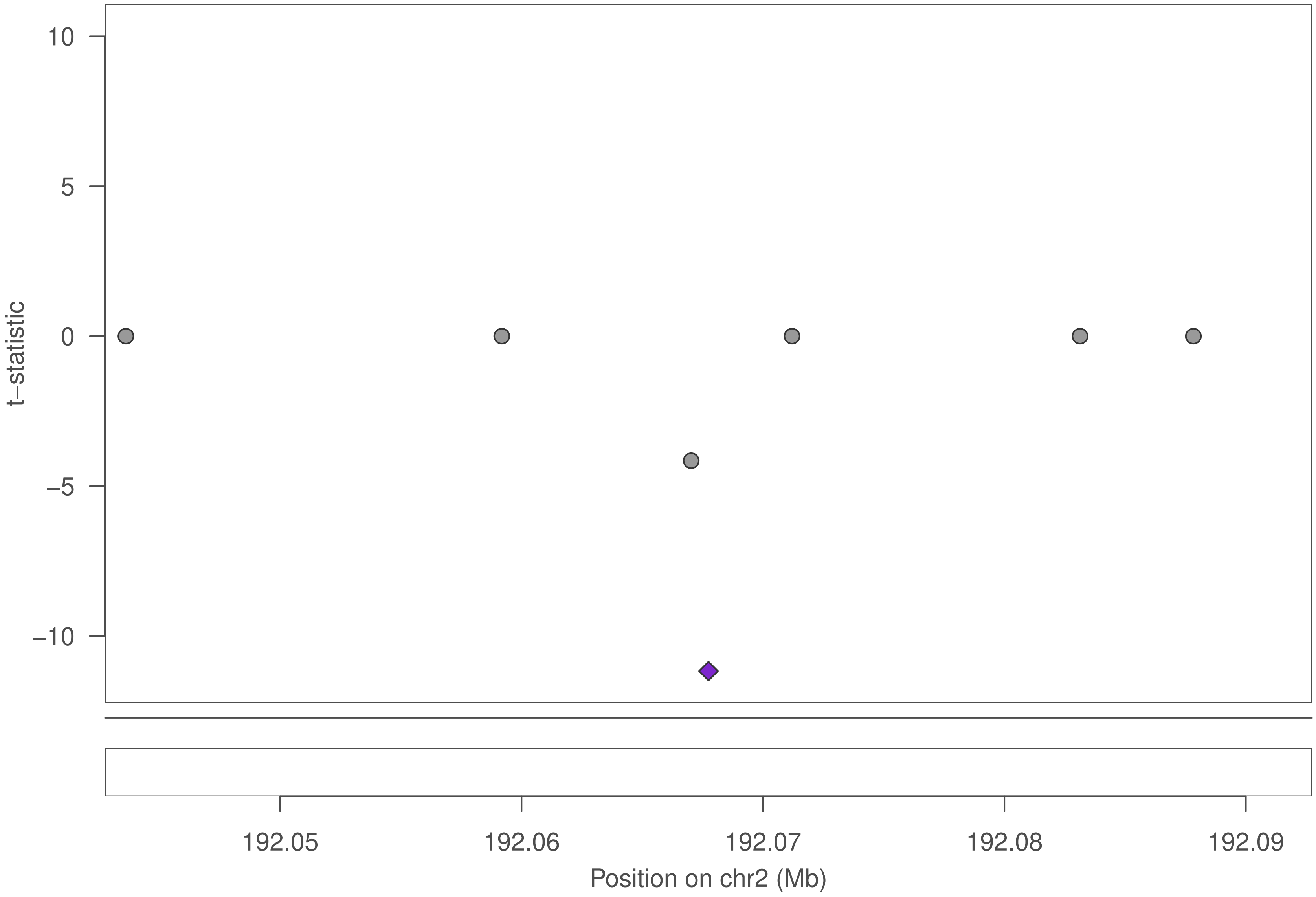

10
5
0
−5
−10
t−statistic
192.05
192.06
192.07
192.08
192.09
Position on chr2 (Mb)

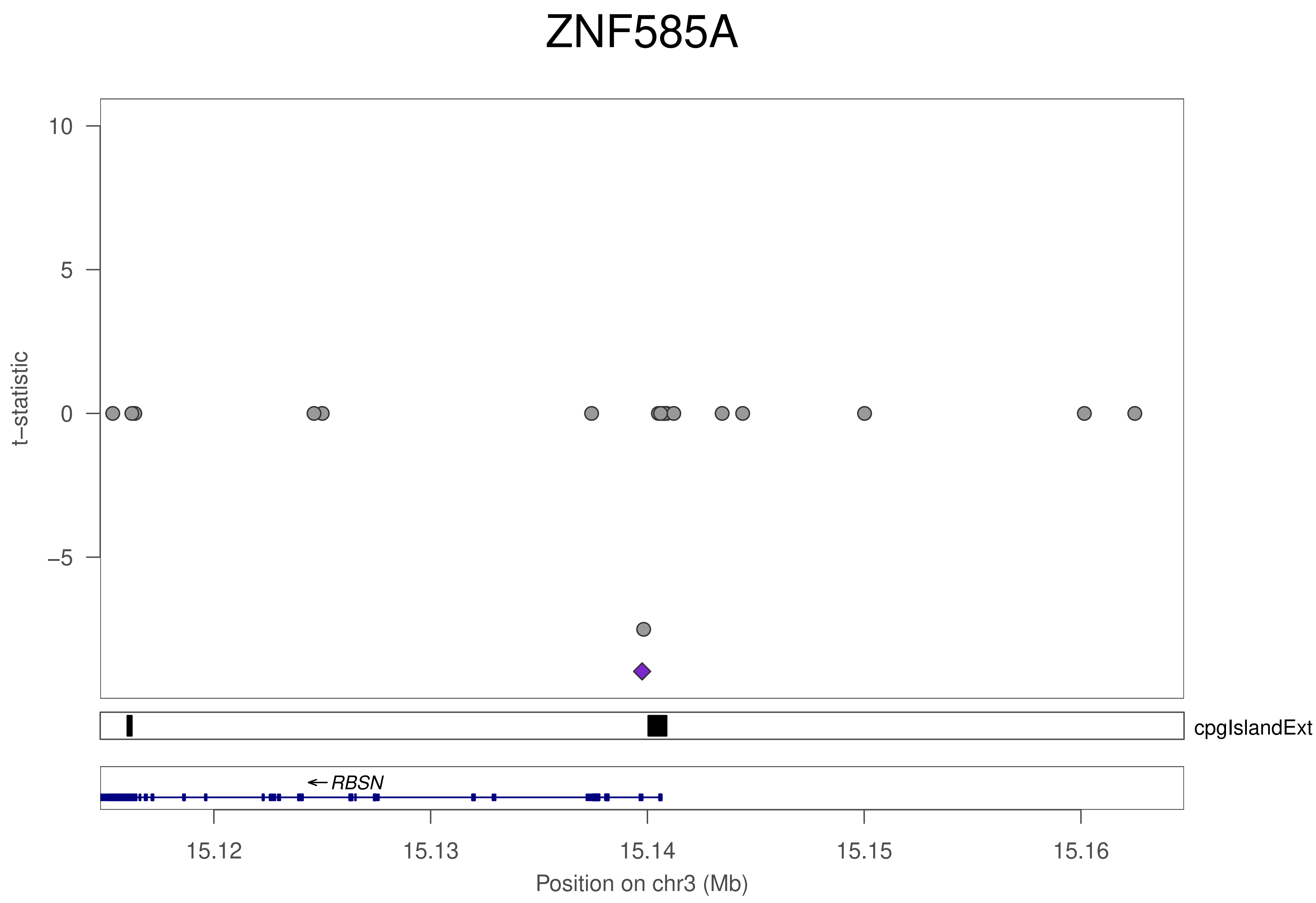

ZNF585A
10
5
0
−5
t−statistic
cpgIslandExt
← RBSN
15.12
15.13
15.14
15.15
15.16
Position on chr3 (Mb)

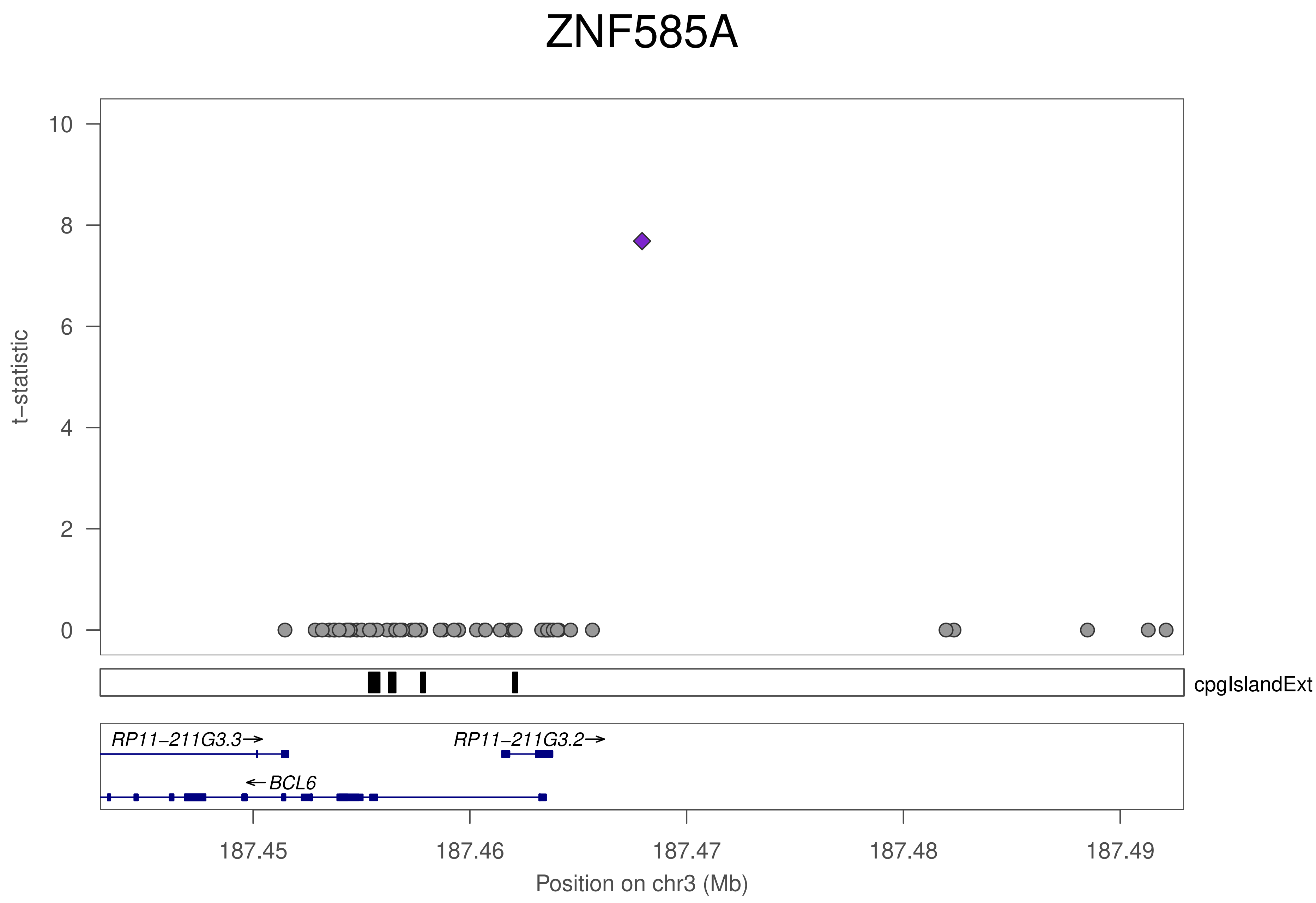

ZNF585A
t−statistic
10
8
6
4
2
0
cpgIslandExt
RP11−211G3.3→
RP11−211G3.2→
←BCL6
187.45
187.46
187.47
187.48
187.49
Position on chr3 (Mb)

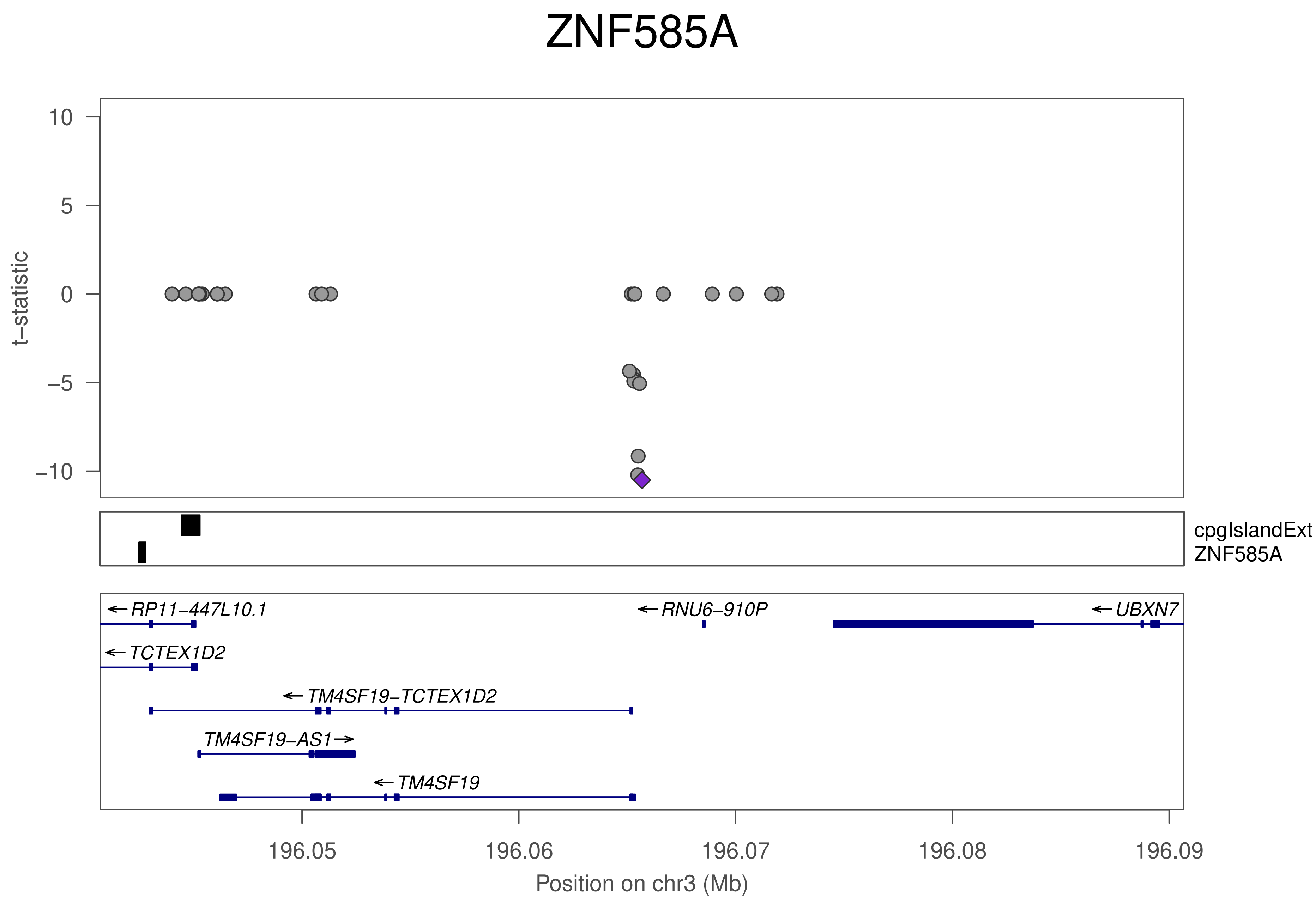

ZNF585A
t−statistic
10
5
0
−5
−10
cpgIslandExt
ZNF585A
← RP11−447L10.1
← RNU6−910P
← UBXN7
← TCTEX1D2
← TM4SF19−TCTEX1D2
TM4SF19−AS1 →
← TM4SF19
196.05
196.06
196.07
196.08
196.09
Position on chr3 (Mb)

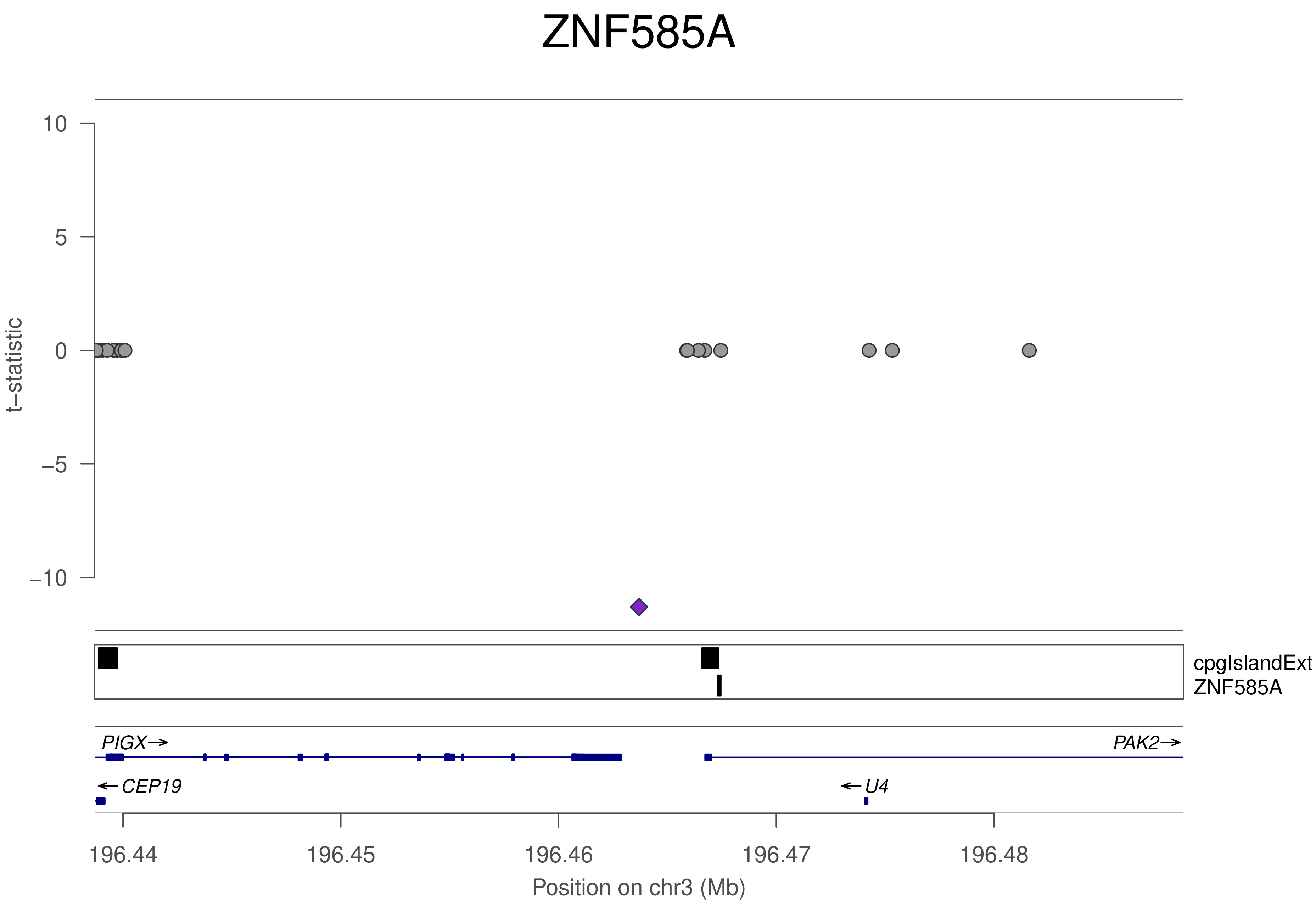
ZNF585A
t−statistic
10
5
0
−5
−10
cpgIslandExt
ZNF585A
PIGX→
PAK2→
←CEP19
←U4
196.44
196.45
196.46
196.47
196.48
Position on chr3 (Mb)

### ZNF585A

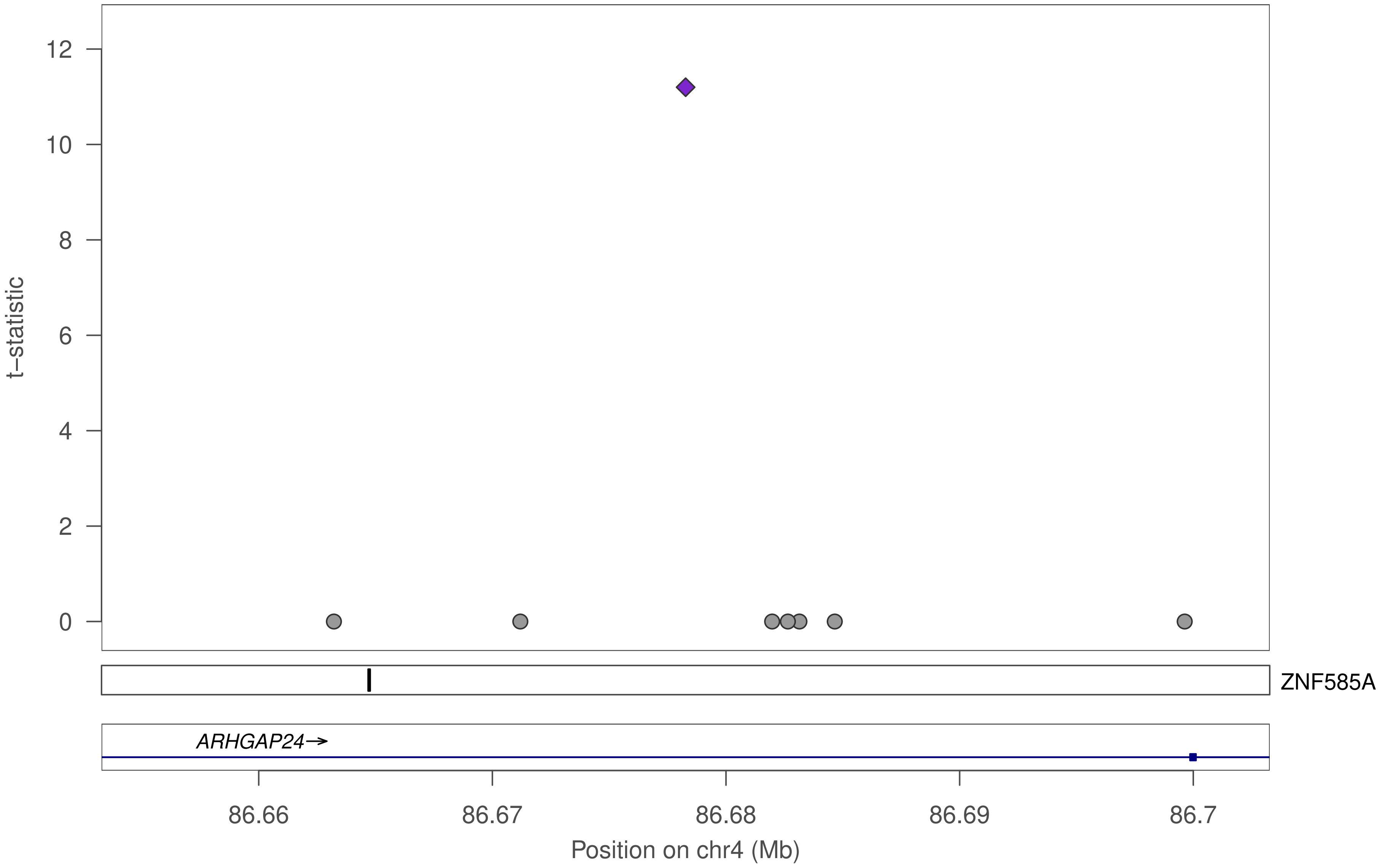

t−statistic
12
10
8
6
4
2
0
ZNF585A
ARHGAP24→
86.66
86.67
86.68
86.69
86.7
Position on chr4 (Mb)

### ZNF585A

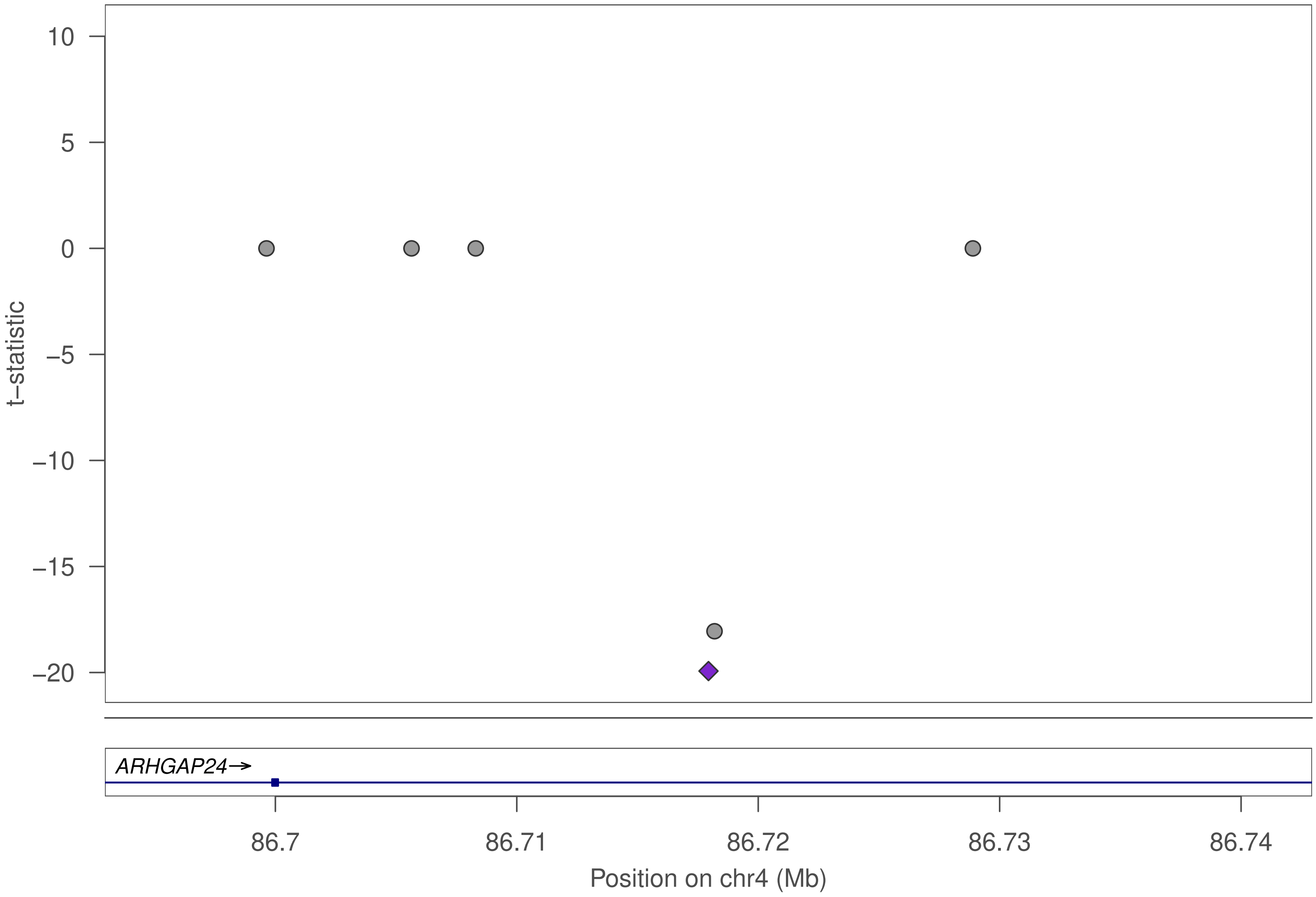

10
5
0
−5
−10
−15
−20
t−statistic
ARHGAP24→
86.7
86.71
86.72
86.73
86.74
Position on chr4 (Mb)

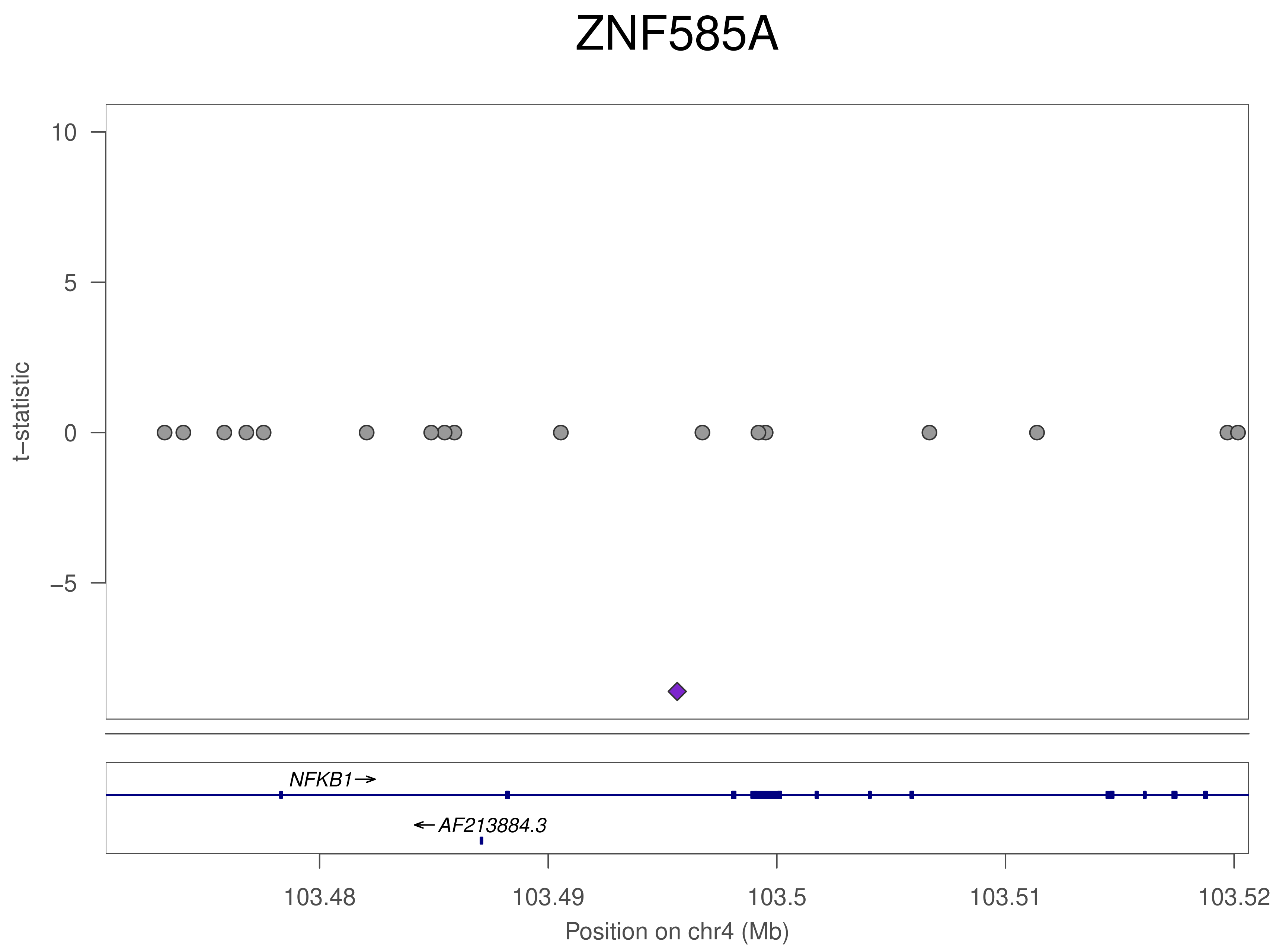
ZNF585A
t−statistic
10
5
0
−5
NFKB1→
←AF213884.3
103.48
103.49
103.5
103.51
103.52
Position on chr4 (Mb)

### ZNF585A

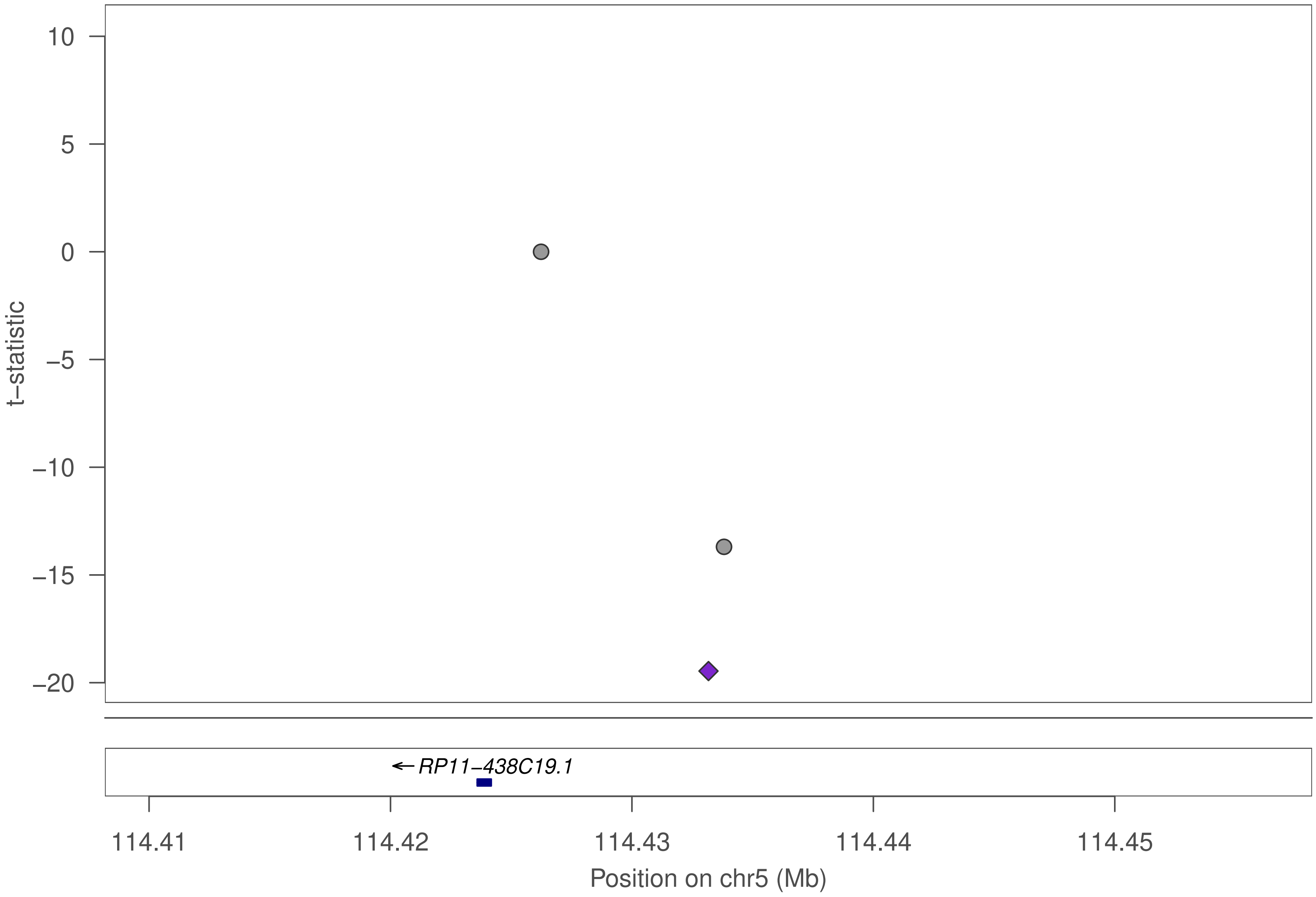

10
5
0
−5
−10
−15
−20
t−statistic
RP11−438C19.1
114.41
114.42
114.43
114.44
114.45
Position on chr5 (Mb)

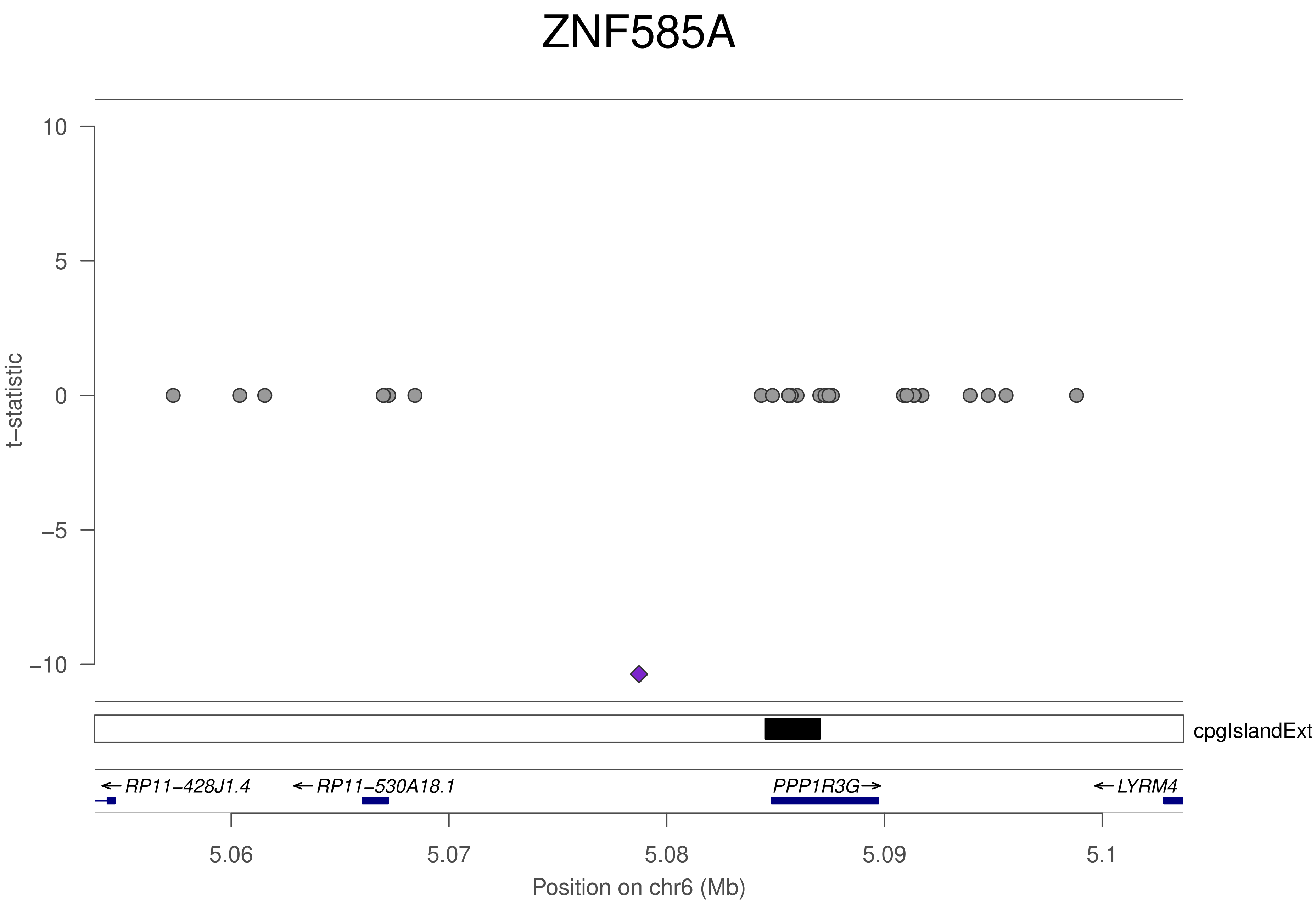

ZNF585A
t–statistic
10
5
0
–5
–10
cpgIslandExt
←RP11–428J1.4
←RP11–530A18.1
PPP1R3G→
←LYRM4
5.06
5.07
5.08
5.09
5.1
Position on chr6 (Mb)

### ZNF585A

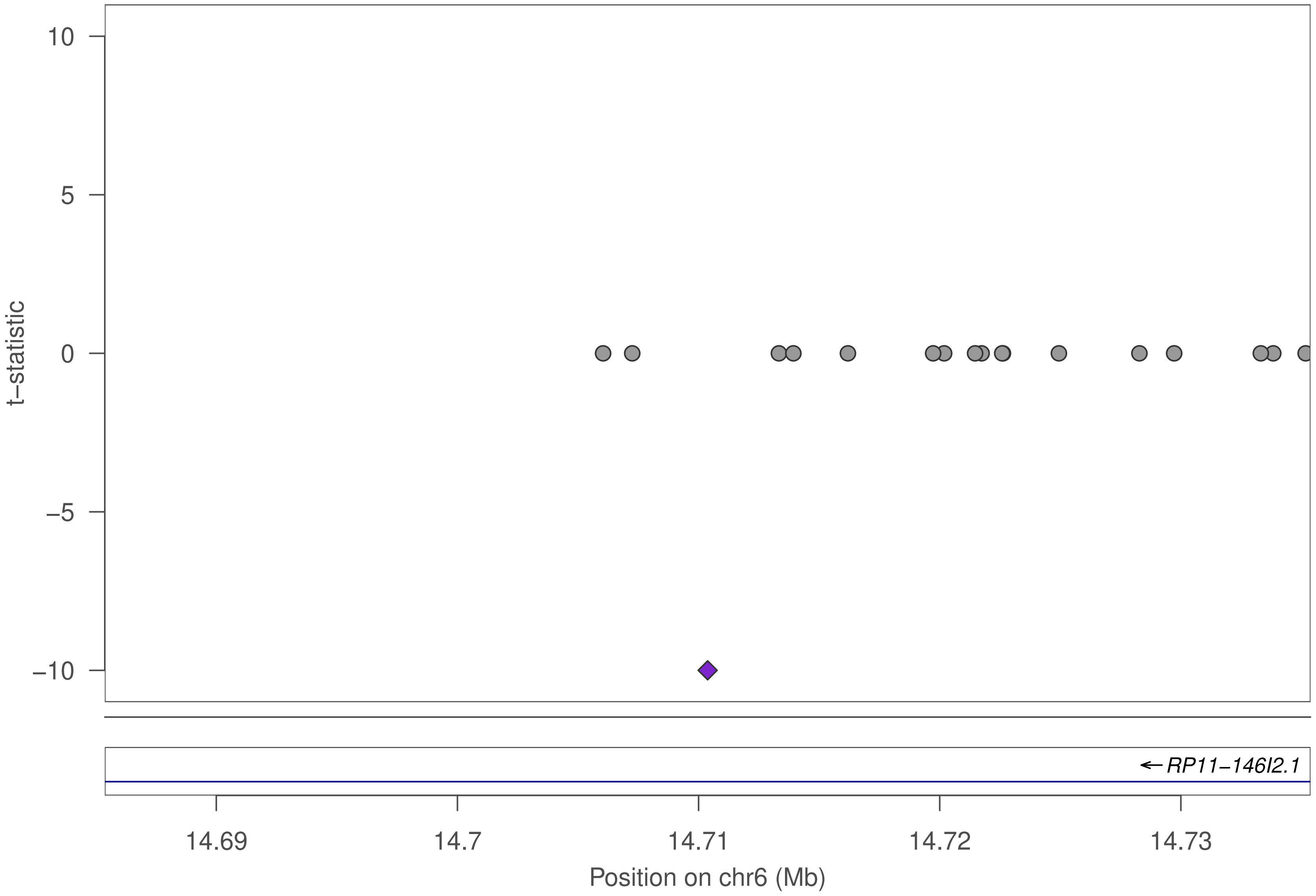

10
5
0
−5
−10
t−statistic
RP11−146I2.1
14.69
14.7
14.71
14.72
14.73
Position on chr6 (Mb)

### ZNF585A

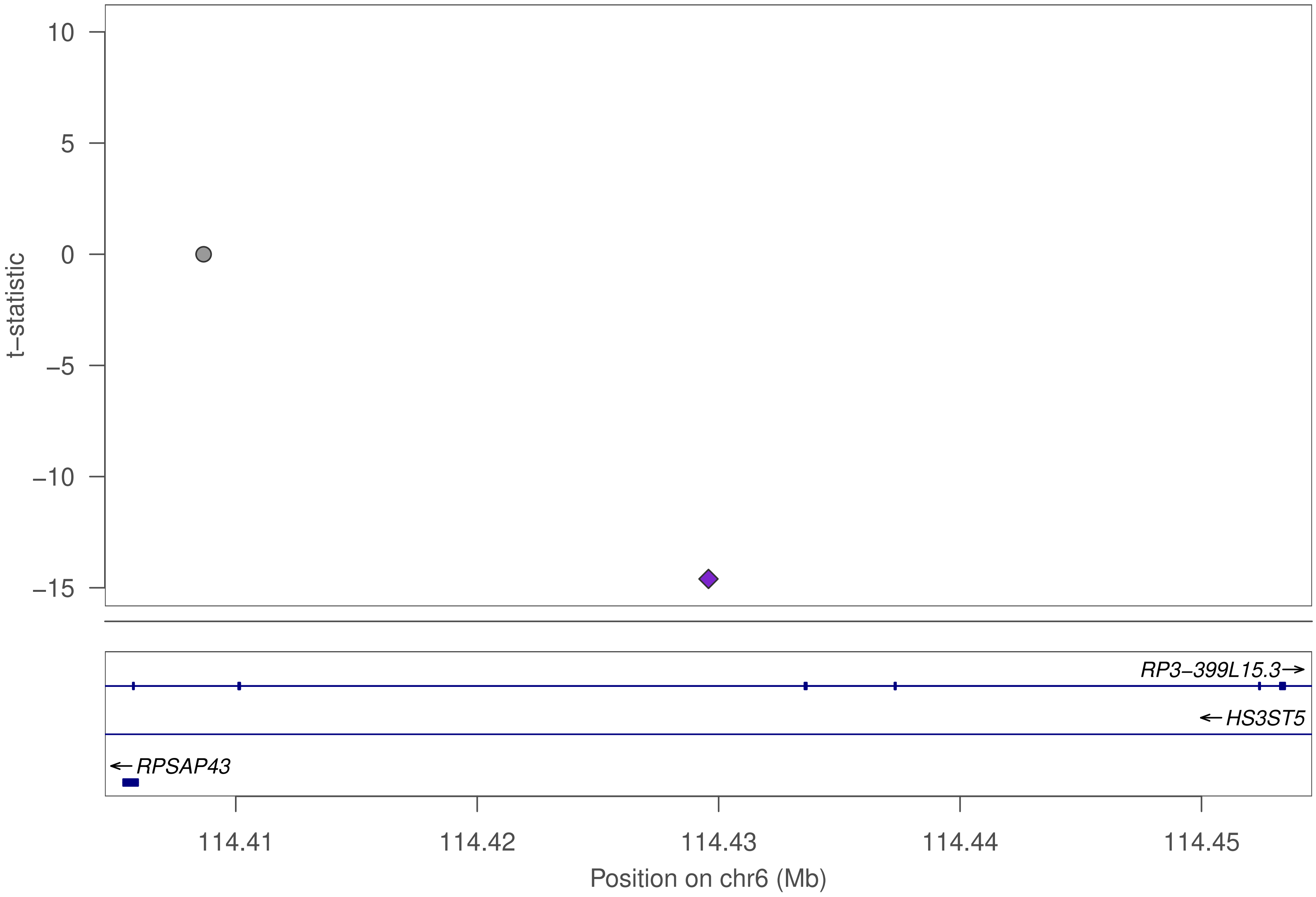

10
5
0
−5
−10
−15
t−statistic
RP3−399L15.3→
←HS3ST5
←RPSAP43
114.41
114.42
114.43
114.44
114.45
Position on chr6 (Mb)

### ZNF585A

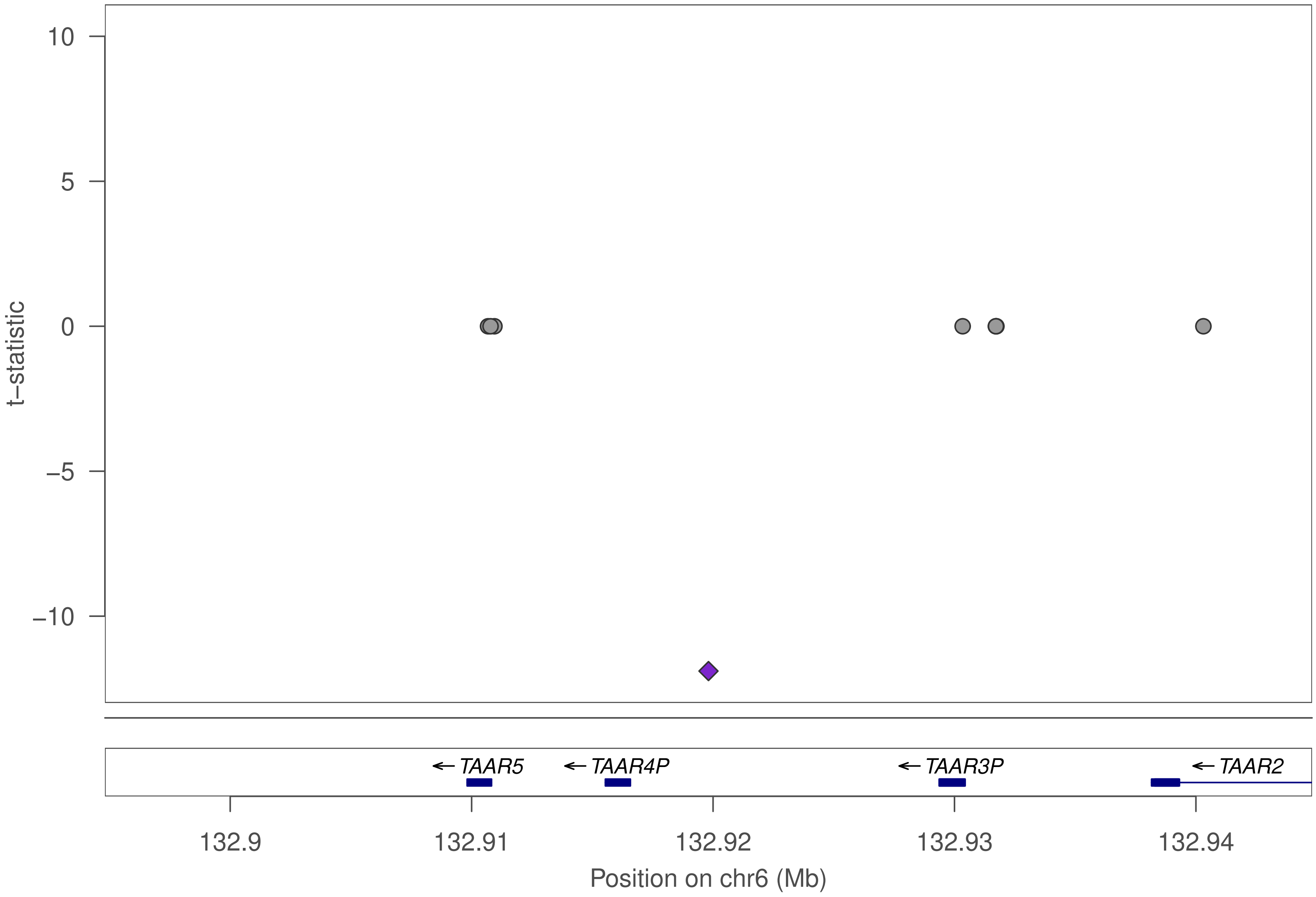

10
5
0
−5
−10
t−statistic
← TAAR5
← TAAR4P
← TAAR3P
← TAAR2
132.9
132.91
132.92
132.93
132.94
Position on chr6 (Mb)

### ZNF585A

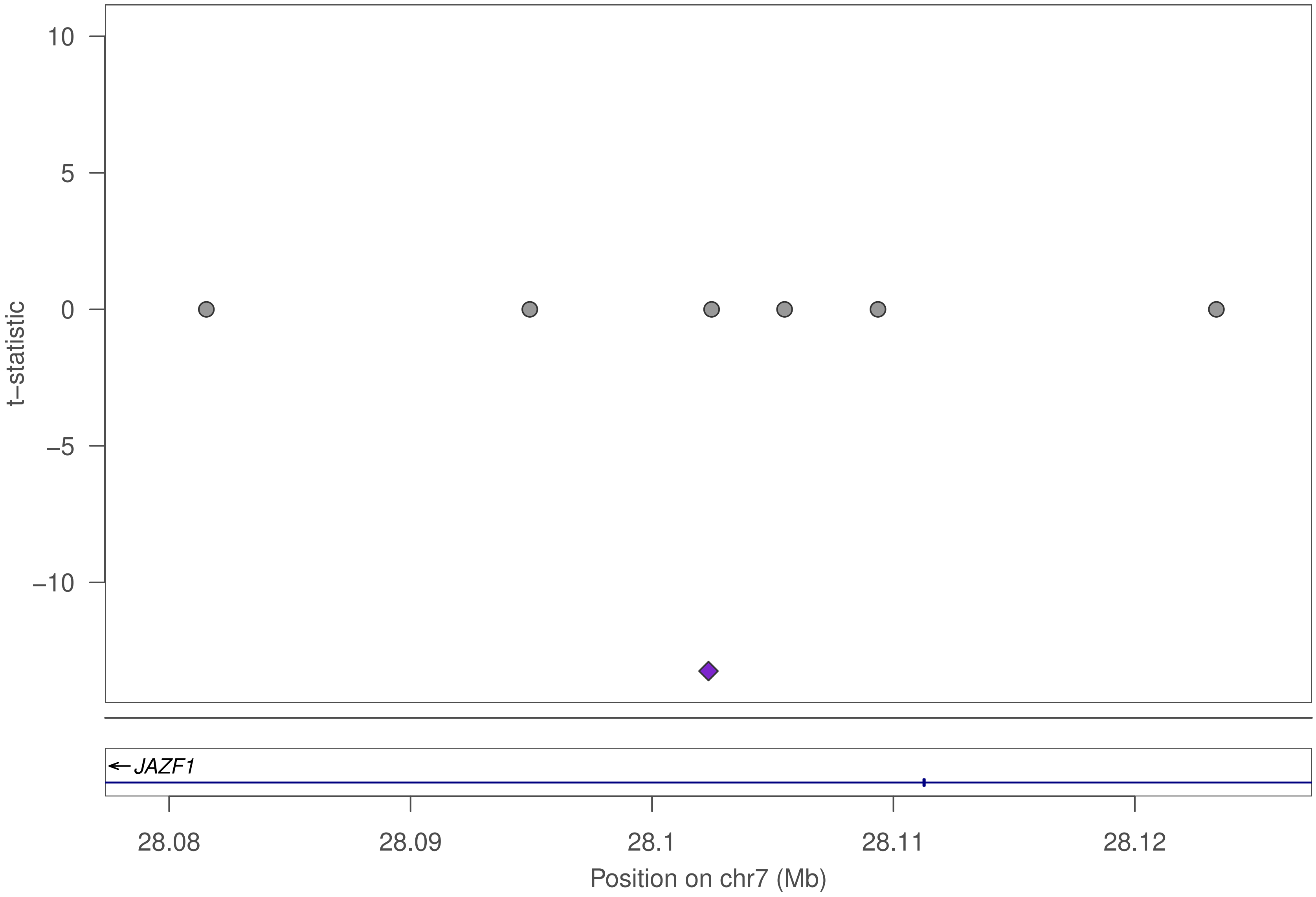

10
5
0
−5
−10
t−statistic
JAZF1
28.08
28.09
28.1
28.11
28.12
Position on chr7 (Mb)

### ZNF585A

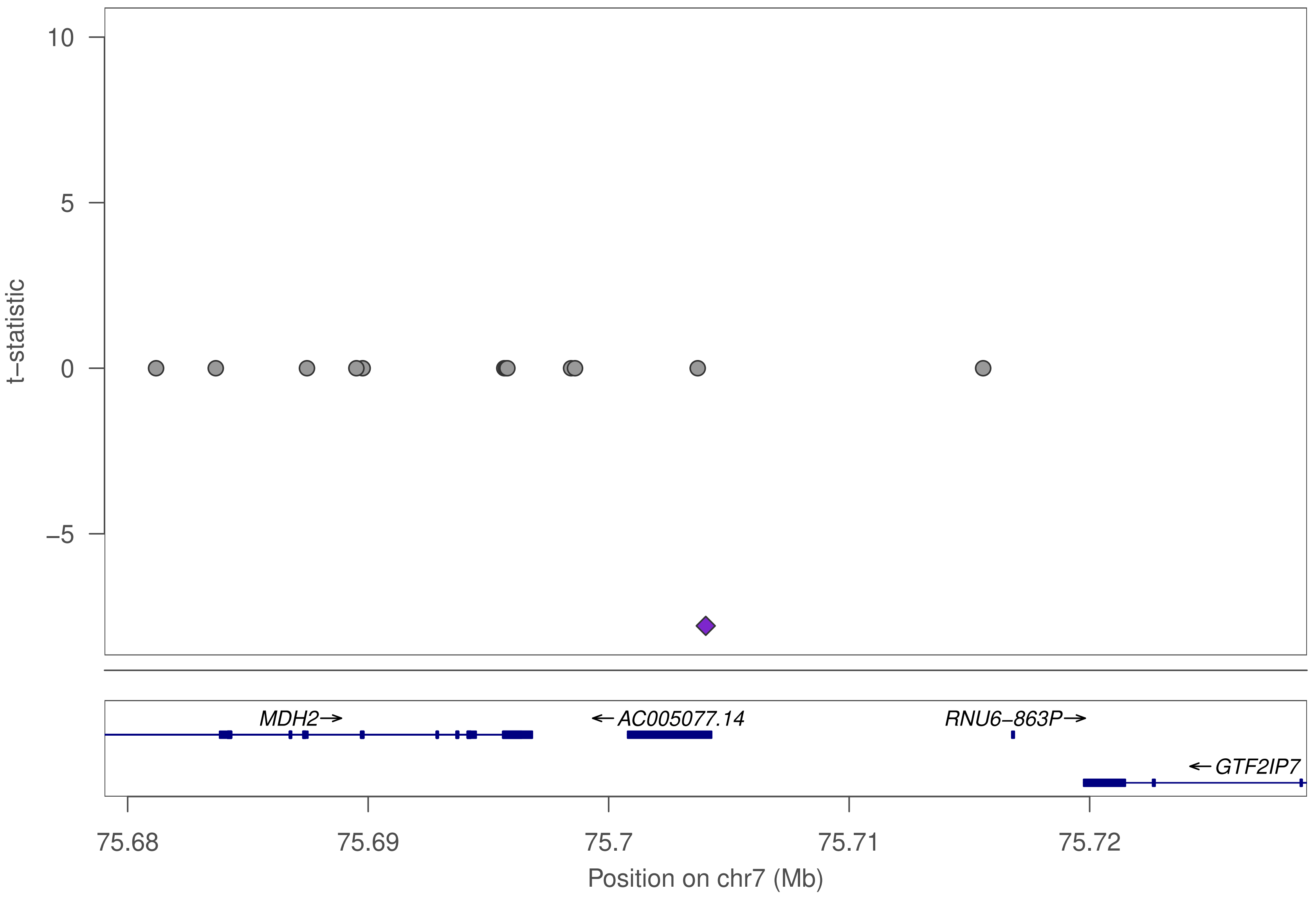

10
5
0
−5
t−statistic
MDH2→
←AC005077.14
RNU6−863P→
←GTF2IP7
75.68
75.69
75.7
75.71
75.72
Position on chr7 (Mb)

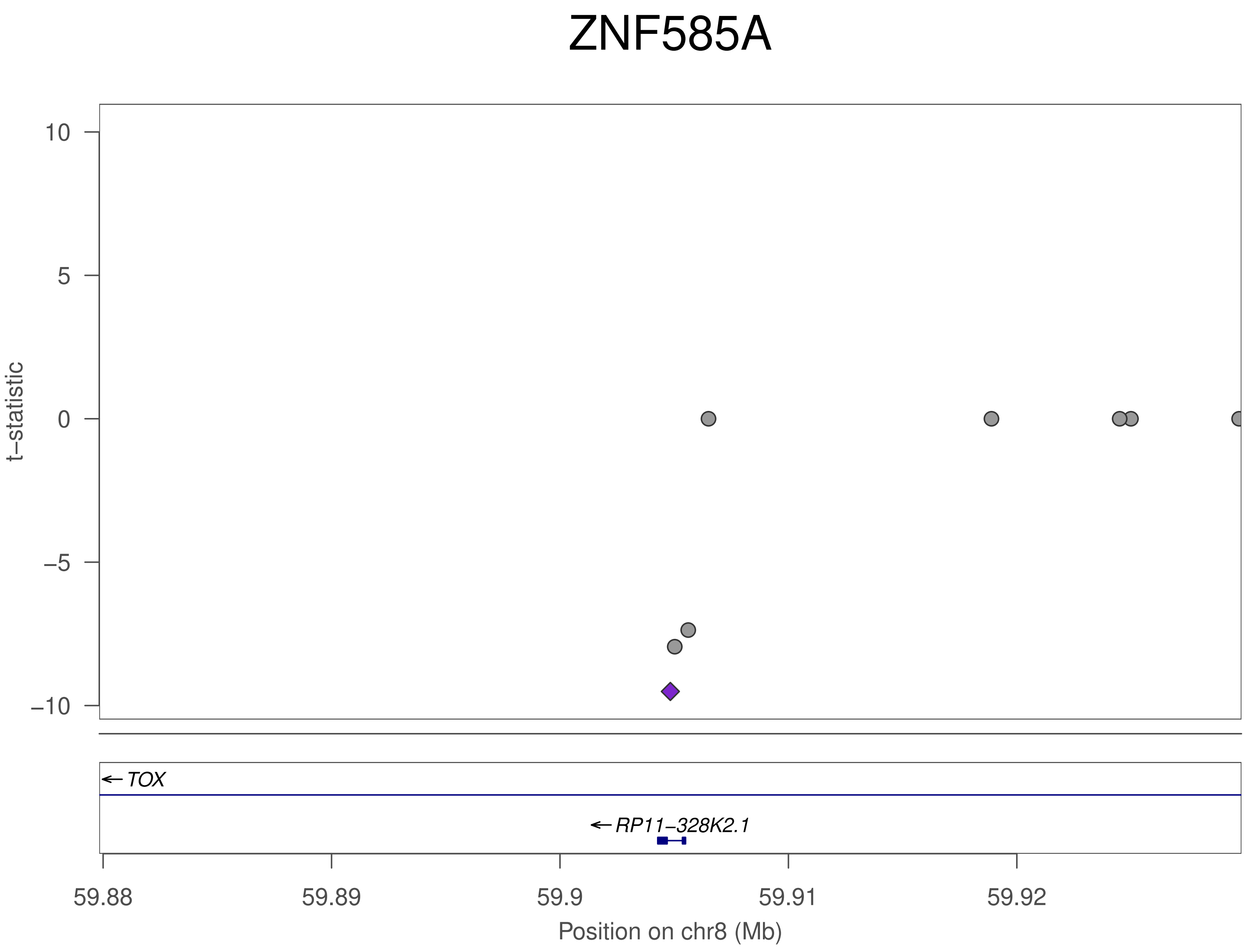

ZNF585A
t−statistic
10
5
0
−5
−10
TOX
RP11−328K2.1
59.88
59.89
59.9
59.91
59.92
Position on chr8 (Mb)

### ZNF585A

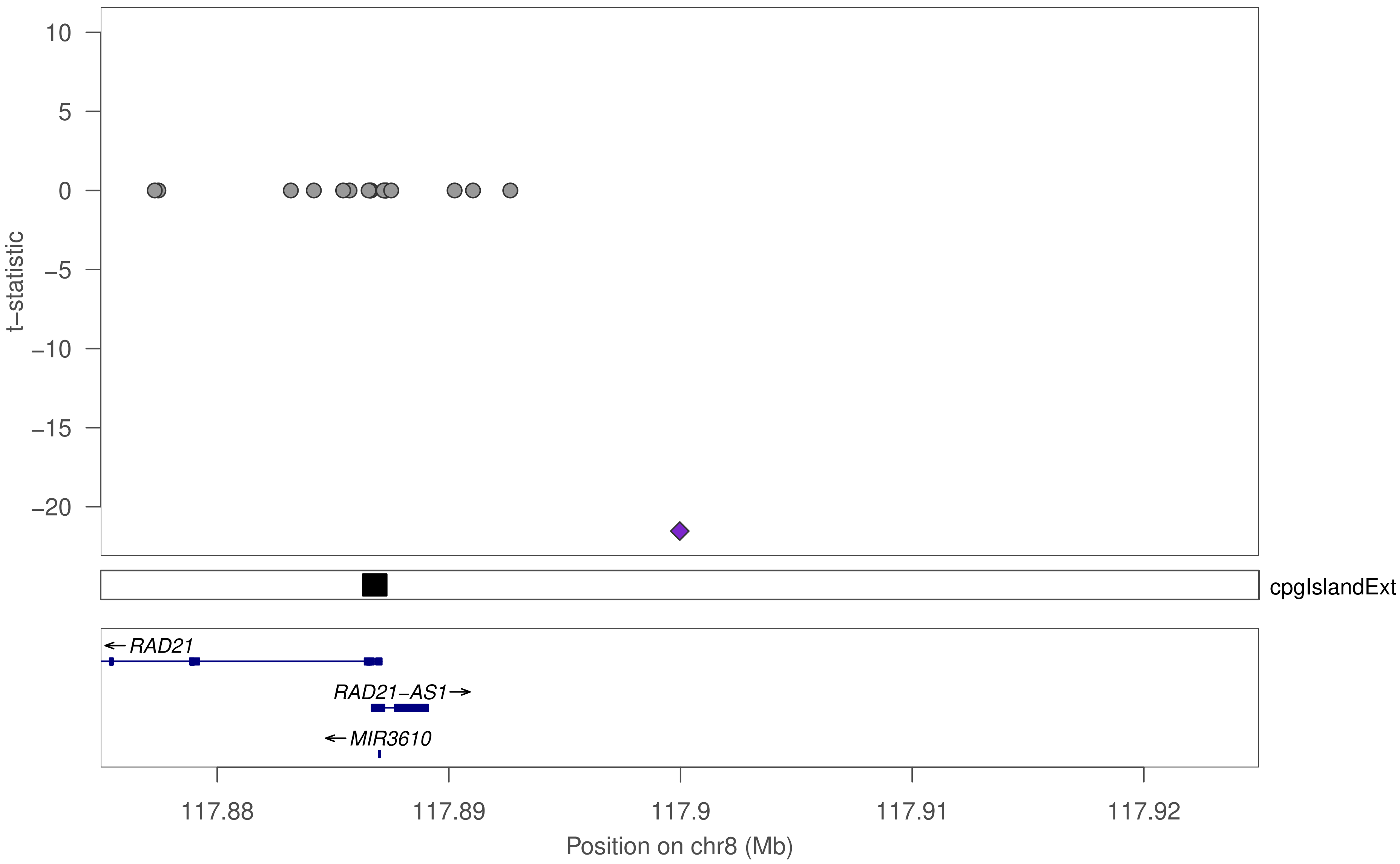

10
5
0
−5
−10
−15
−20
t−statistic
cpgIslandExt
←RAD21
RAD21−AS1→
←MIR3610
117.88
117.89
117.9
117.91
117.92
Position on chr8 (Mb)

### ZNF585A

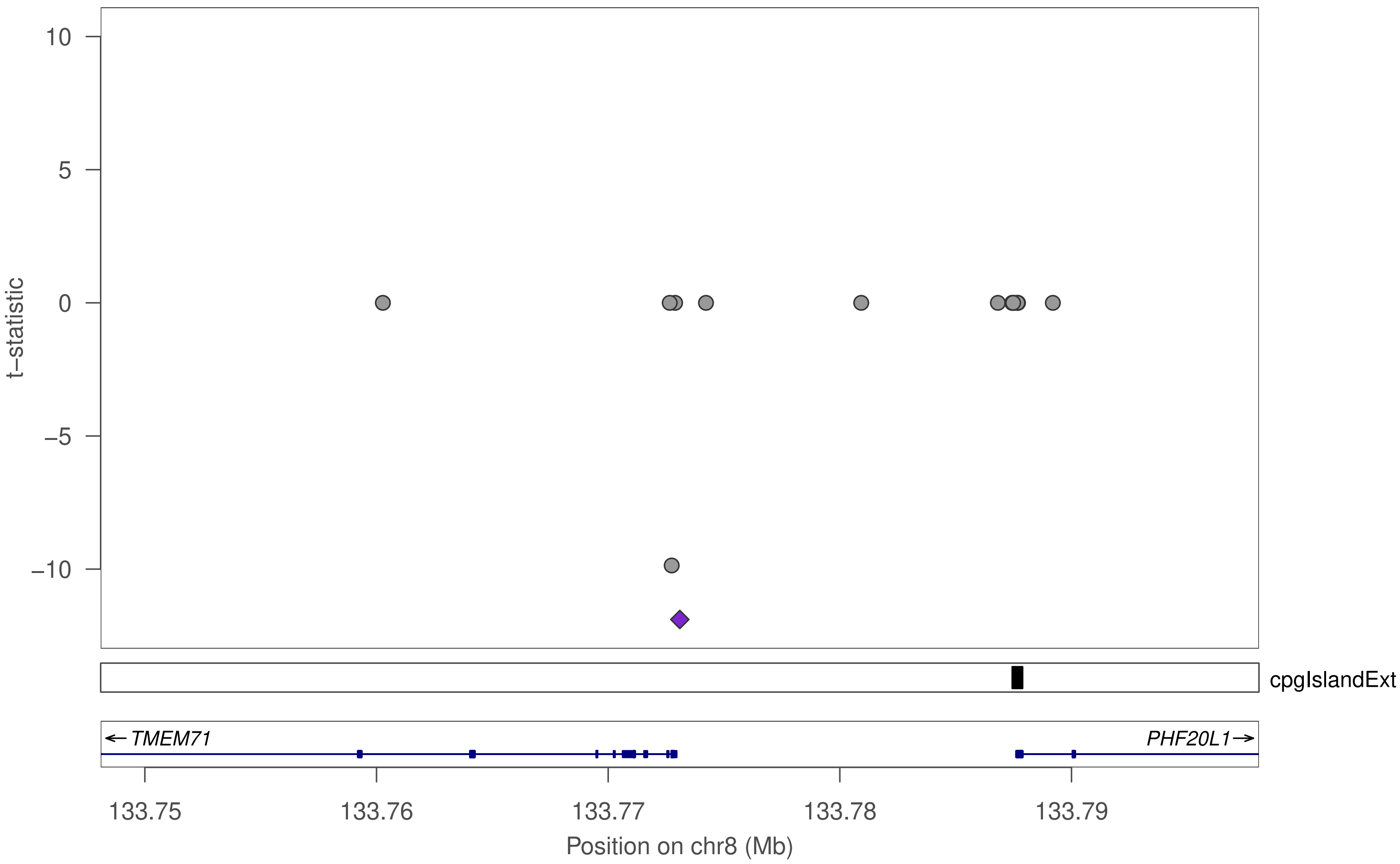

10
5
0
−5
−10
t−statistic
cpgIslandExt
← TMEM71
PHF20L1 →
133.75
133.76
133.77
133.78
133.79
Position on chr8 (Mb)

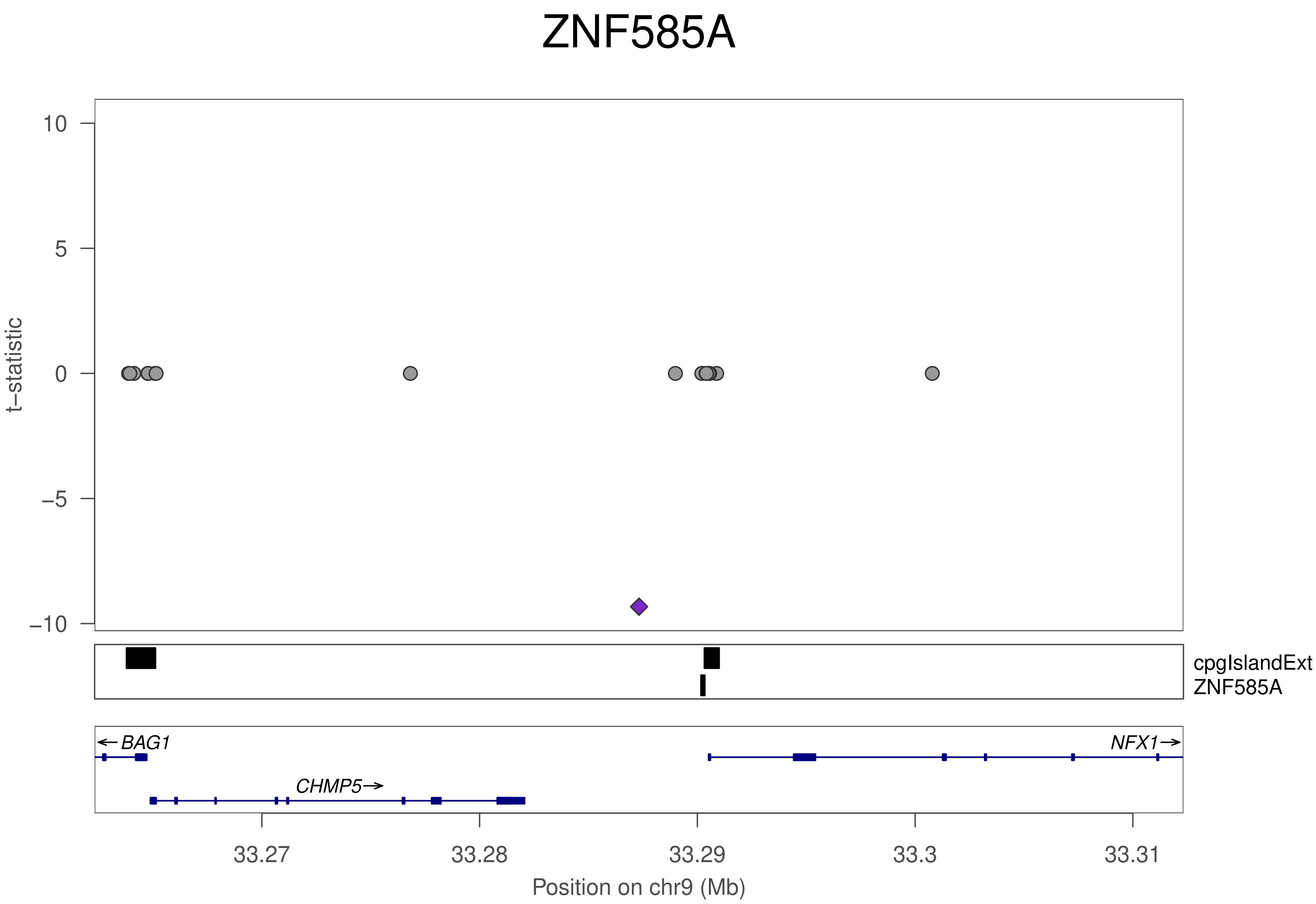
ZNF585A
t–statistic
10
5
0
–5
–10
cpgIslandExt
ZNF585A
BAG1
NFX1
CHMP5
33.27
33.28
33.29
33.3
33.31
Position on chr9 (Mb)

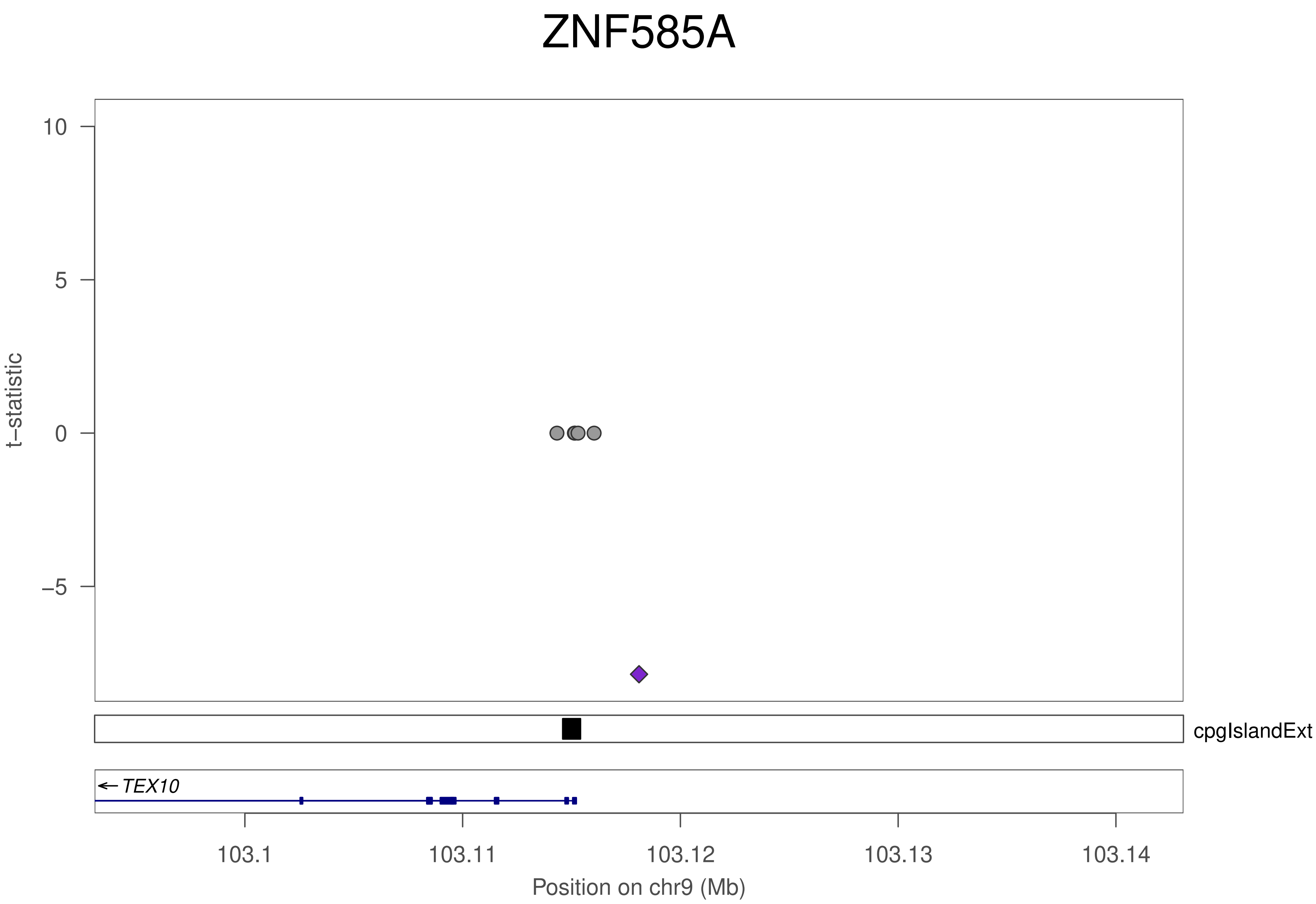

ZNF585A
t–statistic
10
5
0
–5
cpgIslandExt
TEX10
103.1
103.11
103.12
103.13
103.14
Position on chr9 (Mb)

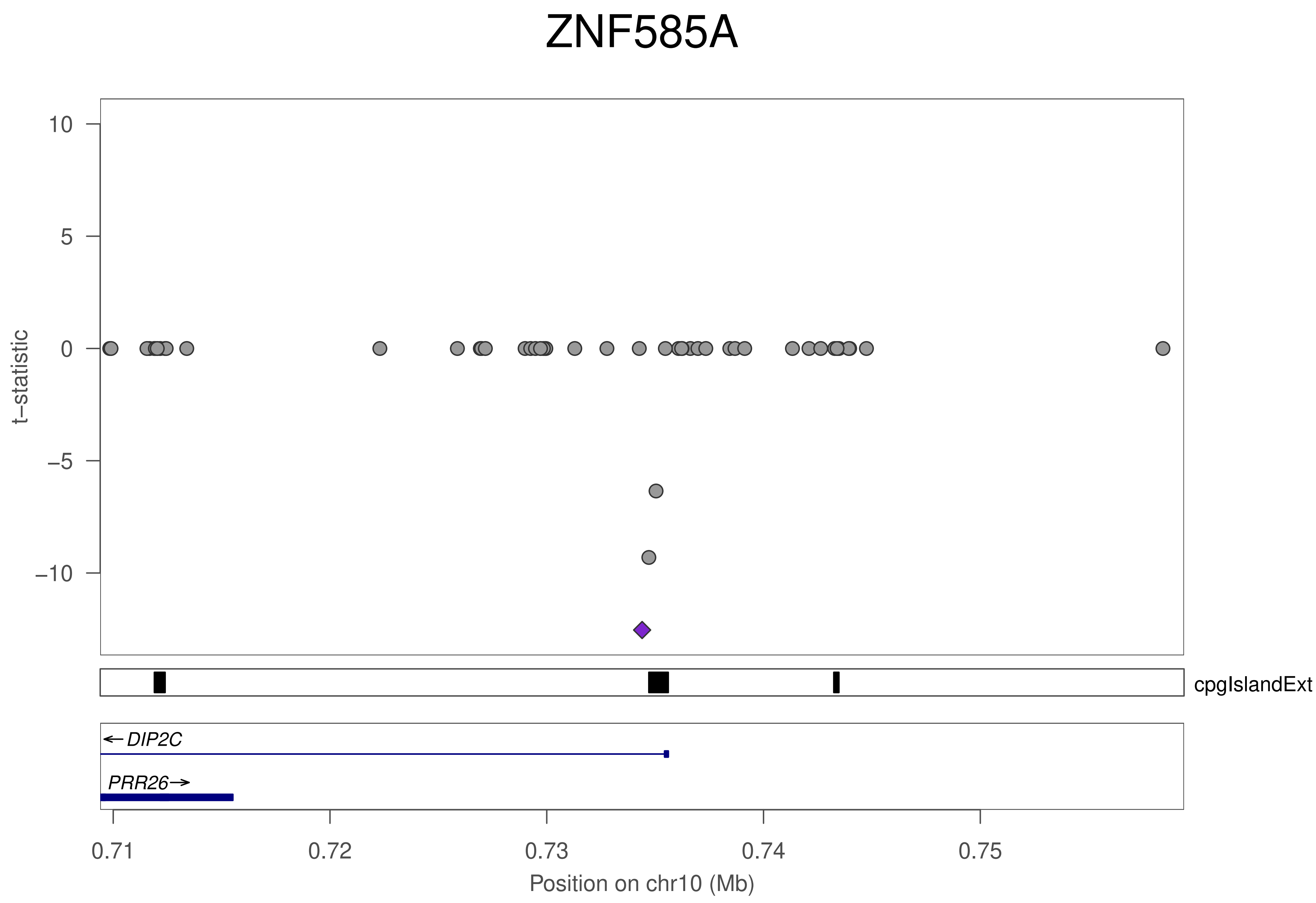
ZNF585A
t–statistic
10
5
0
–5
–10
cpgIslandExt
DIP2C
PRR26
0.71
0.72
0.73
0.74
0.75
Position on chr10 (Mb)

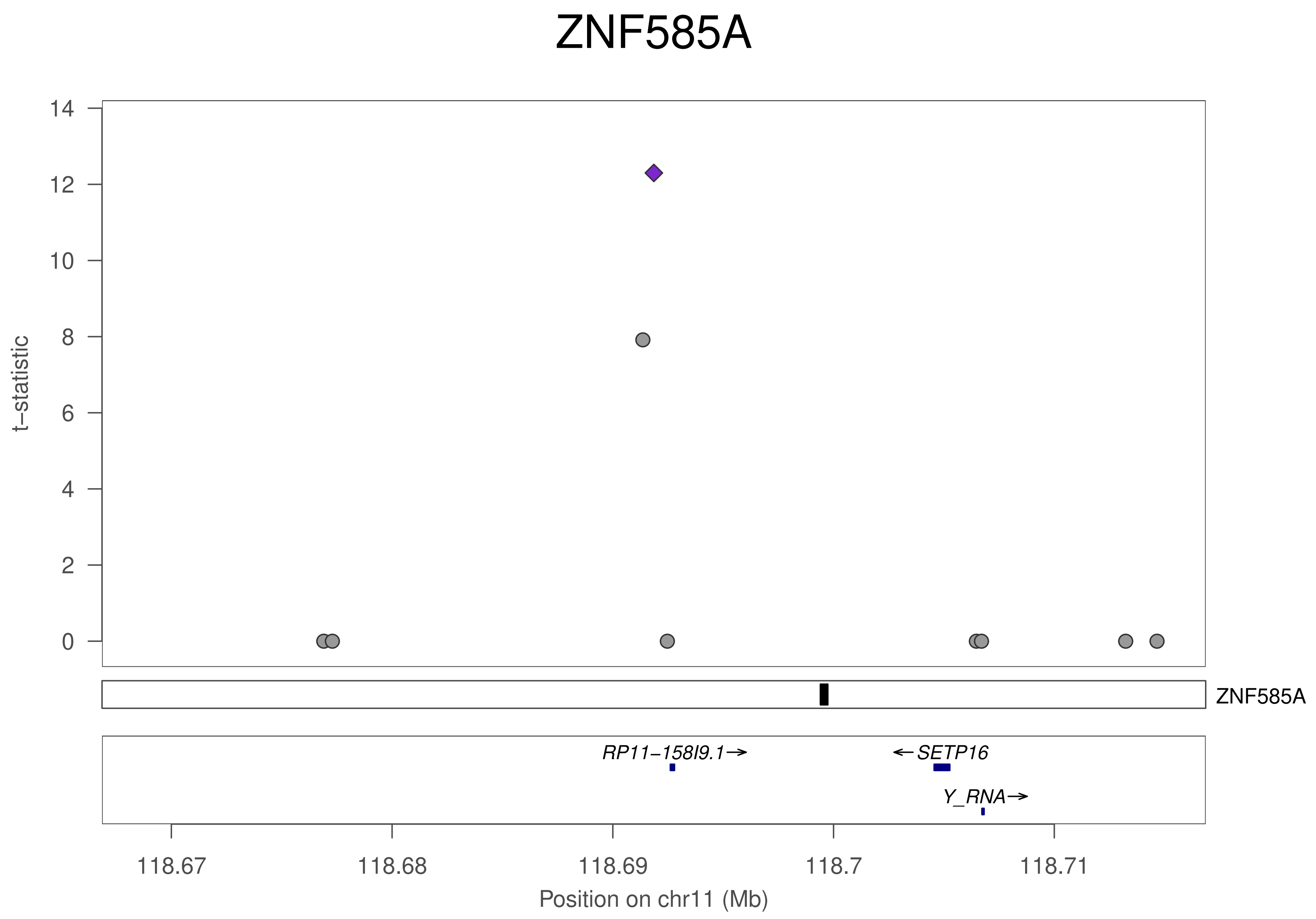

ZNF585A
t−statistic
14
12
10
8
6
4
2
0
ZNF585A
RP11−158I9.1→
←SETP16
Y_RNA→
118.67
118.68
118.69
118.7
118.71
Position on chr11 (Mb)

### ZNF585A

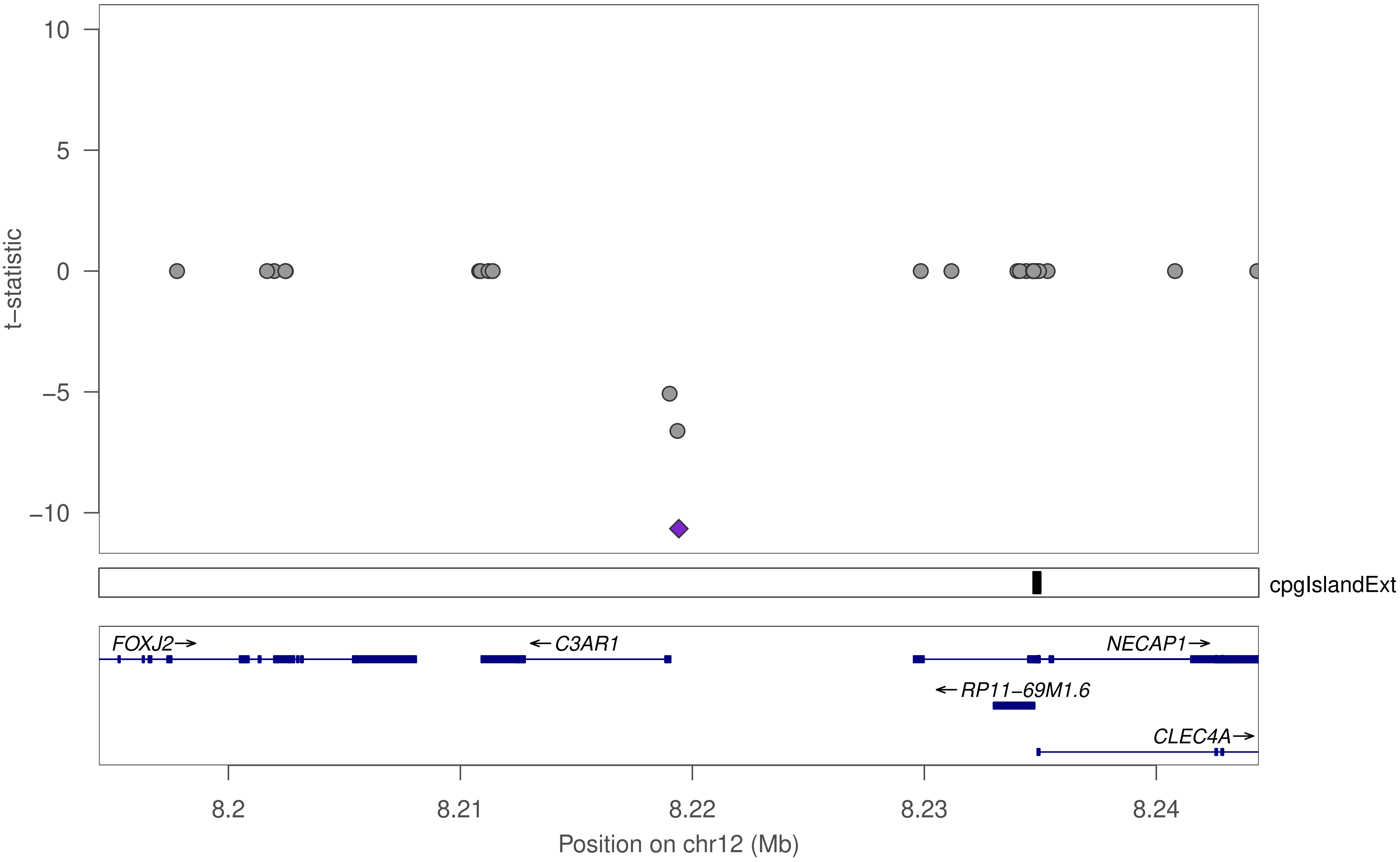

10
5
0
–5
–10
t–statistic
cpgIslandExt
FOXJ2
C3AR1
NECAP1
RP11–69M1.6
CLEC4A
8.2
8.21
8.22
8.23
8.24
Position on chr12 (Mb)

### ZNF585A

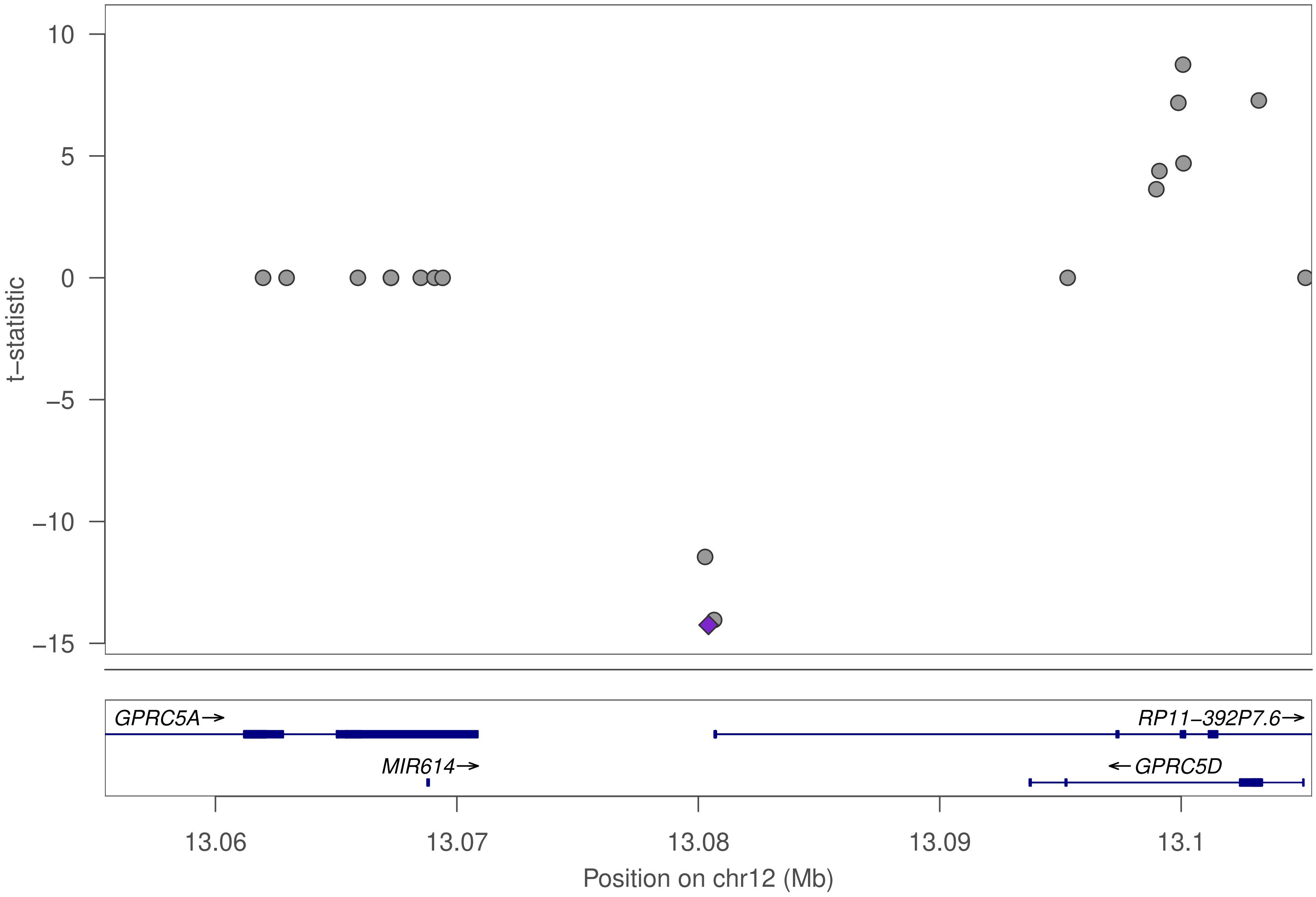

t−statistic
10
5
0
−5
−10
−15
GPRC5A→
MIR614→
RP11−392P7.6→
←GPRC5D
13.06
13.07
13.08
13.09
13.1
Position on chr12 (Mb)

### ZNF585A

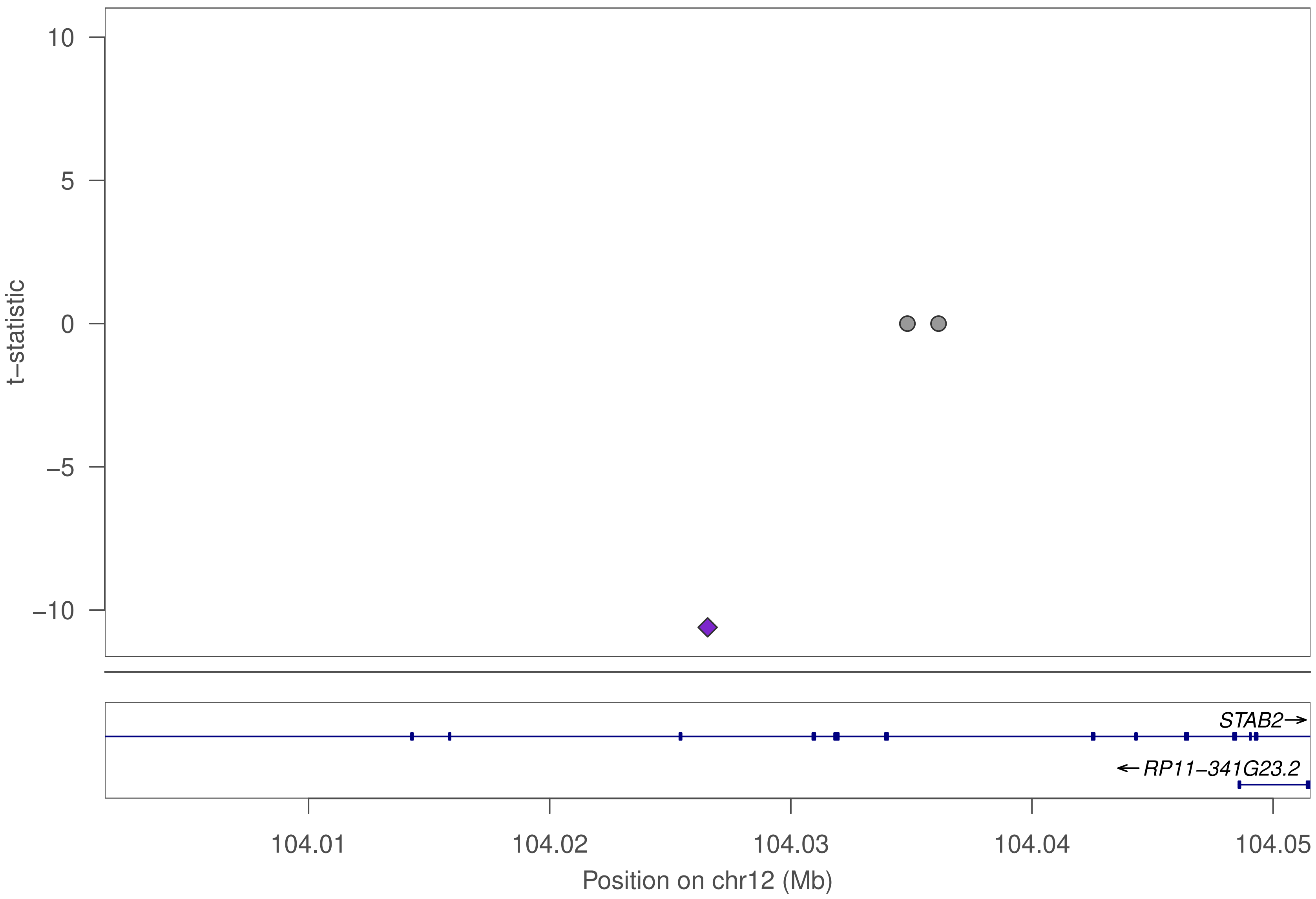

10
5
0
−5
−10
t−statistic
STAB2→
←RP11−341G23.2
104.01
104.02
104.03
104.04
104.05
Position on chr12 (Mb)

### ZNF585A

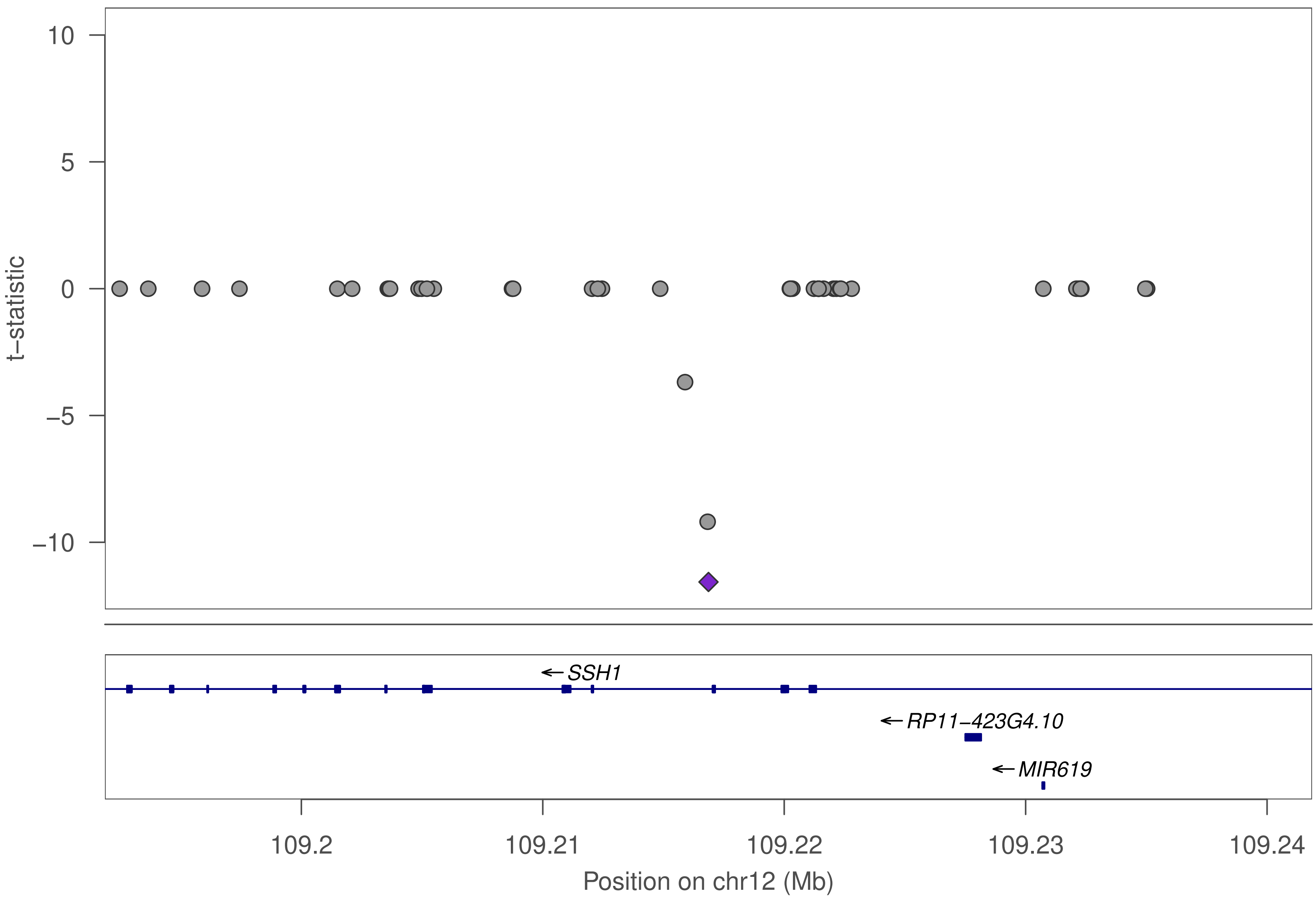

10
5
0
−5
−10
t−statistic
SSH1
RP11−423G4.10
MIR619
109.2
109.21
109.22
109.23
109.24
Position on chr12 (Mb)

### ZNF585A

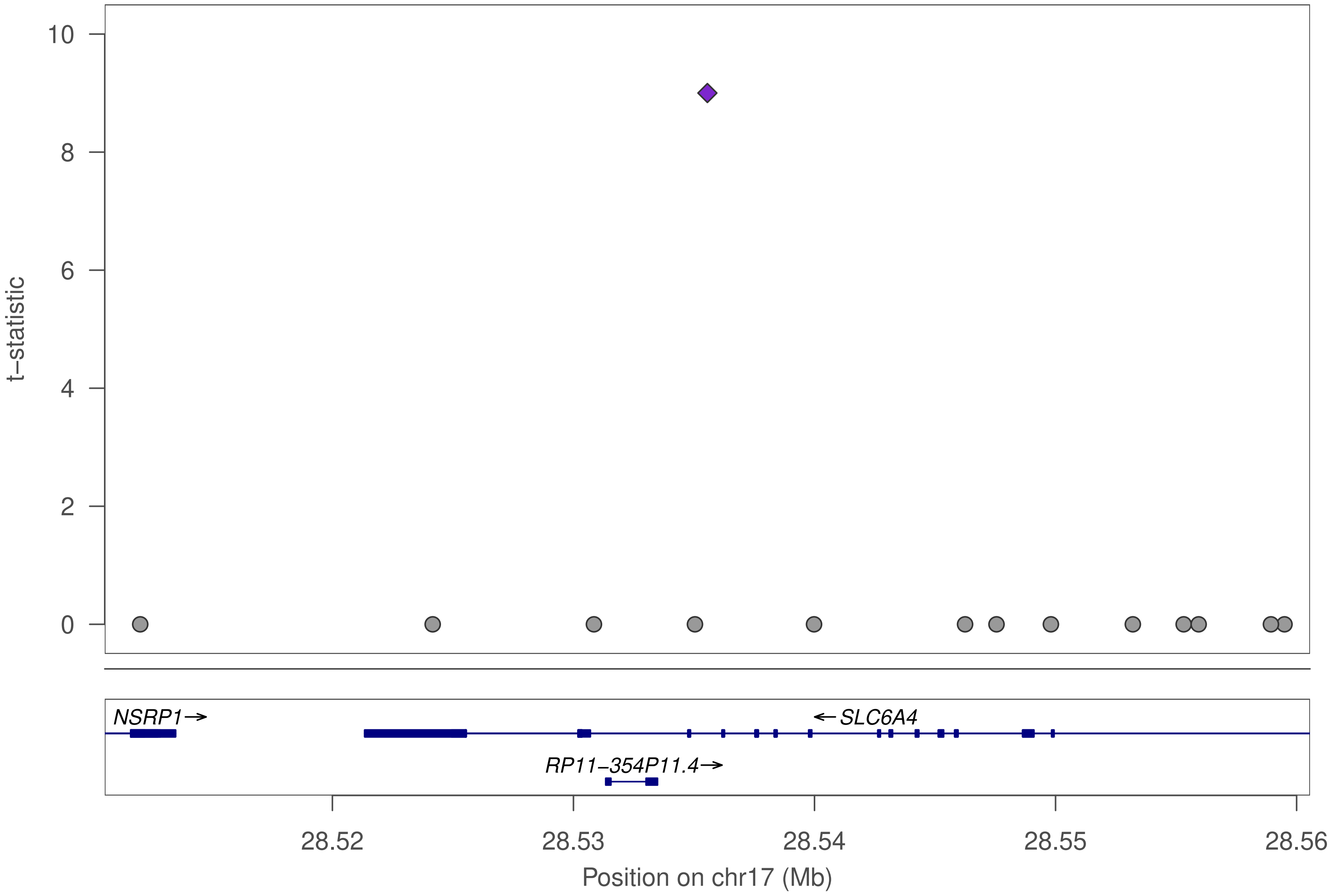
10
8
6
4
2
0
t−statistic
NSRP1→
←SLC6A4
RP11−354P11.4→
28.52
28.53
28.54
28.55
28.56
Position on chr17 (Mb)

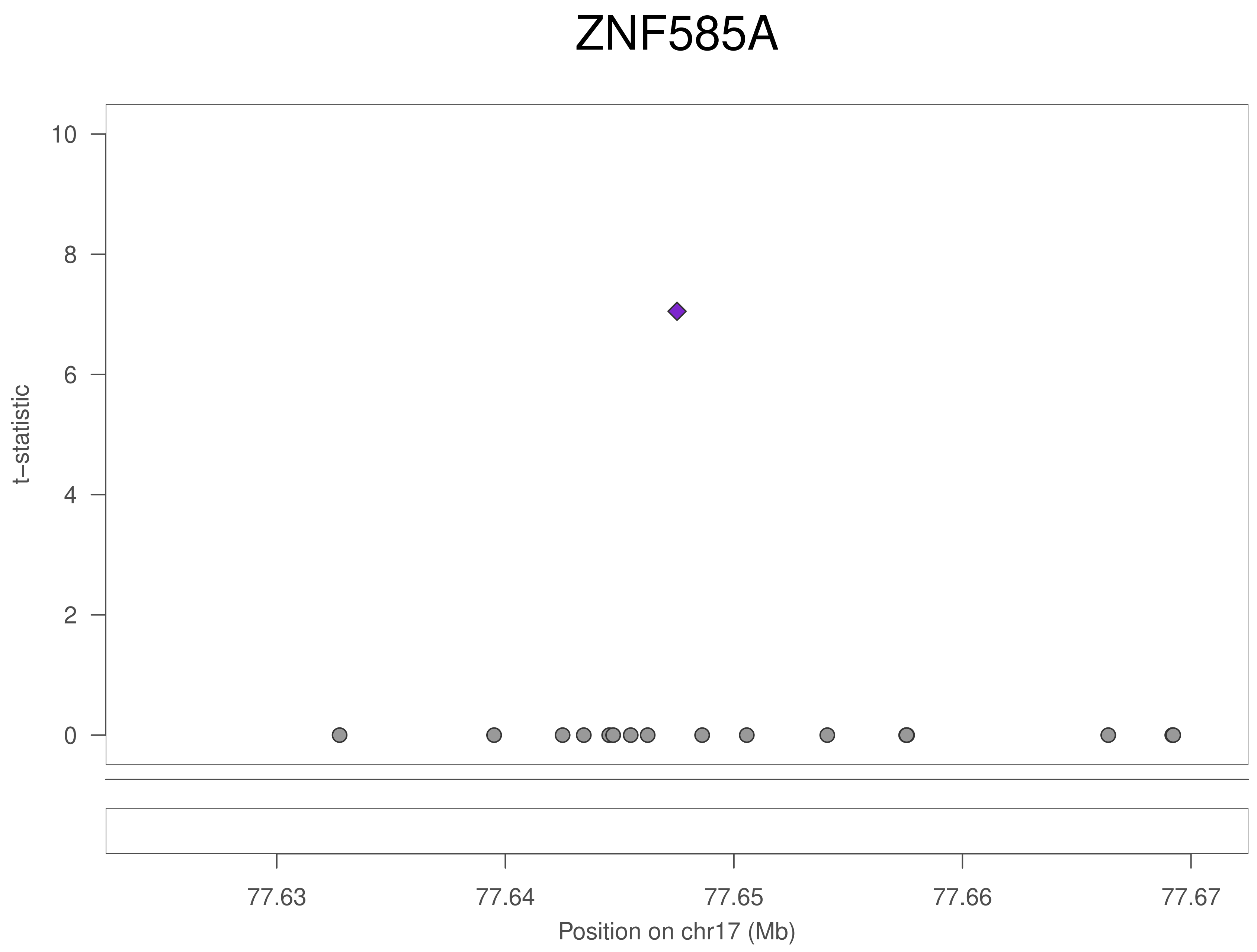
ZNF585A
t−statistic
0
2
4
6
8
10
77.63
77.64
77.65
77.66
77.67
Position on chr17 (Mb)

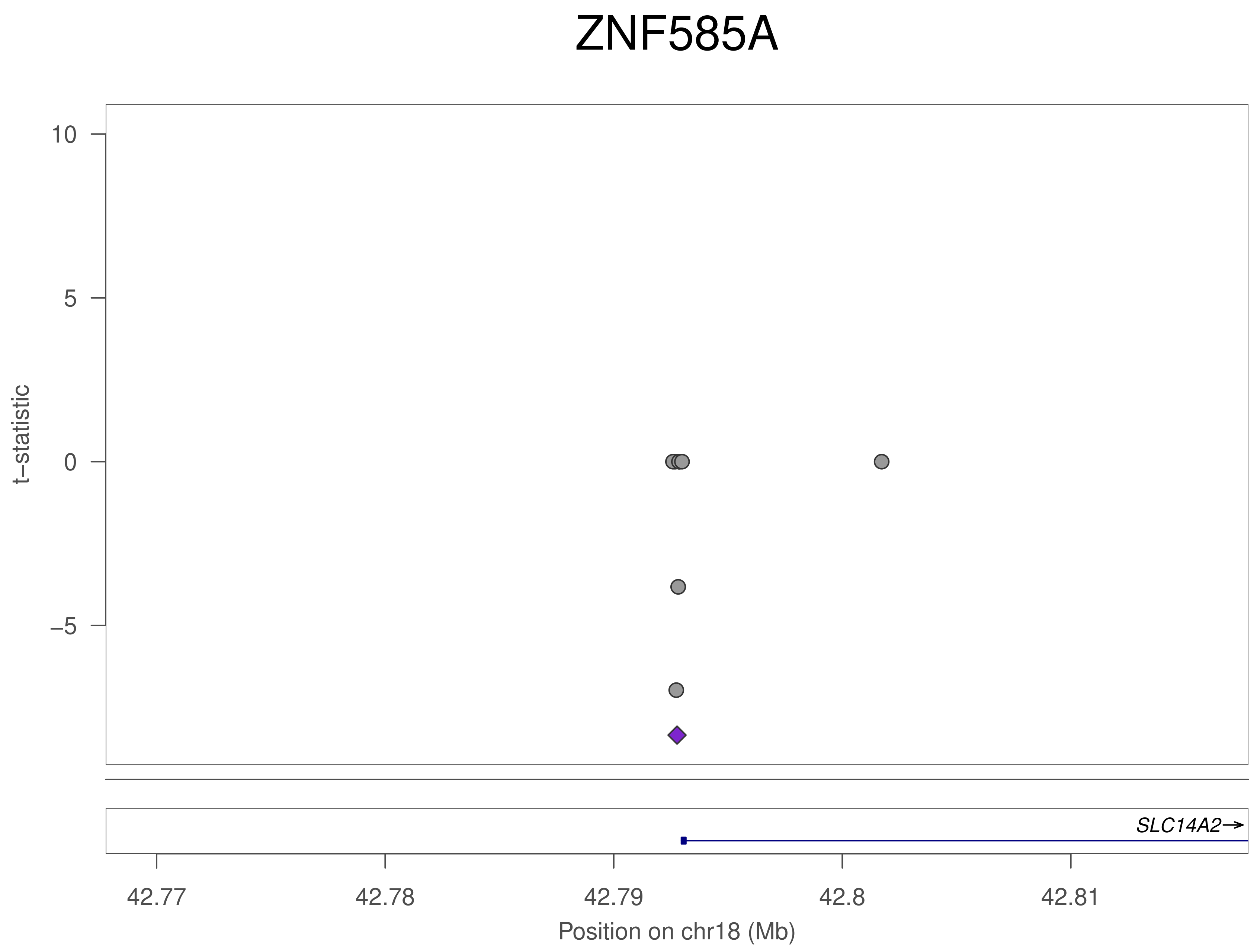

ZNF585A
10
5
0
−5
t−statistic
SLC14A2→
42.77
42.78
42.79
42.8
42.81
Position on chr18 (Mb)

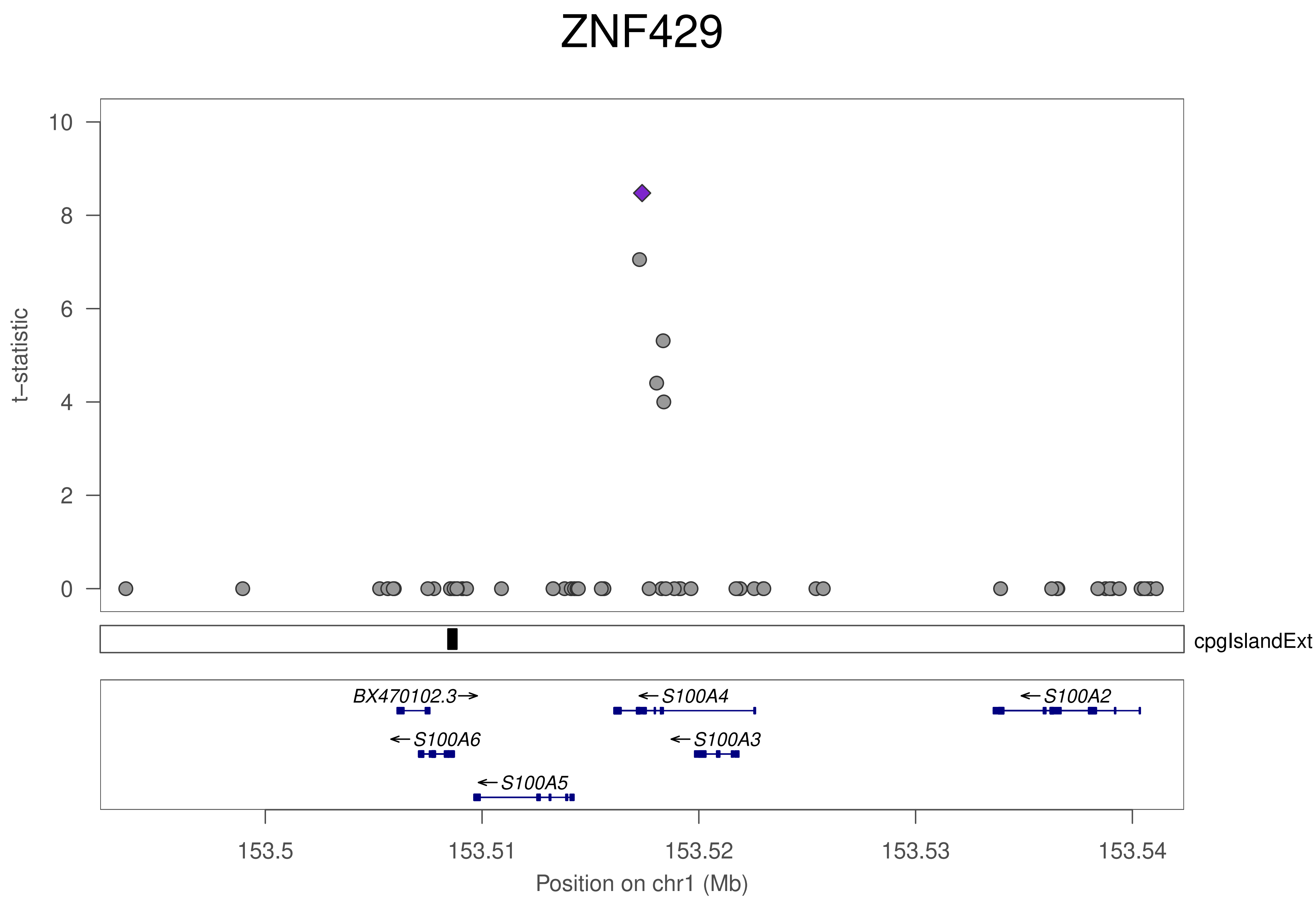

ZNF429
t–statistic
10
8
6
4
2
0
cpgIslandExt
BX470102.3→
←S100A4
←S100A2
←S100A6
←S100A3
←S100A5
153.5
153.51
153.52
153.53
153.54
Position on chr1 (Mb)

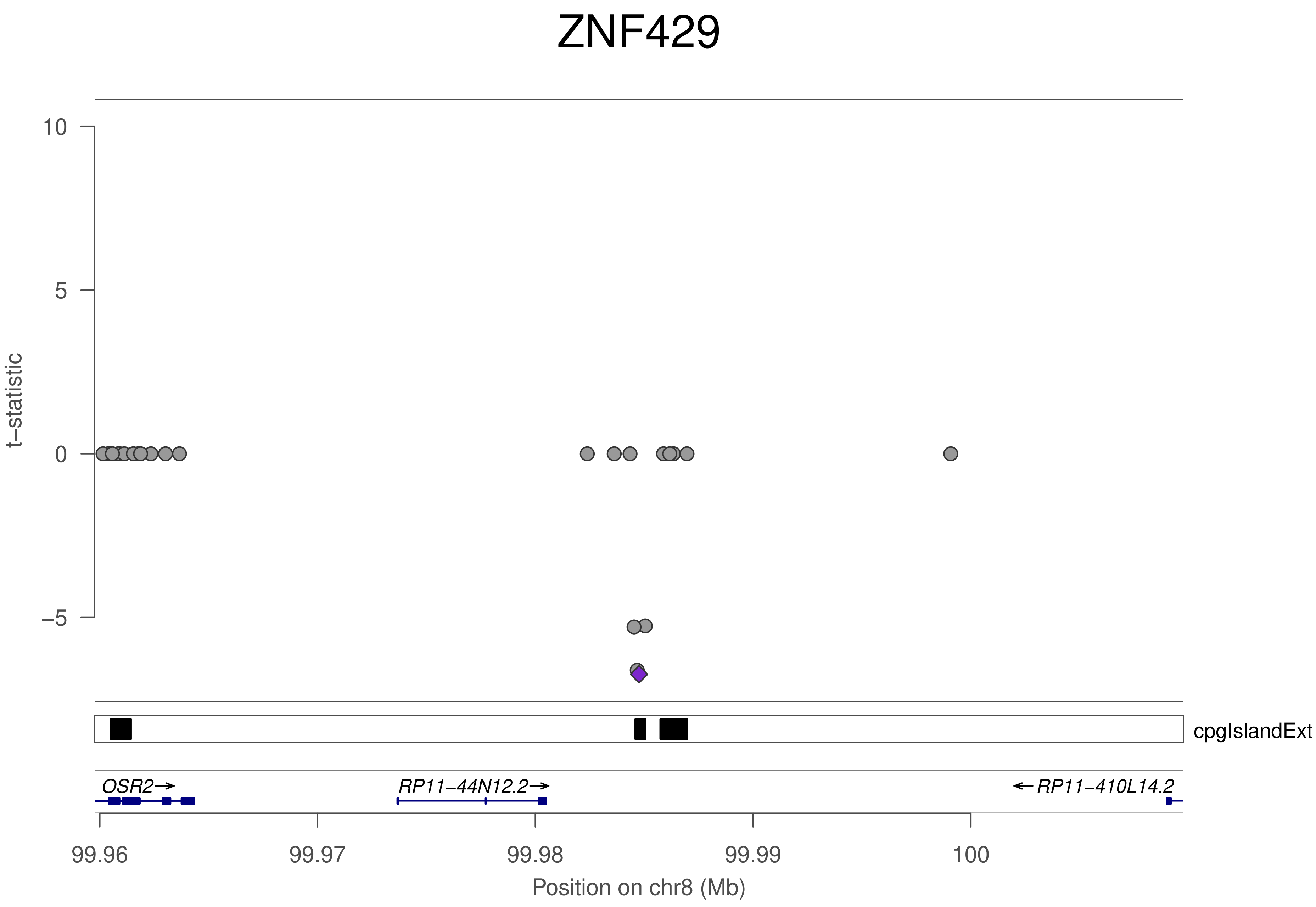
ZNF429
t–statistic
10
5
0
−5
cpgIslandExt
OSR2→
RP11−44N12.2→
←RP11−410L14.2
99.96
99.97
99.98
99.99
100
Position on chr8 (Mb)

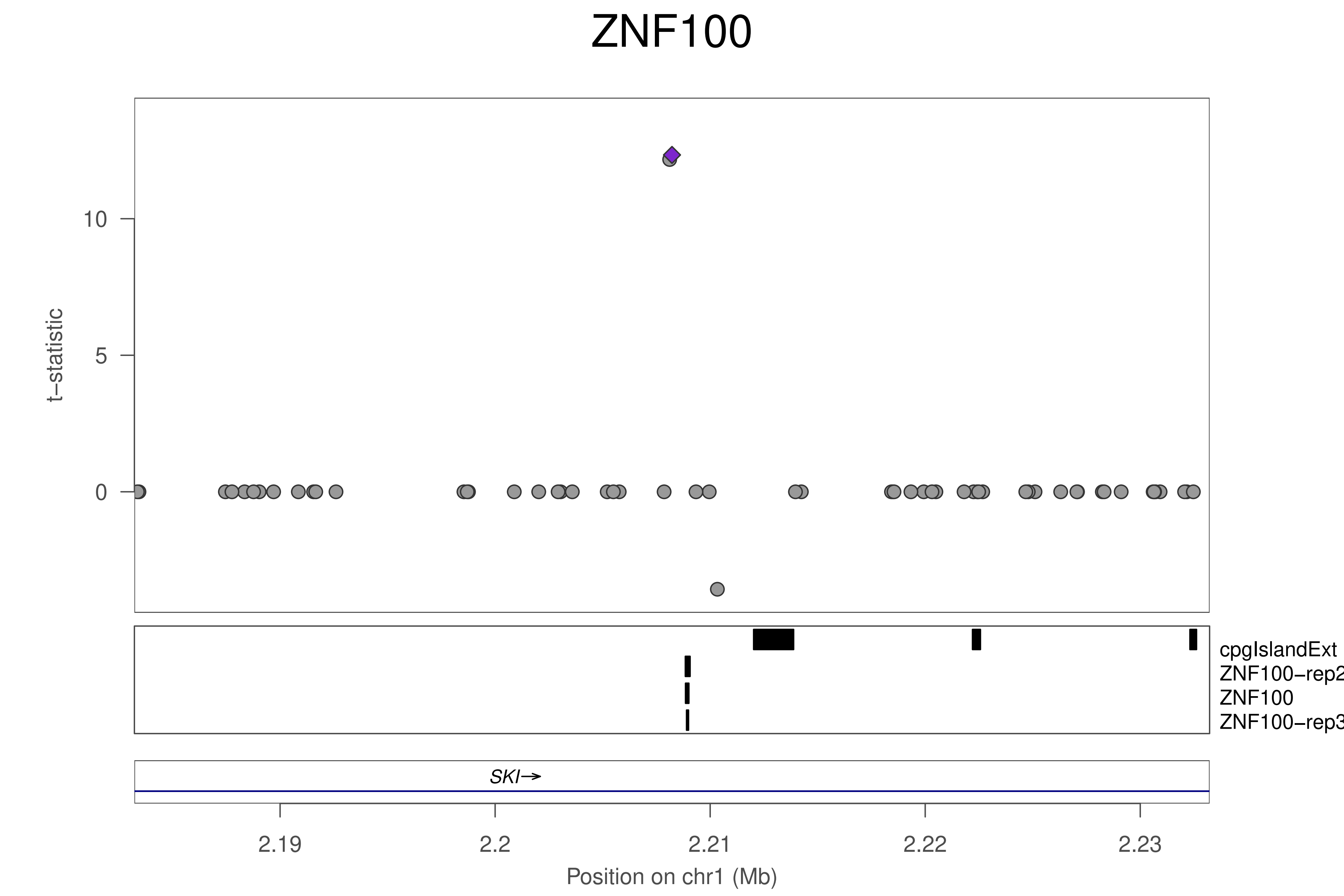

ZNF100
t−statistic
10
5
0
cpgIslandExt
ZNF100−rep2
ZNF100
ZNF100−rep3
SKI→
2.19
2.2
2.21
2.22
2.23
Position on chr1 (Mb)

### ZNF100

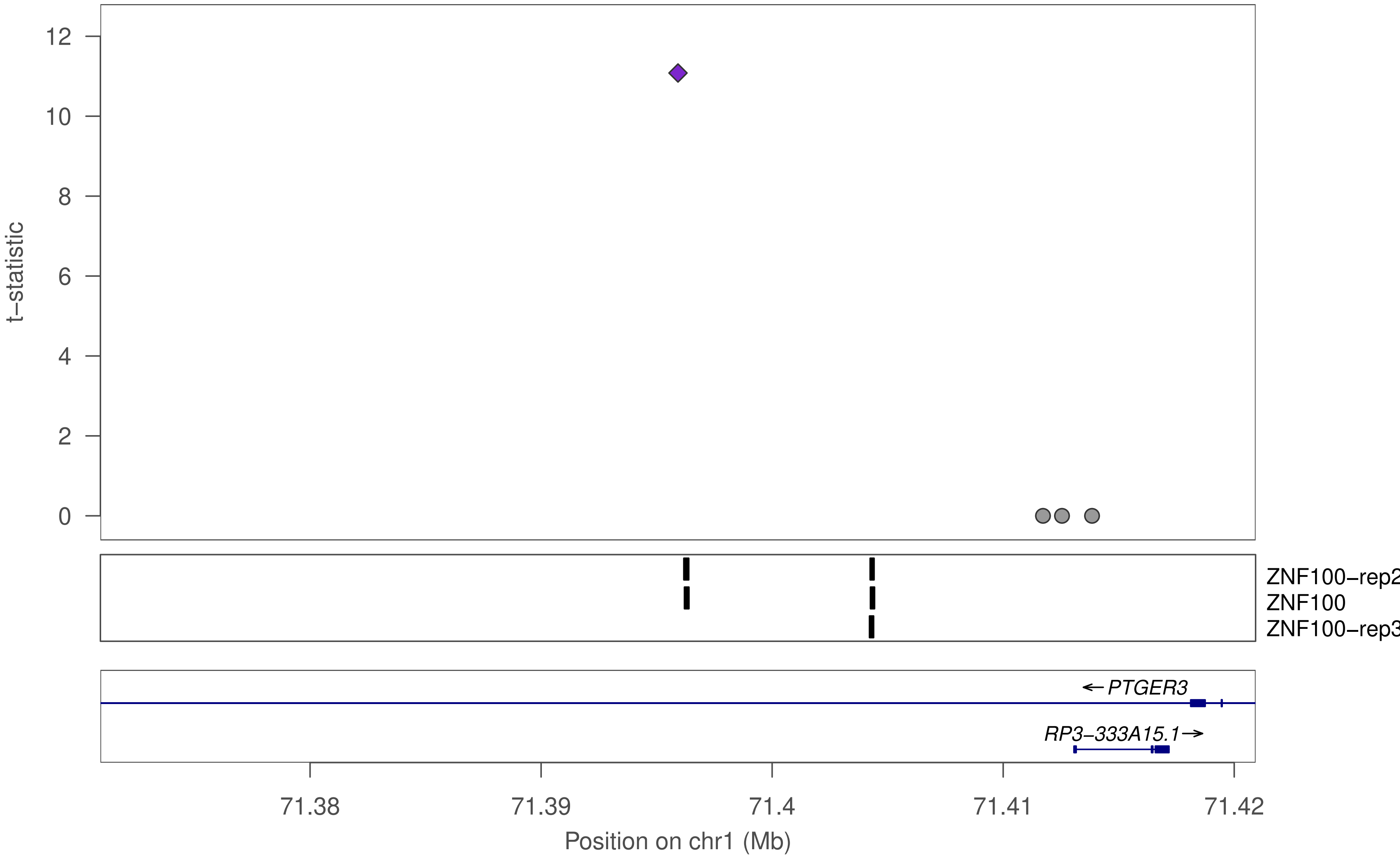

t−statistic
12
10
8
6
4
2
0
ZNF100−rep2
ZNF100
ZNF100−rep3
PTGER3
RP3−333A15.1
71.38
71.39
71.4
71.41
71.42
Position on chr1 (Mb)

### ZNF100

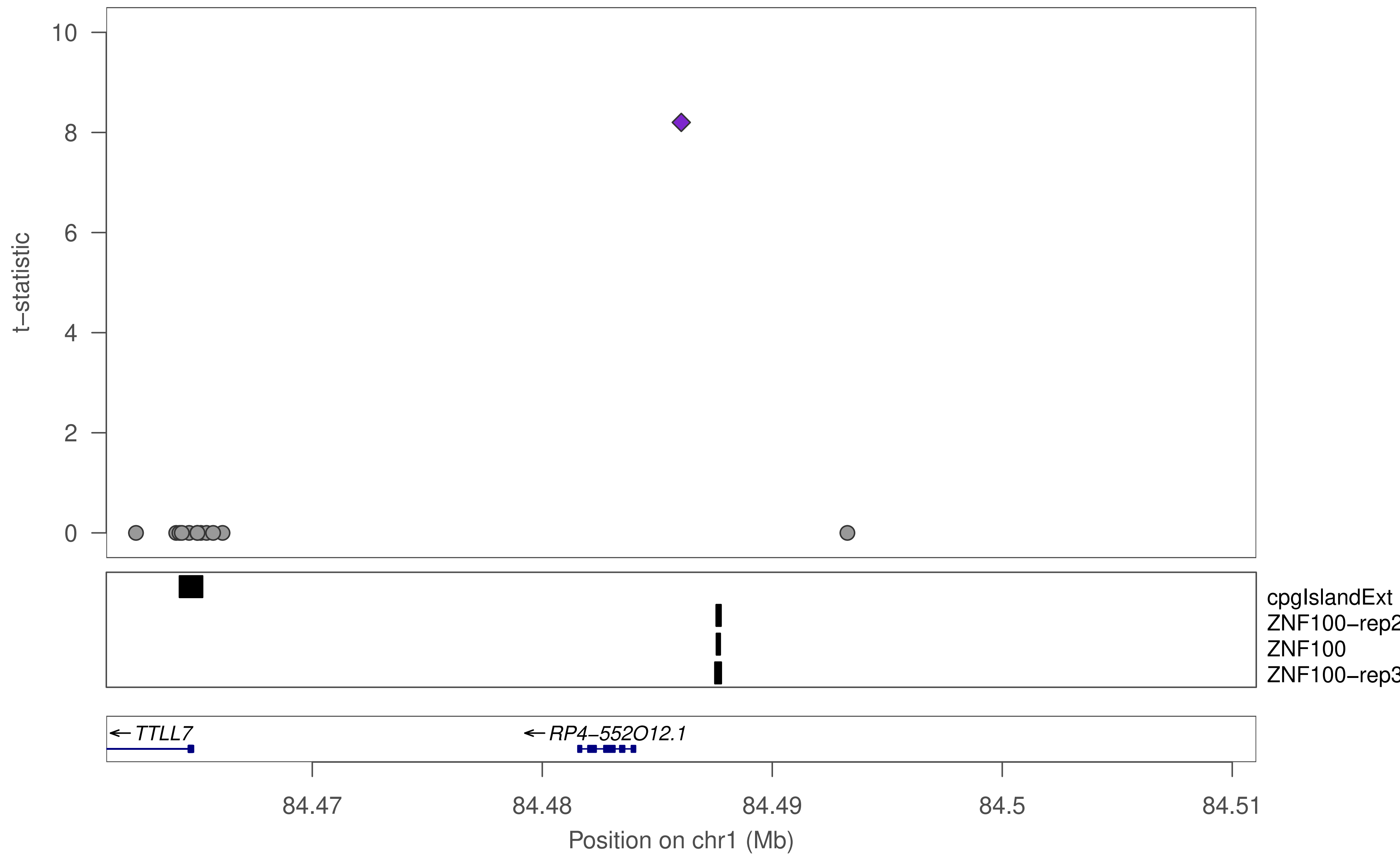

10
8
6
4
2
0
t−statistic
cpgIslandExt
ZNF100−rep2
ZNF100
ZNF100−rep3
← TTLL7
← RP4−552O12.1
84.47
84.48
84.49
84.5
84.51
Position on chr1 (Mb)

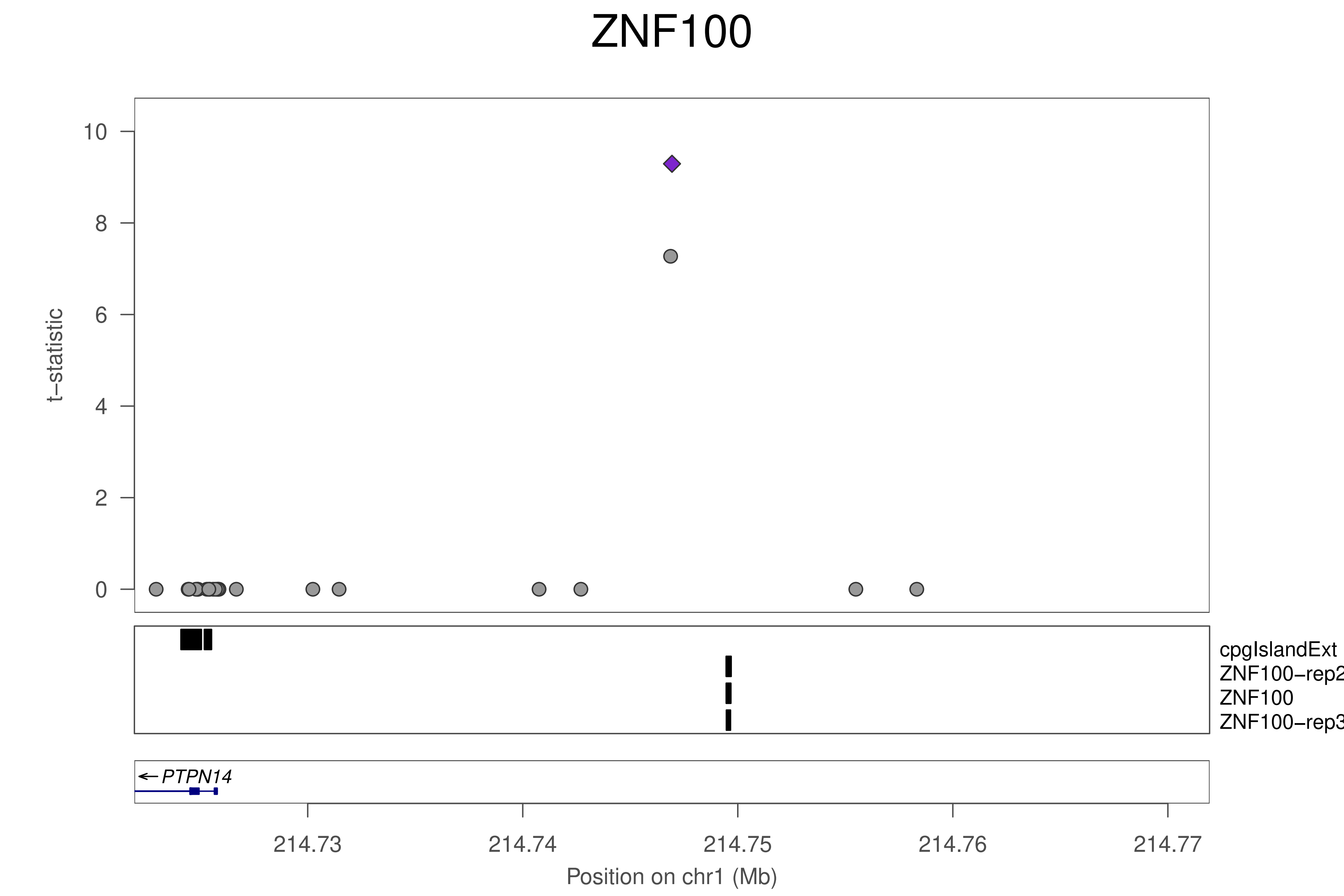

ZNF100
t−statistic
10
8
6
4
2
0
cpgIslandExt
ZNF100−rep2
ZNF100
ZNF100−rep3
← PTPN14
214.73
214.74
214.75
214.76
214.77
Position on chr1 (Mb)

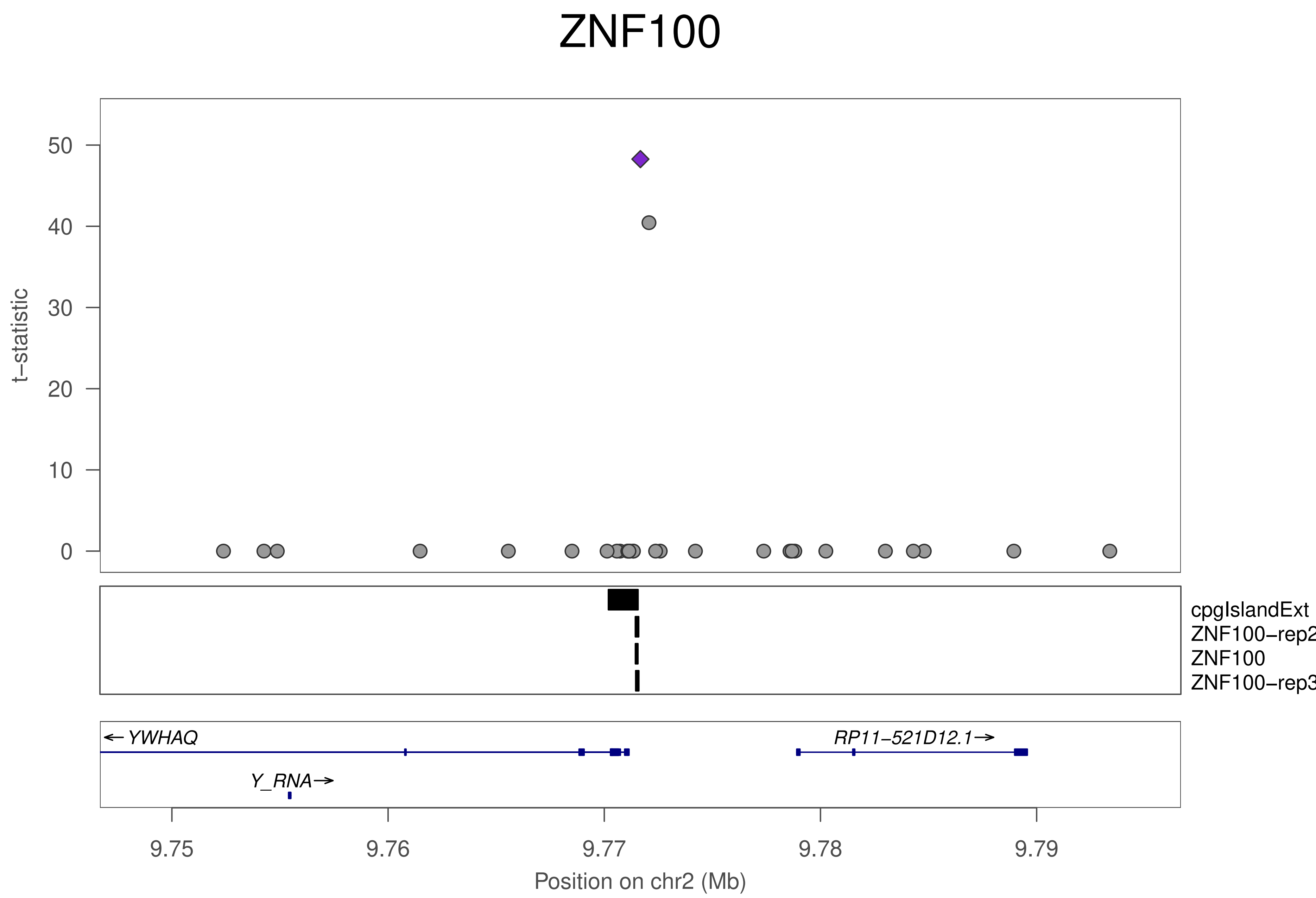

ZNF100
t−statistic
0
10
20
30
40
50
cpgIslandExt
ZNF100−rep2
ZNF100
ZNF100−rep3
← YWHAQ
RP11−521D12.1→
Y_RNA→
9.75
9.76
9.77
9.78
9.79
Position on chr2 (Mb)

### ZNF100

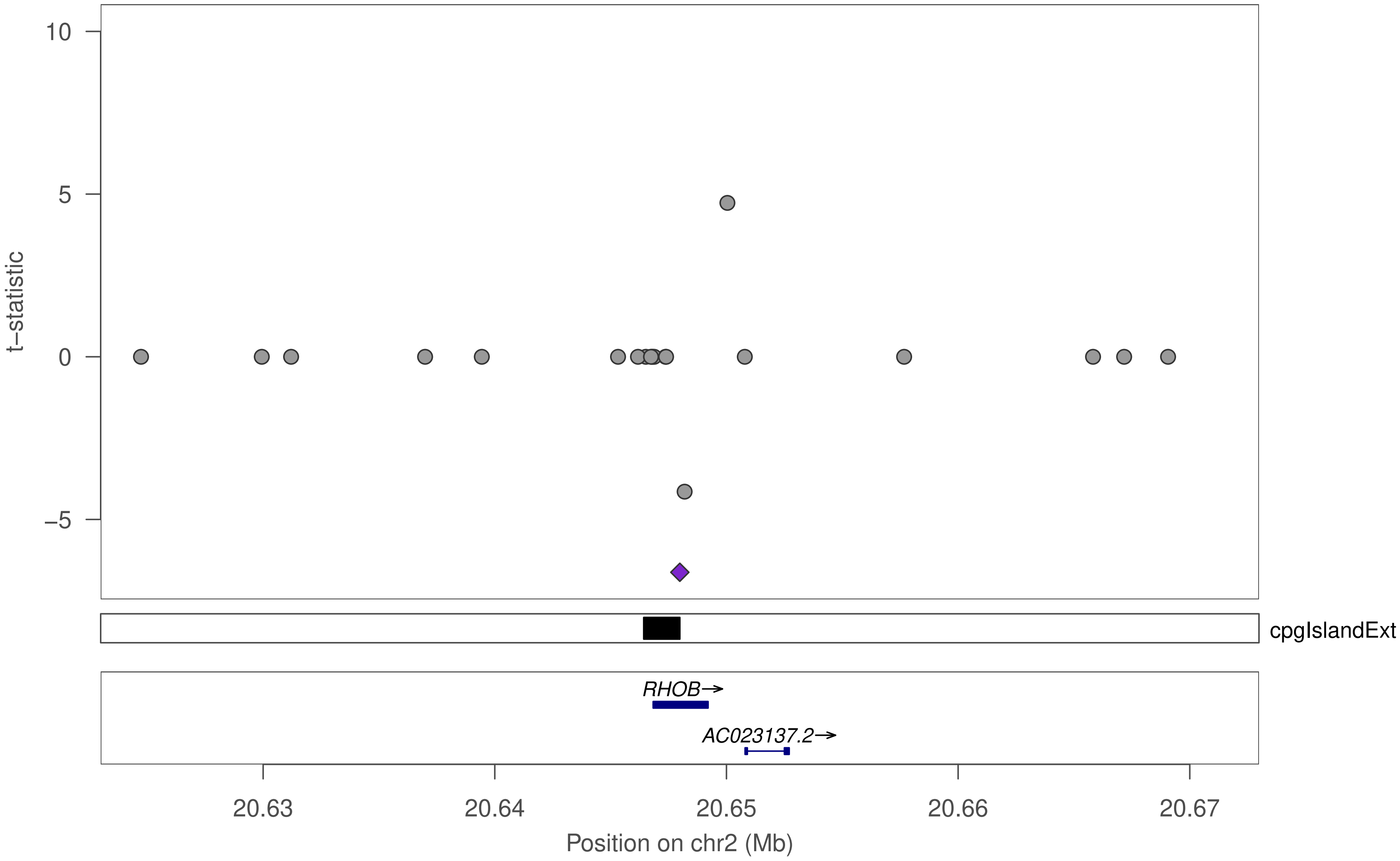

10
5
0
−5
t−statistic
cpgIslandExt
RHOB→
AC023137.2→
20.63
20.64
20.65
20.66
20.67
Position on chr2 (Mb)

### ZNF100

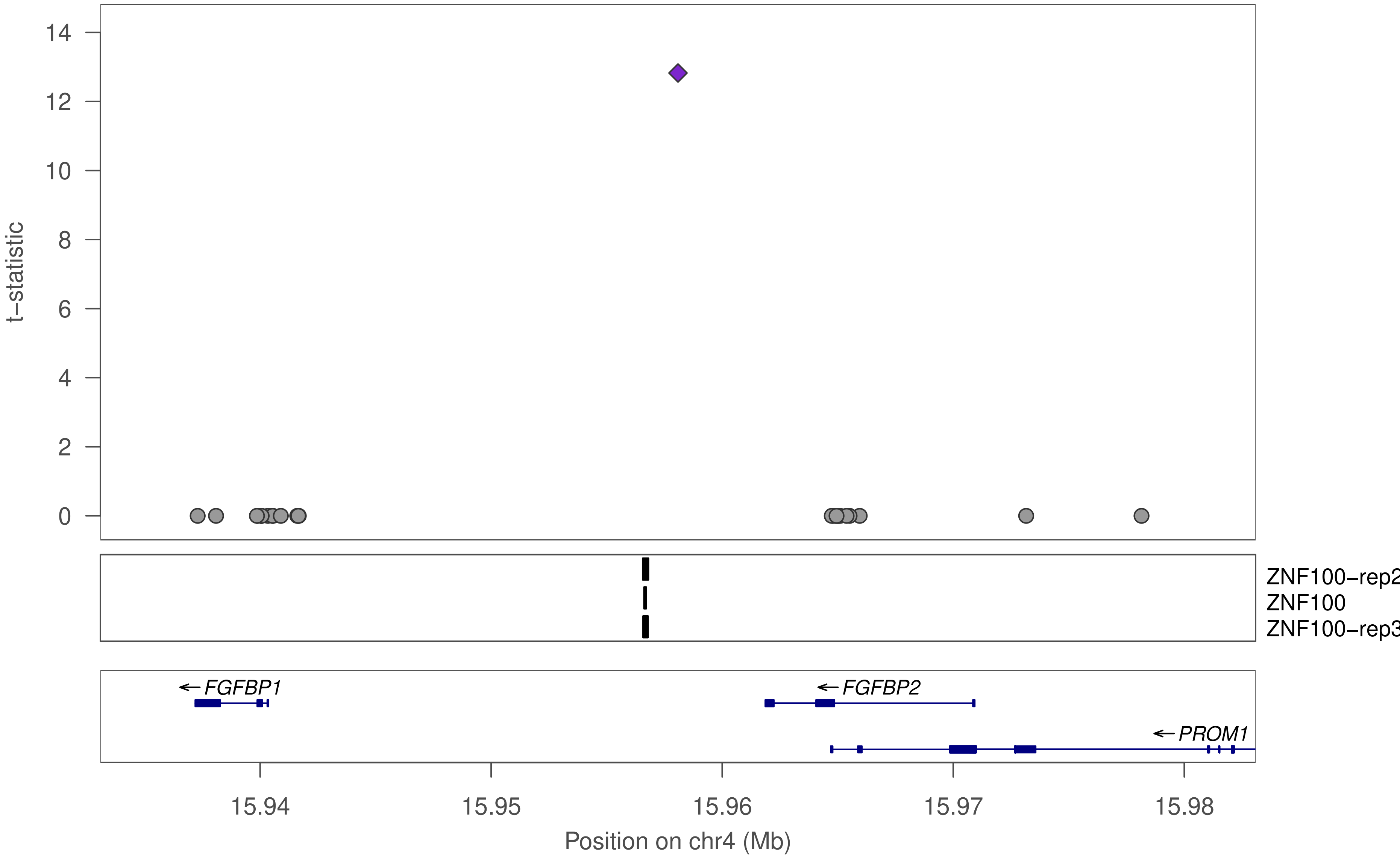

t−statistic
0
2
4
6
8
10
12
14
ZNF100−rep2
ZNF100
ZNF100−rep3
FGFBP1
FGFBP2
PROM1
15.94
15.95
15.96
15.97
15.98
Position on chr4 (Mb)

### ZNF100

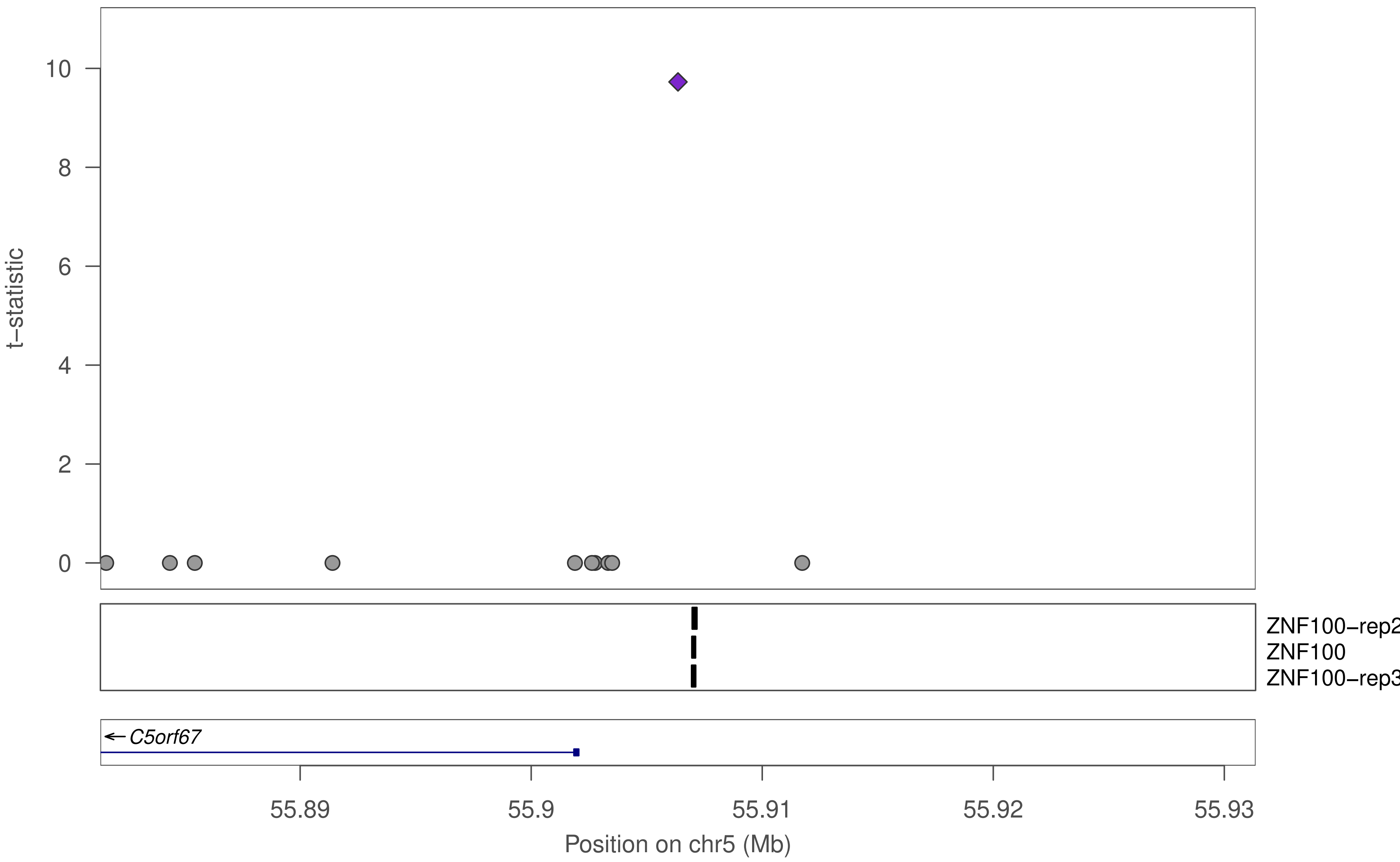

t−statistic
0
2
4
6
8
10
ZNF100−rep2
ZNF100
ZNF100−rep3
← C5orf67
55.89
55.9
55.91
55.92
55.93
Position on chr5 (Mb)

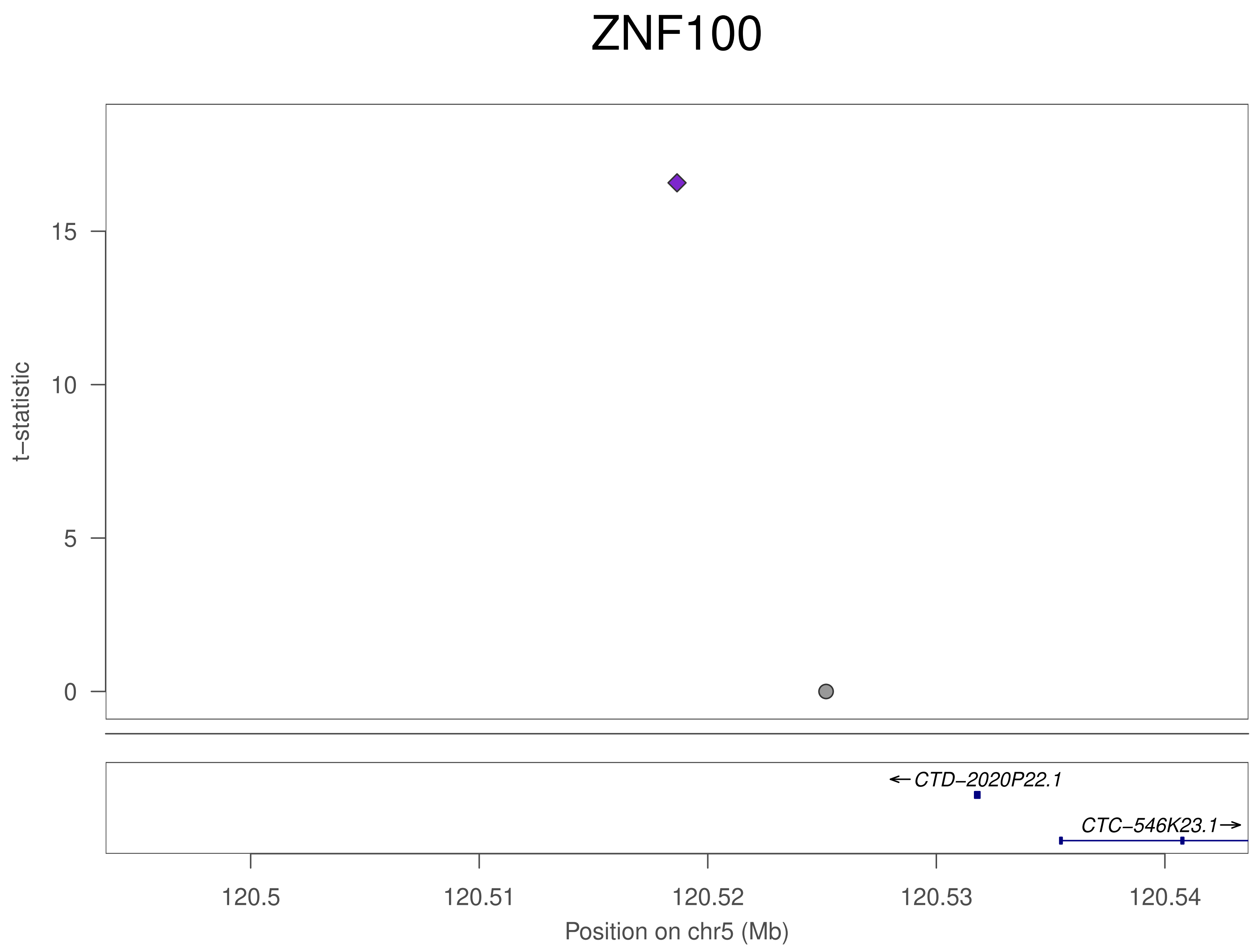
ZNF100
t−statistic
15
10
5
0
←CTD−2020P22.1
CTC−546K23.1→
120.5
120.51
120.52
120.53
120.54
Position on chr5 (Mb)

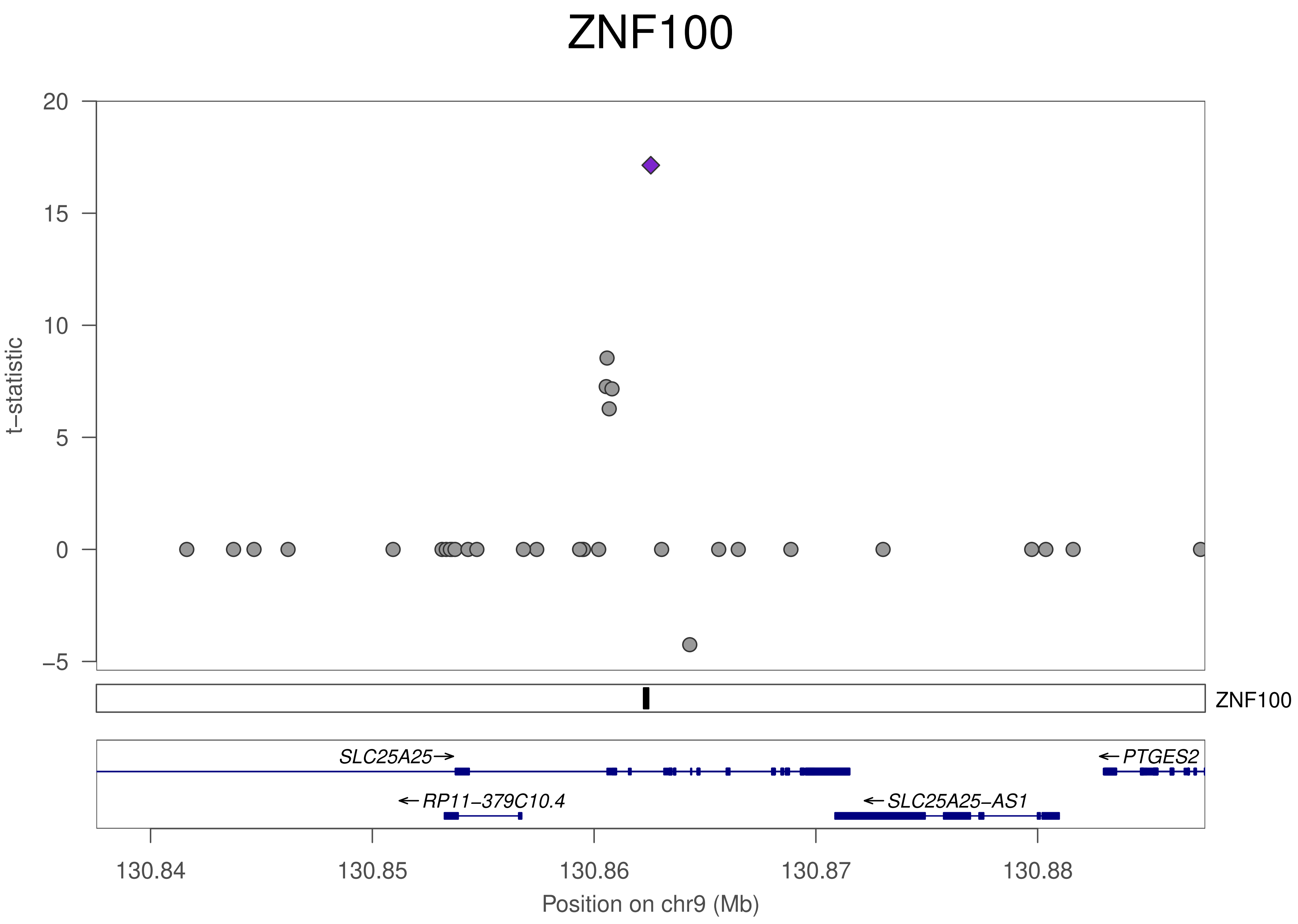

ZNF100
t−statistic
20
15
10
5
0
−5
ZNF100
SLC25A25→
←PTGES2
←RP11−379C10.4
←SLC25A25−AS1
130.84
130.85
130.86
130.87
130.88
Position on chr9 (Mb)

### ZNF100

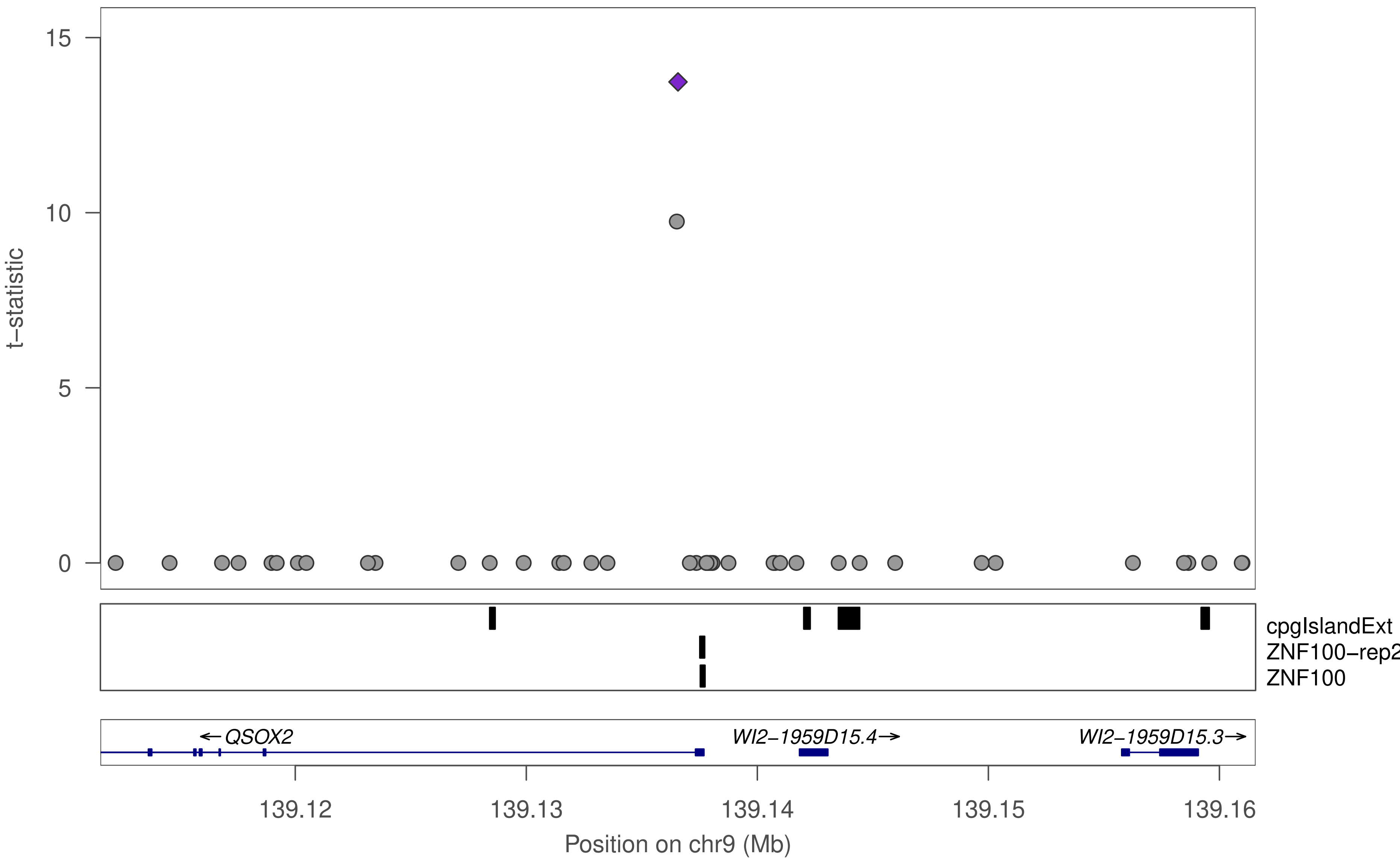

15
10
5
0
t−statistic
cpgIslandExt
ZNF100−rep2
ZNF100
←QSOX2
WI2−1959D15.4→
WI2−1959D15.3→
139.12
139.13
139.14
139.15
139.16
Position on chr9 (Mb)

### ZNF100

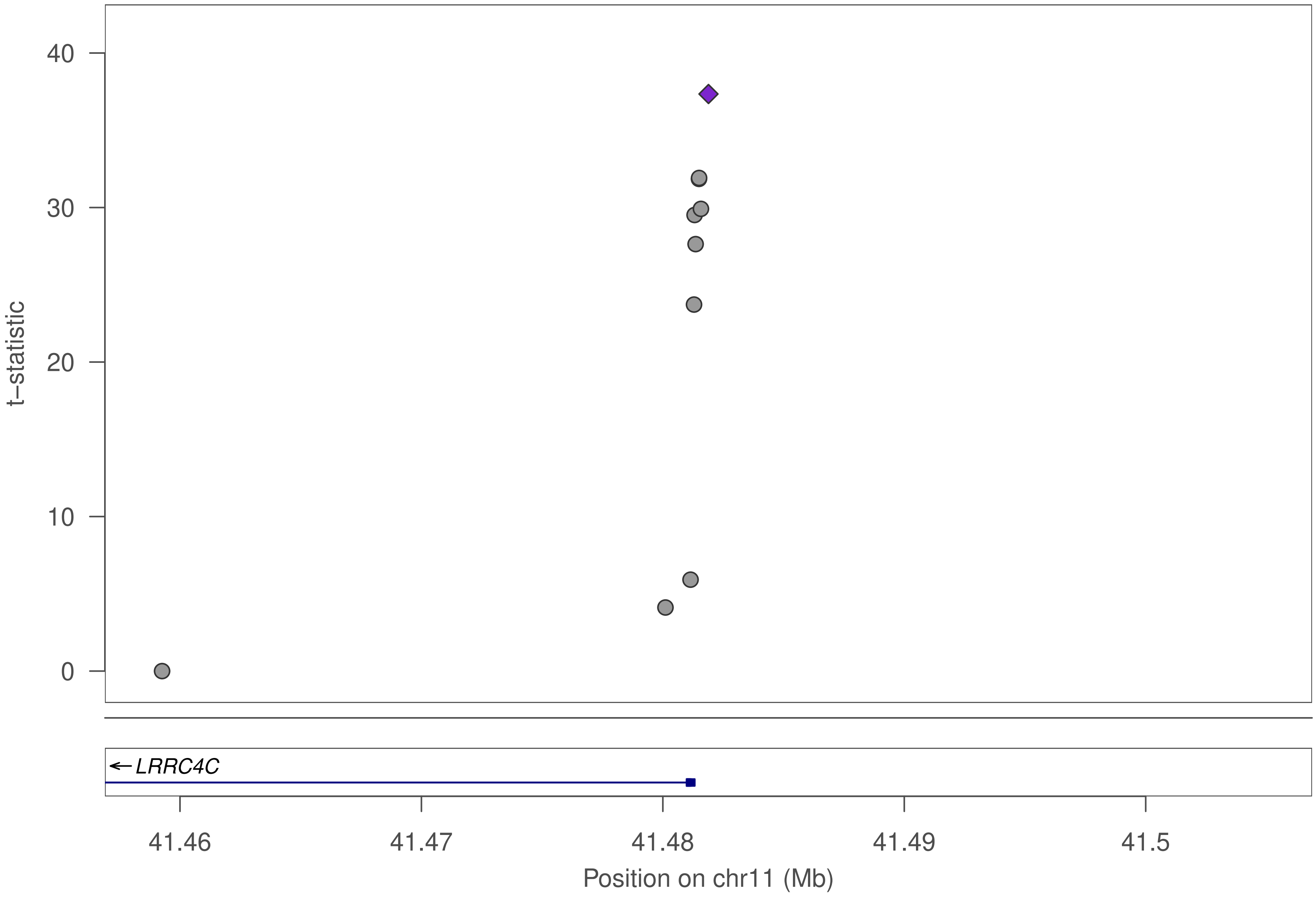

t–statistic
40
30
20
10
0
LRRC4C
41.46
41.47
41.48
41.49
41.5
Position on chr11 (Mb)

### ZNF100

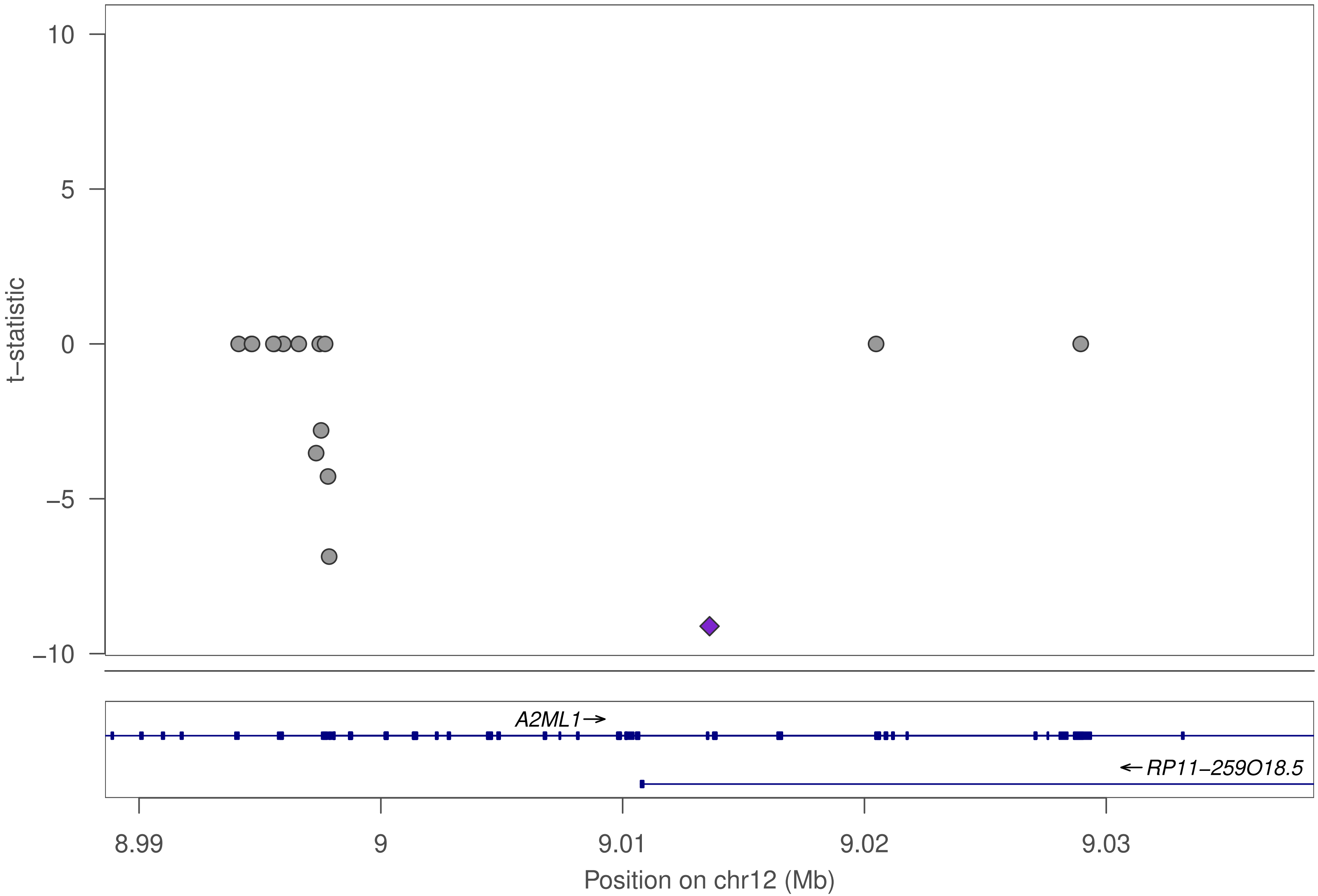

10
5
0
−5
−10
t−statistic
A2ML1→
←RP11−259O18.5
8.99
9
9.01
9.02
9.03
Position on chr12 (Mb)

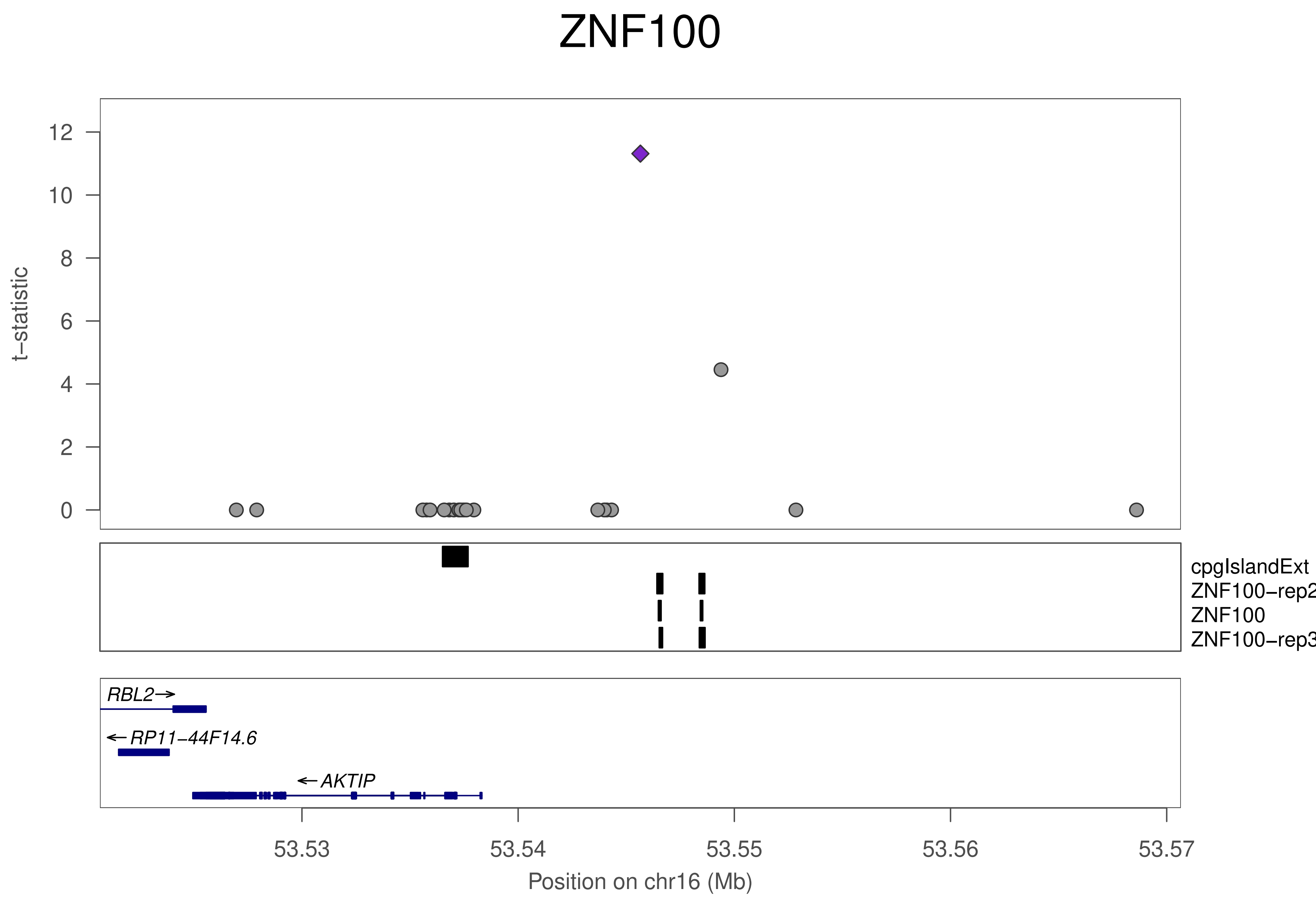
ZNF100
t−statistic
12
10
8
6
4
2
0
cpgIslandExt
ZNF100−rep2
ZNF100
ZNF100−rep3
RBL2
RP11−44F14.6
AKTIP
53.53
53.54
53.55
53.56
53.57
Position on chr16 (Mb)

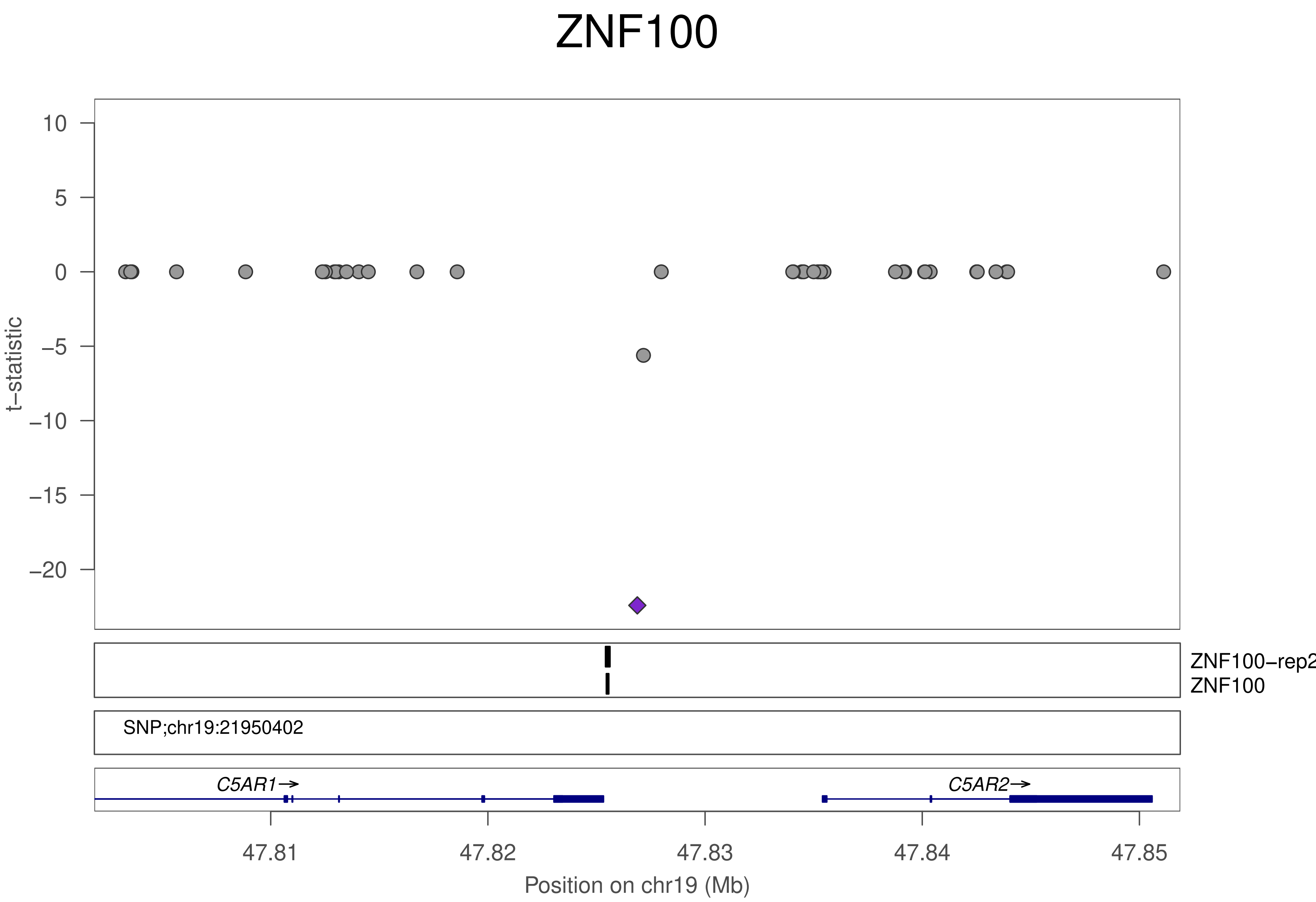
ZNF100
t−statistic
10
5
0
−5
−10
−15
−20
ZNF100−rep2
ZNF100
SNP;chr19:21950402
C5AR1→
C5AR2→
47.81
47.82
47.83
47.84
47.85
Position on chr19 (Mb)

### ZNF398

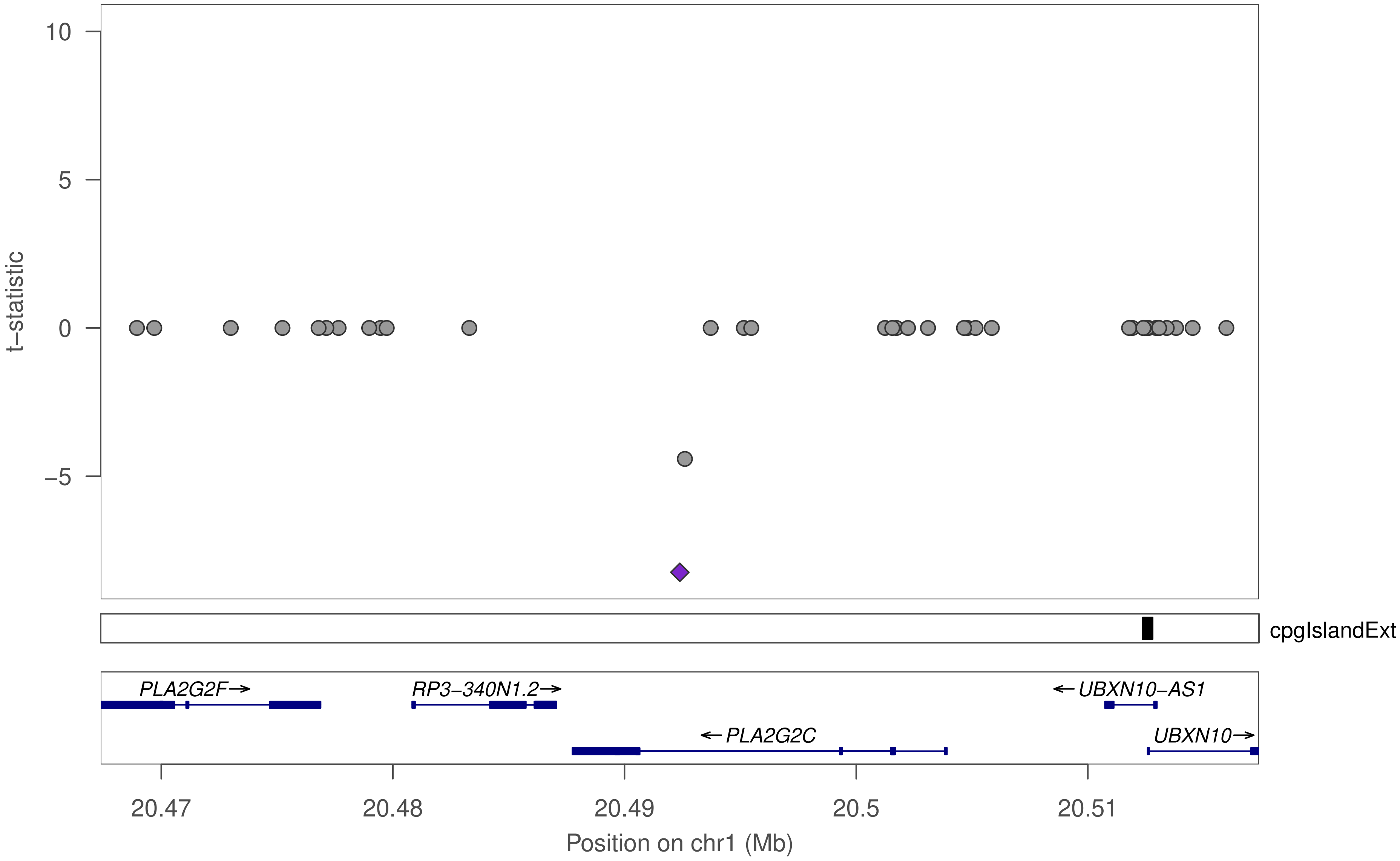

10
5
0
−5
t−statistic
cpgIslandExt
PLA2G2F→
RP3−340N1.2→
←UBXN10−AS1
←PLA2G2C
UBXN10→
20.47
20.48
20.49
20.5
20.51
Position on chr1 (Mb)

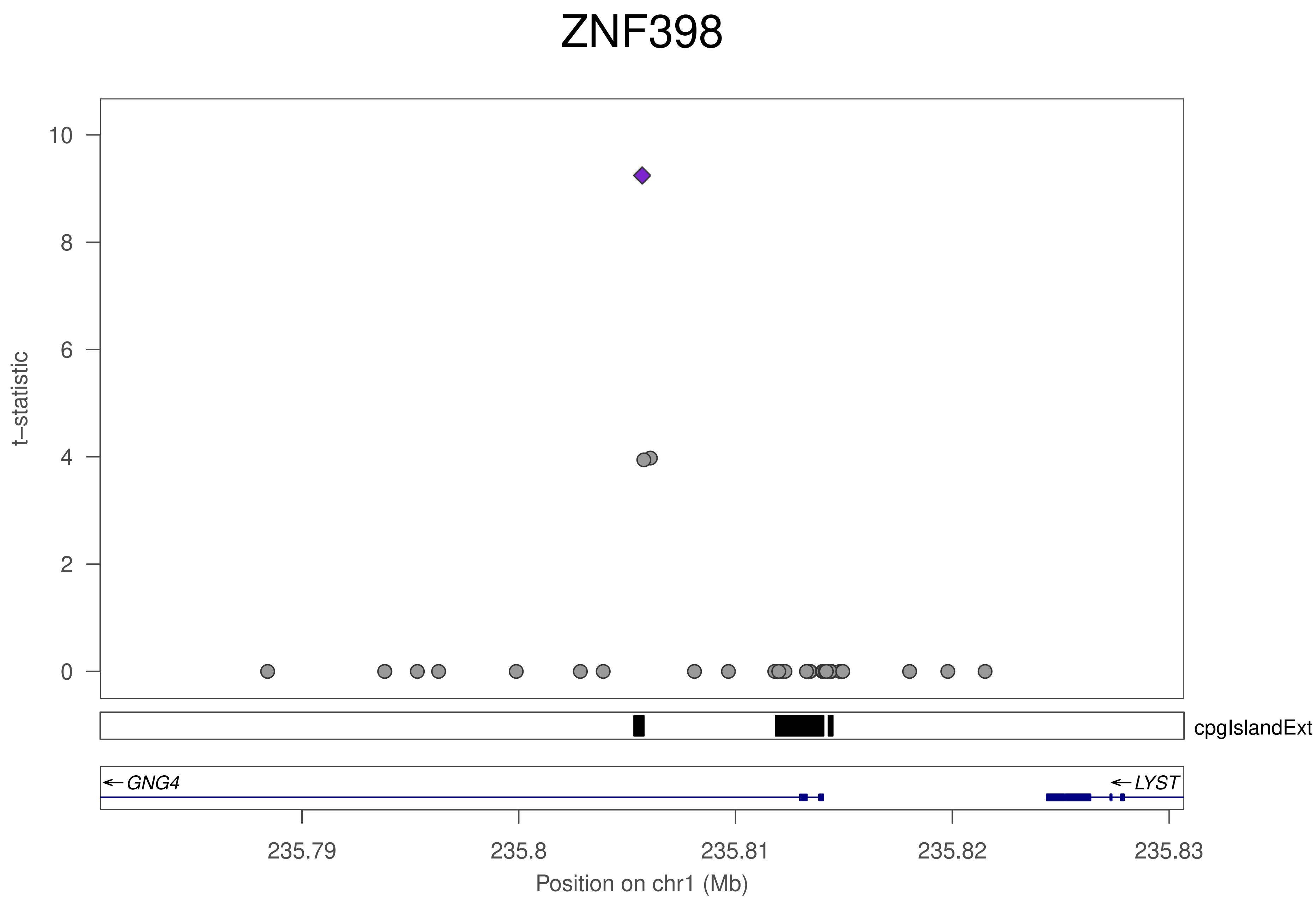
ZNF398
t−statistic
10
8
6
4
2
0
cpgIslandExt
GNG4
LYST
235.79
235.8
235.81
235.82
235.83
Position on chr1 (Mb)

### ZNF398

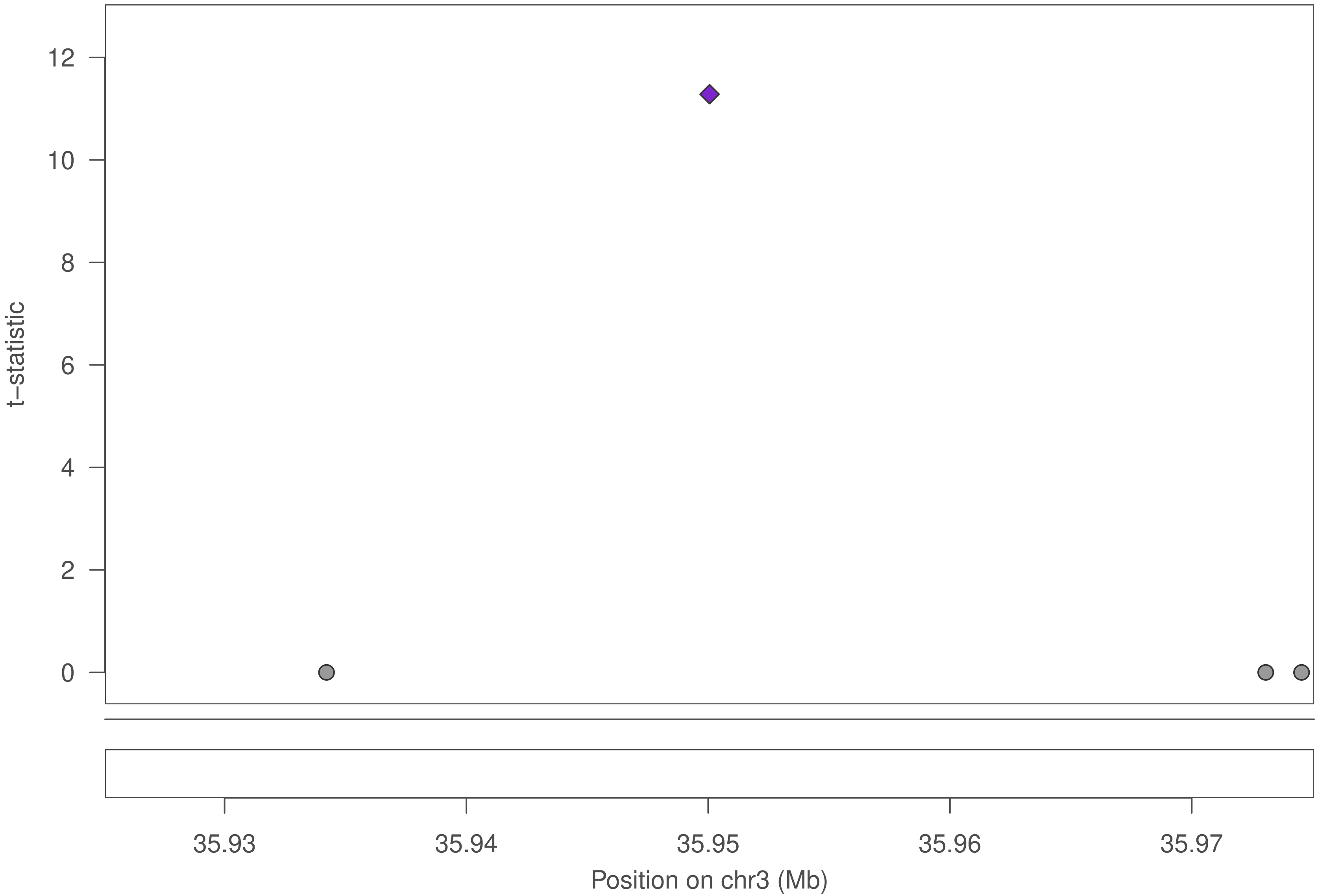

t–statistic
0
2
4
6
8
10
12
35.93
35.94
35.95
35.96
35.97
Position on chr3 (Mb)

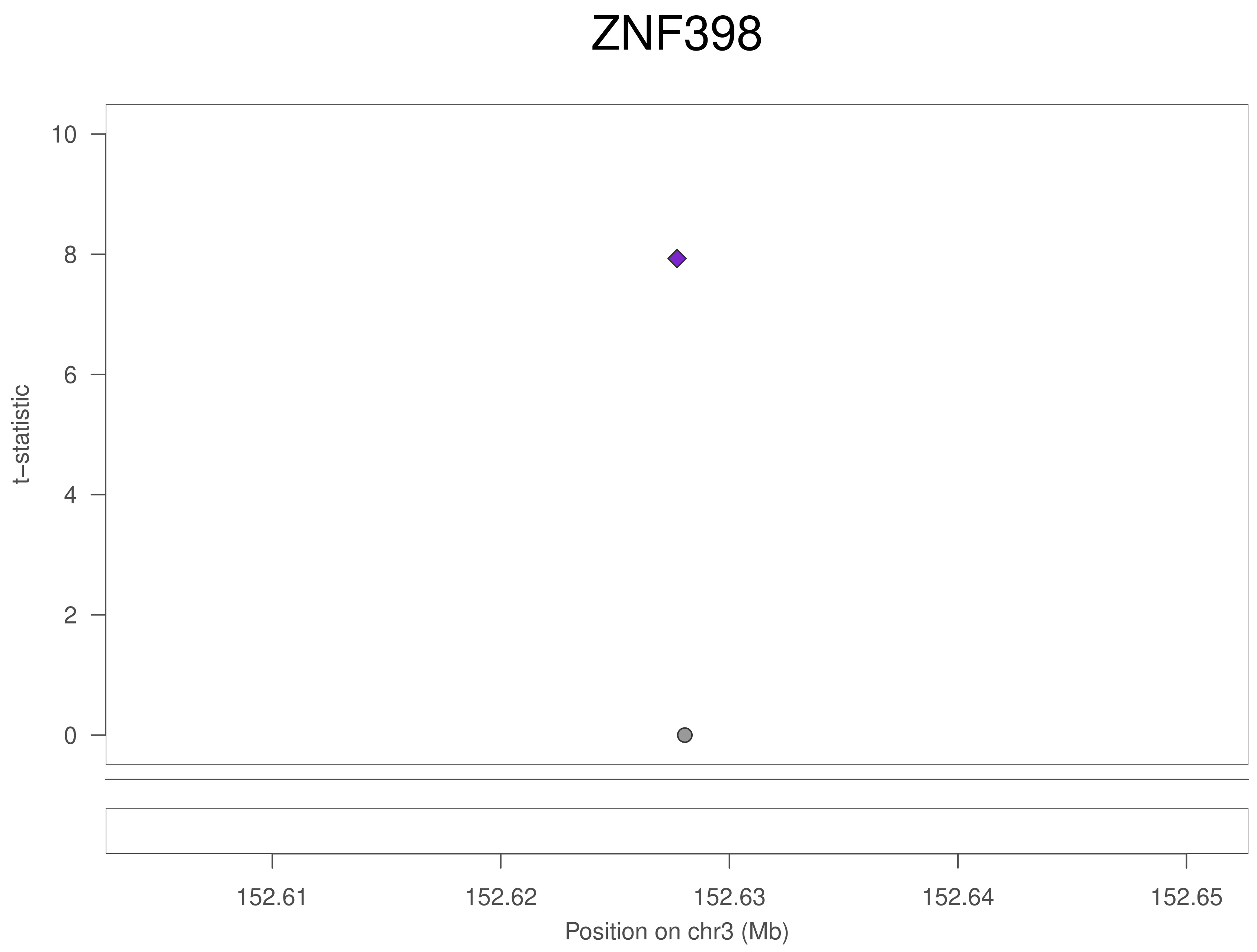
ZNF398
t–statistic
10
8
6
4
2
0
152.61
152.62
152.63
152.64
152.65
Position on chr3 (Mb)

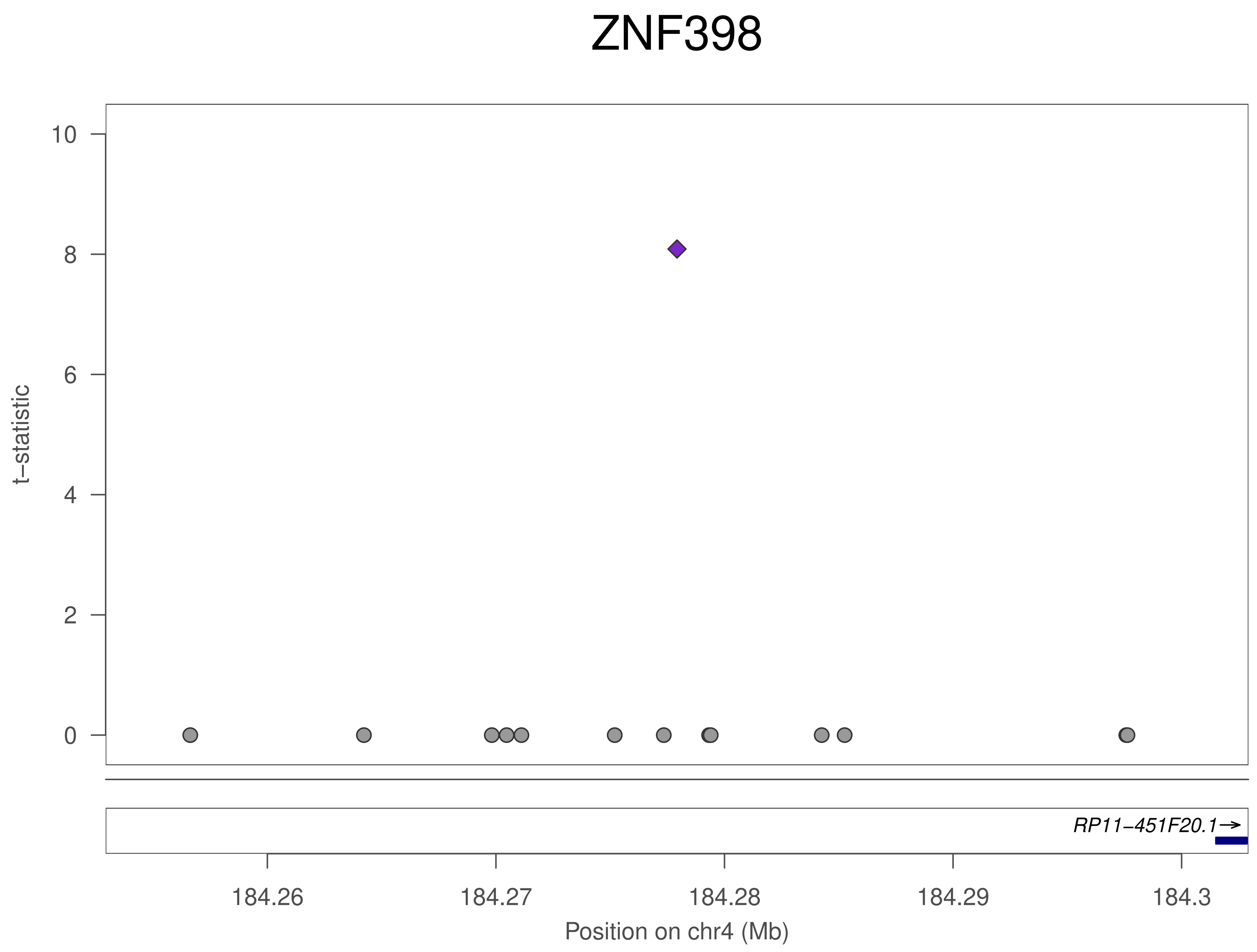

ZNF398
t−statistic
10
8
6
4
2
0
RP11−451F20.1→
184.26
184.27
184.28
184.29
184.3
Position on chr4 (Mb)

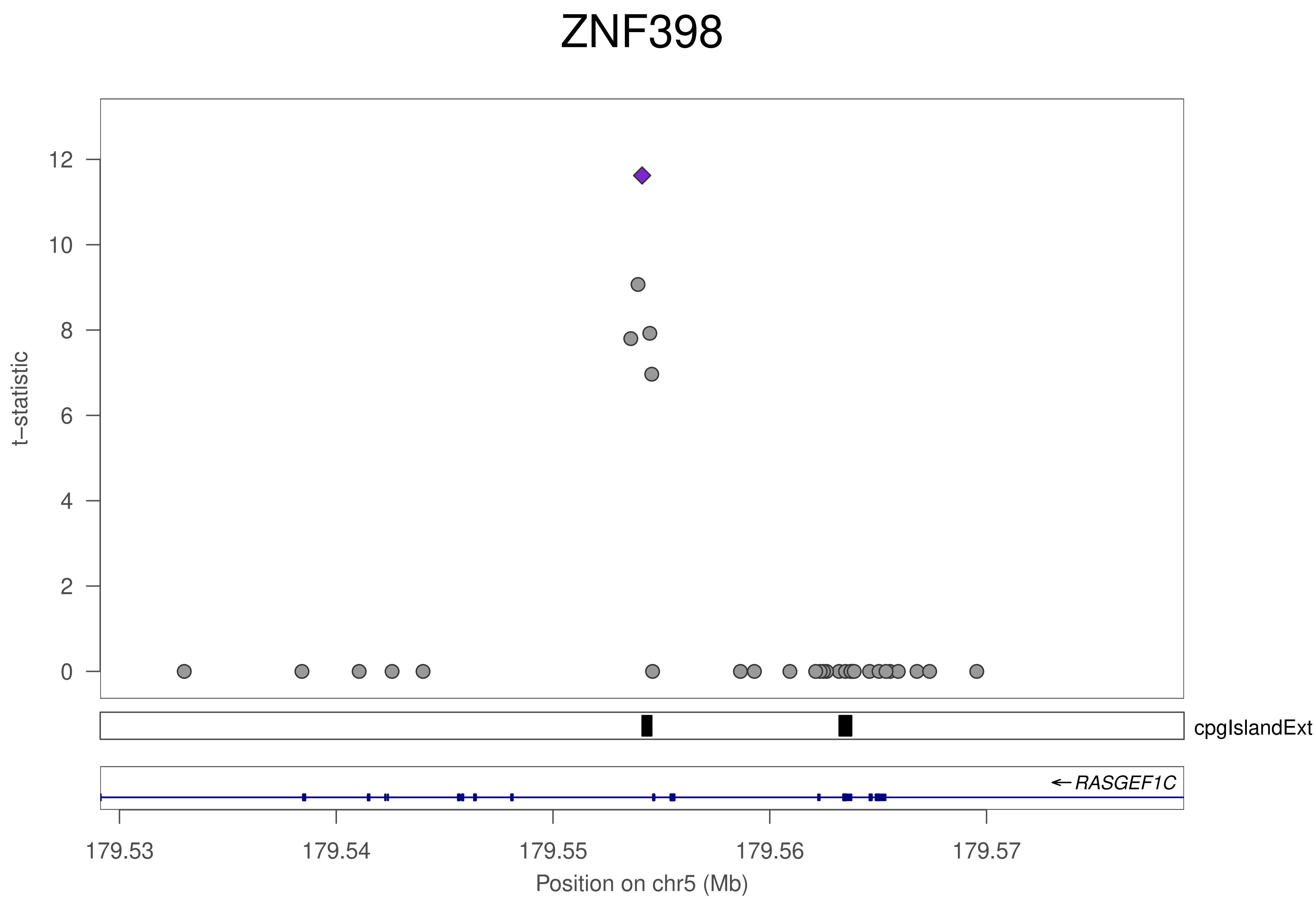

ZNF398
t–statistic
12
10
8
6
4
2
0
cpgIslandExt
← RASGEF1C
179.53
179.54
179.55
179.56
179.57
Position on chr5 (Mb)

### ZNF398

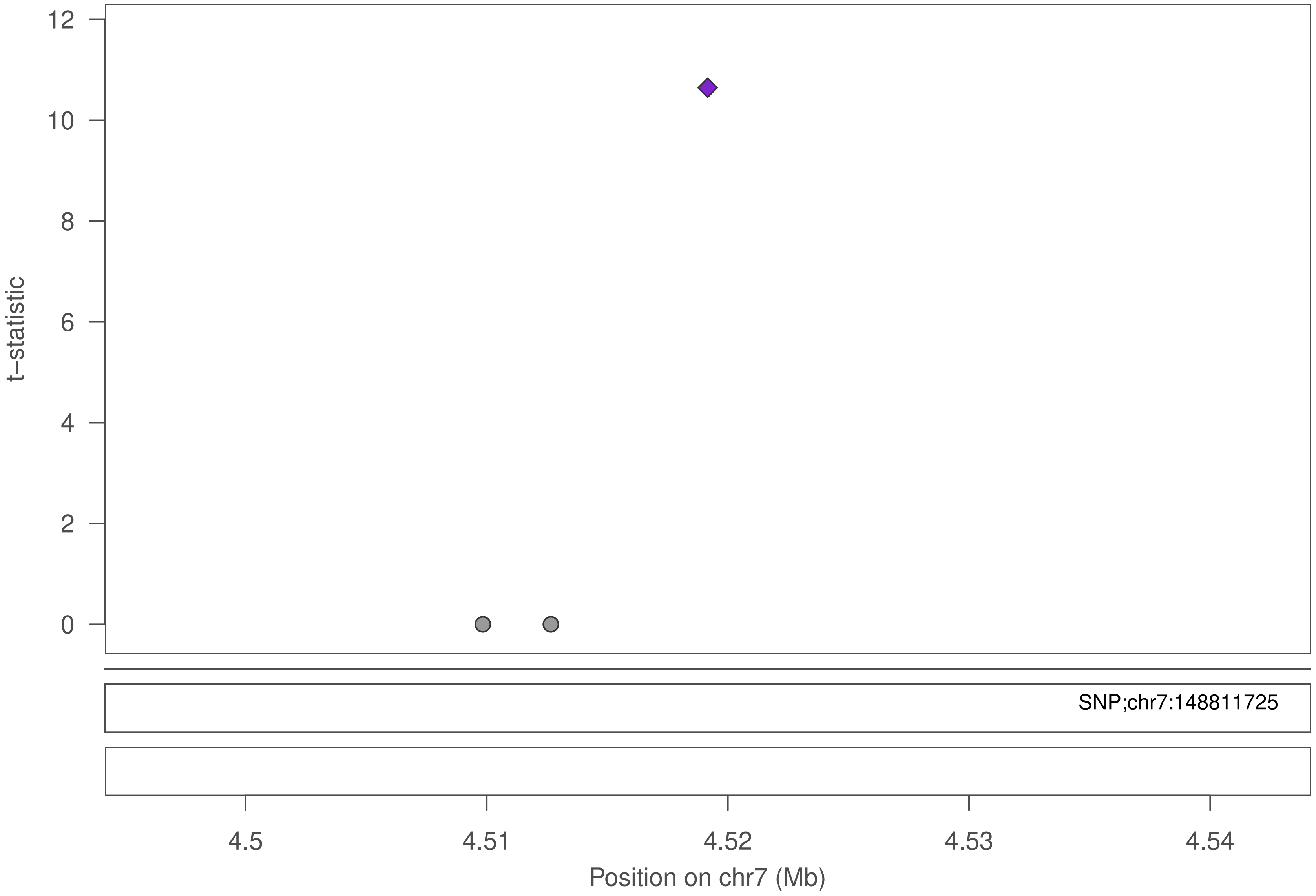

12
10
8
6
4
2
0
t−statistic
SNP;chr7:148811725
4.5
4.51
4.52
4.53
4.54
Position on chr7 (Mb)

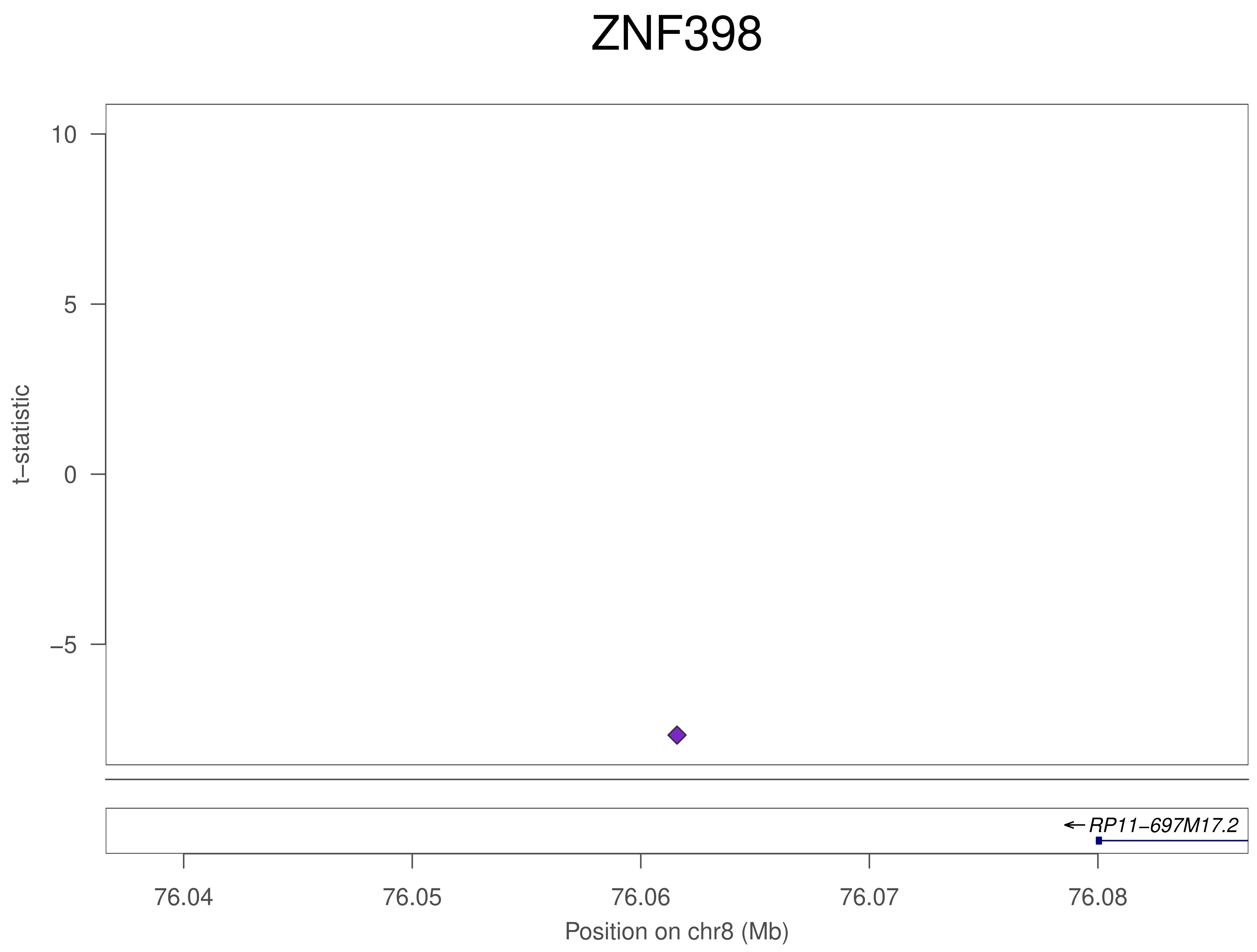
ZNF398
10
5
0
−5
t−statistic
RP11−697M17.2
76.04
76.05
76.06
76.07
76.08
Position on chr8 (Mb)

### ZNF398

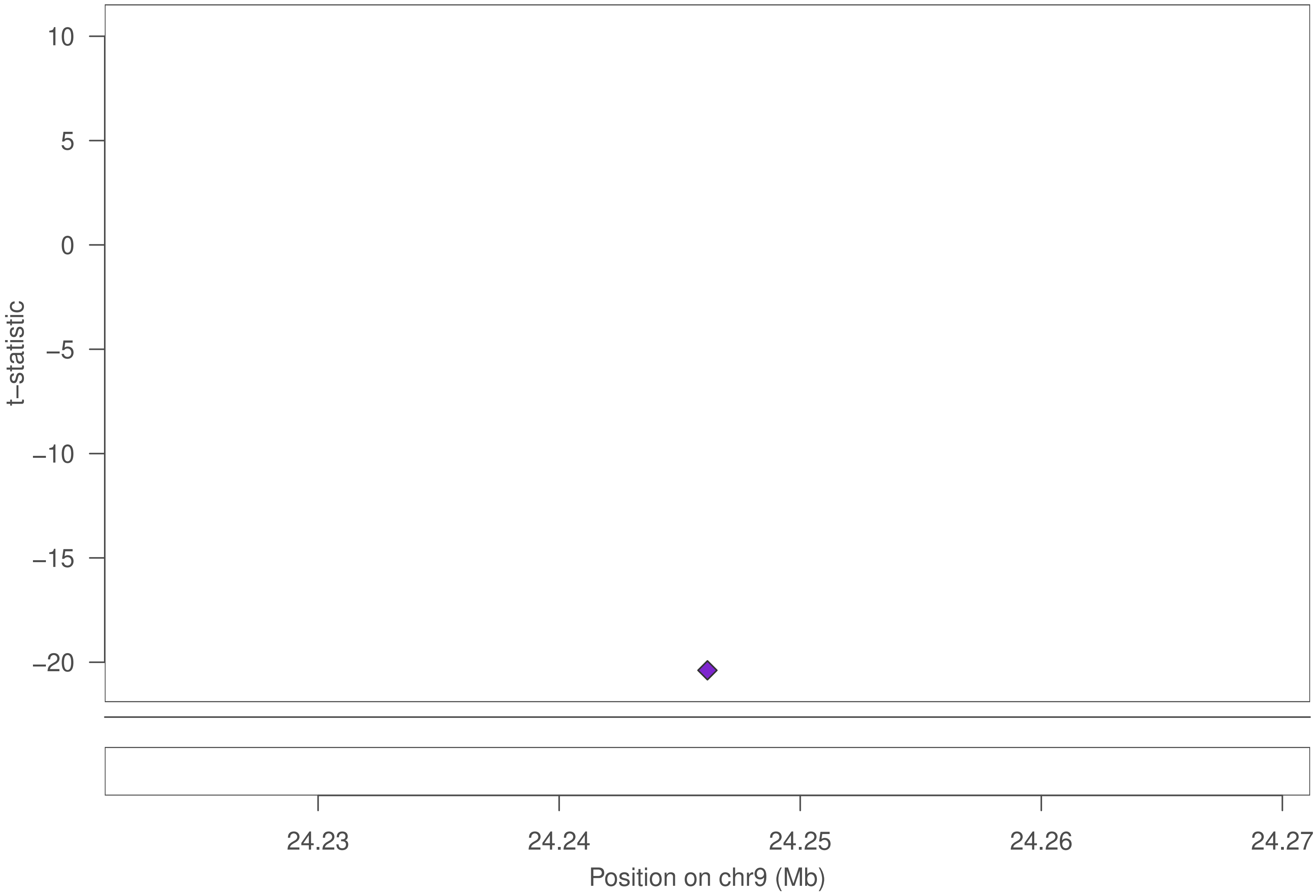

10
5
0
−5
−10
−15
−20
t−statistic
24.23
24.24
24.25
24.26
24.27
Position on chr9 (Mb)

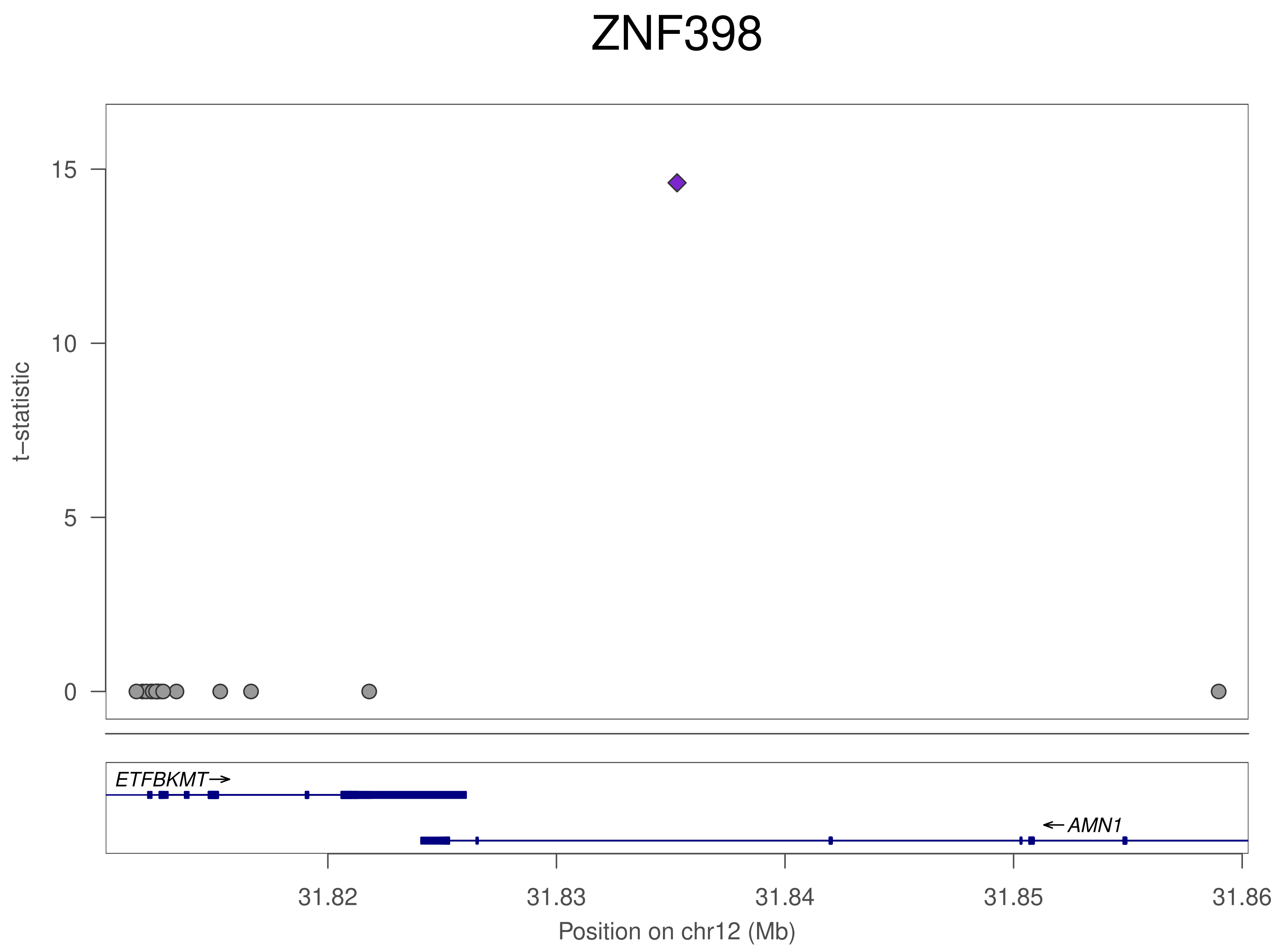
ZNF398
t−statistic
15
10
5
0
ETFBKMT
AMN1
31.82
31.83
31.84
31.85
31.86
Position on chr12 (Mb)

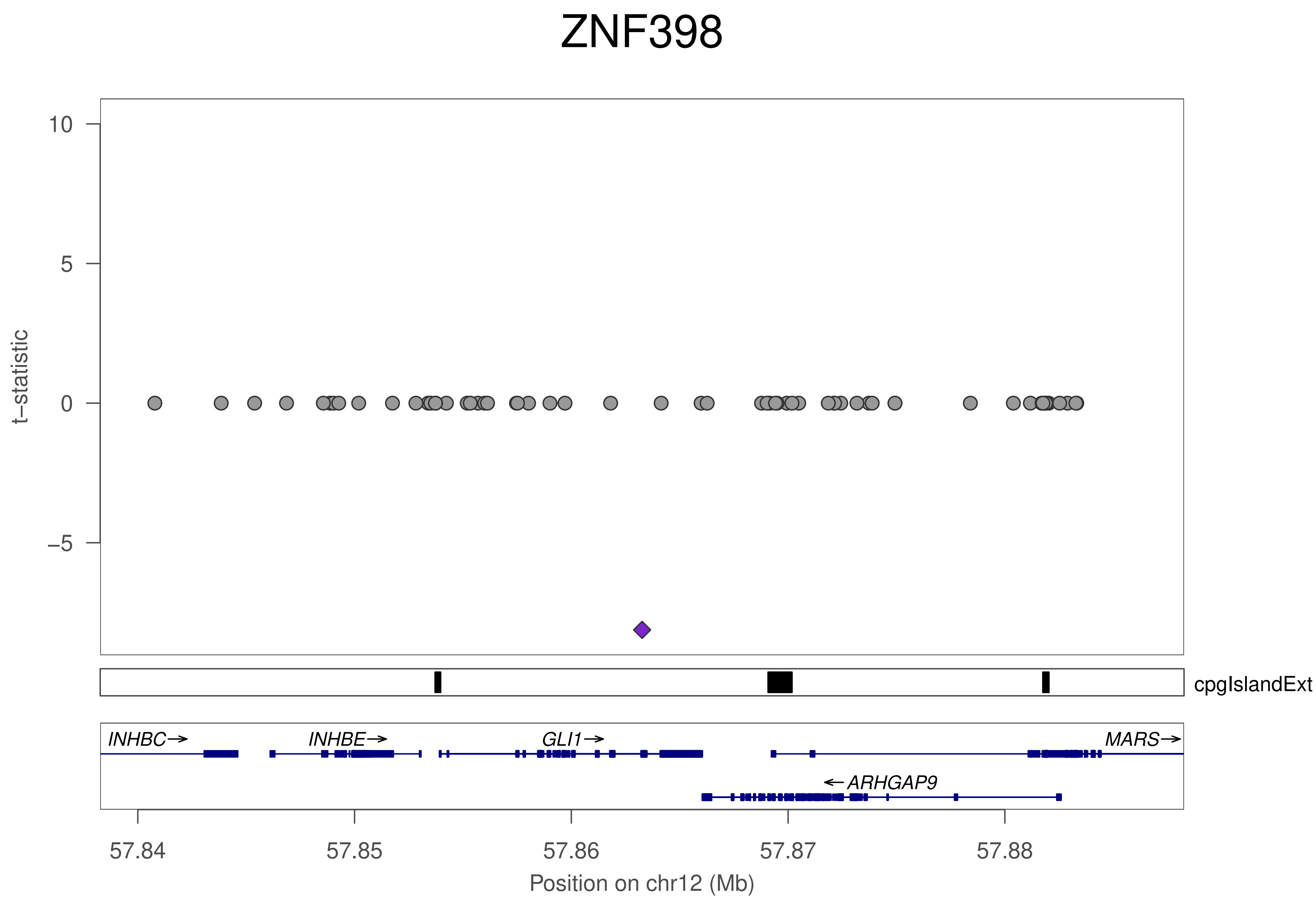
ZNF398
t−statistic
10
5
0
−5
cpgIslandExt
INHBC→
INHBE→
GLI1→
MARS→
←ARHGAP9
57.84
57.85
57.86
57.87
57.88
Position on chr12 (Mb)

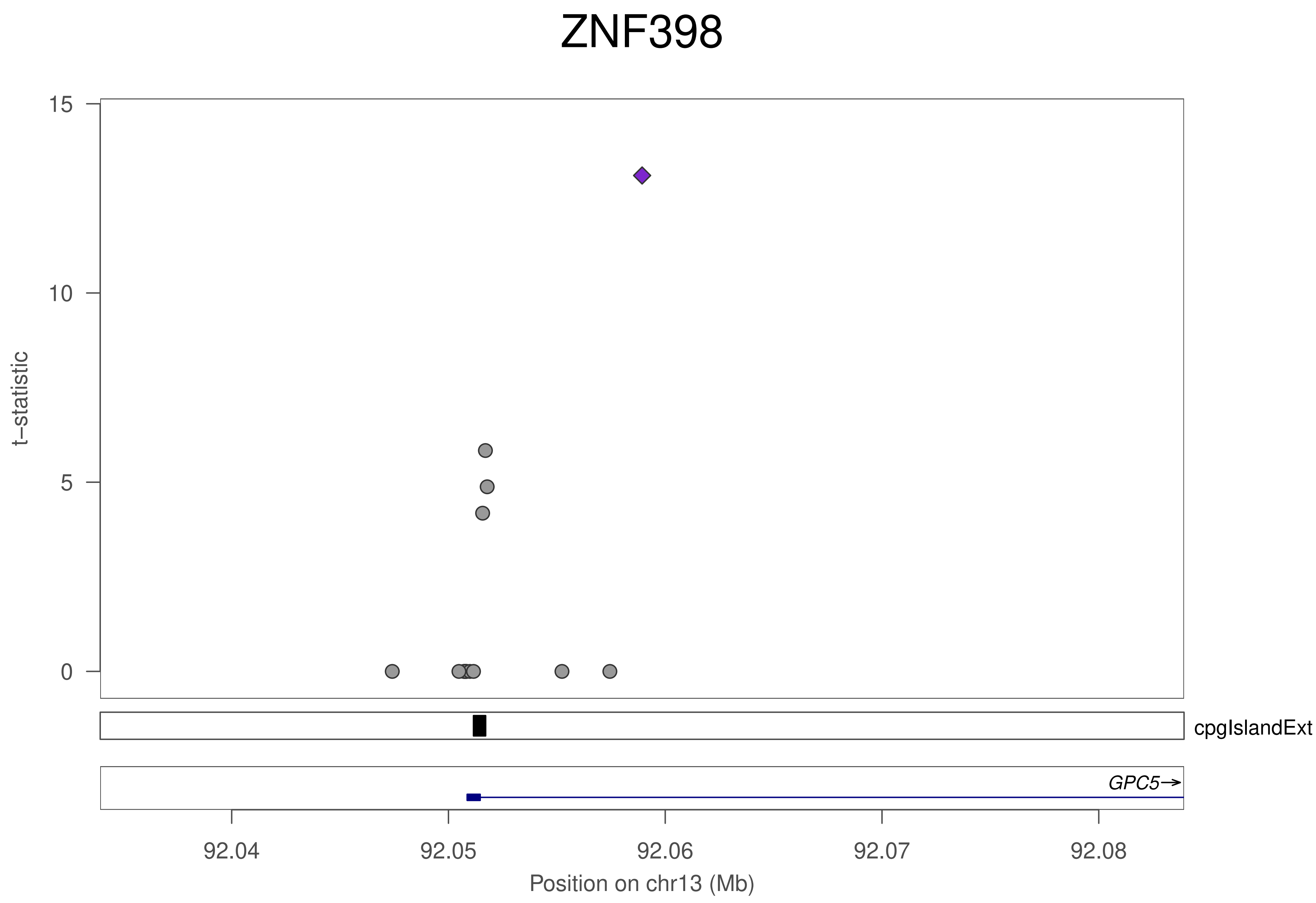
ZNF398
t−statistic
15
10
5
0
cpgIslandExt
GPC5→
92.04
92.05
92.06
92.07
92.08
Position on chr13 (Mb)

### ZNF398

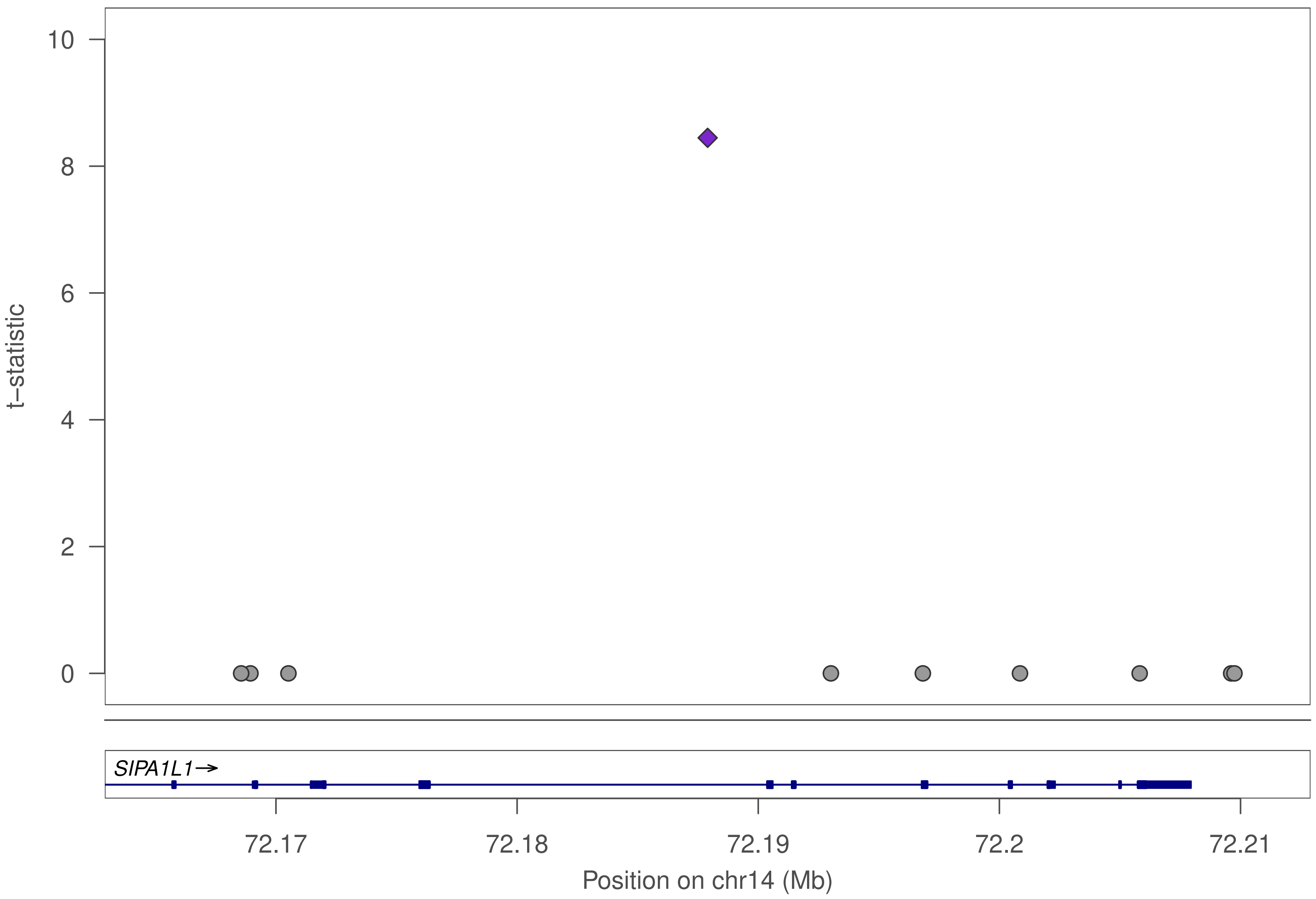

t−statistic
10
8
6
4
2
0
SIPA1L1→
72.17
72.18
72.19
72.2
72.21
Position on chr14 (Mb)

### ZNF398

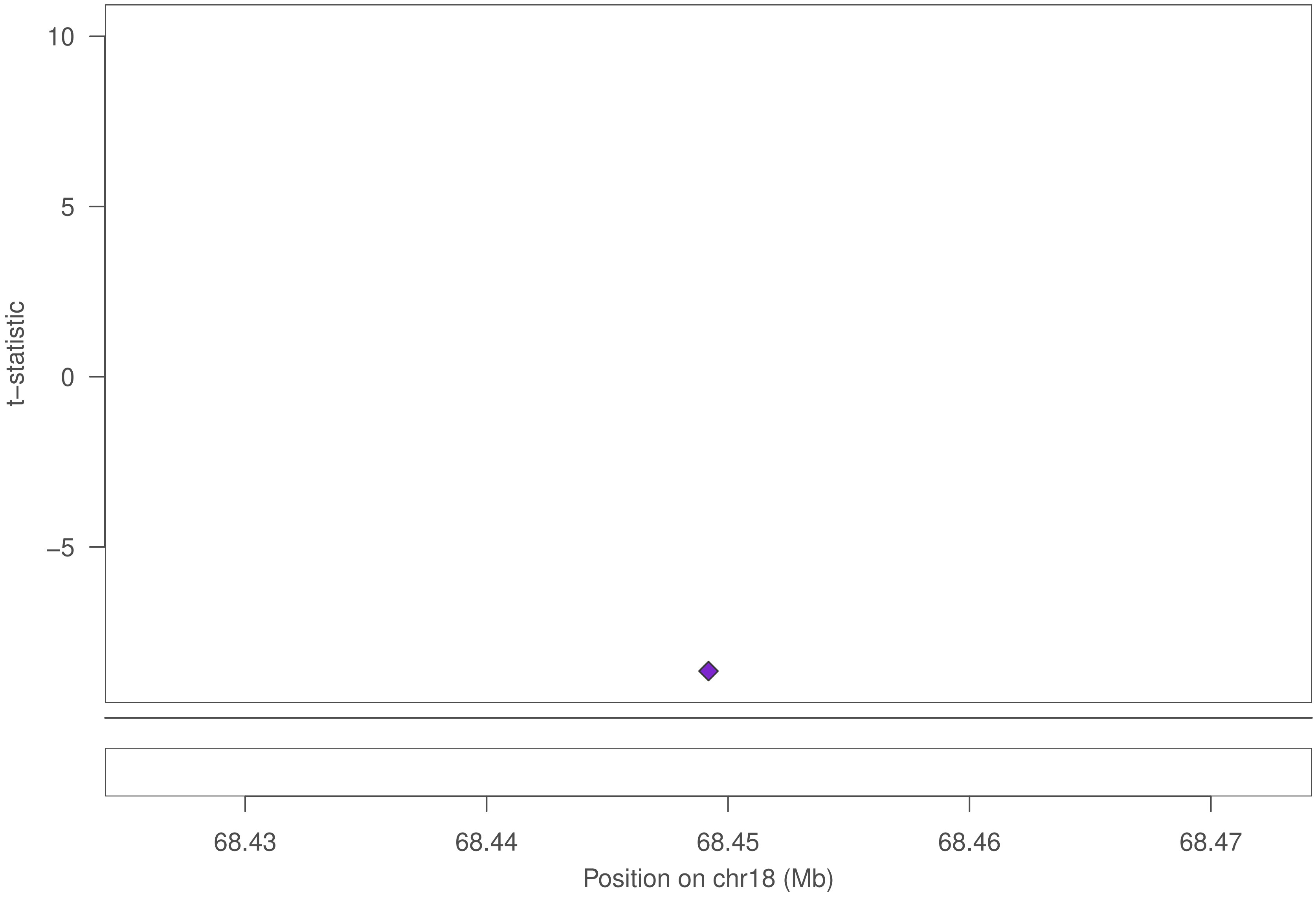

10
5
0
−5
t−statistic
68.43
68.44
68.45
68.46
68.47
Position on chr18 (Mb)

### ZNF257

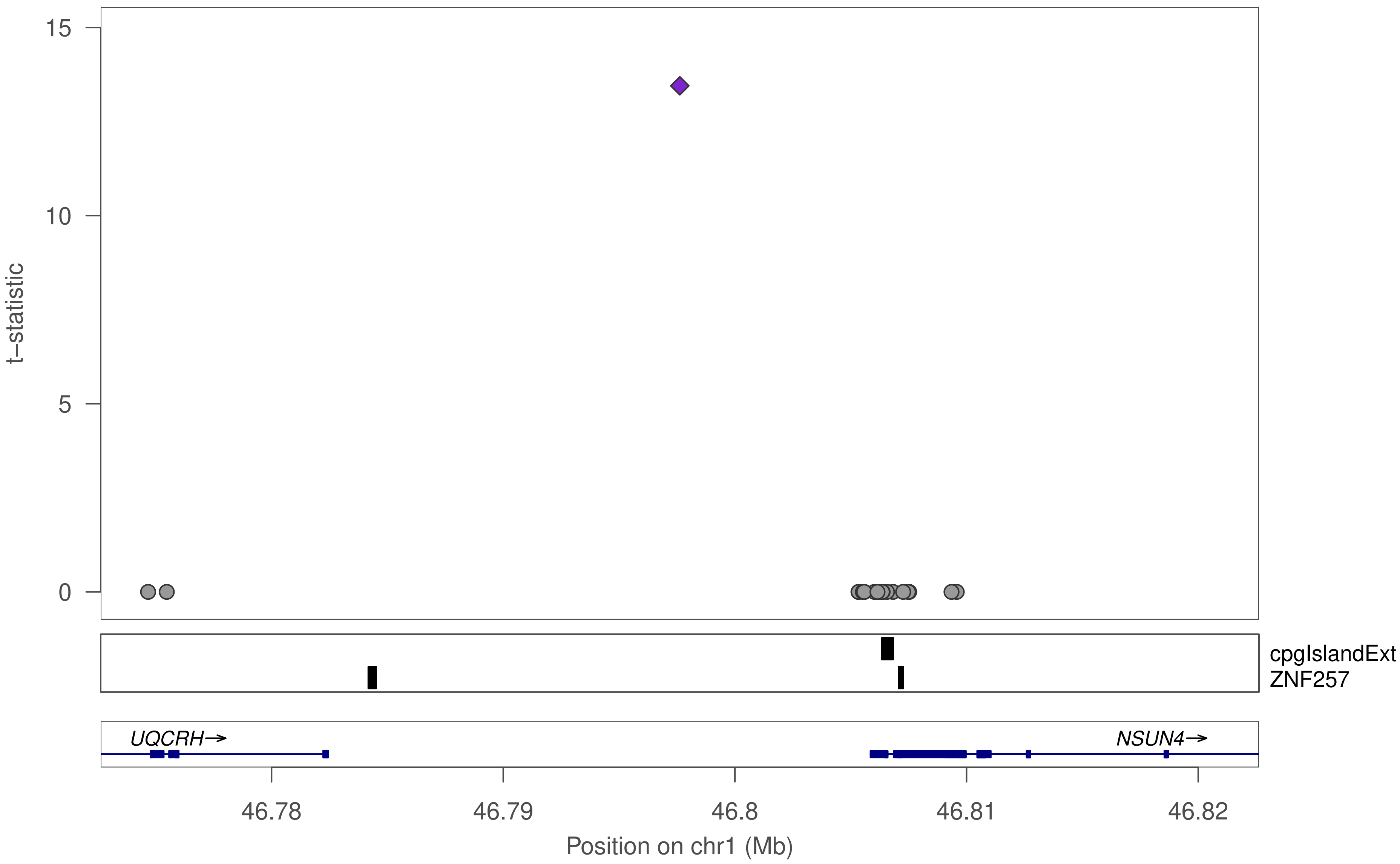

15
10
5
0
t−statistic
cpgIslandExt
ZNF257
UQCRH→
NSUN4→
46.78
46.79
46.8
46.81
46.82
Position on chr1 (Mb)

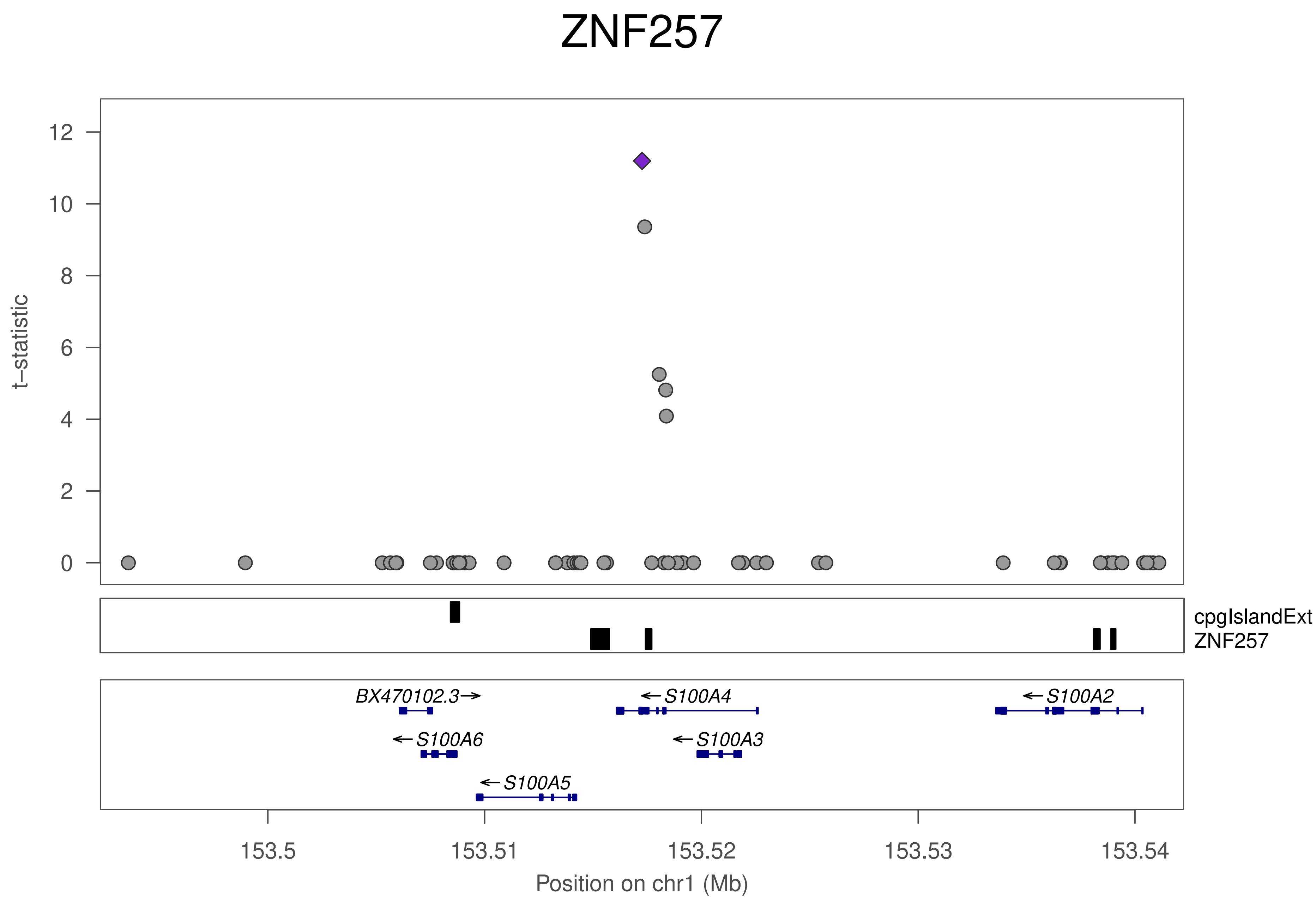
ZNF257
t−statistic
0
2
4
6
8
10
12
cpgIslandExt
ZNF257
BX470102.3
S100A4
S100A2
S100A6
S100A3
S100A5
153.5
153.51
153.52
153.53
153.54
Position on chr1 (Mb)

### ZNF257

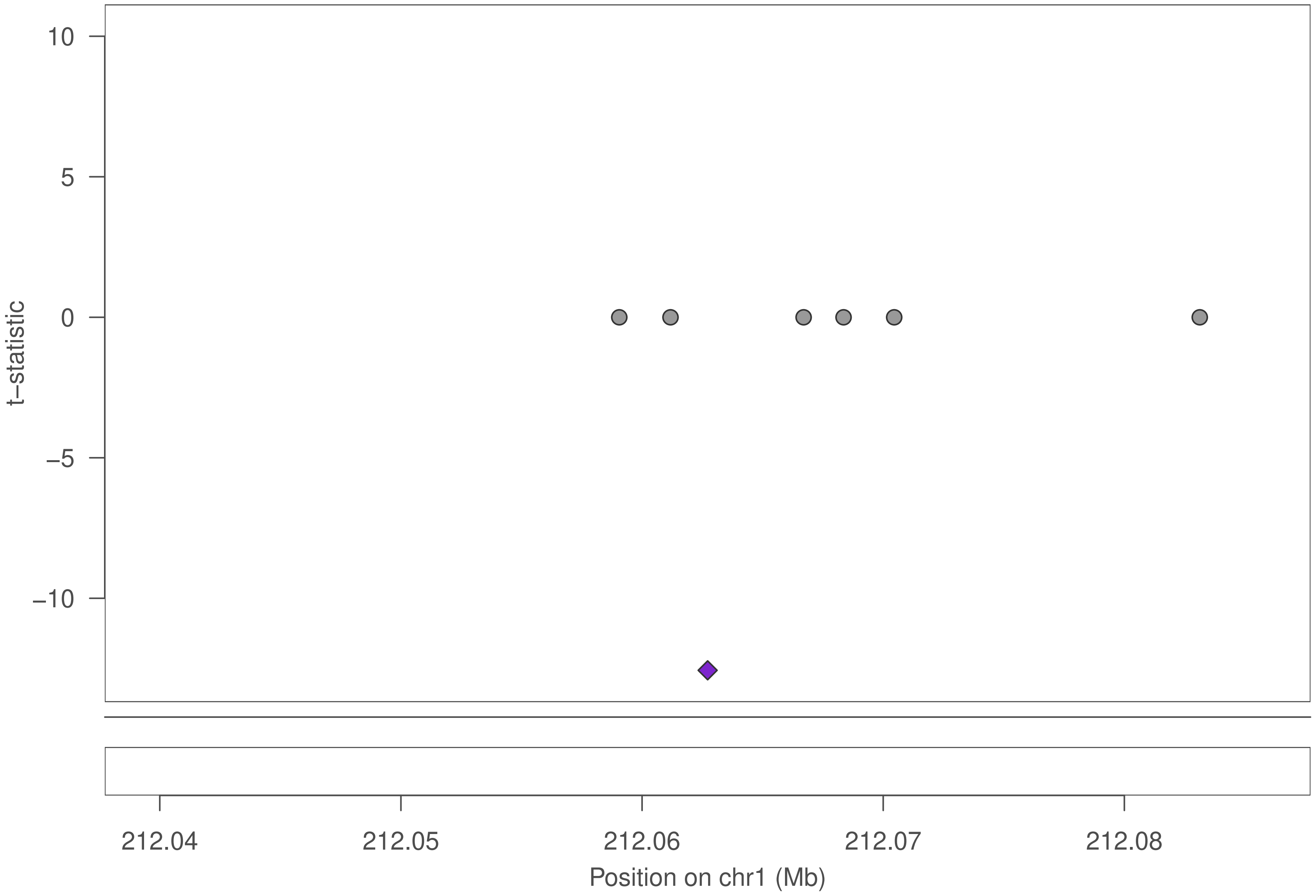
10
5
0
−5
−10
t−statistic
212.04
212.05
212.06
212.07
212.08
Position on chr1 (Mb)

### ZNF257

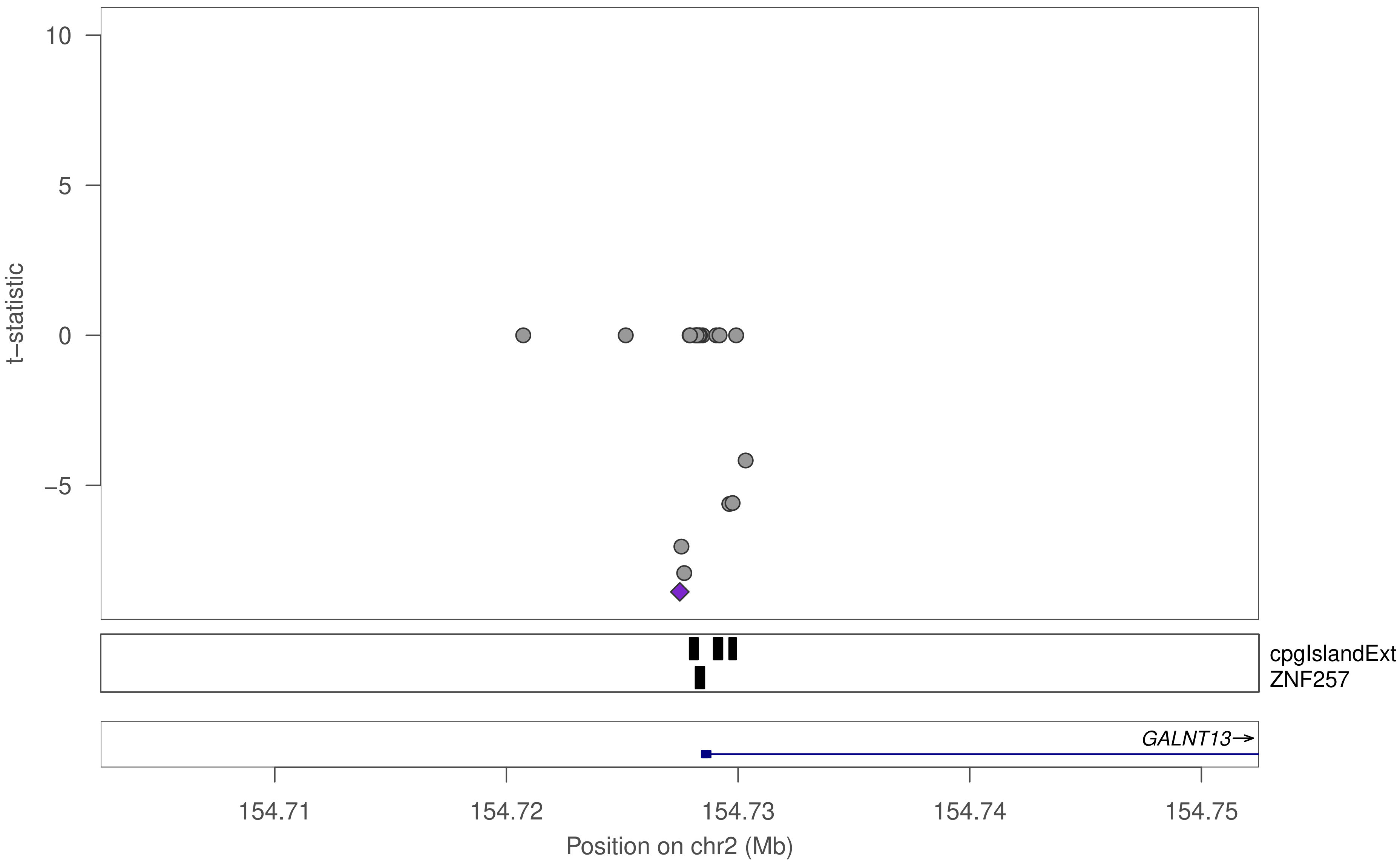

10
5
0
−5
t−statistic
cpgIslandExt
ZNF257
GALNT13→
154.71
154.72
154.73
154.74
154.75
Position on chr2 (Mb)

### ZNF257

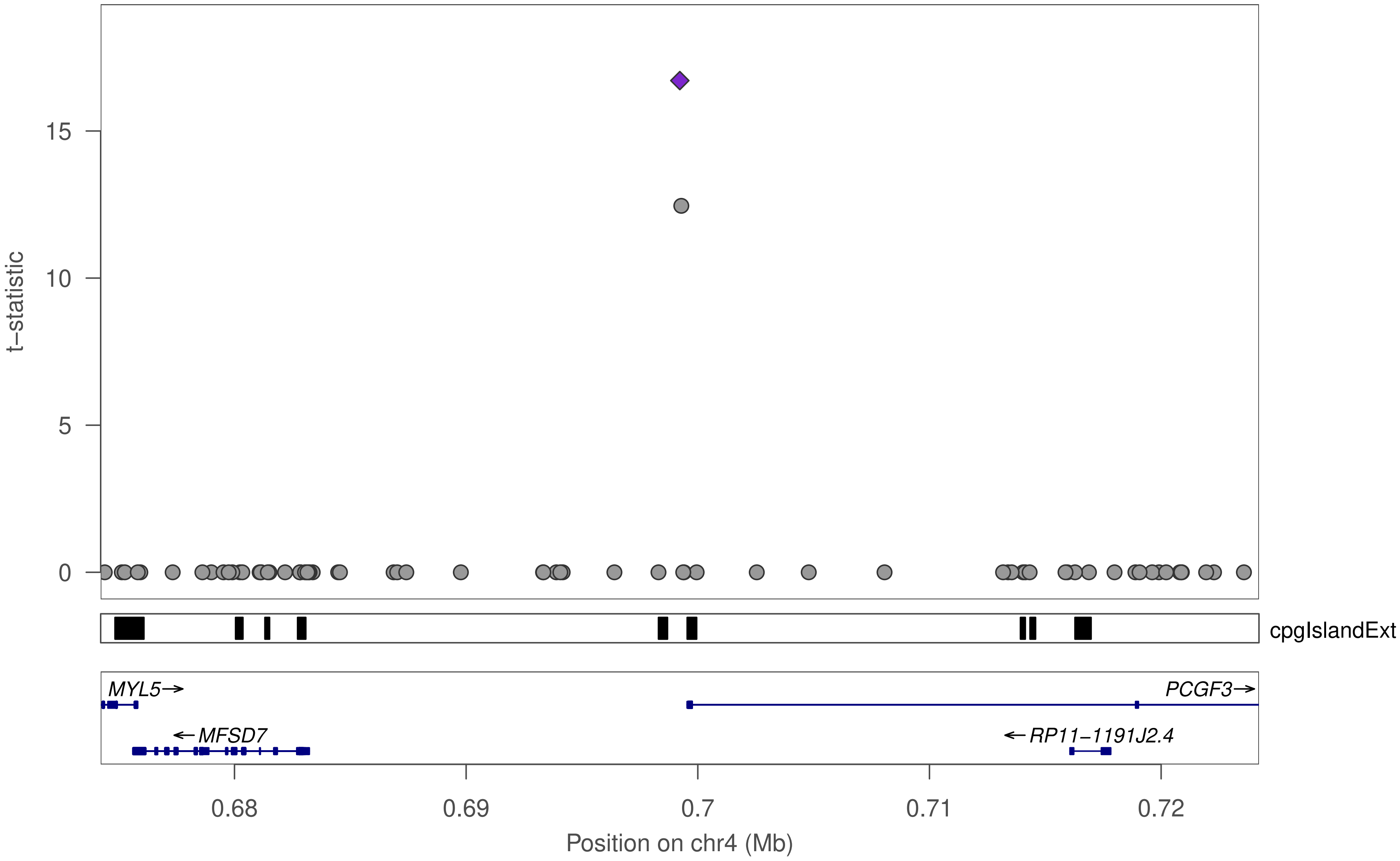

t−statistic
0
5
10
15
cpgIslandExt
MYL5→
←MFSD7
PCGF3→
←RP11−1191J2.4
0.68
0.69
0.7
0.71
0.72
Position on chr4 (Mb)

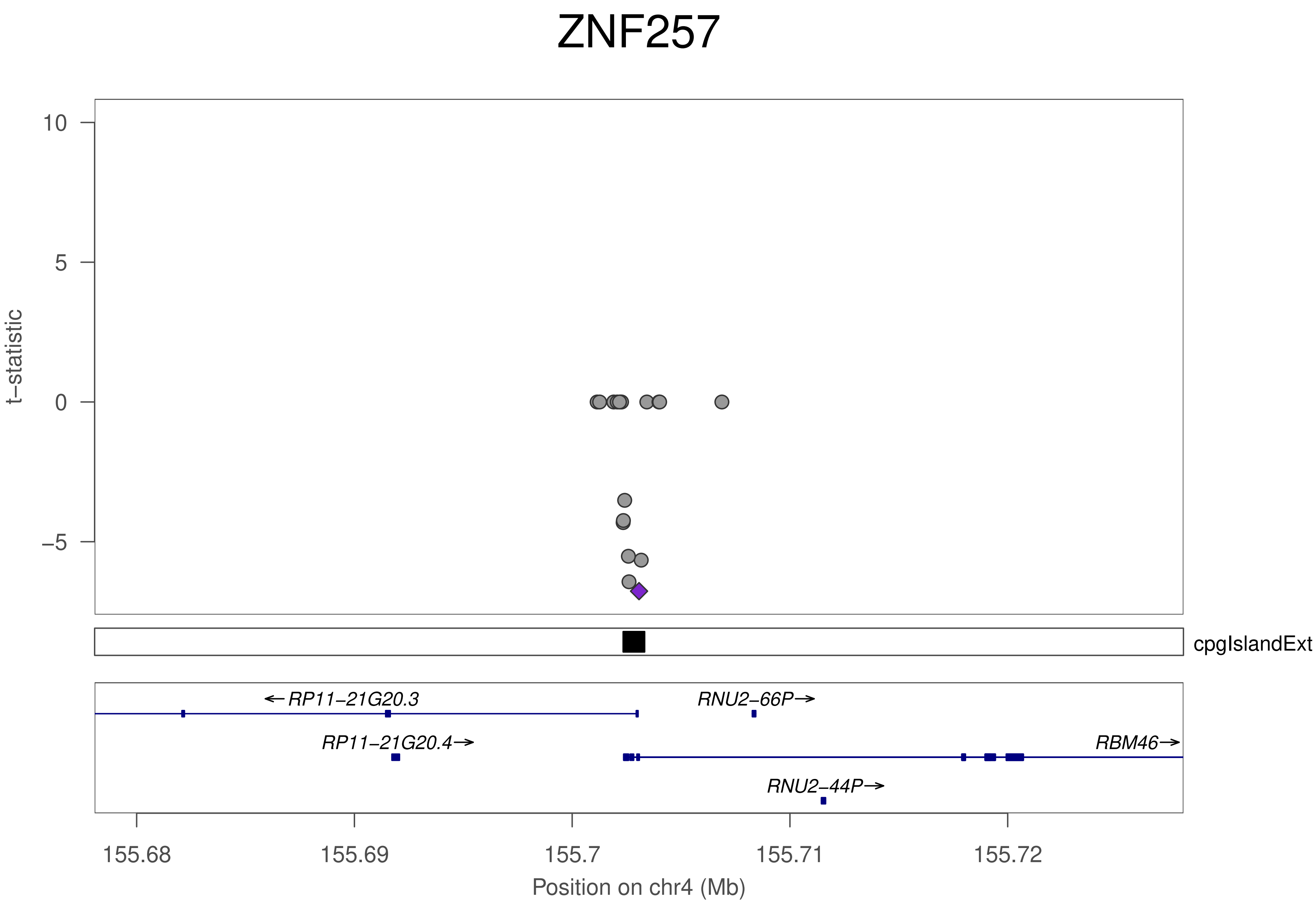
ZNF257
t−statistic
10
5
0
−5
cpgIslandExt
←RP11−21G20.3
RNU2−66P→
RP11−21G20.4→
RBM46→
RNU2−44P→
155.68
155.69
155.7
155.71
155.72
Position on chr4 (Mb)

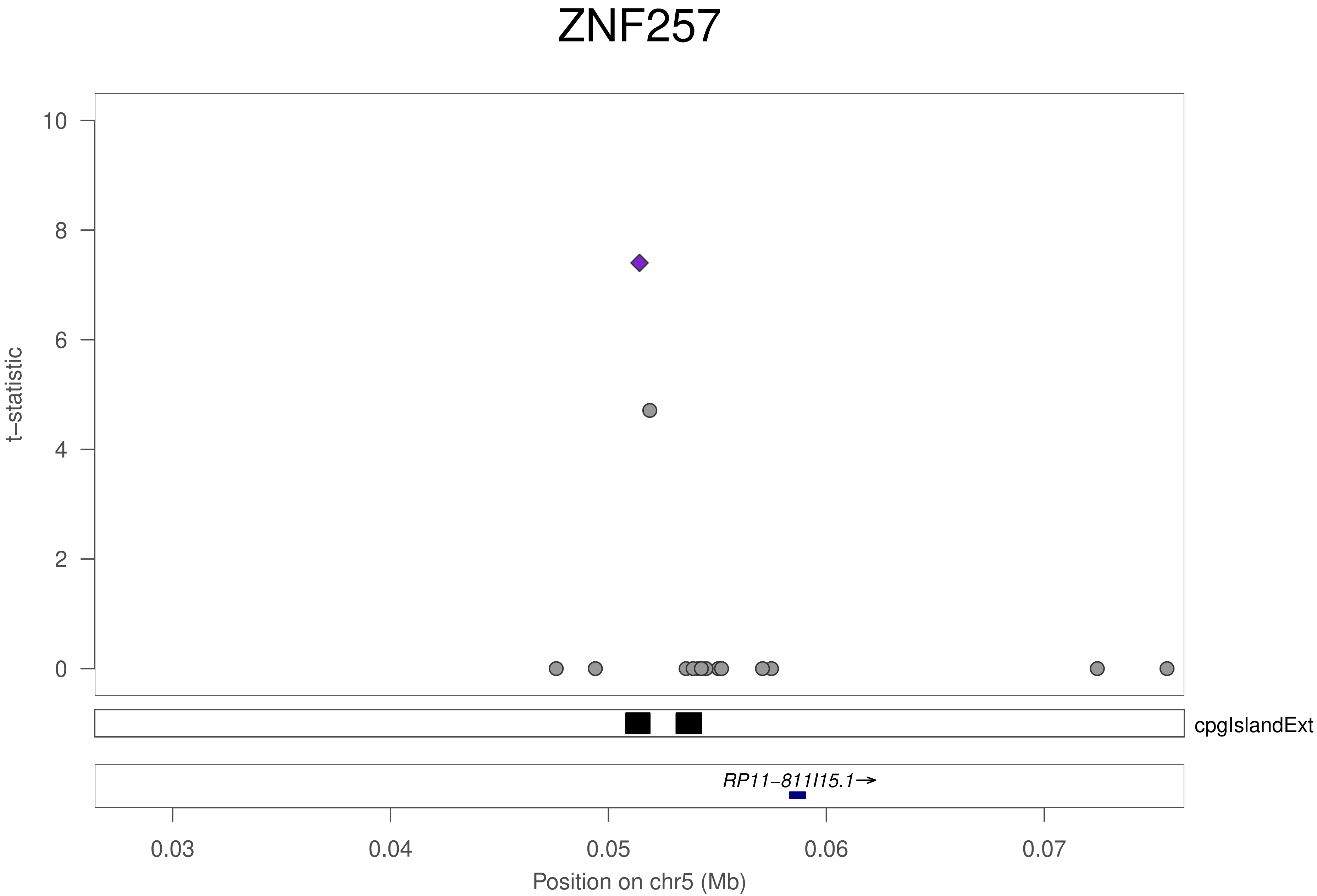

ZNF257
t−statistic
10
8
6
4
2
0
cpgIslandExt
RP11−811I15.1→
0.03
0.04
0.05
0.06
0.07
Position on chr5 (Mb)

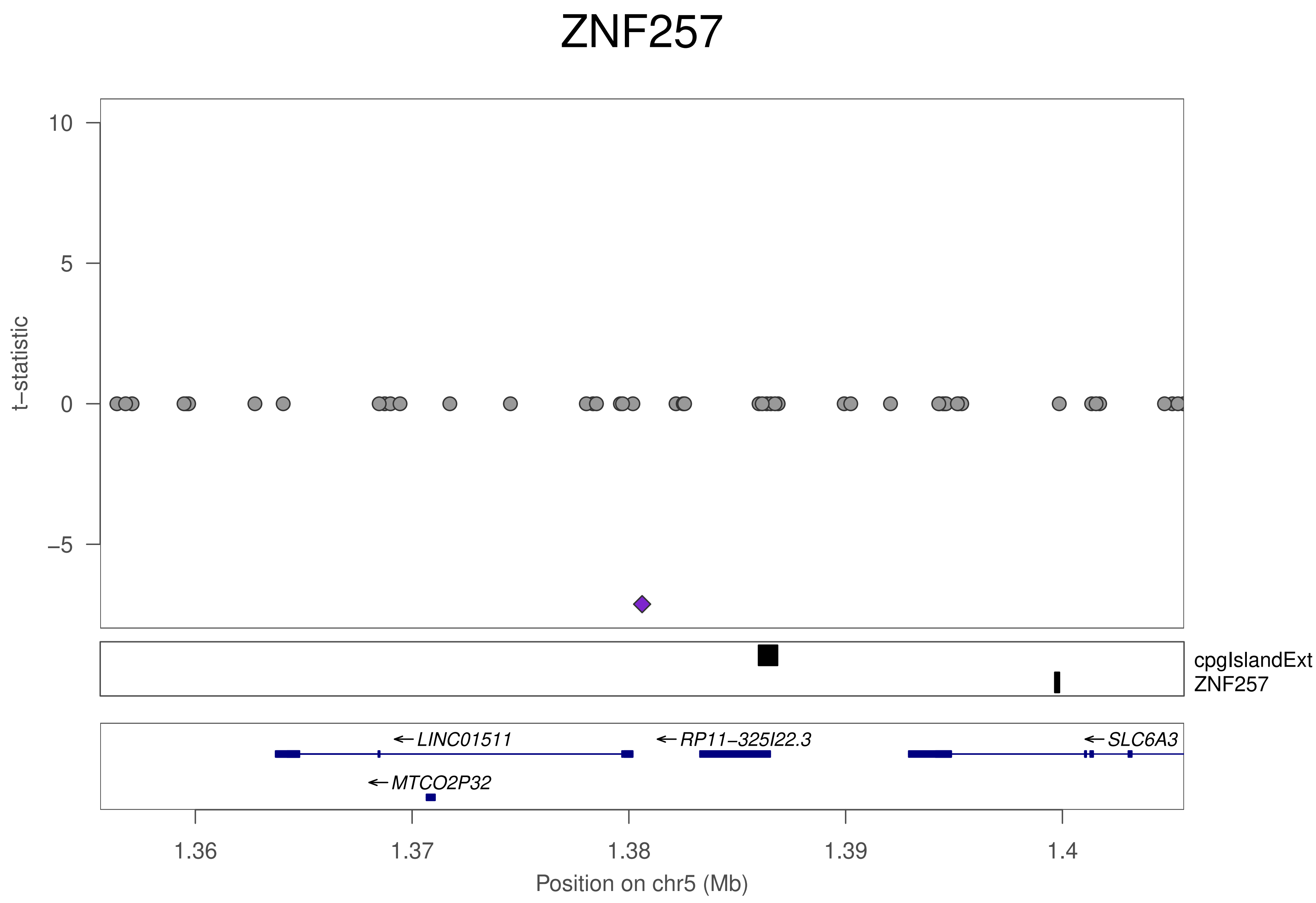
ZNF257
t−statistic
10
5
0
−5
cpgIslandExt
ZNF257
LINC01511
RP11−325I22.3
SLC6A3
MTCO2P32
1.36
1.37
1.38
1.39
1.4
Position on chr5 (Mb)

### ZNF257

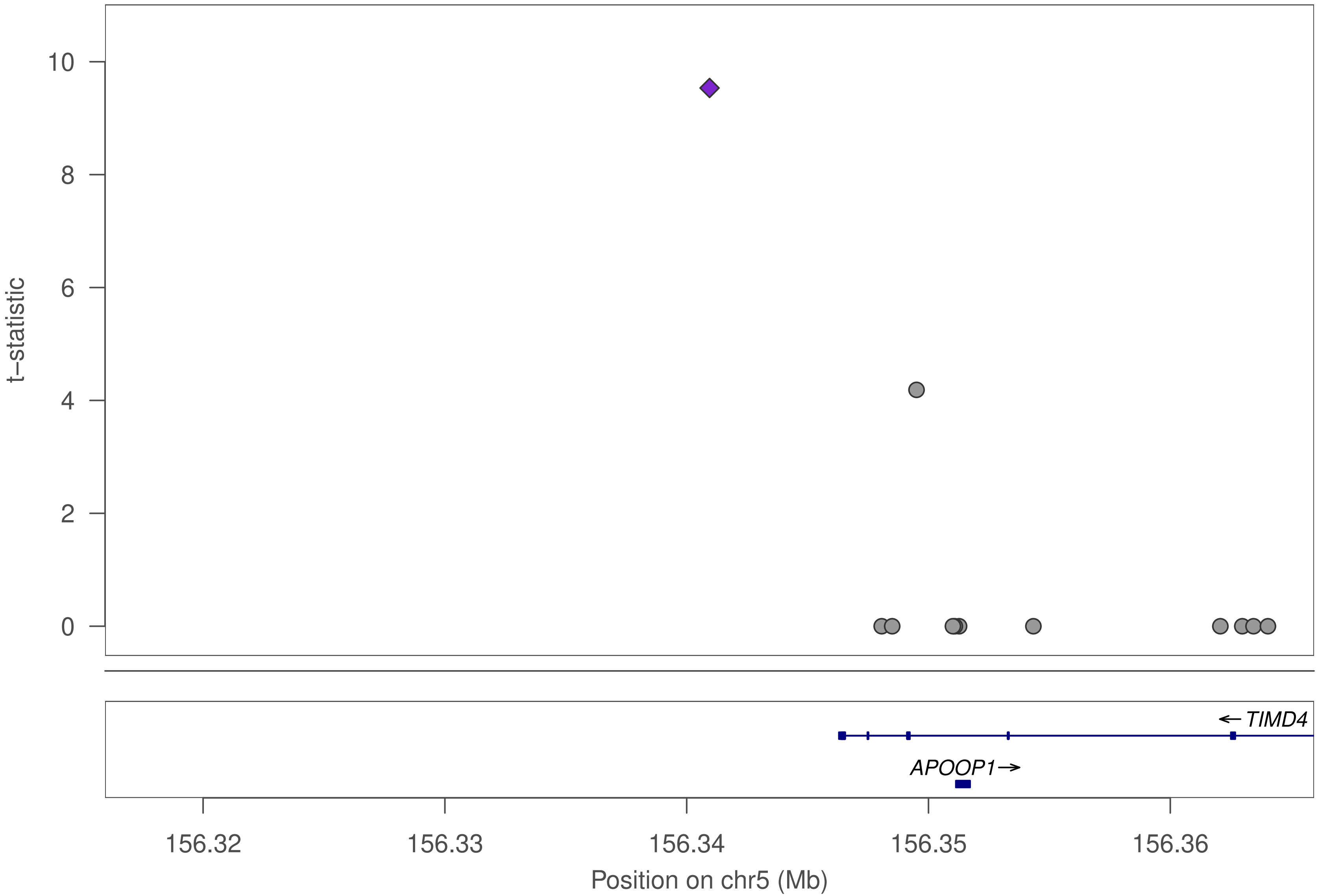

10
8
6
4
2
0
t–statistic
← TIMD4
APOOP1→
156.32
156.33
156.34
156.35
156.36
Position on chr5 (Mb)

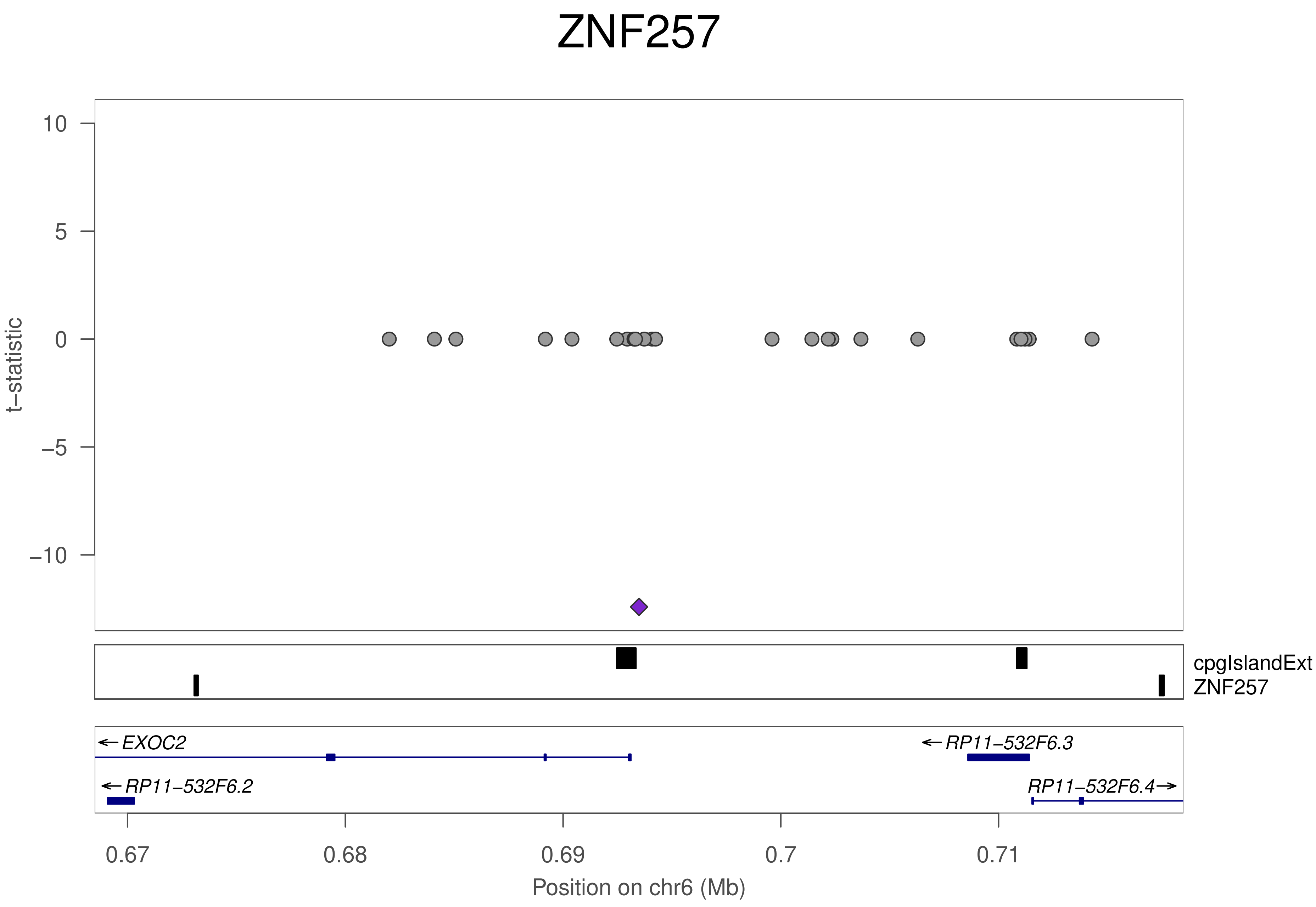

ZNF257
t−statistic
10
5
0
−5
−10
cpgIslandExt
ZNF257
EXOC2
RP11−532F6.3
RP11−532F6.2
RP11−532F6.4
0.67
0.68
0.69
0.7
0.71
Position on chr6 (Mb)

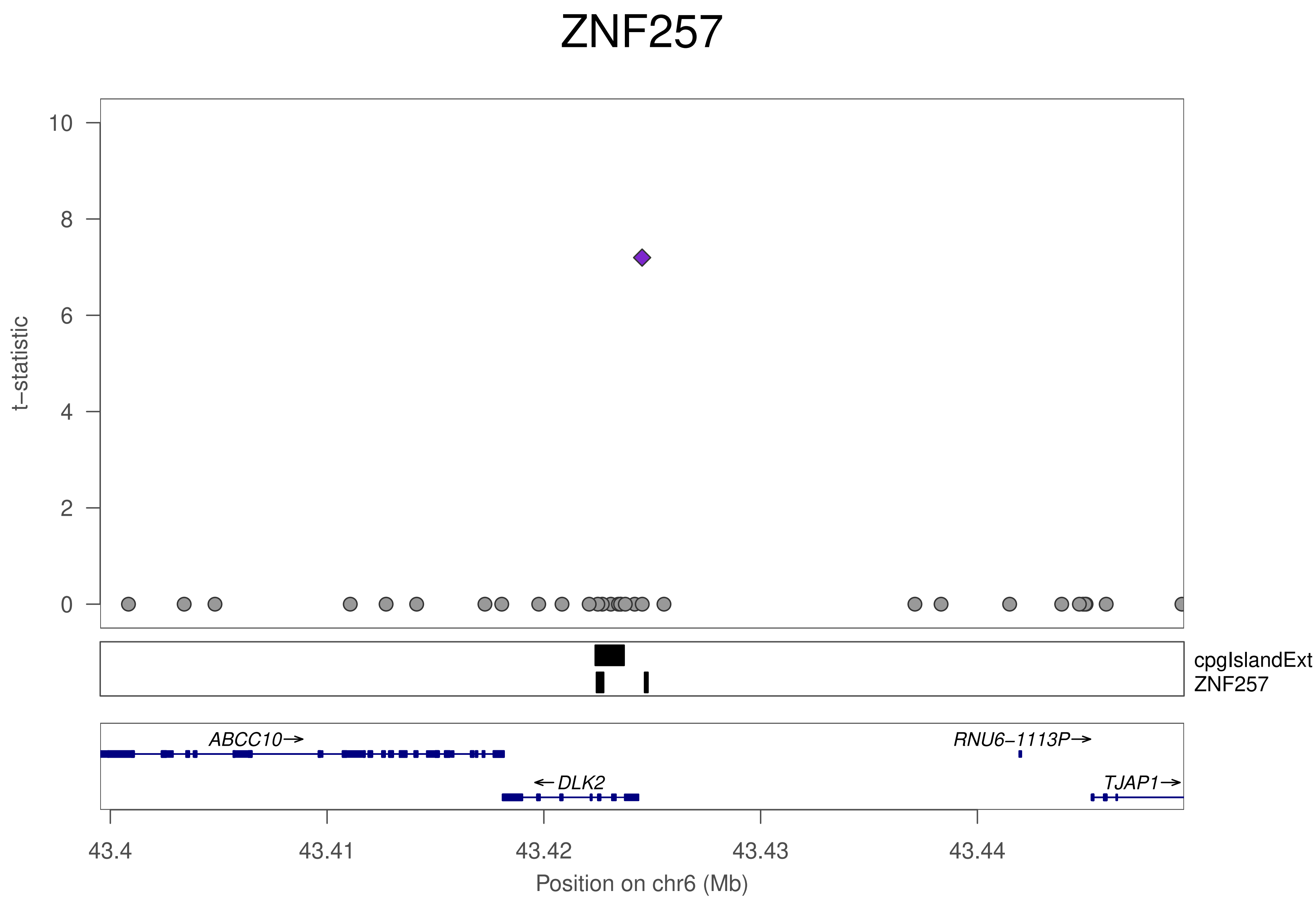

ZNF257
10
8
6
4
2
0
t−statistic
cpgIslandExt
ZNF257
ABCC10→
RNU6−1113P→
←DLK2
TJAP1→
43.4
43.41
43.42
43.43
43.44
Position on chr6 (Mb)

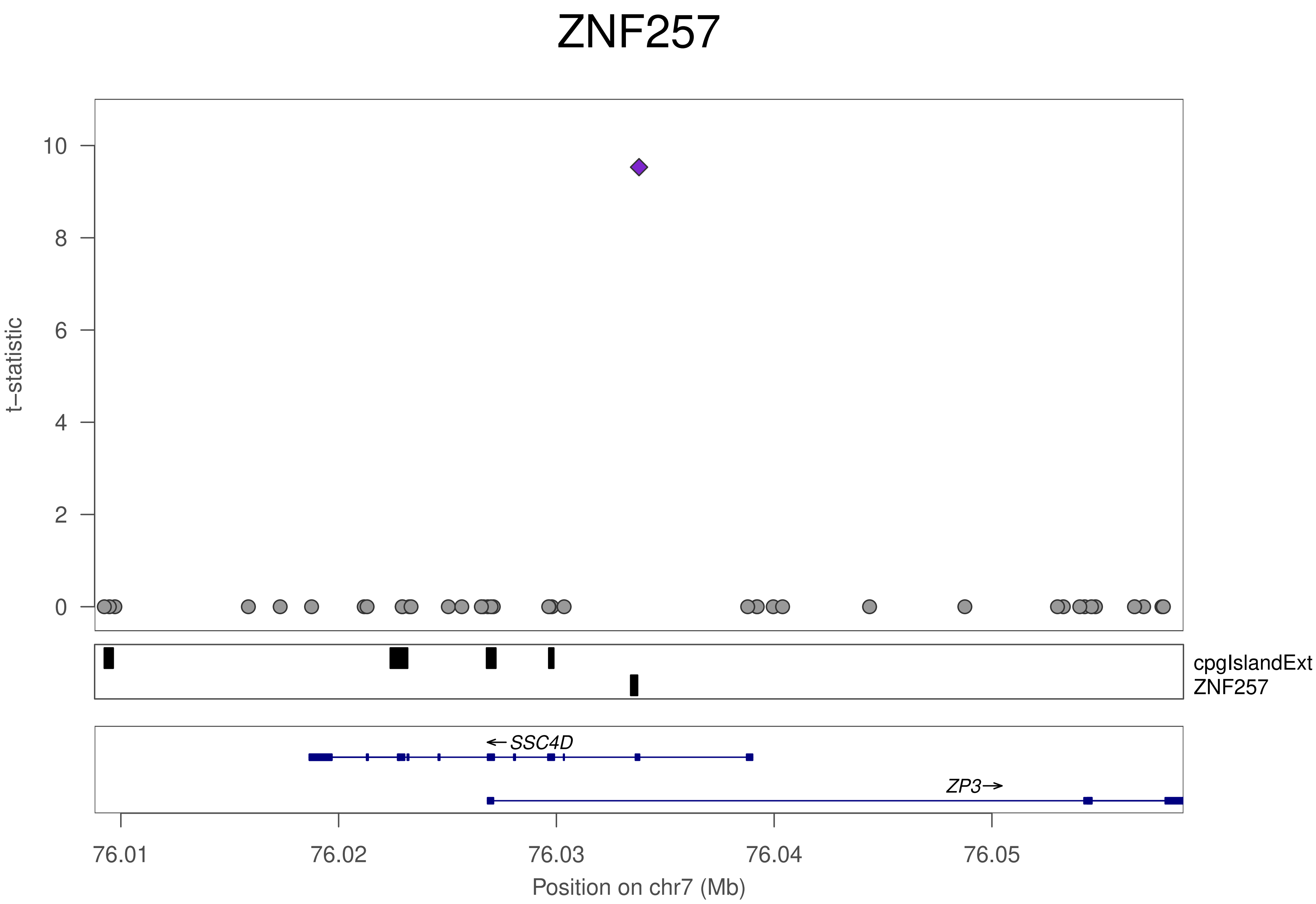

ZNF257
t−statistic
10
8
6
4
2
0
cpgIslandExt
ZNF257
←SSC4D
ZP3→
76.01
76.02
76.03
76.04
76.05
Position on chr7 (Mb)

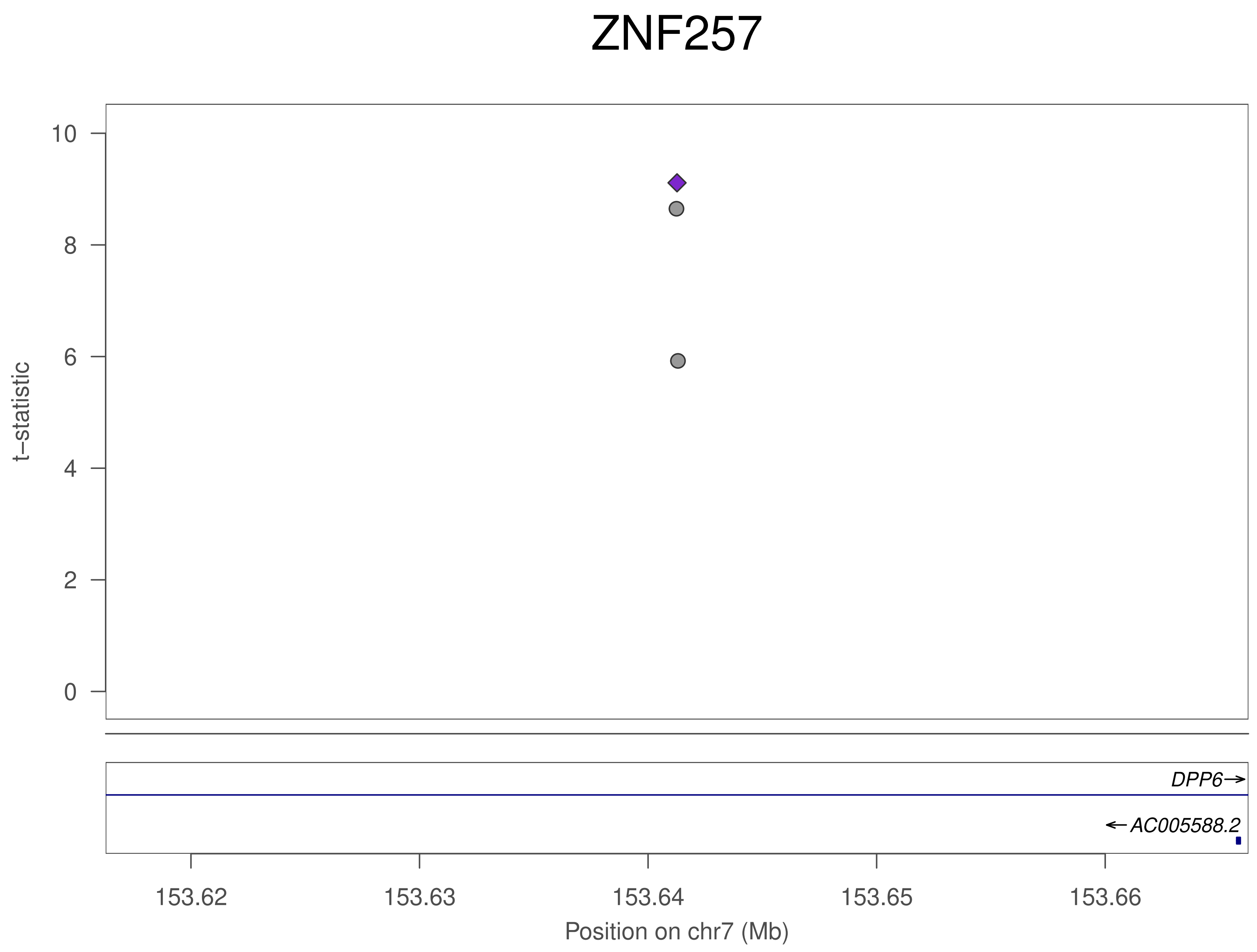

ZNF257
t−statistic
0
2
4
6
8
10
DPP6→
←AC005588.2
153.62
153.63
153.64
153.65
153.66
Position on chr7 (Mb)

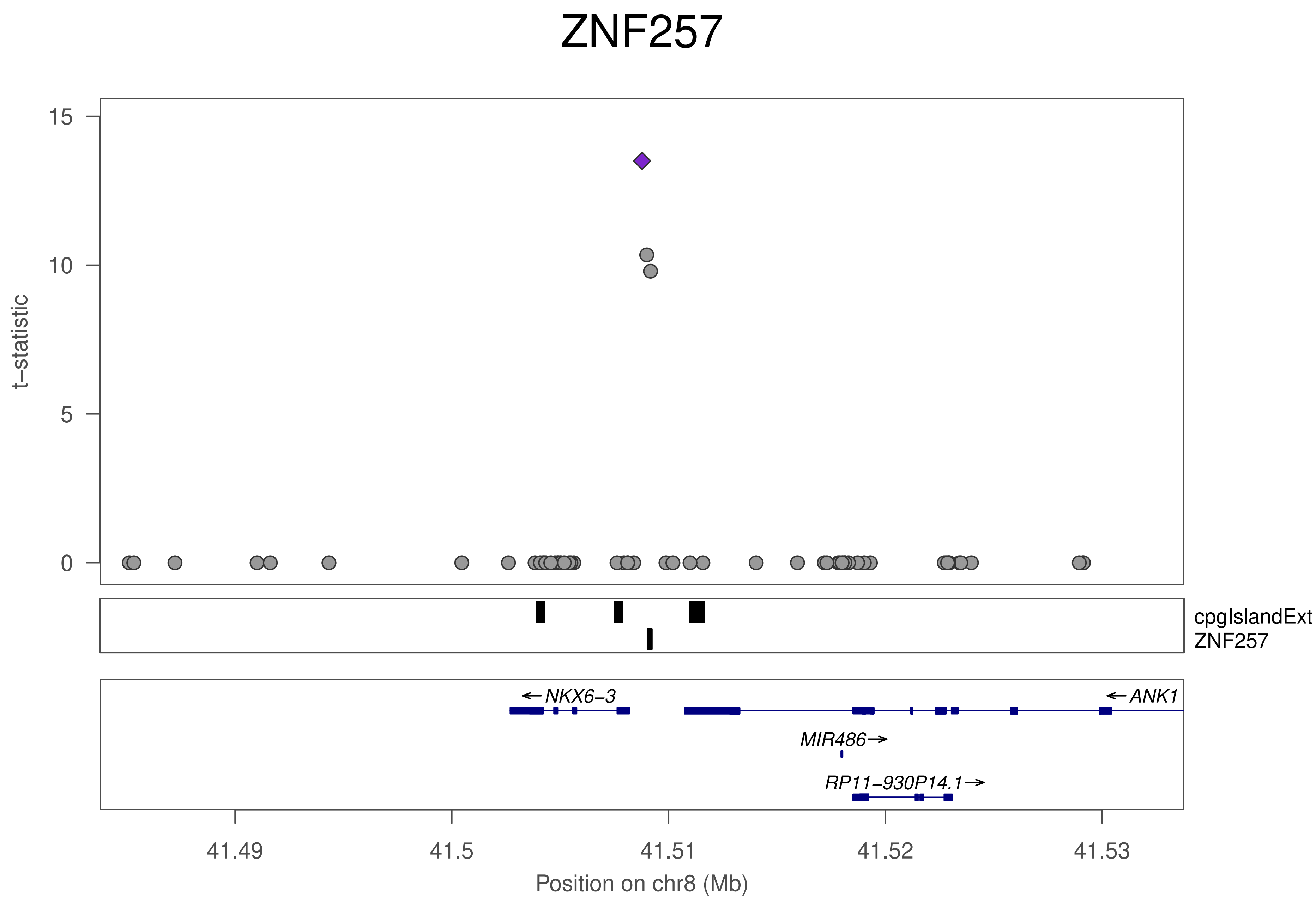

ZNF257
15
10
5
0
t–statistic
cpgIslandExt
ZNF257
←NKX6–3
←ANK1
MIR486→
RP11–930P14.1→
41.49
41.5
41.51
41.52
41.53
Position on chr8 (Mb)

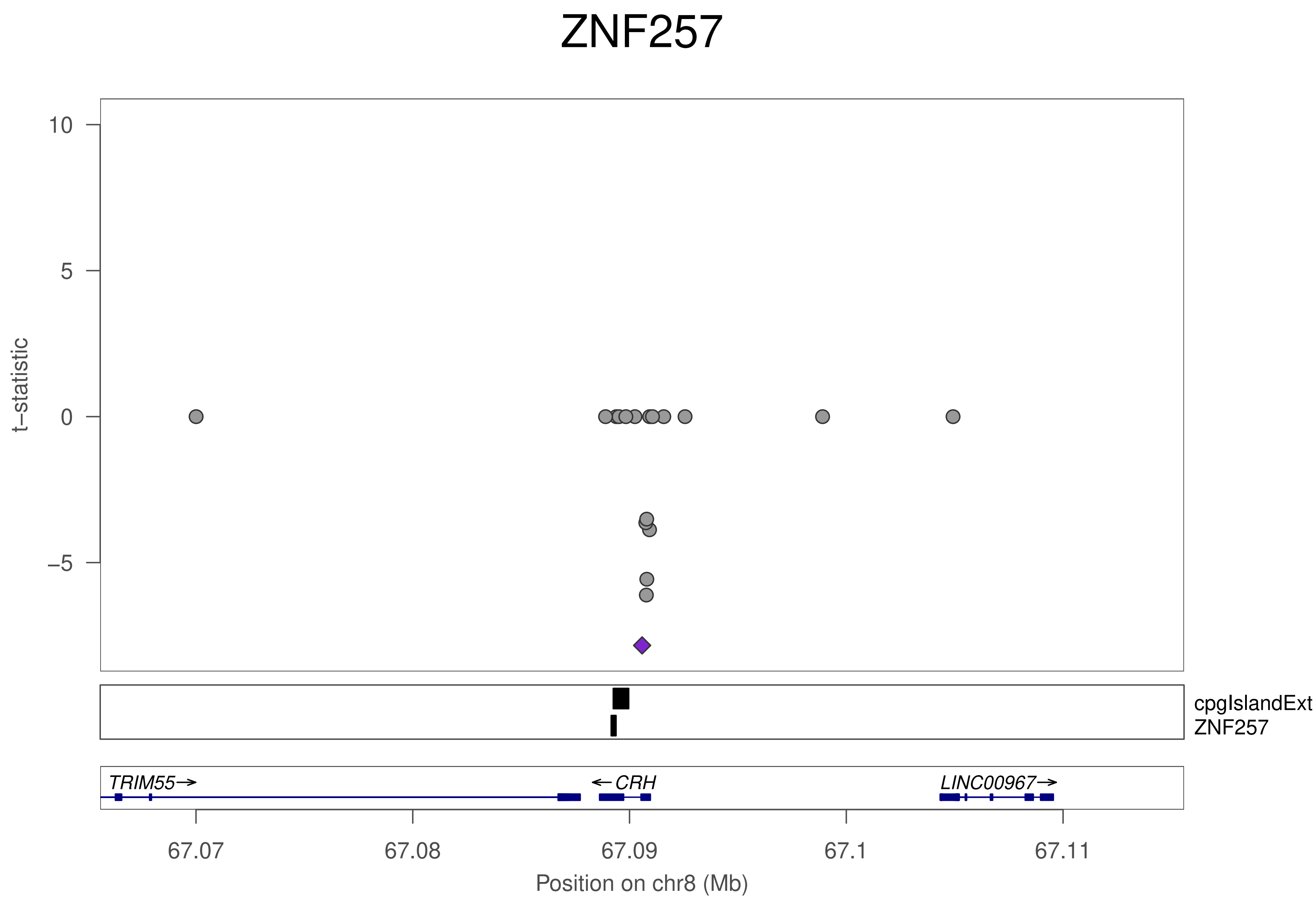
ZNF257
10
5
0
−5
t−statistic
cpgIslandExt
ZNF257
TRIM55→
←CRH
LINC00967→
67.07
67.08
67.09
67.1
67.11
Position on chr8 (Mb)

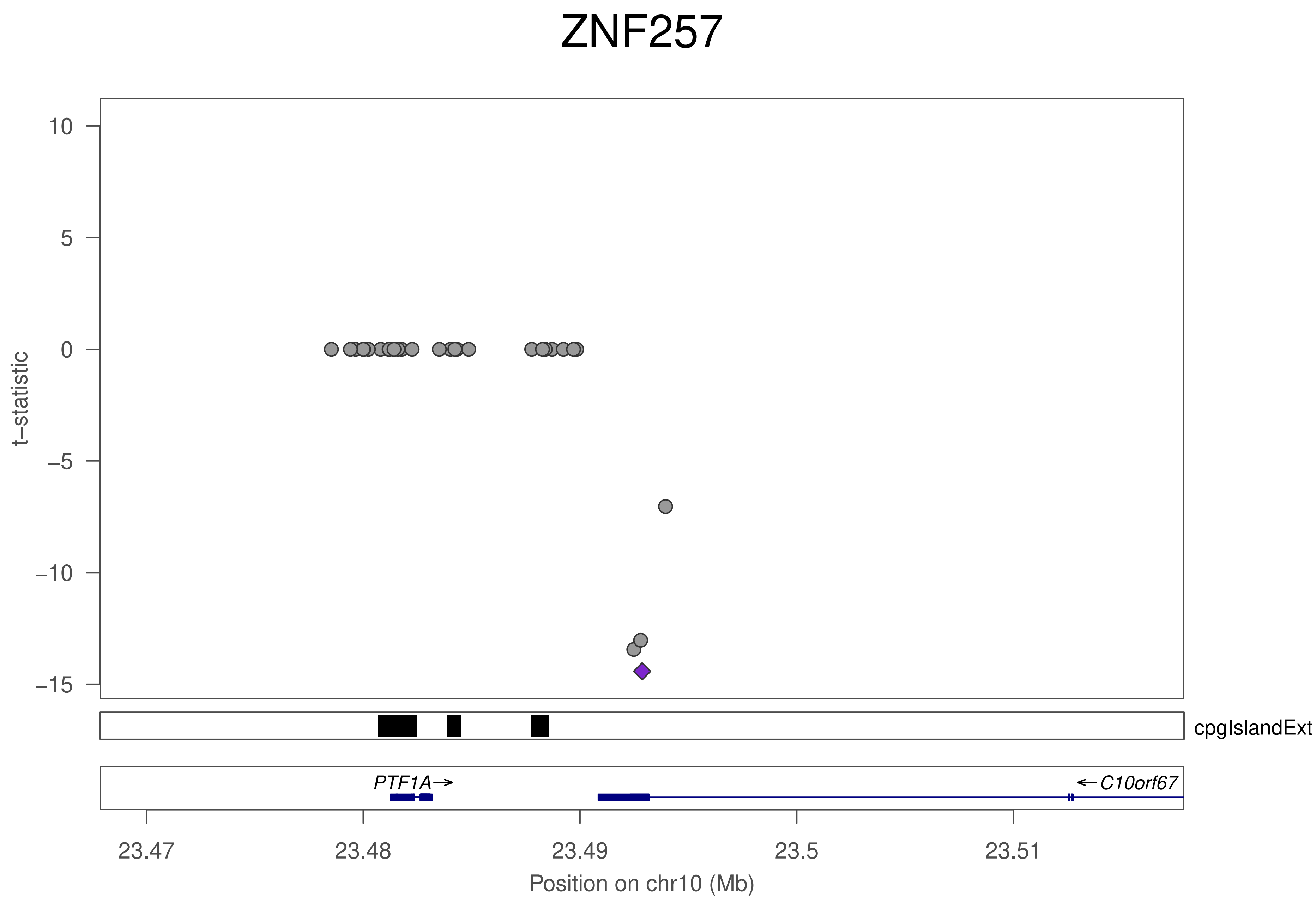

ZNF257
t−statistic
10
5
0
−5
−10
−15
cpgIslandExt
PTF1A→
←C10orf67
23.47
23.48
23.49
23.5
23.51
Position on chr10 (Mb)

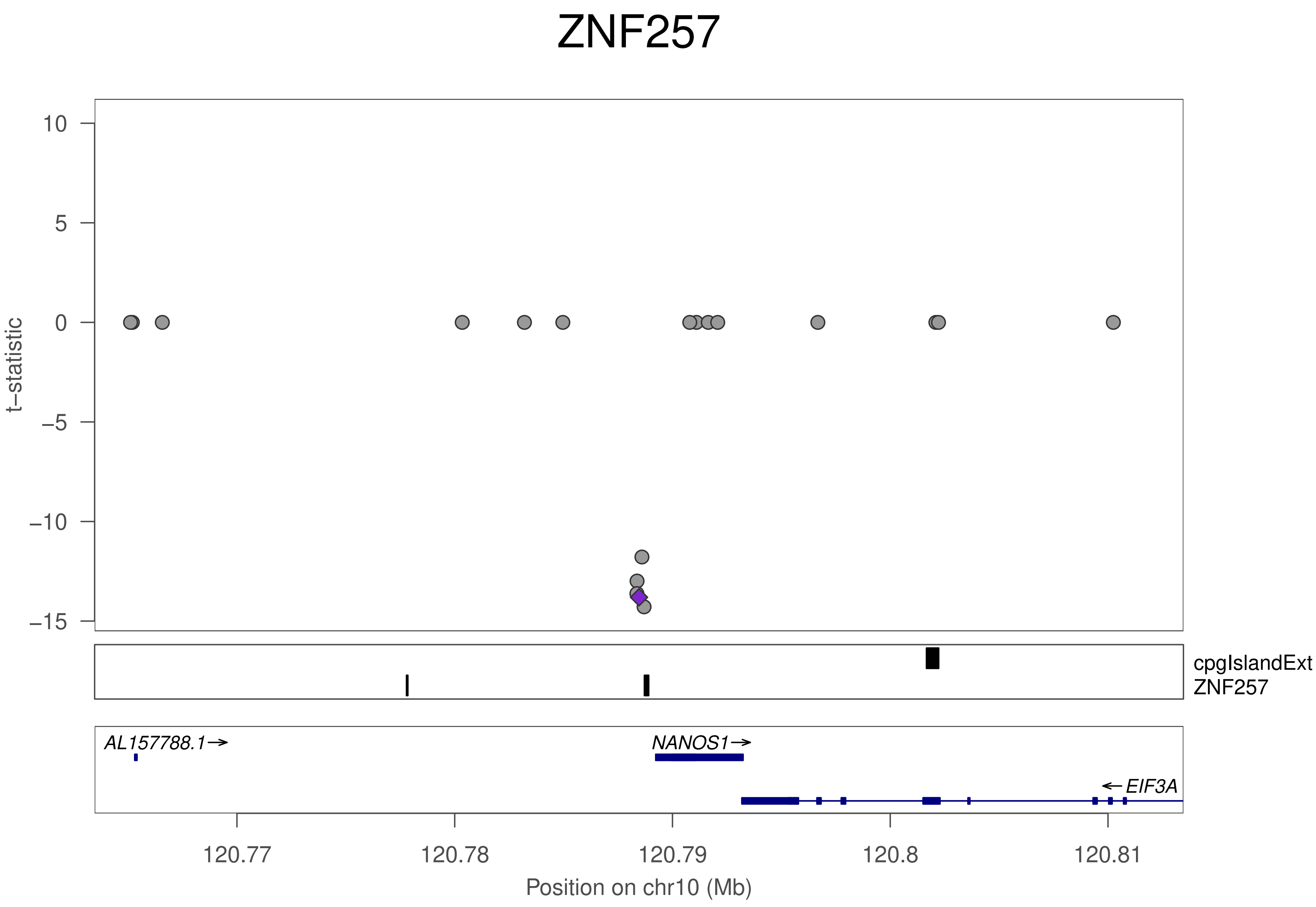
ZNF257
t−statistic
10
5
0
−5
−10
−15
cpgIslandExt
ZNF257
AL157788.1→
NANOS1→
← EIF3A
120.77
120.78
120.79
120.8
120.81
Position on chr10 (Mb)

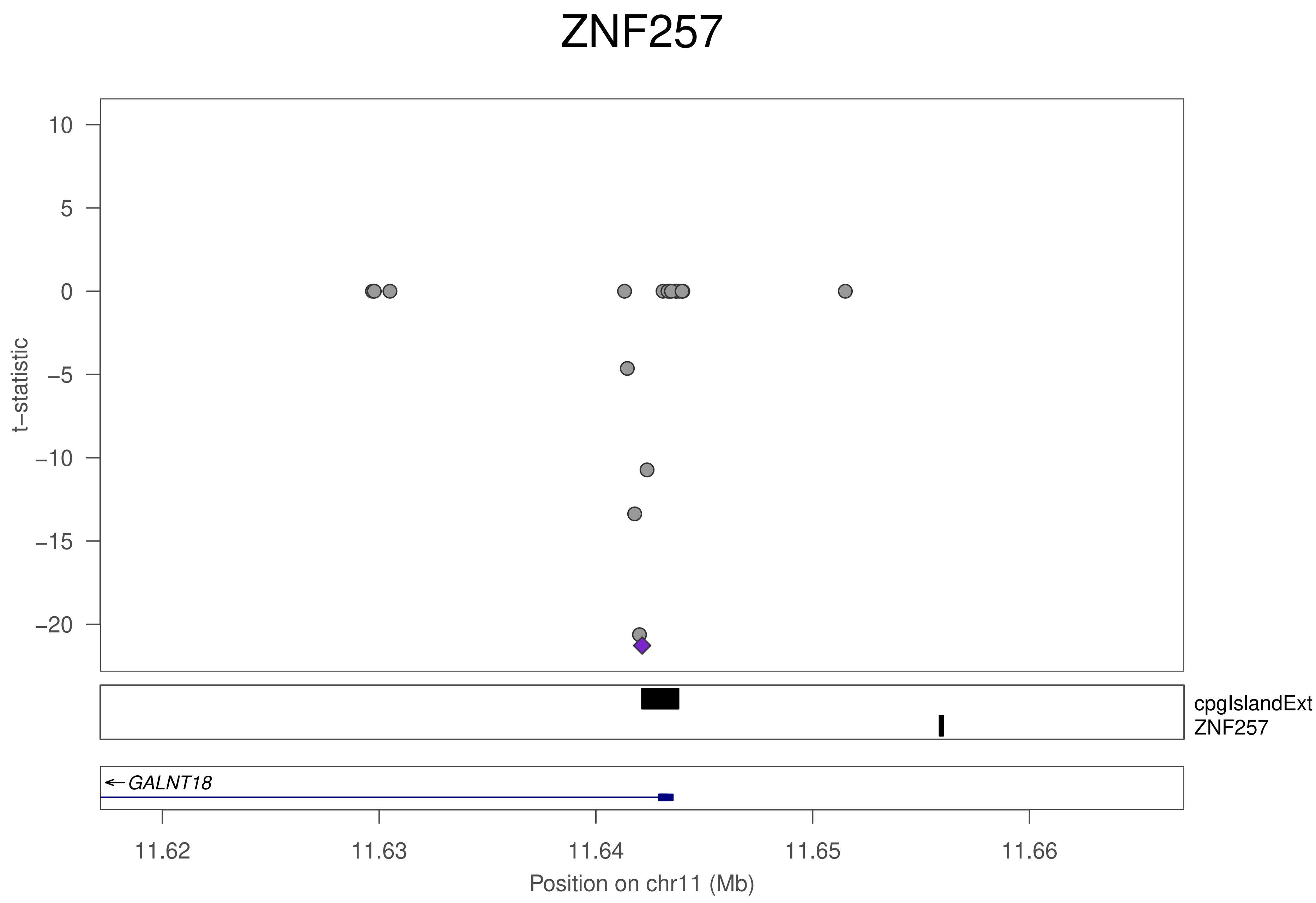
ZNF257
10
5
0
−5
−10
−15
−20
t−statistic
cpgIslandExt
ZNF257
GALNT18
11.62
11.63
11.64
11.65
11.66
Position on chr11 (Mb)

### ZNF257

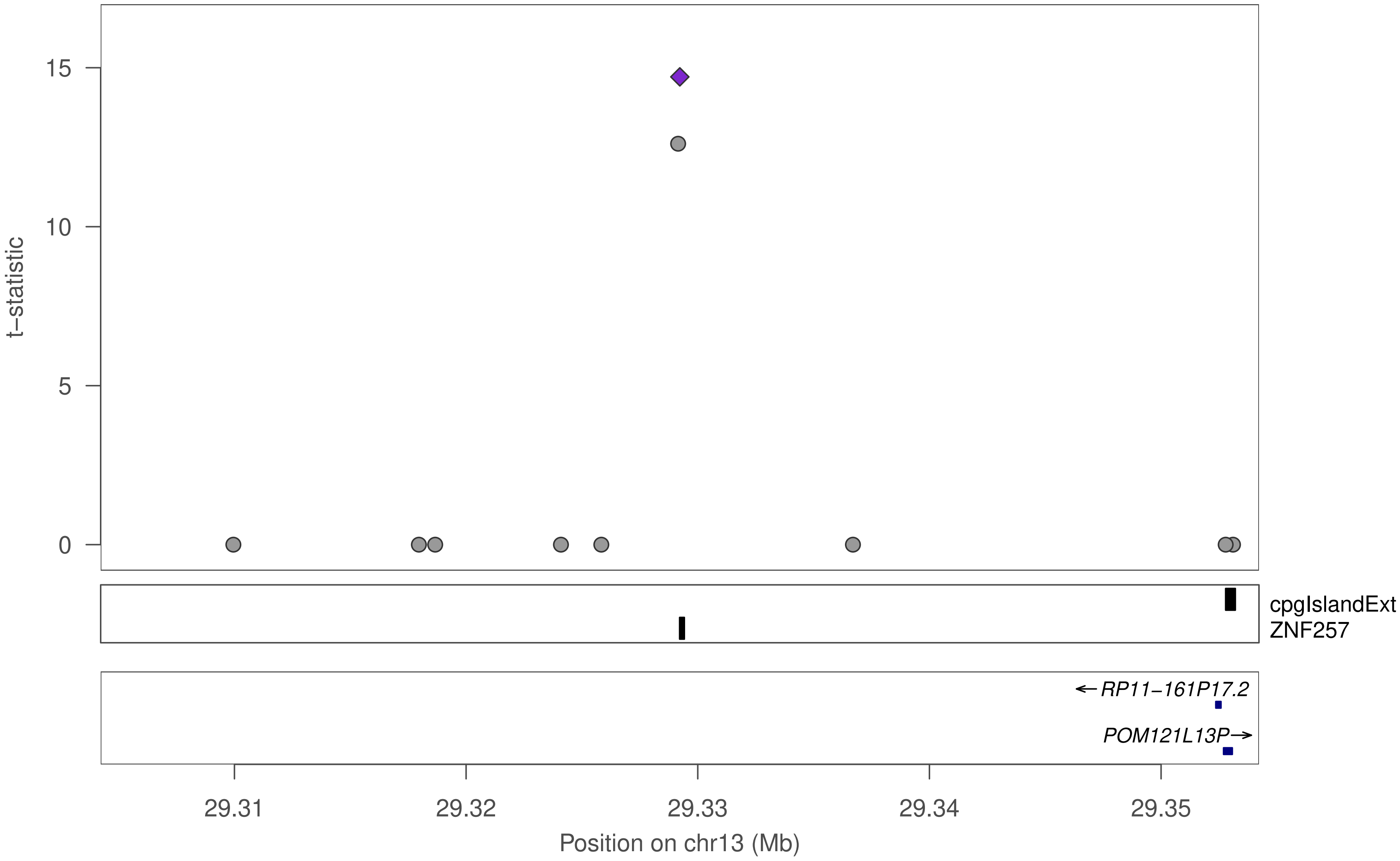
15
10
5
0
t−statistic
cpgIslandExt
ZNF257
←RP11−161P17.2
POM121L13P→
29.31
29.32
29.33
29.34
29.35
Position on chr13 (Mb)

### ZNF257

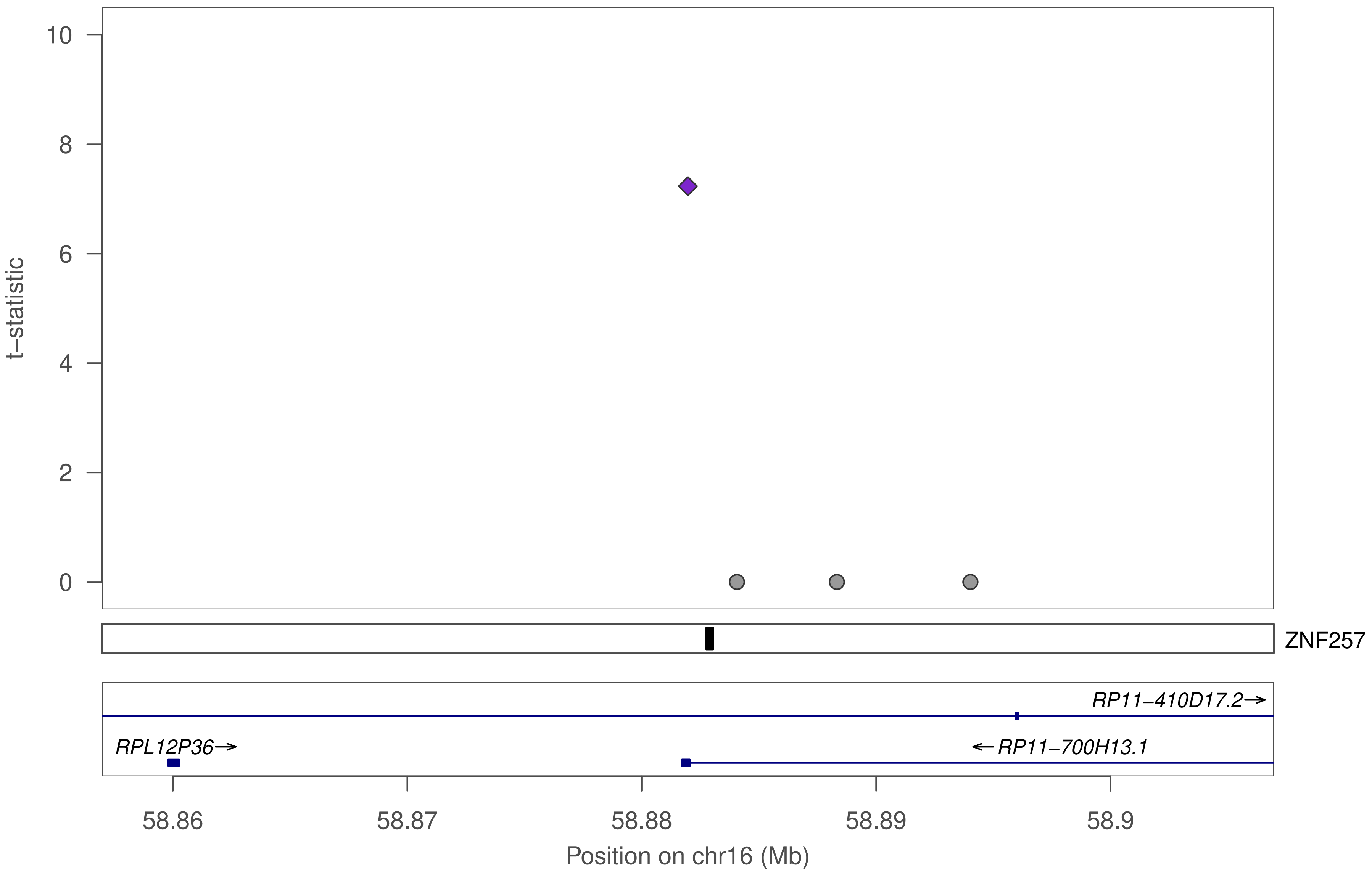

10
8
6
4
2
0
t−statistic
ZNF257
RP11−410D17.2→
RPL12P36→
←RP11−700H13.1
58.86
58.87
58.88
58.89
58.9
Position on chr16 (Mb)

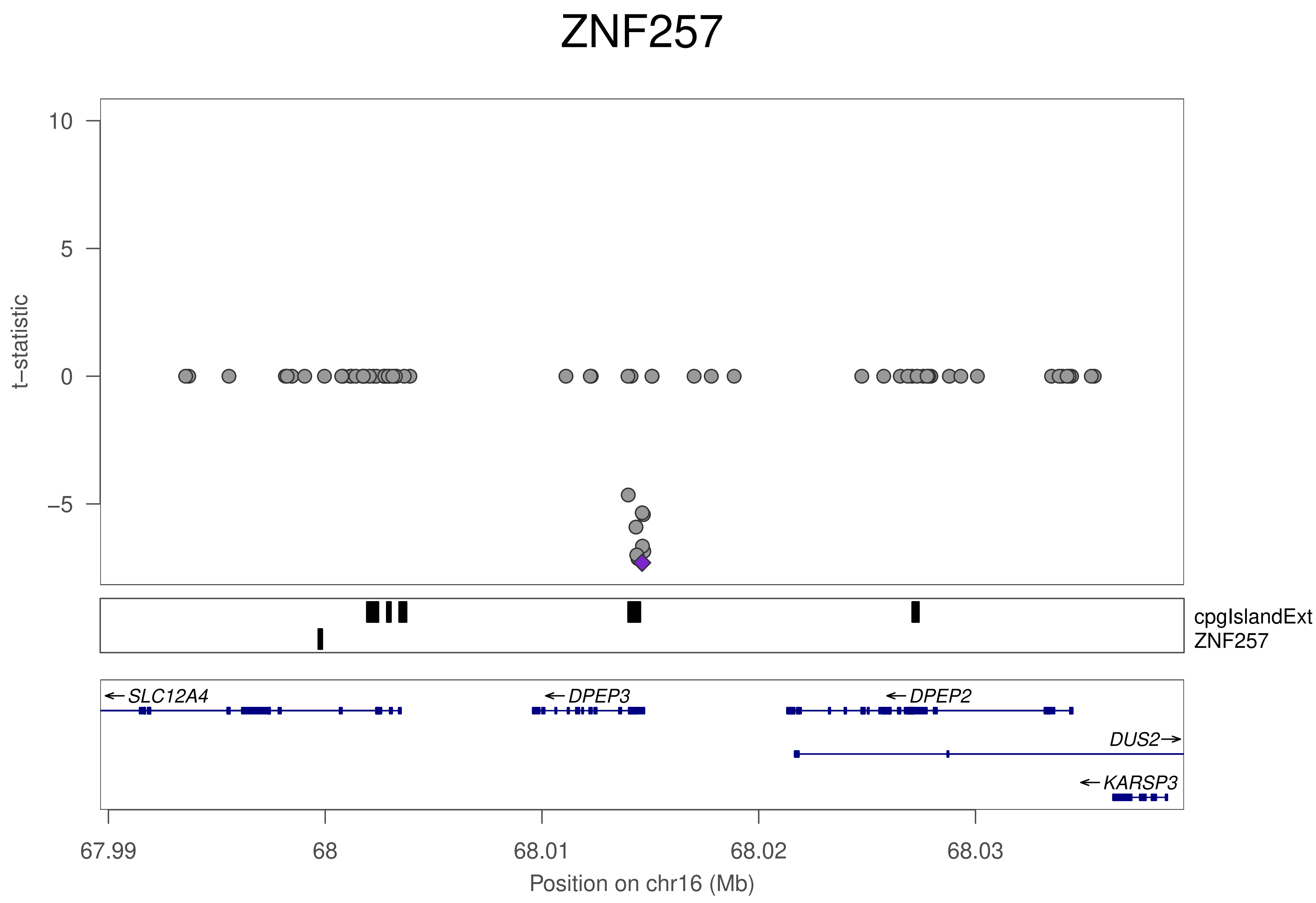
ZNF257
t−statistic
10
5
0
−5
cpgIslandExt
ZNF257
SLC12A4
DPEP3
DPEP2
DUS2
KARSP3
67.99
68
68.01
68.02
68.03
Position on chr16 (Mb)

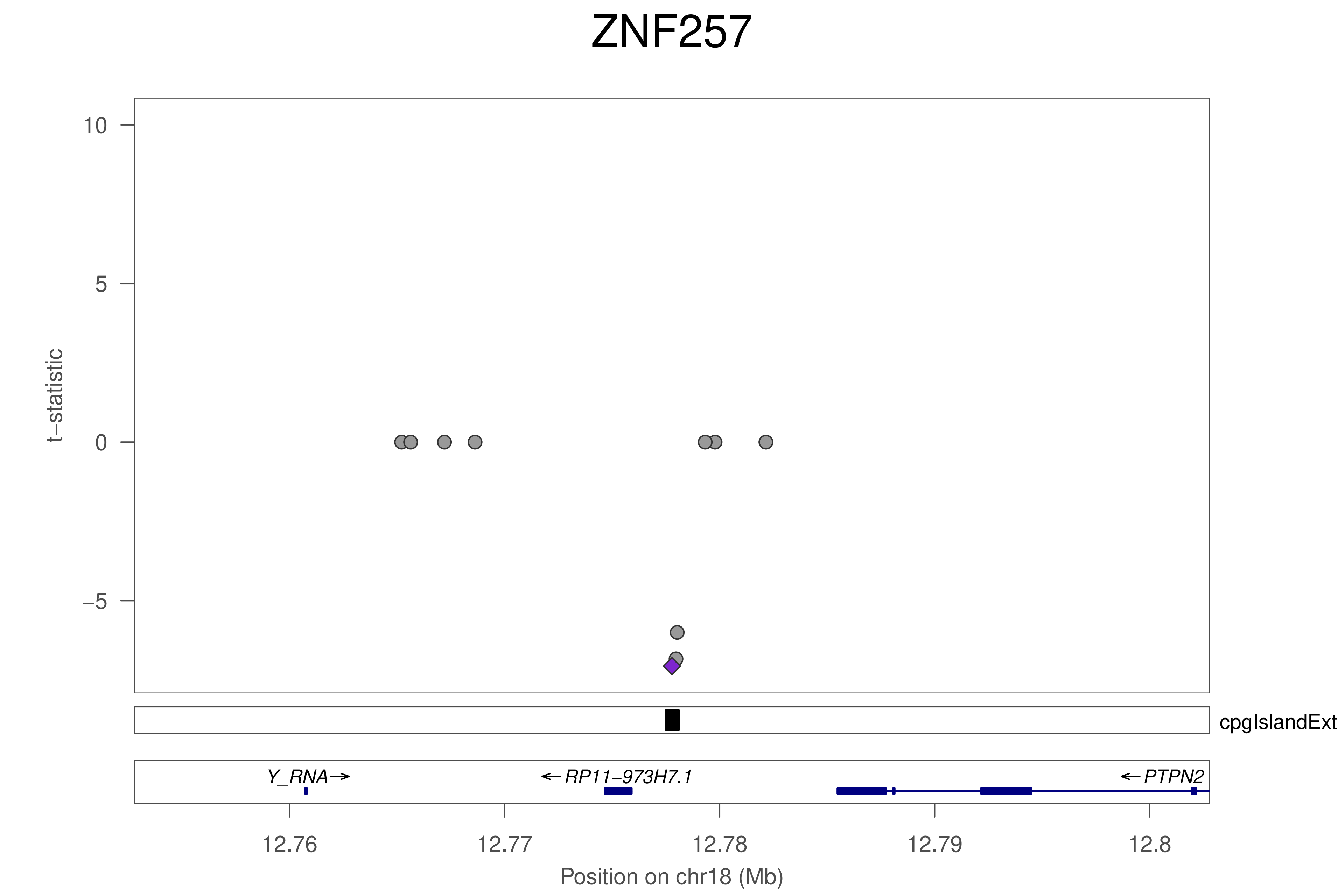
ZNF257
t–statistic
10
5
0
–5
cpgIslandExt
Y_RNA→
←RP11–973H7.1
←PTPN2
12.76
12.77
12.78
12.79
12.8
Position on chr18 (Mb)

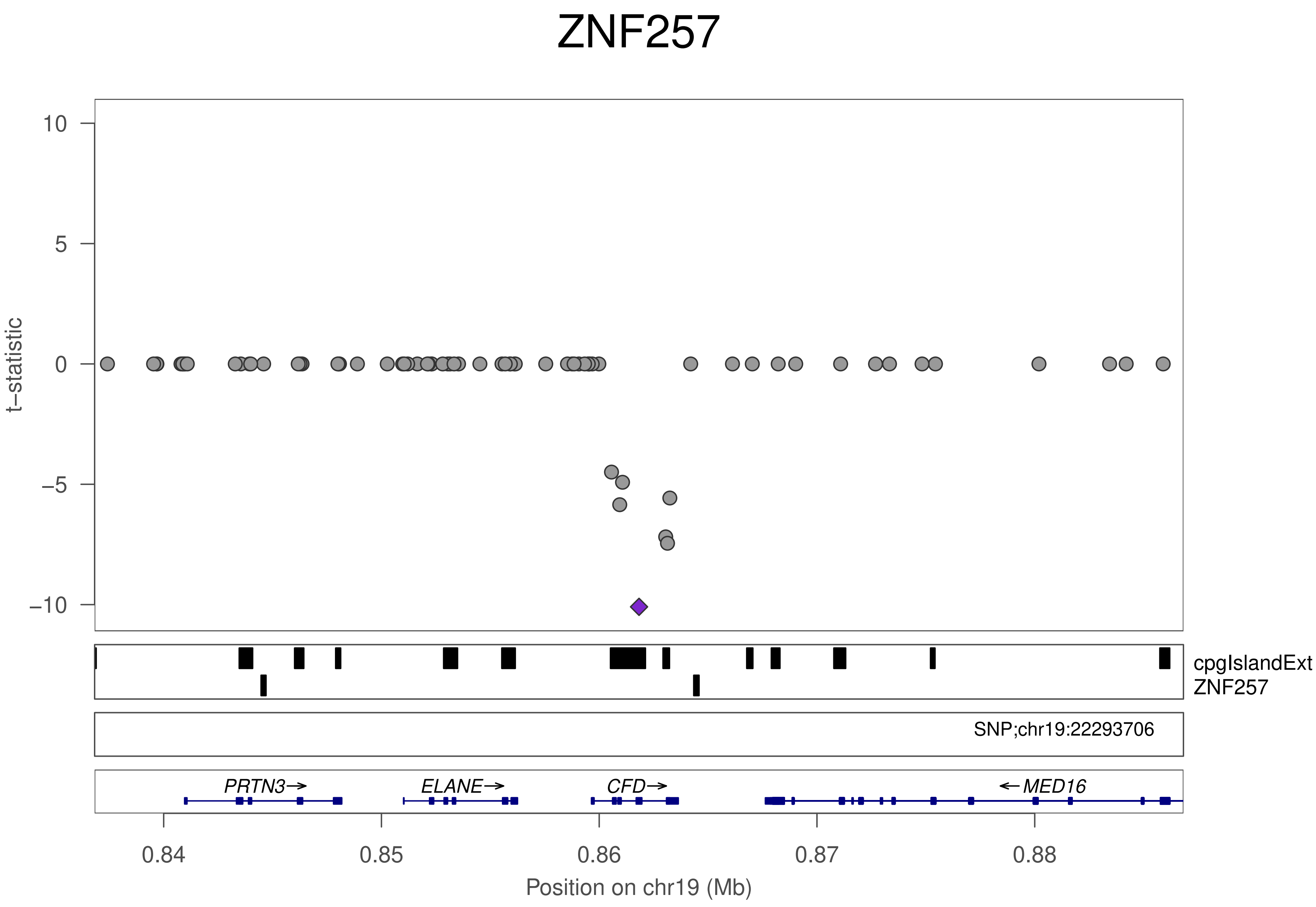
ZNF257
t−statistic
10
5
0
−5
−10
cpgIslandExt
ZNF257
SNP;chr19:22293706
PRTN3→
ELANE→
CFD→
←MED16
0.84
0.85
0.86
0.87
0.88
Position on chr19 (Mb)

### ZNF786

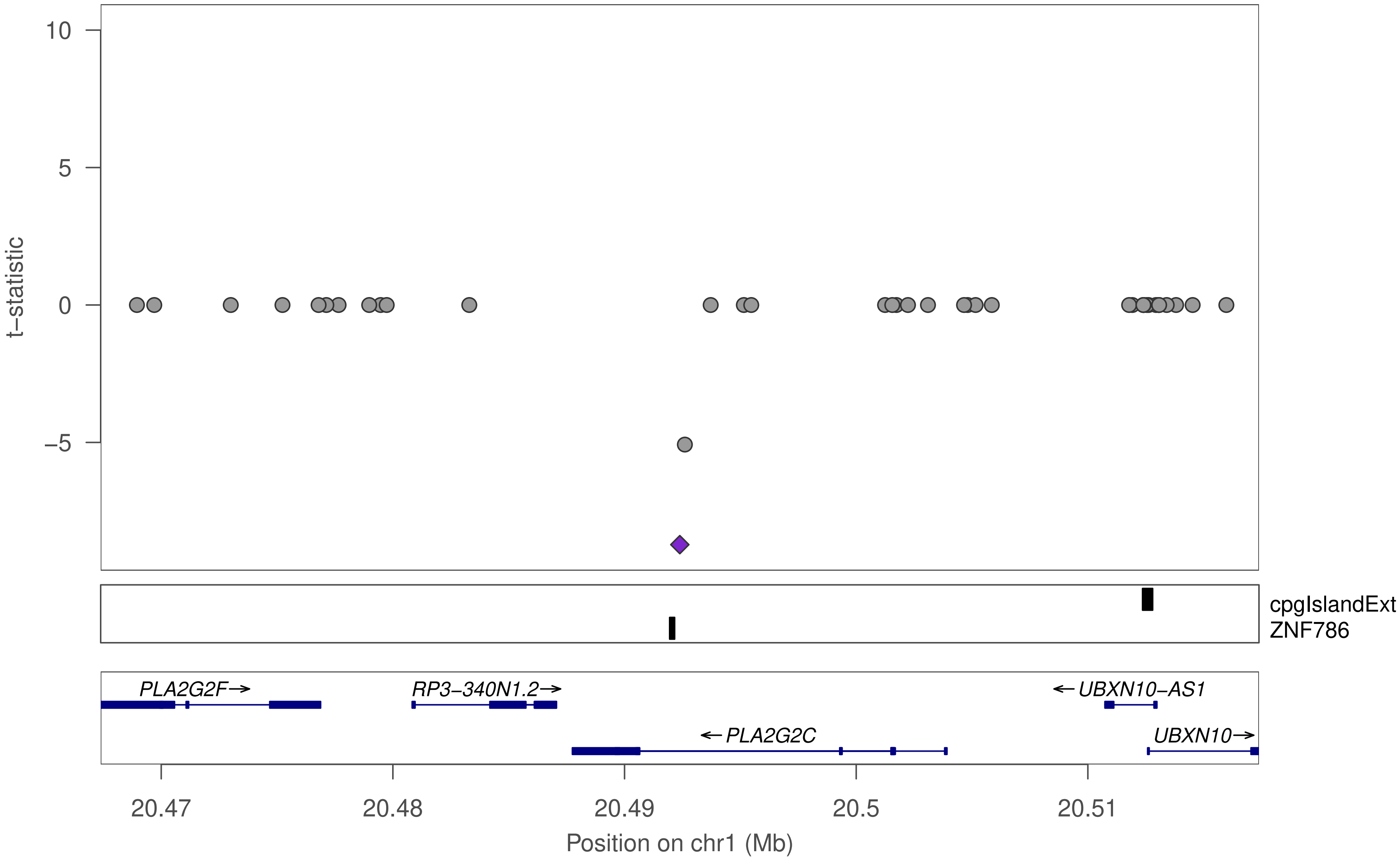
10
5
0
−5
t−statistic
cpgIslandExt
ZNF786
PLA2G2F→
RP3−340N1.2→
←UBXN10−AS1
←PLA2G2C
UBXN10→
20.47
20.48
20.49
20.5
20.51
Position on chr1 (Mb)

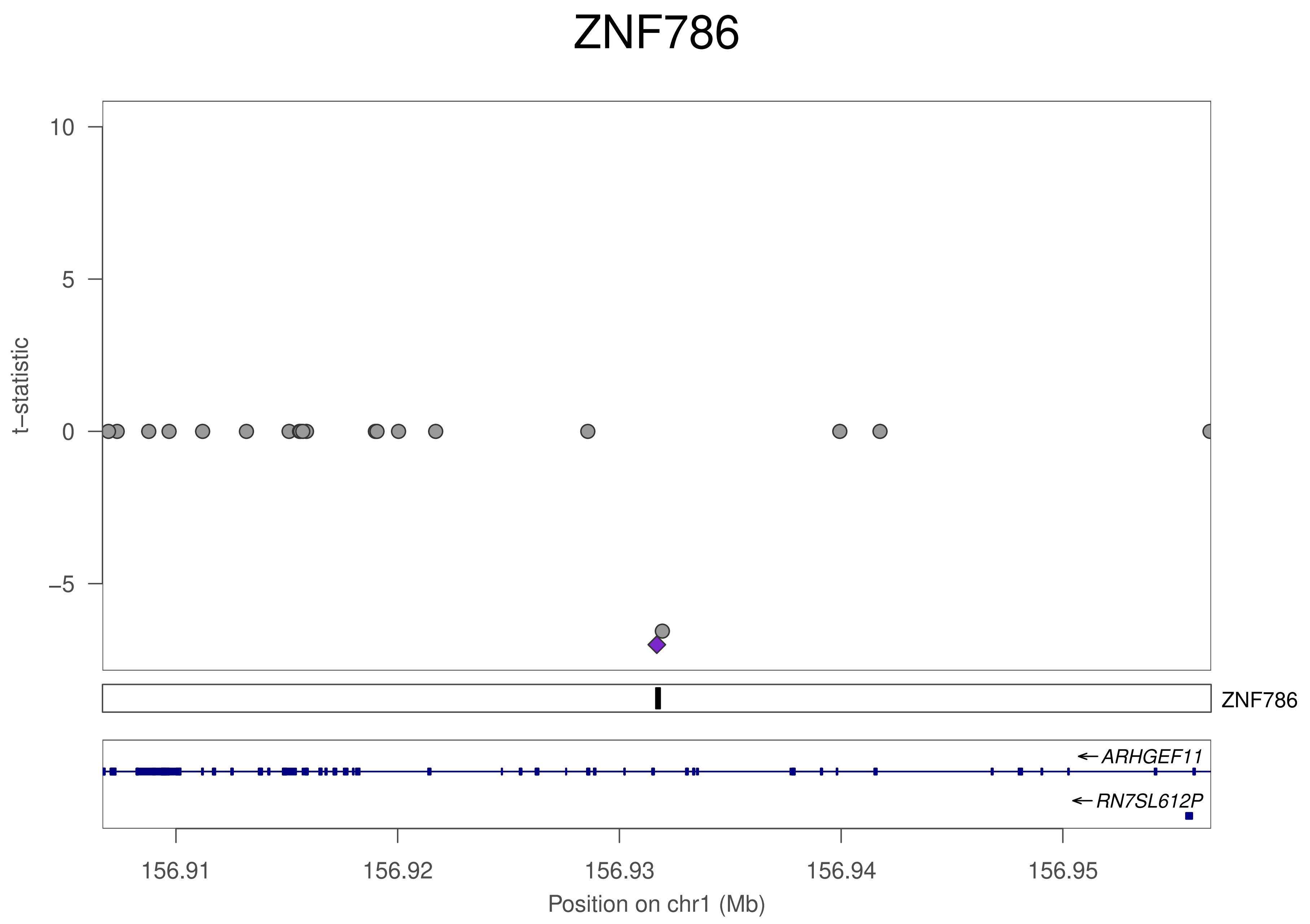
ZNF786
t−statistic
10
5
0
−5
ZNF786
ARHGEF11
RN7SL612P
156.91
156.92
156.93
156.94
156.95
Position on chr1 (Mb)

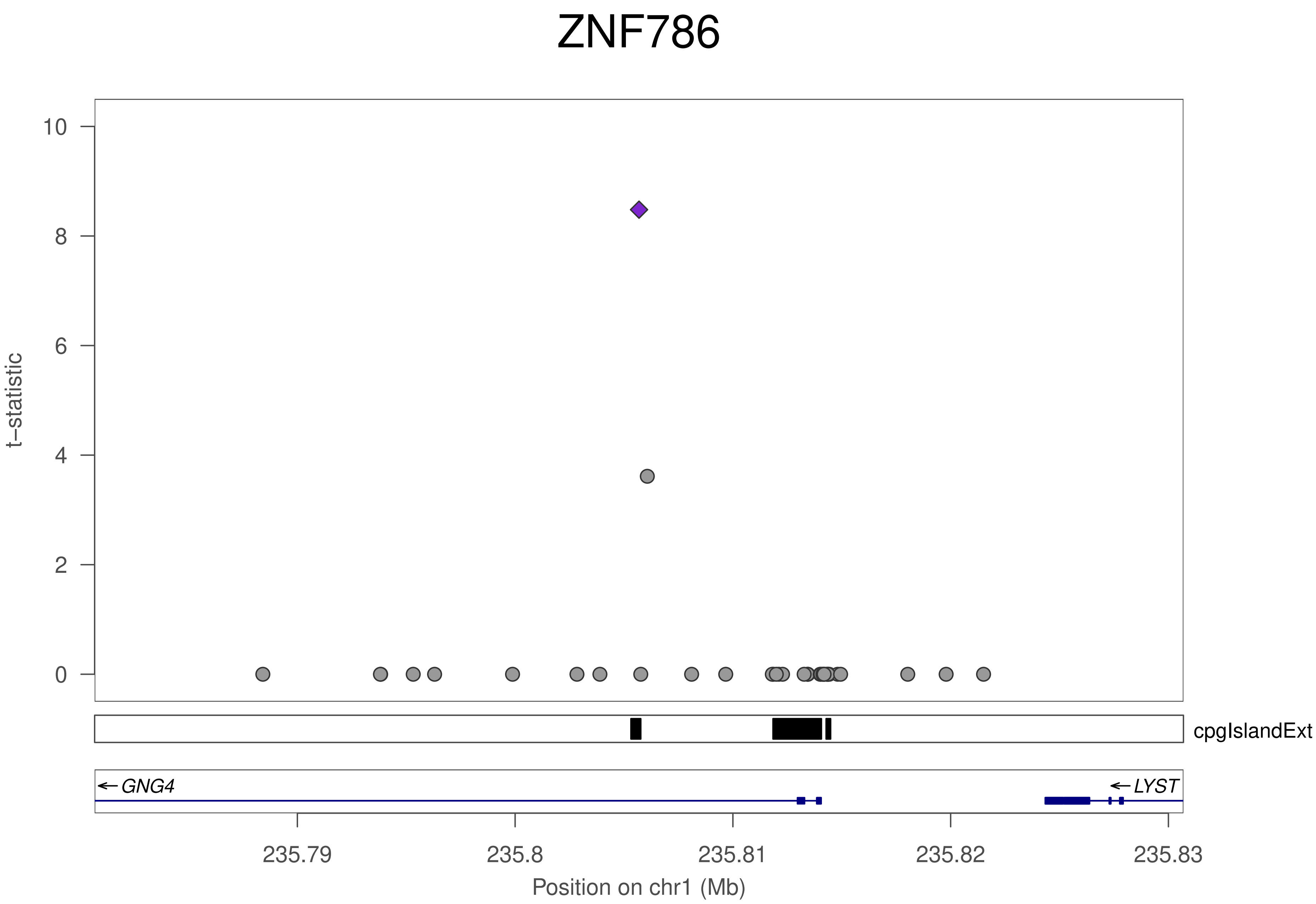
ZNF786
t–statistic
0
2
4
6
8
10
cpgIslandExt
GNG4
LYST
235.79
235.8
235.81
235.82
235.83
Position on chr1 (Mb)

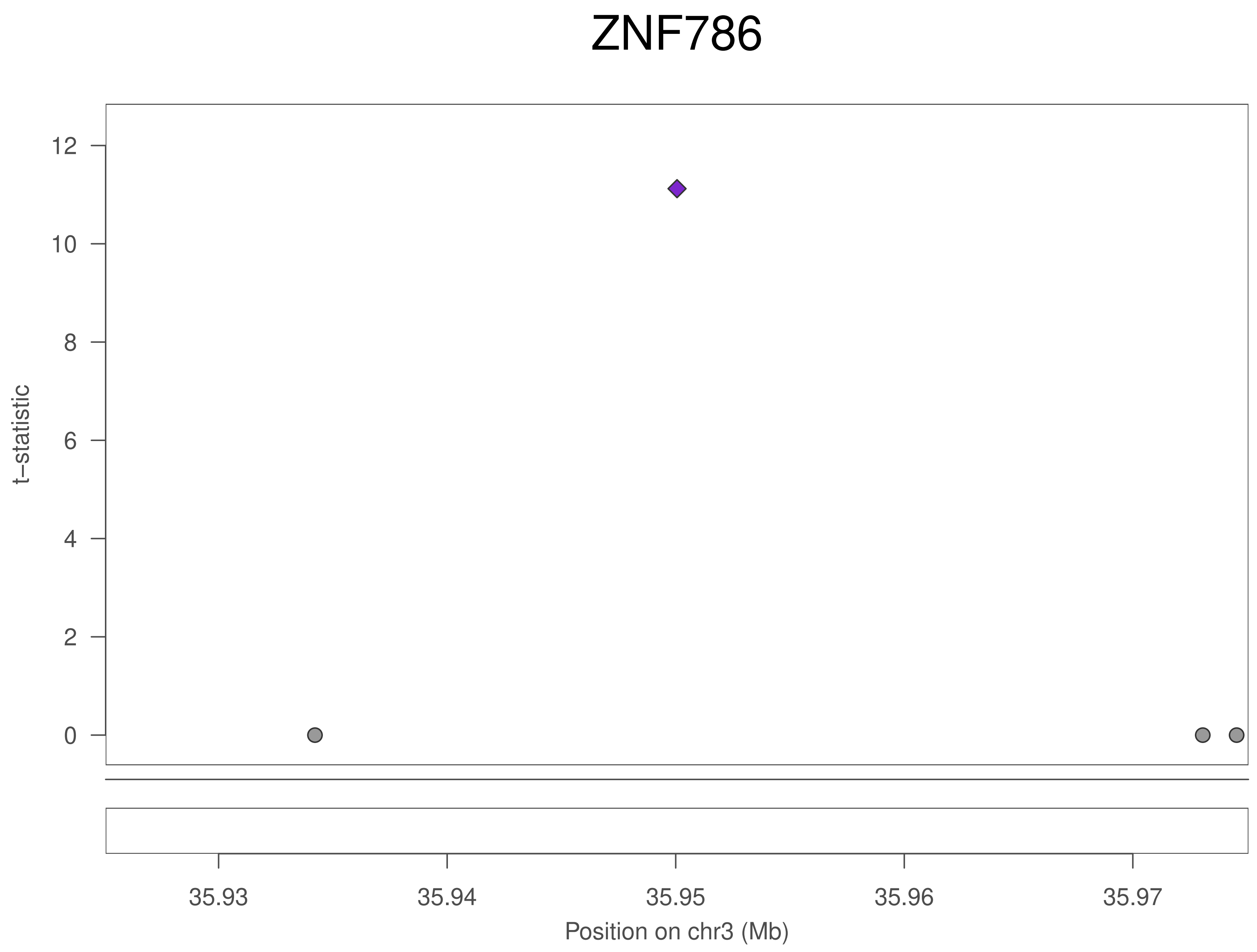

ZNF786
t–statistic
12
10
8
6
4
2
0
35.93
35.94
35.95
35.96
35.97
Position on chr3 (Mb)

### ZNF786

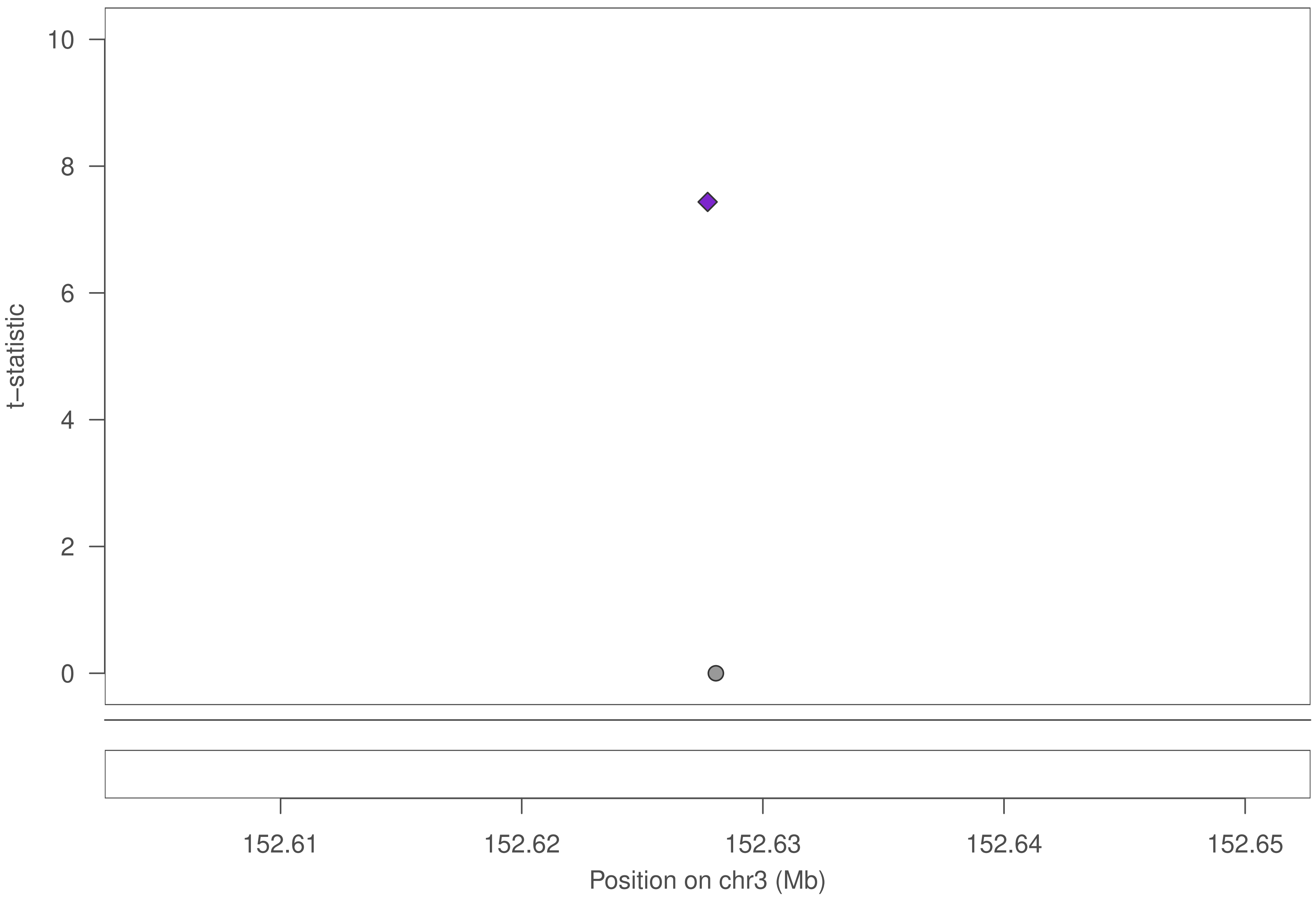
10
8
6
4
2
0
t–statistic
152.61
152.62
152.63
152.64
152.65
Position on chr3 (Mb)

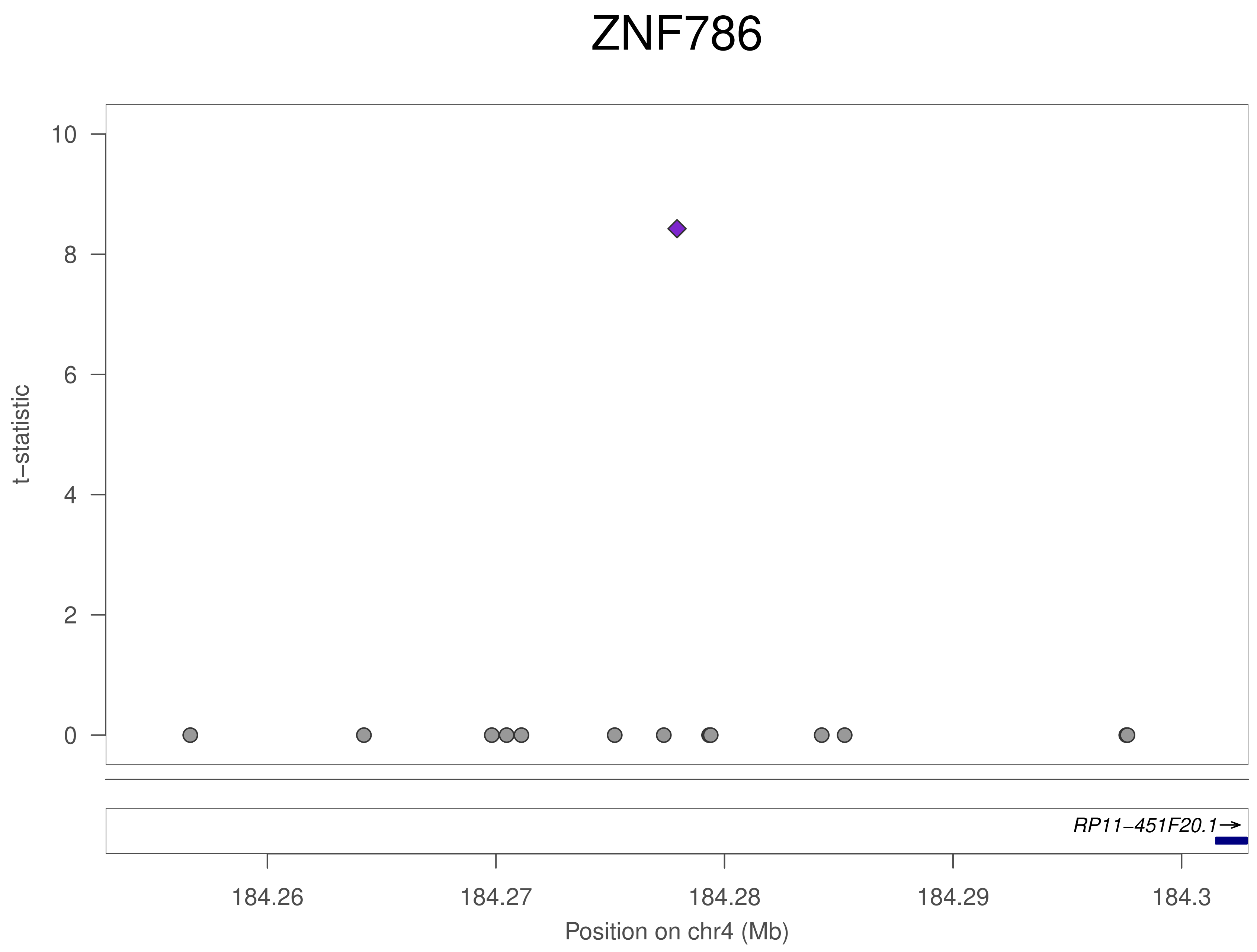
ZNF786
t−statistic
10
8
6
4
2
0
RP11−451F20.1→
184.26
184.27
184.28
184.29
184.3
Position on chr4 (Mb)

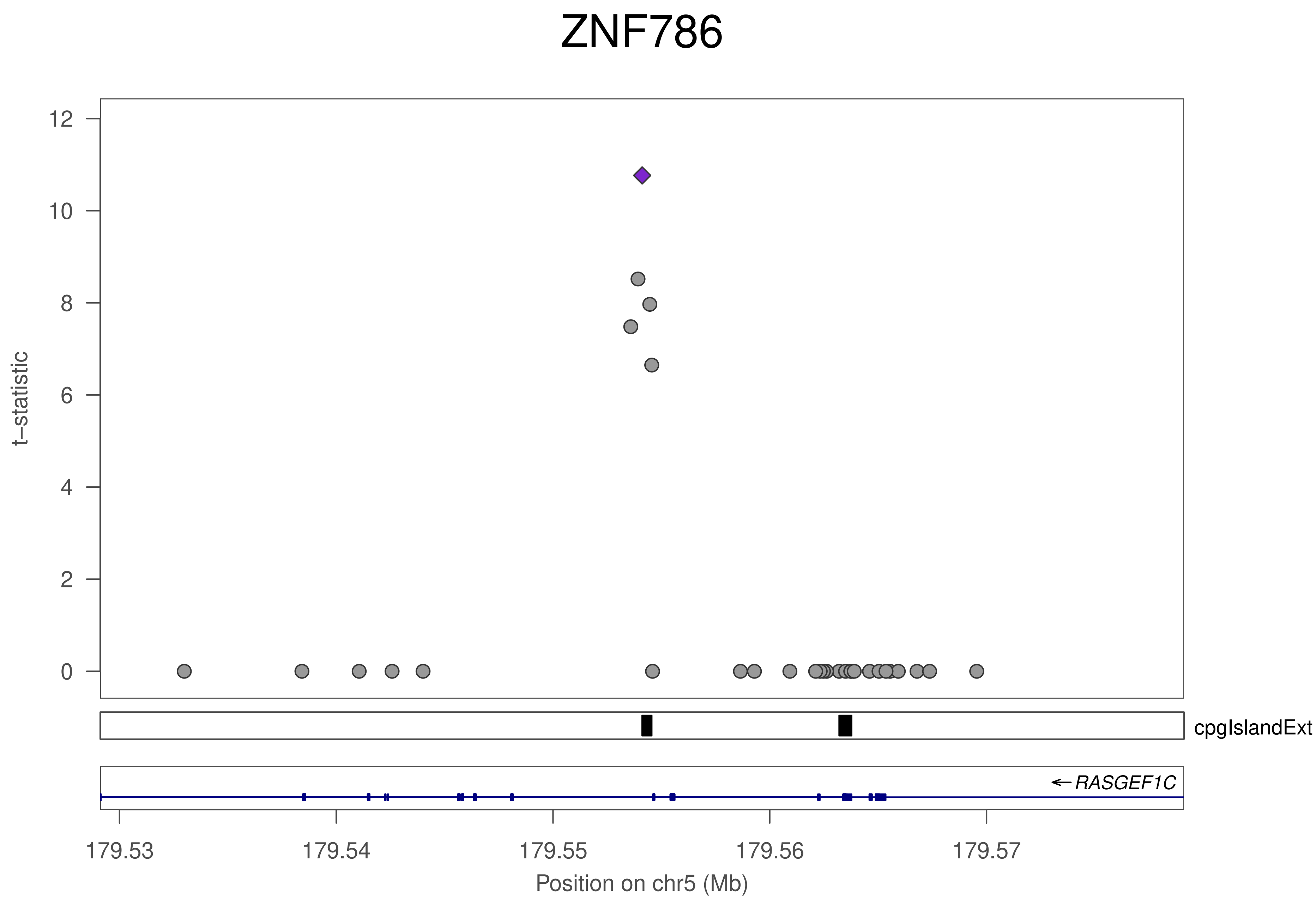
ZNF786
t–statistic
0
2
4
6
8
10
12
cpgIslandExt
← RASGEF1C
179.53
179.54
179.55
179.56
179.57
Position on chr5 (Mb)

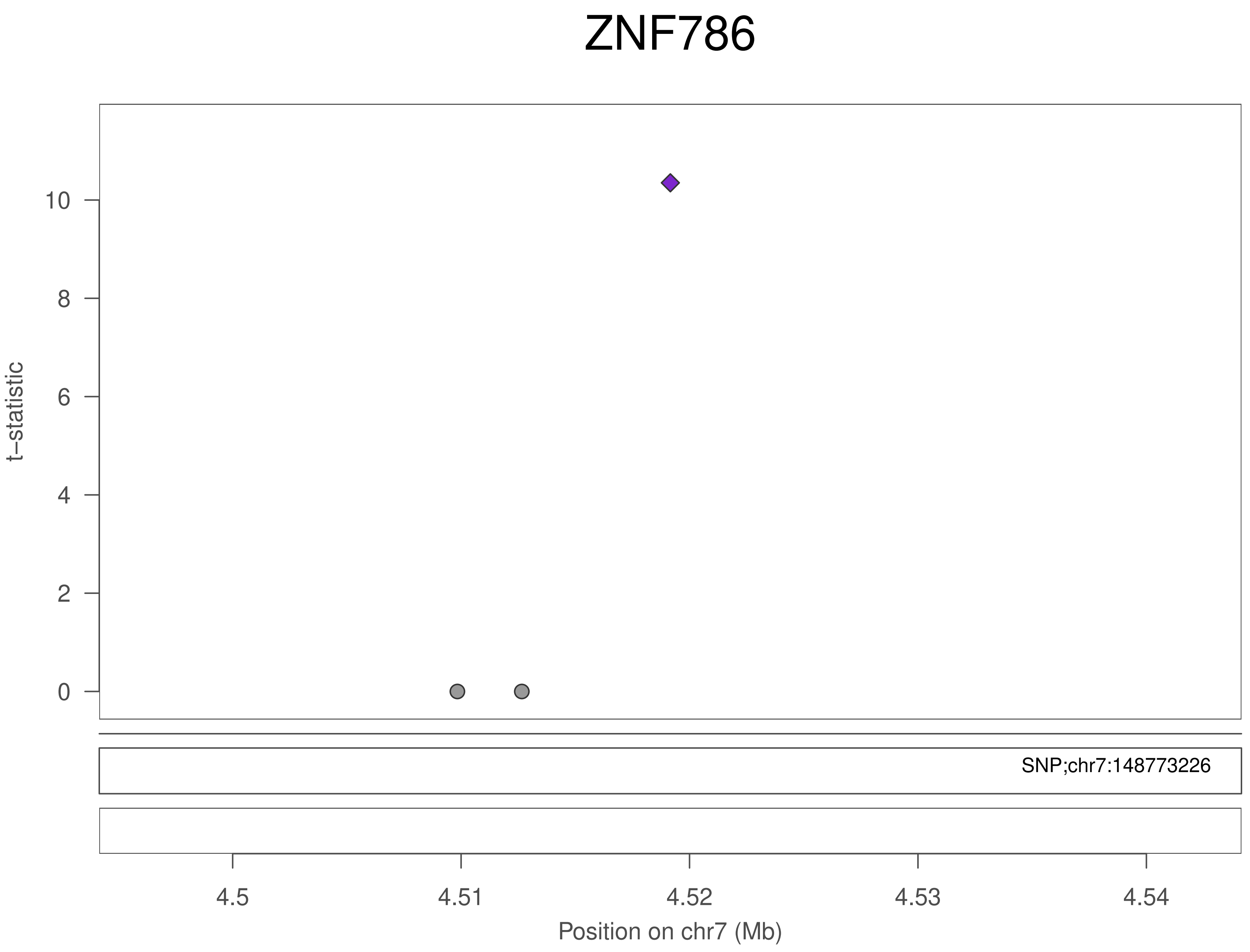
ZNF786
t–statistic
10
8
6
4
2
0
SNP;chr7:148773226
4.5
4.51
4.52
4.53
4.54
Position on chr7 (Mb)

### ZNF786

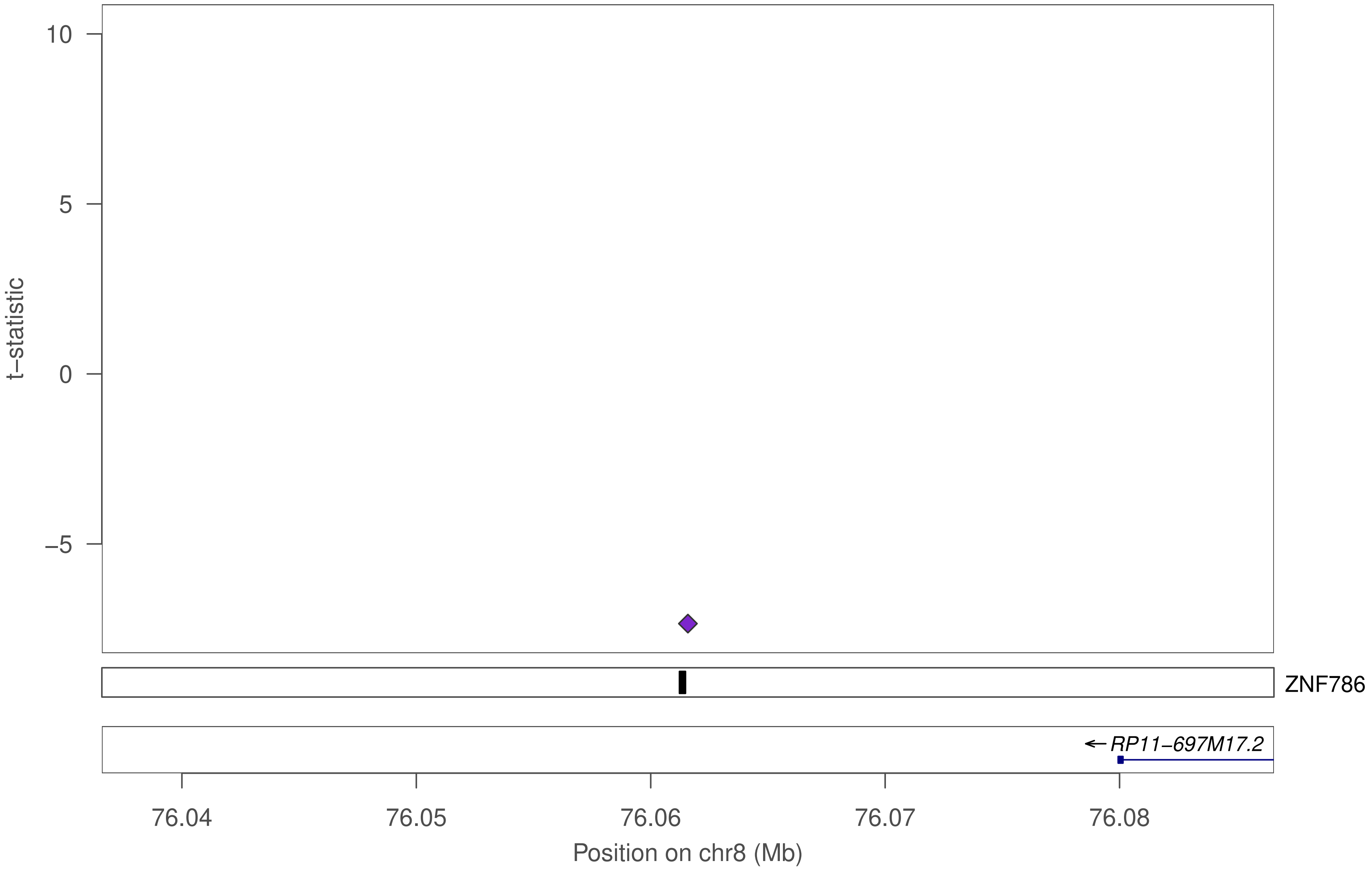

10
5
0
−5
t−statistic
ZNF786
←RP11−697M17.2
76.04
76.05
76.06
76.07
76.08
Position on chr8 (Mb)

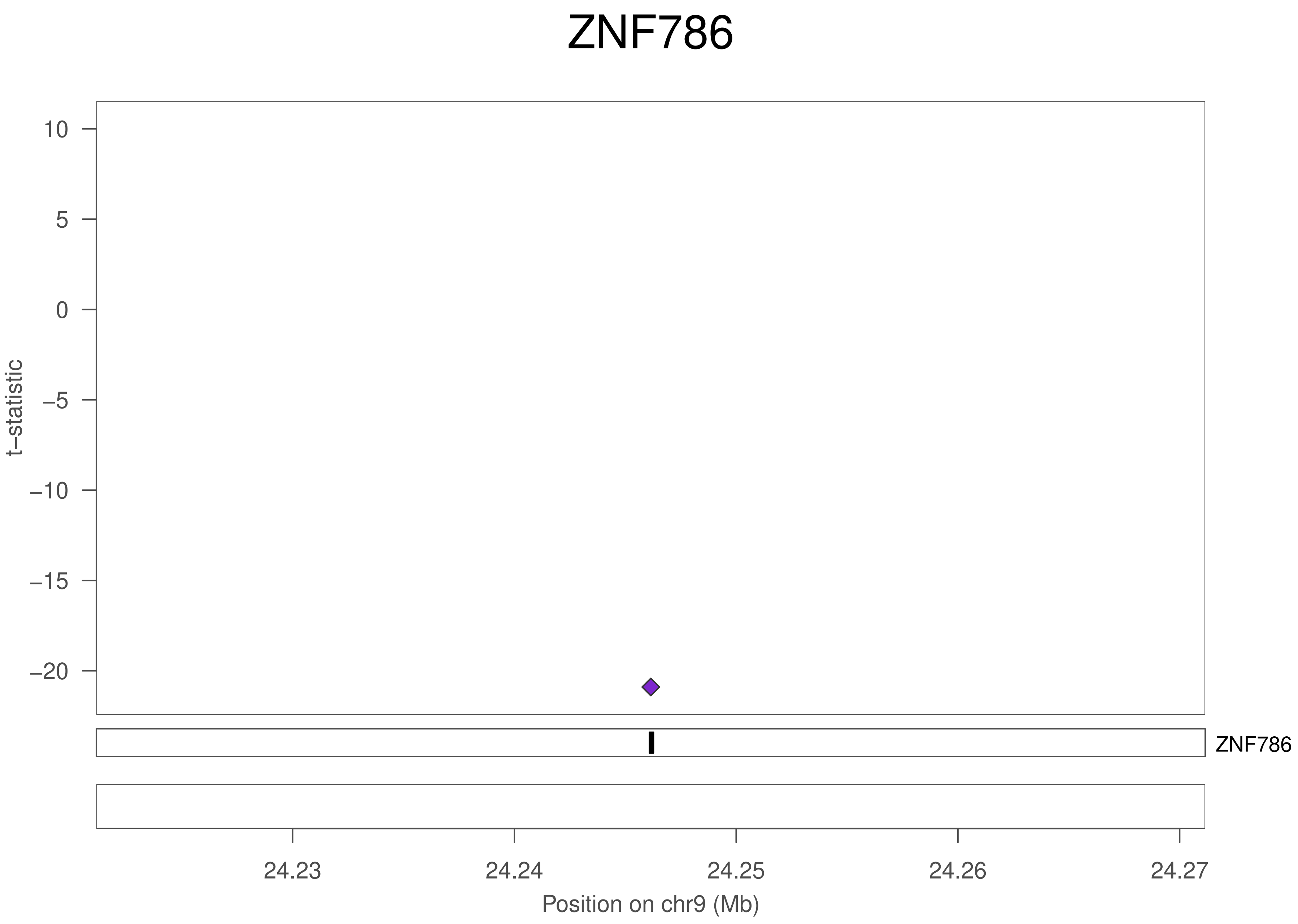
ZNF786
10
5
0
−5
−10
−15
−20
t−statistic
ZNF786
24.23
24.24
24.25
24.26
24.27
Position on chr9 (Mb)

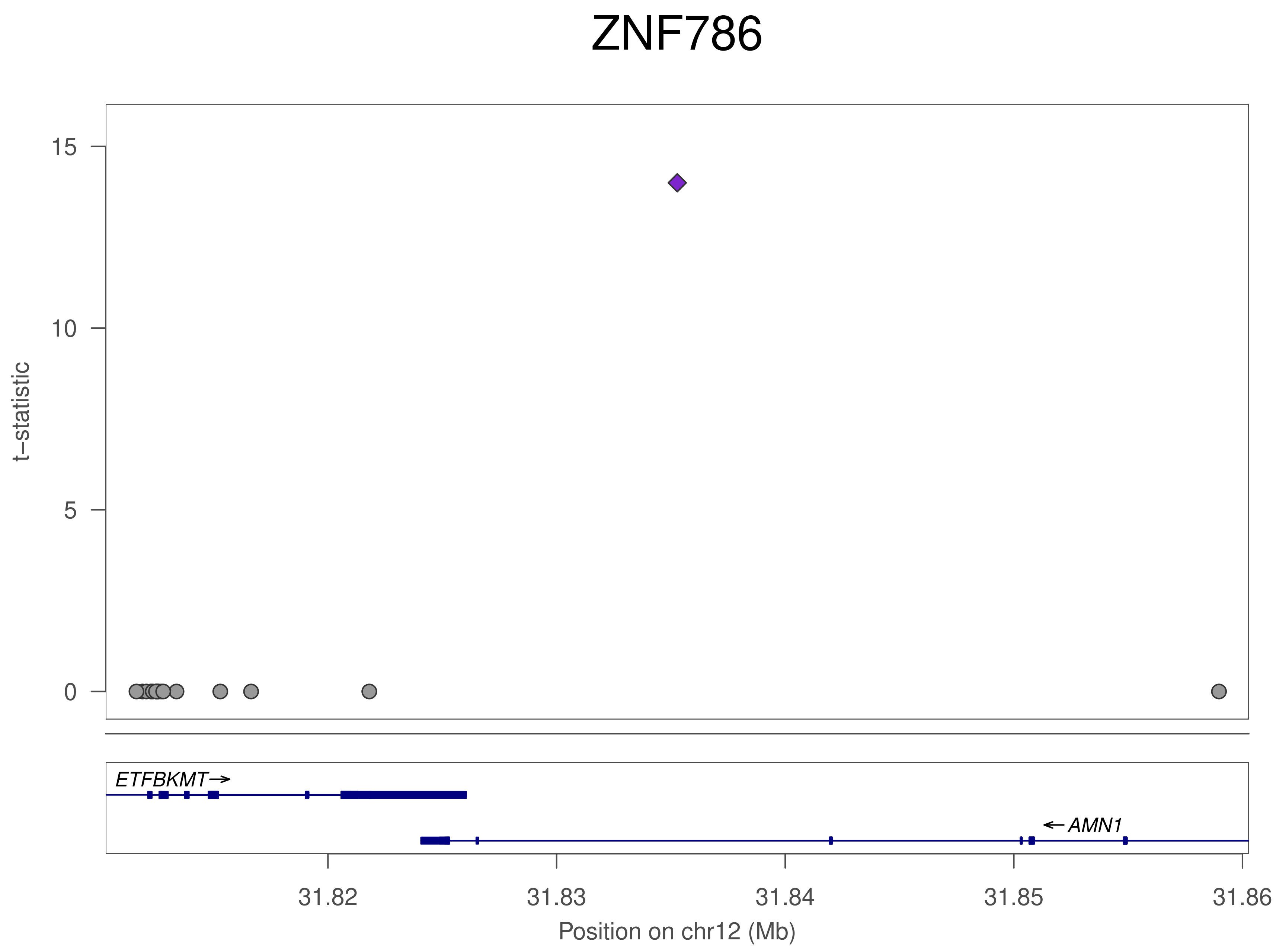
ZNF786
t−statistic
15
10
5
0
ETFBKMT
AMN1
31.82
31.83
31.84
31.85
31.86
Position on chr12 (Mb)

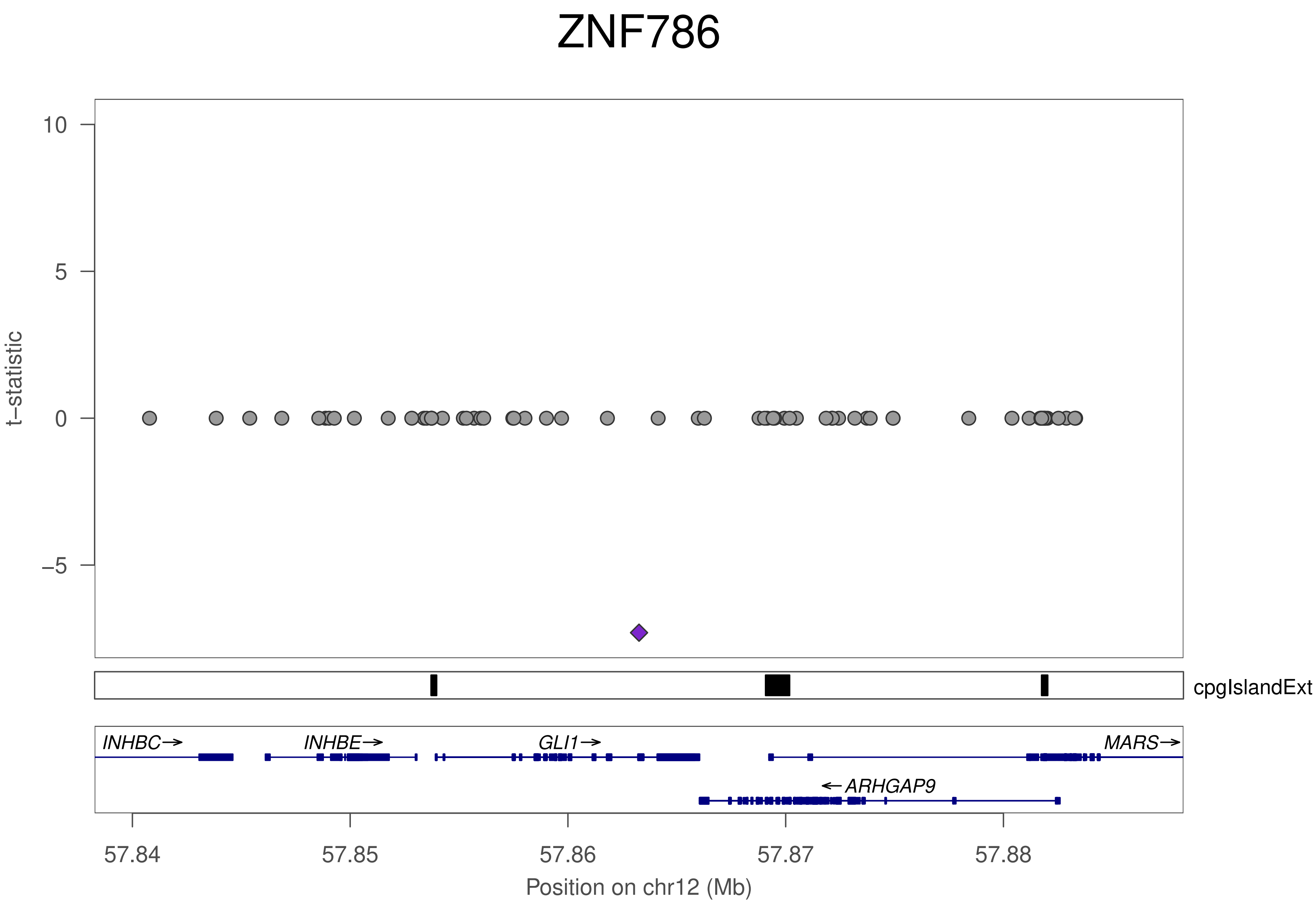

ZNF786
t−statistic
10
5
0
−5
cpgIslandExt
INHBC→
INHBE→
GLI1→
MARS→
←ARHGAP9
57.84
57.85
57.86
57.87
57.88
Position on chr12 (Mb)

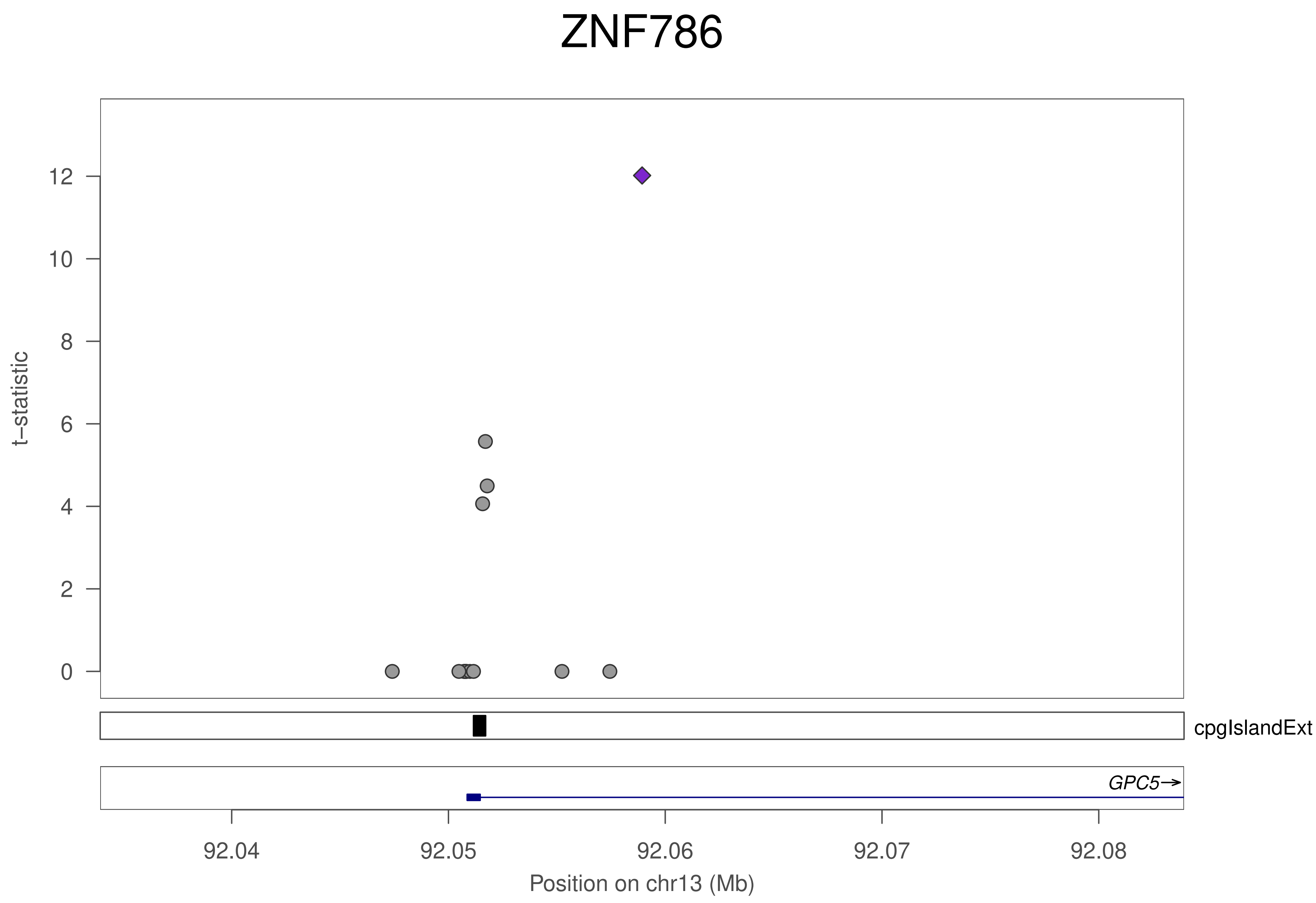
ZNF786
t−statistic
0
2
4
6
8
10
12
cpgIslandExt
GPC5
92.04
92.05
92.06
92.07
92.08
Position on chr13 (Mb)

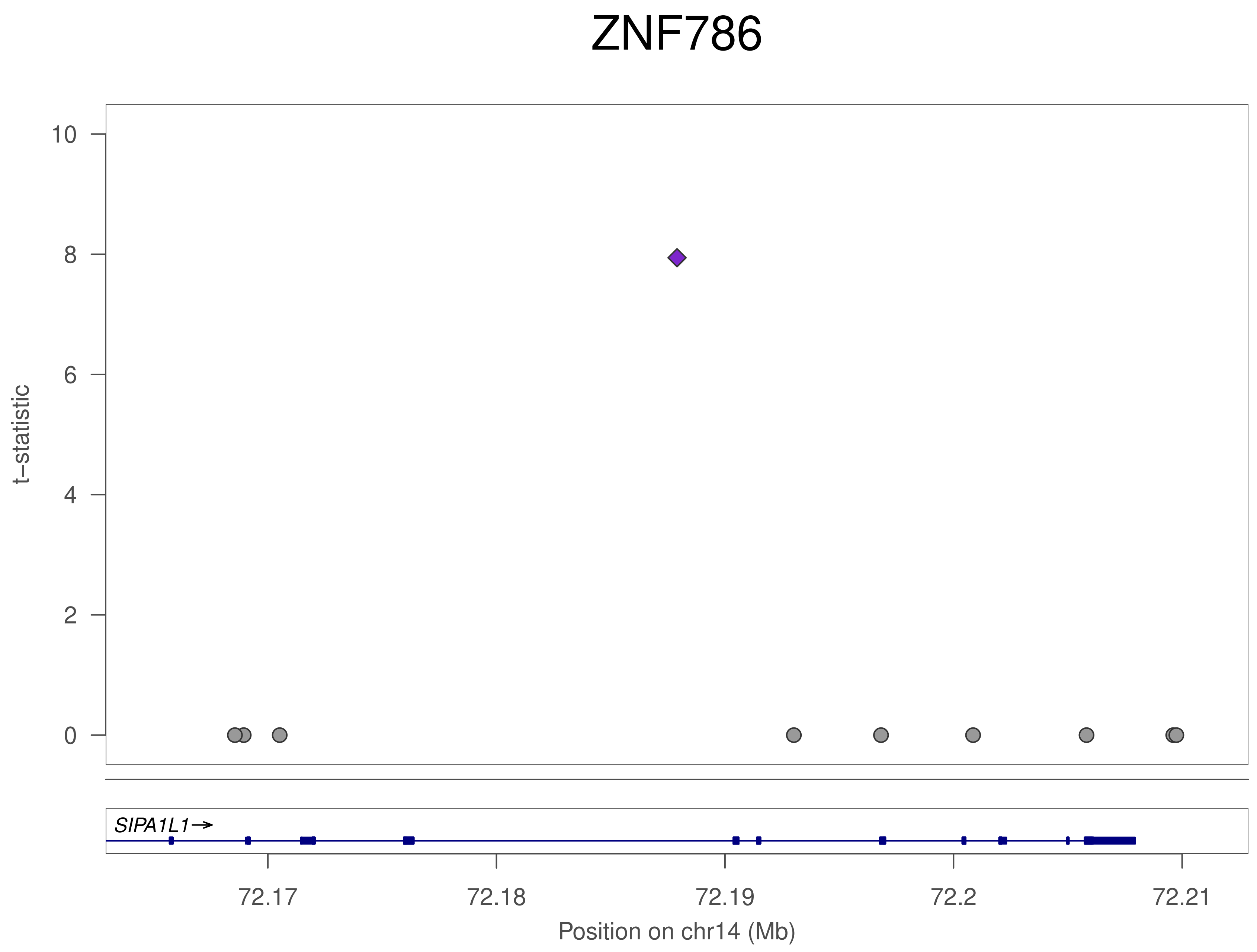
ZNF786
t−statistic
10
8
6
4
2
0
SIPA1L1→
72.17
72.18
72.19
72.2
72.21
Position on chr14 (Mb)

### ZNF786

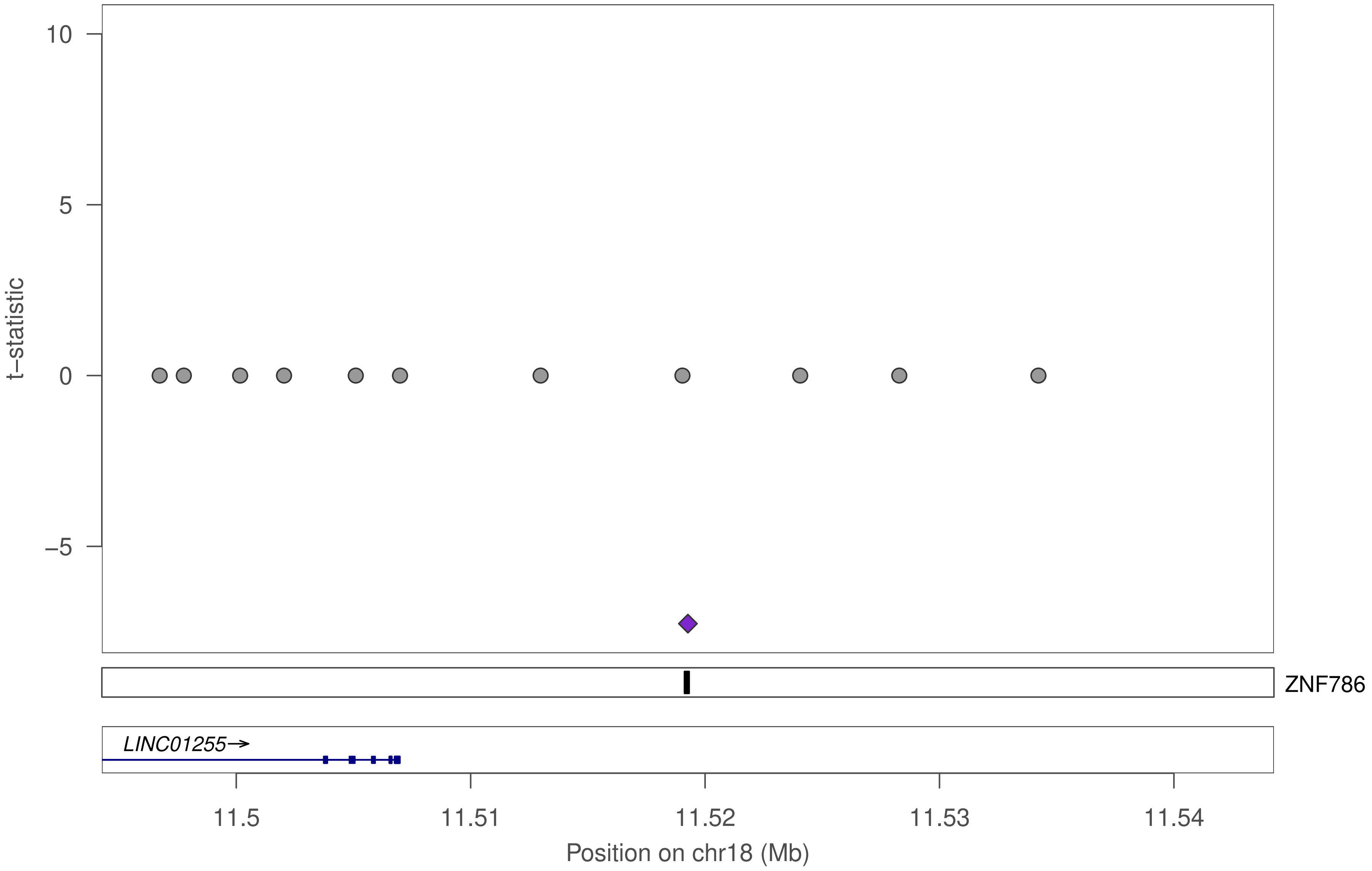

10
5
0
−5
t−statistic
ZNF786
LINC01255→
11.5
11.51
11.52
11.53
11.54
Position on chr18 (Mb)

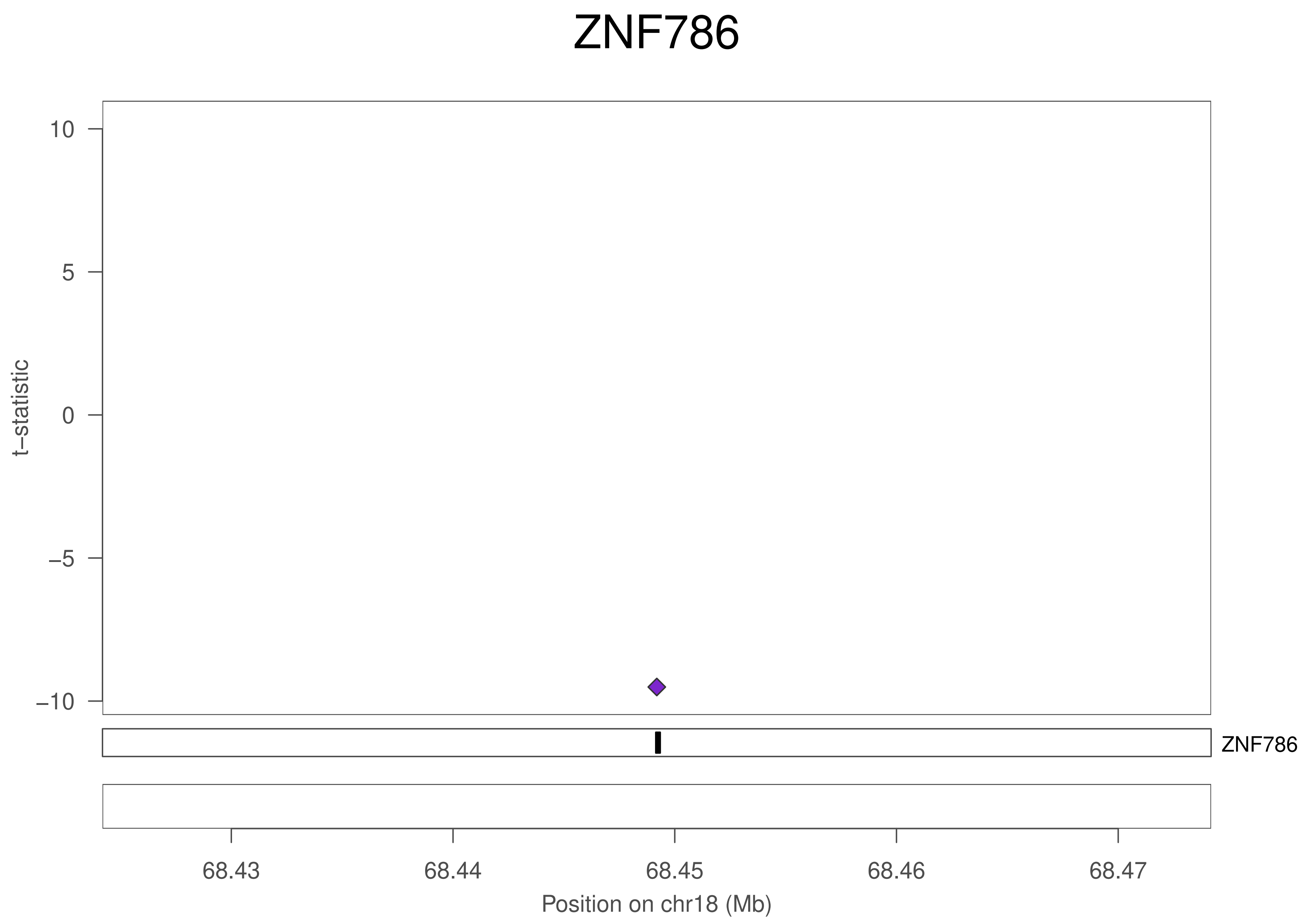
ZNF786
10
5
0
−5
−10
t−statistic
ZNF786
68.43
68.44
68.45
68.46
68.47
Position on chr18 (Mb)

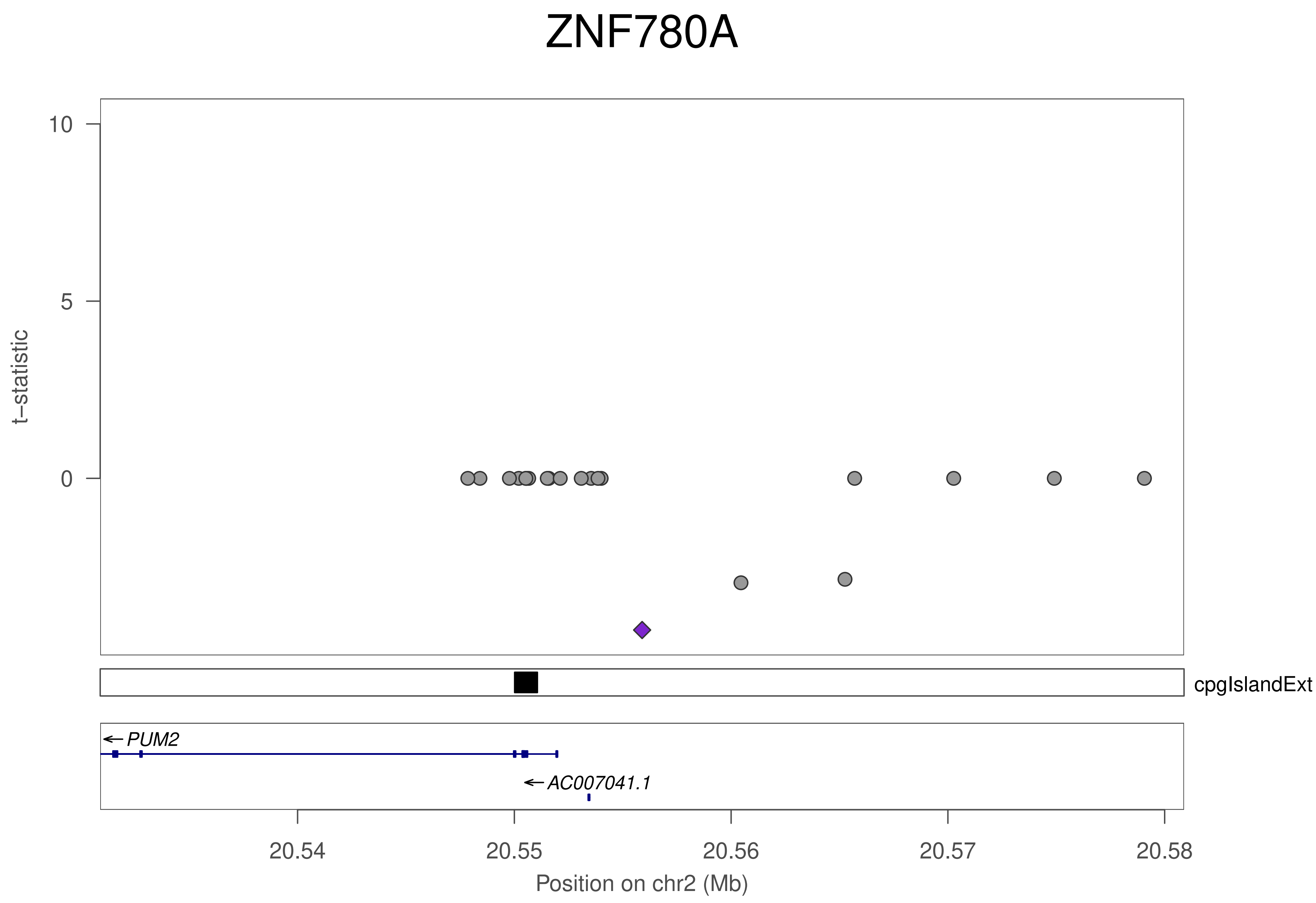

ZNF780A
10
5
0
t−statistic
cpgIslandExt
PUM2
AC007041.1
20.54
20.55
20.56
20.57
20.58
Position on chr2 (Mb)

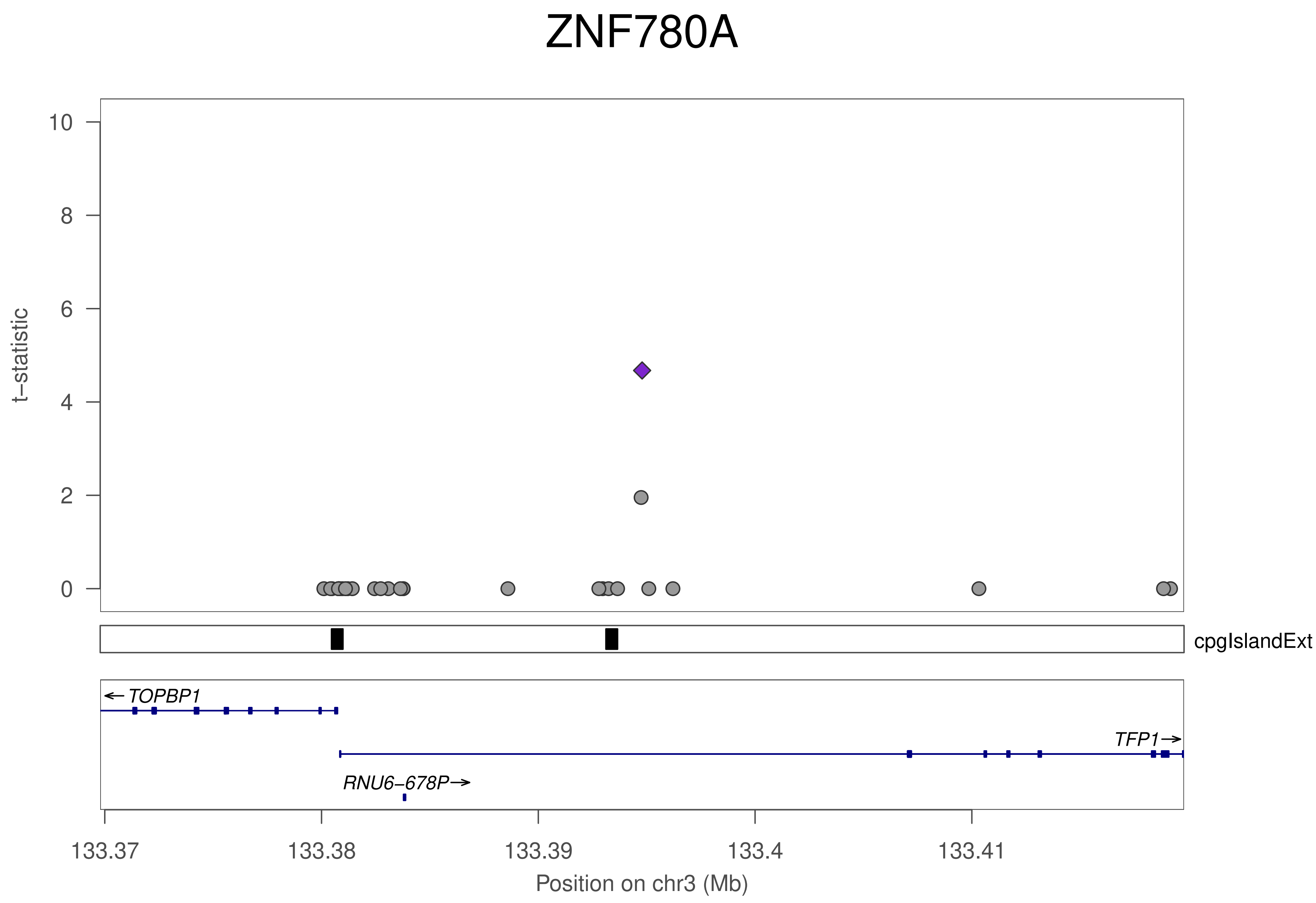
ZNF780A
t–statistic
10
8
6
4
2
0
cpgIslandExt
← TOPBP1
TFP1→
RNU6–678P→
133.37
133.38
133.39
133.4
133.41
Position on chr3 (Mb)

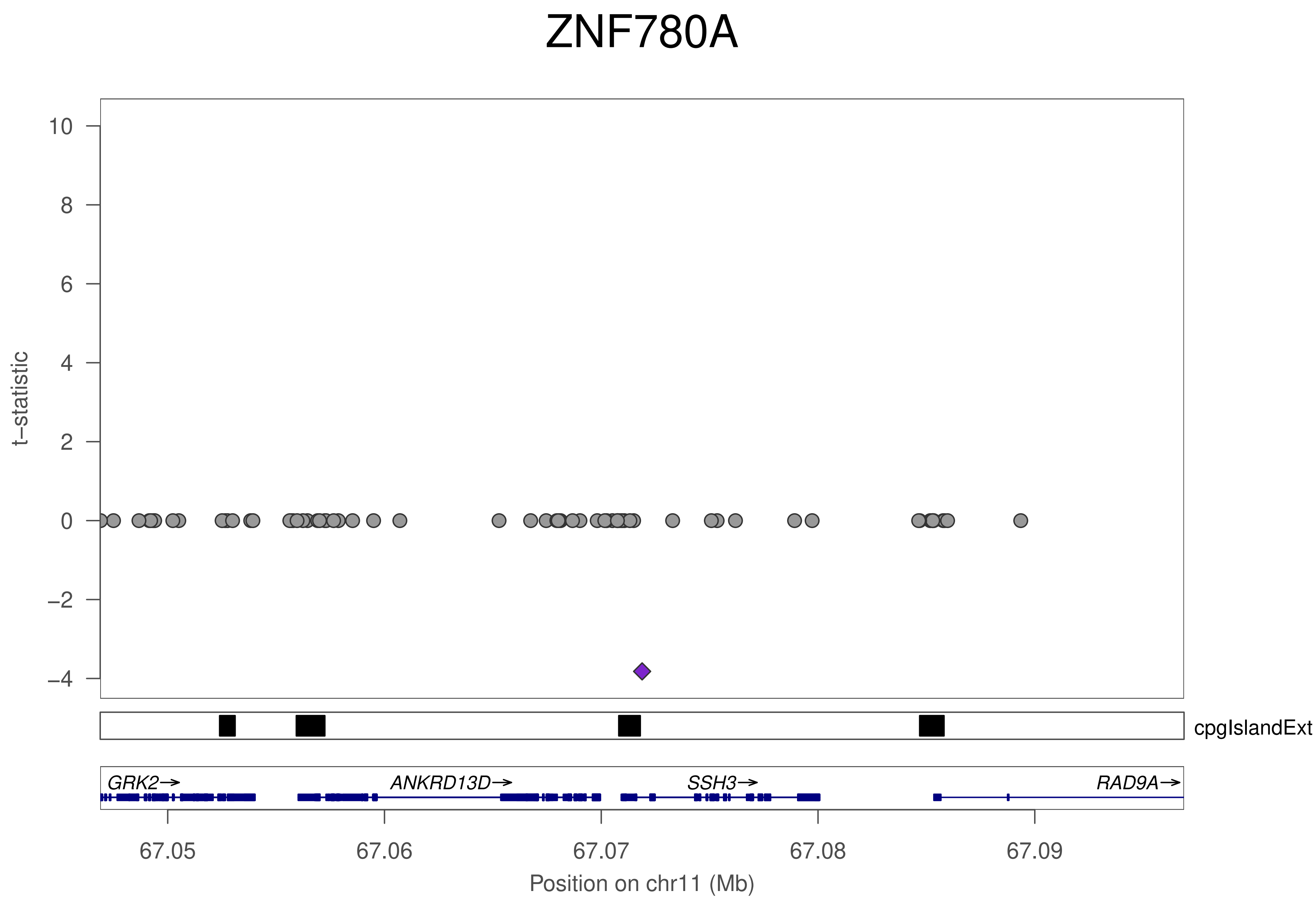

ZNF780A
t−statistic
10
8
6
4
2
0
−2
−4
cpgIslandExt
GRK2→
ANKRD13D→
SSH3→
RAD9A→
67.05
67.06
67.07
67.08
67.09
Position on chr11 (Mb)

### ZNF44

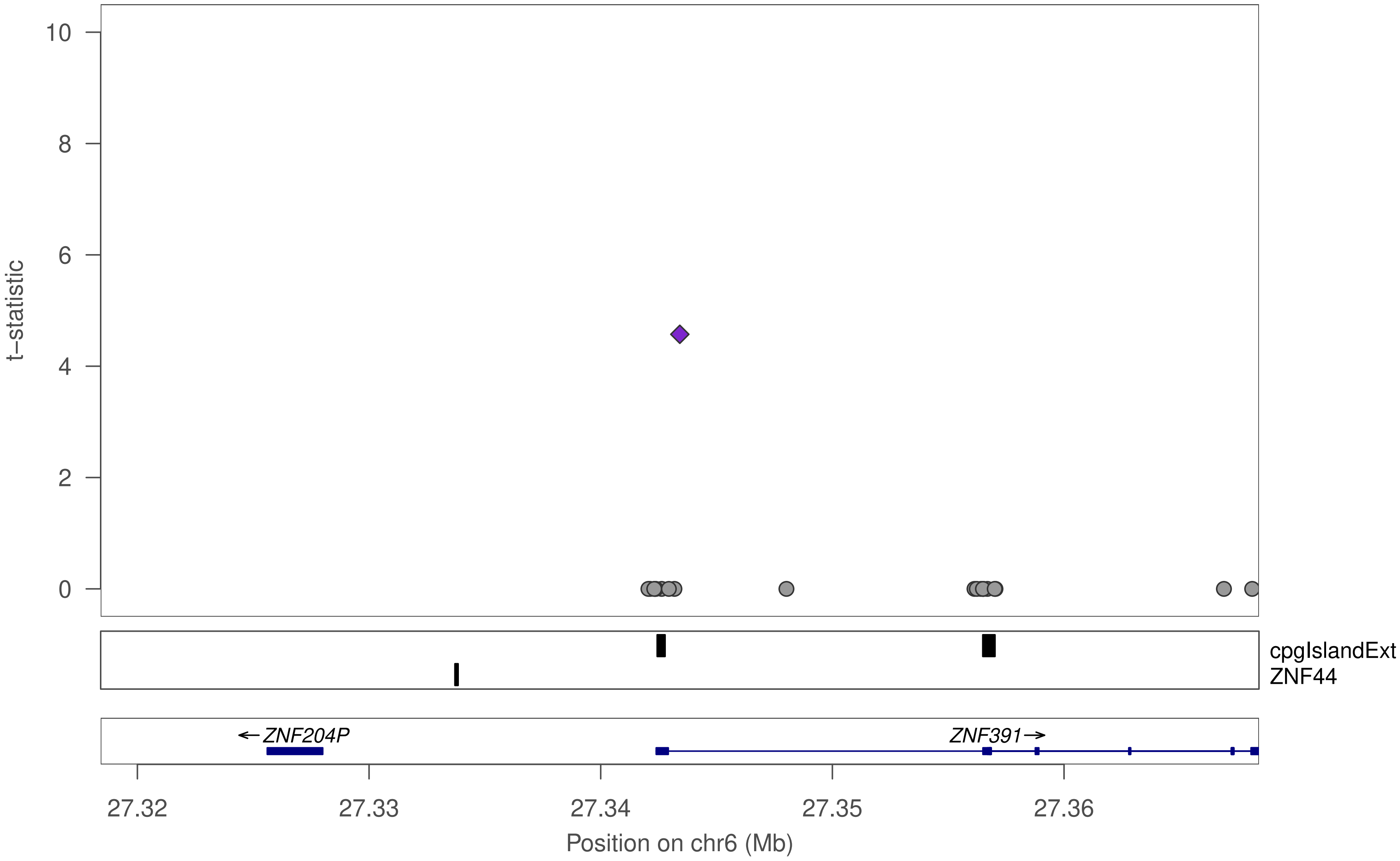

10
8
6
4
2
0
t−statistic
cpgIslandExt
ZNF44
←ZNF204P
ZNF391→
27.32
27.33
27.34
27.35
27.36
Position on chr6 (Mb)

### ZNF44

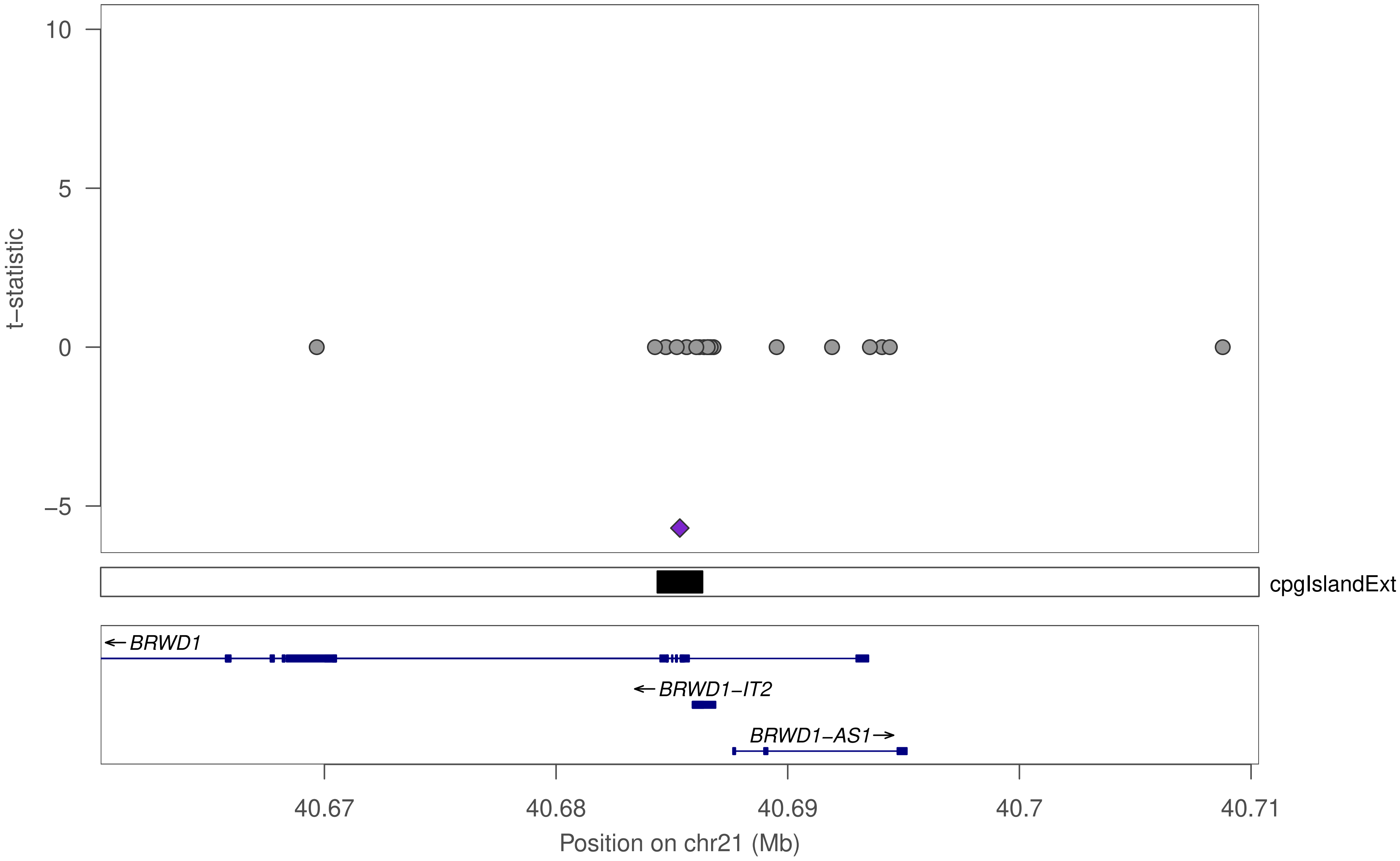
10
5
0
−5
t−statistic
cpgIslandExt
←BRWD1
←BRWD1−IT2
BRWD1−AS1→
40.67
40.68
40.69
40.7
40.71
Position on chr21 (Mb)

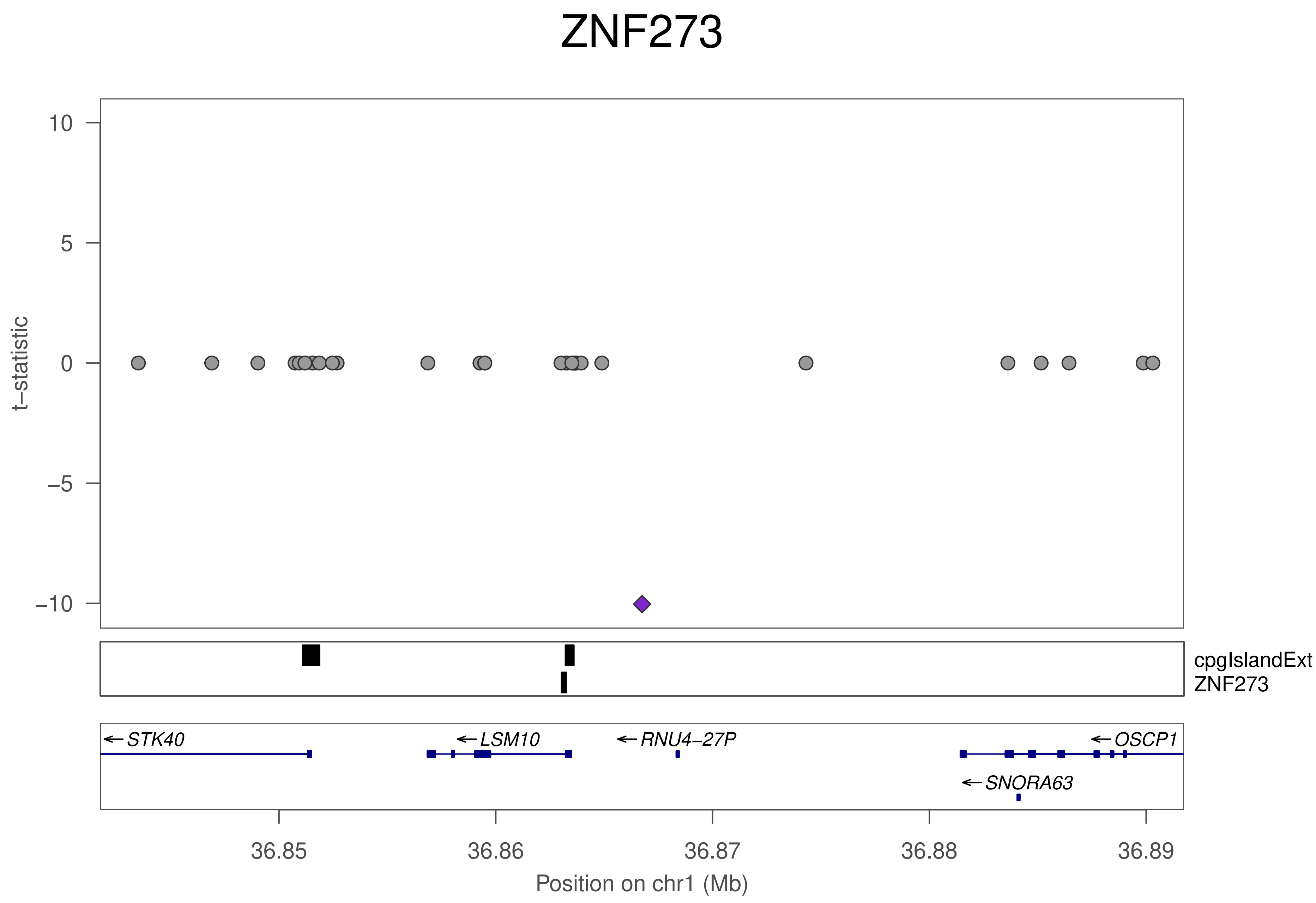

ZNF273
t−statistic
10
5
0
−5
−10
cpgIslandExt
ZNF273
← STK40
← LSM10
← RNU4−27P
← OSCP1
← SNORA63
36.85
36.86
36.87
36.88
36.89
Position on chr1 (Mb)

### ZNF273

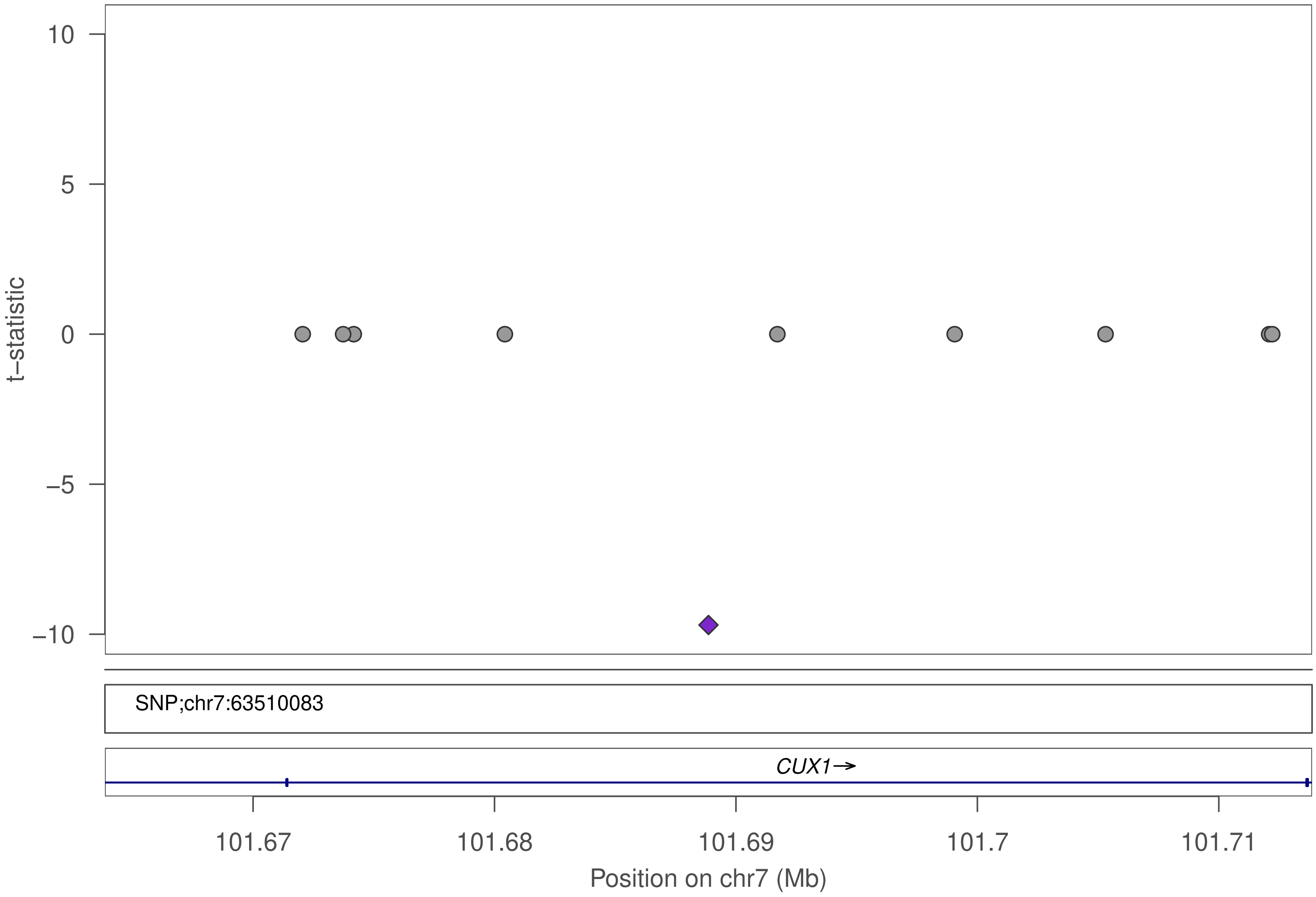

10
5
0
−5
−10
t−statistic
SNP;chr7:63510083
CUX1→
101.67
101.68
101.69
101.7
101.71
Position on chr7 (Mb)

### ZNF273

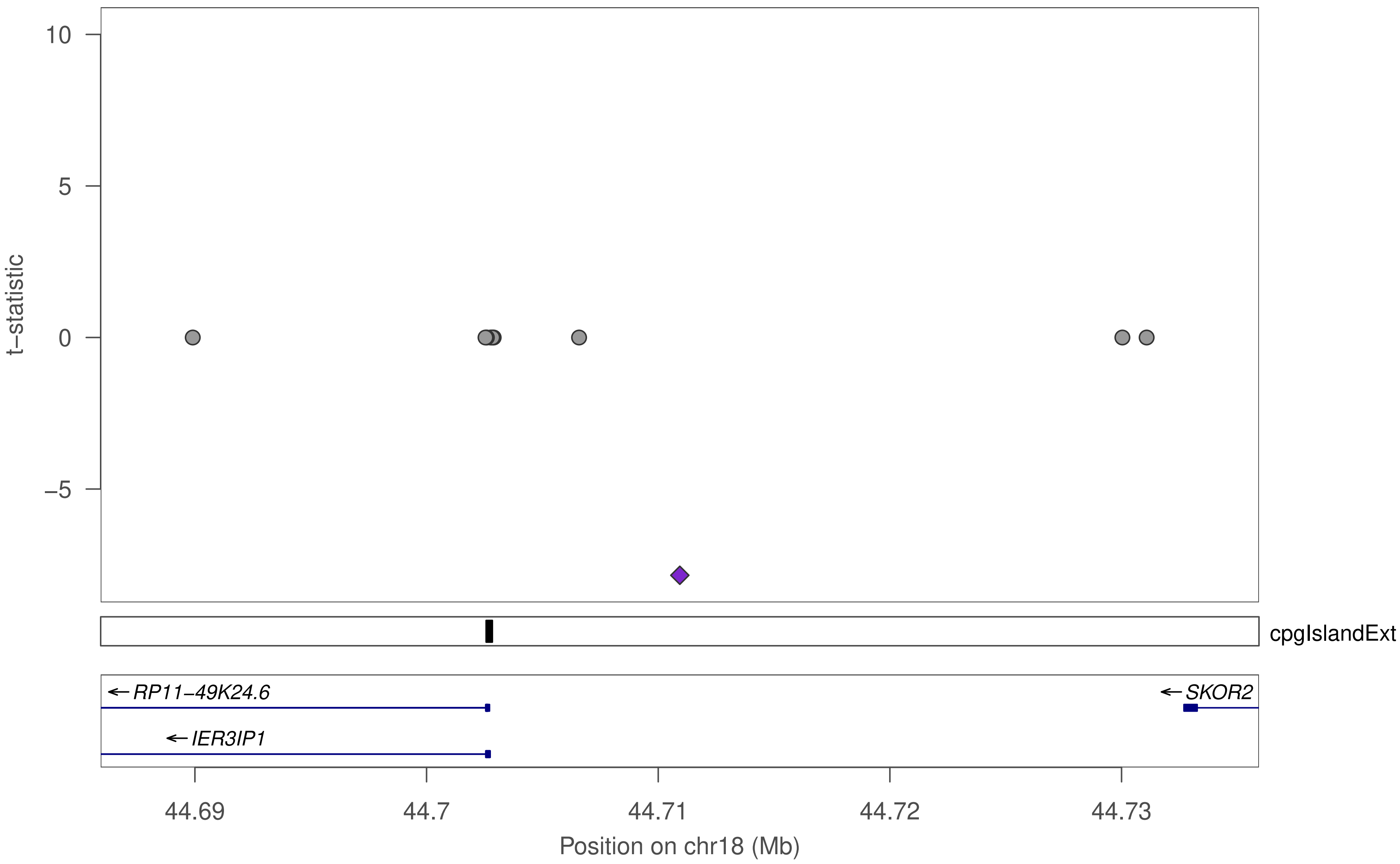

10
5
0
−5
t−statistic
cpgIslandExt
RP11−49K24.6
SKOR2
IER3IP1
44.69
44.7
44.71
44.72
44.73
Position on chr18 (Mb)

### ZNF273

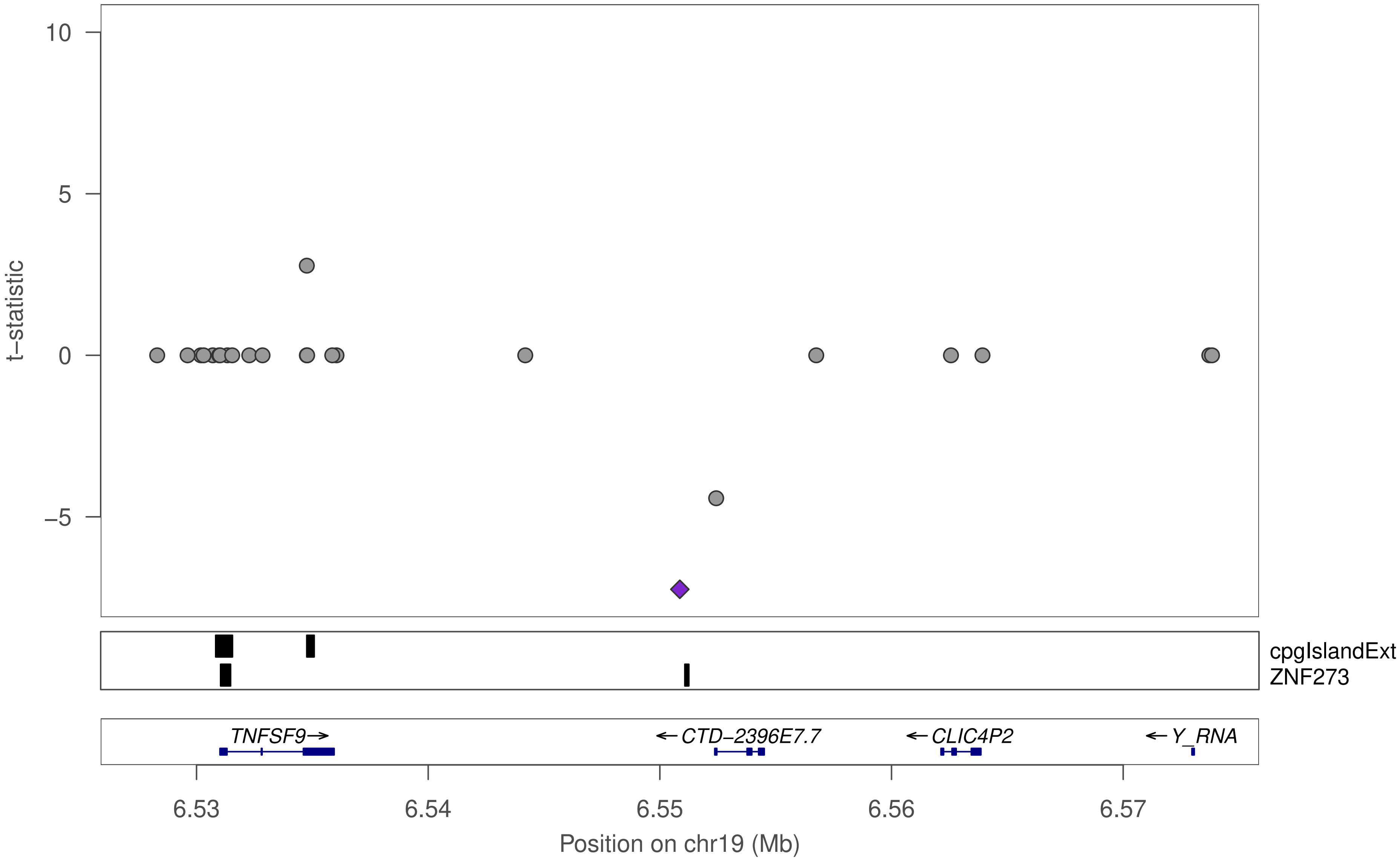

10
5
0
−5
t−statistic
cpgIslandExt
ZNF273
TNFSF9→
←CTD−2396E7.7
←CLIC4P2
←Y_RNA
6.53
6.54
6.55
6.56
6.57
Position on chr19 (Mb)

### ZNF248

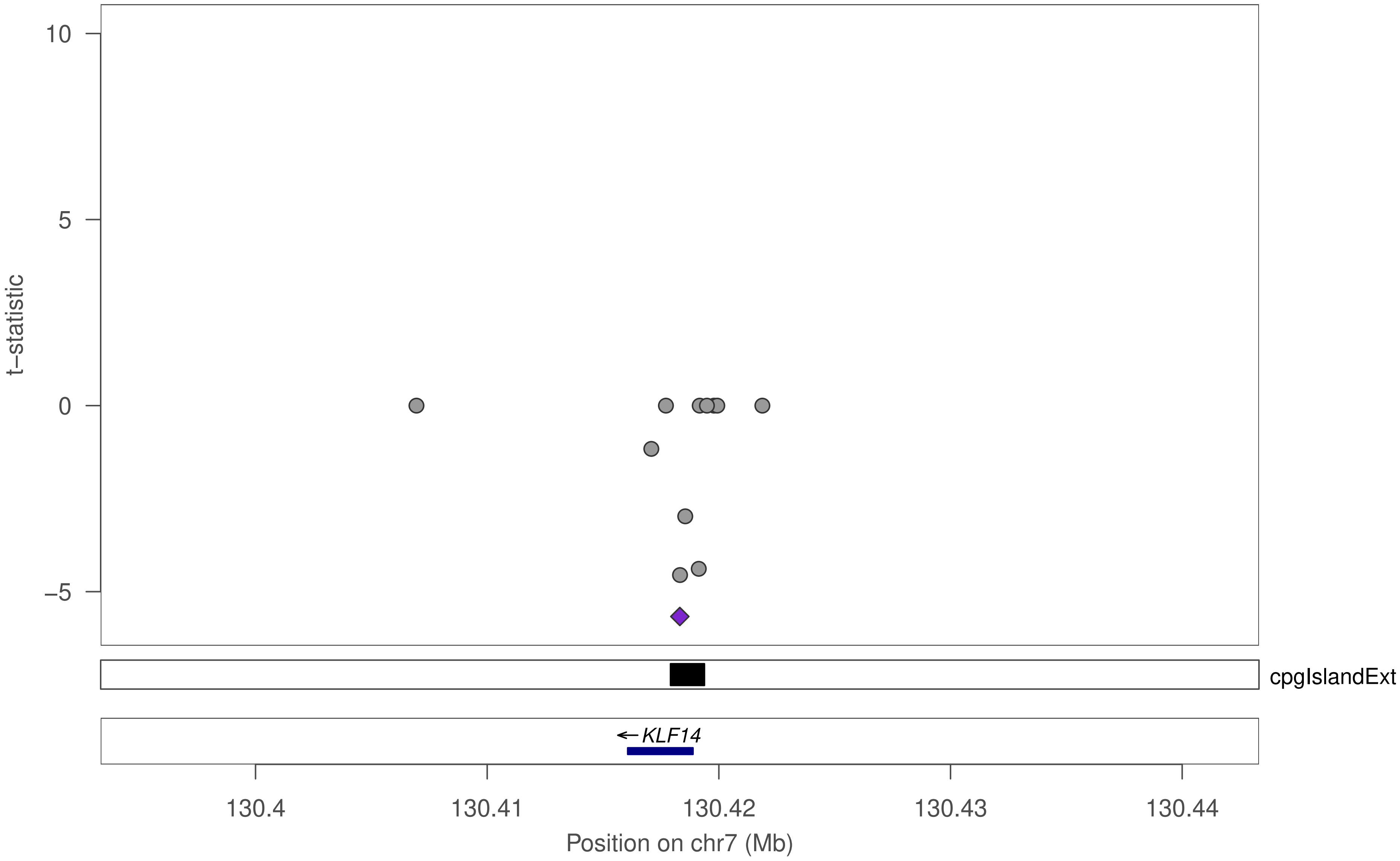

t–statistic
10
5
0
−5
cpgIslandExt
← KLF14
130.4
130.41
130.42
130.43
130.44
Position on chr7 (Mb)

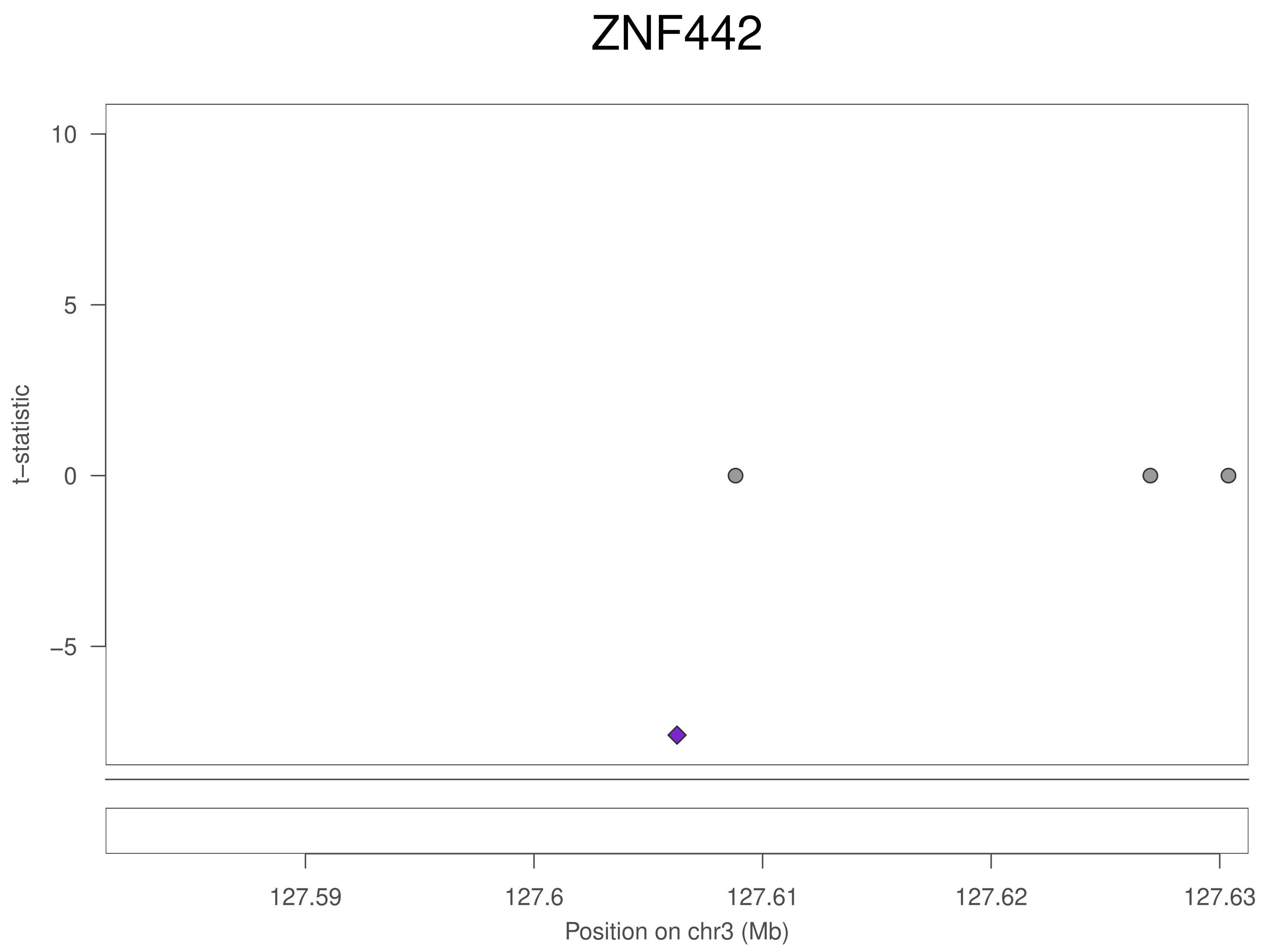
ZNF442
10
5
0
−5
t−statistic
127.59
127.6
127.61
127.62
127.63
Position on chr3 (Mb)

### ZNF442

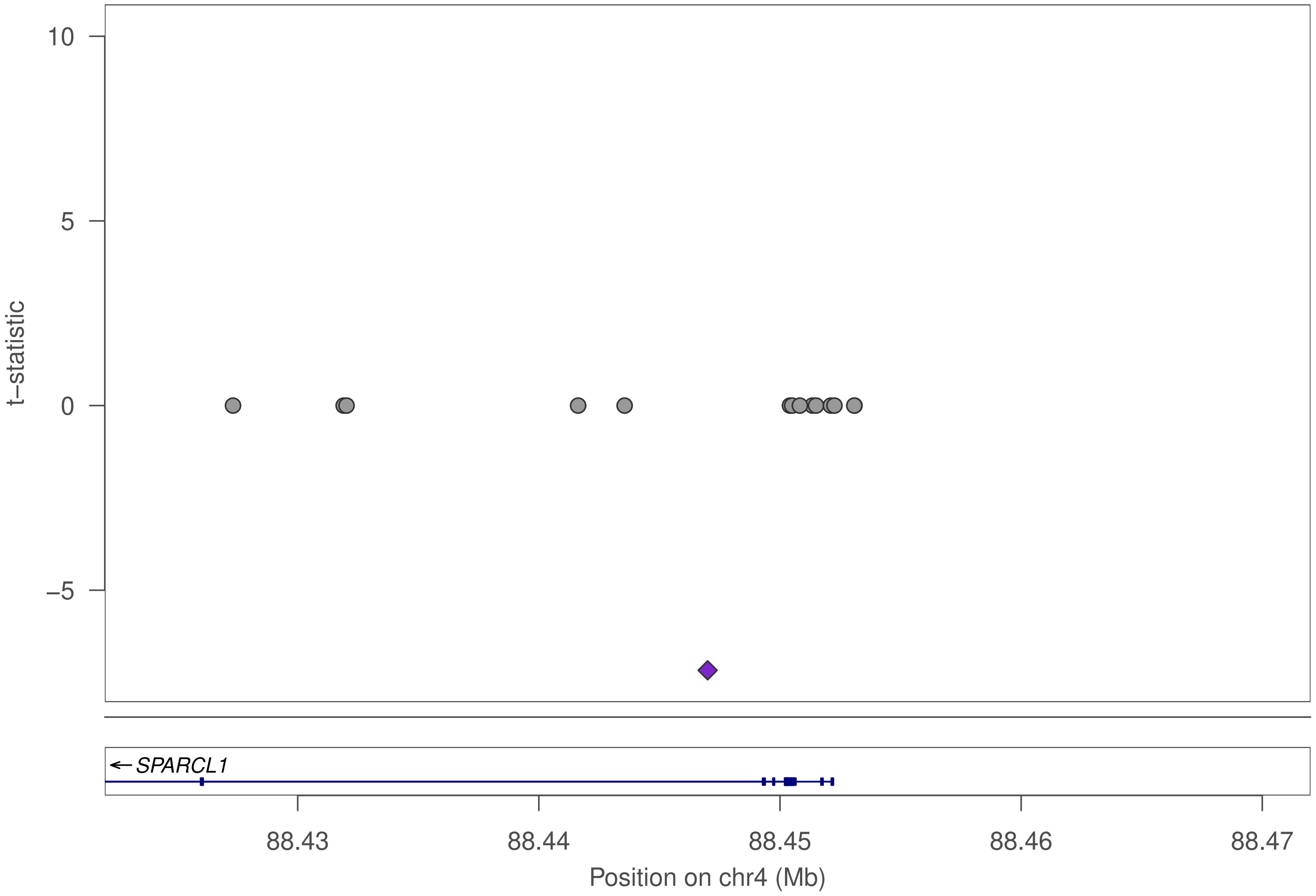

10
5
0
−5
t−statistic
SPARCL1
88.43
88.44
88.45
88.46
88.47
Position on chr4 (Mb)

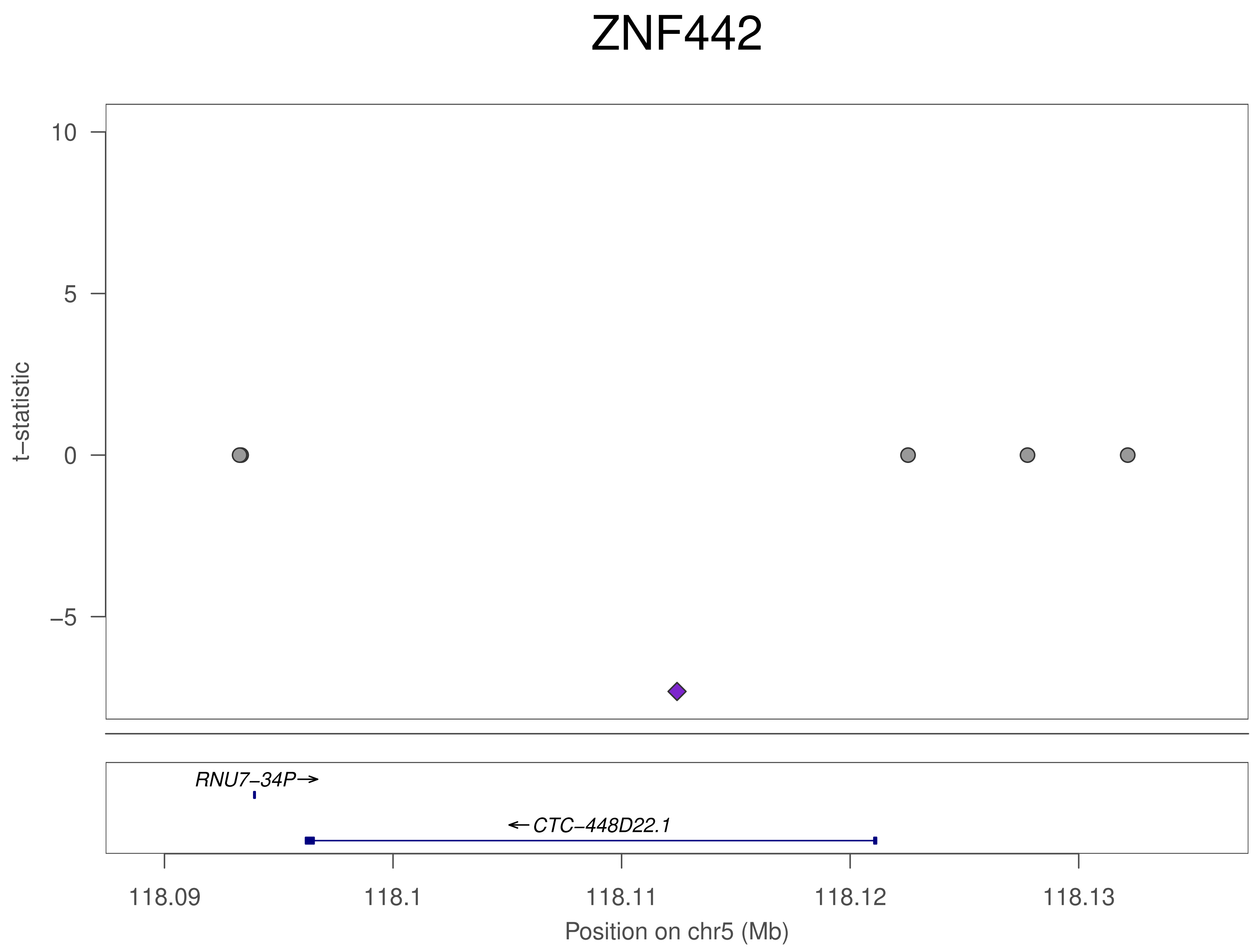

ZNF442
t−statistic
10
5
0
−5
RNU7−34P→
←CTC−448D22.1
118.09
118.1
118.11
118.12
118.13
Position on chr5 (Mb)

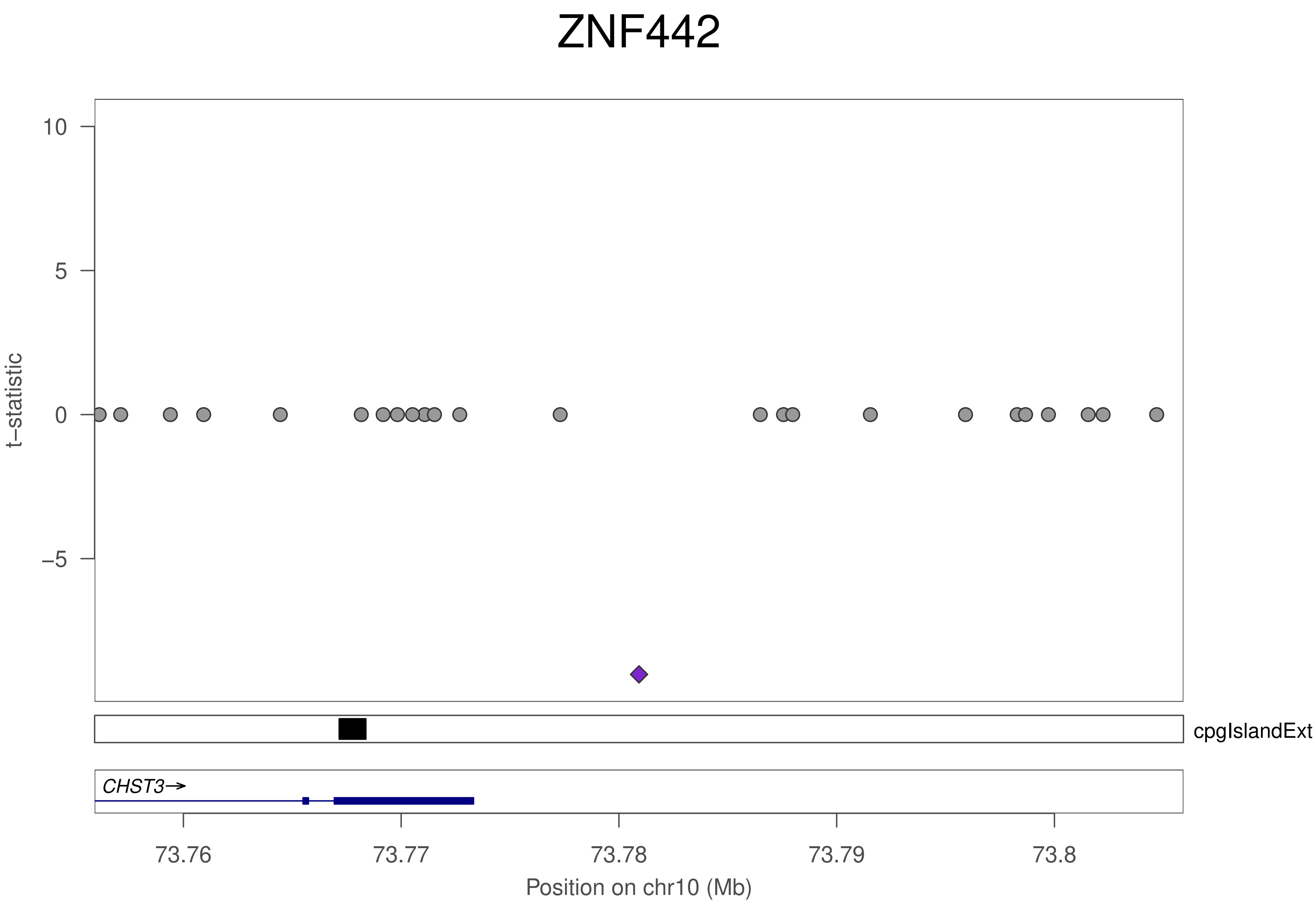
ZNF442
10
5
0
−5
t−statistic
cpgIslandExt
CHST3→
73.76
73.77
73.78
73.79
73.8
Position on chr10 (Mb)

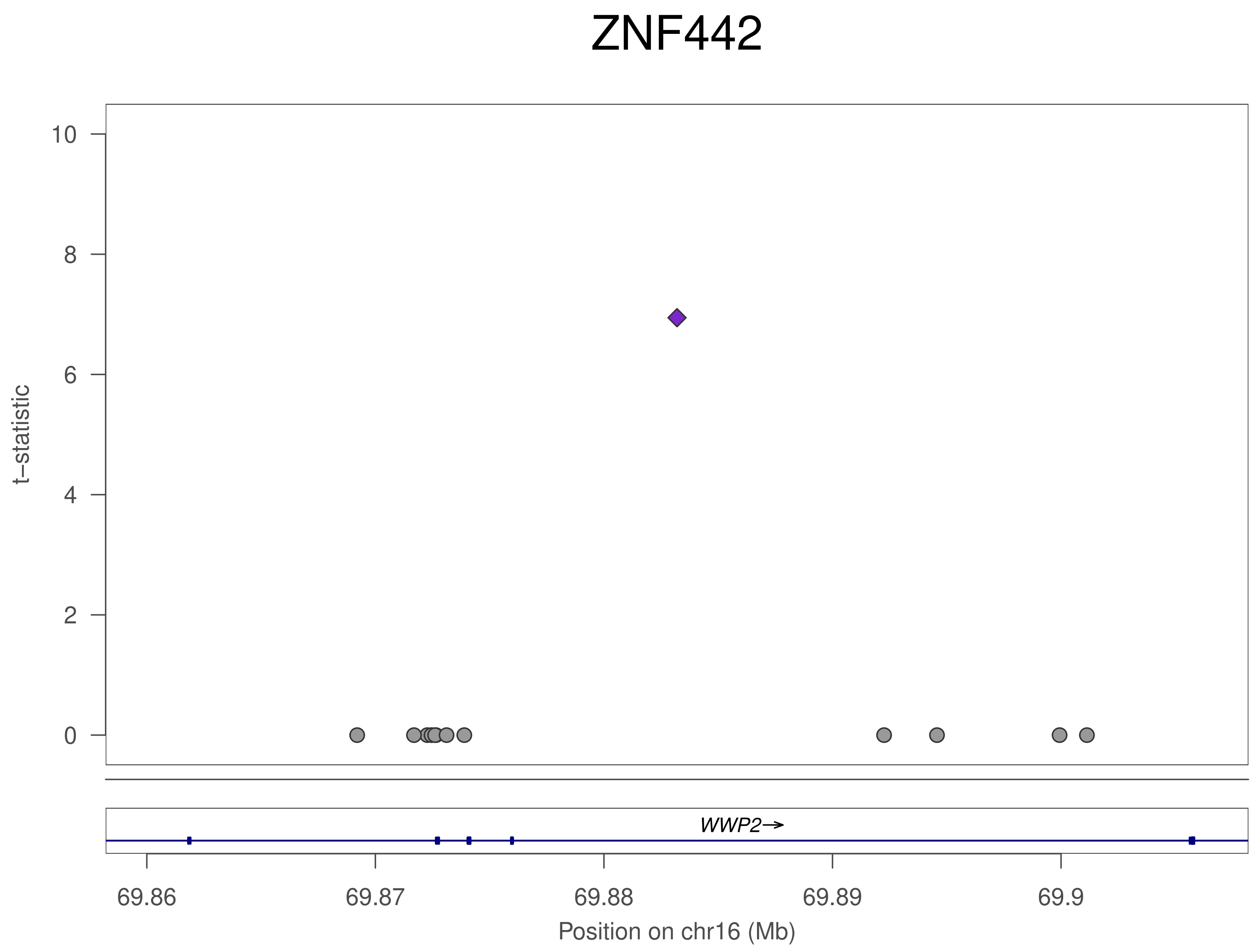

ZNF442
t–statistic
10
8
6
4
2
0
WWP2→
69.86
69.87
69.88
69.89
69.9
Position on chr16 (Mb)

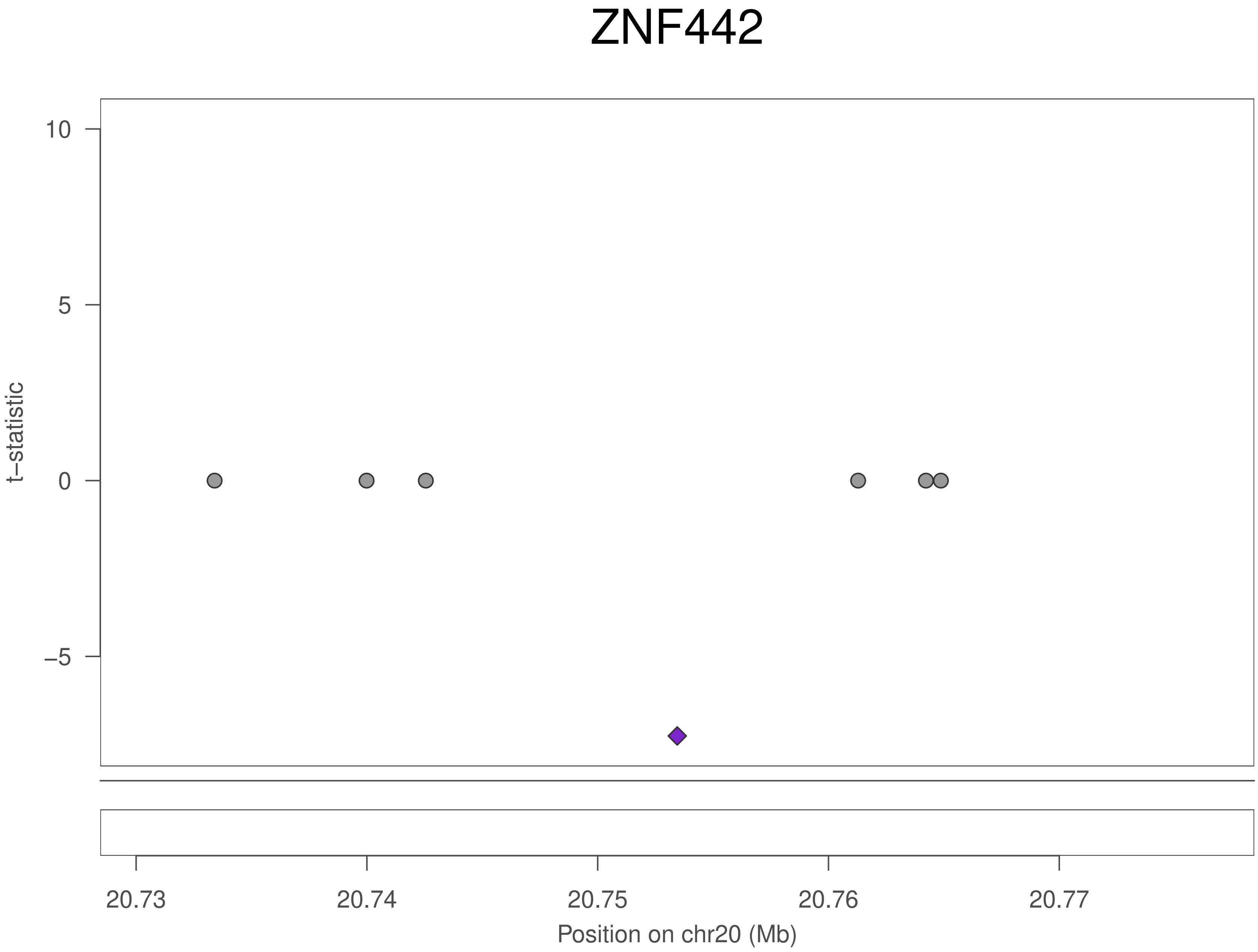
ZNF442
t–statistic
10
5
0
–5
20.73
20.74
20.75
20.76
20.77
Position on chr20 (Mb)

### ZNF442

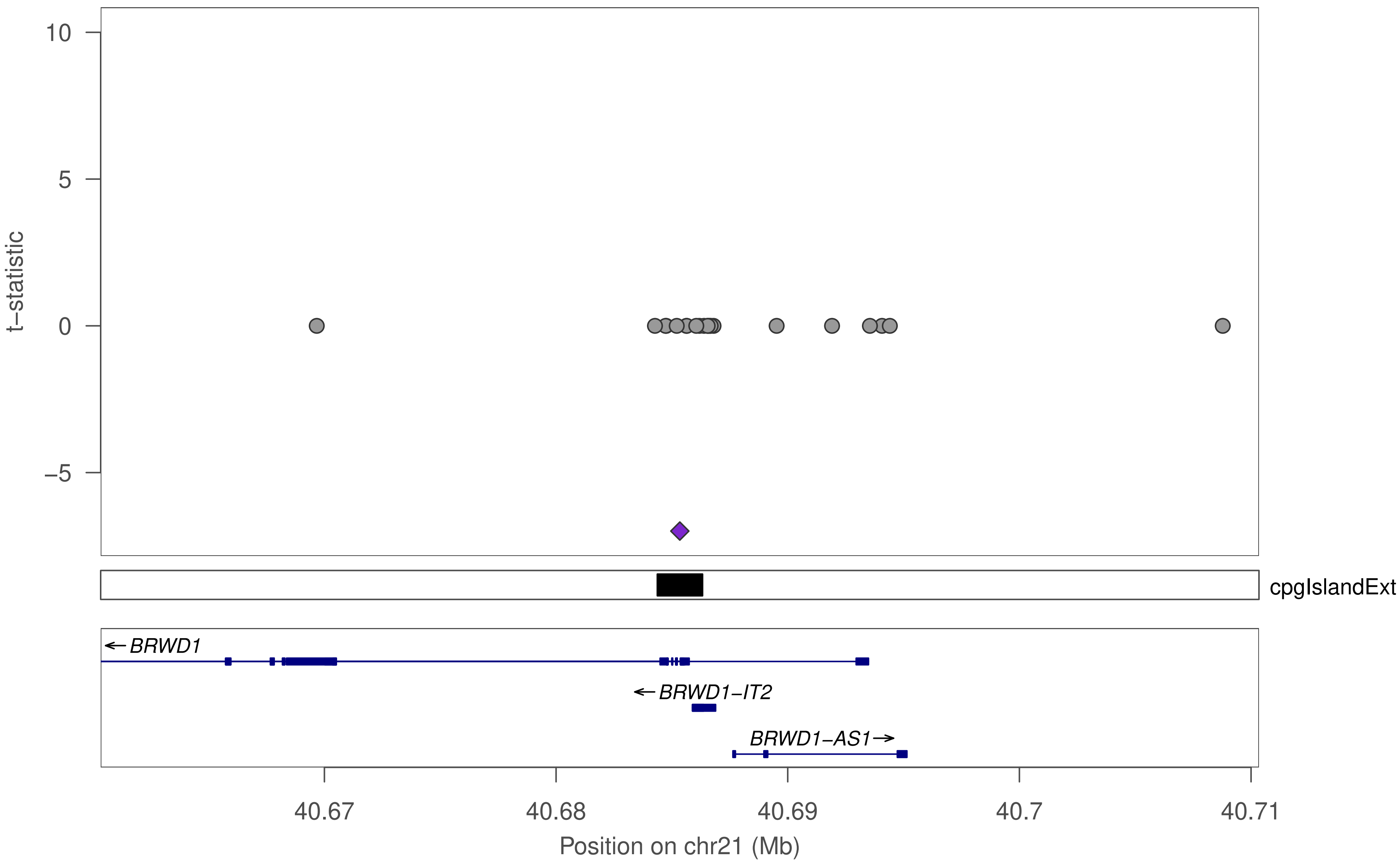
t−statistic
10
5
0
−5
cpgIslandExt
←BRWD1
←BRWD1−IT2
BRWD1−AS1→
40.67
40.68
40.69
40.7
40.71
Position on chr21 (Mb)

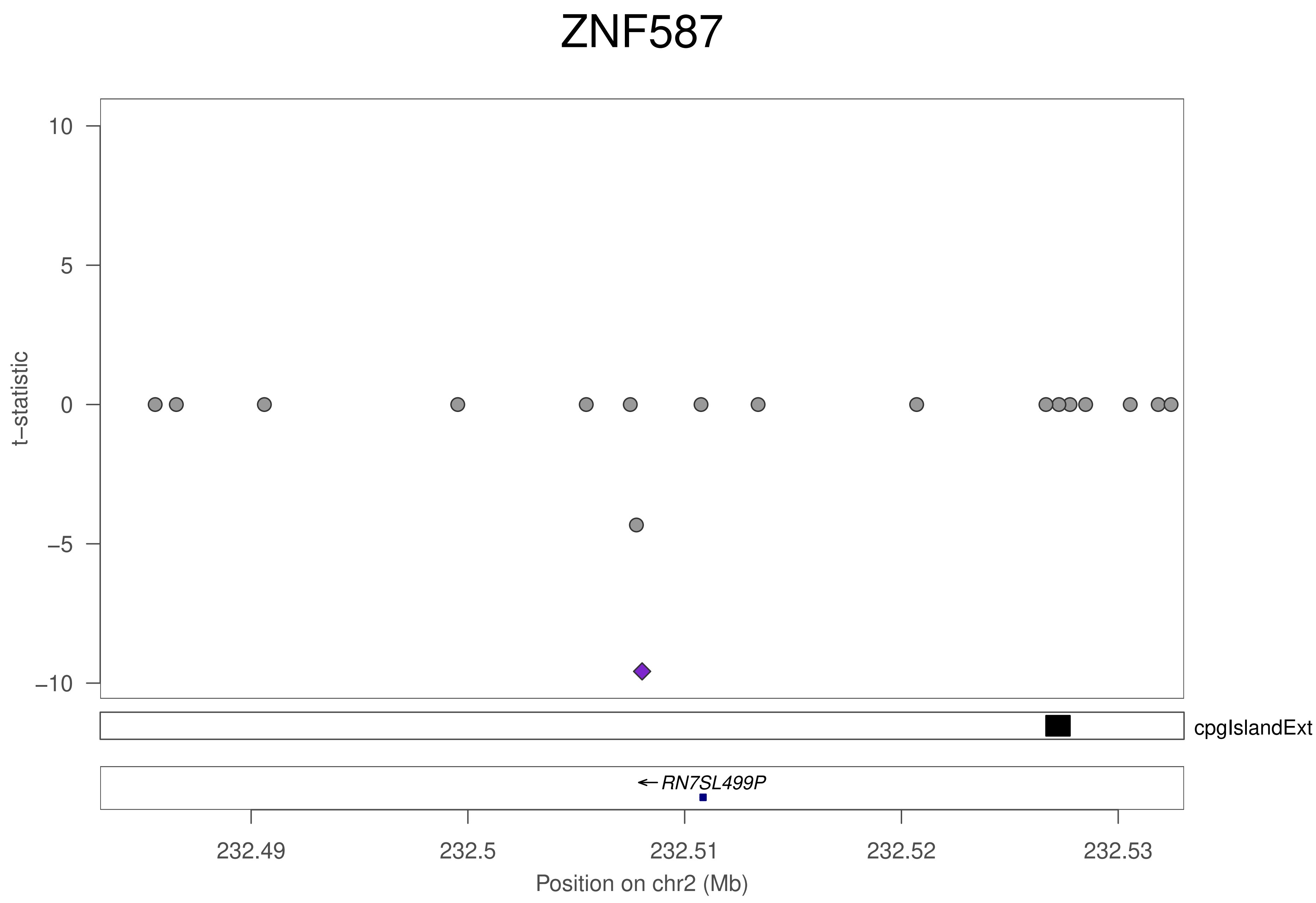

ZNF587
10
5
0
−5
−10
t−statistic
cpgIslandExt
RN7SL499P
232.49
232.5
232.51
232.52
232.53
Position on chr2 (Mb)

### ZNF587

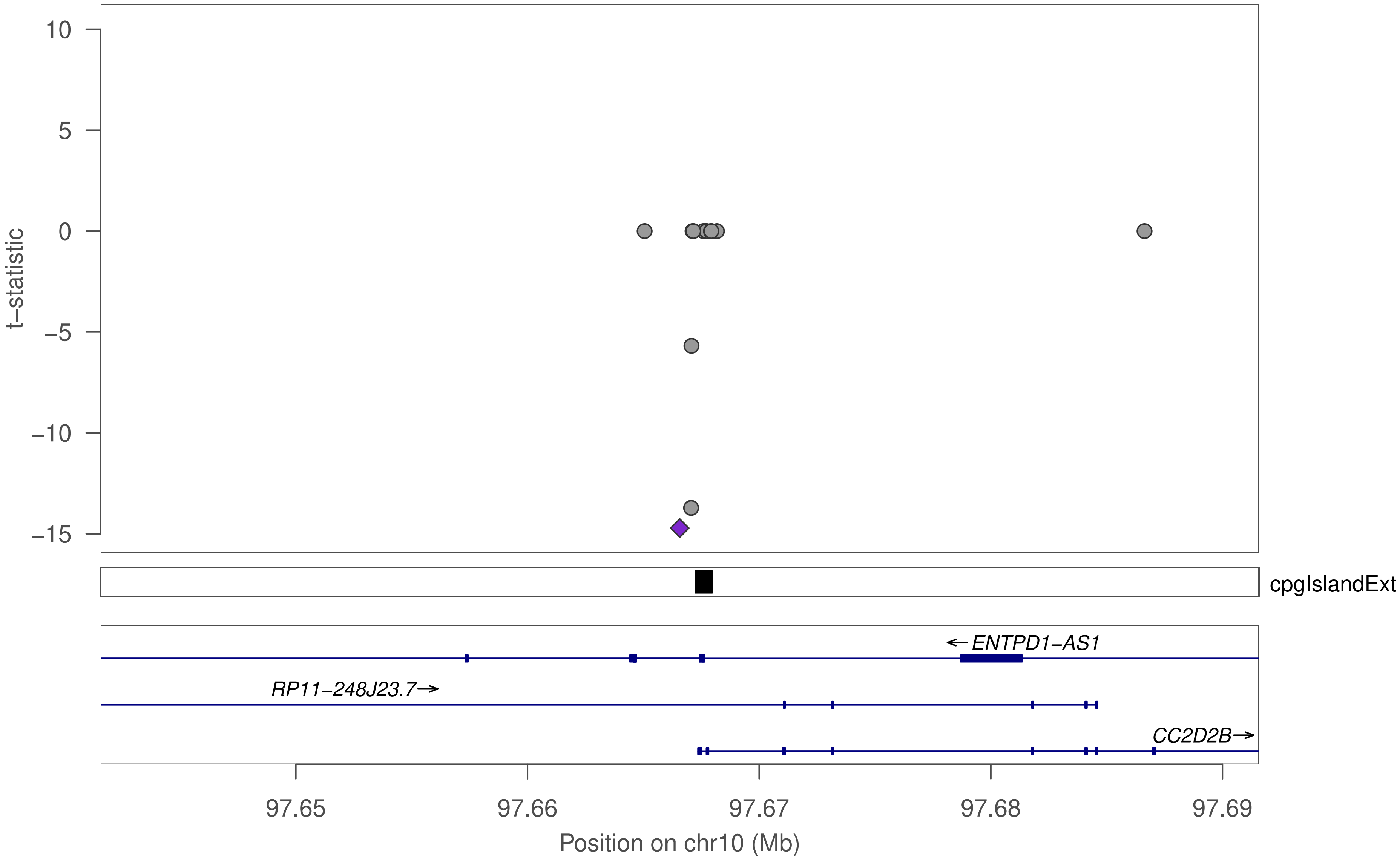

t−statistic
10
5
0
−5
−10
−15
cpgIslandExt
←ENTPD1−AS1
RP11−248J23.7→
CC2D2B→
97.65
97.66
97.67
97.68
97.69
Position on chr10 (Mb)

### ZNF468

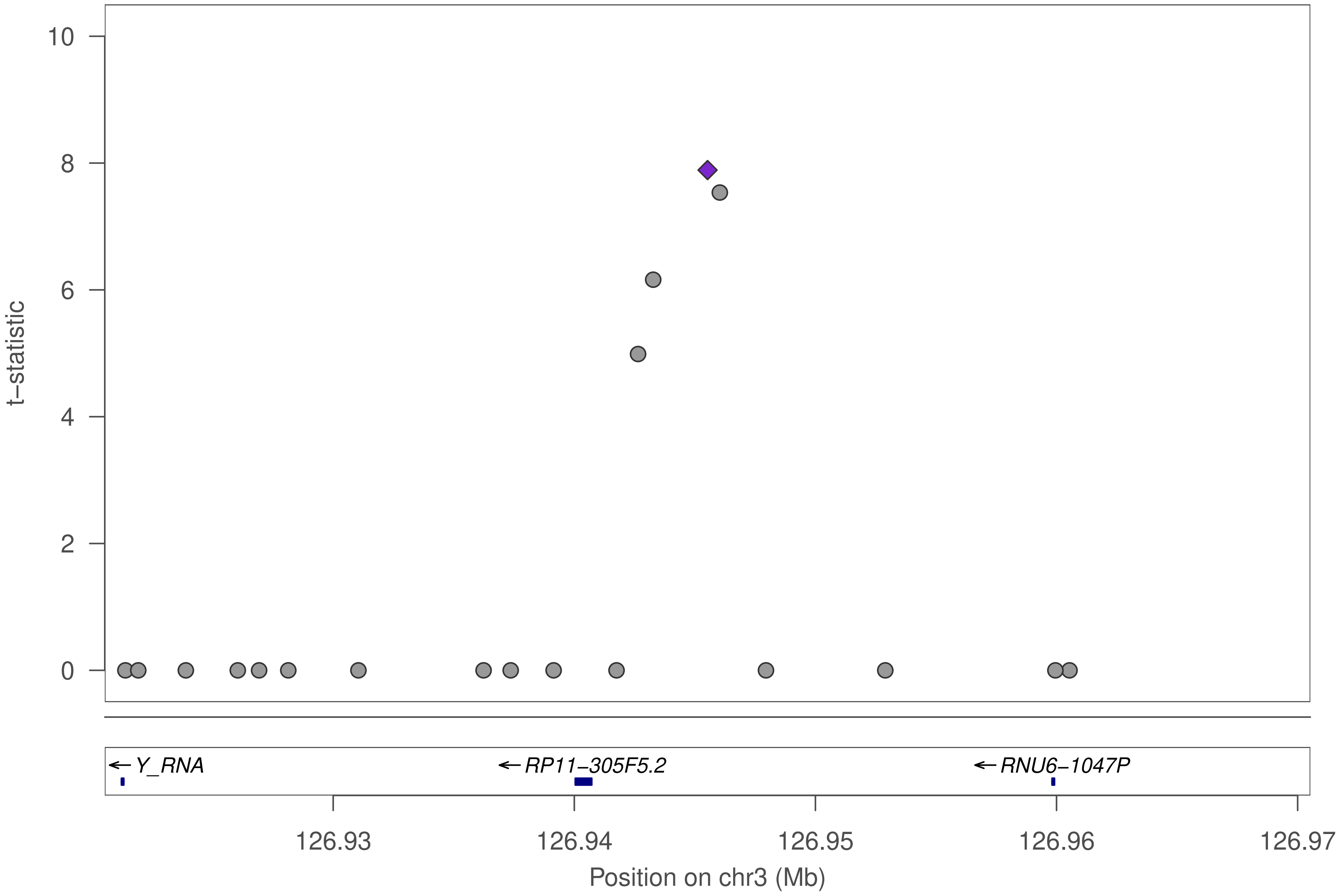

t−statistic
10
8
6
4
2
0
Y_RNA
RP11−305F5.2
RNU6−1047P
126.93
126.94
126.95
126.96
126.97
Position on chr3 (Mb)

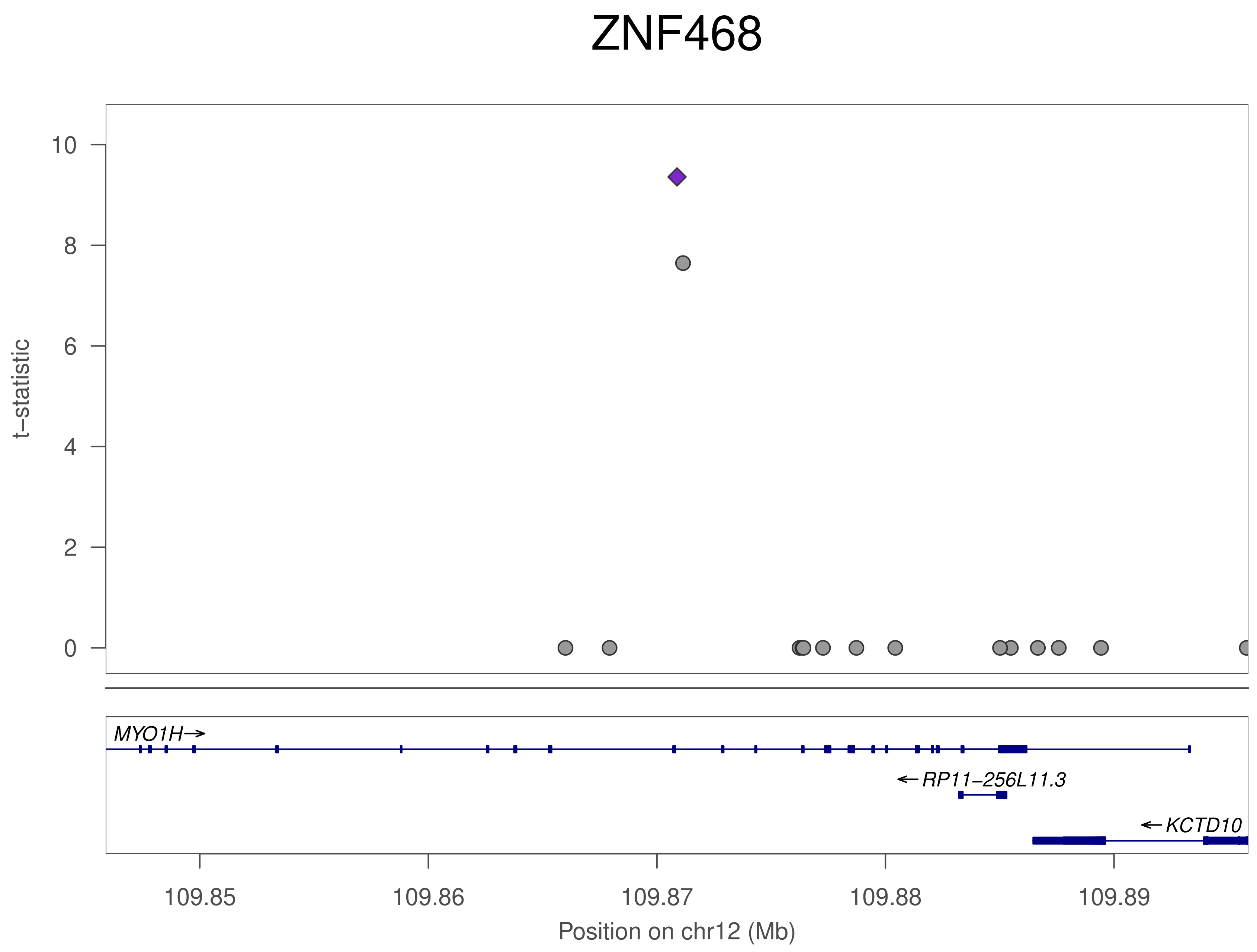
ZNF468
t−statistic
10
8
6
4
2
0
MYO1H→
←RP11−256L11.3
←KCTD10
109.85
109.86
109.87
109.88
109.89
Position on chr12 (Mb)

### ZFP57

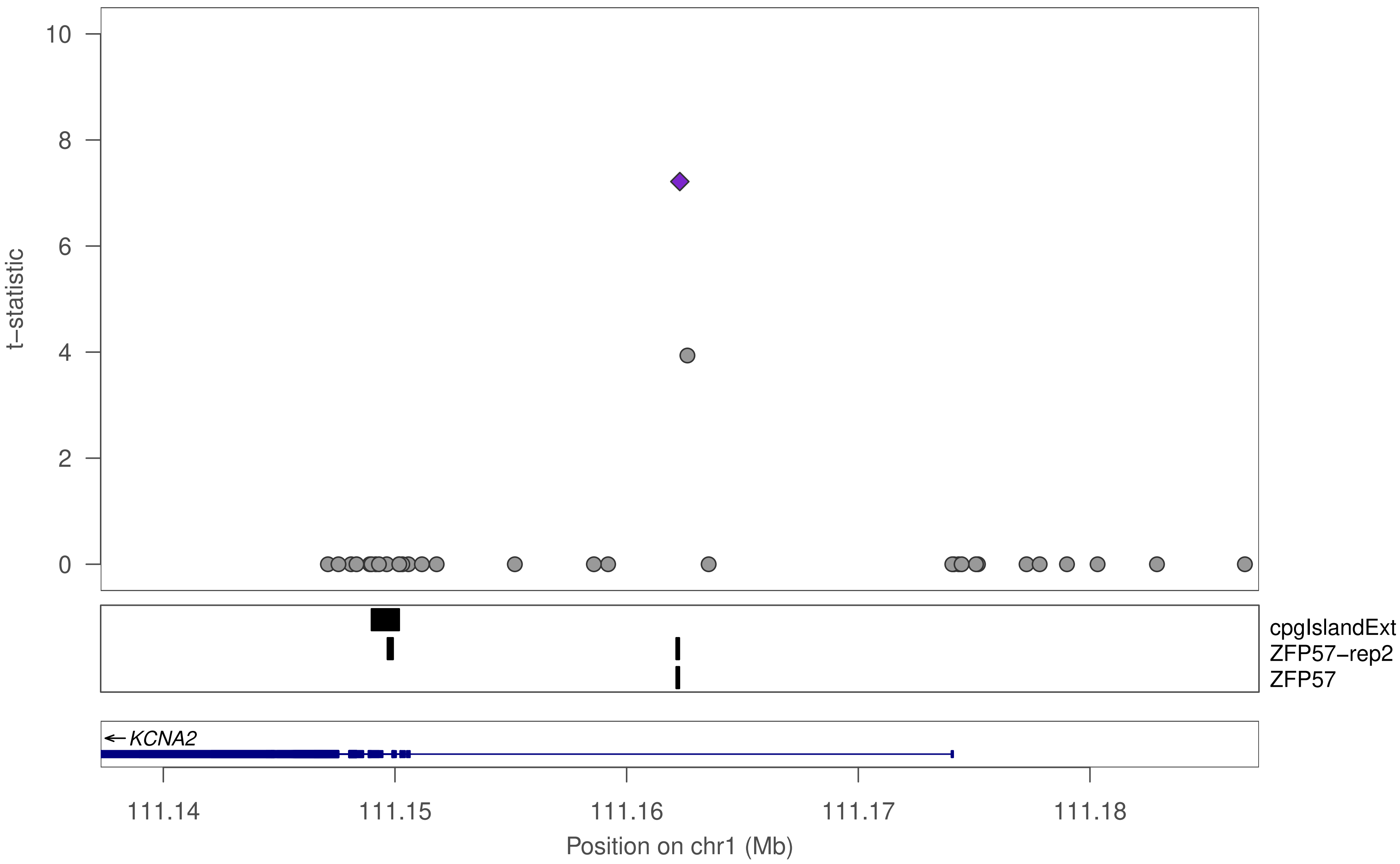
t−statistic
10
8
6
4
2
0
cpgIslandExt
ZFP57−rep2
ZFP57
KCNA2
111.14
111.15
111.16
111.17
111.18
Position on chr1 (Mb)

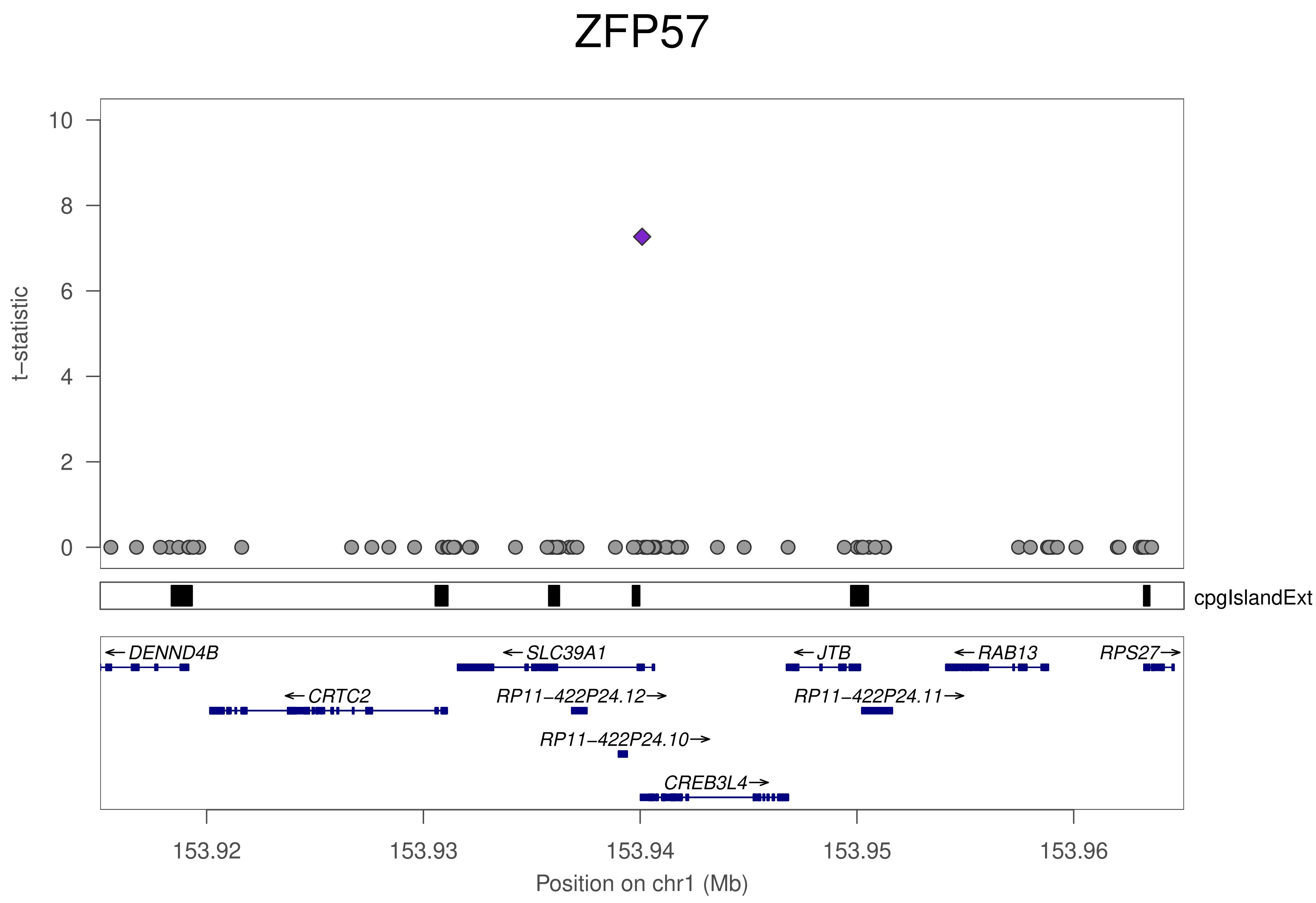
ZFP57
t−statistic
10
8
6
4
2
0
cpgIslandExt
←DENND4B
←SLC39A1
←JTB
←RAB13
RPS27→
←CRTC2
RP11−422P24.12→
RP11−422P24.11→
RP11−422P24.10→
CREB3L4→
153.92
153.93
153.94
153.95
153.96
Position on chr1 (Mb)

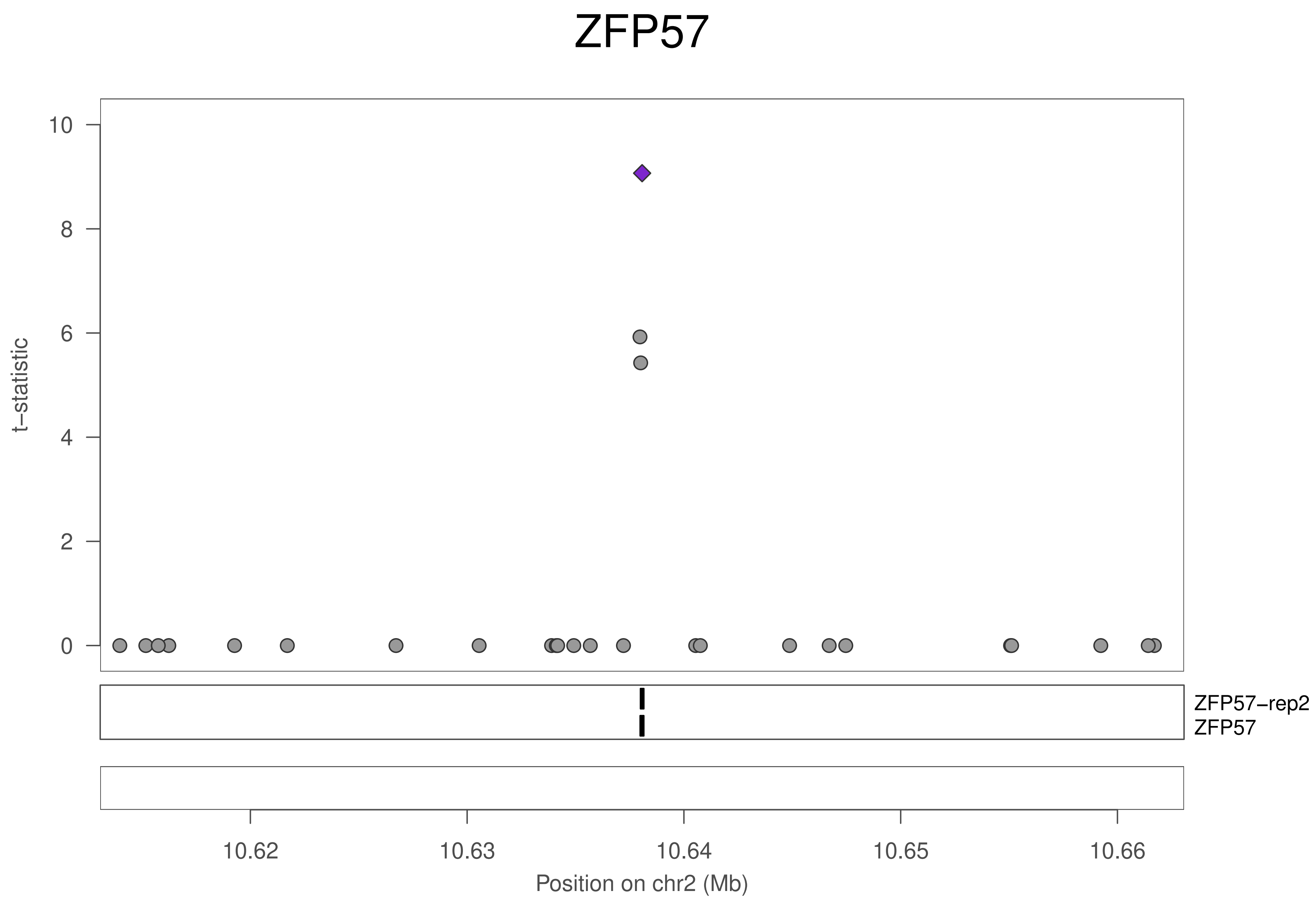

ZFP57
t−statistic
10
8
6
4
2
0
ZFP57−rep2
ZFP57
10.62
10.63
10.64
10.65
10.66
Position on chr2 (Mb)

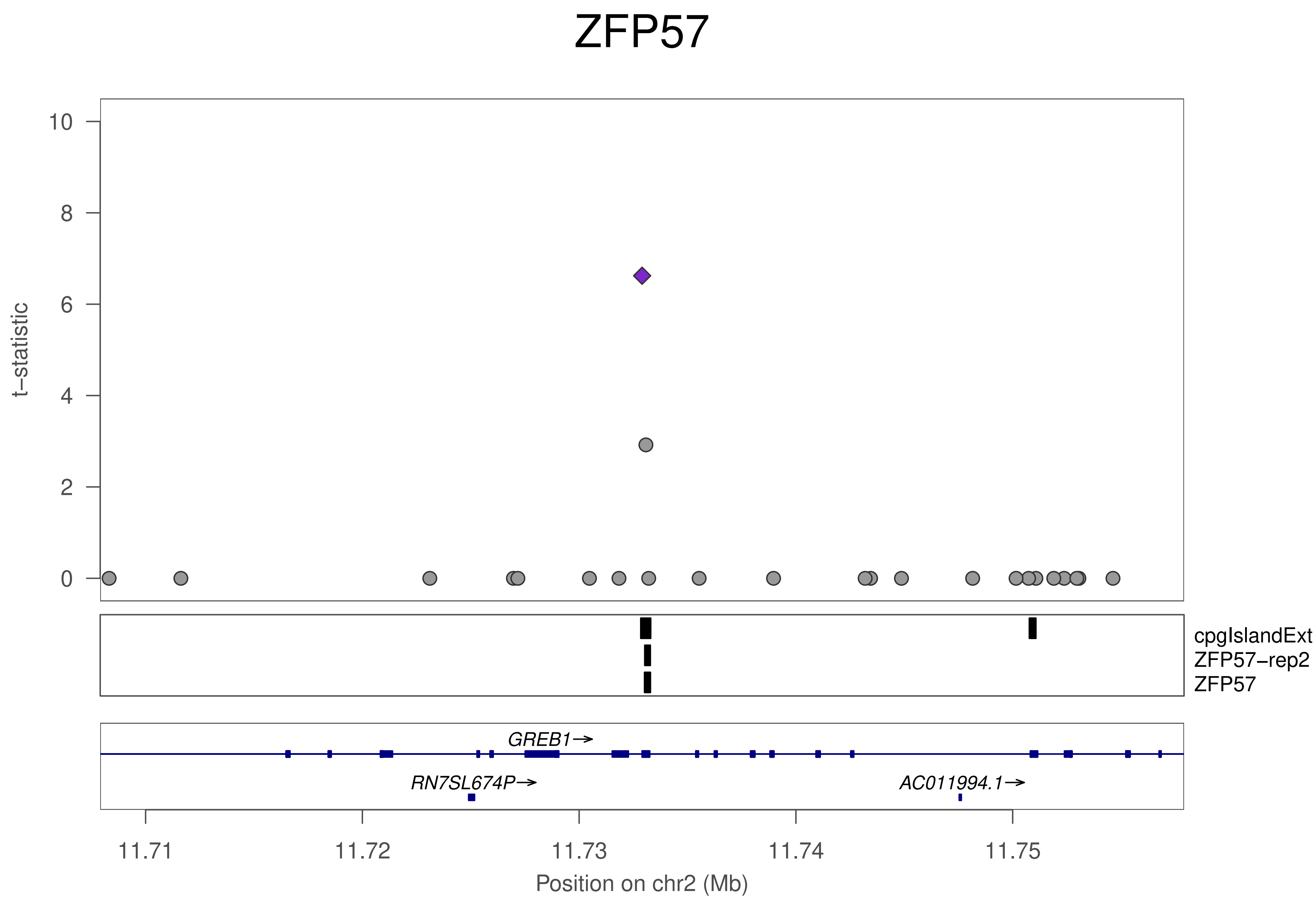

ZFP57
t−statistic
0
2
4
6
8
10
cpgIslandExt
ZFP57−rep2
ZFP57
GREB1→
RN7SL674P→
AC011994.1→
11.71
11.72
11.73
11.74
11.75
Position on chr2 (Mb)

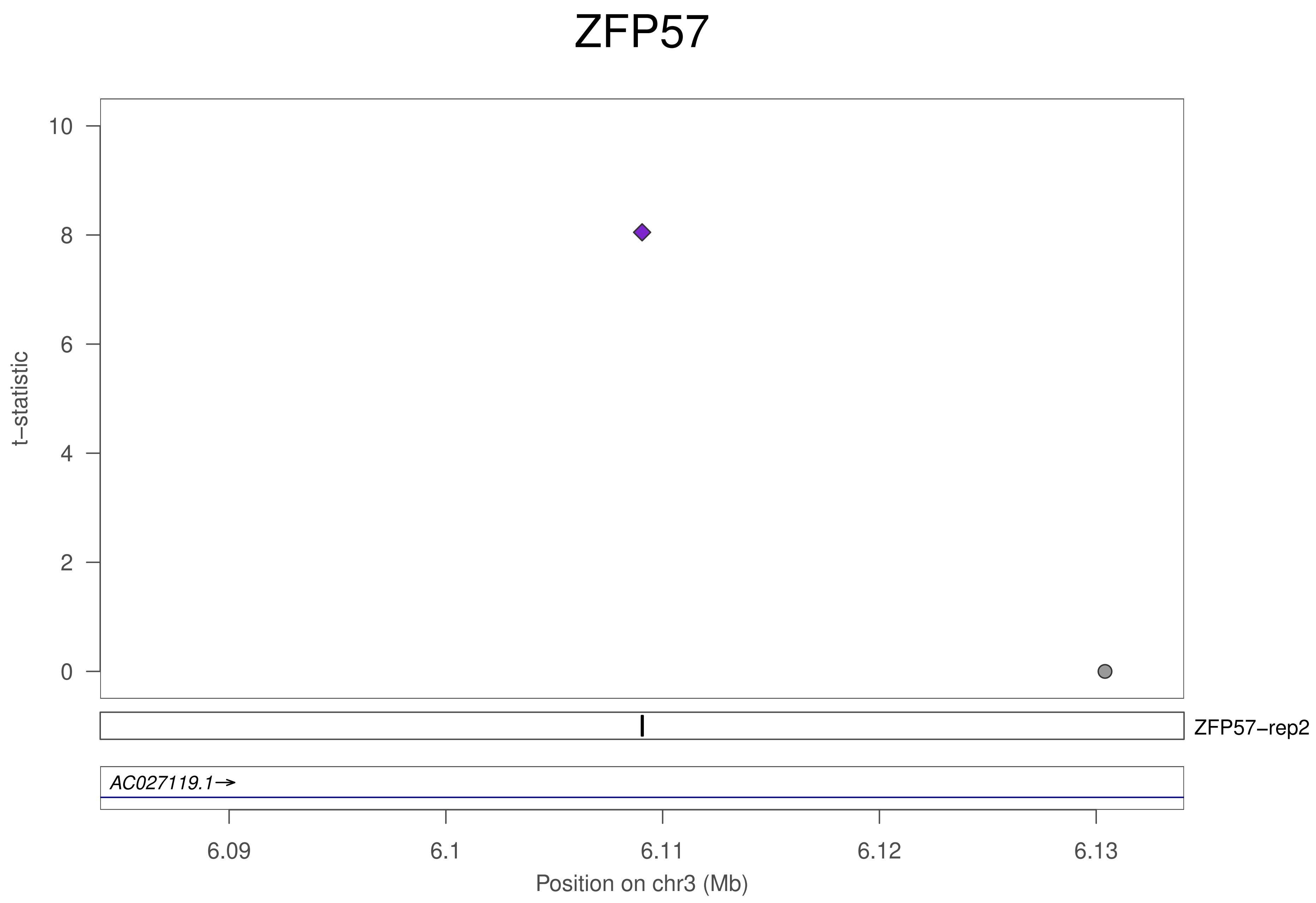

ZFP57
t−statistic
10
8
6
4
2
0
ZFP57−rep2
AC027119.1→
6.09
6.1
6.11
6.12
6.13
Position on chr3 (Mb)

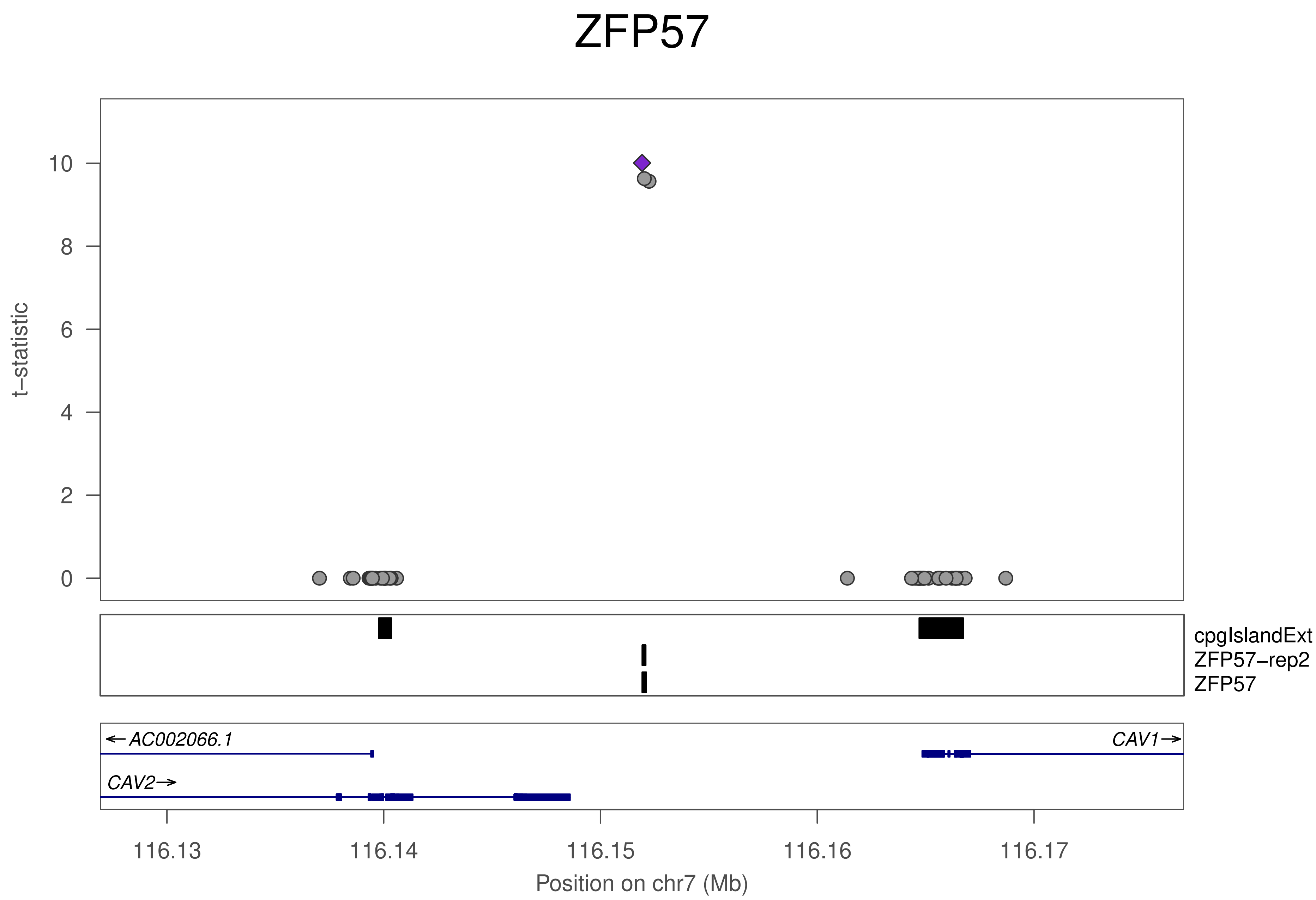

ZFP57
t−statistic
0
2
4
6
8
10
cpgIslandExt
ZFP57−rep2
ZFP57
←AC002066.1
CAV1→
CAV2→
116.13
116.14
116.15
116.16
116.17
Position on chr7 (Mb)

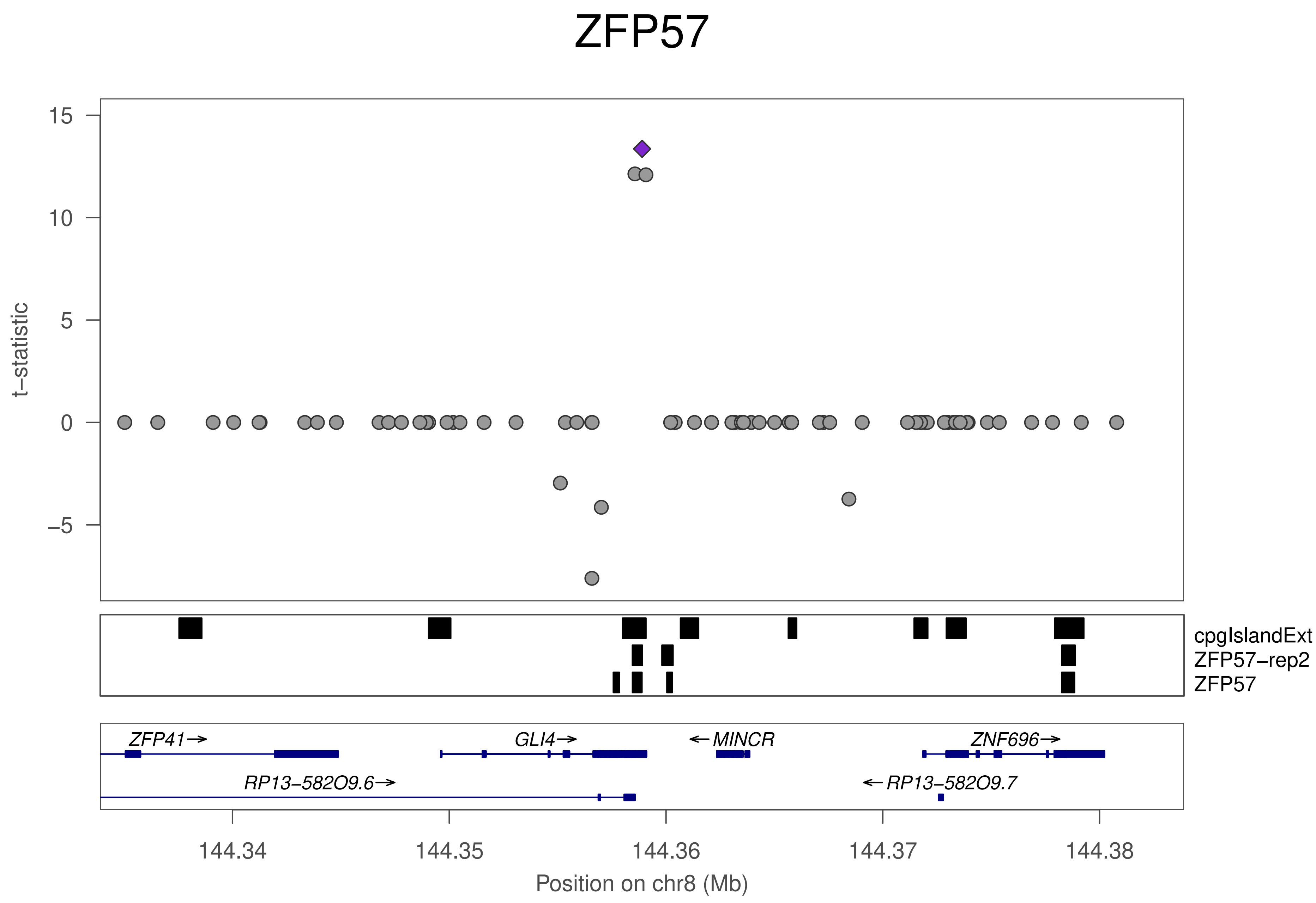

ZFP57
t−statistic
15
10
5
0
−5
cpgIslandExt
ZFP57−rep2
ZFP57
ZFP41→
GLI4→
←MINCR
ZNF696→
RP13−582O9.6→
←RP13−582O9.7
144.34
144.35
144.36
144.37
144.38
Position on chr8 (Mb)

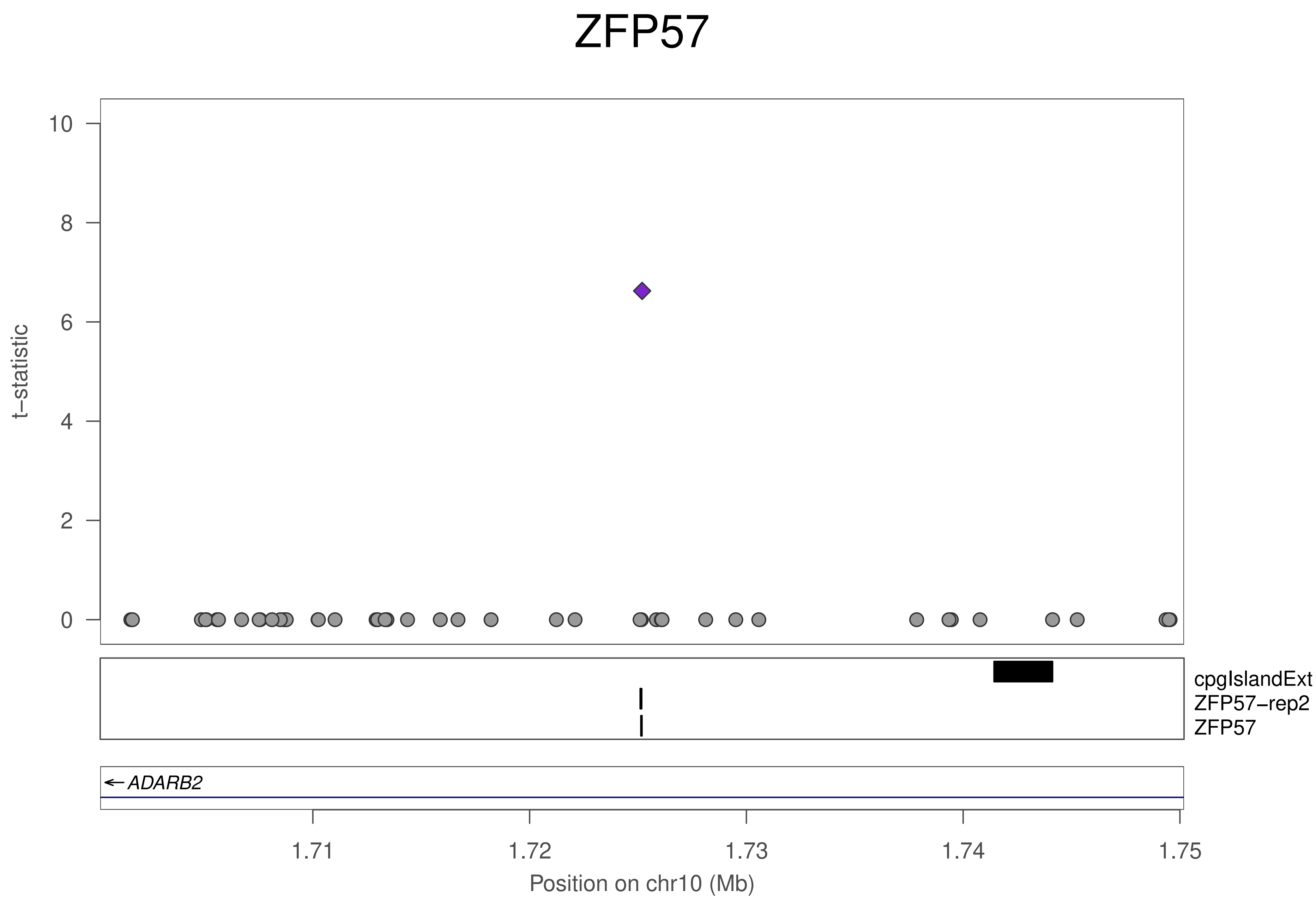
ZFP57
t−statistic
10
8
6
4
2
0
cpgIslandExt
ZFP57−rep2
ZFP57
ADARB2
1.71
1.72
1.73
1.74
1.75
Position on chr10 (Mb)

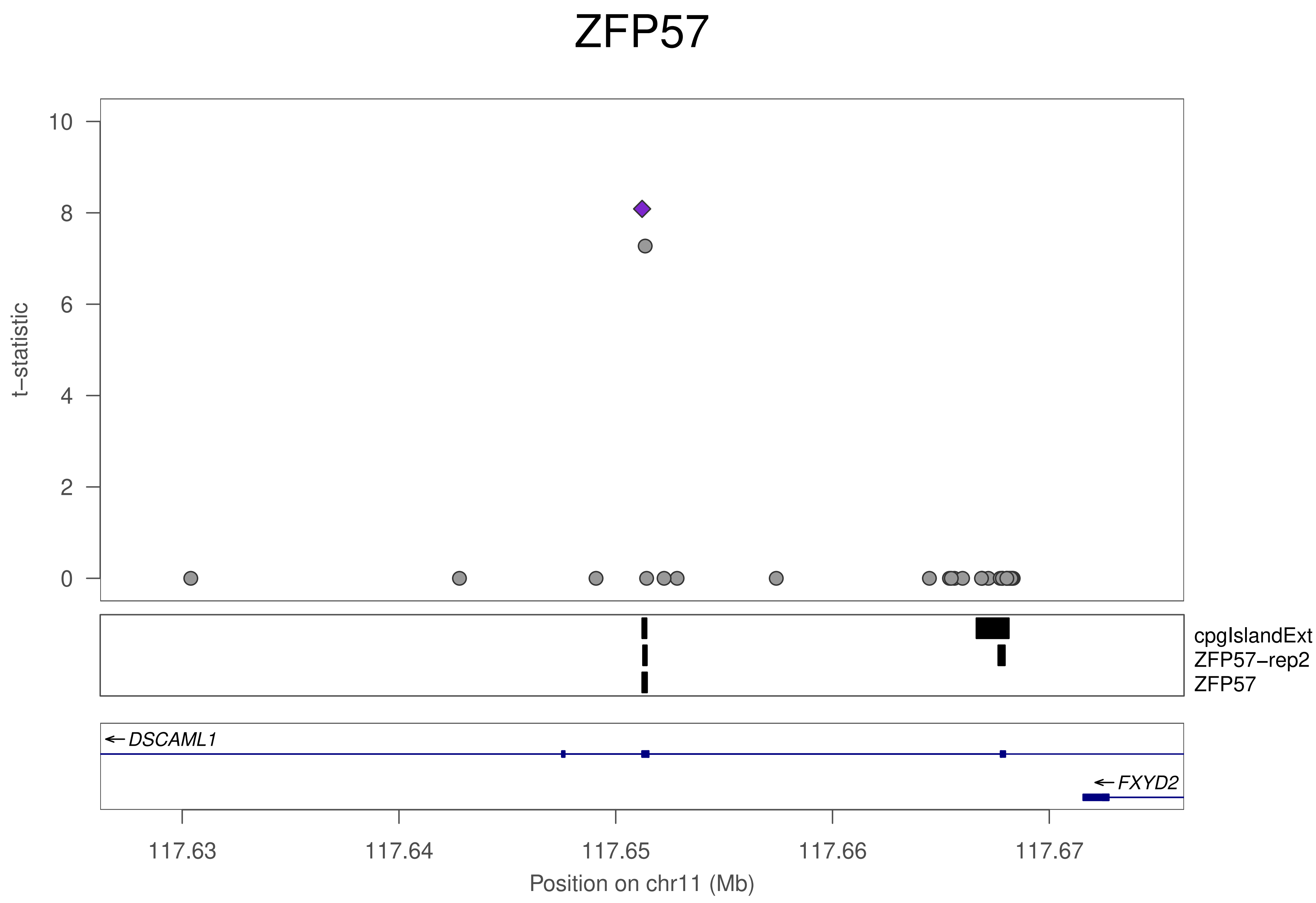
ZFP57
t−statistic
10
8
6
4
2
0
cpgIslandExt
ZFP57−rep2
ZFP57
DSCAML1
FXYD2
117.63
117.64
117.65
117.66
117.67
Position on chr11 (Mb)

### ZFP57

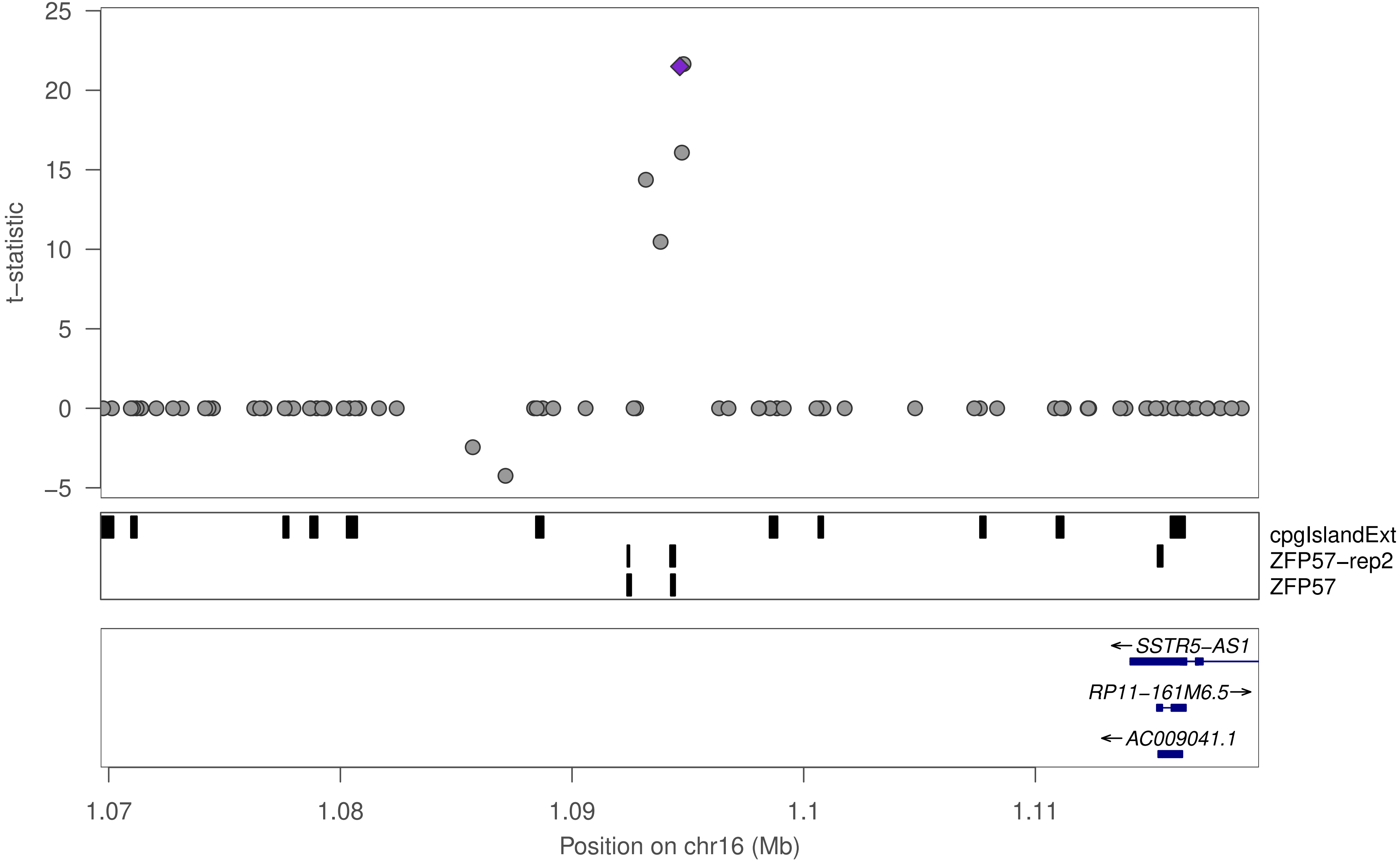

25
20
15
10
5
0
−5
t−statistic
cpgIslandExt
ZFP57−rep2
ZFP57
← SSTR5−AS1
RP11−161M6.5→
← AC009041.1
1.07
1.08
1.09
1.1
1.11
Position on chr16 (Mb)

### ZFP57

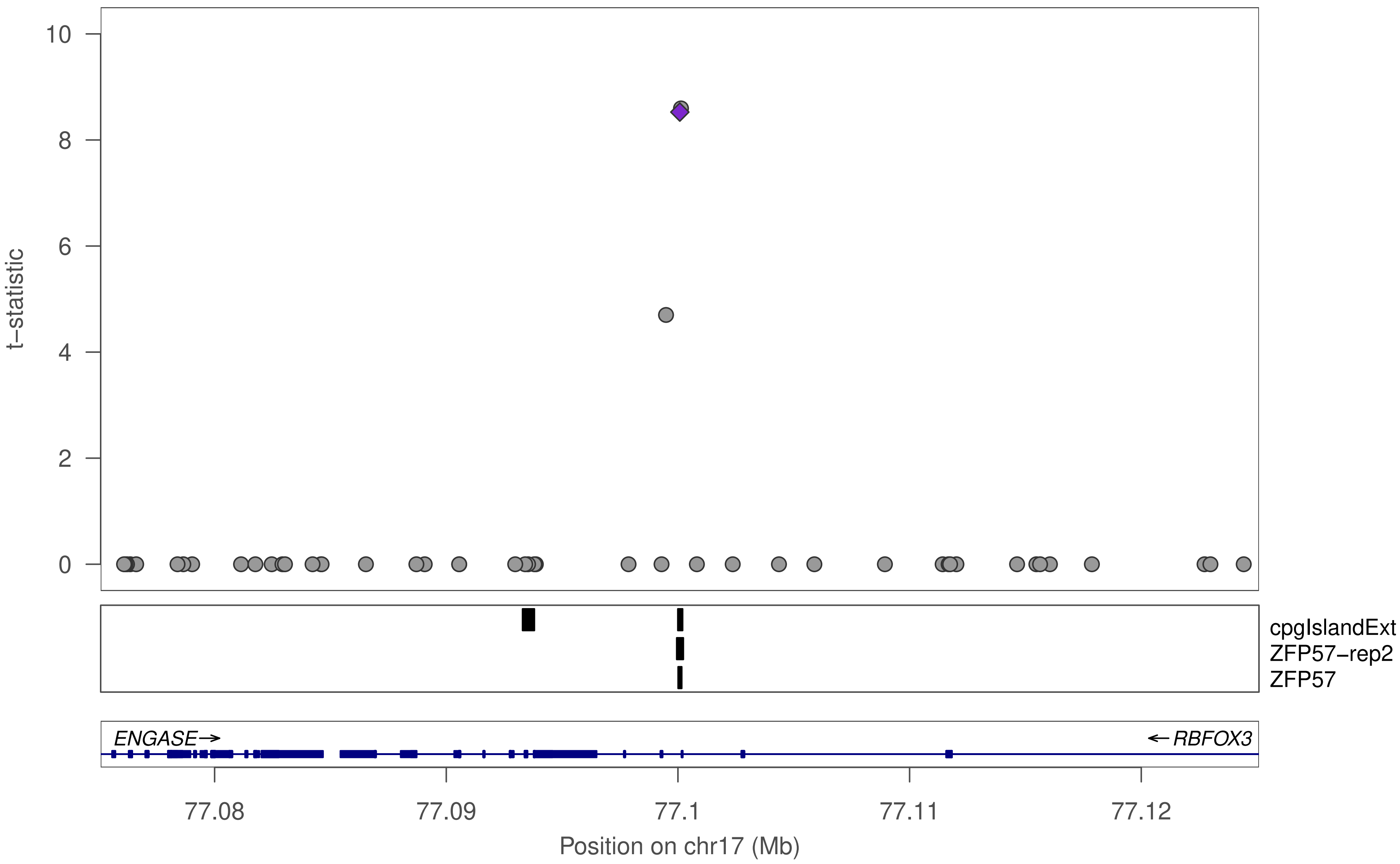

t−statistic
10
8
6
4
2
0
cpgIslandExt
ZFP57−rep2
ZFP57
ENGASE→
←RBFOX3
77.08
77.09
77.1
77.11
77.12
Position on chr17 (Mb)

### ZFP57

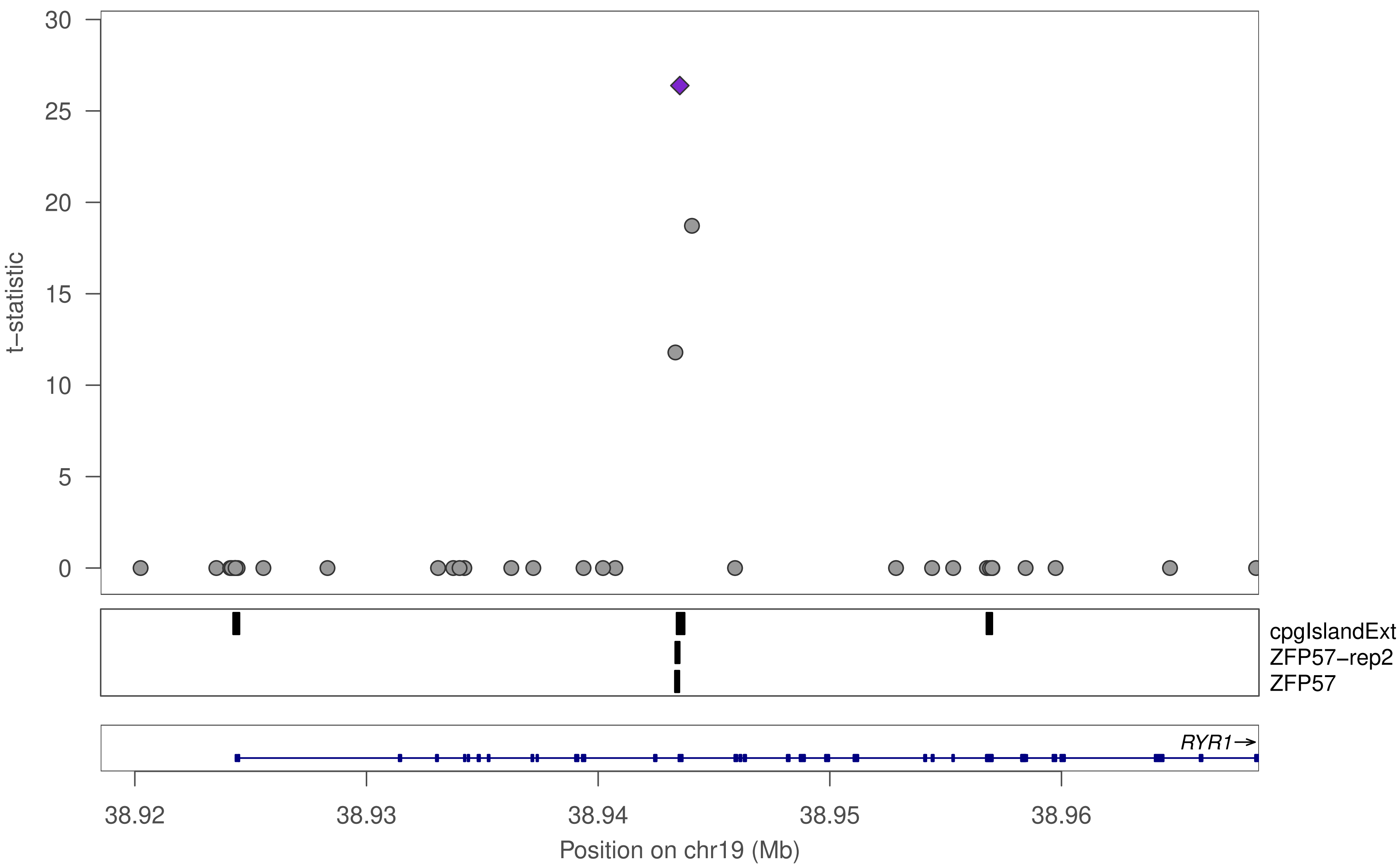

30
25
20
15
10
5
0
t−statistic
cpgIslandExt
ZFP57−rep2
ZFP57
RYR1→
38.92
38.93
38.94
38.95
38.96
Position on chr19 (Mb)

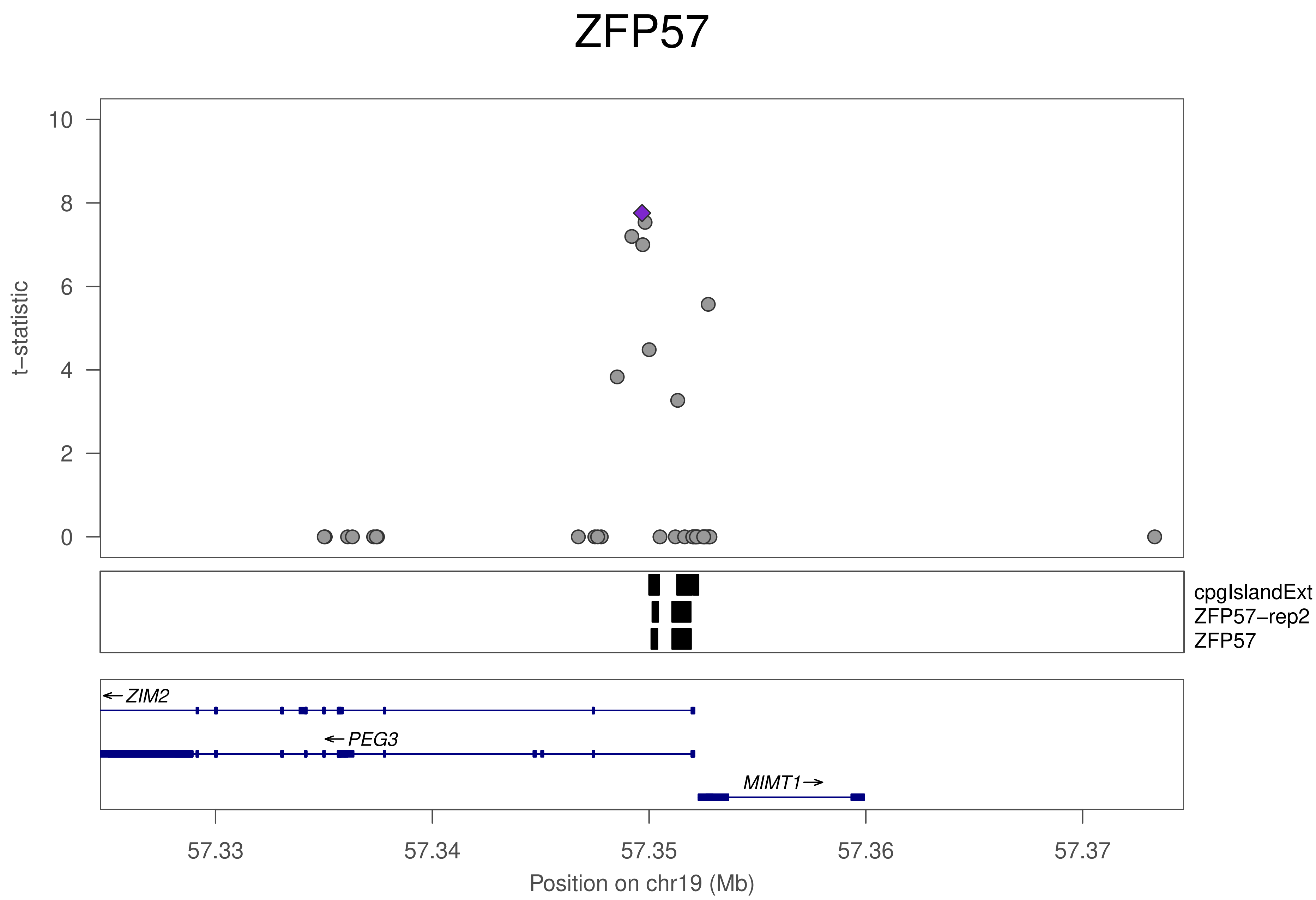
ZFP57
t−statistic
10
8
6
4
2
0
cpgIslandExt
ZFP57−rep2
ZFP57
ZIM2
PEG3
MIMT1
57.33
57.34
57.35
57.36
57.37
Position on chr19 (Mb)

### ZFP57

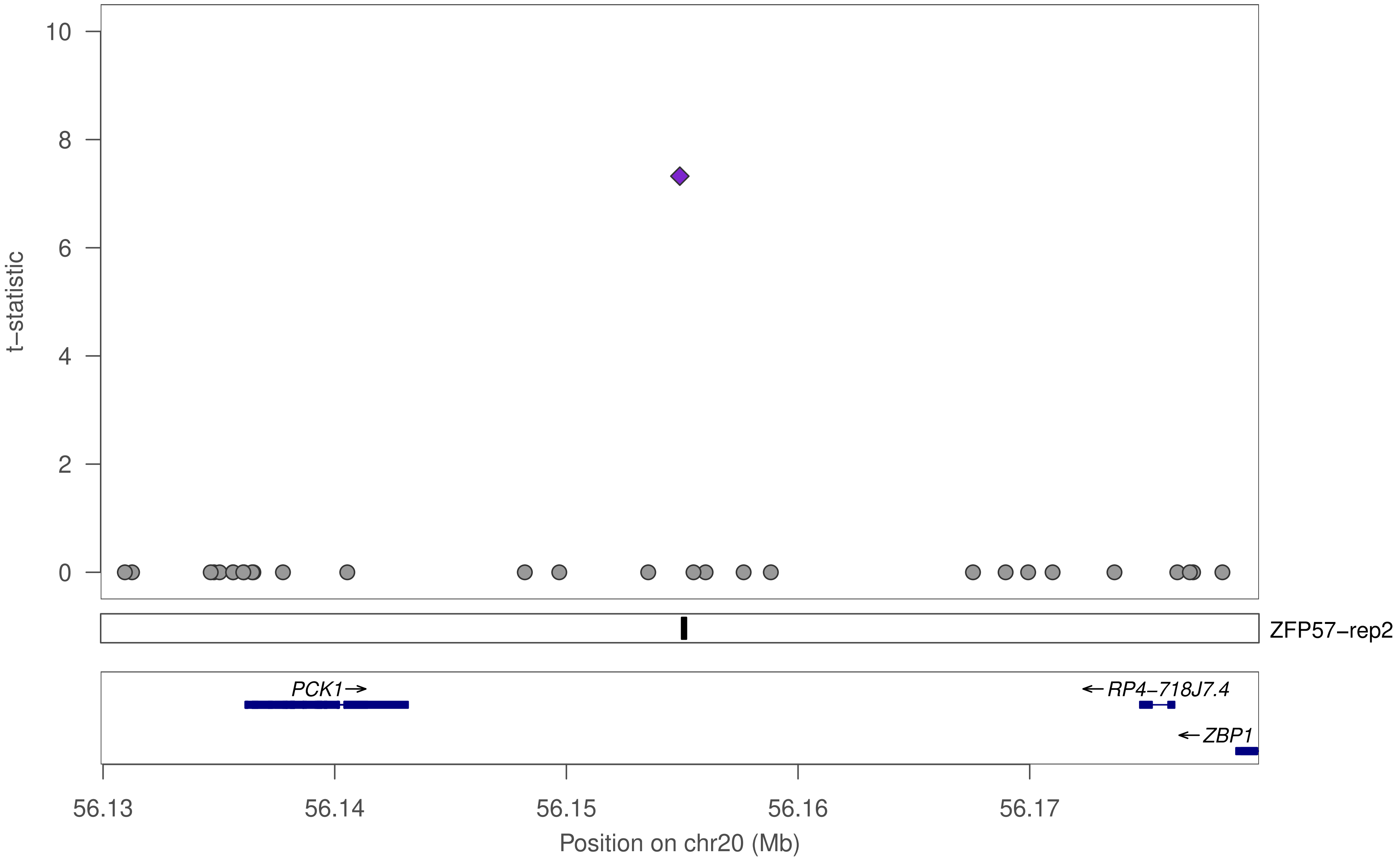

t−statistic
10
8
6
4
2
0
ZFP57−rep2
PCK1→
←RP4−718J7.4
←ZBP1
56.13
56.14
56.15
56.16
56.17
Position on chr20 (Mb)

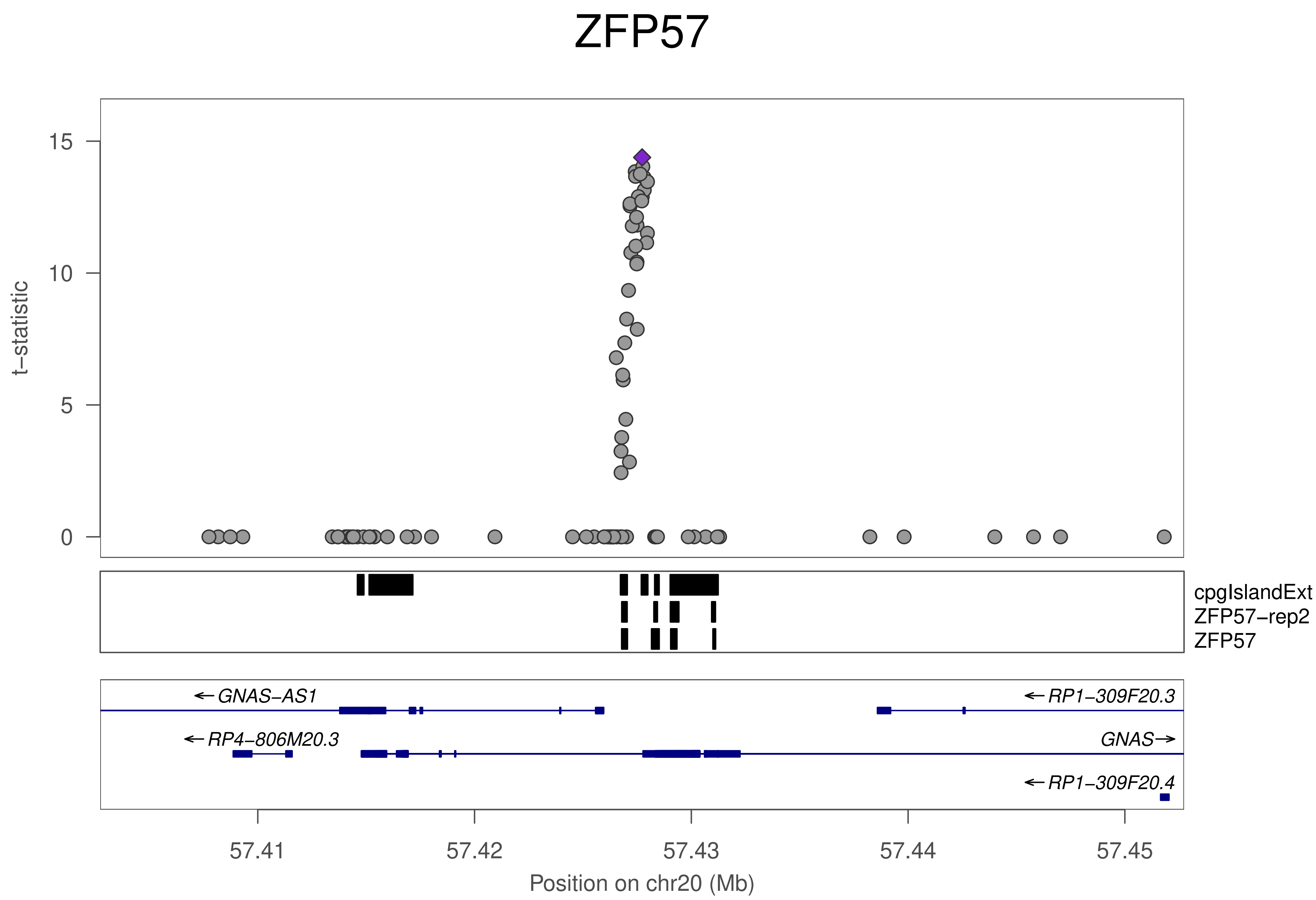
ZFP57
t−statistic
15
10
5
0
cpgIslandExt
ZFP57−rep2
ZFP57
← GNAS−AS1
← RP1−309F20.3
← RP4−806M20.3
GNAS →
← RP1−309F20.4
57.41
57.42
57.43
57.44
57.45
Position on chr20 (Mb)

### ZNF783

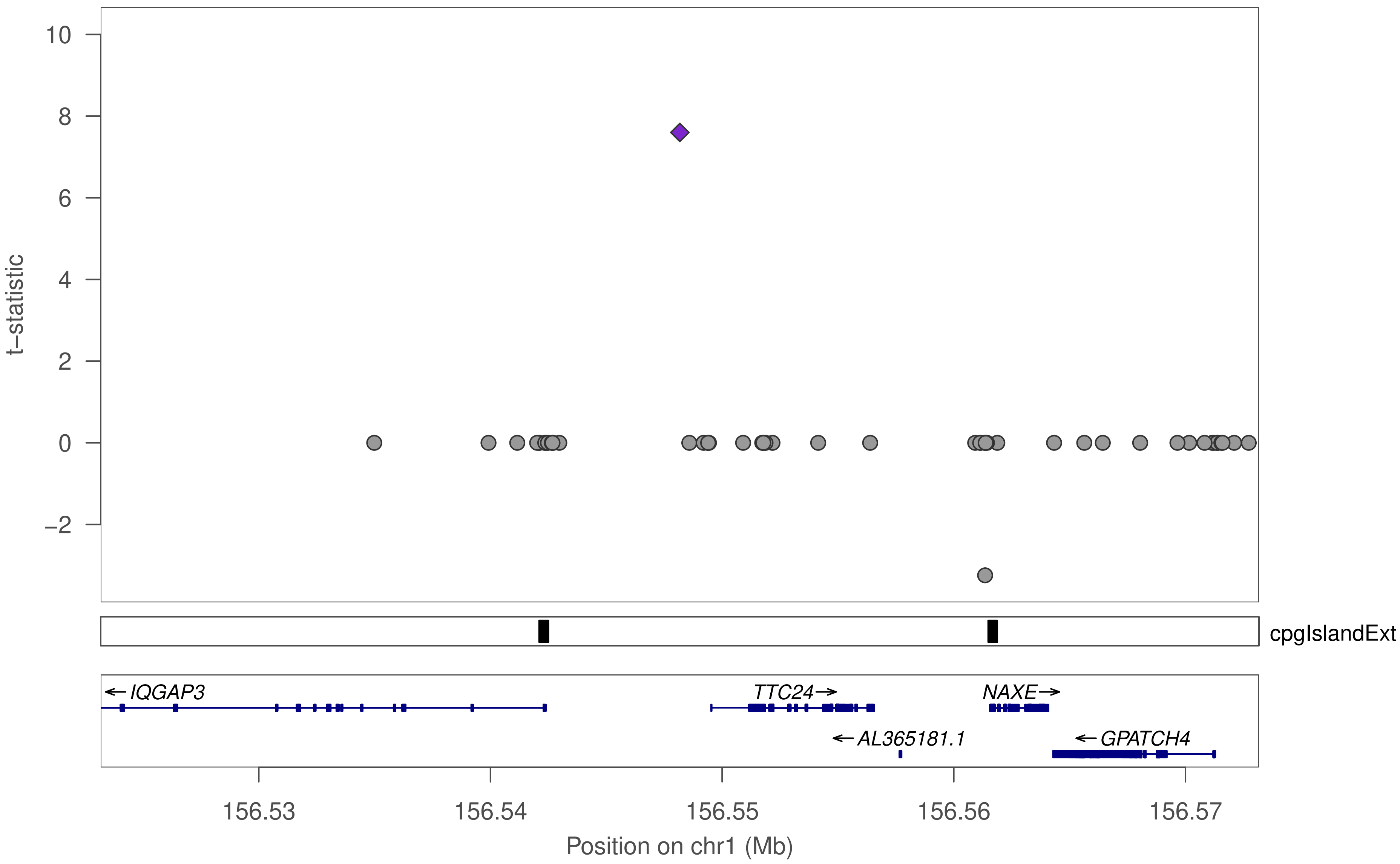

10
8
6
4
2
0
−2
t−statistic
cpgIslandExt
← IQGAP3
TTC24→
NAXE→
← AL365181.1
← GPATCH4
156.53
156.54
156.55
156.56
156.57
Position on chr1 (Mb)

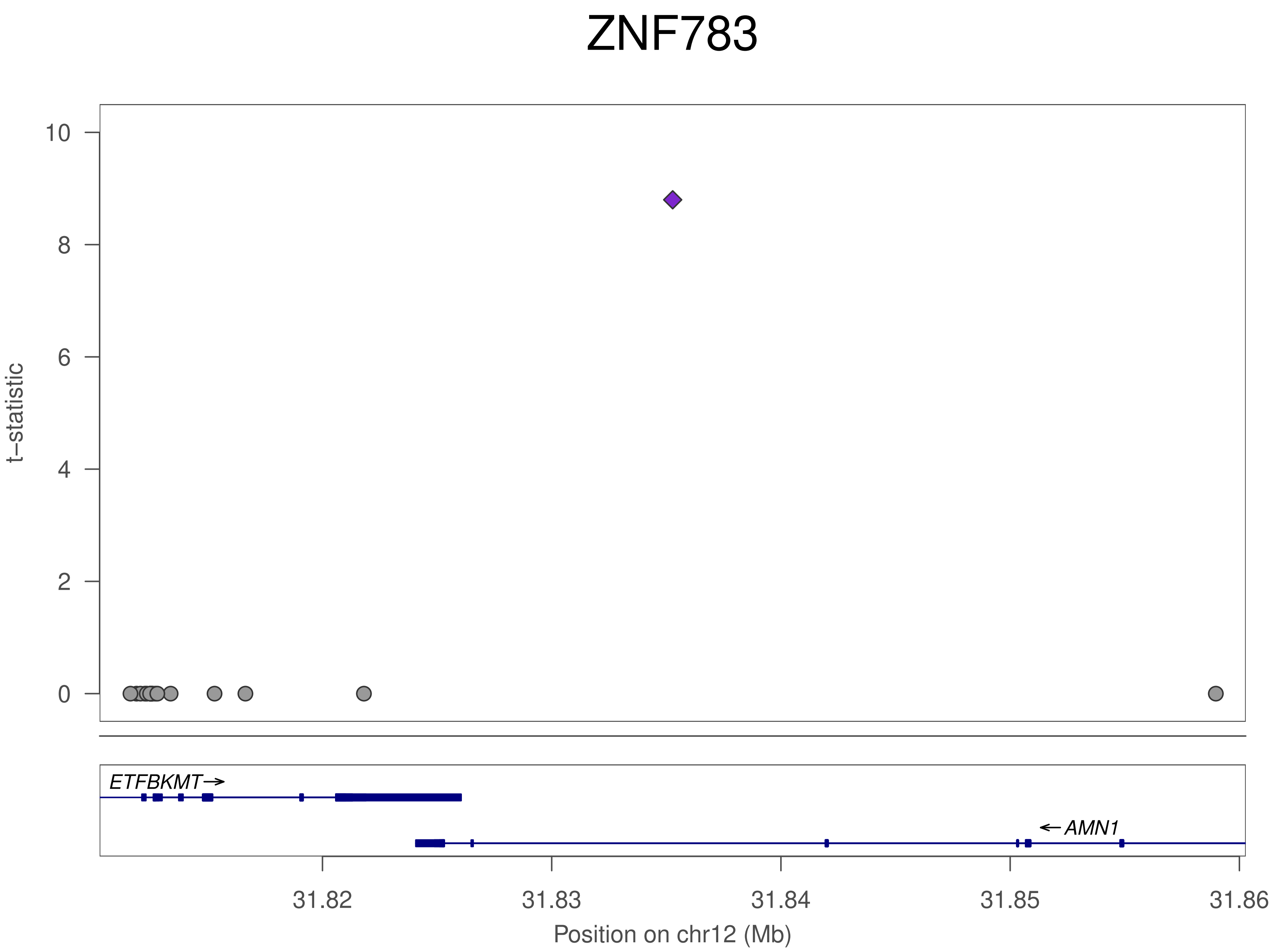
ZNF783
t−statistic
10
8
6
4
2
0
ETFBKMT
AMN1
31.82
31.83
31.84
31.85
31.86
Position on chr12 (Mb)

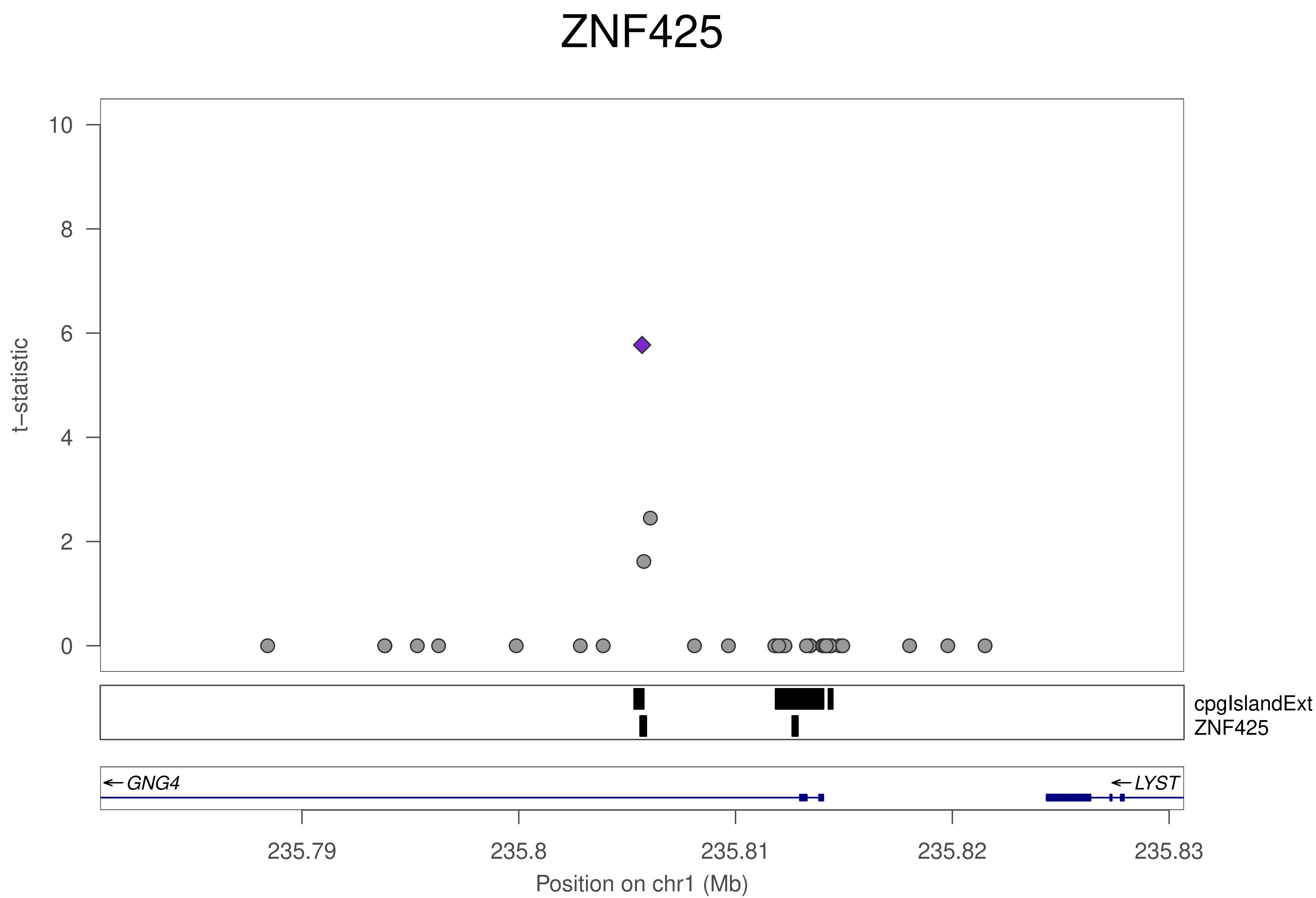
ZNF425
t−statistic
10
8
6
4
2
0
cpgIslandExt
ZNF425
GNG4
LYST
235.79
235.8
235.81
235.82
235.83
Position on chr1 (Mb)

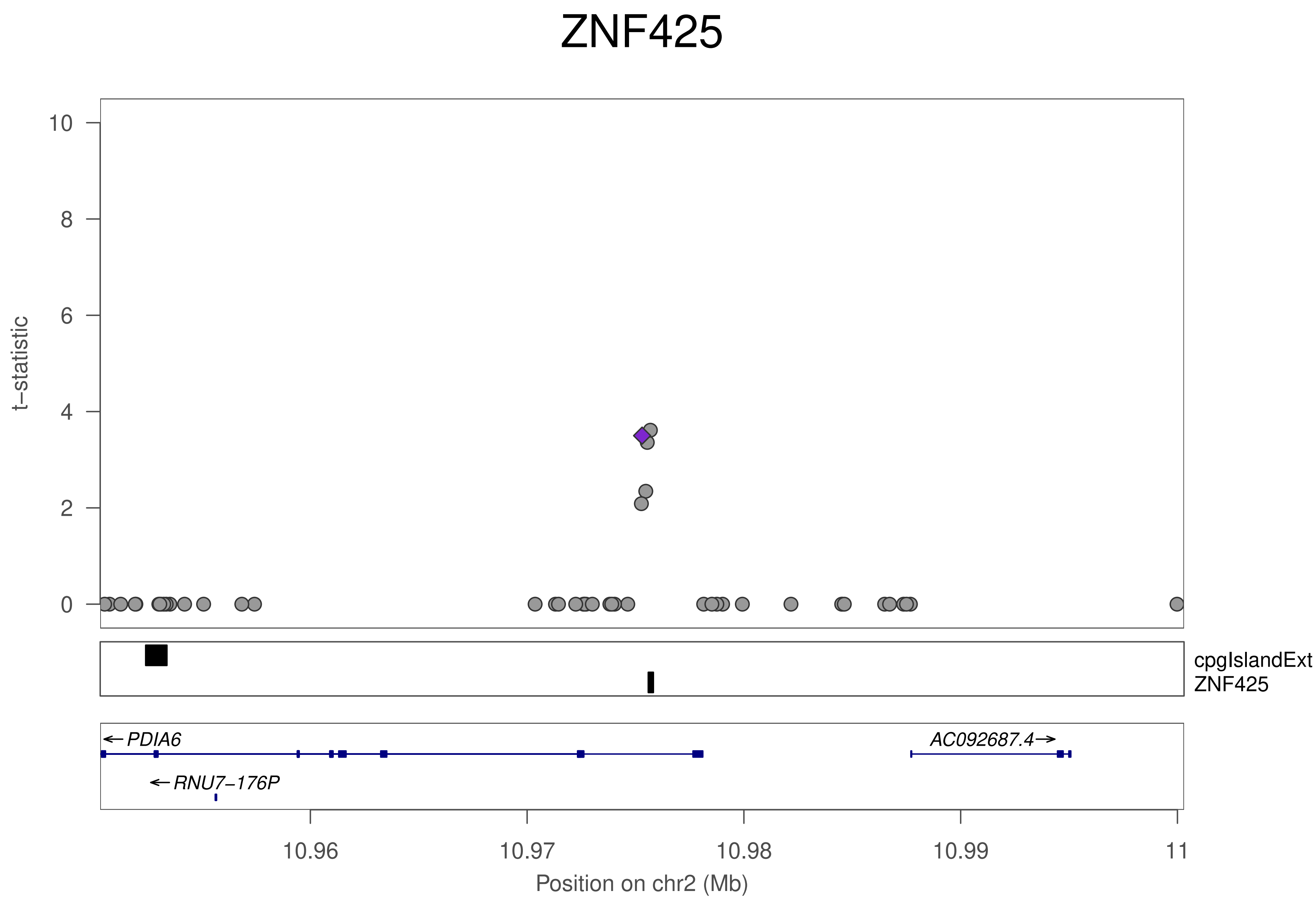
ZNF425
t−statistic
0
2
4
6
8
10
cpgIslandExt
ZNF425
← PDIA6
AC092687.4→
← RNU7−176P
10.96
10.97
10.98
10.99
11
Position on chr2 (Mb)

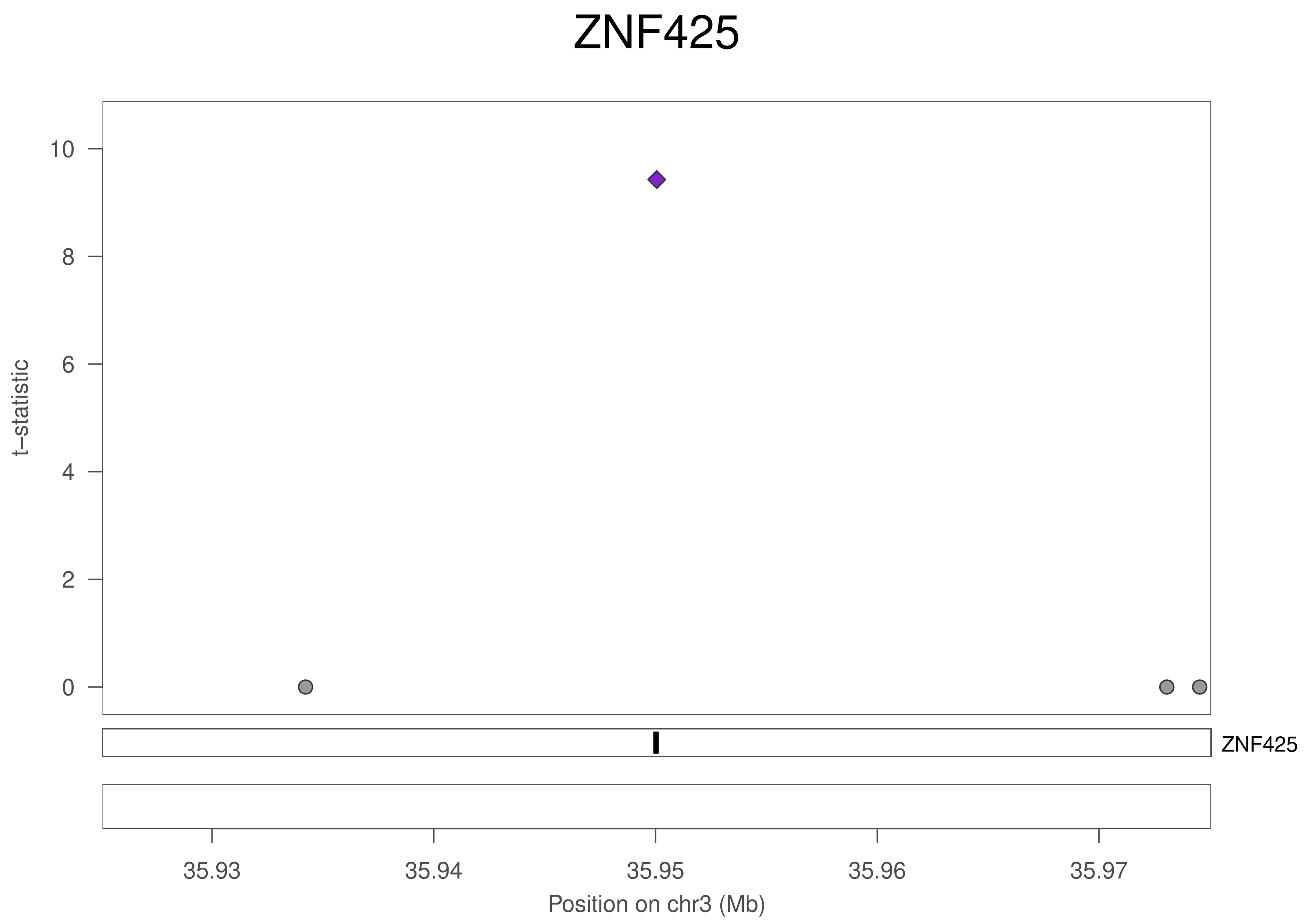

ZNF425
t–statistic
10
8
6
4
2
0
ZNF425
35.93
35.94
35.95
35.96
35.97
Position on chr3 (Mb)

### ZNF425

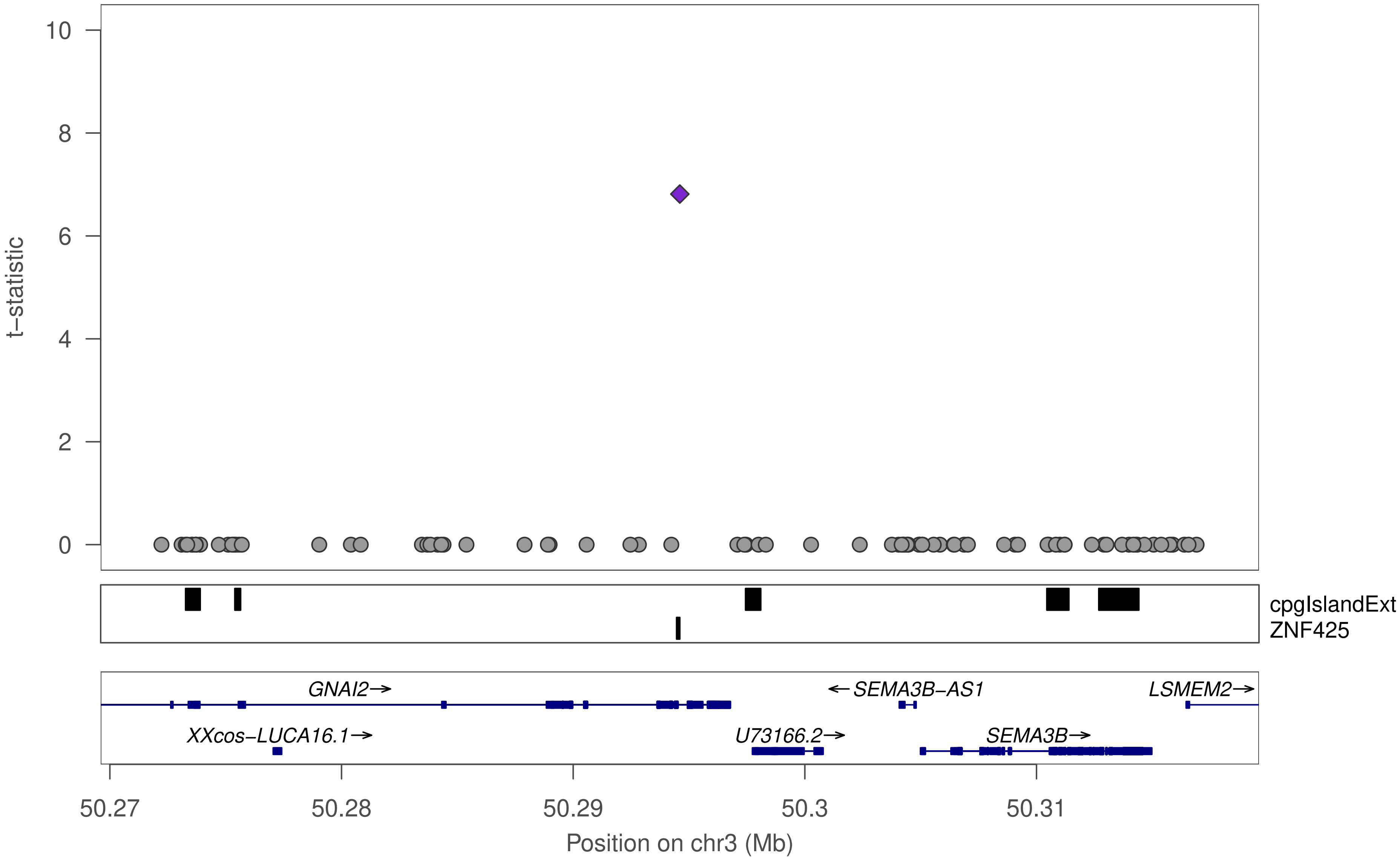

10
8
6
4
2
0
t−statistic
cpgIslandExt
ZNF425
GNAI2→
←SEMA3B−AS1
LSMEM2→
XXcos−LUCA16.1→
U73166.2→
SEMA3B→
50.27
50.28
50.29
50.3
50.31
Position on chr3 (Mb)

### ZNF425

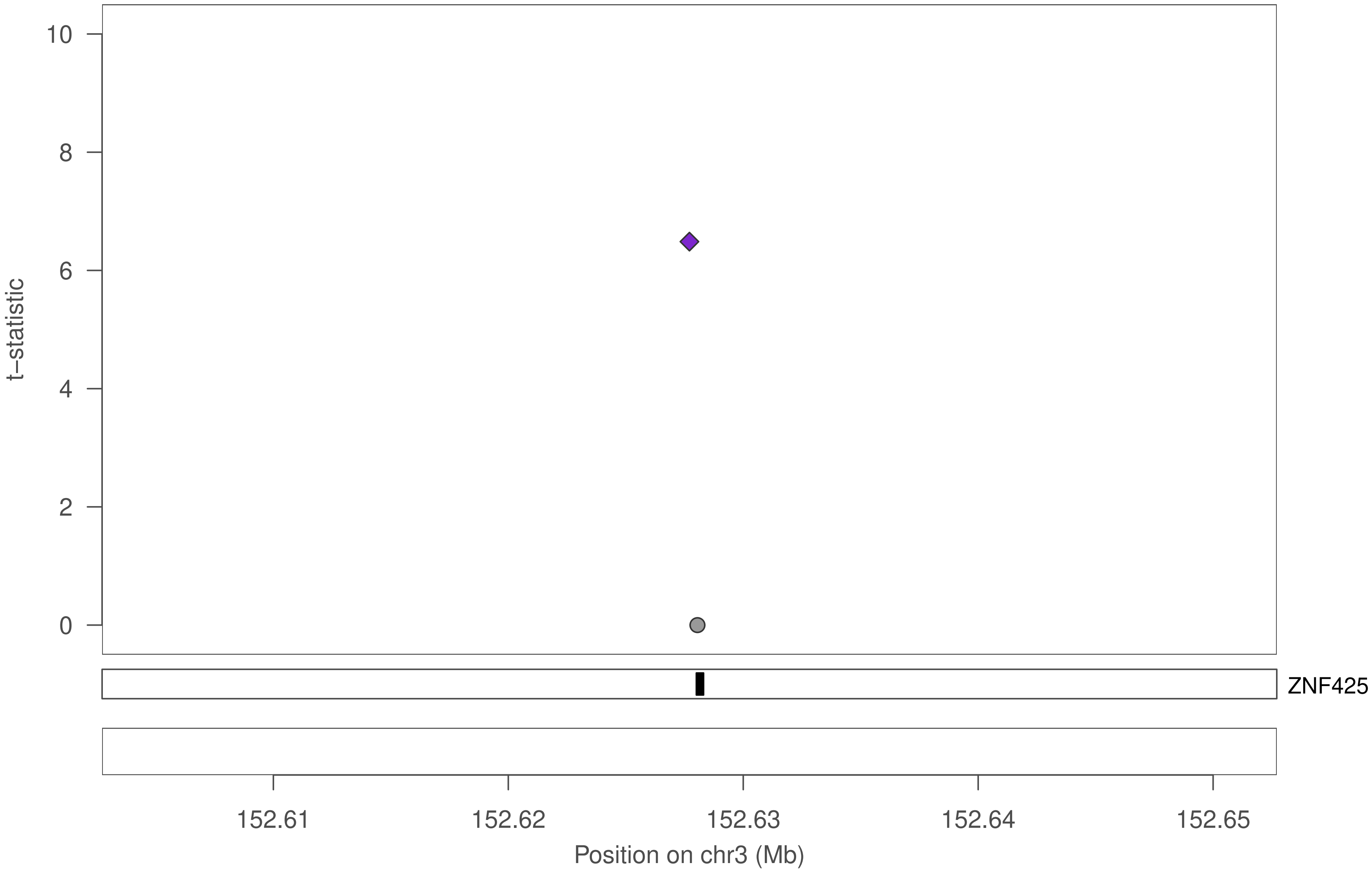

10
8
6
4
2
0
t−statistic
ZNF425
152.61
152.62
152.63
152.64
152.65
Position on chr3 (Mb)

### ZNF425

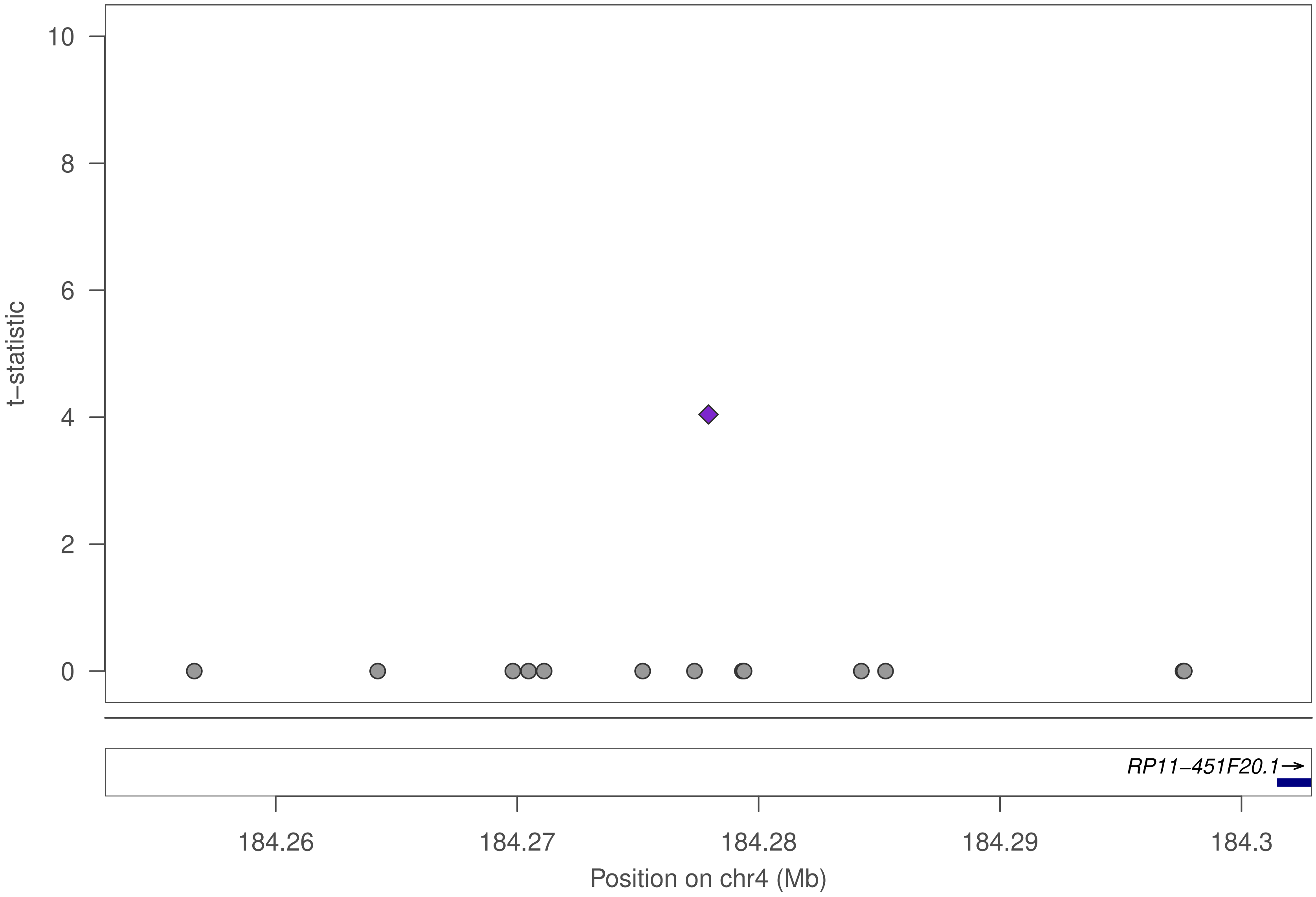

t–statistic
10
8
6
4
2
0
RP11–451F20.1→
184.26
184.27
184.28
184.29
184.3
Position on chr4 (Mb)

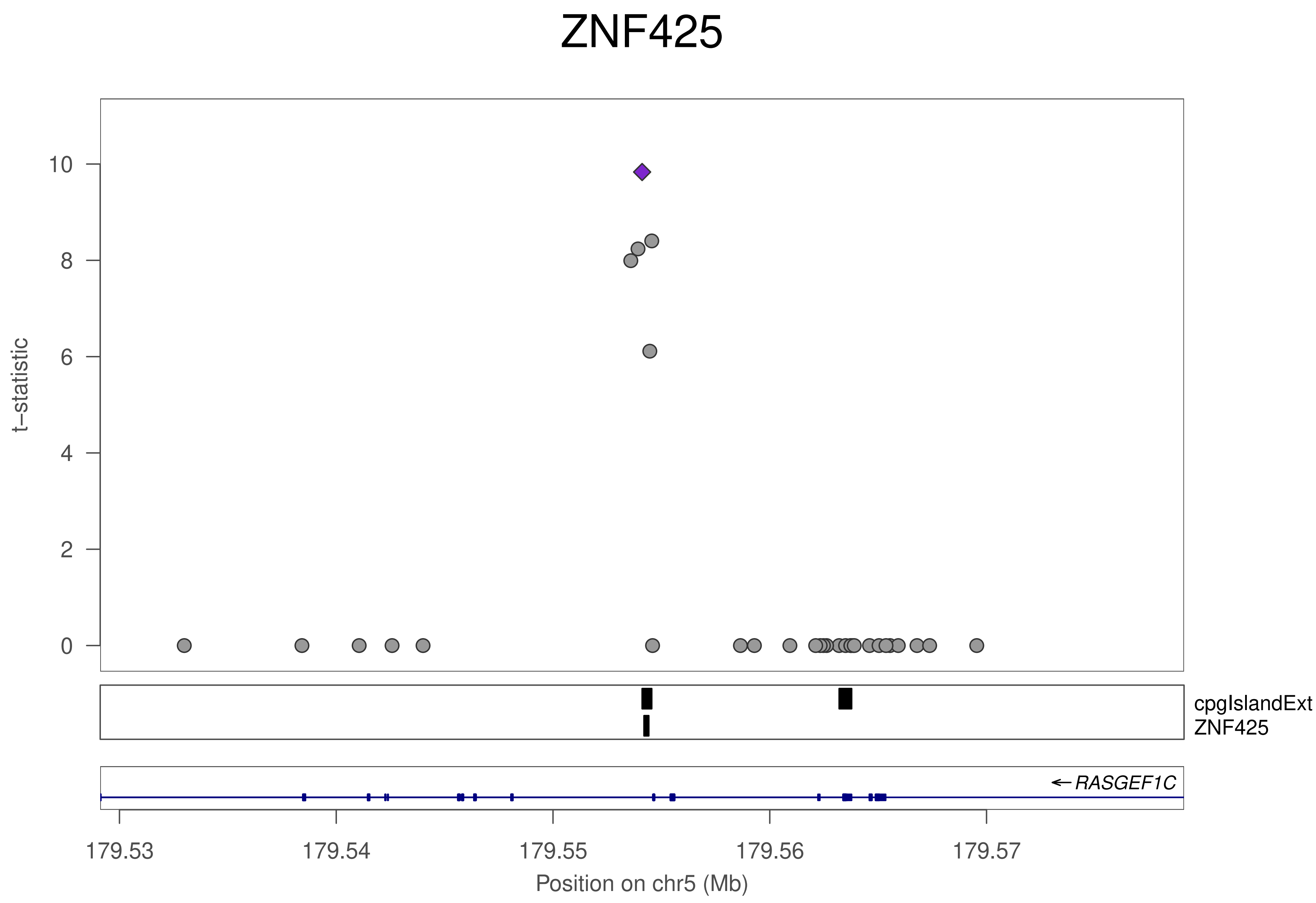

ZNF425
t−statistic
10
8
6
4
2
0
cpgIslandExt
ZNF425
← RASGEF1C
179.53
179.54
179.55
179.56
179.57
Position on chr5 (Mb)

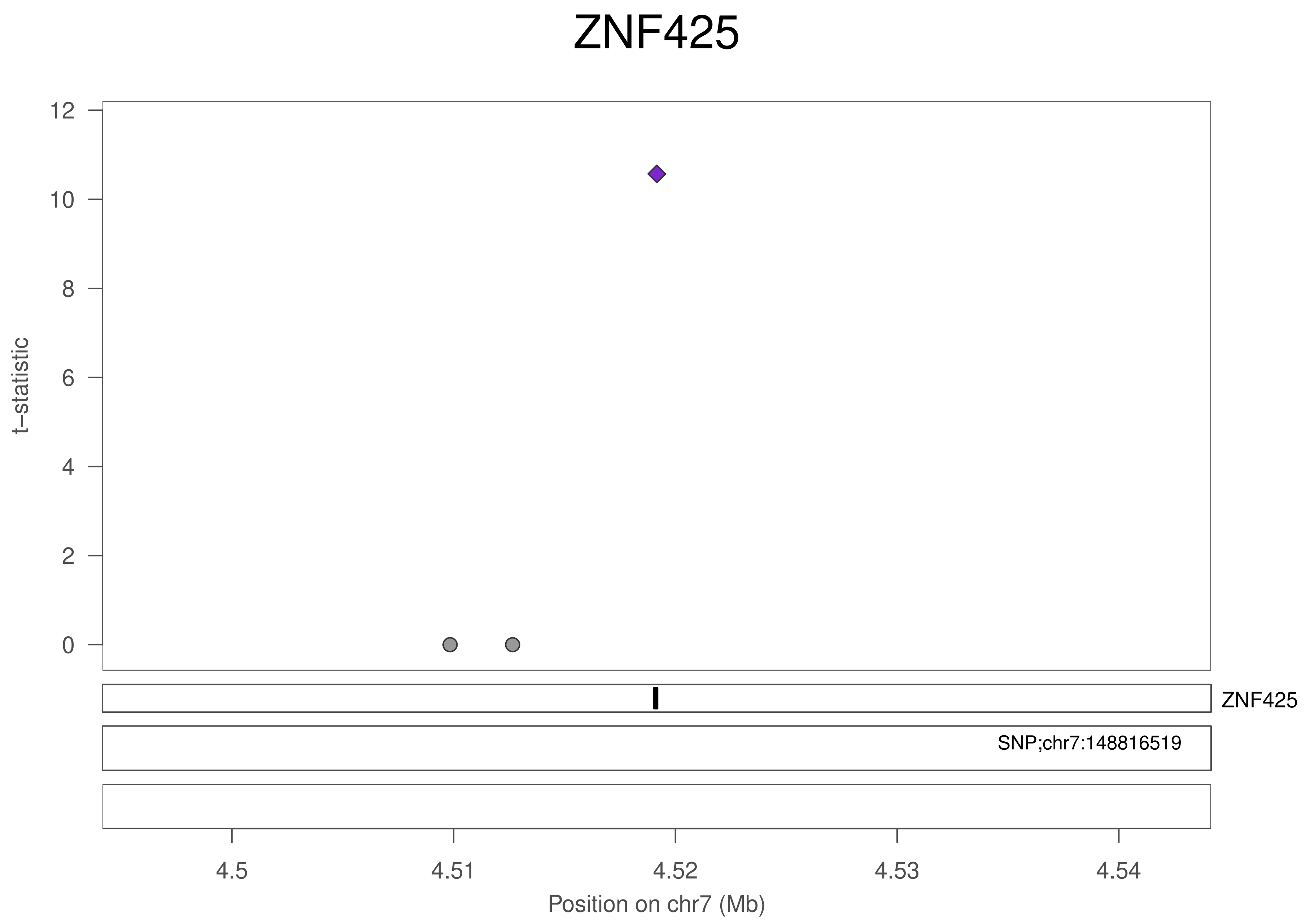
ZNF425
t−statistic
12
10
8
6
4
2
0
ZNF425
SNP;chr7:148816519
4.5
4.51
4.52
4.53
4.54
Position on chr7 (Mb)

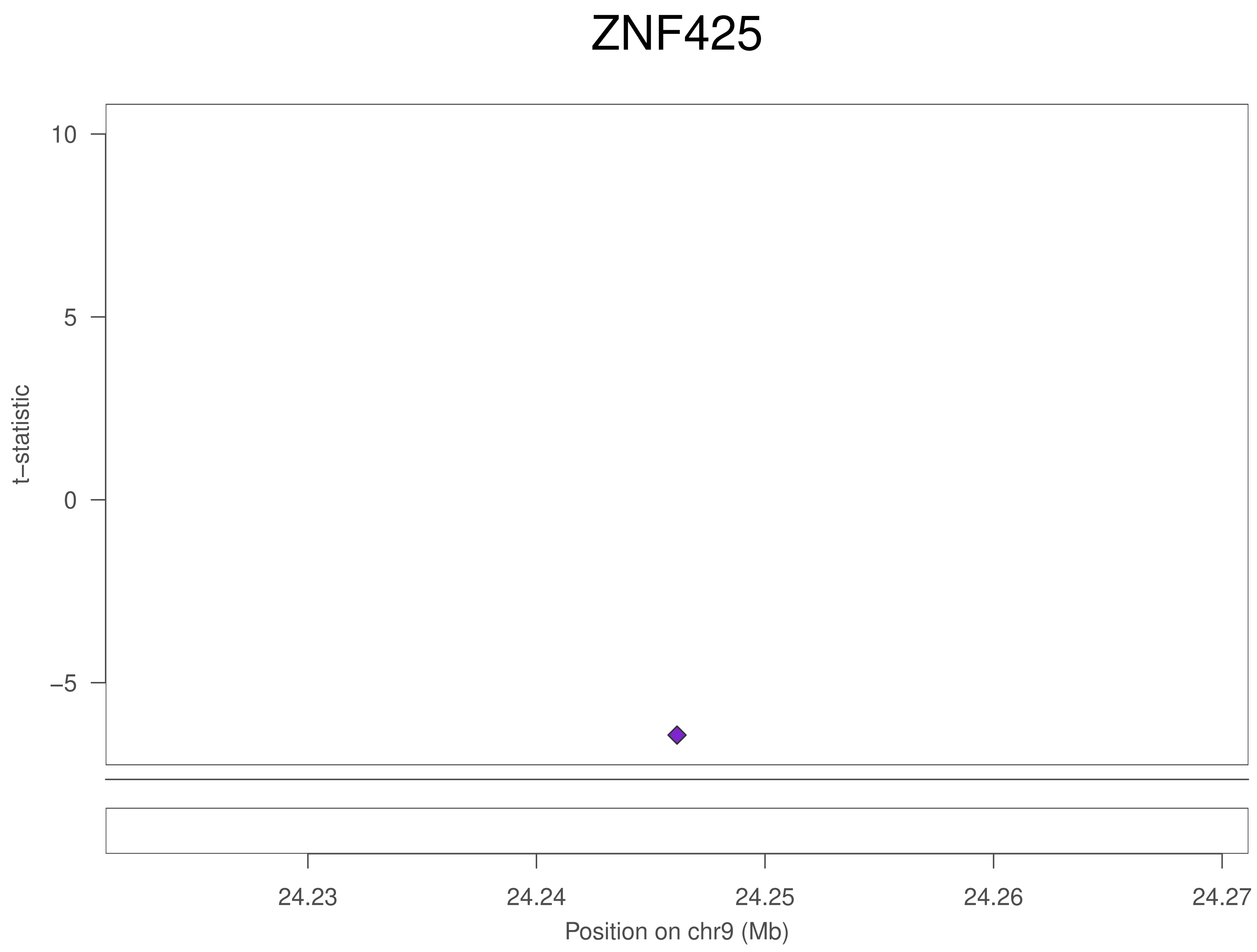
ZNF425
t−statistic
10
5
0
−5
24.23
24.24
24.25
24.26
24.27
Position on chr9 (Mb)

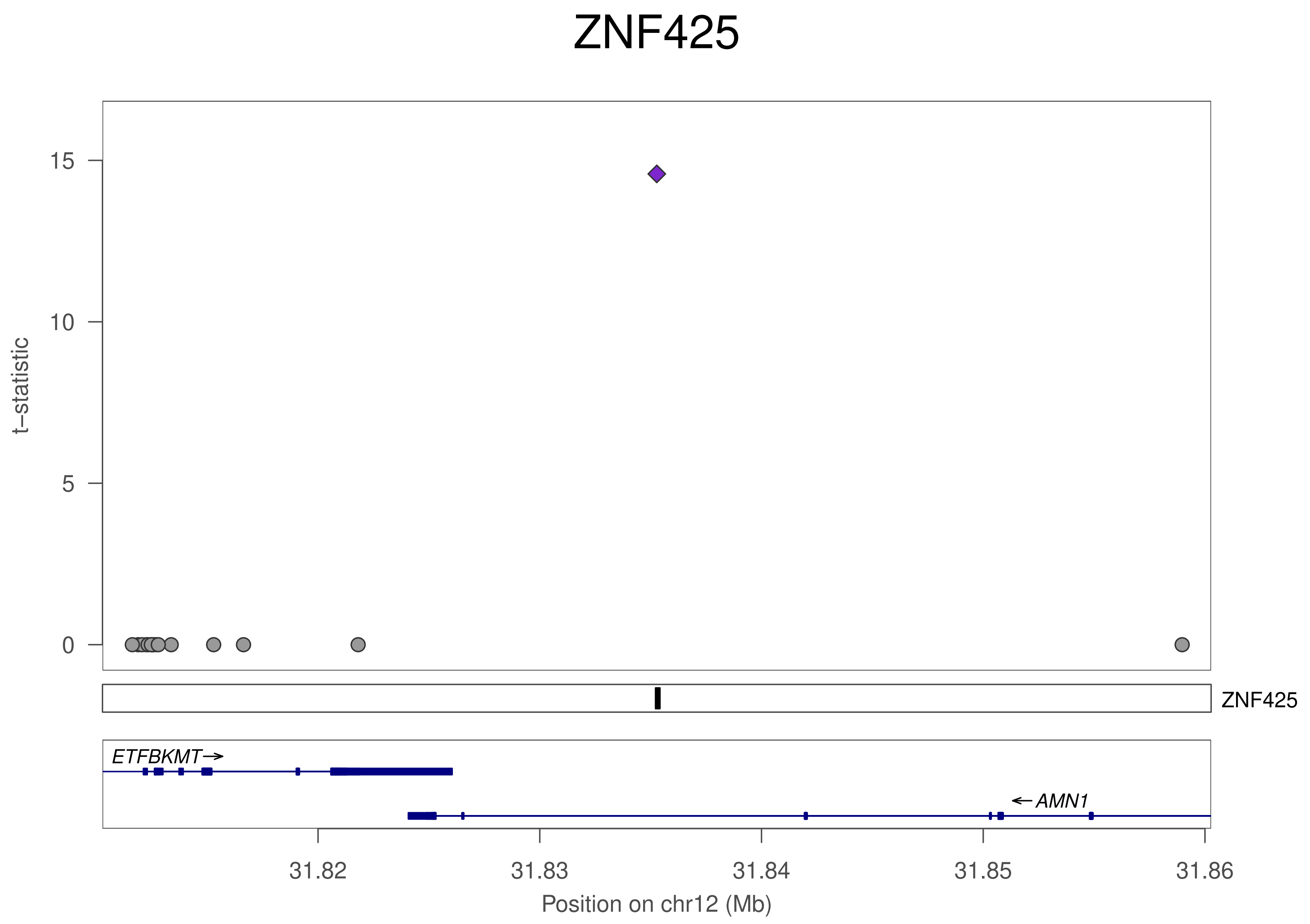
ZNF425
t−statistic
15
10
5
0
ZNF425
ETFBKMT
AMN1
31.82
31.83
31.84
31.85
31.86
Position on chr12 (Mb)

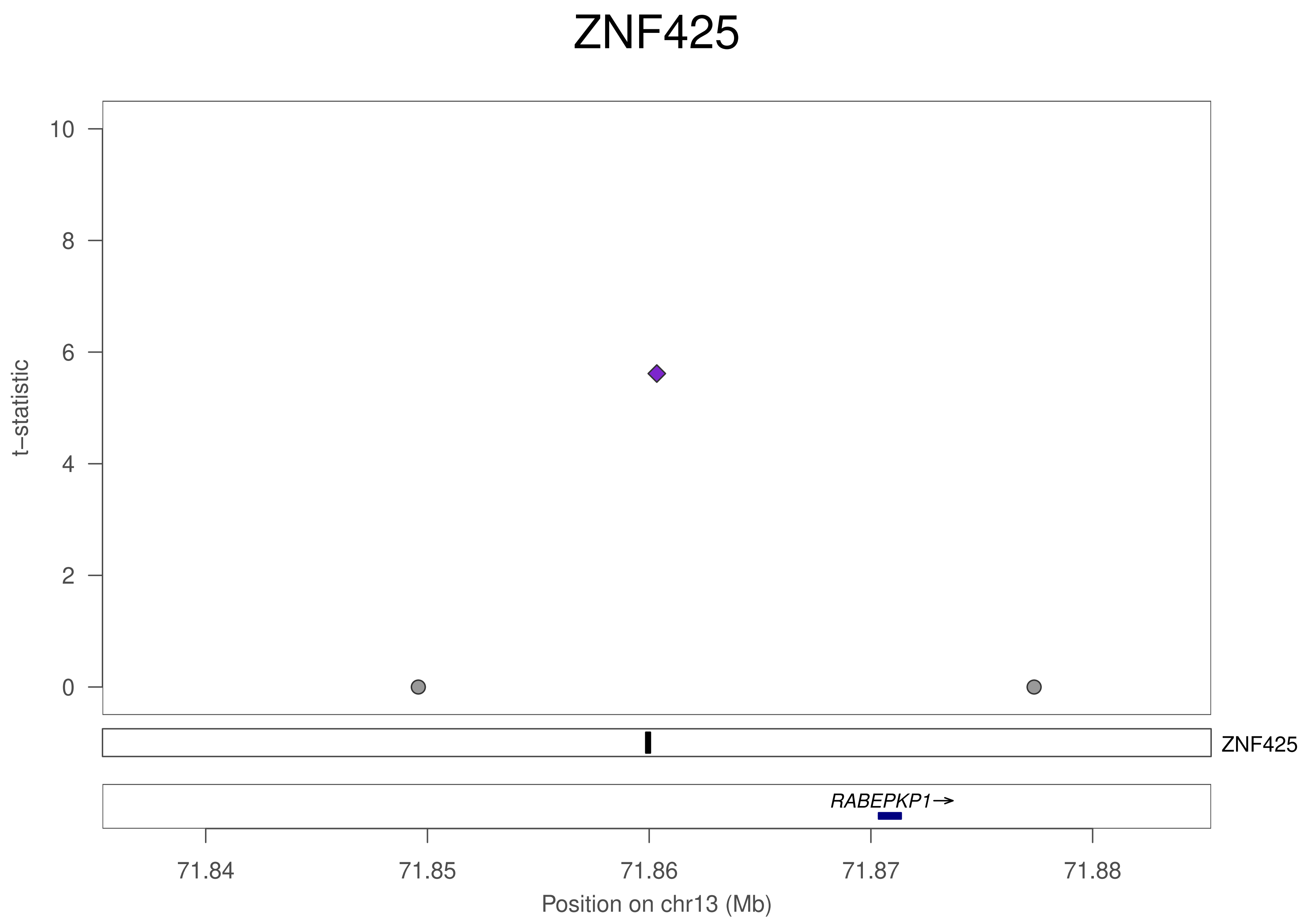
ZNF425
10
8
6
4
2
0
t−statistic
ZNF425
RABEPKP1→
71.84
71.85
71.86
71.87
71.88
Position on chr13 (Mb)

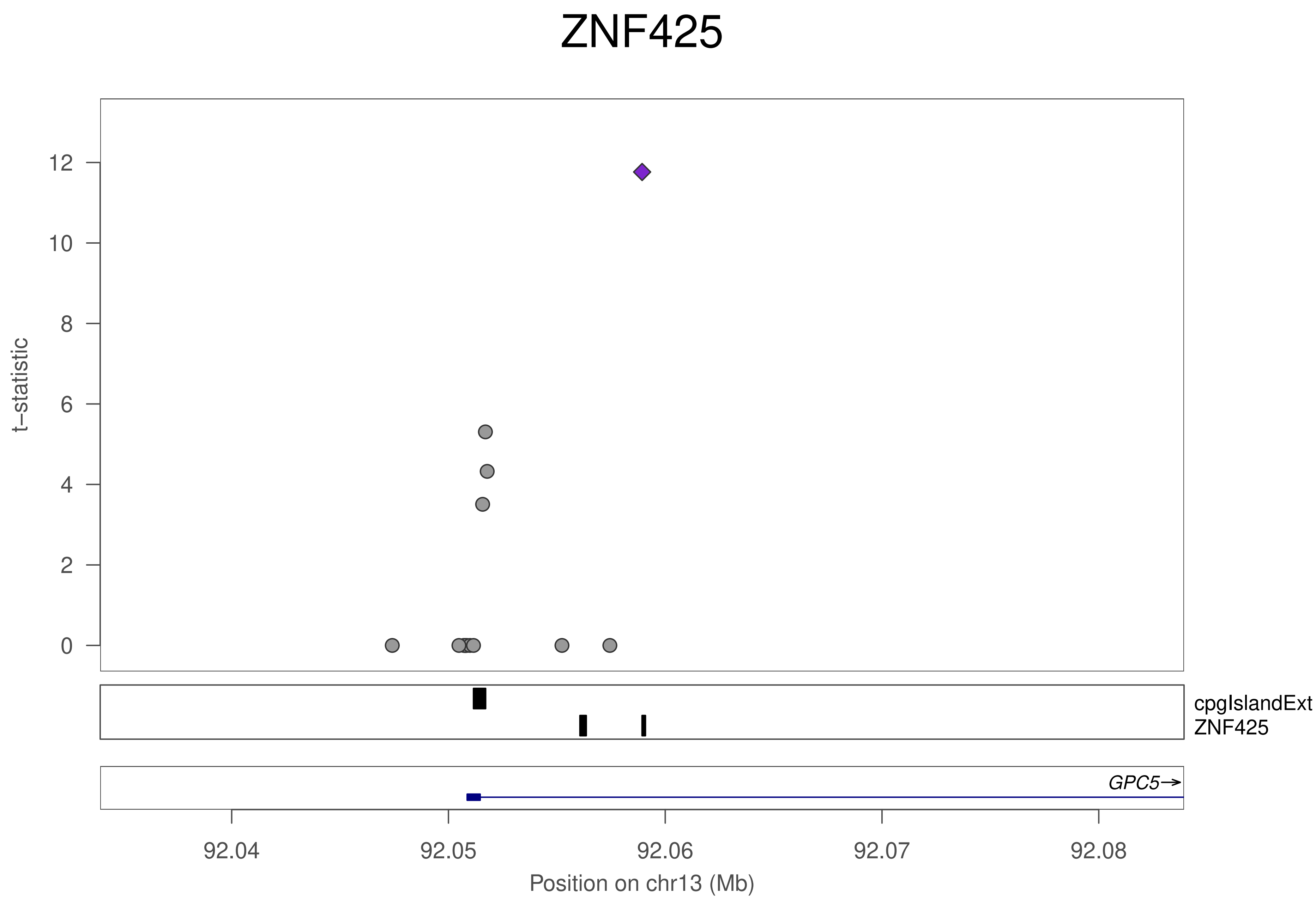
ZNF425
t−statistic
12
10
8
6
4
2
0
cpgIslandExt
ZNF425
GPC5→
92.04
92.05
92.06
92.07
92.08
Position on chr13 (Mb)

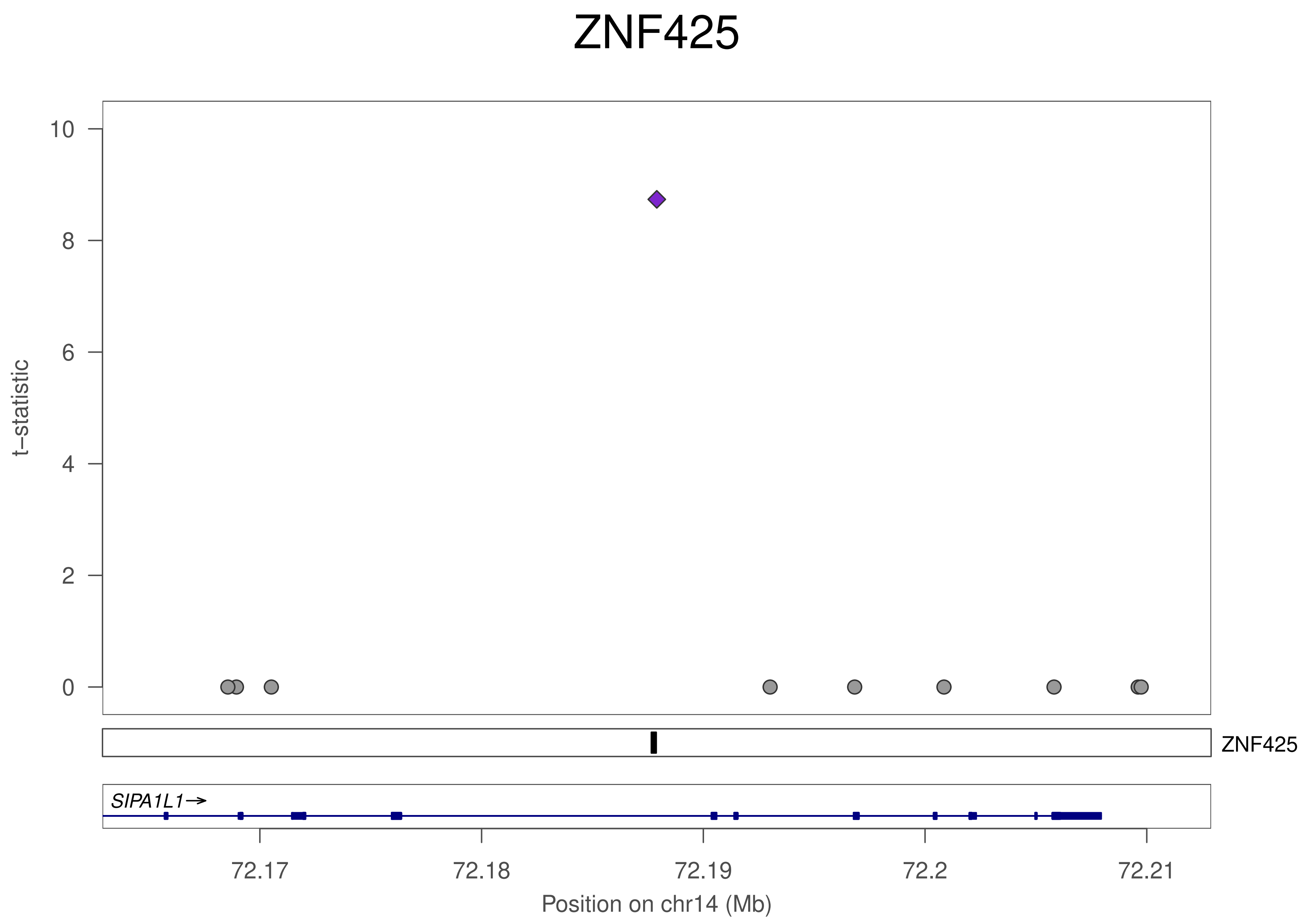

ZNF425
10
8
6
4
2
0
t−statistic
ZNF425
SIPA1L1→
72.17
72.18
72.19
72.2
72.21
Position on chr14 (Mb)

### ZNF425

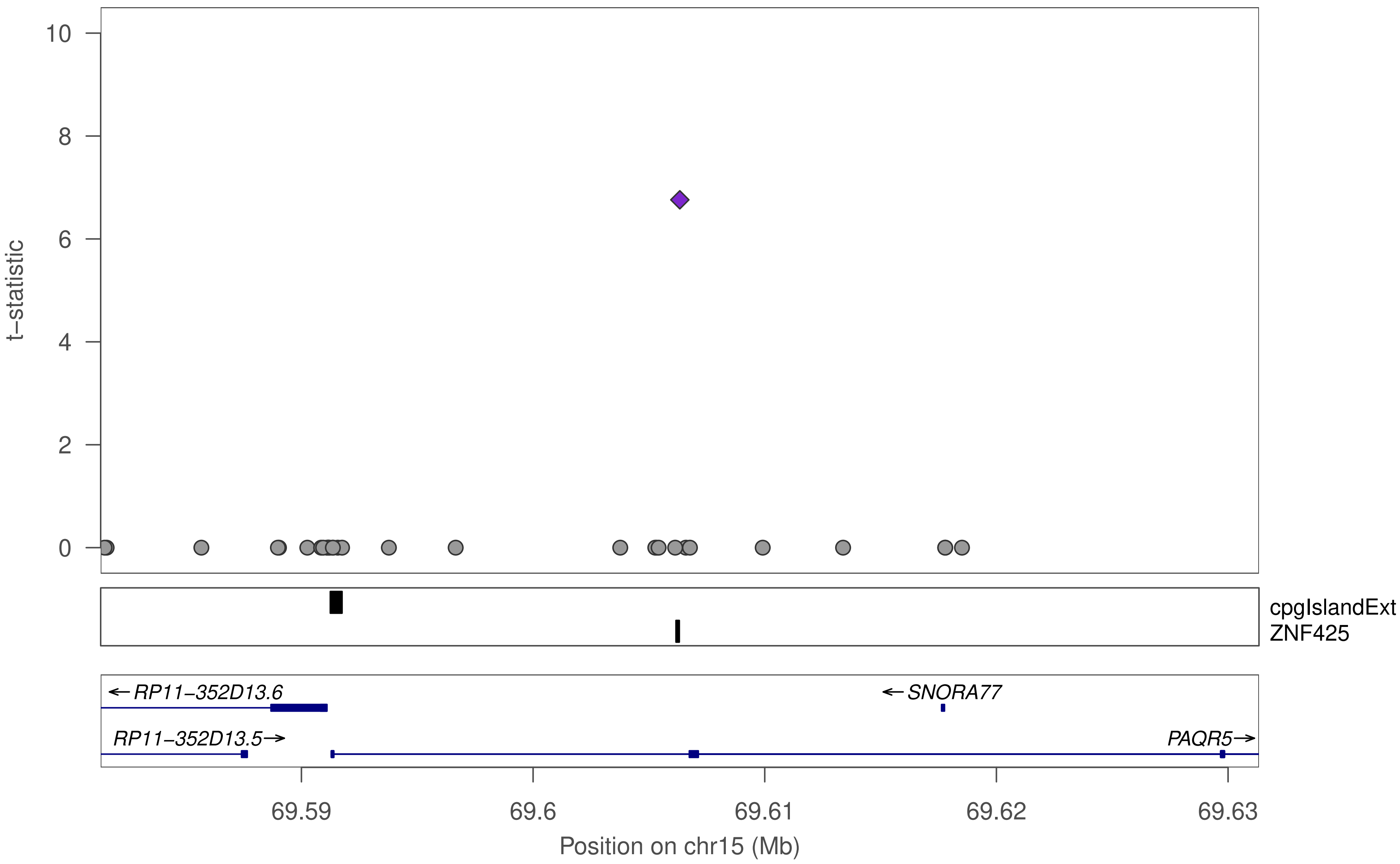

10
8
6
4
2
0
t−statistic
cpgIslandExt
ZNF425
←RP11−352D13.6
←SNORA77
RP11−352D13.5→
PAQR5→
69.59
69.6
69.61
69.62
69.63
Position on chr15 (Mb)

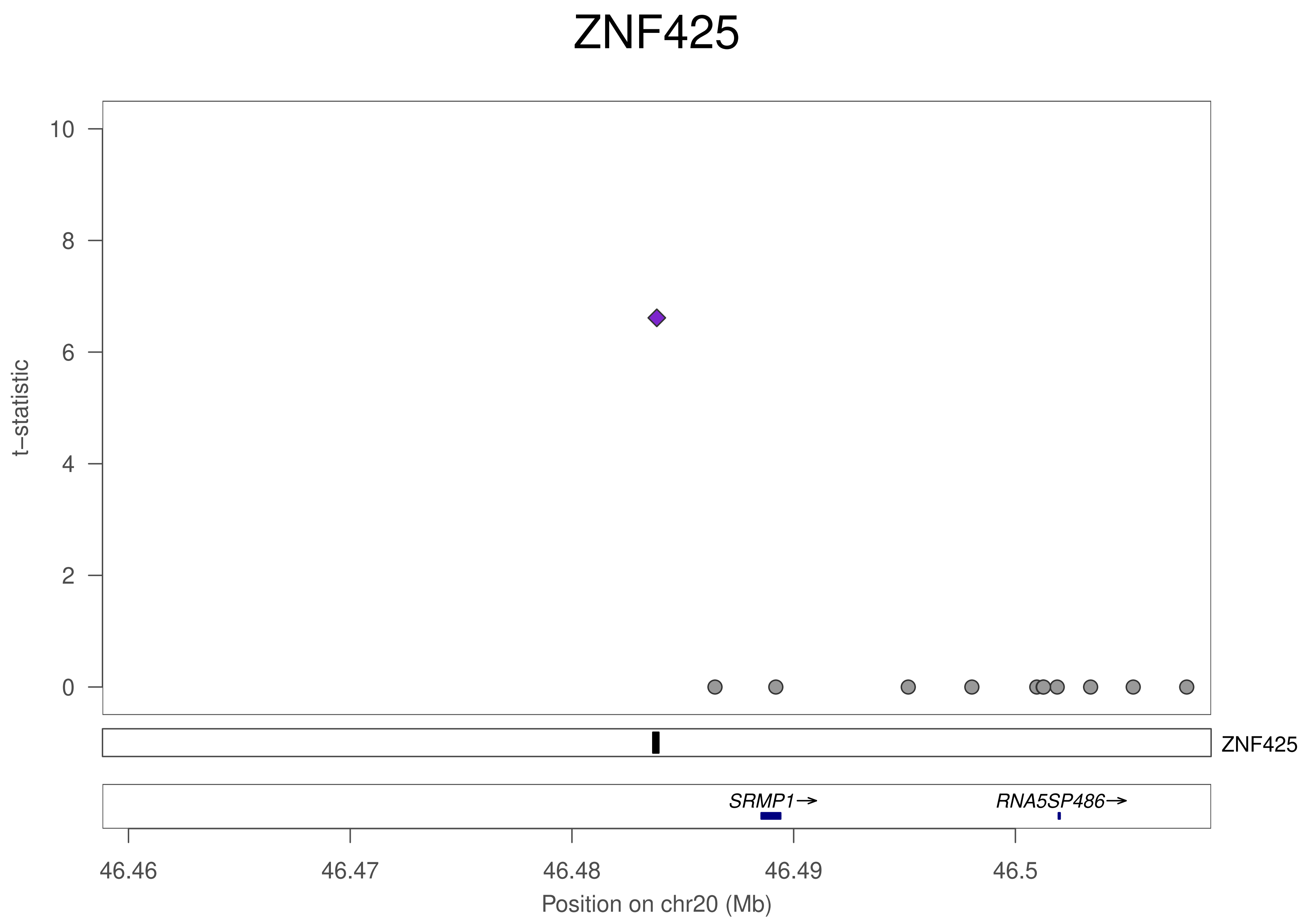

ZNF425
10
8
6
4
2
0
t−statistic
ZNF425
SRMP1→
RNA5SP486→
46.46
46.47
46.48
46.49
46.5
Position on chr20 (Mb)

### ZNF736

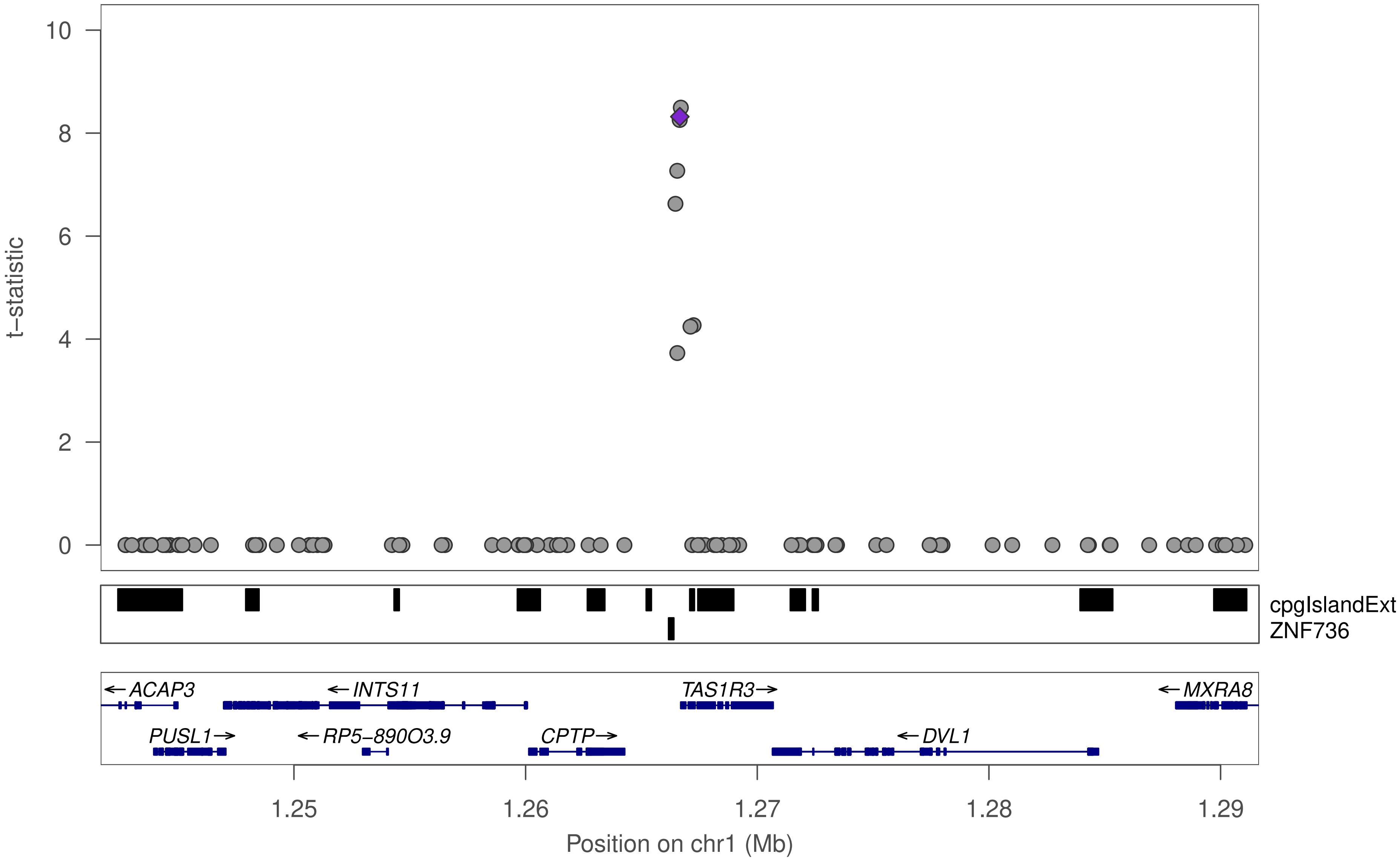

t−statistic
10
8
6
4
2
0
cpgIslandExt
ZNF736
←ACAP3
←INTS11
TAS1R3→
←MXRA8
PUSL1→
←RP5−890O3.9
CPTP→
←DVL1
1.25
1.26
1.27
1.28
1.29
Position on chr1 (Mb)

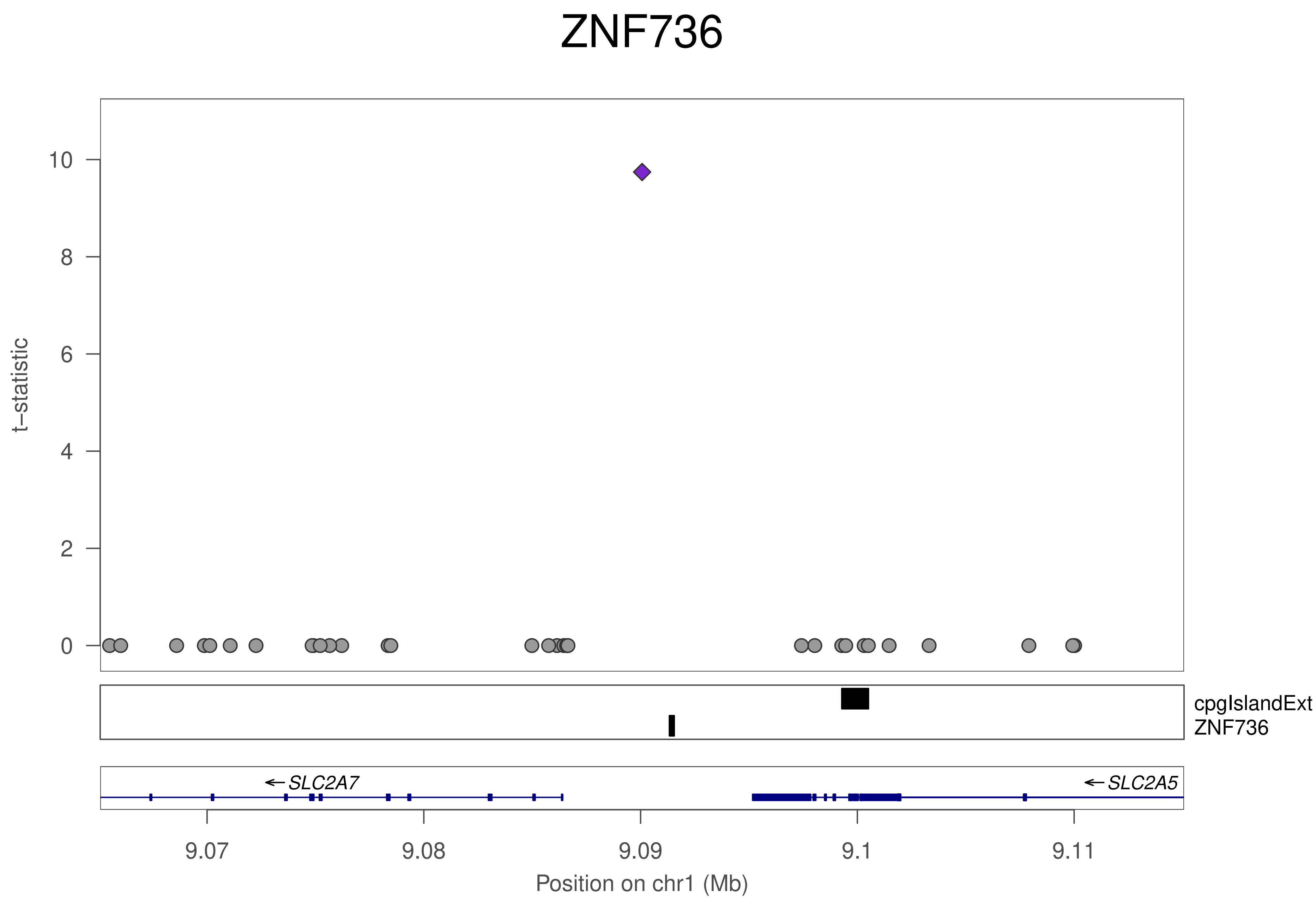

ZNF736
t−statistic
0
2
4
6
8
10
cpgIslandExt
ZNF736
←SLC2A7
←SLC2A5
9.07
9.08
9.09
9.1
9.11
Position on chr1 (Mb)

### ZNF736

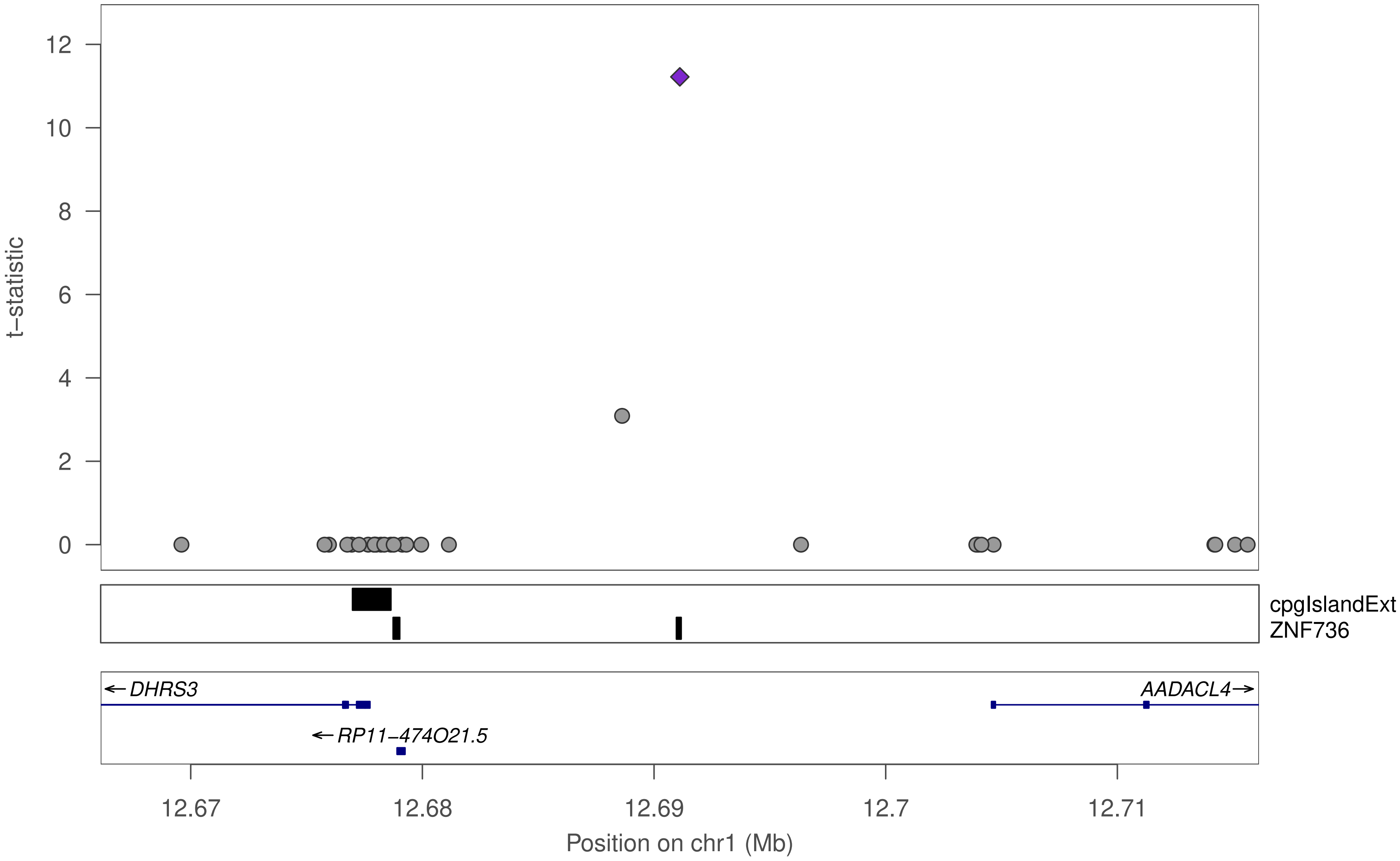

t−statistic
0
2
4
6
8
10
12
cpgIslandExt
ZNF736
← DHRS3
AADACL4→
← RP11−474O21.5
12.67
12.68
12.69
12.7
12.71
Position on chr1 (Mb)

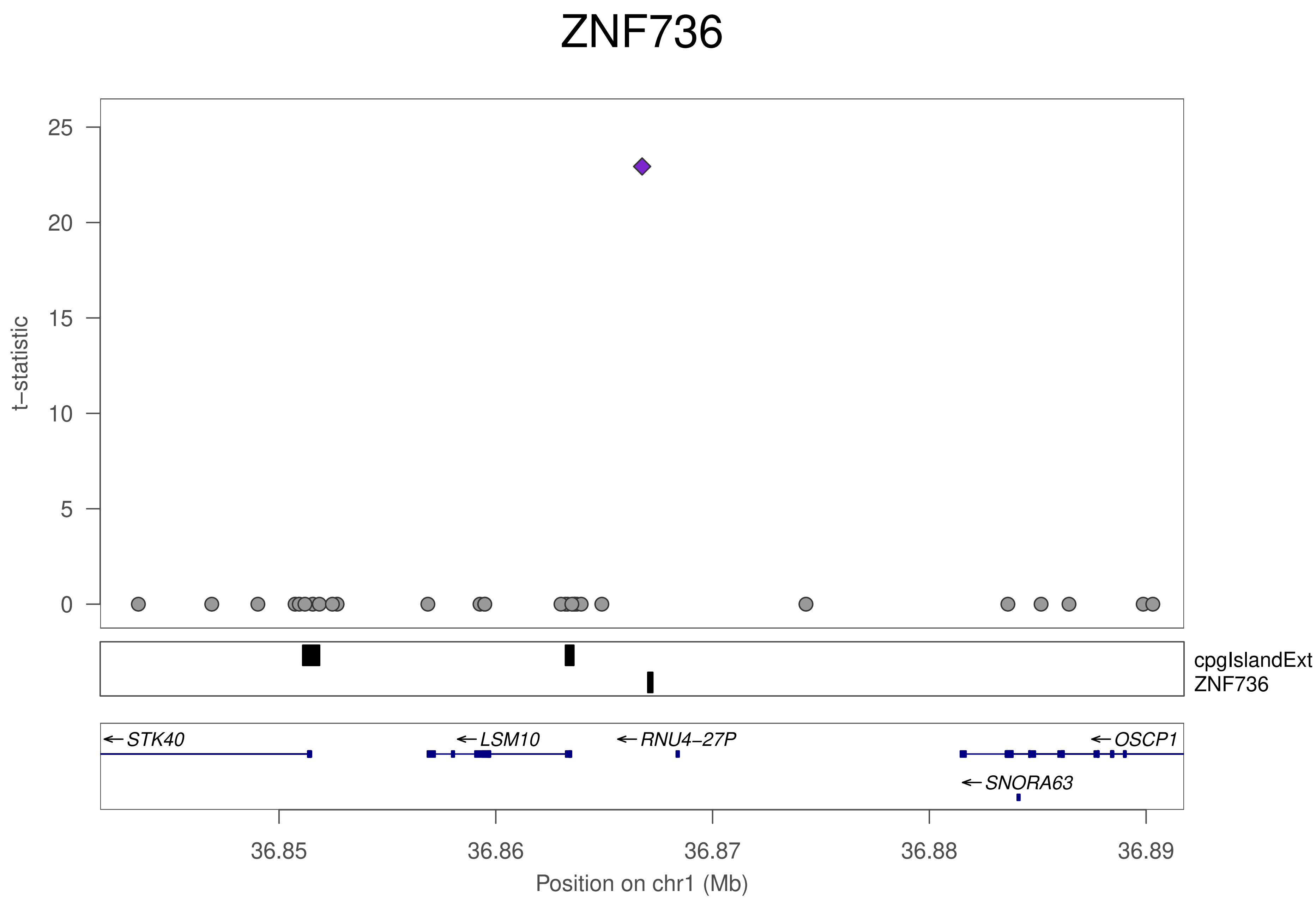
ZNF736
t−statistic
25
20
15
10
5
0
cpgIslandExt
ZNF736
STK40
LSM10
RNU4−27P
OSCP1
SNORA63
36.85
36.86
36.87
36.88
36.89
Position on chr1 (Mb)

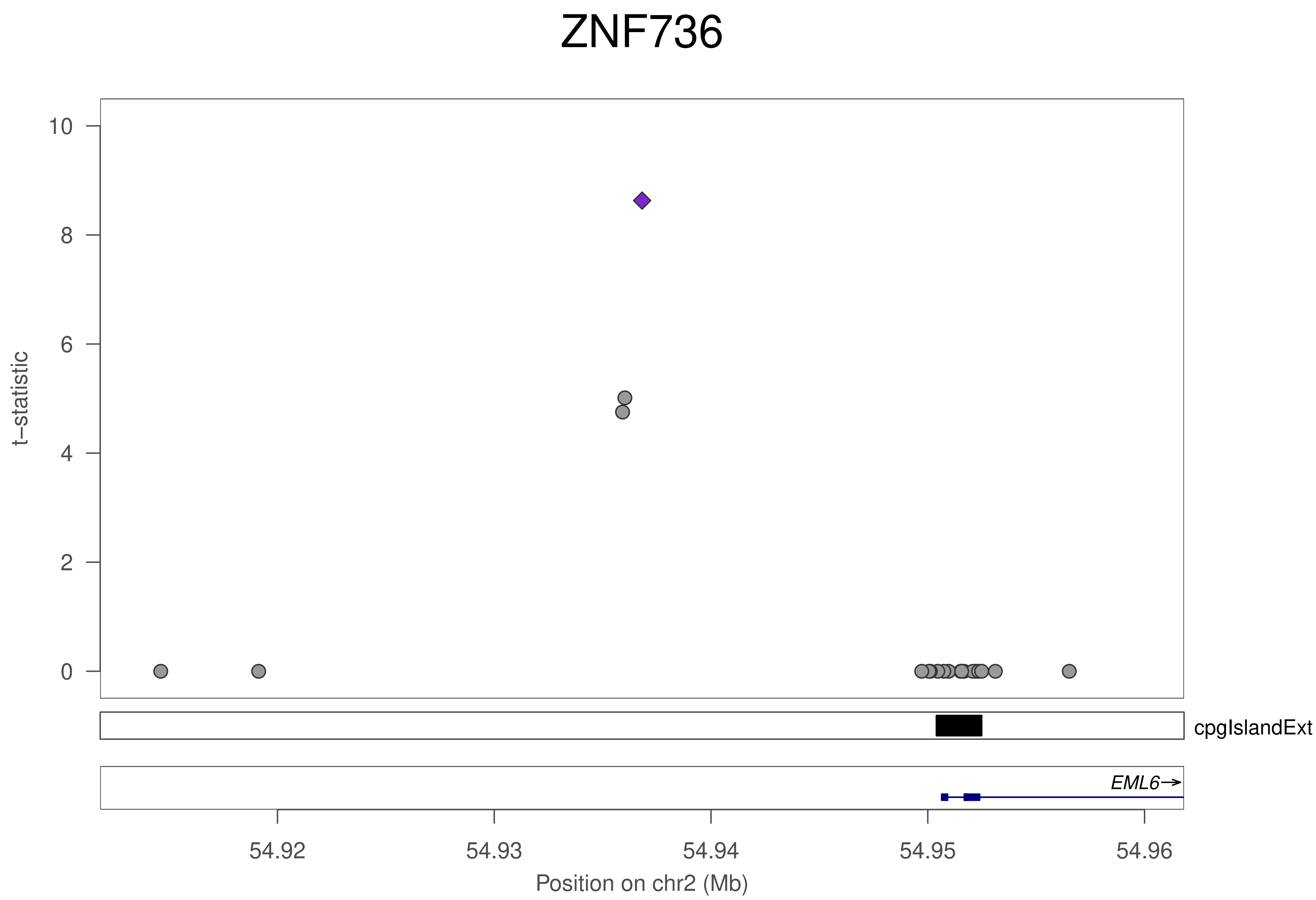
ZNF736
t–statistic
10
8
6
4
2
0
cpgIslandExt
EML6→
54.92
54.93
54.94
54.95
54.96
Position on chr2 (Mb)

### ZNF736

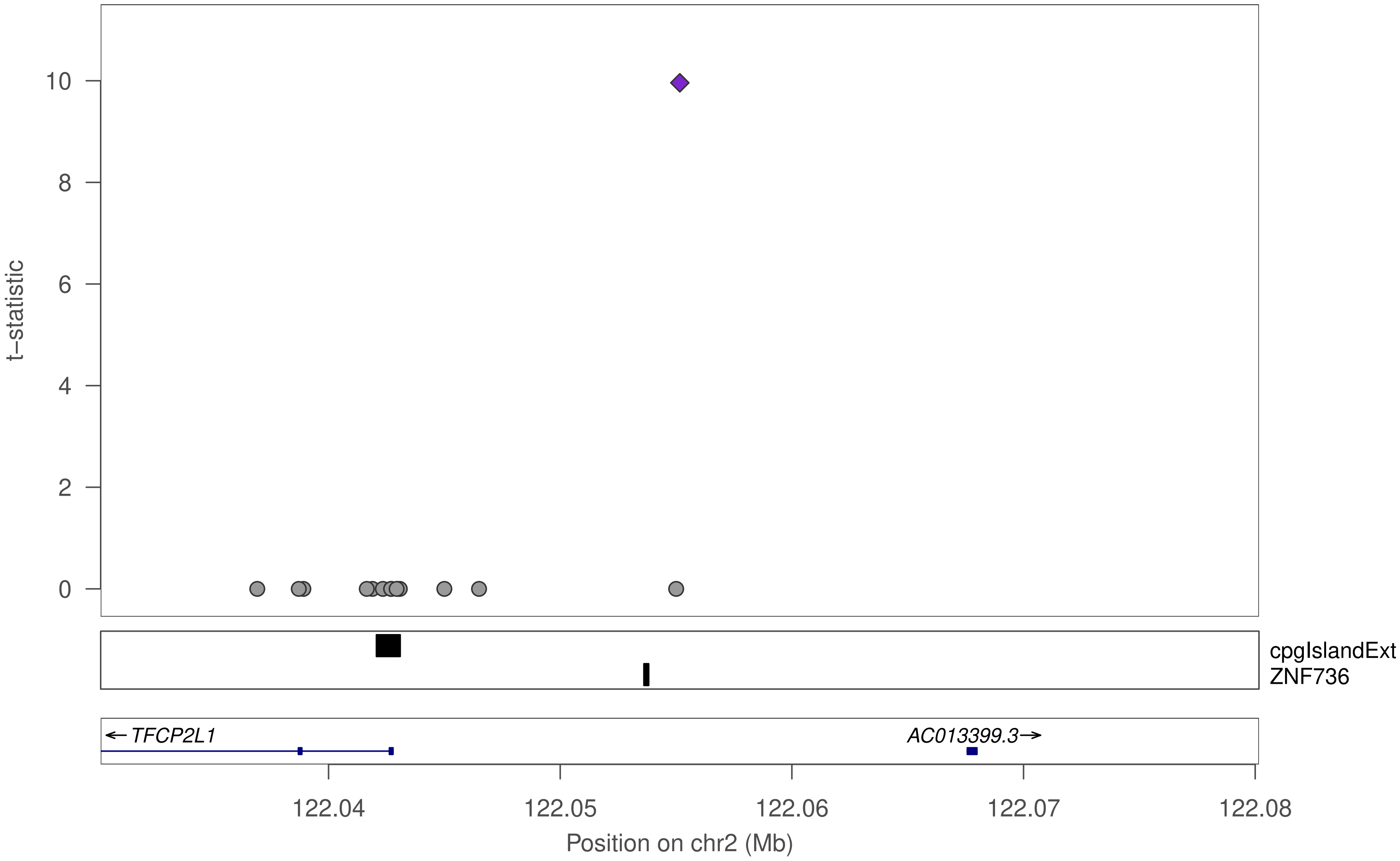

10
8
6
4
2
0
t−statistic
cpgIslandExt
ZNF736
← TFCP2L1
AC013399.3→
122.04
122.05
122.06
122.07
122.08
Position on chr2 (Mb)

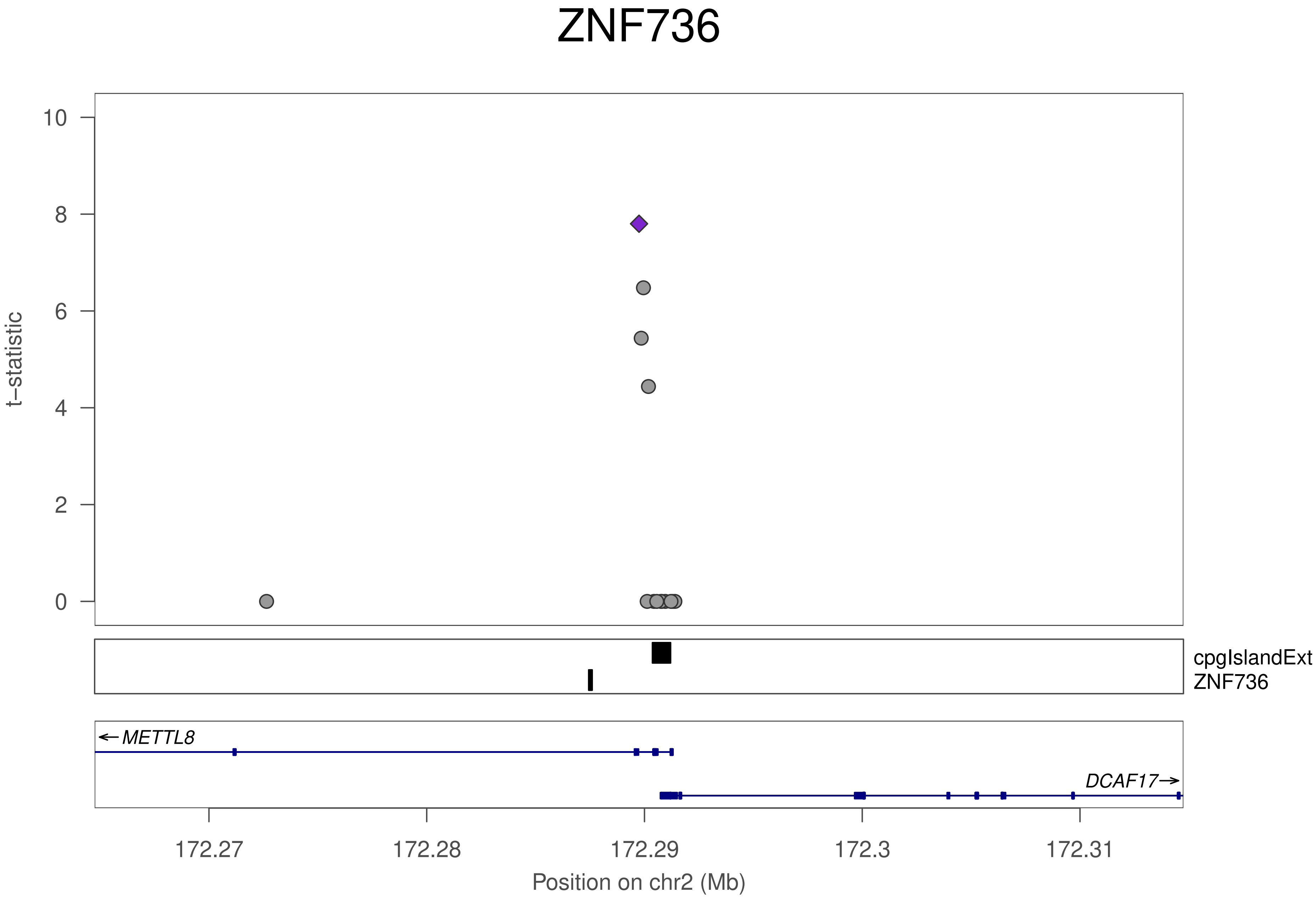
ZNF736
t−statistic
10
8
6
4
2
0
cpgIslandExt
ZNF736
← METTL8
DCAF17 →
172.27
172.28
172.29
172.3
172.31
Position on chr2 (Mb)

### ZNF736

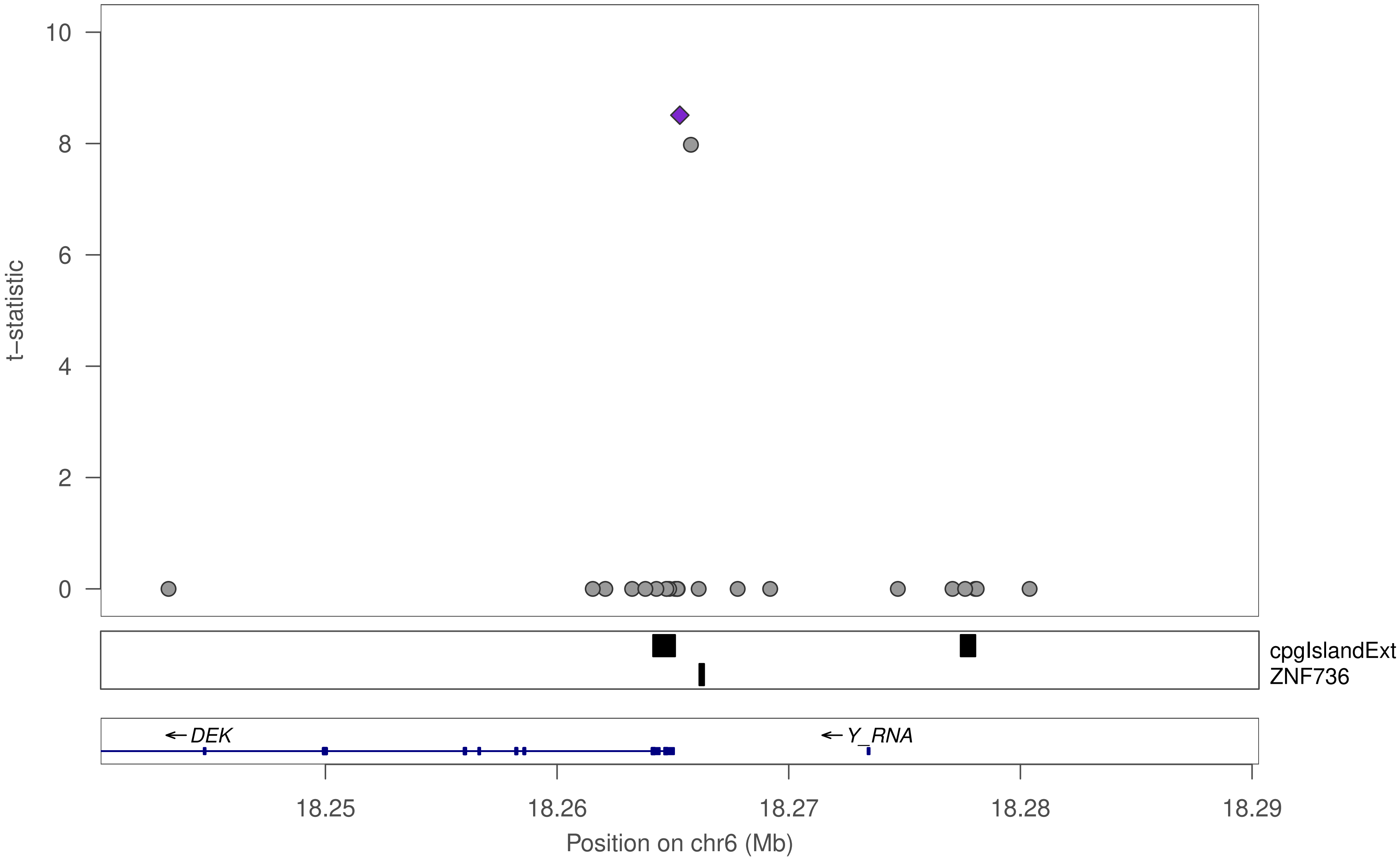

t−statistic
10
8
6
4
2
0
cpgIslandExt
ZNF736
← DEK
← Y_RNA
18.25
18.26
18.27
18.28
18.29
Position on chr6 (Mb)

### ZNF736

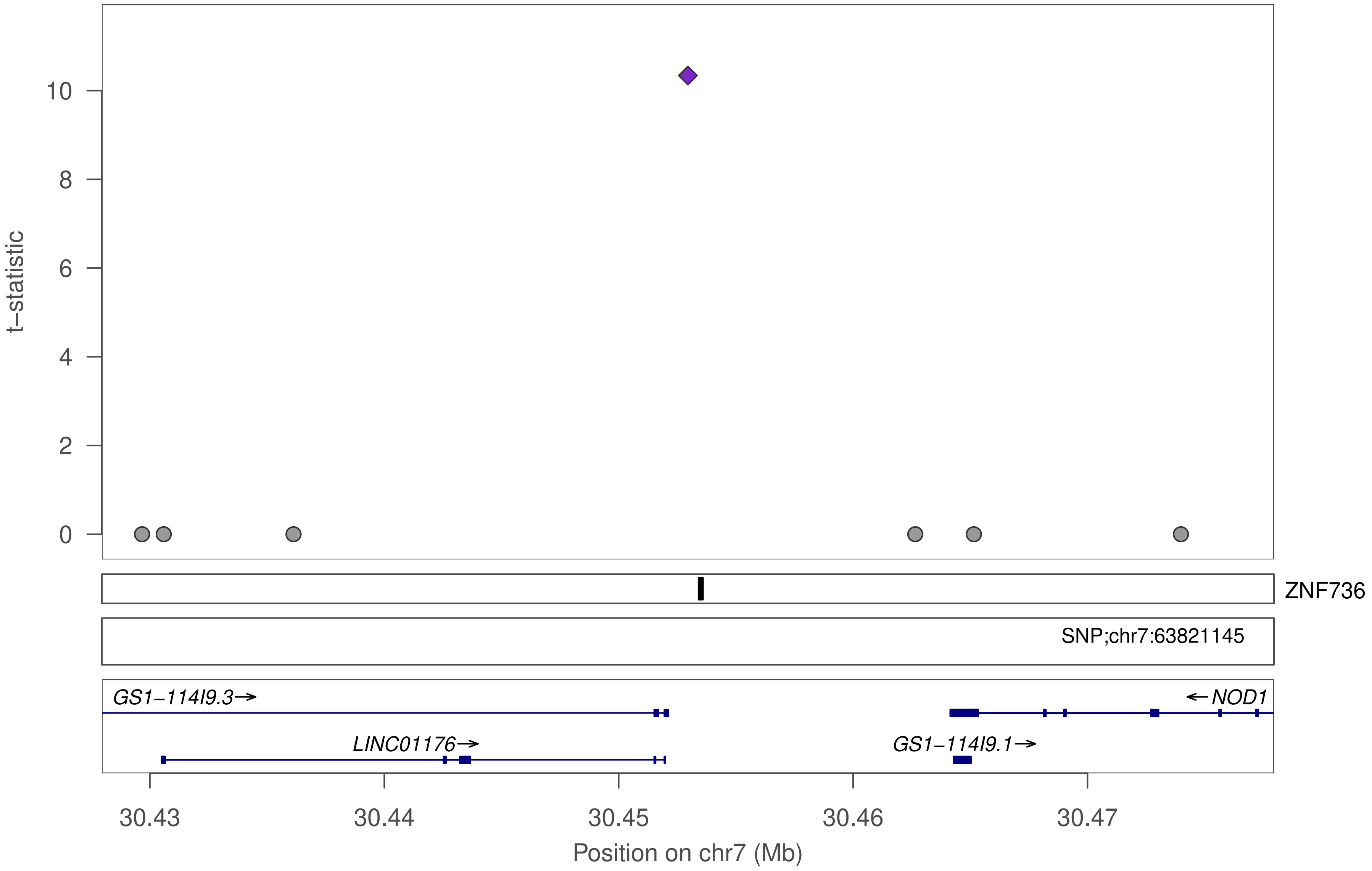

t−statistic
0
2
4
6
8
10
ZNF736
SNP;chr7:63821145
GS1−114I9.3→
←NOD1
LINC01176→
GS1−114I9.1→
30.43
30.44
30.45
30.46
30.47
Position on chr7 (Mb)

### ZNF736

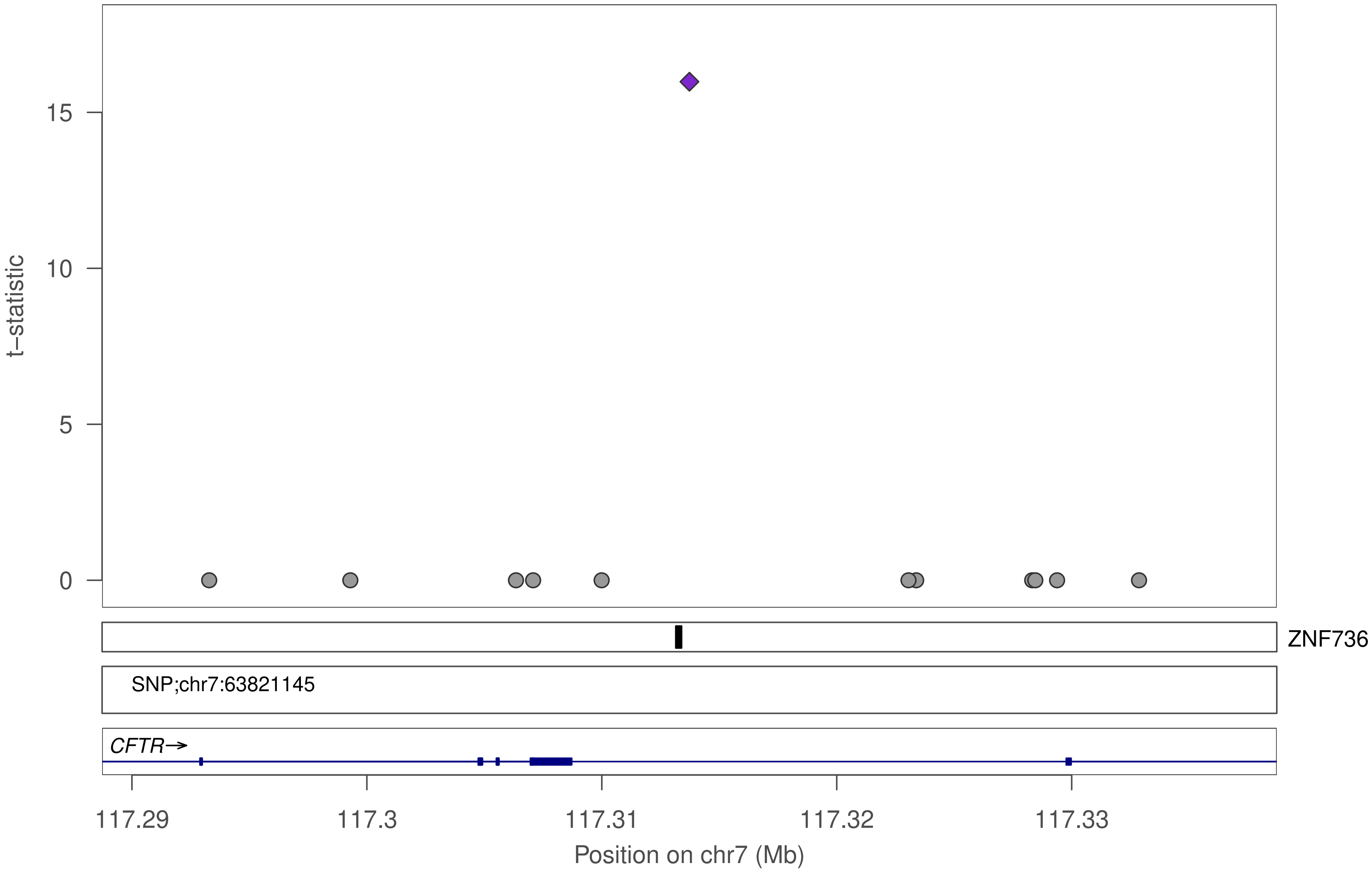

15
10
5
0
t−statistic
ZNF736
SNP;chr7:63821145
CFTR→
117.29
117.3
117.31
117.32
117.33
Position on chr7 (Mb)

### ZNF736

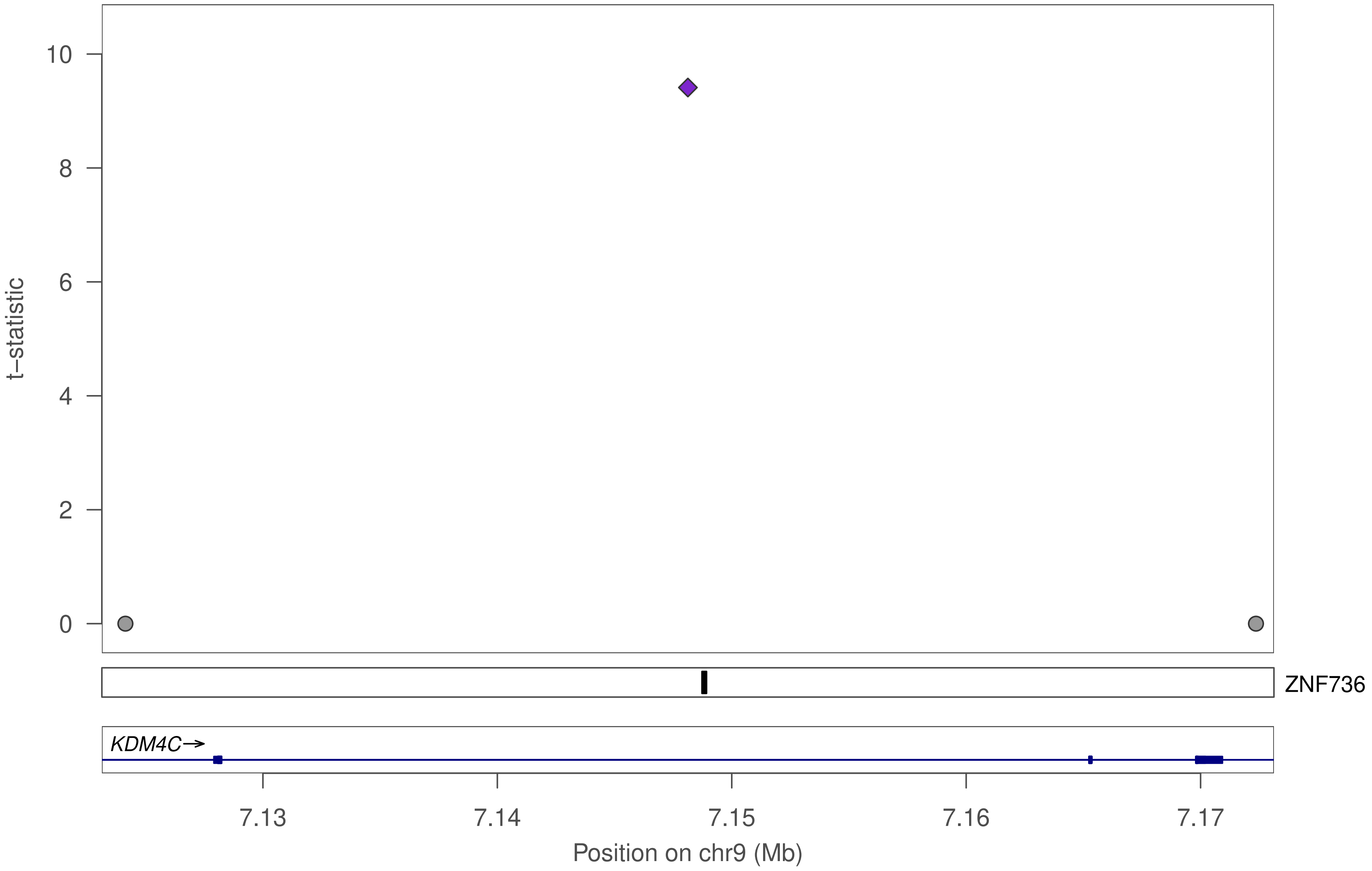

10
8
6
4
2
0
t−statistic
ZNF736
KDM4C→
7.13
7.14
7.15
7.16
7.17
Position on chr9 (Mb)

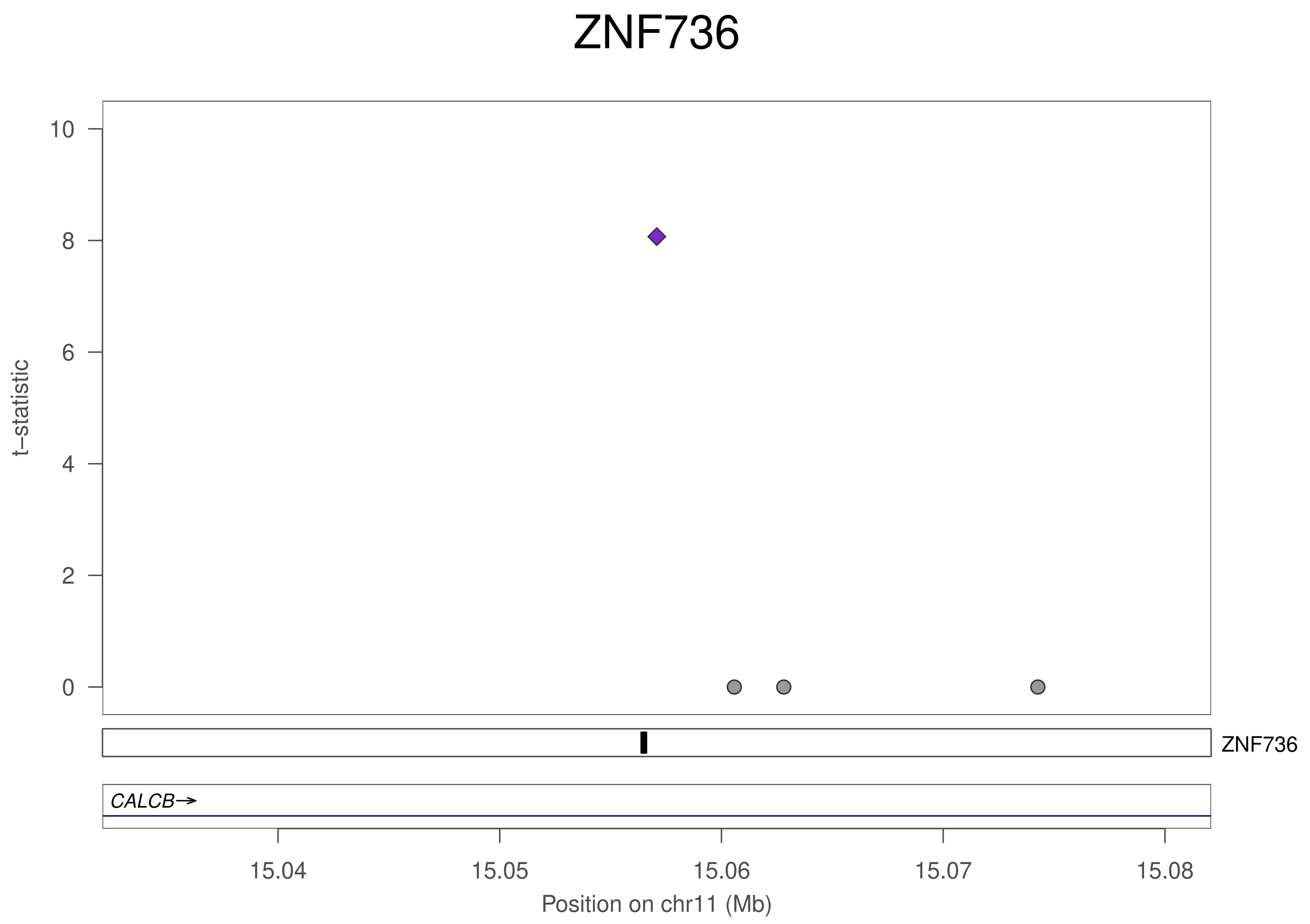
ZNF736
10
8
6
4
2
0
t−statistic
ZNF736
CALCB→
15.04
15.05
15.06
15.07
15.08
Position on chr11 (Mb)

### ZNF736

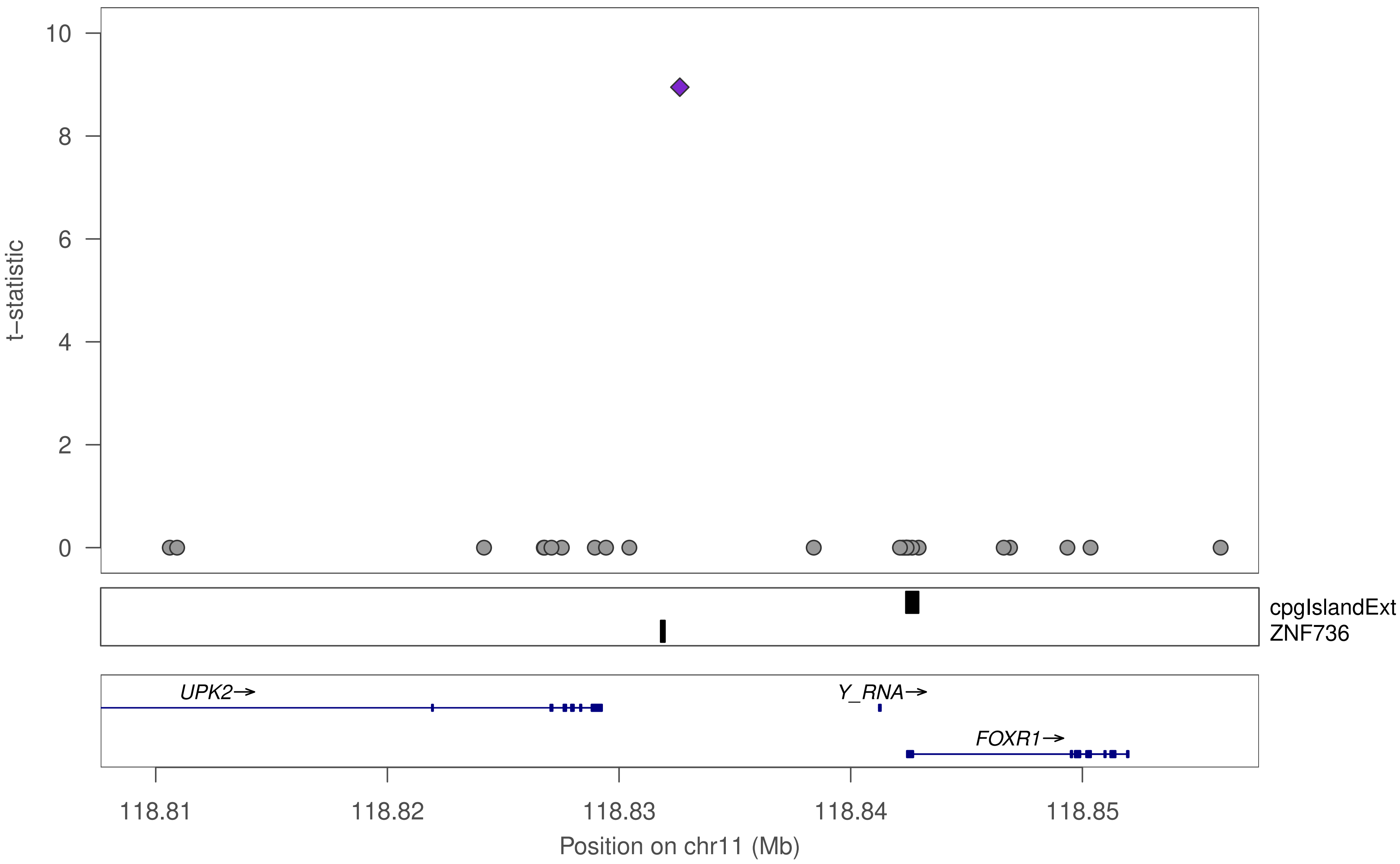

10
8
6
4
2
0
t−statistic
cpgIslandExt
ZNF736
UPK2→
Y_RNA→
FOXR1→
118.81
118.82
118.83
118.84
118.85
Position on chr11 (Mb)

### ZNF736

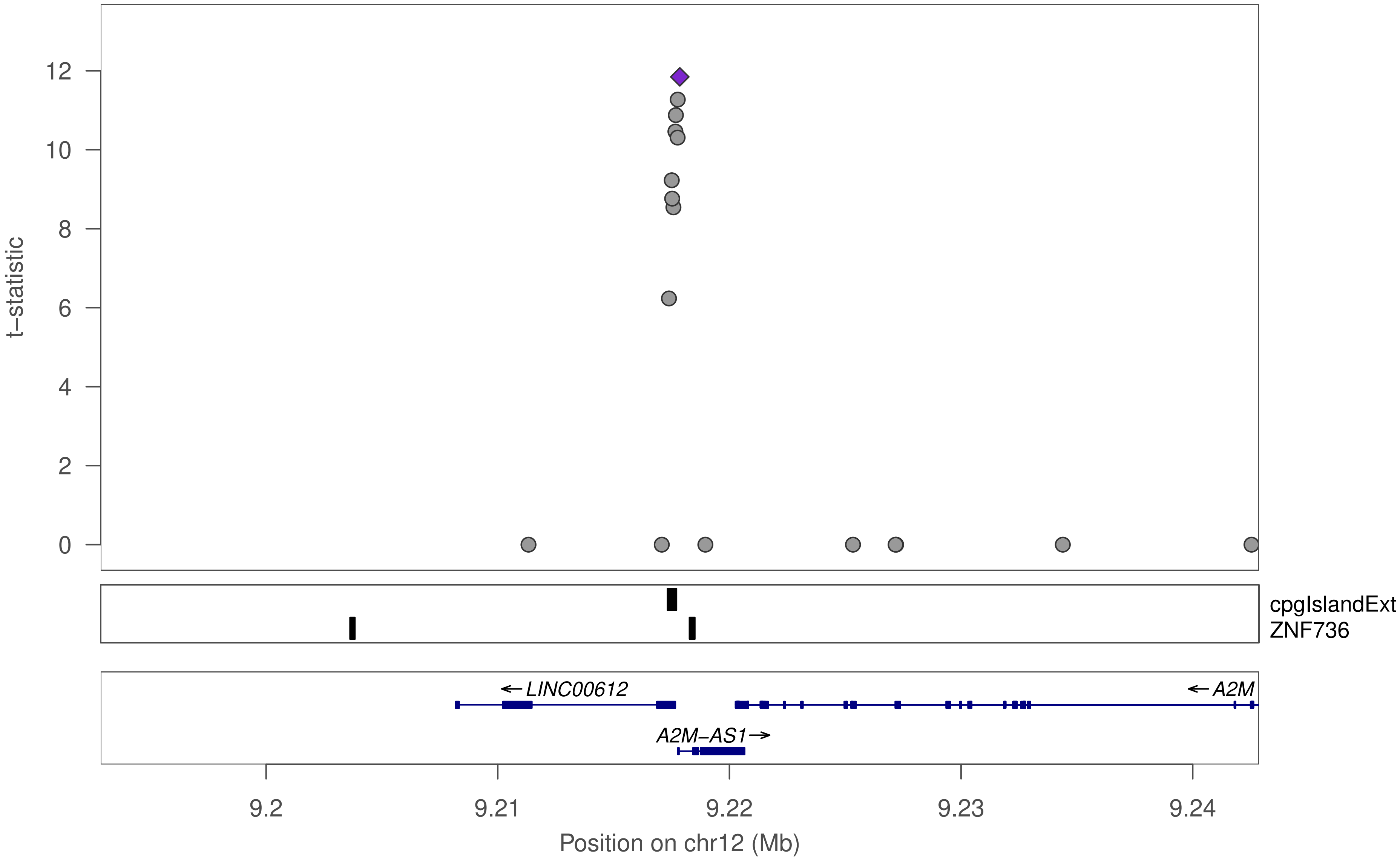

t−statistic
0
2
4
6
8
10
12
cpgIslandExt
ZNF736
←LINC00612
←A2M
A2M−AS1→
9.2
9.21
9.22
9.23
9.24
Position on chr12 (Mb)

### ZNF736

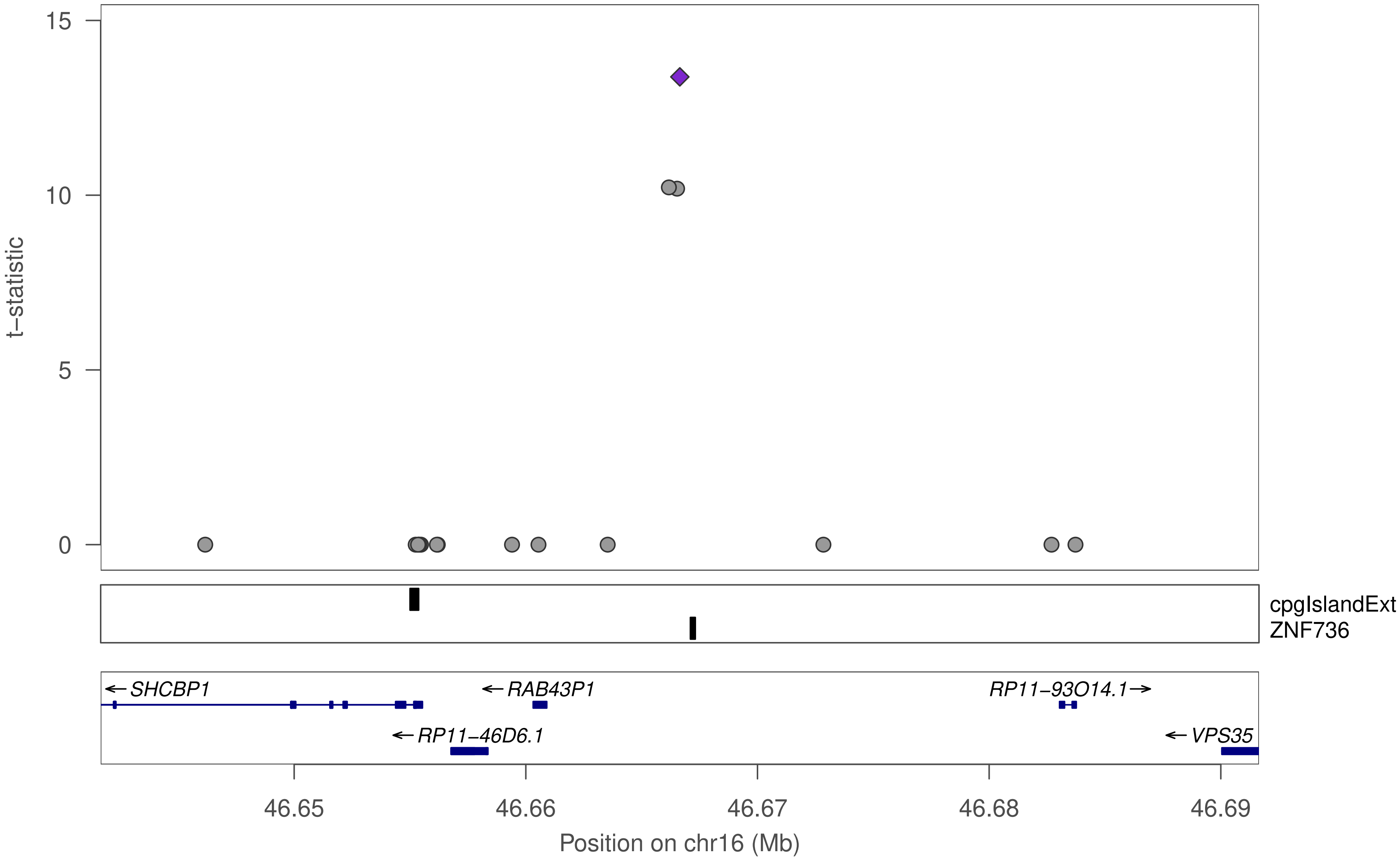

15
10
5
0
t−statistic
cpgIslandExt
ZNF736
SHCBP1
RAB43P1
RP11−93O14.1
RP11−46D6.1
VPS35
46.65
46.66
46.67
46.68
46.69
Position on chr16 (Mb)

### ZNF736

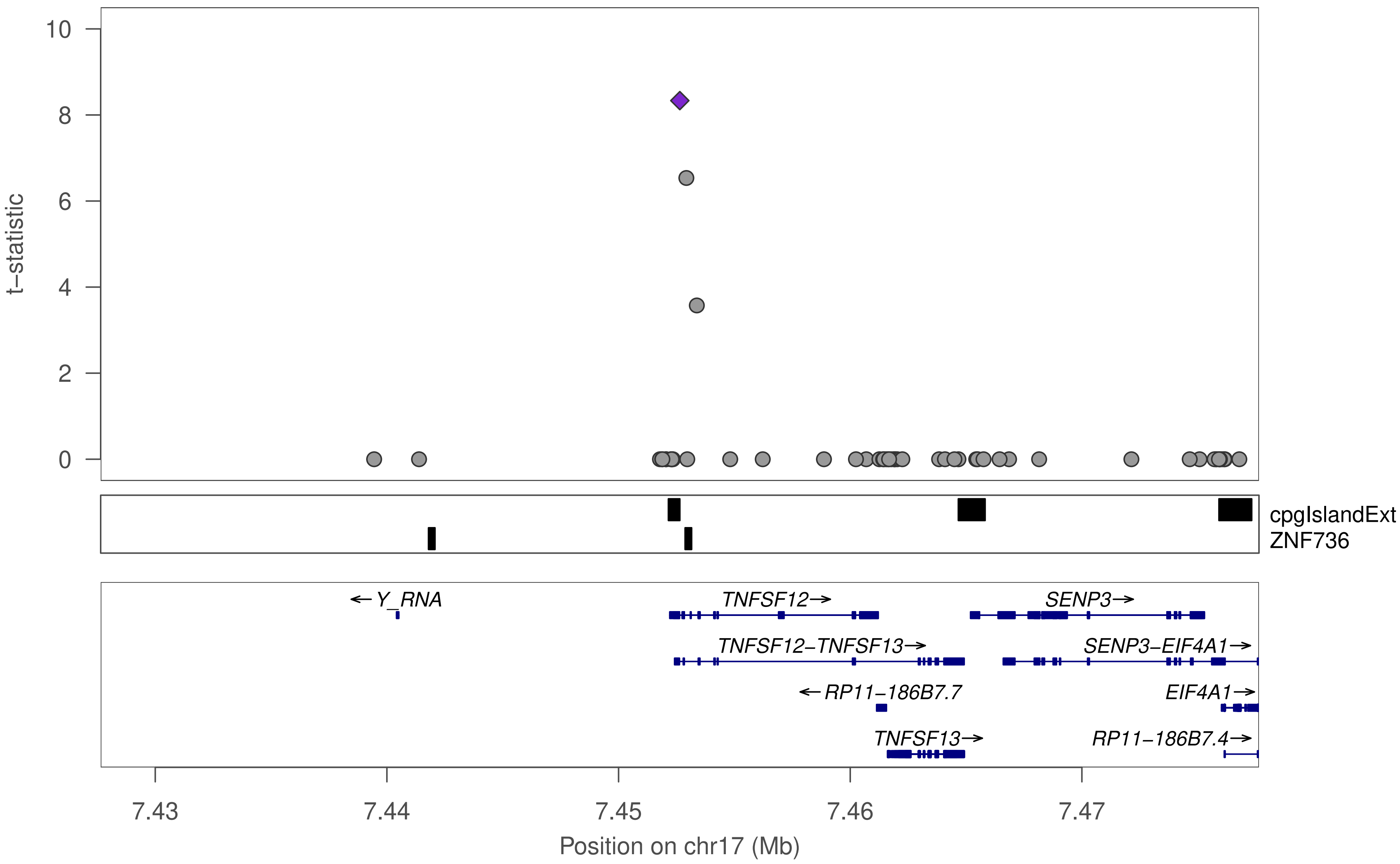
t−statistic
10
8
6
4
2
0
cpgIslandExt
ZNF736
← Y_RNA
TNFSF12→
SENP3→
TNFSF12−TNFSF13→
SENP3−EIF4A1→
← RP11−186B7.7
EIF4A1→
TNFSF13→
RP11−186B7.4→
7.43
7.44
7.45
7.46
7.47
Position on chr17 (Mb)

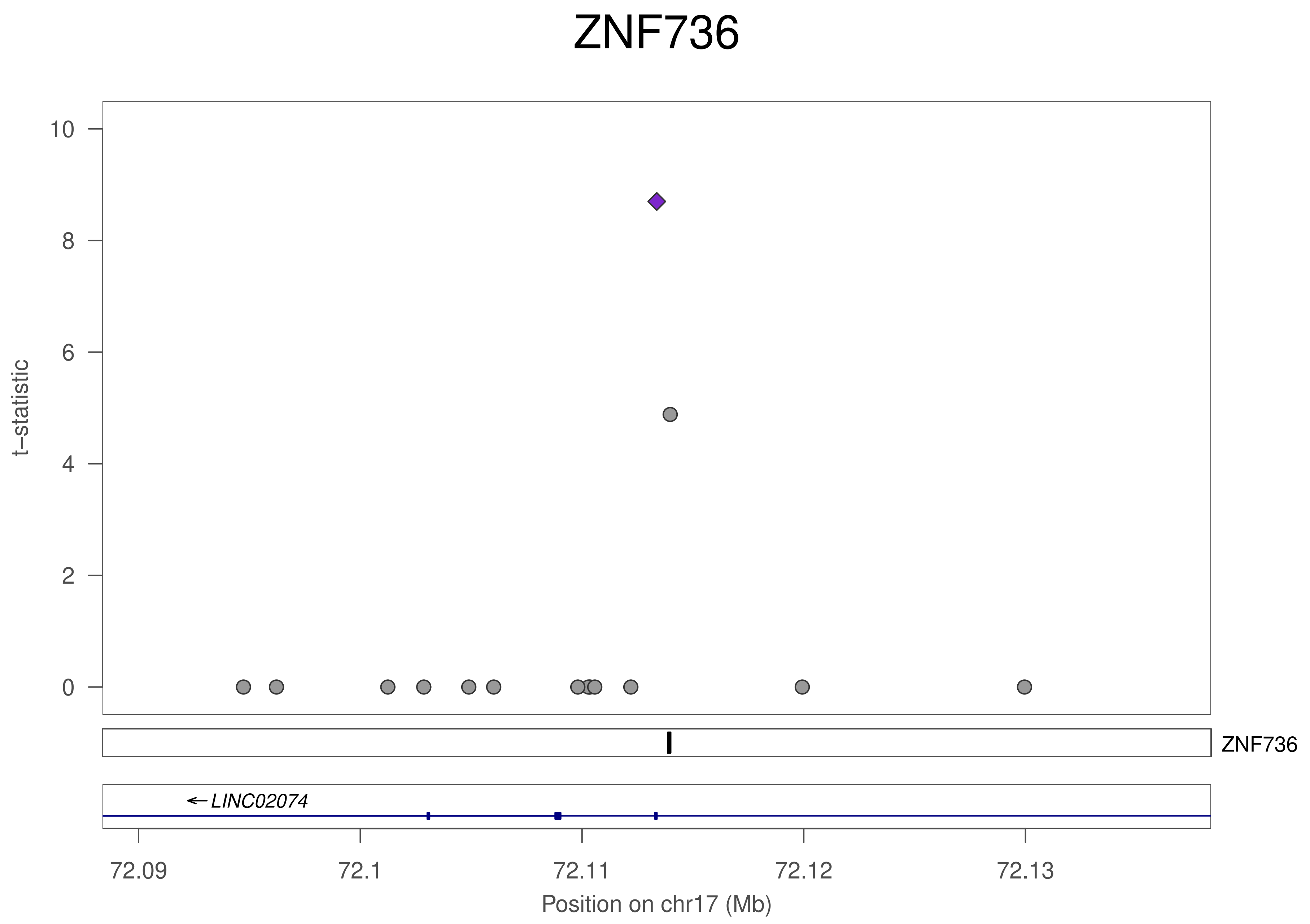
ZNF736
t−statistic
10
8
6
4
2
0
ZNF736
←LINC02074
72.09
72.1
72.11
72.12
72.13
Position on chr17 (Mb)

### ZNF736

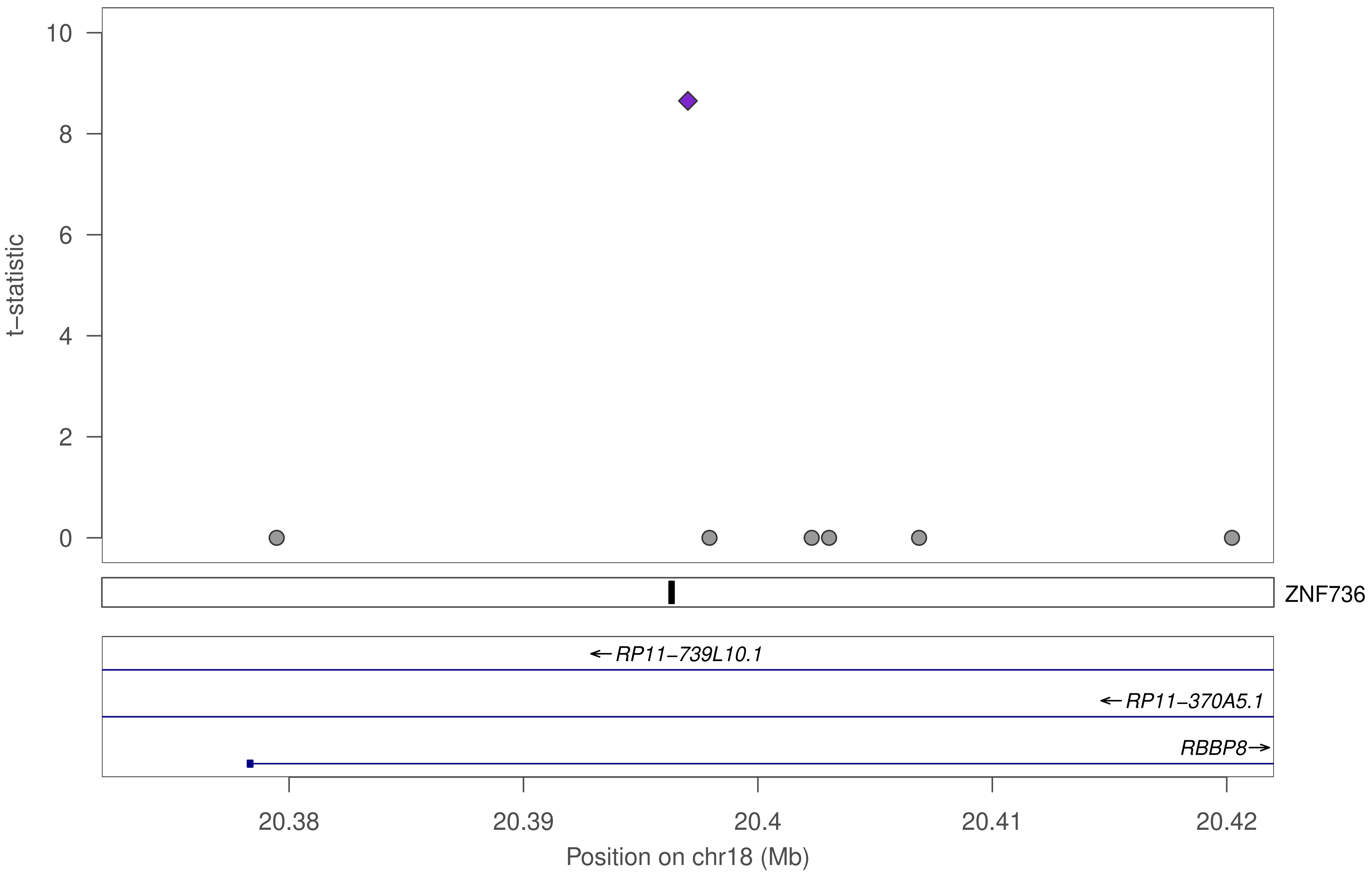

10
8
6
4
2
0
t−statistic
ZNF736
←RP11−739L10.1
←RP11−370A5.1
RBBP8→
20.38
20.39
20.4
20.41
20.42
Position on chr18 (Mb)

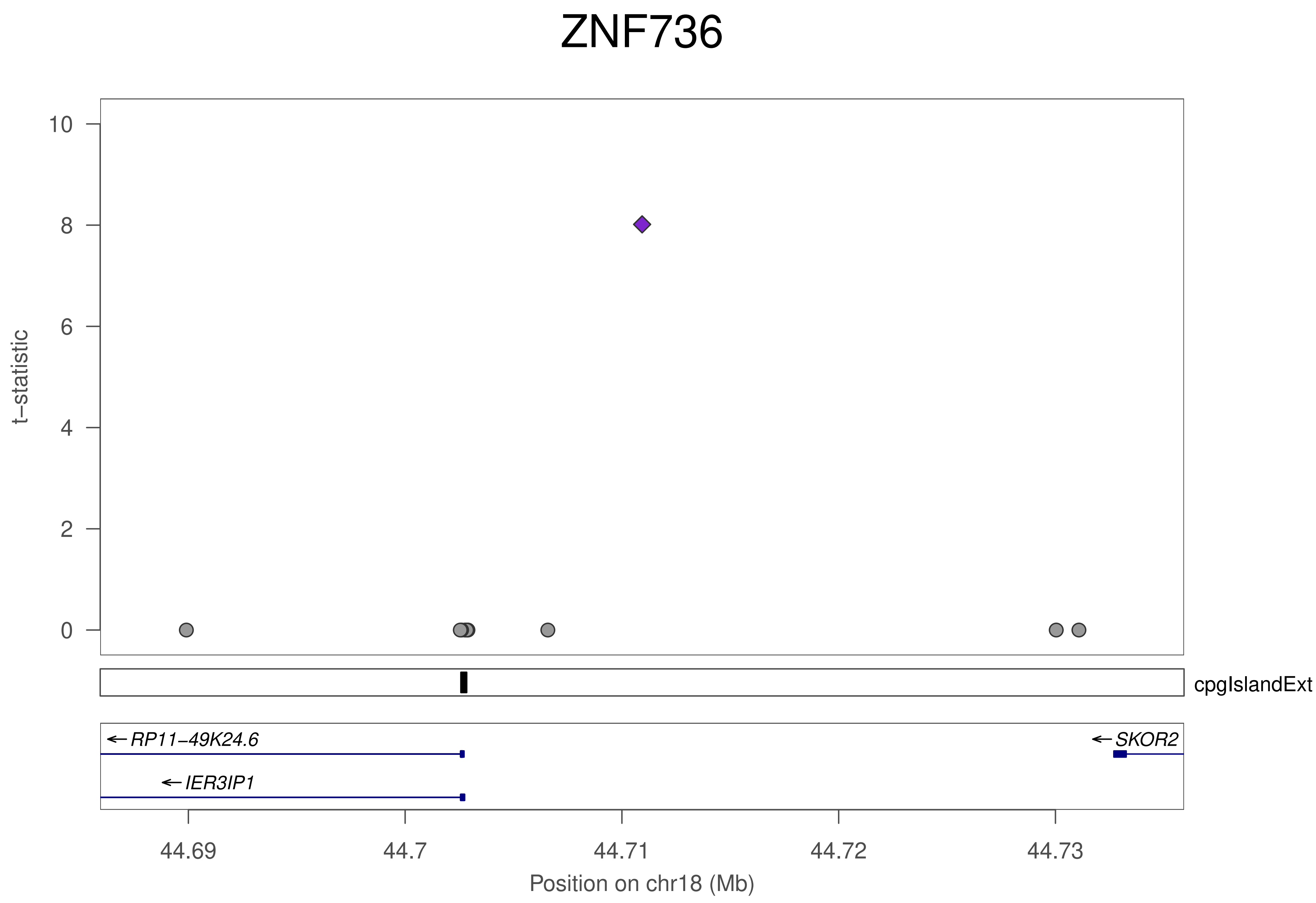
ZNF736
t−statistic
10
8
6
4
2
0
cpgIslandExt
RP11−49K24.6
IER3IP1
SKOR2
44.69
44.7
44.71
44.72
44.73
Position on chr18 (Mb)

### ZNF736

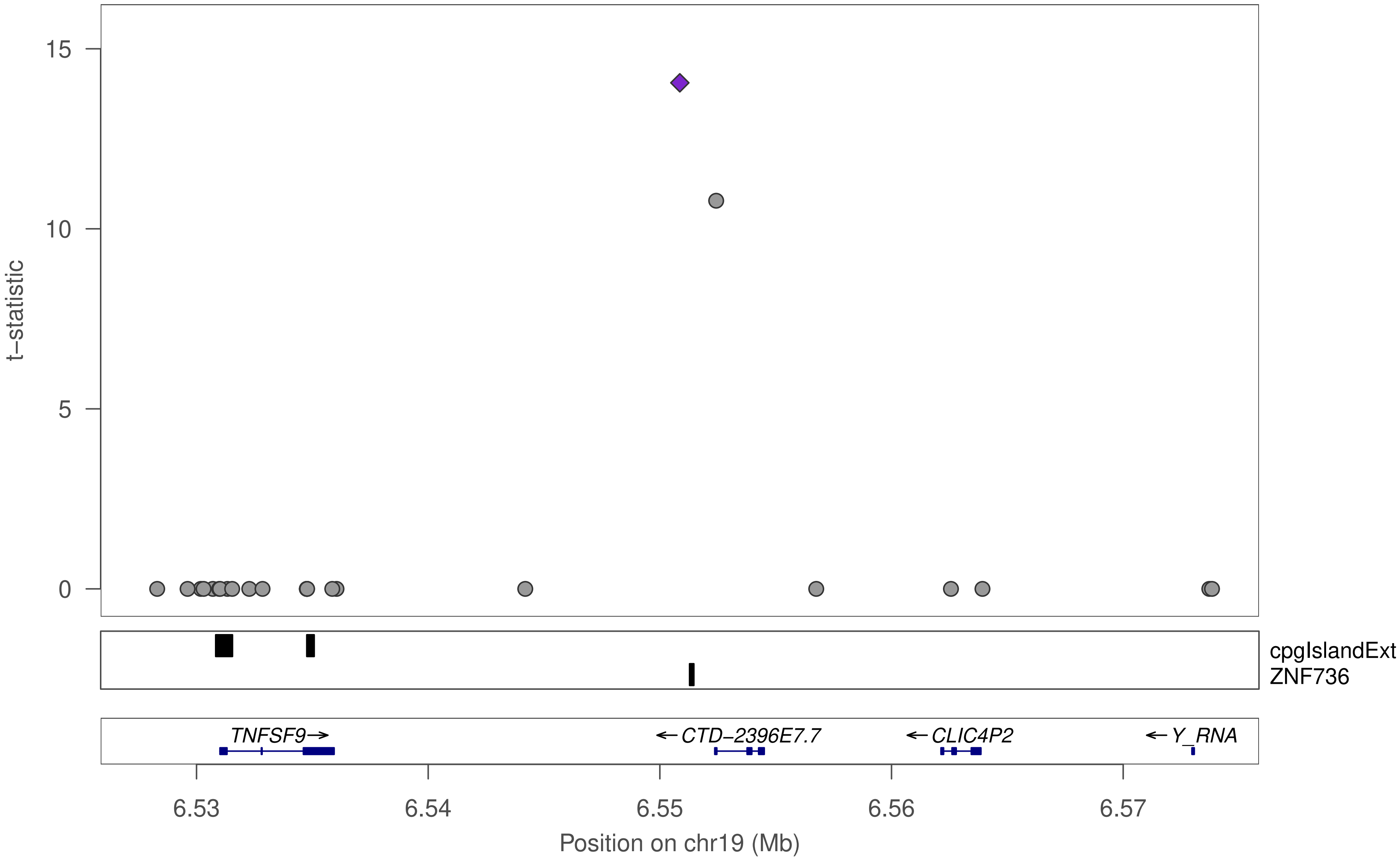

15
10
5
0
t−statistic
cpgIslandExt
ZNF736
TNFSF9→
←CTD−2396E7.7
←CLIC4P2
←Y_RNA
6.53
6.54
6.55
6.56
6.57
Position on chr19 (Mb)

### ZNF736

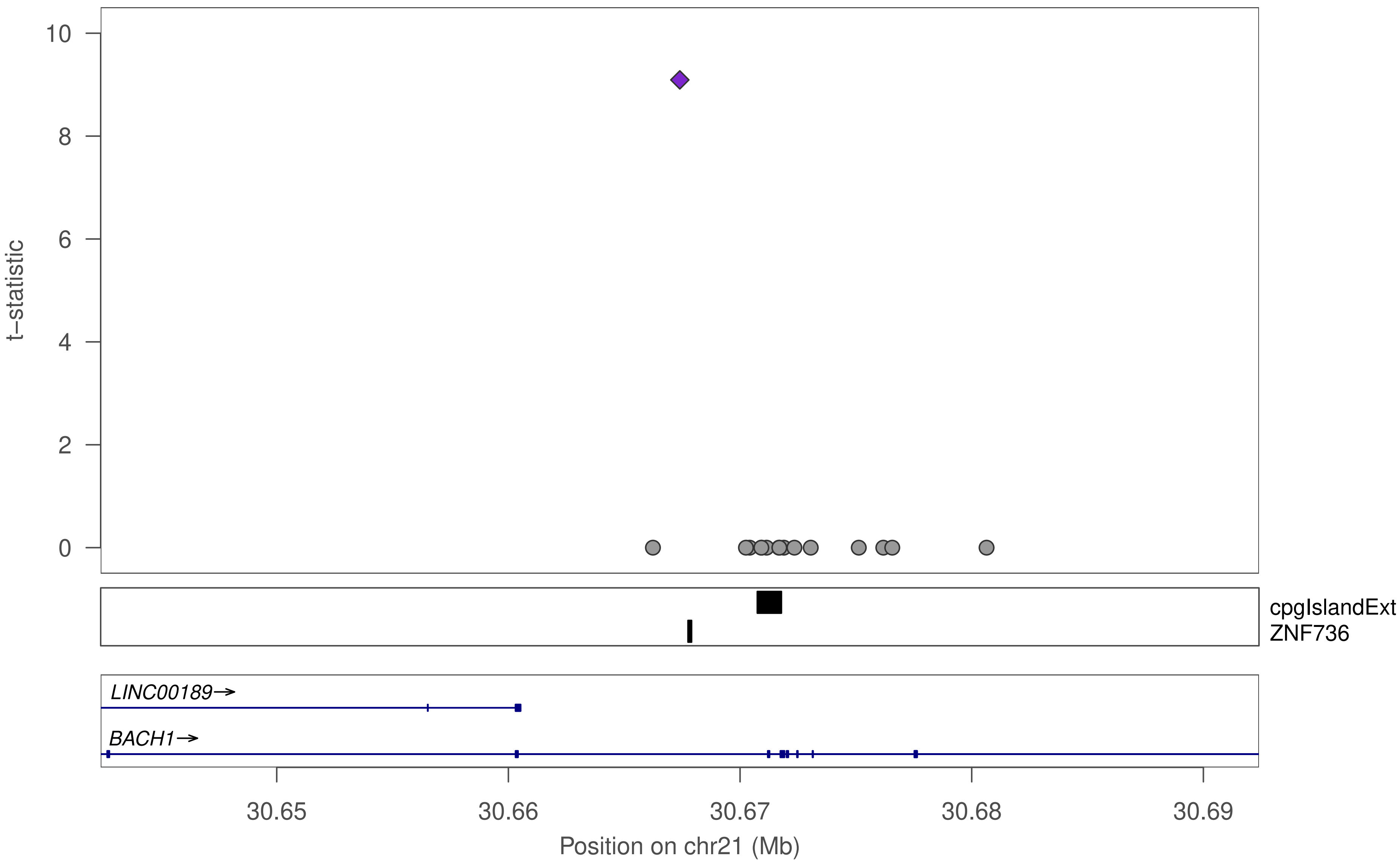

10
8
6
4
2
0
t−statistic
cpgIslandExt
ZNF736
LINC00189→
BACH1→
30.65
30.66
30.67
30.68
30.69
Position on chr21 (Mb)

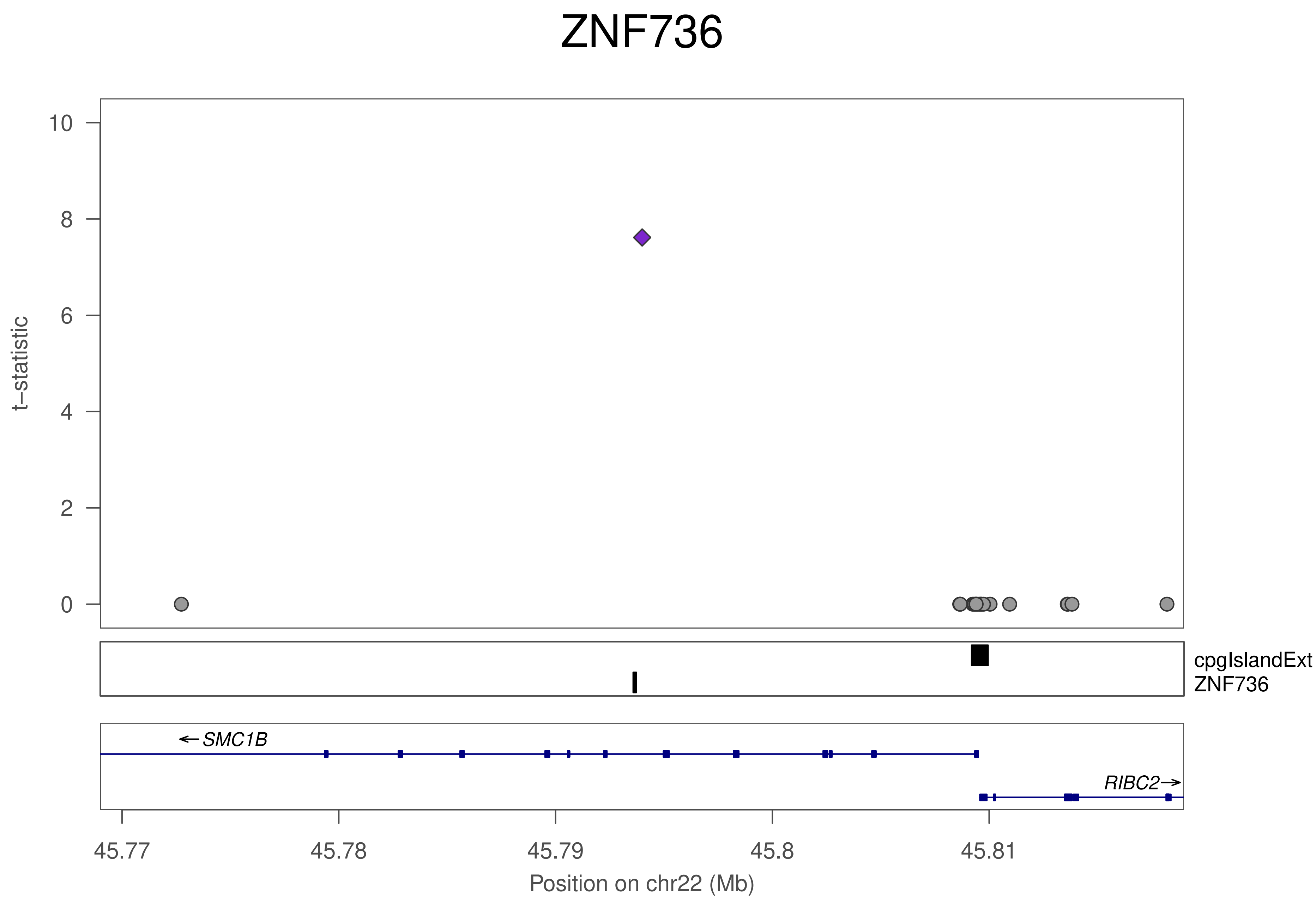
ZNF736
t−statistic
10
8
6
4
2
0
cpgIslandExt
ZNF736
SMC1B
RIBC2
45.77
45.78
45.79
45.8
45.81
Position on chr22 (Mb)

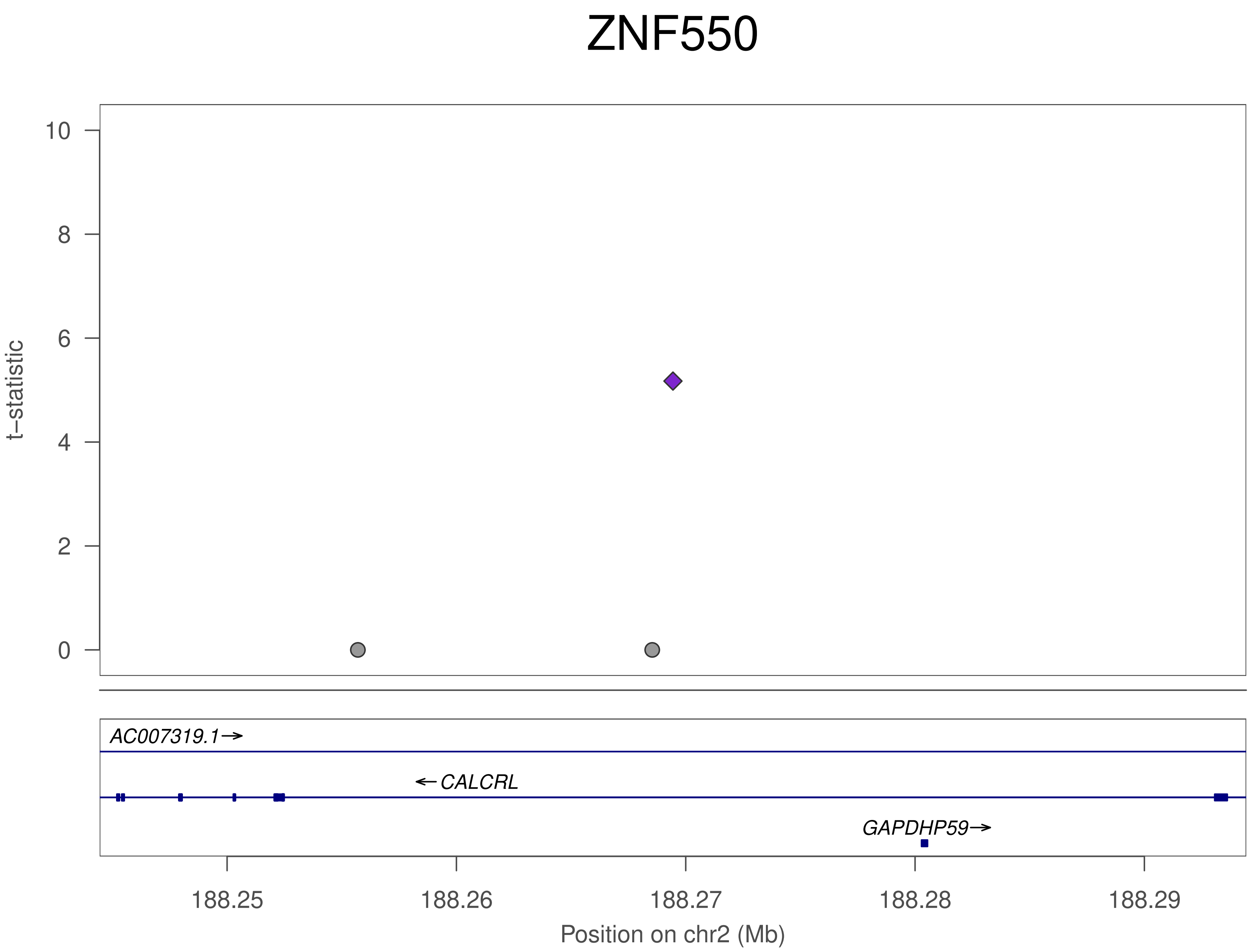
ZNF550
t−statistic
10
8
6
4
2
0
AC007319.1→
←CALCRL
GAPDHP59→
188.25
188.26
188.27
188.28
188.29
Position on chr2 (Mb)

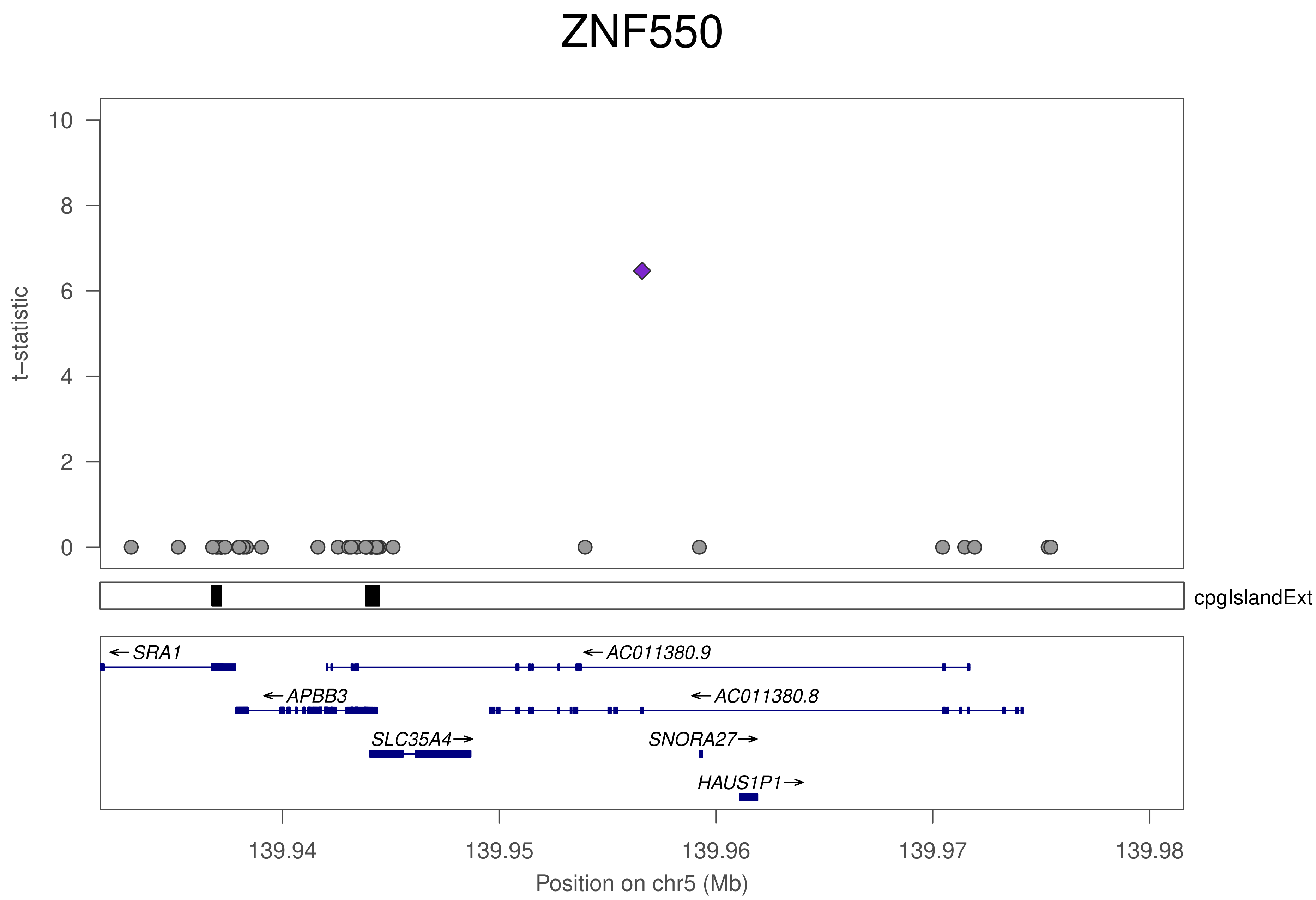
ZNF550
t–statistic
10
8
6
4
2
0
cpgIslandExt
SRA1
AC011380.9
APBB3
AC011380.8
SLC35A4
SNORA27
HAUS1P1
139.94
139.95
139.96
139.97
139.98
Position on chr5 (Mb)

### ZNF550

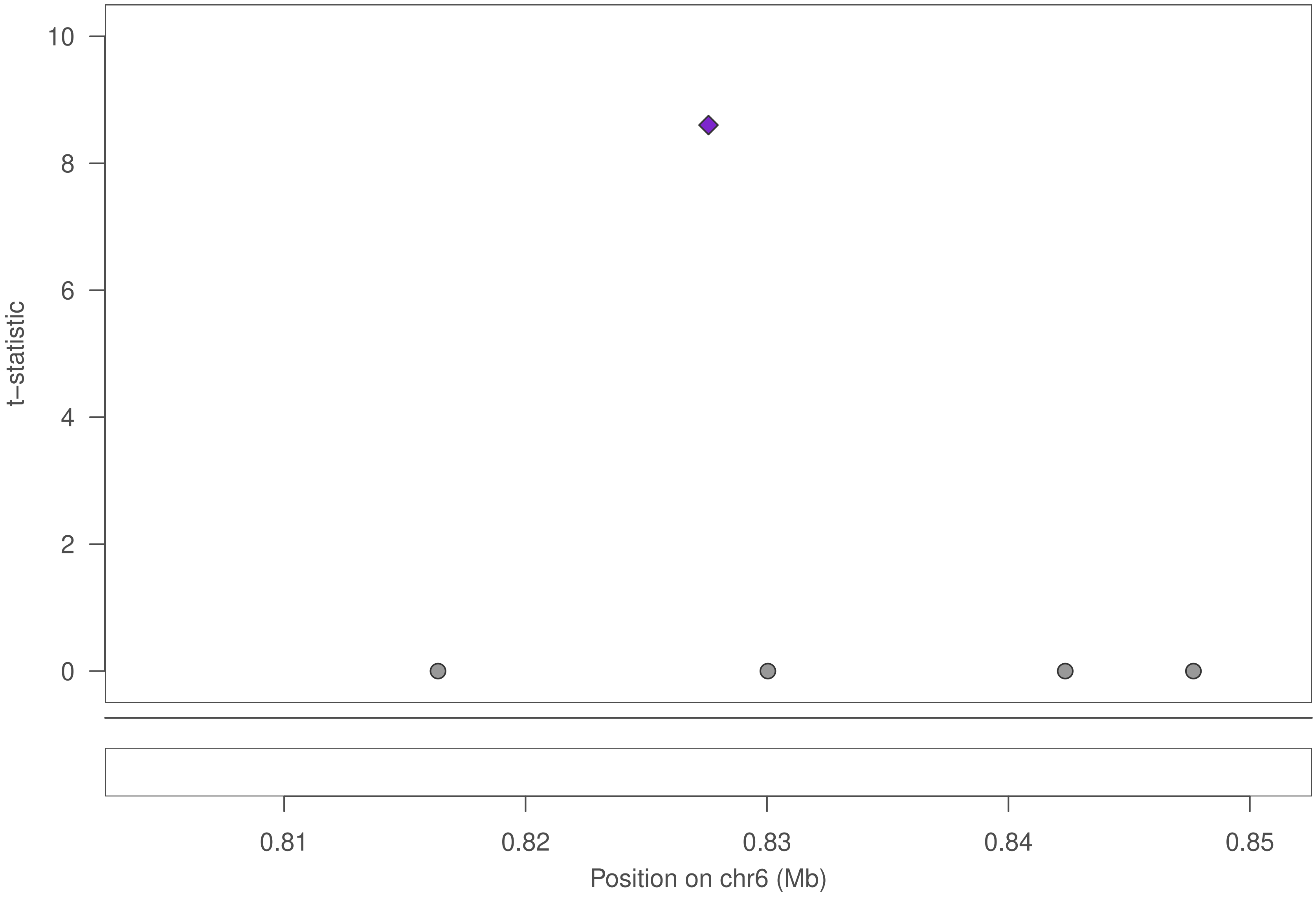

### ZNF550

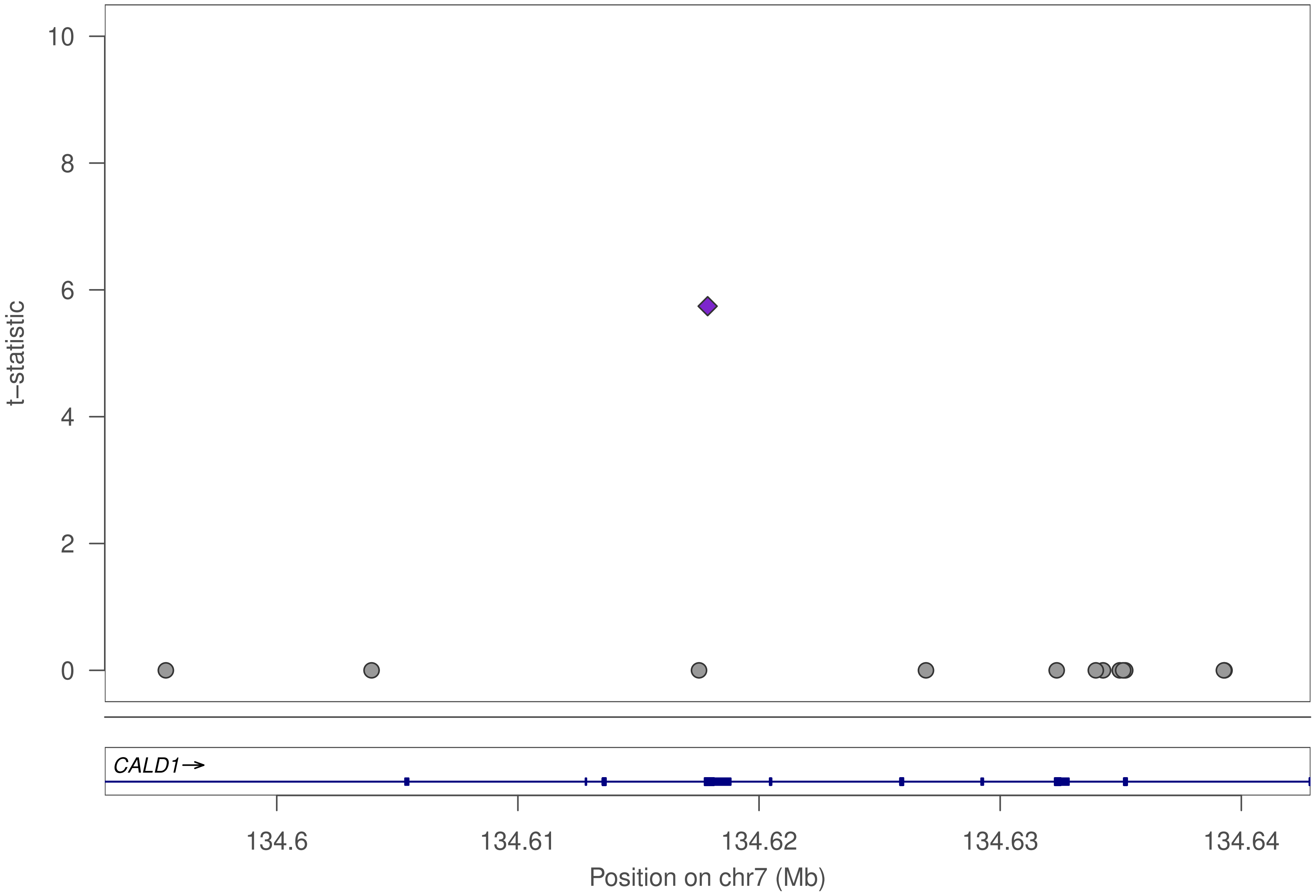

10
8
6
4
2
0
t−statistic
CALD1→
134.6
134.61
134.62
134.63
134.64
Position on chr7 (Mb)

### ZNF550

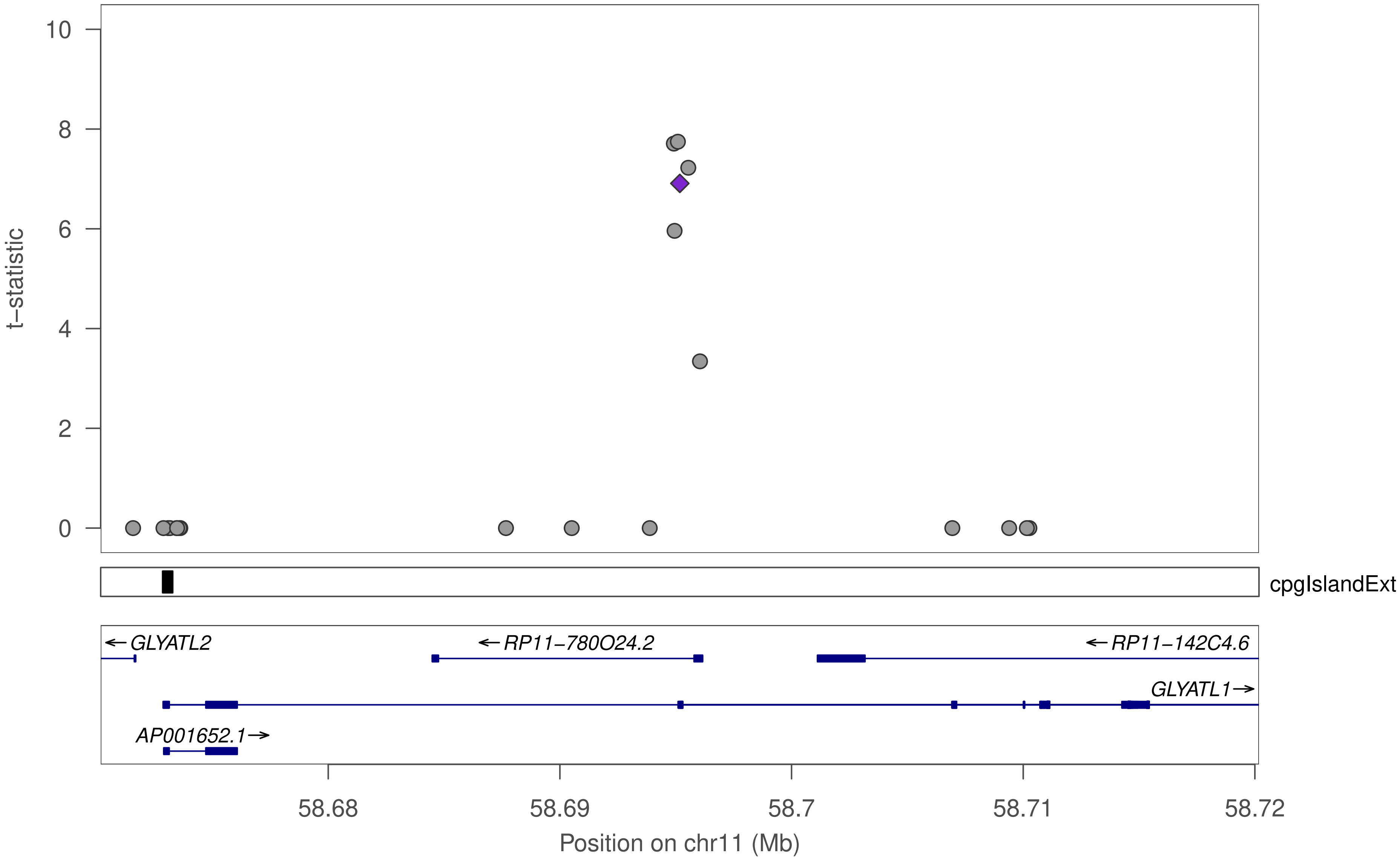

10
8
6
4
2
0
t–statistic
cpgIslandExt
←GLYATL2
←RP11–780O24.2
←RP11–142C4.6
GLYATL1→
AP001652.1→
58.68
58.69
58.7
58.71
58.72
Position on chr11 (Mb)

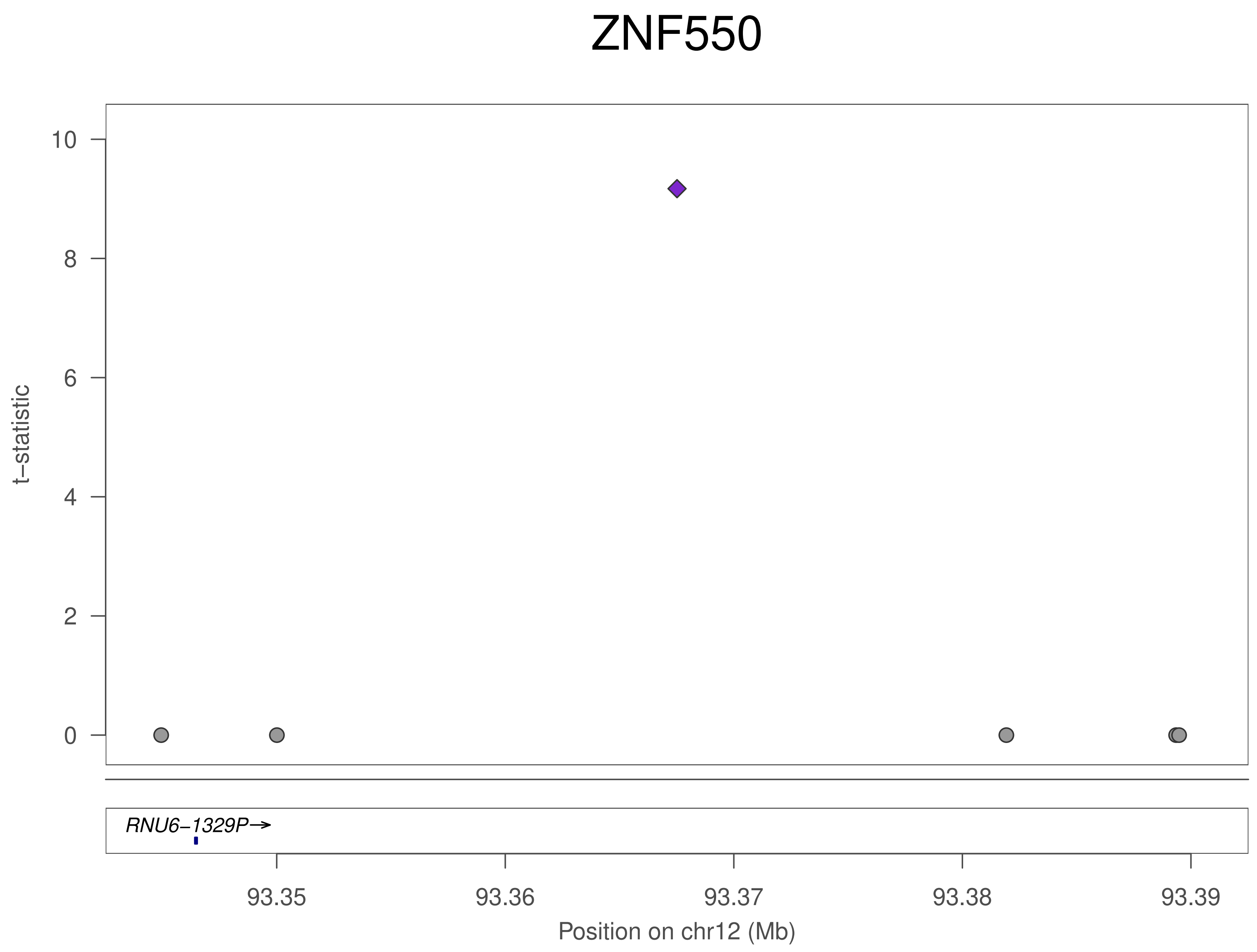
ZNF550
t−statistic
10
8
6
4
2
0
RNU6−1329P→
93.35
93.36
93.37
93.38
93.39
Position on chr12 (Mb)

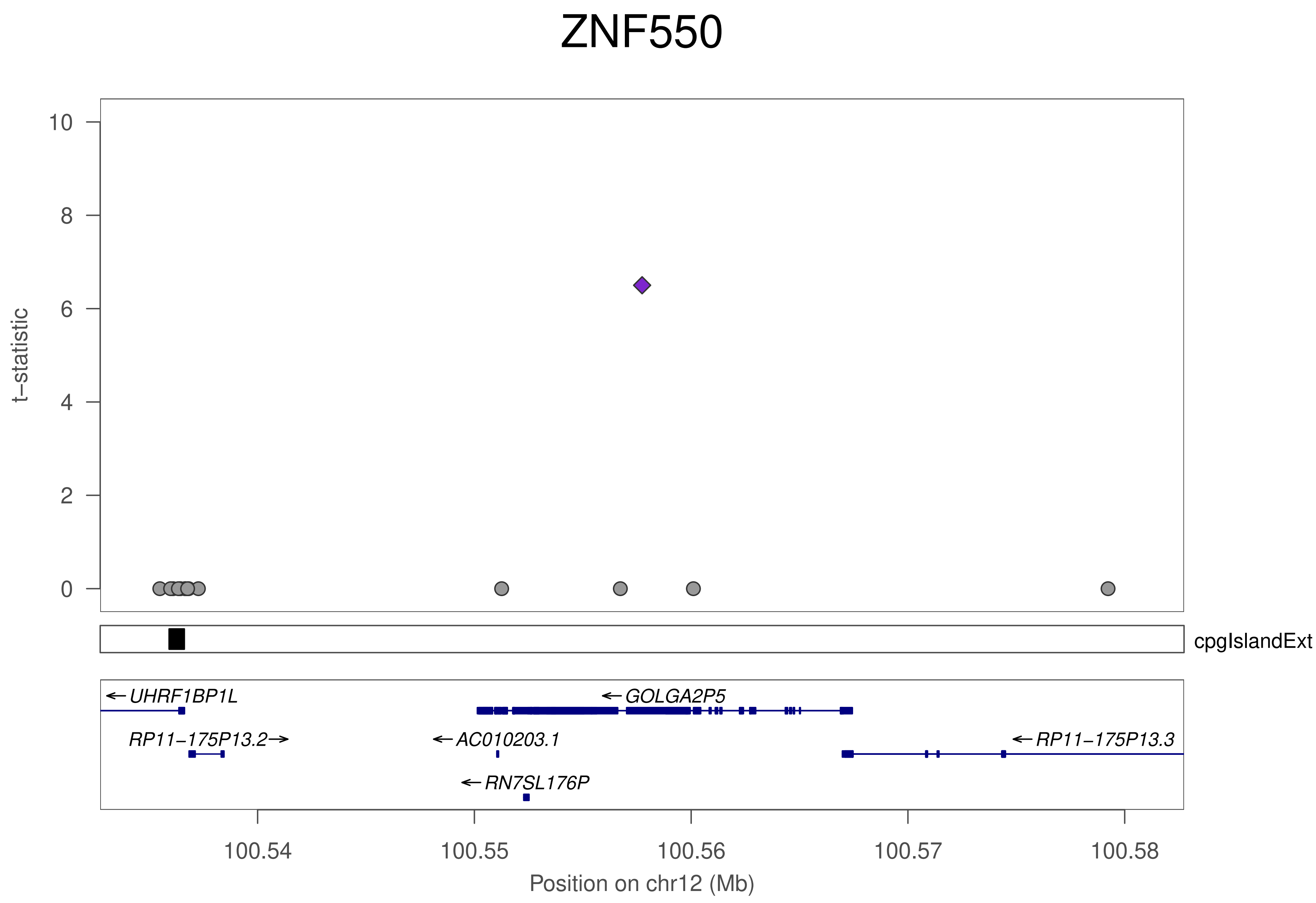

ZNF550
t−statistic
10
8
6
4
2
0
cpgIslandExt
UHRF1BP1L
GOLGA2P5
RP11−175P13.2
AC010203.1
RP11−175P13.3
RN7SL176P
100.54
100.55
100.56
100.57
100.58
Position on chr12 (Mb)
