## Supplementary File 6 for "Genetic control of KRAB-ZFP genes explains distal CpG-site methylation which associates with human disease phenotypes"

### ZNF195

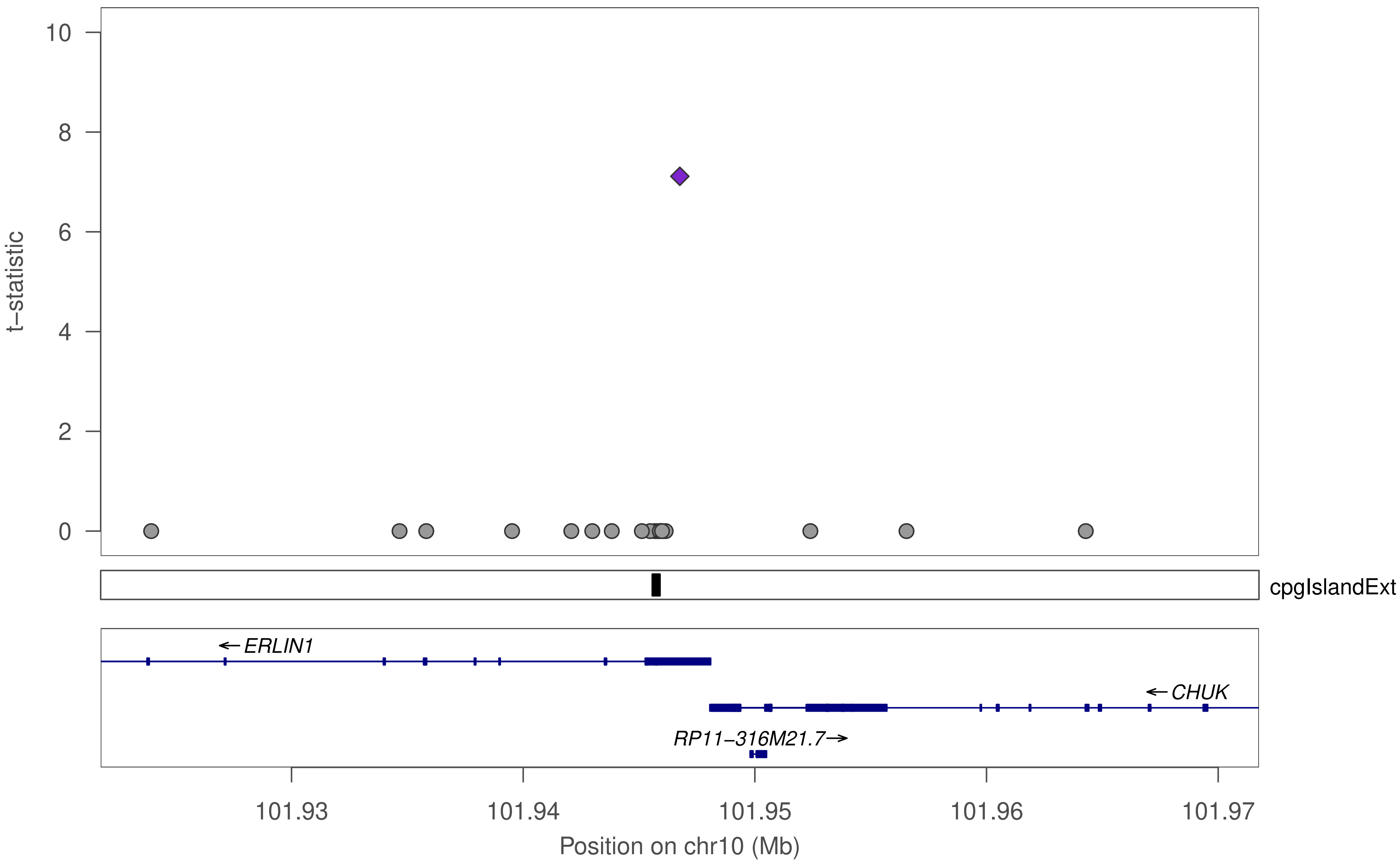

10
8
6
4
2
0
t–statistic
cpgIslandExt
← ERLIN1
← CHUK
RP11–316M21.7→
101.93
101.94
101.95
101.96
101.97
Position on chr10 (Mb)

### ZNF37A

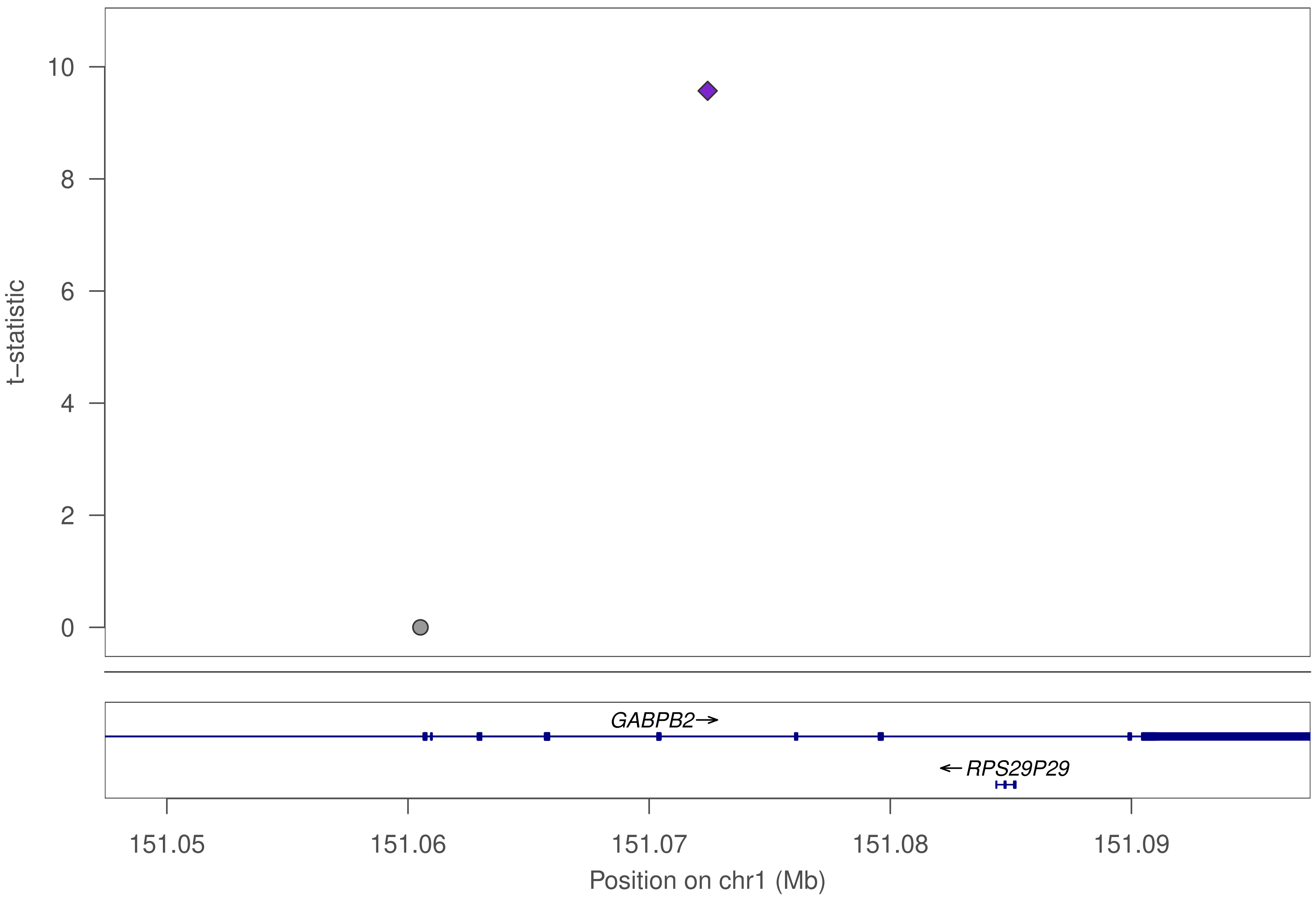

10
8
6
4
2
0
t–statistic
GABPB2
RPS29P29
151.05
151.06
151.07
151.08
151.09
Position on chr1 (Mb)

### ZNF37A

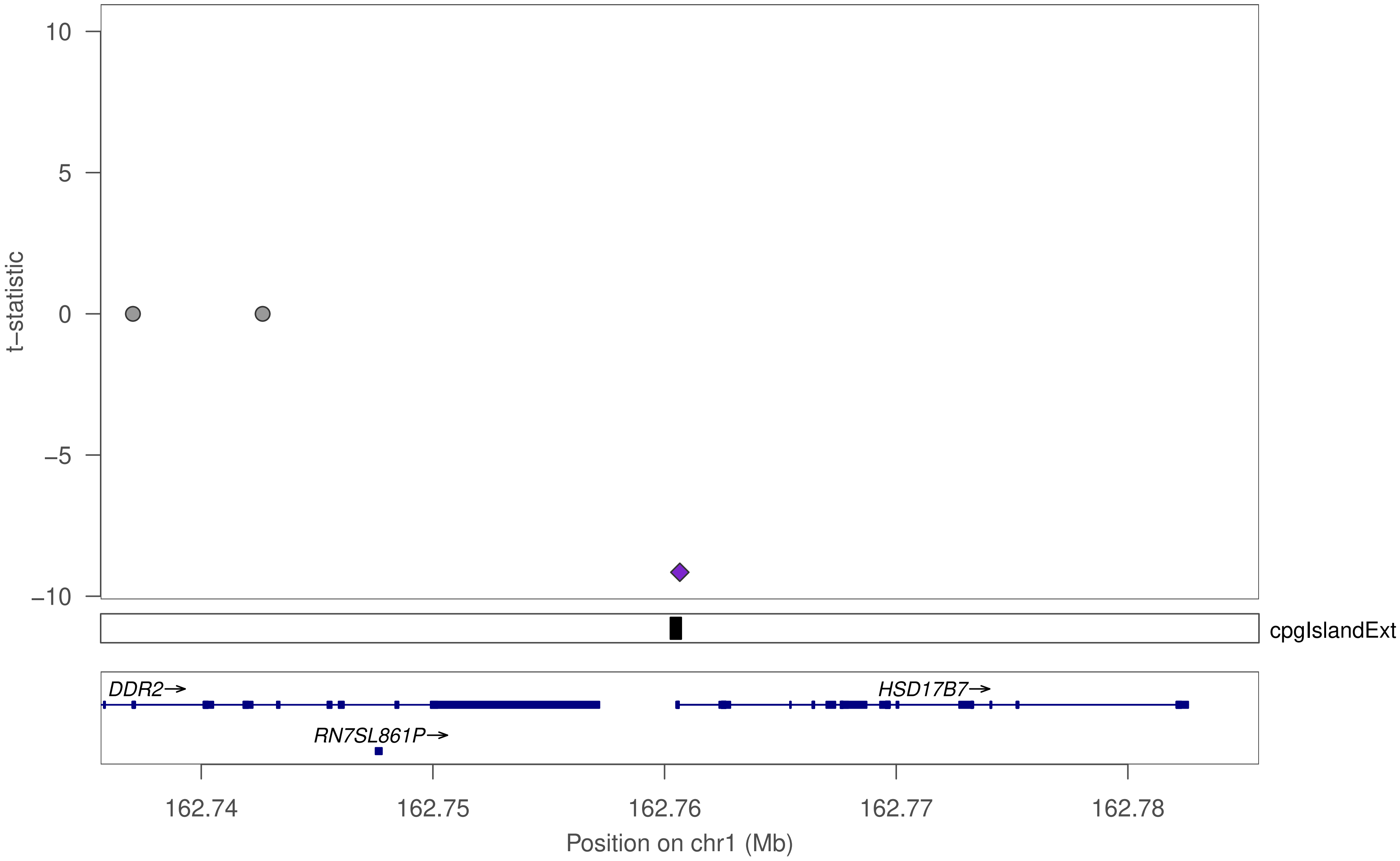

10
5
0
−5
−10
t−statistic
cpgIslandExt
DDR2→
HSD17B7→
RN7SL861P→
162.74
162.75
162.76
162.77
162.78
Position on chr1 (Mb)

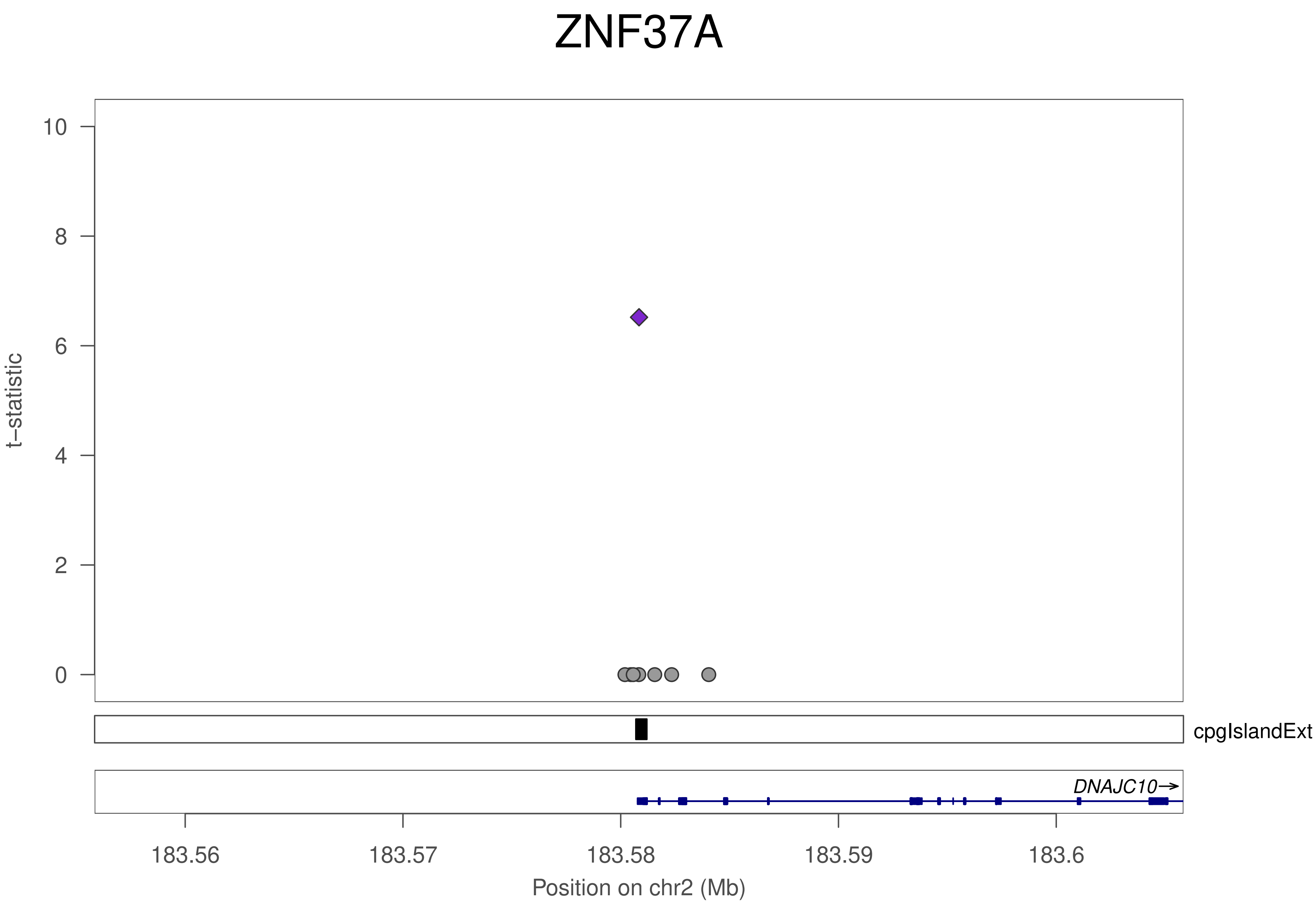
ZNF37A
10
8
6
4
2
0
t−statistic
cpgIslandExt
DNAJC10→
183.56
183.57
183.58
183.59
183.6
Position on chr2 (Mb)

### ZNF37A

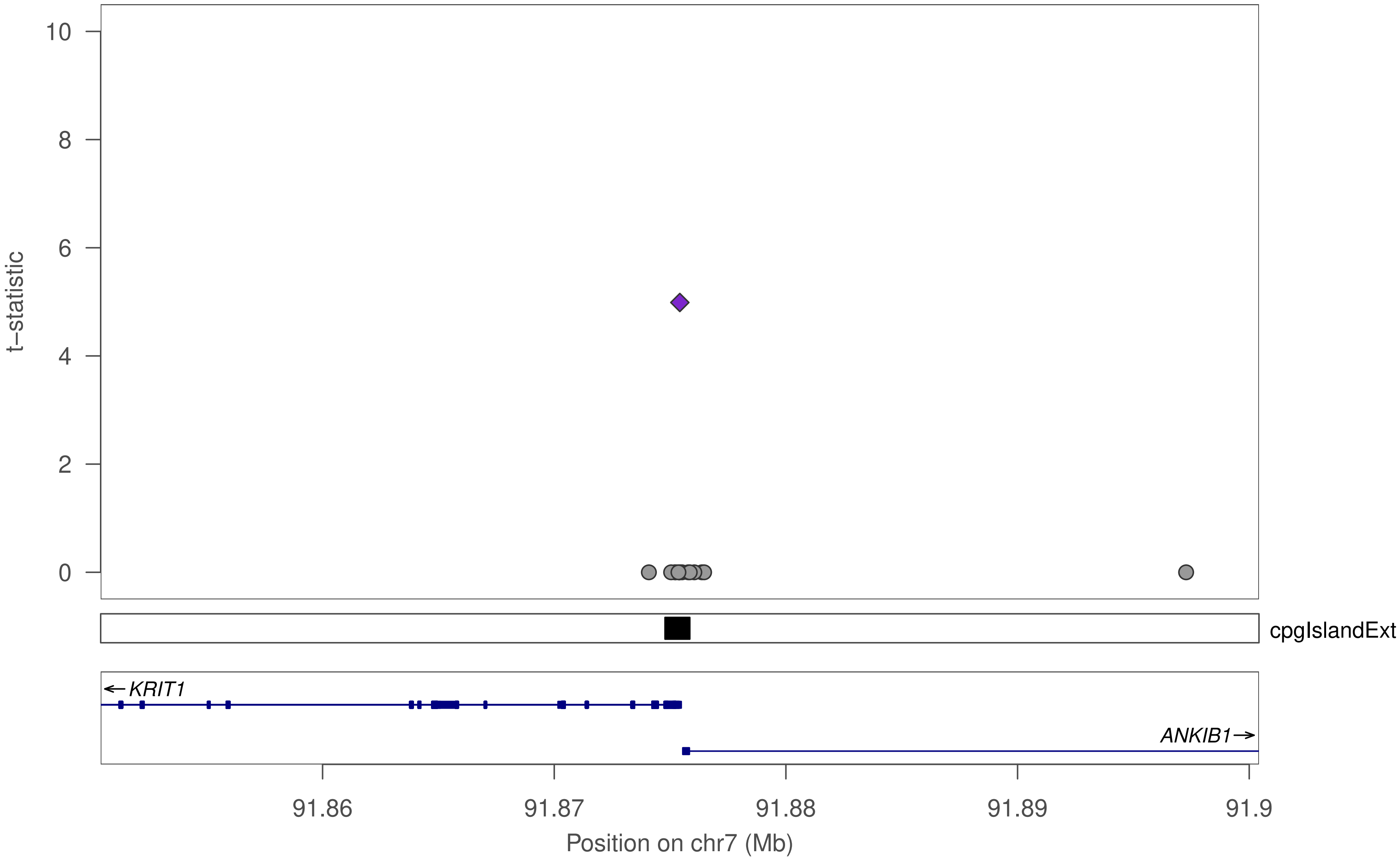
t−statistic
10
8
6
4
2
0
cpgIslandExt
KRIT1
ANKIB1
91.86
91.87
91.88
91.89
91.9
Position on chr7 (Mb)

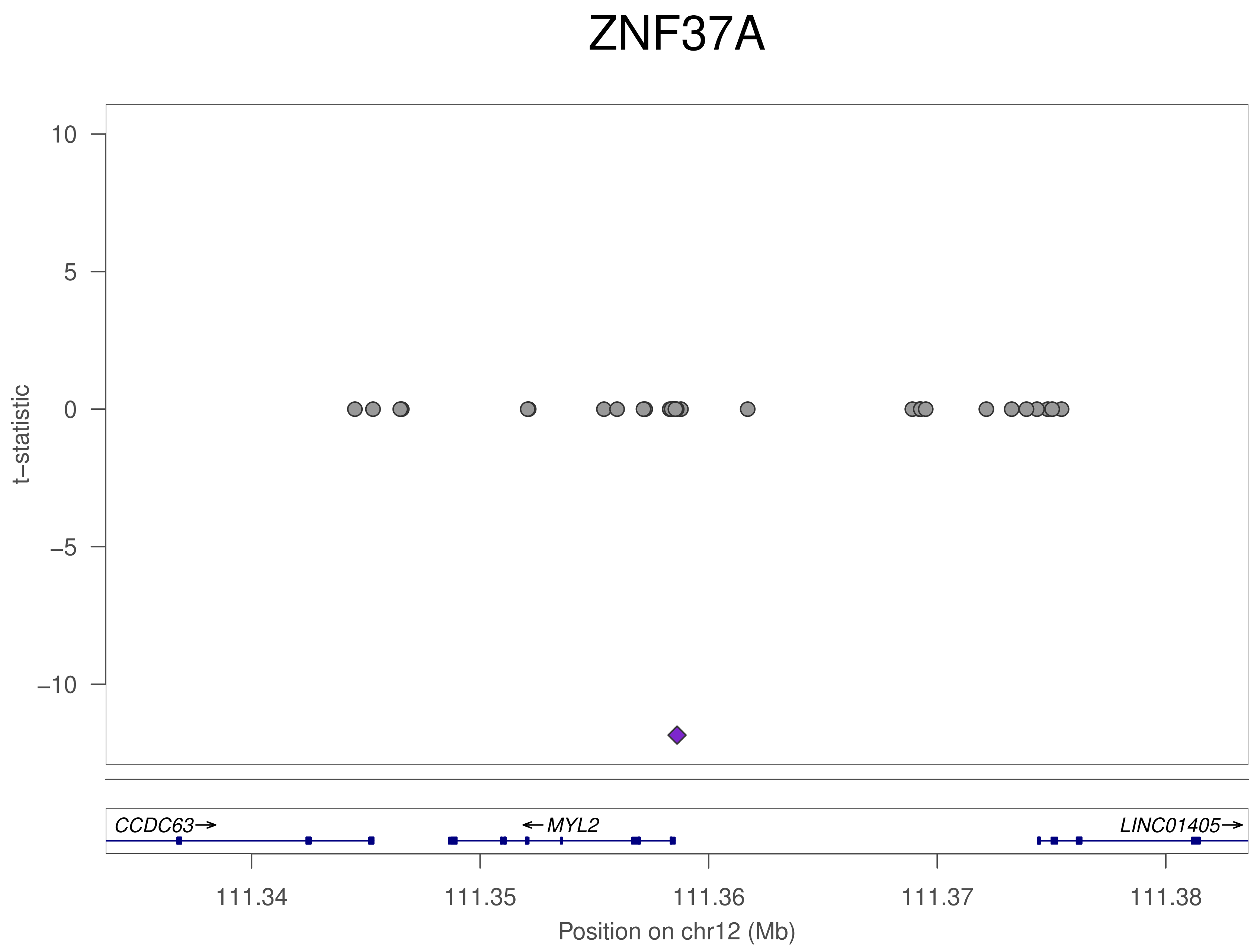
ZNF37A
t−statistic
10
5
0
−5
−10
CCDC63→
←MYL2
LINC01405→
111.34
111.35
111.36
111.37
111.38
Position on chr12 (Mb)

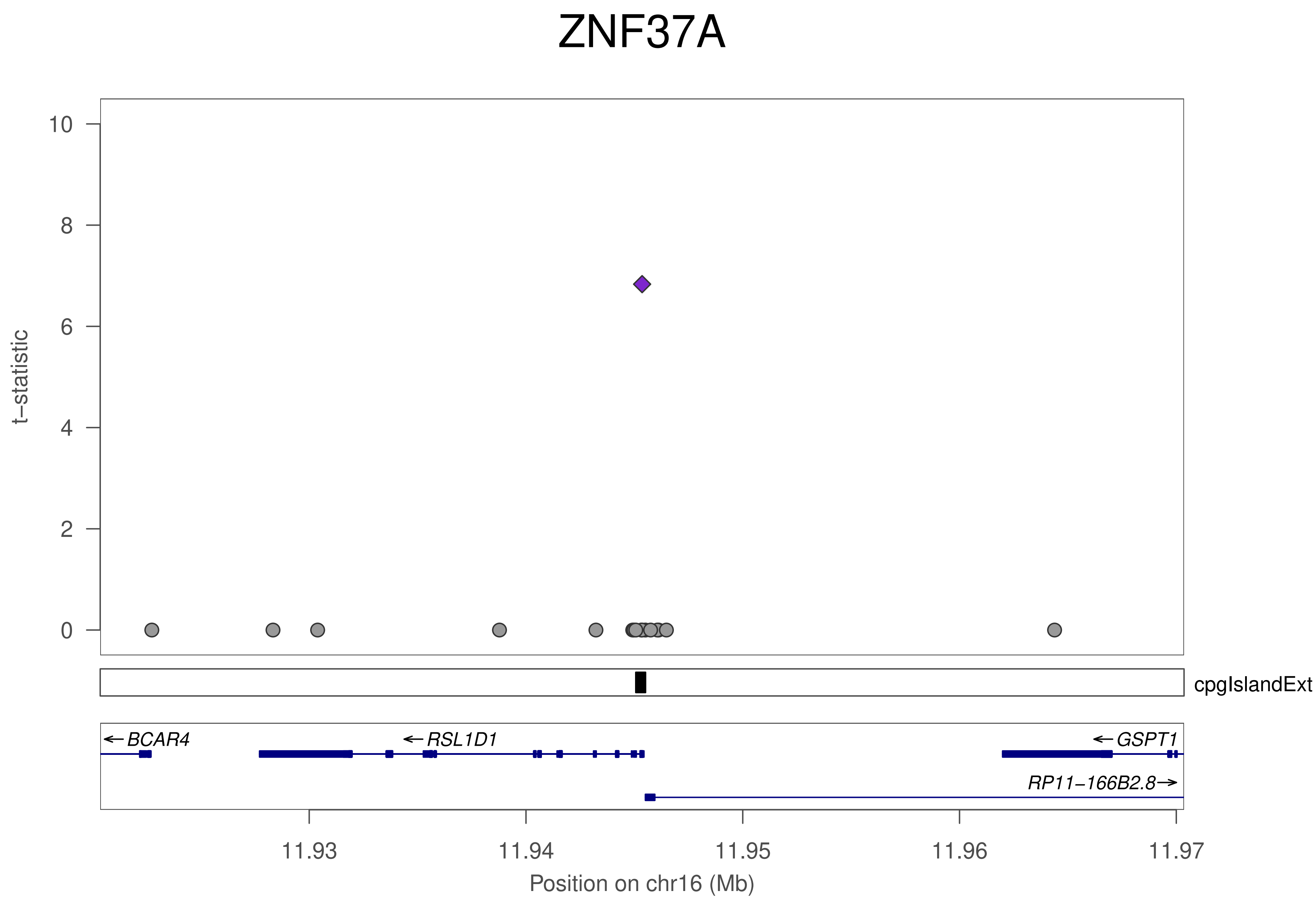
ZNF37A
t−statistic
10
8
6
4
2
0
cpgIslandExt
BCAR4
RSL1D1
GSPT1
RP11−166B2.8
11.93
11.94
11.95
11.96
11.97
Position on chr16 (Mb)

### ZNF37A

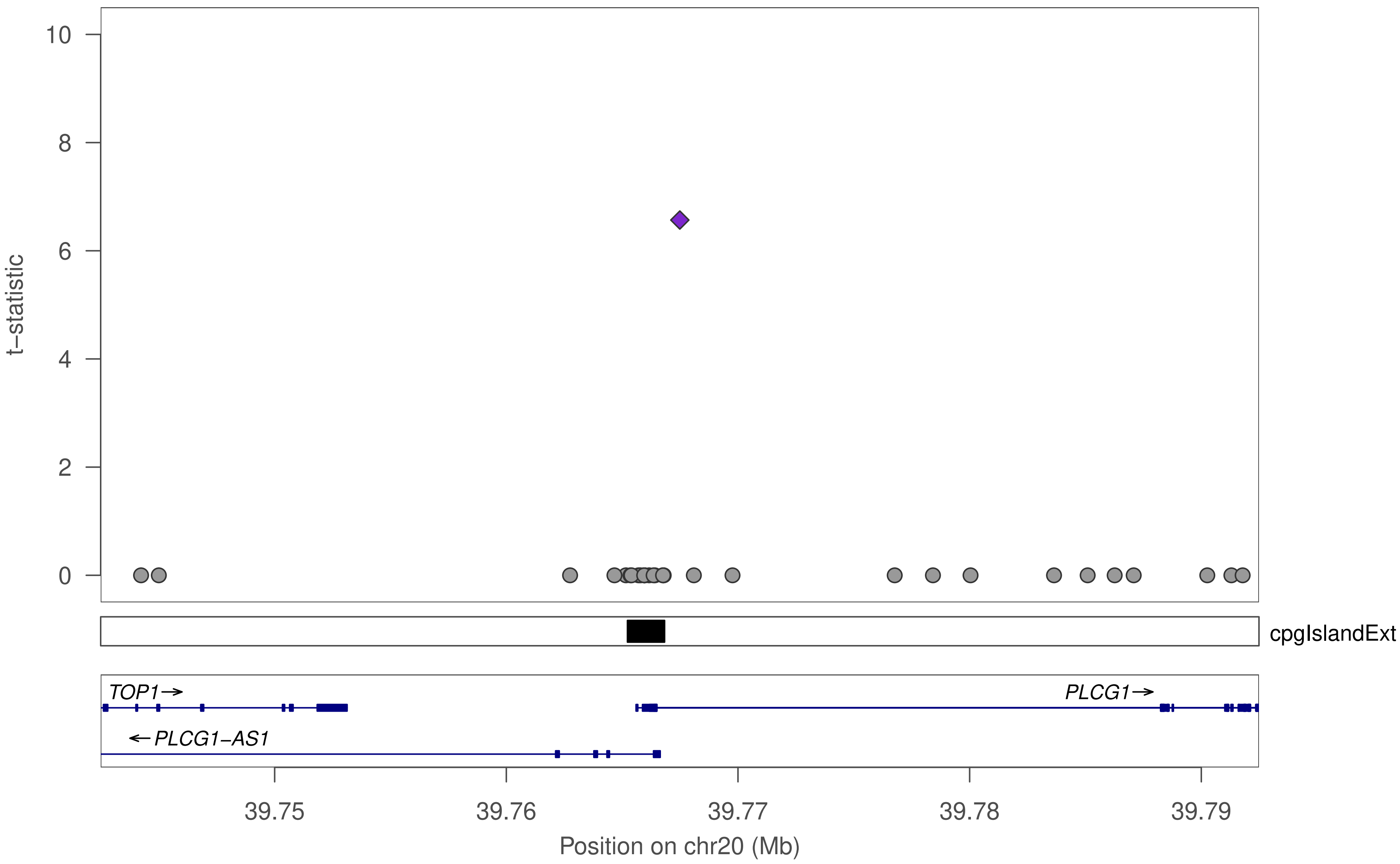

t–statistic
10
8
6
4
2
0
cpgIslandExt
TOP1→
PLCG1→
←PLCG1–AS1
39.75
39.76
39.77
39.78
39.79
Position on chr20 (Mb)

### ZNF586

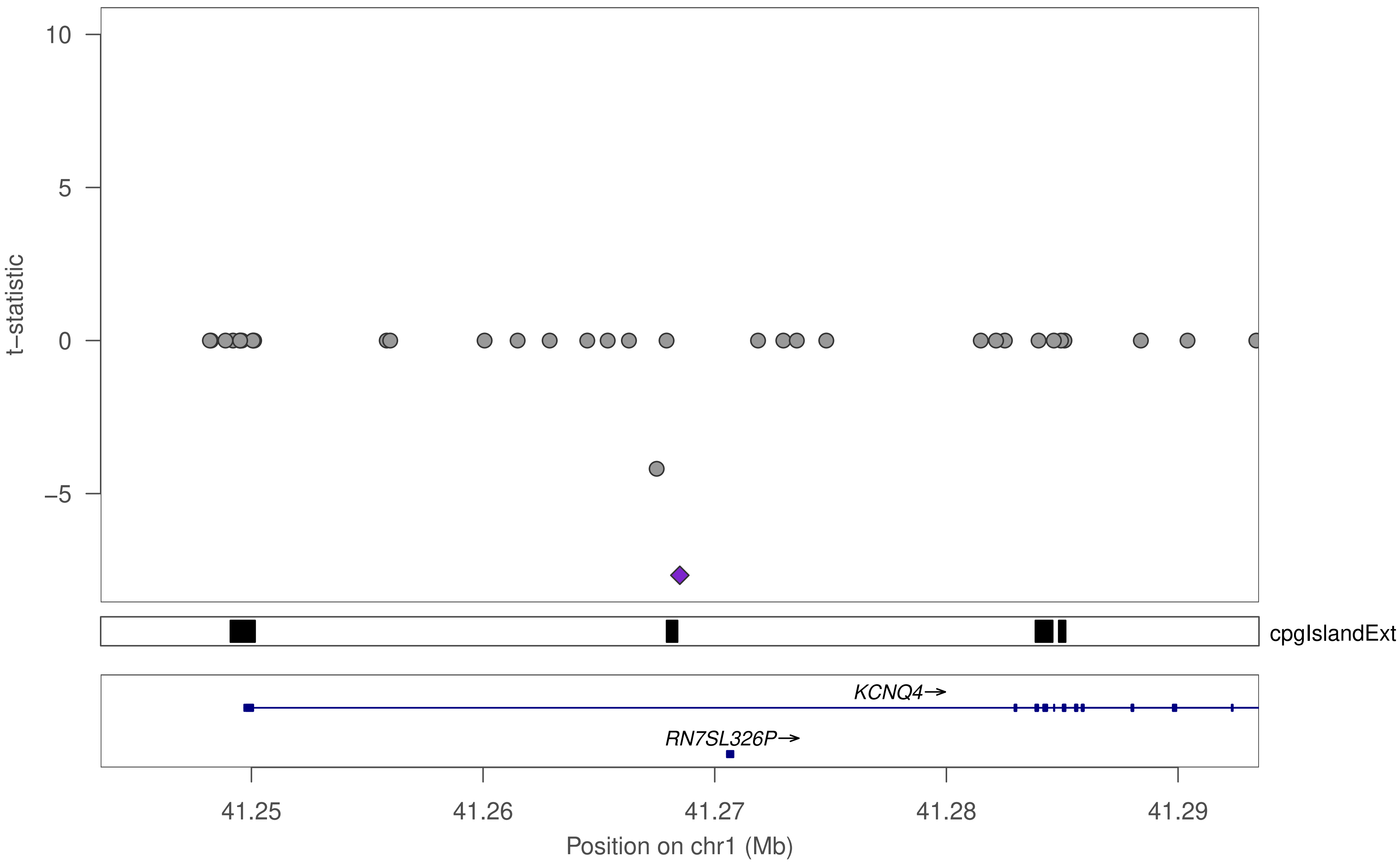

10
5
0
−5
t−statistic
cpgIslandExt
KCNQ4→
RN7SL326P→
41.25
41.26
41.27
41.28
41.29
Position on chr1 (Mb)

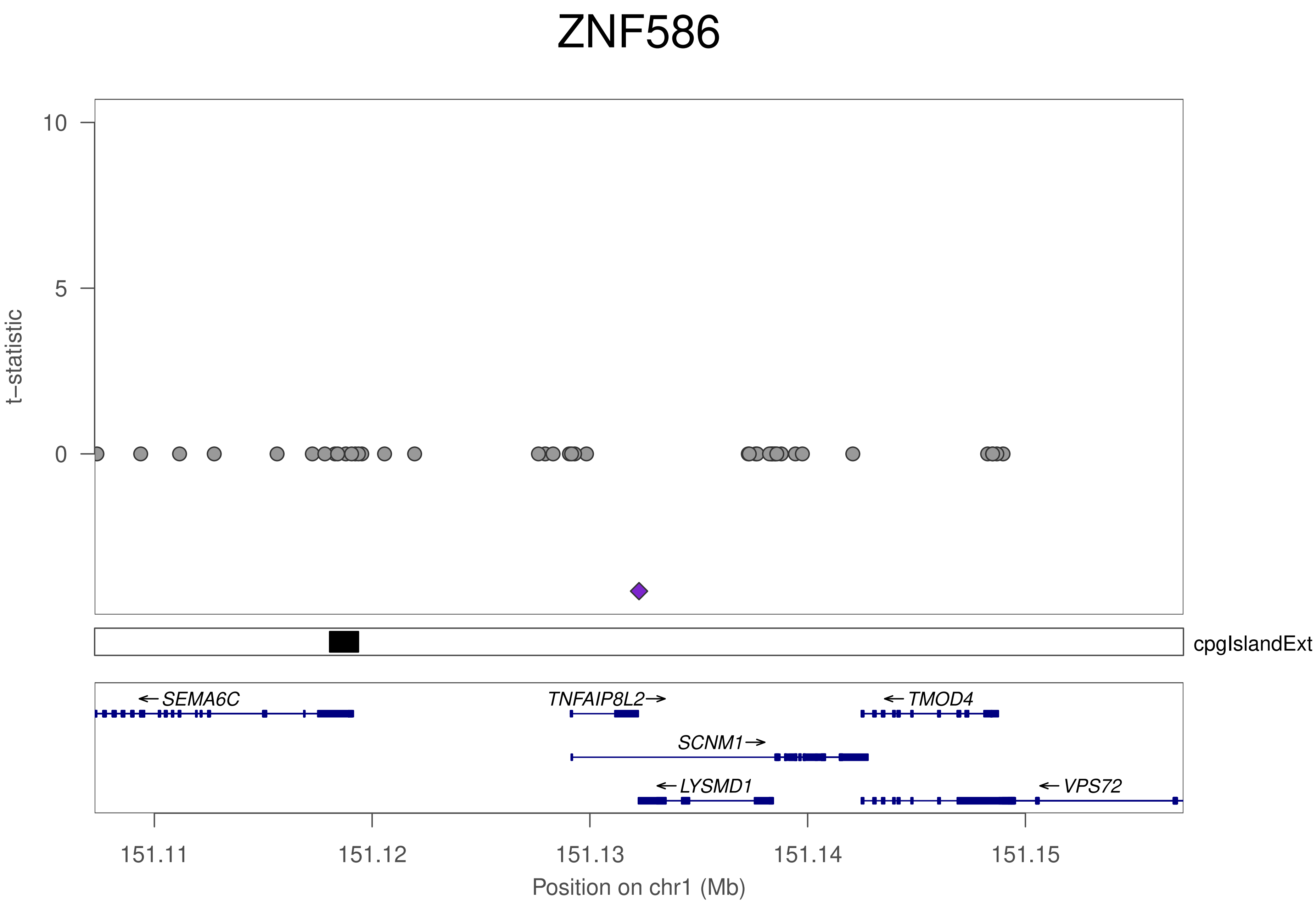
ZNF586
t–statistic
10
5
0
cpgIslandExt
←SEMA6C
TNFAIP8L2→
←TMOD4
SCNM1→
←LYSMD1
←VPS72
151.11
151.12
151.13
151.14
151.15
Position on chr1 (Mb)

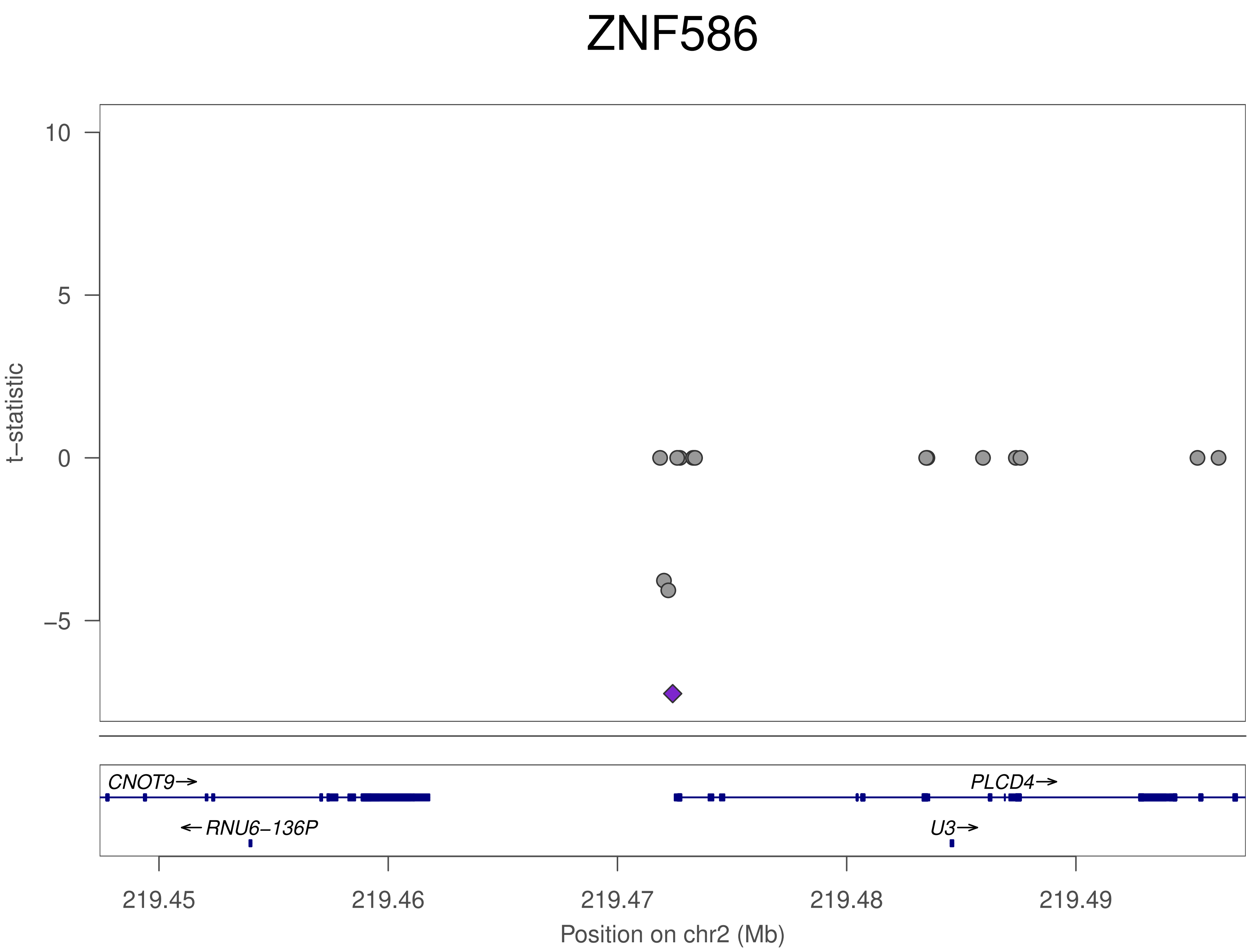

ZNF586
t−statistic
10
5
0
−5
CNOT9→
←RNU6−136P
PLCD4→
U3→
219.45
219.46
219.47
219.48
219.49
Position on chr2 (Mb)

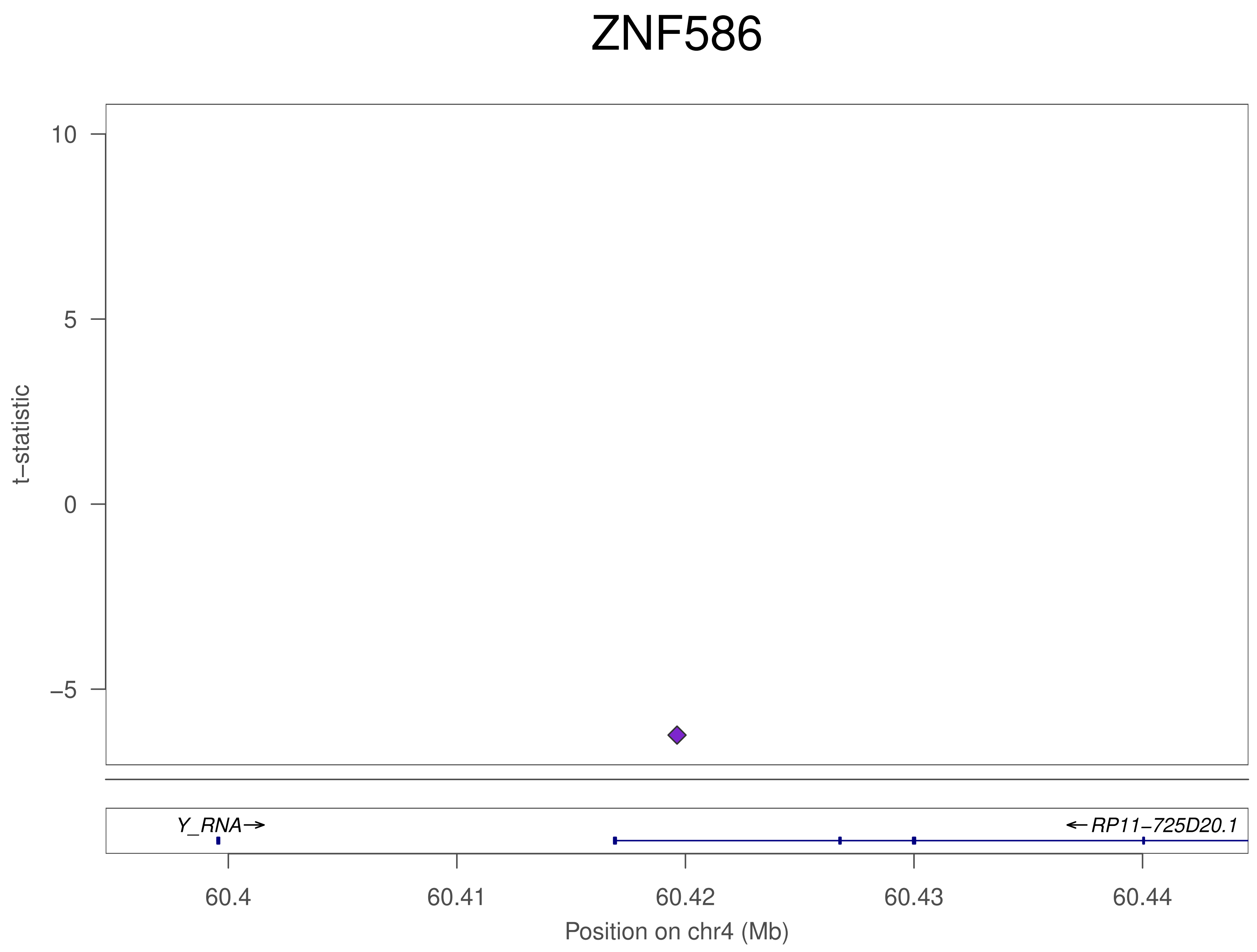

ZNF586
10
5
0
−5
t−statistic
Y_RNA
RP11−725D20.1
60.4
60.41
60.42
60.43
60.44
Position on chr4 (Mb)

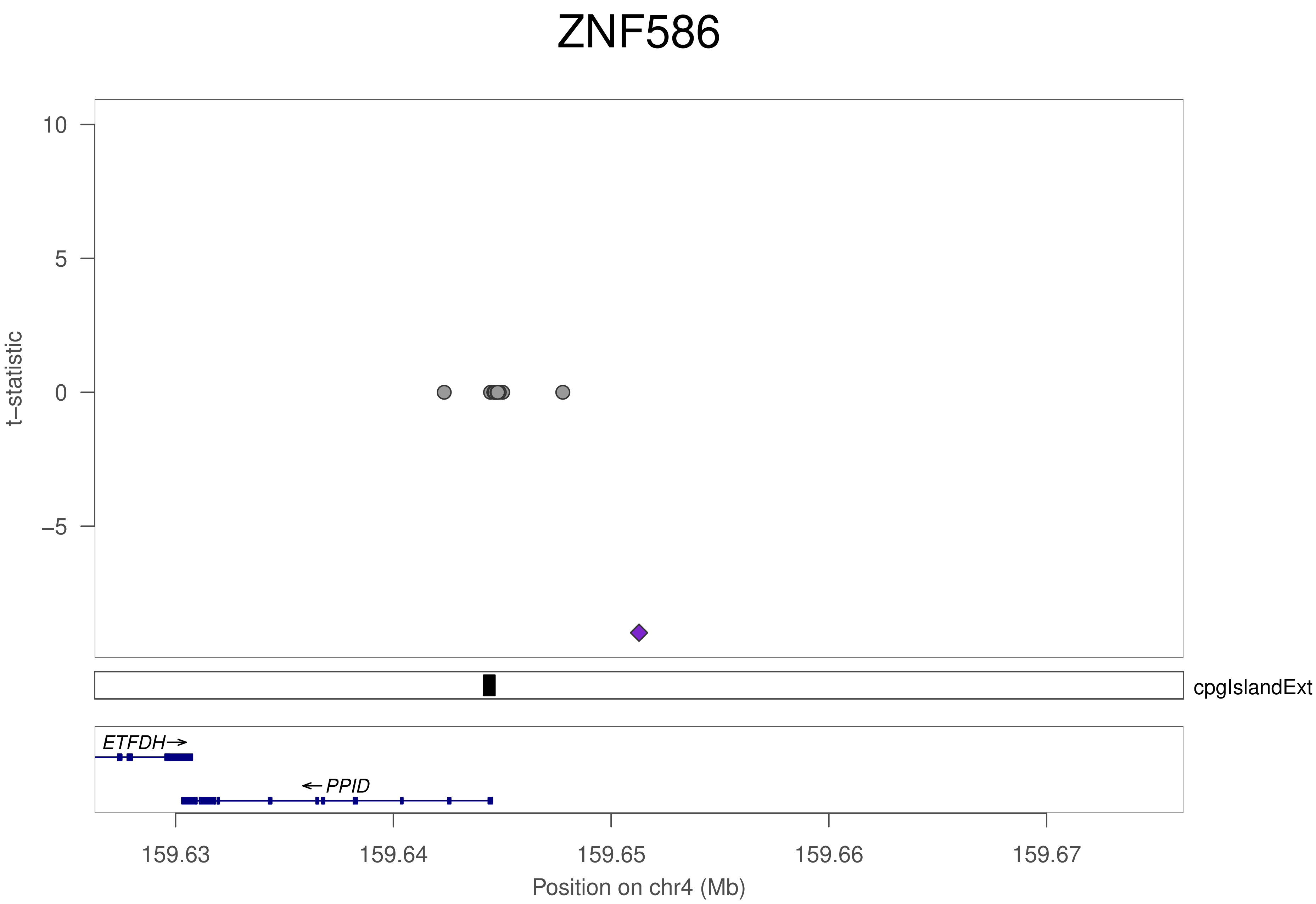

ZNF586
t−statistic
10
5
0
−5
cpgIslandExt
ETFDH
PPID
159.63
159.64
159.65
159.66
159.67
Position on chr4 (Mb)

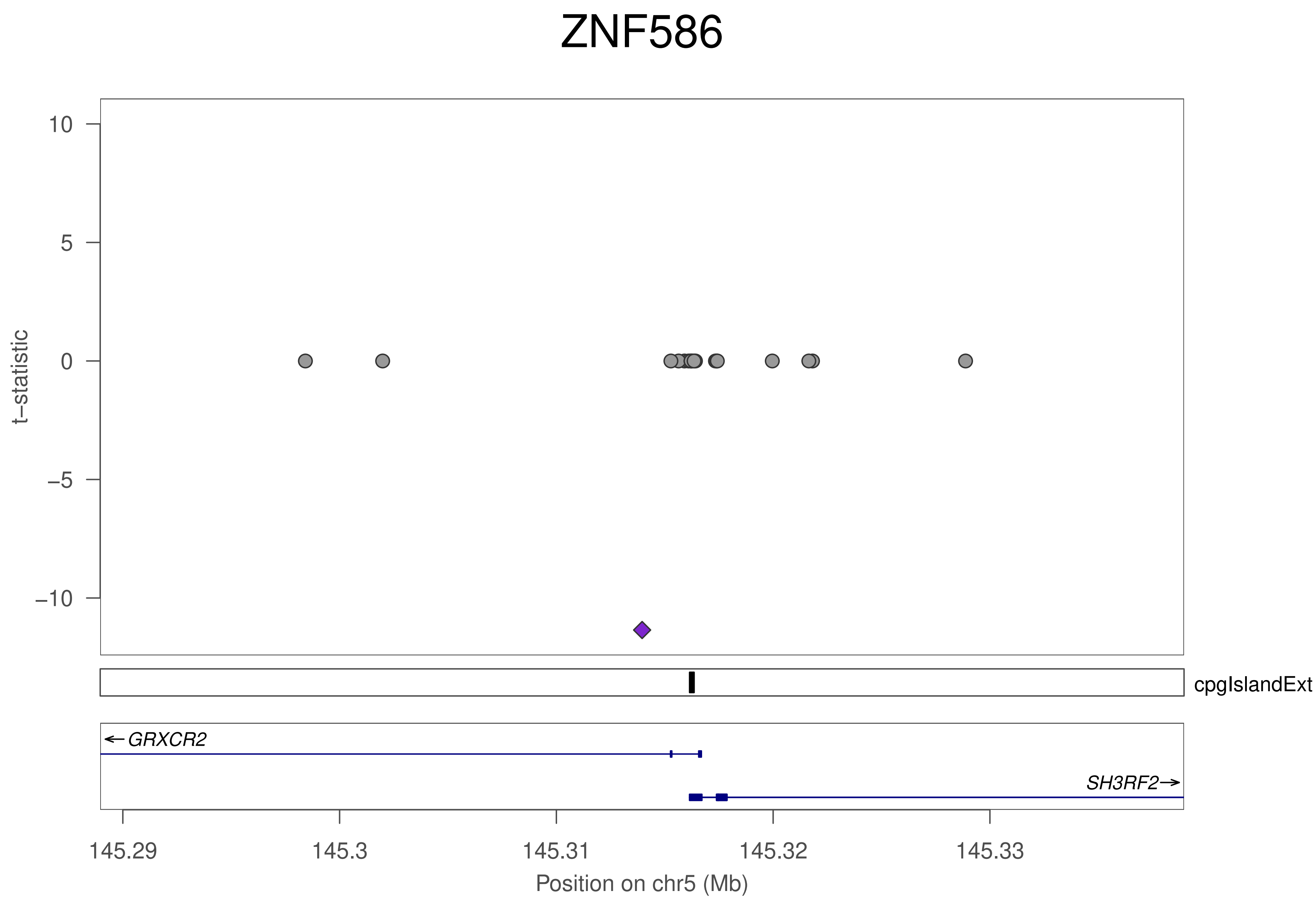
ZNF586
t−statistic
10
5
0
−5
−10
cpgIslandExt
GRXCR2
SH3RF2
145.29
145.3
145.31
145.32
145.33
Position on chr5 (Mb)

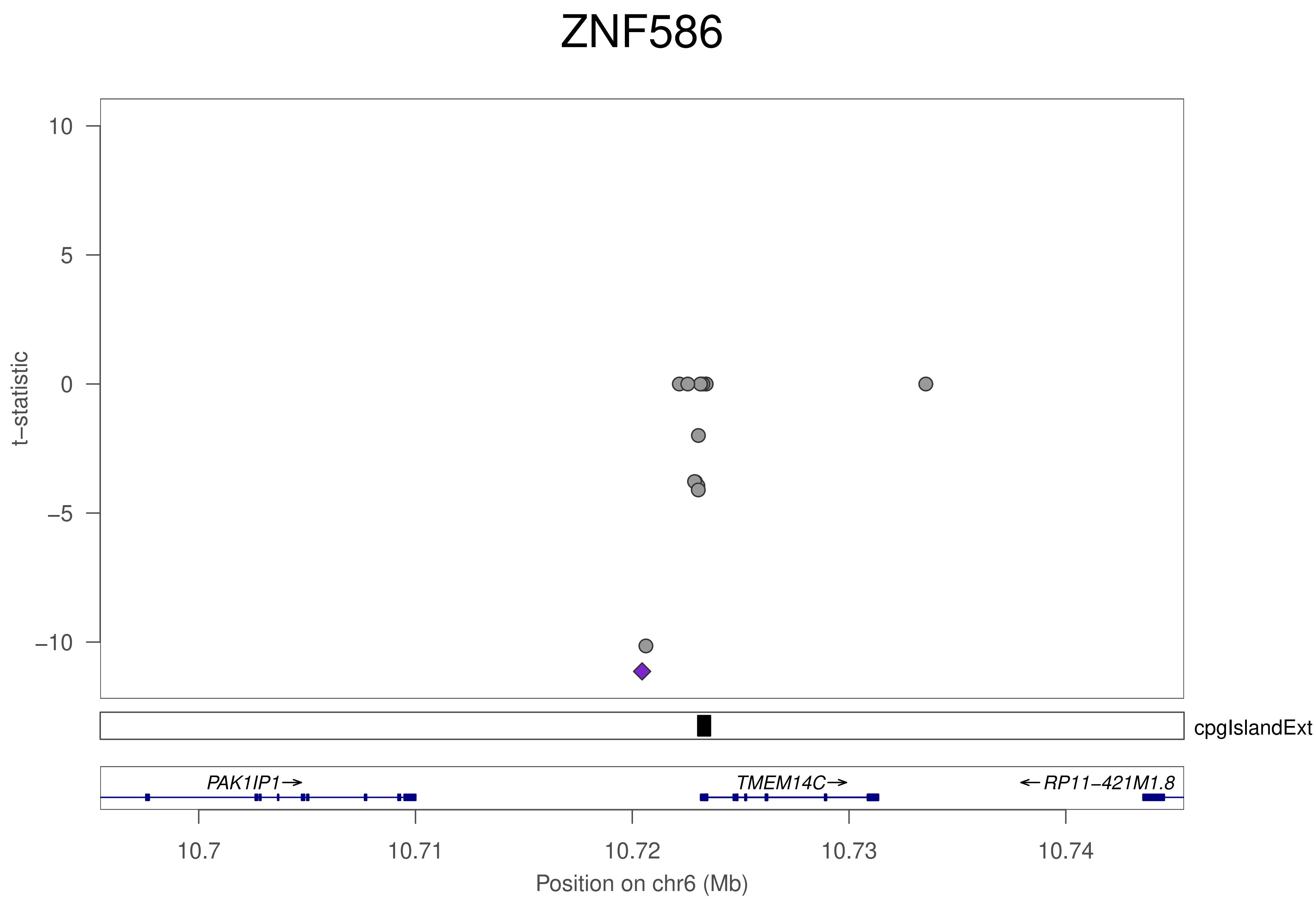

ZNF586
t−statistic
10
5
0
−5
−10
cpgIslandExt
PAK1IP1→
TMEM14C→
←RP11−421M1.8
10.7
10.71
10.72
10.73
10.74
Position on chr6 (Mb)

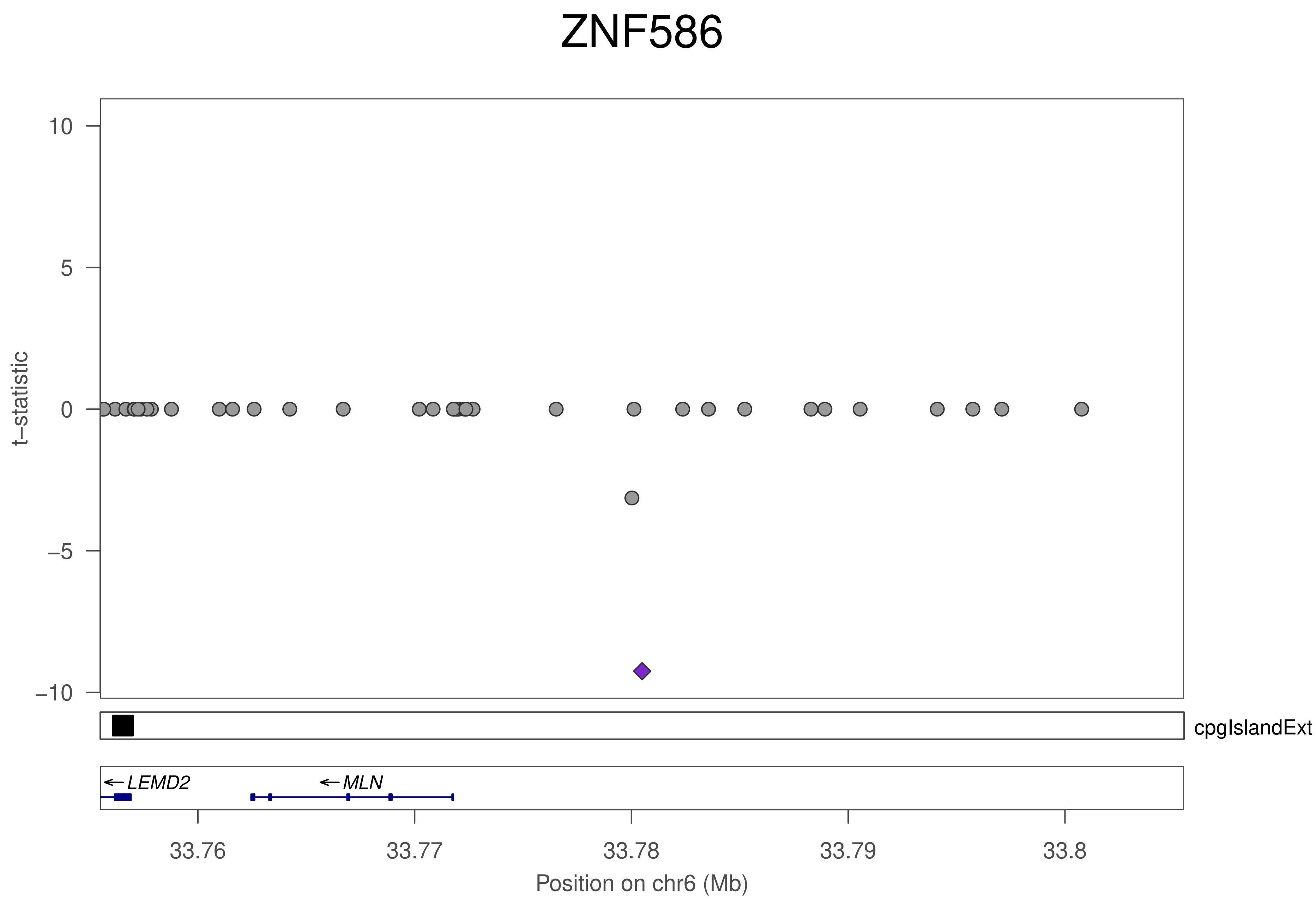

ZNF586
t−statistic
10
5
0
−5
−10
cpgIslandExt
LEMD2
MLN
33.76
33.77
33.78
33.79
33.8
Position on chr6 (Mb)

### ZNF586

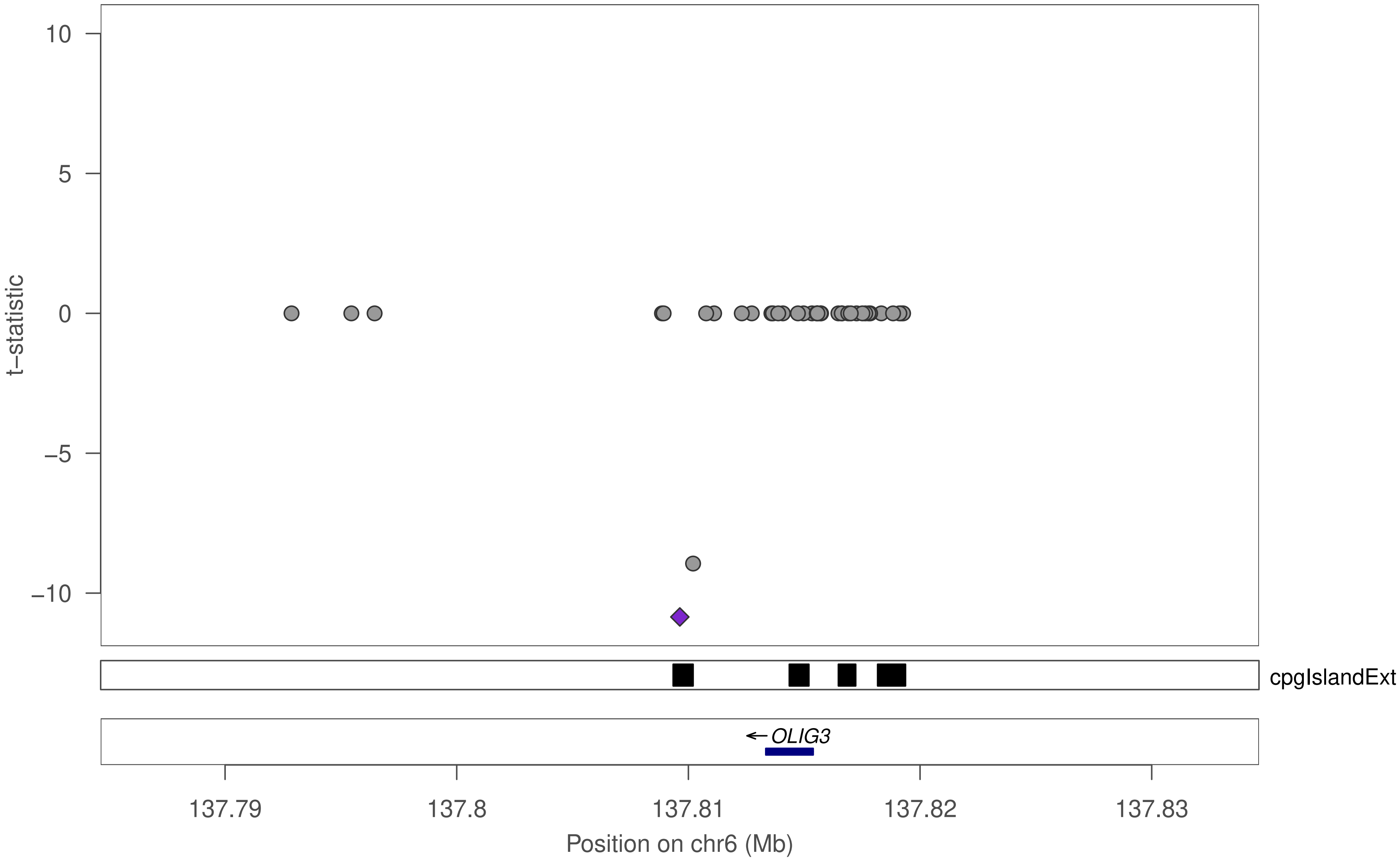

10
5
0
−5
−10
t−statistic
cpgIslandExt
←OLIG3
137.79
137.8
137.81
137.82
137.83
Position on chr6 (Mb)

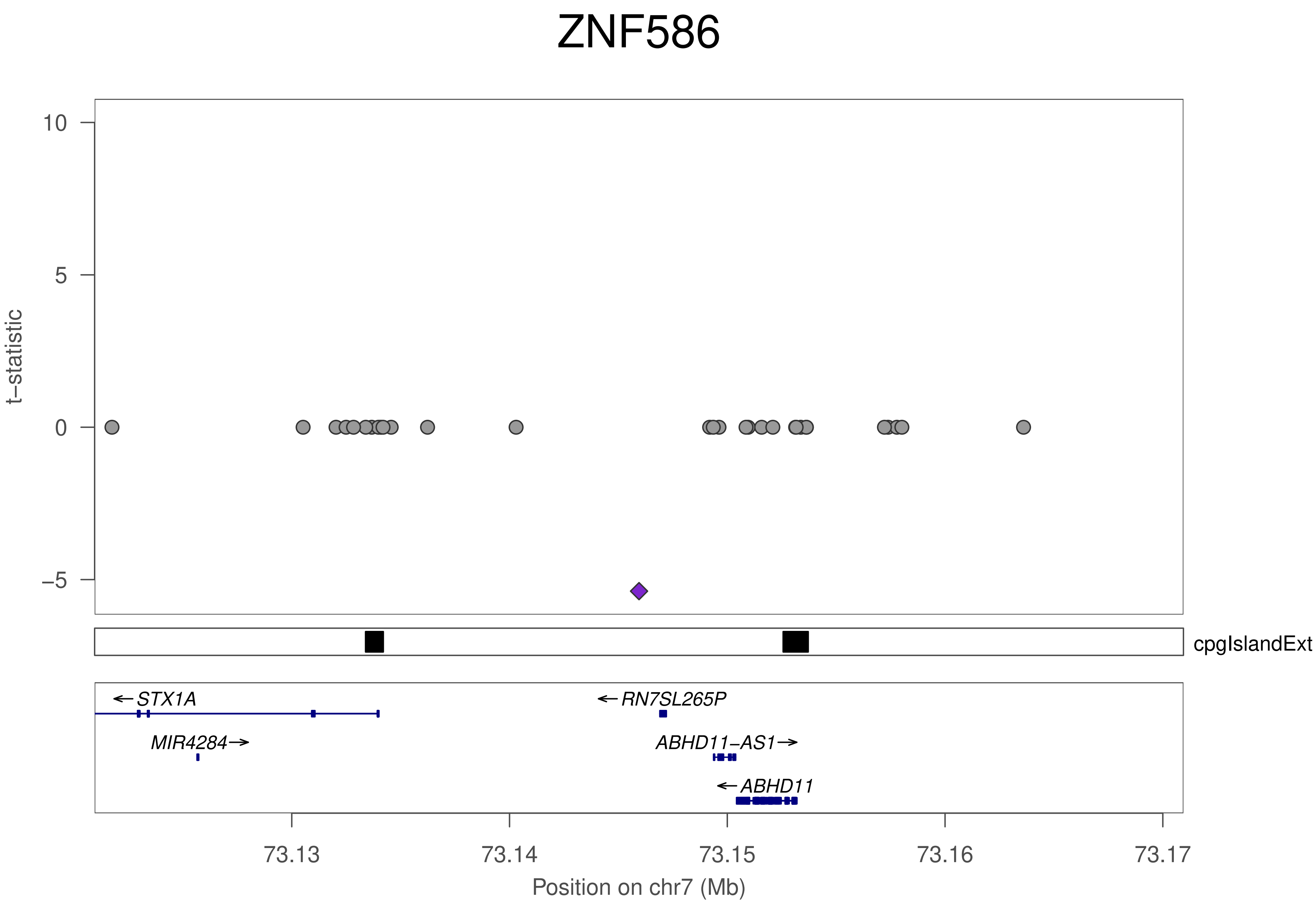
ZNF586
t–statistic
10
5
0
–5
cpgIslandExt
←STX1A
←RN7SL265P
MIR4284→
ABHD11–AS1→
←ABHD11
73.13
73.14
73.15
73.16
73.17
Position on chr7 (Mb)

### ZNF586

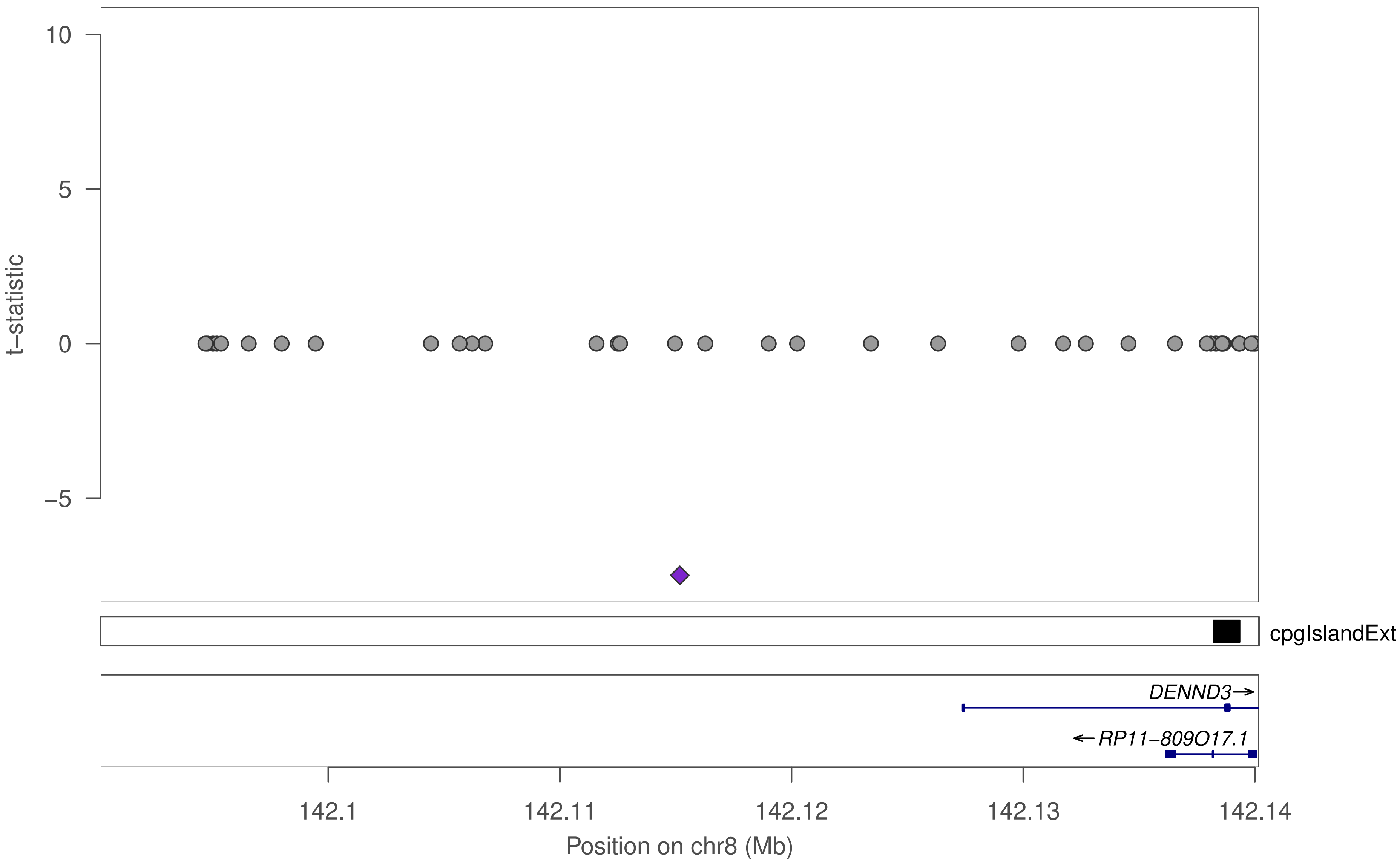

10
5
0
−5
t−statistic
cpgIslandExt
DENND3→
←RP11−809O17.1
142.1
142.11
142.12
142.13
142.14
Position on chr8 (Mb)

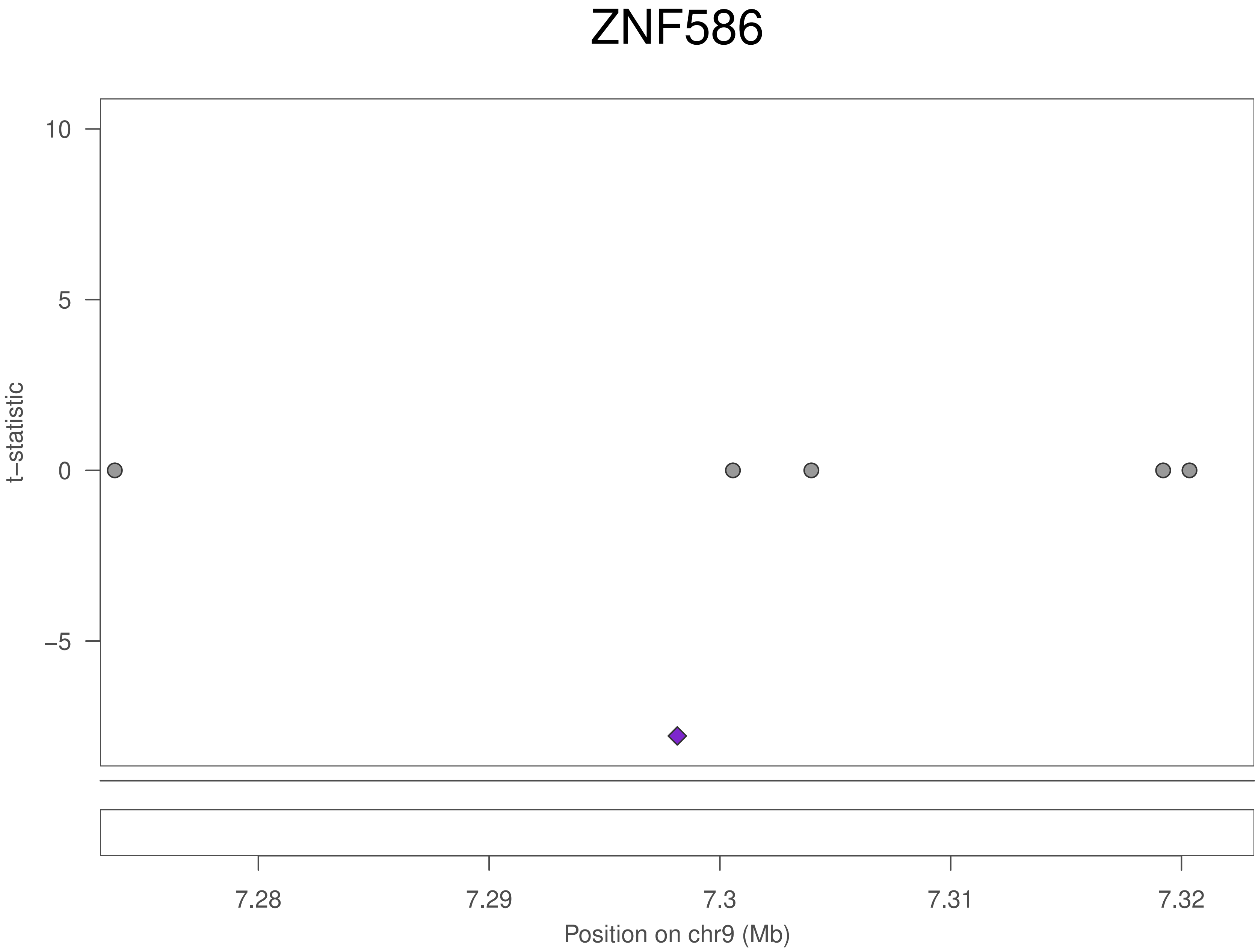
ZNF586
10
5
0
−5
t−statistic
7.28
7.29
7.3
7.31
7.32
Position on chr9 (Mb)

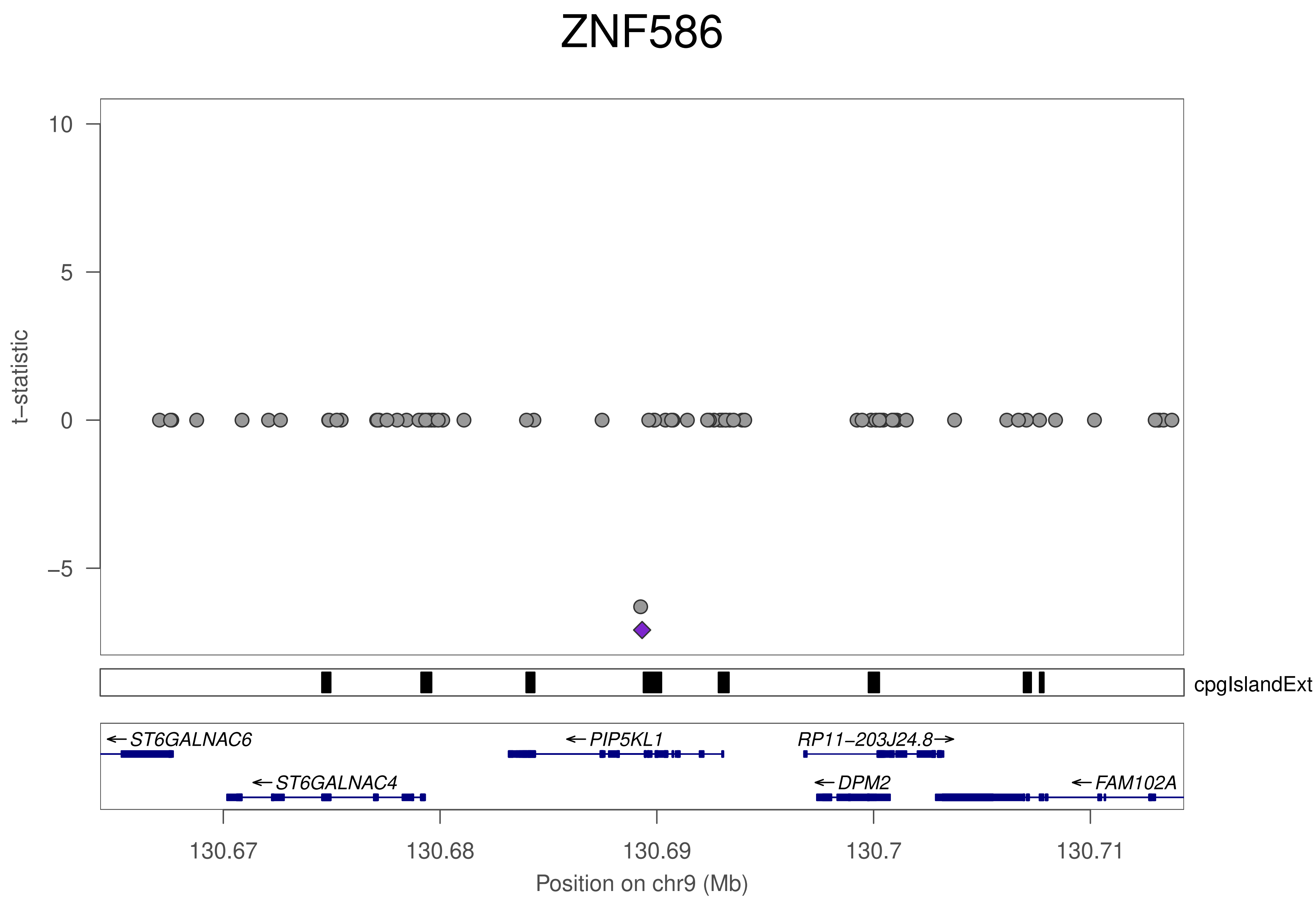
ZNF586
t−statistic
10
5
0
−5
cpgIslandExt
←ST6GALNAC6
←ST6GALNAC4
←PIP5KL1
RP11−203J24.8→
←DPM2
←FAM102A
130.67
130.68
130.69
130.7
130.71
Position on chr9 (Mb)

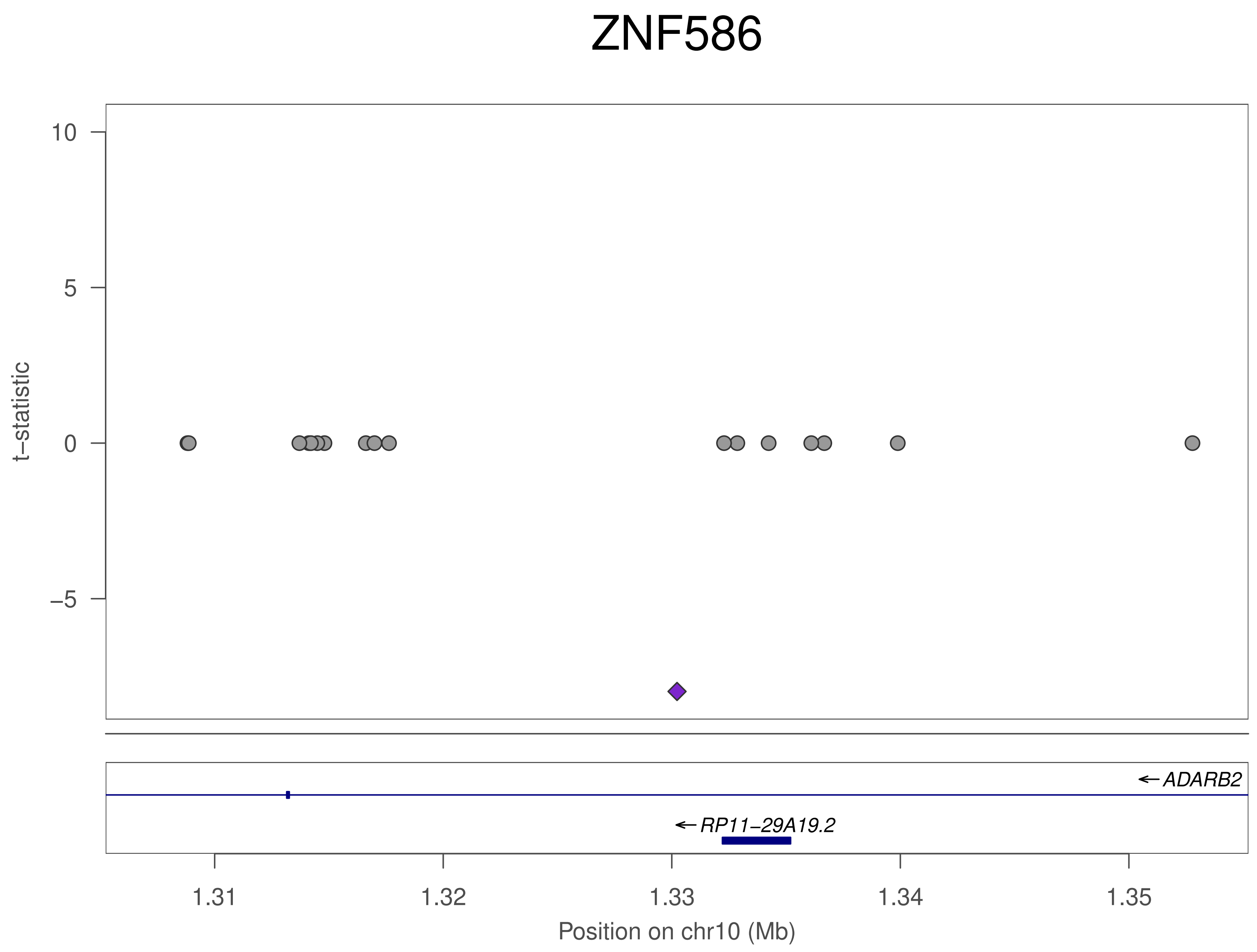

ZNF586
10
5
0
−5
t−statistic
ADARB2
RP11−29A19.2
1.31
1.32
1.33
1.34
1.35
Position on chr10 (Mb)

### ZNF586

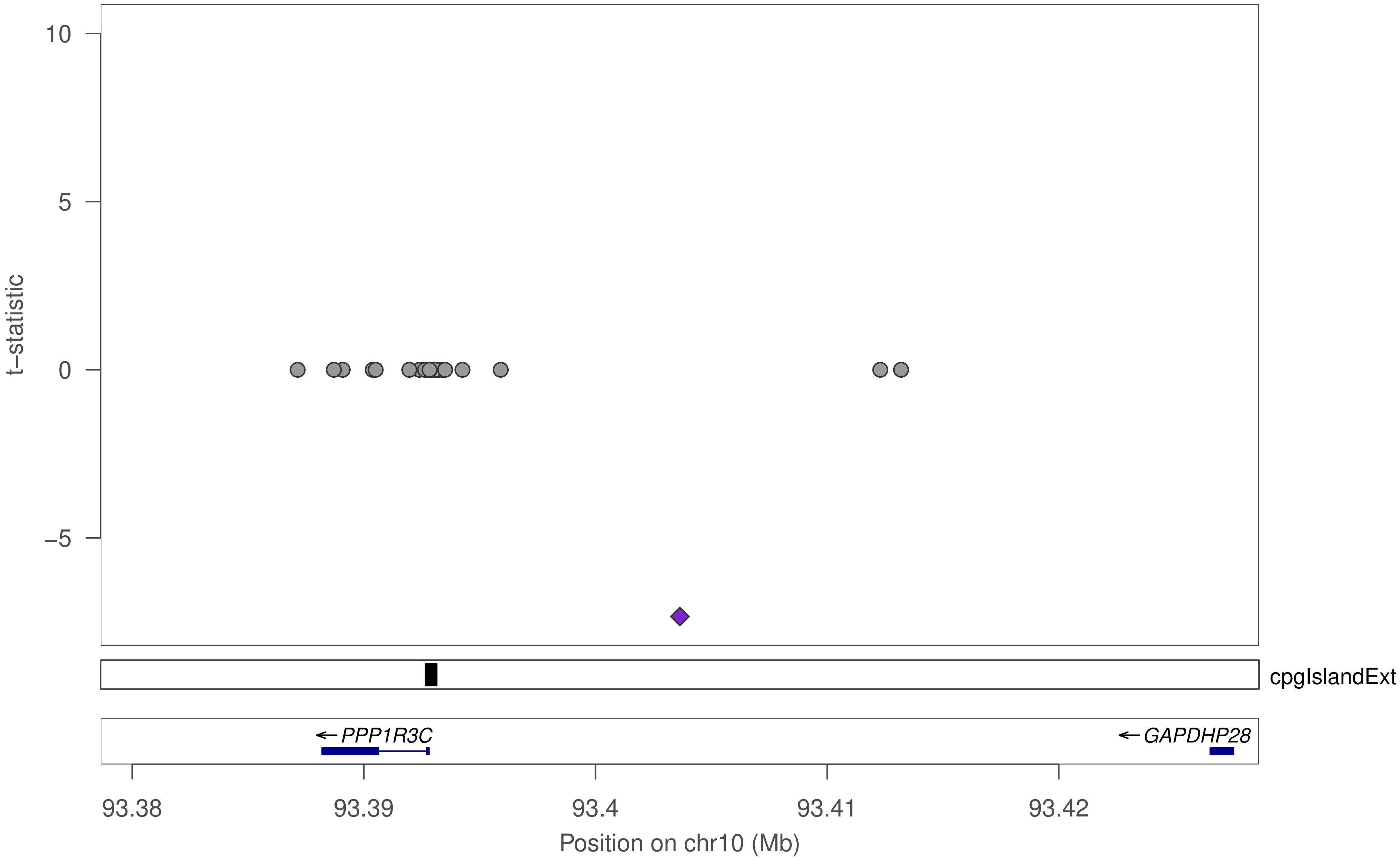

10
5
0
−5
t−statistic
cpgIslandExt
← PPP1R3C
← GAPDHP28
93.38
93.39
93.4
93.41
93.42
Position on chr10 (Mb)

### ZNF586

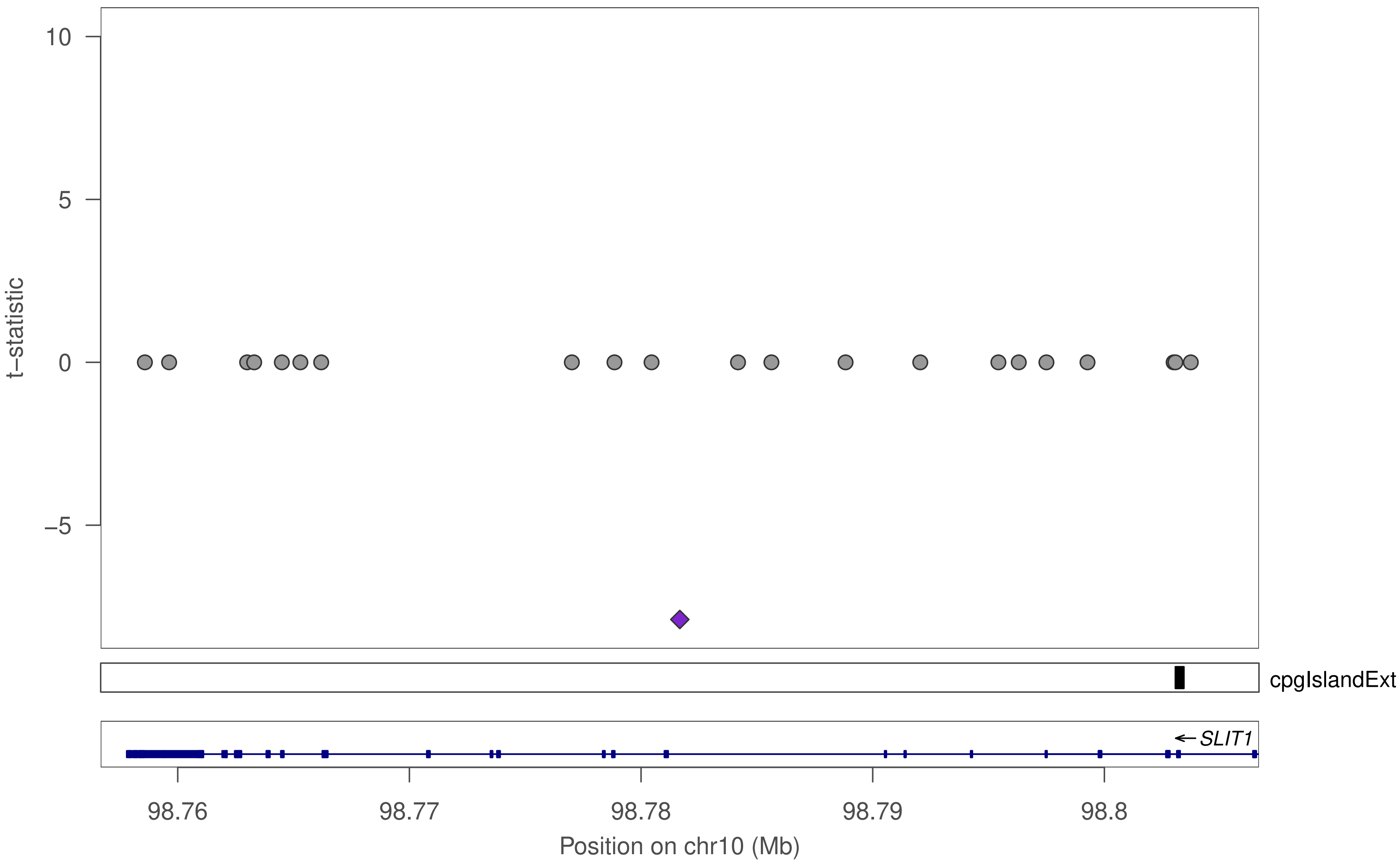

10
5
0
−5
t−statistic
cpgIslandExt
← SLIT1
98.76
98.77
98.78
98.79
98.8
Position on chr10 (Mb)

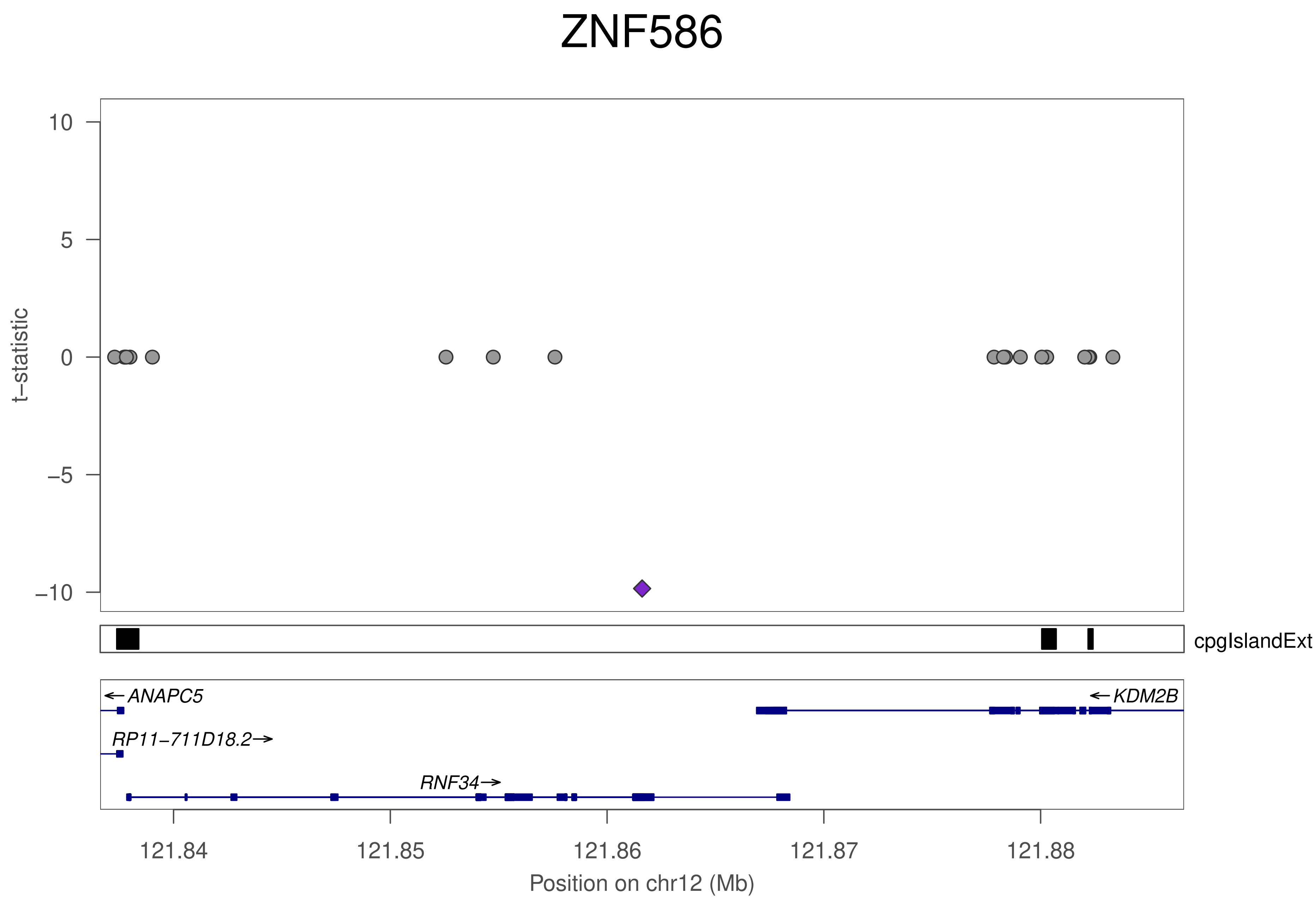

ZNF586
10
5
0
−5
−10
t−statistic
cpgIslandExt
ANAPC5
KDM2B
RP11−711D18.2
RNF34
121.84
121.85
121.86
121.87
121.88
Position on chr12 (Mb)

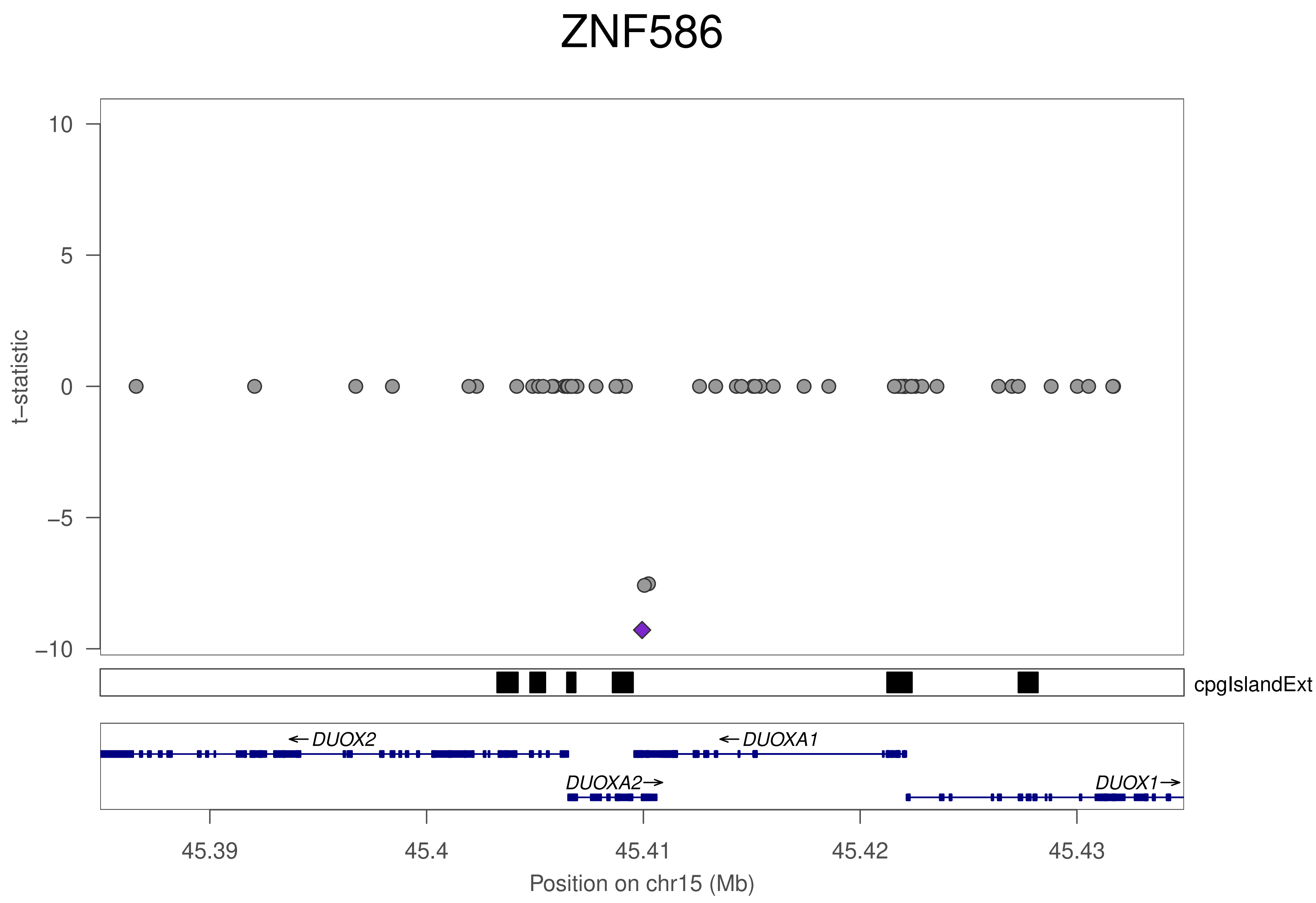

ZNF586
t−statistic
10
5
0
−5
−10
cpgIslandExt
← DUOX2
← DUOXA1
DUOXA2 →
DUOX1 →
45.39
45.4
45.41
45.42
45.43
Position on chr15 (Mb)

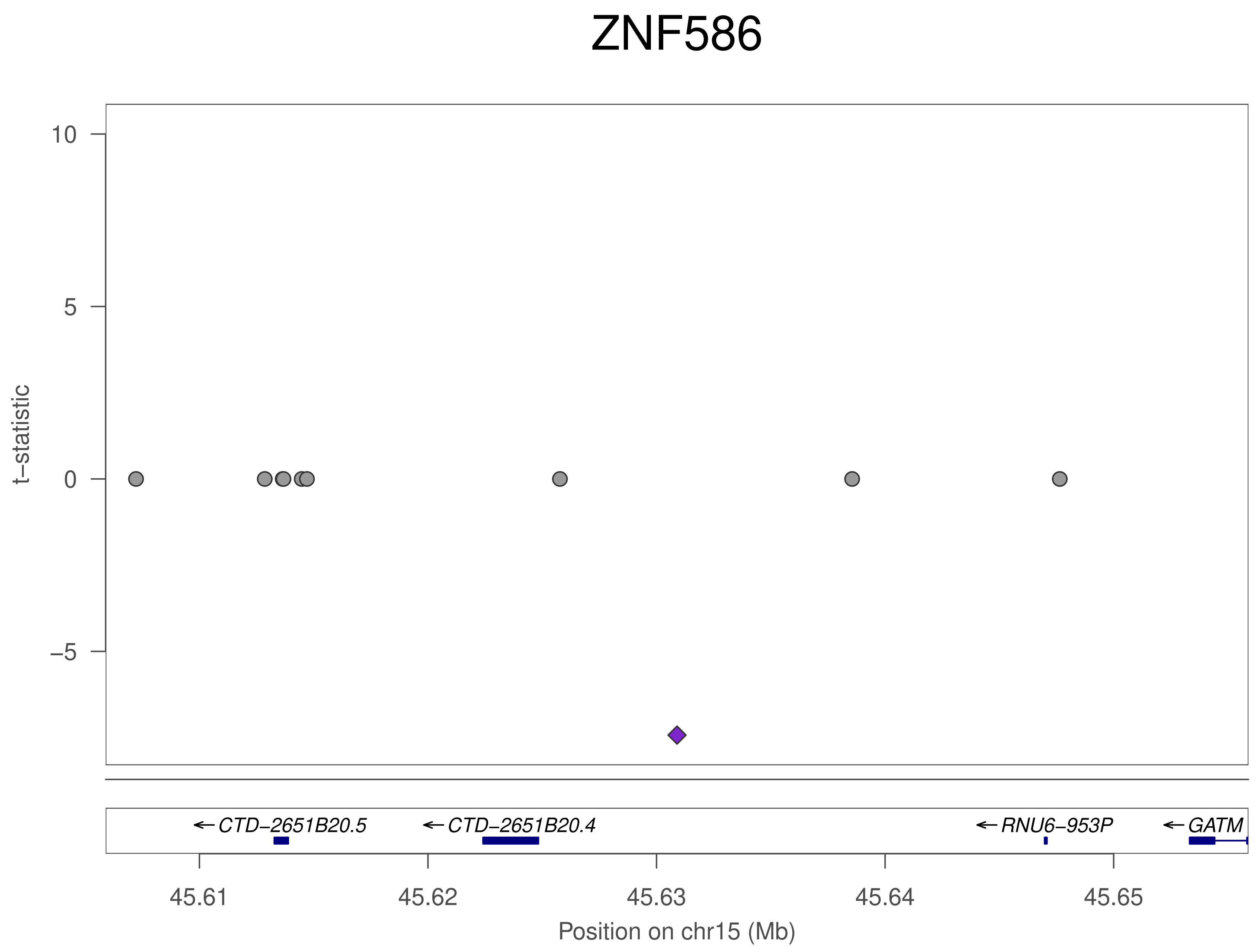
ZNF586
t−statistic
10
5
0
−5
CTD−2651B20.5
CTD−2651B20.4
RNU6−953P
GATM
45.61
45.62
45.63
45.64
45.65
Position on chr15 (Mb)

### ZNF586

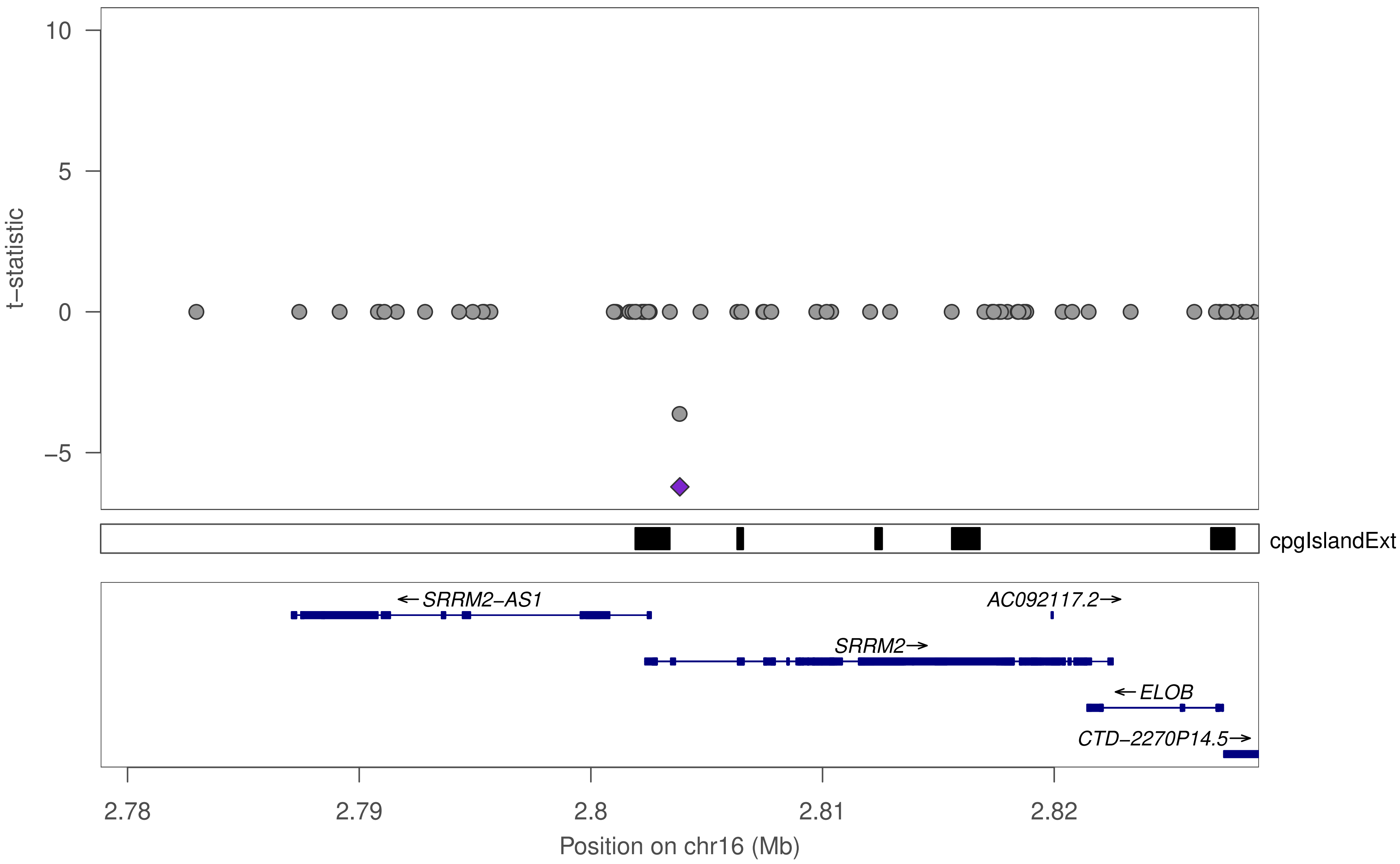

t−statistic
10
5
0
−5
cpgIslandExt
←SRRM2−AS1
AC092117.2→
SRRM2→
←ELOB
CTD−2270P14.5→
2.78
2.79
2.8
2.81
2.82
Position on chr16 (Mb)

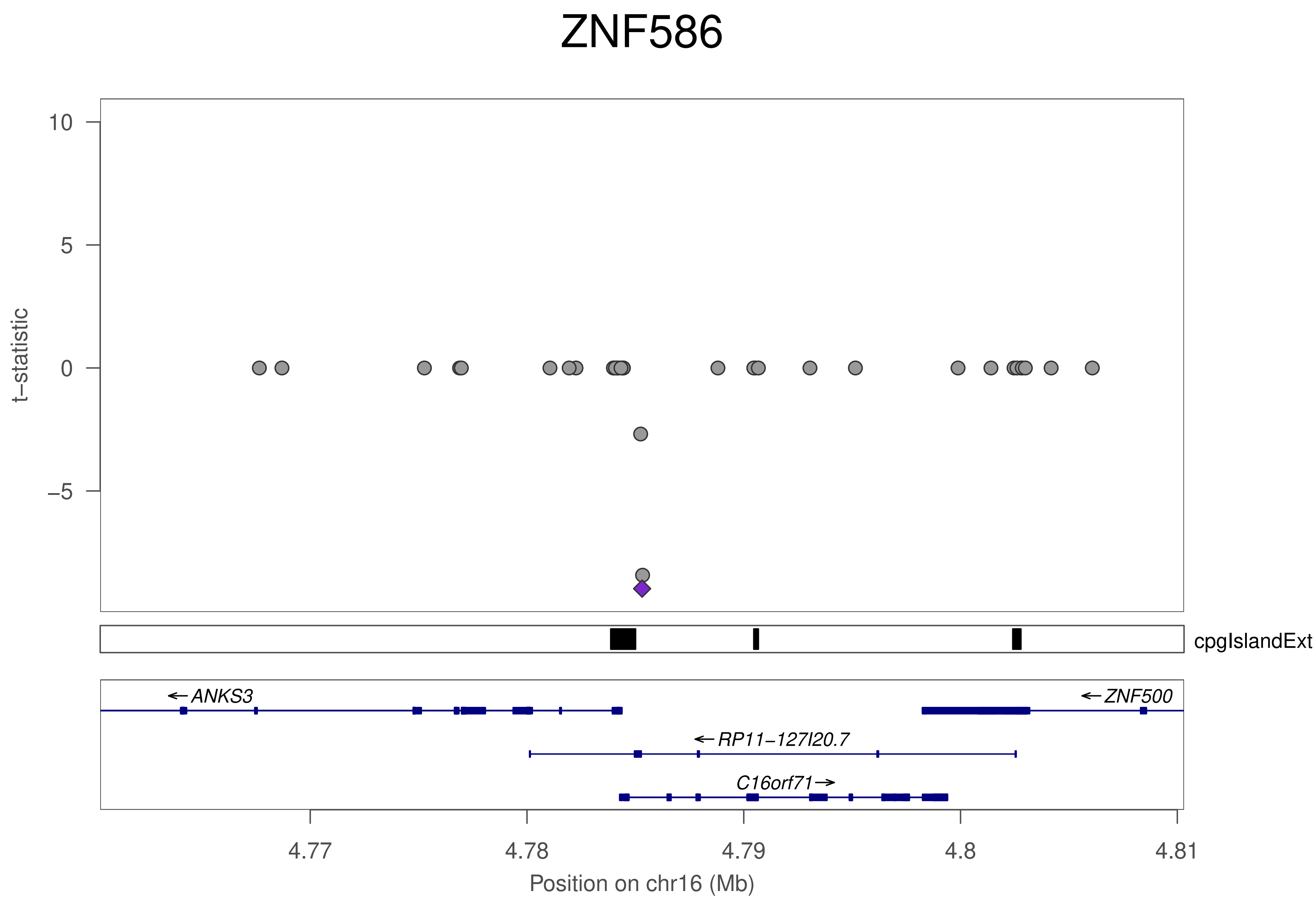

ZNF586
t−statistic
10
5
0
−5
cpgIslandExt
←ANKS3
←ZNF500
←RP11−127I20.7
C16orf71→
4.77
4.78
4.79
4.8
4.81
Position on chr16 (Mb)

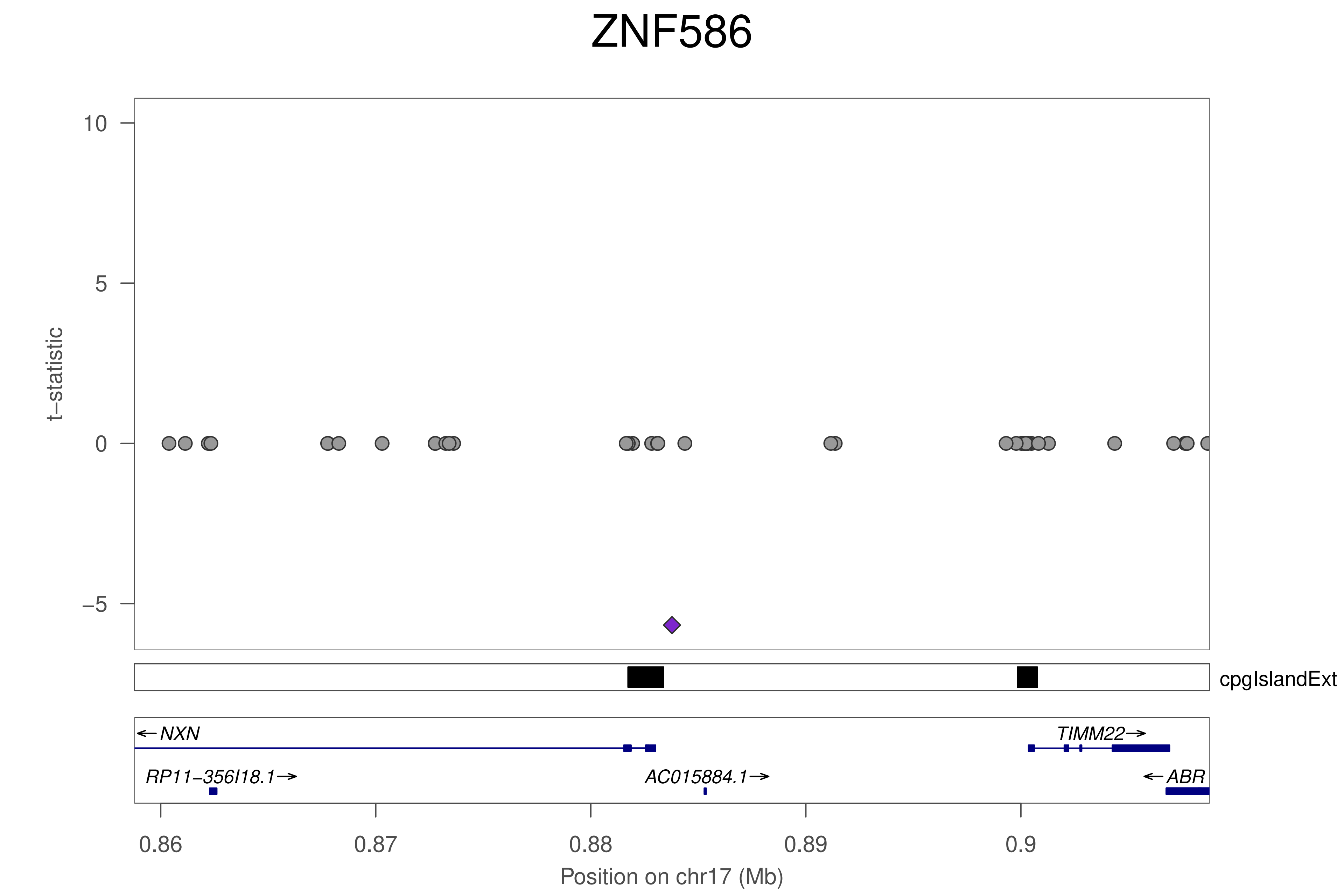
ZNF586
t−statistic
10
5
0
−5
cpgIslandExt
←NXN
TIMM22→
RP11−356I18.1→
AC015884.1→
←ABR
0.86
0.87
0.88
0.89
0.9
Position on chr17 (Mb)

### ZNF586

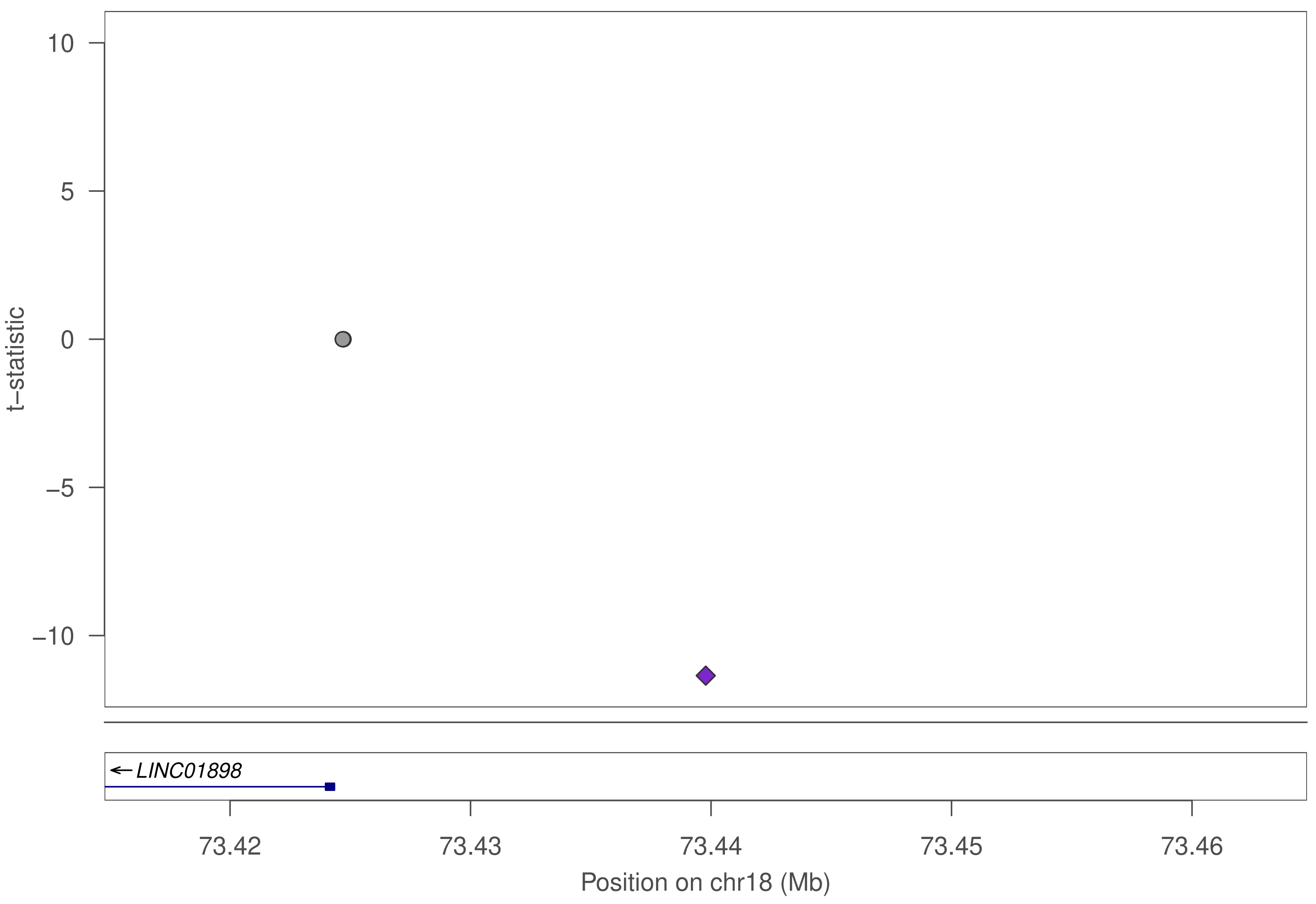

10
5
0
−5
−10
t−statistic
LINC01898
73.42
73.43
73.44
73.45
73.46
Position on chr18 (Mb)

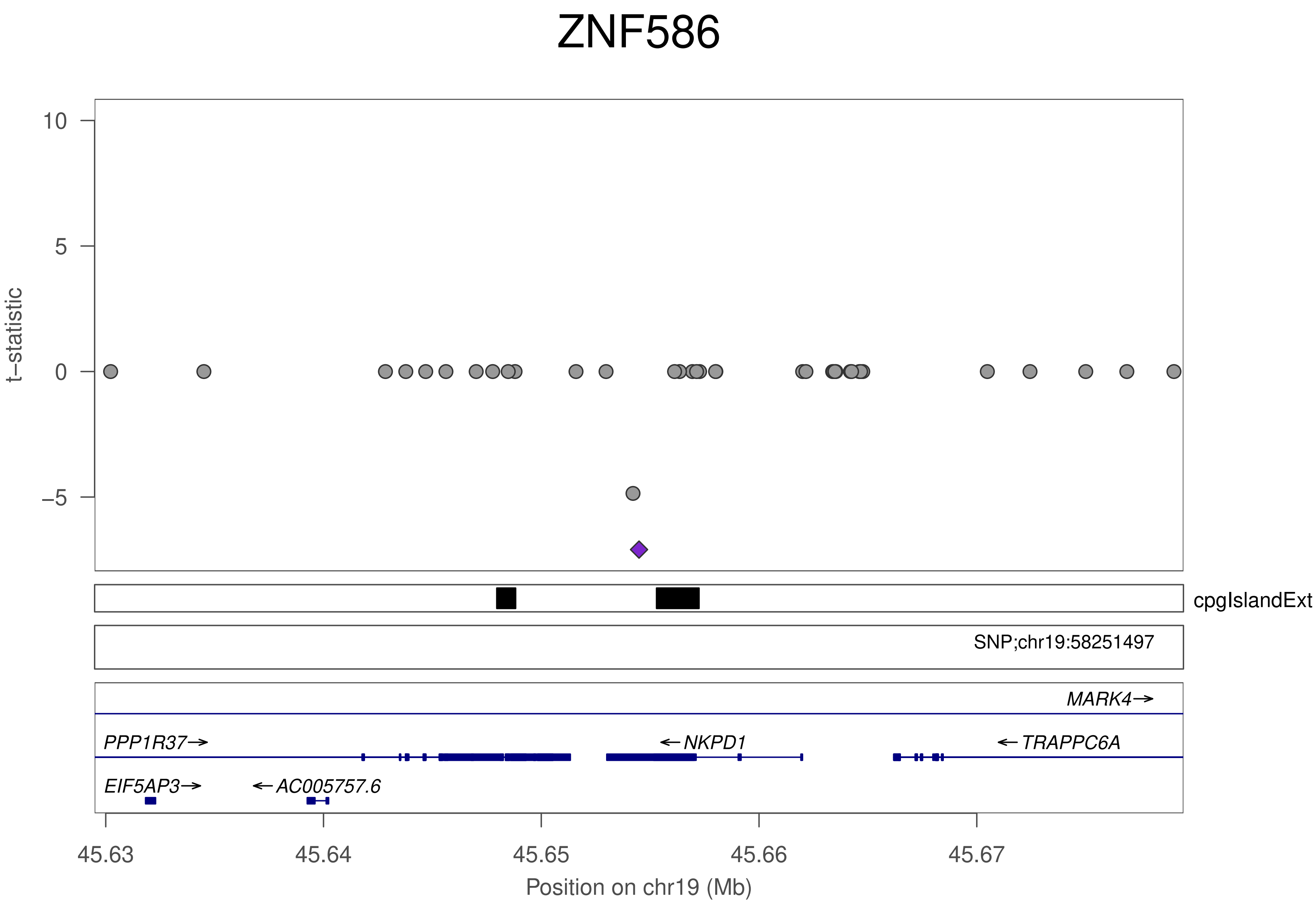
ZNF586
t−statistic
10
5
0
−5
cpgIslandExt
SNP;chr19:58251497
MARK4→
PPP1R37→
←NKPD1
←TRAPPC6A
EIF5AP3→
←AC005757.6
45.63
45.64
45.65
45.66
45.67
Position on chr19 (Mb)

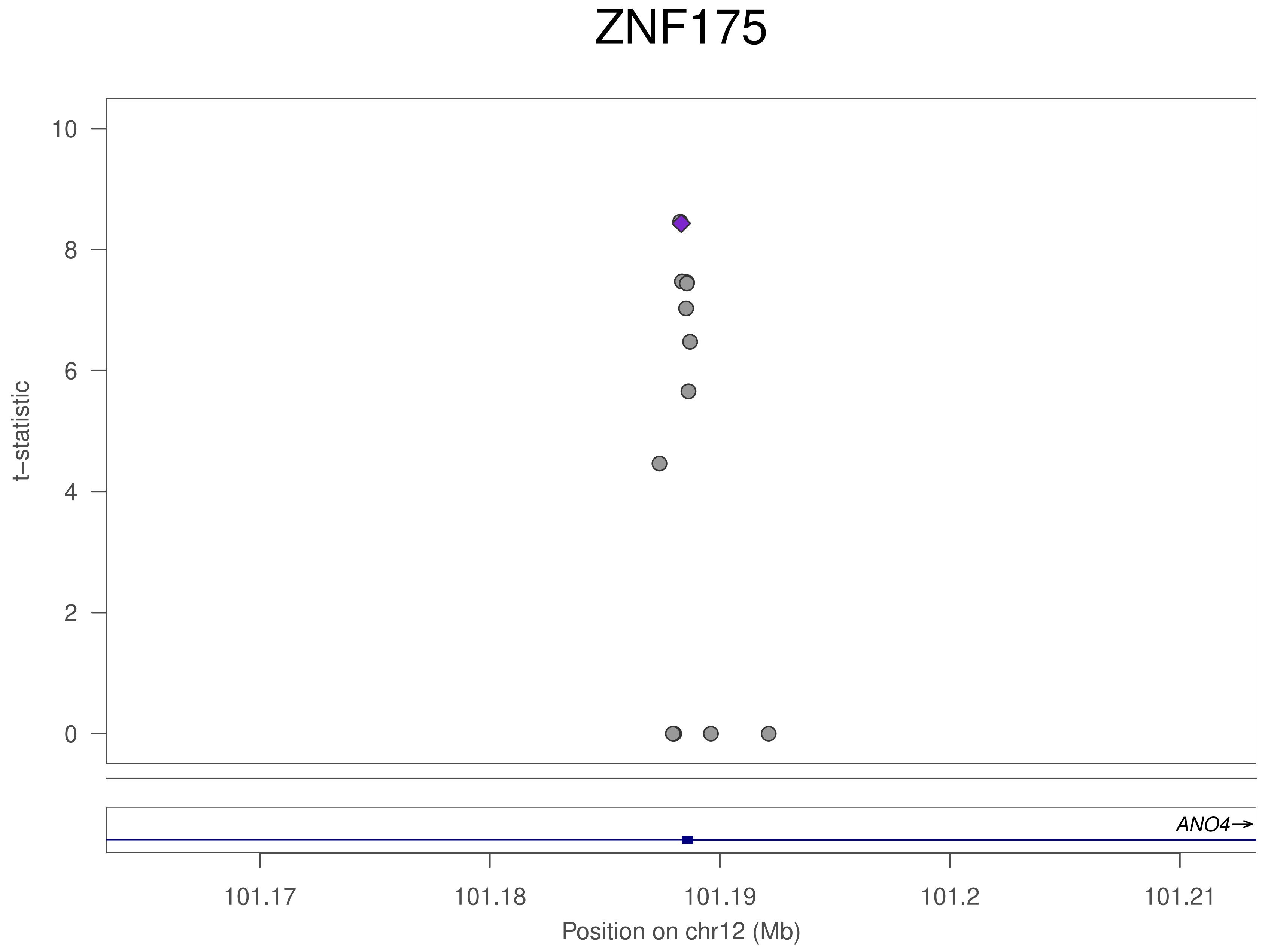
ZNF175
t−statistic
0
2
4
6
8
10
ANO4→
101.17
101.18
101.19
101.2
101.21
Position on chr12 (Mb)

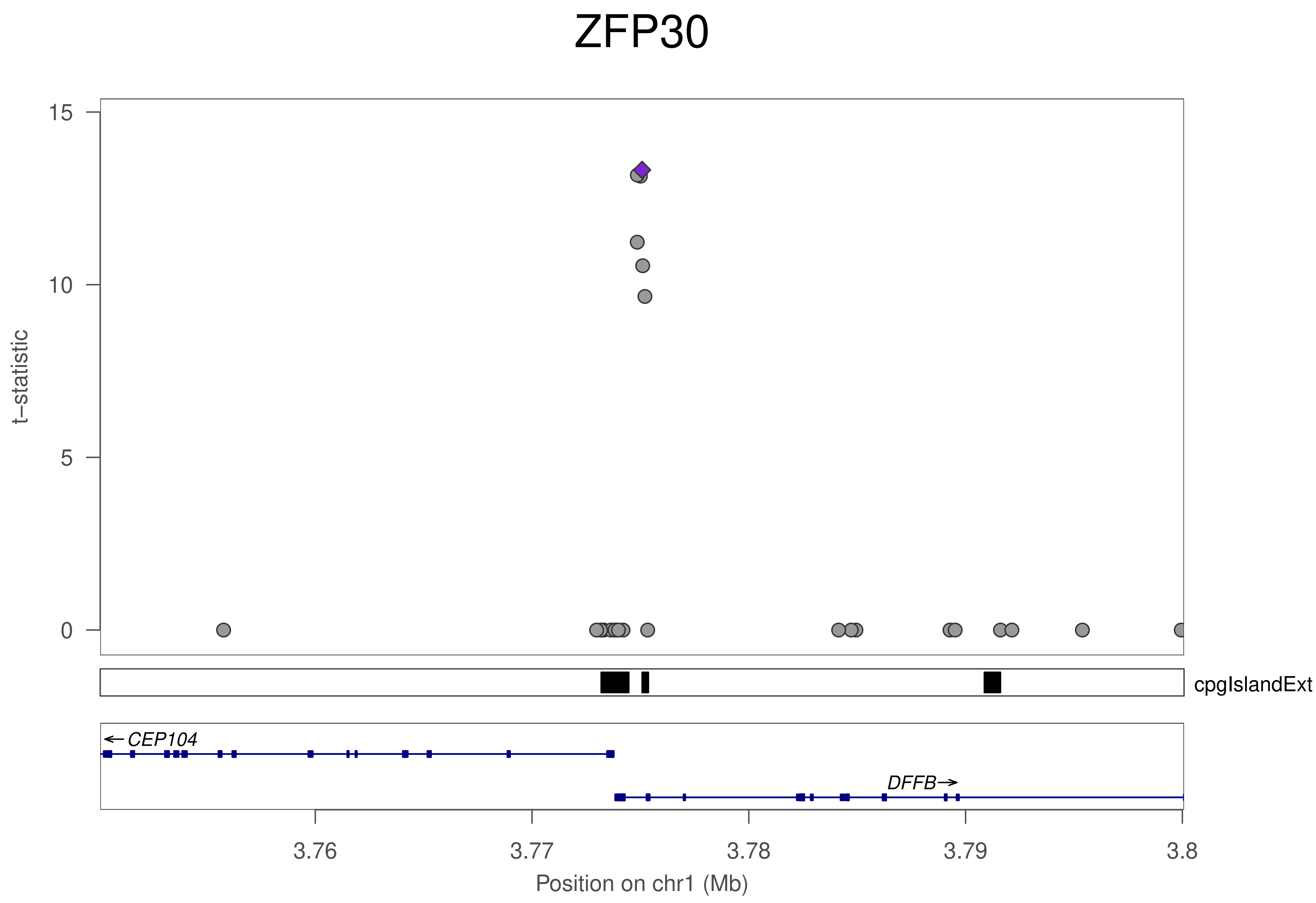
ZFP30
t−statistic
15
10
5
0
cpgIslandExt
CEP104
DFFB
3.76
3.77
3.78
3.79
3.8
Position on chr1 (Mb)

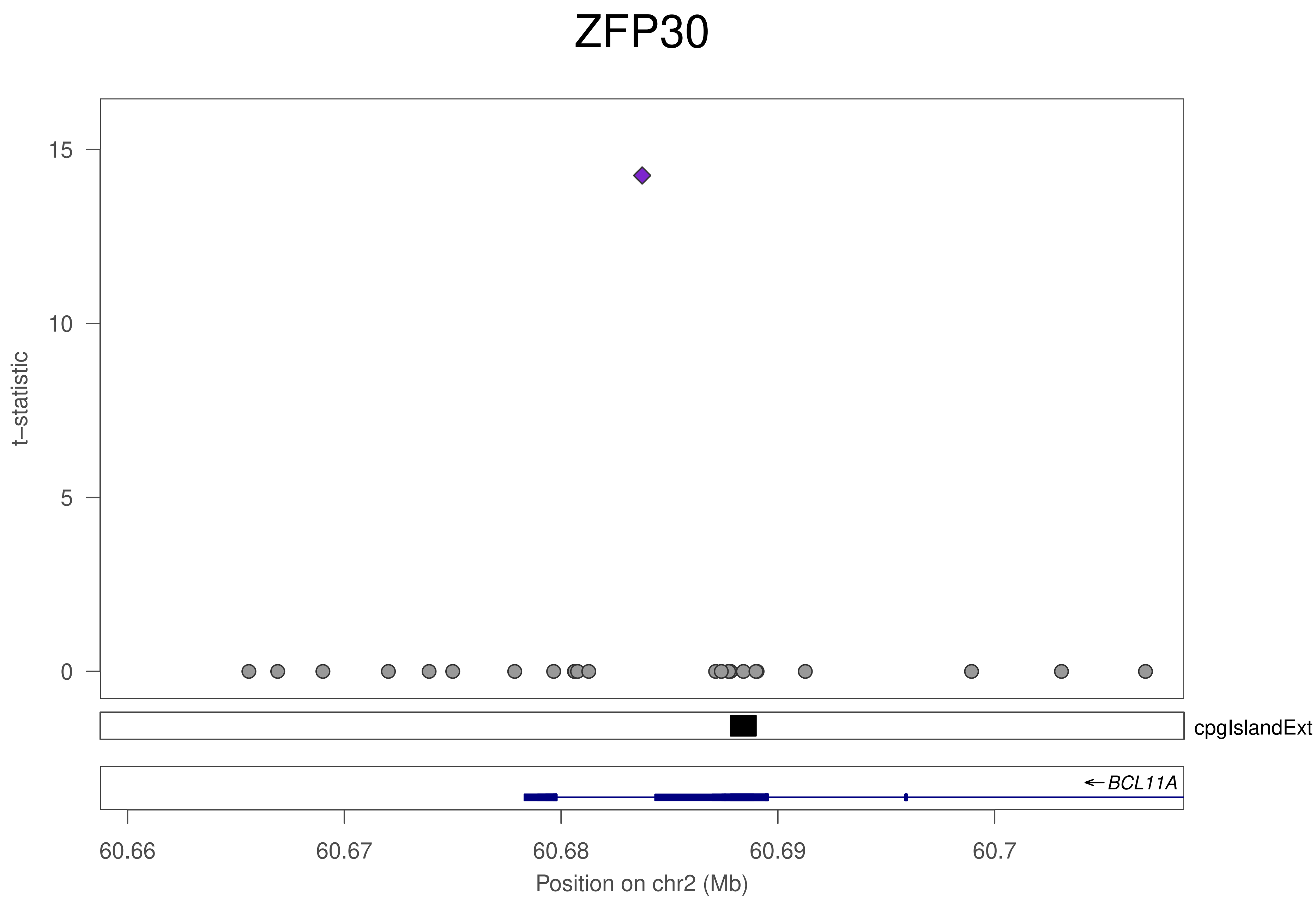
ZFP30
t–statistic
15
10
5
0
cpgIslandExt
←BCL11A
60.66
60.67
60.68
60.69
60.7
Position on chr2 (Mb)

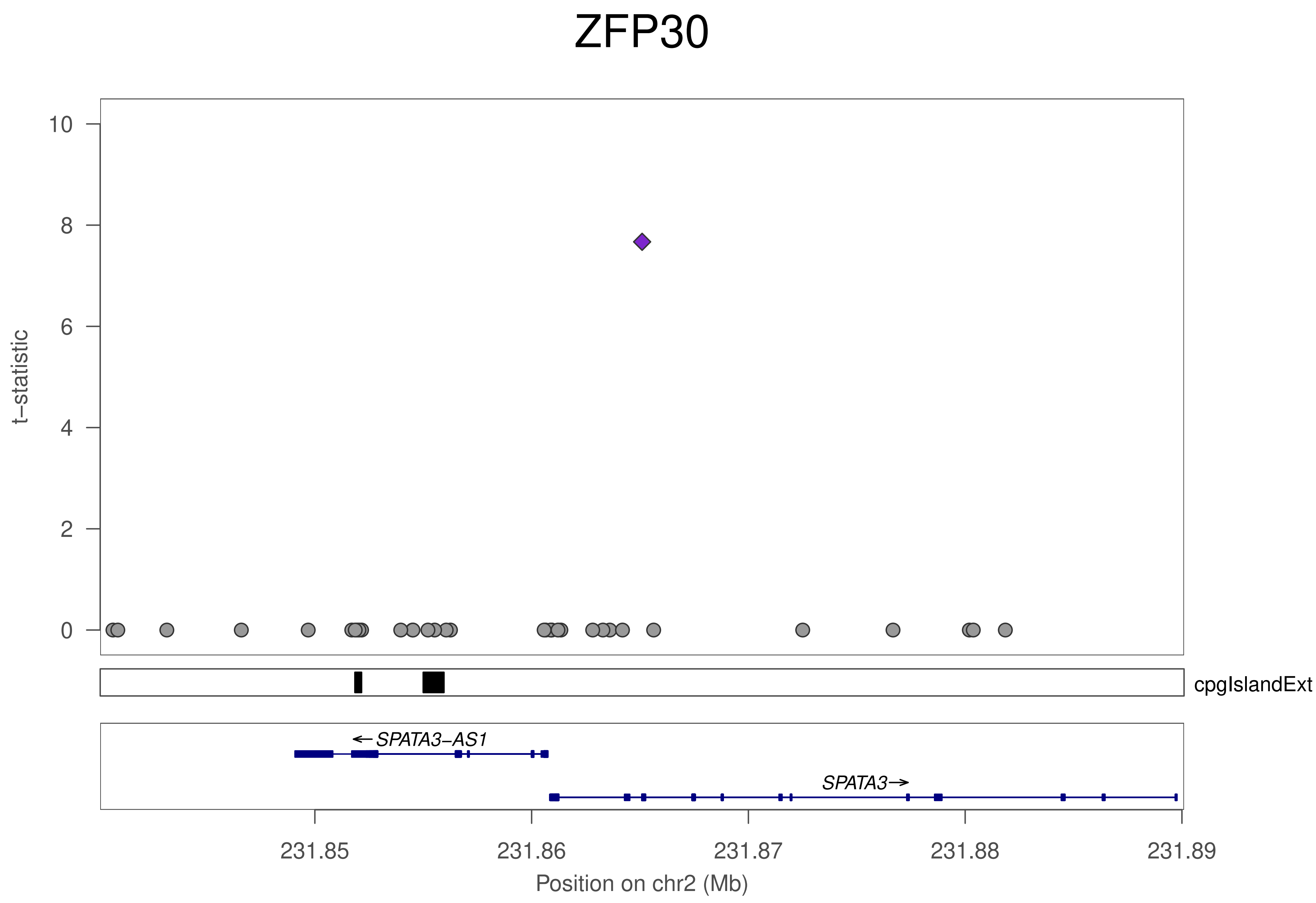

ZFP30
t–statistic
10
8
6
4
2
0
cpgIslandExt
←SPATA3–AS1
SPATA3→
231.85
231.86
231.87
231.88
231.89
Position on chr2 (Mb)

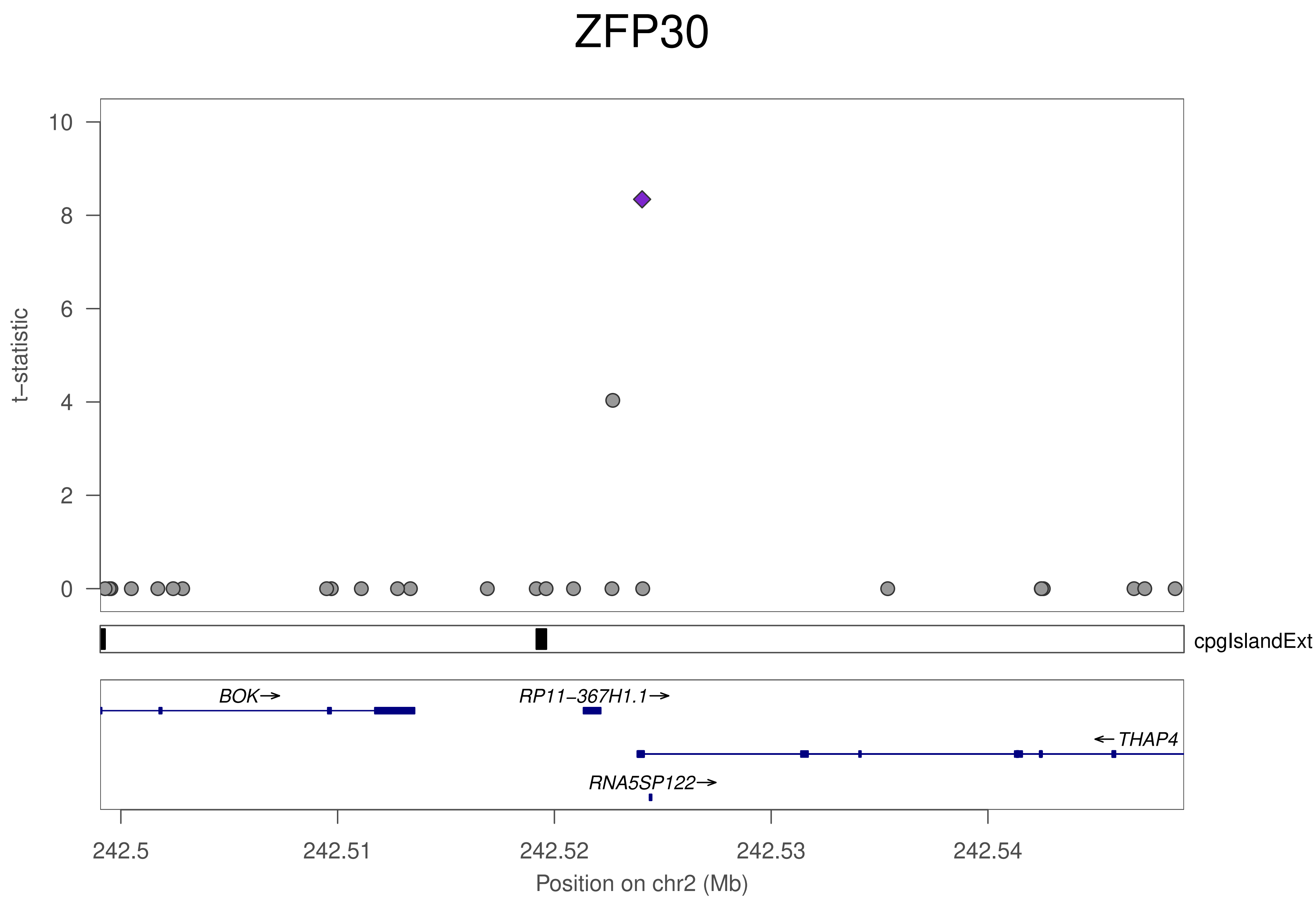

ZFP30
10
8
6
4
2
0
t−statistic
cpgIslandExt
BOK→
RP11−367H1.1→
←THAP4
RNA5SP122→
242.5
242.51
242.52
242.53
242.54
Position on chr2 (Mb)

### ZFP30

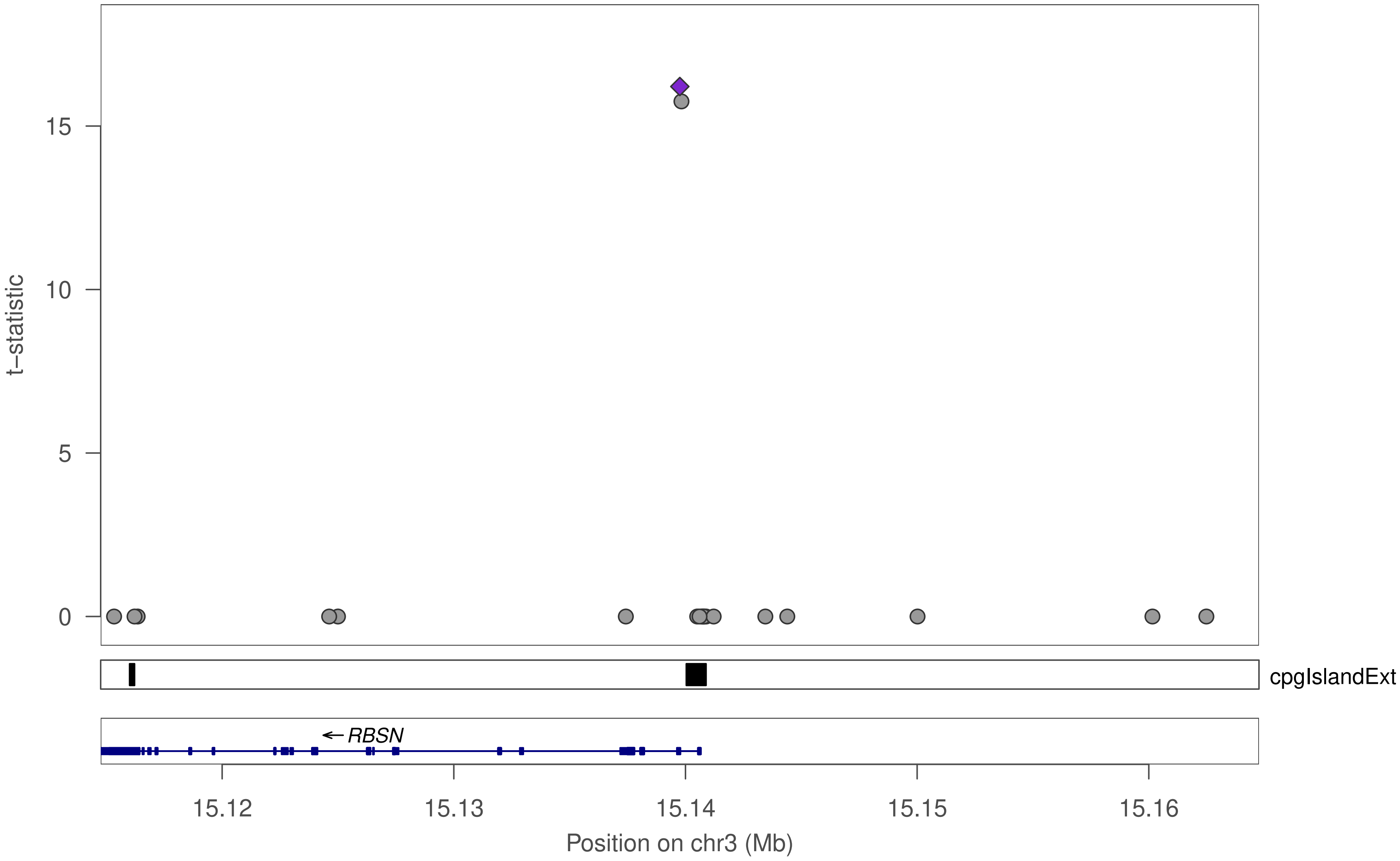

t−statistic
15
10
5
0
cpgIslandExt
← RBSN
15.12
15.13
15.14
15.15
15.16
Position on chr3 (Mb)

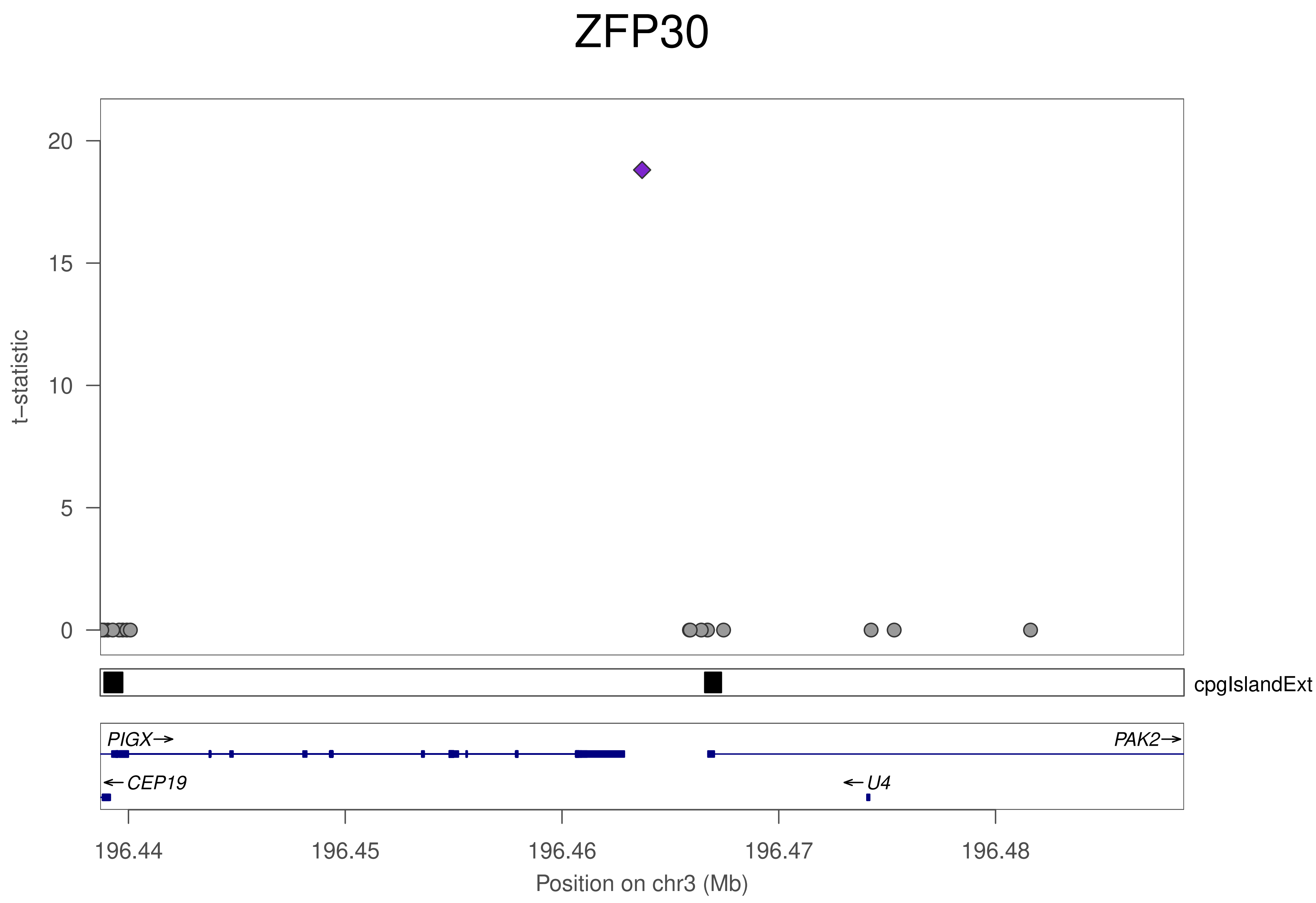
ZFP30
t−statistic
20
15
10
5
0
cpgIslandExt
PIGX→
PAK2→
←CEP19
←U4
196.44
196.45
196.46
196.47
196.48
Position on chr3 (Mb)

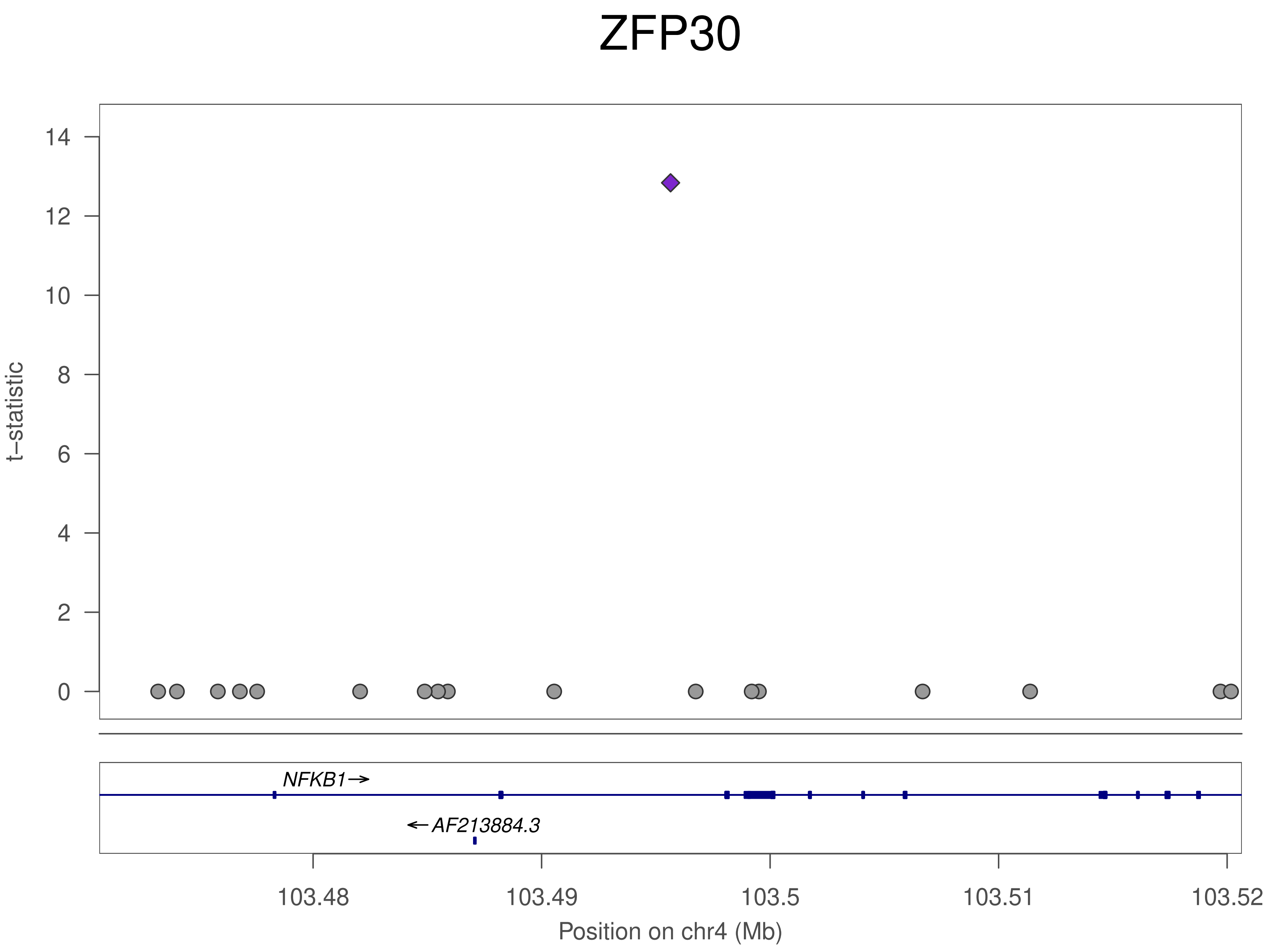
ZFP30
t–statistic
0
2
4
6
8
10
12
14
NFKB1→
←AF213884.3
103.48
103.49
103.5
103.51
103.52
Position on chr4 (Mb)

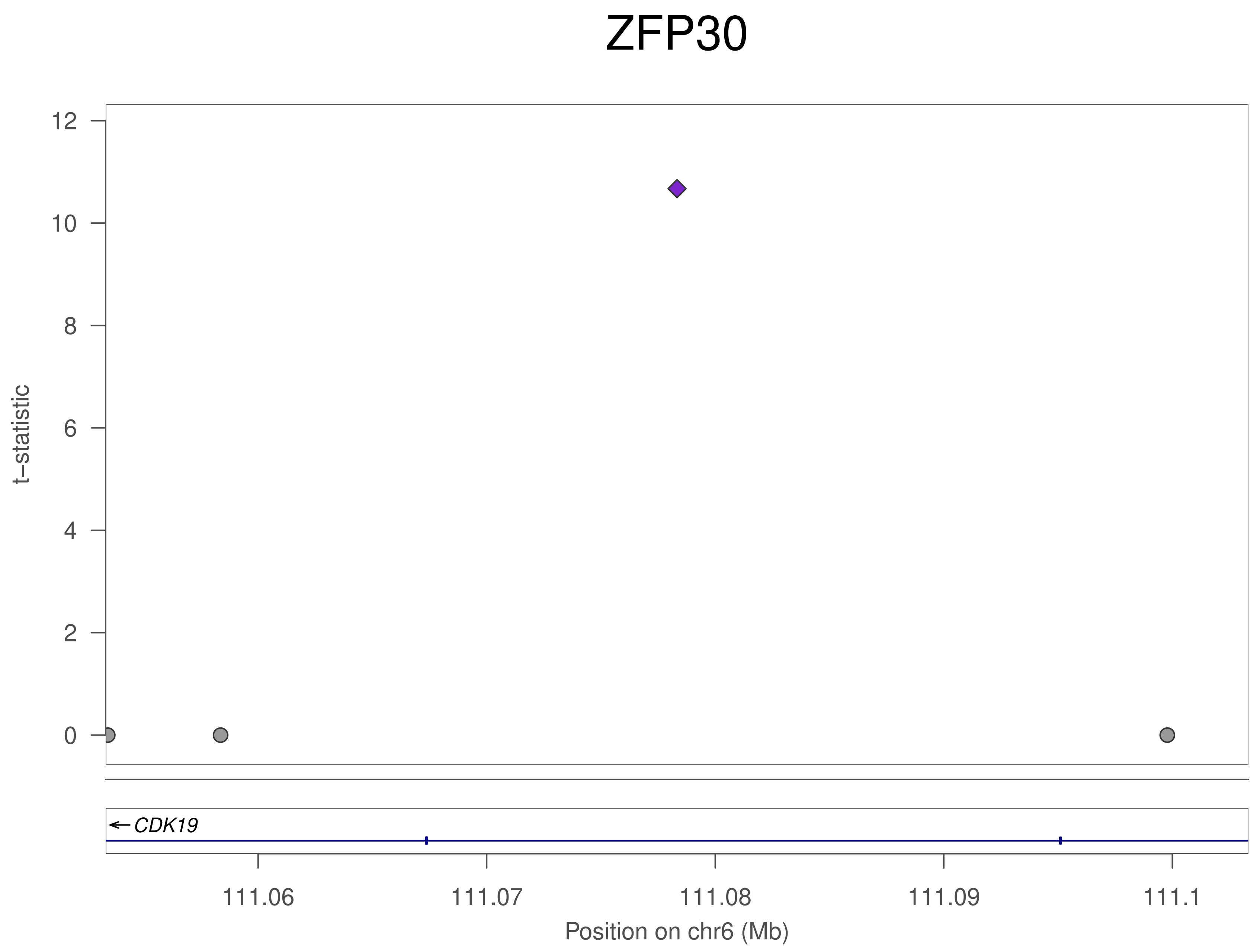

ZFP30
t–statistic
12
10
8
6
4
2
0
← CDK19
111.06
111.07
111.08
111.09
111.1
Position on chr6 (Mb)

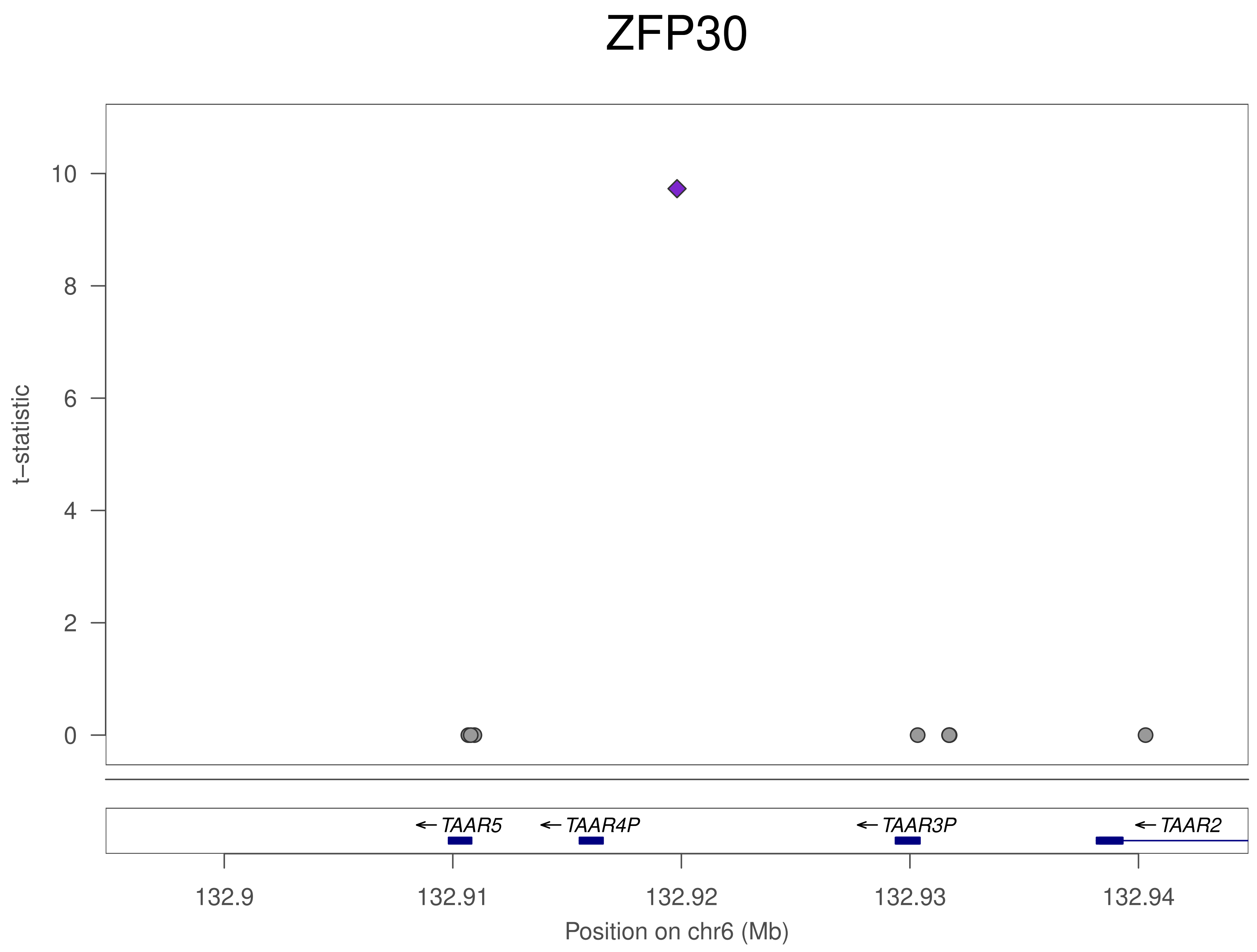
ZFP30
t–statistic
10
8
6
4
2
0
TAAR5
TAAR4P
TAAR3P
TAAR2
132.9
132.91
132.92
132.93
132.94
Position on chr6 (Mb)

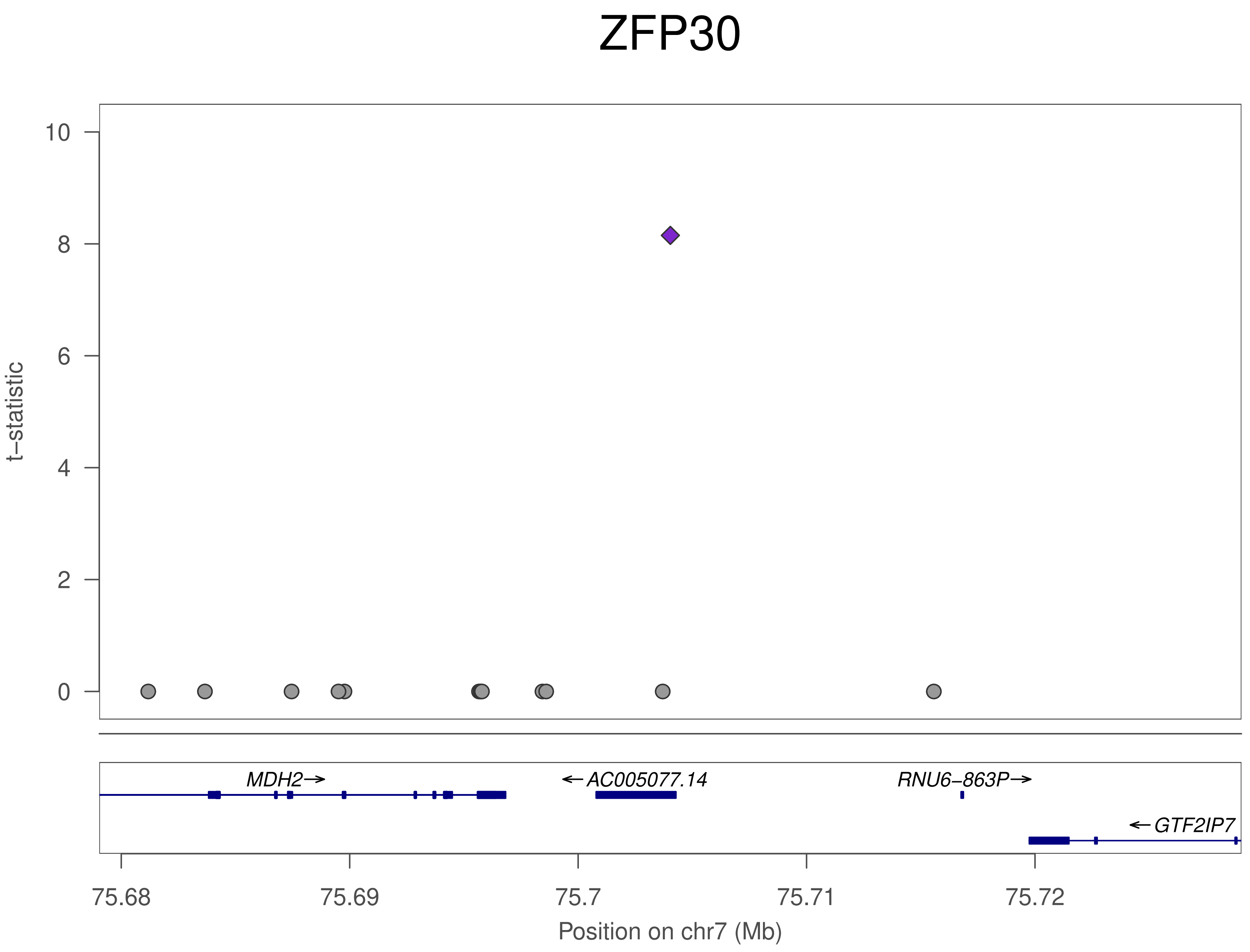
ZFP30
t−statistic
10
8
6
4
2
0
MDH2→
←AC005077.14
RNU6−863P→
←GTF2IP7
75.68
75.69
75.7
75.71
75.72
Position on chr7 (Mb)

### ZFP30

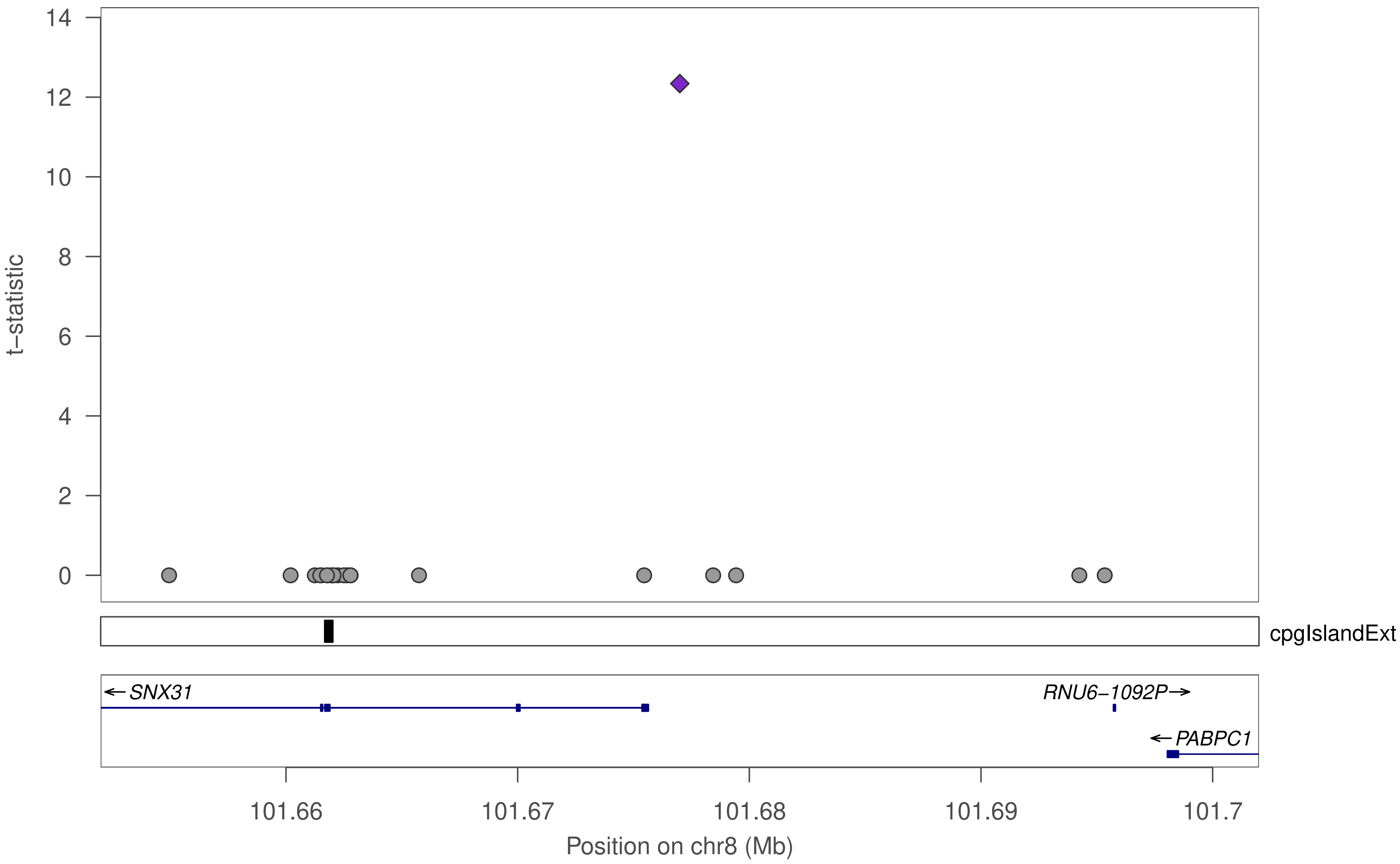

t−statistic
0
2
4
6
8
10
12
14
cpgIslandExt
←SNX31
RNU6−1092P→
←PABPC1
101.66
101.67
101.68
101.69
101.7
Position on chr8 (Mb)

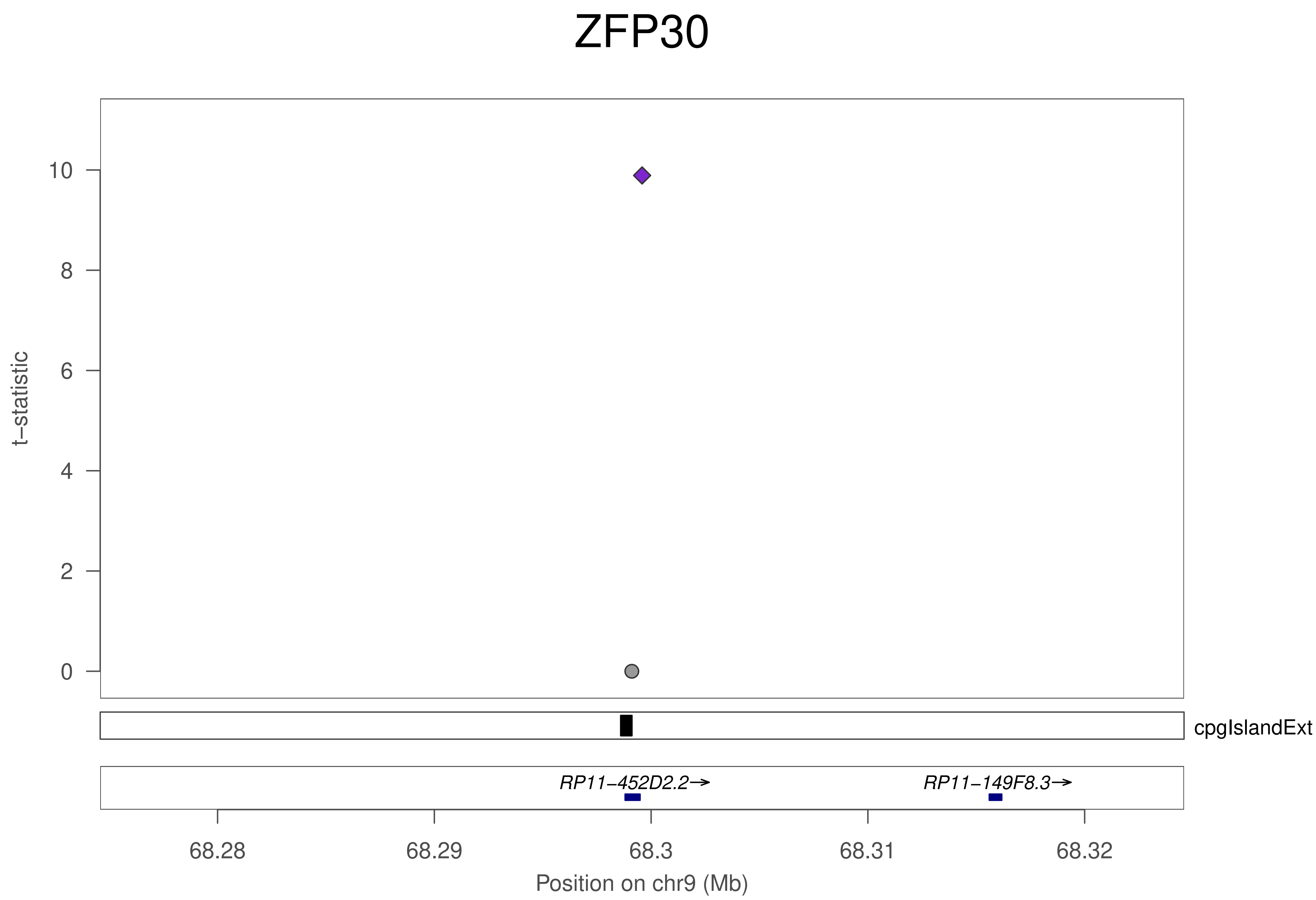
ZFP30
t−statistic
10
8
6
4
2
0
cpgIslandExt
RP11−452D2.2→
RP11−149F8.3→
68.28
68.29
68.3
68.31
68.32
Position on chr9 (Mb)

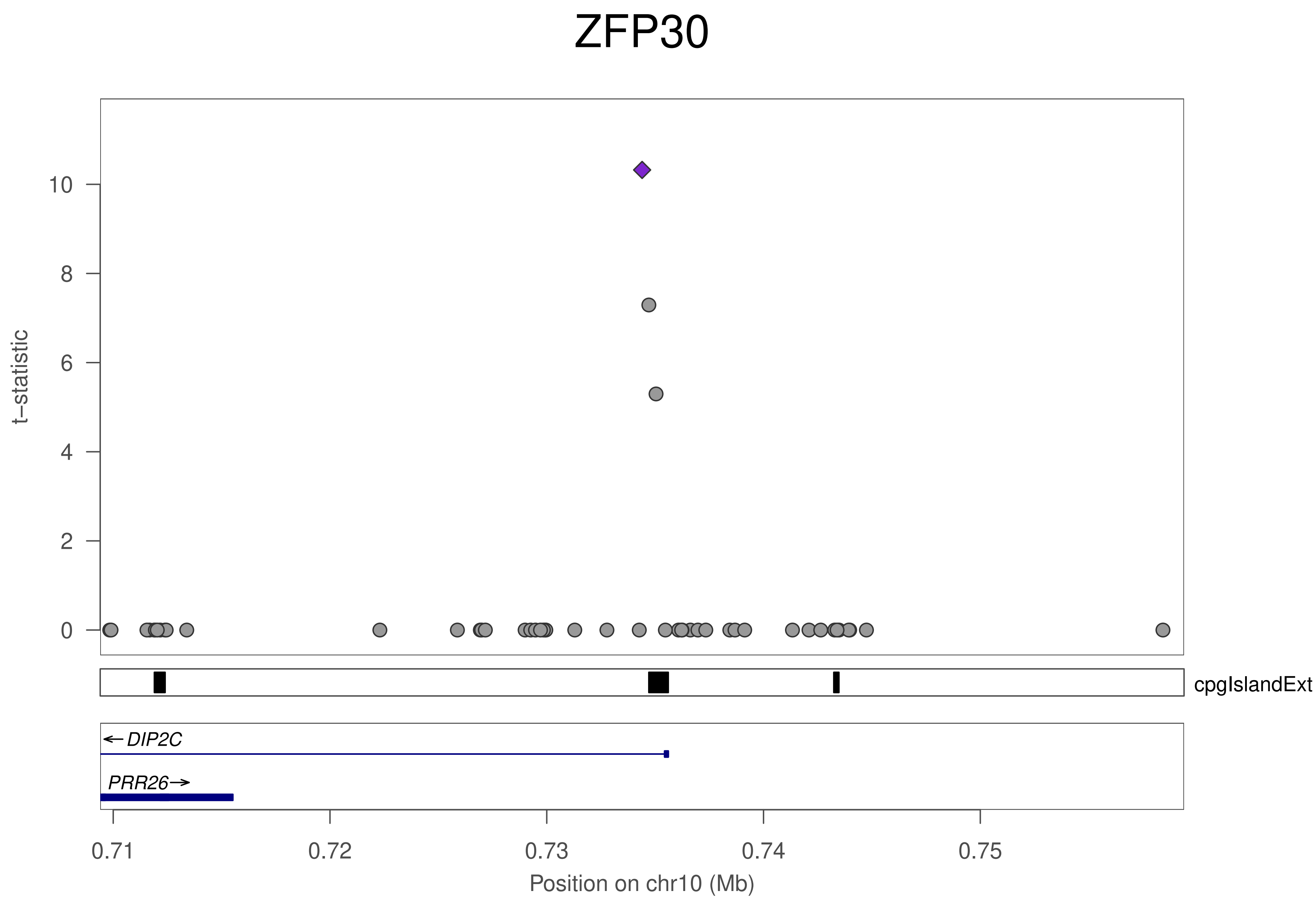

ZFP30
t–statistic
10
8
6
4
2
0
cpgIslandExt
DIP2C
PRR26
0.71
0.72
0.73
0.74
0.75
Position on chr10 (Mb)

### ZFP30

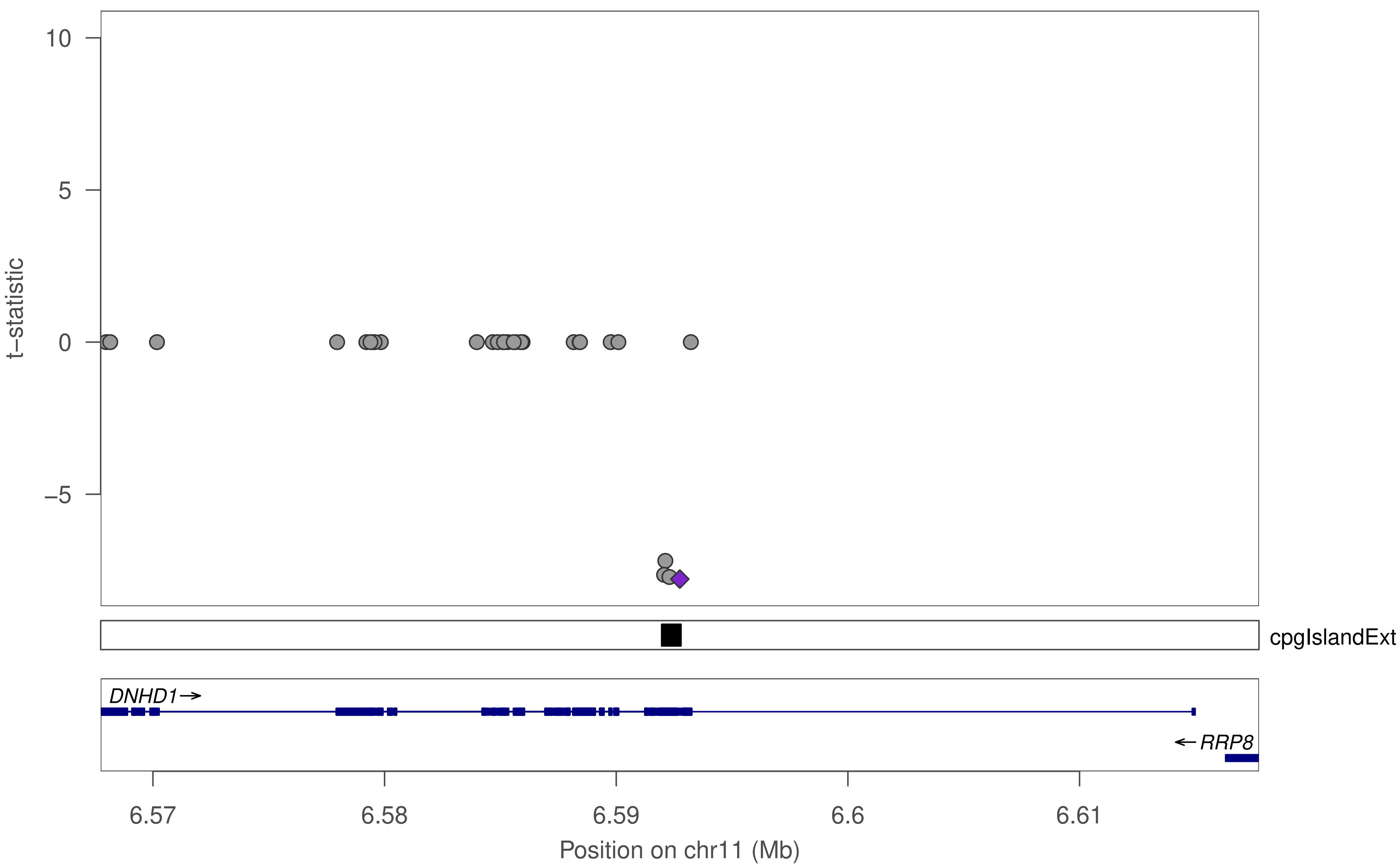

10
5
0
−5
t−statistic
cpgIslandExt
DNHD1→
←RRP8
6.57
6.58
6.59
6.6
6.61
Position on chr11 (Mb)

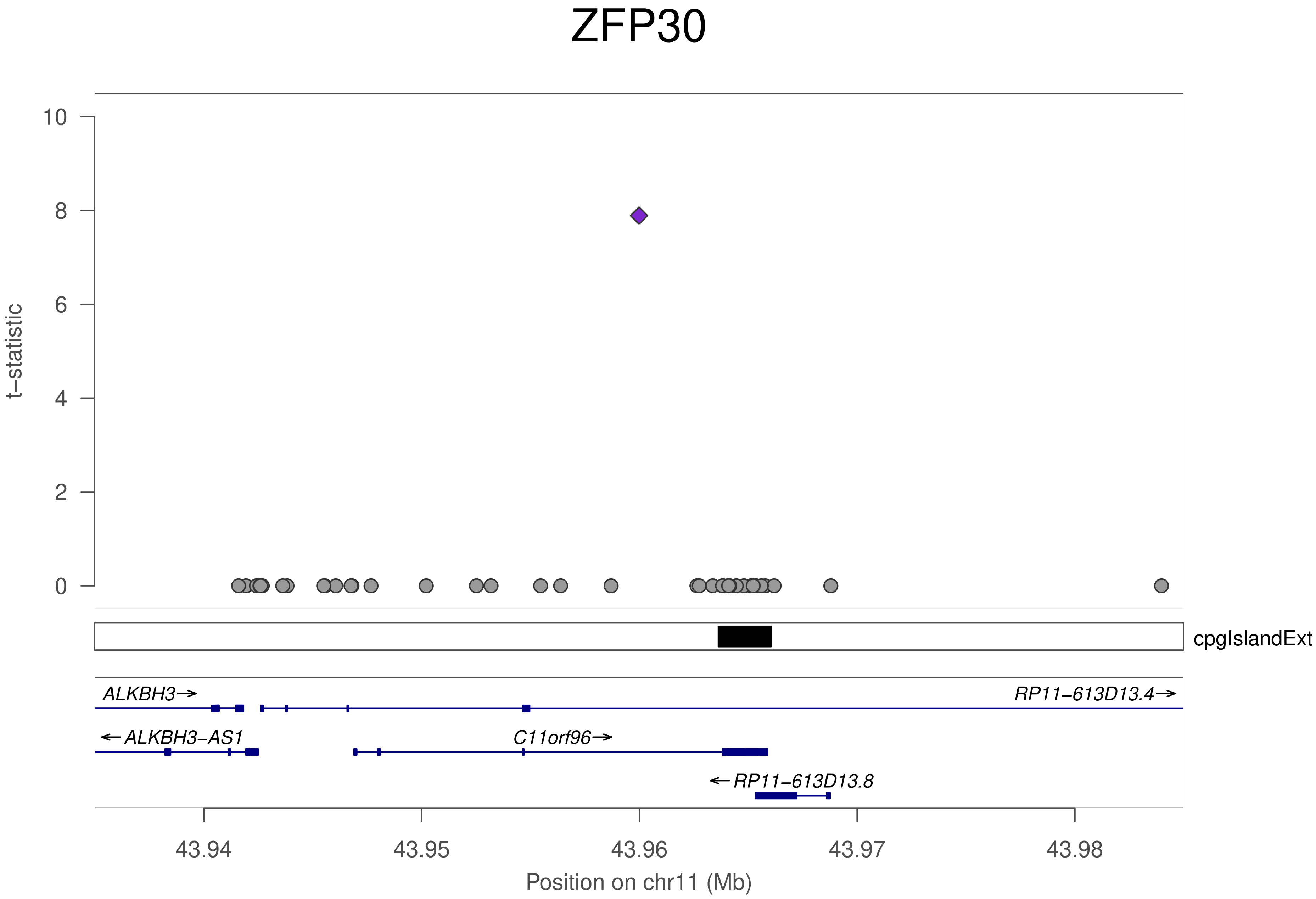
ZFP30
t−statistic
10
8
6
4
2
0
cpgIslandExt
ALKBH3→
RP11−613D13.4→
←ALKBH3−AS1
C11orf96→
←RP11−613D13.8
43.94
43.95
43.96
43.97
43.98
Position on chr11 (Mb)

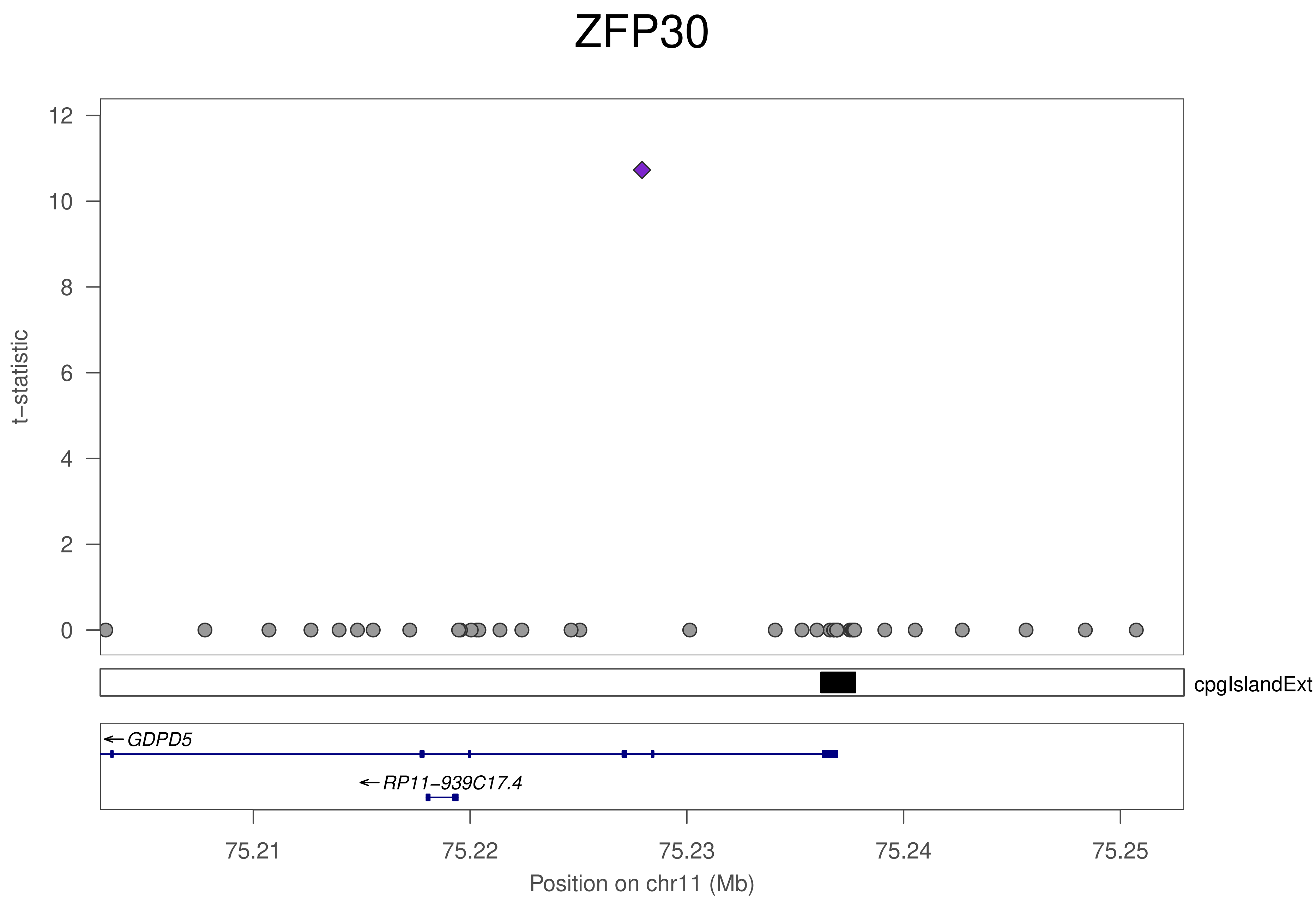
ZFP30
t−statistic
12
10
8
6
4
2
0
cpgIslandExt
GDPD5
RP11−939C17.4
75.21
75.22
75.23
75.24
75.25
Position on chr11 (Mb)

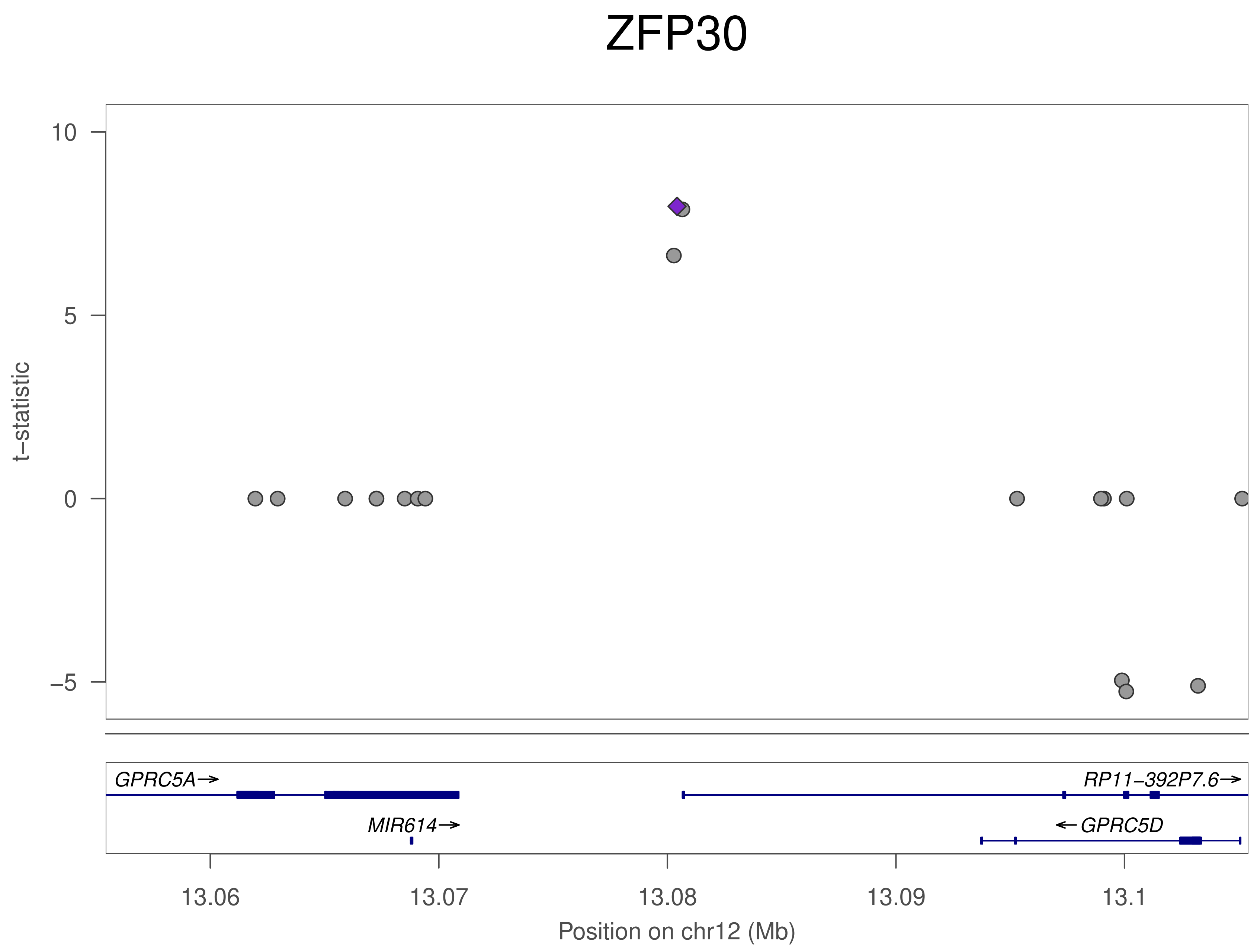
ZFP30
10
5
0
−5
t−statistic
GPRC5A→
RP11−392P7.6→
MIR614→
←GPRC5D
13.06
13.07
13.08
13.09
13.1
Position on chr12 (Mb)

### ZFP30

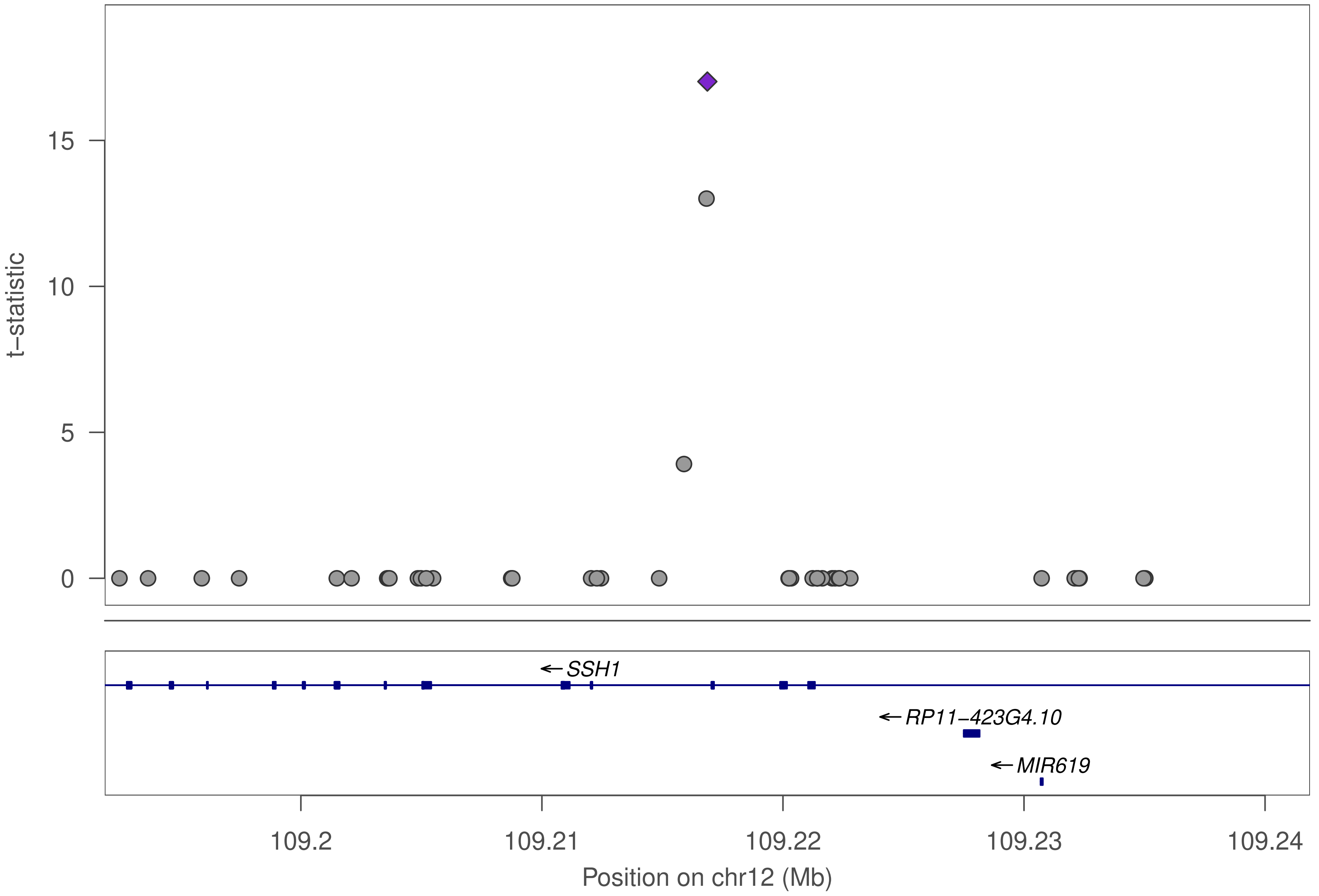

t−statistic
0
5
10
15
←SSH1
←RP11−423G4.10
←MIR619
109.2
109.21
109.22
109.23
109.24
Position on chr12 (Mb)

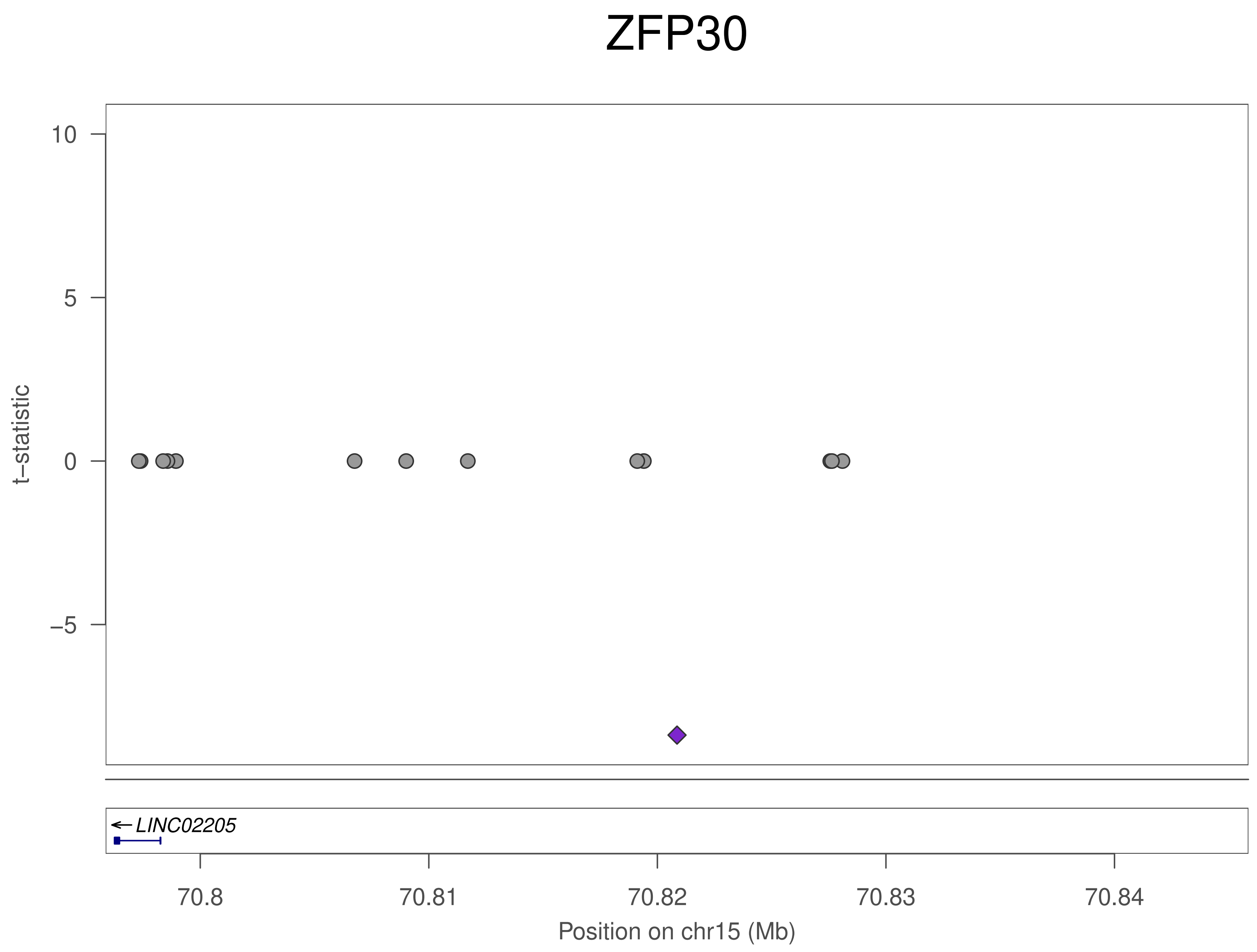
ZFP30
10
5
0
−5
t−statistic
LINC02205
70.8
70.81
70.82
70.83
70.84
Position on chr15 (Mb)

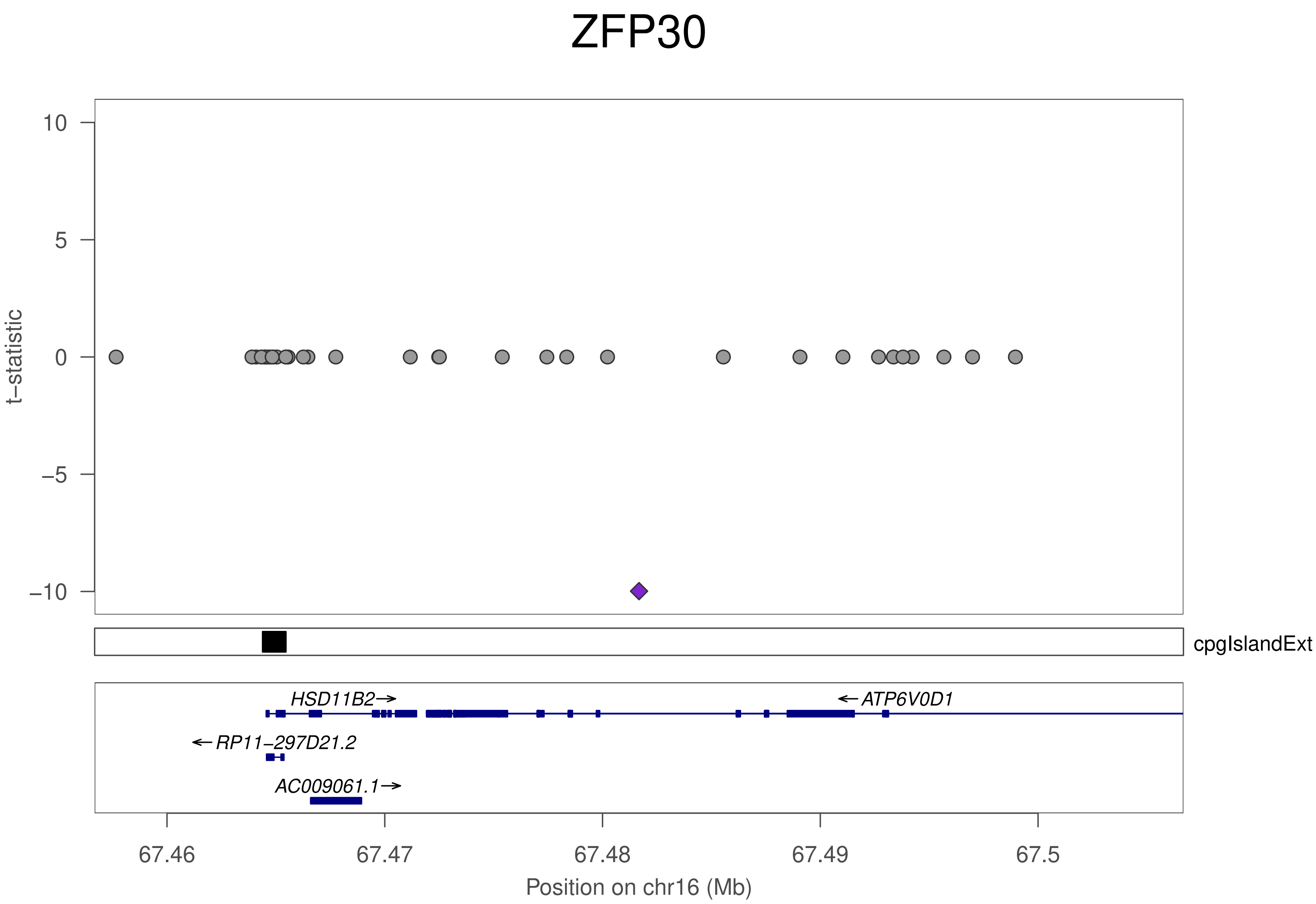
ZFP30
t−statistic
10
5
0
−5
−10
cpgIslandExt
HSD11B2→
←ATP6V0D1
←RP11−297D21.2
AC009061.1→
67.46
67.47
67.48
67.49
67.5
Position on chr16 (Mb)

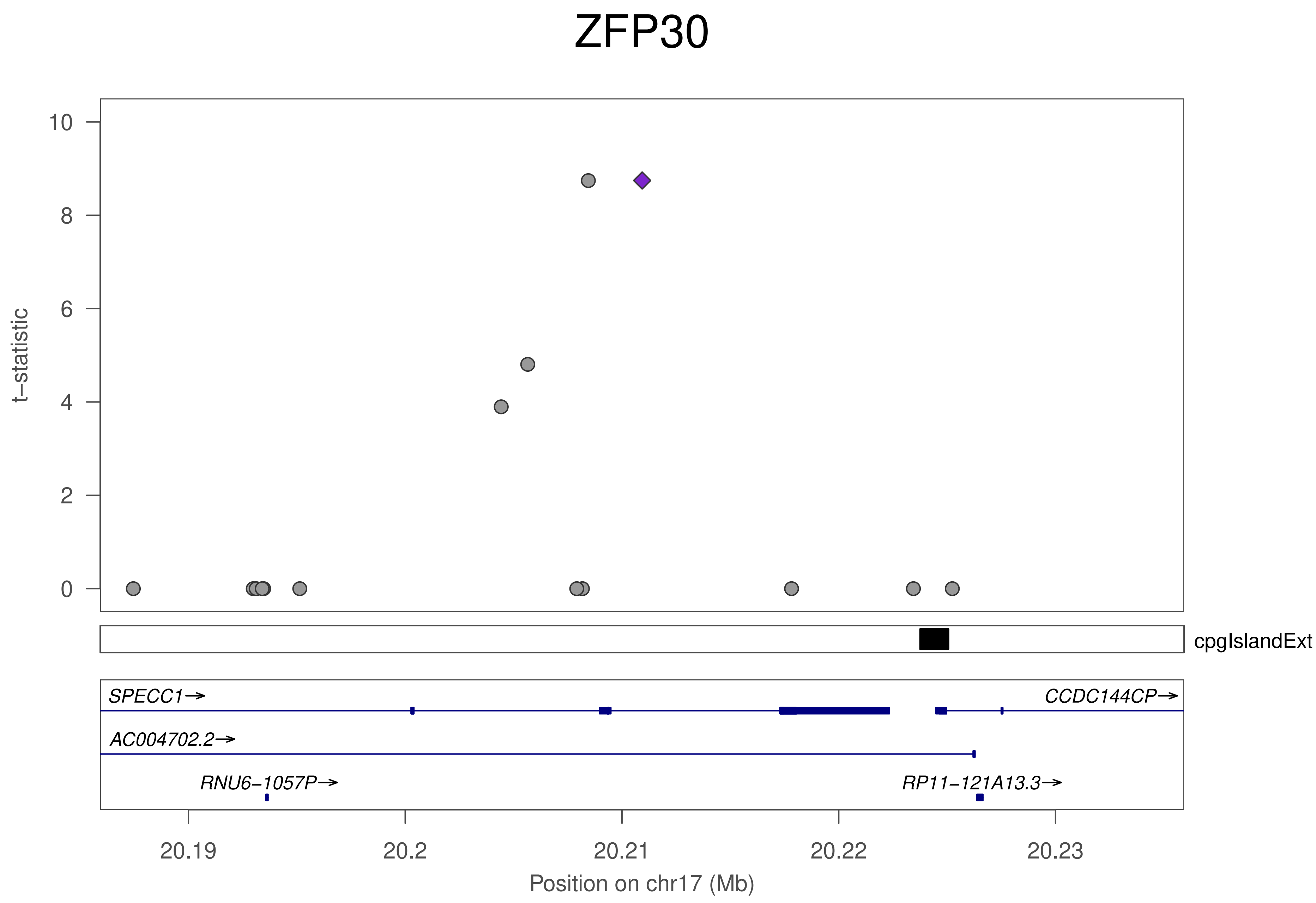
ZFP30
t−statistic
10
8
6
4
2
0
cpgIslandExt
SPECC1→
CCDC144CP→
AC004702.2→
RNU6−1057P→
RP11−121A13.3→
20.19
20.2
20.21
20.22
20.23
Position on chr17 (Mb)

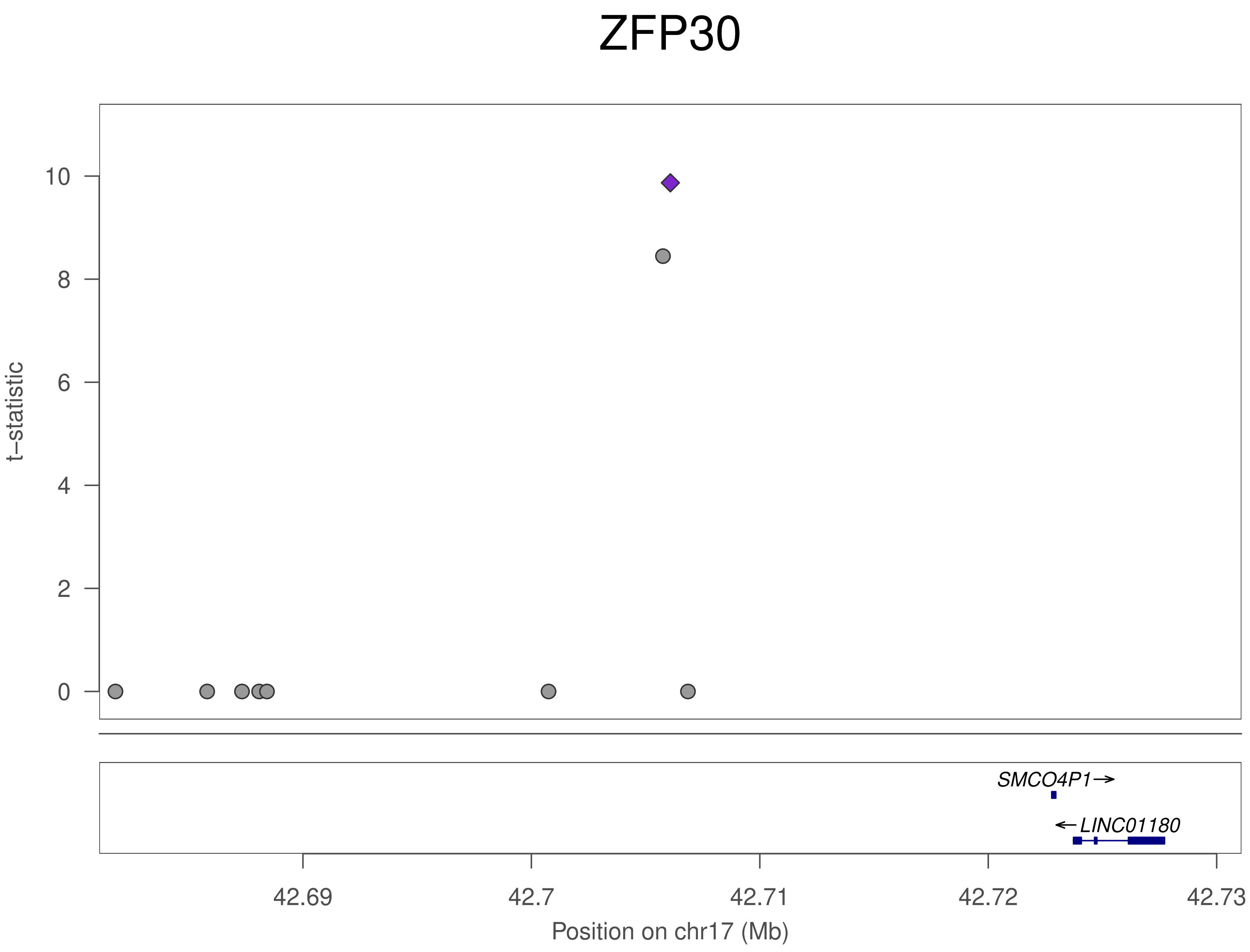

ZFP30
t–statistic
10
8
6
4
2
0
SMCO4P1→
←LINC01180
42.69
42.7
42.71
42.72
42.73
Position on chr17 (Mb)

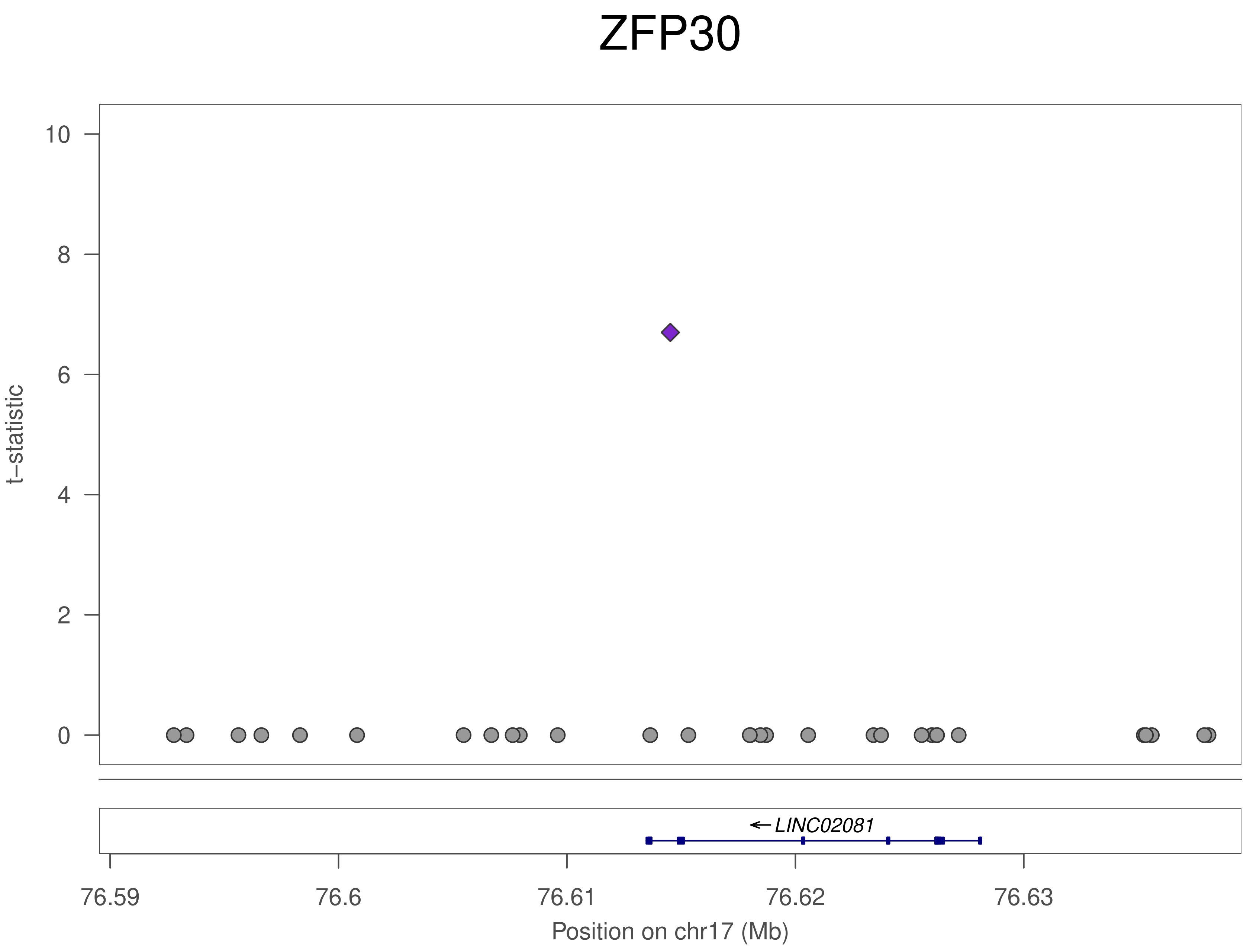

ZFP30
t–statistic
10
8
6
4
2
0
LINC02081
76.59
76.6
76.61
76.62
76.63
Position on chr17 (Mb)

### ZFP30

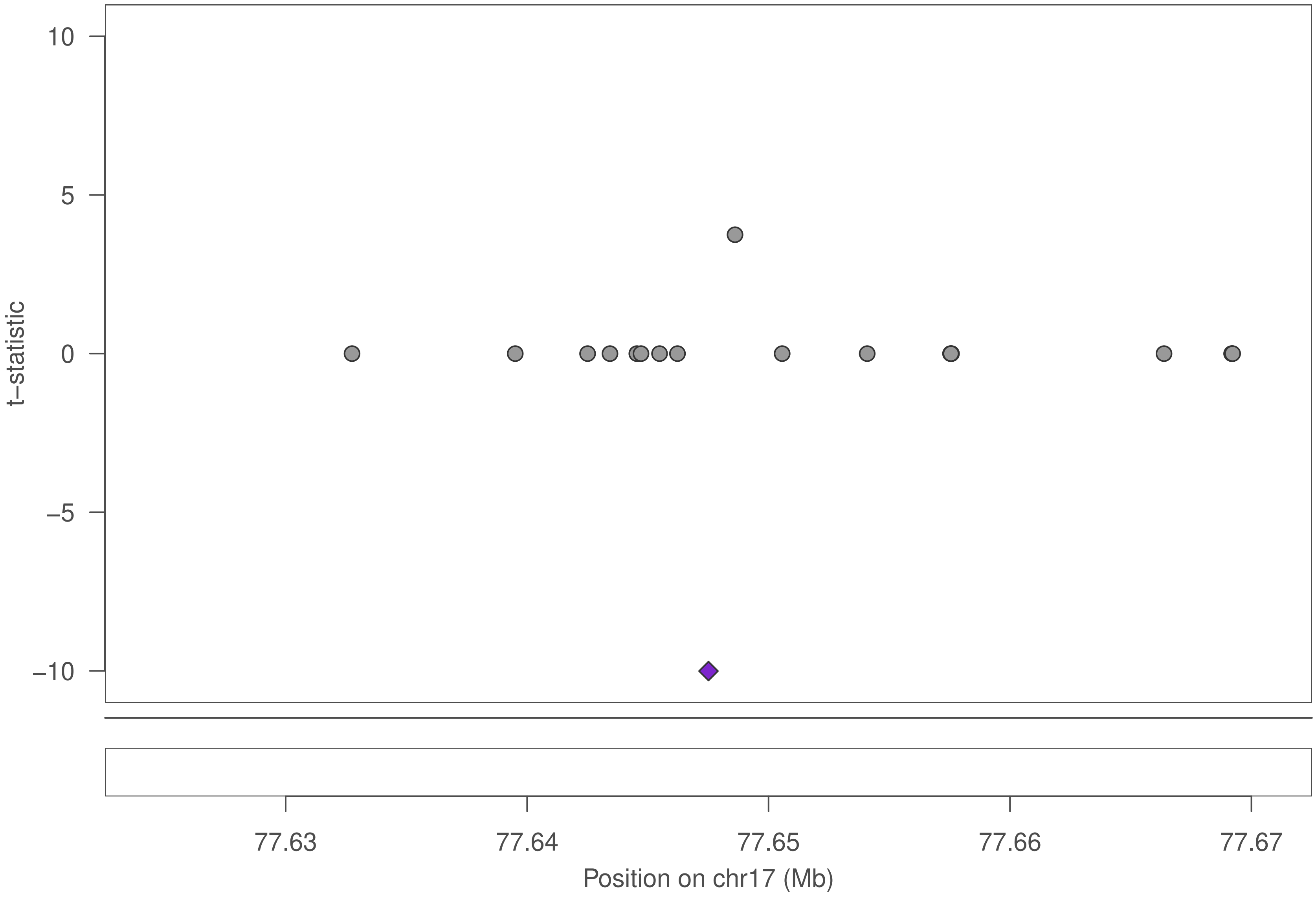

10
5
0
−5
−10
t−statistic
77.63
77.64
77.65
77.66
77.67
Position on chr17 (Mb)

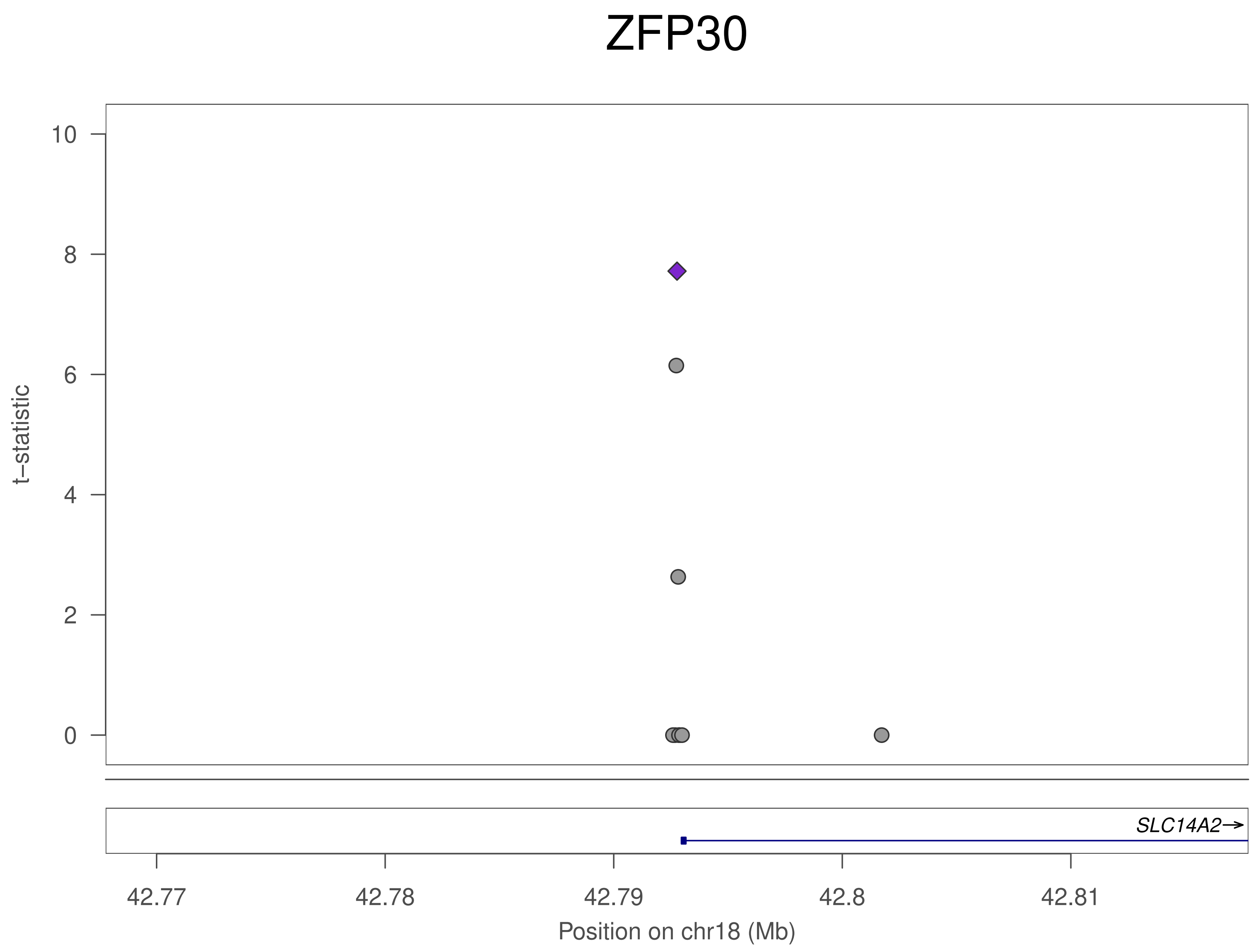
ZFP30
t–statistic
10
8
6
4
2
0
SLC14A2→
42.77
42.78
42.79
42.8
42.81
Position on chr18 (Mb)

### ZFP30

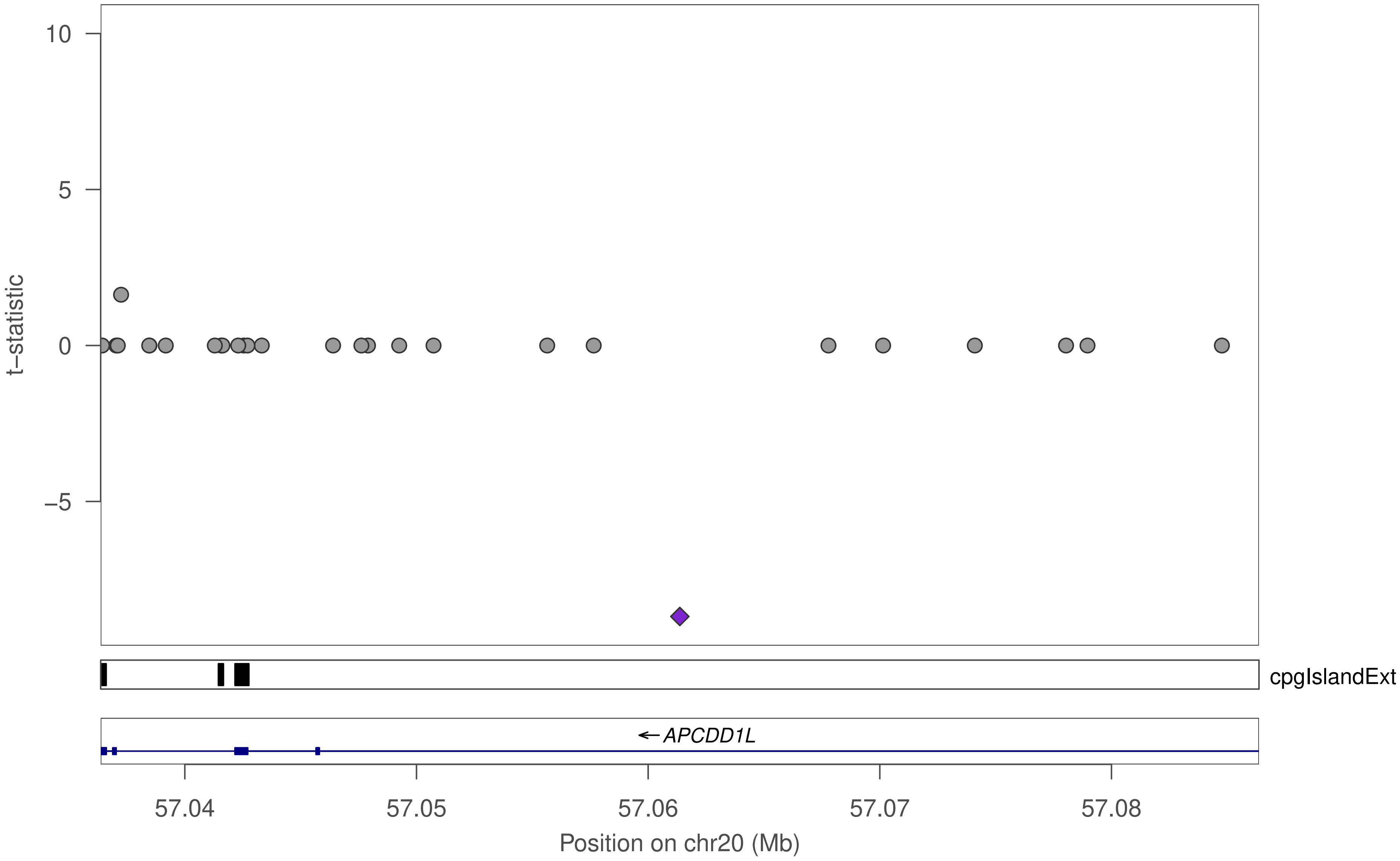

10
5
0
−5
t−statistic
cpgIslandExt
← APCDD1L
57.04
57.05
57.06
57.07
57.08
Position on chr20 (Mb)

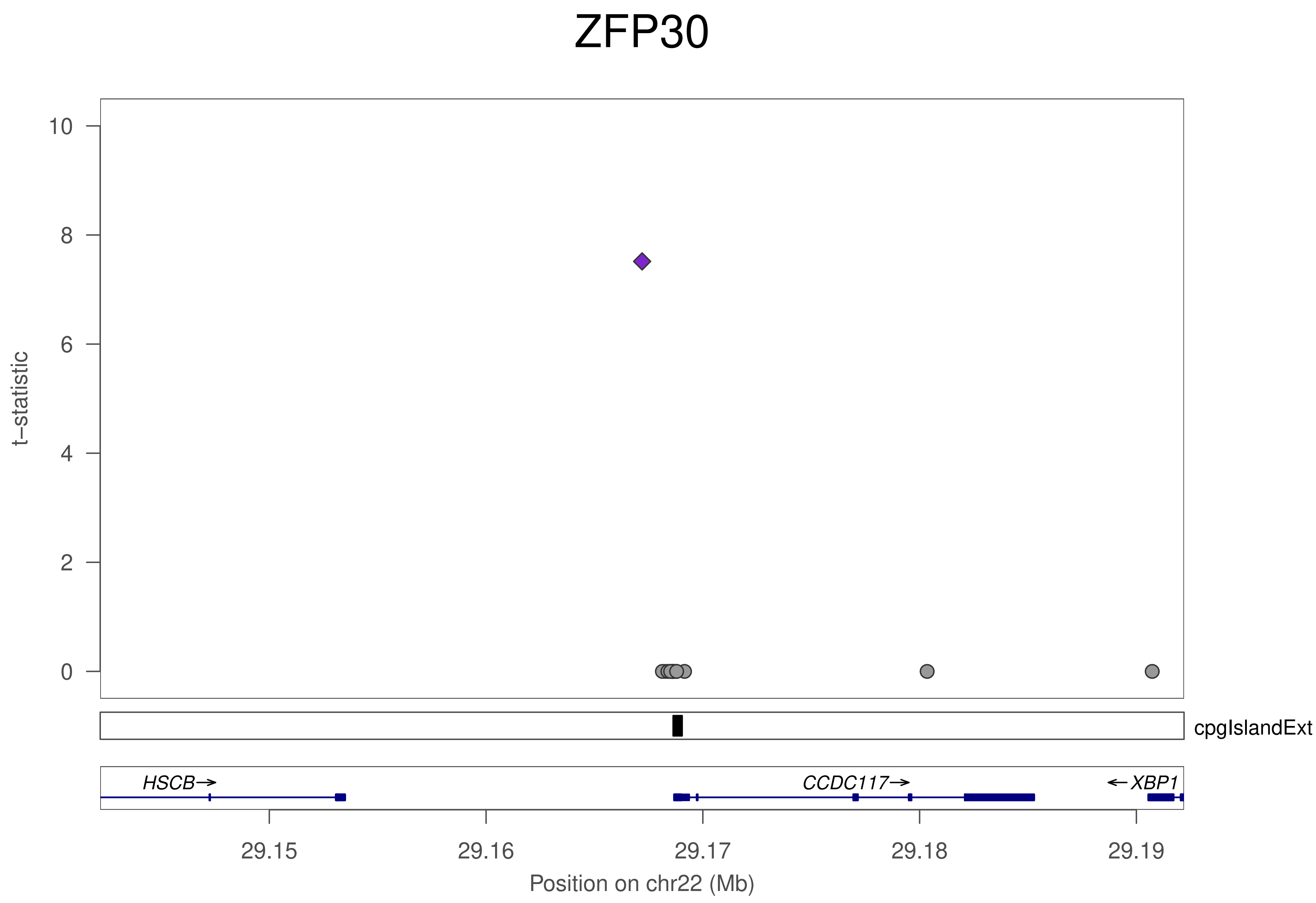
ZFP30
t−statistic
10
8
6
4
2
0
cpgIslandExt
HSCB
CCDC117
XBP1
29.15
29.16
29.17
29.18
29.19
Position on chr22 (Mb)

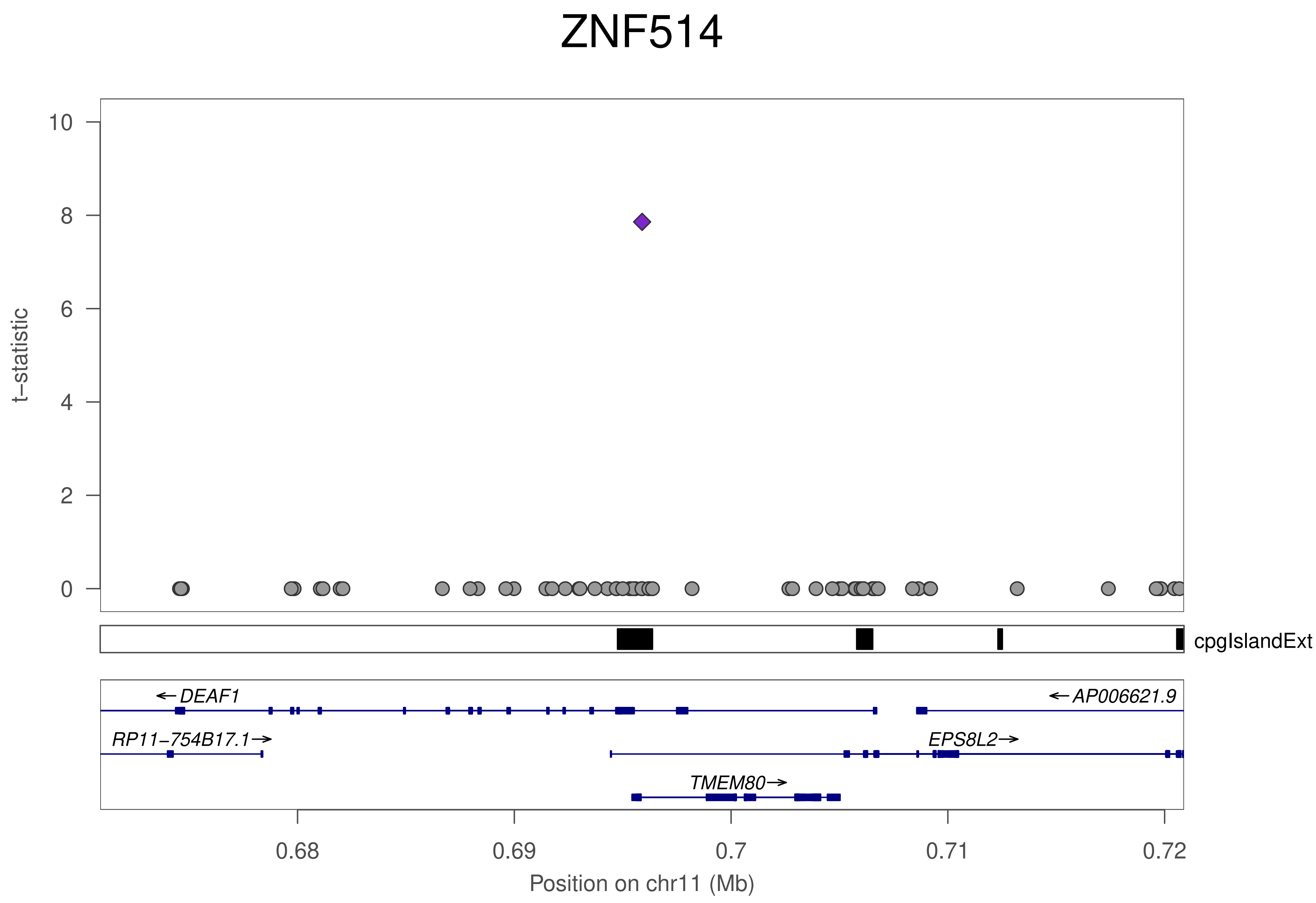

ZNF514
t−statistic
10
8
6
4
2
0
cpgIslandExt
←DEAF1
←AP006621.9
RP11−754B17.1→
EPS8L2→
TMEM80→
0.68
0.69
0.7
0.71
0.72
Position on chr11 (Mb)

### ZNF773

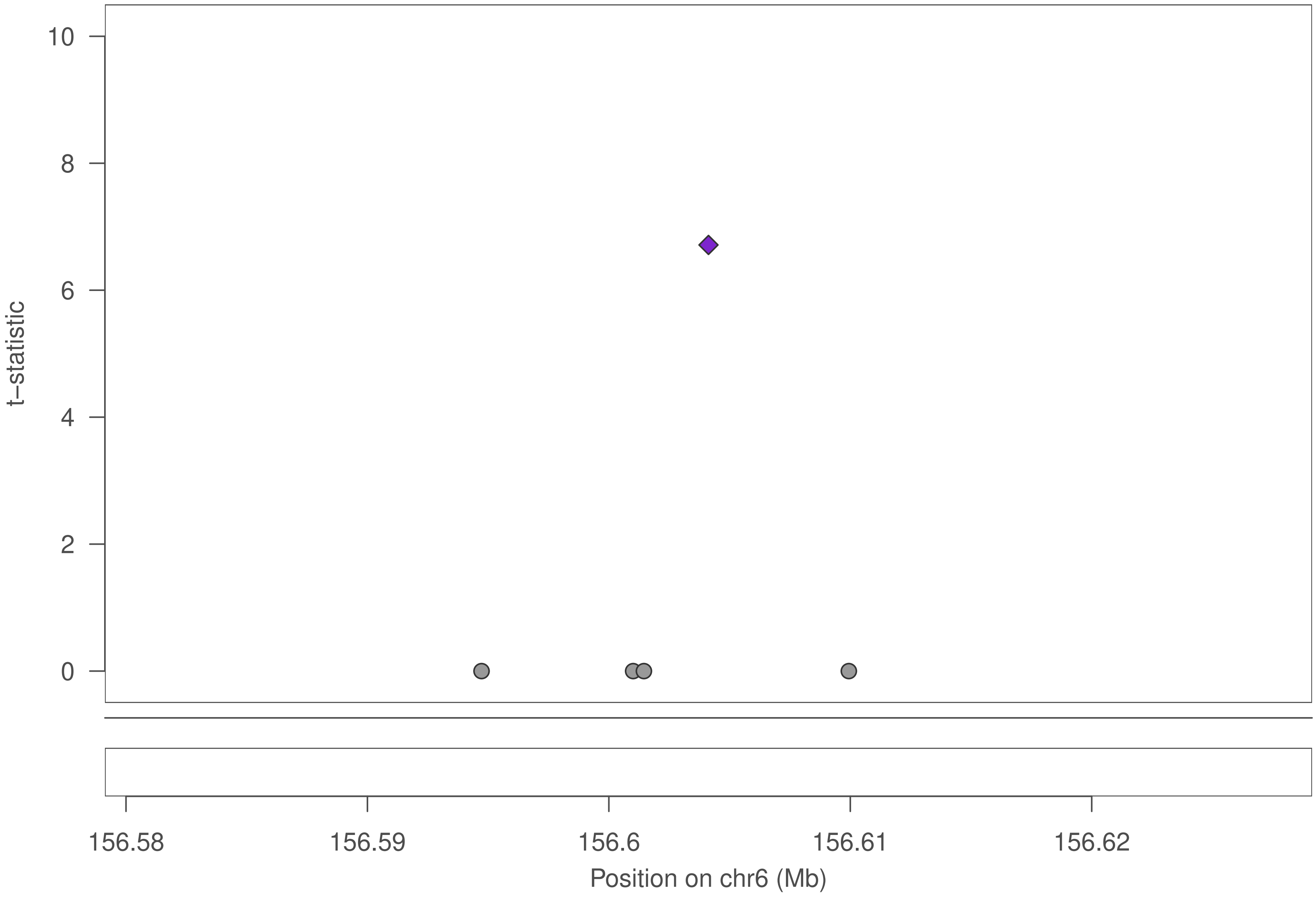

10
8
6
4
2
0
t–statistic
156.58
156.59
156.6
156.61
156.62
Position on chr6 (Mb)

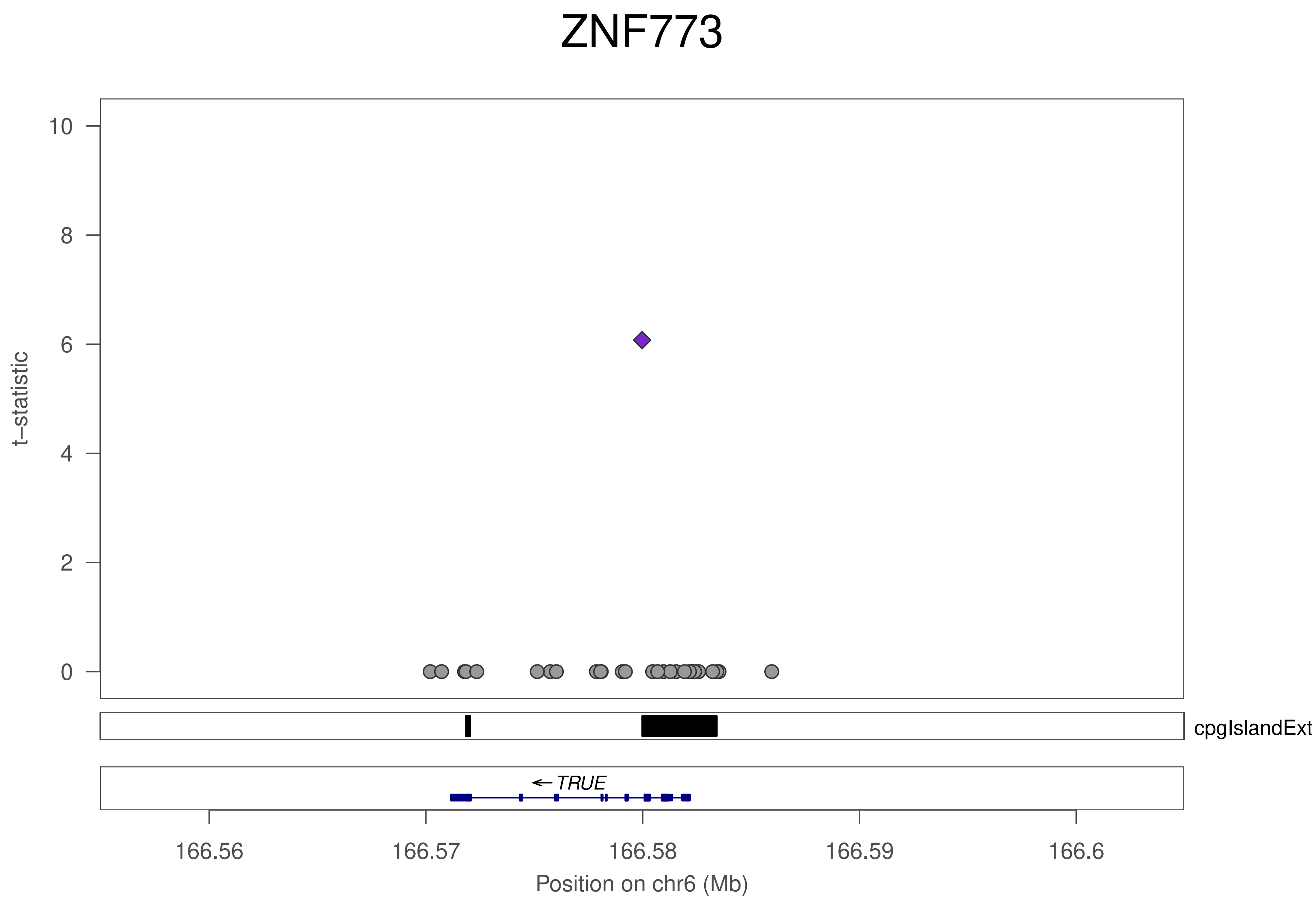
ZNF773
t−statistic
10
8
6
4
2
0
cpgIslandExt
← TRUE
166.56
166.57
166.58
166.59
166.6
Position on chr6 (Mb)

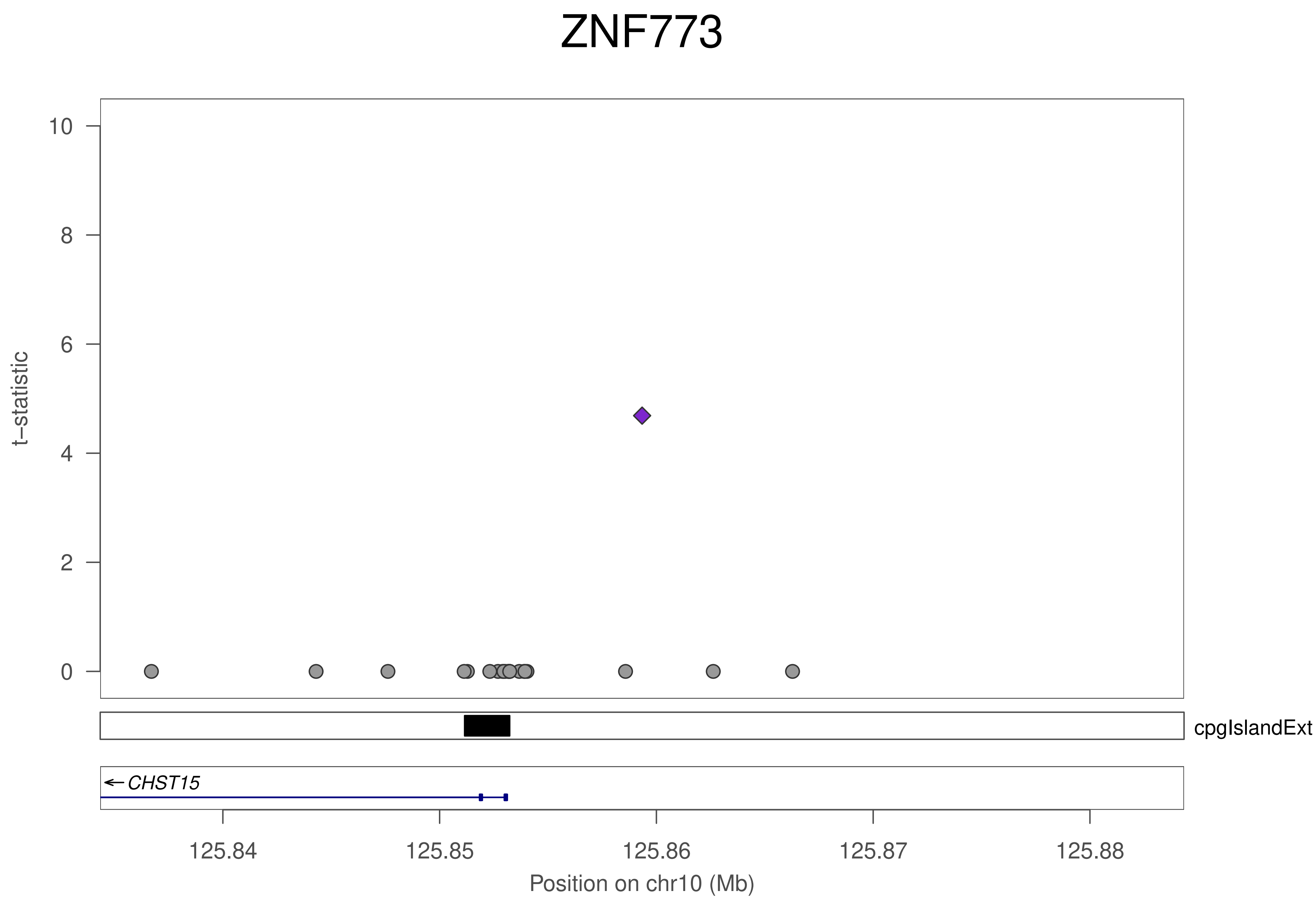
ZNF773
t−statistic
0
2
4
6
8
10
cpgIslandExt
CHST15
125.84
125.85
125.86
125.87
125.88
Position on chr10 (Mb)

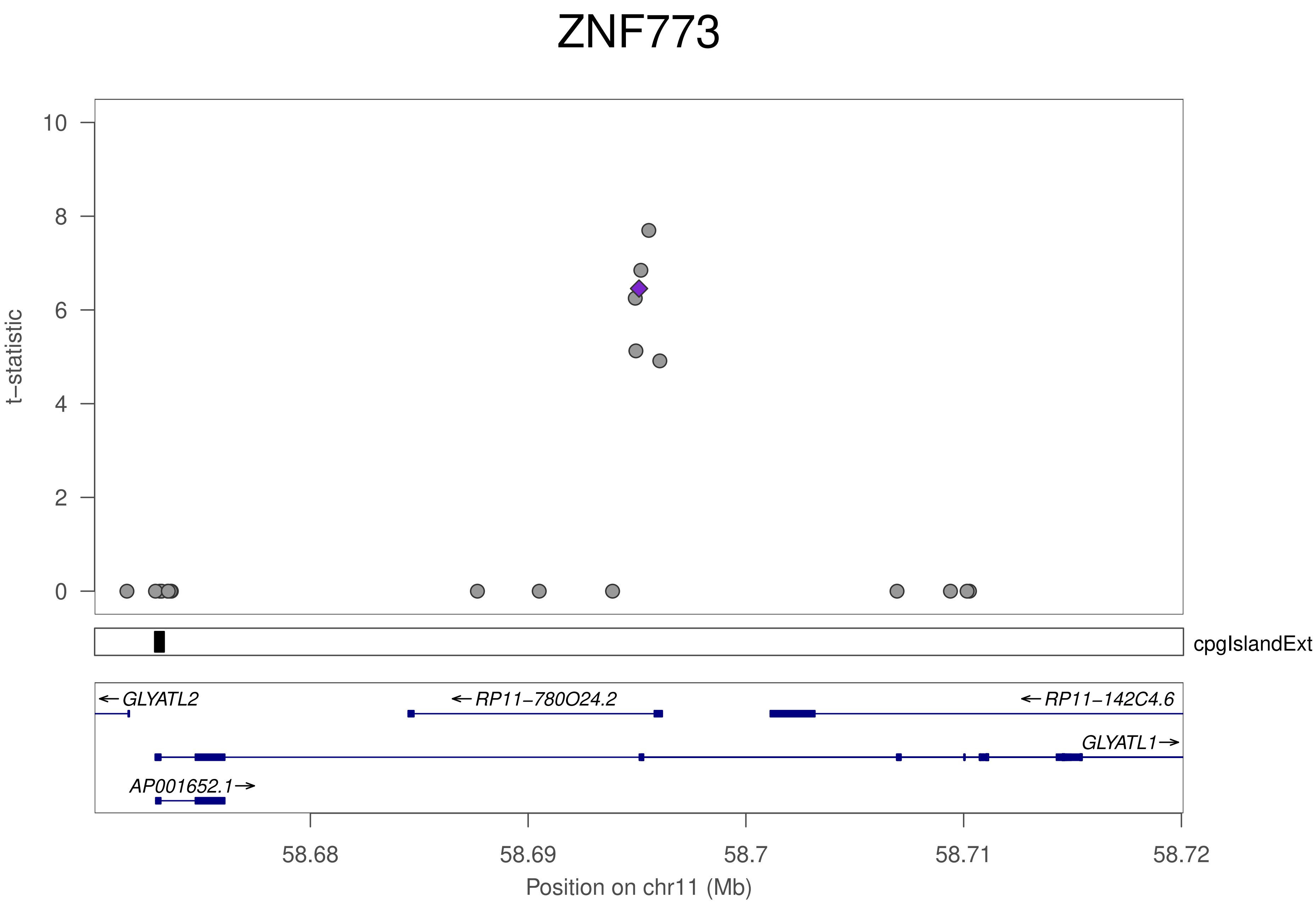

ZNF773
t−statistic
0
2
4
6
8
10
cpgIslandExt
←GLYATL2
←RP11−780O24.2
←RP11−142C4.6
GLYATL1→
AP001652.1→
58.68
58.69
58.7
58.71
58.72
Position on chr11 (Mb)

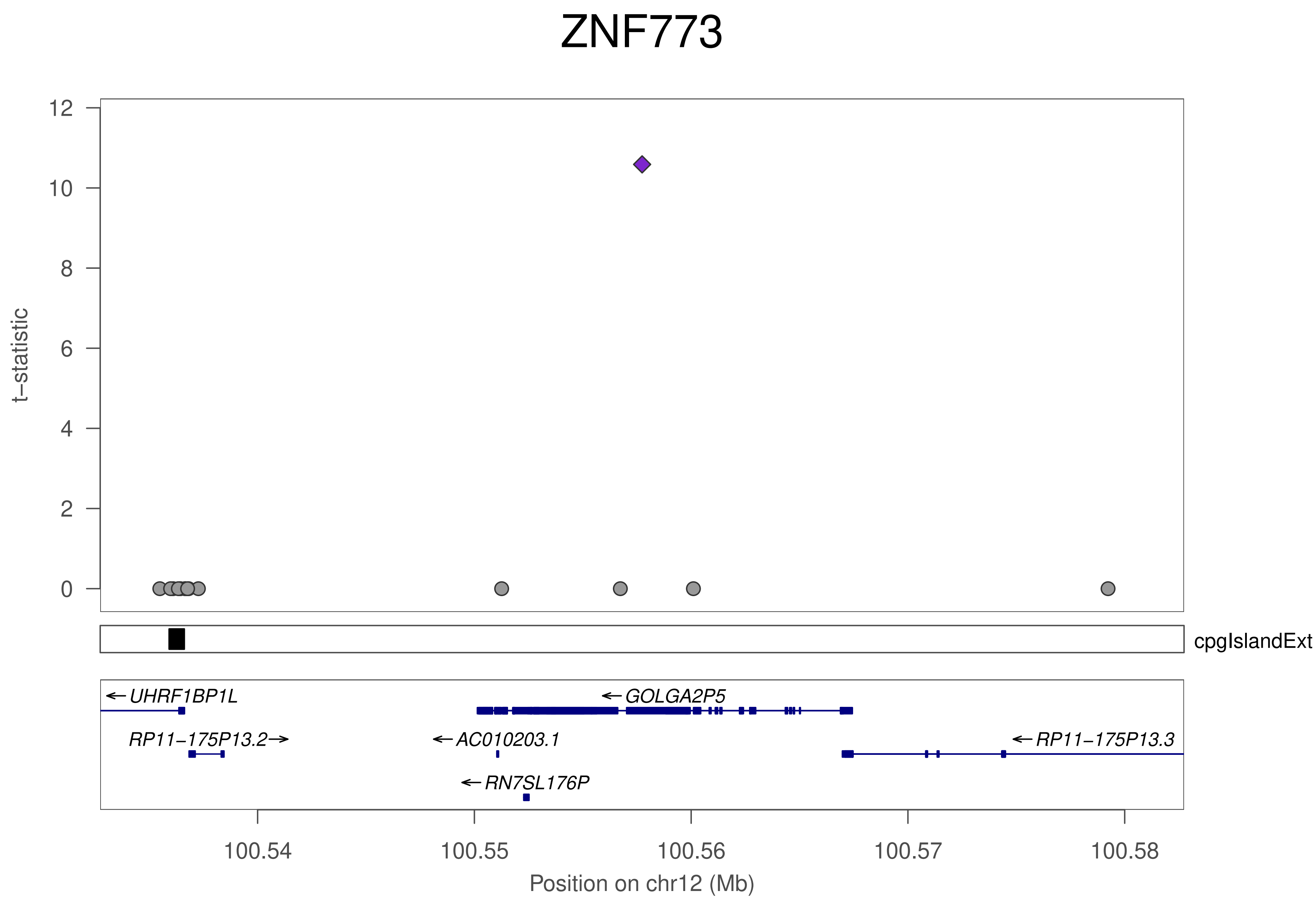

ZNF773
t−statistic
12
10
8
6
4
2
0
cpgIslandExt
←UHRF1BP1L
←GOLGA2P5
RP11−175P13.2→
←AC010203.1
←RP11−175P13.3
←RN7SL176P
100.54
100.55
100.56
100.57
100.58
Position on chr12 (Mb)

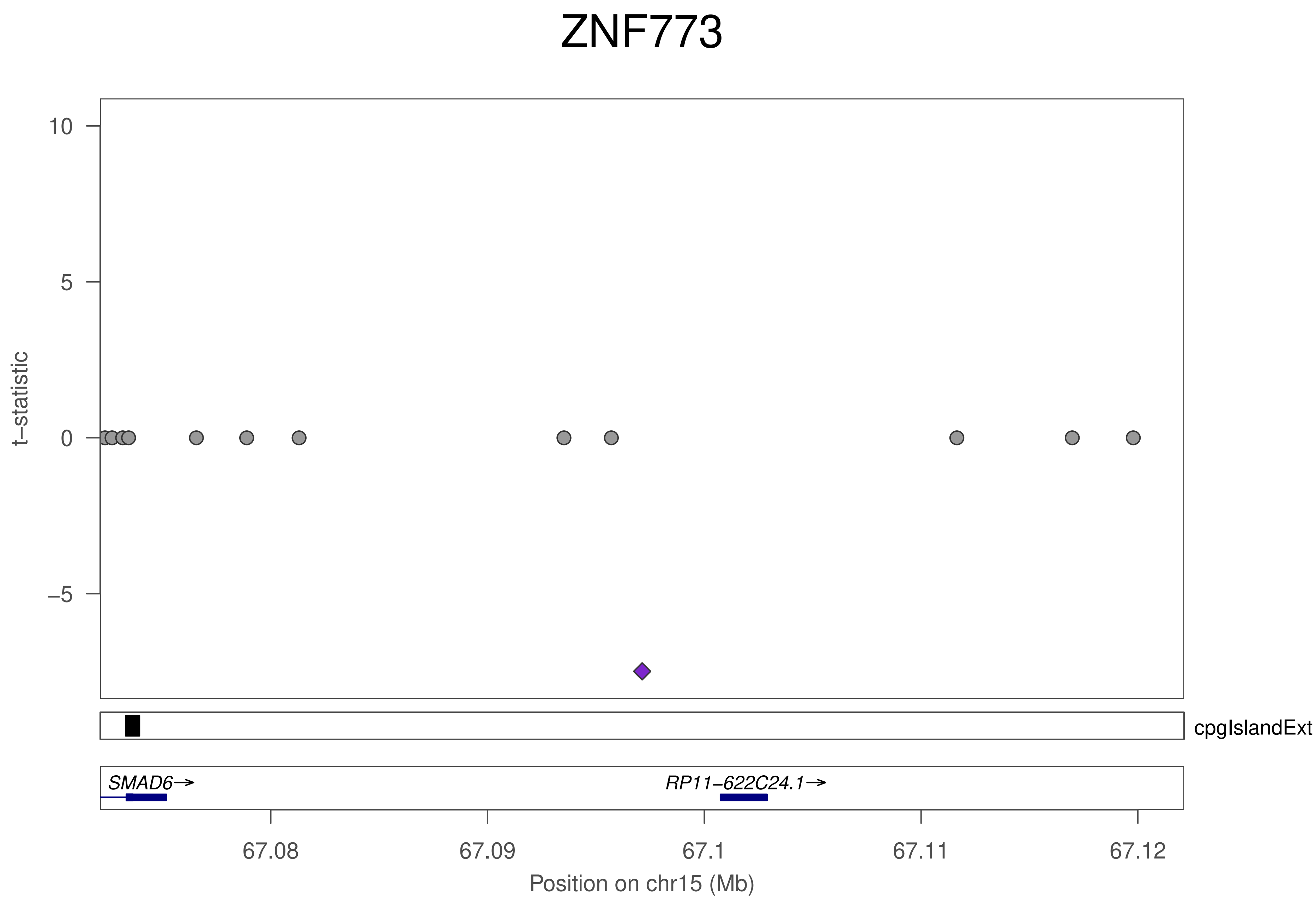

ZNF773
10
5
0
−5
t−statistic
cpgIslandExt
SMAD6→
RP11−622C24.1→
67.08
67.09
67.1
67.11
67.12
Position on chr15 (Mb)

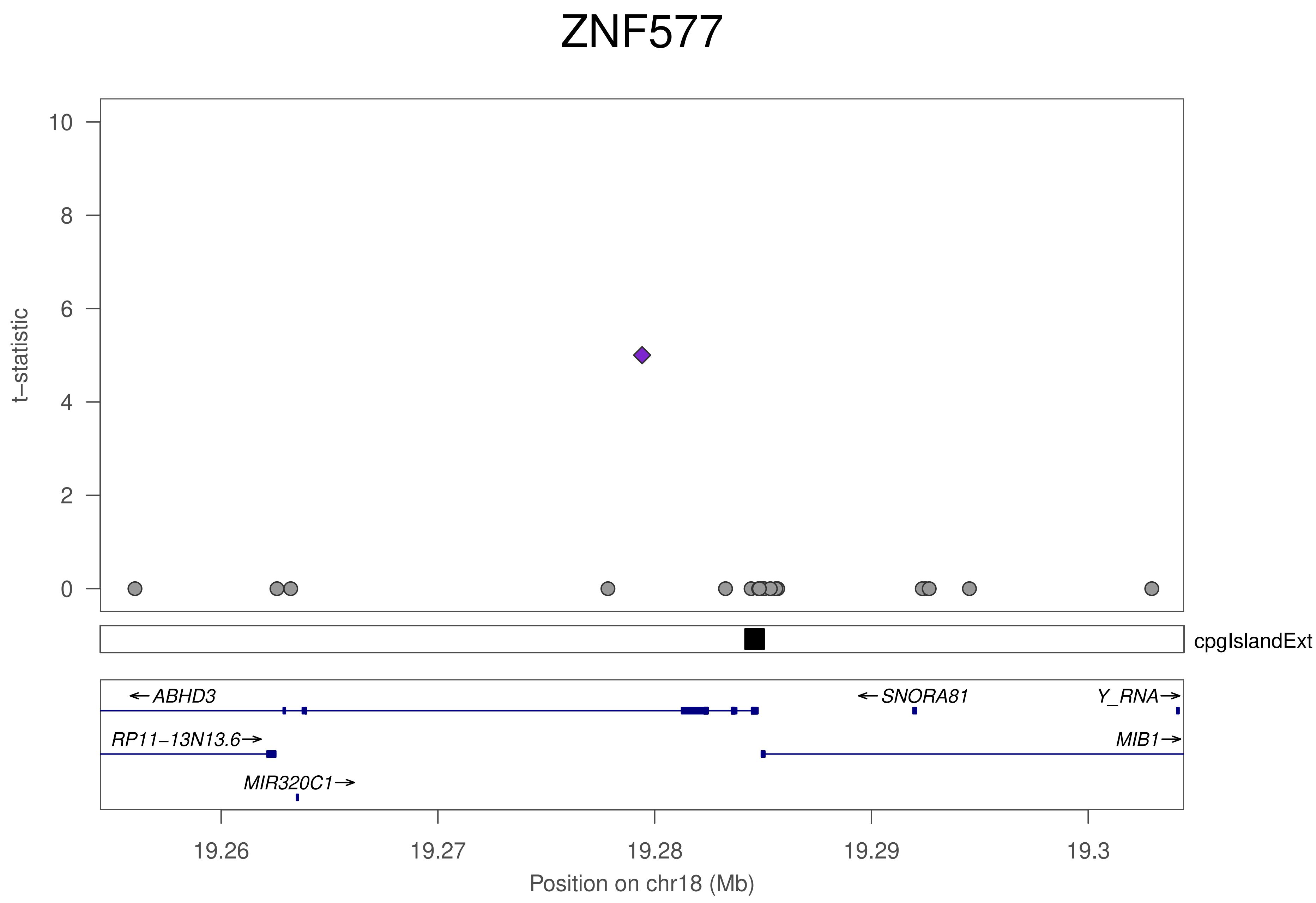

ZNF577
t−statistic
10
8
6
4
2
0
cpgIslandExt
←ABHD3
←SNORA81
Y_RNA→
RP11−13N13.6→
MIB1→
MIR320C1→
19.26
19.27
19.28
19.29
19.3
Position on chr18 (Mb)

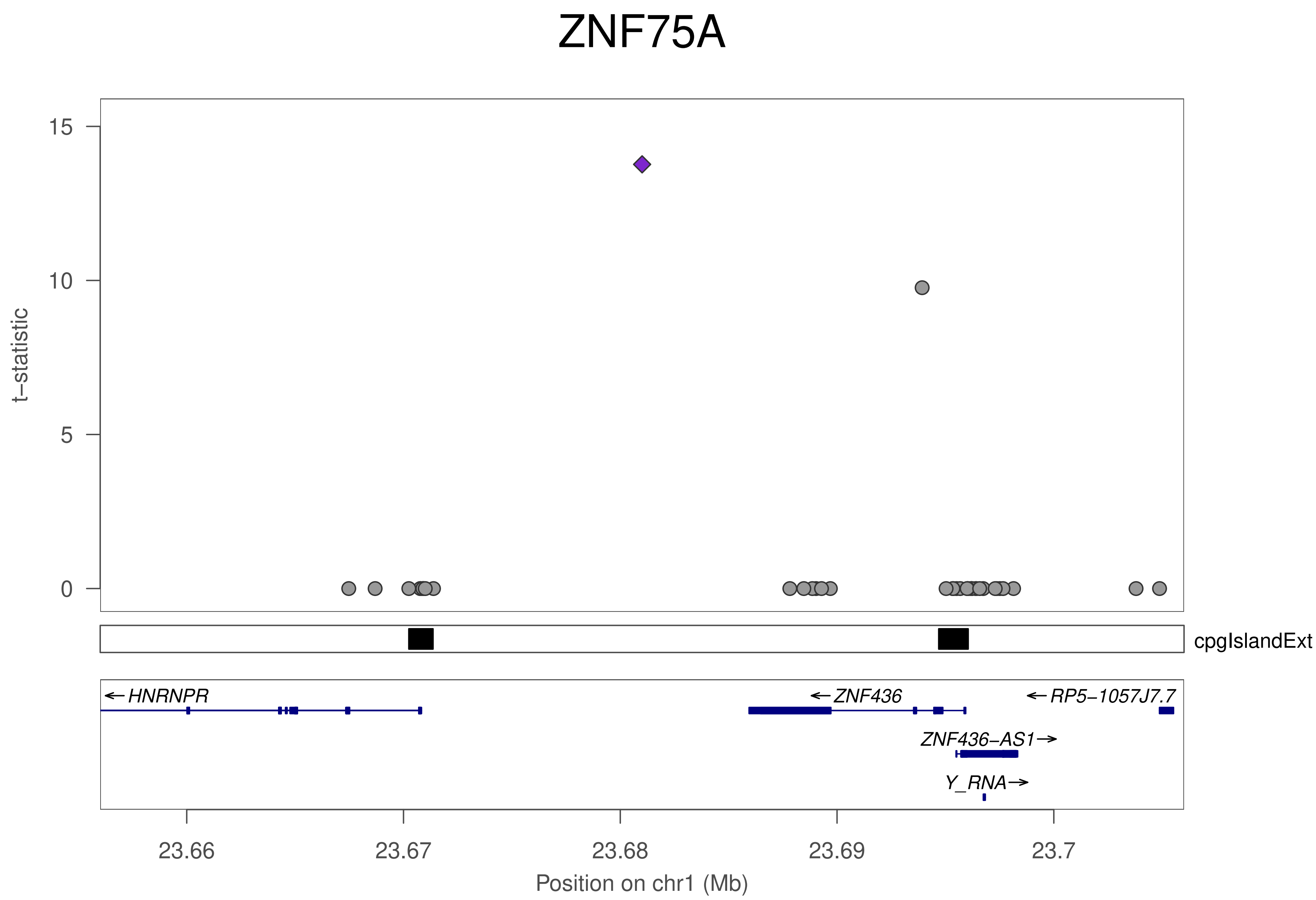

ZNF75A
t−statistic
15
10
5
0
cpgIslandExt
←HNRNPR
←ZNF436
←RP5−1057J7.7
ZNF436−AS1→
Y_RNA→
23.66
23.67
23.68
23.69
23.7
Position on chr1 (Mb)

### ZNF75A

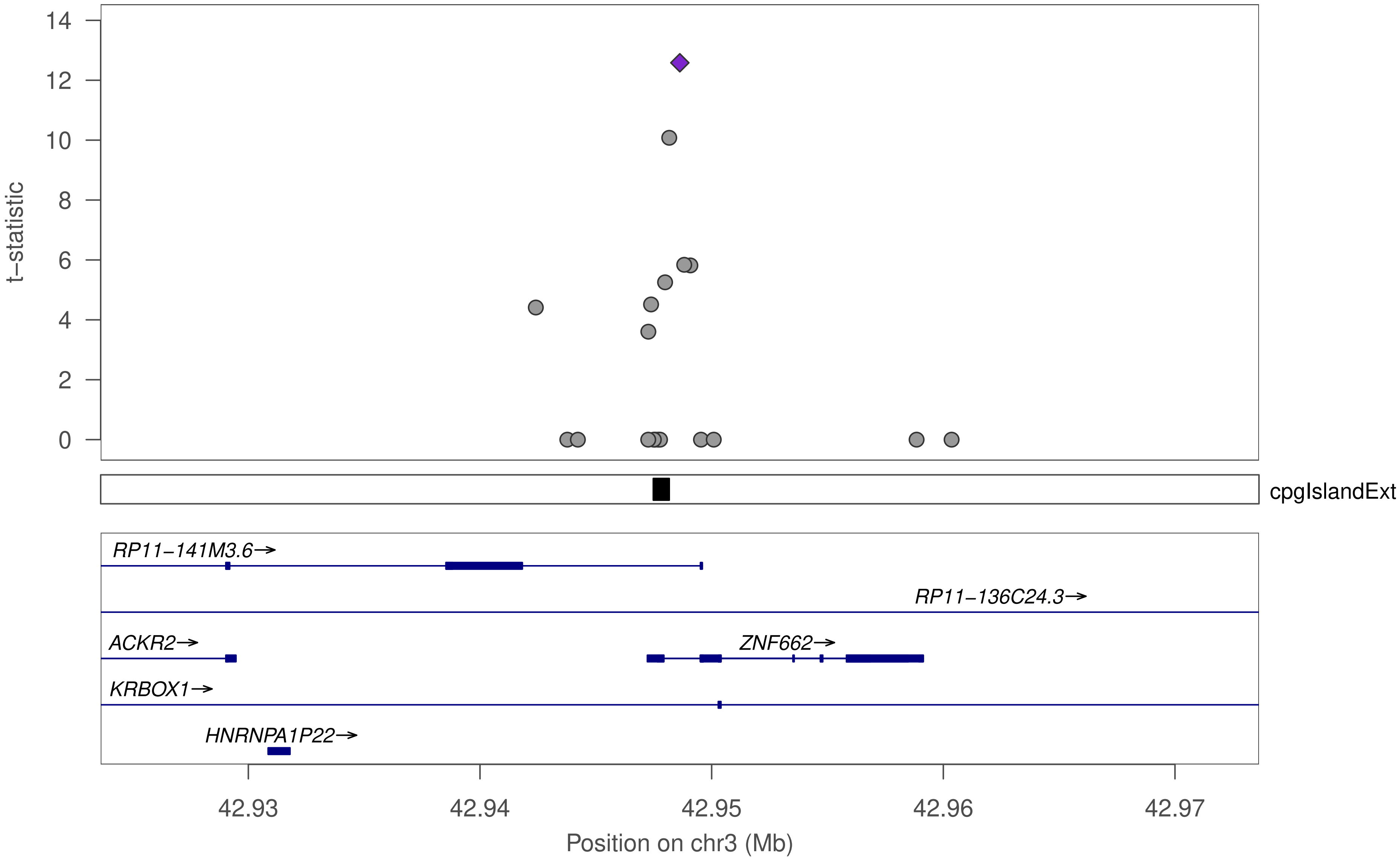

t−statistic
14
12
10
8
6
4
2
0
cpgIslandExt
RP11−141M3.6→
RP11−136C24.3→
ACKR2→
ZNF662→
KRBOX1→
HNRNPA1P22→
42.93
42.94
42.95
42.96
42.97
Position on chr3 (Mb)

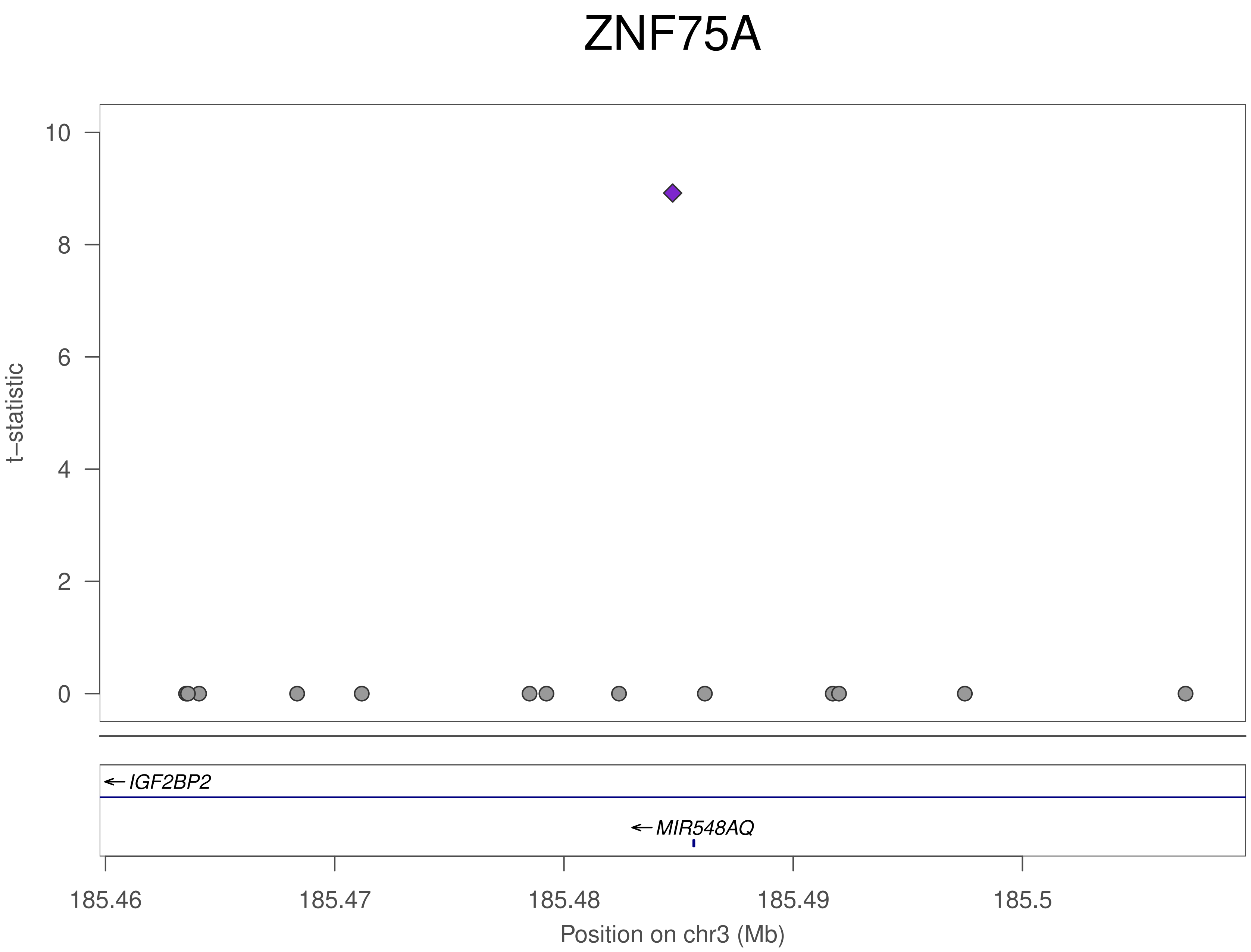
ZNF75A
t–statistic
10
8
6
4
2
0
IGF2BP2
MIR548AQ
185.46
185.47
185.48
185.49
185.5
Position on chr3 (Mb)

### ZNF75A

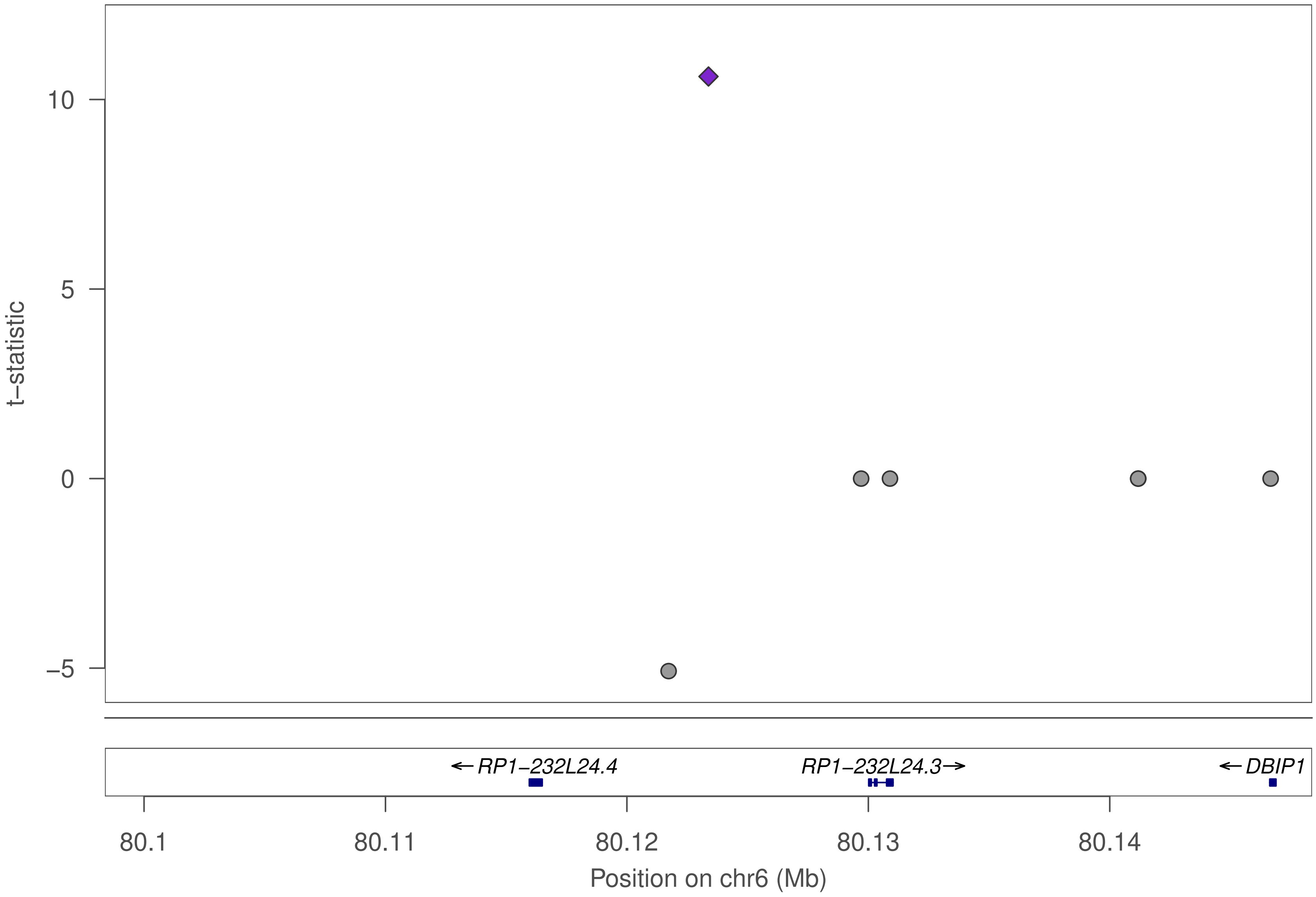

t–statistic
10
5
0
–5
←RP1–232L24.4
RP1–232L24.3→
←DBIP1
80.1
80.11
80.12
80.13
80.14
Position on chr6 (Mb)

### ZNF75A

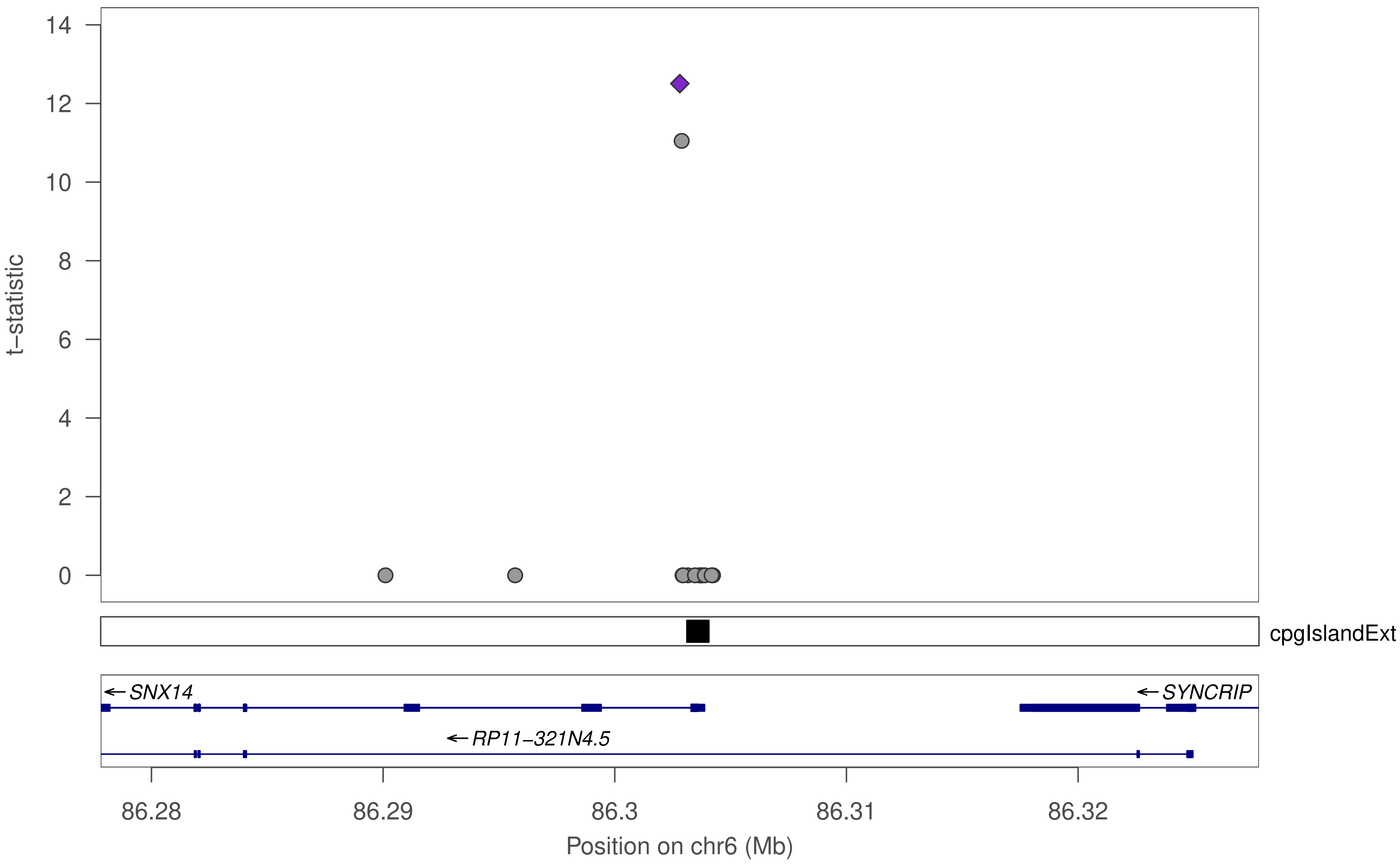
t−statistic
0
2
4
6
8
10
12
14
cpgIslandExt
SNX14
SYNCRIP
RP11−321N4.5
86.28
86.29
86.3
86.31
86.32
Position on chr6 (Mb)

### ZNF75A

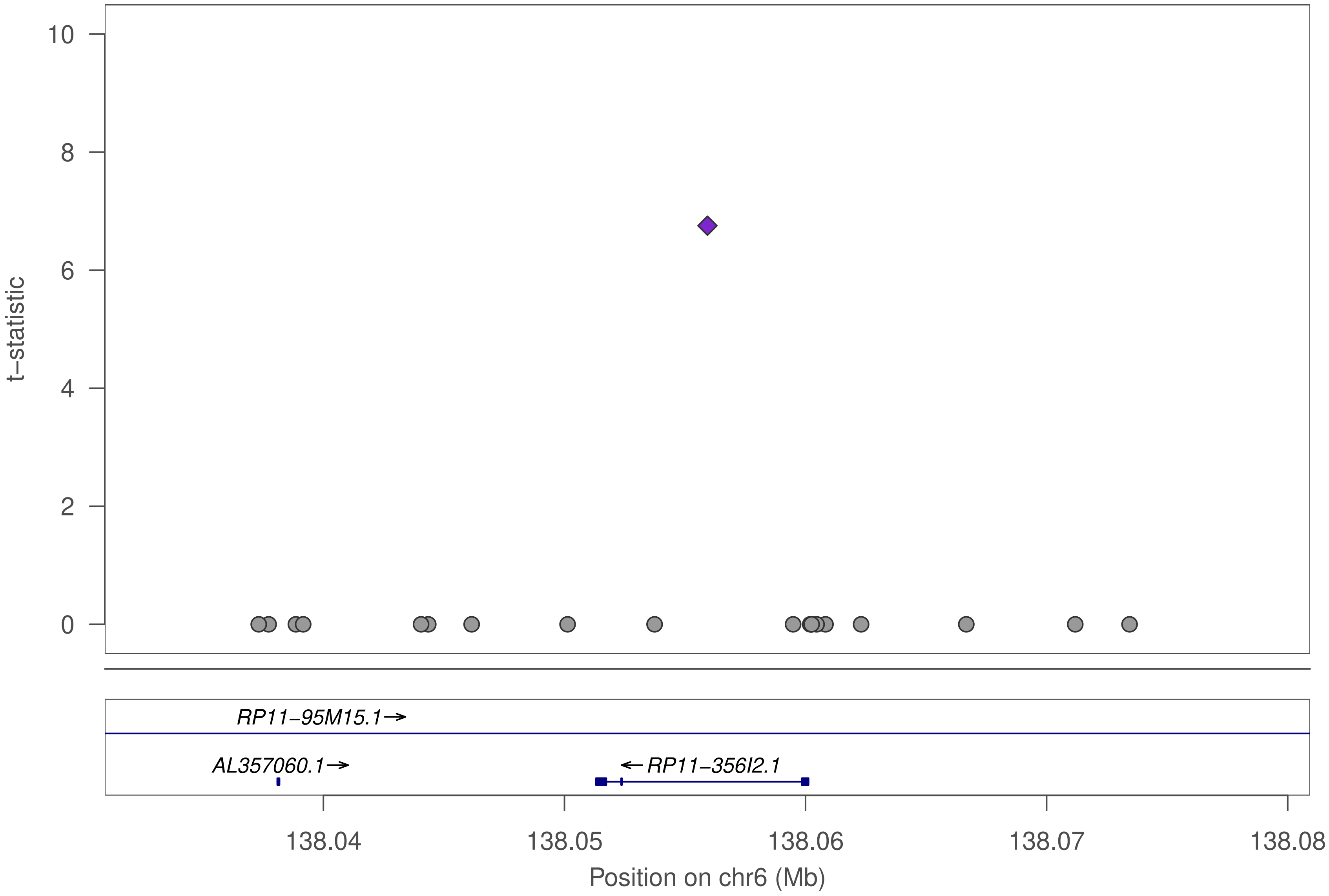

t−statistic
10
8
6
4
2
0
RP11−95M15.1→
AL357060.1→
←RP11−356I2.1
138.04
138.05
138.06
138.07
138.08
Position on chr6 (Mb)

### ZNF75A

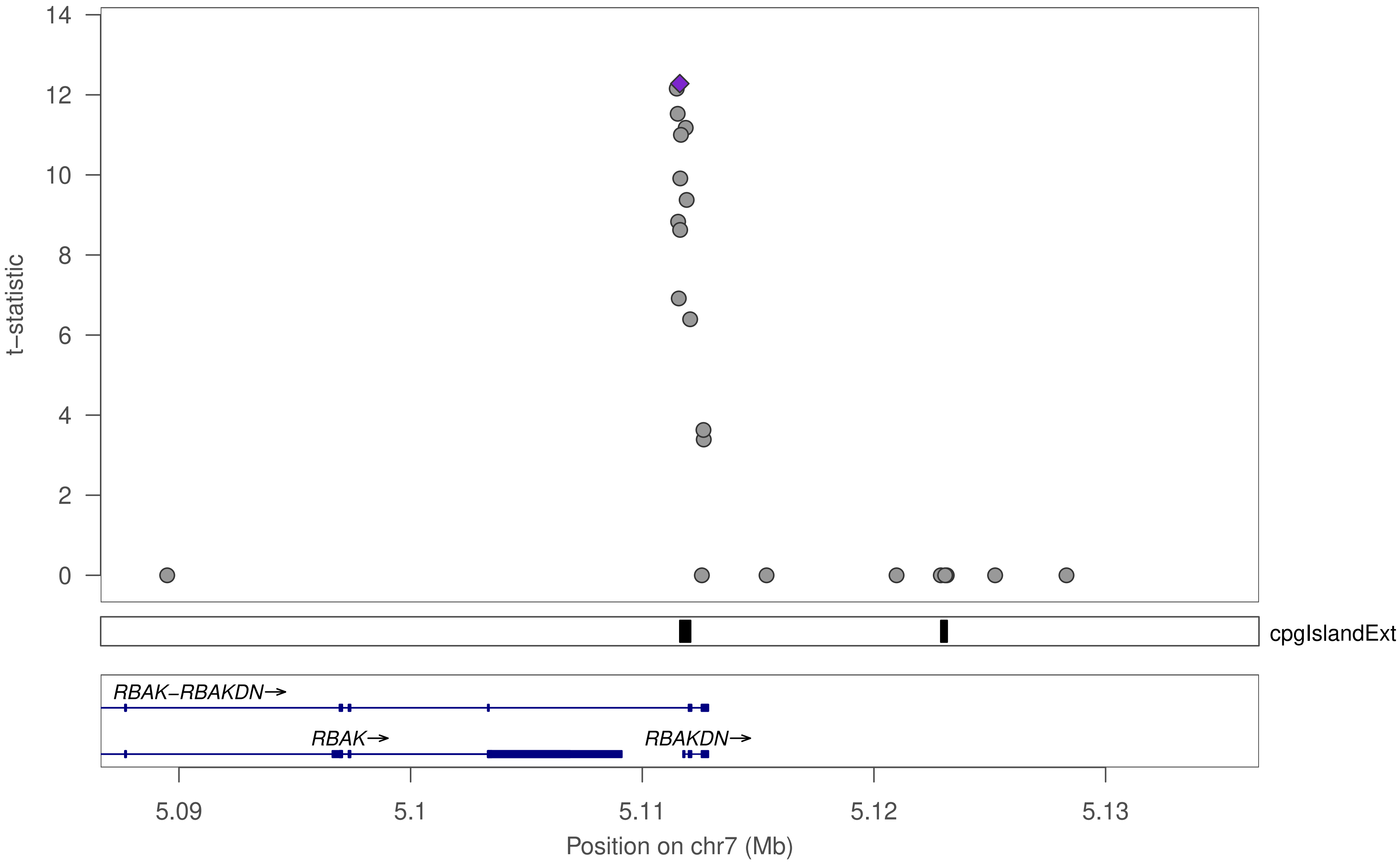

14
12
10
8
6
4
2
0
t−statistic
cpgIslandExt
RBAK−RBAKDN→
RBAK→
RBAKDN→
5.09
5.1
5.11
5.12
5.13
Position on chr7 (Mb)

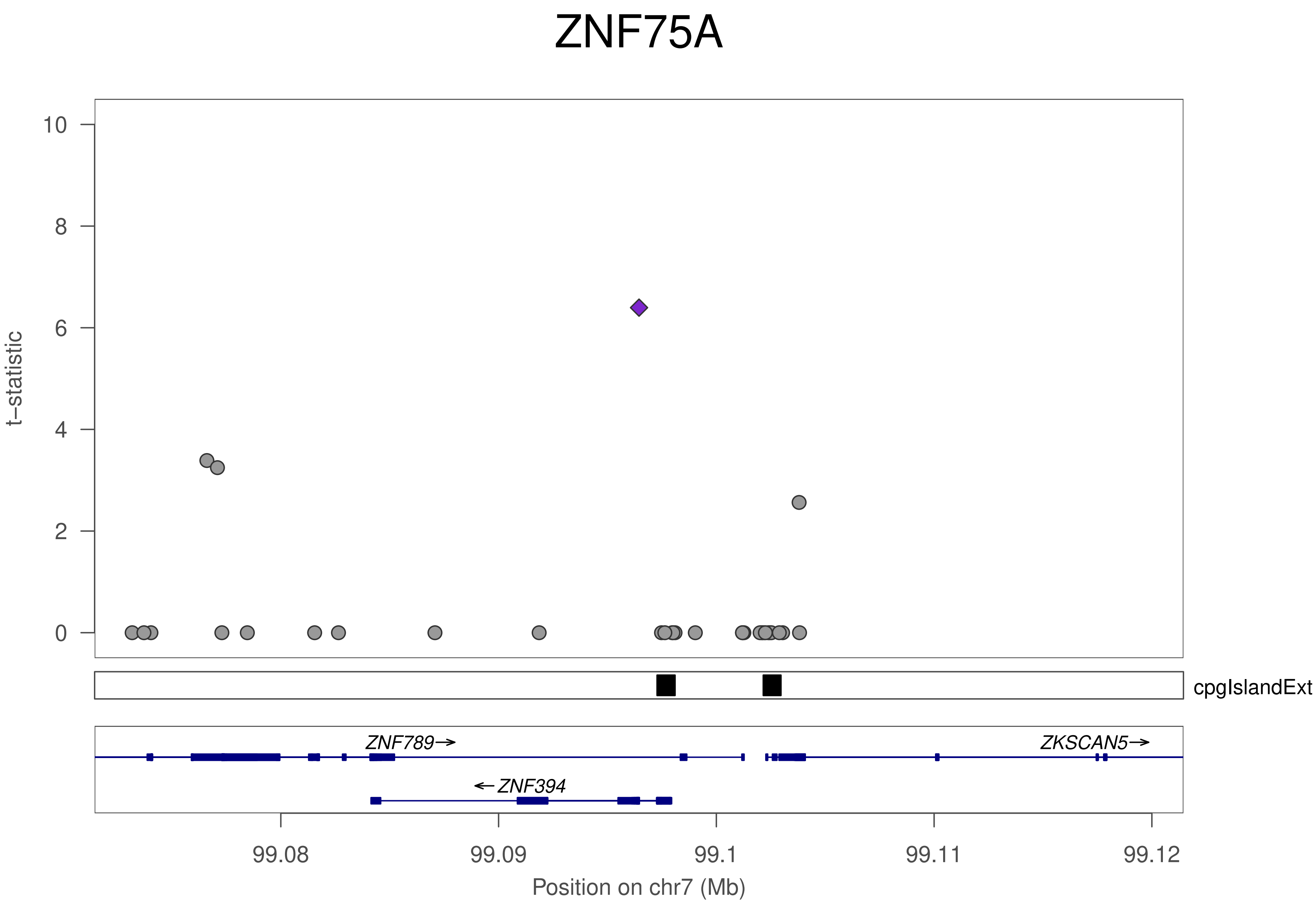
ZNF75A
t−statistic
10
8
6
4
2
0
cpgIslandExt
ZNF789→
ZKSCAN5→
←ZNF394
99.08
99.09
99.1
99.11
99.12
Position on chr7 (Mb)

### ZNF75A

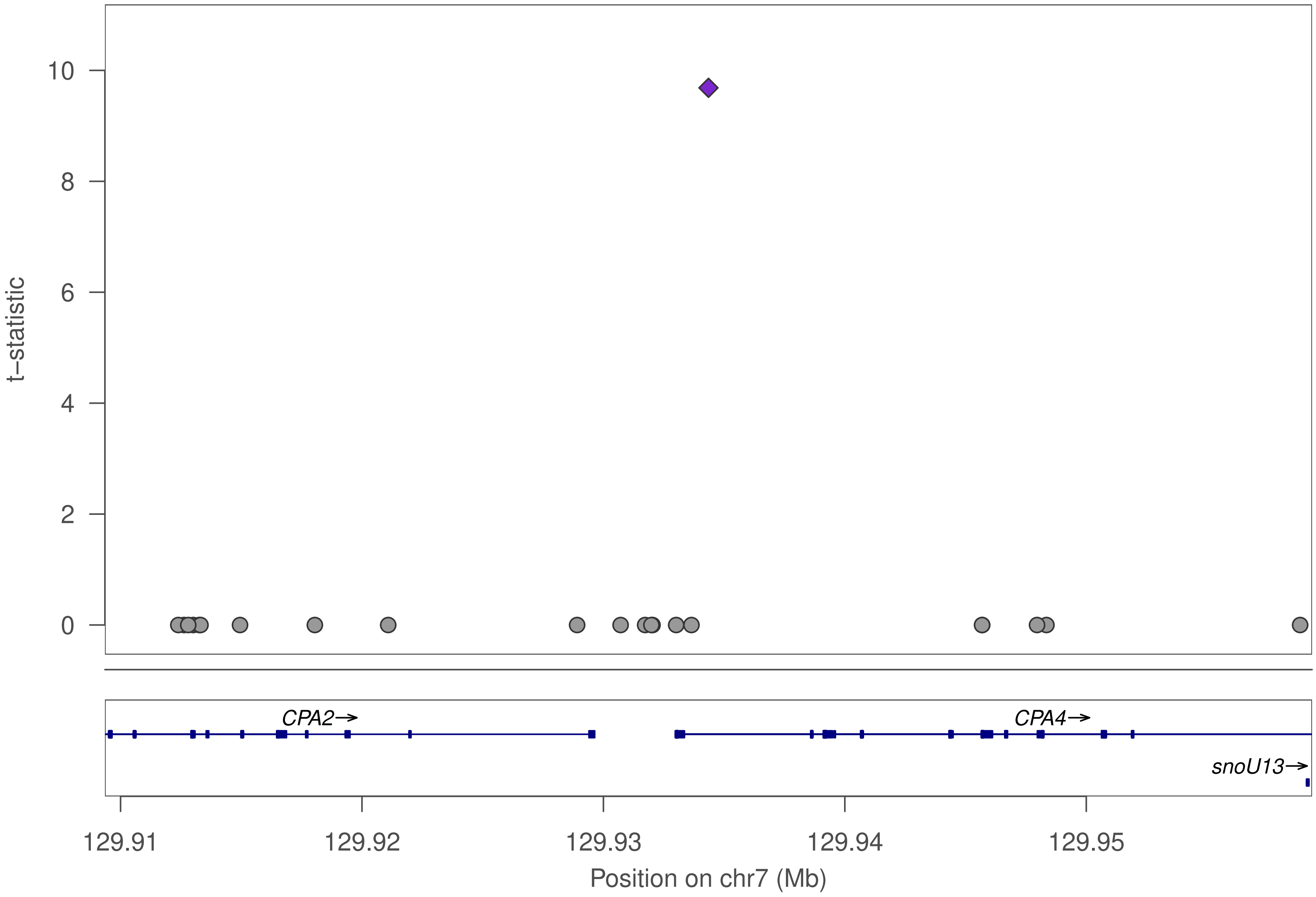

t–statistic
0
2
4
6
8
10
CPA2→
CPA4→
snoU13→
129.91
129.92
129.93
129.94
129.95
Position on chr7 (Mb)

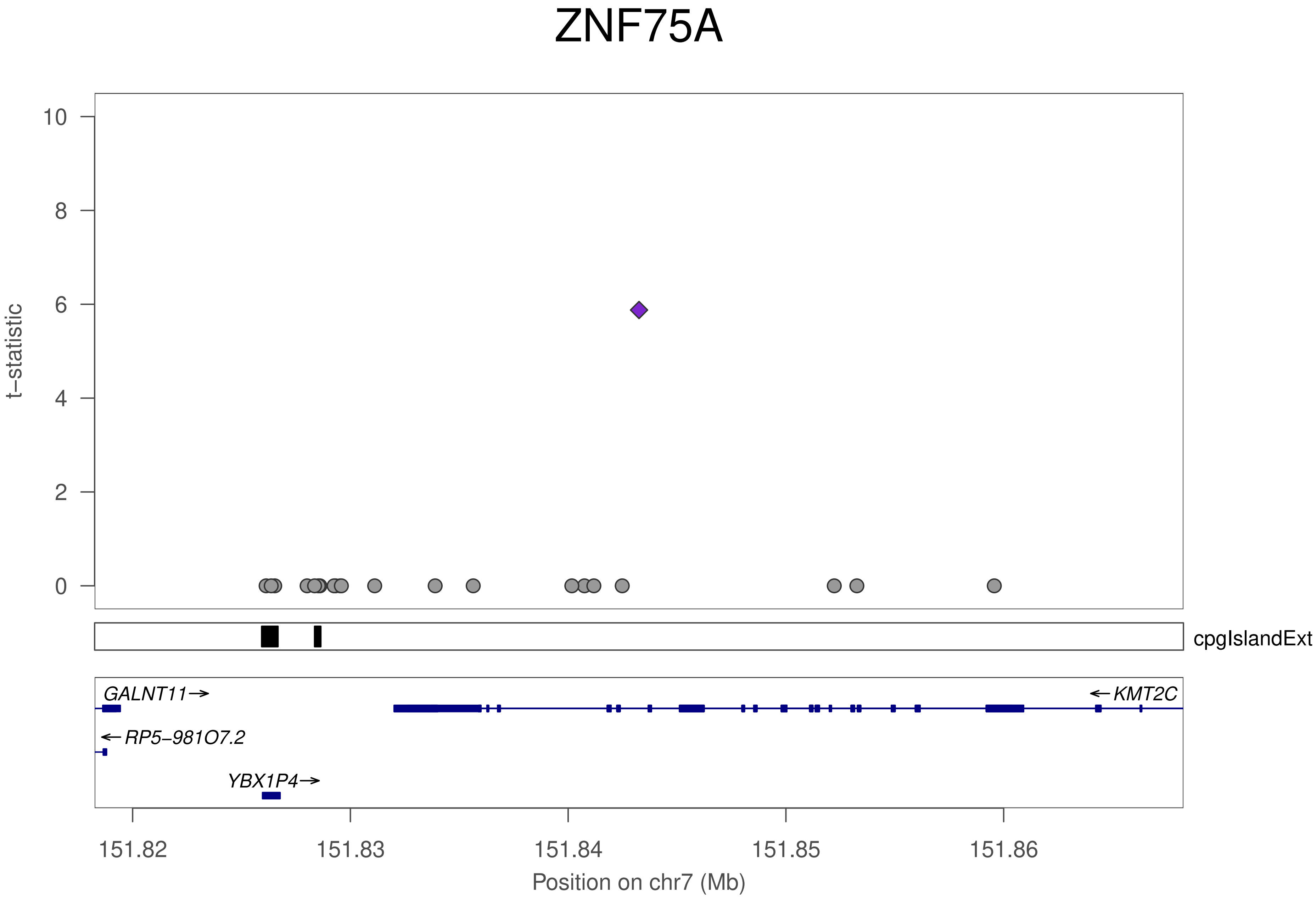
ZNF75A
t−statistic
10
8
6
4
2
0
cpgIslandExt
GALNT11→
←KMT2C
←RP5−981O7.2
YBX1P4→
151.82
151.83
151.84
151.85
151.86
Position on chr7 (Mb)

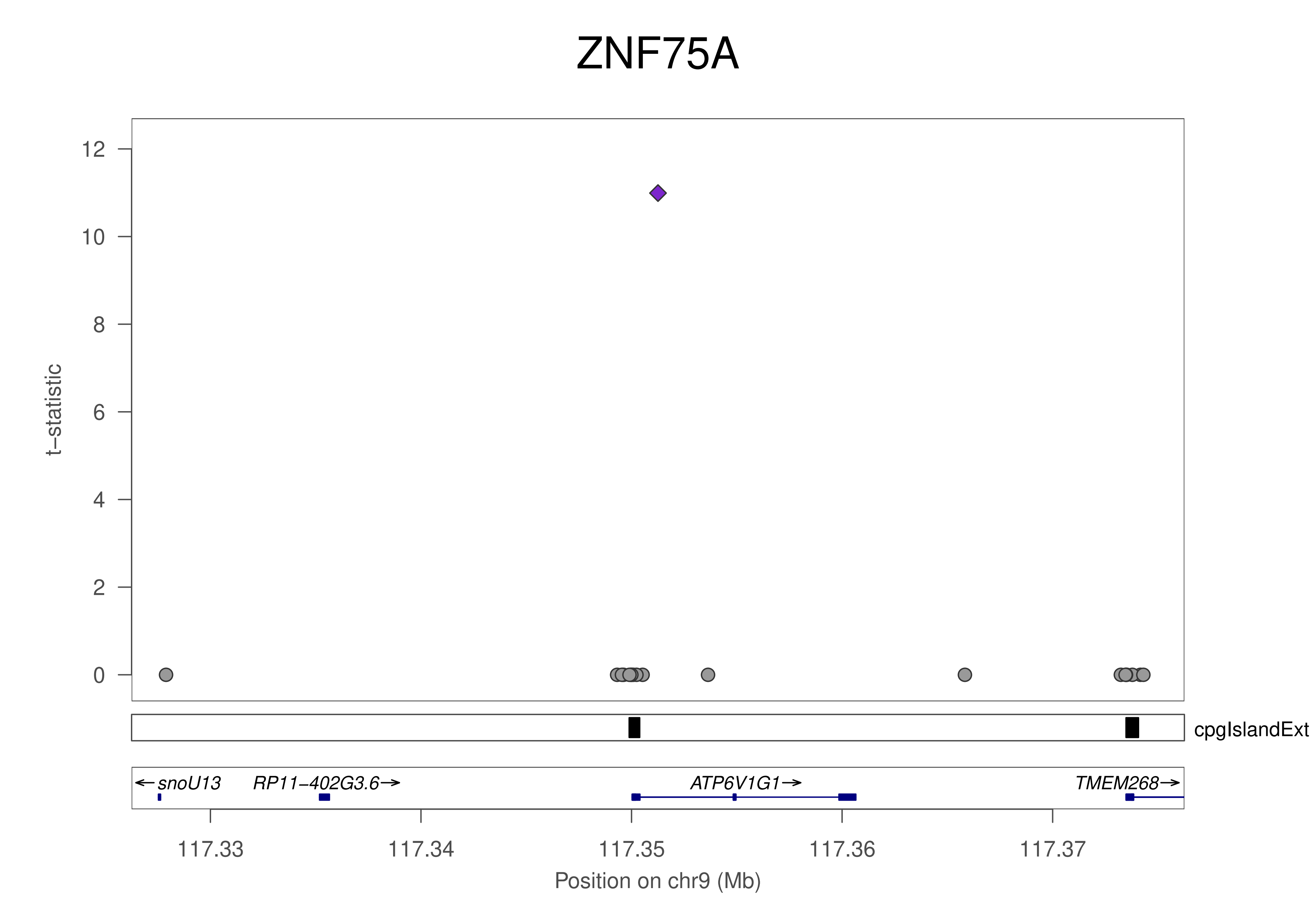
ZNF75A
t−statistic
12
10
8
6
4
2
0
cpgIslandExt
←snoU13
RP11−402G3.6→
ATP6V1G1→
TMEM268→
117.33
117.34
117.35
117.36
117.37
Position on chr9 (Mb)

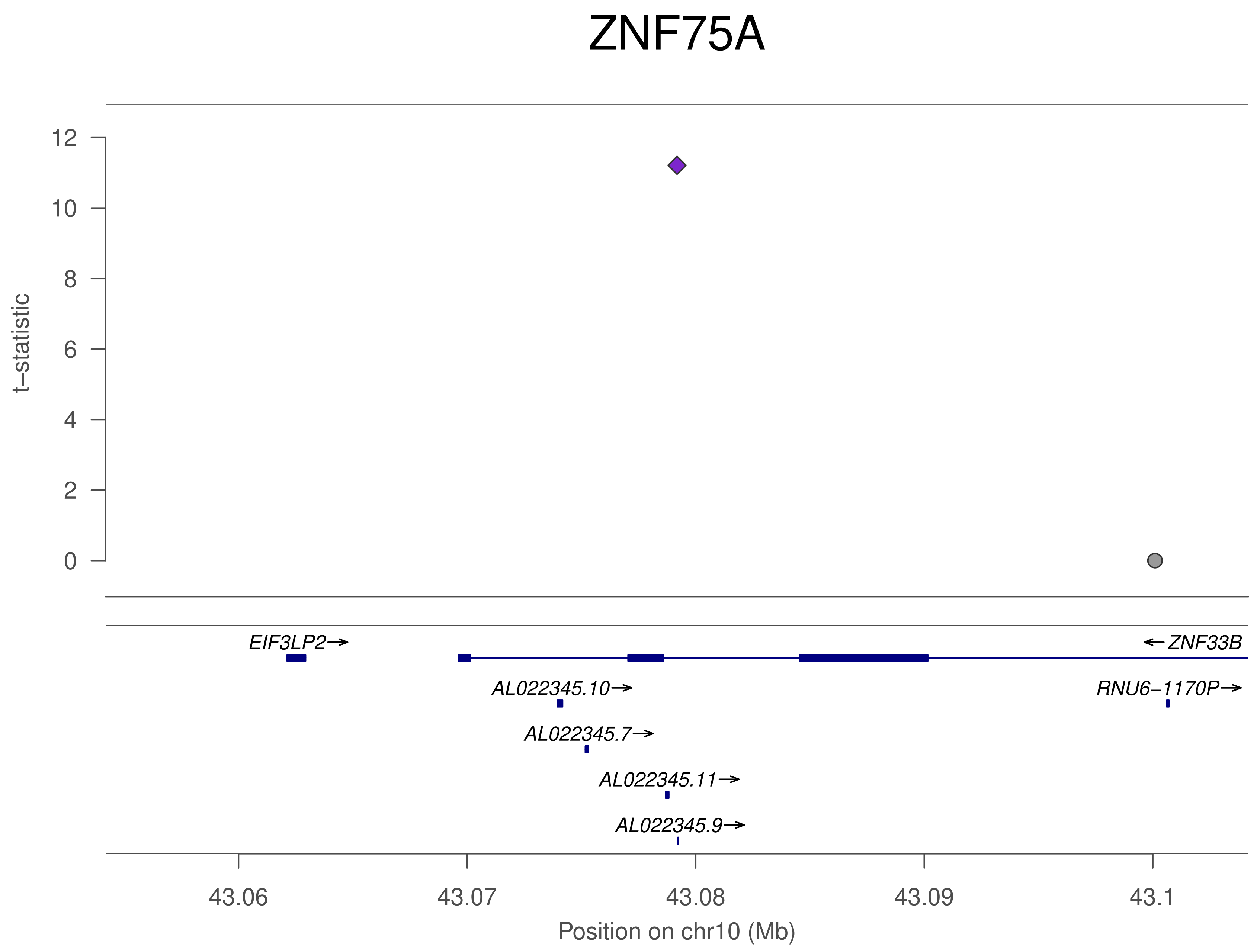
ZNF75A
t–statistic
12
10
8
6
4
2
0
EIF3LP2→
←ZNF33B
AL022345.10→
RNU6–1170P→
AL022345.7→
AL022345.11→
AL022345.9→
43.06
43.07
43.08
43.09
43.1
Position on chr10 (Mb)

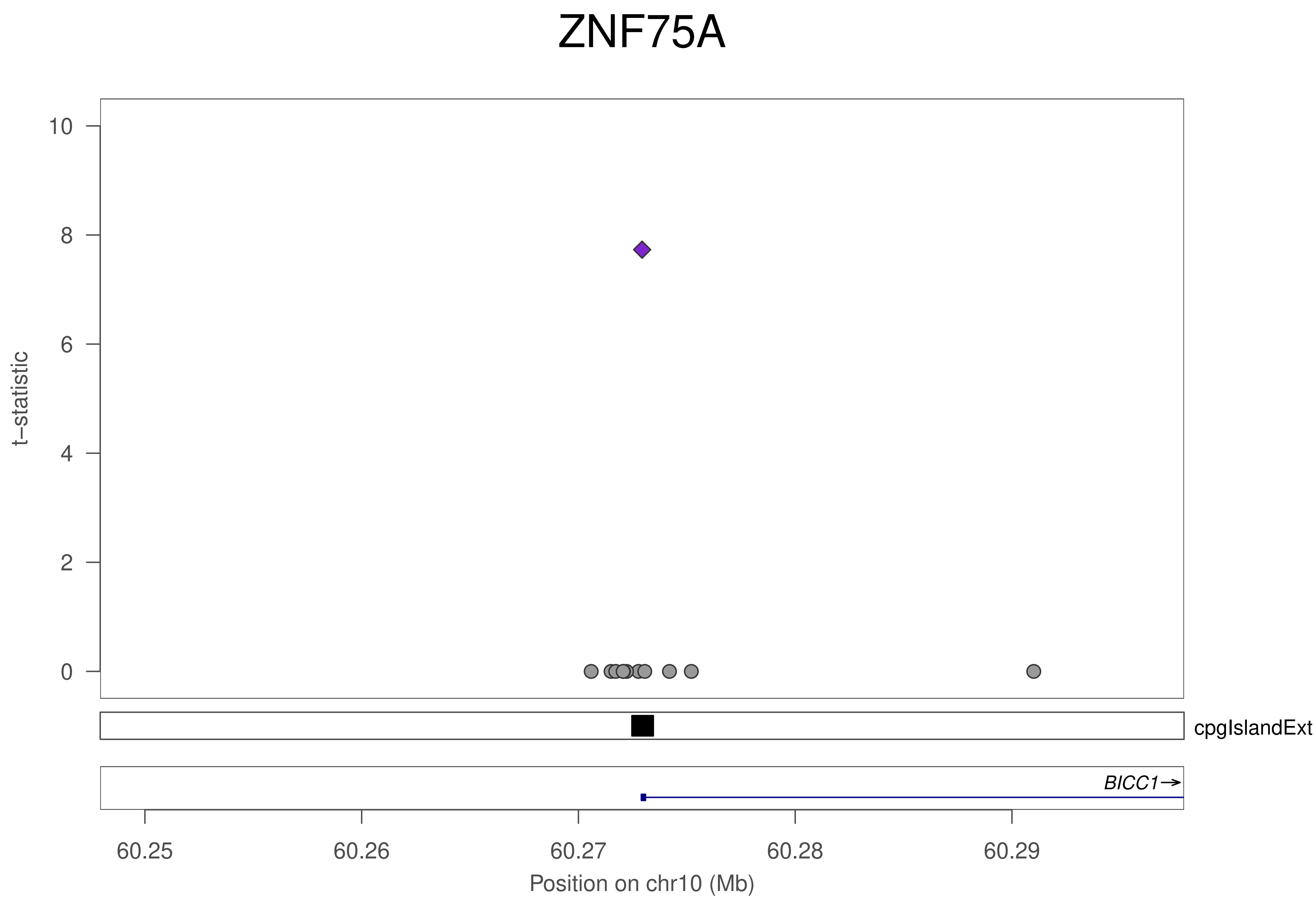
ZNF75A
t–statistic
0
2
4
6
8
10
cpgIslandExt
BICC1→
60.25
60.26
60.27
60.28
60.29
Position on chr10 (Mb)

### ZNF75A

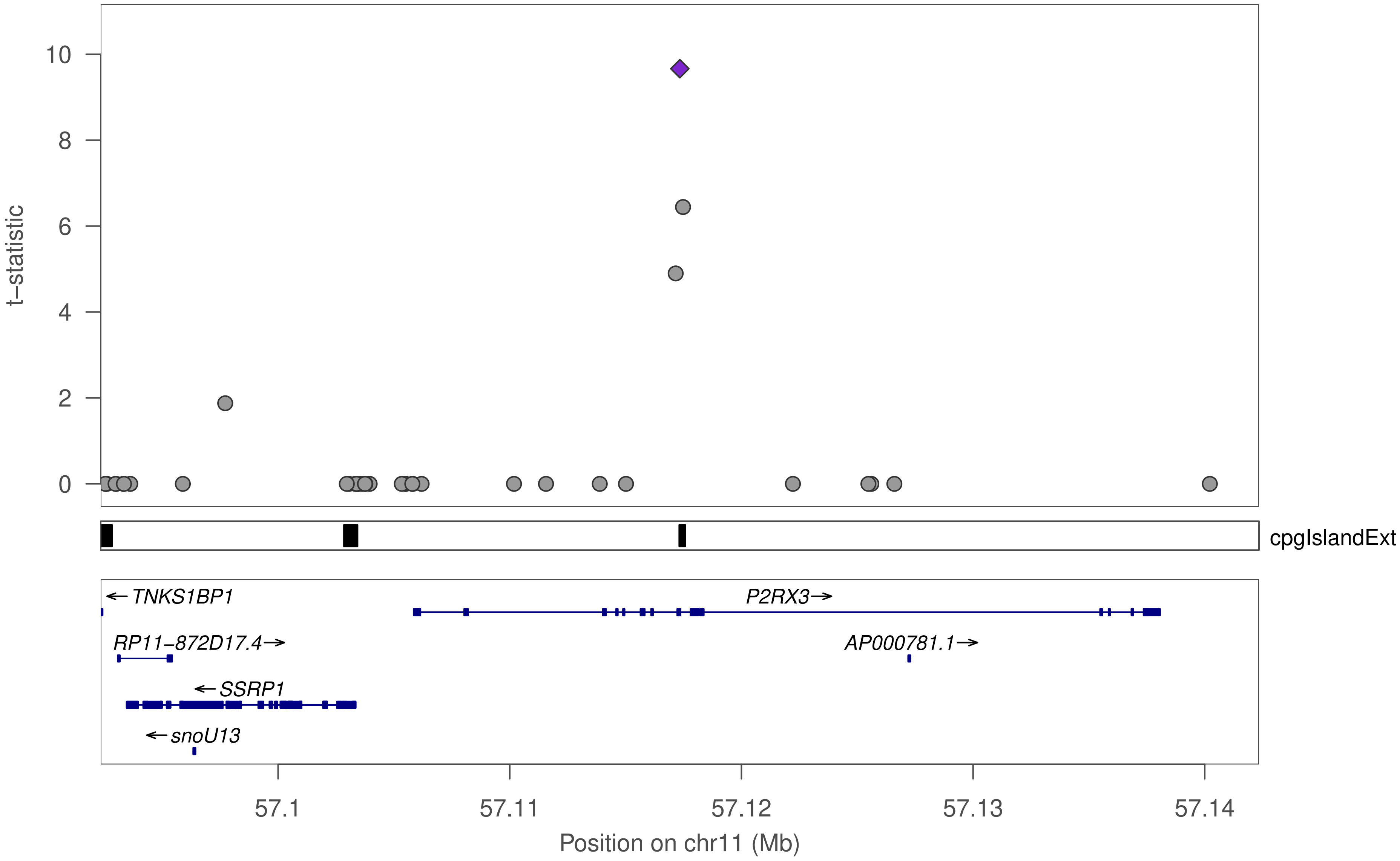

t−statistic
10
8
6
4
2
0
cpgIslandExt
←TNKS1BP1
P2RX3→
RP11−872D17.4→
AP000781.1→
←SSRP1
←snoU13
57.1
57.11
57.12
57.13
57.14
Position on chr11 (Mb)

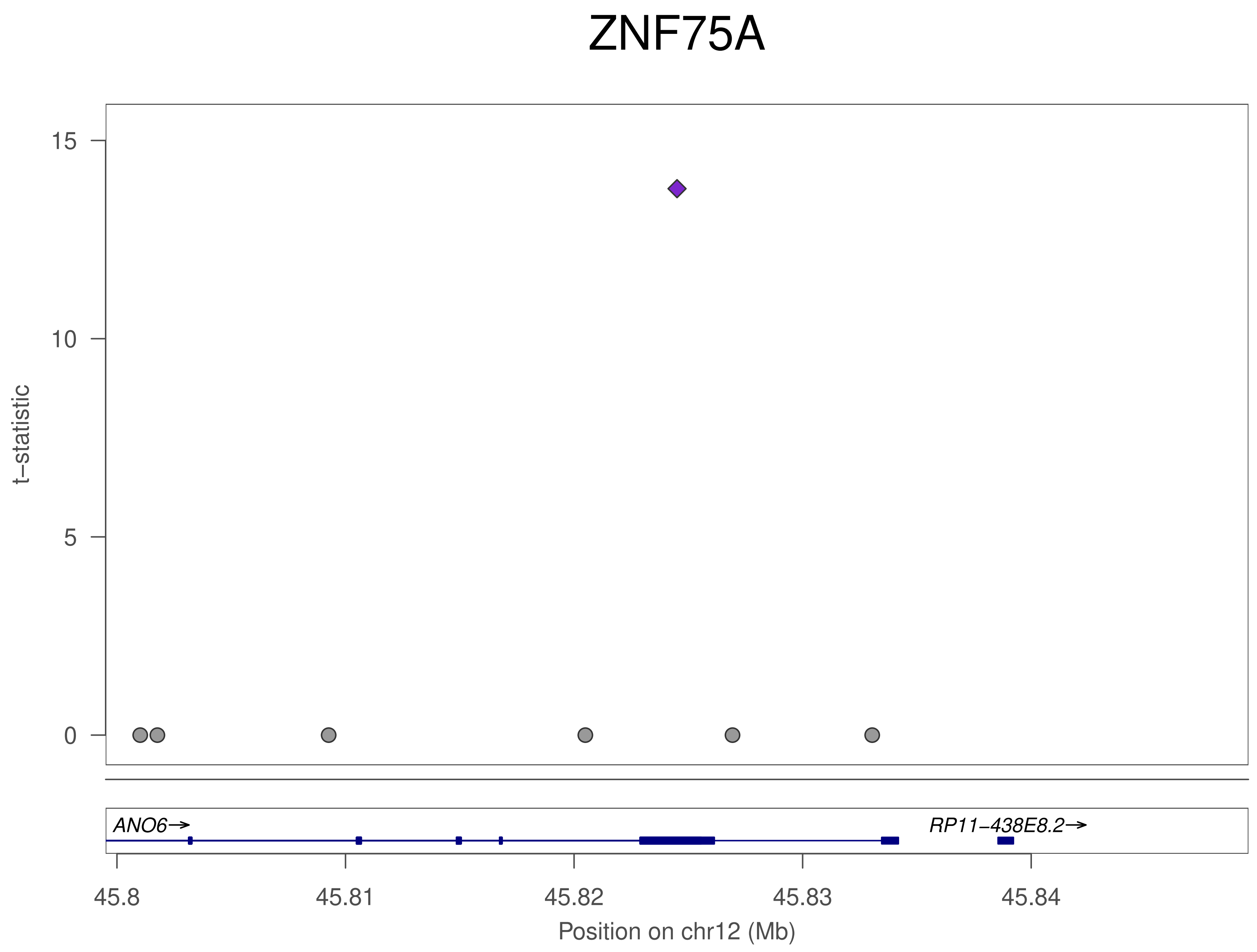

ZNF75A
15
10
5
0
t−statistic
ANO6→
RP11−438E8.2→
45.8
45.81
45.82
45.83
45.84
Position on chr12 (Mb)

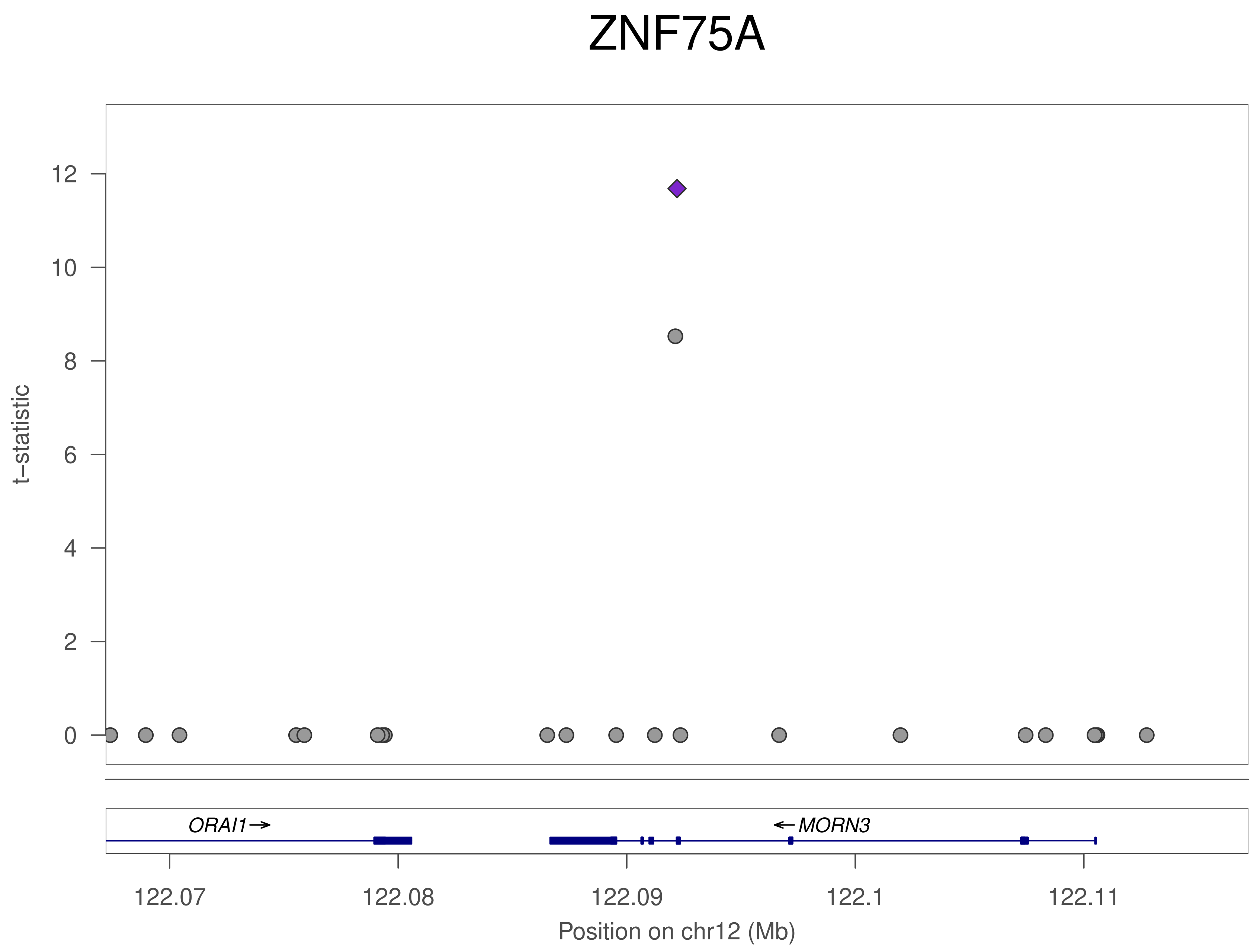
ZNF75A
t−statistic
12
10
8
6
4
2
0
ORAI1→
←MORN3
122.07
122.08
122.09
122.1
122.11
Position on chr12 (Mb)

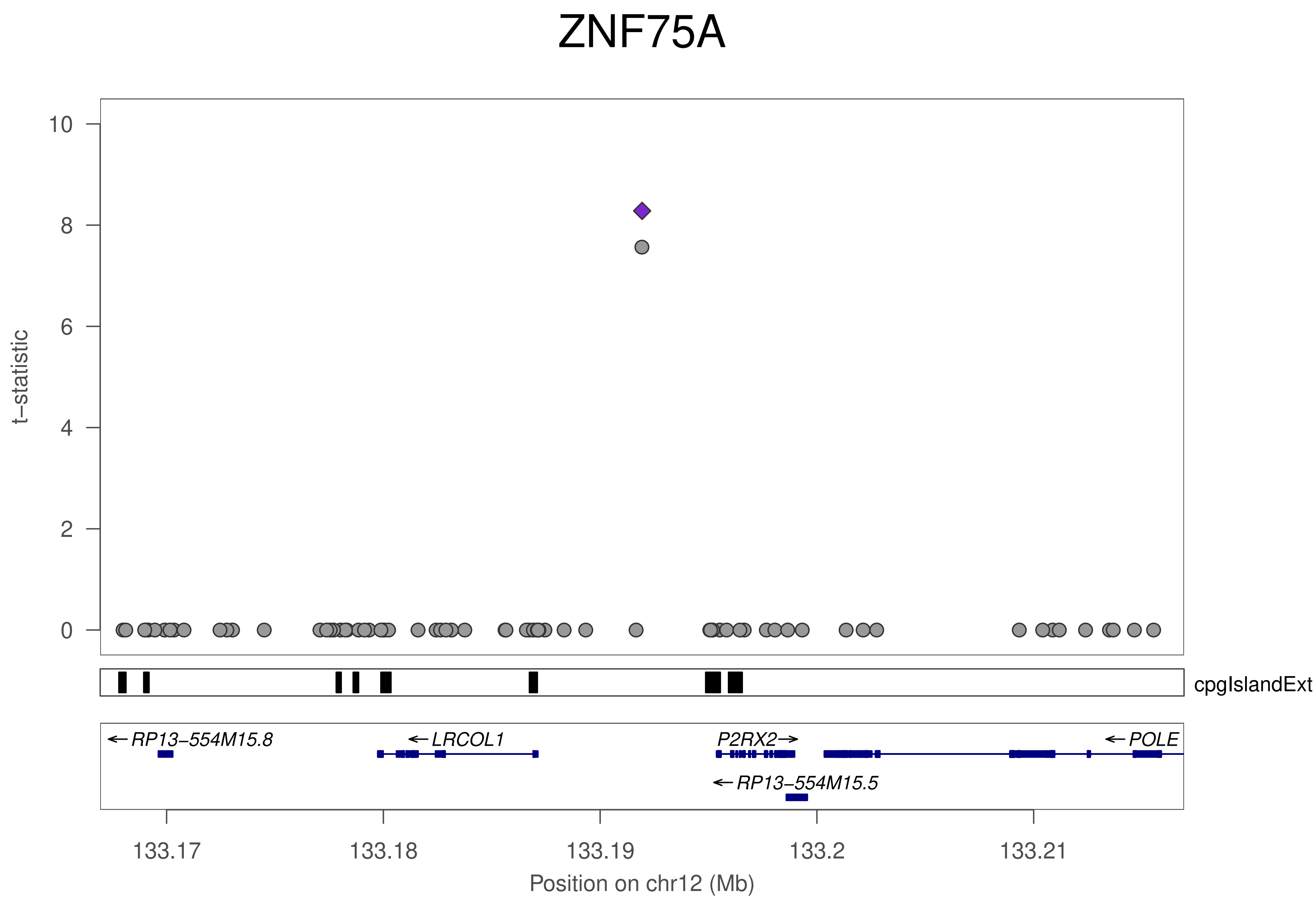
ZNF75A
t–statistic
10
8
6
4
2
0
cpgIslandExt
←RP13–554M15.8
←LRCOL1
P2RX2→
←POLE
←RP13–554M15.5
133.17
133.18
133.19
133.2
133.21
Position on chr12 (Mb)

### ZNF75A

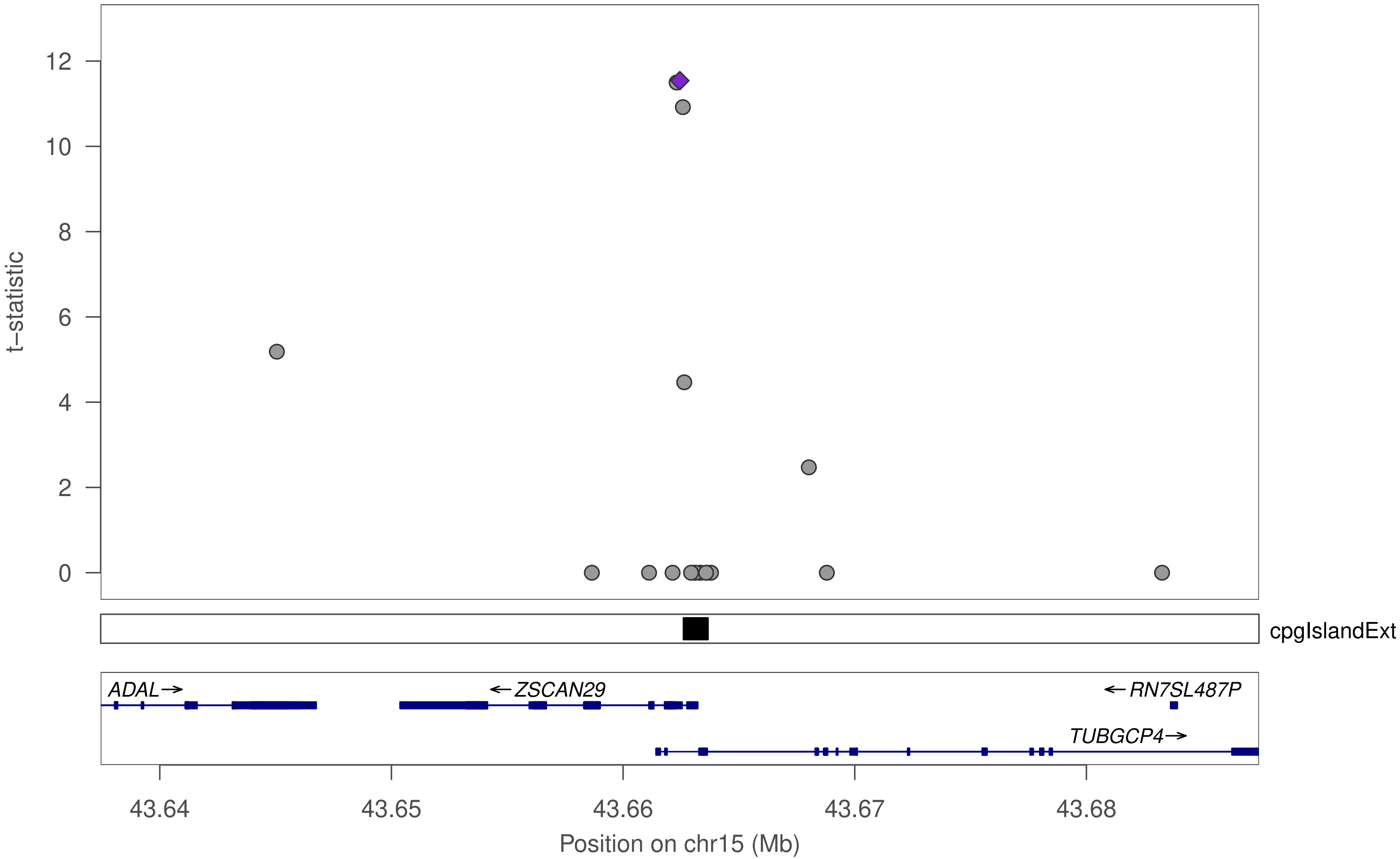

t−statistic
12
10
8
6
4
2
0
cpgIslandExt
ADAL→
←ZSCAN29
←RN7SL487P
TUBGCP4→
43.64
43.65
43.66
43.67
43.68
Position on chr15 (Mb)

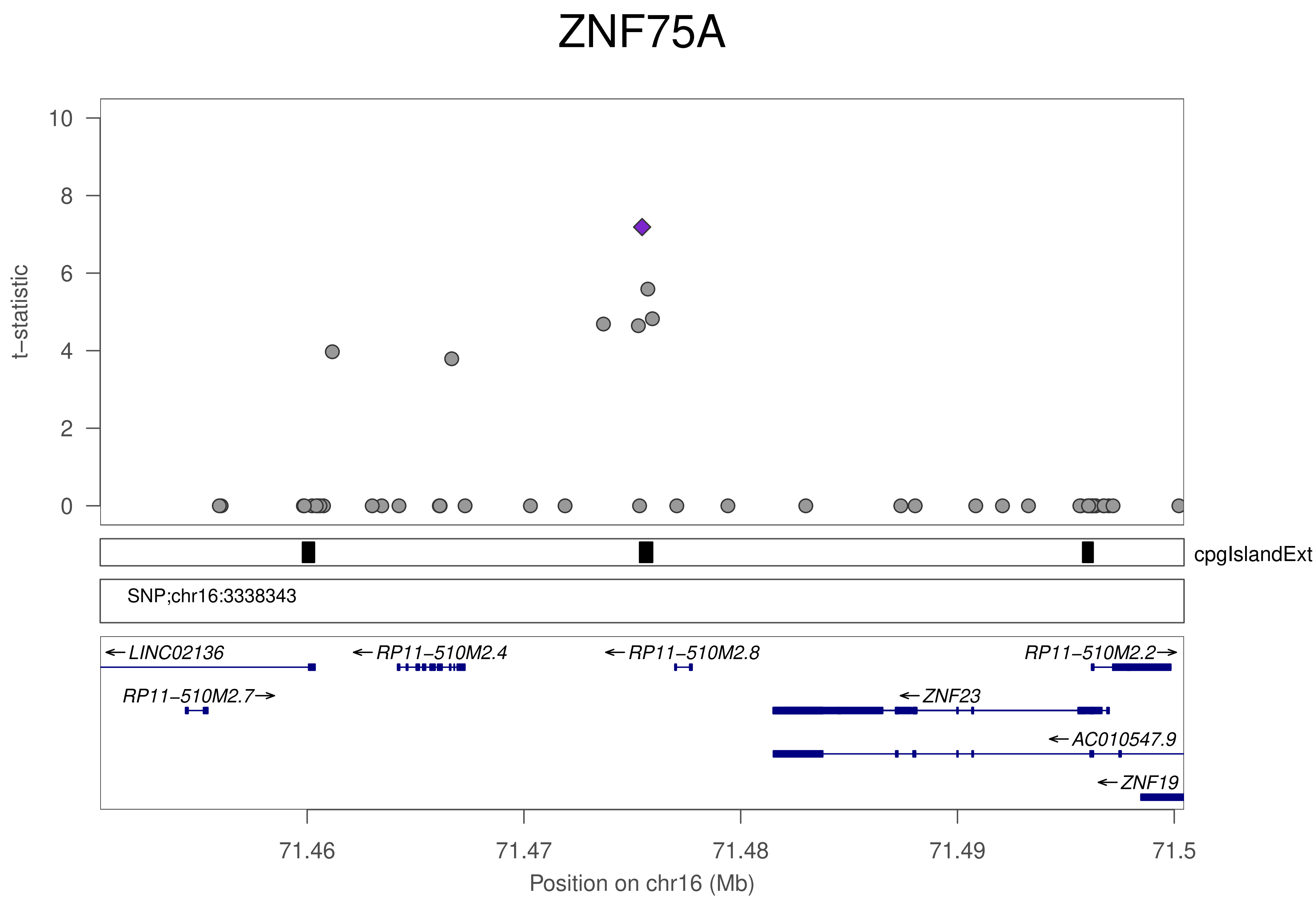
ZNF75A
t–statistic
10
8
6
4
2
0
cpgIslandExt
SNP;chr16:3338343
←LINC02136
←RP11–510M2.4
←RP11–510M2.8
RP11–510M2.2→
RP11–510M2.7→
←ZNF23
←AC010547.9
←ZNF19
71.46
71.47
71.48
71.49
71.5
Position on chr16 (Mb)

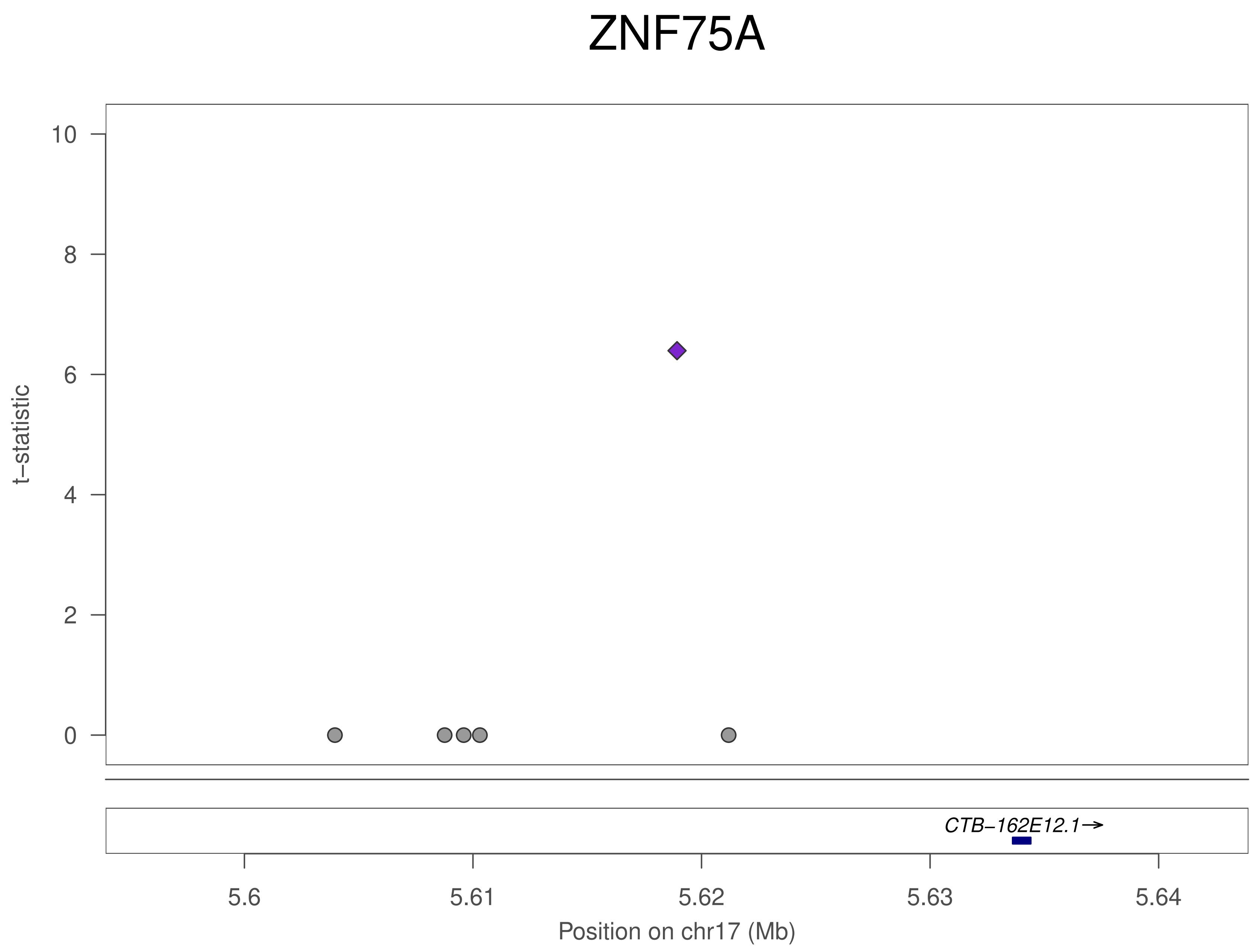
ZNF75A
t−statistic
10
8
6
4
2
0
CTB−162E12.1→
5.6
5.61
5.62
5.63
5.64
Position on chr17 (Mb)

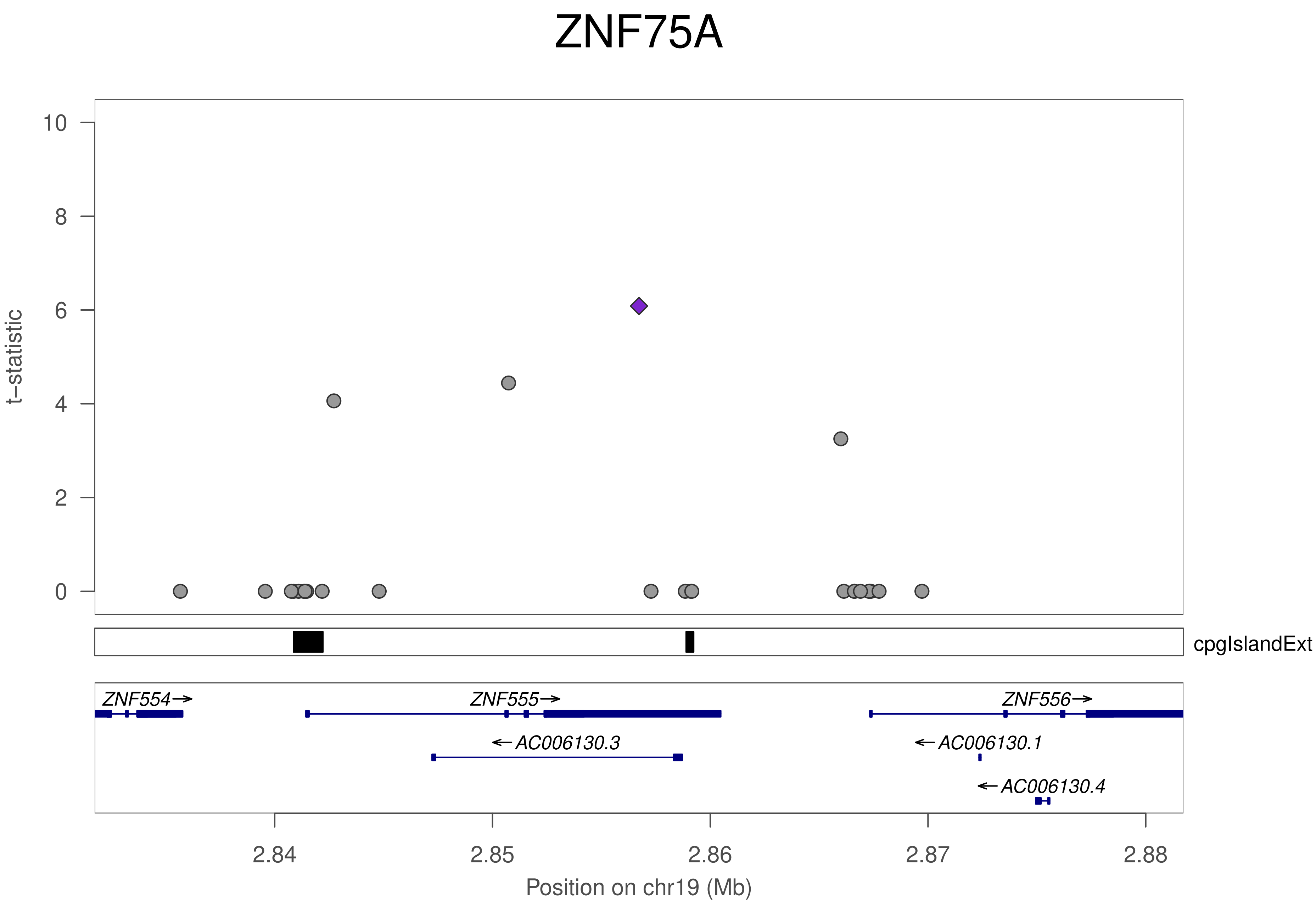

ZNF75A
t−statistic
10
8
6
4
2
0
cpgIslandExt
ZNF554→
ZNF555→
ZNF556→
←AC006130.3
←AC006130.1
←AC006130.4
2.84
2.85
2.86
2.87
2.88
Position on chr19 (Mb)

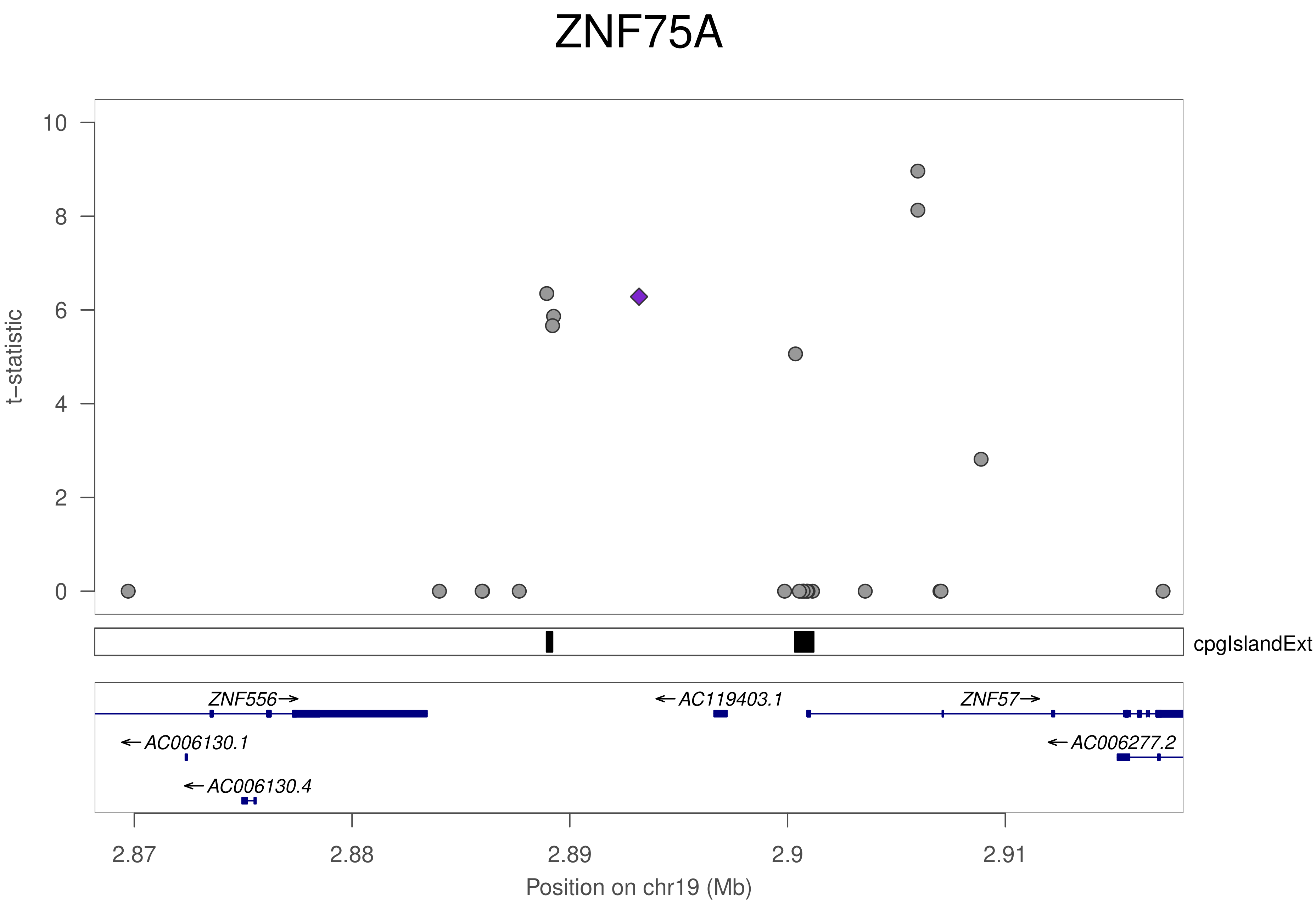

ZNF75A
t–statistic
10
8
6
4
2
0
cpgIslandExt
ZNF556
AC119403.1
ZNF57
AC006130.1
AC006277.2
AC006130.4
2.87
2.88
2.89
2.9
2.91
Position on chr19 (Mb)

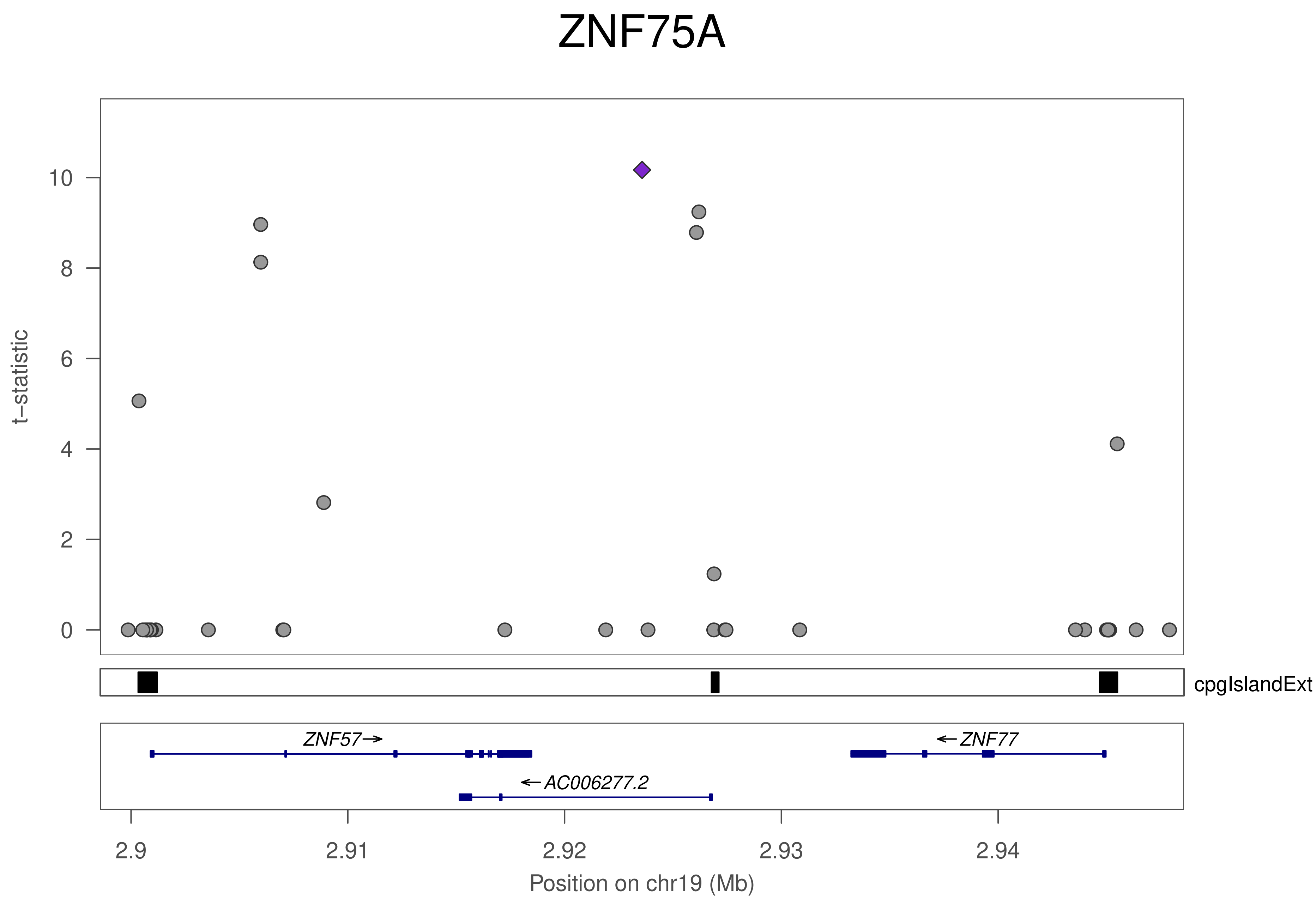
ZNF75A
t–statistic
0
2
4
6
8
10
cpgIslandExt
ZNF57→
←ZNF77
←AC006277.2
2.9
2.91
2.92
2.93
2.94
Position on chr19 (Mb)

### ZNF75A

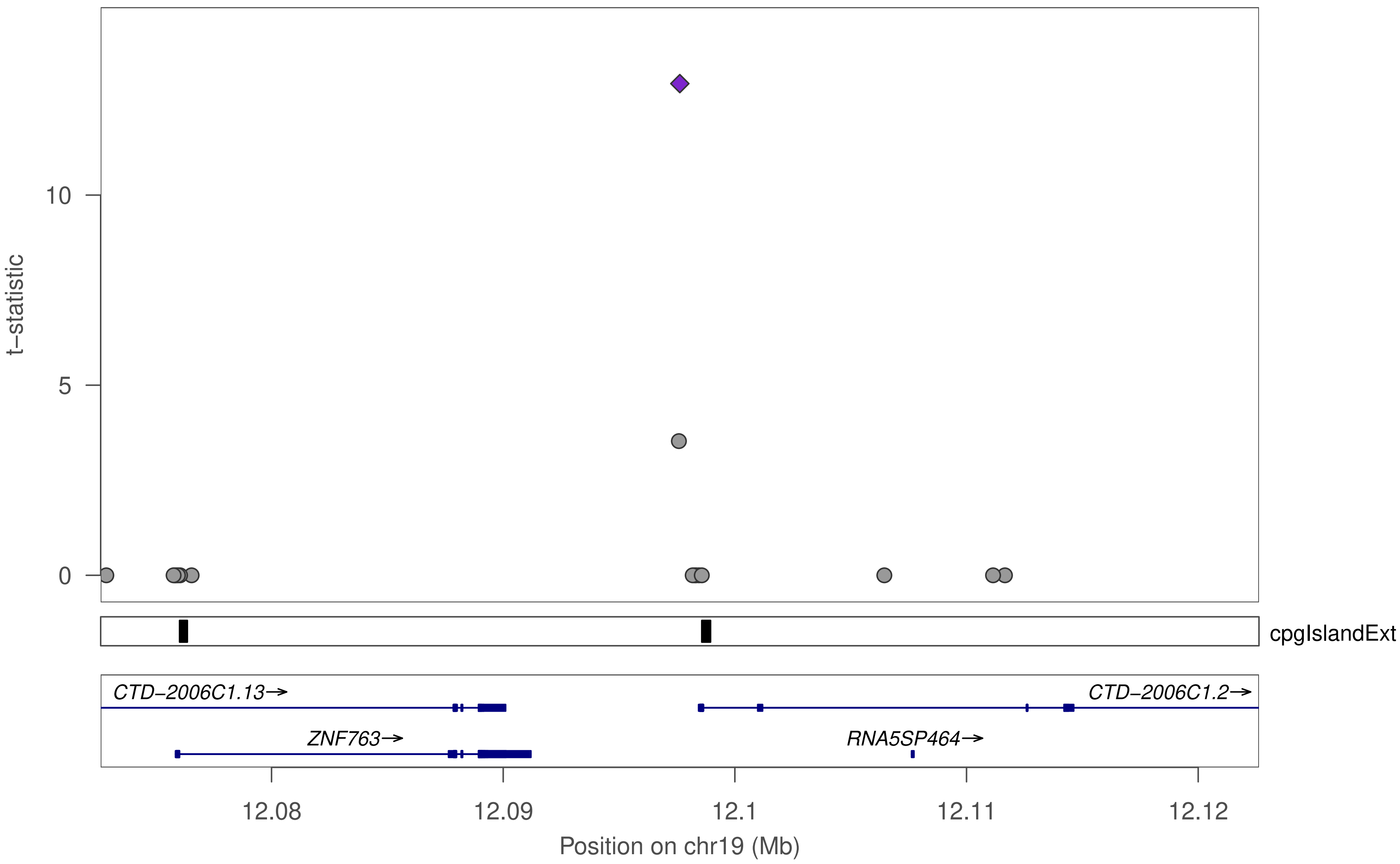

t−statistic
10
5
0
cpgIslandExt
CTD−2006C1.13→
CTD−2006C1.2→
ZNF763→
RNA5SP464→
12.08
12.09
12.1
12.11
12.12
Position on chr19 (Mb)

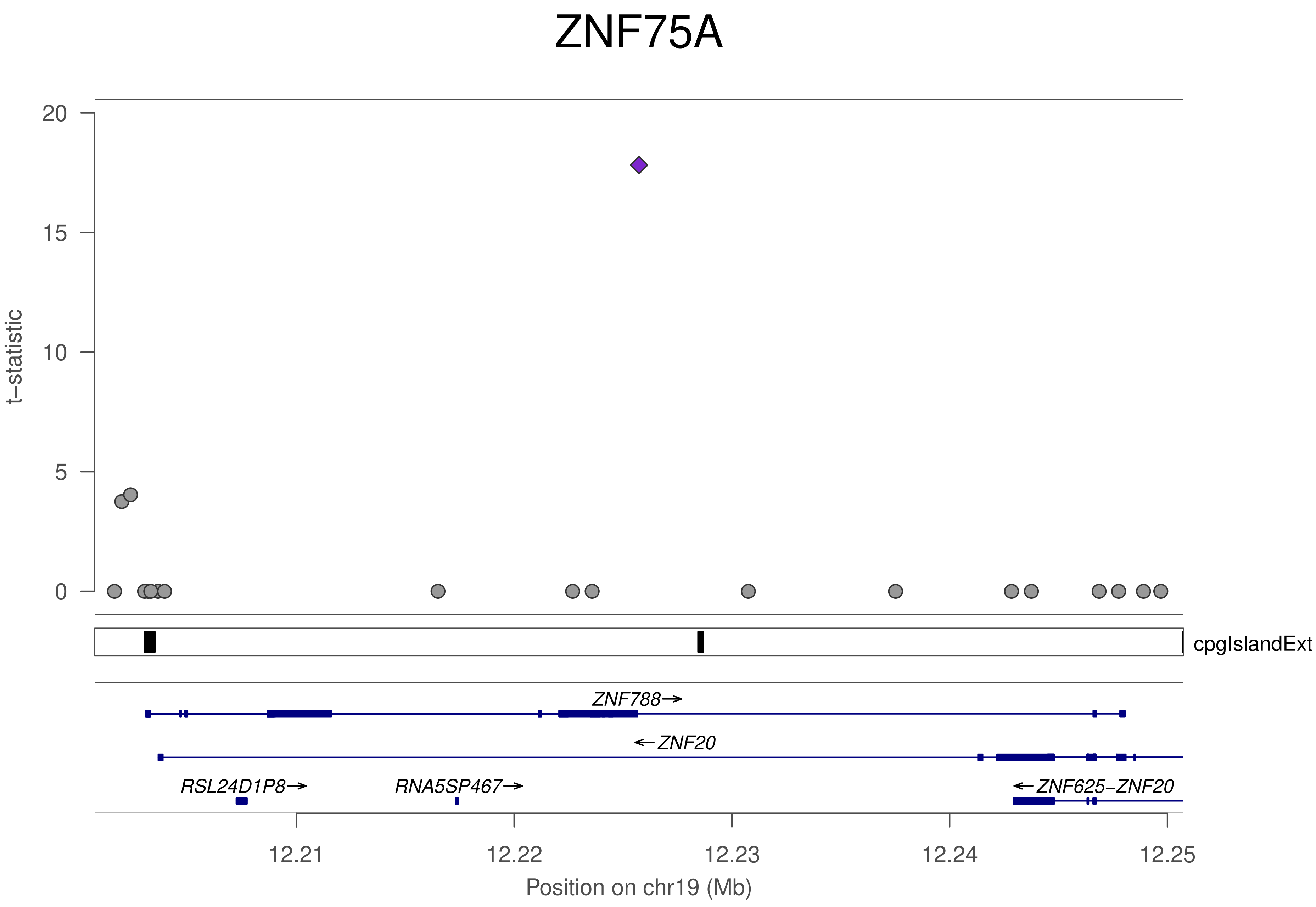

ZNF75A
t−statistic
20
15
10
5
0
cpgIslandExt
ZNF788→
←ZNF20
RSL24D1P8→
RNA5SP467→
←ZNF625−ZNF20
12.21
12.22
12.23
12.24
12.25
Position on chr19 (Mb)

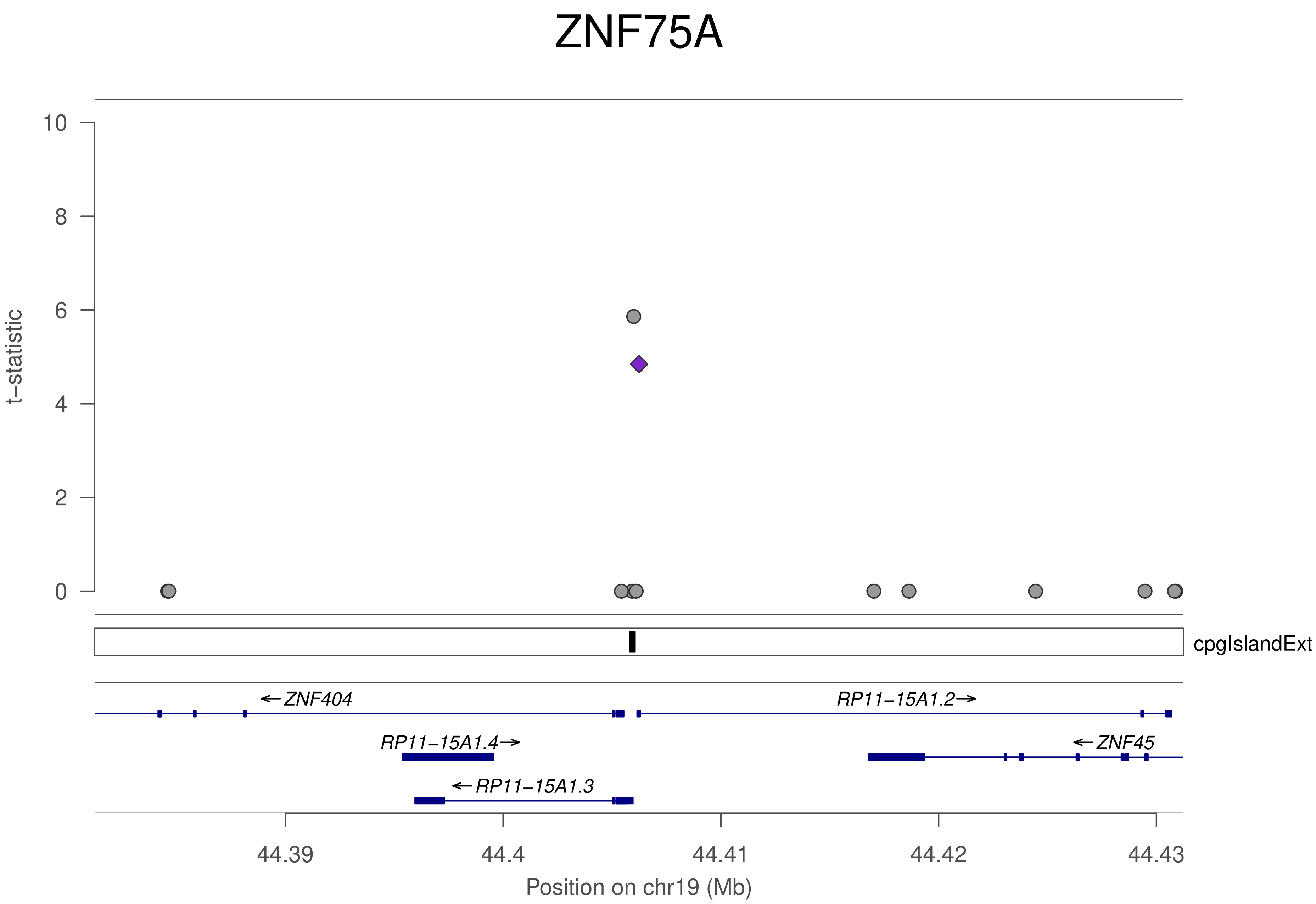
ZNF75A
t−statistic
10
8
6
4
2
0
cpgIslandExt
←ZNF404
RP11−15A1.2→
RP11−15A1.4→
←ZNF45
←RP11−15A1.3
44.39
44.4
44.41
44.42
44.43
Position on chr19 (Mb)

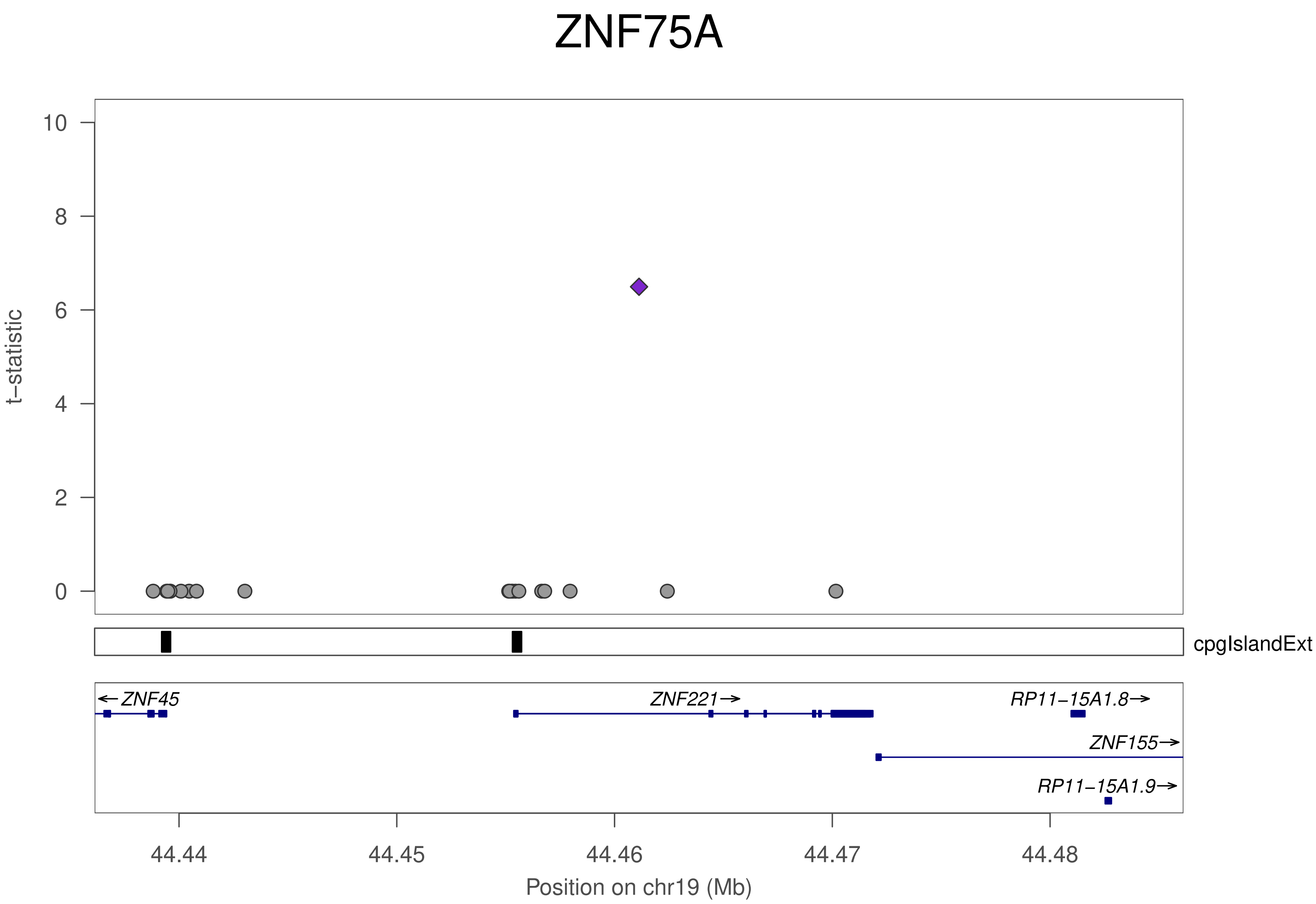
ZNF75A
t−statistic
10
8
6
4
2
0
cpgIslandExt
←ZNF45
ZNF221→
RP11−15A1.8→
ZNF155→
RP11−15A1.9→
44.44
44.45
44.46
44.47
44.48
Position on chr19 (Mb)

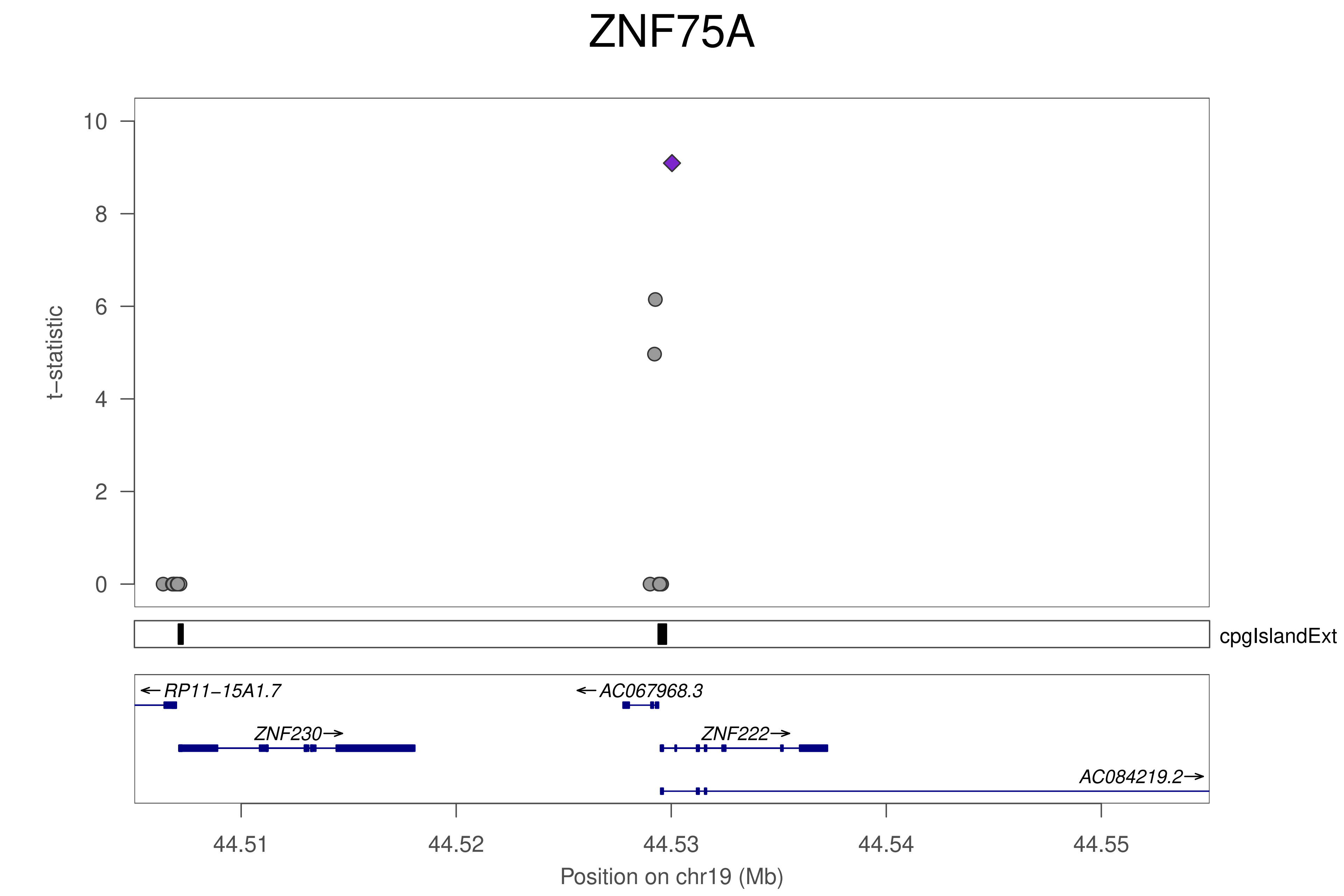
ZNF75A
t−statistic
10
8
6
4
2
0
cpgIslandExt
←RP11−15A1.7
←AC067968.3
ZNF230→
ZNF222→
AC084219.2→
44.51
44.52
44.53
44.54
44.55
Position on chr19 (Mb)

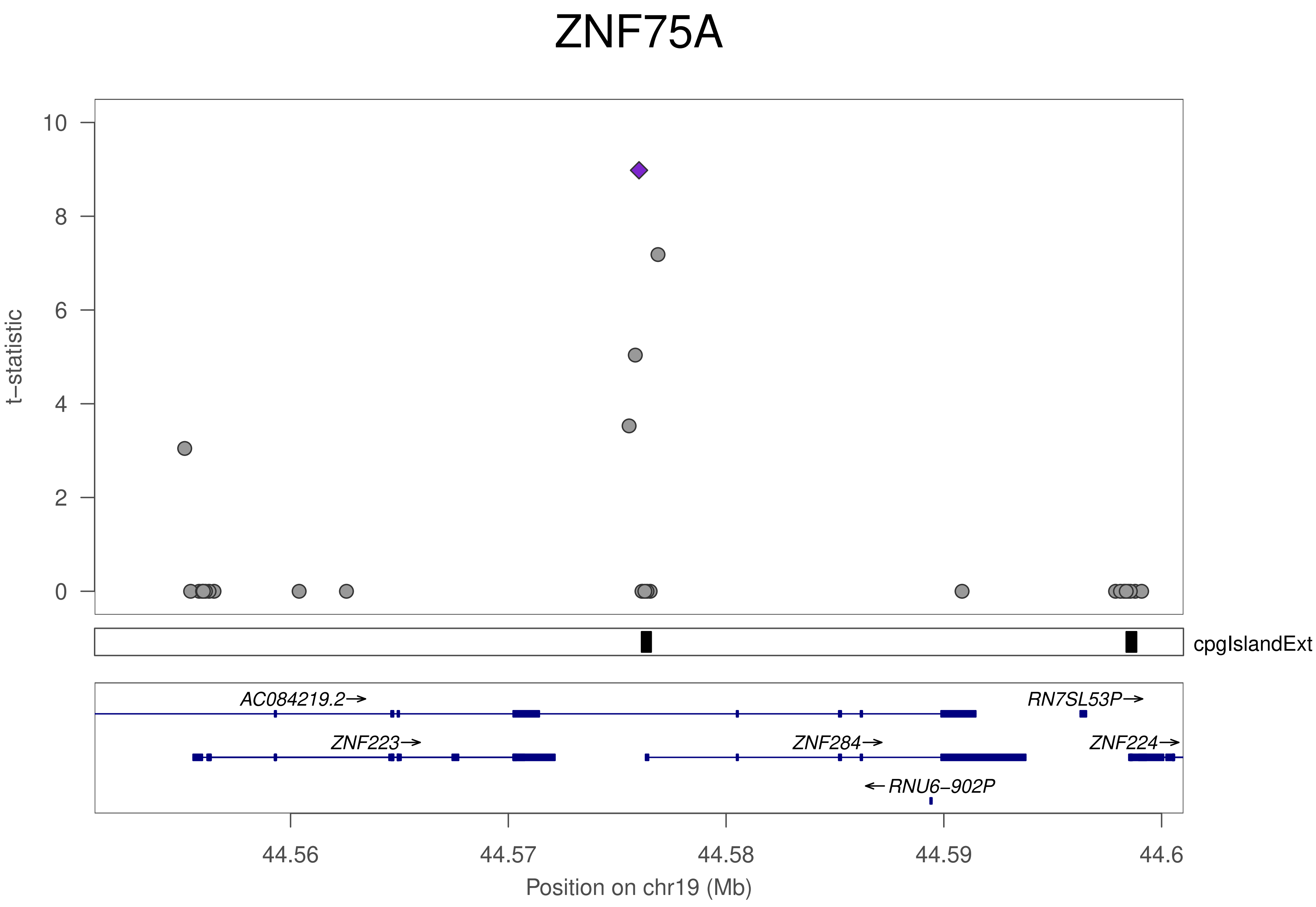
ZNF75A
10
8
6
4
2
0
t−statistic
cpgIslandExt
AC084219.2→
RN7SL53P→
ZNF223→
ZNF284→
ZNF224→
←RNU6−902P
44.56
44.57
44.58
44.59
44.6
Position on chr19 (Mb)

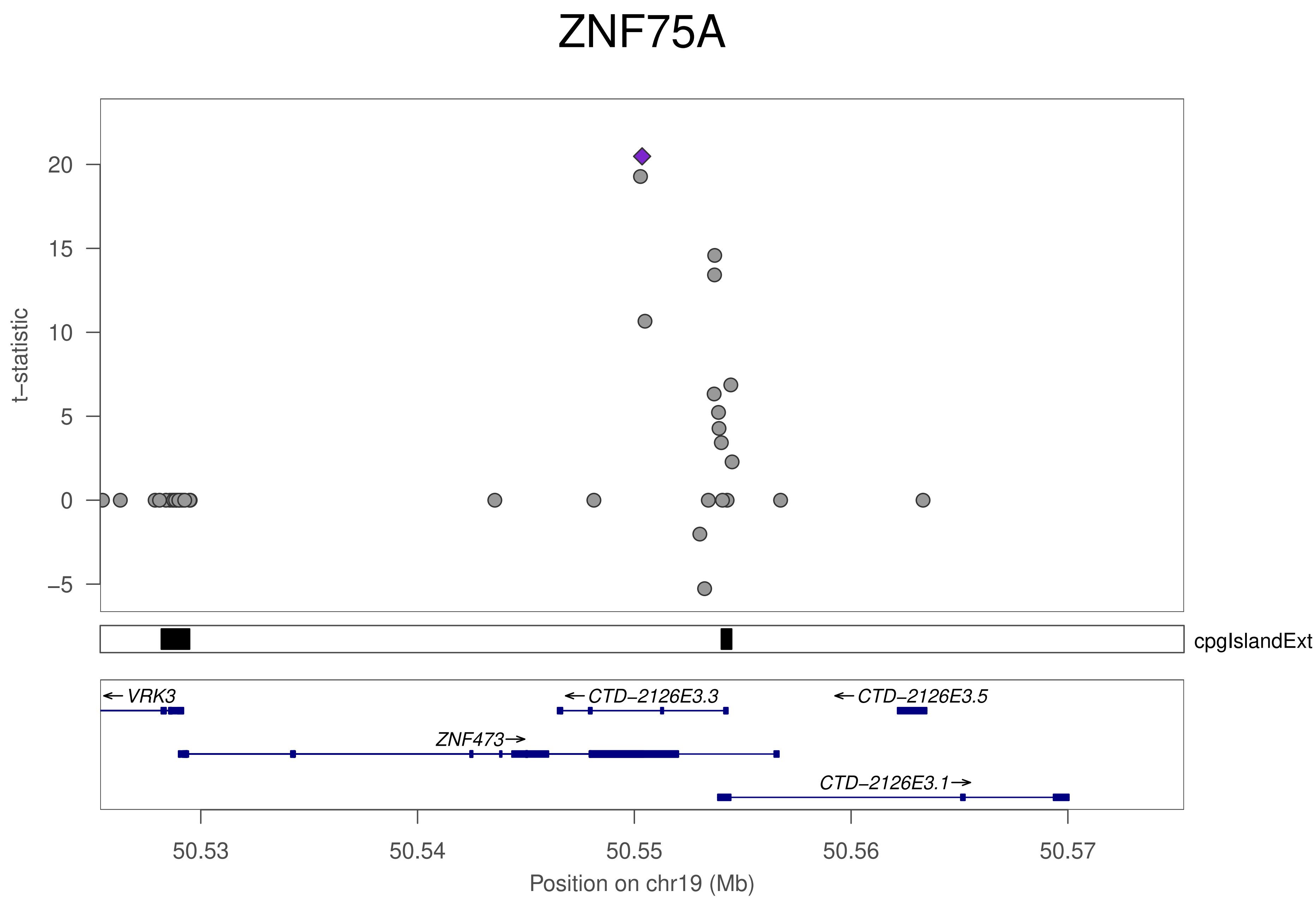

ZNF75A
t−statistic
20
15
10
5
0
−5
cpgIslandExt
← VRK3
← CTD−2126E3.3
← CTD−2126E3.5
ZNF473 →
CTD−2126E3.1 →
50.53
50.54
50.55
50.56
50.57
Position on chr19 (Mb)

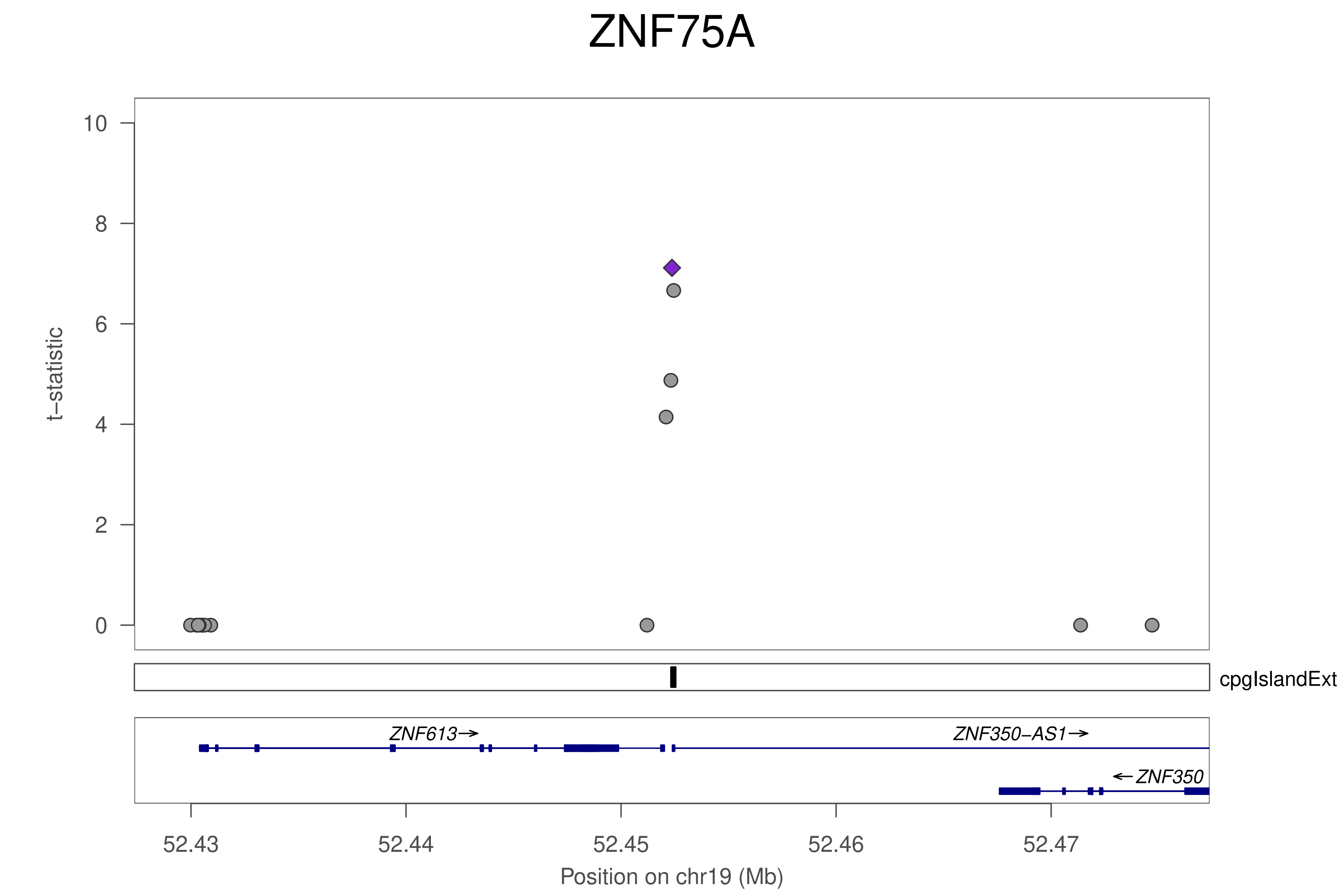
ZNF75A
t−statistic
10
8
6
4
2
0
cpgIslandExt
ZNF613→
ZNF350−AS1→
←ZNF350
52.43
52.44
52.45
52.46
52.47
Position on chr19 (Mb)

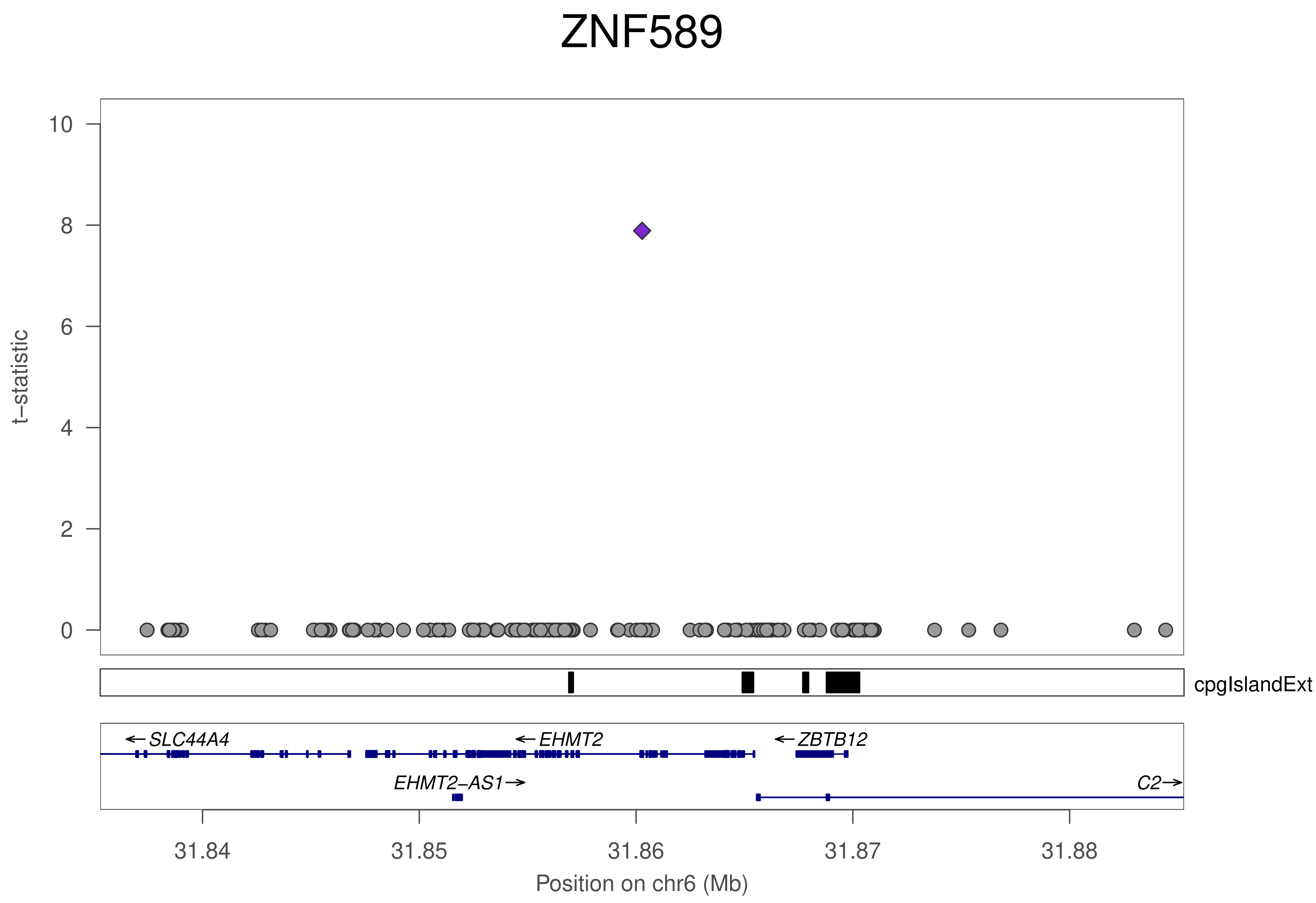
ZNF589
t−statistic
10
8
6
4
2
0
cpgIslandExt
←SLC44A4
←EHMT2
←ZBTB12
EHMT2−AS1→
C2→
31.84
31.85
31.86
31.87
31.88
Position on chr6 (Mb)

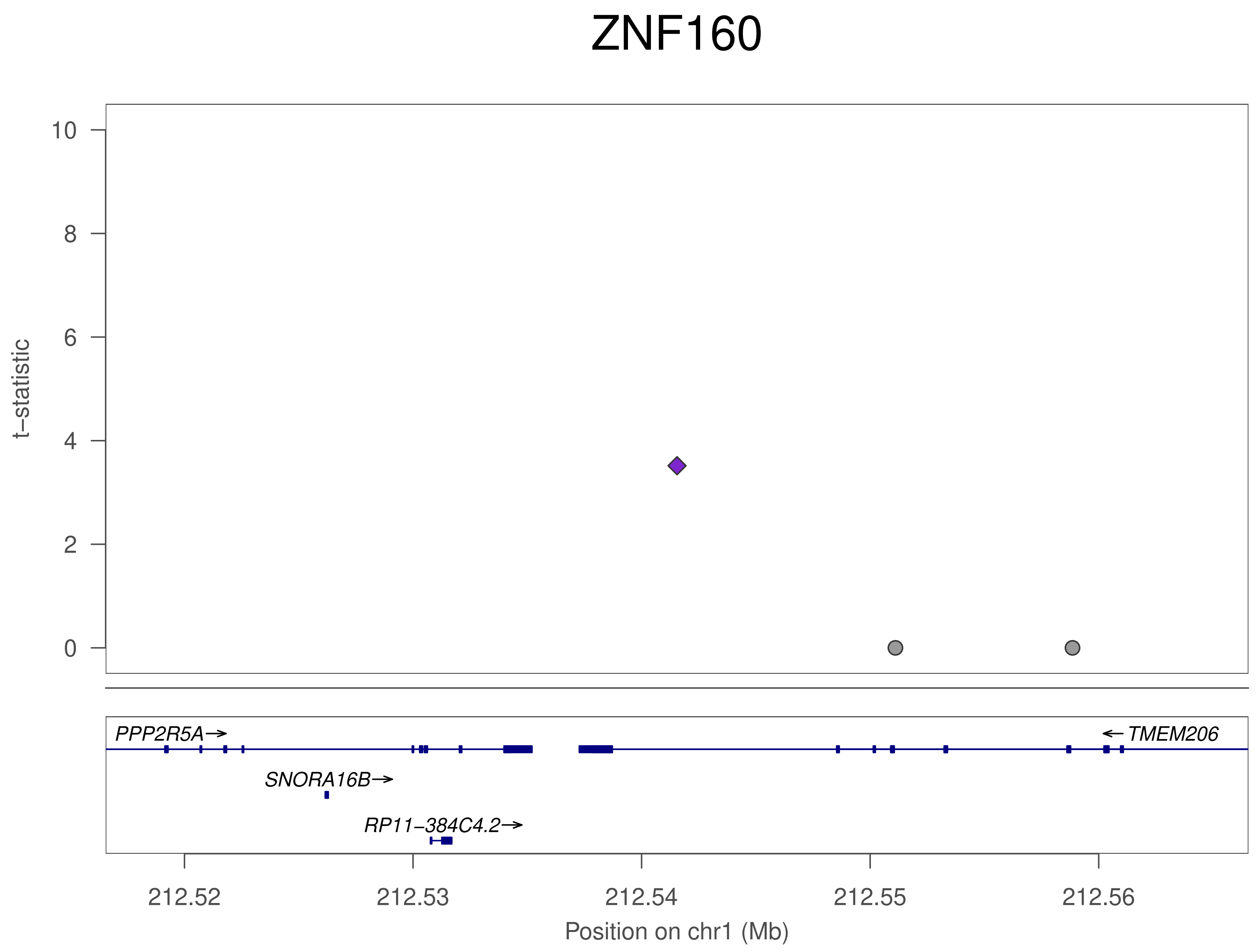
ZNF160
t−statistic
10
8
6
4
2
0
PPP2R5A→
SNORA16B→
RP11−384C4.2→
← TMEM206
212.52
212.53
212.54
212.55
212.56
Position on chr1 (Mb)

### ZNF554

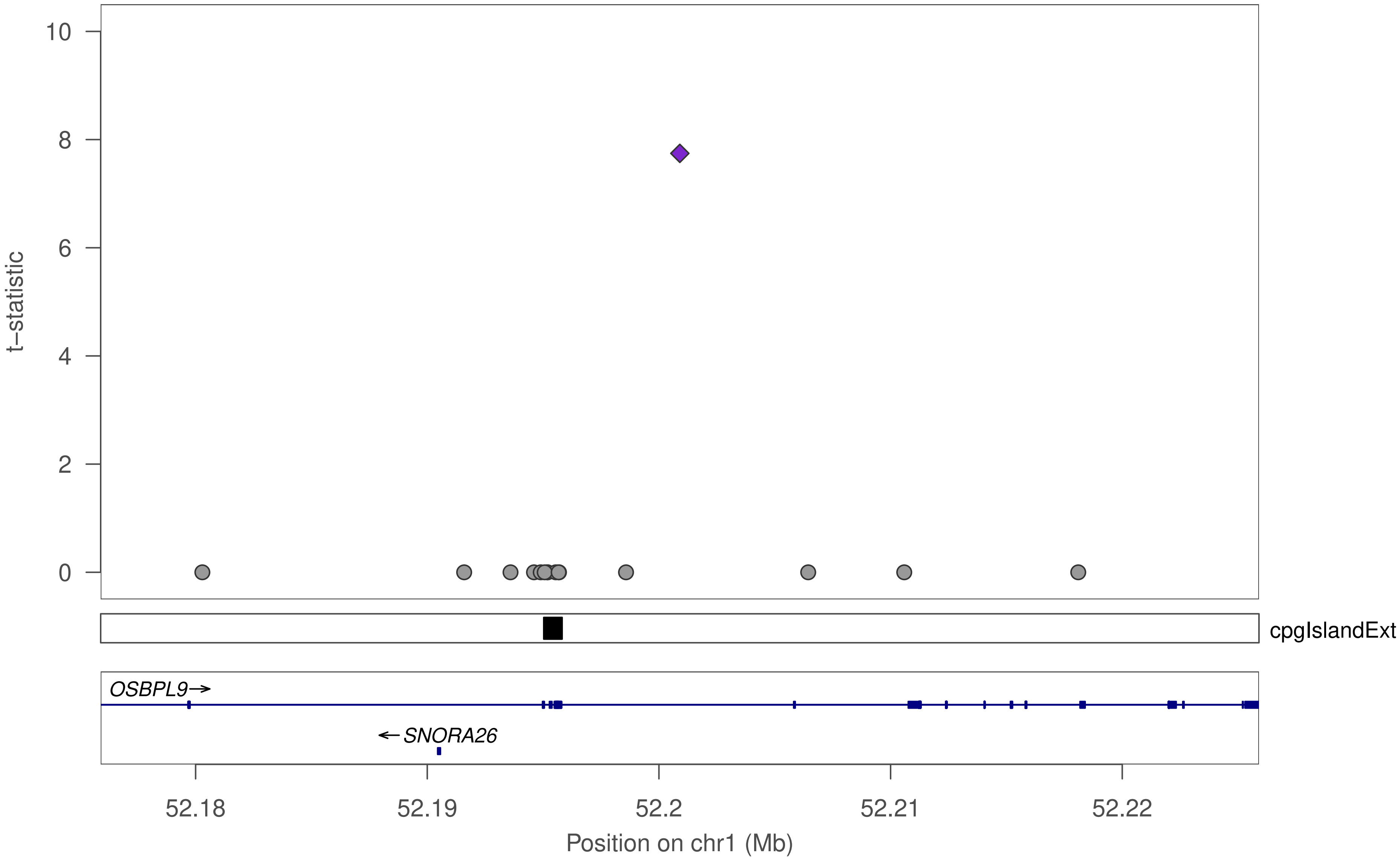

t–statistic
10
8
6
4
2
0
cpgIslandExt
OSBPL9→
←SNORA26
52.18
52.19
52.2
52.21
52.22
Position on chr1 (Mb)

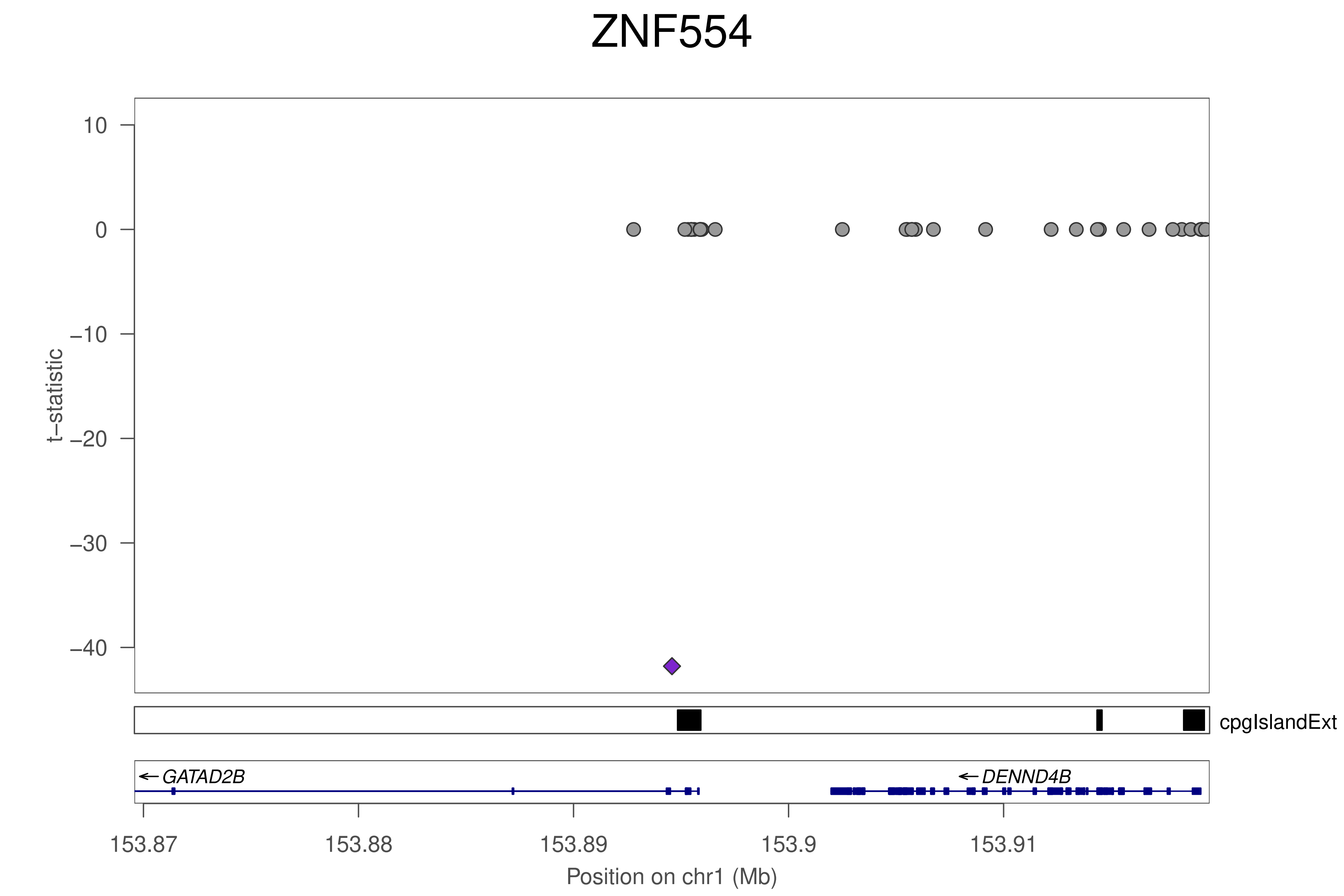

ZNF554
t–statistic
10
0
−10
−20
−30
−40
cpgIslandExt
GATAD2B
DENND4B
153.87
153.88
153.89
153.9
153.91
Position on chr1 (Mb)

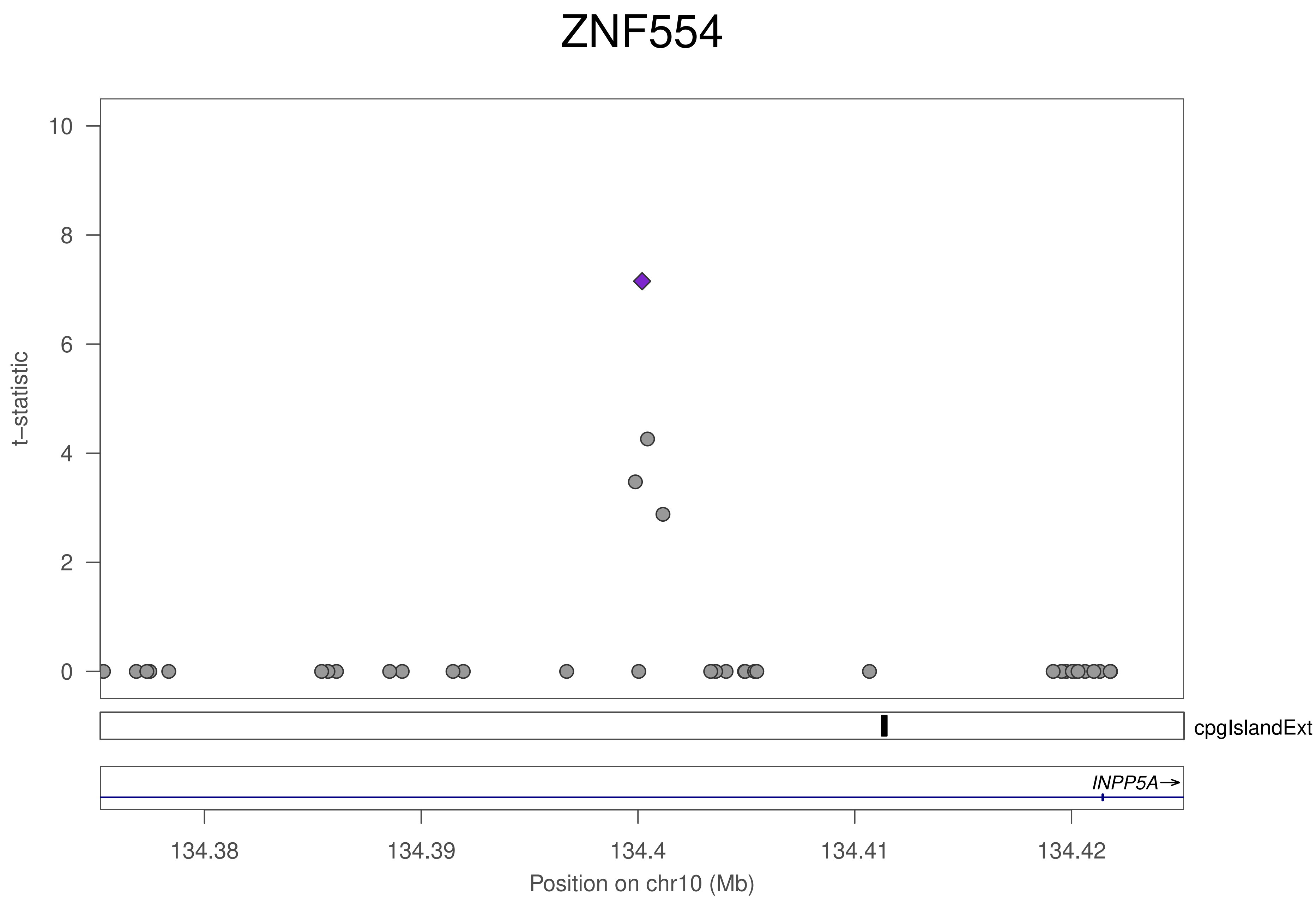

ZNF554
t−statistic
10
8
6
4
2
0
cpgIslandExt
INPP5A→
134.38
134.39
134.4
134.41
134.42
Position on chr10 (Mb)

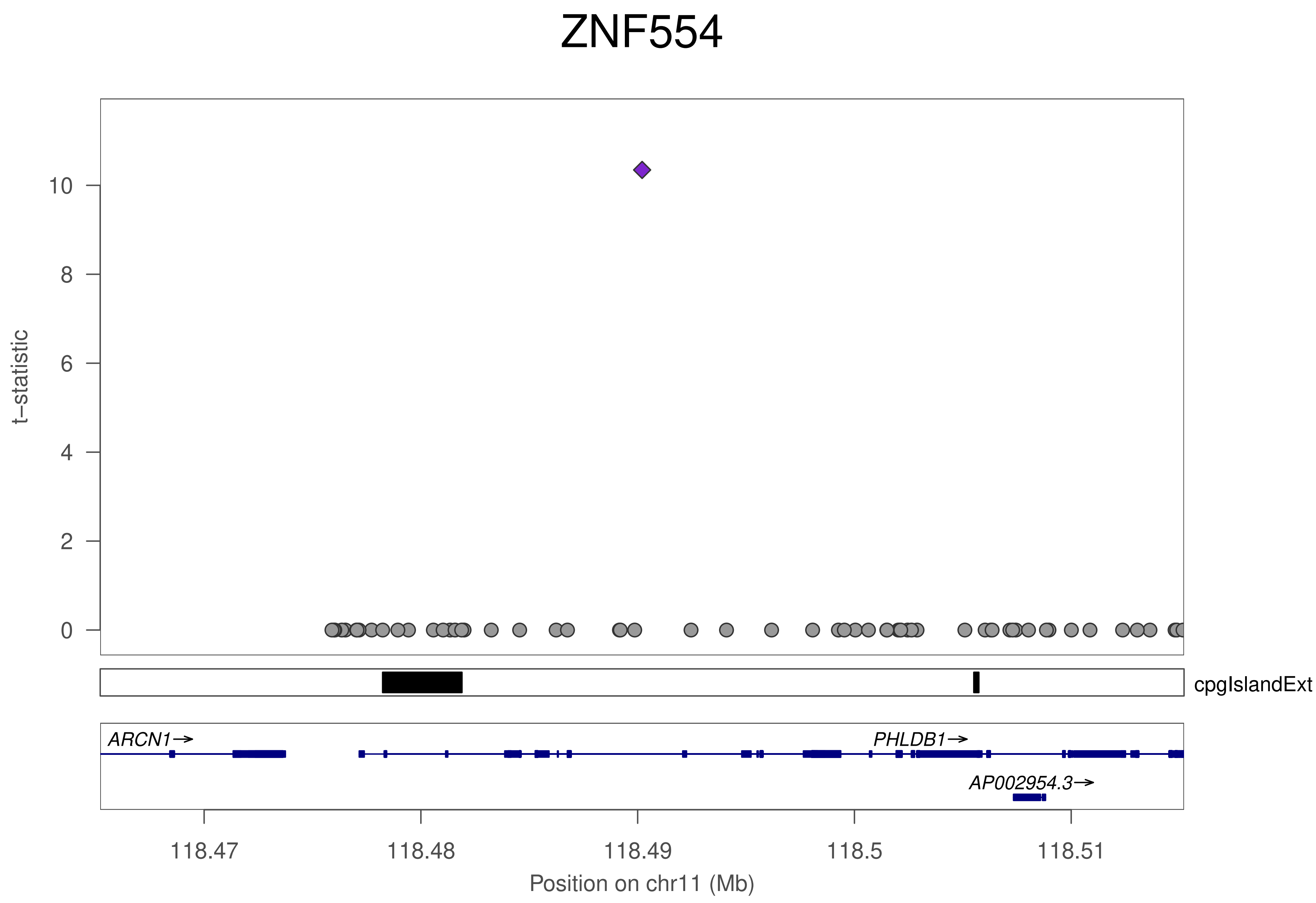
ZNF554
t−statistic
0
2
4
6
8
10
cpgIslandExt
ARCN1→
PHLDB1→
AP002954.3→
118.47
118.48
118.49
118.5
118.51
Position on chr11 (Mb)

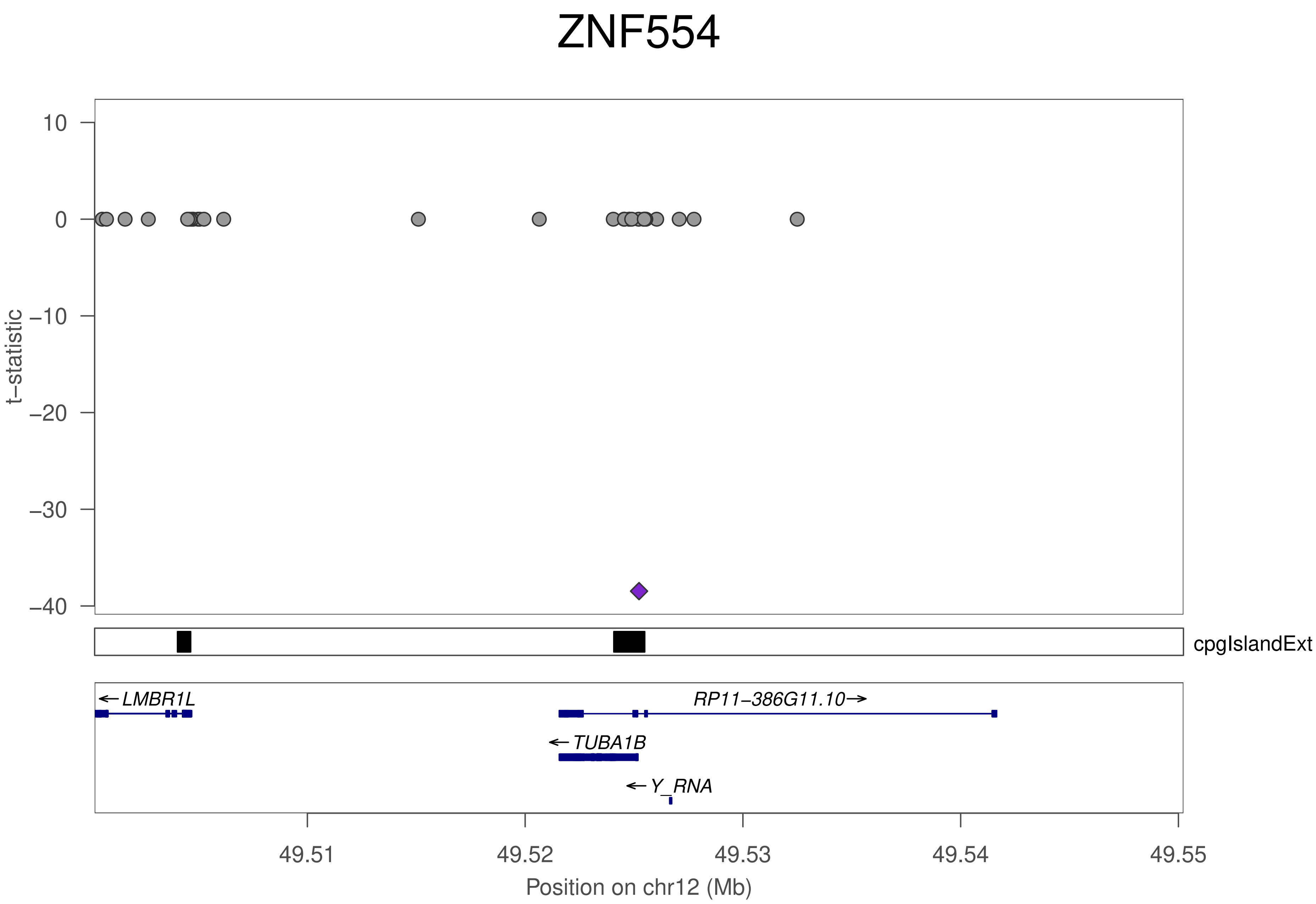
ZNF554
t−statistic
10
0
−10
−20
−30
−40
cpgIslandExt
LMBR1L
RP11−386G11.10
TUBA1B
Y_RNA
49.51
49.52
49.53
49.54
49.55
Position on chr12 (Mb)

### ZNF554

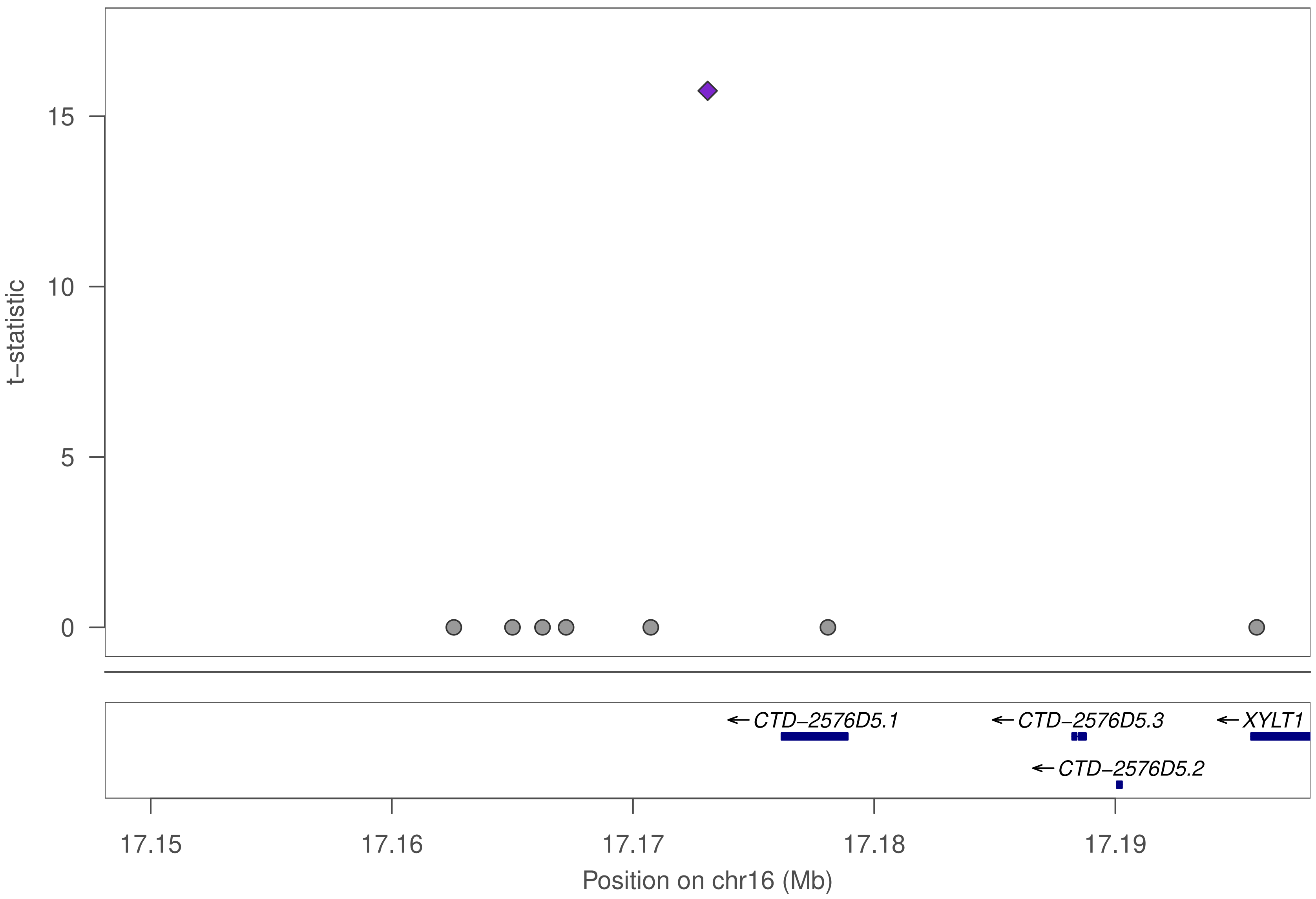

t−statistic
15
10
5
0
CTD−2576D5.1
CTD−2576D5.3
XYLT1
CTD−2576D5.2
17.15
17.16
17.17
17.18
17.19
Position on chr16 (Mb)

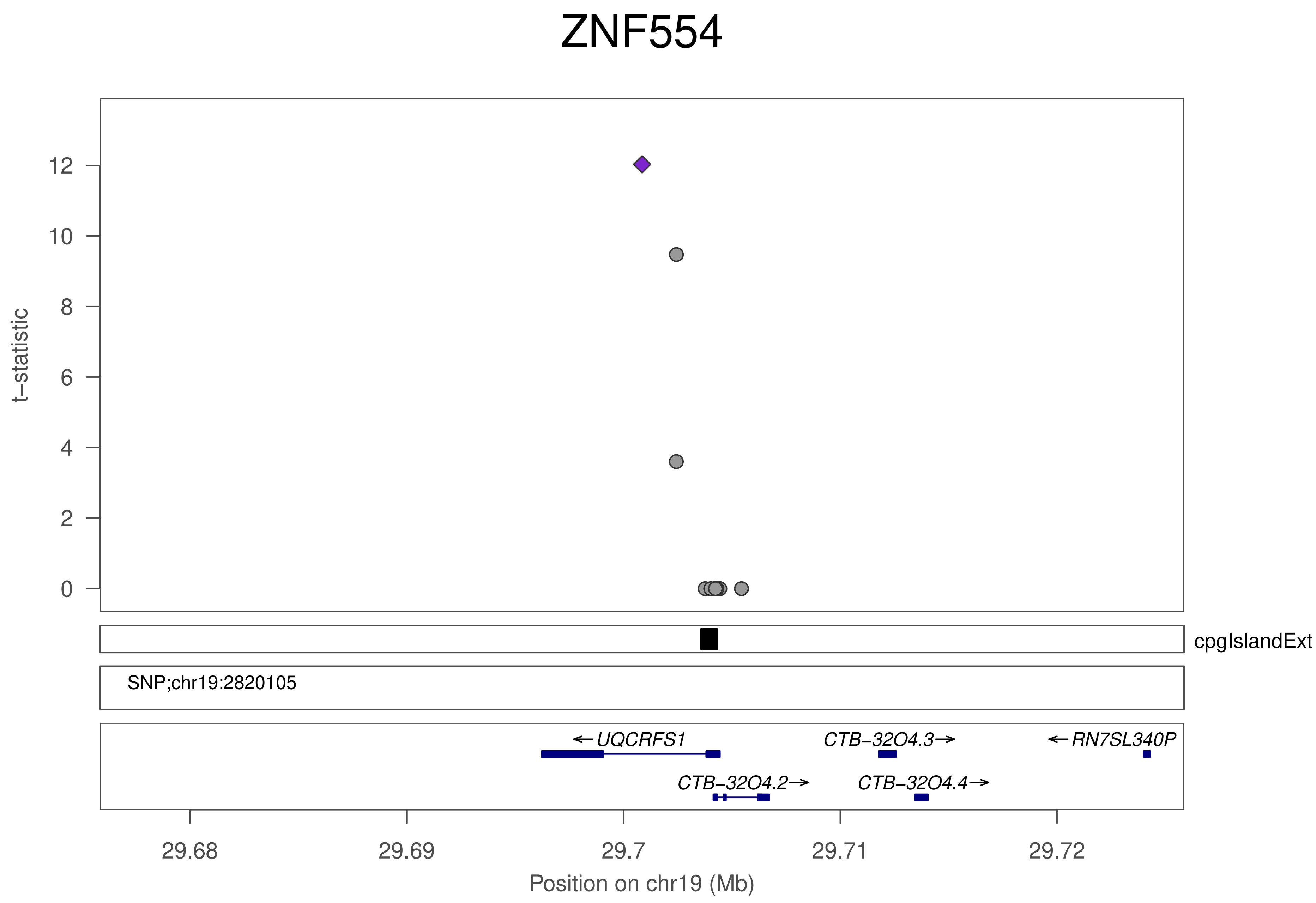
ZNF554
t−statistic
12
10
8
6
4
2
0
cpgIslandExt
SNP;chr19:2820105
← UQCRFS1
CTB−32O4.3→
← RN7SL340P
CTB−32O4.2→
CTB−32O4.4→
29.68
29.69
29.7
29.71
29.72
Position on chr19 (Mb)

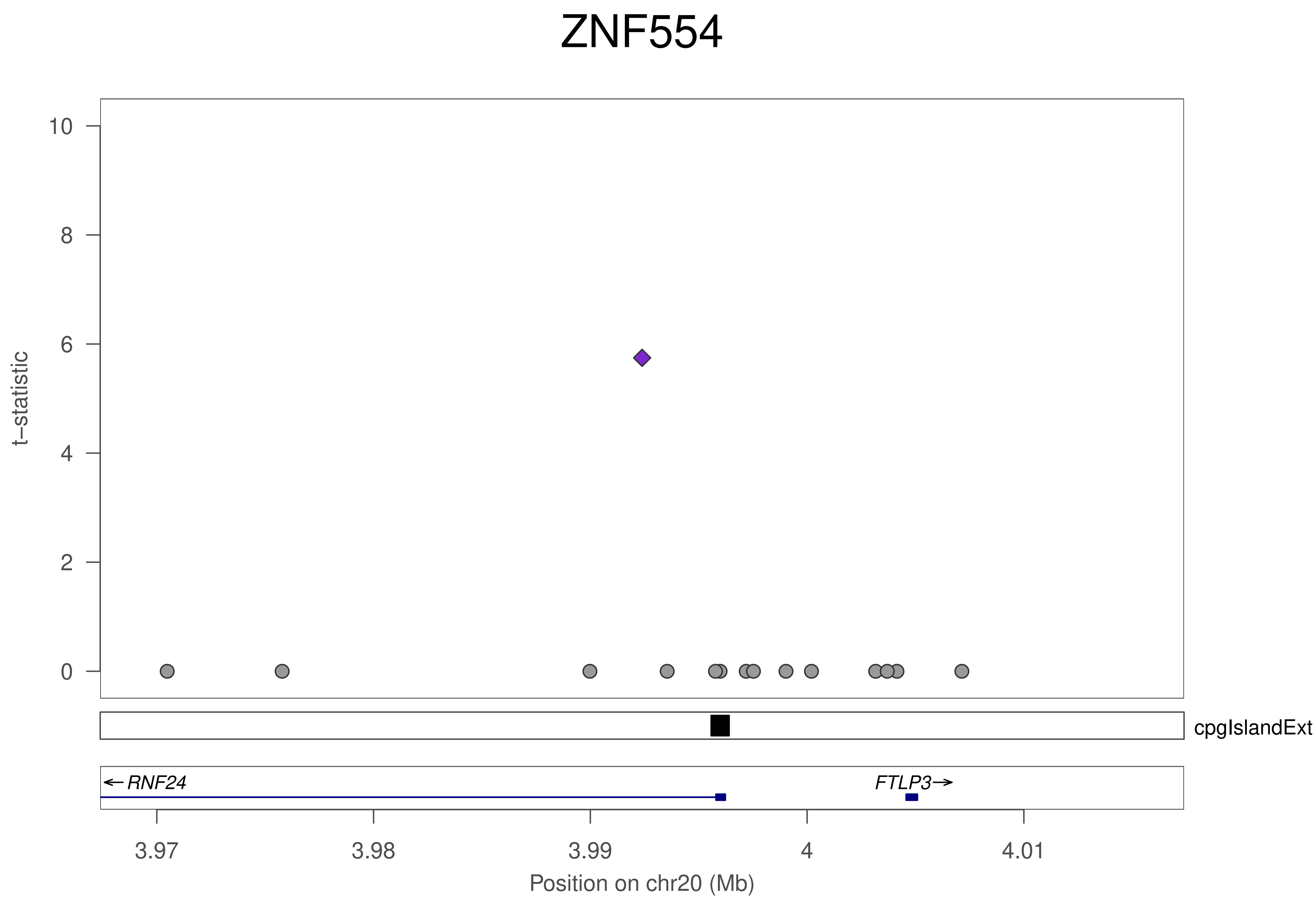

ZNF554
t−statistic
0
2
4
6
8
10
cpgIslandExt
← RNF24
FTLP3 →
3.97
3.98
3.99
4
4.01
Position on chr20 (Mb)

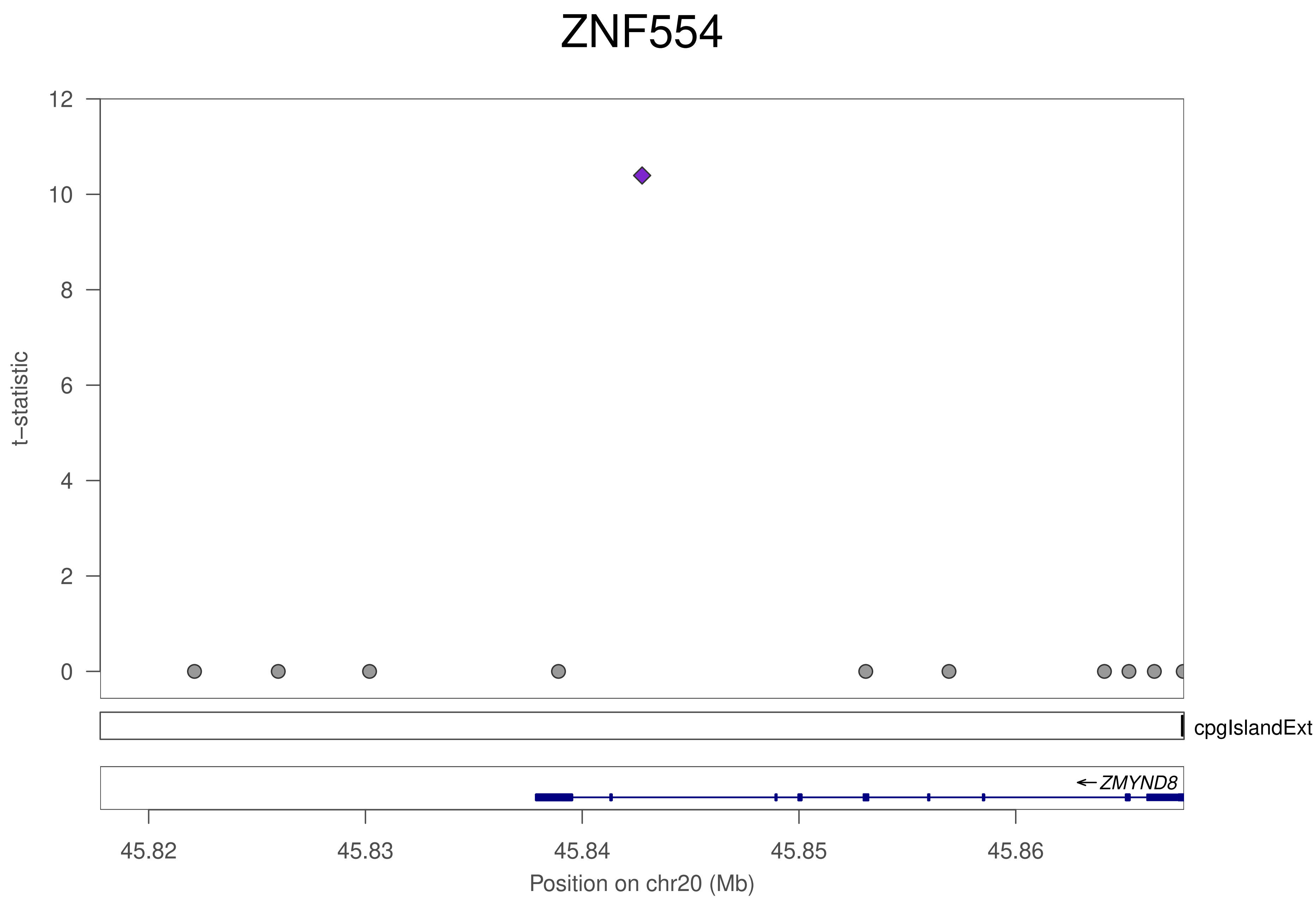
ZNF554
t−statistic
12
10
8
6
4
2
0
cpgIslandExt
ZMYND8
45.82
45.83
45.84
45.85
45.86
Position on chr20 (Mb)

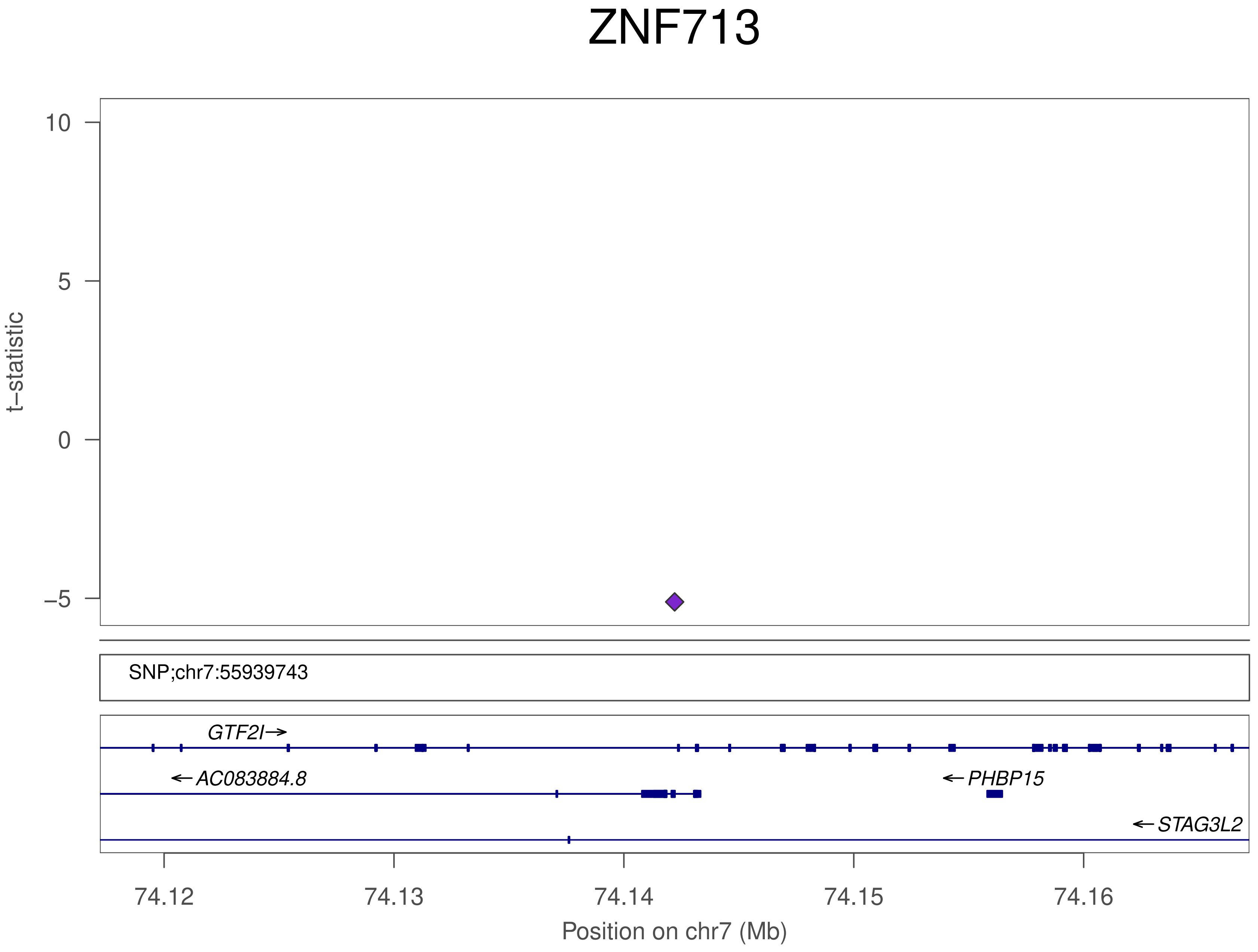
ZNF713
t−statistic
10
5
0
−5
SNP;chr7:55939743
GTF2I
AC083884.8
PHBP15
STAG3L2
74.12
74.13
74.14
74.15
74.16
Position on chr7 (Mb)

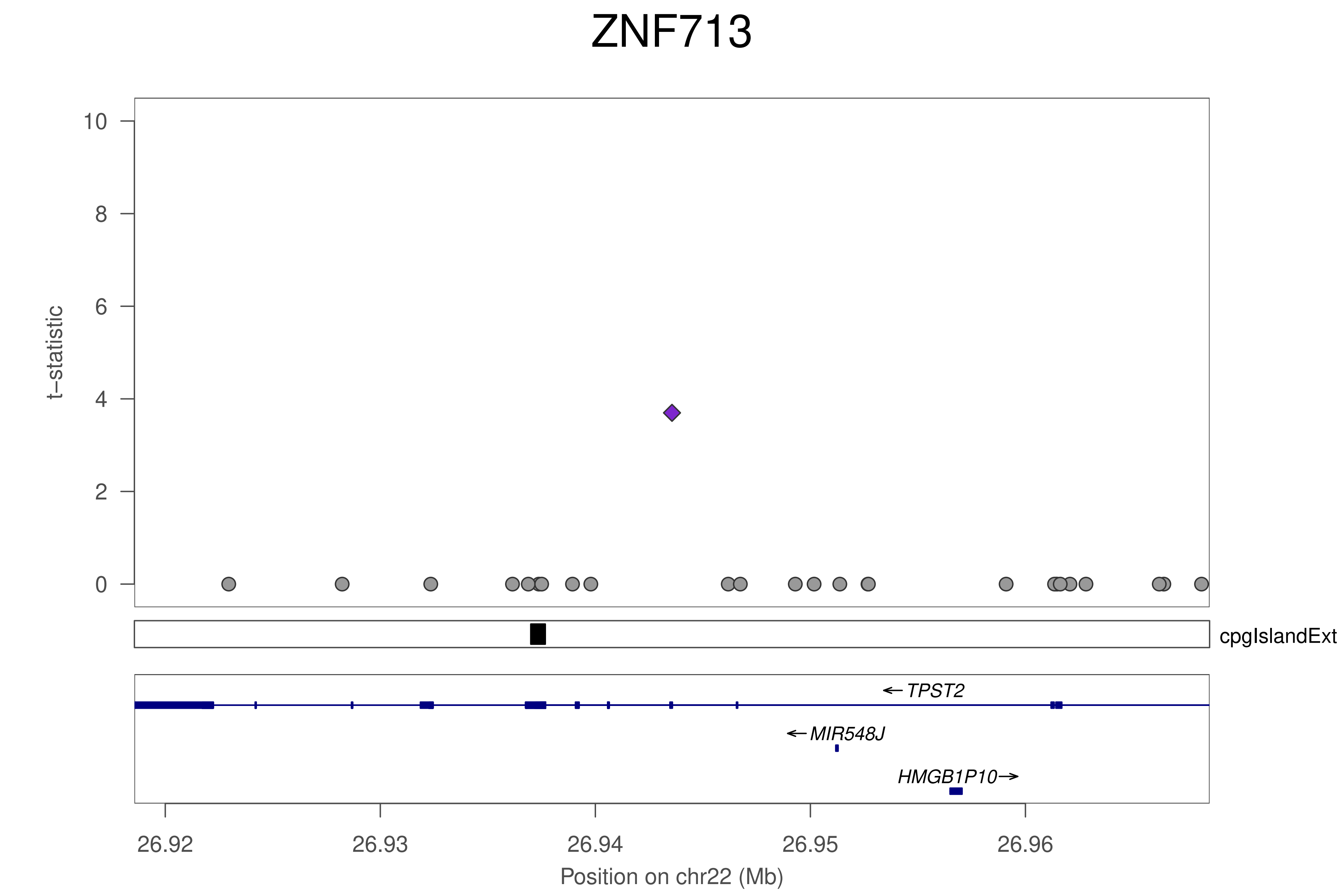
ZNF713
t−statistic
10
8
6
4
2
0
cpgIslandExt
TPST2
MIR548J
HMGB1P10
26.92
26.93
26.94
26.95
26.96
Position on chr22 (Mb)

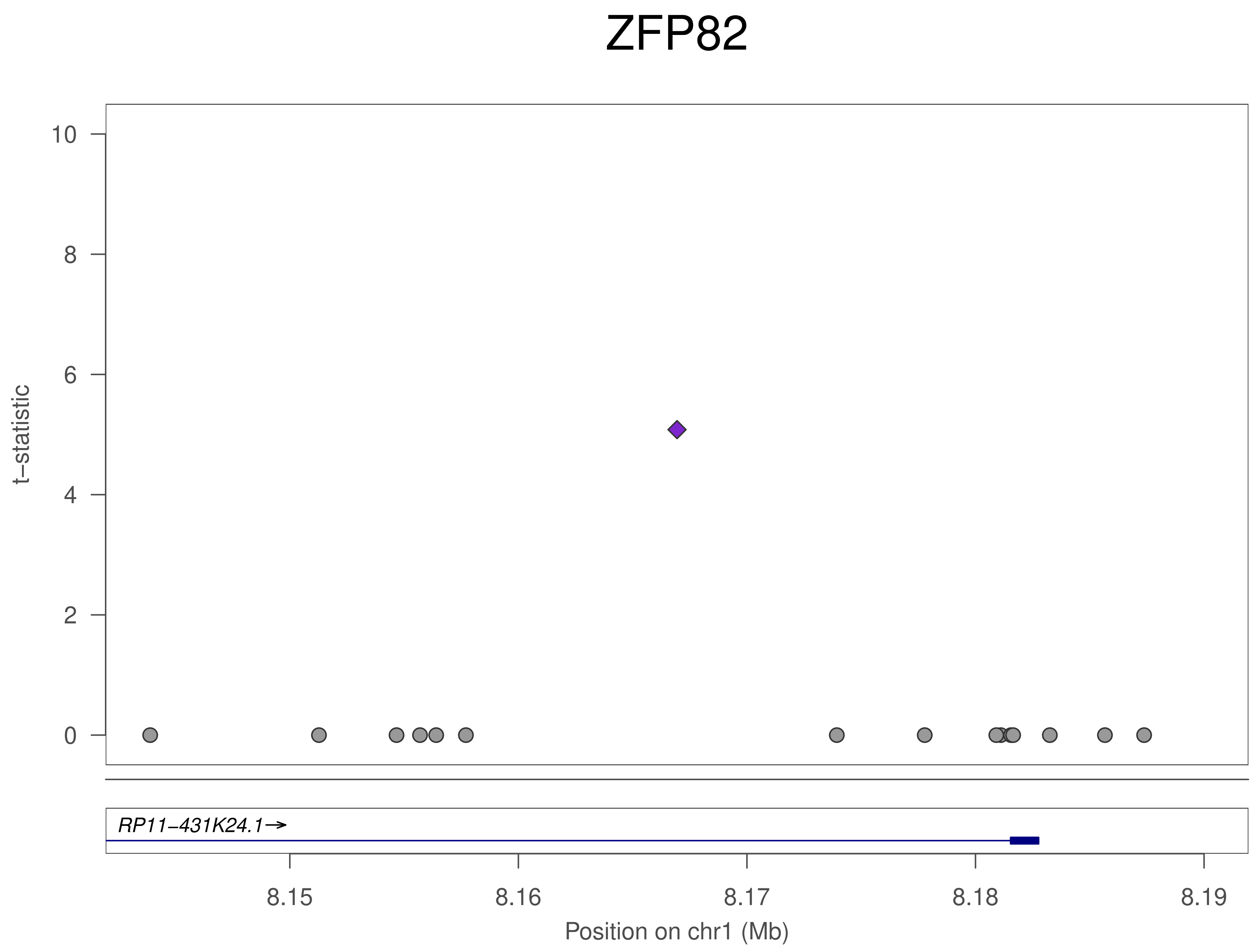
ZFP82
t−statistic
10
8
6
4
2
0
RP11−431K24.1→
8.15
8.16
8.17
8.18
8.19
Position on chr1 (Mb)

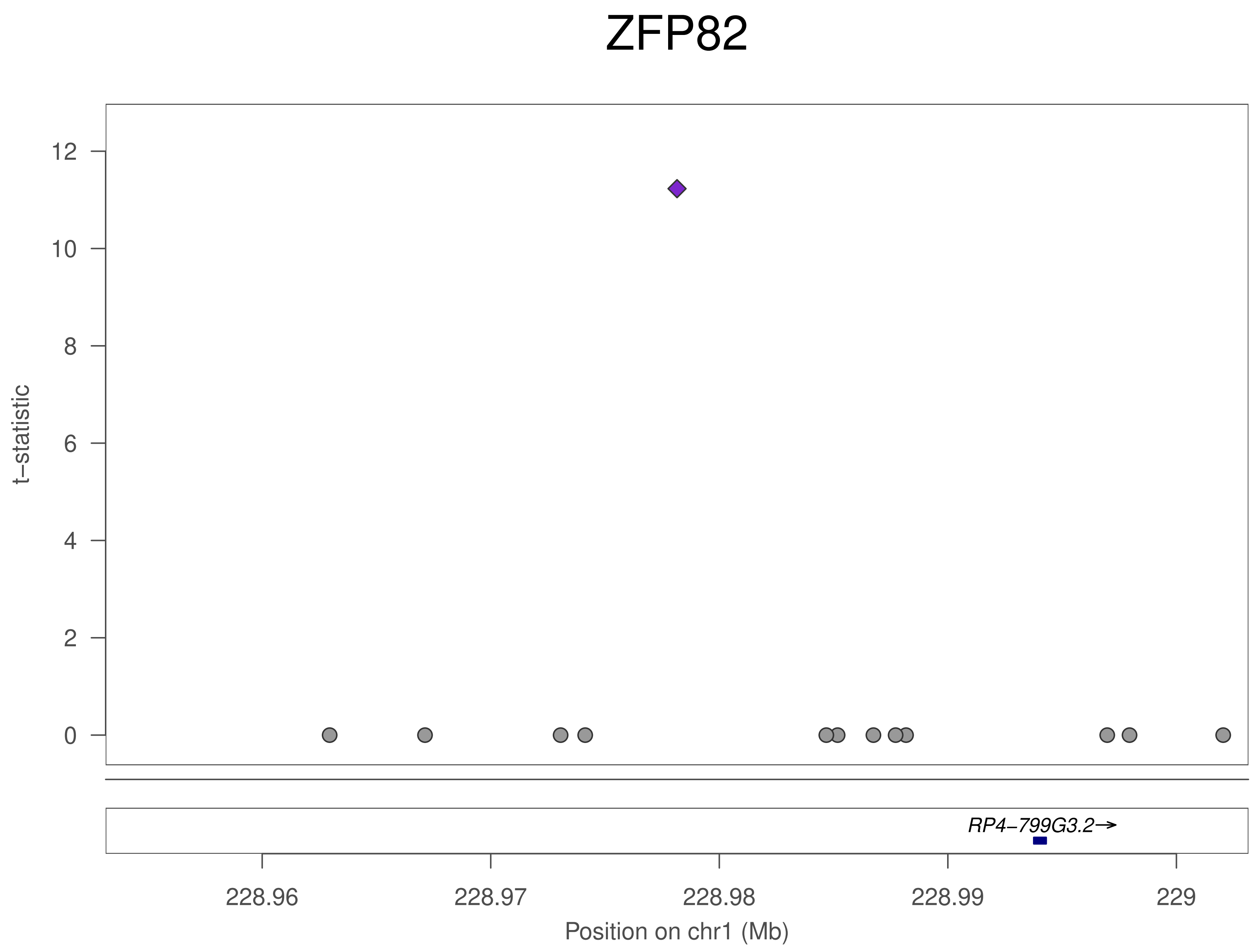

ZFP82
t−statistic
12
10
8
6
4
2
0
RP4−799G3.2→
228.96
228.97
228.98
228.99
229
Position on chr1 (Mb)

### ZFP82

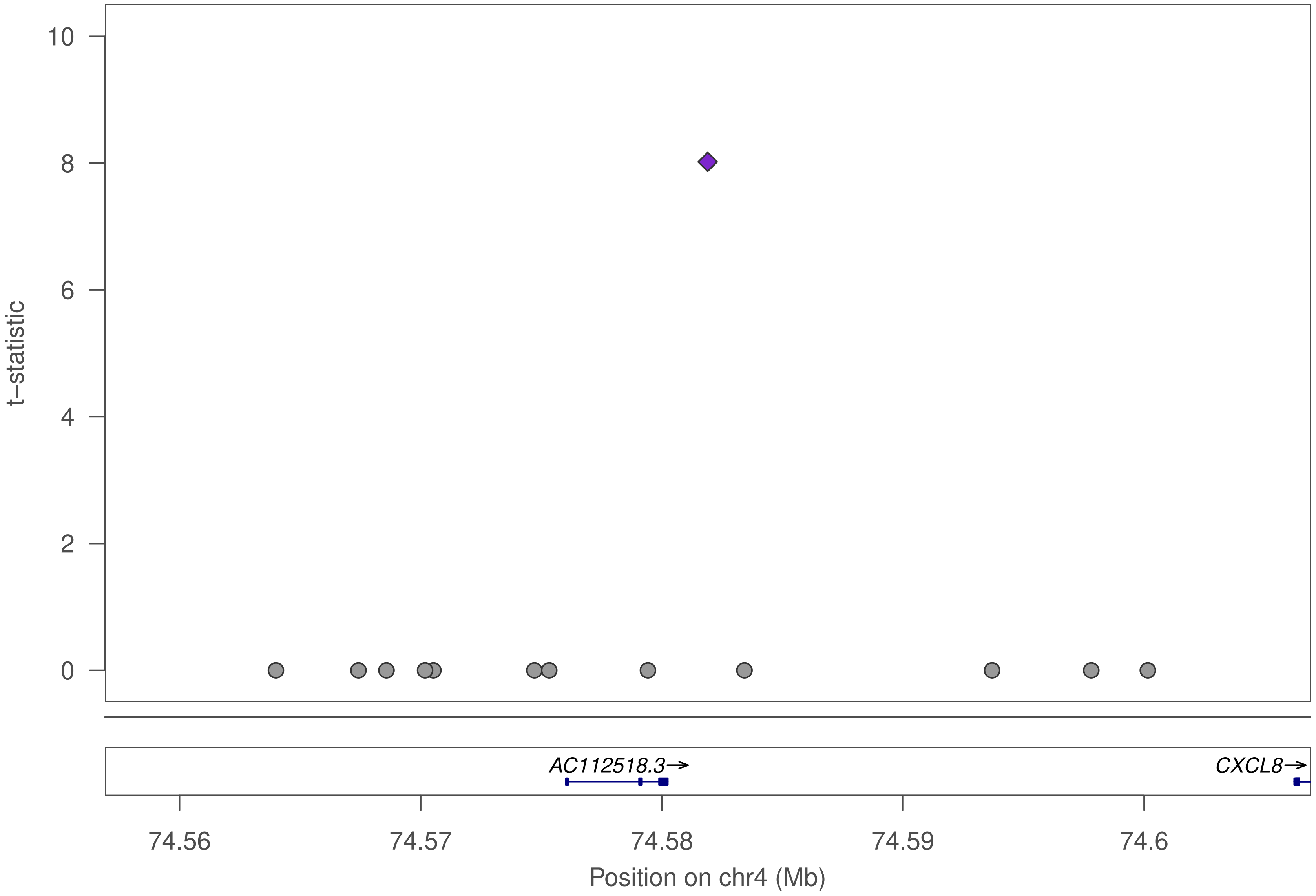
t–statistic
10
8
6
4
2
0
AC112518.3→
CXCL8→
74.56
74.57
74.58
74.59
74.6
Position on chr4 (Mb)

### ZFP82

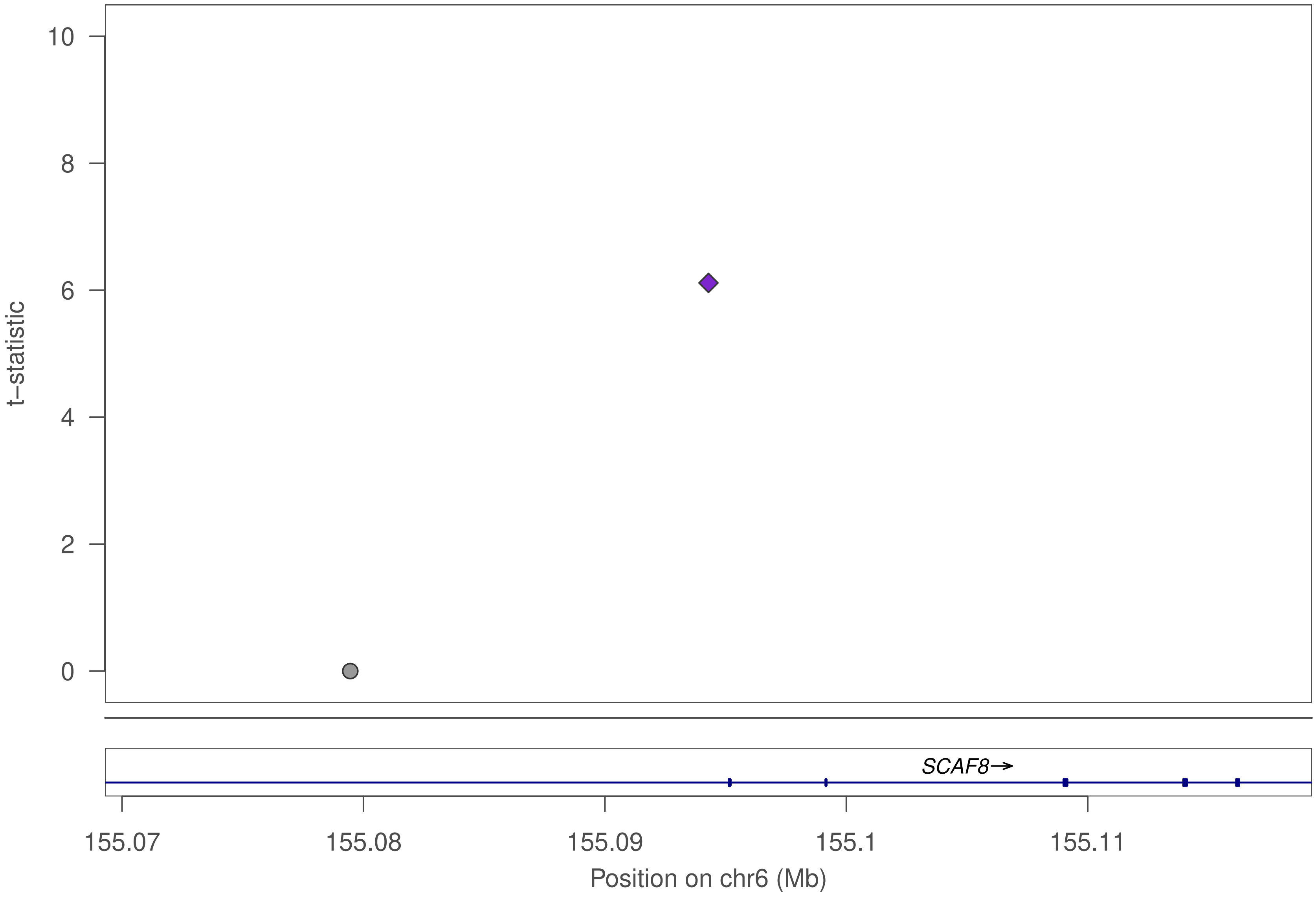

10
8
6
4
2
0
t–statistic
SCAF8→
155.07
155.08
155.09
155.1
155.11
Position on chr6 (Mb)

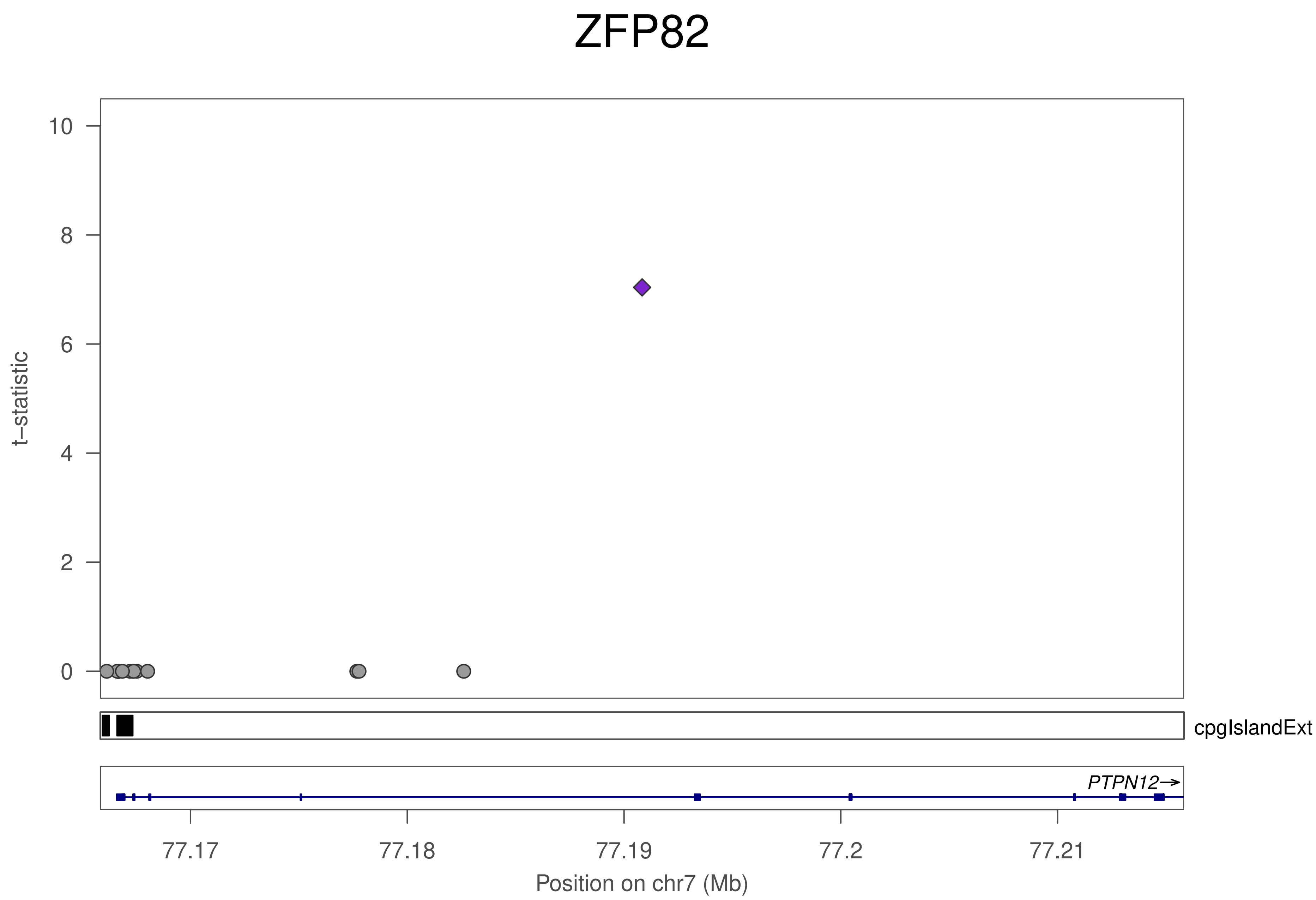
ZFP82
t–statistic
10
8
6
4
2
0
cpgIslandExt
PTPN12→
77.17
77.18
77.19
77.2
77.21
Position on chr7 (Mb)

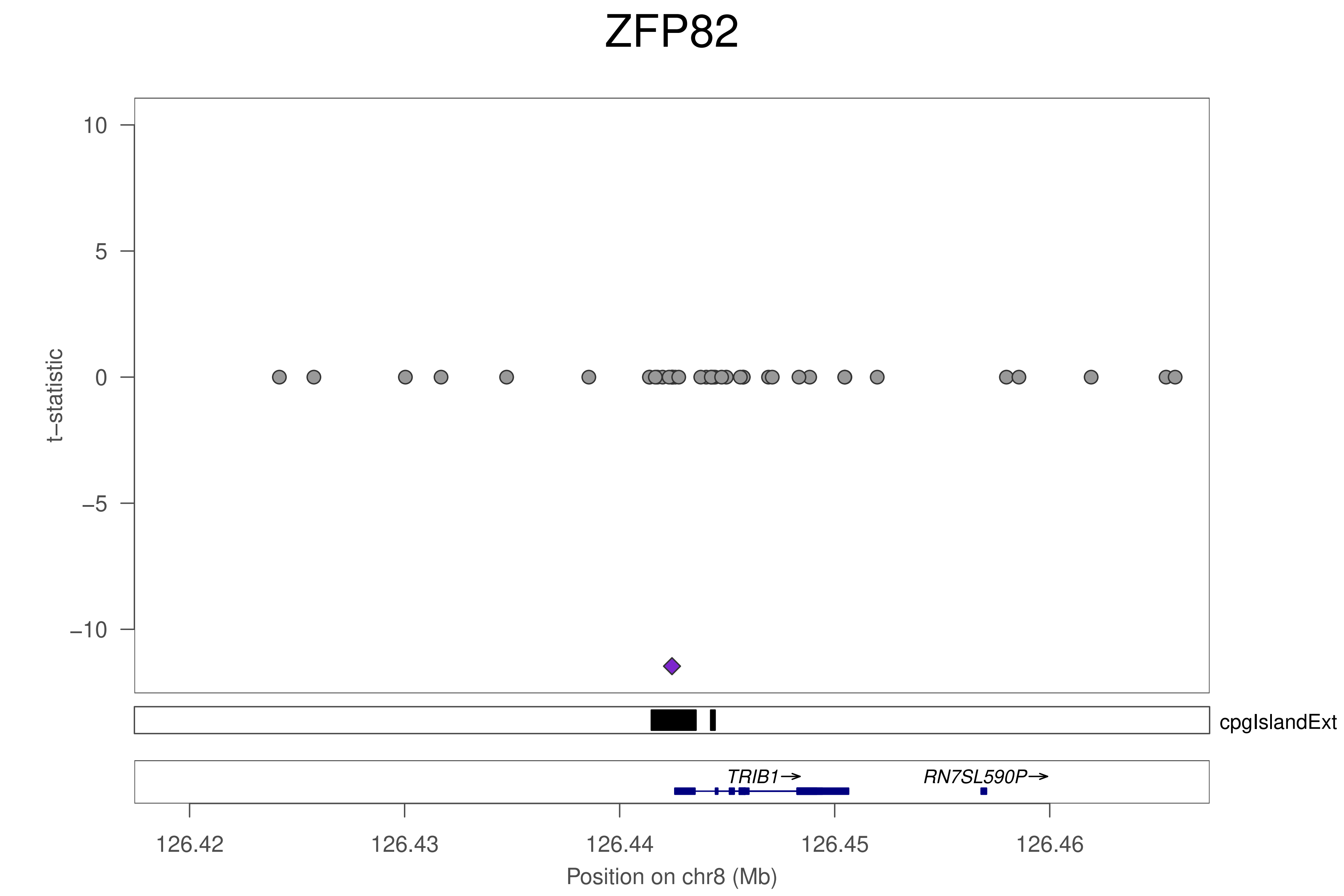
ZFP82
t−statistic
10
5
0
−5
−10
cpgIslandExt
TRIB1→
RN7SL590P→
126.42
126.43
126.44
126.45
126.46
Position on chr8 (Mb)

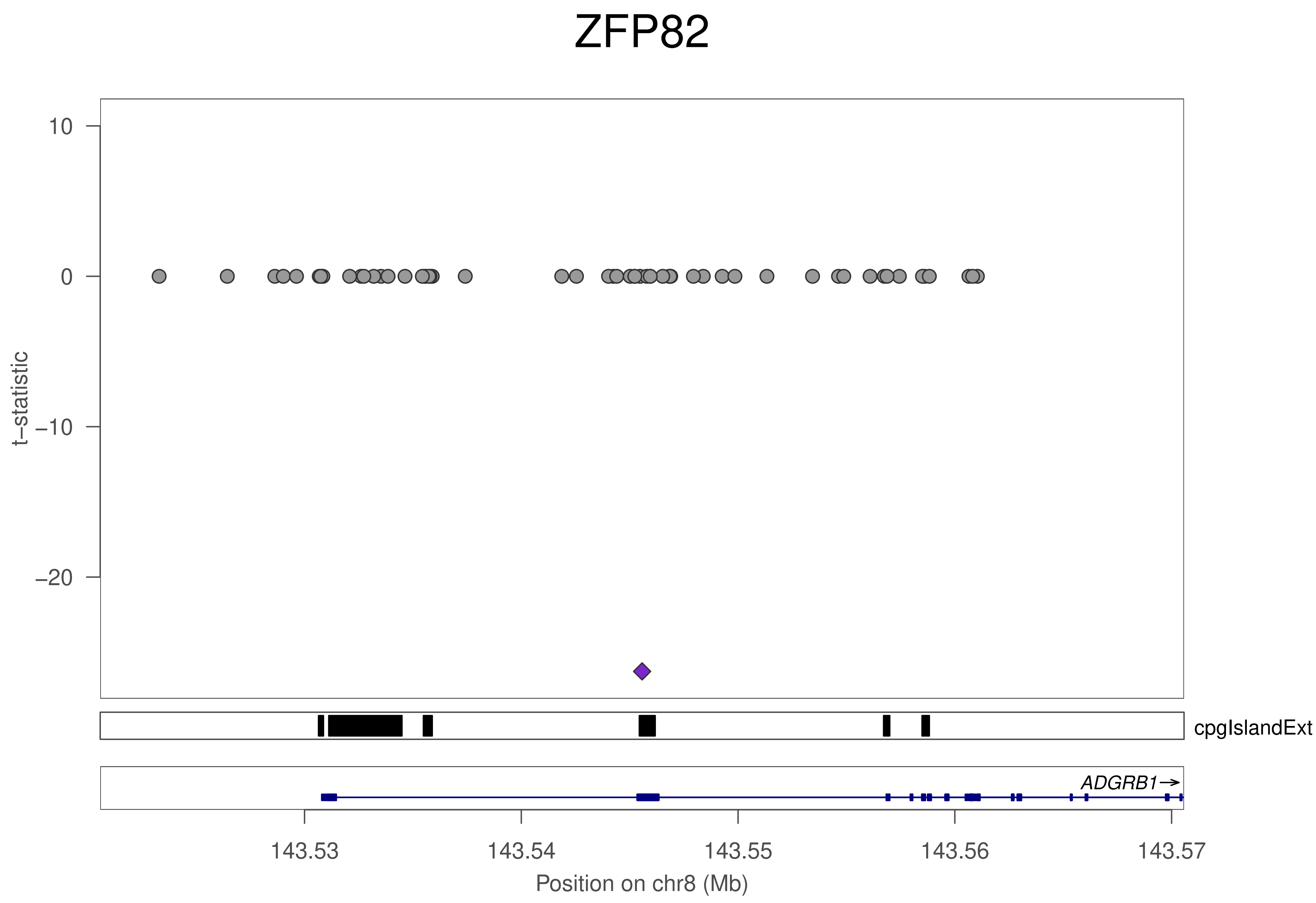

ZFP82
10
0
−10
−20
t−statistic
cpgIslandExt
ADGRB1→
143.53
143.54
143.55
143.56
143.57
Position on chr8 (Mb)

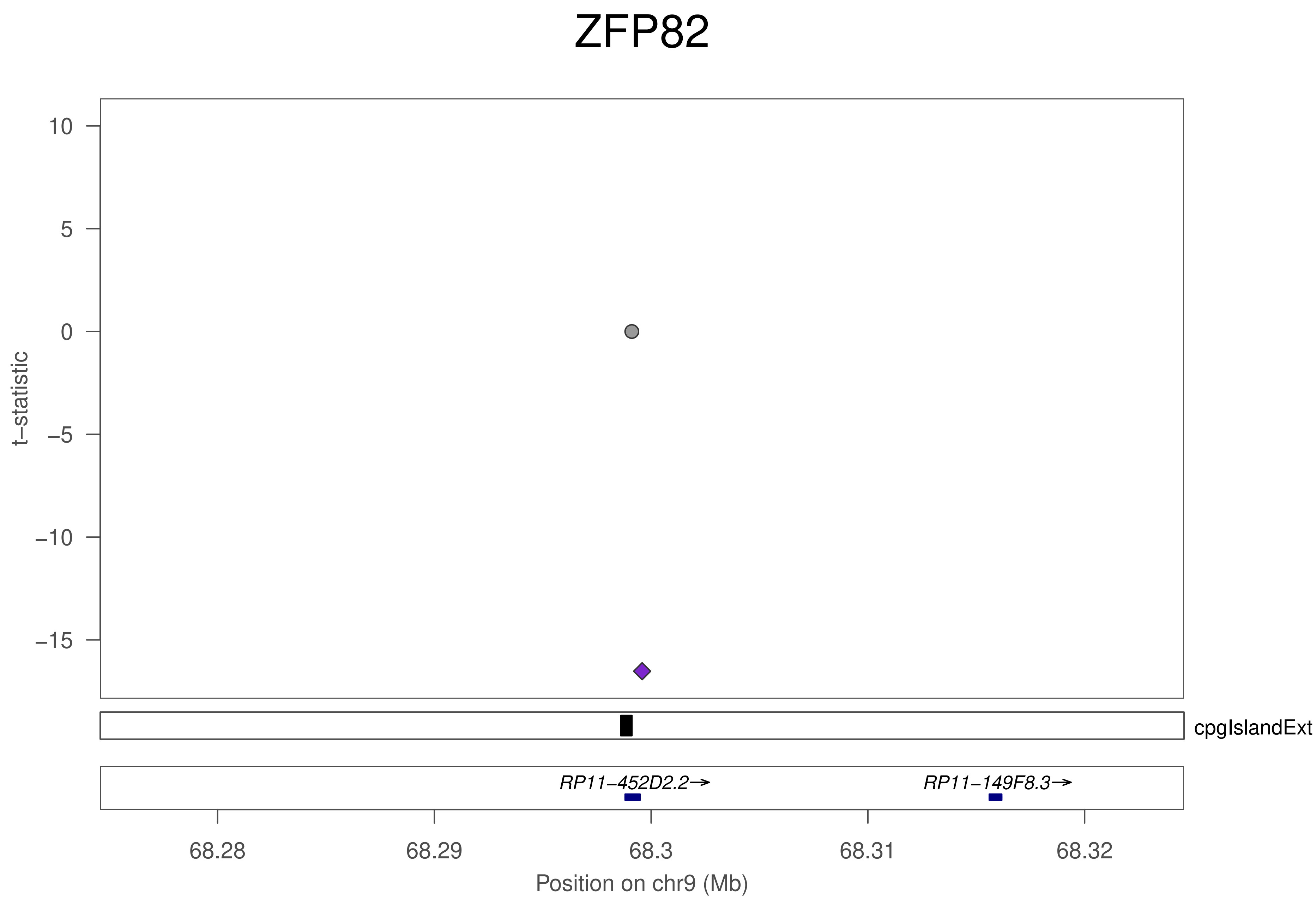
ZFP82
t−statistic
10
5
0
−5
−10
−15
cpgIslandExt
RP11−452D2.2→
RP11−149F8.3→
68.28
68.29
68.3
68.31
68.32
Position on chr9 (Mb)

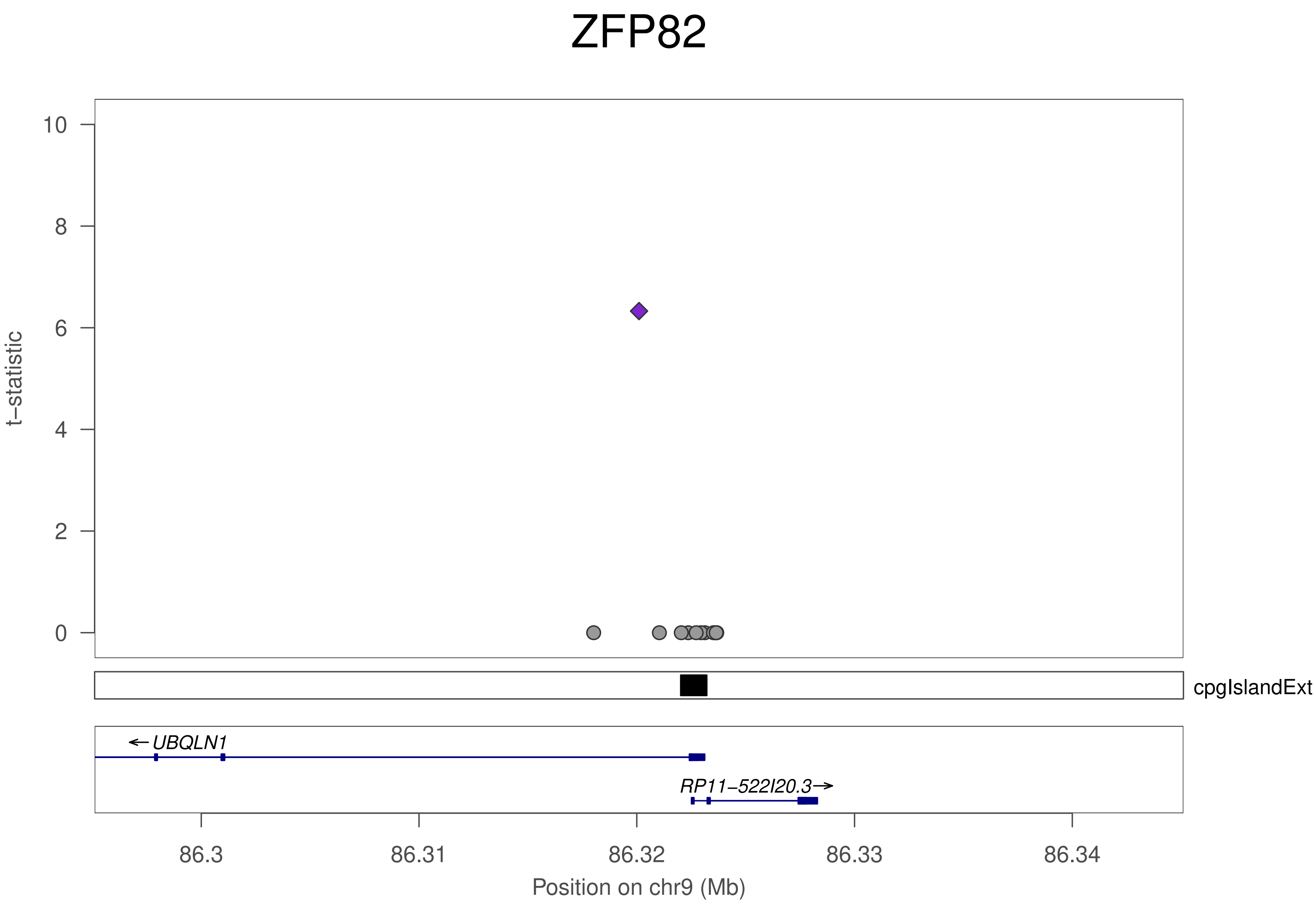

ZFP82
10
8
6
4
2
0
t−statistic
cpgIslandExt
←UBQLN1
RP11−522I20.3→
86.3
86.31
86.32
86.33
86.34
Position on chr9 (Mb)

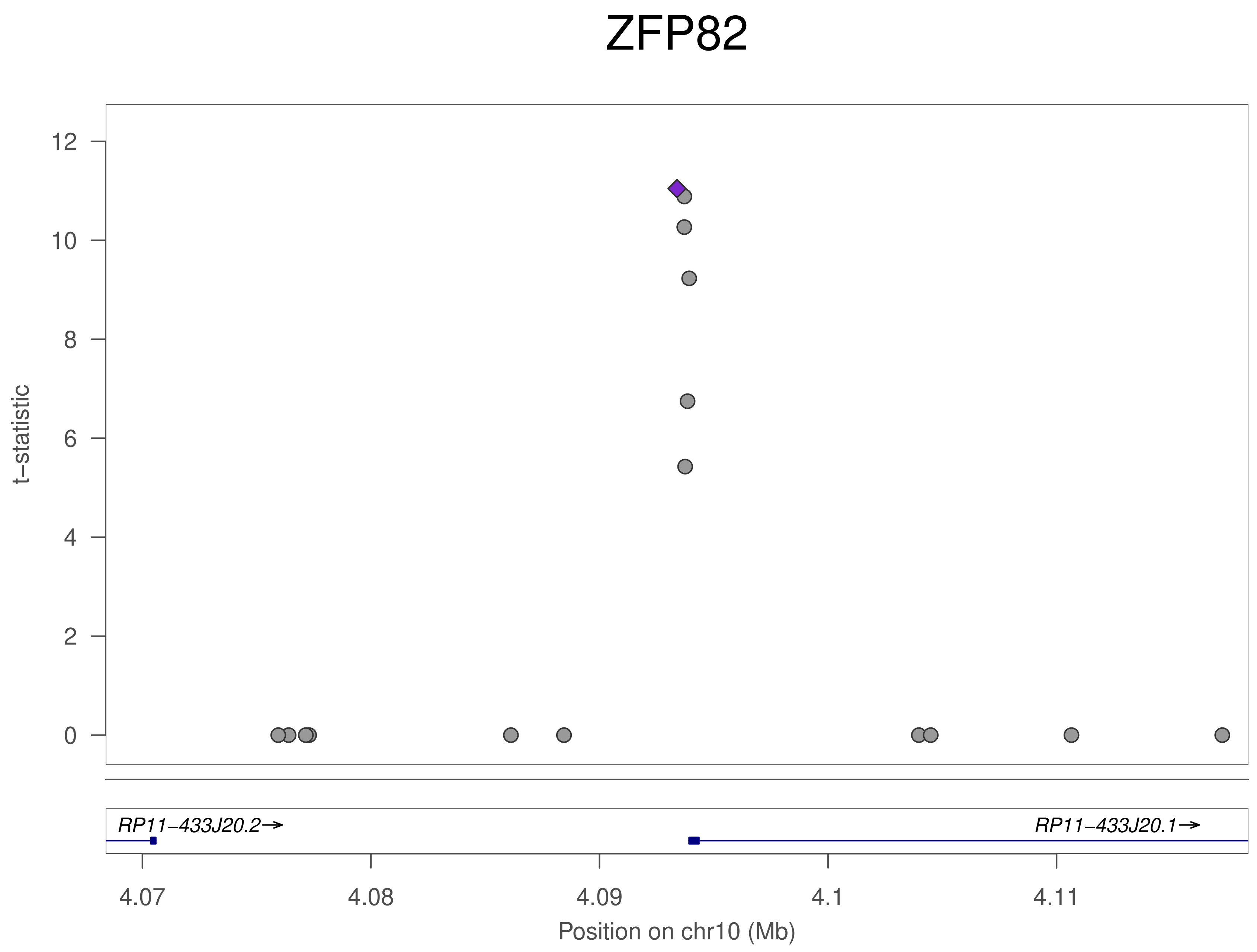
ZFP82
t−statistic
12
10
8
6
4
2
0
RP11−433J20.2→
RP11−433J20.1→
4.07
4.08
4.09
4.1
4.11
Position on chr10 (Mb)

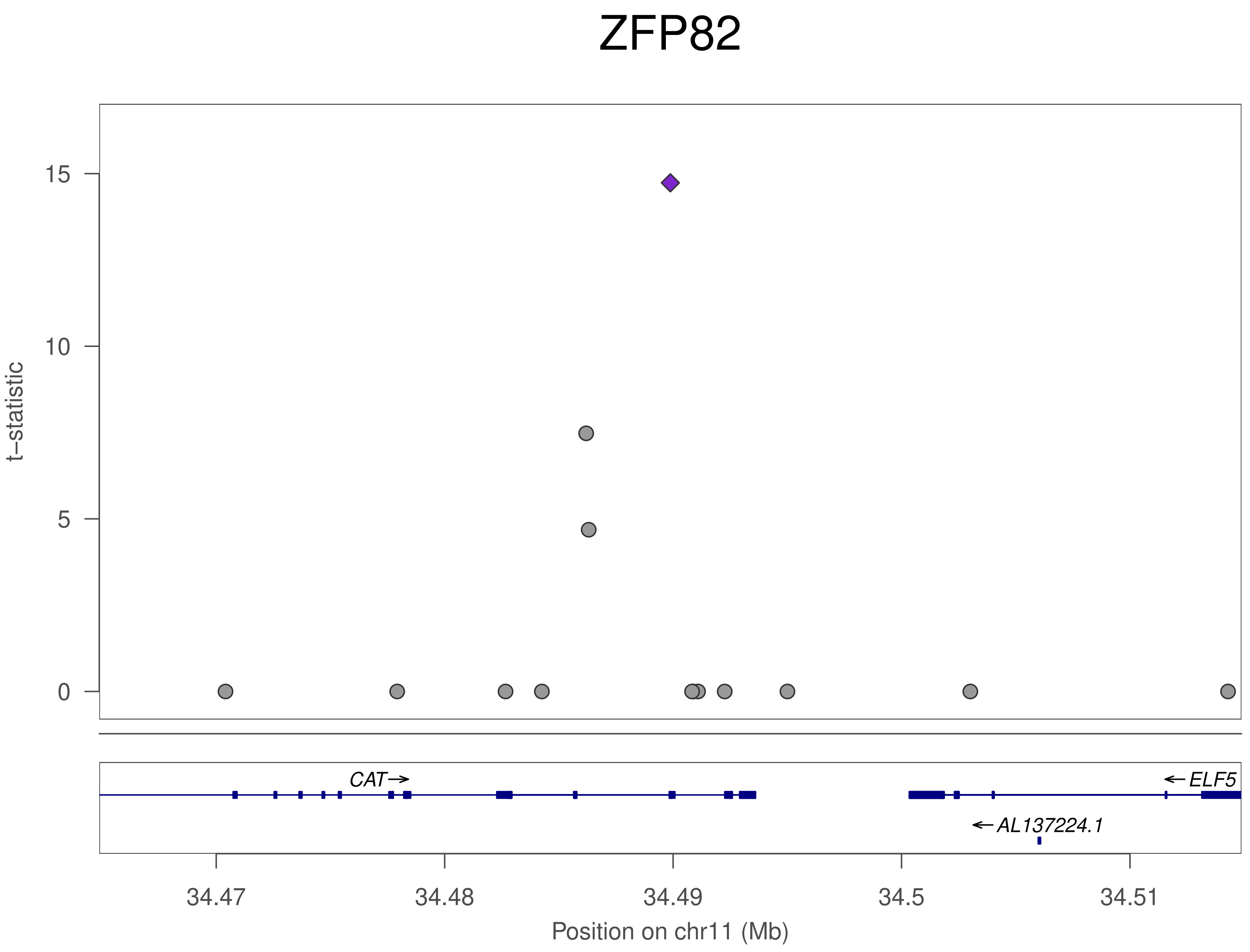
ZFP82
t−statistic
15
10
5
0
CAT→
←ELF5
←AL137224.1
34.47
34.48
34.49
34.5
34.51
Position on chr11 (Mb)

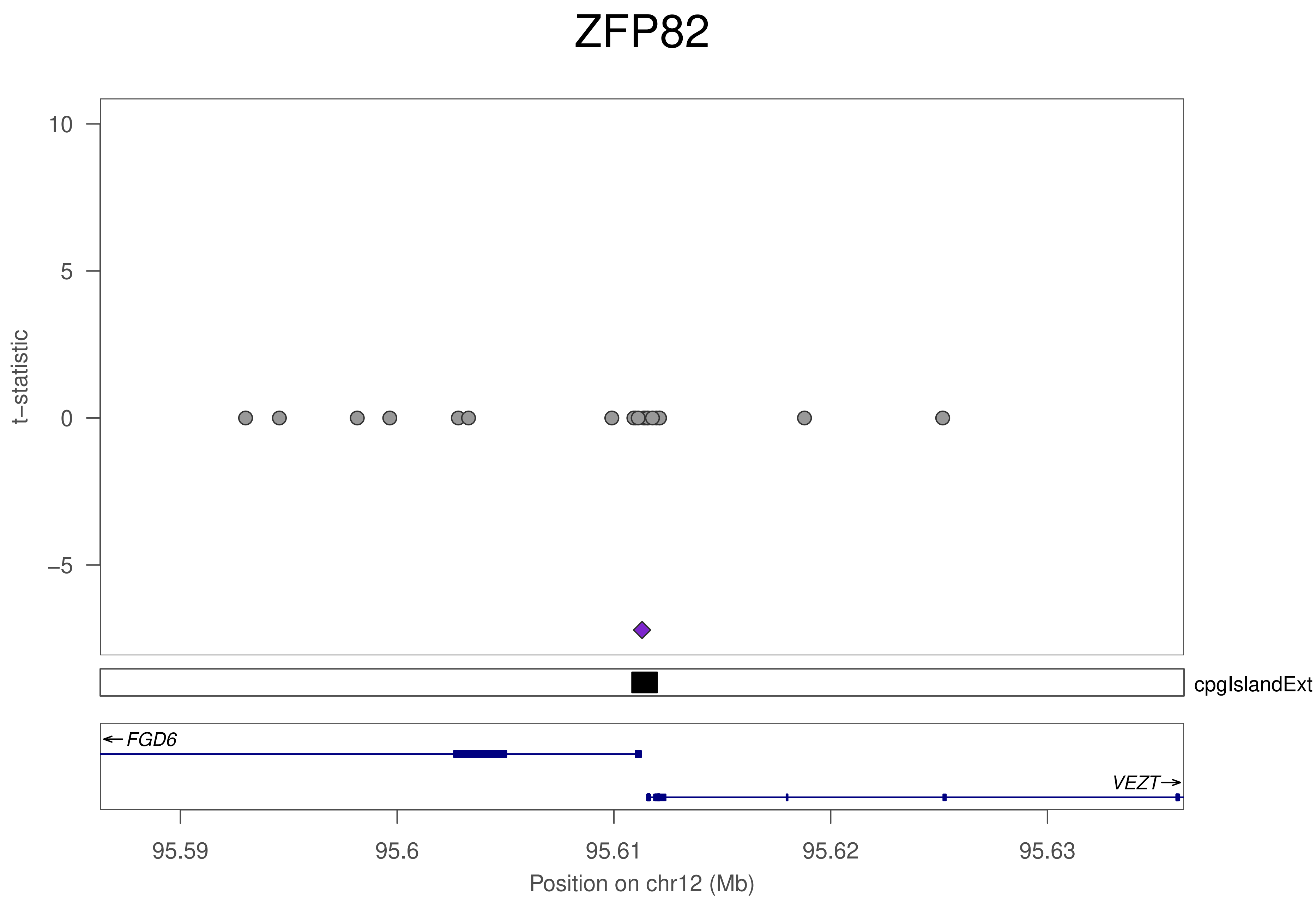
ZFP82
t–statistic
10
5
0
–5
cpgIslandExt
FGD6
VEZT
95.59
95.6
95.61
95.62
95.63
Position on chr12 (Mb)

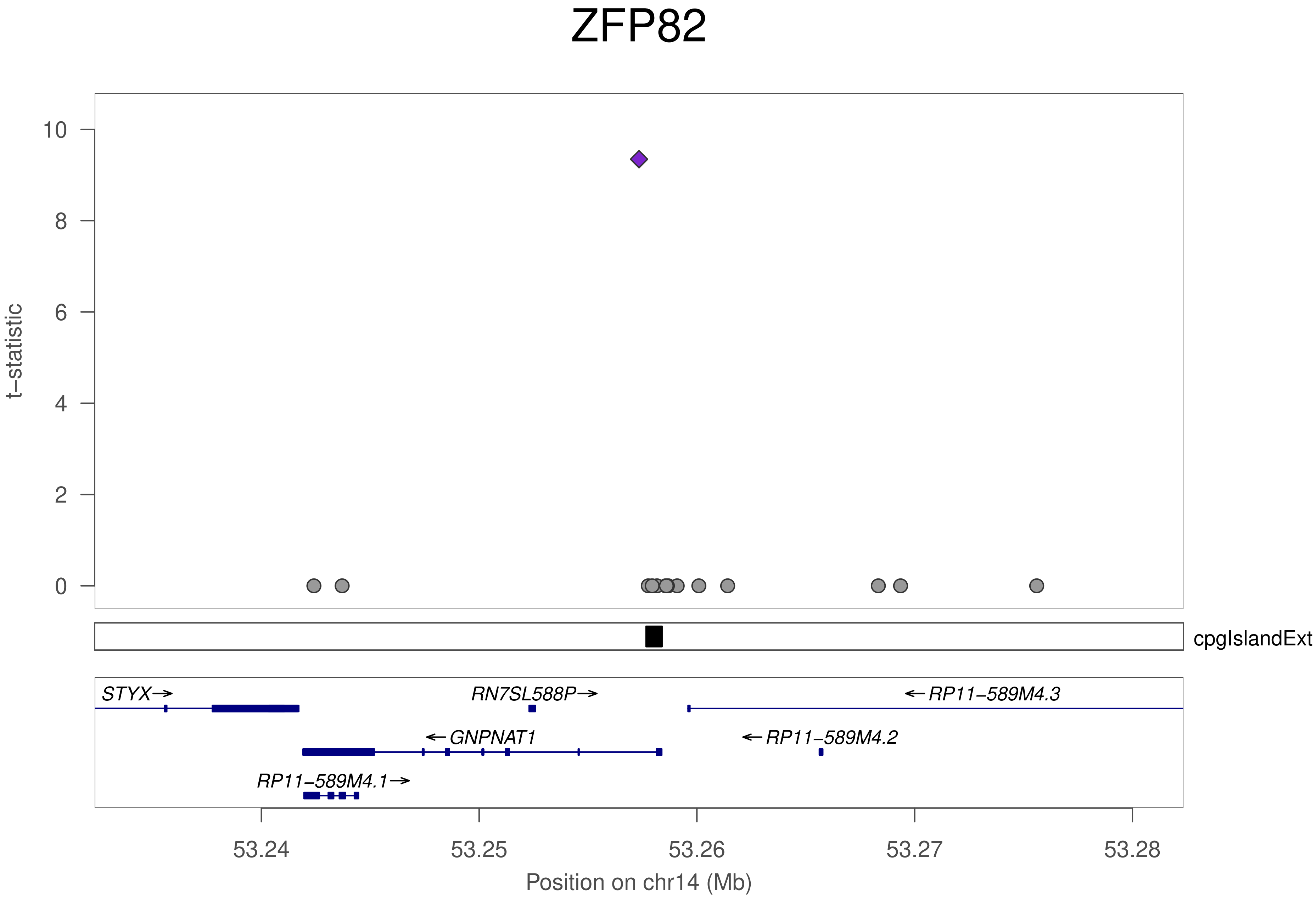
ZFP82
t−statistic
10
8
6
4
2
0
cpgIslandExt
STYX→
RN7SL588P→
←RP11−589M4.3
←GNPNAT1
←RP11−589M4.2
RP11−589M4.1→
53.24
53.25
53.26
53.27
53.28
Position on chr14 (Mb)

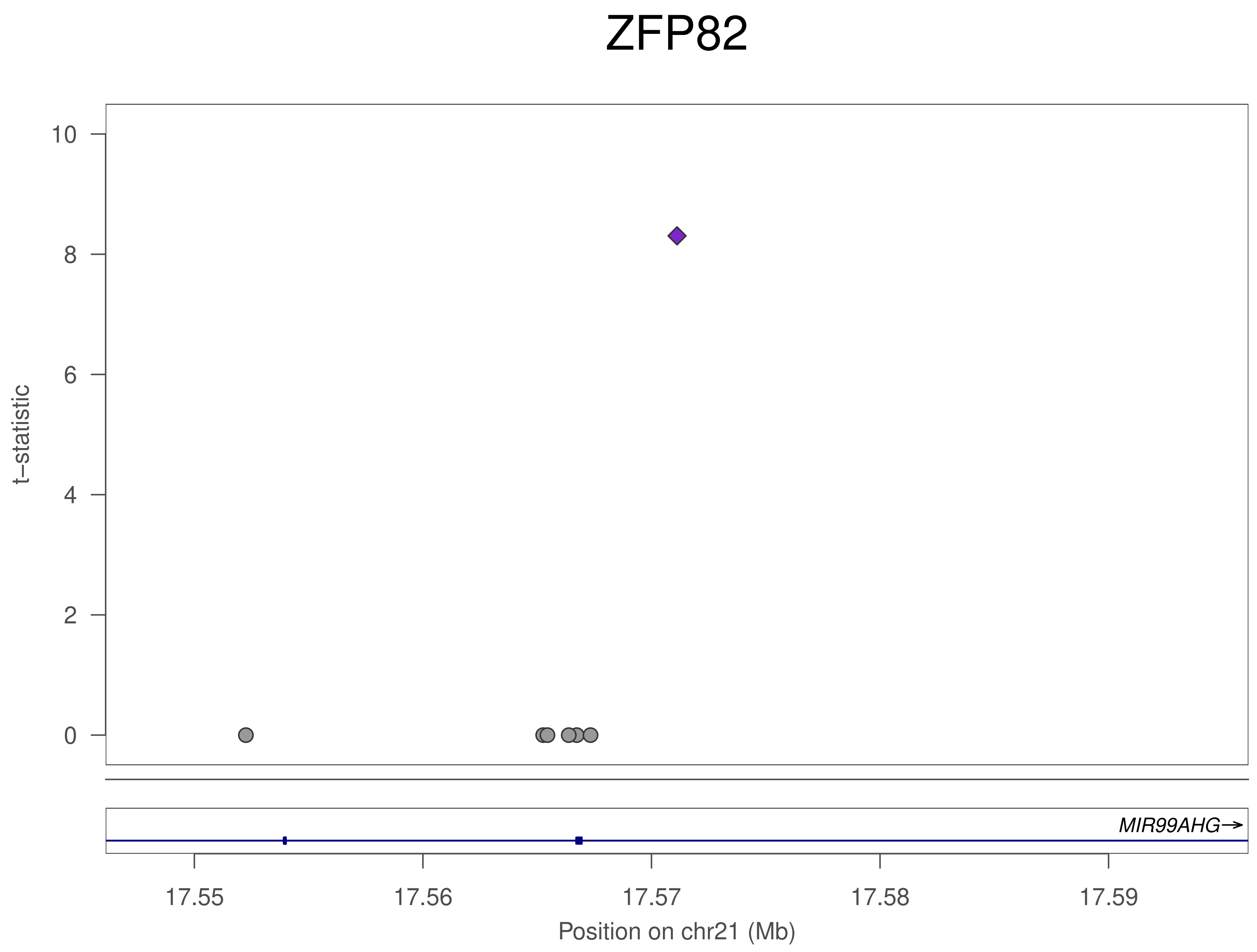
ZFP82
t−statistic
10
8
6
4
2
0
MIR99AHG→
17.55
17.56
17.57
17.58
17.59
Position on chr21 (Mb)

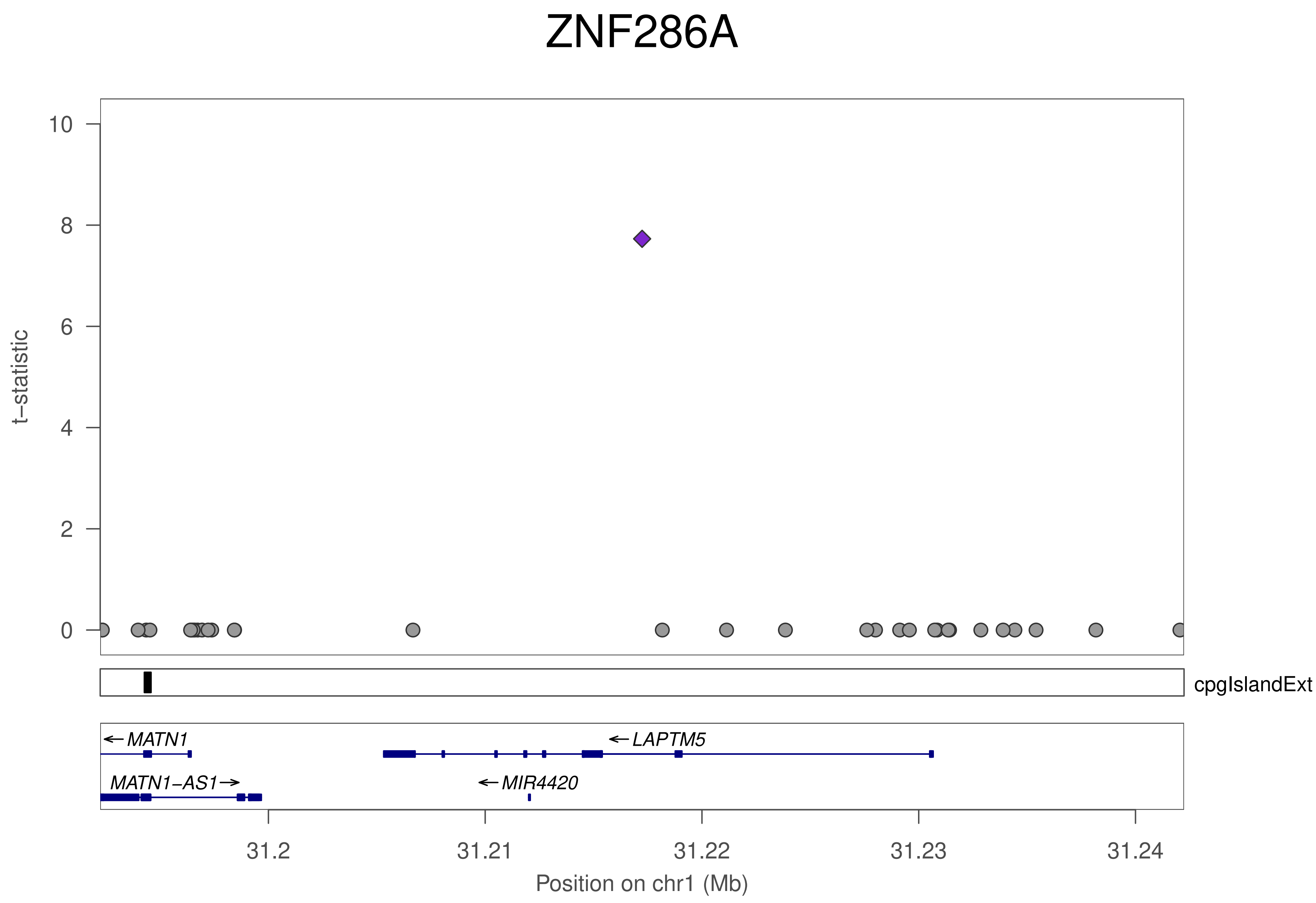
ZNF286A
t−statistic
10
8
6
4
2
0
cpgIslandExt
MATN1
LAPTM5
MATN1−AS1
MIR4420
31.2
31.21
31.22
31.23
31.24
Position on chr1 (Mb)

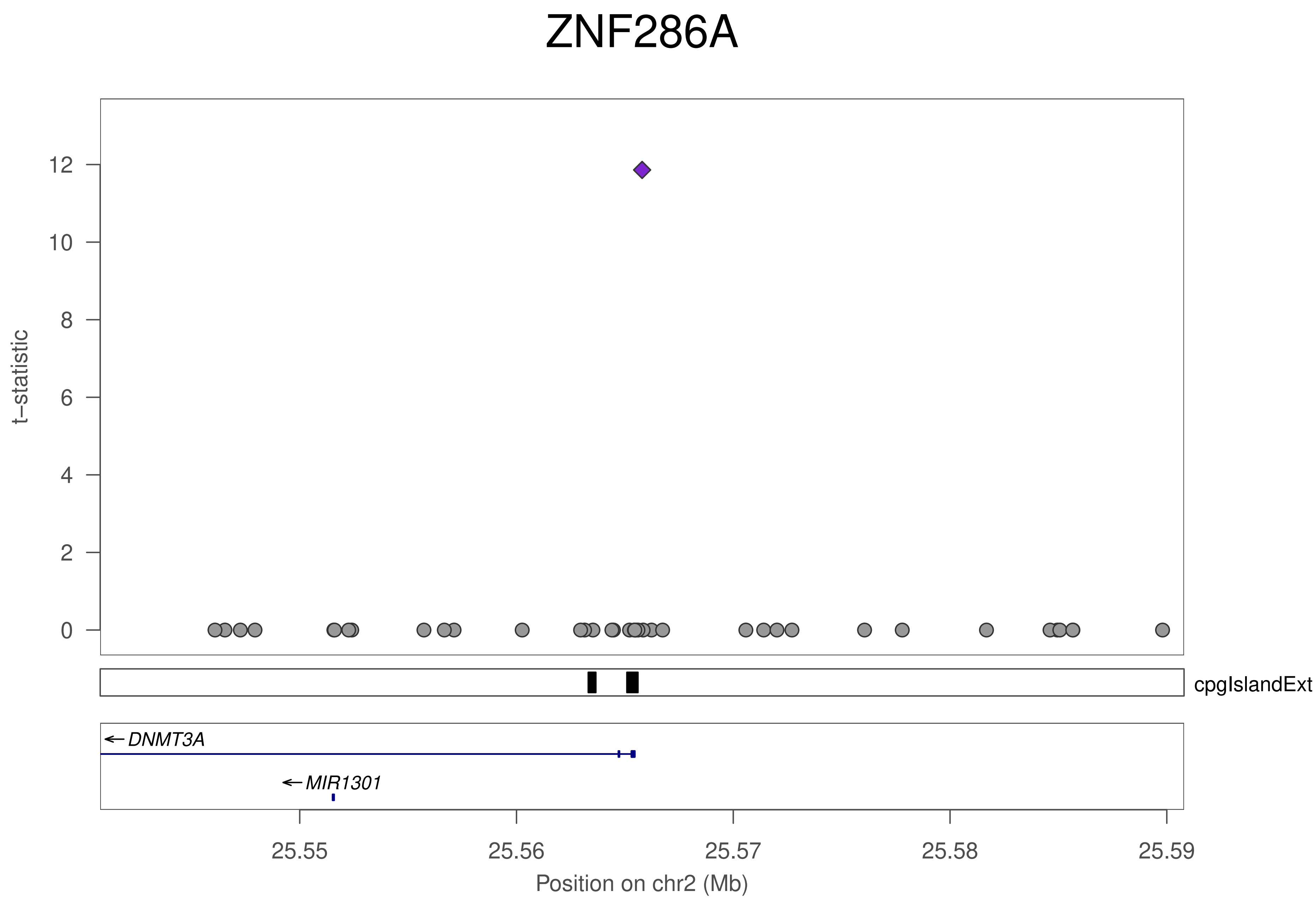
ZNF286A
t−statistic
0
2
4
6
8
10
12
cpgIslandExt
DNMT3A
MIR1301
25.55
25.56
25.57
25.58
25.59
Position on chr2 (Mb)

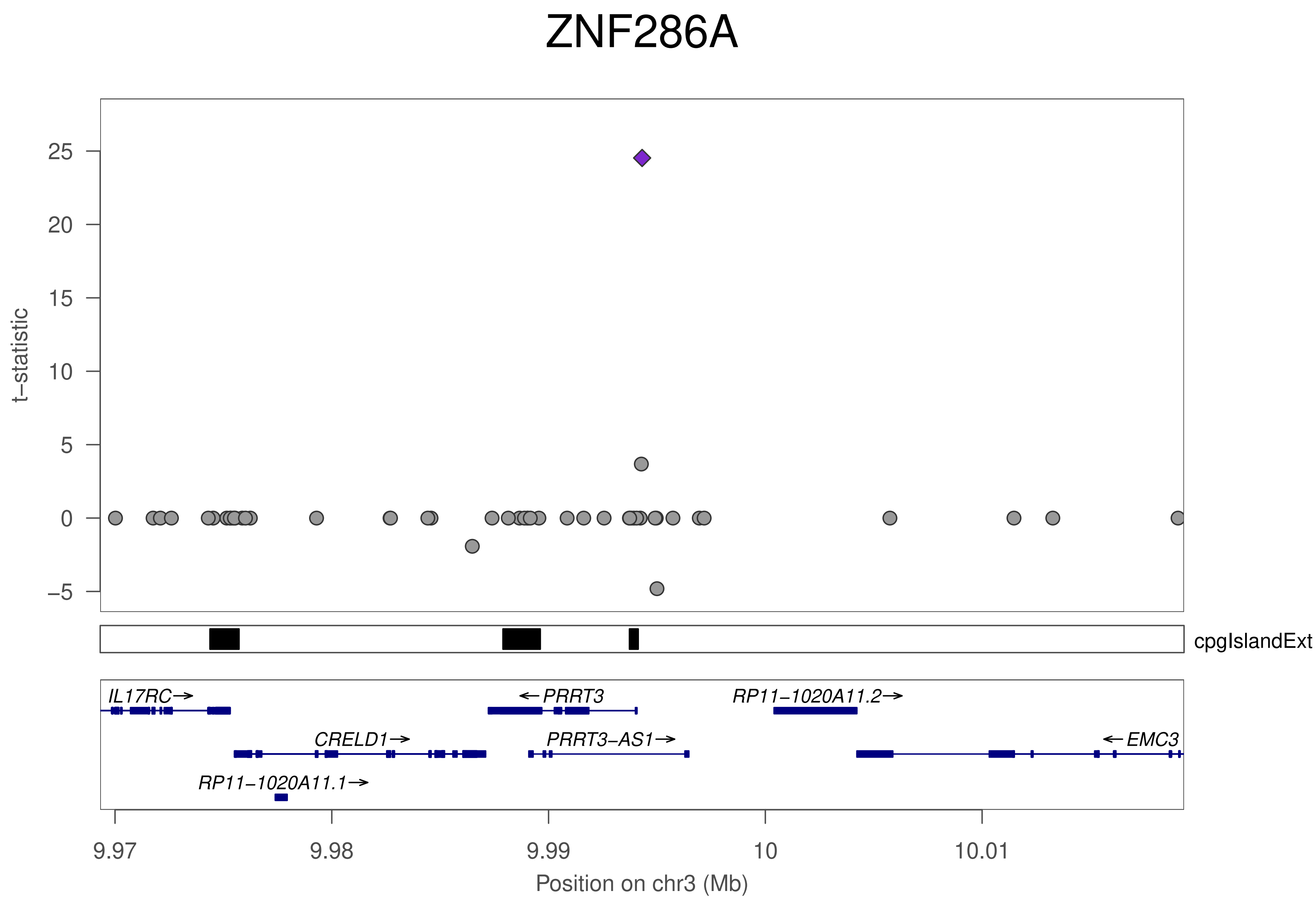

ZNF286A
t−statistic
25
20
15
10
5
0
−5
cpgIslandExt
IL17RC→
←PRRT3
RP11−1020A11.2→
CRELD1→
PRRT3−AS1→
←EMC3
RP11−1020A11.1→
9.97
9.98
9.99
10
10.01
Position on chr3 (Mb)

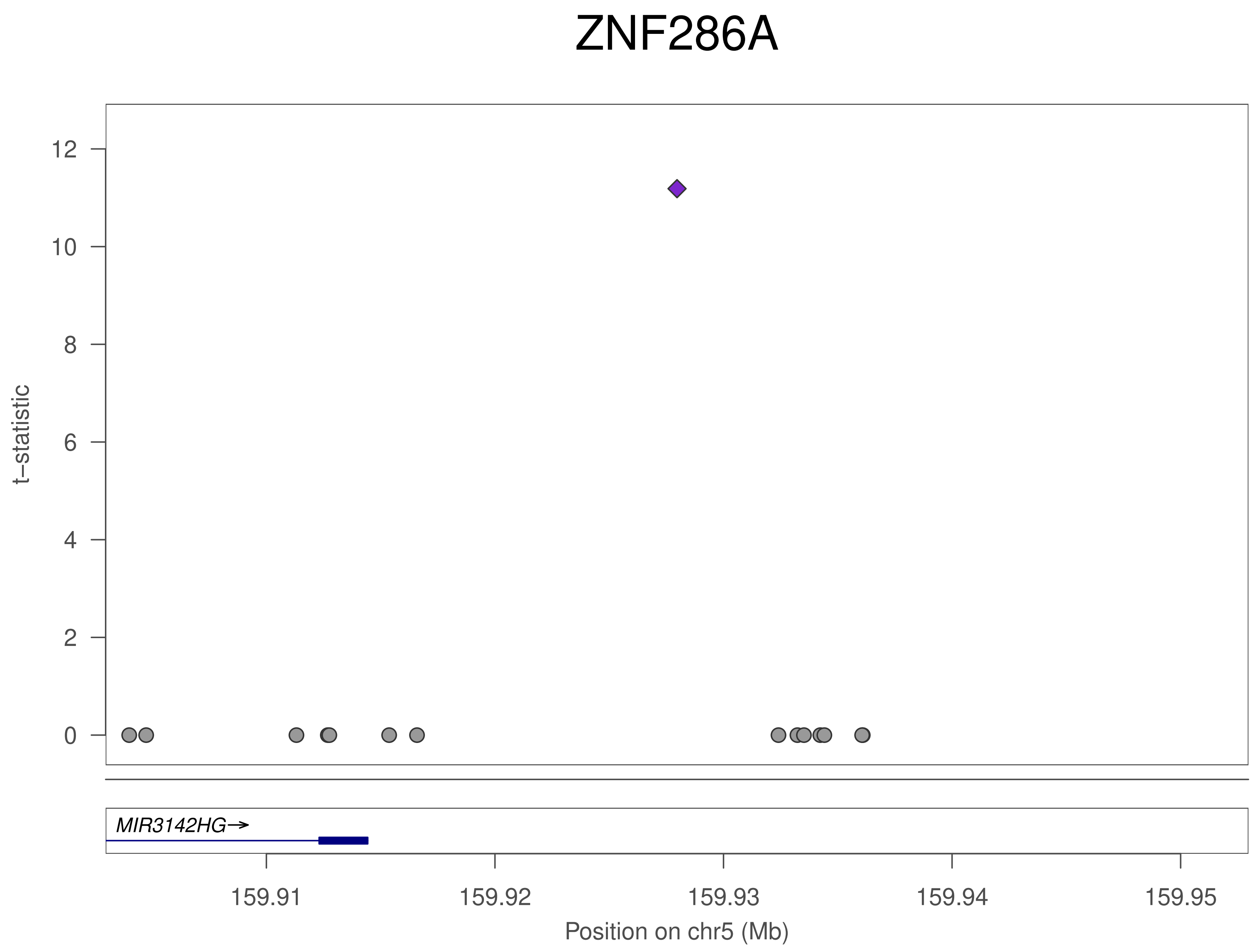

ZNF286A
t–statistic
12
10
8
6
4
2
0
MIR3142HG→
159.91
159.92
159.93
159.94
159.95
Position on chr5 (Mb)

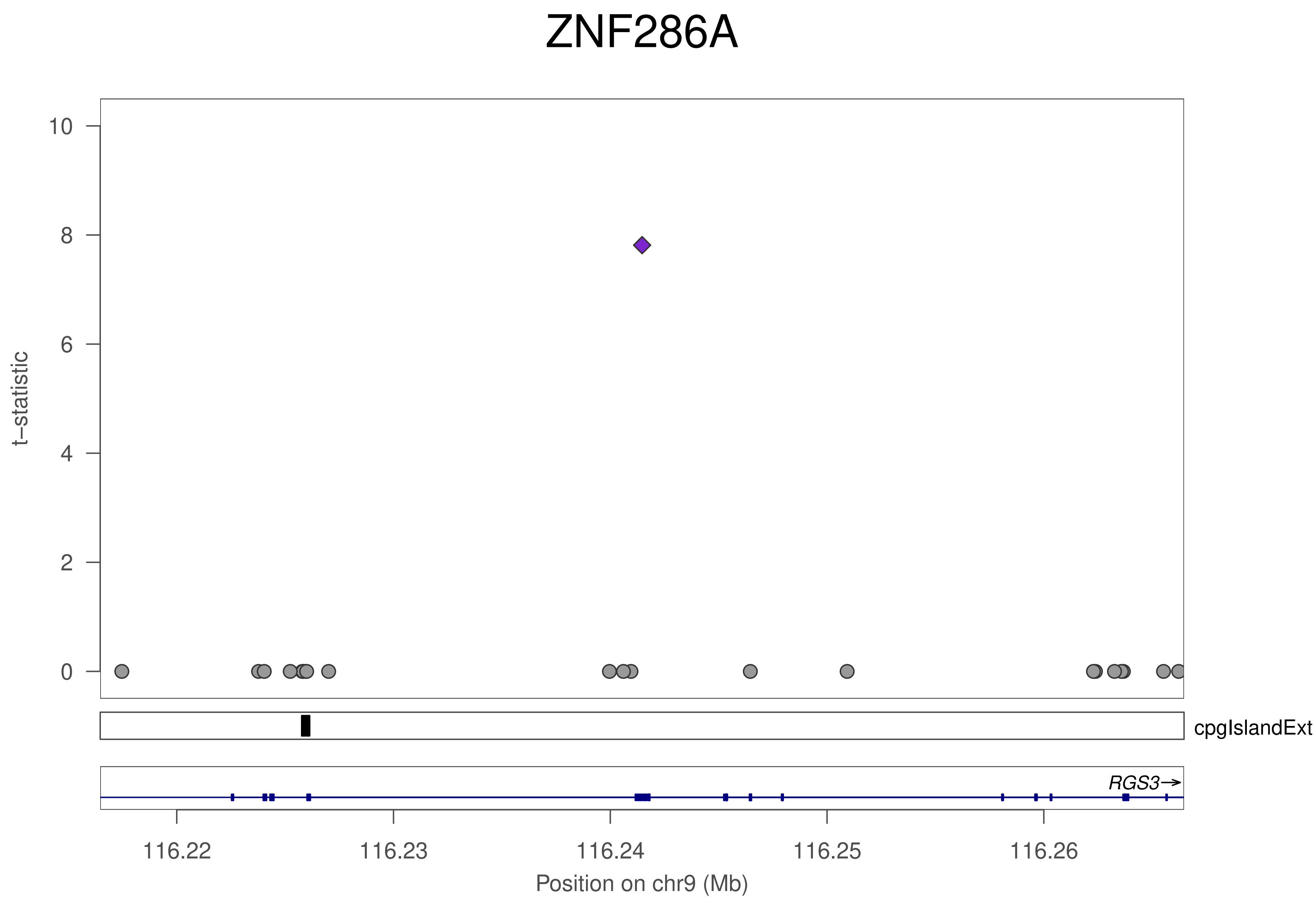
ZNF286A
t−statistic
10
8
6
4
2
0
cpgIslandExt
RGS3→
116.22
116.23
116.24
116.25
116.26
Position on chr9 (Mb)

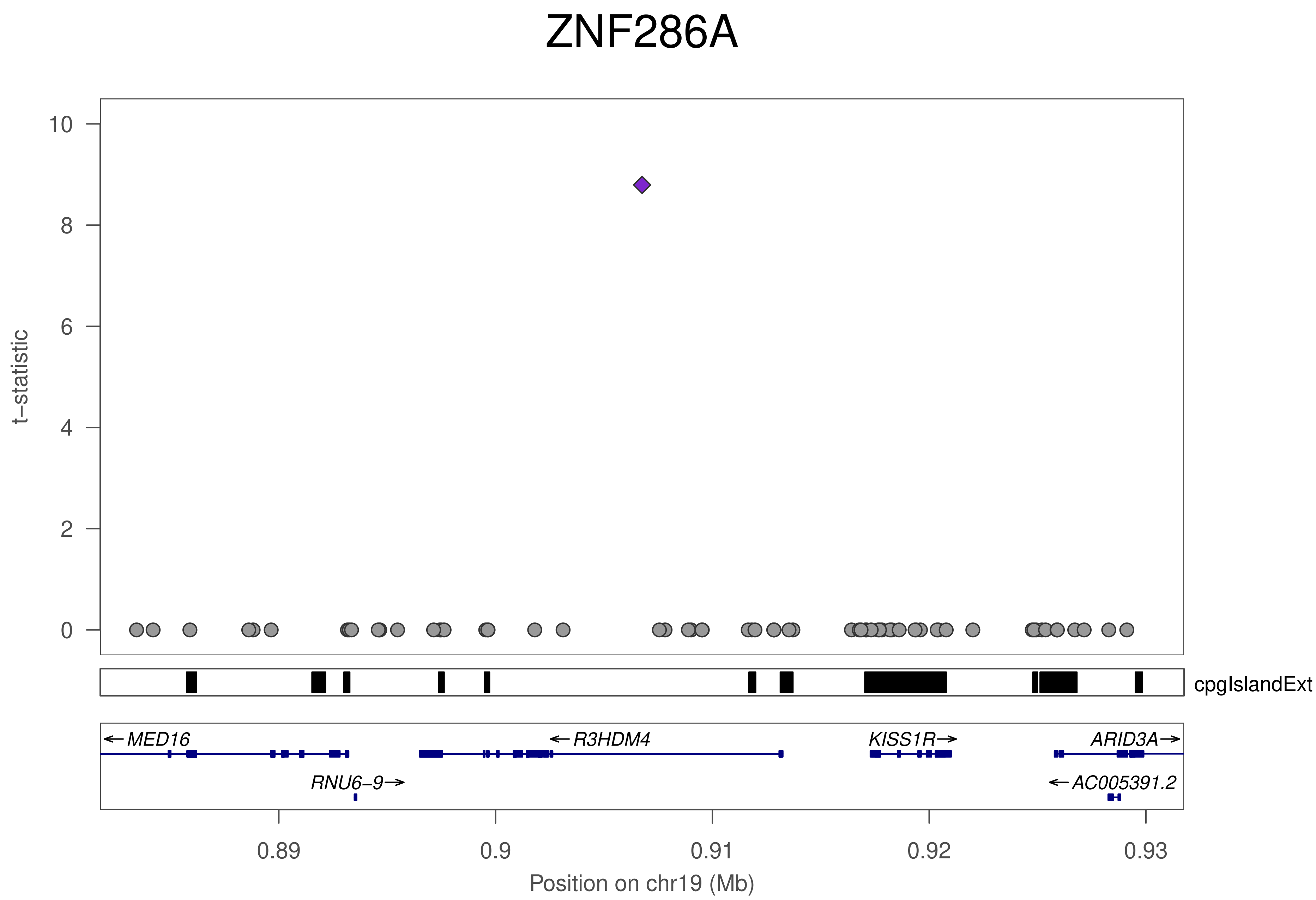

ZNF286A
t−statistic
10
8
6
4
2
0
cpgIslandExt
←MED16
←R3HDM4
KISS1R→
ARID3A→
RNU6−9→
←AC005391.2
0.89
0.9
0.91
0.92
0.93
Position on chr19 (Mb)

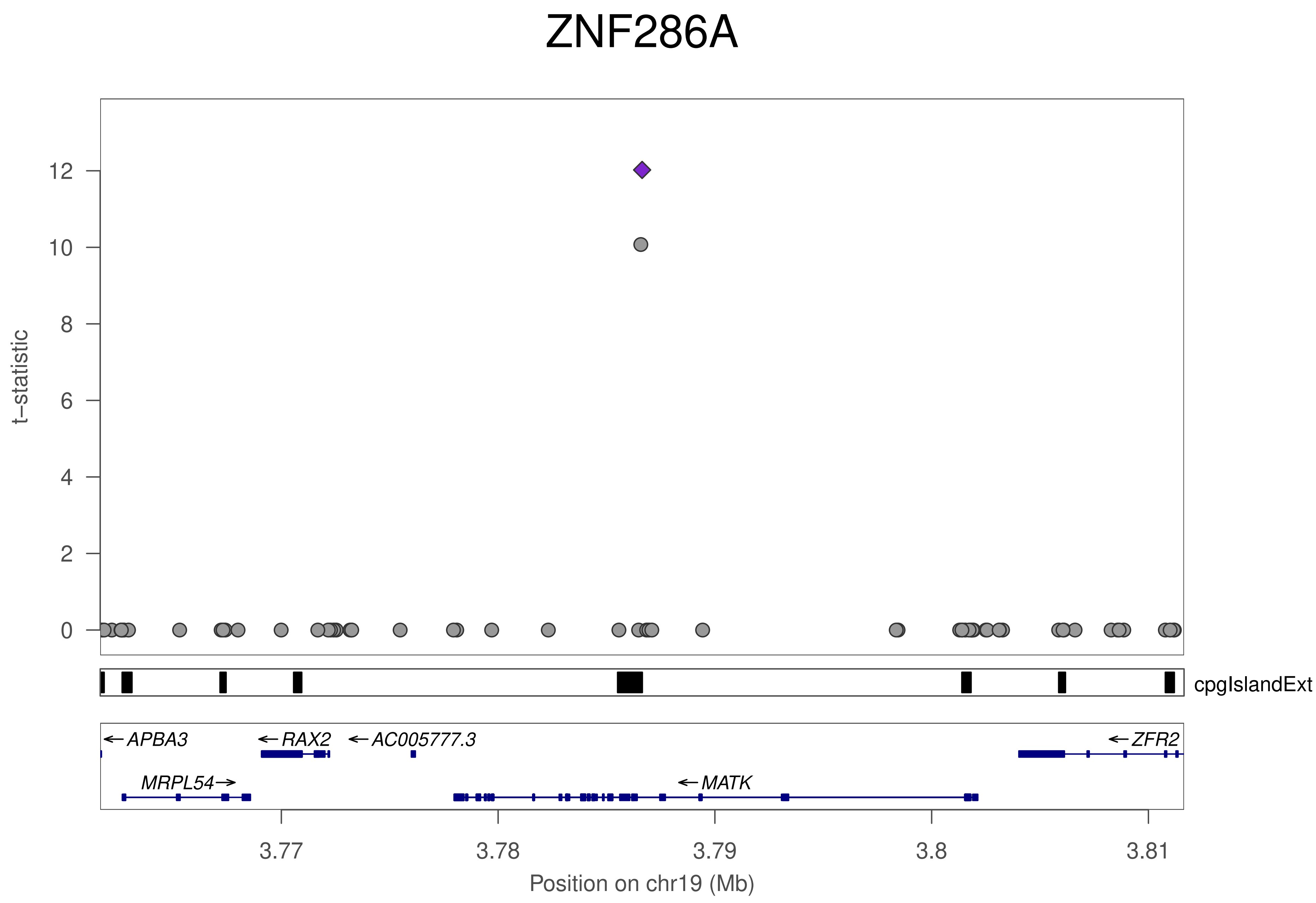

ZNF286A
t−statistic
12
10
8
6
4
2
0
cpgIslandExt
APBA3
RAX2
AC005777.3
ZFR2
MRPL54
MATK
3.77
3.78
3.79
3.8
3.81
Position on chr19 (Mb)

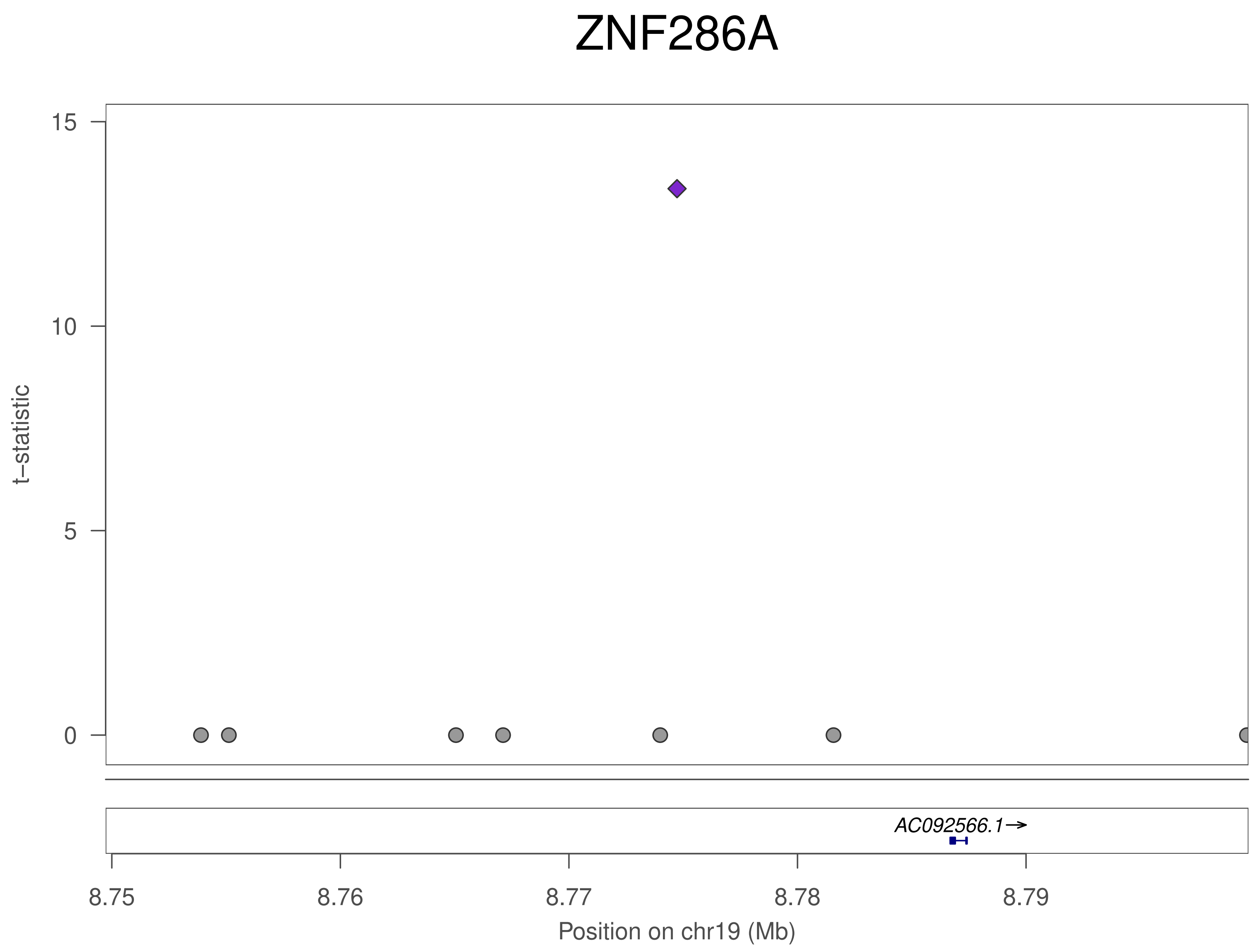
ZNF286A
t–statistic
15
10
5
0
AC092566.1→
8.75
8.76
8.77
8.78
8.79
Position on chr19 (Mb)

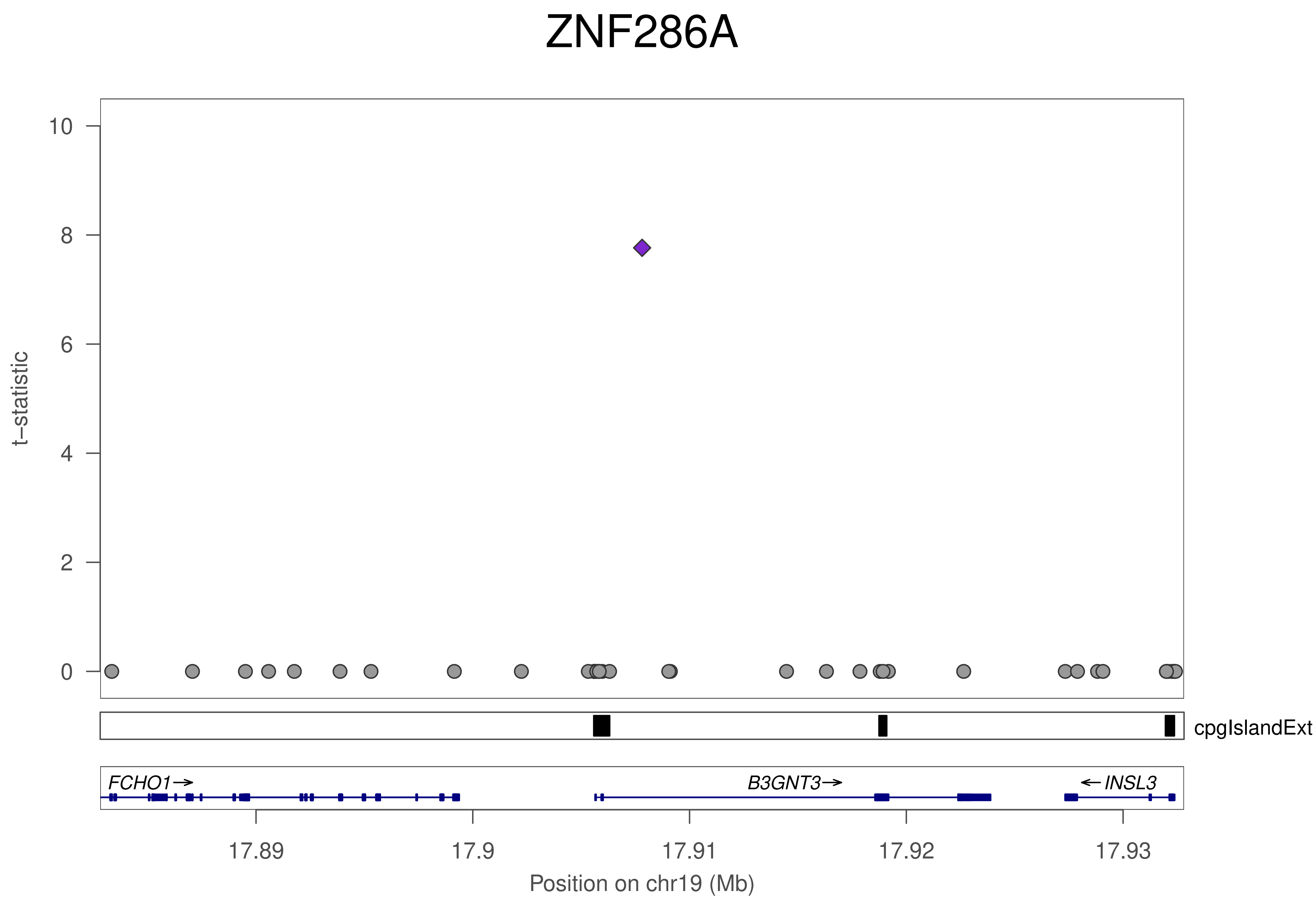
ZNF286A
t−statistic
10
8
6
4
2
0
cpgIslandExt
FCHO1→
B3GNT3→
←INSL3
17.89
17.9
17.91
17.92
17.93
Position on chr19 (Mb)

### ZNF286A

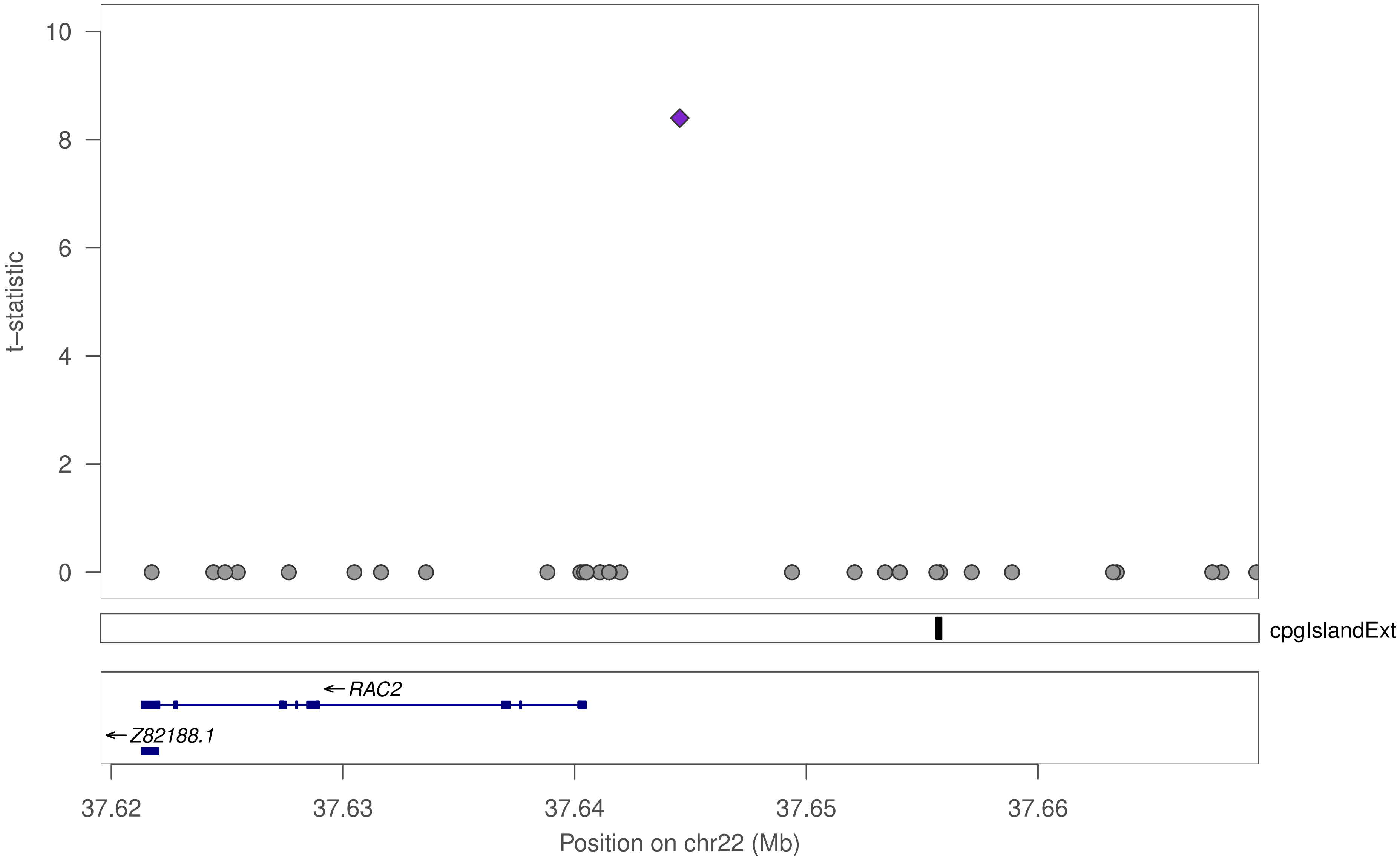

10
8
6
4
2
0
t−statistic
cpgIslandExt
RAC2
Z82188.1
37.62
37.63
37.64
37.65
37.66
Position on chr22 (Mb)

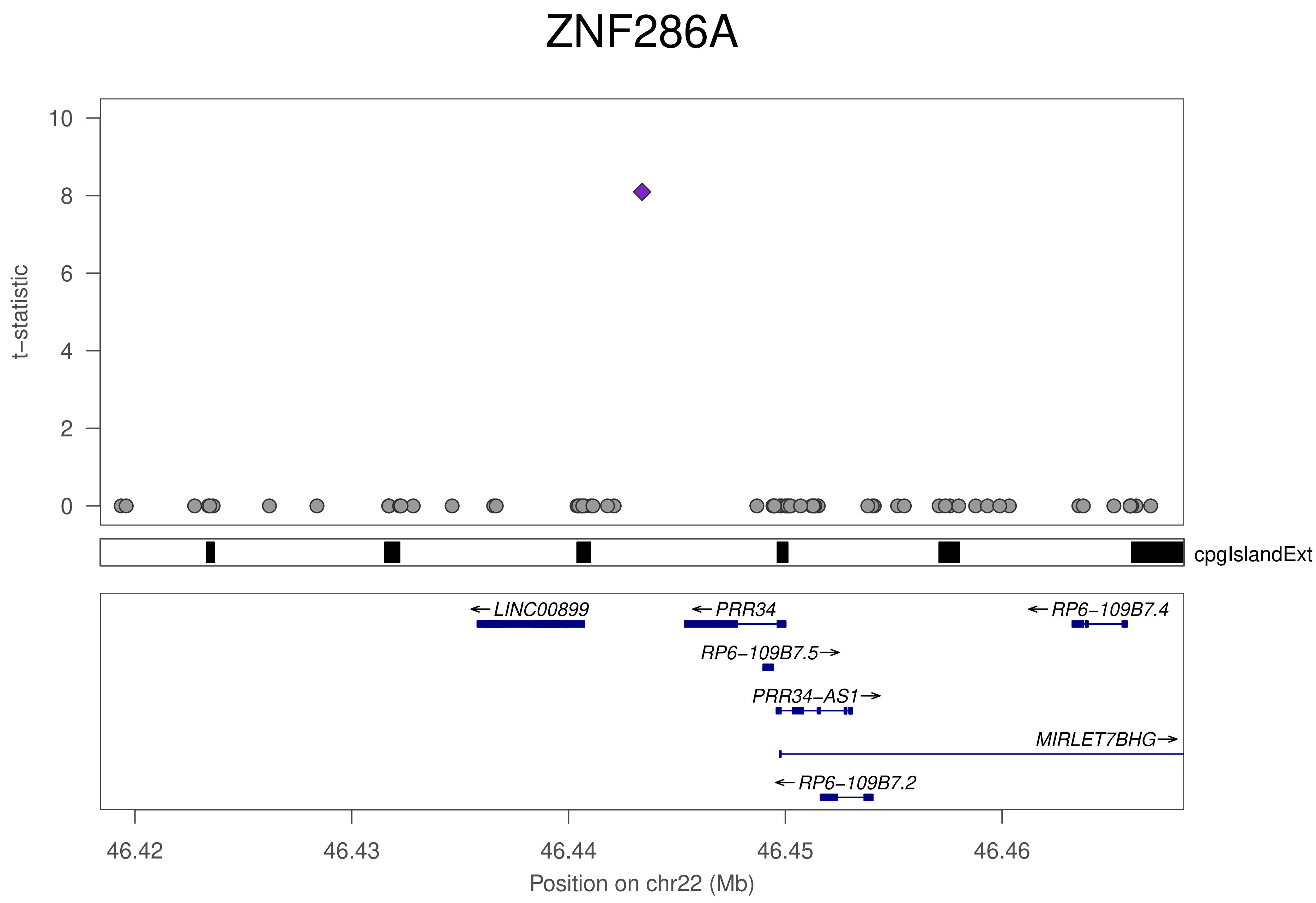

ZNF286A
t–statistic
10
8
6
4
2
0
cpgIslandExt
←LINC00899
←PRR34
←RP6–109B7.4
RP6–109B7.5→
PRR34–AS1→
MIRLET7BHG→
←RP6–109B7.2
46.42
46.43
46.44
46.45
46.46
Position on chr22 (Mb)

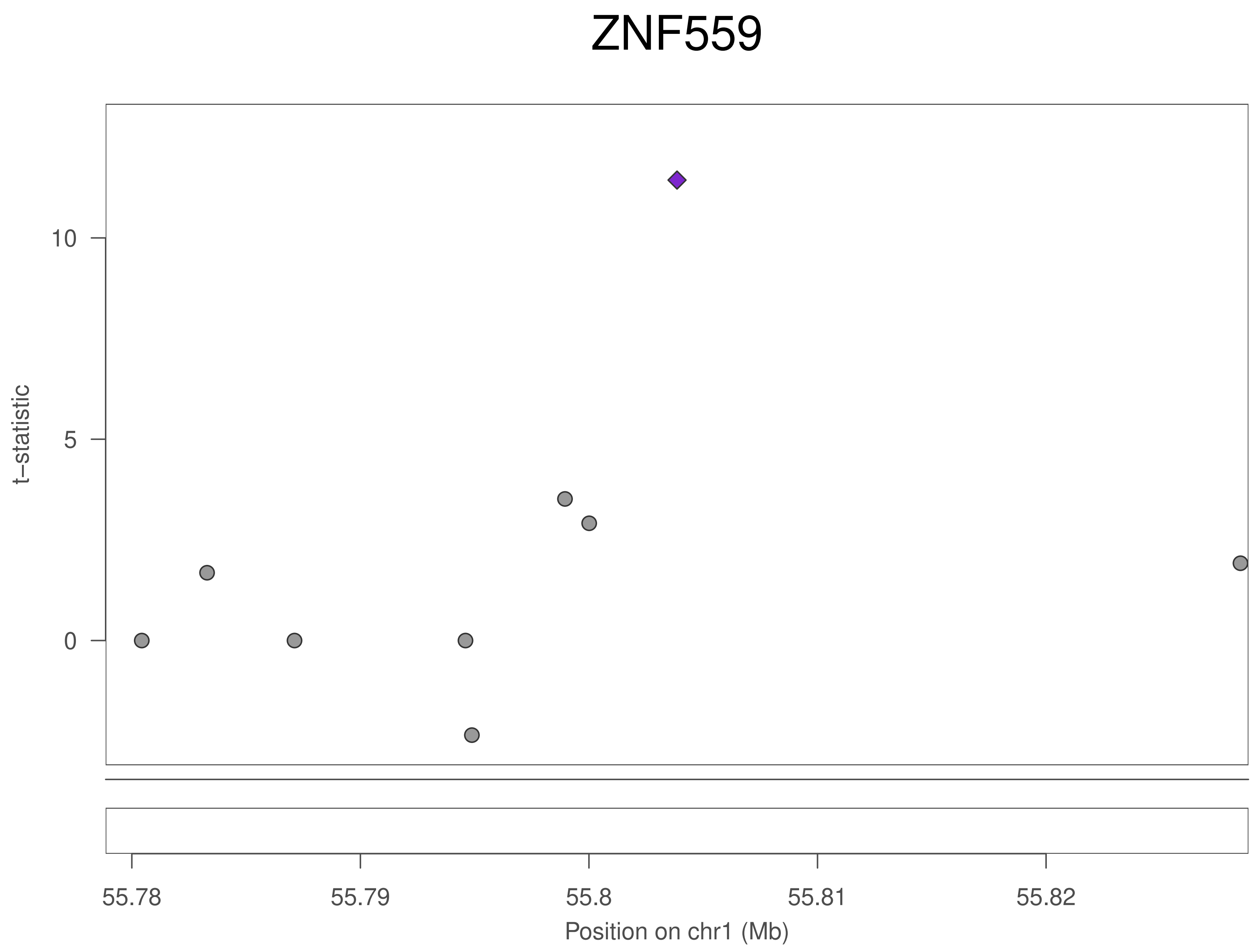

ZNF559
t–statistic
10
5
0
55.78
55.79
55.8
55.81
55.82
Position on chr1 (Mb)

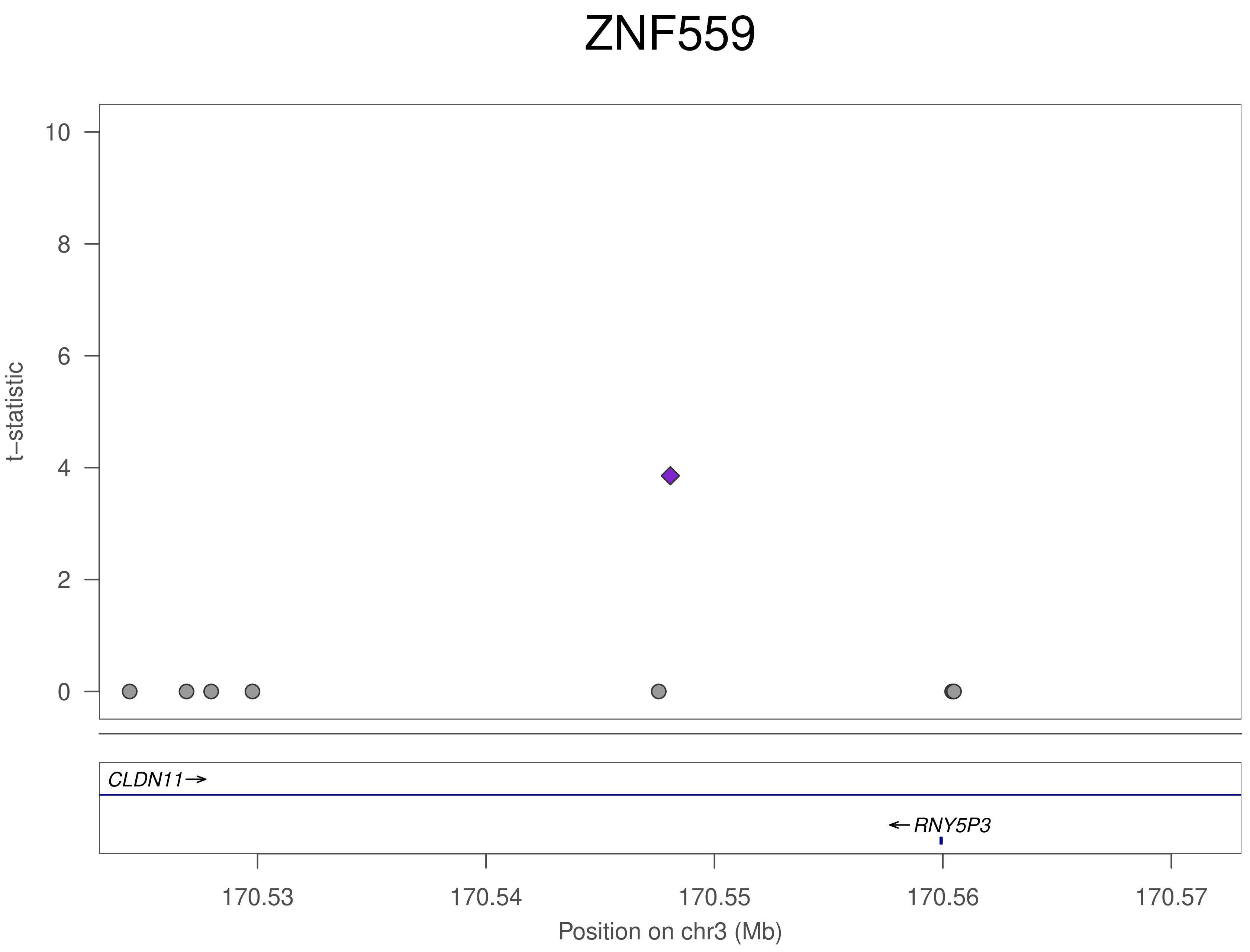

ZNF559
t–statistic
10
8
6
4
2
0
CLDN11→
←RNY5P3
170.53
170.54
170.55
170.56
170.57
Position on chr3 (Mb)

### ZNF559

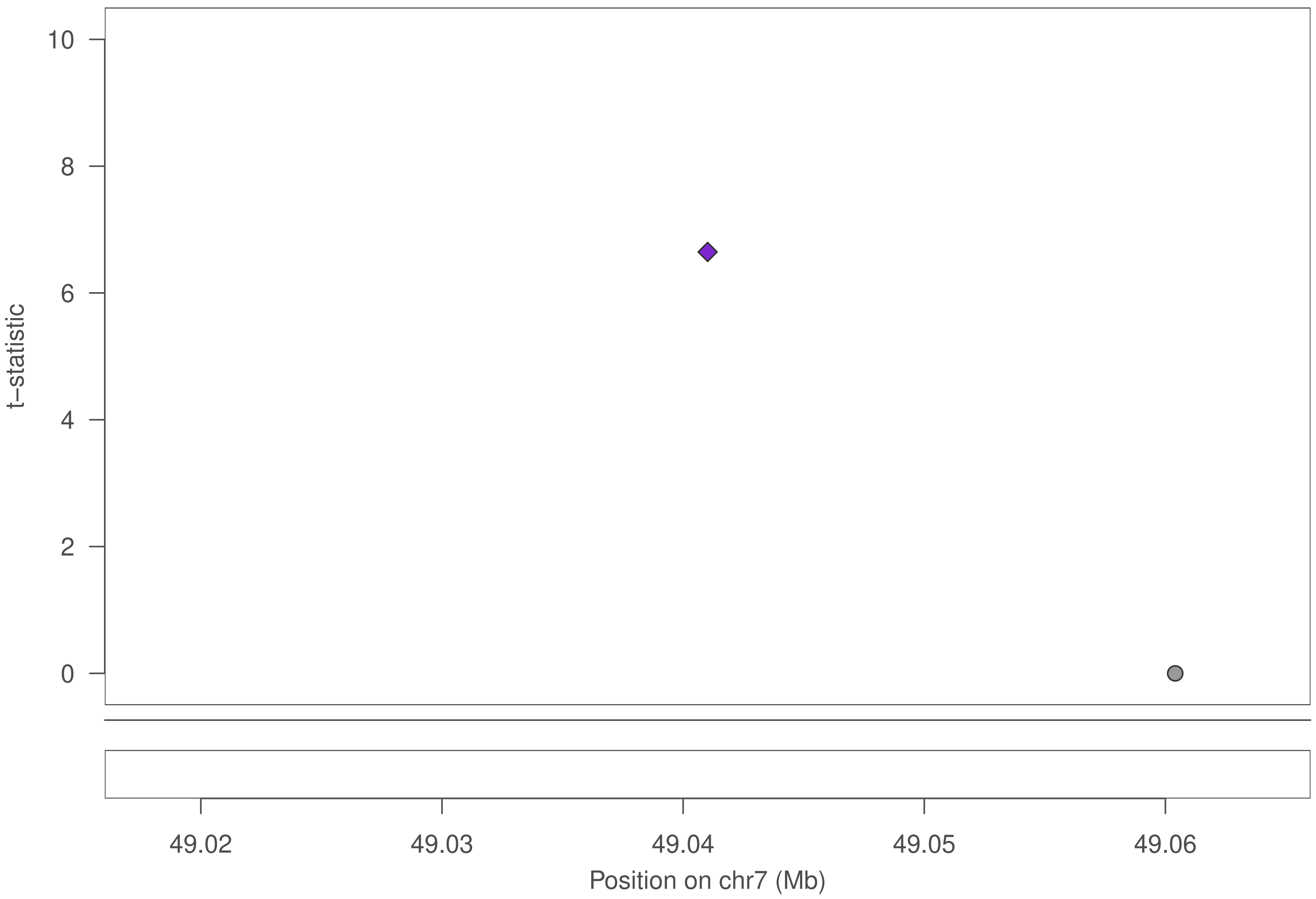

10
8
6
4
2
0
t–statistic
49.02
49.03
49.04
49.05
49.06
Position on chr7 (Mb)

### ZNF559

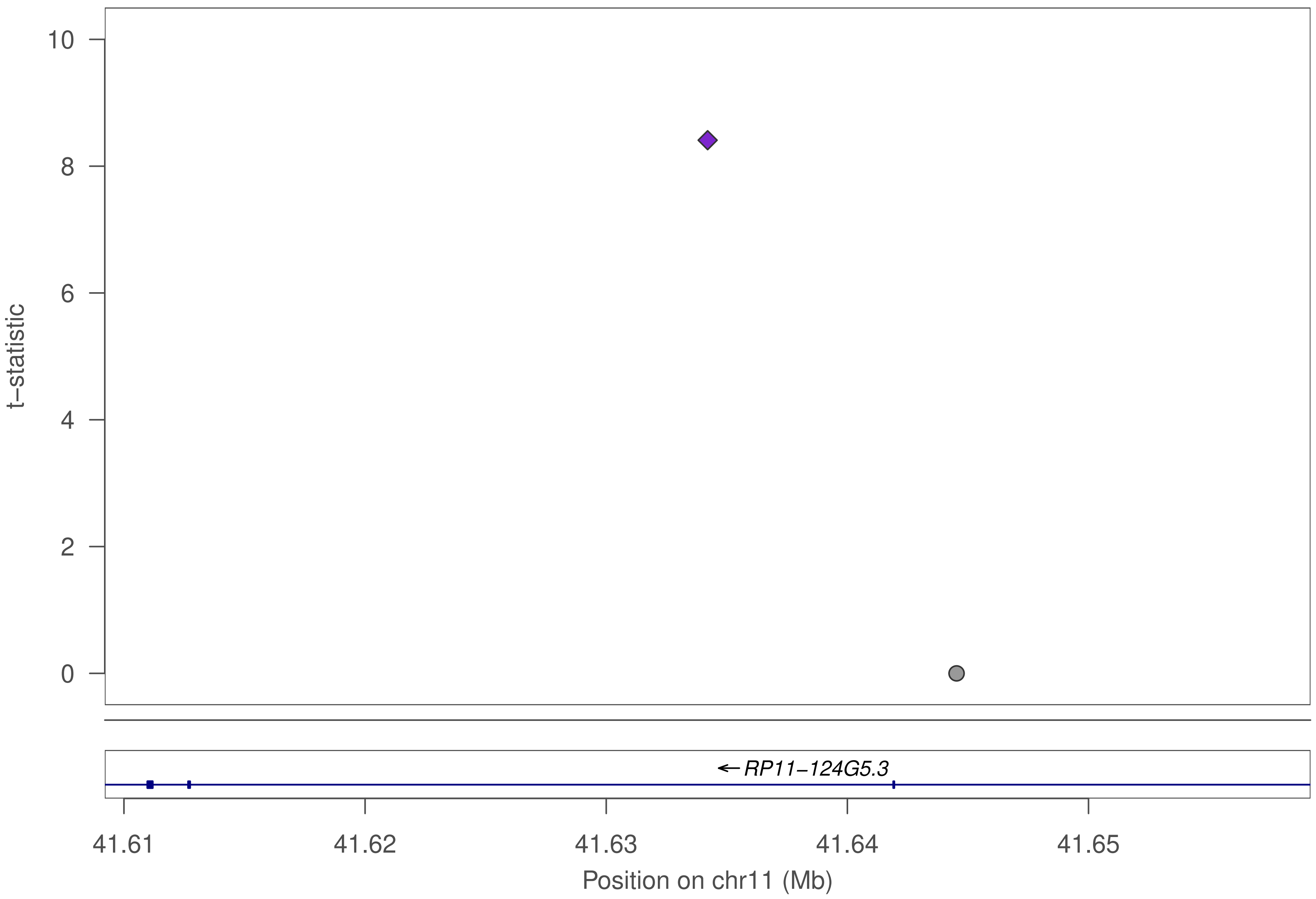

10
8
6
4
2
0
t–statistic
RP11–124G5.3
41.61
41.62
41.63
41.64
41.65
Position on chr11 (Mb)

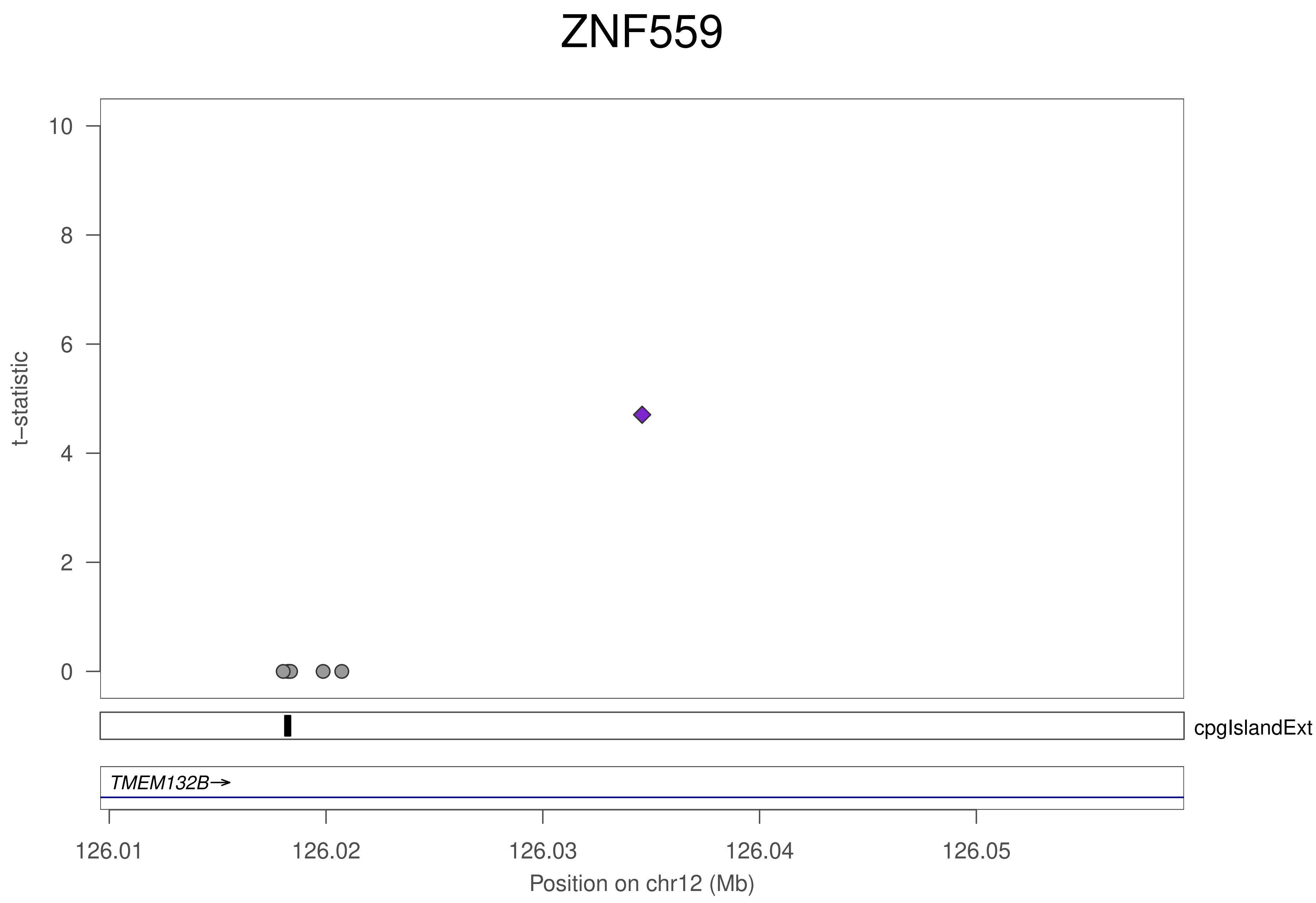
ZNF559
t−statistic
10
8
6
4
2
0
cpgIslandExt
TMEM132B→
126.01
126.02
126.03
126.04
126.05
Position on chr12 (Mb)

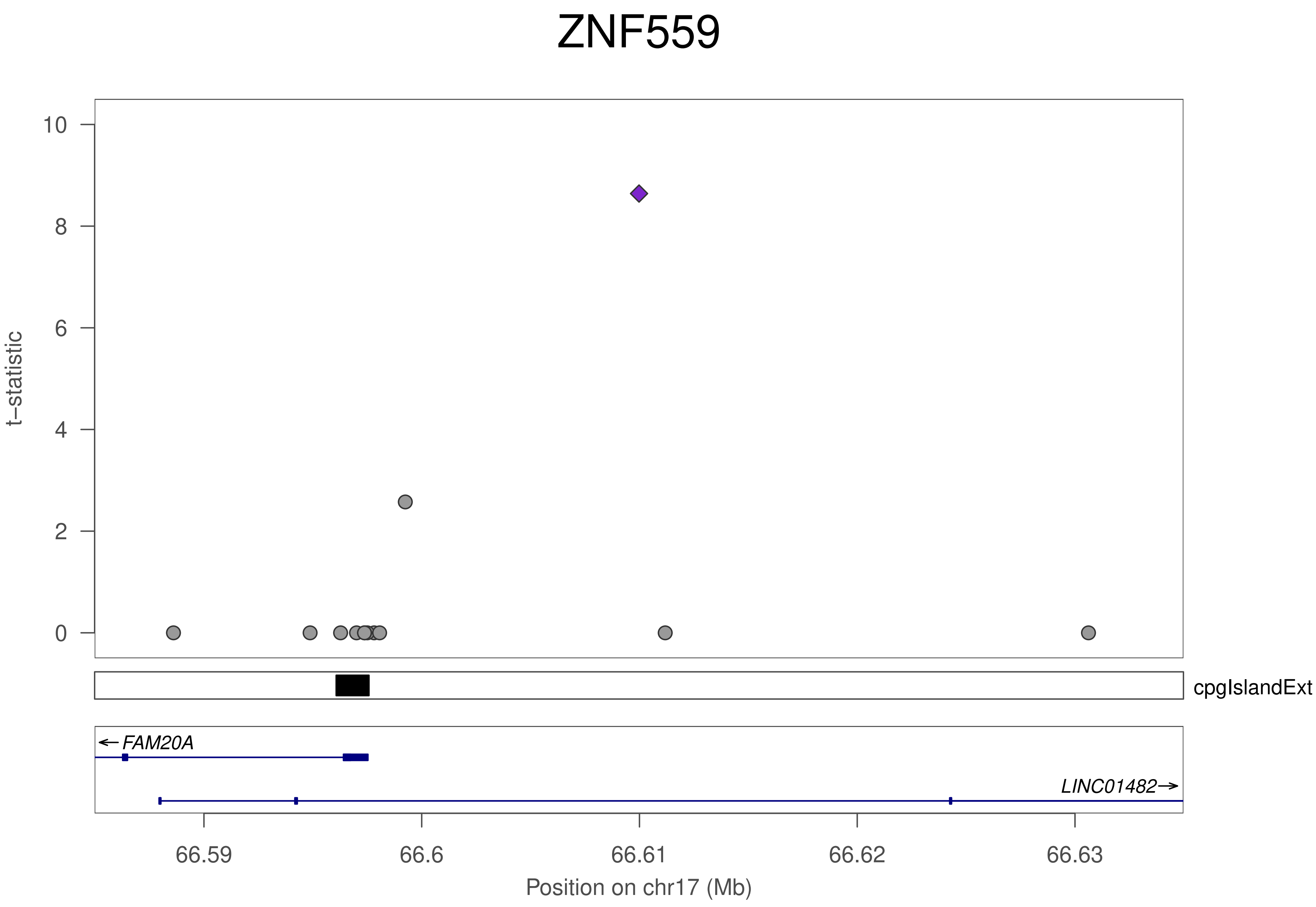

ZNF559
t−statistic
10
8
6
4
2
0
cpgIslandExt
FAM20A
LINC01482
66.59
66.6
66.61
66.62
66.63
Position on chr17 (Mb)

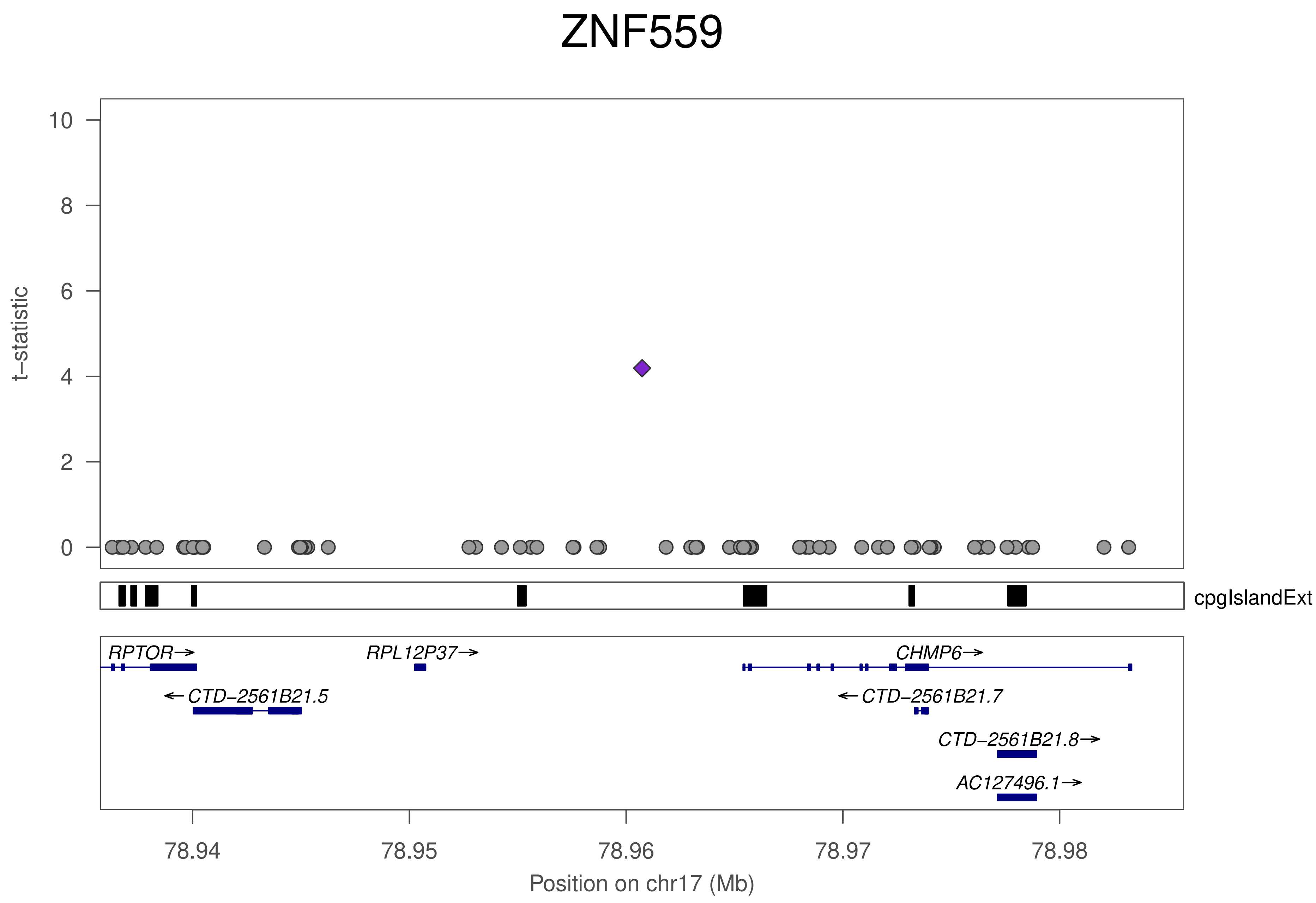
ZNF559
t−statistic
10
8
6
4
2
0
cpgIslandExt
RPTOR→
RPL12P37→
CHMP6→
←CTD−2561B21.5
←CTD−2561B21.7
CTD−2561B21.8→
AC127496.1→
78.94
78.95
78.96
78.97
78.98
Position on chr17 (Mb)

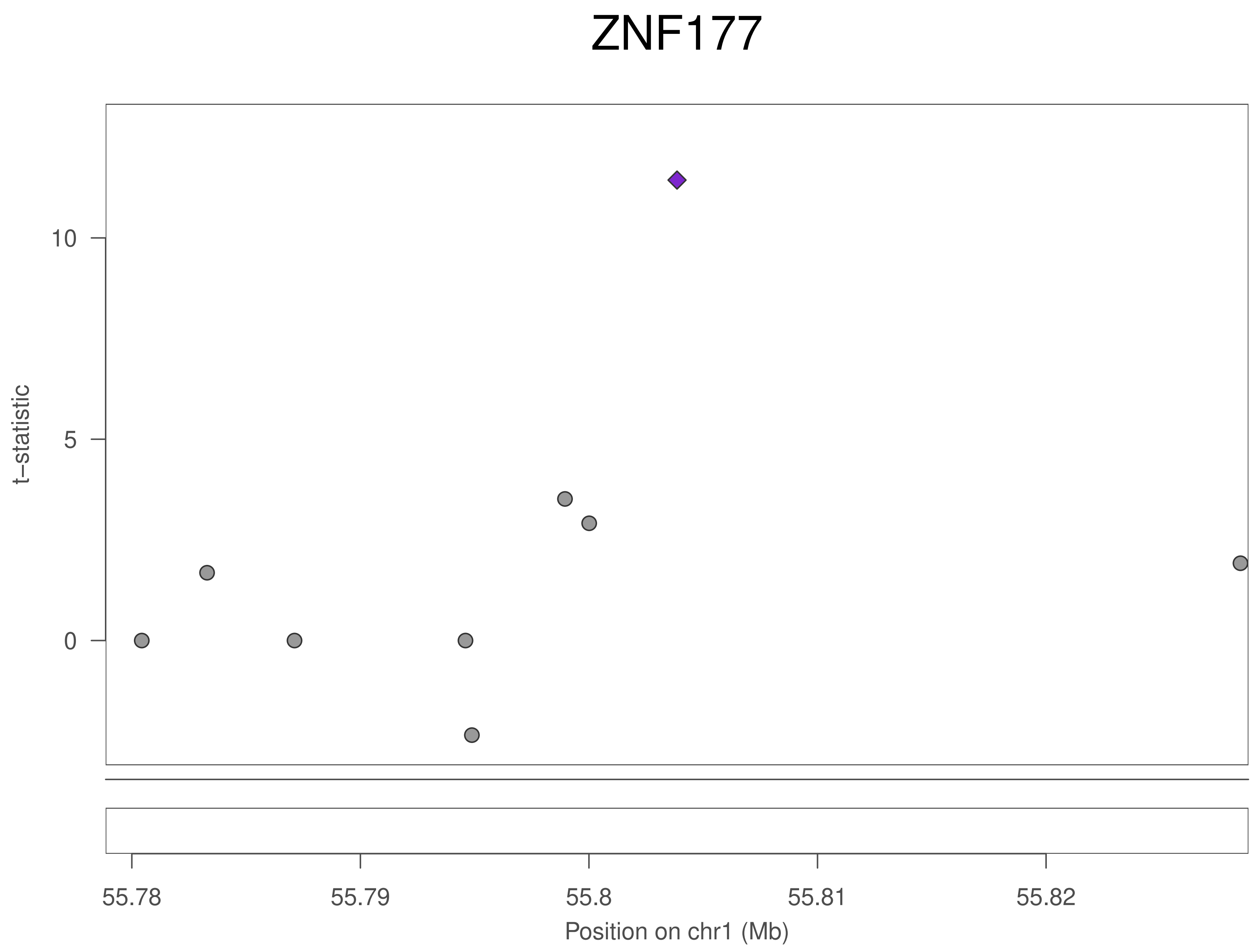
ZNF177
t−statistic
10
5
0
55.78
55.79
55.8
55.81
55.82
Position on chr1 (Mb)

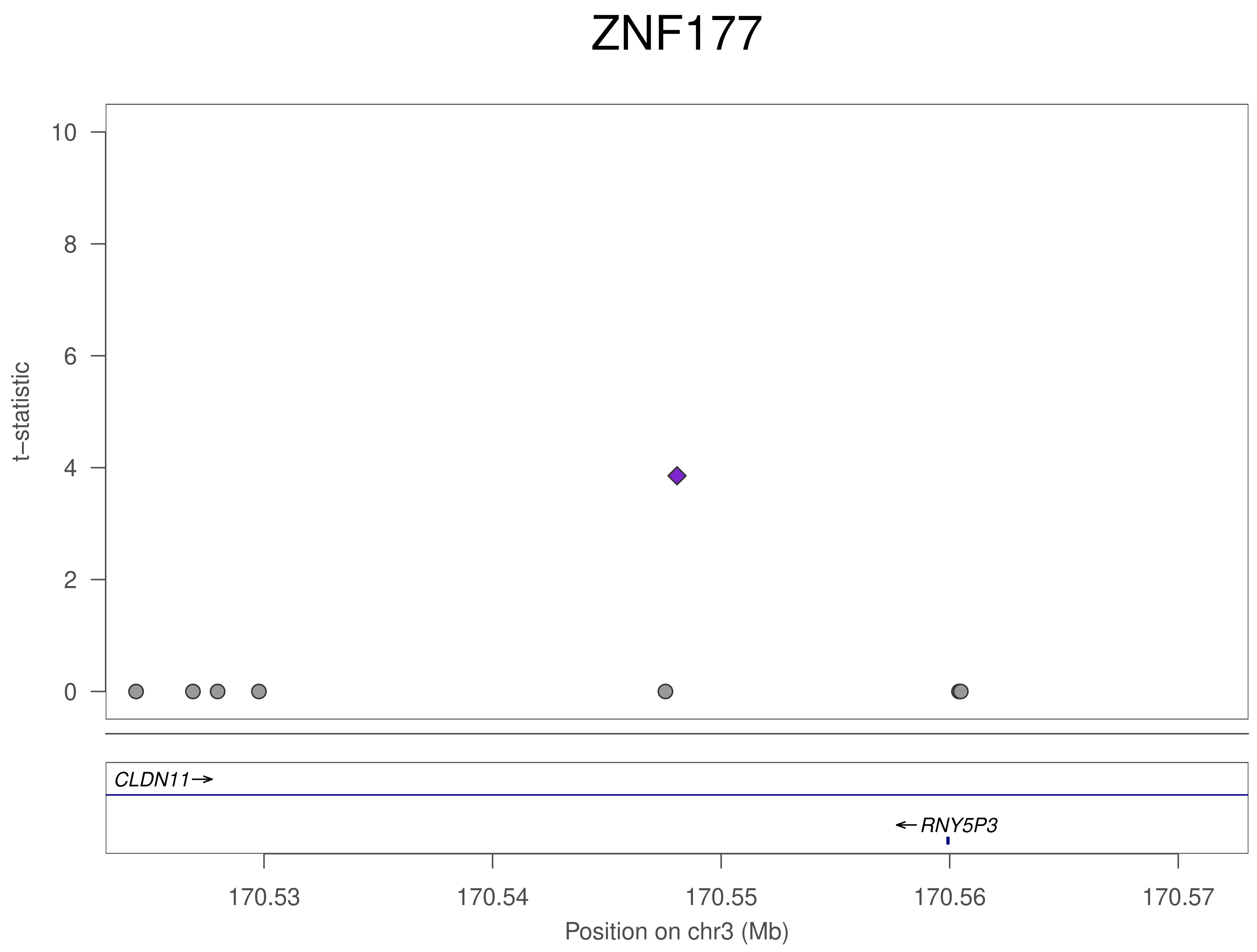

ZNF177
t−statistic
10
8
6
4
2
0
CLDN11→
←RNY5P3
170.53
170.54
170.55
170.56
170.57
Position on chr3 (Mb)

### ZNF177

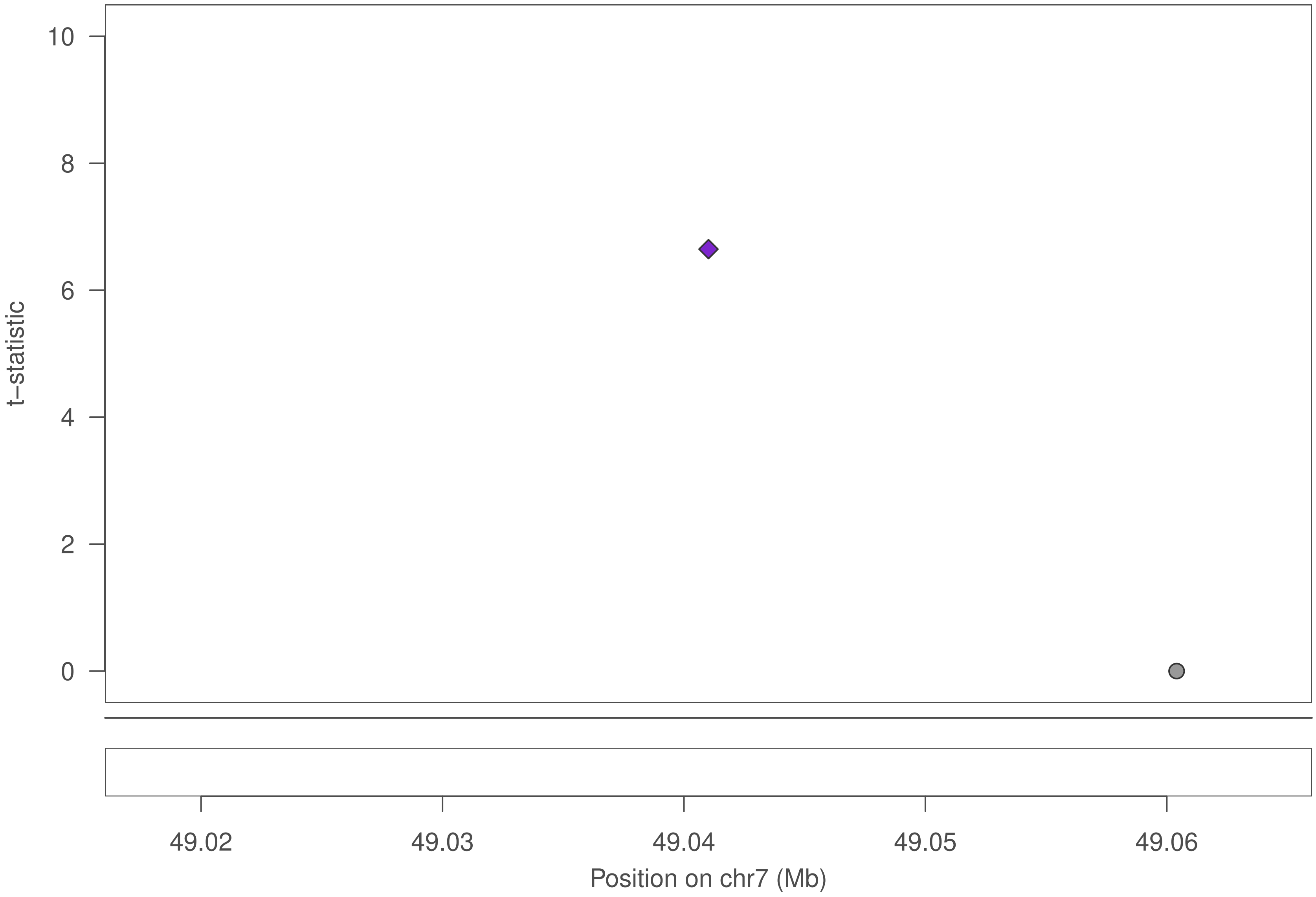

10
8
6
4
2
0
t–statistic
49.02
49.03
49.04
49.05
49.06
Position on chr7 (Mb)

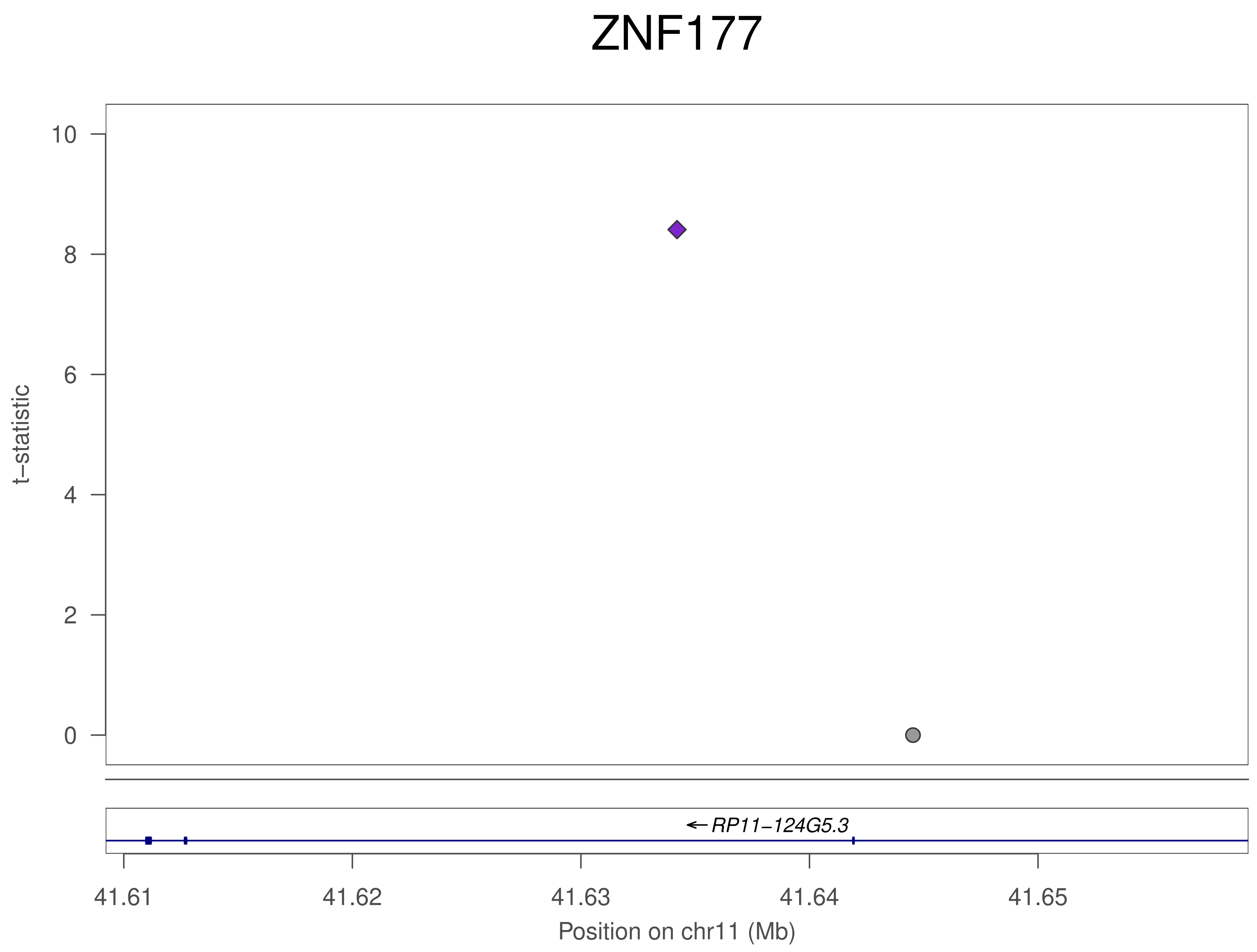
ZNF177
t−statistic
10
8
6
4
2
0
RP11−124G5.3
41.61
41.62
41.63
41.64
41.65
Position on chr11 (Mb)

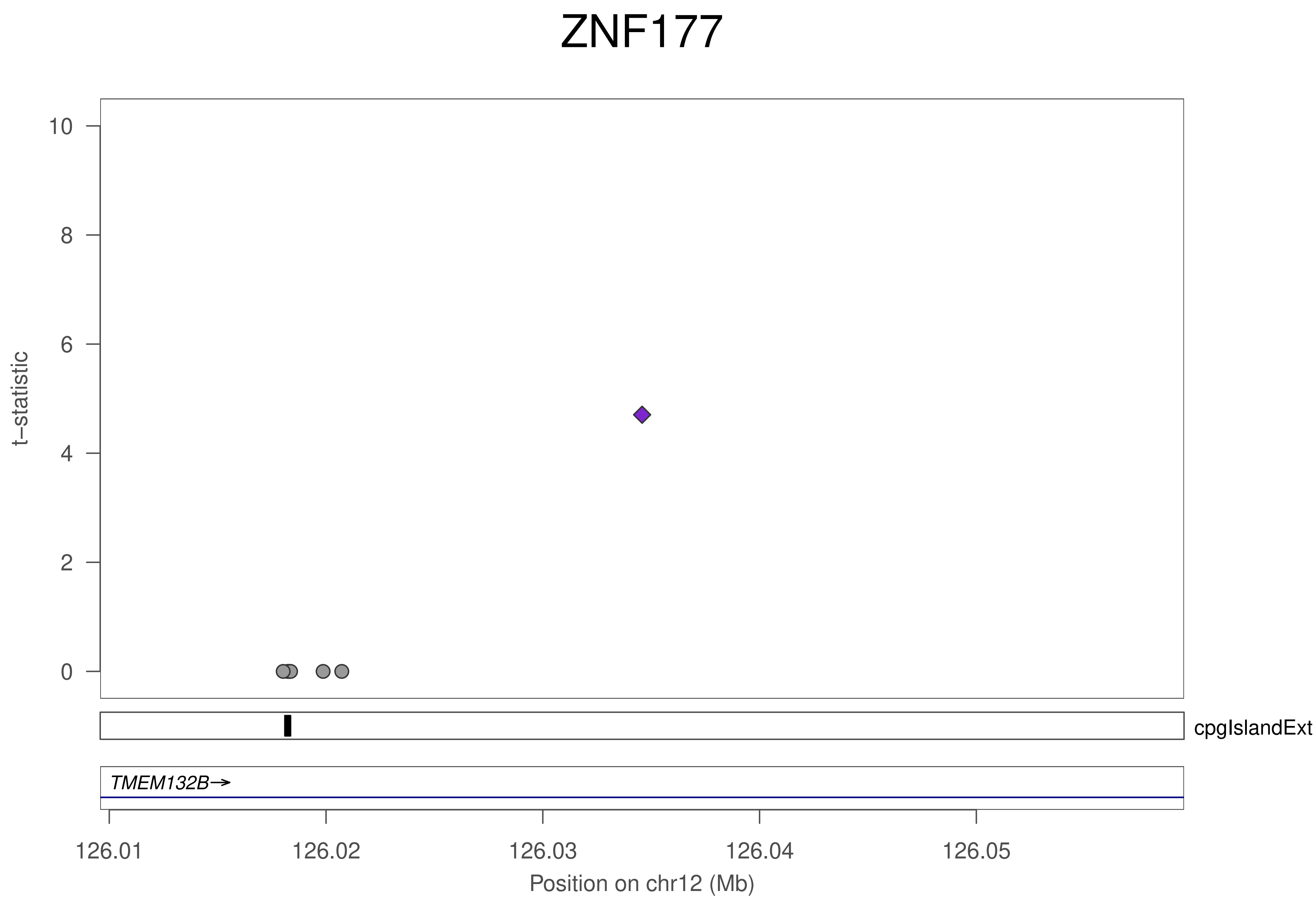
ZNF177
t−statistic
10
8
6
4
2
0
cpgIslandExt
TMEM132B→
126.01
126.02
126.03
126.04
126.05
Position on chr12 (Mb)

### ZNF177

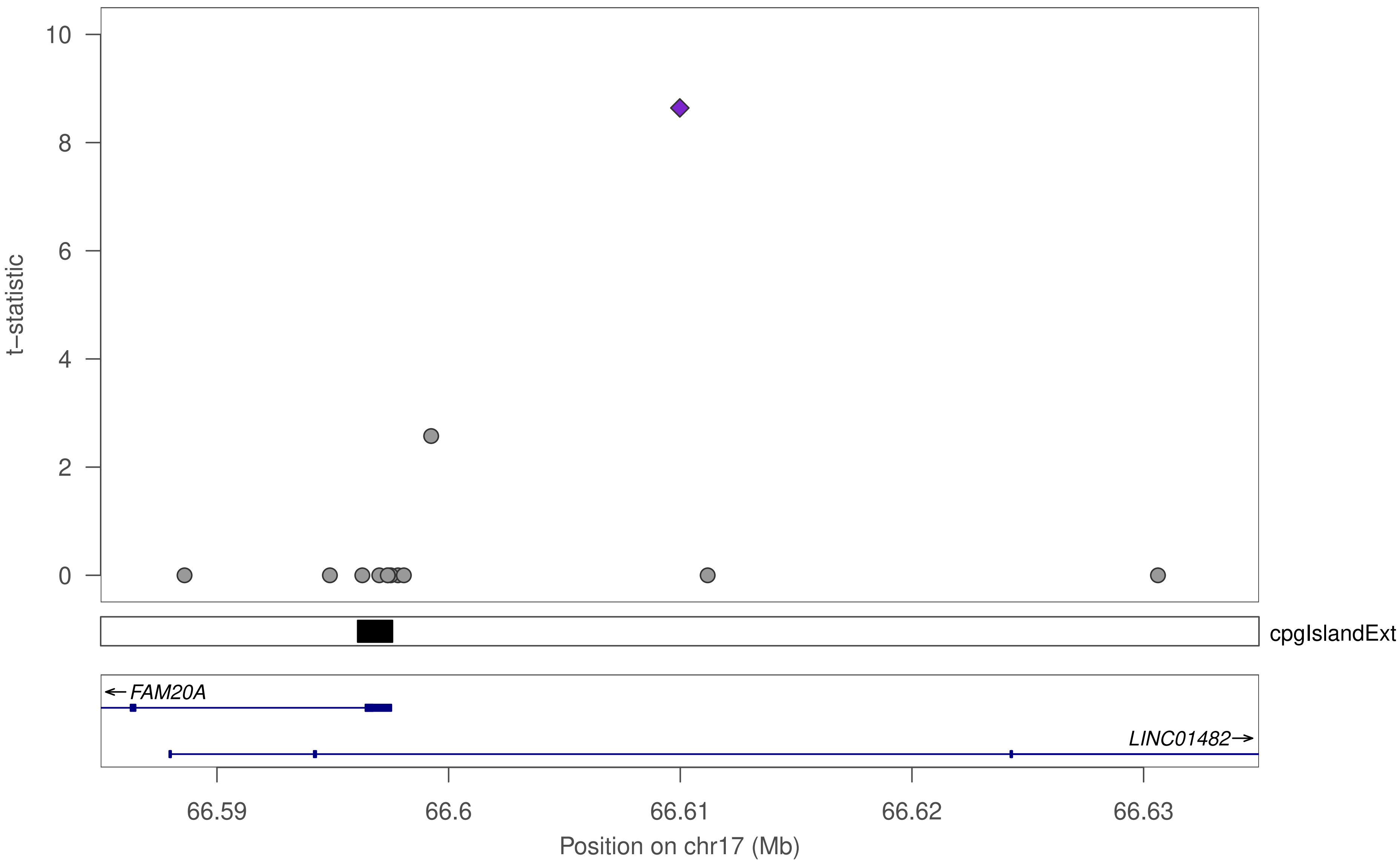

10
8
6
4
2
0
t−statistic
cpgIslandExt
FAM20A
LINC01482
66.59
66.6
66.61
66.62
66.63
Position on chr17 (Mb)

### ZNF177

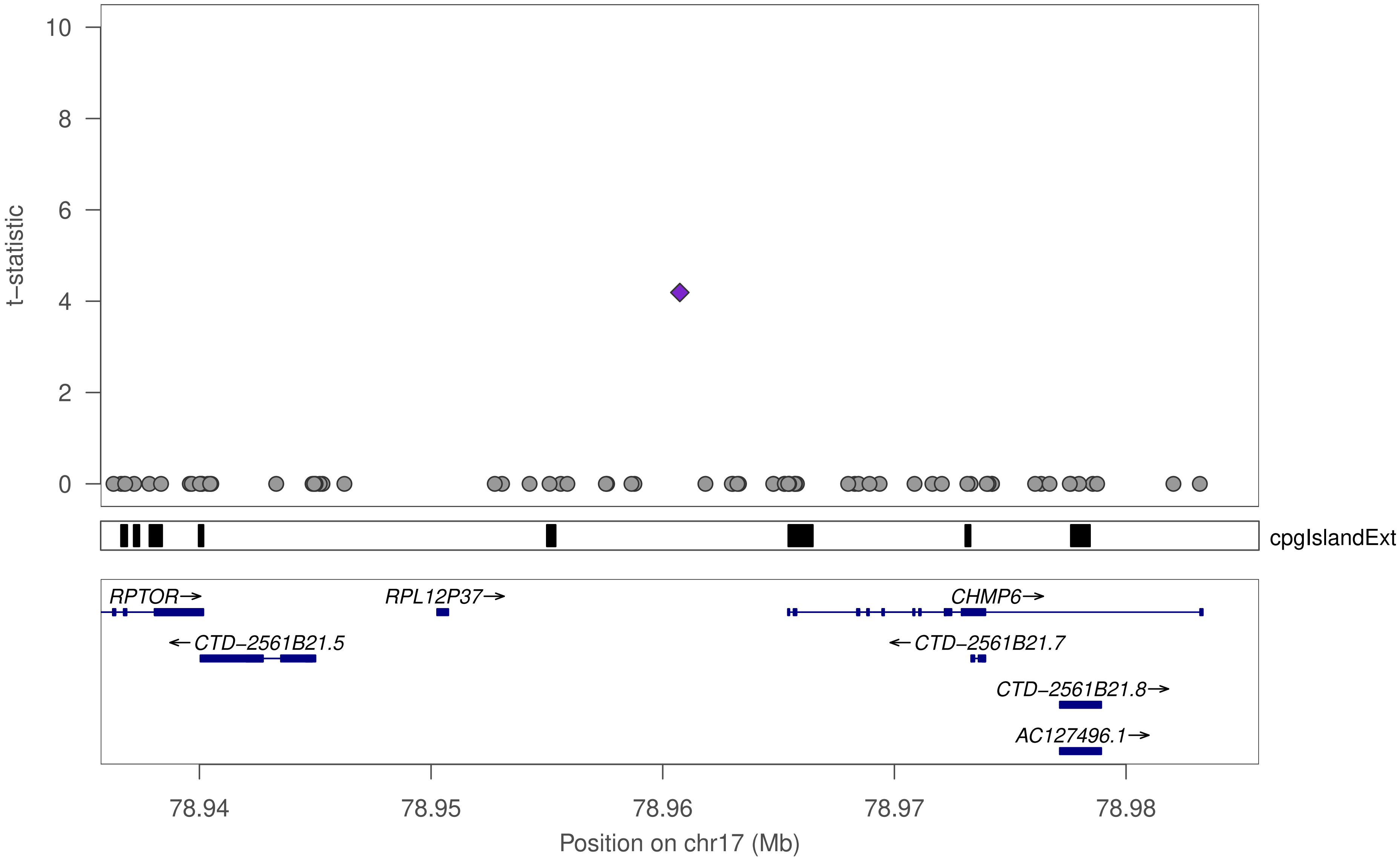

t−statistic
0
2
4
6
8
10
cpgIslandExt
RPTOR→
RPL12P37→
CHMP6→
←CTD−2561B21.5
←CTD−2561B21.7
CTD−2561B21.8→
AC127496.1→
78.94
78.95
78.96
78.97
78.98
Position on chr17 (Mb)

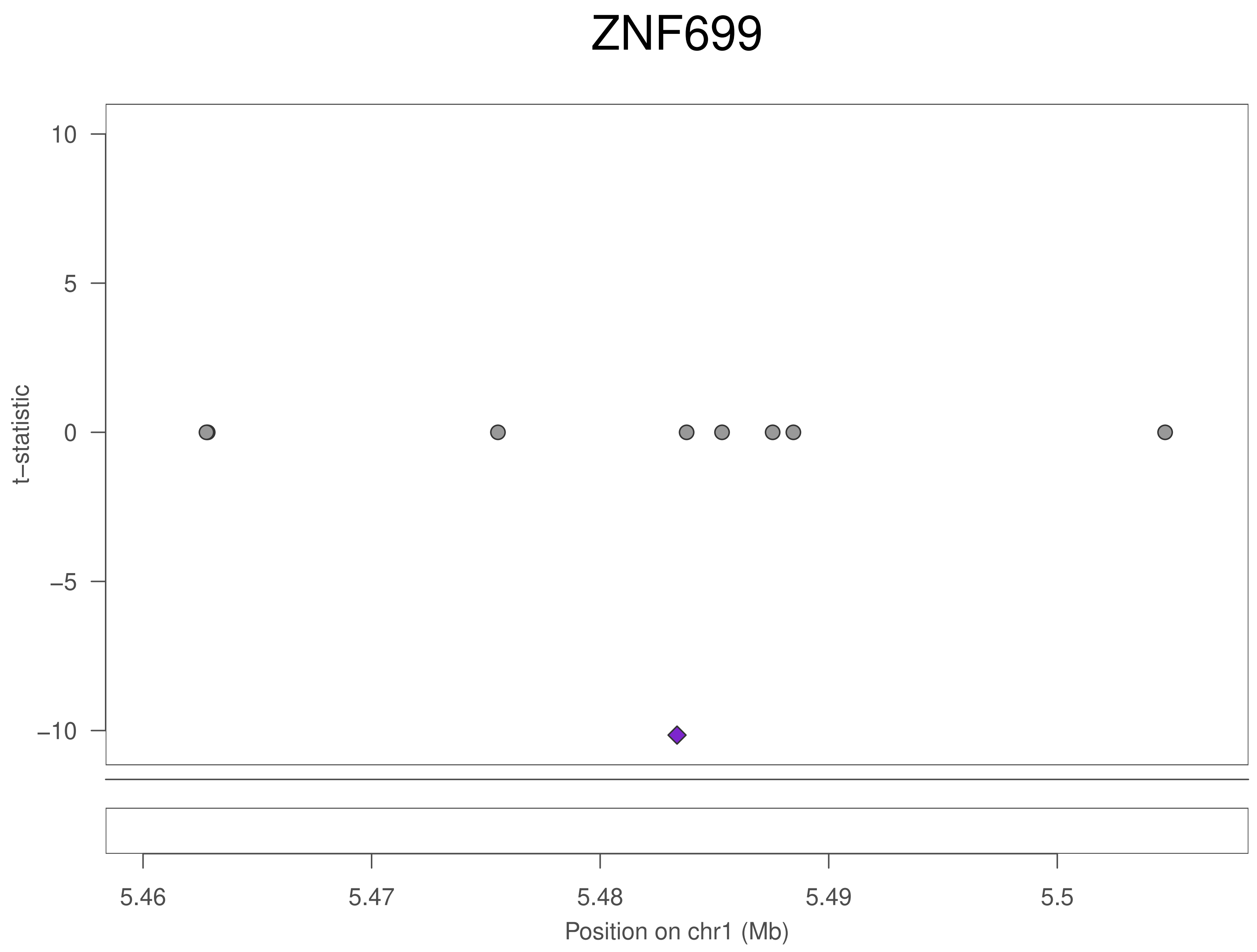
ZNF699
10
5
0
−5
−10
t−statistic
5.46
5.47
5.48
5.49
5.5
Position on chr1 (Mb)

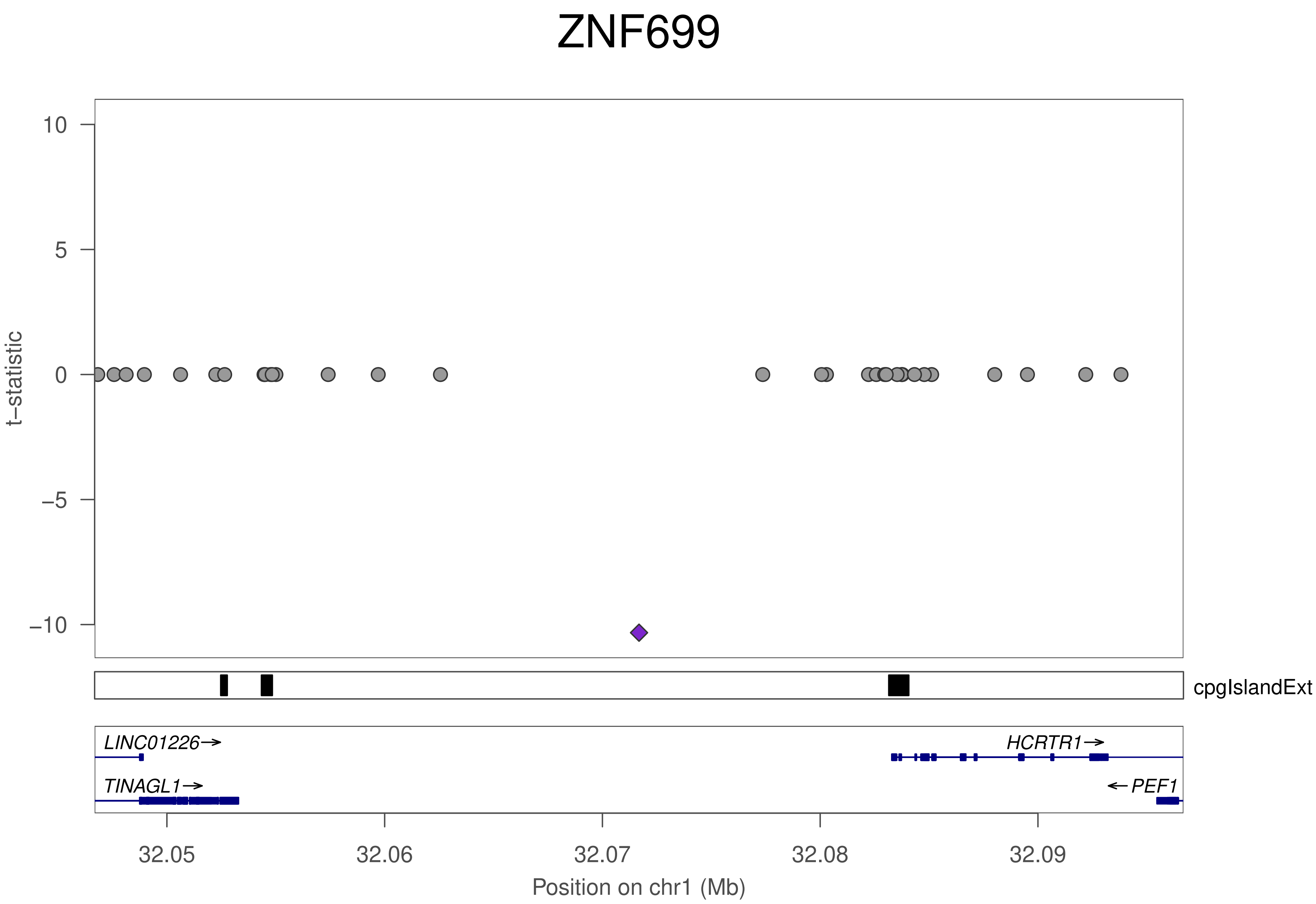
ZNF699
t−statistic
10
5
0
−5
−10
cpgIslandExt
LINC01226→
HCRTR1→
TINAGL1→
←PEF1
32.05
32.06
32.07
32.08
32.09
Position on chr1 (Mb)

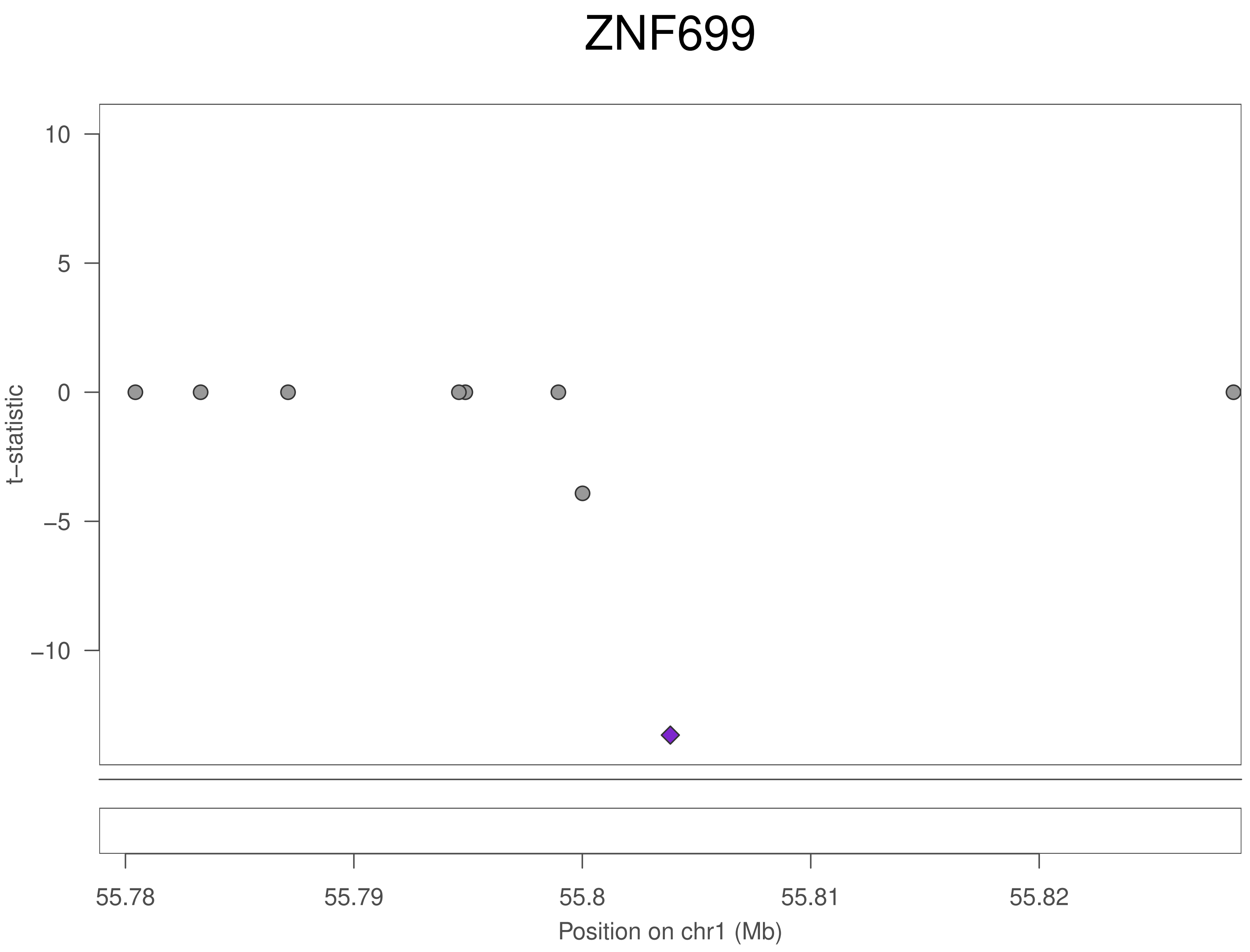
ZNF699
10
5
0
−5
−10
t−statistic
55.78
55.79
55.8
55.81
55.82
Position on chr1 (Mb)

### ZNF699

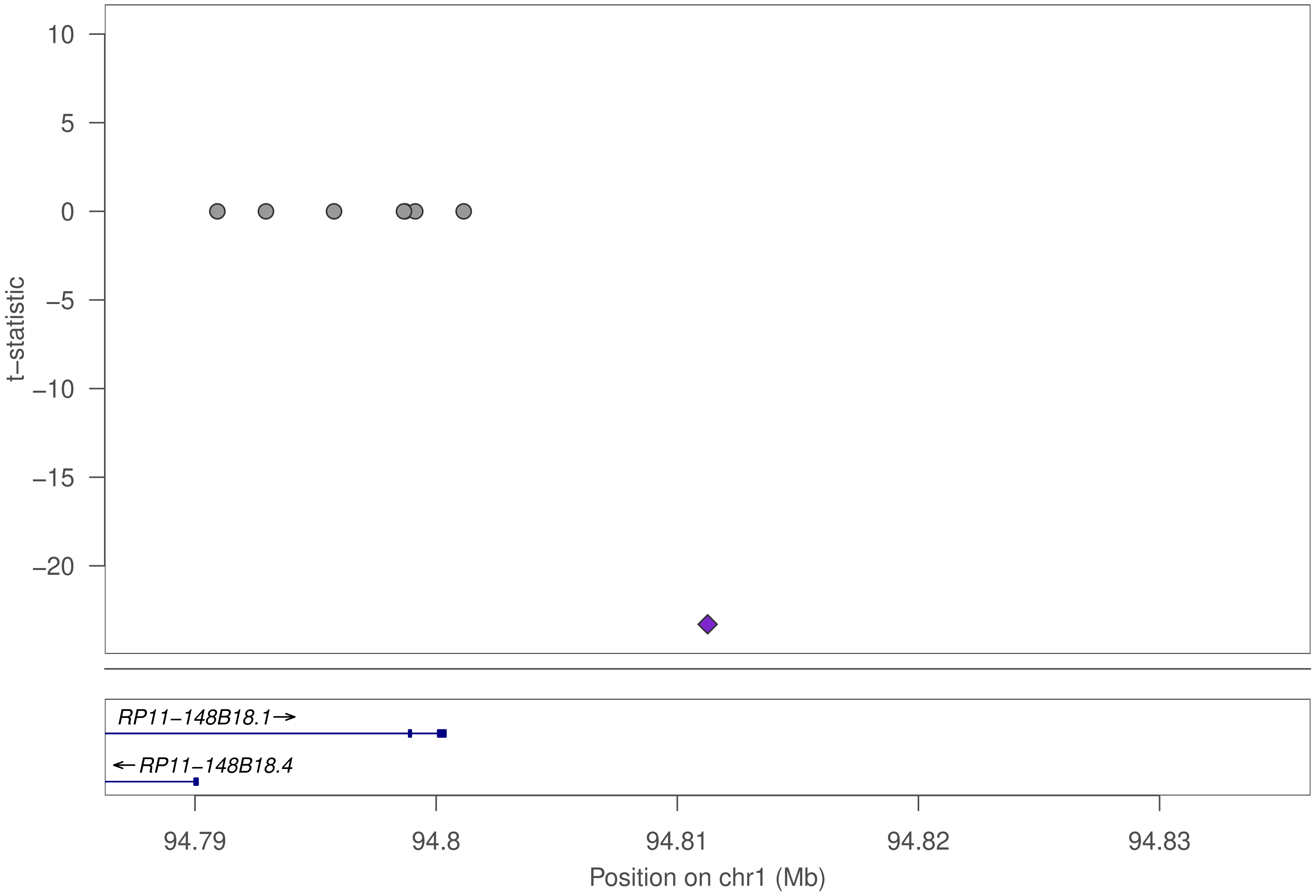

t−statistic
10
5
0
−5
−10
−15
−20
RP11−148B18.1→
←RP11−148B18.4
94.79
94.8
94.81
94.82
94.83
Position on chr1 (Mb)

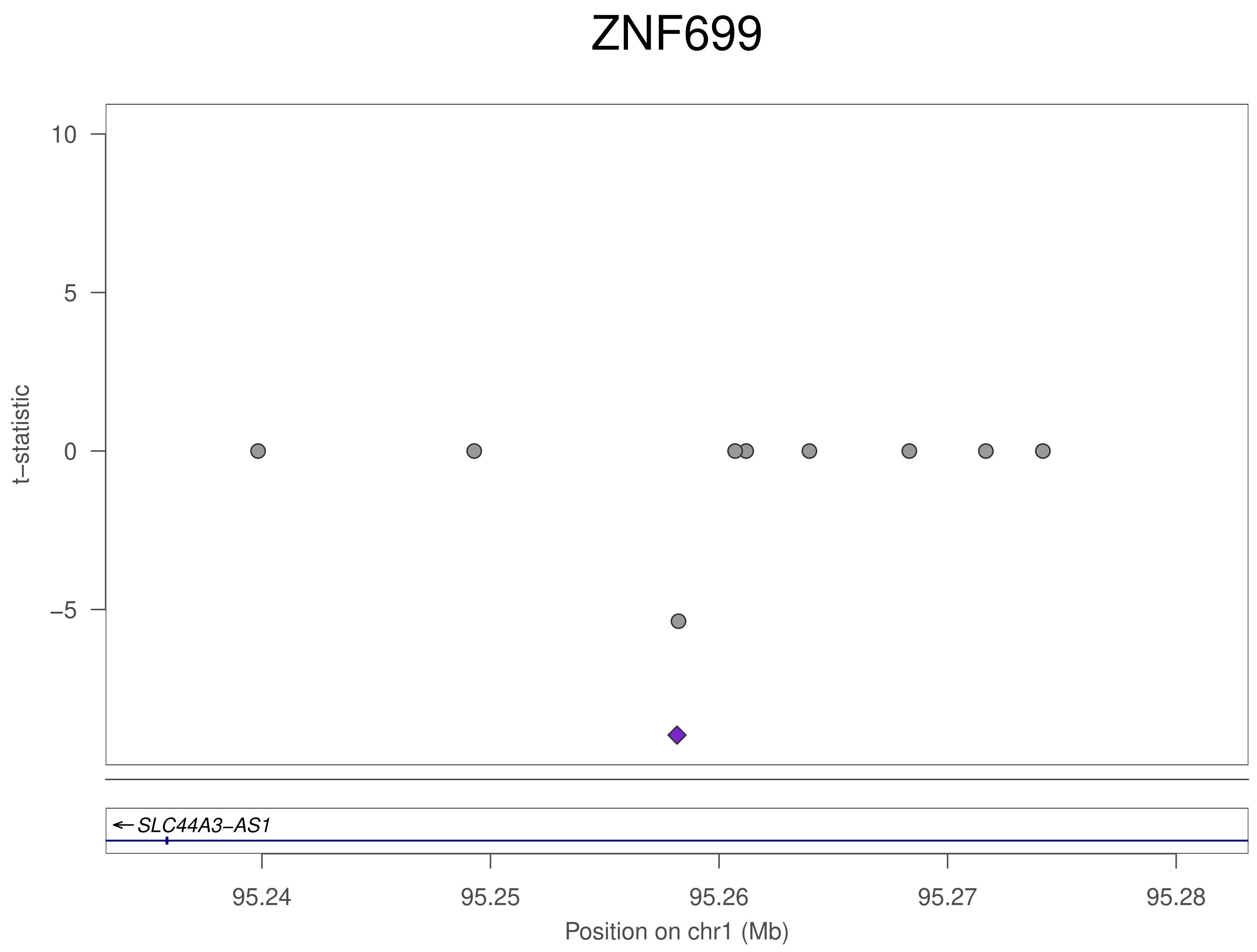
ZNF699
10
5
0
−5
t−statistic
SLC44A3−AS1
95.24
95.25
95.26
95.27
95.28
Position on chr1 (Mb)

### ZNF699

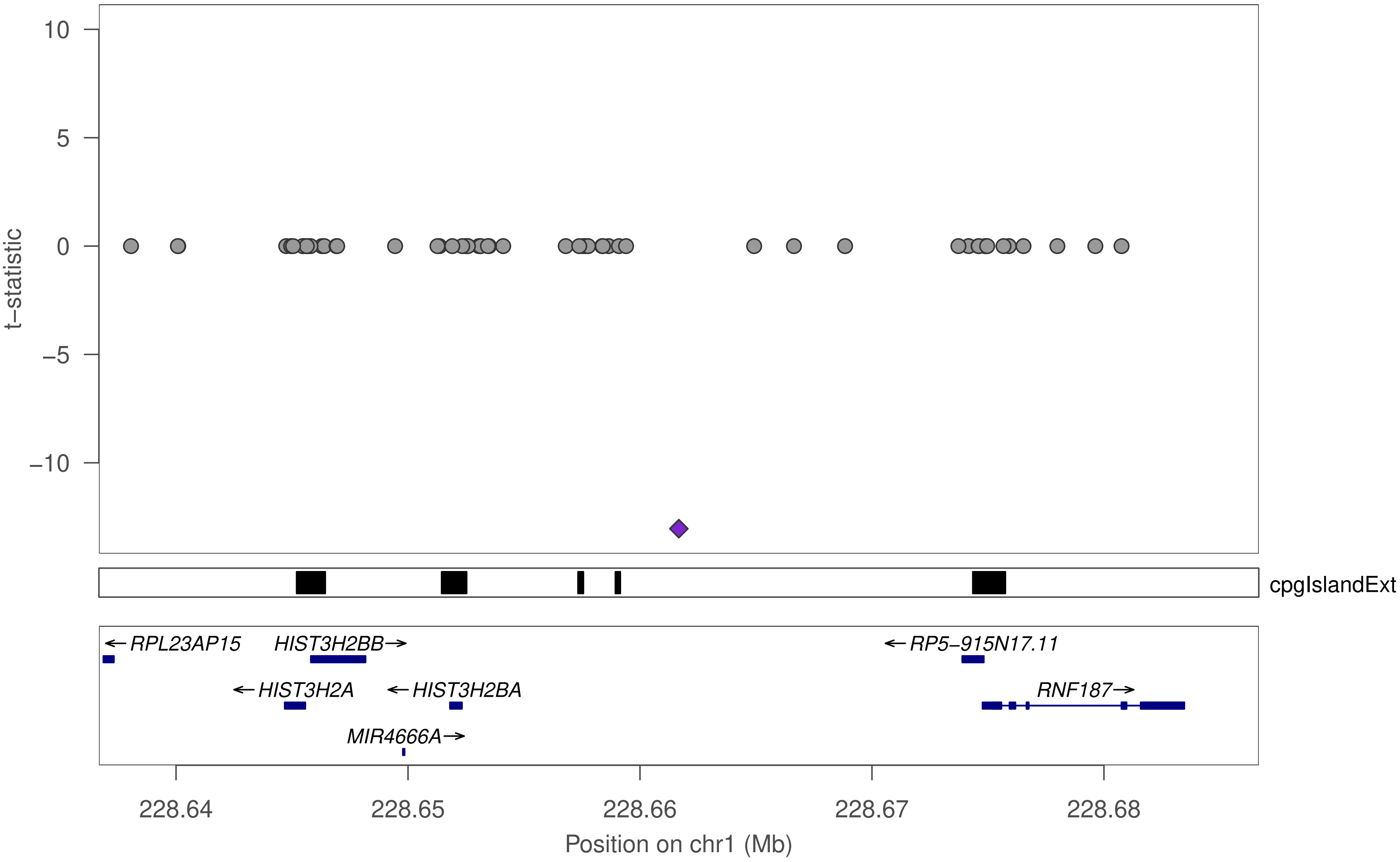

10
5
0
−5
−10
t−statistic
cpgIslandExt
←RPL23AP15
HIST3H2BB→
←RP5−915N17.11
←HIST3H2A
←HIST3H2BA
RNF187→
MIR4666A→
228.64
228.65
228.66
228.67
228.68
Position on chr1 (Mb)

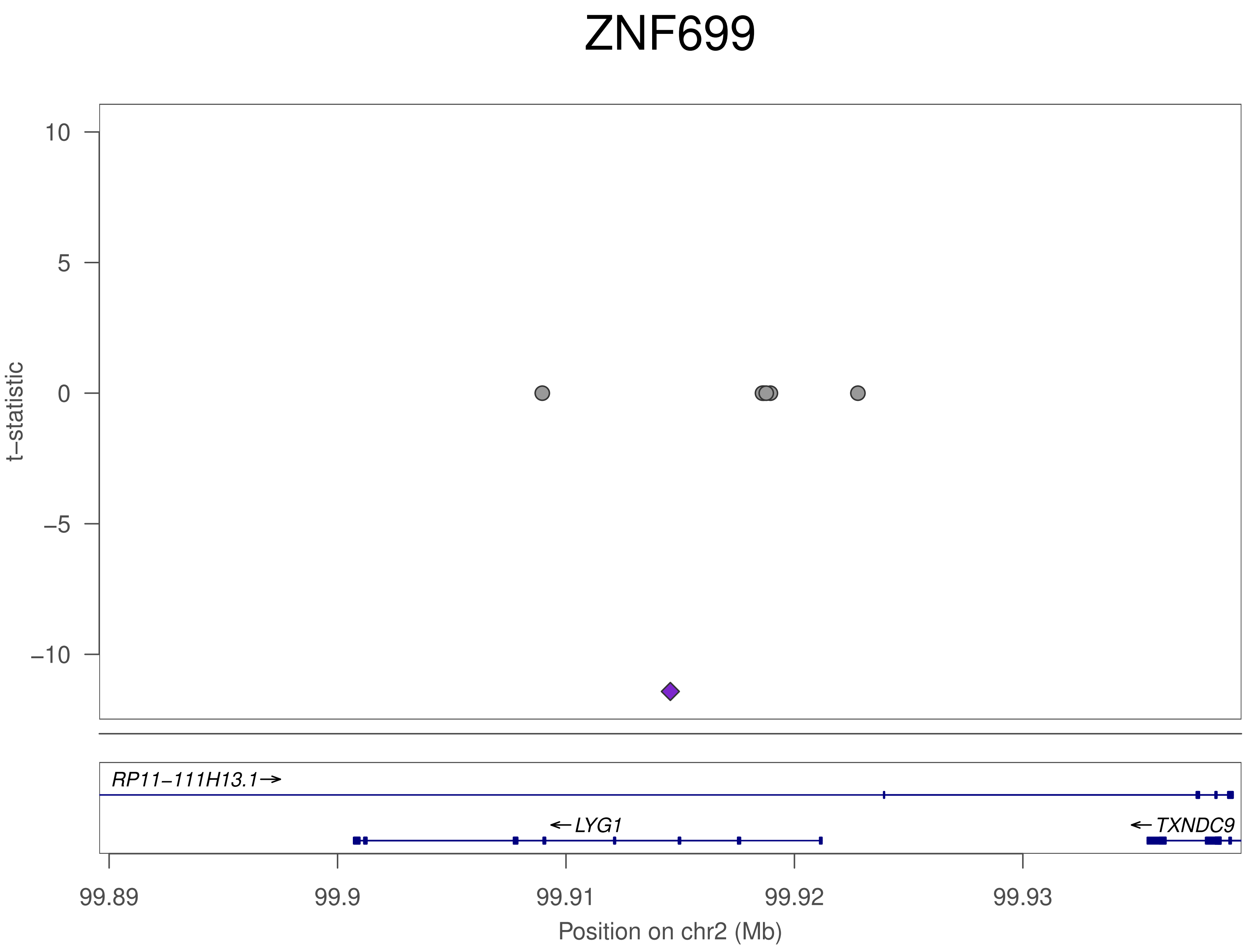

ZNF699
10
5
0
−5
−10
t−statistic
RP11−111H13.1→
←LYG1
←TXNDC9
99.89
99.9
99.91
99.92
99.93
Position on chr2 (Mb)

### ZNF699

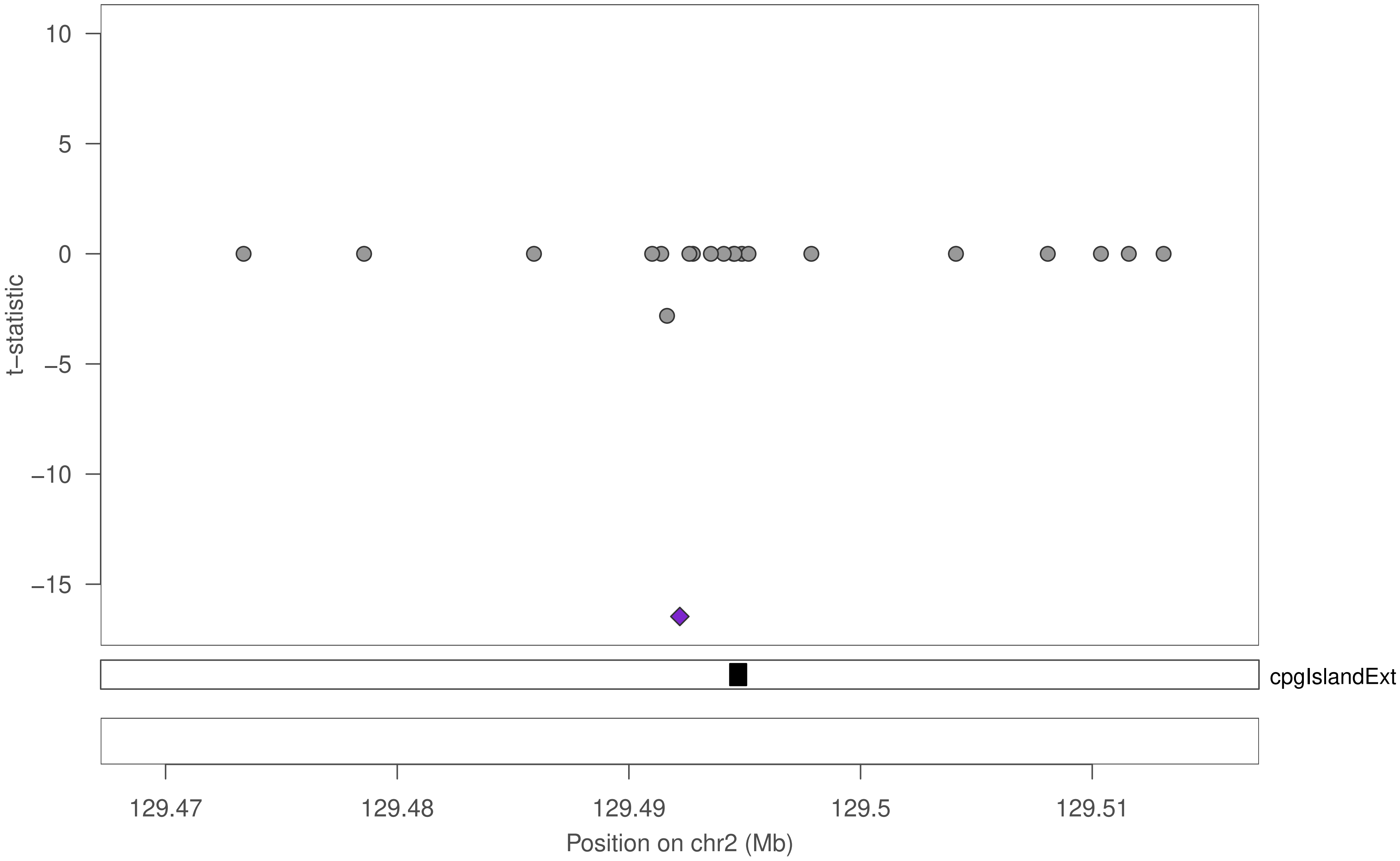

10
5
0
−5
−10
−15
t−statistic
cpgIslandExt
129.47
129.48
129.49
129.5
129.51
Position on chr2 (Mb)

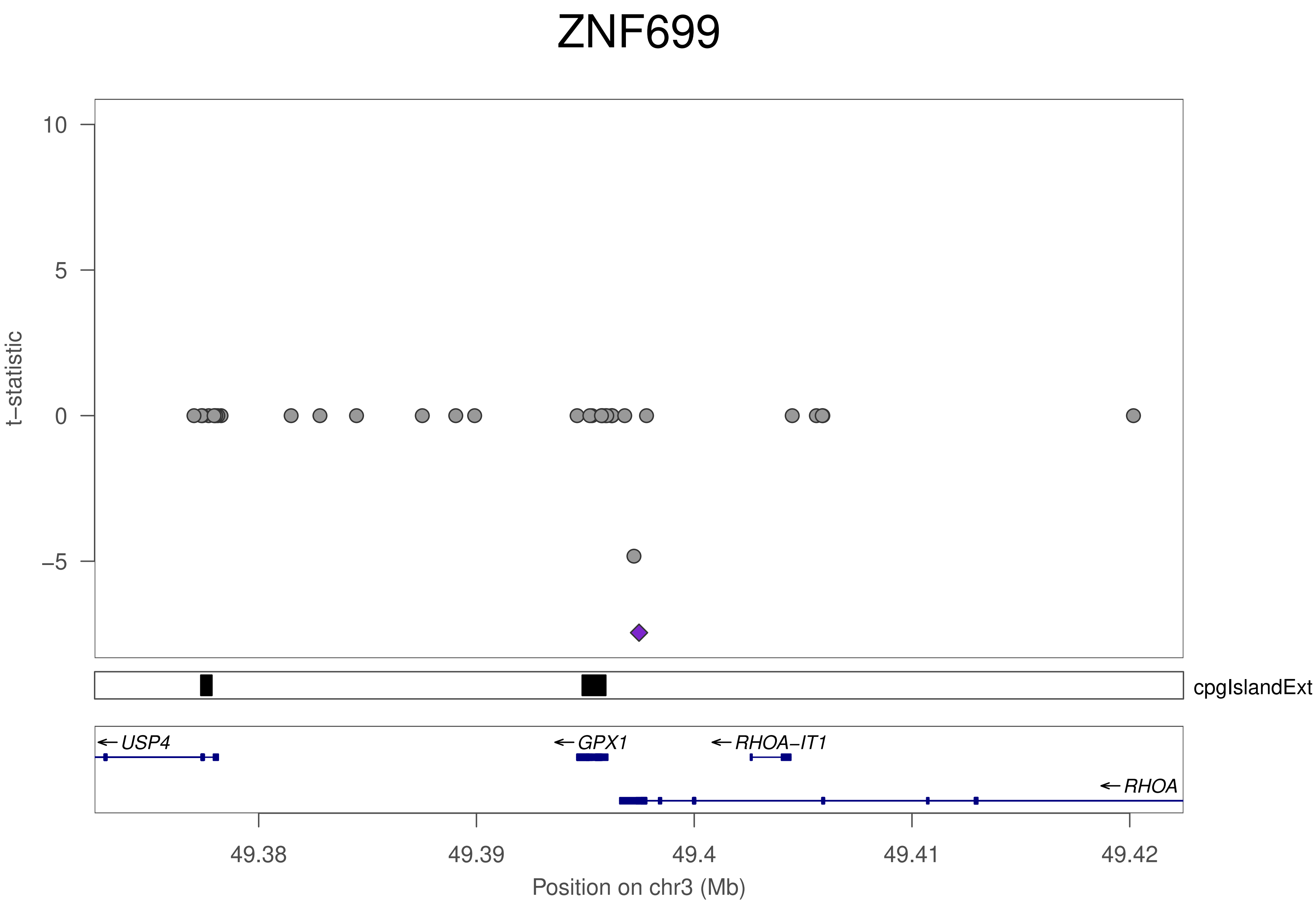
ZNF699
t–statistic
10
5
0
–5
cpgIslandExt
USP4
GPX1
RHOA–IT1
RHOA
49.38
49.39
49.4
49.41
49.42
Position on chr3 (Mb)

### ZNF699

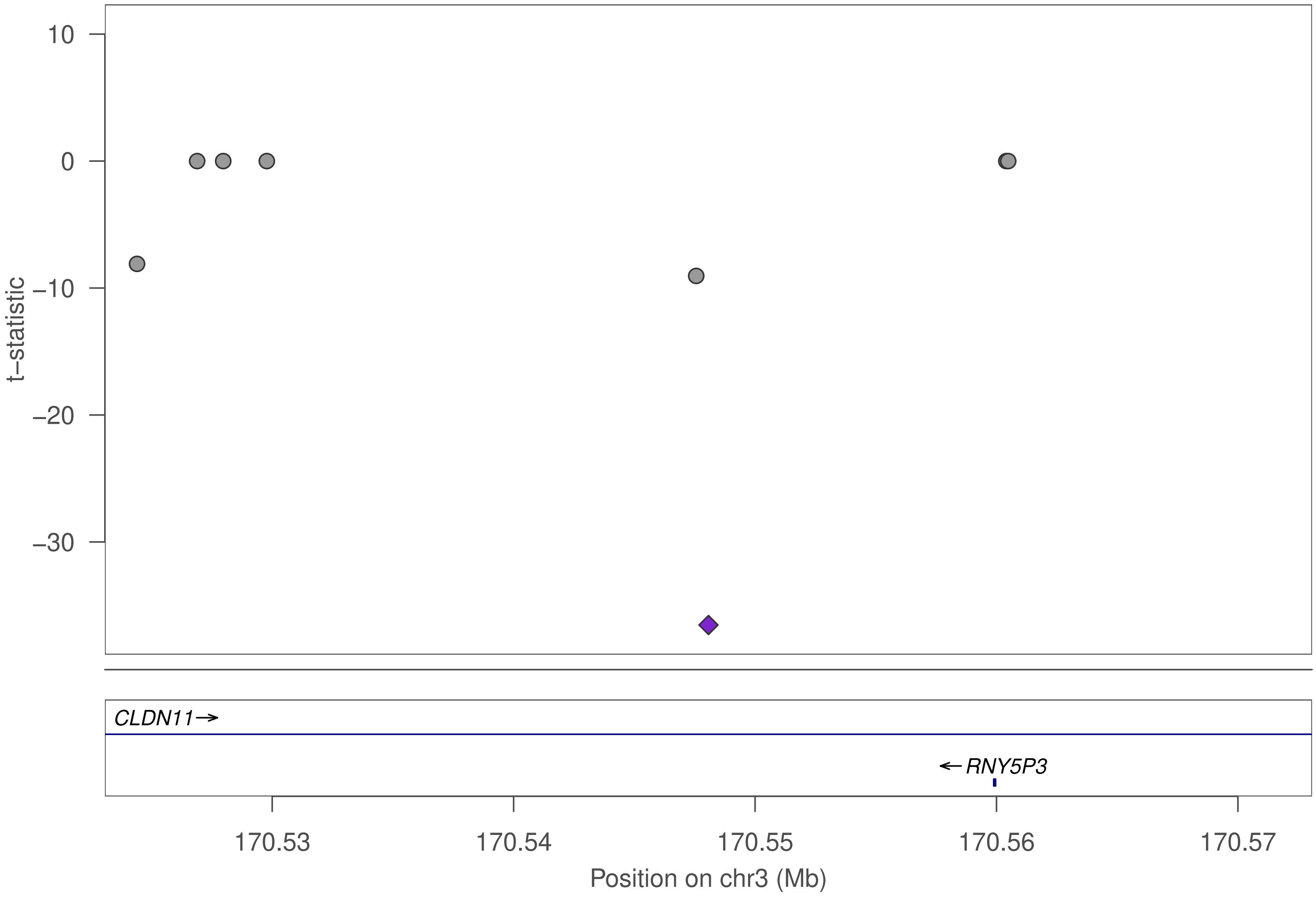

10
0
−10
−20
−30
t−statistic
CLDN11→
←RNY5P3
170.53
170.54
170.55
170.56
170.57
Position on chr3 (Mb)

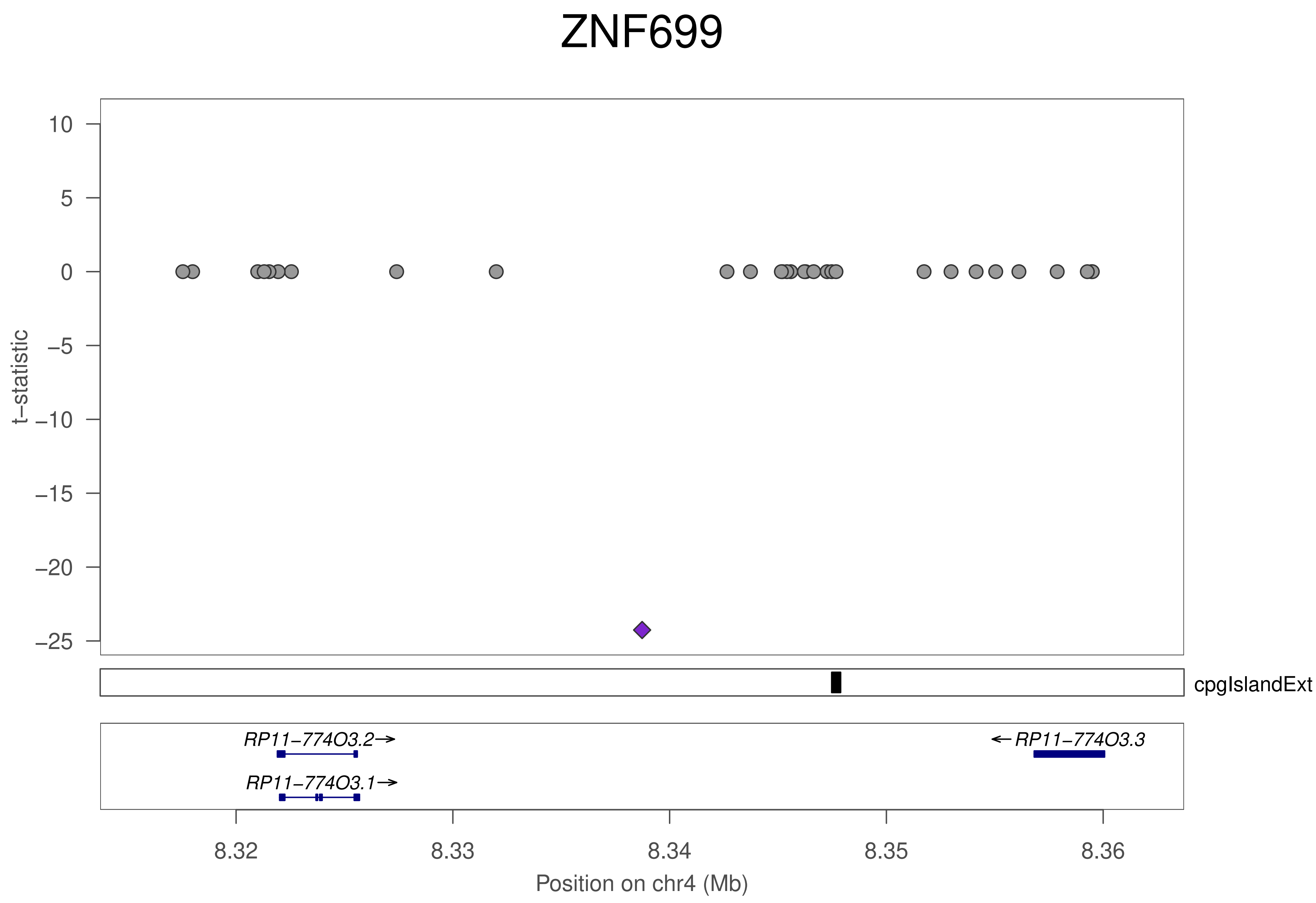
ZNF699
t−statistic
10
5
0
−5
−10
−15
−20
−25
cpgIslandExt
RP11−774O3.2→
RP11−774O3.1→
←RP11−774O3.3
8.32
8.33
8.34
8.35
8.36
Position on chr4 (Mb)

### ZNF699

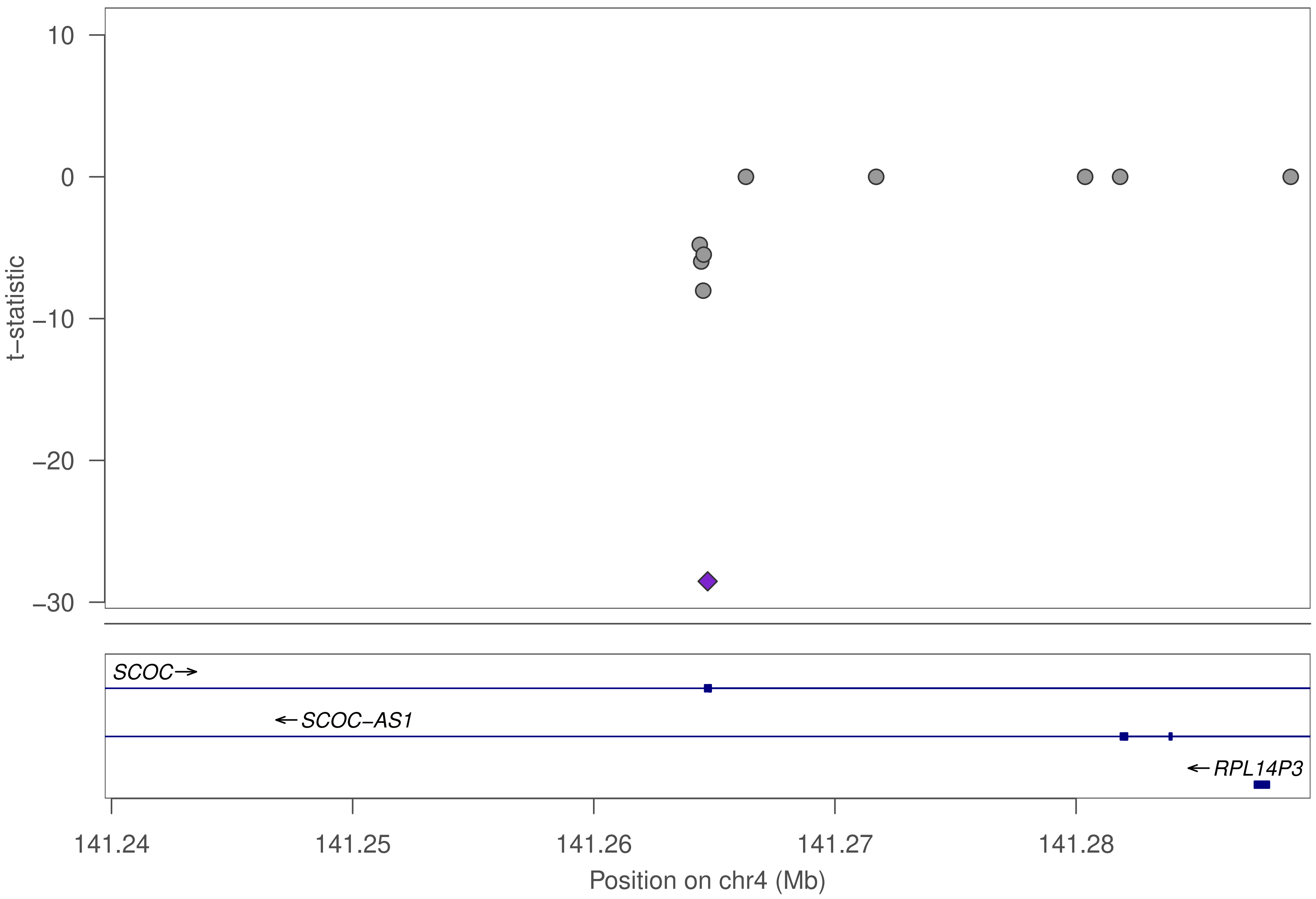

10
0
−10
−20
−30
t−statistic
SCOC→
←SCOC−AS1
←RPL14P3
141.24
141.25
141.26
141.27
141.28
Position on chr4 (Mb)

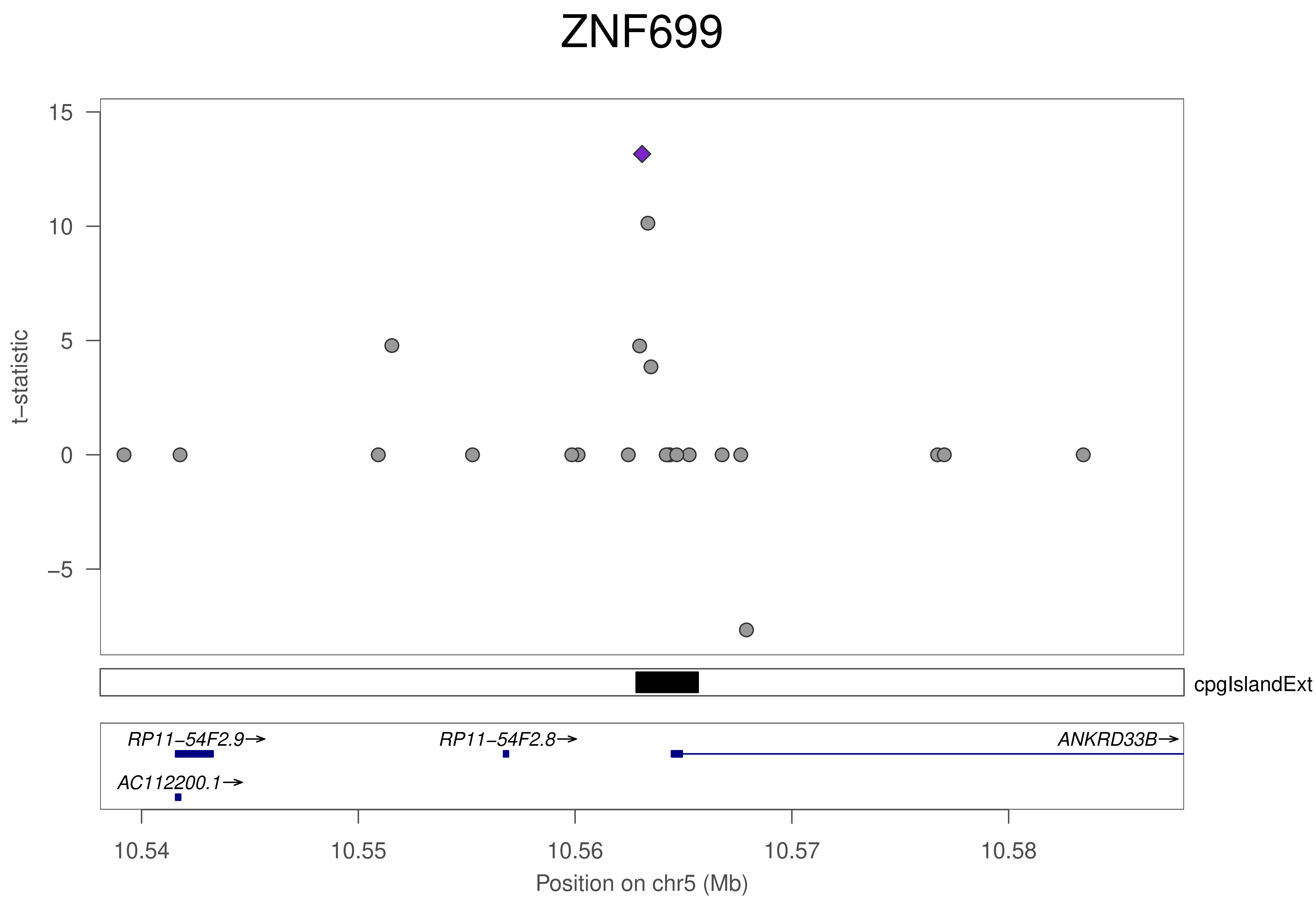

ZNF699
t−statistic
15
10
5
0
−5
cpgIslandExt
RP11−54F2.9→
RP11−54F2.8→
ANKRD33B→
AC112200.1→
10.54
10.55
10.56
10.57
10.58
Position on chr5 (Mb)

### ZNF699

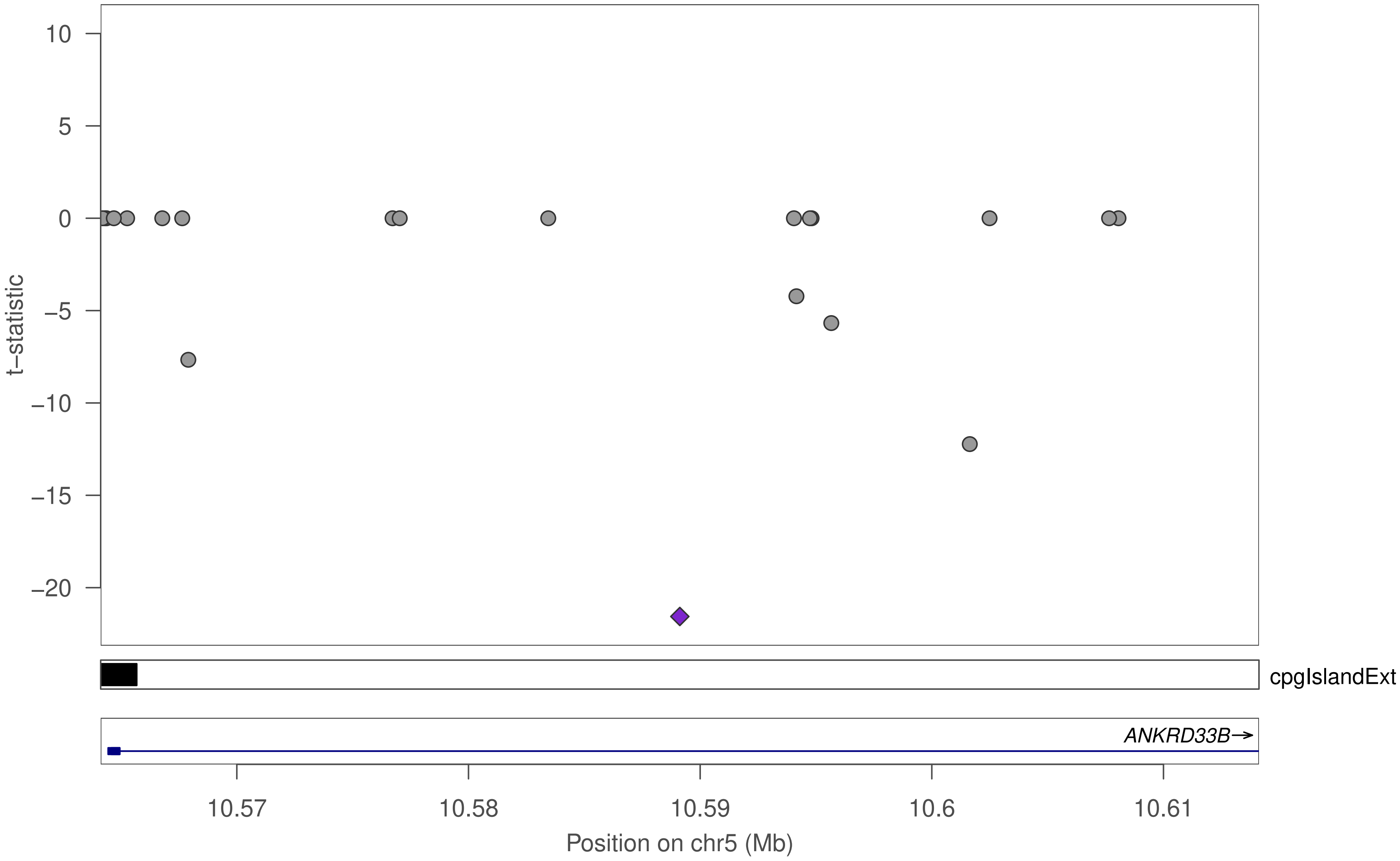

10
5
0
−5
−10
−15
−20
t−statistic
cpgIslandExt
ANKRD33B→
10.57
10.58
10.59
10.6
10.61
Position on chr5 (Mb)

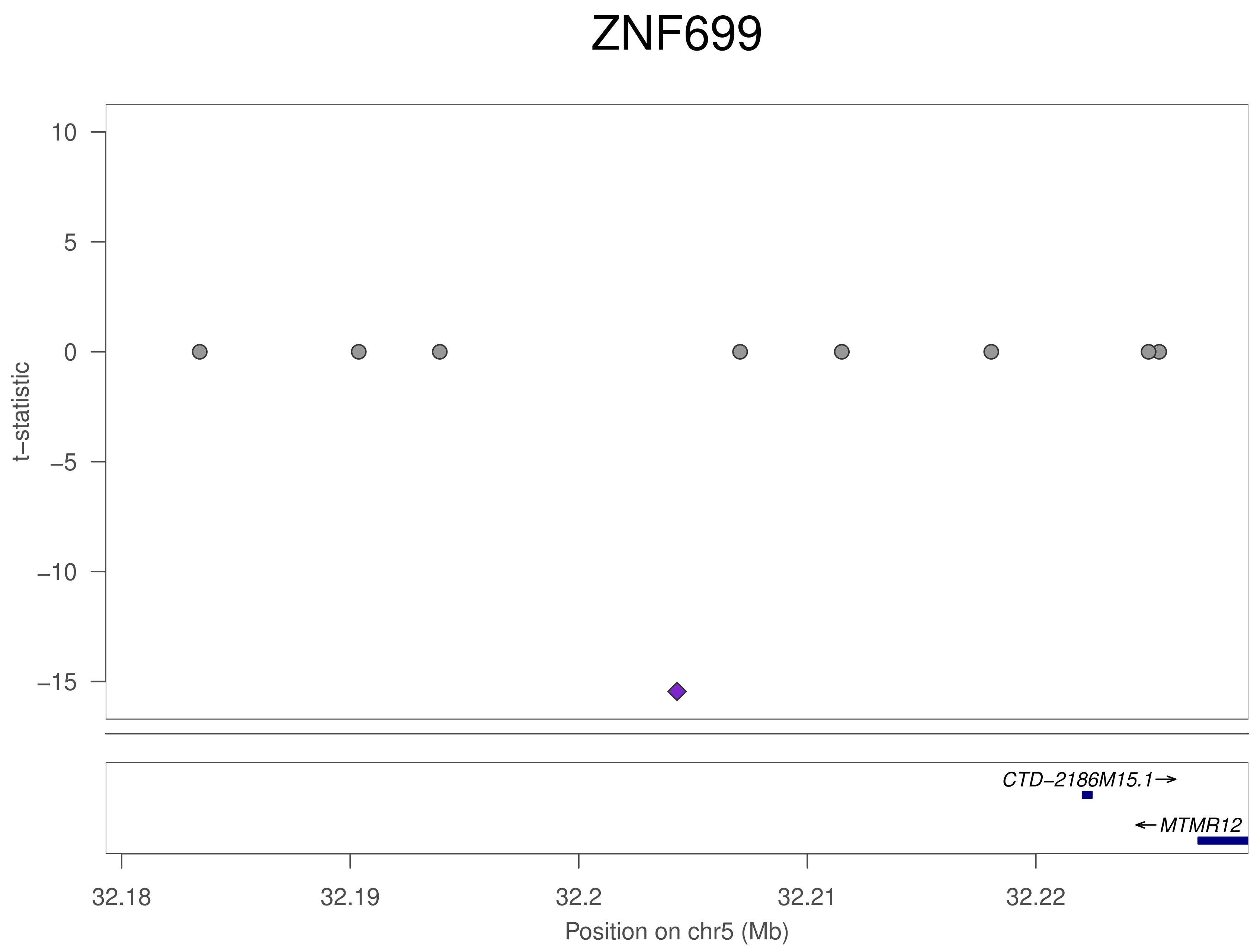
ZNF699
10
5
0
−5
−10
−15
t−statistic
CTD−2186M15.1→
←MTMR12
32.18
32.19
32.2
32.21
32.22
Position on chr5 (Mb)

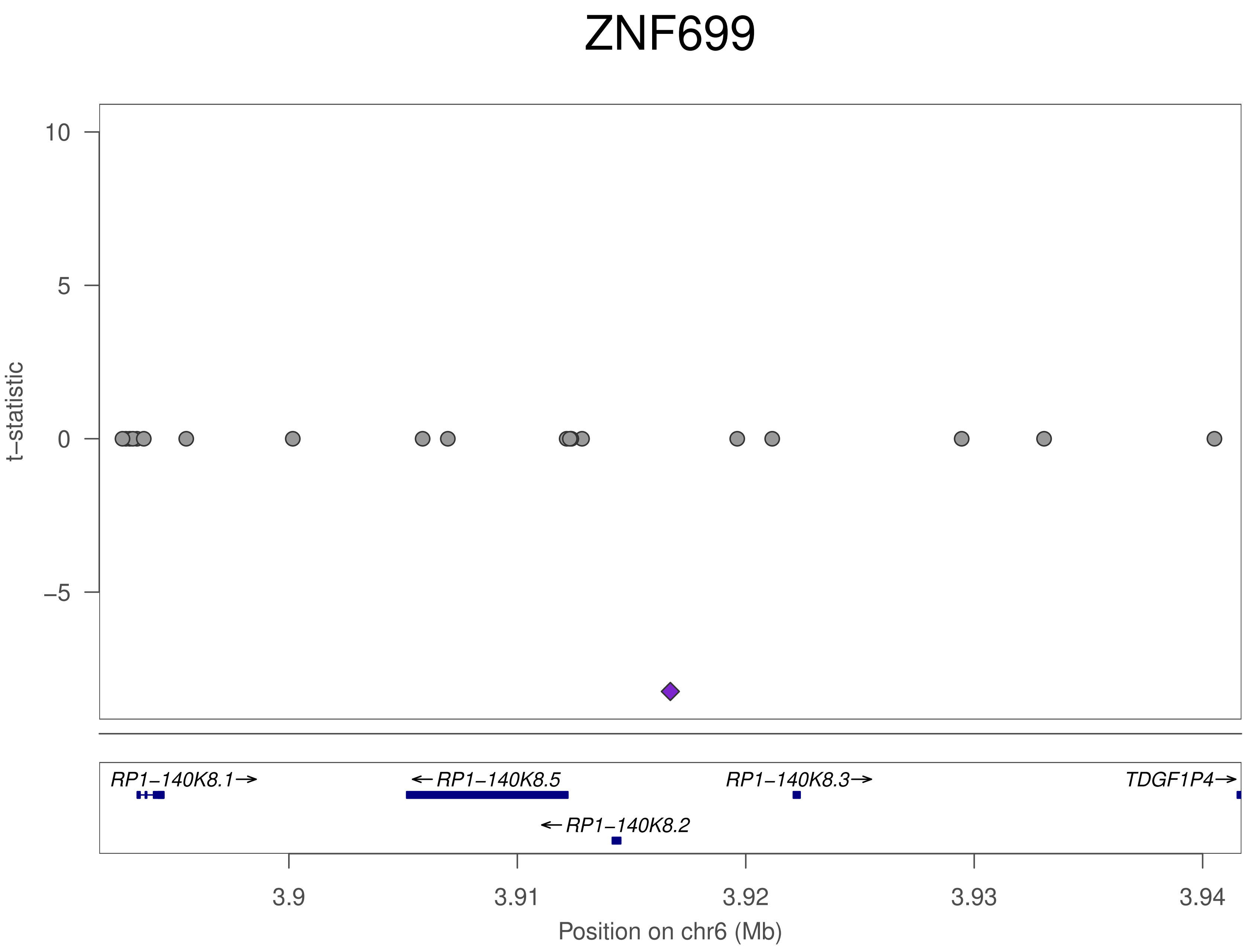
ZNF699
t–statistic
10
5
0
−5
RP1−140K8.1→
←RP1−140K8.5
RP1−140K8.3→
TDGF1P4→
←RP1−140K8.2
3.9
3.91
3.92
3.93
3.94
Position on chr6 (Mb)

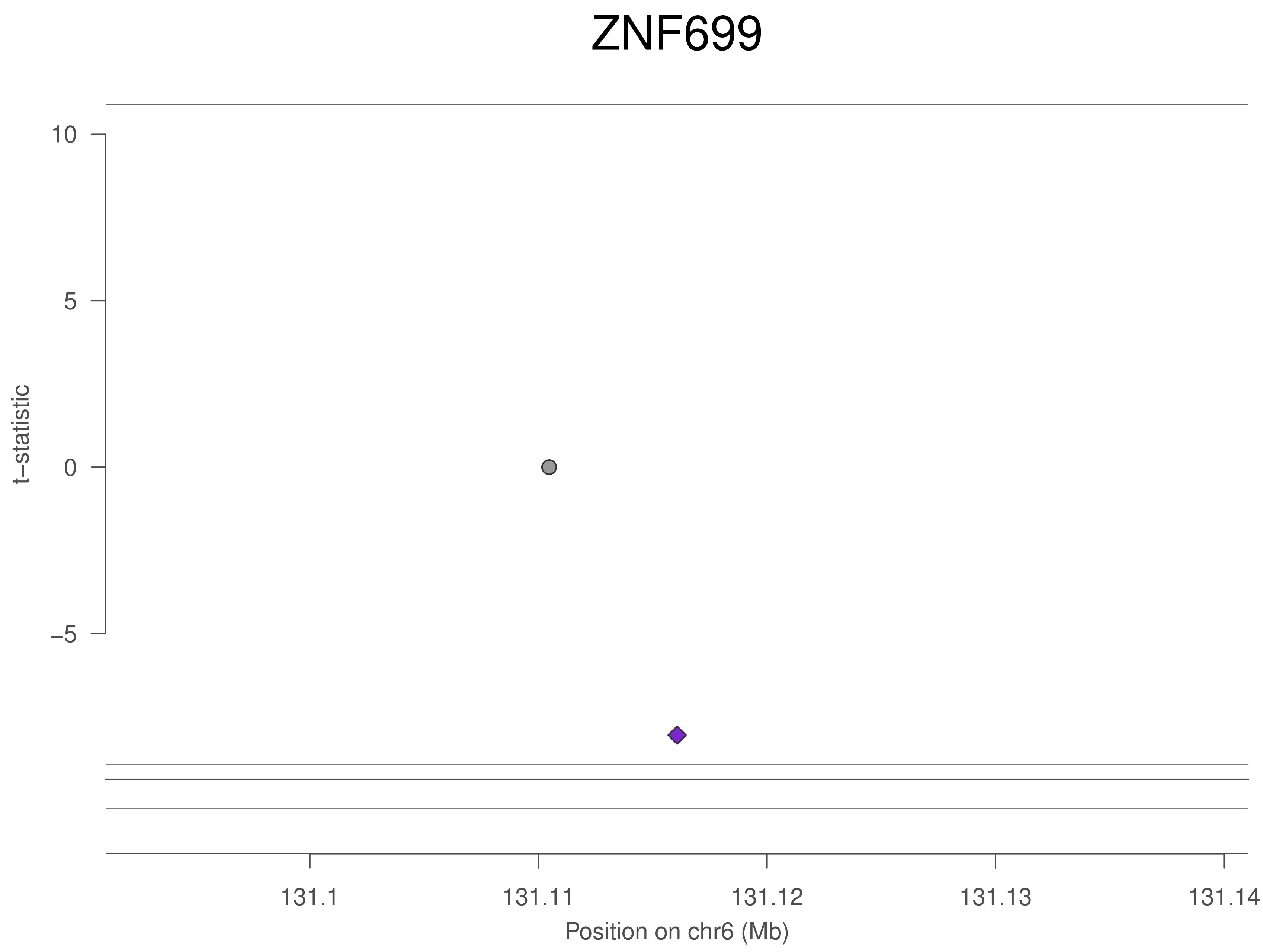
ZNF699
10
5
0
−5
t−statistic
131.1
131.11
131.12
131.13
131.14
Position on chr6 (Mb)

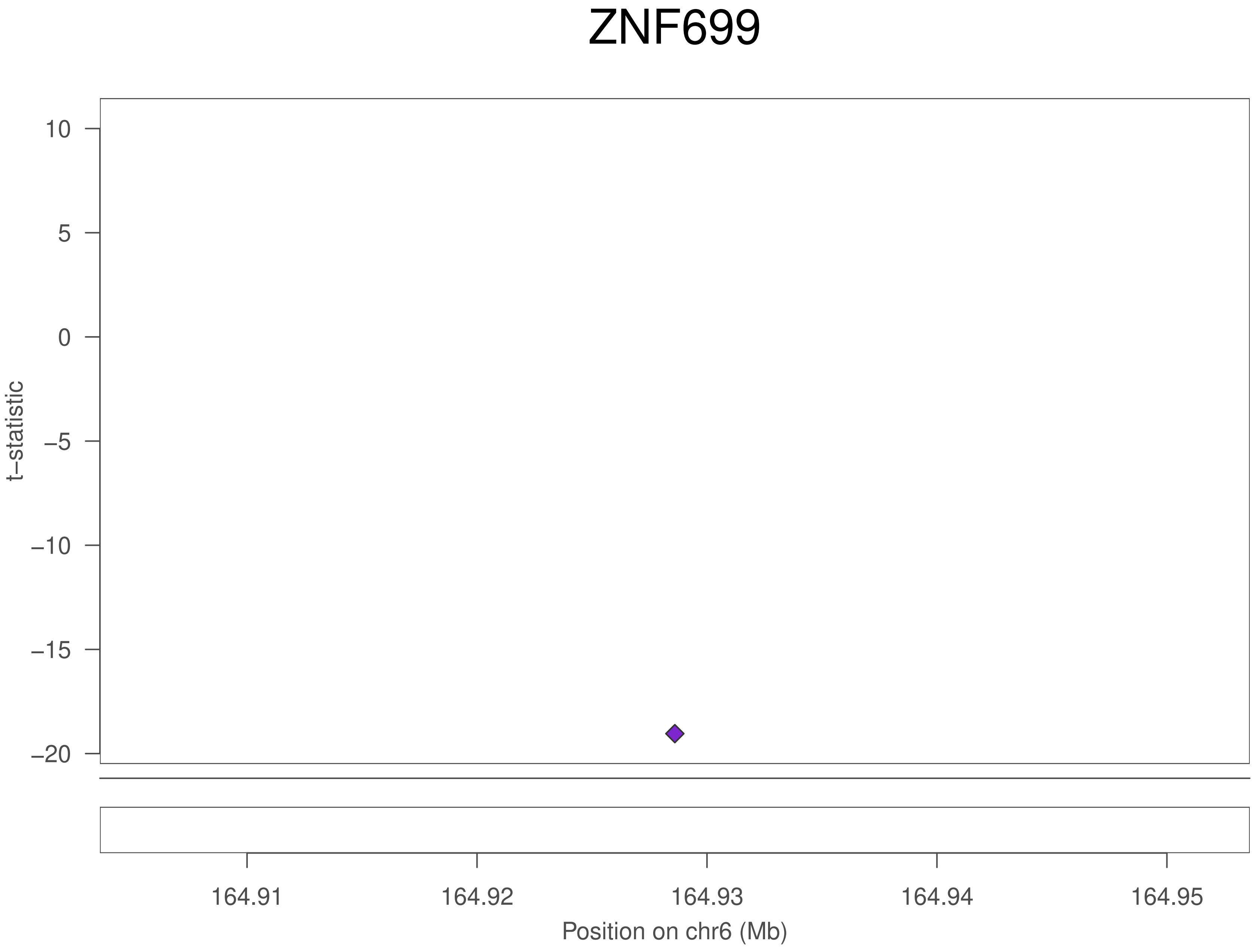
ZNF699
10
5
0
−5
−10
−15
−20
t−statistic
164.91
164.92
164.93
164.94
164.95
Position on chr6 (Mb)

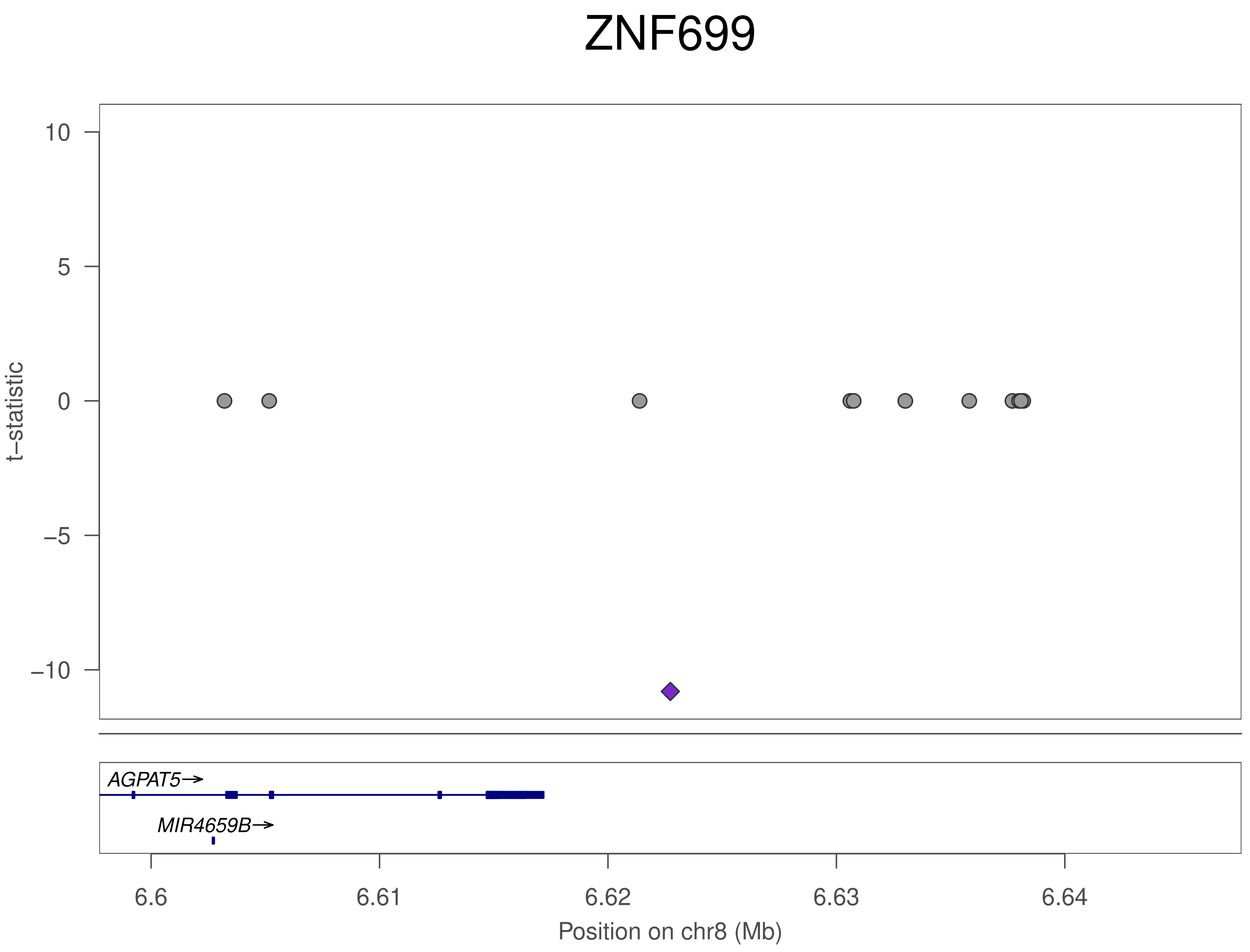
ZNF699
10
5
0
−5
−10
t−statistic
AGPAT5→
MIR4659B→
6.6
6.61
6.62
6.63
6.64
Position on chr8 (Mb)

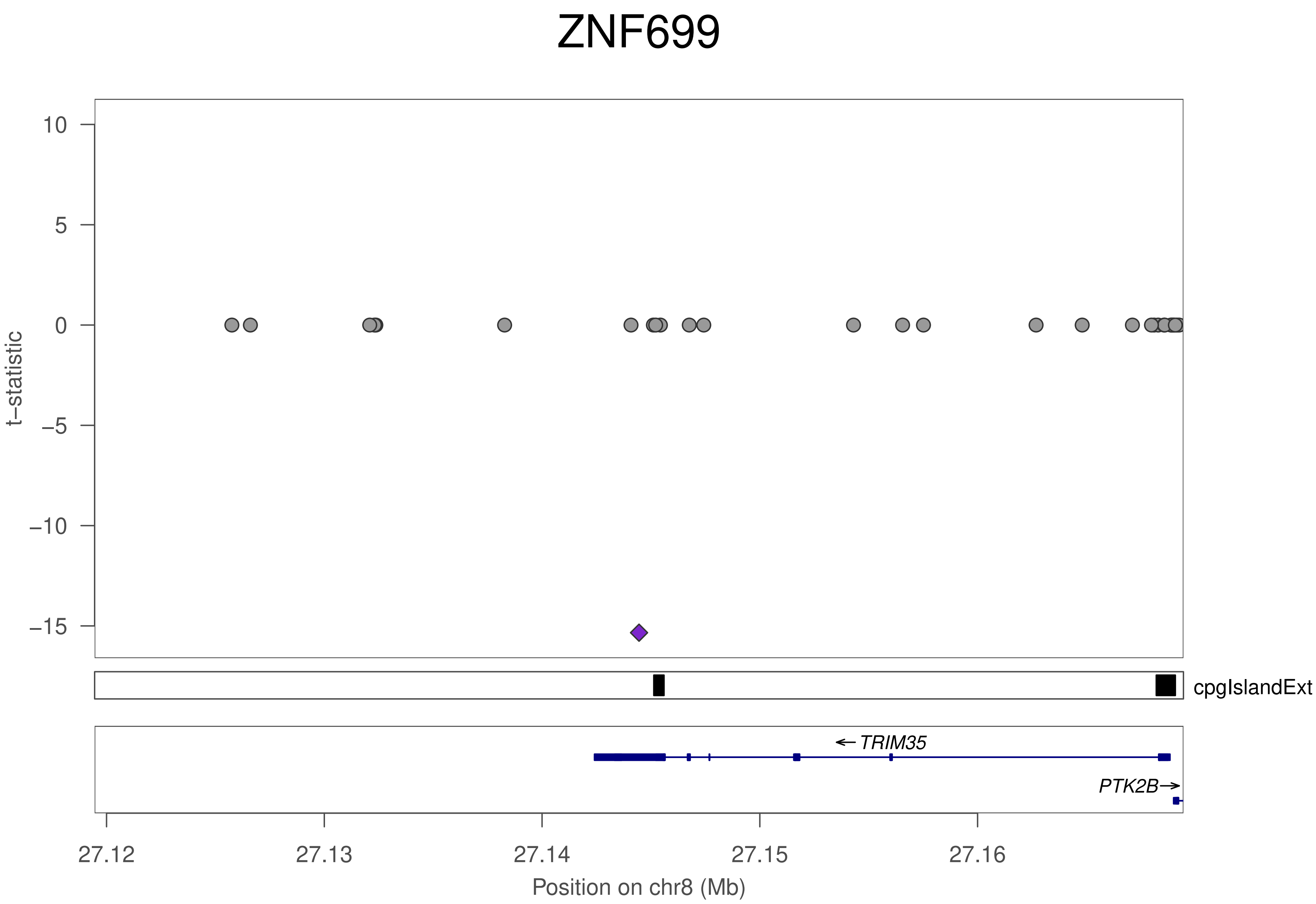

ZNF699
t−statistic
10
5
0
−5
−10
−15
cpgIslandExt
← TRIM35
PTK2B→
27.12
27.13
27.14
27.15
27.16
Position on chr8 (Mb)

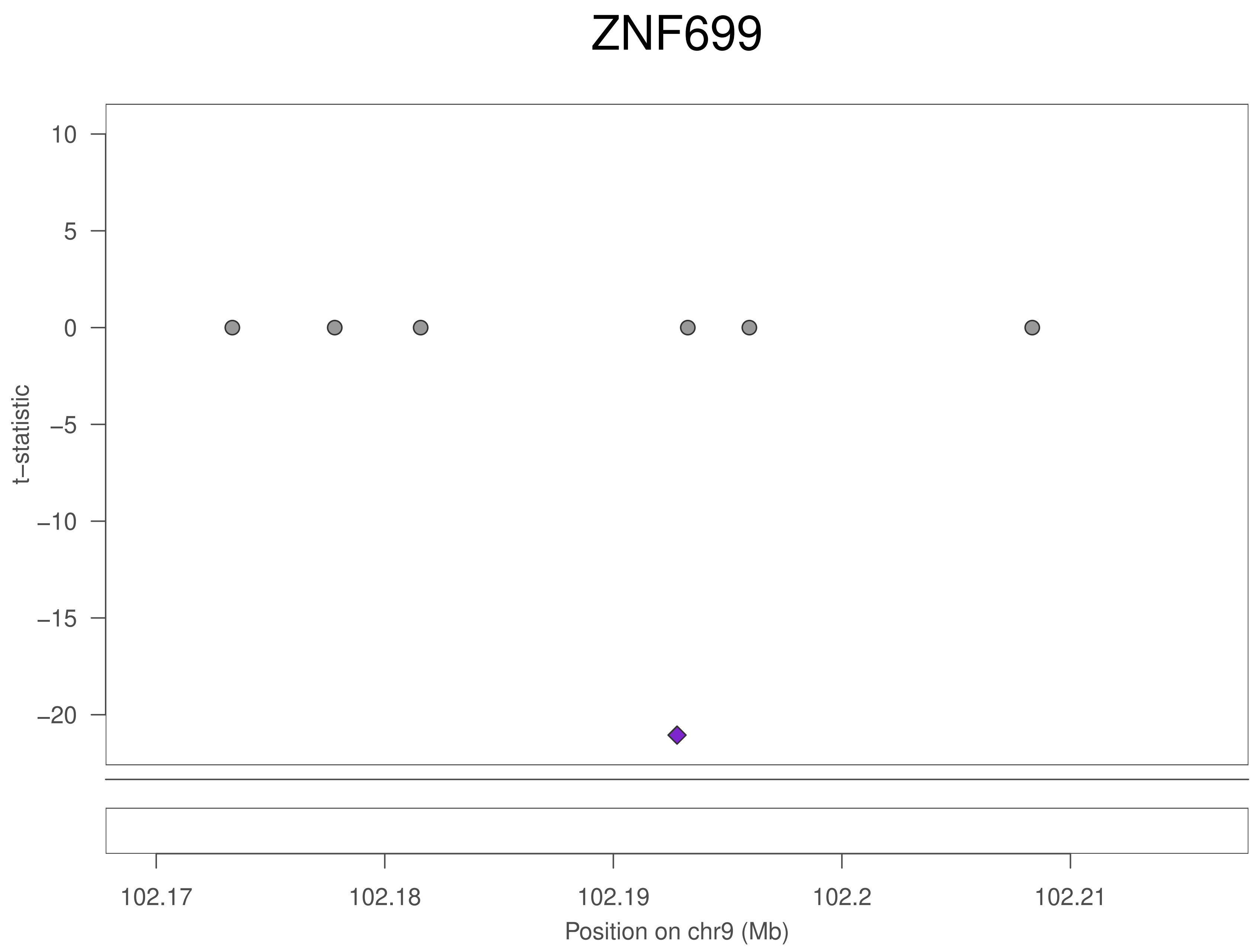
ZNF699
10
5
0
−5
−10
−15
−20
t−statistic
102.17
102.18
102.19
102.2
102.21
Position on chr9 (Mb)

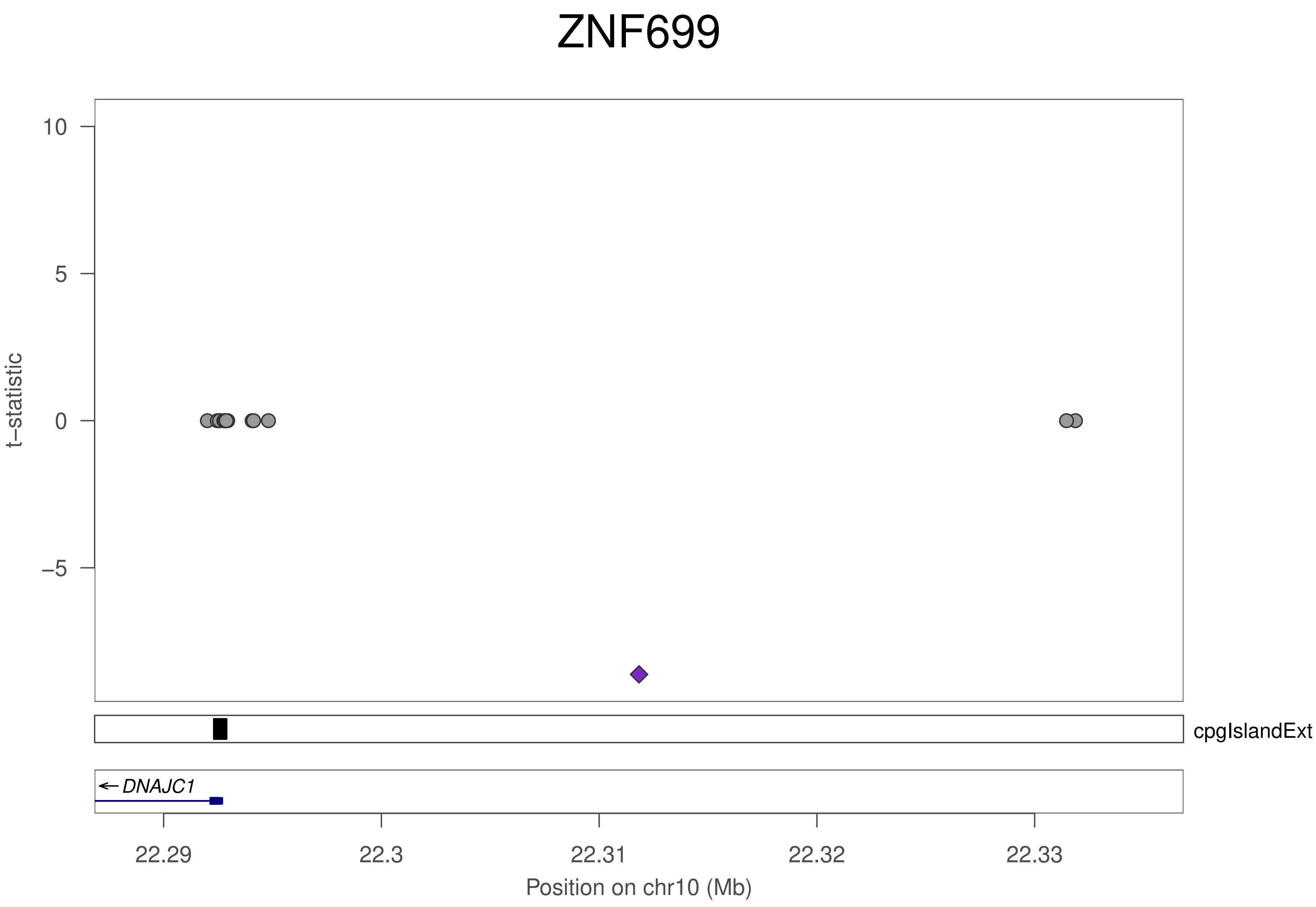

ZNF699
10
5
0
−5
t−statistic
cpgIslandExt
DNAJC1
22.29
22.3
22.31
22.32
22.33
Position on chr10 (Mb)

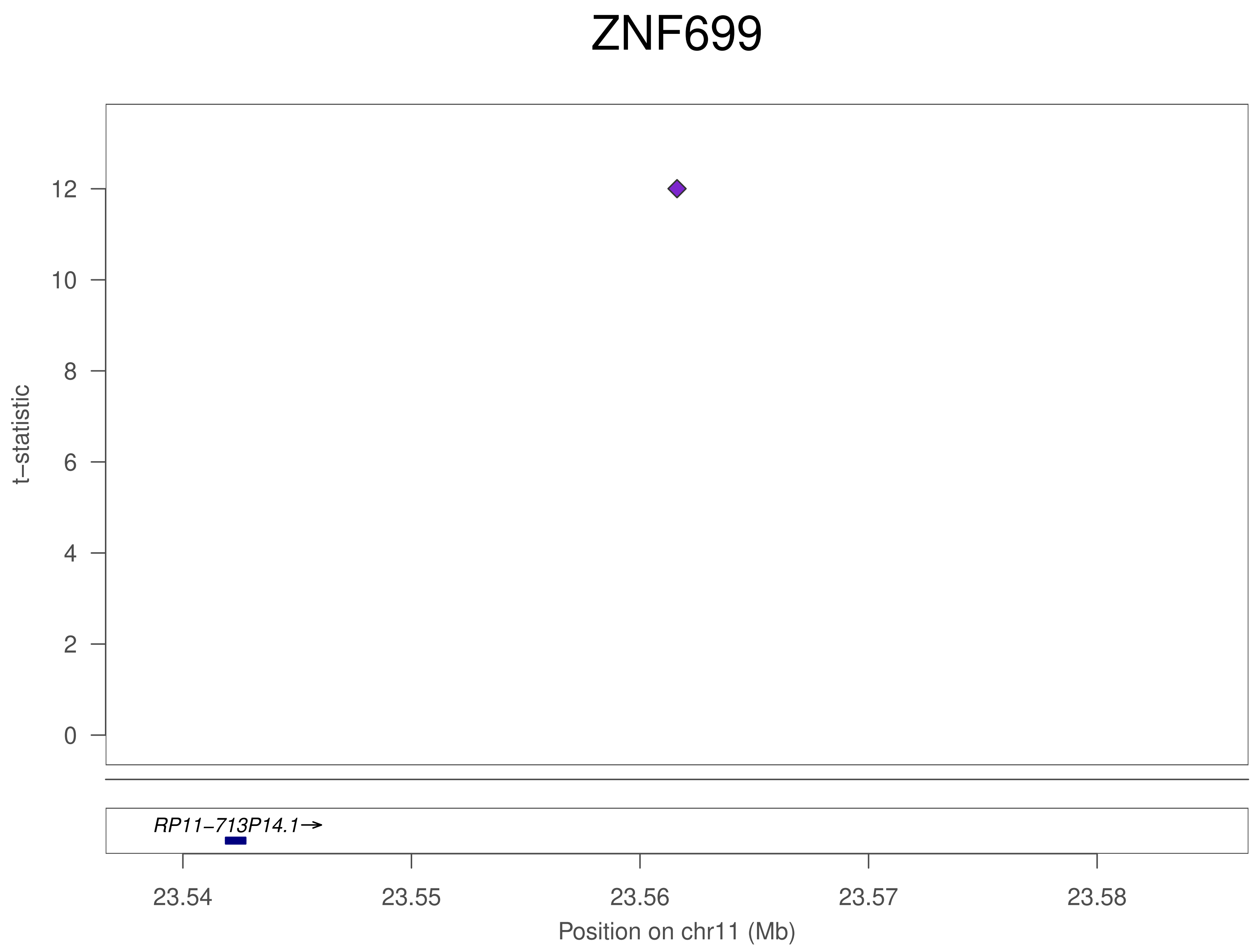
ZNF699
t–statistic
0
2
4
6
8
10
12
RP11–713P14.1→
23.54
23.55
23.56
23.57
23.58
Position on chr11 (Mb)

### ZNF699

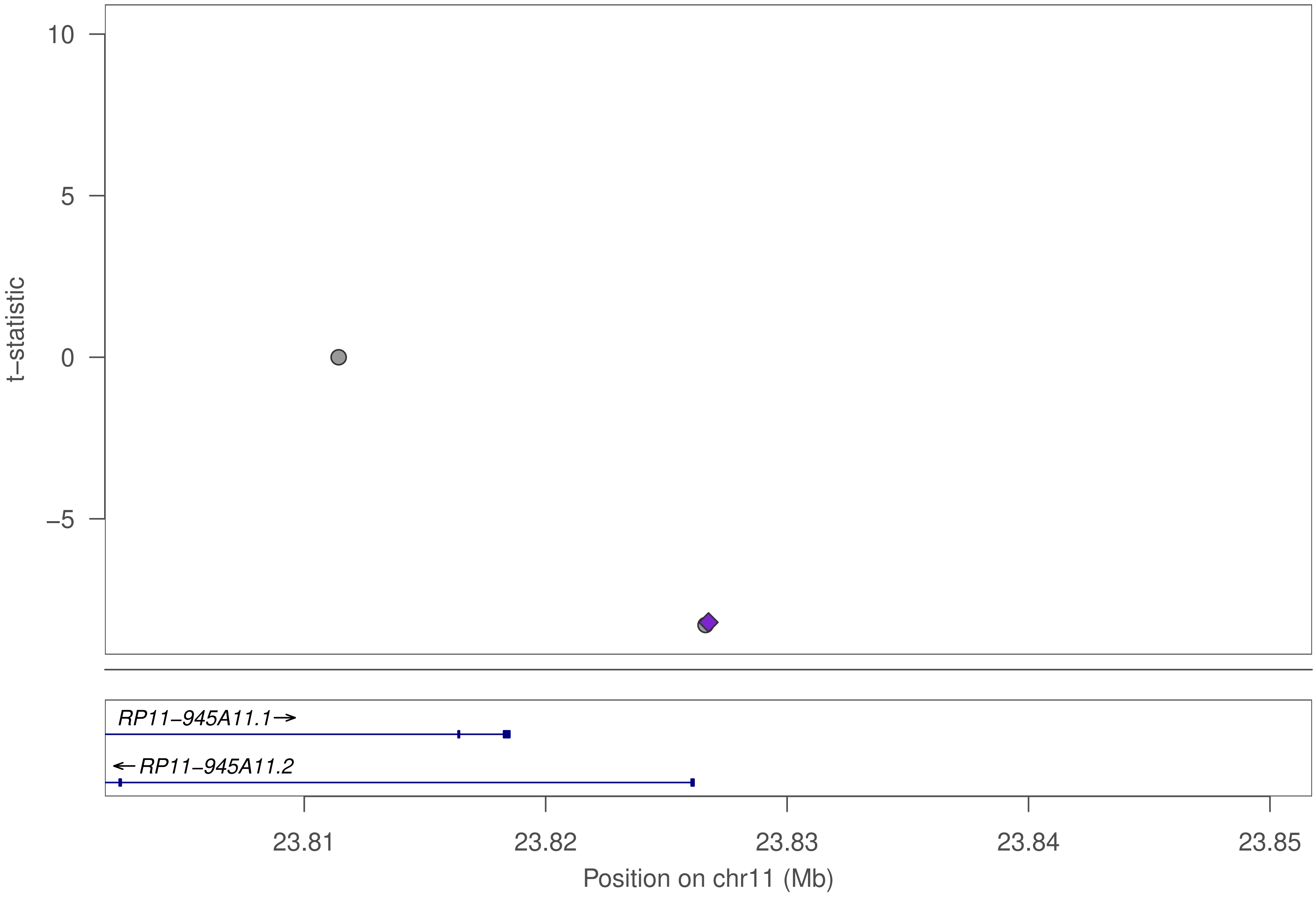

10
5
0
−5
t−statistic
RP11−945A11.1→
←RP11−945A11.2
23.81
23.82
23.83
23.84
23.85
Position on chr11 (Mb)

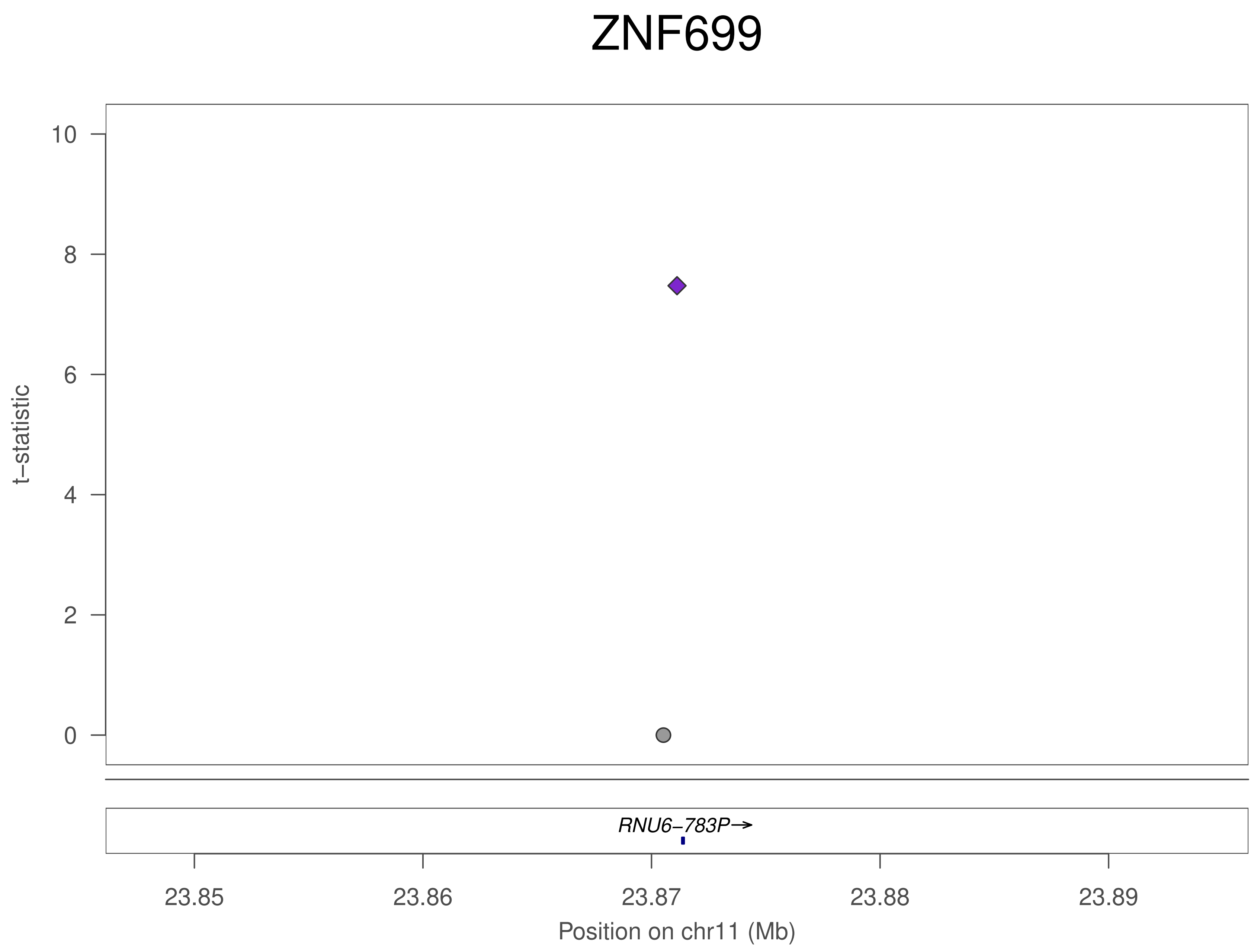

ZNF699
t−statistic
0
2
4
6
8
10
RNU6−783P→
23.85
23.86
23.87
23.88
23.89
Position on chr11 (Mb)

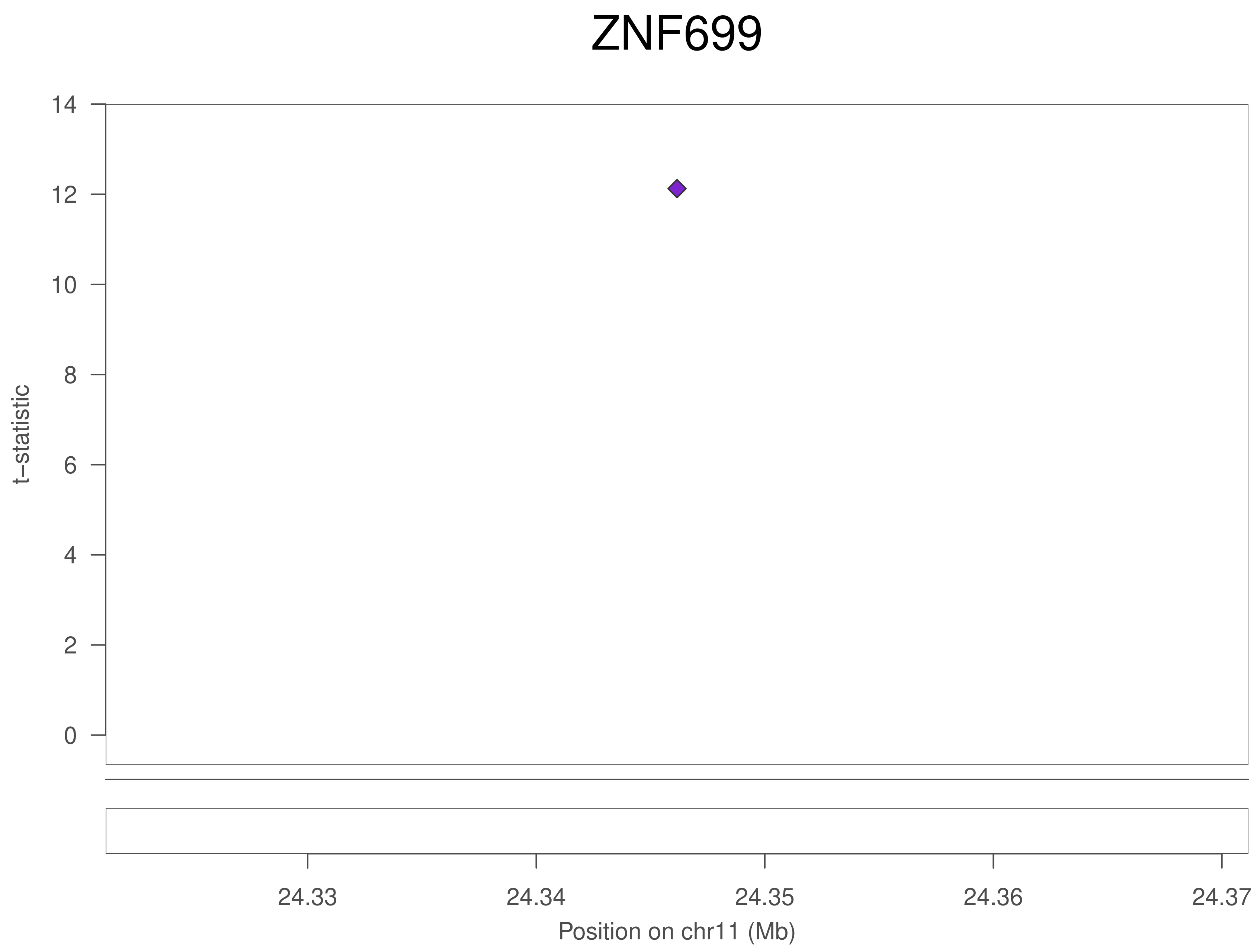

ZNF699
t–statistic
0
2
4
6
8
10
12
14
24.33
24.34
24.35
24.36
24.37
Position on chr11 (Mb)

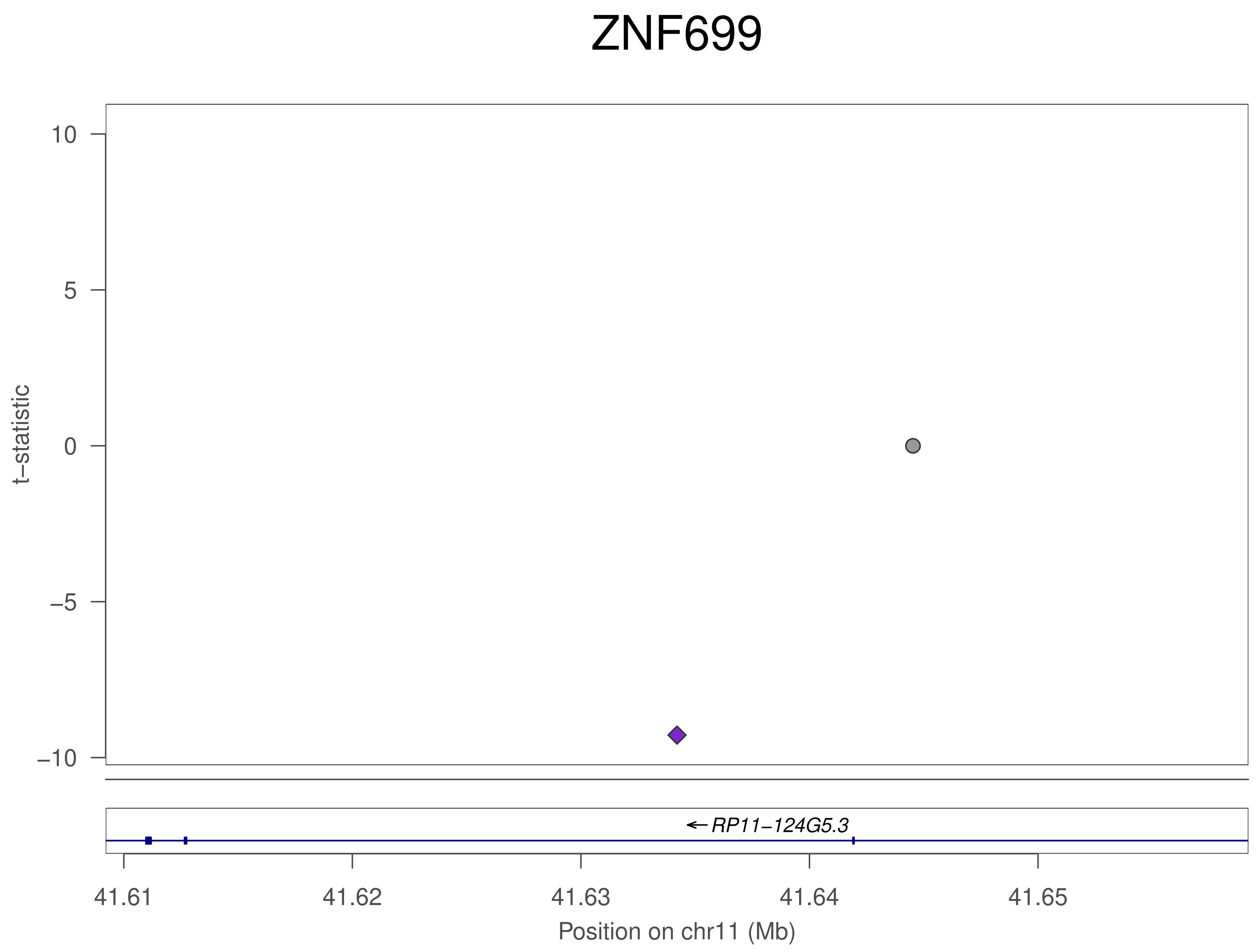
ZNF699
10
5
0
−5
−10
t−statistic
RP11−124G5.3
41.61
41.62
41.63
41.64
41.65
Position on chr11 (Mb)

### ZNF699

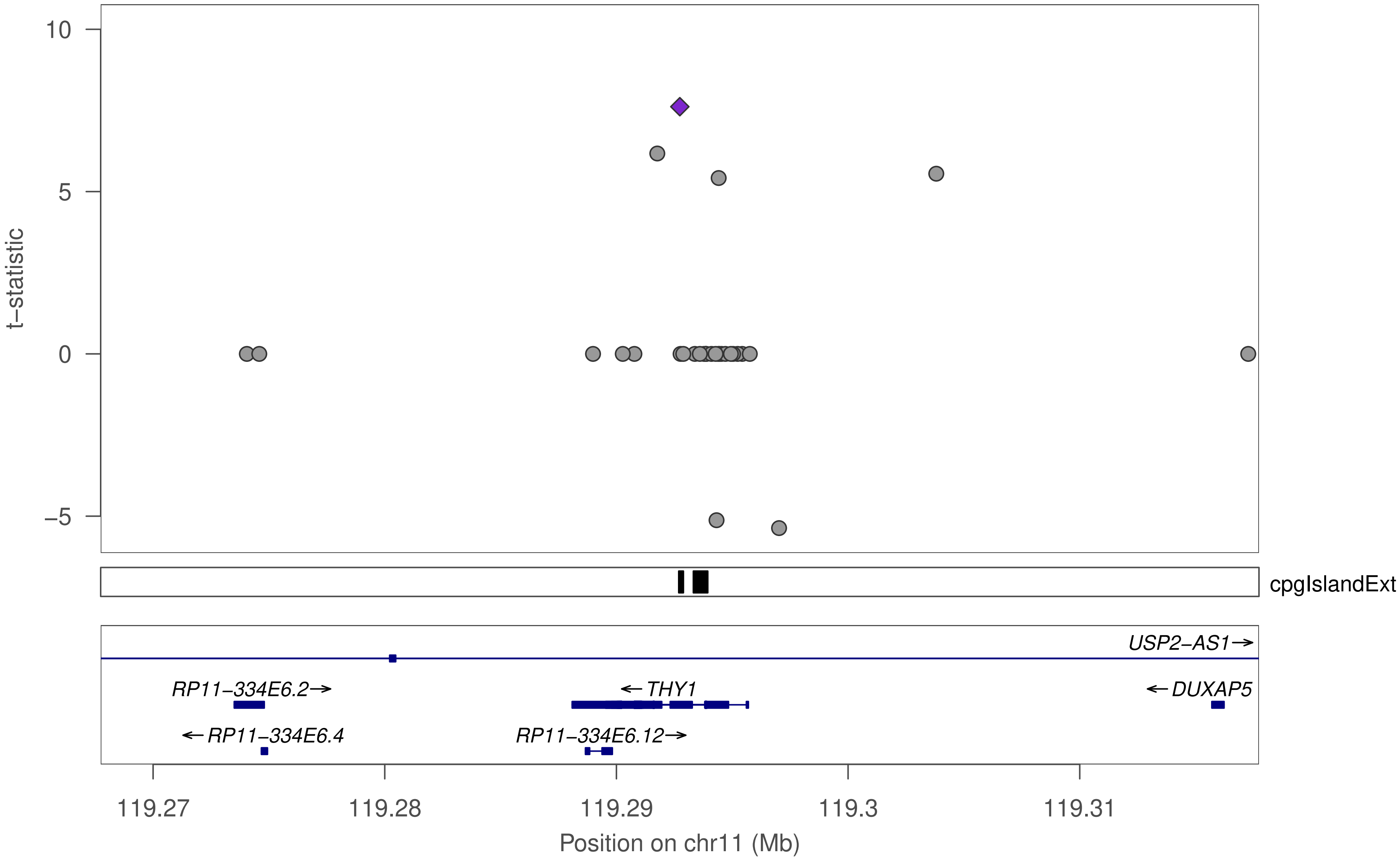

10
5
0
−5
t−statistic
cpgIslandExt
USP2−AS1→
RP11−334E6.2→
←THY1
←DUXAP5
←RP11−334E6.4
RP11−334E6.12→
119.27
119.28
119.29
119.3
119.31
Position on chr11 (Mb)

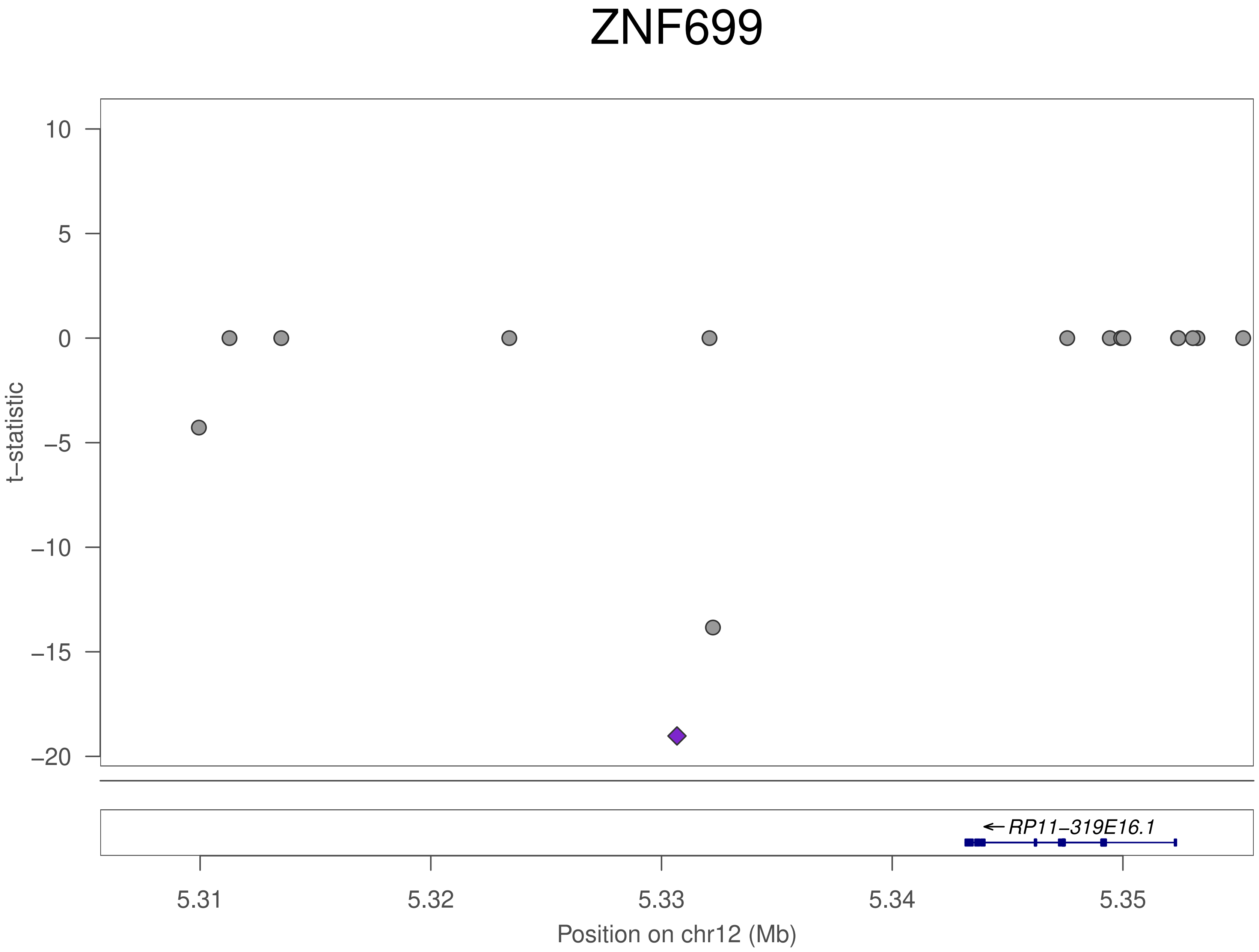

ZNF699
t–statistic
10
5
0
–5
–10
–15
–20
←RP11–319E16.1
5.31
5.32
5.33
5.34
5.35
Position on chr12 (Mb)

### ZNF699

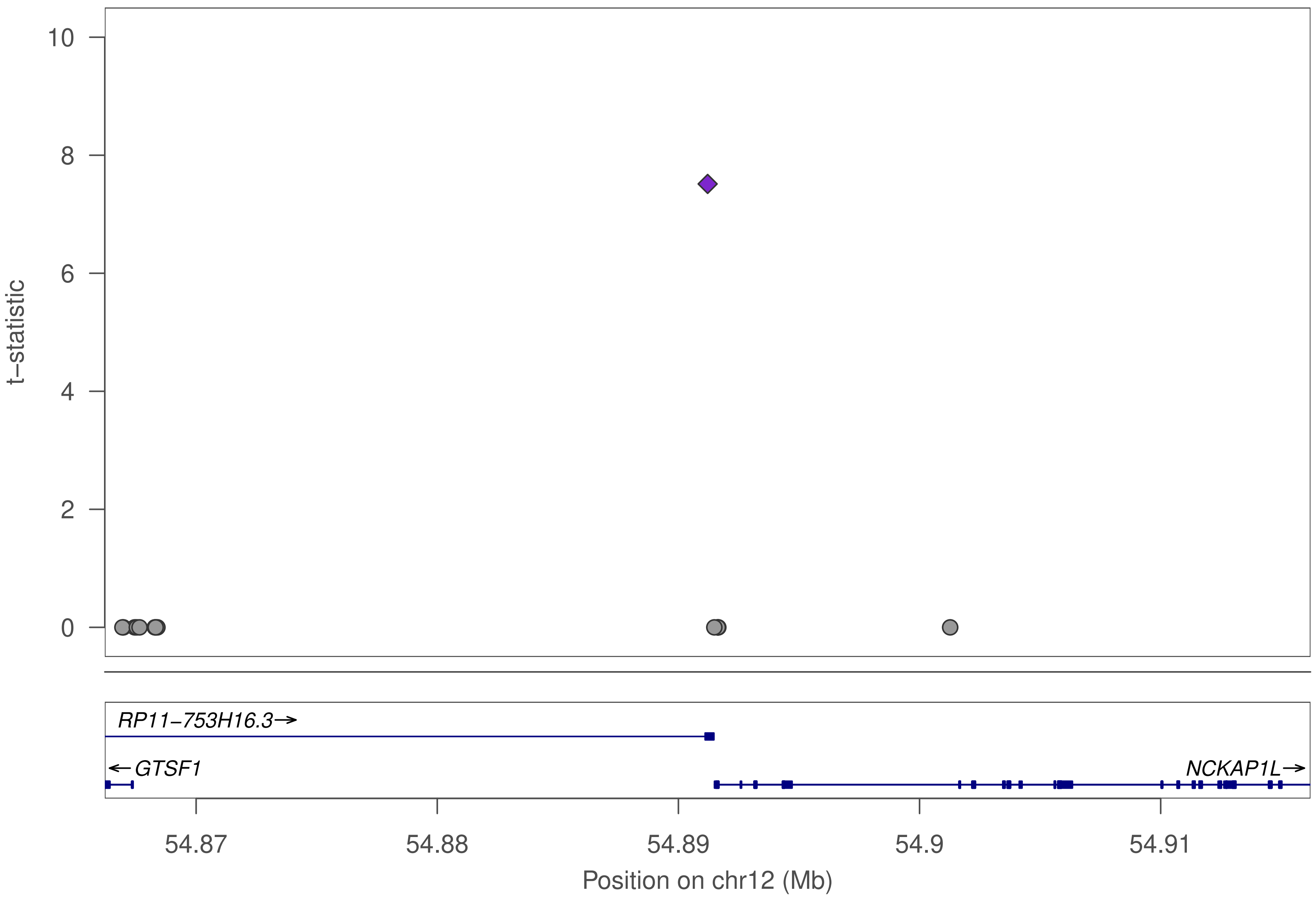

t−statistic
10
8
6
4
2
0
RP11−753H16.3→
←GTSF1
NCKAP1L→
54.87
54.88
54.89
54.9
54.91
Position on chr12 (Mb)

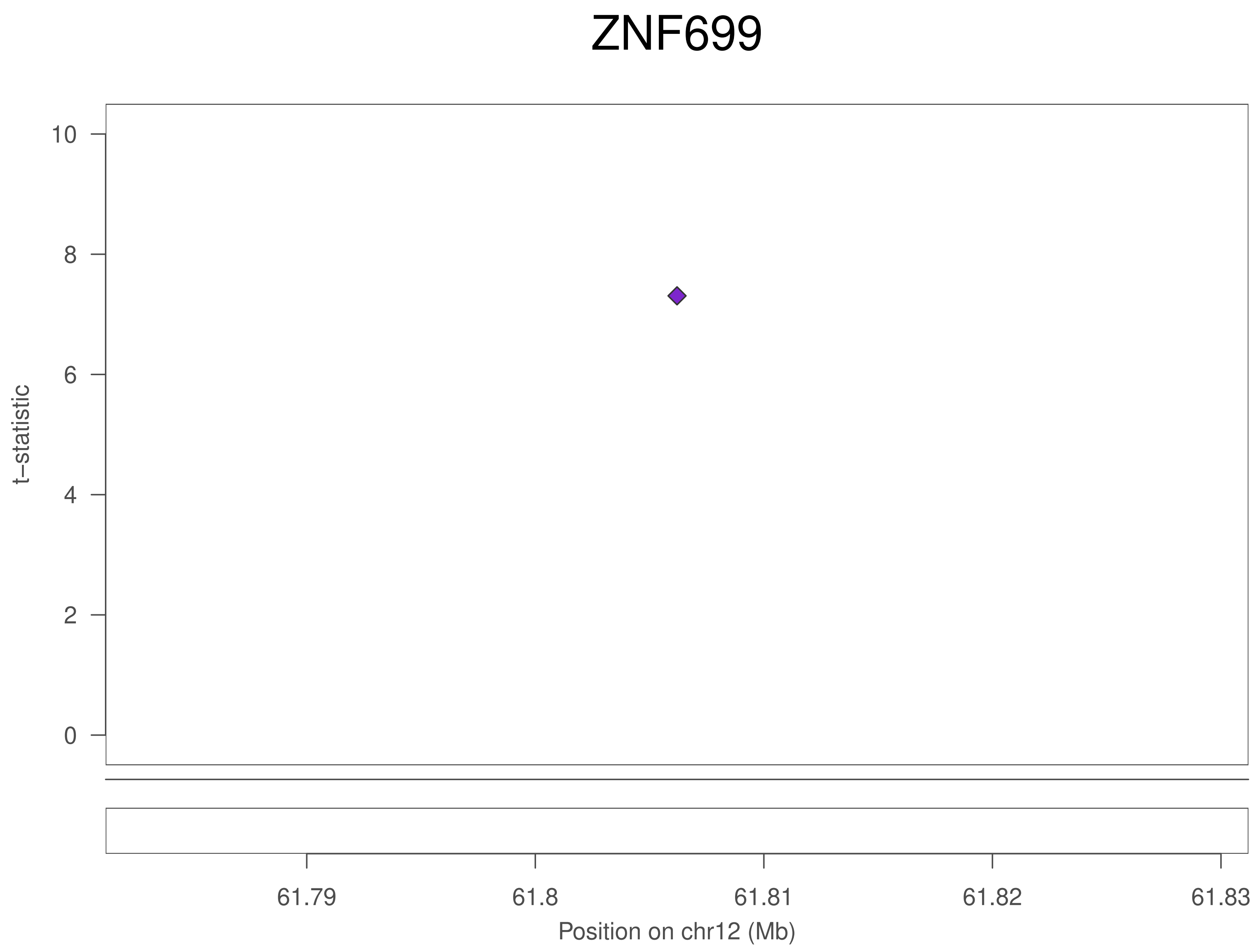
ZNF699
10
8
6
4
2
0
t−statistic
61.79
61.8
61.81
61.82
61.83
Position on chr12 (Mb)

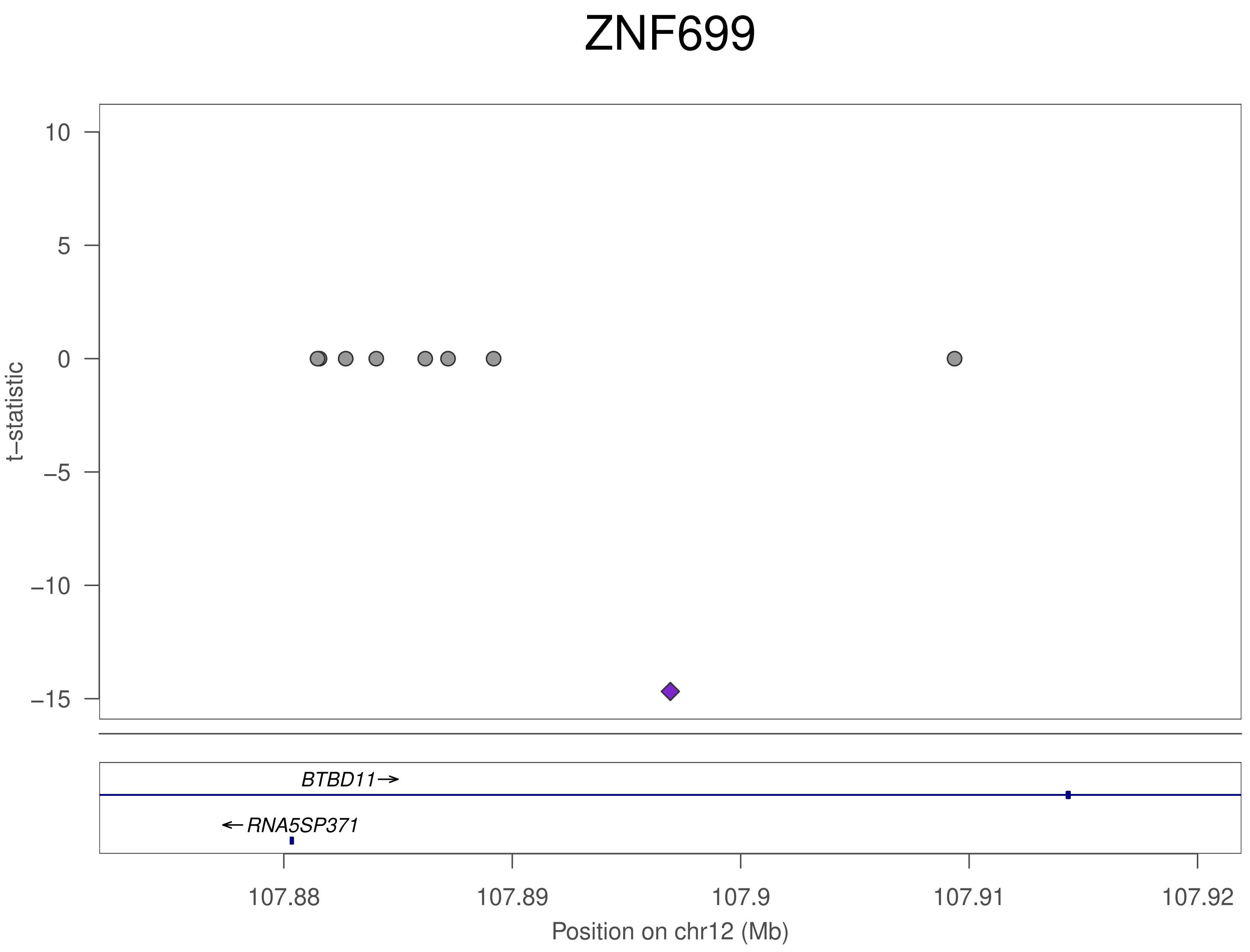

ZNF699
t–statistic
10
5
0
–5
–10
–15
BTBD11
RNA5SP371
107.88
107.89
107.9
107.91
107.92
Position on chr12 (Mb)

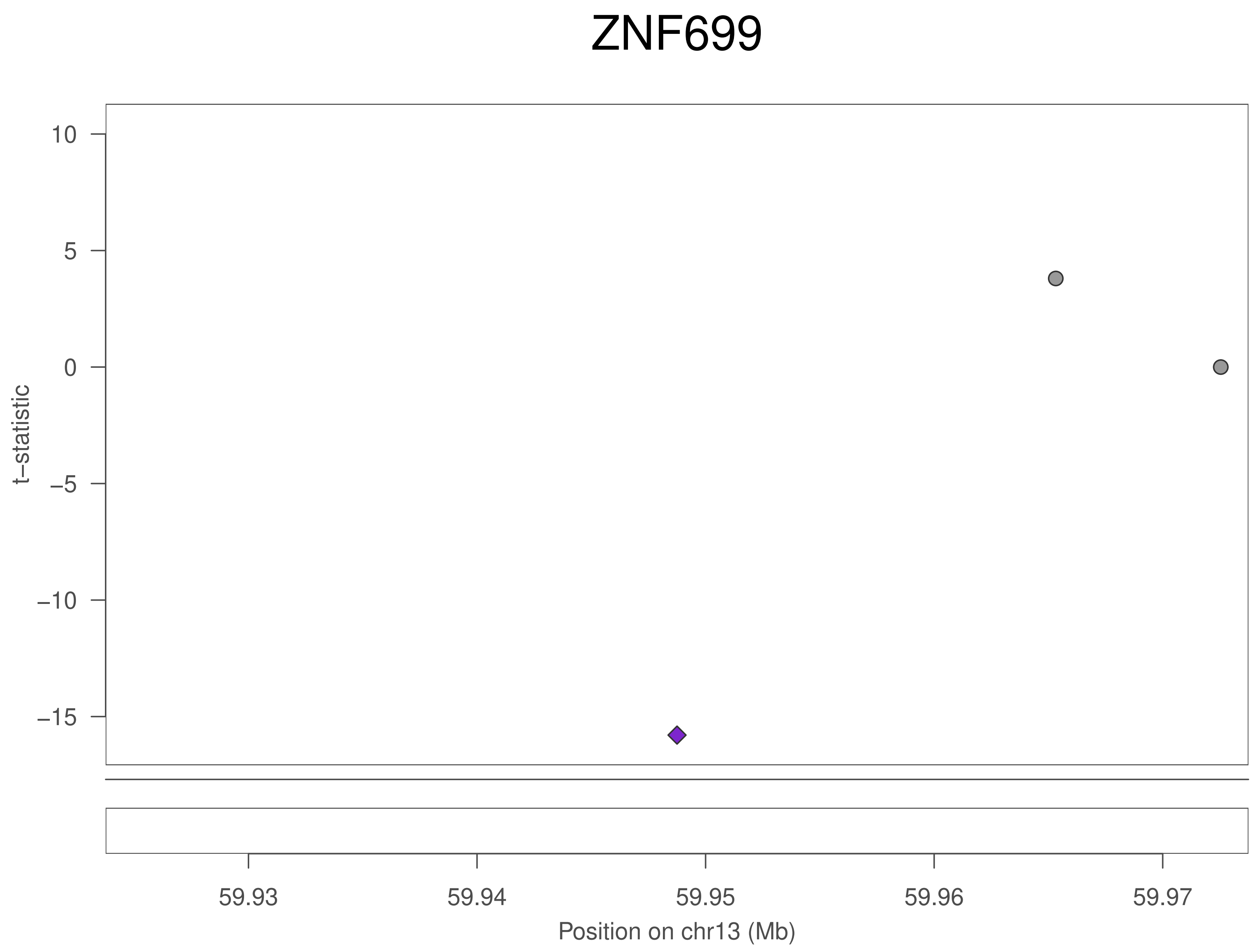
ZNF699
t−statistic
10
5
0
−5
−10
−15
59.93
59.94
59.95
59.96
59.97
Position on chr13 (Mb)

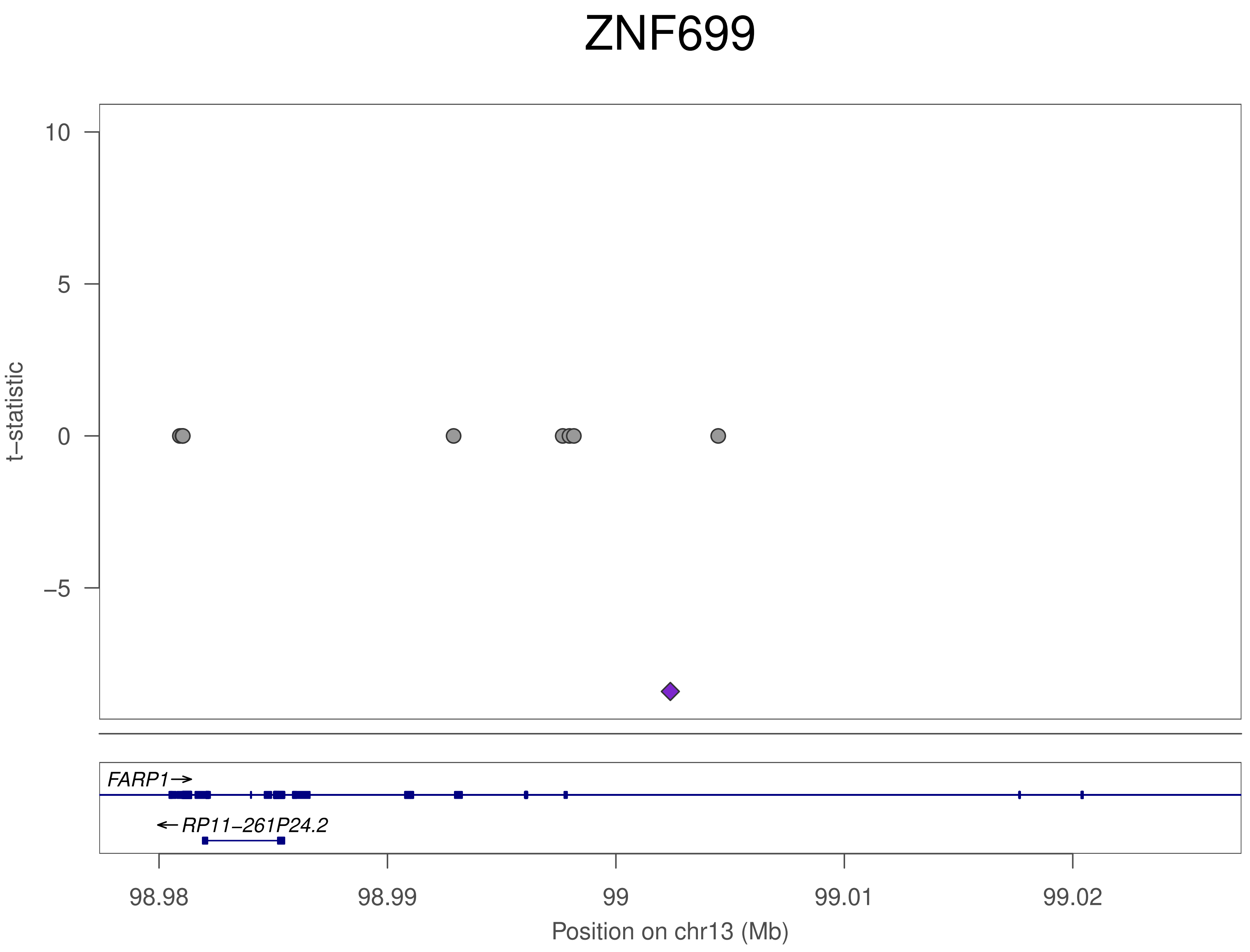
ZNF699
t−statistic
10
5
0
−5
FARP1→
←RP11−261P24.2
98.98
98.99
99
99.01
99.02
Position on chr13 (Mb)

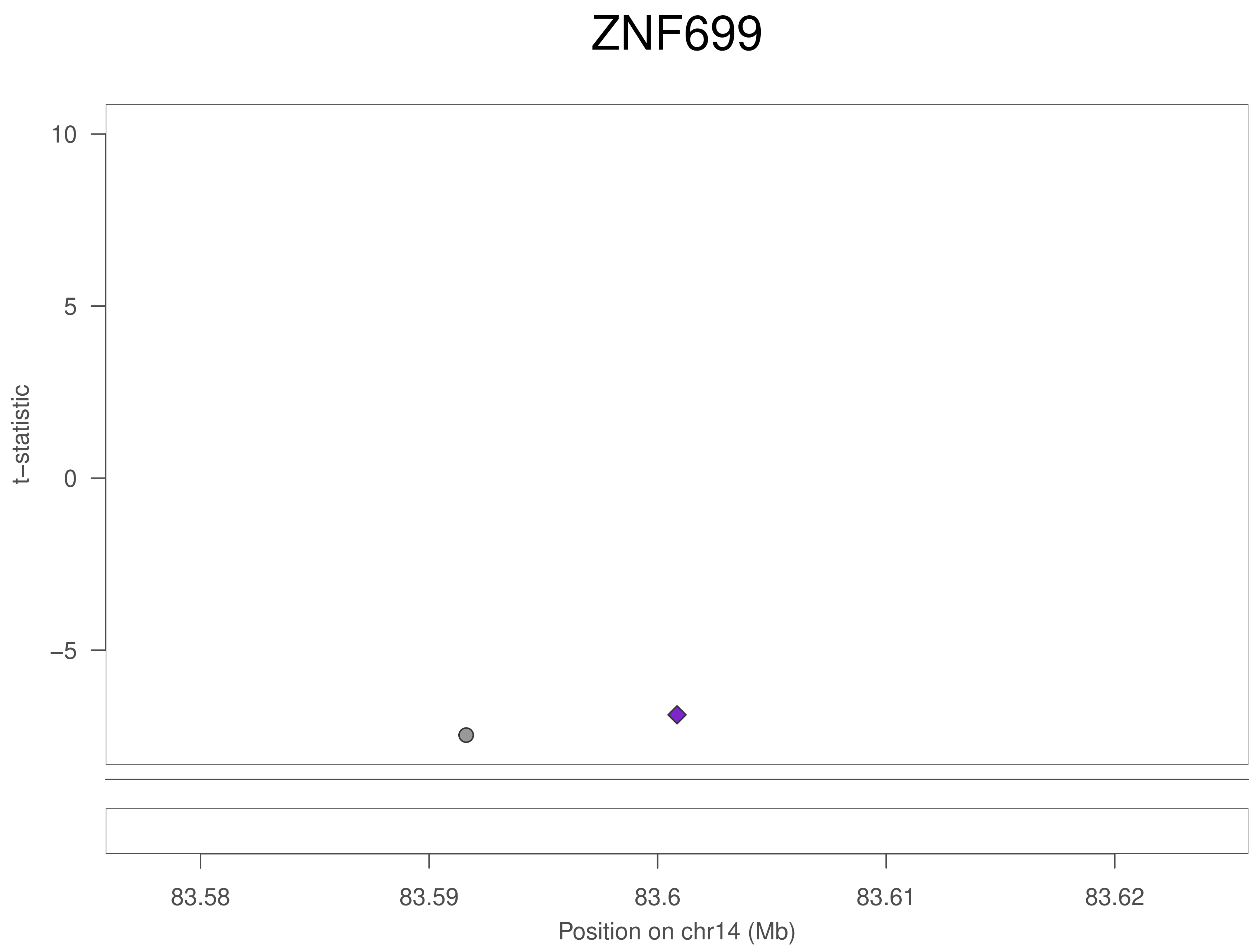
ZNF699
t−statistic
10
5
0
−5
83.58
83.59
83.6
83.61
83.62
Position on chr14 (Mb)

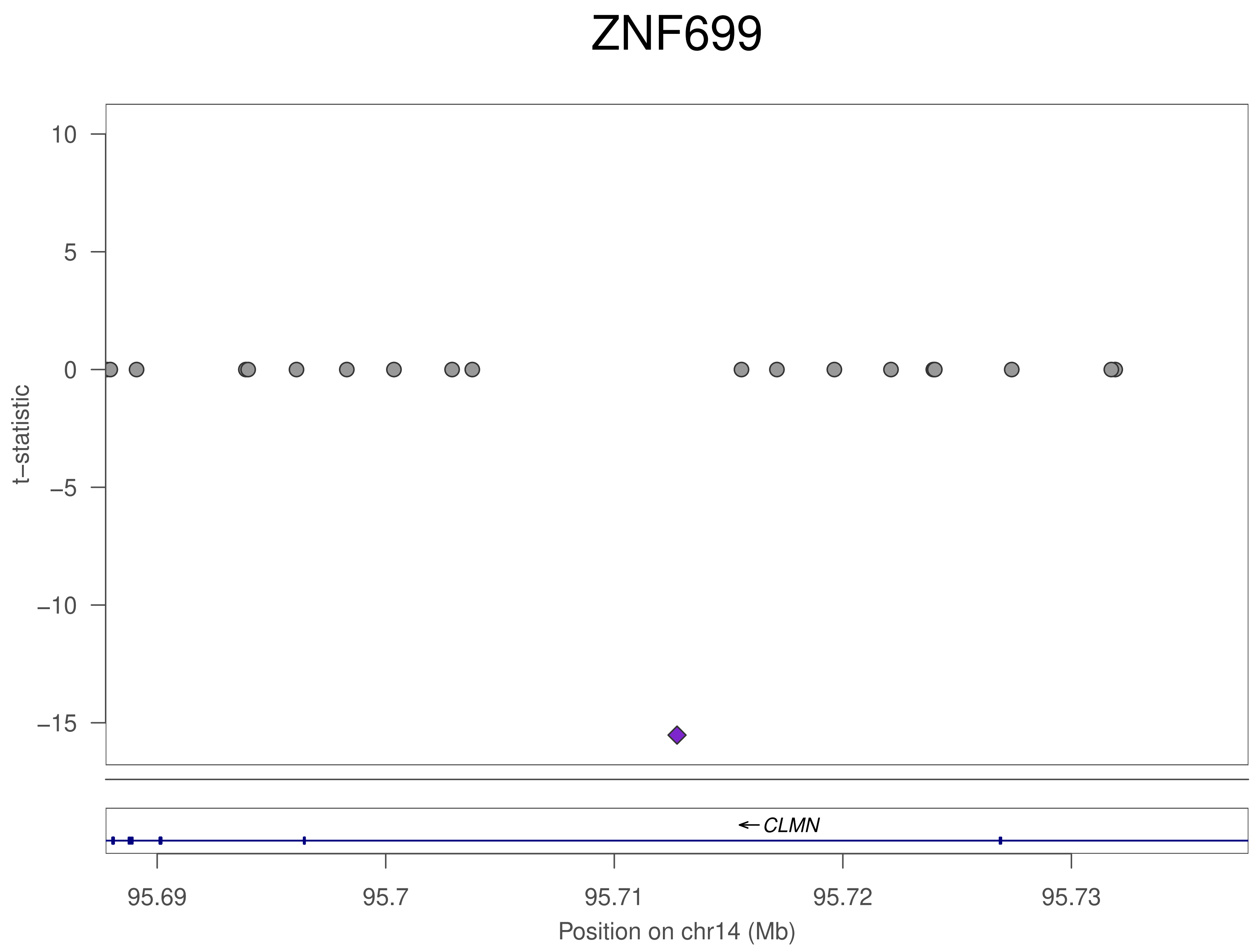
ZNF699
t−statistic
10
5
0
−5
−10
−15
CLMN
95.69
95.7
95.71
95.72
95.73
Position on chr14 (Mb)

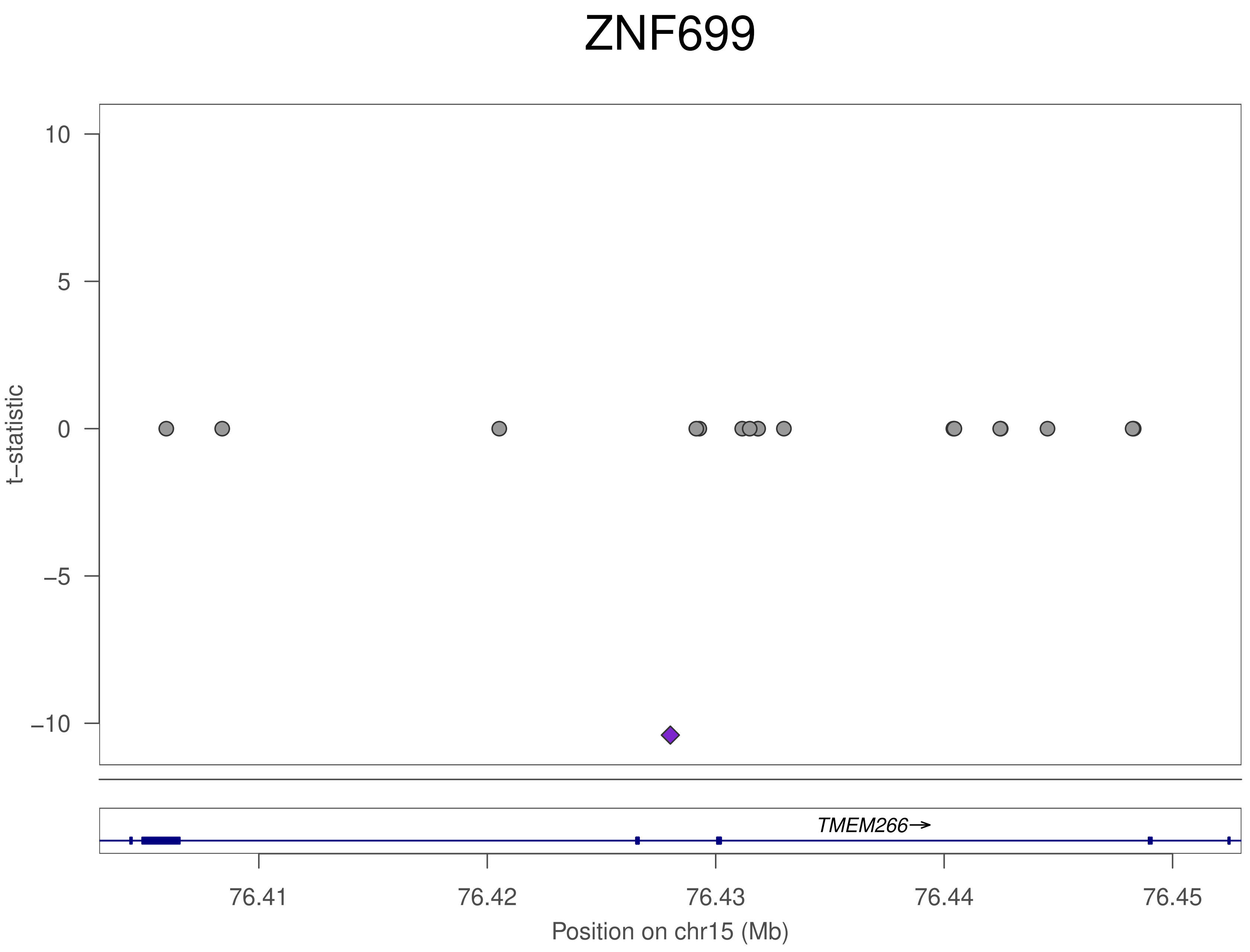

ZNF699
t–statistic
10
5
0
–5
–10
TMEM266→
76.41
76.42
76.43
76.44
76.45
Position on chr15 (Mb)

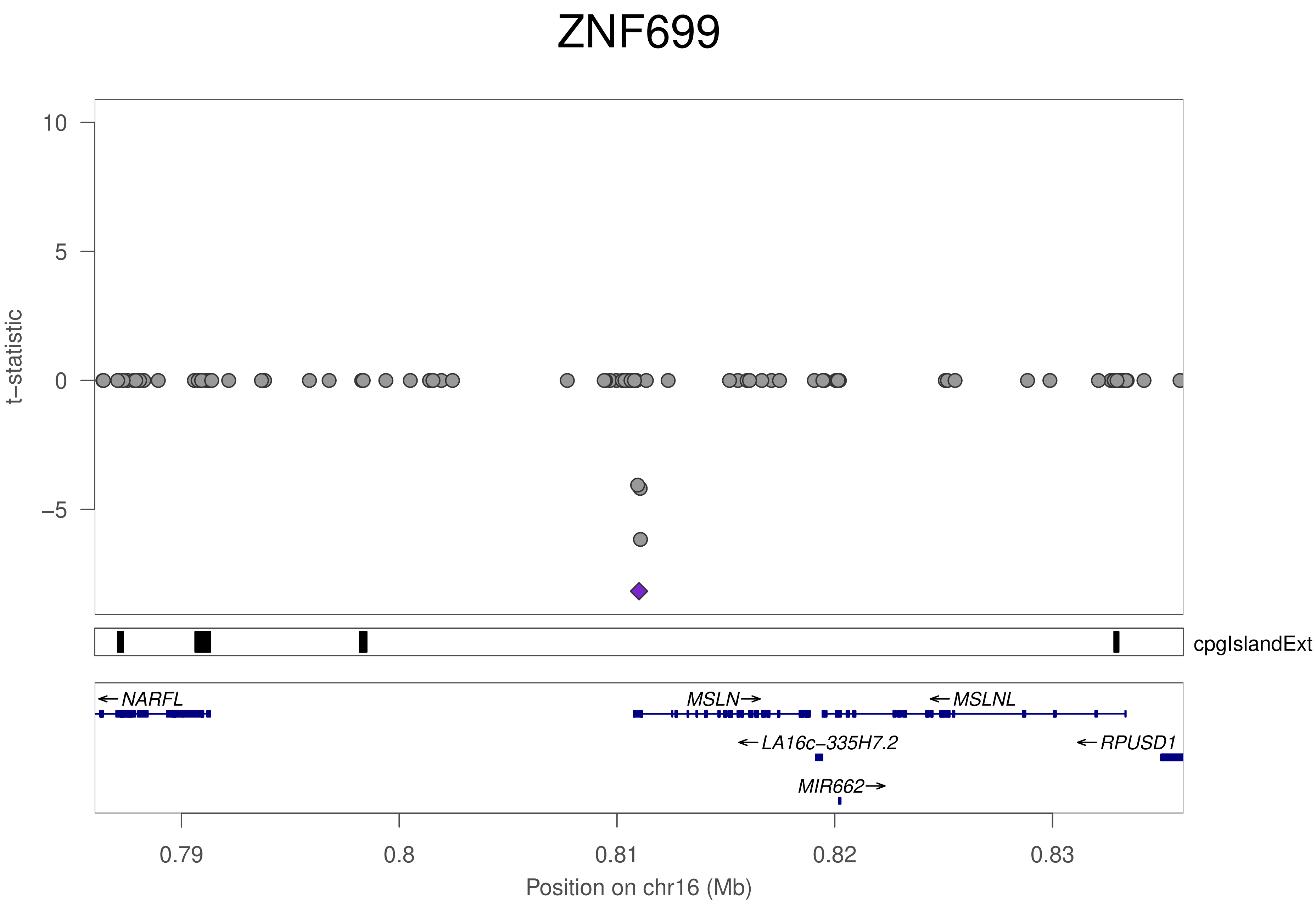

ZNF699
10
5
0
−5
t−statistic
cpgIslandExt
NARFL
MSLN
MSLNL
LA16c−335H7.2
RPUSD1
MIR662
0.79
0.8
0.81
0.82
0.83
Position on chr16 (Mb)

### ZNF699

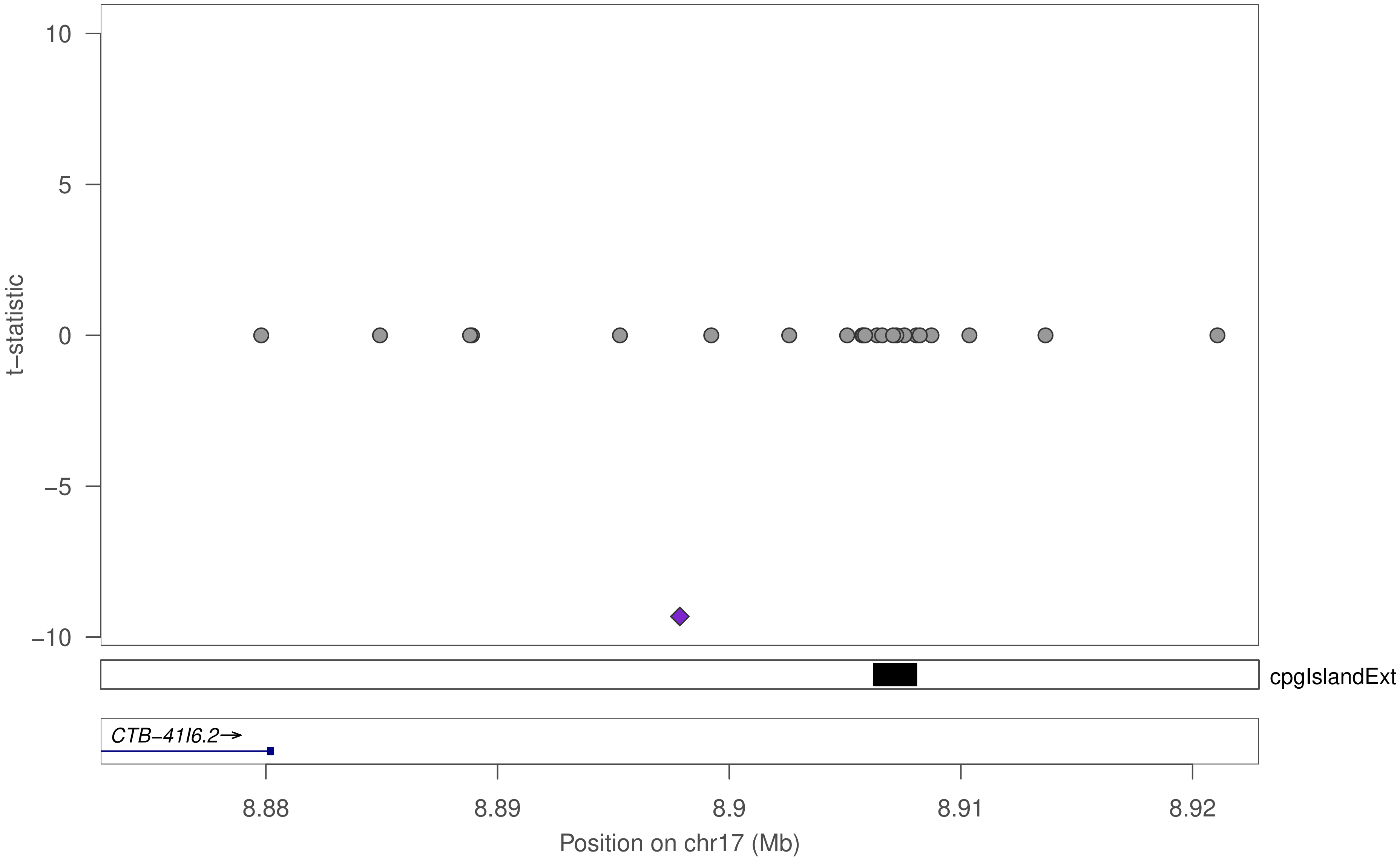

10
5
0
−5
−10
t−statistic
cpgIslandExt
CTB−41I6.2→
8.88
8.89
8.9
8.91
8.92
Position on chr17 (Mb)

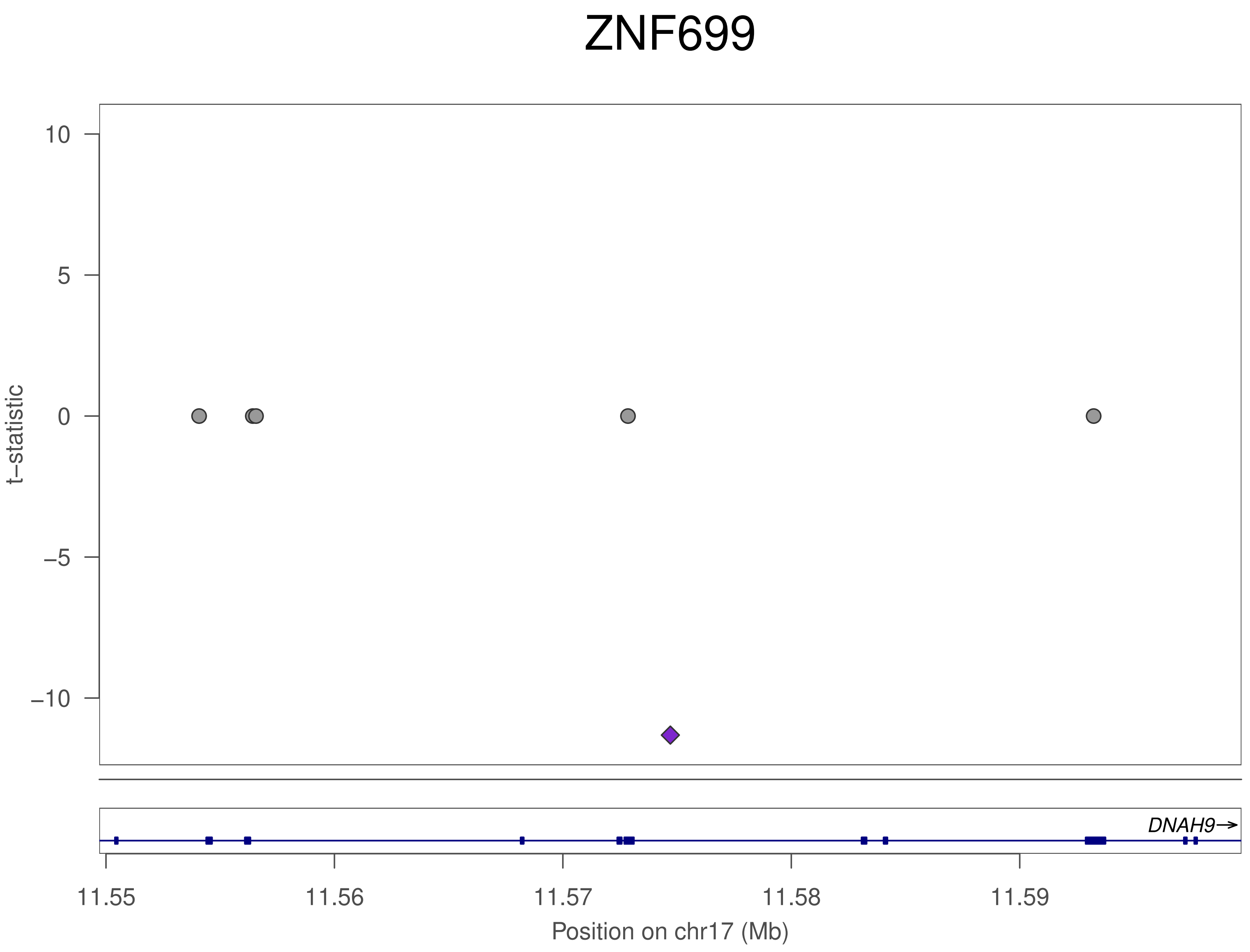

ZNF699
10
5
0
−5
−10
t−statistic
DNAH9→
11.55
11.56
11.57
11.58
11.59
Position on chr17 (Mb)

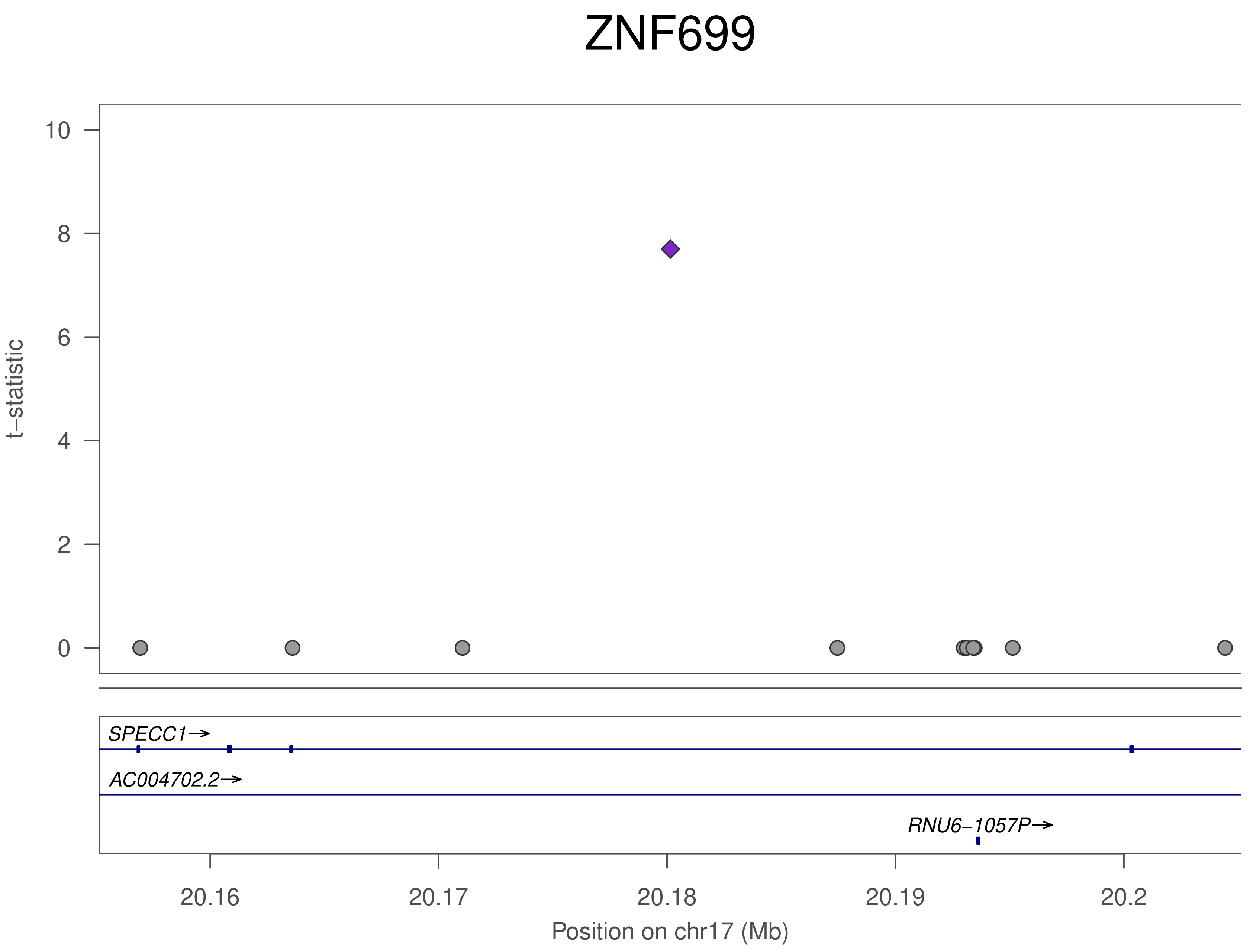

ZNF699
t−statistic
10
8
6
4
2
0
SPECC1→
AC004702.2→
RNU6−1057P→
20.16
20.17
20.18
20.19
20.2
Position on chr17 (Mb)

### ZNF699

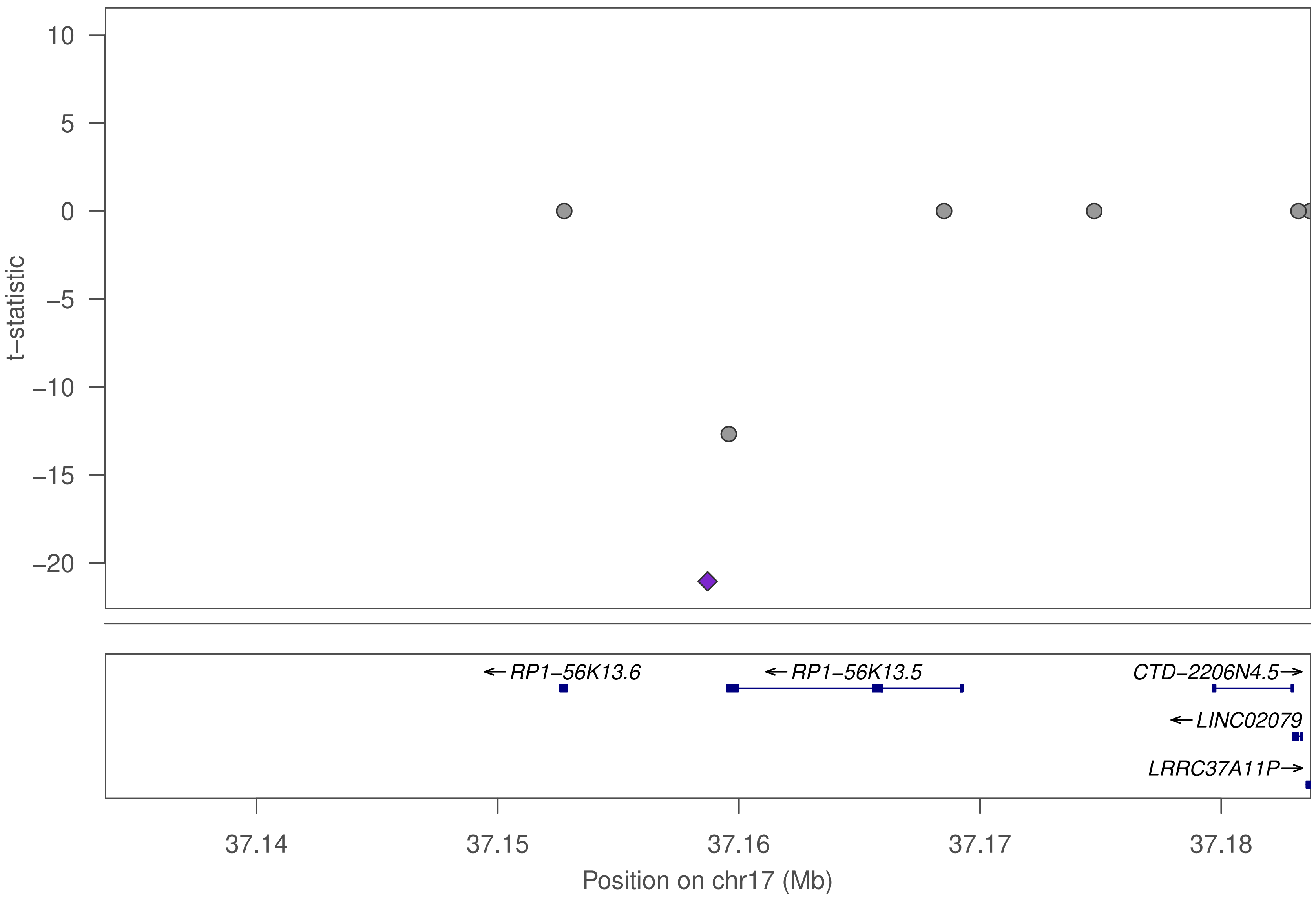

10
5
0
−5
−10
−15
−20
t−statistic
←RP1−56K13.6
←RP1−56K13.5
CTD−2206N4.5→
←LINC02079
LRRC37A11P→
37.14
37.15
37.16
37.17
37.18
Position on chr17 (Mb)

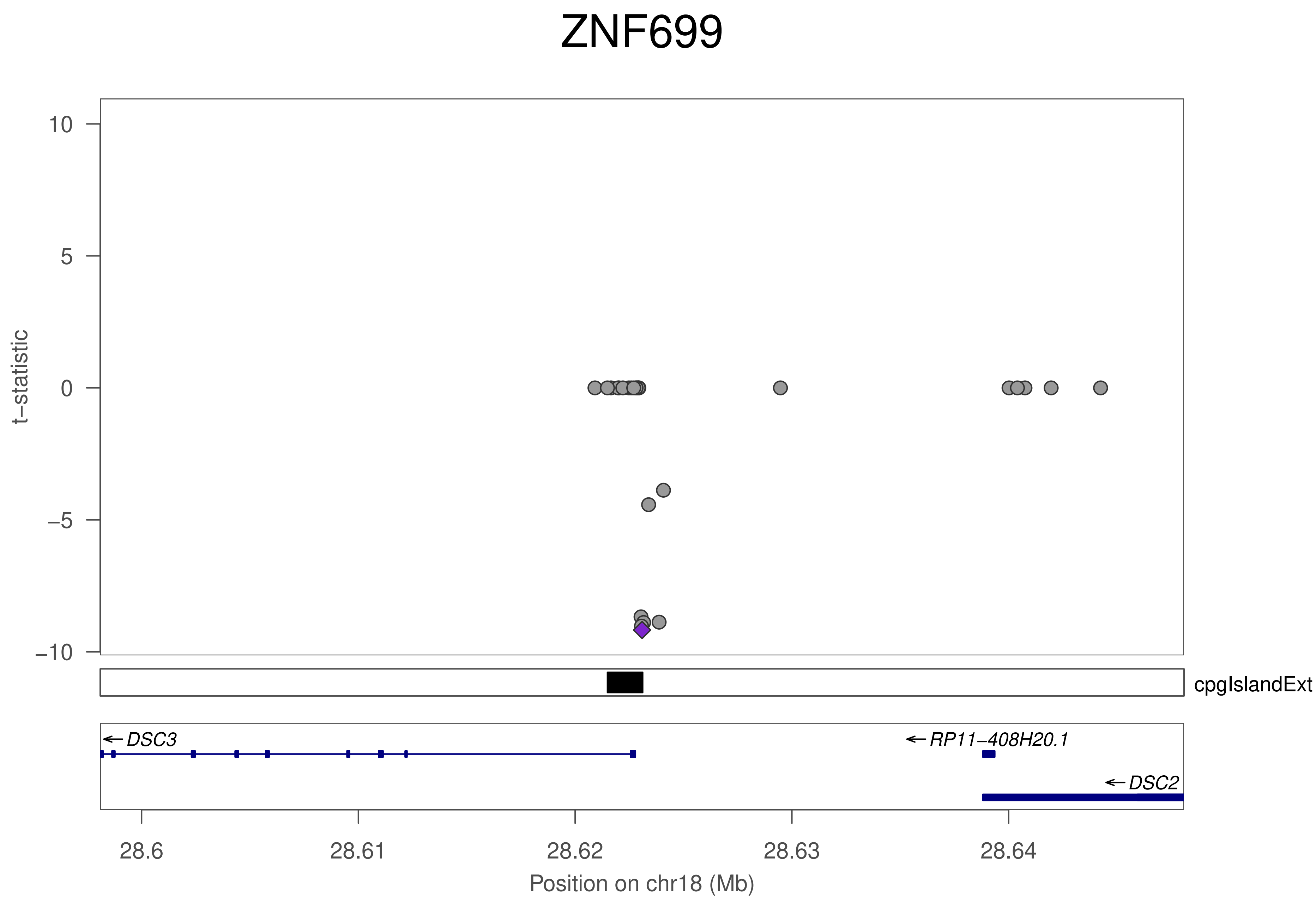
ZNF699
t−statistic
10
5
0
−5
−10
cpgIslandExt
DSC3
RP11−408H20.1
DSC2
28.6
28.61
28.62
28.63
28.64
Position on chr18 (Mb)

### ZNF699

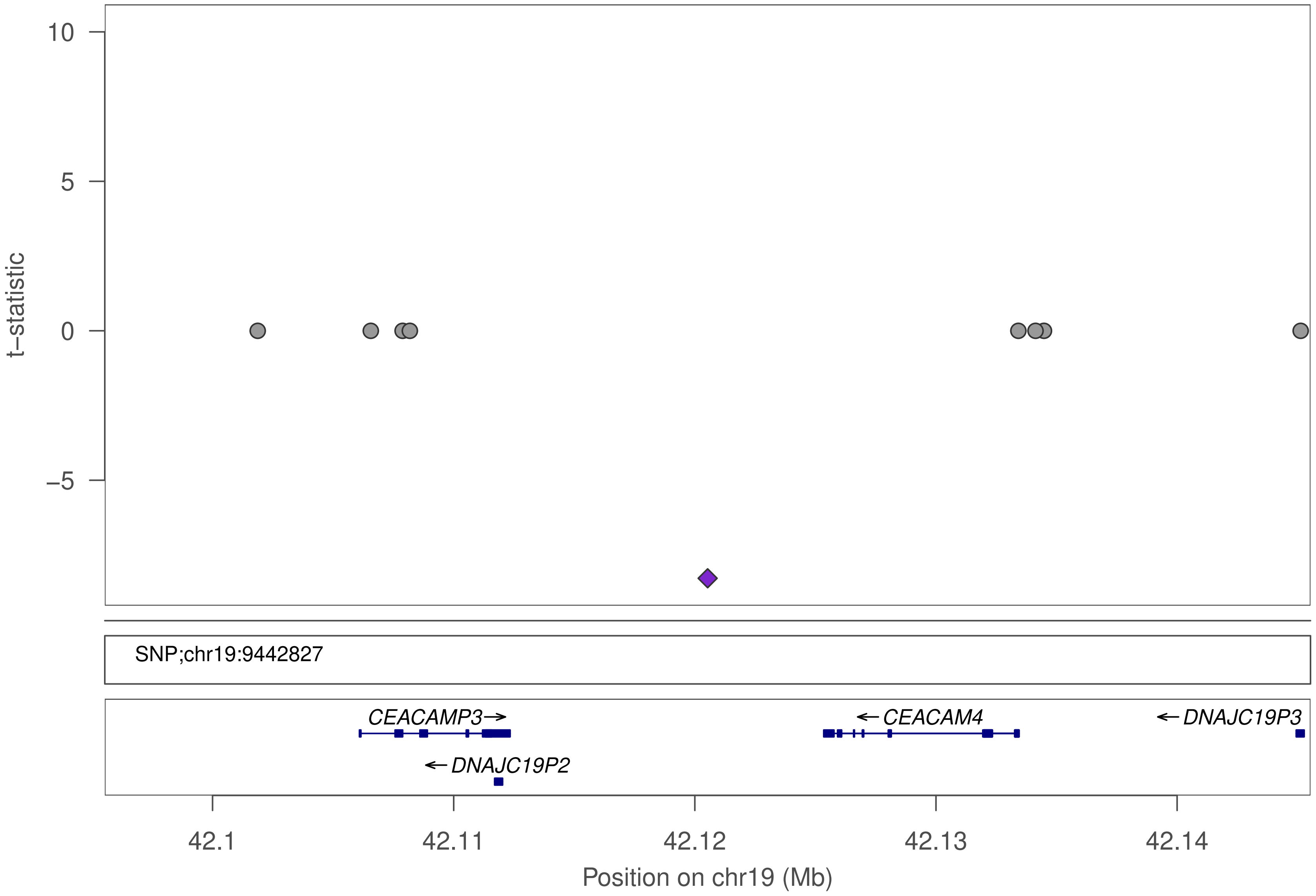

10
5
0
−5
t−statistic
SNP;chr19:9442827
CEACAMP3→
←CEACAM4
←DNAJC19P3
←DNAJC19P2
42.1
42.11
42.12
42.13
42.14
Position on chr19 (Mb)

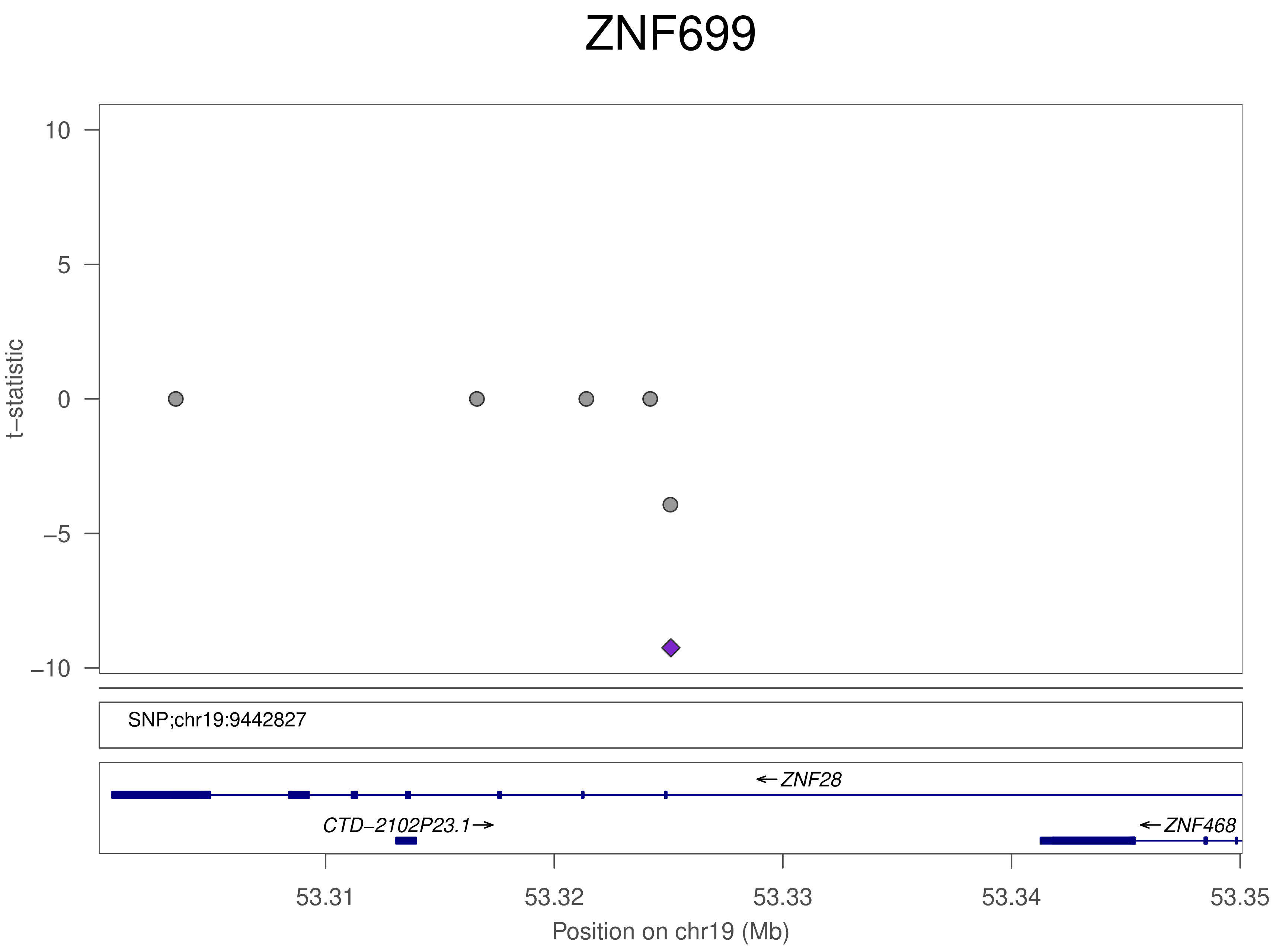

ZNF699
10
5
0
−5
−10
t−statistic
SNP;chr19:9442827
←ZNF28
CTD−2102P23.1→
←ZNF468
53.31
53.32
53.33
53.34
53.35
Position on chr19 (Mb)

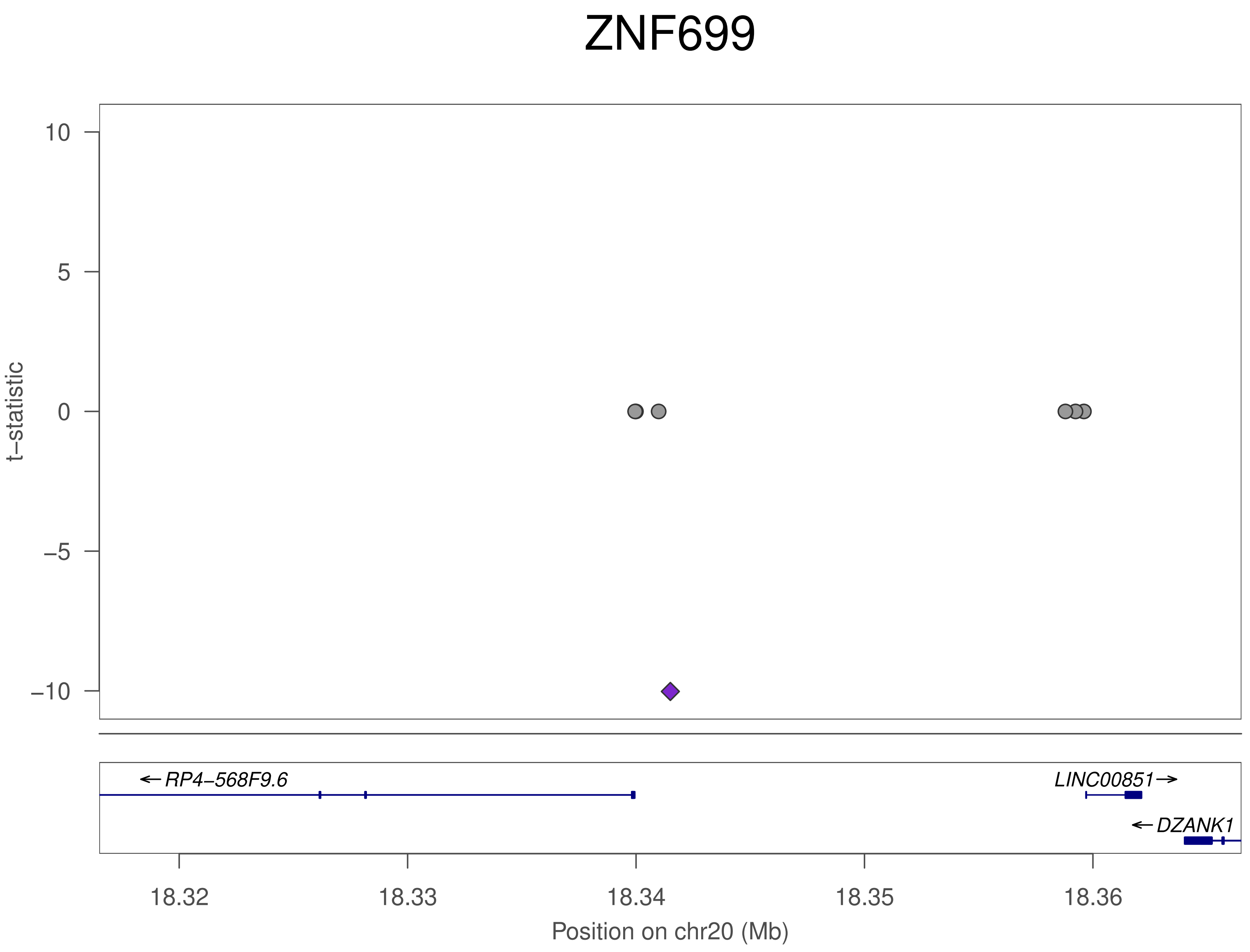

ZNF699
10
5
0
−5
−10
t−statistic
←RP4−568F9.6
LINC00851→
←DZANK1
18.32
18.33
18.34
18.35
18.36
Position on chr20 (Mb)

### ZNF699

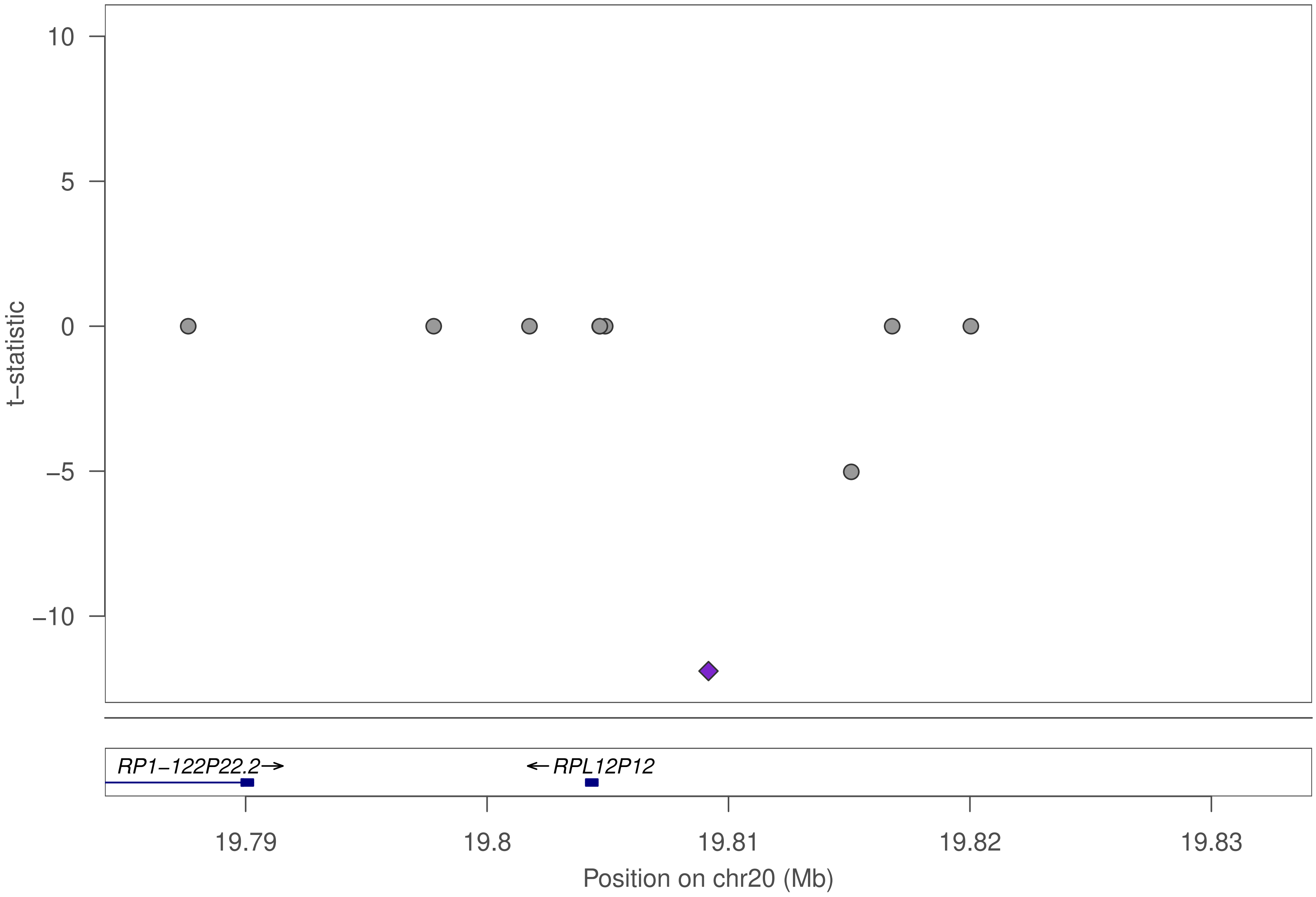

10
5
0
−5
−10
t−statistic
RP1−122P22.2→
←RPL12P12
19.79
19.8
19.81
19.82
19.83
Position on chr20 (Mb)

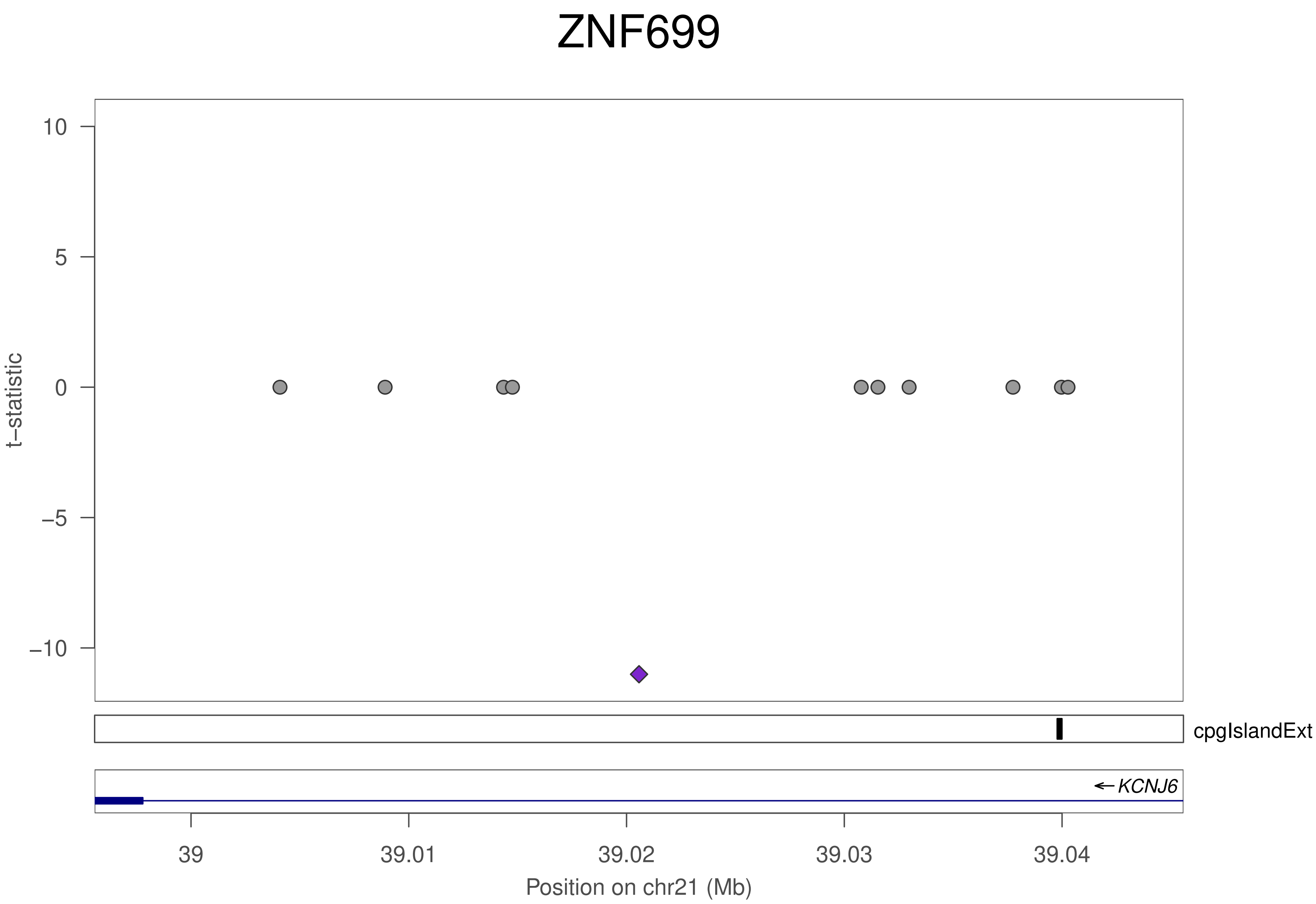
ZNF699
10
5
0
−5
−10
t−statistic
cpgIslandExt
← KCNJ6
39
39.01
39.02
39.03
39.04
Position on chr21 (Mb)

### ZNF699

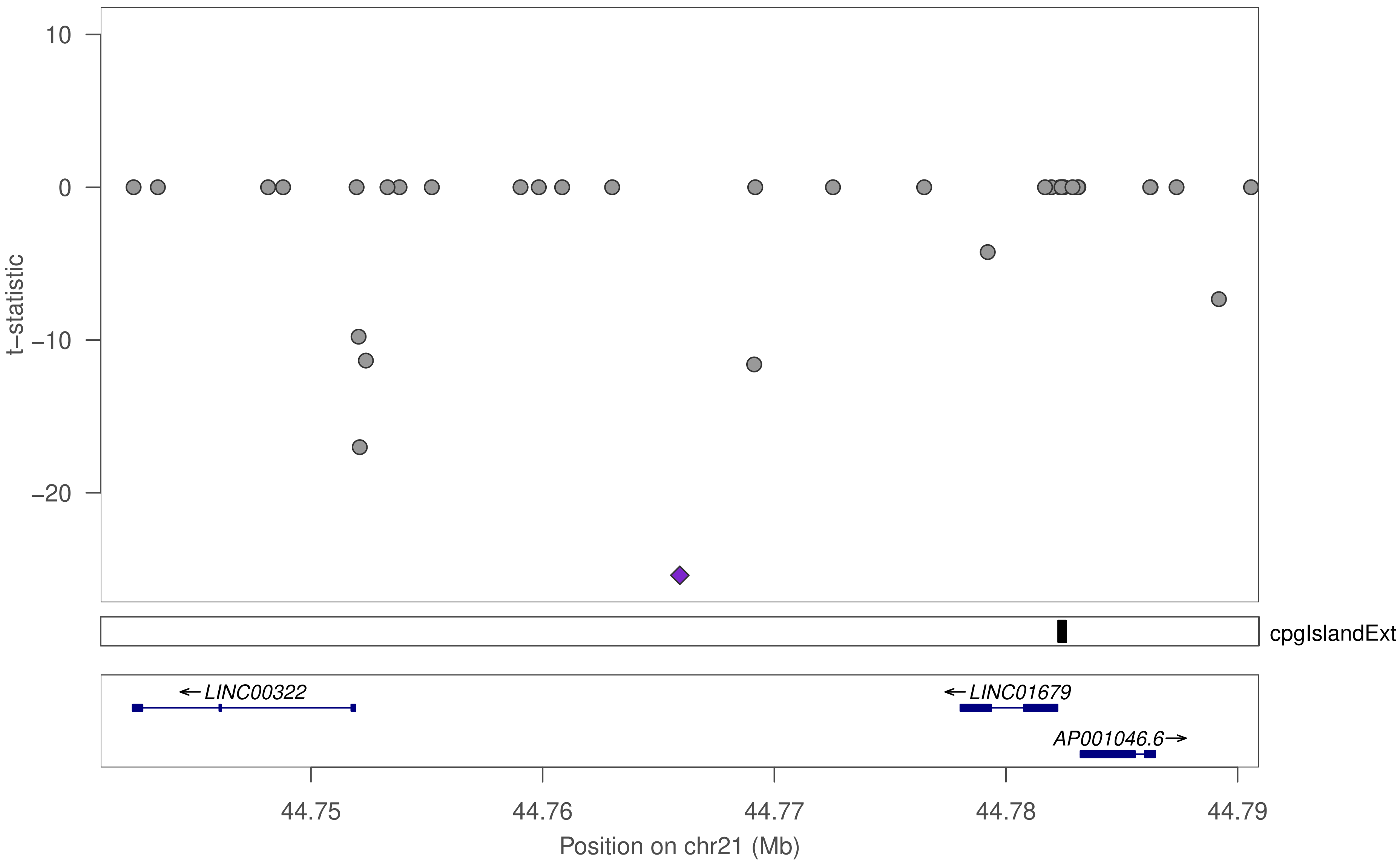
10
0
−10
−20
t−statistic
cpgIslandExt
←LINC00322
←LINC01679
AP001046.6→
44.75
44.76
44.77
44.78
44.79
Position on chr21 (Mb)

### ZNF699

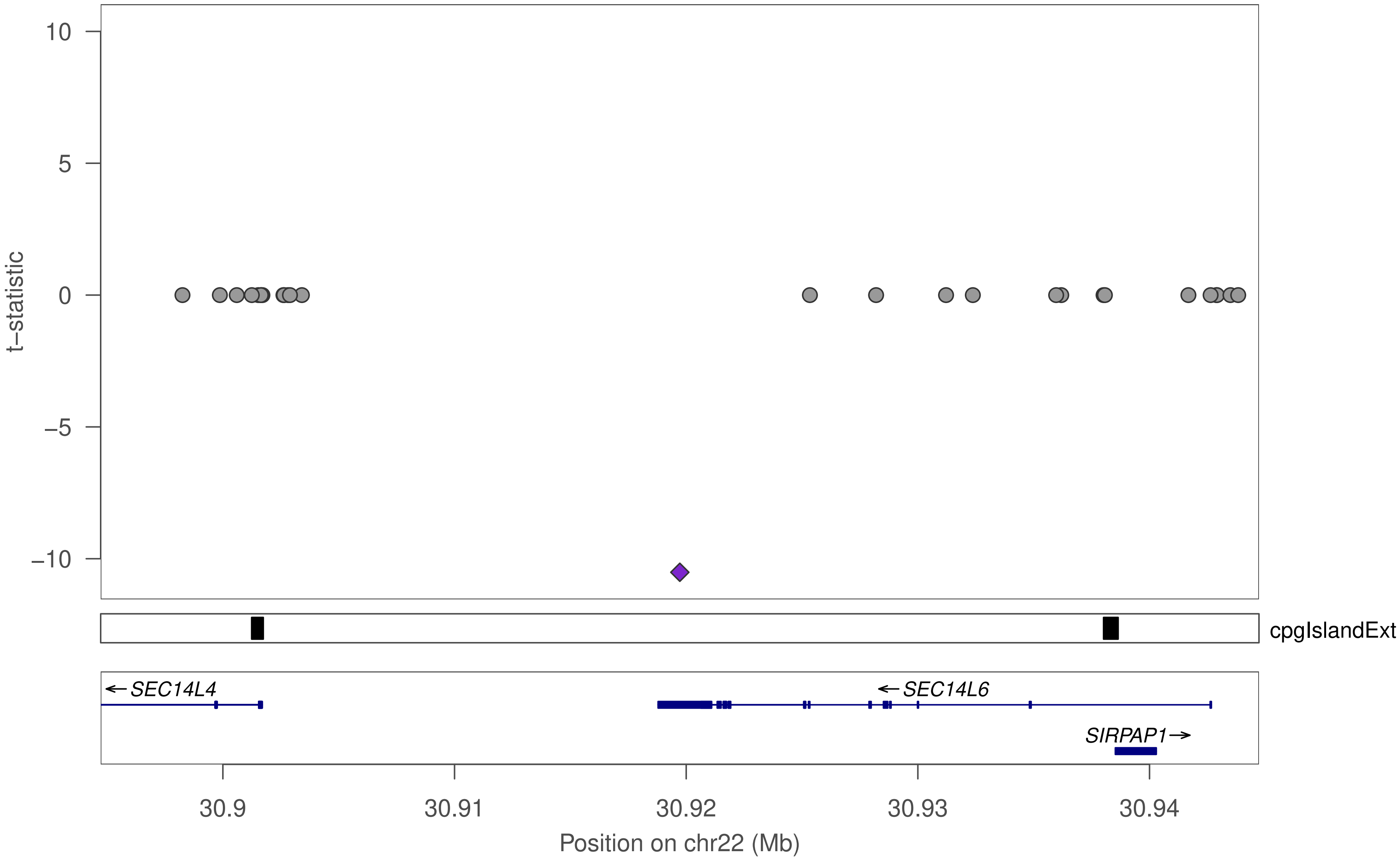

10
5
0
−5
−10
t−statistic
cpgIslandExt
←SEC14L4
←SEC14L6
SIRPAP1→
30.9
30.91
30.92
30.93
30.94
Position on chr22 (Mb)

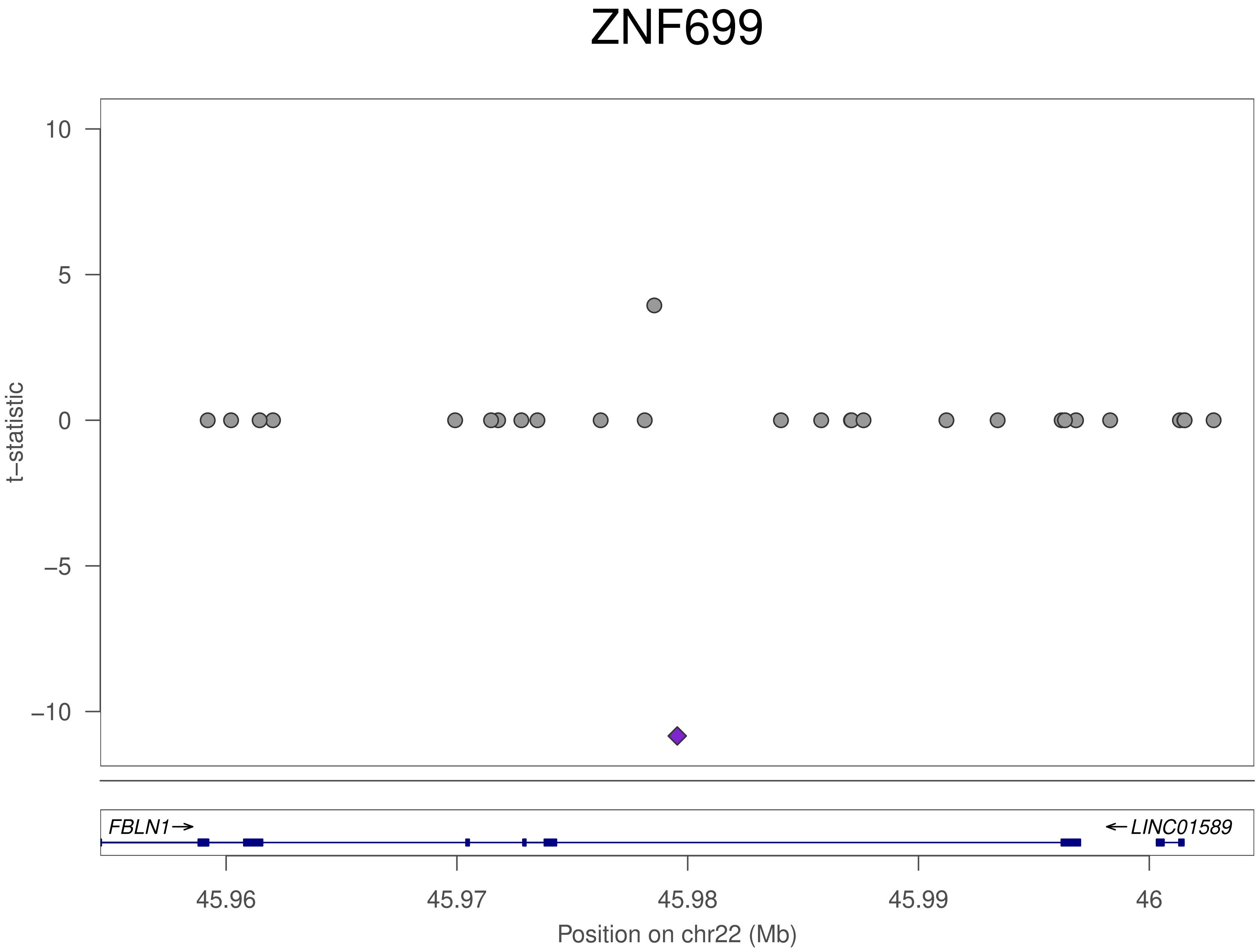
ZNF699
t−statistic
10
5
0
−5
−10
FBLN1→
←LINC01589
45.96
45.97
45.98
45.99
46
Position on chr22 (Mb)

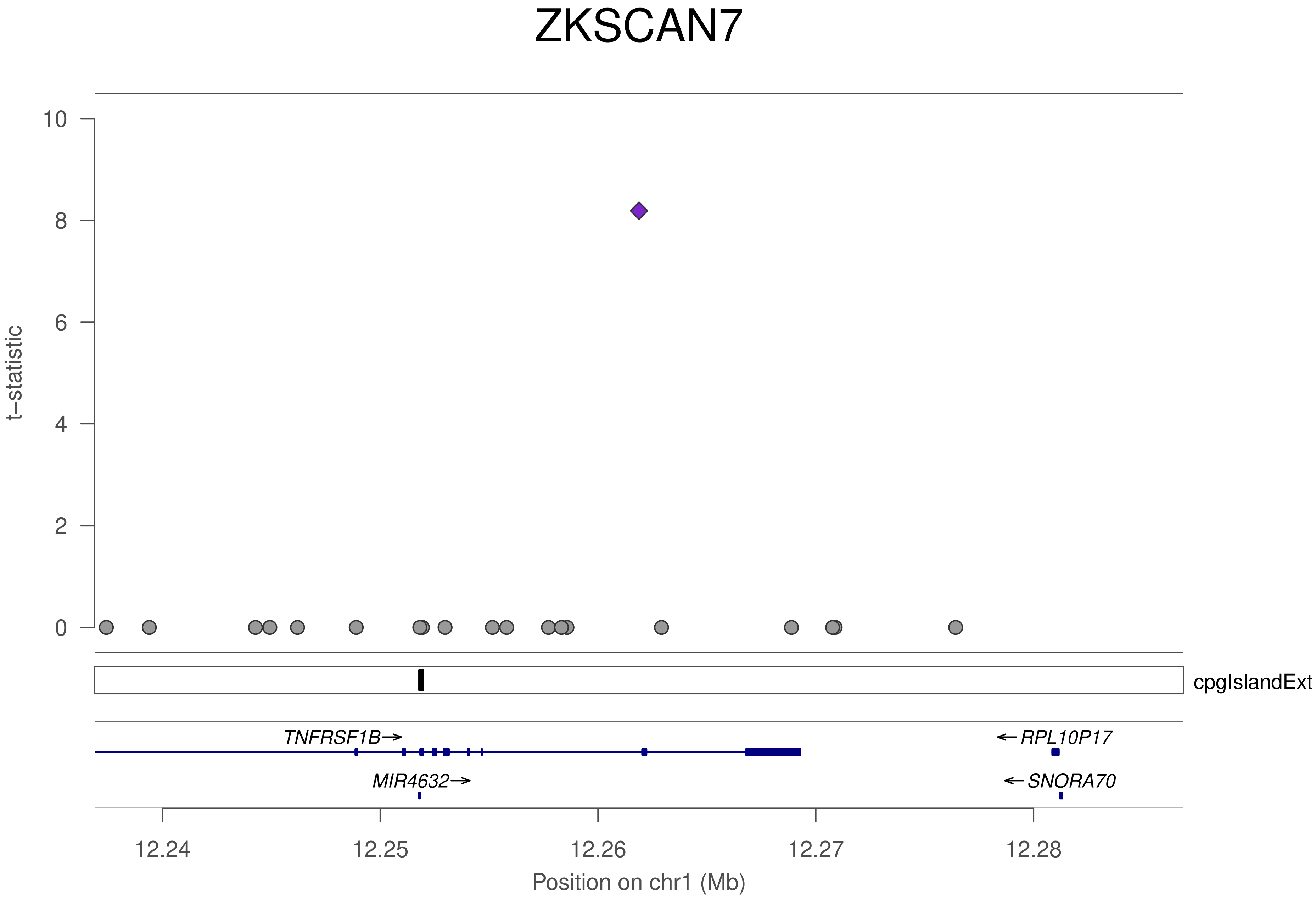
ZKSCAN7
t−statistic
10
8
6
4
2
0
cpgIslandExt
TNFRSF1B→
MIR4632→
←RPL10P17
←SNORA70
12.24
12.25
12.26
12.27
12.28
Position on chr1 (Mb)

### ZKSCAN7

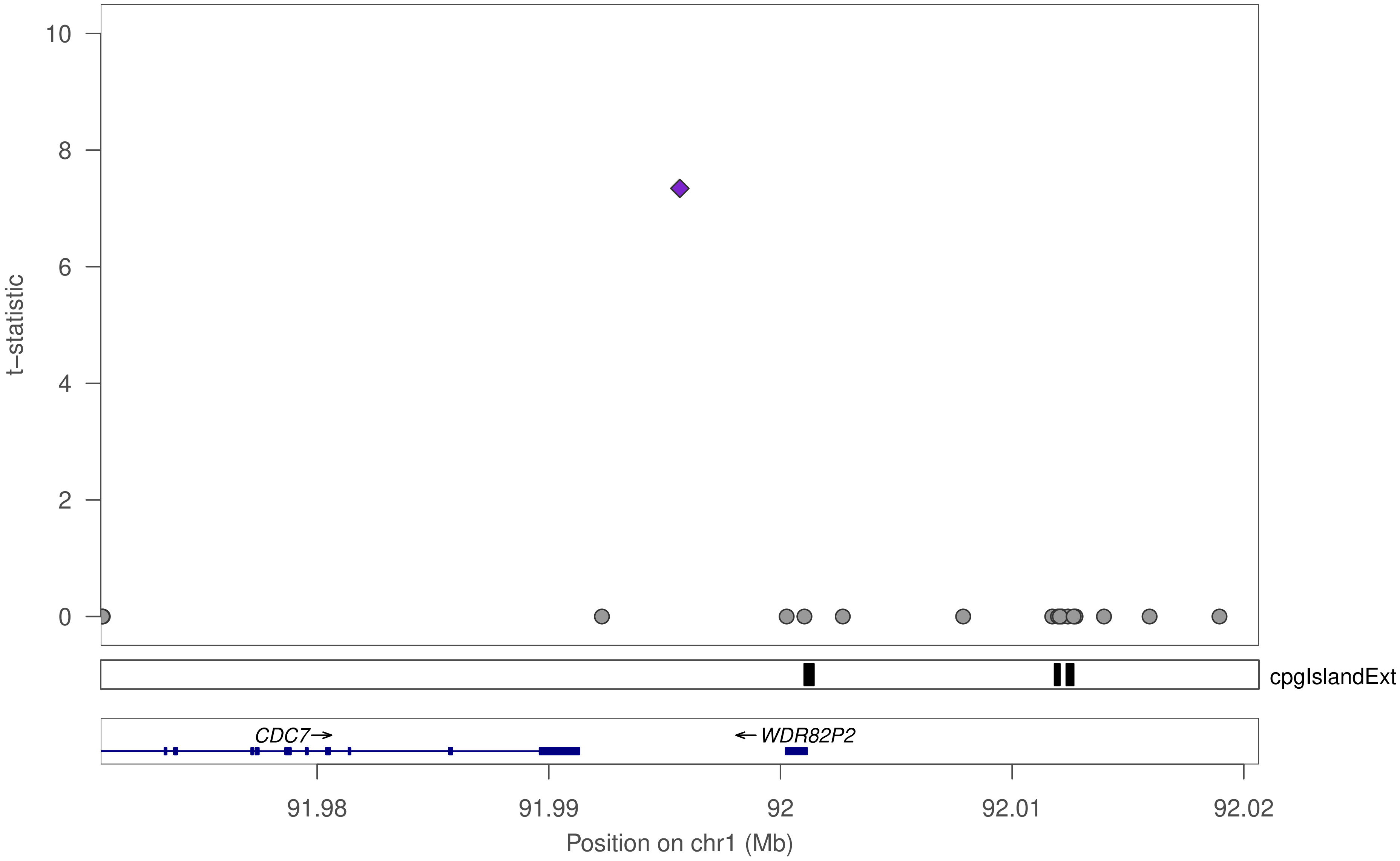

t–statistic
10
8
6
4
2
0
cpgIslandExt
CDC7→
←WDR82P2
91.98
91.99
92
92.01
92.02
Position on chr1 (Mb)

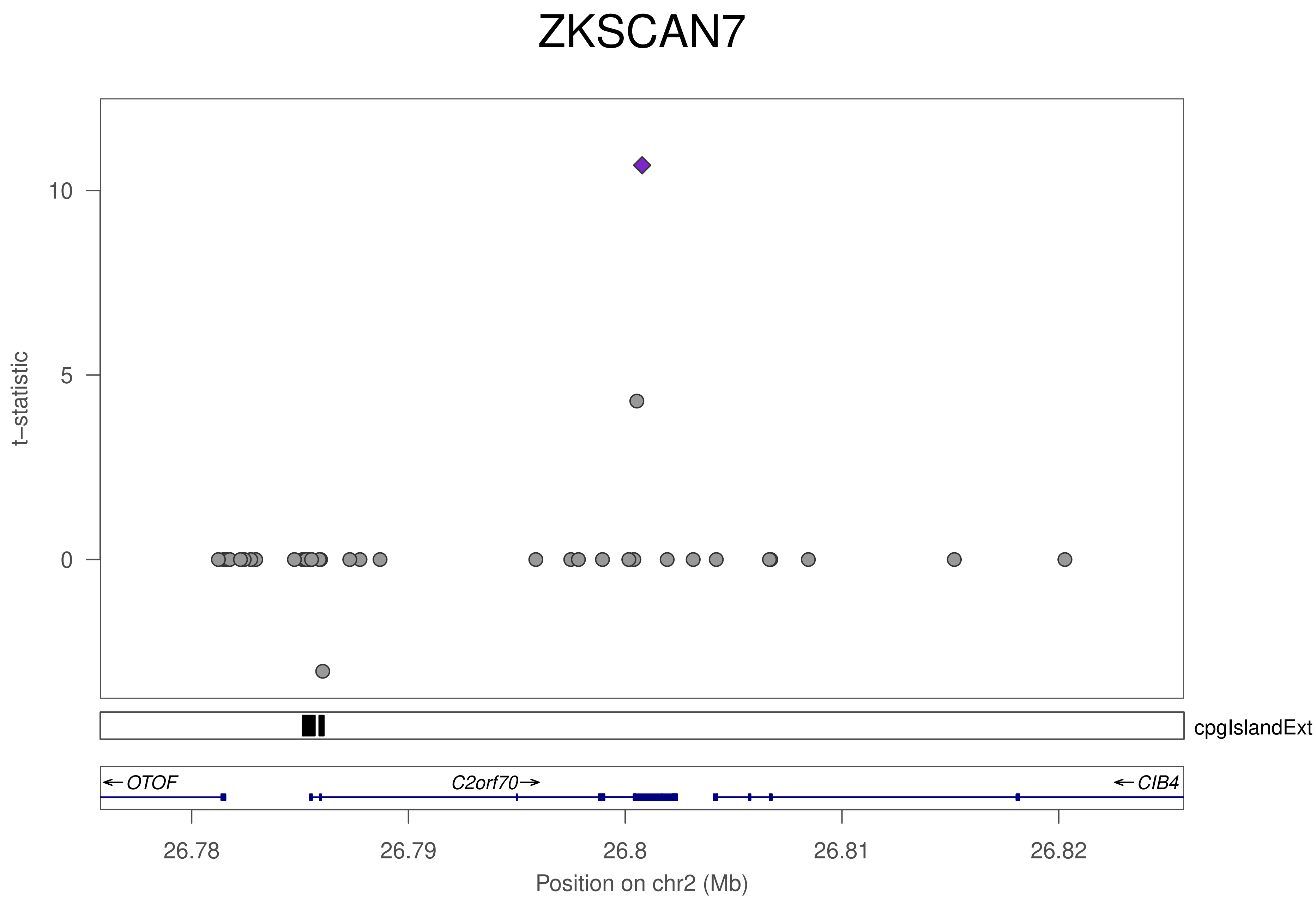
ZKSCAN7
t–statistic
10
5
0
cpgIslandExt
OTOF
C2orf70
CIB4
26.78
26.79
26.8
26.81
26.82
Position on chr2 (Mb)

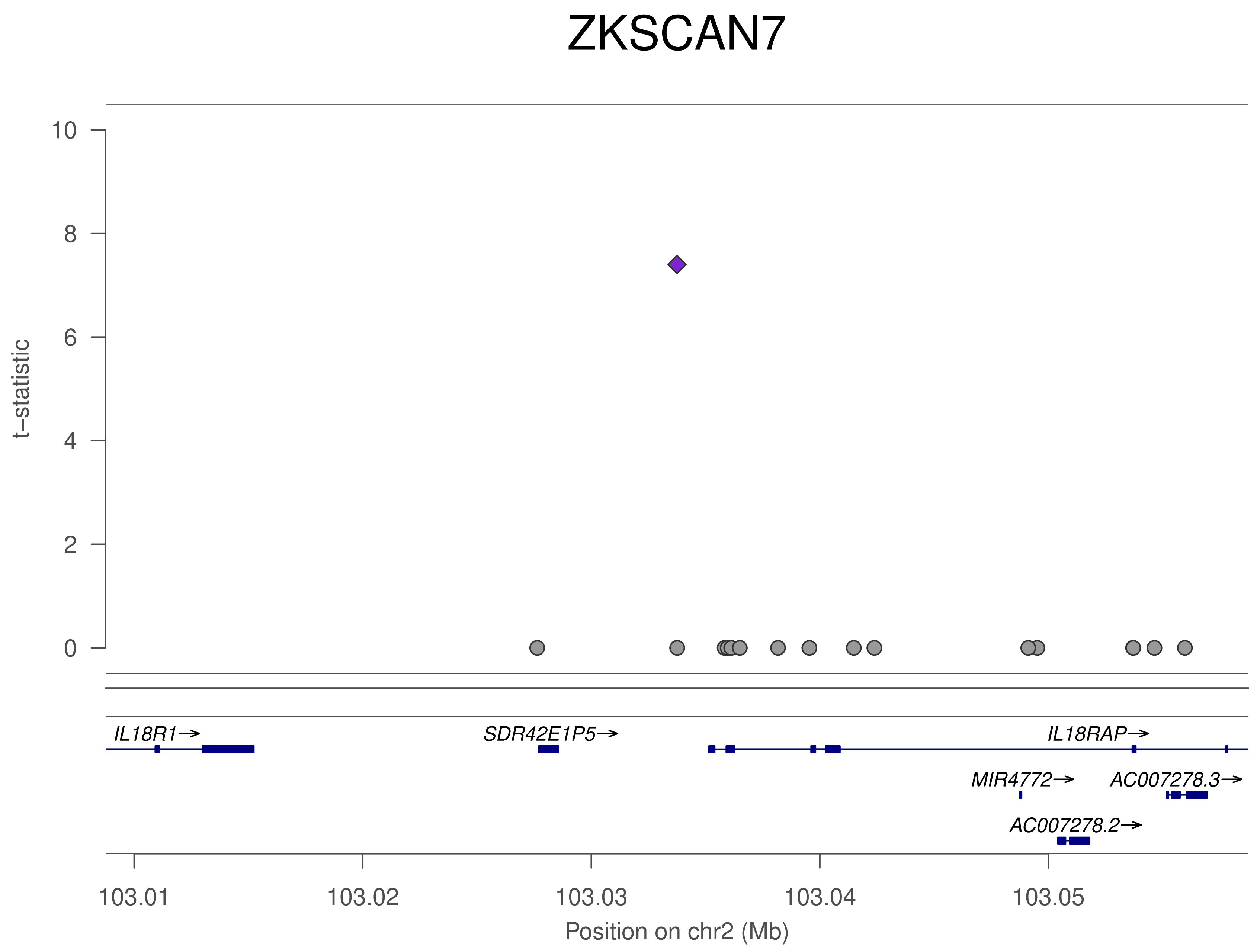
ZKSCAN7
t–statistic
10
8
6
4
2
0
IL18R1→
SDR42E1P5→
IL18RAP→
MIR4772→
AC007278.3→
AC007278.2→
103.01
103.02
103.03
103.04
103.05
Position on chr2 (Mb)

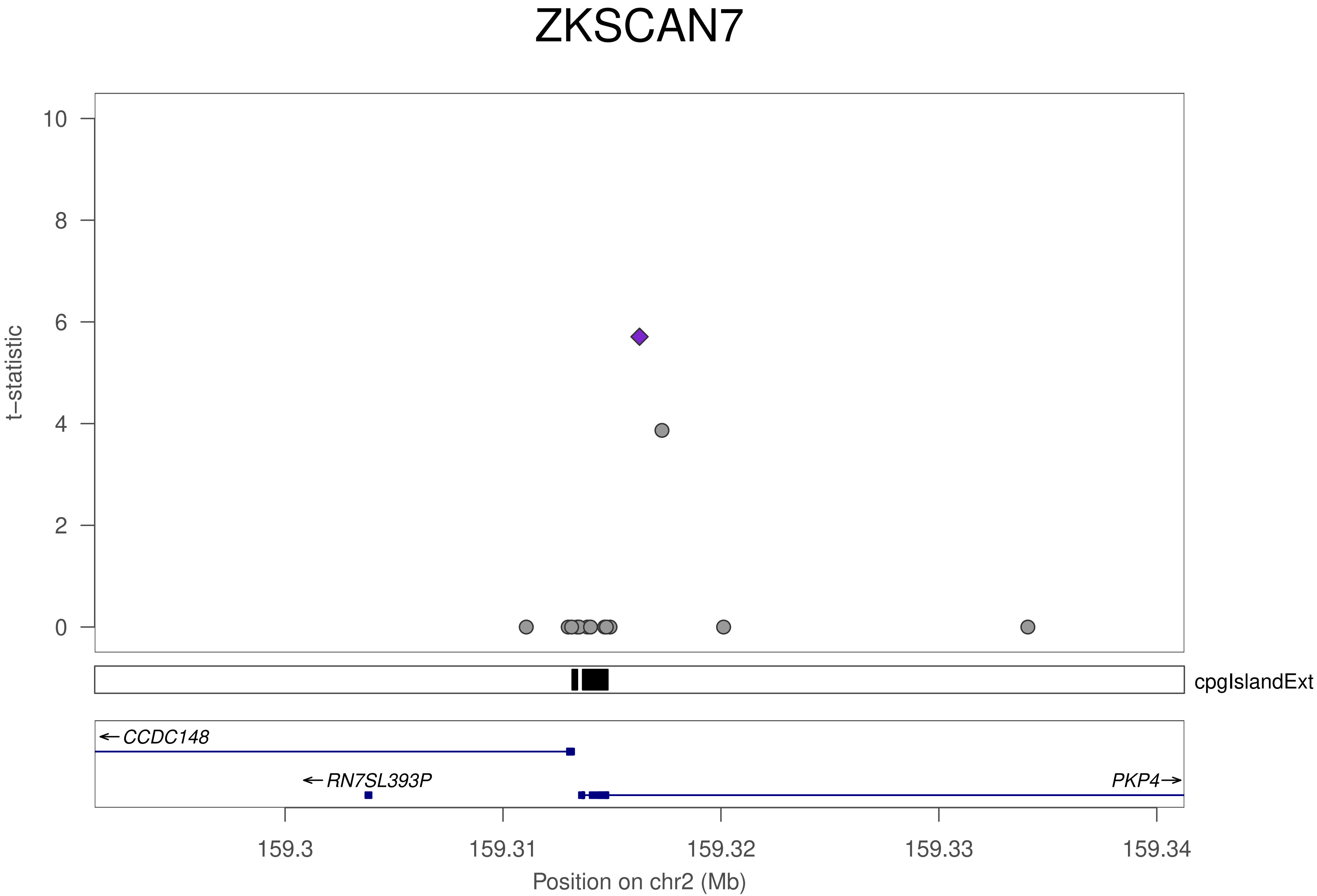

ZKSCAN7
t−statistic
0
2
4
6
8
10
cpgIslandExt
← CCDC148
← RN7SL393P
PKP4 →
159.3
159.31
159.32
159.33
159.34
Position on chr2 (Mb)

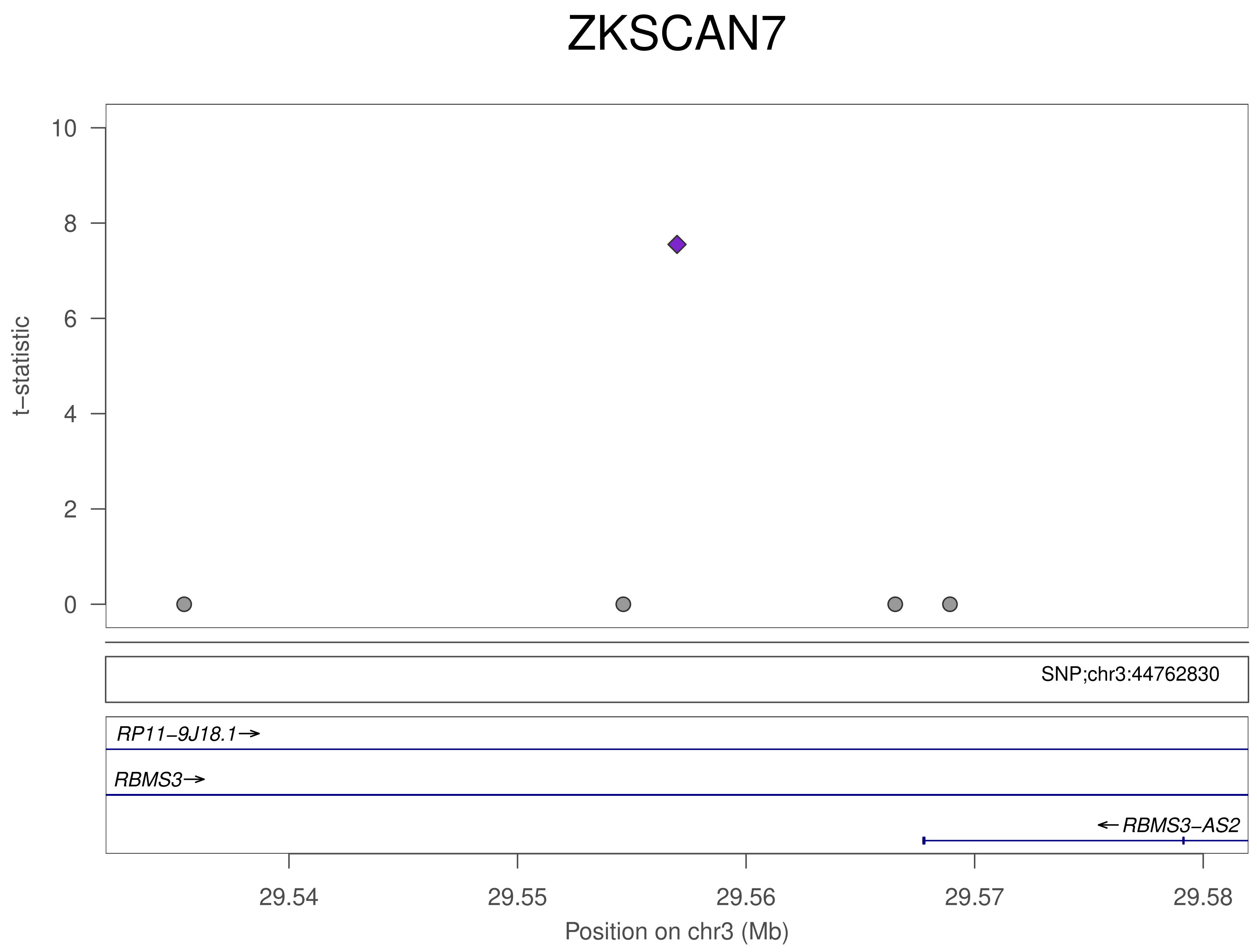
ZKSCAN7
t−statistic
10
8
6
4
2
0
SNP;chr3:44762830
RP11−9J18.1→
RBMS3→
←RBMS3−AS2
29.54
29.55
29.56
29.57
29.58
Position on chr3 (Mb)

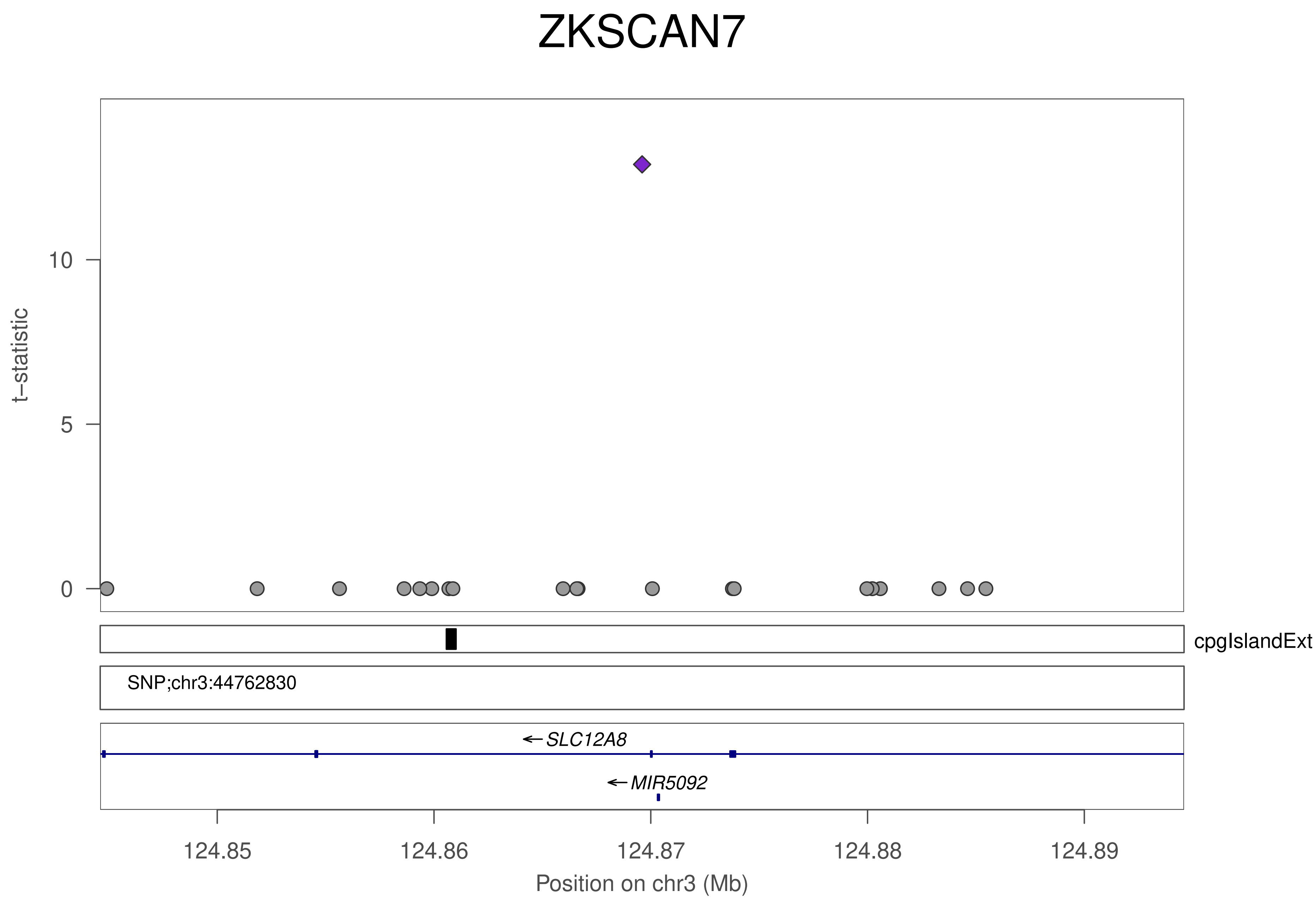

ZKSCAN7
t−statistic
10
5
0
cpgIslandExt
SNP;chr3:44762830
SLC12A8
MIR5092
124.85
124.86
124.87
124.88
124.89
Position on chr3 (Mb)

### ZKSCAN7

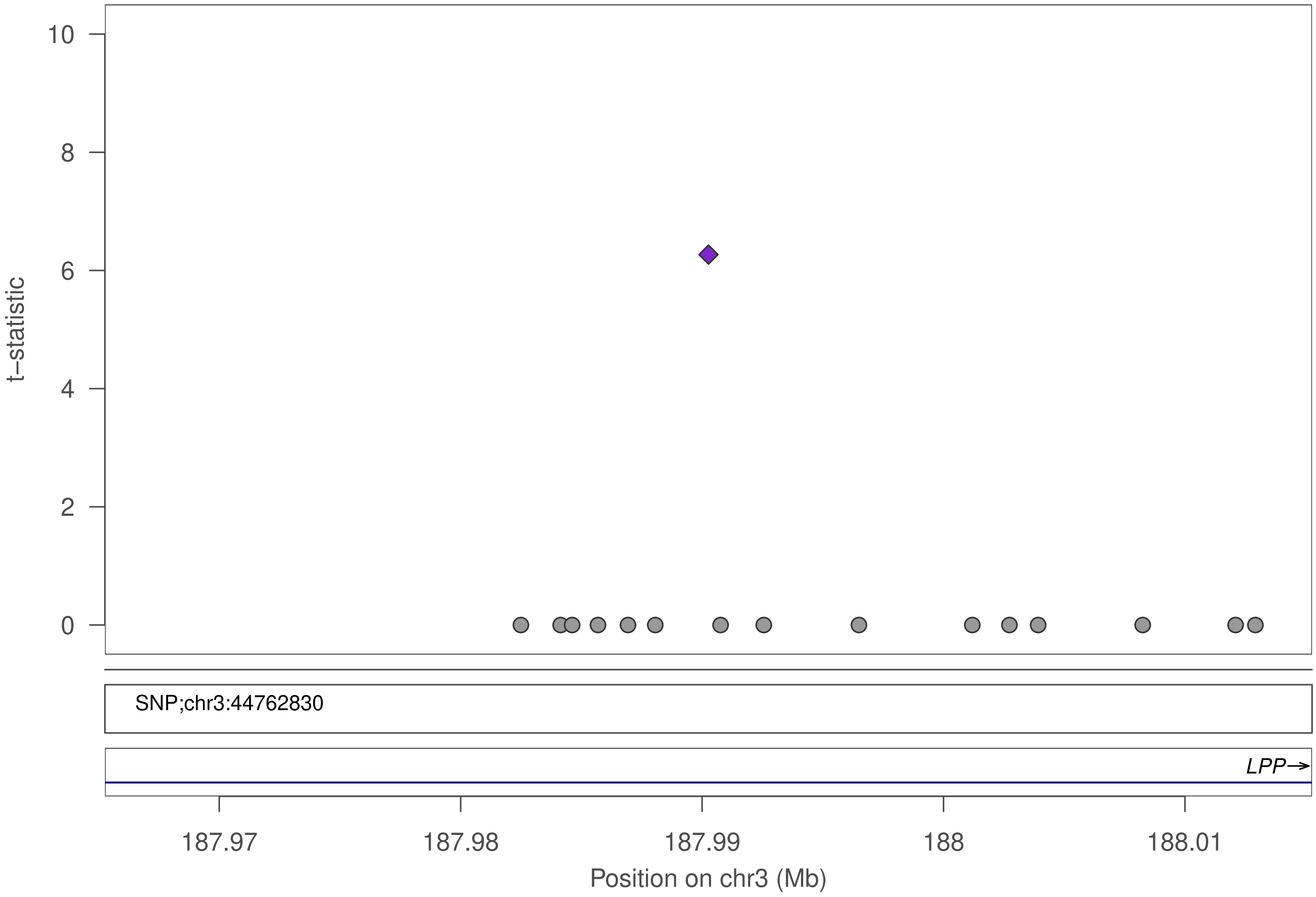

10
8
6
4
2
0
t–statistic
SNP;chr3:44762830
LPP→
187.97
187.98
187.99
188
188.01
Position on chr3 (Mb)

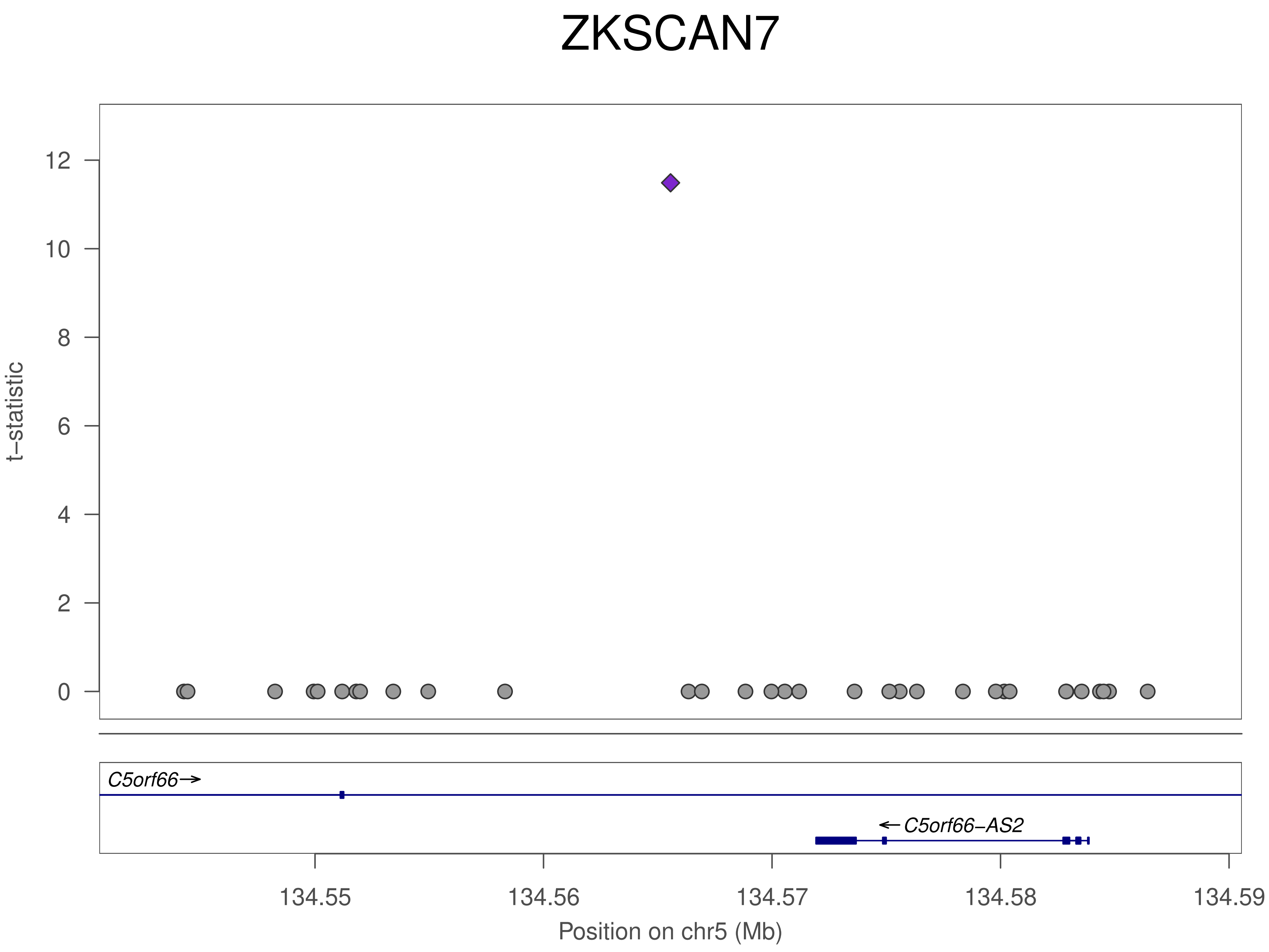

ZKSCAN7
t−statistic
12
10
8
6
4
2
0
C5orf66→
← C5orf66−AS2
134.55
134.56
134.57
134.58
134.59
Position on chr5 (Mb)

### ZKSCAN7

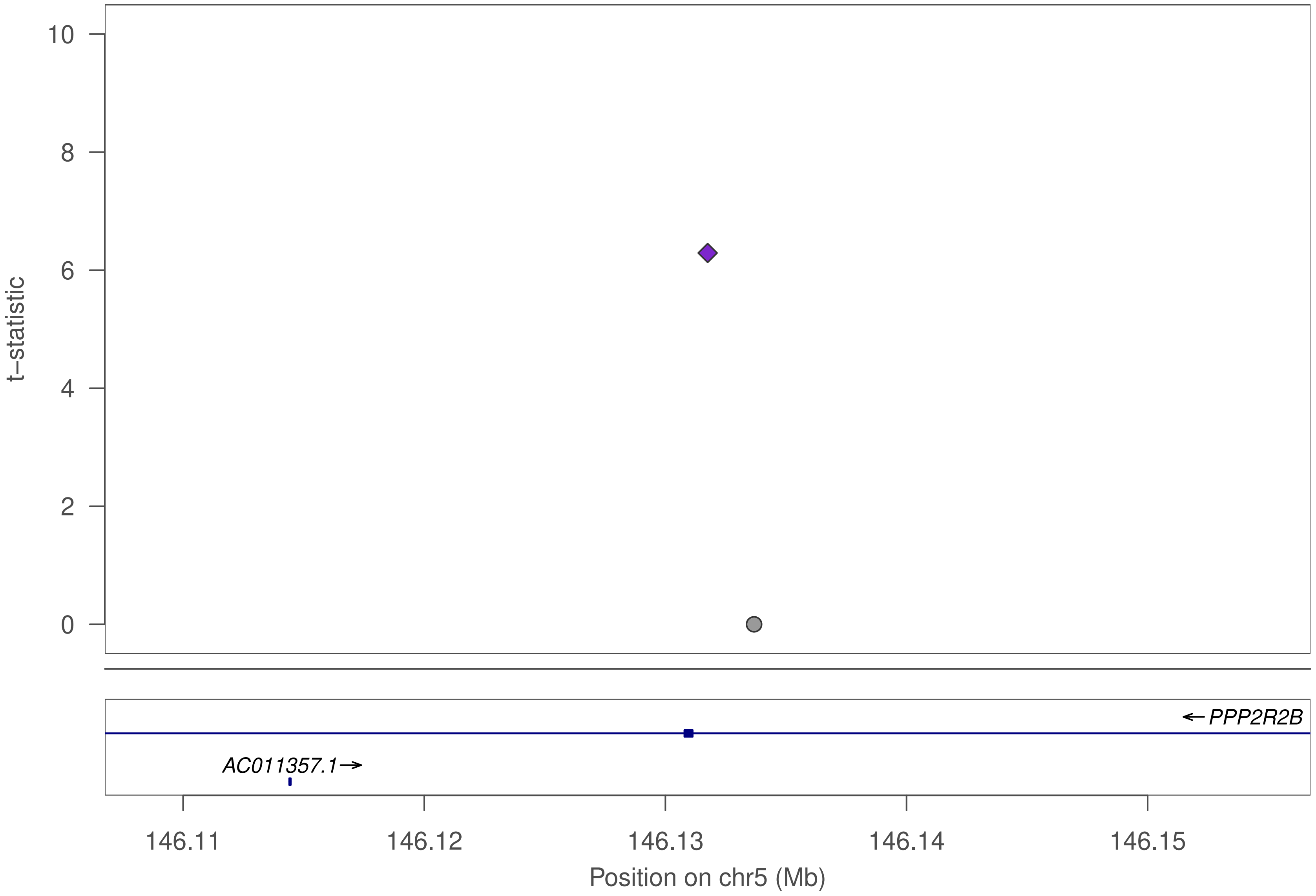

10
8
6
4
2
0
t–statistic
← PPP2R2B
AC011357.1→
146.11
146.12
146.13
146.14
146.15
Position on chr5 (Mb)

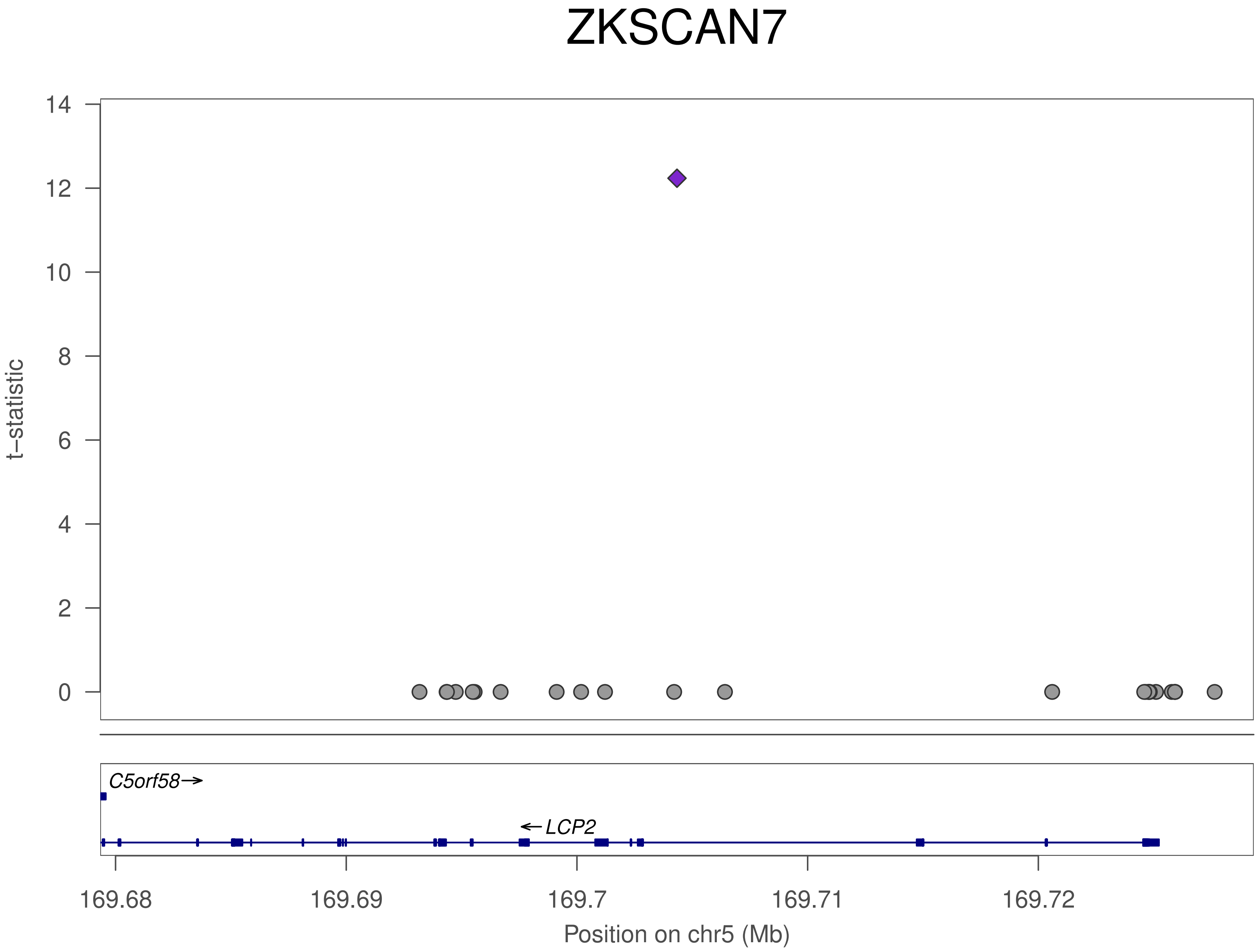
ZKSCAN7
t–statistic
14
12
10
8
6
4
2
0
C5orf58→
←LCP2
169.68
169.69
169.7
169.71
169.72
Position on chr5 (Mb)

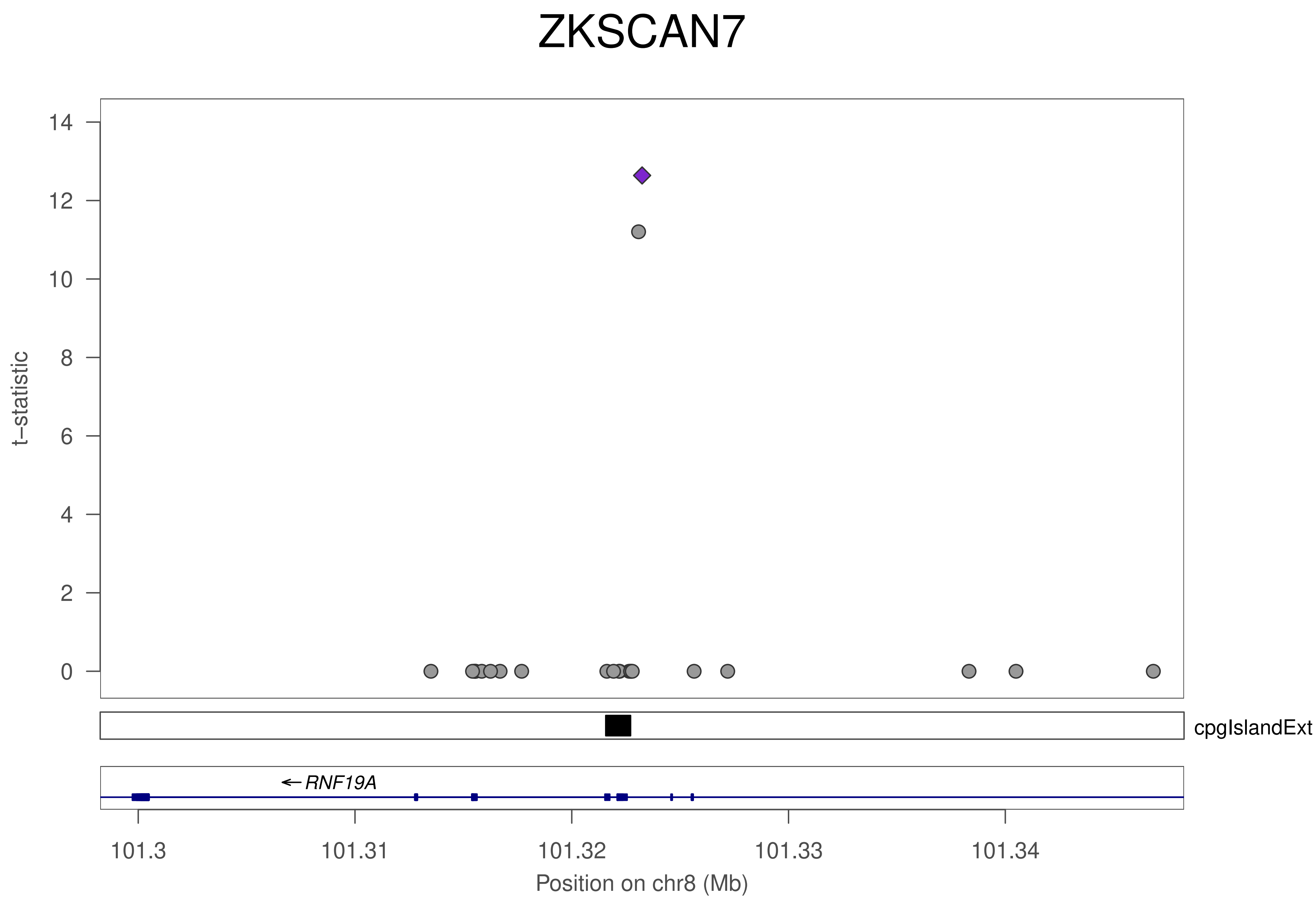

ZKSCAN7
t−statistic
0
2
4
6
8
10
12
14
cpgIslandExt
← RNF19A
101.3
101.31
101.32
101.33
101.34
Position on chr8 (Mb)

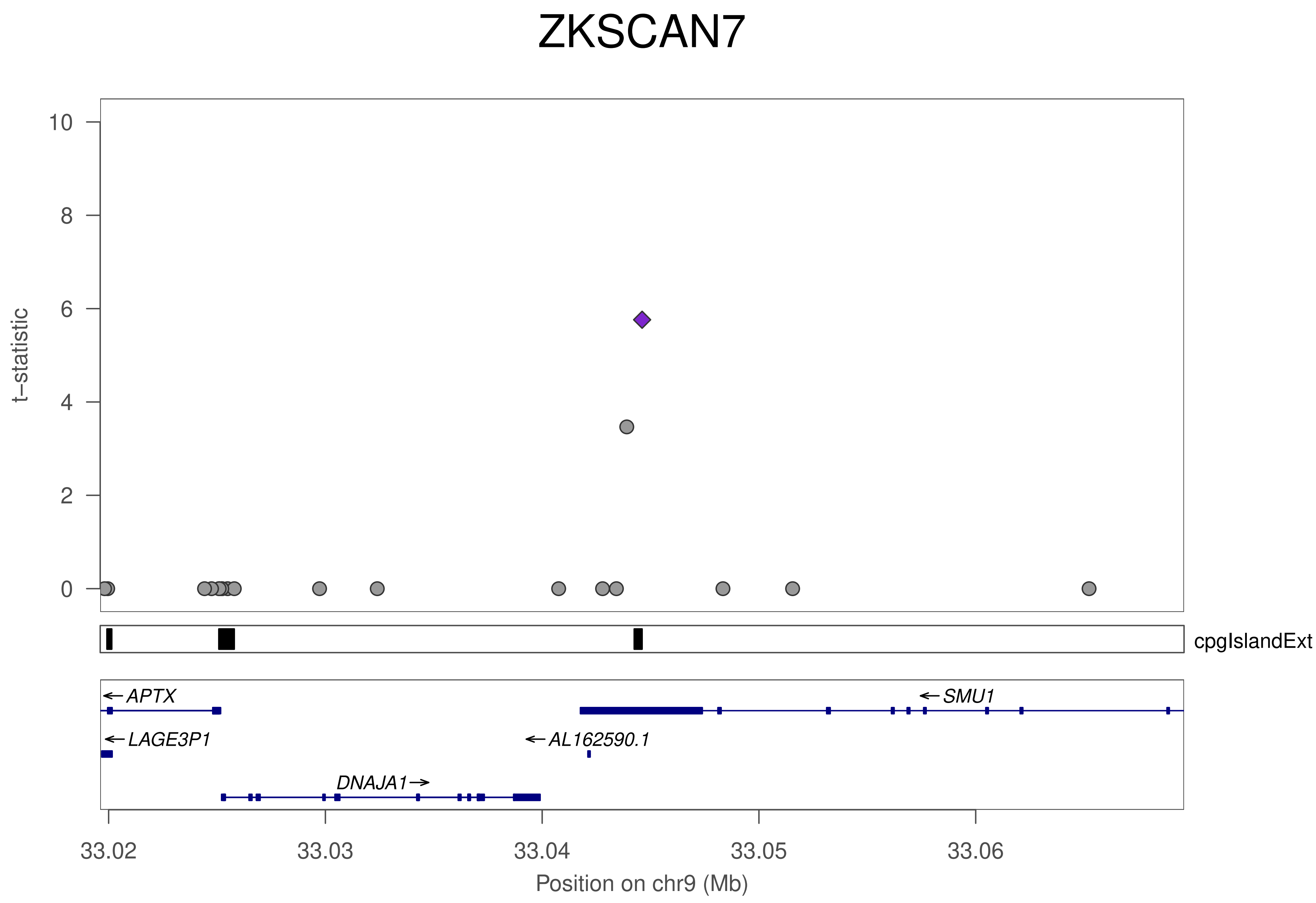
ZKSCAN7
t–statistic
10
8
6
4
2
0
cpgIslandExt
APTX
SMU1
LAGE3P1
AL162590.1
DNAJA1
33.02
33.03
33.04
33.05
33.06
Position on chr9 (Mb)

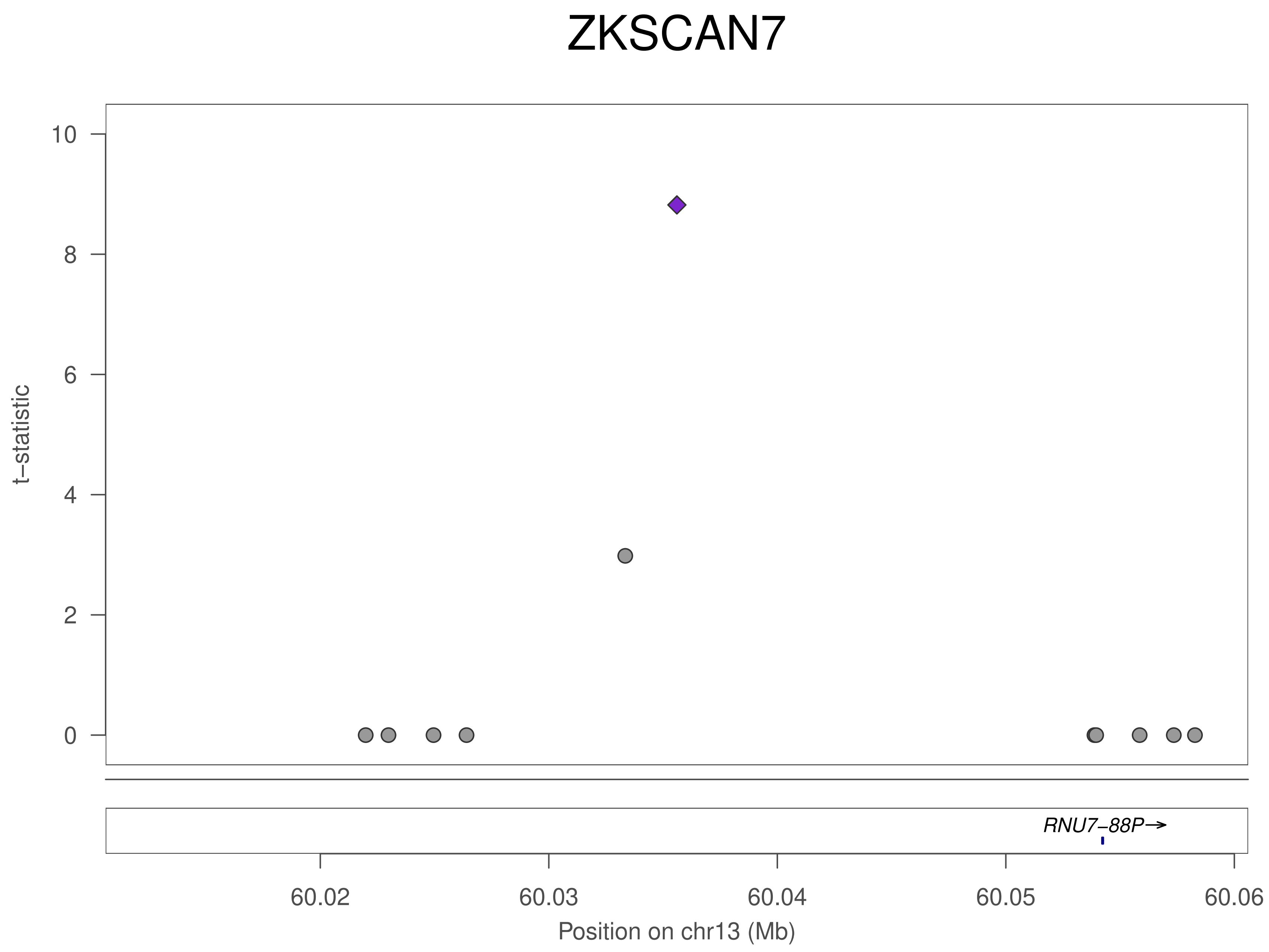

ZKSCAN7
t−statistic
10
8
6
4
2
0
RNU7−88P→
60.02
60.03
60.04
60.05
60.06
Position on chr13 (Mb)

### ZKSCAN7

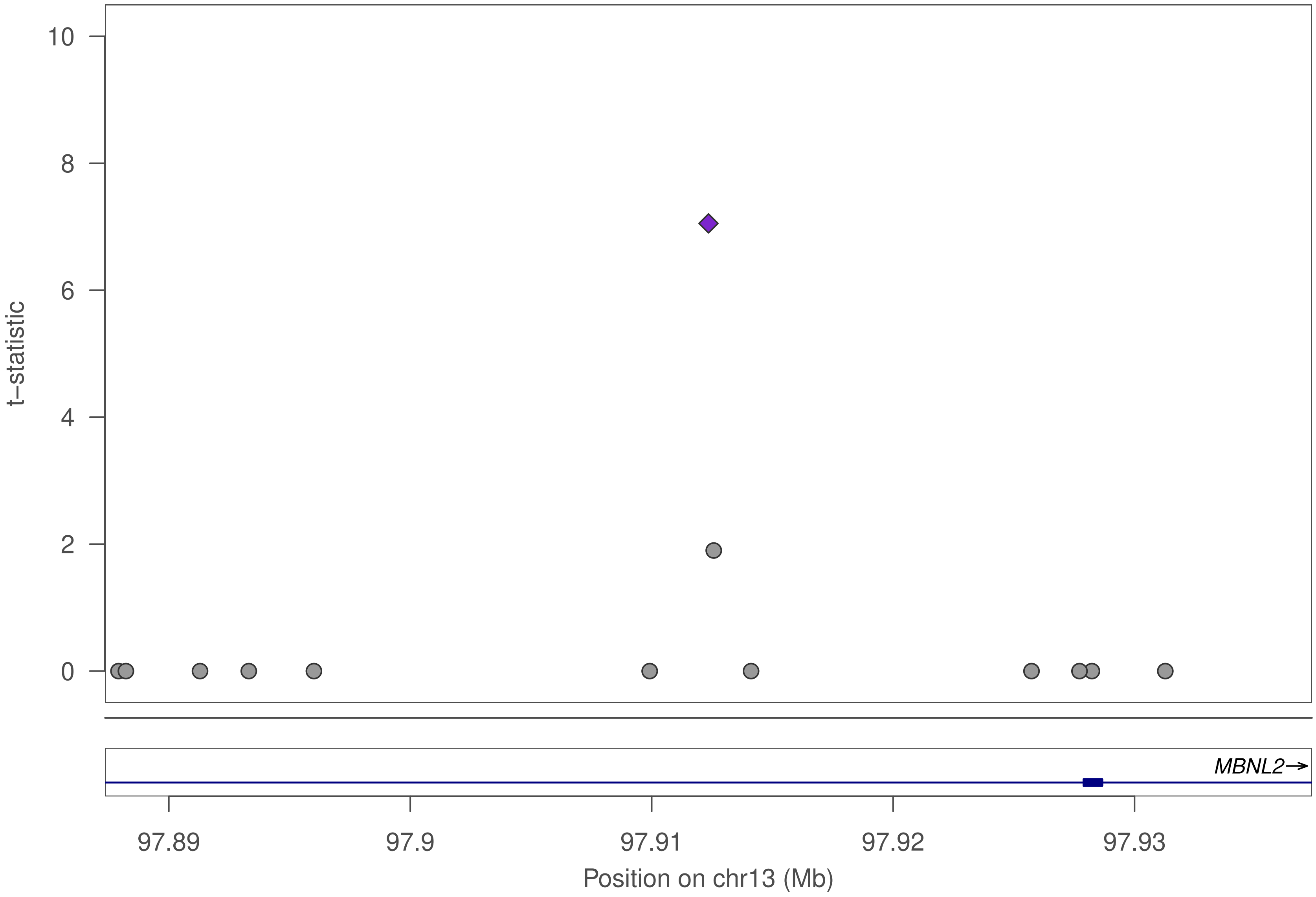

10
8
6
4
2
0
t–statistic
MBNL2→
97.89
97.9
97.91
97.92
97.93
Position on chr13 (Mb)

### ZKSCAN7

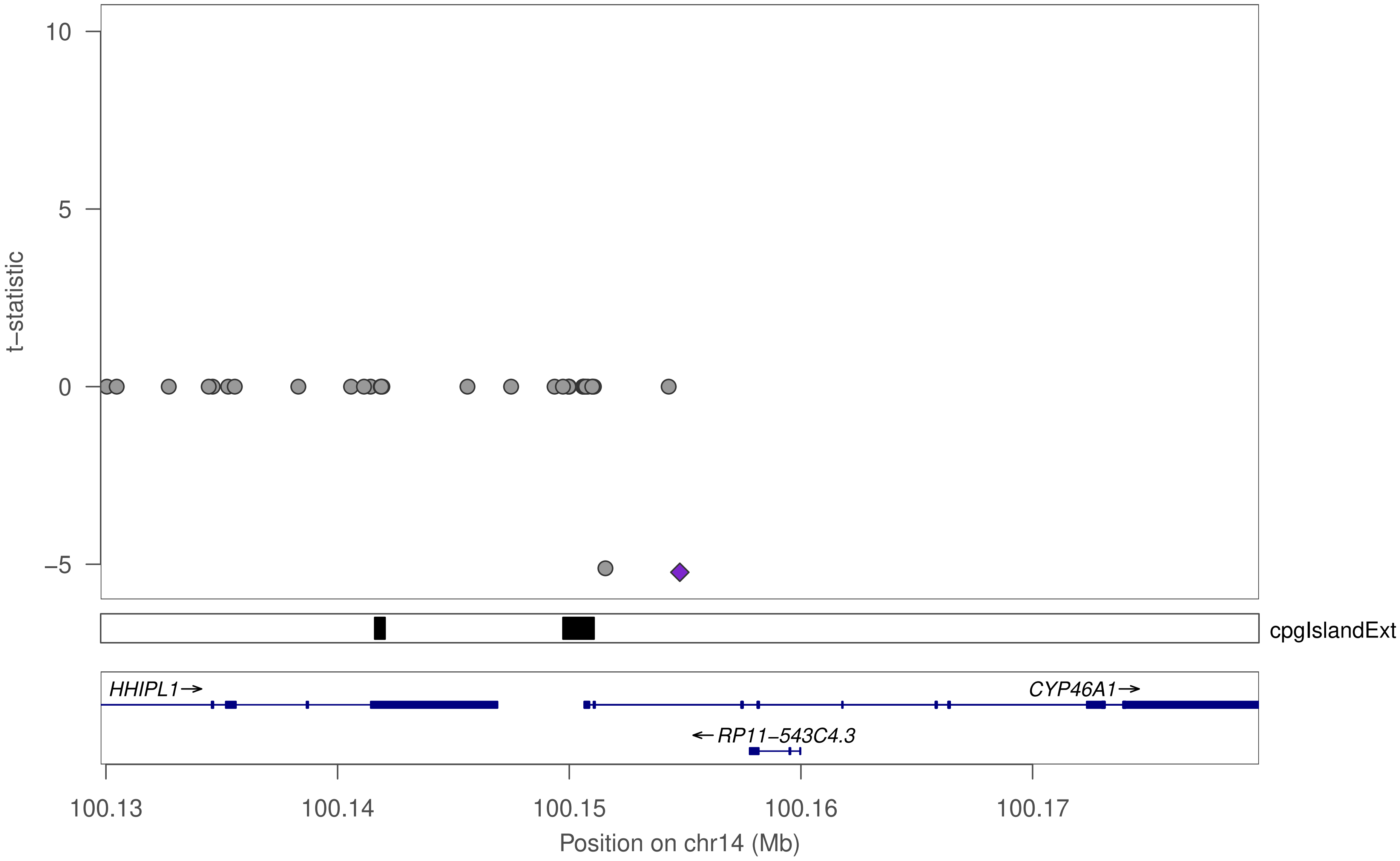

10
5
0
−5
t−statistic
cpgIslandExt
HHIPL1→
CYP46A1→
←RP11−543C4.3
100.13
100.14
100.15
100.16
100.17
Position on chr14 (Mb)

### ZKSCAN7

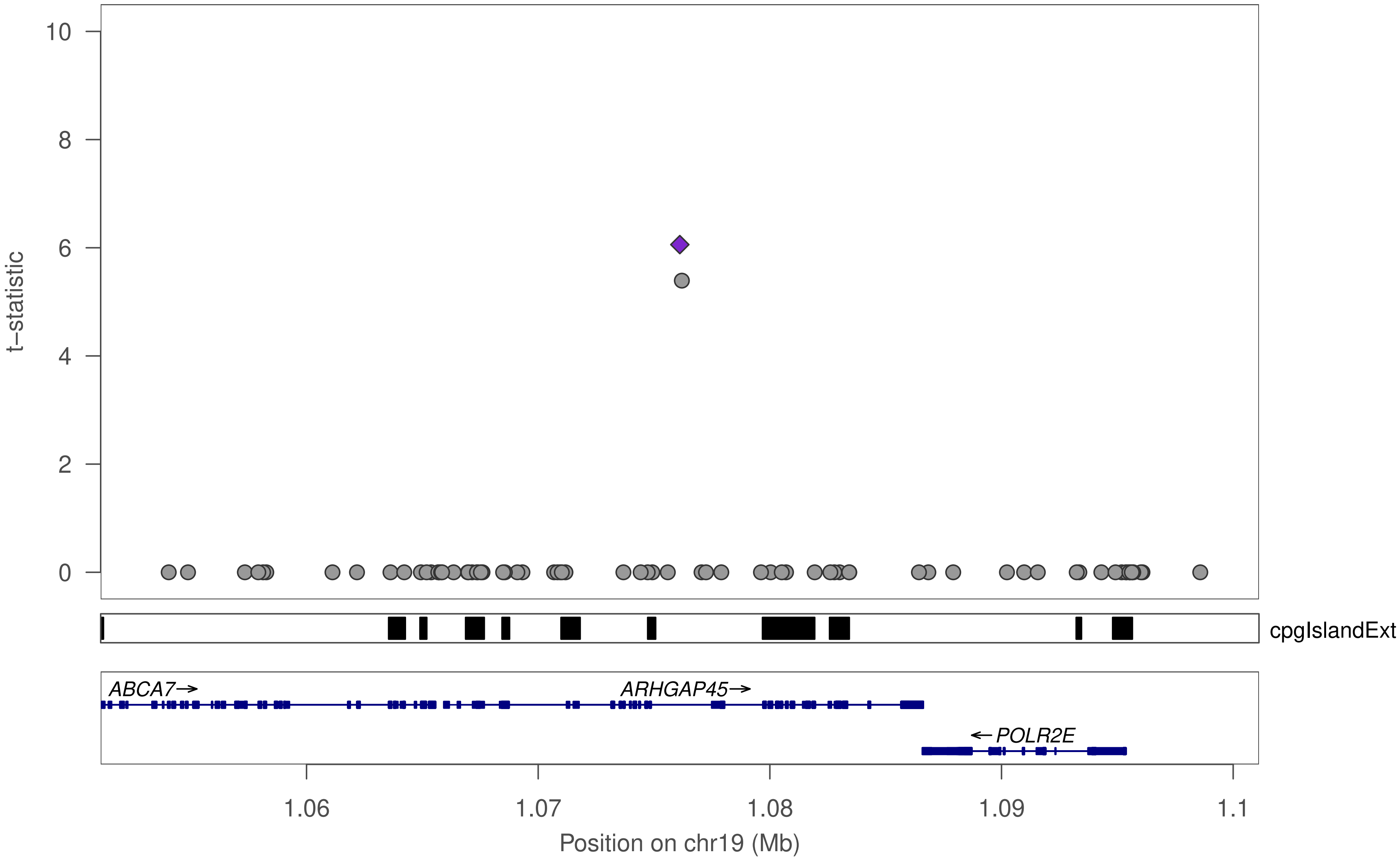

t−statistic
10
8
6
4
2
0
cpgIslandExt
ABCA7→
ARHGAP45→
←POLR2E
1.06
1.07
1.08
1.09
1.1
Position on chr19 (Mb)

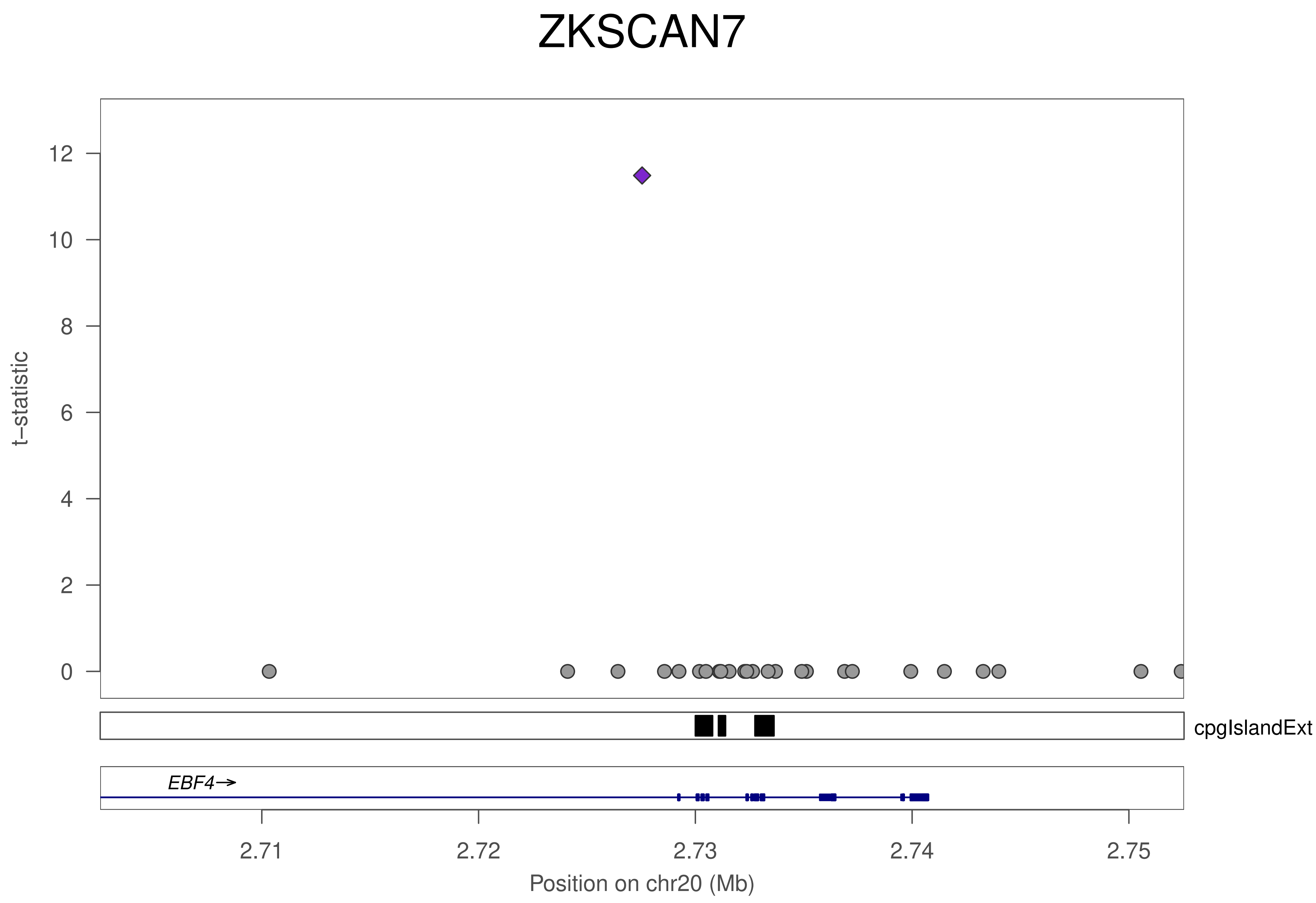

ZKSCAN7
t–statistic
0
2
4
6
8
10
12
cpgIslandExt
EBF4→
2.71
2.72
2.73
2.74
2.75
Position on chr20 (Mb)

### ZNF34

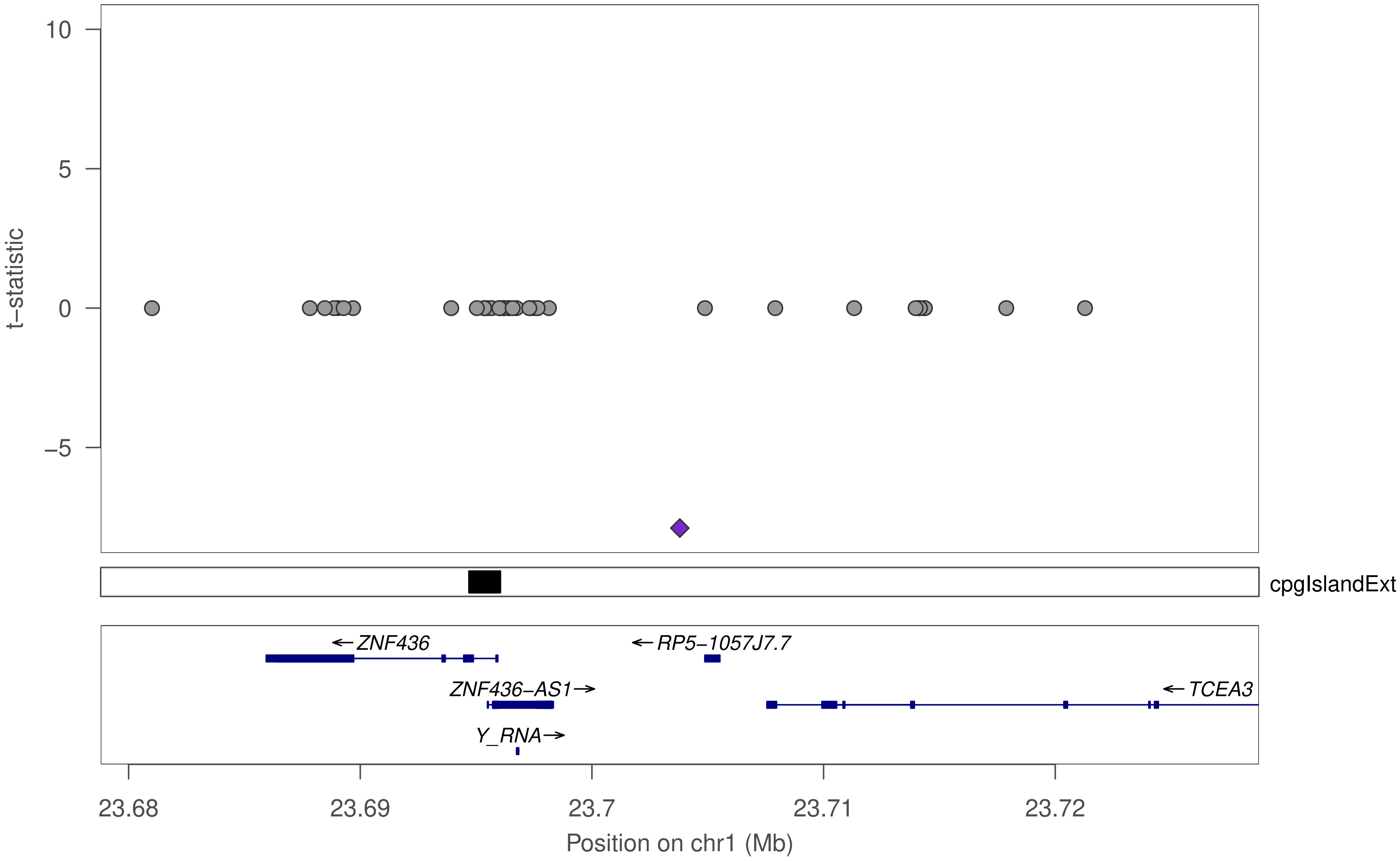

t–statistic
10
5
0
–5
cpgIslandExt
←ZNF436
←RP5–1057J7.7
ZNF436–AS1→
←TCEA3
Y_RNA→
23.68
23.69
23.7
23.71
23.72
Position on chr1 (Mb)

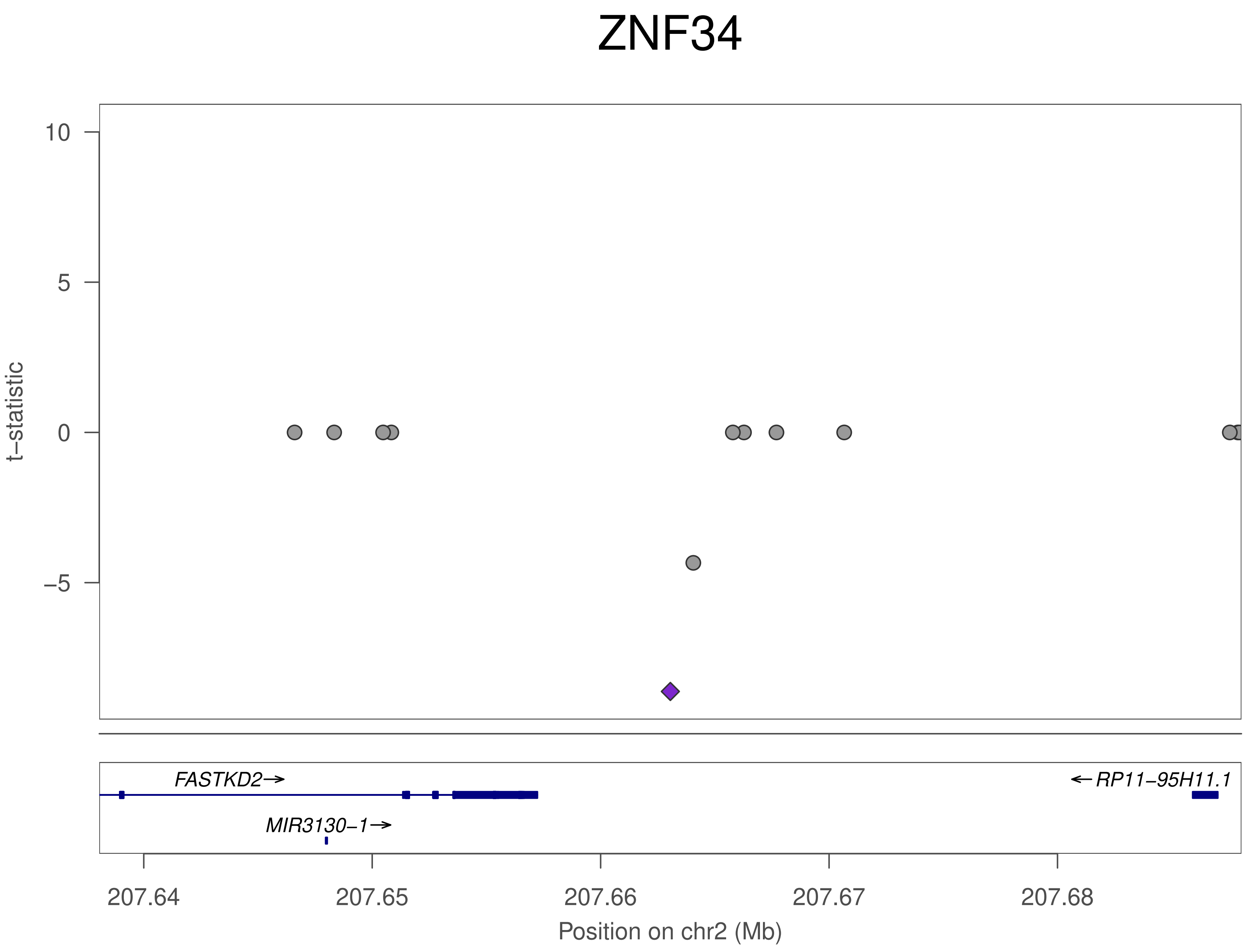
ZNF34
t−statistic
10
5
0
−5
FASTKD2→
←RP11−95H11.1
MIR3130−1→
207.64
207.65
207.66
207.67
207.68
Position on chr2 (Mb)

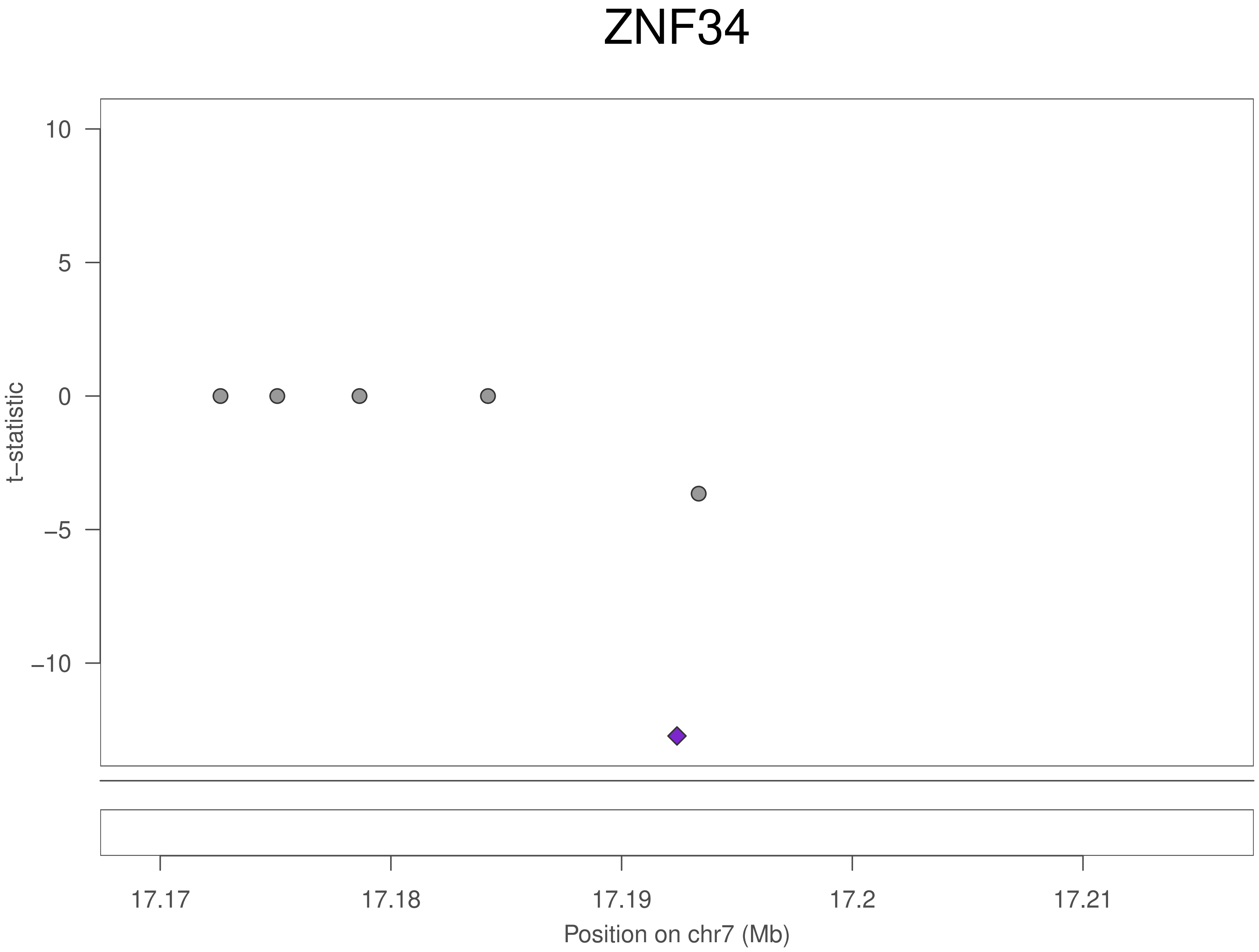

ZNF34
10
5
0
−5
−10
t−statistic
17.17
17.18
17.19
17.2
17.21
Position on chr7 (Mb)

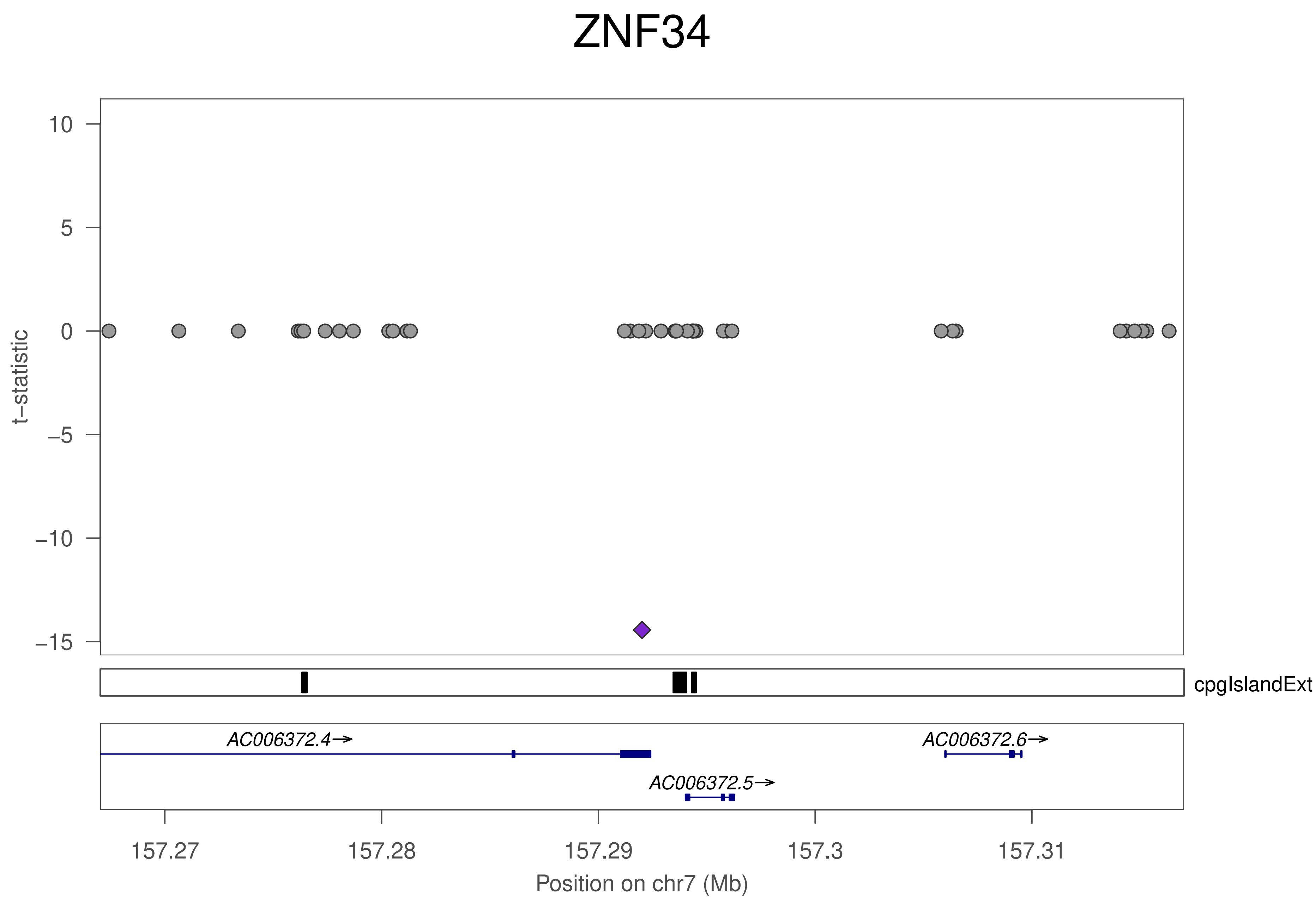
ZNF34
t−statistic
10
5
0
−5
−10
−15
cpgIslandExt
AC006372.4→
AC006372.6→
AC006372.5→
157.27
157.28
157.29
157.3
157.31
Position on chr7 (Mb)

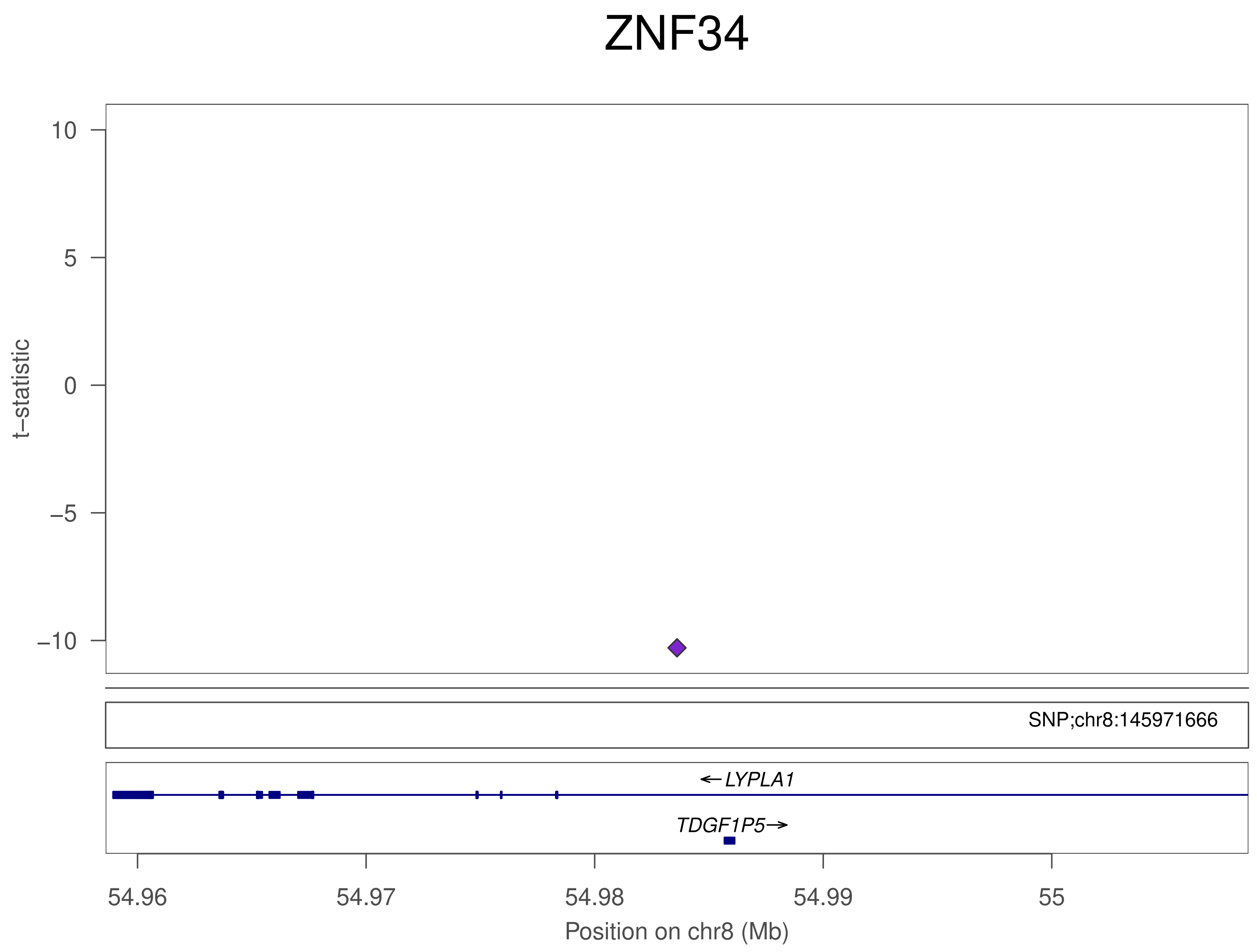

ZNF34
t−statistic
10
5
0
−5
−10
SNP;chr8:145971666
LYPLA1
TDGF1P5
54.96
54.97
54.98
54.99
55
Position on chr8 (Mb)

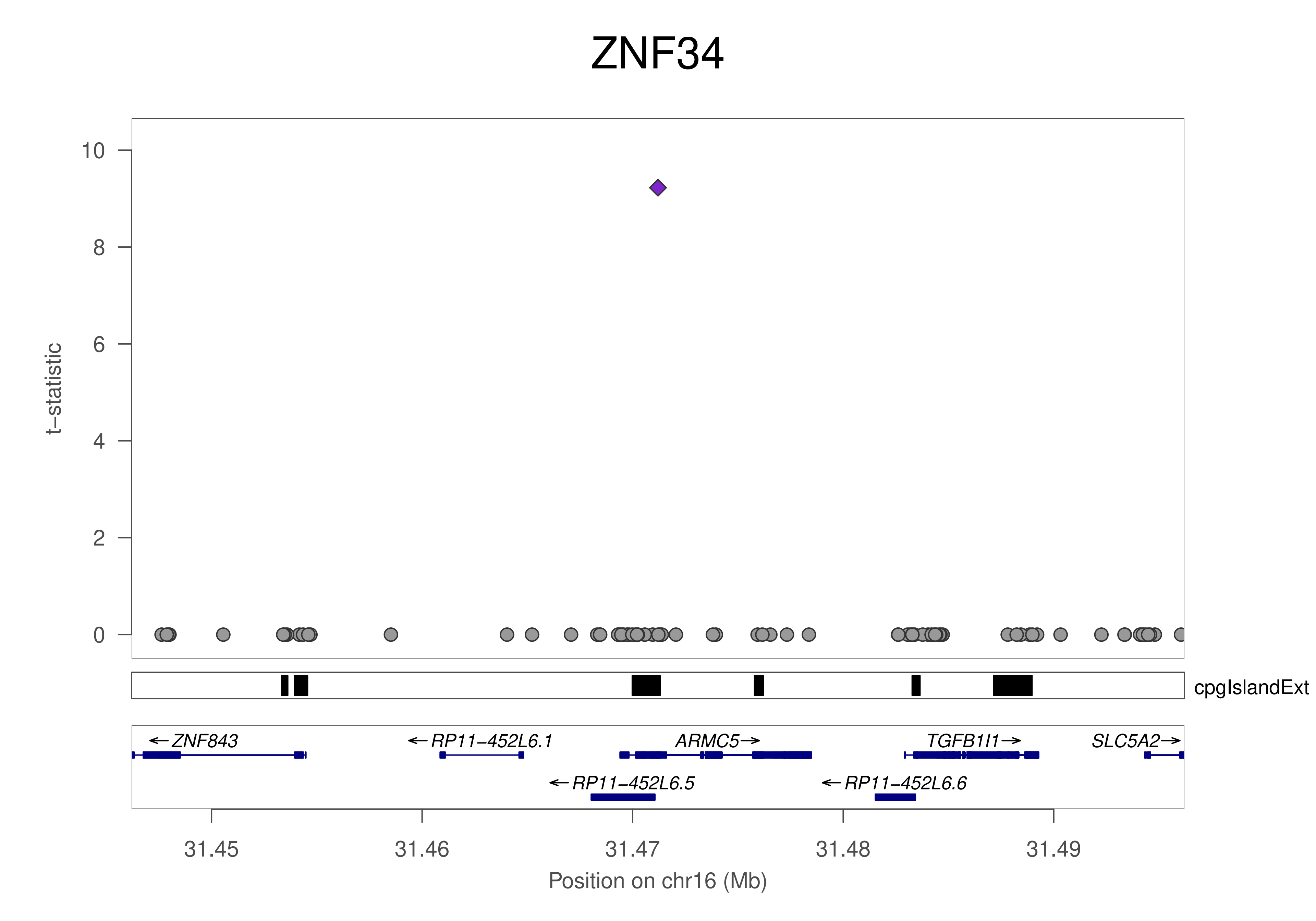
ZNF34
t−statistic
0
2
4
6
8
10
cpgIslandExt
←ZNF843
←RP11−452L6.1
ARMC5→
TGFB1I1→
SLC5A2→
←RP11−452L6.5
←RP11−452L6.6
31.45
31.46
31.47
31.48
31.49
Position on chr16 (Mb)

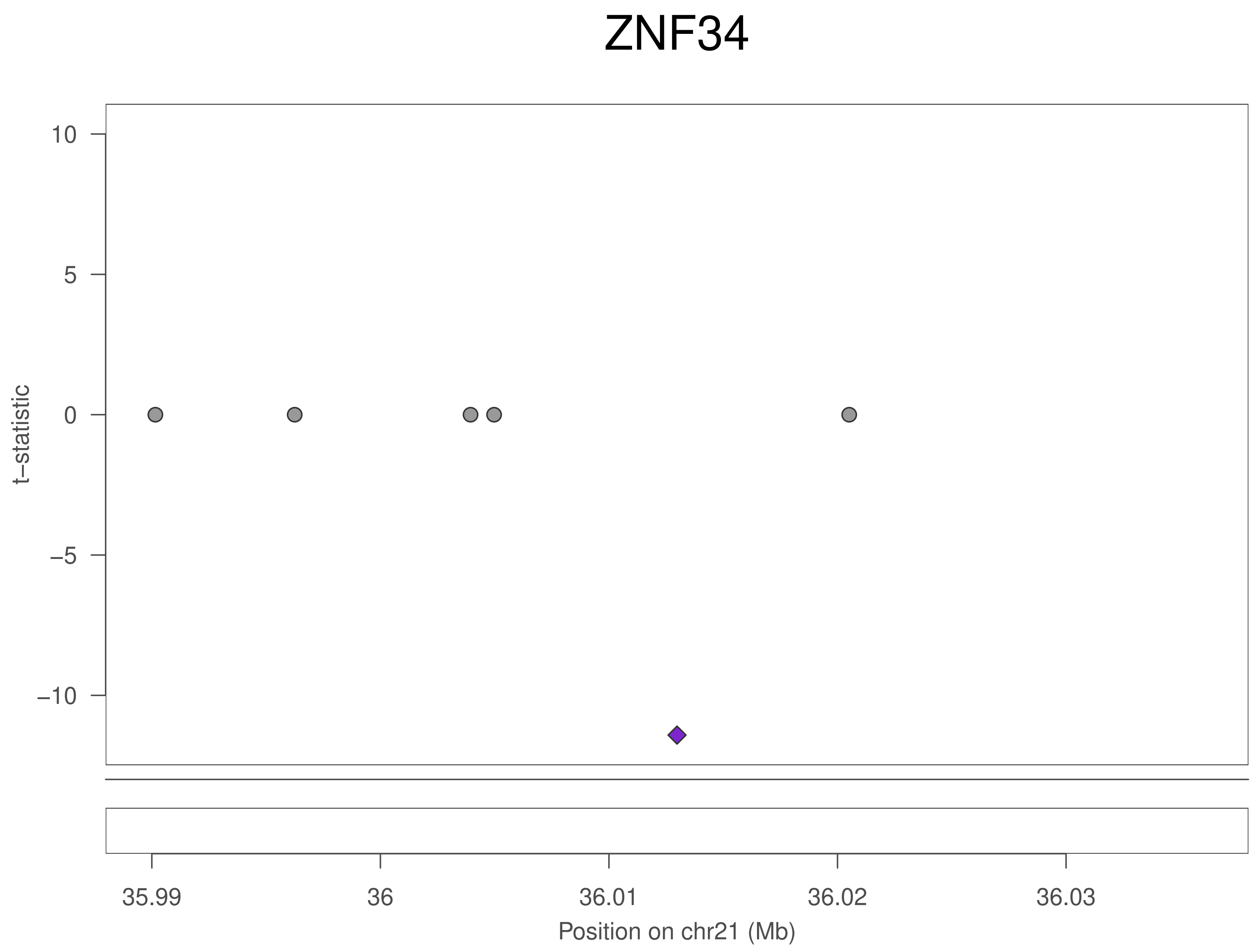

ZNF34
t−statistic
10
5
0
−5
−10
35.99
36
36.01
36.02
36.03
Position on chr21 (Mb)

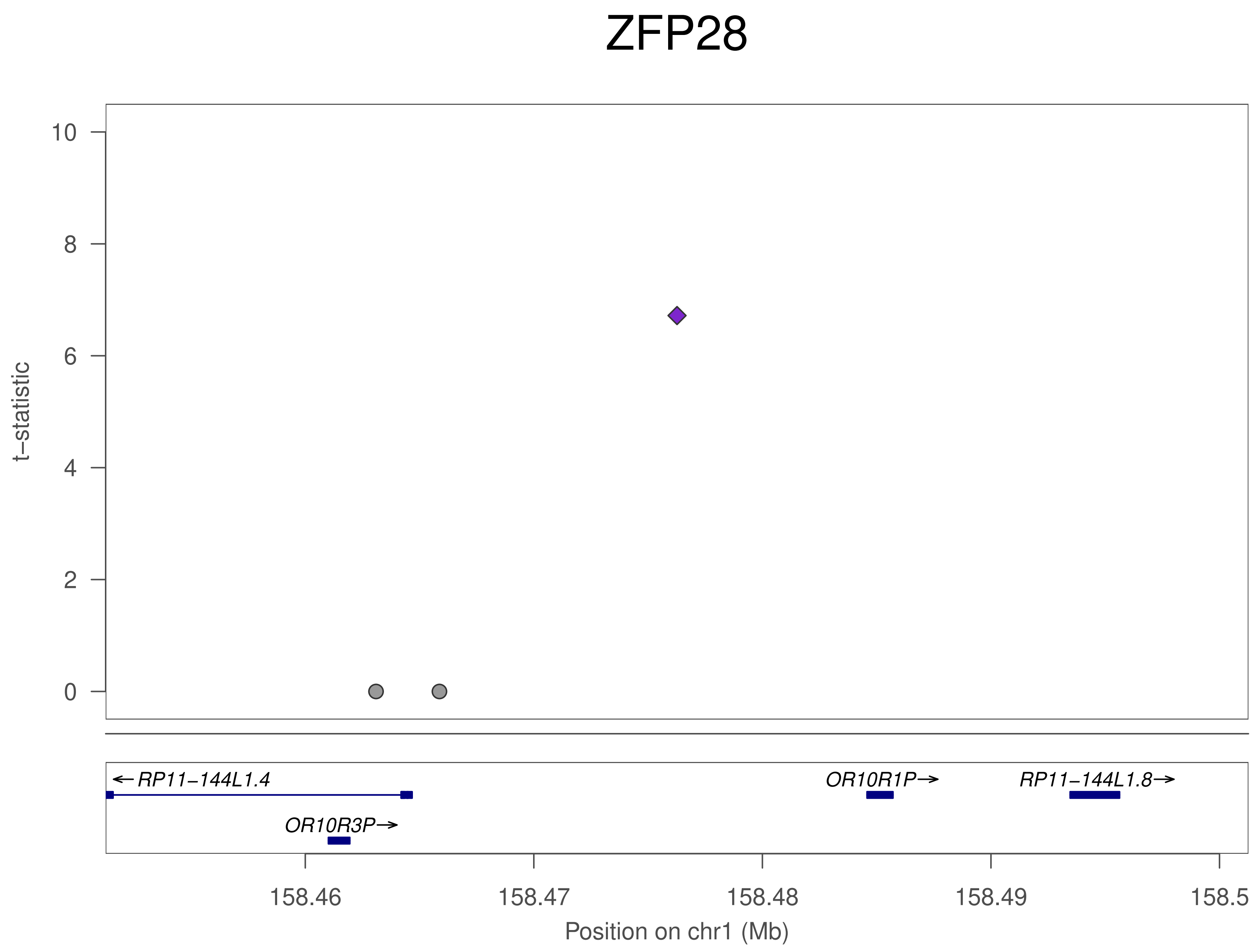
ZFP28
t−statistic
10
8
6
4
2
0
←RP11−144L1.4
OR10R3P→
OR10R1P→
RP11−144L1.8→
158.46
158.47
158.48
158.49
158.5
Position on chr1 (Mb)

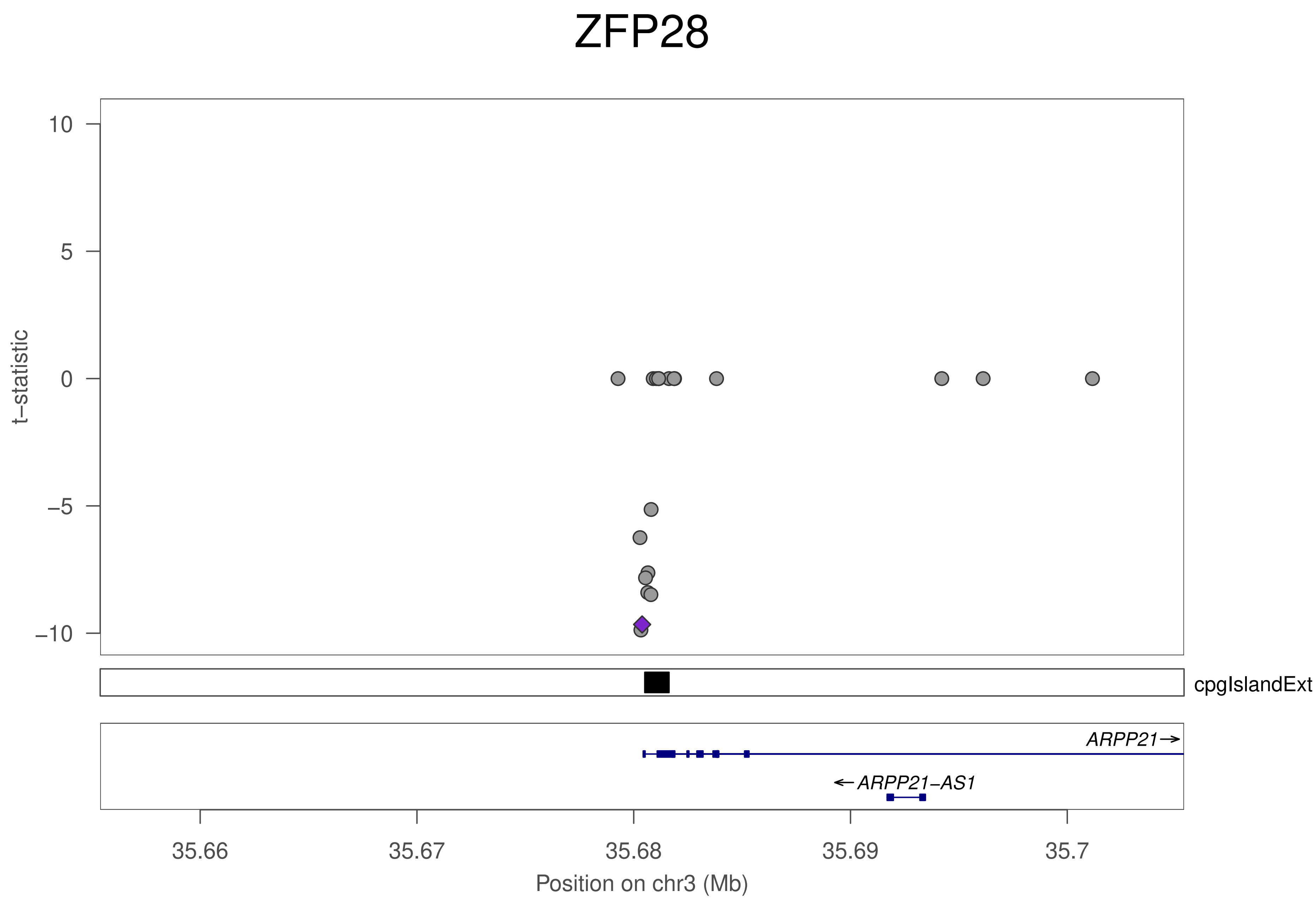

ZFP28
10
5
0
−5
−10
t−statistic
cpgIslandExt
ARPP21→
←ARPP21−AS1
35.66
35.67
35.68
35.69
35.7
Position on chr3 (Mb)

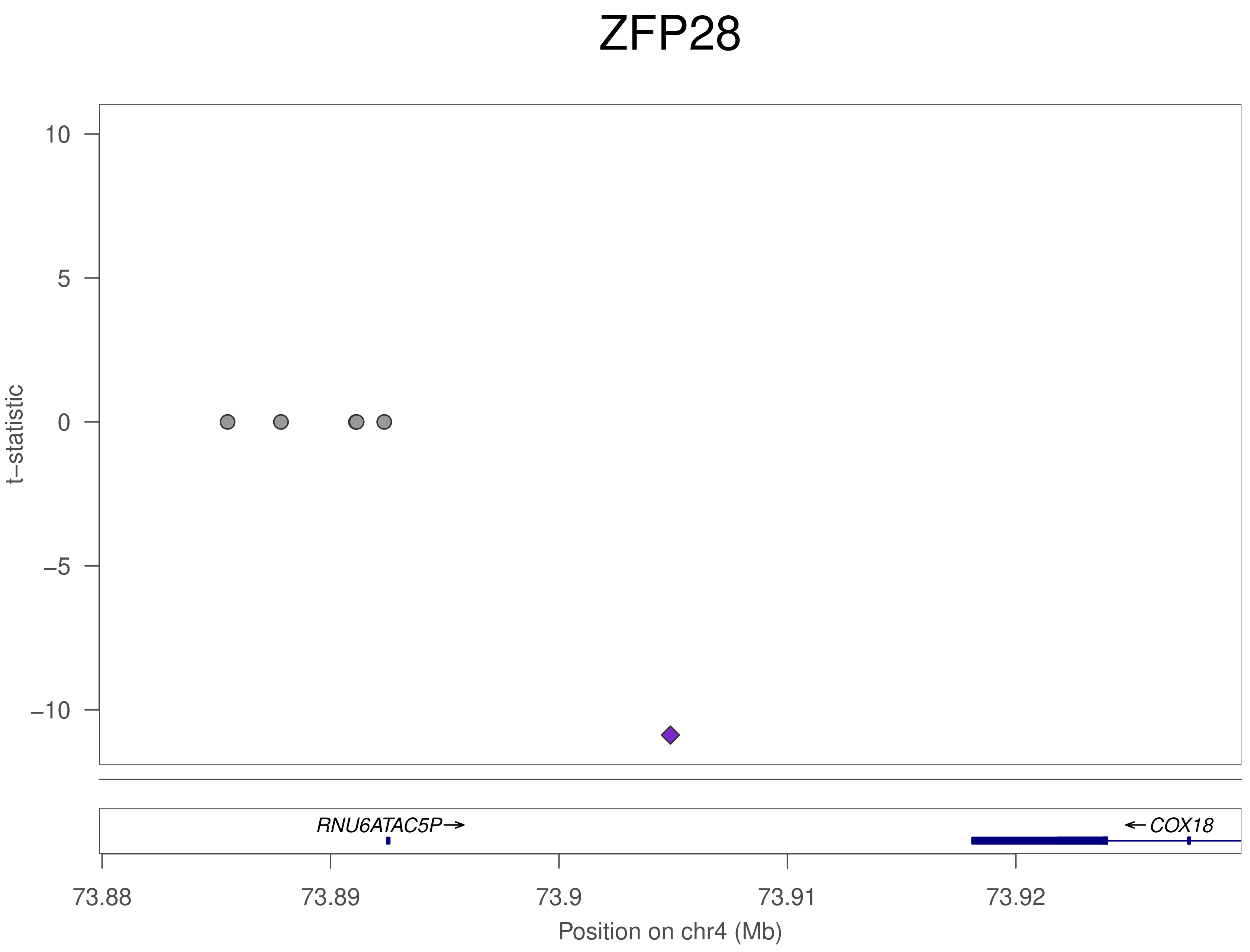
ZFP28
t–statistic
10
5
0
–5
–10
RNU6ATAC5P→
←COX18
73.88
73.89
73.9
73.91
73.92
Position on chr4 (Mb)

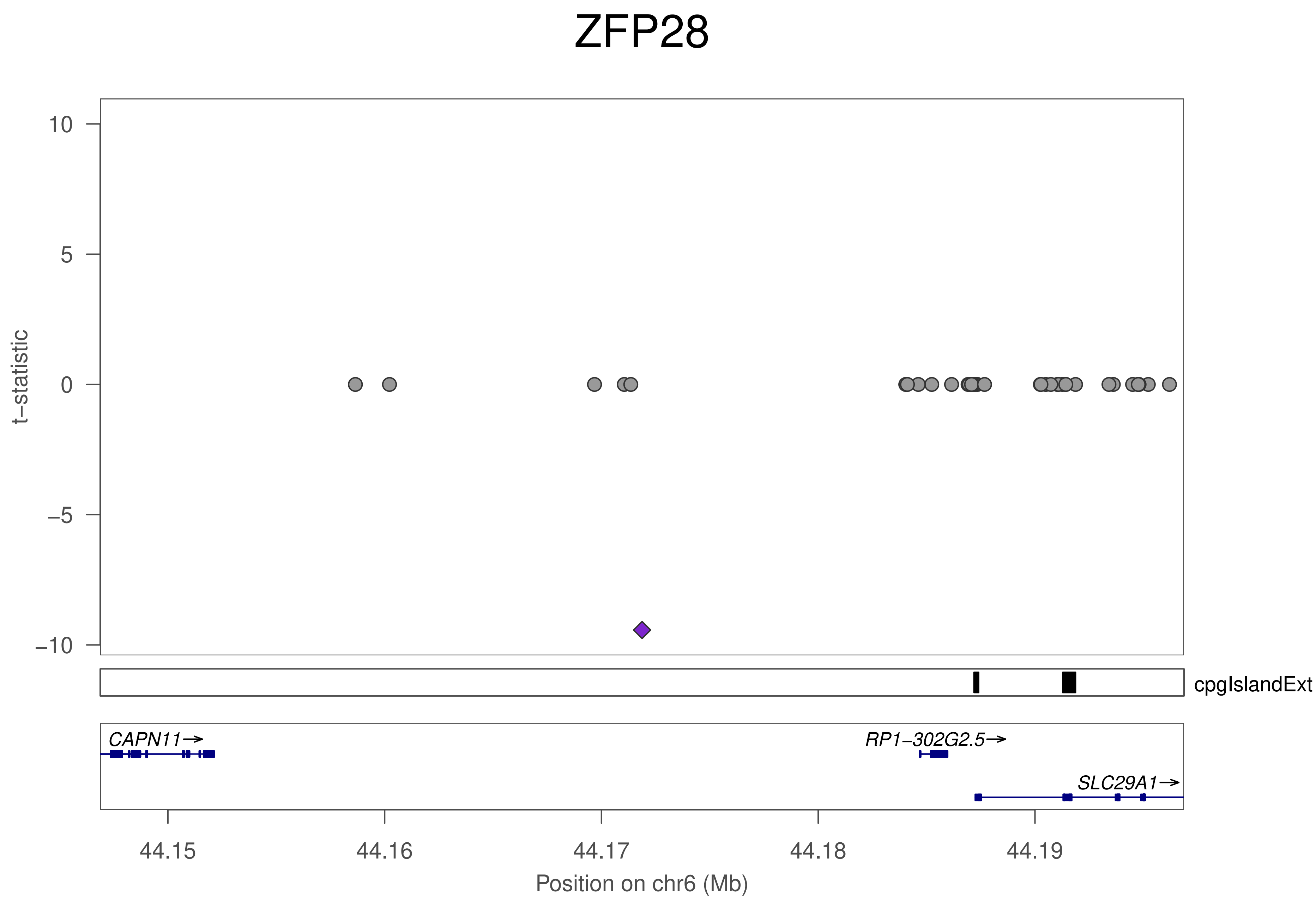
ZFP28
t−statistic
10
5
0
−5
−10
cpgIslandExt
CAPN11→
RP1−302G2.5→
SLC29A1→
44.15
44.16
44.17
44.18
44.19
Position on chr6 (Mb)

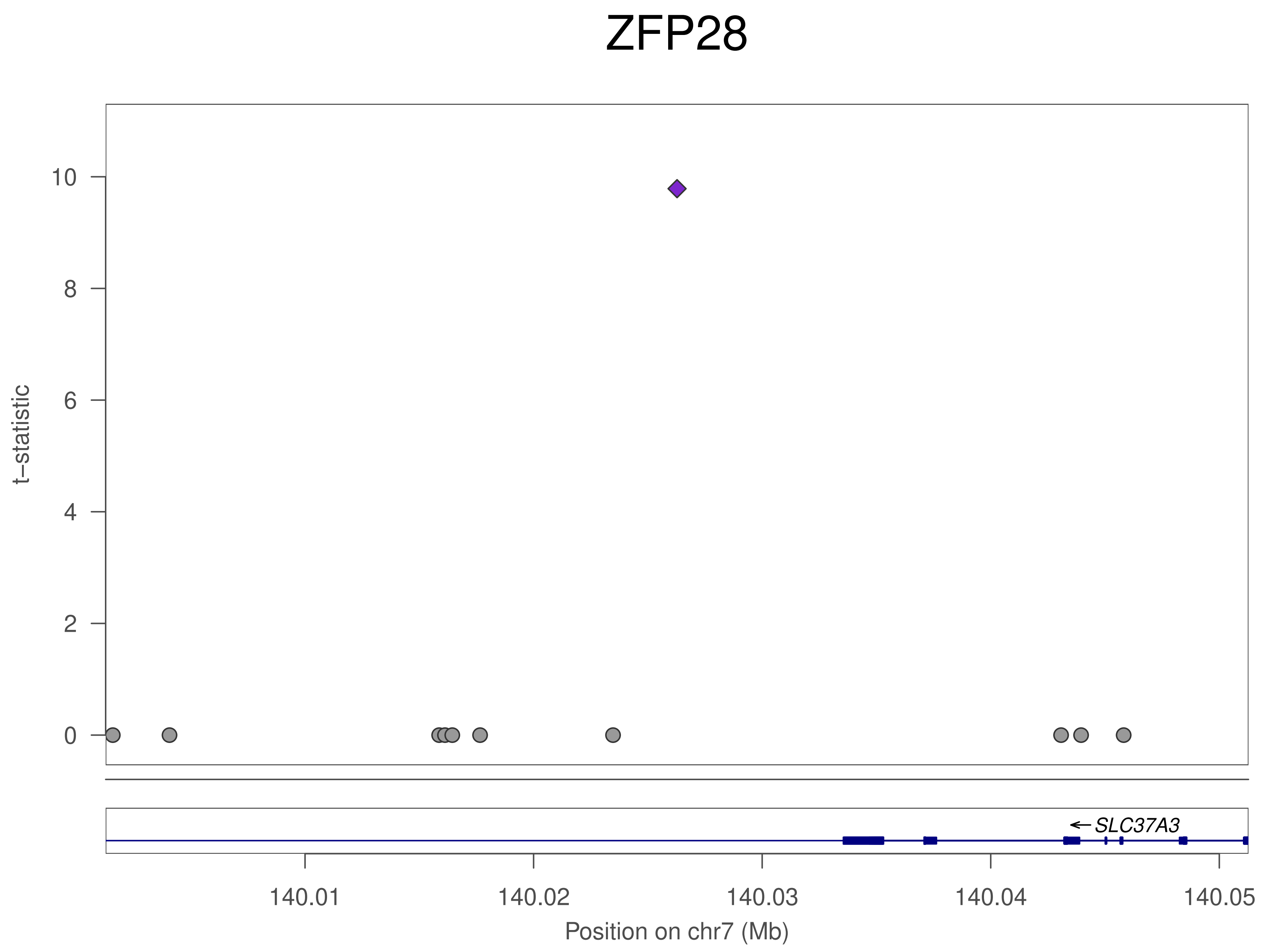

ZFP28
t–statistic
0
2
4
6
8
10
SLC37A3
140.01
140.02
140.03
140.04
140.05
Position on chr7 (Mb)

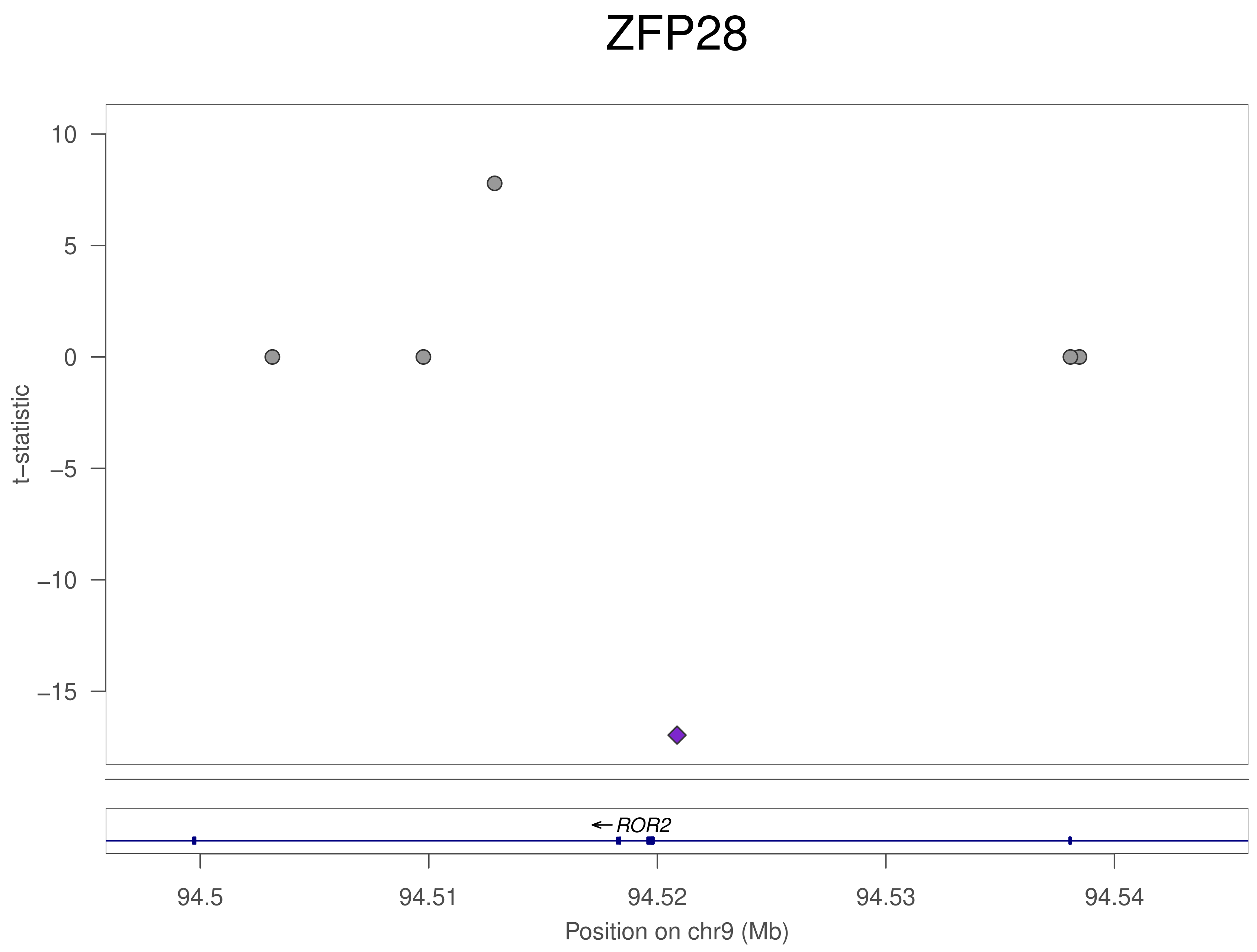

ZFP28
10
5
0
−5
−10
−15
t−statistic
ROR2
94.5
94.51
94.52
94.53
94.54
Position on chr9 (Mb)

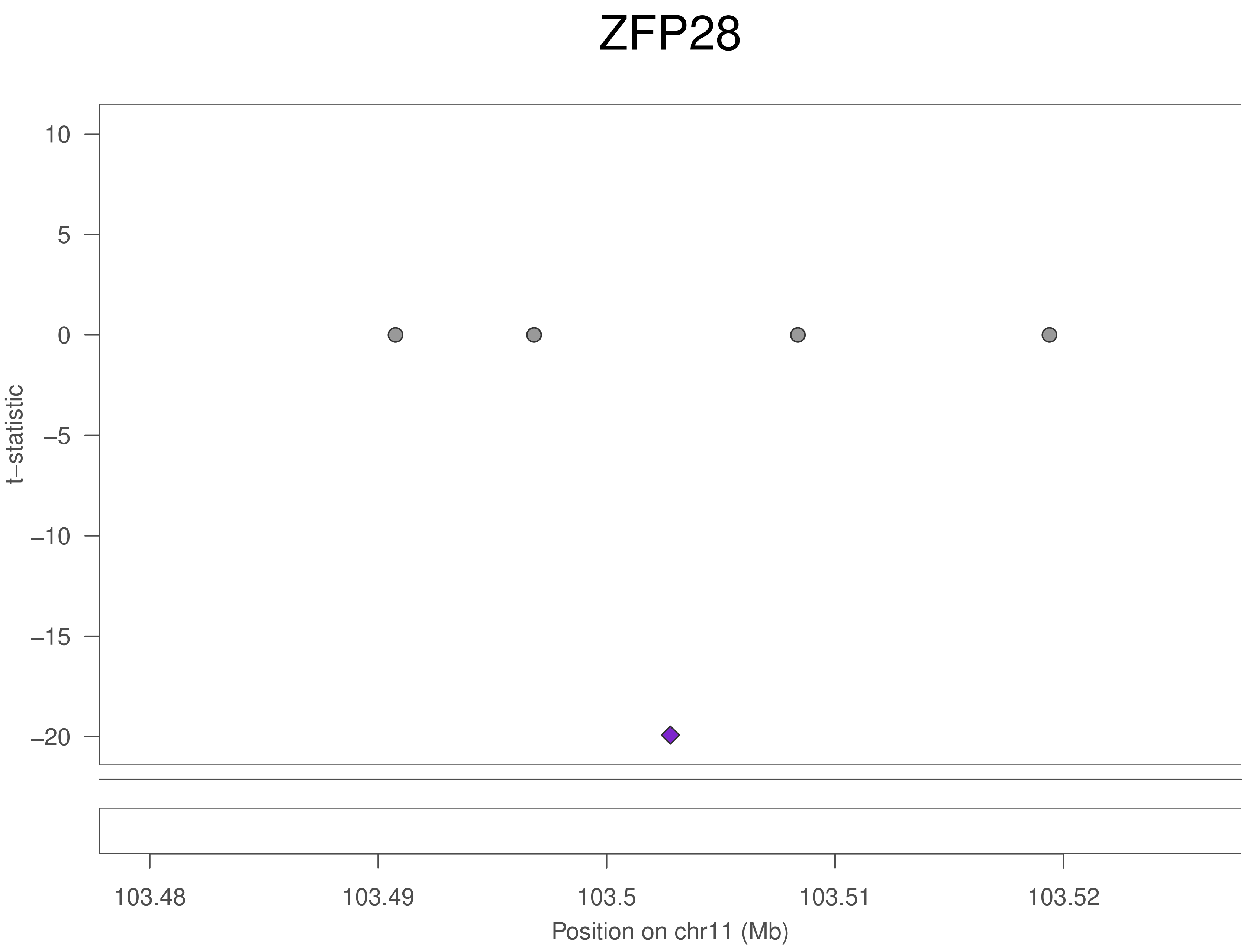
ZFP28
t−statistic
10
5
0
−5
−10
−15
−20
103.48
103.49
103.5
103.51
103.52
Position on chr11 (Mb)

### ZFP28

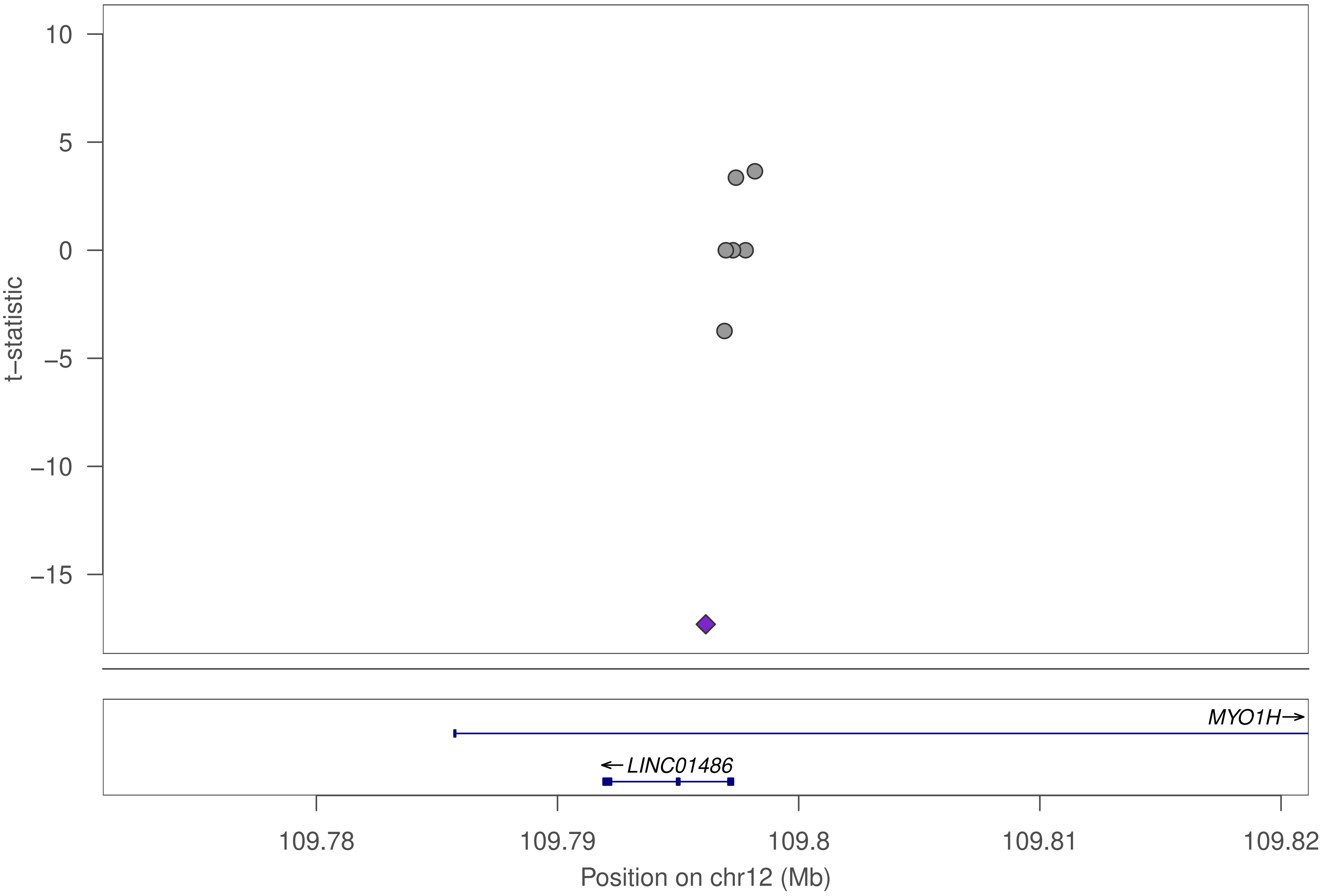

10
5
0
−5
−10
−15
t−statistic
MYO1H→
←LINC01486
109.78
109.79
109.8
109.81
109.82
Position on chr12 (Mb)

### ZFP28

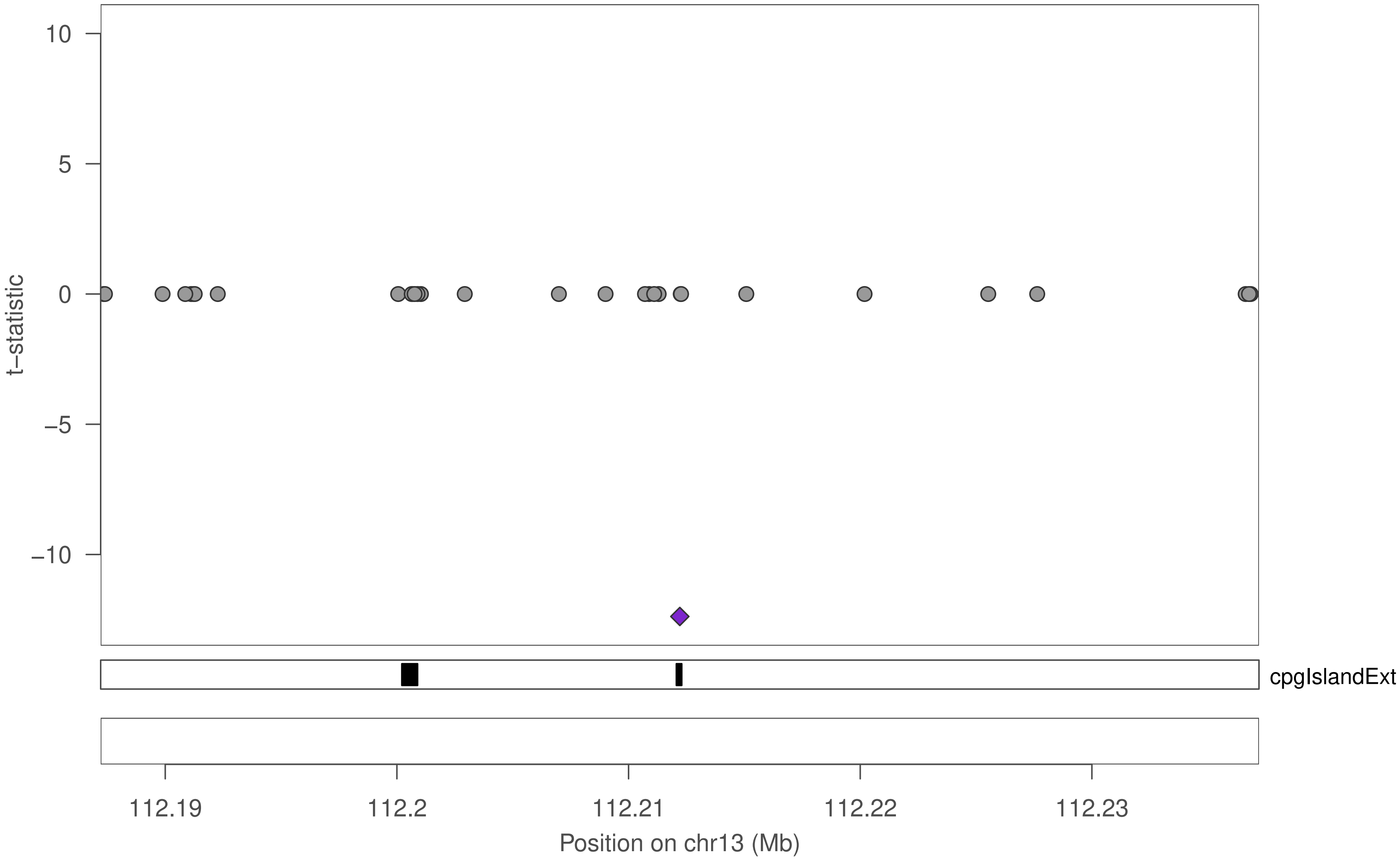

10
5
0
−5
−10
t−statistic
cpgIslandExt
112.19
112.2
112.21
112.22
112.23
Position on chr13 (Mb)

### ZFP28

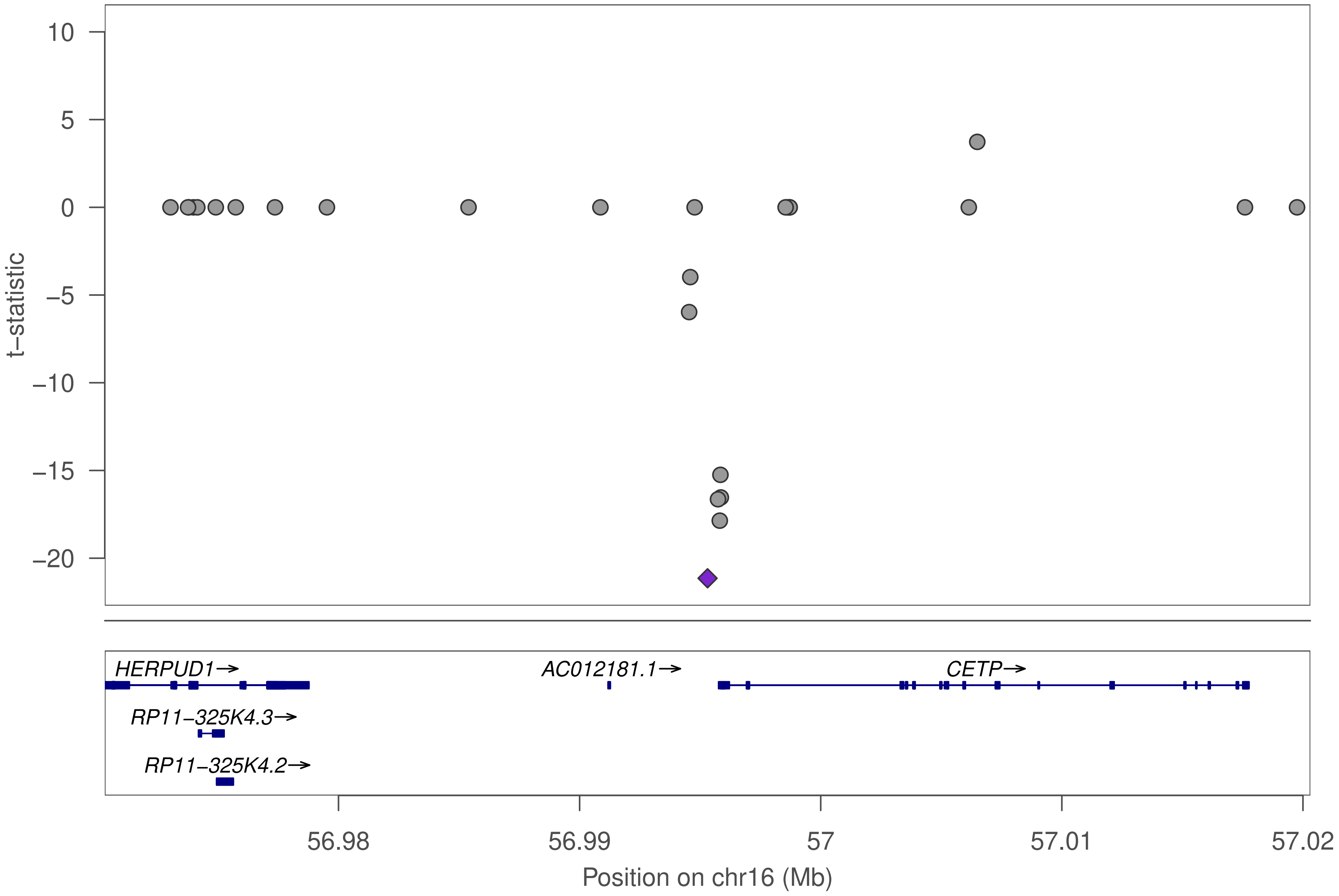

10
5
0
−5
−10
−15
−20
t−statistic
HERPUD1→
AC012181.1→
CETP→
RP11−325K4.3→
RP11−325K4.2→
56.98
56.99
57
57.01
57.02
Position on chr16 (Mb)

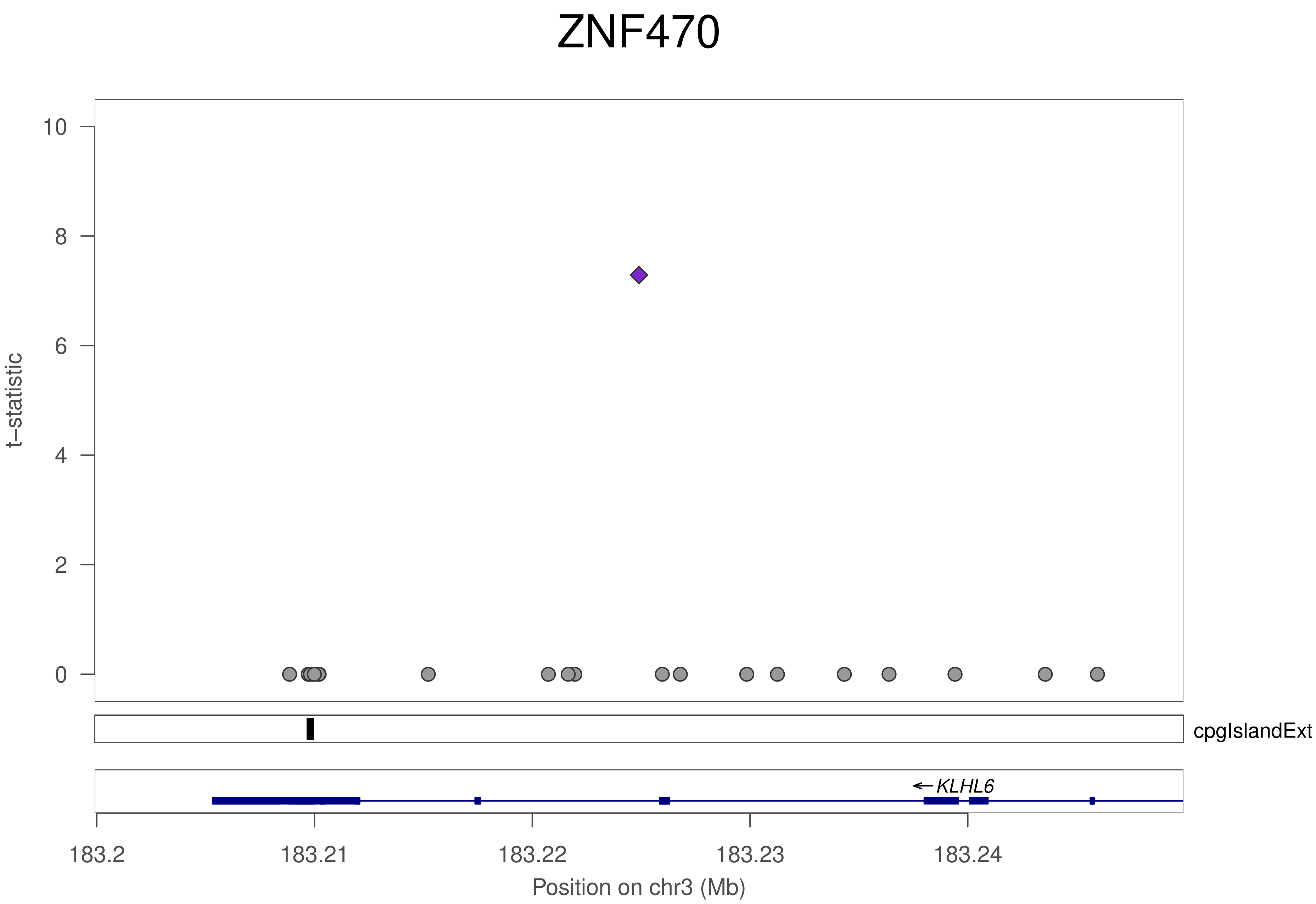

ZNF470
10
8
6
4
2
0
t−statistic
cpgIslandExt
← KLHL6
183.2
183.21
183.22
183.23
183.24
Position on chr3 (Mb)

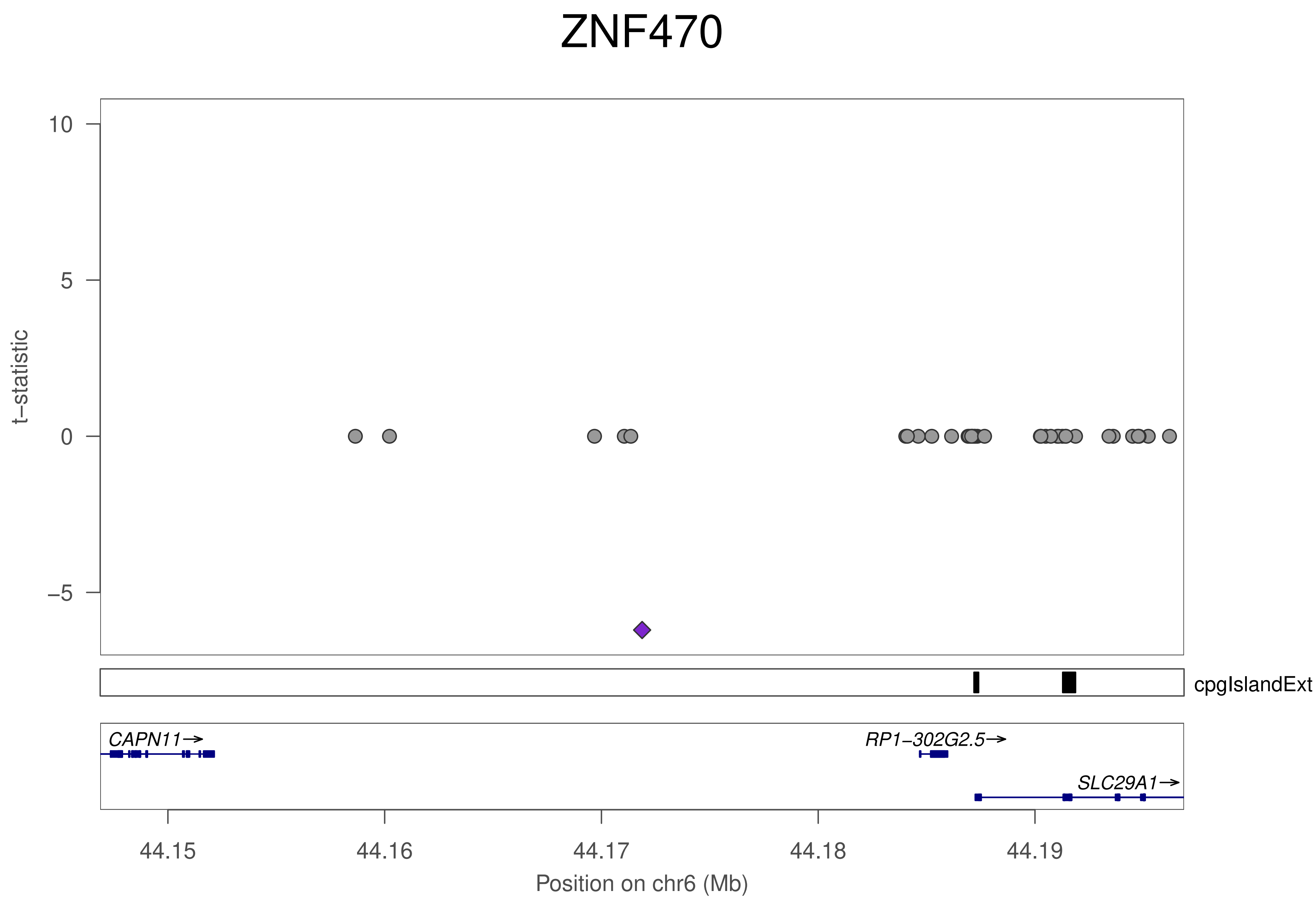

ZNF470
t−statistic
10
5
0
−5
cpgIslandExt
CAPN11→
RP1−302G2.5→
SLC29A1→
44.15
44.16
44.17
44.18
44.19
Position on chr6 (Mb)

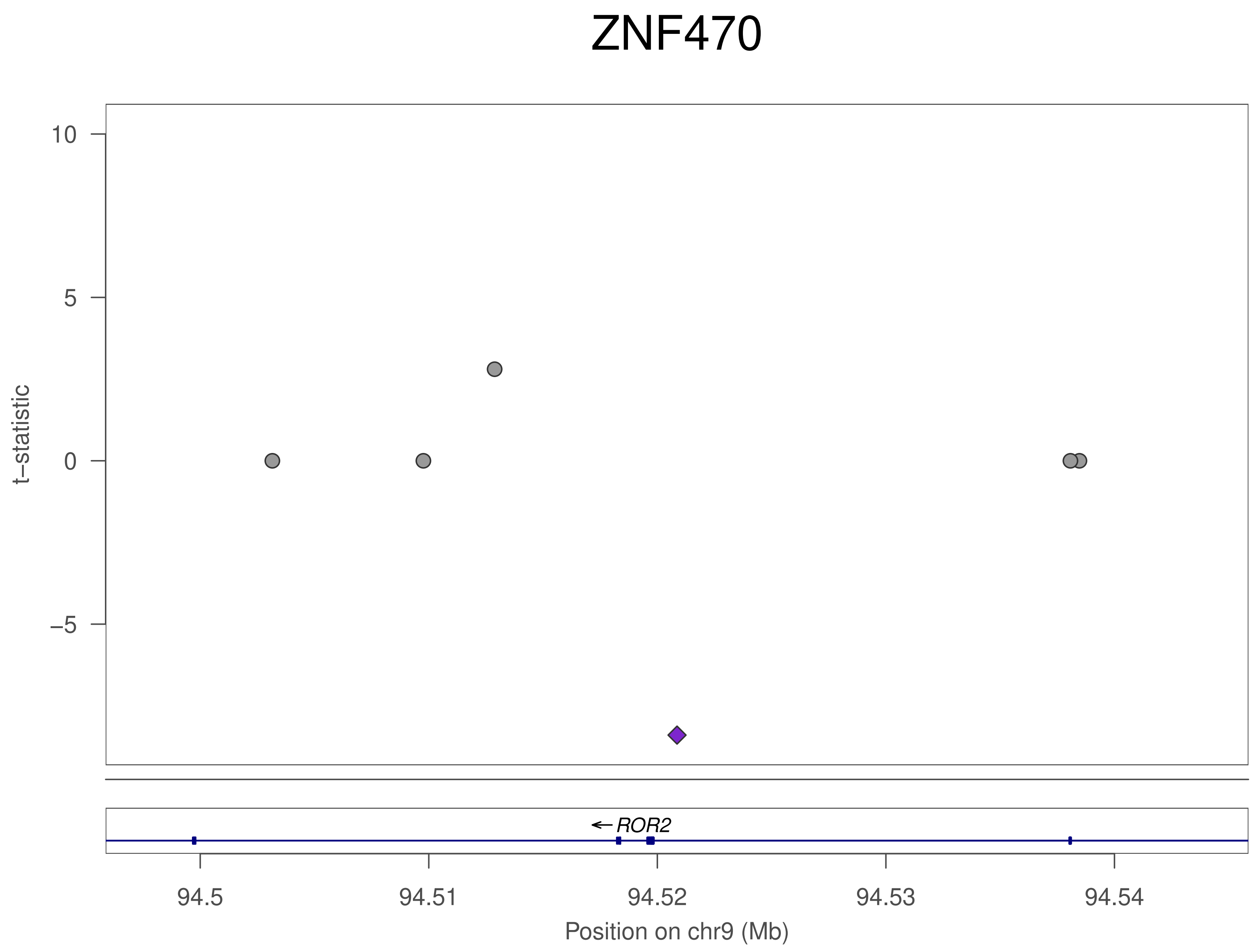

ZNF470
t–statistic
10
5
0
–5
ROR2
94.5
94.51
94.52
94.53
94.54
Position on chr9 (Mb)

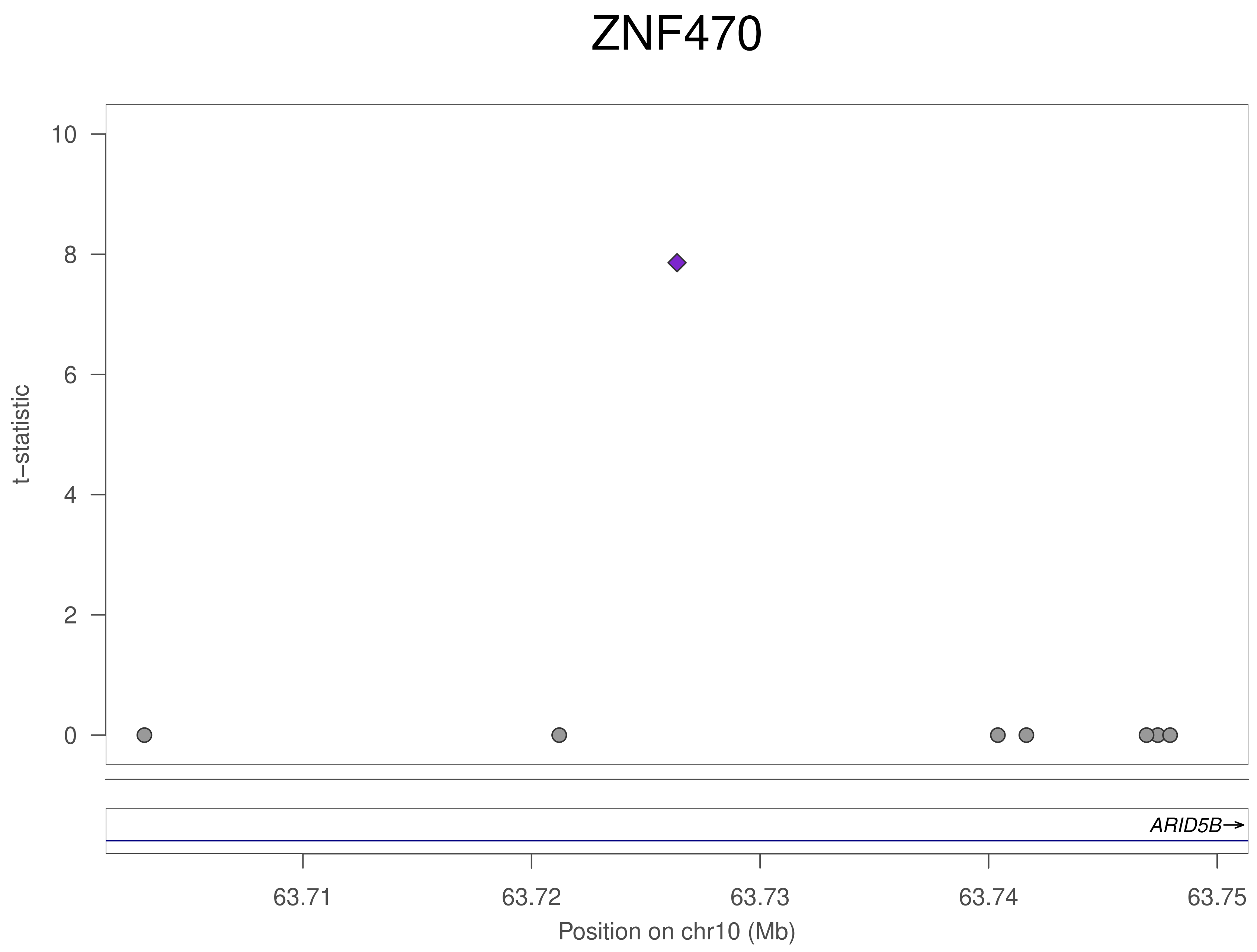
ZNF470
t–statistic
10
8
6
4
2
0
ARID5B→
63.71
63.72
63.73
63.74
63.75
Position on chr10 (Mb)

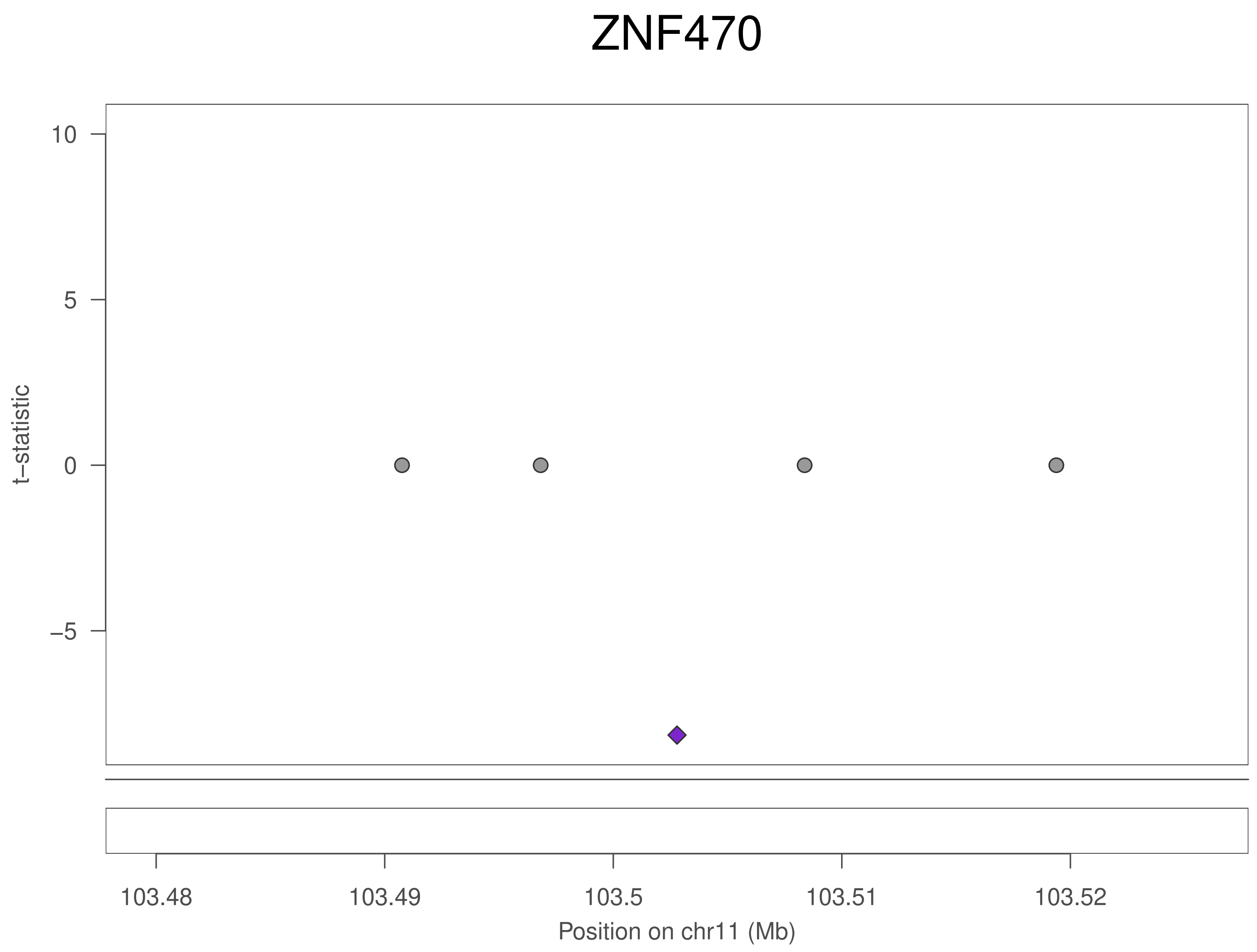
ZNF470
t–statistic
10
5
0
–5
103.48
103.49
103.5
103.51
103.52
Position on chr11 (Mb)

### ZNF470

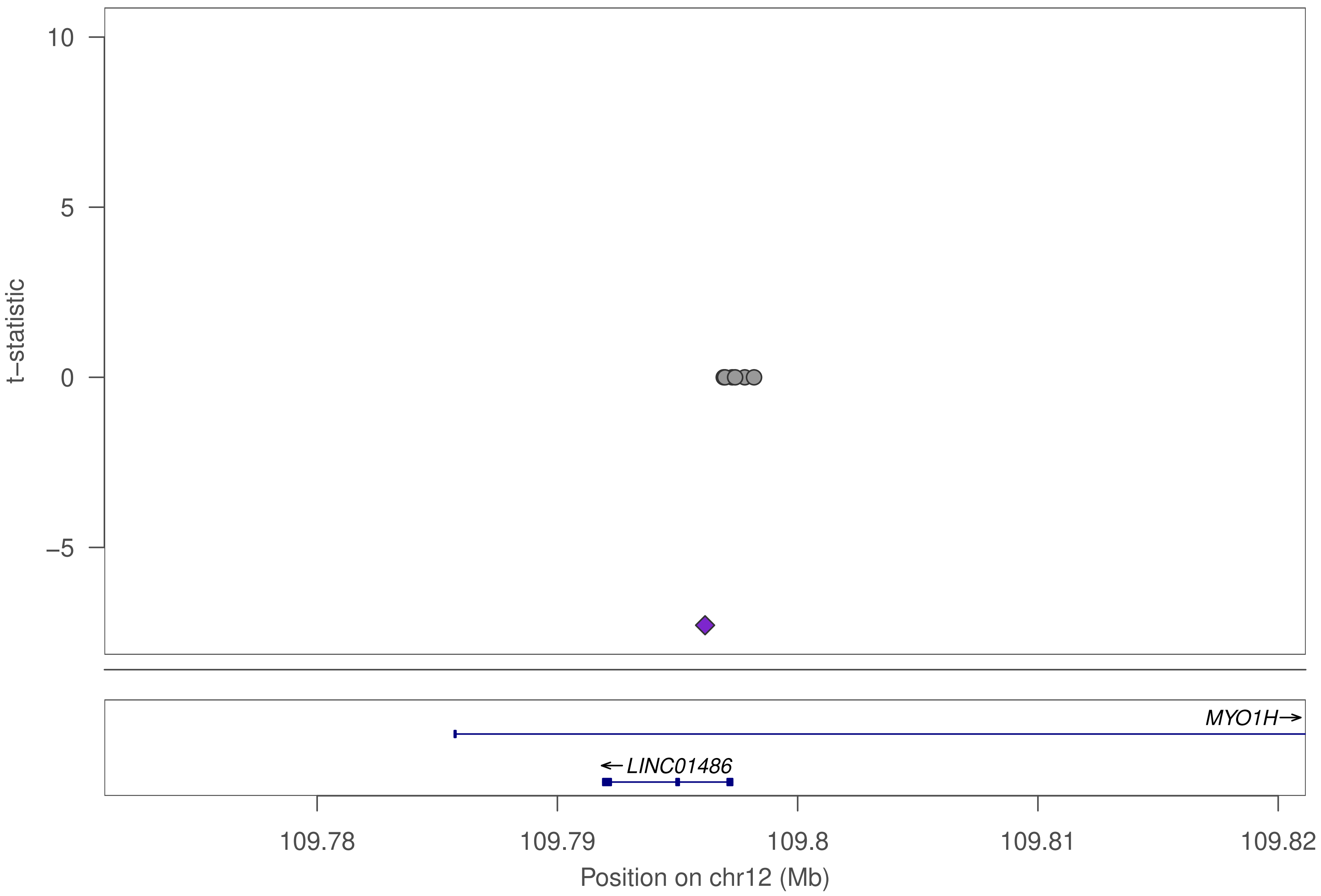

10
5
0
−5
t−statistic
MYO1H→
←LINC01486
109.78
109.79
109.8
109.81
109.82
Position on chr12 (Mb)

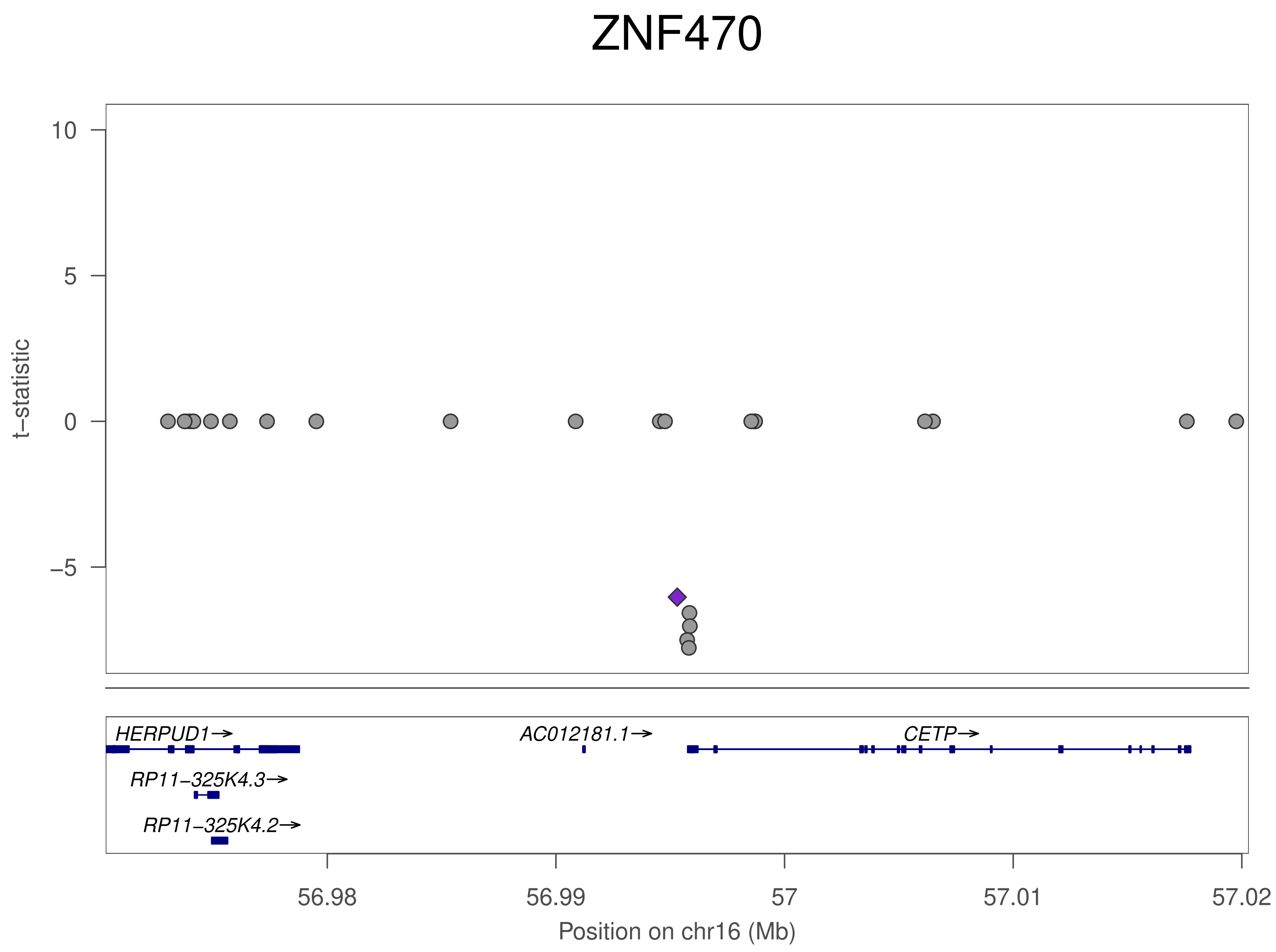

ZNF470
t−statistic
10
5
0
−5
HERPUD1→
AC012181.1→
CETP→
RP11−325K4.3→
RP11−325K4.2→
56.98
56.99
57
57.01
57.02
Position on chr16 (Mb)

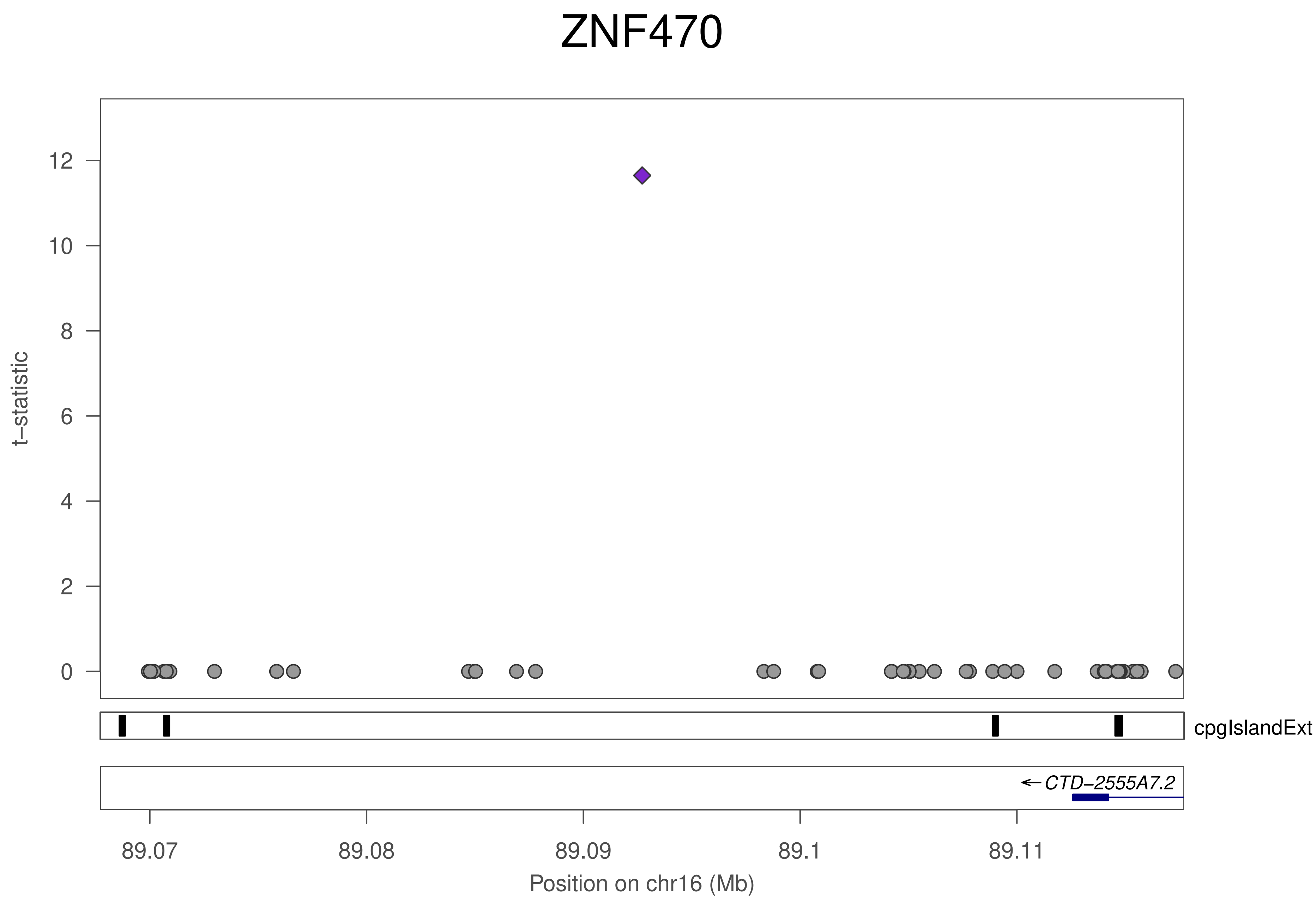

ZNF470
t−statistic
12
10
8
6
4
2
0
cpgIslandExt
← CTD−2555A7.2
89.07
89.08
89.09
89.1
89.11
Position on chr16 (Mb)

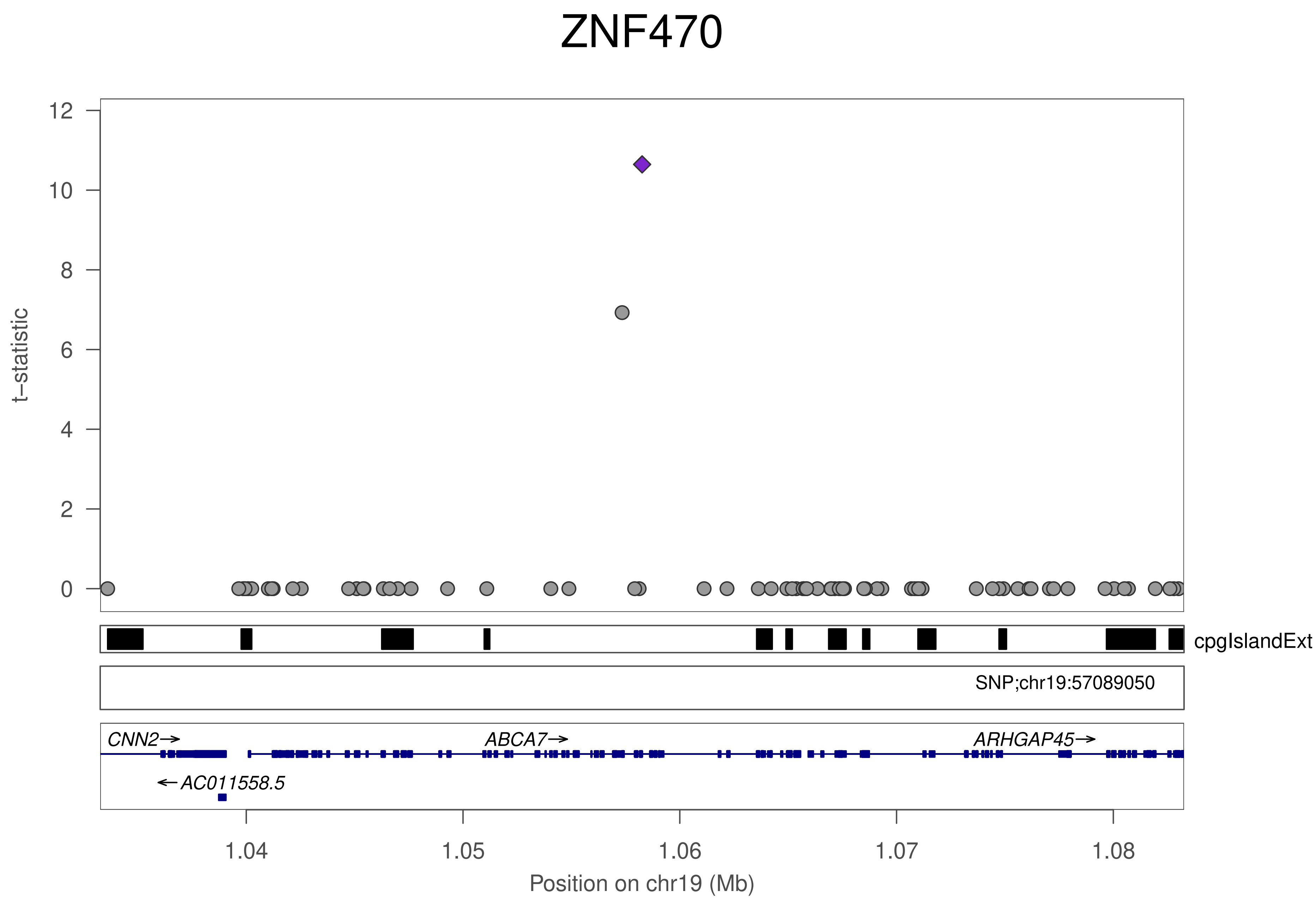

ZNF470
t−statistic
0
2
4
6
8
10
12
cpgIslandExt
SNP;chr19:57089050
CNN2→
ABCA7→
ARHGAP45→
←AC011558.5
1.04
1.05
1.06
1.07
1.08
Position on chr19 (Mb)

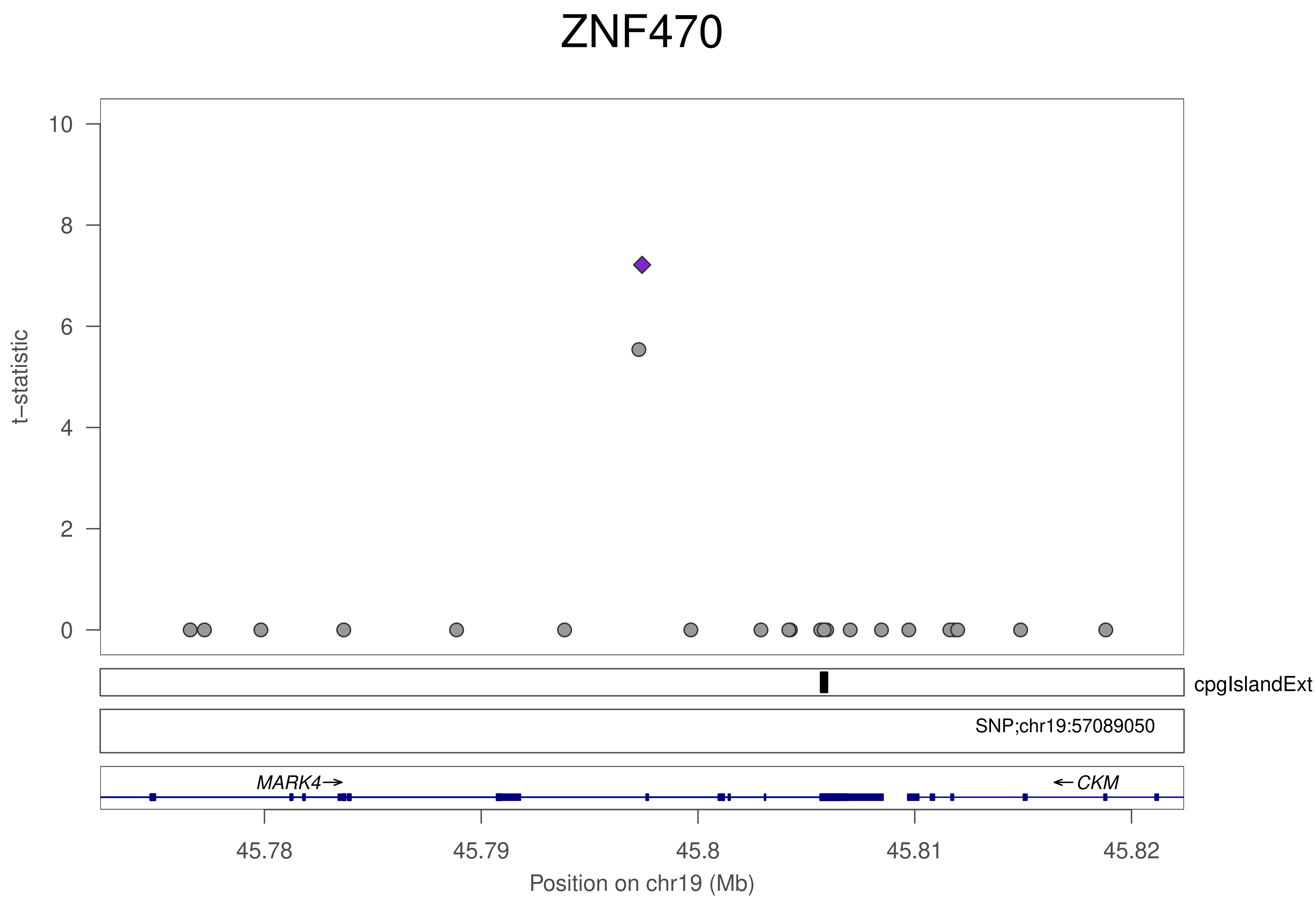
ZNF470
t−statistic
10
8
6
4
2
0
cpgIslandExt
SNP;chr19:57089050
MARK4→
←CKM
45.78
45.79
45.8
45.81
45.82
Position on chr19 (Mb)

### ZNF763

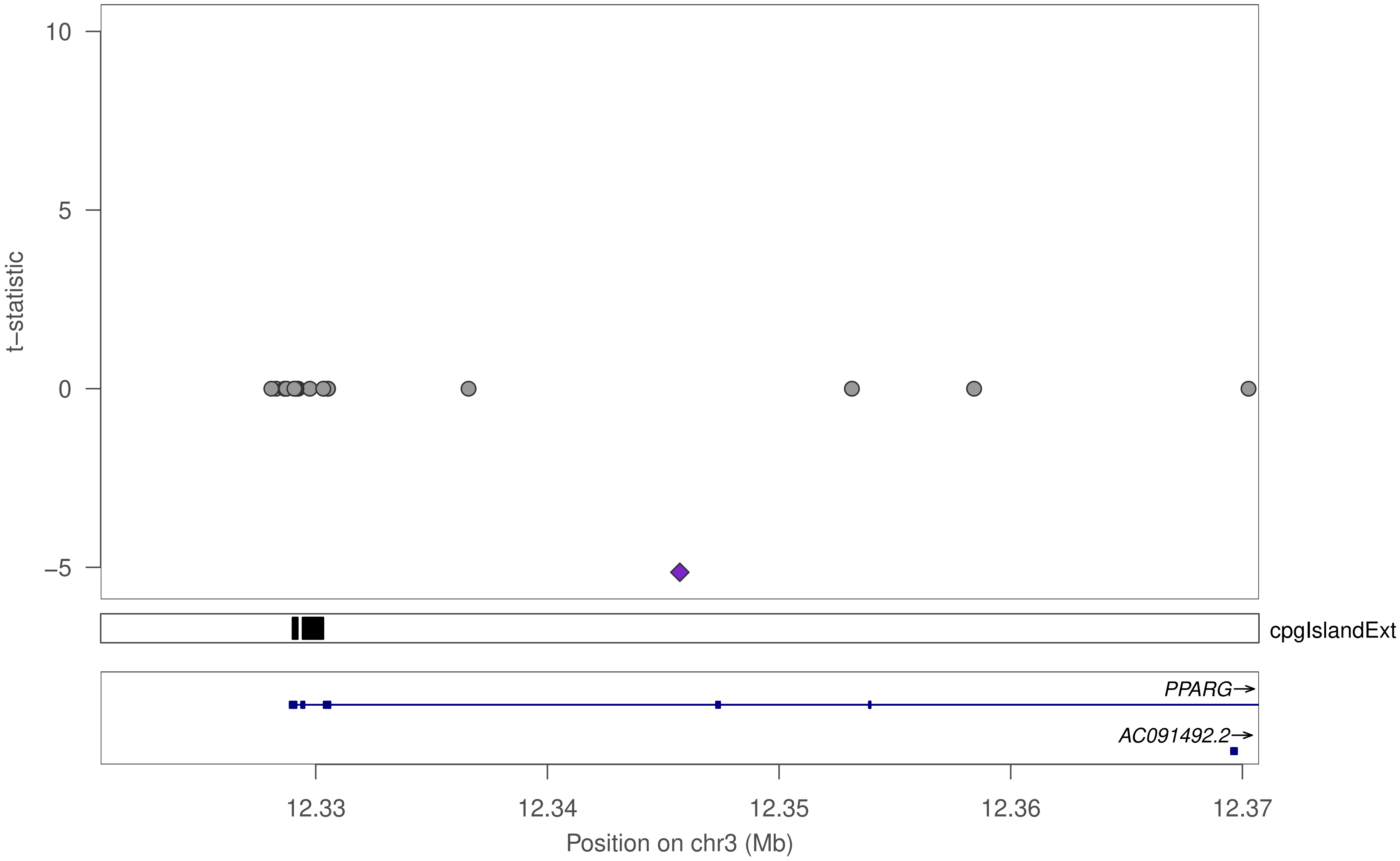

10
5
0
−5
t−statistic
cpgIslandExt
PPARG→
AC091492.2→
12.33
12.34
12.35
12.36
12.37
Position on chr3 (Mb)

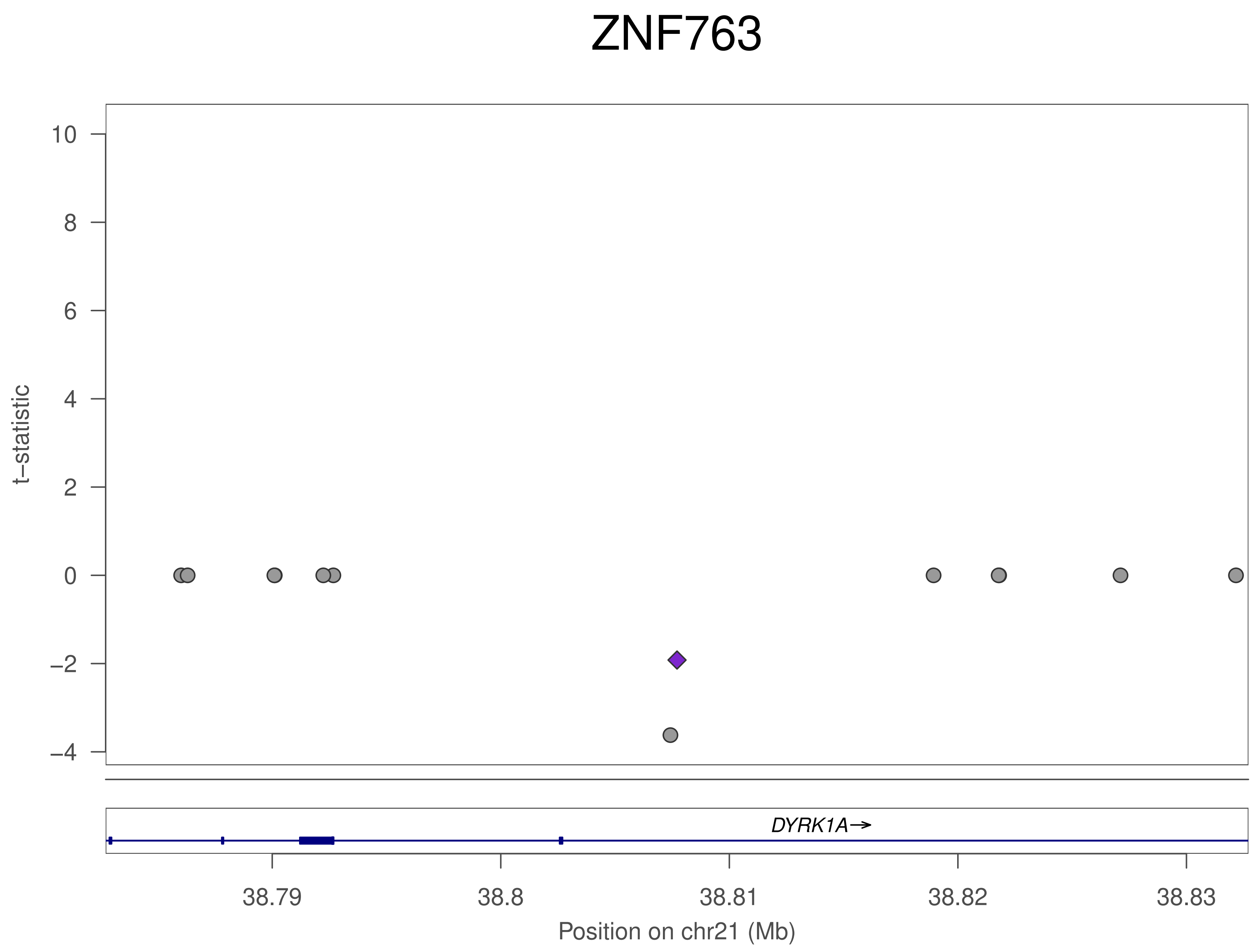

ZNF763
t−statistic
10
8
6
4
2
0
−2
−4
DYRK1A→
38.79
38.8
38.81
38.82
38.83
Position on chr21 (Mb)

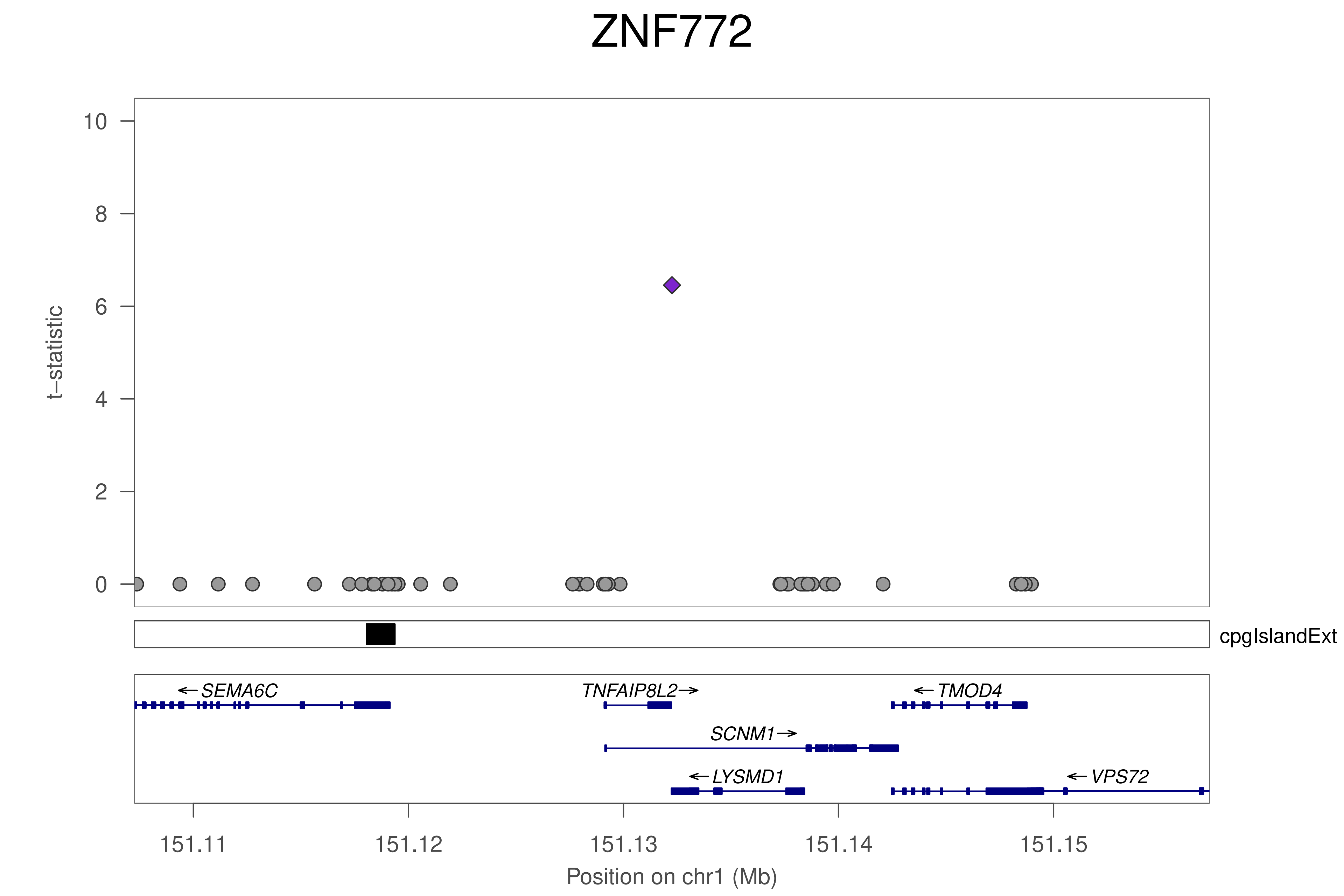
ZNF772
t−statistic
0
2
4
6
8
10
cpgIslandExt
SEMA6C
TNFAIP8L2
TMOD4
SCNM1
LYSMD1
VPS72
151.11
151.12
151.13
151.14
151.15
Position on chr1 (Mb)

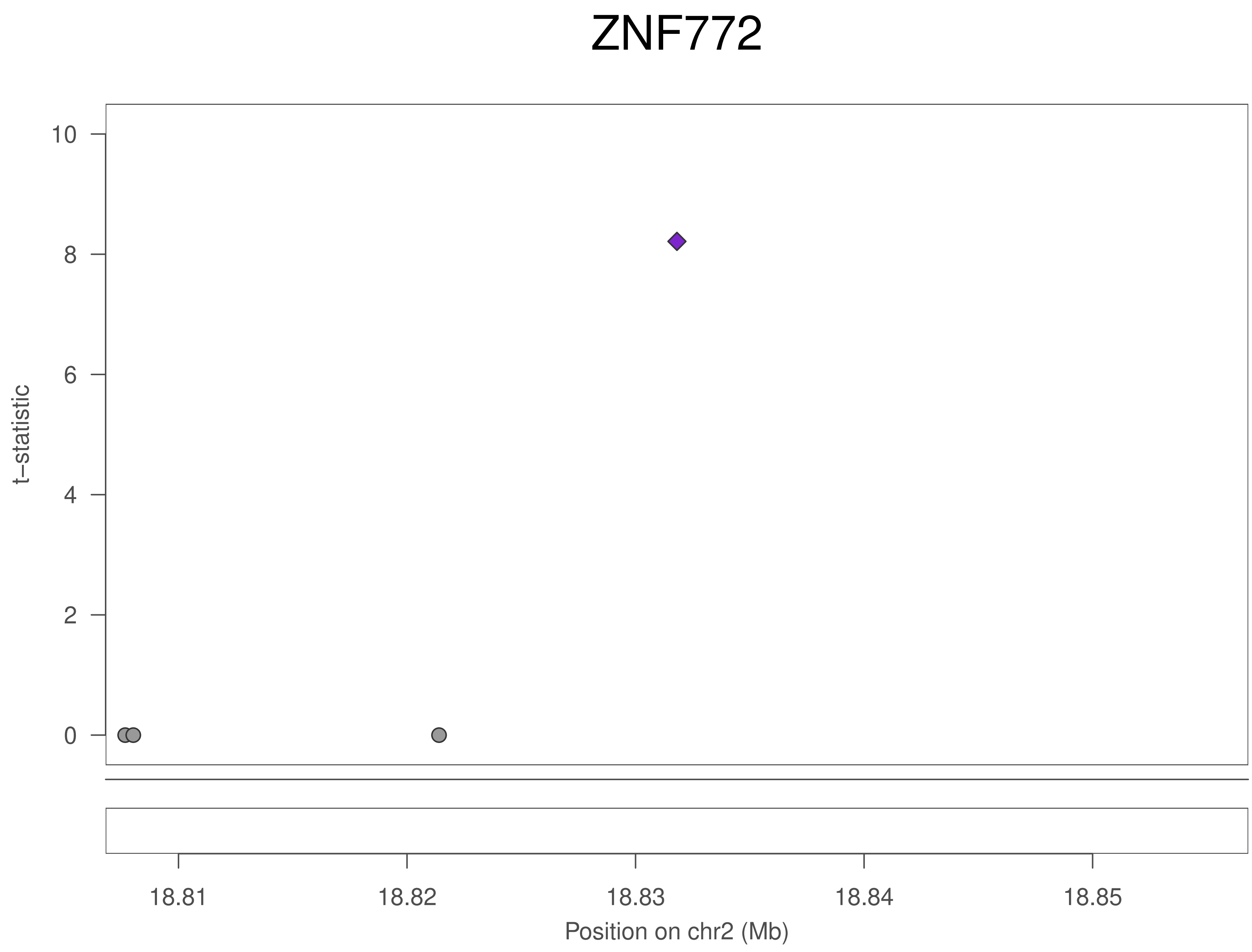
ZNF772
t–statistic
0
2
4
6
8
10
18.81
18.82
18.83
18.84
18.85
Position on chr2 (Mb)

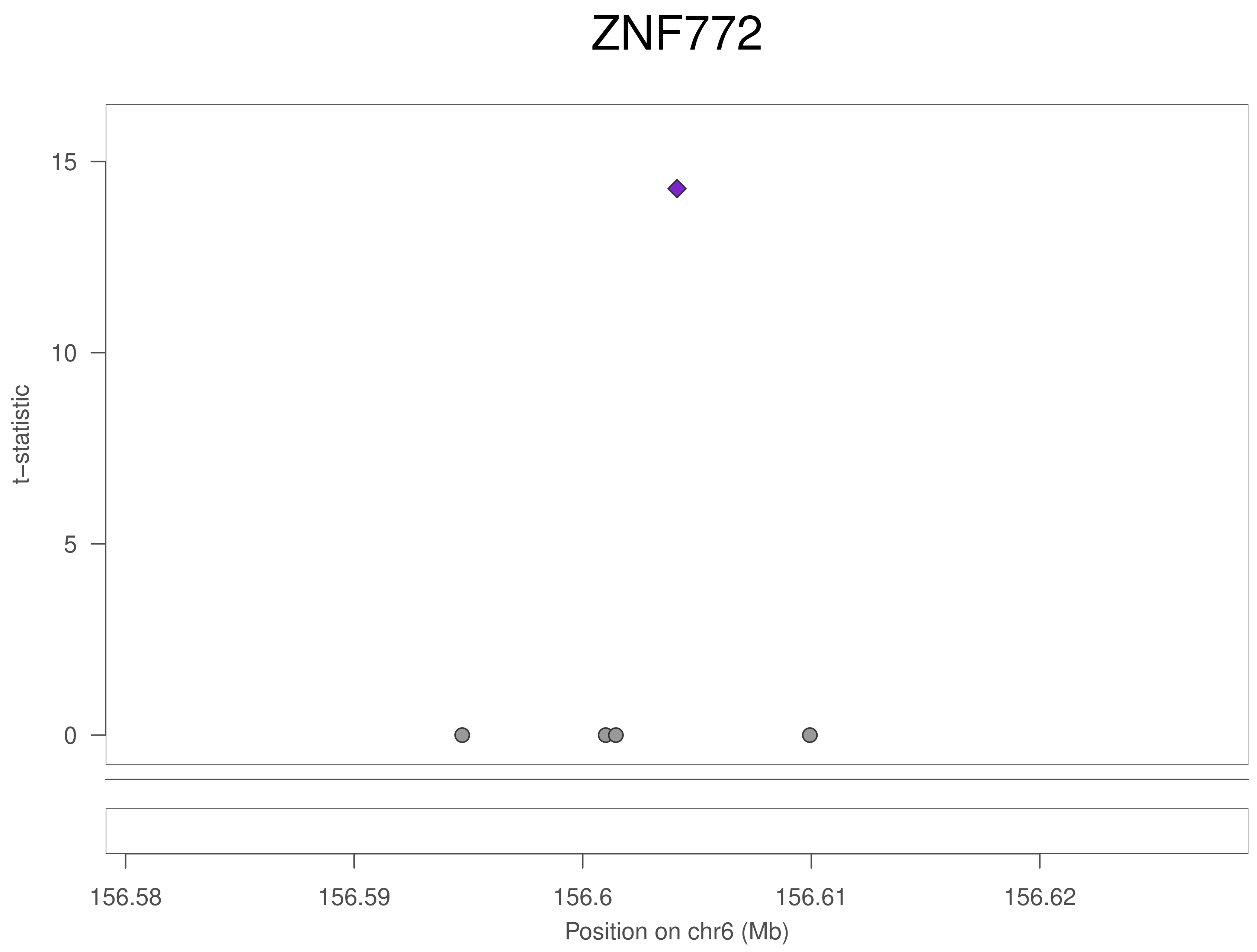
ZNF772
t–statistic
15
10
5
0
156.58
156.59
156.6
156.61
156.62
Position on chr6 (Mb)

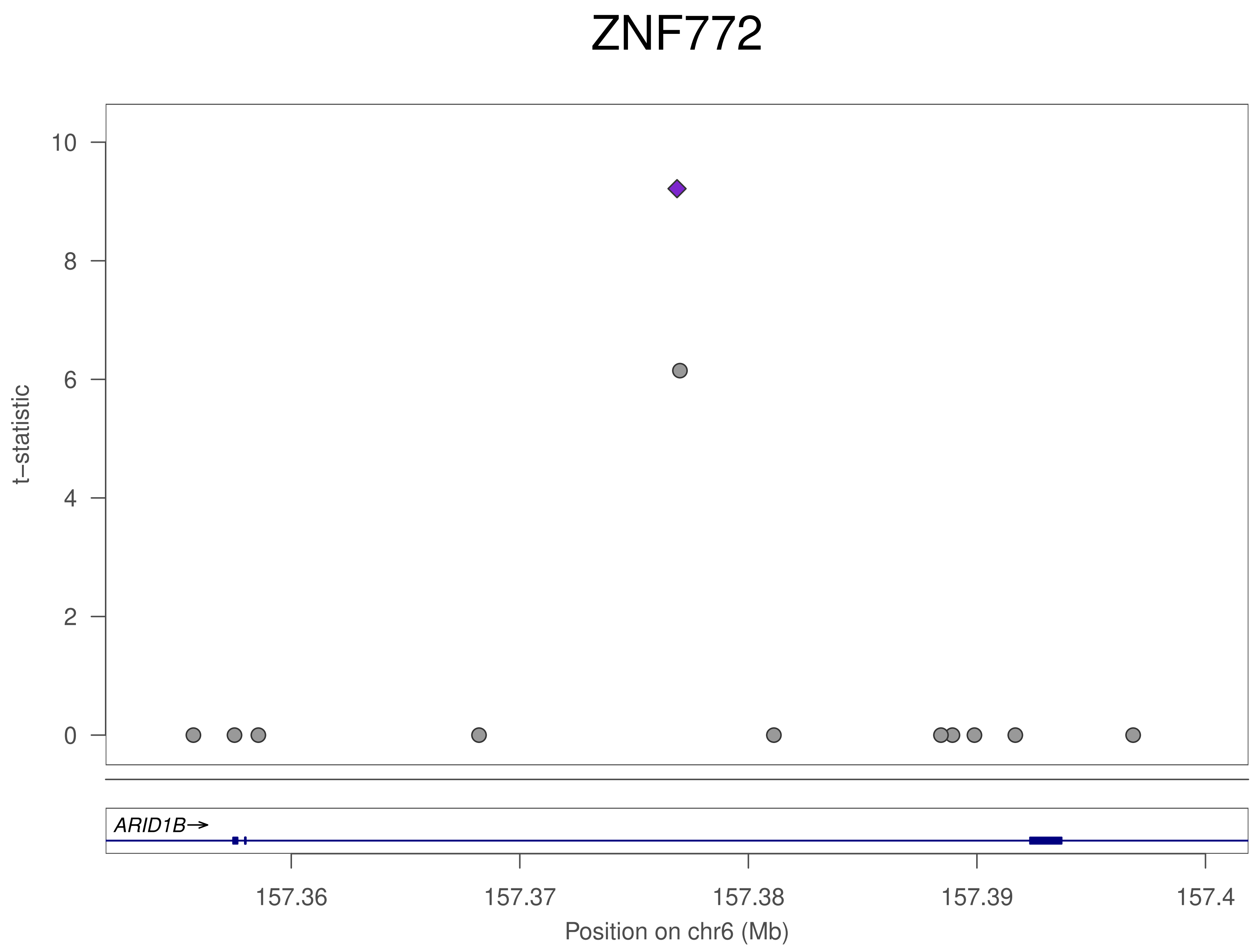
ZNF772
t–statistic
0
2
4
6
8
10
ARID1B→
157.36
157.37
157.38
157.39
157.4
Position on chr6 (Mb)

ZNF772

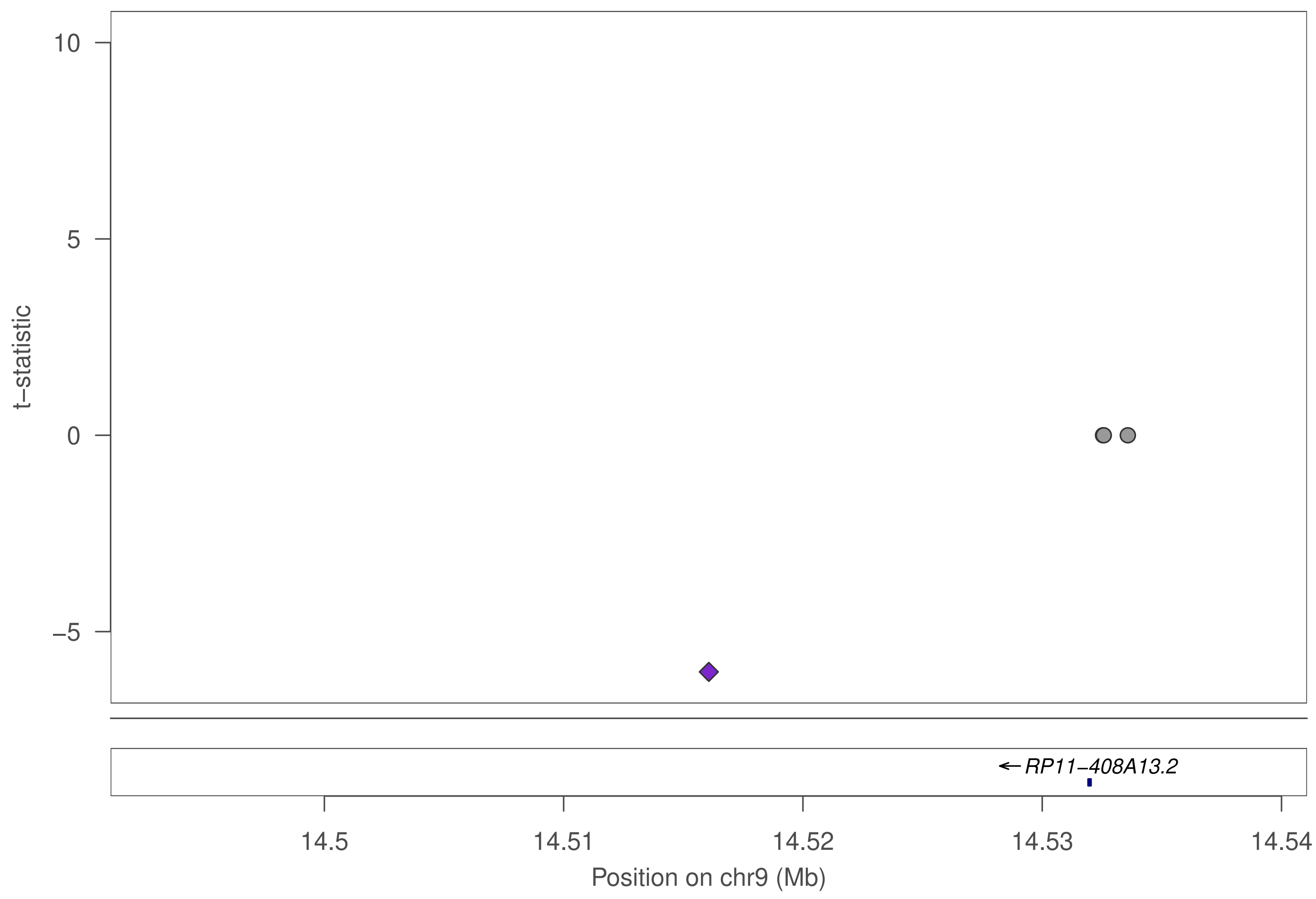

10
5
0
−5
t−statistic
RP11−408A13.2
14.5
14.51
14.52
14.53
14.54
Position on chr9 (Mb)

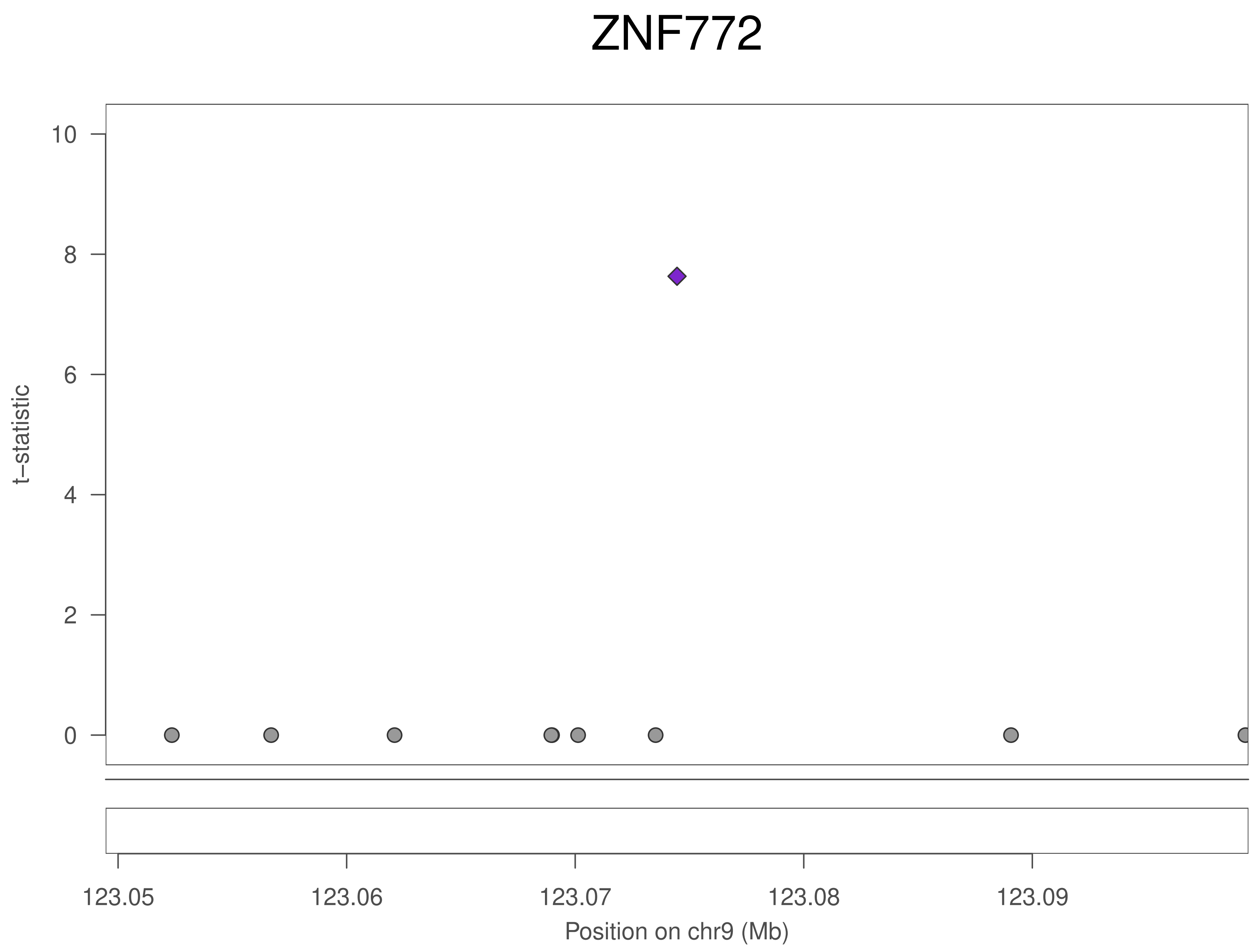
ZNF772
t−statistic
10
8
6
4
2
0
123.05
123.06
123.07
123.08
123.09
Position on chr9 (Mb)

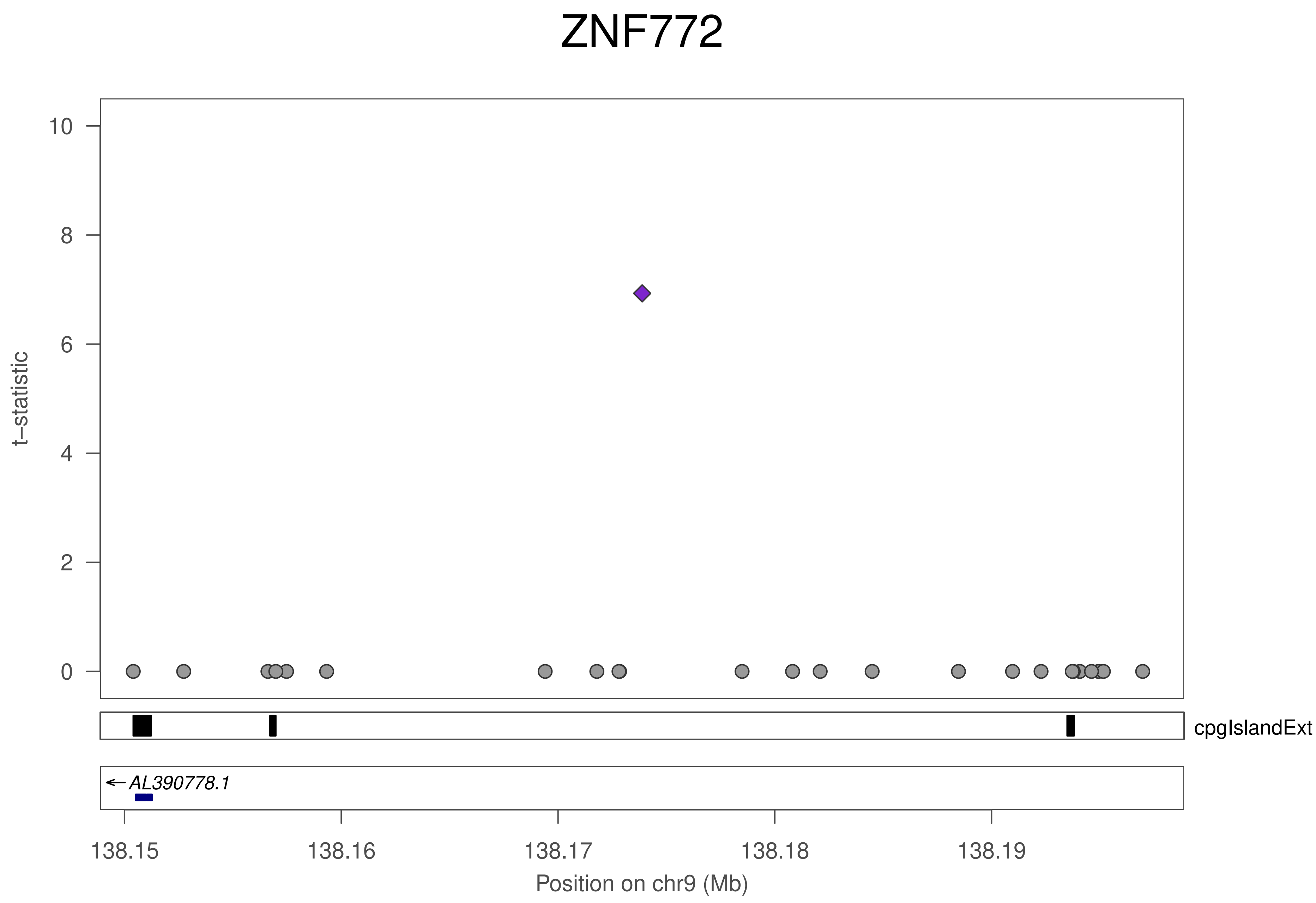

ZNF772
t–statistic
0
2
4
6
8
10
cpgIslandExt
AL390778.1
138.15
138.16
138.17
138.18
138.19
Position on chr9 (Mb)

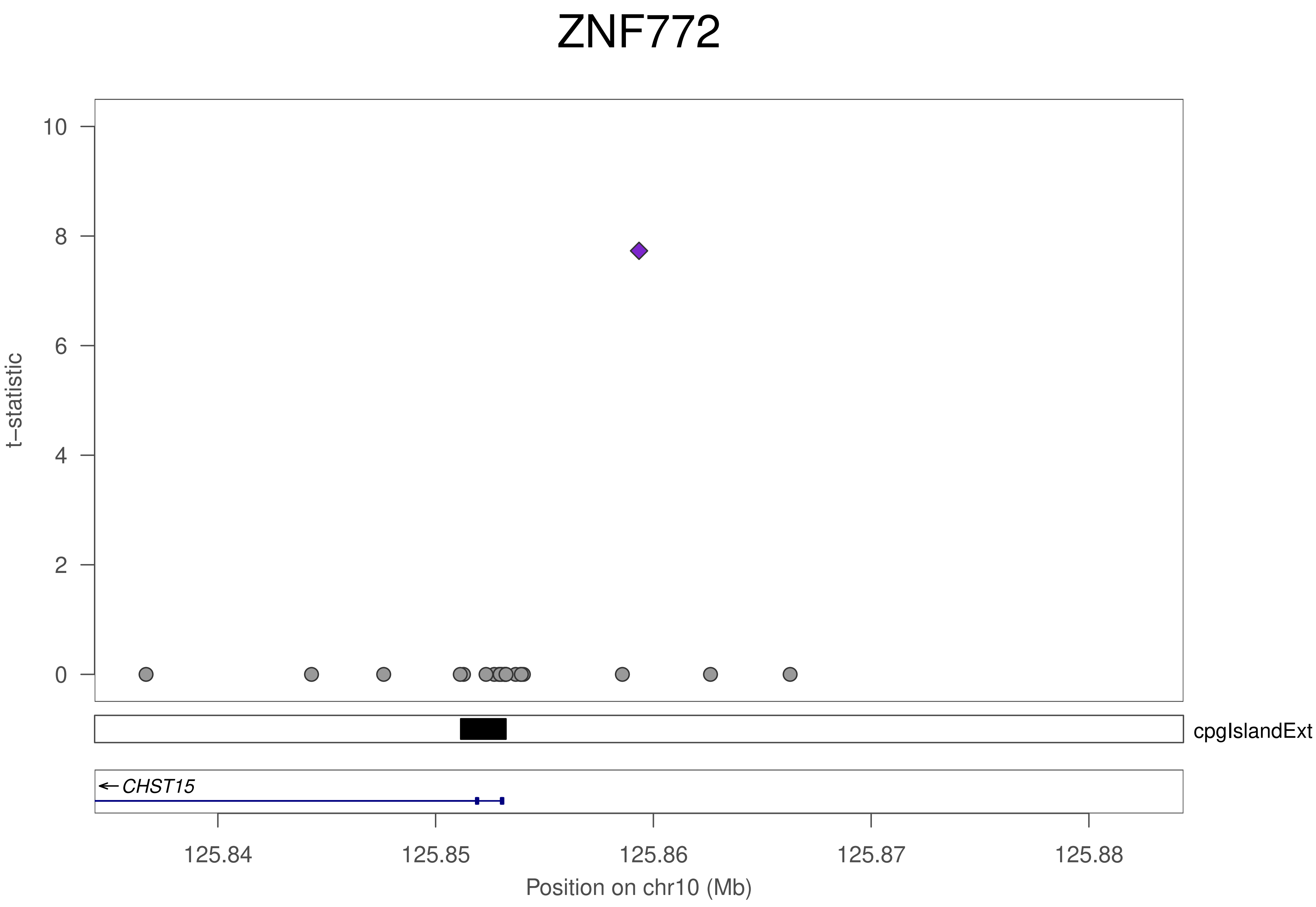
ZNF772
t–statistic
10
8
6
4
2
0
cpgIslandExt
CHST15
125.84
125.85
125.86
125.87
125.88
Position on chr10 (Mb)

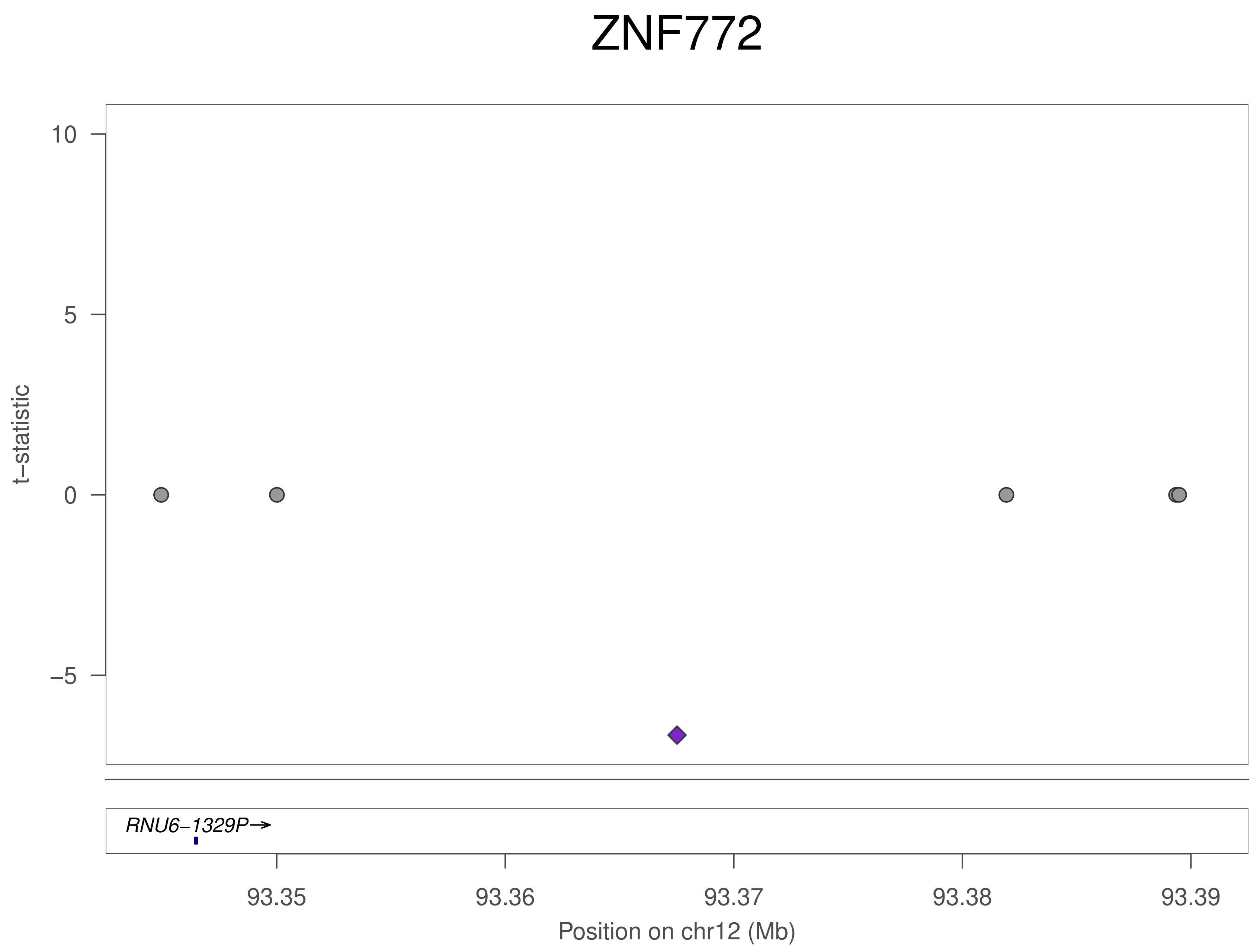
ZNF772
t−statistic
10
5
0
−5
RNU6−1329P→
93.35
93.36
93.37
93.38
93.39
Position on chr12 (Mb)

### ZNF772

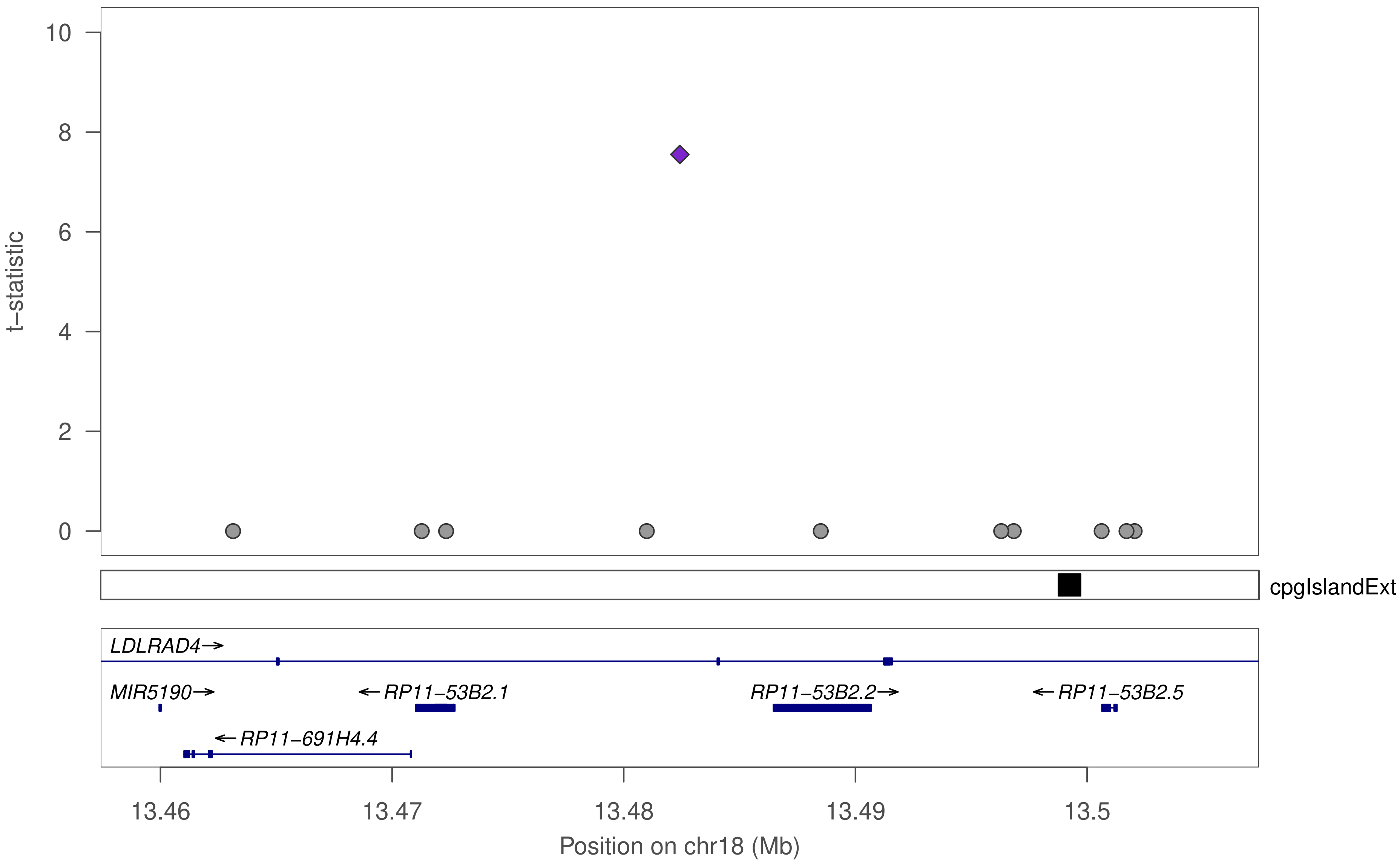

t–statistic
10
8
6
4
2
0
cpgIslandExt
LDLRAD4→
MIR5190→
←RP11–53B2.1
RP11–53B2.2→
←RP11–53B2.5
←RP11–691H4.4
13.46
13.47
13.48
13.49
13.5
Position on chr18 (Mb)

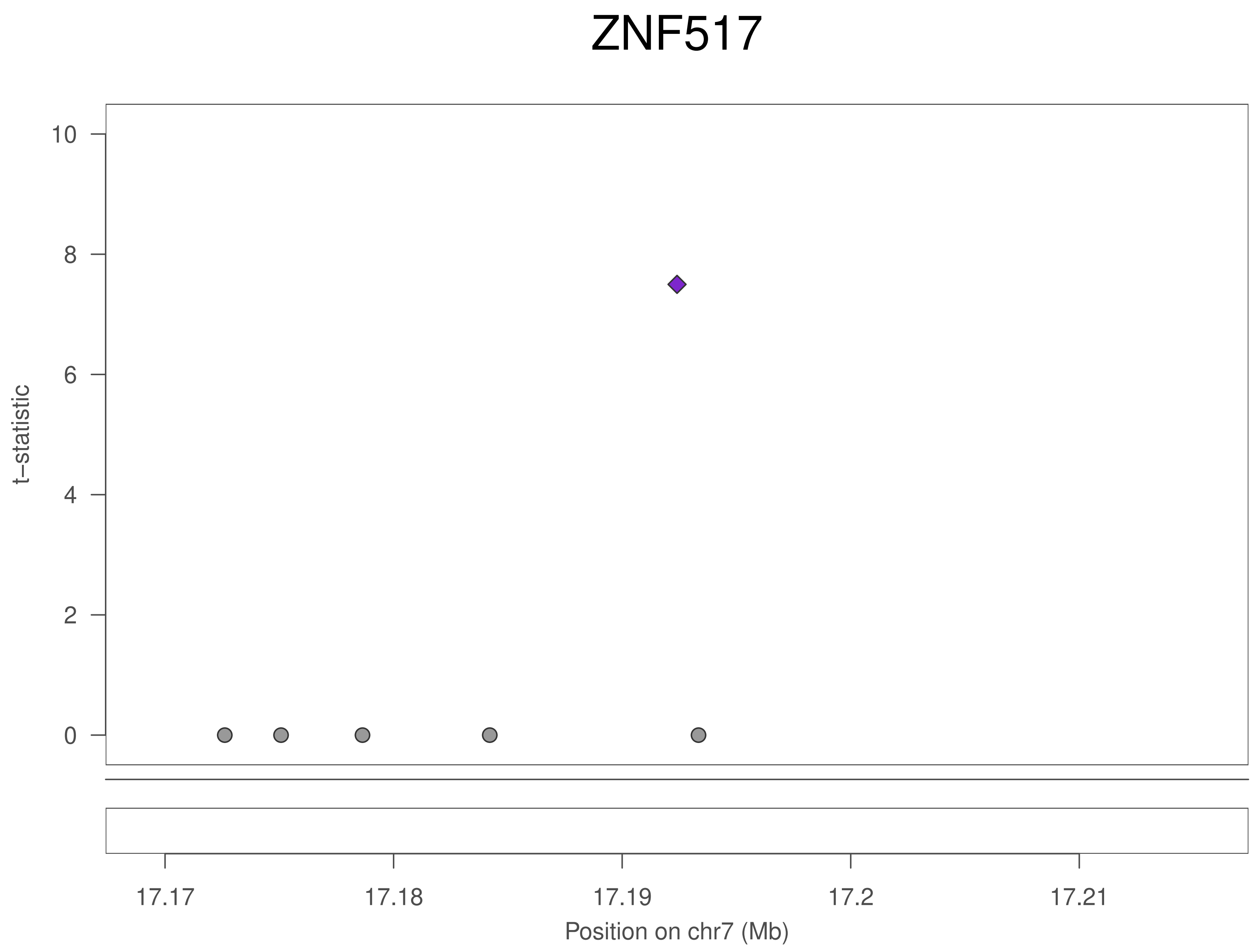
ZNF517
t–statistic
10
8
6
4
2
0
17.17
17.18
17.19
17.2
17.21
Position on chr7 (Mb)

### ZNF517

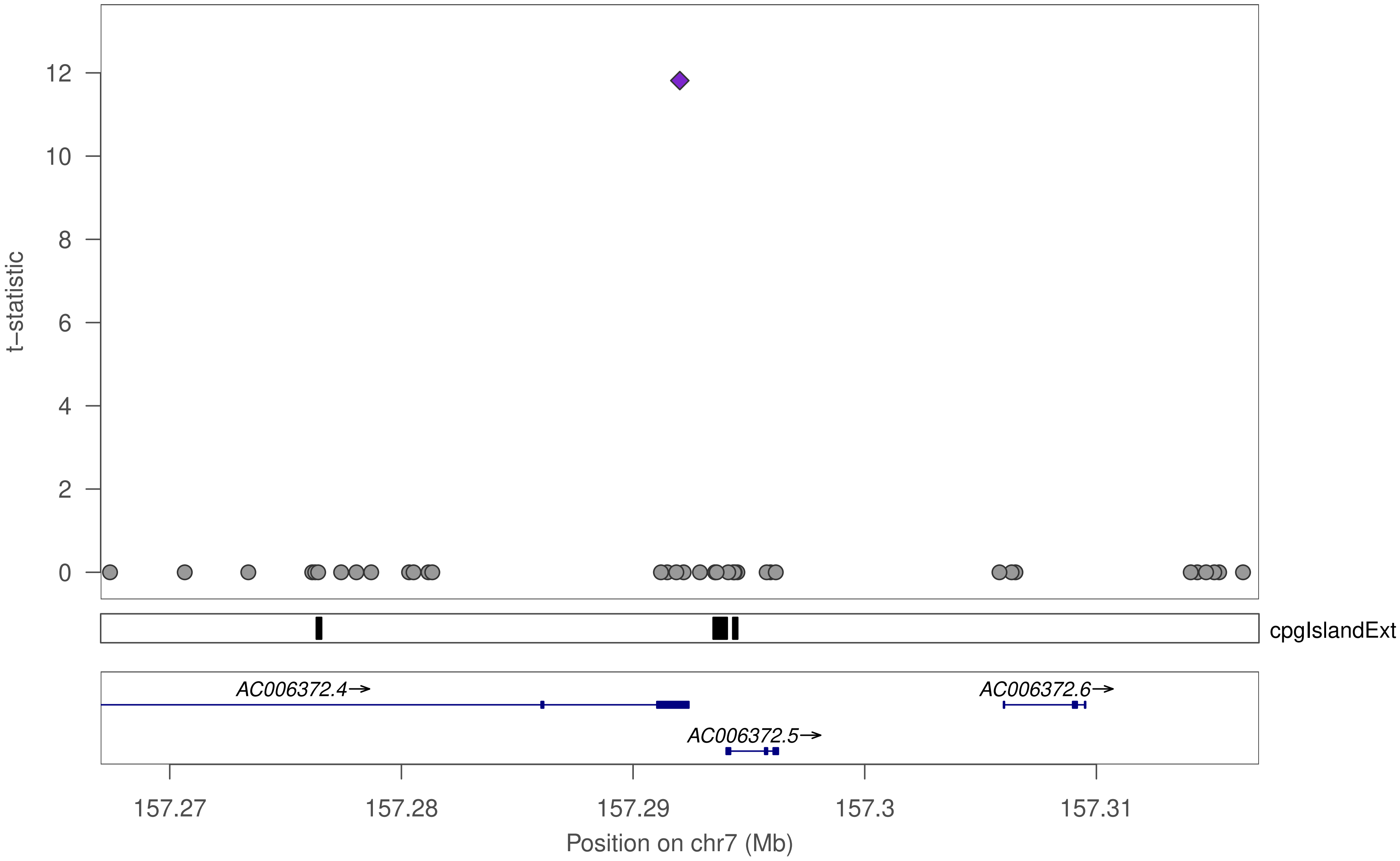

t−statistic
0
2
4
6
8
10
12
cpgIslandExt
AC006372.4→
AC006372.6→
AC006372.5→
157.27
157.28
157.29
157.3
157.31
Position on chr7 (Mb)

### ZNF517

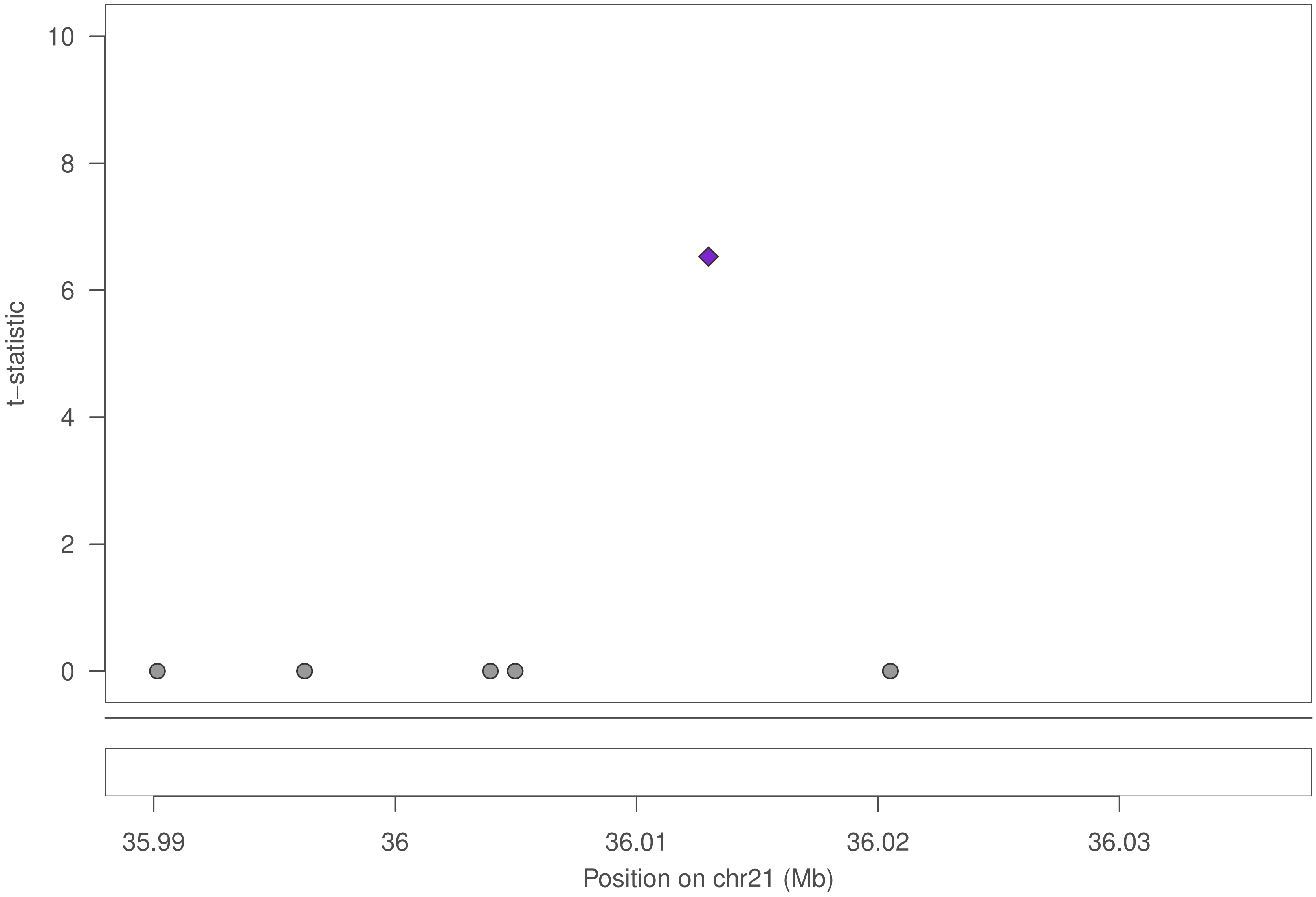

10
8
6
4
2
0
t−statistic
35.99
36
36.01
36.02
36.03
Position on chr21 (Mb)

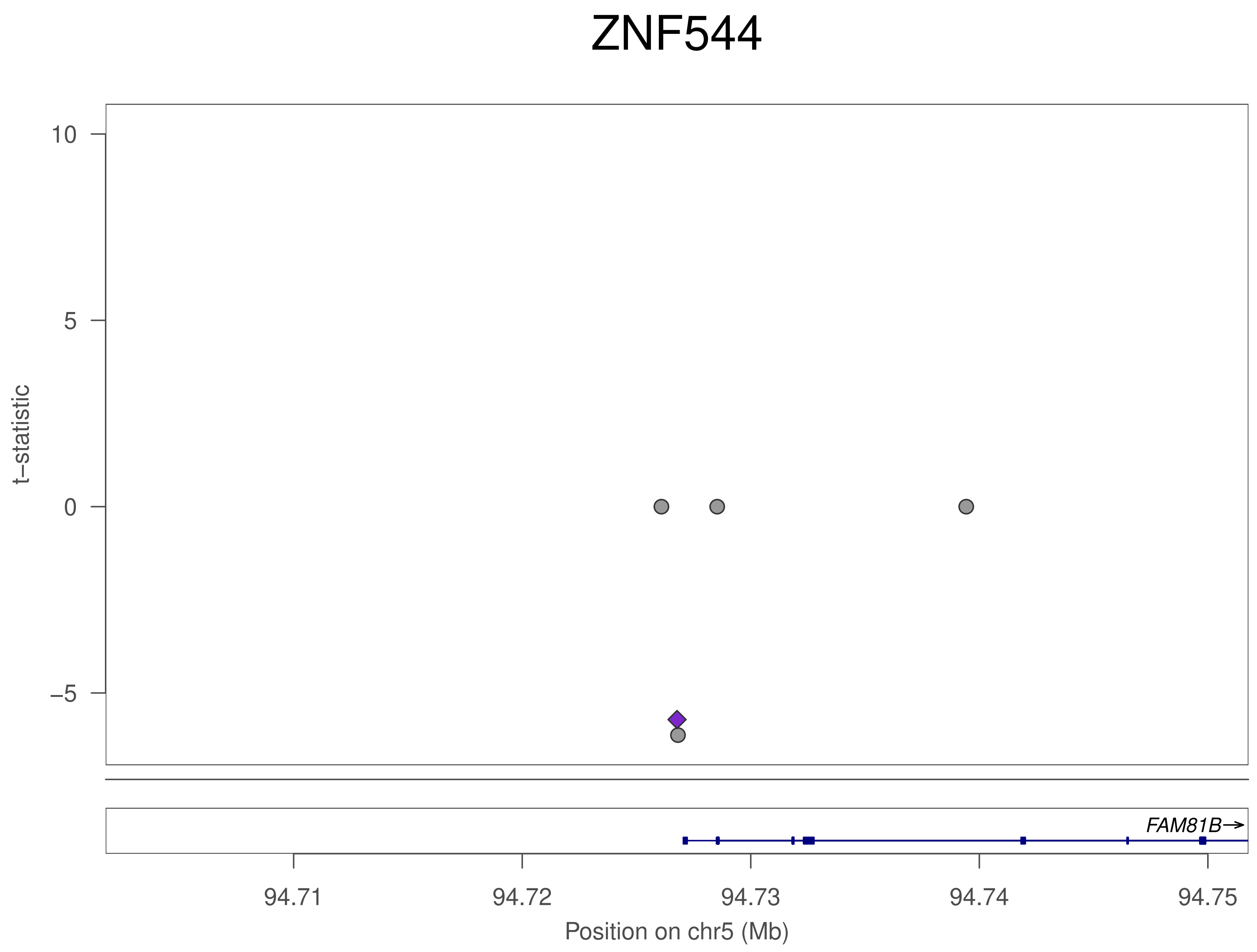

ZNF544
t–statistic
10
5
0
–5
FAM81B→
94.71
94.72
94.73
94.74
94.75
Position on chr5 (Mb)

### ZNF544

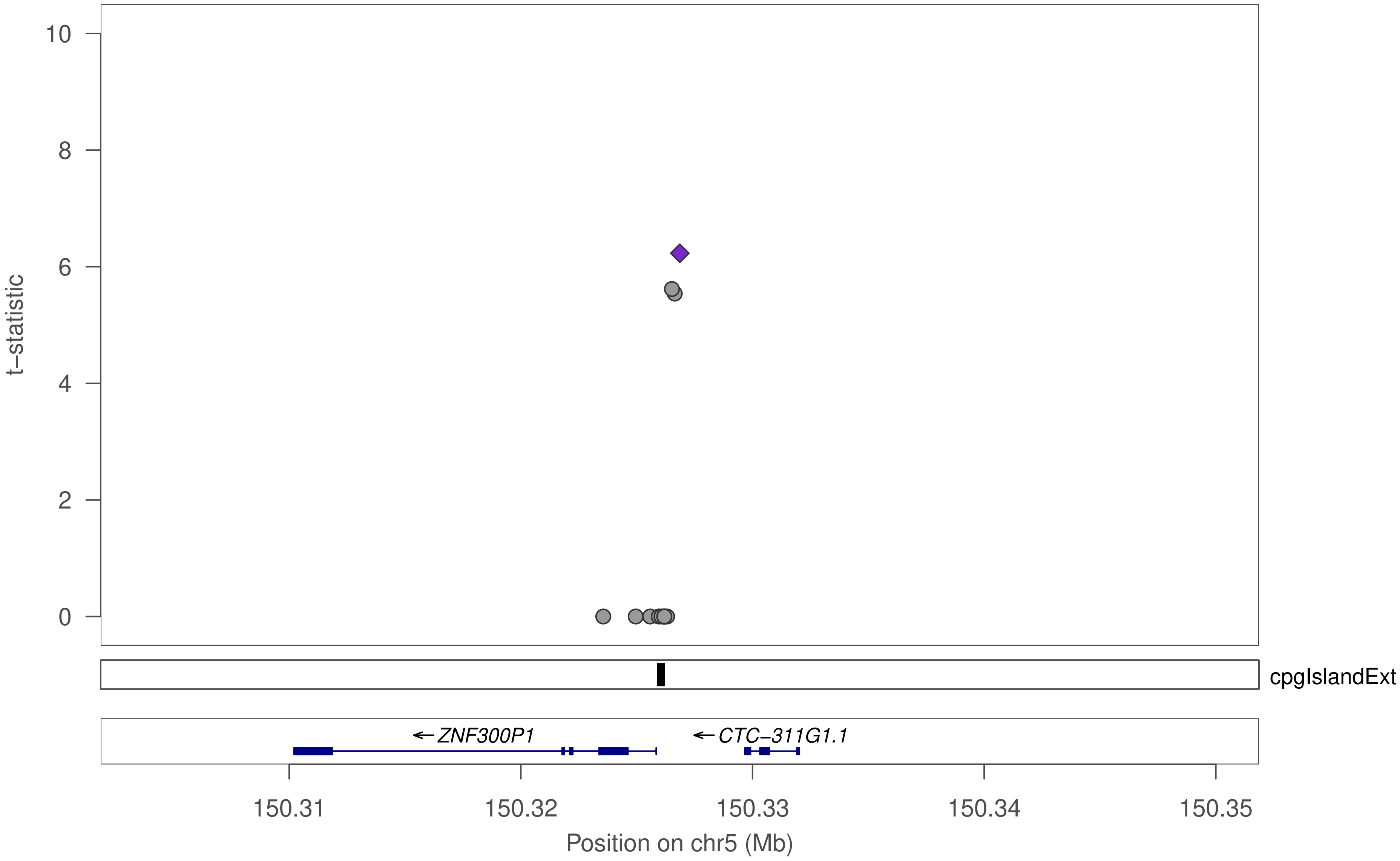

10
8
6
4
2
0
t−statistic
cpgIslandExt
←ZNF300P1
←CTC−311G1.1
150.31
150.32
150.33
150.34
150.35
Position on chr5 (Mb)

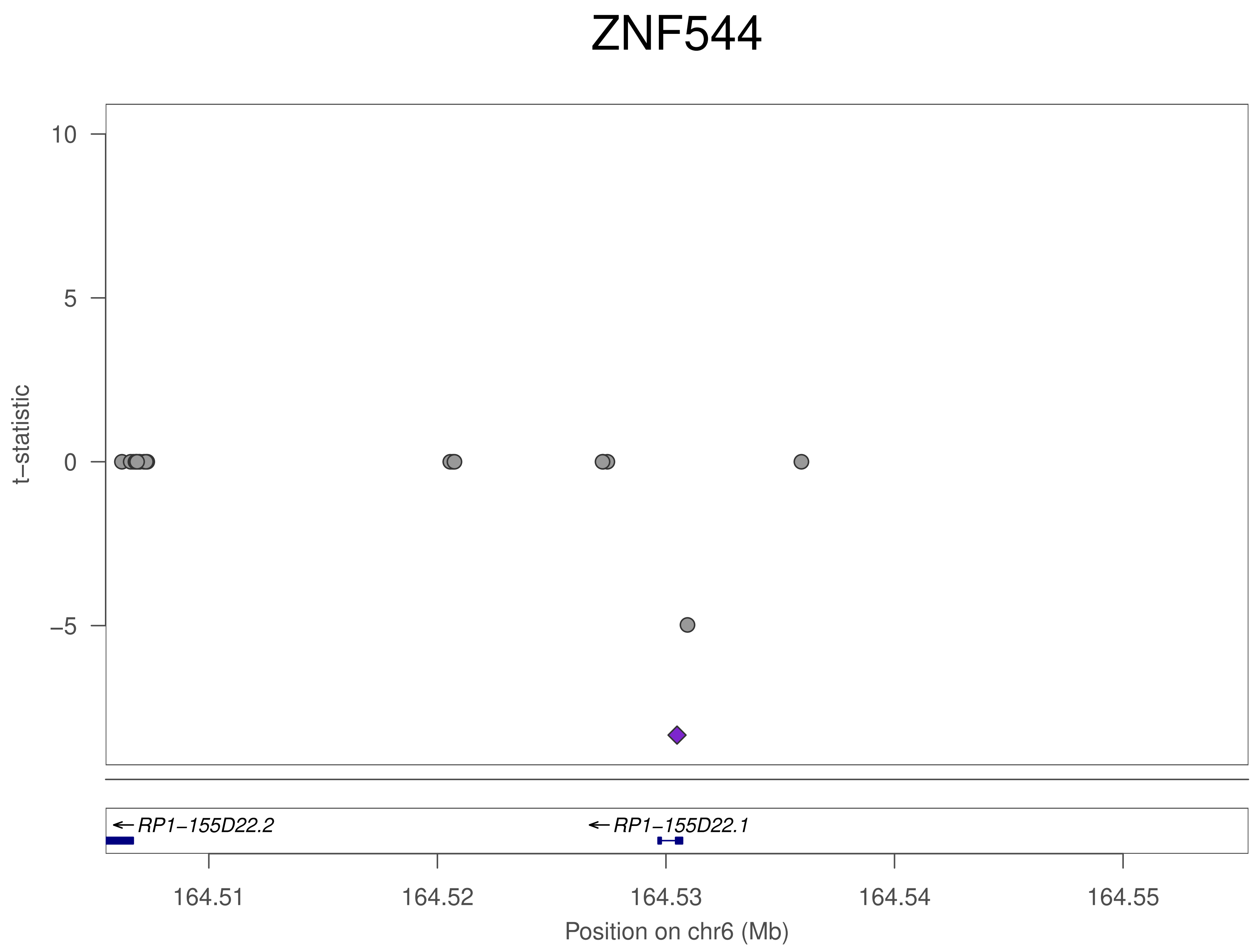

ZNF544
t–statistic
10
5
0
–5
RP1–155D22.2
RP1–155D22.1
164.51
164.52
164.53
164.54
164.55
Position on chr6 (Mb)

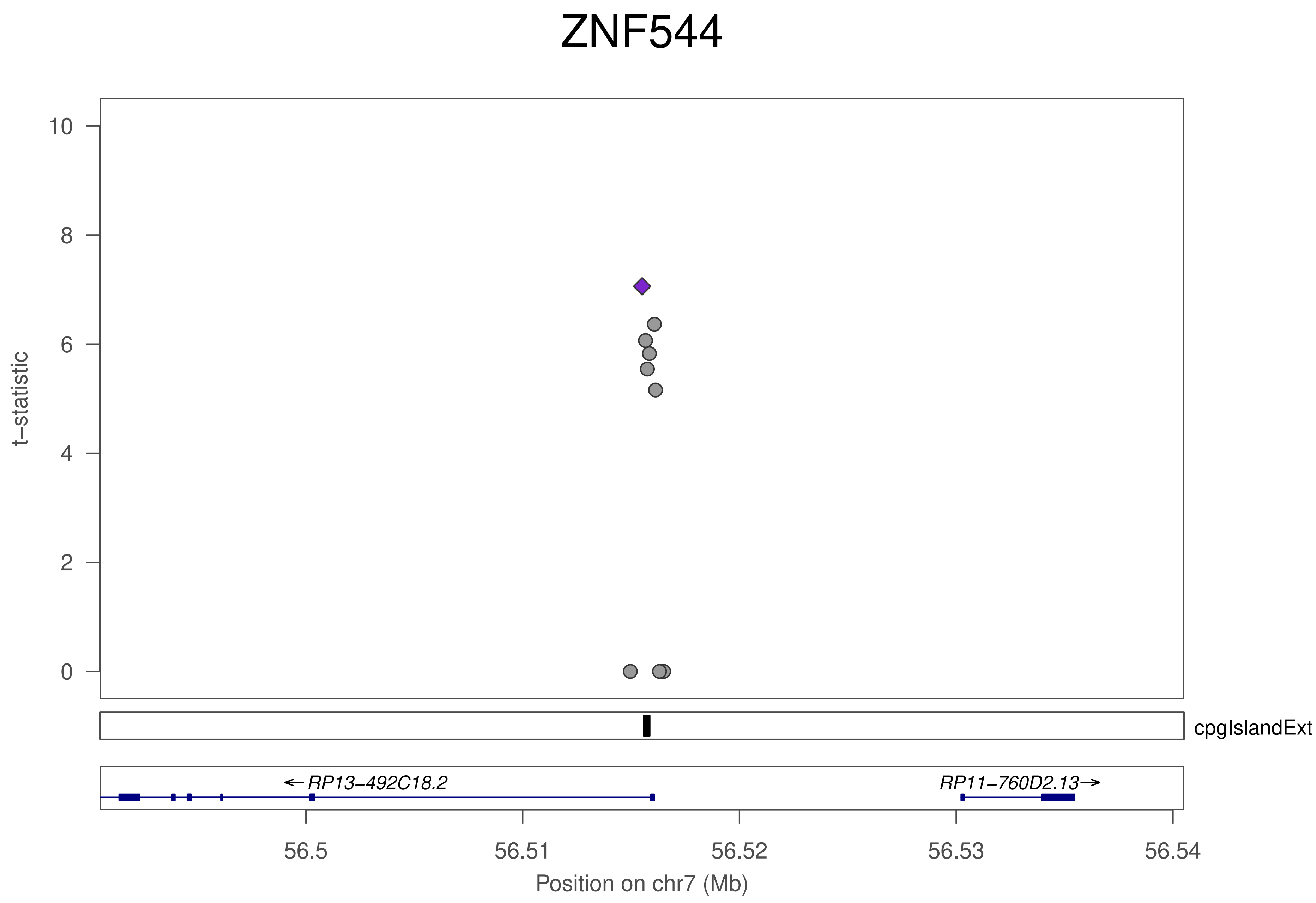
ZNF544
10
8
6
4
2
0
t−statistic
cpgIslandExt
←RP13−492C18.2
RP11−760D2.13→
56.5
56.51
56.52
56.53
56.54
Position on chr7 (Mb)

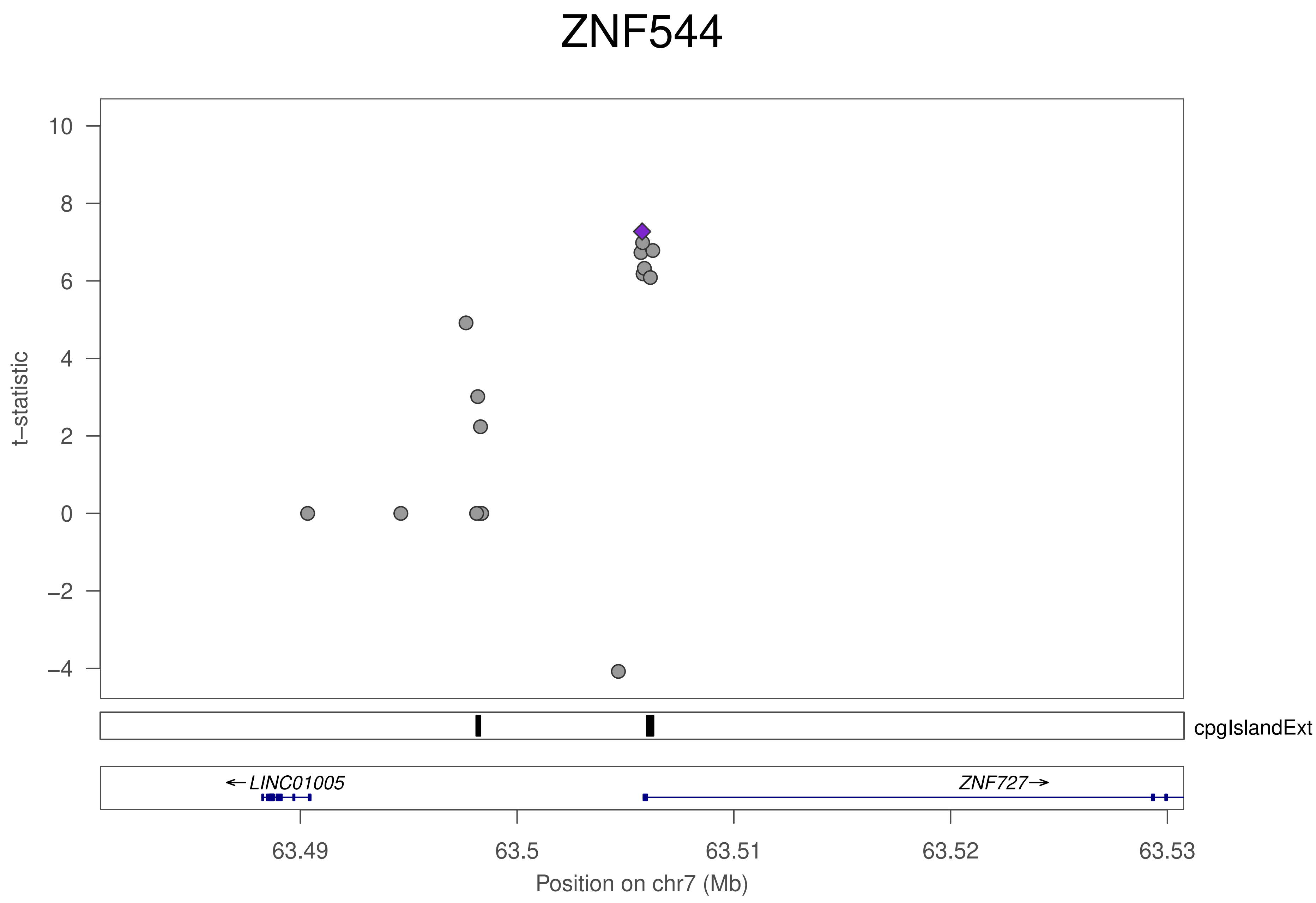
ZNF544
t–statistic
10
8
6
4
2
0
–2
–4
cpgIslandExt
←LINC01005
ZNF727→
63.49
63.5
63.51
63.52
63.53
Position on chr7 (Mb)

### ZNF544

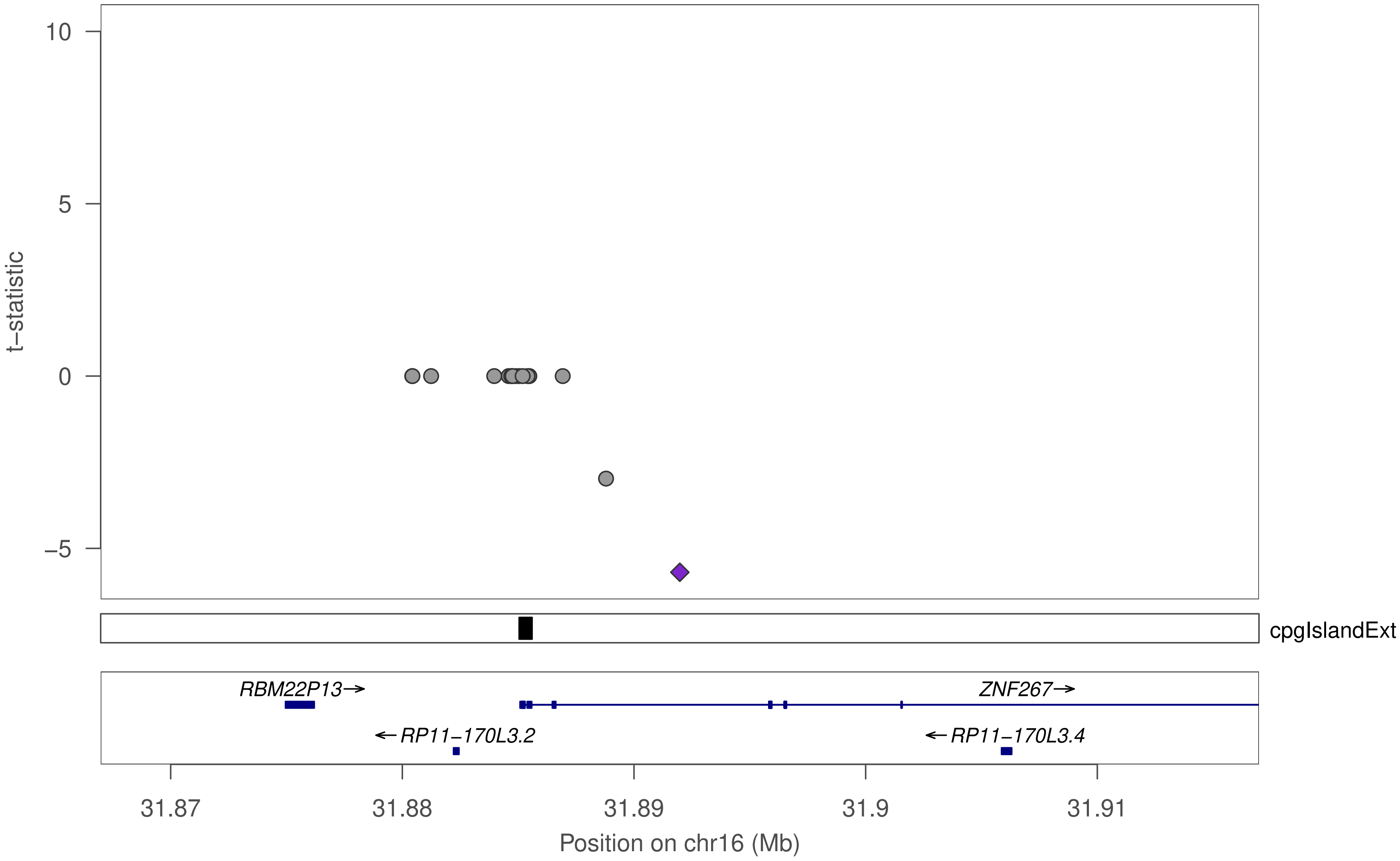

10
5
0
−5
t−statistic
cpgIslandExt
RBM22P13→
ZNF267→
←RP11−170L3.2
←RP11−170L3.4
31.87
31.88
31.89
31.9
31.91
Position on chr16 (Mb)

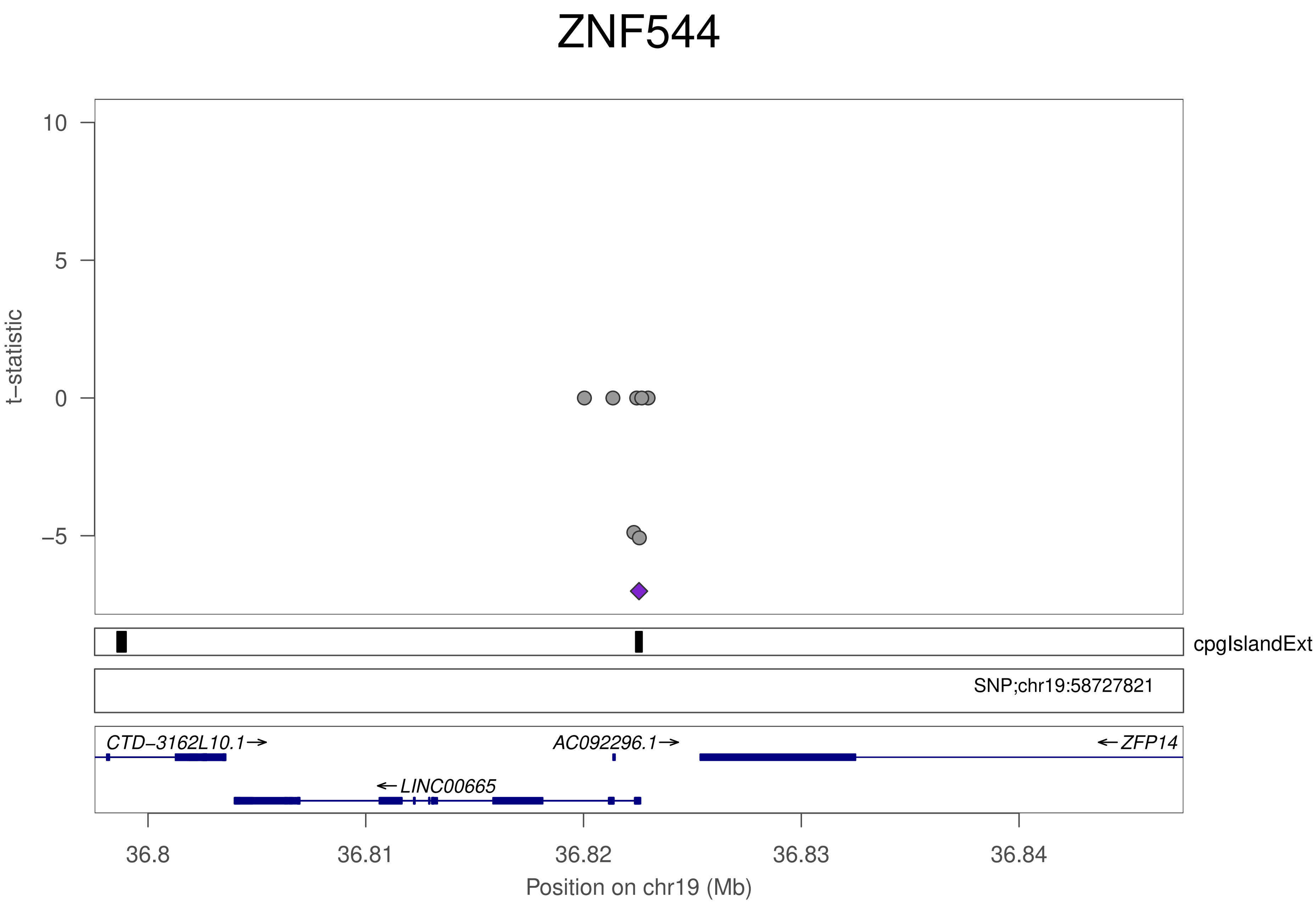

ZNF544
t−statistic
10
5
0
−5
cpgIslandExt
SNP;chr19:58727821
CTD−3162L10.1→
AC092296.1→
←ZFP14
←LINC00665
36.8
36.81
36.82
36.83
36.84
Position on chr19 (Mb)

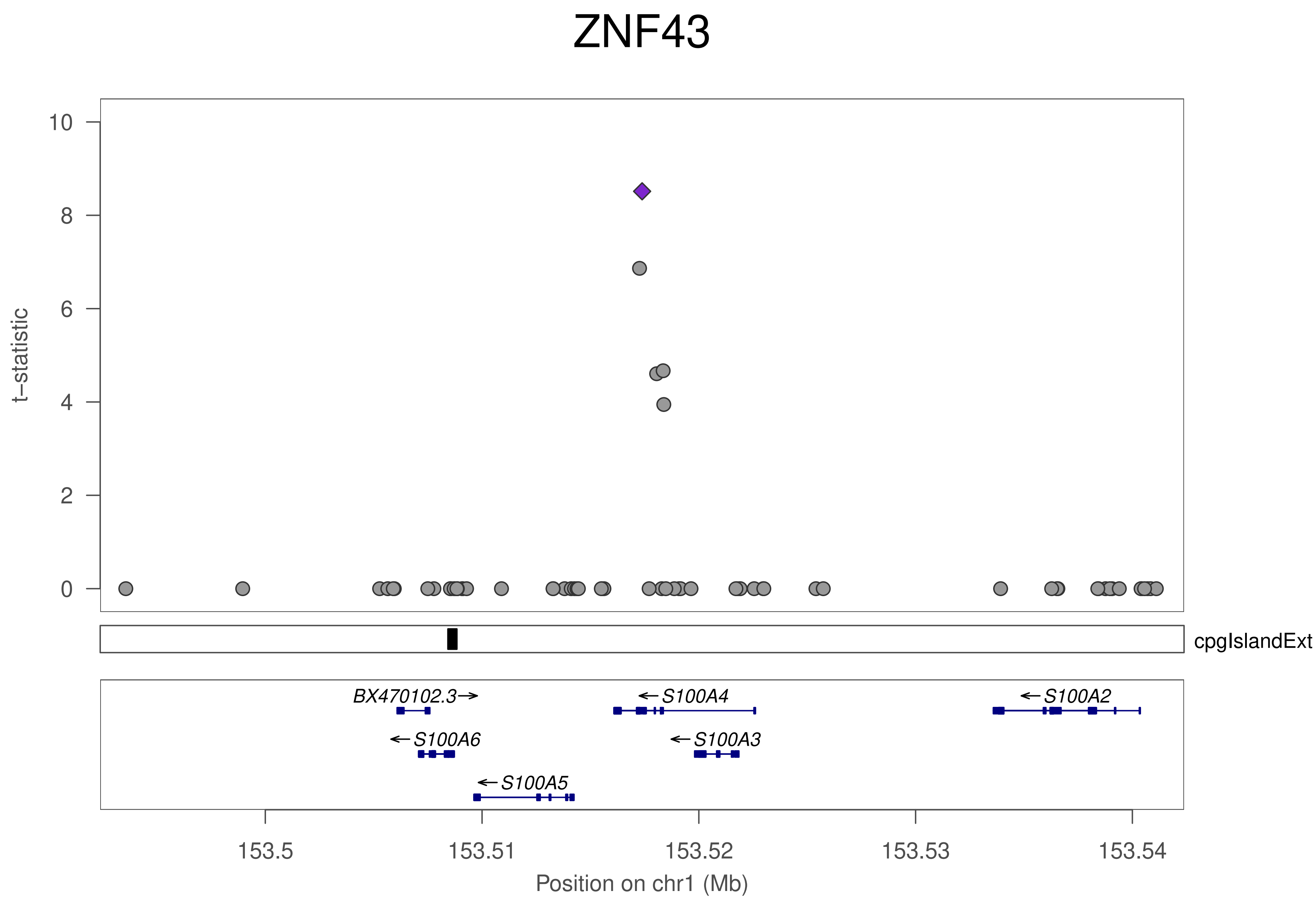

ZNF43
t–statistic
10
8
6
4
2
0
cpgIslandExt
BX470102.3→
←S100A4
←S100A2
←S100A6
←S100A3
←S100A5
153.5
153.51
153.52
153.53
153.54
Position on chr1 (Mb)

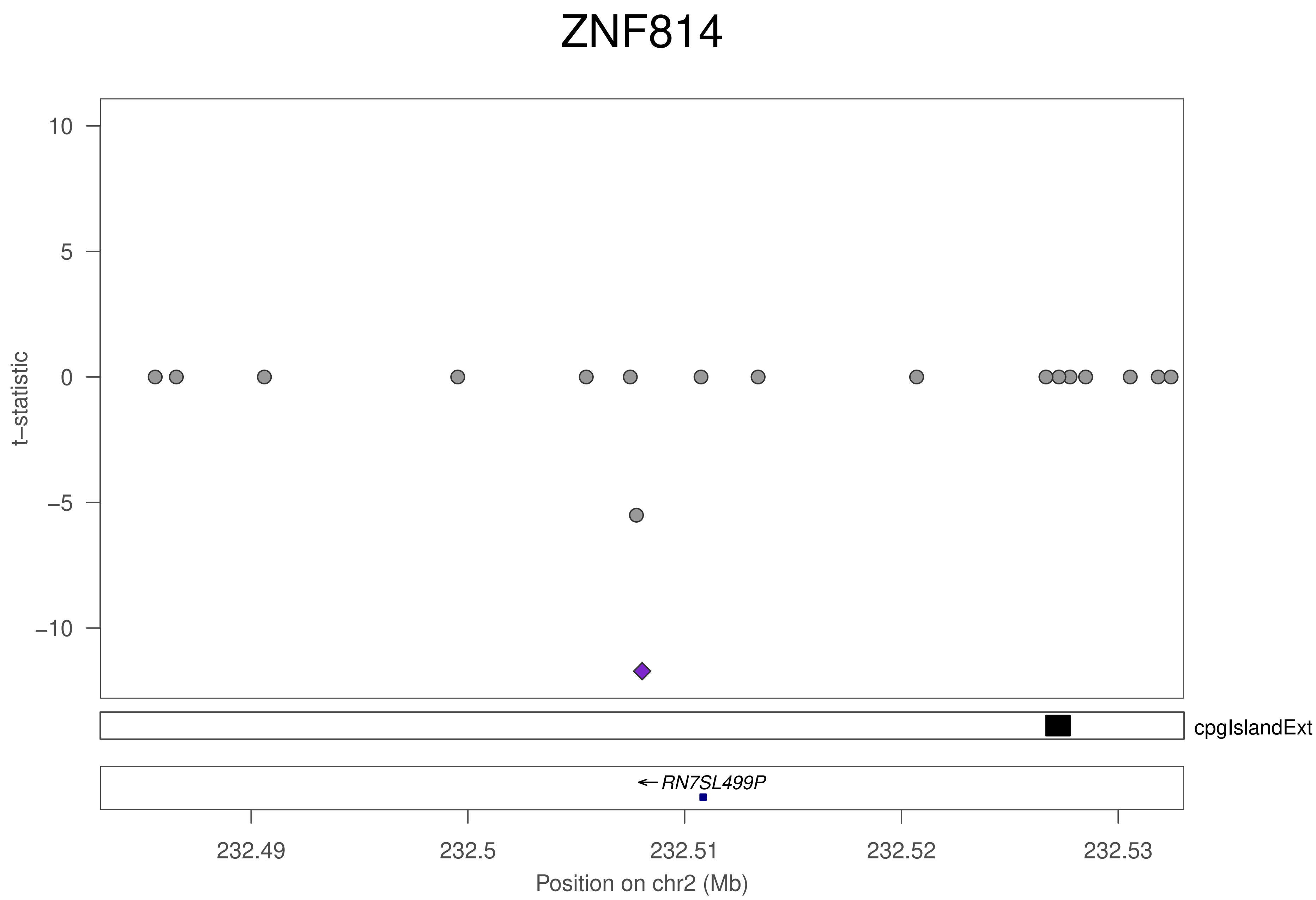
ZNF814
t−statistic
10
5
0
−5
−10
cpgIslandExt
RN7SL499P
232.49
232.5
232.51
232.52
232.53
Position on chr2 (Mb)

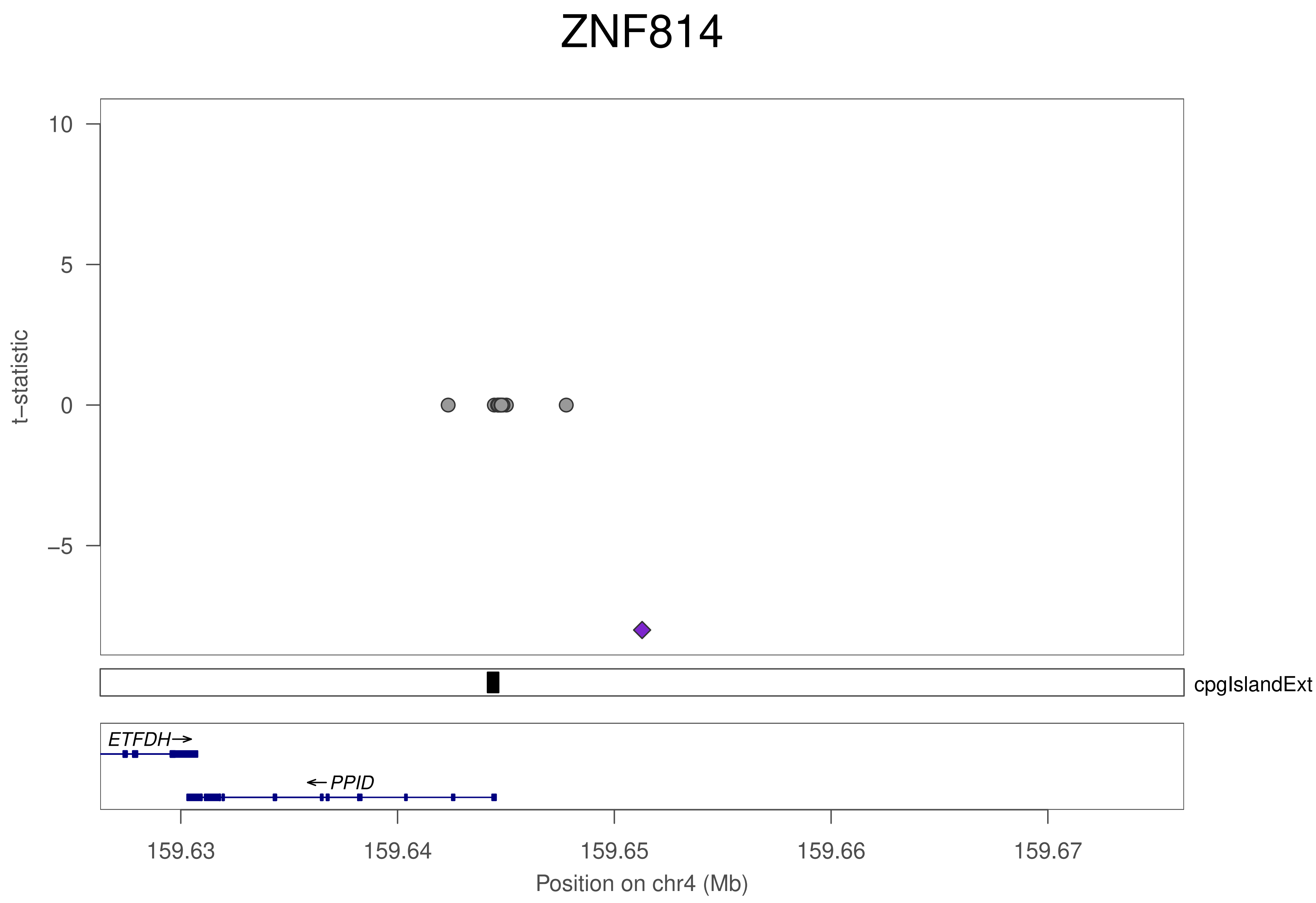
ZNF814
t−statistic
10
5
0
−5
cpgIslandExt
ETFDH
PPID
159.63
159.64
159.65
159.66
159.67
Position on chr4 (Mb)

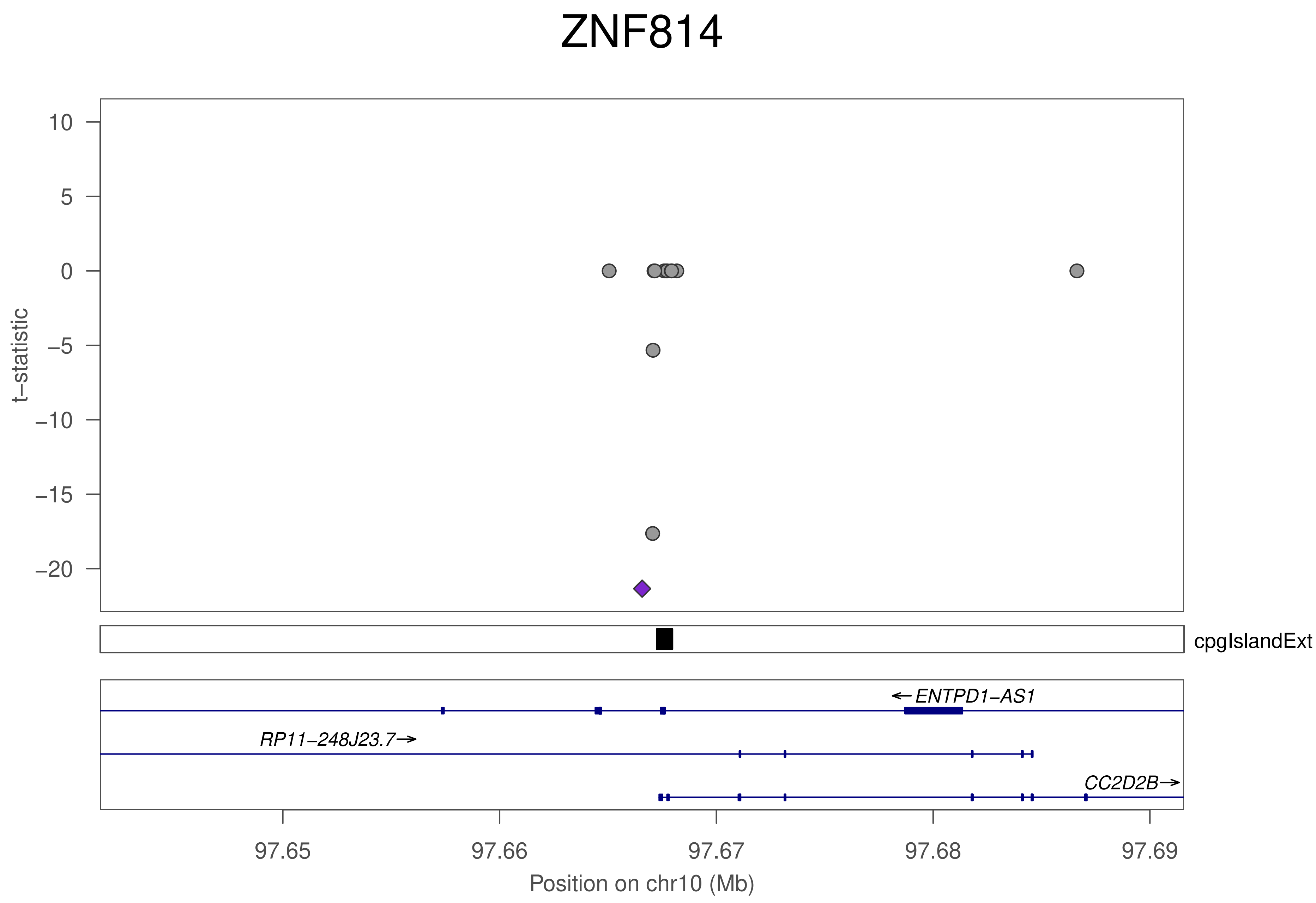
ZNF814
t−statistic
10
5
0
−5
−10
−15
−20
cpgIslandExt
←ENTPD1−AS1
RP11−248J23.7→
CC2D2B→
97.65
97.66
97.67
97.68
97.69
Position on chr10 (Mb)

### ZNF814

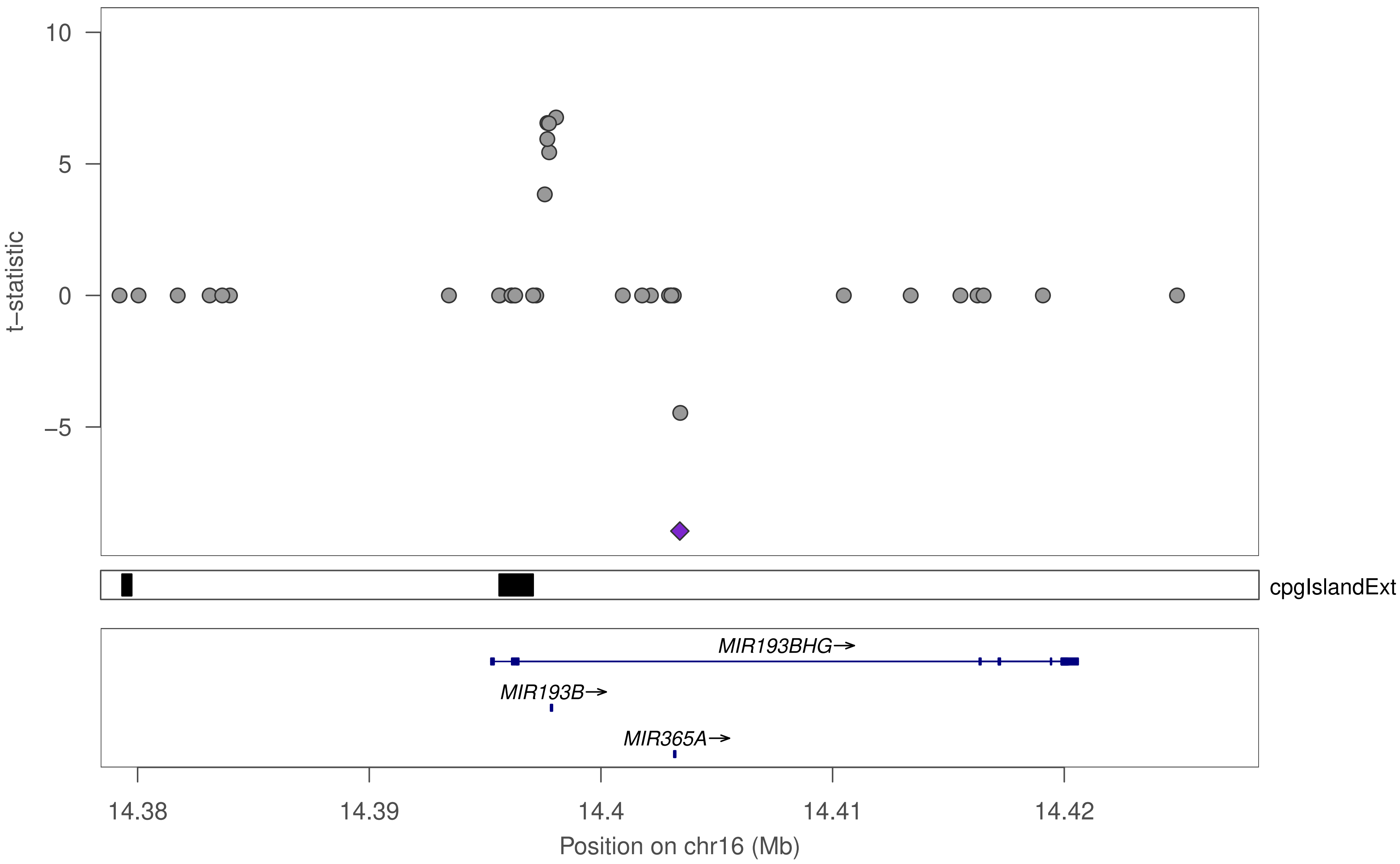
10
5
0
−5
t−statistic
cpgIslandExt
MIR193BHG→
MIR193B→
MIR365A→
14.38
14.39
14.4
14.41
14.42
Position on chr16 (Mb)

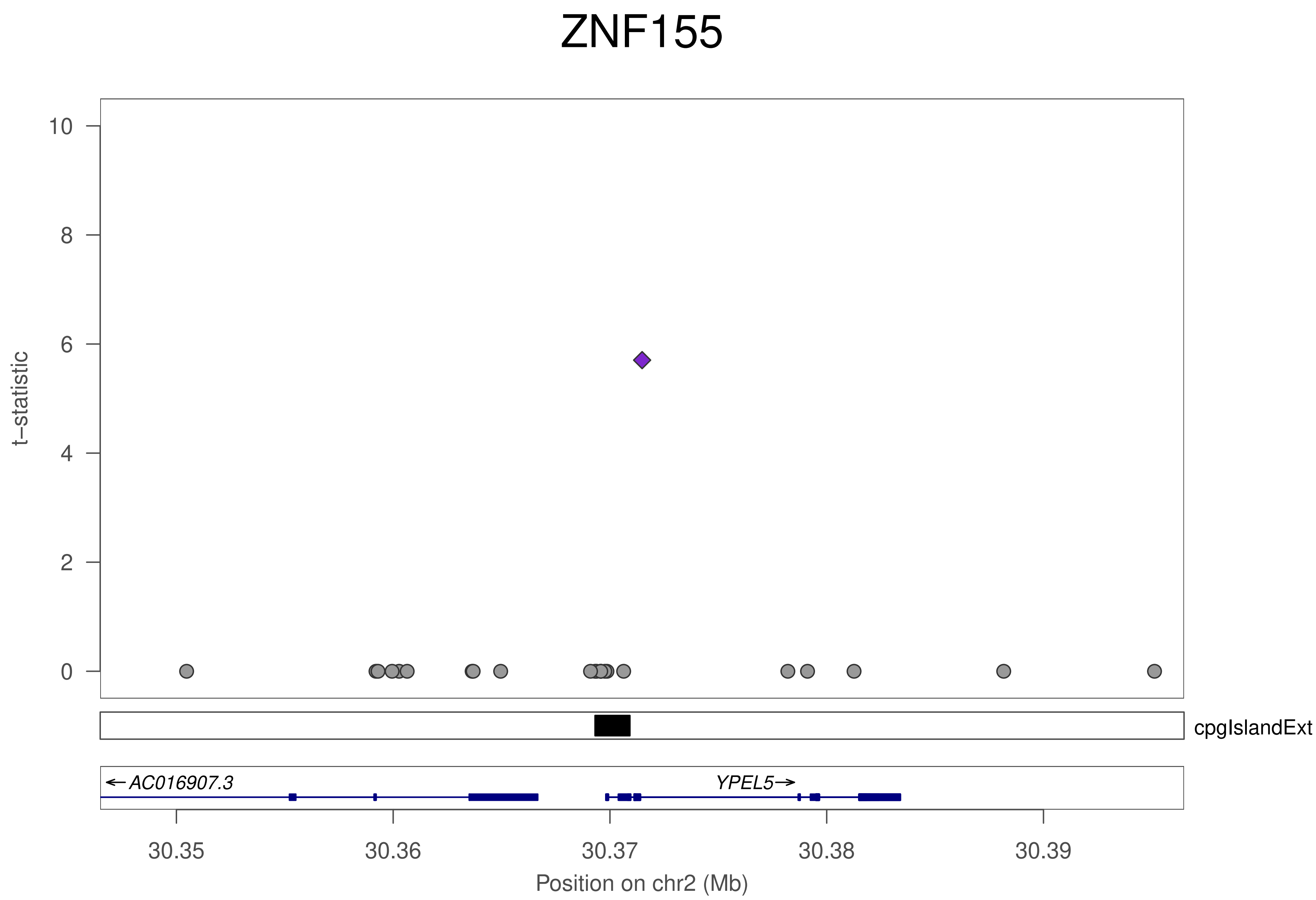

ZNF155
t–statistic
10
8
6
4
2
0
cpgIslandExt
←AC016907.3
YPEL5→
30.35
30.36
30.37
30.38
30.39
Position on chr2 (Mb)

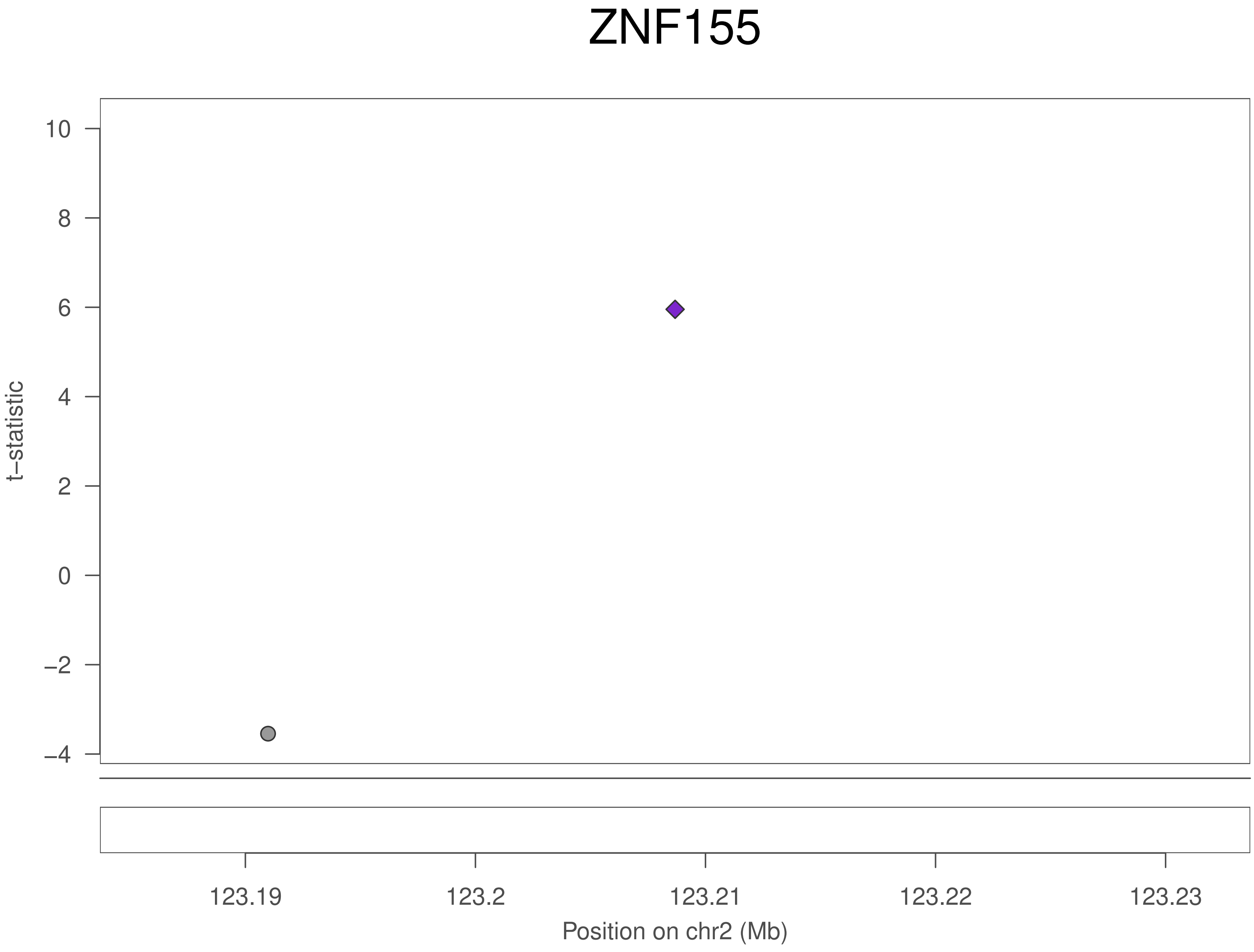
ZNF155
t−statistic
10
8
6
4
2
0
−2
−4
123.19
123.2
123.21
123.22
123.23
Position on chr2 (Mb)

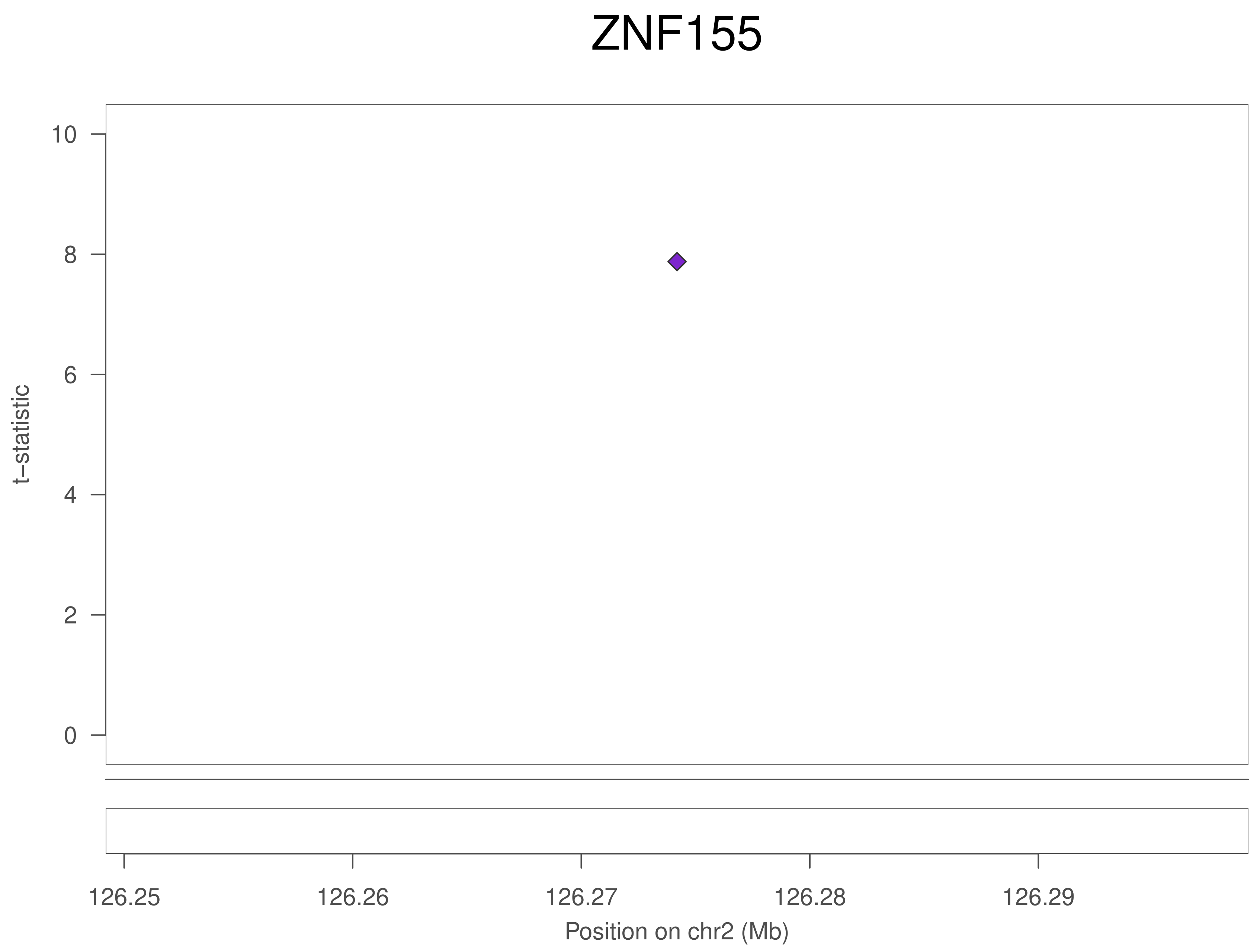
ZNF155
t−statistic
10
8
6
4
2
0
126.25
126.26
126.27
126.28
126.29
Position on chr2 (Mb)

### ZNF155

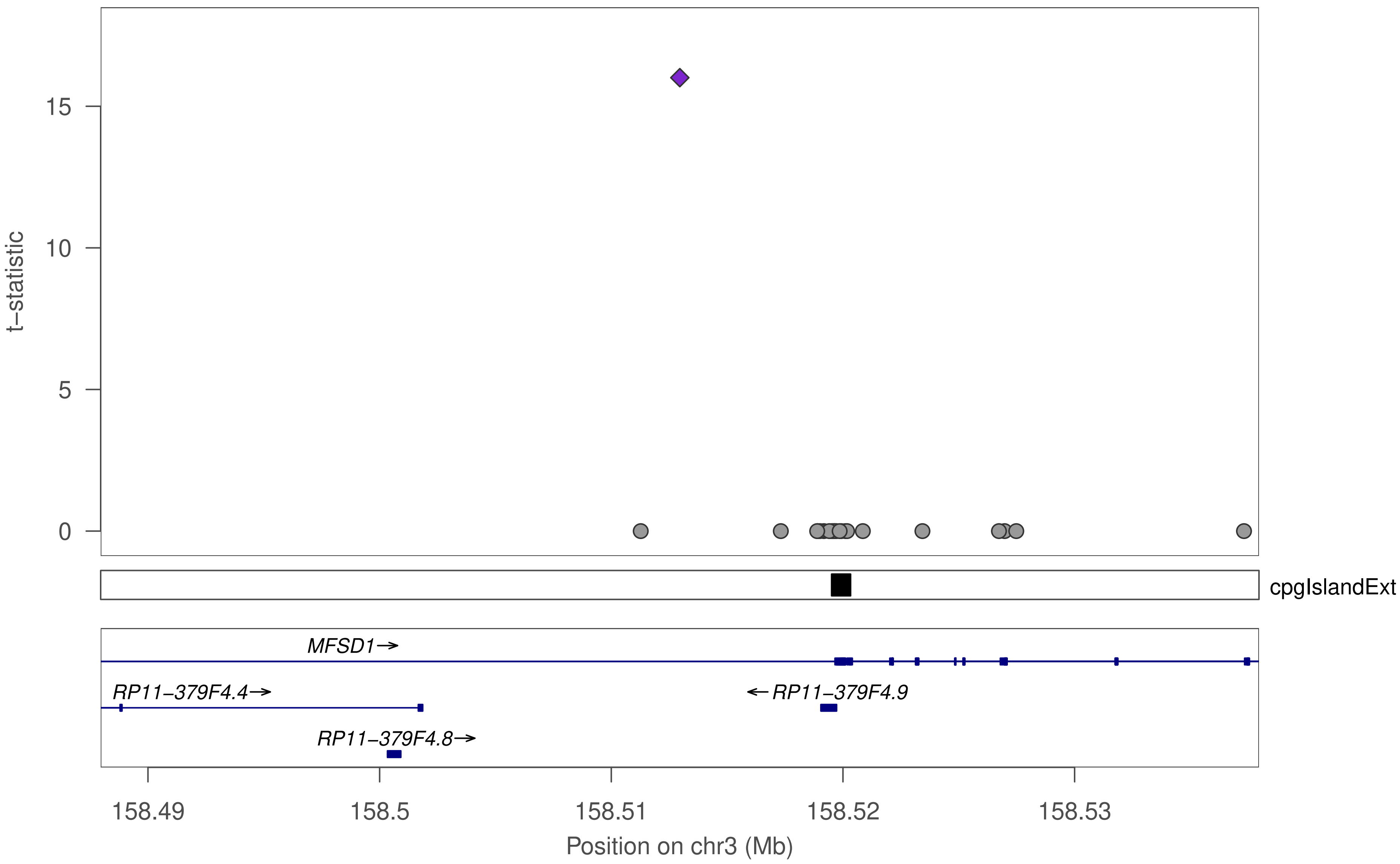

15
10
5
0
t–statistic
cpgIslandExt
MFSD1→
RP11–379F4.4→
←RP11–379F4.9
RP11–379F4.8→
158.49
158.5
158.51
158.52
158.53
Position on chr3 (Mb)

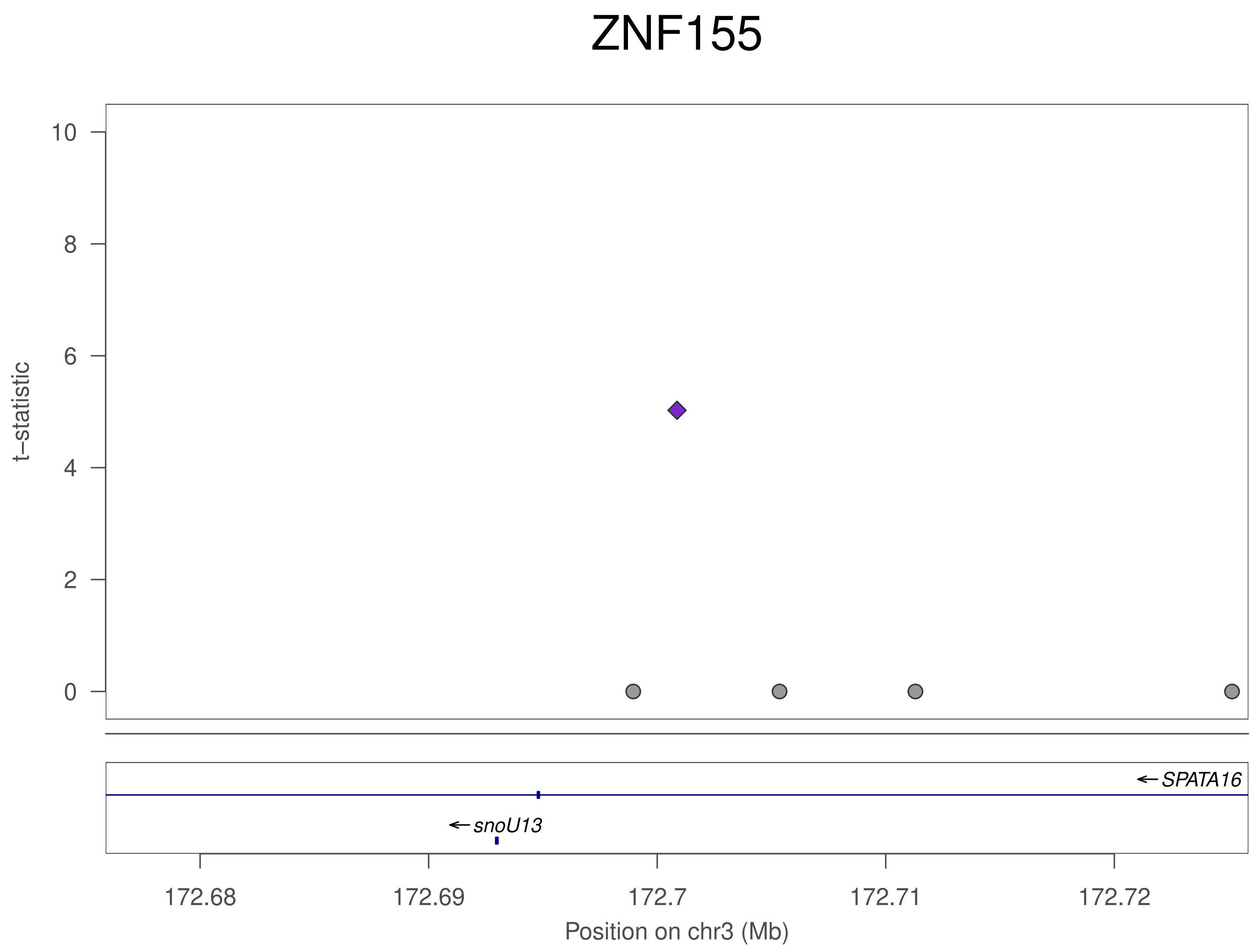

ZNF155
t−statistic
10
8
6
4
2
0
SPATA16
snoU13
172.68
172.69
172.7
172.71
172.72
Position on chr3 (Mb)

### ZNF155

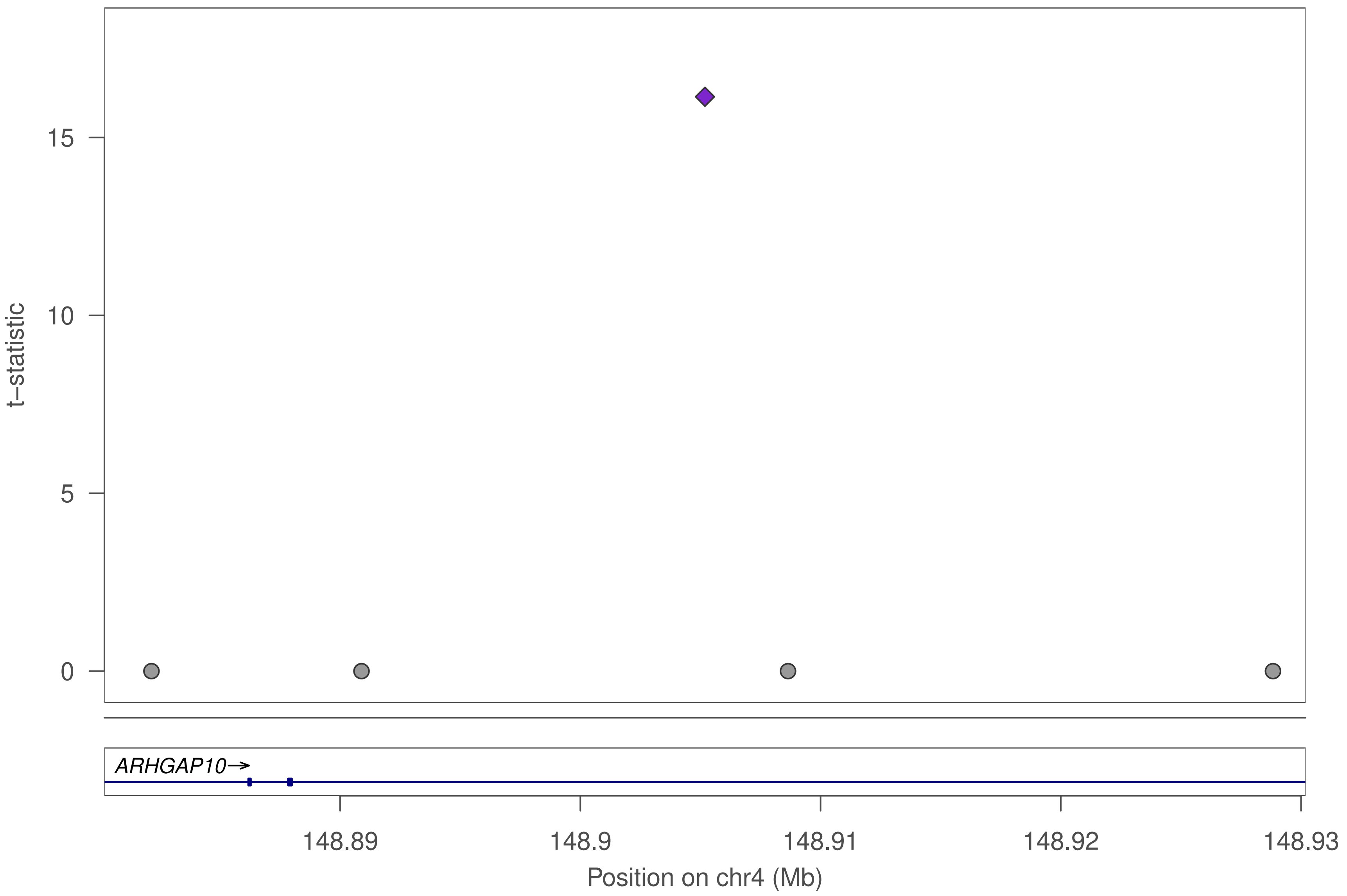

t–statistic
0
5
10
15
ARHGAP10→
148.89
148.9
148.91
148.92
148.93
Position on chr4 (Mb)

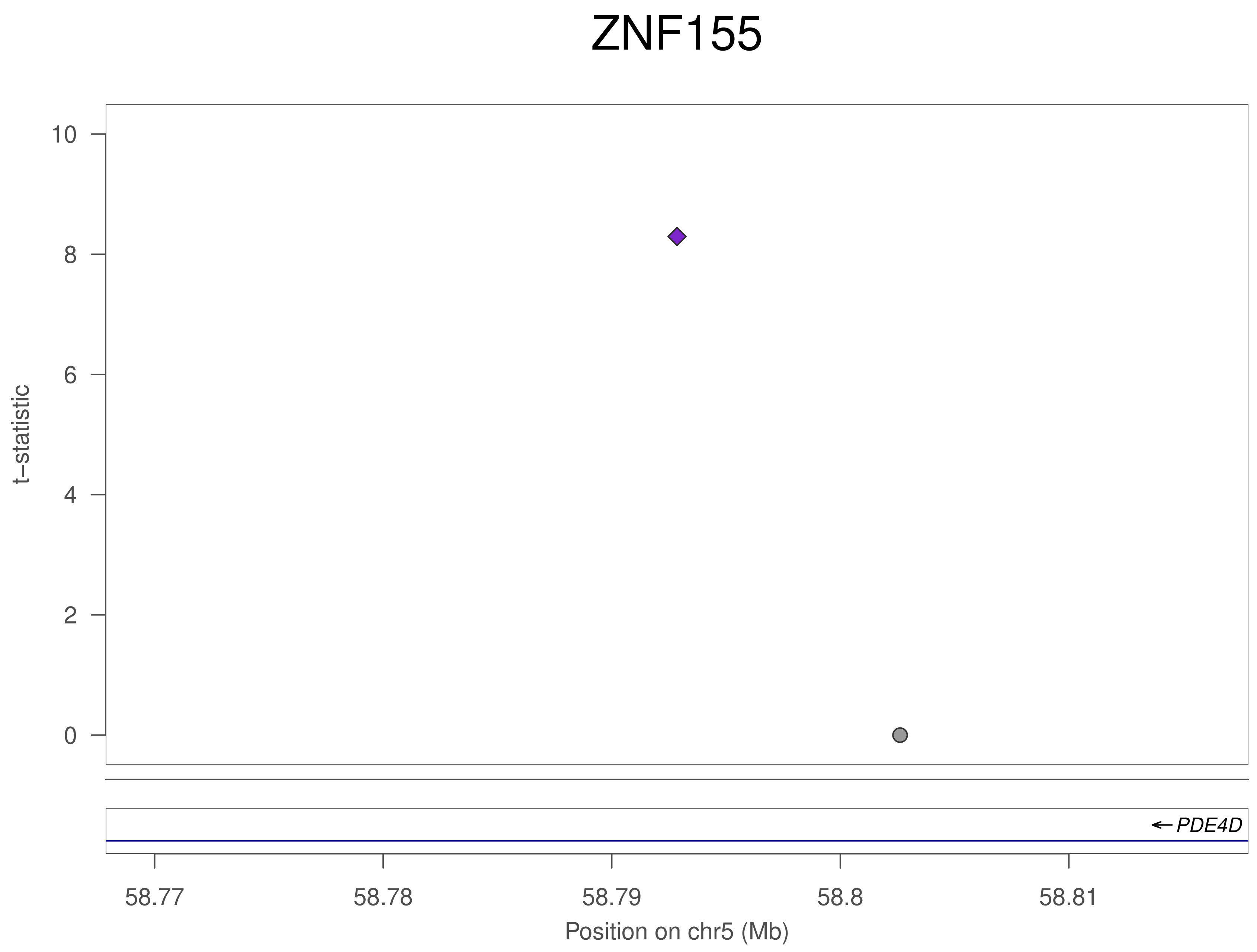
ZNF155
t–statistic
10
8
6
4
2
0
PDE4D
58.77
58.78
58.79
58.8
58.81
Position on chr5 (Mb)

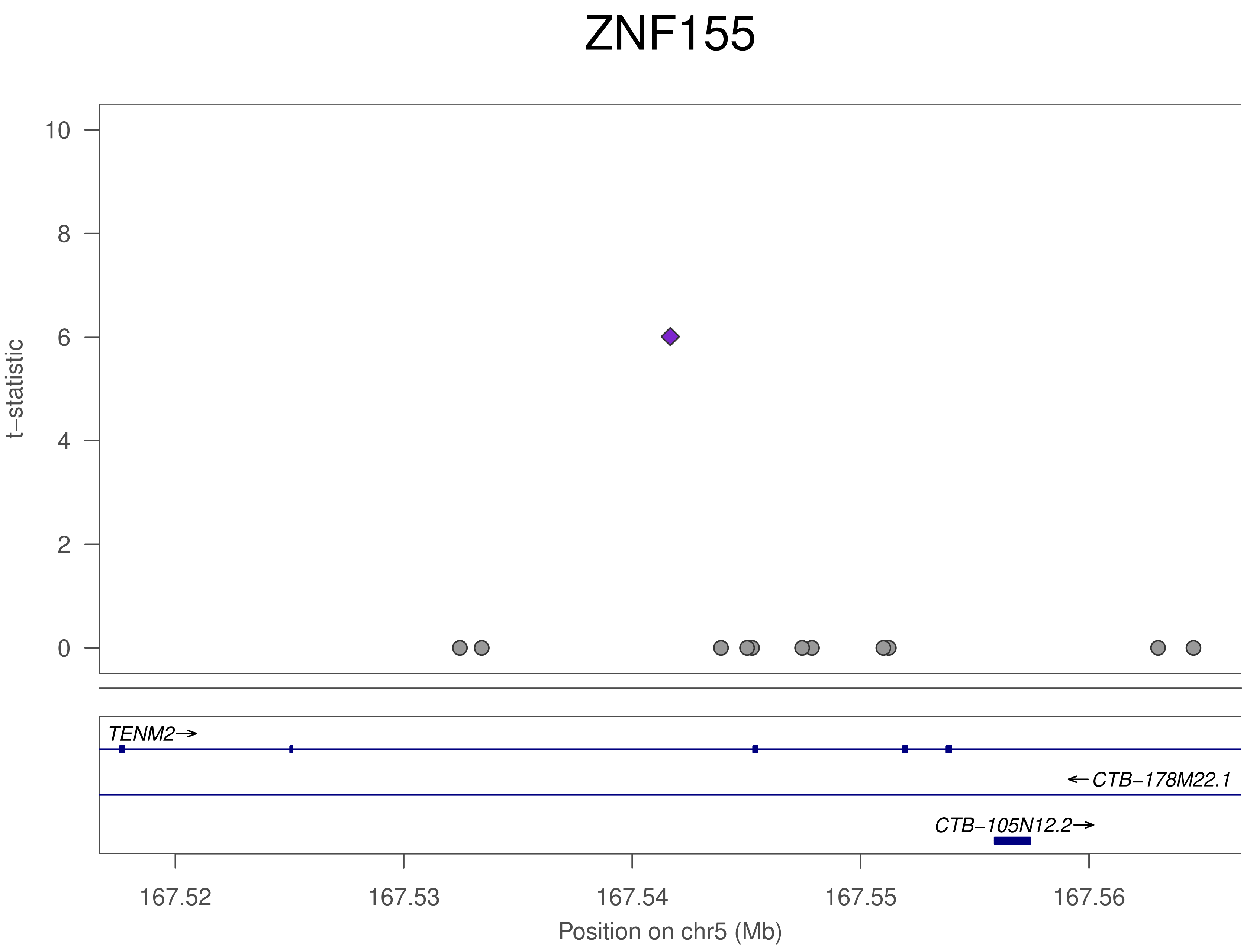

ZNF155
t−statistic
0
2
4
6
8
10
TENM2→
←CTB−178M22.1
CTB−105N12.2→
167.52
167.53
167.54
167.55
167.56
Position on chr5 (Mb)

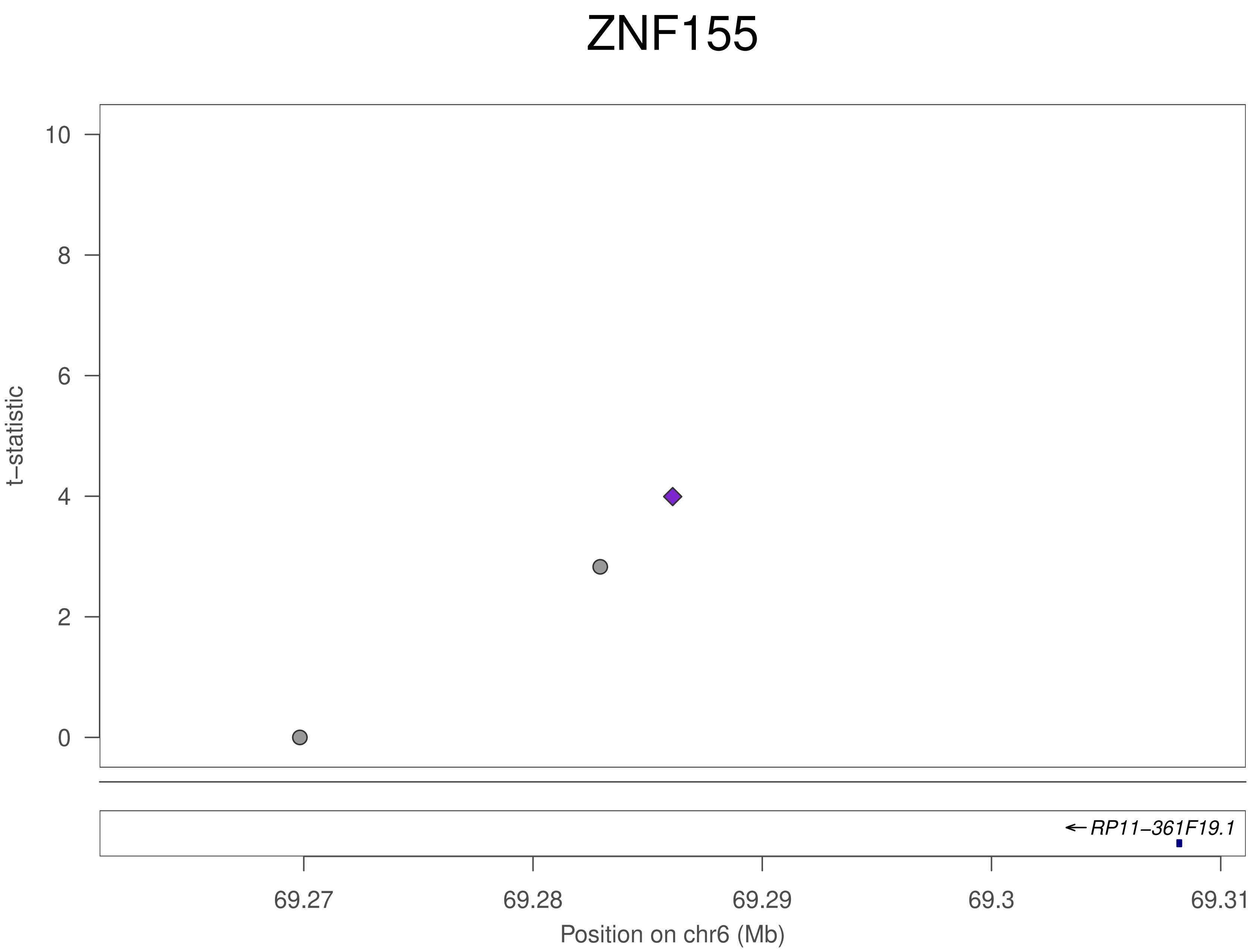
ZNF155
t–statistic
10
8
6
4
2
0
RP11−361F19.1
69.27
69.28
69.29
69.3
69.31
Position on chr6 (Mb)

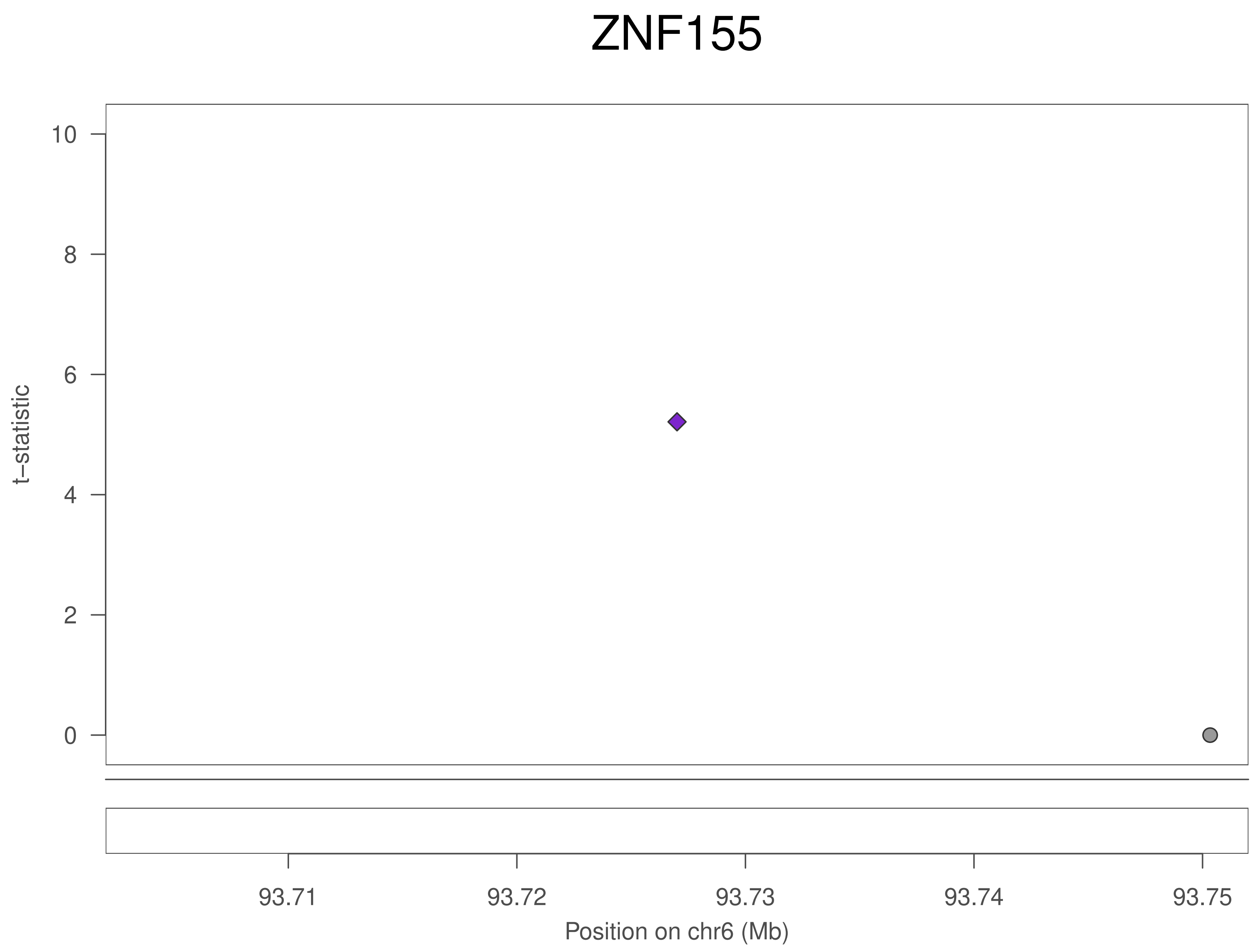

ZNF155
t–statistic
10
8
6
4
2
0
93.71
93.72
93.73
93.74
93.75
Position on chr6 (Mb)

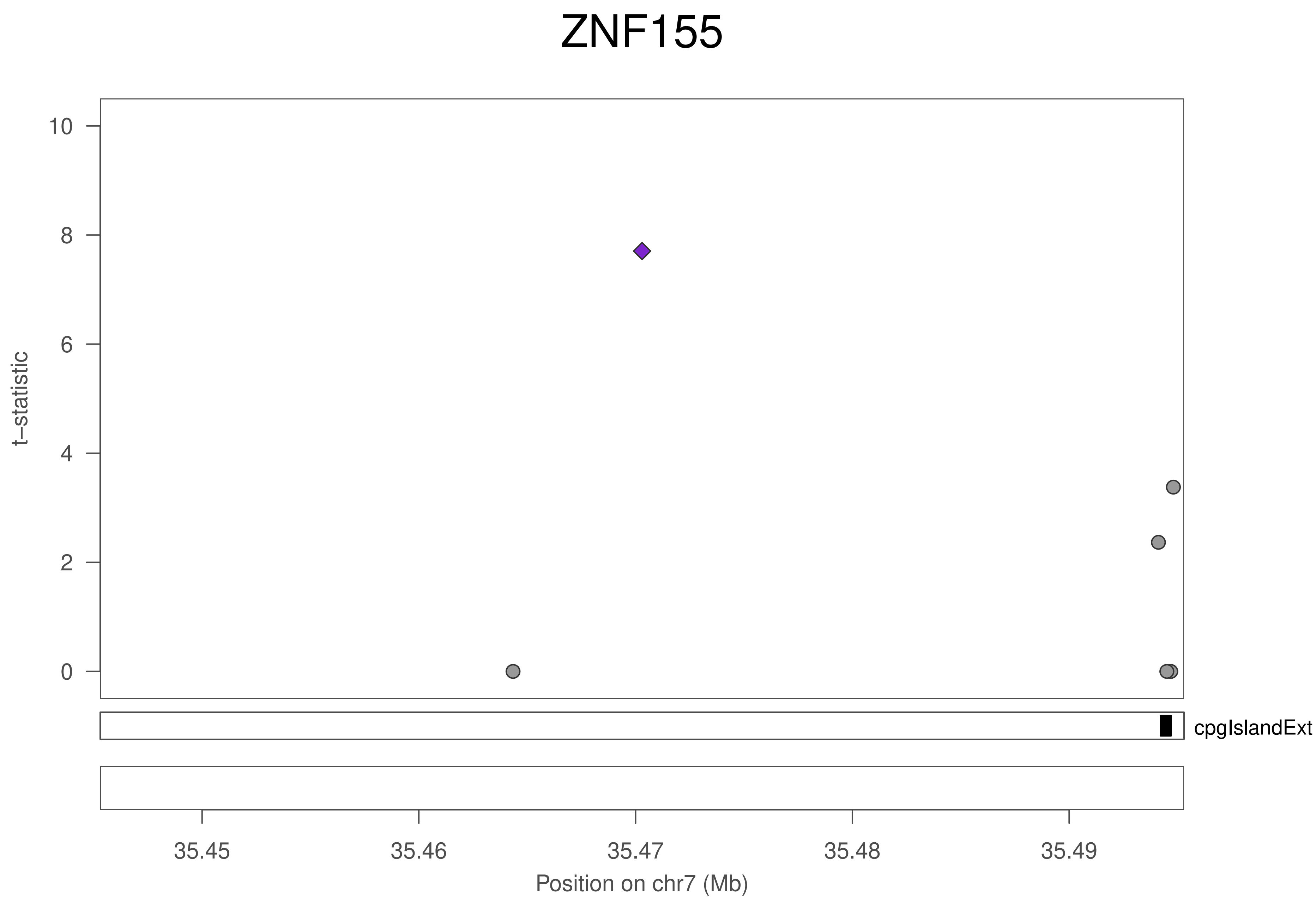

ZNF155
t–statistic
10
8
6
4
2
0
cpgIslandExt
35.45
35.46
35.47
35.48
35.49
Position on chr7 (Mb)

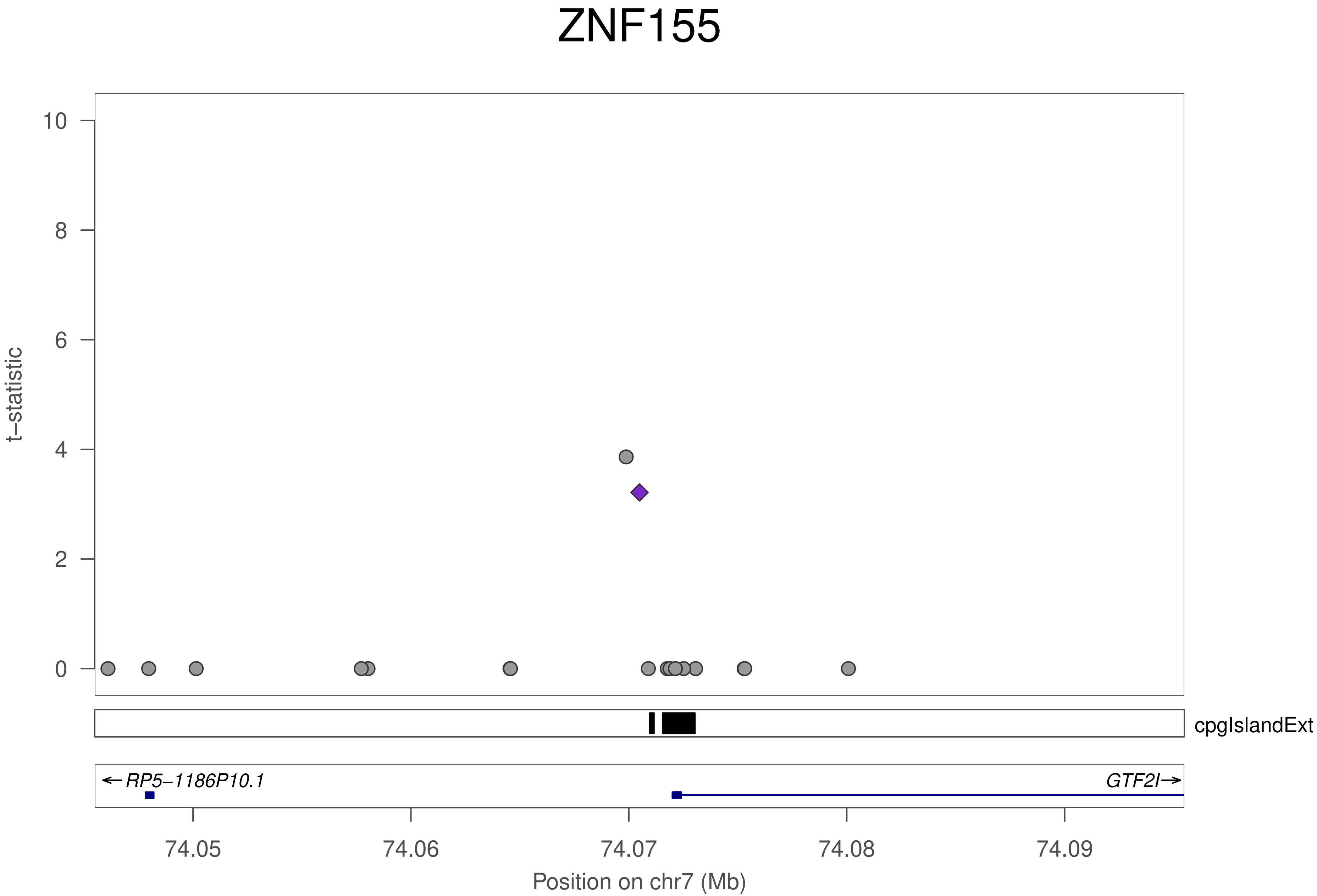
ZNF155
t–statistic
10
8
6
4
2
0
cpgIslandExt
←RP5–1186P10.1
GTF2I→
74.05
74.06
74.07
74.08
74.09
Position on chr7 (Mb)

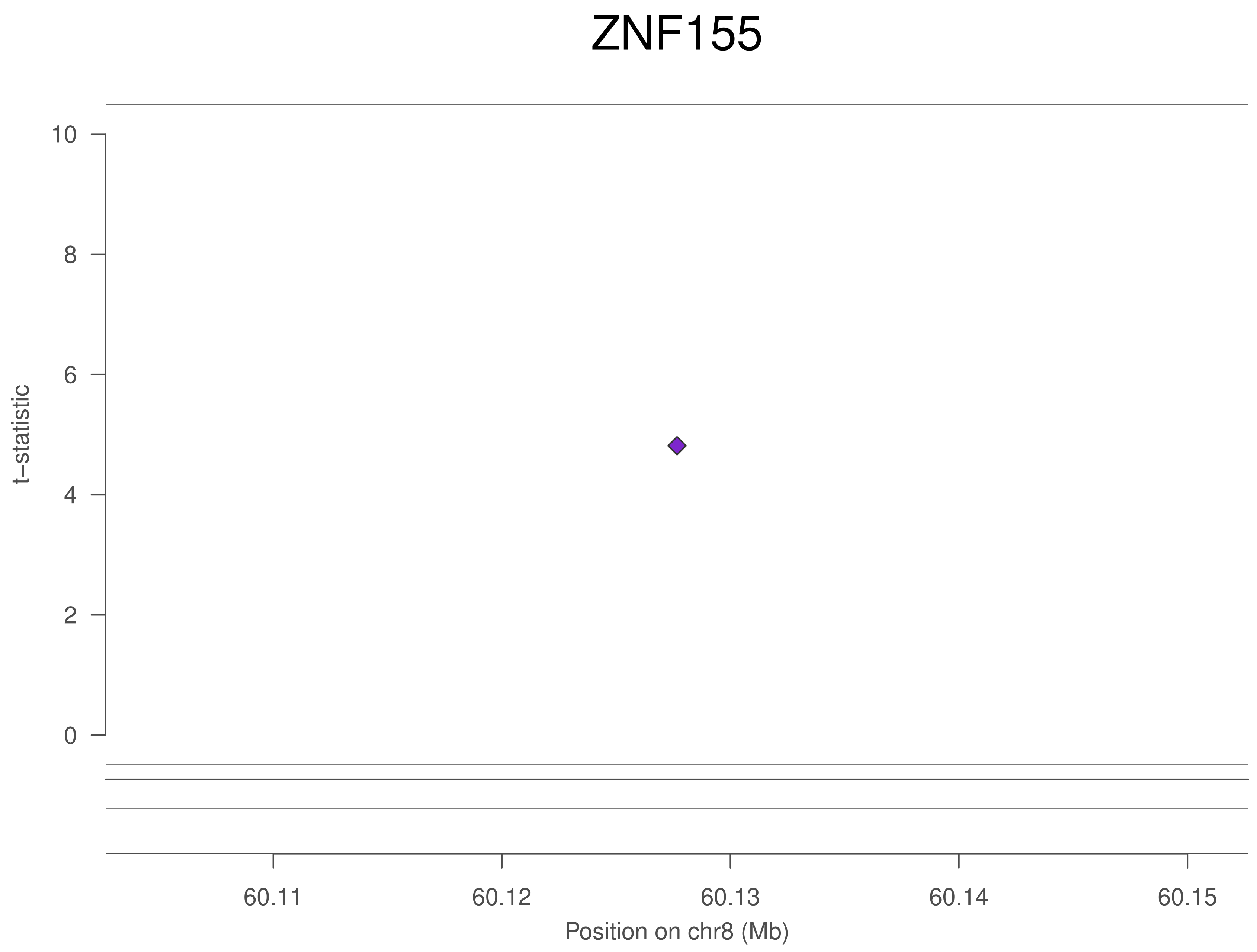
ZNF155
t−statistic
10
8
6
4
2
0
60.11
60.12
60.13
60.14
60.15
Position on chr8 (Mb)

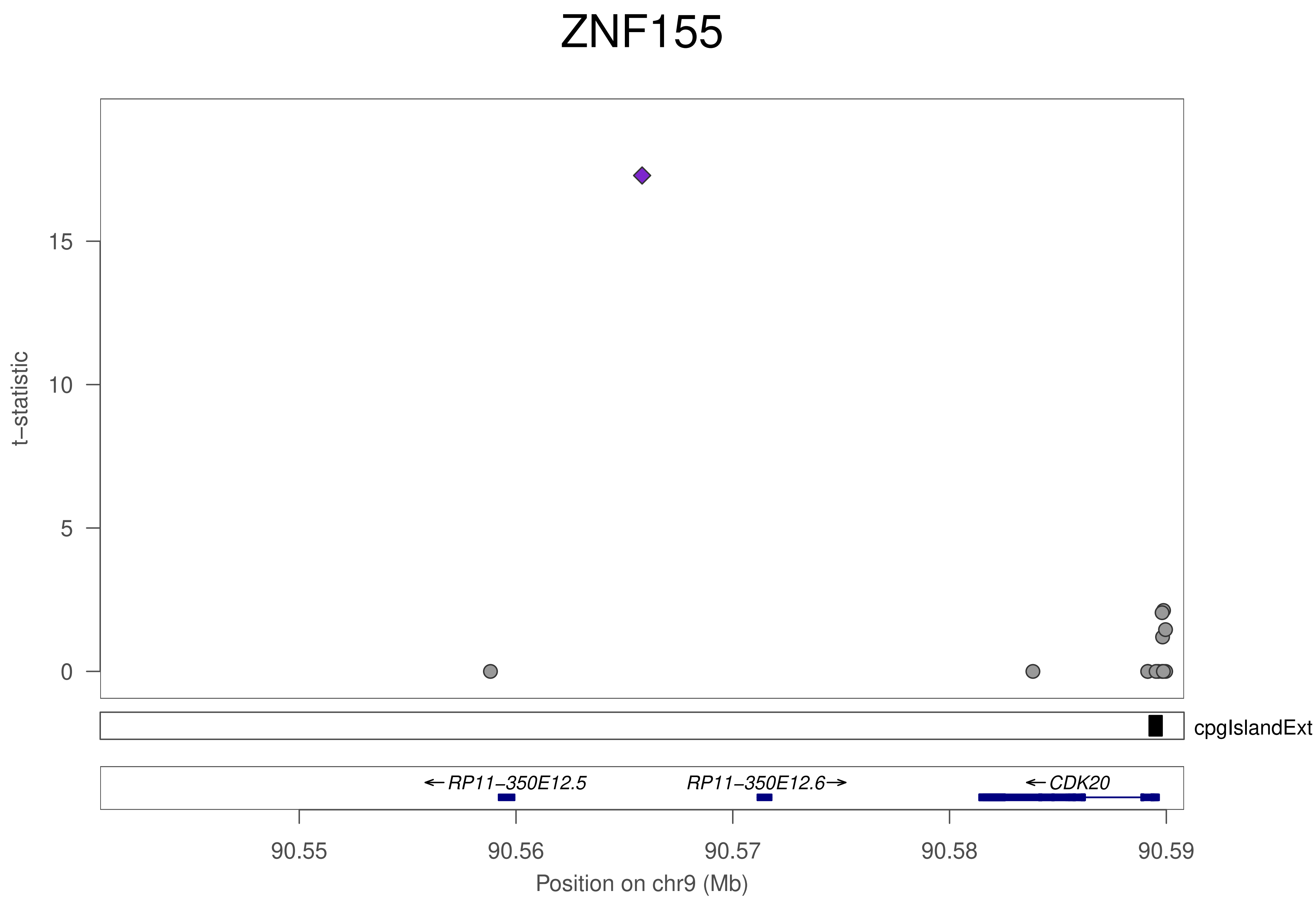
ZNF155
t−statistic
15
10
5
0
cpgIslandExt
←RP11−350E12.5
RP11−350E12.6→
←CDK20
90.55
90.56
90.57
90.58
90.59
Position on chr9 (Mb)

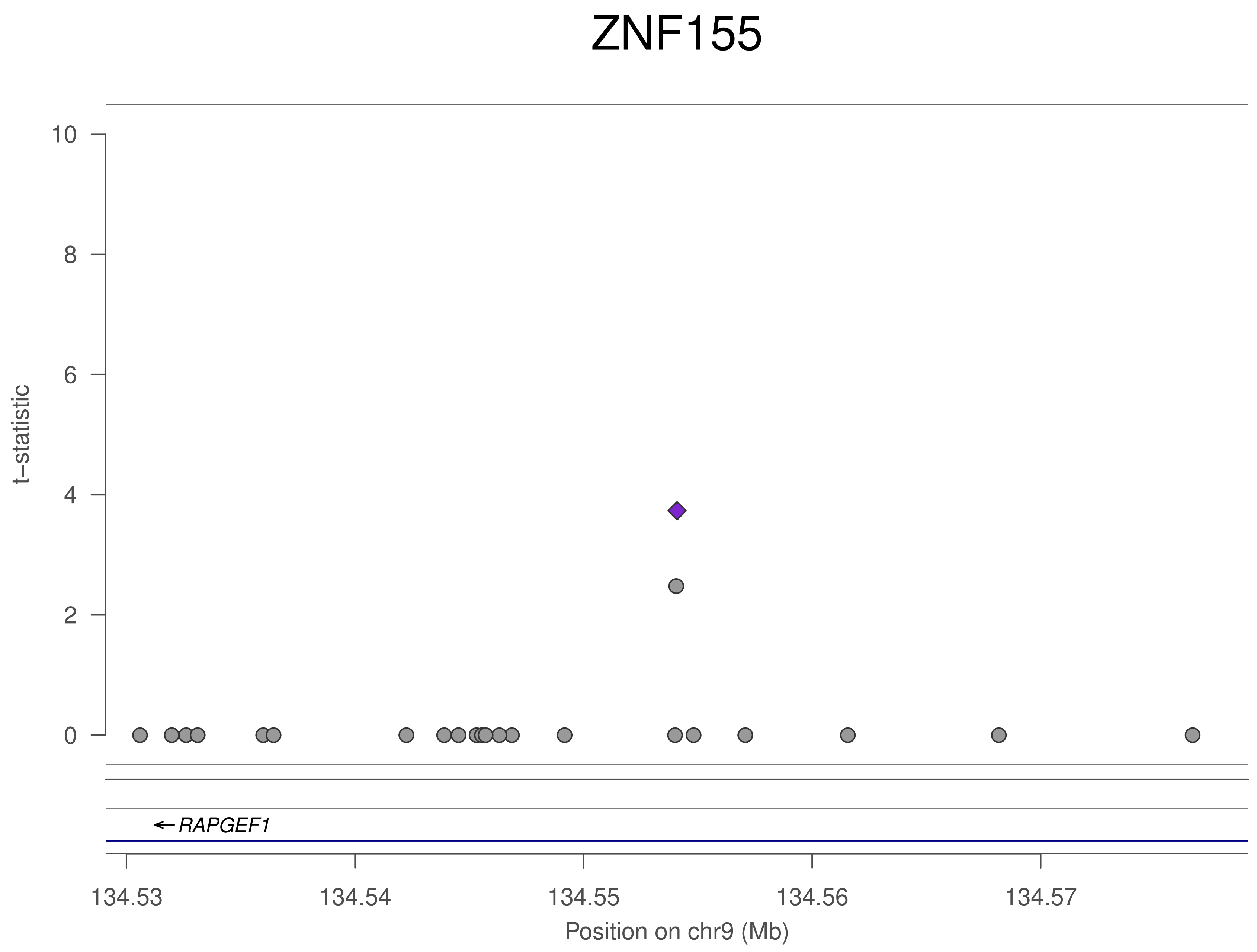

ZNF155
t–statistic
0
2
4
6
8
10
RAPGEF1
134.53
134.54
134.55
134.56
134.57
Position on chr9 (Mb)

### ZNF155

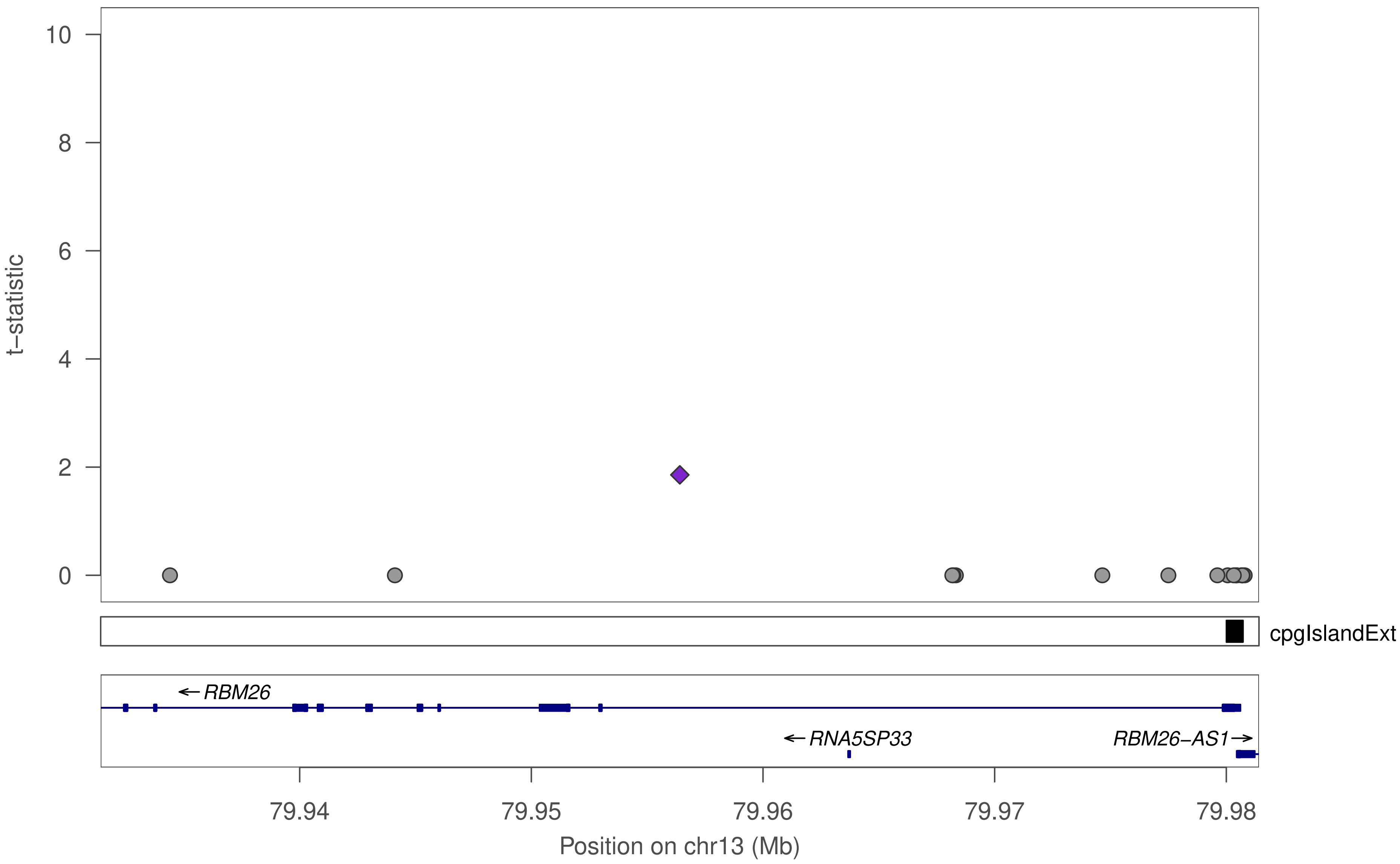

10
8
6
4
2
0
t−statistic
cpgIslandExt
← RBM26
← RNA5SP33
RBM26−AS1 →
79.94
79.95
79.96
79.97
79.98
Position on chr13 (Mb)

### ZNF155

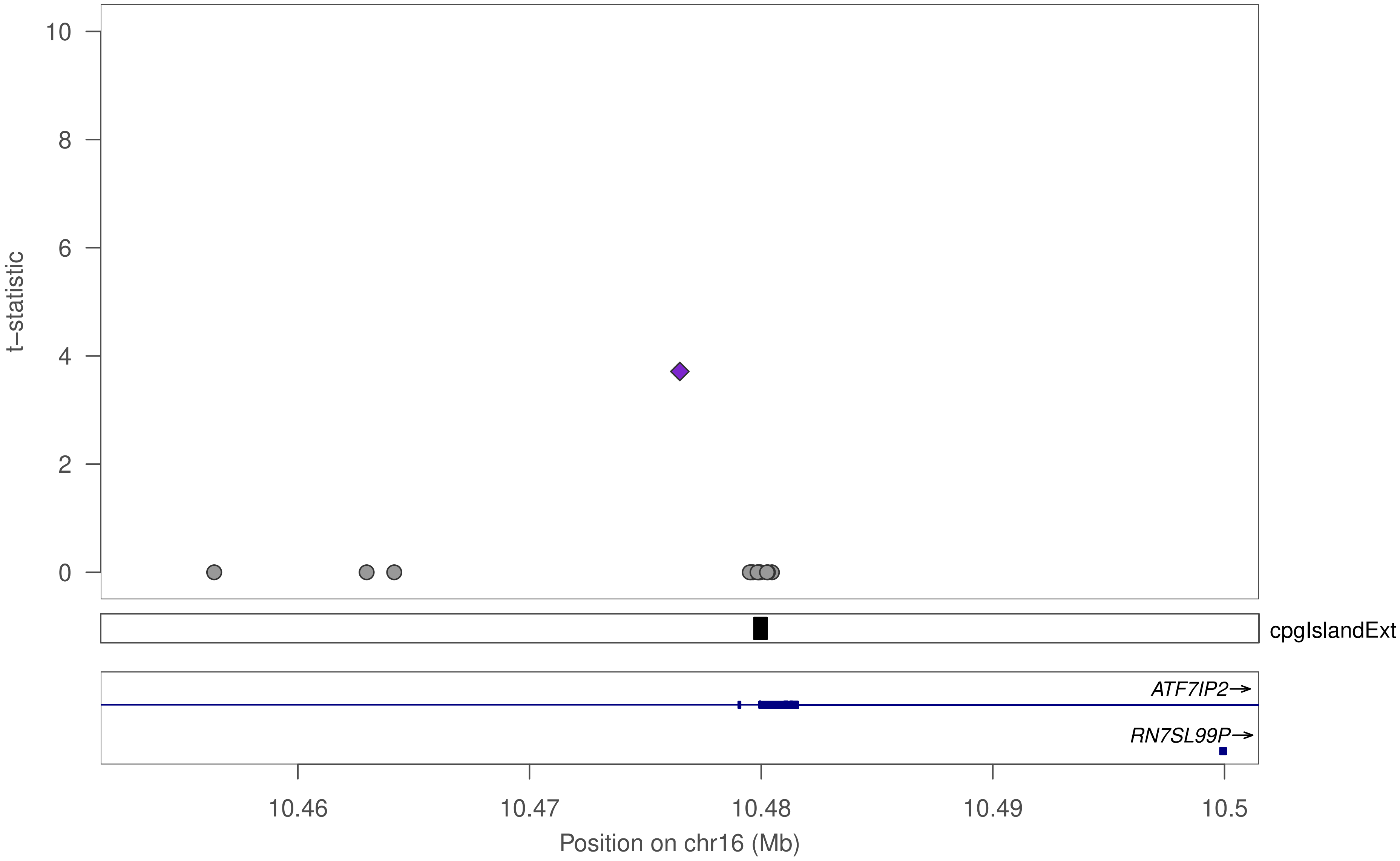

t–statistic
10
8
6
4
2
0
cpgIslandExt
ATF7IP2→
RN7SL99P→
10.46
10.47
10.48
10.49
10.5
Position on chr16 (Mb)

### ZNF155

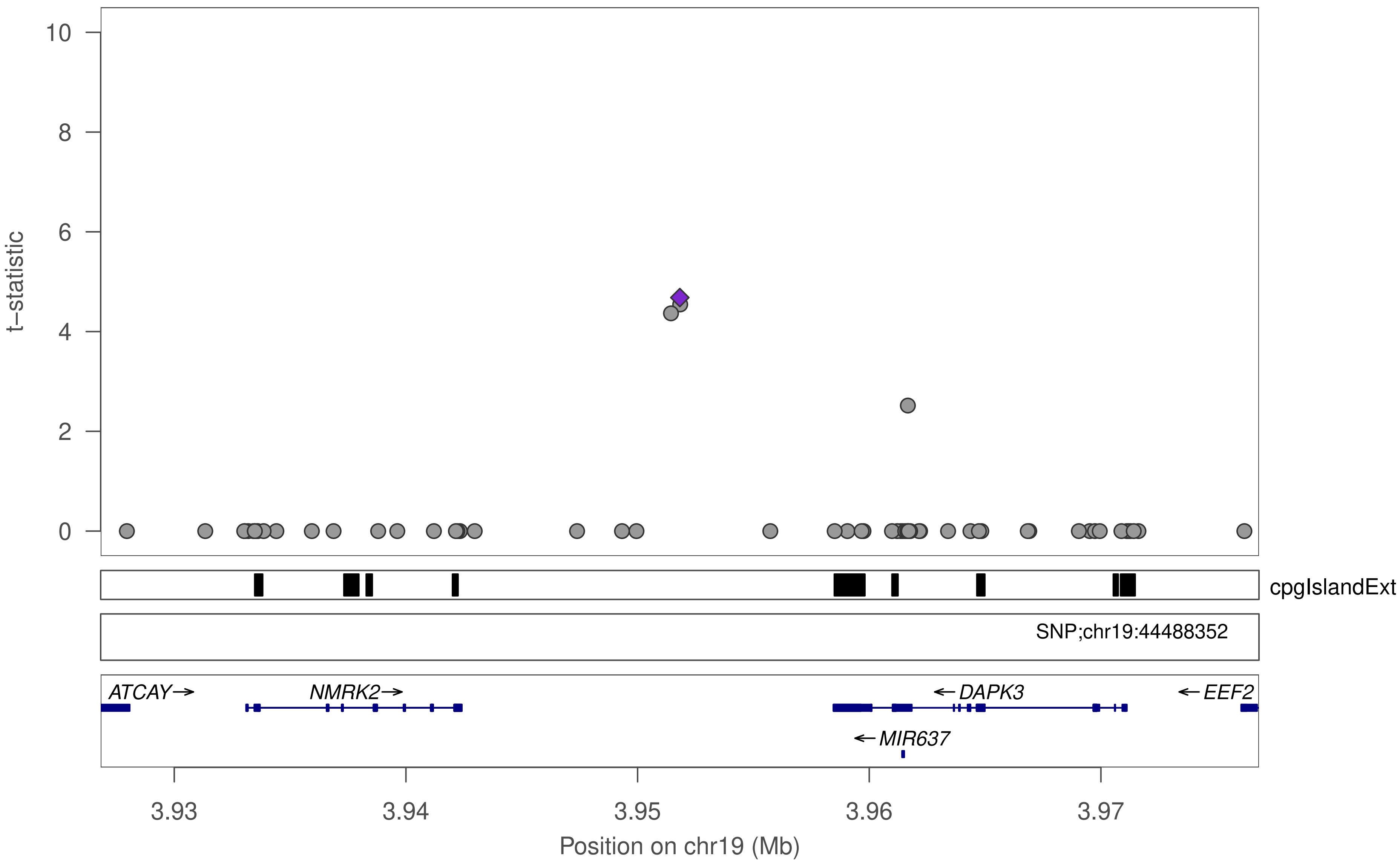

10
8
6
4
2
0
t−statistic
cpgIslandExt
SNP;chr19:44488352
ATCAY→
NMRK2→
←DAPK3
←EEF2
←MIR637
3.93
3.94
3.95
3.96
3.97
Position on chr19 (Mb)

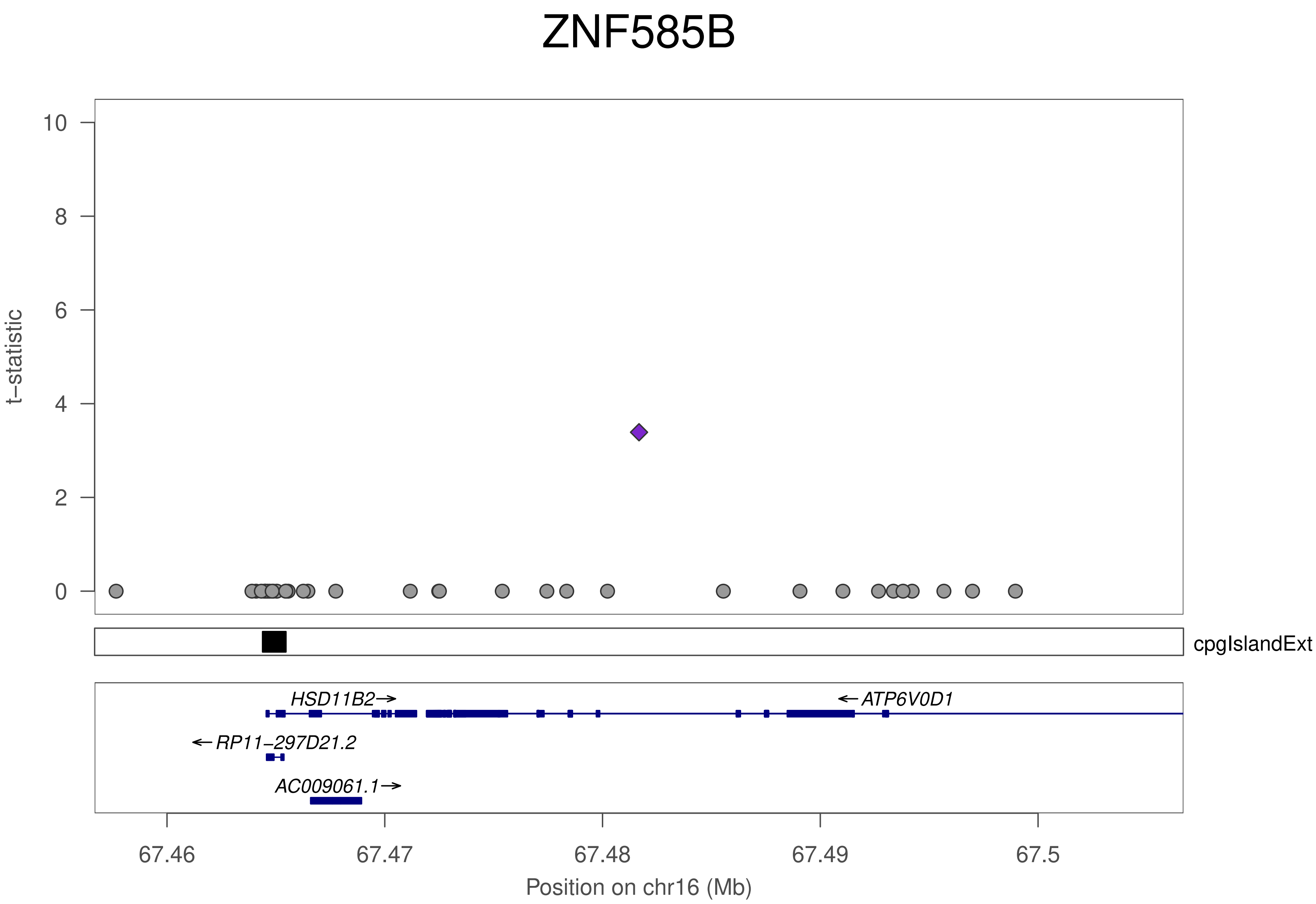
ZNF585B
t–statistic
10
8
6
4
2
0
cpgIslandExt
HSD11B2→
←ATP6V0D1
←RP11–297D21.2
AC009061.1→
67.46
67.47
67.48
67.49
67.5
Position on chr16 (Mb)

### ZNF585B

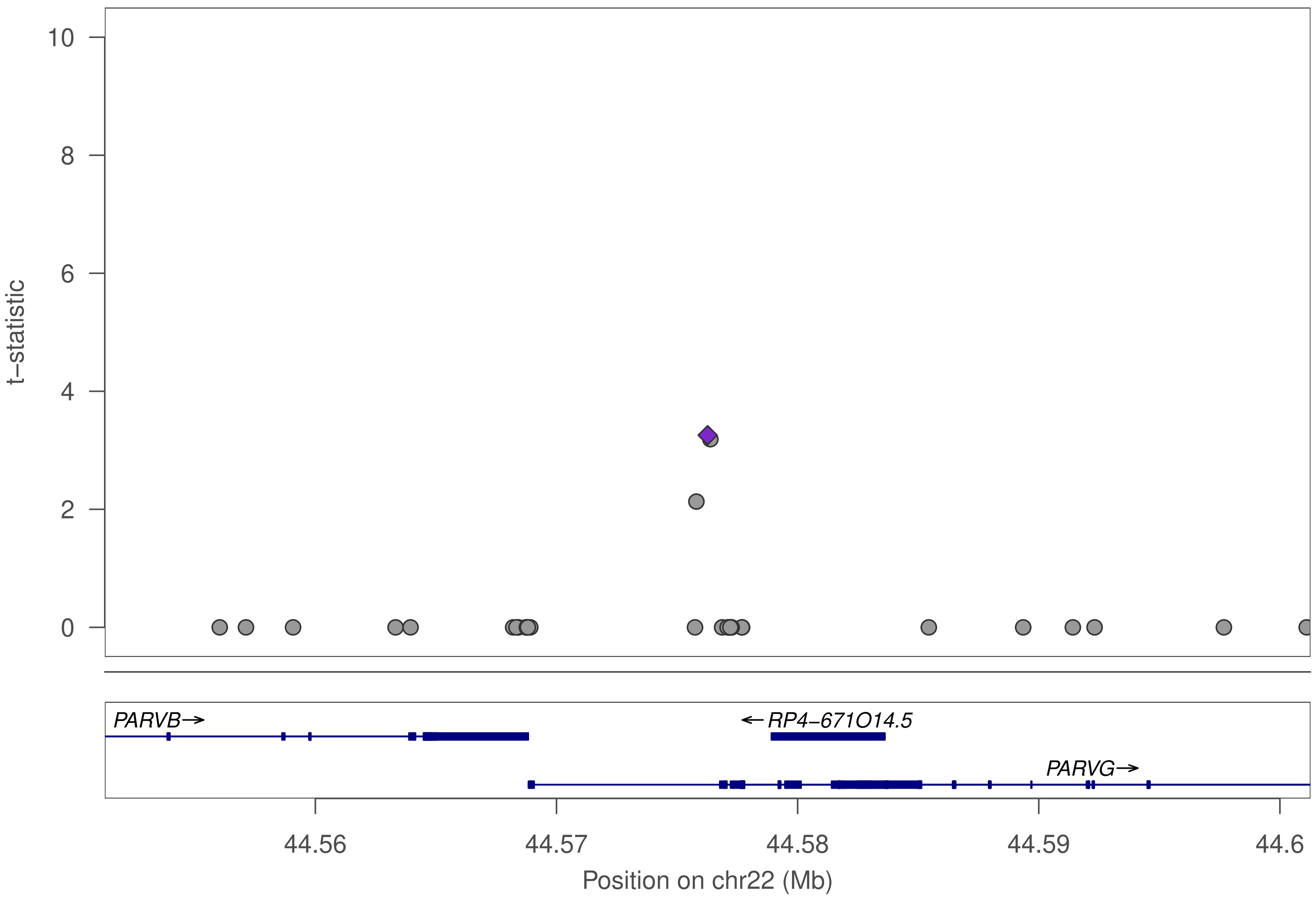

10
8
6
4
2
0
t–statistic
PARVB→
←RP4–671O14.5
PARVG→
44.56
44.57
44.58
44.59
44.6
Position on chr22 (Mb)

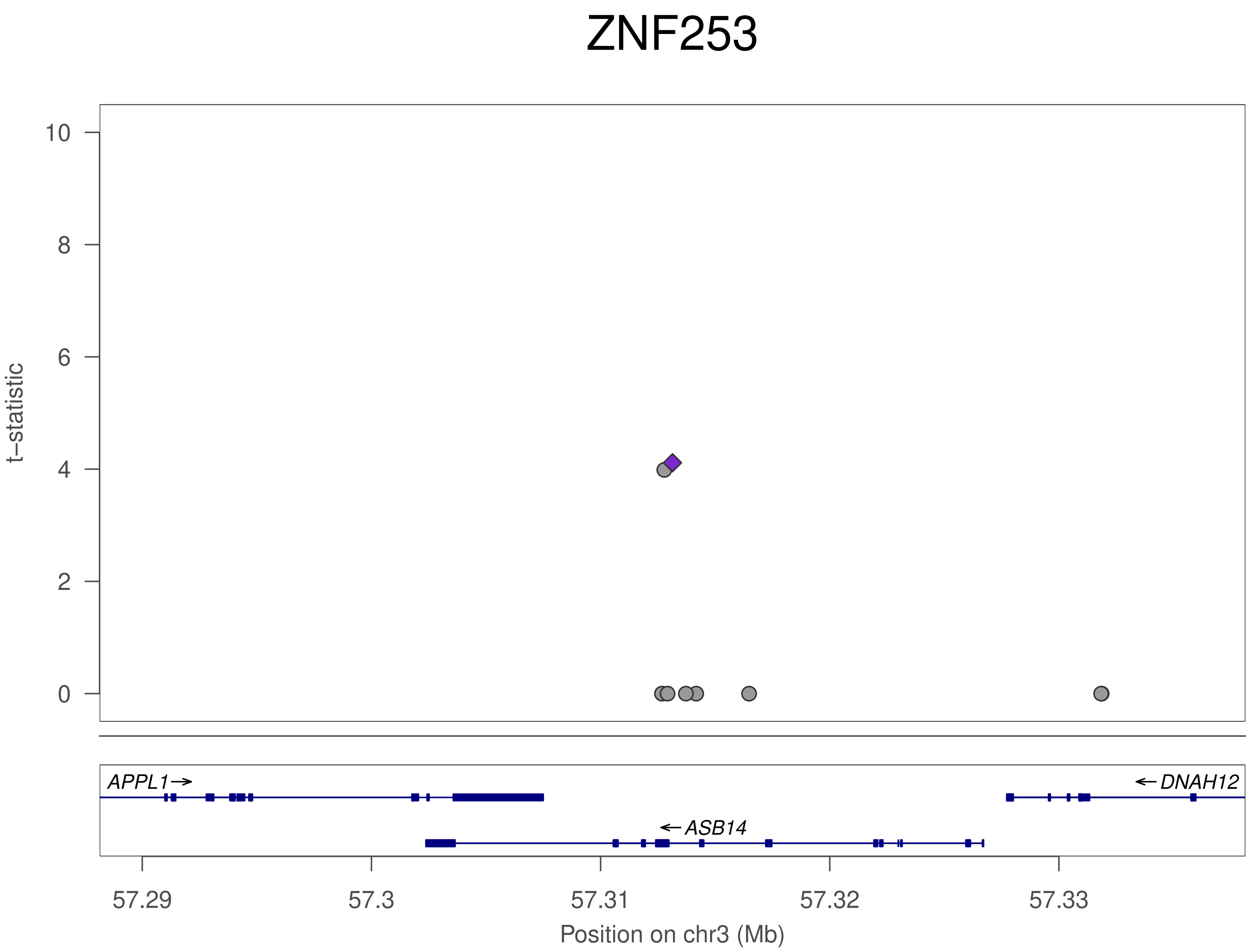

ZNF253
t−statistic
10
8
6
4
2
0
APPL1→
←DNAH12
←ASB14
57.29
57.3
57.31
57.32
57.33
Position on chr3 (Mb)

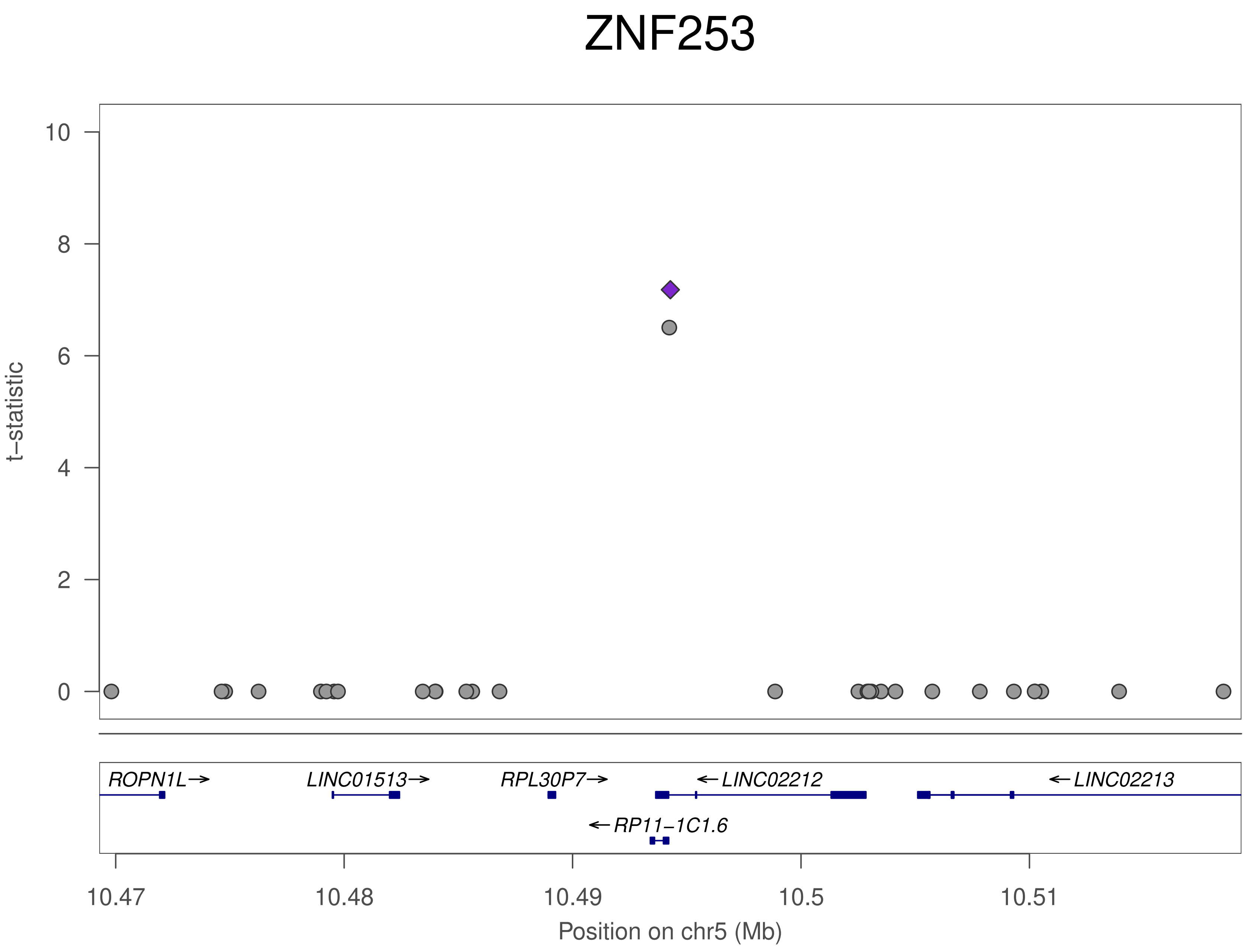
ZNF253
t−statistic
10
8
6
4
2
0
ROPN1L→
LINC01513→
RPL30P7→
←LINC02212
←LINC02213
←RP11−1C1.6
10.47
10.48
10.49
10.5
10.51
Position on chr5 (Mb)

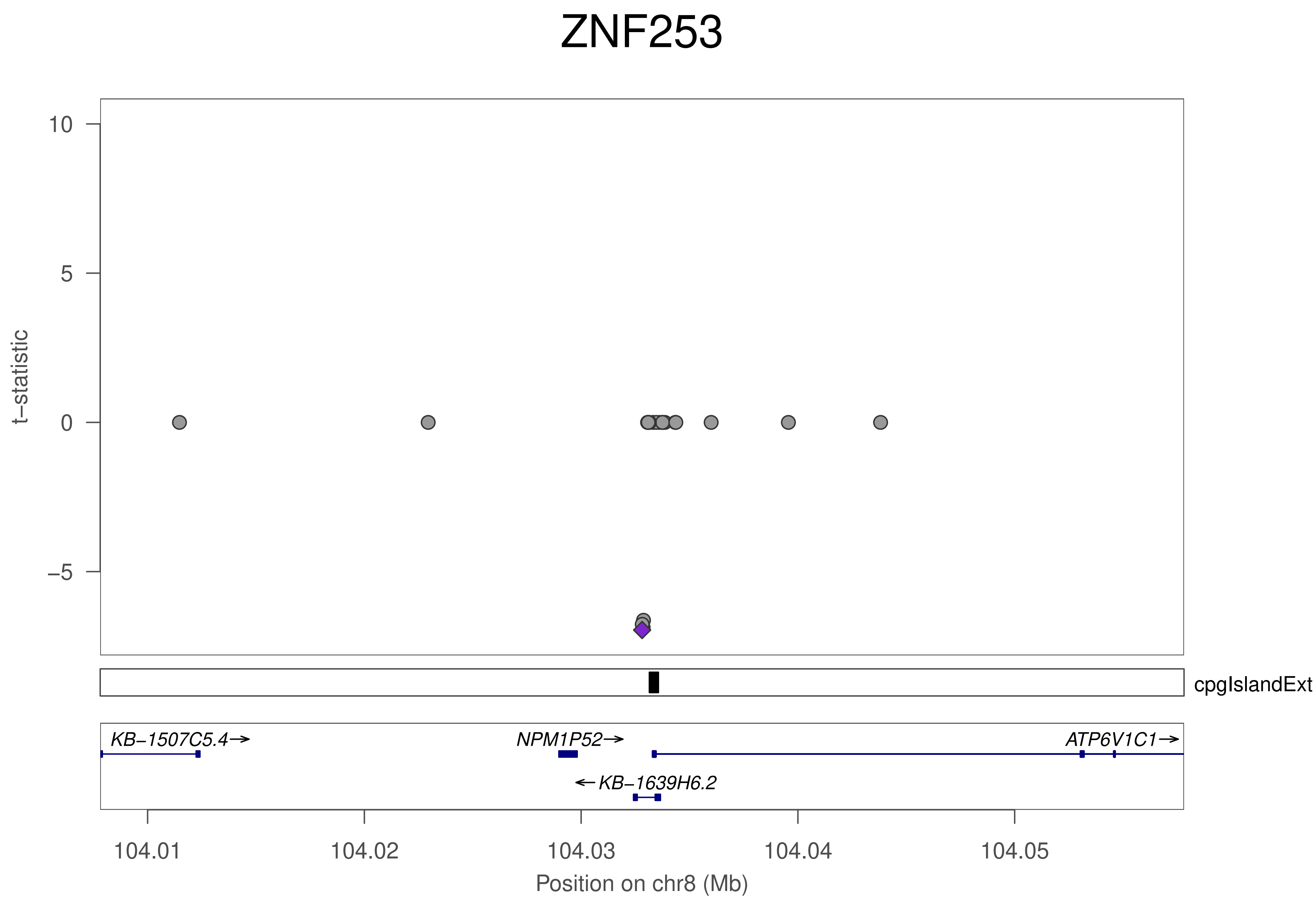
ZNF253
t−statistic
10
5
0
−5
cpgIslandExt
KB−1507C5.4→
NPM1P52→
ATP6V1C1→
←KB−1639H6.2
104.01
104.02
104.03
104.04
104.05
Position on chr8 (Mb)

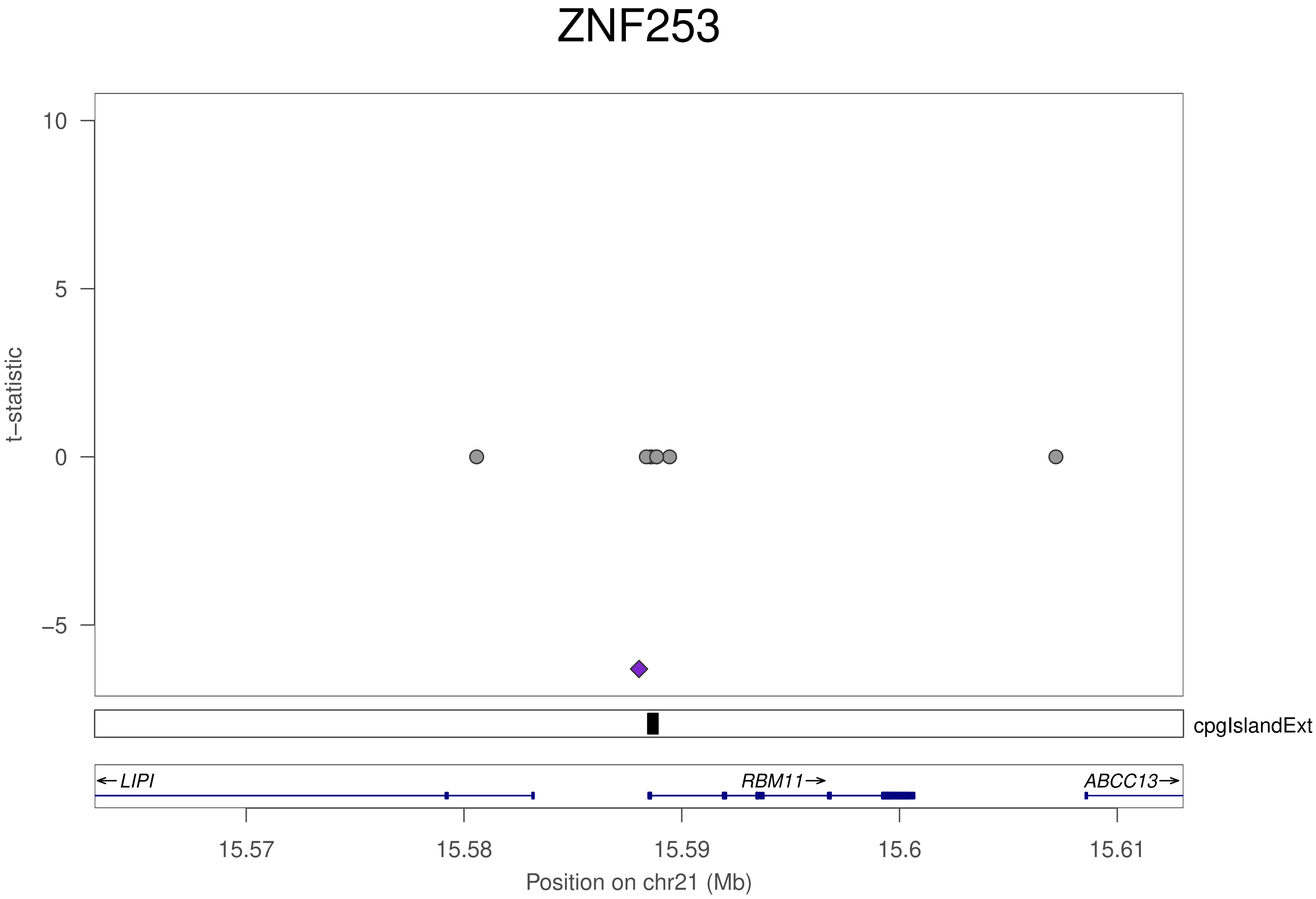

ZNF253
10
5
0
−5
t−statistic
cpgIslandExt
←LIPI
RBM11→
ABCC13→
15.57
15.58
15.59
15.6
15.61
Position on chr21 (Mb)

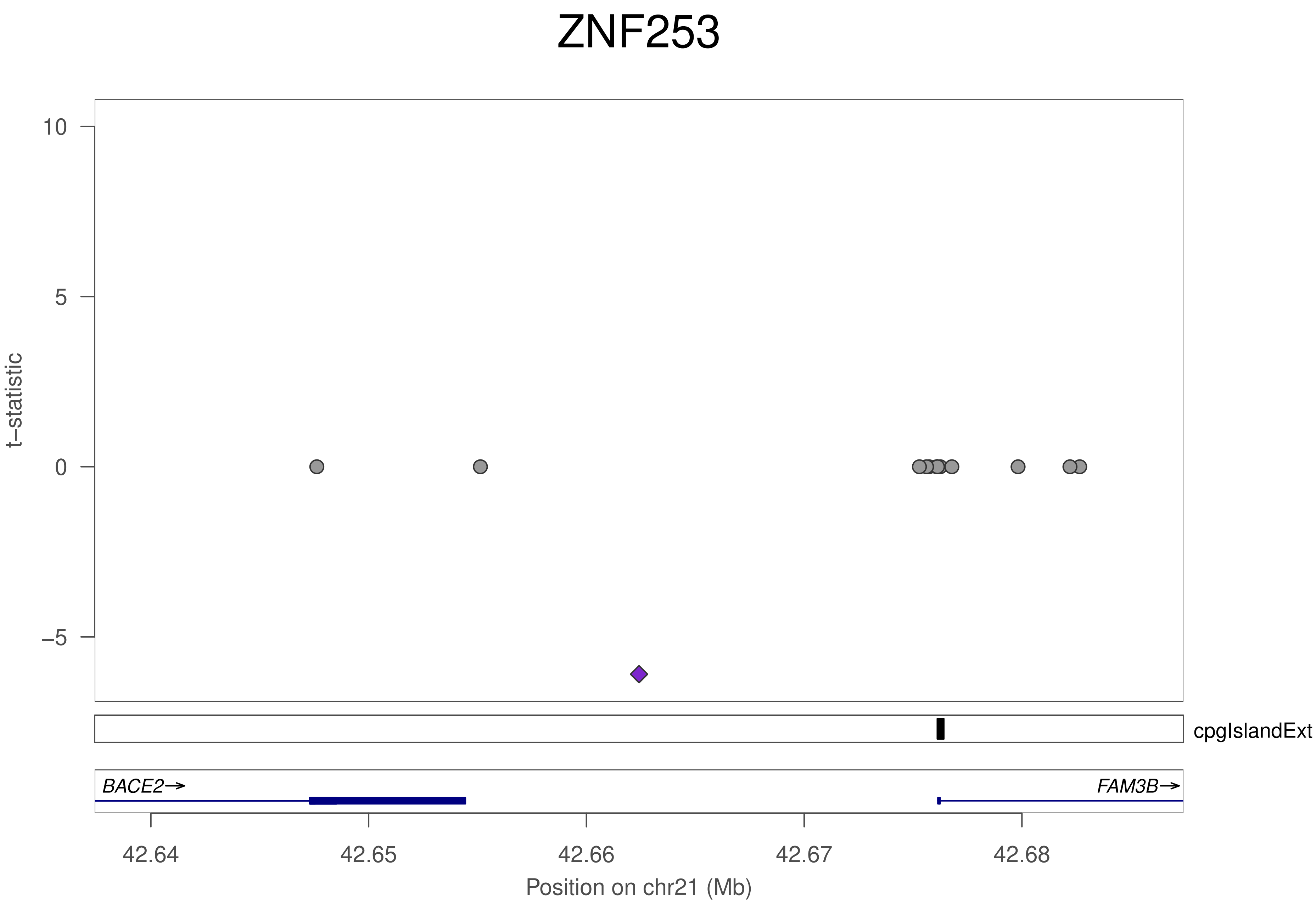

ZNF253
10
5
0
−5
t−statistic
cpgIslandExt
BACE2→
FAM3B→
42.64
42.65
42.66
42.67
42.68
Position on chr21 (Mb)

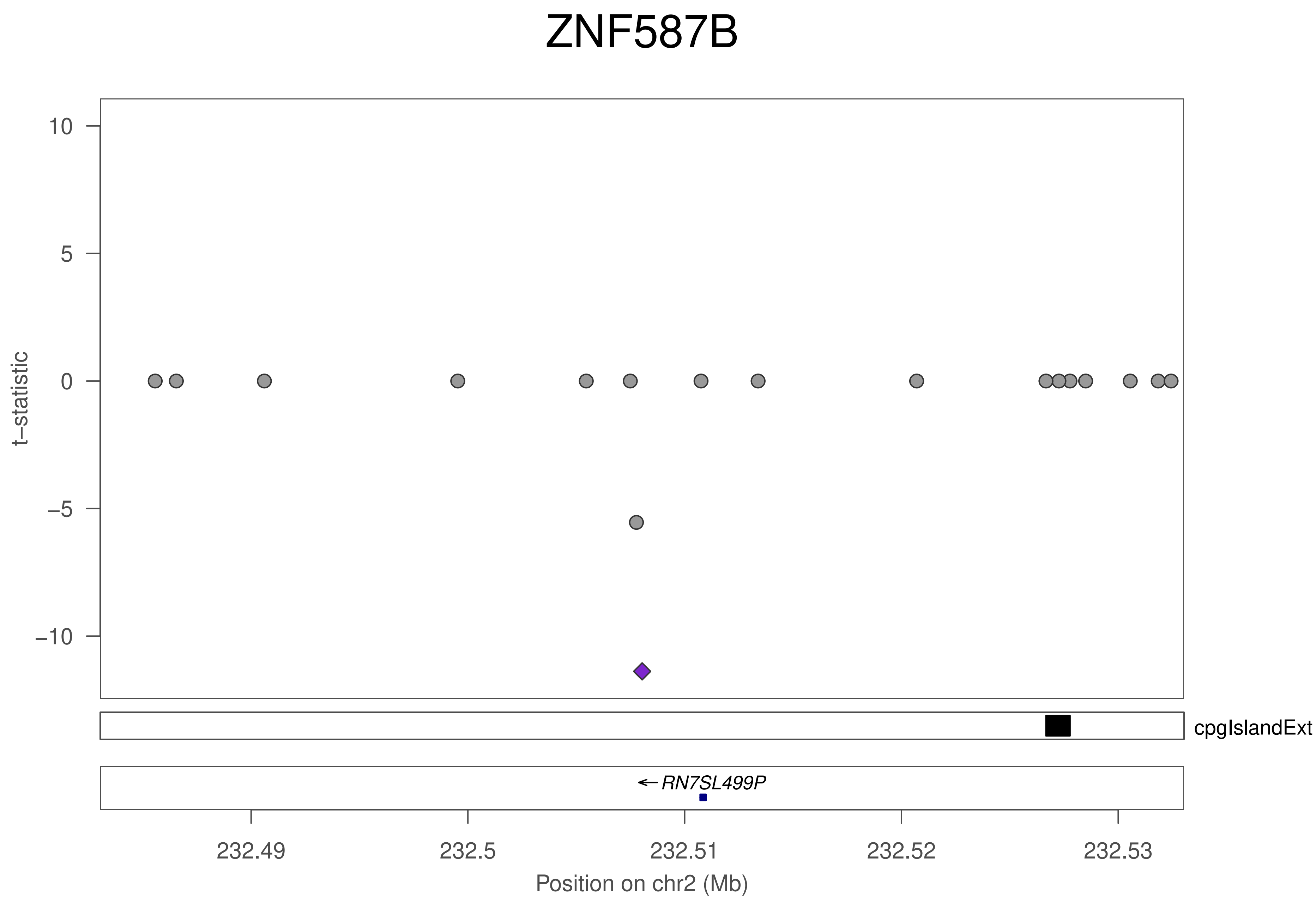
ZNF587B
t–statistic
10
5
0
−5
−10
cpgIslandExt
RN7SL499P
232.49
232.5
232.51
232.52
232.53
Position on chr2 (Mb)

### ZNF587B

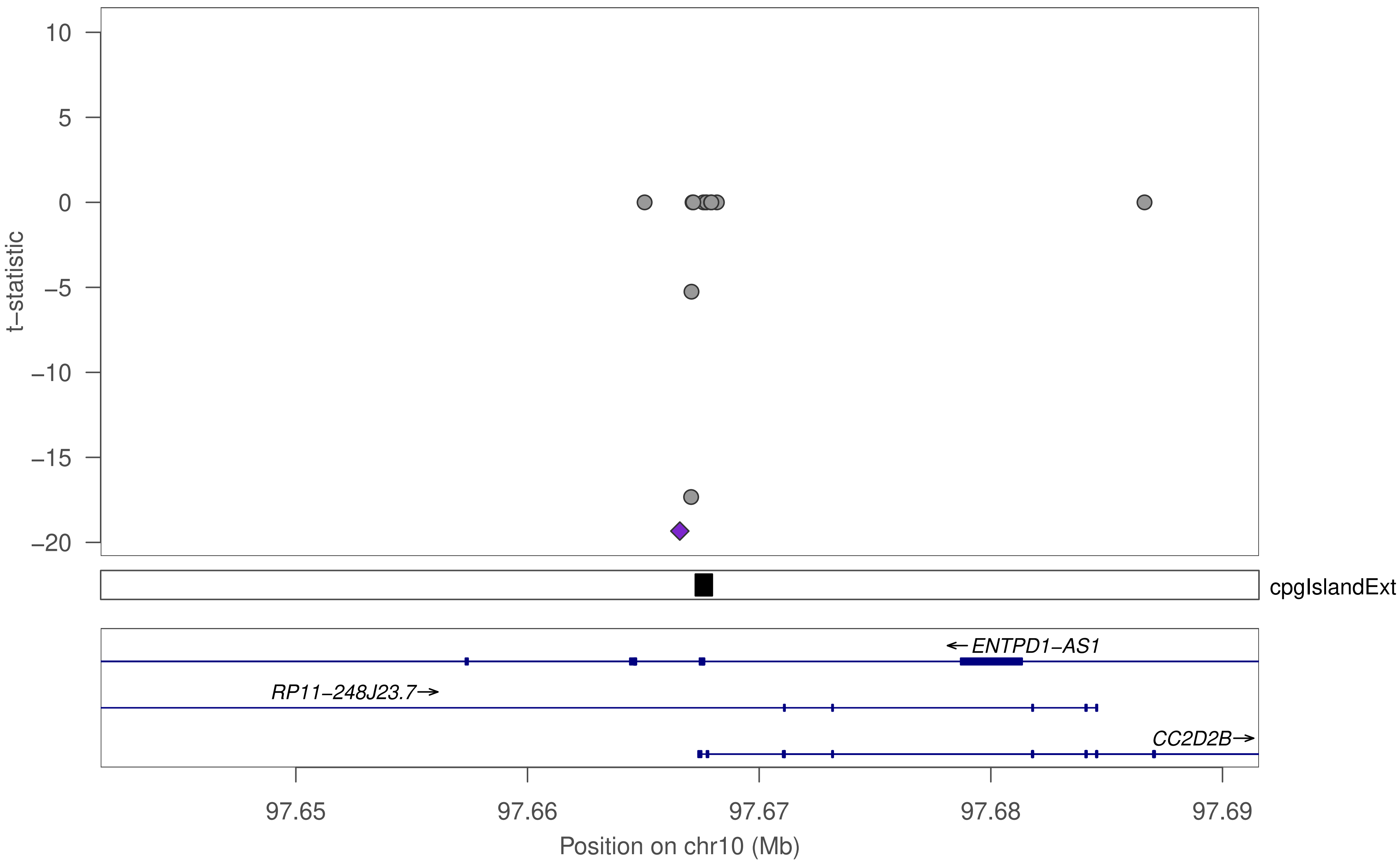

10
5
0
−5
−10
−15
−20
t−statistic
cpgIslandExt
←ENTPD1−AS1
RP11−248J23.7→
CC2D2B→
97.65
97.66
97.67
97.68
97.69
Position on chr10 (Mb)

### ZNF587B

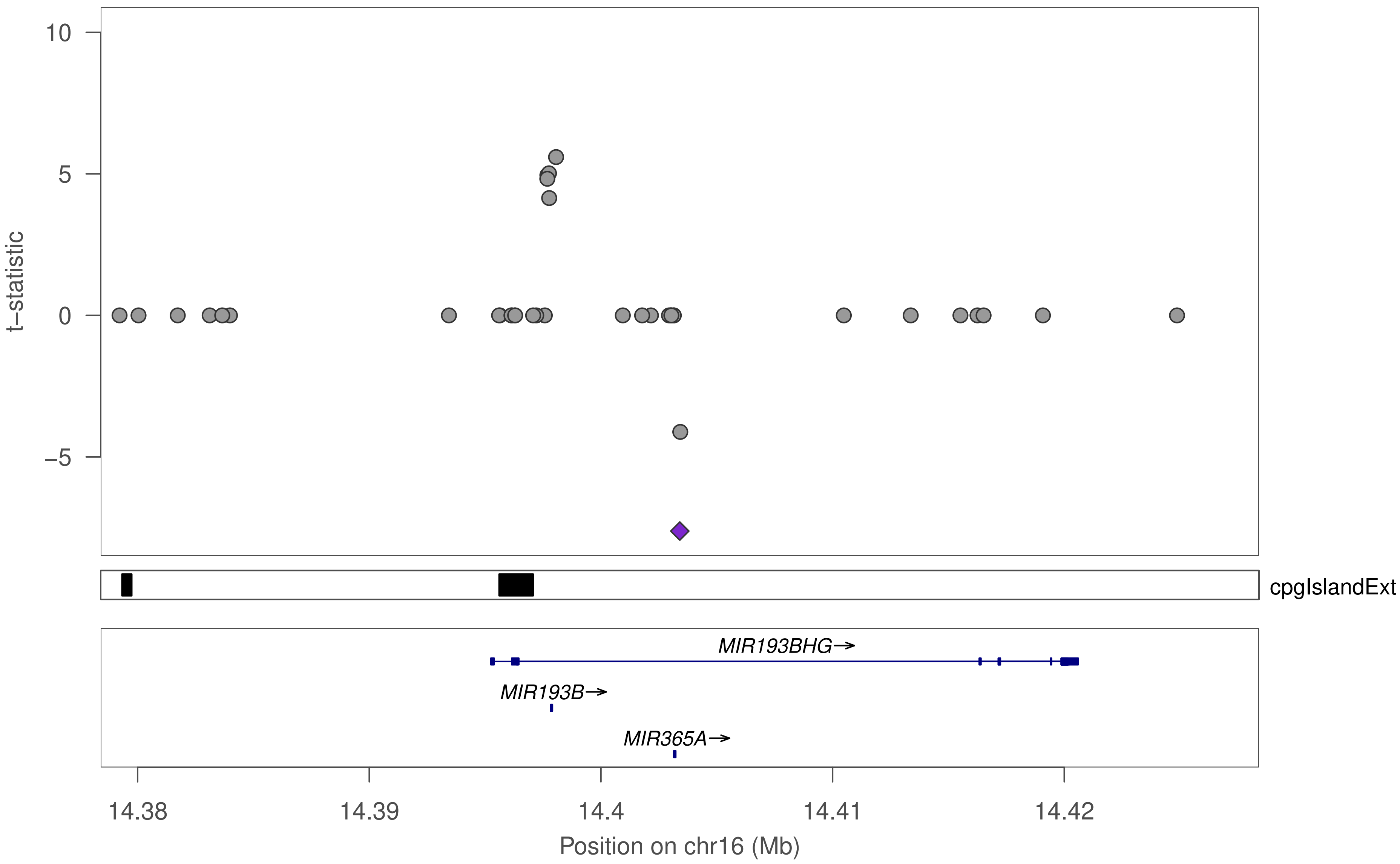

10
5
0
−5
t−statistic
cpgIslandExt
MIR193BHG→
MIR193B→
MIR365A→
14.38
14.39
14.4
14.41
14.42
Position on chr16 (Mb)
